# The genetic architecture of the human cerebral cortex

**DOI:** 10.1101/399402

**Authors:** Katrina L. Grasby, Neda Jahanshad, Jodie N. Painter, Lucía Colodro-Conde, Janita Bralten, Derrek P. Hibar, Penelope A. Lind, Fabrizio Pizzagalli, Christopher R.K. Ching, Mary Agnes B. McMahon, Natalia Shatokhina, Leo C.P. Zsembik, Ingrid Agartz, Saud Alhusaini, Marcio A.A. Almeida, Dag Alnæs, Inge K. Amlien, Micael Andersson, Tyler Ard, Nicola J. Armstrong, Allison Ashley-Koch, Joshua R. Atkins, Manon Bernard, Rachel M. Brouwer, Elizabeth E.L. Buimer, Robin Bülow, Christian Bürger, Dara M. Cannon, Mallar Chakravarty, Qiang Chen, Joshua W. Cheung, Baptiste Couvy-Duchesne, Anders M. Dale, Shareefa Dalvie, Tânia K. de Araujo, Greig I. de Zubicaray, Sonja M.C. de Zwarte, Anouk den Braber, Nhat Trung Doan, Katharina Dohm, Stefan Ehrlich, Hannah-Ruth Engelbrecht, Susanne Erk, Chun Chieh Fan, Iryna O. Fedko, Sonya F. Foley, Judith M. Ford, Masaki Fukunaga, Melanie E. Garrett, Tian Ge, Sudheer Giddaluru, Aaron L. Goldman, Melissa J. Green, Nynke A. Groenewold, Dominik Grotegerd, Tiril P. Gurholt, Boris A. Gutman, Narelle K. Hansell, Mathew A. Harris, Marc B. Harrison, Courtney C. Haswell, Michael Hauser, Stefan Herms, Dirk J. Heslenfeld, New Fei Ho, David Hoehn, Per Hoffmann, Laurena Holleran, Martine Hoogman, Jouke-Jan Hottenga, Masashi Ikeda, Deborah Janowitz, Iris E. Jansen, Tianye Jia, Christiane Jockwitz, Ryota Kanai, Sherif Karama, Dalia Kasperaviciute, Tobias Kaufmann, Sinead Kelly, Masataka Kikuchi, Marieke Klein, Michael Knapp, Annchen R. Knodt, Bernd Krämer, Max Lam, Thomas M. Lancaster, Phil H. Lee, Tristram A. Lett, Lindsay B. Lewis, Iscia Lopes-Cendes, Michelle Luciano, Fabio Macciardi, Andre F. Marquand, Samuel R. Mathias, Tracy R. Melzer, Yuri Milaneschi, Nazanin Mirza-Schreiber, Jose C.V. Moreira, Thomas W. Mühleisen, Bertram Müller-Myhsok, Pablo Najt, Soichiro Nakahara, Kwangsik Nho, Loes M. Olde Loohuis, Dimitri Papadopoulos Orfanos, John F. Pearson, Toni L. Pitcher, Benno Pütz, Yann Quidé, Anjanibhargavi Ragothaman, Faisal M. Rashid, William R. Reay, Ronny Redlich, Céline S. Reinbold, Jonathan Repple, Geneviève Richard, Brandalyn C. Riedel, Shannon L. Risacher, Cristiane S. Rocha, Nina Roth Mota, Lauren Salminen, Arvin Saremi, Andrew J. Saykin, Fenja Schlag, Lianne Schmaal, Peter R. Schofield, Rodrigo Secolin, Chin Yang Shapland, Li Shen, Jean Shin, Elena Shumskaya, Ida E. Sønderby, Emma Sprooten, Lachlan T. Strike, Katherine E. Tansey, Alexander Teumer, Anbupalam Thalamuthu, Sophia I. Thomopoulos, Diana Tordesillas-Gutiérrez, Jessica A. Turner, Anne Uhlmann, Costanza Ludovica Vallerga, Dennis van der Meer, Marjolein M.J. van Donkelaar, Liza van Eijk, Theo G.M. van Erp, Neeltje E.M. van Haren, Daan van Rooij, Marie-José van Tol, Jan H. Veldink, Ellen Verhoef, Esther Walton, Mingyuan Wang, Yunpeng Wang, Joanna M. Wardlaw, Wei Wen, Lars T. Westlye, Christopher D. Whelan, Stephanie H. Witt, Katharina Wittfeld, Christiane Wolf, Thomas Wolfers, Jing Qin Wu, Clarissa L. Yasuda, Dario Zaremba, Zuo Zhang, Alyssa H. Zhu, Marcel P. Zwiers, Eric Artiges, Amelia A. Assareh, Rosa Ayesa-Arriola, Aysenil Belger, Christine L. Brandt, Gregory G. Brown, Sven Cichon, Joanne E. Curran, Gareth E. Davies, Franziska Degenhardt, Michelle F. Dennis, Bruno Dietsche, Srdjan Djurovic, Colin P. Doherty, Ryan Espiritu, Daniel Garijo, Yolanda Gil, Penny A. Gowland, Robert C. Green, Alexander N. Häusler, Walter Heindel, Beng-Choon Ho, Wolfgang U. Hoffmann, Florian Holsboer, Georg Homuth, Norbert Hosten, Clifford R. Jack, MiHyun Jang, Andreas Jansen, Nathan A. Kimbrel, Knut Kolskår, Sanne Koops, Axel Krug, Kelvin O. Lim, Jurjen J. Luykx, Daniel H. Mathalon, Karen A. Mather, Venkata S. Mattay, Sarah Matthews, Jaqueline Mayoral Van Son, Sarah C. McEwen, Ingrid Melle, Derek W. Morris, Bryon A. Mueller, Matthias Nauck, Jan E. Nordvik, Markus M. Nöthen, Daniel S. O’Leary, Nils Opel, Marie - Laure Paillère Martinot, G. Bruce Pike, Adrian Preda, Erin B. Quinlan, Paul E. Rasser, Varun Ratnakar, Simone Reppermund, Vidar M. Steen, Paul A. Tooney, Fábio R. Torres, Dick J. Veltman, James T. Voyvodic, Robert Whelan, Tonya White, Hidenaga Yamamori, Oscar L. Lopez, Hieab H.H. Adams, Joshua C. Bis, Stephanie Debette, Charles Decarli, Myriam Fornage, Vilmundur Gudnason, Edith Hofer, M. Arfan Ikram, Lenore Launer, W. T. Longstreth, Bernard Mazoyer, Thomas H. Mosley, Gennady V. Roshchupkin, Claudia L. Satizabal, Reinhold Schmidt, Sudha Seshadri, Qiong Yang, The Alzheimer’s Disease Neuroimaging Initiative, CHARGE consortium, EPIGEN consortium, IMAGEN consortium, SYS consortium, The Parkinson’s Progression Markers Initiative, Marina K.M. Alvim, David Ames, Tim J. Anderson, Ole A. Andreassen, Alejandro Arias-Vasquez, Mark E. Bastin, Bernhard T. Baune, John Blangero, Dorret I. Boomsma, Henry Brodaty, Han G. Brunner, Randy L. Buckner, Jan K. Buitelaar, Juan R. Bustillo, Wiepke Cahn, Murray J. Cairns, Vince Calhoun, Vaughan J. Carr, Xavier Caseras, Svenja Caspers, Gianpiero L. Cavalleri, Fernando Cendes, Benedicto Crespo-Facorro, John C. Dalrymple-Alford, Udo Dannlowski, Eco J.C. de Geus, Ian J. Deary, Chantal Depondt, Sylvane Desrivières, Gary Donohoe, Thomas Espeseth, Guillén Fernández, Simon E. Fisher, Herta Flor, Andreas J. Forstner, Clyde Francks, Barbara Franke, David C. Glahn, Randy L. Gollub, Hans J. Grabe, Oliver Gruber, Asta K. Håberg, Ahmad R. Hariri, Catharina A. Hartman, Ryota Hashimoto, Andreas Heinz, Frans A. Henskens, Manon H.J. Hillegers, Pieter J. Hoekstra, Avram J. Holmes, L. Elliot Hong, William D. Hopkins, Hilleke E. Hulshoff Pol, Terry L. Jernigan, Erik G. Jönsson, René S. Kahn, Martin A. Kennedy, Tilo T.J. Kircher, Peter Kochunov, John B.J. Kwok, Stephanie Le Hellard, Carmel M. Loughland, Nicholas G. Martin, Jean-Luc Martinot, Colm McDonald, Katie L. McMahon, Andreas Meyer-Lindenberg, Patricia T. Michie, Rajendra A. Morey, Bryan Mowry, Lars Nyberg, Jaap Oosterlaan, Roel A. Ophoff, Christos Pantelis, Tomas Paus, Zdenka Pausova, Brenda W.J.H. Penninx, Tinca J.C. Polderman, Danielle Posthuma, Marcella Rietschel, Joshua L. Roffman, Laura M. Rowland, Perminder S. Sachdev, Philipp G. Sämann, Ulrich Schall, Gunter Schumann, Rodney J. Scott, Kang Sim, Sanjay M. Sisodiya, Jordan W. Smoller, Iris E. Sommer, Beate St Pourcain, Dan J. Stein, Arthur W. Toga, Julian N. Trollor, Nic J.A. Van der Wee, Dennis van ‘t Ent, Henry Völzke, Henrik Walter, Bernd Weber, Daniel R. Weinberger, Margaret J. Wright, Juan Zhou, Jason L. Stein, Paul M. Thompson, Sarah E. Medland, on behalf of the Enhancing NeuroImaging Genetics through Meta-Analysis Consortium - Genetics working group

## Abstract

The cerebral cortex underlies our complex cognitive capabilities, yet we know little about the specific genetic loci influencing human cortical structure. To identify genetic variants, including structural variants, impacting cortical structure, we conducted a genome-wide association meta-analysis of brain MRI data from 51,662 individuals. We analysed the surface area and average thickness of the whole cortex and 34 regions with known functional specialisations. We identified 255 nominally significant loci (*P* ≤ 5 × 10^−8^); 199 survived multiple testing correction (*P* ≤ 8.3 × 10^−10^; 187 surface area; 12 thickness). We found significant enrichment for loci influencing total surface area within regulatory elements active during prenatal cortical development, supporting the radial unit hypothesis. Loci impacting regional surface area cluster near genes in Wnt signalling pathways, known to influence progenitor expansion and areal identity. Variation in cortical structure is genetically correlated with cognitive function, Parkinson’s disease, insomnia, depression and ADHD.

**One Sentence Summary:** Common genetic variation is associated with inter-individual variation in the structure of the human cortex, both globally and within specific regions, and is shared with genetic risk factors for some neuropsychiatric disorders.

---

The human cerebral cortex is the outer grey matter layer of the brain, which is implicated in multiple aspects of higher cognitive function. Its distinct folding pattern is characterised by convex (*gyral*) and concave (*sulcal*) regions. Computational brain mapping approaches use the consistent folding patterns across individual cortices to label brain regions(*1*). During fetal development excitatory neurons, the predominant neuronal cell-type in the cortex, are generated from neural progenitor cells in the developing germinal zone(*2*). The radial unit hypothesis(*3*) posits that the expansion of cortical surface area (SA) is driven by the proliferation of these neural progenitor cells, whereas thickness (TH) is determined by the number of neurogenic divisions. Variation in global and regional measures of cortical SA and TH are associated with neuropsychiatric disorders and psychological traits(*4*) (Table S1). Twin and family-based brain imaging studies show that SA and TH measurements are highly heritable and are largely influenced by independent genetic factors(*5*). Despite extensive studies of genes impacting cortical structure in model organisms(*6*), our current understanding of genetic variation impacting human cortical size and patterning is limited to rare, highly penetrant variants(*7, 8*). These variants often disrupt cortical development, leading to altered post-natal structure. However, little is known about how common genetic variants impact human cortical SA and TH.

To address this, we conducted genome-wide association meta-analyses of cortical SA and TH measures in 51,662 individuals from 60 cohorts from around the world (Tables S2–S4). Cortical measures were extracted from structural brain MRI scans in regions defined by gyral anatomy using the Desikan-Killiany atlas(*9*). We analysed two global measures, total SA and average TH, and SA and TH for 34 regions averaged across both hemispheres, yielding 70 distinct phenotypes (Fig. 1A; Table S1).

**Fig. 1.**
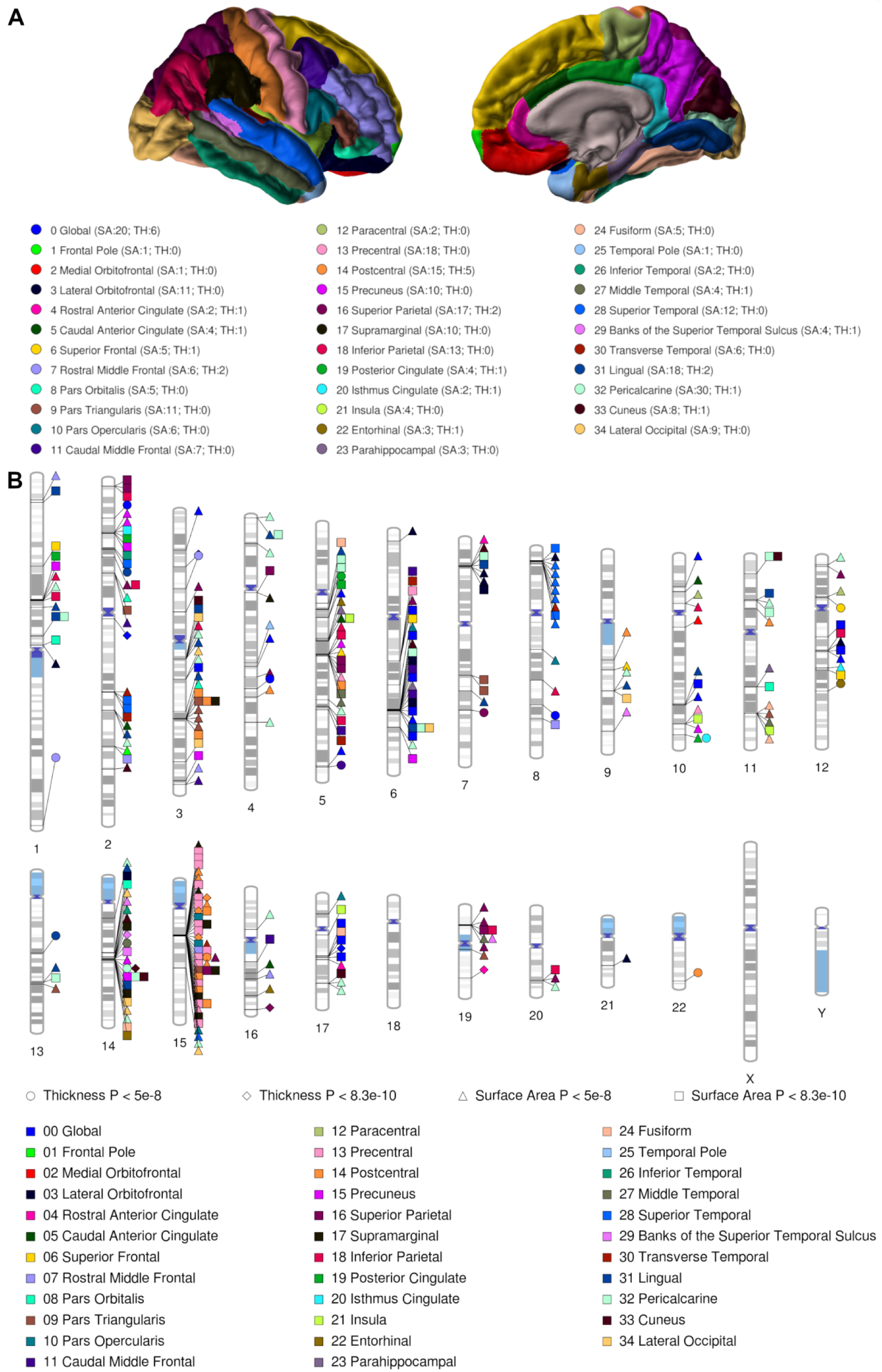
Regions of the human cortex and associated genetic loci. **A**, The 34 cortical regions defined by the Desikan-Killiany atlas; **B**, Ideogram of loci influencing cortical SA and TH.

Within each cohort genome-wide association (GWAS) for each of the 70 phenotypes was conducted using an additive model. To identify genetic influences specific to each region, the primary GWAS of regional measures included the global measure of SA or TH as a covariate. To better localise the global findings, regional GWAS were also run without controlling for global measures. To estimate the multiple testing burden associated with analysing 70 phenotypes we used matrix spectral decomposition(*10*), which yielded 60 independent traits. Therefore, we adopted a significance threshold of *P* ≤ 8.3 × 10^−10^.

The principal meta-analysis comprised results from 49 ENIGMA cohorts of European ancestry (23,909 participants) and the UK Biobank(*11*) (10,083 participants of European ancestry). We sought replication for loci reaching *P* ≤ 5 × 10^−8^ in an additional ENIGMA cohort (777 participants) and with the CHARGE consortium(*12*) (13,950 participants, excluding UK Biobank). In addition, we meta-analysed eight cohorts of non-European ancestry (2,943 participants) to examine the generalization of these effects. High genetic correlations were observed between the meta-analysed ENIGMA European cohorts (excluding UK Biobank) and the UK Biobank cohort using LD-score regression (total SA *r*_G_ = 1.00, *P* = 2.7 ×10^−27^, average TH *r*_G_ = 0.91, *P* = 1.7 × 10^−19^), indicating consistent genetic architecture between the 49 ENIGMA cohorts and the single-site, single-scanner UK Biobank cohort.

Across the 70 cortical phenotypes we identified 306 loci that were nominally genome-wide significant in the principal meta-analysis (*P* ≤ 5 × 10^−8^; Fig. 1B; Table S5). Of these 118 are novel, neither they nor their proxies have been associated with cortical SA or TH or volume in previous studies(*12-14*). Twenty of these were insertions or deletions (INDELs), which were not available in the replication data set. Eleven INDELs could be replicated with a proxy SNP; however, for six INDELs and one single nucleotide polymorphism (SNP) there were no proxies available to assess replication. Of the 299 loci, 255 remained genome-wide significant when the replication data were included in the meta-analysis (241 influencing SA and 14 influencing TH), with 199 passing multiple testing correction (*P* ≤ 8.3 × 10^−10^; 187 influencing SA and 12 influencing TH). Of the 255 loci that replicated in Europeans, eleven SNPs were not available or did not pass quality control in the meta-analysis of non-European cohorts. Of the remaining 244 loci, 241 were supported in the meta-analysis with the non-European cohorts, such that the beta from the principal meta-analysis was contained within the 95% confidence intervals from the non-European meta-analysis. While most effects generalized across ancestry groups, some loci showed evidence of substantial heterogeneity. Table S5 details these results and Figure S1 sumarises these meta-analytic steps and results. Significant gene-based association was observed for 253 genes across the 70 cortical phenotypes (Table S6). Figures summarising the meta-analytic results (Manhattan, QQ, Forest, and Locus Zoom plots) are provided in the additional online materials.

## Genetics of total SA and average TH

Common variants explained 34% (*SE* = 3%) of the variation in total SA and 26% (*SE* = 2%) in average TH, which approaches a third of the heritability estimated from twin and family studies(*5*) (Table S7). We observed a significant negative genetic correlation between total SA and average TH (*r*_G_ = -.32, *SE* = .05, *P* = 6.5 × 10^−12^; Fig. 2A), which persisted after excluding the chromosome 17 inversion region known to influence brain size(*14*) (*r*_G_ = -.31, *SE* = .05, *P* = 3.3 × 10^−12^). The direction of this correlation suggests that opposing genetic influences may constrain the total cortical size. The small magnitude of this correlation is consistent with the radial unit hypothesis(*3*), whereby different developmental mechanisms promote SA and TH expansion.

**Fig. 2.**
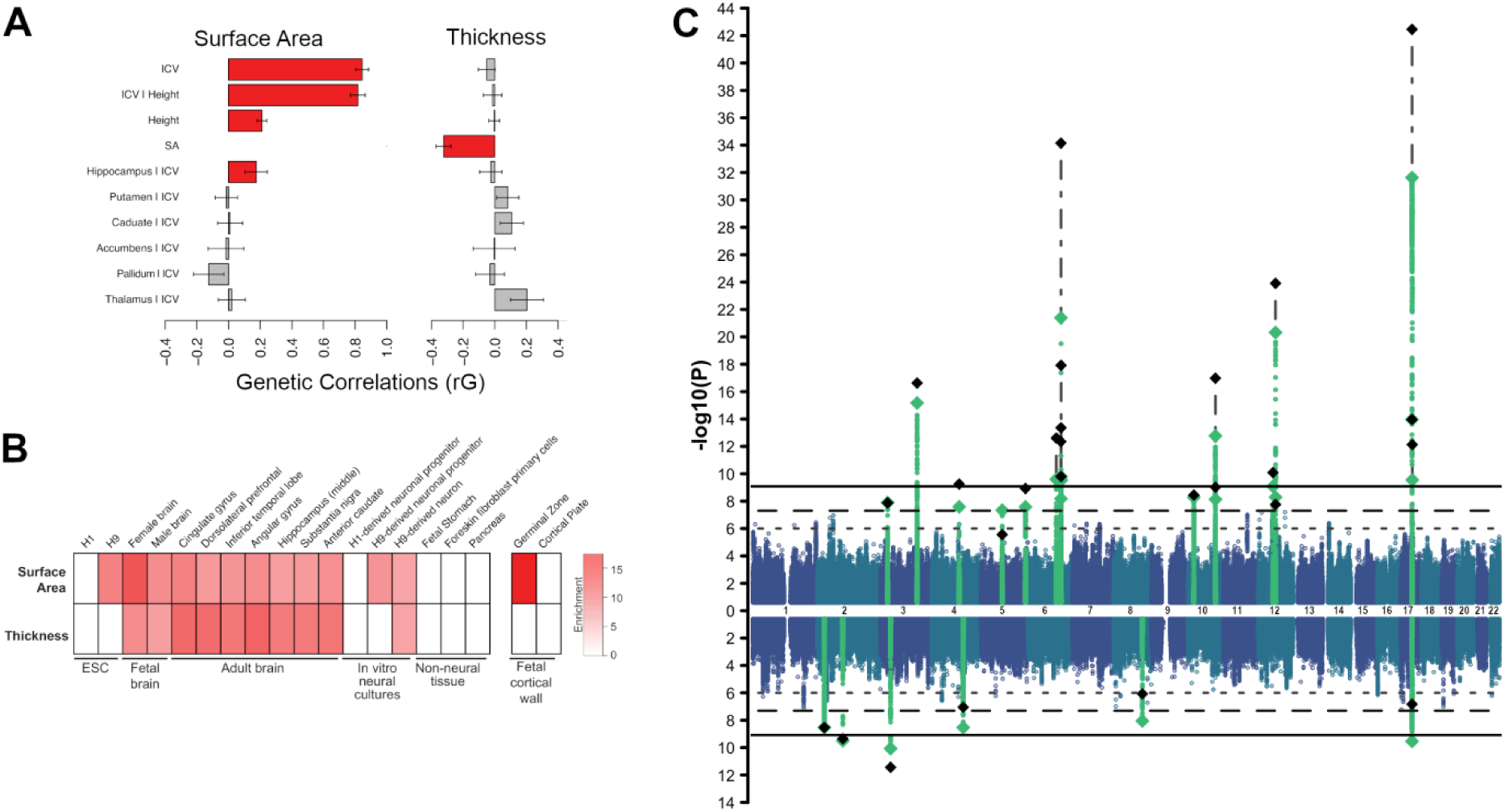
Genetics of Global Measures. **A**, Genetic correlations between global measures and selected traits (red indicates significant correlation, FDR<0.05); **B**, Partitioned heritability enrichment values (significant enrichments are coloured, FDR<0.05); **C**, Manhattan plot of loci associated with global SA (top) and TH (bottom), green diamonds indicate lead SNP in the principal meta-analysis, black diamonds indicate change in *P*-value after replication, dashed horizontal line is genome-wide significance, solid horizontal like is multiple-testing correction threshold.

As expected, total SA showed a positive genetic correlation with intracranial volume (ICV); this correlation remained after controlling for height demonstrating that this relationship is not solely driven by body size (Fig. 2A; Table S8). The global cortical measures did not show significant genetic correlations with the volumes of major subcortical structures (Fig. 2A) except for total SA and the hippocampus, consistent with their shared telencephalic developmental origin. This indicates variation in cortical and subcortical structures have predominantly independent genetic influences, consistent with known differences in cell-type composition between these structures.

To identify if common variation associated with cortical structure perturbs gene regulation during a specific developmental time period or within a given cell-type, we performed partitioned heritability analyses(*15*) using sets of gene regulatory annotations from adult and fetal brain tissues(*16, 17*). The strongest enrichment of the heritability for global SA was seen within areas of active gene regulation (promoters and enhancers) in the mid-fetal human brain (Fig. 2B). We further identified a stronger enrichment in regions of the fetal cortex with more accessible chromatin in the neural progenitor-enriched germinal zone than in the neuron-enriched cortical plate(*16*). There was also enrichment of active regulatory elements within embryonic stem cells differentiated to neural progenitors(*17*). We conducted pathway analyses to determine if there was enrichment of association near genes in known biological pathways. Among the 998 significant gene-sets a number were involved in chromatin modification, a process guiding neurodevelopmental fate decisions(*18*) (Fig. 3C, Table S9). These findings suggest that total SA in adults is influenced by common genetic variants that may alter gene regulatory activity in neural progenitor cells during fetal development, supporting the radial unit hypothesis(*3*). In contrast, the strongest evidence of enrichment for average TH was found in active regulatory elements in the adult brain samples, which may reflect processes occurring after mid-fetal development, such as myelination, branching, or pruning(*19*).

**Fig. 3.**
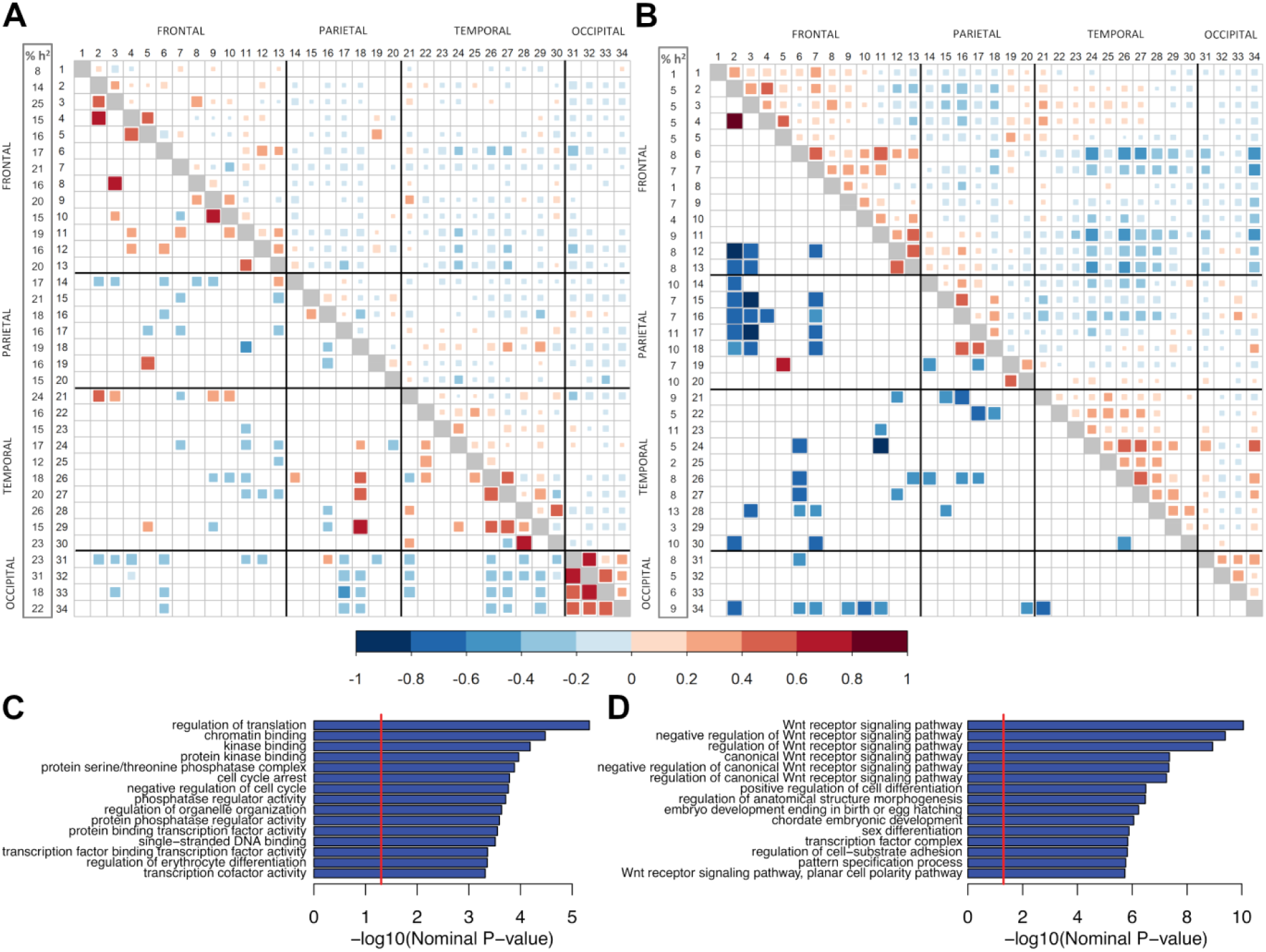
Genetic and Phenotypic Correlations Between Cortical Regions. **A**, Surface Area; **B**, Thickness. The regions are numbered according to the legend of Fig. 1A. The proportion of variance accounted for by common genetic variants is shown in the first column (*h*^2^_SNP_). Phenotypic correlations from the UK Biobank are in the upper triangle. Genetic correlations from the principal meta-analysis are in the lower triangle. Only significant correlations are shown. **C**, Enrichment of gene ontology annotations for total surface area; **D**, Enrichment of gene ontology annotations for regional surface area. The horizontal red lines (**C** and **D**) indicate nominal significance.

## Loci influencing total SA and average TH

Of the replicated loci, 17 loci were nominally associated with total SA; 12 survived correction for multiple testing (Fig. 2C, Table S5). Eight loci influencing total SA have been previously associated with ICV(*14*). Of these, rs79600142 (*P* = 2.3 × 10^−32^; *P*_*rep*_ = 3.5 × 10^−43^), in the highly pleiotropic chromosome 17q21.31 inversion region, has been associated with Parkinson’s disease(*20*), educational attainment(*21*), and neuroticism(*22*). On 10q24.33, rs1628768 (*P* = 1.7 × 10^−13^; *P*_*rep*_ = 1.0 × 10^−17^) is a cortical expression quantitative trait locus (eQTL)(*23*) in adult cortex for *INA*, and schizophrenia candidate genes *AS3MT, NT5C2* and *WBP1L(24) (P*_*ADULT*_ = 9.0 × 10^−3^; Tables S10–S11). This region has been associated with schizophrenia, however, rs1628768 is in low LD with the schizophrenia-associated SNP rs11191419 (*r*^2^ = 0.15). The 6q21 locus influencing total SA is intronic to *FOXO3* (which also showed a significant gene-based association with total SA, Table S6). The minor allele of the lead variant rs2802295 is associated with decreased total SA (*P* = 2.5 × 10^−10^; *P*_*rep*_ = 2.5 × 10^−13^) and has previously been associated with lower general cognitive function(*25*) (rs2490272: *P*_*Cognition*_ = 9.9 × 10-14; *r2*_rs2802295:rs2490272_ = 1).

Of the loci not previously associated with ICV, our novel loci include rs11171739 (*P* = 8.4 × 10^−10^; *P*_*rep*_ = 8.1 × 10^−11^) on 12q13.2. In high LD with SNPs associated with educational attainment(*21*), rs11171739 is an eQTL for *RPS26* in fetal(*26*) and adult cortex (*P*_*FETAL*_ = 6.1 × 10^−27^, P_ADULT_ = 8.8 × 10^−49^; Tables S10–S11). This eQTL association was recently highlighted in a brain expression GWAS including subjects with Alzheimer’s disease and other brain pathologies(*27*). On 2q24.2, rs13021985 (*P* = 3.4 × 10^−9^; *P*_*rep*_ = 8.1 × 10^−12^) is a fetal cortex eQTL for *TBR1* (*P*_*FETAL*_ = 1.4 × 10^−4^; Tables S10–S11), a transcription factor specifically expressed in postmitotic projection neurons and part of the *Pax6*-*Tbr2*-*Tbr1* cascade that modulates numerous neurodevelopmental processes(*28*). On 3p24.1, rs12630663 (*P* = 1.3 × 10^−8^; *P*_*rep*_ = 1.4 × 10^−8^) is of interest due to its proximity (∼200kb) to *EOMES* (also known as *TBR2*), which is expressed specifically in intermediate progenitor cells(*29*) in the developing fetal cortex(*2*). rs12630663 is located in a chromosomal region with chromatin accessibility specific to the human fetal cortex germinal zone of human(*16*). This region shows significant chromatin interaction with the *EOMES* promoter(*29*) and contains numerous regulatory elements that when excised via CRISPR/Cas9 in differentiating neural progenitor cells significantly reduced *EOMES* expression(*16*). A rare homozygous chromosomal translocation in the region separating the regulatory elements from *EOMES* (Fig. S2) silences its expression and causes microcephaly(*30*) demonstrating that rare and common non-coding variation can have similar phenotypic consequences, but to different degrees.

The two replicated loci associated with average TH, both of which are novel, survived correction for multiple testing (Fig. 2C; Table S5). On 3p22.1 rs533577 (*P* = 8.4 × 10^−11^; *P*_*rep*_ = 3.7 × 10^−12^) is a fetal cortex eQTL (*P*_*FETAL*_ = 8.9 × 10^−6^) for *RPSA*, encoding a 40S ribosomal protein with a potential role as a laminin receptor(*31*). Laminins are major constituents of extracellular matrix, and have critical roles in neurogenesis, neuronal differentiation and migration(*32*). On 2q11.2, rs11692435 (*P* = 3.2 × 10^−10^; *P*_*rep*_ = 4.5 × 10^−10^) encodes a missense variant (p.A143V) predicted to impact ACTR1B protein function, and is an *ACTR1B* eQTL in fetal cortex (*P*_*FETAL*_ = 6.5 × 10^−3^) (Tables S10–S11). *ACTR1B* is a subunit of the dynactin complex involved in microtubule remodeling, which is important for neuronal migration(*33*).

## Genetics of regional SA and TH

Within individual cortical regions the amount of phenotypic variance explained by common variants was higher for SA (8–31%) than for TH (1–13%) (Fig. 3A–B; Table S7). With few exceptions, the genetic correlations between SA and TH within the same region were moderate and negative (Tables S12–S13), suggesting that genetic variants contributing to the expansion of SA tend to decrease TH. Most genetic correlations between regional surface areas did not survive multiple testing correction, and those that did implied a general pattern of positive correlations between physically adjacent regions and negative correlations with more distal regions (Fig. 3A). This pattern mirrored the phenotypic correlations between regions and was also observed for TH (Fig. 3A–B). The positive genetic correlations were typically between SA of regions surrounding the major, early forming sulci (e.g., pericalcarine, lingual, cuneus, and lateral occipital regions surrounding the calcarine sulcus), which may potentially reflect genetic effects acting on the development of the sulci. However, the general pattern of correlations may, in part, depend on the regional partitioning by the Desikan-Killiany atlas(*9*) (supplementary text). Hierarchical clustering of the genetic correlations resulted in a general grouping by physical proximity (Fig. S3).

To further investigate biological pathways influencing areal identity, we summarised the individual regional results using multivariate GWAS analyses(*34*) separately for SA and TH that modelled the phenotypic correlations between regions. Pathway analyses of the multivariate SA results showed significant enrichment for 903 gene sets (Fig. 3D; Table S9), many of which are involved in Wnt signalling, with the canonical Wnt signalling pathway showing the strongest enrichment (*P* = 8.8 × 10^−11^). Wnt proteins regulate neural progenitor fate decisions(*35, 36*) and are expressed in spatially specific manners influencing areal identity(*6*). Pathway analyses of the multivariate TH results did not yield any findings that survived multiple testing.

## Loci influencing regional SA and TH

A total of 224 loci were nominally associated with regional SA and 12 with TH; of these 175 SA and 10 TH loci survived multiple testing correction (Table S5). As shown in Fig. 1C, most loci were associated with a single cortical region. Of the loci influencing regional measures, few were also associated with global measures, and those that were showed effects in the same direction, implying that the significant regional loci were not due to collider bias(*37*) (Fig. S4).

The strongest regional association was observed on chromosome 15q14 with the precentral SA (rs1080066, *P* = 1.8 × 10^−137^; *P*_*rep*_ = 4.6 × 10^−189^; variance explained = 1.03%; Fig 4A). Across 11 traits we observed 41 independent significant associations from 18 LD blocks (*r*^2^ threshold ≤ .02; see Fig. 4D, Table S5). As we observed strong association with the SA of both pre- and post-central gyri, we localised the association within the central sulcus in 5,993 unrelated individuals from the UK Biobank. The maximal association between rs1080066 and sulcal depth was observed around the *pli de passage fronto-pariétal moyen (P* = 7.9 × 10^− 21^), a region associated with hand fine-motor function in humans(*38*) and shows distinct depth patterns across different species of primates(*39*) (Fig. 4C). Variants in the rs1080066 LD block are fetal cortex eQTLs for an upsteam lncRNA *RP11-275I4.2* (*P*_*FETAL*_ = 4.0 × 10^−4^) and a downstream gene *EIF2AK4* (*P*_*FETAL*_ = 7.4 × 10^−3^) encoding the GCN2 protein, a negative regulator of synaptic plasticity, memory and neuritogenesis(*40*). The functional data also highlight *THBS1*, with roles in synaptogenesis and the maintenance of synaptic integrity(*41*), with chromatin interaction between the rs1080066 region and the *THBS1* promoter in neural progenitor cells and an eQTL effect in whole blood (*P*_*BIOSgenelevel*_ = 1.5 × 10^−9^). There was evidence of heterogeneity in the effect of rs1080066 across the non-European cohorts (Table S5), which might be due in part to the strength of the effect and the disparate power across ancestry groups.

**Fig. 4.**
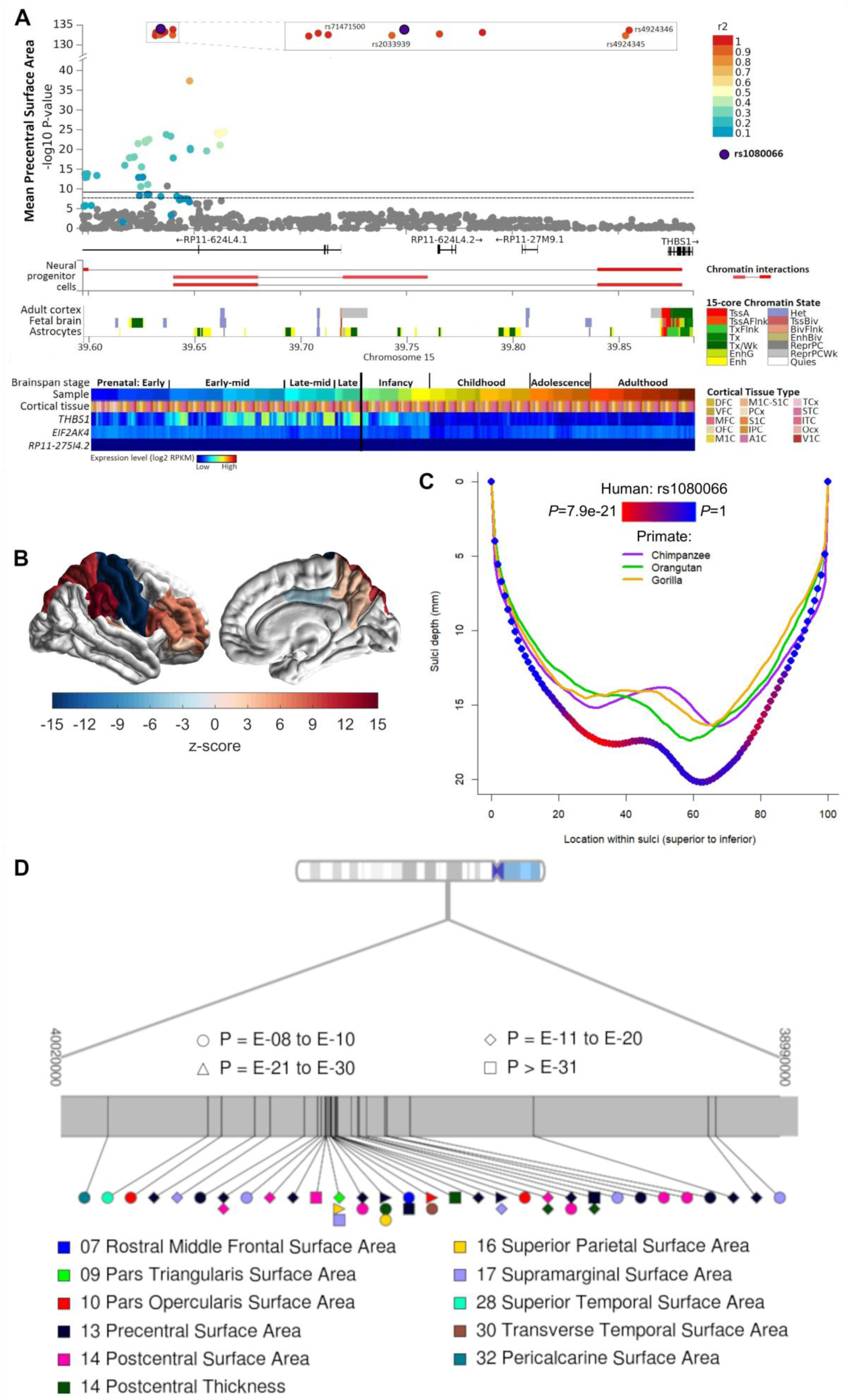
Genetics of Regional Measures. **A**, Regional plot for rs1080066, including additional lead SNPs within the LD block and surrounding genes, chromatin interactions in neural progenitor cells, chromatin state in RoadMap brain tissues*, and BRAINSPAN candidate gene expression in brain tissue** (abbreviations detailed below); **B**, rs1080066 (G allele) association with SA of regions; C, rs1080066 association with central sulcus depth and depth of several primate species; **D** ideogram of 15q14, detailing the significant independent loci and cortical regions. *TssA:Active Transcription Start Site (TSS); TssAFlnk:Flanking Active TSS; TxFlnk:Transcription at gene 5’ and 3’; Tx:Strong transcription; TxWk:Weak transcription; EnhG:Genic enhancers; Enh:Enhancers; Het:Heterochromatin; TssBiv:Bivalent/Poised TSS; BivFlnk:Flanking Bivalent TSS/Enhancer; EnhBiv:Bivalent Enhancer; ReprPC:Repressed; PolyComb; ReprPCWk:Weak Repressed PolyComb; Quies:Quiescent/Low; **DFC:dorsolateral prefrontal cortex; VFC:ventrolateral prefrontal cortex; MFC:anterior cingulate cortex; OFC:orbital frontal cortex; M1C:primary motor cortex; M1C-S1C:primary motor-sensory cortex; PCx:parietal neocortex; S1C:primary somatosensory cortex; IPC:posteroventral parietal cortex; A1C:primary auditory cortex; TCx:temporal neocortex; STC:posterior superior temporal cortex; ITC:inferolateral temporal cortex; Ocx:occipital neocortex; V1C:primary visual cortex.

At another region containing multiple regional hits, on 14q23.1, we observed 20 significant loci (Table S5) from four LD blocks. Our strongest association here was for the precuneus SA (rs73313052: *P* = 1.1 × 10^−24^; *P*_*rep*_ = 2.2 × 10^−35^; variance explained = 0.18%). These loci are located near *DACT1* and *DAAM1*, both involved in synapse formation and critical members of the Wnt signalling cascade(*42, 43*). rs73313052 and high LD proxies are eQTLs for *DAAM1 (P*_*ADULT*_ = 9.0 × 10^−3^) in adult cortex and for *LRRC9* (*P*_*FETAL*_ = 3.9 × 10^−3^) in fetal cortex, *LRRC9* is primarily expressed in brain tissue but is of unknown function (Tables S10– S11).

Consistent with enrichment in the pathway analyses, a number of other loci were located in regions with functional links to genes involved in Wnt signalling, including 1p13.2, where rs2999158 (lingual SA, *P* = 1.9 × 10^−11^, *P*_*rep*_ = 3.0 × 10^−11^; pericalcarine SA, *P* = 1.9 × 10^−11^; *P*_*rep*_ = 9.9 × 10^−16^) is an eQTL for for *ST7L* and *WNT2B* (minimum *P*_*ADULT*_ = 9.0 × 10^−3^) in adult cortex (Tables S10–S11). A number of our novel regional associations occur near genes with known roles in brain development. For example, on chromosome 1p22.2, rs1413536 (inferior parietal SA: *P* = 1.6 × 10^−10^; *P*_*rep*_ = 3.1 × 10^−14^) is an eQTL in adult cortex for *LMO4* (*P*_*ADULT*_ = 9.0 × 10^−3^), with chromatin interactions between the region housing both this SNP and rs59373415 (precuneus SA: *P*_*rep*_ = 5.3 x× 10^−12^) and the *LMO4* promoter in neural progenitor cells (Table S10–S11). *Lmo4* is one of the few genes already known to be involved in areal identity specification in mammalian brain(*44*).

## Genetic correlations with other traits

To examine shared genetic effects between cortical structure and other traits, we performed genetic correlation analyses with GWAS summary statistics from 23 selected traits. We observed significant positive genetic correlations between total SA and general cognitive function(*45*), educational attainment(*21*), and Parkinson’s disease(*46*). For total SA, significant negative genetic correlations were detected with insomnia(*47*), attention deficit hyperactivity disorder (ADHD)(*48*), depressive symptoms(*49*), major depressive disorder(*50*), and neuroticism(*51*)(Fig. 5A; Table S14). Genetic correlations with average TH did not survive multiple testing correction due to the weaker genetic association seen in the TH analyses. We mapped genetic correlation patterns across the cortical regions without correction for the global measures to map the magnitude of these effects across the brain (Fig. 5B). No additional neuropsychiatric or psychological traits were significant at a regional level.

**Fig. 5.**
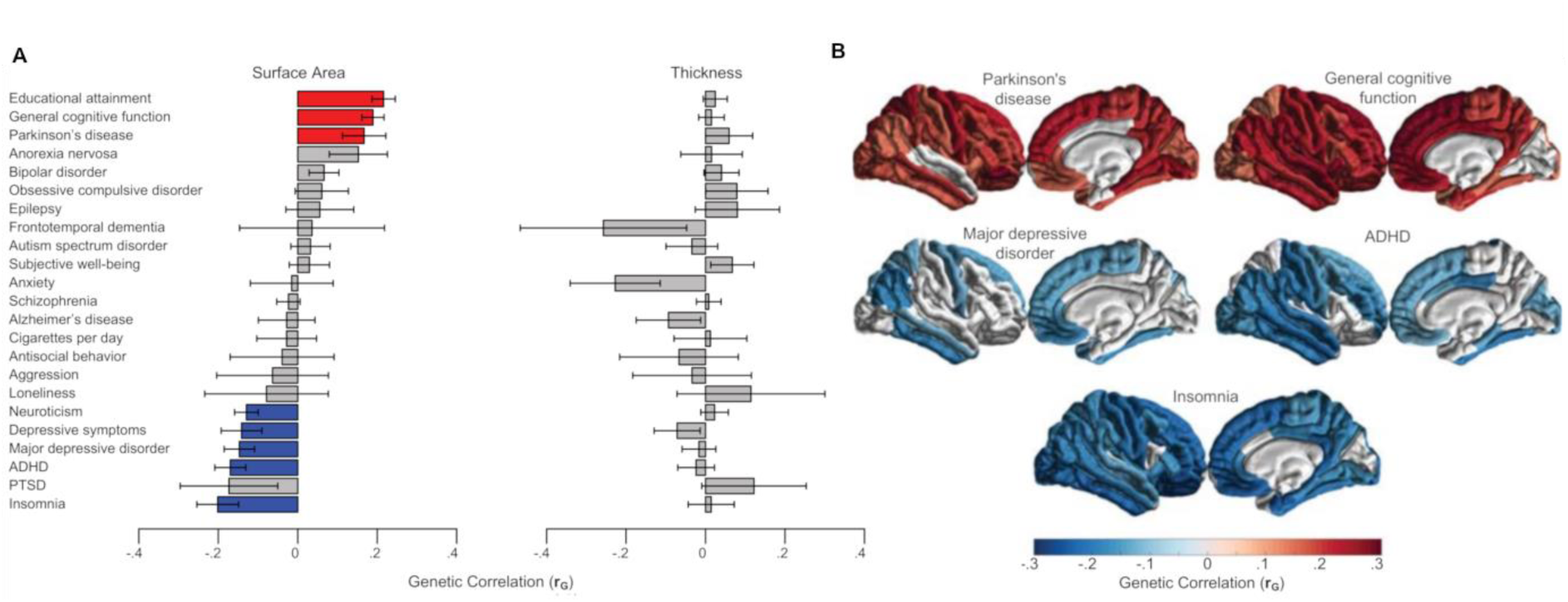
Genetic correlations with neuropsychiatric and psychological traits. **A**, genetic correlations with total SA and average TH positive correlations are shown in red, while negative correlations are shown in blue; **B**, regional variation in the strength of genetic correlations between regional surface area (without correction for total surface area) and traits showing significant genetic correlations with total surface area.

## Discussion

Here we present a large-scale collaborative investigation of the effects of common genetic variation on human cortical structure using data from 51,662 individuals from 60 cohorts from around the world. We identify specific loci influencing cortical surface area (187 loci surviving multiple testing) and thickness (12 loci), implicating genes involved in areal patterning and cortical development. Our results support the radial unit hypothesis of surface area expansion in humans(*3*): genetic variation within regulatory elements in fetal neural progenitor cells(*16*) is associated with variability in adult cortical surface area. We also find that Wnt signalling genes influence areal expansion in humans, as has been reported in model organisms such as mice(*6*). Cortical thickness was associated with loci near genes implicated in cell differentiation, migration, adhesion, and myelination. Consequently, molecular studies in the appropriate tissues, such as neural progenitor cells and their differentiated neurons, will be critical to map the involvement of specific genes. Genetic variation associated with brain structure is functionally relevant, as evidenced by genetic correlations with a range of neuropsychiatric disorders and psychological traits, including general cognitive function, Parkinson’s disease, depression, ADHD and insomnia. This work identifies novel genome-wide significant loci associated with cortical surface area and thickness based on the largest imaging genetics study to date, providing a deeper understanding of the genetic architecture of the human cerebral cortex and its patterning.

## Supporting information

Supplemental_Manhattan_Plots

Supplemental_QQ_Plots

Supplemental_Forest_Plots

Supplemental_LocusZoom_Plots

## Supplementary Materials

### Materials and Methods

#### Ethical approval and data availability

Participants in all cohorts in this study gave written informed consent and sites involved obtained approval from local research ethics committees or Institutional Review Boards. Ethics approval for the meta-analysis was granted by the QIMR Berghofer Medical Research Institute Human Research Ethics Committee (approval: *P2204*).

#### Imaging

Measures of cortical surface area (SA) and thickness (TH) were derived from *in-vivo* whole brain T1-weighted magnetic resonance imaging (MRI) scans using FreeSurfer MRI processing software(*1*) (Table S3). SA and TH were quantified for each subject within 34 distinct gyral-defined regions in each brain hemisphere according to the Desikan-Killiany atlas(*9*) (Fig. 1A). SA was measured at the grey-white matter boundary. TH was measured as the average distance between the white matter and pial surfaces. The total SA and average TH of each hemisphere was computed separately. High test-retest correlations have been reported for all measures with the exception of the frontal and temporal poles(*5*). Image processing and quality control were implemented at the cohort level following detailed, harmonized protocols (see http://enigma.ini.usc.edu/protocols/imaging-protocols/ for protocols); phenotype distributions for all traits in all cohorts were inspected centrally prior to meta-analysis. Any cohort where the phenotypic distribution for a given trait showed deviation from expectations that could not be resolved through reanalysis or outlier inspection were excluded from analyses of that trait.

#### Genome-wide association analyses

At each site, genotypes were imputed using either the 1000 Genomes Project(*52*) or Haplotype Reference Consortium(*53*) references (Table S4). To ensure consistency in the correction for ancestry and stability of the correction given the relatively small sample sizes, each cohort also ran the same multidimensional scaling (MDS) analysis protocol in which the data from the HapMap 3 populations were merged with the site level data and MDS components were calculated across this combined data set. Within each cohort, genome-wide association (GWAS) was conducted using an additive model including covariates to control for the effects of age, sex, ancestry (the first four MDS components), diagnostic status (when the cohort followed a case-control design), and scanner (when multiple scanners were used at the same site).

The primary GWAS of regional measures included the global measure of SA or TH as an additional covariate, to test for genetic influences specific to each region. However, to aid interpretation, the regional GWAS were also run without controlling for global measures. Cohort level GWAS results underwent quality control (excluding variants with an imputation *R*^2^ ≤ .5 and MAF ≤ .005). Across all cohorts, for each phenotype, GWAS summary plots (Manhattan and QQ plots) were visually inspected by the central analysis group, if a given trait showed deviation from expectations that could not be resolved through reanalysis that cohort was excluded from analyses of that trait.

#### Multiple testing correction

We analysed 70 traits (total SA, average TH, and the SA and TH of 34 cortical regions averaged across right and left hemispheres). However, after accounting for the correlation between the traits in the UK Biobank (residuals correcting for sex, age, ancestry and global measures) using matrix spectral decomposition (matSpD(*10*)) the effective number of traits was estimated to be 60. Therefore, we applied the significance threshold of P ≤ 8.3 × 10^−10^ to correct for multiple testing in the GWAS meta-analysis results. Multiple testing corrections applied to each of the follow-up analyses are described below.

#### Meta-analysis

The initial meta-analysis was conducted on all of the ENIGMA European cohorts with genome-wide imputed data, which were then meta-anlysed with the UK Biobank European participants to give the principal results. We took the significant principal results and meta-analysed them with an additional ENIGMA cohort and results from the CHARGE consortium. In an additional replication we took these results and meta-analysed them with the ENIGMA non-European cohorts. Cohort information is provided in Table S2. All meta-analyses were conducted using METAL(*54*). The results of the meta-analysis are summarized in Table S5. For the intial and principal meta-analyses we used standard error weighted meta-analyses. In the replication steps we used sample size weighted meta-analyses, in order to include results from the CHARGE consortium for which only sample size weighted results were available. For each meta-analysis, the results were quality controlled, removing strand ambiguous SNPs and INDELs where the effect allele frequency crossed .5, and (for the initial meta-analysis) variants where the total sample size was < 10,000. Independent loci were identified by clumping significant loci in PLINK(*55*), with thresholds of 1 Mb and *r*^2^ < .2. For the chromosome 17 inversion region this was increased to 10 Mb. For clumping, a random sample of 5,000 unrelated individuals from the UK BioBank were used as an LD reference.

Following Rietveld et al(*56*), we estimated the variance explained *R*^*2*^ by each variant *j* as:

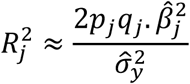

where *p*_*j*_ and *q*_*j*_ are the minor and major allele frequencies, 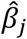 is the estimated effect of the variant within the meta-analysis and 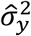 is the estimated variance of the trait (for which we used the pooled variance of the trait across all ENIGMA cohorts and UK Biobank; see Table S1). To obtain beta and standard error estimates from the results from the sample size weighted meta-analyses reported in Table S5 we used the following equations from Rietveld et al(*56*):

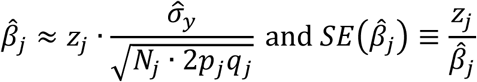

Where *z*_*j*_ is the Z-score and SE 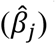 is the estimated standard effect of the variant within the meta-analysis and *N* is the number of contributing alleles.

#### Analyses of UK Biobank data

Analyses of the UK Biobank cohort were conducted on the 2018 (version 3) imputed genotypes, imputed to the Haplotype Reference Consortium and merged UK10K and 1000 Genomes (phase 3) panels. UK Biobank bulk imaging data were made available for 12,962 individuals under application #11559 in July 2017. We processed the raw MRI data using the ENIGMA protocols. Following processing, all images were visually inspected. Analyses of UK Biobank participants within .02 on the first and second MDS components of the European centroid were included in the meta-analyses of the European ancestry cohorts. Analyses of participants beyond this threshold were included in the meta-analysis of non-European ancestry cohorts.

#### Gene-based association analyses

We conducted genome-wide gene-based association analysis using the principal meta-analytic results. We used the 19,427 protein-coding genes from the NCBI 37.3 gene definitions as the basis for the gene-based association analysis using MAGMA(*57*). For each gene we selected all SNPs within exonic, intronic and untranslated regions as well as SNPs within 50 kb upstream and downstream of the gene. After SNP annotation, there were 18,048 genes that were covered by at least one SNP. Gene-based association tests were performed taking LD between SNPs into account. We applied a Bonferroni correction to account for multiple testing, adjusting for the number of genes tested as well as the number of traits tested (60 independent traits), setting the genome-wide threshold for significance at 4.5 × 10^−8^. These results are shown in Table S6.

#### Heritability due to common variants, genetic correlations and partitioned heritability

We used LD score regression(*58, 59*) to estimate the proportion of variance accounted for by common SNPs or SNP heritability (*h*^*2*^_SNP_) of the global measures (total SA and average TH) and the SA and TH of each of the 34 cortical regions. These results are shown in Table S7. LD score regression(*59*) was also used to estimate genetic correlations between regions and with global measures. These results are shown in Table S12−13. We used a threshold of P ≤ 8.3 × 10^−4^ (.05/60) to correct for multiple testing in the genetic and phenotypic correlations shown in Fig. 3. To identify patterns of genetic correlations of SA and TH (both with and without correction for global measures), we used Mclust(*60*) for hierarchical cluster analysis, which uses expectation-maximisation to fit parameterized Gaussian mixture models to the data. The best-fitting model for number and shape of clusters was selected as the one with the largest Bayesian Information Criterion. These results are shown in Supplementary Fig. 3.

Partitioned heritability analysis was used to estimate the percentage of heritability explained by annotated regions of the genome(*61*). Annotations were derived from either Epigenomics Roadmap(*17*) or a study of chromatin accessibility in mid-fetal brains(*16*). For analyses using Epigenomics Roadmap data, chromatin states (15 state model) were downloaded for available tissue types (http://egg2.wustl.edu/roadmap/web_portal/chr_state_learning.html). For each tissue, genomic regions comprising all active regulatory elements (TssA, TssAflnk, Enh, EnhG) within each tissue type were added as an additional annotation to the baseline model provided with the LDSC package (https://github.com/bulik/ldsc). For analyses using chromatin accessibility in mid-fetal brains, the genomic coordinates of peaks more accessible in the germinal zone than the cortical plate (GZ > CP) and peaks more accessible in the cortical plate than the germinal zone (CP > GZ) were added separately to the baseline annotations. Partitioned heritability and the enrichment of heritability explained in these annotations was run using LD score regression(*61*). The significance of enrichment was corrected across all annotations used (including those not displayed) using false discovery rate (FDR) and the enrichment scores were plotted as a heatmap for those that survived significance (Fig. 2b).

Genetic correlations were calculated to determine if shared genetic influences contributed to both cortical structure and neuropsychiatric disorders or psychological traits. Summary statistics were downloaded from the following published genome-wide association studies: general cognitive function(*45*), insomnia(*47*), antisocial behavior(*62*), educational attainment(*21*), subjective well-being(*49*), depressive symptoms(*49*), neuroticism(*51*), attention deficit hyperactivity disorder (ADHD)(*48*), autism(*63*), bipolar disorder(*64*), anorexia nervosa(*65*), major depressive disorder(*50*), obsessive compulsive disorder(*66*), post-traumatic stress disorder (PTSD)(*67*), schizophrenia(*68*), anxiety disorders(*69*), aggression(*70*), Alzheimer’s disease(*71*), loneliness(*72*), cigarettes smoked per day(*73*), epilepsy(*74*), Parkinson’s disease(*46*), and frontotemporal dementia(*75*). LD score regression was used to calculate genetic correlations(*58*). Significance was corrected for multiple comparisons using FDR across all genetic correlations with average TH and total SA, and significant associations were highlighted in Fig. 5A. To explore regional variability in those significant genetic correlations, genetic correlations were conducted between the trait and the cortical regions (without correcting for global measures) are depicted in Fig. 5B.

#### Multivariate GWAS analysis

We used TATES(*34*) to conduct two multivariate analyses: one for the 34 regional SA measures, and one for the 34 regional TH measures. These analyses were run on the meta-analytic results from the second phase of meta-analysis. Briefly, TATES combines the *p*-values from univariate GWAS while correcting for the phenotypic correlations between traits and does not require access to raw genotypic data(*34*). The power of TATES has been shown to be similar or greater than that of multivariate tests using raw data across a range of scenarios for analyses of 20 or more traits(*76*). For these analyses, we used phenotypic correlations calculated from the UK Biobank cohort (residuals correcting for sex, age, ancestry, and global brain measures).

#### Gene-set enrichment analyses

Gene-set enrichment analyses were performed on total SA and average TH as well as the multivariate GWAS results for SA and TH using DEPICT(*77*). Within DEPICT, groups of SNPs were assessed for enrichment in 14,462 gene-sets. These analyses were run using variants with *P* ≤ 1.0 × 10^−5^. Gene-set enrichment analyses were considered significant if they survived FDR correction (q ≤ 0.05)(*77*). These results are shown in Table S9.

#### Functional annotation

Potential functional impact was investigated for lead variants and their proxies (defined here as *r*^2^ > 0.6 to the lead SNP) at each of the 306 loci nominally associated with global and regional SA and TH using a number of publicly available data sources. The majority of the SNP annotations were as provided by FUMA(*23*) which annotates:

- SNP location (e.g., genic/intergenic)
- the potential for functional effects through predicted effects as determined by CADD(*78*) and Regulome(*79*)
- expression quantitative trait (eQTL) effects. We considered eQTLs within cortical structures from GTEx v7, the UK Brain Expression Consortium (http://www.braineac.org/), and the CommonMind Consortium(*80*), and PsychENCODE(*81*) (http://resource.psychencode.org)
- chromatin state
- the presence of enhancers and promoters in SNP regions (RoadMap tissues E053, E073, E081, E082, E125)
- chromatin state (see below) and interactions in numerous brain tissues (GEO GSE87112). We included data for dorsolateral prefrontal cortex and neural progenitor cells, PsychENCODE, and adult and fetal cortex(*82*).

In the main text we provide *P*-values for adult cortical eQTLs as calculated by FUMA across tissues in which significant eQTLs were observed (Table S11). These data were used by FUMA to map coding and non-coding (e.g. lncRNA) genes to each lead SNP based on an eQTL effect with an FDR correction *P* ≤ .05 in cortical tissue, and/or chromatin interactions between the region harbouring the lead SNP and a gene promoter in a second chromosomal region (including interactions with an FDR correction *P* ≤ 1 × 10^−6^)(*23*). HaploReg(*83*) was used to annotate transcription factor binding across multiple tissues, and whether SNPs modified transcription factor binding motifs. The potential for a detrimental effect on protein function due to lead or proxy SNPs located within gene exons was investigated using SIFT and PolyPhen as reported by SNPNexus(*80*). Fetal eQTL data were taken from O’Brien et al(*26*): we have noted only those eQTLs passing our FDR correction *(P* ≤ .05) of the nomimal P-values provided in the original publication. In the main text we provide the nominal *P*-values as reported by O’Brien et al.

In Fig. 4 we annotate the genomic context of rs1080066 and high LD proxies associated with additional traits, chromatin state in relevant tissues, and gene expression in pre- and post-natal brains. Chromatin state represents the degree to which 200 bp genomic regions are accessible for transcription. Around each of our associated loci chromatin state was annotated by FUMA(*23*) utilising the core 15-state model (Table S10). In Fig. 4, genomic regions in three tissues/cells most relevant to our study (RoadMap E073 dorsolateral prefrontal cortex [Adult cortex], E081 female fetal brain [Fetal brain], and E125 NH-A Astrocytes Primary Cells [Astrocytes]) are indicated as one of the 15 possible chromatin states as predicted by Roadmap Epignomics using ChromHMM, based on data for 5 chromatin marks (H3K4me3, H3K4me1, H3K36me3, H3K27me3, H3K9me3) in 127 epigenomes(*17*). Chromatin states are as follows: TssA:Active Transcription Start Site (TSS); TssAFlnk:Flanking Active TSS; TxFlnk:Transcription at gene 5’ and 3’; Tx:Strong transcription; TxWk:Weak transcription; EnhG:Genic enhancers; Enh:Enhancers; ZNF/Rpts:ZNF genes & repeats; Het:Heterochromatin; TssBiv:Bivalent/Poised TSS; BivFlnk:Flanking Bivalent TSS/Enhancer; EnhBiv:Bivalent Enhancer; ReprPC:Repressed; PolyComb; ReprPCWk:Weak Repressed PolyComb; Quies:Quiescent/Low. Pre- and post-natal gene expression data across multiple brain regions was obtained from the BrainSpan Atlas of the Developing Human Brain (http://www.brainspan.org/). These data include gene expression information for cortical tissues indicated on a scale from low (dark blue) to high (dark red) expression on a log_2_ RPKM scale (RPKM = Reads Per Kilobase [of transcript per] Million [mapped reads], which normalises expression levels to account for sequencing depth and gene length). The BRAINSPAN cortical tissues, organised in ontological order, are as follows: DFC:dorsolateral prefrontal cortex; VFC:ventrolateral prefrontal cortex; MFC:anterior (rostral) cingulate (medial prefrontal) cortex; OFC:orbital frontal cortex; M1C:primary motor cortex (area M1, area 4); M1C-S1C:primary motor-sensory cortex (samples); PCx:parietal neocortex; S1C:primary somatosensory cortex (area S1, areas 3,1,2); IPC:posteroventral (inferior) parietal cortex; A1C:primary auditory cortex (core); TCx:temporal neocortex; STC:posterior (caudal) superior temporal cortex (area 22c); ITC:inferolateral temporal cortex (area TEv, area 20); Ocx:occipital neocortex; V1C:primary visual cortex (striate cortex, area V1/17).

For each locus, we evaluated functional annotations for the lead SNP and for additional SNPs considered to be credible causal variants (CCVs) if they were either i) in reasonable LD (*r*^2^ ≥ 0.6 in individuals of European ancestry) with the lead SNP and/or ii) had *P*-values within 2 orders of magnitude of the lead SNP. As lincRNAs show considerable cell/tissue specificity, in the main text we detail SNP location based on neighbouring coding genes, but detail lincRNAs when our lead SNPs show eQTL effects and/or chromatin interactions to these non-coding transcripts. Genes at each associated locus were determined to be potential candidates by considering whether the lead SNP (or a proxy) was an eQTL for a particular gene in adult cortical tissue (e.g. BRAINEAC, CMC or GTEx cortical tissues) and/or when chromatin interactions were observed to occur between the region harbouring the lead/proxy SNPs and a gene promoter in relevant brain tissues (dorsolateral prefrontal cortex and/or neural progenitor cells).

#### Analysis of the central sulcus

To follow-up the precentral surface area association with rs1080066, 10,557 UK Biobank MRI scans were further analyzed using BrainVISA-4.5 Morphologist pipeline for the extraction and parameterization of the central sulcus. Quality controlled FreeSurfer outputs (orig.mgz, ribbon.mgz and talairach.auto) were directly imported into the pipeline to use the same gray and white matter segmentations. Sulci were automatically labeled according to a predefined anatomical nomenclature of 60 sulcal labels per hemisphere(*84, 85*). Extracted meshes for the left and right central sulcus were visually quality checked; subjects with mislabelled central sulcus were discarded from further analysis; 6,045 individuals had good quality extractions for both the left and right hemispheres. The central sulcus depth profile was measured by extending the method introduced in(*38, 86*). The ridges at the fundus of the sulcus and at the convex hull, along with the two extremities, were automatically extracted. Using these landmarks, two coordinate fields (x and y) were extrapolated over the entire mesh surface(*87*). Sulcal depth was defined as the distance between paired points at the sulcal fundus and brain envelope that shared the same y coordinate(*88*). For each individual, the parametrized surface was divided into 100 equally spaced points along the length of the sulcus, and the depth at each point was recorded for comparison. We averaged the corresponding depth measurements across the left and right sulcus and calculated the effect of the rs1080066 G allele on the bilaterally averaged depth at each point. These results are shown in Fig. 4C.

#### Estimating linkage disequilibrium with the 5-HTTLPR variable number tandem repeat

Using PLINK(*55*), we estimated the LD between rs4291964 and the 5-HTTLPR variable number tandem repeat using data from 807 unrelated founders from the QTIM sample who are genotyped for 5-HTTLPR and have rs4291964 imputed (imputation accuracy *r*^2^ = 0.96). These analyses showed the two genotypes to be unlinked, *r*^2^ = 0.03, *D’* = 0.267.

### Supplementary Text

#### Sulcal development

Positive genetic correlations between the SA of neighbouring regions may also be driven by the development of the sulcus, separating the regions. The pre- and post-central regions (also known as the primary motor and sensorimotor cortices, respectively) are consistently labelled across many cortical atlases as the regions directly anterior and posterior to the central sulcus (which appears early in development(*89*)). The SA of all four regions surrounding the calcarine sulcus (the pericalcarine, lingual, cuneus, and lateral occipital region) show positive genetic correlations. The same is also true for the SA of the insula and superior temporal gyri surrounding the lateral sulcus (or Sylvian fissure). These major, early-forming sulci show positive genetic correlations between the regions that directly surround them for SA, but not TH. These observations may imply that part of the genetic influences we observe to be underlying regional SA, may actually be driving the formation of the separating folds, or sulci, during fetal development.

#### The Desikan-Killiany atlas

The Desikan-Killiany atlas(*9*) used here to define the 34 regions of interest is one of many possible atlases. It is one of the coarser atlases, yielding larger, more consistent regions, defined by the common folding patterns visible on standard MRI. More recent efforts partitioning the cortex into 180 regions have used high-resolution multimodal assessments (MMPC)(*90*). It is possible that positive correlations between adjacent structures may reflect suboptimal partitioning of the cortex by the Desikan-Killiany atlas into distinct functional brain regions; for example, we see a positive genetic correlation between the inferior parietal and the superior parietal gyri, whereas in the MMPC atlas, a portion of each of these two regions is included under the *intraparietal* labels. Portions of these genetically correlated regions may in future be re-assigned based on other advanced imaging data, such as multimodal myelin mapping, which may better define cortical cellular architecture.

## Supplementary Tables

**Table S1.** Phenotype descriptions

**Table S2.** Study descriptions

**Table S3.** Description of the imaging data

**Table S4.** Description of the genotype data

**Table S5.** Meta-analytic GWAS results for the 306 loci taken forward for replication

**Table S6.** Results from MAGMA gene based tests

**Table S7.** Variance explained by variants tagged in the GWAS (LDscore h^2^_SNP_)

**Table S8.** Genetic correlations (LDscore *r*_G_) calculated between global cortical measures and selected morphological traits

**Table S9.** Results from DEPICT pathway based tests

**Table S10.** Summary of bioinformatic functional follow-ups

**Table S11.** Functional annotations from FUMA

**Table S12.** Genetic correlations (LDscore *r*_G_) calculated from the GWAS of regional measures corrected for global measures

**Table S13.** Genetic correlations (LDscore *r*_G_) calculated from the GWAS of regional measures not corrected for global measures

**Table S14.** Genetic correlations (LDscore r_G_) calculated between the imaging phenotypes and selected neuropsychiatric disorders and psychological traits

## Supplementary Figures

**Figure S1.**
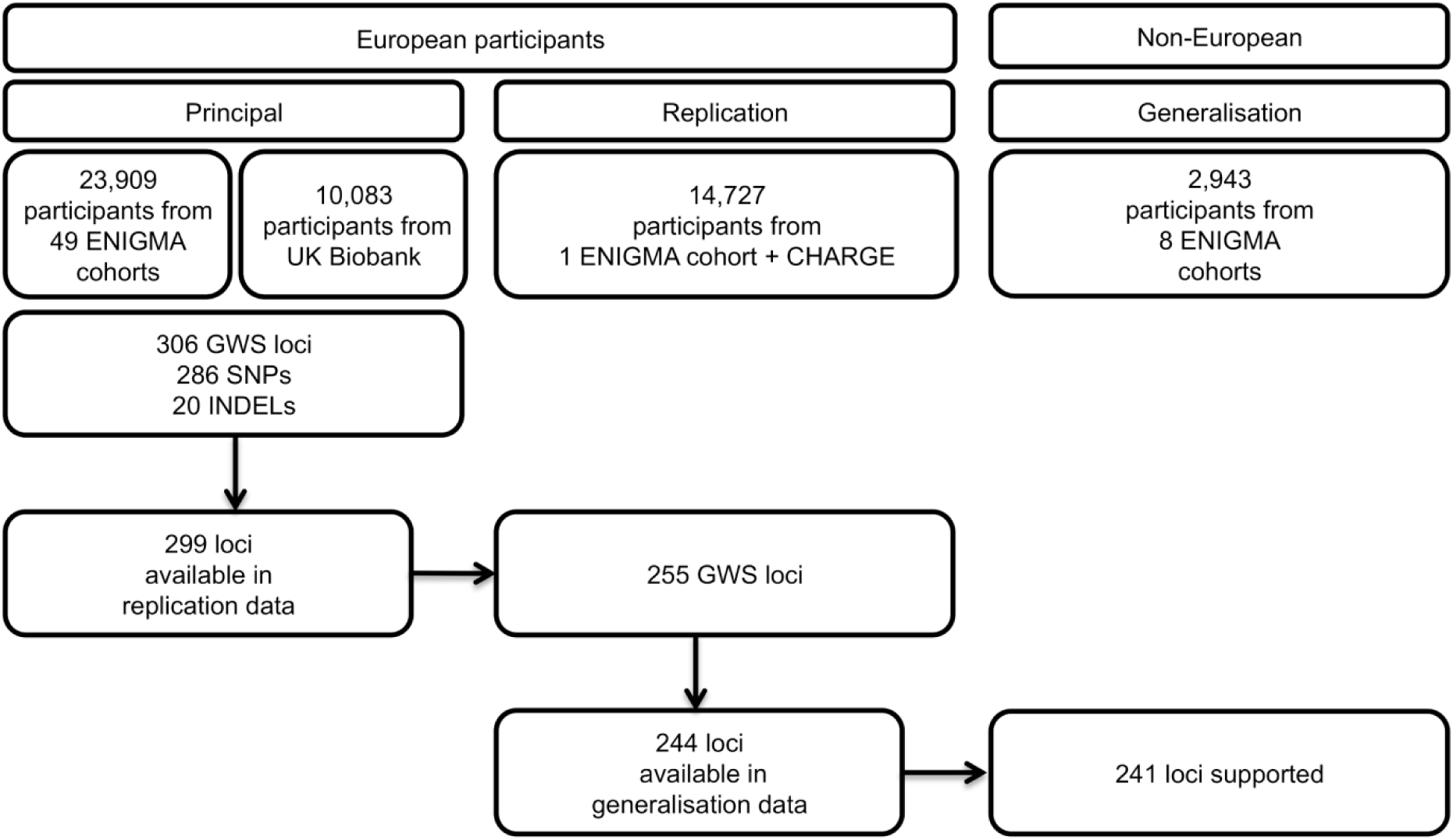
Flow chart summarising the phases of meta-analysis.

**Figure S2.**
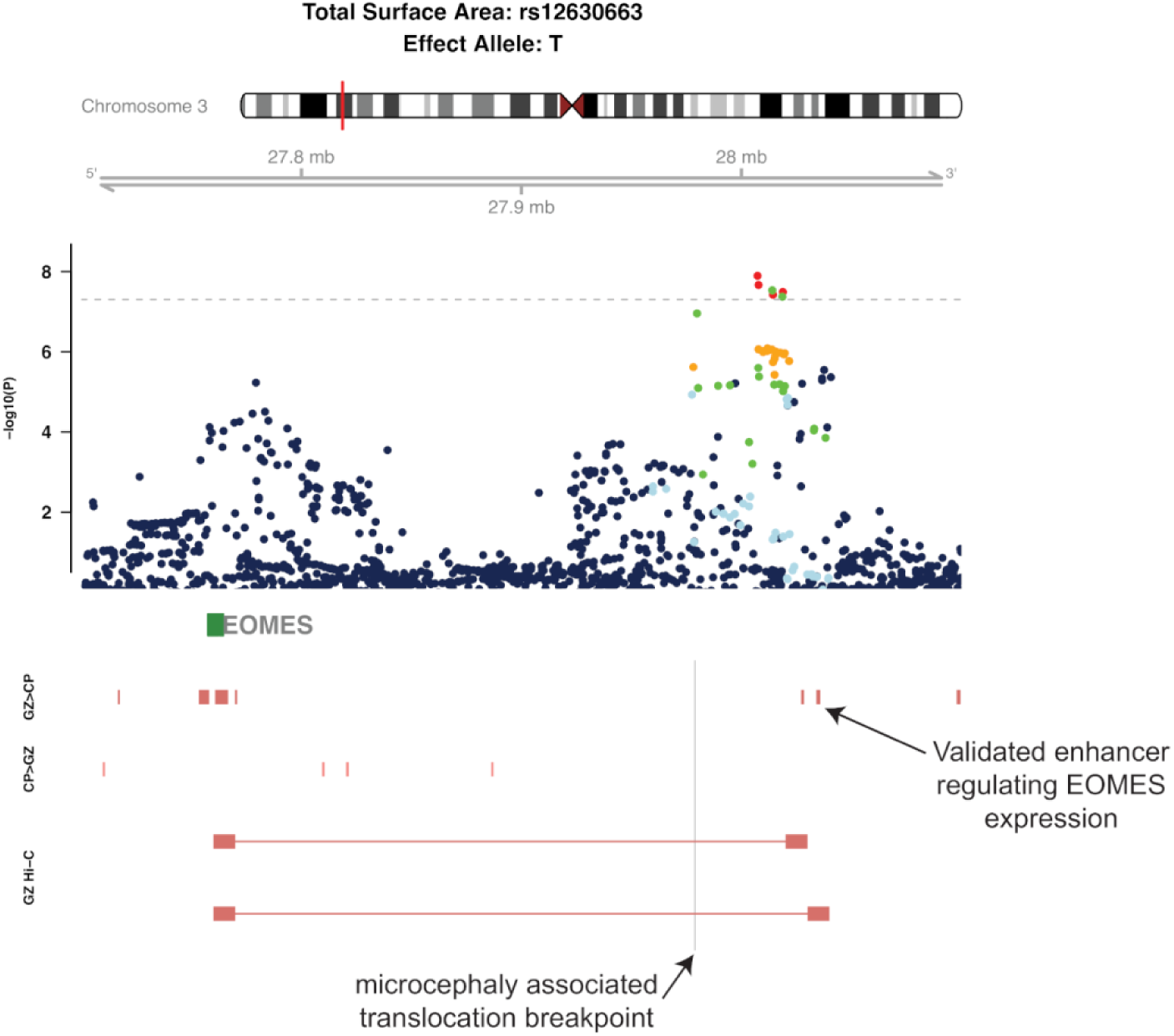
Regional association plot for the 3p24.1 locus (rs12630663).

**Figure S3.**
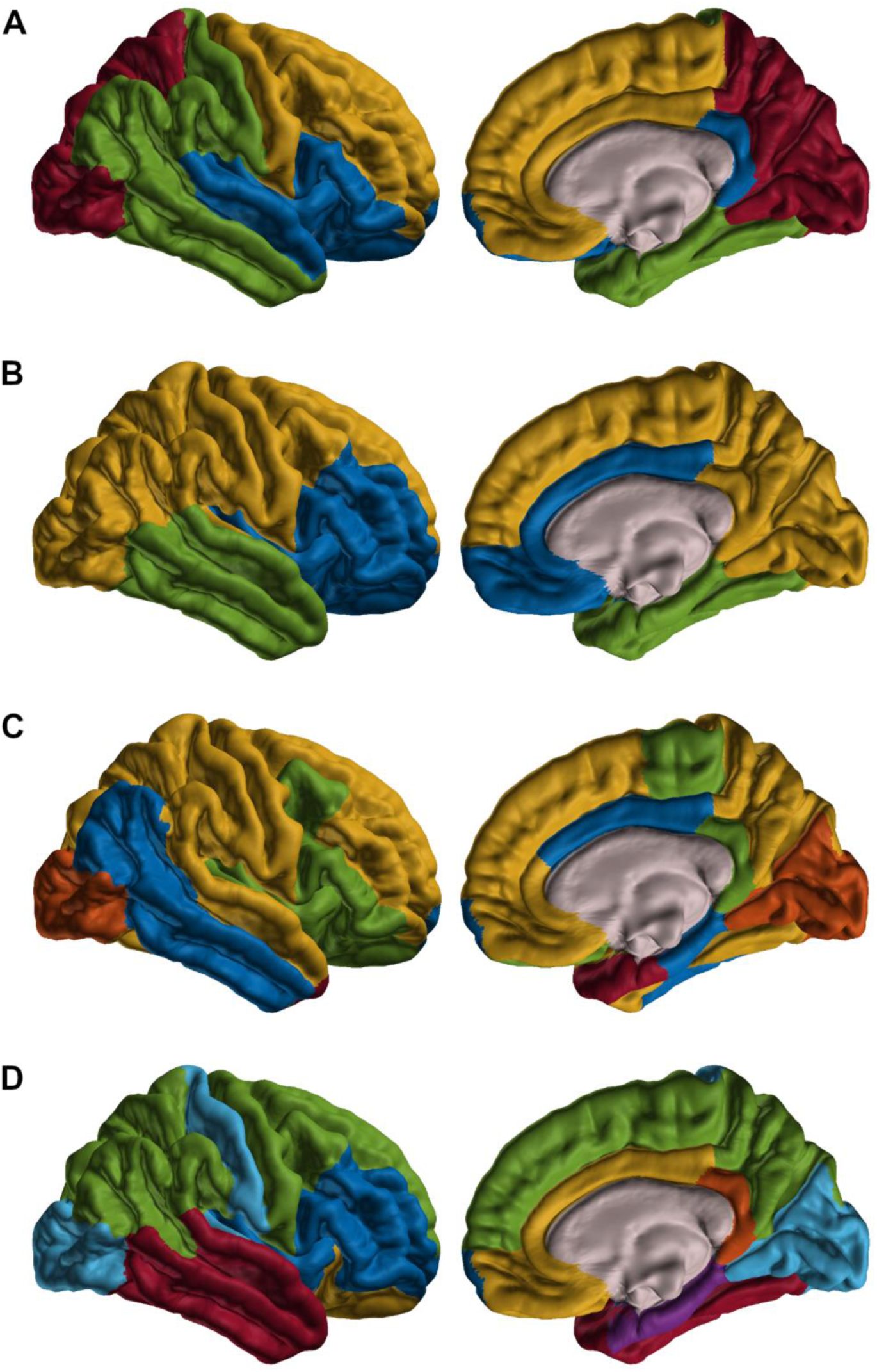
Clustering of genetic correlations among A) surface area and B) thickness regions after correcting for global measures. Clustering of genetic correlations among C) surface area and D) thickness regions without correcting for global measures. The best-fitting model for surface area with global correction was 4 diagonal components with varying volume and shape, and for thickness was 3 spherical components with equal volume. The best-fitting model for surface area without global correction was 5 spherical components with varying volume, and for thickness was 7 diagonal components with equal volume and shape.

**Figure S4.**
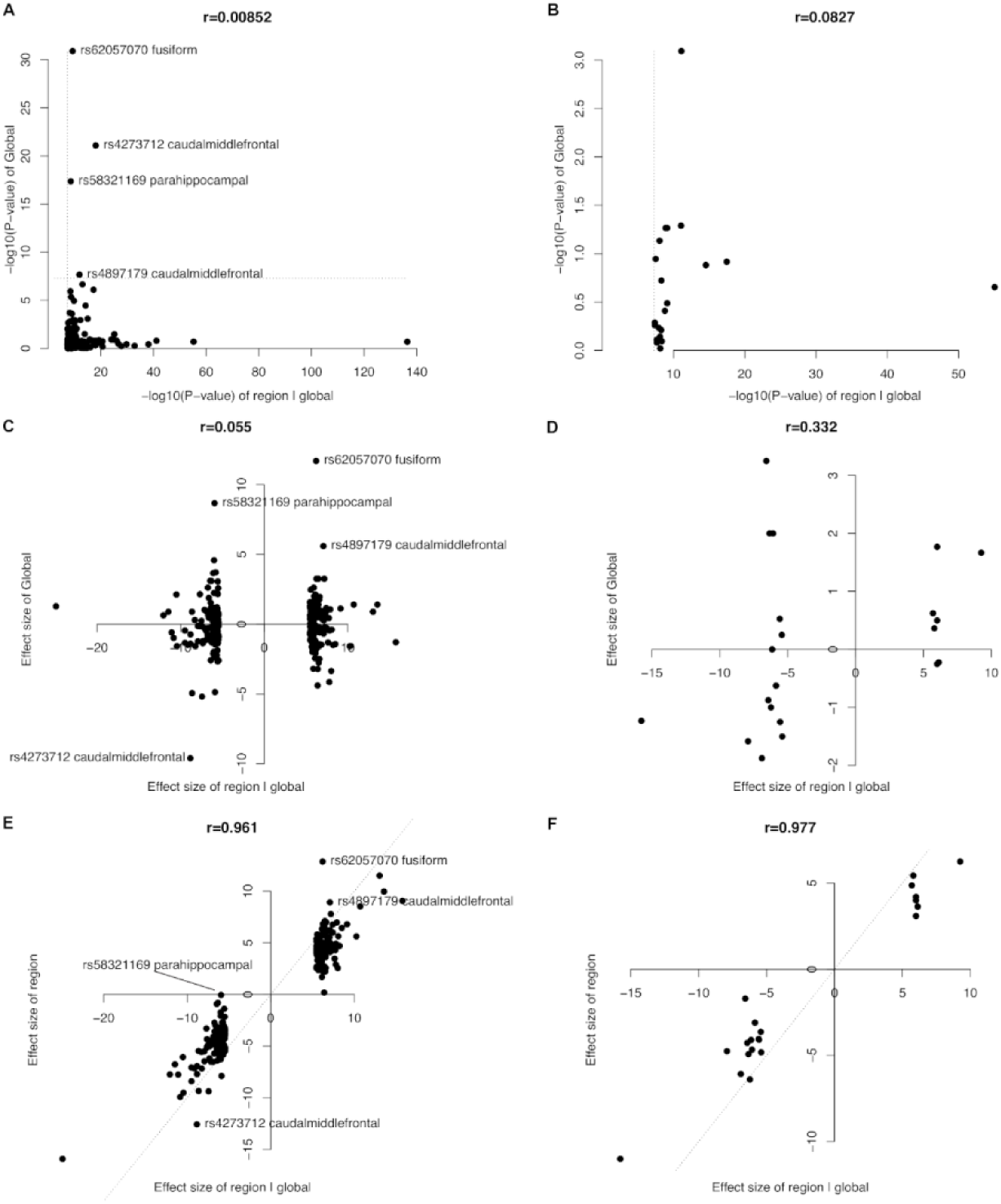
*P*-value of genome-wide significant regional SNPs with global control compared to their *P*-value in the global measure for A) surface area and B) thickness. Effect size of genome-wide significant regional SNPs with global control compared to their effect size in global measures for C) surface area and D) thickness. Effect size of genome-wide significant regional SNPs with global control compared to regional SNPs without global control in E) surface area and F) thickness.

## Consortium Authors

### Alzheimer’s Disease Neuroimaging Initiative (ADNI)

Data used in preparing this article were obtained from the Alzheimer’s Disease Neuroimaging Initiative (ADNI) database (adni.loni.usc.edu). As such, many investigators within the ADNI contributed to the design and implementation of ADNI and/or provided data but did not participate in analysis or writing of this report. A complete listing of ADNI investigators may be found at: http://adni.loni.usc.edu/wp-content/uploads/how_to_apply/ADNI_Acknowledgement_List.pdf *ADNI Infrastructure Investigators* Michael Weiner (UC San Francisco), Paul Aisen (University of Southern California), Ronald Petersen (Mayo Clinic, Rochester), Clifford R. Jack, Jr. (Mayo Clinic, Rochester), William Jagust (UC Berkeley), John Q. Trojanowki (U Pennsylvania), Arthur W. Toga (USC), Laurel Beckett (UC), Davis Robert C. Green (Brigham and Women’s Hospital/Harvard Medical School), Andrew J. Saykin (Indiana University), John Morris (Washington University St. Louis), Leslie M. Shaw (University of Pennsylvania). *ADNI External Advisory Board (ESAB):* Zaven Khachaturian (Prevent Alzheimer’s Disease 2020 (Chair)), Greg Sorensen (Siemens), Maria Carrillo (Alzheimer’s Association), Lew Kuller (University of Pittsburgh), Marc Raichle (Washington University St. Louis), Steven Paul (Cornell University), Peter Davies (Albert Einstein College of Medicine of Yeshiva University), Howard Fillit (AD Drug Discovery Foundation), Franz Hefti (Acumen Pharmaceuticals), David Holtzman (Washington University St. Louis), M. Marcel Mesulam (Northwestern University), William Potter (National Institute of Mental Health), Peter Snyder (Brown University). *ADNI 3 Private Partner Scientific Board (PPSB):* Veronika Logovinsky, (Eli Lilly (Chair)). *Data and Publications Committee:* Robert C. Green (BWH/HMS (Chair)). *Resource Allocation Review Committee:* Tom Montine (University of Washington (Chair)). *Clinical Core Leaders:* Ronald Petersen (Mayo Clinic, Rochester (Core PI)), Paul Aisen (University of Southern California). *Clinical Informatics and Operations:* Gustavo Jimenez (USC), Michael Donohue (USC), Devon Gessert (USC), Kelly Harless (USC), Jennifer Salazar (USC), Yuliana Cabrera (USC), Sarah Walter (USC), Lindsey Hergesheimer (USC). *Biostatistics Core Leaders and Key Personnel:* Laurel Beckett (UC Davis (Core PI)), Danielle Harvey (UC Davis), Michael Donohue (UC San Diego). *MRI Core Leaders and Key Personnel:* Clifford R. Jack, Jr. (Mayo Clinic, Rochester (Core PI)), Matthew Bernstein (Mayo Clinic, Rochester), Nick Fox (University of London), Paul Thompson (UCLA School of Medicine), Norbert Schuff (UCSF MRI), Charles DeCArli (UC Davis), Bret Borowski (RT Mayo Clinic), Jeff Gunter (Mayo Clinic), Matt Senjem (Mayo Clinic), Prashanthi Vemuri (Mayo Clinic), David Jones (Mayo Clinic), Kejal Kantarci (Mayo Clinic), Chad Ward (Mayo Clinic). *PET Core Leaders and Key Personnel:* William Jagust (UC Berkeley (Core PI)), Robert A. Koeppe (University of Michigan), Norm Foster (University of Utah), Eric M. Reiman (Banner Alzheimer’s Institute), Kewei Chen (Banner Alzheimer’s Institute), Chet Mathis (University of Pittsburgh), Susan Landau (UC Berkeley). *Neuropathology Core Leaders:* John C. Morris (Washington University St. Louis), Nigel J. Cairns (Washington University St. Louis), Erin Franklin (Washington University St. Louis), Lisa Taylor-Reinwald (Washington University St. Louis – Past Investigator). *Biomarkers Core Leaders and Key Personnel:* Leslie M. Shaw (UPenn School of Medicine), John Q. Trojanowki (UPenn School of Medicine), Virginia Lee (UPenn School of Medicine), Magdalena Korecka (UPenn School of Medicine), Michal Figurski (UPenn School of Medicine). *Informatics Core Leaders and Key Personnel:* Arthur W. Toga (USC (Core PI)), Karen Crawford (USC), Scott Neu (USC). *Genetics Core Leaders and Key Personnel:* Andrew J. Saykin (Indiana University), Tatiana M. Foroud (Indiana University), Steven Potkin (UC Irvine), Li Shen (Indiana University), Kelley Faber (Indiana University), Sungeun Kim (Indiana University), Kwangsik Nho (Indiana University). *Initial Concept Planning & Development:* Michael W. Weiner (UC San Francisco), Lean Thal (UC San Diego), Zaven Khachaturian (Prevent Alzheimer’s Disease 2020). *Early Project Proposal Development:* Leon Thal (UC San Diego), Neil Buckholtz (National Institute on Aging), Michael W. Weiner (UC San Francisco), Peter J. Snyder (Brown University), William Potter (National Institute of Mental Health), Steven Paul (Cornell University), Marilyn Albert (Johns Hopkins University), Richard Frank (Richard Frank Consulting), Zaven Khachaturian (Prevent Alzheimer’s Disease 2020). *NIA:* John Hsiao (National Institute on Aging). *ADNI Investigators by Site: Oregon Health & Science University*: Joseph Quinn, Lisa C. Silbert, Betty Lind, Jeffrey A. Kaye – Past Investigator, Raina Carter – Past Investigator, Sara Dolen – Past Investigator. *University of Southern California:* Lon S. Schneider, Sonia Pawluczyk, Mauricio Becerra, Liberty Teodoro, Bryan M. Spann – Past Investigator. *University of California – San Diego:* James Brewer, Helen Vanderswag, Adam Fleisher – Past Investigator. *University of Michigan:* Jaimie Ziolkowski, Judith L. Heidebrink, Joanne L. Lord – Past Investigator. *Mayo Clinic, Rochester:* Ronald Petersen, Sara S. Mason, Colleen S. Albers, David Knopman, Kris Johnson – Past Investigator. *Baylor College of Medicine:* Javier Villanueva-Meyer, Valory Pavlik, Nathaniel Pacini, Ashley Lamb, Joseph S. Kass, Rachelle S. Doody – Past Investigator, Victoria Shibley – Past Investigator, Munir Chowdhury – Past Investigator, Susan Rountree – Past Investigator, Mimi Dang – Past Investigator. *Columbia University Medical Center:* Yaakov Stern, Lawrence S. Honig, Karen L. Bell, Randy Yeh. *Washington University, St. Louis:* Beau Ances, John C. Morris, David Winkfield, Maria Carroll, Angela Oliver, Mary L. Creech – Past Investigator, Mark A. Mintun – Past Investigator, Stacy Schneider – Past Investigator. *University of Alabama - Birmingham:* Daniel Marson, David Geldmacher, Marissa Natelson Love, Randall Griffith – Past Investigator, David Clark – Past Investigator, John Brockington – Past Investigator. *Mount Sinai School of Medicine:* Hillel Grossman, Effie Mitsis – Past Investigator. *Rush University Medical Center:* Raj C. Shah, Melissa Lamar, Patricia Samuels. *Wien Center:* Ranjan Duara, Maria T. Greig-Custo, Rosemarie Rodriguez. *Johns Hopkins University:* Marilyn Albert, Chiadi Onyike, Daniel D’Agostino II, Stephanie Kielb – Past Investigator. *New York University:* Martin Sadowski, Mohammed O. Sheikh, Jamika Singleton-Garvin, Anaztasia Ulysse, Mrunalini Gaikwad. *Duke University Medical Center:* P. Murali Doraiswamy, Jeffrey R. Petrella, Olga James, Salvador Borges-Neto, Terence Z. Wong – Past Investigator, Edward Coleman – Past Investigator. *University of Pennsylvania:* Jason H. Karlawish, David A. Wolk, Sanjeev Vaishnavi, Christopher M. Clark – Past Investigator, Steven E. Arnold – Past Investigator. *University of Kentucky:* Charles D. Smith, Greg Jicha, Peter Hardy, Riham El Khouli, Elizabeth Oates, Gary Conrad. *University of Pittsburgh:* Oscar L. Lopez, MaryAnn Oakley, Donna M. Simpson. *University of Rochester Medical Center:* Anton P. Porsteinsson, Kim Martin, Nancy Kowalksi, Melanie Keltz, Bonnie S. Goldstein – Past Investigator, Kelly M. Makino – Past Investigator, M. Saleem Ismail – Past Investigator, Connie Brand – Past Investigator. *University of California Irvine IMIND:* Gaby Thai, Aimee Pierce, Beatriz Yanez, Elizabeth Sosa, Megan Witbracht. *University of Texas Southwestern Medical School:* Kyle Womack, Dana Mathews, Mary Quiceno. *Emory University:* Allan I. Levey, James J. Lah, Janet S. Cellar. *University of Kansas, Medical Center:* Jeffrey M. Burns, Russell H. Swerdlow, William M. Brooks. *University of California, Los Angeles:* Ellen Woo, Daniel H.S. Silverman, Edmond Teng, Sarah Kremen, Liana Apostolova – Past Investigator, Kathleen Tingus – Past Investigator, Po H. Lu – Past Investigator, George Bartzokis – Past Investigator. *Mayo Clinic, Jacksonville:* Neill R Graff-Radford (London), Francine Parfitt, Kim Poki-Walker. *Indiana University:* Martin R. Farlow, Ann Marie Hake, Brandy R. Matthews – Past Investigator, Jared R. Brosch, Scott Herring. *Yale University School of Medicine:* Christopher H. van Dyck, Richard E. Carson, Pradeep Varma. *McGill Univ., Montreal-Jewish General Hospital:* Howard Chertkow, Howard Bergman, Chris Hosein. *Sunnybrook Health Sciences, Ontario:* Sandra Black, Bojana Stefanovic, Chris (Chinthaka) Heyn. *U.B.C. Clinic for AD & Related Disorders:* Ging-Yuek Robin Hsiung, Benita Mudge, Vesna Sossi, Howard Feldman – Past Investigator, Michele Assaly – Past Investigator. *Cognitive Neurology - St. Joseph’s, Ontario:* Elizabeth Finger, Stephen Pasternack, William Pavlosky, Irina Rachinsky – Past Investigator, Dick Drost – Past Investigator, Andrew Kertesz – Past Investigator. *Cleveland Clinic Lou Ruvo Center for Brain Health:* Charles Bernick, Donna Muni. *Northwestern University:* Marek-Marsel Mesulam, Emily Rogalski, Kristine Lipowski, Sandra Weintraub, Borna Bonakdarpour, Diana Kerwin – Past Investigator, Chuang-Kuo Wu,– Past Investigator, Nancy Johnson – Past Investigator. *Premiere Research Inst (Palm Beach Neurology):* Carl Sadowsky, Teresa Villena. *Georgetown University Medical Center:* Raymond Scott Turner, Kathleen Johnson, Brigid Reynolds. *Brigham and Women’s Hospital*: Reisa A. Sperling, Keith A. Johnson, Gad A. Marshall. *Stanford University:* Jerome Yesavage, Joy L. Taylor, Steven Chao, Barton Lane – Past Investigator, Allyson Rosen – Past Investigator, Jared Tinklenberg – Past Investigator. *Banner Sun Health Research Institute:* Edward Zamrini, Christine M. Belden, Sherye A. Sirrel. *Boston University:* Neil Kowall, Ronald Killiany, Andrew E. Budson, Alexander Norbash – Past Investigator, Patricia Lynn Johnson – Past Investigator. *Howard University:* Thomas O. Obisesan, Ntekim E. Oyonumo, Joanne Allard, Olu Ogunlana. *Case Western Reserve University:* Alan Lerner, Paula Ogrocki, Curtis Tatsuoka, Parianne Fatica. *University of California, Davis – Sacramento:* Evan Fletcher, Pauline Maillard, John Olichney, Charles DeCarli, Owen Carmichael – Past Investigator. *Neurological Care of CNY:* Smita Kittur – Past Investigator. *Parkwood Institute:* Michael Borrie, T-Y Lee, Dr Rob Bartha. *University of Wisconsin:* Sterling Johnson, Sanjay Asthana, Cynthia M. Carlsson. *Banner Alzheimer’s Institute:* Pierre Tariot, Anna Burke, Joel Hetelle, Kathryn DeMarco, Nadira Trncic – Past Investigator, Adam Fleisher – Past Investigator, Stephanie Reeder – Past Investigator. *Dent Neurologic Institute:* Vernice Bates, Horacio Capote, Michelle Rainka. *Ohio State University:* Douglas W. Scharre, Maria Kataki, Rawan Tarawneh. *Albany Medical College:* Earl A. Zimmerman, Dzintra Celmins, David Hart. *Hartford Hospital, Olin Neuropsychiatry Research Center:* Godfrey D. Pearlson, Karen Blank, Karen Anderson. *Dartmouth-Hitchcock Medical Center:* Laura A. Flashman, Marc Seltzer, Mary L. Hynes, Robert B. Santulli – Past Investigator. *Wake Forest University Health Sciences:* Kaycee M. Sink, Mia Yang, Akiva Mintz. *Rhode Island Hospital:* Brian R. Ott, Geoffrey Tremont, Lori A. Daiello. *Butler Hospital:* Courtney Bodge, Stephen Salloway, Paul Malloy, Stephen Correia, Athena Lee. *UC San Francisco:* Howard J. Rosen, Bruce L. Miller, David Perry. *Medical University South Carolina:* Jacobo Mintzer, Kenneth Spicer, David Bachman. *St. Joseph’s Health Care:* Elizabeth Finger, Stephen Pasternak, Irina Rachinsky, John Rogers, Andrew Kertesz – Past Investigator, Dick Drost – Past Investigator. *Nathan Kline Institute:* Nunzio Pomara, Raymundo Hernando, Antero Sarrael. *University of Iowa College of Medicine:* Delwyn D. Miller, Karen Ekstam Smith, Hristina Koleva, Ki Won Nam, Hyungsub Shim, Susan K. Schultz – Past Investigator. *Cornell University:* Norman Relkin, Gloria Chiang, Michael Lin, Lisa Ravdin. *University of South Florida: USF Health Byrd Alzheimer’s Institute:* Amanda Smith, Christi Leach, Balebail Ashok Raj – Past Investigator, Kristin Fargher – Past Investigator.

### CHARGE Consortium

Edith Hofer (Clinical Division of Neurogeriatrics, Department of Neurology, Medical University of Graz, Graz, Austria), Gennady V. Roshchupkin (Department of Radiology and Nuclear Medicine, Erasmus MC, Rotterdam, The Netherlands), Hieab H. H. Adams (Department of Radiology and Nuclear Medicine, Erasmus MC, Rotterdam, The Netherlands), Maria J. Knol (Department of Epidemiology, Erasmus MC, Rotterdam, The Netherlands), Honghuang Lin (Section of Computational Biomedicine, Department of Medicine, Boston University School of Medicine, Boston, MA, USA), Shuo Li (Department of Biostatistics, Boston University School of Public Health, Boston, MA, USA), Habil Zare (Glenn Biggs Institute for Alzheimer’s and Neurodegenerative Diseases, UT Health San Antonio, San Antonio, USA), Shahzad Ahmad (Department of Epidemiology, Erasmus MC, Rotterdam, The Netherlands), Nicola J. Armstrong (Mathematics and Statistics, Murdoch University, Perth, Australia), Claudia L. Satizabal (Department of Epidemiology and Biostatistics, Glenn Biggs Institute for Alzheimer’s and Neurodegenerative Diseases, UT Health San Antonio, San Antonio, USA), Manon Bernard (Hospital for Sick Children, Toronto, Canada), Joshua C. Bis (Cardiovascular Health Research Unit, Department of Medicine, University of Washington, Seattle, WA, USA), Nathan A. Gillespie (Virginia Institute for Psychiatric and Behavior Genetics, Virginia Commonwealth University, VA, USA), Michelle Luciano (Centre for Cognitive Epidemiology and Cognitive Ageing, University of Edinburgh, Edinburgh, UK), Aniket Mishra (University of Bordeaux, Bordeaux Population Health Research Center, INSERM UMR 1219, Bordeaux, France), Markus Scholz (Institute for Medical Informatics, Statistics and Epidemiology, University of Leipzig, Leipzig, Germany), Alexander Teumer (Institute for Community Medicine, University Medicine Greifswald, Greifswald, Germany), Rui Xia (Institute of Molecular Medicine and Human Genetics Center, University of Texas Health Science Center at Houston, Houston, TX, USA), Xueqiu Jian (Institute of Molecular Medicine and Human Genetics Center, University of Texas Health Science Center at Houston, Houston, TX, USA), Thomas H. Mosley (Department of Medicine, University of Mississippi Medical Center, Jackson, MS, USA), Yasaman Saba (Gottfried Schatz Research Center for Cell Signaling, Metabolism and Aging, Medical University of Graz, Graz, Austria), Lukas Pirpamer (Clinical Division of Neurogeriatrics, Department of Neurology, Medical University of Graz, Graz, Austria), Stephan Seiler (Imaging of Dementia and Aging (IDeA) Laboratory, Department of Neurology, University of California-Davis, Davis, CA, USA), James T. Becker (Departments of Psychiatry, Neurology, and Psychology, University of Pittsburgh, Pittsburgh, PA, USA), Owen Carmichael (Pennington Biomedical Research Center, Baton Rouge, LA, USA), Jerome I. Rotter (Institute for Translational Genomics and Population Sciences, Los Angeles Biomedical Research Institute and Pediatrics at Harbor-UCLA Medical Center, Torrance, CA, USA), Bruce M. Psaty (Cardiovascular Health Research Unit, Departments of Medicine, Epidemiology and Health Services, University of Washington, Seattle, WA, USA), Oscar L. Lopez (Departments of Psychiatry, Neurology, and Psychology, University of Pittsburgh, Pittsburgh, PA, USA), Najaf Amin (Department of Epidemiology, Erasmus MC, Rotterdam, The Netherlands), Sven J. van der Lee (Department of Epidemiology, Erasmus MC, Rotterdam, The Netherlands), Qiong Yang (Department of Biostatistics, Boston University School of Public Health, Boston, MA, USA), Jayandra J. Himali (Department of Biostatistics, Boston University School of Public Health, Boston, MA, USA), Pauline Maillard (Imaging of Dementia and Aging (IDeA) Laboratory, Department of Neurology, University of California-Davis, Davis, CA, USA), Alexa S. Beiser (Department of Neurology, Boston University School of Medicine, Boston, MA, USA), Charles DeCarli (Imaging of Dementia and Aging (IDeA) Laboratory, Department of Neurology, University of California-Davis, Davis, CA, USA), Sherif Karama (McGill University, Montreal Neurological Institute, Montreal, Canada), Lindsay Lewis (McGill University, Montreal Neurological Institute, Montreal, Canada), Mark Bastin (Centre for Cognitive Epidemiology and Cognitive Ageing, University of Edinburgh, Edinburgh, UK), Ian J. Deary (Centre for Cognitive Epidemiology and Cognitive Ageing, University of Edinburgh, Edinburgh, UK), Veronica Witte (Department of Neurology, Max Planck Institute of Cognitive and Brain Sciences, Leipzig, Germany), Frauke Beyer (Department of Neurology, Max Planck Institute of Cognitive and Brain Sciences, Leipzig, Germany), Markus Loeffler (Institute for Medical Informatics, Statistics and Epidemiology, University of Leipzig, Leipzig, Germany), Karen A. Mather (Centre for Healthy Brain Ageing, School of Psychiatry, University of New South Wales, Sydney, Australia), Peter R. Schofield (Neuroscience Research Australia, Sydney, Australia), Anbupalam Thalamuthu (Centre for Healthy Brain Ageing, School of Psychiatry, University of New South Wales, Sydney, Australia), John B. Kwok (Brain and Mind Centre - The University of Sydney, Camperdown, NSW, Australia), Margaret J. Wright (Queensland Brain Institute, The University of Queensland, St Lucia, QLD, Australia), David Ames (National Ageing Research Institute, Royal Melbourne Hospital, Victoria, Australia), Julian Trollor (Centre for Healthy Brain Ageing, School of Psychiatry, University of New South Wales, Sydney, Australia), Jiyang Jiang (Centre for Healthy Brain Ageing, School of Psychiatry, University of New South Wales, Sydney, Australia), Henry Brodaty (Dementia Centre for Research Collaboration, University of New South Wales, Sydney, NSW, Australia), Wei Wen (Centre for Healthy Brain Ageing, School of Psychiatry, University of New South Wales, Sydney, Australia), Meike W Vernooij (Department of Radiology and Nuclear Medicine, Erasmus MC, Rotterdam, The Netherlands), Albert Hofman (Department of Epidemiology, Harvard T.H. Chan School of Public Health, Boston, MA, USA), André G. Uitterlinden (Department of Epidemiology, Erasmus MC, Rotterdam, The Netherlands), Wiro J. Niessen (Imaging Physics, Faculty of Applied Sciences, Delft University of Technology, The Netherlands), Katharina Wittfeld (German Center for Neurodegenerative Diseases (DZNE), Site Rostock/ Greifswald, Germany), Robin Bülow (Institute for Diagnostic Radiology and Neuroradiology, University Medicine Greifswald, Greifswald, Germany), Uwe Völker (Interfaculty Institute for Genetics and Functional Genomics, University Medicine Greifswald, Greifswald, Germany), Zdenka Pausova (Hospital for Sick Children, Toronto, Canada), G. Bruce Pike (Departments of Radiology and Clinial Neurosciences, University of Calgary, Calgary, Canada), Sophie Maingault (University of Bordeaux, Institut des Maladies NeurodégénrativesUMR5293, CEA, CNRS, Ubordeaux, Bordeaux, France), Fabrice Crivello (University of Bordeaux, Institut des Maladies NeurodégénrativesUMR5293, CEA, CNRS, Ubordeaux, Bordeaux, France), Bernard Mazoyer (Neurodegeneratives Diseases Institute UMR 5293, CNRS, CEA, University of Bordeaux, Bordeaux, France), Michael C. Neale (Virginia Institute for Psychiatric and Behavior Genetics, Virginia Commonwealth University, VA, USA), Carol E. Franz (Department of Psychiatry, University of California San Diego, CA, USA), Michael J. Lyons (Department of Psychological and Brain Sciences, Boston University, Boston, MA, USA), Matthew S. Panizzon (Department of Psychiatry, University of California San Diego, CA, USA), Ole A. Andreassen (ORMENT, KG Jebsen Centre for Psychosis Research, Institute of Clinical Medicine, University of Oslo and Division of Mental Health and Addiction, Oslo University Hospital, Oslo, Norway), Anders M. Dale (Departments of Radiology and Neurosciences, University of California, San Diego, La Jolla, CA, USA), Mark Logue (National Center for PTSD at Boston VA Healthcare System, Boston, MA, USA), Perminder S. Sachdev (Centre for Healthy Brain Ageing, School of Psychiatry, University of New South Wales, Sydney, Australia), William S. Kremen (Department of Psychiatry, University of California San Diego, CA, USA), Joanna A. Wardlaw (Centre for Cognitive Epidemiology and Cognitive Ageing, University of Edinburgh, Edinburgh, UK), Arno Villringer (Department of Neurology, Max Planck Institute of Cognitive and Brain Sciences, Leipzig, Germany), Cornelia M. van Duijn (Department of Epidemiology, Erasmus MC, Rotterdam, The Netherlands), Hans Jörgen Grabe (Department of Psychiatry and Psychotherapy, University Medicine Greifswald, Germany), William T. Longstreth Jr (Departments of Neurology and Epidemiology, University of Washington, Seattle, WA, USA), Myriam Fornage (Institute of Molecular Medicine and Human Genetics Center, University of Texas Health Science Center at Houston, Houston, TX, USA), Tomas Paus (Bloorview Research Institute, Holland Bloorview Kids Rehabilitation Hospital, Toronto, Ontario, Canada), Stephanie Debette (University of Bordeaux, Bordeaux Population Health Research Center, INSERM UMR 1219, Bordeaux, France), M. Arfan Ikram (Department of Radiology and Nuclear Medicine, Erasmus MC, Rotterdam, The Netherlands), Helena Schmidt (Gottfried Schatz Research Center for Cell Signaling, Metabolism and Aging, Medical University of Graz, Graz, Austria), Reinhold Schmidt (Clinical Division of Neurogeriatrics, Department of Neurology, Medical University of Graz, Graz, Austria), Sudha Seshadri (Department of Epidemiology and Biostatistics, Glenn Biggs Institute for Alzheimer’s and Neurodegenerative Diseases, UT Health San Antonio, San Antonio, USA.

### EPIGEN Consortium

David B. Goldstein (The Centre for Genomics and Population Genetics, Duke University Institute for Genome Sciences and Policy, Durham, North Carolina, USA), Erin L. Heinzen (The Centre for Genomics and Population Genetics, Duke University Institute for Genome Sciences and Policy, Durham, North Carolina, USA), Kevin Shianna (The Centre for Genomics and Population Genetics, Duke University Institute for Genome Sciences and Policy, Durham, North Carolina, USA), Rodney Radtke (Department of Medicine, Duke University Medical Center, Durham, North Carolina, USA) and Ruth Ottmann (Departments of Epidemiology, Neurology, and the G.H. Sergievsky Center, Columbia University, New York, NY).

### IMAGEN Consortium

Dr. Eric Artiges (INSERM), Semiha Aydin (Physikalisch-Technische Bundesanstalt), Prof. Dr. Dr. Tobias Banaschewski (Central Institute of Mental Health), Alexis Barbot (Commissariat à l’Energie Atomique), Prof. Dr. Gareth Barker (King’s College London), Andreas Becker (Georg-August-Universität Göttingen), Pauline Bezivin-Frere (INSERM), Dr. Francesca Biondo (King’s College London), Dr. Arun Bokde (Trinity College Dublin), Uli Bromberg (University of Hamburg), Dr. Ruediger Bruehl, Prof. Dr. Christian Büchel (University of Hamburg), Dr. Congying Chu (King’s College London), Dr. Patricia Conrod (King’s College London), Laura Daedelow (Charité Universitätsmedizin Berlin), Dr. Jeffrey Dalley (Cambridge University), Dr. Sylvane Desrivieres (King’s College London), Eoin Dooley (Trinity College Dublin), Irina Filippi (INSERM), Dr Ariane Fillmer (Physikalisch-Technische Bundesanstalt), Prof. Dr. Herta Flor (Central Institute of Mental Health), Juliane Fröhner (Technische Universität Dresden), Vincent Frouin (Commissariat à l’Energie Atomique), Dr. Hugh Garavan (University of Vermont), Prof. Penny Gowland (University of Nottingham), Yvonne Grimmer (Central Institute of Mental Health), Prof. Dr. Andreas Heinz (Charité Universitätsmedizin Berlin), Dr. Sarah Hohmann (Central Institute of Mental Health), Albrecht Ihlenfeld (Physikalisch-Technische Bundesanstalt), Alex Ing (King’s College London), Corinna Isensee (University Medical Center Göttingen), Dr. Bernd Ittermann (Physikalisch-Technische Bundesanstalt), Dr. Tianye Jia (King’s College London), Dr. Hervé Lemaitre (INSERM), Emma Lethbridge (University of Nottingham), Prof. Dr. Jean-Luc Martinot (INSERM), Sabina Millenet (Central Institute of Mental Health), Sarah Miller (Charité Universitätsmedizin Berlin), Ruben Miranda (INSERM), PD Dr. Frauke Nees (Central Institute of Mental Health), Dr. Marie-Laure Paillere (INSERM), Dimitri Papadopoulos (INSERM), Prof. Dr. Tomáš Paus (Bloorview Research Institute, Holland Bloorview Kids Rehabilitation Hospital and Departments of Psychology and Psychatry, University of Toronto), Dr. Zdenka Pausova (University of Toronto), Dr. Dr. Jani Pentilla (INSERM), Dr. Jean-Baptiste Poline (Commissariat à l’Energie Atomique), Prof. Dr. Luise Poustka (University Medical Center Göttingen), Dr. Erin Burke Quinlan (King’s College London), Dr. Michael Rapp (Charité Universitätsmedizin Berlin), Prof. Dr. Trevor Robbins (Cambridge University), Dr. Gabriel Robert (King’s College London), John Rogers (Delosis), Dr. Barbara Ruggeri (King’s College London), Prof. Dr. Gunter Schumann (King’s College London), Prof. Dr. Michael Smolka (Technische Universität Dresden), Argyris Stringaris (National Institute of Mental Health), Betteke van Noort (Charité Universitätsmedizin Berlin), Dr. Henrik Walter (Charité Universitätsmedizin Berlin), Dr. Robert Whelan (Trinity College Dublin), Prof. Dr. Steve Williams (King’s College London).

### Parkinson’s Progression Markers Initiative (PPMI)

Data used in preparing this article were obtained from the PPMI database (http://www.ppmi-info.org/). As such, many investigators within the PPMI contributed to the design and implementation of PPMI and/or provided data but did not participate in analysis or writing of this report. A complete listing of PPMI investigators may be found at: http://www.ppmi-info.org/authorslist/. Kenneth Marek (Institute for Neurodegenerative Disorders, New Haven), Danna Jennings (Institute for Neurodegenerative Disorders, New Haven), Shirley Lasch (Institute for Neurodegenerative Disorders, New Haven), Caroline Tanner (University of California, San Francisco), Tanya Simuni (Northwestern University, Chicago), Christopher Coffey (University of Iowa, Iowa City), Karl Kieburtz (Clinical Trials Coordination Center, University of Rochester), Renee Wilson (Clinical Trials Coordination Center, University of Rochester), Werner Poewe (Innsbruck Medical University, Innsbruck), Brit Mollenhauer (Paracelsus-Elena Klinik, Kassel), Douglas Galasko (University of California, San Diego), Tatiana Foroud (Indiana University, Indianapolis), Todd Sherer (The Michael J. Fox Foundation for Parkinson’s Research, New York), Sohini Chowdhury (The Michael J. Fox Foundation for Parkinson’s Research, New York), Mark Frasier (The Michael J. Fox Foundation for Parkinson’s Research, New York), Catherine Kopil (The Michael J. Fox Foundation for Parkinson’s Research, New York), Vanessa Arnedo (The Michael J. Fox Foundation for Parkinson’s Research, New York), Alice Rudolph (Clinical Trials Coordination Center, University of Rochester), Cynthia Casaceli (Clinical Trials Coordination Center, University of Rochester), John Seibyl (Institute for Neurodegenerative Disorders, New Haven), Susan Mendick (Institute for Neurodegenerative Disorders, New Haven), Norbert Schuff (University of California, San Francisco), Chelsea Caspell (University of Iowa, Iowa City), Liz Uribe (University of Iowa, Iowa City), Eric Foster (University of Iowa, Iowa City), Katherine Gloer (University of Iowa, Iowa City), Jon Yankey (University of Iowa, Iowa City), Arthur Toga (Laboratory of Neuroimaging (LONI), University of Southern California), Karen Crawford (Laboratory of Neuroimaging (LONI), University of Southern California), Paola Casalin (BioRep, Milan), Giulia Malferrari (BioRep, Milan), Andrew Singleton (National Institute on Aging, NIH, Bethesda), Keith A. Hawkins (Yale University, New Haven), David Russell (Institute for Neurodegenerative Disorders, New Haven), Stewart Factor (Emory University of Medicine, Atlanta), Penelope Hogarth (Oregon Health and Science University, Portland), David Standaert (University of Alabama at Birmingham, Birmingham), Robert Hauser (University of South Florida, Tampa), Joseph Jankovic (Baylor College of Medicine, Houston), Matthew Stern (University of Pennsylvania, Philadelphia), Lama Chahine (University of Pennsylvania, Philadelphia), James Leverenz (University of Washington, Seattle), Samuel Frank (Boston University, Boston), Irene Richard (University of Rochester, Rochester), Klaus Seppi (Innsbruck Medical University, Innsbruck), Holly Shill (Banner Research Institute, Sun City), Hubert Fernandez (Cleveland Clinic, Cleveland), Daniela Berg (University of Tuebingen, Tuebingen), Isabel Wurster (University of Tuebingen, Tuebingen), Zoltan Mari (Johns Hopkins University, Baltimore), David Brooks (Imperial College of London, London), Nicola Pavese (Imperial College of London, London), Paolo Barone (University of Salerno, Salerno), Stuart Isaacson (Parkinson’s Disease and Movement Disorders Center, Boca Raton), Alberto Espay (University of Cincinnati, Cincinnati), Dominic Rowe (Macquarie University, Sydney), Melanie Brandabur (The Parkinson’s Institute, Sunnyvale), James Tetrud (The Parkinson’s Institute, Sunnyvale), Grace Liang (The Parkinson’s Institute, Sunnyvale), Alex Iranzo (Hospital Clinic of Barcelona, Barcelona), Eduardo Tolosa (Hospital Clinic of Barcelona, Barcelona), Shu-Ching Hu (University of Washington, Seattle), Gretchen Todd (University of Washington, Seattle), Laura Leary (Institute for Neurodegenerative Disorders, New Haven), Cheryl Riordan (Institute for Neurodegenerative Disorders, New Haven), Linda Rees (The Parkinson’s Institute, Sunnyvale), Alicia Portillo (Oregon Health and Science University, Portland), Art Lenahan (Oregon Health and Science University, Portland), Karen Williams (Northwestern University, Chicago), Stephanie Guthrie (University of Alabama at Birmingham, Birmingham), Ashlee Rawlins (University of Alabama at Birmingham, Birmingham), Sherry Harlan (University of South Florida, Tampa), Christine Hunter (Baylor College of Medicine, Houston), Baochan Tran (University of Pennsylvania, Philadelphia), Abigail Darin (University of Pennsylvania, Philadelphia), Carly Linder (University of Pennsylvania, Philadelphia), Marne Baca (University of Washington, Seattle), Heli Venkov (University of Washington, Seattle), Cathi-Ann Thomas (Boston University, Boston), Raymond James (Boston University, Boston), Cheryl Deeley (University of Rochester, Rochester), Courtney Bishop (University of Rochester, Rochester), Fabienne Sprenger (Innsbruck Medical University, Innsbruck), Diana Willeke (Paracelsus-Elena Klinik, Kassel), Sanja Obradov (Banner Research Institute, Sun City), Jennifer Mule (Cleveland Clinic, Cleveland), Nancy Monahan (Cleveland Clinic, Cleveland), Katharina Gauss (University of Tuebingen, Tuebingen), Deborah Fontaine (University of California, San Diego), Christina Gigliotti (University of California, San Diego), Arita McCoy (Johns Hopkins University, Baltimore), Becky Dunlop (Johns Hopkins University, Baltimore), Bina Shah (Imperial College of London, London), Susan Ainscough (University of Salerno, Salerno), Angela James (Parkinson’s Disease and Movement Disorders Center, Boca Raton), Rebecca Silverstein (Parkinson’s Disease and Movement Disorders Center, Boca Raton), Kristy Espay (University of Cincinnati, Cincinnati), Madelaine Ranola (Macquarie University, Sydney), Thomas Comery (Pfizer, Inc., Groton), Jesse Cedarbaum (Biogen Idec, Cambridge), Bernard Ravina (Biogen Idec, Cambridge), Igor D. Grachev (GE Healthcare, Princeton), Jordan S. Dubow (AbbVie, Abbot Park), Michael Ahlijanian (Bristol-Myers Squibb Company), Holly Soares (Bristol-Myers Squibb Company), Suzanne Ostrowizki (F.Hoffmann La-Roche, Basel), Paulo Fontoura (F.Hoffmann La-Roche, Basel), Alison Chalker (Merck & Co., North Wales), David L. Hewitt (Merck & Co., North Wales), Marcel van der Brug (Genentech, Inc., South San Francisco), Alastair D. Reith (GlaxoSmithKline, Stevenage), Peggy Taylor (Covance, Dedham), Jan Egebjerg (H. Lundbeck), Mark Minton (Avid Radiopharmaceuticals, Philadelphia), Andrew Siderowf (Avid Radiopharmaceuticals, Philadelphia), Pierandrea Muglia (UCB Pharma S.A., Brussels), Robert Umek (Meso Scale Discovery), Ana Catafau (Meso Scale Discovery), Vera Kiyasova (Servier), Barbara Saba (Servier).

### SYS Consortium

Tomáš Paus MD PhD (Bloorview Research Institute, University of Toronto, Canada), Zdenka Pausova, MD (The Hospital for Sick Children, University of Toronto, Canada), G. Bruce Pike PhD (Department of Radiology, University of Calgary, Canada), Louis Richer PhD (Department of Health Sciences, University of Quebec in Chicoutimi, Canada), Gabriel Leonard PhD (Montreal Neurological Institute, McGill University, Canada), Michel Perron PhD (CEGEP Jonquiere, Canada), Suzanne Veillette PhD (CEGEP Jonquiere, Canada) and Manon Bernard BComp (The Hospital for Sick Children, University of Toronto, Canada).

### Additional cohort information

#### ADNI

Data used in the preparation of this article were obtained from the Alzheimer’s Disease Neuroimaging Initiative database. The ADNI was launched in 2003 as a 5-year public– private partnership to assess and optimize biomarkers for clinical trials in Alzheimer’s disease. The initial sample included older adults who were cognitive normal (CN) as well as meeting criteria for MCI and clinical AD. In 2011, ADNI-2 began to recruit an additional CN group as well as individuals with significant memory concerns (SMC), early MCI and late MCI, and AD. These subjects, and others carried forward from ADNI-1, were scanned with an updated neuroimaging protocol. Participants were recruited from over 60 sites across the U.S. and Canada. For up-to-date information, please see www.adni-info.org.

#### ALSPAC

Pregnant women resident in Avon, UK with expected dates of delivery 1st April 1991 to 31st December 1992 were invited to take part in the study. The initial number of pregnancies enrolled is 14,541 (for these at least one questionnaire has been returned or a “Children in Focus” clinic had been attended by 19/07/99). Of these initial pregnancies, there was a total of 14,676 fetuses, resulting in 14,062 live births and 13,988 children who were alive at 1 year of age. When the oldest children were approximately 7 years of age, an attempt was made to bolster the initial sample with eligible cases who had failed to join the study originally. As a result, when considering variables collected from the age of seven onwards (and potentially abstracted from obstetric notes) there are data available for more than the 14,541 pregnancies mentioned above. The number of new pregnancies not in the initial sample (known as Phase I enrolment) that are currently represented on the built files and reflecting enrolment status at the age of 18 is 706 (452 and 254 recruited during Phases II and III respectively), resulting in an additional 713 children being enrolled. The phases of enrolment are described in more detail in the cohort profile paper (see footnote 4 below). The total sample size for analyses using any data collected after the age of seven is therefore 15,247 pregnancies, resulting in 15,458 fetuses. Of this total sample of 15,458 fetuses, 14,775 were live births and 14,701 were alive at 1 year of age. A 10% sample of the ALSPAC cohort, known as the Children in Focus (CiF) group, attended clinics at the University of Bristol at various time intervals between 4 to 61 months of age. The CiF group were chosen at random from the last 6 months of ALSPAC births (1432 families attended at least one clinic). Excluded were those mothers who had moved out of the area or were lost to follow-up, and those partaking in another study of infant development in Avon. The data used in the present study were collected from 391 males and further description of this subset and the variables used in this study are provided in Supplementary Tables 2–4.

The study website contains details of all the data that is available through a fully searchable data dictionary (http://www.bris.ac.uk/alspac/researchers/data-access/data-dictionary/).

Further information can be found in the following papers:

Boyd A, Golding J, Macleod J, Lawlor DA, Fraser A, Henderson J, Molloy L, Ness A, Ring S, Davey Smith G. Cohort Profile: The ‘Children of the 90s’; the index offspring of The Avon Longitudinal Study of Parents and Children (ALSPAC). International Journal of Epidemiology 2013; 42: 111-127;

Fraser A, Macdonald-Wallis C, Tilling K, Boyd A, Golding J, Davey Smith G, Henderson J, Macleod J, Molloy L, Ness A, Ring S, Nelson SM, Lawlor DA. Cohort Profile: The Avon Longitudinal Study of Parents and Children: ALSPAC mothers cohort. International Journal of Epidemiology 2013; 42:97-110.

## ENIGMA

The study was supported in part by grant U54 EB020403 from the NIH Big Data to Knowledge (BD2K) Initiative, a cross-NIH partnership. Additional support was provided by R01MH116147, P41 EB015922, RF1AG051710, RF1 AG041915 (to P.T.), by P01 AG026572, R01 AG059874 and by R01 MH117601 (to N.J. and L.S.). S.E.M. was funded by an NHMRC Senior Research Fellowship (APP1103623). L.C.-C. was supported by a QIMR Berghofer Fellowship. J.L.S was supported by R01MH118349 and R00MH102357.

## 1000BRAINS

Is a population-based cohort based on the Heinz-Nixdorf Recall Study and is supported in part by the German National Cohort. We thank the Heinz Nixdorf Foundation (Germany) for their generous support in terms of the Heinz Nixdorf Study. The HNR study is also supported by the German Ministry of Education and Science (FKZ 01EG940), and the German Research Council (DFG, ER 155/6-1). The authors are supported by the Initiative and Networking Fund of the Helmholtz Association (Svenja Caspers) and the European Union’s Horizon 2020 Research and Innovation Programme under Grant Agreements 720270 (Human Brain Project SGA1; Sven Cichon) and 785907 (Human Brain Project SGA2; Svenja Caspers, Sven Cichon). This work was further supported by the German Federal Ministry of Education and Research (BMBF) through the Integrated Network IntegraMent (Integrated Understanding of Causes and Mechanisms in Mental Disorders) under the auspices of the e:Med Program (grant 01ZX1314A; Sven Cichon), and by the Swiss National Science Foundation (SNSF, grant 156791; Sven Cichon).

## ADNI1 and ADNI2GO

Data used in the preparation of this article were obtained from the Alzheimer’s Disease Neuroimaging Initiative (ADNI) database (adni.loni.usc.edu). The ADNI was launched in 2003 as a public-private partnership, led by Principal Investigator Michael W. Weiner, MD. The primary goal of ADNI has been to test whether serial magnetic resonance imaging (MRI), positron emission tomography (PET), other biological markers, and clinical and neuropsychological assessment can be combined to measure the progression of mild cognitive impairment (MCI) and early Alzheimer’s disease (AD). Data collection and sharing for this project was funded by the Alzheimer’s Disease Neuroimaging Initiative (ADNI) (National Institutes of Health Grant U01 AG024904) and DOD ADNI (Department of Defense award number W81XWH-12-2-0012). ADNI is funded by the National Institute on Aging, the National Institute of Biomedical Imaging and Bioengineering, and through generous contributions from the following: AbbVie, Alzheimer’s Association; Alzheimer’s Drug Discovery Foundation; Araclon Biotech; BioClinica, Inc.; Biogen; Bristol-Myers Squibb Company; CereSpir, Inc.; Cogstate; Eisai Inc.; Elan Pharmaceuticals, Inc.; Eli Lilly and Company; EuroImmun; F. Hoffmann-La Roche Ltd and its affiliated company Genentech, Inc.; Fujirebio; GE Healthcare; IXICO Ltd.; Janssen Alzheimer Immunotherapy Research & Development, LLC.; Johnson & Johnson Pharmaceutical Research & Development LLC.; Lumosity; Lundbeck; Merck & Co., Inc.; Meso Scale Diagnostics, LLC.; NeuroRx Research; Neurotrack Technologies; Novartis Pharmaceuticals Corporation; Pfizer Inc.; Piramal Imaging; Servier; Takeda Pharmaceutical Company; and Transition Therapeutics. The Canadian Institutes of Health Research is providing funds to support ADNI clinical sites in Canada. Private sector contributions are facilitated by the Foundation for the National Institutes of Health (www.fnih.org). The grantee organization is the Northern California Institute for Research and Education, and the study is coordinated by the Alzheimer’s Therapeutic Research Institute at the University of Southern California. ADNI data are disseminated by the Laboratory for Neuro Imaging at the University of Southern California. Samples from the National Centralized Repository for Alzheimer’s Disease and Related Dementias (NCRAD), which receives government support under a cooperative agreement grant (U24 AG21886) awarded by the National Institute on Aging (NIA), were used in this study. We thank contributors who collected samples used in this study, as well as patients and their families, whose help and participation made this work possible. Additional support for data analysis was provided by NLM R01 LM012535 and NIA R03 AG054936 (to K.N.).

## ALSPAC

We are extremely grateful to all the families who took part in this study, the midwives for their help in recruiting them, and the whole ALSPAC team, which includes interviewers, computer and laboratory technicians, clerical workers, research scientists, volunteers, managers, receptionists and nurses. Ethical approval for the study was obtained from the ALSPAC Ethics and Law Committee and the Local Research Ethics Committees. The UK Medical Research Council and Wellcome (Grant ref: 102215/2/13/2) and the University of Bristol provide core support for ALSPAC. This publication is the work of the authors and they will serve as guarantors for the contents of this paper. *A comprehensive list of grants funding is available on the ALSPAC website (http://www.bristol.ac.uk/alspac/external/documents/grant-acknowledgements.pdf).* ALSPAC neuroimaging data was specifically funded by RO1 MH085772 (Axon, Testosterone and Mental Health during Adolescence; PI: T. Paus). GWAS data was generated by Sample Logistics and Genotyping Facilities at Wellcome Sanger Institute and LabCorp (Laboratory Corporation of America) using support from 23andMe. We would like to acknowledge the help of Lara B Clauss during the quality control process of the ALSPAC neuroimaging data.

## ASRB

Data and samples were collected by the Australian Schizophrenia Research Bank (ASRB), supported by the Australian NHMRC, the Pratt Foundation, Ramsay Health Care, and the Viertel Charitable Foundation. The ASRB were also supported by the Schizophrenia Research Institute (Australia), utilizing infrastructure funding from NSW Health and the Macquarie Group Foundation. DNA analysis was supported by the Neurobehavioral Genetics Unit, utilising funding from NSW Health and the National Health and Medical Research Council (NHMRC) Project Grants (1067137, 1147644, 1051672). MC was supported by an NHMRC Senior Research Fellowship (1121474). CP was supported by a NHMRC Senior Principal Research Fellowship (628386 & 1105825).

## BETULA

This work was supported by a Wallenberg Scholar grant from the Knut and Alice Wallenberg (KAW) Foundation and a grant from Torsten and Ragnar Söderbergs Foundation to LN, a grant from HelseVest RHF (Grant 911554) to SLH, grants from the Bergen Research Foundation and the University of Bergen to SLH, grants from the Dr Einar Martens Fund and the K.G. Jebsen Foundation to SLH and VMS, the Research Council of Norway to TE (Grant 177458/V 50) and LTW (Grant 204966/F 20). We thank the Centre for Advanced Study (CAS) at the Norwegian Academy of Science and Letters in Oslo for hosting collaborative projects and workshops between Norway and Sweden in 2011–2012. Image analyses were performed on resources provided by the Swedish National Infrastructure for Computing (SNIC) at HPC2N in Umeå.

## BIG

The Brain Imaging Genetics (BIG) database was established in Nijmegen, the Netherlands in 2007. This resource is now part of Cognomics, a joint initiative by researchers of the Donders Centre for Cognitive Neuroimaging, the Human Genetics and Cognitive Neuroscience departments of the Radboud University Medical Center, and the Max Planck Institute for Psycholinguistics. The present study includes two subsamples of BIG, from successive waves of genotyping on Affymetrix (BIG-Affy) and PsychChip (BIG-PsychChip) arrays. Analyses for this project were carried out on the Dutch national e-infrastructure with the support of SURF Cooperative. Nijmegen’s BIG resource is part of Cognomics, a joint initiative by researchers of the Donders Centre for Cognitive Neuroimaging, the Human Genetics and Cognitive Neuroscience departments of the Radboud University Medical Center, and the Max Planck Institute for Psycholinguistics (funded by the Max Planck Society). Support for the Cognomics Initiative, including phenotyping and genotyping of BIG cohorts, comes from funds of the participating departments and centres and from external national grants, i.e. the Biobanking and Biomolecular Resources Research Infrastructure (Netherlands) (BBMRI-NL), the Hersenstichting Nederland, and the Netherlands Organisation for Scientific Research (NWO), including the NWO Brain & Cognition Excellence Program (grant 433-09-229) and the Vici Innovation Program (grant 016-130-669 to BF). Additional support was received from the European Community’s Seventh Framework Programme (FP7/2007–2013) under grant agreements n◦ 602805 (Aggressotype), n◦ 602450 (IMAGEMEND), and n◦ 278948 (TACTICS) as well as from the European Community’s Horizon 2020 Programme (H2020/2014–2020) under grant agreements n◦ 643051 (MiND) and n◦ 667302 (CoCA) and and from the Innovative Medicines Initiative (IMI) 2 Joint Undertaking (H2020/EFPIA) under grant agreement no. 115916 (PRISM). The work was also supported by grants for the ENIGMA Consortium (Foundation for the National Institutes of Health (NIH); grant number U54 EB020403) from the BD2K Initiative of a cross-NIH partnership.

## BONN

The authors would like to thank (in alphabetical order) Marcel Bartling, Ulrike Broicher, Laura Ehrmantraut, Anna Maaser, Bettina Mahlow, Stephanie Mentges, Karolina Raczka, Laura Schinabeck, and Peter Trautner for their support and help. The study was partly funded by the Frankfurt Institute for Risk Management and Regulation (FIRM) and BW was supported by a Heisenberg Grant of the German Research Foundation ((Deutsche Forschungsgemeinschaft (DFG)), WE 4427 (3-2)).

## BrainScale

This work was supported by Nederlandse Organisatie voor Wetenschappelijk Onderzoek (NWO 51.02.061 to H.H., NWO 51.02.062 to D.B., NWO-NIHC Programs of excellence 433-09-220 to H.H., NWO-MagW 480-04-004 to D.B., and NWO/SPI 56-464-14192 to D.B.); FP7 Ideas: European Research Council (ERC-230374 to D.B.); and Universiteit Utrecht (High Potential Grant to H.H.).

## CARDIFF

We are grateful to all researchers within Cardiff University who contributed to the MBBrains panel and to NCMH for their support with genotyping. NCMH supported genotyping.

## CHARGE

Infrastructure for the CHARGE Consortium is supported in part by the National Heart, Lung, and Blood Institute grant HL105756 and for the neuroCHARGE phenotype working group through the National Institute on Aging grant AG033193. *Atherosclerosis Risk in Communities Study (ARIC):* The Atherosclerosis Risk in Communities study was performed as a collaborative study supported by National Heart, Lung, and Blood Institute (NHLBI) contracts (HHSN268201100005C, HSN268201100006C, HSN268201100007C, HHSN268201100008C, HHSN268201100009C, HHSN268201100010C, HHSN268201100011C, and HHSN268201100012C), R01HL70825, R01HL087641, R01HL59367, and R01HL086694; National Human Genome Research Institute contract U01HG004402; and National Institutes of Health (NIH) contract HHSN268200625226C. Infrastructure was partly supported by grant No. UL1RR025005, a component of the NIH and NIH Roadmap for Medical Research. This project was partially supported by National Institutes of Health R01 grants HL084099 and NS087541 to MF. *Austrian Stroke Prevention Family (ASPS) / Austrian Stroke Prevention Family Study:* The authors thank the staff and the participants for their valuable contributions. We thank Birgit Reinhart for her long-term administrative commitment, Elfi Hofer for the technical assistance at creating the DNA bank, Ing. Johann Semmler and Anita Harb for DNA sequencing and DNA analyses by TaqMan assays and Irmgard Poelzl for supervising the quality management processes after ISO9001 at the biobanking and DNA analyses. The Medical University of Graz and the Steiermärkische Krankenanstaltengesellschaft support the databank of the ASPS/ASPS-Fam. The research reported in this article was funded by the Austrian Science Fund (FWF) grant numbers PI904, P20545-P05 and P13180 and supported by the Austrian National Bank Anniversary Fund, P15435 and the Austrian Ministry of Science under the aegis of the EU Joint Programme-Neurodegenerative Disease Research (JPND)-www.jpnd.eu. *Cardiovascular Health Study (CHS):* This CHS research was supported by NHLBI contracts HHSN268201200036C, HHSN268200800007C, N01HC55222, N01HC85079, N01HC85080, N01HC85081, N01HC85082, N01HC85083, N01HC85086; and NHLBI grants U01HL080295, R01HL087652, R01HL105756, R01HL103612, R01HL120393, and R01HL130114 with additional contribution from the National Institute of Neurological Disorders and Stroke (NINDS). Additional support was provided through R01AG023629, R01AG15928, and R01AG033193 from the National Institute on Aging (NIA). A full list of principal CHS investigators and institutions can be found at CHS-NHLBI.org. The provision of genotyping data was supported in part by the National Center for Advancing Translational Sciences, CTSI grant UL1TR000124, and the National Institute of Diabetes and Digestive and Kidney Disease Diabetes Research Center (DRC) grant DK063491 to the Southern California Diabetes Endocrinology Research Center. The content is solely the responsibility of the authors and does not necessarily represent the official views of the National Institutes of Health. *Erasmus Rucphen Family Study (ERF):* Erasmus Rucphen Family (ERF) was supported by the Consortium for Systems Biology (NCSB), both within the framework of the Netherlands Genomics Initiative (NGI)/Netherlands Organisation for Scientific Research (NWO). ERF study as a part of EUROSPAN (European Special Populations Research Network) was supported by European Commission FP6 STRP grant number 018947 (LSHG-CT-2006-01947) and also received funding from the European Community’s Seventh Framework Programme (FP7/2007-2013)/grant agreement HEALTH-F4-2007-201413 by the European Commission under the programme “Quality of Life and Management of the Living Resources” of 5th Framework Programme (No. QLG2-CT-2002-01254) as well as FP7 project EUROHEADPAIN (nr602633). High-throughput analysis of the ERF data was supported by joint grant from Netherlands Organisation for Scientific Research and the Russian Foundation for Basic Research (NWO-RFBR 047.017.043). High throughput metabolomics measurements of the ERF study has been supported by BBMRI-NL (Biobanking and Biomolecular Resources Research Infrastructure Netherlands). *Framingham Heart Study (FHS):* This work was supported by the National Heart, Lung and Blood Institute’s Framingham Heart Study (Contract No. N01-HC-25195 and No. HHSN268201500001I) and its contract with Affymetrix, Inc. for genotyping services (Contract No. N02-HL-6-4278). A portion of this research utilized the Linux Cluster for Genetic Analysis (LinGA-II) funded by the Robert Dawson Evans Endowment of the Department of Medicine at Boston University School of Medicine and Boston Medical Center. This study was also supported by grants from the National Institute of Aging (R01s AG033040, AG033193, AG054076, AG049607, AG008122, AG016495; and U01-AG049505) and the National Institute of Neurological Disorders and Stroke (R01-NS017950). We would like to thank the dedication of the Framingham Study participants, as well as the Framingham Study team, especially investigators and staff from the Neurology group, for their contributions to data collection. Dr. DeCarli is supported by the Alzheimer’s Disease Center (P30 AG 010129). The views expressed in this manuscript are those of the authors and do not necessarily represent the views of the National Heart, Lung, and Blood Institute; the National Institutes of Health; or the U.S. Department of Health and Human Services. *Lothian Birth Cohort 1936 (LBC1936):* This project is funded by the Age UK’s Disconnected Mind programme (http://www.disconnectedmind.ed.ac.uk) and also by Research Into Ageing (Refs. 251 and 285). The whole genome association part of the study was funded by the Biotechnology and Biological Sciences Research Council (BBSRC; Ref. BB/F019394/1). Analysis of the brain images was funded by the Medical Research Council Grants G1001401 and 8200. The imaging was performed at the Brain Research Imaging Centre, The University of Edinburgh (http://www.bric.ed.ac.uk), a centre in the SINAPSE Collaboration (http://www.sinapse.ac.uk). The work was undertaken by The University of Edinburgh Centre for Cognitive Ageing and Cognitive Epidemiology (http://www.ccace.ed.ac.uk), part of the cross council Lifelong Health and Wellbeing Initiative (Ref. G0700704/84698). Funding from the BBSRC, Engineering and Physical Sciences Research Council (EPSRC), Economic and Social Research Council (ESRC), Medical Research Council (MRC) and Scottish Funding Council through the SINAPSE Collaboration is gratefully acknowledged. We thank the LBC1936 participants and research team members. We also thank the nurses and staff at the Wellcome Trust Clinical Research Facility (http://www.wtcrf.ed.ac.uk), where subjects were tested and the genotyping was performed. *LIFE-Adult:* LIFE-Adult is funded by the Leipzig Research Center for Civilization Diseases (LIFE). LIFE is an organizational unit affiliated to the Medical Faculty of the University of Leipzig. LIFE is funded by means of the European Union, by the European Regional Development Fund (ERDF) and by funds of the Free State of Saxony within the framework of the excellence initiative. This work was also funded by the Deutsche Forschungsgemeinschaft (Grant Number: CRC 1052 “Obesity mechanisms” project A1 to AV) and by the Max Planck Society. *Sydney Memory and Ageing Study (MAS):* MAS is funded by the Australian National Health and Medical Research Council (NHMRC)/Australian Research Council Strategic Award (Grant 401162), NHMRC Project grant 1405325. We would like to gratefully acknowledge and thank the Sydney MAS participants and supporters and the Sydney MAS Research Team. *Older Australian Twin Study (OATS):* OATS is funded by the Australian National Health and Medical Research Council (NHMRC)/Australian Research Council Strategic Award (Grant 401162), NHMRC Program Grants (350833, 568969, 109308) We would like thank and gratefully acknowledge the OATS participants, their supporters and the OATS Research Team. *Rotterdam Study (RSI, RSII, RSIII):* The Rotterdam Study is funded by Erasmus Medical Center and Erasmus University, Rotterdam, Netherlands Organization for the Health Research and Development (ZonMw), the Research Institute for Diseases in the Elderly (RIDE), the Ministry of Education, Culture and Science, the Ministry for Health, Welfare and Sports, the European Commission (DG XII), and the Municipality of Rotterdam. The authors are grateful to the study participants, the staff from the Rotterdam Study and the participating general practitioners and pharmacists. The generation and management of GWAS genotype data for the Rotterdam Study (RS I, RS II, RS III) were executed by the Human Genotyping Facility of the Genetic Laboratory of the Department of Internal Medicine, Erasmus MC, Rotterdam, The Netherlands. The GWAS datasets are supported by the Netherlands Organisation of Scientific Research NWO Investments (nr. 175.010.2005.011, 911-03-012), the Genetic Laboratory of the Department of Internal Medicine, Erasmus MC, the Research Institute for Diseases in the Elderly (014-93-015; RIDE2), the Netherlands Genomics Initiative (NGI)/Netherlands Organisation for Scientific Research (NWO) Netherlands Consortium for Healthy Aging (NCHA), project nr. 050-060-810. We thank Pascal Arp, Mila Jhamai, Marijn Verkerk, Lizbeth Herrera and Marjolein Peters, and Carolina Medina-Gomez, for their help in creating the GWAS database, and Karol Estrada, Yurii Aulchenko, and Carolina Medina-Gomez, for the creation and analysis of imputed data. This work has been performed as part of the CoSTREAM project (www.costream.eu) and has received funding from the European Union’s Horizon 2020 research and innovation programme under grant agreement No 667375. *Study of Health in Pomerania (SHIP) / Study of Health in Pomerania Trend (SHIP-Trend):* SHIP is part of the Community Medicine Research net of the University of Greifswald, Germany, which is funded by the Federal Ministry of Education and Research (grants no. 01ZZ9603, 01ZZ0103, and 01ZZ0403), the Ministry of Cultural Affairs as well as the Social Ministry of the Federal State of Mecklenburg-West Pomerania, and the network ‘Greifswald Approach to Individualized Medicine (GANI_MED)’ funded by the Federal Ministry of Education and Research (grant 03IS2061A). Genome-wide data have been supported by the Federal Ministry of Education and Research (grant no. 03ZIK012) and a joint grant from Siemens Healthineers, Erlangen, Germany and the Federal State of Mecklenburg-West Pomerania. Whole-body MR imaging was supported by a joint grant from Siemens Healthineers, Erlangen, Germany and the Federal State of Mecklenburg West Pomerania. The University of Greifswald is a member of the Caché Campus program of the InterSystems GmbH. *Saguenay Youth Study (SYS):* The Saguenay Youth Study has been funded by the Canadian Institutes of Health Research (TP, ZP), Heart and Stroke Foundation of Canada (ZP), and the Canadian Foundation for Innovation (ZP). We thank all families who took part in the Saguenay Youth Study. SYS is supported by the Canadian Institutes of Health Research: NET54015, NRF86678, TMH109788. *Three-City Dijon (3C-Dijon):* The Three City (3C) Study is conducted under a partnership agreement among the Institut National de la Santé et de la Recherche Médicale (INSERM), the University of Bordeaux, and Sanofi-Aventis. The Fondation pour la Recherche Médicale funded the preparation and initiation of the study. The 3C Study is also supported by the Caisse Nationale Maladie des Travailleurs Salariés, Direction Générale de la Santé, Mutuelle Générale de l’Education Nationale (MGEN), Institut de la Longévité, Conseils Régionaux of Aquitaine and Bourgogne, Fondation de France, and Ministry of Research–INSERM Programme “Cohortes et collections de données biologiques.” Christophe Tzourio and Stéphanie Debette have received investigator-initiated research funding from the French National Research Agency (ANR) and from the Fondation Leducq. Stéphanie Debette is supported by a starting grant from the European Research Council (SEGWAY) and a grant from the Joint Programme of Neurodegenerative Disease research (BRIDGET), from the European Union’s Horizon 2020 research and innovation programme under grant agreements No 643417 & No 640643, and by the Initiative of Excellence of Bordeaux University. We thank Dr. Anne Boland (CNG) for her technical help in preparing the DNA samples for analyses. This work was supported by the National Foundation for Alzheimer’s disease and related disorders, the Institut Pasteur de Lille, the labex DISTALZ and the Centre National de Génotypage. *Vietnam Era Twin Study of Aging (VETSA):* United States National Institute of Health VA San Diego Center of Excellence for Stress and Mental Health R00DA023549; DA-18673; NIA R01 AG018384; R01 AG018386; R01 AG022381; R01 AG022982; R01 DA025109 05; R01 HD050735; K08 AG047903; R03 AG 046413; U54 EB020403; and R01 HD050735-01A2

## DNS

We thank the Duke Neurogenetics Study participants and the staff of the Laboratory of NeuroGenetics. The Duke Neurogenetics Study received support from Duke University as well as US-National Institutes of Health grants R01DA033369 and R01DA031579.

## EPIGEN

Work from the London Cohort was supported by research grants from the Wellcome Trust (grant 084730 to S.M.S.), University College London (UCL)/University College London Hospitals (UCLH) NIHR Biomedical Research Centre/Specialist Biomedical Research Centres (CBRC/SBRC) (grant 114 to S.M.S.), the Comprehensive Local Research Network (CLRN) Flexibility and Sustainability Funding (FSF) (grant CEL1300 to S.M.S.), The Big Lottery Fund, the Wolfson Trust and the Epilepsy Society. This work was partly undertaken at UCLH/UCL, which received a proportion of funding from the UK Department of Health’s NIHR Biomedical Research Centres funding scheme.

## FBIRN

We are thankful to Mrs. Liv McMillan, BS for overall study coordination, Harry Mangalam, PhD, Joseph Farran, BS, and Adam Brenner, BS, for administering the University of California, Irvine High-Performance Computing cluster, and to the research subjects for their participation. This work was supported by the National Center for Research Resources at the National Institutes of Health [grant numbers: NIH 1 U24 RR021992 (Function Biomedical Informatics Research Network), NIH 1 U24 RR025736-01 (Biomedical Informatics Research Network Coordinating Center)], the National Center for Research Resources and the National Center for Advancing Translational Sciences, National Institutes of Health, through Grant UL1 TR000153, and the National Institutes of Health through 5R01MH094524, and P20GM103472.

## FOR2107

This work was funded by the German Research Foundation (DFG, grant FOR2107 DA1151/5-1 and DA1151/5-2 to UD; JA1890/7-1, JA1890/7-2 to AJ; KI 588/14-1, KI 588/14-2 to TK; KR 3822/7-1, KR 3822/7-2 to AK; NO246/10-1, NO246/10-2 to MMN).

## Frontal-temporal dementia GWAS

(utlised to calculate the genetic correlations). We acknowledge the investigators of the original study (Ferrari et al.(*75*)): Raffaele Ferrari, Dena G Hernandez, Michael A Nalls, Jonathan D Rohrer, Adaikalavan Ramasamy, John BJ Kwok, Carol Dobson-Stone, William S Brooks, Peter R Schofield, Glenda M Halliday, John R Hodges, Olivier Piguet, Lauren Bartley, Elizabeth Thompson, Eric Haan, Isabel Hernández, Agustín Ruiz, Mercè Boada, Barbara Borroni, Alessandro Padovani, Carlos Cruchaga, Nigel J Cairns, Luisa Benussi, Giuliano Binetti, Roberta Ghidoni, Gianluigi Forloni, Diego Albani, Daniela Galimberti, Chiara Fenoglio, Maria Serpente, Elio Scarpini, Jordi Clarimón, Alberto Lleó, Rafael Blesa, Maria Landqvist Waldö, Karin Nilsson, Christer Nilsson, Ian RA Mackenzie, Ging-Yuek R Hsiung, David MA Mann, Jordan Grafman, Christopher M Morris, Johannes Attems, Ian G McKeith, Alan J Thomas, Pietro Pietrini, Edward D Huey, Eric M Wassermann, Atik Baborie, Evelyn Jaros, Michael C Tierney, Pau Pastor, Cristina Razquin, Sara Ortega-Cubero, Elena Alonso, Robert Perneczky, Janine Diehl-Schmid, Panagiotis Alexopoulos, Alexander Kurz, Innocenzo Rainero, Elisa Rubino, Lorenzo Pinessi, Ekaterina Rogaeva, Peter St George-Hyslop, Giacomina Rossi, Fabrizio Tagliavini, Giorgio Giaccone, James B Rowe, Johannes CM Schlachetzki, James Uphill, John Collinge, Simon Mead, Adrian Danek, Vivianna M Van Deerlin, Murray Grossman, John Q Trojanowski, Julie van der Zee, Marc Cruts, Christine Van Broeckhoven, Stefano F Cappa, Isabelle Leber, Didier Hannequin, Véronique Golfier, Martine Vercelletto, Alexis Brice, Benedetta Nacmias, Sandro Sorbi, Silvia Bagnoli, Irene Piaceri, Jørgen E Nielsen, Lena E Hjermind, Matthias Riemenschneider, Manuel Mayhaus, Bernd Ibach, Gilles Gasparoni, Sabrina Pichler, Wei Gu, Martin N Rossor, Nick C Fox, Jason D Warren, Maria Grazia Spillantini, Huw R Morris, Patrizia Rizzu, Peter Heutink, Julie S Snowden, Sara Rollinson, Anna Richardson, Alexander Gerhard, Amalia C Bruni, Raffaele Maletta, Francesca Frangipane, Chiara Cupidi, Livia Bernardi, Maria Anfossi, Maura Gallo, Maria Elena Conidi, Nicoletta Smirne, Rosa Rademakers, Matt Baker, Dennis W Dickson, Neill R Graff-Radford, Ronald C Petersen, David Knopman, Keith A Josephs, Bradley F Boeve, Joseph E Parisi, William W Seeley, Bruce L Miller, Anna M Karydas, Howard Rosen, John C van Swieten, Elise GP Dopper, Harro Seelaar, Yolande AL Pijnenburg, Philip Scheltens, Giancarlo Logroscino, Rosa Capozzo, Valeria Novelli, Annibale A Puca, Massimo Franceschi, Alfredo Postiglione, Graziella Milan, Paolo Sorrentino, Mark Kristiansen, Huei-Hsin Chiang, Caroline Graff, Florence Pasquier, Adeline Rollin, Vincent Deramecourt, Thibaud Lebouvier, Dimitrios Kapogiannis, Luigi Ferrucci, Stuart Pickering-Brown, Andrew B Singleton, John Hardy, Parastoo Momeni. Acknowledgement: Intramural funding from the National Institute of Neurological Disorders and Stroke (NINDS) and National Institute on Aging (NIA), the Wellcome/MRC Centre on Parkinson’s disease, Alzheimer’s Research UK (ARUK, Grant ARUK-PG2012-18) and by the office of the Dean of the School of Medicine, Department of Internal Medicine, at Texas Tech University Health Sciences Center. We thank Mike Hubank and Kerra Pearce at the Genomic core facility at the Institute of Child Health (ICH), University College of London (UCL), for assisting RF in performing Illumina genotyping experiments (FTD-GWAS genotyping). This study utilized the high-performance computational capabilities of the Biowulf Linux cluster at the National Institutes of Health, Bethesda, Md. (http://biowulf.nih.gov). North American Brain Expression Consortium (NABEC) - The work performed by the North American Brain Expression Consortium (NABEC) was supported in part by the Intramural Research Program of the National Institute on Aging, National Institutes of Health, part of the US Department of Health and Human Services; project number ZIA AG000932-04. In addition this work was supported by a Research Grant from the Department of Defense, W81XWH-09-2-0128. UK Brain Expression Consortium (UKBEC) - This work performed by the UK Brain Expression Consortium (UKBEC) was supported by the MRC through the MRC Sudden Death Brain Bank (C.S.), by a Project Grant (G0901254 to J.H. and M.W.) and by a Fellowship award (G0802462 to M.R.). D.T. was supported by the King Faisal Specialist Hospital and Research Centre, Saudi Arabia. Computing facilities used at King’s College London were supported by the National Institute for Health Research (NIHR) Biomedical Research Centre based at Guy’s and St Thomas’ NHS Foundation Trust and King’s College London. We would like to thank AROS Applied Biotechnology AS company laboratories and Affymetrix for their valuable input. RF’s work is supported by Alzheimer’s Society (grant number 284), UK; JBJK was supported by the National Health and Medical Resarch Council (NHMRC) Australia, Project Grants 510217 and 1005769; CDS was supported by NHMRC Project Grants 630428 and 1005769; PRS was supported by NHMRC Project Grants 510217 and 1005769 and acknowledges that DNA samples were prepared by Genetic Repositories Australia, supported by NHMRC Enabling Grant 401184; GMH was supported by NHMRC Research Fellowship 630434, Project Grant 1029538, Program Grant 1037746; JRH was supported by the Australian Research Council Federation Fellowship, NHMRC Project Grant 1029538, NHMRC Program Grant 1037746; OP was supported by NHMRC Career Development Fellowship 1022684, Project Grant 1003139. IH, AR and MB acknowledge the patients and controls who participated in this project and the Trinitat Port-Carbó and her family who are supporting Fundació ACE research programs. CC was supported by Grant P30- NS069329-01 and acknowledges that the recruitment and clinical characterization of research participants at Washington University were supported by NIH P50 AG05681, P01 AG03991, and P01 AG026276. LB and GB were supported by the Ricerca Corrente, Italian Ministry of Health; RG was supported by Fondazione CARIPLO 2009-2633, Ricerca Corrente, Italian Ministry of Health; GF was supported by Fondazione CARIPLO 2009-2633. ES was supported by the Italian Ministry of Health; CF was supported by Fondazione Cariplo; MS was supported from the Italian Ministry of Health (Ricerca Corrente); MLW was supported by Government funding of clinical research within NHS Sweden (ALF); KN was supported by Thure Carlsson Foundation; CN was supported by Swedish Alzheimer Fund. IRAM and GYRH were supported by CIHR (grant 74580) PARF (grant C06-01). JG was supported by the NINDS intramural research funds for FTD research. CMM was supported by Medical Research Council UK, Brains for Dementia Research, Alzheimer’s Society, Alzheimer’s Research UK, National Institutes for Health Research, Department of Health, Yvonne Mairy Bequest and acknowledges that tissue made available for this study was provided by the Newcastle Brain Tissue Resource, which was funded in part by grants G0400074 and G1100540 from the UK MRC, the Alzheimer’s Research Trust and Alzheimer’s Society through the Brains for Dementia Research Initiative and an NIHR Biomedical Research Centre Grant in Ageing and Health, and NIHR Biomedical Research Unit in Lewy Body Disorders. CMM was supported by the UK Department of Health and Medical Research Council and the Research was supported by the National Institute for Health Research Newcastle Biomedical Research Centre based at Newcastle Hospitals Foundation Trust and Newcastle University and acknowledges that the views expressed are those of the authors and not necessarily those of the NHS, the NIHR or the Department of Health; JA was supported by MRC, Dunhill Medical Trust, Alzheimer’s Research UK; TDG was supported by Wellcome Trust Senior Clinical Fellow; IGM was supported by NIHR Biomedical Research Centre and Unit on Ageing Grants and acknowledges the National Institute for Health Research Newcastle Biomedical Research Centre based at Newcastle Hospitals Foundation Trust and Newcastle University. The views expressed are those of the author (s) and not necessarily those of the NHS, the NIHR or the Department of Health; AJT was supported by Medical Research Council, Alzheimer’s Society, Alzheimer’s Research UK, National Institutes for Health Research. EJ was supported by NIHR, Newcastle Biomedical Research Centre. PP, CR, SOC and EA were supported partially by FIMA (Foundation for Applied Medical Research); PP acknowledges Manuel Seijo-Martínez (Department of Neurology, Hospital do Salnés, Pontevedra, Spain), Ramon Rene, Jordi Gascon and Jaume Campdelacreu (Department of Neurology, Hospital de Bellvitge, Barcelona, Spain) for providing FTD DNA samples. RP, JDS, PA and AK were supported by German Federal Ministry of Education and Research (BMBF; grant number FKZ 01GI1007A – German FTLD consortium). IR was supported by Ministero dell’Istruzione, dell’Università e della Ricerca (MIUR) of Italy. PStGH was supported by the Canadian Institutes of Health Research, Wellcome Trust, Ontario Research Fund. FT was supported by the Italian Ministry of Health (ricerca corrente) and MIUR grant RBAP11FRE9; GR and GG were supported by the Italian Ministry of Health (ricerca corrente). JBR was supported by Camrbidge NIHR Biomedical Research Centre and Wellcome Trust (088324). JU, JC, SM were supported by the MRC Prion Unit core funding and acknowledge MRC UK, UCLH Biomedical Research Centre, Queen Square Dementia BRU; SM acknowledges the work of John Beck, Tracy Campbell, Gary Adamson, Ron Druyeh, Jessica Lowe, Mark Poulter. AD acknowledges the work of Benedikt Bader and of Manuela Neumann, Sigrun Roeber, Thomas Arzberger and Hans Kretzschmar†; VMVD and JQT were supported by Grants AG032953, AG017586 and AG010124; MG was supported by Grants AG032953, AG017586, AG010124 and NS044266; VMVD acknowledges EunRan Suh, PhD for assistance with sample handling and Elisabeth McCarty-Wood for help in selection of cases; JQT acknowledges Terry Schuck, John Robinson and Kevin Raible for assistance with neuropathological evaluation of cases. CVB and the Antwerp site were in part funded by the MetLife Foundation for Medical Research Award (to CVB), the Belgian Science Policy Office (BELSPO) Interuniversity Attraction Poles program; the Alzheimer Research Foundation (SAO-FRA); the Medical Foundation Queen Elisabeth (GSKE); the Flemish Government initiated Methusalem Excellence Program (to CVB); the Research Foundation Flanders (FWO) and the University of Antwerp Research Fund.. CVB, MC and JvdZ acknowledge the neurologists S Engelborghs, PP De Deyn, A Sieben, R Vandenberghe and the neuropathologist JJ Martin for the clinical and pathological diagnoses. CVB, MC and JvdZ further thank the personnel of the Genetic Service Facility of the VIB Department of Molecular Genetics (http://www.vibgeneticservicefacility.be) and the Antwerp Biobank of the Institute Born-Bunge for their expert support. IL and AB were supported by the program “Investissements d’avenir” ANR-10-IAIHU-06 and acknowledges the contribution of The French research network on FTLD/FTLD-ALS for the contribution in samples collection. BN is founded by Fondazione Cassa di Risparmio di Pistoia e Pescia (grant 2014.0365), SS is founded by the Cassa di Risparmio di Firenze (grant 2014.0310) and a grant from Ministry of Health n° RF-2010-2319722. JEN was supported by the Novo Nordisk Foundation, Denmark. MR was supported by the German National Genome Network (NGFN); German Ministry for Education and Research Grant Number 01GS0465. JDR, MNR, NCF and JDW were supported by an MRC programme grant and the Dementia Platform UK, the NIHR Queen Square Dementia Biomedical Research Unit (BRU) and the Leonard Wolfson Experimental Neurology Centre. MGS was supported by MRC grant n G0301152, Cambridge Biomedical Research Centre and acknowledges Mrs K Westmore for extracting DNA. HM was supported by the Motor Neuron Disease Association (Grant 6057). RR was supported by P50 AG016574, R01 NS080882, R01 NS065782, P50 NS72187 and the Consortium for Frontotemporal Dementia; DWD was supported by P50NS072187, P50AG016574, State of Florida Alzheimer Disease Initiative, & CurePSP, Inc.; NRGR, JEP, RCP, DK, BFB were supported by P50 AG016574; KAJ was supported by R01 AG037491; WWS was supported by NIH AG023501, AG019724, Consortium for Frontotemporal Dementia Research; BLM was supported by P50AG023501, P01AG019724, Consortium for FTD Research; HR was supported by AG032306. JCvS was supported by Stichting Dioraphte Foundation (11 02 03 00), Nuts Ohra Foundation (0801-69), Hersenstichting Nederland (BG 2010-02) and Alzheimer Nederland. CG and HHC acknowledge families, patients, clinicians including Dr Inger Nennesmo and Dr Vesna Jelic, Professor Laura Fratiglioni for control samples and Jenny Björkström, Håkan Thonberg, Charlotte Forsell, Anna-Karin Lindström and Lena Lilius for sample handling. CG was supported by Swedish Brain Power (SBP), the Strategic Research Programme in Neuroscience at Karolinska Institutet (StratNeuro), the regional agreement on medical training and clinical research (ALF) between Stockholm County Council and Karolinska Institutet, Swedish Alzheimer Foundation, Swedish Research Council, Karolinska Institutet PhD-student funding, King Gustaf V and Queen Victoria’s Free Mason Foundation. FP, AR, VD and FL acknowledge Labex DISTALZ. RF acknowledges the help and support of Mrs. June Howard at the Texas Tech University Health Sciences Center Office of Sponsored Programs for tremendous help in managing Material Transfer Agreement at TTUHSC.

## GIG

The GIG (Genomic Imaging Göttingen) sample was established at the Center for Translational Research in Systems Neuroscience and Psychiatry (Head: Prof. Dr. O. Gruber) at Göttingen University. We thank Maria Keil, Esther Diekhof, Tobias Melcher and Ilona Henseler for assistance in data acquisition. We are grateful to all persons who kindly participated in the GIG study.

## GOBS

The GOBS study (PI DG and JB) was supported by the National Institute of Mental Health Grants MH0708143 (Principal Investigator [PI]: DCG), MH078111 (PI: JB), and MH083824 (PI: DCG & JB).

## GSP

Brain Genomics Superstruct Project (GSP): Data were provided [in part] by the Brain GSP of Harvard University and the Massachusetts General Hospital, with support from the Center for BrainScience Neuroinformatics Research Group, the Athinoula A. Martinos Center for Biomedical Imaging and the Center for Human Genetic Research. Twenty individual investigators at Harvard and Massachusetts General Hospital generously contributed data to GSP. This work was made possible by the resources provided through Shared Instrumentation Grants 1S10RR023043 and 1S10RR023401 and was supported by funding from the Simons Foundation (RLB), the Howard Hughes Medical Institute (RLB), NIMH grants R01-MH079799 (JWS), K24MH094614 (JWS), K01MH099232 (AJH), and the Massachusetts General Hospital-University of Southern California Human Connectome Project (U54MH091665).

## HUBIN

This work was supported by the Swedish Research Council (2006-2992, 2006-986, K2007-62X-15077-04-1, K2008-62P-20597-01-3, 2008-2167, 2008-7573, K2010-62X-15078-07-2, K2012-61X-15078-09-3, 14266-01A,02-03, 2017-949), the regional agreement on medical training and clinical research between Stockholm County Council and the Karolinska Institutet, the Knut and Alice Wallenberg Foundation, and the HUBIN project.

## HUNT

The HUNT Study is a collaboration between HUNT Research Centre (Faculty of Medicine and Movement Sciences, NTNU – Norwegian University of Science and Technology), Nord-Trøndelag County Council, Central Norway Health Authority, and the Norwegian Institute of Public Health. HUNT-MRI was funded by the Liaison Committee between the Central Norway Regional Health Authority and the Norwegian University of Science and Technology, and the Norwegian National Advisory Unit for functional MRI.

## IMAGEN

This work received support from the following sources: the European Union-funded FP6 Integrated Project IMAGEN (Reinforcement-related behaviour in normal brain function and psychopathology) (LSHM-CT-2007-037286), the Horizon 2020 funded ERC Advanced Grant ‘STRATIFY’ (Brain network based stratification of reinforcement-related disorders) (695313), ERANID (Understanding the Interplay between Cultural, Biological and Subjective Factors in Drug Use Pathways) (PR-ST-0416-10004), BRIDGET (JPND: BRain Imaging, cognition Dementia and next generation GEnomics) (MR/N027558/1), the FP7 projects IMAGEMEND (602450; IMAging GEnetics for MENtal Disorders) and MATRICS (603016), the Innovative Medicine Initiative Project EU-AIMS (115300-2), the Medical Research Foundation and Medical Research Council grant MR/R00465X/1; the Medical Research Council Grant ‘c-VEDA’ (Consortium on Vulnerability to Externalizing Disorders and Addictions) (MR/N000390/1), the Swedish Research Council FORMAS, the Medical Research Council, the National Institute for Health Research (NIHR) Biomedical Research Centre at South London and Maudsley NHS Foundation Trust and King’s College London, the Bundesministeriumfür Bildung und Forschung (BMBF grants 01GS08152; 01EV0711; eMED SysAlc01ZX1311A; Forschungsnetz AERIAL), the Deutsche Forschungsgemeinschaft (DFG grants SM 80/7-1, SM 80/7-2, SFB 940/1). Further support was provided by grants from: ANR (project AF12-NEUR0008-01 – WM2NA, and ANR-12-SAMA-0004), the Fondation de France, the Fondation pour la Recherche Médicale, the Mission Interministérielle de Lutte-contre-les-Drogues-et-les-Conduites-Addictives (MILDECA), the Fondation pour la Recherche Médicale (DPA20140629802), the Fondation de l’Avenir, Paris Sud University IDEX 2012; the National Institutes of Health, Science Foundation Ireland (16/ERCD/3797), U.S.A. (Axon, Testosterone and Mental Health during Adolescence; RO1 MH085772-01A1), and by NIH Consortium grant U54 EB020403, supported by a cross-NIH alliance that funds Big Data to Knowledge Centres of Excellence.

## IMH

We thank the participants, their carers and research support staff of this study. This work was supported by research grants from the National Healthcare Group, Singapore (SIG/05004; SIG/05028), and the Singapore Bioimaging Consortium (RP C-009/2006) research grants awarded to KS. ML is supported by an National Medical Research Council Research Training Fellowship (MH095: 003/008-1014); Singapore Ministry of Health National Medical Research Council Center Grant (NMRC/CG/004/2013)

## IMpACT

The International Multi-centre persistent ADHD CollaboraTion (IMpACT), is a consortium of clinical and basic researchers from several European countries (The Netherlands, Germany, Spain, Norway, The United Kingdom, Sweden), from the United States of America, and from Brazil. In the current study, the samples from the Netherlands node of IMpACT were used. This work was carried out on the Dutch national e-infrastructure with the support of SURF Cooperative. This works was supported by the Netherlands Organization for Scientific Research (Nederlandse Organisatie voor Wetenschappelijk Onderzoek, NWO), i.e., the NWO Brain & Cognition Excellence Program (grant 433-09-229) and the Vici Innovation Program (grant 016-130-669 to BF). Additional support was received from the European Community’s Seventh Framework Programme (FP7/2007–2013) under grant agreements n◦ 602805 (Aggressotype), n◦ 602450 (IMAGEMEND), and n◦ 278948 (TACTICS) as well as from the European Community’s Horizon 2020 Programme (H2020/2014–2020) under grant agreements n◦ 643051 (MiND), n◦ 667302 (CoCA), and n◦ 728018 (Eat2beNICE). The work was also supported by grants for the ENIGMA Consortium (Foundation for the National Institutes of Health (NIH); grant number U54 EB020403) from the BD2K Initiative of a cross-NIH partnership.

## LBC1936

We thank the participants and research support staff of this study. The work was undertaken as part of the Cross Council and University of Edinburgh Centre for Cognitive Ageing and Cognitive Epidemiology (CCACE; http://www.ccace.ed.ac.uk). This work was supported by a Research into Ageing programme grant (to I.J.D.) and the Age UK-funded Disconnected Mind project (http://www.disconnectedmind.ed.ac.uk; to I.J.D. and J.M.W.), with additional funding from the UK Medical Research Council (MRC Mr/M01311/1, G1001245/96077, G0701120/79365 to I.J.D., J.M.W. and M.E.B.). The whole genome association part of this study was funded by the Biotechnology and Biological Sciences Research Council (BBSRC; Ref. BB/F019394/1). J.M.W. is supported by the Scottish Funding Council through the SINAPSE Collaboration (http://www.sinapse.ac.uk). CCACE (MRC MR/K026992/1) is funded by the BBSRC and MRC. The image acquisition and analysis was performed at the Brain Research Imaging Centre, University of Edinburgh (http://www.bric.ed.ac.uk). MVH is supported by the Row Fogo Charitable Trust.

## LIBD

This work was supported by direct funding from the NIMH intramural research program of the NIH to the Weinberger Lab and by support from the LIeber Institute for Brain Development and the Maltz Research Laboratories.

## MCIC

The authors wish to thank our many colleagues who served as mentors, advisors and supporters during the inception and conduct of the study including Donald Goff, Gina Kuperberg, Jill Goldstein, Martha Shenton, Robert McCarley, Stephan Heckers, Cynthia Wible, Raquelle Mesholam-Gately, and Mark Vangel. We thank the study staff and clinicians at each site that were responsible for the data acquisition. These include: Stuart Wallace, Ann Cousins, Raquelle Mesholam-Gately, Steven Stufflebeam, Oliver Freudenreich, Daphne Holt, Laura Kunkel, Frank Fleming, George He, Hans Johnson, Ron Pierson, Arvind Caprihan, Phyllis Somers, Christine Portal, Kaila Norman, Diana South, Michael Doty and Haley Milner. We would also like to acknowledge the expert guidance on image and other types of data acquisition we obtained from Lee Friedman, Stephan Posse, Jorge Jovicich, and Tom Wassink. We would also like to acknowledge the many research assistants, students and colleagues who assisted in data curation over the years since data acquisition was completed. These include: Stuart Wallace, Carolyn Zyloney, Komal Sawlani, Jill Fries, Adam Scott, Dylan Wood, Runtang Wang, William Courtney, Angie Guimaraes, Lisa Shenkman, Mustafa Kendi, Aysa Tuba Karagulle Kendi, Ryan Muetzel, Tara Biehl, and Marcus Schmidt. This work was supported primarily by the Department of Energy DE-FG02-99ER62764 through its support of the Mind Research Network (MRN, formerly known as the MIND Institute) and the consortium as well as by the National Association for Research in Schizophrenia and Affective Disorders (NARSAD) Young Investigator Award (to SE) as well as through the Blowitz-Ridgeway and Essel Foundations and a ZonMw TOP 91211021 (to TW), the DFG research fellowship (to SE), the Mind Research Network, National Institutes of Health through NCRR 5MO1-RR001066 (MGH General Clinical Research Center), NIMH K08 MH068540, the Biomedical Informatics Research Network with NCRR Supplements to P41 RR14075 (MGH), M01 RR 01066 (MGH), NIBIB R01EB006841 (MRN), R01EB005846 (MRN), 2R01 EB000840 (MRN), 1RC1MH089257 (MRN), as well as grant U24 RR021992.

## Meth-CT

This study was supported by the Medical Research Council, South Africa.

## MIRECC

We thank the US military veterans who participated in this research. The Study was supported by National Institute of Mental Health (1R01MH111671) and the US Department of Veterans Affairs (VISN6 MIRECC).

## MooDS

This work was supported by the German Ministry for Education and Research (BMBF) grants (BMBF National Genome Research Network: NGFN-Plus MooDS “Systematic Investigation of the Molecular Causes of Major Mood Disorders and Schizophrenia”; see under http://www.ngfn.de/en/schizophrenie.html; e:Med Programme: Integrated Network IntegraMent, Integrated Understanding of Causes and Mechanisms in Mental Disorders, grant 01ZX1314A to MMN, grant 01ZX1614A to FD and MMN), and supported by grants from the German Research Foundation the German Research Foundation (Deutsche Forschungsgemeinschaft, DFG, FOR 1617 as well as Excellence Cluster Exc 257).

## MPIP

The MPIP Munich Morphometry Sample comprises images acquired as part of the Munich Antidepressant Response Signature (MARS) Study and the Recurrent Unipolar Depression (RUD) Case-Control study performed at the MPIP, and control subjects acquired at the Ludwig-Maximilians-University, Munich, Department of Psychiatry. We would like to acknowledge all patients and control subjects who have participated in these studies. We are grateful to Rosa Schirmer, Elke Schreiter, Reinhold Borschke, Ines Eidner and Anna Olynyik for supporting MR acquisition and data management. We thank the staff of the Center of Applied Genotyping for generating the genotypes of the MARS cohort. We further thank Dorothee P. Auer for initiating the RUD-MR substudy and Elisabeth Binder for supporting participation in ENIGMA. We are grateful to GlaxoSmithKline for providing the genotypes of the Recurrent Unipolar Depression Case-Control Sample. The study was supported by a grant of the Exzellenz-Stiftung of the Max Planck Society. This work has also been funded by the Federal Ministry of Education and Research (BMBF) in the framework of the National Genome Research Network (NGFN), FKZ 01GS0481.

## MPRC

Support was received from NIH grants U01MH108148, 2R01EB015611, R01DA027680, R01MH085646, P50MH103222, U54 EB020403, and T32MH067533, NSF grants IIS-1302755 and MRI-1531491, a State of Maryland contract (M00B6400091), and a Pfizer research grant.

## MÜNSTER

This work was funded by the German Research Foundation (SFB-TRR58, Projects C09 and Z02 to UD) and the Interdisciplinary Center for Clinical Research (IZKF) of the medical faculty of Münster (grant Dan3/012/17 to UD).

## NCNG

The study was supported by the Bergen Research Foundation (BFS), the University of Bergen, the Research Council of Norway (RCN) (including FUGE grant nos. 151904 and 183327, Psykisk Helse grant no. 175345, RCN grants 154313/V50 to I.R. and 177458/V50 to T.E.), Helse Sørøst RHF to T.E. (grant 2012086), and Dr Einar Martens Fund.

## NESDA

The infrastructure for the NESDA study (www.nesda.nl) is funded through the Geestkracht program of the Netherlands Organisation for Health Research and Development (Zon-Mw, grant number 10-000-1002) and is supported by participating universities (VU University Medical Center, GGZ inGeest, Arkin, Leiden University Medical Center, GGZ Rivierduinen, University Medical Center Groningen) and mental health care organizations.Funding was obtained from the Netherlands Organization for Scientific Research (Geestkracht program grant 10-000-1002); the Center for Medical Systems Biology (CSMB, NWO Genomics), Biobanking and Biomolecular Resources Research Infrastructure (BBMRI-NL), VU University’s Institutes for Health and Care Research (EMGO+) and Neuroscience Campus Amsterdam, University Medical Center Groningen, Leiden University Medical Center, National Institutes of Health (NIH, R01D0042157-01A, MH081802, Grand Opportunity grants 1RC2 MH089951 and 1RC2 MH089995). Part of the genotyping and analyses were funded by the Genetic Association Information Network (GAIN) of the Foundation for the National Institutes of Health.Computing was supported by BiG Grid, the Dutch e-Science Grid, which is financially supported by NWO.

## NeuroIMAGE

The NeuroIMAGE study was supported by NIH Grant R01MH62873 (to Stephen V. Faraone), NWO Large Investment Grant 1750102007010 (to Jan Buitelaar), ZonMW grant 60-60600-97-193, NWO grants 056-13-015 and 433-09-242, and matching grants from Radboud University Nijmegen Medical Center, University Medical Center Groningen and Accare, and Vrije Universiteit Amsterdam. Further support was received from the European Union FP7 programmes TACTICS (grant agreement 278948), IMAGEMEND (grant agreement 602450), Aggressotype (grant agreement 602805), CoCA (grant agreement 667302), and Eat2beNICE (grant agreement 728018).

## NTR

Netherlands Twin Register: Funding was obtained from the Netherlands Organization for Scientific Research (NWO) and The Netherlands Organisation for Health Research and Development (ZonMW) grants 904-61-090, 985-10-002, 912-10-020, 904-61-193,480-04-004, 463-06-001, 451-04-034, 400-05-717, Addiction-31160008, 016-115-035, 481-08-011, 056-32-010, Middelgroot-911-09-032, OCW_NWO Gravity program –024.001.003, NWO-Groot 480-15-001/674, Center for Medical Systems Biology (CSMB, NWO Genomics), NBIC/BioAssist/RK (2008.024), Biobanking and Biomolecular Resources Research Infrastructure (BBMRI –NL, 184.021.007 an 184.033.111); Spinozapremie (NWO-56-464-14192), KNAW Academy Professor Award (PAH/6635) and University Research Fellow grant (URF) to DIB; Amsterdam Public Health research institute (former EMGO+), Neuroscience Amsterdam research institute (former NCA); the European Science Foundation (ESF, EU/QLRT-2001-01254), the European Community’s Seventh Framework Program (FP7-HEALTH-F4-2007-2013, grant 01413: ENGAGE and grant 602768: ACTION); the European Research Council (ERC Advanced, 230374, ERC Starting grant 284167), Rutgers University Cell and DNA Repository (NIMH U24 MH068457-06), the National Institutes of Health (NIH, R01D0042157-01A1, R01MH58799-03, MH081802, DA018673, R01 DK092127-04, Grand Opportunity grants 1RC2 MH089951, and 1RC2 MH089995); the Avera Institute for Human Genetics, Sioux Falls, South Dakota (USA). Part of the genotyping and analyses were funded by the Genetic Association Information Network (GAIN) of the Foundation for the National Institutes of Health. Computing was supported by NWO through grant 2018/EW/00408559, BiG Grid, the Dutch e-Science Grid and SURFSARA.

## OATS

We would like to thank the participants and their supporters for their time and generosity. We acknowledge and thank the contributions of the OATS research team. OATS is supported by the Australian National Health and Medical Research Council (NHMRC)/Australian Research Council Strategic Award (Grant 401162) and the NHMRC Project grant 1405325. This study was facilitated through Twins Research Australia, a national resource in part supported by a Centre for Research Excellence from the NHMRC. DNA was extracted by Genetic Repositories Australia (NHMRC Grant 401184). Genome-wide genotyping was performed by the Diamantina Institute, University of Queensland. A CSIRO Flagship Collaboration Fund Grant partly funded the genotyping.

## OSAKA

This research was supported by AMED under Grant Number JP18dm0307002, JP18dm0207006 (Brain/MINDS) and JSPS KAKENHI Grant Number J16H05375.

## PAFIP

The authors wish to thank all PAFIP research team and all patients and family members who participated in the study. We wish to acknowledge IDIVAL Neuroimaging Unit for imaging acquirement and analysis. We thank Valdecilla Biobank for its help in the technical execution of this work. This work was supported by the Instituto de Salud Carlos III (PI14/00639 and PI14/00918), MINECO (SAF2010-20840-C02-02 and SAF2013-46292-R) and Fundación Instituto de Investigación Marqués de Valdecilla (NCT0235832 and NCT02534363). No pharmaceutical company has financially supported the study.

## PDNZ

We thank the patients who participated in this study, staff at the New Zealand Brain Research Institute and Pacific Radiology Christchurch for study co-ordination and image acquisition, and Ms Allison Miller for DNA preparation and banking. Neurological Foundation of New Zealand; Canterbury Medical Research Foundation; University of Otago Research Grant; Jim and Mary Carney Charitable Trust (Whangarei, New Zealand).

## PING

PING was supported by the National Institute on Drug Abuse (RC2DA029475) and the National Institute of Child Health and Human Development (R01HD061414) in the U.S.

## PPMI

Data used in the preparation of this article were obtained from the Parkinson’s Progression Markers Initiative (PPMI) database (www.ppmi-info.org/data). For up-to-date information on the study, visit www.ppmi-info.org. Parkinson’s Progression Markers Initiative, a public-private partnership, is funded by the Michael J. Fox Foundation for Parkinson’s Research and funding partners, including AbbVie, Allegran, Avid Radiopharmaceuticals, Biogen Idec, BioLegend, Bristol-Meyers Squibb, Denali Therapeutics, GE Healthcare, Genentech, GSK-GlaxoSmithKline, Eli Lilly & Co., F. Hoffman-La Roche Ltd., Lundbeck Pharmaceuticals, Merck and Company, MSD-Meso Scale Discovery, Pfizer, Piramal, Sanofi Genzyme, Servier, Takeda Pharmaceutical Company, TEVA Pharmaceutical Industries, UCB Pharma SA, and Golub Capital (http://www.ppmi-info.org/about-ppmi/who-we-are/study-sponsors/). Support for the image and genetic data analysis of PPMI was provided in part by MJFF grant number 14848 (to N.J.)

## QTIM

We thank the twins and singleton siblings who gave generously of their time to participate in the QTIM study. We also thank the many research assistants, radiographers, and IT support staff for data acquisition and DNA sample preparation. National Institute of Child Health & Human Development (R01 HD050735); National Institute of Biomedical Imaging and Bioengineering (Award 1U54EB020403-01, Subaward 56929223); National Health and Medical Research Council (Project Grants 496682, 1009064 and Medical Bioinformatics Genomics Proteomics Program 389891).

## SHIP and SHIP/TREND

SHIP is part of the Community Medicine Research net of the University of Greifswald, Germany, which is funded by the Federal Ministry of Education and Research (grants no. 01ZZ9603, 01ZZ0103, and 01ZZ0403), the Ministry of Cultural Affairs as well as the Social Ministry of the Federal State of Mecklenburg-West Pomerania, and the network ‘Greifswald Approach to Individualized Medicine (GANI_MED)’ funded by the Federal Ministry of Education and Research (grant 03IS2061A). Whole-body MR imaging was supported by a joint grant from Siemens Healthineers, Erlangen, Germany and the Federal State of Mecklenburg West Pomerania. Genome-wide data have been supported by the Federal Ministry of Education and Research (grant no. 03ZIK012) and a joint grant from Siemens Healthineers, Erlangen, Germany and the Federal State of Mecklenburg-West Pomerania. The University of Greifswald is a member of the Caché Campus program of the InterSystems GmbH. The SHIP authors are grateful to Mario Stanke for the opportunity to use his Server Cluster for the SNP imputation as well as to Holger Prokisch and Thomas Meitinger (Helmholtz Zentrum München) for the genotyping of the SHIP-Trend cohort.

## Sydney MAS

We would like to thank and acknowledge the generosity of our participants and their supporters in contributing to this study. We would also like to thank the Sydney MAS Research Team. Sydney MAS is supported by the National Health and Medical Research Council (NHMRC)/Australian Research Council Strategic Award (Grant 401162) and NHMRC Program Grants (350833, 568969). DNA was extracted by Genetic Repositories Australia (NHMRC Grant 401184). Genome-wide genotyping was performed by the Ramaciotti Centre, University of New South Wales.

## SYS

The SYS has been funded by the Canadian Institutes of Health Research and the Heart and Stroke Foundation of Canada. Computations were performed on the GPC supercomputer at the SciNet HPC Consortium. SciNet is funded by: the Canada Foundation for Innovation under the auspices of Compute Canada; the Government of Ontario; Ontario Research Fund - Research Excellence; and the University of Toronto.

## TCD-NUIG

Included data from two sites. *NUI Galway:* data collection was supported by the Health Research Board (HRA_POR/2011/100). *Trinity College Dublin:* this research was supported by The Science Foundation Ireland Research Investigator project, awarded to Gary Donohoe (SFI: 12.IP.1359).

## TOP and TOP3T

are part of TOP, which is supported by the Research Council of Norway (223273, 213837, 249711, 226971, 262656), the South East Norway Health Authority (2017-112), the Kristian Gerhard Jebsen Stiftelsen (SKGJ-MED-008) and the European Community’s Seventh Framework Programme (FP7/2007–2013), grant agreement no. 602450 (IMAGEMEND).

## UiO2016 and UiO2017

are part of TOP and STROKEMRI, which is supported by the Norwegian ExtraFoundation for Health and Rehabilitation (2015/FO5146), the Research Council of Norway (249795, 248238), and the South-Eastern Norway Regional Health Authority (2014097, 2015044, 2015073).

## UK Biobank

This research has been conducted using the UK Biobank Resource under Application Number ‘11559’.

## UMCU

The UMCU cohort consists of several independent studies, which were supported by The Netherlands Organisation for Health Research and Development (ZonMw) TOP 40-008-12-98-13009, Geestkracht programme of the Netherlands Organisation for Health Research and Development (ZonMw, grant number 10-000-1001), the Stanley Medical Research Institute (Dr. Nolen), the Brain and Behavior Research Foundation (2013-2015 NARSAD Independent Investigator grant number 20244 to M.H.J.H.), The Netherlands Organisation for Scientific Research (2012-2017 VIDI grant number 452-11-014 to N.E.M.H.), The Netherlands Organisation for Health Research and Development (ZonMw grant number 908-02-123 to H.E.H.), The Netherlands Organisation for Scientific Research (VIDI grant number 917-46-370 to H.E.H.).

## UNICAMP

Supported by FAPESP (São Paulo Research Foundation) grant #2013/07559-3: The Brazilian Institute of Neuroscience and Neurotechnology (BRAINN).

## Central Analysis and Coordination Group

C.R.K.C., D.P.H., F.P., J.Br., J.L.S., J.N.P., K.L.G., L.C.-C., L.C.P.Z., M.A.B.M., N.J., N.S., P.A.L., P.M.T., S.E.M.

## Manuscript Writing and Preparation

D.P.H., J.Br., J.L.S., J.N.P., K.L.G., L.C.-C., N.J., P.A.L., P.M.T., S.E.M.

## Project Support

D.Ga., M.A.B.M., M.J., N.S., R.E., V.R., Y.G.

## Cohort Principal Investigator

A.A.V., A.C., A.H., A.J.F., A.J.H., A.K.H., A.M.D., A.M.-L., A.R.H., A.W.T., B.C.-F., B.F., B.Mo., B.S.P., B.T.B., B.W., B.W.J.H.P., C.A.H., C.Dep., C.F., C.M., C.M.L., C.P., D.Am., D.C.G., D.I.B., D.J.S., D.P., D.R.W., D.v.E., E.G.J., E.J.C.d.G., E.L.H., F.A.H., F.C., G.D., G.F., G.G.B., G.L.C., G.S., H.B., H.E.H.P., H.F., H.G.B., H.J.G., H.V., H.W., I.A., I.E.Som., I.J.D., I.M., J.B.J.K., J.Bl., J.C.D.-A., J.K.B., J.-L.M., J.L.R., J.N.T., J.O., J.R.B., J.W.S., J.Z., K.L.M., K.S., L.M.R., L.N., L.R., L.T.W., M.E.B., M.H.J.H., M.J.C., M.J.W., M.K.M.A., M.R., N.D., N.J., N.J.A.v.d.W., O.A.A., O.G., P.G.S., P.J.H., P.K., P.M.T., P.S.S., P.T.M., R.A.M., R.A.O., R.H., R.J.S., R.L.B., R.L.G., R.S.K., S.Ca., S.Des., S.E.F., S.L.H., S.M.S., S.R., T.E., T.J.A., T.J.C.P., T.L.J., T.P., T.T.J.K., U.D., V.C., V.J.C., W.C., W.U.H., X.C., Z.P.

## Imaging Data Collection

A.B., A.d.B., A.F.M., A.J., A.J.H., A.K., A.K.H., A.L.G., A.M.D., A.N.H., A.P., A.R.H., A.R.K., A.U., B.A.M., B.-C.H., B.D., B.F., B.Pi., B.W., B.W.J.H.P., C.B., C.D.W., C.J., C.L.B., C.L.Y., C.M., C.P., C.R.J., C.S.Re., D.Am., D.C.G., D.Gr., D.H.M., D.J., D.J.H., D.J.V., D.M.C., D.P.O., D.R.W., D.S.O., D.T.-G., D.v.E., D.v.R., D.Z., E.A., E.B.Q., E.J.C.d.G., E.L.H., E.Sh., G.B.P., G.D., G.F., G.I.d.Z., G.L.C., G.R., G.S., H.V., H.Y., I.A., I.E.Som., J.A.T., J.E.C., J.E.N., J.K.B., J.-L.M., J.-L.M., J.L.R., J.M.F., J.M.W., J.N.T., J.R., J.T.V., K.D., K.K., K.L.M., K.O.L., K.S., L.M.R., L.R., L.T.W., M.B.H., M.E.B., M.Fu., M.H.J.H., M.Ho., M.-J.v.T., M.J.W., M.-L.P.M., N.E.M.v.H., N.F.H., N.H., N.J.A.v.d.W., N.K.H., N.O., O.G., P.A.G., P.E.R., P.G.S., P.K., P.N., P.S.S., R.A.O., R.B., R.H., R.L.B., R.L.G., R.R., R.S.K., R.W., S.A., S.C.M., S.Ca., S.Er., S.Ko., S.M., S.M.S., T.G.M.v.E., T.R.M., T.Wh., T.W.M., U.D., U.S., V.C., V.J.C., V.S.M., W.D.H., W.H., W.W., X.C.

## Imaging Data Analysis

A.F.M., A.H.Z., A.J.H., A.J.S., A.L.G., A.M.D., A.R., A.R.K., A.S., A.Th., A.U., B.A.G., B.C.R., B.F., B.K., B.S.P., C.B., C.C.F., C.C.H., C.D.W., C.J., C.L.Y., C.R.K.C., C.S.Ro., D.Al., D.C.G., D.Gr., D.H., D.J., D.J.H., D.M.C., D.P.H., D.P.O., D.T.-G., D.v.d.M., D.v.E., D.v.R., D.Z., E.E.L.B., E.Sh., E.Sp., E.W., F.M.R., F.P., F.S., G.I.d.Z., G.R., H.J.G., I.A.,I.E.Som., I.K.A., J.A.T., J.B.J.K., J.C.V.M., J.-L.M., J.L.R., J.L.S., J.M.W., J.R., J.Z., K.D., K.L.M., K.N., K.S., K.W., L.B.L., L.H., L.Sa., L.Sc., L.Sh., L.T.S., L.T.W., L.v.E., L.C.P.Z., M.A., M.A.H., M.B.H., M.C., M.E.B., M.Fu., M.Ho., M.J.G., M.-J.v.T., M.J.W., M.Ki., M.La., M.P.Z., M.W., N.E.M.v.H., N.F.H., N.J., N.O., N.T.D., O.G., P.G.S., P.K., P.M.T., P.N., R.B., R.K., R.L.G., R.M.B., R.R., R.R.-S., S.A., S.Ca., S.Des., S.Eh., S.Er., S.F.F., S.I.T., S.Ka., S.Ke., S.L.R., S.M.C.d.Z., S.R.M., T.A., T.A.L., T.G., T.G.M.v.E., T.J., T.K., T.L.P., T.P.G., T.R.M., T.Wh., T.Wo., T.W.M., U.D., W.W., X.C., Y.Q., Z.Z.

### Genetic Data collection

A.A.A., A.A.-K., A.d.B., A.J.F., A.J.H., A.J.S., A.K.H., A.M.D., A.P., A.R.H., A.R.K., B.-C.H., B.F., B.Mo., B.T.B., B.W., B.W.J.H.P., C.B., C.D.W., C.F., C.M., C.P., C.P.D.,C.S.Re., D.C.G., D.H.M., D.R.W., D.W.M., D.Z., E.A., E.B.Q., E.G.J., E.J.C.d.G., E.L.H., F.D., F.M., F.R.T., G.D., G.E.D., G.F., G.H., G.L.C., G.S., H.V., H.Y., I.E.Som., I.L.-C., J.A.T., J.B.J.K., J.Bl., J.E.C., J.E.N., J.-J.H., J.J.L., J.K.B., J.-L.M., J.-L.M., J.L.R., J.M.F., J.Q.W., J.R., J.W.S., K.A.M., K.D., K.O.L., K.S., L.M.R., L.R., L.Sh., M.A.K., M.F.D., M.H.J.H., M.Ha., M.Ho., M.J.C., M.J.W., M.La., M.-L.P.M., M.M.N., M.N., N.A.K., N.E.M.v.H., N.G.M., N.J.A.v.d.W., N.K.H., N.O., O.G., P.A.T., P.H., P.K., P.R.S., P.S.S., R.A.O., R.C.G., R.H., R.L.B., R.R., R.Se., R.S.K., R.W., S.A., S.Ci., S.Dj., S.E.F., S.Eh., S.Er., S.H., S.L.H., S.M.S., T.G.M.v.E., T.J.A., T.K.d.A., T.L.P., T.W.M., U.D., V.C., V.J.C., V.M.S., X.C.

### Genetic Data Analysis

A.A.-K., A.J.F., A.J.H., A.J.S., A.M.D., A.R.K., A.Te., A.Th., B.C.-D., B.F., B.K., B.M.-M., B.Pü., B.S.P., B.T.B., C.C.F., C.D.W., C.L.V., C.S.Re., C.S.Ro., C.W., C.Y.S., D.C.G., D.K., D.P.H., D.v.d.M., D.v.E., E.G.J., E.L.H., E.V., E.W., F.M., H.-R.E., I.E.J., I.E.Som., I.E.Søn., I.L.-C., I.O.F., J.Bl., J.Br., J.F.P., J.H.V., J.-J.H., J.L.R., J.L.S., J.N.P., J.Q.W., J.R.A., J.S., J.W.C., J.W.S., K.E.T., K.L.G., K.N., L.C.-C., L.M.O.L., L.Sh., L.C.P.Z.,M.A.A.A., M.B., M.E.G., M.Fu., M.Ha., M.I., M.J., M.J.C., M.J.W., M.Ki., M.Kl., M.Kn., M.La., M.Lu., M.M.J.v.D., N.A.G., N.G.M., N.J., N.J.A., N.K.H., N.M.-S., N.R.M., O.G.,P.A.L., P.G.S., P.H., P.H.L., P.K., P.M.T., P.R.S., Q.C., R.A.O., R.M.B., R.R., R.Se., S.Da., S.Des., S.E.M., S.Eh., S.G., S.H., S.H.W., S.L.H., S.M.C.d.Z., S.N., S.R.M., T.A.L., T.G., T.G.M.v.E., T.J., T.K.d.A., T.M.L., W.R.R., Y.M., Y.W.

### CHARGE Study Design

B.Ma., C.Dec., C.L.S., E.H., G.V.R., H.H.H.A., H.J.G., J.C.B., L.J.L., M.A.I., M.Fo., O.L.L., Q.Y., R.Sc., S.Deb., S.S., T.H.M., V.G., W.T.L.

## Data and materials availability

The meta-analytic summary results will be available to download from the ENIGMA consortium webpage upon publication http://enigma.ini.usc.edu/research/download-enigma-gwas-results.

