## Supplemental_Manhattan_Plots for "The genetic architecture of the human cerebral cortex"

**Manhattan plots - within each plot, the Manhattan plot for surface area is shown above the x axis and the Manhattan plot for thickness is shown below.**

| Region# | Phenotype | Page |
|---|---|---|
| 0 | Global | 2 |
| 1 | Frontal Pole | 3 |
| 2 | Medial Orbitofrontal | 4 |
| 3 | Lateral Orbitofrontal | 5 |
| 4 | Rostral Anterior Cingulate | 6 |
| 5 | Caudal Anterior Cingulate | 7 |
| 6 | Superior Frontal | 8 |
| 7 | Rostral Middle Frontal | 9 |
| 8 | Pars Orbitalis | 10 |
| 9 | Pars Triangularis | 11 |
| 10 | Pars Opercularis | 12 |
| 11 | Caudal Middle Frontal | 13 |
| 12 | Paracentral | 14 |
| 13 | Precentral | 15 |
| 14 | Postcentral | 16 |
| 15 | Precuneus | 17 |
| 16 | Superior Parietal | 18 |
| 17 | Supramarginal | 19 |
| 18 | Inferior Parietal | 20 |
| 19 | Posterior Cingulate | 21 |
| 20 | Isthmus Cingulate | 22 |
| 21 | Insula | 23 |
| 22 | Entorhinal | 24 |
| 23 | Parahippocampal | 25 |
| 24 | Fusiform | 26 |
| 25 | Temporal Pole | 27 |
| 26 | Inferior Temporal | 28 |
| 27 | Middle Temporal | 29 |
| 28 | Superior Temporal | 30 |
| 29 | Banks of the Superior Temporal Sulcus | 31 |
| 30 | Transverse Temporal | 32 |
| 31 | Lingual | 33 |
| 32 | Pericalcarine | 34 |
| 33 | Cuneus | 35 |
| 34 | Lateral Occipital | 36 |

#### Global

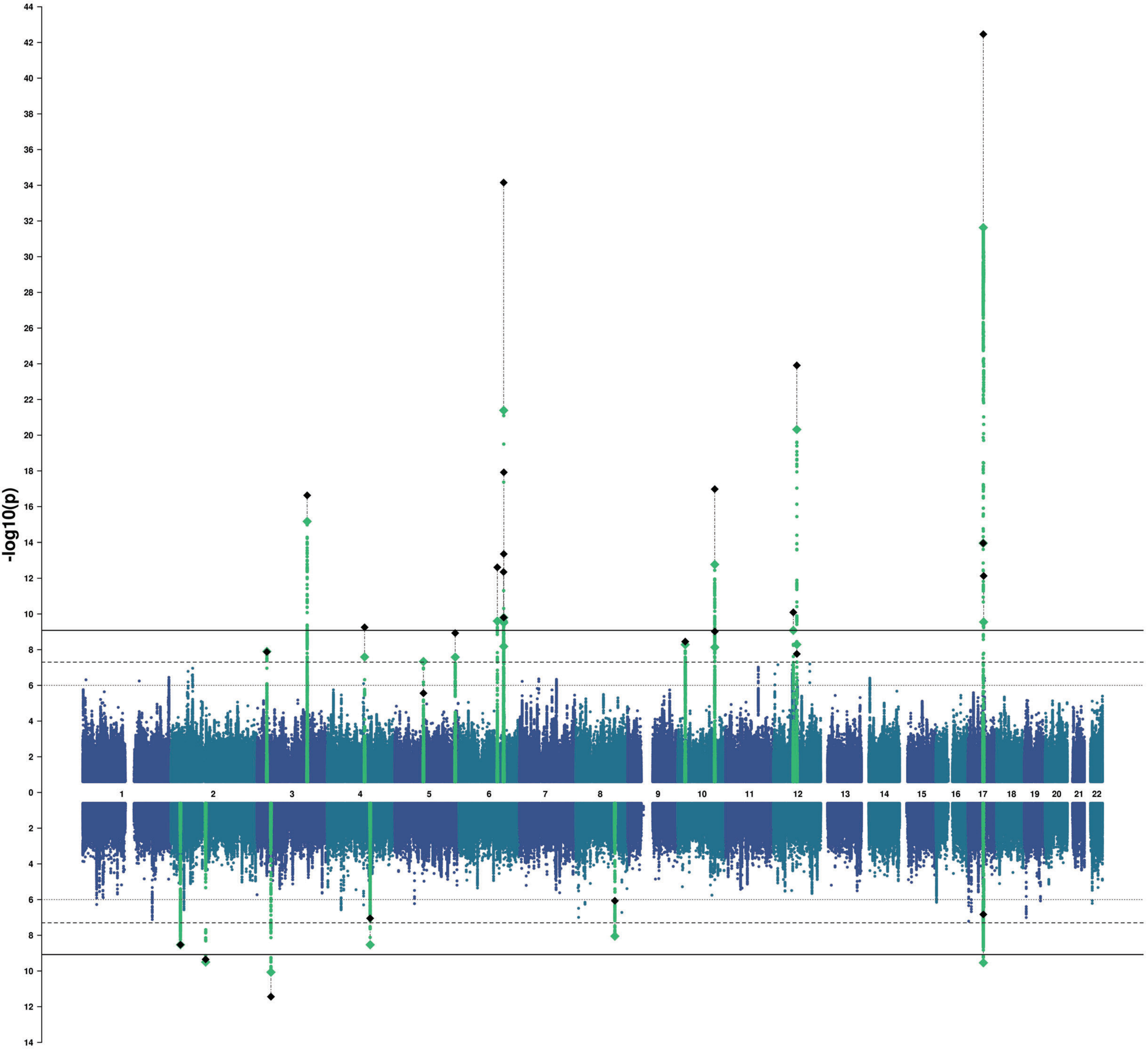

-log10(p)
44
42
40
38
36
34
32
30
28
26
24
22
20
18
16
14
12
10
8
6
4
2
0
2
4
6
8
10
12
14

### Frontal Pole

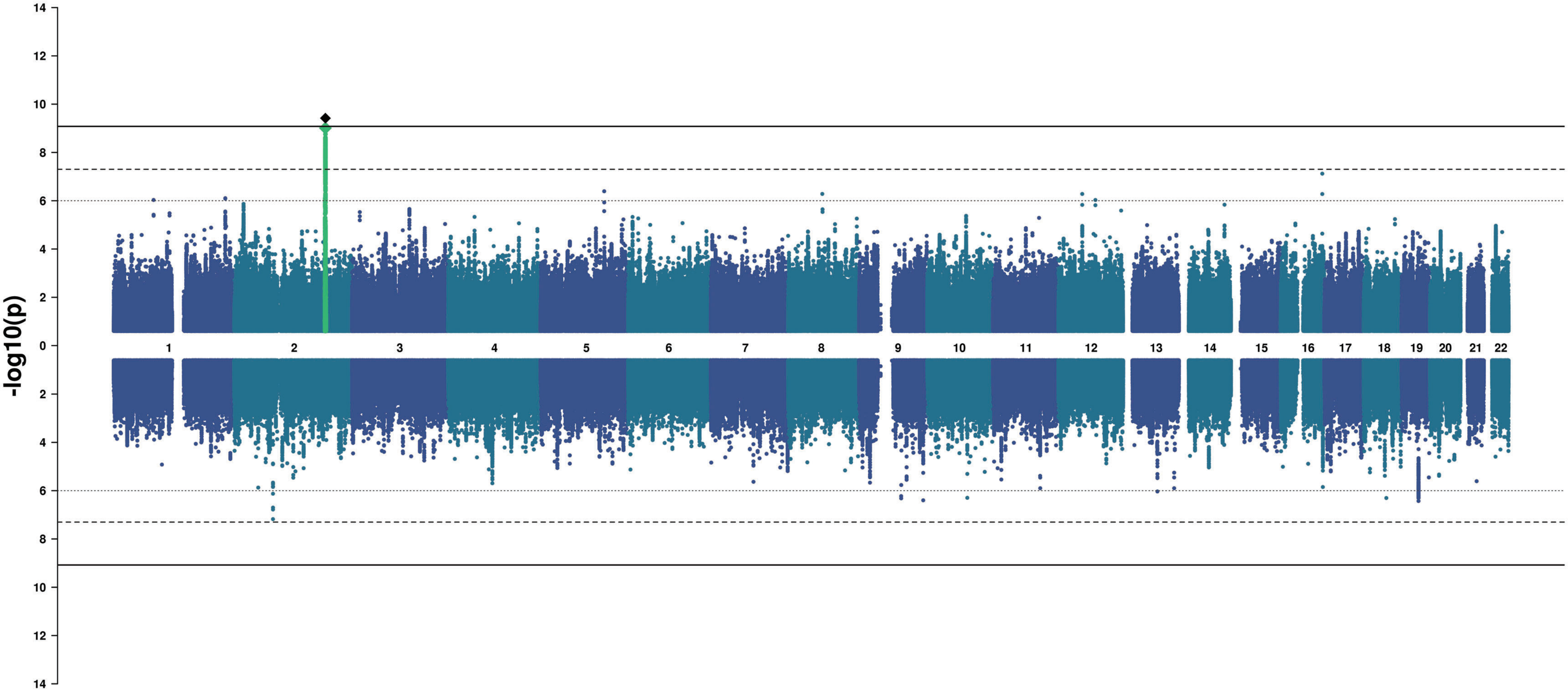

#### Medial Orbitofrontal

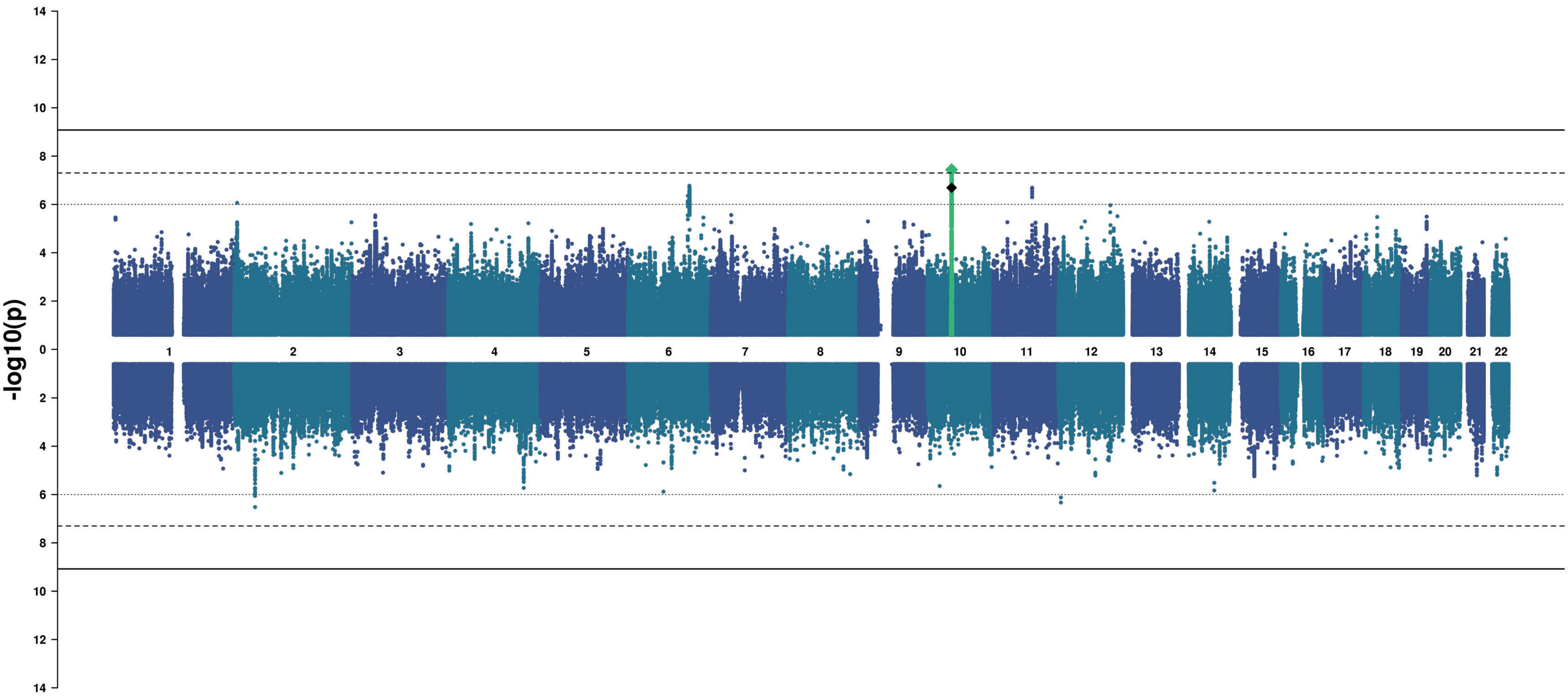

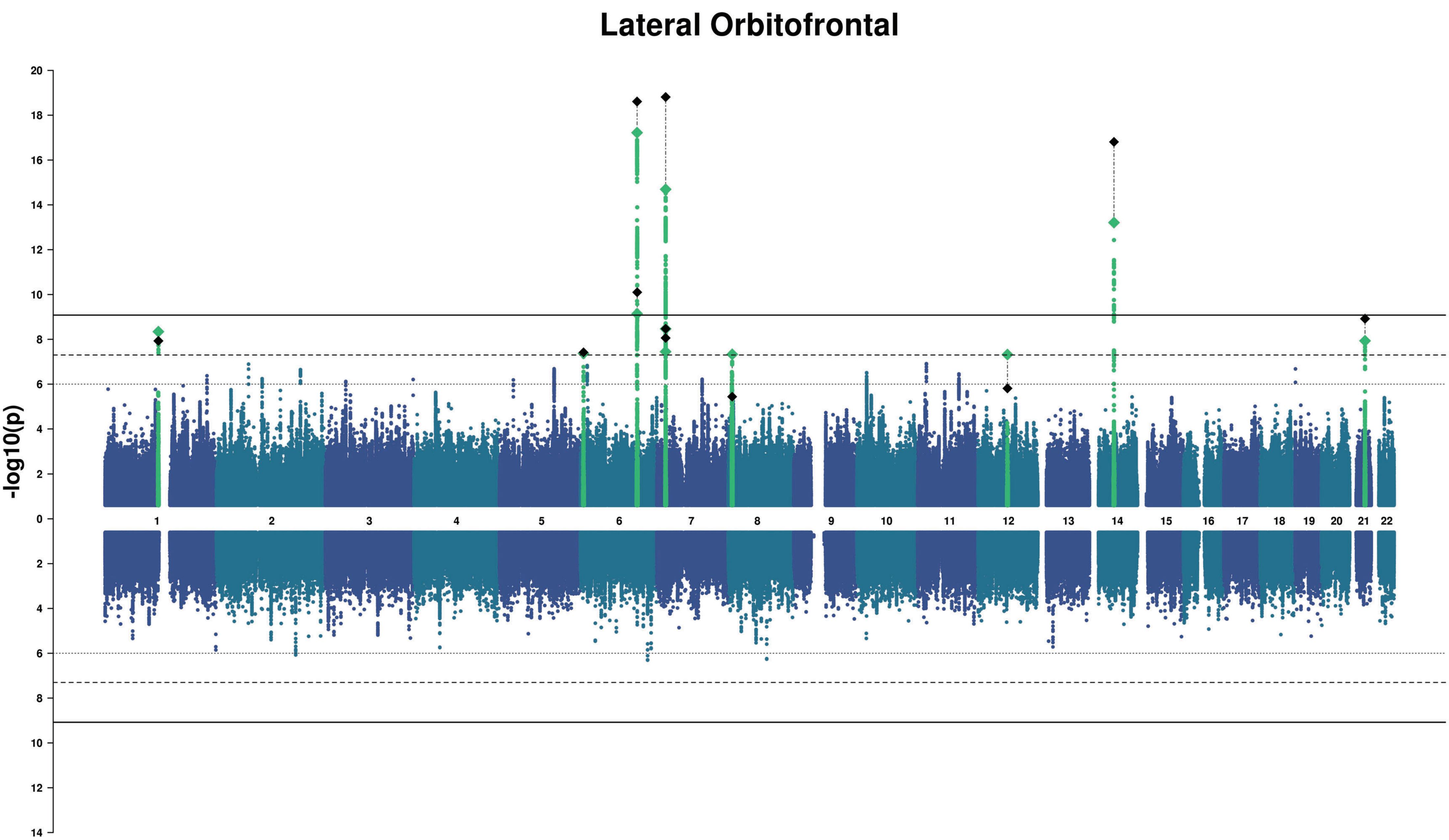
Lateral Orbitofrontal
-log10(p)
20
18
16
14
12
10
8
6
4
2
0
2
4
6
8
10
12
14

### Rostral Anterior Cingulate

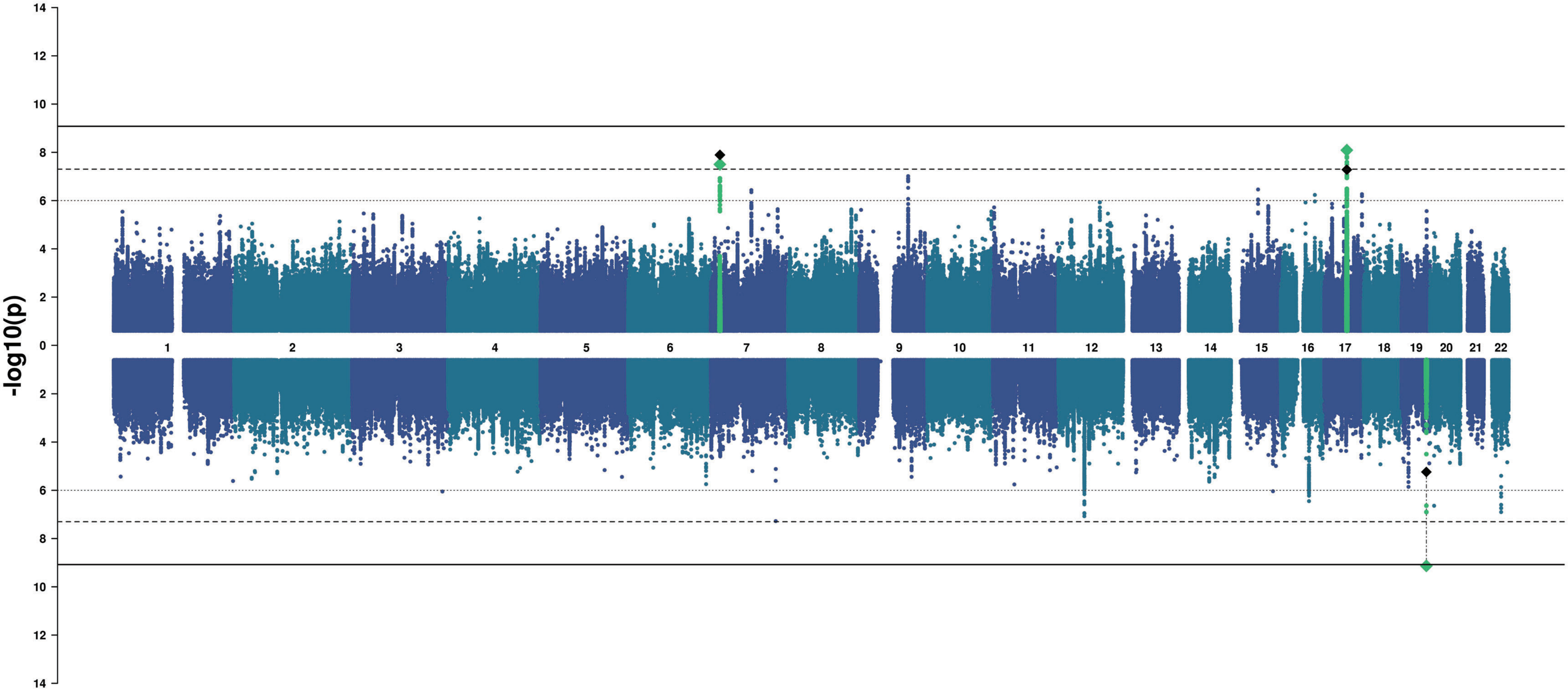

### Caudal Anterior Cingulate

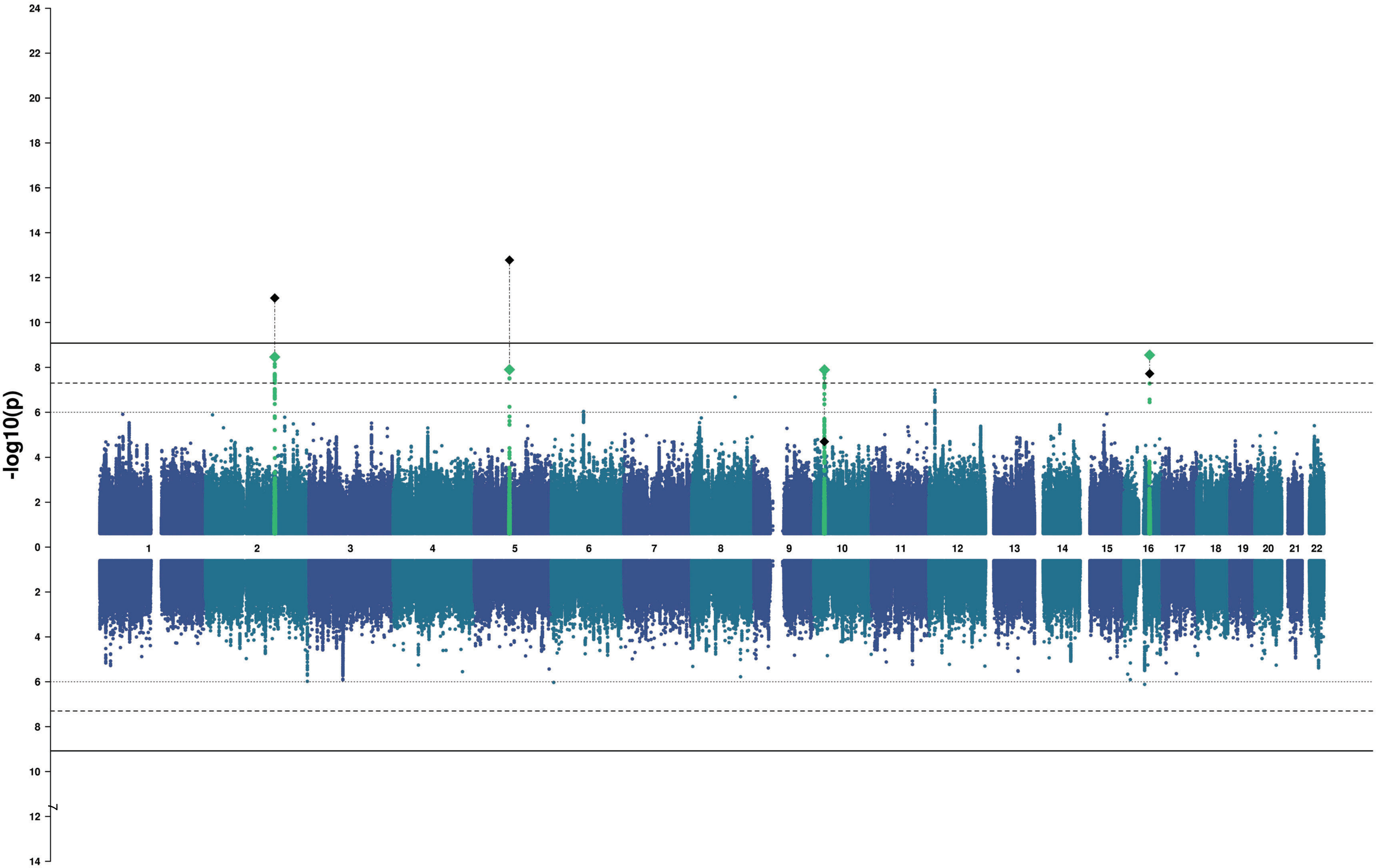

#### Superior Frontal

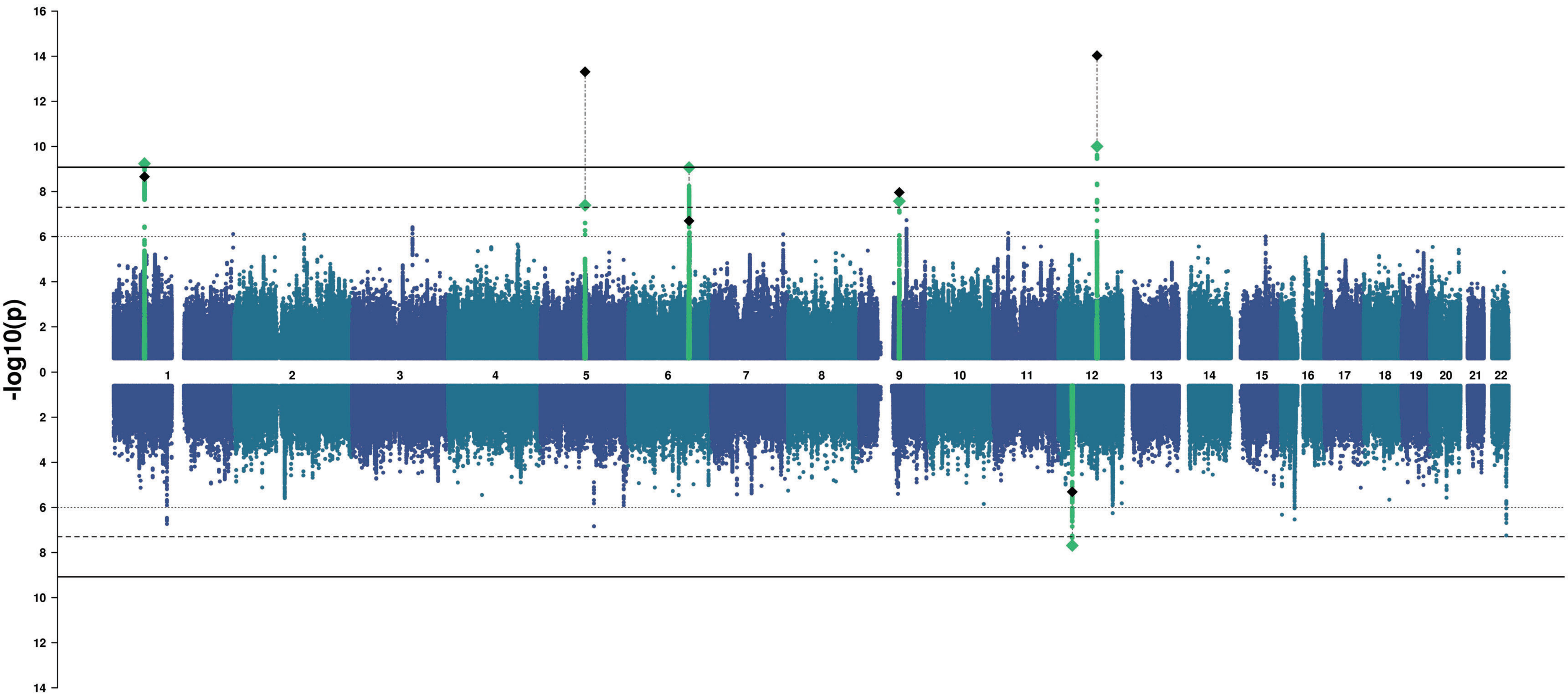

### Rostral Middle Frontal

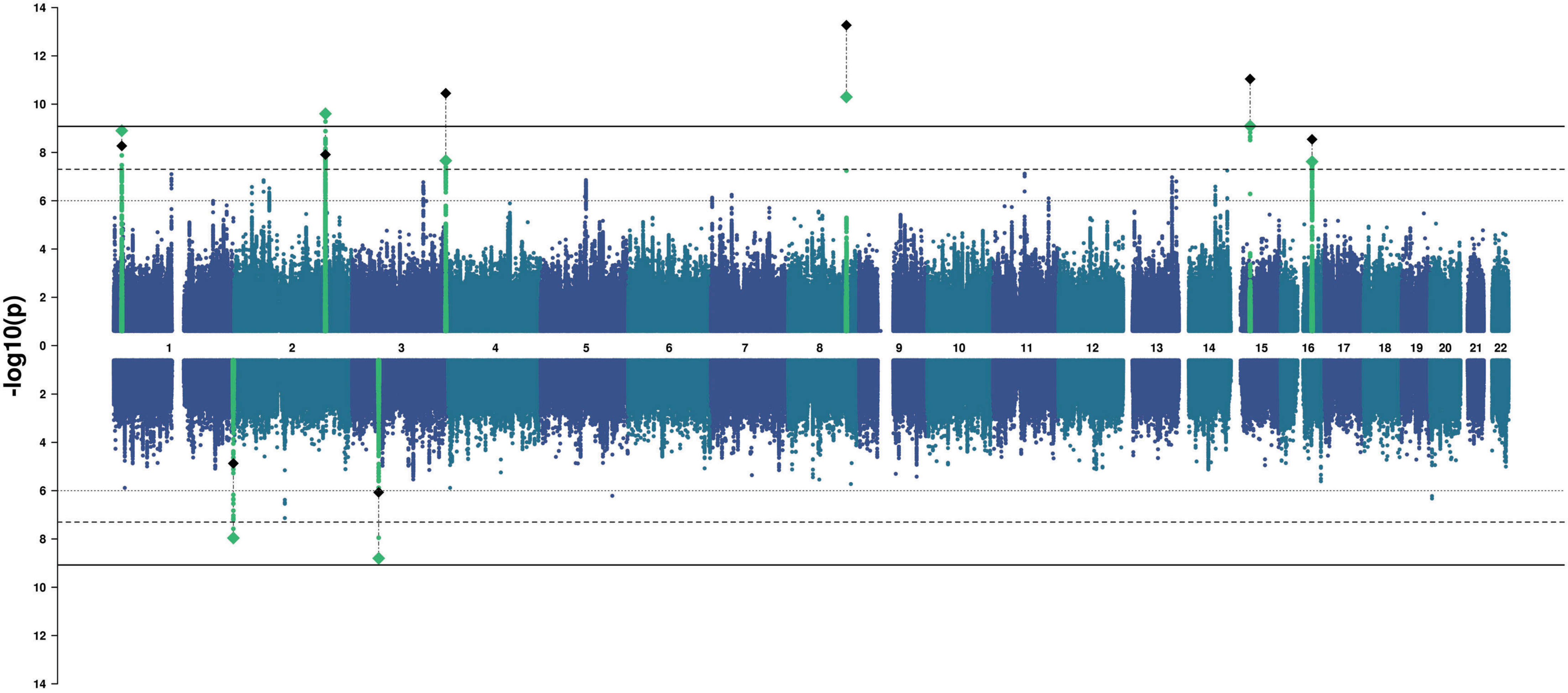

### Pars Orbitalis

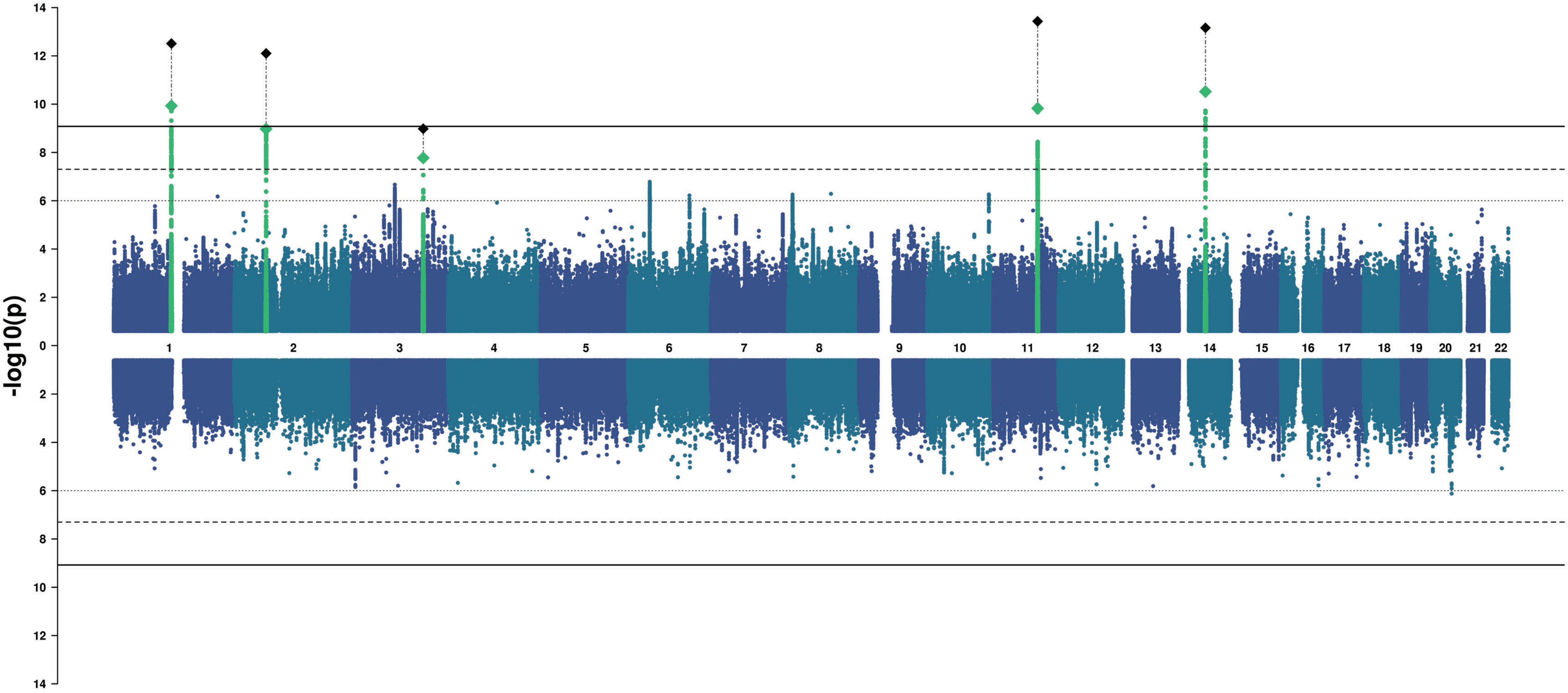

### Pars Triangularis

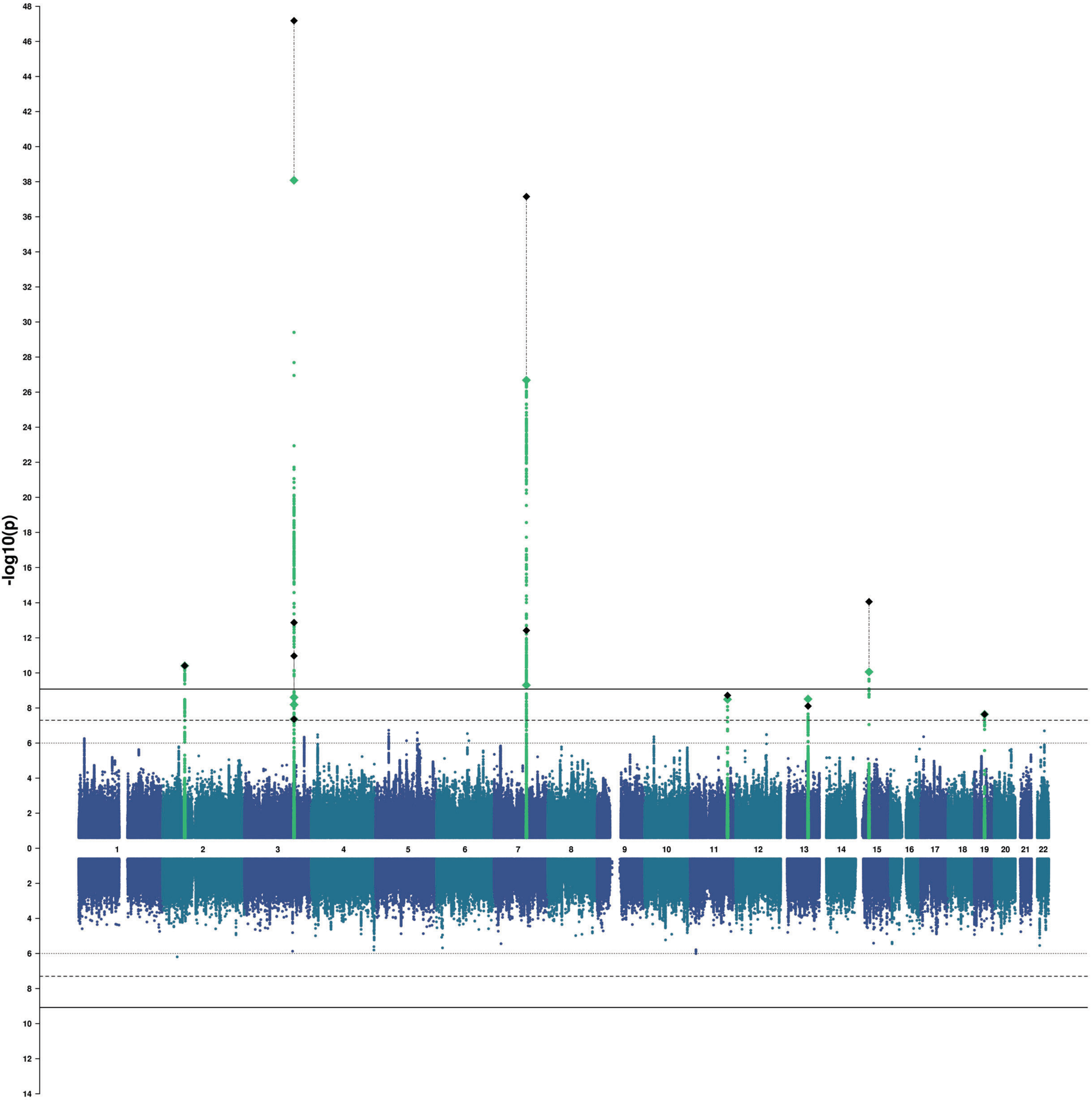

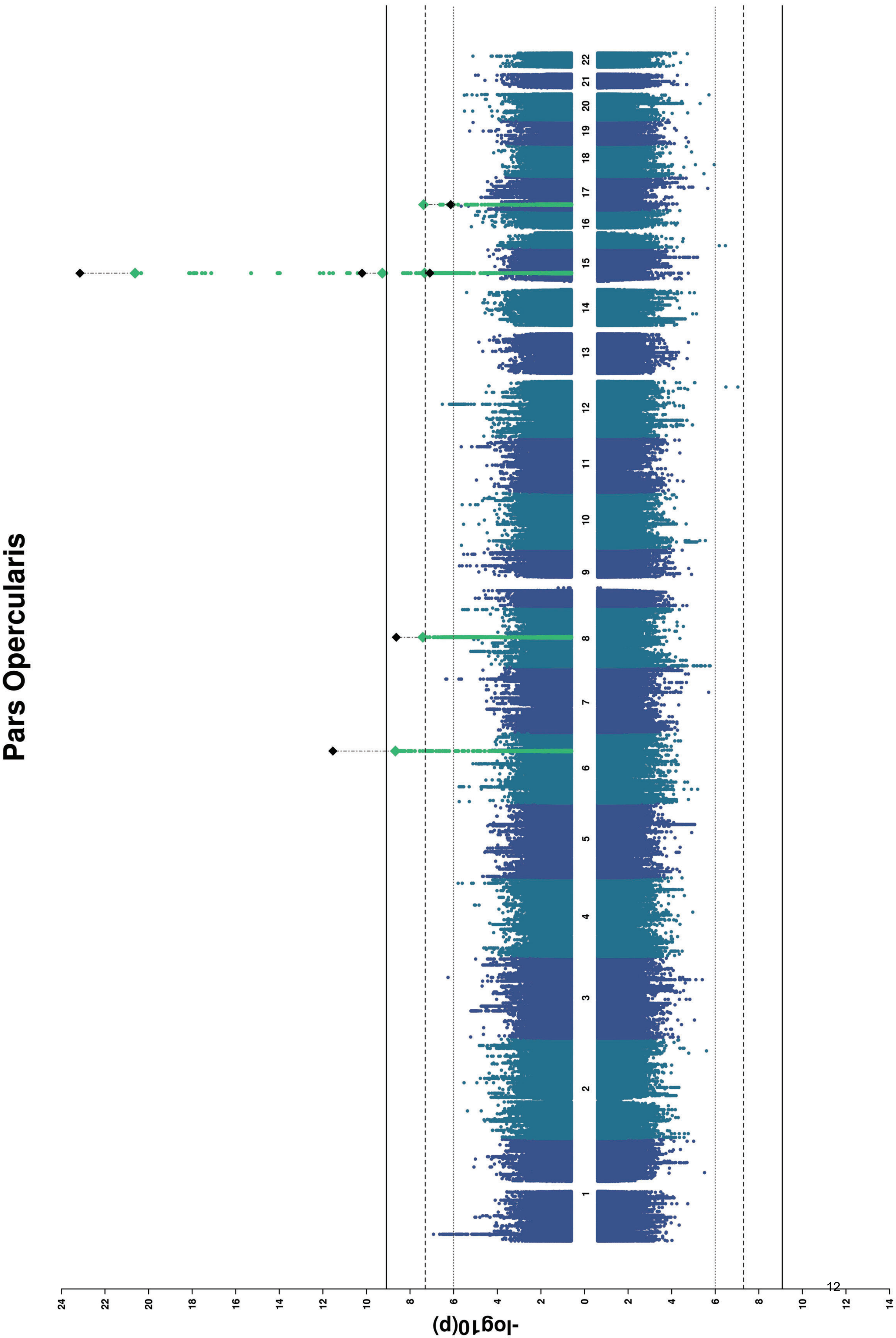
Pars Opercularis
-log10(p)
1 2 3 4 5 6 7 8 9 10 11 12 13 14 15 16 17 18 19 20 21 22
24 22 20 18 16 14 12 10 8 6 4 2 0 2 4 6 8 10 12 14

### Caudal Middle Frontal

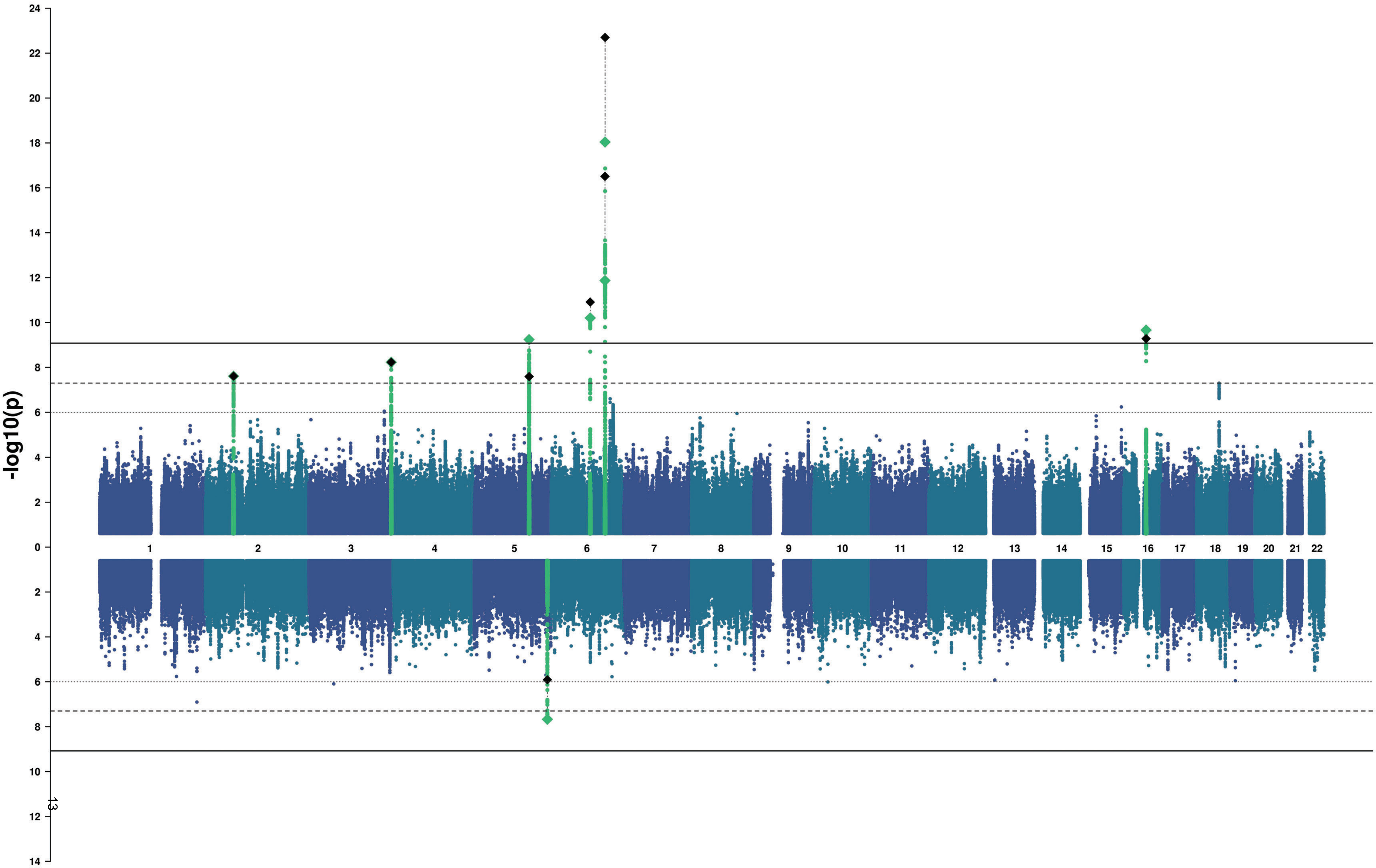

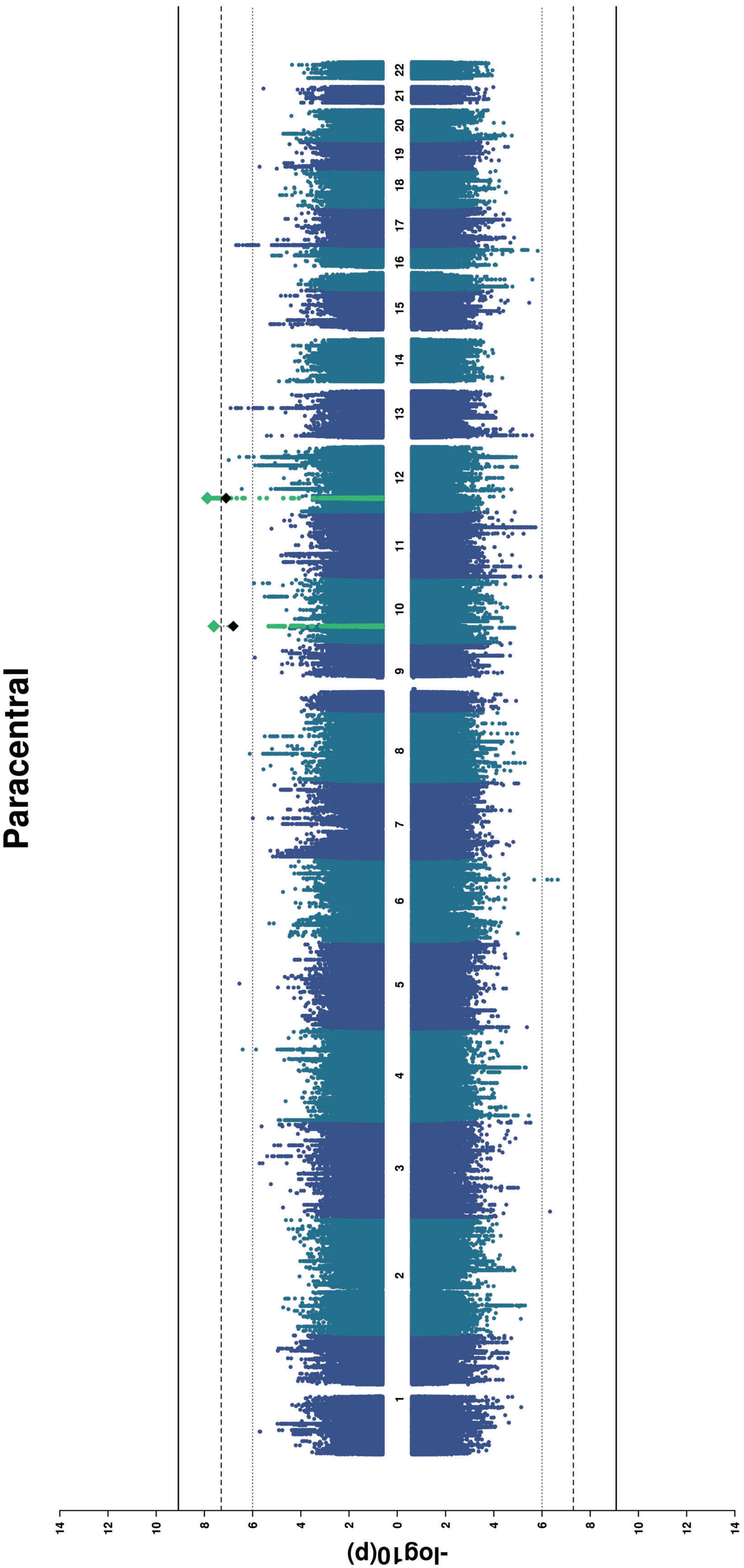
Paracentral
-log10(p)
1 2 3 4 5 6 7 8 9 10 11 12 13 14 15 16 17 18 19 20 21 22
14 12 10 8 6 4 2 0 2 4 6 8 10 12 14

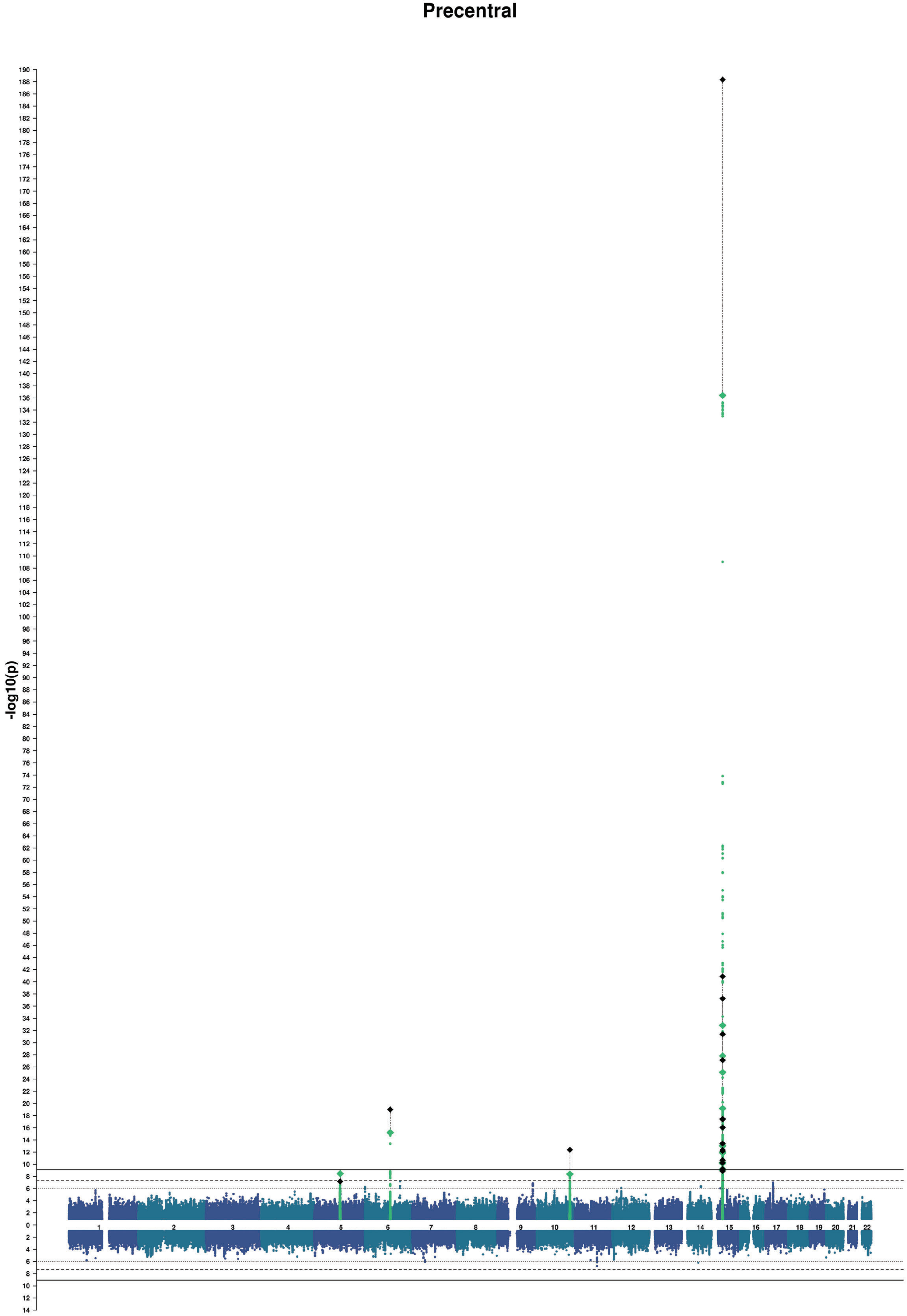
Precentral
-log10(p)

### Postcentral

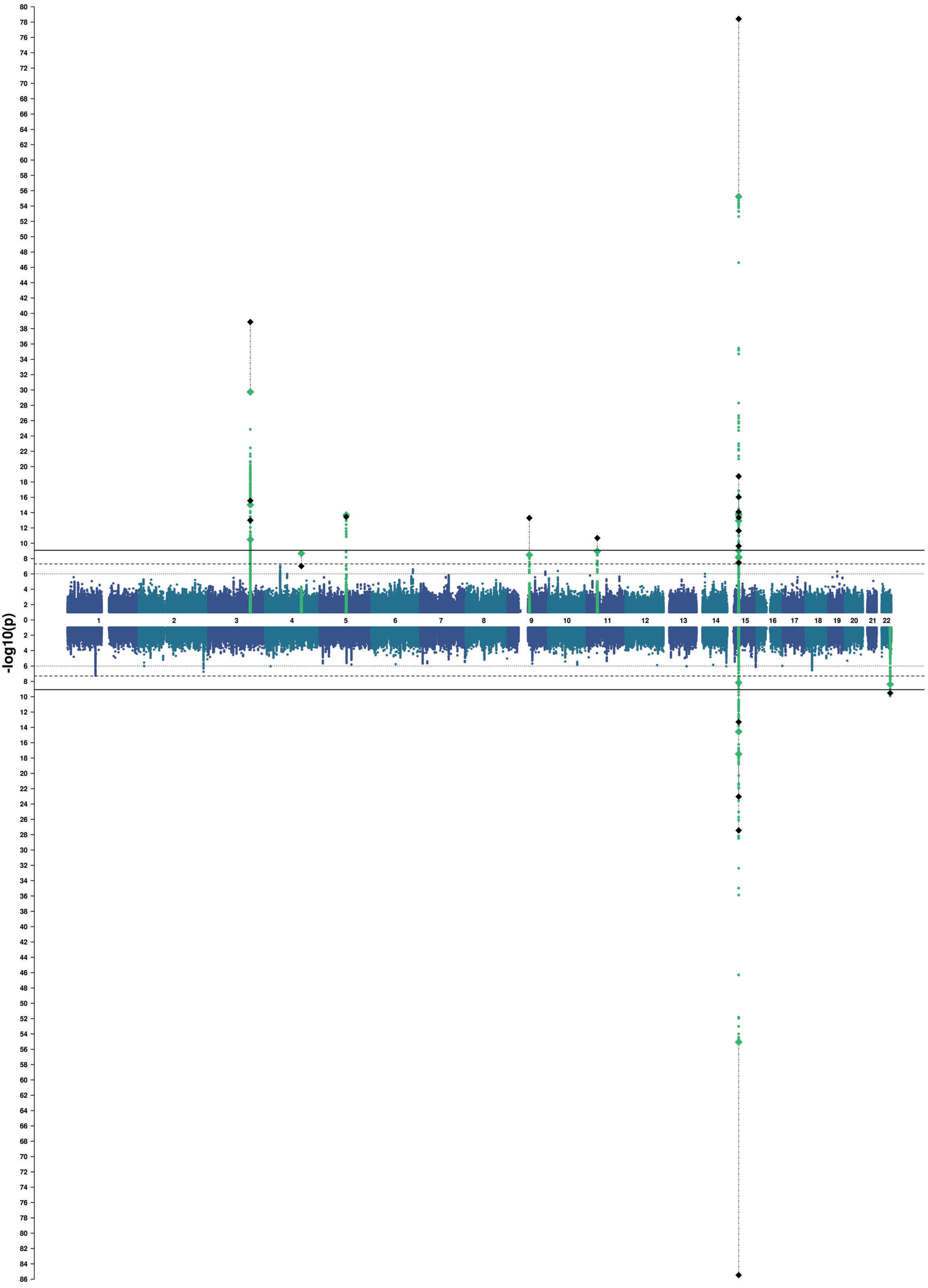

### Precuneus

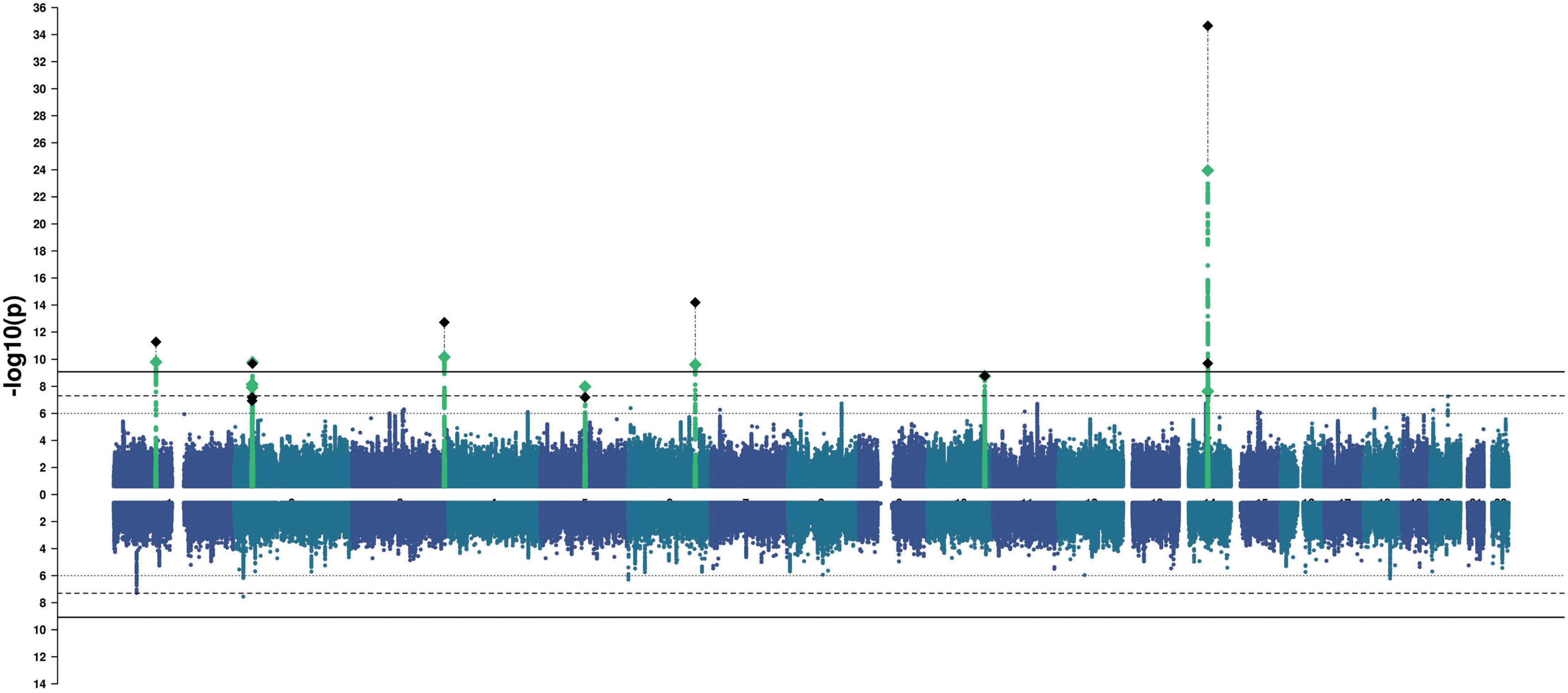

### Superior Parietal

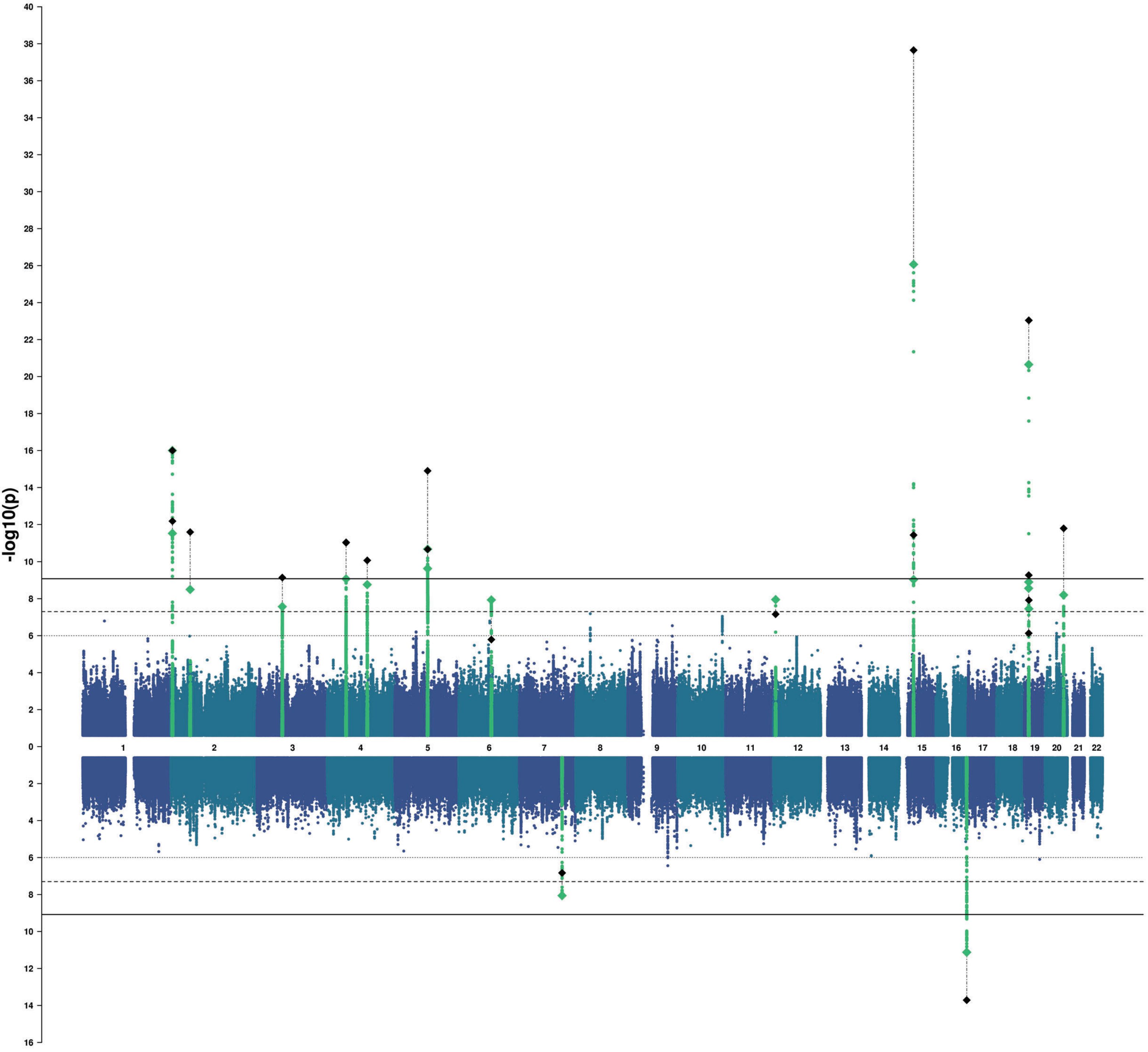

### Supramarginal

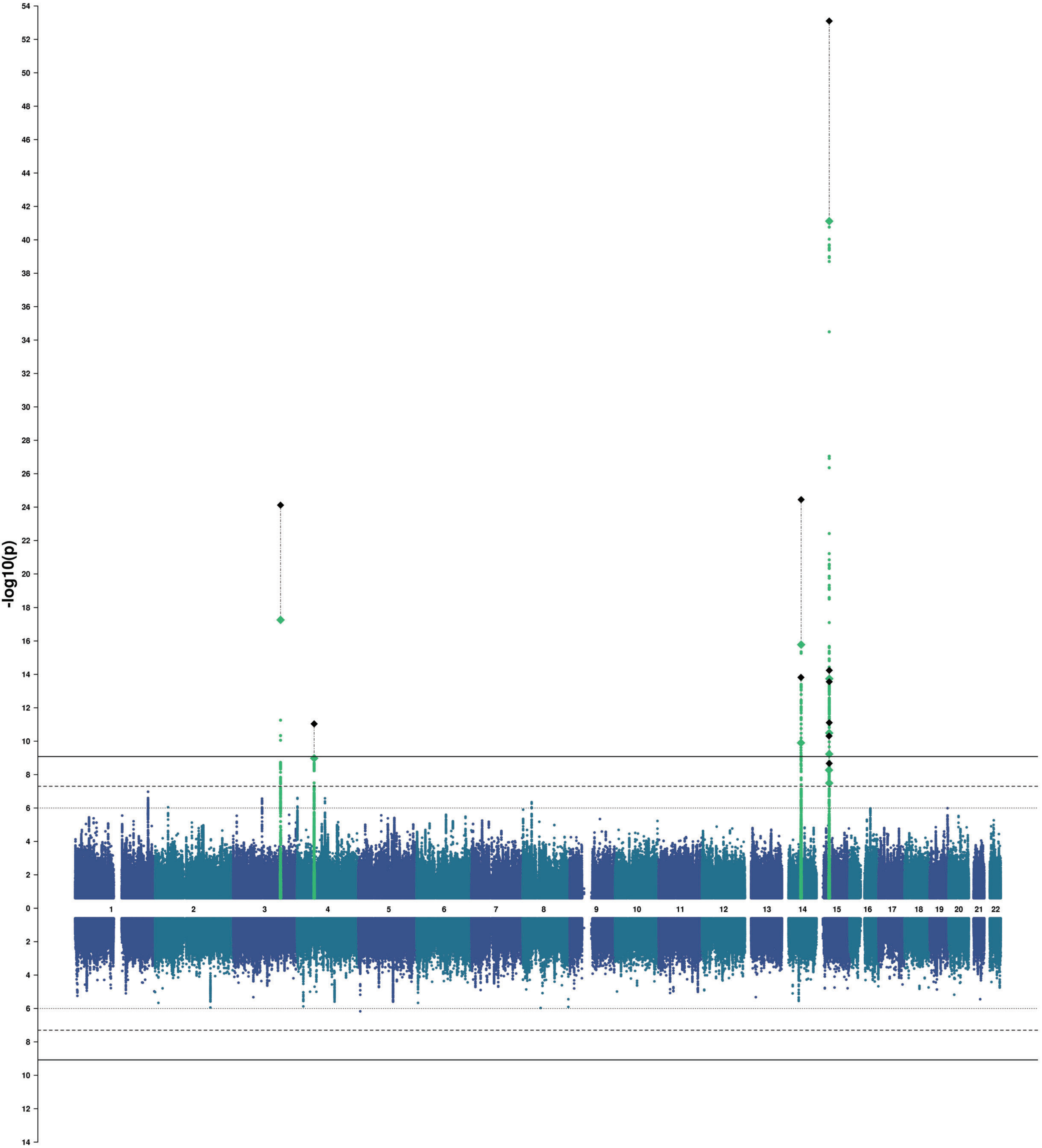

### Inferior Parietal

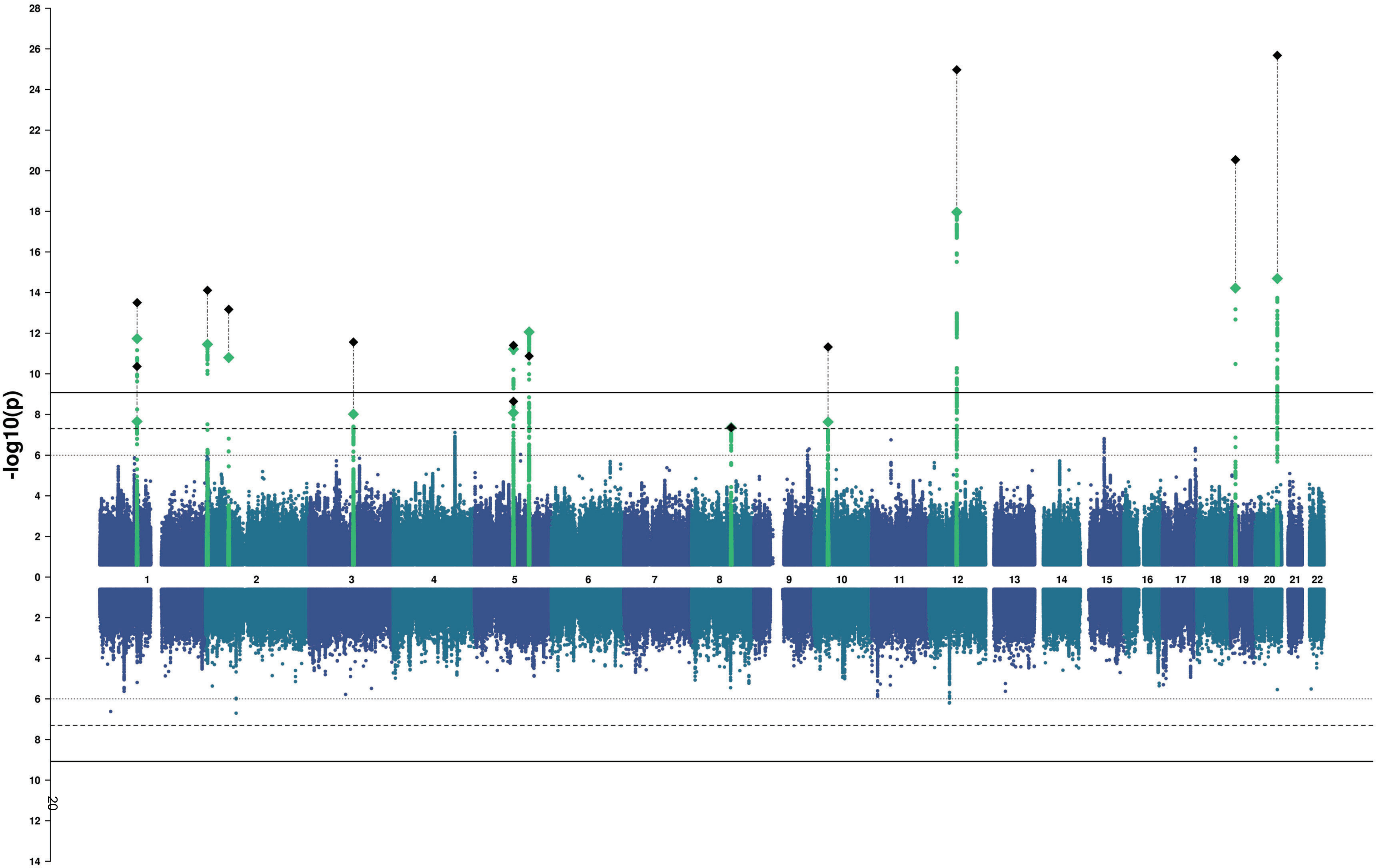

### Posterior Cingulate

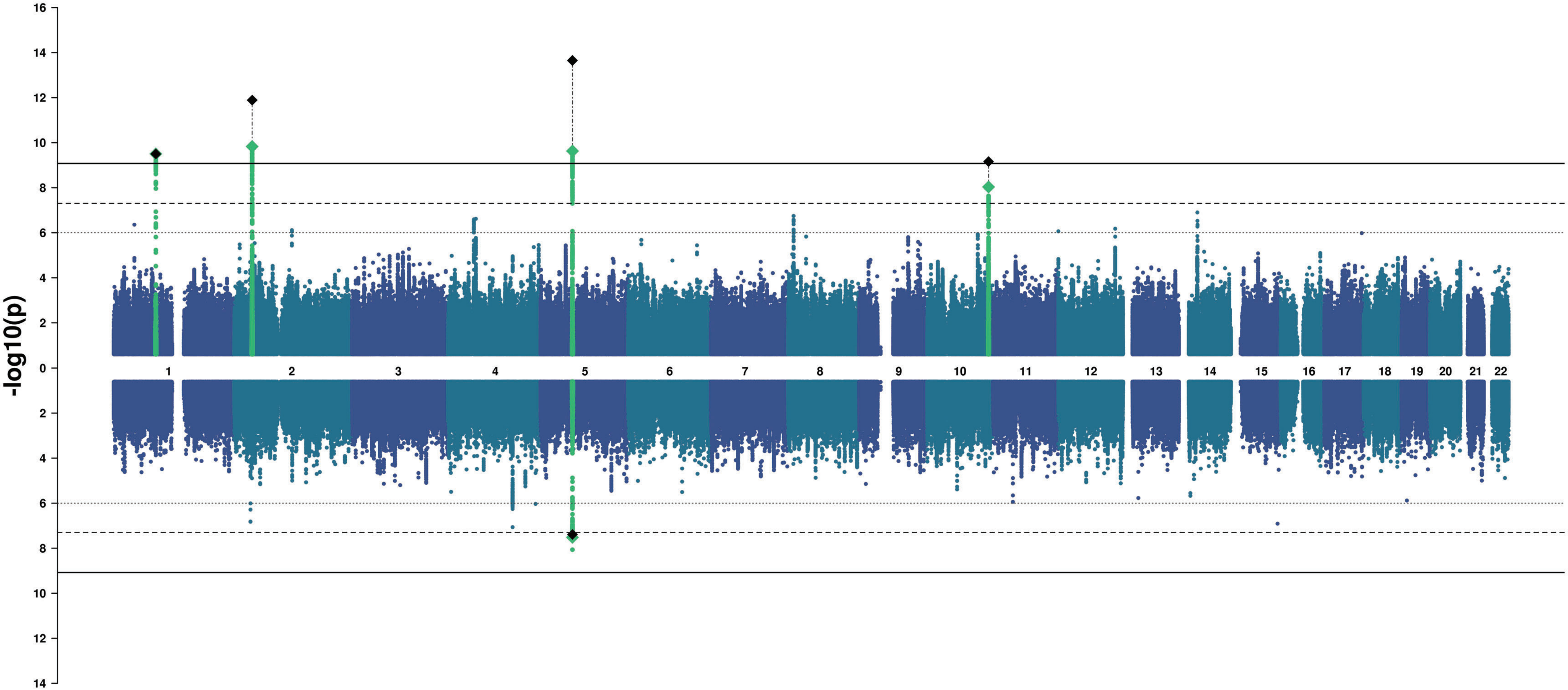

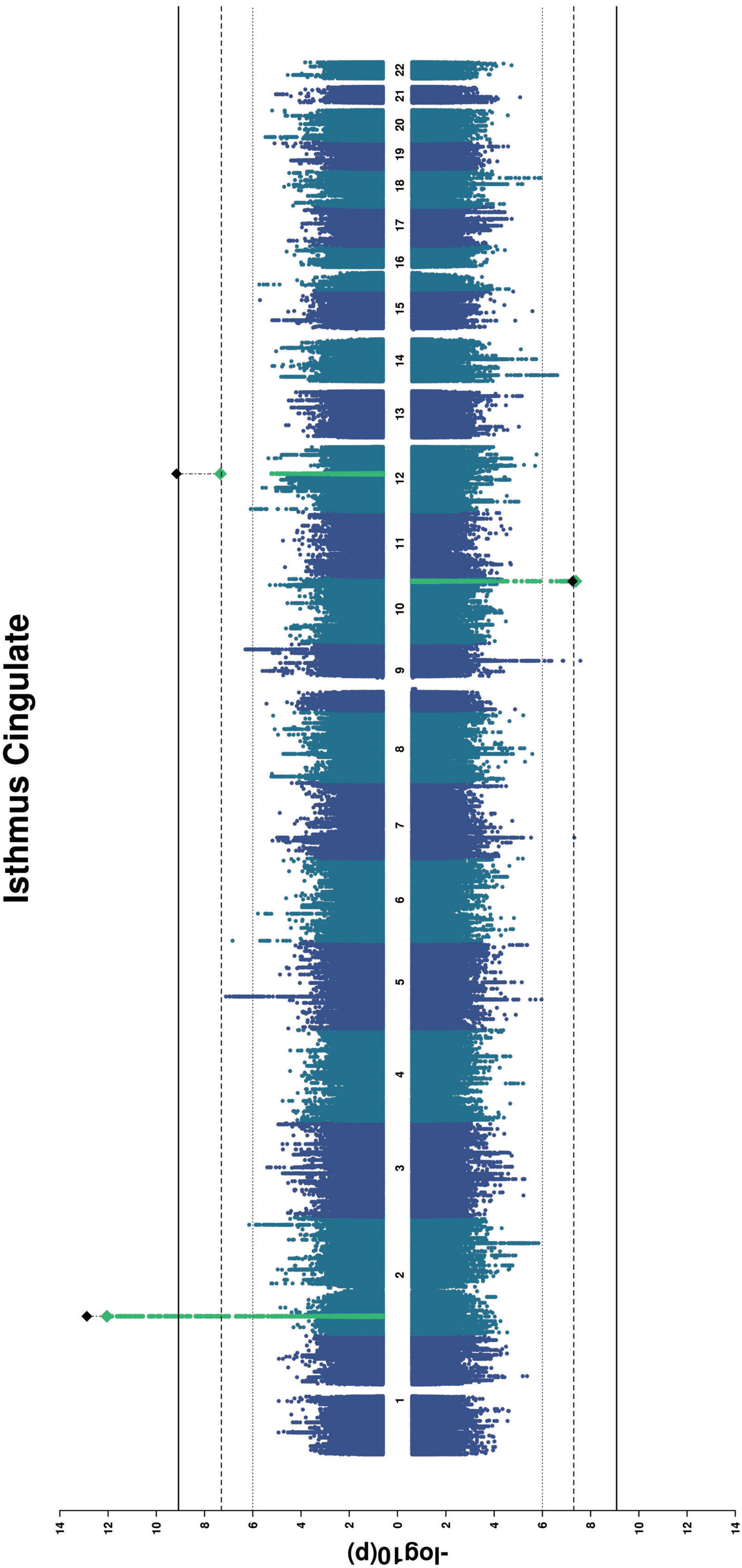
Isthmus Cingulate
-log10(p)
14
12
10
8
6
4
2
0
2
4
6
8
10
12
14

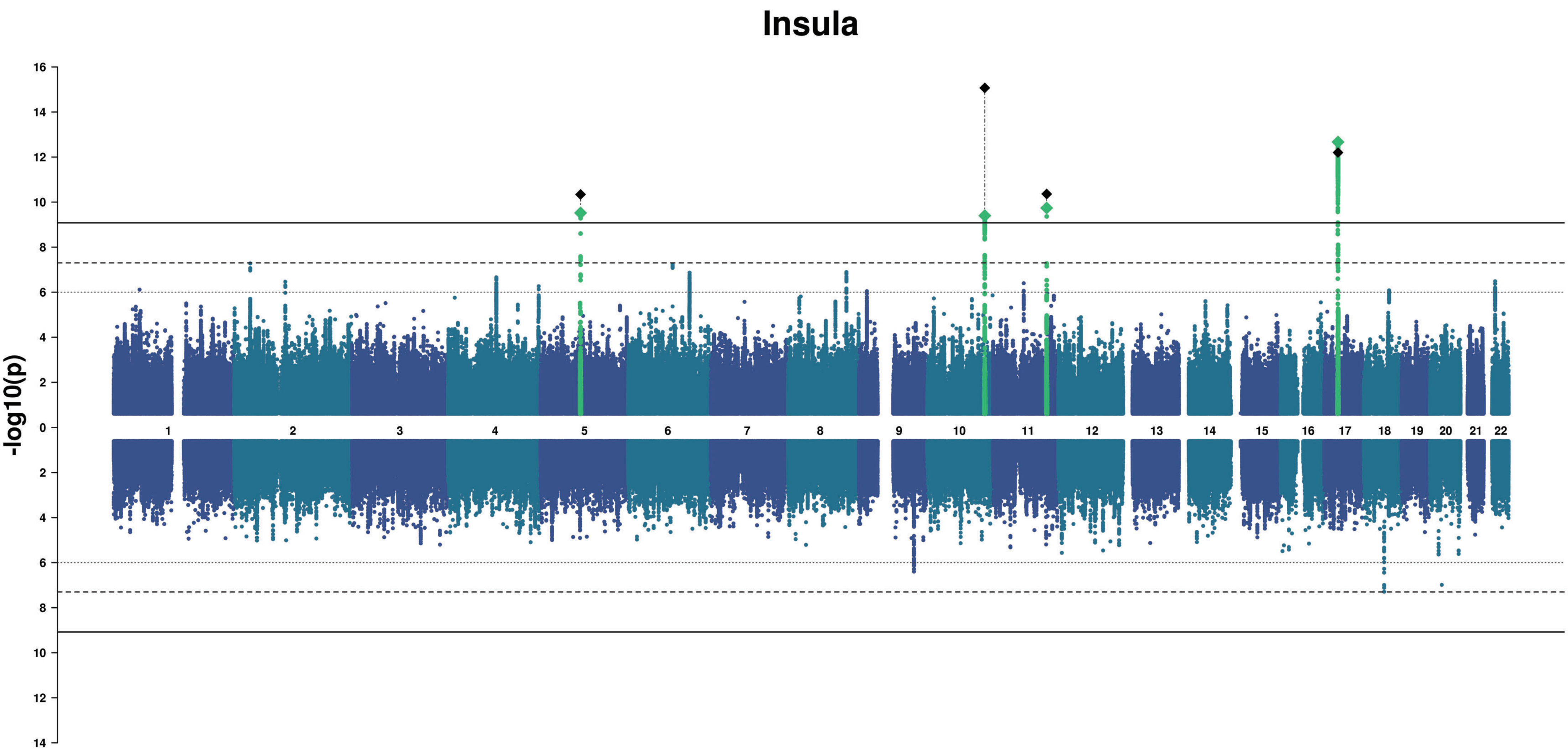
Insula
-log10(p)
16
14
12
10
8
6
4
2
0
2
4
6
8
10
12
14

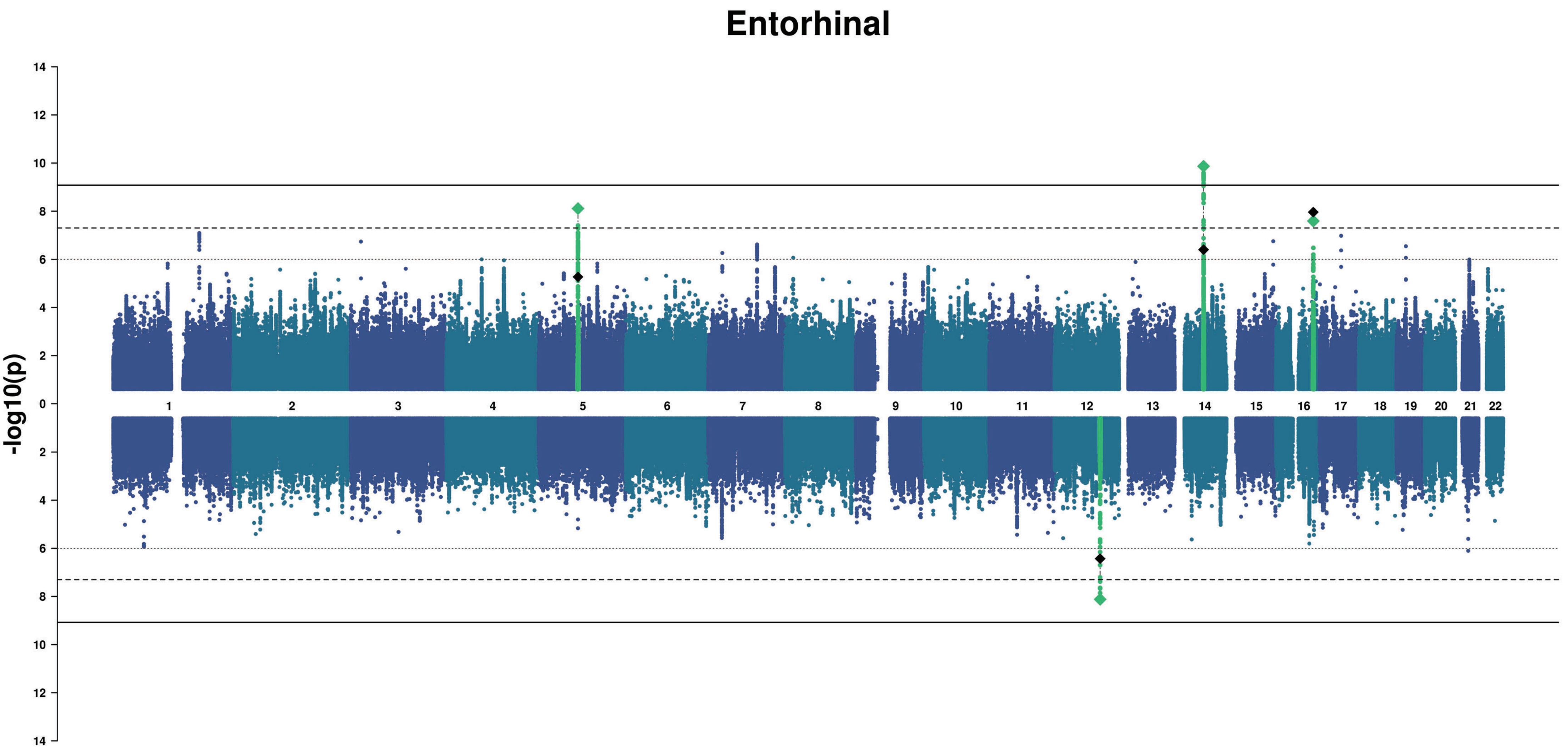
Entorhinal
-log10(p)
14
12
10
8
6
4
2

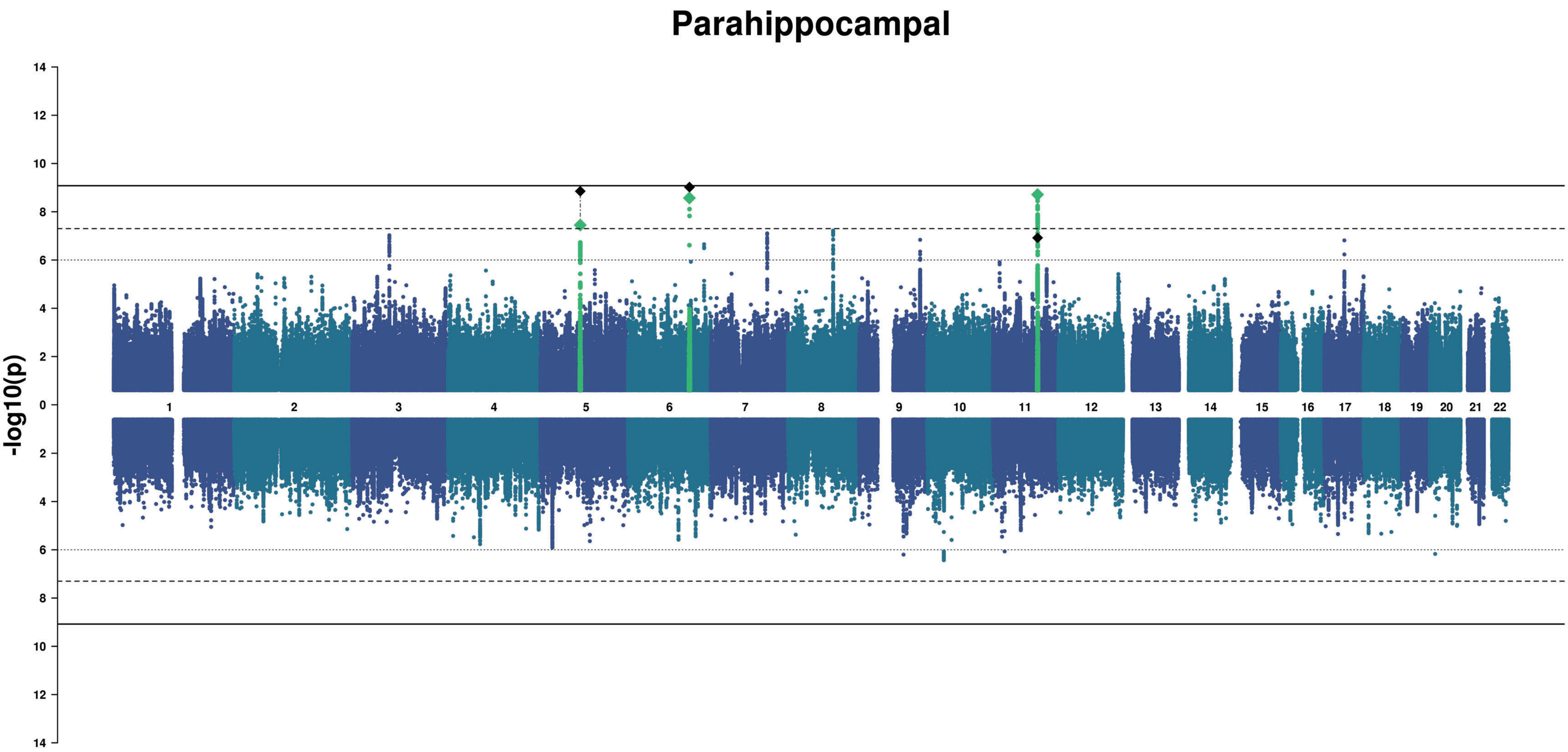
Parahippocampal
-log10(p)
14
12
10
8
6
4
2

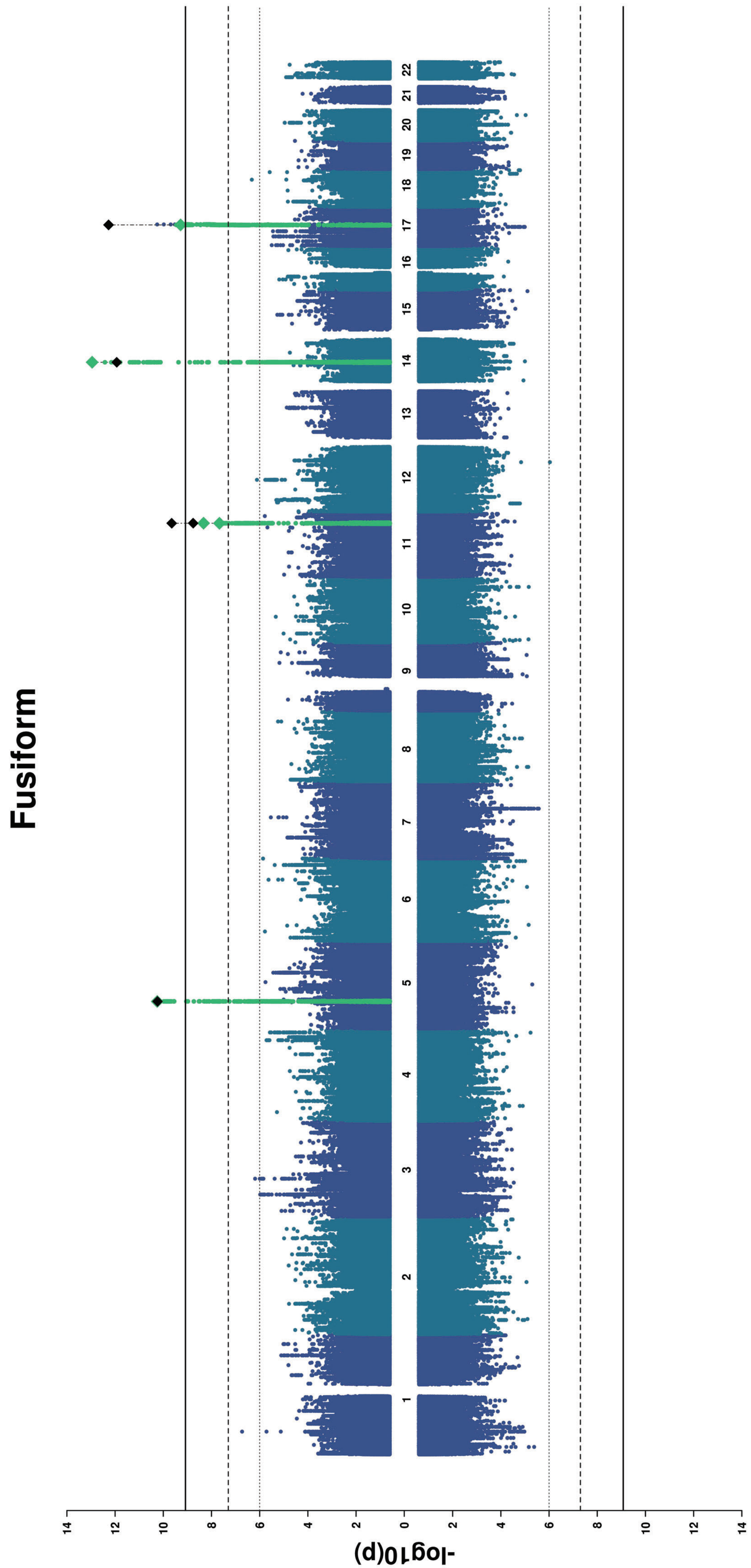

Fusiform
-log10(p)
14 12 10 8 6 4 2 0 2 4 6 8 10 12 14
1 2 3 4 5 6 7 8 9 10 11 12 13 14 15 16 17 18 19 20 21 22

### Temporal Pole

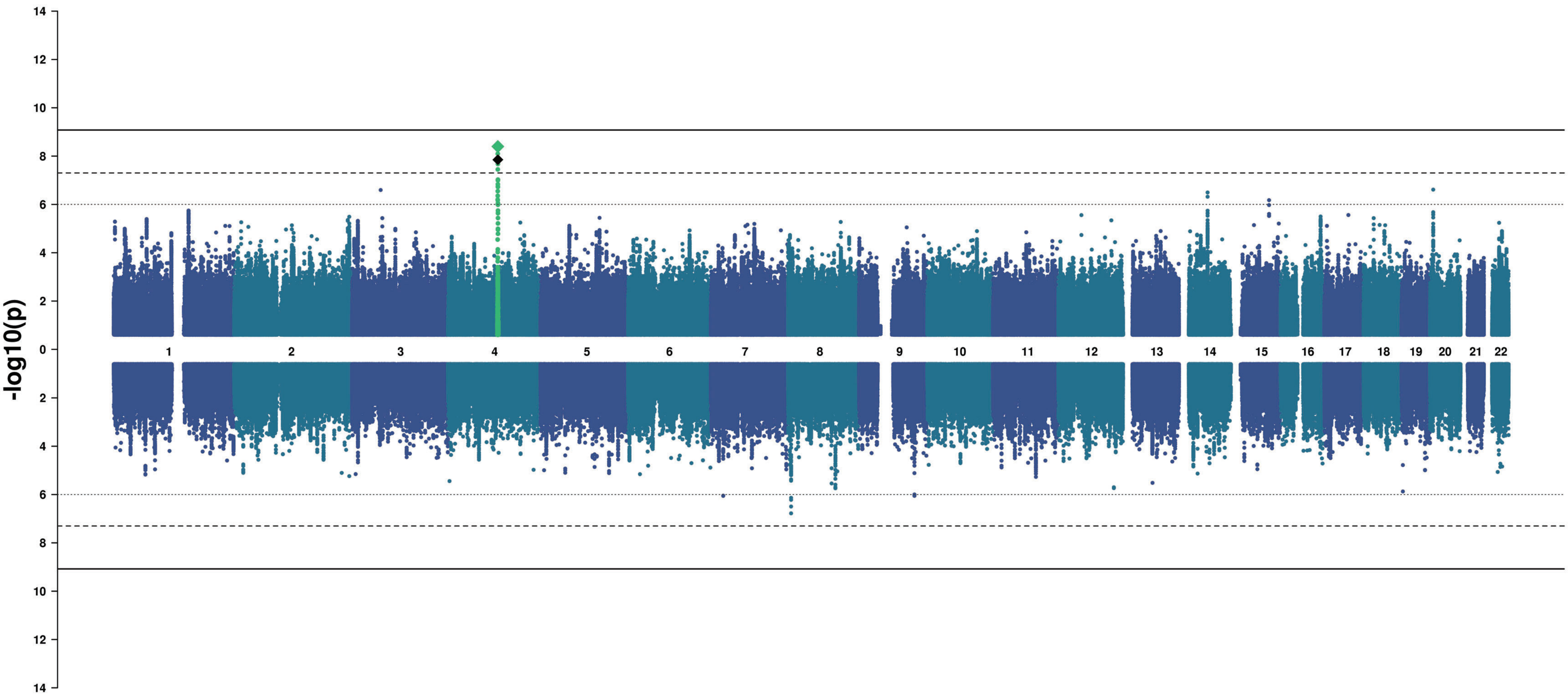

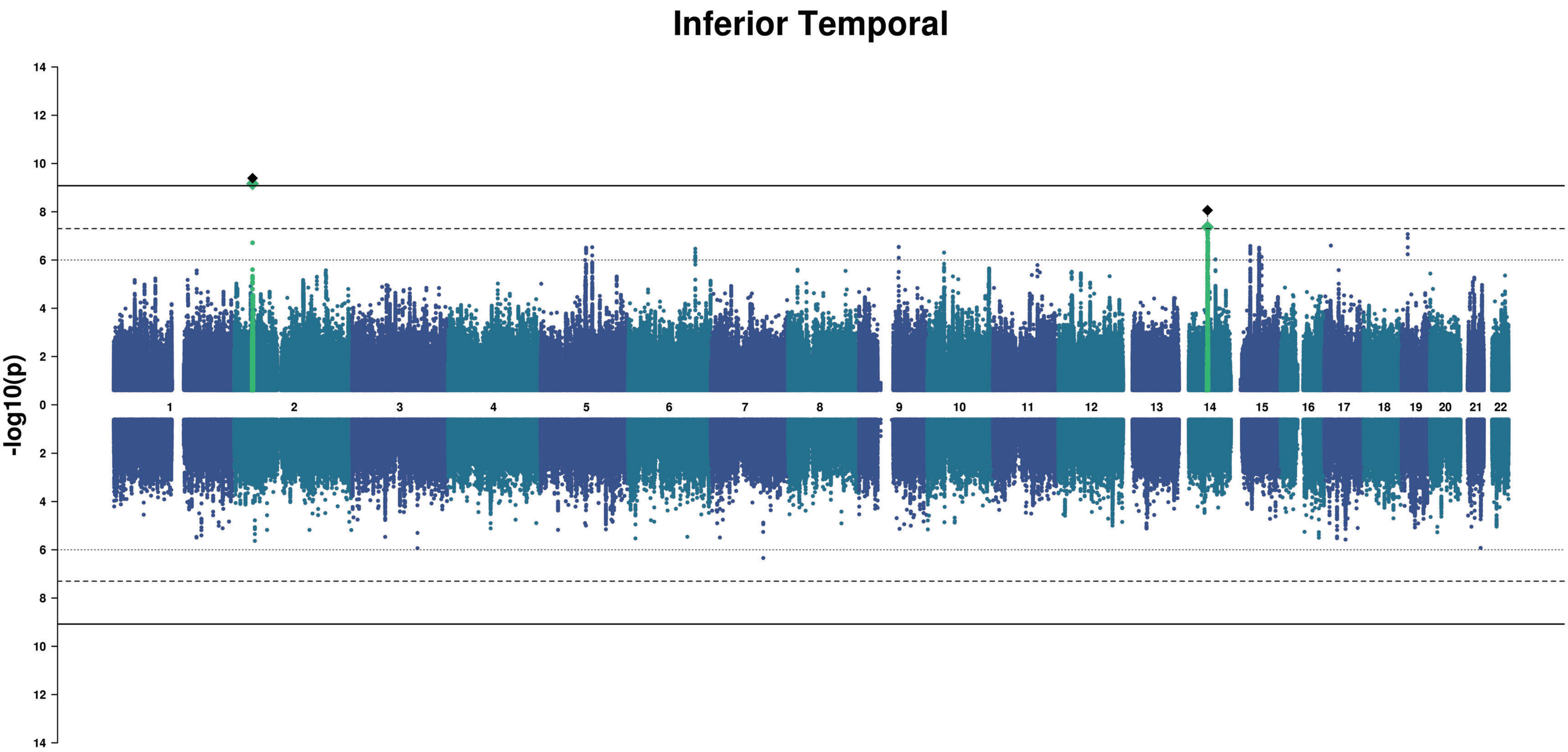

Inferior Temporal
-log10(p)
14
12
10
8
6
4
2

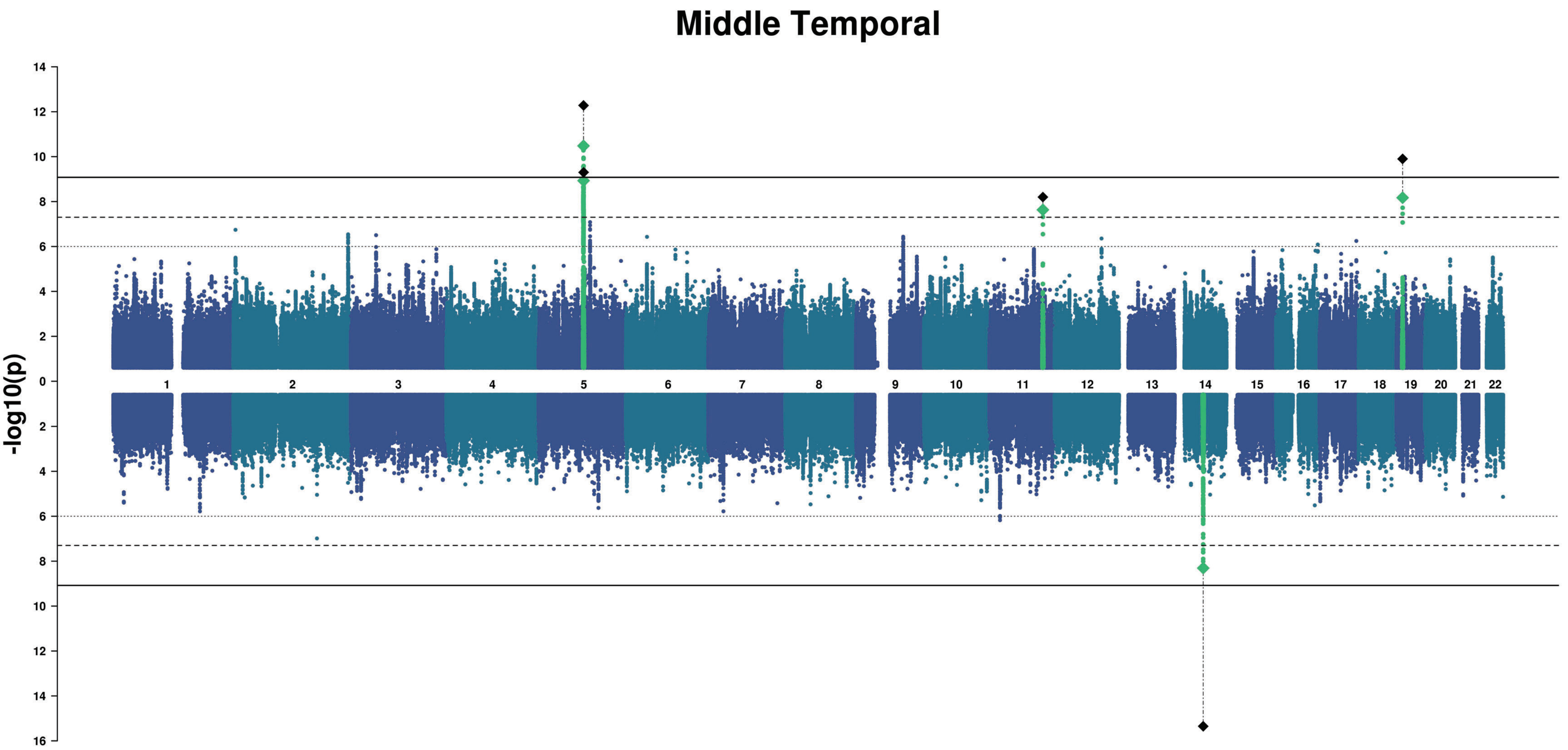
Middle Temporal
-log10(p)
14
12
10
8
6
4
2
0
2
4
6
8
10
12
14
16

### Superior Temporal

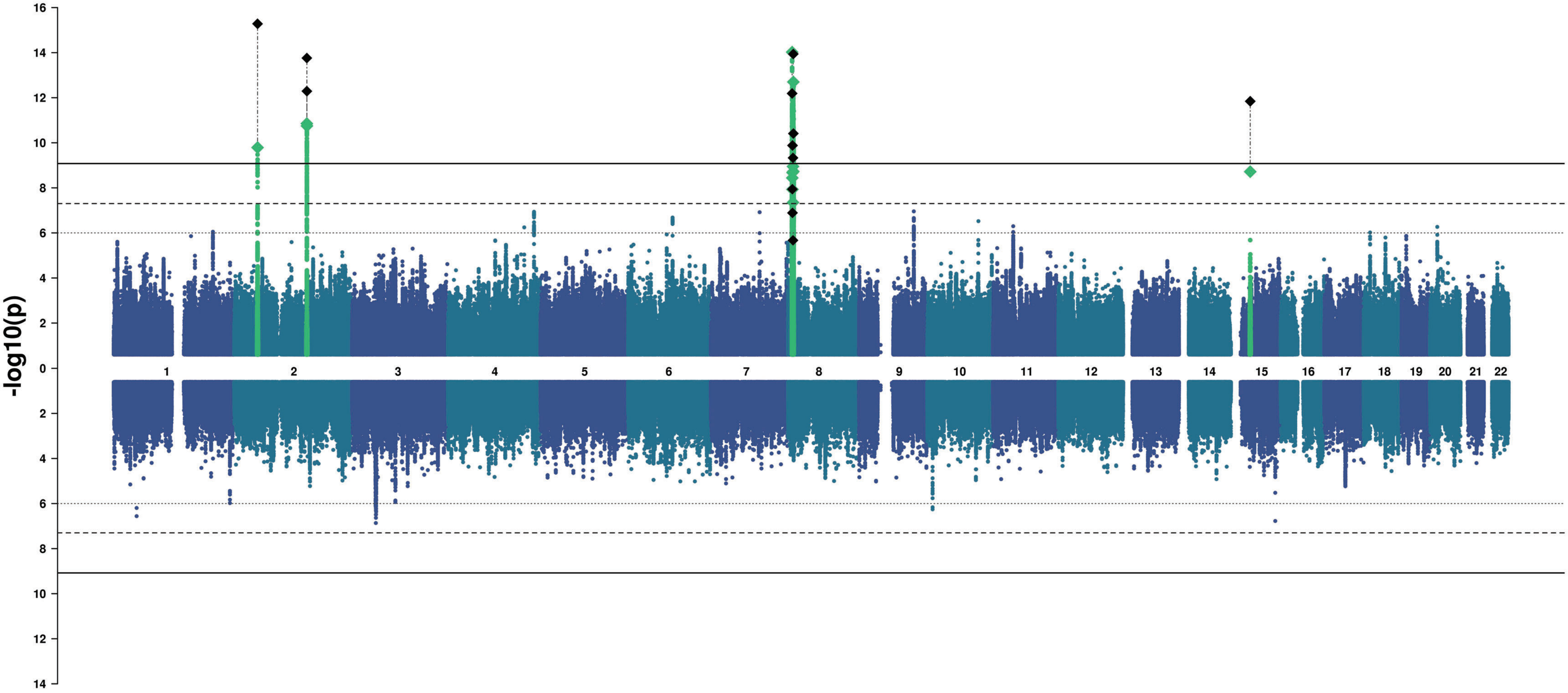

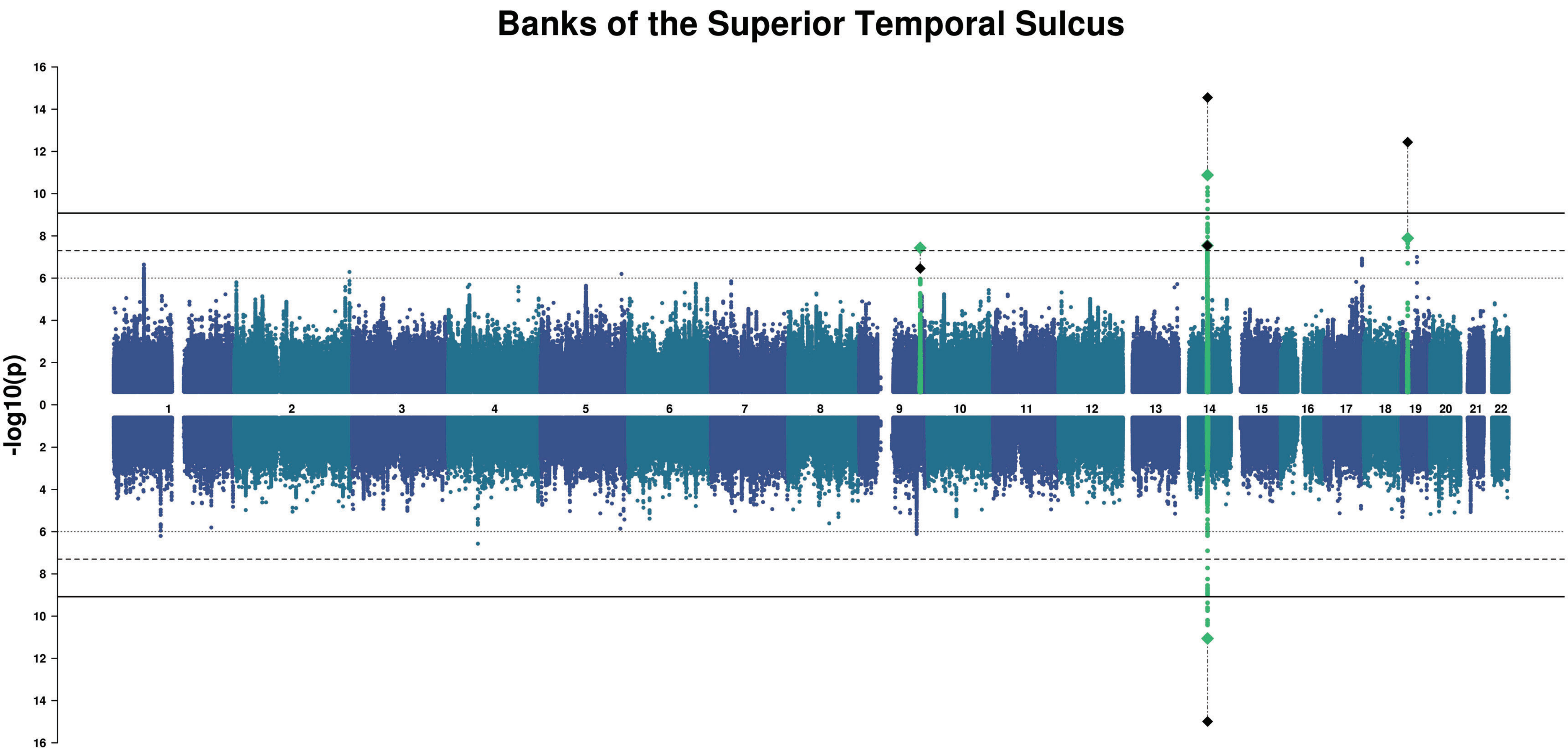

Banks of the Superior Temporal Sulcus
-log10(p)
16
14
12
10
8
6
4
2
0
2
4
6
8
10
12
14
16

#### Transverse Temporal

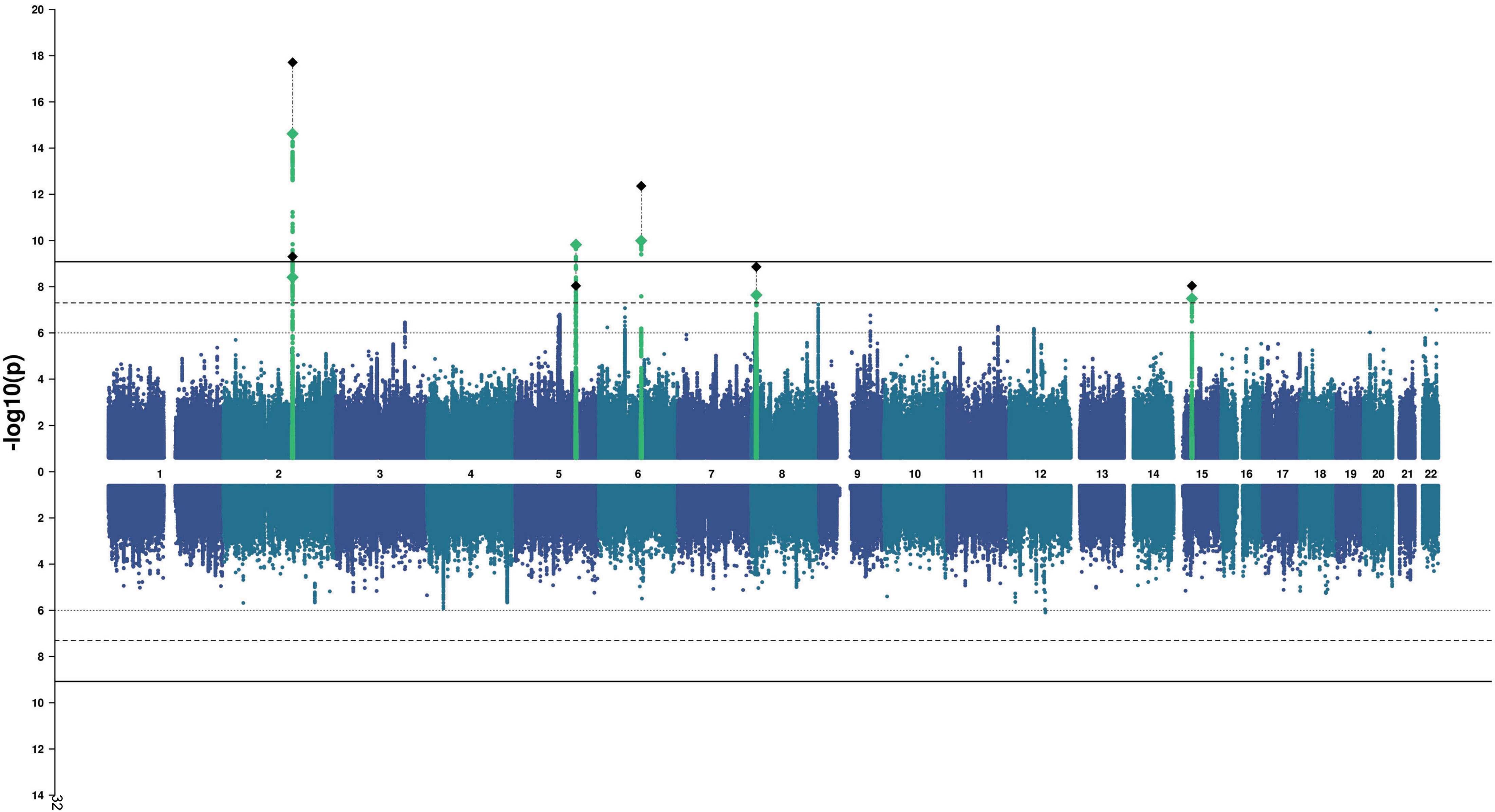

### Lingual

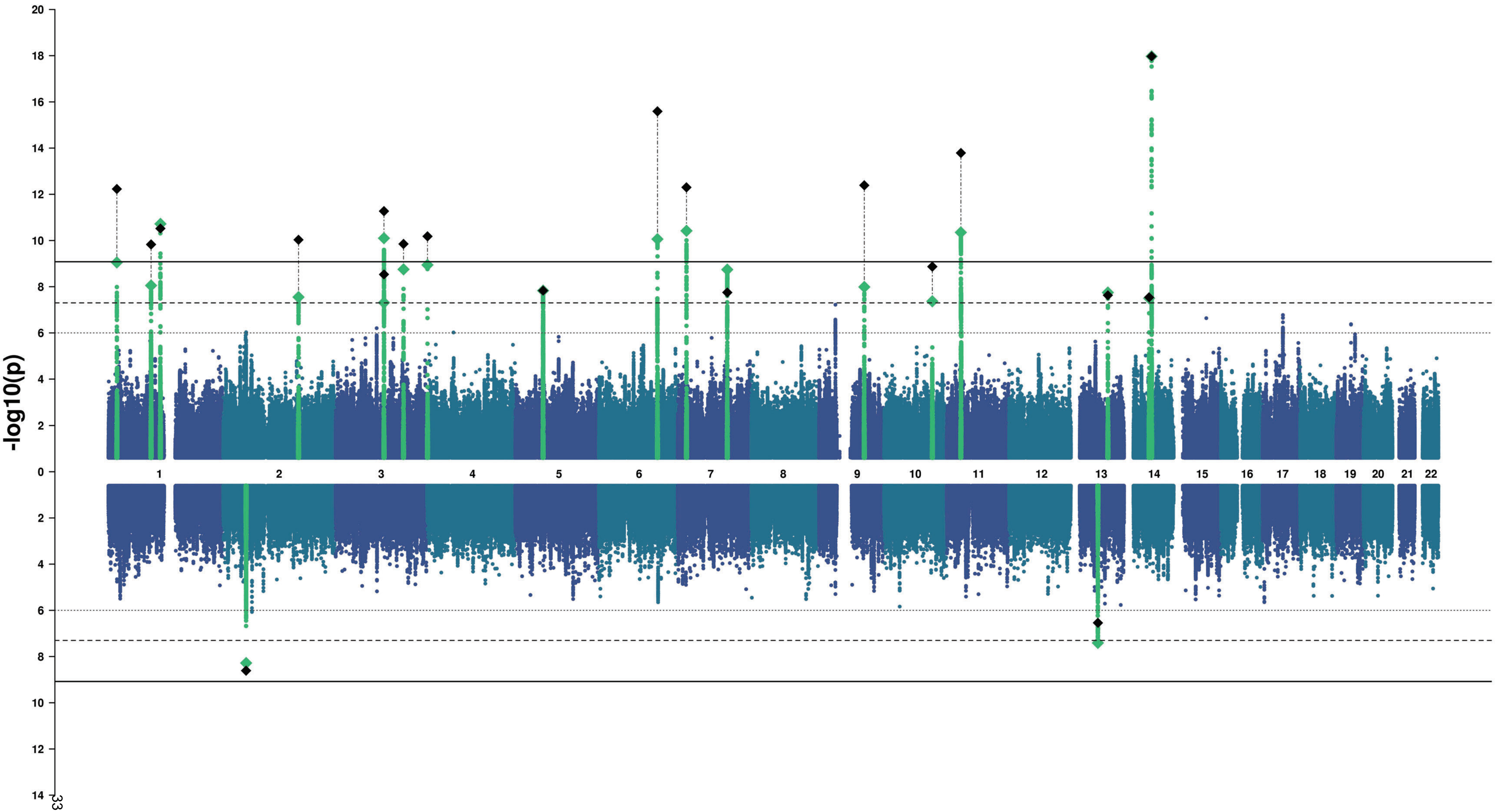

-log10(p)
20
18
16
14
12
10
8
6
4
2
0
2
4
6
8
10
12
14

### Pericalcarine

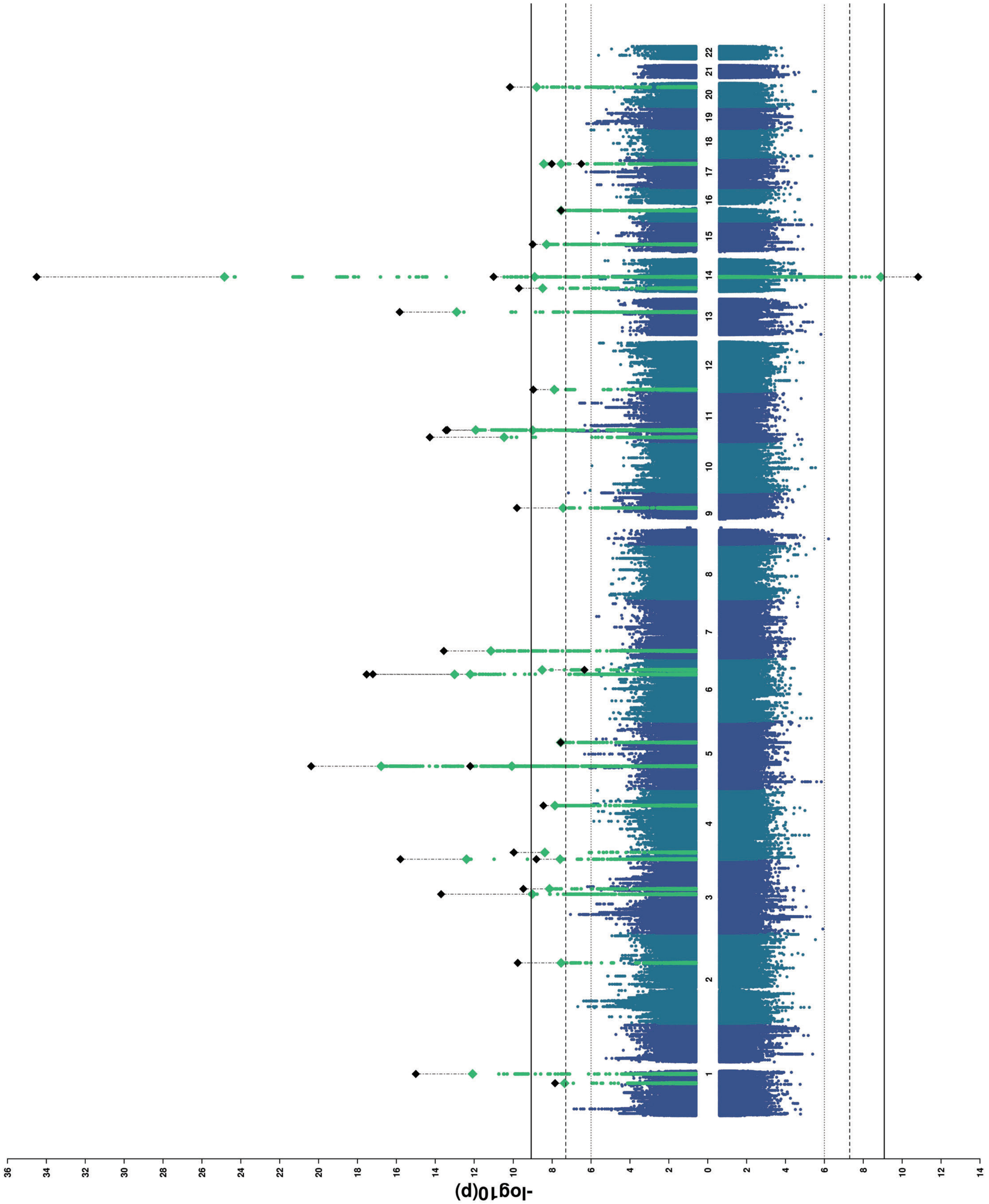

### Cuneus

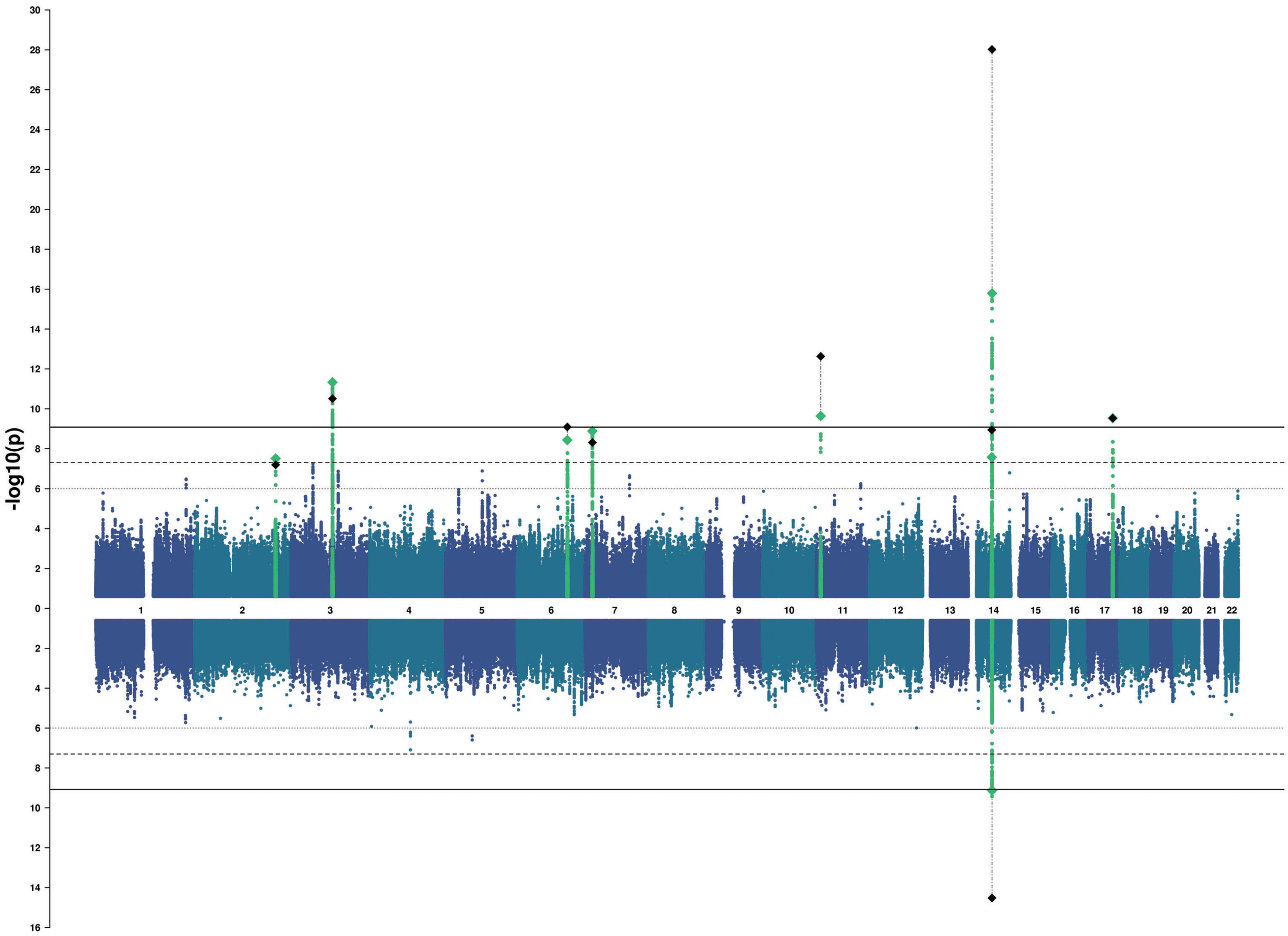

-log10(p)
30
28
26
24
22
20
18
16
14
12
10
8
6
4
2
0
2
4
6
8
10
12
14
16

### Lateral Occipital

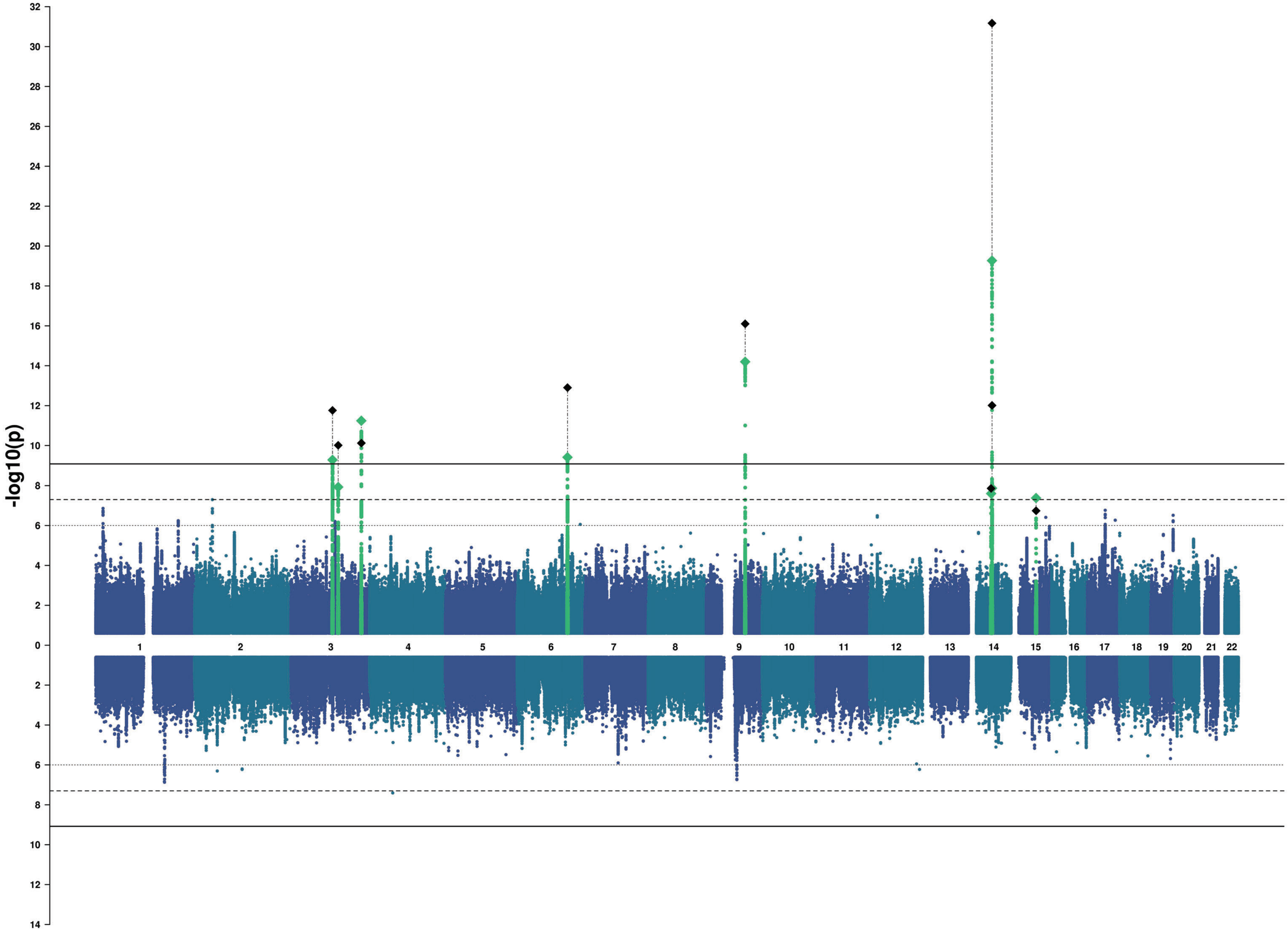
