## Supplemental_QQ_Plots for "The genetic architecture of the human cerebral cortex"


| Region# | Phenotype | Page |
|---|---|---|
| 0 | Global | 2 |
| 1 | Frontal Pole | 3 |
| 2 | Medial Orbitofrontal | 4 |
| 3 | Lateral Orbitofrontal | 5 |
| 4 | Rostral Anterior Cingulate | 6 |
| 5 | Caudal Anterior Cingulate | 7 |
| 6 | Superior Frontal | 8 |
| 7 | Rostral Middle Frontal | 9 |
| 8 | Pars Orbitalis | 10 |
| 9 | Pars Triangularis | 11 |
| 10 | Pars Opercularis | 12 |
| 11 | Caudal Middle Frontal | 13 |
| 12 | Paracentral | 14 |
| 13 | Precentral | 15 |
| 14 | Postcentral | 16 |
| 15 | Precuneus | 17 |
| 16 | Superior Parietal | 18 |
| 17 | Supramarginal | 19 |
| 18 | Inferior Parietal | 20 |
| 19 | Posterior Cingulate | 21 |
| 20 | Isthmus Cingulate | 22 |
| 21 | Insula | 23 |
| 22 | Entorhinal | 24 |
| 23 | Parahippocampal | 25 |
| 24 | Fusiform | 26 |
| 25 | Temporal Pole | 27 |
| 26 | Inferior Temporal | 28 |
| 27 | Middle Temporal | 29 |
| 28 | Superior Temporal | 30 |
| 29 | Banks of the Superior Temporal Sulcus | 31 |
| 30 | Transverse Temporal | 32 |
| 31 | Lingual | 33 |
| 32 | Pericalcarine | 34 |
| 33 | Cuneus | 35 |
| 34 | Lateral Occipital | 36 |

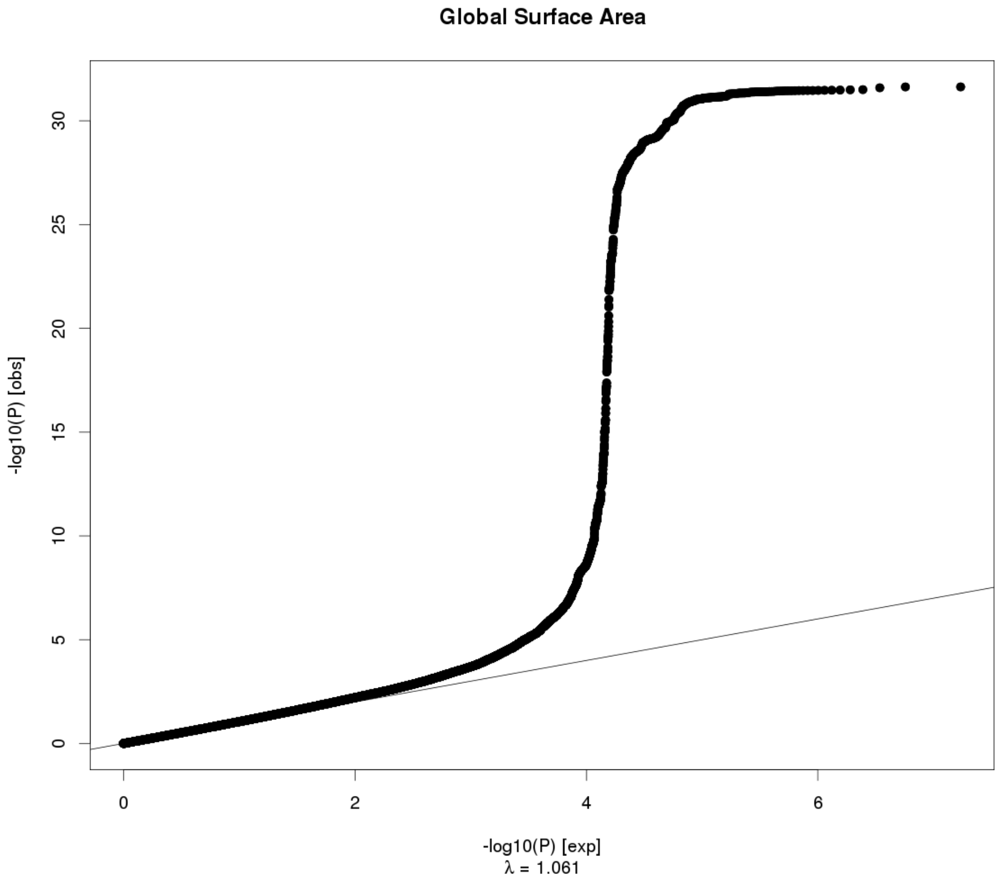
Global Surface Area
-log10(P) [obs]
0
5
10
15
20
25
30
0
2
4
6
-log10(P) [exp]
λ = 1.061

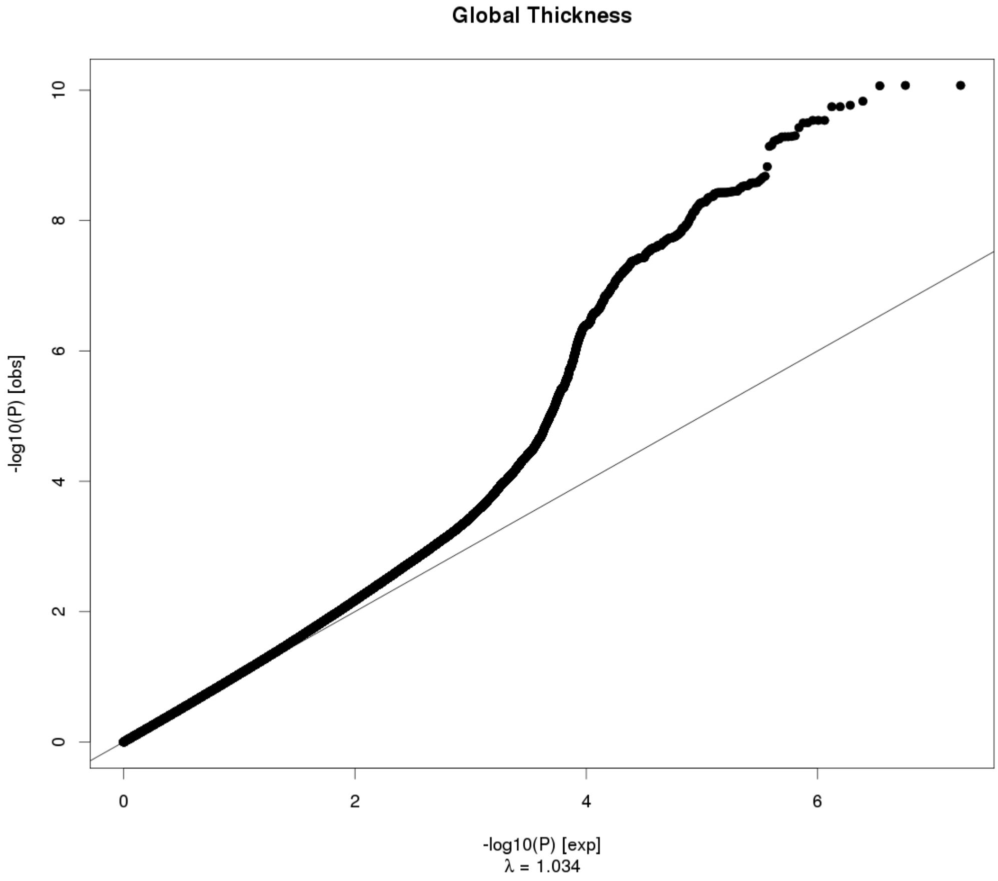
Global Thickness
-log10(P) [obs]
0
2
4
6
8
10
0
2
4
6
-log10(P) [exp]
λ = 1.034

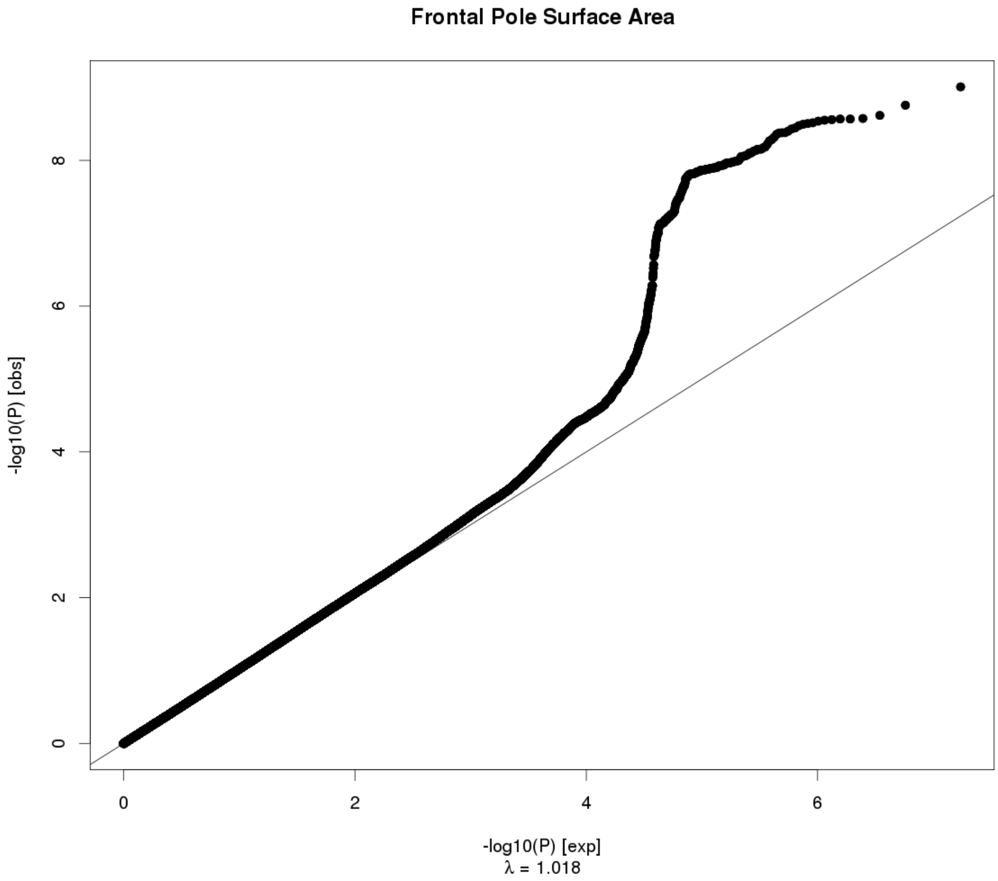
Frontal Pole Surface Area
-log10(P) [obs]
0
2
4
6
8
0
2
4
6
-log10(P) [exp]
λ = 1.018

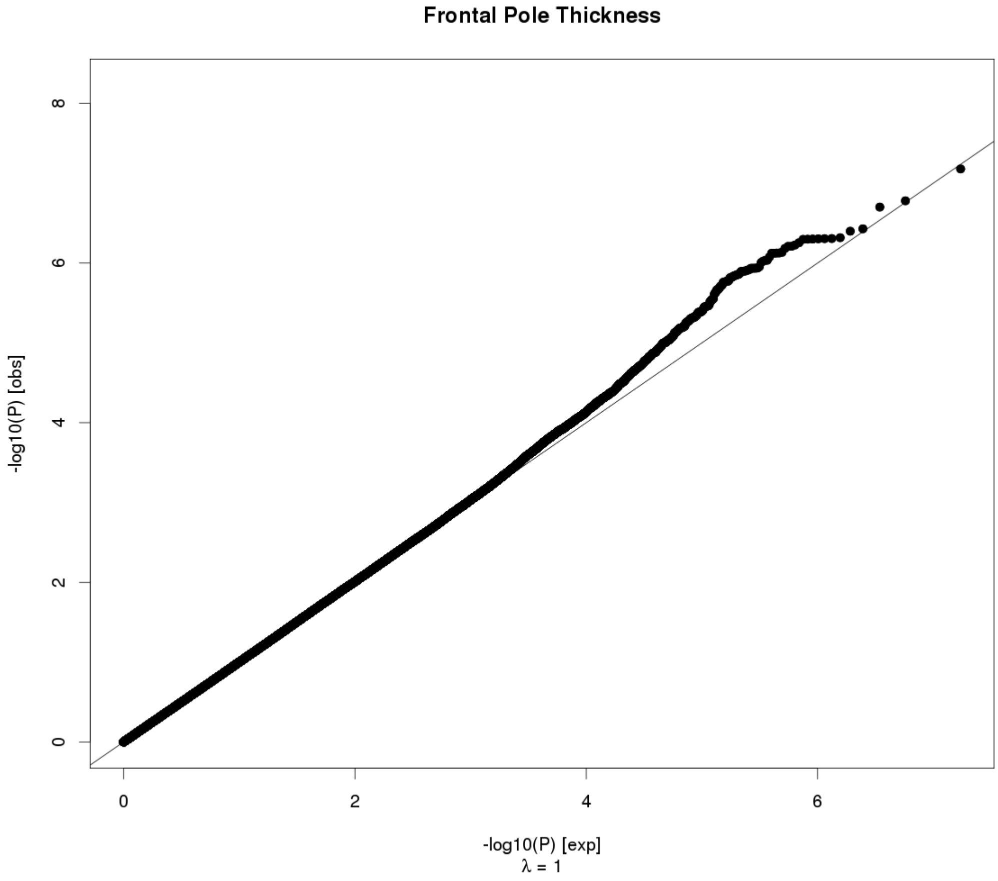
Frontal Pole Thickness
-log10(P) [obs]
0
2
4
6
8
0
2
4
6
-log10(P) [exp]
λ = 1

**Medial Orbitofrontal Surface Area**

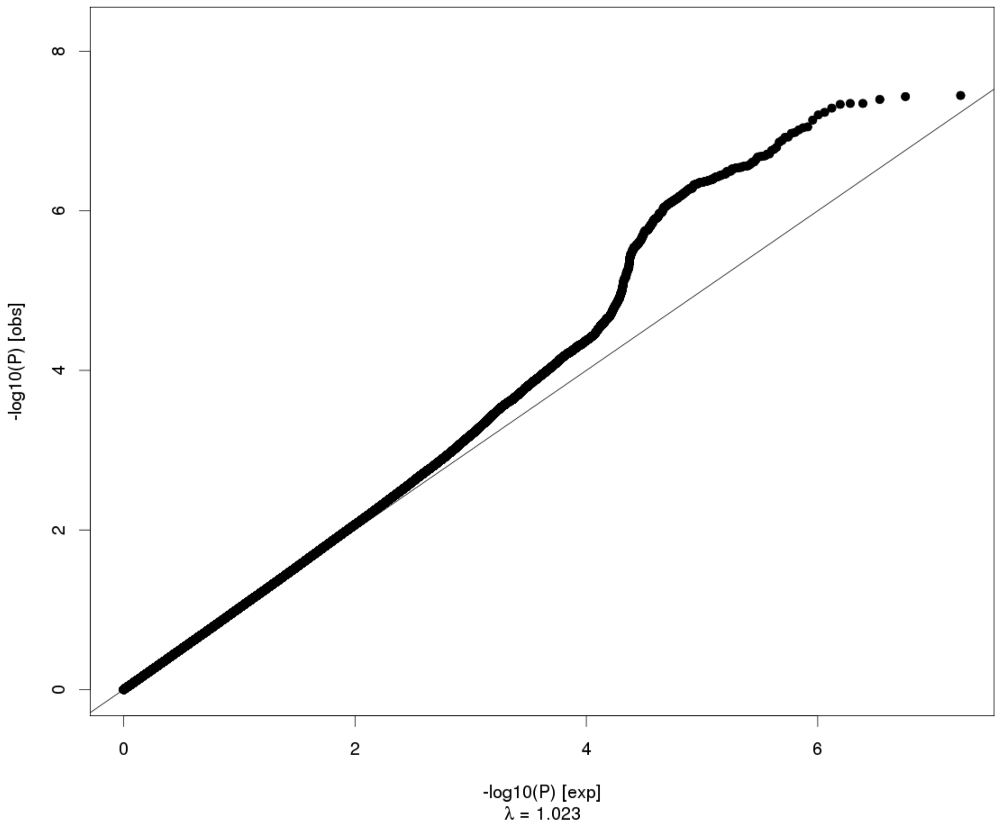


**Medial Orbitofrontal Thickness**

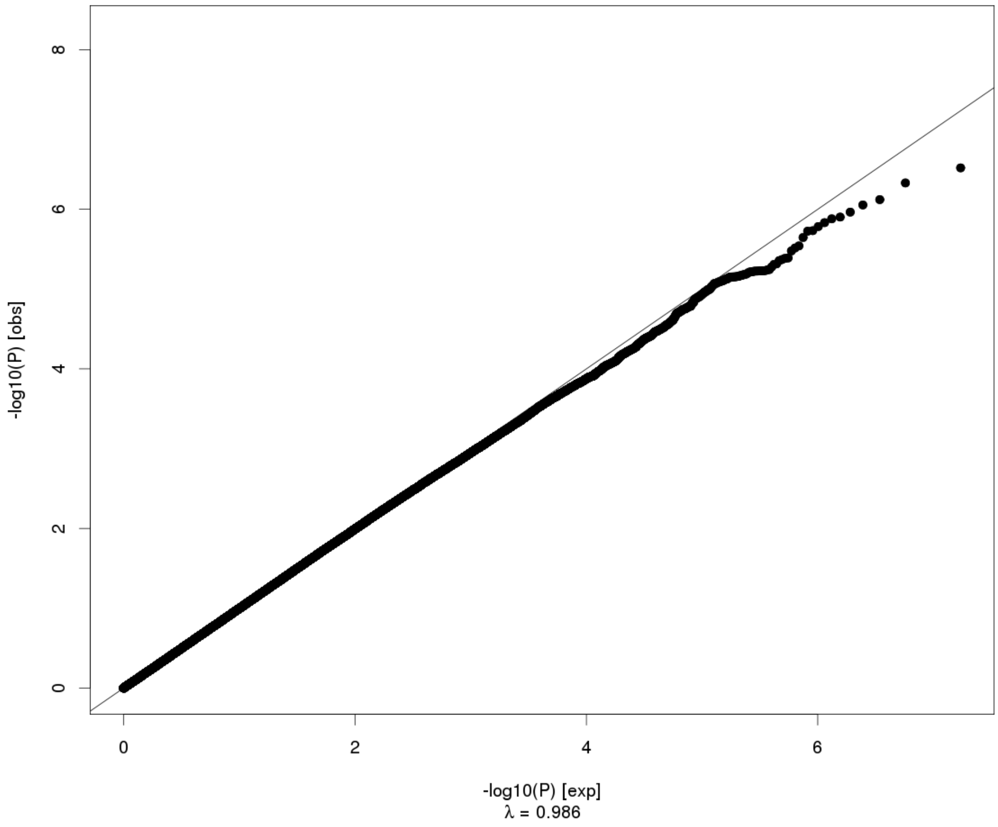

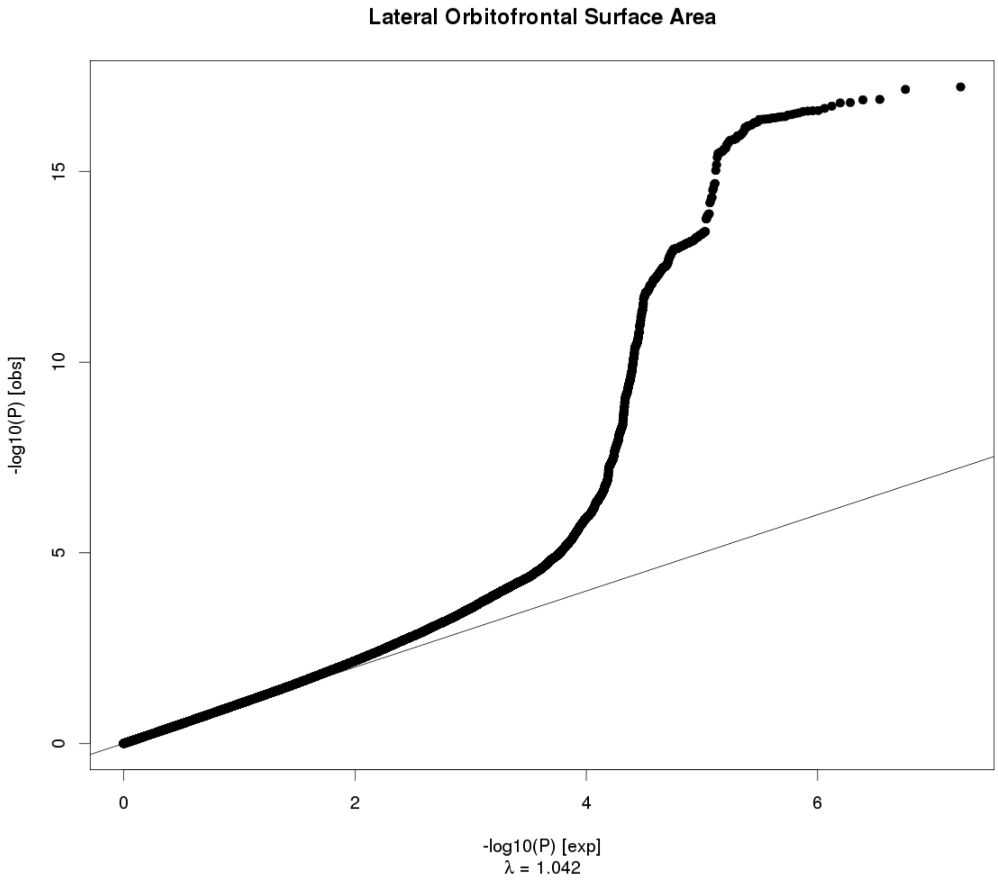

Lateral Orbitofrontal Surface Area
-log10(P) [obs]
0
5
10
15
0
2
4
6
-log10(P) [exp]
λ = 1.042


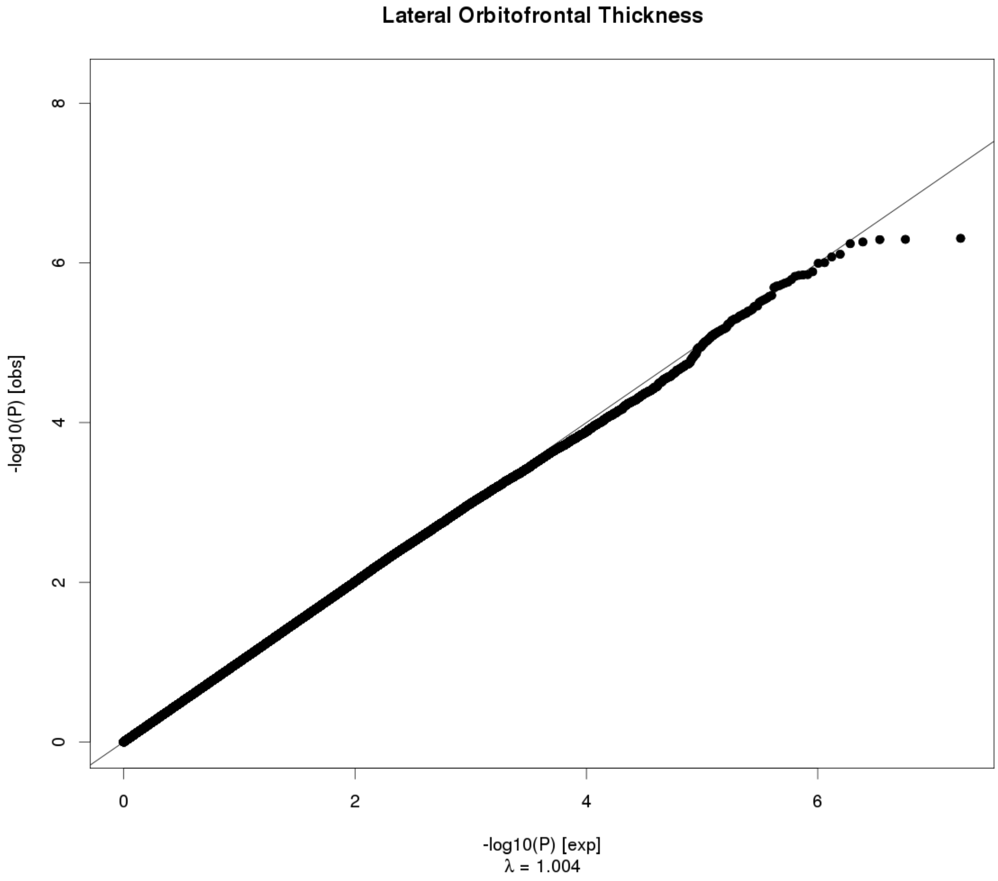

Lateral Orbitofrontal Thickness
-log10(P) [obs]
0
2
4
6
8
0
2
4
6
-log10(P) [exp]
λ = 1.004

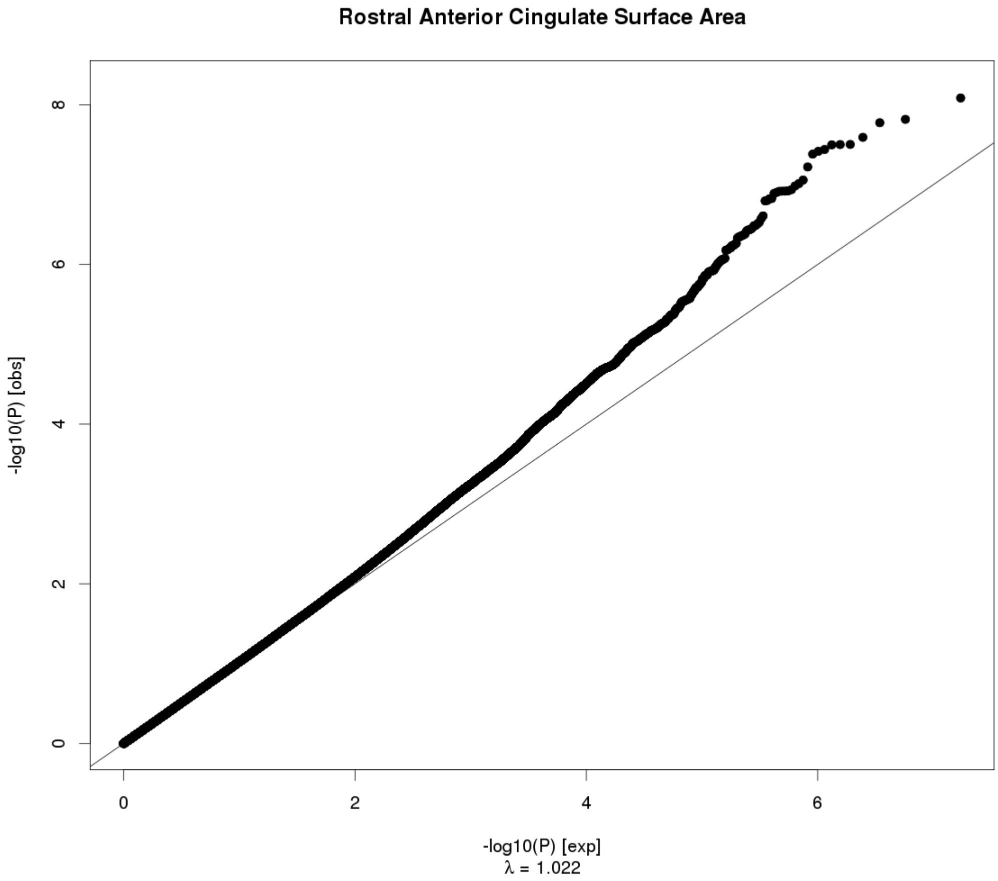
Rostral Anterior Cingulate Surface Area
-log10(P) [obs]
-log10(P) [exp]
λ = 1.022
0
2
4
6
8

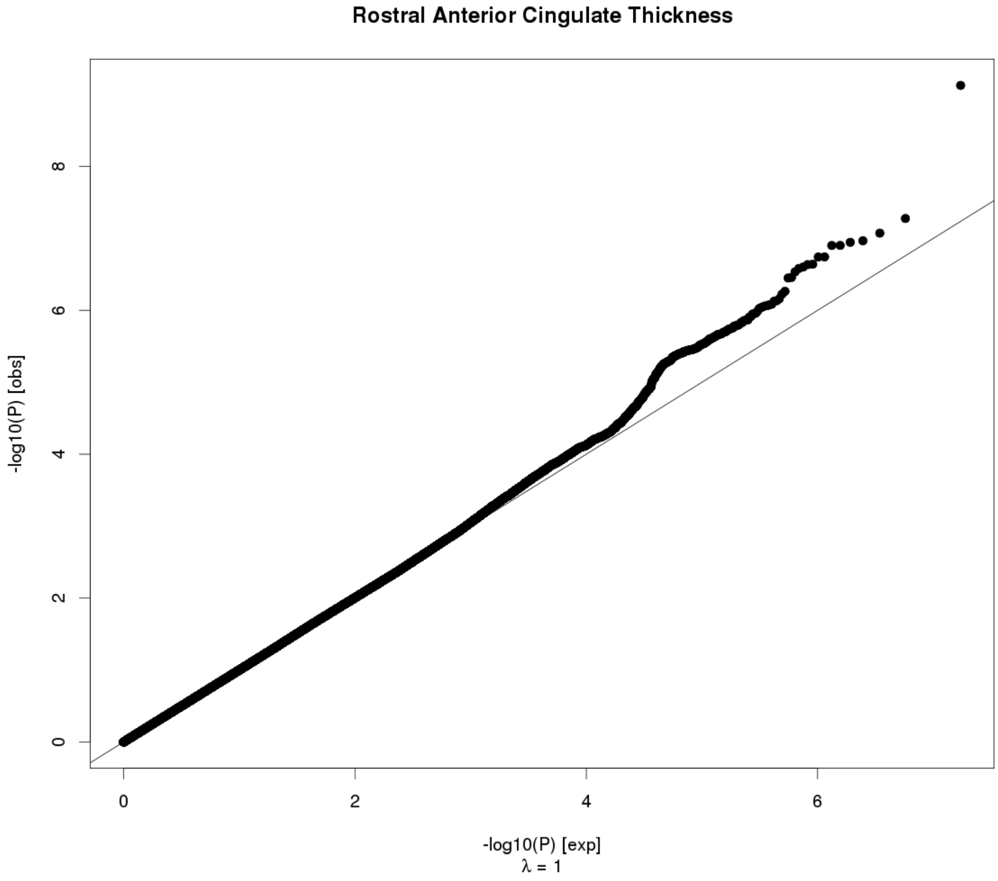
Rostral Anterior Cingulate Thickness
-log10(P) [obs]
-log10(P) [exp]
λ = 1
0
2
4
6
8

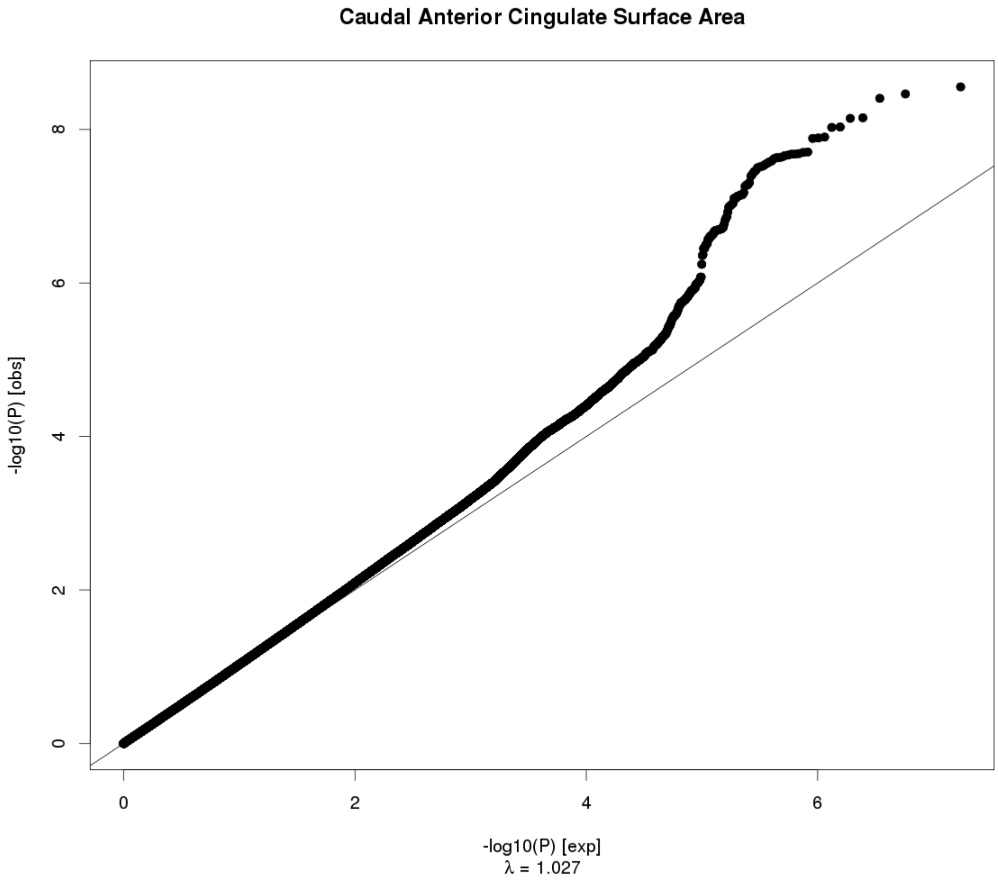
Caudal Anterior Cingulate Surface Area
-log10(P) [obs]
0
2
4
6
8
0
2
4
6
-log10(P) [exp]
λ = 1.027

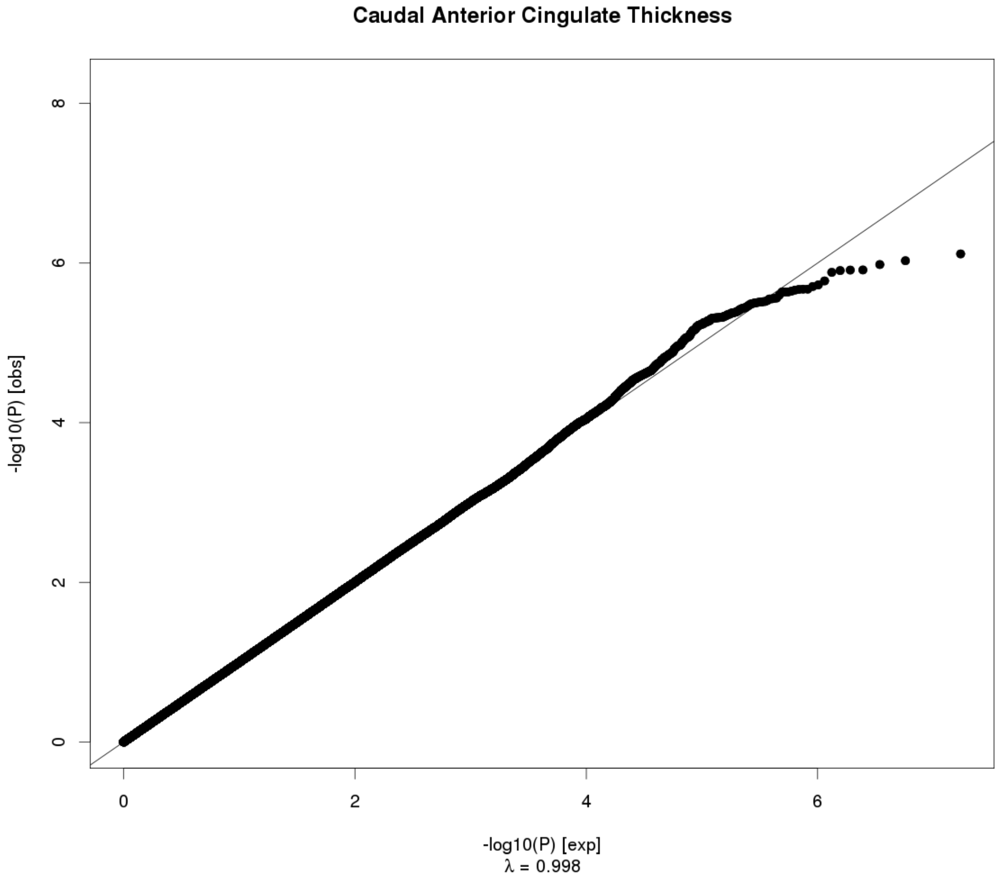
Caudal Anterior Cingulate Thickness
-log10(P) [obs]
0
2
4
6
8
0
2
4
6
-log10(P) [exp]
λ = 0.998

### Superior Frontal Surface Area

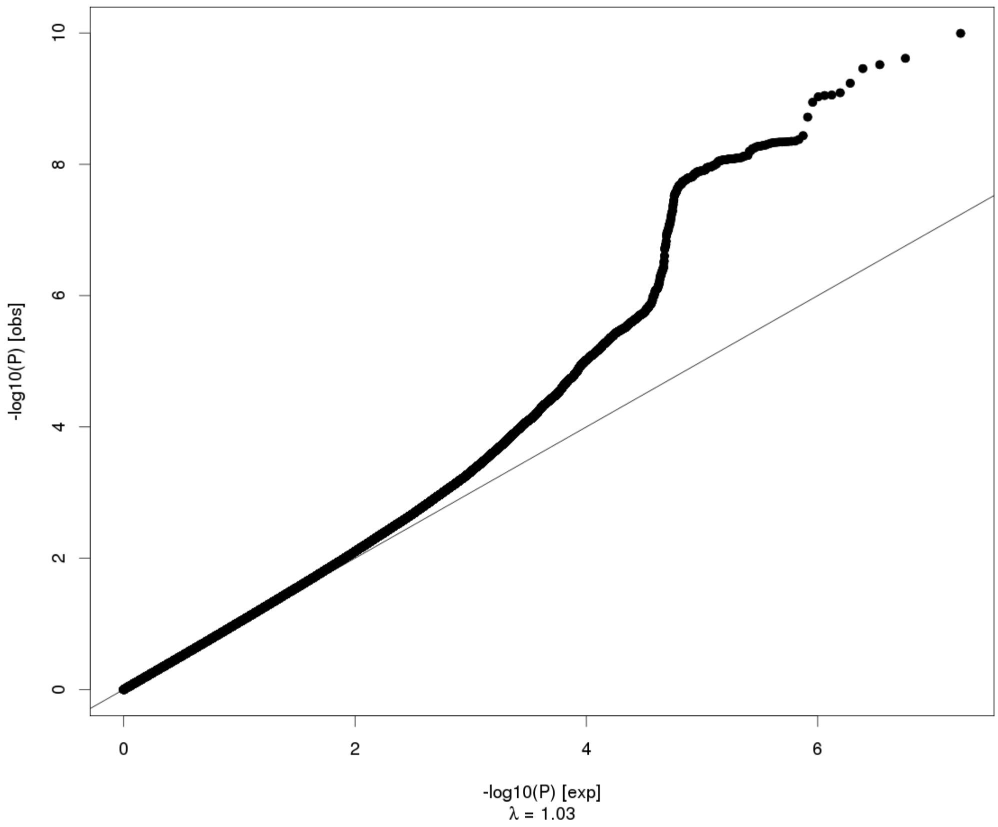


### Superior Frontal Thickness

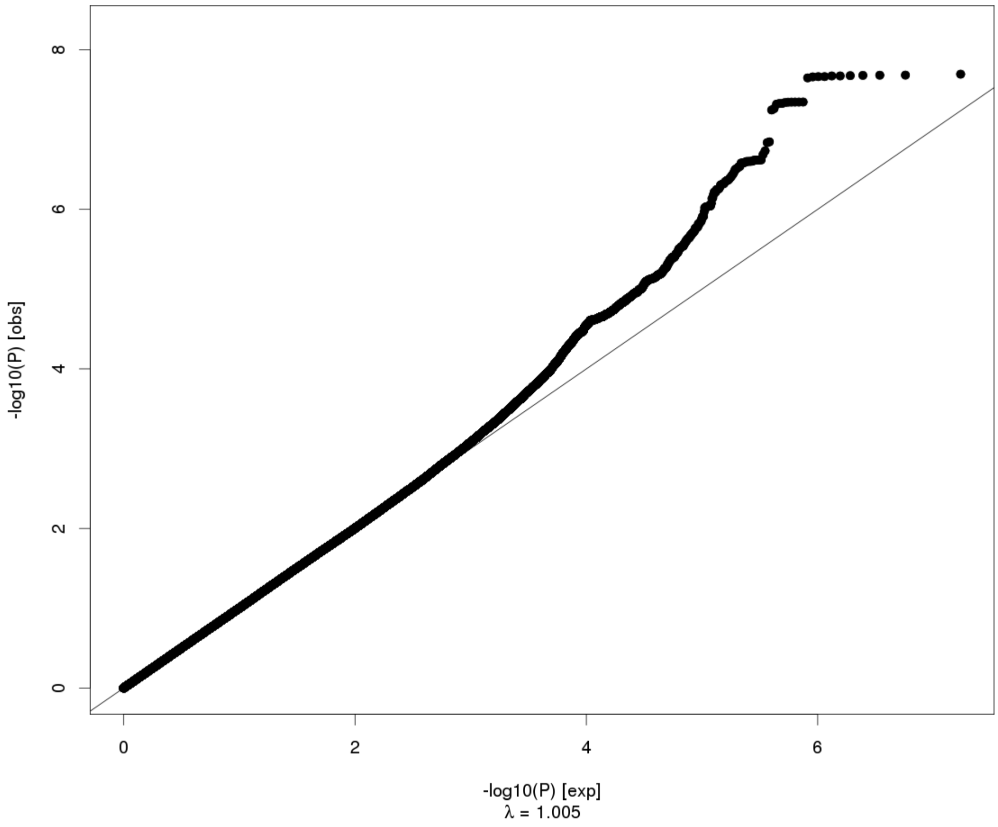

### Rostral Middle Frontal Surface Area

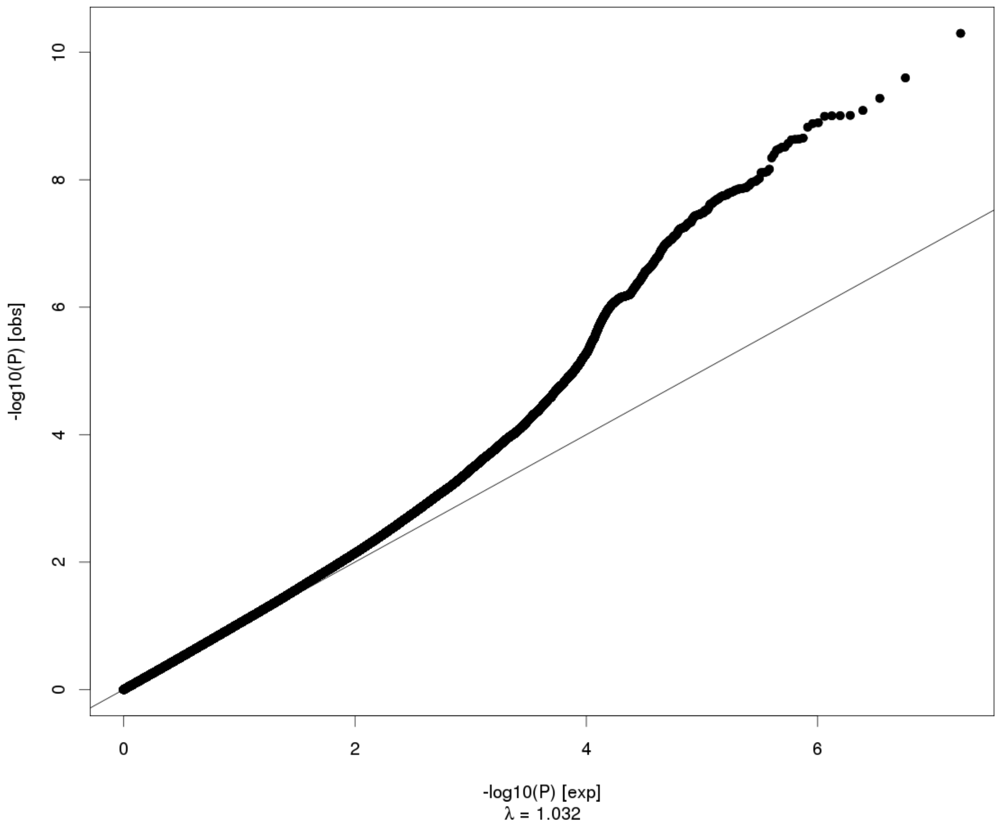


### Rostral Middle Frontal Thickness

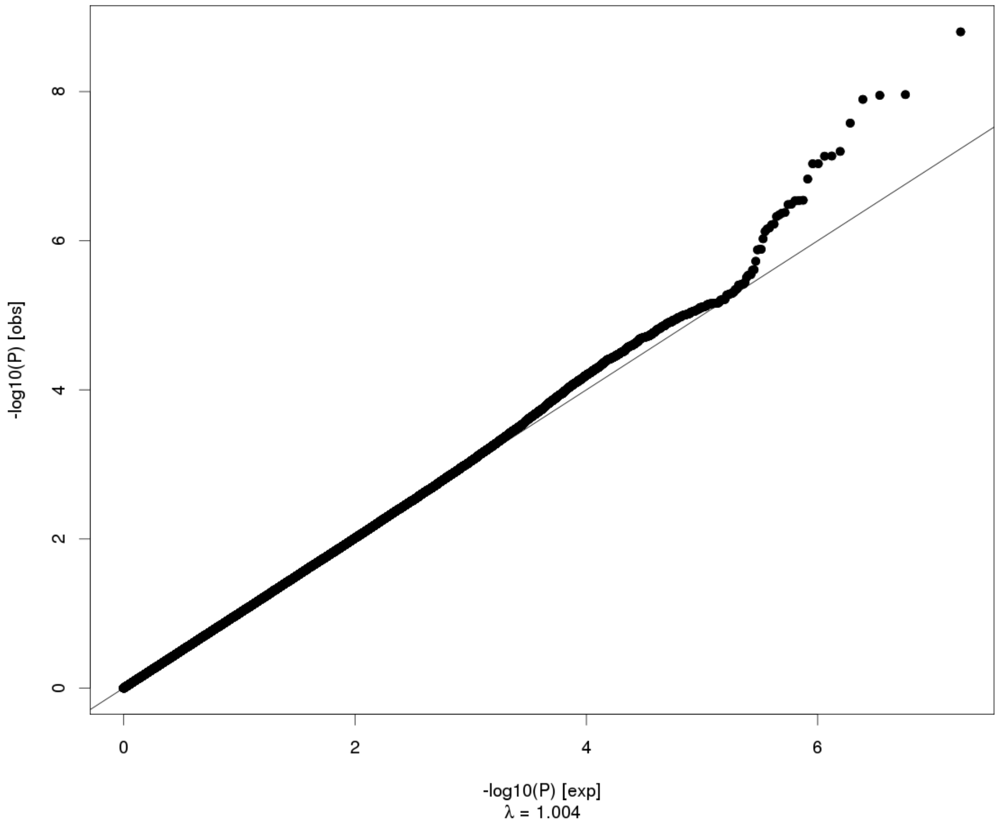

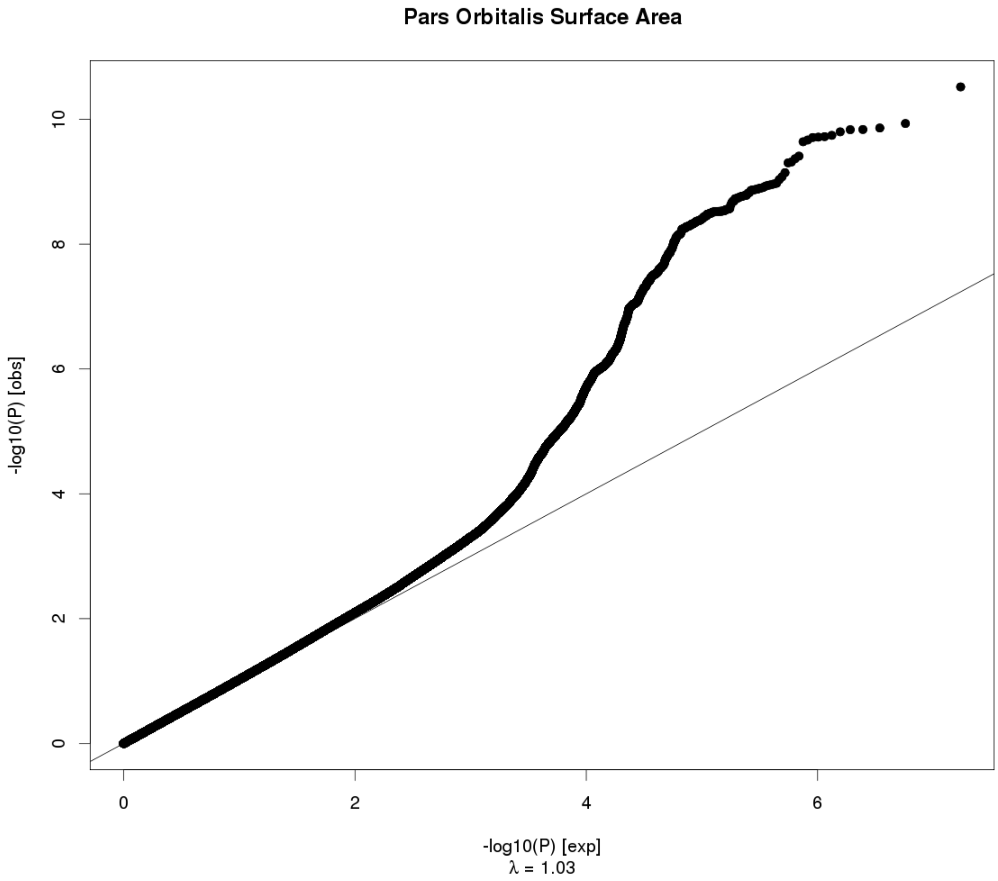

Pars Orbitalis Surface Area
-log10(P) [obs]
0
2
4
6
8
10
0
2
4
6
-log10(P) [exp]
λ = 1.03


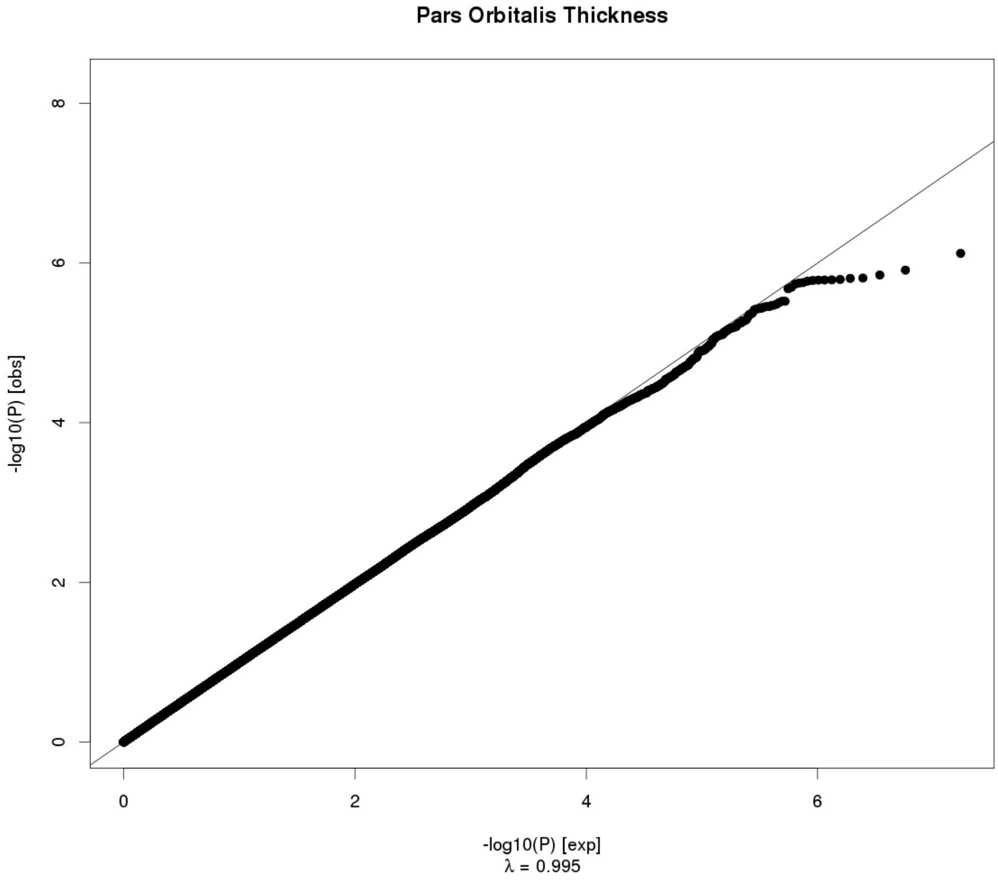

Pars Orbitalis Thickness
-log10(P) [obs]
0
2
4
6
8
0
2
4
6
-log10(P) [exp]
λ = 0.995

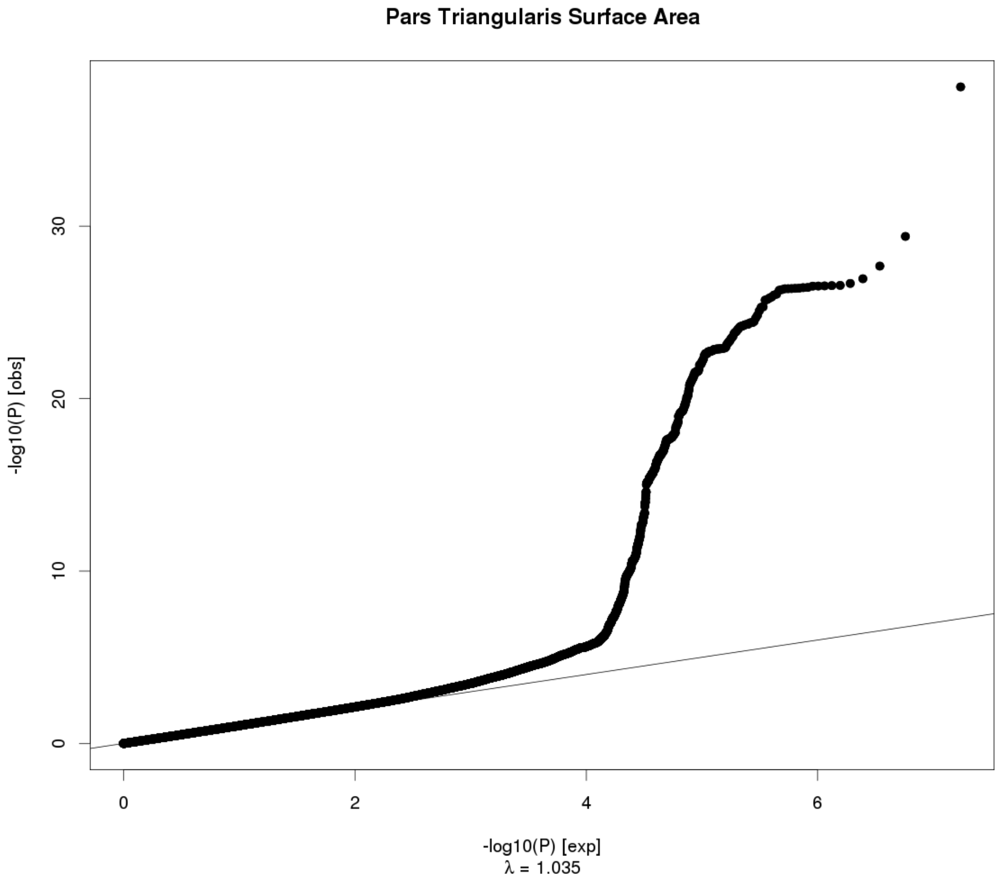
Pars Triangularis Surface Area
-log10(P) [obs]
0
10
20
30
0
2
4
6
-log10(P) [exp]
λ = 1.035

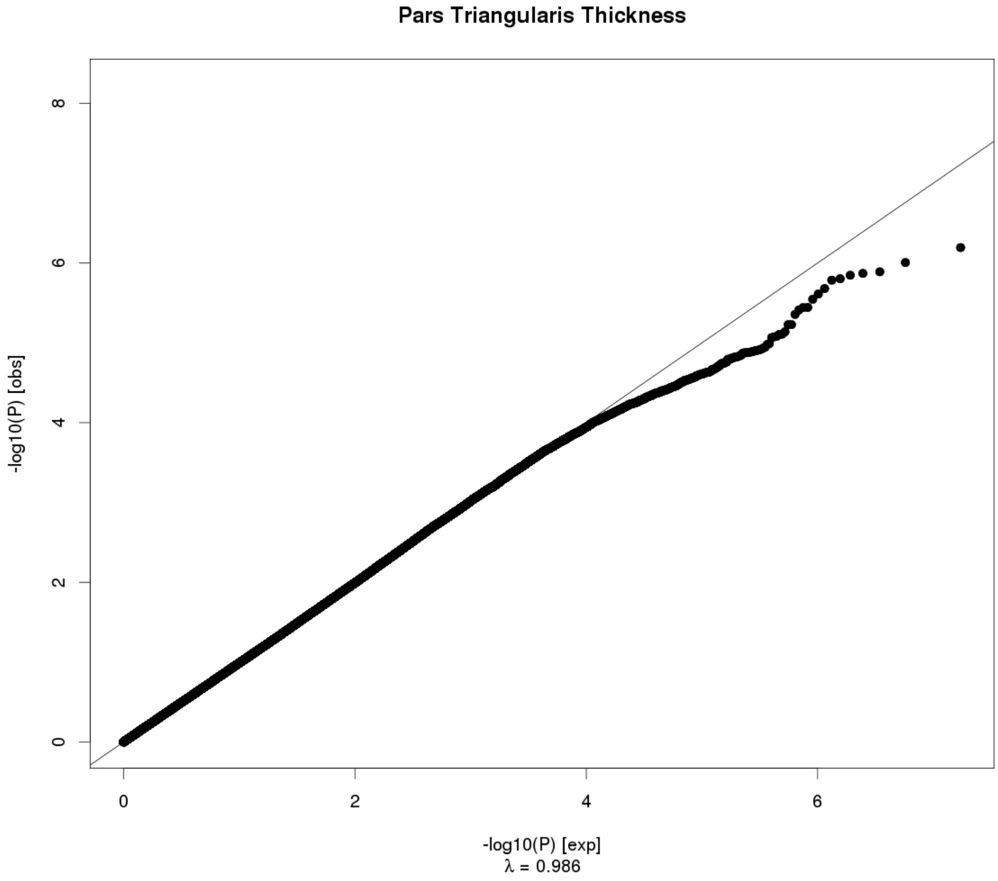
Pars Triangularis Thickness
-log10(P) [obs]
0
2
4
6
8
0
2
4
6
-log10(P) [exp]
λ = 0.986

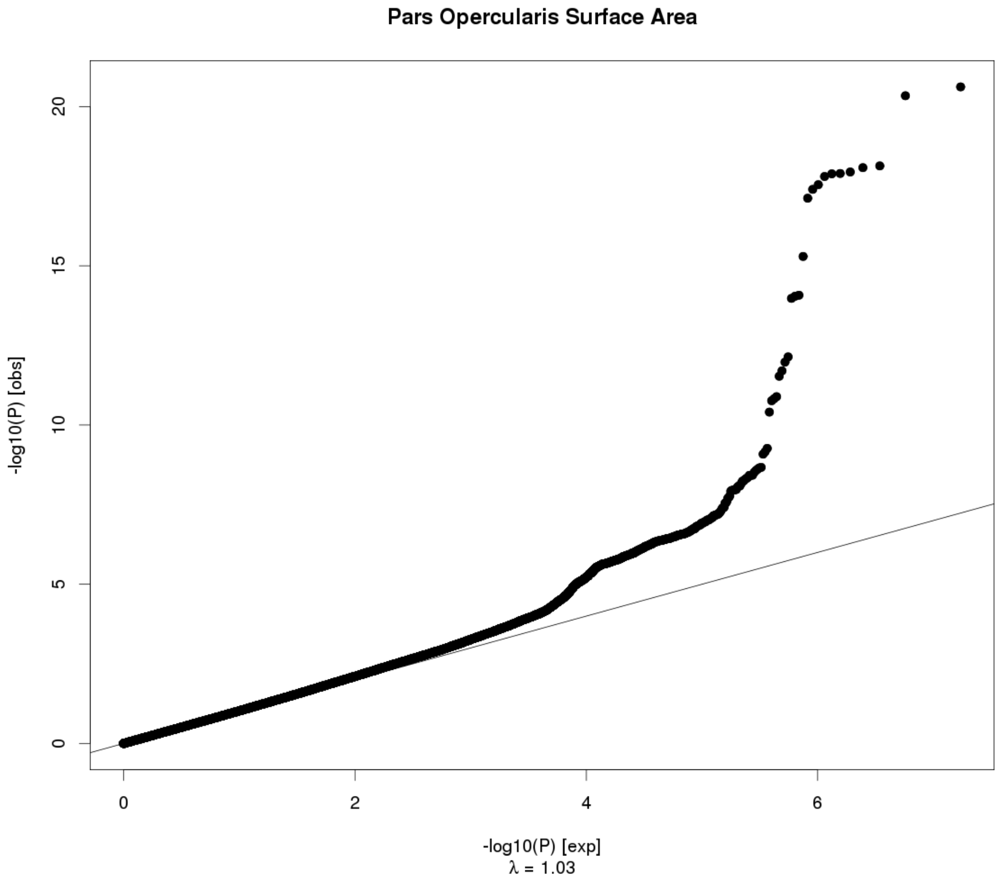
Pars Opercularis Surface Area
-log10(P) [obs]
-log10(P) [exp]
λ = 1.03

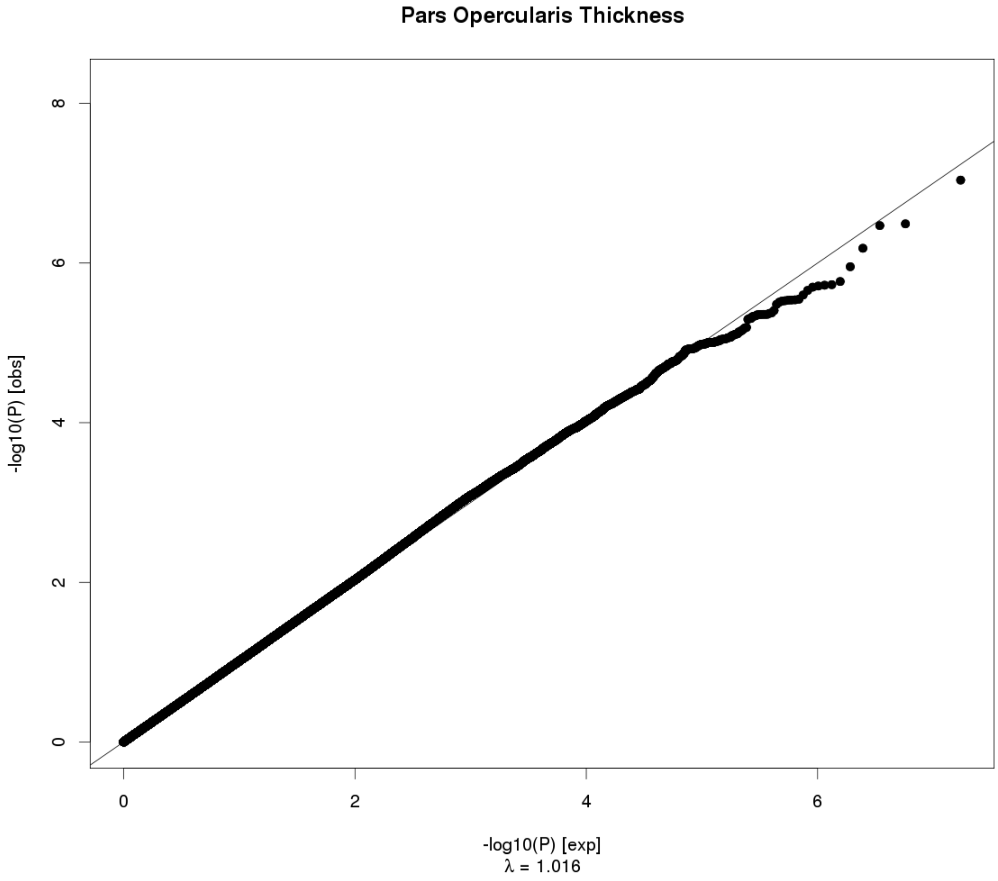
Pars Opercularis Thickness
-log10(P) [obs]
-log10(P) [exp]
λ = 1.016

**Caudal Middle Frontal Surface Area**

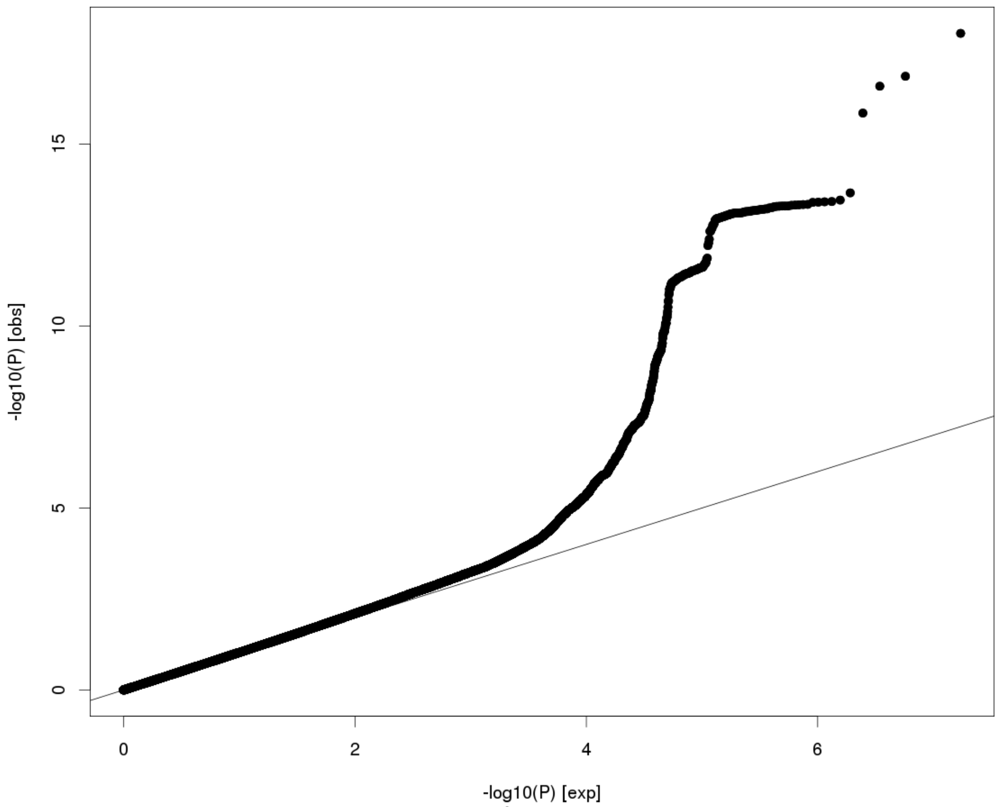


$\lambda$ = 1.034

**Caudal Middle Frontal Thickness**

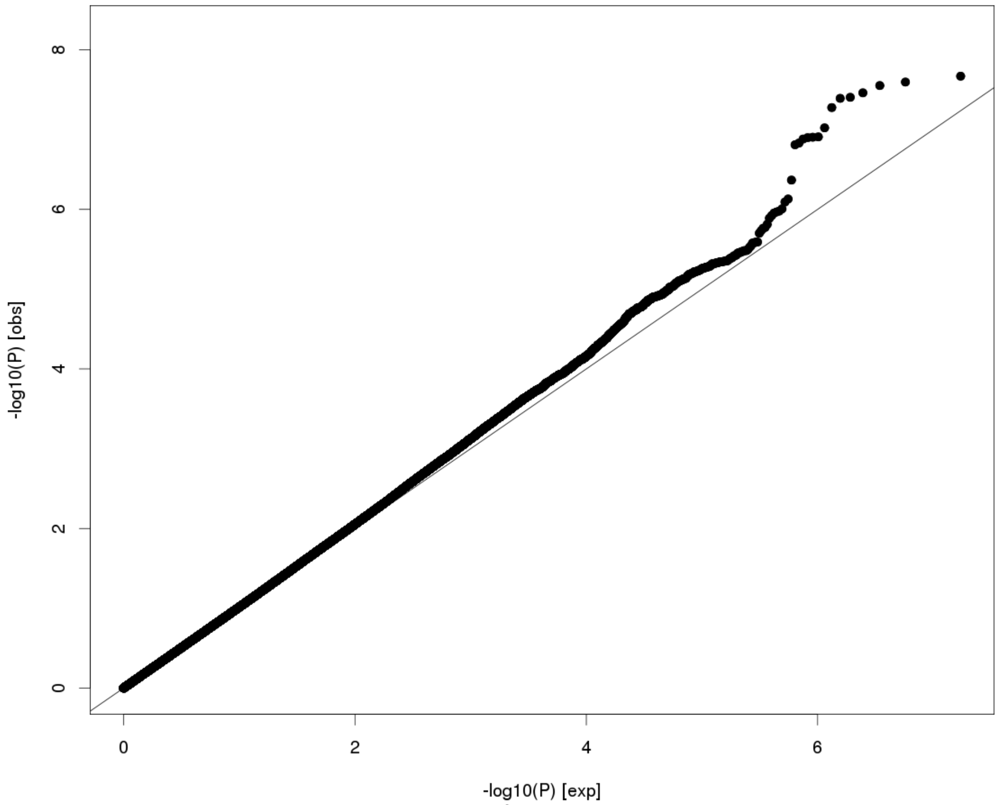


$\lambda$ = 1.019

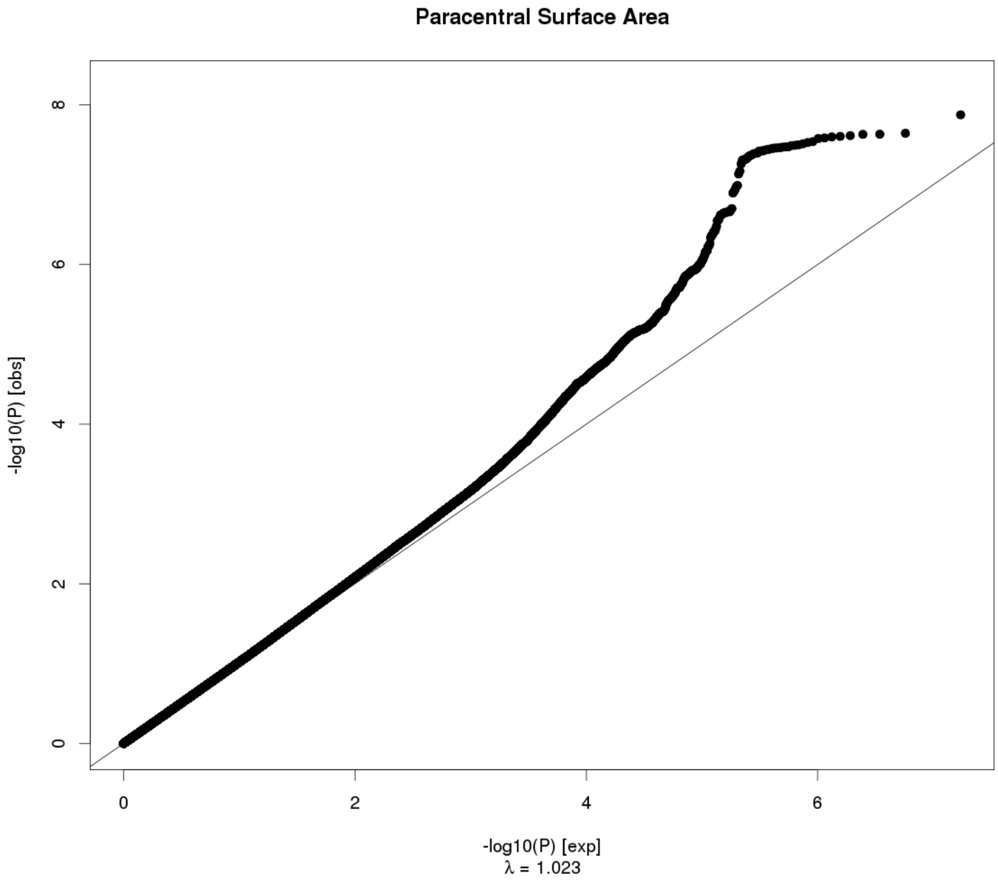

Paracentral Surface Area
-log10(P) [obs]
-log10(P) [exp]
λ = 1.023


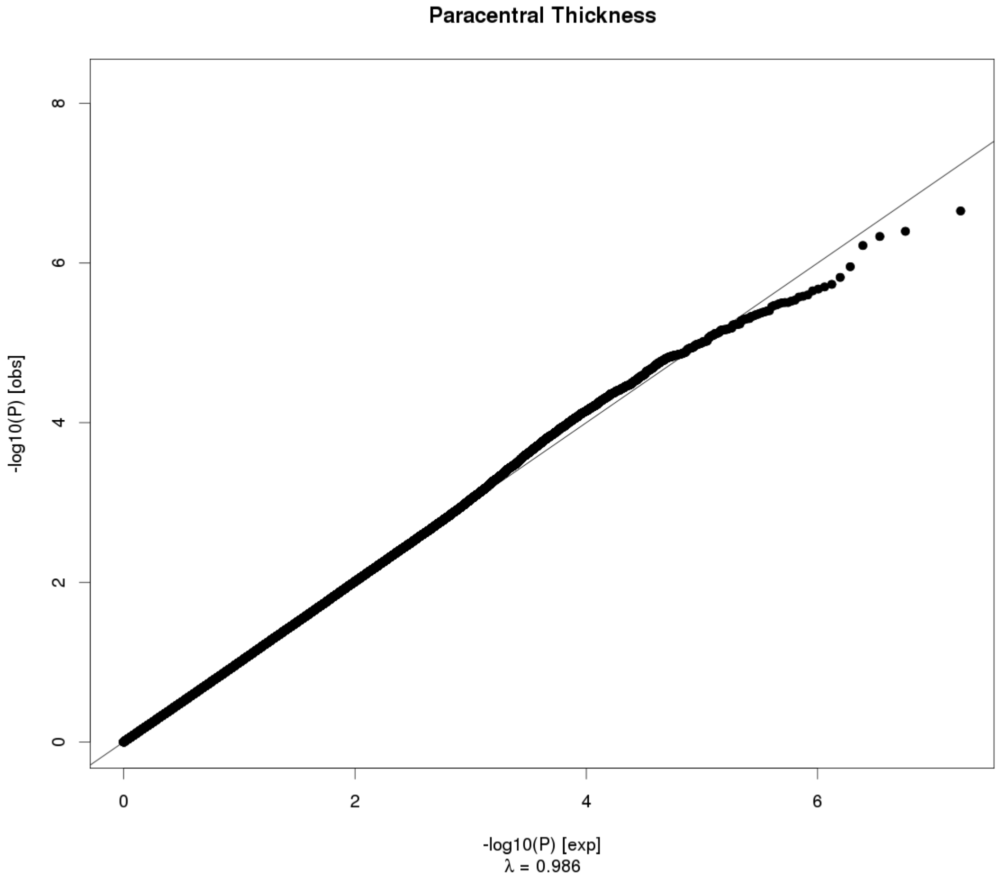

Paracentral Thickness
-log10(P) [obs]
-log10(P) [exp]
λ = 0.986

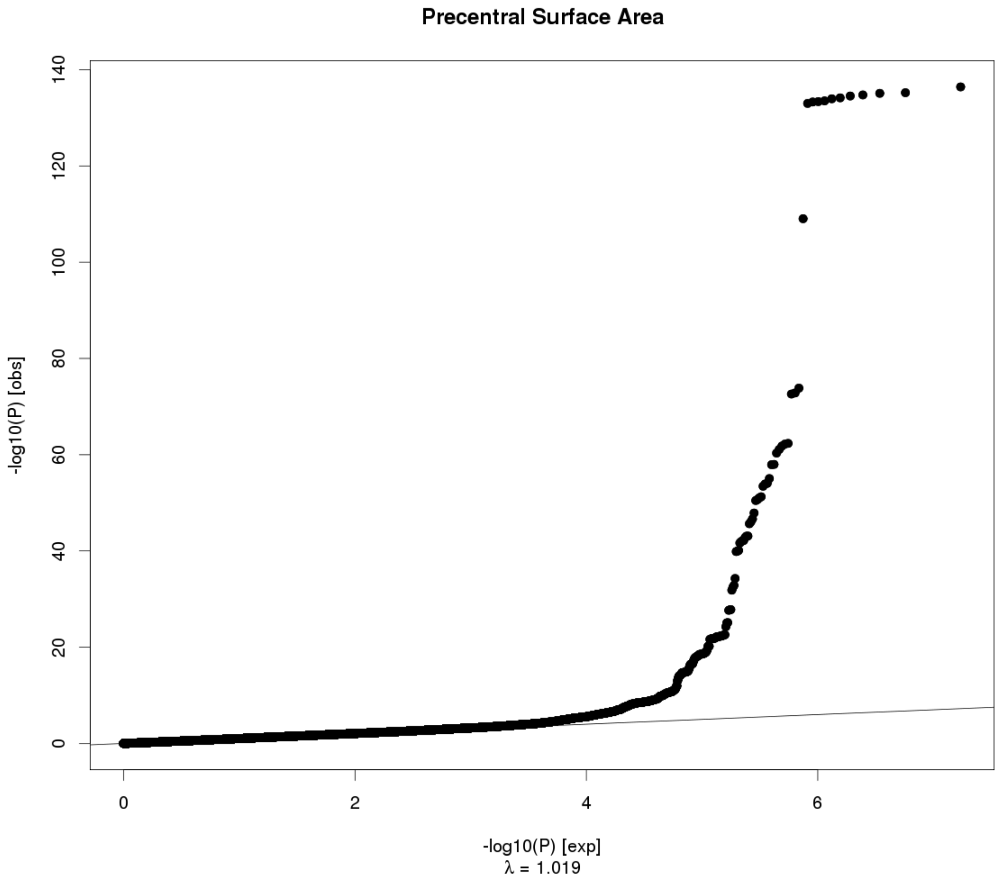
Precentral Surface Area
-log10(P) [obs]
-log10(P) [exp]
λ = 1.019
0
20
40
60
80
100
120
140
0
2
4
6

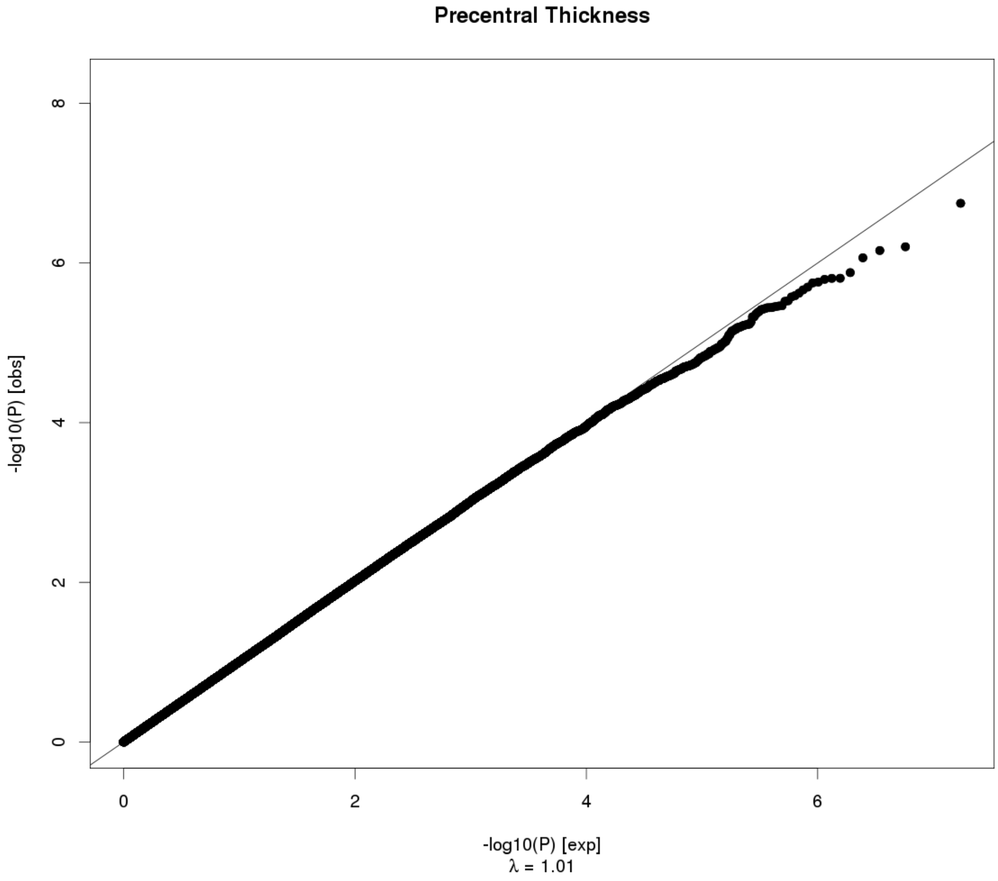
Precentral Thickness
-log10(P) [obs]
-log10(P) [exp]
λ = 1.01
0
2
4
6
8
0
2
4
6

**Postcentral Surface Area**

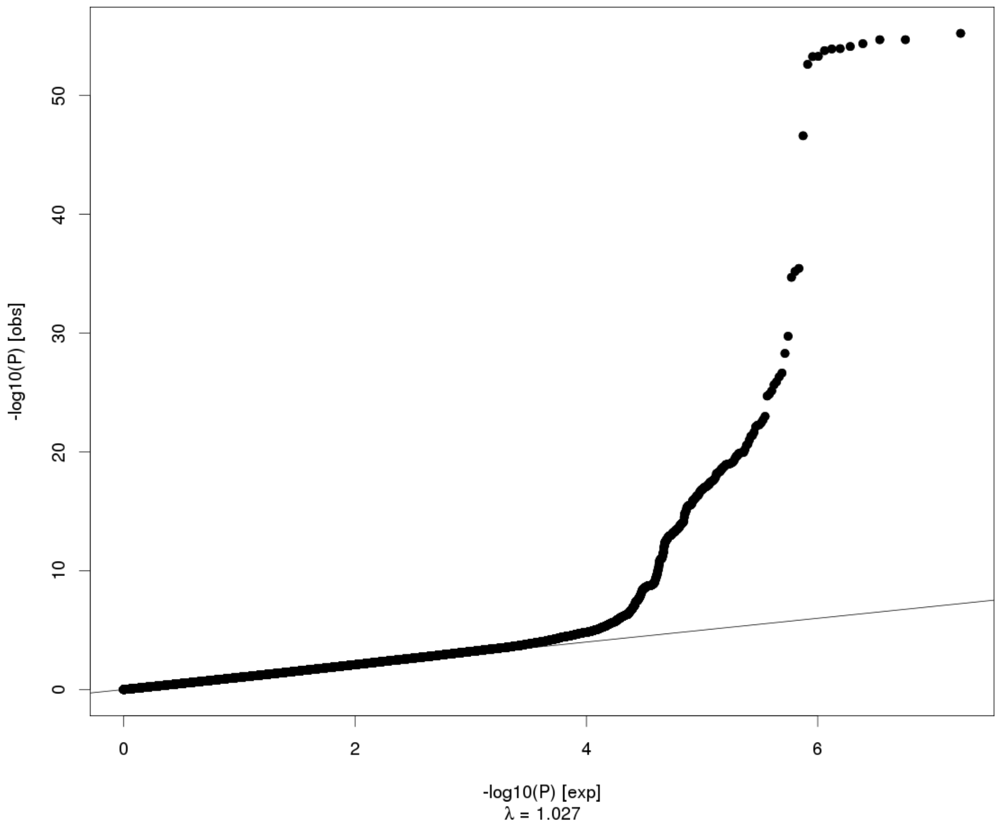


**Postcentral Thickness**

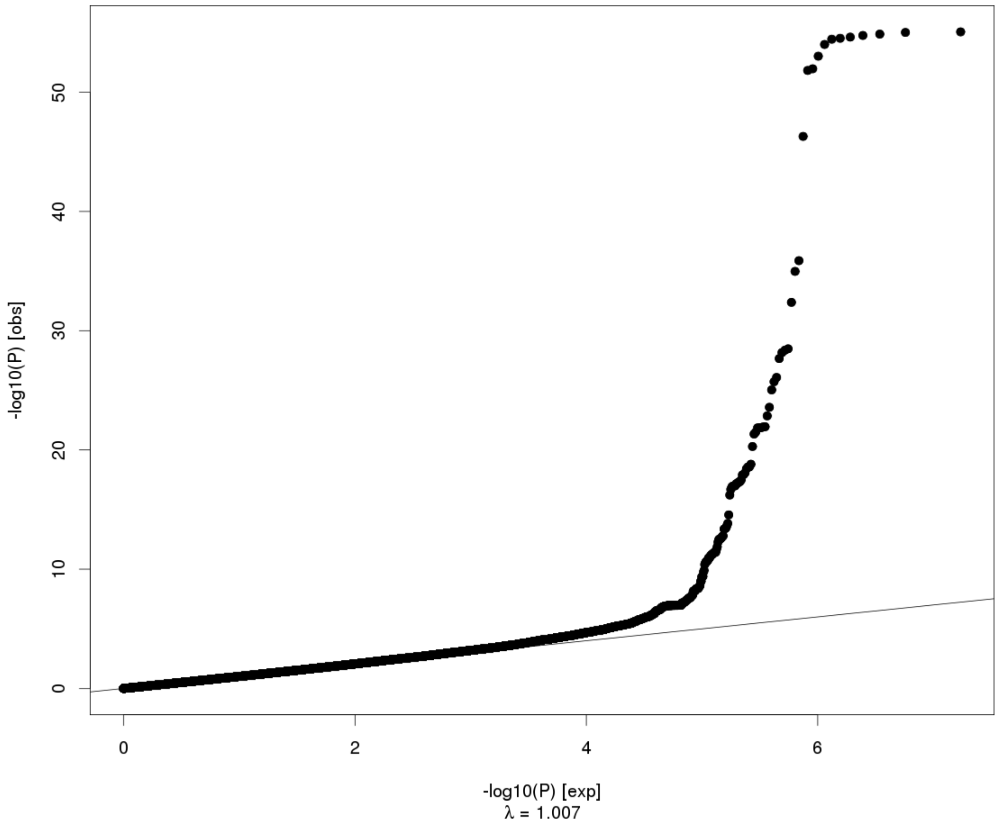

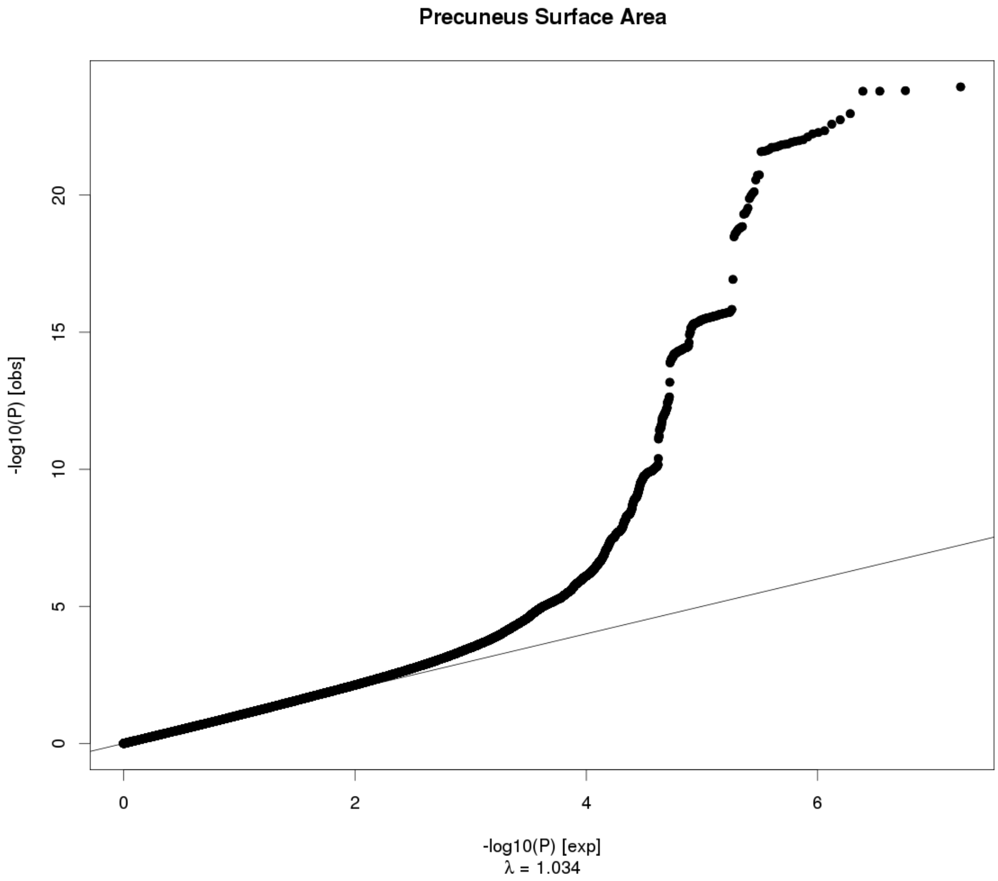

Precuneus Surface Area
-log10(P) [obs]
-log10(P) [exp]
λ = 1.034


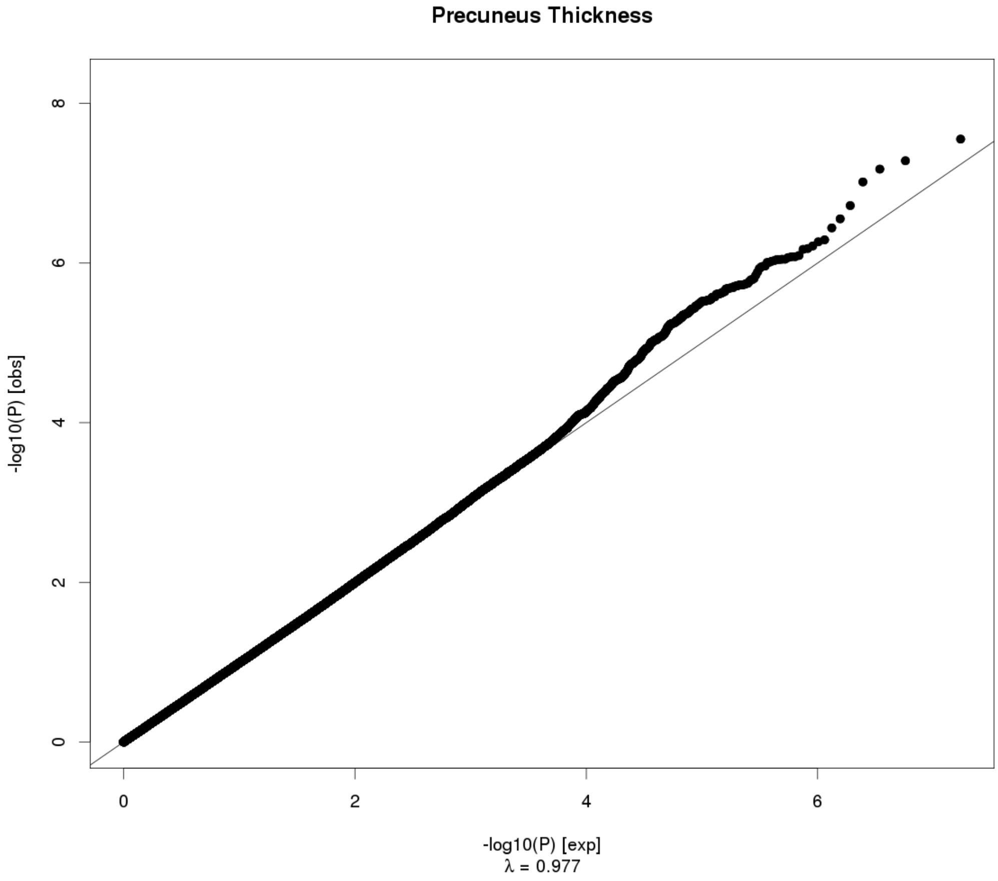

Precuneus Thickness
-log10(P) [obs]
-log10(P) [exp]
λ = 0.977

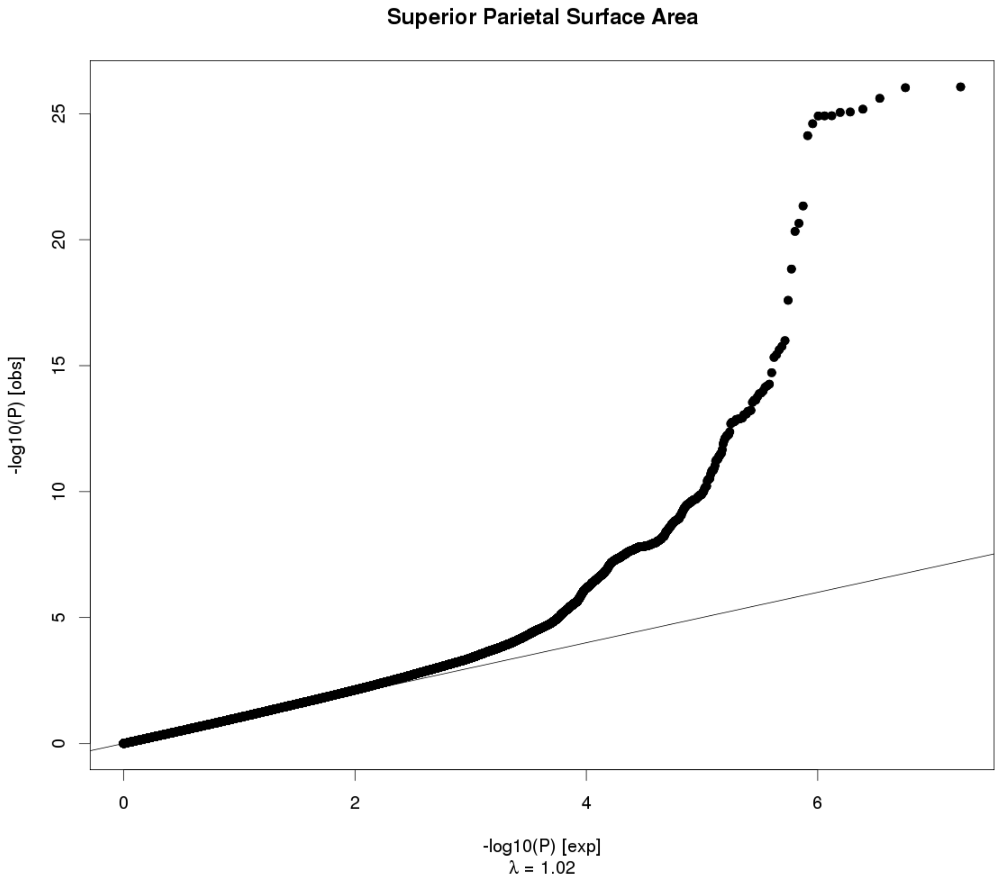
Superior Parietal Surface Area
-log10(P) [obs]
0
5
10
15
20
25
0
2
4
6
-log10(P) [exp]
λ = 1.02

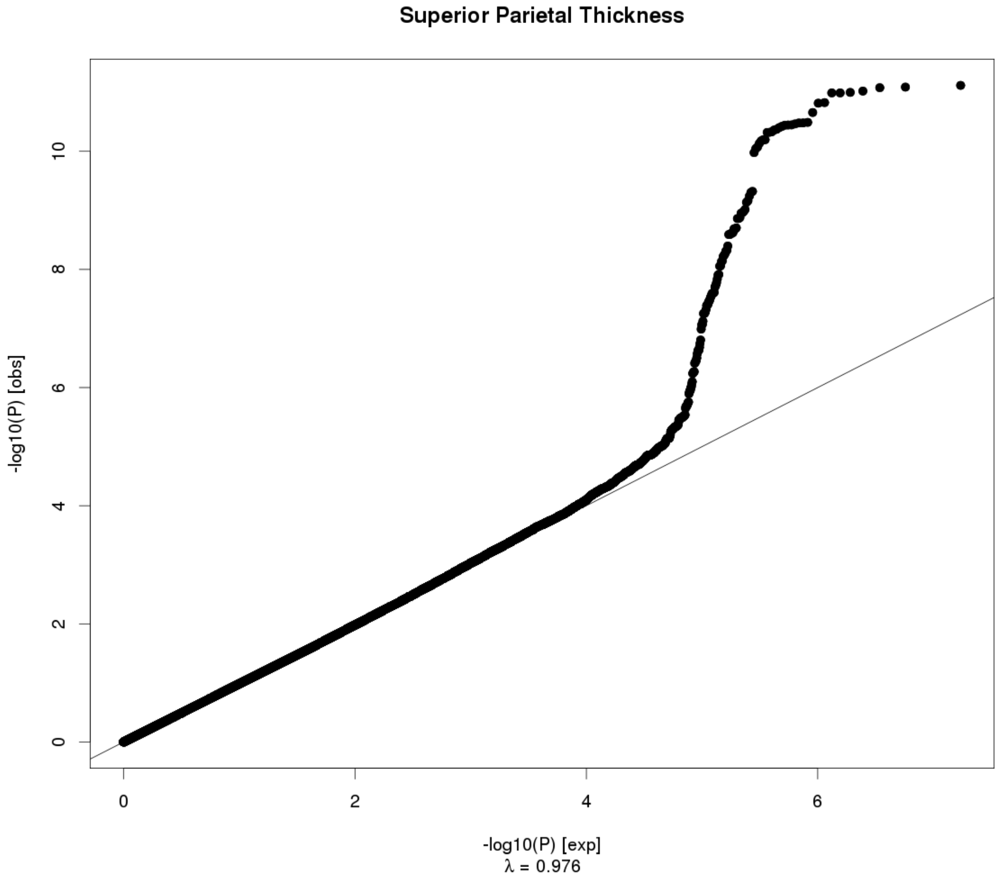
Superior Parietal Thickness
-log10(P) [obs]
0
2
4
6
8
10
0
2
4
6
-log10(P) [exp]
λ = 0.976

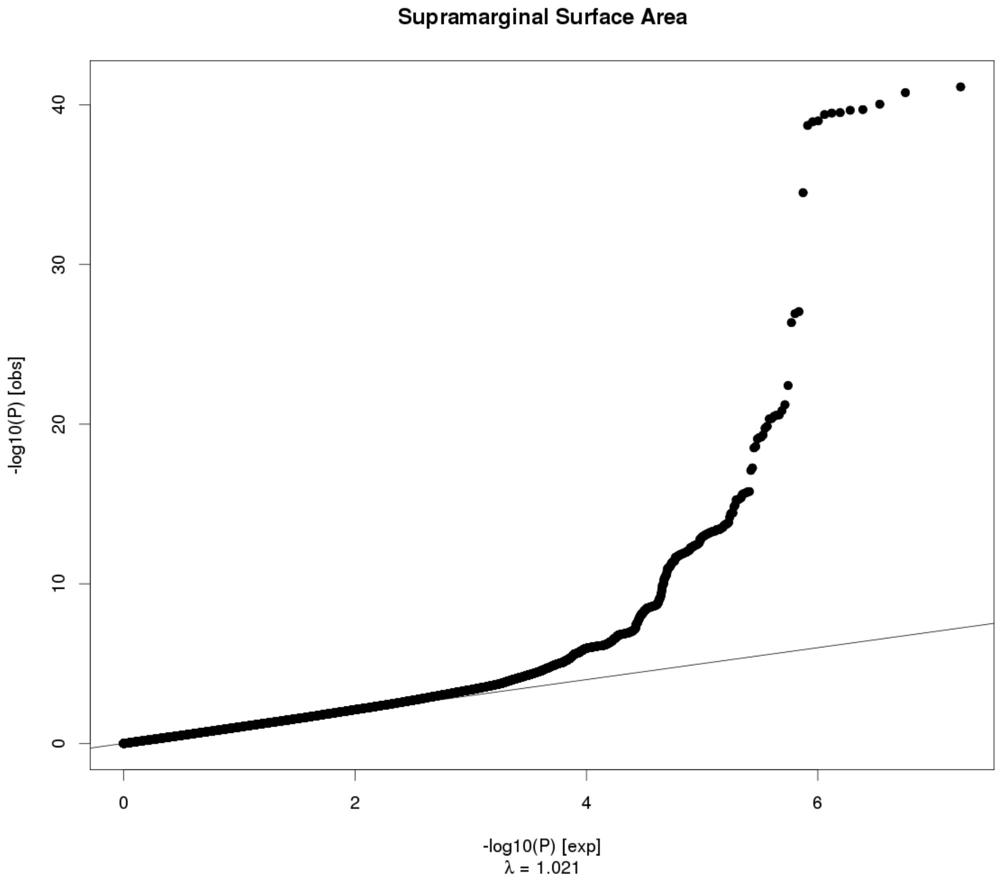
Supramarginal Surface Area
-log10(P) [obs]
-log10(P) [exp]
λ = 1.021

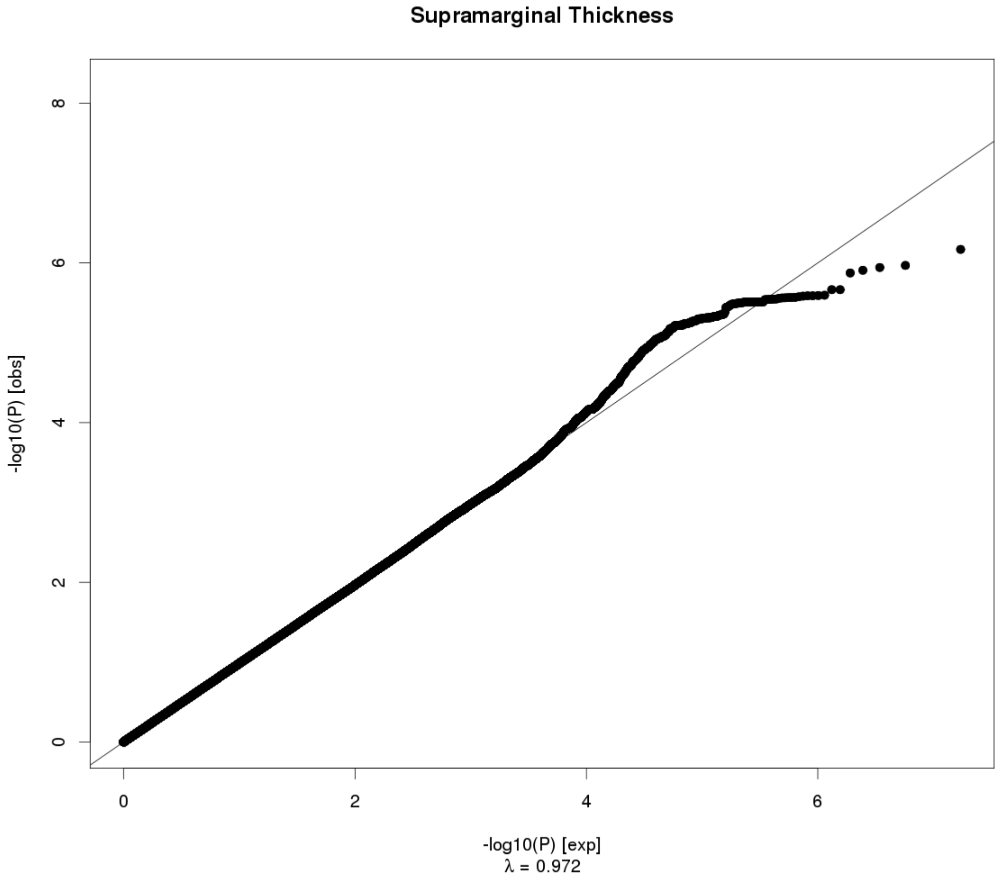
Supramarginal Thickness
-log10(P) [obs]
-log10(P) [exp]
λ = 0.972

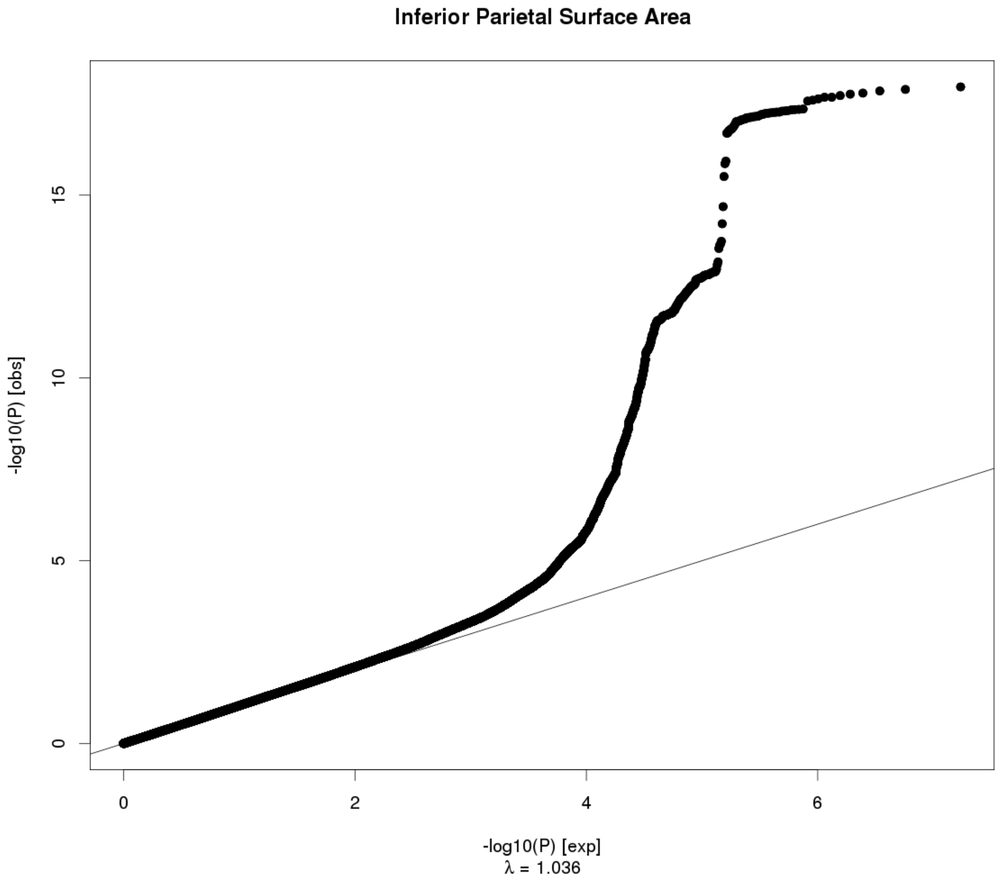
Inferior Parietal Surface Area
-log10(P) [obs]
0
5
10
15
0
2
4
6
-log10(P) [exp]
λ = 1.036

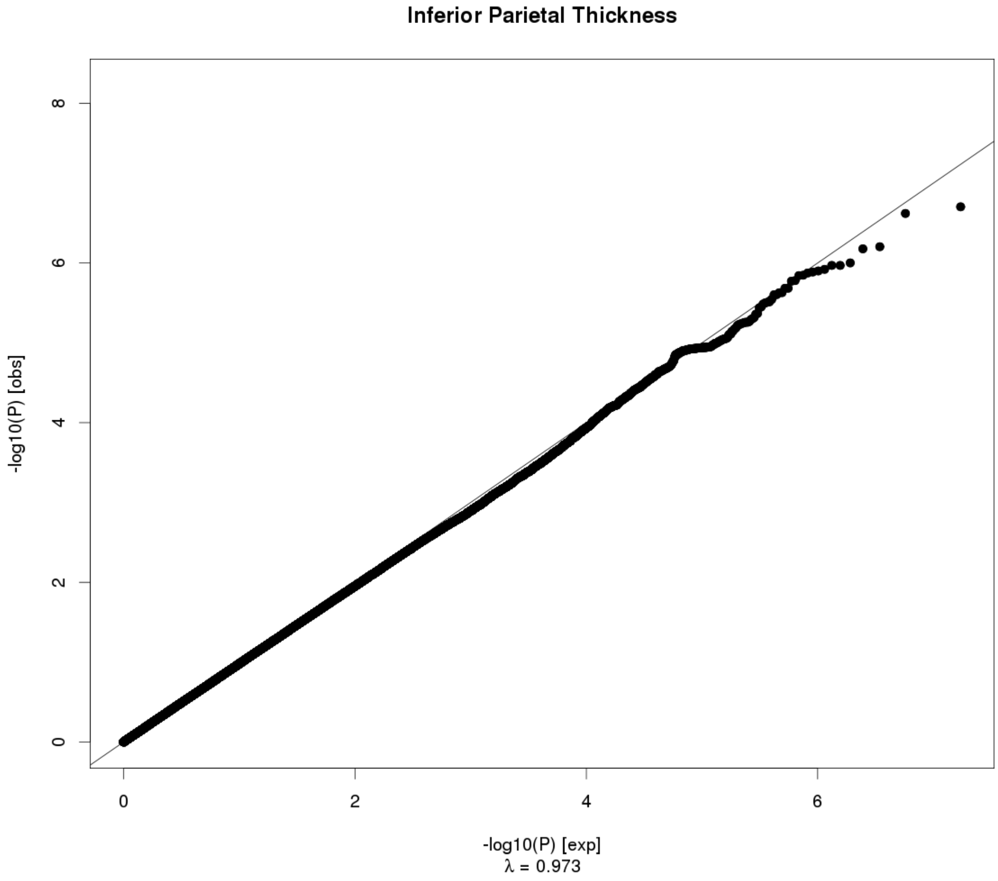
Inferior Parietal Thickness
-log10(P) [obs]
0
2
4
6
8
0
2
4
6
-log10(P) [exp]
λ = 0.973

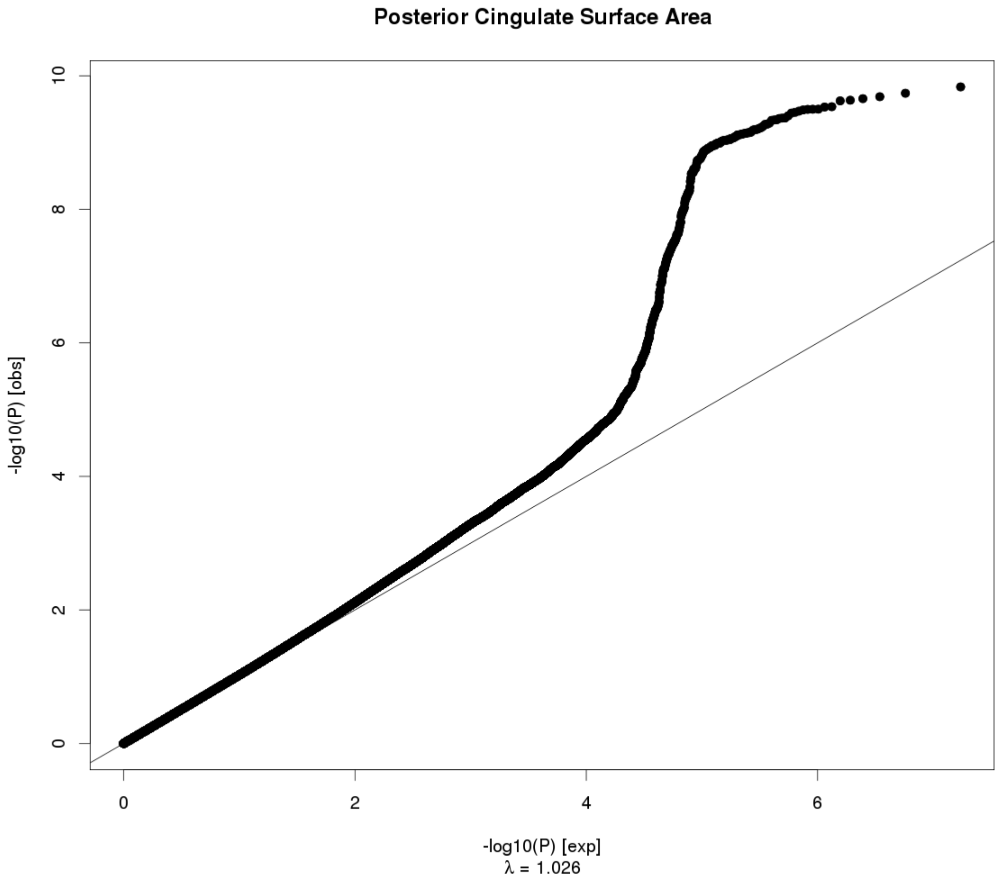
Posterior Cingulate Surface Area
-log10(P) [obs]
-log10(P) [exp]
λ = 1.026

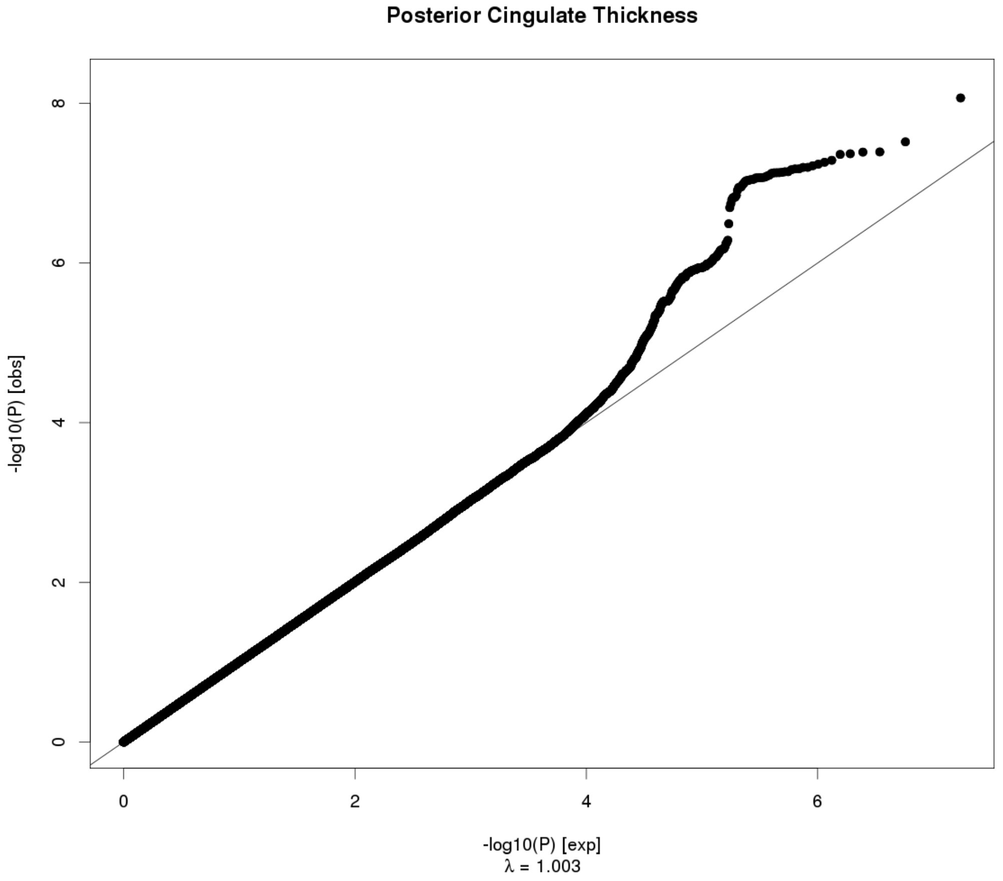
Posterior Cingulate Thickness
-log10(P) [obs]
-log10(P) [exp]
λ = 1.003

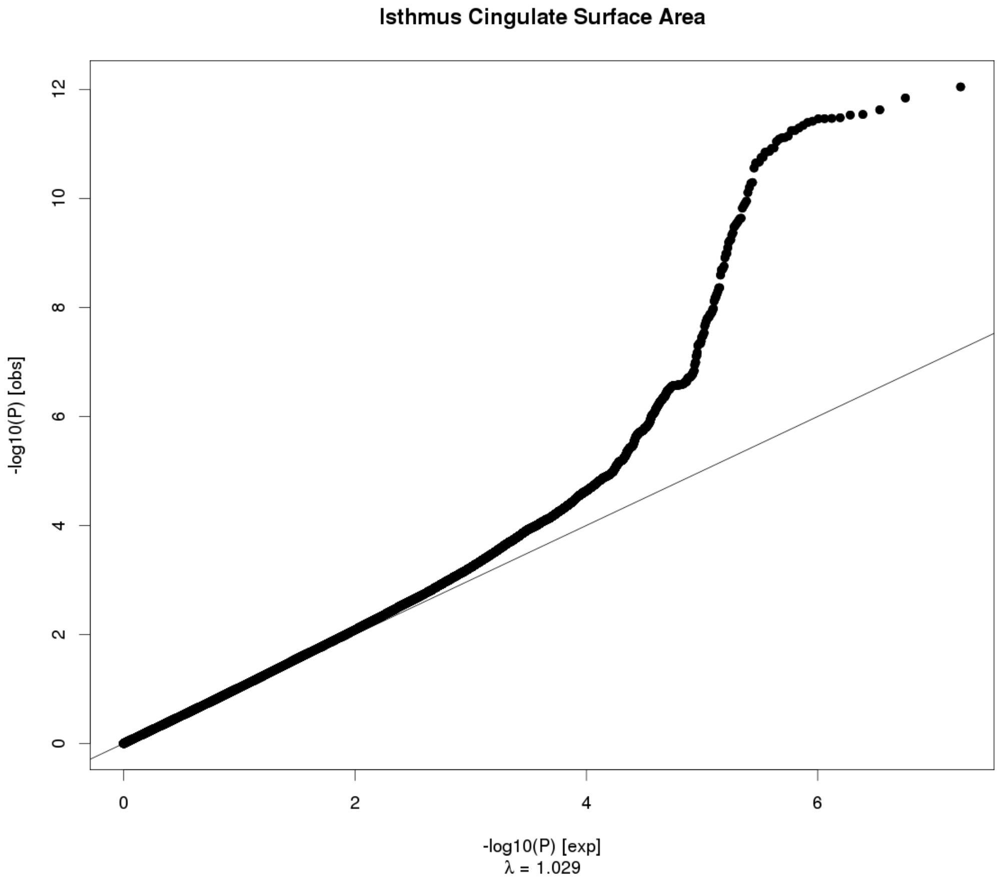

Isthmus Cingulate Surface Area
-log10(P) [obs]
0
2
4
6
8
10
12
-log10(P) [exp]
λ = 1.029


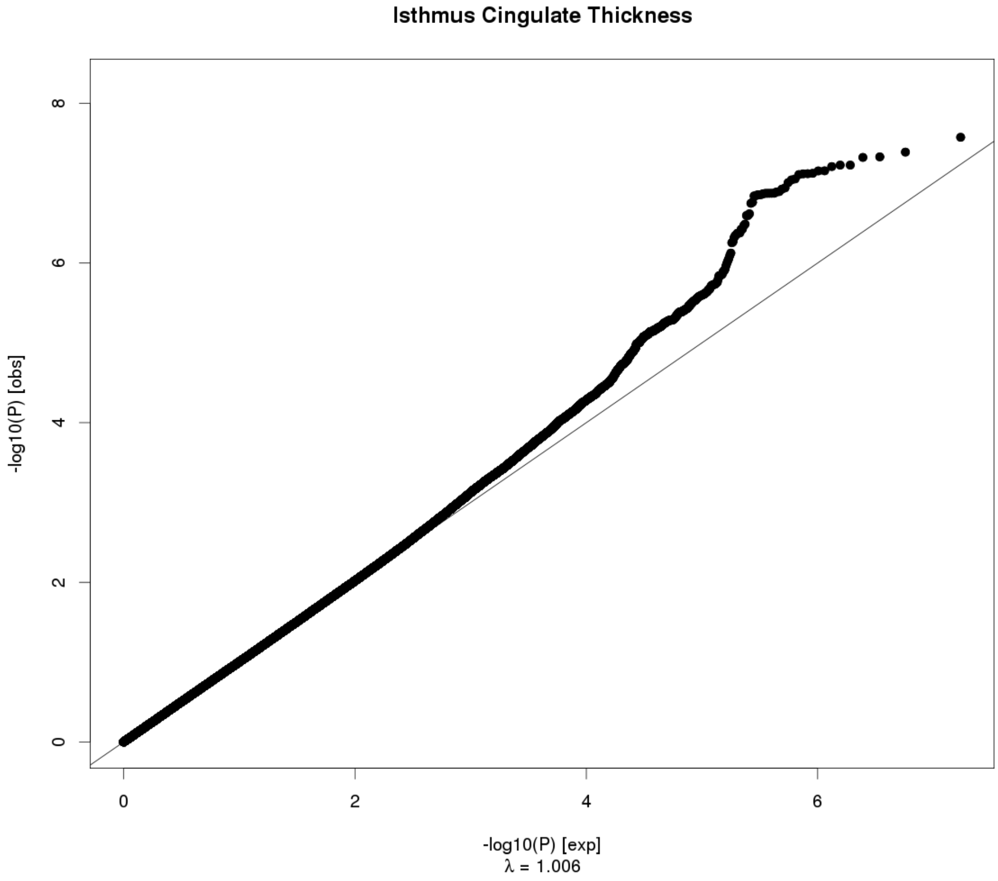

Isthmus Cingulate Thickness
-log10(P) [obs]
0
2
4
6
8
-log10(P) [exp]
λ = 1.006

### Insula Surface Area

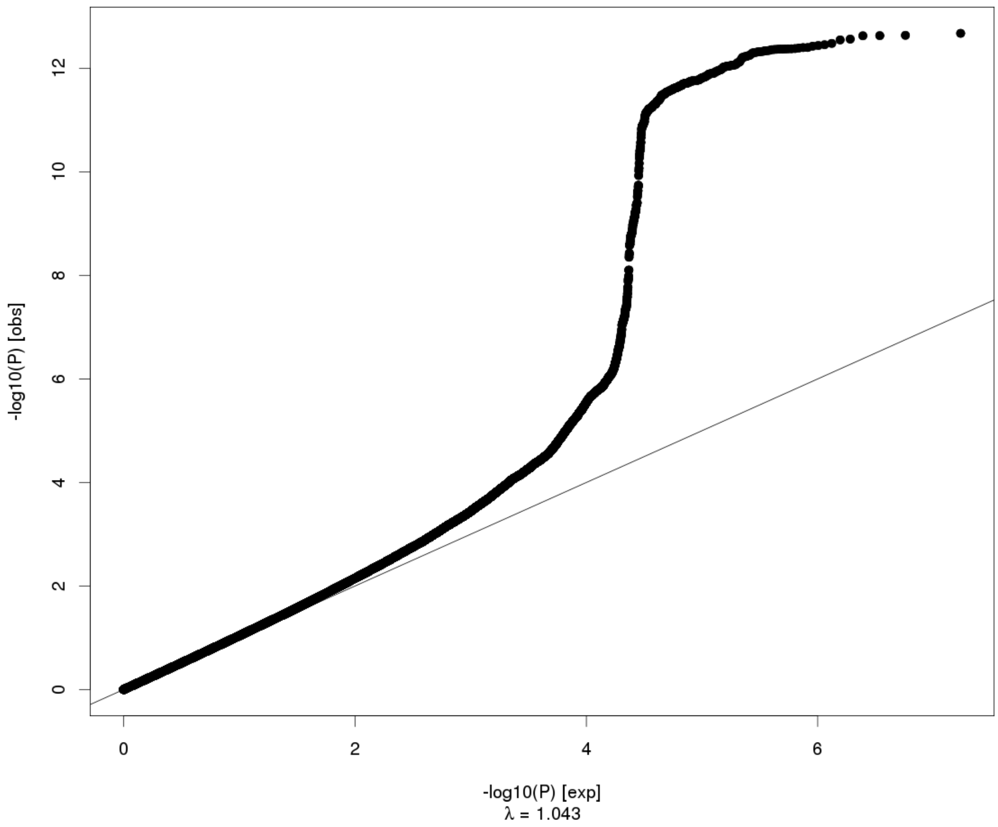


### Insula Thickness

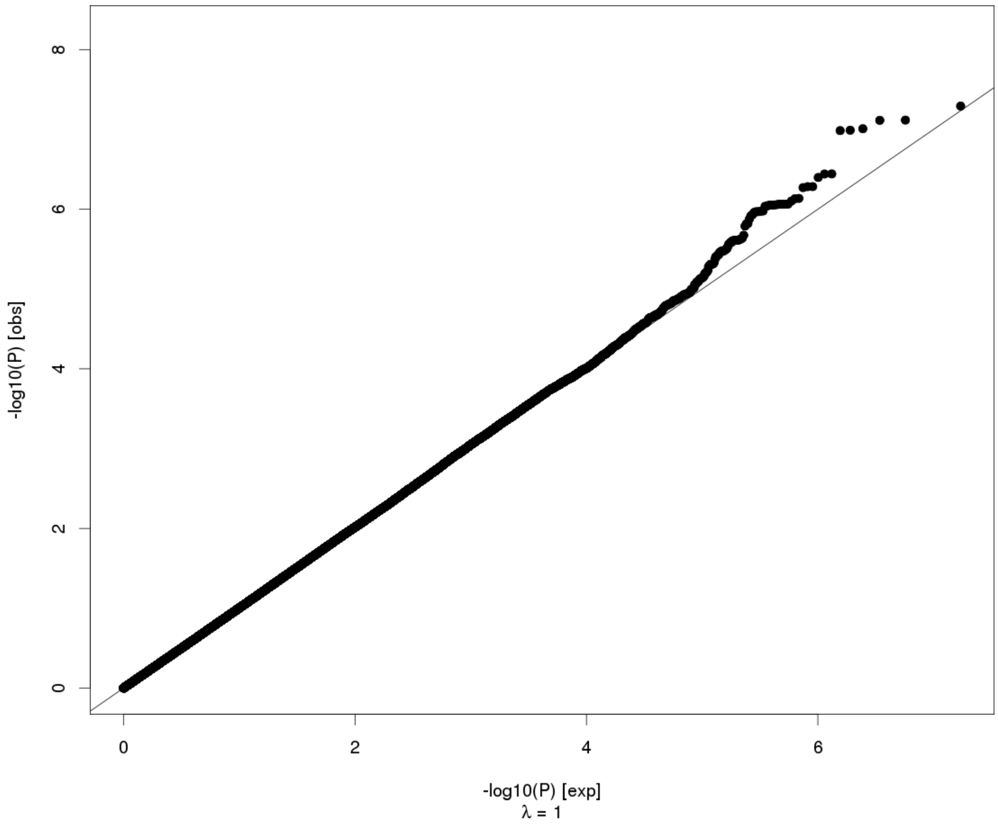

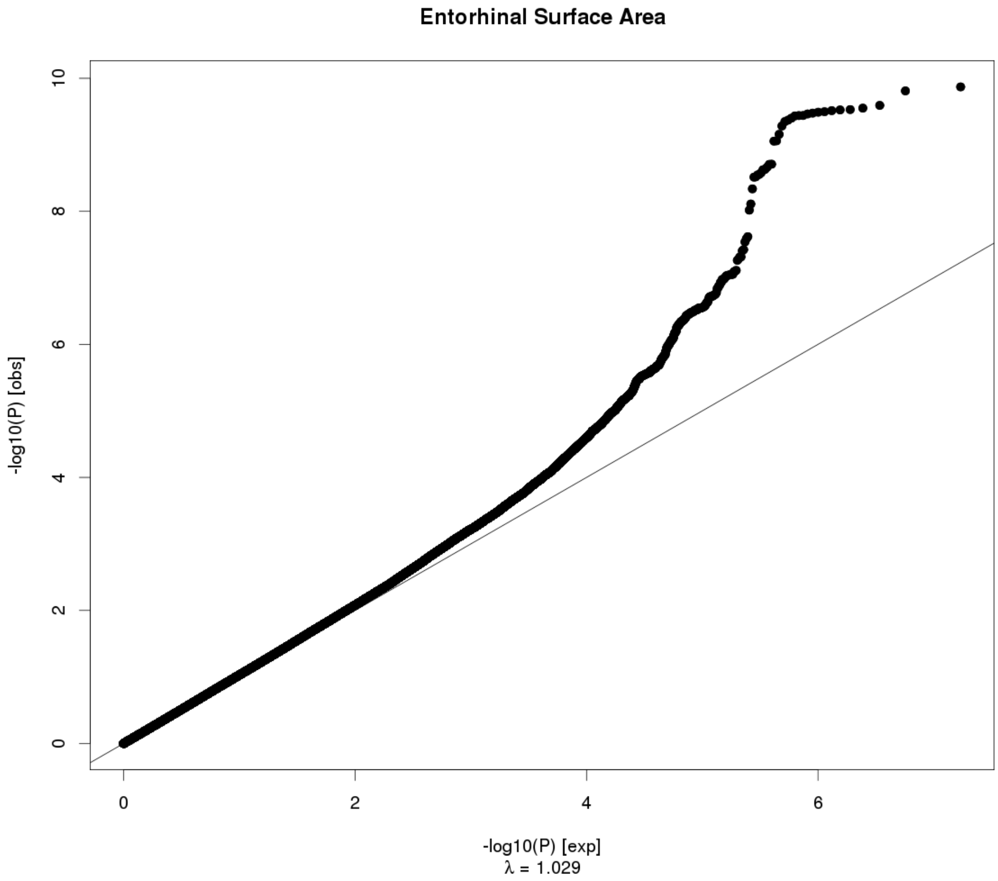
Entorhinal Surface Area
-log10(P) [obs]
0
2
4
6
8
10
0
2
4
6
-log10(P) [exp]
λ = 1.029

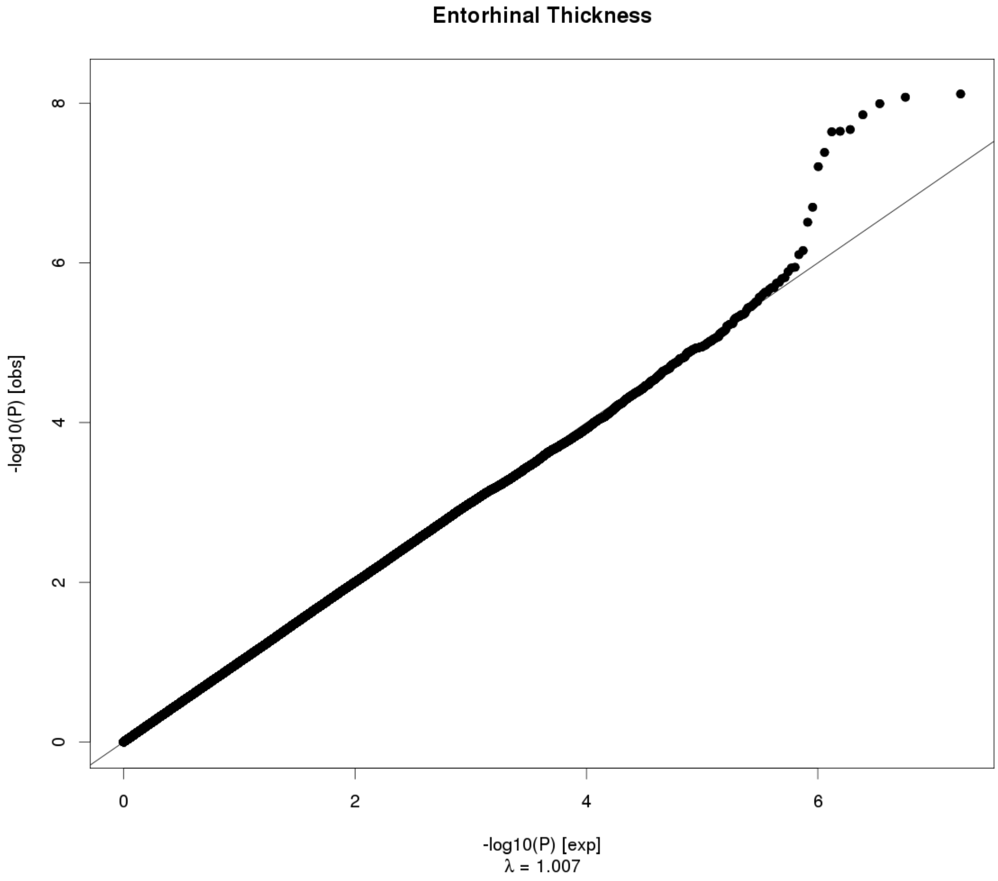
Entorhinal Thickness
-log10(P) [obs]
0
2
4
6
8
0
2
4
6
-log10(P) [exp]
λ = 1.007

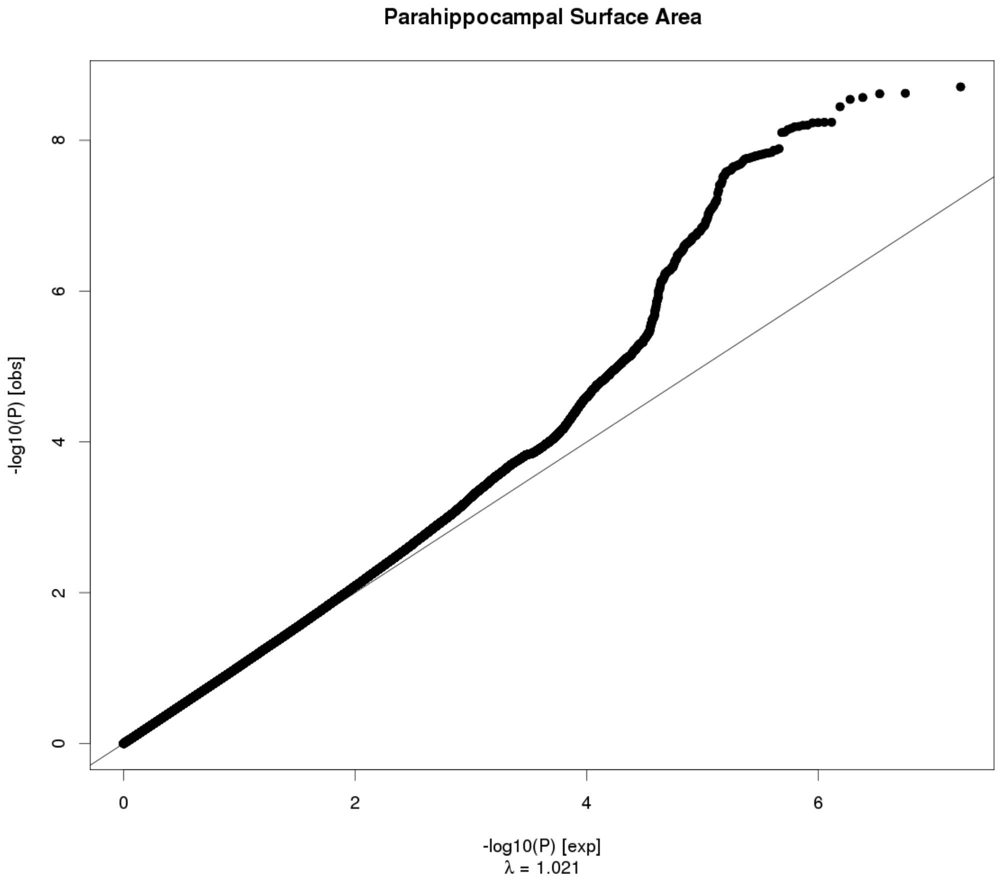
Parahippocampal Surface Area
-log10(P) [obs]
-log10(P) [exp]
λ = 1.021
0
2
4
6
8

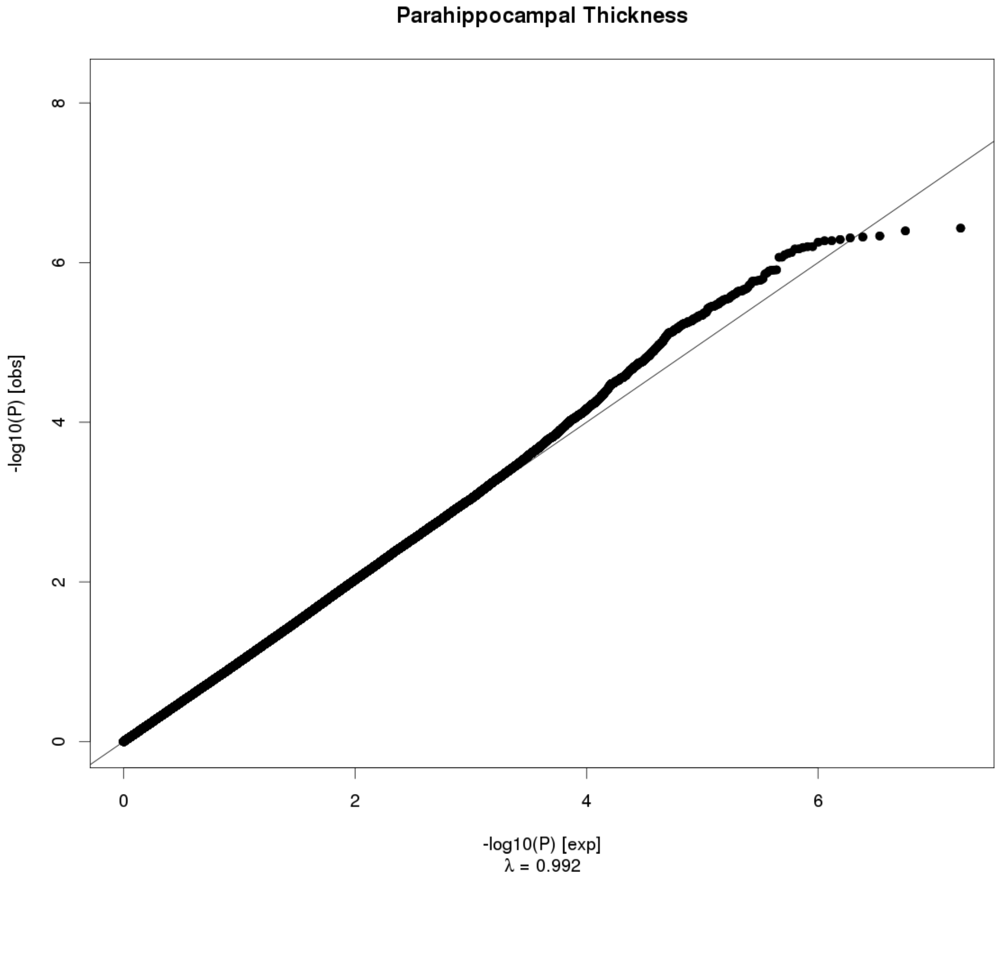
Parahippocampal Thickness
-log10(P) [obs]
-log10(P) [exp]
λ = 0.992
0
2
4
6
8

**Fusiform Surface Area**

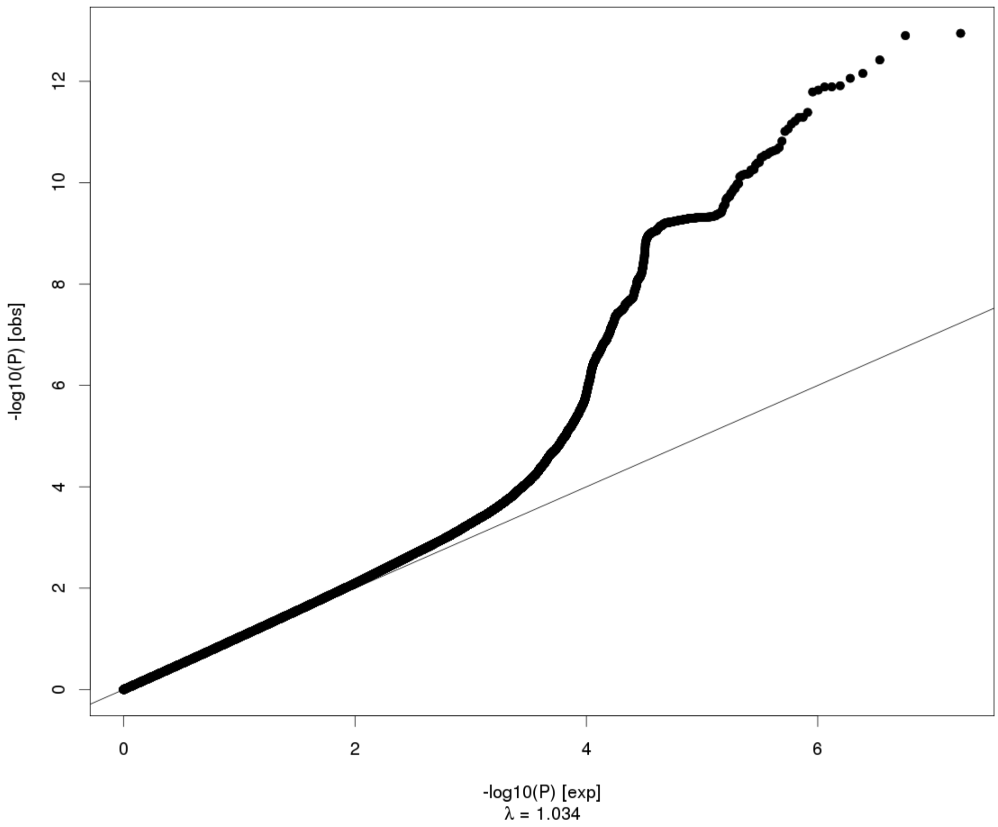


**Fusiform Thickness**

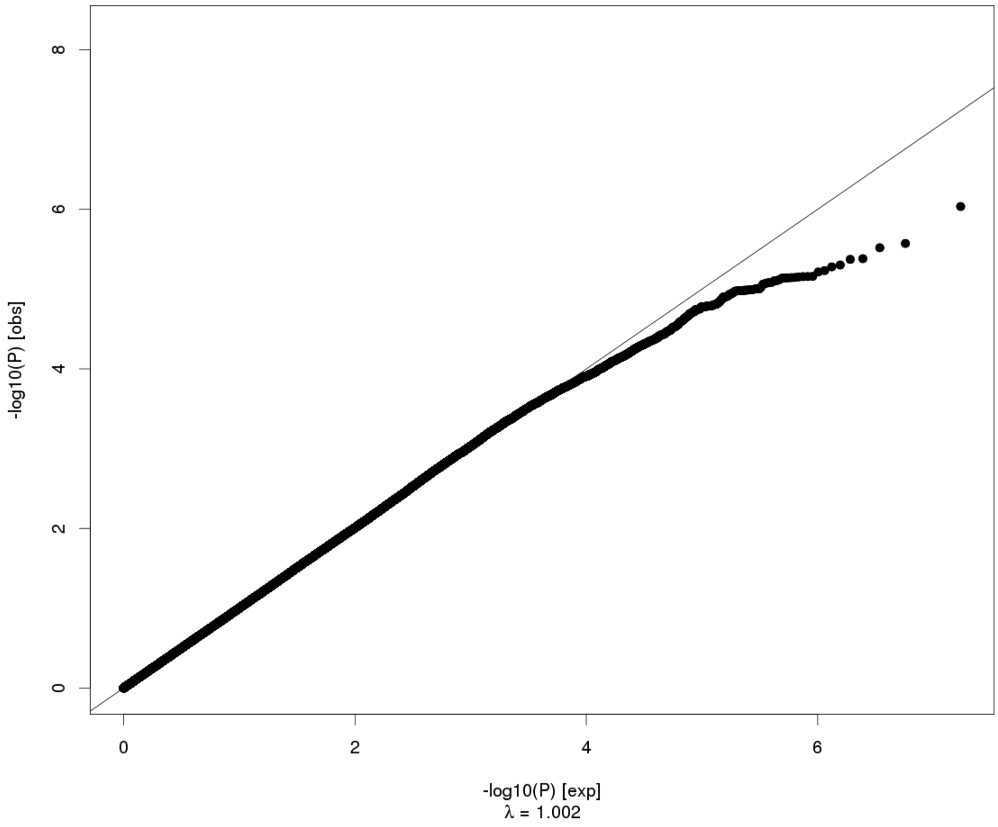

Temporal Pole Surface Area

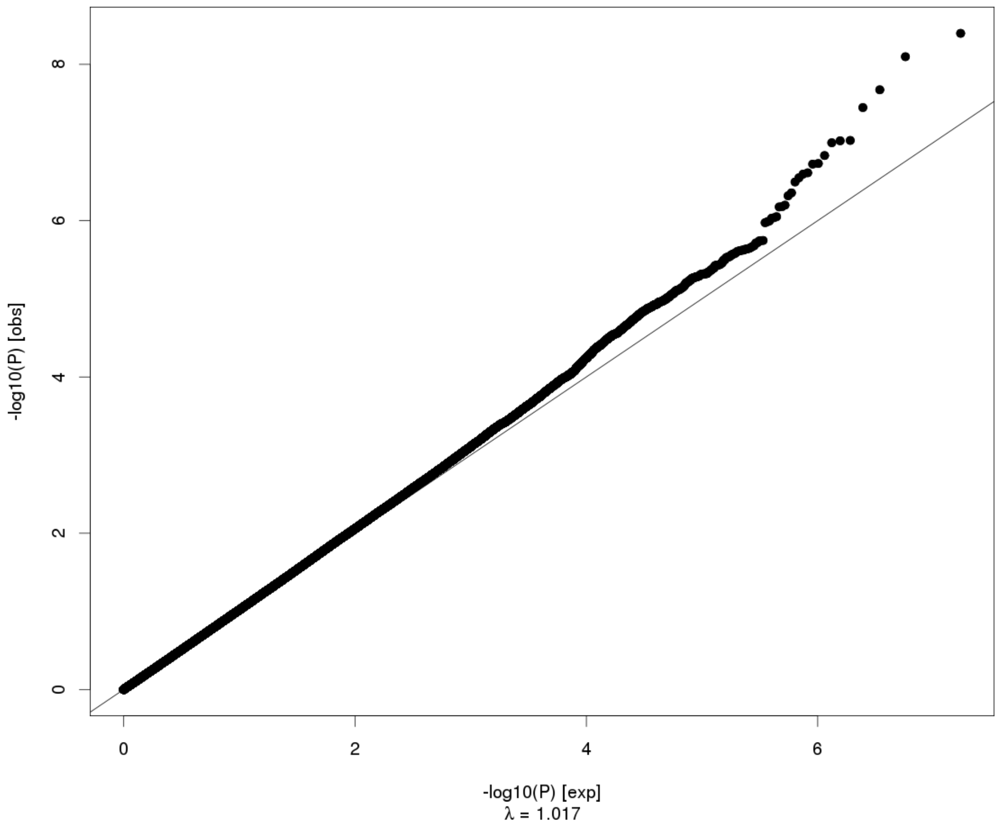


Temporal Pole Thickness

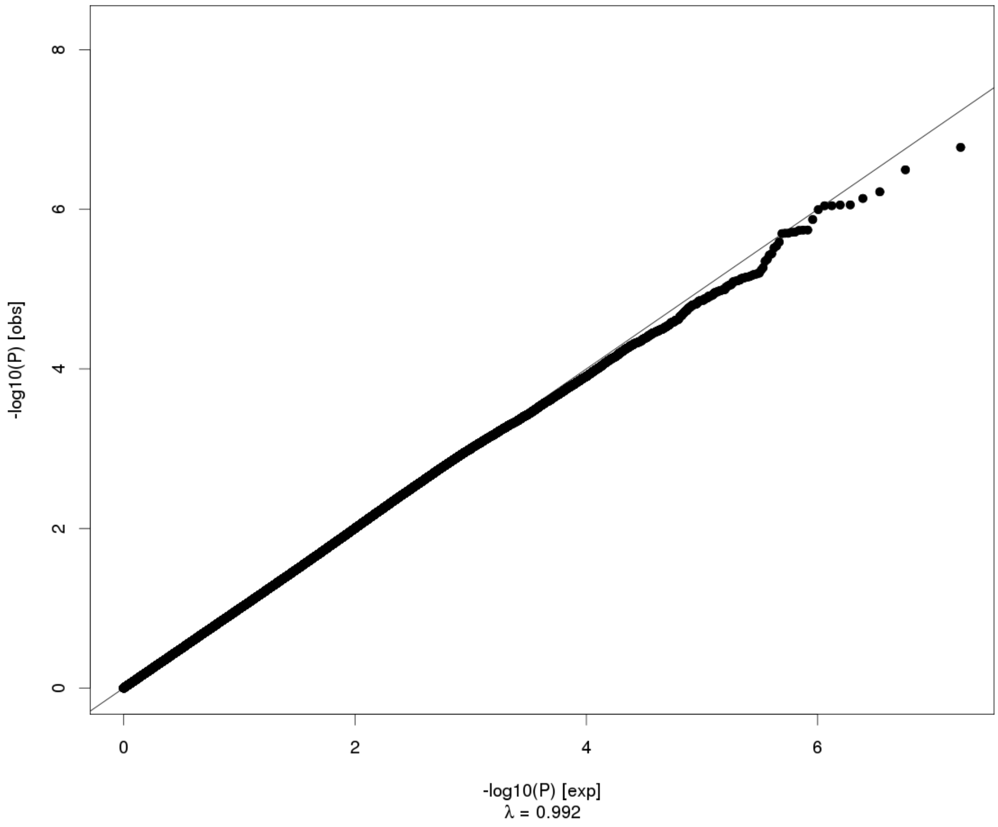

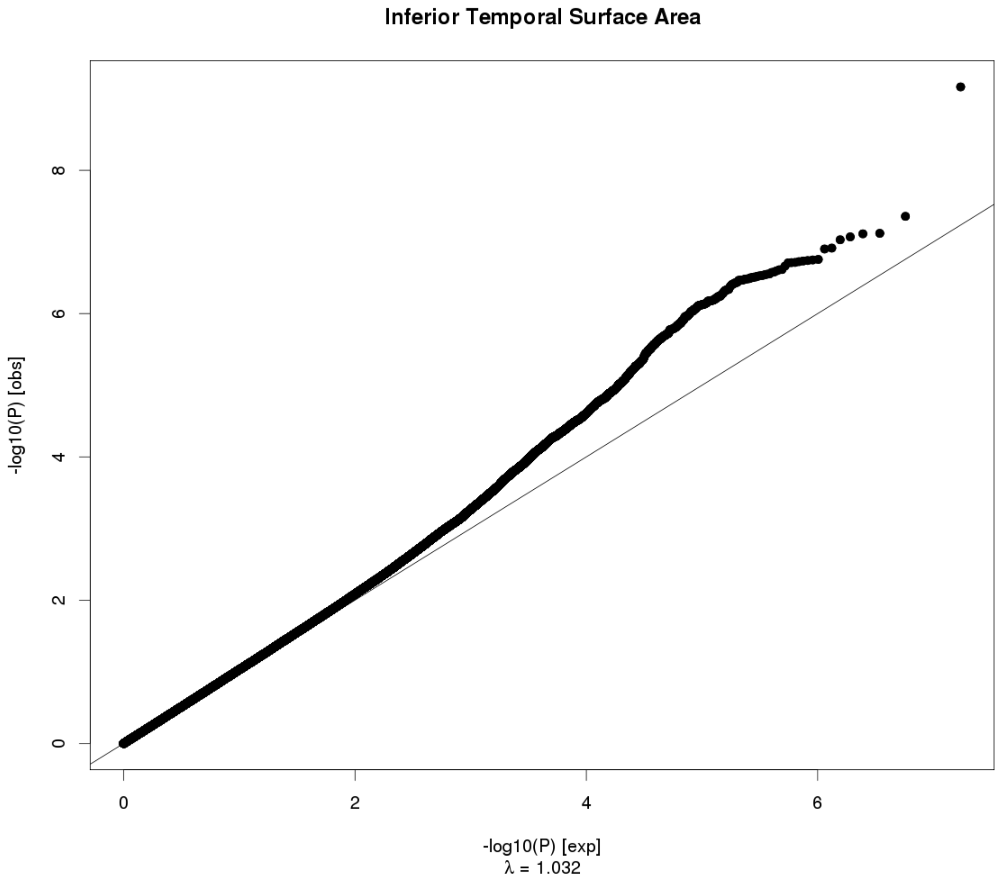
Inferior Temporal Surface Area
-log10(P) [obs]
-log10(P) [exp]
λ = 1.032
0
2
4
6
8

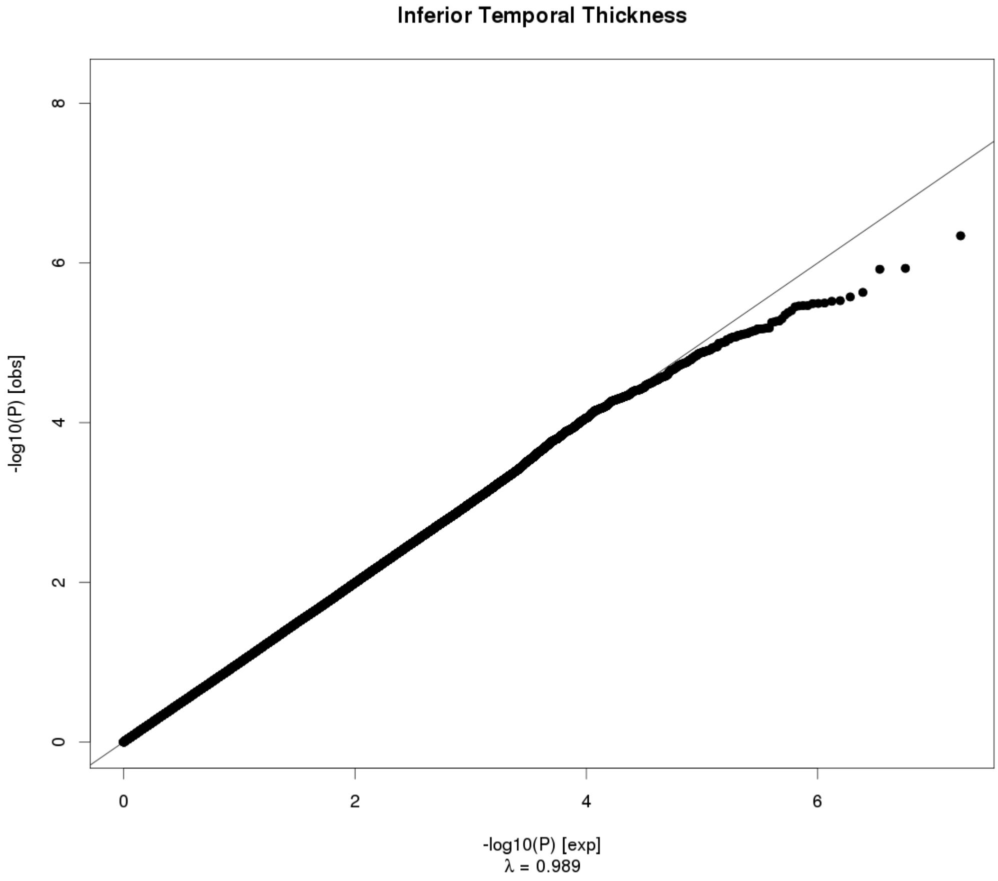
Inferior Temporal Thickness
-log10(P) [obs]
-log10(P) [exp]
λ = 0.989
0
2
4
6
8

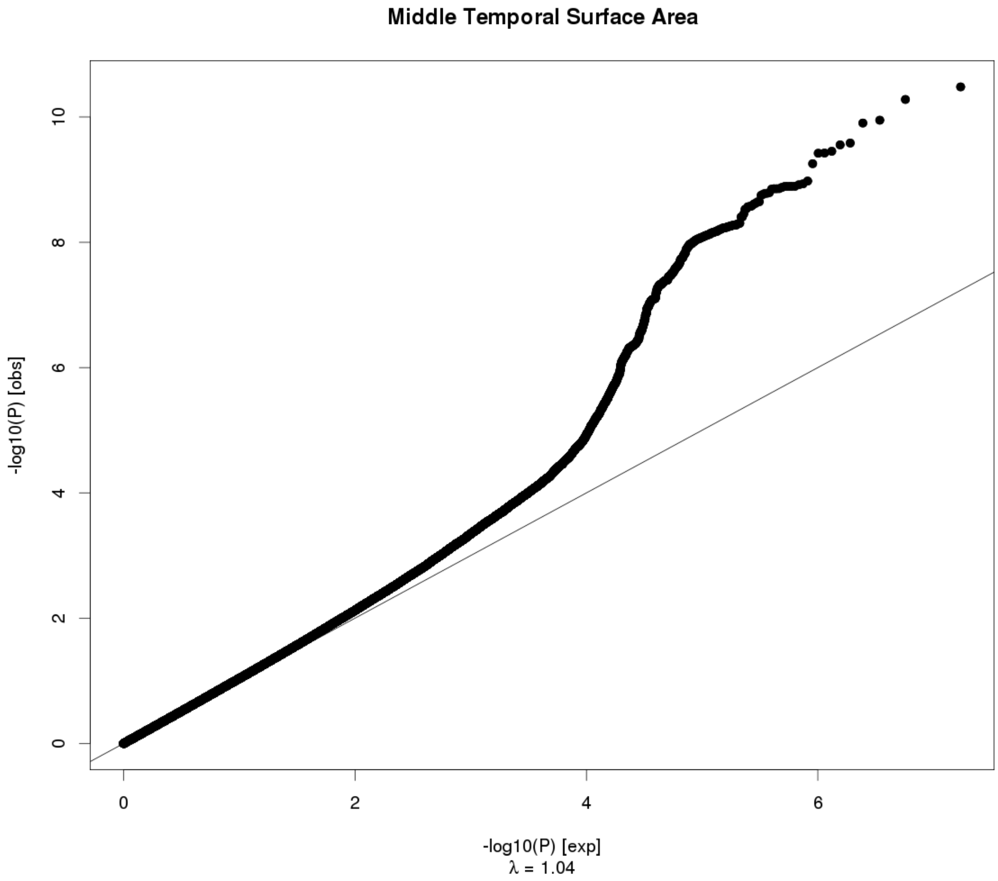
Middle Temporal Surface Area
-log10(P) [obs]
-log10(P) [exp]
λ = 1.04
0
2
4
6
8
10

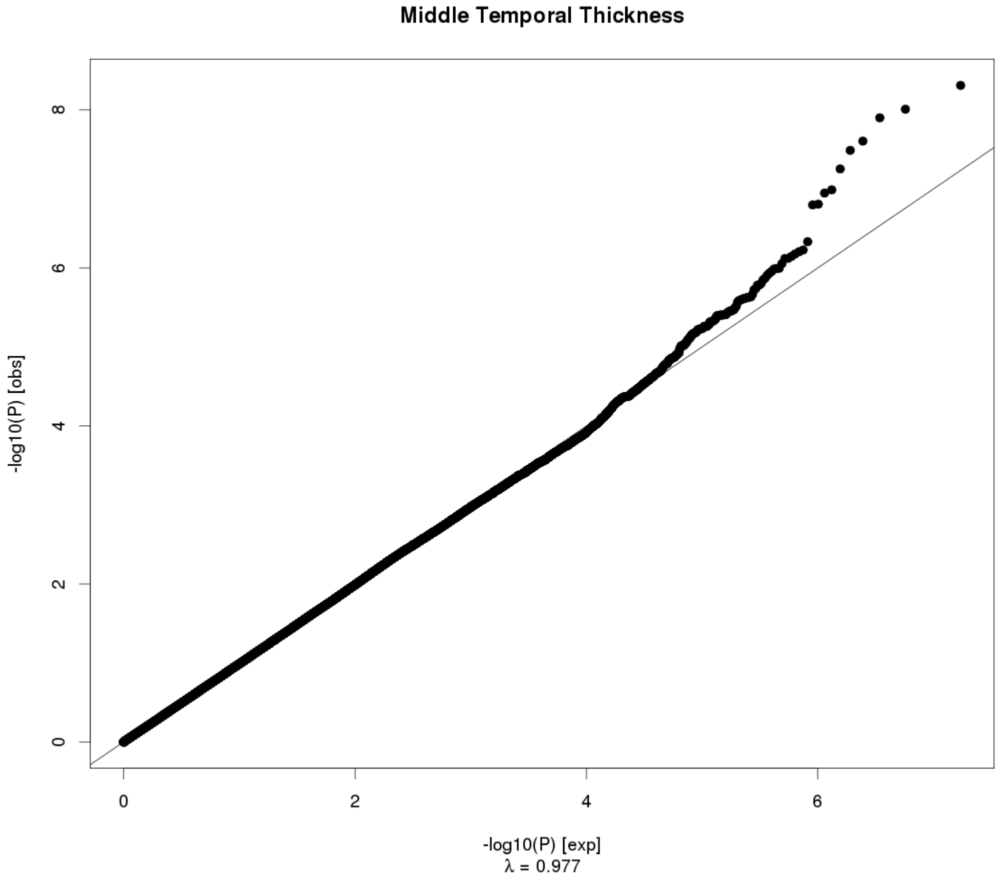
Middle Temporal Thickness
-log10(P) [obs]
-log10(P) [exp]
λ = 0.977
0
2
4
6
8

Superior Temporal Surface Area

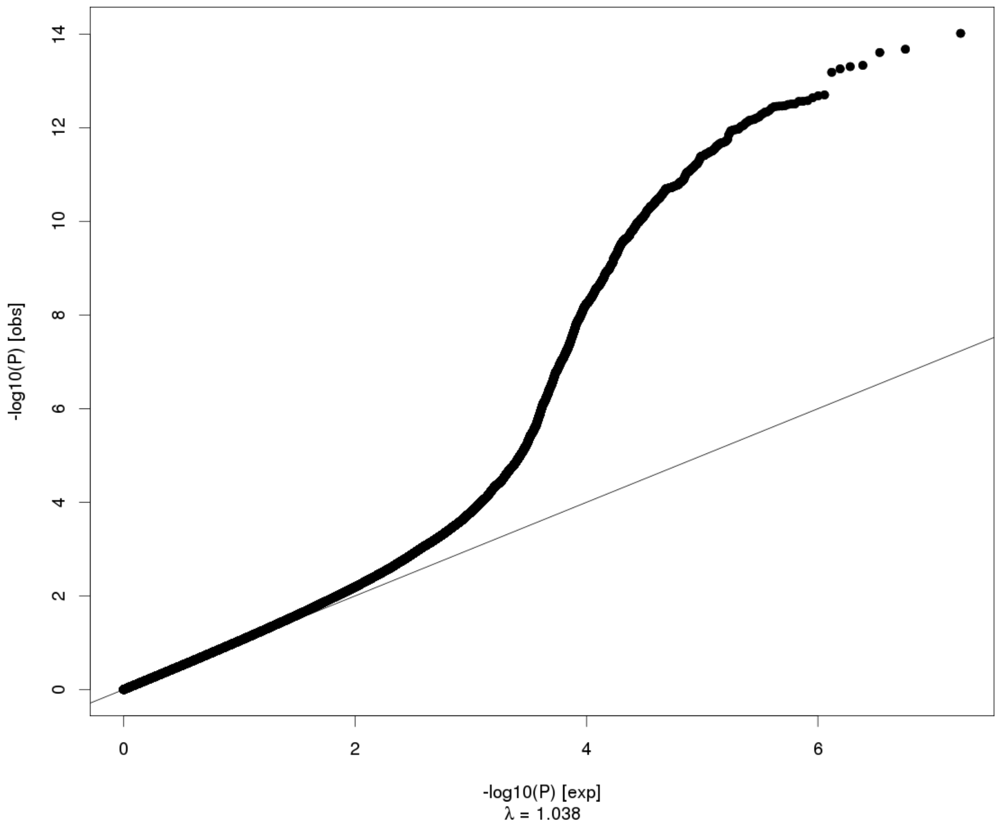


Superior Temporal Thickness

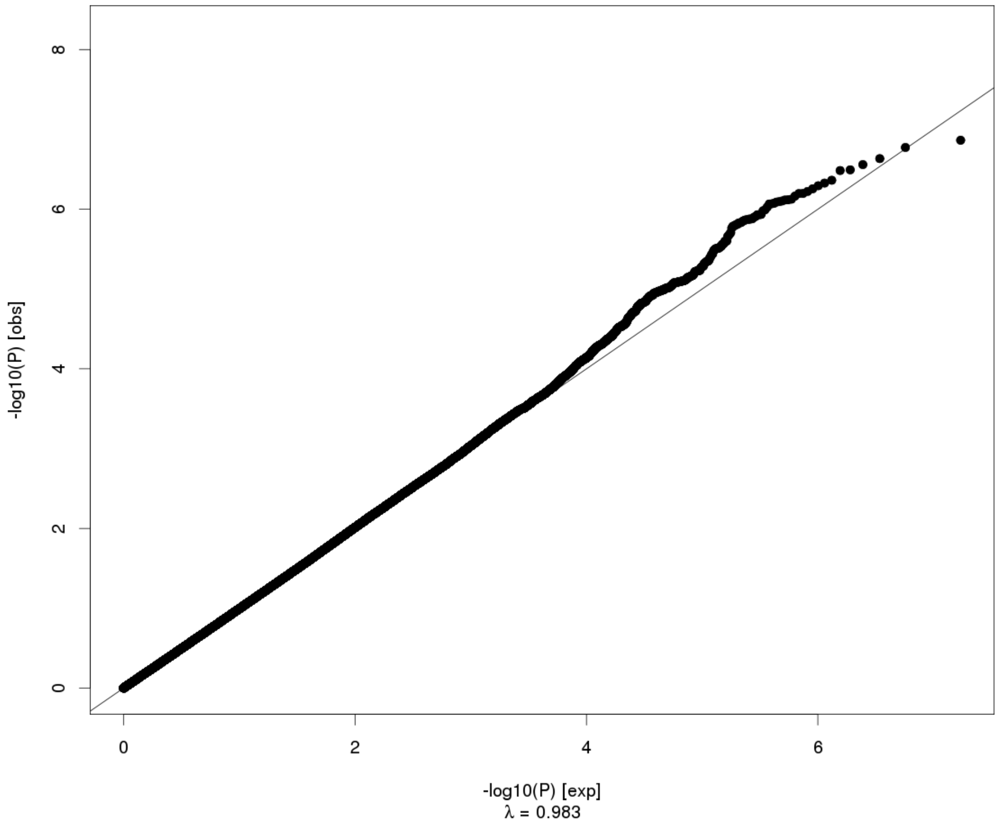

### Banks of the Superior Temporal Sulcus Surface Area

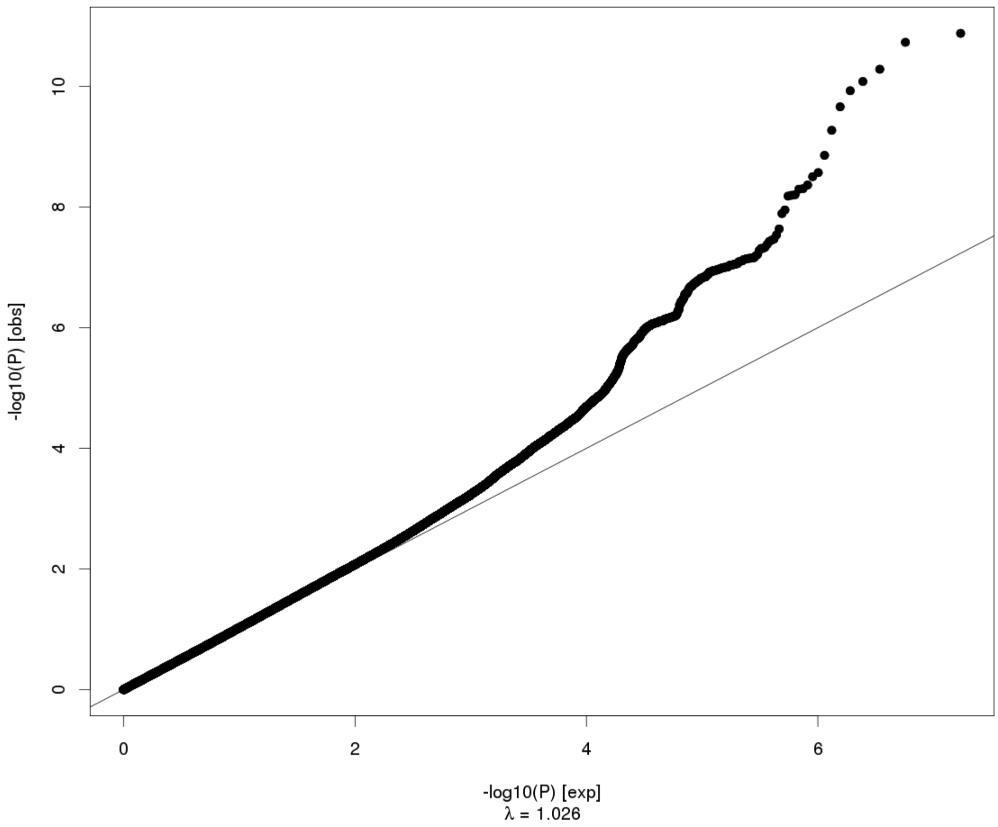


### Banks of the Superior Temporal Sulcus Thickness

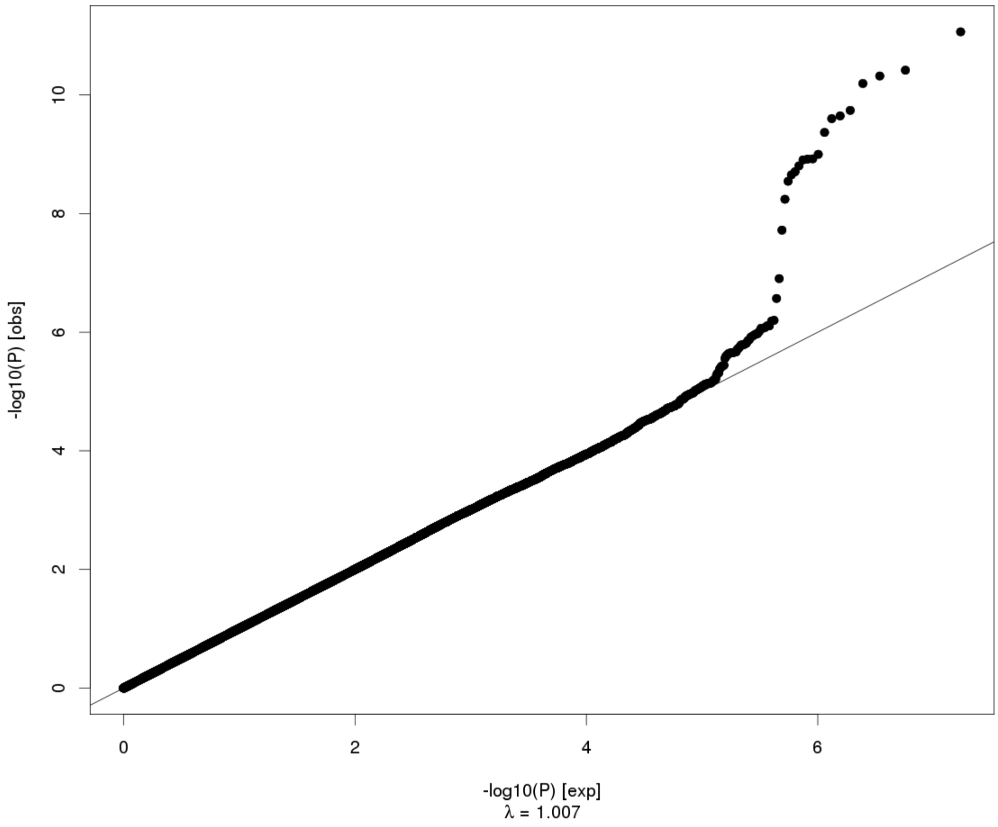

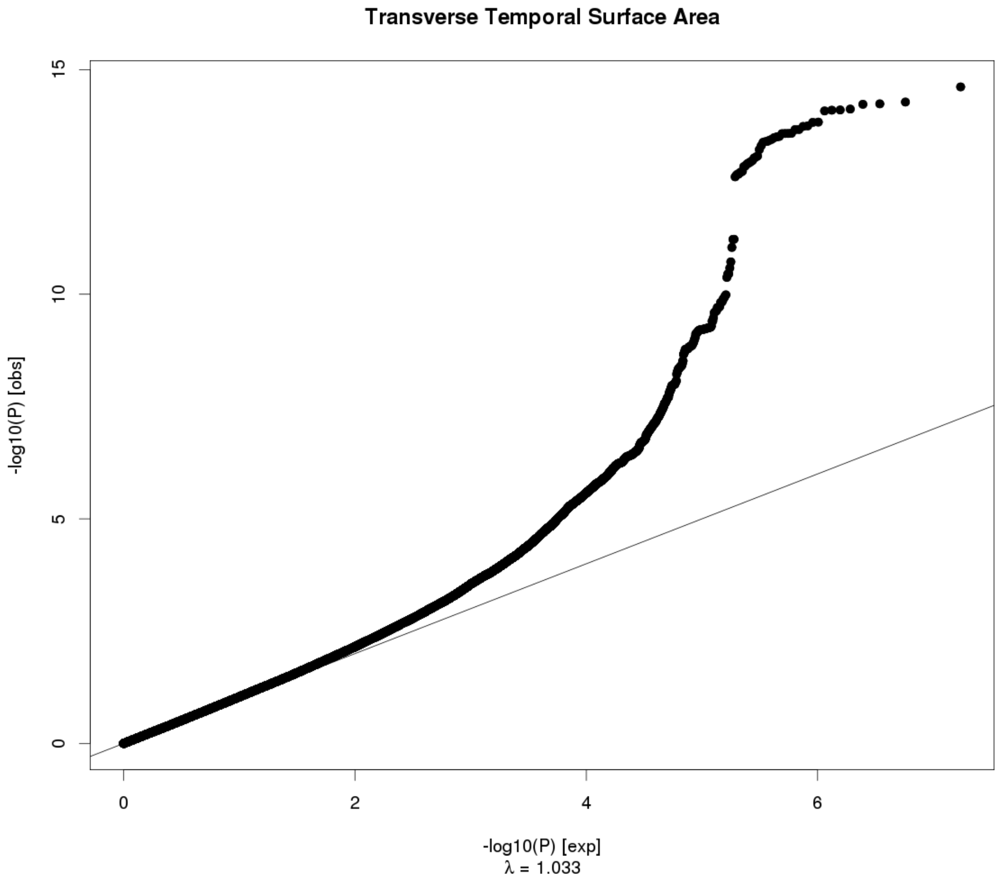
Transverse Temporal Surface Area
15
10
5
0
-log10(P) [obs]
0
2
4
6
-log10(P) [exp]
λ = 1.033

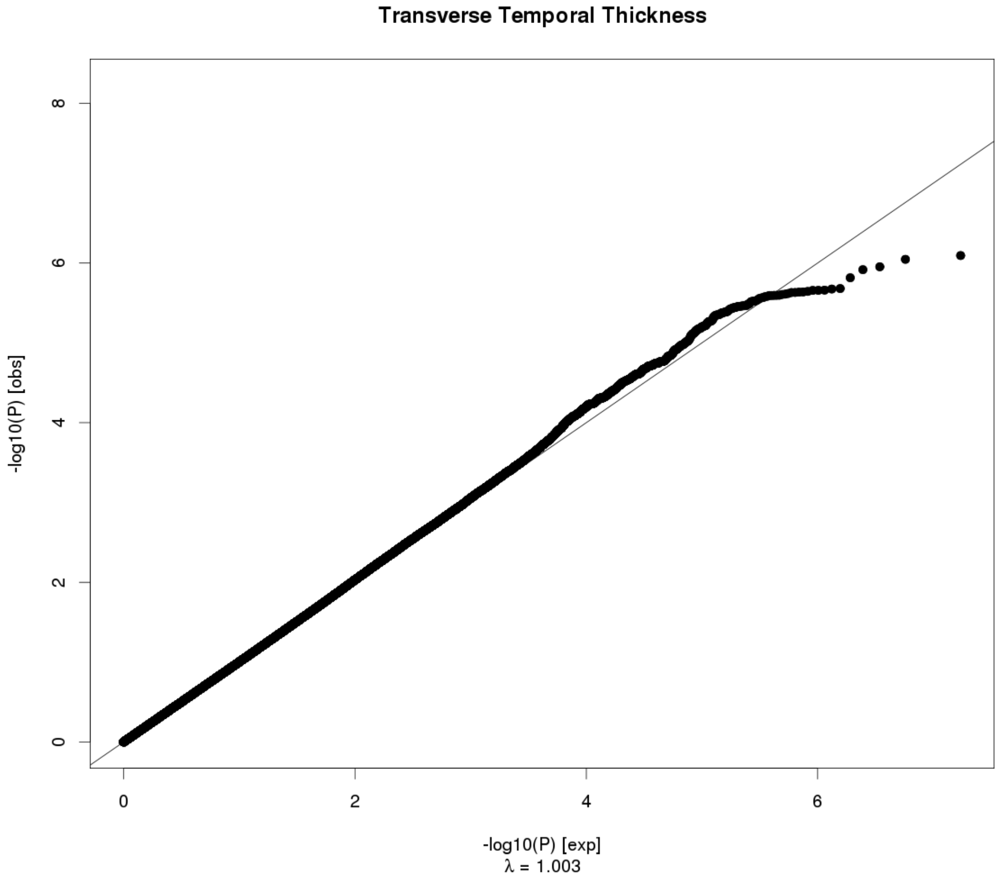
Transverse Temporal Thickness
8
6
4
2
0
-log10(P) [obs]
0
2
4
6
-log10(P) [exp]
λ = 1.003

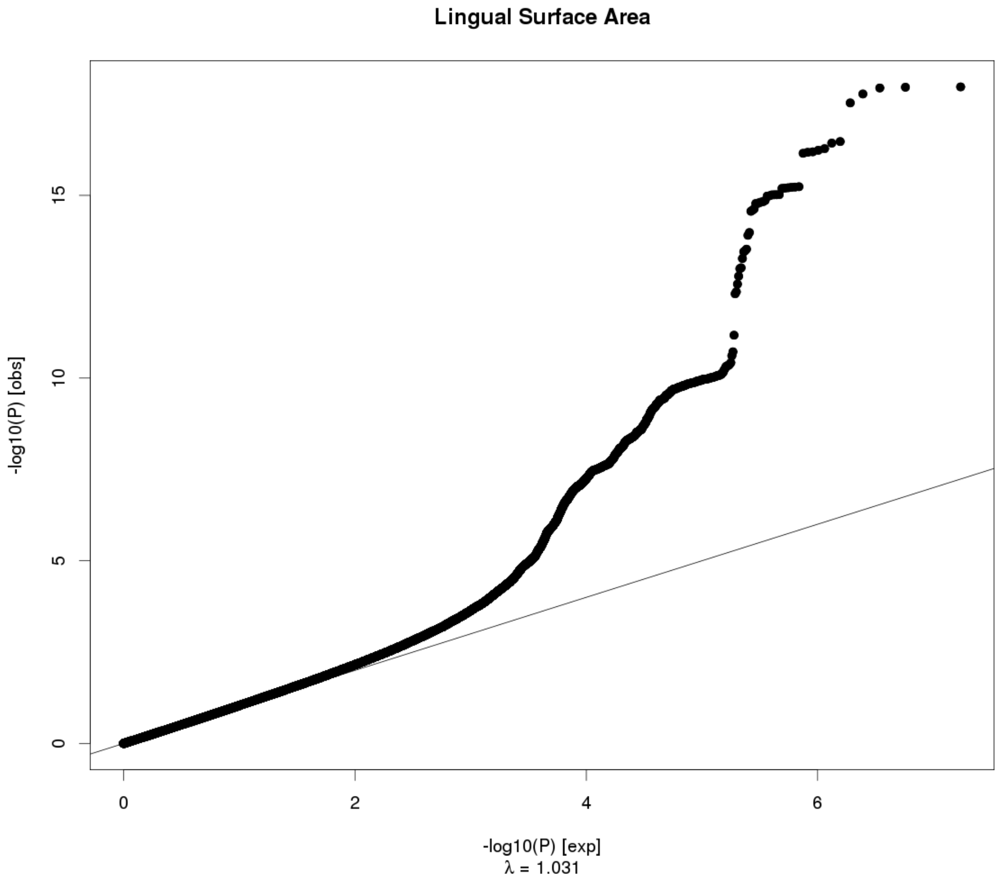
Lingual Surface Area
-log10(P) [obs]
0
5
10
15
0
2
4
6
-log10(P) [exp]
λ = 1.031

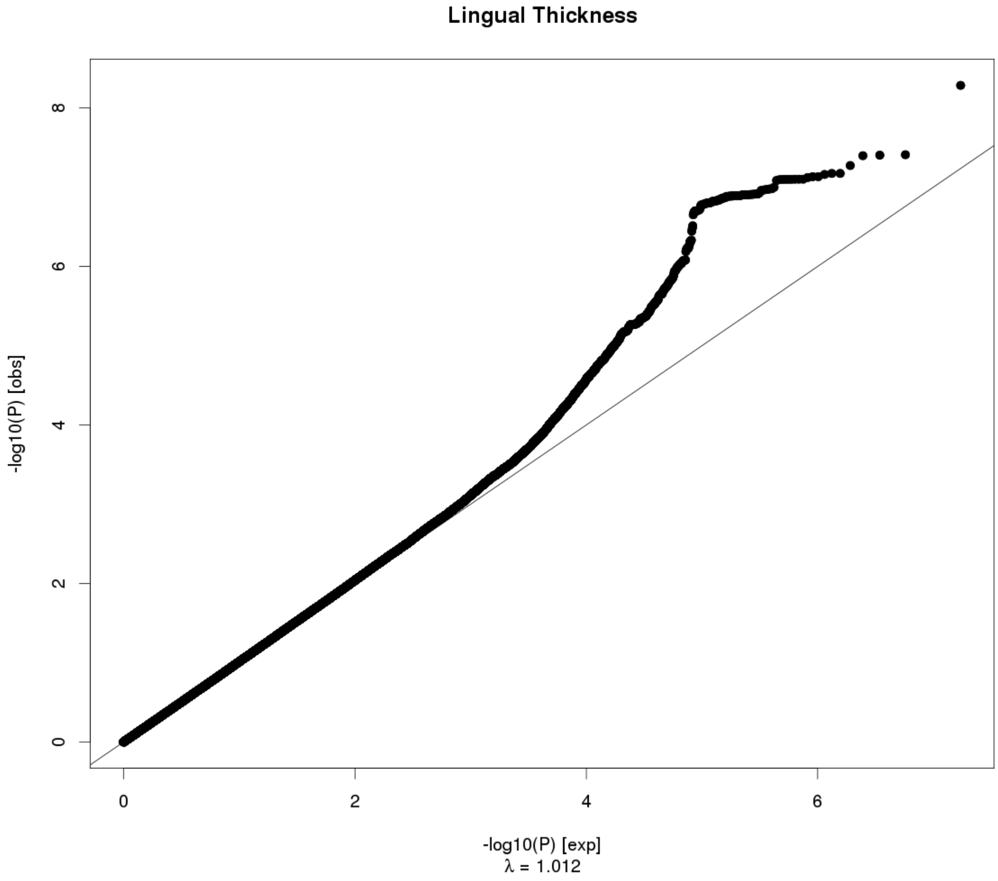
Lingual Thickness
-log10(P) [obs]
0
2
4
6
8
0
2
4
6
-log10(P) [exp]
λ = 1.012

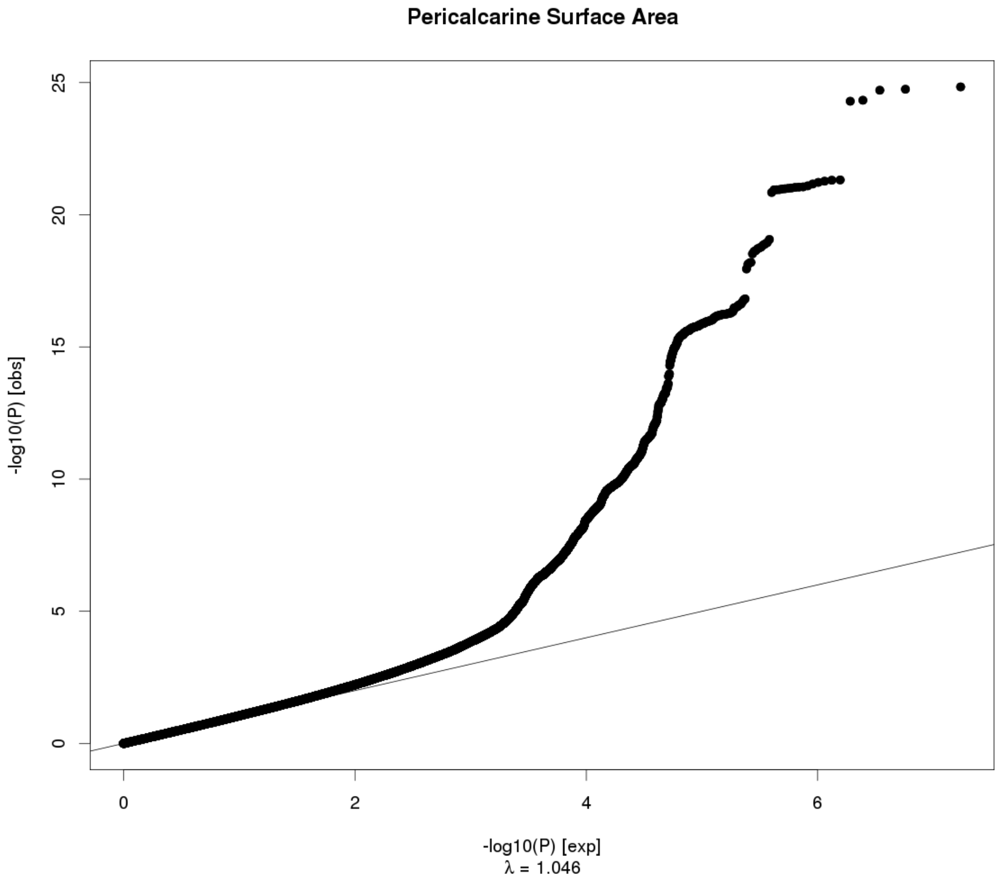

Pericalcarine Surface Area
-log10(P) [obs]
-log10(P) [exp]
λ = 1.046
0
5
10
15
20
25
0
2
4
6


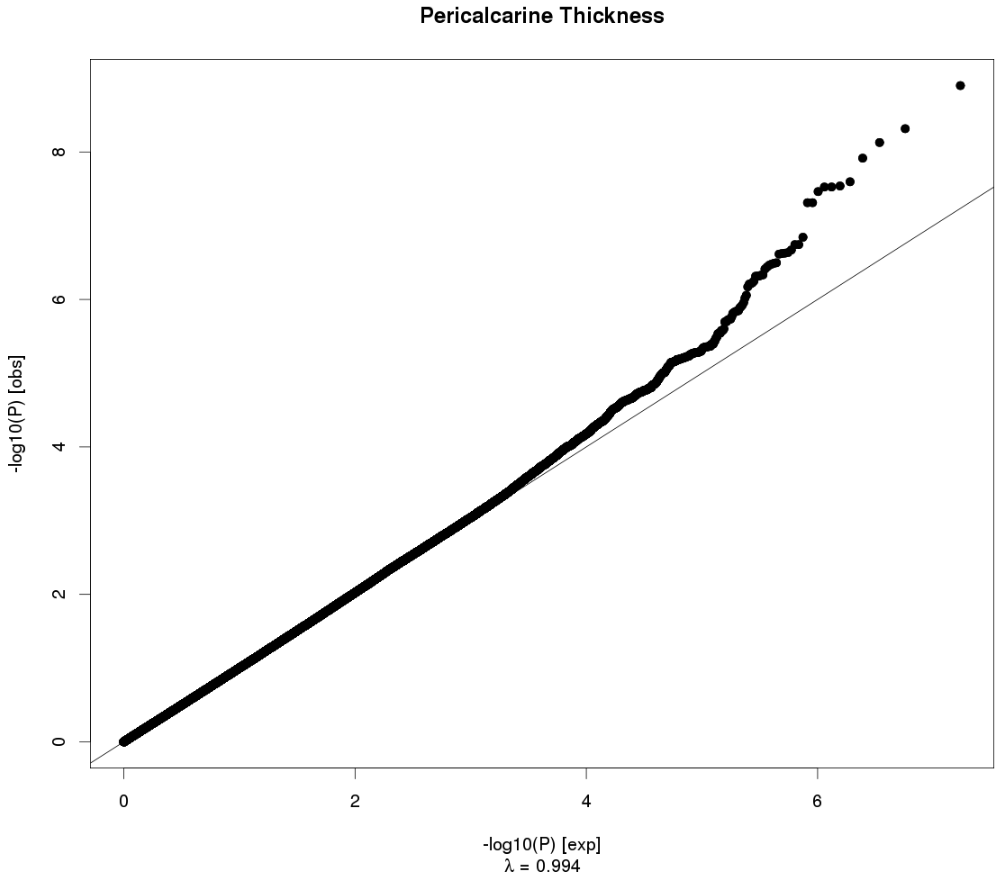

Pericalcarine Thickness
-log10(P) [obs]
-log10(P) [exp]
λ = 0.994
0
2
4
6
8
0
2
4
6

**Cuneus Surface Area**

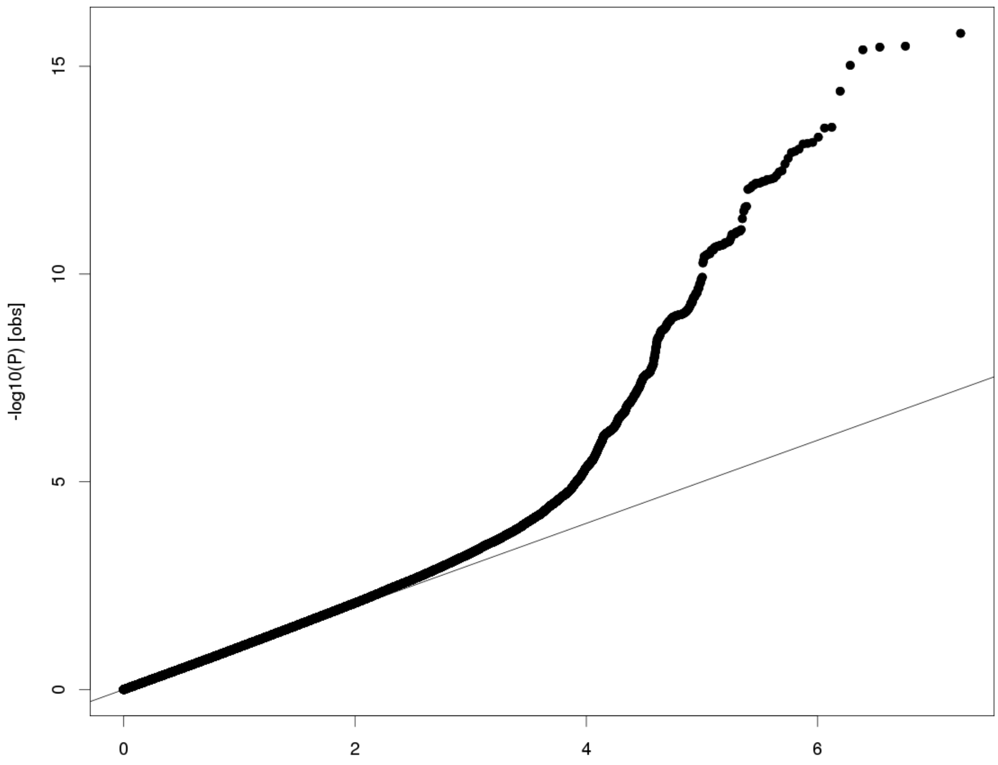


-log10(P) [exp]
λ = 1.02

**Cuneus Thickness**

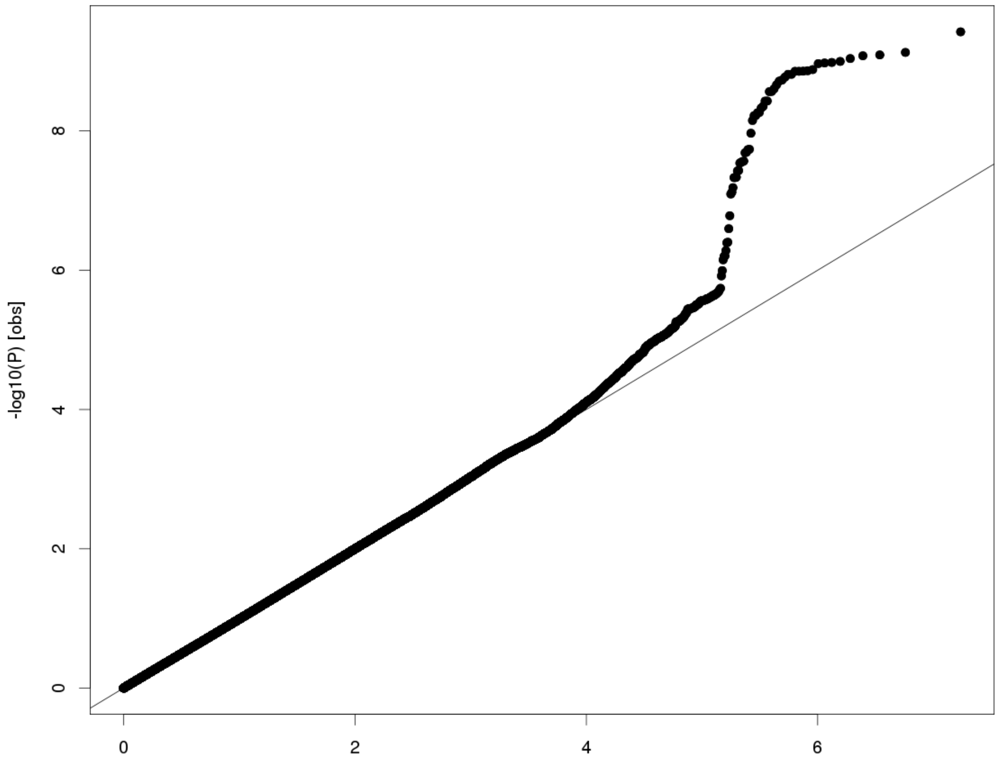


-log10(P) [exp]
λ = 0.986

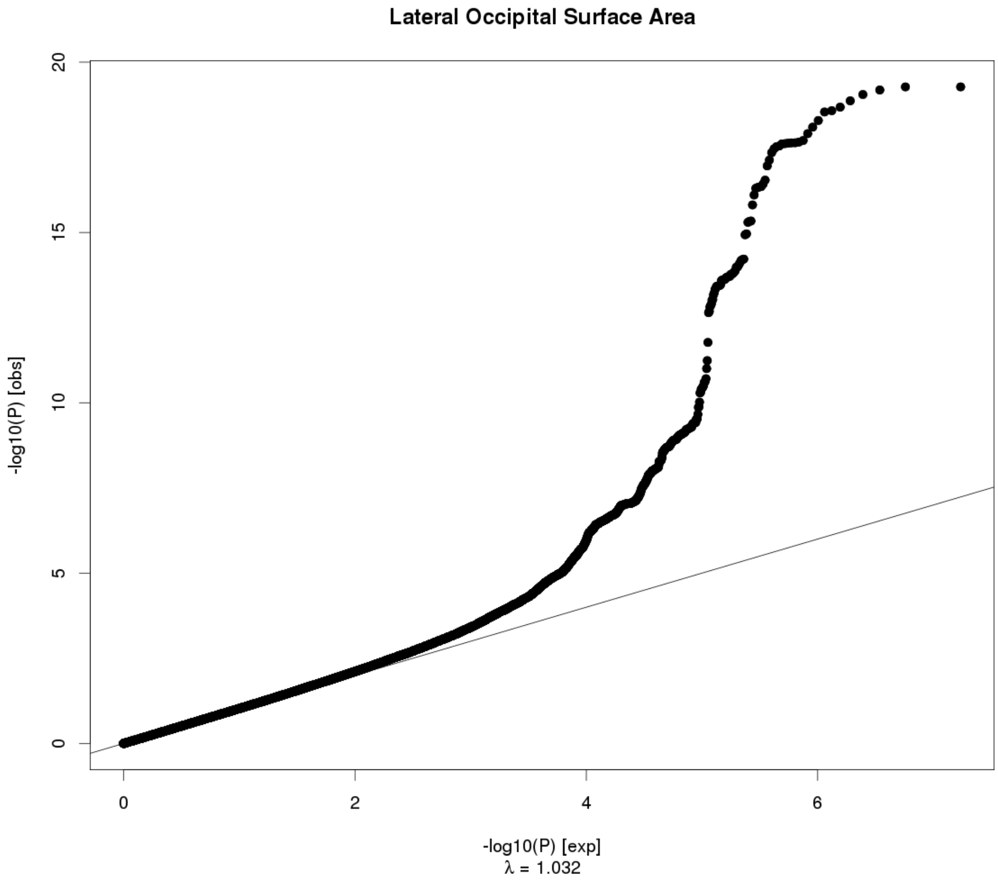
Lateral Occipital Surface Area
-log10(P) [obs]
0
5
10
15
20
0
2
4
6
-log10(P) [exp]
λ = 1.032

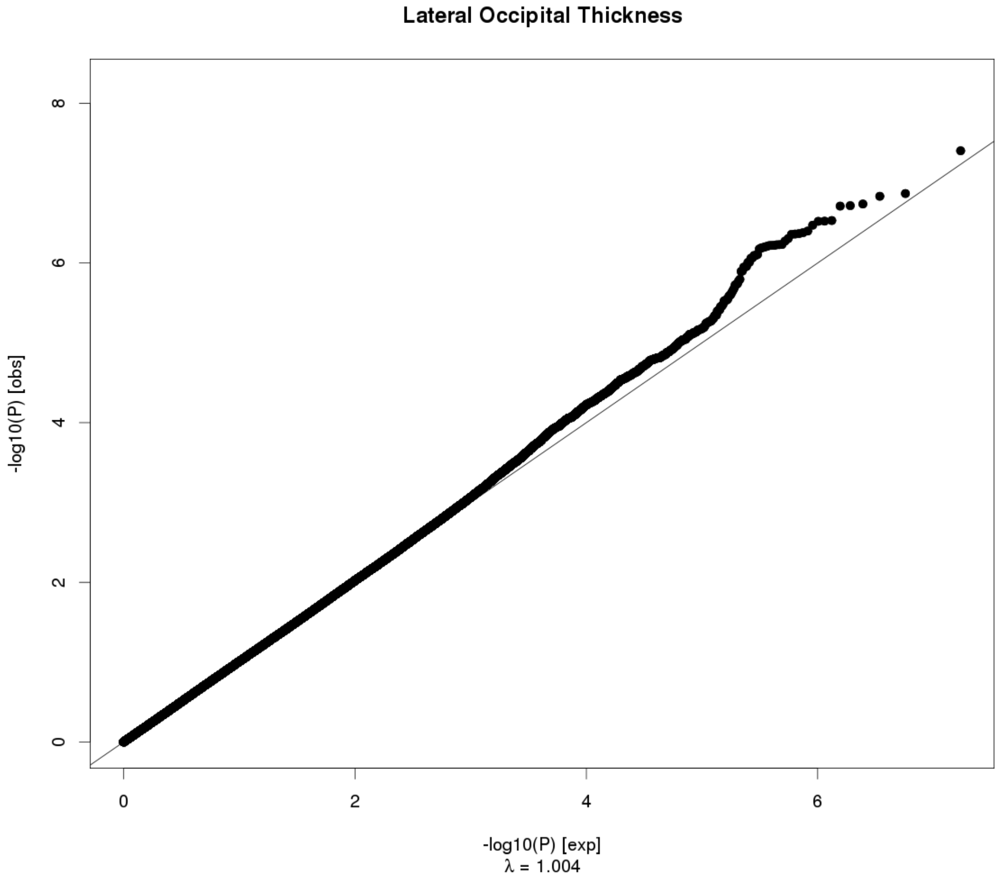
Lateral Occipital Thickness
-log10(P) [obs]
0
2
4
6
8
0
2
4
6
-log10(P) [exp]
λ = 1.004
