## Supplemental_Forest_Plots for "The genetic architecture of the human cerebral cortex"

| Locus # | Area | Trait | Marker | SNP | Page |
|---|---|---|---|---|---|
| 1 | Rostral Middle Frontal | SA | 1:17311882 | rs6682671 | 8 |
| 2 | Lingual | SA | 1:18962095 | rs1934057 | 9 |
| 3 | Superior Frontal | SA | 1:64235727 | rs4915928 | 10 |
| 4 | Posterior Cingulate | SA | 1:87633483:T_TGA | rs113198643 | 11 |
| 5 | Precuneus | SA | 1:87873700 | rs59373415 | 12 |
| 6 | Inferior Parietal | SA | 1:88216936 | rs305437 | 13 |
| 7 | Pericalcarine | SA | 1:88269969 | rs147753572 | 14 |
| 8 | Inferior Parietal | SA | 1:88423397 | rs1413536 | 15 |
| 9 | Lingual | SA | 1:93032583 | rs6603991 | 16 |
| 10 | Lingual | SA | 1:113239478 | rs2999158 | 17 |
| 11 | Pericalcarine | SA | 1:113239478 | rs2999158 | 18 |
| 12 | Pars Orbitalis | SA | 1:119762175 | rs72691108 | 19 |
| 13 | Lateral Orbitofrontal | SA | 1:119799099 | rs7529542 | 20 |
| 14 | Rostral Middle Frontal | TH | 1:247527273 | rs12058942 | 21 |
| 15 | Superior Parietal | SA | 2:4550112:CTG_C | rs769344141 | 22 |
| 16 | Superior Parietal | SA | 2:4550411 | rs688409 | 23 |
| 17 | Inferior Parietal | SA | 2:4554304 | rs639016 | 24 |
| 18 | Average Thickness | TH | 2:27148654:C_CT | rs11386753 | 25 |
| 19 | Precuneus | SA | 2:37076594 | rs62132521 | 26 |
| 20 | Precuneus | SA | 2:37079583 | rs62132522 | 27 |
| 21 | Isthmus Cingulate | SA | 2:37150793 | rs3770776 | 28 |
| 22 | Posterior Cingulate | SA | 2:37175444 | rs11695609 | 29 |
| 23 | Precuneus | SA | 2:37748367 | rs7559976 | 30 |
| 24 | Inferior Temporal | SA | 2:37899310 | rs9309013 | 31 |
| 25 | Superior Temporal | SA | 2:48277490 | rs7601767 | 32 |
| 26 | Lingual | TH | 2:49391013 | rs10495963 | 33 |
| 27 | Inferior Parietal | SA | 2:54615325 | rs79272390 | 34 |
| 28 | Superior Parietal | SA | 2:54615325 | rs79272390 | 35 |
| 29 | Pars Orbitalis | SA | 2:65971597 | rs2287283 | 36 |
| 30 | Pars Triangularis | SA | 2:65971789:C_CAG | rs142706617 | 37 |
| 31 | Caudal Middle Frontal | SA | 2:65974661:AG_A | rs770408932 | 38 |
| 32 | Average Thickness | TH | 2:98275354 | rs11692435 | 39 |
| 33 | Transverse Temporal | SA | 2:149940486 | rs2889657 | 40 |
| 34 | Superior Temporal | SA | 2:149994571 | rs13011264 | 41 |
| 35 | Superior Temporal | SA | 2:150007758 | rs389020 | 42 |
| 36 | Transverse Temporal | SA | 2:150037201 | rs11684511 | 43 |
| 37 | Caudal Anterior Cingulate | SA | 2:162822709 | rs13021985 | 44 |
| 38 | Lingual | SA | 2:162868858 | rs1014444 | 45 |
| 39 | Pericalcarine | SA | 2:162901327 | rs16822665 | 46 |
| 40 | Frontal Pole | SA | 2:188278203 | rs17464221 | 47 |
| 41 | Rostral Middle Frontal | SA | 2:188379320 | rs35612915 | 48 |
| 42 | Cuneus | SA | 2:204517033 | rs71427711 | 49 |
| 43 | Total Surface Area | SA | 3:28007315 | rs12630663 | 50 |
| 44 | Average Thickness | TH | 3:39489651 | rs533577 | 51 |
| 45 | Rostral Middle Frontal | TH | 3:55068255 | rs4955920 | 52 |
| 46 | Superior Parietal | SA | 3:71557865 | rs17718831 | 53 |
| 47 | Cuneus | SA | 3:104647183 | rs4895120 | 54 |
| 48 | Lingual | SA | 3:104662433 | rs13318870 | 55 |
| 49 | Lateral Occipital | SA | 3:104678133 | rs9863836 | 56 |
| 50 | Inferior Parietal | SA | 3:104724219 | rs9856782 | 57 |
| 51 | Pericalcarine | SA | 3:104724787 | rs971550 | 58 |
| 52 | Lingual | SA | 3:104735633 | rs28551708 | 59 |

| | | | | | |
|---|---|---|---|---|---|
| 53 | Lateral Occipital | SA | 3:118994959 | rs56007616 | 60 |
| 54 | Pericalcarine | SA | 3:118995379 | rs16829649 | 61 |
| 55 | Total Surface Area | SA | 3:141721762 | rs34464850 | 62 |
| 56 | Lingual | SA | 3:146993779 | rs61508189 | 63 |
| 57 | Pars Orbitalis | SA | 3:147093600 | rs1503738 | 64 |
| 58 | Postcentral | SA | 3:147105790 | rs2279830 | 65 |
| 59 | Pars Triangularis | SA | 3:147106319 | rs2279829 | 66 |
| 60 | Postcentral | SA | 3:147106319 | rs2279829 | 67 |
| 61 | Supramarginal | SA | 3:147106319 | rs2279829 | 68 |
| 62 | Pars Triangularis | SA | 3:147121751 | rs6766244 | 69 |
| 63 | Pars Triangularis | SA | 3:147171352 | rs9881533 | 70 |
| 64 | Pars Triangularis | SA | 3:147218844:CCAT_C | 3:147218844_CCAT_C | 71 |
| 65 | Postcentral | SA | 3:147227163 | rs78155705 | 72 |
| 66 | Lateral Occipital | SA | 3:177295215 | rs552305 | 73 |
| 67 | Precuneus | SA | 3:190657360 | rs905124 | 74 |
| 68 | Rostral Middle Frontal | SA | 3:193536820 | rs1165645 | 75 |
| 69 | Caudal Middle Frontal | SA | 3:193547618:T_TG | rs140045876 | 76 |
| 70 | Pericalcarine | SA | 4:980896 | rs11248061 | 77 |
| 71 | Lingual | SA | 4:1002448 | rs6812278 | 78 |
| 72 | Pericalcarine | SA | 4:1002448 | rs6812278 | 79 |
| 73 | Pericalcarine | SA | 4:19095652 | rs13115025 | 80 |
| 74 | Superior Parietal | SA | 4:53702875 | rs6554054 | 81 |
| 75 | Supramarginal | SA | 4:53738411 | rs2200225 | 82 |
| 76 | Temporal Pole | SA | 4:103112470 | rs6855246 | 83 |
| 77 | Total Surface Area | SA | 4:106009763 | rs2301718 | 84 |
| 78 | Superior Parietal | SA | 4:113416783 | rs6840242 | 85 |
| 79 | Average Thickness | TH | 4:121643239 | rs35021943 | 86 |
| 80 | Postcentral | SA | 4:127537734 | rs313135 | 87 |
| 81 | Pericalcarine | SA | 4:145284208 | rs7378179 | 88 |
| 82 | Fusiform | SA | 5:56082094 | rs10940512 | 89 |
| 83 | Lingual | SA | 5:60299840:C_CT | rs75625671 | 90 |
| 84 | Pericalcarine | SA | 5:60318973 | rs62367903 | 91 |
| 85 | Pericalcarine | SA | 5:60484660 | rs159540 | 92 |
| 86 | Posterior Cingulate | TH | 5:66144113 | rs12110247 | 93 |
| 87 | Posterior Cingulate | SA | 5:66155020 | rs72761270 | 94 |
| 88 | Total Surface Area | SA | 5:81092787 | rs386424 | 95 |
| 89 | Entorhinal | SA | 5:81882571 | rs4147321 | 96 |
| 90 | Parahippocampal | SA | 5:82350191 | rs27493 | 97 |
| 91 | Caudal Anterior Cingulate | SA | 5:82843609 | rs7728751 | 98 |
| 92 | Insula | SA | 5:82843609 | rs7728751 | 99 |
| 93 | Inferior Parietal | SA | 5:92090343 | rs34969 | 100 |
| 94 | Inferior Parietal | SA | 5:92134273 | rs27540 | 101 |
| 95 | Precuneus | SA | 5:92186429 | rs888814 | 102 |
| 96 | Superior Frontal | SA | 5:92187932 | rs17669337 | 103 |
| 97 | Superior Parietal | SA | 5:92797166 | rs115877304 | 104 |
| 98 | Superior Parietal | SA | 5:92948485 | rs114489117 | 105 |
| 99 | Precentral | SA | 5:92950673 | rs10064431 | 106 |
| 100 | Postcentral | SA | 5:93347836 | rs34322452 | 107 |
| 101 | Middle Temporal | SA | 5:93385558 | rs141834426 | 108 |
| 102 | Middle Temporal | SA | 5:93557702 | rs17376456 | 109 |
| 103 | Pericalcarine | SA | 5:124283820:T_TA | rs149998495 | 110 |
| 104 | Inferior Parietal | SA | 5:128971797 | rs62399042 | 111 |
| 105 | Caudal Middle Frontal | SA | 5:129018314 | rs30641 | 112 |
| 106 | Transverse Temporal | SA | 5:131341541 | rs7714191 | 113 |
| 107 | Total Surface Area | SA | 5:170778824 | rs7715167 | 114 |

| | | | | | |
|---|---|---|---|---|---|
| 108 | Caudal Middle Frontal | TH | 5:171927246 | rs11745941 | 115 |
| 109 | Lateral Orbitofrontal | SA | 6:7119134 | rs13208234 | 116 |
| 110 | Caudal Middle Frontal | SA | 6:92002653 | rs9345125 | 117 |
| 111 | Transverse Temporal | SA | 6:92009564 | rs4706391 | 118 |
| 112 | Precentral | SA | 6:92009596 | rs4706392 | 119 |
| 113 | Superior Parietal | SA | 6:92010245 | rs2144366 | 120 |
| 114 | Total Surface Area | SA | 6:108926496 | rs2802295 | 121 |
| 115 | Superior Frontal | SA | 6:126024301 | rs142301939 | 122 |
| 116 | Pars Opercularis | SA | 6:126070789 | rs7764016 | 123 |
| 117 | Total Surface Area | SA | 6:126412953 | rs11154343 | 124 |
| 118 | Cuneus | SA | 6:126613946 | rs76470478 | 125 |
| 119 | Pericalcarine | SA | 6:126666648 | rs117892760 | 126 |
| 120 | Lateral Orbitofrontal | SA | 6:126727908 | rs4897178 | 127 |
| 121 | Caudal Middle Frontal | SA | 6:126727930 | rs4897179 | 128 |
| 122 | Total Surface Area | SA | 6:126792095 | rs11759026 | 129 |
| 123 | Parahippocampal | SA | 6:126868567 | rs58321169 | 130 |
| 124 | Caudal Middle Frontal | SA | 6:126964510 | rs4273712 | 131 |
| 125 | Lateral Orbitofrontal | SA | 6:126995044 | rs1262478 | 132 |
| 126 | Total Surface Area | SA | 6:127000881 | rs74580701 | 133 |
| 127 | Total Surface Area | SA | 6:127025661 | rs13212044 | 134 |
| 128 | Lateral Occipital | SA | 6:127096181 | rs9401907 | 135 |
| 129 | Lingual | SA | 6:127096181 | rs9401907 | 136 |
| 130 | Pericalcarine | SA | 6:127096181 | rs9401907 | 137 |
| 131 | Total Surface Area | SA | 6:127204623 | rs9375477 | 138 |
| 132 | Pericalcarine | SA | 6:138865324 | rs4895532 | 139 |
| 133 | Precuneus | SA | 6:138868516 | rs9399245 | 140 |
| 134 | Rostral Anterior Cingulate | SA | 7:18737197 | rs1178101 | 141 |
| 135 | Cuneus | SA | 7:18881075 | rs12536836 | 142 |
| 136 | Pericalcarine | SA | 7:18883690 | rs6461386 | 143 |
| 137 | Lingual | SA | 7:18915666 | rs10237280 | 144 |
| 138 | Lateral Orbitofrontal | SA | 7:19520977 | rs73685918 | 145 |
| 139 | Lateral Orbitofrontal | SA | 7:19619484:T_TAG | rs371426453 | 146 |
| 140 | Lateral Orbitofrontal | SA | 7:19626622 | rs4721802 | 147 |
| 141 | Pars Triangularis | SA | 7:96202718 | rs10278627 | 148 |
| 142 | Pars Triangularis | SA | 7:96227720 | rs56290730 | 149 |
| 143 | Lingual | SA | 7:107237807 | rs7809950 | 150 |
| 144 | Superior Parietal | TH | 7:120721087 | rs73215353 | 151 |
| 145 | Superior Temporal | SA | 8:8605632 | rs4841029 | 152 |
| 146 | Lateral Orbitofrontal | SA | 8:8975103 | rs1822951 | 153 |
| 147 | Superior Temporal | SA | 8:9055772 | rs10094141 | 154 |
| 148 | Superior Temporal | SA | 8:9370559 | rs10100760 | 155 |
| 149 | Superior Temporal | SA | 8:9408271 | rs4840425 | 156 |
| 150 | Superior Temporal | SA | 8:10431422 | rs11250033 | 157 |
| 151 | Superior Temporal | SA | 8:10431486 | rs12548232 | 158 |
| 152 | Transverse Temporal | SA | 8:10943276 | rs2409691 | 159 |
| 153 | Superior Temporal | SA | 8:10962800 | rs1057626 | 160 |
| 154 | Superior Temporal | SA | 8:11109691 | rs2736373 | 161 |
| 155 | Pars Opercularis | SA | 8:71564667 | rs441890 | 162 |
| 156 | Inferior Parietal | SA | 8:93204400:T_TA | rs35576477 | 163 |
| 157 | Average Thickness | TH | 8:110585288 | rs7824177 | 164 |
| 158 | Rostral Middle Frontal | SA | 8:120596023 | rs10283100 | 165 |
| 159 | Postcentral | SA | 9:76157130 | rs11789773 | 166 |
| 160 | Superior Frontal | SA | 9:83480392 | rs76696867 | 167 |
| 161 | Pericalcarine | SA | 9:98257305 | rs28633576 | 168 |
| 162 | Lingual | SA | 9:98276371 | rs28410513 | 169 |

| | | | | | |
|---|---|---|---|---|---|
| 163 | Lateral Occipital | SA | 9:98278332 | rs28496034 | 170 |
| 164 | Banks of the Superior Temporal Sulcus | SA | 9:126603989 | rs7862092 | 171 |
| 165 | Total Surface Area | SA | 10:21790476 | rs12357321 | 172 |
| 166 | Caudal Anterior Cingulate | SA | 10:25123182 | rs4747503 | 173 |
| 167 | Paracentral | SA | 10:33539487 | rs2269084 | 174 |
| 168 | Inferior Parietal | SA | 10:33613967 | rs75921753 | 175 |
| 169 | Medial Orbitofrontal | SA | 10:50302895 | rs7097933 | 176 |
| 170 | Lingual | SA | 10:104422321 | rs7914158 | 177 |
| 171 | Total Surface Area | SA | 10:105012994 | rs1628768 | 178 |
| 172 | Total Surface Area | SA | 10:105218254 | rs4918016 | 179 |
| 173 | Precentral | SA | 10:118696266 | rs4751614 | 180 |
| 174 | Insula | SA | 10:118752250 | rs1122688 | 181 |
| 175 | Precuneus | SA | 10:118777998 | rs10749233 | 182 |
| 176 | Isthmus Cingulate | TH | 10:126486443 | rs12764880 | 183 |
| 177 | Posterior Cingulate | SA | 10:126486443 | rs12764880 | 184 |
| 178 | Cuneus | SA | 11:12071855 | rs10765918 | 185 |
| 179 | Pericalcarine | SA | 11:12071855 | rs10765918 | 186 |
| 180 | Lingual | SA | 11:30819811 | rs2022130 | 187 |
| 181 | Pericalcarine | SA | 11:30883876 | rs273587 | 188 |
| 182 | Pericalcarine | SA | 11:31499705 | rs1223090 | 189 |
| 183 | Postcentral | SA | 11:37001542 | rs11033898 | 190 |
| 184 | Parahippocampal | SA | 11:92319654 | rs1792354 | 191 |
| 185 | Pars Orbitalis | SA | 11:92348702 | rs61901866 | 192 |
| 186 | Fusiform | SA | 11:110962056 | rs7123402 | 193 |
| 187 | Pars Triangularis | SA | 11:111050526 | rs59614433 | 194 |
| 188 | Middle Temporal | SA | 11:111055661 | rs12794347 | 195 |
| 189 | Insula | SA | 11:111056158 | rs58066679 | 196 |
| 190 | Fusiform | SA | 11:111072779 | rs949279 | 197 |
| 191 | Pericalcarine | SA | 12:5314508 | rs17179798 | 198 |
| 192 | Superior Parietal | SA | 12:6545417 | rs7980991 | 199 |
| 193 | Paracentral | SA | 12:27790516 | rs2346756 | 200 |
| 194 | Superior Frontal | TH | 12:28833981 | rs4572176 | 201 |
| 195 | Total Surface Area | SA | 12:56470625 | rs11171739 | 202 |
| 196 | Inferior Parietal | SA | 12:65797106 | rs2336714 | 203 |
| 197 | Lateral Orbitofrontal | SA | 12:65905126 | rs79487293 | 204 |
| 198 | Total Surface Area | SA | 12:66327632 | rs10878349 | 205 |
| 199 | Total Surface Area | SA | 12:66367070 | rs12826248 | 206 |
| 200 | Isthmus Cingulate | SA | 12:78241285 | rs1123680 | 207 |
| 201 | Superior Frontal | SA | 12:79951566 | rs4842266 | 208 |
| 202 | Entorhinal | TH | 12:94558576 | rs3847803 | 209 |
| 203 | Lingual | TH | 13:58591051 | rs9537915 | 210 |
| 204 | Lingual | SA | 13:80173760 | rs9545145 | 211 |
| 205 | Pericalcarine | SA | 13:80192906 | rs9545158 | 212 |
| 206 | Pars Triangularis | SA | 13:81198667 | rs7996803 | 213 |
| 207 | Pericalcarine | SA | 14:29228427 | rs35342371 | 214 |
| 208 | Lingual | SA | 14:54316607 | rs7148896 | 215 |
| 209 | Lateral Orbitofrontal | SA | 14:54724054 | rs2358483 | 216 |
| 210 | Pars Orbitalis | SA | 14:54777745 | rs7147119 | 217 |
| 211 | Lateral Occipital | SA | 14:57337018 | rs76398229 | 218 |
| 212 | Banks of the Superior Temporal Sulcus | SA | 14:58724817:C_CTT | rs35486640 | 219 |
| 213 | Inferior Temporal | SA | 14:59063842 | rs7155669 | 220 |
| 214 | Cuneus | SA | 14:59064739 | rs170239 | 221 |
| 215 | Supramarginal | SA | 14:59069053 | rs221326 | 222 |

| | | | | | |
|---|---|---|---|---|---|
| 216 | Banks of the Superior Temporal Sulcus | TH | 14:59071725 | rs149142 | 223 |
| 217 | Middle Temporal | TH | 14:59074136 | rs160459 | 224 |
| 218 | Banks of the Superior Temporal Sulcus | SA | 14:59074878 | rs160458 | 225 |
| 219 | Precuneus | SA | 14:59598286 | rs7143623 | 226 |
| 220 | Cuneus | TH | 14:59605116 | rs117461235 | 227 |
| 221 | Pericalcarine | TH | 14:59605116 | rs117461235 | 228 |
| 222 | Cuneus | SA | 14:59625997 | rs73313052 | 229 |
| 223 | Pericalcarine | SA | 14:59625997 | rs73313052 | 230 |
| 224 | Precuneus | SA | 14:59625997 | rs73313052 | 231 |
| 225 | Lingual | SA | 14:59627434:AC_A | 14:59627434_AC_A | 232 |
| 226 | Supramarginal | SA | 14:59627631 | rs2164950 | 233 |
| 227 | Lateral Occipital | SA | 14:59628679 | rs76341705 | 234 |
| 228 | Lateral Occipital | SA | 14:59683086 | rs78445564 | 235 |
| 229 | Pericalcarine | SA | 14:59805480 | rs75341124 | 236 |
| 230 | Fusiform | SA | 14:59827306 | rs17834032 | 237 |
| 231 | Entorhinal | SA | 14:59860705 | rs7141150 | 238 |
| 232 | Supramarginal | SA | 15:39081795 | rs62007727 | 239 |
| 233 | Precentral | SA | 15:39092485:AG_A | 15:39092485_AG_A | 240 |
| 234 | Precentral | SA | 15:39343032 | rs11070172 | 241 |
| 235 | Precentral | SA | 15:39519925 | rs156795 | 242 |
| 236 | Postcentral | SA | 15:39553167 | rs77470370 | 243 |
| 237 | Postcentral | SA | 15:39556139:TAC_T | 15:39556139_TAC_T | 244 |
| 238 | Precentral | SA | 15:39582377:TA_T | 15:39582377_TA_T | 245 |
| 239 | Supramarginal | SA | 15:39593078 | rs11631253 | 246 |
| 240 | Postcentral | TH | 15:39595340 | rs62002282 | 247 |
| 241 | Precentral | SA | 15:39595340 | rs62002282 | 248 |
| 242 | Postcentral | SA | 15:39604340 | rs3862145 | 249 |
| 243 | Precentral | SA | 15:39604340 | rs3862145 | 250 |
| 244 | Postcentral | SA | 15:39623847 | rs10851383 | 251 |
| 245 | Postcentral | TH | 15:39623847 | rs10851383 | 252 |
| 246 | Pars Opercularis | SA | 15:39625618 | rs11070185 | 253 |
| 247 | Precentral | SA | 15:39625778 | rs10851385 | 254 |
| 248 | Supramarginal | SA | 15:39625778 | rs10851385 | 255 |
| 249 | Precentral | SA | 15:39627855 | rs76465453 | 256 |
| 250 | Postcentral | TH | 15:39632013 | rs71471500 | 257 |
| 251 | Pars Opercularis | SA | 15:39633904 | rs2033939 | 258 |
| 252 | Transverse Temporal | SA | 15:39633904 | rs2033939 | 259 |
| 253 | Precentral | SA | 15:39634222 | rs1080066 | 260 |
| 254 | Rostral Middle Frontal | SA | 15:39634222 | rs1080066 | 261 |
| 255 | Postcentral | TH | 15:39634303 | rs1822105 | 262 |
| 256 | Precentral | SA | 15:39634303 | rs1822105 | 263 |
| 257 | Superior Parietal | SA | 15:39634303 | rs1822105 | 264 |
| 258 | Postcentral | SA | 15:39637355 | rs117193619 | 265 |
| 259 | Precentral | SA | 15:39637355 | rs117193619 | 266 |
| 260 | Pars Triangularis | SA | 15:39639898 | rs4924345 | 267 |
| 261 | Superior Parietal | SA | 15:39639898 | rs4924345 | 268 |
| 262 | Supramarginal | SA | 15:39639898 | rs4924345 | 269 |
| 263 | Postcentral | SA | 15:39639992 | rs4924346 | 270 |
| 264 | Precentral | SA | 15:39640301 | rs148182077 | 271 |
| 265 | Postcentral | SA | 15:39641885 | rs10520129 | 272 |
| 266 | Supramarginal | SA | 15:39647278 | rs35391898 | 273 |
| 267 | Postcentral | SA | 15:39652450 | rs72722993 | 274 |
| 268 | Precentral | SA | 15:39652450 | rs72722993 | 275 |

| | | | | | |
|---|---|---|---|---|---|
| 269 | Precentral | SA | 15:39671514 | rs11070197 | 276 |
| 270 | Supramarginal | SA | 15:39720600 | rs78502100 | 277 |
| 271 | Precentral | SA | 15:39742570 | rs62005276 | 278 |
| 272 | Pars Opercularis | SA | 15:39790176 | rs10459586 | 279 |
| 273 | Superior Temporal | SA | 15:39809640 | rs115241741 | 280 |
| 274 | Pericalcarine | SA | 15:39953061 | rs8034885 | 281 |
| 275 | Lateral Occipital | SA | 15:63297719 | rs28514429 | 282 |
| 276 | Pericalcarine | SA | 16:28919583:A_ATT | rs10653411 | 283 |
| 277 | Caudal Middle Frontal | SA | 16:52538337 | rs7184835 | 284 |
| 278 | Caudal Anterior Cingulate | SA | 16:60827388 | rs80241863 | 285 |
| 279 | Rostral Middle Frontal | SA | 16:65014909 | rs40115 | 286 |
| 280 | Entorhinal | SA | 16:77320605 | rs12921392 | 287 |
| 281 | Superior Parietal | TH | 16:87229344 | rs56023709 | 288 |
| 282 | Pars Opercularis | SA | 17:13054525 | rs12938190 | 289 |
| 283 | Insula | SA | 17:28338304 | rs4291964 | 290 |
| 284 | Total Surface Area | SA | 17:43543075:C_CA | rs200291097 | 291 |
| 285 | Fusiform | SA | 17:43859640 | rs62057070 | 292 |
| 286 | Total Surface Area | SA | 17:43897722 | rs79600142 | 293 |
| 287 | Average Thickness | TH | 17:43919068 | rs2316766 | 294 |
| 288 | Total Surface Area | SA | 17:44808360 | rs35937770 | 295 |
| 289 | Rostral Anterior Cingulate | SA | 17:46584657 | rs2202895 | 296 |
| 290 | Cuneus | SA | 17:63906955:C_CAA | rs143442432 | 297 |
| 291 | Pericalcarine | SA | 17:63914750 | rs1420791 | 298 |
| 292 | Pericalcarine | SA | 17:63990193 | rs150476910 | 299 |
| 293 | Superior Parietal | SA | 19:12904400 | rs17884482 | 300 |
| 294 | Superior Parietal | SA | 19:13096285 | rs4334415 | 301 |
| 295 | Inferior Parietal | SA | 19:13109763 | rs68175985 | 302 |
| 296 | Superior Parietal | SA | 19:13109763 | rs68175985 | 303 |
| 297 | Banks of the Superior Temporal Sulcus | SA | 19:13109955 | rs73006822 | 304 |
| 298 | Middle Temporal | SA | 19:13109955 | rs73006822 | 305 |
| 299 | Superior Parietal | SA | 19:13292745 | rs8103974 | 306 |
| 300 | Pars Triangularis | SA | 19:30852938:CTACT_G_C | 19:30852938_CTACTG_C | 307 |
| 301 | Rostral Anterior Cingulate | TH | 19:51621561 | rs62115964 | 308 |
| 302 | Inferior Parietal | SA | 20:52432451 | rs4437022 | 309 |
| 303 | Superior Parietal | SA | 20:52447303 | rs6022786 | 310 |
| 304 | Pericalcarine | SA | 20:52468966 | rs4811476 | 311 |
| 305 | Lateral Orbitofrontal | SA | 21:34298974 | rs12626790 | 312 |
| 306 | Postcentral | TH | 22:47200354 | rs4823878 | 313 |

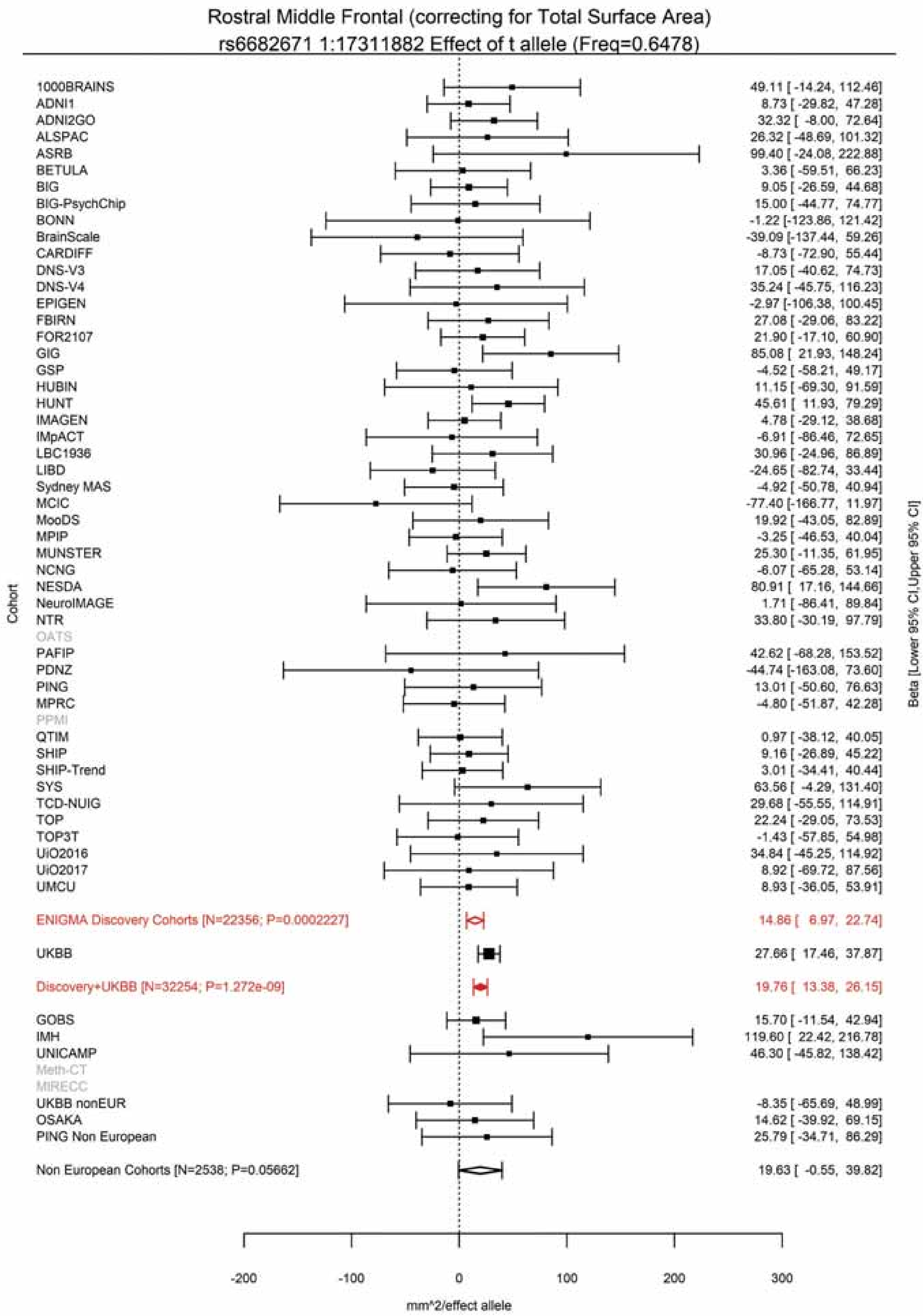
Rostral Middle Frontal (correcting for Total Surface Area)
rs6682671 1:17311882 Effect of t allele (Freq=0.6478)
1000BRAINS 49.11 [ -14.24, 112.46]
ADNI1 8.73 [ -29.82, 47.28]
ADNI2GO 32.32 [ -8.00, 72.64]
ALSPAC 26.32 [ -48.69, 101.32]
ASRB 99.40 [ -24.08, 222.88]
BETULA 3.36 [ -59.51, 66.23]
BIG 9.05 [ -26.59, 44.68]
BIG-PsychChip 15.00 [ -44.77, 74.77]
BONN -1.22 [-123.86, 121.42]
BrainScale -39.09 [-137.44, 59.26]
CARDIFF -8.73 [ -72.90, 55.44]
DNS-V3 17.05 [ -40.62, 74.73]
DNS-V4 35.24 [ -45.75, 116.23]
EPIGEN -2.97 [-106.38, 100.45]
FBIRN 27.08 [ -29.06, 83.22]
FOR2107 21.90 [ -17.10, 60.90]
GIG 85.08 [ 21.93, 148.24]
GSP -4.52 [ -58.21, 49.17]
HUBIN 11.15 [ -69.30, 91.59]
HUNT 45.61 [ 11.93, 79.29]
IMAGEN 4.78 [ -29.12, 38.68]
IMpACT -6.91 [ -86.46, 72.65]
LBC1936 30.96 [ -24.96, 86.89]
LIBD -24.65 [ -82.74, 33.44]
Sydney MAS -4.92 [ -50.78, 40.94]
MCIC -77.40 [-166.77, 11.97]
MooDS 19.92 [ -43.05, 82.89]
MPIP -3.25 [ -46.53, 40.04]
MUNSTER 25.30 [ -11.35, 61.95]
NCNG -6.07 [ -65.28, 53.14]
NESDA 80.91 [ 17.16, 144.66]
NeuroIMAGE 1.71 [ -86.41, 89.84]
NTR 33.80 [ -30.19, 97.79]
OATS
PAFIP 42.62 [ -68.28, 153.52]
PDNZ -44.74 [-163.08, 73.60]
PING 13.01 [ -50.60, 76.63]
MPRC -4.80 [ -51.87, 42.28]
PPMI
QTIM 0.97 [ -38.12, 40.05]
SHIP 9.16 [ -26.89, 45.22]
SHIP-Trend 3.01 [ -34.41, 40.44]
SYS 63.56 [ -4.29, 131.40]
TCD-NUIG 29.68 [ -55.55, 114.91]
TOP 22.24 [ -29.05, 73.53]
TOP3T -1.43 [ -57.85, 54.98]
UiO2016 34.84 [ -45.25, 114.92]
UiO2017 8.92 [ -69.72, 87.56]
UMCU 8.93 [ -36.05, 53.91]
ENIGMA Discovery Cohorts [N=22356; P=0.0002227] 14.86 [ 6.97, 22.74]
UKBB 27.66 [ 17.46, 37.87]
Discovery+UKBB [N=32254; P=1.272e-09] 19.76 [ 13.38, 26.15]
GOBS 15.70 [ -11.54, 42.94]
IMH 119.60 [ 22.42, 216.78]
UNICAMP 46.30 [ -45.82, 138.42]
Meth-CT
MIRECC
UKBB nonEUR -8.35 [ -65.69, 48.99]
OSAKA 14.62 [ -39.92, 69.15]
PING Non European 25.79 [ -34.71, 86.29]
Non European Cohorts [N=2538; P=0.05662] 19.63 [ -0.55, 39.82]
Cohort
Beta [Lower 95% CI,Upper 95% CI]
-200
-100
0
100
200
300
mm^2/effect allele

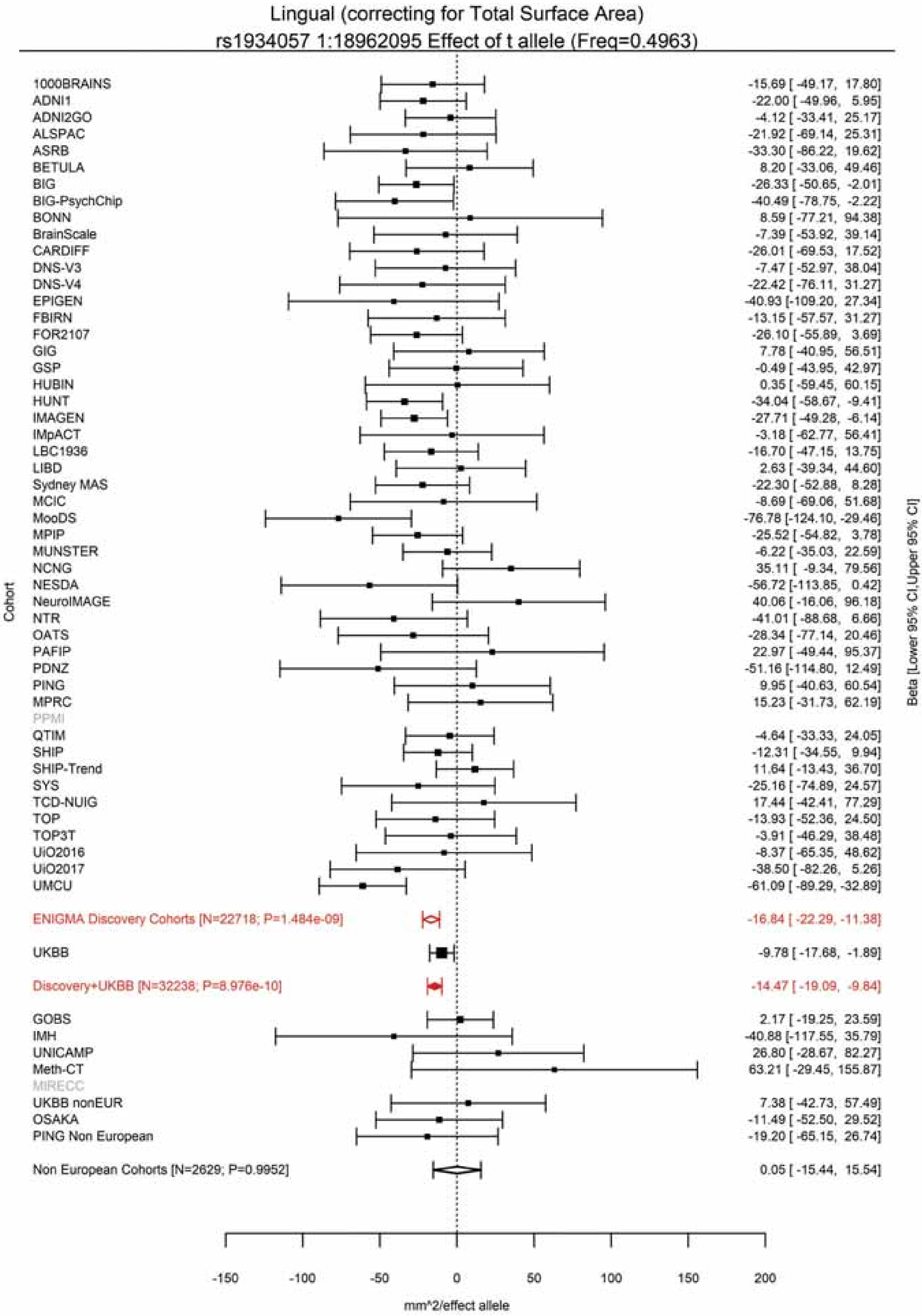

Lingual (correcting for Total Surface Area)
rs1934057 1:18962095 Effect of t allele (Freq=0.4963)
Cohort
1000BRAINS -15.69 [ -49.17, 17.80]
ADNI1 -22.00 [ -49.96, 5.95]
ADNI2GO -4.12 [ -33.41, 25.17]
ALSPAC -21.92 [ -69.14, 25.31]
ASRB -33.30 [ -86.22, 19.62]
BETULA 8.20 [ -33.06, 49.46]
BIG -26.33 [ -50.65, -2.01]
BIG-PsychChip -40.49 [ -78.75, -2.22]
BONN 8.59 [ -77.21, 94.38]
BrainScale -7.39 [ -53.92, 39.14]
CARDIFF -26.01 [ -69.53, 17.52]
DNS-V3 -7.47 [ -52.97, 38.04]
DNS-V4 -22.42 [ -76.11, 31.27]
EPIGEN -40.93 [-109.20, 27.34]
FBIRN -13.15 [ -57.57, 31.27]
FOR2107 -26.10 [ -55.89, 3.69]
GIG 7.78 [ -40.95, 56.51]
GSP -0.49 [ -43.95, 42.97]
HUBIN 0.35 [ -59.45, 60.15]
HUNT -34.04 [ -58.67, -9.41]
IMAGEN -27.71 [ -49.28, -6.14]
IMpACT -3.18 [ -62.77, 56.41]
LBC1936 -16.70 [ -47.15, 13.75]
LIBD 2.63 [ -39.34, 44.60]
Sydney MAS -22.30 [ -52.88, 8.28]
MCIC -8.69 [ -69.06, 51.68]
MooDS -76.78 [-124.10, -29.46]
MPIP -25.52 [ -54.82, 3.78]
MUNSTER -6.22 [ -35.03, 22.59]
NCNG 35.11 [ -9.34, 79.56]
NESDA -56.72 [-113.85, 0.42]
NeuroIMAGE 40.06 [ -16.06, 96.18]
NTR -41.01 [ -88.68, 6.66]
OATS -28.34 [ -77.14, 20.46]
PAFIP 22.97 [ -49.44, 95.37]
PDNZ -51.16 [-114.80, 12.49]
PING 9.95 [ -40.63, 60.54]
MPRC 15.23 [ -31.73, 62.19]
PPMI
QTIM -4.64 [ -33.33, 24.05]
SHIP -12.31 [ -34.55, 9.94]
SHIP-Trend 11.64 [ -13.43, 36.70]
SYS -25.16 [ -74.89, 24.57]
TCD-NUIG 17.44 [ -42.41, 77.29]
TOP -13.93 [ -52.36, 24.50]
TOP3T -3.91 [ -46.29, 38.48]
UiO2016 -8.37 [ -65.35, 48.62]
UiO2017 -38.50 [ -82.26, 5.26]
UMCU -61.09 [ -89.29, -32.89]
ENIGMA Discovery Cohorts [N=22718; P=1.484e-09] -16.84 [ -22.29, -11.38]
UKBB -9.78 [ -17.68, -1.89]
Discovery+UKBB [N=32238; P=8.976e-10] -14.47 [ -19.09, -9.84]
GOBS 2.17 [ -19.25, 23.59]
IMH -40.88 [-117.55, 35.79]
UNICAMP 26.80 [ -28.67, 82.27]
Meth-CT 63.21 [ -29.45, 155.87]
MIRECC
UKBB nonEUR 7.38 [ -42.73, 57.49]
OSAKA -11.49 [ -52.50, 29.52]
PING Non European -19.20 [ -65.15, 26.74]
Non European Cohorts [N=2629; P=0.9952] 0.05 [ -15.44, 15.54]
Beta [Lower 95% CI,Upper 95% CI]
-150
-100
-50
0
50
100
150
200
mm^2/effect allele

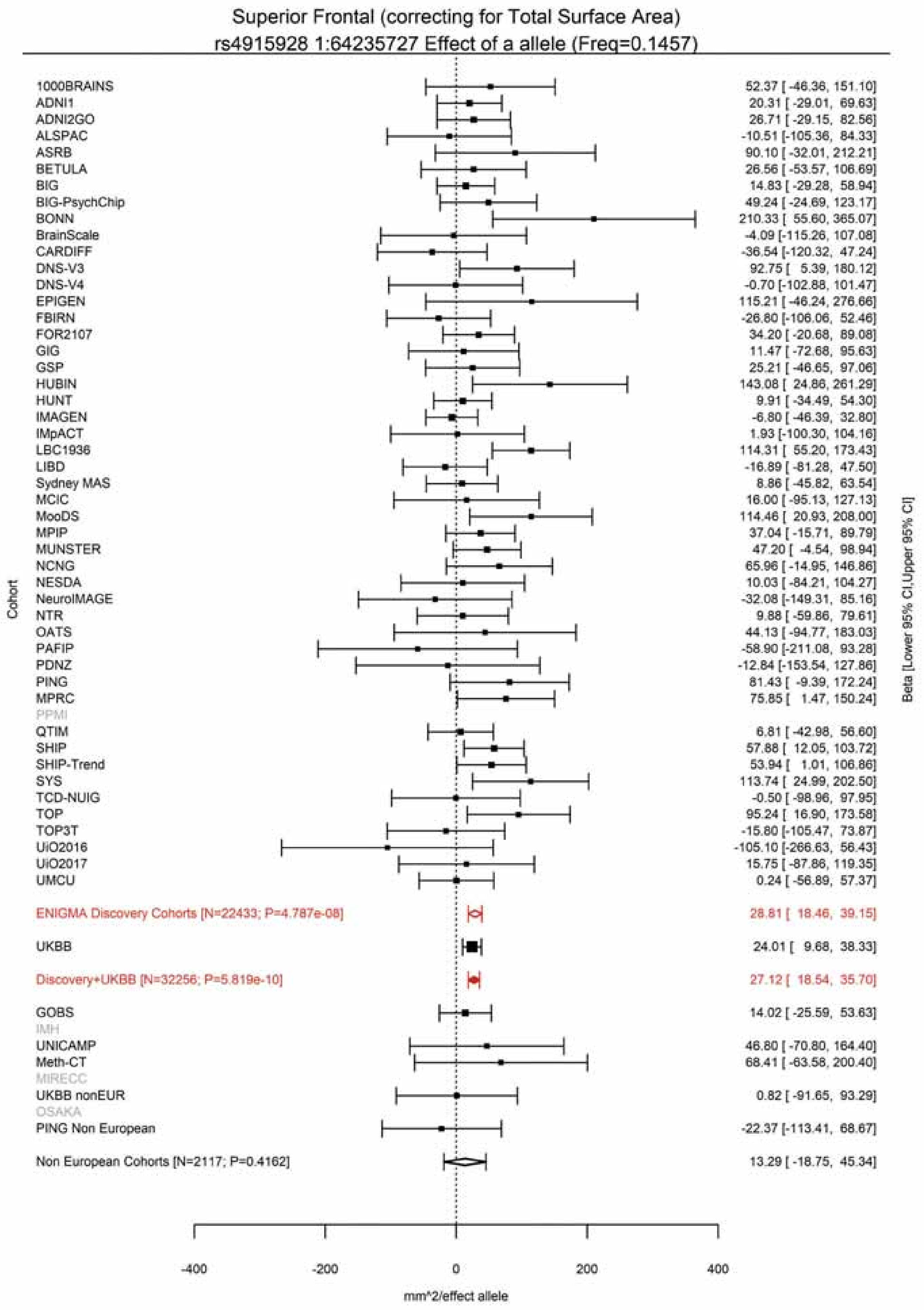

Superior Frontal (correcting for Total Surface Area)
rs4915928 1:64235727 Effect of a allele (Freq=0.1457)
Cohort
1000BRAINS 52.37 [ -46.36, 151.10]
ADNI1 20.31 [ -29.01, 69.63]
ADNI2GO 26.71 [ -29.15, 82.56]
ALSPAC -10.51 [-105.36, 84.33]
ASRB 90.10 [ -32.01, 212.21]
BETULA 26.56 [ -53.57, 106.69]
BIG 14.83 [ -29.28, 58.94]
BIG-PsychChip 49.24 [ -24.69, 123.17]
BONN 210.33 [ 55.60, 365.07]
BrainScale -4.09 [-115.26, 107.08]
CARDIFF -36.54 [-120.32, 47.24]
DNS-V3 92.75 [ 5.39, 180.12]
DNS-V4 -0.70 [-102.88, 101.47]
EPIGEN 115.21 [ -46.24, 276.66]
FBIRN -26.80 [-106.06, 52.46]
FOR2107 34.20 [ -20.68, 89.08]
GIG 11.47 [ -72.68, 95.63]
GSP 25.21 [ -46.65, 97.06]
HUBIN 143.08 [ 24.86, 261.29]
HUNT 9.91 [ -34.49, 54.30]
IMAGEN -6.80 [ -46.39, 32.80]
IMpACT 1.93 [-100.30, 104.16]
LBC1936 114.31 [ 55.20, 173.43]
LIBD -16.89 [ -81.28, 47.50]
Sydney MAS 8.86 [ -45.82, 63.54]
MCIC 16.00 [ -95.13, 127.13]
MooDS 114.46 [ 20.93, 208.00]
MPIP 37.04 [ -15.71, 89.79]
MUNSTER 47.20 [ -4.54, 98.94]
NCNG 65.96 [ -14.95, 146.86]
NESDA 10.03 [ -84.21, 104.27]
NeuroIMAGE -32.08 [-149.31, 85.16]
NTR 9.88 [ -59.86, 79.61]
OATS 44.13 [ -94.77, 183.03]
PAFIP -58.90 [-211.08, 93.28]
PDNZ -12.84 [-153.54, 127.86]
PING 81.43 [ -9.39, 172.24]
MPRC 75.85 [ 1.47, 150.24]
PPMI
QTIM 6.81 [ -42.98, 56.60]
SHIP 57.88 [ 12.05, 103.72]
SHIP-Trend 53.94 [ 1.01, 106.86]
SYS 113.74 [ 24.99, 202.50]
TCD-NUIG -0.50 [ -98.96, 97.95]
TOP 95.24 [ 16.90, 173.58]
TOP3T -15.80 [-105.47, 73.87]
UiO2016 -105.10 [-266.63, 56.43]
UiO2017 15.75 [ -87.86, 119.35]
UMCU 0.24 [ -56.89, 57.37]
ENIGMA Discovery Cohorts [N=22433; P=4.787e-08] 28.81 [ 18.46, 39.15]
UKBB 24.01 [ 9.68, 38.33]
Discovery+UKBB [N=32256; P=5.819e-10] 27.12 [ 18.54, 35.70]
GOBS 14.02 [ -25.59, 53.63]
IMH
UNICAMP 46.80 [ -70.80, 164.40]
Meth-CT 68.41 [ -63.58, 200.40]
MIRECC
UKBB nonEUR 0.82 [ -91.65, 93.29]
OSAKA
PING Non European -22.37 [-113.41, 68.67]
Non European Cohorts [N=2117; P=0.4162] 13.29 [ -18.75, 45.34]
-400
-200
0
200
400
mm^2/effect allele
Beta [Lower 95% CI,Upper 95% CI]

Posterior Cingulate (correcting for Total Surface Area)
rs113198643 1:87633483:T_TGA Effect of TGA allele (Freq=0.1325)

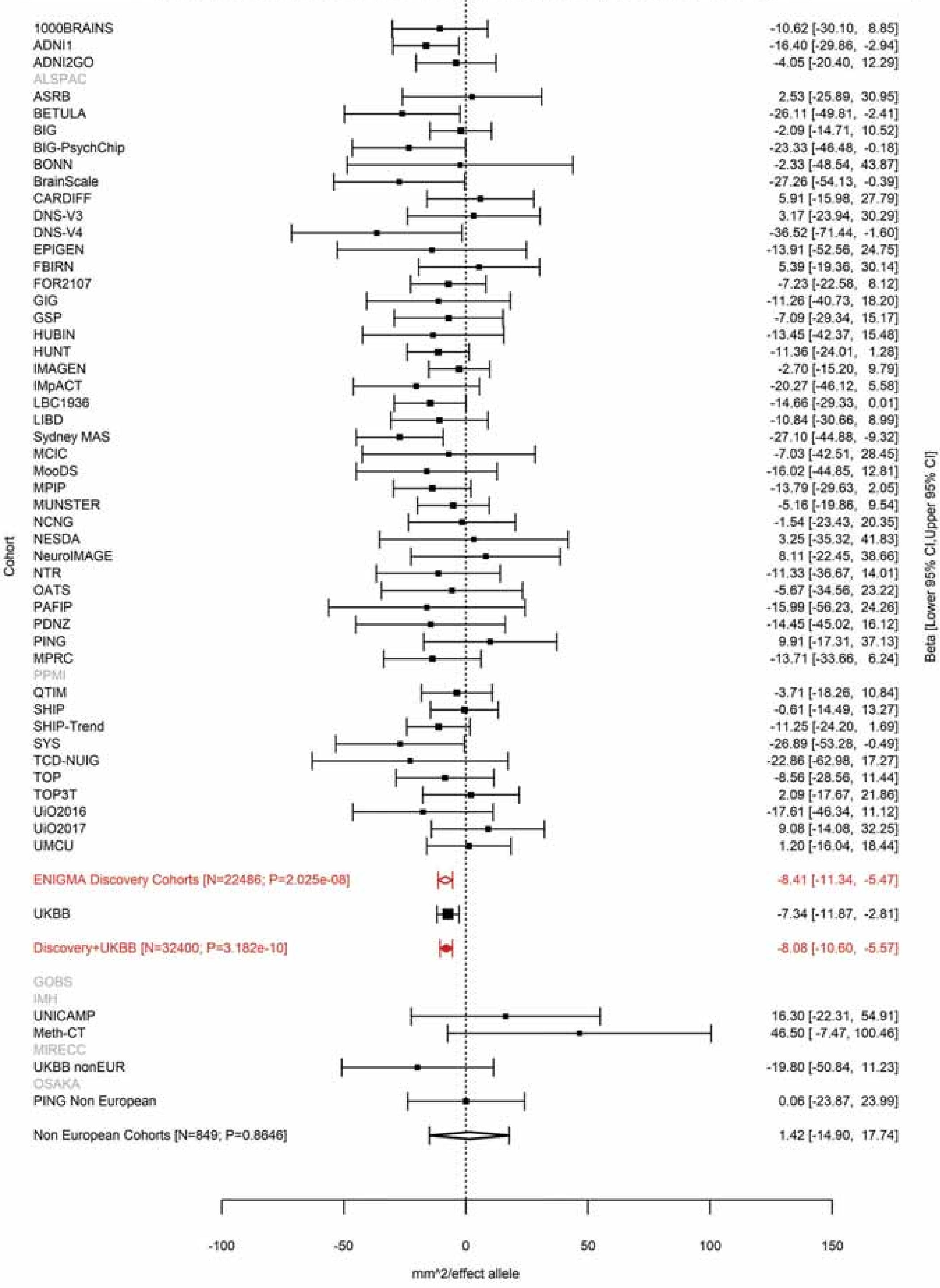

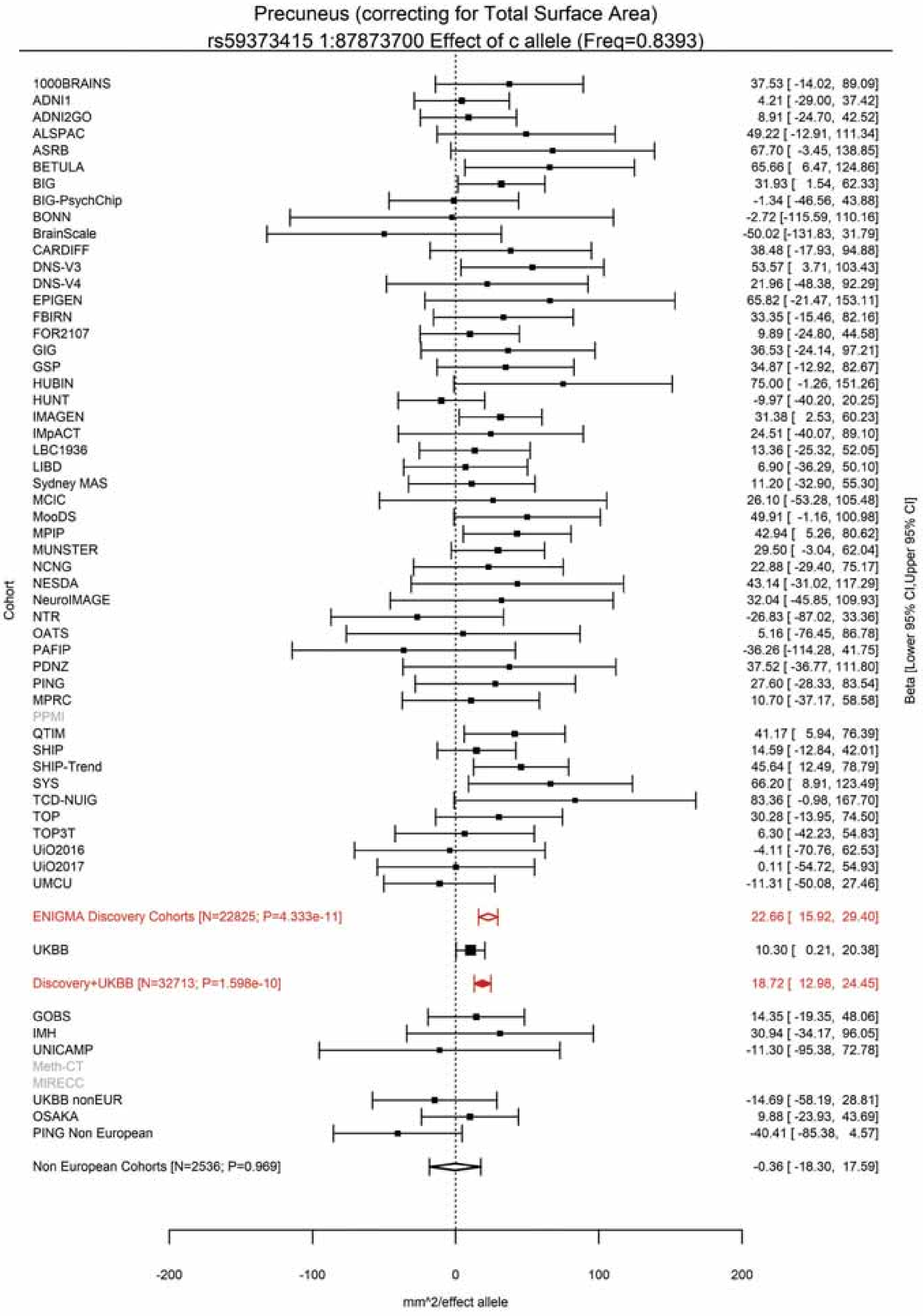

Precuneus (correcting for Total Surface Area)
rs59373415 1:87873700 Effect of c allele (Freq=0.8393)
Cohort
1000BRAINS 37.53 [ -14.02, 89.09]
ADNI1 4.21 [ -29.00, 37.42]
ADNI2GO 8.91 [ -24.70, 42.52]
ALSPAC 49.22 [ -12.91, 111.34]
ASRB 67.70 [ -3.45, 138.85]
BETULA 65.66 [ 6.47, 124.86]
BIG 31.93 [ 1.54, 62.33]
BIG-PsychChip -1.34 [ -46.56, 43.88]
BONN -2.72 [-115.59, 110.16]
BrainScale -50.02 [-131.83, 31.79]
CARDIFF 38.48 [ -17.93, 94.88]
DNS-V3 53.57 [ 3.71, 103.43]
DNS-V4 21.96 [ -48.38, 92.29]
EPIGEN 65.82 [ -21.47, 153.11]
FBIRN 33.35 [ -15.46, 82.16]
FOR2107 9.89 [ -24.80, 44.58]
GIG 36.53 [ -24.14, 97.21]
GSP 34.87 [ -12.92, 82.67]
HUBIN 75.00 [ -1.26, 151.26]
HUNT -9.97 [ -40.20, 20.25]
IMAGEN 31.38 [ 2.53, 60.23]
IMpACT 24.51 [ -40.07, 89.10]
LBC1936 13.36 [ -25.32, 52.05]
LIBD 6.90 [ -36.29, 50.10]
Sydney MAS 11.20 [ -32.90, 55.30]
MCIC 26.10 [ -53.28, 105.48]
MooDS 49.91 [ -1.16, 100.98]
MPIP 42.94 [ 5.26, 80.62]
MUNSTER 29.50 [ -3.04, 62.04]
NCNG 22.88 [ -29.40, 75.17]
NESDA 43.14 [ -31.02, 117.29]
NeuroIMAGE 32.04 [ -45.85, 109.93]
NTR -26.83 [ -87.02, 33.36]
OATS 5.16 [ -76.45, 86.78]
PAFIP -36.26 [-114.28, 41.75]
PDNZ 37.52 [ -36.77, 111.80]
PING 27.60 [ -28.33, 83.54]
MPRC 10.70 [ -37.17, 58.58]
PPMI
QTIM 41.17 [ 5.94, 76.39]
SHIP 14.59 [ -12.84, 42.01]
SHIP-Trend 45.64 [ 12.49, 78.79]
SYS 66.20 [ 8.91, 123.49]
TCD-NUIG 83.36 [ -0.98, 167.70]
TOP 30.28 [ -13.95, 74.50]
TOP3T 6.30 [ -42.23, 54.83]
UiO2016 -4.11 [ -70.76, 62.53]
UiO2017 0.11 [ -54.72, 54.93]
UMCU -11.31 [ -50.08, 27.46]
ENIGMA Discovery Cohorts [N=22825; P=4.333e-11] 22.66 [ 15.92, 29.40]
UKBB 10.30 [ 0.21, 20.38]
Discovery+UKBB [N=32713; P=1.598e-10] 18.72 [ 12.98, 24.45]
GOBS 14.35 [ -19.35, 48.06]
IMH 30.94 [ -34.17, 96.05]
UNICAMP -11.30 [ -95.38, 72.78]
Meth-CT
MIRECC
UKBB nonEUR -14.69 [ -58.19, 28.81]
OSAKA 9.88 [ -23.93, 43.69]
PING Non European -40.41 [ -85.38, 4.57]
Non European Cohorts [N=2536; P=0.969] -0.36 [ -18.30, 17.59]
-200
-100
0
100
200
mm^2/effect allele
Beta [Lower 95% CI,Upper 95% CI]

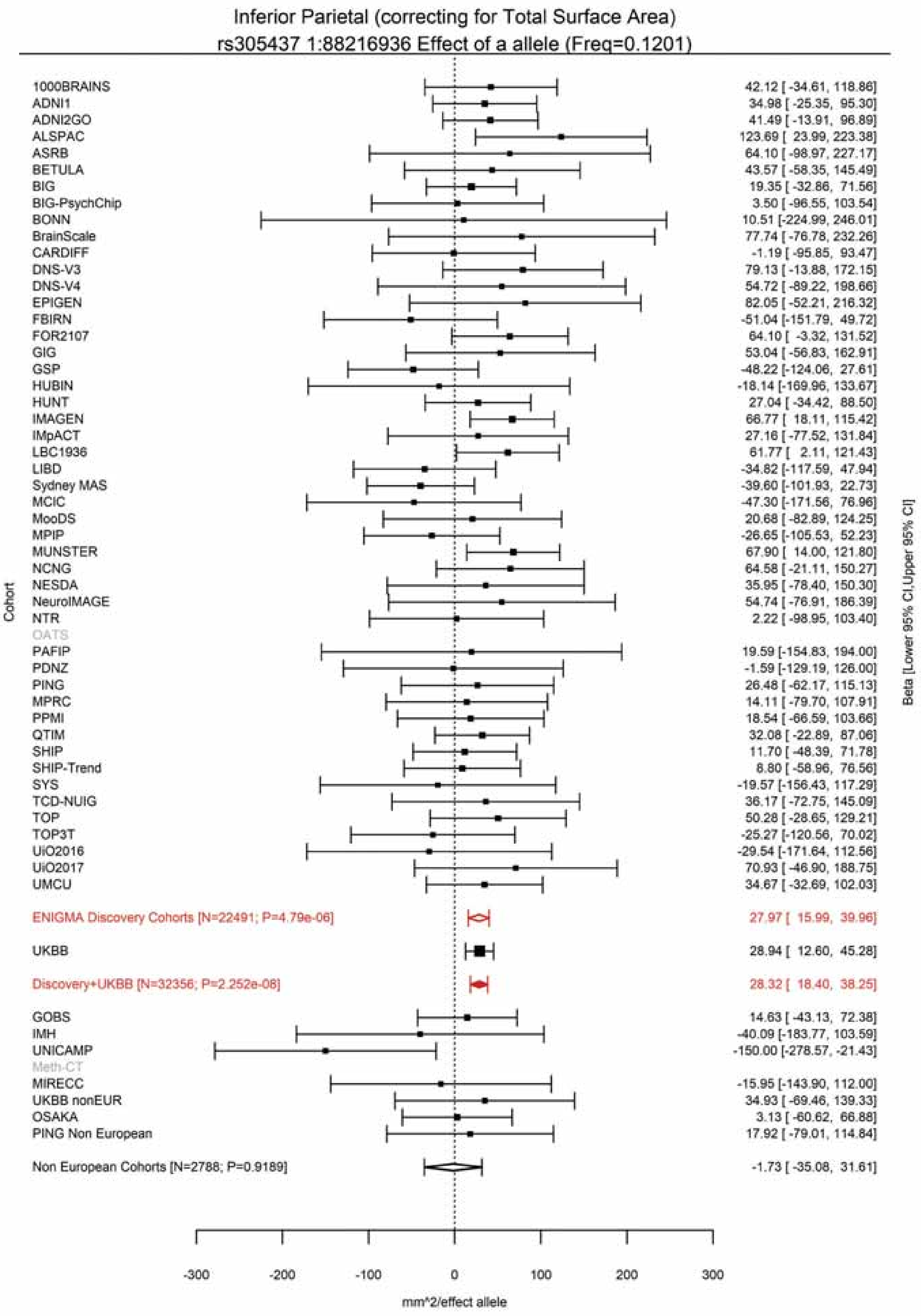
Inferior Parietal (correcting for Total Surface Area)
rs305437 1:88216936 Effect of a allele (Freq=0.1201)
1000BRAINS
ADNI1
ADNI2GO
ALSPAC
ASRB
BETULA
BIG
BIG-PsychChip
BONN
BrainScale
CARDIFF
DNS-V3
DNS-V4
EPIGEN
FBIRN
FOR2107
GIG
GSP
HUBIN
HUNT
IMAGEN
IMpACT
LBC1936
LIBD
Sydney MAS
MCIC
MooDS
MPIP
MUNSTER
NCNG
NESDA
NeuroIMAGE
NTR
OATS
PAFIP
PDNZ
PING
MPRC
PPMI
QTIM
SHIP
SHIP-Trend
SYS
TCD-NUIG
TOP
TOP3T
UiO2016
UiO2017
UMCU
ENIGMA Discovery Cohorts [N=22491; P=4.79e-06]
UKBB
Discovery+UKBB [N=32356; P=2.252e-08]
GOBS
IMH
UNICAMP
Meth-CT
MIRECC
UKBB nonEUR
OSAKA
PING Non European
Non European Cohorts [N=2788; P=0.9189]
42.12 [ -34.61, 118.86]
34.98 [ -25.35, 95.30]
41.49 [ -13.91, 96.89]
123.69 [ 23.99, 223.38]
64.10 [ -98.97, 227.17]
43.57 [ -58.35, 145.49]
19.35 [ -32.86, 71.56]
3.50 [ -96.55, 103.54]
10.51 [-224.99, 246.01]
77.74 [ -76.78, 232.26]
-1.19 [ -95.85, 93.47]
79.13 [ -13.88, 172.15]
54.72 [ -89.22, 198.66]
82.05 [ -52.21, 216.32]
-51.04 [-151.79, 49.72]
64.10 [ -3.32, 131.52]
53.04 [ -56.83, 162.91]
-48.22 [-124.06, 27.61]
-18.14 [-169.96, 133.67]
27.04 [ -34.42, 88.50]
66.77 [ 18.11, 115.42]
27.16 [ -77.52, 131.84]
61.77 [ 2.11, 121.43]
-34.82 [-117.59, 47.94]
-39.60 [-101.93, 22.73]
-47.30 [-171.56, 76.96]
20.68 [ -82.89, 124.25]
-26.65 [-105.53, 52.23]
67.90 [ 14.00, 121.80]
64.58 [ -21.11, 150.27]
35.95 [ -78.40, 150.30]
54.74 [ -76.91, 186.39]
2.22 [ -98.95, 103.40]
19.59 [-154.83, 194.00]
-1.59 [-129.19, 126.00]
26.48 [ -62.17, 115.13]
14.11 [ -79.70, 107.91]
18.54 [ -66.59, 103.66]
32.08 [ -22.89, 87.06]
11.70 [ -48.39, 71.78]
8.80 [ -58.96, 76.56]
-19.57 [-156.43, 117.29]
36.17 [ -72.75, 145.09]
50.28 [ -28.65, 129.21]
-25.27 [-120.56, 70.02]
-29.54 [-171.64, 112.56]
70.93 [ -46.90, 188.75]
34.67 [ -32.69, 102.03]
27.97 [ 15.99, 39.96]
28.94 [ 12.60, 45.28]
28.32 [ 18.40, 38.25]
14.63 [ -43.13, 72.38]
-40.09 [-183.77, 103.59]
-150.00 [-278.57, -21.43]
-15.95 [-143.90, 112.00]
34.93 [ -69.46, 139.33]
3.13 [ -60.62, 66.88]
17.92 [ -79.01, 114.84]
-1.73 [ -35.08, 31.61]
Cohort
Beta [Lower 95% CI,Upper 95% CI]
-300
-200
-100
0
100
200
300
mm^2/effect allele

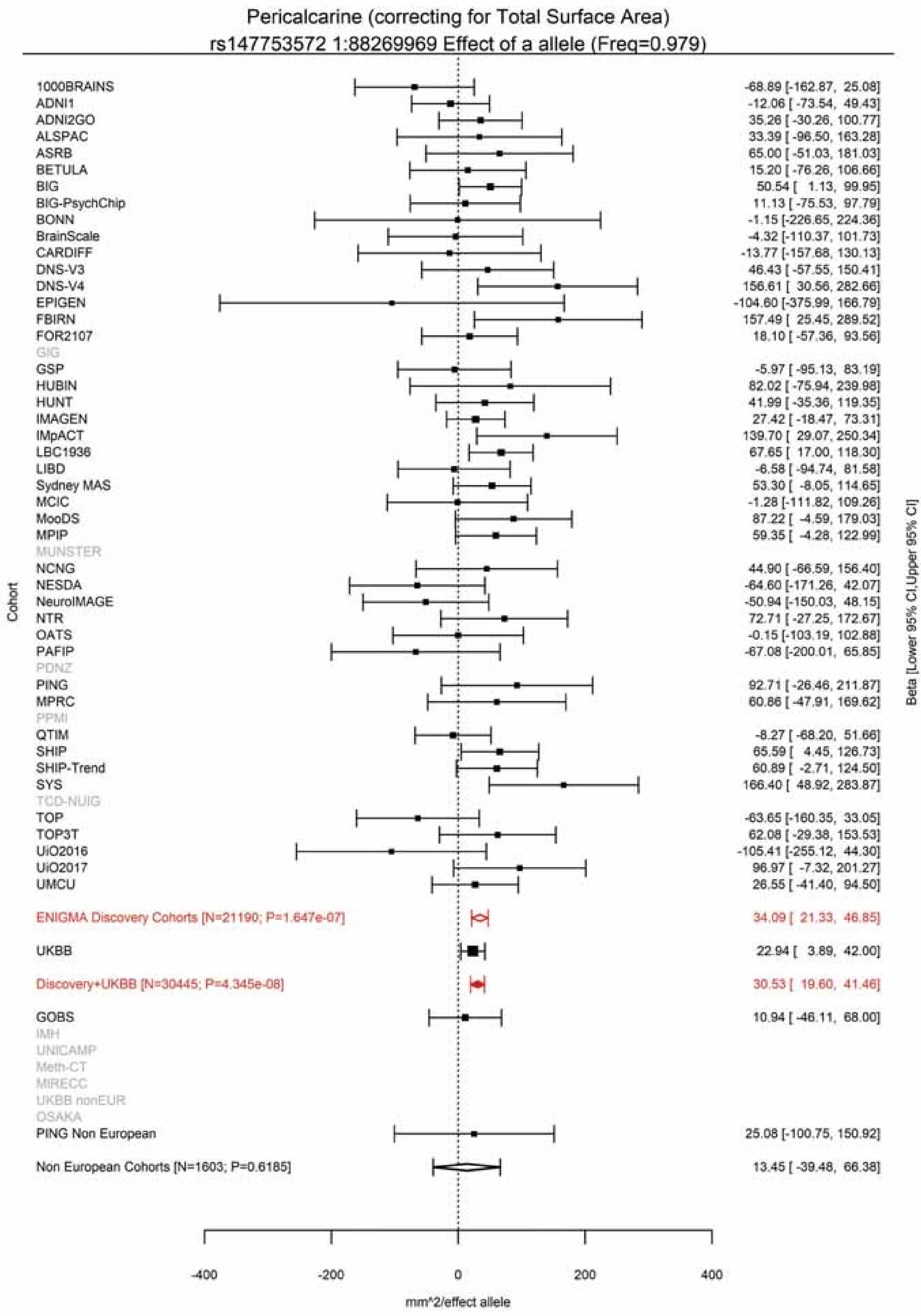

Pericalcarine (correcting for Total Surface Area)
rs147753572 1:88269969 Effect of a allele (Freq=0.979)
1000BRAINS -68.89 [-162.87, 25.08]
ADNI1 -12.06 [-73.54, 49.43]
ADNI2GO 35.26 [-30.26, 100.77]
ALSPAC 33.39 [-96.50, 163.28]
ASRB 65.00 [-51.03, 181.03]
BETULA 15.20 [-76.26, 106.66]
BIG 50.54 [1.13, 99.95]
BIG-PsychChip 11.13 [-75.53, 97.79]
BONN -1.15 [-226.65, 224.36]
BrainScale -4.32 [-110.37, 101.73]
CARDIFF -13.77 [-157.68, 130.13]
DNS-V3 46.43 [-57.55, 150.41]
DNS-V4 156.61 [30.56, 282.66]
EPIGEN -104.60 [-375.99, 166.79]
FBIRN 157.49 [25.45, 289.52]
FOR2107 18.10 [-57.36, 93.56]
GIG
GSP -5.97 [-95.13, 83.19]
HUBIN 82.02 [-75.94, 239.98]
HUNT 41.99 [-35.36, 119.35]
IMAGEN 27.42 [-18.47, 73.31]
IMpACT 139.70 [29.07, 250.34]
LBC1936 67.65 [17.00, 118.30]
LIBD -6.58 [-94.74, 81.58]
Sydney MAS 53.30 [-8.05, 114.65]
MCIC -1.28 [-111.82, 109.26]
MooDS 87.22 [-4.59, 179.03]
MPIP 59.35 [-4.28, 122.99]
MUNSTER
NCNG 44.90 [-66.59, 156.40]
NESDA -64.60 [-171.26, 42.07]
NeuroIMAGE -50.94 [-150.03, 48.15]
NTR 72.71 [-27.25, 172.67]
OATS -0.15 [-103.19, 102.88]
PAFIP -67.08 [-200.01, 65.85]
PDNZ
PING 92.71 [-26.46, 211.87]
MPRC 60.86 [-47.91, 169.62]
PPMI
QTIM -8.27 [-68.20, 51.66]
SHIP 65.59 [4.45, 126.73]
SHIP-Trend 60.89 [-2.71, 124.50]
SYS 166.40 [48.92, 283.87]
TCD-NUIG
TOP -63.65 [-160.35, 33.05]
TOP3T 62.08 [-29.38, 153.53]
UiO2016 -105.41 [-255.12, 44.30]
UiO2017 96.97 [-7.32, 201.27]
UMCU 26.55 [-41.40, 94.50]
ENIGMA Discovery Cohorts [N=21190; P=1.647e-07] 34.09 [21.33, 46.85]
UKBB 22.94 [3.89, 42.00]
Discovery+UKBB [N=30445; P=4.345e-08] 30.53 [19.60, 41.46]
GOBS 10.94 [-46.11, 68.00]
IMH
UNICAMP
Meth-CT
MIRECC
UKBB nonEUR
OSAKA
PING Non European 25.08 [-100.75, 150.92]
Non European Cohorts [N=1603; P=0.6185] 13.45 [-39.48, 66.38]
Cohort
Beta [Lower 95% CI,Upper 95% CI]
-400
-200
0
200
400
mm^2/effect allele

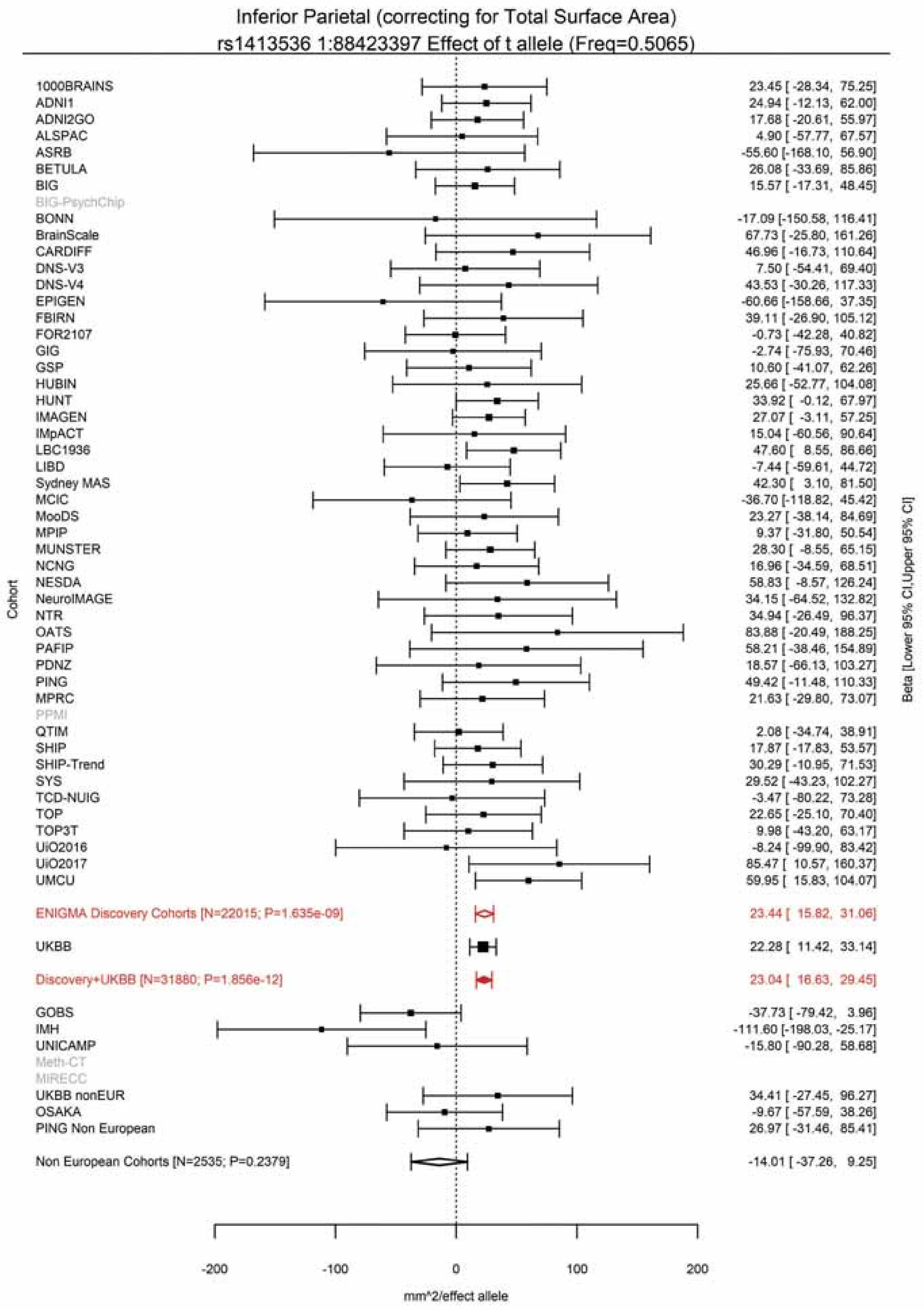

Inferior Parietal (correcting for Total Surface Area)
rs1413536 1:88423397 Effect of t allele (Freq=0.5065)
Cohort
1000BRAINS 23.45 [ -28.34, 75.25]
ADNI1 24.94 [ -12.13, 62.00]
ADNI2GO 17.68 [ -20.61, 55.97]
ALSPAC 4.90 [ -57.77, 67.57]
ASRB -55.60 [-168.10, 56.90]
BETULA 26.08 [ -33.69, 85.86]
BIG 15.57 [ -17.31, 48.45]
BIG-PsychChip
BONN -17.09 [-150.58, 116.41]
BrainScale 67.73 [ -25.80, 161.26]
CARDIFF 46.96 [ -16.73, 110.64]
DNS-V3 7.50 [ -54.41, 69.40]
DNS-V4 43.53 [ -30.26, 117.33]
EPIGEN -60.66 [-158.66, 37.35]
FBIRN 39.11 [ -26.90, 105.12]
FOR2107 -0.73 [ -42.28, 40.82]
GIG -2.74 [ -75.93, 70.46]
GSP 10.60 [ -41.07, 62.26]
HUBIN 25.66 [ -52.77, 104.08]
HUNT 33.92 [ -0.12, 67.97]
IMAGEN 27.07 [ -3.11, 57.25]
IMpACT 15.04 [ -60.56, 90.64]
LBC1936 47.60 [ 8.55, 86.66]
LIBD -7.44 [ -59.61, 44.72]
Sydney MAS 42.30 [ 3.10, 81.50]
MCIC -36.70 [-118.82, 45.42]
MooDS 23.27 [ -38.14, 84.69]
MPIP 9.37 [ -31.80, 50.54]
MUNSTER 28.30 [ -8.55, 65.15]
NCNG 16.96 [ -34.59, 68.51]
NESDA 58.83 [ -8.57, 126.24]
NeuroIMAGE 34.15 [ -64.52, 132.82]
NTR 34.94 [ -26.49, 96.37]
OATS 83.88 [ -20.49, 188.25]
PAFIP 58.21 [ -38.46, 154.89]
PDNZ 18.57 [ -66.13, 103.27]
PING 49.42 [ -11.48, 110.33]
MPRC 21.63 [ -29.80, 73.07]
PPMI
QTIM 2.08 [ -34.74, 38.91]
SHIP 17.87 [ -17.83, 53.57]
SHIP-Trend 30.29 [ -10.95, 71.53]
SYS 29.52 [ -43.23, 102.27]
TCD-NUIG -3.47 [ -80.22, 73.28]
TOP 22.65 [ -25.10, 70.40]
TOP3T 9.98 [ -43.20, 63.17]
UiO2016 -8.24 [ -99.90, 83.42]
UiO2017 85.47 [ 10.57, 160.37]
UMCU 59.95 [ 15.83, 104.07]
ENIGMA Discovery Cohorts [N=22015; P=1.635e-09] 23.44 [ 15.82, 31.06]
UKBB 22.28 [ 11.42, 33.14]
Discovery+UKBB [N=31880; P=1.856e-12] 23.04 [ 16.63, 29.45]
GOBS -37.73 [ -79.42, 3.96]
IMH -111.60 [-198.03, -25.17]
UNICAMP -15.80 [ -90.28, 58.68]
Meth-CT
MIRECC
UKBB nonEUR 34.41 [ -27.45, 96.27]
OSAKA -9.67 [ -57.59, 38.26]
PING Non European 26.97 [ -31.46, 85.41]
Non European Cohorts [N=2535; P=0.2379] -14.01 [ -37.26, 9.25]
-200
-100
0
100
200
mm^2/effect allele
Beta [Lower 95% CI,Upper 95% CI]

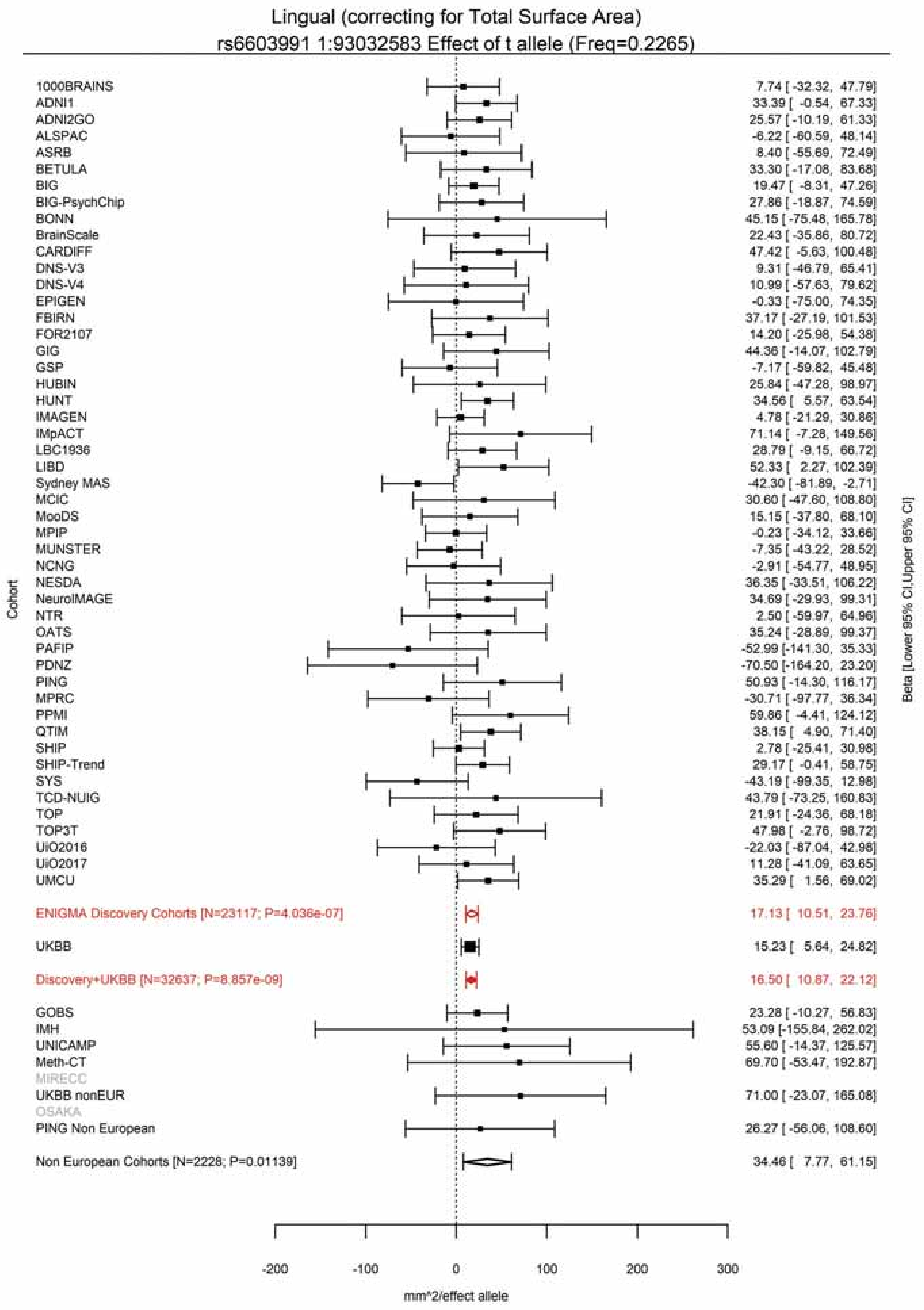
Lingual (correcting for Total Surface Area)
rs6603991 1:93032583 Effect of t allele (Freq=0.2265)
Cohort
1000BRAINS 7.74 [ -32.32, 47.79]
ADNI1 33.39 [ -0.54, 67.33]
ADNI2GO 25.57 [ -10.19, 61.33]
ALSPAC -6.22 [ -60.59, 48.14]
ASRB 8.40 [ -55.69, 72.49]
BETULA 33.30 [ -17.08, 83.68]
BIG 19.47 [ -8.31, 47.26]
BIG-PsychChip 27.86 [ -18.87, 74.59]
BONN 45.15 [ -75.48, 165.78]
BrainScale 22.43 [ -35.86, 80.72]
CARDIFF 47.42 [ -5.63, 100.48]
DNS-V3 9.31 [ -46.79, 65.41]
DNS-V4 10.99 [ -57.63, 79.62]
EPIGEN -0.33 [ -75.00, 74.35]
FBIRN 37.17 [ -27.19, 101.53]
FOR2107 14.20 [ -25.98, 54.38]
GIG 44.36 [ -14.07, 102.79]
GSP -7.17 [ -59.82, 45.48]
HUBIN 25.84 [ -47.28, 98.97]
HUNT 34.56 [ 5.57, 63.54]
IMAGEN 4.78 [ -21.29, 30.86]
IMpACT 71.14 [ -7.28, 149.56]
LBC1936 28.79 [ -9.15, 66.72]
LIBD 52.33 [ 2.27, 102.39]
Sydney MAS -42.30 [ -81.89, -2.71]
MCIC 30.60 [ -47.60, 108.80]
MooDS 15.15 [ -37.80, 68.10]
MPIP -0.23 [ -34.12, 33.66]
MUNSTER -7.35 [ -43.22, 28.52]
NCNG -2.91 [ -54.77, 48.95]
NESDA 36.35 [ -33.51, 106.22]
NeuroIMAGE 34.69 [ -29.93, 99.31]
NTR 2.50 [ -59.97, 64.96]
OATS 35.24 [ -28.89, 99.37]
PAFIP -52.99 [-141.30, 35.33]
PDNZ -70.50 [-164.20, 23.20]
PING 50.93 [ -14.30, 116.17]
MPRC -30.71 [ -97.77, 36.34]
PPMI 59.86 [ -4.41, 124.12]
QTIM 38.15 [ 4.90, 71.40]
SHIP 2.78 [ -25.41, 30.98]
SHIP-Trend 29.17 [ -0.41, 58.75]
SYS -43.19 [ -99.35, 12.98]
TCD-NUIG 43.79 [ -73.25, 160.83]
TOP 21.91 [ -24.36, 68.18]
TOP3T 47.98 [ -2.76, 98.72]
UiO2016 -22.03 [ -87.04, 42.98]
UiO2017 11.28 [ -41.09, 63.65]
UMCU 35.29 [ 1.56, 69.02]
ENIGMA Discovery Cohorts [N=23117; P=4.036e-07] 17.13 [ 10.51, 23.76]
UKBB 15.23 [ 5.64, 24.82]
Discovery+UKBB [N=32637; P=8.857e-09] 16.50 [ 10.87, 22.12]
GOBS 23.28 [ -10.27, 56.83]
IMH 53.09 [-155.84, 262.02]
UNICAMP 55.60 [ -14.37, 125.57]
Meth-CT 69.70 [ -53.47, 192.87]
MIRECC
UKBB nonEUR 71.00 [ -23.07, 165.08]
OSAKA
PING Non European 26.27 [ -56.06, 108.60]
Non European Cohorts [N=2228; P=0.01139] 34.46 [ 7.77, 61.15]
-200
-100
0
100
200
300
mm^2/effect allele
Beta [Lower 95% CI,Upper 95% CI]

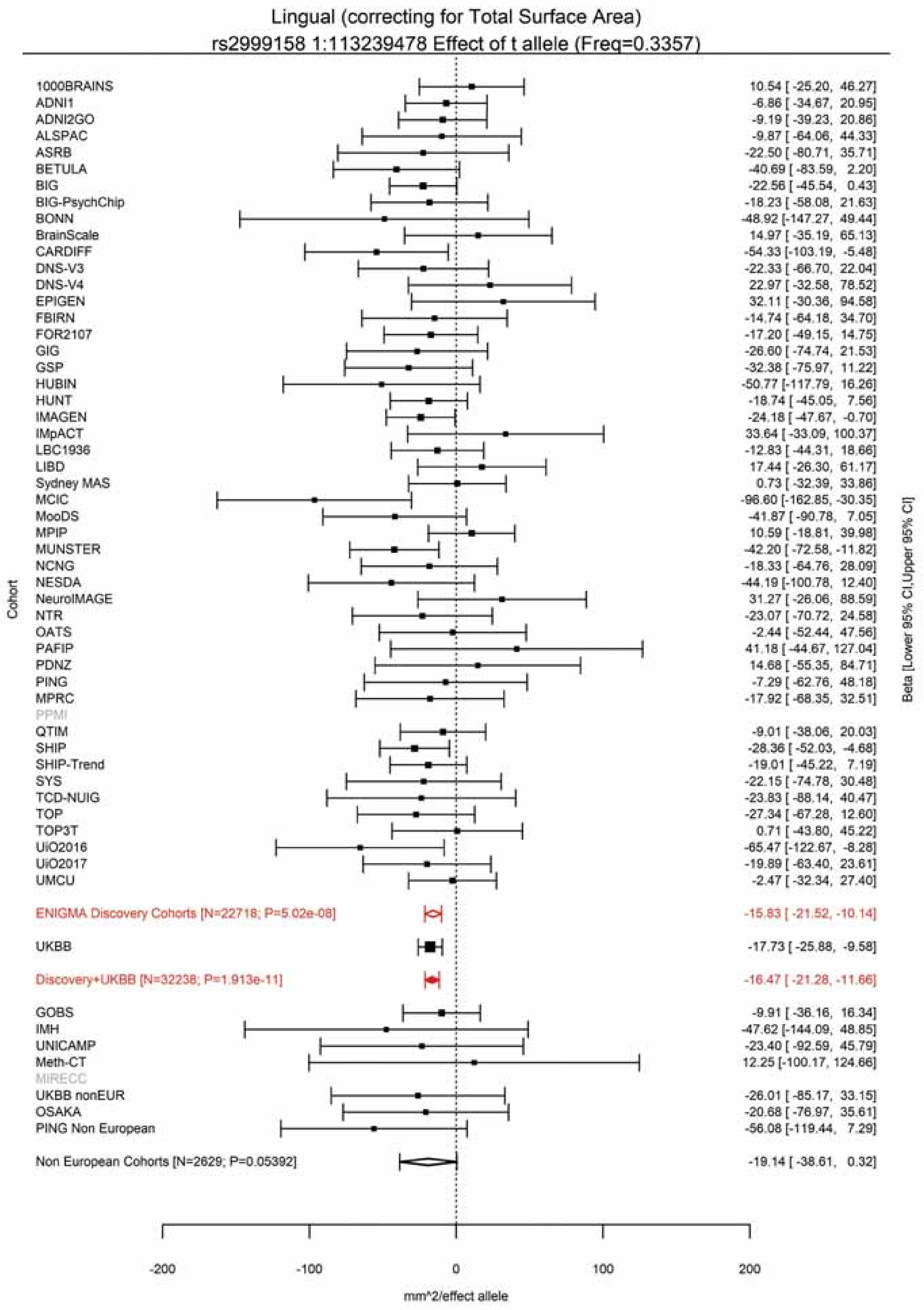
Lingual (correcting for Total Surface Area)
rs2999158 1:113239478 Effect of t allele (Freq=0.3357)
Cohort
1000BRAINS 10.54 [ -25.20, 46.27]
ADNI1 -6.86 [ -34.67, 20.95]
ADNI2GO -9.19 [ -39.23, 20.86]
ALSPAC -9.87 [ -64.06, 44.33]
ASRB -22.50 [ -80.71, 35.71]
BETULA -40.69 [ -83.59, 2.20]
BIG -22.56 [ -45.54, 0.43]
BIG-PsychChip -18.23 [ -58.08, 21.63]
BONN -48.92 [-147.27, 49.44]
BrainScale 14.97 [ -35.19, 65.13]
CARDIFF -54.33 [-103.19, -5.48]
DNS-V3 -22.33 [ -66.70, 22.04]
DNS-V4 22.97 [ -32.58, 78.52]
EPIGEN 32.11 [ -30.36, 94.58]
FBIRN -14.74 [ -64.18, 34.70]
FOR2107 -17.20 [ -49.15, 14.75]
GIG -26.60 [ -74.74, 21.53]
GSP -32.38 [ -75.97, 11.22]
HUBIN -50.77 [-117.79, 16.26]
HUNT -18.74 [ -45.05, 7.56]
IMAGEN -24.18 [ -47.67, -0.70]
IMpACT 33.64 [ -33.09, 100.37]
LBC1936 -12.83 [ -44.31, 18.66]
LIBD 17.44 [ -26.30, 61.17]
Sydney MAS 0.73 [ -32.39, 33.86]
MCIC -96.60 [-162.85, -30.35]
MooDS -41.87 [ -90.78, 7.05]
MPIP 10.59 [ -18.81, 39.98]
MUNSTER -42.20 [ -72.58, -11.82]
NCNG -18.33 [ -64.76, 28.09]
NESDA -44.19 [-100.78, 12.40]
NeuroIMAGE 31.27 [ -26.06, 88.59]
NTR -23.07 [ -70.72, 24.58]
OATS -2.44 [ -52.44, 47.56]
PAFIP 41.18 [ -44.67, 127.04]
PDNZ 14.68 [ -55.35, 84.71]
PING -7.29 [ -62.76, 48.18]
MPRC -17.92 [ -68.35, 32.51]
PPMI
QTIM -9.01 [ -38.06, 20.03]
SHIP -28.36 [ -52.03, -4.68]
SHIP-Trend -19.01 [ -45.22, 7.19]
SYS -22.15 [ -74.78, 30.48]
TCD-NUIG -23.83 [ -88.14, 40.47]
TOP -27.34 [ -67.28, 12.60]
TOP3T 0.71 [ -43.80, 45.22]
UiO2016 -65.47 [-122.67, -8.28]
UiO2017 -19.89 [ -63.40, 23.61]
UMCU -2.47 [ -32.34, 27.40]
ENIGMA Discovery Cohorts [N=22718; P=5.02e-08] -15.83 [ -21.52, -10.14]
UKBB -17.73 [ -25.88, -9.58]
Discovery+UKBB [N=32238; P=1.913e-11] -16.47 [ -21.28, -11.66]
GOBS -9.91 [ -36.16, 16.34]
IMH -47.62 [-144.09, 48.85]
UNICAMP -23.40 [ -92.59, 45.79]
Meth-CT 12.25 [-100.17, 124.66]
MIRECC
UKBB nonEUR -26.01 [ -85.17, 33.15]
OSAKA -20.68 [ -76.97, 35.61]
PING Non European -56.08 [-119.44, 7.29]
Non European Cohorts [N=2629; P=0.05392] -19.14 [ -38.61, 0.32]
Beta [Lower 95% CI,Upper 95% CI]
-200
-100
0
100
200
mm^2/effect allele

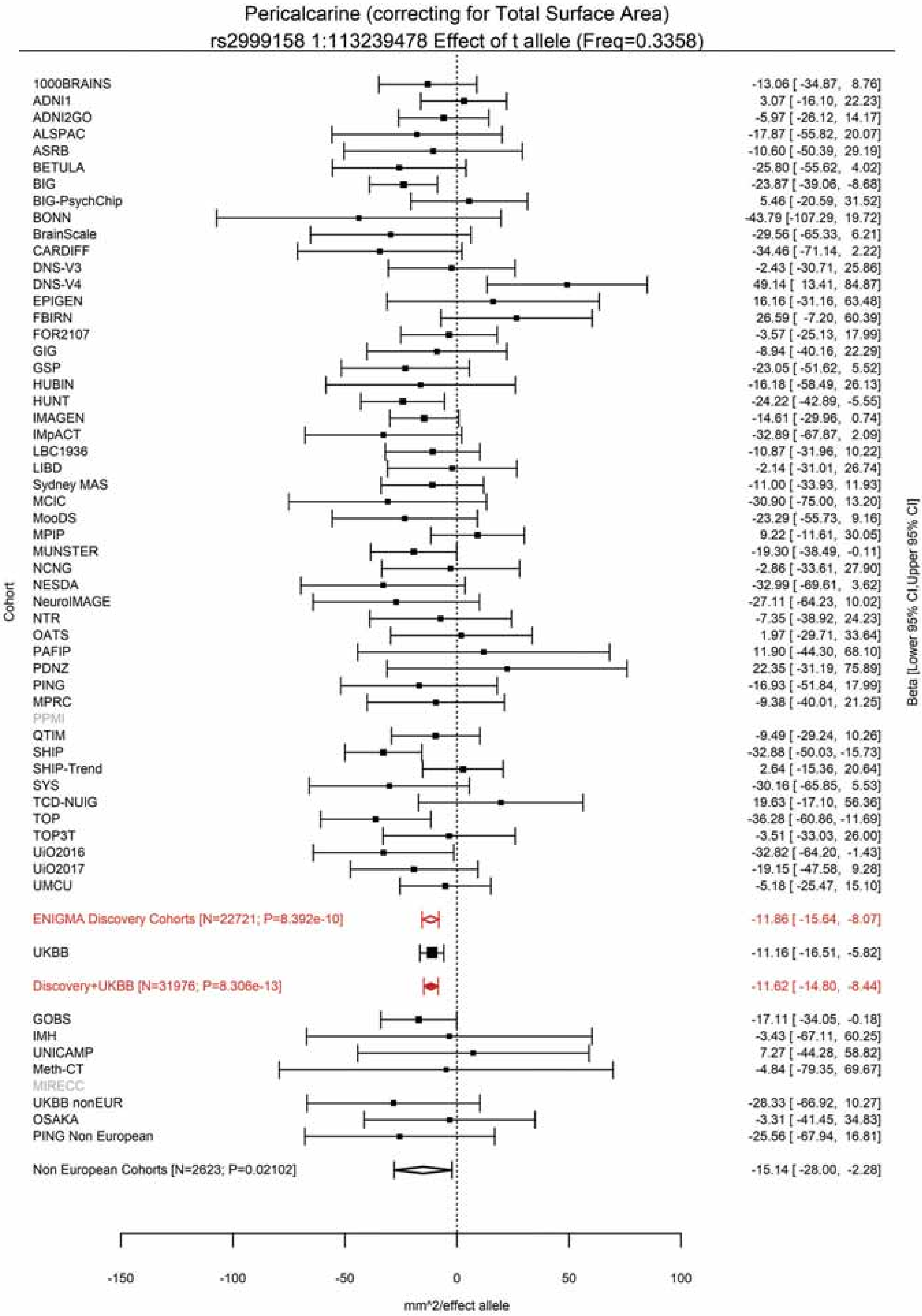
Pericalcarine (correcting for Total Surface Area)
rs2999158 1:113239478 Effect of t allele (Freq=0.3358)
Cohort
1000BRAINS -13.06 [ -34.87, 8.76]
ADNI1 3.07 [ -16.10, 22.23]
ADNI2GO -5.97 [ -26.12, 14.17]
ALSPAC -17.87 [ -55.82, 20.07]
ASRB -10.60 [ -50.39, 29.19]
BETULA -25.80 [ -55.62, 4.02]
BIG -23.87 [ -39.06, -8.68]
BIG-PsychChip 5.46 [ -20.59, 31.52]
BONN -43.79 [-107.29, 19.72]
BrainScale -29.56 [ -65.33, 6.21]
CARDIFF -34.46 [ -71.14, 2.22]
DNS-V3 -2.43 [ -30.71, 25.86]
DNS-V4 49.14 [ 13.41, 84.87]
EPIGEN 16.16 [ -31.16, 63.48]
FBIRN 26.59 [ -7.20, 60.39]
FOR2107 -3.57 [ -25.13, 17.99]
GIG -8.94 [ -40.16, 22.29]
GSP -23.05 [ -51.62, 5.52]
HUBIN -16.18 [ -58.49, 26.13]
HUNT -24.22 [ -42.89, -5.55]
IMAGEN -14.61 [ -29.96, 0.74]
IMpACT -32.89 [ -67.87, 2.09]
LBC1936 -10.87 [ -31.96, 10.22]
LIBD -2.14 [ -31.01, 26.74]
Sydney MAS -11.00 [ -33.93, 11.93]
MCIC -30.90 [ -75.00, 13.20]
MooDS -23.29 [ -55.73, 9.16]
MPIP 9.22 [ -11.61, 30.05]
MUNSTER -19.30 [ -38.49, -0.11]
NCNG -2.86 [ -33.61, 27.90]
NESDA -32.99 [ -69.61, 3.62]
NeuroIMAGE -27.11 [ -64.23, 10.02]
NTR -7.35 [ -38.92, 24.23]
OATS 1.97 [ -29.71, 33.64]
PAFIP 11.90 [ -44.30, 68.10]
PDNZ 22.35 [ -31.19, 75.89]
PING -16.93 [ -51.84, 17.99]
MPRC -9.38 [ -40.01, 21.25]
PPMI
QTIM -9.49 [ -29.24, 10.26]
SHIP -32.88 [ -50.03, -15.73]
SHIP-Trend 2.64 [ -15.36, 20.64]
SYS -30.16 [ -65.85, 5.53]
TCD-NUIG 19.63 [ -17.10, 56.36]
TOP -36.28 [ -60.86, -11.69]
TOP3T -3.51 [ -33.03, 26.00]
UiO2016 -32.82 [ -64.20, -1.43]
UiO2017 -19.15 [ -47.58, 9.28]
UMCU -5.18 [ -25.47, 15.10]
ENIGMA Discovery Cohorts [N=22721; P=8.392e-10] -11.86 [ -15.64, -8.07]
UKBB -11.16 [ -16.51, -5.82]
Discovery+UKBB [N=31976; P=8.306e-13] -11.62 [ -14.80, -8.44]
GOBS -17.11 [ -34.05, -0.18]
IMH -3.43 [ -67.11, 60.25]
UNICAMP 7.27 [ -44.28, 58.82]
Meth-CT -4.84 [ -79.35, 69.67]
MIRECC
UKBB nonEUR -28.33 [ -66.92, 10.27]
OSAKA -3.31 [ -41.45, 34.83]
PING Non European -25.56 [ -67.94, 16.81]
Non European Cohorts [N=2623; P=0.02102] -15.14 [ -28.00, -2.28]
Beta [Lower 95% CI,Upper 95% CI]
-150
-100
-50
0
50
100
mm^2/effect allele

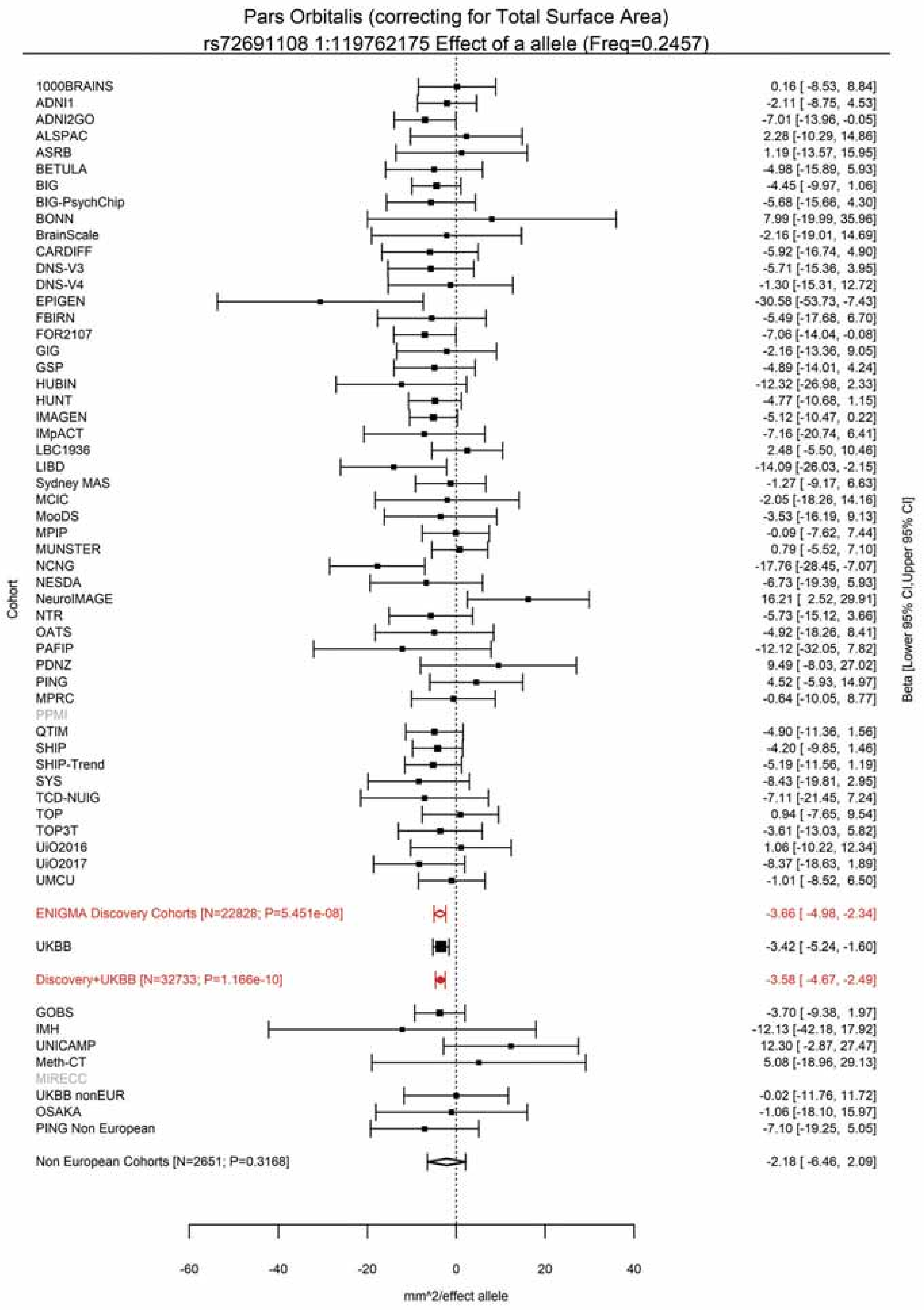

Pars Orbitalis (correcting for Total Surface Area)
rs72691108 1:119762175 Effect of a allele (Freq=0.2457)
Cohort
1000BRAINS 0.16 [ -8.53, 8.84]
ADNI1 -2.11 [ -8.75, 4.53]
ADNI2GO -7.01 [-13.96, -0.05]
ALSPAC 2.28 [-10.29, 14.86]
ASRB 1.19 [-13.57, 15.95]
BETULA -4.98 [-15.89, 5.93]
BIG -4.45 [ -9.97, 1.06]
BIG-PsychChip -5.68 [-15.66, 4.30]
BONN 7.99 [-19.99, 35.96]
BrainScale -2.16 [-19.01, 14.69]
CARDIFF -5.92 [-16.74, 4.90]
DNS-V3 -5.71 [-15.36, 3.95]
DNS-V4 -1.30 [-15.31, 12.72]
EPIGEN -30.58 [-53.73, -7.43]
FBIRN -5.49 [-17.68, 6.70]
FOR2107 -7.06 [-14.04, -0.08]
GIG -2.16 [-13.36, 9.05]
GSP -4.89 [-14.01, 4.24]
HUBIN -12.32 [-26.98, 2.33]
HUNT -4.77 [-10.68, 1.15]
IMAGEN -5.12 [-10.47, 0.22]
IMpACT -7.16 [-20.74, 6.41]
LBC1936 2.48 [ -5.50, 10.46]
LIBD -14.09 [-26.03, -2.15]
Sydney MAS -1.27 [ -9.17, 6.63]
MCIC -2.05 [-18.26, 14.16]
MooDS -3.53 [-16.19, 9.13]
MPIP -0.09 [ -7.62, 7.44]
MUNSTER 0.79 [ -5.52, 7.10]
NCNG -17.76 [-28.45, -7.07]
NESDA -6.73 [-19.39, 5.93]
NeuroIMAGE 16.21 [ 2.52, 29.91]
NTR -5.73 [-15.12, 3.66]
OATS -4.92 [-18.26, 8.41]
PAFIP -12.12 [-32.05, 7.82]
PDNZ 9.49 [ -8.03, 27.02]
PING 4.52 [ -5.93, 14.97]
MPRC -0.64 [-10.05, 8.77]
PPMI
QTIM -4.90 [-11.36, 1.56]
SHIP -4.20 [ -9.85, 1.46]
SHIP-Trend -5.19 [-11.56, 1.19]
SYS -8.43 [-19.81, 2.95]
TCD-NUIG -7.11 [-21.45, 7.24]
TOP 0.94 [ -7.65, 9.54]
TOP3T -3.61 [-13.03, 5.82]
UiO2016 1.06 [-10.22, 12.34]
UiO2017 -8.37 [-18.63, 1.89]
UMCU -1.01 [ -8.52, 6.50]
ENIGMA Discovery Cohorts [N=22828; P=5.451e-08] -3.66 [ -4.98, -2.34]
UKBB -3.42 [ -5.24, -1.60]
Discovery+UKBB [N=32733; P=1.166e-10] -3.58 [ -4.67, -2.49]
GOBS -3.70 [ -9.38, 1.97]
IMH -12.13 [-42.18, 17.92]
UNICAMP 12.30 [ -2.87, 27.47]
Meth-CT 5.08 [-18.96, 29.13]
MIRECC
UKBB nonEUR -0.02 [-11.76, 11.72]
OSAKA -1.06 [-18.10, 15.97]
PING Non European -7.10 [-19.25, 5.05]
Non European Cohorts [N=2651; P=0.3168] -2.18 [ -6.46, 2.09]
Beta [Lower 95% CI,Upper 95% CI]
-60 -40 -20 0 20 40
mm^2/effect allele

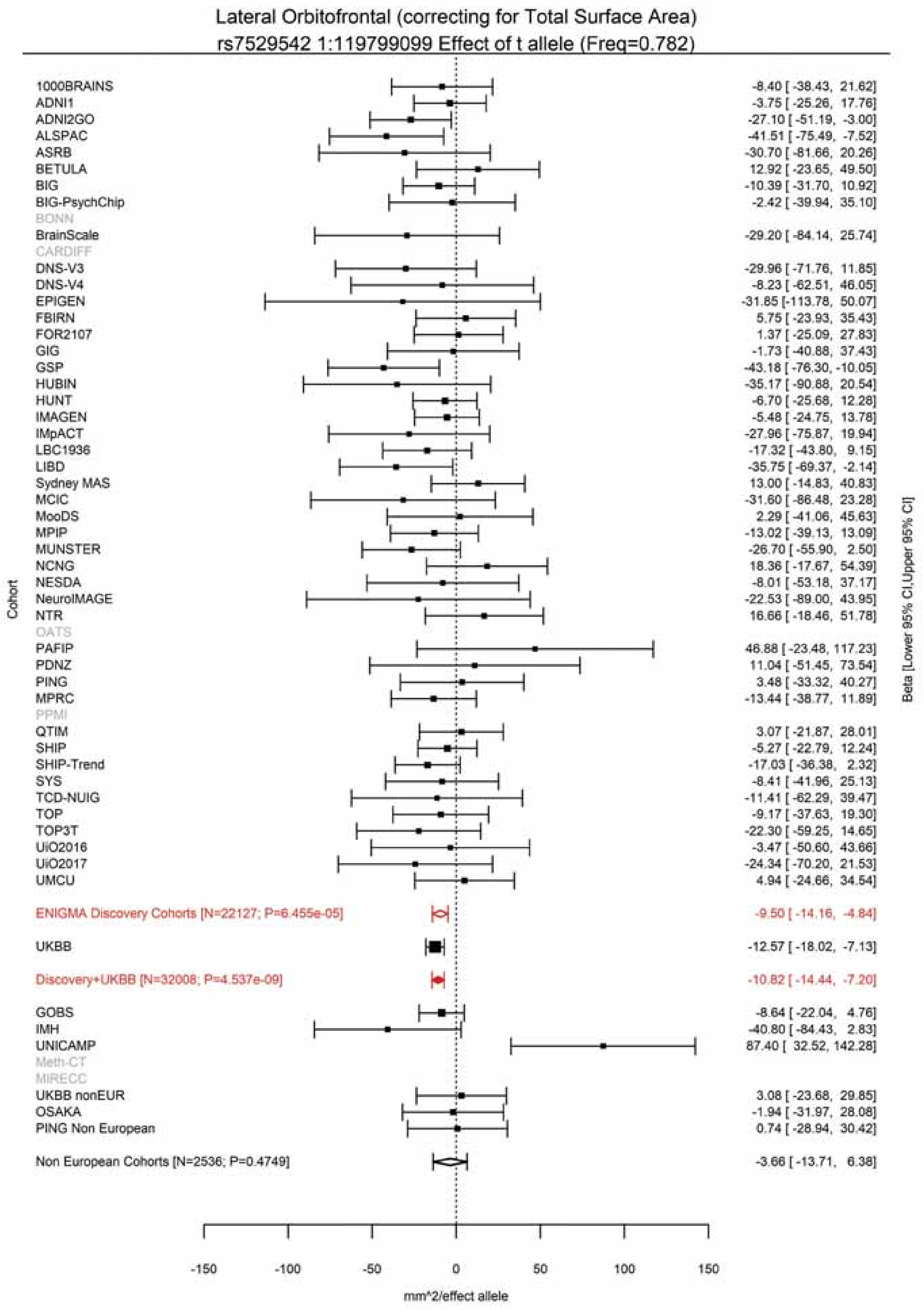
Lateral Orbitofrontal (correcting for Total Surface Area)
rs7529542 1:119799099 Effect of t allele (Freq=0.782)
Cohort
Beta [Lower 95% CI,Upper 95% CI]
1000BRAINS -8.40 [ -38.43, 21.62]
ADNI1 -3.75 [ -25.26, 17.76]
ADNI2GO -27.10 [ -51.19, -3.00]
ALSPAC -41.51 [ -75.49, -7.52]
ASRB -30.70 [ -81.66, 20.26]
BETULA 12.92 [ -23.65, 49.50]
BIG -10.39 [ -31.70, 10.92]
BIG-PsychChip -2.42 [ -39.94, 35.10]
BONN
BrainScale -29.20 [ -84.14, 25.74]
CARDIFF
DNS-V3 -29.96 [ -71.76, 11.85]
DNS-V4 -8.23 [ -62.51, 46.05]
EPIGEN -31.85 [-113.78, 50.07]
FBIRN 5.75 [ -23.93, 35.43]
FOR2107 1.37 [ -25.09, 27.83]
GIG -1.73 [ -40.88, 37.43]
GSP -43.18 [ -76.30, -10.05]
HUBIN -35.17 [ -90.88, 20.54]
HUNT -6.70 [ -25.68, 12.28]
IMAGEN -5.48 [ -24.75, 13.78]
IMpACT -27.96 [ -75.87, 19.94]
LBC1936 -17.32 [ -43.80, 9.15]
LIBD -35.75 [ -69.37, -2.14]
Sydney MAS 13.00 [ -14.83, 40.83]
MCIC -31.60 [ -86.48, 23.28]
MooDS 2.29 [ -41.06, 45.63]
MPIP -13.02 [ -39.13, 13.09]
MUNSTER -26.70 [ -55.90, 2.50]
NCNG 18.36 [ -17.67, 54.39]
NESDA -8.01 [ -53.18, 37.17]
NeuroIMAGE -22.53 [ -89.00, 43.95]
NTR 16.66 [ -18.46, 51.78]
OATS
PAFIP 46.88 [ -23.48, 117.23]
PDNZ 11.04 [ -51.45, 73.54]
PING 3.48 [ -33.32, 40.27]
MPRC -13.44 [ -38.77, 11.89]
PPMI
QTIM 3.07 [ -21.87, 28.01]
SHIP -5.27 [ -22.79, 12.24]
SHIP-Trend -17.03 [ -36.38, 2.32]
SYS -8.41 [ -41.96, 25.13]
TCD-NUIG -11.41 [ -62.29, 39.47]
TOP -9.17 [ -37.63, 19.30]
TOP3T -22.30 [ -59.25, 14.65]
UiO2016 -3.47 [ -50.60, 43.66]
UiO2017 -24.34 [ -70.20, 21.53]
UMCU 4.94 [ -24.66, 34.54]
ENIGMA Discovery Cohorts [N=22127; P=6.455e-05] -9.50 [ -14.16, -4.84]
UKBB -12.57 [ -18.02, -7.13]
Discovery+UKBB [N=32008; P=4.537e-09] -10.82 [ -14.44, -7.20]
GOBS -8.64 [ -22.04, 4.76]
IMH -40.80 [ -84.43, 2.83]
UNICAMP 87.40 [ 32.52, 142.28]
Meth-CT
MIRECC
UKBB nonEUR 3.08 [ -23.68, 29.85]
OSAKA -1.94 [ -31.97, 28.08]
PING Non European 0.74 [ -28.94, 30.42]
Non European Cohorts [N=2536; P=0.4749] -3.66 [ -13.71, 6.38]
-150 -100 -50 0 50 100 150
mm^2/effect allele

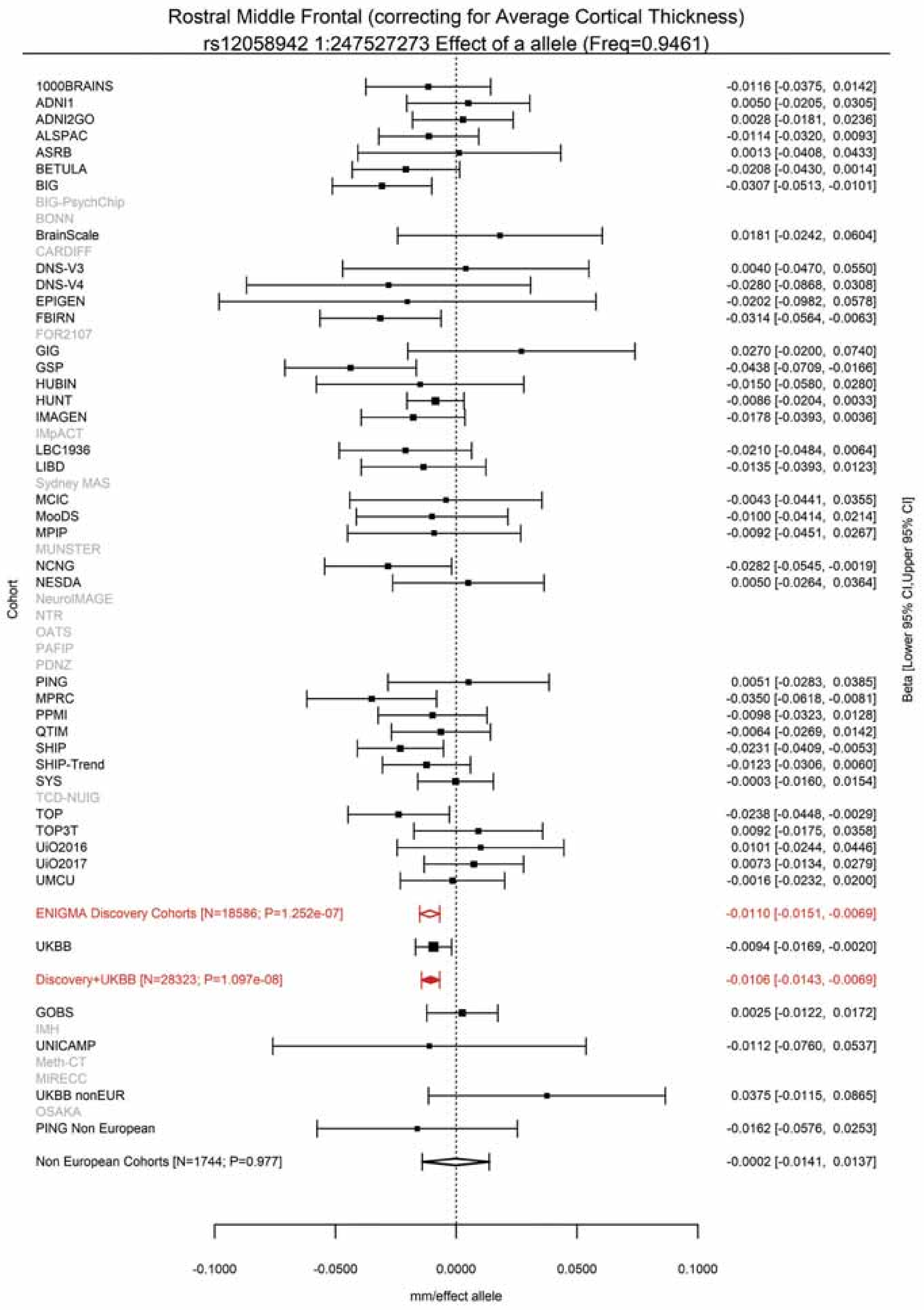

Rostral Middle Frontal (correcting for Average Cortical Thickness)
rs12058942 1:247527273 Effect of a allele (Freq=0.9461)
Cohort
1000BRAINS -0.0116 [-0.0375, 0.0142]
ADNI1 0.0050 [-0.0205, 0.0305]
ADNI2GO 0.0028 [-0.0181, 0.0236]
ALSPAC -0.0114 [-0.0320, 0.0093]
ASRB 0.0013 [-0.0408, 0.0433]
BETULA -0.0208 [-0.0430, 0.0014]
BIG -0.0307 [-0.0513, -0.0101]
BIG-PsychChip
BONN
BrainScale 0.0181 [-0.0242, 0.0604]
CARDIFF
DNS-V3 0.0040 [-0.0470, 0.0550]
DNS-V4 -0.0280 [-0.0868, 0.0308]
EPIGEN -0.0202 [-0.0982, 0.0578]
FBIRN -0.0314 [-0.0564, -0.0063]
FOR2107
GIG 0.0270 [-0.0200, 0.0740]
GSP -0.0438 [-0.0709, -0.0166]
HUBIN -0.0150 [-0.0580, 0.0280]
HUNT -0.0086 [-0.0204, 0.0033]
IMAGEN -0.0178 [-0.0393, 0.0036]
IMpACT
LBC1936 -0.0210 [-0.0484, 0.0064]
LIBD -0.0135 [-0.0393, 0.0123]
Sydney MAS
MCIC -0.0043 [-0.0441, 0.0355]
MooDS -0.0100 [-0.0414, 0.0214]
MPIP -0.0092 [-0.0451, 0.0267]
MUNSTER
NCNG -0.0282 [-0.0545, -0.0019]
NESDA 0.0050 [-0.0264, 0.0364]
NeuroIMAGE
NTR
OATS
PAFIP
PDNZ
PING 0.0051 [-0.0283, 0.0385]
MPRC -0.0350 [-0.0618, -0.0081]
PPMI -0.0098 [-0.0323, 0.0128]
QTIM -0.0064 [-0.0269, 0.0142]
SHIP -0.0231 [-0.0409, -0.0053]
SHIP-Trend -0.0123 [-0.0306, 0.0060]
SYS -0.0003 [-0.0160, 0.0154]
TCD-NUIG
TOP -0.0238 [-0.0448, -0.0029]
TOP3T 0.0092 [-0.0175, 0.0358]
UiO2016 0.0101 [-0.0244, 0.0446]
UiO2017 0.0073 [-0.0134, 0.0279]
UMCU -0.0016 [-0.0232, 0.0200]
ENIGMA Discovery Cohorts [N=18586; P=1.252e-07] -0.0110 [-0.0151, -0.0069]
UKBB -0.0094 [-0.0169, -0.0020]
Discovery+UKBB [N=28323; P=1.097e-08] -0.0106 [-0.0143, -0.0069]
GOBS 0.0025 [-0.0122, 0.0172]
IMH
UNICAMP -0.0112 [-0.0760, 0.0537]
Meth-CT
MIRECC
UKBB nonEUR 0.0375 [-0.0115, 0.0865]
OSAKA
PING Non European -0.0162 [-0.0576, 0.0253]
Non European Cohorts [N=1744; P=0.977] -0.0002 [-0.0141, 0.0137]
-0.1000
-0.0500
0.0000
0.0500
0.1000
mm/effect allele
Beta [Lower 95% CI,Upper 95% CI]

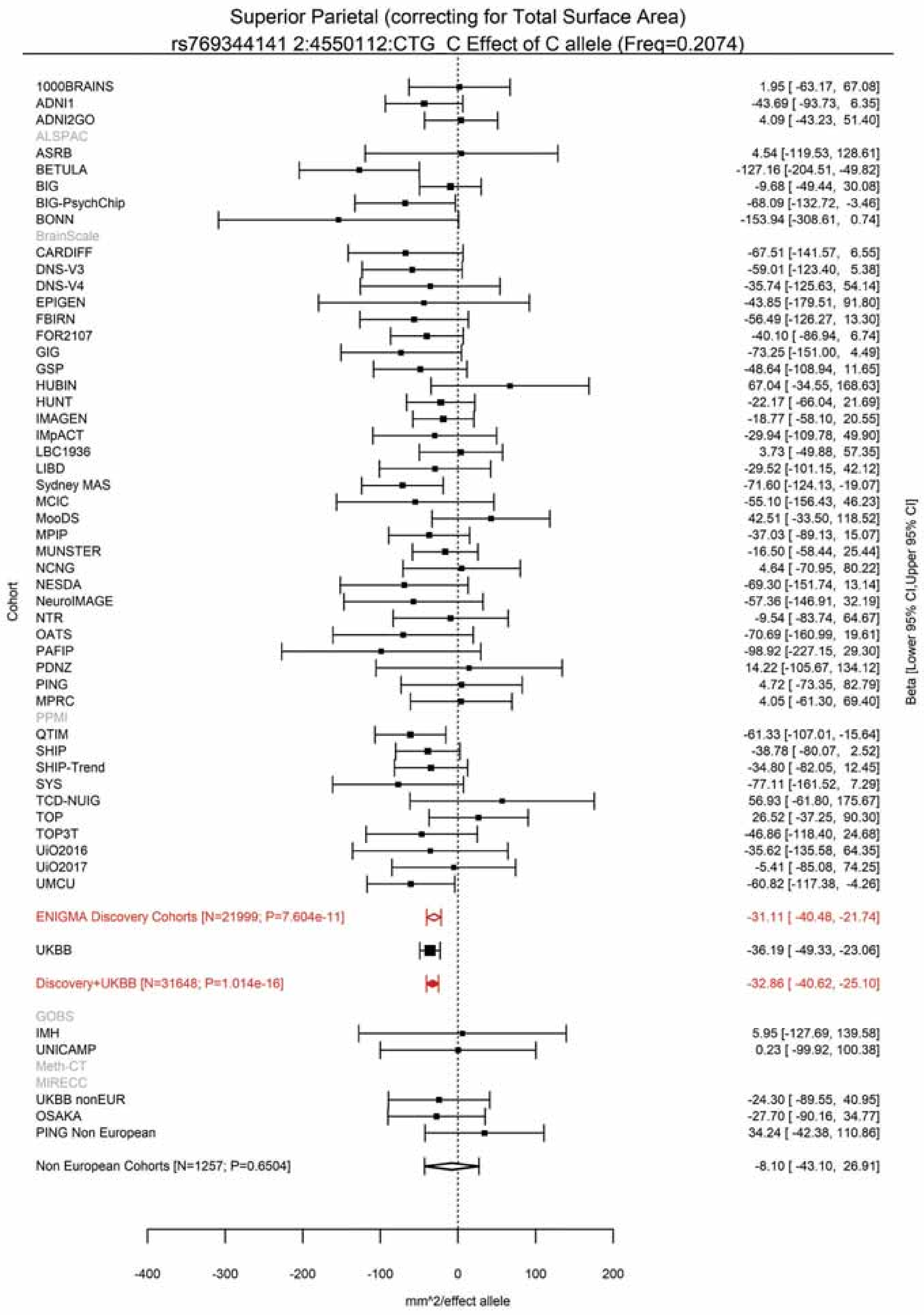
Superior Parietal (correcting for Total Surface Area)
rs769344141 2:4550112:CTG_C Effect of C allele (Freq=0.2074)
Cohort
1000BRAINS 1.95 [ -63.17, 67.08]
ADNI1 -43.69 [ -93.73, 6.35]
ADNI2GO 4.09 [ -43.23, 51.40]
ALSPAC
ASRB 4.54 [-119.53, 128.61]
BETULA -127.16 [-204.51, -49.82]
BIG -9.68 [ -49.44, 30.08]
BIG-PsychChip -68.09 [-132.72, -3.46]
BONN -153.94 [-308.61, 0.74]
BrainScale
CARDIFF -67.51 [-141.57, 6.55]
DNS-V3 -59.01 [-123.40, 5.38]
DNS-V4 -35.74 [-125.63, 54.14]
EPIGEN -43.85 [-179.51, 91.80]
FBIRN -56.49 [-126.27, 13.30]
FOR2107 -40.10 [ -86.94, 6.74]
GIG -73.25 [-151.00, 4.49]
GSP -48.64 [-108.94, 11.65]
HUBIN 67.04 [ -34.55, 168.63]
HUNT -22.17 [ -66.04, 21.69]
IMAGEN -18.77 [ -58.10, 20.55]
IMpACT -29.94 [-109.78, 49.90]
LBC1936 3.73 [ -49.88, 57.35]
LIBD -29.52 [-101.15, 42.12]
Sydney MAS -71.60 [-124.13, -19.07]
MCIC -55.10 [-156.43, 46.23]
MooDS 42.51 [ -33.50, 118.52]
MPIP -37.03 [ -89.13, 15.07]
MUNSTER -16.50 [ -58.44, 25.44]
NCNG 4.64 [ -70.95, 80.22]
NESDA -69.30 [-151.74, 13.14]
NeuroIMAGE -57.36 [-146.91, 32.19]
NTR -9.54 [ -83.74, 64.67]
OATS -70.69 [-160.99, 19.61]
PAFIP -98.92 [-227.15, 29.30]
PDNZ 14.22 [-105.67, 134.12]
PING 4.72 [ -73.35, 82.79]
MPRC 4.05 [ -61.30, 69.40]
PPMI
QTIM -61.33 [-107.01, -15.64]
SHIP -38.78 [ -80.07, 2.52]
SHIP-Trend -34.80 [ -82.05, 12.45]
SYS -77.11 [-161.52, 7.29]
TCD-NUIG 56.93 [ -61.80, 175.67]
TOP 26.52 [ -37.25, 90.30]
TOP3T -46.86 [-118.40, 24.68]
UiO2016 -35.62 [-135.58, 64.35]
UiO2017 -5.41 [ -85.08, 74.25]
UMCU -60.82 [-117.38, -4.26]
ENIGMA Discovery Cohorts [N=21999; P=7.604e-11] -31.11 [ -40.48, -21.74]
UKBB -36.19 [ -49.33, -23.06]
Discovery+UKBB [N=31648; P=1.014e-16] -32.86 [ -40.62, -25.10]
GOBS
IMH 5.95 [-127.69, 139.58]
UNICAMP 0.23 [ -99.92, 100.38]
Meth-CT
MIRECC
UKBB nonEUR -24.30 [ -89.55, 40.95]
OSAKA -27.70 [ -90.16, 34.77]
PING Non European 34.24 [ -42.38, 110.86]
Non European Cohorts [N=1257; P=0.6504] -8.10 [ -43.10, 26.91]
-400
-300
-200
-100
0
100
200
mm^2/effect allele
Beta [Lower 95% CI,Upper 95% CI]

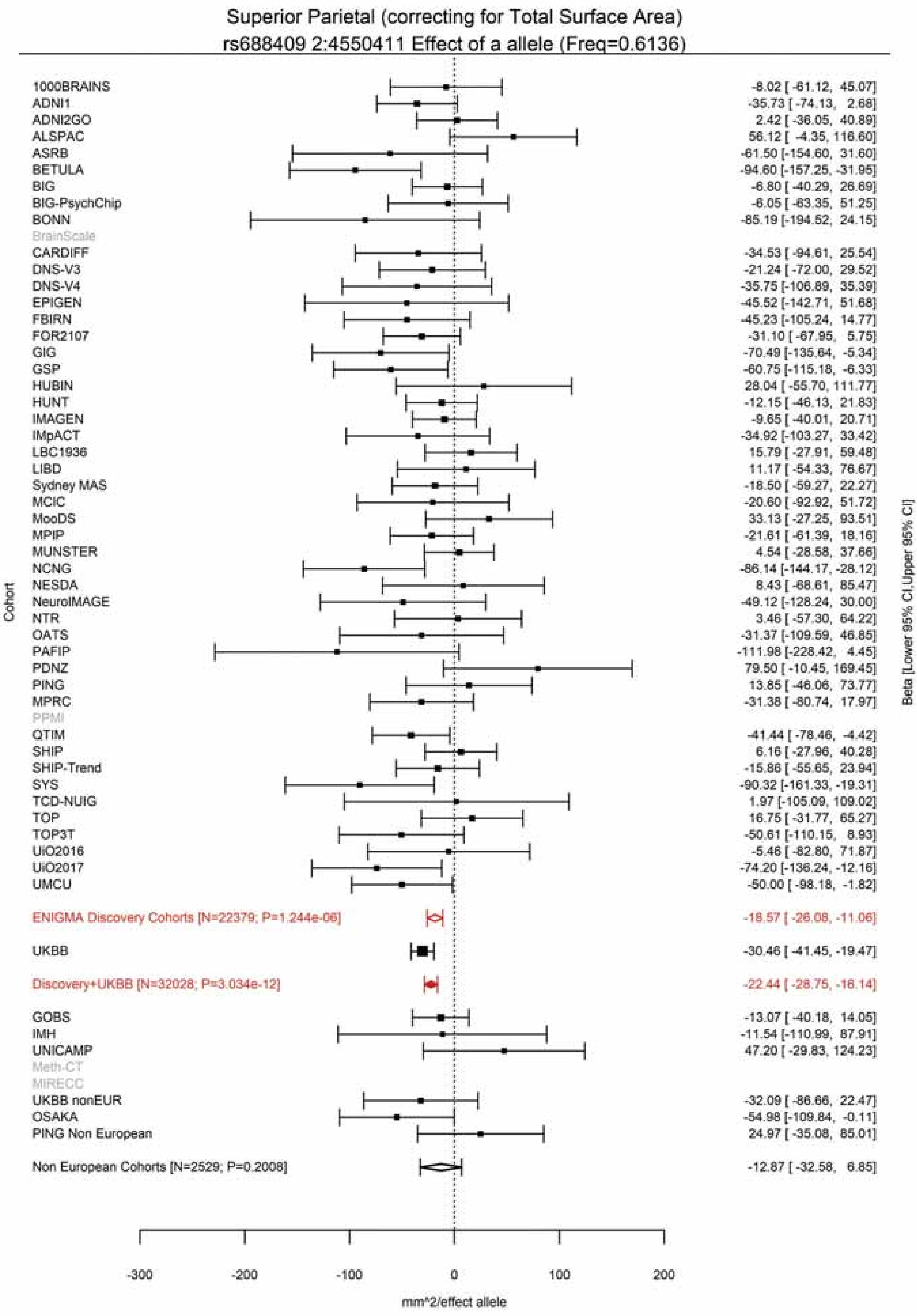
Superior Parietal (correcting for Total Surface Area)
rs688409 2:4550411 Effect of a allele (Freq=0.6136)
Cohort
1000BRAINS -8.02 [ -61.12, 45.07]
ADNI1 -35.73 [ -74.13, 2.68]
ADNI2GO 2.42 [ -36.05, 40.89]
ALSPAC 56.12 [ -4.35, 116.60]
ASRB -61.50 [-154.60, 31.60]
BETULA -94.60 [-157.25, -31.95]
BIG -6.80 [ -40.29, 26.69]
BIG-PsychChip -6.05 [ -63.35, 51.25]
BONN -85.19 [-194.52, 24.15]
BrainScale
CARDIFF -34.53 [ -94.61, 25.54]
DNS-V3 -21.24 [ -72.00, 29.52]
DNS-V4 -35.75 [-106.89, 35.39]
EPIGEN -45.52 [-142.71, 51.68]
FBIRN -45.23 [-105.24, 14.77]
FOR2107 -31.10 [ -67.95, 5.75]
GIG -70.49 [-135.64, -5.34]
GSP -60.75 [-115.18, -6.33]
HUBIN 28.04 [ -55.70, 111.77]
HUNT -12.15 [ -46.13, 21.83]
IMAGEN -9.65 [ -40.01, 20.71]
IMpACT -34.92 [-103.27, 33.42]
LBC1936 15.79 [ -27.91, 59.48]
LIBD 11.17 [ -54.33, 76.67]
Sydney MAS -18.50 [ -59.27, 22.27]
MCIC -20.60 [ -92.92, 51.72]
MooDS 33.13 [ -27.25, 93.51]
MPIP -21.61 [ -61.39, 18.16]
MUNSTER 4.54 [ -28.58, 37.66]
NCNG -86.14 [-144.17, -28.12]
NESDA 8.43 [ -68.61, 85.47]
NeuroIMAGE -49.12 [-128.24, 30.00]
NTR 3.46 [ -57.30, 64.22]
OATS -31.37 [-109.59, 46.85]
PAFIP -111.98 [-228.42, 4.45]
PDNZ 79.50 [ -10.45, 169.45]
PING 13.85 [ -46.06, 73.77]
MPRC -31.38 [ -80.74, 17.97]
PPMI
QTIM -41.44 [ -78.46, -4.42]
SHIP 6.16 [ -27.96, 40.28]
SHIP-Trend -15.86 [ -55.65, 23.94]
SYS -90.32 [-161.33, -19.31]
TCD-NUIG 1.97 [-105.09, 109.02]
TOP 16.75 [ -31.77, 65.27]
TOP3T -50.61 [-110.15, 8.93]
UiO2016 -5.46 [ -82.80, 71.87]
UiO2017 -74.20 [-136.24, -12.16]
UMCU -50.00 [ -98.18, -1.82]
ENIGMA Discovery Cohorts [N=22379; P=1.244e-06] -18.57 [ -26.08, -11.06]
UKBB -30.46 [ -41.45, -19.47]
Discovery+UKBB [N=32028; P=3.034e-12] -22.44 [ -28.75, -16.14]
GOBS -13.07 [ -40.18, 14.05]
IMH -11.54 [-110.99, 87.91]
UNICAMP 47.20 [ -29.83, 124.23]
Meth-CT
MIRECC
UKBB nonEUR -32.09 [ -86.66, 22.47]
OSAKA -54.98 [-109.84, -0.11]
PING Non European 24.97 [ -35.08, 85.01]
Non European Cohorts [N=2529; P=0.2008] -12.87 [ -32.58, 6.85]
Beta [Lower 95% CI,Upper 95% CI]
-300
-200
-100
0
100
200
mm^2/effect allele

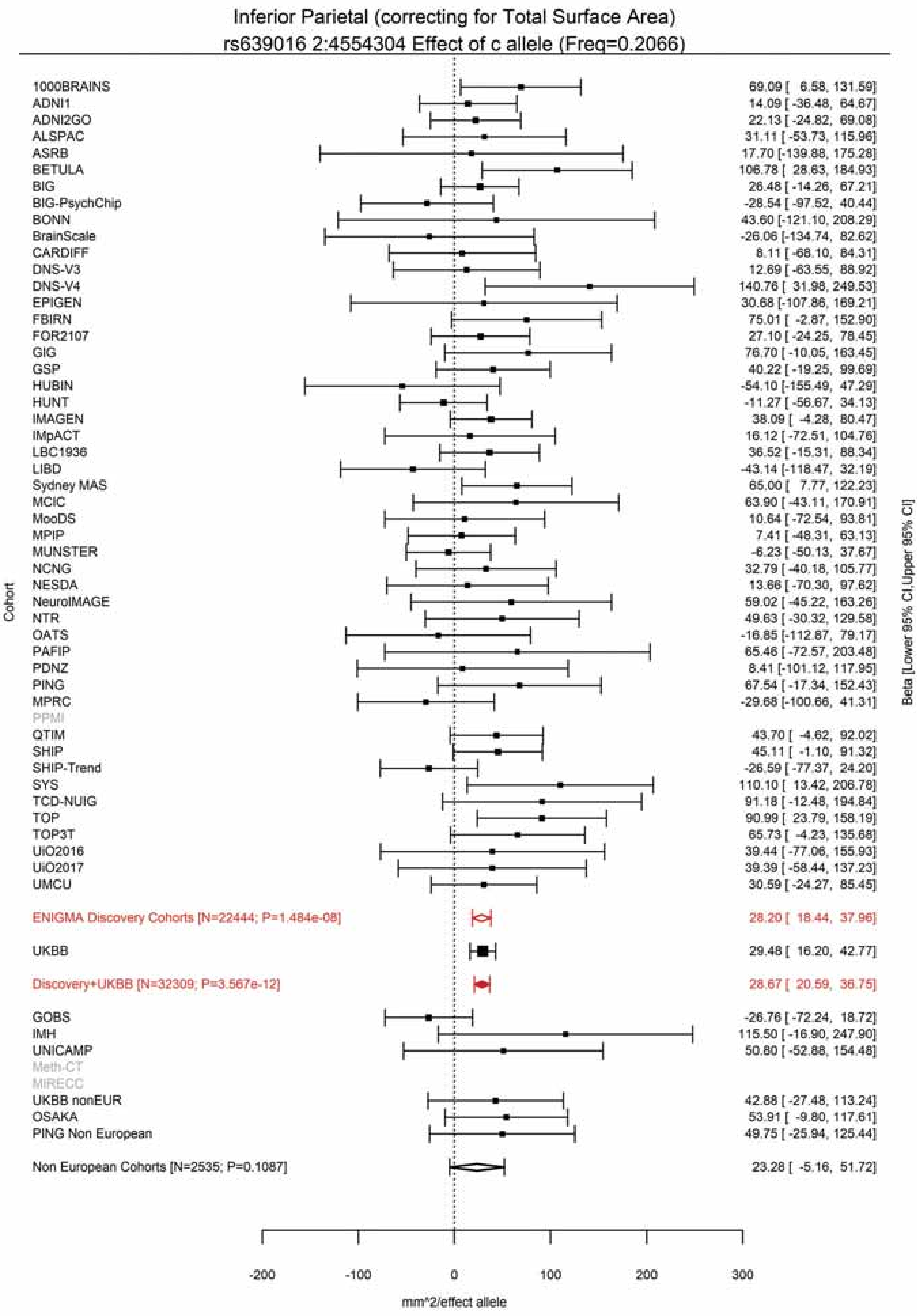

Inferior Parietal (correcting for Total Surface Area)
rs639016 2:4554304 Effect of c allele (Freq=0.2066)
Cohort
1000BRAINS 69.09 [ 6.58, 131.59]
ADNI1 14.09 [ -36.48, 64.67]
ADNI2GO 22.13 [ -24.82, 69.08]
ALSPAC 31.11 [ -53.73, 115.96]
ASRB 17.70 [-139.88, 175.28]
BETULA 106.78 [ 28.63, 184.93]
BIG 26.48 [ -14.26, 67.21]
BIG-PsychChip -28.54 [ -97.52, 40.44]
BONN 43.60 [-121.10, 208.29]
BrainScale -26.06 [-134.74, 82.62]
CARDIFF 8.11 [ -68.10, 84.31]
DNS-V3 12.69 [ -63.55, 88.92]
DNS-V4 140.76 [ 31.98, 249.53]
EPIGEN 30.68 [-107.86, 169.21]
FBIRN 75.01 [ -2.87, 152.90]
FOR2107 27.10 [ -24.25, 78.45]
GIG 76.70 [ -10.05, 163.45]
GSP 40.22 [ -19.25, 99.69]
HUBIN -54.10 [-155.49, 47.29]
HUNT -11.27 [ -56.67, 34.13]
IMAGEN 38.09 [ -4.28, 80.47]
IMpACT 16.12 [ -72.51, 104.76]
LBC1936 36.52 [ -15.31, 88.34]
LIBD -43.14 [-118.47, 32.19]
Sydney MAS 65.00 [ 7.77, 122.23]
MCIC 63.90 [ -43.11, 170.91]
MooDS 10.64 [ -72.54, 93.81]
MPIP 7.41 [ -48.31, 63.13]
MUNSTER -6.23 [ -50.13, 37.67]
NCNG 32.79 [ -40.18, 105.77]
NESDA 13.66 [ -70.30, 97.62]
NeuroIMAGE 59.02 [ -45.22, 163.26]
NTR 49.63 [ -30.32, 129.58]
OATS -16.85 [-112.87, 79.17]
PAFIP 65.46 [ -72.57, 203.48]
PDNZ 8.41 [-101.12, 117.95]
PING 67.54 [ -17.34, 152.43]
MPRC -29.68 [-100.66, 41.31]
PPMI
QTIM 43.70 [ -4.62, 92.02]
SHIP 45.11 [ -1.10, 91.32]
SHIP-Trend -26.59 [ -77.37, 24.20]
SYS 110.10 [ 13.42, 206.78]
TCD-NUIG 91.18 [ -12.48, 194.84]
TOP 90.99 [ 23.79, 158.19]
TOP3T 65.73 [ -4.23, 135.68]
UiO2016 39.44 [ -77.06, 155.93]
UiO2017 39.39 [ -58.44, 137.23]
UMCU 30.59 [ -24.27, 85.45]
ENIGMA Discovery Cohorts [N=22444; P=1.484e-08] 28.20 [ 18.44, 37.96]
UKBB 29.48 [ 16.20, 42.77]
Discovery+UKBB [N=32309; P=3.567e-12] 28.67 [ 20.59, 36.75]
GOBS -26.76 [ -72.24, 18.72]
IMH 115.50 [ -16.90, 247.90]
UNICAMP 50.80 [ -52.88, 154.48]
Meth-CT
MIRECC
UKBB nonEUR 42.88 [ -27.48, 113.24]
OSAKA 53.91 [ -9.80, 117.61]
PING Non European 49.75 [ -25.94, 125.44]
Non European Cohorts [N=2535; P=0.1087] 23.28 [ -5.16, 51.72]
Beta [Lower 95% CI,Upper 95% CI]
-200
-100
0
100
200
300
mm^2/effect allele

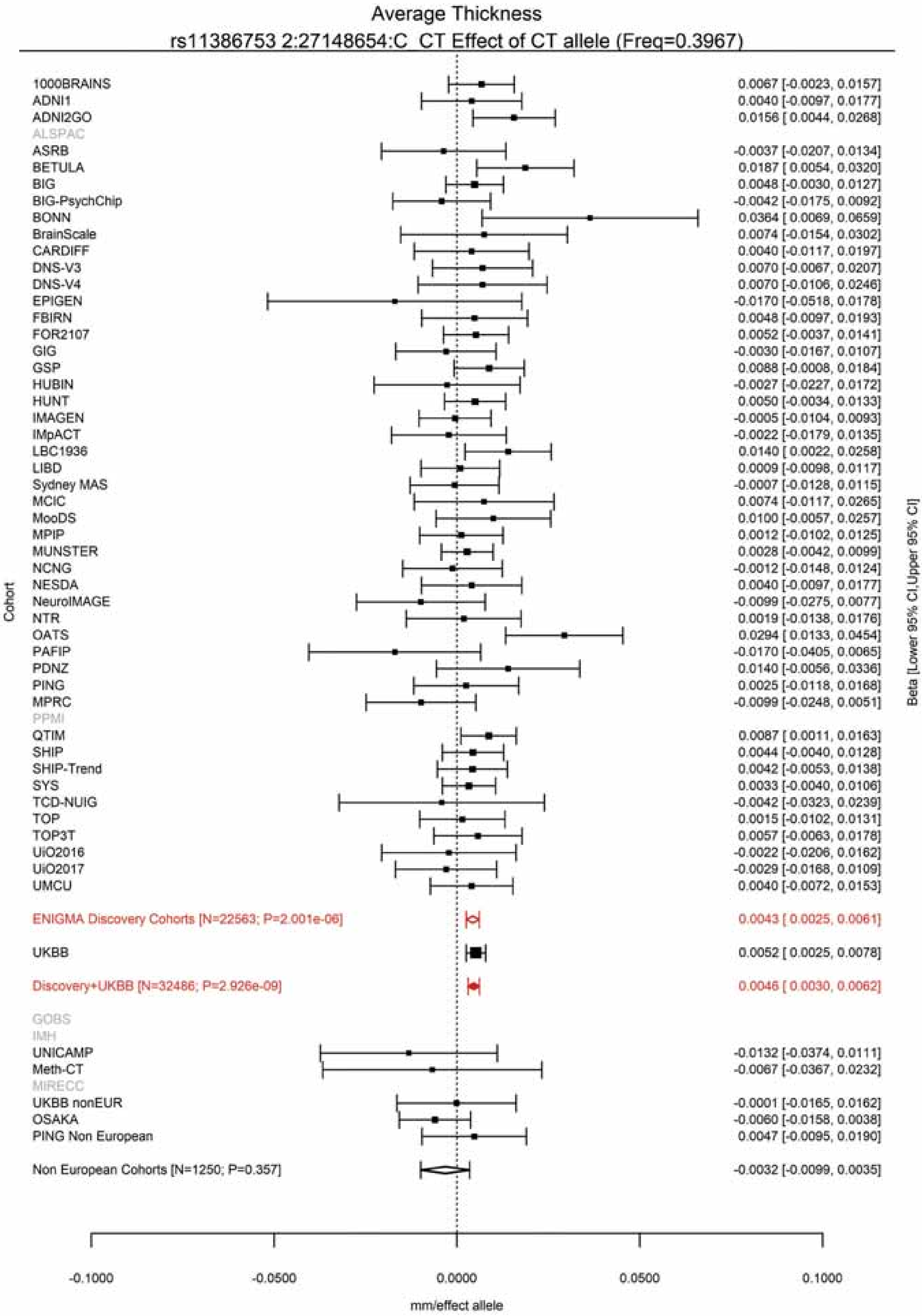
Average Thickness
rs11386753 2:27148654:C_CT Effect of CT allele (Freq=0.3967)
1000BRAINS 0.0067 [-0.0023, 0.0157]
ADNI1 0.0040 [-0.0097, 0.0177]
ADNI2GO 0.0156 [ 0.0044, 0.0268]
ALSPAC
ASRB -0.0037 [-0.0207, 0.0134]
BETULA 0.0187 [ 0.0054, 0.0320]
BIG 0.0048 [-0.0030, 0.0127]
BIG-PsychChip -0.0042 [-0.0175, 0.0092]
BONN 0.0364 [ 0.0069, 0.0659]
BrainScale 0.0074 [-0.0154, 0.0302]
CARDIFF 0.0040 [-0.0117, 0.0197]
DNS-V3 0.0070 [-0.0067, 0.0207]
DNS-V4 0.0070 [-0.0106, 0.0246]
EPIGEN -0.0170 [-0.0518, 0.0178]
FBIRN 0.0048 [-0.0097, 0.0193]
FOR2107 0.0052 [-0.0037, 0.0141]
GIG -0.0030 [-0.0167, 0.0107]
GSP 0.0088 [-0.0008, 0.0184]
HUBIN -0.0027 [-0.0227, 0.0172]
HUNT 0.0050 [-0.0034, 0.0133]
IMAGEN -0.0005 [-0.0104, 0.0093]
IMpACT -0.0022 [-0.0179, 0.0135]
LBC1936 0.0140 [ 0.0022, 0.0258]
LIBD 0.0009 [-0.0098, 0.0117]
Sydney MAS -0.0007 [-0.0128, 0.0115]
MCIC 0.0074 [-0.0117, 0.0265]
MooDS 0.0100 [-0.0057, 0.0257]
MPIP 0.0012 [-0.0102, 0.0125]
MUNSTER 0.0028 [-0.0042, 0.0099]
NCNG -0.0012 [-0.0148, 0.0124]
NESDA 0.0040 [-0.0097, 0.0177]
NeuroIMAGE -0.0099 [-0.0275, 0.0077]
NTR 0.0019 [-0.0138, 0.0176]
OATS 0.0294 [ 0.0133, 0.0454]
PAFIP -0.0170 [-0.0405, 0.0065]
PDNZ 0.0140 [-0.0056, 0.0336]
PING 0.0025 [-0.0118, 0.0168]
MPRC -0.0099 [-0.0248, 0.0051]
PPMI
QTIM 0.0087 [ 0.0011, 0.0163]
SHIP 0.0044 [-0.0040, 0.0128]
SHIP-Trend 0.0042 [-0.0053, 0.0138]
SYS 0.0033 [-0.0040, 0.0106]
TCD-NUIG -0.0042 [-0.0323, 0.0239]
TOP 0.0015 [-0.0102, 0.0131]
TOP3T 0.0057 [-0.0063, 0.0178]
UiO2016 -0.0022 [-0.0206, 0.0162]
UiO2017 -0.0029 [-0.0168, 0.0109]
UMCU 0.0040 [-0.0072, 0.0153]
ENIGMA Discovery Cohorts [N=22563; P=2.001e-06] 0.0043 [ 0.0025, 0.0061]
UKBB 0.0052 [ 0.0025, 0.0078]
Discovery+UKBB [N=32486; P=2.926e-09] 0.0046 [ 0.0030, 0.0062]
GOBS
IMH
UNICAMP -0.0132 [-0.0374, 0.0111]
Meth-CT -0.0067 [-0.0367, 0.0232]
MIRECC
UKBB nonEUR -0.0001 [-0.0165, 0.0162]
OSAKA -0.0060 [-0.0158, 0.0038]
PING Non European 0.0047 [-0.0095, 0.0190]
Non European Cohorts [N=1250; P=0.357] -0.0032 [-0.0099, 0.0035]
Cohort
Beta [Lower 95% CI,Upper 95% CI]
-0.1000
-0.0500
0.0000
0.0500
0.1000
mm/effect allele

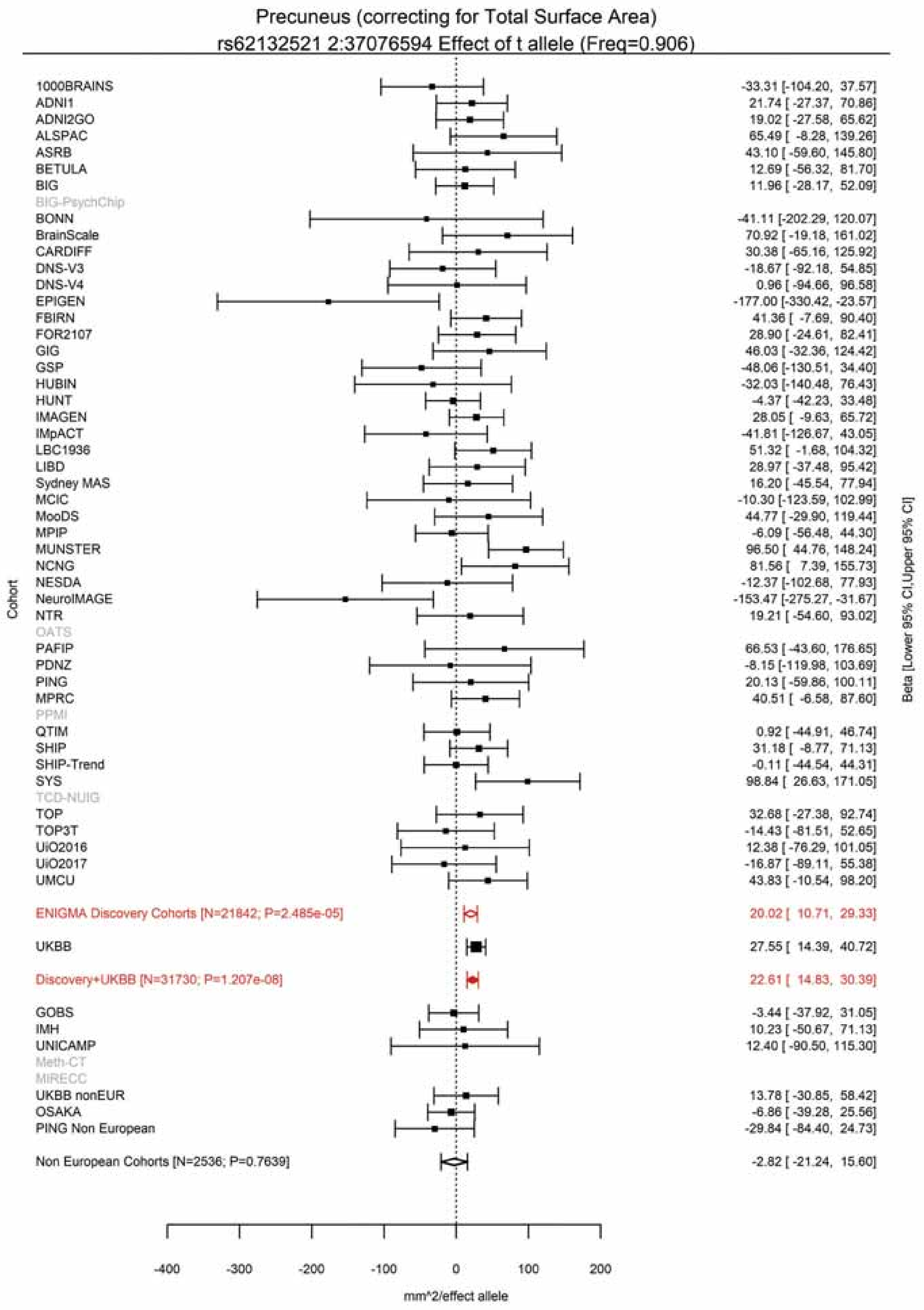
Precuneus (correcting for Total Surface Area)
rs62132521 2:37076594 Effect of t allele (Freq=0.906)
1000BRAINS -33.31 [-104.20, 37.57]
ADNI1 21.74 [ -27.37, 70.86]
ADNI2GO 19.02 [ -27.58, 65.62]
ALSPAC 65.49 [ -8.28, 139.26]
ASRB 43.10 [ -59.60, 145.80]
BETULA 12.69 [ -56.32, 81.70]
BIG 11.96 [ -28.17, 52.09]
BIG-PsychChip
BONN -41.11 [-202.29, 120.07]
BrainScale 70.92 [ -19.18, 161.02]
CARDIFF 30.38 [ -65.16, 125.92]
DNS-V3 -18.67 [ -92.18, 54.85]
DNS-V4 0.96 [ -94.66, 96.58]
EPIGEN -177.00 [-330.42, -23.57]
FBIRN 41.36 [ -7.69, 90.40]
FOR2107 28.90 [ -24.61, 82.41]
GIG 46.03 [ -32.36, 124.42]
GSP -48.06 [-130.51, 34.40]
HUBIN -32.03 [-140.48, 76.43]
HUNT -4.37 [ -42.23, 33.48]
IMAGEN 28.05 [ -9.63, 65.72]
IMpACT -41.81 [-126.67, 43.05]
LBC1936 51.32 [ -1.68, 104.32]
LIBD 28.97 [ -37.48, 95.42]
Sydney MAS 16.20 [ -45.54, 77.94]
MCIC -10.30 [-123.59, 102.99]
MooDS 44.77 [ -29.90, 119.44]
MPIP -6.09 [ -56.48, 44.30]
MUNSTER 96.50 [ 44.76, 148.24]
NCNG 81.56 [ 7.39, 155.73]
NESDA -12.37 [-102.68, 77.93]
NeuroIMAGE -153.47 [-275.27, -31.67]
NTR 19.21 [ -54.60, 93.02]
OATS
PAFIP 66.53 [ -43.60, 176.65]
PDNZ -8.15 [-119.98, 103.69]
PING 20.13 [ -59.86, 100.11]
MPRC 40.51 [ -6.58, 87.60]
PPMI
QTIM 0.92 [ -44.91, 46.74]
SHIP 31.18 [ -8.77, 71.13]
SHIP-Trend -0.11 [ -44.54, 44.31]
SYS 98.84 [ 26.63, 171.05]
TCD-NUIG
TOP 32.68 [ -27.38, 92.74]
TOP3T -14.43 [ -81.51, 52.65]
UiO2016 12.38 [ -76.29, 101.05]
UiO2017 -16.87 [ -89.11, 55.38]
UMCU 43.83 [ -10.54, 98.20]
ENIGMA Discovery Cohorts [N=21842; P=2.485e-05] 20.02 [ 10.71, 29.33]
UKBB 27.55 [ 14.39, 40.72]
Discovery+UKBB [N=31730; P=1.207e-08] 22.61 [ 14.83, 30.39]
GOBS -3.44 [ -37.92, 31.05]
IMH 10.23 [ -50.67, 71.13]
UNICAMP 12.40 [ -90.50, 115.30]
Meth-CT
MIRECC
UKBB nonEUR 13.78 [ -30.85, 58.42]
OSAKA -6.86 [ -39.28, 25.56]
PING Non European -29.84 [ -84.40, 24.73]
Non European Cohorts [N=2536; P=0.7639] -2.82 [ -21.24, 15.60]
Cohort
Beta [Lower 95% CI,Upper 95% CI]
-400
-300
-200
-100
0
100
200
mm^2/effect allele

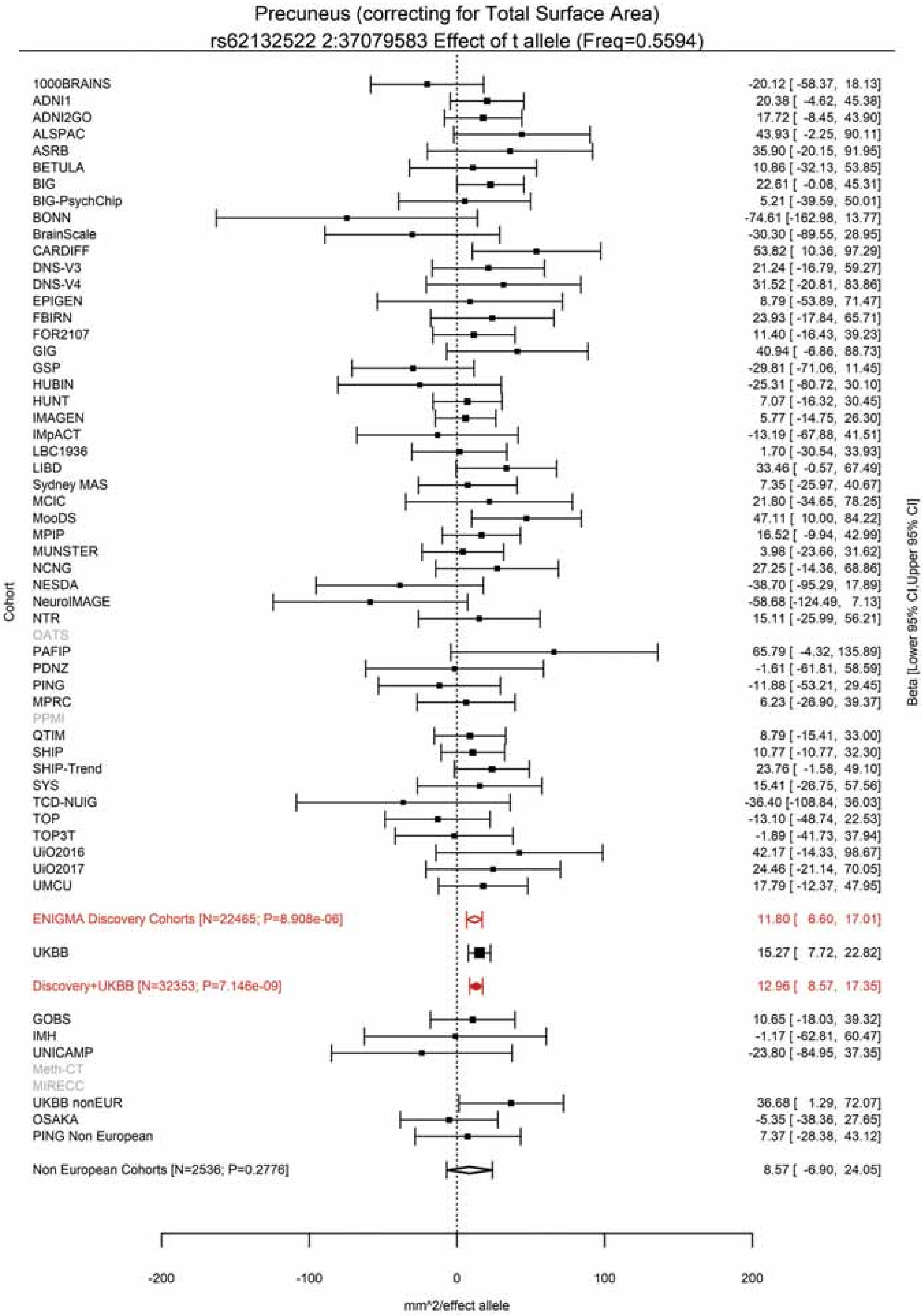
Precuneus (correcting for Total Surface Area)
rs62132522 2:37079583 Effect of t allele (Freq=0.5594)
Cohort
1000BRAINS -20.12 [ -58.37, 18.13]
ADNI1 20.38 [ -4.62, 45.38]
ADNI2GO 17.72 [ -8.45, 43.90]
ALSPAC 43.93 [ -2.25, 90.11]
ASRB 35.90 [ -20.15, 91.95]
BETULA 10.86 [ -32.13, 53.85]
BIG 22.61 [ -0.08, 45.31]
BIG-PsychChip 5.21 [ -39.59, 50.01]
BONN -74.61 [-162.98, 13.77]
BrainScale -30.30 [ -89.55, 28.95]
CARDIFF 53.82 [ 10.36, 97.29]
DNS-V3 21.24 [ -16.79, 59.27]
DNS-V4 31.52 [ -20.81, 83.86]
EPIGEN 8.79 [ -53.89, 71.47]
FBIRN 23.93 [ -17.84, 65.71]
FOR2107 11.40 [ -16.43, 39.23]
GIG 40.94 [ -6.86, 88.73]
GSP -29.81 [ -71.06, 11.45]
HUBIN -25.31 [ -80.72, 30.10]
HUNT 7.07 [ -16.32, 30.45]
IMAGEN 5.77 [ -14.75, 26.30]
IMpACT -13.19 [ -67.88, 41.51]
LBC1936 1.70 [ -30.54, 33.93]
LIBD 33.46 [ -0.57, 67.49]
Sydney MAS 7.35 [ -25.97, 40.67]
MCIC 21.80 [ -34.65, 78.25]
MooDS 47.11 [ 10.00, 84.22]
MPIP 16.52 [ -9.94, 42.99]
MUNSTER 3.98 [ -23.66, 31.62]
NCNG 27.25 [ -14.36, 68.86]
NESDA -38.70 [ -95.29, 17.89]
NeuroIMAGE -58.68 [-124.49, 7.13]
NTR 15.11 [ -25.99, 56.21]
OATS
PAFIP 65.79 [ -4.32, 135.89]
PDNZ -1.61 [ -61.81, 58.59]
PING -11.88 [ -53.21, 29.45]
MPRC 6.23 [ -26.90, 39.37]
PPMI
QTIM 8.79 [ -15.41, 33.00]
SHIP 10.77 [ -10.77, 32.30]
SHIP-Trend 23.76 [ -1.58, 49.10]
SYS 15.41 [ -26.75, 57.56]
TCD-NUIG -36.40 [-108.84, 36.03]
TOP -13.10 [ -48.74, 22.53]
TOP3T -1.89 [ -41.73, 37.94]
UiO2016 42.17 [ -14.33, 98.67]
UiO2017 24.46 [ -21.14, 70.05]
UMCU 17.79 [ -12.37, 47.95]
ENIGMA Discovery Cohorts [N=22465; P=8.908e-06] 11.80 [ 6.60, 17.01]
UKBB 15.27 [ 7.72, 22.82]
Discovery+UKBB [N=32353; P=7.146e-09] 12.96 [ 8.57, 17.35]
GOBS 10.65 [ -18.03, 39.32]
IMH -1.17 [ -62.81, 60.47]
UNICAMP -23.80 [ -84.95, 37.35]
Meth-CT
MIRECC
UKBB nonEUR 36.68 [ 1.29, 72.07]
OSAKA -5.35 [ -38.36, 27.65]
PING Non European 7.37 [ -28.38, 43.12]
Non European Cohorts [N=2536; P=0.2776] 8.57 [ -6.90, 24.05]
Beta [Lower 95% CI,Upper 95% CI]
-200
-100
0
100
200
mm^2/effect allele

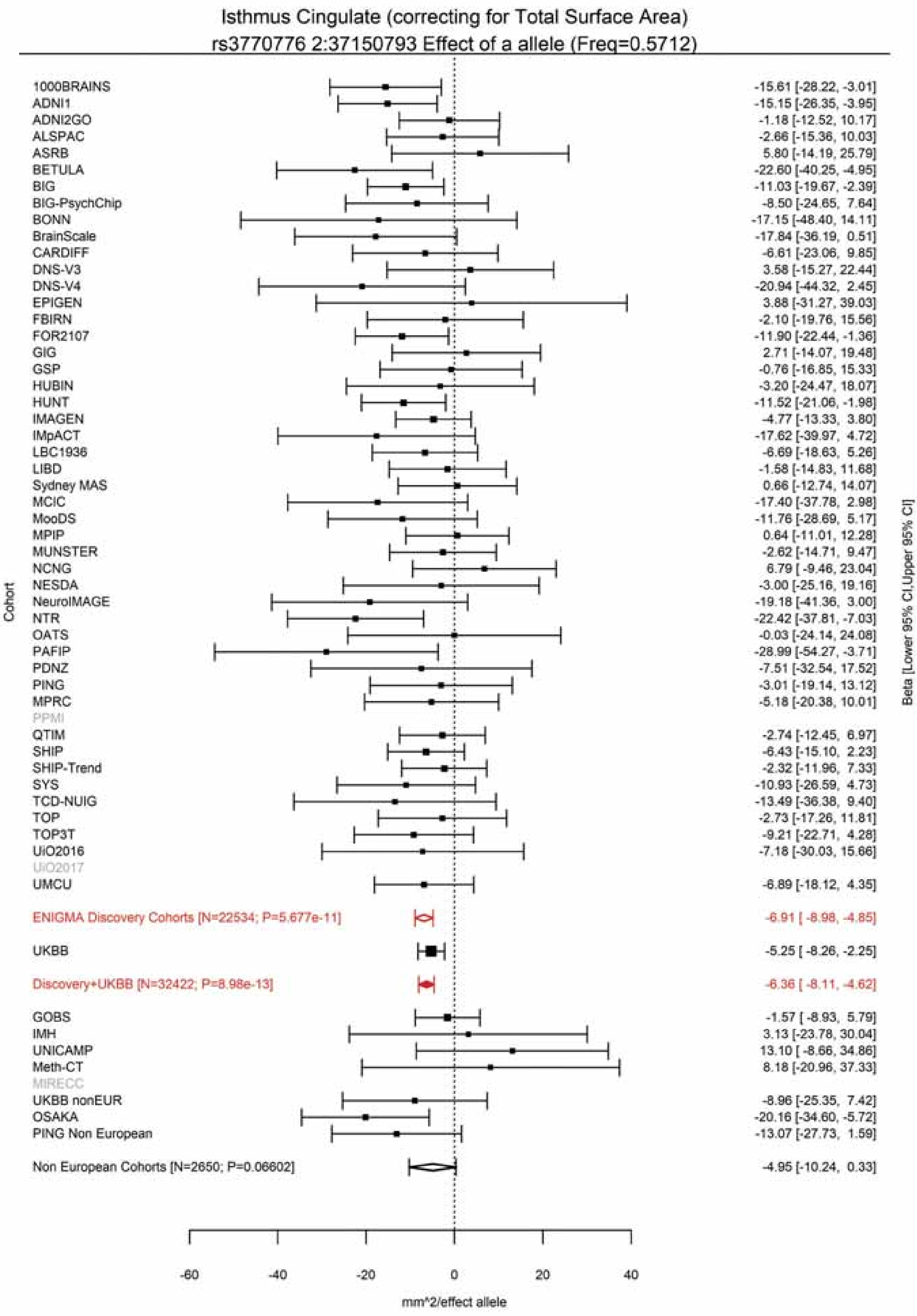
Isthmus Cingulate (correcting for Total Surface Area)
rs3770776 2:37150793 Effect of a allele (Freq=0.5712)
1000BRAINS -15.61 [-28.22, -3.01]
ADNI1 -15.15 [-26.35, -3.95]
ADNI2GO -1.18 [-12.52, 10.17]
ALSPAC -2.66 [-15.36, 10.03]
ASRB 5.80 [-14.19, 25.79]
BETULA -22.60 [-40.25, -4.95]
BIG -11.03 [-19.67, -2.39]
BIG-PsychChip -8.50 [-24.65, 7.64]
BONN -17.15 [-48.40, 14.11]
BrainScale -17.84 [-36.19, 0.51]
CARDIFF -6.61 [-23.06, 9.85]
DNS-V3 3.58 [-15.27, 22.44]
DNS-V4 -20.94 [-44.32, 2.45]
EPIGEN 3.88 [-31.27, 39.03]
FBIRN -2.10 [-19.76, 15.56]
FOR2107 -11.90 [-22.44, -1.36]
GIG 2.71 [-14.07, 19.48]
GSP -0.76 [-16.85, 15.33]
HUBIN -3.20 [-24.47, 18.07]
HUNT -11.52 [-21.06, -1.98]
IMAGEN -4.77 [-13.33, 3.80]
IMpACT -17.62 [-39.97, 4.72]
LBC1936 -6.69 [-18.63, 5.26]
LIBD -1.58 [-14.83, 11.68]
Sydney MAS 0.66 [-12.74, 14.07]
MCIC -17.40 [-37.78, 2.98]
MooDS -11.76 [-28.69, 5.17]
MPIP 0.64 [-11.01, 12.28]
MUNSTER -2.62 [-14.71, 9.47]
NCNG 6.79 [-9.46, 23.04]
NESDA -3.00 [-25.16, 19.16]
NeuroIMAGE -19.18 [-41.36, 3.00]
NTR -22.42 [-37.81, -7.03]
OATS -0.03 [-24.14, 24.08]
PAFIP -28.99 [-54.27, -3.71]
PDNZ -7.51 [-32.54, 17.52]
PING -3.01 [-19.14, 13.12]
MPRC -5.18 [-20.38, 10.01]
PPMI
QTIM -2.74 [-12.45, 6.97]
SHIP -6.43 [-15.10, 2.23]
SHIP-Trend -2.32 [-11.96, 7.33]
SYS -10.93 [-26.59, 4.73]
TCD-NUIG -13.49 [-36.38, 9.40]
TOP -2.73 [-17.26, 11.81]
TOP3T -9.21 [-22.71, 4.28]
UiO2016 -7.18 [-30.03, 15.66]
UiO2017
UMCU -6.89 [-18.12, 4.35]
ENIGMA Discovery Cohorts [N=22534; P=5.677e-11] -6.91 [-8.98, -4.85]
UKBB -5.25 [-8.26, -2.25]
Discovery+UKBB [N=32422; P=8.98e-13] -6.36 [-8.11, -4.62]
GOBS -1.57 [-8.93, 5.79]
IMH 3.13 [-23.78, 30.04]
UNICAMP 13.10 [-8.66, 34.86]
Meth-CT 8.18 [-20.96, 37.33]
MIRECC
UKBB nonEUR -8.96 [-25.35, 7.42]
OSAKA -20.16 [-34.60, -5.72]
PING Non European -13.07 [-27.73, 1.59]
Non European Cohorts [N=2650; P=0.06602] -4.95 [-10.24, 0.33]
Cohort
Beta [Lower 95% CI,Upper 95% CI]
-60
-40
-20
0
20
40
mm^2/effect allele

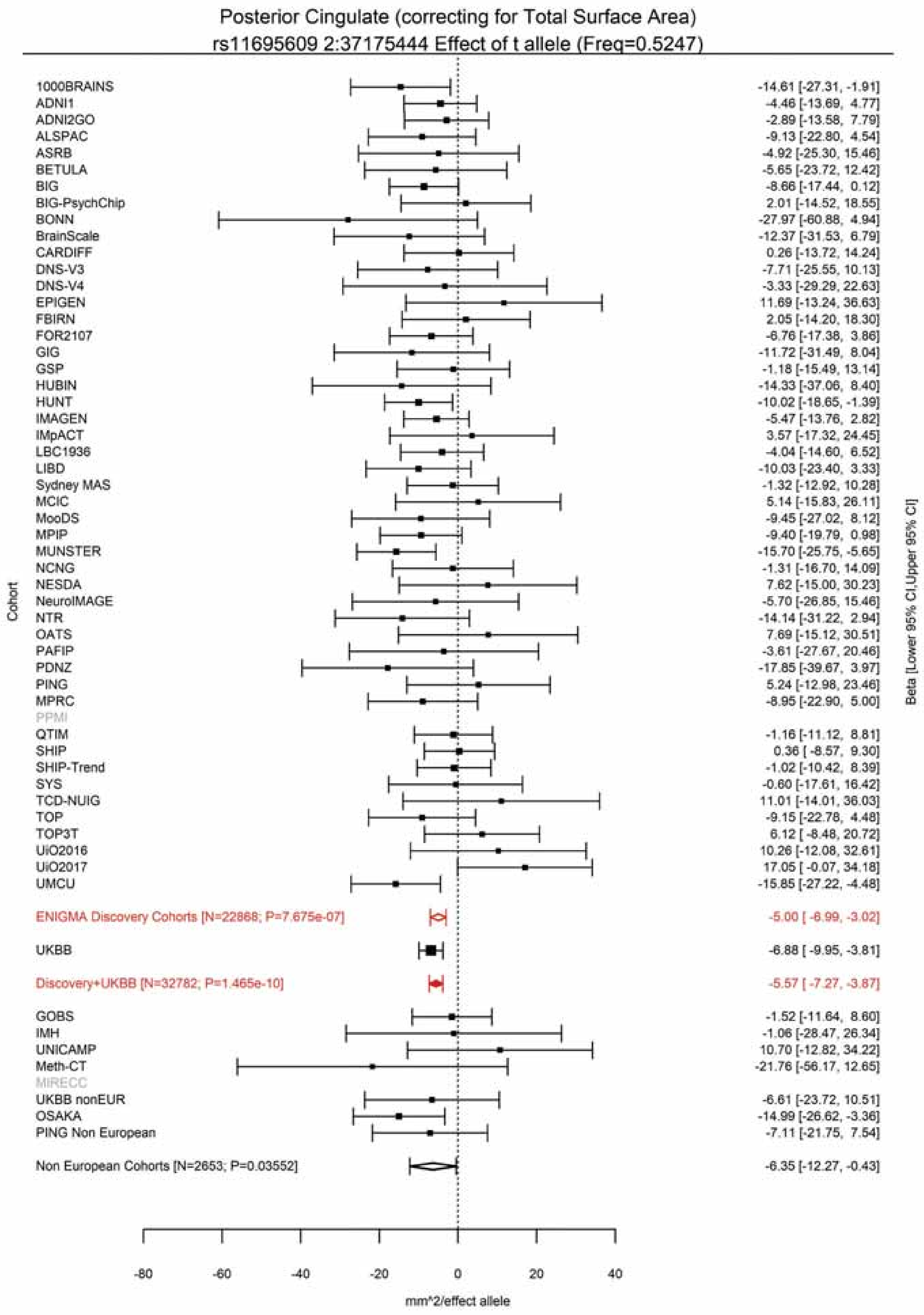
Posterior Cingulate (correcting for Total Surface Area)
rs11695609 2:37175444 Effect of t allele (Freq=0.5247)
Cohort
1000BRAINS -14.61 [-27.31, -1.91]
ADNI1 -4.46 [-13.69, 4.77]
ADNI2GO -2.89 [-13.58, 7.79]
ALSPAC -9.13 [-22.80, 4.54]
ASRB -4.92 [-25.30, 15.46]
BETULA -5.65 [-23.72, 12.42]
BIG -8.66 [-17.44, 0.12]
BIG-PsychChip 2.01 [-14.52, 18.55]
BONN -27.97 [-60.88, 4.94]
BrainScale -12.37 [-31.53, 6.79]
CARDIFF 0.26 [-13.72, 14.24]
DNS-V3 -7.71 [-25.55, 10.13]
DNS-V4 -3.33 [-29.29, 22.63]
EPIGEN 11.69 [-13.24, 36.63]
FBIRN 2.05 [-14.20, 18.30]
FOR2107 -6.76 [-17.38, 3.86]
GIG -11.72 [-31.49, 8.04]
GSP -1.18 [-15.49, 13.14]
HUBIN -14.33 [-37.06, 8.40]
HUNT -10.02 [-18.65, -1.39]
IMAGEN -5.47 [-13.76, 2.82]
IMpACT 3.57 [-17.32, 24.45]
LBC1936 -4.04 [-14.60, 6.52]
LIBD -10.03 [-23.40, 3.33]
Sydney MAS -1.32 [-12.92, 10.28]
MCIC 5.14 [-15.83, 26.11]
MooDS -9.45 [-27.02, 8.12]
MPIP -9.40 [-19.79, 0.98]
MUNSTER -15.70 [-25.75, -5.65]
NCNG -1.31 [-16.70, 14.09]
NESDA 7.62 [-15.00, 30.23]
NeuroIMAGE -5.70 [-26.85, 15.46]
NTR -14.14 [-31.22, 2.94]
OATS 7.69 [-15.12, 30.51]
PAFIP -3.61 [-27.67, 20.46]
PDNZ -17.85 [-39.67, 3.97]
PING 5.24 [-12.98, 23.46]
MPRC -8.95 [-22.90, 5.00]
PPMI
QTIM -1.16 [-11.12, 8.81]
SHIP 0.36 [ -8.57, 9.30]
SHIP-Trend -1.02 [-10.42, 8.39]
SYS -0.60 [-17.61, 16.42]
TCD-NUIG 11.01 [-14.01, 36.03]
TOP -9.15 [-22.78, 4.48]
TOP3T 6.12 [ -8.48, 20.72]
UiO2016 10.26 [-12.08, 32.61]
UiO2017 17.05 [ -0.07, 34.18]
UMCU -15.85 [-27.22, -4.48]
ENIGMA Discovery Cohorts [N=22868; P=7.675e-07] -5.00 [ -6.99, -3.02]
UKBB -6.88 [ -9.95, -3.81]
Discovery+UKBB [N=32782; P=1.465e-10] -5.57 [ -7.27, -3.87]
GOBS -1.52 [-11.64, 8.60]
IMH -1.06 [-28.47, 26.34]
UNICAMP 10.70 [-12.82, 34.22]
Meth-CT -21.76 [-56.17, 12.65]
MIRECC
UKBB nonEUR -6.61 [-23.72, 10.51]
OSAKA -14.99 [-26.62, -3.36]
PING Non European -7.11 [-21.75, 7.54]
Non European Cohorts [N=2653; P=0.03552] -6.35 [-12.27, -0.43]
Beta [Lower 95% CI,Upper 95% CI]
-80 -60 -40 -20 0 20 40
mm^2/effect allele

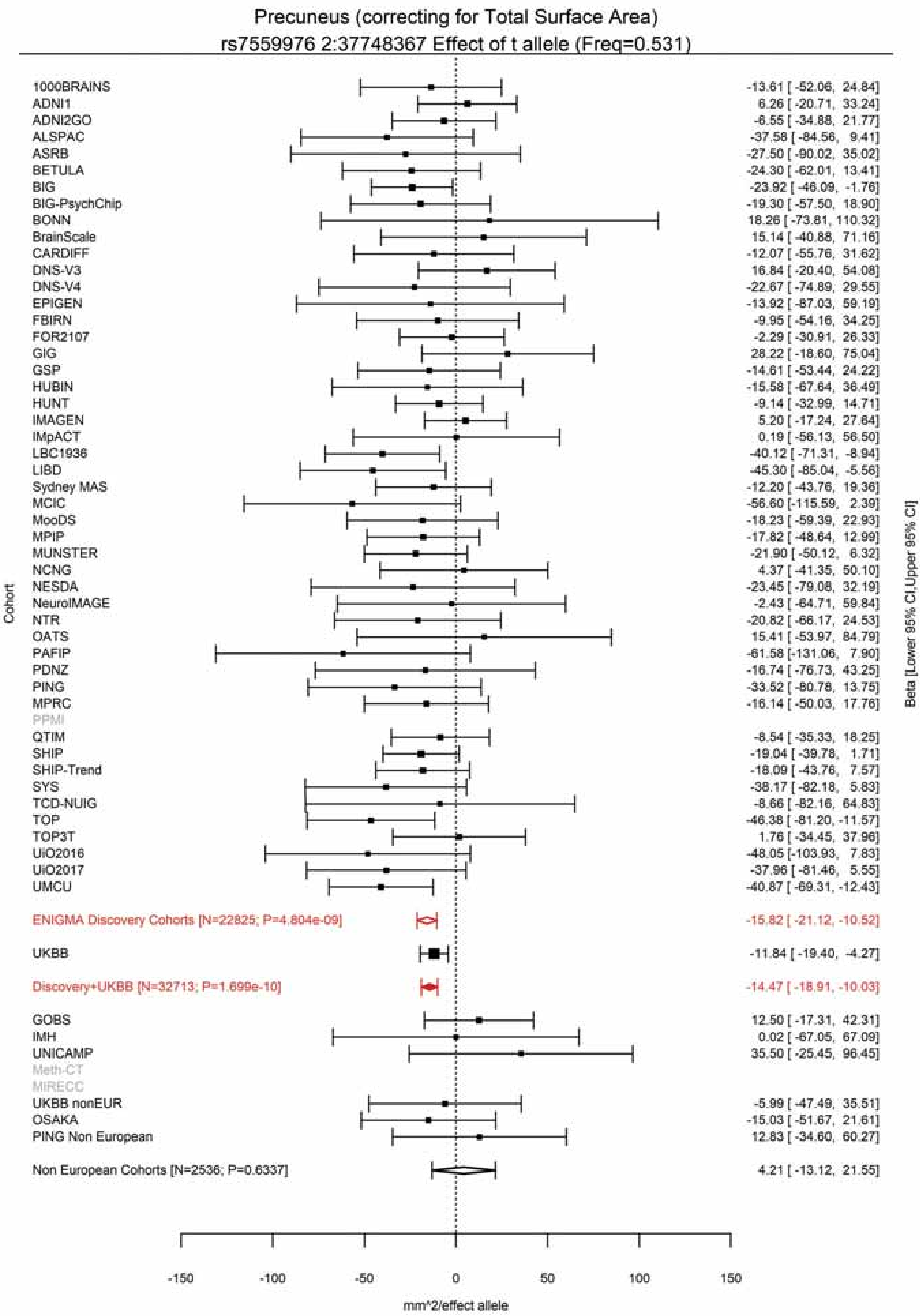

Precuneus (correcting for Total Surface Area)
rs7559976 2:37748367 Effect of t allele (Freq=0.531)
Cohort
Beta [Lower 95% CI,Upper 95% CI]
1000BRAINS -13.61 [ -52.06, 24.84]
ADNI1 6.26 [ -20.71, 33.24]
ADNI2GO -6.55 [ -34.88, 21.77]
ALSPAC -37.58 [ -84.56, 9.41]
ASRB -27.50 [ -90.02, 35.02]
BETULA -24.30 [ -62.01, 13.41]
BIG -23.92 [ -46.09, -1.76]
BIG-PsychChip -19.30 [ -57.50, 18.90]
BONN 18.26 [ -73.81, 110.32]
BrainScale 15.14 [ -40.88, 71.16]
CARDIFF -12.07 [ -55.76, 31.62]
DNS-V3 16.84 [ -20.40, 54.08]
DNS-V4 -22.67 [ -74.89, 29.55]
EPIGEN -13.92 [ -87.03, 59.19]
FBIRN -9.95 [ -54.16, 34.25]
FOR2107 -2.29 [ -30.91, 26.33]
GIG 28.22 [ -18.60, 75.04]
GSP -14.61 [ -53.44, 24.22]
HUBIN -15.58 [ -67.64, 36.49]
HUNT -9.14 [ -32.99, 14.71]
IMAGEN 5.20 [ -17.24, 27.64]
IMpACT 0.19 [ -56.13, 56.50]
LBC1936 -40.12 [ -71.31, -8.94]
LIBD -45.30 [ -85.04, -5.56]
Sydney MAS -12.20 [ -43.76, 19.36]
MCIC -56.60 [-115.59, 2.39]
MooDS -18.23 [ -59.39, 22.93]
MPIP -17.82 [ -48.64, 12.99]
MUNSTER -21.90 [ -50.12, 6.32]
NCNG 4.37 [ -41.35, 50.10]
NESDA -23.45 [ -79.08, 32.19]
NeuroIMAGE -2.43 [ -64.71, 59.84]
NTR -20.82 [ -66.17, 24.53]
OATS 15.41 [ -53.97, 84.79]
PAFIP -61.58 [-131.06, 7.90]
PDNZ -16.74 [ -76.73, 43.25]
PING -33.52 [ -80.78, 13.75]
MPRC -16.14 [ -50.03, 17.76]
PPMI
QTIM -8.54 [ -35.33, 18.25]
SHIP -19.04 [ -39.78, 1.71]
SHIP-Trend -18.09 [ -43.76, 7.57]
SYS -38.17 [ -82.18, 5.83]
TCD-NUIG -8.66 [ -82.16, 64.83]
TOP -46.38 [ -81.20, -11.57]
TOP3T 1.76 [ -34.45, 37.96]
UiO2016 -48.05 [-103.93, 7.83]
UiO2017 -37.96 [ -81.46, 5.55]
UMCU -40.87 [ -69.31, -12.43]
ENIGMA Discovery Cohorts [N=22825; P=4.804e-09] -15.82 [ -21.12, -10.52]
UKBB -11.84 [ -19.40, -4.27]
Discovery+UKBB [N=32713; P=1.699e-10] -14.47 [ -18.91, -10.03]
GOBS 12.50 [ -17.31, 42.31]
IMH 0.02 [ -67.05, 67.09]
UNICAMP 35.50 [ -25.45, 96.45]
Meth-CT
MIRECC
UKBB nonEUR -5.99 [ -47.49, 35.51]
OSAKA -15.03 [ -51.67, 21.61]
PING Non European 12.83 [ -34.60, 60.27]
Non European Cohorts [N=2536; P=0.6337] 4.21 [ -13.12, 21.55]
-150
-100
-50
0
50
100
150
mm^2/effect allele

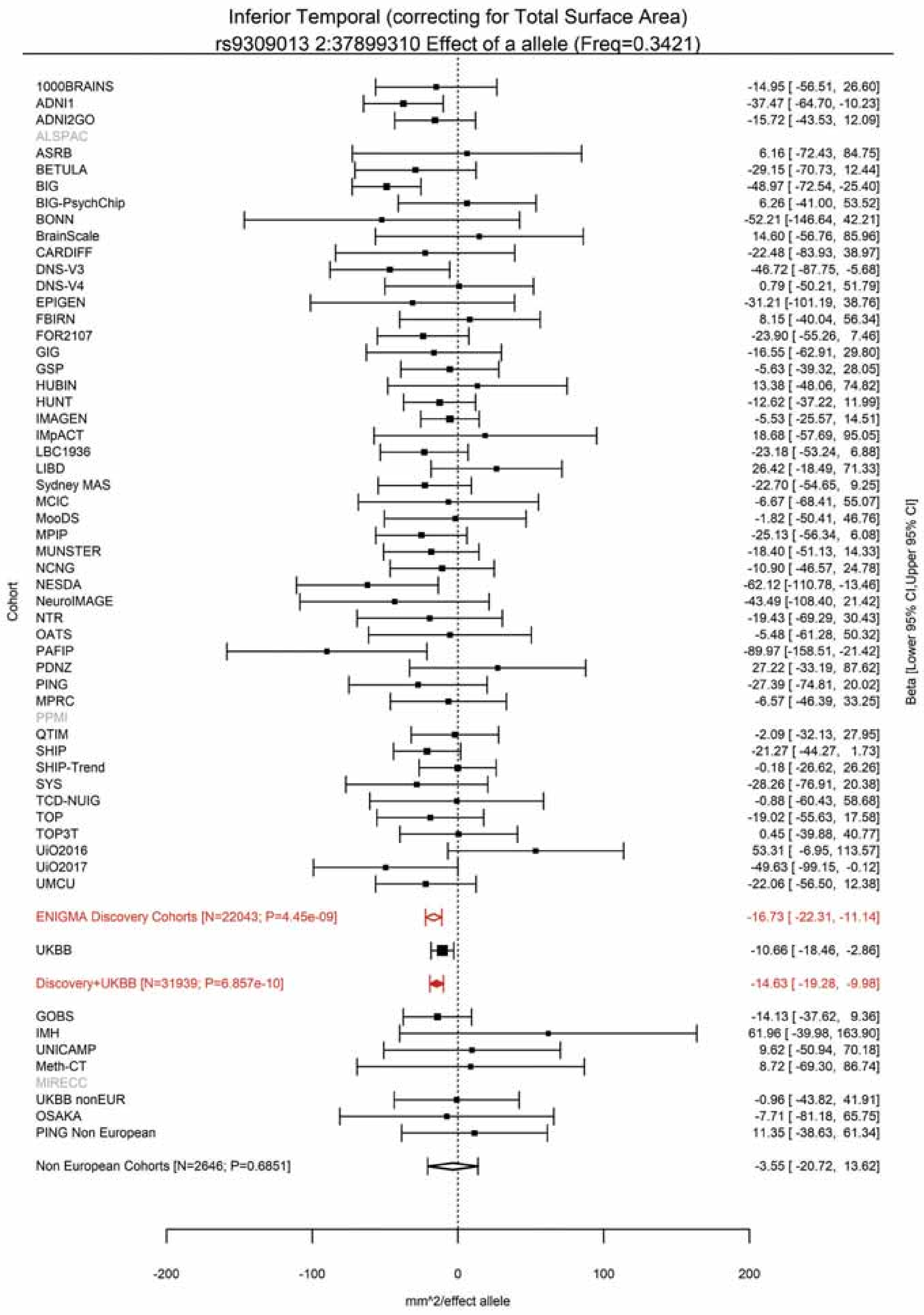

Inferior Temporal (correcting for Total Surface Area)
rs9309013 2:37899310 Effect of a allele (Freq=0.3421)
1000BRAINS -14.95 [ -56.51, 26.60]
ADNI1 -37.47 [ -64.70, -10.23]
ADNI2GO -15.72 [ -43.53, 12.09]
ALSPAC
ASRB 6.16 [ -72.43, 84.75]
BETULA -29.15 [ -70.73, 12.44]
BIG -48.97 [ -72.54, -25.40]
BIG-PsychChip 6.26 [ -41.00, 53.52]
BONN -52.21 [-146.64, 42.21]
BrainScale 14.60 [ -56.76, 85.96]
CARDIFF -22.48 [ -83.93, 38.97]
DNS-V3 -46.72 [ -87.75, -5.68]
DNS-V4 0.79 [ -50.21, 51.79]
EPIGEN -31.21 [-101.19, 38.76]
FBIRN 8.15 [ -40.04, 56.34]
FOR2107 -23.90 [ -55.26, 7.46]
GIG -16.55 [ -62.91, 29.80]
GSP -5.63 [ -39.32, 28.05]
HUBIN 13.38 [ -48.06, 74.82]
HUNT -12.62 [ -37.22, 11.99]
IMAGEN -5.53 [ -25.57, 14.51]
IMpACT 18.68 [ -57.69, 95.05]
LBC1936 -23.18 [ -53.24, 6.88]
LIBD 26.42 [ -18.49, 71.33]
Sydney MAS -22.70 [ -54.65, 9.25]
MCIC -6.67 [ -68.41, 55.07]
MooDS -1.82 [ -50.41, 46.76]
MPIP -25.13 [ -56.34, 6.08]
MUNSTER -18.40 [ -51.13, 14.33]
NCNG -10.90 [ -46.57, 24.78]
NESDA -62.12 [-110.78, -13.46]
NeuroIMAGE -43.49 [-108.40, 21.42]
NTR -19.43 [ -69.29, 30.43]
OATS -5.48 [ -61.28, 50.32]
PAFIP -89.97 [-158.51, -21.42]
PDNZ 27.22 [ -33.19, 87.62]
PING -27.39 [ -74.81, 20.02]
MPRC -6.57 [ -46.39, 33.25]
PPMI
QTIM -2.09 [ -32.13, 27.95]
SHIP -21.27 [ -44.27, 1.73]
SHIP-Trend -0.18 [ -26.62, 26.26]
SYS -28.26 [ -76.91, 20.38]
TCD-NUIG -0.88 [ -60.43, 58.68]
TOP -19.02 [ -55.63, 17.58]
TOP3T 0.45 [ -39.88, 40.77]
UiO2016 53.31 [ -6.95, 113.57]
UiO2017 -49.63 [ -99.15, -0.12]
UMCU -22.06 [ -56.50, 12.38]
ENIGMA Discovery Cohorts [N=22043; P=4.45e-09] -16.73 [ -22.31, -11.14]
UKBB -10.66 [ -18.46, -2.86]
Discovery+UKBB [N=31939; P=6.857e-10] -14.63 [ -19.28, -9.98]
GOBS -14.13 [ -37.62, 9.36]
IMH 61.96 [ -39.98, 163.90]
UNICAMP 9.62 [ -50.94, 70.18]
Meth-CT 8.72 [ -69.30, 86.74]
MIRECC
UKBB nonEUR -0.96 [ -43.82, 41.91]
OSAKA -7.71 [ -81.18, 65.75]
PING Non European 11.35 [ -38.63, 61.34]
Non European Cohorts [N=2646; P=0.6851] -3.55 [ -20.72, 13.62]
Cohort
Beta [Lower 95% CI,Upper 95% CI]
-200
-100
0
100
200
mm^2/effect allele

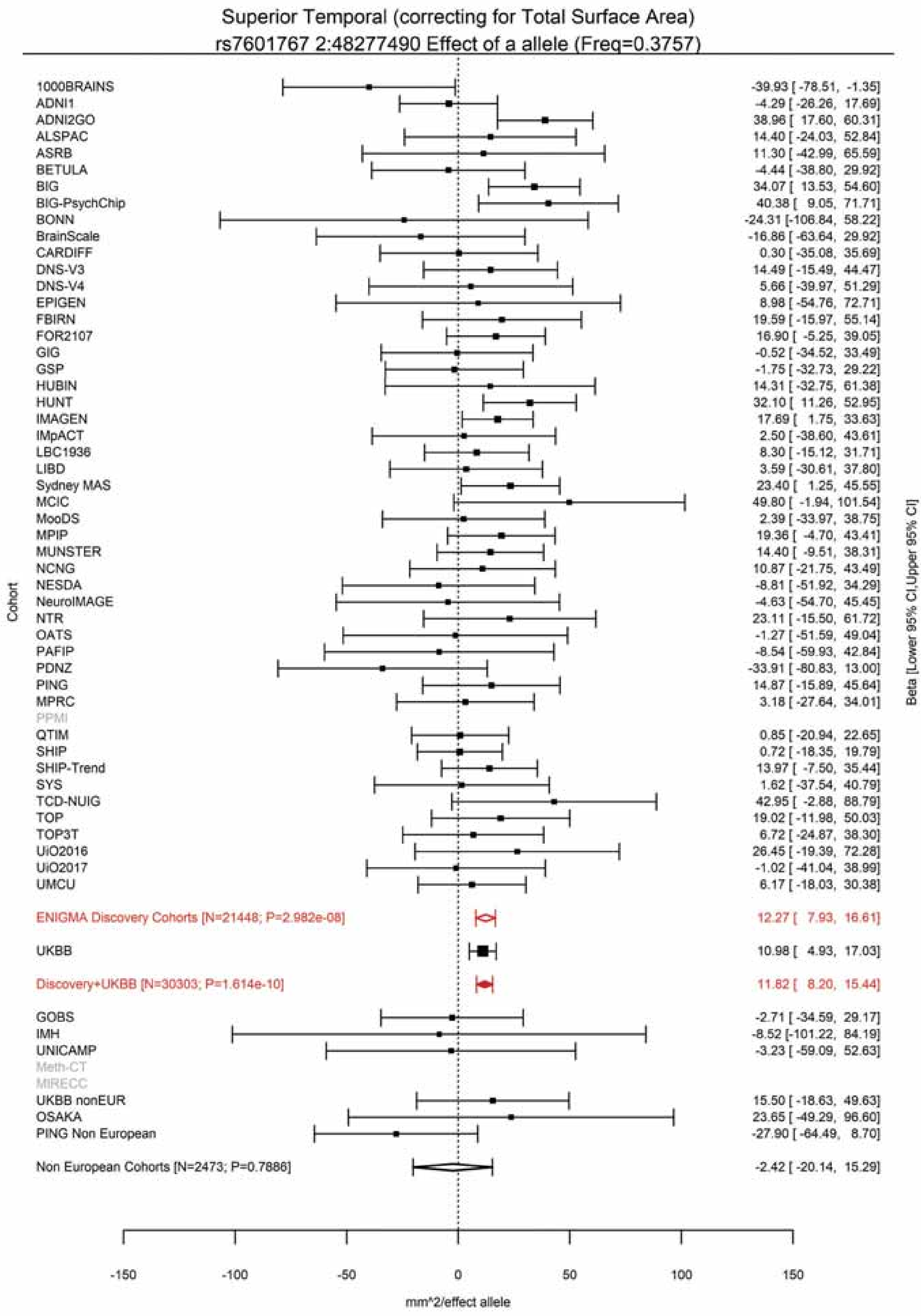

Superior Temporal (correcting for Total Surface Area)
rs7601767 2:48277490 Effect of a allele (Freq=0.3757)
Cohort
1000BRAINS -39.93 [ -78.51, -1.35]
ADNI1 -4.29 [ -26.26, 17.69]
ADNI2GO 38.96 [ 17.60, 60.31]
ALSPAC 14.40 [ -24.03, 52.84]
ASRB 11.30 [ -42.99, 65.59]
BETULA -4.44 [ -38.80, 29.92]
BIG 34.07 [ 13.53, 54.60]
BIG-PsychChip 40.38 [ 9.05, 71.71]
BONN -24.31 [-106.84, 58.22]
BrainScale -16.86 [ -63.64, 29.92]
CARDIFF 0.30 [ -35.08, 35.69]
DNS-V3 14.49 [ -15.49, 44.47]
DNS-V4 5.66 [ -39.97, 51.29]
EPIGEN 8.98 [ -54.76, 72.71]
FBIRN 19.59 [ -15.97, 55.14]
FOR2107 16.90 [ -5.25, 39.05]
GIG -0.52 [ -34.52, 33.49]
GSP -1.75 [ -32.73, 29.22]
HUBIN 14.31 [ -32.75, 61.38]
HUNT 32.10 [ 11.26, 52.95]
IMAGEN 17.69 [ 1.75, 33.63]
IMpACT 2.50 [ -38.60, 43.61]
LBC1936 8.30 [ -15.12, 31.71]
LIBD 3.59 [ -30.61, 37.80]
Sydney MAS 23.40 [ 1.25, 45.55]
MCIC 49.80 [ -1.94, 101.54]
MooDS 2.39 [ -33.97, 38.75]
MPIP 19.36 [ -4.70, 43.41]
MUNSTER 14.40 [ -9.51, 38.31]
NCNG 10.87 [ -21.75, 43.49]
NESDA -8.81 [ -51.92, 34.29]
NeuroIMAGE -4.63 [ -54.70, 45.45]
NTR 23.11 [ -15.50, 61.72]
OATS -1.27 [ -51.59, 49.04]
PAFIP -8.54 [ -59.93, 42.84]
PDNZ -33.91 [ -80.83, 13.00]
PING 14.87 [ -15.89, 45.64]
MPRC 3.18 [ -27.64, 34.01]
PPMI
QTIM 0.85 [ -20.94, 22.65]
SHIP 0.72 [ -18.35, 19.79]
SHIP-Trend 13.97 [ -7.50, 35.44]
SYS 1.62 [ -37.54, 40.79]
TCD-NUIG 42.95 [ -2.88, 88.79]
TOP 19.02 [ -11.98, 50.03]
TOP3T 6.72 [ -24.87, 38.30]
UiO2016 26.45 [ -19.39, 72.28]
UiO2017 -1.02 [ -41.04, 38.99]
UMCU 6.17 [ -18.03, 30.38]
ENIGMA Discovery Cohorts [N=21448; P=2.982e-08] 12.27 [ 7.93, 16.61]
UKBB 10.98 [ 4.93, 17.03]
Discovery+UKBB [N=30303; P=1.614e-10] 11.82 [ 8.20, 15.44]
GOBS -2.71 [ -34.59, 29.17]
IMH -8.52 [-101.22, 84.19]
UNICAMP -3.23 [ -59.09, 52.63]
Meth-CT
MIRECC
UKBB nonEUR 15.50 [ -18.63, 49.63]
OSAKA 23.65 [ -49.29, 96.60]
PING Non European -27.90 [ -64.49, 8.70]
Non European Cohorts [N=2473; P=0.7886] -2.42 [ -20.14, 15.29]
Beta [Lower 95% CI,Upper 95% CI]
-150 -100 -50 0 50 100 150
mm^2/effect allele

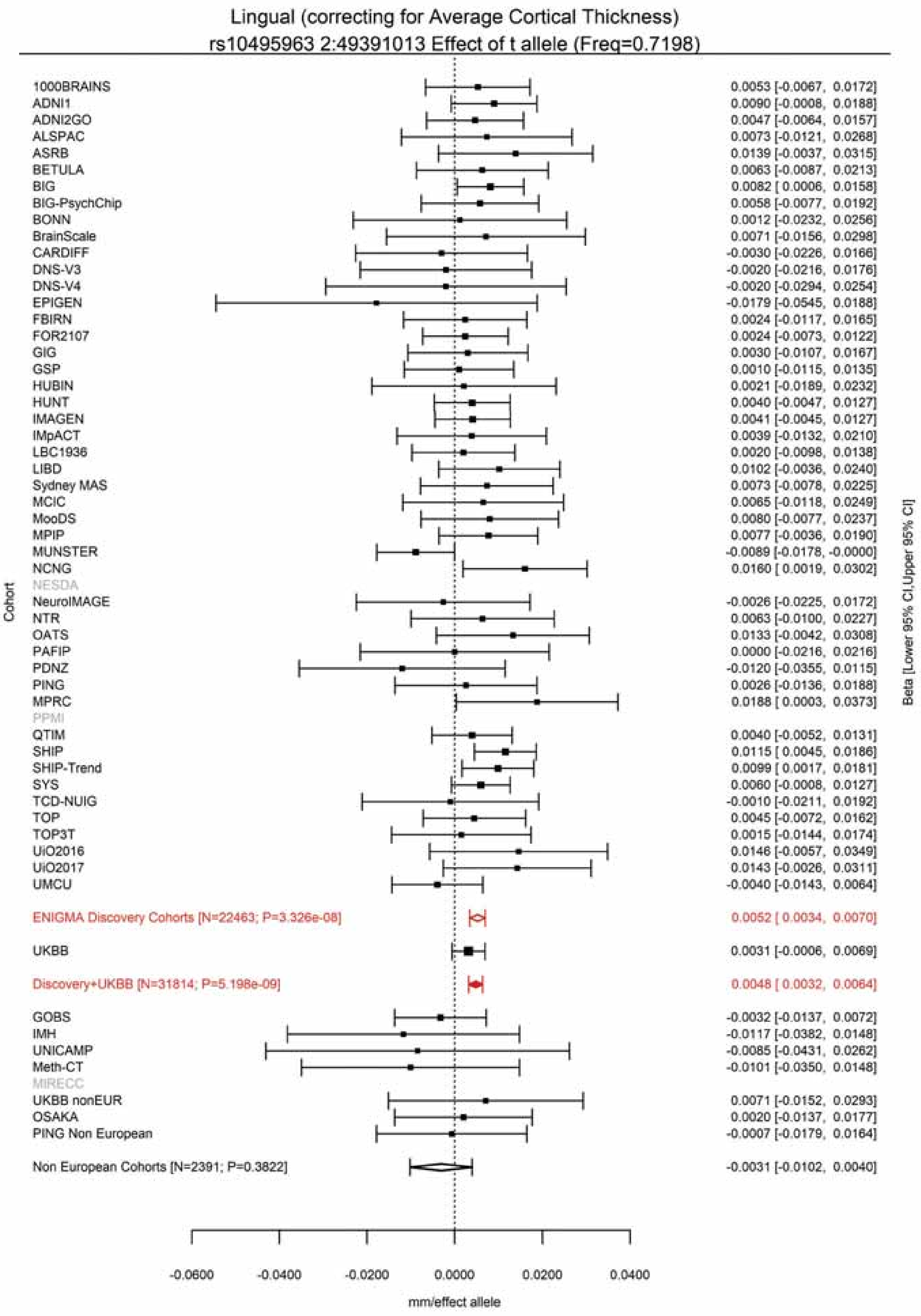
Lingual (correcting for Average Cortical Thickness)
rs10495963 2:49391013 Effect of t allele (Freq=0.7198)
Cohort
1000BRAINS 0.0053 [-0.0067, 0.0172]
ADNI1 0.0090 [-0.0008, 0.0188]
ADNI2GO 0.0047 [-0.0064, 0.0157]
ALSPAC 0.0073 [-0.0121, 0.0268]
ASRB 0.0139 [-0.0037, 0.0315]
BETULA 0.0063 [-0.0087, 0.0213]
BIG 0.0082 [ 0.0006, 0.0158]
BIG-PsychChip 0.0058 [-0.0077, 0.0192]
BONN 0.0012 [-0.0232, 0.0256]
BrainScale 0.0071 [-0.0156, 0.0298]
CARDIFF -0.0030 [-0.0226, 0.0166]
DNS-V3 -0.0020 [-0.0216, 0.0176]
DNS-V4 -0.0020 [-0.0294, 0.0254]
EPIGEN -0.0179 [-0.0545, 0.0188]
FBIRN 0.0024 [-0.0117, 0.0165]
FOR2107 0.0024 [-0.0073, 0.0122]
GIG 0.0030 [-0.0107, 0.0167]
GSP 0.0010 [-0.0115, 0.0135]
HUBIN 0.0021 [-0.0189, 0.0232]
HUNT 0.0040 [-0.0047, 0.0127]
IMAGEN 0.0041 [-0.0045, 0.0127]
IMpACT 0.0039 [-0.0132, 0.0210]
LBC1936 0.0020 [-0.0098, 0.0138]
LIBD 0.0102 [-0.0036, 0.0240]
Sydney MAS 0.0073 [-0.0078, 0.0225]
MCIC 0.0065 [-0.0118, 0.0249]
MooDS 0.0080 [-0.0077, 0.0237]
MPIP 0.0077 [-0.0036, 0.0190]
MUNSTER -0.0089 [-0.0178, -0.0000]
NCNG 0.0160 [ 0.0019, 0.0302]
NESDA
NeuroIMAGE -0.0026 [-0.0225, 0.0172]
NTR 0.0063 [-0.0100, 0.0227]
OATS 0.0133 [-0.0042, 0.0308]
PAFIP 0.0000 [-0.0216, 0.0216]
PDNZ -0.0120 [-0.0355, 0.0115]
PING 0.0026 [-0.0136, 0.0188]
MPRC 0.0188 [ 0.0003, 0.0373]
PPMI
QTIM 0.0040 [-0.0052, 0.0131]
SHIP 0.0115 [ 0.0045, 0.0186]
SHIP-Trend 0.0099 [ 0.0017, 0.0181]
SYS 0.0060 [-0.0008, 0.0127]
TCD-NUIG -0.0010 [-0.0211, 0.0192]
TOP 0.0045 [-0.0072, 0.0162]
TOP3T 0.0015 [-0.0144, 0.0174]
UiO2016 0.0146 [-0.0057, 0.0349]
UiO2017 0.0143 [-0.0026, 0.0311]
UMCU -0.0040 [-0.0143, 0.0064]
ENIGMA Discovery Cohorts [N=22463; P=3.326e-08] 0.0052 [ 0.0034, 0.0070]
UKBB 0.0031 [-0.0006, 0.0069]
Discovery+UKBB [N=31814; P=5.198e-09] 0.0048 [ 0.0032, 0.0064]
GOBS -0.0032 [-0.0137, 0.0072]
IMH -0.0117 [-0.0382, 0.0148]
UNICAMP -0.0085 [-0.0431, 0.0262]
Meth-CT -0.0101 [-0.0350, 0.0148]
MIRECC
UKBB nonEUR 0.0071 [-0.0152, 0.0293]
OSAKA 0.0020 [-0.0137, 0.0177]
PING Non European -0.0007 [-0.0179, 0.0164]
Non European Cohorts [N=2391; P=0.3822] -0.0031 [-0.0102, 0.0040]
-0.0600
-0.0400
-0.0200
0.0000
0.0200
0.0400
mm/effect allele
Beta [Lower 95% CI,Upper 95% CI]

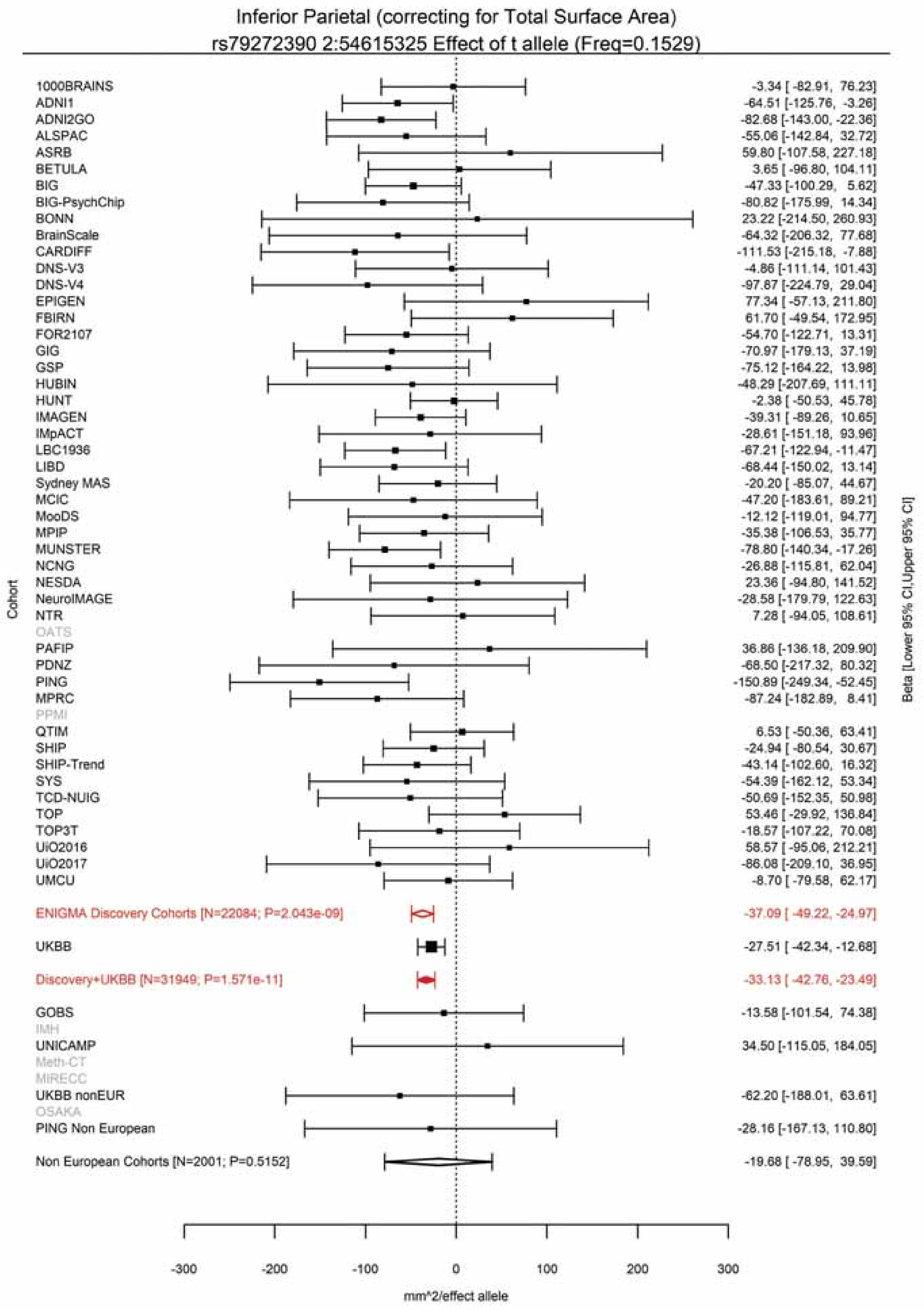

Inferior Parietal (correcting for Total Surface Area)
rs79272390 2:54615325 Effect of t allele (Freq=0.1529)
Cohort
1000BRAINS -3.34 [ -82.91, 76.23]
ADNI1 -64.51 [-125.76, -3.26]
ADNI2GO -82.68 [-143.00, -22.36]
ALSPAC -55.06 [-142.84, 32.72]
ASRB 59.80 [-107.58, 227.18]
BETULA 3.65 [ -96.80, 104.11]
BIG -47.33 [-100.29, 5.62]
BIG-PsychChip -80.82 [-175.99, 14.34]
BONN 23.22 [-214.50, 260.93]
BrainScale -64.32 [-206.32, 77.68]
CARDIFF -111.53 [-215.18, -7.88]
DNS-V3 -4.86 [-111.14, 101.43]
DNS-V4 -97.87 [-224.79, 29.04]
EPIGEN 77.34 [ -57.13, 211.80]
FBIRN 61.70 [ -49.54, 172.95]
FOR2107 -54.70 [-122.71, 13.31]
GIG -70.97 [-179.13, 37.19]
GSP -75.12 [-164.22, 13.98]
HUBIN -48.29 [-207.69, 111.11]
HUNT -2.38 [ -50.53, 45.78]
IMAGEN -39.31 [ -89.26, 10.65]
IMpACT -28.61 [-151.18, 93.96]
LBC1936 -67.21 [-122.94, -11.47]
LIBD -68.44 [-150.02, 13.14]
Sydney MAS -20.20 [ -85.07, 44.67]
MCIC -47.20 [-183.61, 89.21]
MooDS -12.12 [-119.01, 94.77]
MPIP -35.38 [-106.53, 35.77]
MUNSTER -78.80 [-140.34, -17.26]
NCNG -26.88 [-115.81, 62.04]
NESDA 23.36 [ -94.80, 141.52]
NeuroIMAGE -28.58 [-179.79, 122.63]
NTR 7.28 [ -94.05, 108.61]
OATS
PAFIP 36.86 [-136.18, 209.90]
PDNZ -68.50 [-217.32, 80.32]
PING -150.89 [-249.34, -52.45]
MPRC -87.24 [-182.89, 8.41]
PPMI
QTIM 6.53 [ -50.36, 63.41]
SHIP -24.94 [ -80.54, 30.67]
SHIP-Trend -43.14 [-102.60, 16.32]
SYS -54.39 [-162.12, 53.34]
TCD-NUIG -50.69 [-152.35, 50.98]
TOP 53.46 [ -29.92, 136.84]
TOP3T -18.57 [-107.22, 70.08]
UiO2016 58.57 [ -95.06, 212.21]
UiO2017 -86.08 [-209.10, 36.95]
UMCU -8.70 [ -79.58, 62.17]
ENIGMA Discovery Cohorts [N=22084; P=2.043e-09] -37.09 [ -49.22, -24.97]
UKBB -27.51 [ -42.34, -12.68]
Discovery+UKBB [N=31949; P=1.571e-11] -33.13 [ -42.76, -23.49]
GOBS -13.58 [-101.54, 74.38]
IMH
UNICAMP 34.50 [-115.05, 184.05]
Meth-CT
MIRECC
UKBB nonEUR -62.20 [-188.01, 63.61]
OSAKA
PING Non European -28.16 [-167.13, 110.80]
Non European Cohorts [N=2001; P=0.5152] -19.68 [ -78.95, 39.59]
-300
-200
-100
0
100
200
300
mm^2/effect allele
Beta [Lower 95% CI,Upper 95% CI]

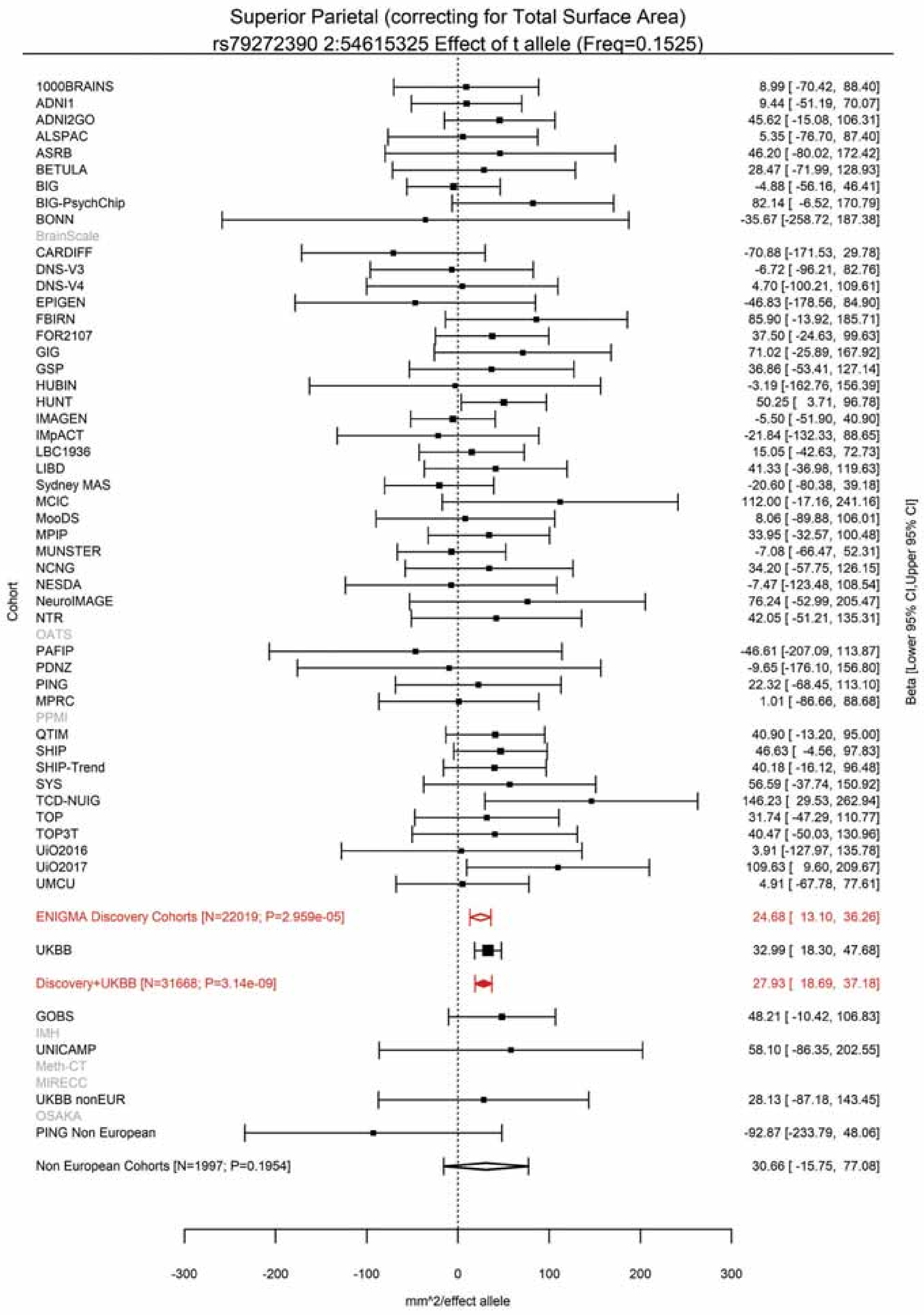

Superior Parietal (correcting for Total Surface Area)
rs79272390 2:54615325 Effect of t allele (Freq=0.1525)
Cohort
1000BRAINS 8.99 [ -70.42, 88.40]
ADNI1 9.44 [ -51.19, 70.07]
ADNI2GO 45.62 [ -15.08, 106.31]
ALSPAC 5.35 [ -76.70, 87.40]
ASRB 46.20 [ -80.02, 172.42]
BETULA 28.47 [ -71.99, 128.93]
BIG -4.88 [ -56.16, 46.41]
BIG-PsychChip 82.14 [ -6.52, 170.79]
BONN -35.67 [-258.72, 187.38]
BrainScale
CARDIFF -70.88 [-171.53, 29.78]
DNS-V3 -6.72 [ -96.21, 82.76]
DNS-V4 4.70 [-100.21, 109.61]
EPIGEN -46.83 [-178.56, 84.90]
FBIRN 85.90 [ -13.92, 185.71]
FOR2107 37.50 [ -24.63, 99.63]
GIG 71.02 [ -25.89, 167.92]
GSP 36.86 [ -53.41, 127.14]
HUBIN -3.19 [-162.76, 156.39]
HUNT 50.25 [ 3.71, 96.78]
IMAGEN -5.50 [ -51.90, 40.90]
IMpACT -21.84 [-132.33, 88.65]
LBC1936 15.05 [ -42.63, 72.73]
LIBD 41.33 [ -36.98, 119.63]
Sydney MAS -20.60 [ -80.38, 39.18]
MCIC 112.00 [ -17.16, 241.16]
MooDS 8.06 [ -89.88, 106.01]
MPIP 33.95 [ -32.57, 100.48]
MUNSTER -7.08 [ -66.47, 52.31]
NCNG 34.20 [ -57.75, 126.15]
NESDA -7.47 [-123.48, 108.54]
NeuroIMAGE 76.24 [ -52.99, 205.47]
NTR 42.05 [ -51.21, 135.31]
OATS
PAFIP -46.61 [-207.09, 113.87]
PDNZ -9.65 [-176.10, 156.80]
PING 22.32 [ -68.45, 113.10]
MPRC 1.01 [ -86.66, 88.68]
PPMI
QTIM 40.90 [ -13.20, 95.00]
SHIP 46.63 [ -4.56, 97.83]
SHIP-Trend 40.18 [ -16.12, 96.48]
SYS 56.59 [ -37.74, 150.92]
TCD-NUIG 146.23 [ 29.53, 262.94]
TOP 31.74 [ -47.29, 110.77]
TOP3T 40.47 [ -50.03, 130.96]
UiO2016 3.91 [-127.97, 135.78]
UiO2017 109.63 [ 9.60, 209.67]
UMCU 4.91 [ -67.78, 77.61]
ENIGMA Discovery Cohorts [N=22019; P=2.959e-05] 24.68 [ 13.10, 36.26]
UKBB 32.99 [ 18.30, 47.68]
Discovery+UKBB [N=31668; P=3.14e-09] 27.93 [ 18.69, 37.18]
GOBS 48.21 [ -10.42, 106.83]
IMH
UNICAMP 58.10 [ -86.35, 202.55]
Meth-CT
MIRECC
UKBB nonEUR 28.13 [ -87.18, 143.45]
OSAKA
PING Non European -92.87 [-233.79, 48.06]
Non European Cohorts [N=1997; P=0.1954] 30.66 [ -15.75, 77.08]
Beta [Lower 95% CI,Upper 95% CI]
-300
-200
-100
0
100
200
300
mm^2/effect allele

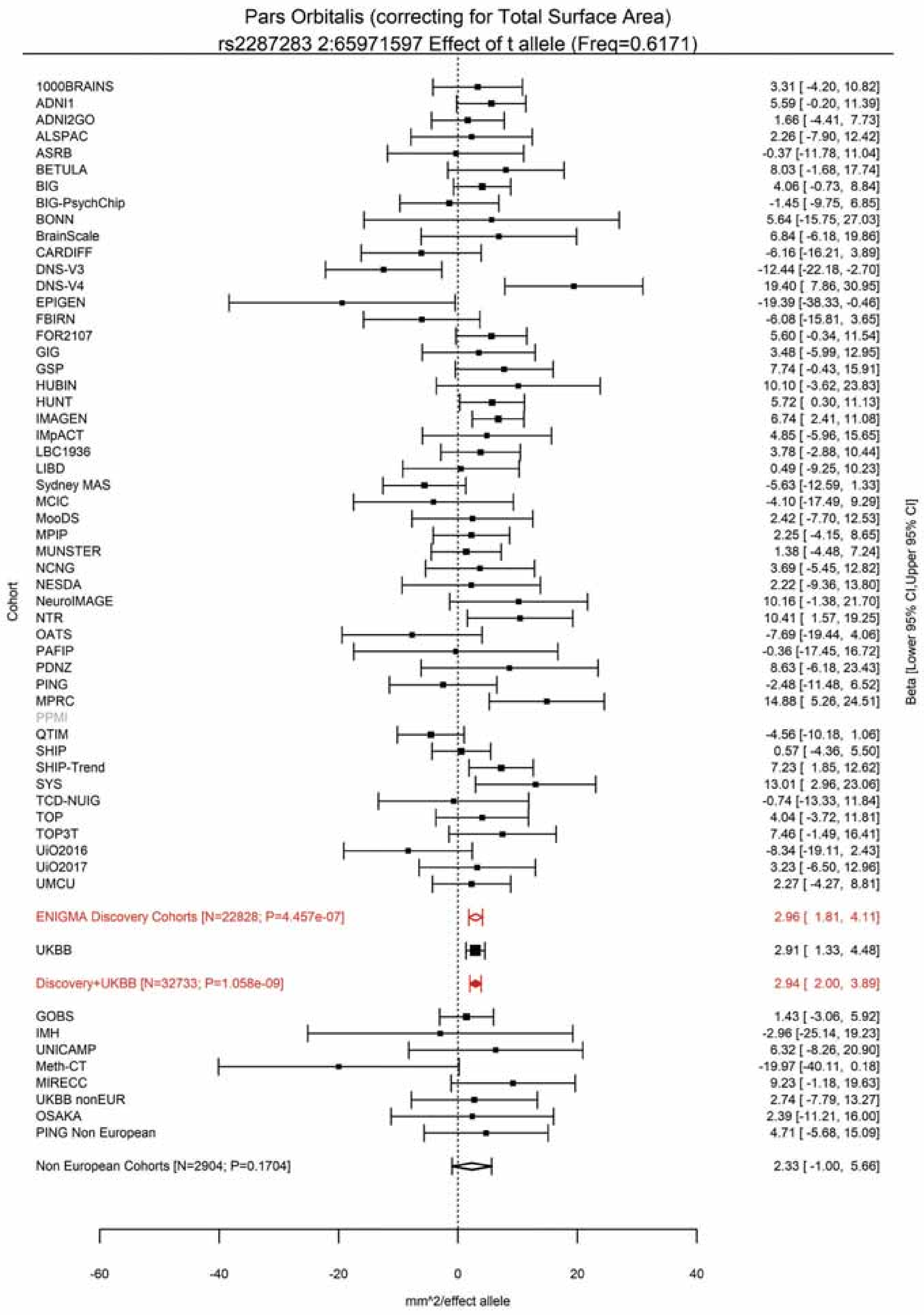
Pars Orbitalis (correcting for Total Surface Area)
rs2287283 2:65971597 Effect of t allele (Freq=0.6171)
Cohort
Beta [Lower 95% CI,Upper 95% CI]
1000BRAINS 3.31 [ -4.20, 10.82]
ADNI1 5.59 [ -0.20, 11.39]
ADNI2GO 1.66 [ -4.41, 7.73]
ALSPAC 2.26 [ -7.90, 12.42]
ASRB -0.37 [-11.78, 11.04]
BETULA 8.03 [ -1.68, 17.74]
BIG 4.06 [ -0.73, 8.84]
BIG-PsychChip -1.45 [ -9.75, 6.85]
BONN 5.64 [-15.75, 27.03]
BrainScale 6.84 [ -6.18, 19.86]
CARDIFF -6.16 [-16.21, 3.89]
DNS-V3 -12.44 [-22.18, -2.70]
DNS-V4 19.40 [ 7.86, 30.95]
EPIGEN -19.39 [-38.33, -0.46]
FBIRN -6.08 [-15.81, 3.65]
FOR2107 5.60 [ -0.34, 11.54]
GIG 3.48 [ -5.99, 12.95]
GSP 7.74 [ -0.43, 15.91]
HUBIN 10.10 [ -3.62, 23.83]
HUNT 5.72 [ 0.30, 11.13]
IMAGEN 6.74 [ 2.41, 11.08]
IMpACT 4.85 [ -5.96, 15.65]
LBC1936 3.78 [ -2.88, 10.44]
LIBD 0.49 [ -9.25, 10.23]
Sydney MAS -5.63 [-12.59, 1.33]
MCIC -4.10 [-17.49, 9.29]
MooDS 2.42 [ -7.70, 12.53]
MPIP 2.25 [ -4.15, 8.65]
MUNSTER 1.38 [ -4.48, 7.24]
NCNG 3.69 [ -5.45, 12.82]
NESDA 2.22 [ -9.36, 13.80]
NeuroIMAGE 10.16 [ -1.38, 21.70]
NTR 10.41 [ 1.57, 19.25]
OATS -7.69 [-19.44, 4.06]
PAFIP -0.36 [-17.45, 16.72]
PDNZ 8.63 [ -6.18, 23.43]
PING -2.48 [-11.48, 6.52]
MPRC 14.88 [ 5.26, 24.51]
PPMI
QTIM -4.56 [-10.18, 1.06]
SHIP 0.57 [ -4.36, 5.50]
SHIP-Trend 7.23 [ 1.85, 12.62]
SYS 13.01 [ 2.96, 23.06]
TCD-NUIG -0.74 [-13.33, 11.84]
TOP 4.04 [ -3.72, 11.81]
TOP3T 7.46 [ -1.49, 16.41]
UiO2016 -8.34 [-19.11, 2.43]
UiO2017 3.23 [ -6.50, 12.96]
UMCU 2.27 [ -4.27, 8.81]
ENIGMA Discovery Cohorts [N=22828; P=4.457e-07] 2.96 [ 1.81, 4.11]
UKBB 2.91 [ 1.33, 4.48]
Discovery+UKBB [N=32733; P=1.058e-09] 2.94 [ 2.00, 3.89]
GOBS 1.43 [ -3.06, 5.92]
IMH -2.96 [-25.14, 19.23]
UNICAMP 6.32 [ -8.26, 20.90]
Meth-CT -19.97 [-40.11, 0.18]
MIRECC 9.23 [ -1.18, 19.63]
UKBB nonEUR 2.74 [ -7.79, 13.27]
OSAKA 2.39 [-11.21, 16.00]
PING Non European 4.71 [ -5.68, 15.09]
Non European Cohorts [N=2904; P=0.1704] 2.33 [ -1.00, 5.66]
-60
-40
-20
0
20
40
mm^2/effect allele

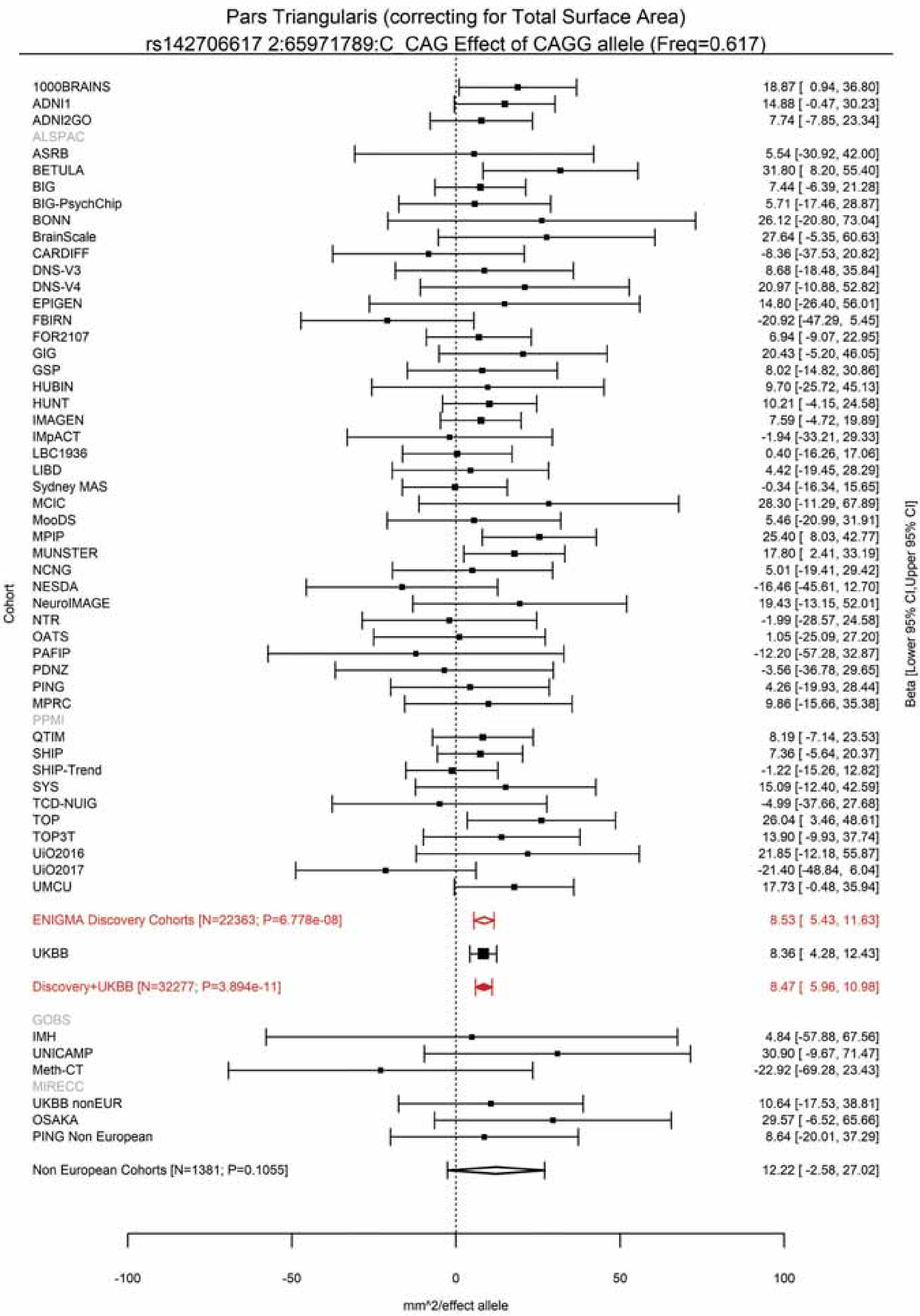
Pars Triangularis (correcting for Total Surface Area)
rs142706617 2:65971789:C_CAG Effect of CAGG allele (Freq=0.617)
Cohort
1000BRAINS 18.87 [ 0.94, 36.80]
ADNI1 14.88 [ -0.47, 30.23]
ADNI2GO 7.74 [ -7.85, 23.34]
ALSPAC
ASRB 5.54 [-30.92, 42.00]
BETULA 31.80 [ 8.20, 55.40]
BIG 7.44 [ -6.39, 21.28]
BIG-PsychChip 5.71 [-17.46, 28.87]
BONN 26.12 [-20.80, 73.04]
BrainScale 27.64 [ -5.35, 60.63]
CARDIFF -8.36 [-37.53, 20.82]
DNS-V3 8.68 [-18.48, 35.84]
DNS-V4 20.97 [-10.88, 52.82]
EPIGEN 14.80 [-26.40, 56.01]
FBIRN -20.92 [-47.29, 5.45]
FOR2107 6.94 [ -9.07, 22.95]
GIG 20.43 [ -5.20, 46.05]
GSP 8.02 [-14.82, 30.86]
HUBIN 9.70 [-25.72, 45.13]
HUNT 10.21 [ -4.15, 24.58]
IMAGEN 7.59 [ -4.72, 19.89]
IMpACT -1.94 [-33.21, 29.33]
LBC1936 0.40 [-16.26, 17.06]
LIBD 4.42 [-19.45, 28.29]
Sydney MAS -0.34 [-16.34, 15.65]
MCIC 28.30 [-11.29, 67.89]
MooDS 5.46 [-20.99, 31.91]
MPIP 25.40 [ 8.03, 42.77]
MUNSTER 17.80 [ 2.41, 33.19]
NCNG 5.01 [-19.41, 29.42]
NESDA -16.46 [-45.61, 12.70]
NeuroIMAGE 19.43 [-13.15, 52.01]
NTR -1.99 [-28.57, 24.58]
OATS 1.05 [-25.09, 27.20]
PAFIP -12.20 [-57.28, 32.87]
PDNZ -3.56 [-36.78, 29.65]
PING 4.26 [-19.93, 28.44]
MPRC 9.86 [-15.66, 35.38]
PPMI
QTIM 8.19 [ -7.14, 23.53]
SHIP 7.36 [ -5.64, 20.37]
SHIP-Trend -1.22 [-15.26, 12.82]
SYS 15.09 [-12.40, 42.59]
TCD-NUIG -4.99 [-37.66, 27.68]
TOP 26.04 [ 3.46, 48.61]
TOP3T 13.90 [ -9.93, 37.74]
UiO2016 21.85 [-12.18, 55.87]
UiO2017 -21.40 [-48.84, 6.04]
UMCU 17.73 [ -0.48, 35.94]
ENIGMA Discovery Cohorts [N=22363; P=6.778e-08] 8.53 [ 5.43, 11.63]
UKBB 8.36 [ 4.28, 12.43]
Discovery+UKBB [N=32277; P=3.894e-11] 8.47 [ 5.96, 10.98]
GOBS
IMH 4.84 [-57.88, 67.56]
UNICAMP 30.90 [ -9.67, 71.47]
Meth-CT -22.92 [-69.28, 23.43]
MIRECC
UKBB nonEUR 10.64 [-17.53, 38.81]
OSAKA 29.57 [ -6.52, 65.66]
PING Non European 8.64 [-20.01, 37.29]
Non European Cohorts [N=1381; P=0.1055] 12.22 [ -2.58, 27.02]
Beta [Lower 95% CI,Upper 95% CI]
-100
-50
0
50
100
mm^2/effect allele

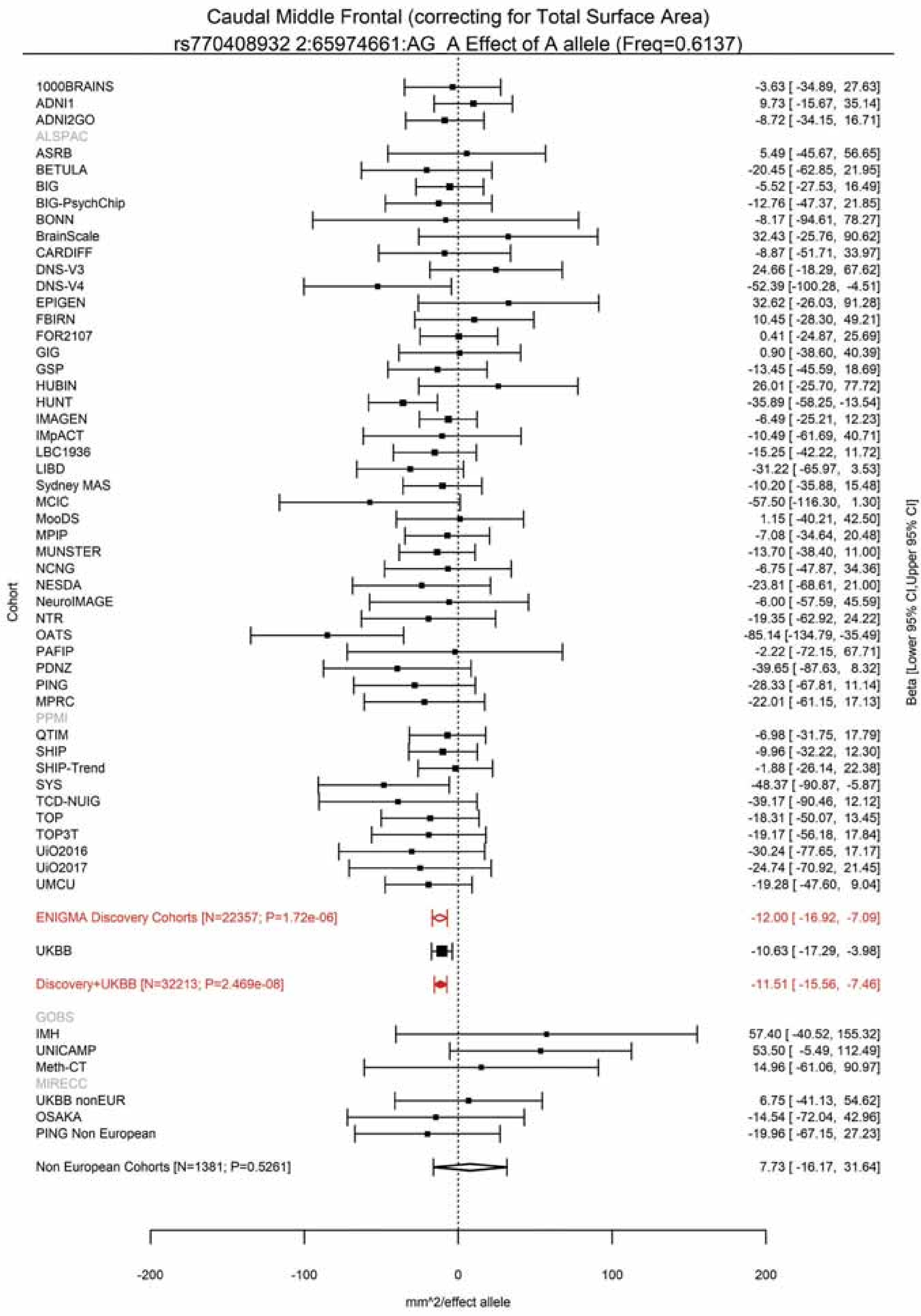
Caudal Middle Frontal (correcting for Total Surface Area)
rs770408932 2:65974661:AG A Effect of A allele (Freq=0.6137)
Cohort
Beta [Lower 95% CI,Upper 95% CI]
1000BRAINS -3.63 [ -34.89, 27.63]
ADNI1 9.73 [ -15.67, 35.14]
ADNI2GO -8.72 [ -34.15, 16.71]
ALSPAC
ASRB 5.49 [ -45.67, 56.65]
BETULA -20.45 [ -62.85, 21.95]
BIG -5.52 [ -27.53, 16.49]
BIG-PsychChip -12.76 [ -47.37, 21.85]
BONN -8.17 [ -94.61, 78.27]
BrainScale 32.43 [ -25.76, 90.62]
CARDIFF -8.87 [ -51.71, 33.97]
DNS-V3 24.66 [ -18.29, 67.62]
DNS-V4 -52.39 [-100.28, -4.51]
EPIGEN 32.62 [ -26.03, 91.28]
FBIRN 10.45 [ -28.30, 49.21]
FOR2107 0.41 [ -24.87, 25.69]
GIG 0.90 [ -38.60, 40.39]
GSP -13.45 [ -45.59, 18.69]
HUBIN 26.01 [ -25.70, 77.72]
HUNT -35.89 [ -58.25, -13.54]
IMAGEN -6.49 [ -25.21, 12.23]
IMpACT -10.49 [ -61.69, 40.71]
LBC1936 -15.25 [ -42.22, 11.72]
LIBD -31.22 [ -65.97, 3.53]
Sydney MAS -10.20 [ -35.88, 15.48]
MCIC -57.50 [-116.30, 1.30]
MooDS 1.15 [ -40.21, 42.50]
MPIP -7.08 [ -34.64, 20.48]
MUNSTER -13.70 [ -38.40, 11.00]
NCNG -6.75 [ -47.87, 34.36]
NESDA -23.81 [ -68.61, 21.00]
NeuroIMAGE -6.00 [ -57.59, 45.59]
NTR -19.35 [ -62.92, 24.22]
OATS -85.14 [-134.79, -35.49]
PAFIP -2.22 [ -72.15, 67.71]
PDNZ -39.65 [ -87.63, 8.32]
PING -28.33 [ -67.81, 11.14]
MPRC -22.01 [ -61.15, 17.13]
PPMI
QTIM -6.98 [ -31.75, 17.79]
SHIP -9.96 [ -32.22, 12.30]
SHIP-Trend -1.88 [ -26.14, 22.38]
SYS -48.37 [ -90.87, -5.87]
TCD-NUIG -39.17 [ -90.46, 12.12]
TOP -18.31 [ -50.07, 13.45]
TOP3T -19.17 [ -56.18, 17.84]
UiO2016 -30.24 [ -77.65, 17.17]
UiO2017 -24.74 [ -70.92, 21.45]
UMCU -19.28 [ -47.60, 9.04]
ENIGMA Discovery Cohorts [N=22357; P=1.72e-06] -12.00 [ -16.92, -7.09]
UKBB -10.63 [ -17.29, -3.98]
Discovery+UKBB [N=32213; P=2.469e-08] -11.51 [ -15.56, -7.46]
GOBS
IMH 57.40 [ -40.52, 155.32]
UNICAMP 53.50 [ -5.49, 112.49]
Meth-CT 14.96 [ -61.06, 90.97]
MIRECC
UKBB nonEUR 6.75 [ -41.13, 54.62]
OSAKA -14.54 [ -72.04, 42.96]
PING Non European -19.96 [ -67.15, 27.23]
Non European Cohorts [N=1381; P=0.5261] 7.73 [ -16.17, 31.64]
-200
-100
0
100
200
mm^2/effect allele

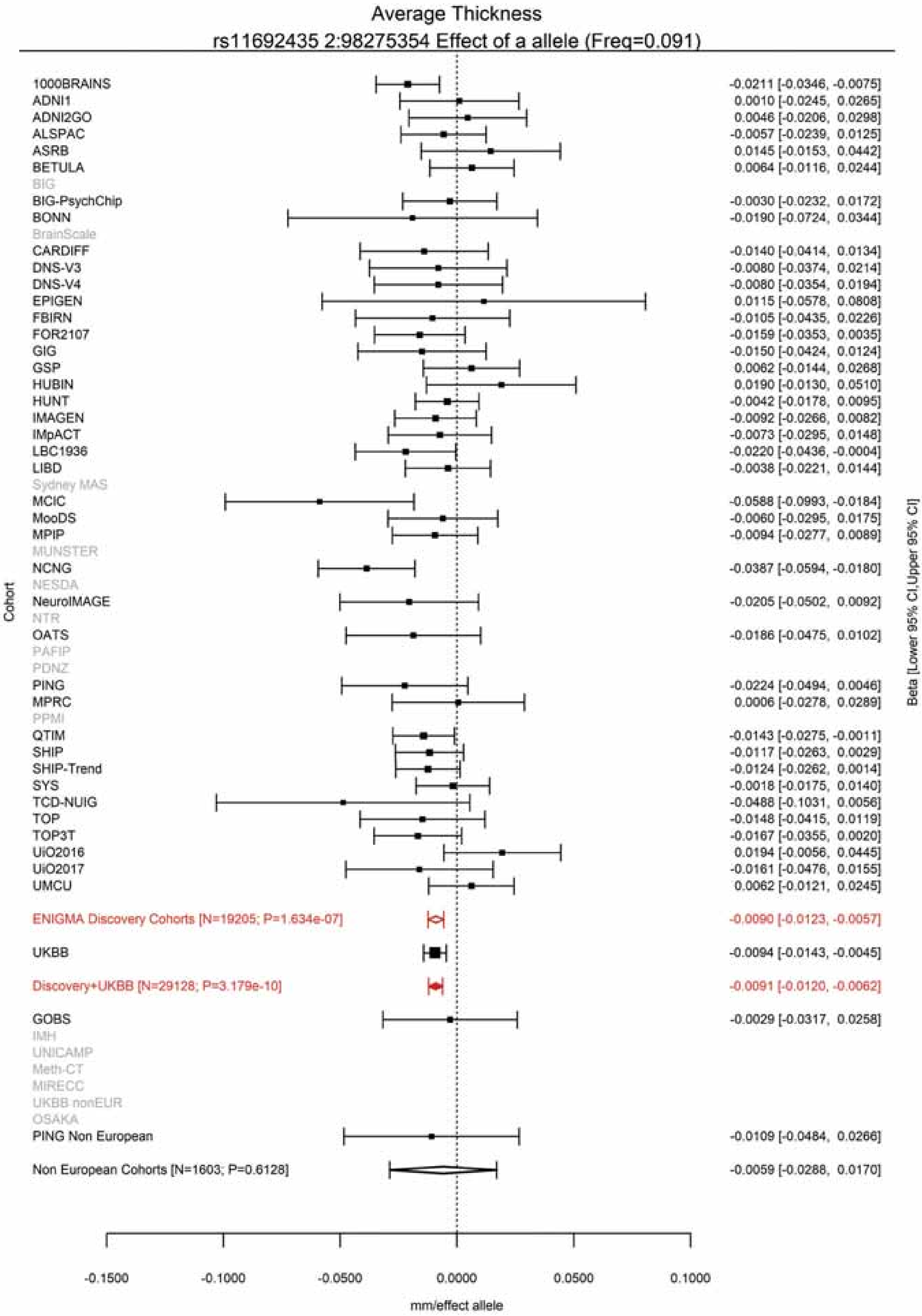

Average Thickness
rs11692435 2:98275354 Effect of a allele (Freq=0.091)
1000BRAINS -0.0211 [-0.0346, -0.0075]
ADNI1 0.0010 [-0.0245, 0.0265]
ADNI2GO 0.0046 [-0.0206, 0.0298]
ALSPAC -0.0057 [-0.0239, 0.0125]
ASRB 0.0145 [-0.0153, 0.0442]
BETULA 0.0064 [-0.0116, 0.0244]
BIG
BIG-PsychChip -0.0030 [-0.0232, 0.0172]
BONN -0.0190 [-0.0724, 0.0344]
BrainScale
CARDIFF -0.0140 [-0.0414, 0.0134]
DNS-V3 -0.0080 [-0.0374, 0.0214]
DNS-V4 -0.0080 [-0.0354, 0.0194]
EPIGEN 0.0115 [-0.0578, 0.0808]
FBIRN -0.0105 [-0.0435, 0.0226]
FOR2107 -0.0159 [-0.0353, 0.0035]
GIG -0.0150 [-0.0424, 0.0124]
GSP 0.0062 [-0.0144, 0.0268]
HUBIN 0.0190 [-0.0130, 0.0510]
HUNT -0.0042 [-0.0178, 0.0095]
IMAGEN -0.0092 [-0.0266, 0.0082]
IMpACT -0.0073 [-0.0295, 0.0148]
LBC1936 -0.0220 [-0.0436, -0.0004]
LIBD -0.0038 [-0.0221, 0.0144]
Sydney MAS
MCIC -0.0588 [-0.0993, -0.0184]
MooDS -0.0060 [-0.0295, 0.0175]
MPIP -0.0094 [-0.0277, 0.0089]
MUNSTER
NCNG -0.0387 [-0.0594, -0.0180]
NESDA
NeuroIMAGE -0.0205 [-0.0502, 0.0092]
NTR
OATS -0.0186 [-0.0475, 0.0102]
PAFIP
PDNZ
PING -0.0224 [-0.0494, 0.0046]
MPRC 0.0006 [-0.0278, 0.0289]
PPMI
QTIM -0.0143 [-0.0275, -0.0011]
SHIP -0.0117 [-0.0263, 0.0029]
SHIP-Trend -0.0124 [-0.0262, 0.0014]
SYS -0.0018 [-0.0175, 0.0140]
TCD-NUIG -0.0488 [-0.1031, 0.0056]
TOP -0.0148 [-0.0415, 0.0119]
TOP3T -0.0167 [-0.0355, 0.0020]
UiO2016 0.0194 [-0.0056, 0.0445]
UiO2017 -0.0161 [-0.0476, 0.0155]
UMCU 0.0062 [-0.0121, 0.0245]
ENIGMA Discovery Cohorts [N=19205; P=1.634e-07] -0.0090 [-0.0123, -0.0057]
UKBB -0.0094 [-0.0143, -0.0045]
Discovery+UKBB [N=29128; P=3.179e-10] -0.0091 [-0.0120, -0.0062]
GOBS -0.0029 [-0.0317, 0.0258]
IMH
UNICAMP
Meth-CT
MIRECC
UKBB nonEUR
OSAKA
PING Non European -0.0109 [-0.0484, 0.0266]
Non European Cohorts [N=1603; P=0.6128] -0.0059 [-0.0288, 0.0170]
Cohort
Beta [Lower 95% CI,Upper 95% CI]
-0.1500
-0.1000
-0.0500
0.0000
0.0500
0.1000
mm/effect allele

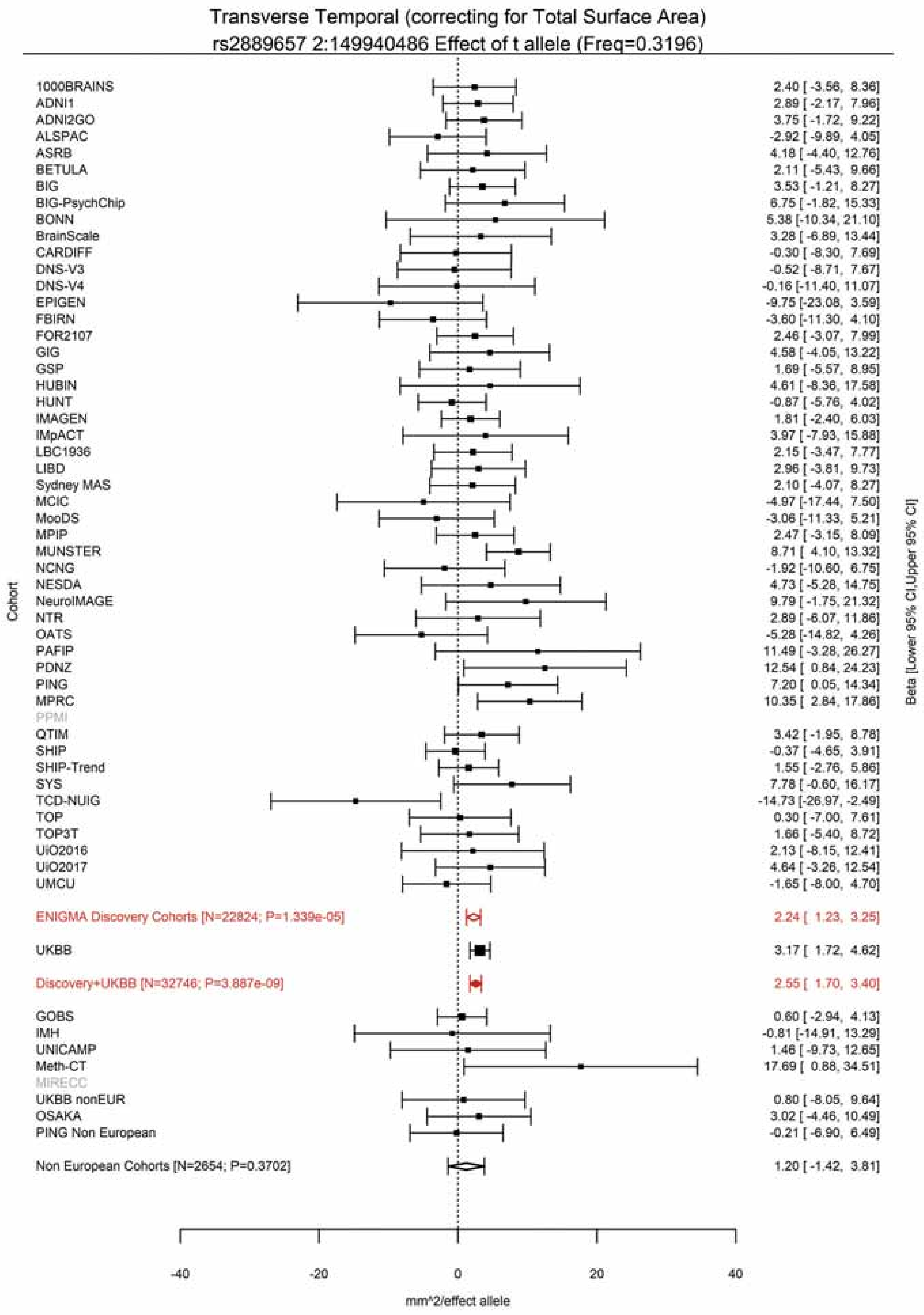

Transverse Temporal (correcting for Total Surface Area)
rs2889657 2:149940486 Effect of t allele (Freq=0.3196)
Cohort
1000BRAINS 2.40 [ -3.56, 8.36]
ADNI1 2.89 [ -2.17, 7.96]
ADNI2GO 3.75 [ -1.72, 9.22]
ALSPAC -2.92 [ -9.89, 4.05]
ASRB 4.18 [ -4.40, 12.76]
BETULA 2.11 [ -5.43, 9.66]
BIG 3.53 [ -1.21, 8.27]
BIG-PsychChip 6.75 [ -1.82, 15.33]
BONN 5.38 [-10.34, 21.10]
BrainScale 3.28 [ -6.89, 13.44]
CARDIFF -0.30 [ -8.30, 7.69]
DNS-V3 -0.52 [ -8.71, 7.67]
DNS-V4 -0.16 [-11.40, 11.07]
EPIGEN -9.75 [-23.08, 3.59]
FBIRN -3.60 [-11.30, 4.10]
FOR2107 2.46 [ -3.07, 7.99]
GIG 4.58 [ -4.05, 13.22]
GSP 1.69 [ -5.57, 8.95]
HUBIN 4.61 [ -8.36, 17.58]
HUNT -0.87 [ -5.76, 4.02]
IMAGEN 1.81 [ -2.40, 6.03]
IMpACT 3.97 [ -7.93, 15.88]
LBC1936 2.15 [ -3.47, 7.77]
LIBD 2.96 [ -3.81, 9.73]
Sydney MAS 2.10 [ -4.07, 8.27]
MCIC -4.97 [-17.44, 7.50]
MooDS -3.06 [-11.33, 5.21]
MPIP 2.47 [ -3.15, 8.09]
MUNSTER 8.71 [ 4.10, 13.32]
NCNG -1.92 [-10.60, 6.75]
NESDA 4.73 [ -5.28, 14.75]
NeuroIMAGE 9.79 [ -1.75, 21.32]
NTR 2.89 [ -6.07, 11.86]
OATS -5.28 [-14.82, 4.26]
PAFIP 11.49 [ -3.28, 26.27]
PDNZ 12.54 [ 0.84, 24.23]
PING 7.20 [ 0.05, 14.34]
MPRC 10.35 [ 2.84, 17.86]
PPMI
QTIM 3.42 [ -1.95, 8.78]
SHIP -0.37 [ -4.65, 3.91]
SHIP-Trend 1.55 [ -2.76, 5.86]
SYS 7.78 [ -0.60, 16.17]
TCD-NUIG -14.73 [-26.97, -2.49]
TOP 0.30 [ -7.00, 7.61]
TOP3T 1.66 [ -5.40, 8.72]
UiO2016 2.13 [ -8.15, 12.41]
UiO2017 4.64 [ -3.26, 12.54]
UMCU -1.65 [ -8.00, 4.70]
ENIGMA Discovery Cohorts [N=22824; P=1.339e-05] 2.24 [ 1.23, 3.25]
UKBB 3.17 [ 1.72, 4.62]
Discovery+UKBB [N=32746; P=3.887e-09] 2.55 [ 1.70, 3.40]
GOBS 0.60 [ -2.94, 4.13]
IMH -0.81 [-14.91, 13.29]
UNICAMP 1.46 [ -9.73, 12.65]
Meth-CT 17.69 [ 0.88, 34.51]
MIRECC
UKBB nonEUR 0.80 [ -8.05, 9.64]
OSAKA 3.02 [ -4.46, 10.49]
PING Non European -0.21 [ -6.90, 6.49]
Non European Cohorts [N=2654; P=0.3702] 1.20 [ -1.42, 3.81]
Beta [Lower 95% CI,Upper 95% CI]
-40
-20
0
20
40
mm^2/effect allele

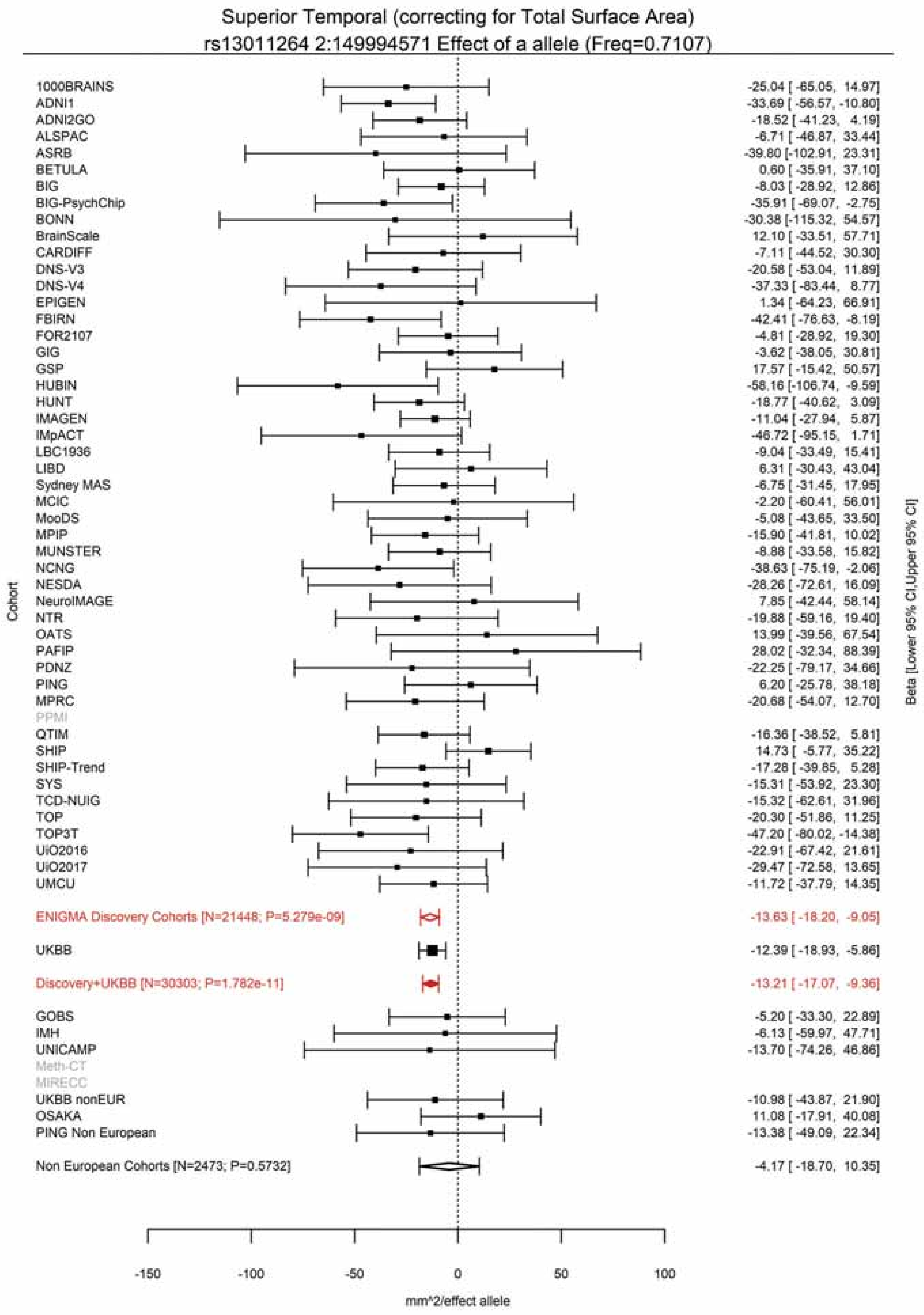
Superior Temporal (correcting for Total Surface Area)
rs13011264 2:149994571 Effect of a allele (Freq=0.7107)
Cohort
1000BRAINS -25.04 [ -65.05, 14.97]
ADNI1 -33.69 [ -56.57, -10.80]
ADNI2GO -18.52 [ -41.23, 4.19]
ALSPAC -6.71 [ -46.87, 33.44]
ASRB -39.80 [-102.91, 23.31]
BETULA 0.60 [ -35.91, 37.10]
BIG -8.03 [ -28.92, 12.86]
BIG-PsychChip -35.91 [ -69.07, -2.75]
BONN -30.38 [-115.32, 54.57]
BrainScale 12.10 [ -33.51, 57.71]
CARDIFF -7.11 [ -44.52, 30.30]
DNS-V3 -20.58 [ -53.04, 11.89]
DNS-V4 -37.33 [ -83.44, 8.77]
EPIGEN 1.34 [ -64.23, 66.91]
FBIRN -42.41 [ -76.63, -8.19]
FOR2107 -4.81 [ -28.92, 19.30]
GIG -3.62 [ -38.05, 30.81]
GSP 17.57 [ -15.42, 50.57]
HUBIN -58.16 [-106.74, -9.59]
HUNT -18.77 [ -40.62, 3.09]
IMAGEN -11.04 [ -27.94, 5.87]
IMpACT -46.72 [ -95.15, 1.71]
LBC1936 -9.04 [ -33.49, 15.41]
LIBD 6.31 [ -30.43, 43.04]
Sydney MAS -6.75 [ -31.45, 17.95]
MCIC -2.20 [ -60.41, 56.01]
MooDS -5.08 [ -43.65, 33.50]
MPIP -15.90 [ -41.81, 10.02]
MUNSTER -8.88 [ -33.58, 15.82]
NCNG -38.63 [ -75.19, -2.06]
NESDA -28.26 [ -72.61, 16.09]
NeuroIMAGE 7.85 [ -42.44, 58.14]
NTR -19.88 [ -59.16, 19.40]
OATS 13.99 [ -39.56, 67.54]
PAFIP 28.02 [ -32.34, 88.39]
PDNZ -22.25 [ -79.17, 34.66]
PING 6.20 [ -25.78, 38.18]
MPRC -20.68 [ -54.07, 12.70]
PPMI
QTIM -16.36 [ -38.52, 5.81]
SHIP 14.73 [ -5.77, 35.22]
SHIP-Trend -17.28 [ -39.85, 5.28]
SYS -15.31 [ -53.92, 23.30]
TCD-NUIG -15.32 [ -62.61, 31.96]
TOP -20.30 [ -51.86, 11.25]
TOP3T -47.20 [ -80.02, -14.38]
UiO2016 -22.91 [ -67.42, 21.61]
UiO2017 -29.47 [ -72.58, 13.65]
UMCU -11.72 [ -37.79, 14.35]
ENIGMA Discovery Cohorts [N=21448; P=5.279e-09] -13.63 [ -18.20, -9.05]
UKBB -12.39 [ -18.93, -5.86]
Discovery+UKBB [N=30303; P=1.782e-11] -13.21 [ -17.07, -9.36]
GOBS -5.20 [ -33.30, 22.89]
IMH -6.13 [ -59.97, 47.71]
UNICAMP -13.70 [ -74.26, 46.86]
Meth-CT
MIRECC
UKBB nonEUR -10.98 [ -43.87, 21.90]
OSAKA 11.08 [ -17.91, 40.08]
PING Non European -13.38 [ -49.09, 22.34]
Non European Cohorts [N=2473; P=0.5732] -4.17 [ -18.70, 10.35]
Beta [Lower 95% CI,Upper 95% CI]
-150 -100 -50 0 50 100
mm^2/effect allele

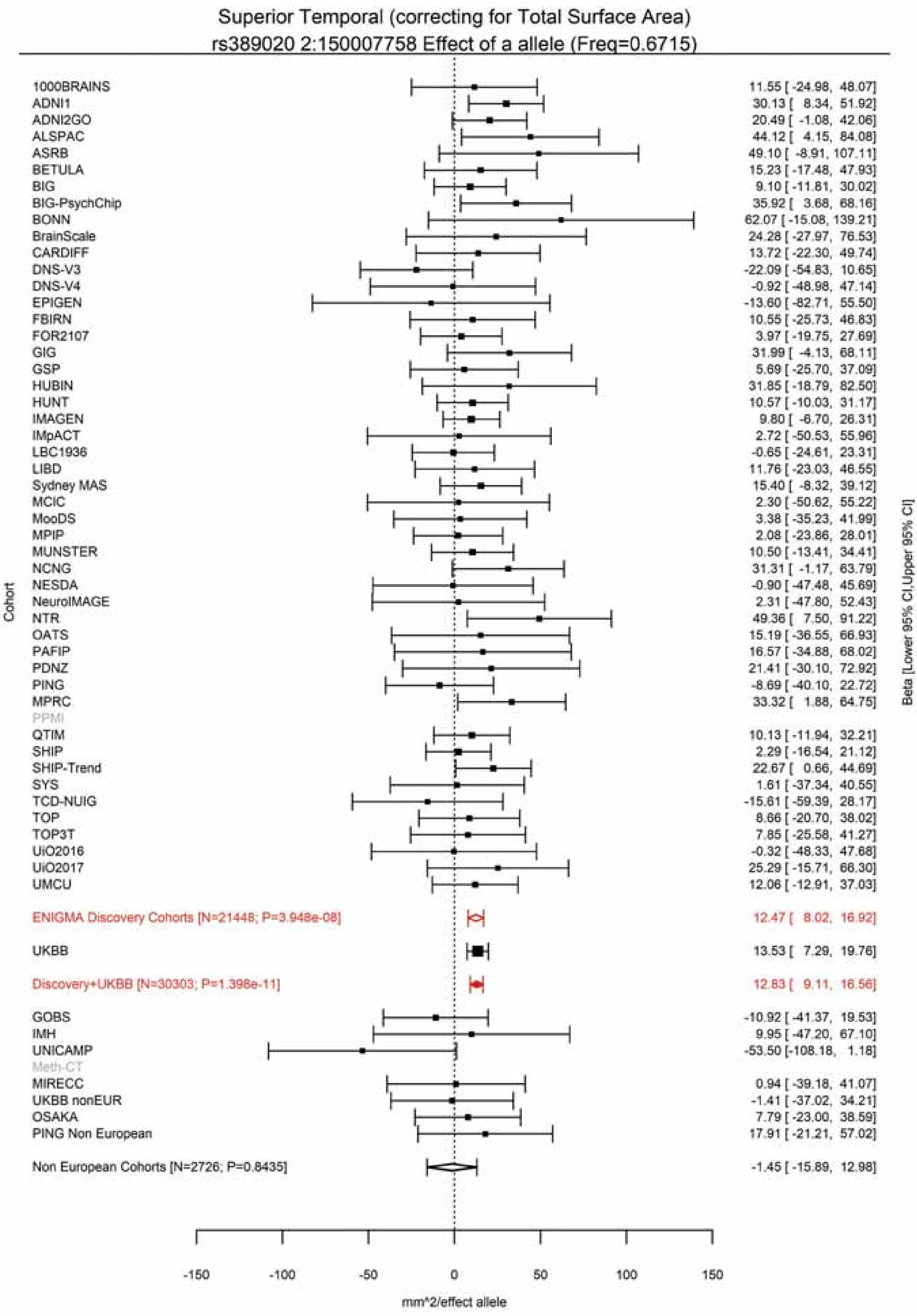
Superior Temporal (correcting for Total Surface Area)
rs389020 2:150007758 Effect of a allele (Freq=0.6715)
Cohort
Beta [Lower 95% CI,Upper 95% CI]
1000BRAINS 11.55 [ -24.98, 48.07]
ADNI1 30.13 [ 8.34, 51.92]
ADNI2GO 20.49 [ -1.08, 42.06]
ALSPAC 44.12 [ 4.15, 84.08]
ASRB 49.10 [ -8.91, 107.11]
BETULA 15.23 [ -17.48, 47.93]
BIG 9.10 [ -11.81, 30.02]
BIG-PsychChip 35.92 [ 3.68, 68.16]
BONN 62.07 [ -15.08, 139.21]
BrainScale 24.28 [ -27.97, 76.53]
CARDIFF 13.72 [ -22.30, 49.74]
DNS-V3 -22.09 [ -54.83, 10.65]
DNS-V4 -0.92 [ -48.98, 47.14]
EPIGEN -13.60 [ -82.71, 55.50]
FBIRN 10.55 [ -25.73, 46.83]
FOR2107 3.97 [ -19.75, 27.69]
GIG 31.99 [ -4.13, 68.11]
GSP 5.69 [ -25.70, 37.09]
HUBIN 31.85 [ -18.79, 82.50]
HUNT 10.57 [ -10.03, 31.17]
IMAGEN 9.80 [ -6.70, 26.31]
IMpACT 2.72 [ -50.53, 55.96]
LBC1936 -0.65 [ -24.61, 23.31]
LIBD 11.76 [ -23.03, 46.55]
Sydney MAS 15.40 [ -8.32, 39.12]
MCIC 2.30 [ -50.62, 55.22]
MooDS 3.38 [ -35.23, 41.99]
MPIP 2.08 [ -23.86, 28.01]
MUNSTER 10.50 [ -13.41, 34.41]
NCNG 31.31 [ -1.17, 63.79]
NESDA -0.90 [ -47.48, 45.69]
NeuroIMAGE 2.31 [ -47.80, 52.43]
NTR 49.36 [ 7.50, 91.22]
OATS 15.19 [ -36.55, 66.93]
PAFIP 16.57 [ -34.88, 68.02]
PDNZ 21.41 [ -30.10, 72.92]
PING -8.69 [ -40.10, 22.72]
MPRC 33.32 [ 1.88, 64.75]
PPMI
QTIM 10.13 [ -11.94, 32.21]
SHIP 2.29 [ -16.54, 21.12]
SHIP-Trend 22.67 [ 0.66, 44.69]
SYS 1.61 [ -37.34, 40.55]
TCD-NUIG -15.61 [ -59.39, 28.17]
TOP 8.66 [ -20.70, 38.02]
TOP3T 7.85 [ -25.58, 41.27]
UiO2016 -0.32 [ -48.33, 47.68]
UiO2017 25.29 [ -15.71, 66.30]
UMCU 12.06 [ -12.91, 37.03]
ENIGMA Discovery Cohorts [N=21448; P=3.948e-08] 12.47 [ 8.02, 16.92]
UKBB 13.53 [ 7.29, 19.76]
Discovery+UKBB [N=30303; P=1.398e-11] 12.83 [ 9.11, 16.56]
GOBS -10.92 [ -41.37, 19.53]
IMH 9.95 [ -47.20, 67.10]
UNICAMP -53.50 [ -108.18, 1.18]
Meth-CT
MIRECC 0.94 [ -39.18, 41.07]
UKBB nonEUR -1.41 [ -37.02, 34.21]
OSAKA 7.79 [ -23.00, 38.59]
PING Non European 17.91 [ -21.21, 57.02]
Non European Cohorts [N=2726; P=0.8435] -1.45 [ -15.89, 12.98]
-150
-100
-50
0
50
100
150
mm^2/effect allele

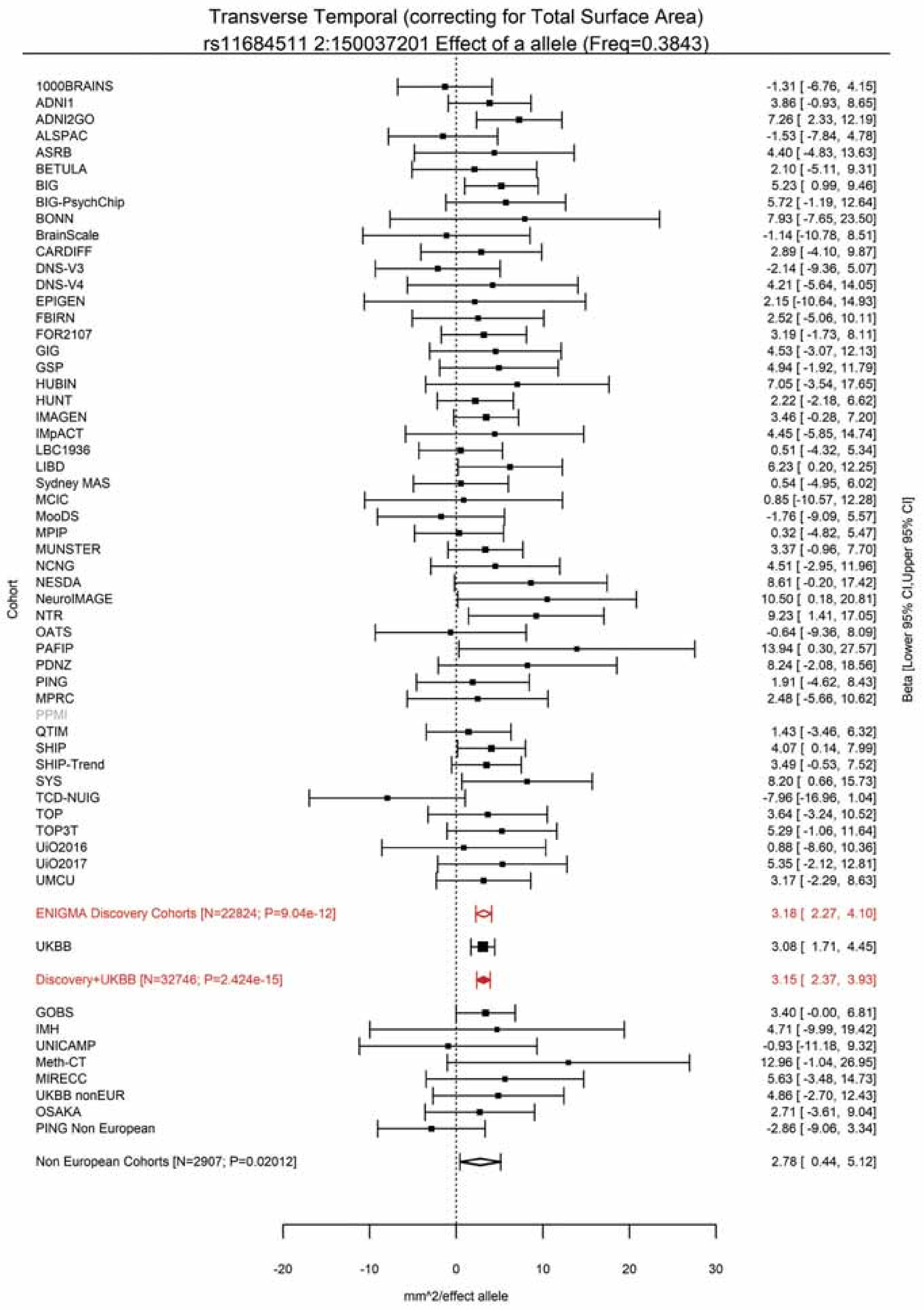

Transverse Temporal (correcting for Total Surface Area)
rs11684511 2:150037201 Effect of a allele (Freq=0.3843)
Cohort
1000BRAINS -1.31 [ -6.76, 4.15]
ADNI1 3.86 [ -0.93, 8.65]
ADNI2GO 7.26 [ 2.33, 12.19]
ALSPAC -1.53 [ -7.84, 4.78]
ASRB 4.40 [ -4.83, 13.63]
BETULA 2.10 [ -5.11, 9.31]
BIG 5.23 [ 0.99, 9.46]
BIG-PsychChip 5.72 [ -1.19, 12.64]
BONN 7.93 [ -7.65, 23.50]
BrainScale -1.14 [-10.78, 8.51]
CARDIFF 2.89 [ -4.10, 9.87]
DNS-V3 -2.14 [ -9.36, 5.07]
DNS-V4 4.21 [ -5.64, 14.05]
EPIGEN 2.15 [-10.64, 14.93]
FBIRN 2.52 [ -5.06, 10.11]
FOR2107 3.19 [ -1.73, 8.11]
GIG 4.53 [ -3.07, 12.13]
GSP 4.94 [ -1.92, 11.79]
HUBIN 7.05 [ -3.54, 17.65]
HUNT 2.22 [ -2.18, 6.62]
IMAGEN 3.46 [ -0.28, 7.20]
IMpACT 4.45 [ -5.85, 14.74]
LBC1936 0.51 [ -4.32, 5.34]
LIBD 6.23 [ 0.20, 12.25]
Sydney MAS 0.54 [ -4.95, 6.02]
MCIC 0.85 [-10.57, 12.28]
MooDS -1.76 [ -9.09, 5.57]
MPIP 0.32 [ -4.82, 5.47]
MUNSTER 3.37 [ -0.96, 7.70]
NCNG 4.51 [ -2.95, 11.96]
NESDA 8.61 [ -0.20, 17.42]
NeuroIMAGE 10.50 [ 0.18, 20.81]
NTR 9.23 [ 1.41, 17.05]
OATS -0.64 [ -9.36, 8.09]
PAFIP 13.94 [ 0.30, 27.57]
PDNZ 8.24 [ -2.08, 18.56]
PING 1.91 [ -4.62, 8.43]
MPRC 2.48 [ -5.66, 10.62]
PPMI
QTIM 1.43 [ -3.46, 6.32]
SHIP 4.07 [ 0.14, 7.99]
SHIP-Trend 3.49 [ -0.53, 7.52]
SYS 8.20 [ 0.66, 15.73]
TCD-NUIG -7.96 [-16.96, 1.04]
TOP 3.64 [ -3.24, 10.52]
TOP3T 5.29 [ -1.06, 11.64]
UiO2016 0.88 [ -8.60, 10.36]
UiO2017 5.35 [ -2.12, 12.81]
UMCU 3.17 [ -2.29, 8.63]
ENIGMA Discovery Cohorts [N=22824; P=9.04e-12] 3.18 [ 2.27, 4.10]
UKBB 3.08 [ 1.71, 4.45]
Discovery+UKBB [N=32746; P=2.424e-15] 3.15 [ 2.37, 3.93]
GOBS 3.40 [ -0.00, 6.81]
IMH 4.71 [ -9.99, 19.42]
UNICAMP -0.93 [-11.18, 9.32]
Meth-CT 12.96 [ -1.04, 26.95]
MIRECC 5.63 [ -3.48, 14.73]
UKBB nonEUR 4.86 [ -2.70, 12.43]
OSAKA 2.71 [ -3.61, 9.04]
PING Non European -2.86 [ -9.06, 3.34]
Non European Cohorts [N=2907; P=0.02012] 2.78 [ 0.44, 5.12]
Beta [Lower 95% CI,Upper 95% CI]
-20
-10
0
10
20
30
mm^2/effect allele

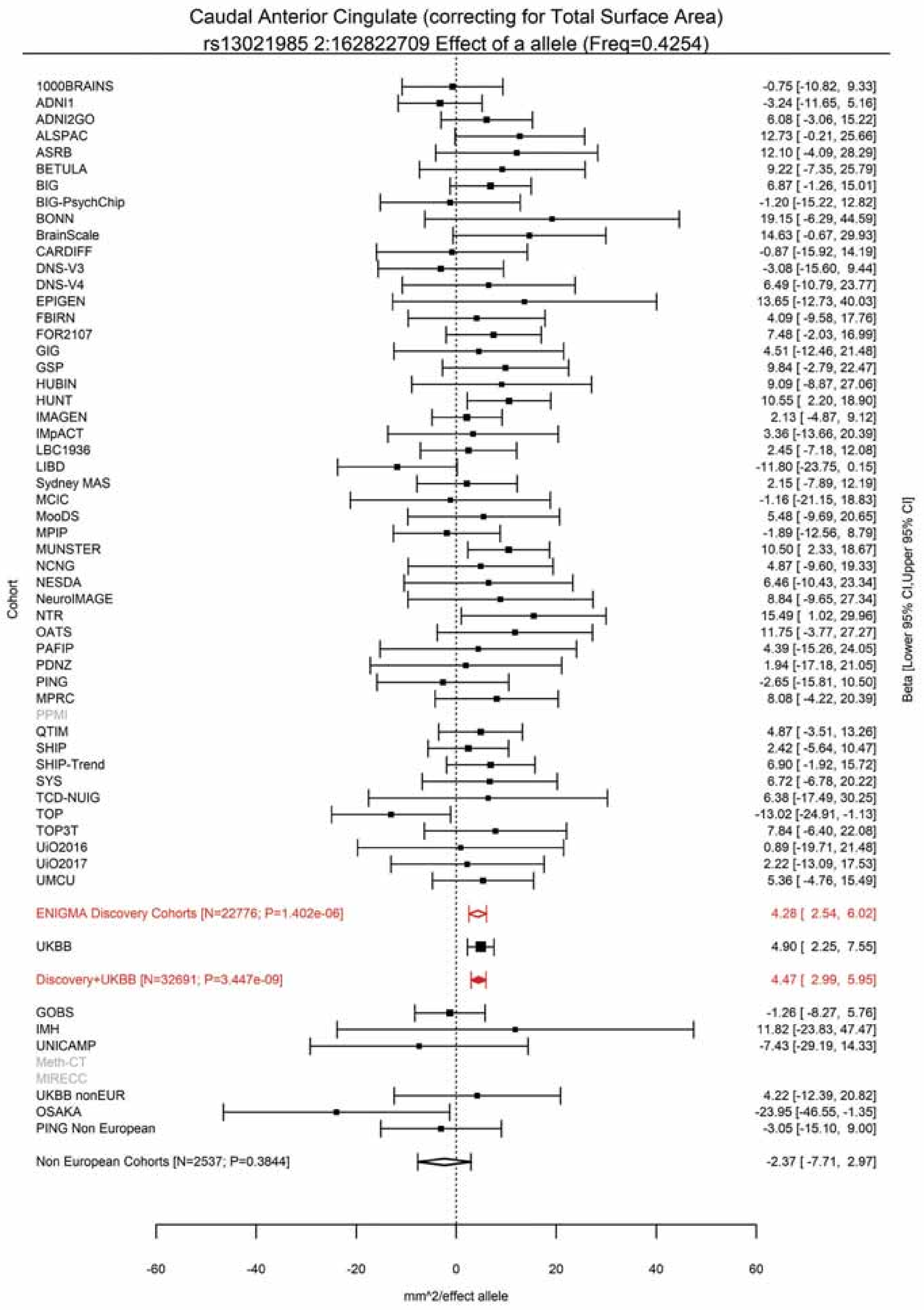

Caudal Anterior Cingulate (correcting for Total Surface Area)
rs13021985 2:162822709 Effect of a allele (Freq=0.4254)
1000BRAINS -0.75 [-10.82, 9.33]
ADNI1 -3.24 [-11.65, 5.16]
ADNI2GO 6.08 [ -3.06, 15.22]
ALSPAC 12.73 [ -0.21, 25.66]
ASRB 12.10 [ -4.09, 28.29]
BETULA 9.22 [ -7.35, 25.79]
BIG 6.87 [ -1.26, 15.01]
BIG-PsychChip -1.20 [-15.22, 12.82]
BONN 19.15 [ -6.29, 44.59]
BrainScale 14.63 [ -0.67, 29.93]
CARDIFF -0.87 [-15.92, 14.19]
DNS-V3 -3.08 [-15.60, 9.44]
DNS-V4 6.49 [-10.79, 23.77]
EPIGEN 13.65 [-12.73, 40.03]
FBIRN 4.09 [ -9.58, 17.76]
FOR2107 7.48 [ -2.03, 16.99]
GIG 4.51 [-12.46, 21.48]
GSP 9.84 [ -2.79, 22.47]
HUBIN 9.09 [ -8.87, 27.06]
HUNT 10.55 [ 2.20, 18.90]
IMAGEN 2.13 [ -4.87, 9.12]
IMpACT 3.36 [-13.66, 20.39]
LBC1936 2.45 [ -7.18, 12.08]
LIBD -11.80 [-23.75, 0.15]
Sydney MAS 2.15 [ -7.89, 12.19]
MCIC -1.16 [-21.15, 18.83]
MooDS 5.48 [ -9.69, 20.65]
MPIP -1.89 [-12.56, 8.79]
MUNSTER 10.50 [ 2.33, 18.67]
NCNG 4.87 [ -9.60, 19.33]
NESDA 6.46 [-10.43, 23.34]
NeuroIMAGE 8.84 [ -9.65, 27.34]
NTR 15.49 [ 1.02, 29.96]
OATS 11.75 [ -3.77, 27.27]
PAFIP 4.39 [-15.26, 24.05]
PDNZ 1.94 [-17.18, 21.05]
PING -2.65 [-15.81, 10.50]
MPRC 8.08 [ -4.22, 20.39]
PPMI
QTIM 4.87 [ -3.51, 13.26]
SHIP 2.42 [ -5.64, 10.47]
SHIP-Trend 6.90 [ -1.92, 15.72]
SYS 6.72 [ -6.78, 20.22]
TCD-NUIG 6.38 [-17.49, 30.25]
TOP -13.02 [-24.91, -1.13]
TOP3T 7.84 [ -6.40, 22.08]
UiO2016 0.89 [-19.71, 21.48]
UiO2017 2.22 [-13.09, 17.53]
UMCU 5.36 [ -4.76, 15.49]
ENIGMA Discovery Cohorts [N=22776; P=1.402e-06] 4.28 [ 2.54, 6.02]
UKBB 4.90 [ 2.25, 7.55]
Discovery+UKBB [N=32691; P=3.447e-09] 4.47 [ 2.99, 5.95]
GOBS -1.26 [ -8.27, 5.76]
IMH 11.82 [-23.83, 47.47]
UNICAMP -7.43 [-29.19, 14.33]
Meth-CT
MIRECC
UKBB nonEUR 4.22 [-12.39, 20.82]
OSAKA -23.95 [-46.55, -1.35]
PING Non European -3.05 [-15.10, 9.00]
Non European Cohorts [N=2537; P=0.3844] -2.37 [ -7.71, 2.97]
Cohort
Beta [Lower 95% CI,Upper 95% CI]
-60 -40 -20 0 20 40 60
mm^2/effect allele

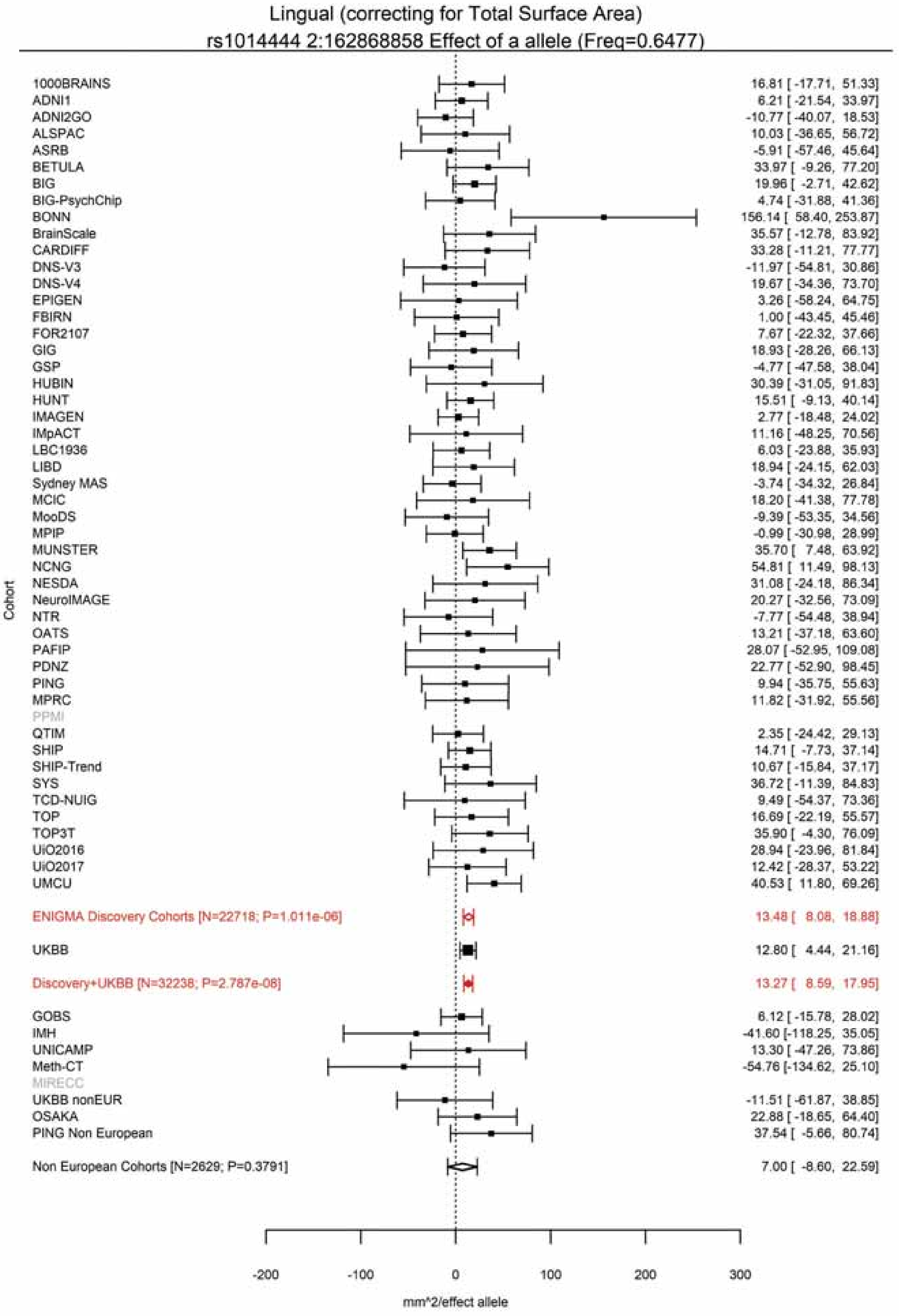

Lingual (correcting for Total Surface Area)
rs1014444 2:162868858 Effect of a allele (Freq=0.6477)
Cohort
1000BRAINS 16.81 [ -17.71, 51.33]
ADNI1 6.21 [ -21.54, 33.97]
ADNI2GO -10.77 [ -40.07, 18.53]
ALSPAC 10.03 [ -36.65, 56.72]
ASRB -5.91 [ -57.46, 45.64]
BETULA 33.97 [ -9.26, 77.20]
BIG 19.96 [ -2.71, 42.62]
BIG-PsychChip 4.74 [ -31.88, 41.36]
BONN 156.14 [ 58.40, 253.87]
BrainScale 35.57 [ -12.78, 83.92]
CARDIFF 33.28 [ -11.21, 77.77]
DNS-V3 -11.97 [ -54.81, 30.86]
DNS-V4 19.67 [ -34.36, 73.70]
EPIGEN 3.26 [ -58.24, 64.75]
FBIRN 1.00 [ -43.45, 45.46]
FOR2107 7.67 [ -22.32, 37.66]
GIG 18.93 [ -28.26, 66.13]
GSP -4.77 [ -47.58, 38.04]
HUBIN 30.39 [ -31.05, 91.83]
HUNT 15.51 [ -9.13, 40.14]
IMAGEN 2.77 [ -18.48, 24.02]
IMpACT 11.16 [ -48.25, 70.56]
LBC1936 6.03 [ -23.88, 35.93]
LIBD 18.94 [ -24.15, 62.03]
Sydney MAS -3.74 [ -34.32, 26.84]
MCIC 18.20 [ -41.38, 77.78]
MooDS -9.39 [ -53.35, 34.56]
MPIP -0.99 [ -30.98, 28.99]
MUNSTER 35.70 [ 7.48, 63.92]
NCNG 54.81 [ 11.49, 98.13]
NESDA 31.08 [ -24.18, 86.34]
NeuroIMAGE 20.27 [ -32.56, 73.09]
NTR -7.77 [ -54.48, 38.94]
OATS 13.21 [ -37.18, 63.60]
PAFIP 28.07 [ -52.95, 109.08]
PDNZ 22.77 [ -52.90, 98.45]
PING 9.94 [ -35.75, 55.63]
MPRC 11.82 [ -31.92, 55.56]
PPMI
QTIM 2.35 [ -24.42, 29.13]
SHIP 14.71 [ -7.73, 37.14]
SHIP-Trend 10.67 [ -15.84, 37.17]
SYS 36.72 [ -11.39, 84.83]
TCD-NUIG 9.49 [ -54.37, 73.36]
TOP 16.69 [ -22.19, 55.57]
TOP3T 35.90 [ -4.30, 76.09]
UiO2016 28.94 [ -23.96, 81.84]
UiO2017 12.42 [ -28.37, 53.22]
UMCU 40.53 [ 11.80, 69.26]
ENIGMA Discovery Cohorts [N=22718; P=1.011e-06] 13.48 [ 8.08, 18.88]
UKBB 12.80 [ 4.44, 21.16]
Discovery+UKBB [N=32238; P=2.787e-08] 13.27 [ 8.59, 17.95]
GOBS 6.12 [ -15.78, 28.02]
IMH -41.60 [-118.25, 35.05]
UNICAMP 13.30 [ -47.26, 73.86]
Meth-CT -54.76 [-134.62, 25.10]
MIRECC
UKBB nonEUR -11.51 [ -61.87, 38.85]
OSAKA 22.88 [ -18.65, 64.40]
PING Non European 37.54 [ -5.66, 80.74]
Non European Cohorts [N=2629; P=0.3791] 7.00 [ -8.60, 22.59]
Beta [Lower 95% CI,Upper 95% CI]
-200
-100
0
100
200
300
mm^2/effect allele

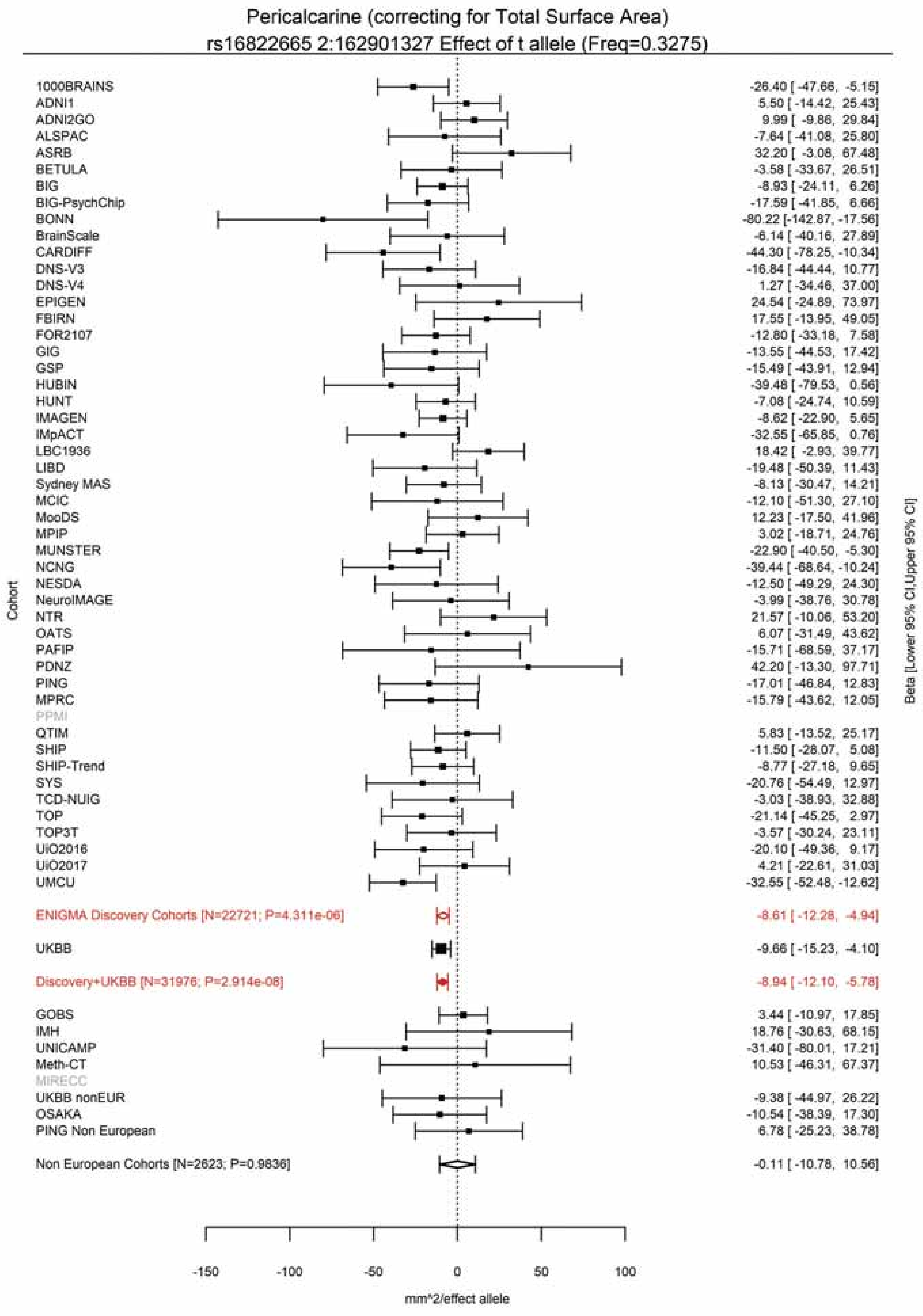

Pericalcarine (correcting for Total Surface Area)
rs16822665 2:162901327 Effect of t allele (Freq=0.3275)
Cohort
1000BRAINS -26.40 [ -47.66, -5.15]
ADNI1 5.50 [ -14.42, 25.43]
ADNI2GO 9.99 [ -9.86, 29.84]
ALSPAC -7.64 [ -41.08, 25.80]
ASRB 32.20 [ -3.08, 67.48]
BETULA -3.58 [ -33.67, 26.51]
BIG -8.93 [ -24.11, 6.26]
BIG-PsychChip -17.59 [ -41.85, 6.66]
BONN -80.22 [-142.87, -17.56]
BrainScale -6.14 [ -40.16, 27.89]
CARDIFF -44.30 [ -78.25, -10.34]
DNS-V3 -16.84 [ -44.44, 10.77]
DNS-V4 1.27 [ -34.46, 37.00]
EPIGEN 24.54 [ -24.89, 73.97]
FBIRN 17.55 [ -13.95, 49.05]
FOR2107 -12.80 [ -33.18, 7.58]
GIG -13.55 [ -44.53, 17.42]
GSP -15.49 [ -43.91, 12.94]
HUBIN -39.48 [ -79.53, 0.56]
HUNT -7.08 [ -24.74, 10.59]
IMAGEN -8.62 [ -22.90, 5.65]
IMpACT -32.55 [ -65.85, 0.76]
LBC1936 18.42 [ -2.93, 39.77]
LIBD -19.48 [ -50.39, 11.43]
Sydney MAS -8.13 [ -30.47, 14.21]
MCIC -12.10 [ -51.30, 27.10]
MooDS 12.23 [ -17.50, 41.96]
MPIP 3.02 [ -18.71, 24.76]
MUNSTER -22.90 [ -40.50, -5.30]
NCNG -39.44 [ -68.64, -10.24]
NESDA -12.50 [ -49.29, 24.30]
NeuroIMAGE -3.99 [ -38.76, 30.78]
NTR 21.57 [ -10.06, 53.20]
OATS 6.07 [ -31.49, 43.62]
PAFIP -15.71 [ -68.59, 37.17]
PDNZ 42.20 [ -13.30, 97.71]
PING -17.01 [ -46.84, 12.83]
MPRC -15.79 [ -43.62, 12.05]
PPMI
QTIM 5.83 [ -13.52, 25.17]
SHIP -11.50 [ -28.07, 5.08]
SHIP-Trend -8.77 [ -27.18, 9.65]
SYS -20.76 [ -54.49, 12.97]
TCD-NUIG -3.03 [ -38.93, 32.88]
TOP -21.14 [ -45.25, 2.97]
TOP3T -3.57 [ -30.24, 23.11]
UiO2016 -20.10 [ -49.36, 9.17]
UiO2017 4.21 [ -22.61, 31.03]
UMCU -32.55 [ -52.48, -12.62]
ENIGMA Discovery Cohorts [N=22721; P=4.311e-06] -8.61 [ -12.28, -4.94]
UKBB -9.66 [ -15.23, -4.10]
Discovery+UKBB [N=31976; P=2.914e-08] -8.94 [ -12.10, -5.78]
GOBS 3.44 [ -10.97, 17.85]
IMH 18.76 [ -30.63, 68.15]
UNICAMP -31.40 [ -80.01, 17.21]
Meth-CT 10.53 [ -46.31, 67.37]
MIRECC
UKBB nonEUR -9.38 [ -44.97, 26.22]
OSAKA -10.54 [ -38.39, 17.30]
PING Non European 6.78 [ -25.23, 38.78]
Non European Cohorts [N=2623; P=0.9836] -0.11 [ -10.78, 10.56]
Beta [Lower 95% CI,Upper 95% CI]
-150
-100
-50
0
50
100
mm^2/effect allele

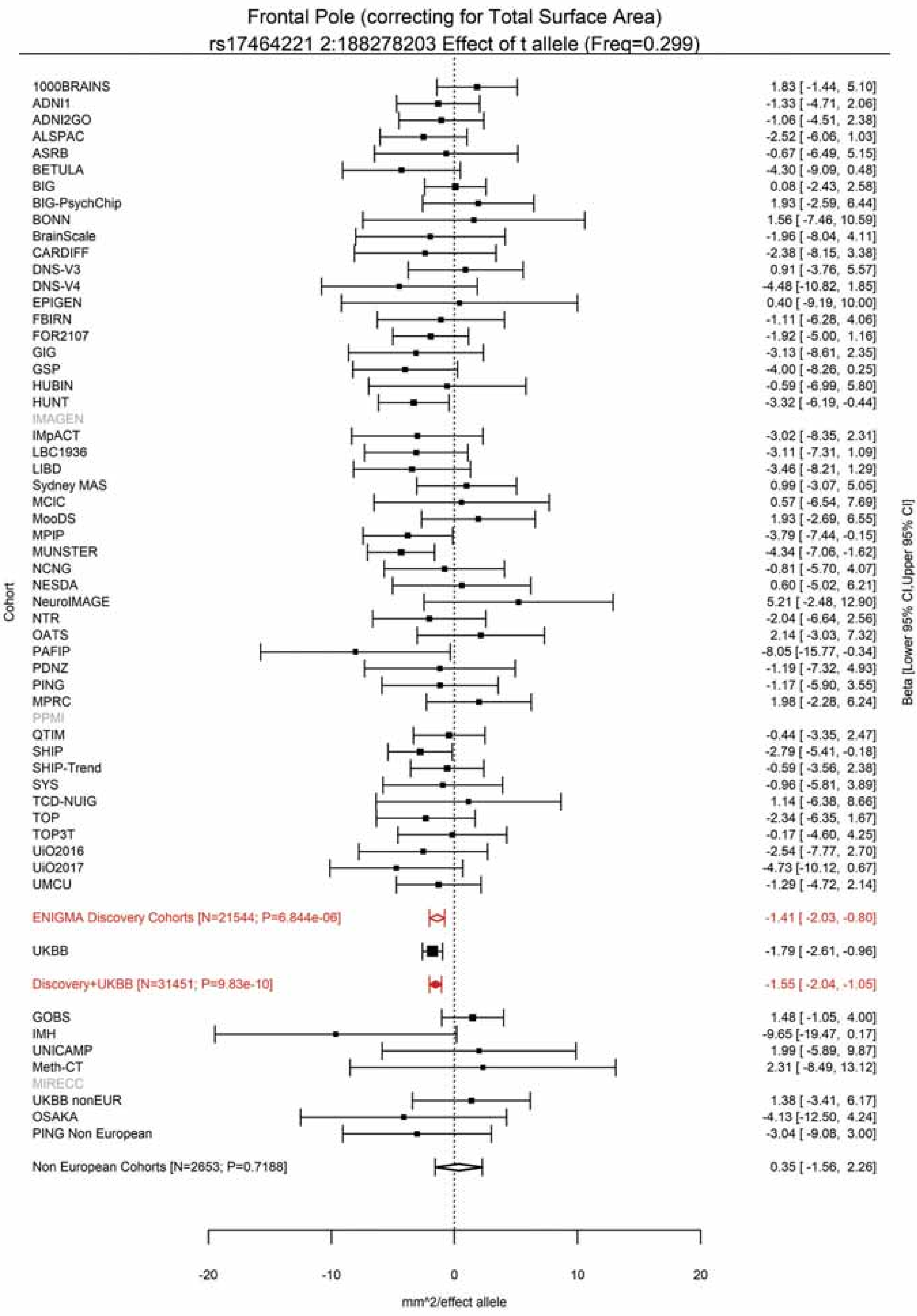
Frontal Pole (correcting for Total Surface Area)
rs17464221 2:188278203 Effect of t allele (Freq=0.299)
1000BRAINS 1.83 [ -1.44, 5.10]
ADNI1 -1.33 [ -4.71, 2.06]
ADNI2GO -1.06 [ -4.51, 2.38]
ALSPAC -2.52 [ -6.06, 1.03]
ASRB -0.67 [ -6.49, 5.15]
BETULA -4.30 [ -9.09, 0.48]
BIG 0.08 [ -2.43, 2.58]
BIG-PsychChip 1.93 [ -2.59, 6.44]
BONN 1.56 [ -7.46, 10.59]
BrainScale -1.96 [ -8.04, 4.11]
CARDIFF -2.38 [ -8.15, 3.38]
DNS-V3 0.91 [ -3.76, 5.57]
DNS-V4 -4.48 [-10.82, 1.85]
EPIGEN 0.40 [ -9.19, 10.00]
FBIRN -1.11 [ -6.28, 4.06]
FOR2107 -1.92 [ -5.00, 1.16]
GIG -3.13 [ -8.61, 2.35]
GSP -4.00 [ -8.26, 0.25]
HUBIN -0.59 [ -6.99, 5.80]
HUNT -3.32 [ -6.19, -0.44]
IMAGEN
IMpACT -3.02 [ -8.35, 2.31]
LBC1936 -3.11 [ -7.31, 1.09]
LIBD -3.46 [ -8.21, 1.29]
Sydney MAS 0.99 [ -3.07, 5.05]
MCIC 0.57 [ -6.54, 7.69]
MooDS 1.93 [ -2.69, 6.55]
MPIP -3.79 [ -7.44, -0.15]
MUNSTER -4.34 [ -7.06, -1.62]
NCNG -0.81 [ -5.70, 4.07]
NESDA 0.60 [ -5.02, 6.21]
NeuroIMAGE 5.21 [ -2.48, 12.90]
NTR -2.04 [ -6.64, 2.56]
OATS 2.14 [ -3.03, 7.32]
PAFIP -8.05 [-15.77, -0.34]
PDNZ -1.19 [ -7.32, 4.93]
PING -1.17 [ -5.90, 3.55]
MPRC 1.98 [ -2.28, 6.24]
PPMI
QTIM -0.44 [ -3.35, 2.47]
SHIP -2.79 [ -5.41, -0.18]
SHIP-Trend -0.59 [ -3.56, 2.38]
SYS -0.96 [ -5.81, 3.89]
TCD-NUIG 1.14 [ -6.38, 8.66]
TOP -2.34 [ -6.35, 1.67]
TOP3T -0.17 [ -4.60, 4.25]
UiO2016 -2.54 [ -7.77, 2.70]
UiO2017 -4.73 [-10.12, 0.67]
UMCU -1.29 [ -4.72, 2.14]
ENIGMA Discovery Cohorts [N=21544; P=6.844e-06] -1.41 [ -2.03, -0.80]
UKBB -1.79 [ -2.61, -0.96]
Discovery+UKBB [N=31451; P=9.83e-10] -1.55 [ -2.04, -1.05]
GOBS 1.48 [ -1.05, 4.00]
IMH -9.65 [-19.47, 0.17]
UNICAMP 1.99 [ -5.89, 9.87]
Meth-CT 2.31 [ -8.49, 13.12]
MIRECC
UKBB nonEUR 1.38 [ -3.41, 6.17]
OSAKA -4.13 [-12.50, 4.24]
PING Non European -3.04 [ -9.08, 3.00]
Non European Cohorts [N=2653; P=0.7188] 0.35 [ -1.56, 2.26]
Cohort
Beta [Lower 95% CI,Upper 95% CI]
-20
-10
0
10
20
mm^2/effect allele

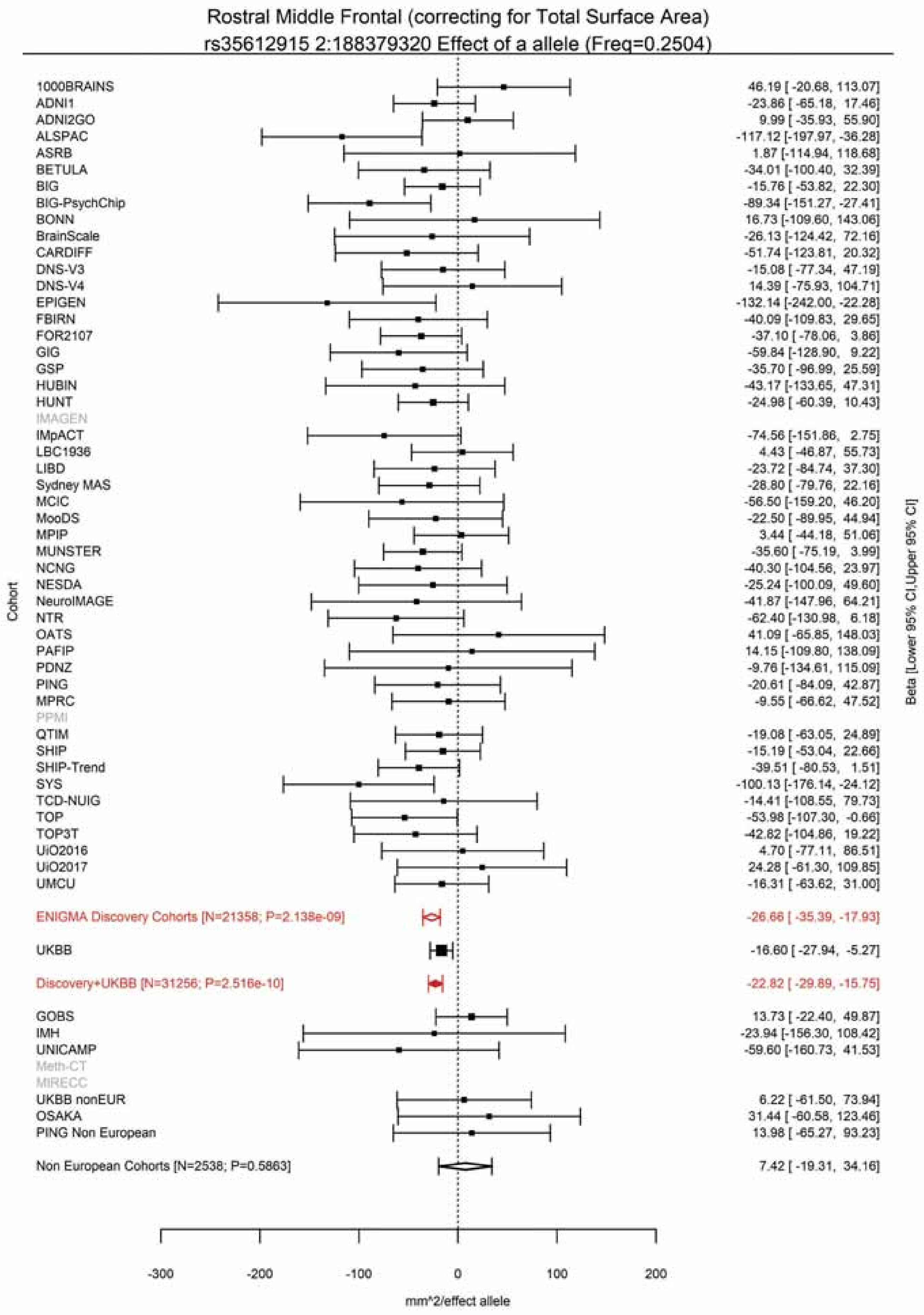

Rostral Middle Frontal (correcting for Total Surface Area)
rs35612915 2:188379320 Effect of a allele (Freq=0.2504)
Cohort
1000BRAINS 46.19 [ -20.68, 113.07]
ADNI1 -23.86 [ -65.18, 17.46]
ADNI2GO 9.99 [ -35.93, 55.90]
ALSPAC -117.12 [-197.97, -36.28]
ASRB 1.87 [-114.94, 118.68]
BETULA -34.01 [-100.40, 32.39]
BIG -15.76 [ -53.82, 22.30]
BIG-PsychChip -89.34 [-151.27, -27.41]
BONN 16.73 [-109.60, 143.06]
BrainScale -26.13 [-124.42, 72.16]
CARDIFF -51.74 [-123.81, 20.32]
DNS-V3 -15.08 [ -77.34, 47.19]
DNS-V4 14.39 [ -75.93, 104.71]
EPIGEN -132.14 [-242.00, -22.28]
FBIRN -40.09 [-109.83, 29.65]
FOR2107 -37.10 [ -78.06, 3.86]
GIG -59.84 [-128.90, 9.22]
GSP -35.70 [ -96.99, 25.59]
HUBIN -43.17 [-133.65, 47.31]
HUNT -24.98 [ -60.39, 10.43]
IMAGEN
IMpACT -74.56 [-151.86, 2.75]
LBC1936 4.43 [ -46.87, 55.73]
LIBD -23.72 [ -84.74, 37.30]
Sydney MAS -28.80 [ -79.76, 22.16]
MCIC -56.50 [-159.20, 46.20]
MooDS -22.50 [ -89.95, 44.94]
MPIP 3.44 [ -44.18, 51.06]
MUNSTER -35.60 [ -75.19, 3.99]
NCNG -40.30 [-104.56, 23.97]
NESDA -25.24 [-100.09, 49.60]
NeuroIMAGE -41.87 [-147.96, 64.21]
NTR -62.40 [-130.98, 6.18]
OATS 41.09 [ -65.85, 148.03]
PAFIP 14.15 [-109.80, 138.09]
PDNZ -9.76 [-134.61, 115.09]
PING -20.61 [ -84.09, 42.87]
MPRC -9.55 [ -66.62, 47.52]
PPMI
QTIM -19.08 [ -63.05, 24.89]
SHIP -15.19 [ -53.04, 22.66]
SHIP-Trend -39.51 [ -80.53, 1.51]
SYS -100.13 [-176.14, -24.12]
TCD-NUIG -14.41 [-108.55, 79.73]
TOP -53.98 [-107.30, -0.66]
TOP3T -42.82 [-104.86, 19.22]
UiO2016 4.70 [ -77.11, 86.51]
UiO2017 24.28 [ -61.30, 109.85]
UMCU -16.31 [ -63.62, 31.00]
ENIGMA Discovery Cohorts [N=21358; P=2.138e-09] -26.66 [ -35.39, -17.93]
UKBB -16.60 [ -27.94, -5.27]
Discovery+UKBB [N=31256; P=2.516e-10] -22.82 [ -29.89, -15.75]
GOBS 13.73 [ -22.40, 49.87]
IMH -23.94 [-156.30, 108.42]
UNICAMP -59.60 [-160.73, 41.53]
Meth-CT
MIRECC
UKBB nonEUR 6.22 [ -61.50, 73.94]
OSAKA 31.44 [ -60.58, 123.46]
PING Non European 13.98 [ -65.27, 93.23]
Non European Cohorts [N=2538; P=0.5863] 7.42 [ -19.31, 34.16]
Beta [Lower 95% CI,Upper 95% CI]
-300
-200
-100
0
100
200
mm^2/effect allele

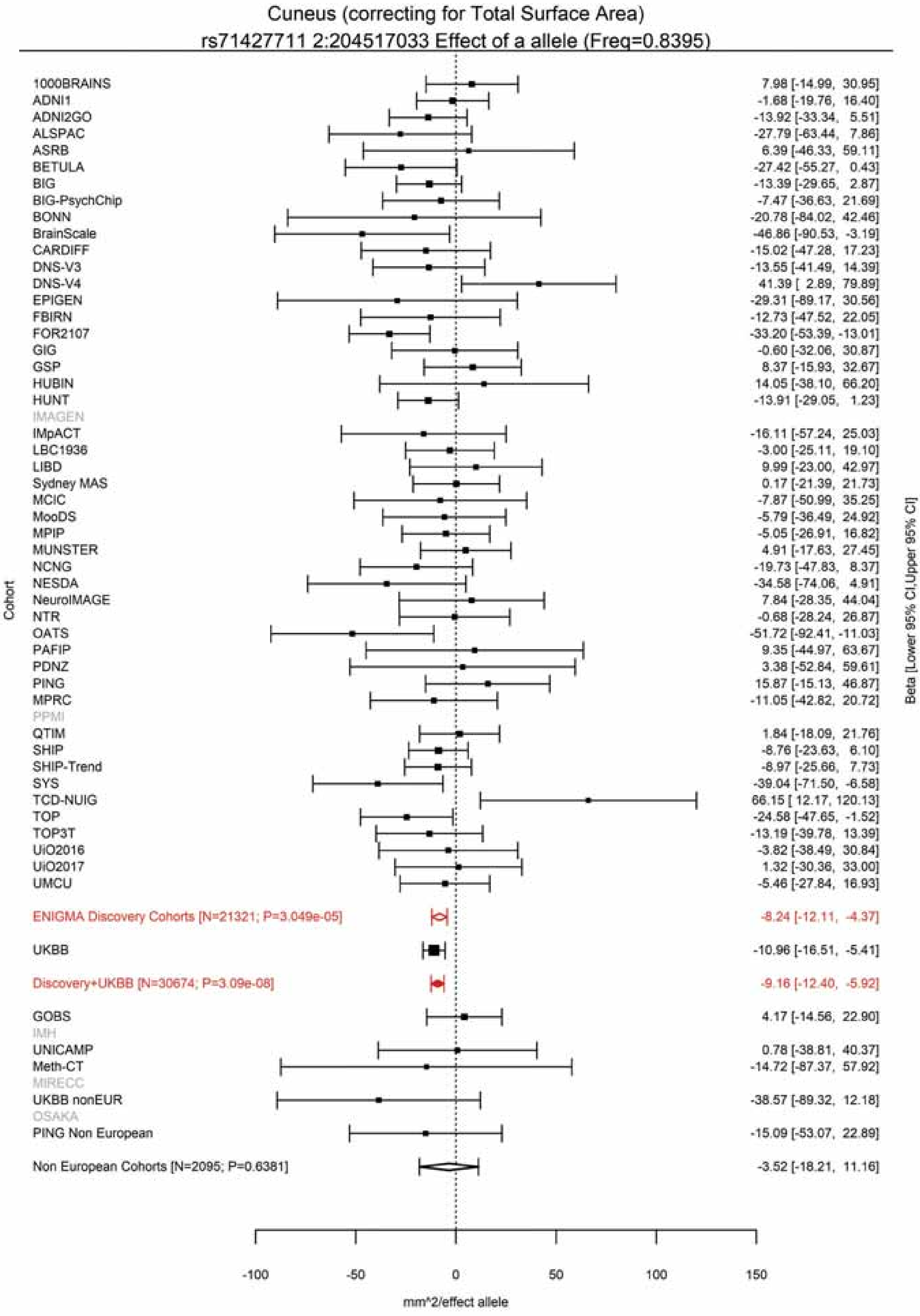
Cuneus (correcting for Total Surface Area)
rs71427711 2:204517033 Effect of a allele (Freq=0.8395)
1000BRAINS 7.98 [-14.99, 30.95]
ADNI1 -1.68 [-19.76, 16.40]
ADNI2GO -13.92 [-33.34, 5.51]
ALSPAC -27.79 [-63.44, 7.86]
ASRB 6.39 [-46.33, 59.11]
BETULA -27.42 [-55.27, 0.43]
BIG -13.39 [-29.65, 2.87]
BIG-PsychChip -7.47 [-36.63, 21.69]
BONN -20.78 [-84.02, 42.46]
BrainScale -46.86 [-90.53, -3.19]
CARDIFF -15.02 [-47.28, 17.23]
DNS-V3 -13.55 [-41.49, 14.39]
DNS-V4 41.39 [ 2.89, 79.89]
EPIGEN -29.31 [-89.17, 30.56]
FBIRN -12.73 [-47.52, 22.05]
FOR2107 -33.20 [-53.39, -13.01]
GIG -0.60 [-32.06, 30.87]
GSP 8.37 [-15.93, 32.67]
HUBIN 14.05 [-38.10, 66.20]
HUNT -13.91 [-29.05, 1.23]
IMAGEN
IMpACT -16.11 [-57.24, 25.03]
LBC1936 -3.00 [-25.11, 19.10]
LIBD 9.99 [-23.00, 42.97]
Sydney MAS 0.17 [-21.39, 21.73]
MCIC -7.87 [-50.99, 35.25]
MooDS -5.79 [-36.49, 24.92]
MPIP -5.05 [-26.91, 16.82]
MUNSTER 4.91 [-17.63, 27.45]
NCNG -19.73 [-47.83, 8.37]
NESDA -34.58 [-74.06, 4.91]
NeuroIMAGE 7.84 [-28.35, 44.04]
NTR -0.68 [-28.24, 26.87]
OATS -51.72 [-92.41, -11.03]
PAFIP 9.35 [-44.97, 63.67]
PDNZ 3.38 [-52.84, 59.61]
PING 15.87 [-15.13, 46.87]
MPRC -11.05 [-42.82, 20.72]
PPMI
QTIM 1.84 [-18.09, 21.76]
SHIP -8.76 [-23.63, 6.10]
SHIP-Trend -8.97 [-25.66, 7.73]
SYS -39.04 [-71.50, -6.58]
TCD-NUIG 66.15 [ 12.17, 120.13]
TOP -24.58 [-47.65, -1.52]
TOP3T -13.19 [-39.78, 13.39]
UiO2016 -3.82 [-38.49, 30.84]
UiO2017 1.32 [-30.36, 33.00]
UMCU -5.46 [-27.84, 16.93]
ENIGMA Discovery Cohorts [N=21321; P=3.049e-05] -8.24 [-12.11, -4.37]
UKBB -10.96 [-16.51, -5.41]
Discovery+UKBB [N=30674; P=3.09e-08] -9.16 [-12.40, -5.92]
GOBS 4.17 [-14.56, 22.90]
IMH
UNICAMP 0.78 [-38.81, 40.37]
Meth-CT -14.72 [-87.37, 57.92]
MIRECC
UKBB nonEUR -38.57 [-89.32, 12.18]
OSAKA
PING Non European -15.09 [-53.07, 22.89]
Non European Cohorts [N=2095; P=0.6381] -3.52 [-18.21, 11.16]
Cohort
Beta [Lower 95% CI,Upper 95% CI]
-100
-50
0
50
100
150
mm^2/effect allele

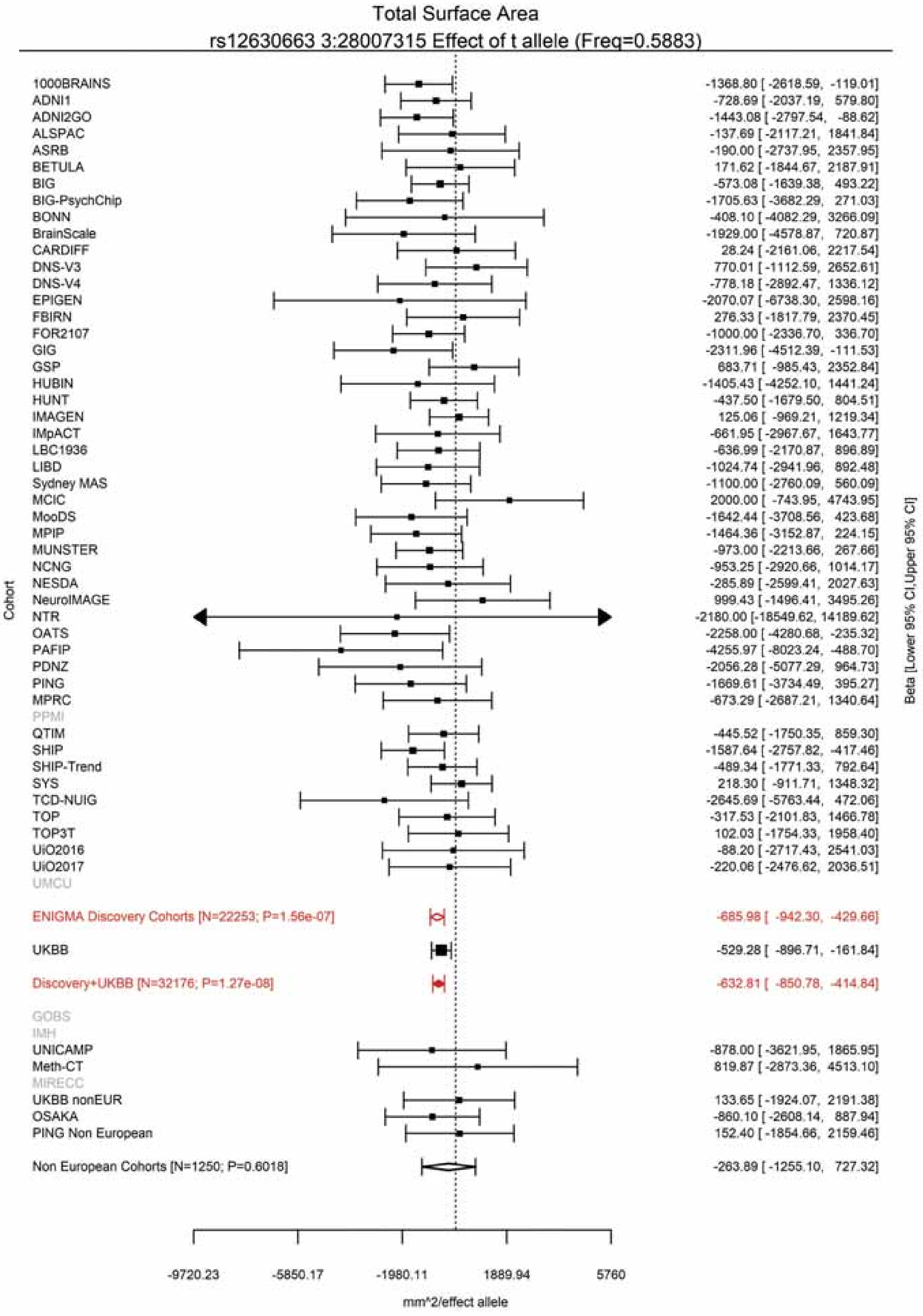

Total Surface Area
rs12630663 3:28007315 Effect of t allele (Freq=0.5883)
Cohort
1000BRAINS -1368.80 [ -2618.59, -119.01]
ADNI1 -728.69 [ -2037.19, 579.80]
ADNI2GO -1443.08 [ -2797.54, -88.62]
ALSPAC -137.69 [ -2117.21, 1841.84]
ASRB -190.00 [ -2737.95, 2357.95]
BETULA 171.62 [ -1844.67, 2187.91]
BIG -573.08 [ -1639.38, 493.22]
BIG-PsychChip -1705.63 [ -3682.29, 271.03]
BONN -408.10 [ -4082.29, 3266.09]
BrainScale -1929.00 [ -4578.87, 720.87]
CARDIFF 28.24 [ -2161.06, 2217.54]
DNS-V3 770.01 [ -1112.59, 2652.61]
DNS-V4 -778.18 [ -2892.47, 1336.12]
EPIGEN -2070.07 [ -6738.30, 2598.16]
FBIRN 276.33 [ -1817.79, 2370.45]
FOR2107 -1000.00 [ -2336.70, 336.70]
GIG -2311.96 [ -4512.39, -111.53]
GSP 683.71 [ -985.43, 2352.84]
HUBIN -1405.43 [ -4252.10, 1441.24]
HUNT -437.50 [ -1679.50, 804.51]
IMAGEN 125.06 [ -969.21, 1219.34]
IMpACT -661.95 [ -2967.67, 1643.77]
LBC1936 -636.99 [ -2170.87, 896.89]
LIBD -1024.74 [ -2941.96, 892.48]
Sydney MAS -1100.00 [ -2760.09, 560.09]
MCIC 2000.00 [ -743.95, 4743.95]
MooDS -1642.44 [ -3708.56, 423.68]
MPIP -1464.36 [ -3152.87, 224.15]
MUNSTER -973.00 [ -2213.66, 267.66]
NCNG -953.25 [ -2920.66, 1014.17]
NESDA -285.89 [ -2599.41, 2027.63]
NeuroIMAGE 999.43 [ -1496.41, 3495.26]
NTR -2180.00 [-18549.62, 14189.62]
OATS -2258.00 [ -4280.68, -235.32]
PAFIP -4255.97 [ -8023.24, -488.70]
PDNZ -2056.28 [ -5077.29, 964.73]
PING -1669.61 [ -3734.49, 395.27]
MPRC -673.29 [ -2687.21, 1340.64]
PPMI
QTIM -445.52 [ -1750.35, 859.30]
SHIP -1587.64 [ -2757.82, -417.46]
SHIP-Trend -489.34 [ -1771.33, 792.64]
SYS 218.30 [ -911.71, 1348.32]
TCD-NUIG -2645.69 [ -5763.44, 472.06]
TOP -317.53 [ -2101.83, 1466.78]
TOP3T 102.03 [ -1754.33, 1958.40]
UiO2016 -88.20 [ -2717.43, 2541.03]
UiO2017 -220.06 [ -2476.62, 2036.51]
UMCU
ENIGMA Discovery Cohorts [N=22253; P=1.56e-07] -685.98 [ -942.30, -429.66]
UKBB -529.28 [ -896.71, -161.84]
Discovery+UKBB [N=32176; P=1.27e-08] -632.81 [ -850.78, -414.84]
GOBS
IMH
UNICAMP -878.00 [ -3621.95, 1865.95]
Meth-CT 819.87 [ -2873.36, 4513.10]
MIRECC
UKBB nonEUR 133.65 [ -1924.07, 2191.38]
OSAKA -860.10 [ -2608.14, 887.94]
PING Non European 152.40 [ -1854.66, 2159.46]
Non European Cohorts [N=1250; P=0.6018] -263.89 [ -1255.10, 727.32]
Beta [Lower 95% CI,Upper 95% CI]
-9720.23
-5850.17
-1980.11
1889.94
5760
mm^2/effect allele

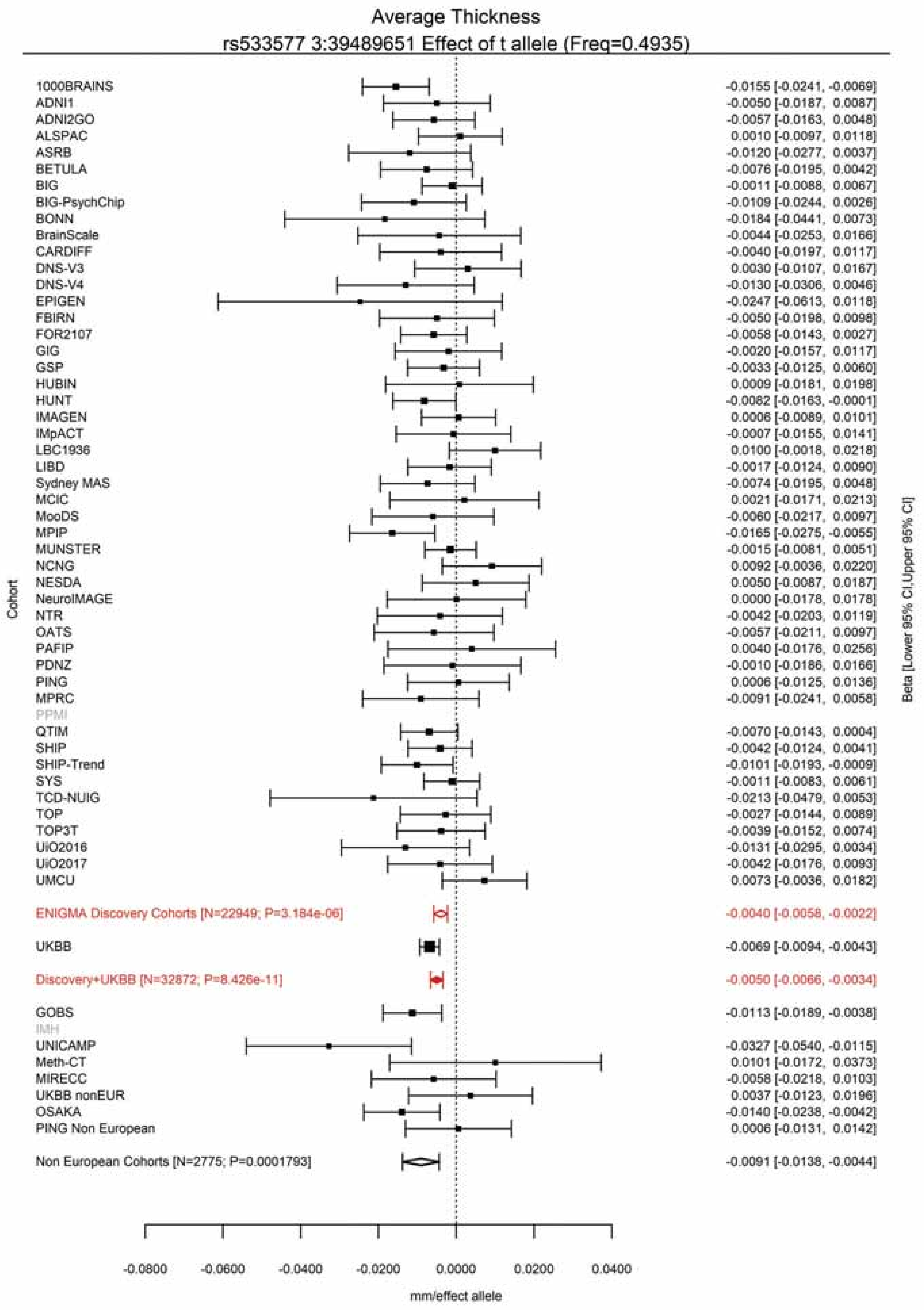
Average Thickness
rs533577 3:39489651 Effect of t allele (Freq=0.4935)
Cohort
1000BRAINS -0.0155 [-0.0241, -0.0069]
ADNI1 -0.0050 [-0.0187, 0.0087]
ADNI2GO -0.0057 [-0.0163, 0.0048]
ALSPAC 0.0010 [-0.0097, 0.0118]
ASRB -0.0120 [-0.0277, 0.0037]
BETULA -0.0076 [-0.0195, 0.0042]
BIG -0.0011 [-0.0088, 0.0067]
BIG-PsychChip -0.0109 [-0.0244, 0.0026]
BONN -0.0184 [-0.0441, 0.0073]
BrainScale -0.0044 [-0.0253, 0.0166]
CARDIFF -0.0040 [-0.0197, 0.0117]
DNS-V3 0.0030 [-0.0107, 0.0167]
DNS-V4 -0.0130 [-0.0306, 0.0046]
EPIGEN -0.0247 [-0.0613, 0.0118]
FBIRN -0.0050 [-0.0198, 0.0098]
FOR2107 -0.0058 [-0.0143, 0.0027]
GIG -0.0020 [-0.0157, 0.0117]
GSP -0.0033 [-0.0125, 0.0060]
HUBIN 0.0009 [-0.0181, 0.0198]
HUNT -0.0082 [-0.0163, -0.0001]
IMAGEN 0.0006 [-0.0089, 0.0101]
IMpACT -0.0007 [-0.0155, 0.0141]
LBC1936 0.0100 [-0.0018, 0.0218]
LIBD -0.0017 [-0.0124, 0.0090]
Sydney MAS -0.0074 [-0.0195, 0.0048]
MCIC 0.0021 [-0.0171, 0.0213]
MooDS -0.0060 [-0.0217, 0.0097]
MPIP -0.0165 [-0.0275, -0.0055]
MUNSTER -0.0015 [-0.0081, 0.0051]
NCNG 0.0092 [-0.0036, 0.0220]
NESDA 0.0050 [-0.0087, 0.0187]
NeuroIMAGE 0.0000 [-0.0178, 0.0178]
NTR -0.0042 [-0.0203, 0.0119]
OATS -0.0057 [-0.0211, 0.0097]
PAFIP 0.0040 [-0.0176, 0.0256]
PDNZ -0.0010 [-0.0186, 0.0166]
PING 0.0006 [-0.0125, 0.0136]
MPRC -0.0091 [-0.0241, 0.0058]
PPMI
QTIM -0.0070 [-0.0143, 0.0004]
SHIP -0.0042 [-0.0124, 0.0041]
SHIP-Trend -0.0101 [-0.0193, -0.0009]
SYS -0.0011 [-0.0083, 0.0061]
TCD-NUIG -0.0213 [-0.0479, 0.0053]
TOP -0.0027 [-0.0144, 0.0089]
TOP3T -0.0039 [-0.0152, 0.0074]
UiO2016 -0.0131 [-0.0295, 0.0034]
UiO2017 -0.0042 [-0.0176, 0.0093]
UMCU 0.0073 [-0.0036, 0.0182]
ENIGMA Discovery Cohorts [N=22949; P=3.184e-06] -0.0040 [-0.0058, -0.0022]
UKBB -0.0069 [-0.0094, -0.0043]
Discovery+UKBB [N=32872; P=8.426e-11] -0.0050 [-0.0066, -0.0034]
GOBS -0.0113 [-0.0189, -0.0038]
IMH
UNICAMP -0.0327 [-0.0540, -0.0115]
Meth-CT 0.0101 [-0.0172, 0.0373]
MIRECC -0.0058 [-0.0218, 0.0103]
UKBB nonEUR 0.0037 [-0.0123, 0.0196]
OSAKA -0.0140 [-0.0238, -0.0042]
PING Non European 0.0006 [-0.0131, 0.0142]
Non European Cohorts [N=2775; P=0.0001793] -0.0091 [-0.0138, -0.0044]
Beta [Lower 95% CI,Upper 95% CI]
-0.0800 -0.0600 -0.0400 -0.0200 0.0000 0.0200 0.0400
mm/effect allele

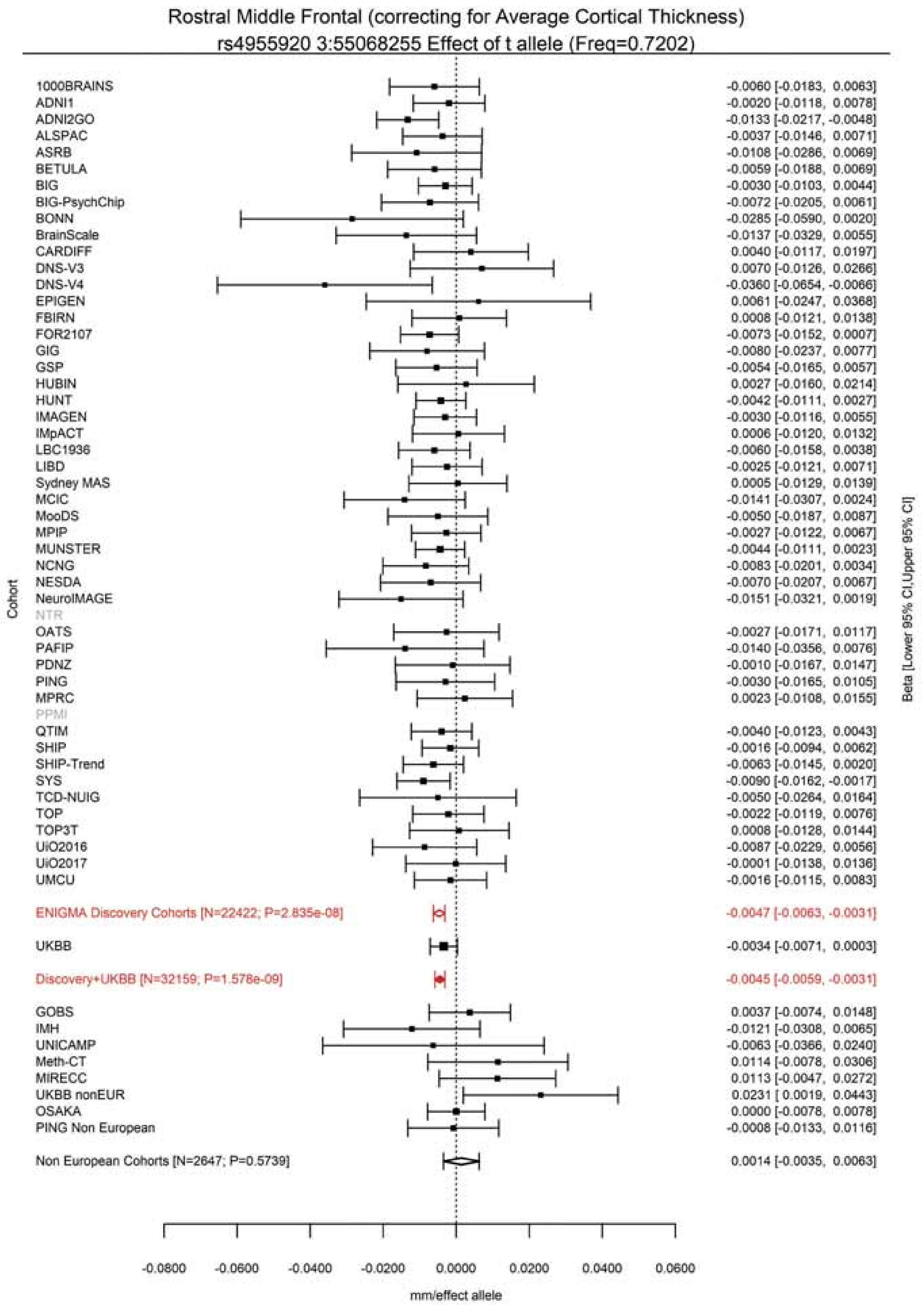
Rostral Middle Frontal (correcting for Average Cortical Thickness)
rs4955920 3:55068255 Effect of t allele (Freq=0.7202)
1000BRAINS -0.0060 [-0.0183, 0.0063]
ADNI1 -0.0020 [-0.0118, 0.0078]
ADNI2GO -0.0133 [-0.0217, -0.0048]
ALSPAC -0.0037 [-0.0146, 0.0071]
ASRB -0.0108 [-0.0286, 0.0069]
BETULA -0.0059 [-0.0188, 0.0069]
BIG -0.0030 [-0.0103, 0.0044]
BIG-PsychChip -0.0072 [-0.0205, 0.0061]
BONN -0.0285 [-0.0590, 0.0020]
BrainScale -0.0137 [-0.0329, 0.0055]
CARDIFF 0.0040 [-0.0117, 0.0197]
DNS-V3 0.0070 [-0.0126, 0.0266]
DNS-V4 -0.0360 [-0.0654, -0.0066]
EPIGEN 0.0061 [-0.0247, 0.0368]
FBIRN 0.0008 [-0.0121, 0.0138]
FOR2107 -0.0073 [-0.0152, 0.0007]
GIG -0.0080 [-0.0237, 0.0077]
GSP -0.0054 [-0.0165, 0.0057]
HUBIN 0.0027 [-0.0160, 0.0214]
HUNT -0.0042 [-0.0111, 0.0027]
IMAGEN -0.0030 [-0.0116, 0.0055]
IMpACT 0.0006 [-0.0120, 0.0132]
LBC1936 -0.0060 [-0.0158, 0.0038]
LIBD -0.0025 [-0.0121, 0.0071]
Sydney MAS 0.0005 [-0.0129, 0.0139]
MCIC -0.0141 [-0.0307, 0.0024]
MooDS -0.0050 [-0.0187, 0.0087]
MPIP -0.0027 [-0.0122, 0.0067]
MUNSTER -0.0044 [-0.0111, 0.0023]
NCNG -0.0083 [-0.0201, 0.0034]
NESDA -0.0070 [-0.0207, 0.0067]
NeuroIMAGE -0.0151 [-0.0321, 0.0019]
NTR
OATS -0.0027 [-0.0171, 0.0117]
PAFIP -0.0140 [-0.0356, 0.0076]
PDNZ -0.0010 [-0.0167, 0.0147]
PING -0.0030 [-0.0165, 0.0105]
MPRC 0.0023 [-0.0108, 0.0155]
PPMI
QTIM -0.0040 [-0.0123, 0.0043]
SHIP -0.0016 [-0.0094, 0.0062]
SHIP-Trend -0.0063 [-0.0145, 0.0020]
SYS -0.0090 [-0.0162, -0.0017]
TCD-NUIG -0.0050 [-0.0264, 0.0164]
TOP -0.0022 [-0.0119, 0.0076]
TOP3T 0.0008 [-0.0128, 0.0144]
UiO2016 -0.0087 [-0.0229, 0.0056]
UiO2017 -0.0001 [-0.0138, 0.0136]
UMCU -0.0016 [-0.0115, 0.0083]
ENIGMA Discovery Cohorts [N=22422; P=2.835e-08] -0.0047 [-0.0063, -0.0031]
UKBB -0.0034 [-0.0071, 0.0003]
Discovery+UKBB [N=32159; P=1.578e-09] -0.0045 [-0.0059, -0.0031]
GOBS 0.0037 [-0.0074, 0.0148]
IMH -0.0121 [-0.0308, 0.0065]
UNICAMP -0.0063 [-0.0366, 0.0240]
Meth-CT 0.0114 [-0.0078, 0.0306]
MIRECC 0.0113 [-0.0047, 0.0272]
UKBB nonEUR 0.0231 [ 0.0019, 0.0443]
OSAKA 0.0000 [-0.0078, 0.0078]
PING Non European -0.0008 [-0.0133, 0.0116]
Non European Cohorts [N=2647; P=0.5739] 0.0014 [-0.0035, 0.0063]
Cohort
Beta [Lower 95% CI,Upper 95% CI]
-0.0800
-0.0600
-0.0400
-0.0200
0.0000
0.0200
0.0400
0.0600
mm/effect allele

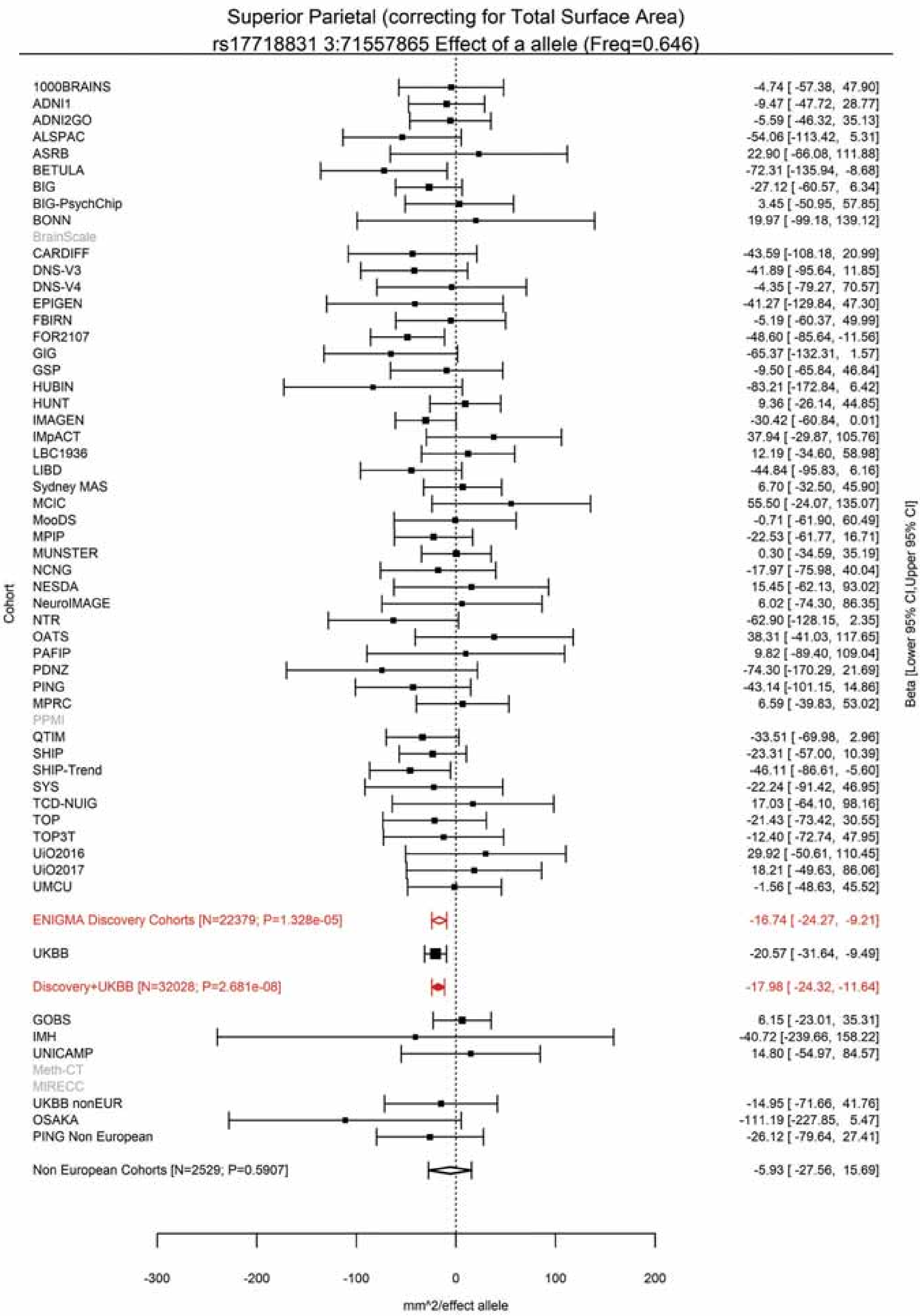
Superior Parietal (correcting for Total Surface Area)
rs17718831 3:71557865 Effect of a allele (Freq=0.646)
Cohort
1000BRAINS -4.74 [ -57.38, 47.90]
ADNI1 -9.47 [ -47.72, 28.77]
ADNI2GO -5.59 [ -46.32, 35.13]
ALSPAC -54.06 [-113.42, 5.31]
ASRB 22.90 [ -66.08, 111.88]
BETULA -72.31 [-135.94, -8.68]
BIG -27.12 [ -60.57, 6.34]
BIG-PsychChip 3.45 [ -50.95, 57.85]
BONN 19.97 [ -99.18, 139.12]
BrainScale
CARDIFF -43.59 [-108.18, 20.99]
DNS-V3 -41.89 [ -95.64, 11.85]
DNS-V4 -4.35 [ -79.27, 70.57]
EPIGEN -41.27 [-129.84, 47.30]
FBIRN -5.19 [ -60.37, 49.99]
FOR2107 -48.60 [ -85.64, -11.56]
GIG -65.37 [-132.31, 1.57]
GSP -9.50 [ -65.84, 46.84]
HUBIN -83.21 [-172.84, 6.42]
HUNT 9.36 [ -26.14, 44.85]
IMAGEN -30.42 [ -60.84, 0.01]
IMpACT 37.94 [ -29.87, 105.76]
LBC1936 12.19 [ -34.60, 58.98]
LIBD -44.84 [ -95.83, 6.16]
Sydney MAS 6.70 [ -32.50, 45.90]
MCIC 55.50 [ -24.07, 135.07]
MooDS -0.71 [ -61.90, 60.49]
MPIP -22.53 [ -61.77, 16.71]
MUNSTER 0.30 [ -34.59, 35.19]
NCNG -17.97 [ -75.98, 40.04]
NESDA 15.45 [ -62.13, 93.02]
NeuroIMAGE 6.02 [ -74.30, 86.35]
NTR -62.90 [-128.15, 2.35]
OATS 38.31 [ -41.03, 117.65]
PAFIP 9.82 [ -89.40, 109.04]
PDNZ -74.30 [-170.29, 21.69]
PING -43.14 [-101.15, 14.86]
MPRC 6.59 [ -39.83, 53.02]
PPMI
QTIM -33.51 [ -69.98, 2.96]
SHIP -23.31 [ -57.00, 10.39]
SHIP-Trend -46.11 [ -86.61, -5.60]
SYS -22.24 [ -91.42, 46.95]
TCD-NUIG 17.03 [ -64.10, 98.16]
TOP -21.43 [ -73.42, 30.55]
TOP3T -12.40 [ -72.74, 47.95]
UiO2016 29.92 [ -50.61, 110.45]
UiO2017 18.21 [ -49.63, 86.06]
UMCU -1.56 [ -48.63, 45.52]
ENIGMA Discovery Cohorts [N=22379; P=1.328e-05] -16.74 [ -24.27, -9.21]
UKBB -20.57 [ -31.64, -9.49]
Discovery+UKBB [N=32028; P=2.681e-08] -17.98 [ -24.32, -11.64]
GOBS 6.15 [ -23.01, 35.31]
IMH -40.72 [-239.66, 158.22]
UNICAMP 14.80 [ -54.97, 84.57]
Meth-CT
MIRECC
UKBB nonEUR -14.95 [ -71.66, 41.76]
OSAKA -111.19 [-227.85, 5.47]
PING Non European -26.12 [ -79.64, 27.41]
Non European Cohorts [N=2529; P=0.5907] -5.93 [ -27.56, 15.69]
Beta [Lower 95% CI,Upper 95% CI]
-300
-200
-100
0
100
200
mm^2/effect allele

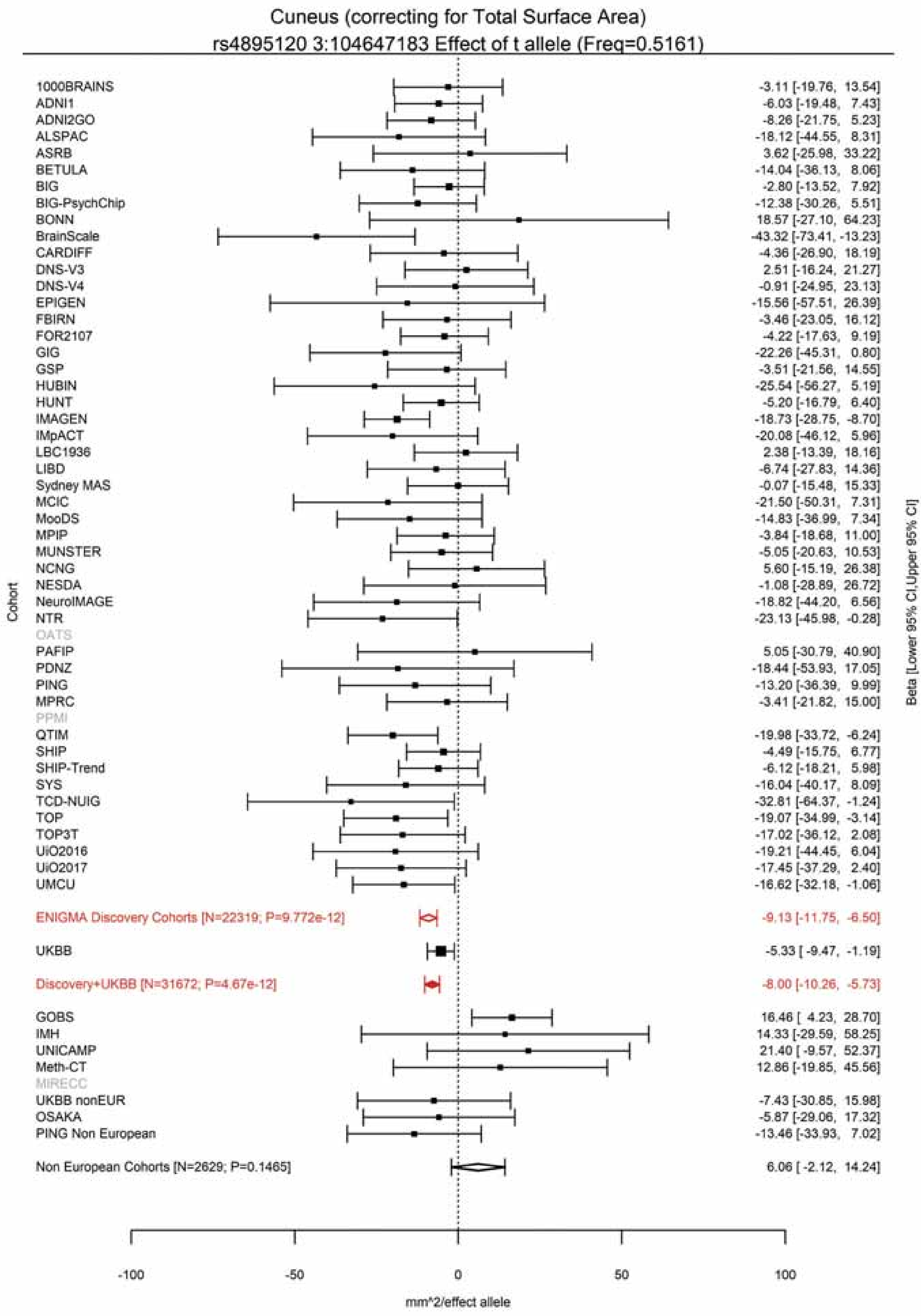

Cuneus (correcting for Total Surface Area)
rs4895120 3:104647183 Effect of t allele (Freq=0.5161)
Cohort
1000BRAINS -3.11 [-19.76, 13.54]
ADNI1 -6.03 [-19.48, 7.43]
ADNI2GO -8.26 [-21.75, 5.23]
ALSPAC -18.12 [-44.55, 8.31]
ASRB 3.62 [-25.98, 33.22]
BETULA -14.04 [-36.13, 8.06]
BIG -2.80 [-13.52, 7.92]
BIG-PsychChip -12.38 [-30.26, 5.51]
BONN 18.57 [-27.10, 64.23]
BrainScale -43.32 [-73.41, -13.23]
CARDIFF -4.36 [-26.90, 18.19]
DNS-V3 2.51 [-16.24, 21.27]
DNS-V4 -0.91 [-24.95, 23.13]
EPIGEN -15.56 [-57.51, 26.39]
FBIRN -3.46 [-23.05, 16.12]
FOR2107 -4.22 [-17.63, 9.19]
GIG -22.26 [-45.31, 0.80]
GSP -3.51 [-21.56, 14.55]
HUBIN -25.54 [-56.27, 5.19]
HUNT -5.20 [-16.79, 6.40]
IMAGEN -18.73 [-28.75, -8.70]
IMpACT -20.08 [-46.12, 5.96]
LBC1936 2.38 [-13.39, 18.16]
LIBD -6.74 [-27.83, 14.36]
Sydney MAS -0.07 [-15.48, 15.33]
MCIC -21.50 [-50.31, 7.31]
MooDS -14.83 [-36.99, 7.34]
MPIP -3.84 [-18.68, 11.00]
MUNSTER -5.05 [-20.63, 10.53]
NCNG 5.60 [-15.19, 26.38]
NESDA -1.08 [-28.89, 26.72]
NeuroIMAGE -18.82 [-44.20, 6.56]
NTR -23.13 [-45.98, -0.28]
OATS
PAFIP 5.05 [-30.79, 40.90]
PDNZ -18.44 [-53.93, 17.05]
PING -13.20 [-36.39, 9.99]
MPRC -3.41 [-21.82, 15.00]
PPMI
QTIM -19.98 [-33.72, -6.24]
SHIP -4.49 [-15.75, 6.77]
SHIP-Trend -6.12 [-18.21, 5.98]
SYS -16.04 [-40.17, 8.09]
TCD-NUIG -32.81 [-64.37, -1.24]
TOP -19.07 [-34.99, -3.14]
TOP3T -17.02 [-36.12, 2.08]
UiO2016 -19.21 [-44.45, 6.04]
UiO2017 -17.45 [-37.29, 2.40]
UMCU -16.62 [-32.18, -1.06]
ENIGMA Discovery Cohorts [N=22319; P=9.772e-12] -9.13 [-11.75, -6.50]
UKBB -5.33 [ -9.47, -1.19]
Discovery+UKBB [N=31672; P=4.67e-12] -8.00 [-10.26, -5.73]
GOBS 16.46 [ 4.23, 28.70]
IMH 14.33 [-29.59, 58.25]
UNICAMP 21.40 [ -9.57, 52.37]
Meth-CT 12.86 [-19.85, 45.56]
MIRECC
UKBB nonEUR -7.43 [-30.85, 15.98]
OSAKA -5.87 [-29.06, 17.32]
PING Non European -13.46 [-33.93, 7.02]
Non European Cohorts [N=2629; P=0.1465] 6.06 [ -2.12, 14.24]
-100
-50
0
50
100
mm^2/effect allele
Beta [Lower 95% CI,Upper 95% CI]

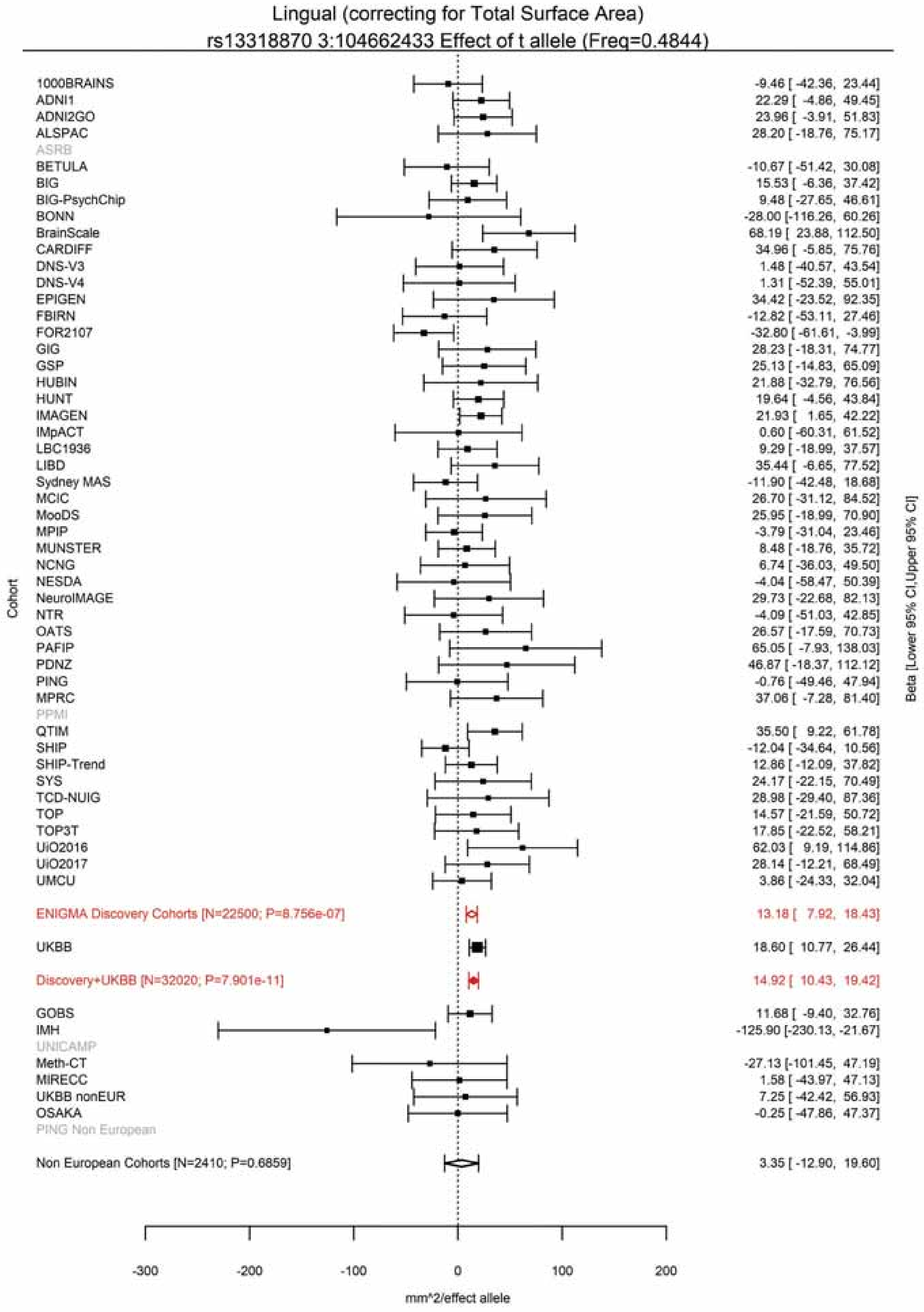
Lingual (correcting for Total Surface Area)
rs13318870 3:104662433 Effect of t allele (Freq=0.4844)
Cohort
1000BRAINS -9.46 [ -42.36, 23.44]
ADNI1 22.29 [ -4.86, 49.45]
ADNI2GO 23.96 [ -3.91, 51.83]
ALSPAC 28.20 [ -18.76, 75.17]
ASRB
BETULA -10.67 [ -51.42, 30.08]
BIG 15.53 [ -6.36, 37.42]
BIG-PsychChip 9.48 [ -27.65, 46.61]
BONN -28.00 [-116.26, 60.26]
BrainScale 68.19 [ 23.88, 112.50]
CARDIFF 34.96 [ -5.85, 75.76]
DNS-V3 1.48 [ -40.57, 43.54]
DNS-V4 1.31 [ -52.39, 55.01]
EPIGEN 34.42 [ -23.52, 92.35]
FBIRN -12.82 [ -53.11, 27.46]
FOR2107 -32.80 [ -61.61, -3.99]
GIG 28.23 [ -18.31, 74.77]
GSP 25.13 [ -14.83, 65.09]
HUBIN 21.88 [ -32.79, 76.56]
HUNT 19.64 [ -4.56, 43.84]
IMAGEN 21.93 [ 1.65, 42.22]
IMpACT 0.60 [ -60.31, 61.52]
LBC1936 9.29 [ -18.99, 37.57]
LIBD 35.44 [ -6.65, 77.52]
Sydney MAS -11.90 [ -42.48, 18.68]
MCIC 26.70 [ -31.12, 84.52]
MooDS 25.95 [ -18.99, 70.90]
MPIP -3.79 [ -31.04, 23.46]
MUNSTER 8.48 [ -18.76, 35.72]
NCNG 6.74 [ -36.03, 49.50]
NESDA -4.04 [ -58.47, 50.39]
NeuroIMAGE 29.73 [ -22.68, 82.13]
NTR -4.09 [ -51.03, 42.85]
OATS 26.57 [ -17.59, 70.73]
PAFIP 65.05 [ -7.93, 138.03]
PDNZ 46.87 [ -18.37, 112.12]
PING -0.76 [ -49.46, 47.94]
MPRC 37.06 [ -7.28, 81.40]
PPMI
QTIM 35.50 [ 9.22, 61.78]
SHIP -12.04 [ -34.64, 10.56]
SHIP-Trend 12.86 [ -12.09, 37.82]
SYS 24.17 [ -22.15, 70.49]
TCD-NUIG 28.98 [ -29.40, 87.36]
TOP 14.57 [ -21.59, 50.72]
TOP3T 17.85 [ -22.52, 58.21]
UiO2016 62.03 [ 9.19, 114.86]
UiO2017 28.14 [ -12.21, 68.49]
UMCU 3.86 [ -24.33, 32.04]
ENIGMA Discovery Cohorts [N=22500; P=8.756e-07] 13.18 [ 7.92, 18.43]
UKBB 18.60 [ 10.77, 26.44]
Discovery+UKBB [N=32020; P=7.901e-11] 14.92 [ 10.43, 19.42]
GOBS 11.68 [ -9.40, 32.76]
IMH -125.90 [-230.13, -21.67]
UNICAMP
Meth-CT -27.13 [-101.45, 47.19]
MIRECC 1.58 [ -43.97, 47.13]
UKBB nonEUR 7.25 [ -42.42, 56.93]
OSAKA -0.25 [ -47.86, 47.37]
PING Non European
Non European Cohorts [N=2410; P=0.6859] 3.35 [ -12.90, 19.60]
Beta [Lower 95% CI,Upper 95% CI]
-300
-200
-100
0
100
200
mm^2/effect allele

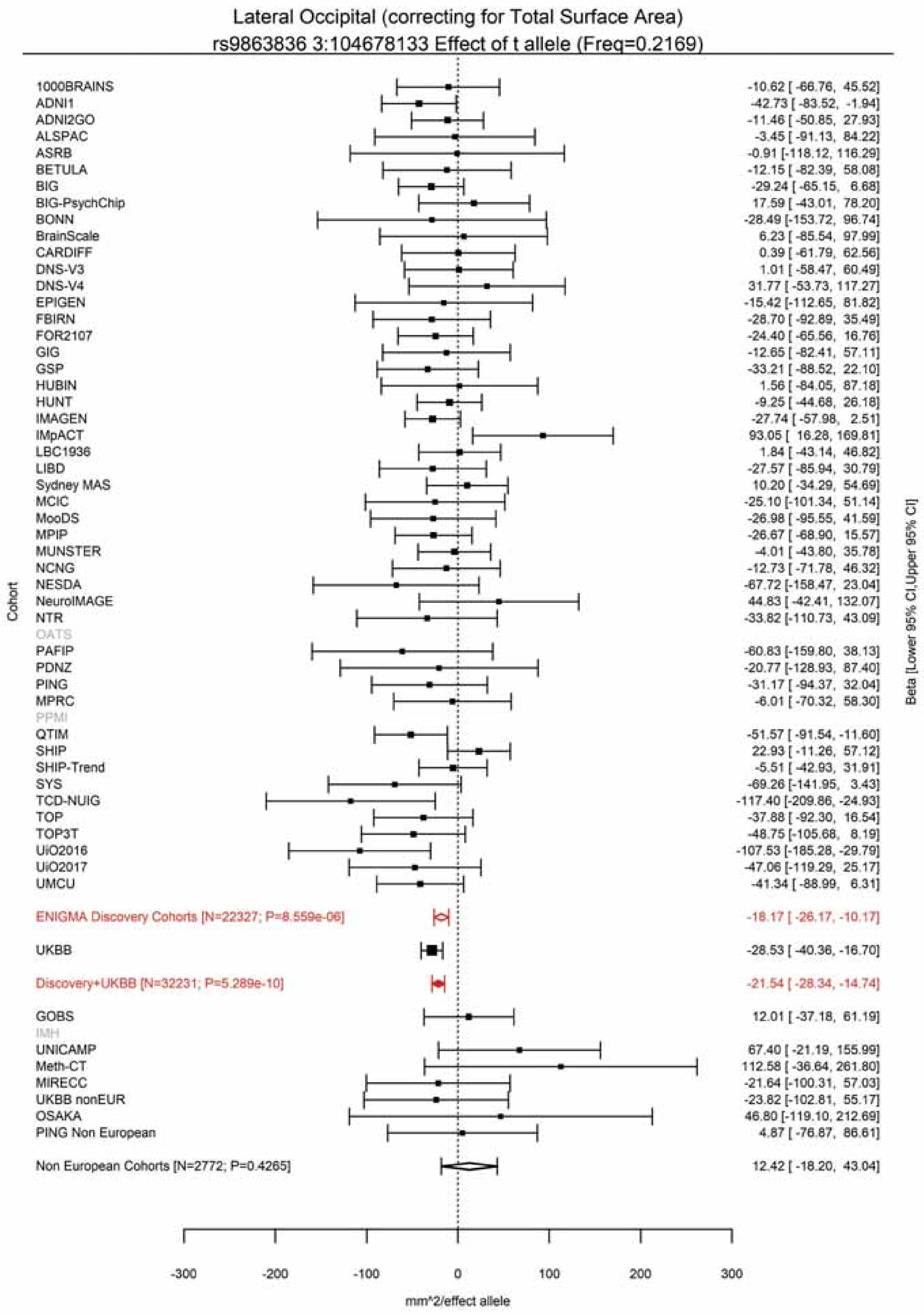

Lateral Occipital (correcting for Total Surface Area)
rs9863836 3:104678133 Effect of t allele (Freq=0.2169)
Cohort
1000BRAINS -10.62 [ -66.76, 45.52]
ADNI1 -42.73 [ -83.52, -1.94]
ADNI2GO -11.46 [ -50.85, 27.93]
ALSPAC -3.45 [ -91.13, 84.22]
ASRB -0.91 [-118.12, 116.29]
BETULA -12.15 [ -82.39, 58.08]
BIG -29.24 [ -65.15, 6.68]
BIG-PsychChip 17.59 [ -43.01, 78.20]
BONN -28.49 [-153.72, 96.74]
BrainScale 6.23 [ -85.54, 97.99]
CARDIFF 0.39 [ -61.79, 62.56]
DNS-V3 1.01 [ -58.47, 60.49]
DNS-V4 31.77 [ -53.73, 117.27]
EPIGEN -15.42 [-112.65, 81.82]
FBIRN -28.70 [ -92.89, 35.49]
FOR2107 -24.40 [ -65.56, 16.76]
GIG -12.65 [ -82.41, 57.11]
GSP -33.21 [ -88.52, 22.10]
HUBIN 1.56 [ -84.05, 87.18]
HUNT -9.25 [ -44.68, 26.18]
IMAGEN -27.74 [ -57.98, 2.51]
IMpACT 93.05 [ 16.28, 169.81]
LBC1936 1.84 [ -43.14, 46.82]
LIBD -27.57 [ -85.94, 30.79]
Sydney MAS 10.20 [ -34.29, 54.69]
MCIC -25.10 [-101.34, 51.14]
MooDS -26.98 [ -95.55, 41.59]
MPIP -26.67 [ -68.90, 15.57]
MUNSTER -4.01 [ -43.80, 35.78]
NCNG -12.73 [ -71.78, 46.32]
NESDA -67.72 [-158.47, 23.04]
NeuroIMAGE 44.83 [ -42.41, 132.07]
NTR -33.82 [-110.73, 43.09]
OATS
PAFIP -60.83 [-159.80, 38.13]
PDNZ -20.77 [-128.93, 87.40]
PING -31.17 [ -94.37, 32.04]
MPRC -6.01 [ -70.32, 58.30]
PPMI
QTIM -51.57 [ -91.54, -11.60]
SHIP 22.93 [ -11.26, 57.12]
SHIP-Trend -5.51 [ -42.93, 31.91]
SYS -69.26 [-141.95, 3.43]
TCD-NUIG -117.40 [-209.86, -24.93]
TOP -37.88 [ -92.30, 16.54]
TOP3T -48.75 [-105.68, 8.19]
UiO2016 -107.53 [-185.28, -29.79]
UiO2017 -47.06 [-119.29, 25.17]
UMCU -41.34 [ -88.99, 6.31]
ENIGMA Discovery Cohorts [N=22327; P=8.559e-06] -18.17 [ -26.17, -10.17]
UKBB -28.53 [ -40.36, -16.70]
Discovery+UKBB [N=32231; P=5.289e-10] -21.54 [ -28.34, -14.74]
GOBS 12.01 [ -37.18, 61.19]
IMH
UNICAMP 67.40 [ -21.19, 155.99]
Meth-CT 112.58 [ -36.64, 261.80]
MIRECC -21.64 [-100.31, 57.03]
UKBB nonEUR -23.82 [-102.81, 55.17]
OSAKA 46.80 [-119.10, 212.69]
PING Non European 4.87 [ -76.87, 86.61]
Non European Cohorts [N=2772; P=0.4265] 12.42 [ -18.20, 43.04]
Beta [Lower 95% CI,Upper 95% CI]
-300
-200
-100
0
100
200
300
mm^2/effect allele

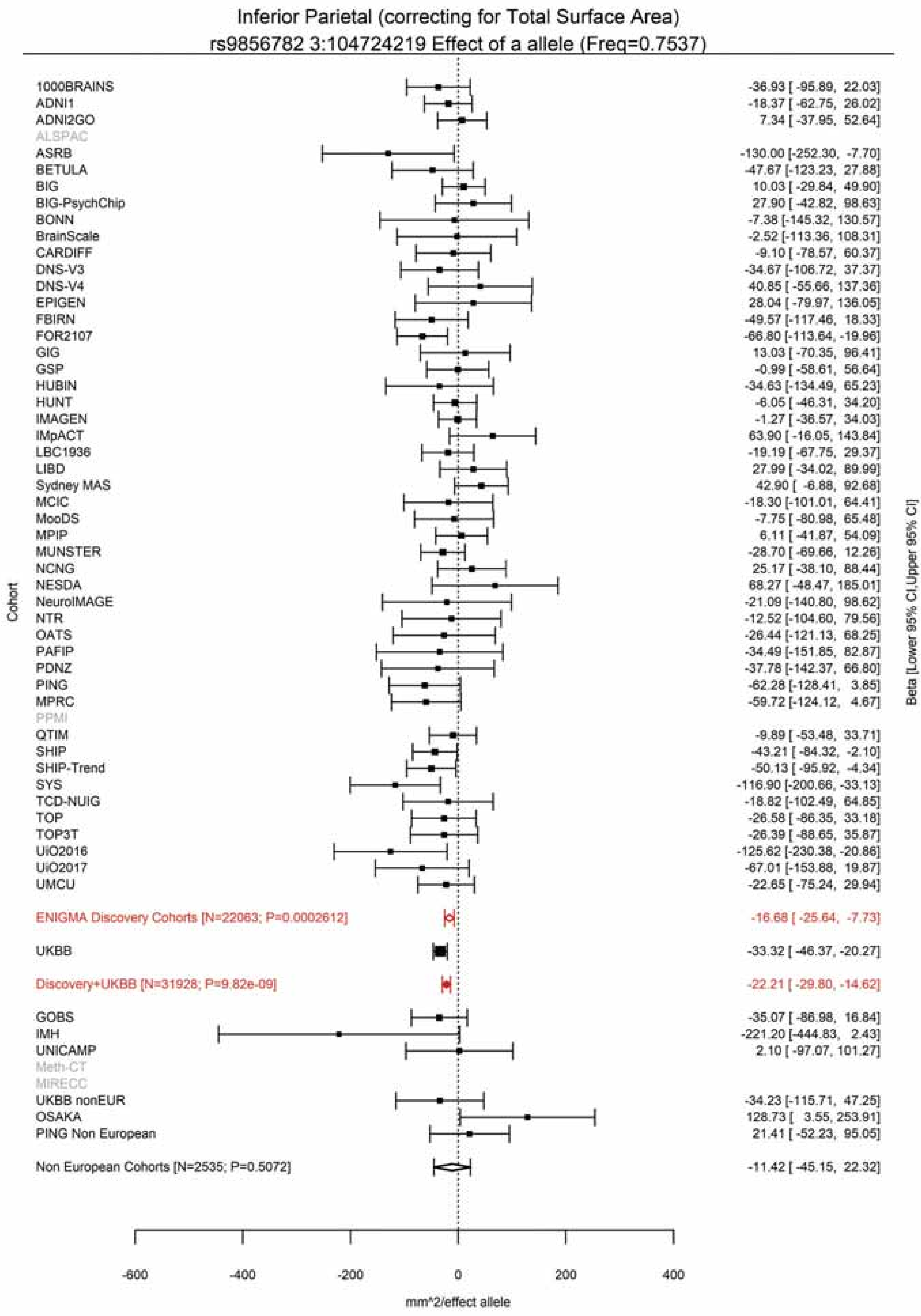

Inferior Parietal (correcting for Total Surface Area)
rs9856782 3:104724219 Effect of a allele (Freq=0.7537)
Cohort
1000BRAINS -36.93 [ -95.89, 22.03]
ADNI1 -18.37 [ -62.75, 26.02]
ADNI2GO 7.34 [ -37.95, 52.64]
ALSPAC
ASRB -130.00 [-252.30, -7.70]
BETULA -47.67 [-123.23, 27.88]
BIG 10.03 [ -29.84, 49.90]
BIG-PsychChip 27.90 [ -42.82, 98.63]
BONN -7.38 [-145.32, 130.57]
BrainScale -2.52 [-113.36, 108.31]
CARDIFF -9.10 [ -78.57, 60.37]
DNS-V3 -34.67 [-106.72, 37.37]
DNS-V4 40.85 [ -55.66, 137.36]
EPIGEN 28.04 [ -79.97, 136.05]
FBIRN -49.57 [-117.46, 18.33]
FOR2107 -66.80 [-113.64, -19.96]
GIG 13.03 [ -70.35, 96.41]
GSP -0.99 [ -58.61, 56.64]
HUBIN -34.63 [-134.49, 65.23]
HUNT -6.05 [ -46.31, 34.20]
IMAGEN -1.27 [ -36.57, 34.03]
IMpACT 63.90 [ -16.05, 143.84]
LBC1936 -19.19 [ -67.75, 29.37]
LIBD 27.99 [ -34.02, 89.99]
Sydney MAS 42.90 [ -6.88, 92.68]
MCIC -18.30 [-101.01, 64.41]
MooDS -7.75 [ -80.98, 65.48]
MPIP 6.11 [ -41.87, 54.09]
MUNSTER -28.70 [ -69.66, 12.26]
NCNG 25.17 [ -38.10, 88.44]
NESDA 68.27 [ -48.47, 185.01]
NeuroIMAGE -21.09 [-140.80, 98.62]
NTR -12.52 [-104.60, 79.56]
OATS -26.44 [-121.13, 68.25]
PAFIP -34.49 [-151.85, 82.87]
PDNZ -37.78 [-142.37, 66.80]
PING -62.28 [-128.41, 3.85]
MPRC -59.72 [-124.12, 4.67]
PPMI
QTIM -9.89 [ -53.48, 33.71]
SHIP -43.21 [ -84.32, -2.10]
SHIP-Trend -50.13 [ -95.92, -4.34]
SYS -116.90 [-200.66, -33.13]
TCD-NUIG -18.82 [-102.49, 64.85]
TOP -26.58 [ -86.35, 33.18]
TOP3T -26.39 [ -88.65, 35.87]
UiO2016 -125.62 [-230.38, -20.86]
UiO2017 -67.01 [-153.88, 19.87]
UMCU -22.65 [ -75.24, 29.94]
ENIGMA Discovery Cohorts [N=22063; P=0.0002612] -16.68 [ -25.64, -7.73]
UKBB -33.32 [ -46.37, -20.27]
Discovery+UKBB [N=31928; P=9.82e-09] -22.21 [ -29.80, -14.62]
GOBS -35.07 [ -86.98, 16.84]
IMH -221.20 [-444.83, 2.43]
UNICAMP 2.10 [ -97.07, 101.27]
Meth-CT
MIRECC
UKBB nonEUR -34.23 [-115.71, 47.25]
OSAKA 128.73 [ 3.55, 253.91]
PING Non European 21.41 [ -52.23, 95.05]
Non European Cohorts [N=2535; P=0.5072] -11.42 [ -45.15, 22.32]
-600
-400
-200
0
200
400
mm^2/effect allele
Beta [Lower 95% CI,Upper 95% CI]

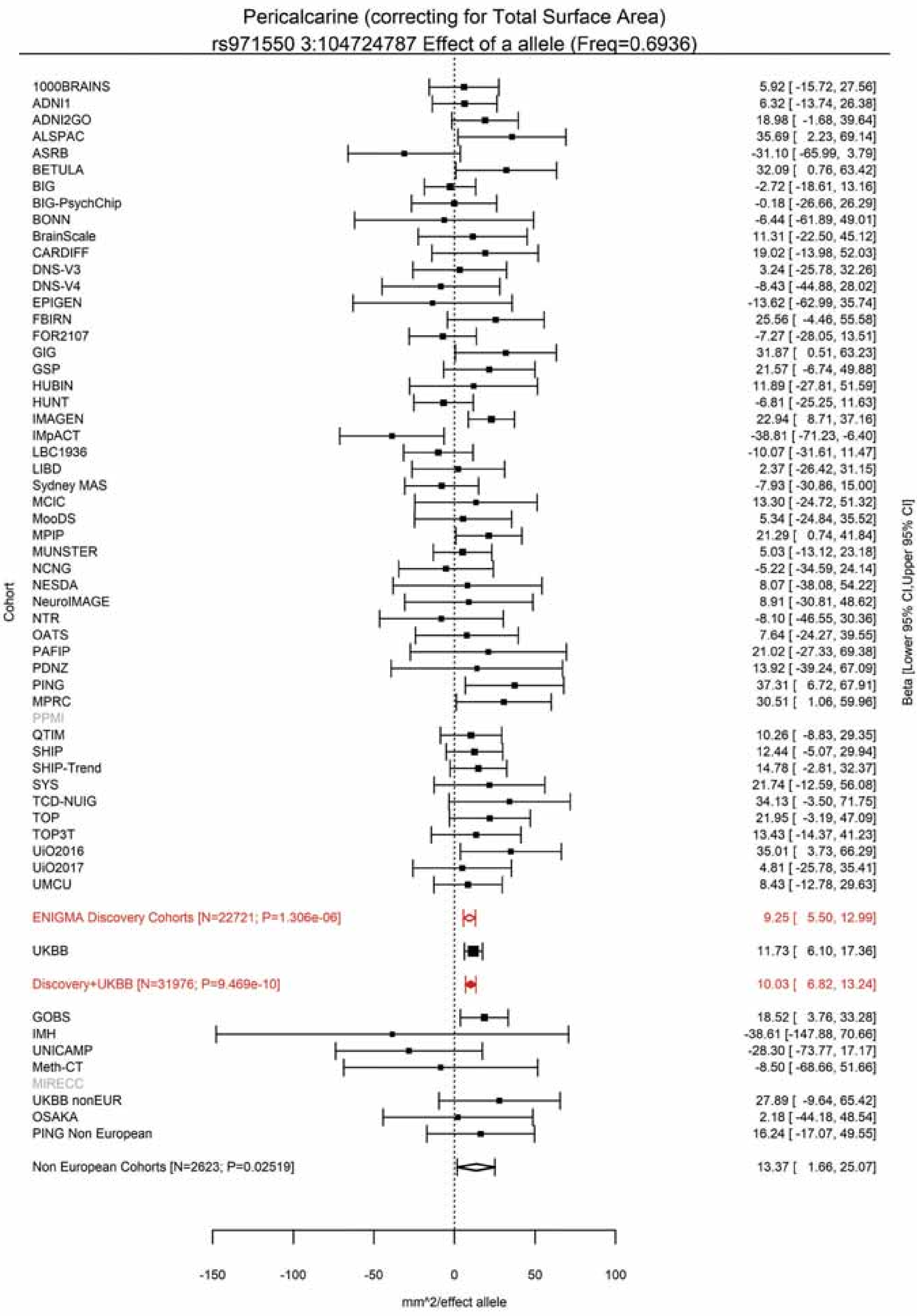

Pericalcarine (correcting for Total Surface Area)
rs971550 3:104724787 Effect of a allele (Freq=0.6936)
Cohort
1000BRAINS 5.92 [ -15.72, 27.56]
ADNI1 6.32 [ -13.74, 26.38]
ADNI2GO 18.98 [ -1.68, 39.64]
ALSPAC 35.69 [ 2.23, 69.14]
ASRB -31.10 [ -65.99, 3.79]
BETULA 32.09 [ 0.76, 63.42]
BIG -2.72 [ -18.61, 13.16]
BIG-PsychChip -0.18 [ -26.66, 26.29]
BONN -6.44 [ -61.89, 49.01]
BrainScale 11.31 [ -22.50, 45.12]
CARDIFF 19.02 [ -13.98, 52.03]
DNS-V3 3.24 [ -25.78, 32.26]
DNS-V4 -8.43 [ -44.88, 28.02]
EPIGEN -13.62 [ -62.99, 35.74]
FBIRN 25.56 [ -4.46, 55.58]
FOR2107 -7.27 [ -28.05, 13.51]
GIG 31.87 [ 0.51, 63.23]
GSP 21.57 [ -6.74, 49.88]
HUBIN 11.89 [ -27.81, 51.59]
HUNT -6.81 [ -25.25, 11.63]
IMAGEN 22.94 [ 8.71, 37.16]
IMpACT -38.81 [ -71.23, -6.40]
LBC1936 -10.07 [ -31.61, 11.47]
LIBD 2.37 [ -26.42, 31.15]
Sydney MAS -7.93 [ -30.86, 15.00]
MCIC 13.30 [ -24.72, 51.32]
MooDS 5.34 [ -24.84, 35.52]
MPIP 21.29 [ 0.74, 41.84]
MUNSTER 5.03 [ -13.12, 23.18]
NCNG -5.22 [ -34.59, 24.14]
NESDA 8.07 [ -38.08, 54.22]
NeuroIMAGE 8.91 [ -30.81, 48.62]
NTR -8.10 [ -46.55, 30.36]
OATS 7.64 [ -24.27, 39.55]
PAFIP 21.02 [ -27.33, 69.38]
PDNZ 13.92 [ -39.24, 67.09]
PING 37.31 [ 6.72, 67.91]
MPRC 30.51 [ 1.06, 59.96]
PPMI
QTIM 10.26 [ -8.83, 29.35]
SHIP 12.44 [ -5.07, 29.94]
SHIP-Trend 14.78 [ -2.81, 32.37]
SYS 21.74 [ -12.59, 56.08]
TCD-NUIG 34.13 [ -3.50, 71.75]
TOP 21.95 [ -3.19, 47.09]
TOP3T 13.43 [ -14.37, 41.23]
UiO2016 35.01 [ 3.73, 66.29]
UiO2017 4.81 [ -25.78, 35.41]
UMCU 8.43 [ -12.78, 29.63]
ENIGMA Discovery Cohorts [N=22721; P=1.306e-06] 9.25 [ 5.50, 12.99]
UKBB 11.73 [ 6.10, 17.36]
Discovery+UKBB [N=31976; P=9.469e-10] 10.03 [ 6.82, 13.24]
GOBS 18.52 [ 3.76, 33.28]
IMH -38.61 [-147.88, 70.66]
UNICAMP -28.30 [ -73.77, 17.17]
Meth-CT -8.50 [ -68.66, 51.66]
MIRECC
UKBB nonEUR 27.89 [ -9.64, 65.42]
OSAKA 2.18 [ -44.18, 48.54]
PING Non European 16.24 [ -17.07, 49.55]
Non European Cohorts [N=2623; P=0.02519] 13.37 [ 1.66, 25.07]
-150
-100
-50
0
50
100
mm^2/effect allele
Beta [Lower 95% CI,Upper 95% CI]

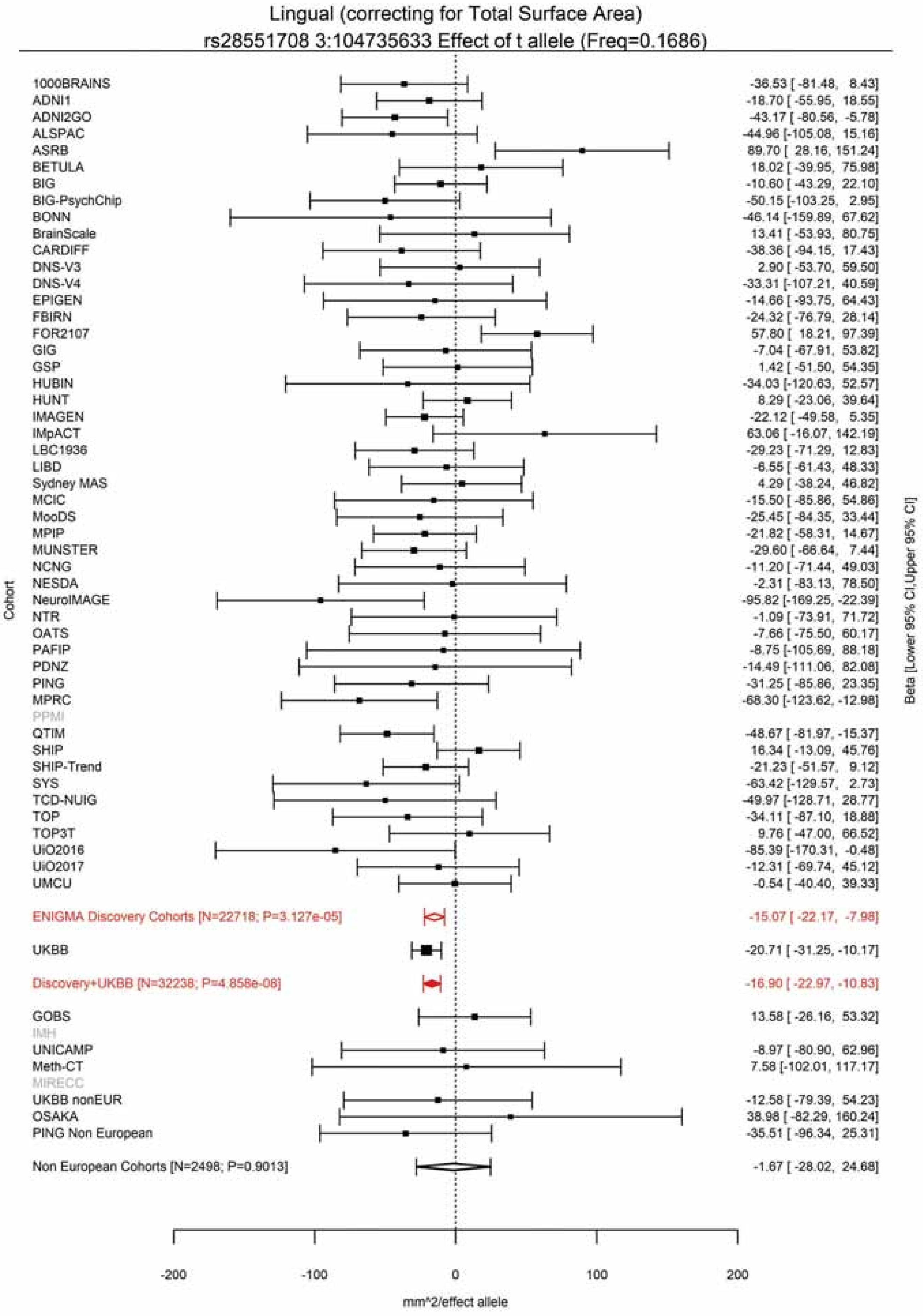
Lingual (correcting for Total Surface Area)
rs28551708 3:104735633 Effect of t allele (Freq=0.1686)
Cohort
1000BRAINS -36.53 [ -81.48, 8.43]
ADNI1 -18.70 [ -55.95, 18.55]
ADNI2GO -43.17 [ -80.56, -5.78]
ALSPAC -44.96 [-105.08, 15.16]
ASRB 89.70 [ 28.16, 151.24]
BETULA 18.02 [ -39.95, 75.98]
BIG -10.60 [ -43.29, 22.10]
BIG-PsychChip -50.15 [-103.25, 2.95]
BONN -46.14 [-159.89, 67.62]
BrainScale 13.41 [ -53.93, 80.75]
CARDIFF -38.36 [ -94.15, 17.43]
DNS-V3 2.90 [ -53.70, 59.50]
DNS-V4 -33.31 [-107.21, 40.59]
EPIGEN -14.66 [ -93.75, 64.43]
FBIRN -24.32 [ -76.79, 28.14]
FOR2107 57.80 [ 18.21, 97.39]
GIG -7.04 [ -67.91, 53.82]
GSP 1.42 [ -51.50, 54.35]
HUBIN -34.03 [-120.63, 52.57]
HUNT 8.29 [ -23.06, 39.64]
IMAGEN -22.12 [ -49.58, 5.35]
IMpACT 63.06 [ -16.07, 142.19]
LBC1936 -29.23 [ -71.29, 12.83]
LIBD -6.55 [ -61.43, 48.33]
Sydney MAS 4.29 [ -38.24, 46.82]
MCIC -15.50 [ -85.86, 54.86]
MooDS -25.45 [ -84.35, 33.44]
MPIP -21.82 [ -58.31, 14.67]
MUNSTER -29.60 [ -66.64, 7.44]
NCNG -11.20 [ -71.44, 49.03]
NESDA -2.31 [ -83.13, 78.50]
NeuroIMAGE -95.82 [-169.25, -22.39]
NTR -1.09 [ -73.91, 71.72]
OATS -7.66 [ -75.50, 60.17]
PAFIP -8.75 [-105.69, 88.18]
PDNZ -14.49 [-111.06, 82.08]
PING -31.25 [ -85.86, 23.35]
MPRC -68.30 [-123.62, -12.98]
PPMI
QTIM -48.67 [ -81.97, -15.37]
SHIP 16.34 [ -13.09, 45.76]
SHIP-Trend -21.23 [ -51.57, 9.12]
SYS -63.42 [-129.57, 2.73]
TCD-NUIG -49.97 [-128.71, 28.77]
TOP -34.11 [ -87.10, 18.88]
TOP3T 9.76 [ -47.00, 66.52]
UiO2016 -85.39 [-170.31, -0.48]
UiO2017 -12.31 [ -69.74, 45.12]
UMCU -0.54 [ -40.40, 39.33]
ENIGMA Discovery Cohorts [N=22718; P=3.127e-05] -15.07 [ -22.17, -7.98]
UKBB -20.71 [ -31.25, -10.17]
Discovery+UKBB [N=32238; P=4.858e-08] -16.90 [ -22.97, -10.83]
GOBS 13.58 [ -26.16, 53.32]
IMH
UNICAMP -8.97 [ -80.90, 62.96]
Meth-CT 7.58 [-102.01, 117.17]
MIRECC
UKBB nonEUR -12.58 [ -79.39, 54.23]
OSAKA 38.98 [ -82.29, 160.24]
PING Non European -35.51 [ -96.34, 25.31]
Non European Cohorts [N=2498; P=0.9013] -1.67 [ -28.02, 24.68]
-200
-100
0
100
200
mm^2/effect allele
Beta [Lower 95% CI,Upper 95% CI]

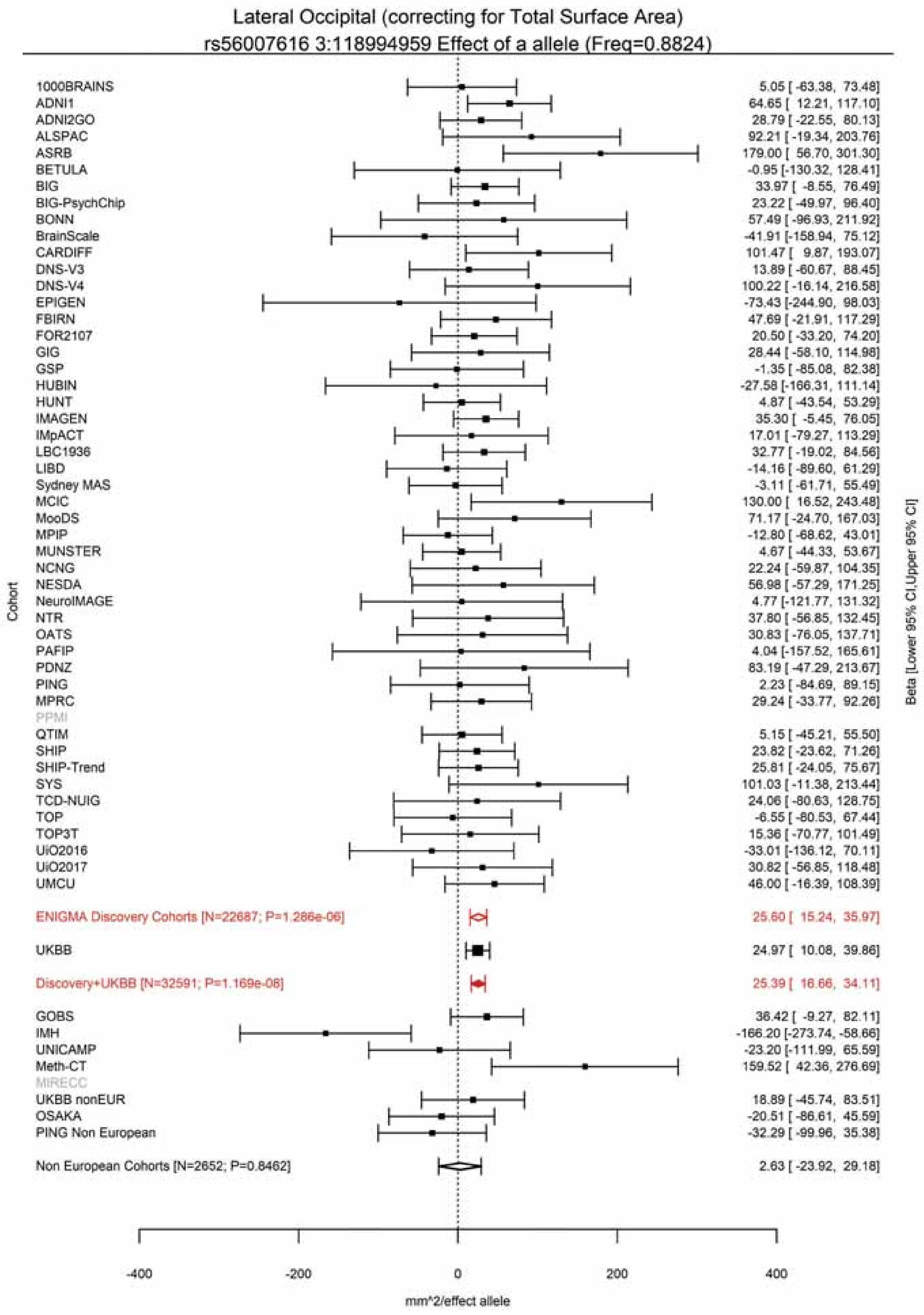
Lateral Occipital (correcting for Total Surface Area)
rs56007616 3:118994959 Effect of a allele (Freq=0.8824)
Cohort
1000BRAINS 5.05 [ -63.38, 73.48]
ADNI1 64.65 [ 12.21, 117.10]
ADNI2GO 28.79 [ -22.55, 80.13]
ALSPAC 92.21 [ -19.34, 203.76]
ASRB 179.00 [ 56.70, 301.30]
BETULA -0.95 [-130.32, 128.41]
BIG 33.97 [ -8.55, 76.49]
BIG-PsychChip 23.22 [ -49.97, 96.40]
BONN 57.49 [ -96.93, 211.92]
BrainScale -41.91 [-158.94, 75.12]
CARDIFF 101.47 [ 9.87, 193.07]
DNS-V3 13.89 [ -60.67, 88.45]
DNS-V4 100.22 [ -16.14, 216.58]
EPIGEN -73.43 [-244.90, 98.03]
FBIRN 47.69 [ -21.91, 117.29]
FOR2107 20.50 [ -33.20, 74.20]
GIG 28.44 [ -58.10, 114.98]
GSP -1.35 [ -85.08, 82.38]
HUBIN -27.58 [-166.31, 111.14]
HUNT 4.87 [ -43.54, 53.29]
IMAGEN 35.30 [ -5.45, 76.05]
IMpACT 17.01 [ -79.27, 113.29]
LBC1936 32.77 [ -19.02, 84.56]
LIBD -14.16 [ -89.60, 61.29]
Sydney MAS -3.11 [ -61.71, 55.49]
MCIC 130.00 [ 16.52, 243.48]
MooDS 71.17 [ -24.70, 167.03]
MPIP -12.80 [ -68.62, 43.01]
MUNSTER 4.67 [ -44.33, 53.67]
NCNG 22.24 [ -59.87, 104.35]
NESDA 56.98 [ -57.29, 171.25]
NeuroIMAGE 4.77 [-121.77, 131.32]
NTR 37.80 [ -56.85, 132.45]
OATS 30.83 [ -76.05, 137.71]
PAFIP 4.04 [-157.52, 165.61]
PDNZ 83.19 [ -47.29, 213.67]
PING 2.23 [ -84.69, 89.15]
MPRC 29.24 [ -33.77, 92.26]
PPMI
QTIM 5.15 [ -45.21, 55.50]
SHIP 23.82 [ -23.62, 71.26]
SHIP-Trend 25.81 [ -24.05, 75.67]
SYS 101.03 [ -11.38, 213.44]
TCD-NUIG 24.06 [ -80.63, 128.75]
TOP -6.55 [ -80.53, 67.44]
TOP3T 15.36 [ -70.77, 101.49]
UiO2016 -33.01 [-136.12, 70.11]
UiO2017 30.82 [ -56.85, 118.48]
UMCU 46.00 [ -16.39, 108.39]
ENIGMA Discovery Cohorts [N=22687; P=1.286e-06] 25.60 [ 15.24, 35.97]
UKBB 24.97 [ 10.08, 39.86]
Discovery+UKBB [N=32591; P=1.169e-08] 25.39 [ 16.66, 34.11]
GOBS 36.42 [ -9.27, 82.11]
IMH -166.20 [-273.74, -58.66]
UNICAMP -23.20 [-111.99, 65.59]
Meth-CT 159.52 [ 42.36, 276.69]
MIRECC
UKBB nonEUR 18.89 [ -45.74, 83.51]
OSAKA -20.51 [ -86.61, 45.59]
PING Non European -32.29 [ -99.96, 35.38]
Non European Cohorts [N=2652; P=0.8462] 2.63 [ -23.92, 29.18]
Beta [Lower 95% CI,Upper 95% CI]
-400
-200
0
200
400
mm^2/effect allele

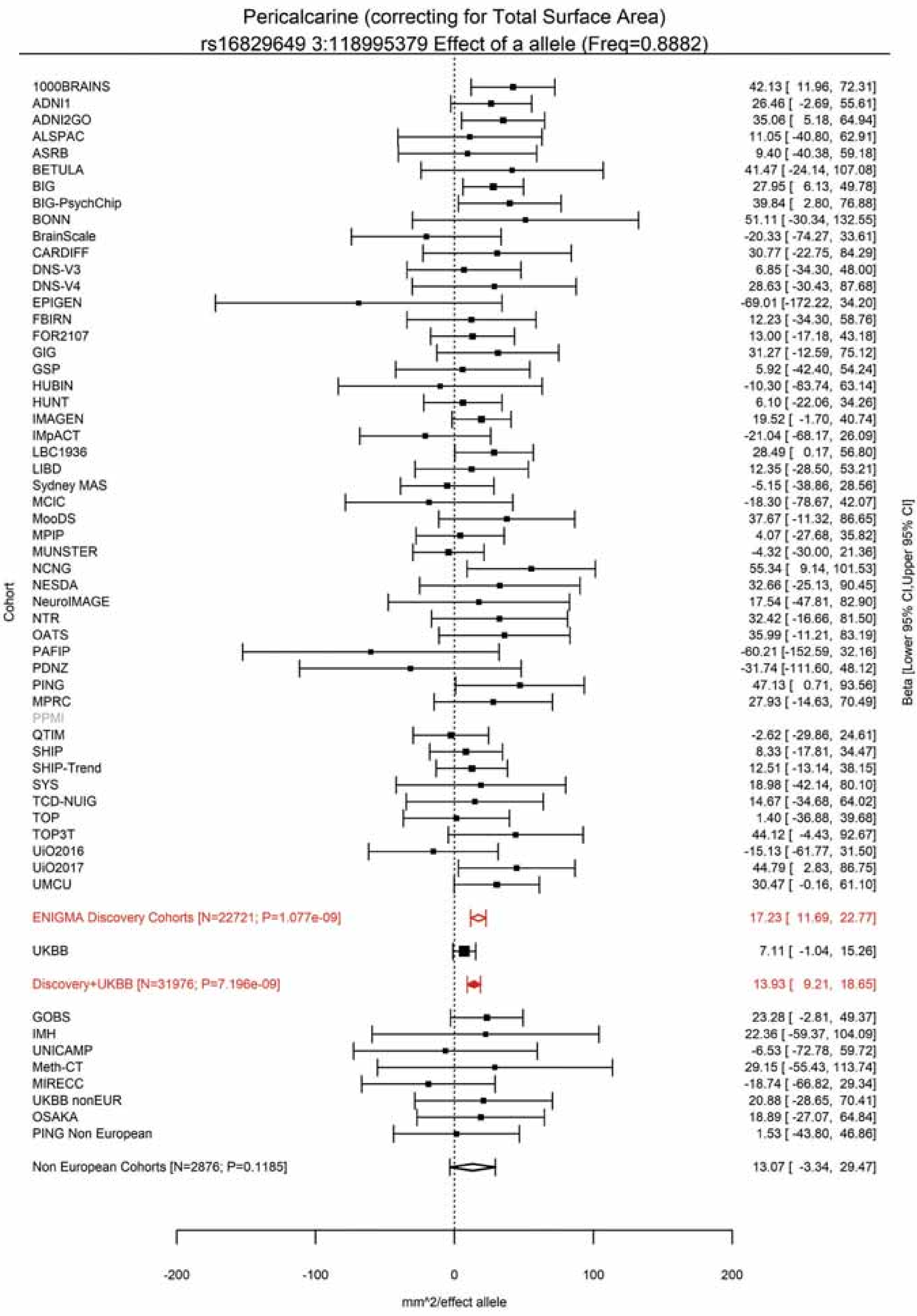

Pericalcarine (correcting for Total Surface Area)
rs16829649 3:118995379 Effect of a allele (Freq=0.8882)
1000BRAINS 42.13 [ 11.96, 72.31]
ADNI1 26.46 [ -2.69, 55.61]
ADNI2GO 35.06 [ 5.18, 64.94]
ALSPAC 11.05 [ -40.80, 62.91]
ASRB 9.40 [ -40.38, 59.18]
BETULA 41.47 [ -24.14, 107.08]
BIG 27.95 [ 6.13, 49.78]
BIG-PsychChip 39.84 [ 2.80, 76.88]
BONN 51.11 [ -30.34, 132.55]
BrainScale -20.33 [ -74.27, 33.61]
CARDIFF 30.77 [ -22.75, 84.29]
DNS-V3 6.85 [ -34.30, 48.00]
DNS-V4 28.63 [ -30.43, 87.68]
EPIGEN -69.01 [-172.22, 34.20]
FBIRN 12.23 [ -34.30, 58.76]
FOR2107 13.00 [ -17.18, 43.18]
GIG 31.27 [ -12.59, 75.12]
GSP 5.92 [ -42.40, 54.24]
HUBIN -10.30 [ -83.74, 63.14]
HUNT 6.10 [ -22.06, 34.26]
IMAGEN 19.52 [ -1.70, 40.74]
IMpACT -21.04 [ -68.17, 26.09]
LBC1936 28.49 [ 0.17, 56.80]
LIBD 12.35 [ -28.50, 53.21]
Sydney MAS -5.15 [ -38.86, 28.56]
MCIC -18.30 [ -78.67, 42.07]
MooDS 37.67 [ -11.32, 86.65]
MPIP 4.07 [ -27.68, 35.82]
MUNSTER -4.32 [ -30.00, 21.36]
NCNG 55.34 [ 9.14, 101.53]
NESDA 32.66 [ -25.13, 90.45]
NeuroIMAGE 17.54 [ -47.81, 82.90]
NTR 32.42 [ -16.66, 81.50]
OATS 35.99 [ -11.21, 83.19]
PAFIP -60.21 [-152.59, 32.16]
PDNZ -31.74 [-111.60, 48.12]
PING 47.13 [ 0.71, 93.56]
MPRC 27.93 [ -14.63, 70.49]
PPMI
QTIM -2.62 [ -29.86, 24.61]
SHIP 8.33 [ -17.81, 34.47]
SHIP-Trend 12.51 [ -13.14, 38.15]
SYS 18.98 [ -42.14, 80.10]
TCD-NUIG 14.67 [ -34.68, 64.02]
TOP 1.40 [ -36.88, 39.68]
TOP3T 44.12 [ -4.43, 92.67]
UiO2016 -15.13 [ -61.77, 31.50]
UiO2017 44.79 [ 2.83, 86.75]
UMCU 30.47 [ -0.16, 61.10]
ENIGMA Discovery Cohorts [N=22721; P=1.077e-09] 17.23 [ 11.69, 22.77]
UKBB 7.11 [ -1.04, 15.26]
Discovery+UKBB [N=31976; P=7.196e-09] 13.93 [ 9.21, 18.65]
GOBS 23.28 [ -2.81, 49.37]
IMH 22.36 [ -59.37, 104.09]
UNICAMP -6.53 [ -72.78, 59.72]
Meth-CT 29.15 [ -55.43, 113.74]
MIRECC -18.74 [ -66.82, 29.34]
UKBB nonEUR 20.88 [ -28.65, 70.41]
OSAKA 18.89 [ -27.07, 64.84]
PING Non European 1.53 [ -43.80, 46.86]
Non European Cohorts [N=2876; P=0.1185] 13.07 [ -3.34, 29.47]
Cohort
Beta [Lower 95% CI,Upper 95% CI]
-200
-100
0
100
200
mm^2/effect allele

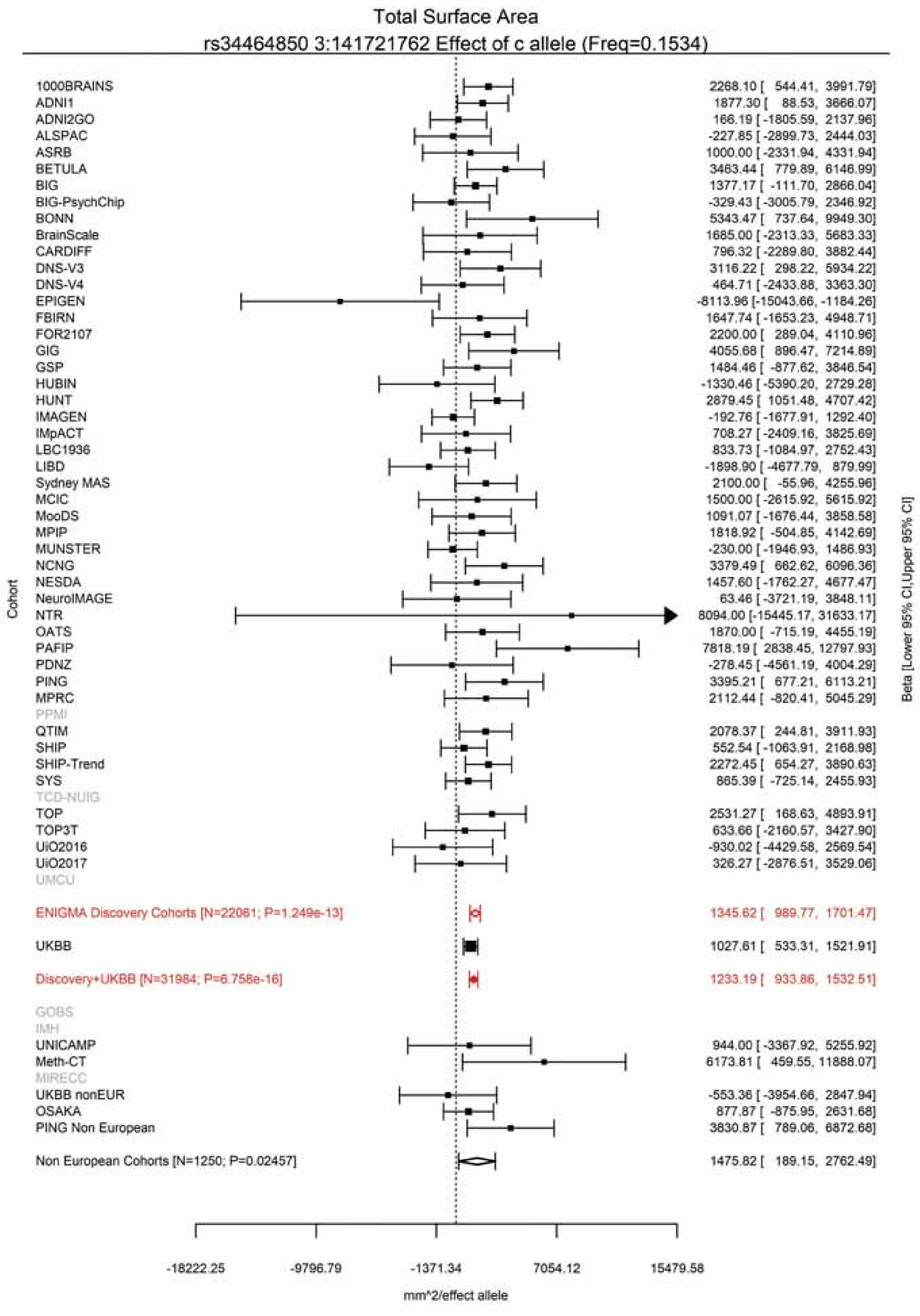
Total Surface Area
rs34464850 3:141721762 Effect of c allele (Freq=0.1534)
1000BRAINS 2268.10 [ 544.41, 3991.79]
ADNI1 1877.30 [ 88.53, 3666.07]
ADNI2GO 166.19 [ -1805.59, 2137.96]
ALSPAC -227.85 [ -2899.73, 2444.03]
ASRB 1000.00 [ -2331.94, 4331.94]
BETULA 3463.44 [ 779.89, 6146.99]
BIG 1377.17 [ -111.70, 2866.04]
BIG-PsychChip -329.43 [ -3005.79, 2346.92]
BONN 5343.47 [ 737.64, 9949.30]
BrainScale 1685.00 [ -2313.33, 5683.33]
CARDIFF 796.32 [ -2289.80, 3882.44]
DNS-V3 3116.22 [ 298.22, 5934.22]
DNS-V4 464.71 [ -2433.88, 3363.30]
EPIGEN -8113.96 [-15043.66, -1184.26]
FBIRN 1647.74 [ -1653.23, 4948.71]
FOR2107 2200.00 [ 289.04, 4110.96]
GIG 4055.68 [ 896.47, 7214.89]
GSP 1484.46 [ -877.62, 3846.54]
HUBIN -1330.46 [ -5390.20, 2729.28]
HUNT 2879.45 [ 1051.48, 4707.42]
IMAGEN -192.76 [ -1677.91, 1292.40]
IMpACT 708.27 [ -2409.16, 3825.69]
LBC1936 833.73 [ -1084.97, 2752.43]
LIBD -1898.90 [ -4677.79, 879.99]
Sydney MAS 2100.00 [ -55.96, 4255.96]
MCIC 1500.00 [ -2615.92, 5615.92]
MooDS 1091.07 [ -1676.44, 3858.58]
MPIP 1818.92 [ -504.85, 4142.69]
MUNSTER -230.00 [ -1946.93, 1486.93]
NCNG 3379.49 [ 662.62, 6096.36]
NESDA 1457.60 [ -1762.27, 4677.47]
NeuroIMAGE 63.46 [ -3721.19, 3848.11]
NTR 8094.00 [-15445.17, 31633.17]
OATS 1870.00 [ -715.19, 4455.19]
PAFIP 7818.19 [ 2838.45, 12797.93]
PDNZ -278.45 [ -4561.19, 4004.29]
PING 3395.21 [ 677.21, 6113.21]
MPRC 2112.44 [ -820.41, 5045.29]
PPMI
QTIM 2078.37 [ 244.81, 3911.93]
SHIP 552.54 [ -1063.91, 2168.98]
SHIP-Trend 2272.45 [ 654.27, 3890.63]
SYS 865.39 [ -725.14, 2455.93]
TCD-NUIG
TOP 2531.27 [ 168.63, 4893.91]
TOP3T 633.66 [ -2160.57, 3427.90]
UiO2016 -930.02 [ -4429.58, 2569.54]
UiO2017 326.27 [ -2876.51, 3529.06]
UMCU
ENIGMA Discovery Cohorts [N=22061; P=1.249e-13] 1345.62 [ 989.77, 1701.47]
UKBB 1027.61 [ 533.31, 1521.91]
Discovery+UKBB [N=31984; P=6.758e-16] 1233.19 [ 933.86, 1532.51]
GOBS
IMH
UNICAMP 944.00 [ -3367.92, 5255.92]
Meth-CT 6173.81 [ 459.55, 11888.07]
MIRECC
UKBB nonEUR -553.36 [ -3954.66, 2847.94]
OSAKA 877.87 [ -875.95, 2631.68]
PING Non European 3830.87 [ 789.06, 6872.68]
Non European Cohorts [N=1250; P=0.02457] 1475.82 [ 189.15, 2762.49]
Cohort
Beta [Lower 95% CI,Upper 95% CI]
-18222.25
-9796.79
-1371.34
7054.12
15479.58
mm^2/effect allele

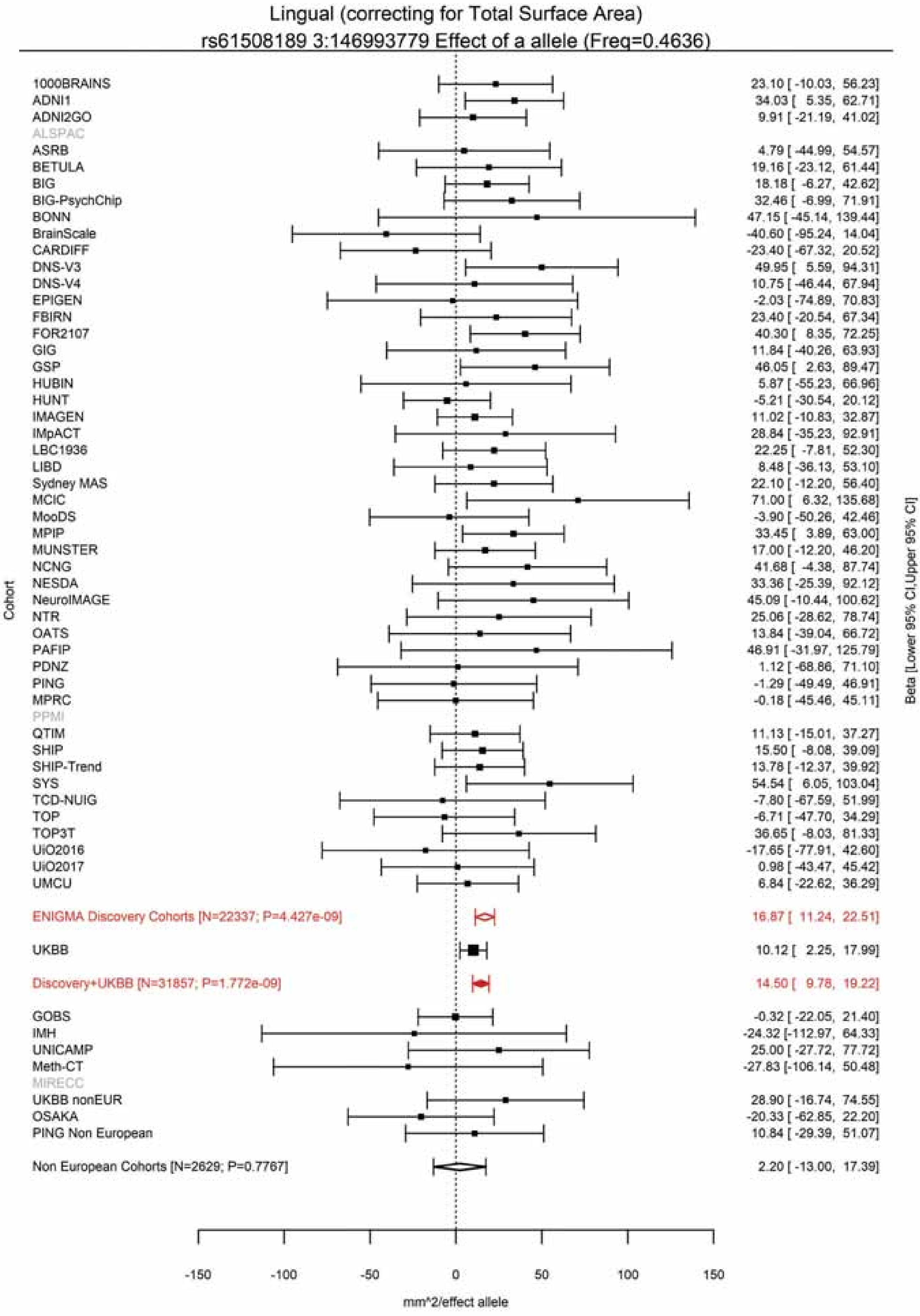
Lingual (correcting for Total Surface Area)
rs61508189 3:146993779 Effect of a allele (Freq=0.4636)
Cohort
1000BRAINS 23.10 [ -10.03, 56.23]
ADNI1 34.03 [ 5.35, 62.71]
ADNI2GO 9.91 [ -21.19, 41.02]
ALSPAC
ASRB 4.79 [ -44.99, 54.57]
BETULA 19.16 [ -23.12, 61.44]
BIG 18.18 [ -6.27, 42.62]
BIG-PsychChip 32.46 [ -6.99, 71.91]
BONN 47.15 [ -45.14, 139.44]
BrainScale -40.60 [ -95.24, 14.04]
CARDIFF -23.40 [ -67.32, 20.52]
DNS-V3 49.95 [ 5.59, 94.31]
DNS-V4 10.75 [ -46.44, 67.94]
EPIGEN -2.03 [ -74.89, 70.83]
FBIRN 23.40 [ -20.54, 67.34]
FOR2107 40.30 [ 8.35, 72.25]
GIG 11.84 [ -40.26, 63.93]
GSP 46.05 [ 2.63, 89.47]
HUBIN 5.87 [ -55.23, 66.96]
HUNT -5.21 [ -30.54, 20.12]
IMAGEN 11.02 [ -10.83, 32.87]
IMpACT 28.84 [ -35.23, 92.91]
LBC1936 22.25 [ -7.81, 52.30]
LIBD 8.48 [ -36.13, 53.10]
Sydney MAS 22.10 [ -12.20, 56.40]
MCIC 71.00 [ 6.32, 135.68]
MooDS -3.90 [ -50.26, 42.46]
MPIP 33.45 [ 3.89, 63.00]
MUNSTER 17.00 [ -12.20, 46.20]
NCNG 41.68 [ -4.38, 87.74]
NESDA 33.36 [ -25.39, 92.12]
NeuroIMAGE 45.09 [ -10.44, 100.62]
NTR 25.06 [ -28.62, 78.74]
OATS 13.84 [ -39.04, 66.72]
PAFIP 46.91 [ -31.97, 125.79]
PDNZ 1.12 [ -68.86, 71.10]
PING -1.29 [ -49.49, 46.91]
MPRC -0.18 [ -45.46, 45.11]
PPMI
QTIM 11.13 [ -15.01, 37.27]
SHIP 15.50 [ -8.08, 39.09]
SHIP-Trend 13.78 [ -12.37, 39.92]
SYS 54.54 [ 6.05, 103.04]
TCD-NUIG -7.80 [ -67.59, 51.99]
TOP -6.71 [ -47.70, 34.29]
TOP3T 36.65 [ -8.03, 81.33]
UiO2016 -17.65 [ -77.91, 42.60]
UiO2017 0.98 [ -43.47, 45.42]
UMCU 6.84 [ -22.62, 36.29]
ENIGMA Discovery Cohorts [N=22337; P=4.427e-09] 16.87 [ 11.24, 22.51]
UKBB 10.12 [ 2.25, 17.99]
Discovery+UKBB [N=31857; P=1.772e-09] 14.50 [ 9.78, 19.22]
GOBS -0.32 [ -22.05, 21.40]
IMH -24.32 [-112.97, 64.33]
UNICAMP 25.00 [ -27.72, 77.72]
Meth-CT -27.83 [-106.14, 50.48]
MIRECC
UKBB nonEUR 28.90 [ -16.74, 74.55]
OSAKA -20.33 [ -62.85, 22.20]
PING Non European 10.84 [ -29.39, 51.07]
Non European Cohorts [N=2629; P=0.7767] 2.20 [ -13.00, 17.39]
Beta [Lower 95% CI,Upper 95% CI]
-150
-100
-50
0
50
100
150
mm^2/effect allele

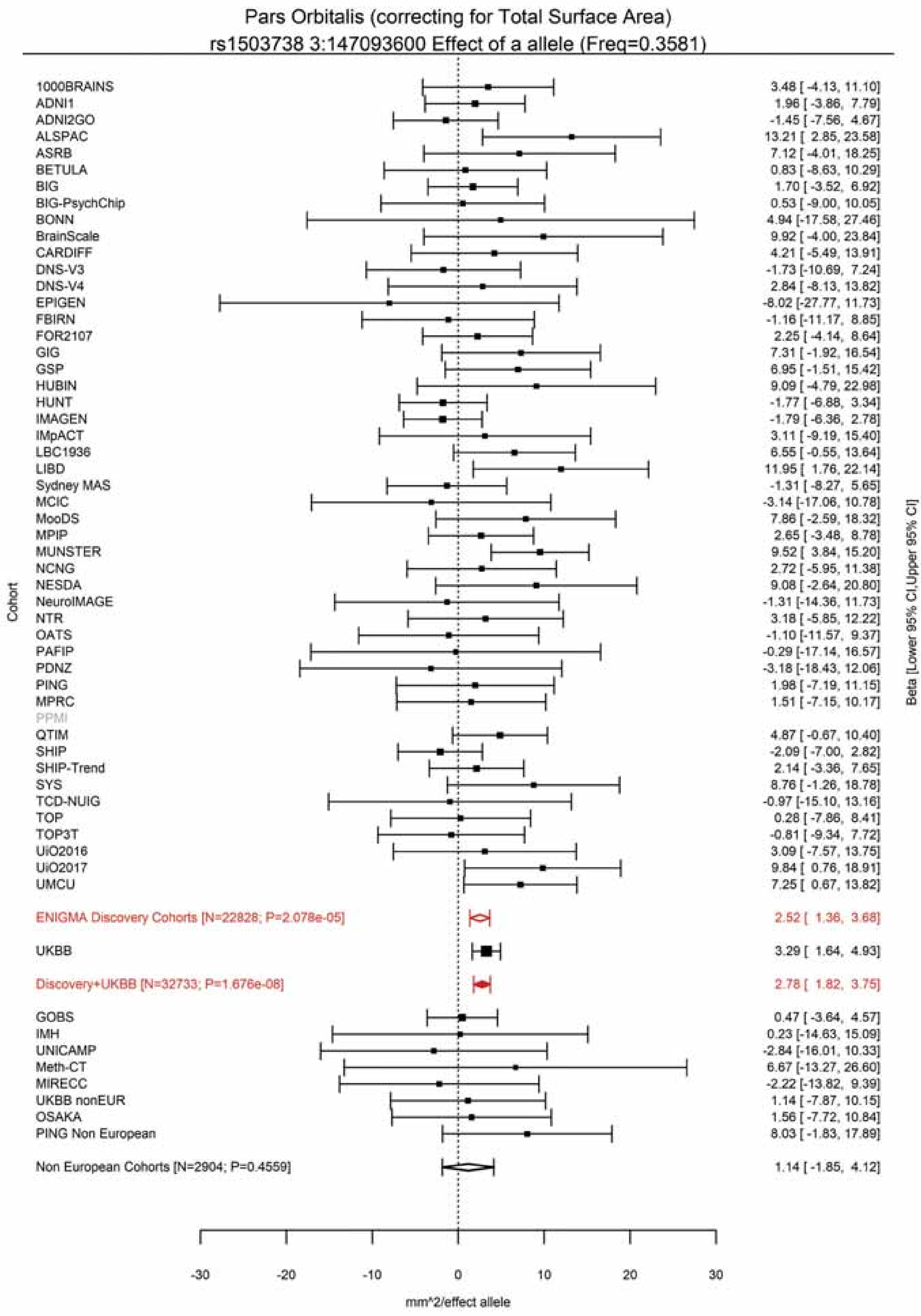

Pars Orbitalis (correcting for Total Surface Area)
rs1503738 3:147093600 Effect of a allele (Freq=0.3581)
Cohort
Beta [Lower 95% CI,Upper 95% CI]
1000BRAINS 3.48 [ -4.13, 11.10]
ADNI1 1.96 [ -3.86, 7.79]
ADNI2GO -1.45 [ -7.56, 4.67]
ALSPAC 13.21 [ 2.85, 23.58]
ASRB 7.12 [ -4.01, 18.25]
BETULA 0.83 [ -8.63, 10.29]
BIG 1.70 [ -3.52, 6.92]
BIG-PsychChip 0.53 [ -9.00, 10.05]
BONN 4.94 [-17.58, 27.46]
BrainScale 9.92 [ -4.00, 23.84]
CARDIFF 4.21 [ -5.49, 13.91]
DNS-V3 -1.73 [-10.69, 7.24]
DNS-V4 2.84 [ -8.13, 13.82]
EPIGEN -8.02 [-27.77, 11.73]
FBIRN -1.16 [-11.17, 8.85]
FOR2107 2.25 [ -4.14, 8.64]
GIG 7.31 [ -1.92, 16.54]
GSP 6.95 [ -1.51, 15.42]
HUBIN 9.09 [ -4.79, 22.98]
HUNT -1.77 [ -6.88, 3.34]
IMAGEN -1.79 [ -6.36, 2.78]
IMpACT 3.11 [ -9.19, 15.40]
LBC1936 6.55 [ -0.55, 13.64]
LIBD 11.95 [ 1.76, 22.14]
Sydney MAS -1.31 [ -8.27, 5.65]
MCIC -3.14 [-17.06, 10.78]
MooDS 7.86 [ -2.59, 18.32]
MPIP 2.65 [ -3.48, 8.78]
MUNSTER 9.52 [ 3.84, 15.20]
NCNG 2.72 [ -5.95, 11.38]
NESDA 9.08 [ -2.64, 20.80]
NeuroIMAGE -1.31 [-14.36, 11.73]
NTR 3.18 [ -5.85, 12.22]
OATS -1.10 [-11.57, 9.37]
PAFIP -0.29 [-17.14, 16.57]
PDNZ -3.18 [-18.43, 12.06]
PING 1.98 [ -7.19, 11.15]
MPRC 1.51 [ -7.15, 10.17]
PPMI
QTIM 4.87 [ -0.67, 10.40]
SHIP -2.09 [ -7.00, 2.82]
SHIP-Trend 2.14 [ -3.36, 7.65]
SYS 8.76 [ -1.26, 18.78]
TCD-NUIG -0.97 [-15.10, 13.16]
TOP 0.28 [ -7.86, 8.41]
TOP3T -0.81 [ -9.34, 7.72]
UiO2016 3.09 [ -7.57, 13.75]
UiO2017 9.84 [ 0.76, 18.91]
UMCU 7.25 [ 0.67, 13.82]
ENIGMA Discovery Cohorts [N=22828; P=2.078e-05] 2.52 [ 1.36, 3.68]
UKBB 3.29 [ 1.64, 4.93]
Discovery+UKBB [N=32733; P=1.676e-08] 2.78 [ 1.82, 3.75]
GOBS 0.47 [ -3.64, 4.57]
IMH 0.23 [-14.63, 15.09]
UNICAMP -2.84 [-16.01, 10.33]
Meth-CT 6.67 [-13.27, 26.60]
MIRECC -2.22 [-13.82, 9.39]
UKBB nonEUR 1.14 [ -7.87, 10.15]
OSAKA 1.56 [ -7.72, 10.84]
PING Non European 8.03 [ -1.83, 17.89]
Non European Cohorts [N=2904; P=0.4559] 1.14 [ -1.85, 4.12]
-30
-20
-10
0
10
20
30
mm^2/effect allele

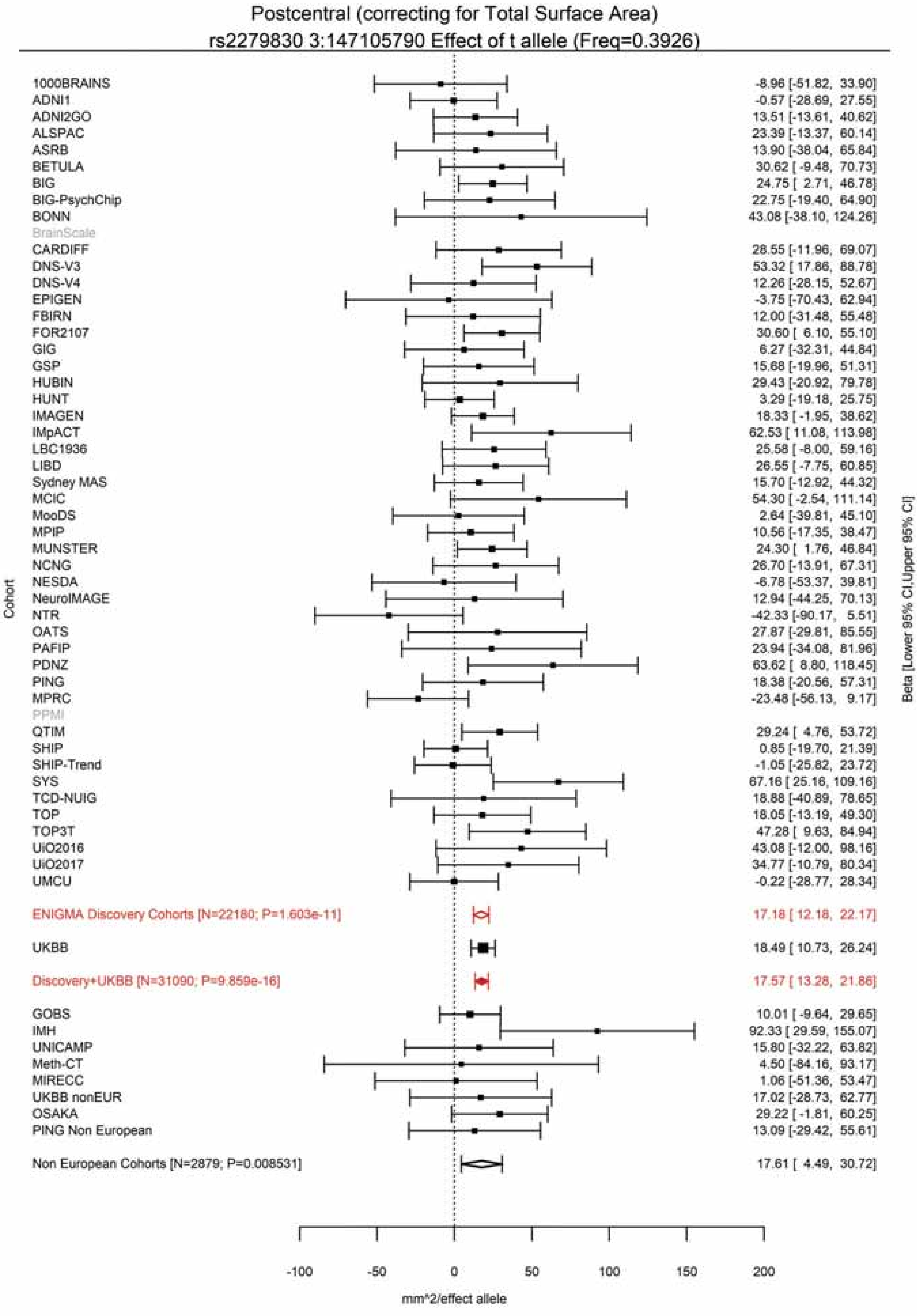

Postcentral (correcting for Total Surface Area)
rs2279830 3:147105790 Effect of t allele (Freq=0.3926)
Cohort
1000BRAINS -8.96 [-51.82, 33.90]
ADNI1 -0.57 [-28.69, 27.55]
ADNI2GO 13.51 [-13.61, 40.62]
ALSPAC 23.39 [-13.37, 60.14]
ASRB 13.90 [-38.04, 65.84]
BETULA 30.62 [ -9.48, 70.73]
BIG 24.75 [ 2.71, 46.78]
BIG-PsychChip 22.75 [-19.40, 64.90]
BONN 43.08 [-38.10, 124.26]
BrainScale
CARDIFF 28.55 [-11.96, 69.07]
DNS-V3 53.32 [ 17.86, 88.78]
DNS-V4 12.26 [-28.15, 52.67]
EPIGEN -3.75 [-70.43, 62.94]
FBIRN 12.00 [-31.48, 55.48]
FOR2107 30.60 [ 6.10, 55.10]
GIG 6.27 [-32.31, 44.84]
GSP 15.68 [-19.96, 51.31]
HUBIN 29.43 [-20.92, 79.78]
HUNT 3.29 [-19.18, 25.75]
IMAGEN 18.33 [ -1.95, 38.62]
IMpACT 62.53 [ 11.08, 113.98]
LBC1936 25.58 [ -8.00, 59.16]
LIBD 26.55 [ -7.75, 60.85]
Sydney MAS 15.70 [-12.92, 44.32]
MCIC 54.30 [ -2.54, 111.14]
MooDS 2.64 [-39.81, 45.10]
MPIP 10.56 [-17.35, 38.47]
MUNSTER 24.30 [ 1.76, 46.84]
NCNG 26.70 [-13.91, 67.31]
NESDA -6.78 [-53.37, 39.81]
NeuroIMAGE 12.94 [-44.25, 70.13]
NTR -42.33 [-90.17, 5.51]
OATS 27.87 [-29.81, 85.55]
PAFIP 23.94 [-34.08, 81.96]
PDNZ 63.62 [ 8.80, 118.45]
PING 18.38 [-20.56, 57.31]
MPRC -23.48 [-56.13, 9.17]
PPMI
QTIM 29.24 [ 4.76, 53.72]
SHIP 0.85 [-19.70, 21.39]
SHIP-Trend -1.05 [-25.82, 23.72]
SYS 67.16 [ 25.16, 109.16]
TCD-NUIG 18.88 [-40.89, 78.65]
TOP 18.05 [-13.19, 49.30]
TOP3T 47.28 [ 9.63, 84.94]
UiO2016 43.08 [-12.00, 98.16]
UiO2017 34.77 [-10.79, 80.34]
UMCU -0.22 [-28.77, 28.34]
ENIGMA Discovery Cohorts [N=22180; P=1.603e-11] 17.18 [ 12.18, 22.17]
UKBB 18.49 [ 10.73, 26.24]
Discovery+UKBB [N=31090; P=9.859e-16] 17.57 [ 13.28, 21.86]
GOBS 10.01 [ -9.64, 29.65]
IMH 92.33 [ 29.59, 155.07]
UNICAMP 15.80 [-32.22, 63.82]
Meth-CT 4.50 [-84.16, 93.17]
MIRECC 1.06 [-51.36, 53.47]
UKBB nonEUR 17.02 [-28.73, 62.77]
OSAKA 29.22 [ -1.81, 60.25]
PING Non European 13.09 [-29.42, 55.61]
Non European Cohorts [N=2879; P=0.008531] 17.61 [ 4.49, 30.72]
Beta [Lower 95% CI,Upper 95% CI]
-100 -50 0 50 100 150 200
mm^2/effect allele

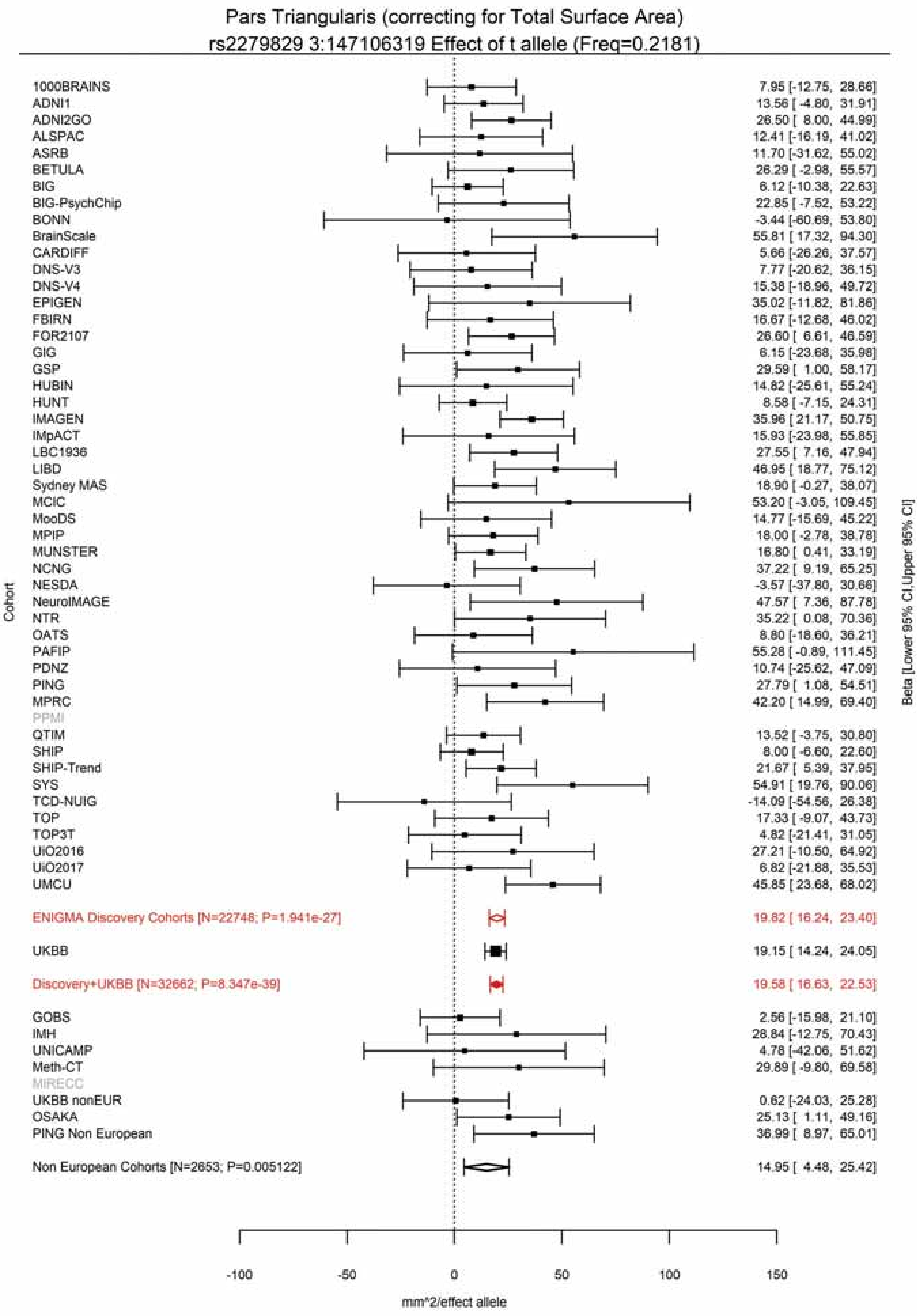
Pars Triangularis (correcting for Total Surface Area)
rs2279829 3:147106319 Effect of t allele (Freq=0.2181)
Cohort
1000BRAINS 7.95 [-12.75, 28.66]
ADNI1 13.56 [ -4.80, 31.91]
ADNI2GO 26.50 [ 8.00, 44.99]
ALSPAC 12.41 [-16.19, 41.02]
ASRB 11.70 [-31.62, 55.02]
BETULA 26.29 [ -2.98, 55.57]
BIG 6.12 [-10.38, 22.63]
BIG-PsychChip 22.85 [ -7.52, 53.22]
BONN -3.44 [-60.69, 53.80]
BrainScale 55.81 [ 17.32, 94.30]
CARDIFF 5.66 [-26.26, 37.57]
DNS-V3 7.77 [-20.62, 36.15]
DNS-V4 15.38 [-18.96, 49.72]
EPIGEN 35.02 [-11.82, 81.86]
FBIRN 16.67 [-12.68, 46.02]
FOR2107 26.60 [ 6.61, 46.59]
GIG 6.15 [-23.68, 35.98]
GSP 29.59 [ 1.00, 58.17]
HUBIN 14.82 [-25.61, 55.24]
HUNT 8.58 [ -7.15, 24.31]
IMAGEN 35.96 [ 21.17, 50.75]
IMpACT 15.93 [-23.98, 55.85]
LBC1936 27.55 [ 7.16, 47.94]
LIBD 46.95 [ 18.77, 75.12]
Sydney MAS 18.90 [ -0.27, 38.07]
MCIC 53.20 [ -3.05, 109.45]
MooDS 14.77 [-15.69, 45.22]
MPIP 18.00 [ -2.78, 38.78]
MUNSTER 16.80 [ 0.41, 33.19]
NCNG 37.22 [ 9.19, 65.25]
NESDA -3.57 [-37.80, 30.66]
NeuroIMAGE 47.57 [ 7.36, 87.78]
NTR 35.22 [ 0.08, 70.36]
OATS 8.80 [-18.60, 36.21]
PAFIP 55.28 [ -0.89, 111.45]
PDNZ 10.74 [-25.62, 47.09]
PING 27.79 [ 1.08, 54.51]
MPRC 42.20 [ 14.99, 69.40]
PPMI
QTIM 13.52 [ -3.75, 30.80]
SHIP 8.00 [ -6.60, 22.60]
SHIP-Trend 21.67 [ 5.39, 37.95]
SYS 54.91 [ 19.76, 90.06]
TCD-NUIG -14.09 [-54.56, 26.38]
TOP 17.33 [ -9.07, 43.73]
TOP3T 4.82 [-21.41, 31.05]
UiO2016 27.21 [-10.50, 64.92]
UiO2017 6.82 [-21.88, 35.53]
UMCU 45.85 [ 23.68, 68.02]
ENIGMA Discovery Cohorts [N=22748; P=1.941e-27] 19.82 [ 16.24, 23.40]
UKBB 19.15 [ 14.24, 24.05]
Discovery+UKBB [N=32662; P=8.347e-39] 19.58 [ 16.63, 22.53]
GOBS 2.56 [-15.98, 21.10]
IMH 28.84 [-12.75, 70.43]
UNICAMP 4.78 [-42.06, 51.62]
Meth-CT 29.89 [ -9.80, 69.58]
MIRECC
UKBB nonEUR 0.62 [-24.03, 25.28]
OSAKA 25.13 [ 1.11, 49.16]
PING Non European 36.99 [ 8.97, 65.01]
Non European Cohorts [N=2653; P=0.005122] 14.95 [ 4.48, 25.42]
Beta [Lower 95% CI,Upper 95% CI]
-100
-50
0
50
100
150
mm^2/effect allele

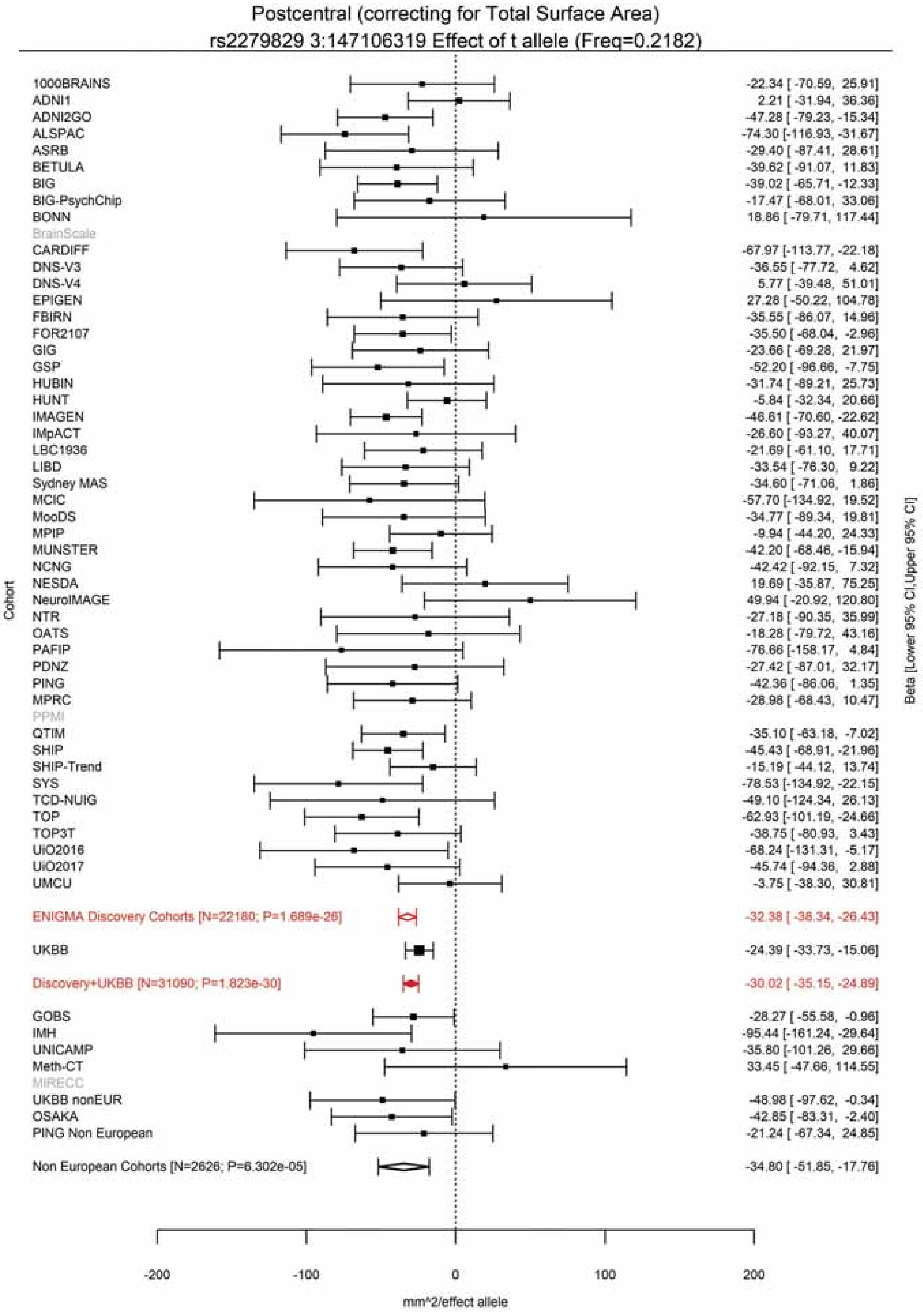
Postcentral (correcting for Total Surface Area)
rs2279829 3:147106319 Effect of t allele (Freq=0.2182)
1000BRAINS -22.34 [ -70.59, 25.91]
ADNI1 2.21 [ -31.94, 36.36]
ADNI2GO -47.28 [ -79.23, -15.34]
ALSPAC -74.30 [-116.93, -31.67]
ASRB -29.40 [ -87.41, 28.61]
BETULA -39.62 [ -91.07, 11.83]
BIG -39.02 [ -65.71, -12.33]
BIG-PsychChip -17.47 [ -68.01, 33.06]
BONN 18.86 [ -79.71, 117.44]
BrainScale
CARDIFF -67.97 [-113.77, -22.18]
DNS-V3 -36.55 [ -77.72, 4.62]
DNS-V4 5.77 [ -39.48, 51.01]
EPIGEN 27.28 [ -50.22, 104.78]
FBIRN -35.55 [ -86.07, 14.96]
FOR2107 -35.50 [ -68.04, -2.96]
GIG -23.66 [ -69.28, 21.97]
GSP -52.20 [ -96.66, -7.75]
HUBIN -31.74 [ -89.21, 25.73]
HUNT -5.84 [ -32.34, 20.66]
IMAGEN -46.61 [ -70.60, -22.62]
IMpACT -26.60 [ -93.27, 40.07]
LBC1936 -21.69 [ -61.10, 17.71]
LIBD -33.54 [ -76.30, 9.22]
Sydney MAS -34.60 [ -71.06, 1.86]
MCIC -57.70 [-134.92, 19.52]
MooDS -34.77 [ -89.34, 19.81]
MPIP -9.94 [ -44.20, 24.33]
MUNSTER -42.20 [ -68.46, -15.94]
NCNG -42.42 [ -92.15, 7.32]
NESDA 19.69 [ -35.87, 75.25]
NeuroIMAGE 49.94 [ -20.92, 120.80]
NTR -27.18 [ -90.35, 35.99]
OATS -18.28 [ -79.72, 43.16]
PAFIP -76.66 [-158.17, 4.84]
PDNZ -27.42 [ -87.01, 32.17]
PING -42.36 [ -86.06, 1.35]
MPRC -28.98 [ -68.43, 10.47]
PPMI
QTIM -35.10 [ -63.18, -7.02]
SHIP -45.43 [ -68.91, -21.96]
SHIP-Trend -15.19 [ -44.12, 13.74]
SYS -78.53 [-134.92, -22.15]
TCD-NUIG -49.10 [-124.34, 26.13]
TOP -62.93 [-101.19, -24.66]
TOP3T -38.75 [ -80.93, 3.43]
UiO2016 -68.24 [-131.31, -5.17]
UiO2017 -45.74 [ -94.36, 2.88]
UMCU -3.75 [ -38.30, 30.81]
ENIGMA Discovery Cohorts [N=22180; P=1.689e-26] -32.38 [ -38.34, -26.43]
UKBB -24.39 [ -33.73, -15.06]
Discovery+UKBB [N=31090; P=1.823e-30] -30.02 [ -35.15, -24.89]
GOBS -28.27 [ -55.58, -0.96]
IMH -95.44 [-161.24, -29.64]
UNICAMP -35.80 [-101.26, 29.66]
Meth-CT 33.45 [ -47.66, 114.55]
MIRECC
UKBB nonEUR -48.98 [ -97.62, -0.34]
OSAKA -42.85 [ -83.31, -2.40]
PING Non European -21.24 [ -67.34, 24.85]
Non European Cohorts [N=2626; P=6.302e-05] -34.80 [ -51.85, -17.76]
Cohort
Beta [Lower 95% CI,Upper 95% CI]
-200
-100
0
100
200
mm^2/effect allele

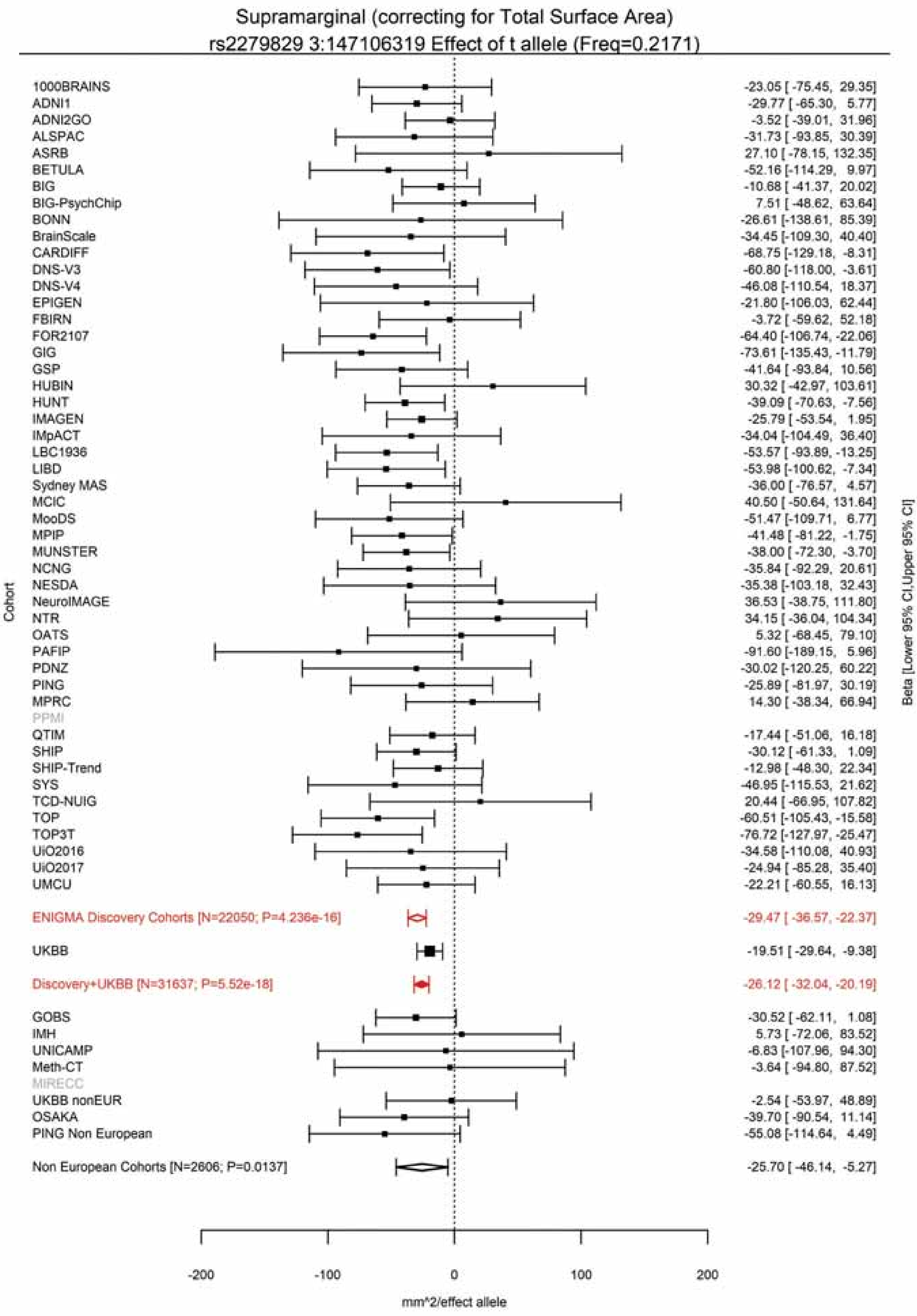
Supramarginal (correcting for Total Surface Area)
rs2279829 3:147106319 Effect of t allele (Freq=0.2171)
1000BRAINS -23.05 [ -75.45, 29.35]
ADNI1 -29.77 [ -65.30, 5.77]
ADNI2GO -3.52 [ -39.01, 31.96]
ALSPAC -31.73 [ -93.85, 30.39]
ASRB 27.10 [ -78.15, 132.35]
BETULA -52.16 [-114.29, 9.97]
BIG -10.68 [ -41.37, 20.02]
BIG-PsychChip 7.51 [ -48.62, 63.64]
BONN -26.61 [-138.61, 85.39]
BrainScale -34.45 [-109.30, 40.40]
CARDIFF -68.75 [-129.18, -8.31]
DNS-V3 -60.80 [-118.00, -3.61]
DNS-V4 -46.08 [-110.54, 18.37]
EPIGEN -21.80 [-106.03, 62.44]
FBIRN -3.72 [ -59.62, 52.18]
FOR2107 -64.40 [-106.74, -22.06]
GIG -73.61 [-135.43, -11.79]
GSP -41.64 [ -93.84, 10.56]
HUBIN 30.32 [ -42.97, 103.61]
HUNT -39.09 [ -70.63, -7.56]
IMAGEN -25.79 [ -53.54, 1.95]
IMpACT -34.04 [-104.49, 36.40]
LBC1936 -53.57 [ -93.89, -13.25]
LIBD -53.98 [-100.62, -7.34]
Sydney MAS -36.00 [ -76.57, 4.57]
MCIC 40.50 [ -50.64, 131.64]
MooDS -51.47 [-109.71, 6.77]
MPIP -41.48 [ -81.22, -1.75]
MUNSTER -38.00 [ -72.30, -3.70]
NCNG -35.84 [ -92.29, 20.61]
NESDA -35.38 [-103.18, 32.43]
NeuroIMAGE 36.53 [ -38.75, 111.80]
NTR 34.15 [ -36.04, 104.34]
OATS 5.32 [ -68.45, 79.10]
PAFIP -91.60 [-189.15, 5.96]
PDNZ -30.02 [-120.25, 60.22]
PING -25.89 [ -81.97, 30.19]
MPRC 14.30 [ -38.34, 66.94]
PPMI
QTIM -17.44 [ -51.06, 16.18]
SHIP -30.12 [ -61.33, 1.09]
SHIP-Trend -12.98 [ -48.30, 22.34]
SYS -46.95 [-115.53, 21.62]
TCD-NUIG 20.44 [ -66.95, 107.82]
TOP -60.51 [-105.43, -15.58]
TOP3T -76.72 [-127.97, -25.47]
UiO2016 -34.58 [-110.08, 40.93]
UiO2017 -24.94 [ -85.28, 35.40]
UMCU -22.21 [ -60.55, 16.13]
ENIGMA Discovery Cohorts [N=22050; P=4.236e-16] -29.47 [ -36.57, -22.37]
UKBB -19.51 [ -29.64, -9.38]
Discovery+UKBB [N=31637; P=5.52e-18] -26.12 [ -32.04, -20.19]
GOBS -30.52 [ -62.11, 1.08]
IMH 5.73 [ -72.06, 83.52]
UNICAMP -6.83 [-107.96, 94.30]
Meth-CT -3.64 [ -94.80, 87.52]
MIRECC
UKBB nonEUR -2.54 [ -53.97, 48.89]
OSAKA -39.70 [ -90.54, 11.14]
PING Non European -55.08 [-114.64, 4.49]
Non European Cohorts [N=2606; P=0.0137] -25.70 [ -46.14, -5.27]
Cohort
Beta [Lower 95% CI,Upper 95% CI]
-200
-100
0
100
200
mm^2/effect allele

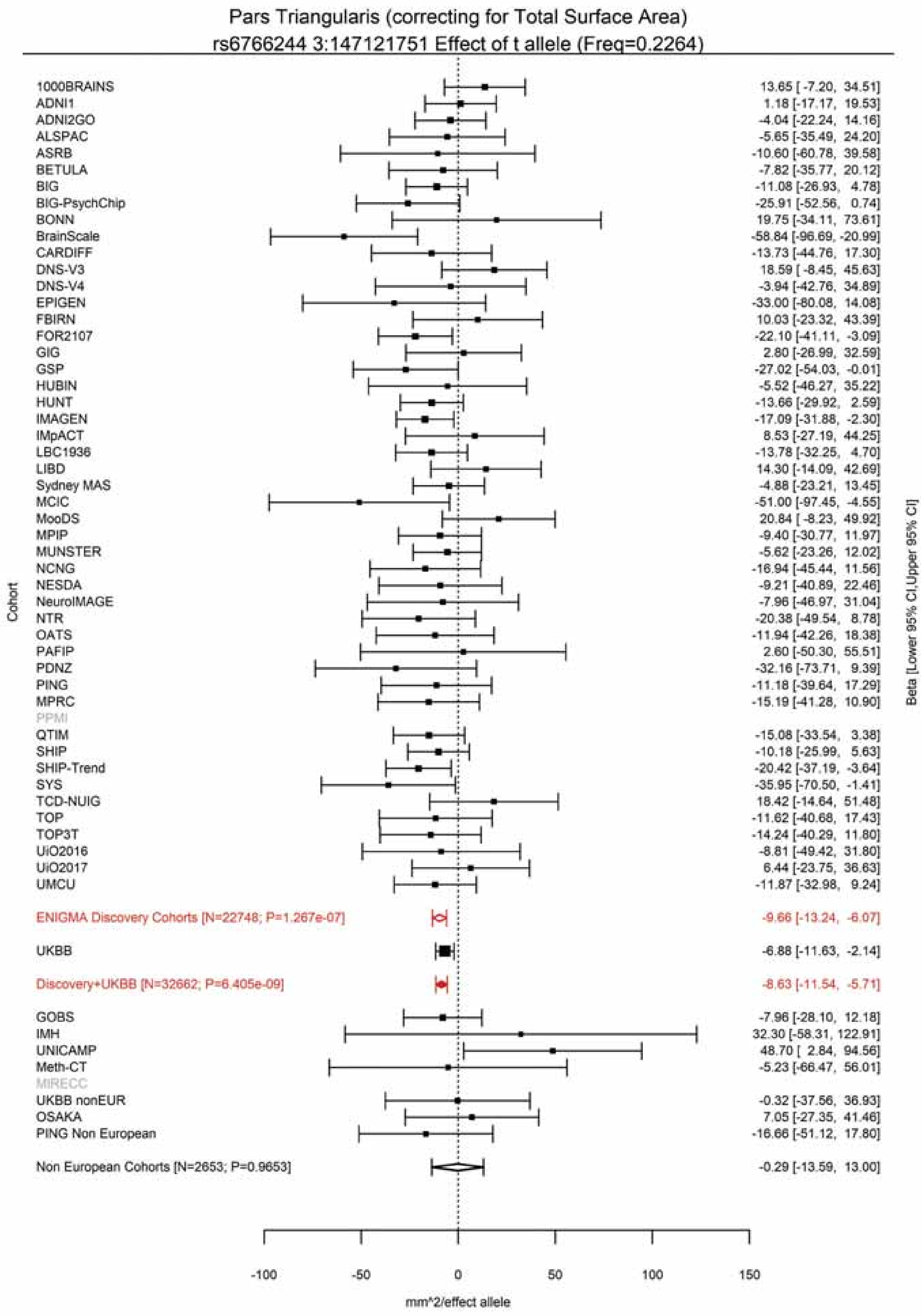
Pars Triangularis (correcting for Total Surface Area)
rs6766244 3:147121751 Effect of t allele (Freq=0.2264)
Cohort
Beta [Lower 95% CI,Upper 95% CI]
1000BRAINS 13.65 [ -7.20, 34.51]
ADNI1 1.18 [-17.17, 19.53]
ADNI2GO -4.04 [-22.24, 14.16]
ALSPAC -5.65 [-35.49, 24.20]
ASRB -10.60 [-60.78, 39.58]
BETULA -7.82 [-35.77, 20.12]
BIG -11.08 [-26.93, 4.78]
BIG-PsychChip -25.91 [-52.56, 0.74]
BONN 19.75 [-34.11, 73.61]
BrainScale -58.84 [-96.69, -20.99]
CARDIFF -13.73 [-44.76, 17.30]
DNS-V3 18.59 [ -8.45, 45.63]
DNS-V4 -3.94 [-42.76, 34.89]
EPIGEN -33.00 [-80.08, 14.08]
FBIRN 10.03 [-23.32, 43.39]
FOR2107 -22.10 [-41.11, -3.09]
GIG 2.80 [-26.99, 32.59]
GSP -27.02 [-54.03, -0.01]
HUBIN -5.52 [-46.27, 35.22]
HUNT -13.66 [-29.92, 2.59]
IMAGEN -17.09 [-31.88, -2.30]
IMpACT 8.53 [-27.19, 44.25]
LBC1936 -13.78 [-32.25, 4.70]
LIBD 14.30 [-14.09, 42.69]
Sydney MAS -4.88 [-23.21, 13.45]
MCIC -51.00 [-97.45, -4.55]
MooDS 20.84 [ -8.23, 49.92]
MPIP -9.40 [-30.77, 11.97]
MUNSTER -5.62 [-23.26, 12.02]
NCNG -16.94 [-45.44, 11.56]
NESDA -9.21 [-40.89, 22.46]
NeuroIMAGE -7.96 [-46.97, 31.04]
NTR -20.38 [-49.54, 8.78]
OATS -11.94 [-42.26, 18.38]
PAFIP 2.60 [-50.30, 55.51]
PDNZ -32.16 [-73.71, 9.39]
PING -11.18 [-39.64, 17.29]
MPRC -15.19 [-41.28, 10.90]
PPMI
QTIM -15.08 [-33.54, 3.38]
SHIP -10.18 [-25.99, 5.63]
SHIP-Trend -20.42 [-37.19, -3.64]
SYS -35.95 [-70.50, -1.41]
TCD-NUIG 18.42 [-14.64, 51.48]
TOP -11.62 [-40.68, 17.43]
TOP3T -14.24 [-40.29, 11.80]
UiO2016 -8.81 [-49.42, 31.80]
UiO2017 6.44 [-23.75, 36.63]
UMCU -11.87 [-32.98, 9.24]
ENIGMA Discovery Cohorts [N=22748; P=1.267e-07] -9.66 [-13.24, -6.07]
UKBB -6.88 [-11.63, -2.14]
Discovery+UKBB [N=32662; P=6.405e-09] -8.63 [-11.54, -5.71]
GOBS -7.96 [-28.10, 12.18]
IMH 32.30 [-58.31, 122.91]
UNICAMP 48.70 [ 2.84, 94.56]
Meth-CT -5.23 [-66.47, 56.01]
MIRECC
UKBB nonEUR -0.32 [-37.56, 36.93]
OSAKA 7.05 [-27.35, 41.46]
PING Non European -16.66 [-51.12, 17.80]
Non European Cohorts [N=2653; P=0.9653] -0.29 [-13.59, 13.00]
-100
-50
0
50
100
150
mm^2/effect allele

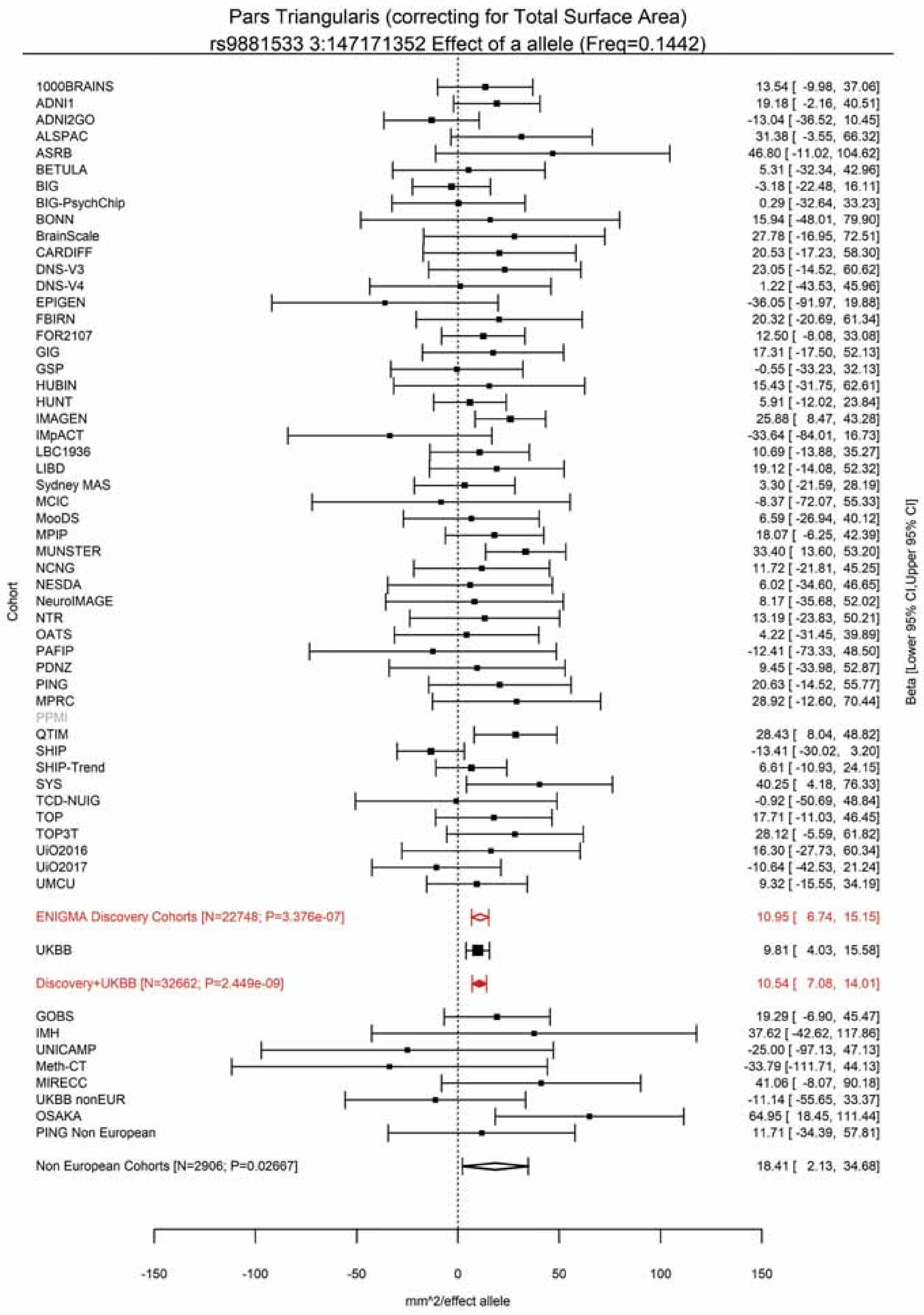
Pars Triangularis (correcting for Total Surface Area)
rs9881533 3:147171352 Effect of a allele (Freq=0.1442)
Cohort
Beta [Lower 95% CI,Upper 95% CI]
1000BRAINS 13.54 [ -9.98, 37.06]
ADNI1 19.18 [ -2.16, 40.51]
ADNI2GO -13.04 [ -36.52, 10.45]
ALSPAC 31.38 [ -3.55, 66.32]
ASRB 46.80 [ -11.02, 104.62]
BETULA 5.31 [ -32.34, 42.96]
BIG -3.18 [ -22.48, 16.11]
BIG-PsychChip 0.29 [ -32.64, 33.23]
BONN 15.94 [ -48.01, 79.90]
BrainScale 27.78 [ -16.95, 72.51]
CARDIFF 20.53 [ -17.23, 58.30]
DNS-V3 23.05 [ -14.52, 60.62]
DNS-V4 1.22 [ -43.53, 45.96]
EPIGEN -36.05 [ -91.97, 19.88]
FBIRN 20.32 [ -20.69, 61.34]
FOR2107 12.50 [ -8.08, 33.08]
GIG 17.31 [ -17.50, 52.13]
GSP -0.55 [ -33.23, 32.13]
HUBIN 15.43 [ -31.75, 62.61]
HUNT 5.91 [ -12.02, 23.84]
IMAGEN 25.88 [ 8.47, 43.28]
IMpACT -33.64 [ -84.01, 16.73]
LBC1936 10.69 [ -13.88, 35.27]
LIBD 19.12 [ -14.08, 52.32]
Sydney MAS 3.30 [ -21.59, 28.19]
MCIC -8.37 [ -72.07, 55.33]
MooDS 6.59 [ -26.94, 40.12]
MPIP 18.07 [ -6.25, 42.39]
MUNSTER 33.40 [ 13.60, 53.20]
NCNG 11.72 [ -21.81, 45.25]
NESDA 6.02 [ -34.60, 46.65]
NeuroIMAGE 8.17 [ -35.68, 52.02]
NTR 13.19 [ -23.83, 50.21]
OATS 4.22 [ -31.45, 39.89]
PAFIP -12.41 [ -73.33, 48.50]
PDNZ 9.45 [ -33.98, 52.87]
PING 20.63 [ -14.52, 55.77]
MPRC 28.92 [ -12.60, 70.44]
PPMI
QTIM 28.43 [ 8.04, 48.82]
SHIP -13.41 [ -30.02, 3.20]
SHIP-Trend 6.61 [ -10.93, 24.15]
SYS 40.25 [ 4.18, 76.33]
TCD-NUIG -0.92 [ -50.69, 48.84]
TOP 17.71 [ -11.03, 46.45]
TOP3T 28.12 [ -5.59, 61.82]
UiO2016 16.30 [ -27.73, 60.34]
UiO2017 -10.64 [ -42.53, 21.24]
UMCU 9.32 [ -15.55, 34.19]
ENIGMA Discovery Cohorts [N=22748; P=3.376e-07] 10.95 [ 6.74, 15.15]
UKBB 9.81 [ 4.03, 15.58]
Discovery+UKBB [N=32662; P=2.449e-09] 10.54 [ 7.08, 14.01]
GOBS 19.29 [ -6.90, 45.47]
IMH 37.62 [ -42.62, 117.86]
UNICAMP -25.00 [ -97.13, 47.13]
Meth-CT -33.79 [ -111.71, 44.13]
MIRECC 41.06 [ -8.07, 90.18]
UKBB nonEUR -11.14 [ -55.65, 33.37]
OSAKA 64.95 [ 18.45, 111.44]
PING Non European 11.71 [ -34.39, 57.81]
Non European Cohorts [N=2906; P=0.02667] 18.41 [ 2.13, 34.68]
-150
-100
-50
0
50
100
150
mm^2/effect allele

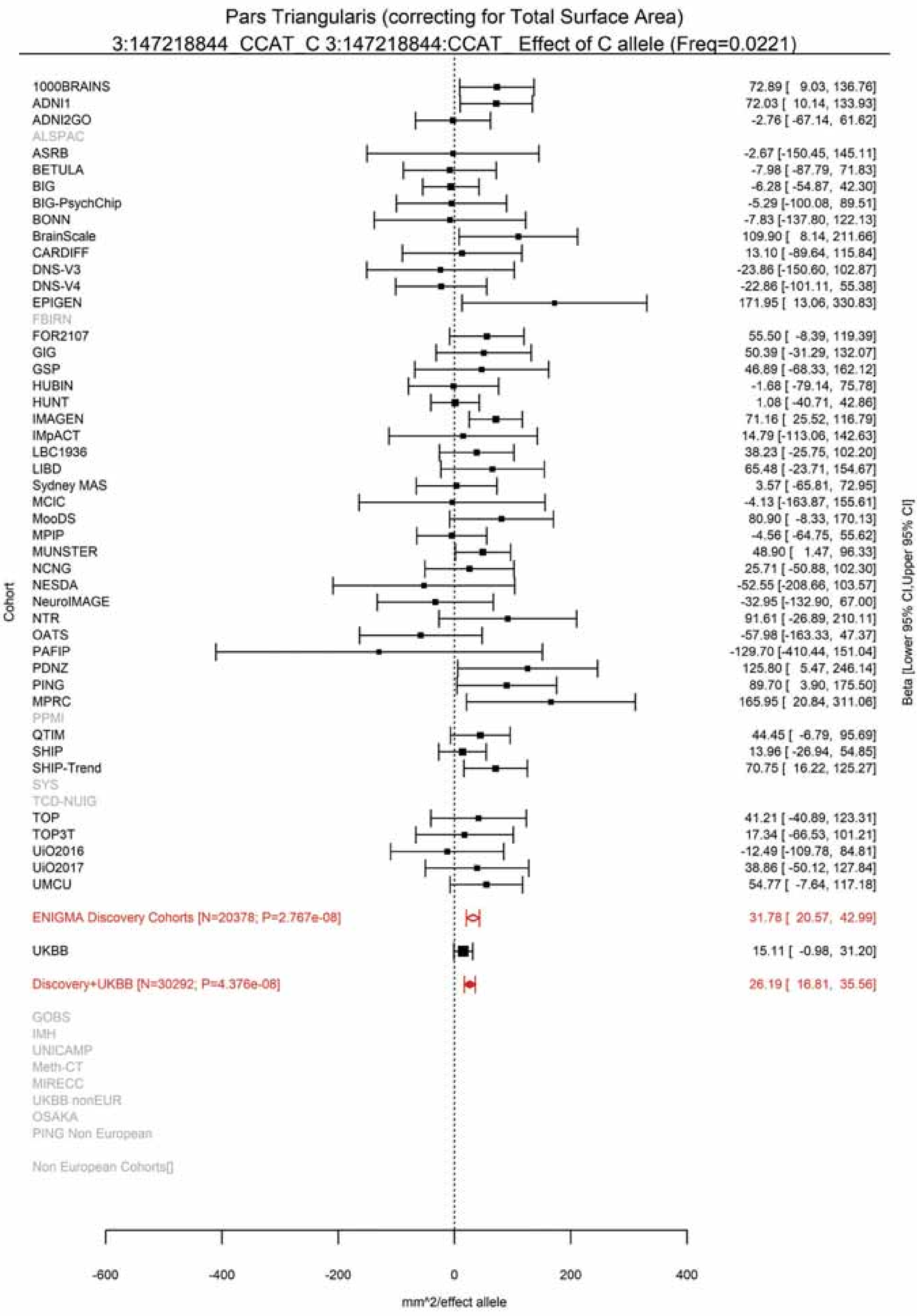
Pars Triangularis (correcting for Total Surface Area)
3:147218844_CCAT_C 3:147218844:CCAT Effect of C allele (Freq=0.0221)
1000BRAINS 72.89 [ 9.03, 136.76]
ADNI1 72.03 [ 10.14, 133.93]
ADNI2GO -2.76 [-67.14, 61.62]
ALSPAC
ASRB -2.67 [-150.45, 145.11]
BETULA -7.98 [-87.79, 71.83]
BIG -6.28 [-54.87, 42.30]
BIG-PsychChip -5.29 [-100.08, 89.51]
BONN -7.83 [-137.80, 122.13]
BrainScale 109.90 [ 8.14, 211.66]
CARDIFF 13.10 [-89.64, 115.84]
DNS-V3 -23.86 [-150.60, 102.87]
DNS-V4 -22.86 [-101.11, 55.38]
EPIGEN 171.95 [ 13.06, 330.83]
FBIRN
FOR2107 55.50 [ -8.39, 119.39]
GIG 50.39 [-31.29, 132.07]
GSP 46.89 [-68.33, 162.12]
HUBIN -1.68 [-79.14, 75.78]
HUNT 1.08 [-40.71, 42.86]
IMAGEN 71.16 [ 25.52, 116.79]
IMpACT 14.79 [-113.06, 142.63]
LBC1936 38.23 [-25.75, 102.20]
LIBD 65.48 [-23.71, 154.67]
Sydney MAS 3.57 [-65.81, 72.95]
MCIC -4.13 [-163.87, 155.61]
MooDS 80.90 [ -8.33, 170.13]
MPIP -4.56 [-64.75, 55.62]
MUNSTER 48.90 [ 1.47, 96.33]
NCNG 25.71 [-50.88, 102.30]
NESDA -52.55 [-208.66, 103.57]
NeuroIMAGE -32.95 [-132.90, 67.00]
NTR 91.61 [-26.89, 210.11]
OATS -57.98 [-163.33, 47.37]
PAFIP -129.70 [-410.44, 151.04]
PDNZ 125.80 [ 5.47, 246.14]
PING 89.70 [ 3.90, 175.50]
MPRC 165.95 [ 20.84, 311.06]
PPMI
QTIM 44.45 [ -6.79, 95.69]
SHIP 13.96 [-26.94, 54.85]
SHIP-Trend 70.75 [ 16.22, 125.27]
SYS
TCD-NUIG
TOP 41.21 [-40.89, 123.31]
TOP3T 17.34 [-66.53, 101.21]
UiO2016 -12.49 [-109.78, 84.81]
UiO2017 38.86 [-50.12, 127.84]
UMCU 54.77 [ -7.64, 117.18]
ENIGMA Discovery Cohorts [N=20378; P=2.767e-08] 31.78 [ 20.57, 42.99]
UKBB 15.11 [ -0.98, 31.20]
Discovery+UKBB [N=30292; P=4.376e-08] 26.19 [ 16.81, 35.56]
GOBS
IMH
UNICAMP
Meth-CT
MIRECC
UKBB nonEUR
OSAKA
PING Non European
Non European Cohorts[]
Cohort
Beta [Lower 95% CI,Upper 95% CI]
-600
-400
-200
0
200
400
mm^2/effect allele

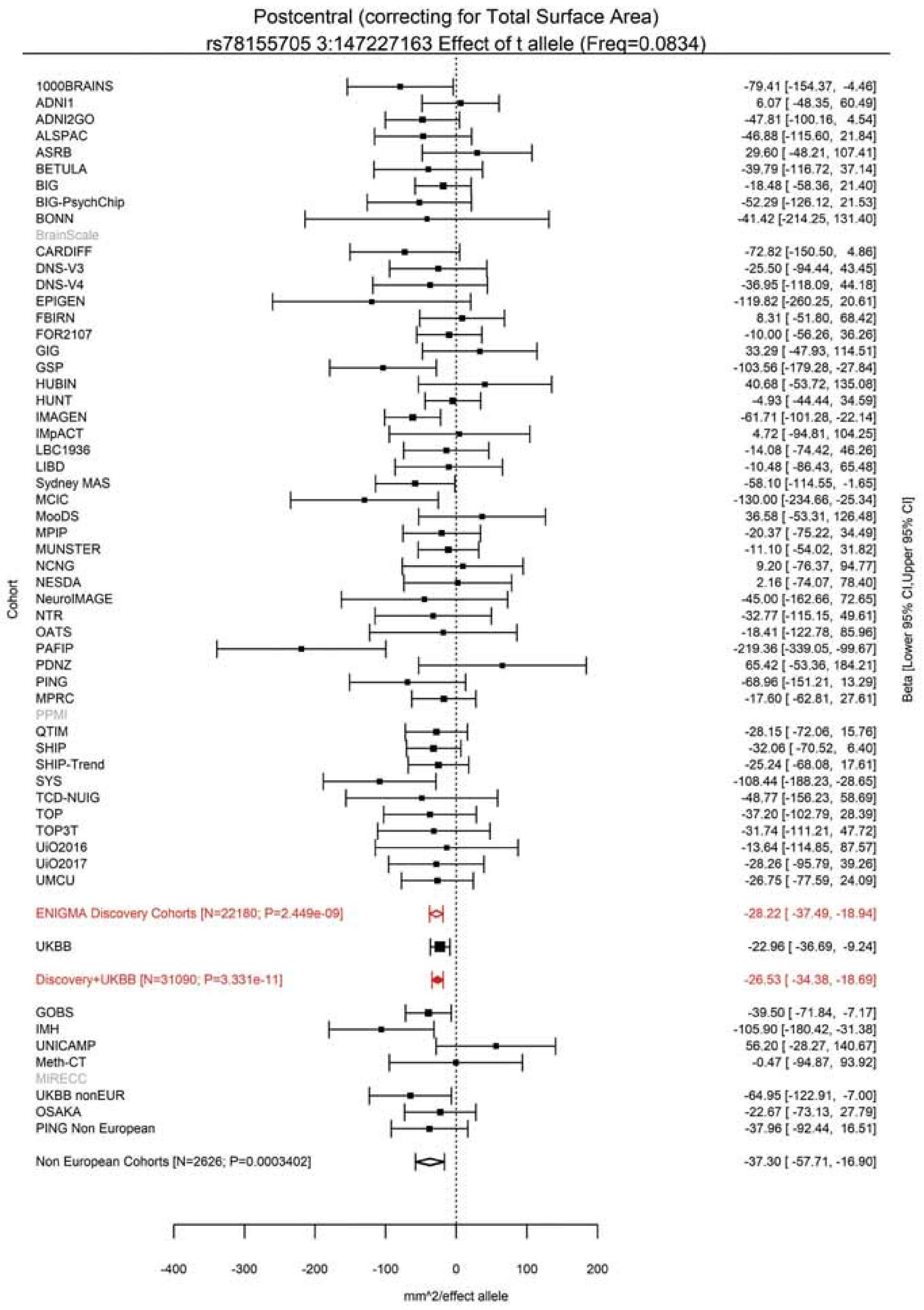

Postcentral (correcting for Total Surface Area)
rs78155705 3:147227163 Effect of t allele (Freq=0.0834)
1000BRAINS -79.41 [-154.37, -4.46]
ADNI1 6.07 [ -48.35, 60.49]
ADNI2GO -47.81 [-100.16, 4.54]
ALSPAC -46.88 [-115.60, 21.84]
ASRB 29.60 [ -48.21, 107.41]
BETULA -39.79 [-116.72, 37.14]
BIG -18.48 [ -58.36, 21.40]
BIG-PsychChip -52.29 [-126.12, 21.53]
BONN -41.42 [-214.25, 131.40]
BrainScale
CARDIFF -72.82 [-150.50, 4.86]
DNS-V3 -25.50 [ -94.44, 43.45]
DNS-V4 -36.95 [-118.09, 44.18]
EPIGEN -119.82 [-260.25, 20.61]
FBIRN 8.31 [ -51.80, 68.42]
FOR2107 -10.00 [ -56.26, 36.26]
GIG 33.29 [ -47.93, 114.51]
GSP -103.56 [-179.28, -27.84]
HUBIN 40.68 [ -53.72, 135.08]
HUNT -4.93 [ -44.44, 34.59]
IMAGEN -61.71 [-101.28, -22.14]
IMpACT 4.72 [ -94.81, 104.25]
LBC1936 -14.08 [ -74.42, 46.26]
LIBD -10.48 [ -86.43, 65.48]
Sydney MAS -58.10 [-114.55, -1.65]
MCIC -130.00 [-234.66, -25.34]
MooDS 36.58 [ -53.31, 126.48]
MPIP -20.37 [ -75.22, 34.49]
MUNSTER -11.10 [ -54.02, 31.82]
NCNG 9.20 [ -76.37, 94.77]
NESDA 2.16 [ -74.07, 78.40]
NeuroIMAGE -45.00 [-162.66, 72.65]
NTR -32.77 [-115.15, 49.61]
OATS -18.41 [-122.78, 85.96]
PAFIP -219.36 [-339.05, -99.67]
PDNZ 65.42 [ -53.36, 184.21]
PING -68.96 [-151.21, 13.29]
MPRC -17.60 [ -62.81, 27.61]
PPMI
QTIM -28.15 [ -72.06, 15.76]
SHIP -32.06 [ -70.52, 6.40]
SHIP-Trend -25.24 [ -68.08, 17.61]
SYS -108.44 [-188.23, -28.65]
TCD-NUIG -48.77 [-156.23, 58.69]
TOP -37.20 [-102.79, 28.39]
TOP3T -31.74 [-111.21, 47.72]
UiO2016 -13.64 [-114.85, 87.57]
UiO2017 -28.26 [ -95.79, 39.26]
UMCU -26.75 [ -77.59, 24.09]
ENIGMA Discovery Cohorts [N=22180; P=2.449e-09] -28.22 [ -37.49, -18.94]
UKBB -22.96 [ -36.69, -9.24]
Discovery+UKBB [N=31090; P=3.331e-11] -26.53 [ -34.38, -18.69]
GOBS -39.50 [ -71.84, -7.17]
IMH -105.90 [-180.42, -31.38]
UNICAMP 56.20 [ -28.27, 140.67]
Meth-CT -0.47 [ -94.87, 93.92]
MIRECC
UKBB nonEUR -64.95 [-122.91, -7.00]
OSAKA -22.67 [ -73.13, 27.79]
PING Non European -37.96 [ -92.44, 16.51]
Non European Cohorts [N=2626; P=0.0003402] -37.30 [ -57.71, -16.90]
Cohort
Beta [Lower 95% CI,Upper 95% CI]
-400
-300
-200
-100
0
100
200
mm^2/effect allele

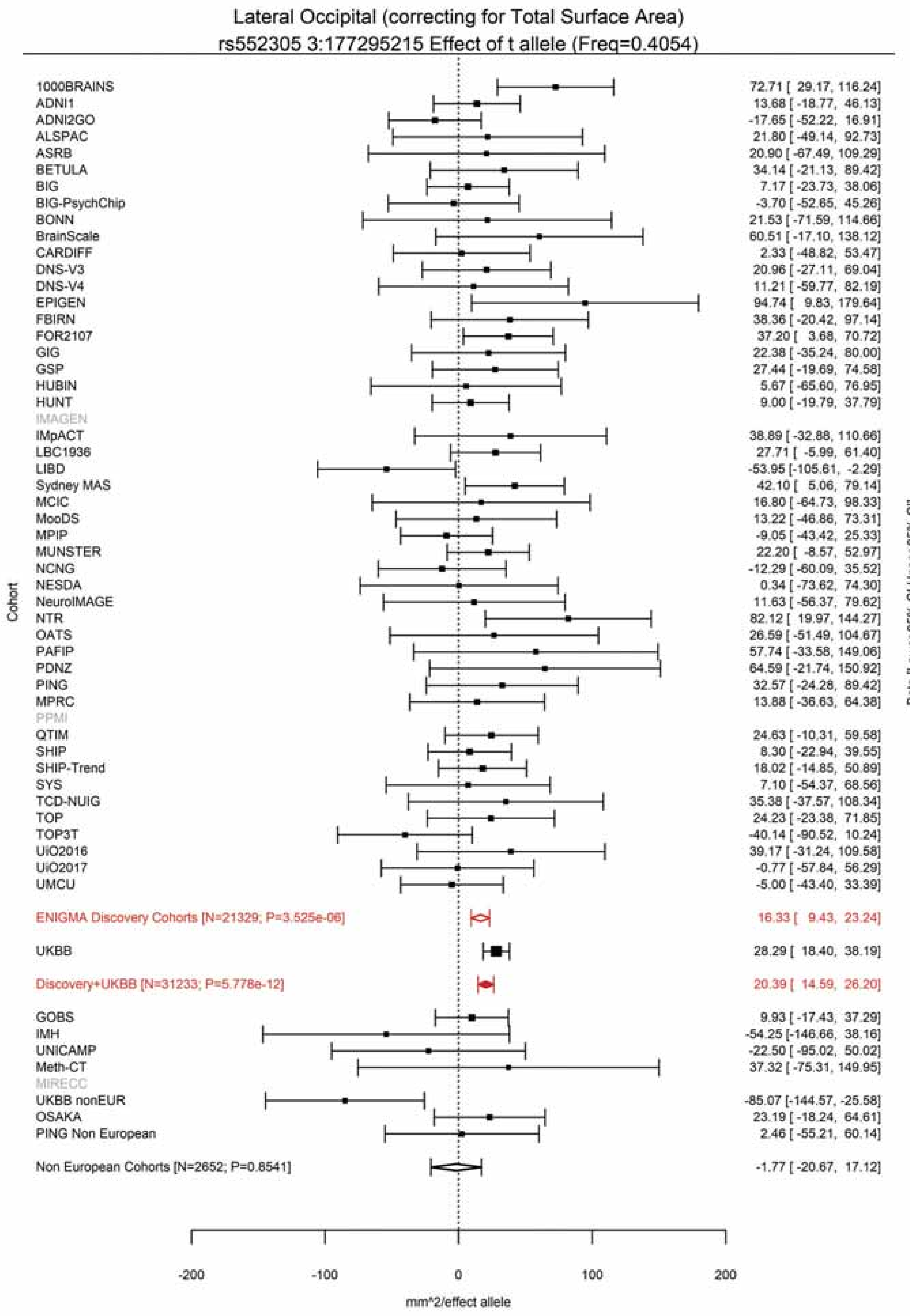
Lateral Occipital (correcting for Total Surface Area)
rs552305 3:177295215 Effect of t allele (Freq=0.4054)
Cohort
1000BRAINS 72.71 [ 29.17, 116.24]
ADNI1 13.68 [ -18.77, 46.13]
ADNI2GO -17.65 [ -52.22, 16.91]
ALSPAC 21.80 [ -49.14, 92.73]
ASRB 20.90 [ -67.49, 109.29]
BETULA 34.14 [ -21.13, 89.42]
BIG 7.17 [ -23.73, 38.06]
BIG-PsychChip -3.70 [ -52.65, 45.26]
BONN 21.53 [ -71.59, 114.66]
BrainScale 60.51 [ -17.10, 138.12]
CARDIFF 2.33 [ -48.82, 53.47]
DNS-V3 20.96 [ -27.11, 69.04]
DNS-V4 11.21 [ -59.77, 82.19]
EPIGEN 94.74 [ 9.83, 179.64]
FBIRN 38.36 [ -20.42, 97.14]
FOR2107 37.20 [ 3.68, 70.72]
GIG 22.38 [ -35.24, 80.00]
GSP 27.44 [ -19.69, 74.58]
HUBIN 5.67 [ -65.60, 76.95]
HUNT 9.00 [ -19.79, 37.79]
IMAGEN
IMpACT 38.89 [ -32.88, 110.66]
LBC1936 27.71 [ -5.99, 61.40]
LIBD -53.95 [-105.61, -2.29]
Sydney MAS 42.10 [ 5.06, 79.14]
MCIC 16.80 [ -64.73, 98.33]
MooDS 13.22 [ -46.86, 73.31]
MPIP -9.05 [ -43.42, 25.33]
MUNSTER 22.20 [ -8.57, 52.97]
NCNG -12.29 [ -60.09, 35.52]
NESDA 0.34 [ -73.62, 74.30]
NeuroIMAGE 11.63 [ -56.37, 79.62]
NTR 82.12 [ 19.97, 144.27]
OATS 26.59 [ -51.49, 104.67]
PAFIP 57.74 [ -33.58, 149.06]
PDNZ 64.59 [ -21.74, 150.92]
PING 32.57 [ -24.28, 89.42]
MPRC 13.88 [ -36.63, 64.38]
PPMI
QTIM 24.63 [ -10.31, 59.58]
SHIP 8.30 [ -22.94, 39.55]
SHIP-Trend 18.02 [ -14.85, 50.89]
SYS 7.10 [ -54.37, 68.56]
TCD-NUIG 35.38 [ -37.57, 108.34]
TOP 24.23 [ -23.38, 71.85]
TOP3T -40.14 [ -90.52, 10.24]
UiO2016 39.17 [ -31.24, 109.58]
UiO2017 -0.77 [ -57.84, 56.29]
UMCU -5.00 [ -43.40, 33.39]
ENIGMA Discovery Cohorts [N=21329; P=3.525e-06] 16.33 [ 9.43, 23.24]
UKBB 28.29 [ 18.40, 38.19]
Discovery+UKBB [N=31233; P=5.778e-12] 20.39 [ 14.59, 26.20]
GOBS 9.93 [ -17.43, 37.29]
IMH -54.25 [-146.66, 38.16]
UNICAMP -22.50 [ -95.02, 50.02]
Meth-CT 37.32 [ -75.31, 149.95]
MIRECC
UKBB nonEUR -85.07 [-144.57, -25.58]
OSAKA 23.19 [ -18.24, 64.61]
PING Non European 2.46 [ -55.21, 60.14]
Non European Cohorts [N=2652; P=0.8541] -1.77 [ -20.67, 17.12]
Beta [Lower 95% CI,Upper 95% CI]
-200
-100
0
100
200
mm^2/effect allele

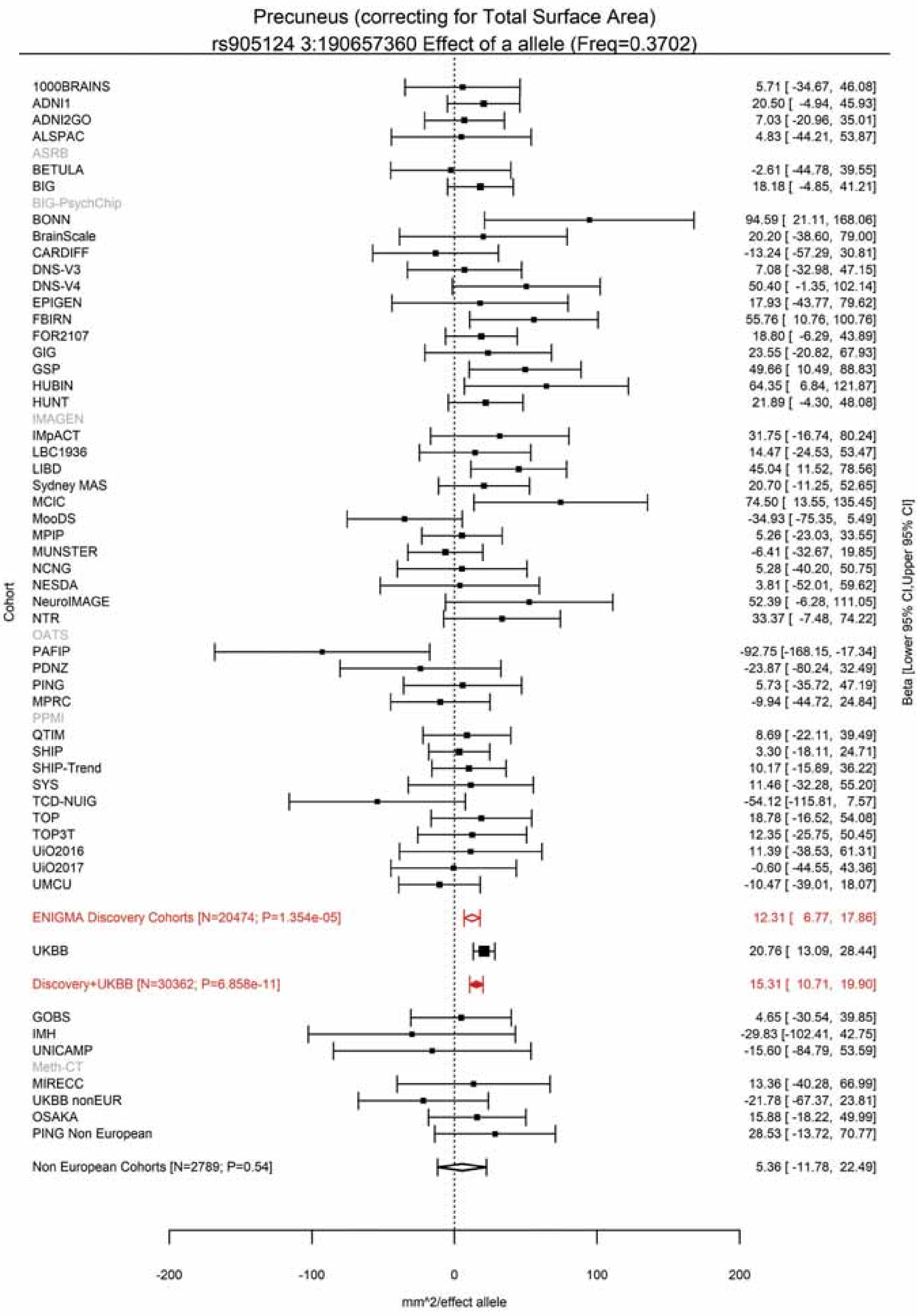

Precuneus (correcting for Total Surface Area)
rs905124 3:190657360 Effect of a allele (Freq=0.3702)
Cohort
1000BRAINS 5.71 [ -34.67, 46.08]
ADNI1 20.50 [ -4.94, 45.93]
ADNI2GO 7.03 [ -20.96, 35.01]
ALSPAC 4.83 [ -44.21, 53.87]
ASRB
BETULA -2.61 [ -44.78, 39.55]
BIG 18.18 [ -4.85, 41.21]
BIG-PsychChip
BONN 94.59 [ 21.11, 168.06]
BrainScale 20.20 [ -38.60, 79.00]
CARDIFF -13.24 [ -57.29, 30.81]
DNS-V3 7.08 [ -32.98, 47.15]
DNS-V4 50.40 [ -1.35, 102.14]
EPIGEN 17.93 [ -43.77, 79.62]
FBIRN 55.76 [ 10.76, 100.76]
FOR2107 18.80 [ -6.29, 43.89]
GIG 23.55 [ -20.82, 67.93]
GSP 49.66 [ 10.49, 88.83]
HUBIN 64.35 [ 6.84, 121.87]
HUNT 21.89 [ -4.30, 48.08]
IMAGEN
IMpACT 31.75 [ -16.74, 80.24]
LBC1936 14.47 [ -24.53, 53.47]
LIBD 45.04 [ 11.52, 78.56]
Sydney MAS 20.70 [ -11.25, 52.65]
MCIC 74.50 [ 13.55, 135.45]
MooDS -34.93 [ -75.35, 5.49]
MPIP 5.26 [ -23.03, 33.55]
MUNSTER -6.41 [ -32.67, 19.85]
NCNG 5.28 [ -40.20, 50.75]
NESDA 3.81 [ -52.01, 59.62]
NeuroIMAGE 52.39 [ -6.28, 111.05]
NTR 33.37 [ -7.48, 74.22]
OATS
PAFIP -92.75 [-168.15, -17.34]
PDNZ -23.87 [ -80.24, 32.49]
PING 5.73 [ -35.72, 47.19]
MPRC -9.94 [ -44.72, 24.84]
PPMI
QTIM 8.69 [ -22.11, 39.49]
SHIP 3.30 [ -18.11, 24.71]
SHIP-Trend 10.17 [ -15.89, 36.22]
SYS 11.46 [ -32.28, 55.20]
TCD-NUIG -54.12 [-115.81, 7.57]
TOP 18.78 [ -16.52, 54.08]
TOP3T 12.35 [ -25.75, 50.45]
UiO2016 11.39 [ -38.53, 61.31]
UiO2017 -0.60 [ -44.55, 43.36]
UMCU -10.47 [ -39.01, 18.07]
ENIGMA Discovery Cohorts [N=20474; P=1.354e-05] 12.31 [ 6.77, 17.86]
UKBB 20.76 [ 13.09, 28.44]
Discovery+UKBB [N=30362; P=6.858e-11] 15.31 [ 10.71, 19.90]
GOBS 4.65 [ -30.54, 39.85]
IMH -29.83 [-102.41, 42.75]
UNICAMP -15.60 [ -84.79, 53.59]
Meth-CT
MIRECC 13.36 [ -40.28, 66.99]
UKBB nonEUR -21.78 [ -67.37, 23.81]
OSAKA 15.88 [ -18.22, 49.99]
PING Non European 28.53 [ -13.72, 70.77]
Non European Cohorts [N=2789; P=0.54] 5.36 [ -11.78, 22.49]
Beta [Lower 95% CI,Upper 95% CI]
-200
-100
0
100
200
mm^2/effect allele

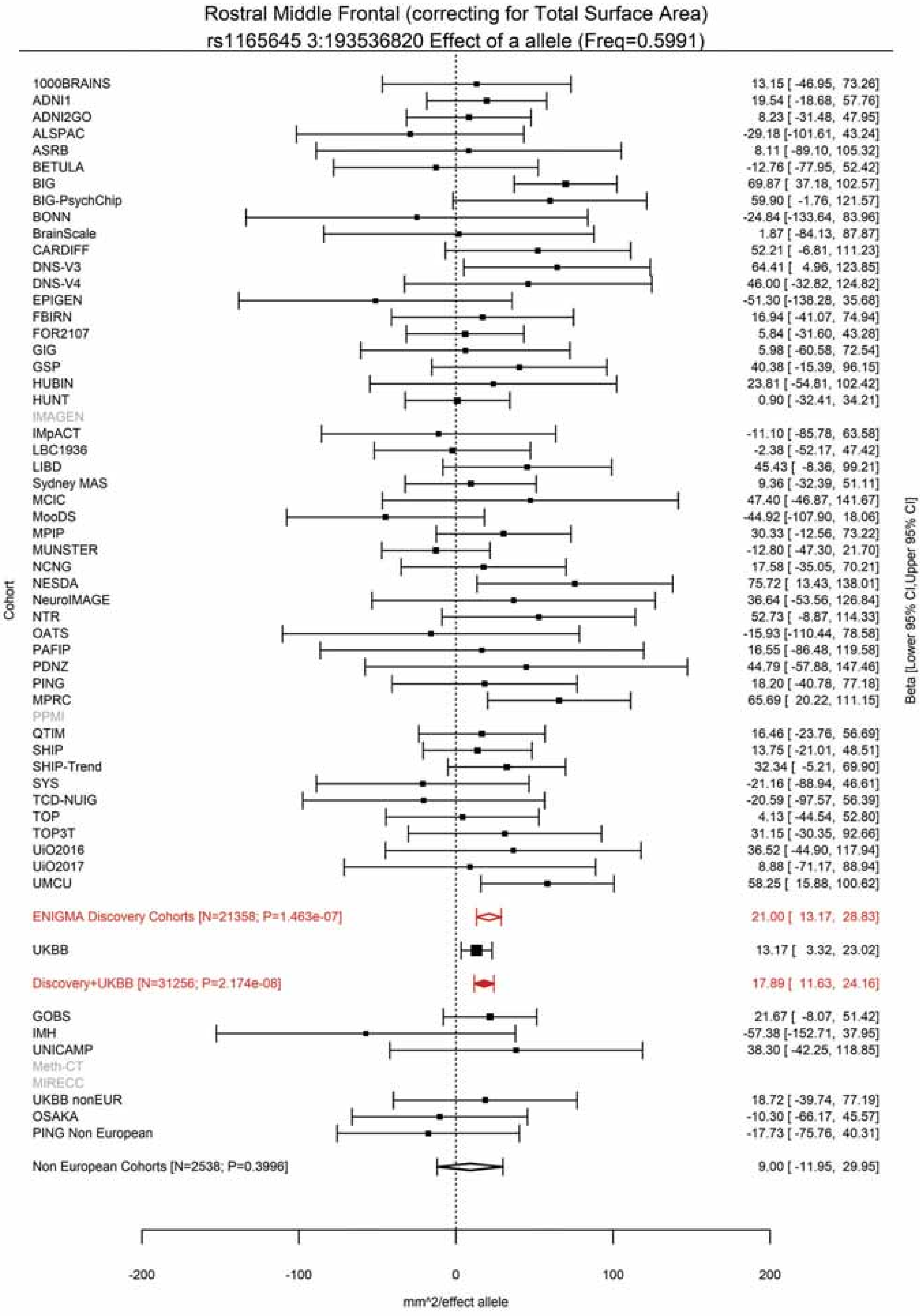
Rostral Middle Frontal (correcting for Total Surface Area)
rs1165645 3:193536820 Effect of a allele (Freq=0.5991)
Cohort
1000BRAINS 13.15 [ -46.95, 73.26]
ADNI1 19.54 [ -18.68, 57.76]
ADNI2GO 8.23 [ -31.48, 47.95]
ALSPAC -29.18 [-101.61, 43.24]
ASRB 8.11 [ -89.10, 105.32]
BETULA -12.76 [ -77.95, 52.42]
BIG 69.87 [ 37.18, 102.57]
BIG-PsychChip 59.90 [ -1.76, 121.57]
BONN -24.84 [-133.64, 83.96]
BrainScale 1.87 [ -84.13, 87.87]
CARDIFF 52.21 [ -6.81, 111.23]
DNS-V3 64.41 [ 4.96, 123.85]
DNS-V4 46.00 [ -32.82, 124.82]
EPIGEN -51.30 [-138.28, 35.68]
FBIRN 16.94 [ -41.07, 74.94]
FOR2107 5.84 [ -31.60, 43.28]
GIG 5.98 [ -60.58, 72.54]
GSP 40.38 [ -15.39, 96.15]
HUBIN 23.81 [ -54.81, 102.42]
HUNT 0.90 [ -32.41, 34.21]
IMAGEN
IMpACT -11.10 [ -85.78, 63.58]
LBC1936 -2.38 [ -52.17, 47.42]
LIBD 45.43 [ -8.36, 99.21]
Sydney MAS 9.36 [ -32.39, 51.11]
MCIC 47.40 [ -46.87, 141.67]
MooDS -44.92 [-107.90, 18.06]
MPIP 30.33 [ -12.56, 73.22]
MUNSTER -12.80 [ -47.30, 21.70]
NCNG 17.58 [ -35.05, 70.21]
NESDA 75.72 [ 13.43, 138.01]
NeuroIMAGE 36.64 [ -53.56, 126.84]
NTR 52.73 [ -8.87, 114.33]
OATS -15.93 [-110.44, 78.58]
PAFIP 16.55 [ -86.48, 119.58]
PDNZ 44.79 [ -57.88, 147.46]
PING 18.20 [ -40.78, 77.18]
MPRC 65.69 [ 20.22, 111.15]
PPMI
QTIM 16.46 [ -23.76, 56.69]
SHIP 13.75 [ -21.01, 48.51]
SHIP-Trend 32.34 [ -5.21, 69.90]
SYS -21.16 [ -88.94, 46.61]
TCD-NUIG -20.59 [ -97.57, 56.39]
TOP 4.13 [ -44.54, 52.80]
TOP3T 31.15 [ -30.35, 92.66]
UiO2016 36.52 [ -44.90, 117.94]
UiO2017 8.88 [ -71.17, 88.94]
UMCU 58.25 [ 15.88, 100.62]
ENIGMA Discovery Cohorts [N=21358; P=1.463e-07] 21.00 [ 13.17, 28.83]
UKBB 13.17 [ 3.32, 23.02]
Discovery+UKBB [N=31256; P=2.174e-08] 17.89 [ 11.63, 24.16]
GOBS 21.67 [ -8.07, 51.42]
IMH -57.38 [-152.71, 37.95]
UNICAMP 38.30 [ -42.25, 118.85]
Meth-CT
MIRECC
UKBB nonEUR 18.72 [ -39.74, 77.19]
OSAKA -10.30 [ -66.17, 45.57]
PING Non European -17.73 [ -75.76, 40.31]
Non European Cohorts [N=2538; P=0.3996] 9.00 [ -11.95, 29.95]
Beta [Lower 95% CI,Upper 95% CI]
-200
-100
0
100
200
mm^2/effect allele

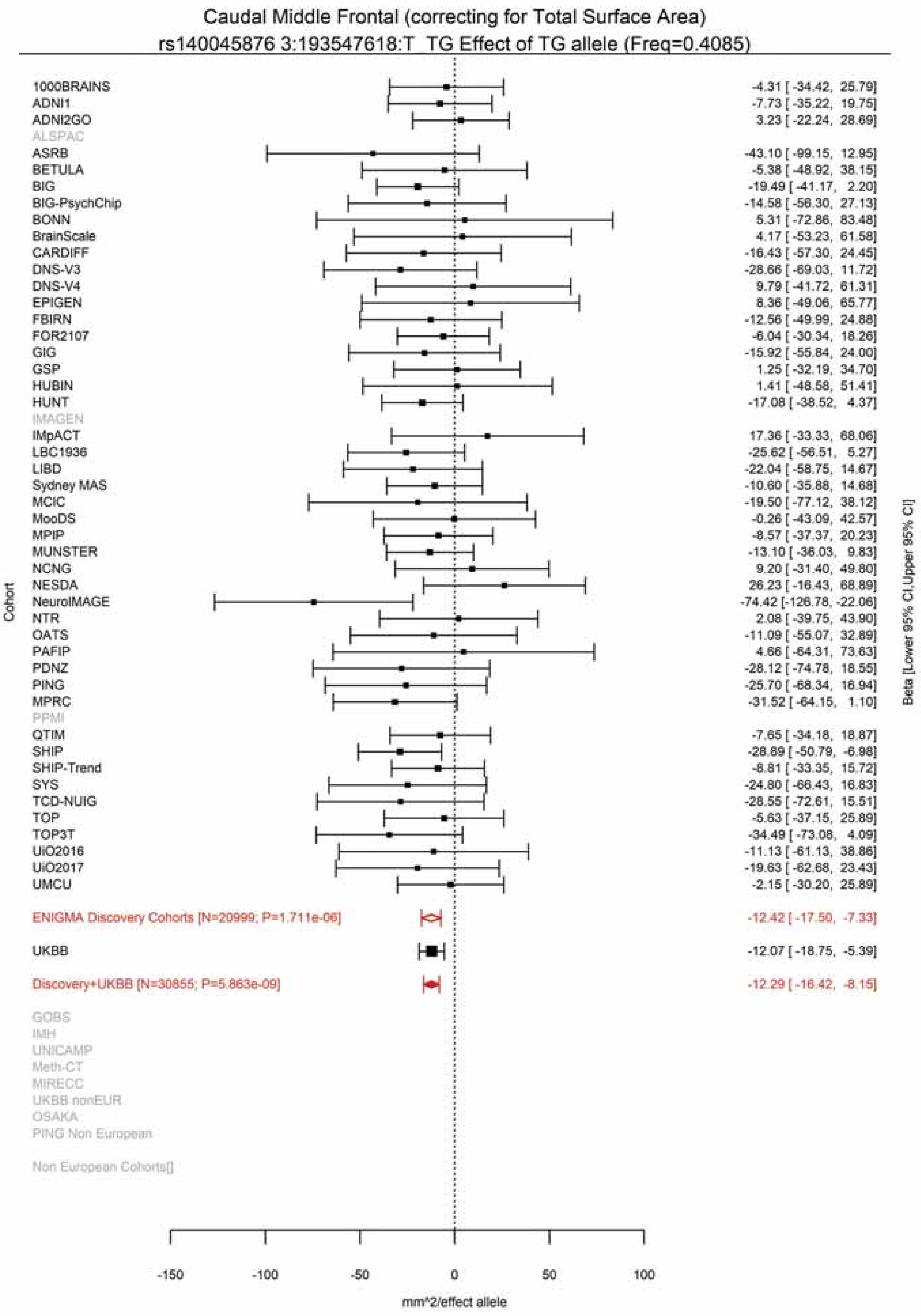

Caudal Middle Frontal (correcting for Total Surface Area)
rs140045876 3:193547618:T_TG Effect of TG allele (Freq=0.4085)
1000BRAINS -4.31 [ -34.42, 25.79]
ADNI1 -7.73 [ -35.22, 19.75]
ADNI2GO 3.23 [ -22.24, 28.69]
ALSPAC
ASRB -43.10 [ -99.15, 12.95]
BETULA -5.38 [ -48.92, 38.15]
BIG -19.49 [ -41.17, 2.20]
BIG-PsychChip -14.58 [ -56.30, 27.13]
BONN 5.31 [ -72.86, 83.48]
BrainScale 4.17 [ -53.23, 61.58]
CARDIFF -16.43 [ -57.30, 24.45]
DNS-V3 -28.66 [ -69.03, 11.72]
DNS-V4 9.79 [ -41.72, 61.31]
EPIGEN 8.36 [ -49.06, 65.77]
FBIRN -12.56 [ -49.99, 24.88]
FOR2107 -6.04 [ -30.34, 18.26]
GIG -15.92 [ -55.84, 24.00]
GSP 1.25 [ -32.19, 34.70]
HUBIN 1.41 [ -48.58, 51.41]
HUNT -17.08 [ -38.52, 4.37]
IMAGEN
IMpACT 17.36 [ -33.33, 68.06]
LBC1936 -25.62 [ -56.51, 5.27]
LIBD -22.04 [ -58.75, 14.67]
Sydney MAS -10.60 [ -35.88, 14.68]
MCIC -19.50 [ -77.12, 38.12]
MooDS -0.26 [ -43.09, 42.57]
MPIP -8.57 [ -37.37, 20.23]
MUNSTER -13.10 [ -36.03, 9.83]
NCNG 9.20 [ -31.40, 49.80]
NESDA 26.23 [ -16.43, 68.89]
NeuroIMAGE -74.42 [-126.78, -22.06]
NTR 2.08 [ -39.75, 43.90]
OATS -11.09 [ -55.07, 32.89]
PAFIP 4.66 [ -64.31, 73.63]
PDNZ -28.12 [ -74.78, 18.55]
PING -25.70 [ -68.34, 16.94]
MPRC -31.52 [ -64.15, 1.10]
PPMI
QTIM -7.65 [ -34.18, 18.87]
SHIP -28.89 [ -50.79, -6.98]
SHIP-Trend -8.81 [ -33.35, 15.72]
SYS -24.80 [ -66.43, 16.83]
TCD-NUIG -28.55 [ -72.61, 15.51]
TOP -5.63 [ -37.15, 25.89]
TOP3T -34.49 [ -73.08, 4.09]
UiO2016 -11.13 [ -61.13, 38.86]
UiO2017 -19.63 [ -62.68, 23.43]
UMCU -2.15 [ -30.20, 25.89]
ENIGMA Discovery Cohorts [N=20999; P=1.711e-06] -12.42 [ -17.50, -7.33]
UKBB -12.07 [ -18.75, -5.39]
Discovery+UKBB [N=30855; P=5.863e-09] -12.29 [ -16.42, -8.15]
GOBS
IMH
UNICAMP
Meth-CT
MIRECC
UKBB nonEUR
OSAKA
PING Non European
Non European Cohorts[]
Cohort
Beta [Lower 95% CI,Upper 95% CI]
-150
-100
-50
0
50
100
mm^2/effect allele

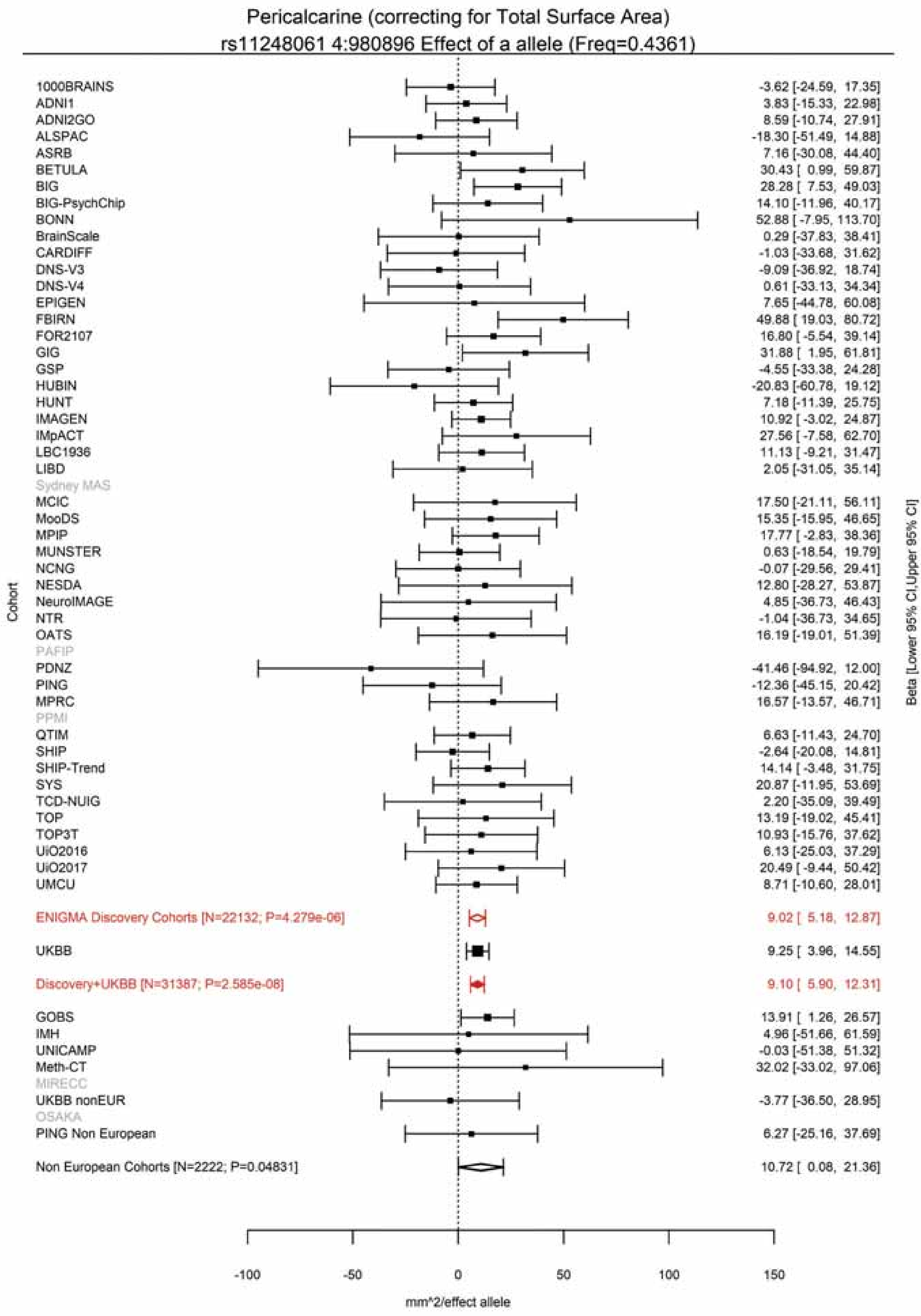
Pericalcarine (correcting for Total Surface Area)
rs11248061 4:980896 Effect of a allele (Freq=0.4361)
Cohort
1000BRAINS -3.62 [-24.59, 17.35]
ADNI1 3.83 [-15.33, 22.98]
ADNI2GO 8.59 [-10.74, 27.91]
ALSPAC -18.30 [-51.49, 14.88]
ASRB 7.16 [-30.08, 44.40]
BETULA 30.43 [ 0.99, 59.87]
BIG 28.28 [ 7.53, 49.03]
BIG-PsychChip 14.10 [-11.96, 40.17]
BONN 52.88 [ -7.95, 113.70]
BrainScale 0.29 [-37.83, 38.41]
CARDIFF -1.03 [-33.68, 31.62]
DNS-V3 -9.09 [-36.92, 18.74]
DNS-V4 0.61 [-33.13, 34.34]
EPIGEN 7.65 [-44.78, 60.08]
FBIRN 49.88 [ 19.03, 80.72]
FOR2107 16.80 [ -5.54, 39.14]
GIG 31.88 [ 1.95, 61.81]
GSP -4.55 [-33.38, 24.28]
HUBIN -20.83 [-60.78, 19.12]
HUNT 7.18 [-11.39, 25.75]
IMAGEN 10.92 [ -3.02, 24.87]
IMpACT 27.56 [ -7.58, 62.70]
LBC1936 11.13 [ -9.21, 31.47]
LIBD 2.05 [-31.05, 35.14]
Sydney MAS
MCIC 17.50 [-21.11, 56.11]
MooDS 15.35 [-15.95, 46.65]
MPIP 17.77 [ -2.83, 38.36]
MUNSTER 0.63 [-18.54, 19.79]
NCNG -0.07 [-29.56, 29.41]
NESDA 12.80 [-28.27, 53.87]
NeuroIMAGE 4.85 [-36.73, 46.43]
NTR -1.04 [-36.73, 34.65]
OATS 16.19 [-19.01, 51.39]
PAFIP
PDNZ -41.46 [-94.92, 12.00]
PING -12.36 [-45.15, 20.42]
MPRC 16.57 [-13.57, 46.71]
PPMI
QTIM 6.63 [-11.43, 24.70]
SHIP -2.64 [-20.08, 14.81]
SHIP-Trend 14.14 [ -3.48, 31.75]
SYS 20.87 [-11.95, 53.69]
TCD-NUIG 2.20 [-35.09, 39.49]
TOP 13.19 [-19.02, 45.41]
TOP3T 10.93 [-15.76, 37.62]
UiO2016 6.13 [-25.03, 37.29]
UiO2017 20.49 [ -9.44, 50.42]
UMCU 8.71 [-10.60, 28.01]
ENIGMA Discovery Cohorts [N=22132; P=4.279e-06] 9.02 [ 5.18, 12.87]
UKBB 9.25 [ 3.96, 14.55]
Discovery+UKBB [N=31387; P=2.585e-08] 9.10 [ 5.90, 12.31]
GOBS 13.91 [ 1.26, 26.57]
IMH 4.96 [-51.66, 61.59]
UNICAMP -0.03 [-51.38, 51.32]
Meth-CT 32.02 [-33.02, 97.06]
MIRECC
UKBB nonEUR -3.77 [-36.50, 28.95]
OSAKA
PING Non European 6.27 [-25.16, 37.69]
Non European Cohorts [N=2222; P=0.04831] 10.72 [ 0.08, 21.36]
Beta [Lower 95% CI,Upper 95% CI]
-100
-50
0
50
100
150
mm^2/effect allele

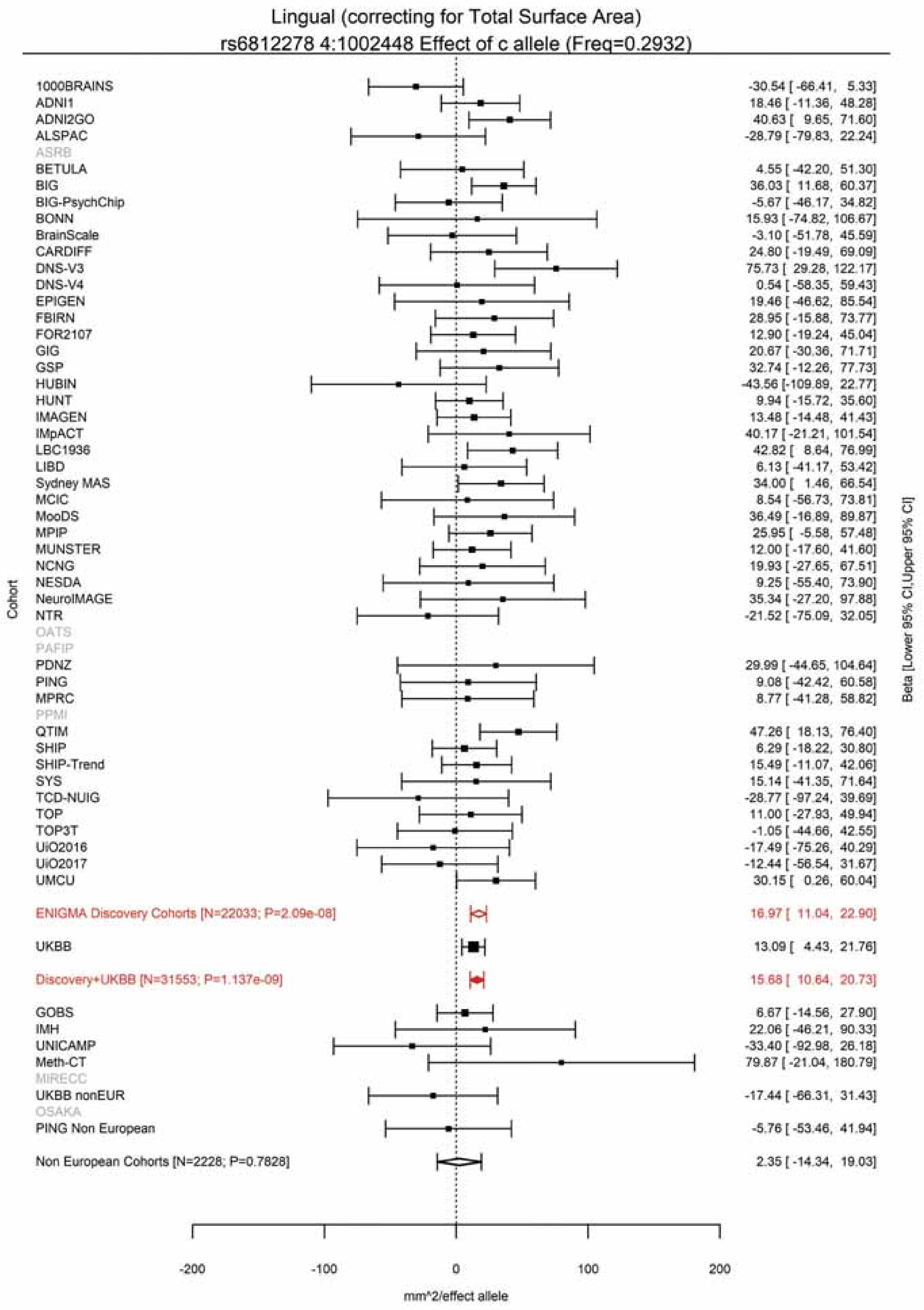

Lingual (correcting for Total Surface Area)
rs6812278 4:1002448 Effect of c allele (Freq=0.2932)
Cohort
1000BRAINS -30.54 [ -66.41, 5.33]
ADNI1 18.46 [ -11.36, 48.28]
ADNI2GO 40.63 [ 9.65, 71.60]
ALSPAC -28.79 [ -79.83, 22.24]
ASRB
BETULA 4.55 [ -42.20, 51.30]
BIG 36.03 [ 11.68, 60.37]
BIG-PsychChip -5.67 [ -46.17, 34.82]
BONN 15.93 [ -74.82, 106.67]
BrainScale -3.10 [ -51.78, 45.59]
CARDIFF 24.80 [ -19.49, 69.09]
DNS-V3 75.73 [ 29.28, 122.17]
DNS-V4 0.54 [ -58.35, 59.43]
EPIGEN 19.46 [ -46.62, 85.54]
FBIRN 28.95 [ -15.88, 73.77]
FOR2107 12.90 [ -19.24, 45.04]
GIG 20.67 [ -30.36, 71.71]
GSP 32.74 [ -12.26, 77.73]
HUBIN -43.56 [-109.89, 22.77]
HUNT 9.94 [ -15.72, 35.60]
IMAGEN 13.48 [ -14.48, 41.43]
IMpACT 40.17 [ -21.21, 101.54]
LBC1936 42.82 [ 8.64, 76.99]
LIBD 6.13 [ -41.17, 53.42]
Sydney MAS 34.00 [ 1.46, 66.54]
MCIC 8.54 [ -56.73, 73.81]
MooDS 36.49 [ -16.89, 89.87]
MPIP 25.95 [ -5.58, 57.48]
MUNSTER 12.00 [ -17.60, 41.60]
NCNG 19.93 [ -27.65, 67.51]
NESDA 9.25 [ -55.40, 73.90]
NeuroIMAGE 35.34 [ -27.20, 97.88]
NTR -21.52 [ -75.09, 32.05]
OATS
PAFIP
PDNZ 29.99 [ -44.65, 104.64]
PING 9.08 [ -42.42, 60.58]
MPRC 8.77 [ -41.28, 58.82]
PPMI
QTIM 47.26 [ 18.13, 76.40]
SHIP 6.29 [ -18.22, 30.80]
SHIP-Trend 15.49 [ -11.07, 42.06]
SYS 15.14 [ -41.35, 71.64]
TCD-NUIG -28.77 [ -97.24, 39.69]
TOP 11.00 [ -27.93, 49.94]
TOP3T -1.05 [ -44.66, 42.55]
UiO2016 -17.49 [ -75.26, 40.29]
UiO2017 -12.44 [ -56.54, 31.67]
UMCU 30.15 [ 0.26, 60.04]
ENIGMA Discovery Cohorts [N=22033; P=2.09e-08] 16.97 [ 11.04, 22.90]
UKBB 13.09 [ 4.43, 21.76]
Discovery+UKBB [N=31553; P=1.137e-09] 15.68 [ 10.64, 20.73]
GOBS 6.67 [ -14.56, 27.90]
IMH 22.06 [ -46.21, 90.33]
UNICAMP -33.40 [ -92.98, 26.18]
Meth-CT 79.87 [ -21.04, 180.79]
MIRECC
UKBB nonEUR -17.44 [ -66.31, 31.43]
OSAKA
PING Non European -5.76 [ -53.46, 41.94]
Non European Cohorts [N=2228; P=0.7828] 2.35 [ -14.34, 19.03]
Beta [Lower 95% CI,Upper 95% CI]
-200
-100
0
100
200
mm^2/effect allele

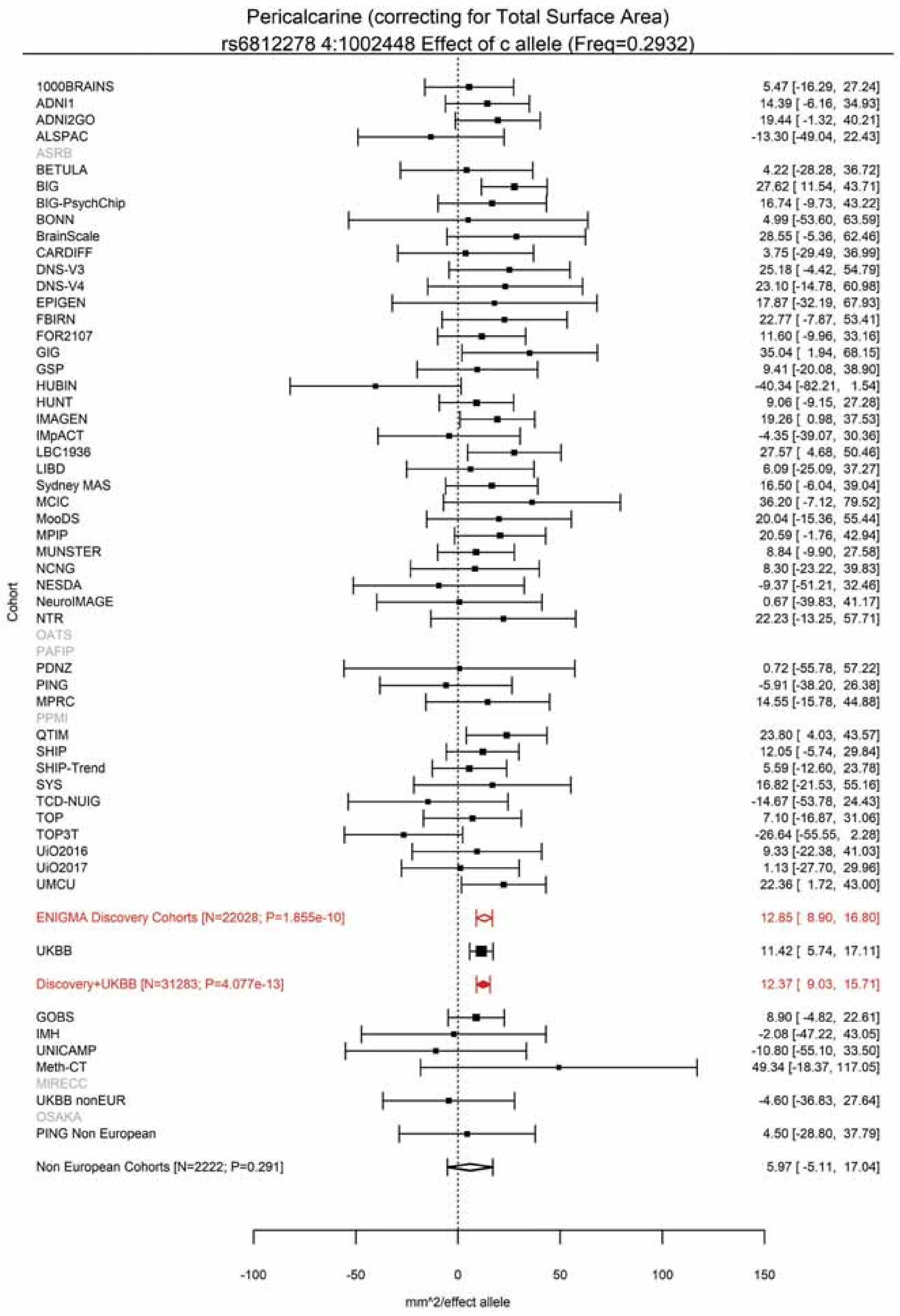

Pericalcarine (correcting for Total Surface Area)
rs6812278 4:1002448 Effect of c allele (Freq=0.2932)
1000BRAINS 5.47 [-16.29, 27.24]
ADNI1 14.39 [ -6.16, 34.93]
ADNI2GO 19.44 [ -1.32, 40.21]
ALSPAC -13.30 [-49.04, 22.43]
ASRB
BETULA 4.22 [-28.28, 36.72]
BIG 27.62 [ 11.54, 43.71]
BIG-PsychChip 16.74 [ -9.73, 43.22]
BONN 4.99 [-53.60, 63.59]
BrainScale 28.55 [ -5.36, 62.46]
CARDIFF 3.75 [-29.49, 36.99]
DNS-V3 25.18 [ -4.42, 54.79]
DNS-V4 23.10 [-14.78, 60.98]
EPIGEN 17.87 [-32.19, 67.93]
FBIRN 22.77 [ -7.87, 53.41]
FOR2107 11.60 [ -9.96, 33.16]
GIG 35.04 [ 1.94, 68.15]
GSP 9.41 [-20.08, 38.90]
HUBIN -40.34 [-82.21, 1.54]
HUNT 9.06 [ -9.15, 27.28]
IMAGEN 19.26 [ 0.98, 37.53]
IMpACT -4.35 [-39.07, 30.36]
LBC1936 27.57 [ 4.68, 50.46]
LIBD 6.09 [-25.09, 37.27]
Sydney MAS 16.50 [ -6.04, 39.04]
MCIC 36.20 [ -7.12, 79.52]
MooDS 20.04 [-15.36, 55.44]
MPIP 20.59 [ -1.76, 42.94]
MUNSTER 8.84 [ -9.90, 27.58]
NCNG 8.30 [-23.22, 39.83]
NESDA -9.37 [-51.21, 32.46]
NeuroIMAGE 0.67 [-39.83, 41.17]
NTR 22.23 [-13.25, 57.71]
OATS
PAFIP
PDNZ 0.72 [-55.78, 57.22]
PING -5.91 [-38.20, 26.38]
MPRC 14.55 [-15.78, 44.88]
PPMI
QTIM 23.80 [ 4.03, 43.57]
SHIP 12.05 [ -5.74, 29.84]
SHIP-Trend 5.59 [-12.60, 23.78]
SYS 16.82 [-21.53, 55.16]
TCD-NUIG -14.67 [-53.78, 24.43]
TOP 7.10 [-16.87, 31.06]
TOP3T -26.64 [-55.55, 2.28]
UiO2016 9.33 [-22.38, 41.03]
UiO2017 1.13 [-27.70, 29.96]
UMCU 22.36 [ 1.72, 43.00]
ENIGMA Discovery Cohorts [N=22028; P=1.855e-10] 12.85 [ 8.90, 16.80]
UKBB 11.42 [ 5.74, 17.11]
Discovery+UKBB [N=31283; P=4.077e-13] 12.37 [ 9.03, 15.71]
GOBS 8.90 [ -4.82, 22.61]
IMH -2.08 [-47.22, 43.05]
UNICAMP -10.80 [-55.10, 33.50]
Meth-CT 49.34 [-18.37, 117.05]
MIRECC
UKBB nonEUR -4.60 [-36.83, 27.64]
OSAKA
PING Non European 4.50 [-28.80, 37.79]
Non European Cohorts [N=2222; P=0.291] 5.97 [ -5.11, 17.04]
Cohort
Beta [Lower 95% CI,Upper 95% CI]
-100
-50
0
50
100
150
mm^2/effect allele

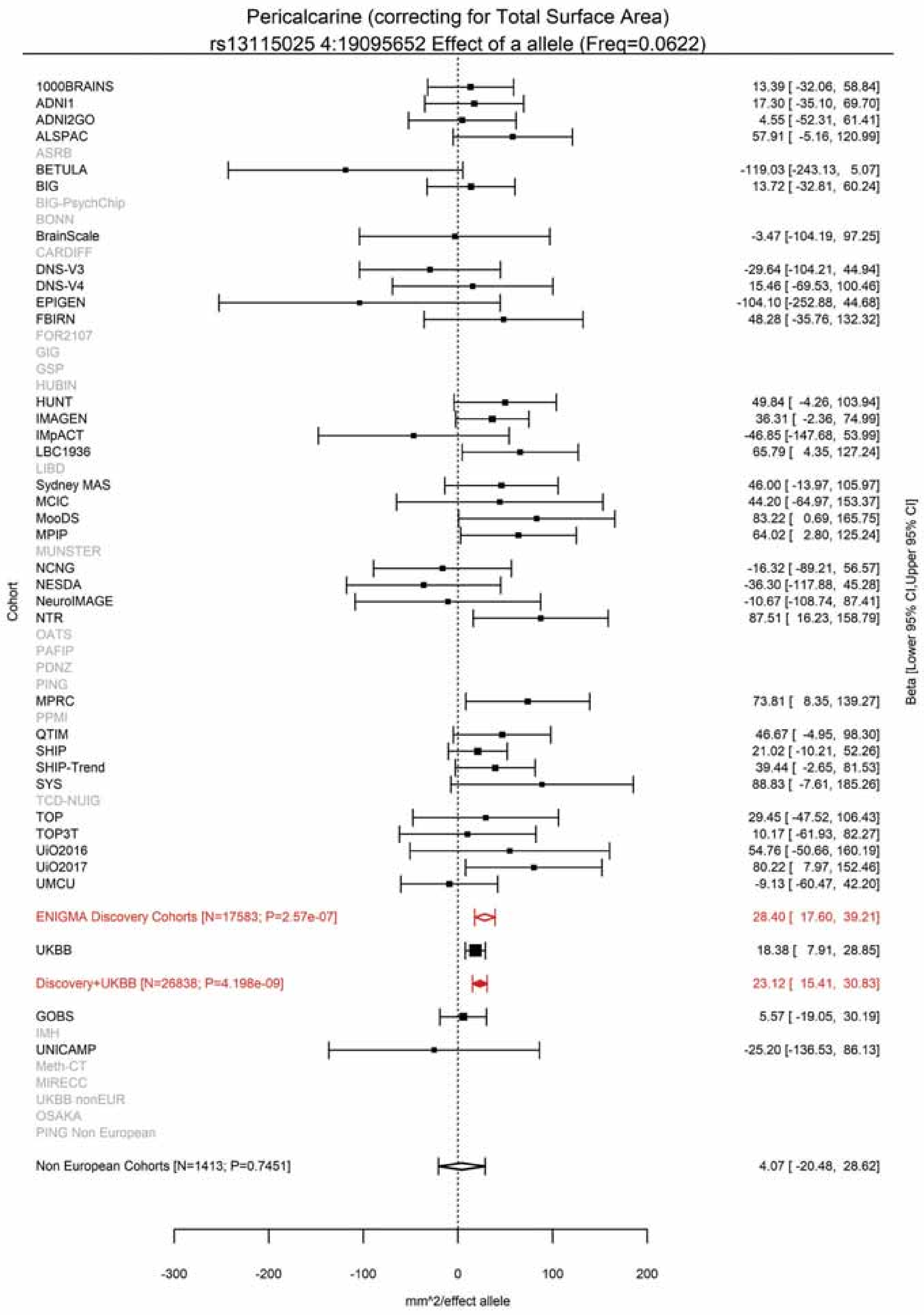
Pericalcarine (correcting for Total Surface Area)
rs13115025 4:19095652 Effect of a allele (Freq=0.0622)
Cohort
1000BRAINS 13.39 [ -32.06, 58.84]
ADNI1 17.30 [ -35.10, 69.70]
ADNI2GO 4.55 [ -52.31, 61.41]
ALSPAC 57.91 [ -5.16, 120.99]
ASRB
BETULA -119.03 [-243.13, 5.07]
BIG 13.72 [ -32.81, 60.24]
BIG-PsychChip
BONN
BrainScale -3.47 [-104.19, 97.25]
CARDIFF
DNS-V3 -29.64 [-104.21, 44.94]
DNS-V4 15.46 [ -69.53, 100.46]
EPIGEN -104.10 [-252.88, 44.68]
FBIRN 48.28 [ -35.76, 132.32]
FOR2107
GIG
GSP
HUBIN
HUNT 49.84 [ -4.26, 103.94]
IMAGEN 36.31 [ -2.36, 74.99]
IMpACT -46.85 [-147.68, 53.99]
LBC1936 65.79 [ 4.35, 127.24]
LIBD
Sydney MAS 46.00 [ -13.97, 105.97]
MCIC 44.20 [ -64.97, 153.37]
MooDS 83.22 [ 0.69, 165.75]
MPIP 64.02 [ 2.80, 125.24]
MUNSTER
NCNG -16.32 [ -89.21, 56.57]
NESDA -36.30 [-117.88, 45.28]
NeuroIMAGE -10.67 [-108.74, 87.41]
NTR 87.51 [ 16.23, 158.79]
OATS
PAFIP
PDNZ
PING
MPRC 73.81 [ 8.35, 139.27]
PPMI
QTIM 46.67 [ -4.95, 98.30]
SHIP 21.02 [ -10.21, 52.26]
SHIP-Trend 39.44 [ -2.65, 81.53]
SYS 88.83 [ -7.61, 185.26]
TCD-NUIG
TOP 29.45 [ -47.52, 106.43]
TOP3T 10.17 [ -61.93, 82.27]
UiO2016 54.76 [ -50.66, 160.19]
UiO2017 80.22 [ 7.97, 152.46]
UMCU -9.13 [ -60.47, 42.20]
ENIGMA Discovery Cohorts [N=17583; P=2.57e-07] 28.40 [ 17.60, 39.21]
UKBB 18.38 [ 7.91, 28.85]
Discovery+UKBB [N=26838; P=4.198e-09] 23.12 [ 15.41, 30.83]
GOBS 5.57 [ -19.05, 30.19]
IMH
UNICAMP -25.20 [-136.53, 86.13]
Meth-CT
MIRECC
UKBB nonEUR
OSAKA
PING Non European
Non European Cohorts [N=1413; P=0.7451] 4.07 [ -20.48, 28.62]
-300
-200
-100
0
100
200
mm^2/effect allele
Beta [Lower 95% CI,Upper 95% CI]

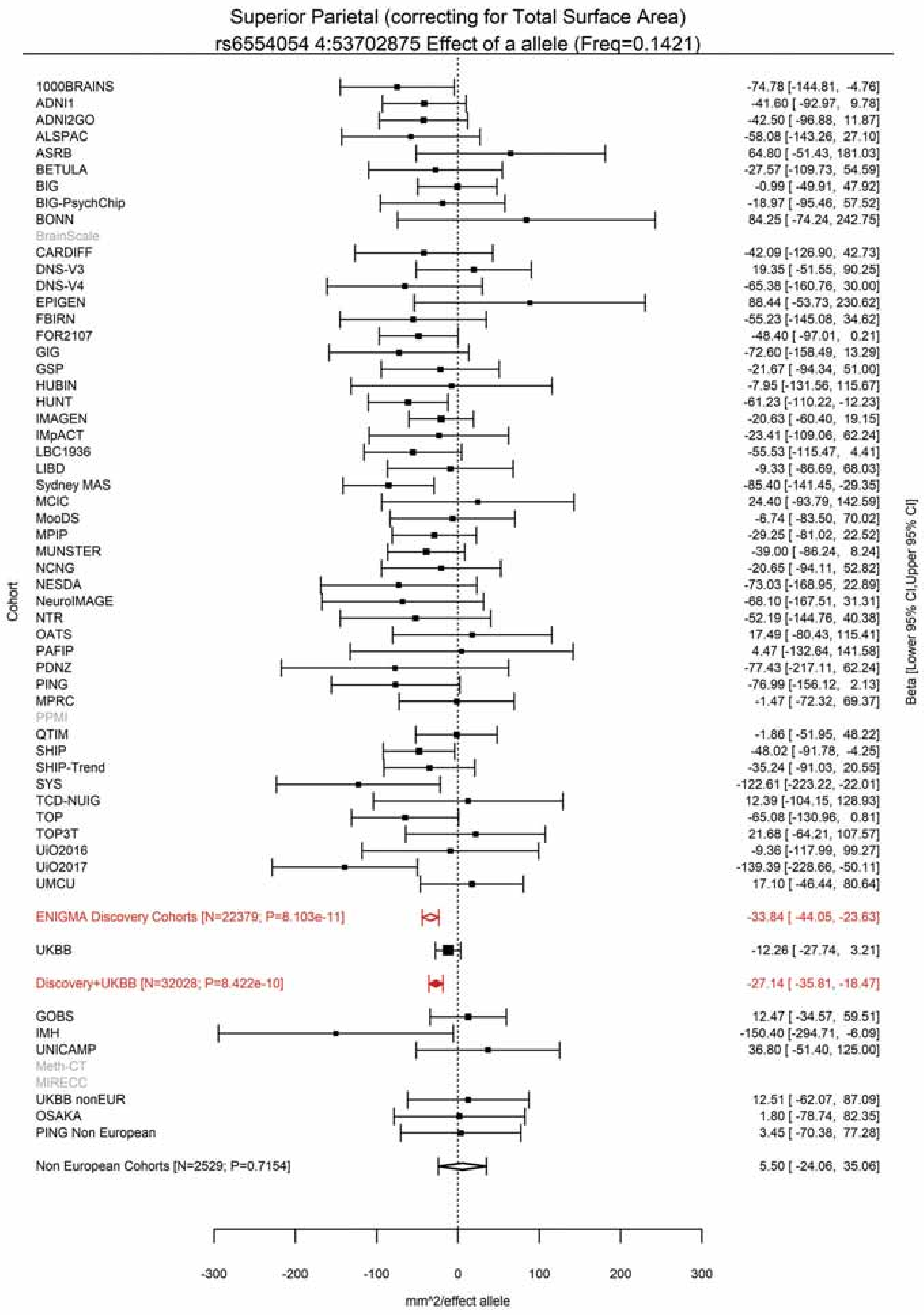
Superior Parietal (correcting for Total Surface Area)
rs6554054 4:53702875 Effect of a allele (Freq=0.1421)
Cohort
1000BRAINS -74.78 [-144.81, -4.76]
ADNI1 -41.60 [ -92.97, 9.78]
ADNI2GO -42.50 [ -96.88, 11.87]
ALSPAC -58.08 [-143.26, 27.10]
ASRB 64.80 [ -51.43, 181.03]
BETULA -27.57 [-109.73, 54.59]
BIG -0.99 [ -49.91, 47.92]
BIG-PsychChip -18.97 [ -95.46, 57.52]
BONN 84.25 [ -74.24, 242.75]
BrainScale
CARDIFF -42.09 [-126.90, 42.73]
DNS-V3 19.35 [ -51.55, 90.25]
DNS-V4 -65.38 [-160.76, 30.00]
EPIGEN 88.44 [ -53.73, 230.62]
FBIRN -55.23 [-145.08, 34.62]
FOR2107 -48.40 [ -97.01, 0.21]
GIG -72.60 [-158.49, 13.29]
GSP -21.67 [ -94.34, 51.00]
HUBIN -7.95 [-131.56, 115.67]
HUNT -61.23 [-110.22, -12.23]
IMAGEN -20.63 [ -60.40, 19.15]
IMpACT -23.41 [-109.06, 62.24]
LBC1936 -55.53 [-115.47, 4.41]
LIBD -9.33 [ -86.69, 68.03]
Sydney MAS -85.40 [-141.45, -29.35]
MCIC 24.40 [ -93.79, 142.59]
MooDS -6.74 [ -83.50, 70.02]
MPIP -29.25 [ -81.02, 22.52]
MUNSTER -39.00 [ -86.24, 8.24]
NCNG -20.65 [ -94.11, 52.82]
NESDA -73.03 [-168.95, 22.89]
NeuroIMAGE -68.10 [-167.51, 31.31]
NTR -52.19 [-144.76, 40.38]
OATS 17.49 [ -80.43, 115.41]
PAFIP 4.47 [-132.64, 141.58]
PDNZ -77.43 [-217.11, 62.24]
PING -76.99 [-156.12, 2.13]
MPRC -1.47 [ -72.32, 69.37]
PPMI
QTIM -1.86 [ -51.95, 48.22]
SHIP -48.02 [ -91.78, -4.25]
SHIP-Trend -35.24 [ -91.03, 20.55]
SYS -122.61 [-223.22, -22.01]
TCD-NUIG 12.39 [-104.15, 128.93]
TOP -65.08 [-130.96, 0.81]
TOP3T 21.68 [ -64.21, 107.57]
UiO2016 -9.36 [-117.99, 99.27]
UiO2017 -139.39 [-228.66, -50.11]
UMCU 17.10 [ -46.44, 80.64]
ENIGMA Discovery Cohorts [N=22379; P=8.103e-11] -33.84 [ -44.05, -23.63]
UKBB -12.26 [ -27.74, 3.21]
Discovery+UKBB [N=32028; P=8.422e-10] -27.14 [ -35.81, -18.47]
GOBS 12.47 [ -34.57, 59.51]
IMH -150.40 [-294.71, -6.09]
UNICAMP 36.80 [ -51.40, 125.00]
Meth-CT
MIRECC
UKBB nonEUR 12.51 [ -62.07, 87.09]
OSAKA 1.80 [ -78.74, 82.35]
PING Non European 3.45 [ -70.38, 77.28]
Non European Cohorts [N=2529; P=0.7154] 5.50 [ -24.06, 35.06]
-300
-200
-100
0
100
200
300
mm^2/effect allele
Beta [Lower 95% CI,Upper 95% CI]

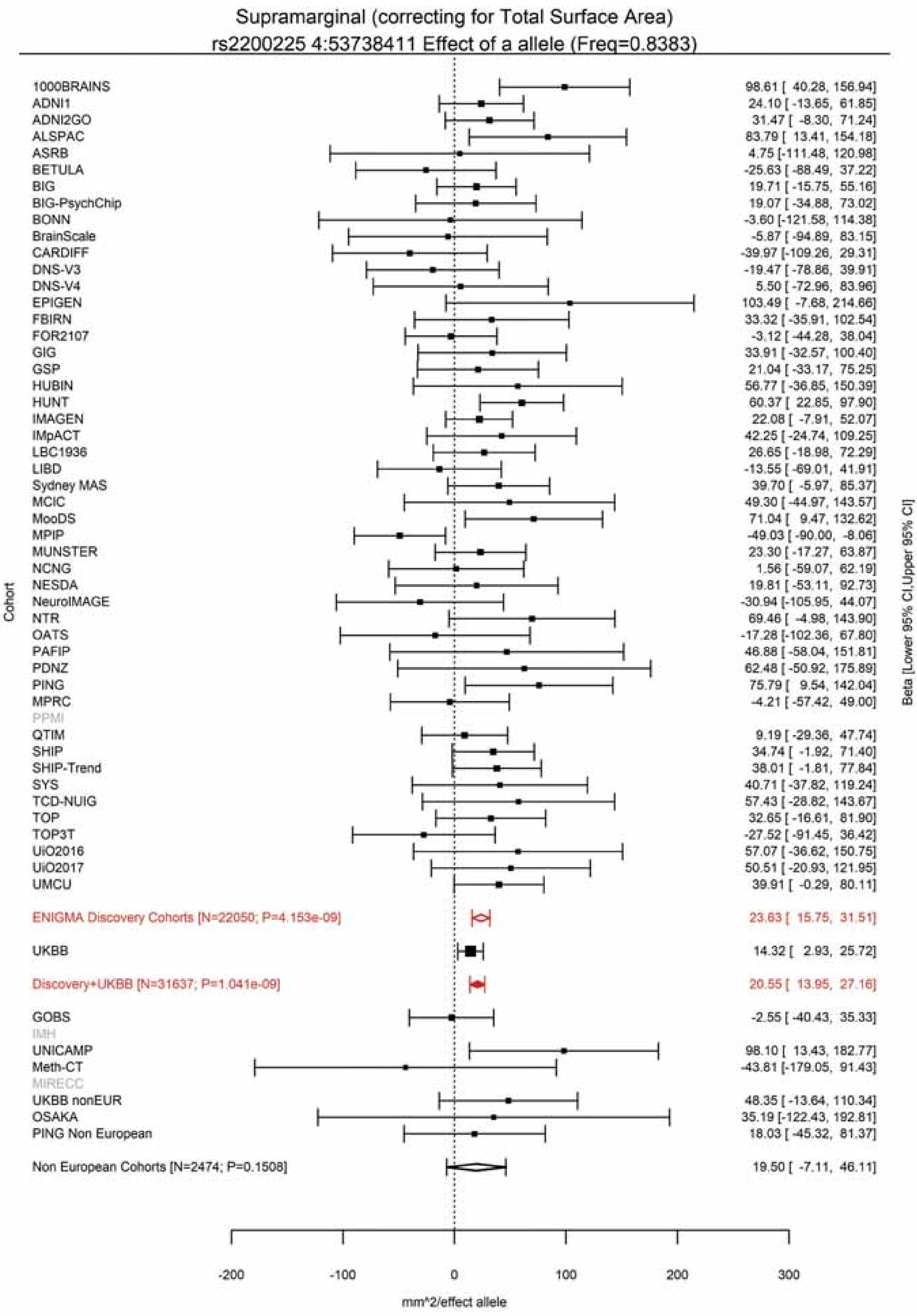
Supramarginal (correcting for Total Surface Area)
rs2200225 4:53738411 Effect of a allele (Freq=0.8383)
Cohort
1000BRAINS 98.61 [ 40.28, 156.94]
ADNI1 24.10 [ -13.65, 61.85]
ADNI2GO 31.47 [ -8.30, 71.24]
ALSPAC 83.79 [ 13.41, 154.18]
ASRB 4.75 [-111.48, 120.98]
BETULA -25.63 [ -88.49, 37.22]
BIG 19.71 [ -15.75, 55.16]
BIG-PsychChip 19.07 [ -34.88, 73.02]
BONN -3.60 [-121.58, 114.38]
BrainScale -5.87 [ -94.89, 83.15]
CARDIFF -39.97 [-109.26, 29.31]
DNS-V3 -19.47 [ -78.86, 39.91]
DNS-V4 5.50 [ -72.96, 83.96]
EPIGEN 103.49 [ -7.68, 214.66]
FBIRN 33.32 [ -35.91, 102.54]
FOR2107 -3.12 [ -44.28, 38.04]
GIG 33.91 [ -32.57, 100.40]
GSP 21.04 [ -33.17, 75.25]
HUBIN 56.77 [ -36.85, 150.39]
HUNT 60.37 [ 22.85, 97.90]
IMAGEN 22.08 [ -7.91, 52.07]
IMpACT 42.25 [ -24.74, 109.25]
LBC1936 26.65 [ -18.98, 72.29]
LIBD -13.55 [ -69.01, 41.91]
Sydney MAS 39.70 [ -5.97, 85.37]
MCIC 49.30 [ -44.97, 143.57]
MooDS 71.04 [ 9.47, 132.62]
MPIP -49.03 [ -90.00, -8.06]
MUNSTER 23.30 [ -17.27, 63.87]
NCNG 1.56 [ -59.07, 62.19]
NESDA 19.81 [ -53.11, 92.73]
NeuroIMAGE -30.94 [-105.95, 44.07]
NTR 69.46 [ -4.98, 143.90]
OATS -17.28 [-102.36, 67.80]
PAFIP 46.88 [ -58.04, 151.81]
PDNZ 62.48 [ -50.92, 175.89]
PING 75.79 [ 9.54, 142.04]
MPRC -4.21 [ -57.42, 49.00]
PPMI
QTIM 9.19 [ -29.36, 47.74]
SHIP 34.74 [ -1.92, 71.40]
SHIP-Trend 38.01 [ -1.81, 77.84]
SYS 40.71 [ -37.82, 119.24]
TCD-NUIG 57.43 [ -28.82, 143.67]
TOP 32.65 [ -16.61, 81.90]
TOP3T -27.52 [ -91.45, 36.42]
UiO2016 57.07 [ -36.62, 150.75]
UiO2017 50.51 [ -20.93, 121.95]
UMCU 39.91 [ -0.29, 80.11]
ENIGMA Discovery Cohorts [N=22050; P=4.153e-09] 23.63 [ 15.75, 31.51]
UKBB 14.32 [ 2.93, 25.72]
Discovery+UKBB [N=31637; P=1.041e-09] 20.55 [ 13.95, 27.16]
GOBS -2.55 [ -40.43, 35.33]
IMH
UNICAMP 98.10 [ 13.43, 182.77]
Meth-CT -43.81 [-179.05, 91.43]
MIRECC
UKBB nonEUR 48.35 [ -13.64, 110.34]
OSAKA 35.19 [-122.43, 192.81]
PING Non European 18.03 [ -45.32, 81.37]
Non European Cohorts [N=2474; P=0.1508] 19.50 [ -7.11, 46.11]
Beta [Lower 95% CI,Upper 95% CI]
-200
-100
0
100
200
300
mm^2/effect allele

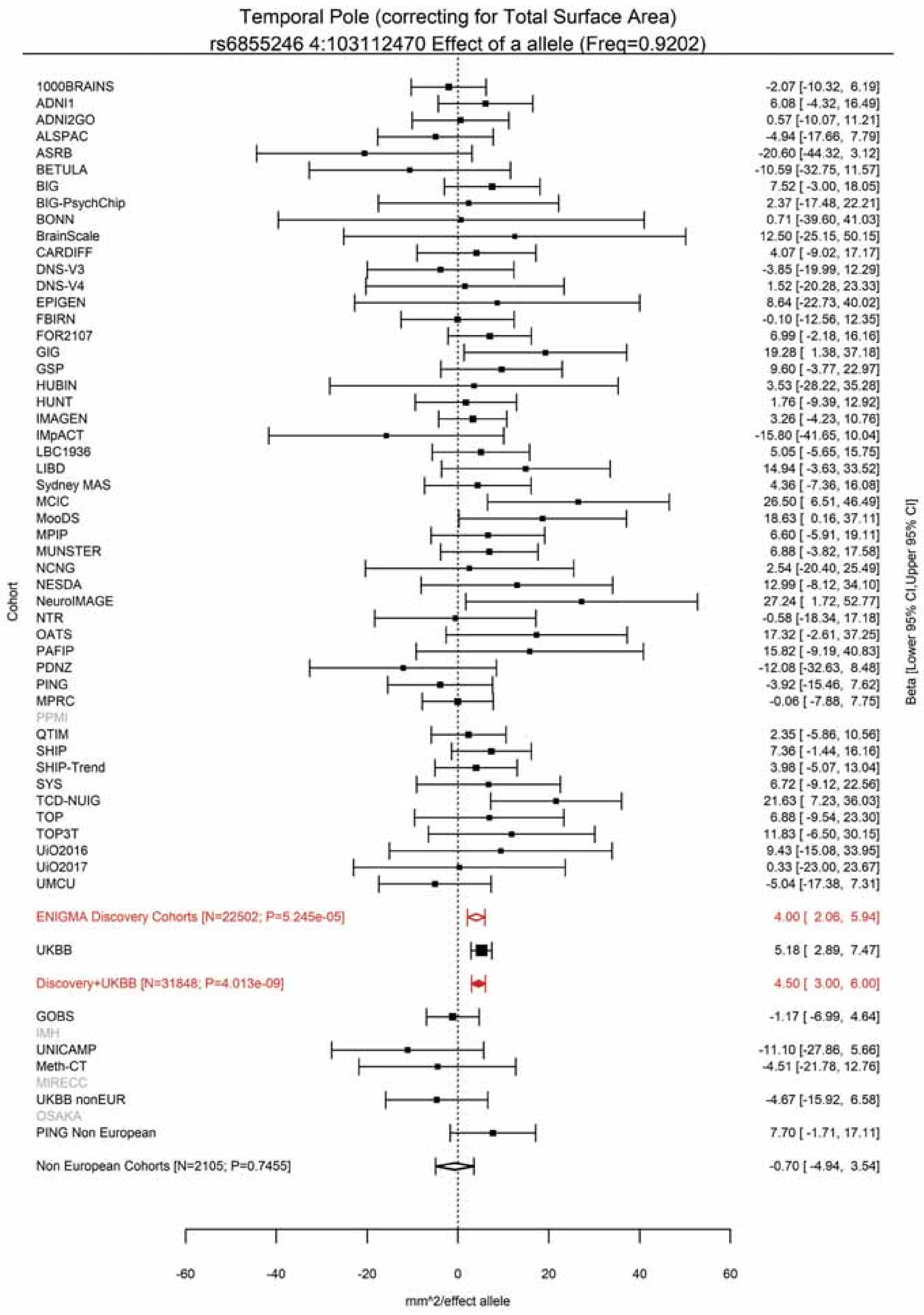
Temporal Pole (correcting for Total Surface Area)
rs6855246 4:103112470 Effect of a allele (Freq=0.9202)
Cohort
1000BRAINS -2.07 [-10.32, 6.19]
ADNI1 6.08 [ -4.32, 16.49]
ADNI2GO 0.57 [-10.07, 11.21]
ALSPAC -4.94 [-17.66, 7.79]
ASRB -20.60 [-44.32, 3.12]
BETULA -10.59 [-32.75, 11.57]
BIG 7.52 [ -3.00, 18.05]
BIG-PsychChip 2.37 [-17.48, 22.21]
BONN 0.71 [-39.60, 41.03]
BrainScale 12.50 [-25.15, 50.15]
CARDIFF 4.07 [ -9.02, 17.17]
DNS-V3 -3.85 [-19.99, 12.29]
DNS-V4 1.52 [-20.28, 23.33]
EPIGEN 8.64 [-22.73, 40.02]
FBIRN -0.10 [-12.56, 12.35]
FOR2107 6.99 [ -2.18, 16.16]
GIG 19.28 [ 1.38, 37.18]
GSP 9.60 [ -3.77, 22.97]
HUBIN 3.53 [-28.22, 35.28]
HUNT 1.76 [ -9.39, 12.92]
IMAGEN 3.26 [ -4.23, 10.76]
IMpACT -15.80 [-41.65, 10.04]
LBC1936 5.05 [ -5.65, 15.75]
LIBD 14.94 [ -3.63, 33.52]
Sydney MAS 4.36 [ -7.36, 16.08]
MCIC 26.50 [ 6.51, 46.49]
MooDS 18.63 [ 0.16, 37.11]
MPIP 6.60 [ -5.91, 19.11]
MUNSTER 6.88 [ -3.82, 17.58]
NCNG 2.54 [-20.40, 25.49]
NESDA 12.99 [ -8.12, 34.10]
NeuroIMAGE 27.24 [ 1.72, 52.77]
NTR -0.58 [-18.34, 17.18]
OATS 17.32 [ -2.61, 37.25]
PAFIP 15.82 [ -9.19, 40.83]
PDNZ -12.08 [-32.63, 8.48]
PING -3.92 [-15.46, 7.62]
MPRC -0.06 [ -7.88, 7.75]
PPMI
QTIM 2.35 [ -5.86, 10.56]
SHIP 7.36 [ -1.44, 16.16]
SHIP-Trend 3.98 [ -5.07, 13.04]
SYS 6.72 [ -9.12, 22.56]
TCD-NUIG 21.63 [ 7.23, 36.03]
TOP 6.88 [ -9.54, 23.30]
TOP3T 11.83 [ -6.50, 30.15]
UiO2016 9.43 [-15.08, 33.95]
UiO2017 0.33 [-23.00, 23.67]
UMCU -5.04 [-17.38, 7.31]
ENIGMA Discovery Cohorts [N=22502; P=5.245e-05] 4.00 [ 2.06, 5.94]
UKBB 5.18 [ 2.89, 7.47]
Discovery+UKBB [N=31848; P=4.013e-09] 4.50 [ 3.00, 6.00]
GOBS -1.17 [ -6.99, 4.64]
IMH
UNICAMP -11.10 [-27.86, 5.66]
Meth-CT -4.51 [-21.78, 12.76]
MIRECC
UKBB nonEUR -4.67 [-15.92, 6.58]
OSAKA
PING Non European 7.70 [ -1.71, 17.11]
Non European Cohorts [N=2105; P=0.7455] -0.70 [ -4.94, 3.54]
-60 -40 -20 0 20 40 60
mm^2/effect allele
Beta [Lower 95% CI,Upper 95% CI]

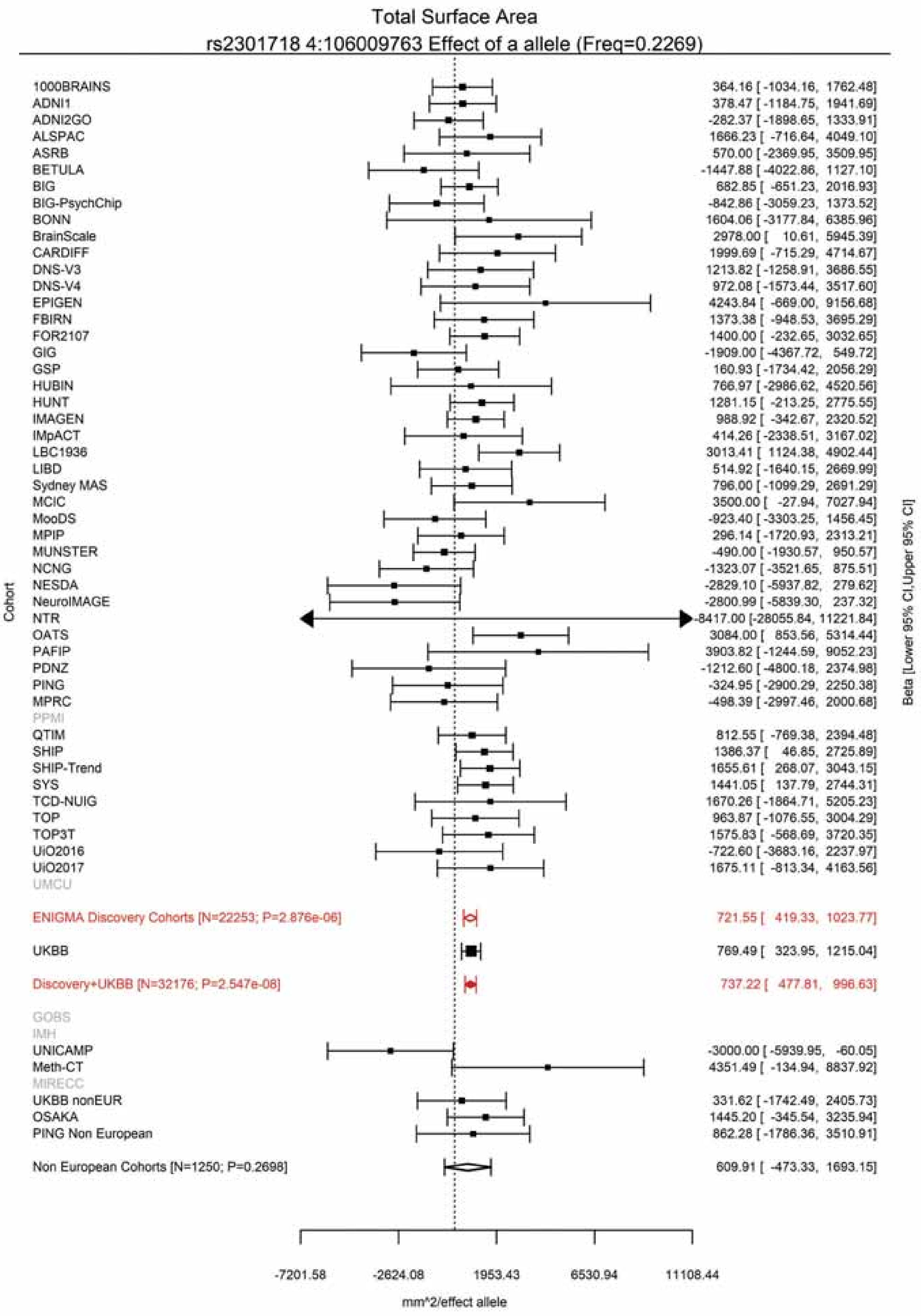

Total Surface Area
rs2301718 4:106009763 Effect of a allele (Freq=0.2269)
1000BRAINS 364.16 [ -1034.16, 1762.48]
ADNI1 378.47 [ -1184.75, 1941.69]
ADNI2GO -282.37 [ -1898.65, 1333.91]
ALSPAC 1666.23 [ -716.64, 4049.10]
ASRB 570.00 [ -2369.95, 3509.95]
BETULA -1447.88 [ -4022.86, 1127.10]
BIG 682.85 [ -651.23, 2016.93]
BIG-PsychChip -842.86 [ -3059.23, 1373.52]
BONN 1604.06 [ -3177.84, 6385.96]
BrainScale 2978.00 [ 10.61, 5945.39]
CARDIFF 1999.69 [ -715.29, 4714.67]
DNS-V3 1213.82 [ -1258.91, 3686.55]
DNS-V4 972.08 [ -1573.44, 3517.60]
EPIGEN 4243.84 [ -669.00, 9156.68]
FBIRN 1373.38 [ -948.53, 3695.29]
FOR2107 1400.00 [ -232.65, 3032.65]
GIG -1909.00 [ -4367.72, 549.72]
GSP 160.93 [ -1734.42, 2056.29]
HUBIN 766.97 [ -2986.62, 4520.56]
HUNT 1281.15 [ -213.25, 2775.55]
IMAGEN 988.92 [ -342.67, 2320.52]
IMpACT 414.26 [ -2338.51, 3167.02]
LBC1936 3013.41 [ 1124.38, 4902.44]
LIBD 514.92 [ -1640.15, 2669.99]
Sydney MAS 796.00 [ -1099.29, 2691.29]
MCIC 3500.00 [ -27.94, 7027.94]
MooDS -923.40 [ -3303.25, 1456.45]
MPIP 296.14 [ -1720.93, 2313.21]
MUNSTER -490.00 [ -1930.57, 950.57]
NCNG -1323.07 [ -3521.65, 875.51]
NESDA -2829.10 [ -5937.82, 279.62]
NeuroIMAGE -2800.99 [ -5839.30, 237.32]
NTR -8417.00 [-28055.84, 11221.84]
OATS 3084.00 [ 853.56, 5314.44]
PAFIP 3903.82 [ -1244.59, 9052.23]
PDNZ -1212.60 [ -4800.18, 2374.98]
PING -324.95 [ -2900.29, 2250.38]
MPRC -498.39 [ -2997.46, 2000.68]
PPMI
QTIM 812.55 [ -769.38, 2394.48]
SHIP 1386.37 [ 46.85, 2725.89]
SHIP-Trend 1655.61 [ 268.07, 3043.15]
SYS 1441.05 [ 137.79, 2744.31]
TCD-NUIG 1670.26 [ -1864.71, 5205.23]
TOP 963.87 [ -1076.55, 3004.29]
TOP3T 1575.83 [ -568.69, 3720.35]
UiO2016 -722.60 [ -3683.16, 2237.97]
UiO2017 1675.11 [ -813.34, 4163.56]
UMCU
ENIGMA Discovery Cohorts [N=22253; P=2.876e-06] 721.55 [ 419.33, 1023.77]
UKBB 769.49 [ 323.95, 1215.04]
Discovery+UKBB [N=32176; P=2.547e-08] 737.22 [ 477.81, 996.63]
GOBS
IMH
UNICAMP -3000.00 [ -5939.95, -60.05]
Meth-CT 4351.49 [ -134.94, 8837.92]
MIRECC
UKBB nonEUR 331.62 [ -1742.49, 2405.73]
OSAKA 1445.20 [ -345.54, 3235.94]
PING Non European 862.28 [ -1786.36, 3510.91]
Non European Cohorts [N=1250; P=0.2698] 609.91 [ -473.33, 1693.15]
Cohort
Beta [Lower 95% CI,Upper 95% CI]
-7201.58
-2624.08
1953.43
6530.94
11108.44
mm^2/effect allele

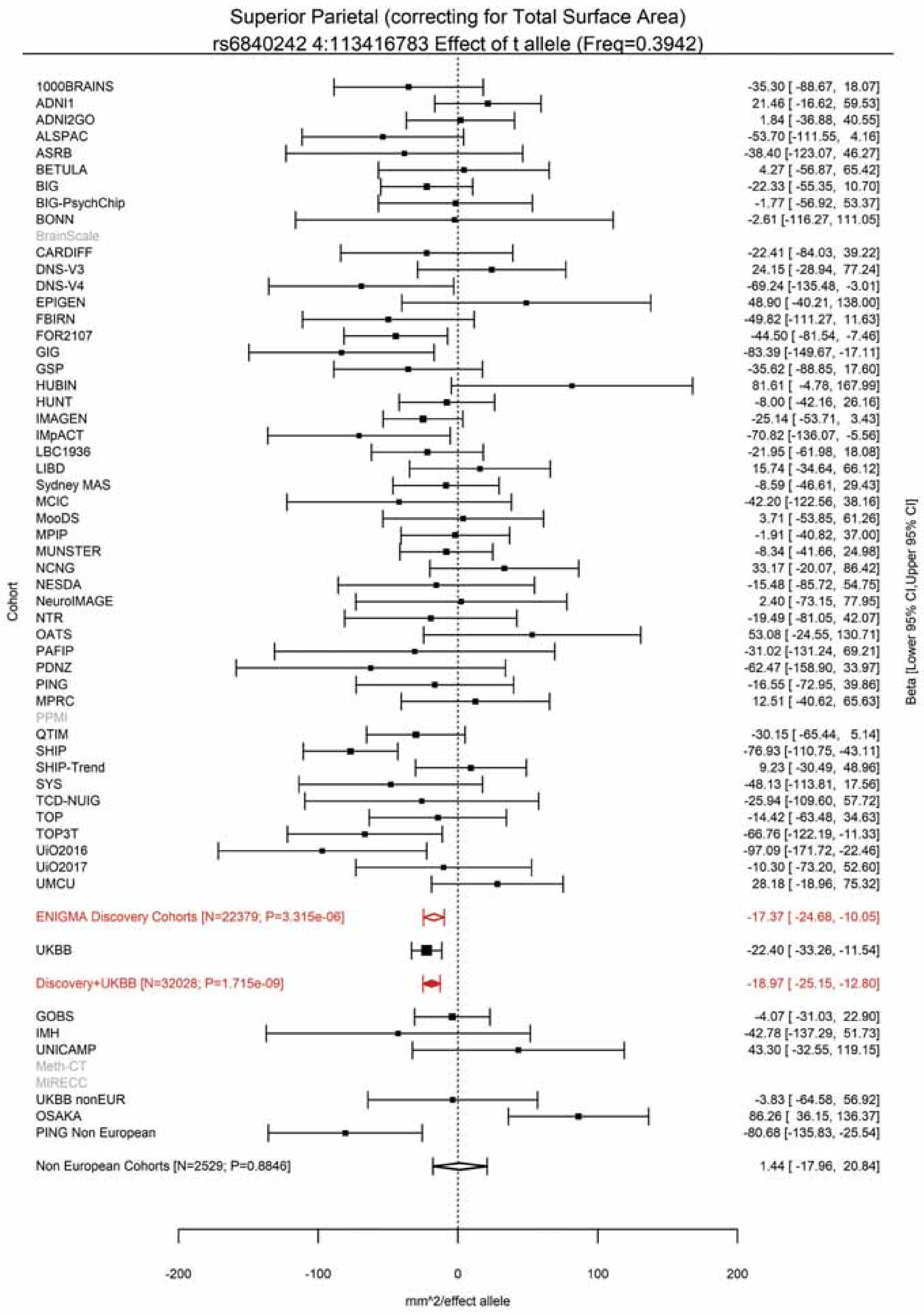
Superior Parietal (correcting for Total Surface Area)
rs6840242 4:113416783 Effect of t allele (Freq=0.3942)
Cohort
1000BRAINS -35.30 [ -88.67, 18.07]
ADNI1 21.46 [ -16.62, 59.53]
ADNI2GO 1.84 [ -36.88, 40.55]
ALSPAC -53.70 [-111.55, 4.16]
ASRB -38.40 [-123.07, 46.27]
BETULA 4.27 [ -56.87, 65.42]
BIG -22.33 [ -55.35, 10.70]
BIG-PsychChip -1.77 [ -56.92, 53.37]
BONN -2.61 [-116.27, 111.05]
BrainScale
CARDIFF -22.41 [ -84.03, 39.22]
DNS-V3 24.15 [ -28.94, 77.24]
DNS-V4 -69.24 [-135.48, -3.01]
EPIGEN 48.90 [ -40.21, 138.00]
FBIRN -49.82 [-111.27, 11.63]
FOR2107 -44.50 [ -81.54, -7.46]
GIG -83.39 [-149.67, -17.11]
GSP -35.62 [ -88.85, 17.60]
HUBIN 81.61 [ -4.78, 167.99]
HUNT -8.00 [ -42.16, 26.16]
IMAGEN -25.14 [ -53.71, 3.43]
IMpACT -70.82 [-136.07, -5.56]
LBC1936 -21.95 [ -61.98, 18.08]
LIBD 15.74 [ -34.64, 66.12]
Sydney MAS -8.59 [ -46.61, 29.43]
MCIC -42.20 [-122.56, 38.16]
MooDS 3.71 [ -53.85, 61.26]
MPIP -1.91 [ -40.82, 37.00]
MUNSTER -8.34 [ -41.66, 24.98]
NCNG 33.17 [ -20.07, 86.42]
NESDA -15.48 [ -85.72, 54.75]
NeuroIMAGE 2.40 [ -73.15, 77.95]
NTR -19.49 [ -81.05, 42.07]
OATS 53.08 [ -24.55, 130.71]
PAFIP -31.02 [-131.24, 69.21]
PDNZ -62.47 [-158.90, 33.97]
PING -16.55 [ -72.95, 39.86]
MPRC 12.51 [ -40.62, 65.63]
PPMI
QTIM -30.15 [ -65.44, 5.14]
SHIP -76.93 [-110.75, -43.11]
SHIP-Trend 9.23 [ -30.49, 48.96]
SYS -48.13 [-113.81, 17.56]
TCD-NUIG -25.94 [-109.60, 57.72]
TOP -14.42 [ -63.48, 34.63]
TOP3T -66.76 [-122.19, -11.33]
UiO2016 -97.09 [-171.72, -22.46]
UiO2017 -10.30 [ -73.20, 52.60]
UMCU 28.18 [ -18.96, 75.32]
ENIGMA Discovery Cohorts [N=22379; P=3.315e-06] -17.37 [ -24.68, -10.05]
UKBB -22.40 [ -33.26, -11.54]
Discovery+UKBB [N=32028; P=1.715e-09] -18.97 [ -25.15, -12.80]
GOBS -4.07 [ -31.03, 22.90]
IMH -42.78 [-137.29, 51.73]
UNICAMP 43.30 [ -32.55, 119.15]
Meth-CT
MIRECC
UKBB nonEUR -3.83 [ -64.58, 56.92]
OSAKA 86.26 [ 36.15, 136.37]
PING Non European -80.68 [-135.83, -25.54]
Non European Cohorts [N=2529; P=0.8846] 1.44 [ -17.96, 20.84]
Beta [Lower 95% CI,Upper 95% CI]
-200
-100
0
100
200
mm^2/effect allele

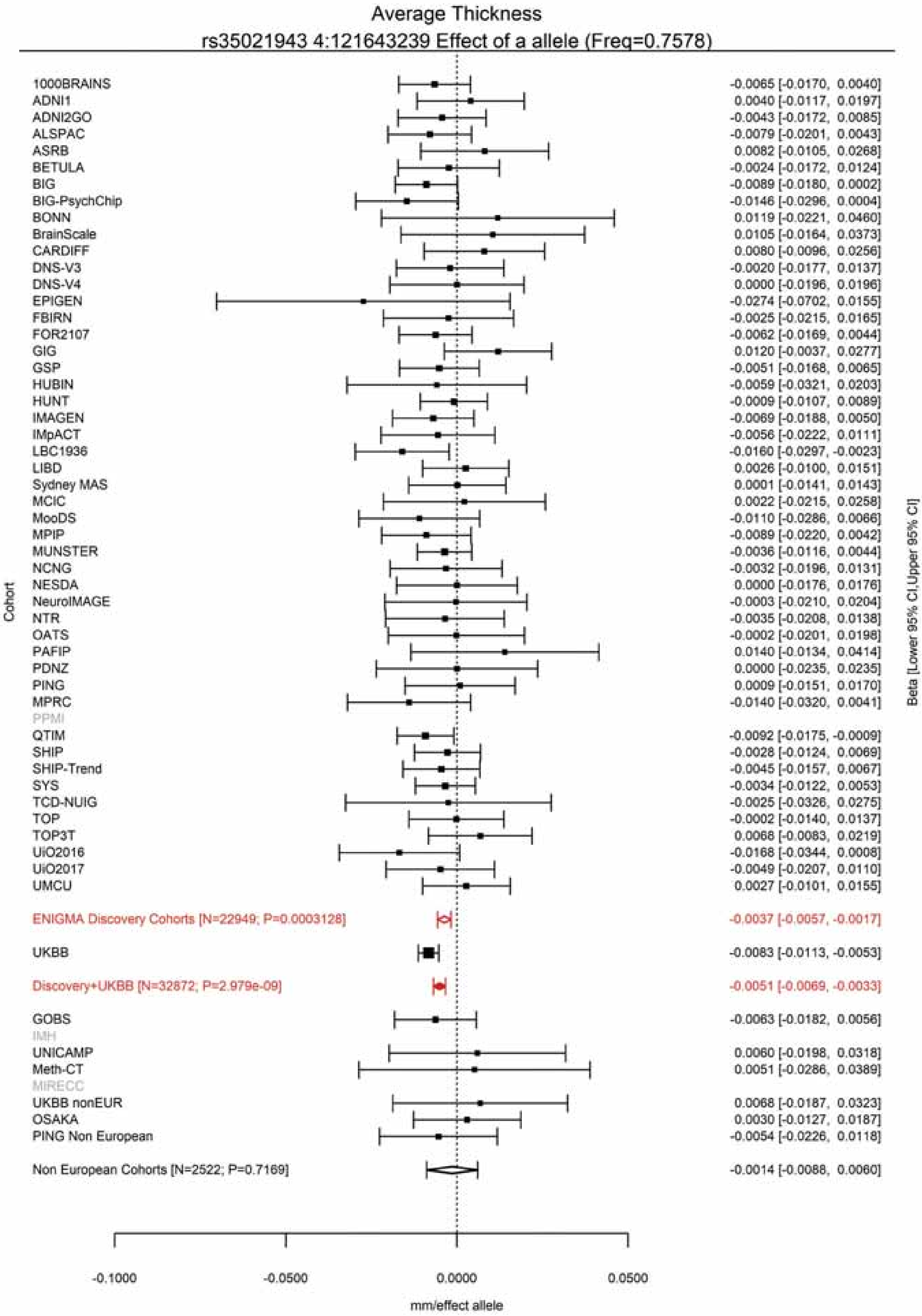
Average Thickness
rs35021943 4:121643239 Effect of a allele (Freq=0.7578)
Cohort
Beta [Lower 95% CI,Upper 95% CI]
1000BRAINS -0.0065 [-0.0170, 0.0040]
ADNI1 0.0040 [-0.0117, 0.0197]
ADNI2GO -0.0043 [-0.0172, 0.0085]
ALSPAC -0.0079 [-0.0201, 0.0043]
ASRB 0.0082 [-0.0105, 0.0268]
BETULA -0.0024 [-0.0172, 0.0124]
BIG -0.0089 [-0.0180, 0.0002]
BIG-PsychChip -0.0146 [-0.0296, 0.0004]
BONN 0.0119 [-0.0221, 0.0460]
BrainScale 0.0105 [-0.0164, 0.0373]
CARDIFF 0.0080 [-0.0096, 0.0256]
DNS-V3 -0.0020 [-0.0177, 0.0137]
DNS-V4 0.0000 [-0.0196, 0.0196]
EPIGEN -0.0274 [-0.0702, 0.0155]
FBIRN -0.0025 [-0.0215, 0.0165]
FOR2107 -0.0062 [-0.0169, 0.0044]
GIG 0.0120 [-0.0037, 0.0277]
GSP -0.0051 [-0.0168, 0.0065]
HUBIN -0.0059 [-0.0321, 0.0203]
HUNT -0.0009 [-0.0107, 0.0089]
IMAGEN -0.0069 [-0.0188, 0.0050]
IMpACT -0.0056 [-0.0222, 0.0111]
LBC1936 -0.0160 [-0.0297, -0.0023]
LIBD 0.0026 [-0.0100, 0.0151]
Sydney MAS 0.0001 [-0.0141, 0.0143]
MCIC 0.0022 [-0.0215, 0.0258]
MooDS -0.0110 [-0.0286, 0.0066]
MPIP -0.0089 [-0.0220, 0.0042]
MUNSTER -0.0036 [-0.0116, 0.0044]
NCNG -0.0032 [-0.0196, 0.0131]
NESDA 0.0000 [-0.0176, 0.0176]
NeuroIMAGE -0.0003 [-0.0210, 0.0204]
NTR -0.0035 [-0.0208, 0.0138]
OATS -0.0002 [-0.0201, 0.0198]
PAFIP 0.0140 [-0.0134, 0.0414]
PDNZ 0.0000 [-0.0235, 0.0235]
PING 0.0009 [-0.0151, 0.0170]
MPRC -0.0140 [-0.0320, 0.0041]
PPMI
QTIM -0.0092 [-0.0175, -0.0009]
SHIP -0.0028 [-0.0124, 0.0069]
SHIP-Trend -0.0045 [-0.0157, 0.0067]
SYS -0.0034 [-0.0122, 0.0053]
TCD-NUIG -0.0025 [-0.0326, 0.0275]
TOP -0.0002 [-0.0140, 0.0137]
TOP3T 0.0068 [-0.0083, 0.0219]
UiO2016 -0.0168 [-0.0344, 0.0008]
UiO2017 -0.0049 [-0.0207, 0.0110]
UMCU 0.0027 [-0.0101, 0.0155]
ENIGMA Discovery Cohorts [N=22949; P=0.0003128] -0.0037 [-0.0057, -0.0017]
UKBB -0.0083 [-0.0113, -0.0053]
Discovery+UKBB [N=32872; P=2.979e-09] -0.0051 [-0.0069, -0.0033]
GOBS -0.0063 [-0.0182, 0.0056]
IMH
UNICAMP 0.0060 [-0.0198, 0.0318]
Meth-CT 0.0051 [-0.0286, 0.0389]
MIRECC
UKBB nonEUR 0.0068 [-0.0187, 0.0323]
OSAKA 0.0030 [-0.0127, 0.0187]
PING Non European -0.0054 [-0.0226, 0.0118]
Non European Cohorts [N=2522; P=0.7169] -0.0014 [-0.0088, 0.0060]
-0.1000
-0.0500
0.0000
0.0500
mm/effect allele

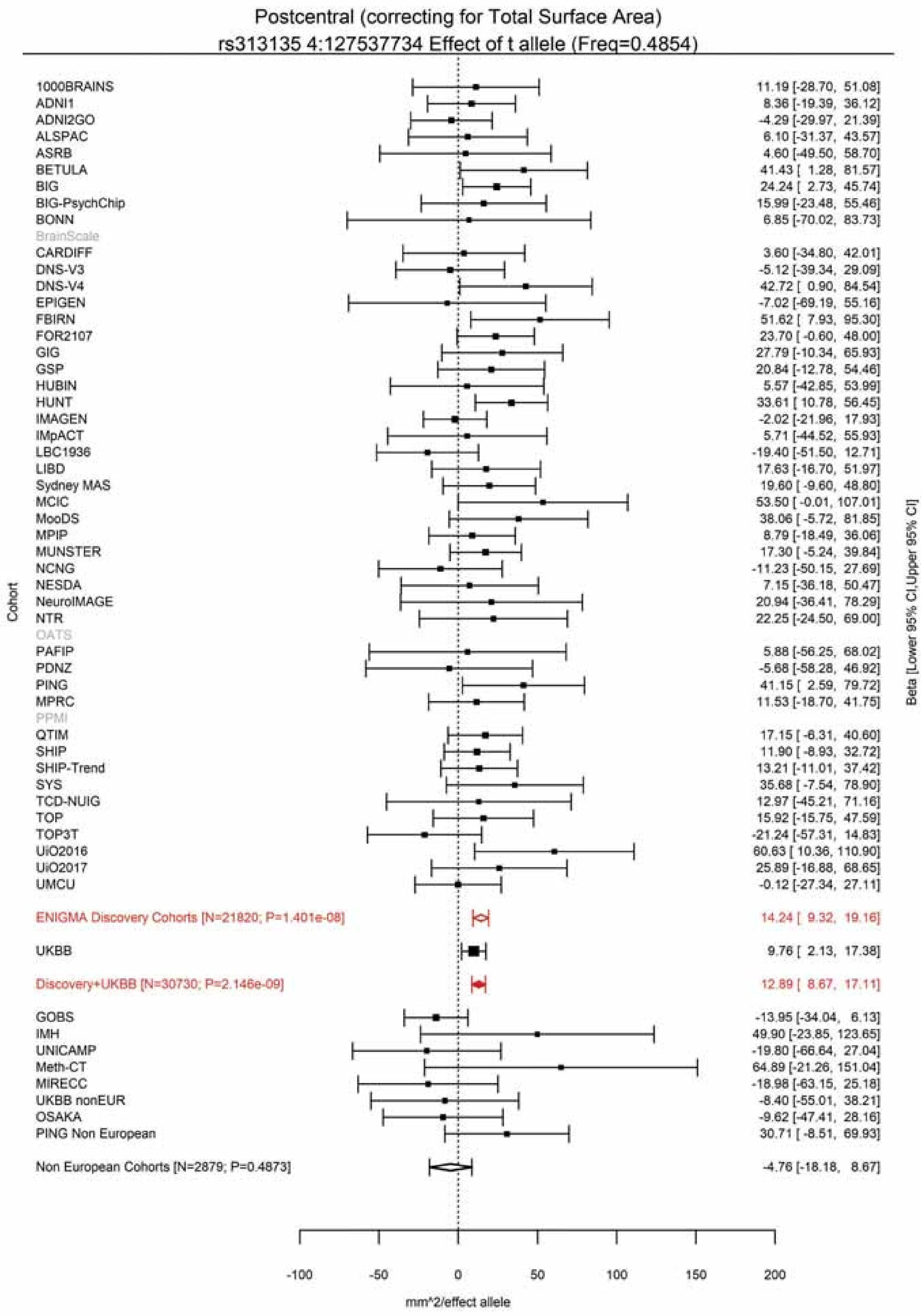
Postcentral (correcting for Total Surface Area)
rs313135 4:127537734 Effect of t allele (Freq=0.4854)
Cohort
1000BRAINS 11.19 [-28.70, 51.08]
ADNI1 8.36 [-19.39, 36.12]
ADNI2GO -4.29 [-29.97, 21.39]
ALSPAC 6.10 [-31.37, 43.57]
ASRB 4.60 [-49.50, 58.70]
BETULA 41.43 [ 1.28, 81.57]
BIG 24.24 [ 2.73, 45.74]
BIG-PsychChip 15.99 [-23.48, 55.46]
BONN 6.85 [-70.02, 83.73]
BrainScale
CARDIFF 3.60 [-34.80, 42.01]
DNS-V3 -5.12 [-39.34, 29.09]
DNS-V4 42.72 [ 0.90, 84.54]
EPIGEN -7.02 [-69.19, 55.16]
FBIRN 51.62 [ 7.93, 95.30]
FOR2107 23.70 [ -0.60, 48.00]
GIG 27.79 [-10.34, 65.93]
GSP 20.84 [-12.78, 54.46]
HUBIN 5.57 [-42.85, 53.99]
HUNT 33.61 [ 10.78, 56.45]
IMAGEN -2.02 [-21.96, 17.93]
IMpACT 5.71 [-44.52, 55.93]
LBC1936 -19.40 [-51.50, 12.71]
LIBD 17.63 [-16.70, 51.97]
Sydney MAS 19.60 [ -9.60, 48.80]
MCIC 53.50 [ -0.01, 107.01]
MooDS 38.06 [ -5.72, 81.85]
MPIP 8.79 [-18.49, 36.06]
MUNSTER 17.30 [ -5.24, 39.84]
NCNG -11.23 [-50.15, 27.69]
NESDA 7.15 [-36.18, 50.47]
NeuroIMAGE 20.94 [-36.41, 78.29]
NTR 22.25 [-24.50, 69.00]
OATS
PAFIP 5.88 [-56.25, 68.02]
PDNZ -5.68 [-58.28, 46.92]
PING 41.15 [ 2.59, 79.72]
MPRC 11.53 [-18.70, 41.75]
PPMI
QTIM 17.15 [ -6.31, 40.60]
SHIP 11.90 [ -8.93, 32.72]
SHIP-Trend 13.21 [-11.01, 37.42]
SYS 35.68 [ -7.54, 78.90]
TCD-NUIG 12.97 [-45.21, 71.16]
TOP 15.92 [-15.75, 47.59]
TOP3T -21.24 [-57.31, 14.83]
UiO2016 60.63 [ 10.36, 110.90]
UiO2017 25.89 [-16.88, 68.65]
UMCU -0.12 [-27.34, 27.11]
ENIGMA Discovery Cohorts [N=21820; P=1.401e-08] 14.24 [ 9.32, 19.16]
UKBB 9.76 [ 2.13, 17.38]
Discovery+UKBB [N=30730; P=2.146e-09] 12.89 [ 8.67, 17.11]
GOBS -13.95 [-34.04, 6.13]
IMH 49.90 [-23.85, 123.65]
UNICAMP -19.80 [-66.64, 27.04]
Meth-CT 64.89 [-21.26, 151.04]
MIRECC -18.98 [-63.15, 25.18]
UKBB nonEUR -8.40 [-55.01, 38.21]
OSAKA -9.62 [-47.41, 28.16]
PING Non European 30.71 [ -8.51, 69.93]
Non European Cohorts [N=2879; P=0.4873] -4.76 [-18.18, 8.67]
Beta [Lower 95% CI,Upper 95% CI]
-100
-50
0
50
100
150
200
mm^2/effect allele

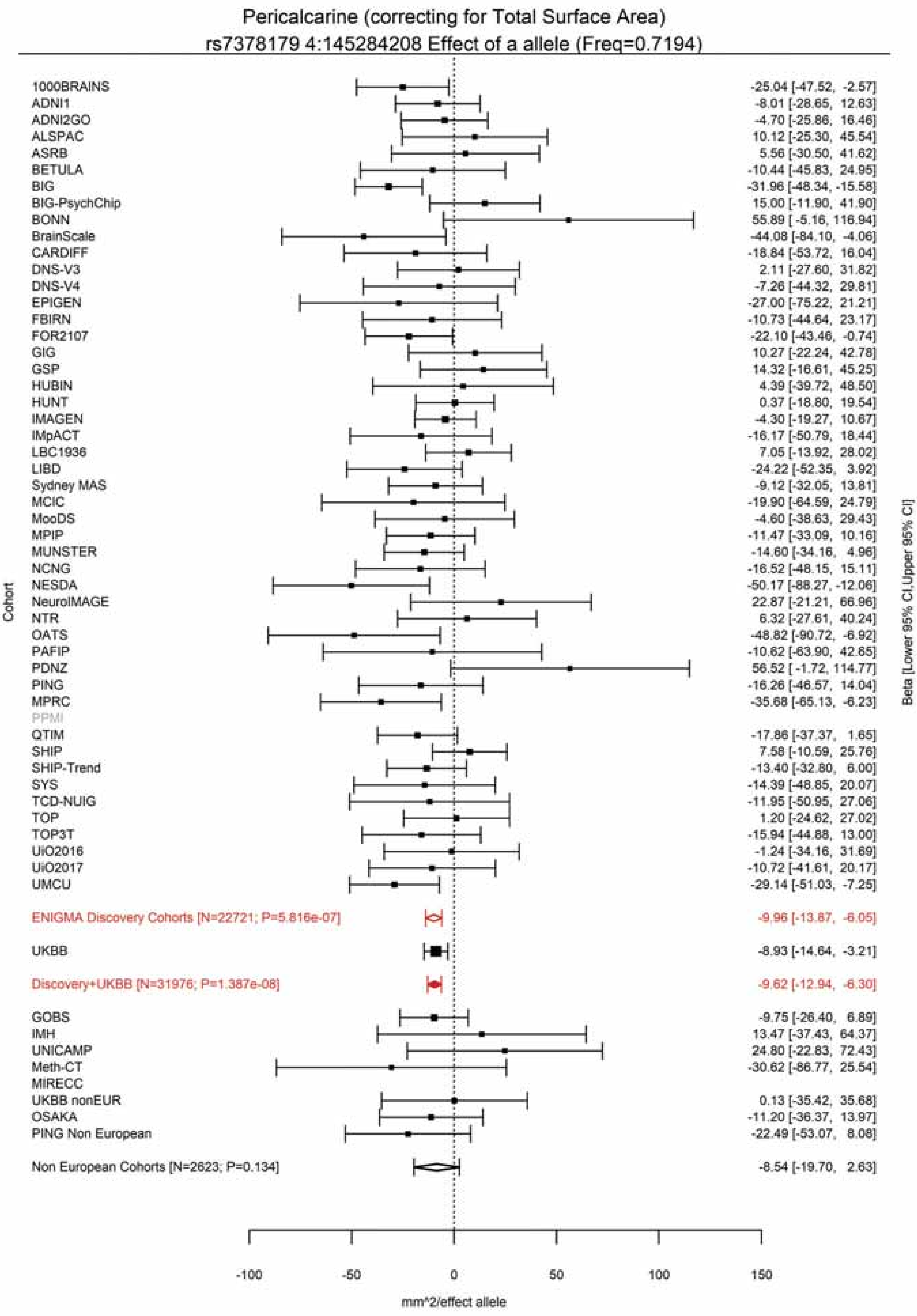
Pericalcarine (correcting for Total Surface Area)
rs7378179 4:145284208 Effect of a allele (Freq=0.7194)
Cohort
1000BRAINS -25.04 [-47.52, -2.57]
ADNI1 -8.01 [-28.65, 12.63]
ADNI2GO -4.70 [-25.86, 16.46]
ALSPAC 10.12 [-25.30, 45.54]
ASRB 5.56 [-30.50, 41.62]
BETULA -10.44 [-45.83, 24.95]
BIG -31.96 [-48.34, -15.58]
BIG-PsychChip 15.00 [-11.90, 41.90]
BONN 55.89 [-5.16, 116.94]
BrainScale -44.08 [-84.10, -4.06]
CARDIFF -18.84 [-53.72, 16.04]
DNS-V3 2.11 [-27.60, 31.82]
DNS-V4 -7.26 [-44.32, 29.81]
EPIGEN -27.00 [-75.22, 21.21]
FBIRN -10.73 [-44.64, 23.17]
FOR2107 -22.10 [-43.46, -0.74]
GIG 10.27 [-22.24, 42.78]
GSP 14.32 [-16.61, 45.25]
HUBIN 4.39 [-39.72, 48.50]
HUNT 0.37 [-18.80, 19.54]
IMAGEN -4.30 [-19.27, 10.67]
IMpACT -16.17 [-50.79, 18.44]
LBC1936 7.05 [-13.92, 28.02]
LIBD -24.22 [-52.35, 3.92]
Sydney MAS -9.12 [-32.05, 13.81]
MCIC -19.90 [-64.59, 24.79]
MooDS -4.60 [-38.63, 29.43]
MPIP -11.47 [-33.09, 10.16]
MUNSTER -14.60 [-34.16, 4.96]
NCNG -16.52 [-48.15, 15.11]
NESDA -50.17 [-88.27, -12.06]
NeuroIMAGE 22.87 [-21.21, 66.96]
NTR 6.32 [-27.61, 40.24]
OATS -48.82 [-90.72, -6.92]
PAFIP -10.62 [-63.90, 42.65]
PDNZ 56.52 [-1.72, 114.77]
PING -16.26 [-46.57, 14.04]
MPRC -35.68 [-65.13, -6.23]
PPMI
QTIM -17.86 [-37.37, 1.65]
SHIP 7.58 [-10.59, 25.76]
SHIP-Trend -13.40 [-32.80, 6.00]
SYS -14.39 [-48.85, 20.07]
TCD-NUIG -11.95 [-50.95, 27.06]
TOP 1.20 [-24.62, 27.02]
TOP3T -15.94 [-44.88, 13.00]
UiO2016 -1.24 [-34.16, 31.69]
UiO2017 -10.72 [-41.61, 20.17]
UMCU -29.14 [-51.03, -7.25]
ENIGMA Discovery Cohorts [N=22721; P=5.816e-07] -9.96 [-13.87, -6.05]
UKBB -8.93 [-14.64, -3.21]
Discovery+UKBB [N=31976; P=1.387e-08] -9.62 [-12.94, -6.30]
GOBS -9.75 [-26.40, 6.89]
IMH 13.47 [-37.43, 64.37]
UNICAMP 24.80 [-22.83, 72.43]
Meth-CT -30.62 [-86.77, 25.54]
MIRECC
UKBB nonEUR 0.13 [-35.42, 35.68]
OSAKA -11.20 [-36.37, 13.97]
PING Non European -22.49 [-53.07, 8.08]
Non European Cohorts [N=2623; P=0.134] -8.54 [-19.70, 2.63]
Beta [Lower 95% CI,Upper 95% CI]
-100
-50
0
50
100
150
mm^2/effect allele

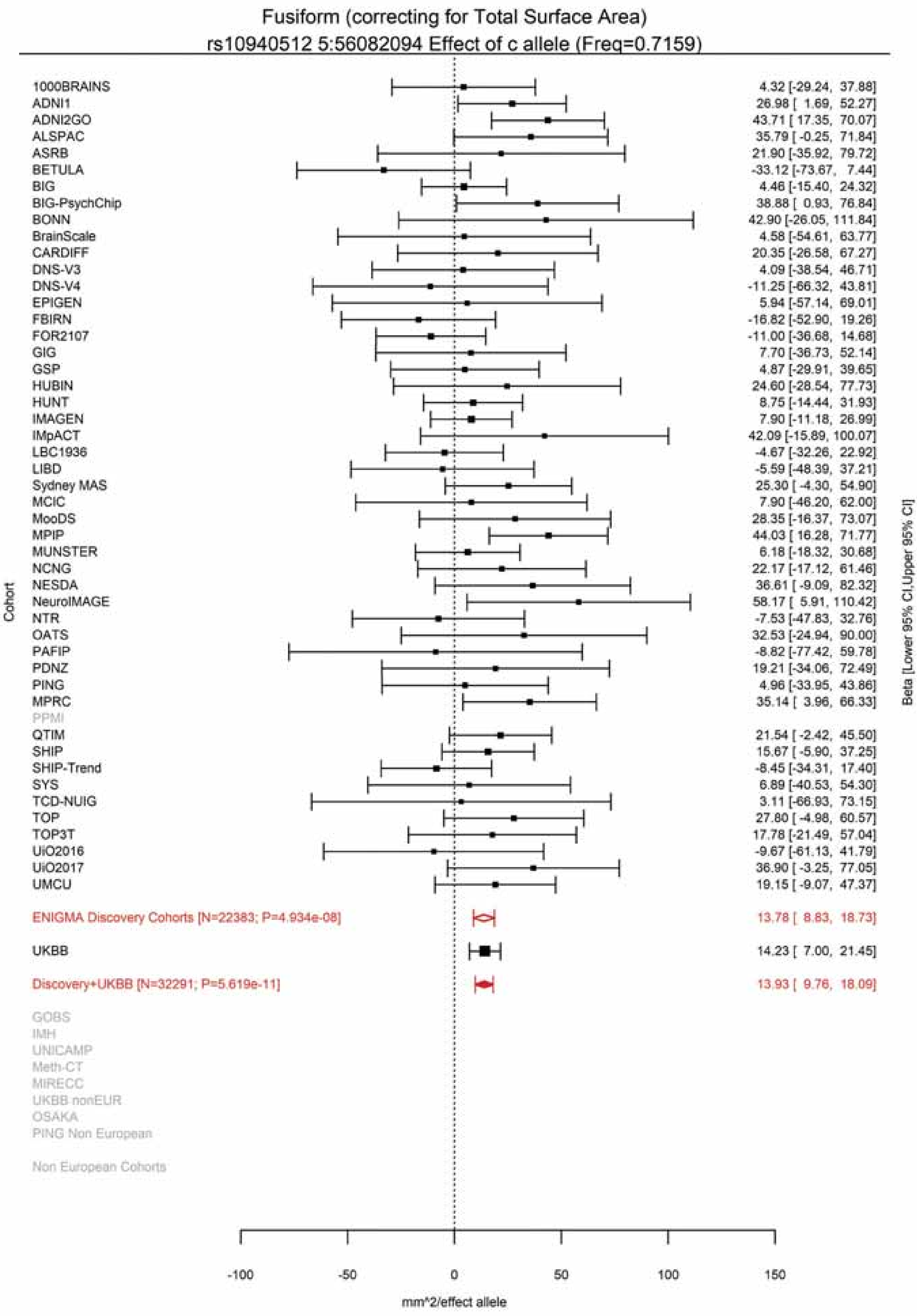
Fusiform (correcting for Total Surface Area)
rs10940512 5:56082094 Effect of c allele (Freq=0.7159)
Cohort
1000BRAINS 4.32 [-29.24, 37.88]
ADNI1 26.98 [ 1.69, 52.27]
ADNI2GO 43.71 [ 17.35, 70.07]
ALSPAC 35.79 [ -0.25, 71.84]
ASRB 21.90 [-35.92, 79.72]
BETULA -33.12 [-73.67, 7.44]
BIG 4.46 [-15.40, 24.32]
BIG-PsychChip 38.88 [ 0.93, 76.84]
BONN 42.90 [-26.05, 111.84]
BrainScale 4.58 [-54.61, 63.77]
CARDIFF 20.35 [-26.58, 67.27]
DNS-V3 4.09 [-38.54, 46.71]
DNS-V4 -11.25 [-66.32, 43.81]
EPIGEN 5.94 [-57.14, 69.01]
FBIRN -16.82 [-52.90, 19.26]
FOR2107 -11.00 [-36.68, 14.68]
GIG 7.70 [-36.73, 52.14]
GSP 4.87 [-29.91, 39.65]
HUBIN 24.60 [-28.54, 77.73]
HUNT 8.75 [-14.44, 31.93]
IMAGEN 7.90 [-11.18, 26.99]
IMpACT 42.09 [-15.89, 100.07]
LBC1936 -4.67 [-32.26, 22.92]
LIBD -5.59 [-48.39, 37.21]
Sydney MAS 25.30 [ -4.30, 54.90]
MCIC 7.90 [-46.20, 62.00]
MooDS 28.35 [-16.37, 73.07]
MPIP 44.03 [ 16.28, 71.77]
MUNSTER 6.18 [-18.32, 30.68]
NCNG 22.17 [-17.12, 61.46]
NESDA 36.61 [ -9.09, 82.32]
NeuroIMAGE 58.17 [ 5.91, 110.42]
NTR -7.53 [-47.83, 32.76]
OATS 32.53 [-24.94, 90.00]
PAFIP -8.82 [-77.42, 59.78]
PDNZ 19.21 [-34.06, 72.49]
PING 4.96 [-33.95, 43.86]
MPRC 35.14 [ 3.96, 66.33]
PPMI
QTIM 21.54 [ -2.42, 45.50]
SHIP 15.67 [ -5.90, 37.25]
SHIP-Trend -8.45 [-34.31, 17.40]
SYS 6.89 [-40.53, 54.30]
TCD-NUIG 3.11 [-66.93, 73.15]
TOP 27.80 [ -4.98, 60.57]
TOP3T 17.78 [-21.49, 57.04]
UiO2016 -9.67 [-61.13, 41.79]
UiO2017 36.90 [ -3.25, 77.05]
UMCU 19.15 [ -9.07, 47.37]
ENIGMA Discovery Cohorts [N=22383; P=4.934e-08] 13.78 [ 8.83, 18.73]
UKBB 14.23 [ 7.00, 21.45]
Discovery+UKBB [N=32291; P=5.619e-11] 13.93 [ 9.76, 18.09]
GOBS
IMH
UNICAMP
Meth-CT
MIRECC
UKBB nonEUR
OSAKA
PING Non European
Non European Cohorts
Beta [Lower 95% CI,Upper 95% CI]
-100
-50
0
50
100
150
mm^2/effect allele

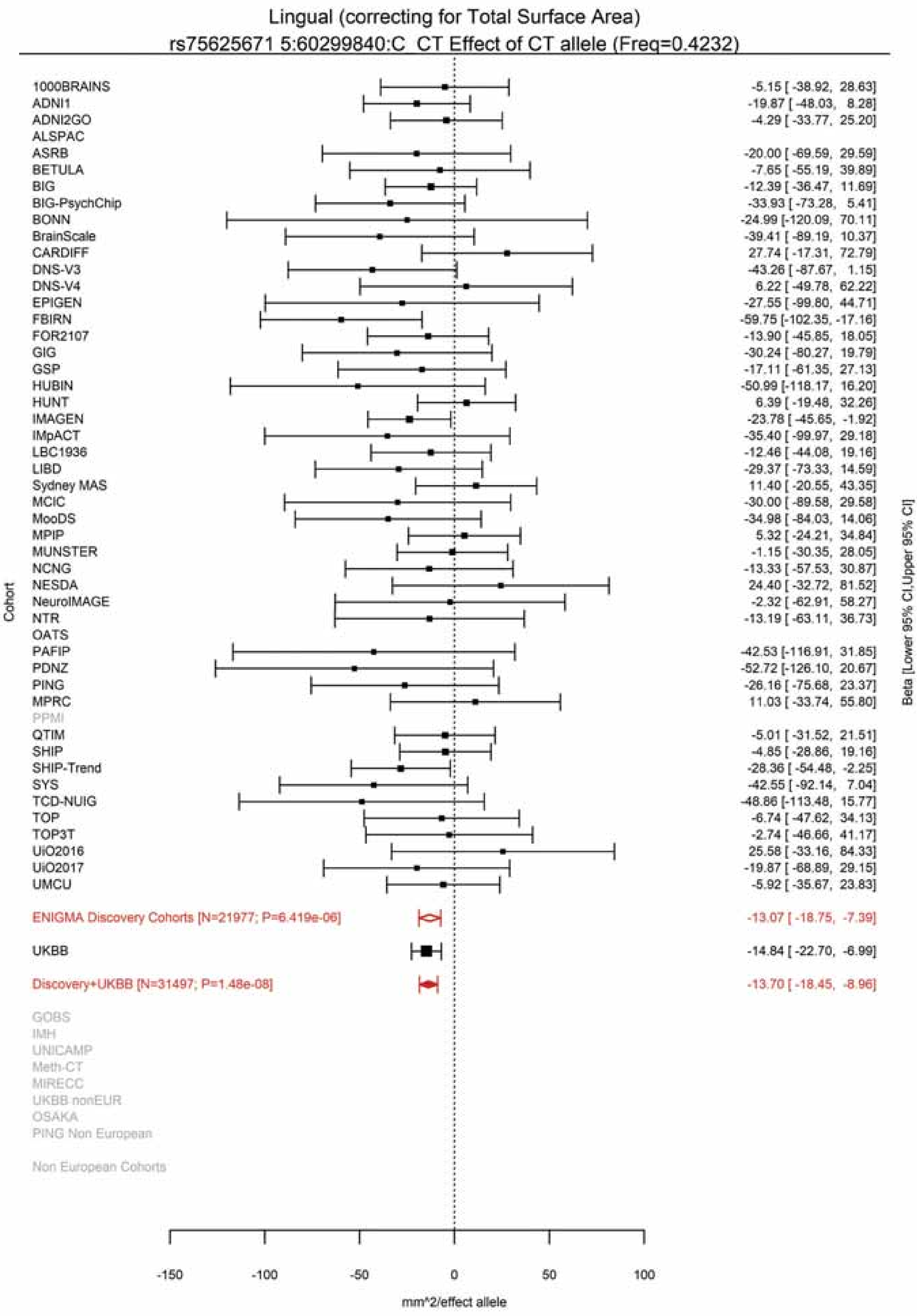
Lingual (correcting for Total Surface Area)
rs75625671 5:60299840:C CT Effect of CT allele (Freq=0.4232)
Cohort
1000BRAINS -5.15 [ -38.92, 28.63]
ADNI1 -19.87 [ -48.03, 8.28]
ADNI2GO -4.29 [ -33.77, 25.20]
ALSPAC
ASRB -20.00 [ -69.59, 29.59]
BETULA -7.65 [ -55.19, 39.89]
BIG -12.39 [ -36.47, 11.69]
BIG-PsychChip -33.93 [ -73.28, 5.41]
BONN -24.99 [-120.09, 70.11]
BrainScale -39.41 [ -89.19, 10.37]
CARDIFF 27.74 [ -17.31, 72.79]
DNS-V3 -43.26 [ -87.67, 1.15]
DNS-V4 6.22 [ -49.78, 62.22]
EPIGEN -27.55 [ -99.80, 44.71]
FBIRN -59.75 [-102.35, -17.16]
FOR2107 -13.90 [ -45.85, 18.05]
GIG -30.24 [ -80.27, 19.79]
GSP -17.11 [ -61.35, 27.13]
HUBIN -50.99 [-118.17, 16.20]
HUNT 6.39 [ -19.48, 32.26]
IMAGEN -23.78 [ -45.65, -1.92]
IMpACT -35.40 [ -99.97, 29.18]
LBC1936 -12.46 [ -44.08, 19.16]
LIBD -29.37 [ -73.33, 14.59]
Sydney MAS 11.40 [ -20.55, 43.35]
MCIC -30.00 [ -89.58, 29.58]
MooDS -34.98 [ -84.03, 14.06]
MPIP 5.32 [ -24.21, 34.84]
MUNSTER -1.15 [ -30.35, 28.05]
NCNG -13.33 [ -57.53, 30.87]
NESDA 24.40 [ -32.72, 81.52]
NeuroIMAGE -2.32 [ -62.91, 58.27]
NTR -13.19 [ -63.11, 36.73]
OATS
PAFIP -42.53 [-116.91, 31.85]
PDNZ -52.72 [-126.10, 20.67]
PING -26.16 [ -75.68, 23.37]
MPRC 11.03 [ -33.74, 55.80]
PPMI
QTIM -5.01 [ -31.52, 21.51]
SHIP -4.85 [ -28.86, 19.16]
SHIP-Trend -28.36 [ -54.48, -2.25]
SYS -42.55 [ -92.14, 7.04]
TCD-NUIG -48.86 [-113.48, 15.77]
TOP -6.74 [ -47.62, 34.13]
TOP3T -2.74 [ -46.66, 41.17]
UiO2016 25.58 [ -33.16, 84.33]
UiO2017 -19.87 [ -68.89, 29.15]
UMCU -5.92 [ -35.67, 23.83]
ENIGMA Discovery Cohorts [N=21977; P=6.419e-06] -13.07 [ -18.75, -7.39]
UKBB -14.84 [ -22.70, -6.99]
Discovery+UKBB [N=31497; P=1.48e-08] -13.70 [ -18.45, -8.96]
GOBS
IMH
UNICAMP
Meth-CT
MIRECC
UKBB nonEUR
OSAKA
PING Non European
Non European Cohorts
Beta [Lower 95% CI,Upper 95% CI]
-150
-100
-50
0
50
100
mm^2/effect allele

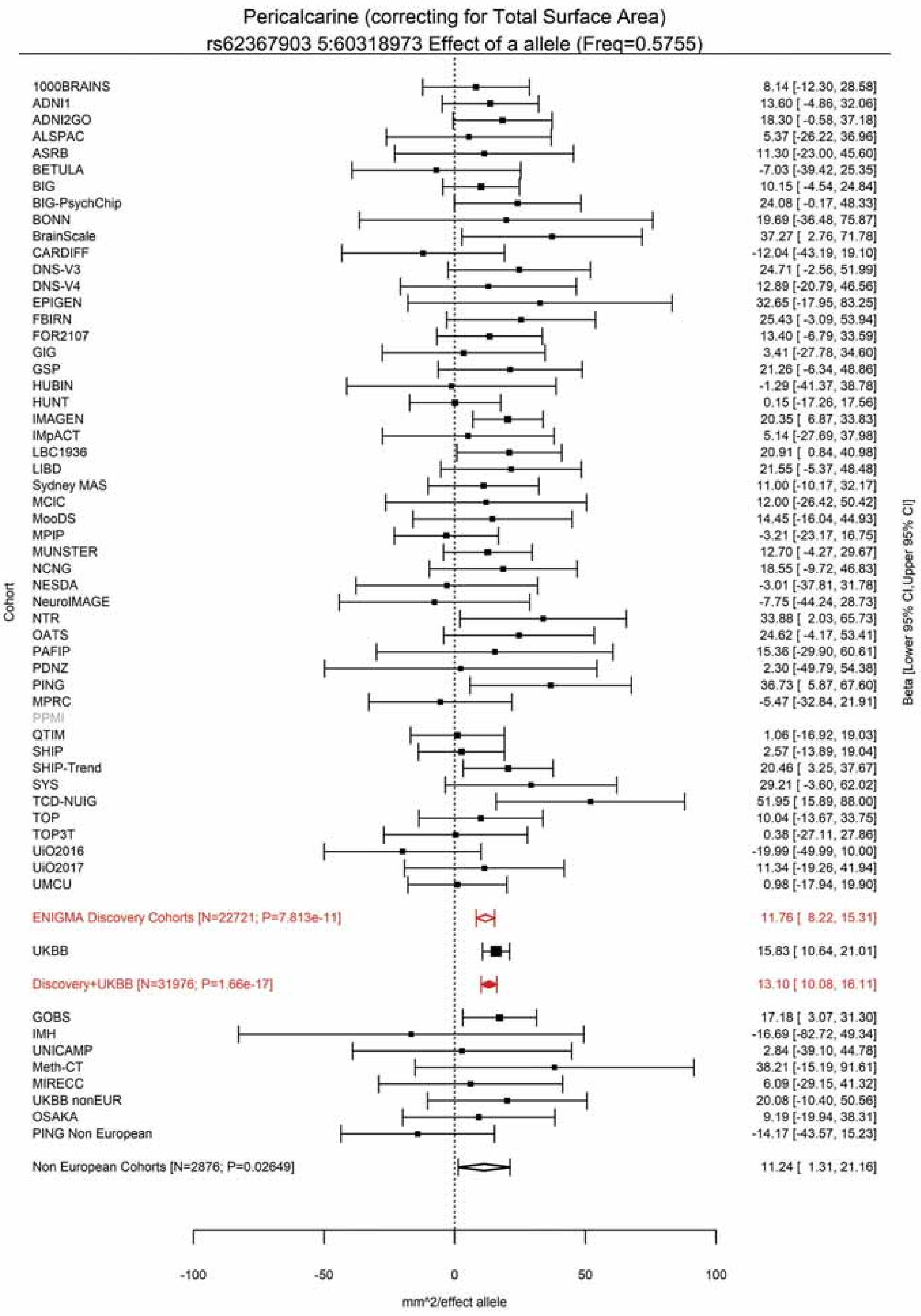
Pericalcarine (correcting for Total Surface Area)
rs62367903 5:60318973 Effect of a allele (Freq=0.5755)
Cohort
1000BRAINS 8.14 [-12.30, 28.58]
ADNI1 13.60 [ -4.86, 32.06]
ADNI2GO 18.30 [ -0.58, 37.18]
ALSPAC 5.37 [-26.22, 36.96]
ASRB 11.30 [-23.00, 45.60]
BETULA -7.03 [-39.42, 25.35]
BIG 10.15 [ -4.54, 24.84]
BIG-PsychChip 24.08 [ -0.17, 48.33]
BONN 19.69 [-36.48, 75.87]
BrainScale 37.27 [ 2.76, 71.78]
CARDIFF -12.04 [-43.19, 19.10]
DNS-V3 24.71 [ -2.56, 51.99]
DNS-V4 12.89 [-20.79, 46.56]
EPIGEN 32.65 [-17.95, 83.25]
FBIRN 25.43 [ -3.09, 53.94]
FOR2107 13.40 [ -6.79, 33.59]
GIG 3.41 [-27.78, 34.60]
GSP 21.26 [ -6.34, 48.86]
HUBIN -1.29 [-41.37, 38.78]
HUNT 0.15 [-17.26, 17.56]
IMAGEN 20.35 [ 6.87, 33.83]
IMpACT 5.14 [-27.69, 37.98]
LBC1936 20.91 [ 0.84, 40.98]
LIBD 21.55 [ -5.37, 48.48]
Sydney MAS 11.00 [-10.17, 32.17]
MCIC 12.00 [-26.42, 50.42]
MooDS 14.45 [-16.04, 44.93]
MPIP -3.21 [-23.17, 16.75]
MUNSTER 12.70 [ -4.27, 29.67]
NCNG 18.55 [ -9.72, 46.83]
NESDA -3.01 [-37.81, 31.78]
NeuroIMAGE -7.75 [-44.24, 28.73]
NTR 33.88 [ 2.03, 65.73]
OATS 24.62 [ -4.17, 53.41]
PAFIP 15.36 [-29.90, 60.61]
PDNZ 2.30 [-49.79, 54.38]
PING 36.73 [ 5.87, 67.60]
MPRC -5.47 [-32.84, 21.91]
PPMI
QTIM 1.06 [-16.92, 19.03]
SHIP 2.57 [-13.89, 19.04]
SHIP-Trend 20.46 [ 3.25, 37.67]
SYS 29.21 [ -3.60, 62.02]
TCD-NUIG 51.95 [ 15.89, 88.00]
TOP 10.04 [-13.67, 33.75]
TOP3T 0.38 [-27.11, 27.86]
UiO2016 -19.99 [-49.99, 10.00]
UiO2017 11.34 [-19.26, 41.94]
UMCU 0.98 [-17.94, 19.90]
ENIGMA Discovery Cohorts [N=22721; P=7.813e-11] 11.76 [ 8.22, 15.31]
UKBB 15.83 [ 10.64, 21.01]
Discovery+UKBB [N=31976; P=1.66e-17] 13.10 [ 10.08, 16.11]
GOBS 17.18 [ 3.07, 31.30]
IMH -16.69 [-82.72, 49.34]
UNICAMP 2.84 [-39.10, 44.78]
Meth-CT 38.21 [-15.19, 91.61]
MIRECC 6.09 [-29.15, 41.32]
UKBB nonEUR 20.08 [-10.40, 50.56]
OSAKA 9.19 [-19.94, 38.31]
PING Non European -14.17 [-43.57, 15.23]
Non European Cohorts [N=2876; P=0.02649] 11.24 [ 1.31, 21.16]
Beta [Lower 95% CI,Upper 95% CI]
-100
-50
0
50
100
mm^2/effect allele

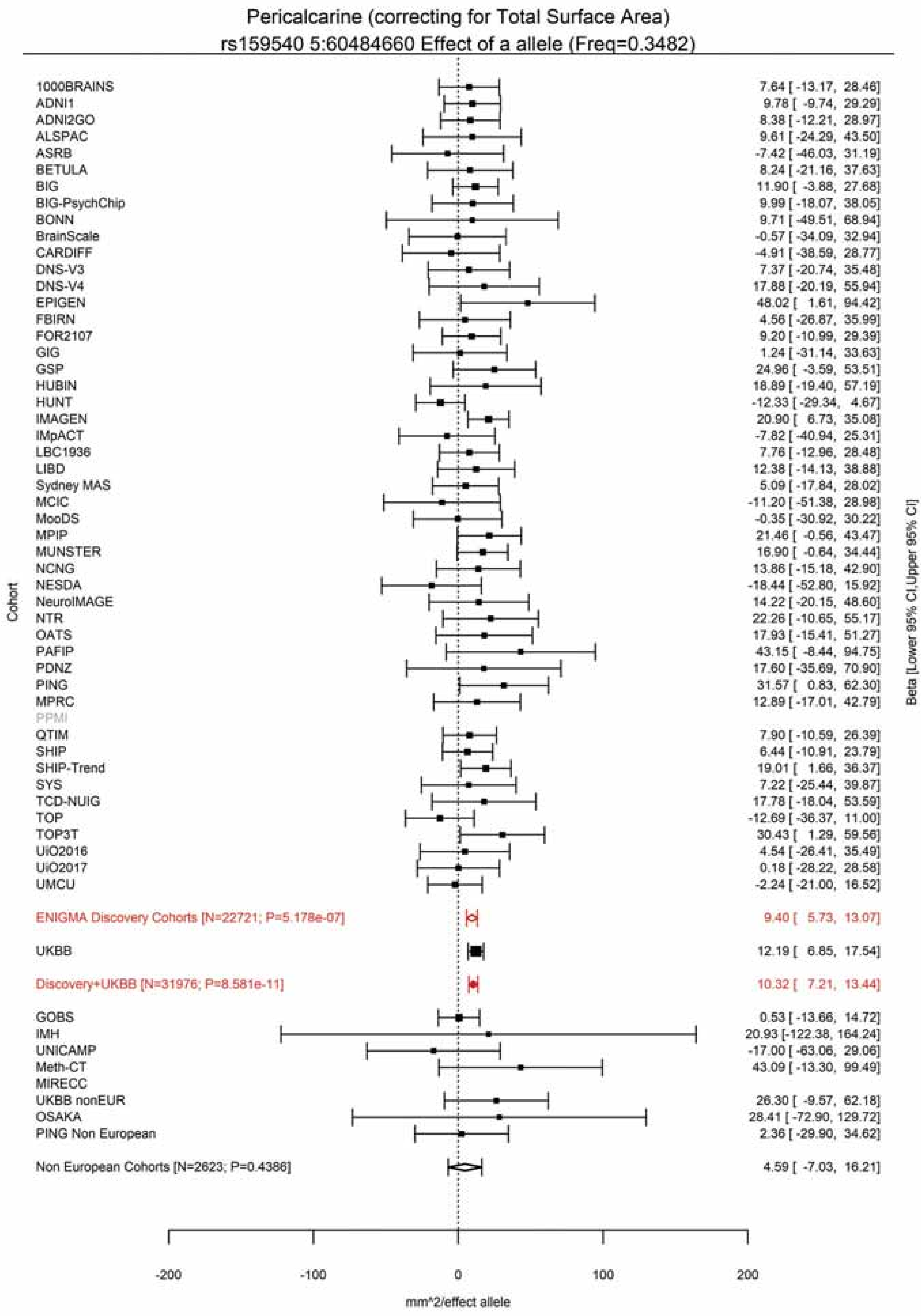

Pericalcarine (correcting for Total Surface Area)
rs159540 5:60484660 Effect of a allele (Freq=0.3482)
Cohort
Beta [Lower 95% CI,Upper 95% CI]
1000BRAINS 7.64 [ -13.17, 28.46]
ADNI1 9.78 [ -9.74, 29.29]
ADNI2GO 8.38 [ -12.21, 28.97]
ALSPAC 9.61 [ -24.29, 43.50]
ASRB -7.42 [ -46.03, 31.19]
BETULA 8.24 [ -21.16, 37.63]
BIG 11.90 [ -3.88, 27.68]
BIG-PsychChip 9.99 [ -18.07, 38.05]
BONN 9.71 [ -49.51, 68.94]
BrainScale -0.57 [ -34.09, 32.94]
CARDIFF -4.91 [ -38.59, 28.77]
DNS-V3 7.37 [ -20.74, 35.48]
DNS-V4 17.88 [ -20.19, 55.94]
EPIGEN 48.02 [ 1.61, 94.42]
FBIRN 4.56 [ -26.87, 35.99]
FOR2107 9.20 [ -10.99, 29.39]
GIG 1.24 [ -31.14, 33.63]
GSP 24.96 [ -3.59, 53.51]
HUBIN 18.89 [ -19.40, 57.19]
HUNT -12.33 [ -29.34, 4.67]
IMAGEN 20.90 [ 6.73, 35.08]
IMpACT -7.82 [ -40.94, 25.31]
LBC1936 7.76 [ -12.96, 28.48]
LIBD 12.38 [ -14.13, 38.88]
Sydney MAS 5.09 [ -17.84, 28.02]
MCIC -11.20 [ -51.38, 28.98]
MooDS -0.35 [ -30.92, 30.22]
MPIP 21.46 [ -0.56, 43.47]
MUNSTER 16.90 [ -0.64, 34.44]
NCNG 13.86 [ -15.18, 42.90]
NESDA -18.44 [ -52.80, 15.92]
NeuroIMAGE 14.22 [ -20.15, 48.60]
NTR 22.26 [ -10.65, 55.17]
OATS 17.93 [ -15.41, 51.27]
PAFIP 43.15 [ -8.44, 94.75]
PDNZ 17.60 [ -35.69, 70.90]
PING 31.57 [ 0.83, 62.30]
MPRC 12.89 [ -17.01, 42.79]
PPMI
QTIM 7.90 [ -10.59, 26.39]
SHIP 6.44 [ -10.91, 23.79]
SHIP-Trend 19.01 [ 1.66, 36.37]
SYS 7.22 [ -25.44, 39.87]
TCD-NUIG 17.78 [ -18.04, 53.59]
TOP -12.69 [ -36.37, 11.00]
TOP3T 30.43 [ 1.29, 59.56]
UiO2016 4.54 [ -26.41, 35.49]
UiO2017 0.18 [ -28.22, 28.58]
UMCU -2.24 [ -21.00, 16.52]
ENIGMA Discovery Cohorts [N=22721; P=5.178e-07] 9.40 [ 5.73, 13.07]
UKBB 12.19 [ 6.85, 17.54]
Discovery+UKBB [N=31976; P=8.581e-11] 10.32 [ 7.21, 13.44]
GOBS 0.53 [ -13.66, 14.72]
IMH 20.93 [-122.38, 164.24]
UNICAMP -17.00 [ -63.06, 29.06]
Meth-CT 43.09 [ -13.30, 99.49]
MIRECC
UKBB nonEUR 26.30 [ -9.57, 62.18]
OSAKA 28.41 [ -72.90, 129.72]
PING Non European 2.36 [ -29.90, 34.62]
Non European Cohorts [N=2623; P=0.4386] 4.59 [ -7.03, 16.21]
-200
-100
0
100
200
mm^2/effect allele

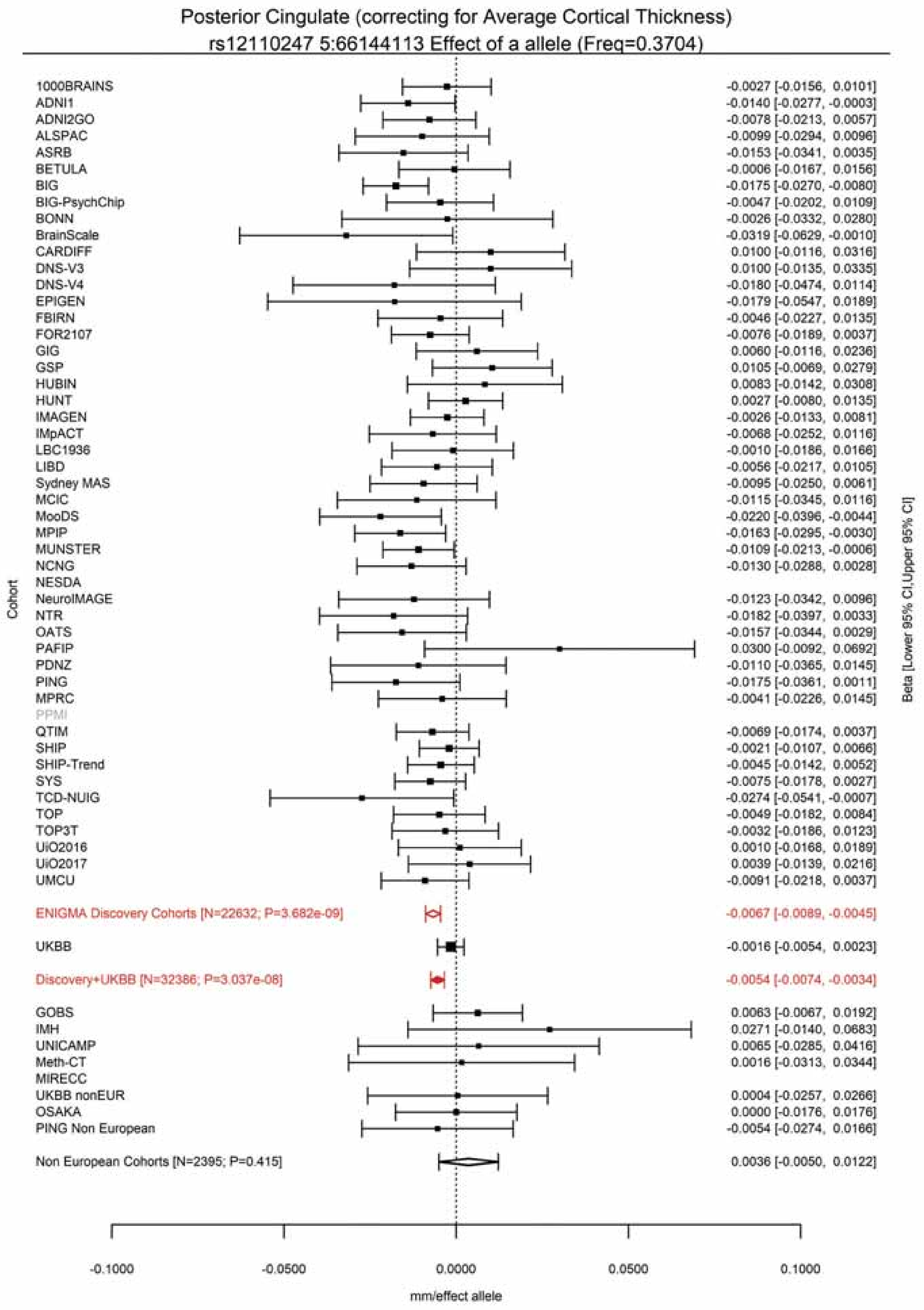

Posterior Cingulate (correcting for Average Cortical Thickness)
rs12110247 5:66144113 Effect of a allele (Freq=0.3704)
Cohort
Beta [Lower 95% CI,Upper 95% CI]
1000BRAINS -0.0027 [-0.0156, 0.0101]
ADNI1 -0.0140 [-0.0277, -0.0003]
ADNI2GO -0.0078 [-0.0213, 0.0057]
ALSPAC -0.0099 [-0.0294, 0.0096]
ASRB -0.0153 [-0.0341, 0.0035]
BETULA -0.0006 [-0.0167, 0.0156]
BIG -0.0175 [-0.0270, -0.0080]
BIG-PsychChip -0.0047 [-0.0202, 0.0109]
BONN -0.0026 [-0.0332, 0.0280]
BrainScale -0.0319 [-0.0629, -0.0010]
CARDIFF 0.0100 [-0.0116, 0.0316]
DNS-V3 0.0100 [-0.0135, 0.0335]
DNS-V4 -0.0180 [-0.0474, 0.0114]
EPIGEN -0.0179 [-0.0547, 0.0189]
FBIRN -0.0046 [-0.0227, 0.0135]
FOR2107 -0.0076 [-0.0189, 0.0037]
GIG 0.0060 [-0.0116, 0.0236]
GSP 0.0105 [-0.0069, 0.0279]
HUBIN 0.0083 [-0.0142, 0.0308]
HUNT 0.0027 [-0.0080, 0.0135]
IMAGEN -0.0026 [-0.0133, 0.0081]
IMpACT -0.0068 [-0.0252, 0.0116]
LBC1936 -0.0010 [-0.0186, 0.0166]
LIBD -0.0056 [-0.0217, 0.0105]
Sydney MAS -0.0095 [-0.0250, 0.0061]
MCIC -0.0115 [-0.0345, 0.0116]
MooDS -0.0220 [-0.0396, -0.0044]
MPIP -0.0163 [-0.0295, -0.0030]
MUNSTER -0.0109 [-0.0213, -0.0006]
NCNG -0.0130 [-0.0288, 0.0028]
NESDA
NeuroIMAGE -0.0123 [-0.0342, 0.0096]
NTR -0.0182 [-0.0397, 0.0033]
OATS -0.0157 [-0.0344, 0.0029]
PAFIP 0.0300 [-0.0092, 0.0692]
PDNZ -0.0110 [-0.0365, 0.0145]
PING -0.0175 [-0.0361, 0.0011]
MPRC -0.0041 [-0.0226, 0.0145]
PPMI
QTIM -0.0069 [-0.0174, 0.0037]
SHIP -0.0021 [-0.0107, 0.0066]
SHIP-Trend -0.0045 [-0.0142, 0.0052]
SYS -0.0075 [-0.0178, 0.0027]
TCD-NUIG -0.0274 [-0.0541, -0.0007]
TOP -0.0049 [-0.0182, 0.0084]
TOP3T -0.0032 [-0.0186, 0.0123]
UiO2016 0.0010 [-0.0168, 0.0189]
UiO2017 0.0039 [-0.0139, 0.0216]
UMCU -0.0091 [-0.0218, 0.0037]
ENIGMA Discovery Cohorts [N=22632; P=3.682e-09] -0.0067 [-0.0089, -0.0045]
UKBB -0.0016 [-0.0054, 0.0023]
Discovery+UKBB [N=32386; P=3.037e-08] -0.0054 [-0.0074, -0.0034]
GOBS 0.0063 [-0.0067, 0.0192]
IMH 0.0271 [-0.0140, 0.0683]
UNICAMP 0.0065 [-0.0285, 0.0416]
Meth-CT 0.0016 [-0.0313, 0.0344]
MIRECC
UKBB nonEUR 0.0004 [-0.0257, 0.0266]
OSAKA 0.0000 [-0.0176, 0.0176]
PING Non European -0.0054 [-0.0274, 0.0166]
Non European Cohorts [N=2395; P=0.415] 0.0036 [-0.0050, 0.0122]
-0.1000
-0.0500
0.0000
0.0500
0.1000
mm/effect allele

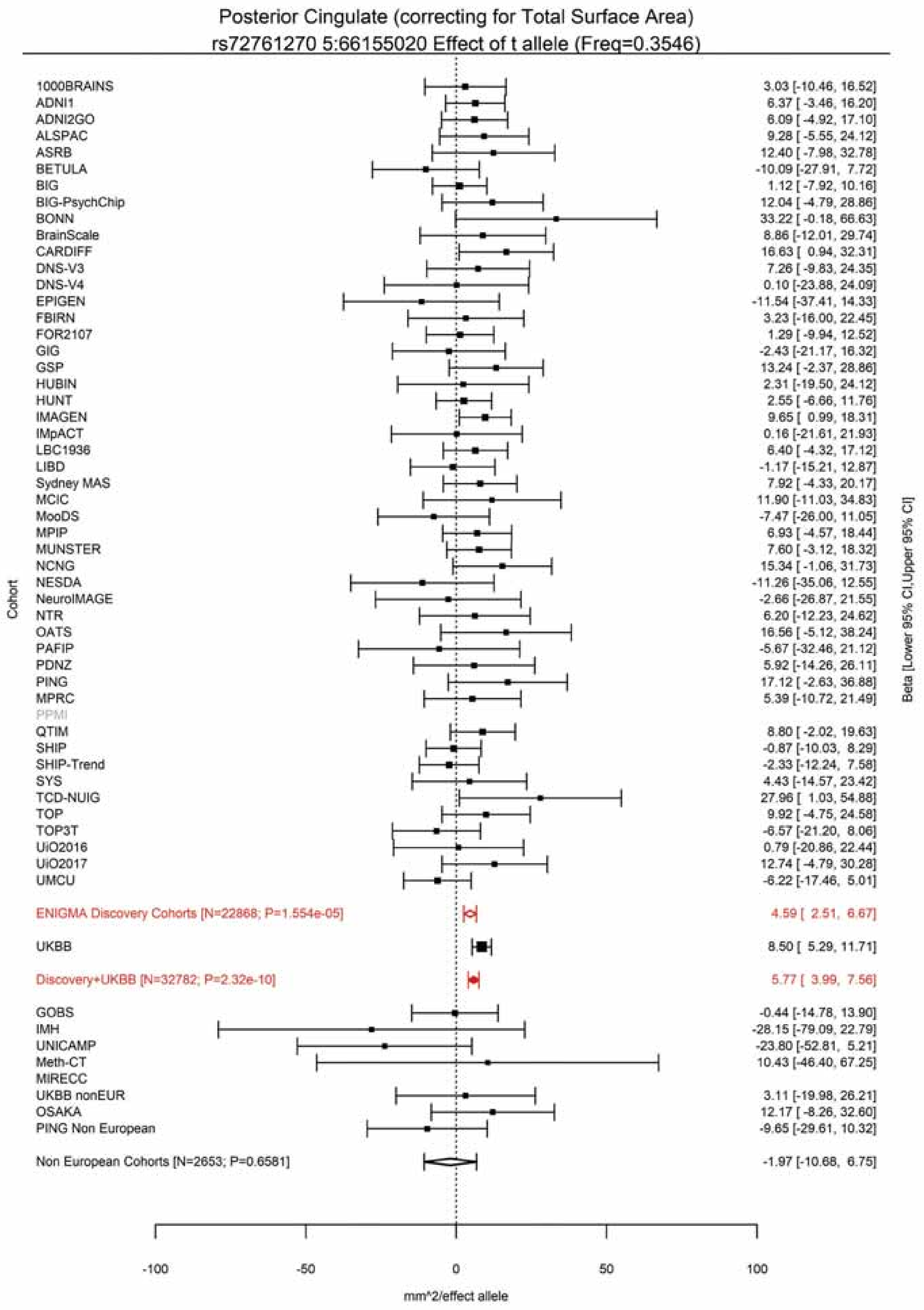
Posterior Cingulate (correcting for Total Surface Area)
rs72761270 5:66155020 Effect of t allele (Freq=0.3546)
Cohort
1000BRAINS 3.03 [-10.46, 16.52]
ADNI1 6.37 [ -3.46, 16.20]
ADNI2GO 6.09 [ -4.92, 17.10]
ALSPAC 9.28 [ -5.55, 24.12]
ASRB 12.40 [ -7.98, 32.78]
BETULA -10.09 [-27.91, 7.72]
BIG 1.12 [ -7.92, 10.16]
BIG-PsychChip 12.04 [ -4.79, 28.86]
BONN 33.22 [ -0.18, 66.63]
BrainScale 8.86 [-12.01, 29.74]
CARDIFF 16.63 [ 0.94, 32.31]
DNS-V3 7.26 [ -9.83, 24.35]
DNS-V4 0.10 [-23.88, 24.09]
EPIGEN -11.54 [-37.41, 14.33]
FBIRN 3.23 [-16.00, 22.45]
FOR2107 1.29 [ -9.94, 12.52]
GIG -2.43 [-21.17, 16.32]
GSP 13.24 [ -2.37, 28.86]
HUBIN 2.31 [-19.50, 24.12]
HUNT 2.55 [ -6.66, 11.76]
IMAGEN 9.65 [ 0.99, 18.31]
IMpACT 0.16 [-21.61, 21.93]
LBC1936 6.40 [ -4.32, 17.12]
LIBD -1.17 [-15.21, 12.87]
Sydney MAS 7.92 [ -4.33, 20.17]
MCIC 11.90 [-11.03, 34.83]
MooDS -7.47 [-26.00, 11.05]
MPIP 6.93 [ -4.57, 18.44]
MUNSTER 7.60 [ -3.12, 18.32]
NCNG 15.34 [ -1.06, 31.73]
NESDA -11.26 [-35.06, 12.55]
NeuroIMAGE -2.66 [-26.87, 21.55]
NTR 6.20 [-12.23, 24.62]
OATS 16.56 [ -5.12, 38.24]
PAFIP -5.67 [-32.46, 21.12]
PDNZ 5.92 [-14.26, 26.11]
PING 17.12 [ -2.63, 36.88]
MPRC 5.39 [-10.72, 21.49]
PPMI
QTIM 8.80 [ -2.02, 19.63]
SHIP -0.87 [-10.03, 8.29]
SHIP-Trend -2.33 [-12.24, 7.58]
SYS 4.43 [-14.57, 23.42]
TCD-NUIG 27.96 [ 1.03, 54.88]
TOP 9.92 [ -4.75, 24.58]
TOP3T -6.57 [-21.20, 8.06]
UiO2016 0.79 [-20.86, 22.44]
UiO2017 12.74 [ -4.79, 30.28]
UMCU -6.22 [-17.46, 5.01]
ENIGMA Discovery Cohorts [N=22868; P=1.554e-05] 4.59 [ 2.51, 6.67]
UKBB 8.50 [ 5.29, 11.71]
Discovery+UKBB [N=32782; P=2.32e-10] 5.77 [ 3.99, 7.56]
GOBS -0.44 [-14.78, 13.90]
IMH -28.15 [-79.09, 22.79]
UNICAMP -23.80 [-52.81, 5.21]
Meth-CT 10.43 [-46.40, 67.25]
MIRECC
UKBB nonEUR 3.11 [-19.98, 26.21]
OSAKA 12.17 [ -8.26, 32.60]
PING Non European -9.65 [-29.61, 10.32]
Non European Cohorts [N=2653; P=0.6581] -1.97 [-10.68, 6.75]
Beta [Lower 95% CI,Upper 95% CI]
-100
-50
0
50
100
mm^2/effect allele

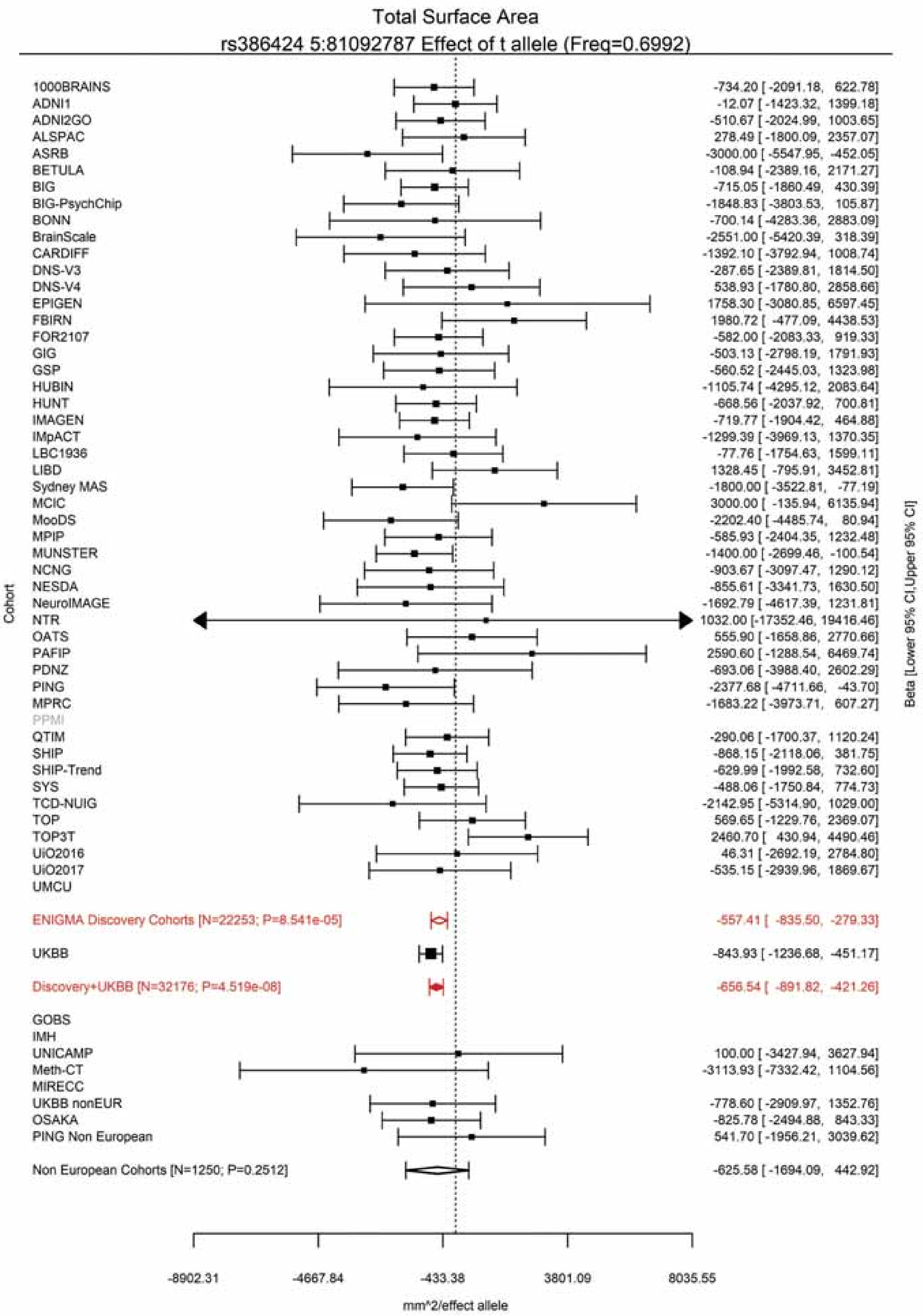

Total Surface Area
rs386424 5:81092787 Effect of t allele (Freq=0.6992)
Cohort
1000BRAINS -734.20 [ -2091.18, 622.78]
ADNI1 -12.07 [ -1423.32, 1399.18]
ADNI2GO -510.67 [ -2024.99, 1003.65]
ALSPAC 278.49 [ -1800.09, 2357.07]
ASRB -3000.00 [ -5547.95, -452.05]
BETULA -108.94 [ -2389.16, 2171.27]
BIG -715.05 [ -1860.49, 430.39]
BIG-PsychChip -1848.83 [ -3803.53, 105.87]
BONN -700.14 [ -4283.36, 2883.09]
BrainScale -2551.00 [ -5420.39, 318.39]
CARDIFF -1392.10 [ -3792.94, 1008.74]
DNS-V3 -287.65 [ -2389.81, 1814.50]
DNS-V4 538.93 [ -1780.80, 2858.66]
EPIGEN 1758.30 [ -3080.85, 6597.45]
FBIRN 1980.72 [ -477.09, 4438.53]
FOR2107 -582.00 [ -2083.33, 919.33]
GIG -503.13 [ -2798.19, 1791.93]
GSP -560.52 [ -2445.03, 1323.98]
HUBIN -1105.74 [ -4295.12, 2083.64]
HUNT -668.56 [ -2037.92, 700.81]
IMAGEN -719.77 [ -1904.42, 464.88]
IMpACT -1299.39 [ -3969.13, 1370.35]
LBC1936 -77.76 [ -1754.63, 1599.11]
LIBD 1328.45 [ -795.91, 3452.81]
Sydney MAS -1800.00 [ -3522.81, -77.19]
MCIC 3000.00 [ -135.94, 6135.94]
MooDS -2202.40 [ -4485.74, 80.94]
MPIP -585.93 [ -2404.35, 1232.48]
MUNSTER -1400.00 [ -2699.46, -100.54]
NCNG -903.67 [ -3097.47, 1290.12]
NESDA -855.61 [ -3341.73, 1630.50]
NeuroIMAGE -1692.79 [ -4617.39, 1231.81]
NTR 1032.00 [-17352.46, 19416.46]
OATS 555.90 [ -1658.86, 2770.66]
PAFIP 2590.60 [ -1288.54, 6469.74]
PDNZ -693.06 [ -3988.40, 2602.29]
PING -2377.68 [ -4711.66, -43.70]
MPRC -1683.22 [ -3973.71, 607.27]
PPMI
QTIM -290.06 [ -1700.37, 1120.24]
SHIP -868.15 [ -2118.06, 381.75]
SHIP-Trend -629.99 [ -1992.58, 732.60]
SYS -488.06 [ -1750.84, 774.73]
TCD-NUIG -2142.95 [ -5314.90, 1029.00]
TOP 569.65 [ -1229.76, 2369.07]
TOP3T 2460.70 [ 430.94, 4490.46]
UiO2016 46.31 [ -2692.19, 2784.80]
UiO2017 -535.15 [ -2939.96, 1869.67]
UMCU
ENIGMA Discovery Cohorts [N=22253; P=8.541e-05] -557.41 [ -835.50, -279.33]
UKBB -843.93 [ -1236.68, -451.17]
Discovery+UKBB [N=32176; P=4.519e-08] -656.54 [ -891.82, -421.26]
GOBS
IMH
UNICAMP 100.00 [ -3427.94, 3627.94]
Meth-CT -3113.93 [ -7332.42, 1104.56]
MIRECC
UKBB nonEUR -778.60 [ -2909.97, 1352.76]
OSAKA -825.78 [ -2494.88, 843.33]
PING Non European 541.70 [ -1956.21, 3039.62]
Non European Cohorts [N=1250; P=0.2512] -625.58 [ -1694.09, 442.92]
Beta [Lower 95% CI,Upper 95% CI]
-8902.31
-4667.84
-433.38
3801.09
8035.55
mm^2/effect allele

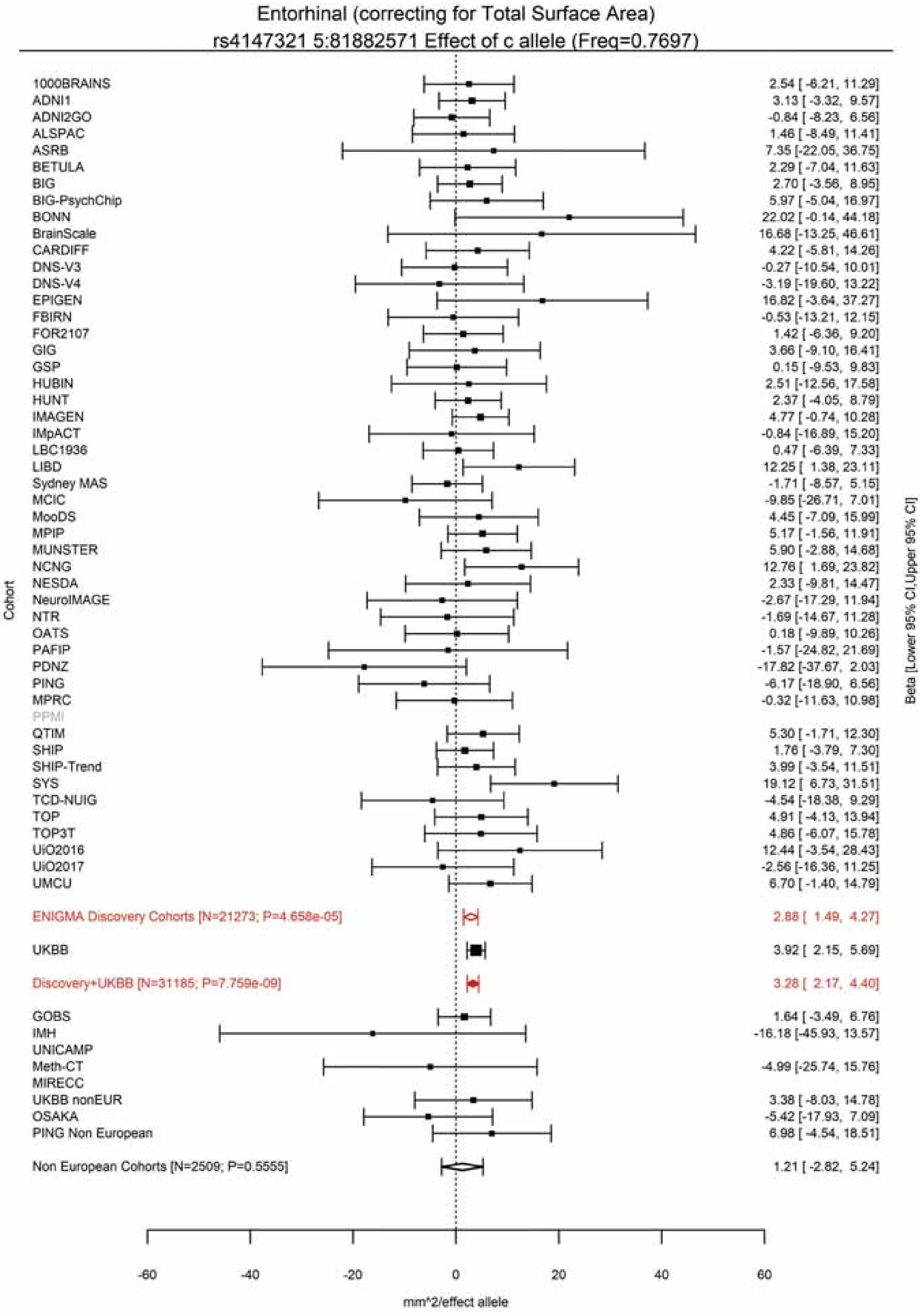

Entorhinal (correcting for Total Surface Area)
rs4147321 5:81882571 Effect of c allele (Freq=0.7697)
1000BRAINS 2.54 [ -6.21, 11.29]
ADNI1 3.13 [ -3.32, 9.57]
ADNI2GO -0.84 [ -8.23, 6.56]
ALSPAC 1.46 [ -8.49, 11.41]
ASRB 7.35 [-22.05, 36.75]
BETULA 2.29 [ -7.04, 11.63]
BIG 2.70 [ -3.56, 8.95]
BIG-PsychChip 5.97 [ -5.04, 16.97]
BONN 22.02 [ -0.14, 44.18]
BrainScale 16.68 [-13.25, 46.61]
CARDIFF 4.22 [ -5.81, 14.26]
DNS-V3 -0.27 [-10.54, 10.01]
DNS-V4 -3.19 [-19.60, 13.22]
EPIGEN 16.82 [ -3.64, 37.27]
FBIRN -0.53 [-13.21, 12.15]
FOR2107 1.42 [ -6.36, 9.20]
GIG 3.66 [ -9.10, 16.41]
GSP 0.15 [ -9.53, 9.83]
HUBIN 2.51 [-12.56, 17.58]
HUNT 2.37 [ -4.05, 8.79]
IMAGEN 4.77 [ -0.74, 10.28]
IMpACT -0.84 [-16.89, 15.20]
LBC1936 0.47 [ -6.39, 7.33]
LIBD 12.25 [ 1.38, 23.11]
Sydney MAS -1.71 [ -8.57, 5.15]
MCIC -9.85 [-26.71, 7.01]
MooDS 4.45 [ -7.09, 15.99]
MPIP 5.17 [ -1.56, 11.91]
MUNSTER 5.90 [ -2.88, 14.68]
NCNG 12.76 [ 1.69, 23.82]
NESDA 2.33 [ -9.81, 14.47]
NeuroIMAGE -2.67 [-17.29, 11.94]
NTR -1.69 [-14.67, 11.28]
OATS 0.18 [ -9.89, 10.26]
PAFIP -1.57 [-24.82, 21.69]
PDNZ -17.82 [-37.67, 2.03]
PING -6.17 [-18.90, 6.56]
MPRC -0.32 [-11.63, 10.98]
PPMI
QTIM 5.30 [ -1.71, 12.30]
SHIP 1.76 [ -3.79, 7.30]
SHIP-Trend 3.99 [ -3.54, 11.51]
SYS 19.12 [ 6.73, 31.51]
TCD-NUIG -4.54 [-18.38, 9.29]
TOP 4.91 [ -4.13, 13.94]
TOP3T 4.86 [ -6.07, 15.78]
UiO2016 12.44 [ -3.54, 28.43]
UiO2017 -2.56 [-16.36, 11.25]
UMCU 6.70 [ -1.40, 14.79]
ENIGMA Discovery Cohorts [N=21273; P=4.658e-05] 2.88 [ 1.49, 4.27]
UKBB 3.92 [ 2.15, 5.69]
Discovery+UKBB [N=31185; P=7.759e-09] 3.28 [ 2.17, 4.40]
GOBS 1.64 [ -3.49, 6.76]
IMH -16.18 [-45.93, 13.57]
UNICAMP
Meth-CT -4.99 [-25.74, 15.76]
MIRECC
UKBB nonEUR 3.38 [ -8.03, 14.78]
OSAKA -5.42 [-17.93, 7.09]
PING Non European 6.98 [ -4.54, 18.51]
Non European Cohorts [N=2509; P=0.5555] 1.21 [ -2.82, 5.24]
Cohort
Beta [Lower 95% CI,Upper 95% CI]
-60 -40 -20 0 20 40 60
mm^2/effect allele

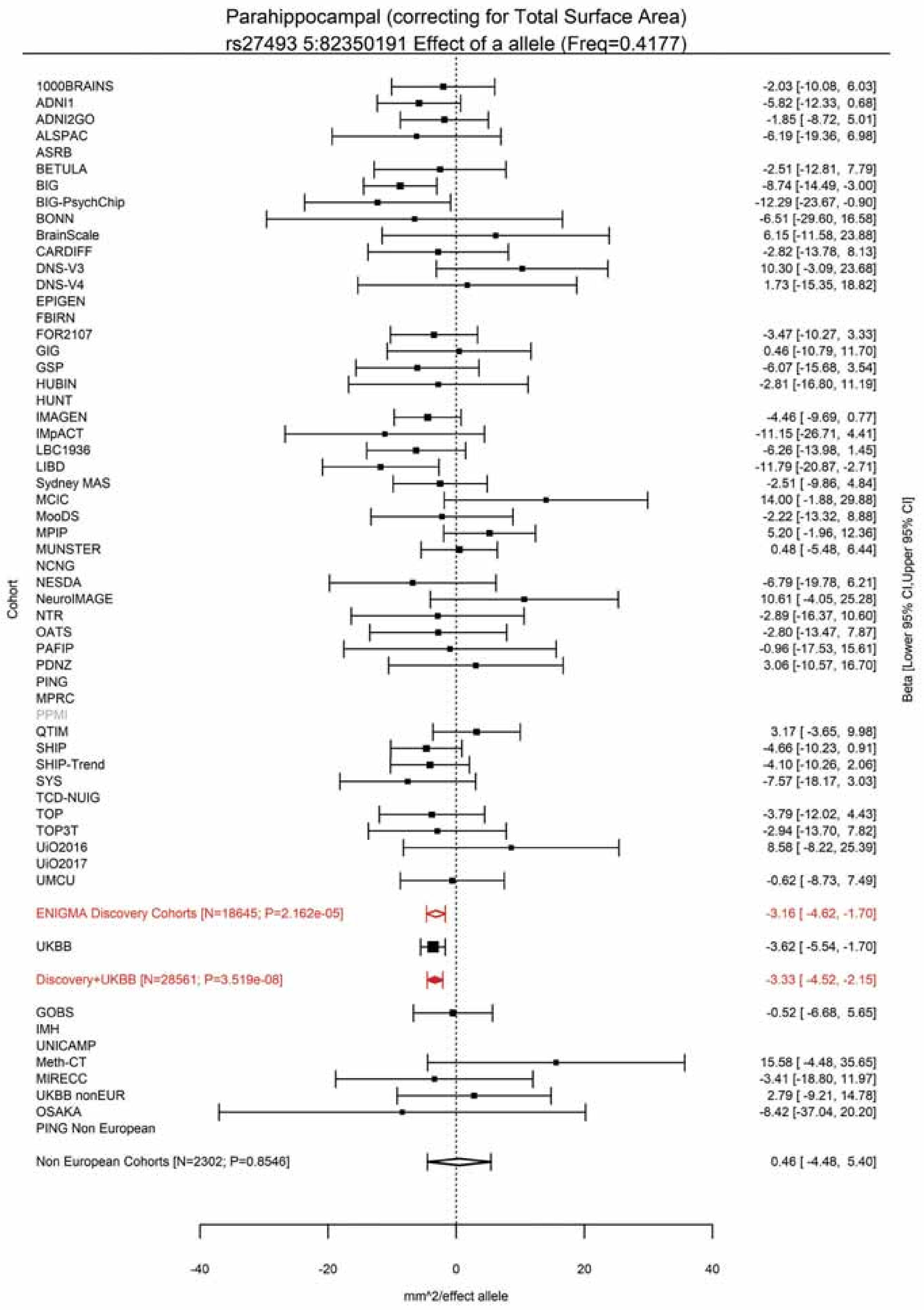
Parahippocampal (correcting for Total Surface Area)
rs27493 5:82350191 Effect of a allele (Freq=0.4177)
Cohort
1000BRAINS -2.03 [-10.08, 6.03]
ADNI1 -5.82 [-12.33, 0.68]
ADNI2GO -1.85 [ -8.72, 5.01]
ALSPAC -6.19 [-19.36, 6.98]
ASRB
BETULA -2.51 [-12.81, 7.79]
BIG -8.74 [-14.49, -3.00]
BIG-PsychChip -12.29 [-23.67, -0.90]
BONN -6.51 [-29.60, 16.58]
BrainScale 6.15 [-11.58, 23.88]
CARDIFF -2.82 [-13.78, 8.13]
DNS-V3 10.30 [ -3.09, 23.68]
DNS-V4 1.73 [-15.35, 18.82]
EPIGEN
FBIRN
FOR2107 -3.47 [-10.27, 3.33]
GIG 0.46 [-10.79, 11.70]
GSP -6.07 [-15.68, 3.54]
HUBIN -2.81 [-16.80, 11.19]
HUNT
IMAGEN -4.46 [ -9.69, 0.77]
IMpACT -11.15 [-26.71, 4.41]
LBC1936 -6.26 [-13.98, 1.45]
LIBD -11.79 [-20.87, -2.71]
Sydney MAS -2.51 [ -9.86, 4.84]
MCIC 14.00 [ -1.88, 29.88]
MooDS -2.22 [-13.32, 8.88]
MPIP 5.20 [ -1.96, 12.36]
MUNSTER 0.48 [ -5.48, 6.44]
NCNG
NESDA -6.79 [-19.78, 6.21]
NeuroIMAGE 10.61 [ -4.05, 25.28]
NTR -2.89 [-16.37, 10.60]
OATS -2.80 [-13.47, 7.87]
PAFIP -0.96 [-17.53, 15.61]
PDNZ 3.06 [-10.57, 16.70]
PING
MPRC
PPMI
QTIM 3.17 [ -3.65, 9.98]
SHIP -4.66 [-10.23, 0.91]
SHIP-Trend -4.10 [-10.26, 2.06]
SYS -7.57 [-18.17, 3.03]
TCD-NUIG
TOP -3.79 [-12.02, 4.43]
TOP3T -2.94 [-13.70, 7.82]
UiO2016 8.58 [ -8.22, 25.39]
UiO2017
UMCU -0.62 [ -8.73, 7.49]
ENIGMA Discovery Cohorts [N=18645; P=2.162e-05] -3.16 [ -4.62, -1.70]
UKBB -3.62 [ -5.54, -1.70]
Discovery+UKBB [N=28561; P=3.519e-08] -3.33 [ -4.52, -2.15]
GOBS -0.52 [ -6.68, 5.65]
IMH
UNICAMP
Meth-CT 15.58 [ -4.48, 35.65]
MIRECC -3.41 [-18.80, 11.97]
UKBB nonEUR 2.79 [ -9.21, 14.78]
OSAKA -8.42 [-37.04, 20.20]
PING Non European
Non European Cohorts [N=2302; P=0.8546] 0.46 [ -4.48, 5.40]
Beta [Lower 95% CI,Upper 95% CI]
-40
-20
0
20
40
mm^2/effect allele

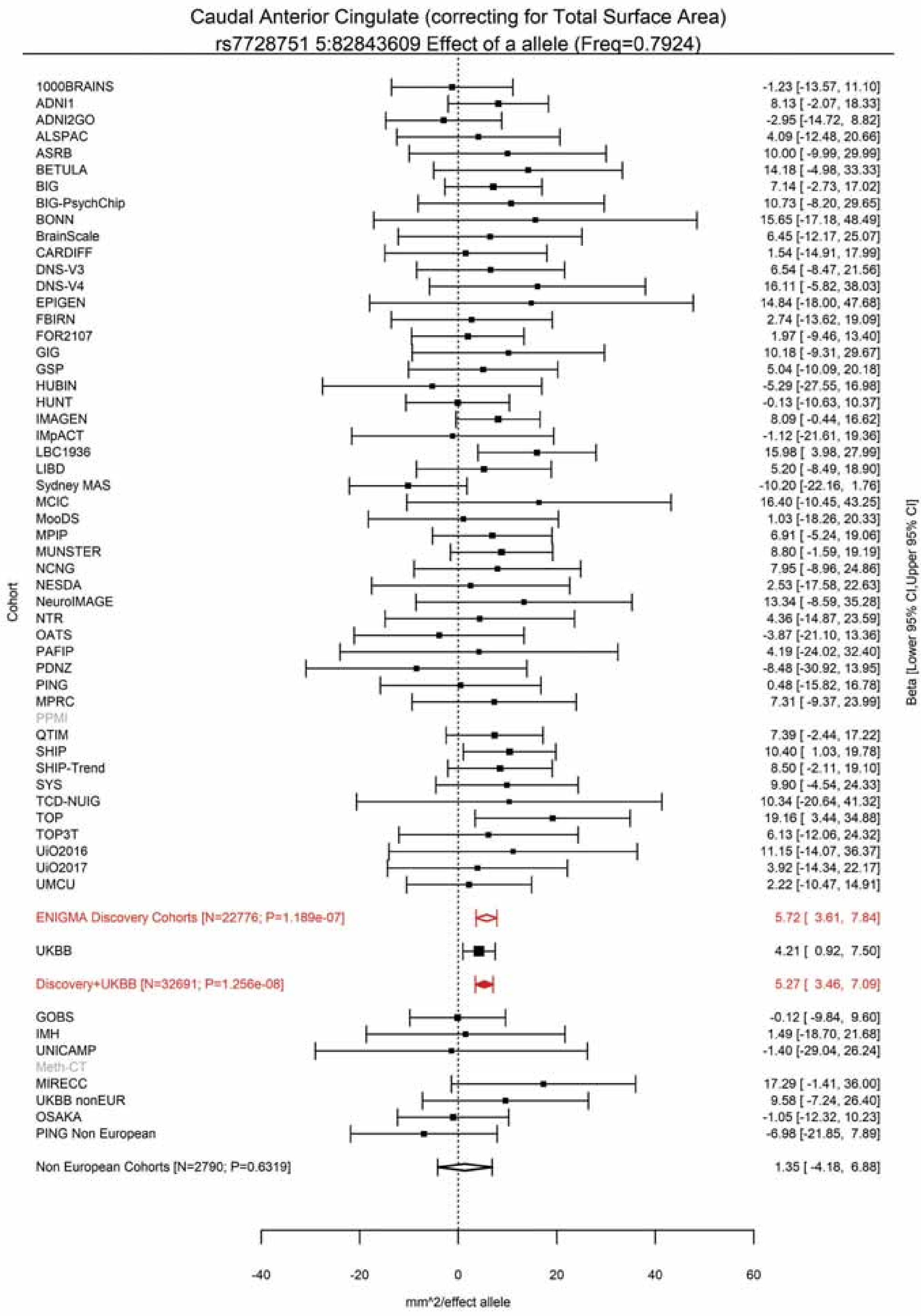

Caudal Anterior Cingulate (correcting for Total Surface Area)
rs7728751 5:82843609 Effect of a allele (Freq=0.7924)
Cohort
1000BRAINS -1.23 [-13.57, 11.10]
ADNI1 8.13 [ -2.07, 18.33]
ADNI2GO -2.95 [-14.72, 8.82]
ALSPAC 4.09 [-12.48, 20.66]
ASRB 10.00 [ -9.99, 29.99]
BETULA 14.18 [ -4.98, 33.33]
BIG 7.14 [ -2.73, 17.02]
BIG-PsychChip 10.73 [ -8.20, 29.65]
BONN 15.65 [-17.18, 48.49]
BrainScale 6.45 [-12.17, 25.07]
CARDIFF 1.54 [-14.91, 17.99]
DNS-V3 6.54 [ -8.47, 21.56]
DNS-V4 16.11 [ -5.82, 38.03]
EPIGEN 14.84 [-18.00, 47.68]
FBIRN 2.74 [-13.62, 19.09]
FOR2107 1.97 [ -9.46, 13.40]
GIG 10.18 [ -9.31, 29.67]
GSP 5.04 [-10.09, 20.18]
HUBIN -5.29 [-27.55, 16.98]
HUNT -0.13 [-10.63, 10.37]
IMAGEN 8.09 [ -0.44, 16.62]
IMpACT -1.12 [-21.61, 19.36]
LBC1936 15.98 [ 3.98, 27.99]
LIBD 5.20 [ -8.49, 18.90]
Sydney MAS -10.20 [-22.16, 1.76]
MCIC 16.40 [-10.45, 43.25]
MooDS 1.03 [-18.26, 20.33]
MPIP 6.91 [ -5.24, 19.06]
MUNSTER 8.80 [ -1.59, 19.19]
NCNG 7.95 [ -8.96, 24.86]
NESDA 2.53 [-17.58, 22.63]
NeuroIMAGE 13.34 [ -8.59, 35.28]
NTR 4.36 [-14.87, 23.59]
OATS -3.87 [-21.10, 13.36]
PAFIP 4.19 [-24.02, 32.40]
PDNZ -8.48 [-30.92, 13.95]
PING 0.48 [-15.82, 16.78]
MPRC 7.31 [ -9.37, 23.99]
PPMI
QTIM 7.39 [ -2.44, 17.22]
SHIP 10.40 [ 1.03, 19.78]
SHIP-Trend 8.50 [ -2.11, 19.10]
SYS 9.90 [ -4.54, 24.33]
TCD-NUIG 10.34 [-20.64, 41.32]
TOP 19.16 [ 3.44, 34.88]
TOP3T 6.13 [-12.06, 24.32]
UiO2016 11.15 [-14.07, 36.37]
UiO2017 3.92 [-14.34, 22.17]
UMCU 2.22 [-10.47, 14.91]
ENIGMA Discovery Cohorts [N=22776; P=1.189e-07] 5.72 [ 3.61, 7.84]
UKBB 4.21 [ 0.92, 7.50]
Discovery+UKBB [N=32691; P=1.256e-08] 5.27 [ 3.46, 7.09]
GOBS -0.12 [ -9.84, 9.60]
IMH 1.49 [-18.70, 21.68]
UNICAMP -1.40 [-29.04, 26.24]
Meth-CT
MIRECC 17.29 [ -1.41, 36.00]
UKBB nonEUR 9.58 [ -7.24, 26.40]
OSAKA -1.05 [-12.32, 10.23]
PING Non European -6.98 [-21.85, 7.89]
Non European Cohorts [N=2790; P=0.6319] 1.35 [ -4.18, 6.88]
Beta [Lower 95% CI,Upper 95% CI]
-40
-20
0
20
40
60
mm^2/effect allele

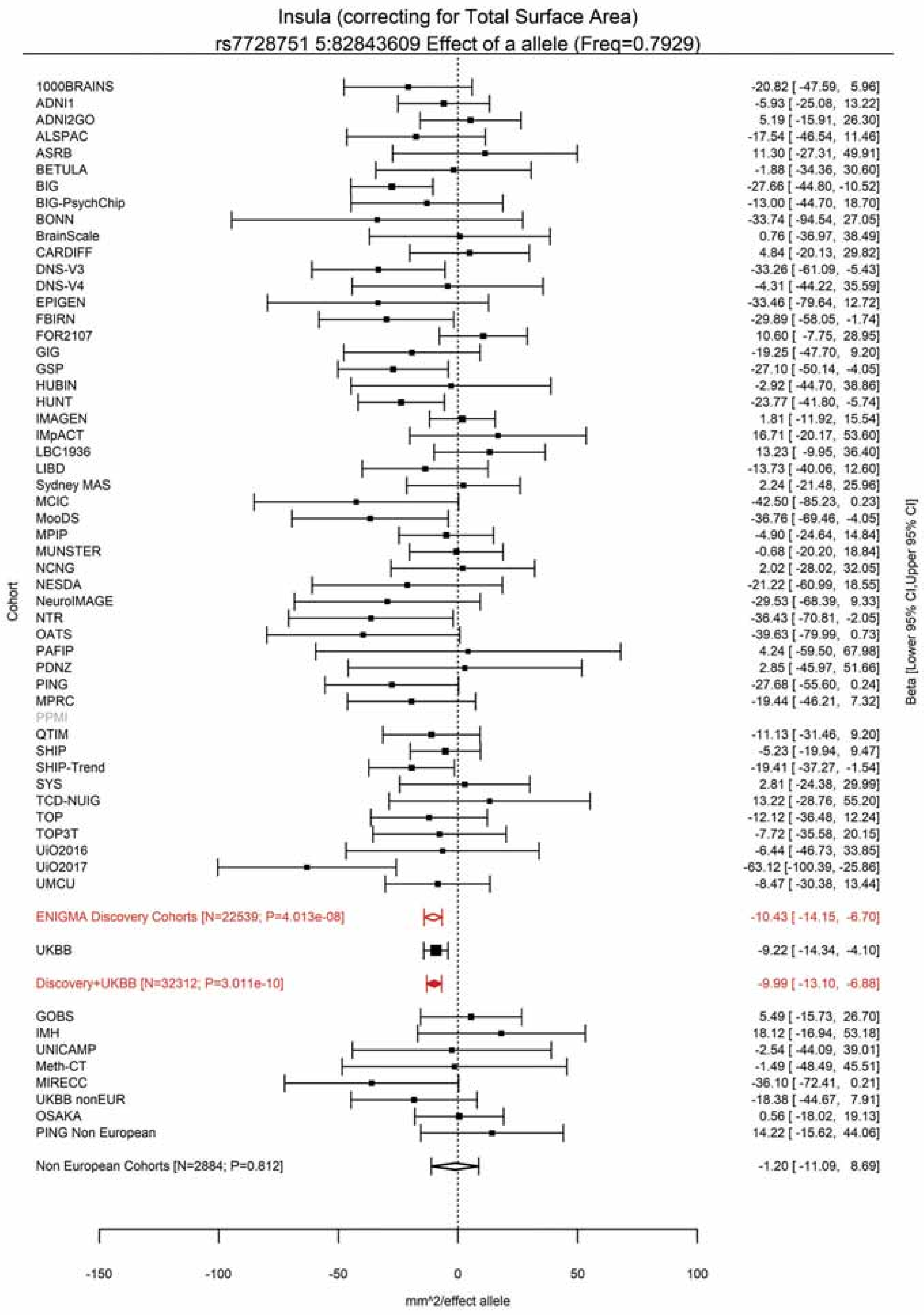
Insula (correcting for Total Surface Area)
rs7728751 5:82843609 Effect of a allele (Freq=0.7929)
Cohort
1000BRAINS -20.82 [ -47.59, 5.96]
ADNI1 -5.93 [ -25.08, 13.22]
ADNI2GO 5.19 [ -15.91, 26.30]
ALSPAC -17.54 [ -46.54, 11.46]
ASRB 11.30 [ -27.31, 49.91]
BETULA -1.88 [ -34.36, 30.60]
BIG -27.66 [ -44.80, -10.52]
BIG-PsychChip -13.00 [ -44.70, 18.70]
BONN -33.74 [ -94.54, 27.05]
BrainScale 0.76 [ -36.97, 38.49]
CARDIFF 4.84 [ -20.13, 29.82]
DNS-V3 -33.26 [ -61.09, -5.43]
DNS-V4 -4.31 [ -44.22, 35.59]
EPIGEN -33.46 [ -79.64, 12.72]
FBIRN -29.89 [ -58.05, -1.74]
FOR2107 10.60 [ -7.75, 28.95]
GIG -19.25 [ -47.70, 9.20]
GSP -27.10 [ -50.14, -4.05]
HUBIN -2.92 [ -44.70, 38.86]
HUNT -23.77 [ -41.80, -5.74]
IMAGEN 1.81 [ -11.92, 15.54]
IMpACT 16.71 [ -20.17, 53.60]
LBC1936 13.23 [ -9.95, 36.40]
LIBD -13.73 [ -40.06, 12.60]
Sydney MAS 2.24 [ -21.48, 25.96]
MCIC -42.50 [ -85.23, 0.23]
MooDS -36.76 [ -69.46, -4.05]
MPIP -4.90 [ -24.64, 14.84]
MUNSTER -0.68 [ -20.20, 18.84]
NCNG 2.02 [ -28.02, 32.05]
NESDA -21.22 [ -60.99, 18.55]
NeuroIMAGE -29.53 [ -68.39, 9.33]
NTR -36.43 [ -70.81, -2.05]
OATS -39.63 [ -79.99, 0.73]
PAFIP 4.24 [ -59.50, 67.98]
PDNZ 2.85 [ -45.97, 51.66]
PING -27.68 [ -55.60, 0.24]
MPRC -19.44 [ -46.21, 7.32]
PPMI
QTIM -11.13 [ -31.46, 9.20]
SHIP -5.23 [ -19.94, 9.47]
SHIP-Trend -19.41 [ -37.27, -1.54]
SYS 2.81 [ -24.38, 29.99]
TCD-NUIG 13.22 [ -28.76, 55.20]
TOP -12.12 [ -36.48, 12.24]
TOP3T -7.72 [ -35.58, 20.15]
UiO2016 -6.44 [ -46.73, 33.85]
UiO2017 -63.12 [-100.39, -25.86]
UMCU -8.47 [ -30.38, 13.44]
ENIGMA Discovery Cohorts [N=22539; P=4.013e-08] -10.43 [ -14.15, -6.70]
UKBB -9.22 [ -14.34, -4.10]
Discovery+UKBB [N=32312; P=3.011e-10] -9.99 [ -13.10, -6.88]
GOBS 5.49 [ -15.73, 26.70]
IMH 18.12 [ -16.94, 53.18]
UNICAMP -2.54 [ -44.09, 39.01]
Meth-CT -1.49 [ -48.49, 45.51]
MIRECC -36.10 [ -72.41, 0.21]
UKBB nonEUR -18.38 [ -44.67, 7.91]
OSAKA 0.56 [ -18.02, 19.13]
PING Non European 14.22 [ -15.62, 44.06]
Non European Cohorts [N=2884; P=0.812] -1.20 [ -11.09, 8.69]
Beta [Lower 95% CI,Upper 95% CI]
-150
-100
-50
0
50
100
mm^2/effect allele

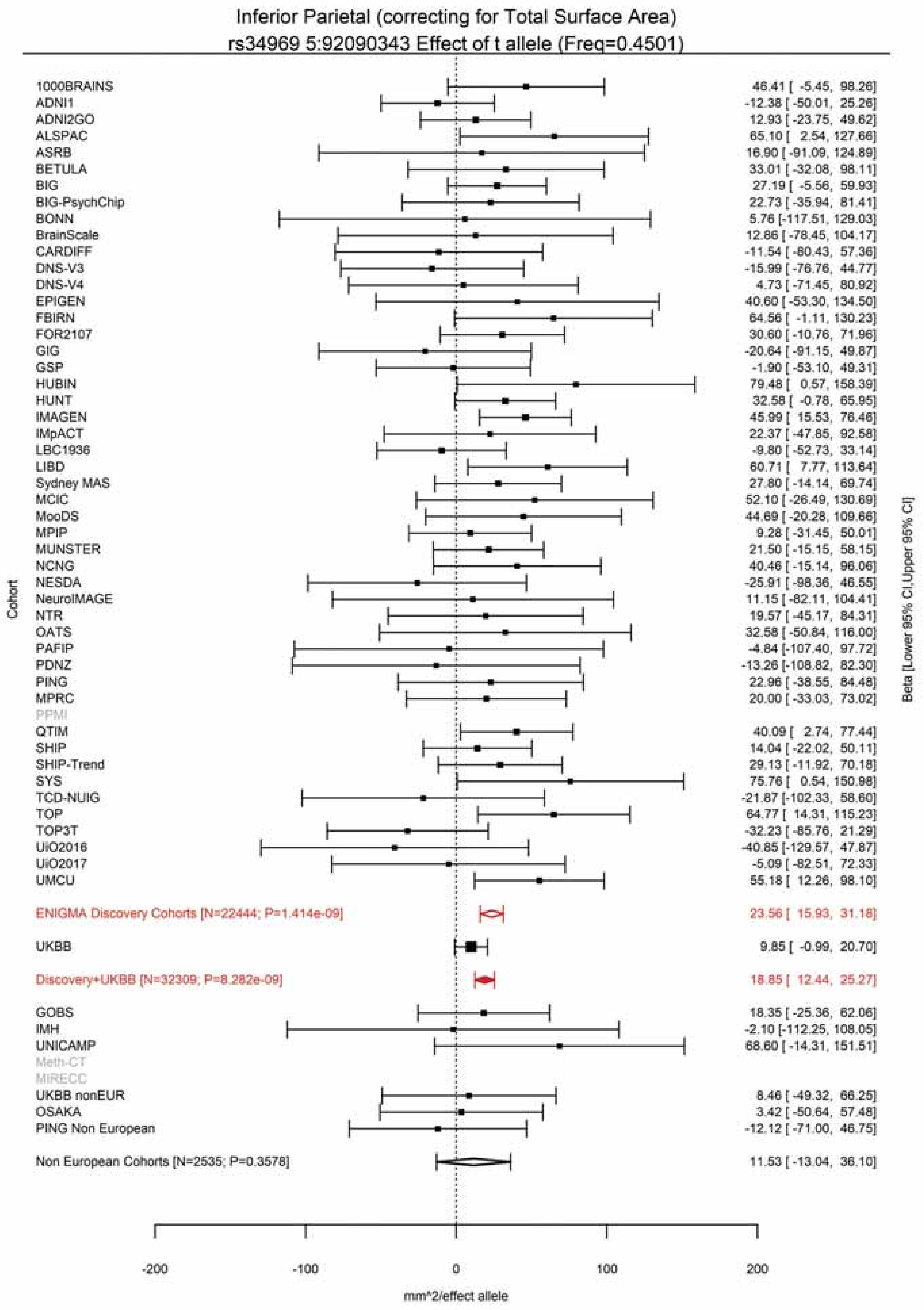
Inferior Parietal (correcting for Total Surface Area)
rs34969 5:92090343 Effect of t allele (Freq=0.4501)
Cohort
1000BRAINS 46.41 [ -5.45, 98.26]
ADNI1 -12.38 [ -50.01, 25.26]
ADNI2GO 12.93 [ -23.75, 49.62]
ALSPAC 65.10 [ 2.54, 127.66]
ASRB 16.90 [ -91.09, 124.89]
BETULA 33.01 [ -32.08, 98.11]
BIG 27.19 [ -5.56, 59.93]
BIG-PsychChip 22.73 [ -35.94, 81.41]
BONN 5.76 [-117.51, 129.03]
BrainScale 12.86 [ -78.45, 104.17]
CARDIFF -11.54 [ -80.43, 57.36]
DNS-V3 -15.99 [ -76.76, 44.77]
DNS-V4 4.73 [ -71.45, 80.92]
EPIGEN 40.60 [ -53.30, 134.50]
FBIRN 64.56 [ -1.11, 130.23]
FOR2107 30.60 [ -10.76, 71.96]
GIG -20.64 [ -91.15, 49.87]
GSP -1.90 [ -53.10, 49.31]
HUBIN 79.48 [ 0.57, 158.39]
HUNT 32.58 [ -0.78, 65.95]
IMAGEN 45.99 [ 15.53, 76.46]
IMpACT 22.37 [ -47.85, 92.58]
LBC1936 -9.80 [ -52.73, 33.14]
LIBD 60.71 [ 7.77, 113.64]
Sydney MAS 27.80 [ -14.14, 69.74]
MCIC 52.10 [ -26.49, 130.69]
MooDS 44.69 [ -20.28, 109.66]
MPIP 9.28 [ -31.45, 50.01]
MUNSTER 21.50 [ -15.15, 58.15]
NCNG 40.46 [ -15.14, 96.06]
NESDA -25.91 [ -98.36, 46.55]
NeuroIMAGE 11.15 [ -82.11, 104.41]
NTR 19.57 [ -45.17, 84.31]
OATS 32.58 [ -50.84, 116.00]
PAFIP -4.84 [-107.40, 97.72]
PDNZ -13.26 [-108.82, 82.30]
PING 22.96 [ -38.55, 84.48]
MPRC 20.00 [ -33.03, 73.02]
PPMI
QTIM 40.09 [ 2.74, 77.44]
SHIP 14.04 [ -22.02, 50.11]
SHIP-Trend 29.13 [ -11.92, 70.18]
SYS 75.76 [ 0.54, 150.98]
TCD-NUIG -21.87 [-102.33, 58.60]
TOP 64.77 [ 14.31, 115.23]
TOP3T -32.23 [ -85.76, 21.29]
UiO2016 -40.85 [-129.57, 47.87]
UiO2017 -5.09 [ -82.51, 72.33]
UMCU 55.18 [ 12.26, 98.10]
ENIGMA Discovery Cohorts [N=22444; P=1.414e-09] 23.56 [ 15.93, 31.18]
UKBB 9.85 [ -0.99, 20.70]
Discovery+UKBB [N=32309; P=8.282e-09] 18.85 [ 12.44, 25.27]
GOBS 18.35 [ -25.36, 62.06]
IMH -2.10 [-112.25, 108.05]
UNICAMP 68.60 [ -14.31, 151.51]
Meth-CT
MIRECC
UKBB nonEUR 8.46 [ -49.32, 66.25]
OSAKA 3.42 [ -50.64, 57.48]
PING Non European -12.12 [ -71.00, 46.75]
Non European Cohorts [N=2535; P=0.3578] 11.53 [ -13.04, 36.10]
Beta [Lower 95% CI,Upper 95% CI]
-200
-100
0
100
200
mm^2/effect allele

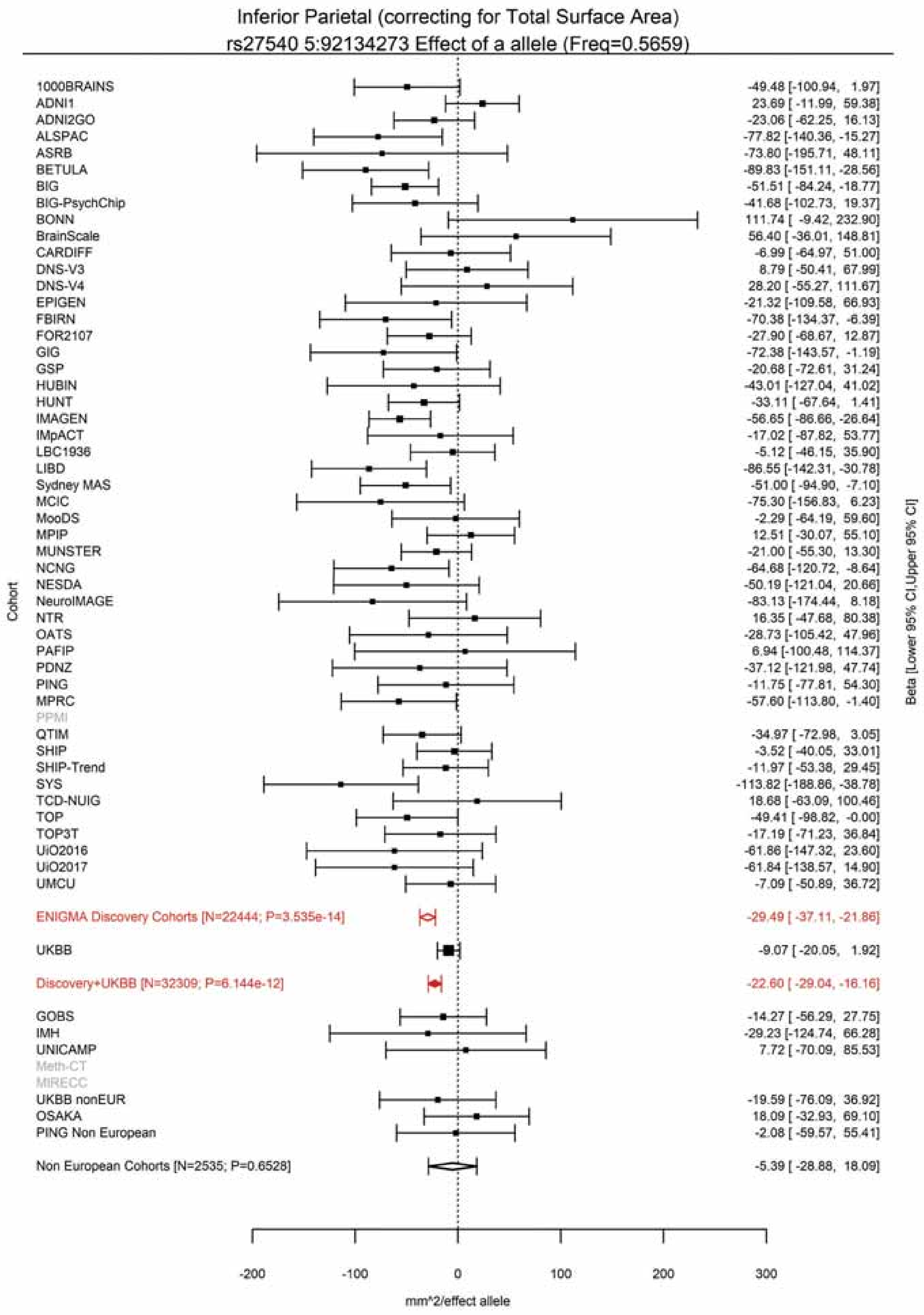
Inferior Parietal (correcting for Total Surface Area)
rs27540 5:92134273 Effect of a allele (Freq=0.5659)
Cohort
Beta [Lower 95% CI,Upper 95% CI]
1000BRAINS -49.48 [-100.94, 1.97]
ADNI1 23.69 [ -11.99, 59.38]
ADNI2GO -23.06 [ -62.25, 16.13]
ALSPAC -77.82 [-140.36, -15.27]
ASRB -73.80 [-195.71, 48.11]
BETULA -89.83 [-151.11, -28.56]
BIG -51.51 [ -84.24, -18.77]
BIG-PsychChip -41.68 [-102.73, 19.37]
BONN 111.74 [ -9.42, 232.90]
BrainScale 56.40 [ -36.01, 148.81]
CARDIFF -6.99 [ -64.97, 51.00]
DNS-V3 8.79 [ -50.41, 67.99]
DNS-V4 28.20 [ -55.27, 111.67]
EPIGEN -21.32 [-109.58, 66.93]
FBIRN -70.38 [-134.37, -6.39]
FOR2107 -27.90 [ -68.67, 12.87]
GIG -72.38 [-143.57, -1.19]
GSP -20.68 [ -72.61, 31.24]
HUBIN -43.01 [-127.04, 41.02]
HUNT -33.11 [ -67.64, 1.41]
IMAGEN -56.65 [ -86.66, -26.64]
IMpACT -17.02 [ -87.82, 53.77]
LBC1936 -5.12 [ -46.15, 35.90]
LIBD -86.55 [-142.31, -30.78]
Sydney MAS -51.00 [ -94.90, -7.10]
MCIC -75.30 [-156.83, 6.23]
MooDS -2.29 [ -64.19, 59.60]
MPIP 12.51 [ -30.07, 55.10]
MUNSTER -21.00 [ -55.30, 13.30]
NCNG -64.68 [-120.72, -8.64]
NESDA -50.19 [-121.04, 20.66]
NeuroIMAGE -83.13 [-174.44, 8.18]
NTR 16.35 [ -47.68, 80.38]
OATS -28.73 [-105.42, 47.96]
PAFIP 6.94 [-100.48, 114.37]
PDNZ -37.12 [-121.98, 47.74]
PING -11.75 [ -77.81, 54.30]
MPRC -57.60 [-113.80, -1.40]
PPMI
QTIM -34.97 [ -72.98, 3.05]
SHIP -3.52 [ -40.05, 33.01]
SHIP-Trend -11.97 [ -53.38, 29.45]
SYS -113.82 [-188.86, -38.78]
TCD-NUIG 18.68 [ -63.09, 100.46]
TOP -49.41 [ -98.82, -0.00]
TOP3T -17.19 [ -71.23, 36.84]
UiO2016 -61.86 [-147.32, 23.60]
UiO2017 -61.84 [-138.57, 14.90]
UMCU -7.09 [ -50.89, 36.72]
ENIGMA Discovery Cohorts [N=22444; P=3.535e-14] -29.49 [ -37.11, -21.86]
UKBB -9.07 [ -20.05, 1.92]
Discovery+UKBB [N=32309; P=6.144e-12] -22.60 [ -29.04, -16.16]
GOBS -14.27 [ -56.29, 27.75]
IMH -29.23 [-124.74, 66.28]
UNICAMP 7.72 [ -70.09, 85.53]
Meth-CT
MIRECC
UKBB nonEUR -19.59 [ -76.09, 36.92]
OSAKA 18.09 [ -32.93, 69.10]
PING Non European -2.08 [ -59.57, 55.41]
Non European Cohorts [N=2535; P=0.6528] -5.39 [ -28.88, 18.09]
-200
-100
0
100
200
300
mm^2/effect allele

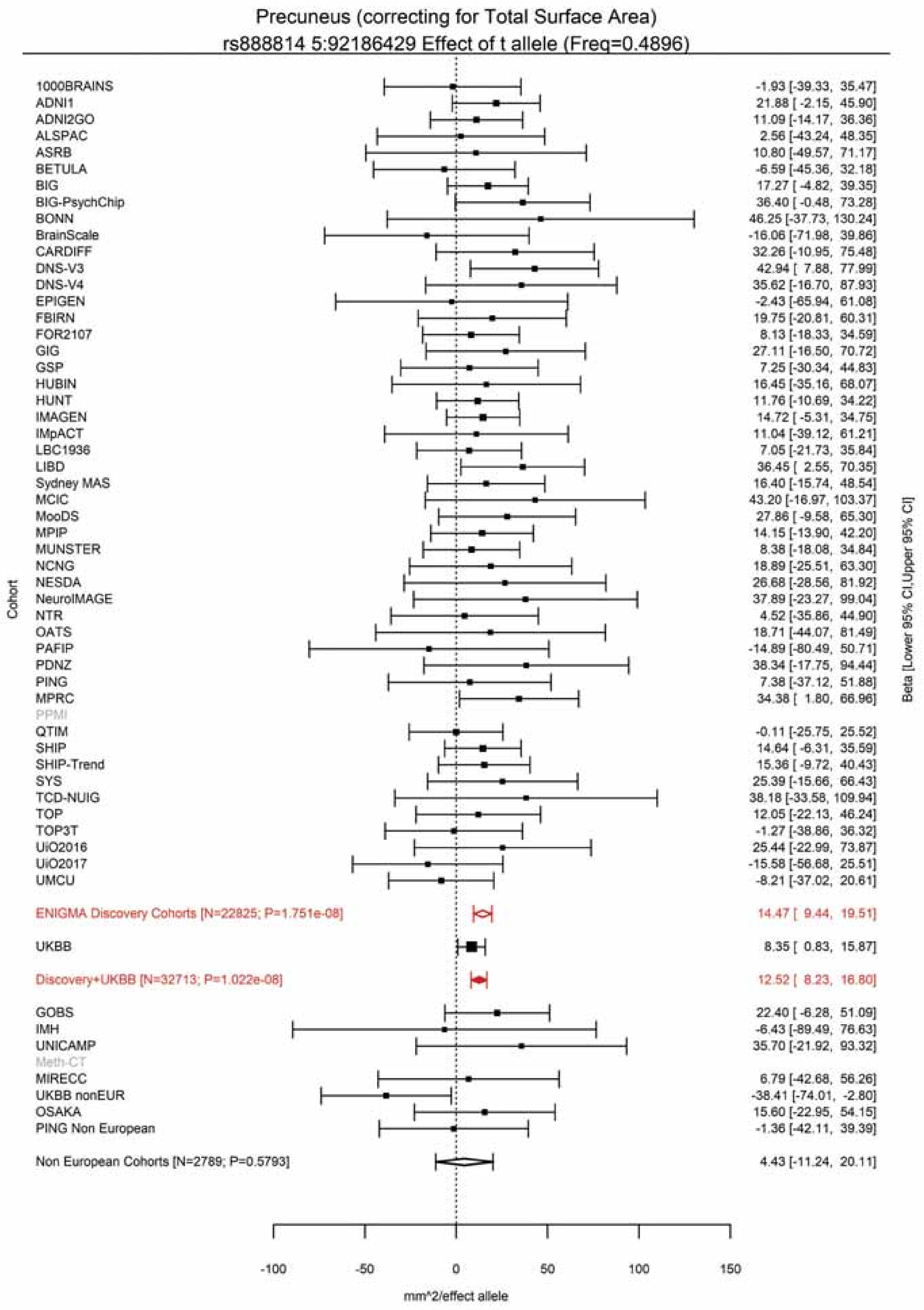

Precuneus (correcting for Total Surface Area)
rs888814 5:92186429 Effect of t allele (Freq=0.4896)
Cohort
1000BRAINS -1.93 [-39.33, 35.47]
ADNI1 21.88 [ -2.15, 45.90]
ADNI2GO 11.09 [-14.17, 36.36]
ALSPAC 2.56 [-43.24, 48.35]
ASRB 10.80 [-49.57, 71.17]
BETULA -6.59 [-45.36, 32.18]
BIG 17.27 [ -4.82, 39.35]
BIG-PsychChip 36.40 [ -0.48, 73.28]
BONN 46.25 [-37.73, 130.24]
BrainScale -16.06 [-71.98, 39.86]
CARDIFF 32.26 [-10.95, 75.48]
DNS-V3 42.94 [ 7.88, 77.99]
DNS-V4 35.62 [-16.70, 87.93]
EPIGEN -2.43 [-65.94, 61.08]
FBIRN 19.75 [-20.81, 60.31]
FOR2107 8.13 [-18.33, 34.59]
GIG 27.11 [-16.50, 70.72]
GSP 7.25 [-30.34, 44.83]
HUBIN 16.45 [-35.16, 68.07]
HUNT 11.76 [-10.69, 34.22]
IMAGEN 14.72 [ -5.31, 34.75]
IMpACT 11.04 [-39.12, 61.21]
LBC1936 7.05 [-21.73, 35.84]
LIBD 36.45 [ 2.55, 70.35]
Sydney MAS 16.40 [-15.74, 48.54]
MCIC 43.20 [-16.97, 103.37]
MooDS 27.86 [ -9.58, 65.30]
MPIP 14.15 [-13.90, 42.20]
MUNSTER 8.38 [-18.08, 34.84]
NCNG 18.89 [-25.51, 63.30]
NESDA 26.68 [-28.56, 81.92]
NeuroIMAGE 37.89 [-23.27, 99.04]
NTR 4.52 [-35.86, 44.90]
OATS 18.71 [-44.07, 81.49]
PAFIP -14.89 [-80.49, 50.71]
PDNZ 38.34 [-17.75, 94.44]
PING 7.38 [-37.12, 51.88]
MPRC 34.38 [ 1.80, 66.96]
PPMI
QTIM -0.11 [-25.75, 25.52]
SHIP 14.64 [ -6.31, 35.59]
SHIP-Trend 15.36 [ -9.72, 40.43]
SYS 25.39 [-15.66, 66.43]
TCD-NUIG 38.18 [-33.58, 109.94]
TOP 12.05 [-22.13, 46.24]
TOP3T -1.27 [-38.86, 36.32]
UiO2016 25.44 [-22.99, 73.87]
UiO2017 -15.58 [-56.68, 25.51]
UMCU -8.21 [-37.02, 20.61]
ENIGMA Discovery Cohorts [N=22825; P=1.751e-08] 14.47 [ 9.44, 19.51]
UKBB 8.35 [ 0.83, 15.87]
Discovery+UKBB [N=32713; P=1.022e-08] 12.52 [ 8.23, 16.80]
GOBS 22.40 [ -6.28, 51.09]
IMH -6.43 [-89.49, 76.63]
UNICAMP 35.70 [-21.92, 93.32]
Meth-CT
MIRECC 6.79 [-42.68, 56.26]
UKBB nonEUR -38.41 [-74.01, -2.80]
OSAKA 15.60 [-22.95, 54.15]
PING Non European -1.36 [-42.11, 39.39]
Non European Cohorts [N=2789; P=0.5793] 4.43 [-11.24, 20.11]
Beta [Lower 95% CI,Upper 95% CI]
-100
-50
0
50
100
150
mm^2/effect allele

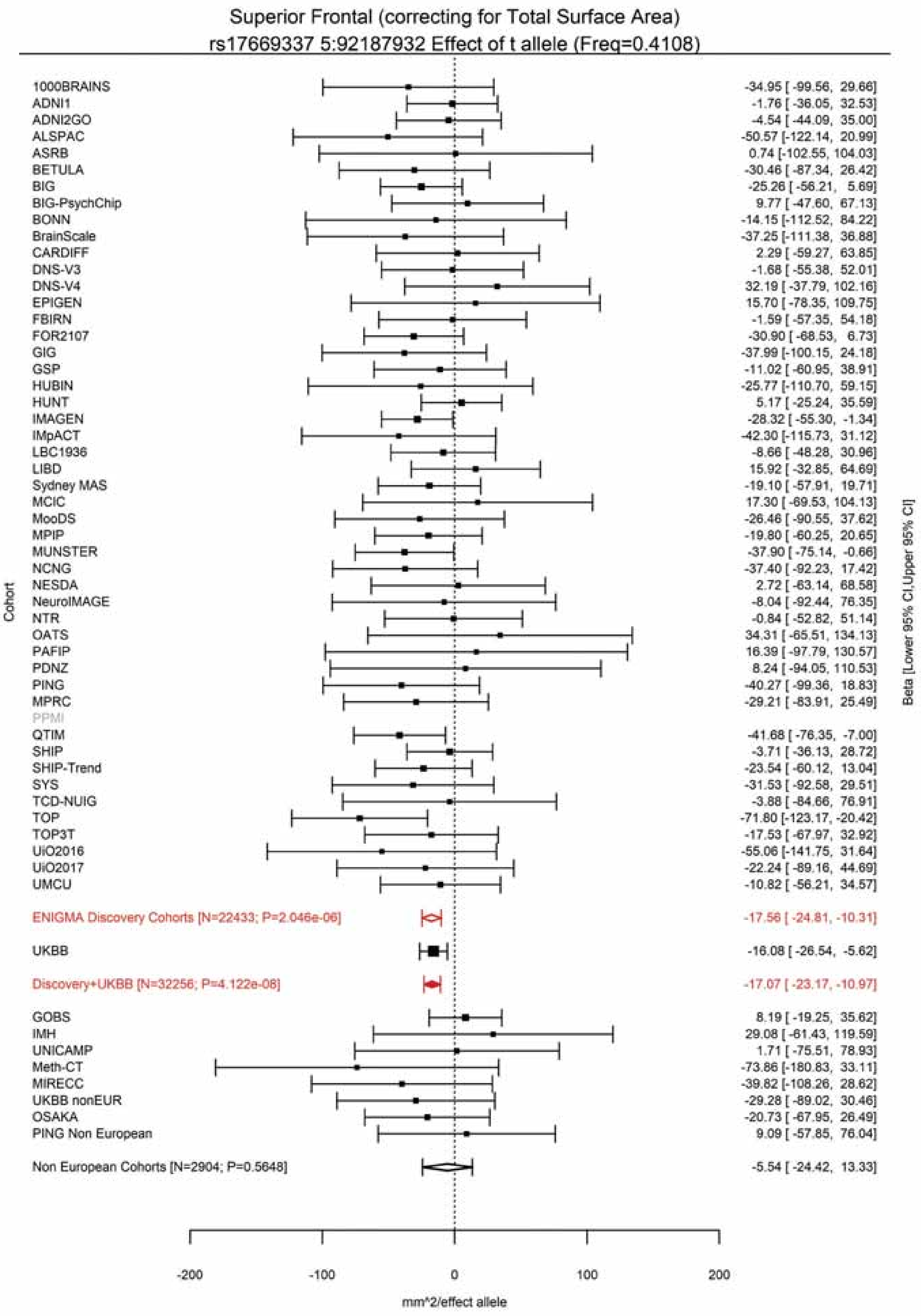

Superior Frontal (correcting for Total Surface Area)
rs17669337 5:92187932 Effect of t allele (Freq=0.4108)
Cohort
Beta [Lower 95% CI,Upper 95% CI]
1000BRAINS -34.95 [ -99.56, 29.66]
ADNI1 -1.76 [ -36.05, 32.53]
ADNI2GO -4.54 [ -44.09, 35.00]
ALSPAC -50.57 [-122.14, 20.99]
ASRB 0.74 [-102.55, 104.03]
BETULA -30.46 [ -87.34, 26.42]
BIG -25.26 [ -56.21, 5.69]
BIG-PsychChip 9.77 [ -47.60, 67.13]
BONN -14.15 [-112.52, 84.22]
BrainScale -37.25 [-111.38, 36.88]
CARDIFF 2.29 [ -59.27, 63.85]
DNS-V3 -1.68 [ -55.38, 52.01]
DNS-V4 32.19 [ -37.79, 102.16]
EPIGEN 15.70 [ -78.35, 109.75]
FBIRN -1.59 [ -57.35, 54.18]
FOR2107 -30.90 [ -68.53, 6.73]
GIG -37.99 [-100.15, 24.18]
GSP -11.02 [ -60.95, 38.91]
HUBIN -25.77 [-110.70, 59.15]
HUNT 5.17 [ -25.24, 35.59]
IMAGEN -28.32 [ -55.30, -1.34]
IMpACT -42.30 [-115.73, 31.12]
LBC1936 -8.66 [ -48.28, 30.96]
LIBD 15.92 [ -32.85, 64.69]
Sydney MAS -19.10 [ -57.91, 19.71]
MCIC 17.30 [ -69.53, 104.13]
MooDS -26.46 [ -90.55, 37.62]
MPIP -19.80 [ -60.25, 20.65]
MUNSTER -37.90 [ -75.14, -0.66]
NCNG -37.40 [ -92.23, 17.42]
NESDA 2.72 [ -63.14, 68.58]
NeuroIMAGE -8.04 [ -92.44, 76.35]
NTR -0.84 [ -52.82, 51.14]
OATS 34.31 [ -65.51, 134.13]
PAFIP 16.39 [ -97.79, 130.57]
PDNZ 8.24 [ -94.05, 110.53]
PING -40.27 [ -99.36, 18.83]
MPRC -29.21 [ -83.91, 25.49]
PPMI
QTIM -41.68 [ -76.35, -7.00]
SHIP -3.71 [ -36.13, 28.72]
SHIP-Trend -23.54 [ -60.12, 13.04]
SYS -31.53 [ -92.58, 29.51]
TCD-NUIG -3.88 [ -84.66, 76.91]
TOP -71.80 [-123.17, -20.42]
TOP3T -17.53 [ -67.97, 32.92]
UiO2016 -55.06 [-141.75, 31.64]
UiO2017 -22.24 [ -89.16, 44.69]
UMCU -10.82 [ -56.21, 34.57]
ENIGMA Discovery Cohorts [N=22433; P=2.046e-06] -17.56 [ -24.81, -10.31]
UKBB -16.08 [ -26.54, -5.62]
Discovery+UKBB [N=32256; P=4.122e-08] -17.07 [ -23.17, -10.97]
GOBS 8.19 [ -19.25, 35.62]
IMH 29.08 [ -61.43, 119.59]
UNICAMP 1.71 [ -75.51, 78.93]
Meth-CT -73.86 [-180.83, 33.11]
MIRECC -39.82 [-108.26, 28.62]
UKBB nonEUR -29.28 [ -89.02, 30.46]
OSAKA -20.73 [ -67.95, 26.49]
PING Non European 9.09 [ -57.85, 76.04]
Non European Cohorts [N=2904; P=0.5648] -5.54 [ -24.42, 13.33]
-200
-100
0
100
200
mm^2/effect allele

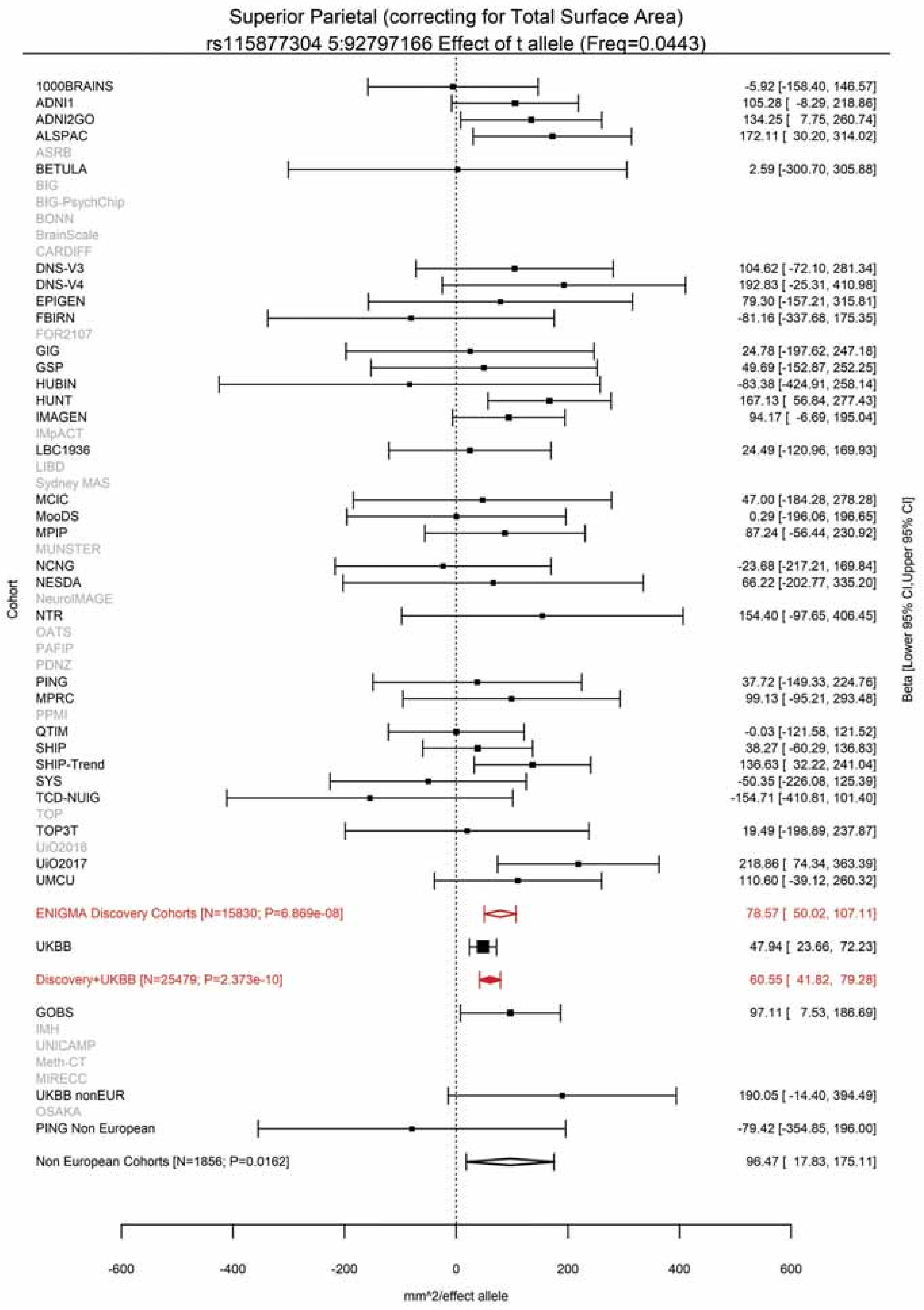

Superior Parietal (correcting for Total Surface Area)
rs115877304 5:92797166 Effect of t allele (Freq=0.0443)
Cohort
1000BRAINS -5.92 [-158.40, 146.57]
ADNI1 105.28 [ -8.29, 218.86]
ADNI2GO 134.25 [ 7.75, 260.74]
ALSPAC 172.11 [ 30.20, 314.02]
ASRB
BETULA 2.59 [-300.70, 305.88]
BIG
BIG-PsychChip
BONN
BrainScale
CARDIFF
DNS-V3 104.62 [ -72.10, 281.34]
DNS-V4 192.83 [ -25.31, 410.98]
EPIGEN 79.30 [-157.21, 315.81]
FBIRN -81.16 [-337.68, 175.35]
FOR2107
GIG 24.78 [-197.62, 247.18]
GSP 49.69 [-152.87, 252.25]
HUBIN -83.38 [-424.91, 258.14]
HUNT 167.13 [ 56.84, 277.43]
IMAGEN 94.17 [ -6.69, 195.04]
IMpACT
LBC1936 24.49 [-120.96, 169.93]
LIBD
Sydney MAS
MCIC 47.00 [-184.28, 278.28]
MooDS 0.29 [-196.06, 196.65]
MPIP 87.24 [ -56.44, 230.92]
MUNSTER
NCNG -23.68 [-217.21, 169.84]
NESDA 66.22 [-202.77, 335.20]
NeuroIMAGE
NTR 154.40 [ -97.65, 406.45]
OATS
PAFIP
PDNZ
PING 37.72 [-149.33, 224.76]
MPRC 99.13 [ -95.21, 293.48]
PPMI
QTIM -0.03 [-121.58, 121.52]
SHIP 38.27 [ -60.29, 136.83]
SHIP-Trend 136.63 [ 32.22, 241.04]
SYS -50.35 [-226.08, 125.39]
TCD-NUIG -154.71 [-410.81, 101.40]
TOP
TOP3T 19.49 [-198.89, 237.87]
UiO2016
UiO2017 218.86 [ 74.34, 363.39]
UMCU 110.60 [ -39.12, 260.32]
ENIGMA Discovery Cohorts [N=15830; P=6.869e-08] 78.57 [ 50.02, 107.11]
UKBB 47.94 [ 23.66, 72.23]
Discovery+UKBB [N=25479; P=2.373e-10] 60.55 [ 41.82, 79.28]
GOBS 97.11 [ 7.53, 186.69]
IMH
UNICAMP
Meth-CT
MIRECC
UKBB nonEUR 190.05 [ -14.40, 394.49]
OSAKA
PING Non European -79.42 [-354.85, 196.00]
Non European Cohorts [N=1856; P=0.0162] 96.47 [ 17.83, 175.11]
-600 -400 -200 0 200 400 600
mm^2/effect allele
Beta [Lower 95% CI,Upper 95% CI]

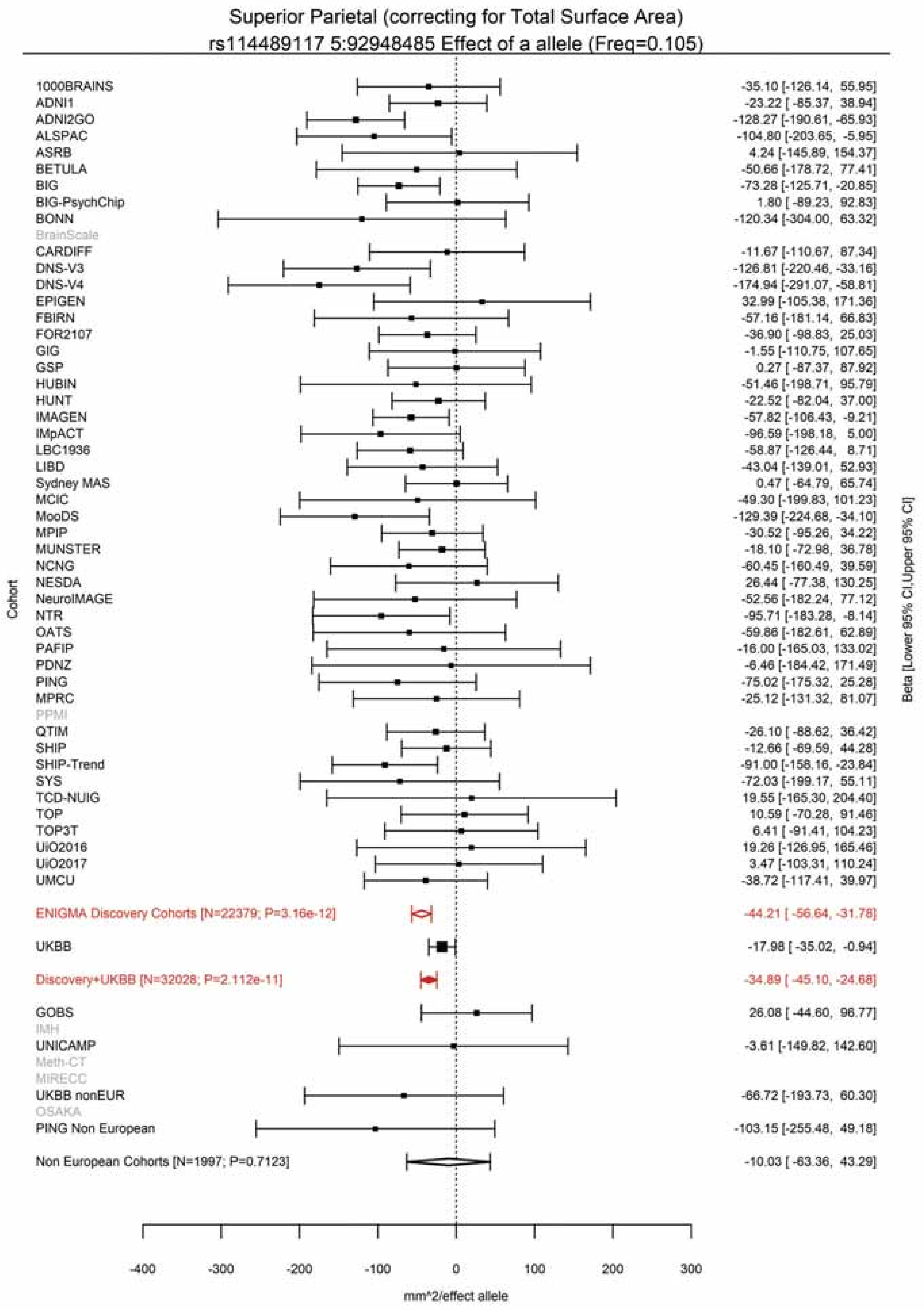

Superior Parietal (correcting for Total Surface Area)
rs114489117 5:92948485 Effect of a allele (Freq=0.105)
Cohort
1000BRAINS -35.10 [-126.14, 55.95]
ADNI1 -23.22 [ -85.37, 38.94]
ADNI2GO -128.27 [-190.61, -65.93]
ALSPAC -104.80 [-203.65, -5.95]
ASRB 4.24 [-145.89, 154.37]
BETULA -50.66 [-178.72, 77.41]
BIG -73.28 [-125.71, -20.85]
BIG-PsychChip 1.80 [ -89.23, 92.83]
BONN -120.34 [-304.00, 63.32]
BrainScale
CARDIFF -11.67 [-110.67, 87.34]
DNS-V3 -126.81 [-220.46, -33.16]
DNS-V4 -174.94 [-291.07, -58.81]
EPIGEN 32.99 [-105.38, 171.36]
FBIRN -57.16 [-181.14, 66.83]
FOR2107 -36.90 [ -98.83, 25.03]
GIG -1.55 [-110.75, 107.65]
GSP 0.27 [ -87.37, 87.92]
HUBIN -51.46 [-198.71, 95.79]
HUNT -22.52 [ -82.04, 37.00]
IMAGEN -57.82 [-106.43, -9.21]
IMpACT -96.59 [-198.18, 5.00]
LBC1936 -58.87 [-126.44, 8.71]
LIBD -43.04 [-139.01, 52.93]
Sydney MAS 0.47 [ -64.79, 65.74]
MCIC -49.30 [-199.83, 101.23]
MooDS -129.39 [-224.68, -34.10]
MPIP -30.52 [ -95.26, 34.22]
MUNSTER -18.10 [ -72.98, 36.78]
NCNG -60.45 [-160.49, 39.59]
NESDA 26.44 [ -77.38, 130.25]
NeuroIMAGE -52.56 [-182.24, 77.12]
NTR -95.71 [-183.28, -8.14]
OATS -59.86 [-182.61, 62.89]
PAFIP -16.00 [-165.03, 133.02]
PDNZ -6.46 [-184.42, 171.49]
PING -75.02 [-175.32, 25.28]
MPRC -25.12 [-131.32, 81.07]
PPMI
QTIM -26.10 [ -88.62, 36.42]
SHIP -12.66 [ -69.59, 44.28]
SHIP-Trend -91.00 [-158.16, -23.84]
SYS -72.03 [-199.17, 55.11]
TCD-NUIG 19.55 [-165.30, 204.40]
TOP 10.59 [ -70.28, 91.46]
TOP3T 6.41 [ -91.41, 104.23]
UiO2016 19.26 [-126.95, 165.46]
UiO2017 3.47 [-103.31, 110.24]
UMCU -38.72 [-117.41, 39.97]
ENIGMA Discovery Cohorts [N=22379; P=3.16e-12] -44.21 [ -56.64, -31.78]
UKBB -17.98 [ -35.02, -0.94]
Discovery+UKBB [N=32028; P=2.112e-11] -34.89 [ -45.10, -24.68]
GOBS 26.08 [ -44.60, 96.77]
IMH
UNICAMP -3.61 [-149.82, 142.60]
Meth-CT
MIRECC
UKBB nonEUR -66.72 [-193.73, 60.30]
OSAKA
PING Non European -103.15 [-255.48, 49.18]
Non European Cohorts [N=1997; P=0.7123] -10.03 [ -63.36, 43.29]
-400 -300 -200 -100 0 100 200 300
mm^2/effect allele
Beta [Lower 95% CI,Upper 95% CI]

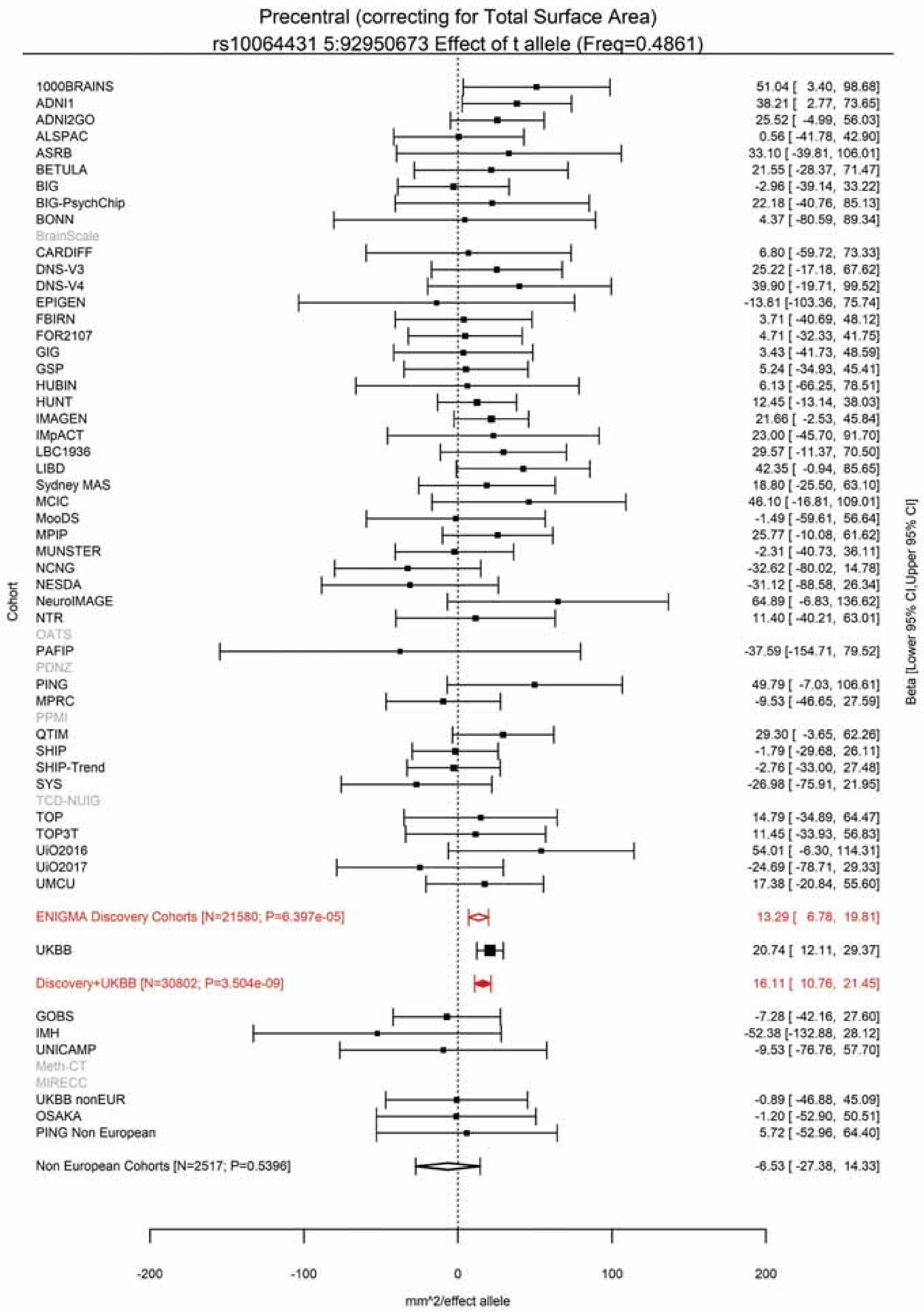
Precentral (correcting for Total Surface Area)
rs10064431 5:92950673 Effect of t allele (Freq=0.4861)
Cohort
1000BRAINS 51.04 [ 3.40, 98.68]
ADNI1 38.21 [ 2.77, 73.65]
ADNI2GO 25.52 [ -4.99, 56.03]
ALSPAC 0.56 [ -41.78, 42.90]
ASRB 33.10 [ -39.81, 106.01]
BETULA 21.55 [ -28.37, 71.47]
BIG -2.96 [ -39.14, 33.22]
BIG-PsychChip 22.18 [ -40.76, 85.13]
BONN 4.37 [ -80.59, 89.34]
BrainScale
CARDIFF 6.80 [ -59.72, 73.33]
DNS-V3 25.22 [ -17.18, 67.62]
DNS-V4 39.90 [ -19.71, 99.52]
EPIGEN -13.81 [-103.36, 75.74]
FBIRN 3.71 [ -40.69, 48.12]
FOR2107 4.71 [ -32.33, 41.75]
GIG 3.43 [ -41.73, 48.59]
GSP 5.24 [ -34.93, 45.41]
HUBIN 6.13 [ -66.25, 78.51]
HUNT 12.45 [ -13.14, 38.03]
IMAGEN 21.66 [ -2.53, 45.84]
IMpACT 23.00 [ -45.70, 91.70]
LBC1936 29.57 [ -11.37, 70.50]
LIBD 42.35 [ -0.94, 85.65]
Sydney MAS 18.80 [ -25.50, 63.10]
MCIC 46.10 [ -16.81, 109.01]
MooDS -1.49 [ -59.61, 56.64]
MPIP 25.77 [ -10.08, 61.62]
MUNSTER -2.31 [ -40.73, 36.11]
NCNG -32.62 [ -80.02, 14.78]
NESDA -31.12 [ -88.58, 26.34]
NeuroIMAGE 64.89 [ -6.83, 136.62]
NTR 11.40 [ -40.21, 63.01]
OATS
PAFIP -37.59 [-154.71, 79.52]
PDNZ
PING 49.79 [ -7.03, 106.61]
MPRC -9.53 [ -46.65, 27.59]
PPMI
QTIM 29.30 [ -3.65, 62.26]
SHIP -1.79 [ -29.68, 26.11]
SHIP-Trend -2.76 [ -33.00, 27.48]
SYS -26.98 [ -75.91, 21.95]
TCD-NUIG
TOP 14.79 [ -34.89, 64.47]
TOP3T 11.45 [ -33.93, 56.83]
UiO2016 54.01 [ -6.30, 114.31]
UiO2017 -24.69 [ -78.71, 29.33]
UMCU 17.38 [ -20.84, 55.60]
ENIGMA Discovery Cohorts [N=21580; P=6.397e-05] 13.29 [ 6.78, 19.81]
UKBB 20.74 [ 12.11, 29.37]
Discovery+UKBB [N=30802; P=3.504e-09] 16.11 [ 10.76, 21.45]
GOBS -7.28 [ -42.16, 27.60]
IMH -52.38 [-132.88, 28.12]
UNICAMP -9.53 [ -76.76, 57.70]
Meth-CT
MIRECC
UKBB nonEUR -0.89 [ -46.88, 45.09]
OSAKA -1.20 [ -52.90, 50.51]
PING Non European 5.72 [ -52.96, 64.40]
Non European Cohorts [N=2517; P=0.5396] -6.53 [ -27.38, 14.33]
Beta [Lower 95% CI,Upper 95% CI]
-200
-100
0
100
200
mm^2/effect allele

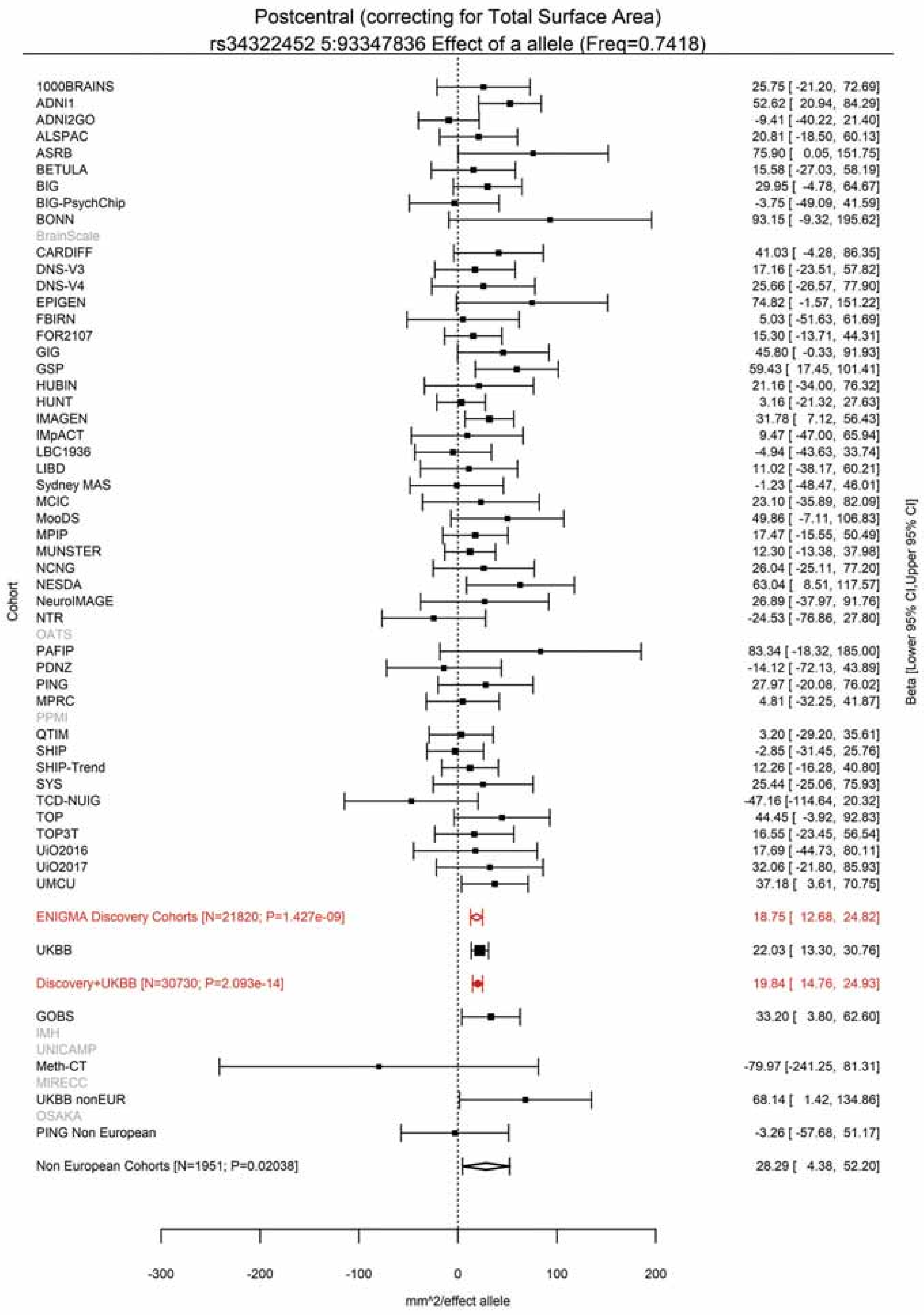

Postcentral (correcting for Total Surface Area)
rs34322452 5:93347836 Effect of a allele (Freq=0.7418)
Cohort
1000BRAINS 25.75 [ -21.20, 72.69]
ADNI1 52.62 [ 20.94, 84.29]
ADNI2GO -9.41 [ -40.22, 21.40]
ALSPAC 20.81 [ -18.50, 60.13]
ASRB 75.90 [ 0.05, 151.75]
BETULA 15.58 [ -27.03, 58.19]
BIG 29.95 [ -4.78, 64.67]
BIG-PsychChip -3.75 [ -49.09, 41.59]
BONN 93.15 [ -9.32, 195.62]
BrainScale
CARDIFF 41.03 [ -4.28, 86.35]
DNS-V3 17.16 [ -23.51, 57.82]
DNS-V4 25.66 [ -26.57, 77.90]
EPIGEN 74.82 [ -1.57, 151.22]
FBIRN 5.03 [ -51.63, 61.69]
FOR2107 15.30 [ -13.71, 44.31]
GIG 45.80 [ -0.33, 91.93]
GSP 59.43 [ 17.45, 101.41]
HUBIN 21.16 [ -34.00, 76.32]
HUNT 3.16 [ -21.32, 27.63]
IMAGEN 31.78 [ 7.12, 56.43]
IMpACT 9.47 [ -47.00, 65.94]
LBC1936 -4.94 [ -43.63, 33.74]
LIBD 11.02 [ -38.17, 60.21]
Sydney MAS -1.23 [ -48.47, 46.01]
MCIC 23.10 [ -35.89, 82.09]
MooDS 49.86 [ -7.11, 106.83]
MPIP 17.47 [ -15.55, 50.49]
MUNSTER 12.30 [ -13.38, 37.98]
NCNG 26.04 [ -25.11, 77.20]
NESDA 63.04 [ 8.51, 117.57]
NeuroIMAGE 26.89 [ -37.97, 91.76]
NTR -24.53 [ -76.86, 27.80]
OATS
PAFIP 83.34 [ -18.32, 185.00]
PDNZ -14.12 [ -72.13, 43.89]
PING 27.97 [ -20.08, 76.02]
MPRC 4.81 [ -32.25, 41.87]
PPMI
QTIM 3.20 [ -29.20, 35.61]
SHIP -2.85 [ -31.45, 25.76]
SHIP-Trend 12.26 [ -16.28, 40.80]
SYS 25.44 [ -25.06, 75.93]
TCD-NUIG -47.16 [-114.64, 20.32]
TOP 44.45 [ -3.92, 92.83]
TOP3T 16.55 [ -23.45, 56.54]
UiO2016 17.69 [ -44.73, 80.11]
UiO2017 32.06 [ -21.80, 85.93]
UMCU 37.18 [ 3.61, 70.75]
ENIGMA Discovery Cohorts [N=21820; P=1.427e-09] 18.75 [ 12.68, 24.82]
UKBB 22.03 [ 13.30, 30.76]
Discovery+UKBB [N=30730; P=2.093e-14] 19.84 [ 14.76, 24.93]
GOBS 33.20 [ 3.80, 62.60]
IMH
UNICAMP
Meth-CT -79.97 [-241.25, 81.31]
MIRECC
UKBB nonEUR 68.14 [ 1.42, 134.86]
OSAKA
PING Non European -3.26 [ -57.68, 51.17]
Non European Cohorts [N=1951; P=0.02038] 28.29 [ 4.38, 52.20]
-300 -200 -100 0 100 200
mm^2/effect allele
Beta [Lower 95% CI,Upper 95% CI]

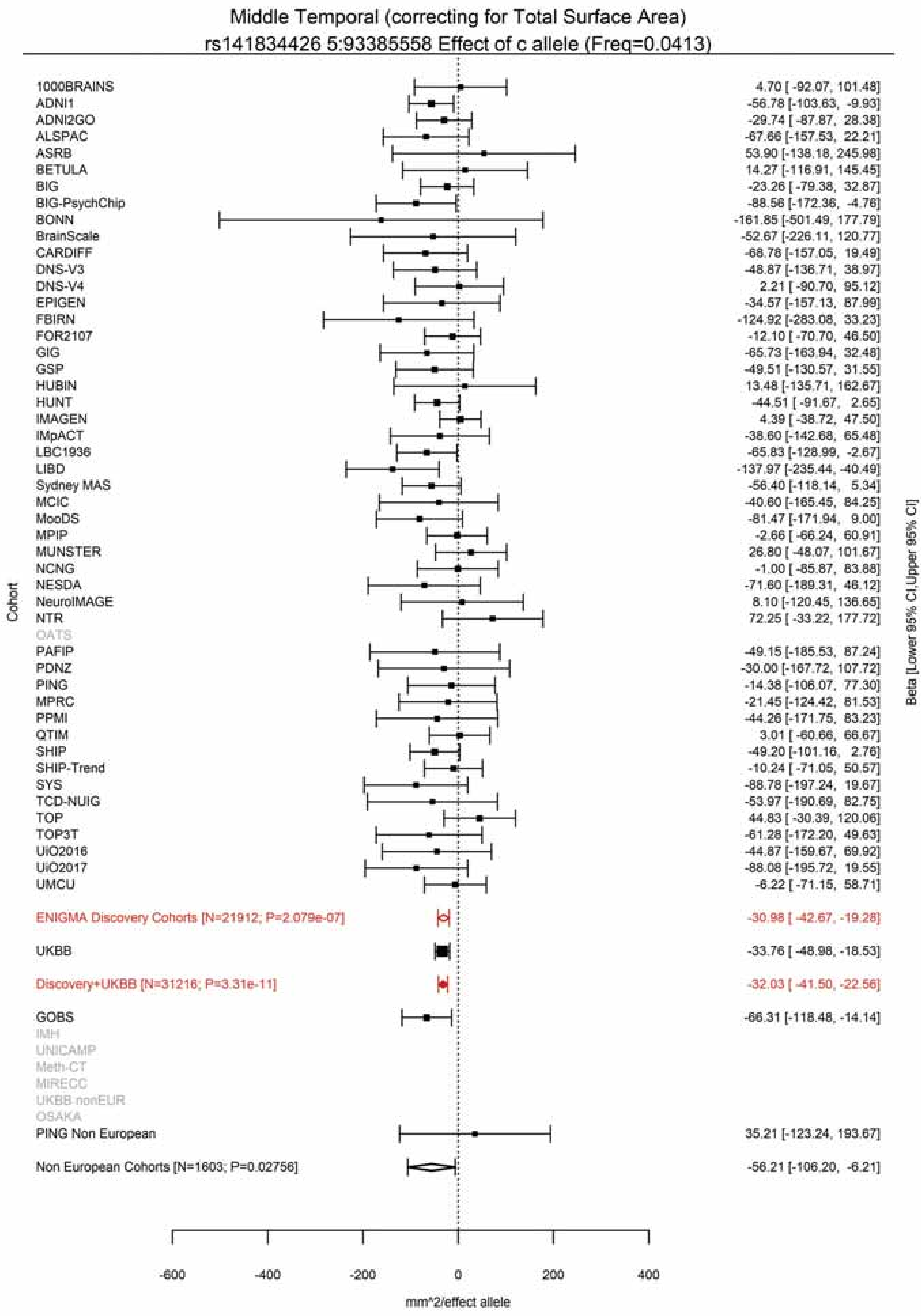

Middle Temporal (correcting for Total Surface Area)
rs141834426 5:93385558 Effect of c allele (Freq=0.0413)
Cohort
1000BRAINS 4.70 [ -92.07, 101.48]
ADNI1 -56.78 [-103.63, -9.93]
ADNI2GO -29.74 [ -87.87, 28.38]
ALSPAC -67.66 [-157.53, 22.21]
ASRB 53.90 [-138.18, 245.98]
BETULA 14.27 [-116.91, 145.45]
BIG -23.26 [ -79.38, 32.87]
BIG-PsychChip -88.56 [-172.36, -4.76]
BONN -161.85 [-501.49, 177.79]
BrainScale -52.67 [-226.11, 120.77]
CARDIFF -68.78 [-157.05, 19.49]
DNS-V3 -48.87 [-136.71, 38.97]
DNS-V4 2.21 [ -90.70, 95.12]
EPIGEN -34.57 [-157.13, 87.99]
FBIRN -124.92 [-283.08, 33.23]
FOR2107 -12.10 [ -70.70, 46.50]
GIG -65.73 [-163.94, 32.48]
GSP -49.51 [-130.57, 31.55]
HUBIN 13.48 [-135.71, 162.67]
HUNT -44.51 [ -91.67, 2.65]
IMAGEN 4.39 [ -38.72, 47.50]
IMpACT -38.60 [-142.68, 65.48]
LBC1936 -65.83 [-128.99, -2.67]
LIBD -137.97 [-235.44, -40.49]
Sydney MAS -56.40 [-118.14, 5.34]
MCIC -40.60 [-165.45, 84.25]
MooDS -81.47 [-171.94, 9.00]
MPIP -2.66 [ -66.24, 60.91]
MUNSTER 26.80 [ -48.07, 101.67]
NCNG -1.00 [ -85.87, 83.88]
NESDA -71.60 [-189.31, 46.12]
NeuroIMAGE 8.10 [-120.45, 136.65]
NTR 72.25 [ -33.22, 177.72]
OATS
PAFIP -49.15 [-185.53, 87.24]
PDNZ -30.00 [-167.72, 107.72]
PING -14.38 [-106.07, 77.30]
MPRC -21.45 [-124.42, 81.53]
PPMI -44.26 [-171.75, 83.23]
QTIM 3.01 [ -60.66, 66.67]
SHIP -49.20 [-101.16, 2.76]
SHIP-Trend -10.24 [ -71.05, 50.57]
SYS -88.78 [-197.24, 19.67]
TCD-NUIG -53.97 [-190.69, 82.75]
TOP 44.83 [ -30.39, 120.06]
TOP3T -61.28 [-172.20, 49.63]
UiO2016 -44.87 [-159.67, 69.92]
UiO2017 -88.08 [-195.72, 19.55]
UMCU -6.22 [ -71.15, 58.71]
ENIGMA Discovery Cohorts [N=21912; P=2.079e-07] -30.98 [ -42.67, -19.28]
UKBB -33.76 [ -48.98, -18.53]
Discovery+UKBB [N=31216; P=3.31e-11] -32.03 [ -41.50, -22.56]
GOBS -66.31 [-118.48, -14.14]
IMH
UNICAMP
Meth-CT
MIRECC
UKBB nonEUR
OSAKA
PING Non European 35.21 [-123.24, 193.67]
Non European Cohorts [N=1603; P=0.02756] -56.21 [-106.20, -6.21]
-600
-400
-200
0
200
400
mm^2/effect allele
Beta [Lower 95% CI,Upper 95% CI]

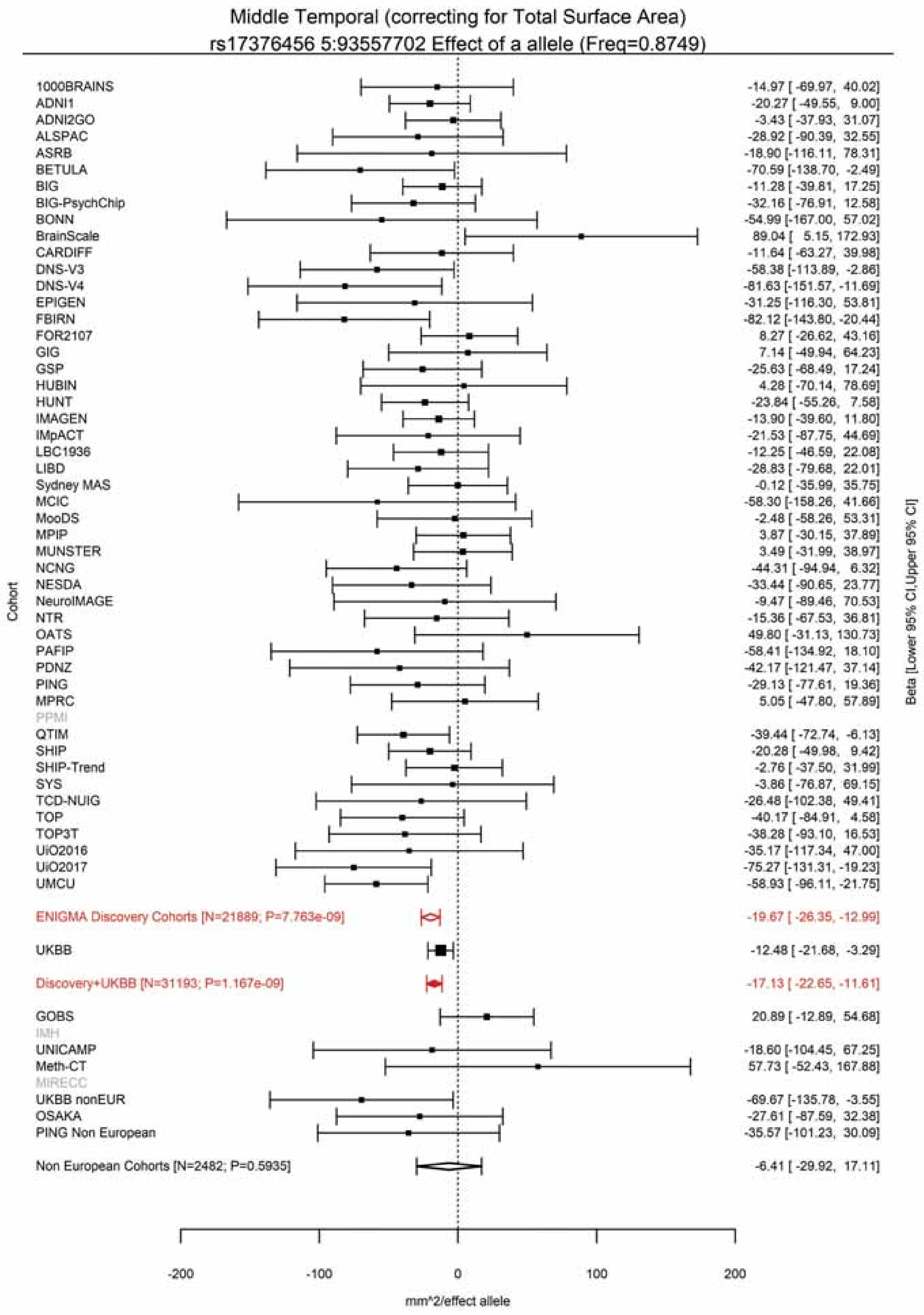

Middle Temporal (correcting for Total Surface Area)
rs17376456 5:93557702 Effect of a allele (Freq=0.8749)
Cohort
Beta [Lower 95% CI,Upper 95% CI]
1000BRAINS -14.97 [ -69.97, 40.02]
ADNI1 -20.27 [ -49.55, 9.00]
ADNI2GO -3.43 [ -37.93, 31.07]
ALSPAC -28.92 [ -90.39, 32.55]
ASRB -18.90 [-116.11, 78.31]
BETULA -70.59 [-138.70, -2.49]
BIG -11.28 [ -39.81, 17.25]
BIG-PsychChip -32.16 [ -76.91, 12.58]
BONN -54.99 [-167.00, 57.02]
BrainScale 89.04 [ 5.15, 172.93]
CARDIFF -11.64 [ -63.27, 39.98]
DNS-V3 -58.38 [-113.89, -2.86]
DNS-V4 -81.63 [-151.57, -11.69]
EPIGEN -31.25 [-116.30, 53.81]
FBIRN -82.12 [-143.80, -20.44]
FOR2107 8.27 [ -26.62, 43.16]
GIG 7.14 [ -49.94, 64.23]
GSP -25.63 [ -68.49, 17.24]
HUBIN 4.28 [ -70.14, 78.69]
HUNT -23.84 [ -55.26, 7.58]
IMAGEN -13.90 [ -39.60, 11.80]
IMpACT -21.53 [ -87.75, 44.69]
LBC1936 -12.25 [ -46.59, 22.08]
LIBD -28.83 [ -79.68, 22.01]
Sydney MAS -0.12 [ -35.99, 35.75]
MCIC -58.30 [-158.26, 41.66]
MooDS -2.48 [ -58.26, 53.31]
MPIP 3.87 [ -30.15, 37.89]
MUNSTER 3.49 [ -31.99, 38.97]
NCNG -44.31 [ -94.94, 6.32]
NESDA -33.44 [ -90.65, 23.77]
NeuroIMAGE -9.47 [ -89.46, 70.53]
NTR -15.36 [ -67.53, 36.81]
OATS 49.80 [ -31.13, 130.73]
PAFIP -58.41 [-134.92, 18.10]
PDNZ -42.17 [-121.47, 37.14]
PING -29.13 [ -77.61, 19.36]
MPRC 5.05 [ -47.80, 57.89]
PPMI
QTIM -39.44 [ -72.74, -6.13]
SHIP -20.28 [ -49.98, 9.42]
SHIP-Trend -2.76 [ -37.50, 31.99]
SYS -3.86 [ -76.87, 69.15]
TCD-NUIG -26.48 [-102.38, 49.41]
TOP -40.17 [ -84.91, 4.58]
TOP3T -38.28 [ -93.10, 16.53]
UiO2016 -35.17 [-117.34, 47.00]
UiO2017 -75.27 [-131.31, -19.23]
UMCU -58.93 [ -96.11, -21.75]
ENIGMA Discovery Cohorts [N=21889; P=7.763e-09] -19.67 [ -26.35, -12.99]
UKBB -12.48 [ -21.68, -3.29]
Discovery+UKBB [N=31193; P=1.167e-09] -17.13 [ -22.65, -11.61]
GOBS 20.89 [ -12.89, 54.68]
IMH
UNICAMP -18.60 [-104.45, 67.25]
Meth-CT 57.73 [ -52.43, 167.88]
MIRECC
UKBB nonEUR -69.67 [-135.78, -3.55]
OSAKA -27.61 [ -87.59, 32.38]
PING Non European -35.57 [-101.23, 30.09]
Non European Cohorts [N=2482; P=0.5935] -6.41 [ -29.92, 17.11]
-200
-100
0
100
200
mm^2/effect allele

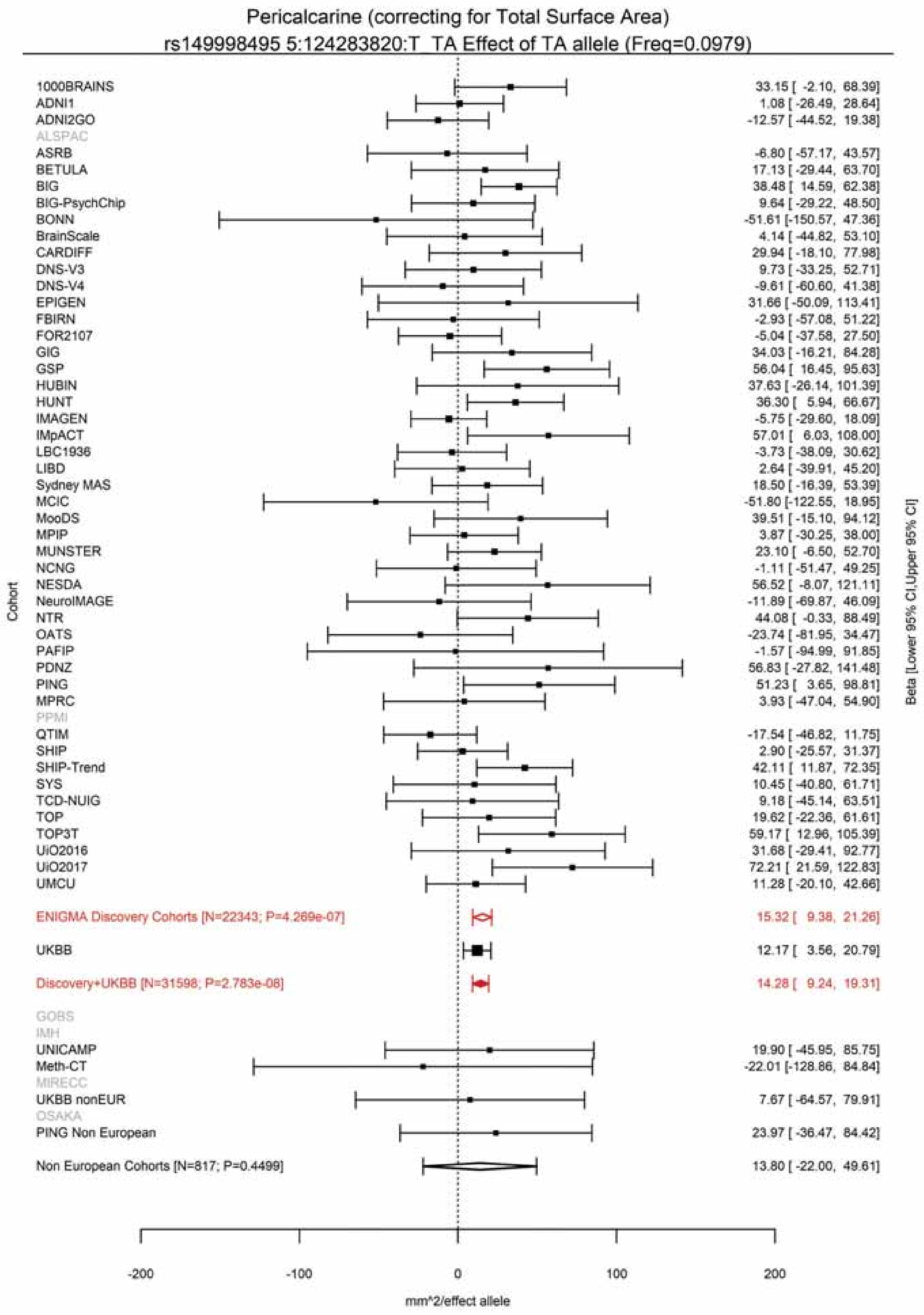
Pericalcarine (correcting for Total Surface Area)
rs149998495 5:124283820:T_TA Effect of TA allele (Freq=0.0979)
1000BRAINS 33.15 [ -2.10, 68.39]
ADNI1 1.08 [ -26.49, 28.64]
ADNI2GO -12.57 [ -44.52, 19.38]
ALSPAC
ASRB -6.80 [ -57.17, 43.57]
BETULA 17.13 [ -29.44, 63.70]
BIG 38.48 [ 14.59, 62.38]
BIG-PsychChip 9.64 [ -29.22, 48.50]
BONN -51.61 [-150.57, 47.36]
BrainScale 4.14 [ -44.82, 53.10]
CARDIFF 29.94 [ -18.10, 77.98]
DNS-V3 9.73 [ -33.25, 52.71]
DNS-V4 -9.61 [ -60.60, 41.38]
EPIGEN 31.66 [ -50.09, 113.41]
FBIRN -2.93 [ -57.08, 51.22]
FOR2107 -5.04 [ -37.58, 27.50]
GIG 34.03 [ -16.21, 84.28]
GSP 56.04 [ 16.45, 95.63]
HUBIN 37.63 [ -26.14, 101.39]
HUNT 36.30 [ 5.94, 66.67]
IMAGEN -5.75 [ -29.60, 18.09]
IMpACT 57.01 [ 6.03, 108.00]
LBC1936 -3.73 [ -38.09, 30.62]
LIBD 2.64 [ -39.91, 45.20]
Sydney MAS 18.50 [ -16.39, 53.39]
MCIC -51.80 [-122.55, 18.95]
MooDS 39.51 [ -15.10, 94.12]
MPIP 3.87 [ -30.25, 38.00]
MUNSTER 23.10 [ -6.50, 52.70]
NCNG -1.11 [ -51.47, 49.25]
NESDA 56.52 [ -8.07, 121.11]
NeuroIMAGE -11.89 [ -69.87, 46.09]
NTR 44.08 [ -0.33, 88.49]
OATS -23.74 [ -81.95, 34.47]
PAFIP -1.57 [ -94.99, 91.85]
PDNZ 56.83 [ -27.82, 141.48]
PING 51.23 [ 3.65, 98.81]
MPRC 3.93 [ -47.04, 54.90]
PPMI
QTIM -17.54 [ -46.82, 11.75]
SHIP 2.90 [ -25.57, 31.37]
SHIP-Trend 42.11 [ 11.87, 72.35]
SYS 10.45 [ -40.80, 61.71]
TCD-NUIG 9.18 [ -45.14, 63.51]
TOP 19.62 [ -22.36, 61.61]
TOP3T 59.17 [ 12.96, 105.39]
UiO2016 31.68 [ -29.41, 92.77]
UiO2017 72.21 [ 21.59, 122.83]
UMCU 11.28 [ -20.10, 42.66]
ENIGMA Discovery Cohorts [N=22343; P=4.269e-07] 15.32 [ 9.38, 21.26]
UKBB 12.17 [ 3.56, 20.79]
Discovery+UKBB [N=31598; P=2.783e-08] 14.28 [ 9.24, 19.31]
GOBS
IMH
UNICAMP 19.90 [ -45.95, 85.75]
Meth-CT -22.01 [-128.86, 84.84]
MIRECC
UKBB nonEUR 7.67 [ -64.57, 79.91]
OSAKA
PING Non European 23.97 [ -36.47, 84.42]
Non European Cohorts [N=817; P=0.4499] 13.80 [ -22.00, 49.61]
Cohort
Beta [Lower 95% CI,Upper 95% CI]
-200
-100
0
100
200
mm^2/effect allele

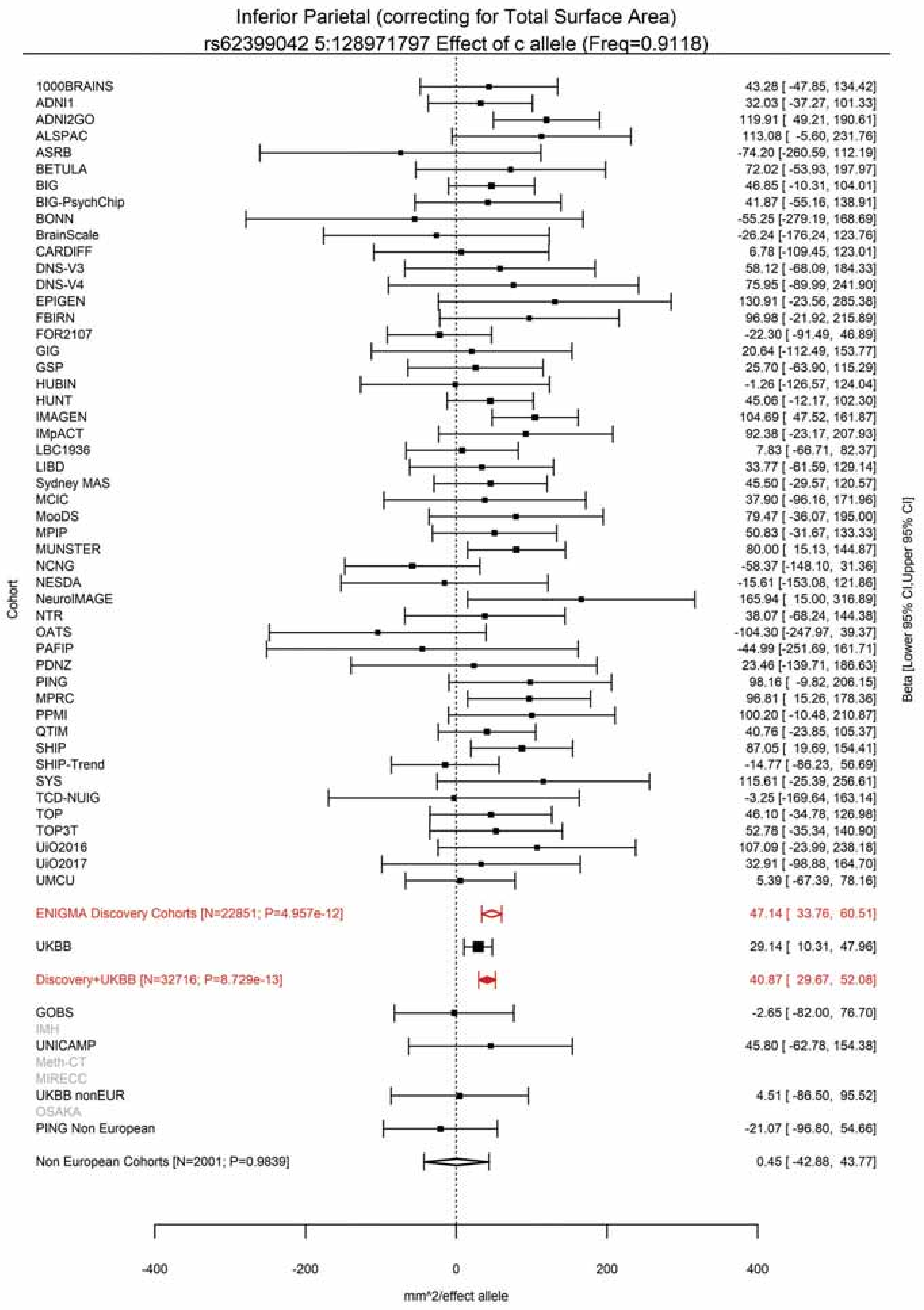

Inferior Parietal (correcting for Total Surface Area)
rs62399042 5:128971797 Effect of c allele (Freq=0.9118)
Cohort
1000BRAINS 43.28 [ -47.85, 134.42]
ADNI1 32.03 [ -37.27, 101.33]
ADNI2GO 119.91 [ 49.21, 190.61]
ALSPAC 113.08 [ -5.60, 231.76]
ASRB -74.20 [-260.59, 112.19]
BETULA 72.02 [ -53.93, 197.97]
BIG 46.85 [ -10.31, 104.01]
BIG-PsychChip 41.87 [ -55.16, 138.91]
BONN -55.25 [-279.19, 168.69]
BrainScale -26.24 [-176.24, 123.76]
CARDIFF 6.78 [-109.45, 123.01]
DNS-V3 58.12 [ -68.09, 184.33]
DNS-V4 75.95 [ -89.99, 241.90]
EPIGEN 130.91 [ -23.56, 285.38]
FBIRN 96.98 [ -21.92, 215.89]
FOR2107 -22.30 [ -91.49, 46.89]
GIG 20.64 [-112.49, 153.77]
GSP 25.70 [ -63.90, 115.29]
HUBIN -1.26 [-126.57, 124.04]
HUNT 45.06 [ -12.17, 102.30]
IMAGEN 104.69 [ 47.52, 161.87]
IMpACT 92.38 [ -23.17, 207.93]
LBC1936 7.83 [ -66.71, 82.37]
LIBD 33.77 [ -61.59, 129.14]
Sydney MAS 45.50 [ -29.57, 120.57]
MCIC 37.90 [ -96.16, 171.96]
MooDS 79.47 [ -36.07, 195.00]
MPIP 50.83 [ -31.67, 133.33]
MUNSTER 80.00 [ 15.13, 144.87]
NCNG -58.37 [-148.10, 31.36]
NESDA -15.61 [-153.08, 121.86]
NeuroIMAGE 165.94 [ 15.00, 316.89]
NTR 38.07 [ -68.24, 144.38]
OATS -104.30 [-247.97, 39.37]
PAFIP -44.99 [-251.69, 161.71]
PDNZ 23.46 [-139.71, 186.63]
PING 98.16 [ -9.82, 206.15]
MPRC 96.81 [ 15.26, 178.36]
PPMI 100.20 [ -10.48, 210.87]
QTIM 40.76 [ -23.85, 105.37]
SHIP 87.05 [ 19.69, 154.41]
SHIP-Trend -14.77 [ -86.23, 56.69]
SYS 115.61 [ -25.39, 256.61]
TCD-NUIG -3.25 [-169.64, 163.14]
TOP 46.10 [ -34.78, 126.98]
TOP3T 52.78 [ -35.34, 140.90]
UiO2016 107.09 [ -23.99, 238.18]
UiO2017 32.91 [ -98.88, 164.70]
UMCU 5.39 [ -67.39, 78.16]
ENIGMA Discovery Cohorts [N=22851; P=4.957e-12] 47.14 [ 33.76, 60.51]
UKBB 29.14 [ 10.31, 47.96]
Discovery+UKBB [N=32716; P=8.729e-13] 40.87 [ 29.67, 52.08]
GOBS -2.65 [ -82.00, 76.70]
IMH
UNICAMP 45.80 [ -62.78, 154.38]
Meth-CT
MIRECC
UKBB nonEUR 4.51 [ -86.50, 95.52]
OSAKA
PING Non European -21.07 [ -96.80, 54.66]
Non European Cohorts [N=2001; P=0.9839] 0.45 [ -42.88, 43.77]
Beta [Lower 95% CI,Upper 95% CI]
-400
-200
0
200
400
mm^2/effect allele

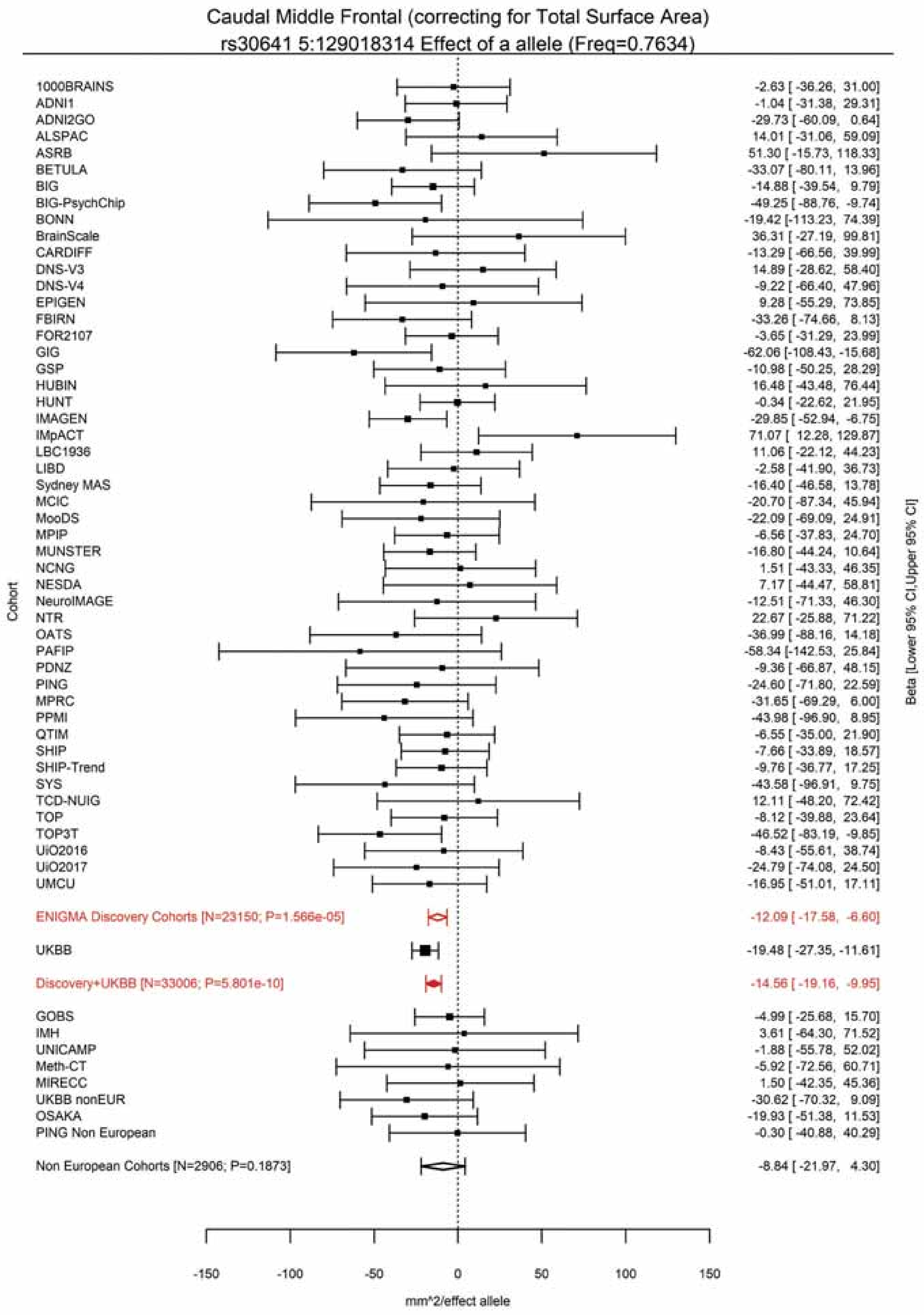

Caudal Middle Frontal (correcting for Total Surface Area)
rs30641 5:129018314 Effect of a allele (Freq=0.7634)
Cohort
Beta [Lower 95% CI,Upper 95% CI]
1000BRAINS -2.63 [ -36.26, 31.00]
ADNI1 -1.04 [ -31.38, 29.31]
ADNI2GO -29.73 [ -60.09, 0.64]
ALSPAC 14.01 [ -31.06, 59.09]
ASRB 51.30 [ -15.73, 118.33]
BETULA -33.07 [ -80.11, 13.96]
BIG -14.88 [ -39.54, 9.79]
BIG-PsychChip -49.25 [ -88.76, -9.74]
BONN -19.42 [-113.23, 74.39]
BrainScale 36.31 [ -27.19, 99.81]
CARDIFF -13.29 [ -66.56, 39.99]
DNS-V3 14.89 [ -28.62, 58.40]
DNS-V4 -9.22 [ -66.40, 47.96]
EPIGEN 9.28 [ -55.29, 73.85]
FBIRN -33.26 [ -74.66, 8.13]
FOR2107 -3.65 [ -31.29, 23.99]
GIG -62.06 [-108.43, -15.68]
GSP -10.98 [ -50.25, 28.29]
HUBIN 16.48 [ -43.48, 76.44]
HUNT -0.34 [ -22.62, 21.95]
IMAGEN -29.85 [ -52.94, -6.75]
IMpACT 71.07 [ 12.28, 129.87]
LBC1936 11.06 [ -22.12, 44.23]
LIBD -2.58 [ -41.90, 36.73]
Sydney MAS -16.40 [ -46.58, 13.78]
MCIC -20.70 [ -87.34, 45.94]
MooDS -22.09 [ -69.09, 24.91]
MPIP -6.56 [ -37.83, 24.70]
MUNSTER -16.80 [ -44.24, 10.64]
NCNG 1.51 [ -43.33, 46.35]
NESDA 7.17 [ -44.47, 58.81]
NeuroIMAGE -12.51 [ -71.33, 46.30]
NTR 22.67 [ -25.88, 71.22]
OATS -36.99 [ -88.16, 14.18]
PAFIP -58.34 [-142.53, 25.84]
PDNZ -9.36 [ -66.87, 48.15]
PING -24.60 [ -71.80, 22.59]
MPRC -31.65 [ -69.29, 6.00]
PPMI -43.98 [ -96.90, 8.95]
QTIM -6.55 [ -35.00, 21.90]
SHIP -7.66 [ -33.89, 18.57]
SHIP-Trend -9.76 [ -36.77, 17.25]
SYS -43.58 [ -96.91, 9.75]
TCD-NUIG 12.11 [ -48.20, 72.42]
TOP -8.12 [ -39.88, 23.64]
TOP3T -46.52 [ -83.19, -9.85]
UiO2016 -8.43 [ -55.61, 38.74]
UiO2017 -24.79 [ -74.08, 24.50]
UMCU -16.95 [ -51.01, 17.11]
ENIGMA Discovery Cohorts [N=23150; P=1.566e-05] -12.09 [ -17.58, -6.60]
UKBB -19.48 [ -27.35, -11.61]
Discovery+UKBB [N=33006; P=5.801e-10] -14.56 [ -19.16, -9.95]
GOBS -4.99 [ -25.68, 15.70]
IMH 3.61 [ -64.30, 71.52]
UNICAMP -1.88 [ -55.78, 52.02]
Meth-CT -5.92 [ -72.56, 60.71]
MIRECC 1.50 [ -42.35, 45.36]
UKBB nonEUR -30.62 [ -70.32, 9.09]
OSAKA -19.93 [ -51.38, 11.53]
PING Non European -0.30 [ -40.88, 40.29]
Non European Cohorts [N=2906; P=0.1873] -8.84 [ -21.97, 4.30]
-150
-100
-50
0
50
100
150
mm^2/effect allele

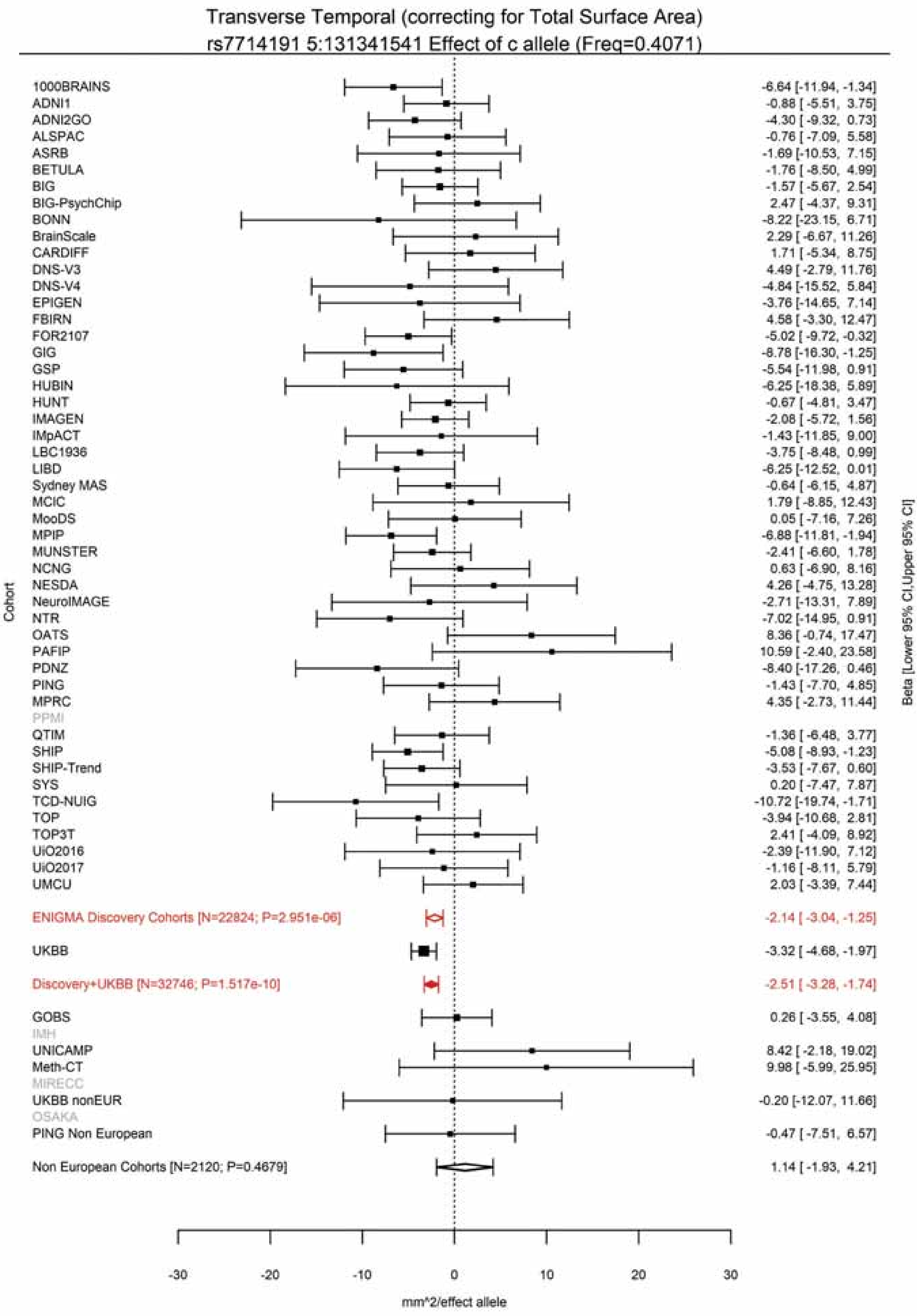
Transverse Temporal (correcting for Total Surface Area)
rs7714191 5:131341541 Effect of c allele (Freq=0.4071)
Cohort
1000BRAINS -6.64 [-11.94, -1.34]
ADNI1 -0.88 [ -5.51, 3.75]
ADNI2GO -4.30 [ -9.32, 0.73]
ALSPAC -0.76 [ -7.09, 5.58]
ASRB -1.69 [-10.53, 7.15]
BETULA -1.76 [ -8.50, 4.99]
BIG -1.57 [ -5.67, 2.54]
BIG-PsychChip 2.47 [ -4.37, 9.31]
BONN -8.22 [-23.15, 6.71]
BrainScale 2.29 [ -6.67, 11.26]
CARDIFF 1.71 [ -5.34, 8.75]
DNS-V3 4.49 [ -2.79, 11.76]
DNS-V4 -4.84 [-15.52, 5.84]
EPIGEN -3.76 [-14.65, 7.14]
FBIRN 4.58 [ -3.30, 12.47]
FOR2107 -5.02 [ -9.72, -0.32]
GIG -8.78 [-16.30, -1.25]
GSP -5.54 [-11.98, 0.91]
HUBIN -6.25 [-18.38, 5.89]
HUNT -0.67 [ -4.81, 3.47]
IMAGEN -2.08 [ -5.72, 1.56]
IMpACT -1.43 [-11.85, 9.00]
LBC1936 -3.75 [ -8.48, 0.99]
LIBD -6.25 [-12.52, 0.01]
Sydney MAS -0.64 [ -6.15, 4.87]
MCIC 1.79 [ -8.85, 12.43]
MooDS 0.05 [ -7.16, 7.26]
MPIP -6.88 [-11.81, -1.94]
MUNSTER -2.41 [ -6.60, 1.78]
NCNG 0.63 [ -6.90, 8.16]
NESDA 4.26 [ -4.75, 13.28]
NeuroIMAGE -2.71 [-13.31, 7.89]
NTR -7.02 [-14.95, 0.91]
OATS 8.36 [ -0.74, 17.47]
PAFIP 10.59 [ -2.40, 23.58]
PDNZ -8.40 [-17.26, 0.46]
PING -1.43 [ -7.70, 4.85]
MPRC 4.35 [ -2.73, 11.44]
PPMI
QTIM -1.36 [ -6.48, 3.77]
SHIP -5.08 [ -8.93, -1.23]
SHIP-Trend -3.53 [ -7.67, 0.60]
SYS 0.20 [ -7.47, 7.87]
TCD-NUIG -10.72 [-19.74, -1.71]
TOP -3.94 [-10.68, 2.81]
TOP3T 2.41 [ -4.09, 8.92]
UiO2016 -2.39 [-11.90, 7.12]
UiO2017 -1.16 [ -8.11, 5.79]
UMCU 2.03 [ -3.39, 7.44]
ENIGMA Discovery Cohorts [N=22824; P=2.951e-06] -2.14 [ -3.04, -1.25]
UKBB -3.32 [ -4.68, -1.97]
Discovery+UKBB [N=32746; P=1.517e-10] -2.51 [ -3.28, -1.74]
GOBS 0.26 [ -3.55, 4.08]
IMH
UNICAMP 8.42 [ -2.18, 19.02]
Meth-CT 9.98 [ -5.99, 25.95]
MIRECC
UKBB nonEUR -0.20 [-12.07, 11.66]
OSAKA
PING Non European -0.47 [ -7.51, 6.57]
Non European Cohorts [N=2120; P=0.4679] 1.14 [ -1.93, 4.21]
Beta [Lower 95% CI,Upper 95% CI]
-30
-20
-10
0
10
20
30
mm^2/effect allele

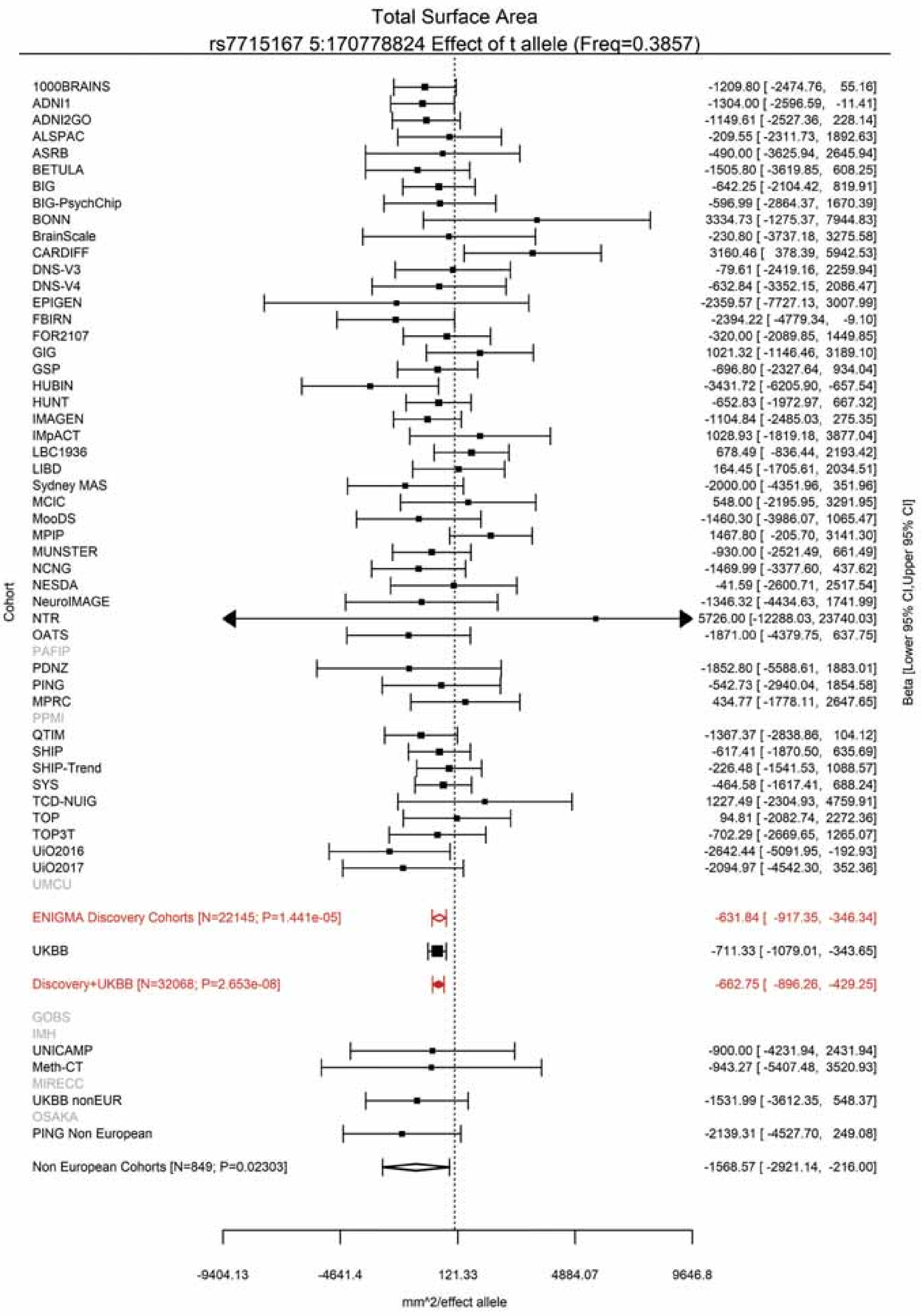
Total Surface Area
rs7715167 5:170778824 Effect of t allele (Freq=0.3857)
1000BRAINS -1209.80 [ -2474.76, 55.16]
ADNI1 -1304.00 [ -2596.59, -11.41]
ADNI2GO -1149.61 [ -2527.36, 228.14]
ALSPAC -209.55 [ -2311.73, 1892.63]
ASRB -490.00 [ -3625.94, 2645.94]
BETULA -1505.80 [ -3619.85, 608.25]
BIG -642.25 [ -2104.42, 819.91]
BIG-PsychChip -596.99 [ -2864.37, 1670.39]
BONN 3334.73 [ -1275.37, 7944.83]
BrainScale -230.80 [ -3737.18, 3275.58]
CARDIFF 3160.46 [ 378.39, 5942.53]
DNS-V3 -79.61 [ -2419.16, 2259.94]
DNS-V4 -632.84 [ -3352.15, 2086.47]
EPIGEN -2359.57 [ -7727.13, 3007.99]
FBIRN -2394.22 [ -4779.34, -9.10]
FOR2107 -320.00 [ -2089.85, 1449.85]
GIG 1021.32 [ -1146.46, 3189.10]
GSP -696.80 [ -2327.64, 934.04]
HUBIN -3431.72 [ -6205.90, -657.54]
HUNT -652.83 [ -1972.97, 667.32]
IMAGEN -1104.84 [ -2485.03, 275.35]
IMpACT 1028.93 [ -1819.18, 3877.04]
LBC1936 678.49 [ -836.44, 2193.42]
LIBD 164.45 [ -1705.61, 2034.51]
Sydney MAS -2000.00 [ -4351.96, 351.96]
MCIC 548.00 [ -2195.95, 3291.95]
MooDS -1460.30 [ -3986.07, 1065.47]
MPIP 1467.80 [ -205.70, 3141.30]
MUNSTER -930.00 [ -2521.49, 661.49]
NCNG -1469.99 [ -3377.60, 437.62]
NESDA -41.59 [ -2600.71, 2517.54]
NeuroIMAGE -1346.32 [ -4434.63, 1741.99]
NTR 5726.00 [-12288.03, 23740.03]
OATS -1871.00 [ -4379.75, 637.75]
PAFIP
PDNZ -1852.80 [ -5588.61, 1883.01]
PING -542.73 [ -2940.04, 1854.58]
MPRC 434.77 [ -1778.11, 2647.65]
PPMI
QTIM -1367.37 [ -2838.86, 104.12]
SHIP -617.41 [ -1870.50, 635.69]
SHIP-Trend -226.48 [ -1541.53, 1088.57]
SYS -464.58 [ -1617.41, 688.24]
TCD-NUIG 1227.49 [ -2304.93, 4759.91]
TOP 94.81 [ -2082.74, 2272.36]
TOP3T -702.29 [ -2669.65, 1265.07]
UiO2016 -2642.44 [ -5091.95, -192.93]
UiO2017 -2094.97 [ -4542.30, 352.36]
UMCU
ENIGMA Discovery Cohorts [N=22145; P=1.441e-05] -631.84 [ -917.35, -346.34]
UKBB -711.33 [ -1079.01, -343.65]
Discovery+UKBB [N=32068; P=2.653e-08] -662.75 [ -896.26, -429.25]
GOBS
IMH
UNICAMP -900.00 [ -4231.94, 2431.94]
Meth-CT -943.27 [ -5407.48, 3520.93]
MIRECC
UKBB nonEUR -1531.99 [ -3612.35, 548.37]
OSAKA
PING Non European -2139.31 [ -4527.70, 249.08]
Non European Cohorts [N=849; P=0.02303] -1568.57 [ -2921.14, -216.00]
Cohort
Beta [Lower 95% CI,Upper 95% CI]
-9404.13
-4641.4
121.33
4884.07
9646.8
mm^2/effect allele

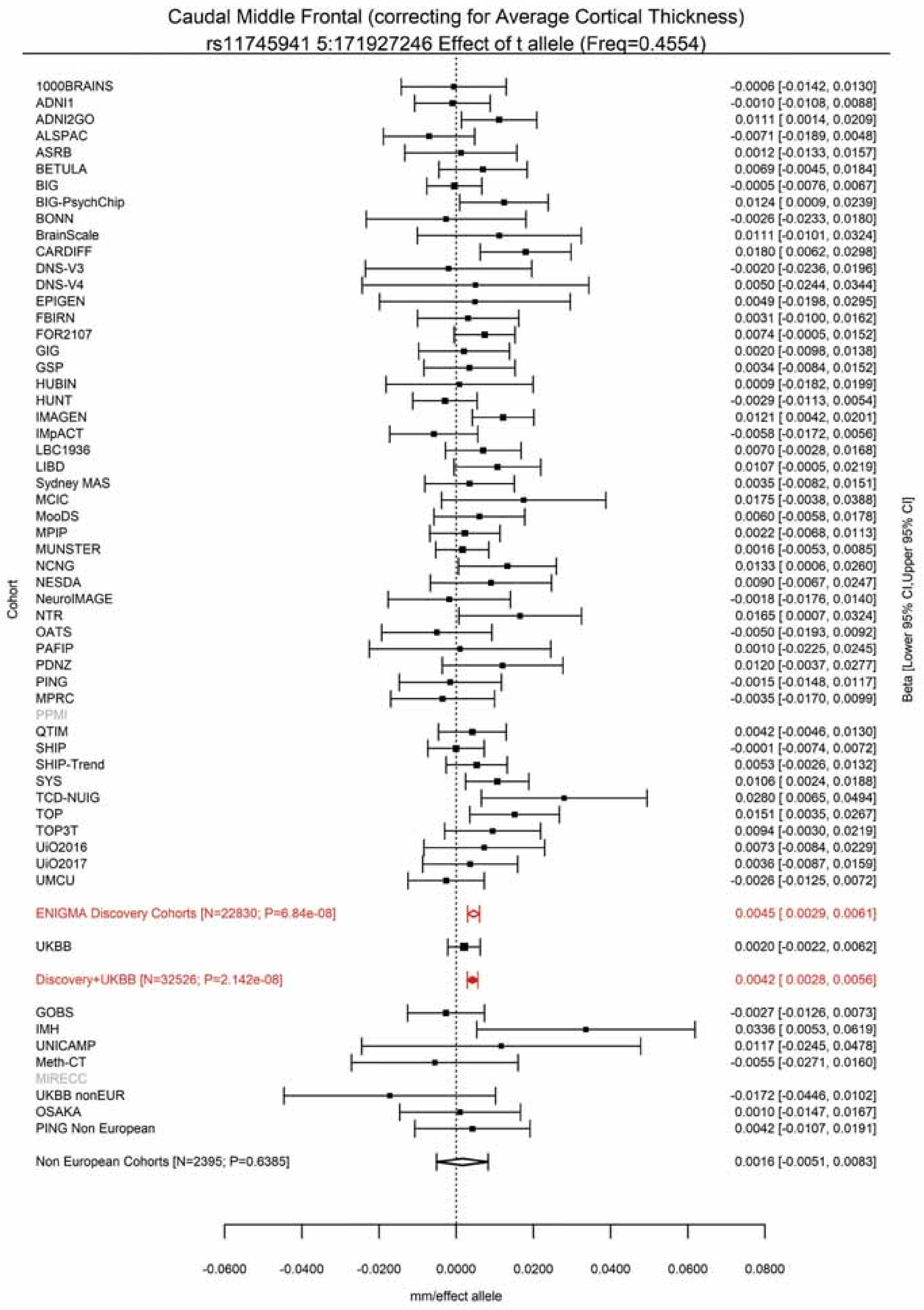
Caudal Middle Frontal (correcting for Average Cortical Thickness)
rs11745941 5:171927246 Effect of t allele (Freq=0.4554)
Cohort
1000BRAINS -0.0006 [-0.0142, 0.0130]
ADNI1 -0.0010 [-0.0108, 0.0088]
ADNI2GO 0.0111 [ 0.0014, 0.0209]
ALSPAC -0.0071 [-0.0189, 0.0048]
ASRB 0.0012 [-0.0133, 0.0157]
BETULA 0.0069 [-0.0045, 0.0184]
BIG -0.0005 [-0.0076, 0.0067]
BIG-PsychChip 0.0124 [ 0.0009, 0.0239]
BONN -0.0026 [-0.0233, 0.0180]
BrainScale 0.0111 [-0.0101, 0.0324]
CARDIFF 0.0180 [ 0.0062, 0.0298]
DNS-V3 -0.0020 [-0.0236, 0.0196]
DNS-V4 0.0050 [-0.0244, 0.0344]
EPIGEN 0.0049 [-0.0198, 0.0295]
FBIRN 0.0031 [-0.0100, 0.0162]
FOR2107 0.0074 [-0.0005, 0.0152]
GIG 0.0020 [-0.0098, 0.0138]
GSP 0.0034 [-0.0084, 0.0152]
HUBIN 0.0009 [-0.0182, 0.0199]
HUNT -0.0029 [-0.0113, 0.0054]
IMAGEN 0.0121 [ 0.0042, 0.0201]
IMpACT -0.0058 [-0.0172, 0.0056]
LBC1936 0.0070 [-0.0028, 0.0168]
LIBD 0.0107 [-0.0005, 0.0219]
Sydney MAS 0.0035 [-0.0082, 0.0151]
MCIC 0.0175 [-0.0038, 0.0388]
MooDS 0.0060 [-0.0058, 0.0178]
MPIP 0.0022 [-0.0068, 0.0113]
MUNSTER 0.0016 [-0.0053, 0.0085]
NCNG 0.0133 [ 0.0006, 0.0260]
NESDA 0.0090 [-0.0067, 0.0247]
NeuroIMAGE -0.0018 [-0.0176, 0.0140]
NTR 0.0165 [ 0.0007, 0.0324]
OATS -0.0050 [-0.0193, 0.0092]
PAFIP 0.0010 [-0.0225, 0.0245]
PDNZ 0.0120 [-0.0037, 0.0277]
PING -0.0015 [-0.0148, 0.0117]
MPRC -0.0035 [-0.0170, 0.0099]
PPMI
QTIM 0.0042 [-0.0046, 0.0130]
SHIP -0.0001 [-0.0074, 0.0072]
SHIP-Trend 0.0053 [-0.0026, 0.0132]
SYS 0.0106 [ 0.0024, 0.0188]
TCD-NUIG 0.0280 [ 0.0065, 0.0494]
TOP 0.0151 [ 0.0035, 0.0267]
TOP3T 0.0094 [-0.0030, 0.0219]
UiO2016 0.0073 [-0.0084, 0.0229]
UiO2017 0.0036 [-0.0087, 0.0159]
UMCU -0.0026 [-0.0125, 0.0072]
ENIGMA Discovery Cohorts [N=22830; P=6.84e-08] 0.0045 [ 0.0029, 0.0061]
UKBB 0.0020 [-0.0022, 0.0062]
Discovery+UKBB [N=32526; P=2.142e-08] 0.0042 [ 0.0028, 0.0056]
GOBS -0.0027 [-0.0126, 0.0073]
IMH 0.0336 [ 0.0053, 0.0619]
UNICAMP 0.0117 [-0.0245, 0.0478]
Meth-CT -0.0055 [-0.0271, 0.0160]
MIRECC
UKBB nonEUR -0.0172 [-0.0446, 0.0102]
OSAKA 0.0010 [-0.0147, 0.0167]
PING Non European 0.0042 [-0.0107, 0.0191]
Non European Cohorts [N=2395; P=0.6385] 0.0016 [-0.0051, 0.0083]
Beta [Lower 95% CI,Upper 95% CI]
-0.0600
-0.0400
-0.0200
0.0000
0.0200
0.0400
0.0600
0.0800
mm/effect allele

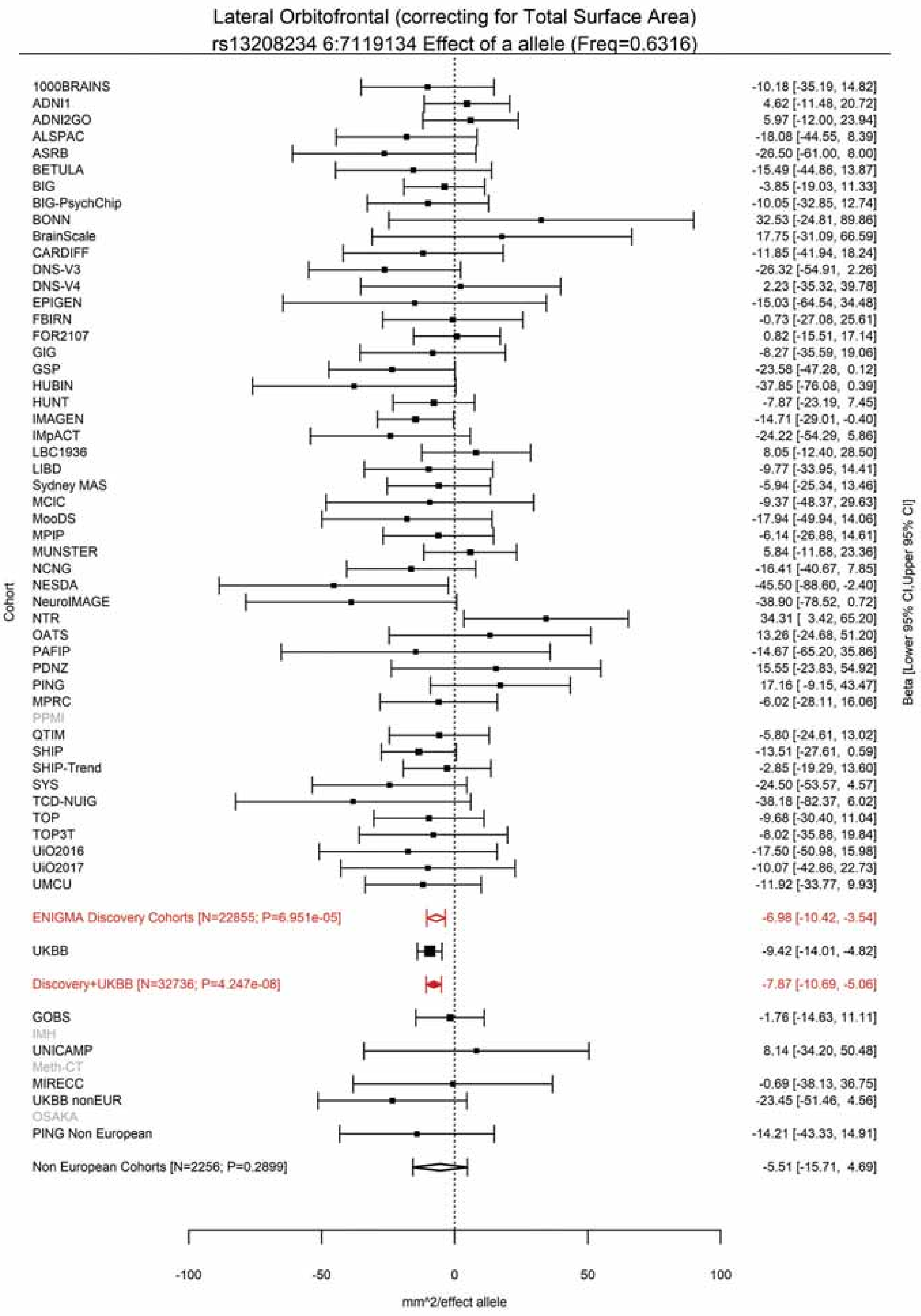
Lateral Orbitofrontal (correcting for Total Surface Area)
rs13208234 6:7119134 Effect of a allele (Freq=0.6316)
Cohort
Beta [Lower 95% CI,Upper 95% CI]
1000BRAINS -10.18 [-35.19, 14.82]
ADNI1 4.62 [-11.48, 20.72]
ADNI2GO 5.97 [-12.00, 23.94]
ALSPAC -18.08 [-44.55, 8.39]
ASRB -26.50 [-61.00, 8.00]
BETULA -15.49 [-44.86, 13.87]
BIG -3.85 [-19.03, 11.33]
BIG-PsychChip -10.05 [-32.85, 12.74]
BONN 32.53 [-24.81, 89.86]
BrainScale 17.75 [-31.09, 66.59]
CARDIFF -11.85 [-41.94, 18.24]
DNS-V3 -26.32 [-54.91, 2.26]
DNS-V4 2.23 [-35.32, 39.78]
EPIGEN -15.03 [-64.54, 34.48]
FBIRN -0.73 [-27.08, 25.61]
FOR2107 0.82 [-15.51, 17.14]
GIG -8.27 [-35.59, 19.06]
GSP -23.58 [-47.28, 0.12]
HUBIN -37.85 [-76.08, 0.39]
HUNT -7.87 [-23.19, 7.45]
IMAGEN -14.71 [-29.01, -0.40]
IMpACT -24.22 [-54.29, 5.86]
LBC1936 8.05 [-12.40, 28.50]
LIBD -9.77 [-33.95, 14.41]
Sydney MAS -5.94 [-25.34, 13.46]
MCIC -9.37 [-48.37, 29.63]
MooDS -17.94 [-49.94, 14.06]
MPIP -6.14 [-26.88, 14.61]
MUNSTER 5.84 [-11.68, 23.36]
NCNG -16.41 [-40.67, 7.85]
NESDA -45.50 [-88.60, -2.40]
NeuroIMAGE -38.90 [-78.52, 0.72]
NTR 34.31 [ 3.42, 65.20]
OATS 13.26 [-24.68, 51.20]
PAFIP -14.67 [-65.20, 35.86]
PDNZ 15.55 [-23.83, 54.92]
PING 17.16 [ -9.15, 43.47]
MPRC -6.02 [-28.11, 16.06]
PPMI
QTIM -5.80 [-24.61, 13.02]
SHIP -13.51 [-27.61, 0.59]
SHIP-Trend -2.85 [-19.29, 13.60]
SYS -24.50 [-53.57, 4.57]
TCD-NUIG -38.18 [-82.37, 6.02]
TOP -9.68 [-30.40, 11.04]
TOP3T -8.02 [-35.88, 19.84]
UiO2016 -17.50 [-50.98, 15.98]
UiO2017 -10.07 [-42.86, 22.73]
UMCU -11.92 [-33.77, 9.93]
ENIGMA Discovery Cohorts [N=22855; P=6.951e-05] -6.98 [-10.42, -3.54]
UKBB -9.42 [-14.01, -4.82]
Discovery+UKBB [N=32736; P=4.247e-08] -7.87 [-10.69, -5.06]
GOBS -1.76 [-14.63, 11.11]
IMH
UNICAMP 8.14 [-34.20, 50.48]
Meth-CT
MIRECC -0.69 [-38.13, 36.75]
UKBB nonEUR -23.45 [-51.46, 4.56]
OSAKA
PING Non European -14.21 [-43.33, 14.91]
Non European Cohorts [N=2256; P=0.2899] -5.51 [-15.71, 4.69]
-100
-50
0
50
100
mm^2/effect allele

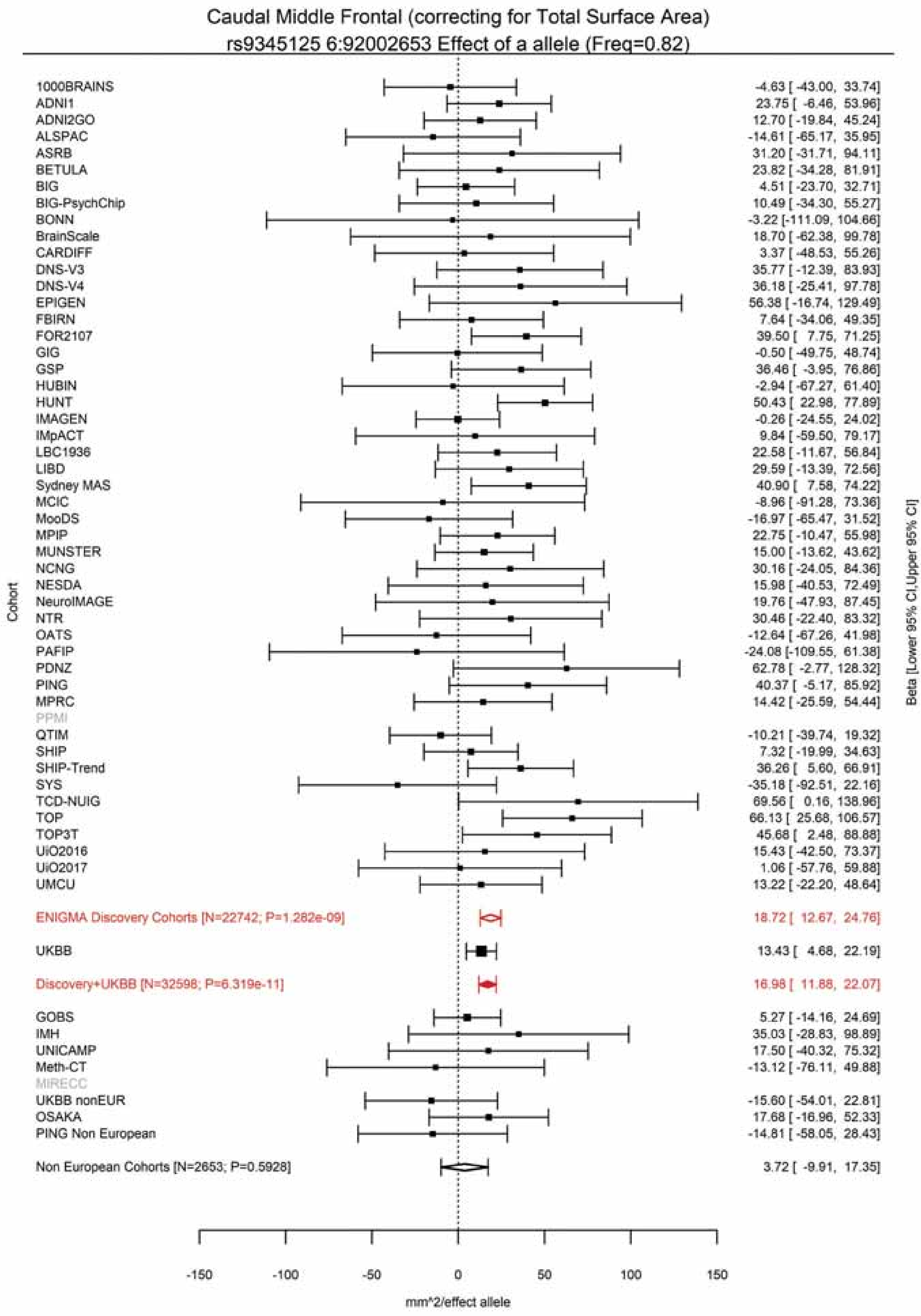

Caudal Middle Frontal (correcting for Total Surface Area)
rs9345125 6:92002653 Effect of a allele (Freq=0.82)
1000BRAINS -4.63 [ -43.00, 33.74]
ADNI1 23.75 [ -6.46, 53.96]
ADNI2GO 12.70 [ -19.84, 45.24]
ALSPAC -14.61 [ -65.17, 35.95]
ASRB 31.20 [ -31.71, 94.11]
BETULA 23.82 [ -34.28, 81.91]
BIG 4.51 [ -23.70, 32.71]
BIG-PsychChip 10.49 [ -34.30, 55.27]
BONN -3.22 [-111.09, 104.66]
BrainScale 18.70 [ -62.38, 99.78]
CARDIFF 3.37 [ -48.53, 55.26]
DNS-V3 35.77 [ -12.39, 83.93]
DNS-V4 36.18 [ -25.41, 97.78]
EPIGEN 56.38 [ -16.74, 129.49]
FBIRN 7.64 [ -34.06, 49.35]
FOR2107 39.50 [ 7.75, 71.25]
GIG -0.50 [ -49.75, 48.74]
GSP 36.46 [ -3.95, 76.86]
HUBIN -2.94 [ -67.27, 61.40]
HUNT 50.43 [ 22.98, 77.89]
IMAGEN -0.26 [ -24.55, 24.02]
IMpACT 9.84 [ -59.50, 79.17]
LBC1936 22.58 [ -11.67, 56.84]
LIBD 29.59 [ -13.39, 72.56]
Sydney MAS 40.90 [ 7.58, 74.22]
MCIC -8.96 [ -91.28, 73.36]
MooDS -16.97 [ -65.47, 31.52]
MPIP 22.75 [ -10.47, 55.98]
MUNSTER 15.00 [ -13.62, 43.62]
NCNG 30.16 [ -24.05, 84.36]
NESDA 15.98 [ -40.53, 72.49]
NeuroIMAGE 19.76 [ -47.93, 87.45]
NTR 30.46 [ -22.40, 83.32]
OATS -12.64 [ -67.26, 41.98]
PAFIP -24.08 [-109.55, 61.38]
PDNZ 62.78 [ -2.77, 128.32]
PING 40.37 [ -5.17, 85.92]
MPRC 14.42 [ -25.59, 54.44]
PPMI
QTIM -10.21 [ -39.74, 19.32]
SHIP 7.32 [ -19.99, 34.63]
SHIP-Trend 36.26 [ 5.60, 66.91]
SYS -35.18 [ -92.51, 22.16]
TCD-NUIG 69.56 [ 0.16, 138.96]
TOP 66.13 [ 25.68, 106.57]
TOP3T 45.68 [ 2.48, 88.88]
UiO2016 15.43 [ -42.50, 73.37]
UiO2017 1.06 [ -57.76, 59.88]
UMCU 13.22 [ -22.20, 48.64]
ENIGMA Discovery Cohorts [N=22742; P=1.282e-09] 18.72 [ 12.67, 24.76]
UKBB 13.43 [ 4.68, 22.19]
Discovery+UKBB [N=32598; P=6.319e-11] 16.98 [ 11.88, 22.07]
GOBS 5.27 [ -14.16, 24.69]
IMH 35.03 [ -28.83, 98.89]
UNICAMP 17.50 [ -40.32, 75.32]
Meth-CT -13.12 [ -76.11, 49.88]
MIRECC
UKBB nonEUR -15.60 [ -54.01, 22.81]
OSAKA 17.68 [ -16.96, 52.33]
PING Non European -14.81 [ -58.05, 28.43]
Non European Cohorts [N=2653; P=0.5928] 3.72 [ -9.91, 17.35]
Cohort
Beta [Lower 95% CI,Upper 95% CI]
-150
-100
-50
0
50
100
150
mm^2/effect allele

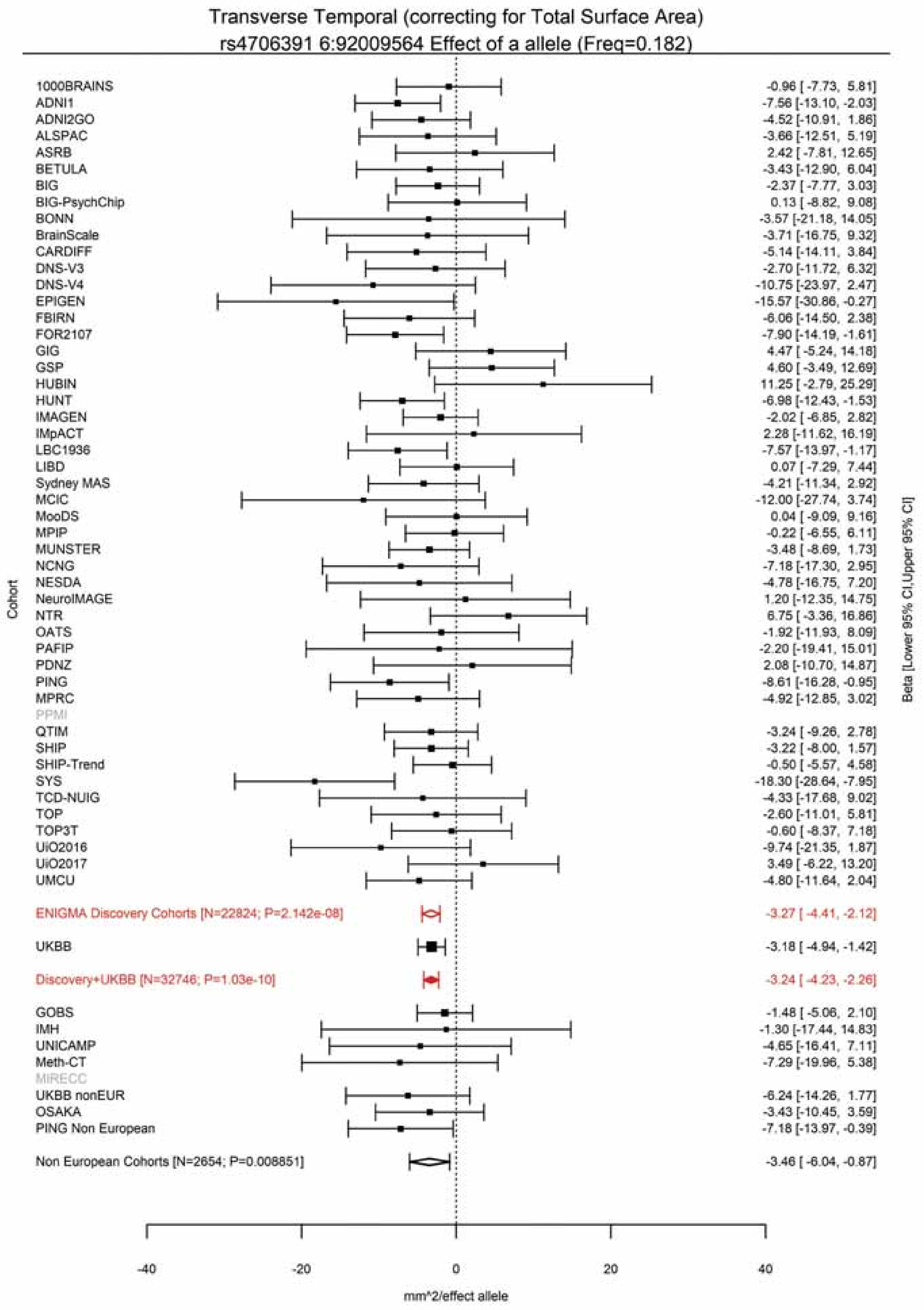
Transverse Temporal (correcting for Total Surface Area)
rs4706391 6:92009564 Effect of a allele (Freq=0.182)
Cohort
Beta [Lower 95% CI,Upper 95% CI]
1000BRAINS -0.96 [ -7.73, 5.81]
ADNI1 -7.56 [-13.10, -2.03]
ADNI2GO -4.52 [-10.91, 1.86]
ALSPAC -3.66 [-12.51, 5.19]
ASRB 2.42 [ -7.81, 12.65]
BETULA -3.43 [-12.90, 6.04]
BIG -2.37 [ -7.77, 3.03]
BIG-PsychChip 0.13 [ -8.82, 9.08]
BONN -3.57 [-21.18, 14.05]
BrainScale -3.71 [-16.75, 9.32]
CARDIFF -5.14 [-14.11, 3.84]
DNS-V3 -2.70 [-11.72, 6.32]
DNS-V4 -10.75 [-23.97, 2.47]
EPIGEN -15.57 [-30.86, -0.27]
FBIRN -6.06 [-14.50, 2.38]
FOR2107 -7.90 [-14.19, -1.61]
GIG 4.47 [ -5.24, 14.18]
GSP 4.60 [ -3.49, 12.69]
HUBIN 11.25 [ -2.79, 25.29]
HUNT -6.98 [-12.43, -1.53]
IMAGEN -2.02 [ -6.85, 2.82]
IMpACT 2.28 [-11.62, 16.19]
LBC1936 -7.57 [-13.97, -1.17]
LIBD 0.07 [ -7.29, 7.44]
Sydney MAS -4.21 [-11.34, 2.92]
MCIC -12.00 [-27.74, 3.74]
MooDS 0.04 [ -9.09, 9.16]
MPIP -0.22 [ -6.55, 6.11]
MUNSTER -3.48 [ -8.69, 1.73]
NCNG -7.18 [-17.30, 2.95]
NESDA -4.78 [-16.75, 7.20]
NeuroIMAGE 1.20 [-12.35, 14.75]
NTR 6.75 [ -3.36, 16.86]
OATS -1.92 [-11.93, 8.09]
PAFIP -2.20 [-19.41, 15.01]
PDNZ 2.08 [-10.70, 14.87]
PING -8.61 [-16.28, -0.95]
MPRC -4.92 [-12.85, 3.02]
PPMI
QTIM -3.24 [ -9.26, 2.78]
SHIP -3.22 [ -8.00, 1.57]
SHIP-Trend -0.50 [ -5.57, 4.58]
SYS -18.30 [-28.64, -7.95]
TCD-NUIG -4.33 [-17.68, 9.02]
TOP -2.60 [-11.01, 5.81]
TOP3T -0.60 [ -8.37, 7.18]
UiO2016 -9.74 [-21.35, 1.87]
UiO2017 3.49 [ -6.22, 13.20]
UMCU -4.80 [-11.64, 2.04]
ENIGMA Discovery Cohorts [N=22824; P=2.142e-08] -3.27 [ -4.41, -2.12]
UKBB -3.18 [ -4.94, -1.42]
Discovery+UKBB [N=32746; P=1.03e-10] -3.24 [ -4.23, -2.26]
GOBS -1.48 [ -5.06, 2.10]
IMH -1.30 [-17.44, 14.83]
UNICAMP -4.65 [-16.41, 7.11]
Meth-CT -7.29 [-19.96, 5.38]
MIRECC
UKBB nonEUR -6.24 [-14.26, 1.77]
OSAKA -3.43 [-10.45, 3.59]
PING Non European -7.18 [-13.97, -0.39]
Non European Cohorts [N=2654; P=0.008851] -3.46 [ -6.04, -0.87]
-40
-20
0
20
40
mm^2/effect allele

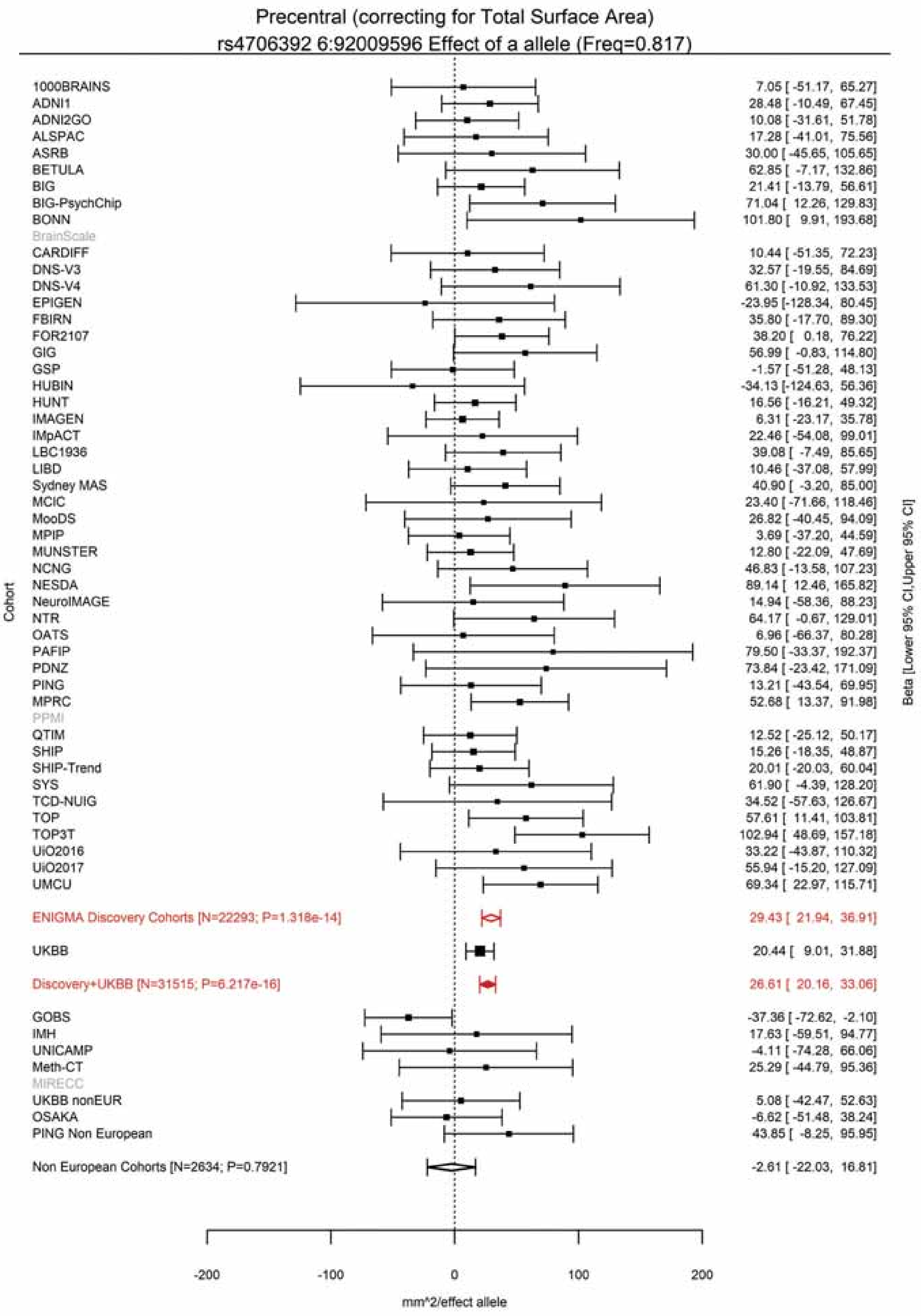
Precentral (correcting for Total Surface Area)
rs4706392 6:92009596 Effect of a allele (Freq=0.817)
Cohort
Beta [Lower 95% CI,Upper 95% CI]
1000BRAINS 7.05 [ -51.17, 65.27]
ADNI1 28.48 [ -10.49, 67.45]
ADNI2GO 10.08 [ -31.61, 51.78]
ALSPAC 17.28 [ -41.01, 75.56]
ASRB 30.00 [ -45.65, 105.65]
BETULA 62.85 [ -7.17, 132.86]
BIG 21.41 [ -13.79, 56.61]
BIG-PsychChip 71.04 [ 12.26, 129.83]
BONN 101.80 [ 9.91, 193.68]
BrainScale
CARDIFF 10.44 [ -51.35, 72.23]
DNS-V3 32.57 [ -19.55, 84.69]
DNS-V4 61.30 [ -10.92, 133.53]
EPIGEN -23.95 [-128.34, 80.45]
FBIRN 35.80 [ -17.70, 89.30]
FOR2107 38.20 [ 0.18, 76.22]
GIG 56.99 [ -0.83, 114.80]
GSP -1.57 [ -51.28, 48.13]
HUBIN -34.13 [-124.63, 56.36]
HUNT 16.56 [ -16.21, 49.32]
IMAGEN 6.31 [ -23.17, 35.78]
IMpACT 22.46 [ -54.08, 99.01]
LBC1936 39.08 [ -7.49, 85.65]
LIBD 10.46 [ -37.08, 57.99]
Sydney MAS 40.90 [ -3.20, 85.00]
MCIC 23.40 [ -71.66, 118.46]
MooDS 26.82 [ -40.45, 94.09]
MPIP 3.69 [ -37.20, 44.59]
MUNSTER 12.80 [ -22.09, 47.69]
NCNG 46.83 [ -13.58, 107.23]
NESDA 89.14 [ 12.46, 165.82]
NeuroIMAGE 14.94 [ -58.36, 88.23]
NTR 64.17 [ -0.67, 129.01]
OATS 6.96 [ -66.37, 80.28]
PAFIP 79.50 [ -33.37, 192.37]
PDNZ 73.84 [ -23.42, 171.09]
PING 13.21 [ -43.54, 69.95]
MPRC 52.68 [ 13.37, 91.98]
PPMI
QTIM 12.52 [ -25.12, 50.17]
SHIP 15.26 [ -18.35, 48.87]
SHIP-Trend 20.01 [ -20.03, 60.04]
SYS 61.90 [ -4.39, 128.20]
TCD-NUIG 34.52 [ -57.63, 126.67]
TOP 57.61 [ 11.41, 103.81]
TOP3T 102.94 [ 48.69, 157.18]
UiO2016 33.22 [ -43.87, 110.32]
UiO2017 55.94 [ -15.20, 127.09]
UMCU 69.34 [ 22.97, 115.71]
ENIGMA Discovery Cohorts [N=22293; P=1.318e-14] 29.43 [ 21.94, 36.91]
UKBB 20.44 [ 9.01, 31.88]
Discovery+UKBB [N=31515; P=6.217e-16] 26.61 [ 20.16, 33.06]
GOBS -37.36 [ -72.62, -2.10]
IMH 17.63 [ -59.51, 94.77]
UNICAMP -4.11 [ -74.28, 66.06]
Meth-CT 25.29 [ -44.79, 95.36]
MIRECC
UKBB nonEUR 5.08 [ -42.47, 52.63]
OSAKA -6.62 [ -51.48, 38.24]
PING Non European 43.85 [ -8.25, 95.95]
Non European Cohorts [N=2634; P=0.7921] -2.61 [ -22.03, 16.81]
-200
-100
0
100
200
mm^2/effect allele

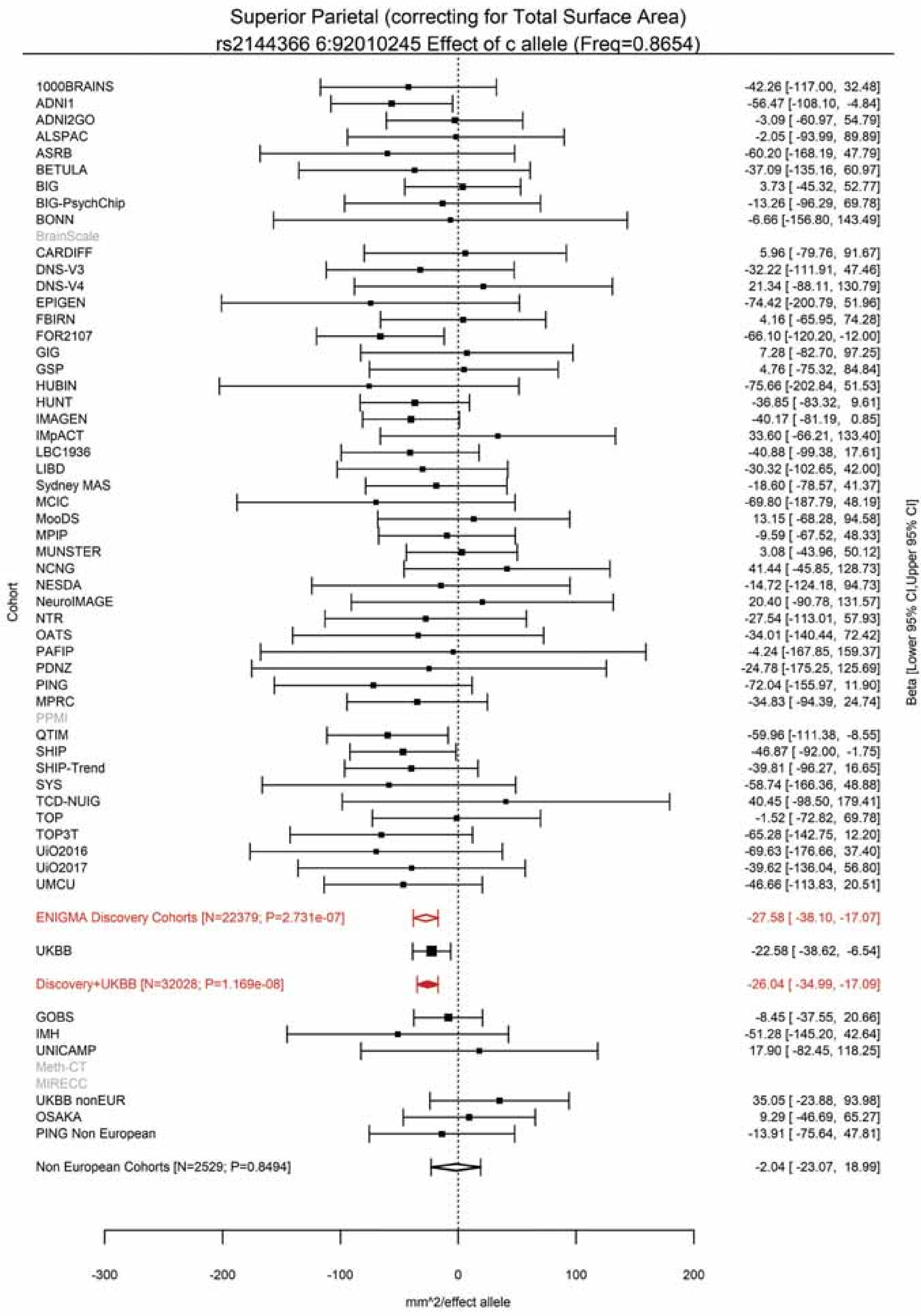
Superior Parietal (correcting for Total Surface Area)
rs2144366 6:92010245 Effect of c allele (Freq=0.8654)
Cohort
1000BRAINS -42.26 [-117.00, 32.48]
ADNI1 -56.47 [-108.10, -4.84]
ADNI2GO -3.09 [ -60.97, 54.79]
ALSPAC -2.05 [ -93.99, 89.89]
ASRB -60.20 [-168.19, 47.79]
BETULA -37.09 [-135.16, 60.97]
BIG 3.73 [ -45.32, 52.77]
BIG-PsychChip -13.26 [ -96.29, 69.78]
BONN -6.66 [-156.80, 143.49]
BrainScale
CARDIFF 5.96 [ -79.76, 91.67]
DNS-V3 -32.22 [-111.91, 47.46]
DNS-V4 21.34 [ -88.11, 130.79]
EPIGEN -74.42 [-200.79, 51.96]
FBIRN 4.16 [ -65.95, 74.28]
FOR2107 -66.10 [-120.20, -12.00]
GIG 7.28 [ -82.70, 97.25]
GSP 4.76 [ -75.32, 84.84]
HUBIN -75.66 [-202.84, 51.53]
HUNT -36.85 [ -83.32, 9.61]
IMAGEN -40.17 [ -81.19, 0.85]
IMpACT 33.60 [ -66.21, 133.40]
LBC1936 -40.88 [ -99.38, 17.61]
LIBD -30.32 [-102.65, 42.00]
Sydney MAS -18.60 [ -78.57, 41.37]
MCIC -69.80 [-187.79, 48.19]
MooDS 13.15 [ -68.28, 94.58]
MPIP -9.59 [ -67.52, 48.33]
MUNSTER 3.08 [ -43.96, 50.12]
NCNG 41.44 [ -45.85, 128.73]
NESDA -14.72 [-124.18, 94.73]
NeuroIMAGE 20.40 [ -90.78, 131.57]
NTR -27.54 [-113.01, 57.93]
OATS -34.01 [-140.44, 72.42]
PAFIP -4.24 [-167.85, 159.37]
PDNZ -24.78 [-175.25, 125.69]
PING -72.04 [-155.97, 11.90]
MPRC -34.83 [ -94.39, 24.74]
PPMI
QTIM -59.96 [-111.38, -8.55]
SHIP -46.87 [ -92.00, -1.75]
SHIP-Trend -39.81 [ -96.27, 16.65]
SYS -58.74 [-166.36, 48.88]
TCD-NUIG 40.45 [ -98.50, 179.41]
TOP -1.52 [ -72.82, 69.78]
TOP3T -65.28 [-142.75, 12.20]
UiO2016 -69.63 [-176.66, 37.40]
UiO2017 -39.62 [-136.04, 56.80]
UMCU -46.66 [-113.83, 20.51]
ENIGMA Discovery Cohorts [N=22379; P=2.731e-07] -27.58 [ -38.10, -17.07]
UKBB -22.58 [ -38.62, -6.54]
Discovery+UKBB [N=32028; P=1.169e-08] -26.04 [ -34.99, -17.09]
GOBS -8.45 [ -37.55, 20.66]
IMH -51.28 [-145.20, 42.64]
UNICAMP 17.90 [ -82.45, 118.25]
Meth-CT
MIRECC
UKBB nonEUR 35.05 [ -23.88, 93.98]
OSAKA 9.29 [ -46.69, 65.27]
PING Non European -13.91 [ -75.64, 47.81]
Non European Cohorts [N=2529; P=0.8494] -2.04 [ -23.07, 18.99]
-300
-200
-100
0
100
200
mm^2/effect allele
Beta [Lower 95% CI,Upper 95% CI]

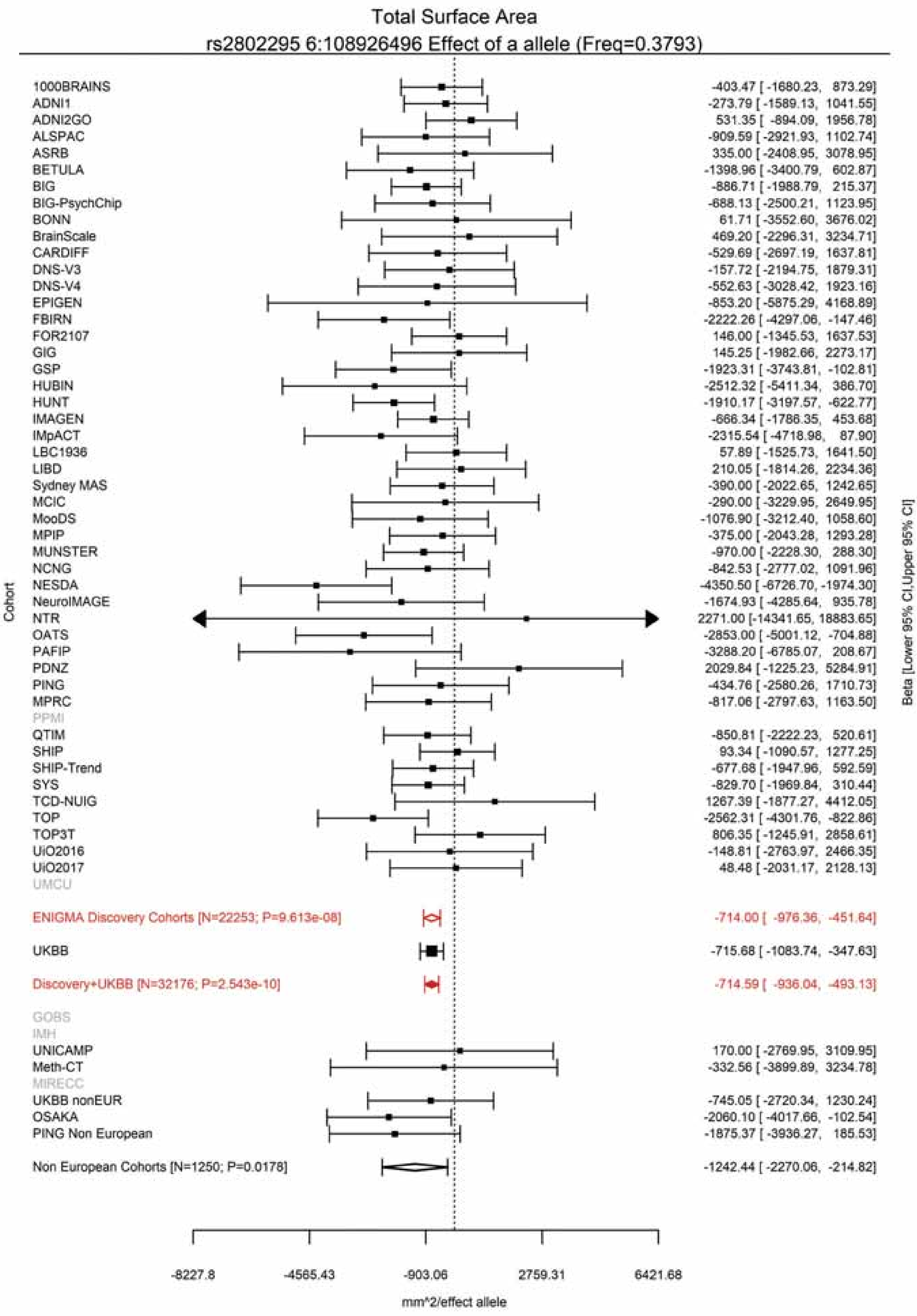
Total Surface Area
rs2802295 6:108926496 Effect of a allele (Freq=0.3793)
1000BRAINS -403.47 [ -1680.23, 873.29]
ADNI1 -273.79 [ -1589.13, 1041.55]
ADNI2GO 531.35 [ -894.09, 1956.78]
ALSPAC -909.59 [ -2921.93, 1102.74]
ASRB 335.00 [ -2408.95, 3078.95]
BETULA -1398.96 [ -3400.79, 602.87]
BIG -886.71 [ -1988.79, 215.37]
BIG-PsychChip -688.13 [ -2500.21, 1123.95]
BONN 61.71 [ -3552.60, 3676.02]
BrainScale 469.20 [ -2296.31, 3234.71]
CARDIFF -529.69 [ -2697.19, 1637.81]
DNS-V3 -157.72 [ -2194.75, 1879.31]
DNS-V4 -552.63 [ -3028.42, 1923.16]
EPIGEN -853.20 [ -5875.29, 4168.89]
FBIRN -2222.26 [ -4297.06, -147.46]
FOR2107 146.00 [ -1345.53, 1637.53]
GIG 145.25 [ -1982.66, 2273.17]
GSP -1923.31 [ -3743.81, -102.81]
HUBIN -2512.32 [ -5411.34, 386.70]
HUNT -1910.17 [ -3197.57, -622.77]
IMAGEN -666.34 [ -1786.35, 453.68]
IMpACT -2315.54 [ -4718.98, 87.90]
LBC1936 57.89 [ -1525.73, 1641.50]
LIBD 210.05 [ -1814.26, 2234.36]
Sydney MAS -390.00 [ -2022.65, 1242.65]
MCIC -290.00 [ -3229.95, 2649.95]
MooDS -1076.90 [ -3212.40, 1058.60]
MPIP -375.00 [ -2043.28, 1293.28]
MUNSTER -970.00 [ -2228.30, 288.30]
NCNG -842.53 [ -2777.02, 1091.96]
NESDA -4350.50 [ -6726.70, -1974.30]
NeuroIMAGE -1674.93 [ -4285.64, 935.78]
NTR 2271.00 [-14341.65, 18883.65]
OATS -2853.00 [ -5001.12, -704.88]
PAFIP -3288.20 [ -6785.07, 208.67]
PDNZ 2029.84 [ -1225.23, 5284.91]
PING -434.76 [ -2580.26, 1710.73]
MPRC -817.06 [ -2797.63, 1163.50]
PPMI
QTIM -850.81 [ -2222.23, 520.61]
SHIP 93.34 [ -1090.57, 1277.25]
SHIP-Trend -677.68 [ -1947.96, 592.59]
SYS -829.70 [ -1969.84, 310.44]
TCD-NUIG 1267.39 [ -1877.27, 4412.05]
TOP -2562.31 [ -4301.76, -822.86]
TOP3T 806.35 [ -1245.91, 2858.61]
UiO2016 -148.81 [ -2763.97, 2466.35]
UiO2017 48.48 [ -2031.17, 2128.13]
UMCU
ENIGMA Discovery Cohorts [N=22253; P=9.613e-08] -714.00 [ -976.36, -451.64]
UKBB -715.68 [ -1083.74, -347.63]
Discovery+UKBB [N=32176; P=2.543e-10] -714.59 [ -936.04, -493.13]
GOBS
IMH
UNICAMP 170.00 [ -2769.95, 3109.95]
Meth-CT -332.56 [ -3899.89, 3234.78]
MIRECC
UKBB nonEUR -745.05 [ -2720.34, 1230.24]
OSAKA -2060.10 [ -4017.66, -102.54]
PING Non European -1875.37 [ -3936.27, 185.53]
Non European Cohorts [N=1250; P=0.0178] -1242.44 [ -2270.06, -214.82]
Cohort
Beta [Lower 95% CI,Upper 95% CI]
-8227.8
-4565.43
-903.06
2759.31
6421.68
mm^2/effect allele

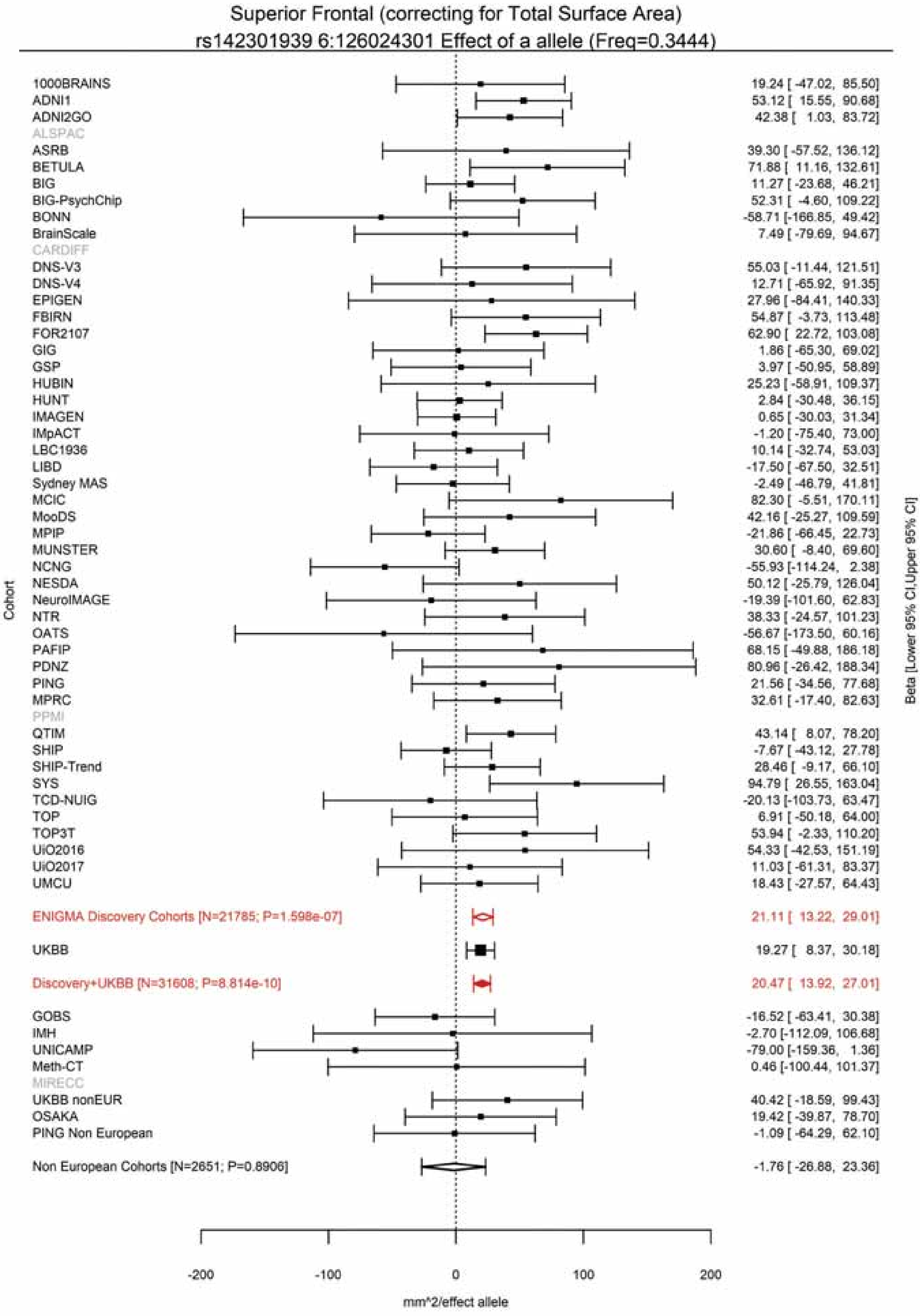

Superior Frontal (correcting for Total Surface Area)
rs142301939 6:126024301 Effect of a allele (Freq=0.3444)
Cohort
1000BRAINS 19.24 [ -47.02, 85.50]
ADNI1 53.12 [ 15.55, 90.68]
ADNI2GO 42.38 [ 1.03, 83.72]
ALSPAC
ASRB 39.30 [ -57.52, 136.12]
BETULA 71.88 [ 11.16, 132.61]
BIG 11.27 [ -23.68, 46.21]
BIG-PsychChip 52.31 [ -4.60, 109.22]
BONN -58.71 [-166.85, 49.42]
BrainScale 7.49 [ -79.69, 94.67]
CARDIFF
DNS-V3 55.03 [ -11.44, 121.51]
DNS-V4 12.71 [ -65.92, 91.35]
EPIGEN 27.96 [ -84.41, 140.33]
FBIRN 54.87 [ -3.73, 113.48]
FOR2107 62.90 [ 22.72, 103.08]
GIG 1.86 [ -65.30, 69.02]
GSP 3.97 [ -50.95, 58.89]
HUBIN 25.23 [ -58.91, 109.37]
HUNT 2.84 [ -30.48, 36.15]
IMAGEN 0.65 [ -30.03, 31.34]
IMpACT -1.20 [ -75.40, 73.00]
LBC1936 10.14 [ -32.74, 53.03]
LIBD -17.50 [ -67.50, 32.51]
Sydney MAS -2.49 [ -46.79, 41.81]
MCIC 82.30 [ -5.51, 170.11]
MooDS 42.16 [ -25.27, 109.59]
MPIP -21.86 [ -66.45, 22.73]
MUNSTER 30.60 [ -8.40, 69.60]
NCNG -55.93 [-114.24, 2.38]
NESDA 50.12 [ -25.79, 126.04]
NeuroIMAGE -19.39 [-101.60, 62.83]
NTR 38.33 [ -24.57, 101.23]
OATS -56.67 [-173.50, 60.16]
PAFIP 68.15 [ -49.88, 186.18]
PDNZ 80.96 [ -26.42, 188.34]
PING 21.56 [ -34.56, 77.68]
MPRC 32.61 [ -17.40, 82.63]
PPMI
QTIM 43.14 [ 8.07, 78.20]
SHIP -7.67 [ -43.12, 27.78]
SHIP-Trend 28.46 [ -9.17, 66.10]
SYS 94.79 [ 26.55, 163.04]
TCD-NUIG -20.13 [-103.73, 63.47]
TOP 6.91 [ -50.18, 64.00]
TOP3T 53.94 [ -2.33, 110.20]
UiO2016 54.33 [ -42.53, 151.19]
UiO2017 11.03 [ -61.31, 83.37]
UMCU 18.43 [ -27.57, 64.43]
ENIGMA Discovery Cohorts [N=21785; P=1.598e-07] 21.11 [ 13.22, 29.01]
UKBB 19.27 [ 8.37, 30.18]
Discovery+UKBB [N=31608; P=8.814e-10] 20.47 [ 13.92, 27.01]
GOBS -16.52 [ -63.41, 30.38]
IMH -2.70 [-112.09, 106.68]
UNICAMP -79.00 [-159.36, 1.36]
Meth-CT 0.46 [-100.44, 101.37]
MIRECC
UKBB nonEUR 40.42 [ -18.59, 99.43]
OSAKA 19.42 [ -39.87, 78.70]
PING Non European -1.09 [ -64.29, 62.10]
Non European Cohorts [N=2651; P=0.8906] -1.76 [ -26.88, 23.36]
Beta [Lower 95% CI,Upper 95% CI]
-200
-100
0
100
200
mm^2/effect allele

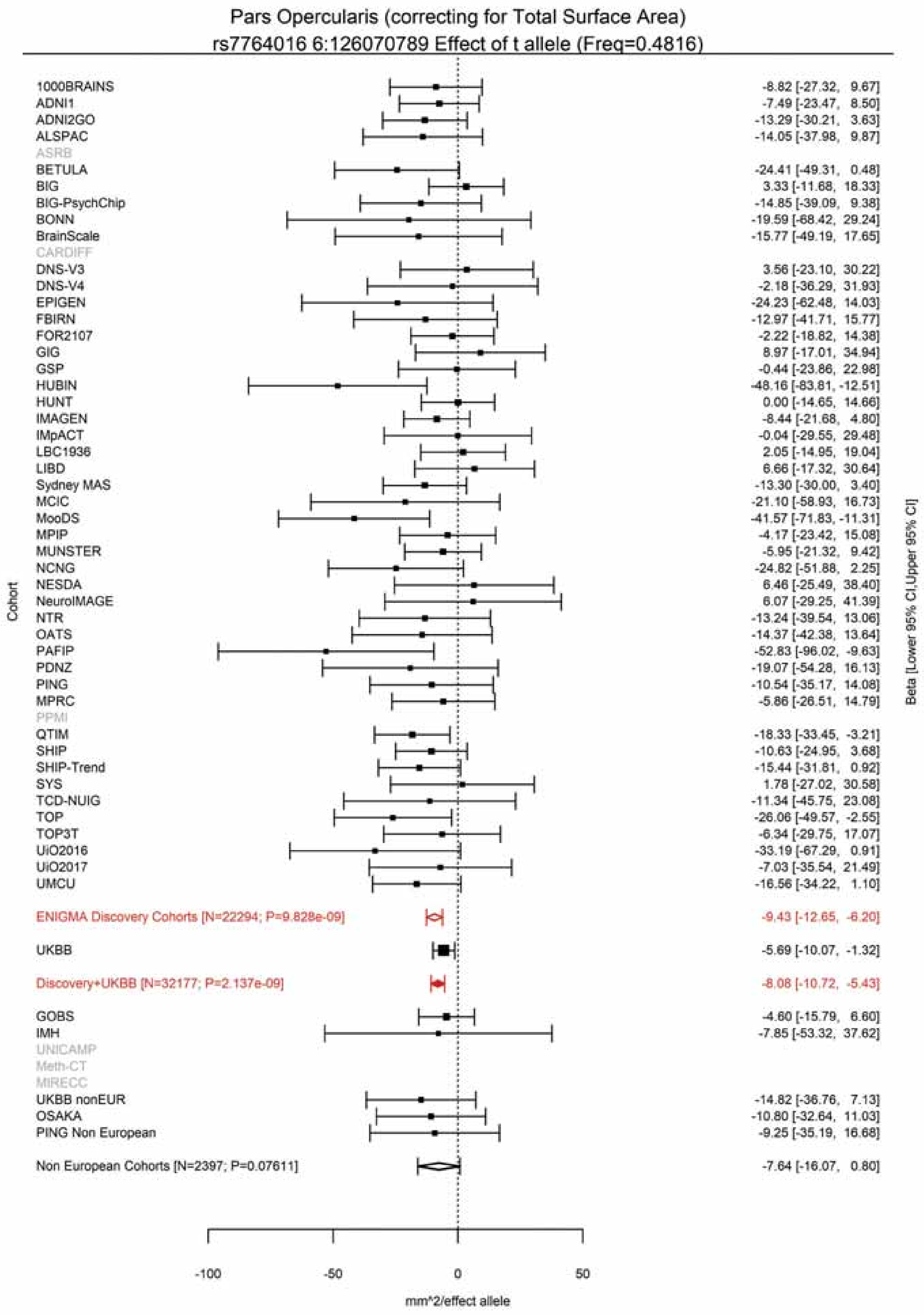
Pars Opercularis (correcting for Total Surface Area)
rs7764016 6:126070789 Effect of t allele (Freq=0.4816)
Cohort
1000BRAINS -8.82 [-27.32, 9.67]
ADNI1 -7.49 [-23.47, 8.50]
ADNI2GO -13.29 [-30.21, 3.63]
ALSPAC -14.05 [-37.98, 9.87]
ASRB
BETULA -24.41 [-49.31, 0.48]
BIG 3.33 [-11.68, 18.33]
BIG-PsychChip -14.85 [-39.09, 9.38]
BONN -19.59 [-68.42, 29.24]
BrainScale -15.77 [-49.19, 17.65]
CARDIFF
DNS-V3 3.56 [-23.10, 30.22]
DNS-V4 -2.18 [-36.29, 31.93]
EPIGEN -24.23 [-62.48, 14.03]
FBIRN -12.97 [-41.71, 15.77]
FOR2107 -2.22 [-18.82, 14.38]
GIG 8.97 [-17.01, 34.94]
GSP -0.44 [-23.86, 22.98]
HUBIN -48.16 [-83.81, -12.51]
HUNT 0.00 [-14.65, 14.66]
IMAGEN -8.44 [-21.68, 4.80]
IMpACT -0.04 [-29.55, 29.48]
LBC1936 2.05 [-14.95, 19.04]
LIBD 6.66 [-17.32, 30.64]
Sydney MAS -13.30 [-30.00, 3.40]
MCIC -21.10 [-58.93, 16.73]
MooDS -41.57 [-71.83, -11.31]
MPIP -4.17 [-23.42, 15.08]
MUNSTER -5.95 [-21.32, 9.42]
NCNG -24.82 [-51.88, 2.25]
NESDA 6.46 [-25.49, 38.40]
NeuroIMAGE 6.07 [-29.25, 41.39]
NTR -13.24 [-39.54, 13.06]
OATS -14.37 [-42.38, 13.64]
PAFIP -52.83 [-96.02, -9.63]
PDNZ -19.07 [-54.28, 16.13]
PING -10.54 [-35.17, 14.08]
MPRC -5.86 [-26.51, 14.79]
PPMI
QTIM -18.33 [-33.45, -3.21]
SHIP -10.63 [-24.95, 3.68]
SHIP-Trend -15.44 [-31.81, 0.92]
SYS 1.78 [-27.02, 30.58]
TCD-NUIG -11.34 [-45.75, 23.08]
TOP -26.06 [-49.57, -2.55]
TOP3T -6.34 [-29.75, 17.07]
UiO2016 -33.19 [-67.29, 0.91]
UiO2017 -7.03 [-35.54, 21.49]
UMCU -16.56 [-34.22, 1.10]
ENIGMA Discovery Cohorts [N=22294; P=9.828e-09] -9.43 [-12.65, -6.20]
UKBB -5.69 [-10.07, -1.32]
Discovery+UKBB [N=32177; P=2.137e-09] -8.08 [-10.72, -5.43]
GOBS -4.60 [-15.79, 6.60]
IMH -7.85 [-53.32, 37.62]
UNICAMP
Meth-CT
MIRECC
UKBB nonEUR -14.82 [-36.76, 7.13]
OSAKA -10.80 [-32.64, 11.03]
PING Non European -9.25 [-35.19, 16.68]
Non European Cohorts [N=2397; P=0.07611] -7.64 [-16.07, 0.80]
-100
-50
0
50
mm^2/effect allele
Beta [Lower 95% CI,Upper 95% CI]

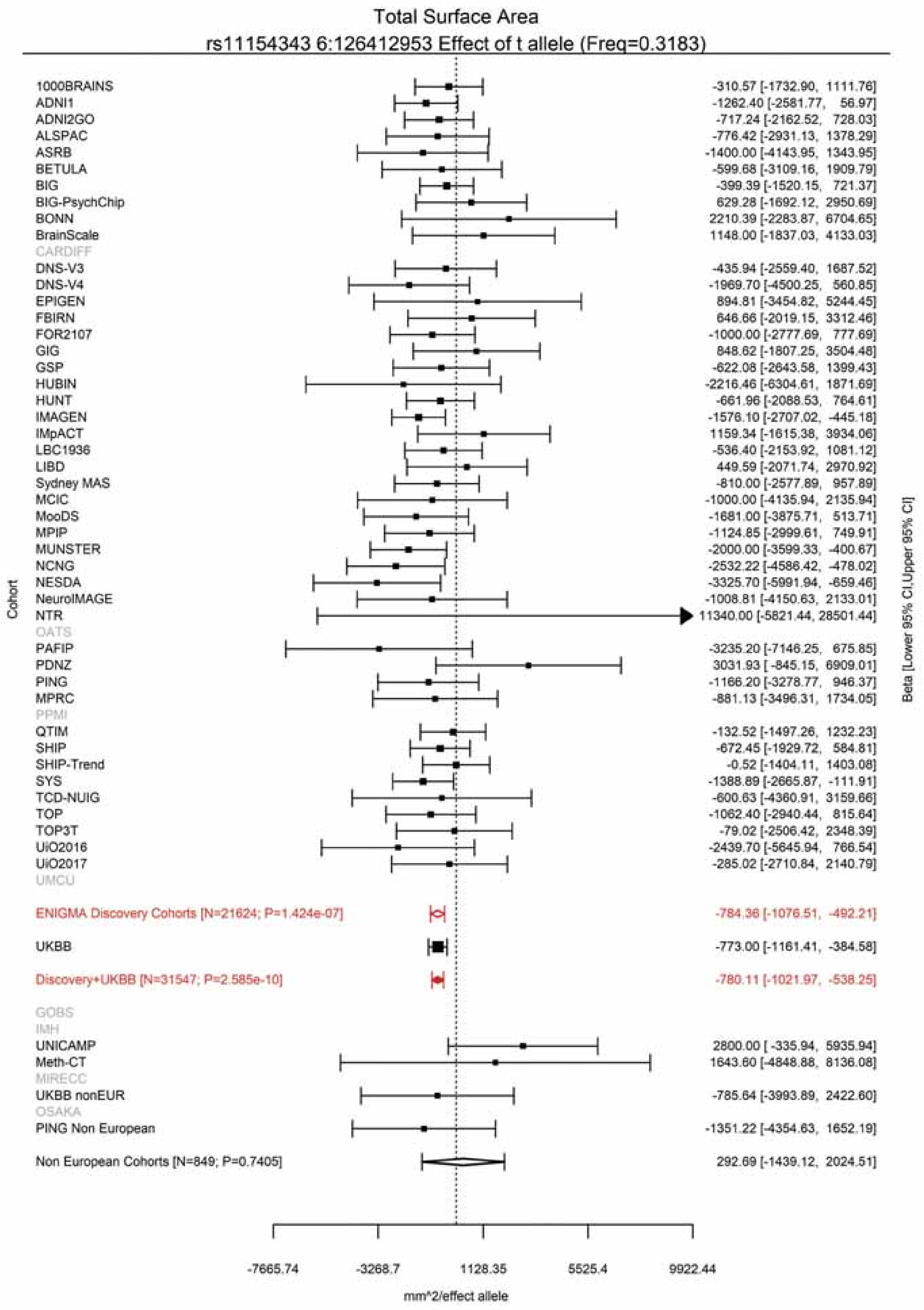

Total Surface Area
rs11154343 6:126412953 Effect of t allele (Freq=0.3183)
Cohort
Beta [Lower 95% CI,Upper 95% CI]
1000BRAINS -310.57 [-1732.90, 1111.76]
ADNI1 -1262.40 [-2581.77, 56.97]
ADNI2GO -717.24 [-2162.52, 728.03]
ALSPAC -776.42 [-2931.13, 1378.29]
ASRB -1400.00 [-4143.95, 1343.95]
BETULA -599.68 [-3109.16, 1909.79]
BIG -399.39 [-1520.15, 721.37]
BIG-PsychChip 629.28 [-1692.12, 2950.69]
BONN 2210.39 [-2283.87, 6704.65]
BrainScale 1148.00 [-1837.03, 4133.03]
CARDIFF
DNS-V3 -435.94 [-2559.40, 1687.52]
DNS-V4 -1969.70 [-4500.25, 560.85]
EPIGEN 894.81 [-3454.82, 5244.45]
FBIRN 646.66 [-2019.15, 3312.46]
FOR2107 -1000.00 [-2777.69, 777.69]
GIG 848.62 [-1807.25, 3504.48]
GSP -622.08 [-2643.58, 1399.43]
HUBIN -2216.46 [-6304.61, 1871.69]
HUNT -661.96 [-2088.53, 764.61]
IMAGEN -1576.10 [-2707.02, -445.18]
IMpACT 1159.34 [-1615.38, 3934.06]
LBC1936 -536.40 [-2153.92, 1081.12]
LIBD 449.59 [-2071.74, 2970.92]
Sydney MAS -810.00 [-2577.89, 957.89]
MCIC -1000.00 [-4135.94, 2135.94]
MooDS -1681.00 [-3875.71, 513.71]
MPIP -1124.85 [-2999.61, 749.91]
MUNSTER -2000.00 [-3599.33, -400.67]
NCNG -2532.22 [-4586.42, -478.02]
NESDA -3325.70 [-5991.94, -659.46]
NeuroIMAGE -1008.81 [-4150.63, 2133.01]
NTR 11340.00 [-5821.44, 28501.44]
OATS
PAFIP -3235.20 [-7146.25, 675.85]
PDNZ 3031.93 [-845.15, 6909.01]
PING -1166.20 [-3278.77, 946.37]
MPRC -881.13 [-3496.31, 1734.05]
PPMI
QTIM -132.52 [-1497.26, 1232.23]
SHIP -672.45 [-1929.72, 584.81]
SHIP-Trend -0.52 [-1404.11, 1403.08]
SYS -1388.89 [-2665.87, -111.91]
TCD-NUIG -600.63 [-4360.91, 3159.66]
TOP -1062.40 [-2940.44, 815.64]
TOP3T -79.02 [-2506.42, 2348.39]
UiO2016 -2439.70 [-5645.94, 766.54]
UiO2017 -285.02 [-2710.84, 2140.79]
UMCU
ENIGMA Discovery Cohorts [N=21624; P=1.424e-07] -784.36 [-1076.51, -492.21]
UKBB -773.00 [-1161.41, -384.58]
Discovery+UKBB [N=31547; P=2.585e-10] -780.11 [-1021.97, -538.25]
GOBS
IMH
UNICAMP 2800.00 [ -335.94, 5935.94]
Meth-CT 1643.60 [-4848.88, 8136.08]
MIRECC
UKBB nonEUR -785.64 [-3993.89, 2422.60]
OSAKA
PING Non European -1351.22 [-4354.63, 1652.19]
Non European Cohorts [N=849; P=0.7405] 292.69 [-1439.12, 2024.51]
-7665.74
-3268.7
1128.35
5525.4
9922.44
mm^2/effect allele

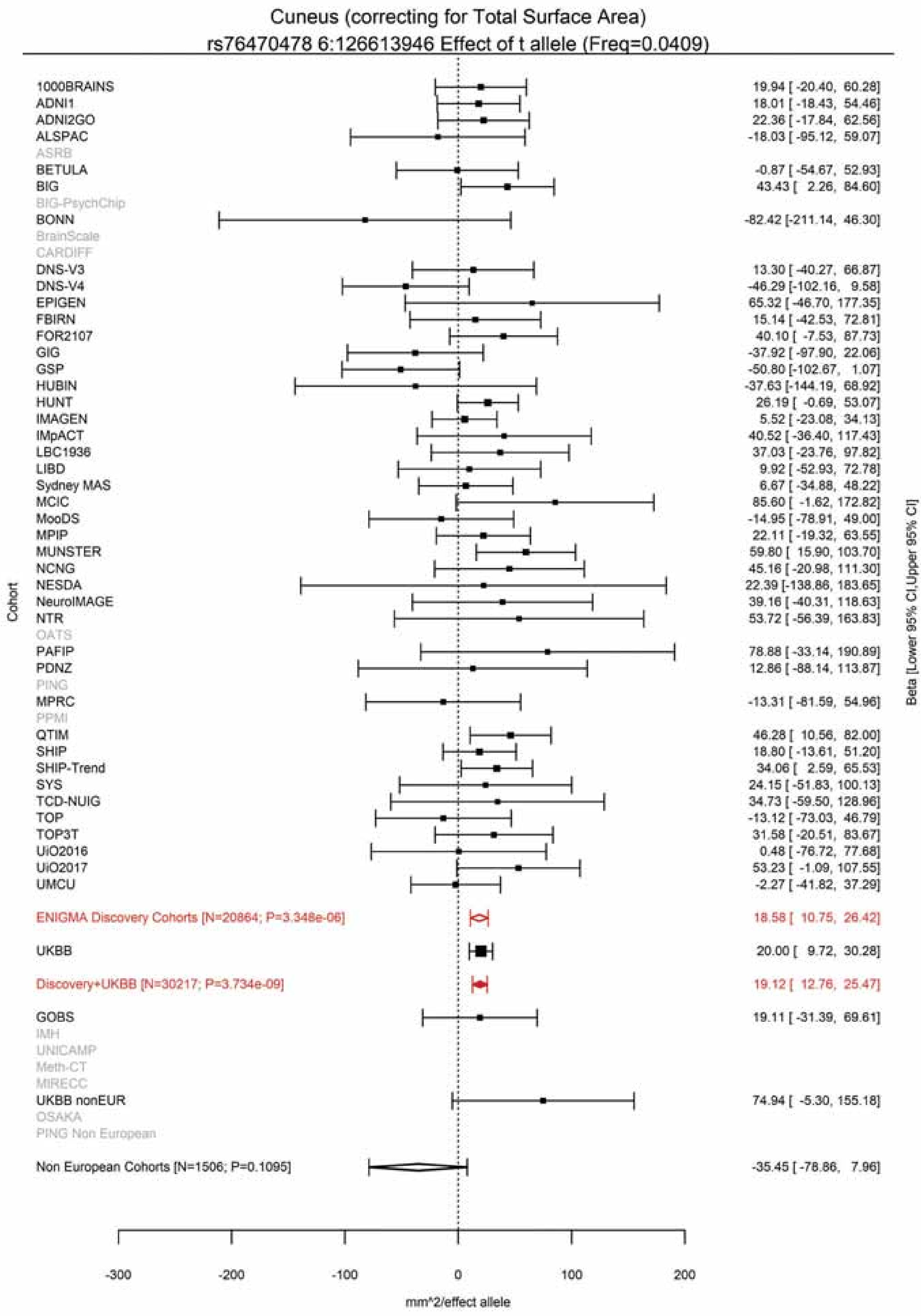

Cuneus (correcting for Total Surface Area)
rs76470478 6:126613946 Effect of t allele (Freq=0.0409)
1000BRAINS 19.94 [ -20.40, 60.28]
ADNI1 18.01 [ -18.43, 54.46]
ADNI2GO 22.36 [ -17.84, 62.56]
ALSPAC -18.03 [ -95.12, 59.07]
ASRB
BETULA -0.87 [ -54.67, 52.93]
BIG 43.43 [ 2.26, 84.60]
BIG-PsychChip
BONN -82.42 [-211.14, 46.30]
BrainScale
CARDIFF
DNS-V3 13.30 [ -40.27, 66.87]
DNS-V4 -46.29 [-102.16, 9.58]
EPIGEN 65.32 [ -46.70, 177.35]
FBIRN 15.14 [ -42.53, 72.81]
FOR2107 40.10 [ -7.53, 87.73]
GIG -37.92 [ -97.90, 22.06]
GSP -50.80 [-102.67, 1.07]
HUBIN -37.63 [-144.19, 68.92]
HUNT 26.19 [ -0.69, 53.07]
IMAGEN 5.52 [ -23.08, 34.13]
IMpACT 40.52 [ -36.40, 117.43]
LBC1936 37.03 [ -23.76, 97.82]
LIBD 9.92 [ -52.93, 72.78]
Sydney MAS 6.67 [ -34.88, 48.22]
MCIC 85.60 [ -1.62, 172.82]
MooDS -14.95 [ -78.91, 49.00]
MPIP 22.11 [ -19.32, 63.55]
MUNSTER 59.80 [ 15.90, 103.70]
NCNG 45.16 [ -20.98, 111.30]
NESDA 22.39 [-138.86, 183.65]
NeuroIMAGE 39.16 [ -40.31, 118.63]
NTR 53.72 [ -56.39, 163.83]
OATS
PAFIP 78.88 [ -33.14, 190.89]
PDNZ 12.86 [ -88.14, 113.87]
PING
MPRC -13.31 [ -81.59, 54.96]
PPMI
QTIM 46.28 [ 10.56, 82.00]
SHIP 18.80 [ -13.61, 51.20]
SHIP-Trend 34.06 [ 2.59, 65.53]
SYS 24.15 [ -51.83, 100.13]
TCD-NUIG 34.73 [ -59.50, 128.96]
TOP -13.12 [ -73.03, 46.79]
TOP3T 31.58 [ -20.51, 83.67]
UiO2016 0.48 [ -76.72, 77.68]
UiO2017 53.23 [ -1.09, 107.55]
UMCU -2.27 [ -41.82, 37.29]
ENIGMA Discovery Cohorts [N=20864; P=3.348e-06] 18.58 [ 10.75, 26.42]
UKBB 20.00 [ 9.72, 30.28]
Discovery+UKBB [N=30217; P=3.734e-09] 19.12 [ 12.76, 25.47]
GOBS 19.11 [ -31.39, 69.61]
IMH
UNICAMP
Meth-CT
MIRECC
UKBB nonEUR 74.94 [ -5.30, 155.18]
OSAKA
PING Non European
Non European Cohorts [N=1506; P=0.1095] -35.45 [ -78.86, 7.96]
Cohort
Beta [Lower 95% CI,Upper 95% CI]
-300
-200
-100
0
100
200
mm^2/effect allele

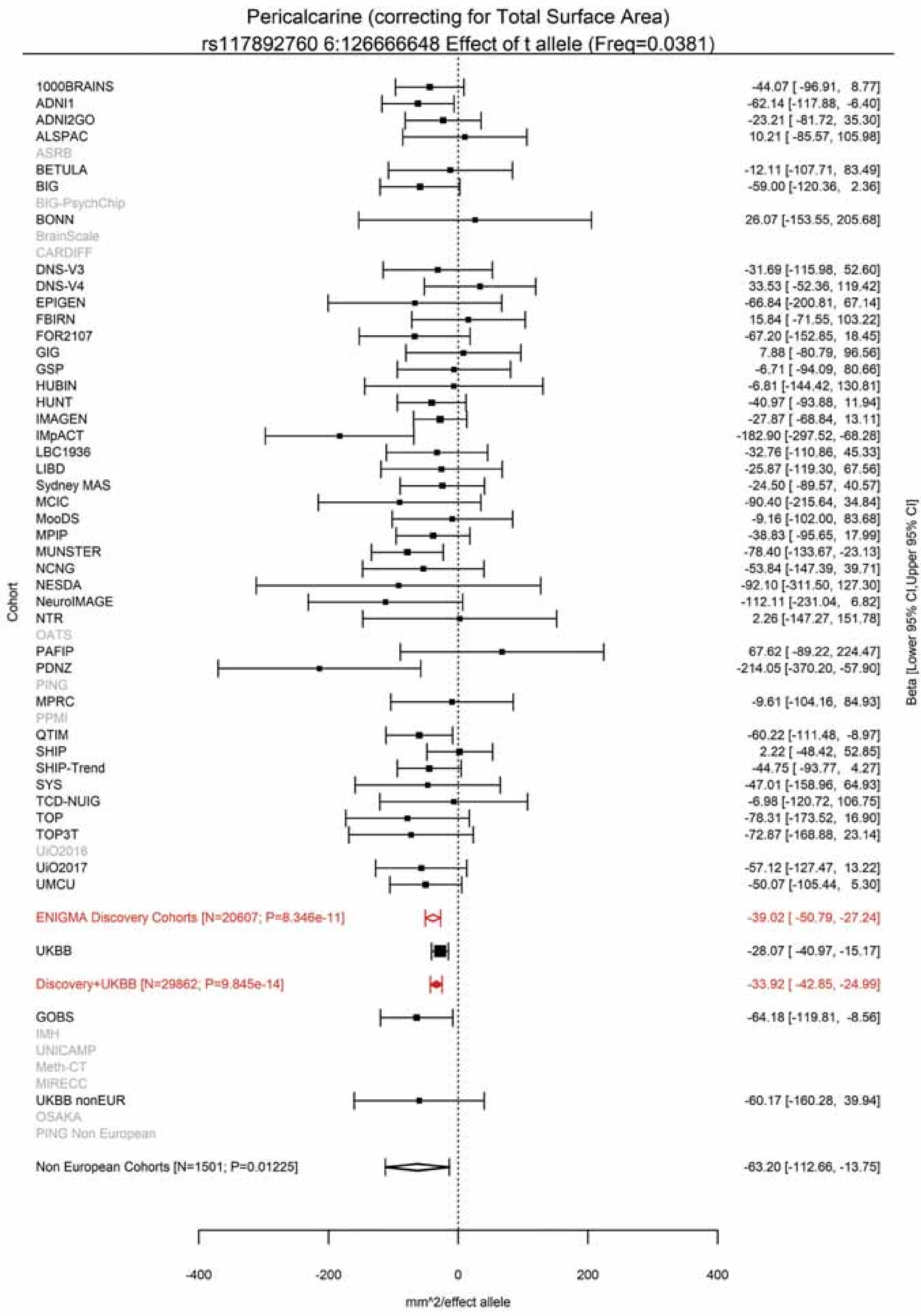

Pericalcarine (correcting for Total Surface Area)
rs117892760 6:126666648 Effect of t allele (Freq=0.0381)
Cohort
Beta [Lower 95% CI,Upper 95% CI]
1000BRAINS -44.07 [-96.91, 8.77]
ADNI1 -62.14 [-117.88, -6.40]
ADNI2GO -23.21 [-81.72, 35.30]
ALSPAC 10.21 [-85.57, 105.98]
ASRB
BETULA -12.11 [-107.71, 83.49]
BIG -59.00 [-120.36, 2.36]
BIG-PsychChip
BONN 26.07 [-153.55, 205.68]
BrainScale
CARDIFF
DNS-V3 -31.69 [-115.98, 52.60]
DNS-V4 33.53 [-52.36, 119.42]
EPIGEN -66.84 [-200.81, 67.14]
FBIRN 15.84 [-71.55, 103.22]
FOR2107 -67.20 [-152.85, 18.45]
GIG 7.88 [-80.79, 96.56]
GSP -6.71 [-94.09, 80.66]
HUBIN -6.81 [-144.42, 130.81]
HUNT -40.97 [-93.88, 11.94]
IMAGEN -27.87 [-68.84, 13.11]
IMpACT -182.90 [-297.52, -68.28]
LBC1936 -32.76 [-110.86, 45.33]
LIBD -25.87 [-119.30, 67.56]
Sydney MAS -24.50 [-89.57, 40.57]
MCIC -90.40 [-215.64, 34.84]
MooDS -9.16 [-102.00, 83.68]
MPIP -38.83 [-95.65, 17.99]
MUNSTER -78.40 [-133.67, -23.13]
NCNG -53.84 [-147.39, 39.71]
NESDA -92.10 [-311.50, 127.30]
NeuroIMAGE -112.11 [-231.04, 6.82]
NTR 2.26 [-147.27, 151.78]
OATS
PAFIP 67.62 [-89.22, 224.47]
PDNZ -214.05 [-370.20, -57.90]
PING
MPRC -9.61 [-104.16, 84.93]
PPMI
QTIM -60.22 [-111.48, -8.97]
SHIP 2.22 [-48.42, 52.85]
SHIP-Trend -44.75 [-93.77, 4.27]
SYS -47.01 [-158.96, 64.93]
TCD-NUIG -6.98 [-120.72, 106.75]
TOP -78.31 [-173.52, 16.90]
TOP3T -72.87 [-168.88, 23.14]
UiO2016
UiO2017 -57.12 [-127.47, 13.22]
UMCU -50.07 [-105.44, 5.30]
ENIGMA Discovery Cohorts [N=20607; P=8.346e-11] -39.02 [-50.79, -27.24]
UKBB -28.07 [-40.97, -15.17]
Discovery+UKBB [N=29862; P=9.845e-14] -33.92 [-42.85, -24.99]
GOBS -64.18 [-119.81, -8.56]
IMH
UNICAMP
Meth-CT
MIRECC
UKBB nonEUR -60.17 [-160.28, 39.94]
OSAKA
PING Non European
Non European Cohorts [N=1501; P=0.01225] -63.20 [-112.66, -13.75]
-400
-200
0
200
400
mm^2/effect allele

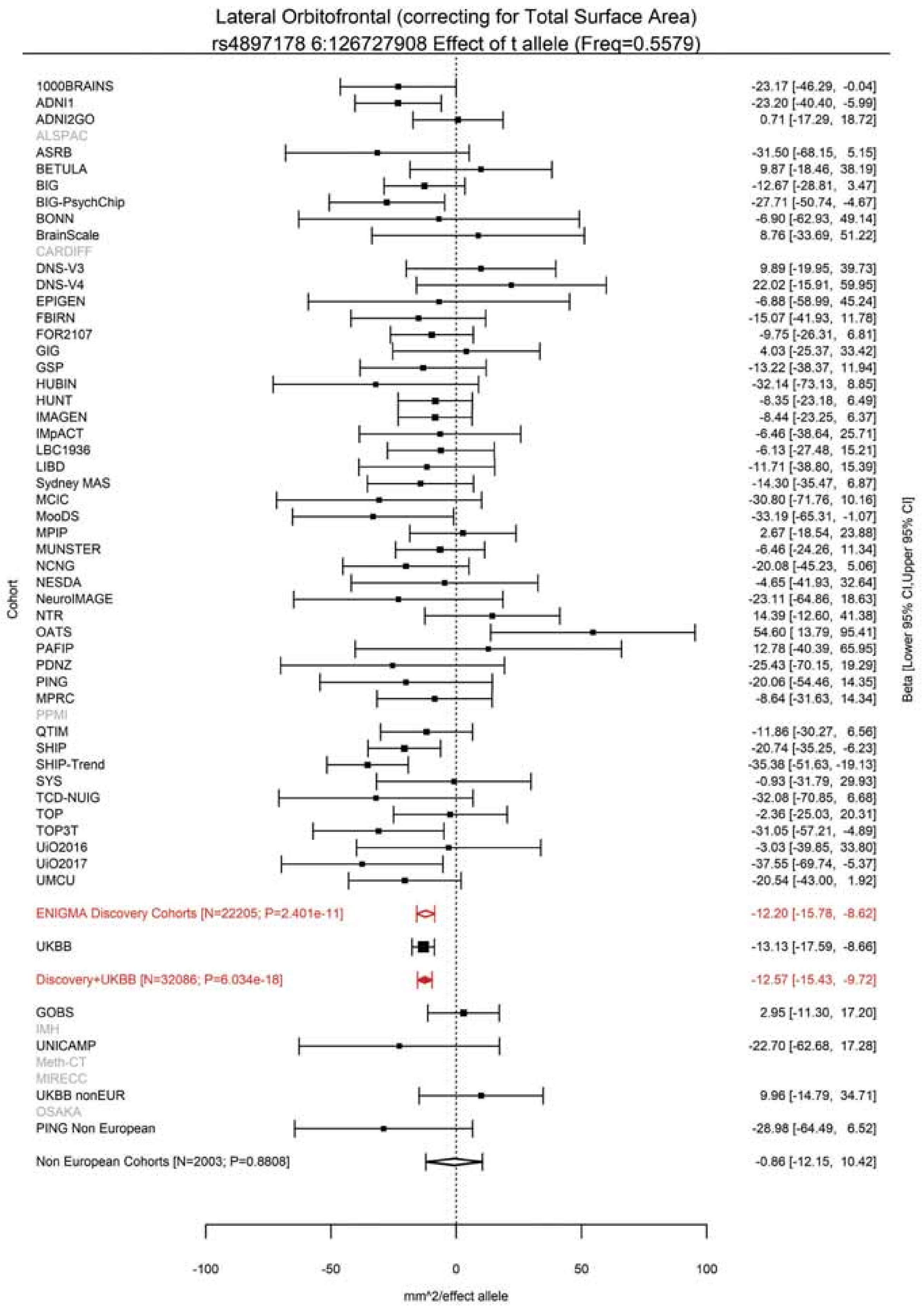
Lateral Orbitofrontal (correcting for Total Surface Area)
rs4897178 6:126727908 Effect of t allele (Freq=0.5579)
1000BRAINS -23.17 [-46.29, -0.04]
ADNI1 -23.20 [-40.40, -5.99]
ADNI2GO 0.71 [-17.29, 18.72]
ALSPAC
ASRB -31.50 [-68.15, 5.15]
BETULA 9.87 [-18.46, 38.19]
BIG -12.67 [-28.81, 3.47]
BIG-PsychChip -27.71 [-50.74, -4.67]
BONN -6.90 [-62.93, 49.14]
BrainScale 8.76 [-33.69, 51.22]
CARDIFF
DNS-V3 9.89 [-19.95, 39.73]
DNS-V4 22.02 [-15.91, 59.95]
EPIGEN -6.88 [-58.99, 45.24]
FBIRN -15.07 [-41.93, 11.78]
FOR2107 -9.75 [-26.31, 6.81]
GIG 4.03 [-25.37, 33.42]
GSP -13.22 [-38.37, 11.94]
HUBIN -32.14 [-73.13, 8.85]
HUNT -8.35 [-23.18, 6.49]
IMAGEN -8.44 [-23.25, 6.37]
IMpACT -6.46 [-38.64, 25.71]
LBC1936 -6.13 [-27.48, 15.21]
LIBD -11.71 [-38.80, 15.39]
Sydney MAS -14.30 [-35.47, 6.87]
MCIC -30.80 [-71.76, 10.16]
MooDS -33.19 [-65.31, -1.07]
MPIP 2.67 [-18.54, 23.88]
MUNSTER -6.46 [-24.26, 11.34]
NCNG -20.08 [-45.23, 5.06]
NESDA -4.65 [-41.93, 32.64]
NeuroIMAGE -23.11 [-64.86, 18.63]
NTR 14.39 [-12.60, 41.38]
OATS 54.60 [ 13.79, 95.41]
PAFIP 12.78 [-40.39, 65.95]
PDNZ -25.43 [-70.15, 19.29]
PING -20.06 [-54.46, 14.35]
MPRC -8.64 [-31.63, 14.34]
PPMI
QTIM -11.86 [-30.27, 6.56]
SHIP -20.74 [-35.25, -6.23]
SHIP-Trend -35.38 [-51.63, -19.13]
SYS -0.93 [-31.79, 29.93]
TCD-NUIG -32.08 [-70.85, 6.68]
TOP -2.36 [-25.03, 20.31]
TOP3T -31.05 [-57.21, -4.89]
UiO2016 -3.03 [-39.85, 33.80]
UiO2017 -37.55 [-69.74, -5.37]
UMCU -20.54 [-43.00, 1.92]
ENIGMA Discovery Cohorts [N=22205; P=2.401e-11] -12.20 [-15.78, -8.62]
UKBB -13.13 [-17.59, -8.66]
Discovery+UKBB [N=32086; P=6.034e-18] -12.57 [-15.43, -9.72]
GOBS 2.95 [-11.30, 17.20]
IMH
UNICAMP -22.70 [-62.68, 17.28]
Meth-CT
MIRECC
UKBB nonEUR 9.96 [-14.79, 34.71]
OSAKA
PING Non European -28.98 [-64.49, 6.52]
Non European Cohorts [N=2003; P=0.8808] -0.86 [-12.15, 10.42]
Cohort
Beta [Lower 95% CI,Upper 95% CI]
-100
-50
0
50
100
mm^2/effect allele

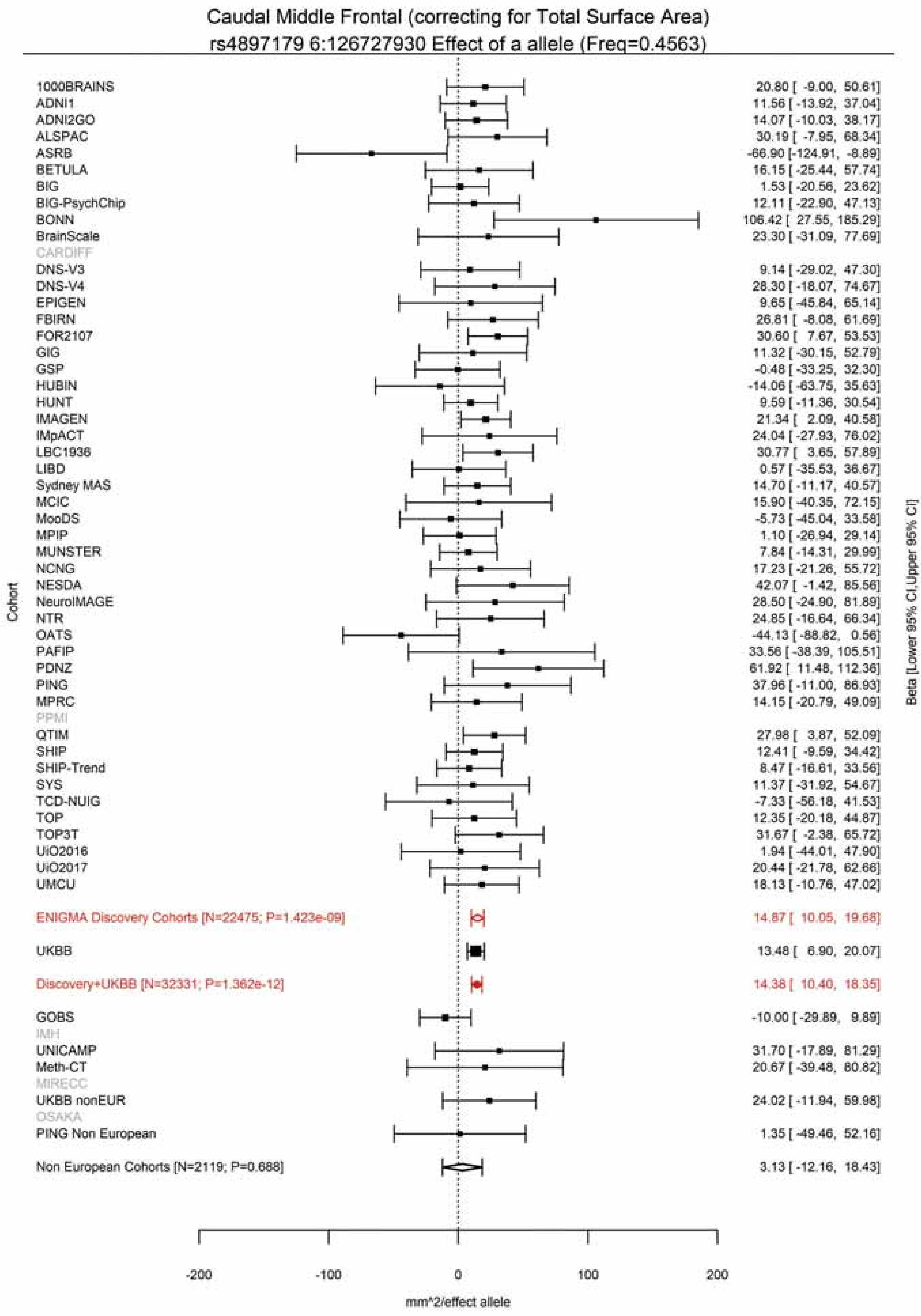
Caudal Middle Frontal (correcting for Total Surface Area)
rs4897179 6:126727930 Effect of a allele (Freq=0.4563)
Cohort
1000BRAINS 20.80 [ -9.00, 50.61]
ADNI1 11.56 [ -13.92, 37.04]
ADNI2GO 14.07 [ -10.03, 38.17]
ALSPAC 30.19 [ -7.95, 68.34]
ASRB -66.90 [-124.91, -8.89]
BETULA 16.15 [ -25.44, 57.74]
BIG 1.53 [ -20.56, 23.62]
BIG-PsychChip 12.11 [ -22.90, 47.13]
BONN 106.42 [ 27.55, 185.29]
BrainScale 23.30 [ -31.09, 77.69]
CARDIFF
DNS-V3 9.14 [ -29.02, 47.30]
DNS-V4 28.30 [ -18.07, 74.67]
EPIGEN 9.65 [ -45.84, 65.14]
FBIRN 26.81 [ -8.08, 61.69]
FOR2107 30.60 [ 7.67, 53.53]
GIG 11.32 [ -30.15, 52.79]
GSP -0.48 [ -33.25, 32.30]
HUBIN -14.06 [ -63.75, 35.63]
HUNT 9.59 [ -11.36, 30.54]
IMAGEN 21.34 [ 2.09, 40.58]
IMpACT 24.04 [ -27.93, 76.02]
LBC1936 30.77 [ 3.65, 57.89]
LIBD 0.57 [ -35.53, 36.67]
Sydney MAS 14.70 [ -11.17, 40.57]
MCIC 15.90 [ -40.35, 72.15]
MooDS -5.73 [ -45.04, 33.58]
MPIP 1.10 [ -26.94, 29.14]
MUNSTER 7.84 [ -14.31, 29.99]
NCNG 17.23 [ -21.26, 55.72]
NESDA 42.07 [ -1.42, 85.56]
NeuroIMAGE 28.50 [ -24.90, 81.89]
NTR 24.85 [ -16.64, 66.34]
OATS -44.13 [ -88.82, 0.56]
PAFIP 33.56 [ -38.39, 105.51]
PDNZ 61.92 [ 11.48, 112.36]
PING 37.96 [ -11.00, 86.93]
MPRC 14.15 [ -20.79, 49.09]
PPMI
QTIM 27.98 [ 3.87, 52.09]
SHIP 12.41 [ -9.59, 34.42]
SHIP-Trend 8.47 [ -16.61, 33.56]
SYS 11.37 [ -31.92, 54.67]
TCD-NUIG -7.33 [ -56.18, 41.53]
TOP 12.35 [ -20.18, 44.87]
TOP3T 31.67 [ -2.38, 65.72]
UiO2016 1.94 [ -44.01, 47.90]
UiO2017 20.44 [ -21.78, 62.66]
UMCU 18.13 [ -10.76, 47.02]
ENIGMA Discovery Cohorts [N=22475; P=1.423e-09] 14.87 [ 10.05, 19.68]
UKBB 13.48 [ 6.90, 20.07]
Discovery+UKBB [N=32331; P=1.362e-12] 14.38 [ 10.40, 18.35]
GOBS -10.00 [ -29.89, 9.89]
IMH
UNICAMP 31.70 [ -17.89, 81.29]
Meth-CT 20.67 [ -39.48, 80.82]
MIRECC
UKBB nonEUR 24.02 [ -11.94, 59.98]
OSAKA
PING Non European 1.35 [ -49.46, 52.16]
Non European Cohorts [N=2119; P=0.688] 3.13 [ -12.16, 18.43]
Beta [Lower 95% CI,Upper 95% CI]
-200
-100
0
100
200
mm^2/effect allele

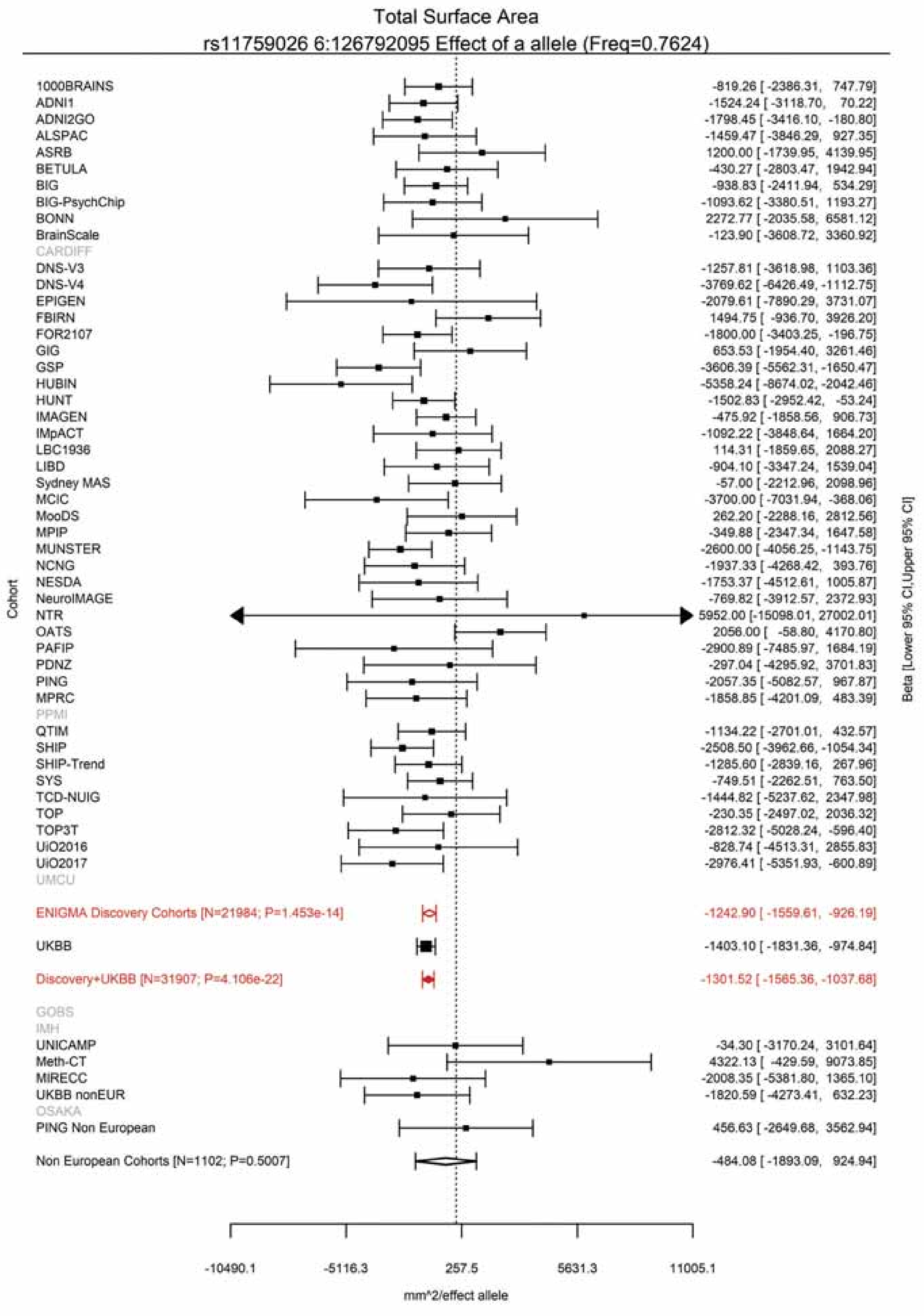

Total Surface Area
rs11759026 6:126792095 Effect of a allele (Freq=0.7624)
1000BRAINS -819.26 [ -2386.31, 747.79]
ADNI1 -1524.24 [ -3118.70, 70.22]
ADNI2GO -1798.45 [ -3416.10, -180.80]
ALSPAC -1459.47 [ -3846.29, 927.35]
ASRB 1200.00 [ -1739.95, 4139.95]
BETULA -430.27 [ -2803.47, 1942.94]
BIG -938.83 [ -2411.94, 534.29]
BIG-PsychChip -1093.62 [ -3380.51, 1193.27]
BONN 2272.77 [ -2035.58, 6581.12]
BrainScale -123.90 [ -3608.72, 3360.92]
CARDIFF
DNS-V3 -1257.81 [ -3618.98, 1103.36]
DNS-V4 -3769.62 [ -6426.49, -1112.75]
EPIGEN -2079.61 [ -7890.29, 3731.07]
FBIRN 1494.75 [ -936.70, 3926.20]
FOR2107 -1800.00 [ -3403.25, -196.75]
GIG 653.53 [ -1954.40, 3261.46]
GSP -3606.39 [ -5562.31, -1650.47]
HUBIN -5358.24 [ -8674.02, -2042.46]
HUNT -1502.83 [ -2952.42, -53.24]
IMAGEN -475.92 [ -1858.56, 906.73]
IMpACT -1092.22 [ -3848.64, 1664.20]
LBC1936 114.31 [ -1859.65, 2088.27]
LIBD -904.10 [ -3347.24, 1539.04]
Sydney MAS -57.00 [ -2212.96, 2098.96]
MCIC -3700.00 [ -7031.94, -368.06]
MooDS 262.20 [ -2288.16, 2812.56]
MPIP -349.88 [ -2347.34, 1647.58]
MUNSTER -2600.00 [ -4056.25, -1143.75]
NCNG -1937.33 [ -4268.42, 393.76]
NESDA -1753.37 [ -4512.61, 1005.87]
NeuroIMAGE -769.82 [ -3912.57, 2372.93]
NTR 5952.00 [-15098.01, 27002.01]
OATS 2056.00 [ -58.80, 4170.80]
PAFIP -2900.89 [ -7485.97, 1684.19]
PDNZ -297.04 [ -4295.92, 3701.83]
PING -2057.35 [ -5082.57, 967.87]
MPRC -1858.85 [ -4201.09, 483.39]
PPMI
QTIM -1134.22 [ -2701.01, 432.57]
SHIP -2508.50 [ -3962.66, -1054.34]
SHIP-Trend -1285.60 [ -2839.16, 267.96]
SYS -749.51 [ -2262.51, 763.50]
TCD-NUIG -1444.82 [ -5237.62, 2347.98]
TOP -230.35 [ -2497.02, 2036.32]
TOP3T -2812.32 [ -5028.24, -596.40]
UiO2016 -828.74 [ -4513.31, 2855.83]
UiO2017 -2976.41 [ -5351.93, -600.89]
UMCU
ENIGMA Discovery Cohorts [N=21984; P=1.453e-14] -1242.90 [ -1559.61, -926.19]
UKBB -1403.10 [ -1831.36, -974.84]
Discovery+UKBB [N=31907; P=4.106e-22] -1301.52 [ -1565.36, -1037.68]
GOBS
IMH
UNICAMP -34.30 [ -3170.24, 3101.64]
Meth-CT 4322.13 [ -429.59, 9073.85]
MIRECC -2008.35 [ -5381.80, 1365.10]
UKBB nonEUR -1820.59 [ -4273.41, 632.23]
OSAKA
PING Non European 456.63 [ -2649.68, 3562.94]
Non European Cohorts [N=1102; P=0.5007] -484.08 [ -1893.09, 924.94]
Cohort
Beta [Lower 95% CI,Upper 95% CI]
-10490.1
-5116.3
257.5
5631.3
11005.1
mm^2/effect allele

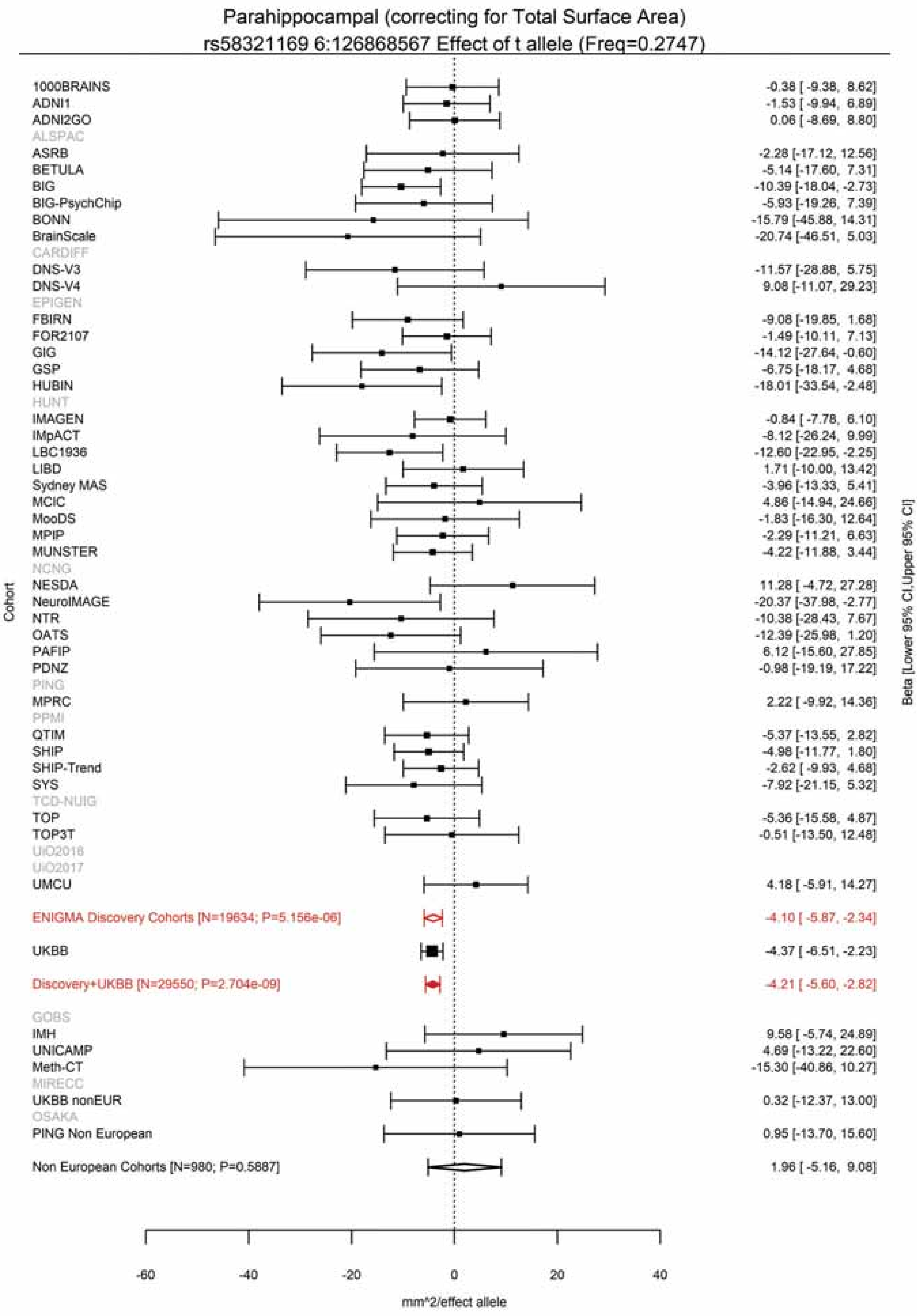
Parahippocampal (correcting for Total Surface Area)
rs58321169 6:126868567 Effect of t allele (Freq=0.2747)
Cohort
1000BRAINS -0.38 [ -9.38, 8.62]
ADNI1 -1.53 [ -9.94, 6.89]
ADNI2GO 0.06 [ -8.69, 8.80]
ALSPAC
ASRB -2.28 [-17.12, 12.56]
BETULA -5.14 [-17.60, 7.31]
BIG -10.39 [-18.04, -2.73]
BIG-PsychChip -5.93 [-19.26, 7.39]
BONN -15.79 [-45.88, 14.31]
BrainScale -20.74 [-46.51, 5.03]
CARDIFF
DNS-V3 -11.57 [-28.88, 5.75]
DNS-V4 9.08 [-11.07, 29.23]
EPIGEN
FBIRN -9.08 [-19.85, 1.68]
FOR2107 -1.49 [-10.11, 7.13]
GIG -14.12 [-27.64, -0.60]
GSP -6.75 [-18.17, 4.68]
HUBIN -18.01 [-33.54, -2.48]
HUNT
IMAGEN -0.84 [ -7.78, 6.10]
IMpACT -8.12 [-26.24, 9.99]
LBC1936 -12.60 [-22.95, -2.25]
LIBD 1.71 [-10.00, 13.42]
Sydney MAS -3.96 [-13.33, 5.41]
MCIC 4.86 [-14.94, 24.66]
MooDS -1.83 [-16.30, 12.64]
MPIP -2.29 [-11.21, 6.63]
MUNSTER -4.22 [-11.88, 3.44]
NCNG
NESDA 11.28 [ -4.72, 27.28]
NeuroIMAGE -20.37 [-37.98, -2.77]
NTR -10.38 [-28.43, 7.67]
OATS -12.39 [-25.98, 1.20]
PAFIP 6.12 [-15.60, 27.85]
PDNZ -0.98 [-19.19, 17.22]
PING
MPRC 2.22 [ -9.92, 14.36]
PPMI
QTIM -5.37 [-13.55, 2.82]
SHIP -4.98 [-11.77, 1.80]
SHIP-Trend -2.62 [ -9.93, 4.68]
SYS -7.92 [-21.15, 5.32]
TCD-NUIG
TOP -5.36 [-15.58, 4.87]
TOP3T -0.51 [-13.50, 12.48]
UiO2016
UiO2017
UMCU 4.18 [ -5.91, 14.27]
ENIGMA Discovery Cohorts [N=19634; P=5.156e-06] -4.10 [ -5.87, -2.34]
UKBB -4.37 [ -6.51, -2.23]
Discovery+UKBB [N=29550; P=2.704e-09] -4.21 [ -5.60, -2.82]
GOBS
IMH 9.58 [ -5.74, 24.89]
UNICAMP 4.69 [-13.22, 22.60]
Meth-CT -15.30 [-40.86, 10.27]
MIRECC
UKBB nonEUR 0.32 [-12.37, 13.00]
OSAKA
PING Non European 0.95 [-13.70, 15.60]
Non European Cohorts [N=980; P=0.5887] 1.96 [ -5.16, 9.08]
-60
-40
-20
0
20
40
mm^2/effect allele
Beta [Lower 95% CI,Upper 95% CI]

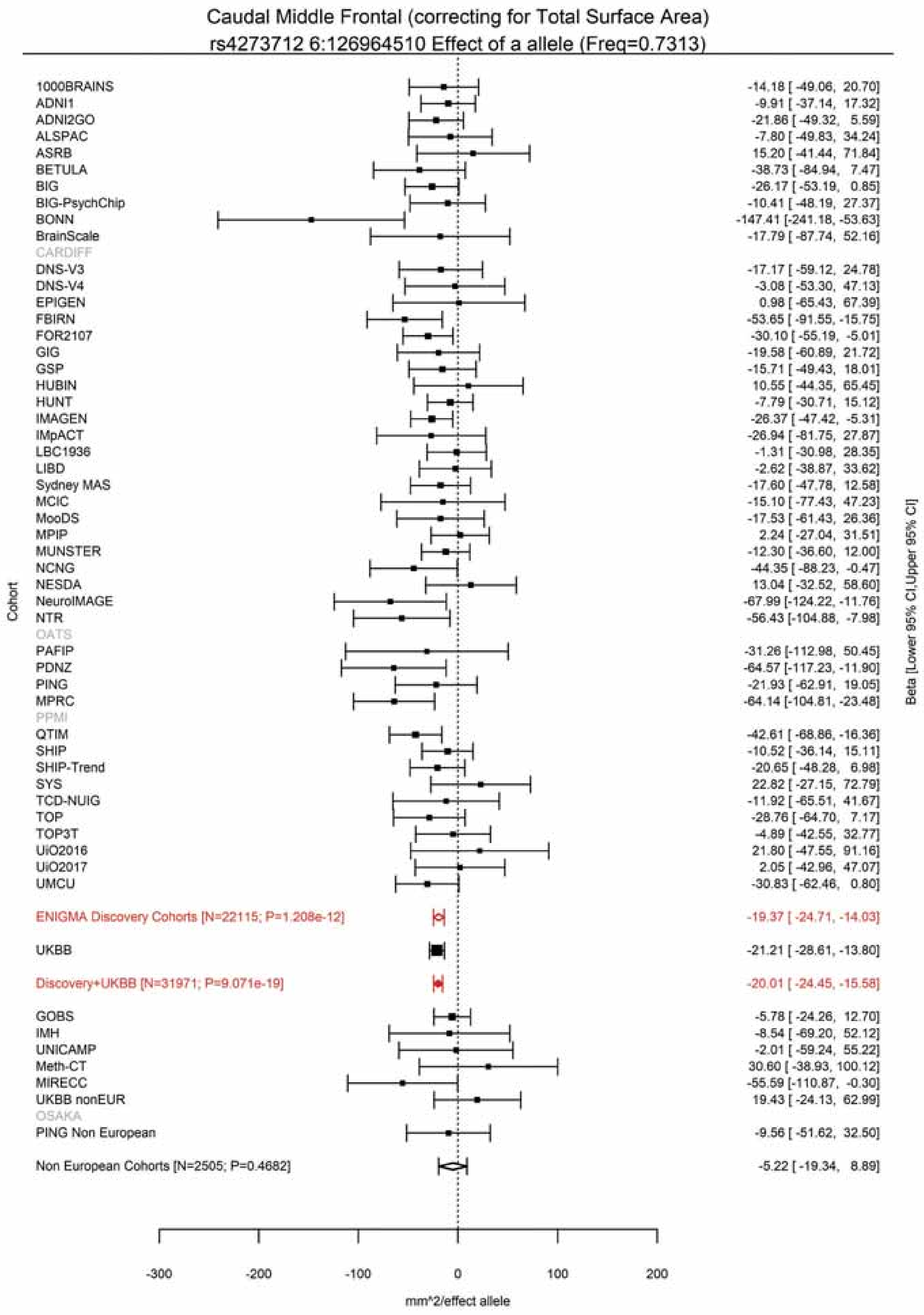
Caudal Middle Frontal (correcting for Total Surface Area)
rs4273712 6:126964510 Effect of a allele (Freq=0.7313)
Cohort
1000BRAINS -14.18 [ -49.06, 20.70]
ADNI1 -9.91 [ -37.14, 17.32]
ADNI2GO -21.86 [ -49.32, 5.59]
ALSPAC -7.80 [ -49.83, 34.24]
ASRB 15.20 [ -41.44, 71.84]
BETULA -38.73 [ -84.94, 7.47]
BIG -26.17 [ -53.19, 0.85]
BIG-PsychChip -10.41 [ -48.19, 27.37]
BONN -147.41 [-241.18, -53.63]
BrainScale -17.79 [ -87.74, 52.16]
CARDIFF
DNS-V3 -17.17 [ -59.12, 24.78]
DNS-V4 -3.08 [ -53.30, 47.13]
EPIGEN 0.98 [ -65.43, 67.39]
FBIRN -53.65 [ -91.55, -15.75]
FOR2107 -30.10 [ -55.19, -5.01]
GIG -19.58 [ -60.89, 21.72]
GSP -15.71 [ -49.43, 18.01]
HUBIN 10.55 [ -44.35, 65.45]
HUNT -7.79 [ -30.71, 15.12]
IMAGEN -26.37 [ -47.42, -5.31]
IMpACT -26.94 [ -81.75, 27.87]
LBC1936 -1.31 [ -30.98, 28.35]
LIBD -2.62 [ -38.87, 33.62]
Sydney MAS -17.60 [ -47.78, 12.58]
MCIC -15.10 [ -77.43, 47.23]
MooDS -17.53 [ -61.43, 26.36]
MPIP 2.24 [ -27.04, 31.51]
MUNSTER -12.30 [ -36.60, 12.00]
NCNG -44.35 [ -88.23, -0.47]
NESDA 13.04 [ -32.52, 58.60]
NeuroIMAGE -67.99 [-124.22, -11.76]
NTR -56.43 [-104.88, -7.98]
OATS
PAFIP -31.26 [-112.98, 50.45]
PDNZ -64.57 [-117.23, -11.90]
PING -21.93 [ -62.91, 19.05]
MPRC -64.14 [-104.81, -23.48]
PPMI
QTIM -42.61 [ -68.86, -16.36]
SHIP -10.52 [ -36.14, 15.11]
SHIP-Trend -20.65 [ -48.28, 6.98]
SYS 22.82 [ -27.15, 72.79]
TCD-NUIG -11.92 [ -65.51, 41.67]
TOP -28.76 [ -64.70, 7.17]
TOP3T -4.89 [ -42.55, 32.77]
UiO2016 21.80 [ -47.55, 91.16]
UiO2017 2.05 [ -42.96, 47.07]
UMCU -30.83 [ -62.46, 0.80]
ENIGMA Discovery Cohorts [N=22115; P=1.208e-12] -19.37 [ -24.71, -14.03]
UKBB -21.21 [ -28.61, -13.80]
Discovery+UKBB [N=31971; P=9.071e-19] -20.01 [ -24.45, -15.58]
GOBS -5.78 [ -24.26, 12.70]
IMH -8.54 [ -69.20, 52.12]
UNICAMP -2.01 [ -59.24, 55.22]
Meth-CT 30.60 [ -38.93, 100.12]
MIRECC -55.59 [-110.87, -0.30]
UKBB nonEUR 19.43 [ -24.13, 62.99]
OSAKA
PING Non European -9.56 [ -51.62, 32.50]
Non European Cohorts [N=2505; P=0.4682] -5.22 [ -19.34, 8.89]
Beta [Lower 95% CI,Upper 95% CI]
-300
-200
-100
0
100
200
mm^2/effect allele

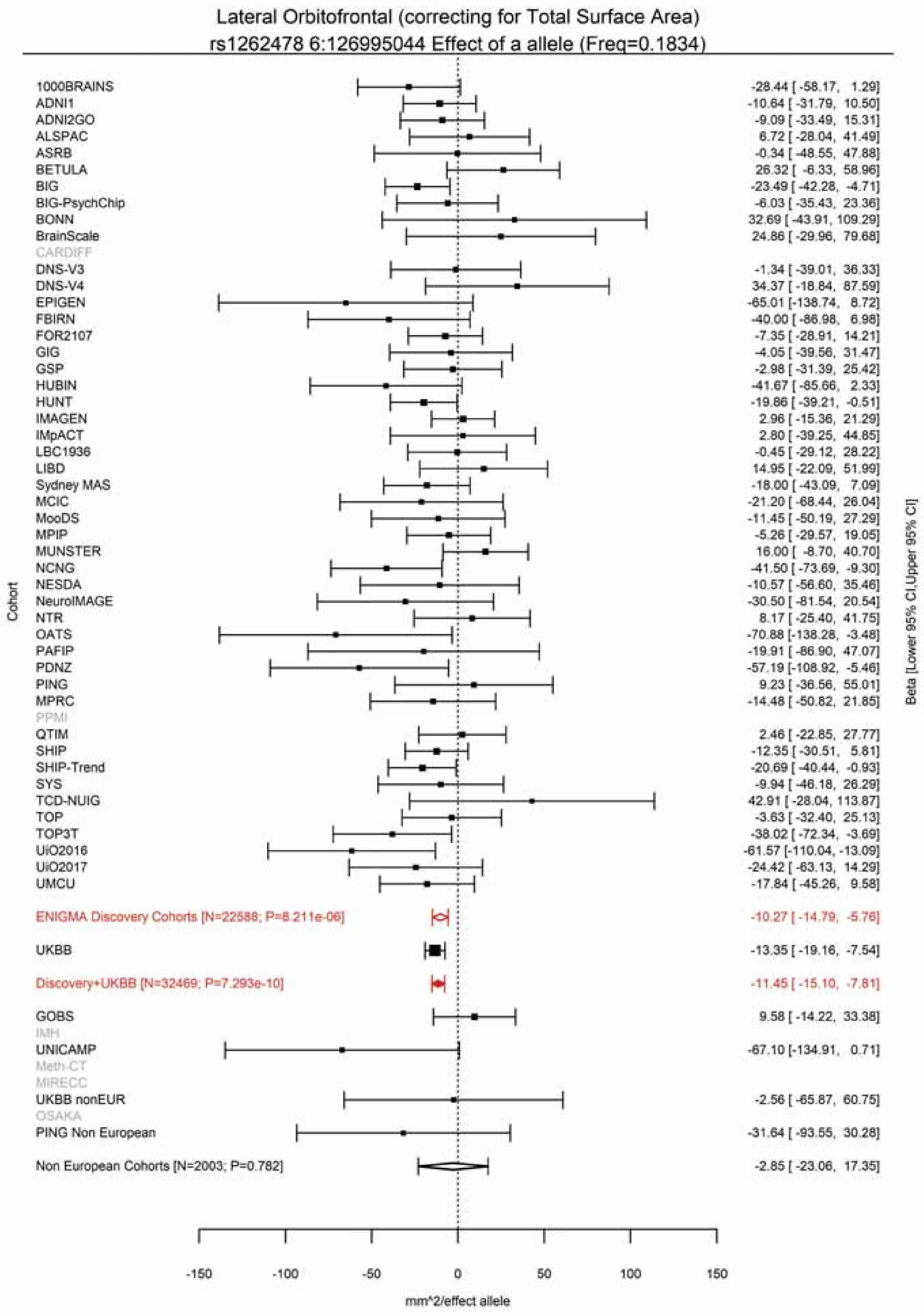
Lateral Orbitofrontal (correcting for Total Surface Area)
rs1262478 6:126995044 Effect of a allele (Freq=0.1834)
Cohort
1000BRAINS -28.44 [ -58.17, 1.29]
ADNI1 -10.64 [ -31.79, 10.50]
ADNI2GO -9.09 [ -33.49, 15.31]
ALSPAC 6.72 [ -28.04, 41.49]
ASRB -0.34 [ -48.55, 47.88]
BETULA 26.32 [ -6.33, 58.96]
BIG -23.49 [ -42.28, -4.71]
BIG-PsychChip -6.03 [ -35.43, 23.36]
BONN 32.69 [ -43.91, 109.29]
BrainScale 24.86 [ -29.96, 79.68]
CARDIFF
DNS-V3 -1.34 [ -39.01, 36.33]
DNS-V4 34.37 [ -18.84, 87.59]
EPIGEN -65.01 [-138.74, 8.72]
FBIRN -40.00 [ -86.98, 6.98]
FOR2107 -7.35 [ -28.91, 14.21]
GIG -4.05 [ -39.56, 31.47]
GSP -2.98 [ -31.39, 25.42]
HUBIN -41.67 [ -85.66, 2.33]
HUNT -19.86 [ -39.21, -0.51]
IMAGEN 2.96 [ -15.36, 21.29]
IMpACT 2.80 [ -39.25, 44.85]
LBC1936 -0.45 [ -29.12, 28.22]
LIBD 14.95 [ -22.09, 51.99]
Sydney MAS -18.00 [ -43.09, 7.09]
MCIC -21.20 [ -68.44, 26.04]
MooDS -11.45 [ -50.19, 27.29]
MPIP -5.26 [ -29.57, 19.05]
MUNSTER 16.00 [ -8.70, 40.70]
NCNG -41.50 [ -73.69, -9.30]
NESDA -10.57 [ -56.60, 35.46]
NeuroIMAGE -30.50 [ -81.54, 20.54]
NTR 8.17 [ -25.40, 41.75]
OATS -70.88 [-138.28, -3.48]
PAFIP -19.91 [ -86.90, 47.07]
PDNZ -57.19 [-108.92, -5.46]
PING 9.23 [ -36.56, 55.01]
MPRC -14.48 [ -50.82, 21.85]
PPMI
QTIM 2.46 [ -22.85, 27.77]
SHIP -12.35 [ -30.51, 5.81]
SHIP-Trend -20.69 [ -40.44, -0.93]
SYS -9.94 [ -46.18, 26.29]
TCD-NUIG 42.91 [ -28.04, 113.87]
TOP -3.63 [ -32.40, 25.13]
TOP3T -38.02 [ -72.34, -3.69]
UiO2016 -61.57 [-110.04, -13.09]
UiO2017 -24.42 [ -63.13, 14.29]
UMCU -17.84 [ -45.26, 9.58]
ENIGMA Discovery Cohorts [N=22588; P=8.211e-06] -10.27 [ -14.79, -5.76]
UKBB -13.35 [ -19.16, -7.54]
Discovery+UKBB [N=32469; P=7.293e-10] -11.45 [ -15.10, -7.81]
GOBS 9.58 [ -14.22, 33.38]
IMH
UNICAMP -67.10 [-134.91, 0.71]
Meth-CT
MIRECC
UKBB nonEUR -2.56 [ -65.87, 60.75]
OSAKA
PING Non European -31.64 [ -93.55, 30.28]
Non European Cohorts [N=2003; P=0.782] -2.85 [ -23.06, 17.35]
-150 -100 -50 0 50 100 150
mm^2/effect allele
Beta [Lower 95% CI,Upper 95% CI]

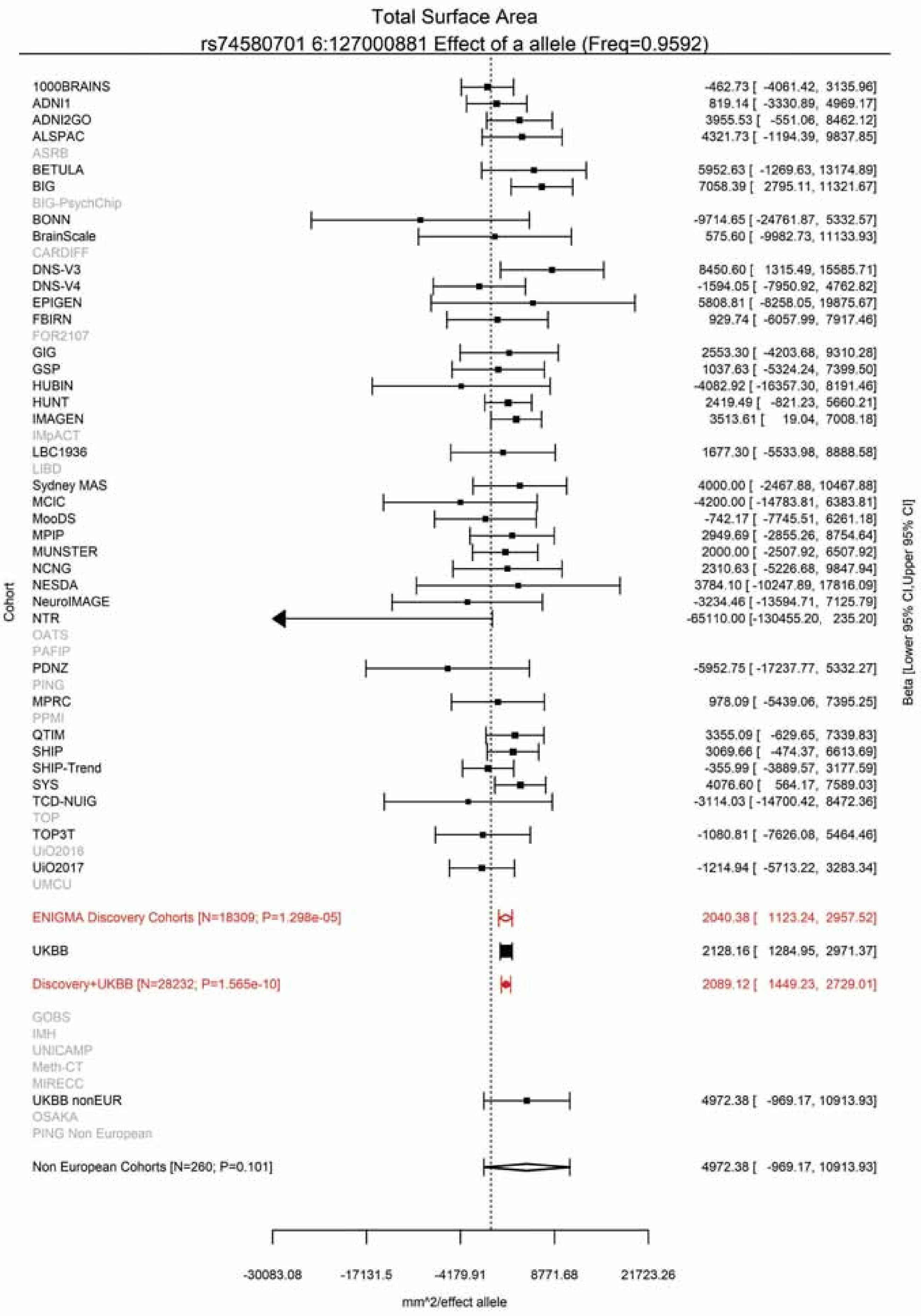

Total Surface Area
rs74580701 6:127000881 Effect of a allele (Freq=0.9592)
1000BRAINS -462.73 [ -4061.42, 3135.96]
ADNI1 819.14 [ -3330.89, 4969.17]
ADNI2GO 3955.53 [ -551.06, 8462.12]
ALSPAC 4321.73 [ -1194.39, 9837.85]
ASRB
BETULA 5952.63 [ -1269.63, 13174.89]
BIG 7058.39 [ 2795.11, 11321.67]
BIG-PsychChip
BONN -9714.65 [ -24761.87, 5332.57]
BrainScale 575.60 [ -9982.73, 11133.93]
CARDIFF
DNS-V3 8450.60 [ 1315.49, 15585.71]
DNS-V4 -1594.05 [ -7950.92, 4762.82]
EPIGEN 5808.81 [ -8258.05, 19875.67]
FBIRN 929.74 [ -6057.99, 7917.46]
FOR2107
GIG 2553.30 [ -4203.68, 9310.28]
GSP 1037.63 [ -5324.24, 7399.50]
HUBIN -4082.92 [ -16357.30, 8191.46]
HUNT 2419.49 [ -821.23, 5660.21]
IMAGEN 3513.61 [ 19.04, 7008.18]
IMpACT
LBC1936 1677.30 [ -5533.98, 8888.58]
LIBD
Sydney MAS 4000.00 [ -2467.88, 10467.88]
MCIC -4200.00 [ -14783.81, 6383.81]
MooDS -742.17 [ -7745.51, 6261.18]
MPIP 2949.69 [ -2855.26, 8754.64]
MUNSTER 2000.00 [ -2507.92, 6507.92]
NCNG 2310.63 [ -5226.68, 9847.94]
NESDA 3784.10 [ -10247.89, 17816.09]
NeuroIMAGE -3234.46 [ -13594.71, 7125.79]
NTR -65110.00 [-130455.20, 235.20]
OATS
PAFIP
PDNZ -5952.75 [ -17237.77, 5332.27]
PING
MPRC 978.09 [ -5439.06, 7395.25]
PPMI
QTIM 3355.09 [ -629.65, 7339.83]
SHIP 3069.66 [ -474.37, 6613.69]
SHIP-Trend -355.99 [ -3889.57, 3177.59]
SYS 4076.60 [ 564.17, 7589.03]
TCD-NUIG -3114.03 [ -14700.42, 8472.36]
TOP
TOP3T -1080.81 [ -7626.08, 5464.46]
UiO2016
UiO2017 -1214.94 [ -5713.22, 3283.34]
UMCU
ENIGMA Discovery Cohorts [N=18309; P=1.298e-05] 2040.38 [ 1123.24, 2957.52]
UKBB 2128.16 [ 1284.95, 2971.37]
Discovery+UKBB [N=28232; P=1.565e-10] 2089.12 [ 1449.23, 2729.01]
GOBS
IMH
UNICAMP
Meth-CT
MIRECC
UKBB nonEUR 4972.38 [ -969.17, 10913.93]
OSAKA
PING Non European
Non European Cohorts [N=260; P=0.101] 4972.38 [ -969.17, 10913.93]
Cohort
Beta [Lower 95% CI,Upper 95% CI]
-30083.08
-17131.5
-4179.91
8771.68
21723.26
mm^2/effect allele

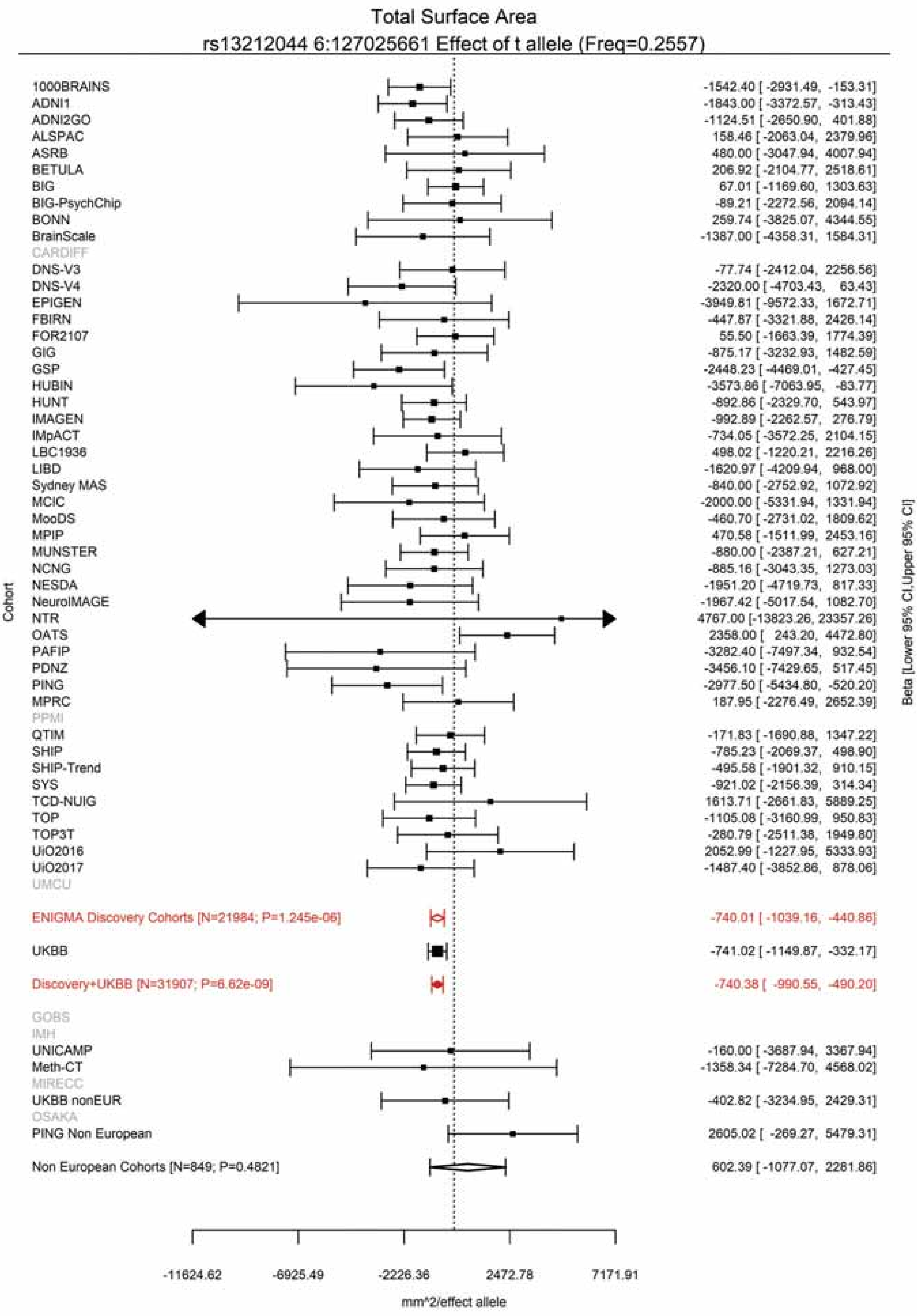
Total Surface Area
rs13212044 6:127025661 Effect of t allele (Freq=0.2557)
1000BRAINS -1542.40 [ -2931.49, -153.31]
ADNI1 -1843.00 [ -3372.57, -313.43]
ADNI2GO -1124.51 [ -2650.90, 401.88]
ALSPAC 158.46 [ -2063.04, 2379.96]
ASRB 480.00 [ -3047.94, 4007.94]
BETULA 206.92 [ -2104.77, 2518.61]
BIG 67.01 [ -1169.60, 1303.63]
BIG-PsychChip -89.21 [ -2272.56, 2094.14]
BONN 259.74 [ -3825.07, 4344.55]
BrainScale -1387.00 [ -4358.31, 1584.31]
CARDIFF
DNS-V3 -77.74 [ -2412.04, 2256.56]
DNS-V4 -2320.00 [ -4703.43, 63.43]
EPIGEN -3949.81 [ -9572.33, 1672.71]
FBIRN -447.87 [ -3321.88, 2426.14]
FOR2107 55.50 [ -1663.39, 1774.39]
GIG -875.17 [ -3232.93, 1482.59]
GSP -2448.23 [ -4469.01, -427.45]
HUBIN -3573.86 [ -7063.95, -83.77]
HUNT -892.86 [ -2329.70, 543.97]
IMAGEN -992.89 [ -2262.57, 276.79]
IMpACT -734.05 [ -3572.25, 2104.15]
LBC1936 498.02 [ -1220.21, 2216.26]
LIBD -1620.97 [ -4209.94, 968.00]
Sydney MAS -840.00 [ -2752.92, 1072.92]
MCIC -2000.00 [ -5331.94, 1331.94]
MooDS -460.70 [ -2731.02, 1809.62]
MPIP 470.58 [ -1511.99, 2453.16]
MUNSTER -880.00 [ -2387.21, 627.21]
NCNG -885.16 [ -3043.35, 1273.03]
NESDA -1951.20 [ -4719.73, 817.33]
NeuroIMAGE -1967.42 [ -5017.54, 1082.70]
NTR 4767.00 [-13823.26, 23357.26]
OATS 2358.00 [ 243.20, 4472.80]
PAFIP -3282.40 [ -7497.34, 932.54]
PDNZ -3456.10 [ -7429.65, 517.45]
PING -2977.50 [ -5434.80, -520.20]
MPRC 187.95 [ -2276.49, 2652.39]
PPMI
QTIM -171.83 [ -1690.88, 1347.22]
SHIP -785.23 [ -2069.37, 498.90]
SHIP-Trend -495.58 [ -1901.32, 910.15]
SYS -921.02 [ -2156.39, 314.34]
TCD-NUIG 1613.71 [ -2661.83, 5889.25]
TOP -1105.08 [ -3160.99, 950.83]
TOP3T -280.79 [ -2511.38, 1949.80]
UiO2016 2052.99 [ -1227.95, 5333.93]
UiO2017 -1487.40 [ -3852.86, 878.06]
UMCU
ENIGMA Discovery Cohorts [N=21984; P=1.245e-06] -740.01 [ -1039.16, -440.86]
UKBB -741.02 [ -1149.87, -332.17]
Discovery+UKBB [N=31907; P=6.62e-09] -740.38 [ -990.55, -490.20]
GOBS
IMH
UNICAMP -160.00 [ -3687.94, 3367.94]
Meth-CT -1358.34 [ -7284.70, 4568.02]
MIRECC
UKBB nonEUR -402.82 [ -3234.95, 2429.31]
OSAKA
PING Non European 2605.02 [ -269.27, 5479.31]
Non European Cohorts [N=849; P=0.4821] 602.39 [ -1077.07, 2281.86]
Cohort
Beta [Lower 95% CI,Upper 95% CI]
-11624.62
-6925.49
-2226.36
2472.78
7171.91
mm^2/effect allele

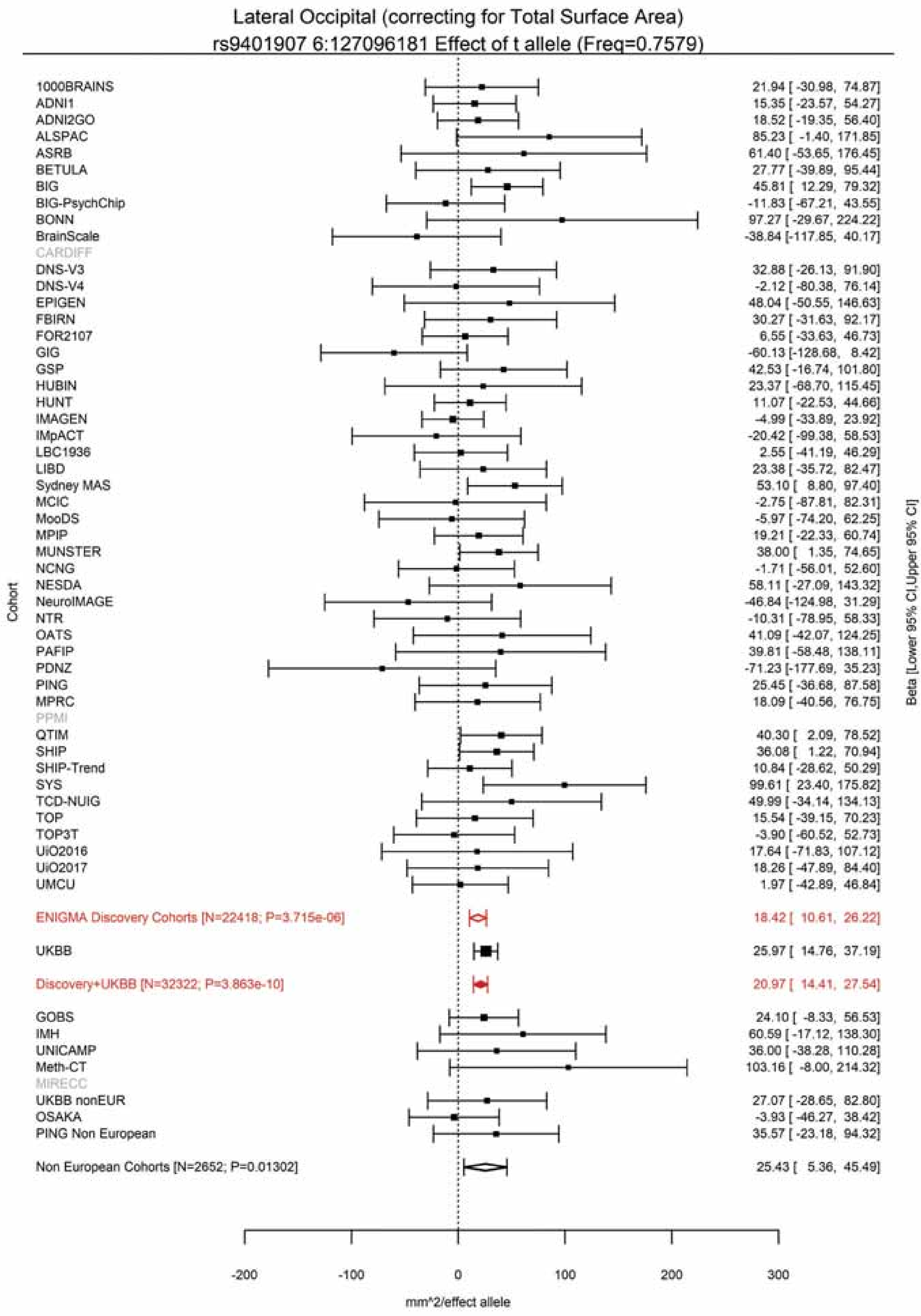

Lateral Occipital (correcting for Total Surface Area)
rs9401907 6:127096181 Effect of t allele (Freq=0.7579)
Cohort
1000BRAINS 21.94 [ -30.98, 74.87]
ADNI1 15.35 [ -23.57, 54.27]
ADNI2GO 18.52 [ -19.35, 56.40]
ALSPAC 85.23 [ -1.40, 171.85]
ASRB 61.40 [ -53.65, 176.45]
BETULA 27.77 [ -39.89, 95.44]
BIG 45.81 [ 12.29, 79.32]
BIG-PsychChip -11.83 [ -67.21, 43.55]
BONN 97.27 [ -29.67, 224.22]
BrainScale -38.84 [-117.85, 40.17]
CARDIFF
DNS-V3 32.88 [ -26.13, 91.90]
DNS-V4 -2.12 [ -80.38, 76.14]
EPIGEN 48.04 [ -50.55, 146.63]
FBIRN 30.27 [ -31.63, 92.17]
FOR2107 6.55 [ -33.63, 46.73]
GIG -60.13 [-128.68, 8.42]
GSP 42.53 [ -16.74, 101.80]
HUBIN 23.37 [ -68.70, 115.45]
HUNT 11.07 [ -22.53, 44.66]
IMAGEN -4.99 [ -33.89, 23.92]
IMpACT -20.42 [ -99.38, 58.53]
LBC1936 2.55 [ -41.19, 46.29]
LIBD 23.38 [ -35.72, 82.47]
Sydney MAS 53.10 [ 8.80, 97.40]
MCIC -2.75 [ -87.81, 82.31]
MooDS -5.97 [ -74.20, 62.25]
MPIP 19.21 [ -22.33, 60.74]
MUNSTER 38.00 [ 1.35, 74.65]
NCNG -1.71 [ -56.01, 52.60]
NESDA 58.11 [ -27.09, 143.32]
NeuroIMAGE -46.84 [-124.98, 31.29]
NTR -10.31 [ -78.95, 58.33]
OATS 41.09 [ -42.07, 124.25]
PAFIP 39.81 [ -58.48, 138.11]
PDNZ -71.23 [-177.69, 35.23]
PING 25.45 [ -36.68, 87.58]
MPRC 18.09 [ -40.56, 76.75]
PPMI
QTIM 40.30 [ 2.09, 78.52]
SHIP 36.08 [ 1.22, 70.94]
SHIP-Trend 10.84 [ -28.62, 50.29]
SYS 99.61 [ 23.40, 175.82]
TCD-NUIG 49.99 [ -34.14, 134.13]
TOP 15.54 [ -39.15, 70.23]
TOP3T -3.90 [ -60.52, 52.73]
UiO2016 17.64 [ -71.83, 107.12]
UiO2017 18.26 [ -47.89, 84.40]
UMCU 1.97 [ -42.89, 46.84]
ENIGMA Discovery Cohorts [N=22418; P=3.715e-06] 18.42 [ 10.61, 26.22]
UKBB 25.97 [ 14.76, 37.19]
Discovery+UKBB [N=32322; P=3.863e-10] 20.97 [ 14.41, 27.54]
GOBS 24.10 [ -8.33, 56.53]
IMH 60.59 [ -17.12, 138.30]
UNICAMP 36.00 [ -38.28, 110.28]
Meth-CT 103.16 [ -8.00, 214.32]
MIRECC
UKBB nonEUR 27.07 [ -28.65, 82.80]
OSAKA -3.93 [ -46.27, 38.42]
PING Non European 35.57 [ -23.18, 94.32]
Non European Cohorts [N=2652; P=0.01302] 25.43 [ 5.36, 45.49]
Beta [Lower 95% CI,Upper 95% CI]
-200
-100
0
100
200
300
mm^2/effect allele

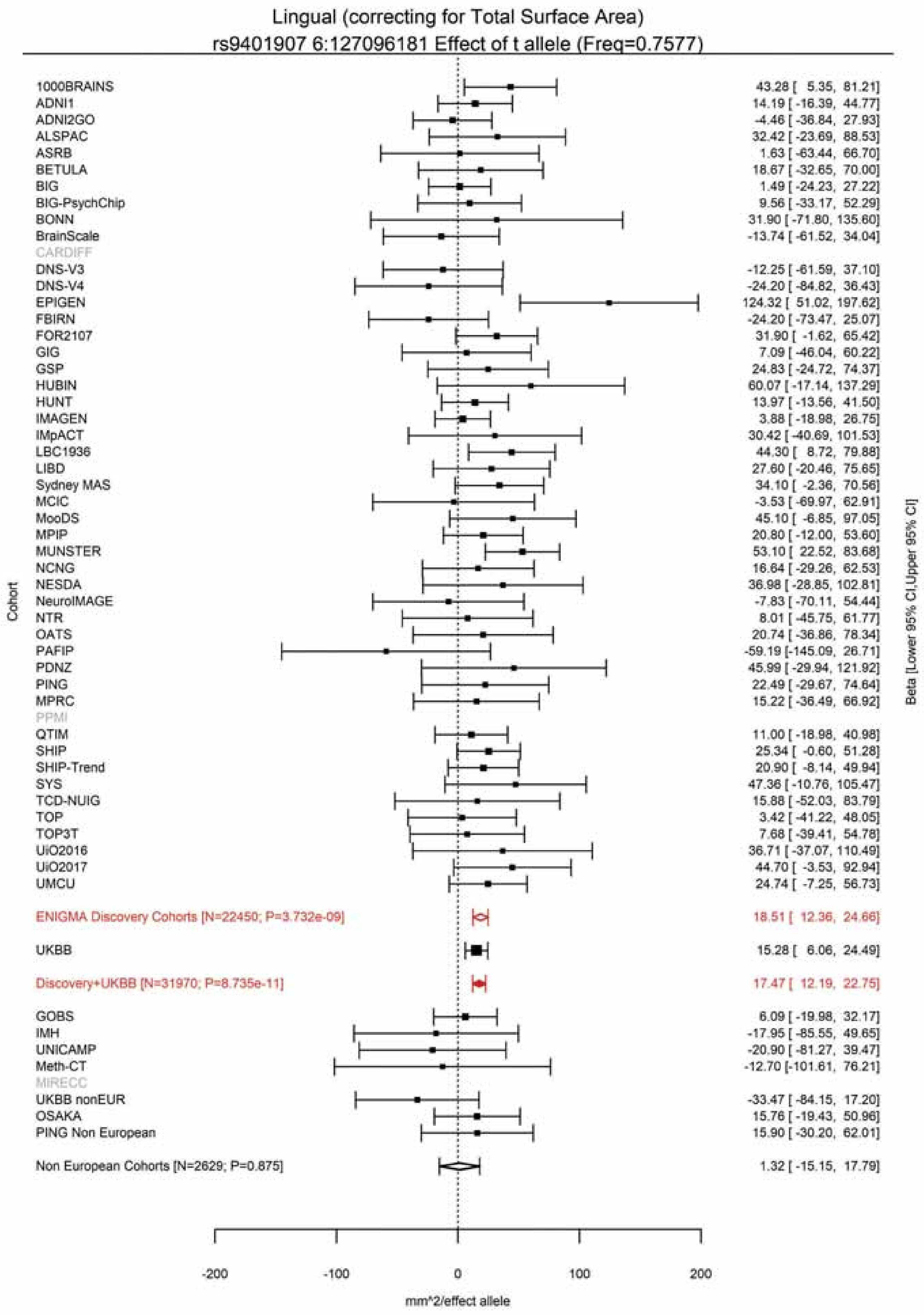

Lingual (correcting for Total Surface Area)
rs9401907 6:127096181 Effect of t allele (Freq=0.7577)
Cohort
Beta [Lower 95% CI,Upper 95% CI]
1000BRAINS 43.28 [ 5.35, 81.21]
ADNI1 14.19 [ -16.39, 44.77]
ADNI2GO -4.46 [ -36.84, 27.93]
ALSPAC 32.42 [ -23.69, 88.53]
ASRB 1.63 [ -63.44, 66.70]
BETULA 18.67 [ -32.65, 70.00]
BIG 1.49 [ -24.23, 27.22]
BIG-PsychChip 9.56 [ -33.17, 52.29]
BONN 31.90 [ -71.80, 135.60]
BrainScale -13.74 [ -61.52, 34.04]
CARDIFF
DNS-V3 -12.25 [ -61.59, 37.10]
DNS-V4 -24.20 [ -84.82, 36.43]
EPIGEN 124.32 [ 51.02, 197.62]
FBIRN -24.20 [ -73.47, 25.07]
FOR2107 31.90 [ -1.62, 65.42]
GIG 7.09 [ -46.04, 60.22]
GSP 24.83 [ -24.72, 74.37]
HUBIN 60.07 [ -17.14, 137.29]
HUNT 13.97 [ -13.56, 41.50]
IMAGEN 3.88 [ -18.98, 26.75]
IMpACT 30.42 [ -40.69, 101.53]
LBC1936 44.30 [ 8.72, 79.88]
LIBD 27.60 [ -20.46, 75.65]
Sydney MAS 34.10 [ -2.36, 70.56]
MCIC -3.53 [ -69.97, 62.91]
MooDS 45.10 [ -6.85, 97.05]
MPIP 20.80 [ -12.00, 53.60]
MUNSTER 53.10 [ 22.52, 83.68]
NCNG 16.64 [ -29.26, 62.53]
NESDA 36.98 [ -28.85, 102.81]
NeuroIMAGE -7.83 [ -70.11, 54.44]
NTR 8.01 [ -45.75, 61.77]
OATS 20.74 [ -36.86, 78.34]
PAFIP -59.19 [-145.09, 26.71]
PDNZ 45.99 [ -29.94, 121.92]
PING 22.49 [ -29.67, 74.64]
MPRC 15.22 [ -36.49, 66.92]
PPMI
QTIM 11.00 [ -18.98, 40.98]
SHIP 25.34 [ -0.60, 51.28]
SHIP-Trend 20.90 [ -8.14, 49.94]
SYS 47.36 [ -10.76, 105.47]
TCD-NUIG 15.88 [ -52.03, 83.79]
TOP 3.42 [ -41.22, 48.05]
TOP3T 7.68 [ -39.41, 54.78]
UiO2016 36.71 [ -37.07, 110.49]
UiO2017 44.70 [ -3.53, 92.94]
UMCU 24.74 [ -7.25, 56.73]
ENIGMA Discovery Cohorts [N=22450; P=3.732e-09] 18.51 [ 12.36, 24.66]
UKBB 15.28 [ 6.06, 24.49]
Discovery+UKBB [N=31970; P=8.735e-11] 17.47 [ 12.19, 22.75]
GOBS 6.09 [ -19.98, 32.17]
IMH -17.95 [ -85.55, 49.65]
UNICAMP -20.90 [ -81.27, 39.47]
Meth-CT -12.70 [-101.61, 76.21]
MIRECC
UKBB nonEUR -33.47 [ -84.15, 17.20]
OSAKA 15.76 [ -19.43, 50.96]
PING Non European 15.90 [ -30.20, 62.01]
Non European Cohorts [N=2629; P=0.875] 1.32 [ -15.15, 17.79]
-200
-100
0
100
200
mm^2/effect allele

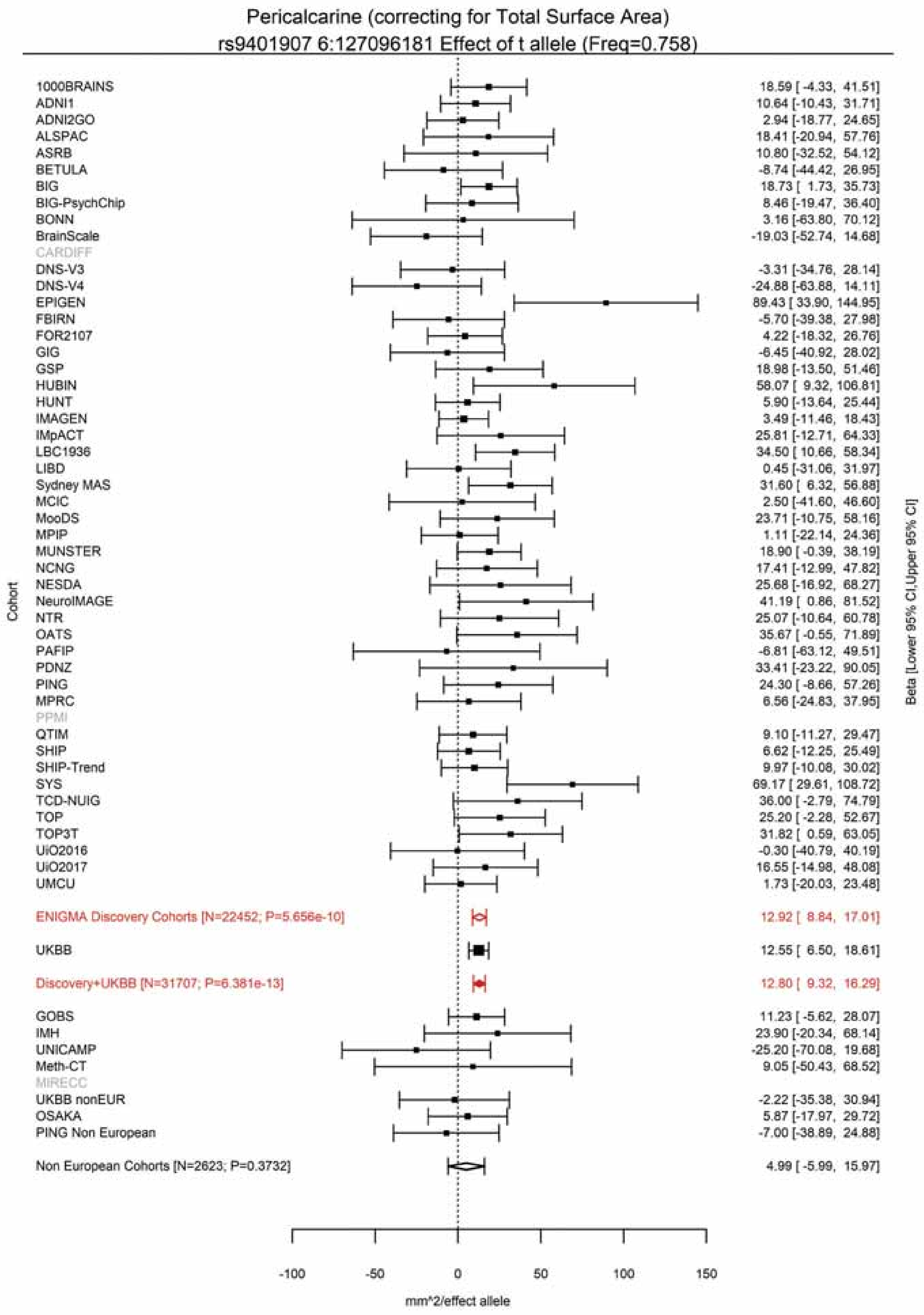

Pericalcarine (correcting for Total Surface Area)
rs9401907 6:127096181 Effect of t allele (Freq=0.758)
Cohort
1000BRAINS 18.59 [ -4.33, 41.51]
ADNI1 10.64 [-10.43, 31.71]
ADNI2GO 2.94 [-18.77, 24.65]
ALSPAC 18.41 [-20.94, 57.76]
ASRB 10.80 [-32.52, 54.12]
BETULA -8.74 [-44.42, 26.95]
BIG 18.73 [ 1.73, 35.73]
BIG-PsychChip 8.46 [-19.47, 36.40]
BONN 3.16 [-63.80, 70.12]
BrainScale -19.03 [-52.74, 14.68]
CARDIFF
DNS-V3 -3.31 [-34.76, 28.14]
DNS-V4 -24.88 [-63.88, 14.11]
EPIGEN 89.43 [ 33.90, 144.95]
FBIRN -5.70 [-39.38, 27.98]
FOR2107 4.22 [-18.32, 26.76]
GIG -6.45 [-40.92, 28.02]
GSP 18.98 [-13.50, 51.46]
HUBIN 58.07 [ 9.32, 106.81]
HUNT 5.90 [-13.64, 25.44]
IMAGEN 3.49 [-11.46, 18.43]
IMpACT 25.81 [-12.71, 64.33]
LBC1936 34.50 [ 10.66, 58.34]
LIBD 0.45 [-31.06, 31.97]
Sydney MAS 31.60 [ 6.32, 56.88]
MCIC 2.50 [-41.60, 46.60]
MooDS 23.71 [-10.75, 58.16]
MPIP 1.11 [-22.14, 24.36]
MUNSTER 18.90 [ -0.39, 38.19]
NCNG 17.41 [-12.99, 47.82]
NESDA 25.68 [-16.92, 68.27]
NeuroIMAGE 41.19 [ 0.86, 81.52]
NTR 25.07 [-10.64, 60.78]
OATS 35.67 [ -0.55, 71.89]
PAFIP -6.81 [-63.12, 49.51]
PDNZ 33.41 [-23.22, 90.05]
PING 24.30 [ -8.66, 57.26]
MPRC 6.56 [-24.83, 37.95]
PPMI
QTIM 9.10 [-11.27, 29.47]
SHIP 6.62 [-12.25, 25.49]
SHIP-Trend 9.97 [-10.08, 30.02]
SYS 69.17 [ 29.61, 108.72]
TCD-NUIG 36.00 [ -2.79, 74.79]
TOP 25.20 [ -2.28, 52.67]
TOP3T 31.82 [ 0.59, 63.05]
UiO2016 -0.30 [-40.79, 40.19]
UiO2017 16.55 [-14.98, 48.08]
UMCU 1.73 [-20.03, 23.48]
ENIGMA Discovery Cohorts [N=22452; P=5.656e-10] 12.92 [ 8.84, 17.01]
UKBB 12.55 [ 6.50, 18.61]
Discovery+UKBB [N=31707; P=6.381e-13] 12.80 [ 9.32, 16.29]
GOBS 11.23 [ -5.62, 28.07]
IMH 23.90 [-20.34, 68.14]
UNICAMP -25.20 [-70.08, 19.68]
Meth-CT 9.05 [-50.43, 68.52]
MIRECC
UKBB nonEUR -2.22 [-35.38, 30.94]
OSAKA 5.87 [-17.97, 29.72]
PING Non European -7.00 [-38.89, 24.88]
Non European Cohorts [N=2623; P=0.3732] 4.99 [ -5.99, 15.97]
-100
-50
0
50
100
150
mm^2/effect allele
Beta [Lower 95% CI,Upper 95% CI]

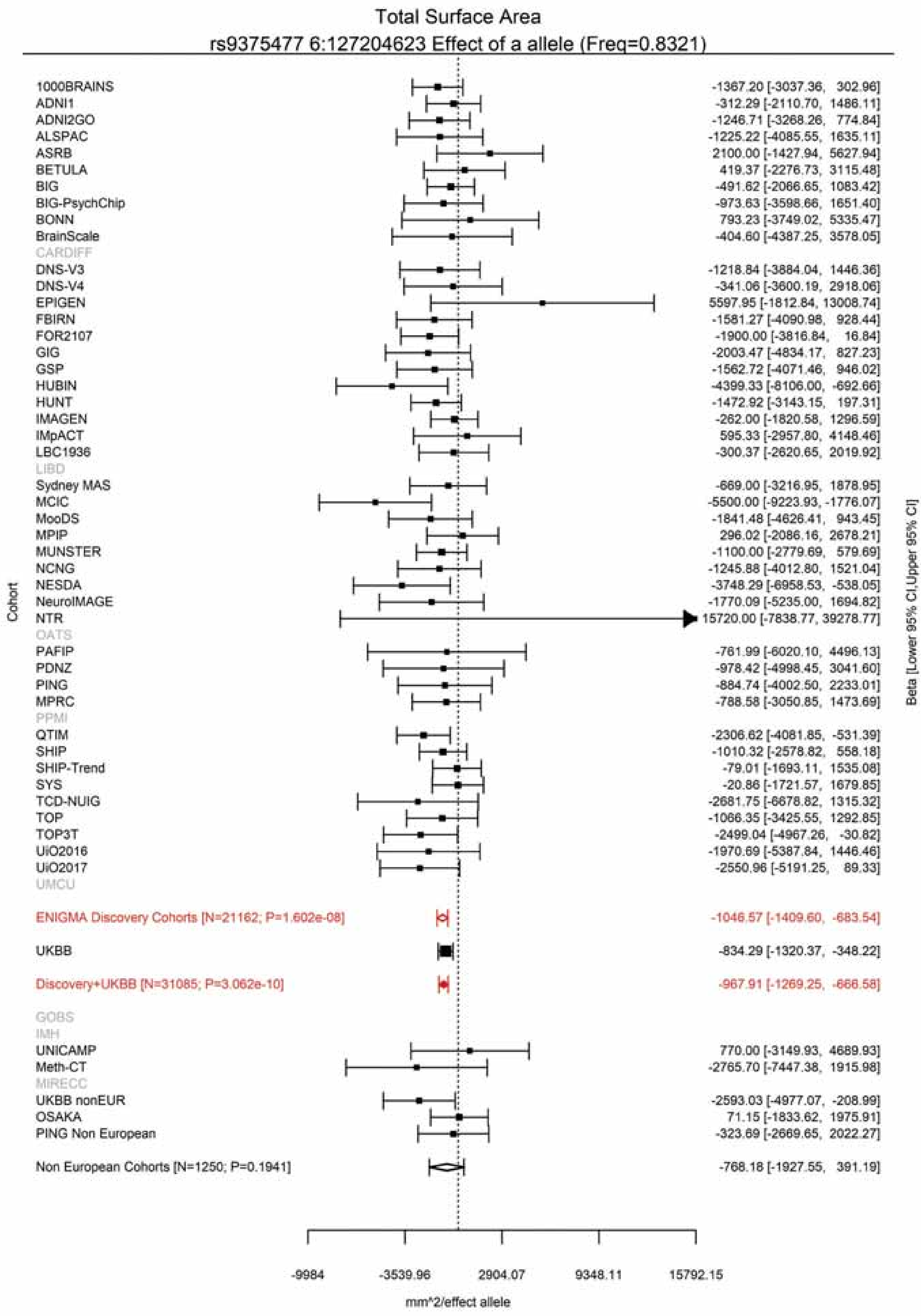
Total Surface Area
rs9375477 6:127204623 Effect of a allele (Freq=0.8321)
1000BRAINS -1367.20 [-3037.36, 302.96]
ADNI1 -312.29 [-2110.70, 1486.11]
ADNI2GO -1246.71 [-3268.26, 774.84]
ALSPAC -1225.22 [-4085.55, 1635.11]
ASRB 2100.00 [-1427.94, 5627.94]
BETULA 419.37 [-2276.73, 3115.48]
BIG -491.62 [-2066.65, 1083.42]
BIG-PsychChip -973.63 [-3598.66, 1651.40]
BONN 793.23 [-3749.02, 5335.47]
BrainScale -404.60 [-4387.25, 3578.05]
CARDIFF
DNS-V3 -1218.84 [-3884.04, 1446.36]
DNS-V4 -341.06 [-3600.19, 2918.06]
EPIGEN 5597.95 [-1812.84, 13008.74]
FBIRN -1581.27 [-4090.98, 928.44]
FOR2107 -1900.00 [-3816.84, 16.84]
GIG -2003.47 [-4834.17, 827.23]
GSP -1562.72 [-4071.46, 946.02]
HUBIN -4399.33 [-8106.00, -692.66]
HUNT -1472.92 [-3143.15, 197.31]
IMAGEN -262.00 [-1820.58, 1296.59]
IMpACT 595.33 [-2957.80, 4148.46]
LBC1936 -300.37 [-2620.65, 2019.92]
LIBD
Sydney MAS -669.00 [-3216.95, 1878.95]
MCIC -5500.00 [-9223.93, -1776.07]
MooDS -1841.48 [-4626.41, 943.45]
MPIP 296.02 [-2086.16, 2678.21]
MUNSTER -1100.00 [-2779.69, 579.69]
NCNG -1245.88 [-4012.80, 1521.04]
NESDA -3748.29 [-6958.53, -538.05]
NeuroIMAGE -1770.09 [-5235.00, 1694.82]
NTR 15720.00 [-7838.77, 39278.77]
OATS
PAFIP -761.99 [-6020.10, 4496.13]
PDNZ -978.42 [-4998.45, 3041.60]
PING -884.74 [-4002.50, 2233.01]
MPRC -788.58 [-3050.85, 1473.69]
PPMI
QTIM -2306.62 [-4081.85, -531.39]
SHIP -1010.32 [-2578.82, 558.18]
SHIP-Trend -79.01 [-1693.11, 1535.08]
SYS -20.86 [-1721.57, 1679.85]
TCD-NUIG -2681.75 [-6678.82, 1315.32]
TOP -1066.35 [-3425.55, 1292.85]
TOP3T -2499.04 [-4967.26, -30.82]
UiO2016 -1970.69 [-5387.84, 1446.46]
UiO2017 -2550.96 [-5191.25, 89.33]
UMCU
ENIGMA Discovery Cohorts [N=21162; P=1.602e-08] -1046.57 [-1409.60, -683.54]
UKBB -834.29 [-1320.37, -348.22]
Discovery+UKBB [N=31085; P=3.062e-10] -967.91 [-1269.25, -666.58]
GOBS
IMH
UNICAMP 770.00 [-3149.93, 4689.93]
Meth-CT -2765.70 [-7447.38, 1915.98]
MIRECC
UKBB nonEUR -2593.03 [-4977.07, -208.99]
OSAKA 71.15 [-1833.62, 1975.91]
PING Non European -323.69 [-2669.65, 2022.27]
Non European Cohorts [N=1250; P=0.1941] -768.18 [-1927.55, 391.19]
Cohort
Beta [Lower 95% CI,Upper 95% CI]
-9984
-3539.96
2904.07
9348.11
15792.15
mm^2/effect allele

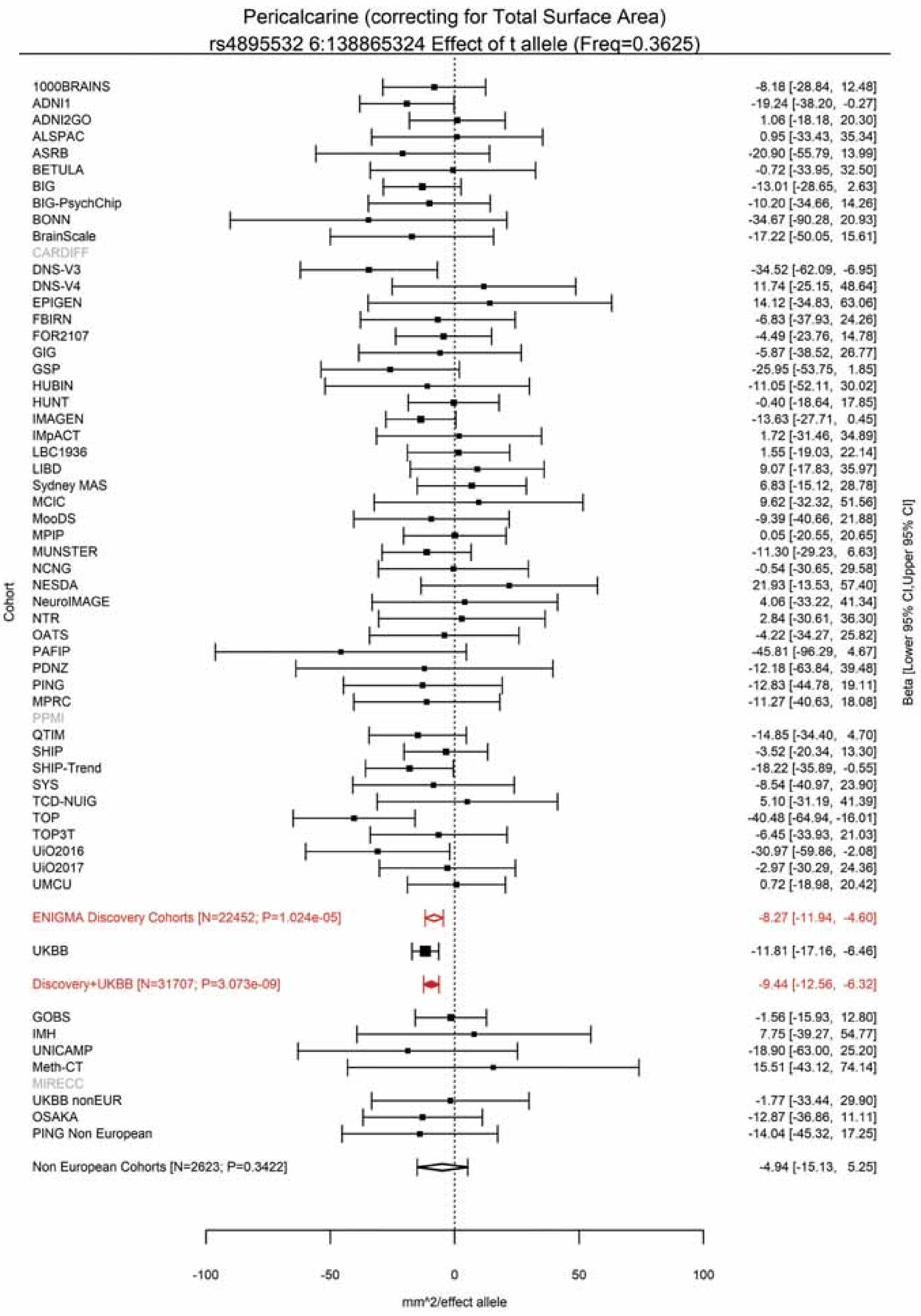

Pericalcarine (correcting for Total Surface Area)
rs4895532 6:138865324 Effect of t allele (Freq=0.3625)
Cohort
1000BRAINS -8.18 [-28.84, 12.48]
ADNI1 -19.24 [-38.20, -0.27]
ADNI2GO 1.06 [-18.18, 20.30]
ALSPAC 0.95 [-33.43, 35.34]
ASRB -20.90 [-55.79, 13.99]
BETULA -0.72 [-33.95, 32.50]
BIG -13.01 [-28.65, 2.63]
BIG-PsychChip -10.20 [-34.66, 14.26]
BONN -34.67 [-90.28, 20.93]
BrainScale -17.22 [-50.05, 15.61]
CARDIFF
DNS-V3 -34.52 [-62.09, -6.95]
DNS-V4 11.74 [-25.15, 48.64]
EPIGEN 14.12 [-34.83, 63.06]
FBIRN -6.83 [-37.93, 24.26]
FOR2107 -4.49 [-23.76, 14.78]
GIG -5.87 [-38.52, 26.77]
GSP -25.95 [-53.75, 1.85]
HUBIN -11.05 [-52.11, 30.02]
HUNT -0.40 [-18.64, 17.85]
IMAGEN -13.63 [-27.71, 0.45]
IMpACT 1.72 [-31.46, 34.89]
LBC1936 1.55 [-19.03, 22.14]
LIBD 9.07 [-17.83, 35.97]
Sydney MAS 6.83 [-15.12, 28.78]
MCIC 9.62 [-32.32, 51.56]
MooDS -9.39 [-40.66, 21.88]
MPIP 0.05 [-20.55, 20.65]
MUNSTER -11.30 [-29.23, 6.63]
NCNG -0.54 [-30.65, 29.58]
NESDA 21.93 [-13.53, 57.40]
NeuroIMAGE 4.06 [-33.22, 41.34]
NTR 2.84 [-30.61, 36.30]
OATS -4.22 [-34.27, 25.82]
PAFIP -45.81 [-96.29, 4.67]
PDNZ -12.18 [-63.84, 39.48]
PING -12.83 [-44.78, 19.11]
MPRC -11.27 [-40.63, 18.08]
PPMI
QTIM -14.85 [-34.40, 4.70]
SHIP -3.52 [-20.34, 13.30]
SHIP-Trend -18.22 [-35.89, -0.55]
SYS -8.54 [-40.97, 23.90]
TCD-NUIG 5.10 [-31.19, 41.39]
TOP -40.48 [-64.94, -16.01]
TOP3T -6.45 [-33.93, 21.03]
UiO2016 -30.97 [-59.86, -2.08]
UiO2017 -2.97 [-30.29, 24.36]
UMCU 0.72 [-18.98, 20.42]
ENIGMA Discovery Cohorts [N=22452; P=1.024e-05] -8.27 [-11.94, -4.60]
UKBB -11.81 [-17.16, -6.46]
Discovery+UKBB [N=31707; P=3.073e-09] -9.44 [-12.56, -6.32]
GOBS -1.56 [-15.93, 12.80]
IMH 7.75 [-39.27, 54.77]
UNICAMP -18.90 [-63.00, 25.20]
Meth-CT 15.51 [-43.12, 74.14]
MIRECC
UKBB nonEUR -1.77 [-33.44, 29.90]
OSAKA -12.87 [-36.86, 11.11]
PING Non European -14.04 [-45.32, 17.25]
Non European Cohorts [N=2623; P=0.3422] -4.94 [-15.13, 5.25]
Beta [Lower 95% CI,Upper 95% CI]
-100
-50
0
50
100
mm^2/effect allele

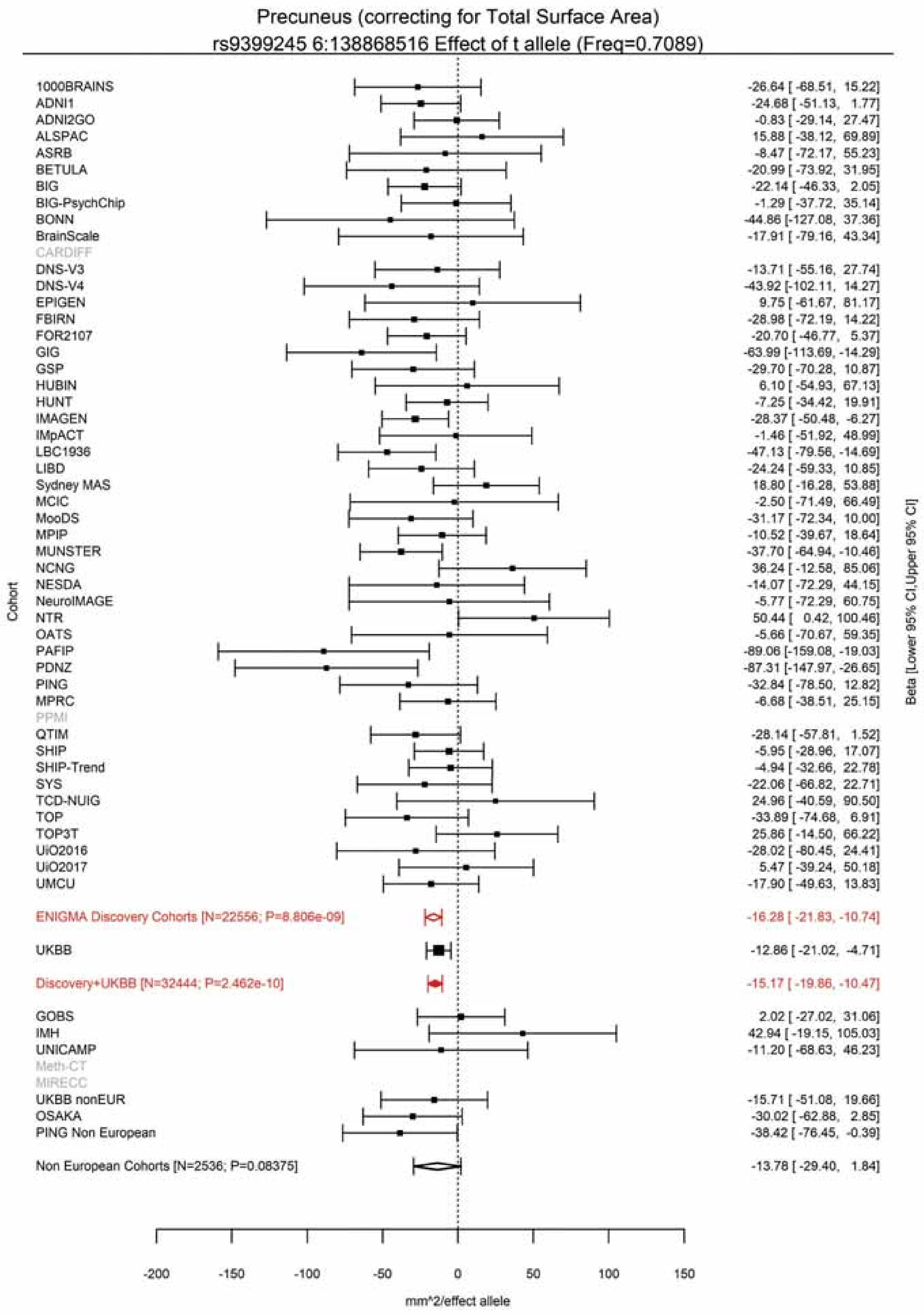
Precuneus (correcting for Total Surface Area)
rs9399245 6:138868516 Effect of t allele (Freq=0.7089)
Cohort
1000BRAINS -26.64 [ -68.51, 15.22]
ADNI1 -24.68 [ -51.13, 1.77]
ADNI2GO -0.83 [ -29.14, 27.47]
ALSPAC 15.88 [ -38.12, 69.89]
ASRB -8.47 [ -72.17, 55.23]
BETULA -20.99 [ -73.92, 31.95]
BIG -22.14 [ -46.33, 2.05]
BIG-PsychChip -1.29 [ -37.72, 35.14]
BONN -44.86 [-127.08, 37.36]
BrainScale -17.91 [ -79.16, 43.34]
CARDIFF
DNS-V3 -13.71 [ -55.16, 27.74]
DNS-V4 -43.92 [-102.11, 14.27]
EPIGEN 9.75 [ -61.67, 81.17]
FBIRN -28.98 [ -72.19, 14.22]
FOR2107 -20.70 [ -46.77, 5.37]
GIG -63.99 [-113.69, -14.29]
GSP -29.70 [ -70.28, 10.87]
HUBIN 6.10 [ -54.93, 67.13]
HUNT -7.25 [ -34.42, 19.91]
IMAGEN -28.37 [ -50.48, -6.27]
IMpACT -1.46 [ -51.92, 48.99]
LBC1936 -47.13 [ -79.56, -14.69]
LIBD -24.24 [ -59.33, 10.85]
Sydney MAS 18.80 [ -16.28, 53.88]
MCIC -2.50 [ -71.49, 66.49]
MooDS -31.17 [ -72.34, 10.00]
MPIP -10.52 [ -39.67, 18.64]
MUNSTER -37.70 [ -64.94, -10.46]
NCNG 36.24 [ -12.58, 85.06]
NESDA -14.07 [ -72.29, 44.15]
NeuroIMAGE -5.77 [ -72.29, 60.75]
NTR 50.44 [ 0.42, 100.46]
OATS -5.66 [ -70.67, 59.35]
PAFIP -89.06 [-159.08, -19.03]
PDNZ -87.31 [-147.97, -26.65]
PING -32.84 [ -78.50, 12.82]
MPRC -6.68 [ -38.51, 25.15]
PPMI
QTIM -28.14 [ -57.81, 1.52]
SHIP -5.95 [ -28.96, 17.07]
SHIP-Trend -4.94 [ -32.66, 22.78]
SYS -22.06 [ -66.82, 22.71]
TCD-NUIG 24.96 [ -40.59, 90.50]
TOP -33.89 [ -74.68, 6.91]
TOP3T 25.86 [ -14.50, 66.22]
UiO2016 -28.02 [ -80.45, 24.41]
UiO2017 5.47 [ -39.24, 50.18]
UMCU -17.90 [ -49.63, 13.83]
ENIGMA Discovery Cohorts [N=22556; P=8.806e-09] -16.28 [ -21.83, -10.74]
UKBB -12.86 [ -21.02, -4.71]
Discovery+UKBB [N=32444; P=2.462e-10] -15.17 [ -19.86, -10.47]
GOBS 2.02 [ -27.02, 31.06]
IMH 42.94 [ -19.15, 105.03]
UNICAMP -11.20 [ -68.63, 46.23]
Meth-CT
MIRECC
UKBB nonEUR -15.71 [ -51.08, 19.66]
OSAKA -30.02 [ -62.88, 2.85]
PING Non European -38.42 [ -76.45, -0.39]
Non European Cohorts [N=2536; P=0.08375] -13.78 [ -29.40, 1.84]
Beta [Lower 95% CI,Upper 95% CI]
-200 -150 -100 -50 0 50 100 150
mm^2/effect allele

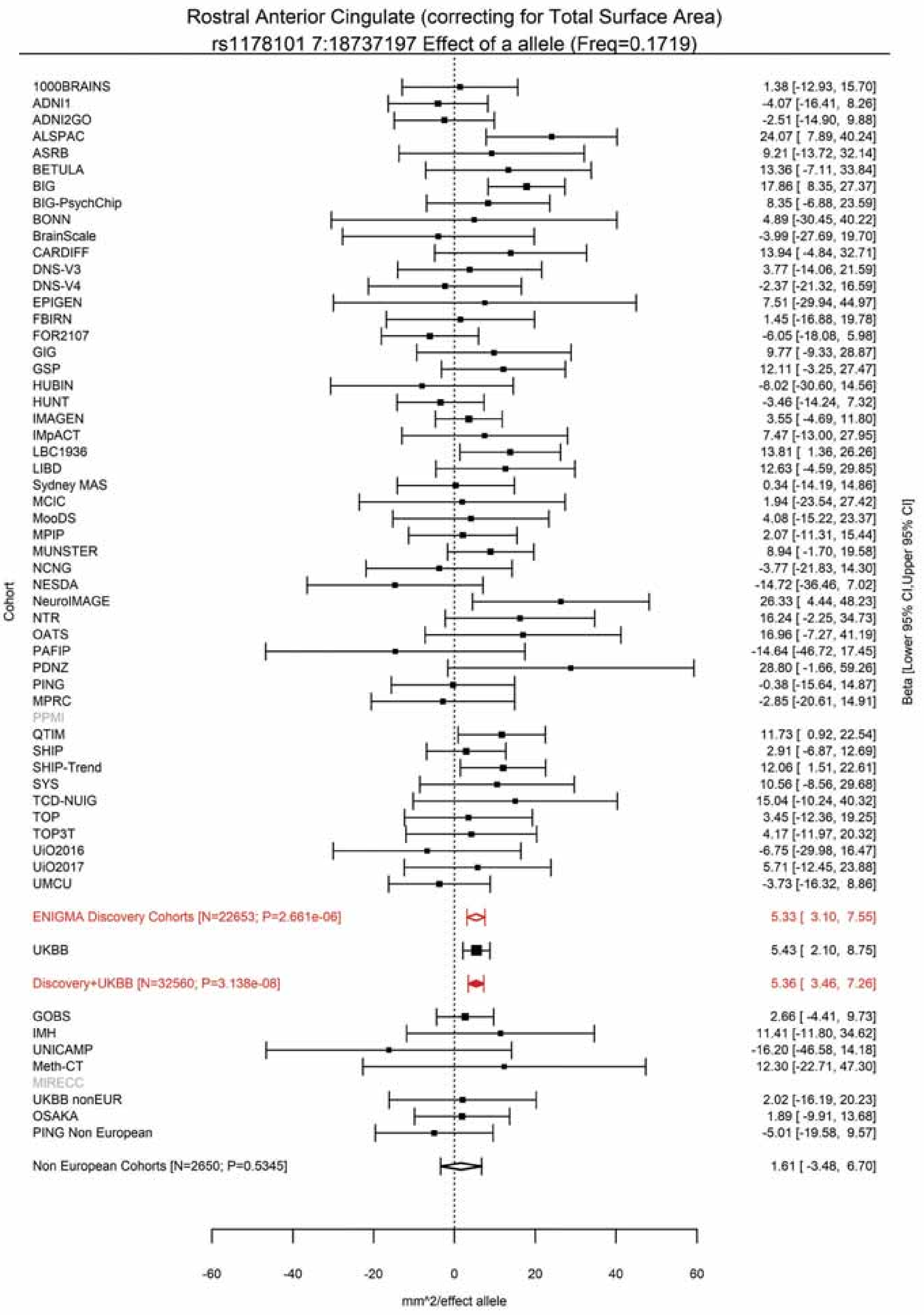
Rostral Anterior Cingulate (correcting for Total Surface Area)
rs1178101 7:18737197 Effect of a allele (Freq=0.1719)
Cohort
Beta [Lower 95% CI,Upper 95% CI]
1000BRAINS 1.38 [-12.93, 15.70]
ADNI1 -4.07 [-16.41, 8.26]
ADNI2GO -2.51 [-14.90, 9.88]
ALSPAC 24.07 [ 7.89, 40.24]
ASRB 9.21 [-13.72, 32.14]
BETULA 13.36 [ -7.11, 33.84]
BIG 17.86 [ 8.35, 27.37]
BIG-PsychChip 8.35 [ -6.88, 23.59]
BONN 4.89 [-30.45, 40.22]
BrainScale -3.99 [-27.69, 19.70]
CARDIFF 13.94 [ -4.84, 32.71]
DNS-V3 3.77 [-14.06, 21.59]
DNS-V4 -2.37 [-21.32, 16.59]
EPIGEN 7.51 [-29.94, 44.97]
FBIRN 1.45 [-16.88, 19.78]
FOR2107 -6.05 [-18.08, 5.98]
GIG 9.77 [ -9.33, 28.87]
GSP 12.11 [ -3.25, 27.47]
HUBIN -8.02 [-30.60, 14.56]
HUNT -3.46 [-14.24, 7.32]
IMAGEN 3.55 [ -4.69, 11.80]
IMpACT 7.47 [-13.00, 27.95]
LBC1936 13.81 [ 1.36, 26.26]
LIBD 12.63 [ -4.59, 29.85]
Sydney MAS 0.34 [-14.19, 14.86]
MCIC 1.94 [-23.54, 27.42]
MooDS 4.08 [-15.22, 23.37]
MPIP 2.07 [-11.31, 15.44]
MUNSTER 8.94 [ -1.70, 19.58]
NCNG -3.77 [-21.83, 14.30]
NESDA -14.72 [-36.46, 7.02]
NeuroIMAGE 26.33 [ 4.44, 48.23]
NTR 16.24 [ -2.25, 34.73]
OATS 16.96 [ -7.27, 41.19]
PAFIP -14.64 [-46.72, 17.45]
PDNZ 28.80 [ -1.66, 59.26]
PING -0.38 [-15.64, 14.87]
MPRC -2.85 [-20.61, 14.91]
PPMI
QTIM 11.73 [ 0.92, 22.54]
SHIP 2.91 [ -6.87, 12.69]
SHIP-Trend 12.06 [ 1.51, 22.61]
SYS 10.56 [ -8.56, 29.68]
TCD-NUIG 15.04 [-10.24, 40.32]
TOP 3.45 [-12.36, 19.25]
TOP3T 4.17 [-11.97, 20.32]
UiO2016 -6.75 [-29.98, 16.47]
UiO2017 5.71 [-12.45, 23.88]
UMCU -3.73 [-16.32, 8.86]
ENIGMA Discovery Cohorts [N=22653; P=2.661e-06] 5.33 [ 3.10, 7.55]
UKBB 5.43 [ 2.10, 8.75]
Discovery+UKBB [N=32560; P=3.138e-08] 5.36 [ 3.46, 7.26]
GOBS 2.66 [ -4.41, 9.73]
IMH 11.41 [-11.80, 34.62]
UNICAMP -16.20 [-46.58, 14.18]
Meth-CT 12.30 [-22.71, 47.30]
MIRECC
UKBB nonEUR 2.02 [-16.19, 20.23]
OSAKA 1.89 [ -9.91, 13.68]
PING Non European -5.01 [-19.58, 9.57]
Non European Cohorts [N=2650; P=0.5345] 1.61 [ -3.48, 6.70]
-60
-40
-20
0
20
40
60
mm^2/effect allele

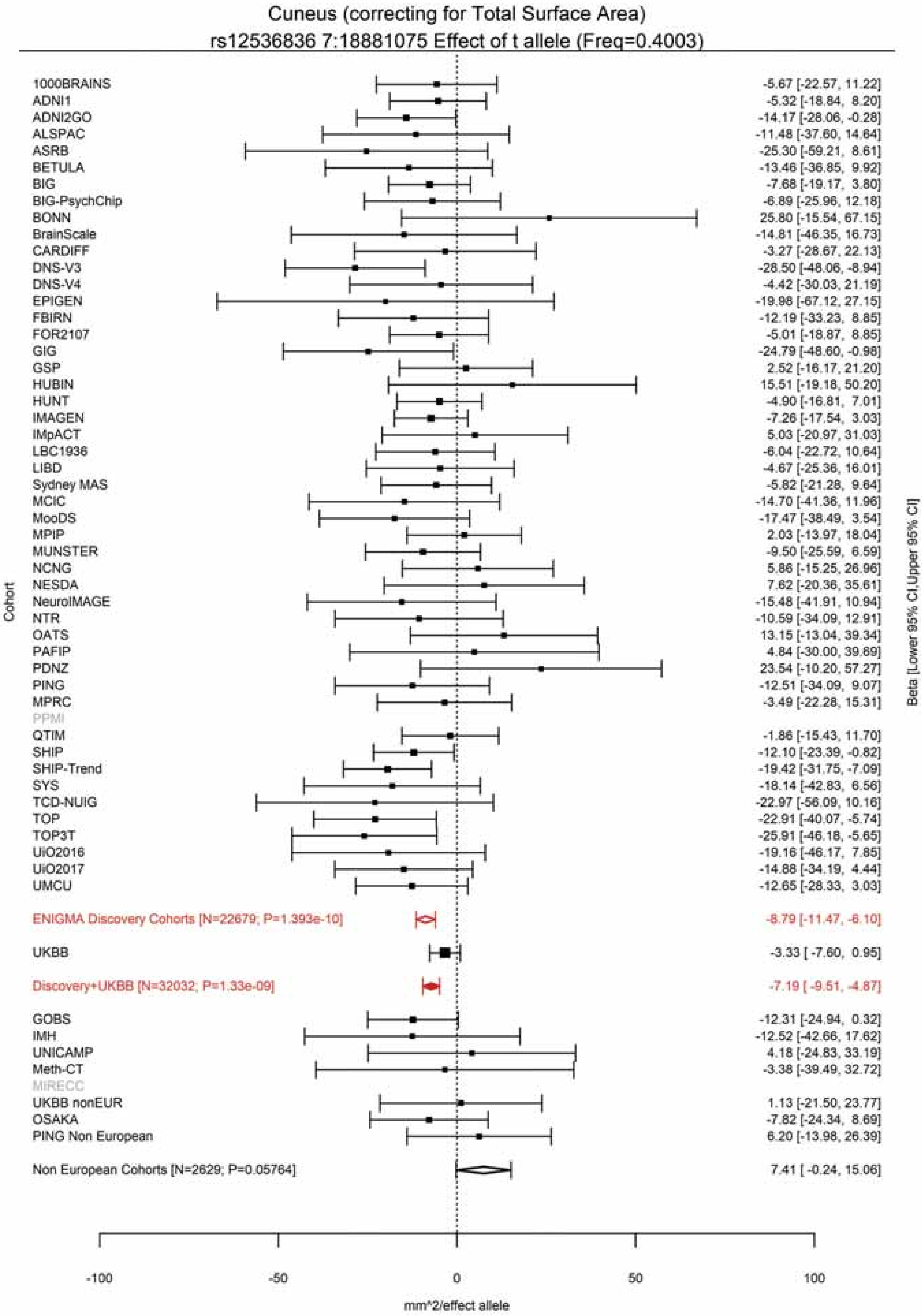

Cuneus (correcting for Total Surface Area)
rs12536836 7:18881075 Effect of t allele (Freq=0.4003)
1000BRAINS -5.67 [-22.57, 11.22]
ADNI1 -5.32 [-18.84, 8.20]
ADNI2GO -14.17 [-28.06, -0.28]
ALSPAC -11.48 [-37.60, 14.64]
ASRB -25.30 [-59.21, 8.61]
BETULA -13.46 [-36.85, 9.92]
BIG -7.68 [-19.17, 3.80]
BIG-PsychChip -6.89 [-25.96, 12.18]
BONN 25.80 [-15.54, 67.15]
BrainScale -14.81 [-46.35, 16.73]
CARDIFF -3.27 [-28.67, 22.13]
DNS-V3 -28.50 [-48.06, -8.94]
DNS-V4 -4.42 [-30.03, 21.19]
EPIGEN -19.98 [-67.12, 27.15]
FBIRN -12.19 [-33.23, 8.85]
FOR2107 -5.01 [-18.87, 8.85]
GIG -24.79 [-48.60, -0.98]
GSP 2.52 [-16.17, 21.20]
HUBIN 15.51 [-19.18, 50.20]
HUNT -4.90 [-16.81, 7.01]
IMAGEN -7.26 [-17.54, 3.03]
IMpACT 5.03 [-20.97, 31.03]
LBC1936 -6.04 [-22.72, 10.64]
LIBD -4.67 [-25.36, 16.01]
Sydney MAS -5.82 [-21.28, 9.64]
MCIC -14.70 [-41.36, 11.96]
MooDS -17.47 [-38.49, 3.54]
MPIP 2.03 [-13.97, 18.04]
MUNSTER -9.50 [-25.59, 6.59]
NCNG 5.86 [-15.25, 26.96]
NESDA 7.62 [-20.36, 35.61]
NeuroIMAGE -15.48 [-41.91, 10.94]
NTR -10.59 [-34.09, 12.91]
OATS 13.15 [-13.04, 39.34]
PAFIP 4.84 [-30.00, 39.69]
PDNZ 23.54 [-10.20, 57.27]
PING -12.51 [-34.09, 9.07]
MPRC -3.49 [-22.28, 15.31]
PPMI
QTIM -1.86 [-15.43, 11.70]
SHIP -12.10 [-23.39, -0.82]
SHIP-Trend -19.42 [-31.75, -7.09]
SYS -18.14 [-42.83, 6.56]
TCD-NUIG -22.97 [-56.09, 10.16]
TOP -22.91 [-40.07, -5.74]
TOP3T -25.91 [-46.18, -5.65]
UiO2016 -19.16 [-46.17, 7.85]
UiO2017 -14.88 [-34.19, 4.44]
UMCU -12.65 [-28.33, 3.03]
ENIGMA Discovery Cohorts [N=22679; P=1.393e-10] -8.79 [-11.47, -6.10]
UKBB -3.33 [-7.60, 0.95]
Discovery+UKBB [N=32032; P=1.33e-09] -7.19 [-9.51, -4.87]
GOBS -12.31 [-24.94, 0.32]
IMH -12.52 [-42.66, 17.62]
UNICAMP 4.18 [-24.83, 33.19]
Meth-CT -3.38 [-39.49, 32.72]
MIRECC
UKBB nonEUR 1.13 [-21.50, 23.77]
OSAKA -7.82 [-24.34, 8.69]
PING Non European 6.20 [-13.98, 26.39]
Non European Cohorts [N=2629; P=0.05764] 7.41 [-0.24, 15.06]
Cohort
Beta [Lower 95% CI,Upper 95% CI]
-100
-50
0
50
100
mm^2/effect allele

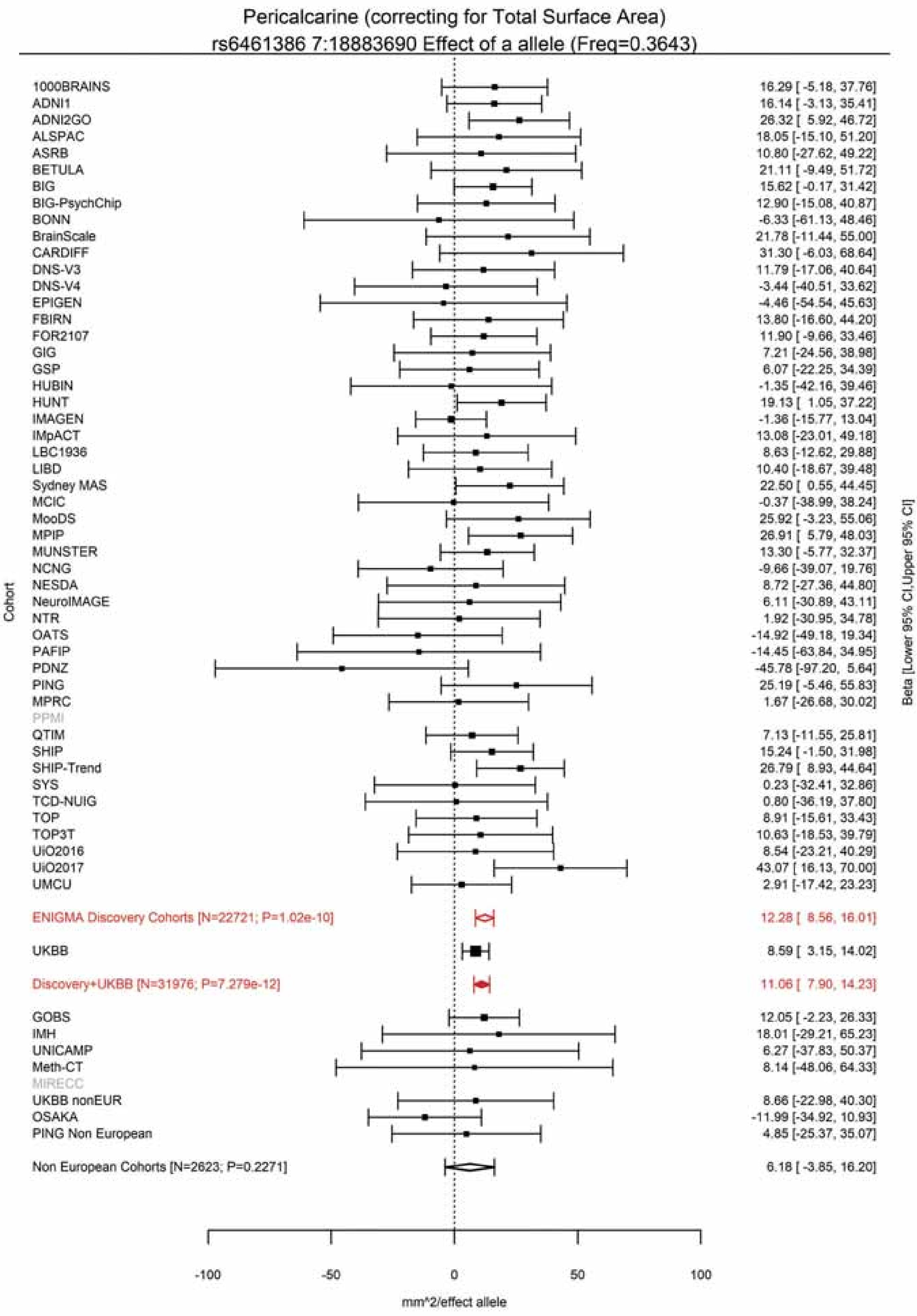
Pericalcarine (correcting for Total Surface Area)
rs6461386 7:18883690 Effect of a allele (Freq=0.3643)
Cohort
1000BRAINS 16.29 [ -5.18, 37.76]
ADNI1 16.14 [ -3.13, 35.41]
ADNI2GO 26.32 [ 5.92, 46.72]
ALSPAC 18.05 [-15.10, 51.20]
ASRB 10.80 [-27.62, 49.22]
BETULA 21.11 [ -9.49, 51.72]
BIG 15.62 [ -0.17, 31.42]
BIG-PsychChip 12.90 [-15.08, 40.87]
BONN -6.33 [-61.13, 48.46]
BrainScale 21.78 [-11.44, 55.00]
CARDIFF 31.30 [ -6.03, 68.64]
DNS-V3 11.79 [-17.06, 40.64]
DNS-V4 -3.44 [-40.51, 33.62]
EPIGEN -4.46 [-54.54, 45.63]
FBIRN 13.80 [-16.60, 44.20]
FOR2107 11.90 [ -9.66, 33.46]
GIG 7.21 [-24.56, 38.98]
GSP 6.07 [-22.25, 34.39]
HUBIN -1.35 [-42.16, 39.46]
HUNT 19.13 [ 1.05, 37.22]
IMAGEN -1.36 [-15.77, 13.04]
IMpACT 13.08 [-23.01, 49.18]
LBC1936 8.63 [-12.62, 29.88]
LIBD 10.40 [-18.67, 39.48]
Sydney MAS 22.50 [ 0.55, 44.45]
MCIC -0.37 [-38.99, 38.24]
MooDS 25.92 [ -3.23, 55.06]
MPIP 26.91 [ 5.79, 48.03]
MUNSTER 13.30 [ -5.77, 32.37]
NCNG -9.66 [-39.07, 19.76]
NESDA 8.72 [-27.36, 44.80]
NeuroIMAGE 6.11 [-30.89, 43.11]
NTR 1.92 [-30.95, 34.78]
OATS -14.92 [-49.18, 19.34]
PAFIP -14.45 [-63.84, 34.95]
PDNZ -45.78 [-97.20, 5.64]
PING 25.19 [ -5.46, 55.83]
MPRC 1.67 [-26.68, 30.02]
PPMI
QTIM 7.13 [-11.55, 25.81]
SHIP 15.24 [ -1.50, 31.98]
SHIP-Trend 26.79 [ 8.93, 44.64]
SYS 0.23 [-32.41, 32.86]
TCD-NUIG 0.80 [-36.19, 37.80]
TOP 8.91 [-15.61, 33.43]
TOP3T 10.63 [-18.53, 39.79]
UiO2016 8.54 [-23.21, 40.29]
UiO2017 43.07 [ 16.13, 70.00]
UMCU 2.91 [-17.42, 23.23]
ENIGMA Discovery Cohorts [N=22721; P=1.02e-10] 12.28 [ 8.56, 16.01]
UKBB 8.59 [ 3.15, 14.02]
Discovery+UKBB [N=31976; P=7.279e-12] 11.06 [ 7.90, 14.23]
GOBS 12.05 [ -2.23, 26.33]
IMH 18.01 [-29.21, 65.23]
UNICAMP 6.27 [-37.83, 50.37]
Meth-CT 8.14 [-48.06, 64.33]
MIRECC
UKBB nonEUR 8.66 [-22.98, 40.30]
OSAKA -11.99 [-34.92, 10.93]
PING Non European 4.85 [-25.37, 35.07]
Non European Cohorts [N=2623; P=0.2271] 6.18 [ -3.85, 16.20]
Beta [Lower 95% CI,Upper 95% CI]
-100
-50
0
50
100
mm^2/effect allele

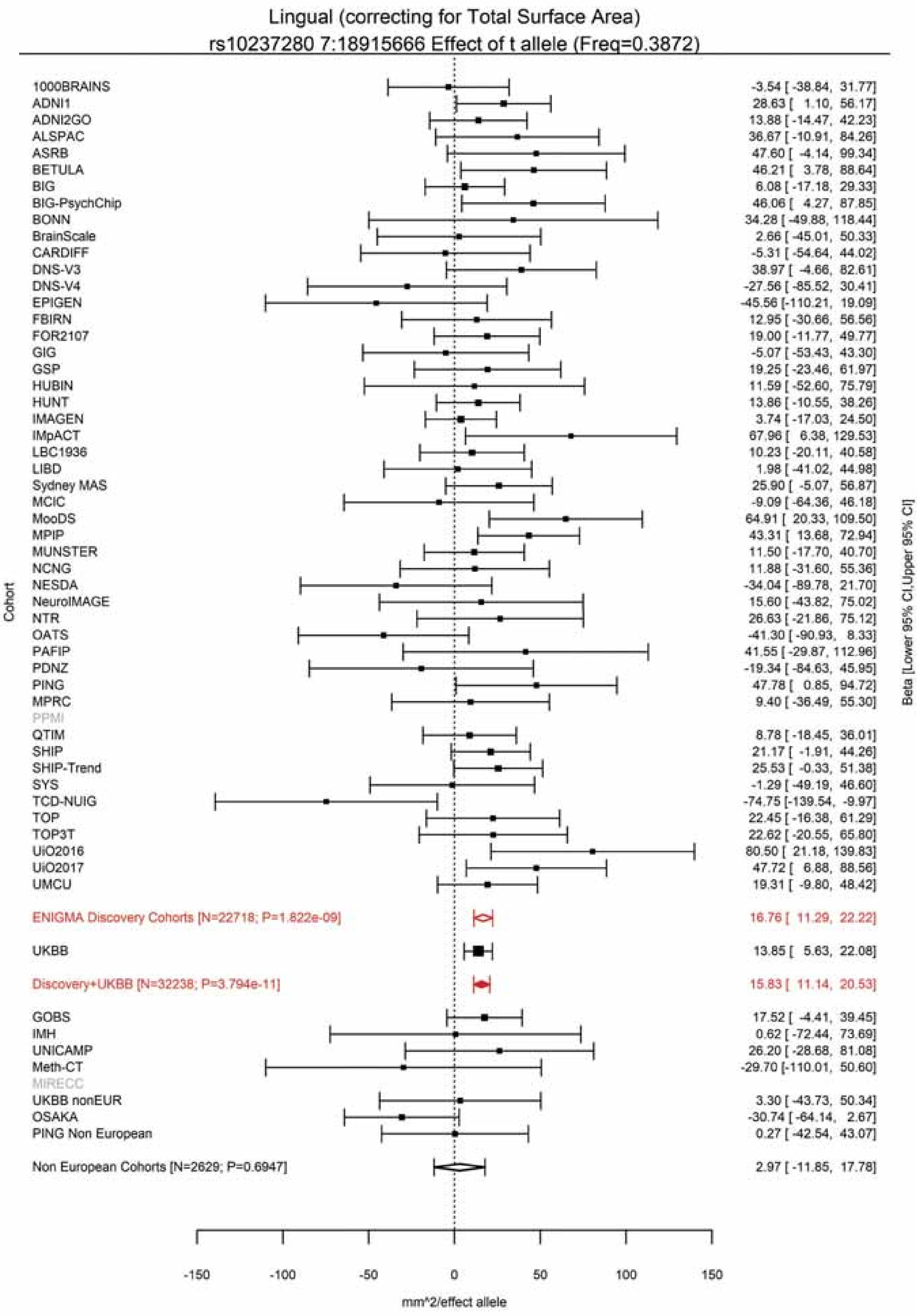
Lingual (correcting for Total Surface Area)
rs10237280 7:18915666 Effect of t allele (Freq=0.3872)
Cohort
1000BRAINS -3.54 [ -38.84, 31.77]
ADNI1 28.63 [ 1.10, 56.17]
ADNI2GO 13.88 [ -14.47, 42.23]
ALSPAC 36.67 [ -10.91, 84.26]
ASRB 47.60 [ -4.14, 99.34]
BETULA 46.21 [ 3.78, 88.64]
BIG 6.08 [ -17.18, 29.33]
BIG-PsychChip 46.06 [ 4.27, 87.85]
BONN 34.28 [ -49.88, 118.44]
BrainScale 2.66 [ -45.01, 50.33]
CARDIFF -5.31 [ -54.64, 44.02]
DNS-V3 38.97 [ -4.66, 82.61]
DNS-V4 -27.56 [ -85.52, 30.41]
EPIGEN -45.56 [-110.21, 19.09]
FBIRN 12.95 [ -30.66, 56.56]
FOR2107 19.00 [ -11.77, 49.77]
GIG -5.07 [ -53.43, 43.30]
GSP 19.25 [ -23.46, 61.97]
HUBIN 11.59 [ -52.60, 75.79]
HUNT 13.86 [ -10.55, 38.26]
IMAGEN 3.74 [ -17.03, 24.50]
IMpACT 67.96 [ 6.38, 129.53]
LBC1936 10.23 [ -20.11, 40.58]
LIBD 1.98 [ -41.02, 44.98]
Sydney MAS 25.90 [ -5.07, 56.87]
MCIC -9.09 [ -64.36, 46.18]
MooDS 64.91 [ 20.33, 109.50]
MPIP 43.31 [ 13.68, 72.94]
MUNSTER 11.50 [ -17.70, 40.70]
NCNG 11.88 [ -31.60, 55.36]
NESDA -34.04 [ -89.78, 21.70]
NeuroIMAGE 15.60 [ -43.82, 75.02]
NTR 26.63 [ -21.86, 75.12]
OATS -41.30 [ -90.93, 8.33]
PAFIP 41.55 [ -29.87, 112.96]
PDNZ -19.34 [ -84.63, 45.95]
PING 47.78 [ 0.85, 94.72]
MPRC 9.40 [ -36.49, 55.30]
PPMI
QTIM 8.78 [ -18.45, 36.01]
SHIP 21.17 [ -1.91, 44.26]
SHIP-Trend 25.53 [ -0.33, 51.38]
SYS -1.29 [ -49.19, 46.60]
TCD-NUIG -74.75 [-139.54, -9.97]
TOP 22.45 [ -16.38, 61.29]
TOP3T 22.62 [ -20.55, 65.80]
UiO2016 80.50 [ 21.18, 139.83]
UiO2017 47.72 [ 6.88, 88.56]
UMCU 19.31 [ -9.80, 48.42]
ENIGMA Discovery Cohorts [N=22718; P=1.822e-09] 16.76 [ 11.29, 22.22]
UKBB 13.85 [ 5.63, 22.08]
Discovery+UKBB [N=32238; P=3.794e-11] 15.83 [ 11.14, 20.53]
GOBS 17.52 [ -4.41, 39.45]
IMH 0.62 [ -72.44, 73.69]
UNICAMP 26.20 [ -28.68, 81.08]
Meth-CT -29.70 [-110.01, 50.60]
MIRECC
UKBB nonEUR 3.30 [ -43.73, 50.34]
OSAKA -30.74 [ -64.14, 2.67]
PING Non European 0.27 [ -42.54, 43.07]
Non European Cohorts [N=2629; P=0.6947] 2.97 [ -11.85, 17.78]
Beta [Lower 95% CI,Upper 95% CI]
-150 -100 -50 0 50 100 150
mm^2/effect allele

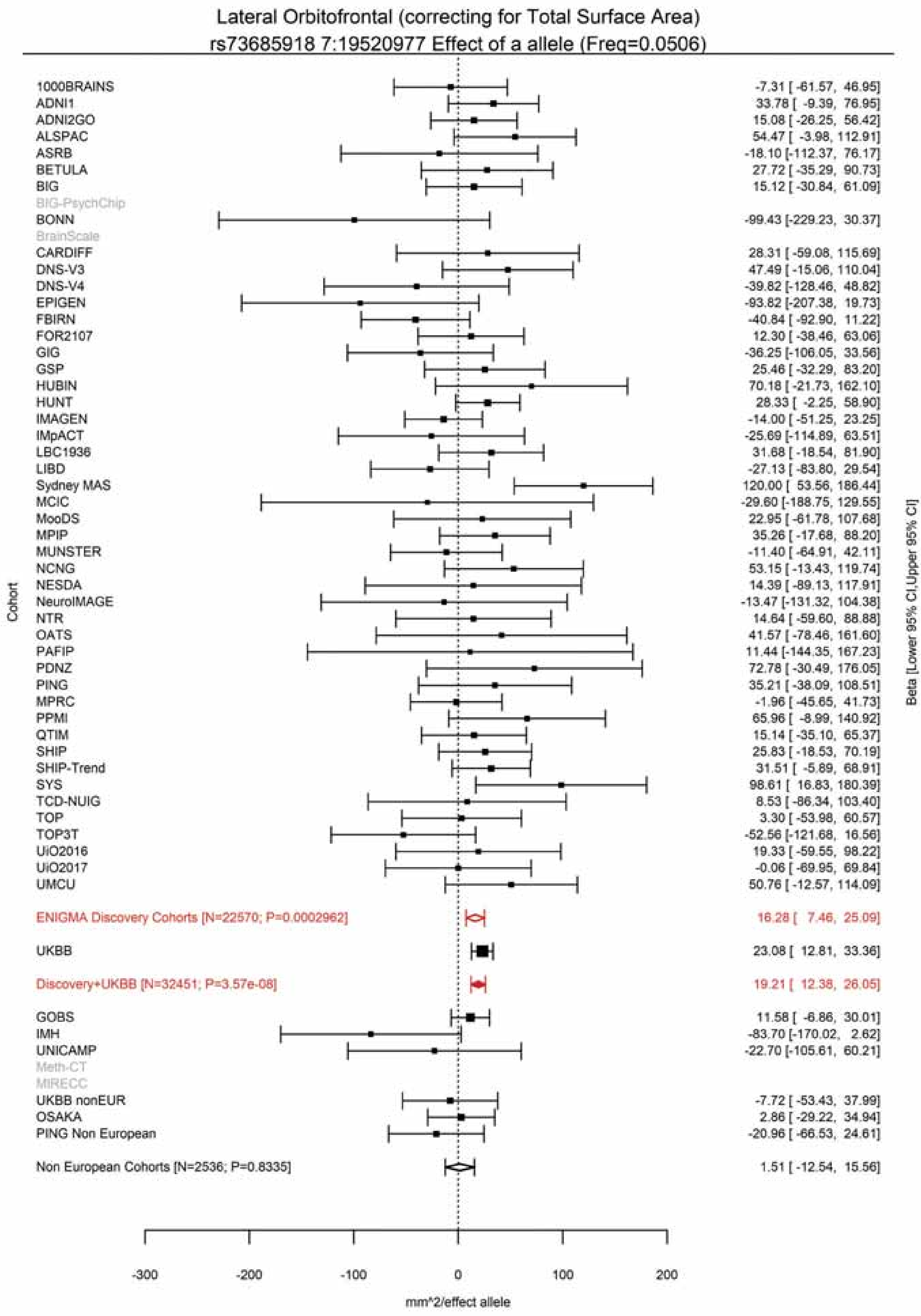
Lateral Orbitofrontal (correcting for Total Surface Area)
rs73685918 7:19520977 Effect of a allele (Freq=0.0506)
Cohort
1000BRAINS -7.31 [ -61.57, 46.95]
ADNI1 33.78 [ -9.39, 76.95]
ADNI2GO 15.08 [ -26.25, 56.42]
ALSPAC 54.47 [ -3.98, 112.91]
ASRB -18.10 [-112.37, 76.17]
BETULA 27.72 [ -35.29, 90.73]
BIG 15.12 [ -30.84, 61.09]
BIG-PsychChip
BONN -99.43 [-229.23, 30.37]
BrainScale
CARDIFF 28.31 [ -59.08, 115.69]
DNS-V3 47.49 [ -15.06, 110.04]
DNS-V4 -39.82 [-128.46, 48.82]
EPIGEN -93.82 [-207.38, 19.73]
FBIRN -40.84 [ -92.90, 11.22]
FOR2107 12.30 [ -38.46, 63.06]
GIG -36.25 [-106.05, 33.56]
GSP 25.46 [ -32.29, 83.20]
HUBIN 70.18 [ -21.73, 162.10]
HUNT 28.33 [ -2.25, 58.90]
IMAGEN -14.00 [ -51.25, 23.25]
IMpACT -25.69 [-114.89, 63.51]
LBC1936 31.68 [ -18.54, 81.90]
LIBD -27.13 [ -83.80, 29.54]
Sydney MAS 120.00 [ 53.56, 186.44]
MCIC -29.60 [-188.75, 129.55]
MooDS 22.95 [ -61.78, 107.68]
MPIP 35.26 [ -17.68, 88.20]
MUNSTER -11.40 [ -64.91, 42.11]
NCNG 53.15 [ -13.43, 119.74]
NESDA 14.39 [ -89.13, 117.91]
NeuroIMAGE -13.47 [-131.32, 104.38]
NTR 14.64 [ -59.60, 88.88]
OATS 41.57 [ -78.46, 161.60]
PAFIP 11.44 [-144.35, 167.23]
PDNZ 72.78 [ -30.49, 176.05]
PING 35.21 [ -38.09, 108.51]
MPRC -1.96 [ -45.65, 41.73]
PPMI 65.96 [ -8.99, 140.92]
QTIM 15.14 [ -35.10, 65.37]
SHIP 25.83 [ -18.53, 70.19]
SHIP-Trend 31.51 [ -5.89, 68.91]
SYS 98.61 [ 16.83, 180.39]
TCD-NUIG 8.53 [ -86.34, 103.40]
TOP 3.30 [ -53.98, 60.57]
TOP3T -52.56 [-121.68, 16.56]
UiO2016 19.33 [ -59.55, 98.22]
UiO2017 -0.06 [ -69.95, 69.84]
UMCU 50.76 [ -12.57, 114.09]
ENIGMA Discovery Cohorts [N=22570; P=0.0002962] 16.28 [ 7.46, 25.09]
UKBB 23.08 [ 12.81, 33.36]
Discovery+UKBB [N=32451; P=3.57e-08] 19.21 [ 12.38, 26.05]
GOBS 11.58 [ -6.86, 30.01]
IMH -83.70 [-170.02, 2.62]
UNICAMP -22.70 [-105.61, 60.21]
Meth-CT
MIRECC
UKBB nonEUR -7.72 [ -53.43, 37.99]
OSAKA 2.86 [ -29.22, 34.94]
PING Non European -20.96 [ -66.53, 24.61]
Non European Cohorts [N=2536; P=0.8335] 1.51 [ -12.54, 15.56]
-300
-200
-100
0
100
200
mm^2/effect allele
Beta [Lower 95% CI,Upper 95% CI]

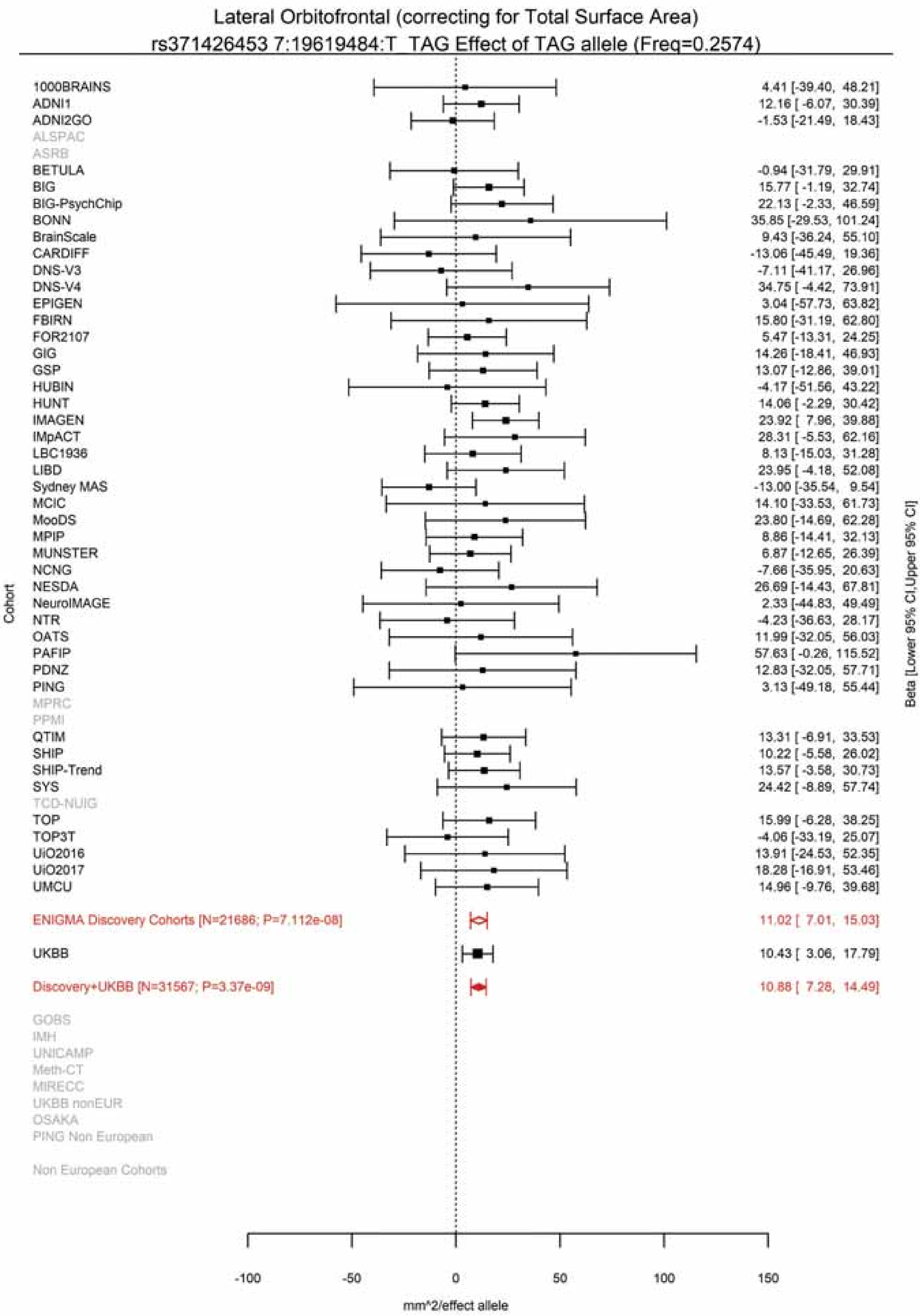
Lateral Orbitofrontal (correcting for Total Surface Area)
rs371426453 7:19619484:T TAG Effect of TAG allele (Freq=0.2574)
1000BRAINS 4.41 [-39.40, 48.21]
ADNI1 12.16 [ -6.07, 30.39]
ADNI2GO -1.53 [-21.49, 18.43]
ALSPAC
ASRB
BETULA -0.94 [-31.79, 29.91]
BIG 15.77 [ -1.19, 32.74]
BIG-PsychChip 22.13 [ -2.33, 46.59]
BONN 35.85 [-29.53, 101.24]
BrainScale 9.43 [-36.24, 55.10]
CARDIFF -13.06 [-45.49, 19.36]
DNS-V3 -7.11 [-41.17, 26.96]
DNS-V4 34.75 [ -4.42, 73.91]
EPIGEN 3.04 [-57.73, 63.82]
FBIRN 15.80 [-31.19, 62.80]
FOR2107 5.47 [-13.31, 24.25]
GIG 14.26 [-18.41, 46.93]
GSP 13.07 [-12.86, 39.01]
HUBIN -4.17 [-51.56, 43.22]
HUNT 14.06 [ -2.29, 30.42]
IMAGEN 23.92 [ 7.96, 39.88]
IMpACT 28.31 [ -5.53, 62.16]
LBC1936 8.13 [-15.03, 31.28]
LIBD 23.95 [ -4.18, 52.08]
Sydney MAS -13.00 [-35.54, 9.54]
MCIC 14.10 [-33.53, 61.73]
MooDS 23.80 [-14.69, 62.28]
MPIP 8.86 [-14.41, 32.13]
MUNSTER 6.87 [-12.65, 26.39]
NCNG -7.66 [-35.95, 20.63]
NESDA 26.69 [-14.43, 67.81]
NeuroIMAGE 2.33 [-44.83, 49.49]
NTR -4.23 [-36.63, 28.17]
OATS 11.99 [-32.05, 56.03]
PAFIP 57.63 [ -0.26, 115.52]
PDNZ 12.83 [-32.05, 57.71]
PING 3.13 [-49.18, 55.44]
MPRC
PPMI
QTIM 13.31 [ -6.91, 33.53]
SHIP 10.22 [ -5.58, 26.02]
SHIP-Trend 13.57 [ -3.58, 30.73]
SYS 24.42 [ -8.89, 57.74]
TCD-NUIG
TOP 15.99 [ -6.28, 38.25]
TOP3T -4.06 [-33.19, 25.07]
UiO2016 13.91 [-24.53, 52.35]
UiO2017 18.28 [-16.91, 53.46]
UMCU 14.96 [ -9.76, 39.68]
ENIGMA Discovery Cohorts [N=21686; P=7.112e-08] 11.02 [ 7.01, 15.03]
UKBB 10.43 [ 3.06, 17.79]
Discovery+UKBB [N=31567; P=3.37e-09] 10.88 [ 7.28, 14.49]
GOBS
IMH
UNICAMP
Meth-CT
MIRECC
UKBB nonEUR
OSAKA
PING Non European
Non European Cohorts
Cohort
Beta [Lower 95% CI,Upper 95% CI]
-100
-50
0
50
100
150
mm^2/effect allele

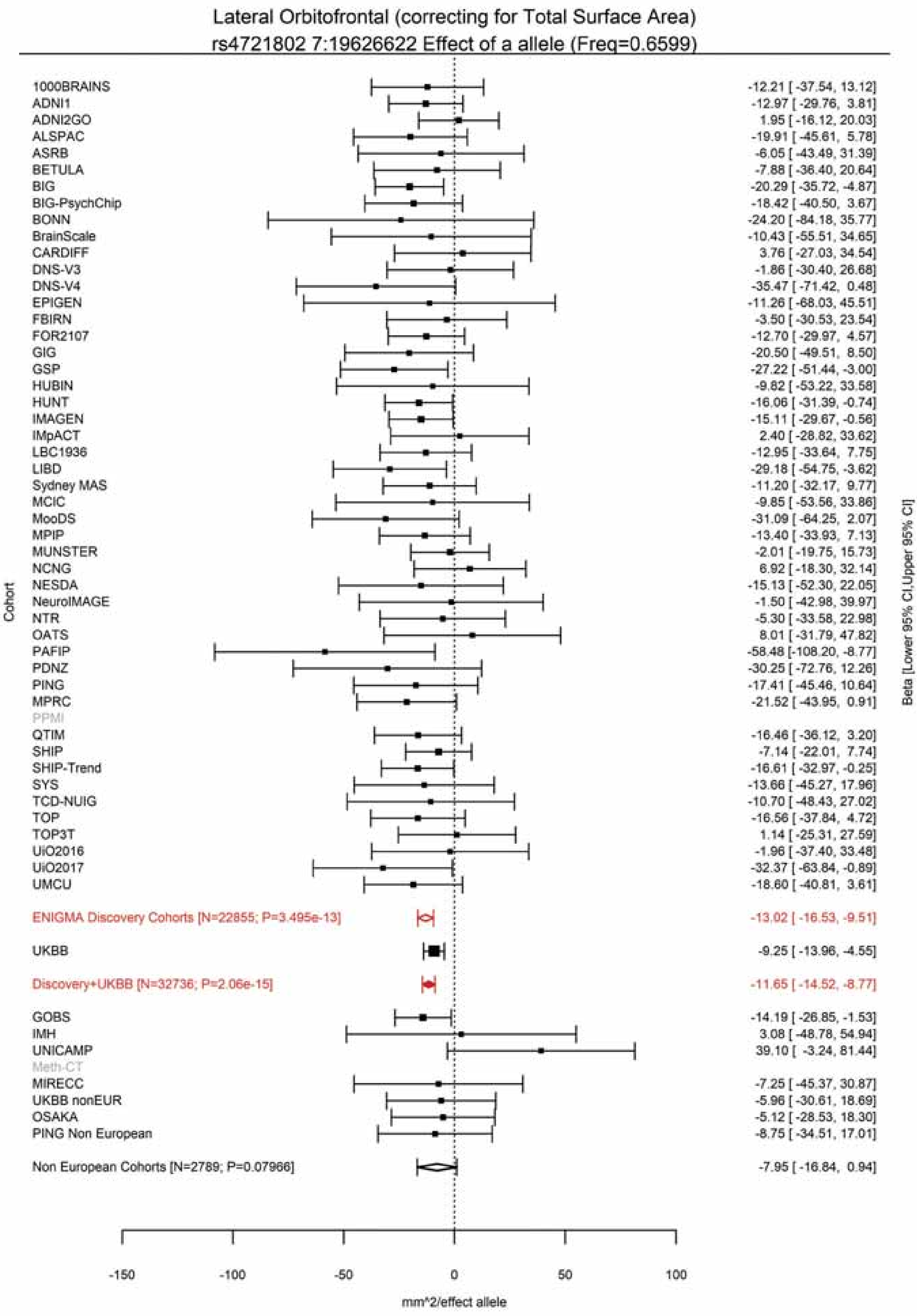
Lateral Orbitofrontal (correcting for Total Surface Area)
rs4721802 7:19626622 Effect of a allele (Freq=0.6599)
Cohort
1000BRAINS -12.21 [ -37.54, 13.12]
ADNI1 -12.97 [ -29.76, 3.81]
ADNI2GO 1.95 [ -16.12, 20.03]
ALSPAC -19.91 [ -45.61, 5.78]
ASRB -6.05 [ -43.49, 31.39]
BETULA -7.88 [ -36.40, 20.64]
BIG -20.29 [ -35.72, -4.87]
BIG-PsychChip -18.42 [ -40.50, 3.67]
BONN -24.20 [ -84.18, 35.77]
BrainScale -10.43 [ -55.51, 34.65]
CARDIFF 3.76 [ -27.03, 34.54]
DNS-V3 -1.86 [ -30.40, 26.68]
DNS-V4 -35.47 [ -71.42, 0.48]
EPIGEN -11.26 [ -68.03, 45.51]
FBIRN -3.50 [ -30.53, 23.54]
FOR2107 -12.70 [ -29.97, 4.57]
GIG -20.50 [ -49.51, 8.50]
GSP -27.22 [ -51.44, -3.00]
HUBIN -9.82 [ -53.22, 33.58]
HUNT -16.06 [ -31.39, -0.74]
IMAGEN -15.11 [ -29.67, -0.56]
IMpACT 2.40 [ -28.82, 33.62]
LBC1936 -12.95 [ -33.64, 7.75]
LIBD -29.18 [ -54.75, -3.62]
Sydney MAS -11.20 [ -32.17, 9.77]
MCIC -9.85 [ -53.56, 33.86]
MooDS -31.09 [ -64.25, 2.07]
MPIP -13.40 [ -33.93, 7.13]
MUNSTER -2.01 [ -19.75, 15.73]
NCNG 6.92 [ -18.30, 32.14]
NESDA -15.13 [ -52.30, 22.05]
NeuroIMAGE -1.50 [ -42.98, 39.97]
NTR -5.30 [ -33.58, 22.98]
OATS 8.01 [ -31.79, 47.82]
PAFIP -58.48 [-108.20, -8.77]
PDNZ -30.25 [ -72.76, 12.26]
PING -17.41 [ -45.46, 10.64]
MPRC -21.52 [ -43.95, 0.91]
PPMI
QTIM -16.46 [ -36.12, 3.20]
SHIP -7.14 [ -22.01, 7.74]
SHIP-Trend -16.61 [ -32.97, -0.25]
SYS -13.66 [ -45.27, 17.96]
TCD-NUIG -10.70 [ -48.43, 27.02]
TOP -16.56 [ -37.84, 4.72]
TOP3T 1.14 [ -25.31, 27.59]
UiO2016 -1.96 [ -37.40, 33.48]
UiO2017 -32.37 [ -63.84, -0.89]
UMCU -18.60 [ -40.81, 3.61]
ENIGMA Discovery Cohorts [N=22855; P=3.495e-13] -13.02 [ -16.53, -9.51]
UKBB -9.25 [ -13.96, -4.55]
Discovery+UKBB [N=32736; P=2.06e-15] -11.65 [ -14.52, -8.77]
GOBS -14.19 [ -26.85, -1.53]
IMH 3.08 [ -48.78, 54.94]
UNICAMP 39.10 [ -3.24, 81.44]
Meth-CT
MIRECC -7.25 [ -45.37, 30.87]
UKBB nonEUR -5.96 [ -30.61, 18.69]
OSAKA -5.12 [ -28.53, 18.30]
PING Non European -8.75 [ -34.51, 17.01]
Non European Cohorts [N=2789; P=0.07966] -7.95 [ -16.84, 0.94]
Beta [Lower 95% CI,Upper 95% CI]
-150
-100
-50
0
50
100
mm^2/effect allele

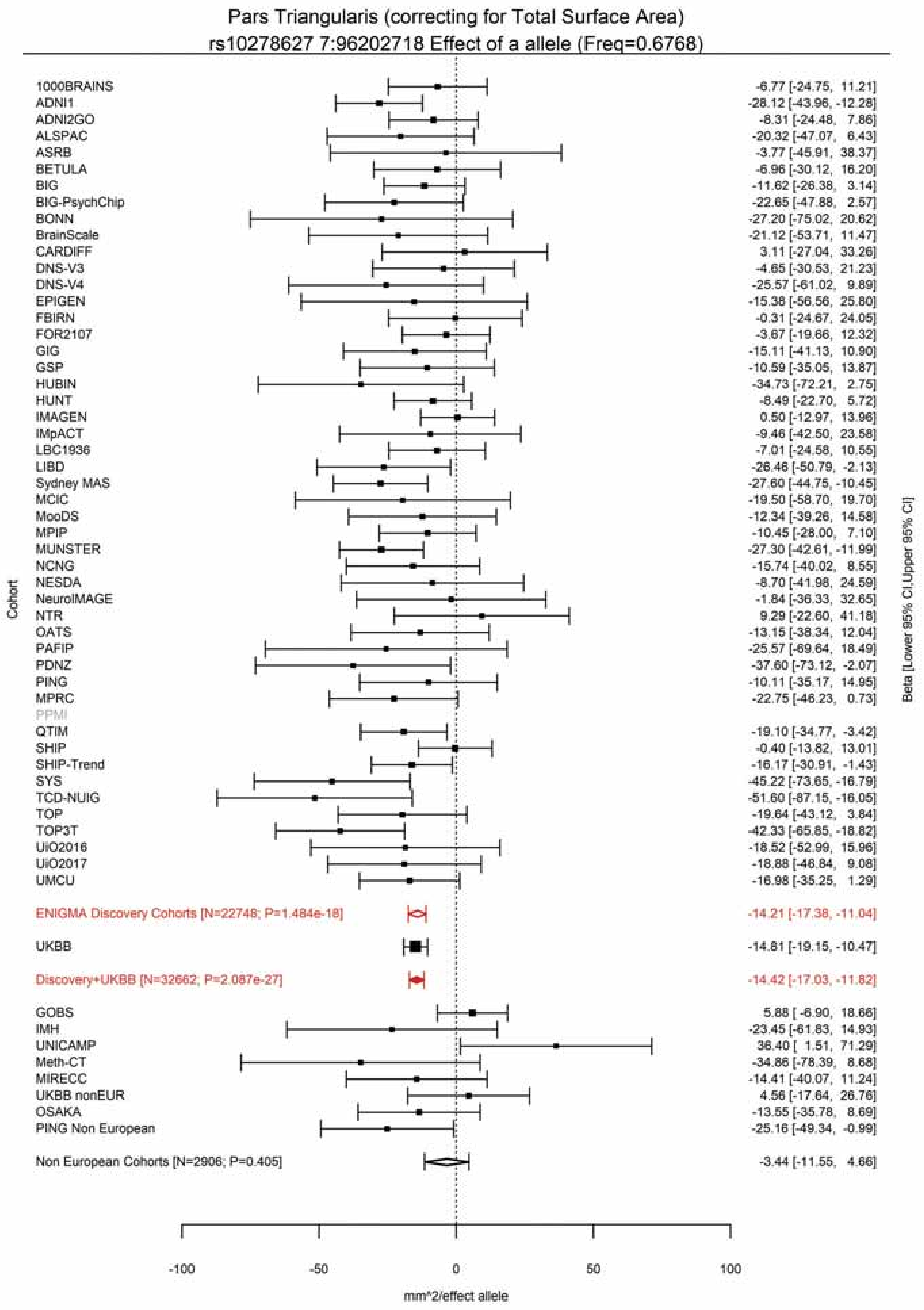

Pars Triangularis (correcting for Total Surface Area)
rs10278627 7:96202718 Effect of a allele (Freq=0.6768)
1000BRAINS -6.77 [-24.75, 11.21]
ADNI1 -28.12 [-43.96, -12.28]
ADNI2GO -8.31 [-24.48, 7.86]
ALSPAC -20.32 [-47.07, 6.43]
ASRB -3.77 [-45.91, 38.37]
BETULA -6.96 [-30.12, 16.20]
BIG -11.62 [-26.38, 3.14]
BIG-PsychChip -22.65 [-47.88, 2.57]
BONN -27.20 [-75.02, 20.62]
BrainScale -21.12 [-53.71, 11.47]
CARDIFF 3.11 [-27.04, 33.26]
DNS-V3 -4.65 [-30.53, 21.23]
DNS-V4 -25.57 [-61.02, 9.89]
EPIGEN -15.38 [-56.56, 25.80]
FBIRN -0.31 [-24.67, 24.05]
FOR2107 -3.67 [-19.66, 12.32]
GIG -15.11 [-41.13, 10.90]
GSP -10.59 [-35.05, 13.87]
HUBIN -34.73 [-72.21, 2.75]
HUNT -8.49 [-22.70, 5.72]
IMAGEN 0.50 [-12.97, 13.96]
IMpACT -9.46 [-42.50, 23.58]
LBC1936 -7.01 [-24.58, 10.55]
LIBD -26.46 [-50.79, -2.13]
Sydney MAS -27.60 [-44.75, -10.45]
MCIC -19.50 [-58.70, 19.70]
MooDS -12.34 [-39.26, 14.58]
MPIP -10.45 [-28.00, 7.10]
MUNSTER -27.30 [-42.61, -11.99]
NCNG -15.74 [-40.02, 8.55]
NESDA -8.70 [-41.98, 24.59]
NeuroIMAGE -1.84 [-36.33, 32.65]
NTR 9.29 [-22.60, 41.18]
OATS -13.15 [-38.34, 12.04]
PAFIP -25.57 [-69.64, 18.49]
PDNZ -37.60 [-73.12, -2.07]
PING -10.11 [-35.17, 14.95]
MPRC -22.75 [-46.23, 0.73]
PPMI
QTIM -19.10 [-34.77, -3.42]
SHIP -0.40 [-13.82, 13.01]
SHIP-Trend -16.17 [-30.91, -1.43]
SYS -45.22 [-73.65, -16.79]
TCD-NUIG -51.60 [-87.15, -16.05]
TOP -19.64 [-43.12, 3.84]
TOP3T -42.33 [-65.85, -18.82]
UiO2016 -18.52 [-52.99, 15.96]
UiO2017 -18.88 [-46.84, 9.08]
UMCU -16.98 [-35.25, 1.29]
ENIGMA Discovery Cohorts [N=22748; P=1.484e-18] -14.21 [-17.38, -11.04]
UKBB -14.81 [-19.15, -10.47]
Discovery+UKBB [N=32662; P=2.087e-27] -14.42 [-17.03, -11.82]
GOBS 5.88 [ -6.90, 18.66]
IMH -23.45 [-61.83, 14.93]
UNICAMP 36.40 [ 1.51, 71.29]
Meth-CT -34.86 [-78.39, 8.68]
MIRECC -14.41 [-40.07, 11.24]
UKBB nonEUR 4.56 [-17.64, 26.76]
OSAKA -13.55 [-35.78, 8.69]
PING Non European -25.16 [-49.34, -0.99]
Non European Cohorts [N=2906; P=0.405] -3.44 [-11.55, 4.66]
Cohort
Beta [Lower 95% CI,Upper 95% CI]
-100
-50
0
50
100
mm^2/effect allele

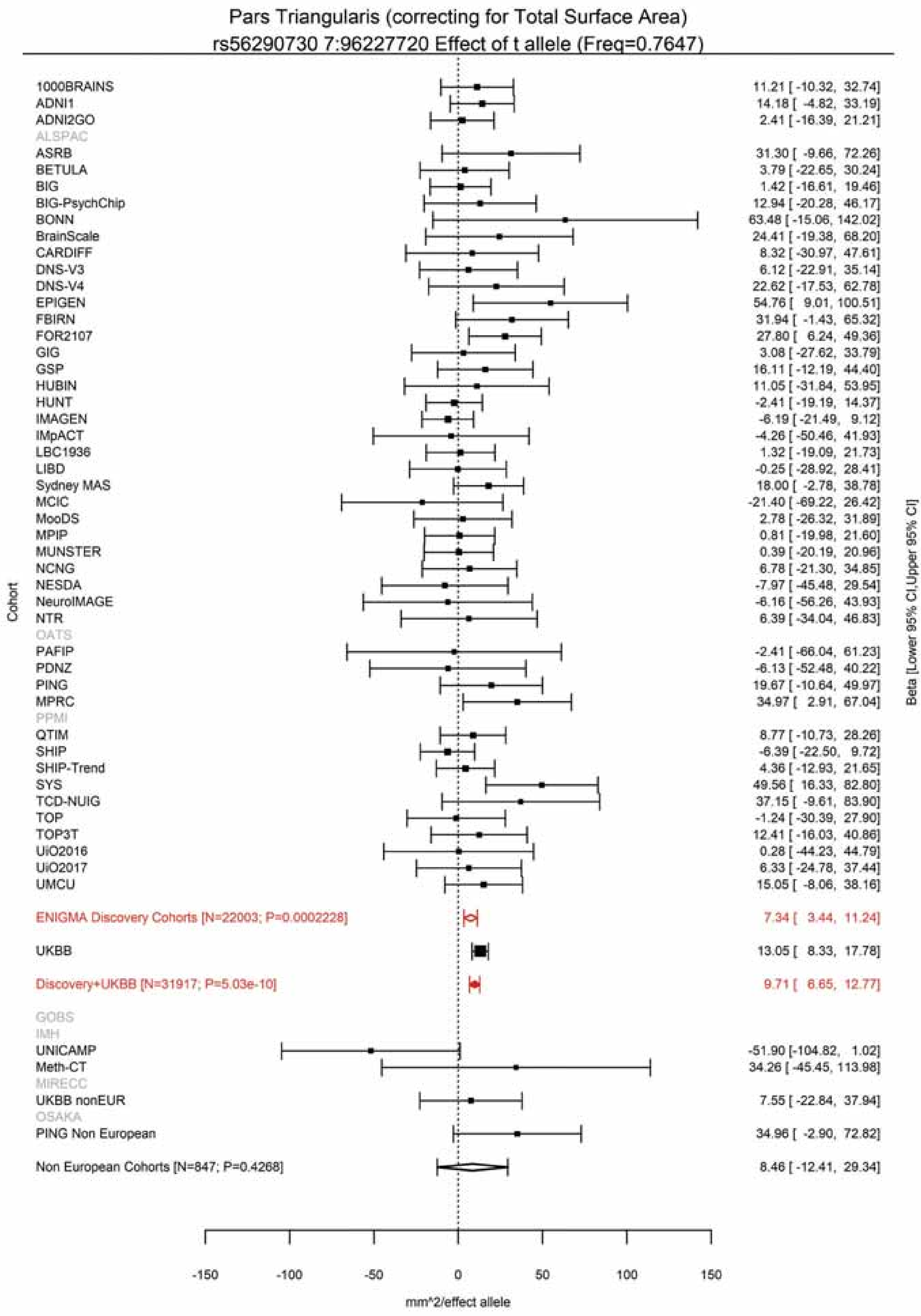
Pars Triangularis (correcting for Total Surface Area)
rs56290730 7:96227720 Effect of t allele (Freq=0.7647)
Cohort
Beta [Lower 95% CI,Upper 95% CI]
1000BRAINS 11.21 [ -10.32, 32.74]
ADNI1 14.18 [ -4.82, 33.19]
ADNI2GO 2.41 [ -16.39, 21.21]
ALSPAC
ASRB 31.30 [ -9.66, 72.26]
BETULA 3.79 [ -22.65, 30.24]
BIG 1.42 [ -16.61, 19.46]
BIG-PsychChip 12.94 [ -20.28, 46.17]
BONN 63.48 [ -15.06, 142.02]
BrainScale 24.41 [ -19.38, 68.20]
CARDIFF 8.32 [ -30.97, 47.61]
DNS-V3 6.12 [ -22.91, 35.14]
DNS-V4 22.62 [ -17.53, 62.78]
EPIGEN 54.76 [ 9.01, 100.51]
FBIRN 31.94 [ -1.43, 65.32]
FOR2107 27.80 [ 6.24, 49.36]
GIG 3.08 [ -27.62, 33.79]
GSP 16.11 [ -12.19, 44.40]
HUBIN 11.05 [ -31.84, 53.95]
HUNT -2.41 [ -19.19, 14.37]
IMAGEN -6.19 [ -21.49, 9.12]
IMpACT -4.26 [ -50.46, 41.93]
LBC1936 1.32 [ -19.09, 21.73]
LIBD -0.25 [ -28.92, 28.41]
Sydney MAS 18.00 [ -2.78, 38.78]
MCIC -21.40 [ -69.22, 26.42]
MooDS 2.78 [ -26.32, 31.89]
MPIP 0.81 [ -19.98, 21.60]
MUNSTER 0.39 [ -20.19, 20.96]
NCNG 6.78 [ -21.30, 34.85]
NESDA -7.97 [ -45.48, 29.54]
NeuroIMAGE -6.16 [ -56.26, 43.93]
NTR 6.39 [ -34.04, 46.83]
OATS
PAFIP -2.41 [ -66.04, 61.23]
PDNZ -6.13 [ -52.48, 40.22]
PING 19.67 [ -10.64, 49.97]
MPRC 34.97 [ 2.91, 67.04]
PPMI
QTIM 8.77 [ -10.73, 28.26]
SHIP -6.39 [ -22.50, 9.72]
SHIP-Trend 4.36 [ -12.93, 21.65]
SYS 49.56 [ 16.33, 82.80]
TCD-NUIG 37.15 [ -9.61, 83.90]
TOP -1.24 [ -30.39, 27.90]
TOP3T 12.41 [ -16.03, 40.86]
UiO2016 0.28 [ -44.23, 44.79]
UiO2017 6.33 [ -24.78, 37.44]
UMCU 15.05 [ -8.06, 38.16]
ENIGMA Discovery Cohorts [N=22003; P=0.0002228] 7.34 [ 3.44, 11.24]
UKBB 13.05 [ 8.33, 17.78]
Discovery+UKBB [N=31917; P=5.03e-10] 9.71 [ 6.65, 12.77]
GOBS
IMH
UNICAMP -51.90 [-104.82, 1.02]
Meth-CT 34.26 [ -45.45, 113.98]
MIRECC
UKBB nonEUR 7.55 [ -22.84, 37.94]
OSAKA
PING Non European 34.96 [ -2.90, 72.82]
Non European Cohorts [N=847; P=0.4268] 8.46 [ -12.41, 29.34]
-150
-100
-50
0
50
100
150
mm^2/effect allele

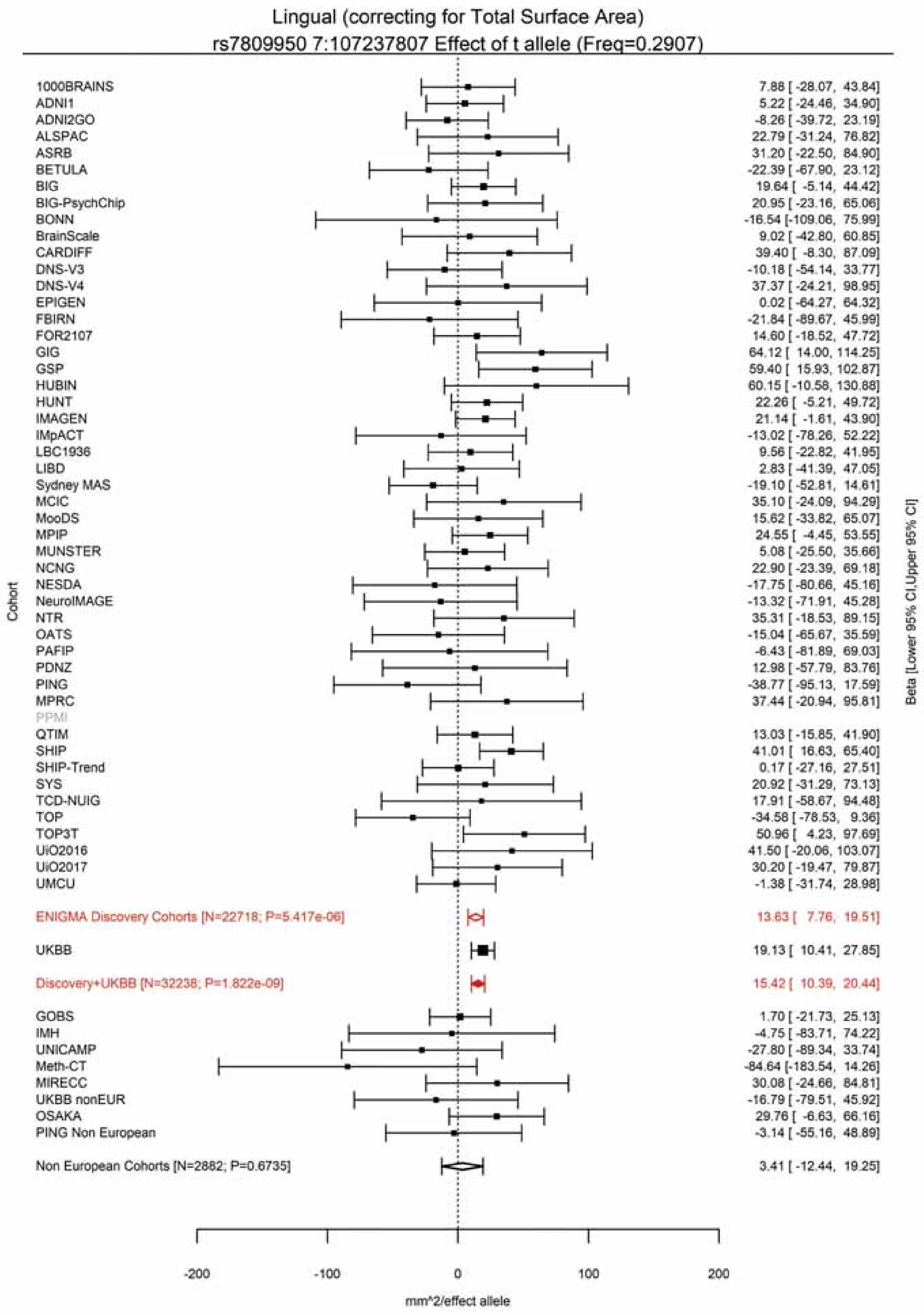
Lingual (correcting for Total Surface Area)
rs7809950 7:107237807 Effect of t allele (Freq=0.2907)
Cohort
1000BRAINS 7.88 [ -28.07, 43.84]
ADNI1 5.22 [ -24.46, 34.90]
ADNI2GO -8.26 [ -39.72, 23.19]
ALSPAC 22.79 [ -31.24, 76.82]
ASRB 31.20 [ -22.50, 84.90]
BETULA -22.39 [ -67.90, 23.12]
BIG 19.64 [ -5.14, 44.42]
BIG-PsychChip 20.95 [ -23.16, 65.06]
BONN -16.54 [-109.06, 75.99]
BrainScale 9.02 [ -42.80, 60.85]
CARDIFF 39.40 [ -8.30, 87.09]
DNS-V3 -10.18 [ -54.14, 33.77]
DNS-V4 37.37 [ -24.21, 98.95]
EPIGEN 0.02 [ -64.27, 64.32]
FBIRN -21.84 [ -89.67, 45.99]
FOR2107 14.60 [ -18.52, 47.72]
GIG 64.12 [ 14.00, 114.25]
GSP 59.40 [ 15.93, 102.87]
HUBIN 60.15 [ -10.58, 130.88]
HUNT 22.26 [ -5.21, 49.72]
IMAGEN 21.14 [ -1.61, 43.90]
IMpACT -13.02 [ -78.26, 52.22]
LBC1936 9.56 [ -22.82, 41.95]
LIBD 2.83 [ -41.39, 47.05]
Sydney MAS -19.10 [ -52.81, 14.61]
MCIC 35.10 [ -24.09, 94.29]
MooDS 15.62 [ -33.82, 65.07]
MPIP 24.55 [ -4.45, 53.55]
MUNSTER 5.08 [ -25.50, 35.66]
NCNG 22.90 [ -23.39, 69.18]
NESDA -17.75 [ -80.66, 45.16]
NeuroIMAGE -13.32 [ -71.91, 45.28]
NTR 35.31 [ -18.53, 89.15]
OATS -15.04 [ -65.67, 35.59]
PAFIP -6.43 [ -81.89, 69.03]
PDNZ 12.98 [ -57.79, 83.76]
PING -38.77 [ -95.13, 17.59]
MPRC 37.44 [ -20.94, 95.81]
PPMI
QTIM 13.03 [ -15.85, 41.90]
SHIP 41.01 [ 16.63, 65.40]
SHIP-Trend 0.17 [ -27.16, 27.51]
SYS 20.92 [ -31.29, 73.13]
TCD-NUIG 17.91 [ -58.67, 94.48]
TOP -34.58 [ -78.53, 9.36]
TOP3T 50.96 [ 4.23, 97.69]
UiO2016 41.50 [ -20.06, 103.07]
UiO2017 30.20 [ -19.47, 79.87]
UMCU -1.38 [ -31.74, 28.98]
ENIGMA Discovery Cohorts [N=22718; P=5.417e-06] 13.63 [ 7.76, 19.51]
UKBB 19.13 [ 10.41, 27.85]
Discovery+UKBB [N=32238; P=1.822e-09] 15.42 [ 10.39, 20.44]
GOBS 1.70 [ -21.73, 25.13]
IMH -4.75 [ -83.71, 74.22]
UNICAMP -27.80 [ -89.34, 33.74]
Meth-CT -84.64 [-183.54, 14.26]
MIRECC 30.08 [ -24.66, 84.81]
UKBB nonEUR -16.79 [ -79.51, 45.92]
OSAKA 29.76 [ -6.63, 66.16]
PING Non European -3.14 [ -55.16, 48.89]
Non European Cohorts [N=2882; P=0.6735] 3.41 [ -12.44, 19.25]
-200
-100
0
100
200
mm^2/effect allele
Beta [Lower 95% CI,Upper 95% CI]

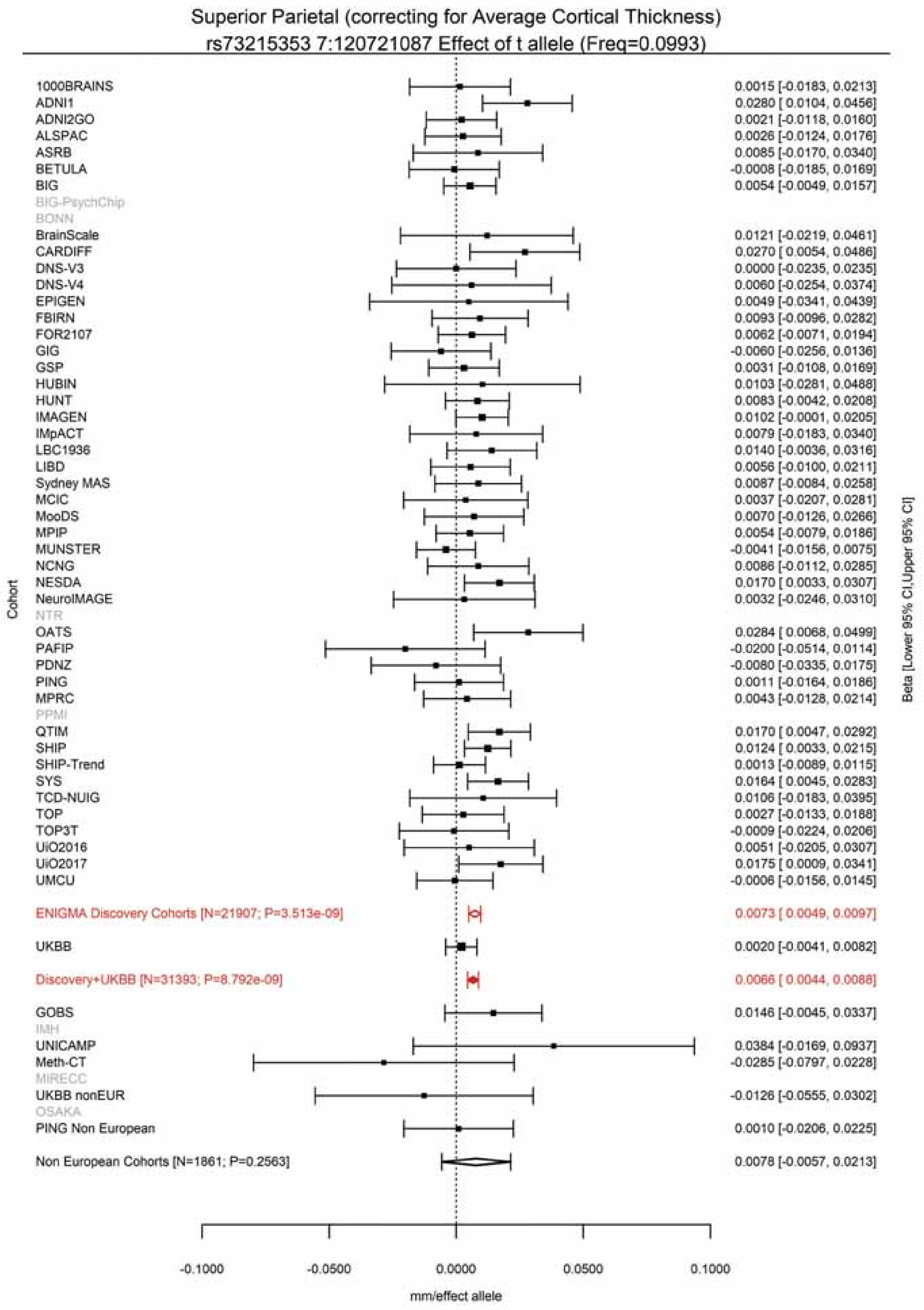
Superior Parietal (correcting for Average Cortical Thickness)
rs73215353 7:120721087 Effect of t allele (Freq=0.0993)
Cohort
1000BRAINS 0.0015 [-0.0183, 0.0213]
ADNI1 0.0280 [ 0.0104, 0.0456]
ADNI2GO 0.0021 [-0.0118, 0.0160]
ALSPAC 0.0026 [-0.0124, 0.0176]
ASRB 0.0085 [-0.0170, 0.0340]
BETULA -0.0008 [-0.0185, 0.0169]
BIG 0.0054 [-0.0049, 0.0157]
BIG-PsychChip
BONN
BrainScale 0.0121 [-0.0219, 0.0461]
CARDIFF 0.0270 [ 0.0054, 0.0486]
DNS-V3 0.0000 [-0.0235, 0.0235]
DNS-V4 0.0060 [-0.0254, 0.0374]
EPIGEN 0.0049 [-0.0341, 0.0439]
FBIRN 0.0093 [-0.0096, 0.0282]
FOR2107 0.0062 [-0.0071, 0.0194]
GIG -0.0060 [-0.0256, 0.0136]
GSP 0.0031 [-0.0108, 0.0169]
HUBIN 0.0103 [-0.0281, 0.0488]
HUNT 0.0083 [-0.0042, 0.0208]
IMAGEN 0.0102 [-0.0001, 0.0205]
IMpACT 0.0079 [-0.0183, 0.0340]
LBC1936 0.0140 [-0.0036, 0.0316]
LIBD 0.0056 [-0.0100, 0.0211]
Sydney MAS 0.0087 [-0.0084, 0.0258]
MCIC 0.0037 [-0.0207, 0.0281]
MooDS 0.0070 [-0.0126, 0.0266]
MPIP 0.0054 [-0.0079, 0.0186]
MUNSTER -0.0041 [-0.0156, 0.0075]
NCNG 0.0086 [-0.0112, 0.0285]
NESDA 0.0170 [ 0.0033, 0.0307]
NeuroIMAGE 0.0032 [-0.0246, 0.0310]
NTR
OATS 0.0284 [ 0.0068, 0.0499]
PAFIP -0.0200 [-0.0514, 0.0114]
PDNZ -0.0080 [-0.0335, 0.0175]
PING 0.0011 [-0.0164, 0.0186]
MPRC 0.0043 [-0.0128, 0.0214]
PPMI
QTIM 0.0170 [ 0.0047, 0.0292]
SHIP 0.0124 [ 0.0033, 0.0215]
SHIP-Trend 0.0013 [-0.0089, 0.0115]
SYS 0.0164 [ 0.0045, 0.0283]
TCD-NUIG 0.0106 [-0.0183, 0.0395]
TOP 0.0027 [-0.0133, 0.0188]
TOP3T -0.0009 [-0.0224, 0.0206]
UiO2016 0.0051 [-0.0205, 0.0307]
UiO2017 0.0175 [ 0.0009, 0.0341]
UMCU -0.0006 [-0.0156, 0.0145]
ENIGMA Discovery Cohorts [N=21907; P=3.513e-09] 0.0073 [ 0.0049, 0.0097]
UKBB 0.0020 [-0.0041, 0.0082]
Discovery+UKBB [N=31393; P=8.792e-09] 0.0066 [ 0.0044, 0.0088]
GOBS 0.0146 [-0.0045, 0.0337]
IMH
UNICAMP 0.0384 [-0.0169, 0.0937]
Meth-CT -0.0285 [-0.0797, 0.0228]
MIRECC
UKBB nonEUR -0.0126 [-0.0555, 0.0302]
OSAKA
PING Non European 0.0010 [-0.0206, 0.0225]
Non European Cohorts [N=1861; P=0.2563] 0.0078 [-0.0057, 0.0213]
-0.1000
-0.0500
0.0000
0.0500
0.1000
mm/effect allele
Beta [Lower 95% CI,Upper 95% CI]

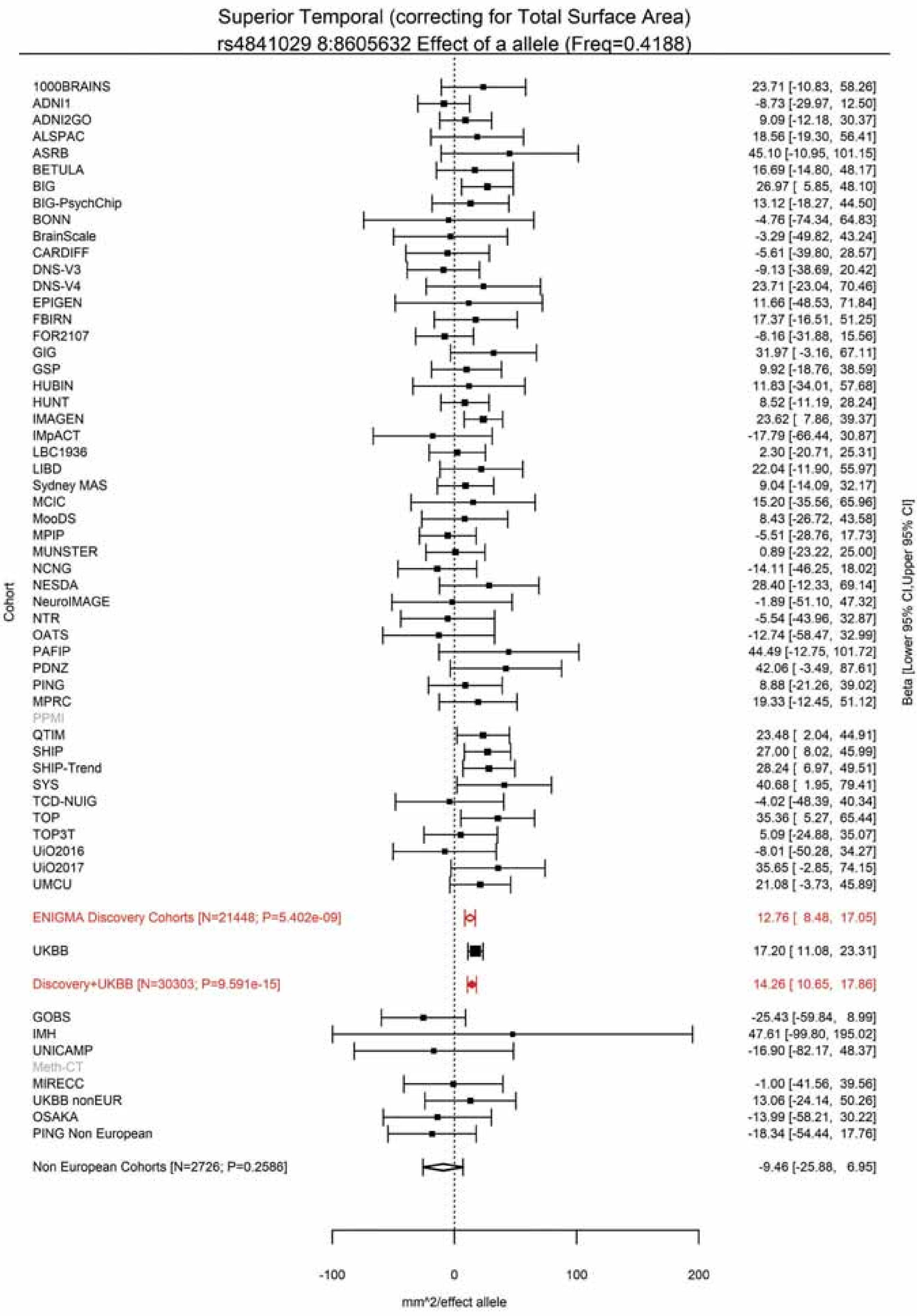
Superior Temporal (correcting for Total Surface Area)
rs4841029 8:8605632 Effect of a allele (Freq=0.4188)
1000BRAINS 23.71 [-10.83, 58.26]
ADNI1 -8.73 [-29.97, 12.50]
ADNI2GO 9.09 [-12.18, 30.37]
ALSPAC 18.56 [-19.30, 56.41]
ASRB 45.10 [-10.95, 101.15]
BETULA 16.69 [-14.80, 48.17]
BIG 26.97 [ 5.85, 48.10]
BIG-PsychChip 13.12 [-18.27, 44.50]
BONN -4.76 [-74.34, 64.83]
BrainScale -3.29 [-49.82, 43.24]
CARDIFF -5.61 [-39.80, 28.57]
DNS-V3 -9.13 [-38.69, 20.42]
DNS-V4 23.71 [-23.04, 70.46]
EPIGEN 11.66 [-48.53, 71.84]
FBIRN 17.37 [-16.51, 51.25]
FOR2107 -8.16 [-31.88, 15.56]
GIG 31.97 [ -3.16, 67.11]
GSP 9.92 [-18.76, 38.59]
HUBIN 11.83 [-34.01, 57.68]
HUNT 8.52 [-11.19, 28.24]
IMAGEN 23.62 [ 7.86, 39.37]
IMpACT -17.79 [-66.44, 30.87]
LBC1936 2.30 [-20.71, 25.31]
LIBD 22.04 [-11.90, 55.97]
Sydney MAS 9.04 [-14.09, 32.17]
MCIC 15.20 [-35.56, 65.96]
MooDS 8.43 [-26.72, 43.58]
MPIP -5.51 [-28.76, 17.73]
MUNSTER 0.89 [-23.22, 25.00]
NCNG -14.11 [-46.25, 18.02]
NESDA 28.40 [-12.33, 69.14]
NeuroIMAGE -1.89 [-51.10, 47.32]
NTR -5.54 [-43.96, 32.87]
OATS -12.74 [-58.47, 32.99]
PAFIP 44.49 [-12.75, 101.72]
PDNZ 42.06 [ -3.49, 87.61]
PING 8.88 [-21.26, 39.02]
MPRC 19.33 [-12.45, 51.12]
PPMI
QTIM 23.48 [ 2.04, 44.91]
SHIP 27.00 [ 8.02, 45.99]
SHIP-Trend 28.24 [ 6.97, 49.51]
SYS 40.68 [ 1.95, 79.41]
TCD-NUIG -4.02 [-48.39, 40.34]
TOP 35.36 [ 5.27, 65.44]
TOP3T 5.09 [-24.88, 35.07]
UiO2016 -8.01 [-50.28, 34.27]
UiO2017 35.65 [ -2.85, 74.15]
UMCU 21.08 [ -3.73, 45.89]
ENIGMA Discovery Cohorts [N=21448; P=5.402e-09] 12.76 [ 8.48, 17.05]
UKBB 17.20 [ 11.08, 23.31]
Discovery+UKBB [N=30303; P=9.591e-15] 14.26 [ 10.65, 17.86]
GOBS -25.43 [-59.84, 8.99]
IMH 47.61 [-99.80, 195.02]
UNICAMP -16.90 [-82.17, 48.37]
Meth-CT
MIRECC -1.00 [-41.56, 39.56]
UKBB nonEUR 13.06 [-24.14, 50.26]
OSAKA -13.99 [-58.21, 30.22]
PING Non European -18.34 [-54.44, 17.76]
Non European Cohorts [N=2726; P=0.2586] -9.46 [-25.88, 6.95]
Cohort
Beta [Lower 95% CI,Upper 95% CI]
-100
0
100
200
mm^2/effect allele

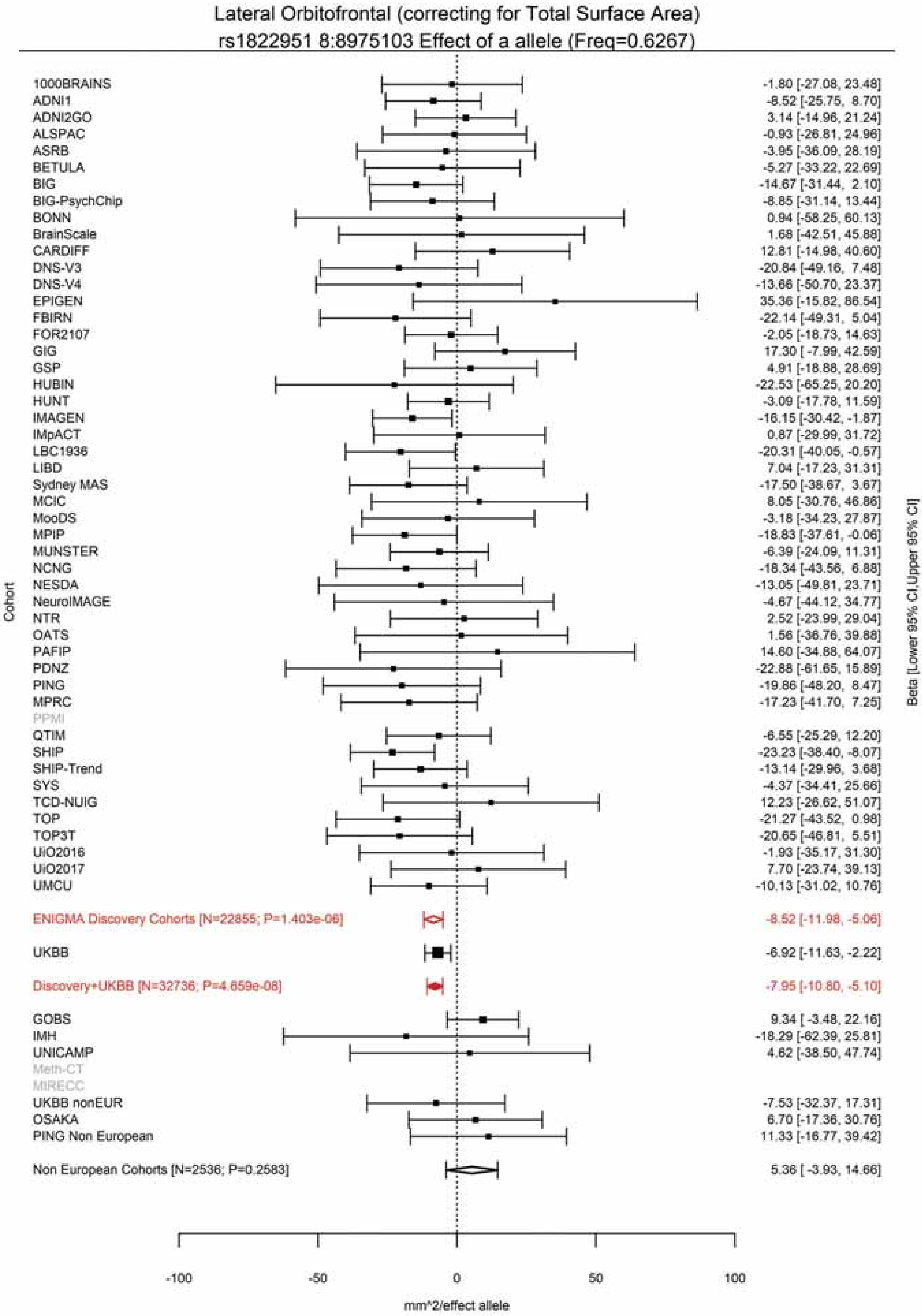
Lateral Orbitofrontal (correcting for Total Surface Area)
rs1822951 8:8975103 Effect of a allele (Freq=0.6267)
Cohort
1000BRAINS -1.80 [-27.08, 23.48]
ADNI1 -8.52 [-25.75, 8.70]
ADNI2GO 3.14 [-14.96, 21.24]
ALSPAC -0.93 [-26.81, 24.96]
ASRB -3.95 [-36.09, 28.19]
BETULA -5.27 [-33.22, 22.69]
BIG -14.67 [-31.44, 2.10]
BIG-PsychChip -8.85 [-31.14, 13.44]
BONN 0.94 [-58.25, 60.13]
BrainScale 1.68 [-42.51, 45.88]
CARDIFF 12.81 [-14.98, 40.60]
DNS-V3 -20.84 [-49.16, 7.48]
DNS-V4 -13.66 [-50.70, 23.37]
EPIGEN 35.36 [-15.82, 86.54]
FBIRN -22.14 [-49.31, 5.04]
FOR2107 -2.05 [-18.73, 14.63]
GIG 17.30 [ -7.99, 42.59]
GSP 4.91 [-18.88, 28.69]
HUBIN -22.53 [-65.25, 20.20]
HUNT -3.09 [-17.78, 11.59]
IMAGEN -16.15 [-30.42, -1.87]
IMpACT 0.87 [-29.99, 31.72]
LBC1936 -20.31 [-40.05, -0.57]
LIBD 7.04 [-17.23, 31.31]
Sydney MAS -17.50 [-38.67, 3.67]
MCIC 8.05 [-30.76, 46.86]
MooDS -3.18 [-34.23, 27.87]
MPIP -18.83 [-37.61, -0.06]
MUNSTER -6.39 [-24.09, 11.31]
NCNG -18.34 [-43.56, 6.88]
NESDA -13.05 [-49.81, 23.71]
NeuroIMAGE -4.67 [-44.12, 34.77]
NTR 2.52 [-23.99, 29.04]
OATS 1.56 [-36.76, 39.88]
PAFIP 14.60 [-34.88, 64.07]
PDNZ -22.88 [-61.65, 15.89]
PING -19.86 [-48.20, 8.47]
MPRC -17.23 [-41.70, 7.25]
PPMI
QTIM -6.55 [-25.29, 12.20]
SHIP -23.23 [-38.40, -8.07]
SHIP-Trend -13.14 [-29.96, 3.68]
SYS -4.37 [-34.41, 25.66]
TCD-NUIG 12.23 [-26.62, 51.07]
TOP -21.27 [-43.52, 0.98]
TOP3T -20.65 [-46.81, 5.51]
UiO2016 -1.93 [-35.17, 31.30]
UiO2017 7.70 [-23.74, 39.13]
UMCU -10.13 [-31.02, 10.76]
ENIGMA Discovery Cohorts [N=22855; P=1.403e-06] -8.52 [-11.98, -5.06]
UKBB -6.92 [-11.63, -2.22]
Discovery+UKBB [N=32736; P=4.659e-08] -7.95 [-10.80, -5.10]
GOBS 9.34 [ -3.48, 22.16]
IMH -18.29 [-62.39, 25.81]
UNICAMP 4.62 [-38.50, 47.74]
Meth-CT
MIRECC
UKBB nonEUR -7.53 [-32.37, 17.31]
OSAKA 6.70 [-17.36, 30.76]
PING Non European 11.33 [-16.77, 39.42]
Non European Cohorts [N=2536; P=0.2583] 5.36 [ -3.93, 14.66]
Beta [Lower 95% CI,Upper 95% CI]
-100
-50
0
50
100
mm^2/effect allele

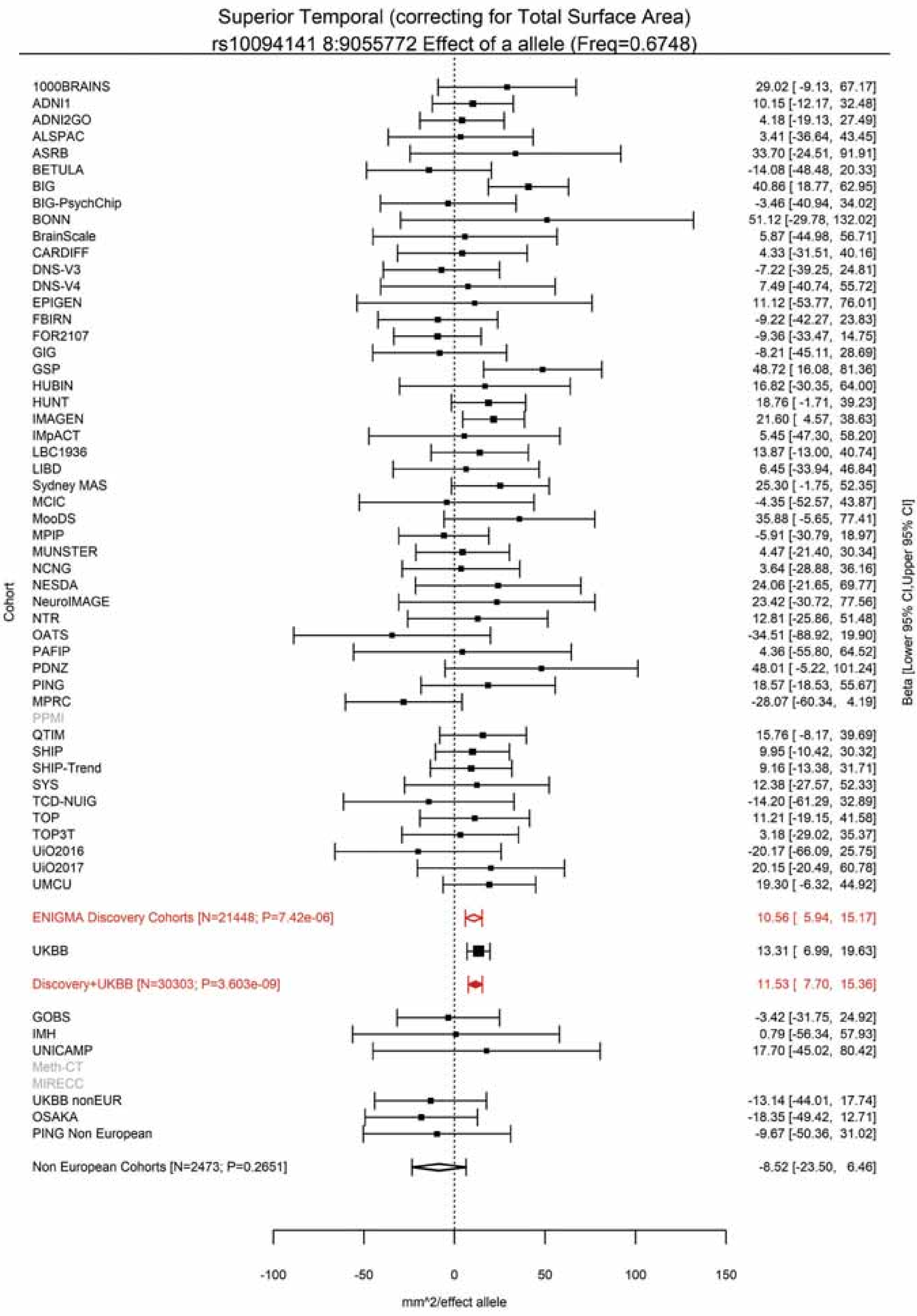
Superior Temporal (correcting for Total Surface Area)
rs10094141 8:9055772 Effect of a allele (Freq=0.6748)
Cohort
Beta [Lower 95% CI,Upper 95% CI]
1000BRAINS 29.02 [ -9.13, 67.17]
ADNI1 10.15 [-12.17, 32.48]
ADNI2GO 4.18 [-19.13, 27.49]
ALSPAC 3.41 [-36.64, 43.45]
ASRB 33.70 [-24.51, 91.91]
BETULA -14.08 [-48.48, 20.33]
BIG 40.86 [ 18.77, 62.95]
BIG-PsychChip -3.46 [-40.94, 34.02]
BONN 51.12 [-29.78, 132.02]
BrainScale 5.87 [-44.98, 56.71]
CARDIFF 4.33 [-31.51, 40.16]
DNS-V3 -7.22 [-39.25, 24.81]
DNS-V4 7.49 [-40.74, 55.72]
EPIGEN 11.12 [-53.77, 76.01]
FBIRN -9.22 [-42.27, 23.83]
FOR2107 -9.36 [-33.47, 14.75]
GIG -8.21 [-45.11, 28.69]
GSP 48.72 [ 16.08, 81.36]
HUBIN 16.82 [-30.35, 64.00]
HUNT 18.76 [ -1.71, 39.23]
IMAGEN 21.60 [ 4.57, 38.63]
IMpACT 5.45 [-47.30, 58.20]
LBC1936 13.87 [-13.00, 40.74]
LIBD 6.45 [-33.94, 46.84]
Sydney MAS 25.30 [ -1.75, 52.35]
MCIC -4.35 [-52.57, 43.87]
MooDS 35.88 [ -5.65, 77.41]
MPIP -5.91 [-30.79, 18.97]
MUNSTER 4.47 [-21.40, 30.34]
NCNG 3.64 [-28.88, 36.16]
NESDA 24.06 [-21.65, 69.77]
NeuroIMAGE 23.42 [-30.72, 77.56]
NTR 12.81 [-25.86, 51.48]
OATS -34.51 [-88.92, 19.90]
PAFIP 4.36 [-55.80, 64.52]
PDNZ 48.01 [ -5.22, 101.24]
PING 18.57 [-18.53, 55.67]
MPRC -28.07 [-60.34, 4.19]
PPMI
QTIM 15.76 [ -8.17, 39.69]
SHIP 9.95 [-10.42, 30.32]
SHIP-Trend 9.16 [-13.38, 31.71]
SYS 12.38 [-27.57, 52.33]
TCD-NUIG -14.20 [-61.29, 32.89]
TOP 11.21 [-19.15, 41.58]
TOP3T 3.18 [-29.02, 35.37]
UiO2016 -20.17 [-66.09, 25.75]
UiO2017 20.15 [-20.49, 60.78]
UMCU 19.30 [ -6.32, 44.92]
ENIGMA Discovery Cohorts [N=21448; P=7.42e-06] 10.56 [ 5.94, 15.17]
UKBB 13.31 [ 6.99, 19.63]
Discovery+UKBB [N=30303; P=3.603e-09] 11.53 [ 7.70, 15.36]
GOBS -3.42 [-31.75, 24.92]
IMH 0.79 [-56.34, 57.93]
UNICAMP 17.70 [-45.02, 80.42]
Meth-CT
MIRECC
UKBB nonEUR -13.14 [-44.01, 17.74]
OSAKA -18.35 [-49.42, 12.71]
PING Non European -9.67 [-50.36, 31.02]
Non European Cohorts [N=2473; P=0.2651] -8.52 [-23.50, 6.46]
-100
-50
0
50
100
150
mm^2/effect allele

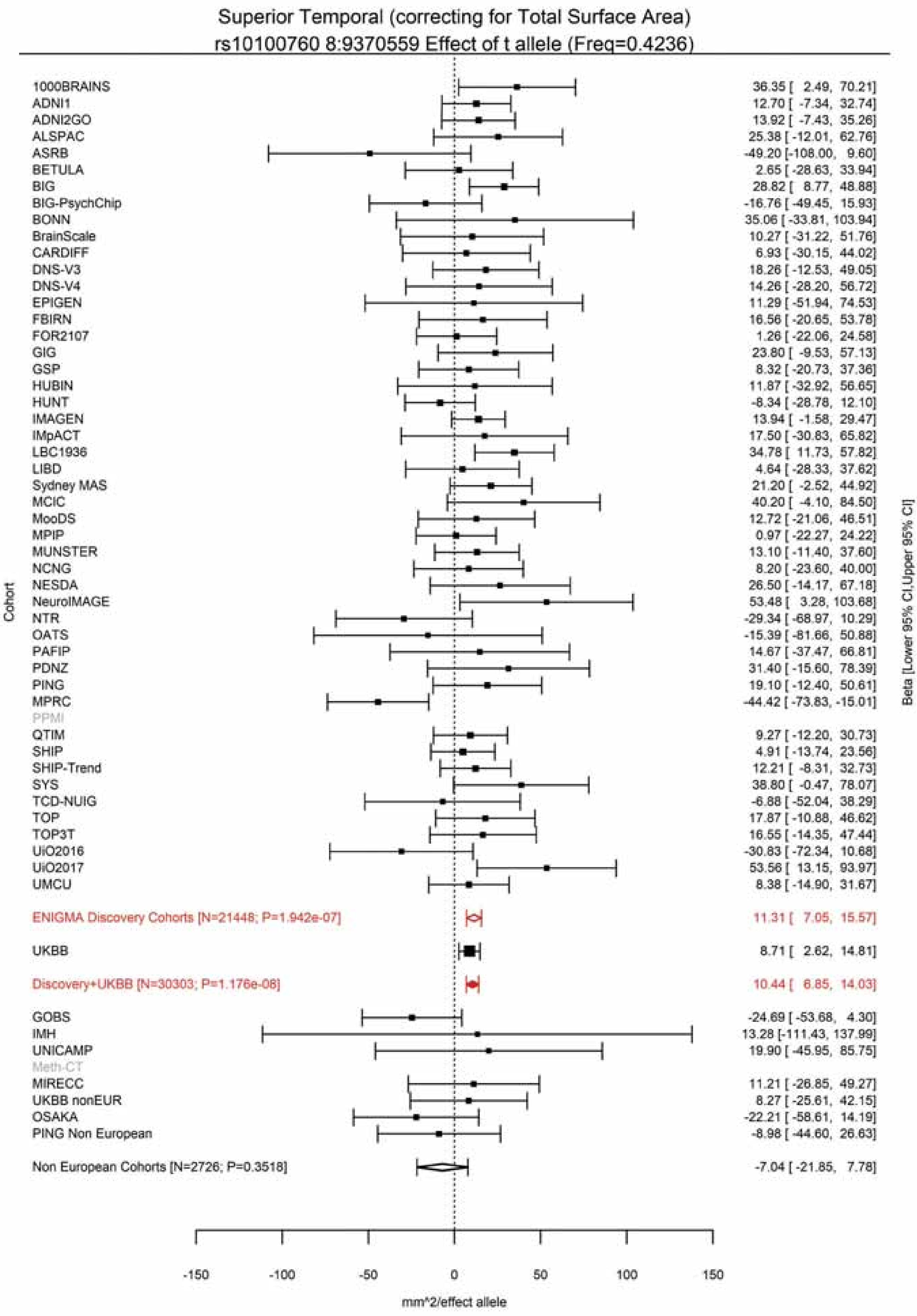
Superior Temporal (correcting for Total Surface Area)
rs10100760 8:9370559 Effect of t allele (Freq=0.4236)
Cohort
1000BRAINS 36.35 [ 2.49, 70.21]
ADNI1 12.70 [ -7.34, 32.74]
ADNI2GO 13.92 [ -7.43, 35.26]
ALSPAC 25.38 [ -12.01, 62.76]
ASRB -49.20 [-108.00, 9.60]
BETULA 2.65 [ -28.63, 33.94]
BIG 28.82 [ 8.77, 48.88]
BIG-PsychChip -16.76 [ -49.45, 15.93]
BONN 35.06 [ -33.81, 103.94]
BrainScale 10.27 [ -31.22, 51.76]
CARDIFF 6.93 [ -30.15, 44.02]
DNS-V3 18.26 [ -12.53, 49.05]
DNS-V4 14.26 [ -28.20, 56.72]
EPIGEN 11.29 [ -51.94, 74.53]
FBIRN 16.56 [ -20.65, 53.78]
FOR2107 1.26 [ -22.06, 24.58]
GIG 23.80 [ -9.53, 57.13]
GSP 8.32 [ -20.73, 37.36]
HUBIN 11.87 [ -32.92, 56.65]
HUNT -8.34 [ -28.78, 12.10]
IMAGEN 13.94 [ -1.58, 29.47]
IMpACT 17.50 [ -30.83, 65.82]
LBC1936 34.78 [ 11.73, 57.82]
LIBD 4.64 [ -28.33, 37.62]
Sydney MAS 21.20 [ -2.52, 44.92]
MCIC 40.20 [ -4.10, 84.50]
MooDS 12.72 [ -21.06, 46.51]
MPIP 0.97 [ -22.27, 24.22]
MUNSTER 13.10 [ -11.40, 37.60]
NCNG 8.20 [ -23.60, 40.00]
NESDA 26.50 [ -14.17, 67.18]
NeuroIMAGE 53.48 [ 3.28, 103.68]
NTR -29.34 [ -68.97, 10.29]
OATS -15.39 [ -81.66, 50.88]
PAFIP 14.67 [ -37.47, 66.81]
PDNZ 31.40 [ -15.60, 78.39]
PING 19.10 [ -12.40, 50.61]
MPRC -44.42 [ -73.83, -15.01]
PPMI
QTIM 9.27 [ -12.20, 30.73]
SHIP 4.91 [ -13.74, 23.56]
SHIP-Trend 12.21 [ -8.31, 32.73]
SYS 38.80 [ -0.47, 78.07]
TCD-NUIG -6.88 [ -52.04, 38.29]
TOP 17.87 [ -10.88, 46.62]
TOP3T 16.55 [ -14.35, 47.44]
UiO2016 -30.83 [ -72.34, 10.68]
UiO2017 53.56 [ 13.15, 93.97]
UMCU 8.38 [ -14.90, 31.67]
ENIGMA Discovery Cohorts [N=21448; P=1.942e-07] 11.31 [ 7.05, 15.57]
UKBB 8.71 [ 2.62, 14.81]
Discovery+UKBB [N=30303; P=1.176e-08] 10.44 [ 6.85, 14.03]
GOBS -24.69 [ -53.68, 4.30]
IMH 13.28 [-111.43, 137.99]
UNICAMP 19.90 [ -45.95, 85.75]
Meth-CT
MIRECC 11.21 [ -26.85, 49.27]
UKBB nonEUR 8.27 [ -25.61, 42.15]
OSAKA -22.21 [ -58.61, 14.19]
PING Non European -8.98 [ -44.60, 26.63]
Non European Cohorts [N=2726; P=0.3518] -7.04 [ -21.85, 7.78]
Beta [Lower 95% CI,Upper 95% CI]
-150
-100
-50
0
50
100
150
mm^2/effect allele

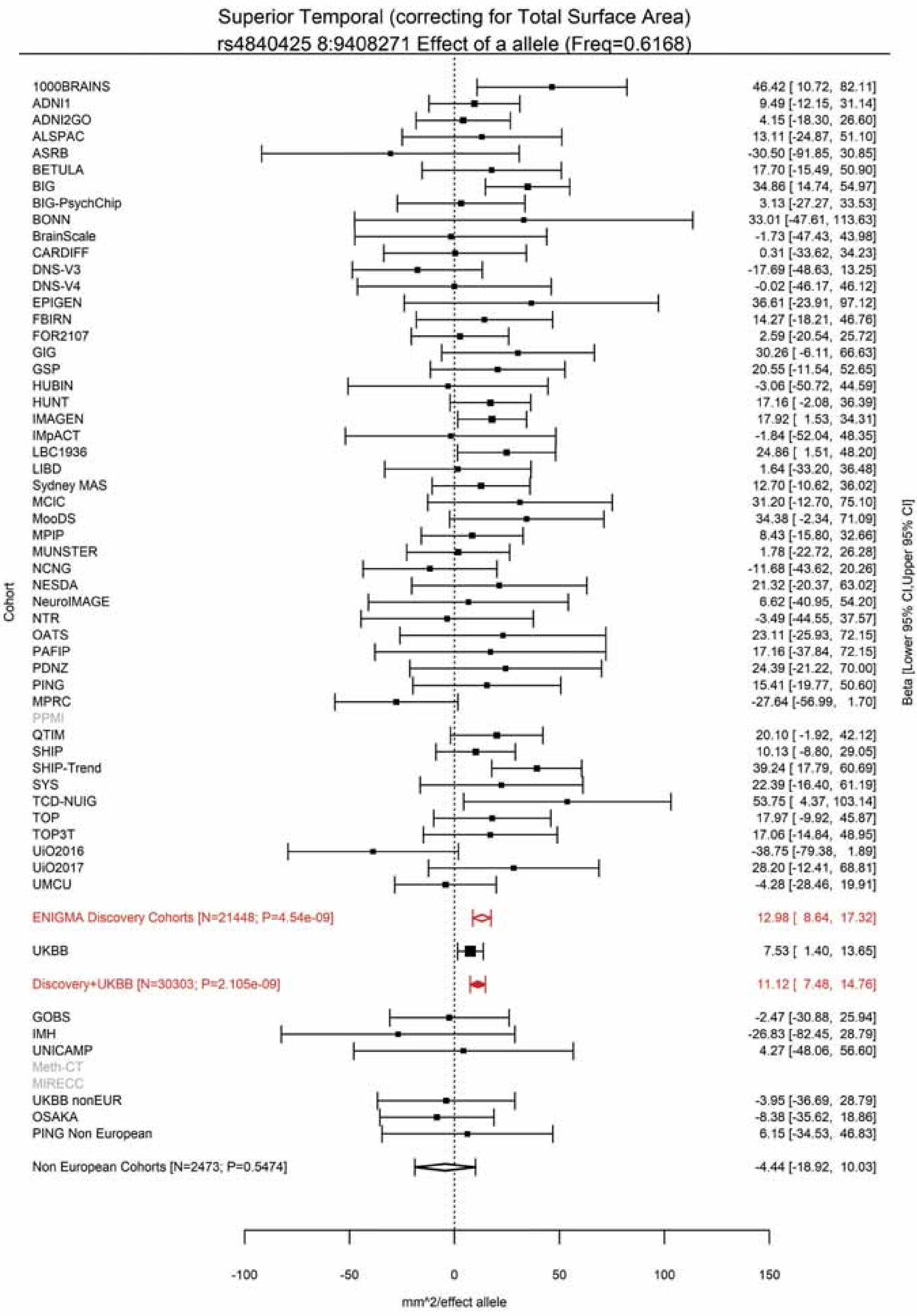

Superior Temporal (correcting for Total Surface Area)
rs4840425 8:9408271 Effect of a allele (Freq=0.6168)
Cohort
1000BRAINS 46.42 [ 10.72, 82.11]
ADNI1 9.49 [-12.15, 31.14]
ADNI2GO 4.15 [-18.30, 26.60]
ALSPAC 13.11 [-24.87, 51.10]
ASRB -30.50 [-91.85, 30.85]
BETULA 17.70 [-15.49, 50.90]
BIG 34.86 [ 14.74, 54.97]
BIG-PsychChip 3.13 [-27.27, 33.53]
BONN 33.01 [-47.61, 113.63]
BrainScale -1.73 [-47.43, 43.98]
CARDIFF 0.31 [-33.62, 34.23]
DNS-V3 -17.69 [-48.63, 13.25]
DNS-V4 -0.02 [-46.17, 46.12]
EPIGEN 36.61 [-23.91, 97.12]
FBIRN 14.27 [-18.21, 46.76]
FOR2107 2.59 [-20.54, 25.72]
GIG 30.26 [ -6.11, 66.63]
GSP 20.55 [-11.54, 52.65]
HUBIN -3.06 [-50.72, 44.59]
HUNT 17.16 [ -2.08, 36.39]
IMAGEN 17.92 [ 1.53, 34.31]
IMpACT -1.84 [-52.04, 48.35]
LBC1936 24.86 [ 1.51, 48.20]
LIBD 1.64 [-33.20, 36.48]
Sydney MAS 12.70 [-10.62, 36.02]
MCIC 31.20 [-12.70, 75.10]
MooDS 34.38 [ -2.34, 71.09]
MPIP 8.43 [-15.80, 32.66]
MUNSTER 1.78 [-22.72, 26.28]
NCNG -11.68 [-43.62, 20.26]
NESDA 21.32 [-20.37, 63.02]
NeuroIMAGE 6.62 [-40.95, 54.20]
NTR -3.49 [-44.55, 37.57]
OATS 23.11 [-25.93, 72.15]
PAFIP 17.16 [-37.84, 72.15]
PDNZ 24.39 [-21.22, 70.00]
PING 15.41 [-19.77, 50.60]
MPRC -27.64 [-56.99, 1.70]
PPMI
QTIM 20.10 [ -1.92, 42.12]
SHIP 10.13 [ -8.80, 29.05]
SHIP-Trend 39.24 [ 17.79, 60.69]
SYS 22.39 [-16.40, 61.19]
TCD-NUIG 53.75 [ 4.37, 103.14]
TOP 17.97 [ -9.92, 45.87]
TOP3T 17.06 [-14.84, 48.95]
UiO2016 -38.75 [-79.38, 1.89]
UiO2017 28.20 [-12.41, 68.81]
UMCU -4.28 [-28.46, 19.91]
ENIGMA Discovery Cohorts [N=21448; P=4.54e-09] 12.98 [ 8.64, 17.32]
UKBB 7.53 [ 1.40, 13.65]
Discovery+UKBB [N=30303; P=2.105e-09] 11.12 [ 7.48, 14.76]
GOBS -2.47 [-30.88, 25.94]
IMH -26.83 [-82.45, 28.79]
UNICAMP 4.27 [-48.06, 56.60]
Meth-CT
MIRECC
UKBB nonEUR -3.95 [-36.69, 28.79]
OSAKA -8.38 [-35.62, 18.86]
PING Non European 6.15 [-34.53, 46.83]
Non European Cohorts [N=2473; P=0.5474] -4.44 [-18.92, 10.03]
-100
-50
0
50
100
150
mm^2/effect allele
Beta [Lower 95% CI,Upper 95% CI]

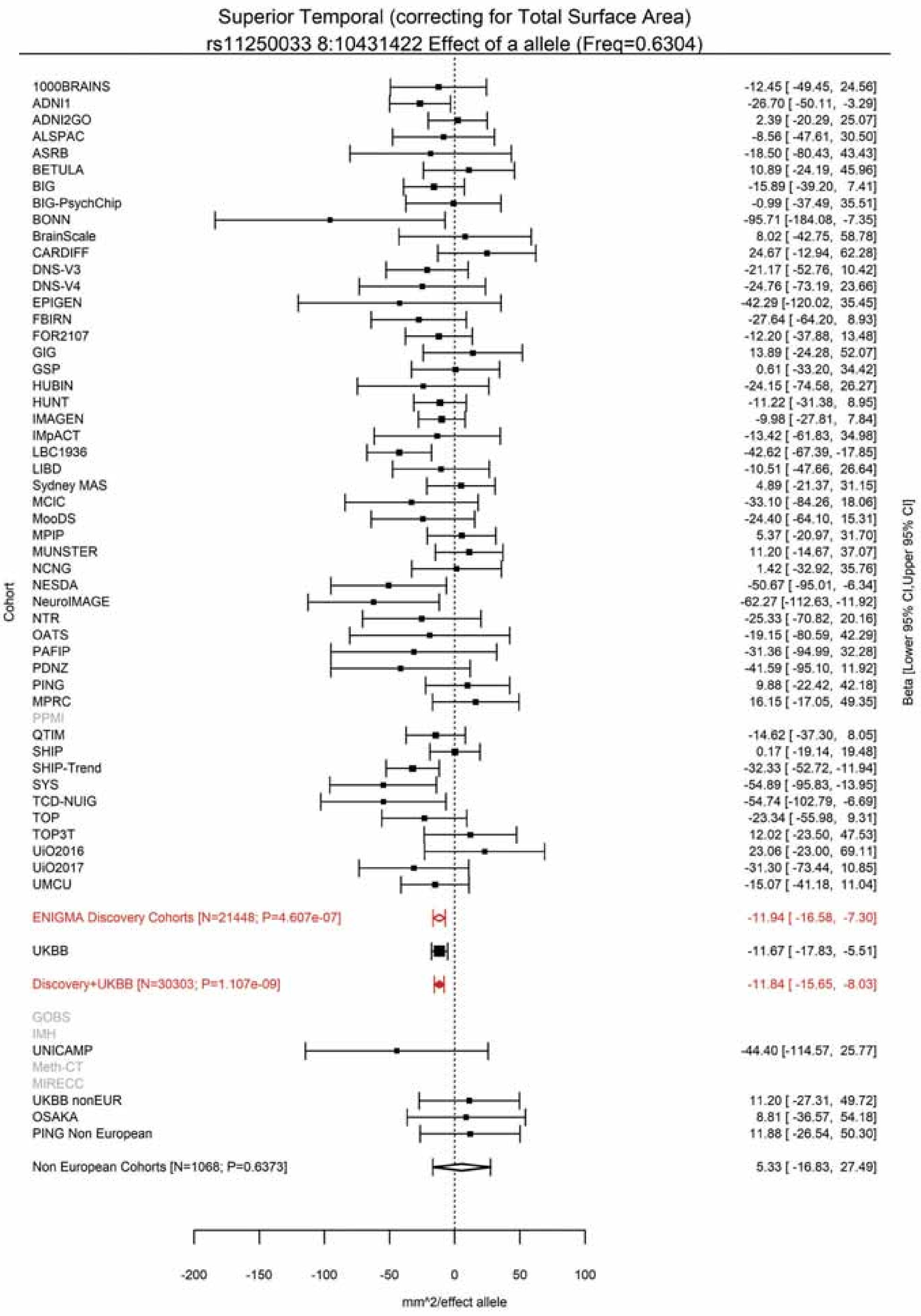
Superior Temporal (correcting for Total Surface Area)
rs11250033 8:10431422 Effect of a allele (Freq=0.6304)
Cohort
Beta [Lower 95% CI,Upper 95% CI]
1000BRAINS -12.45 [ -49.45, 24.56]
ADNI1 -26.70 [ -50.11, -3.29]
ADNI2GO 2.39 [ -20.29, 25.07]
ALSPAC -8.56 [ -47.61, 30.50]
ASRB -18.50 [ -80.43, 43.43]
BETULA 10.89 [ -24.19, 45.96]
BIG -15.89 [ -39.20, 7.41]
BIG-PsychChip -0.99 [ -37.49, 35.51]
BONN -95.71 [-184.08, -7.35]
BrainScale 8.02 [ -42.75, 58.78]
CARDIFF 24.67 [ -12.94, 62.28]
DNS-V3 -21.17 [ -52.76, 10.42]
DNS-V4 -24.76 [ -73.19, 23.66]
EPIGEN -42.29 [-120.02, 35.45]
FBIRN -27.64 [ -64.20, 8.93]
FOR2107 -12.20 [ -37.88, 13.48]
GIG 13.89 [ -24.28, 52.07]
GSP 0.61 [ -33.20, 34.42]
HUBIN -24.15 [ -74.58, 26.27]
HUNT -11.22 [ -31.38, 8.95]
IMAGEN -9.98 [ -27.81, 7.84]
IMpACT -13.42 [ -61.83, 34.98]
LBC1936 -42.62 [ -67.39, -17.85]
LIBD -10.51 [ -47.66, 26.64]
Sydney MAS 4.89 [ -21.37, 31.15]
MCIC -33.10 [ -84.26, 18.06]
MooDS -24.40 [ -64.10, 15.31]
MPIP 5.37 [ -20.97, 31.70]
MUNSTER 11.20 [ -14.67, 37.07]
NCNG 1.42 [ -32.92, 35.76]
NESDA -50.67 [ -95.01, -6.34]
NeuroIMAGE -62.27 [-112.63, -11.92]
NTR -25.33 [ -70.82, 20.16]
OATS -19.15 [ -80.59, 42.29]
PAFIP -31.36 [ -94.99, 32.28]
PDNZ -41.59 [ -95.10, 11.92]
PING 9.88 [ -22.42, 42.18]
MPRC 16.15 [ -17.05, 49.35]
PPMI
QTIM -14.62 [ -37.30, 8.05]
SHIP 0.17 [ -19.14, 19.48]
SHIP-Trend -32.33 [ -52.72, -11.94]
SYS -54.89 [ -95.83, -13.95]
TCD-NUIG -54.74 [-102.79, -6.69]
TOP -23.34 [ -55.98, 9.31]
TOP3T 12.02 [ -23.50, 47.53]
UiO2016 23.06 [ -23.00, 69.11]
UiO2017 -31.30 [ -73.44, 10.85]
UMCU -15.07 [ -41.18, 11.04]
ENIGMA Discovery Cohorts [N=21448; P=4.607e-07] -11.94 [ -16.58, -7.30]
UKBB -11.67 [ -17.83, -5.51]
Discovery+UKBB [N=30303; P=1.107e-09] -11.84 [ -15.65, -8.03]
GOBS
IMH
UNICAMP -44.40 [-114.57, 25.77]
Meth-CT
MIRECC
UKBB nonEUR 11.20 [ -27.31, 49.72]
OSAKA 8.81 [ -36.57, 54.18]
PING Non European 11.88 [ -26.54, 50.30]
Non European Cohorts [N=1068; P=0.6373] 5.33 [ -16.83, 27.49]
-200
-150
-100
-50
0
50
100
mm^2/effect allele

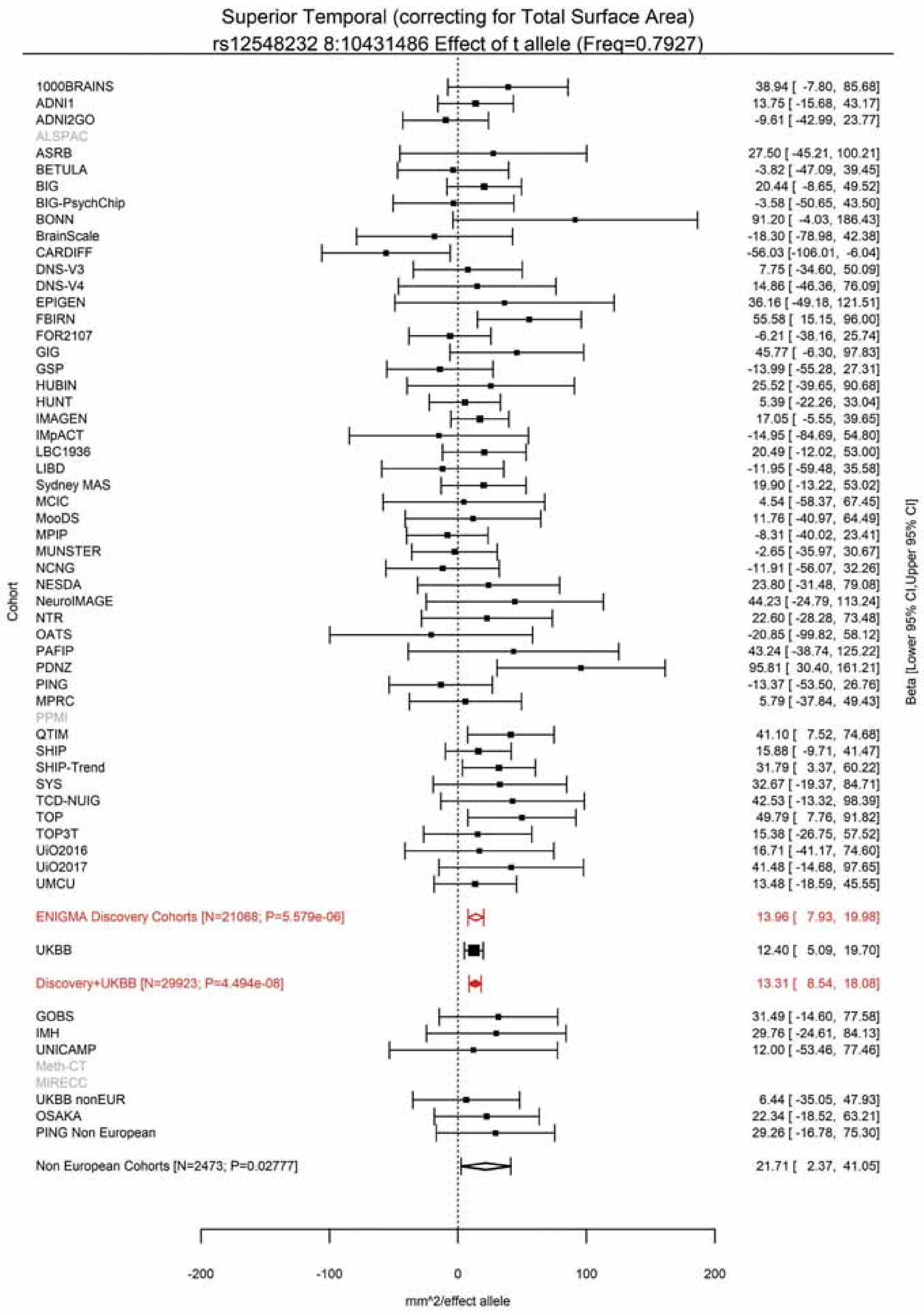
Superior Temporal (correcting for Total Surface Area)
rs12548232 8:10431486 Effect of t allele (Freq=0.7927)
Cohort
1000BRAINS 38.94 [ -7.80, 85.68]
ADNI1 13.75 [ -15.68, 43.17]
ADNI2GO -9.61 [ -42.99, 23.77]
ALSPAC
ASRB 27.50 [ -45.21, 100.21]
BETULA -3.82 [ -47.09, 39.45]
BIG 20.44 [ -8.65, 49.52]
BIG-PsychChip -3.58 [ -50.65, 43.50]
BONN 91.20 [ -4.03, 186.43]
BrainScale -18.30 [ -78.98, 42.38]
CARDIFF -56.03 [-106.01, -6.04]
DNS-V3 7.75 [ -34.60, 50.09]
DNS-V4 14.86 [ -46.36, 76.09]
EPIGEN 36.16 [ -49.18, 121.51]
FBIRN 55.58 [ 15.15, 96.00]
FOR2107 -6.21 [ -38.16, 25.74]
GIG 45.77 [ -6.30, 97.83]
GSP -13.99 [ -55.28, 27.31]
HUBIN 25.52 [ -39.65, 90.68]
HUNT 5.39 [ -22.26, 33.04]
IMAGEN 17.05 [ -5.55, 39.65]
IMpACT -14.95 [ -84.69, 54.80]
LBC1936 20.49 [ -12.02, 53.00]
LIBD -11.95 [ -59.48, 35.58]
Sydney MAS 19.90 [ -13.22, 53.02]
MCIC 4.54 [ -58.37, 67.45]
MooDS 11.76 [ -40.97, 64.49]
MPIP -8.31 [ -40.02, 23.41]
MUNSTER -2.65 [ -35.97, 30.67]
NCNG -11.91 [ -56.07, 32.26]
NESDA 23.80 [ -31.48, 79.08]
NeuroIMAGE 44.23 [ -24.79, 113.24]
NTR 22.60 [ -28.28, 73.48]
OATS -20.85 [ -99.82, 58.12]
PAFIP 43.24 [ -38.74, 125.22]
PDNZ 95.81 [ 30.40, 161.21]
PING -13.37 [ -53.50, 26.76]
MPRC 5.79 [ -37.84, 49.43]
PPMI
QTIM 41.10 [ 7.52, 74.68]
SHIP 15.88 [ -9.71, 41.47]
SHIP-Trend 31.79 [ 3.37, 60.22]
SYS 32.67 [ -19.37, 84.71]
TCD-NUIG 42.53 [ -13.32, 98.39]
TOP 49.79 [ 7.76, 91.82]
TOP3T 15.38 [ -26.75, 57.52]
UiO2016 16.71 [ -41.17, 74.60]
UiO2017 41.48 [ -14.68, 97.65]
UMCU 13.48 [ -18.59, 45.55]
ENIGMA Discovery Cohorts [N=21068; P=5.579e-06] 13.96 [ 7.93, 19.98]
UKBB 12.40 [ 5.09, 19.70]
Discovery+UKBB [N=29923; P=4.494e-08] 13.31 [ 8.54, 18.08]
GOBS 31.49 [ -14.60, 77.58]
IMH 29.76 [ -24.61, 84.13]
UNICAMP 12.00 [ -53.46, 77.46]
Meth-CT
MIRECC
UKBB nonEUR 6.44 [ -35.05, 47.93]
OSAKA 22.34 [ -18.52, 63.21]
PING Non European 29.26 [ -16.78, 75.30]
Non European Cohorts [N=2473; P=0.02777] 21.71 [ 2.37, 41.05]
Beta [Lower 95% CI,Upper 95% CI]
-200
-100
0
100
200
mm^2/effect allele

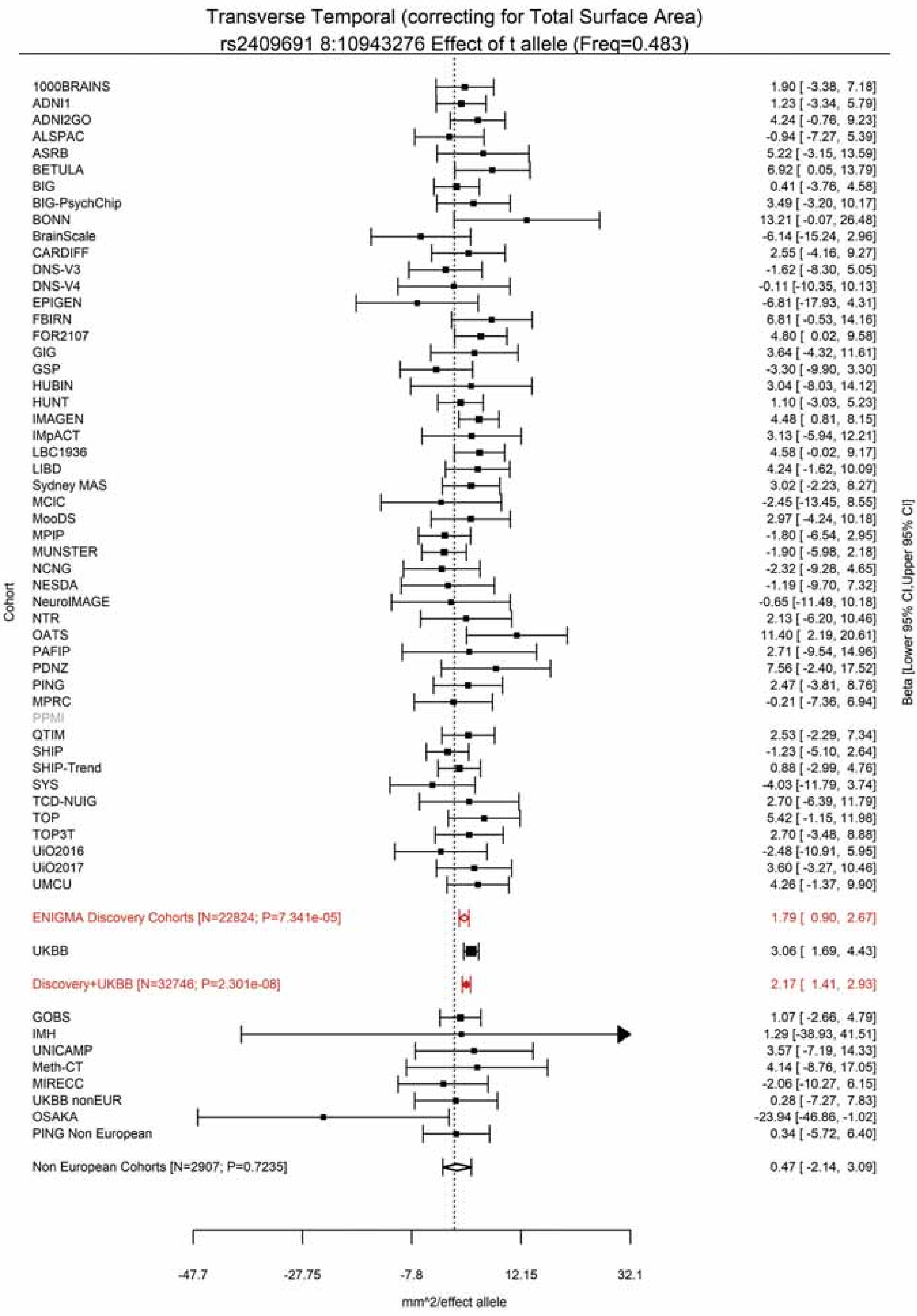

Transverse Temporal (correcting for Total Surface Area)
rs2409691 8:10943276 Effect of t allele (Freq=0.483)
Cohort
Beta [Lower 95% CI,Upper 95% CI]
1000BRAINS 1.90 [ -3.38, 7.18]
ADNI1 1.23 [ -3.34, 5.79]
ADNI2GO 4.24 [ -0.76, 9.23]
ALSPAC -0.94 [ -7.27, 5.39]
ASRB 5.22 [ -3.15, 13.59]
BETULA 6.92 [ 0.05, 13.79]
BIG 0.41 [ -3.76, 4.58]
BIG-PsychChip 3.49 [ -3.20, 10.17]
BONN 13.21 [ -0.07, 26.48]
BrainScale -6.14 [-15.24, 2.96]
CARDIFF 2.55 [ -4.16, 9.27]
DNS-V3 -1.62 [ -8.30, 5.05]
DNS-V4 -0.11 [-10.35, 10.13]
EPIGEN -6.81 [-17.93, 4.31]
FBIRN 6.81 [ -0.53, 14.16]
FOR2107 4.80 [ 0.02, 9.58]
GIG 3.64 [ -4.32, 11.61]
GSP -3.30 [ -9.90, 3.30]
HUBIN 3.04 [ -8.03, 14.12]
HUNT 1.10 [ -3.03, 5.23]
IMAGEN 4.48 [ 0.81, 8.15]
IMpACT 3.13 [ -5.94, 12.21]
LBC1936 4.58 [ -0.02, 9.17]
LIBD 4.24 [ -1.62, 10.09]
Sydney MAS 3.02 [ -2.23, 8.27]
MCIC -2.45 [-13.45, 8.55]
MooDS 2.97 [ -4.24, 10.18]
MPIP -1.80 [ -6.54, 2.95]
MUNSTER -1.90 [ -5.98, 2.18]
NCNG -2.32 [ -9.28, 4.65]
NESDA -1.19 [ -9.70, 7.32]
NeuroIMAGE -0.65 [-11.49, 10.18]
NTR 2.13 [ -6.20, 10.46]
OATS 11.40 [ 2.19, 20.61]
PAFIP 2.71 [ -9.54, 14.96]
PDNZ 7.56 [ -2.40, 17.52]
PING 2.47 [ -3.81, 8.76]
MPRC -0.21 [ -7.36, 6.94]
PPMI
QTIM 2.53 [ -2.29, 7.34]
SHIP -1.23 [ -5.10, 2.64]
SHIP-Trend 0.88 [ -2.99, 4.76]
SYS -4.03 [-11.79, 3.74]
TCD-NUIG 2.70 [ -6.39, 11.79]
TOP 5.42 [ -1.15, 11.98]
TOP3T 2.70 [ -3.48, 8.88]
UiO2016 -2.48 [-10.91, 5.95]
UiO2017 3.60 [ -3.27, 10.46]
UMCU 4.26 [ -1.37, 9.90]
ENIGMA Discovery Cohorts [N=22824; P=7.341e-05] 1.79 [ 0.90, 2.67]
UKBB 3.06 [ 1.69, 4.43]
Discovery+UKBB [N=32746; P=2.301e-08] 2.17 [ 1.41, 2.93]
GOBS 1.07 [ -2.66, 4.79]
IMH 1.29 [-38.93, 41.51]
UNICAMP 3.57 [ -7.19, 14.33]
Meth-CT 4.14 [ -8.76, 17.05]
MIRECC -2.06 [-10.27, 6.15]
UKBB nonEUR 0.28 [ -7.27, 7.83]
OSAKA -23.94 [-46.86, -1.02]
PING Non European 0.34 [ -5.72, 6.40]
Non European Cohorts [N=2907; P=0.7235] 0.47 [ -2.14, 3.09]
-47.7
-27.75
-7.8
12.15
32.1
mm^2/effect allele

### Superior Temporal (correcting for Total Surface Area)
### rs1057626 8:10962800 Effect of t allele (Freq=0.4818)

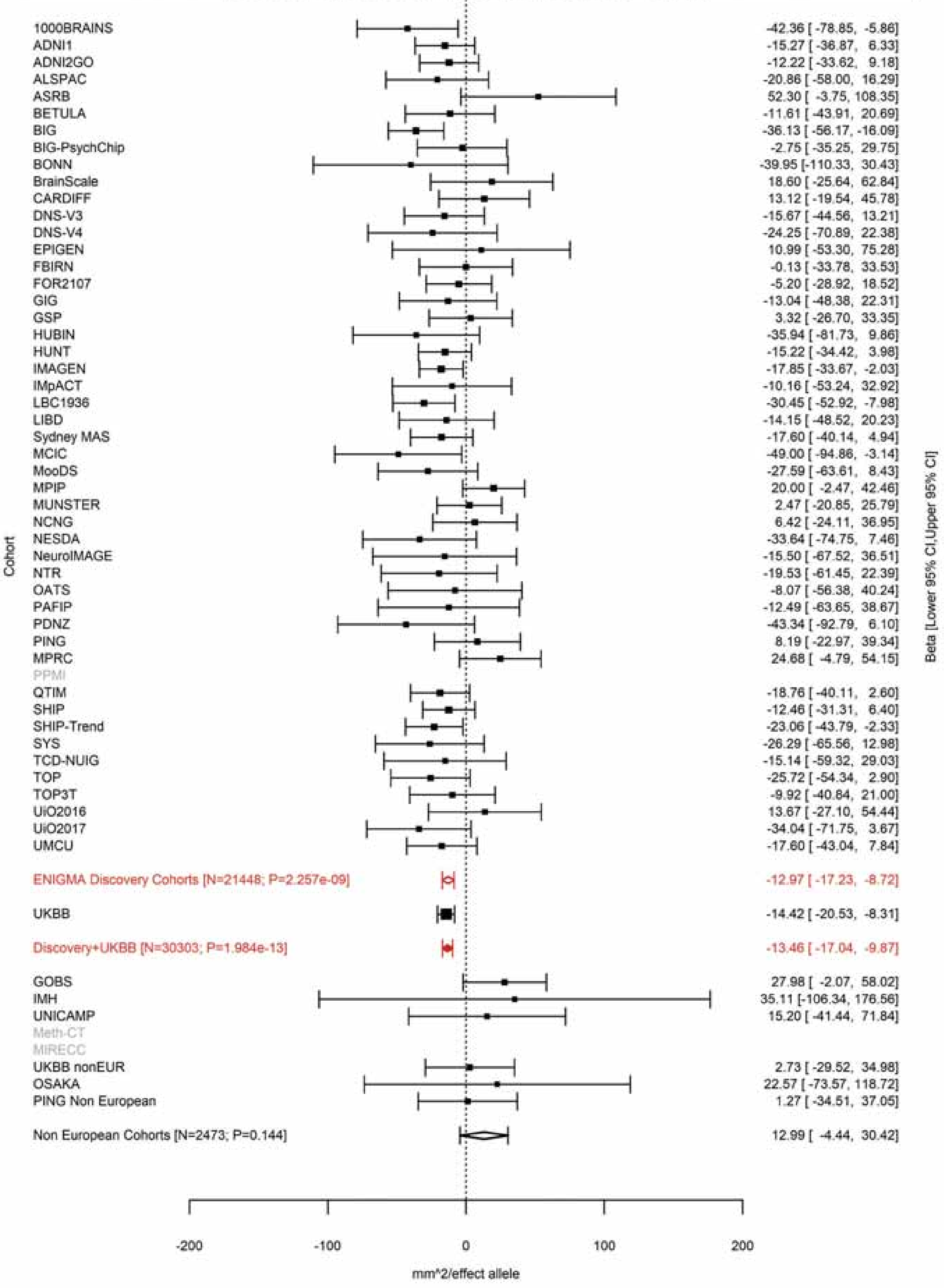

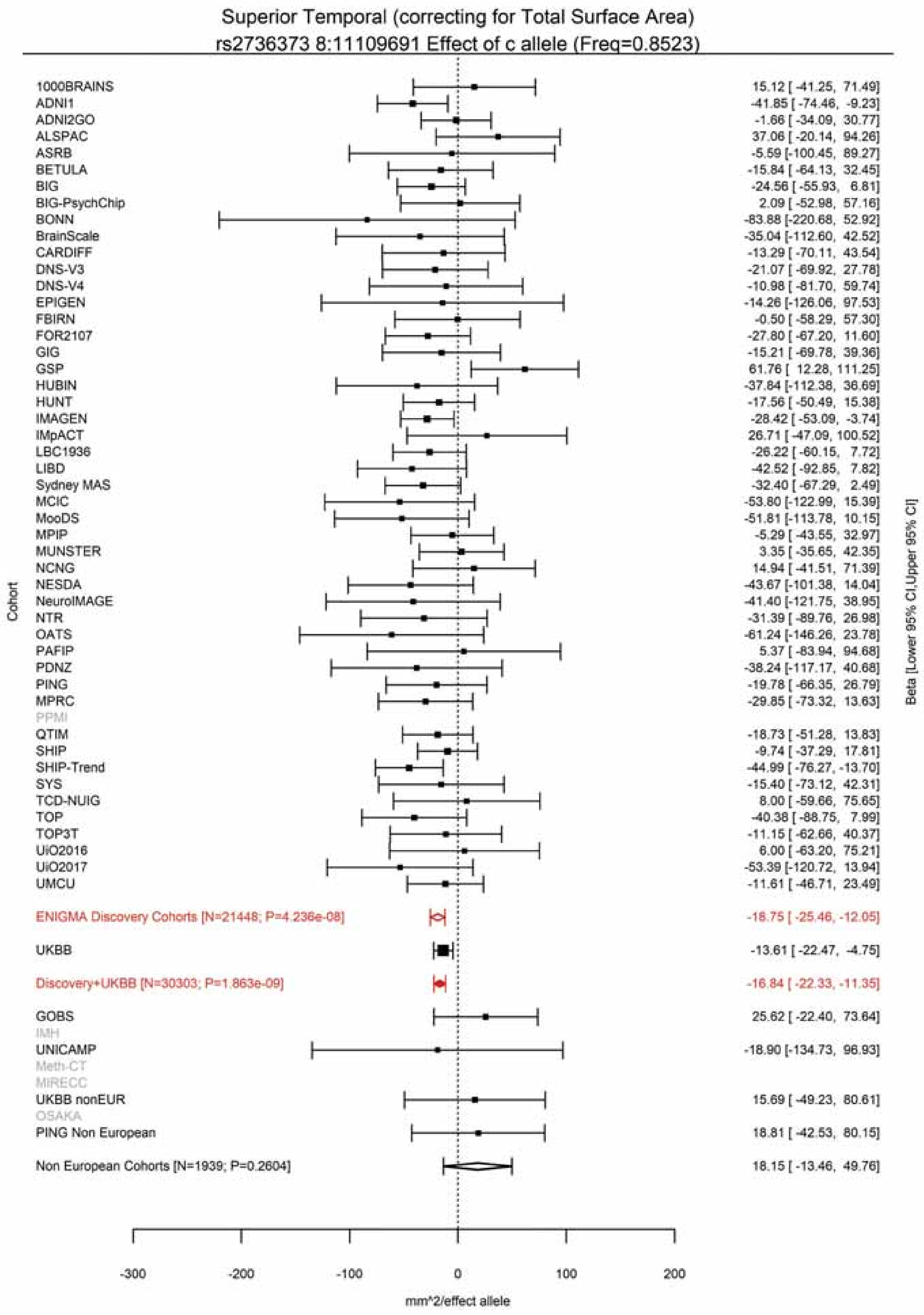
Superior Temporal (correcting for Total Surface Area)
rs2736373 8:11109691 Effect of c allele (Freq=0.8523)
Cohort
Beta [Lower 95% CI,Upper 95% CI]
1000BRAINS 15.12 [ -41.25, 71.49]
ADNI1 -41.85 [ -74.46, -9.23]
ADNI2GO -1.66 [ -34.09, 30.77]
ALSPAC 37.06 [ -20.14, 94.26]
ASRB -5.59 [-100.45, 89.27]
BETULA -15.84 [ -64.13, 32.45]
BIG -24.56 [ -55.93, 6.81]
BIG-PsychChip 2.09 [ -52.98, 57.16]
BONN -83.88 [-220.68, 52.92]
BrainScale -35.04 [-112.60, 42.52]
CARDIFF -13.29 [ -70.11, 43.54]
DNS-V3 -21.07 [ -69.92, 27.78]
DNS-V4 -10.98 [ -81.70, 59.74]
EPIGEN -14.26 [-126.06, 97.53]
FBIRN -0.50 [ -58.29, 57.30]
FOR2107 -27.80 [ -67.20, 11.60]
GIG -15.21 [ -69.78, 39.36]
GSP 61.76 [ 12.28, 111.25]
HUBIN -37.84 [-112.38, 36.69]
HUNT -17.56 [ -50.49, 15.38]
IMAGEN -28.42 [ -53.09, -3.74]
IMpACT 26.71 [ -47.09, 100.52]
LBC1936 -26.22 [ -60.15, 7.72]
LIBD -42.52 [ -92.85, 7.82]
Sydney MAS -32.40 [ -67.29, 2.49]
MCIC -53.80 [-122.99, 15.39]
MooDS -51.81 [-113.78, 10.15]
MPIP -5.29 [ -43.55, 32.97]
MUNSTER 3.35 [ -35.65, 42.35]
NCNG 14.94 [ -41.51, 71.39]
NESDA -43.67 [-101.38, 14.04]
NeuroIMAGE -41.40 [-121.75, 38.95]
NTR -31.39 [ -89.76, 26.98]
OATS -61.24 [-146.26, 23.78]
PAFIP 5.37 [ -83.94, 94.68]
PDNZ -38.24 [-117.17, 40.68]
PING -19.78 [ -66.35, 26.79]
MPRC -29.85 [ -73.32, 13.63]
PPMI
QTIM -18.73 [ -51.28, 13.83]
SHIP -9.74 [ -37.29, 17.81]
SHIP-Trend -44.99 [ -76.27, -13.70]
SYS -15.40 [ -73.12, 42.31]
TCD-NUIG 8.00 [ -59.66, 75.65]
TOP -40.38 [ -88.75, 7.99]
TOP3T -11.15 [ -62.66, 40.37]
UiO2016 6.00 [ -63.20, 75.21]
UiO2017 -53.39 [-120.72, 13.94]
UMCU -11.61 [ -46.71, 23.49]
ENIGMA Discovery Cohorts [N=21448; P=4.236e-08] -18.75 [ -25.46, -12.05]
UKBB -13.61 [ -22.47, -4.75]
Discovery+UKBB [N=30303; P=1.863e-09] -16.84 [ -22.33, -11.35]
GOBS 25.62 [ -22.40, 73.64]
IMH
UNICAMP -18.90 [-134.73, 96.93]
Meth-CT
MIRECC
UKBB nonEUR 15.69 [ -49.23, 80.61]
OSAKA
PING Non European 18.81 [ -42.53, 80.15]
Non European Cohorts [N=1939; P=0.2604] 18.15 [ -13.46, 49.76]
-300
-200
-100
0
100
200
mm^2/effect allele

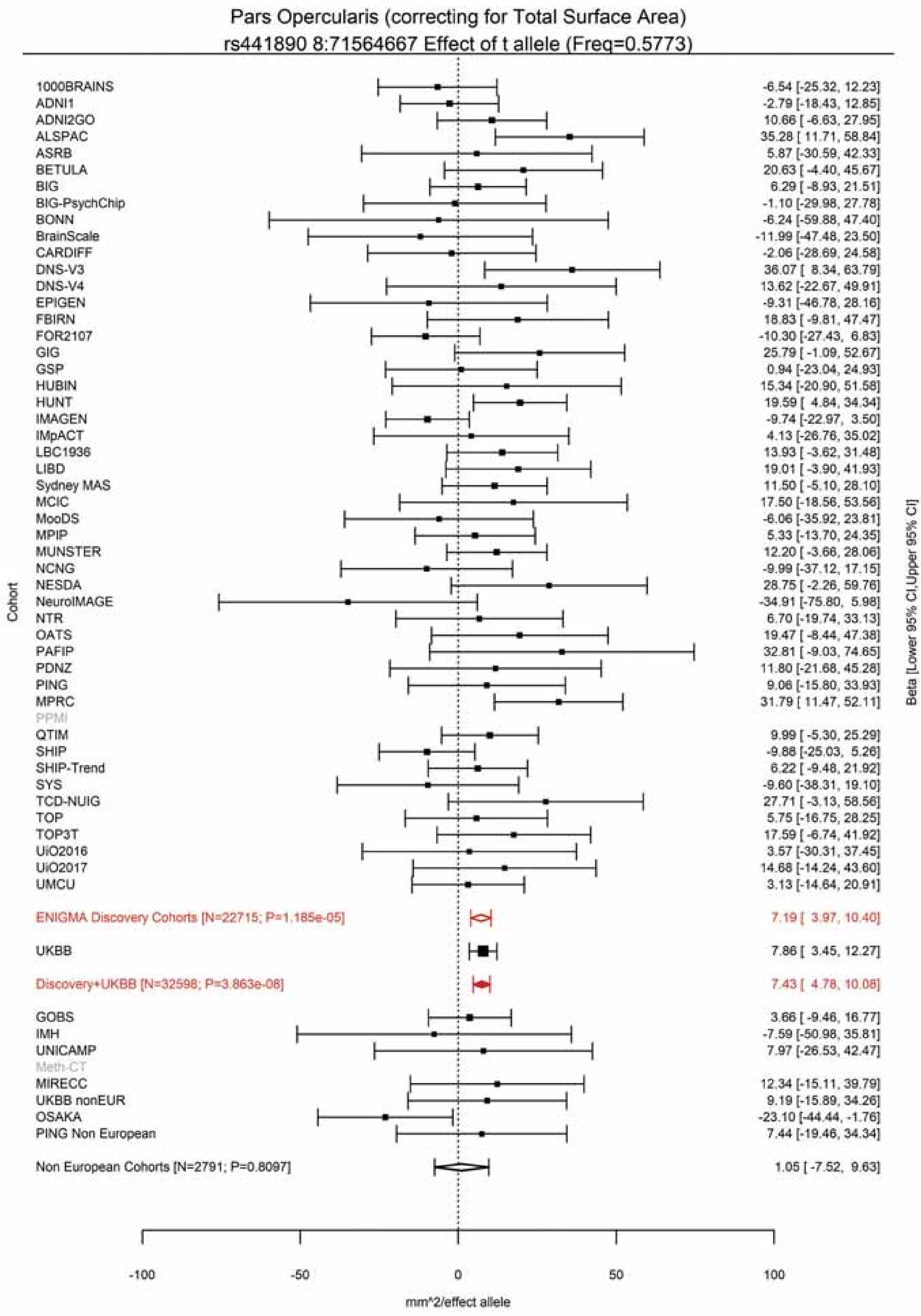
Pars Opercularis (correcting for Total Surface Area)
rs441890 8:71564667 Effect of t allele (Freq=0.5773)
Cohort
1000BRAINS -6.54 [-25.32, 12.23]
ADNI1 -2.79 [-18.43, 12.85]
ADNI2GO 10.66 [ -6.63, 27.95]
ALSPAC 35.28 [ 11.71, 58.84]
ASRB 5.87 [-30.59, 42.33]
BETULA 20.63 [ -4.40, 45.67]
BIG 6.29 [ -8.93, 21.51]
BIG-PsychChip -1.10 [-29.98, 27.78]
BONN -6.24 [-59.88, 47.40]
BrainScale -11.99 [-47.48, 23.50]
CARDIFF -2.06 [-28.69, 24.58]
DNS-V3 36.07 [ 8.34, 63.79]
DNS-V4 13.62 [-22.67, 49.91]
EPIGEN -9.31 [-46.78, 28.16]
FBIRN 18.83 [ -9.81, 47.47]
FOR2107 -10.30 [-27.43, 6.83]
GIG 25.79 [ -1.09, 52.67]
GSP 0.94 [-23.04, 24.93]
HUBIN 15.34 [-20.90, 51.58]
HUNT 19.59 [ 4.84, 34.34]
IMAGEN -9.74 [-22.97, 3.50]
IMpACT 4.13 [-26.76, 35.02]
LBC1936 13.93 [ -3.62, 31.48]
LIBD 19.01 [ -3.90, 41.93]
Sydney MAS 11.50 [ -5.10, 28.10]
MCIC 17.50 [-18.56, 53.56]
MooDS -6.06 [-35.92, 23.81]
MPIP 5.33 [-13.70, 24.35]
MUNSTER 12.20 [ -3.66, 28.06]
NCNG -9.99 [-37.12, 17.15]
NESDA 28.75 [ -2.26, 59.76]
NeuroIMAGE -34.91 [-75.80, 5.98]
NTR 6.70 [-19.74, 33.13]
OATS 19.47 [ -8.44, 47.38]
PAFIP 32.81 [ -9.03, 74.65]
PDNZ 11.80 [-21.68, 45.28]
PING 9.06 [-15.80, 33.93]
MPRC 31.79 [ 11.47, 52.11]
PPMI
QTIM 9.99 [ -5.30, 25.29]
SHIP -9.88 [-25.03, 5.26]
SHIP-Trend 6.22 [ -9.48, 21.92]
SYS -9.60 [-38.31, 19.10]
TCD-NUIG 27.71 [ -3.13, 58.56]
TOP 5.75 [-16.75, 28.25]
TOP3T 17.59 [ -6.74, 41.92]
UiO2016 3.57 [-30.31, 37.45]
UiO2017 14.68 [-14.24, 43.60]
UMCU 3.13 [-14.64, 20.91]
ENIGMA Discovery Cohorts [N=22715; P=1.185e-05] 7.19 [ 3.97, 10.40]
UKBB 7.86 [ 3.45, 12.27]
Discovery+UKBB [N=32598; P=3.863e-08] 7.43 [ 4.78, 10.08]
GOBS 3.66 [ -9.46, 16.77]
IMH -7.59 [-50.98, 35.81]
UNICAMP 7.97 [-26.53, 42.47]
Meth-CT
MIRECC 12.34 [-15.11, 39.79]
UKBB nonEUR 9.19 [-15.89, 34.26]
OSAKA -23.10 [-44.44, -1.76]
PING Non European 7.44 [-19.46, 34.34]
Non European Cohorts [N=2791; P=0.8097] 1.05 [ -7.52, 9.63]
Beta [Lower 95% CI,Upper 95% CI]
-100
-50
0
50
100
mm^2/effect allele

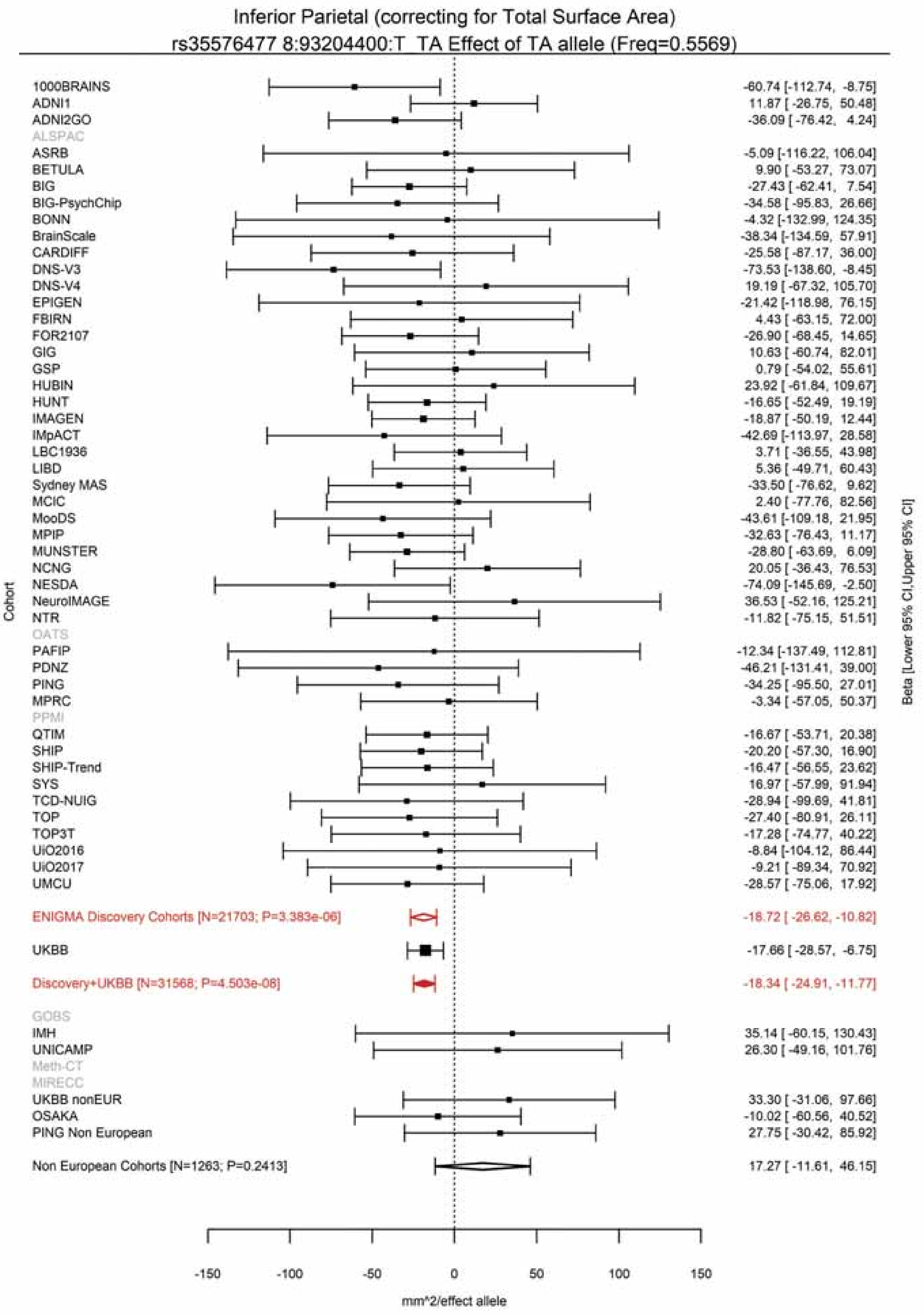
Inferior Parietal (correcting for Total Surface Area)
rs35576477 8:93204400:T_TA Effect of TA allele (Freq=0.5569)
Cohort
1000BRAINS -60.74 [-112.74, -8.75]
ADNI1 11.87 [ -26.75, 50.48]
ADNI2GO -36.09 [ -76.42, 4.24]
ALSPAC
ASRB -5.09 [-116.22, 106.04]
BETULA 9.90 [ -53.27, 73.07]
BIG -27.43 [ -62.41, 7.54]
BIG-PsychChip -34.58 [ -95.83, 26.66]
BONN -4.32 [-132.99, 124.35]
BrainScale -38.34 [-134.59, 57.91]
CARDIFF -25.58 [ -87.17, 36.00]
DNS-V3 -73.53 [-138.60, -8.45]
DNS-V4 19.19 [ -67.32, 105.70]
EPIGEN -21.42 [-118.98, 76.15]
FBIRN 4.43 [ -63.15, 72.00]
FOR2107 -26.90 [ -68.45, 14.65]
GIG 10.63 [ -60.74, 82.01]
GSP 0.79 [ -54.02, 55.61]
HUBIN 23.92 [ -61.84, 109.67]
HUNT -16.65 [ -52.49, 19.19]
IMAGEN -18.87 [ -50.19, 12.44]
IMpACT -42.69 [-113.97, 28.58]
LBC1936 3.71 [ -36.55, 43.98]
LIBD 5.36 [ -49.71, 60.43]
Sydney MAS -33.50 [ -76.62, 9.62]
MCIC 2.40 [ -77.76, 82.56]
MooDS -43.61 [-109.18, 21.95]
MPIP -32.63 [ -76.43, 11.17]
MUNSTER -28.80 [ -63.69, 6.09]
NCNG 20.05 [ -36.43, 76.53]
NESDA -74.09 [-145.69, -2.50]
NeuroIMAGE 36.53 [ -52.16, 125.21]
NTR -11.82 [ -75.15, 51.51]
OATS
PAFIP -12.34 [-137.49, 112.81]
PDNZ -46.21 [-131.41, 39.00]
PING -34.25 [ -95.50, 27.01]
MPRC -3.34 [ -57.05, 50.37]
PPMI
QTIM -16.67 [ -53.71, 20.38]
SHIP -20.20 [ -57.30, 16.90]
SHIP-Trend -16.47 [ -56.55, 23.62]
SYS 16.97 [ -57.99, 91.94]
TCD-NUIG -28.94 [ -99.69, 41.81]
TOP -27.40 [ -80.91, 26.11]
TOP3T -17.28 [ -74.77, 40.22]
UiO2016 -8.84 [-104.12, 86.44]
UiO2017 -9.21 [ -89.34, 70.92]
UMCU -28.57 [ -75.06, 17.92]
ENIGMA Discovery Cohorts [N=21703; P=3.383e-06] -18.72 [ -26.62, -10.82]
UKBB -17.66 [ -28.57, -6.75]
Discovery+UKBB [N=31568; P=4.503e-08] -18.34 [ -24.91, -11.77]
GOBS
IMH 35.14 [ -60.15, 130.43]
UNICAMP 26.30 [ -49.16, 101.76]
Meth-CT
MIRECC
UKBB nonEUR 33.30 [ -31.06, 97.66]
OSAKA -10.02 [ -60.56, 40.52]
PING Non European 27.75 [ -30.42, 85.92]
Non European Cohorts [N=1263; P=0.2413] 17.27 [ -11.61, 46.15]
-150 -100 -50 0 50 100 150
mm^2/effect allele
Beta [Lower 95% CI,Upper 95% CI]

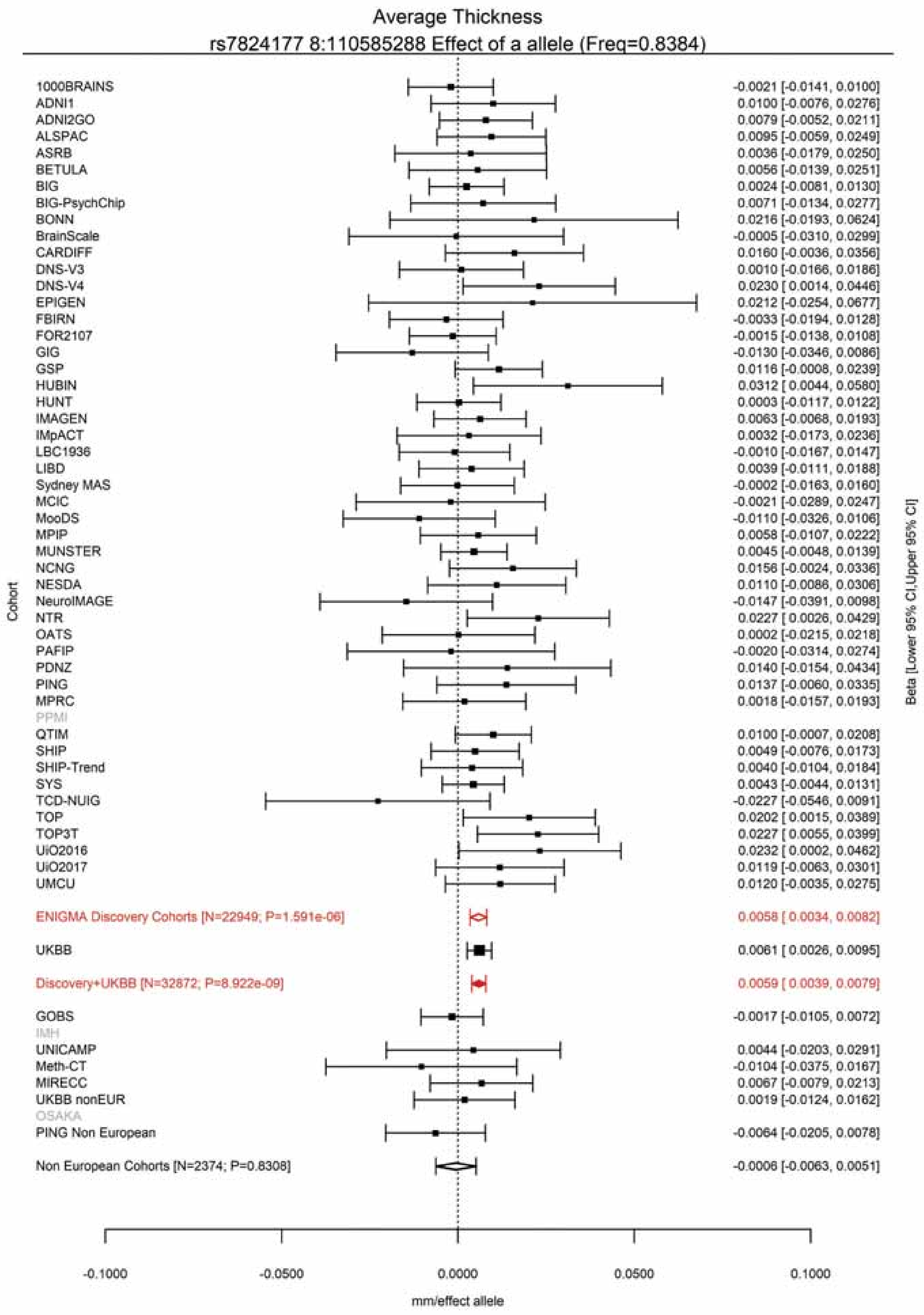
Average Thickness
rs7824177 8:110585288 Effect of a allele (Freq=0.8384)
Cohort
1000BRAINS -0.0021 [-0.0141, 0.0100]
ADNI1 0.0100 [-0.0076, 0.0276]
ADNI2GO 0.0079 [-0.0052, 0.0211]
ALSPAC 0.0095 [-0.0059, 0.0249]
ASRB 0.0036 [-0.0179, 0.0250]
BETULA 0.0056 [-0.0139, 0.0251]
BIG 0.0024 [-0.0081, 0.0130]
BIG-PsychChip 0.0071 [-0.0134, 0.0277]
BONN 0.0216 [-0.0193, 0.0624]
BrainScale -0.0005 [-0.0310, 0.0299]
CARDIFF 0.0160 [-0.0036, 0.0356]
DNS-V3 0.0010 [-0.0166, 0.0186]
DNS-V4 0.0230 [ 0.0014, 0.0446]
EPIGEN 0.0212 [-0.0254, 0.0677]
FBIRN -0.0033 [-0.0194, 0.0128]
FOR2107 -0.0015 [-0.0138, 0.0108]
GIG -0.0130 [-0.0346, 0.0086]
GSP 0.0116 [-0.0008, 0.0239]
HUBIN 0.0312 [ 0.0044, 0.0580]
HUNT 0.0003 [-0.0117, 0.0122]
IMAGEN 0.0063 [-0.0068, 0.0193]
IMpACT 0.0032 [-0.0173, 0.0236]
LBC1936 -0.0010 [-0.0167, 0.0147]
LIBD 0.0039 [-0.0111, 0.0188]
Sydney MAS -0.0002 [-0.0163, 0.0160]
MCIC -0.0021 [-0.0289, 0.0247]
MooDS -0.0110 [-0.0326, 0.0106]
MPIP 0.0058 [-0.0107, 0.0222]
MUNSTER 0.0045 [-0.0048, 0.0139]
NCNG 0.0156 [-0.0024, 0.0336]
NESDA 0.0110 [-0.0086, 0.0306]
NeuroIMAGE -0.0147 [-0.0391, 0.0098]
NTR 0.0227 [ 0.0026, 0.0429]
OATS 0.0002 [-0.0215, 0.0218]
PAFIP -0.0020 [-0.0314, 0.0274]
PDNZ 0.0140 [-0.0154, 0.0434]
PING 0.0137 [-0.0060, 0.0335]
MPRC 0.0018 [-0.0157, 0.0193]
PPMI
QTIM 0.0100 [-0.0007, 0.0208]
SHIP 0.0049 [-0.0076, 0.0173]
SHIP-Trend 0.0040 [-0.0104, 0.0184]
SYS 0.0043 [-0.0044, 0.0131]
TCD-NUIG -0.0227 [-0.0546, 0.0091]
TOP 0.0202 [ 0.0015, 0.0389]
TOP3T 0.0227 [ 0.0055, 0.0399]
UiO2016 0.0232 [ 0.0002, 0.0462]
UiO2017 0.0119 [-0.0063, 0.0301]
UMCU 0.0120 [-0.0035, 0.0275]
ENIGMA Discovery Cohorts [N=22949; P=1.591e-06] 0.0058 [ 0.0034, 0.0082]
UKBB 0.0061 [ 0.0026, 0.0095]
Discovery+UKBB [N=32872; P=8.922e-09] 0.0059 [ 0.0039, 0.0079]
GOBS -0.0017 [-0.0105, 0.0072]
IMH
UNICAMP 0.0044 [-0.0203, 0.0291]
Meth-CT -0.0104 [-0.0375, 0.0167]
MIRECC 0.0067 [-0.0079, 0.0213]
UKBB nonEUR 0.0019 [-0.0124, 0.0162]
OSAKA
PING Non European -0.0064 [-0.0205, 0.0078]
Non European Cohorts [N=2374; P=0.8308] -0.0006 [-0.0063, 0.0051]
Beta [Lower 95% CI,Upper 95% CI]
-0.1000
-0.0500
0.0000
0.0500
0.1000
mm/effect allele

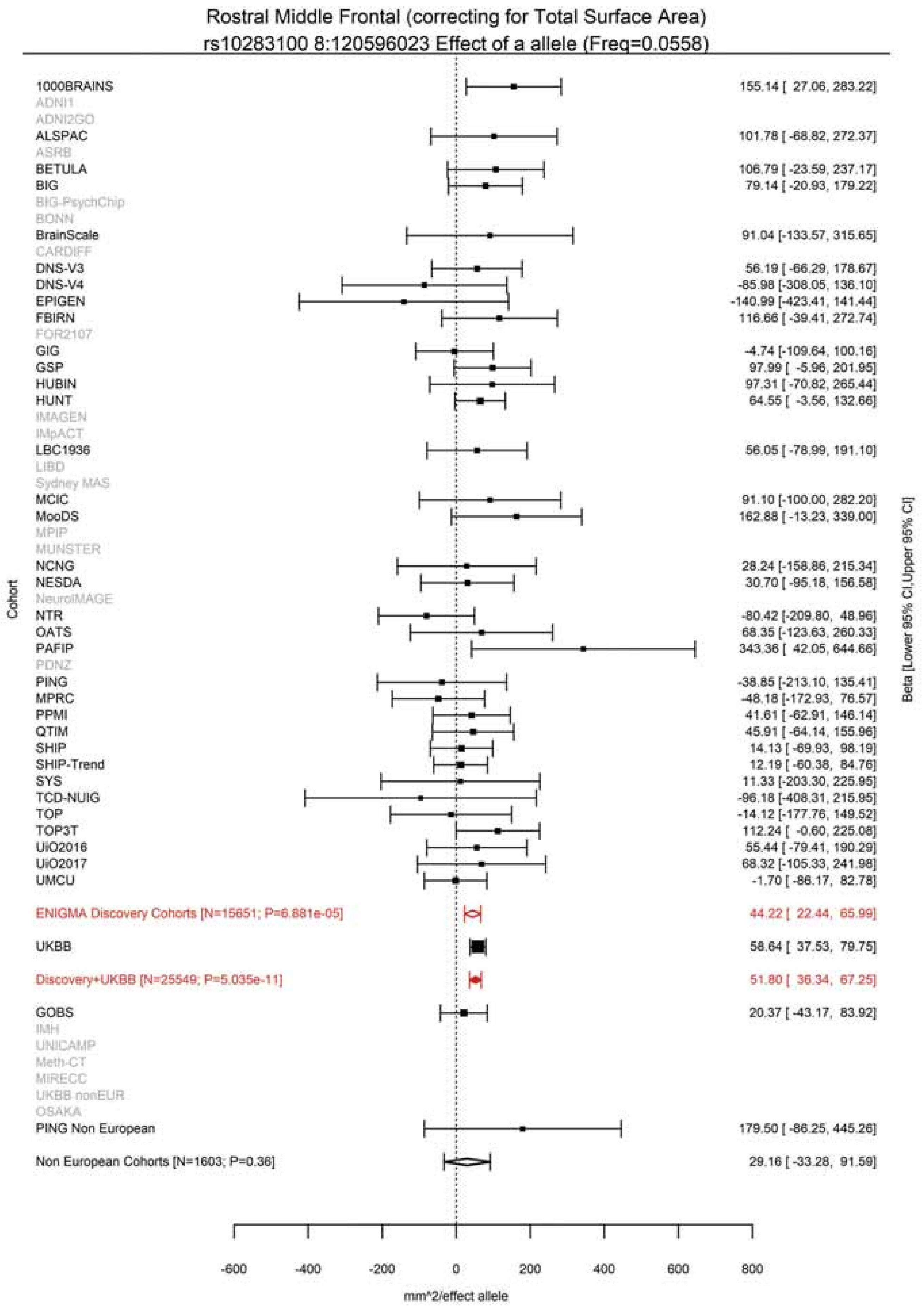

Rostral Middle Frontal (correcting for Total Surface Area)
rs10283100 8:120596023 Effect of a allele (Freq=0.0558)
Cohort
1000BRAINS 155.14 [ 27.06, 283.22]
ADNI1
ADNI2GO
ALSPAC 101.78 [ -68.82, 272.37]
ASRB
BETULA 106.79 [ -23.59, 237.17]
BIG 79.14 [ -20.93, 179.22]
BIG-PsychChip
BONN
BrainScale 91.04 [-133.57, 315.65]
CARDIFF
DNS-V3 56.19 [ -66.29, 178.67]
DNS-V4 -85.98 [-308.05, 136.10]
EPIGEN -140.99 [-423.41, 141.44]
FBIRN 116.66 [ -39.41, 272.74]
FOR2107
GIG -4.74 [-109.64, 100.16]
GSP 97.99 [ -5.96, 201.95]
HUBIN 97.31 [ -70.82, 265.44]
HUNT 64.55 [ -3.56, 132.66]
IMAGEN
IMpACT
LBC1936 56.05 [ -78.99, 191.10]
LIBD
Sydney MAS
MCIC 91.10 [-100.00, 282.20]
MooDS 162.88 [ -13.23, 339.00]
MPIP
MUNSTER
NCNG 28.24 [-158.86, 215.34]
NESDA 30.70 [ -95.18, 156.58]
NeuroIMAGE
NTR -80.42 [-209.80, 48.96]
OATS 68.35 [-123.63, 260.33]
PAFIP 343.36 [ 42.05, 644.66]
PDNZ
PING -38.85 [-213.10, 135.41]
MPRC -48.18 [-172.93, 76.57]
PPMI 41.61 [ -62.91, 146.14]
QTIM 45.91 [ -64.14, 155.96]
SHIP 14.13 [ -69.93, 98.19]
SHIP-Trend 12.19 [ -60.38, 84.76]
SYS 11.33 [-203.30, 225.95]
TCD-NUIG -96.18 [-408.31, 215.95]
TOP -14.12 [-177.76, 149.52]
TOP3T 112.24 [ -0.60, 225.08]
UiO2016 55.44 [ -79.41, 190.29]
UiO2017 68.32 [-105.33, 241.98]
UMCU -1.70 [ -86.17, 82.78]
ENIGMA Discovery Cohorts [N=15651; P=6.881e-05] 44.22 [ 22.44, 65.99]
UKBB 58.64 [ 37.53, 79.75]
Discovery+UKBB [N=25549; P=5.035e-11] 51.80 [ 36.34, 67.25]
GOBS 20.37 [ -43.17, 83.92]
IMH
UNICAMP
Meth-CT
MIRECC
UKBB nonEUR
OSAKA
PING Non European 179.50 [ -86.25, 445.26]
Non European Cohorts [N=1603; P=0.36] 29.16 [ -33.28, 91.59]
Beta [Lower 95% CI,Upper 95% CI]
-600
-400
-200
0
200
400
600
800
mm^2/effect allele

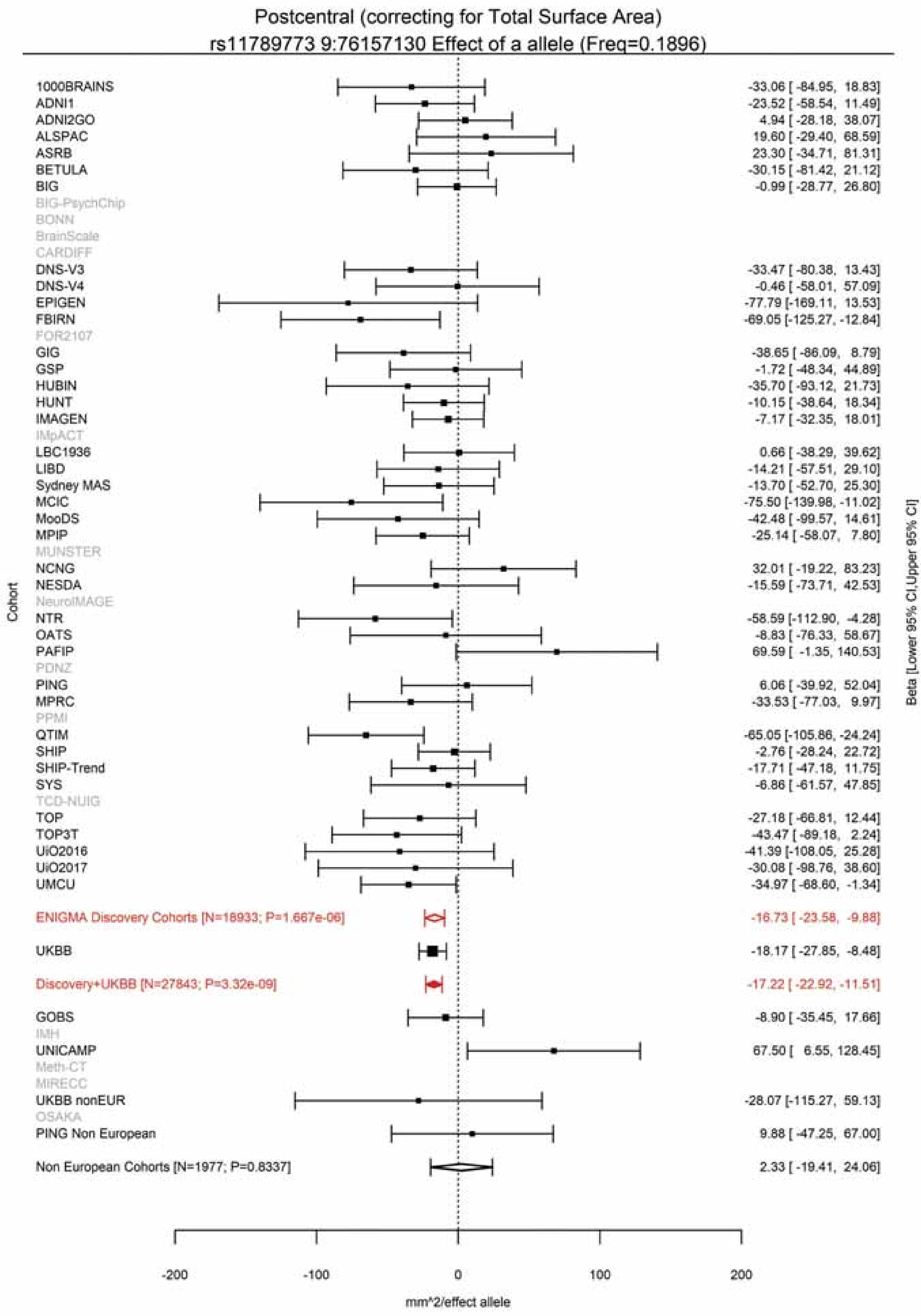

Postcentral (correcting for Total Surface Area)
rs11789773 9:76157130 Effect of a allele (Freq=0.1896)
Cohort
1000BRAINS -33.06 [ -84.95, 18.83]
ADNI1 -23.52 [ -58.54, 11.49]
ADNI2GO 4.94 [ -28.18, 38.07]
ALSPAC 19.60 [ -29.40, 68.59]
ASRB 23.30 [ -34.71, 81.31]
BETULA -30.15 [ -81.42, 21.12]
BIG -0.99 [ -28.77, 26.80]
BIG-PsychChip
BONN
BrainScale
CARDIFF
DNS-V3 -33.47 [ -80.38, 13.43]
DNS-V4 -0.46 [ -58.01, 57.09]
EPIGEN -77.79 [-169.11, 13.53]
FBIRN -69.05 [-125.27, -12.84]
FOR2107
GIG -38.65 [ -86.09, 8.79]
GSP -1.72 [ -48.34, 44.89]
HUBIN -35.70 [ -93.12, 21.73]
HUNT -10.15 [ -38.64, 18.34]
IMAGEN -7.17 [ -32.35, 18.01]
IMpACT
LBC1936 0.66 [ -38.29, 39.62]
LIBD -14.21 [ -57.51, 29.10]
Sydney MAS -13.70 [ -52.70, 25.30]
MCIC -75.50 [-139.98, -11.02]
MooDS -42.48 [ -99.57, 14.61]
MPIP -25.14 [ -58.07, 7.80]
MUNSTER
NCNG 32.01 [ -19.22, 83.23]
NESDA -15.59 [ -73.71, 42.53]
NeuroIMAGE
NTR -58.59 [-112.90, -4.28]
OATS -8.83 [ -76.33, 58.67]
PAFIP 69.59 [ -1.35, 140.53]
PDNZ
PING 6.06 [ -39.92, 52.04]
MPRC -33.53 [ -77.03, 9.97]
PPMI
QTIM -65.05 [-105.86, -24.24]
SHIP -2.76 [ -28.24, 22.72]
SHIP-Trend -17.71 [ -47.18, 11.75]
SYS -6.86 [ -61.57, 47.85]
TCD-NUIG
TOP -27.18 [ -66.81, 12.44]
TOP3T -43.47 [ -89.18, 2.24]
UiO2016 -41.39 [-108.05, 25.28]
UiO2017 -30.08 [ -98.76, 38.60]
UMCU -34.97 [ -68.60, -1.34]
ENIGMA Discovery Cohorts [N=18933; P=1.667e-06] -16.73 [ -23.58, -9.88]
UKBB -18.17 [ -27.85, -8.48]
Discovery+UKBB [N=27843; P=3.32e-09] -17.22 [ -22.92, -11.51]
GOBS -8.90 [ -35.45, 17.66]
IMH
UNICAMP 67.50 [ 6.55, 128.45]
Meth-CT
MIRECC
UKBB nonEUR -28.07 [-115.27, 59.13]
OSAKA
PING Non European 9.88 [ -47.25, 67.00]
Non European Cohorts [N=1977; P=0.8337] 2.33 [ -19.41, 24.06]
-200
-100
0
100
200
mm^2/effect allele
Beta [Lower 95% CI,Upper 95% CI]

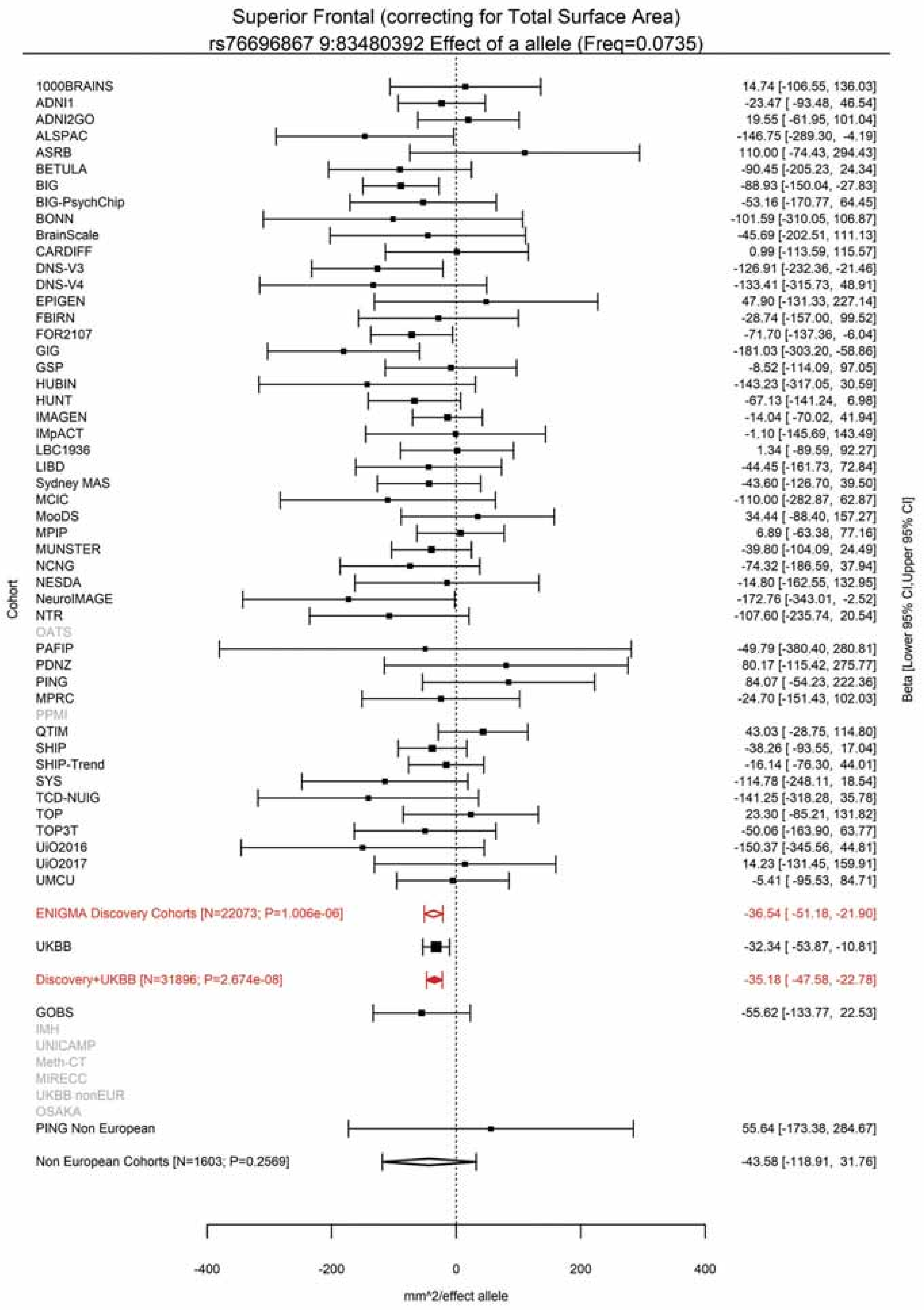
Superior Frontal (correcting for Total Surface Area)
rs76696867 9:83480392 Effect of a allele (Freq=0.0735)
Cohort
Beta [Lower 95% CI,Upper 95% CI]
1000BRAINS 14.74 [-106.55, 136.03]
ADNI1 -23.47 [ -93.48, 46.54]
ADNI2GO 19.55 [ -61.95, 101.04]
ALSPAC -146.75 [-289.30, -4.19]
ASRB 110.00 [ -74.43, 294.43]
BETULA -90.45 [-205.23, 24.34]
BIG -88.93 [-150.04, -27.83]
BIG-PsychChip -53.16 [-170.77, 64.45]
BONN -101.59 [-310.05, 106.87]
BrainScale -45.69 [-202.51, 111.13]
CARDIFF 0.99 [-113.59, 115.57]
DNS-V3 -126.91 [-232.36, -21.46]
DNS-V4 -133.41 [-315.73, 48.91]
EPIGEN 47.90 [-131.33, 227.14]
FBIRN -28.74 [-157.00, 99.52]
FOR2107 -71.70 [-137.36, -6.04]
GIG -181.03 [-303.20, -58.86]
GSP -8.52 [-114.09, 97.05]
HUBIN -143.23 [-317.05, 30.59]
HUNT -67.13 [-141.24, 6.98]
IMAGEN -14.04 [ -70.02, 41.94]
IMpACT -1.10 [-145.69, 143.49]
LBC1936 1.34 [ -89.59, 92.27]
LIBD -44.45 [-161.73, 72.84]
Sydney MAS -43.60 [-126.70, 39.50]
MCIC -110.00 [-282.87, 62.87]
MooDS 34.44 [ -88.40, 157.27]
MPIP 6.89 [ -63.38, 77.16]
MUNSTER -39.80 [-104.09, 24.49]
NCNG -74.32 [-186.59, 37.94]
NESDA -14.80 [-162.55, 132.95]
NeuroIMAGE -172.76 [-343.01, -2.52]
NTR -107.60 [-235.74, 20.54]
OATS
PAFIP -49.79 [-380.40, 280.81]
PDNZ 80.17 [-115.42, 275.77]
PING 84.07 [ -54.23, 222.36]
MPRC -24.70 [-151.43, 102.03]
PPMI
QTIM 43.03 [ -28.75, 114.80]
SHIP -38.26 [ -93.55, 17.04]
SHIP-Trend -16.14 [ -76.30, 44.01]
SYS -114.78 [-248.11, 18.54]
TCD-NUIG -141.25 [-318.28, 35.78]
TOP 23.30 [ -85.21, 131.82]
TOP3T -50.06 [-163.90, 63.77]
UiO2016 -150.37 [-345.56, 44.81]
UiO2017 14.23 [-131.45, 159.91]
UMCU -5.41 [ -95.53, 84.71]
ENIGMA Discovery Cohorts [N=22073; P=1.006e-06] -36.54 [ -51.18, -21.90]
UKBB -32.34 [ -53.87, -10.81]
Discovery+UKBB [N=31896; P=2.674e-08] -35.18 [ -47.58, -22.78]
GOBS -55.62 [-133.77, 22.53]
IMH
UNICAMP
Meth-CT
MIRECC
UKBB nonEUR
OSAKA
PING Non European 55.64 [-173.38, 284.67]
Non European Cohorts [N=1603; P=0.2569] -43.58 [-118.91, 31.76]
-400
-200
0
200
400
mm^2/effect allele

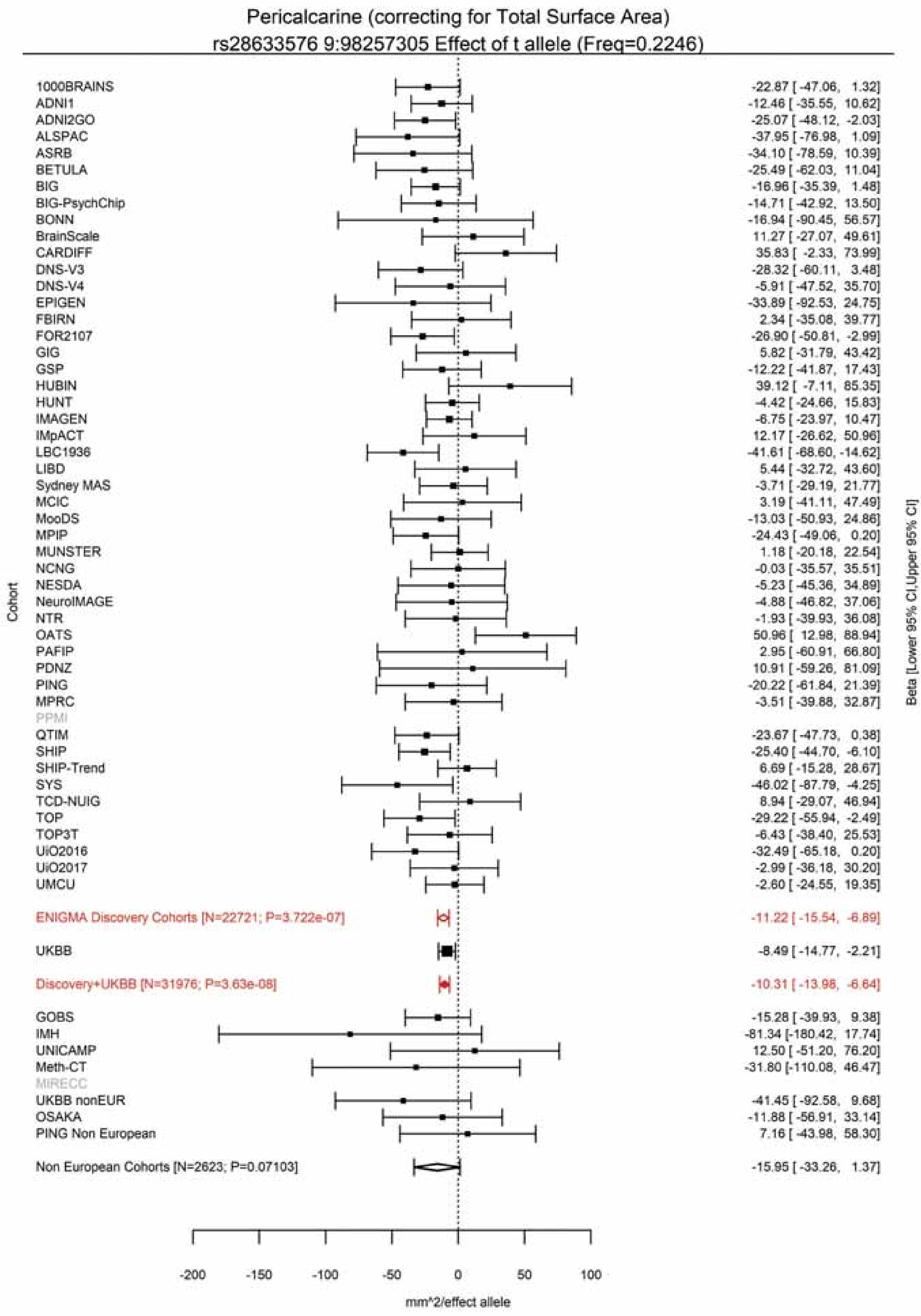

Pericalcarine (correcting for Total Surface Area)
rs28633576 9:98257305 Effect of t allele (Freq=0.2246)
Cohort
1000BRAINS -22.87 [ -47.06, 1.32]
ADNI1 -12.46 [ -35.55, 10.62]
ADNI2GO -25.07 [ -48.12, -2.03]
ALSPAC -37.95 [ -76.98, 1.09]
ASRB -34.10 [ -78.59, 10.39]
BETULA -25.49 [ -62.03, 11.04]
BIG -16.96 [ -35.39, 1.48]
BIG-PsychChip -14.71 [ -42.92, 13.50]
BONN -16.94 [ -90.45, 56.57]
BrainScale 11.27 [ -27.07, 49.61]
CARDIFF 35.83 [ -2.33, 73.99]
DNS-V3 -28.32 [ -60.11, 3.48]
DNS-V4 -5.91 [ -47.52, 35.70]
EPIGEN -33.89 [ -92.53, 24.75]
FBIRN 2.34 [ -35.08, 39.77]
FOR2107 -26.90 [ -50.81, -2.99]
GIG 5.82 [ -31.79, 43.42]
GSP -12.22 [ -41.87, 17.43]
HUBIN 39.12 [ -7.11, 85.35]
HUNT -4.42 [ -24.66, 15.83]
IMAGEN -6.75 [ -23.97, 10.47]
IMpACT 12.17 [ -26.62, 50.96]
LBC1936 -41.61 [ -68.60, -14.62]
LIBD 5.44 [ -32.72, 43.60]
Sydney MAS -3.71 [ -29.19, 21.77]
MCIC 3.19 [ -41.11, 47.49]
MooDS -13.03 [ -50.93, 24.86]
MPIP -24.43 [ -49.06, 0.20]
MUNSTER 1.18 [ -20.18, 22.54]
NCNG -0.03 [ -35.57, 35.51]
NESDA -5.23 [ -45.36, 34.89]
NeuroIMAGE -4.88 [ -46.82, 37.06]
NTR -1.93 [ -39.93, 36.08]
OATS 50.96 [ 12.98, 88.94]
PAFIP 2.95 [ -60.91, 66.80]
PDNZ 10.91 [ -59.26, 81.09]
PING -20.22 [ -61.84, 21.39]
MPRC -3.51 [ -39.88, 32.87]
PPMI
QTIM -23.67 [ -47.73, 0.38]
SHIP -25.40 [ -44.70, -6.10]
SHIP-Trend 6.69 [ -15.28, 28.67]
SYS -46.02 [ -87.79, -4.25]
TCD-NUIG 8.94 [ -29.07, 46.94]
TOP -29.22 [ -55.94, -2.49]
TOP3T -6.43 [ -38.40, 25.53]
UiO2016 -32.49 [ -65.18, 0.20]
UiO2017 -2.99 [ -36.18, 30.20]
UMCU -2.60 [ -24.55, 19.35]
ENIGMA Discovery Cohorts [N=22721; P=3.722e-07] -11.22 [ -15.54, -6.89]
UKBB -8.49 [ -14.77, -2.21]
Discovery+UKBB [N=31976; P=3.63e-08] -10.31 [ -13.98, -6.64]
GOBS -15.28 [ -39.93, 9.38]
IMH -81.34 [-180.42, 17.74]
UNICAMP 12.50 [ -51.20, 76.20]
Meth-CT -31.80 [-110.08, 46.47]
MIRECC
UKBB nonEUR -41.45 [ -92.58, 9.68]
OSAKA -11.88 [ -56.91, 33.14]
PING Non European 7.16 [ -43.98, 58.30]
Non European Cohorts [N=2623; P=0.07103] -15.95 [ -33.26, 1.37]
-200
-150
-100
-50
0
50
100
mm^2/effect allele
Beta [Lower 95% CI,Upper 95% CI]

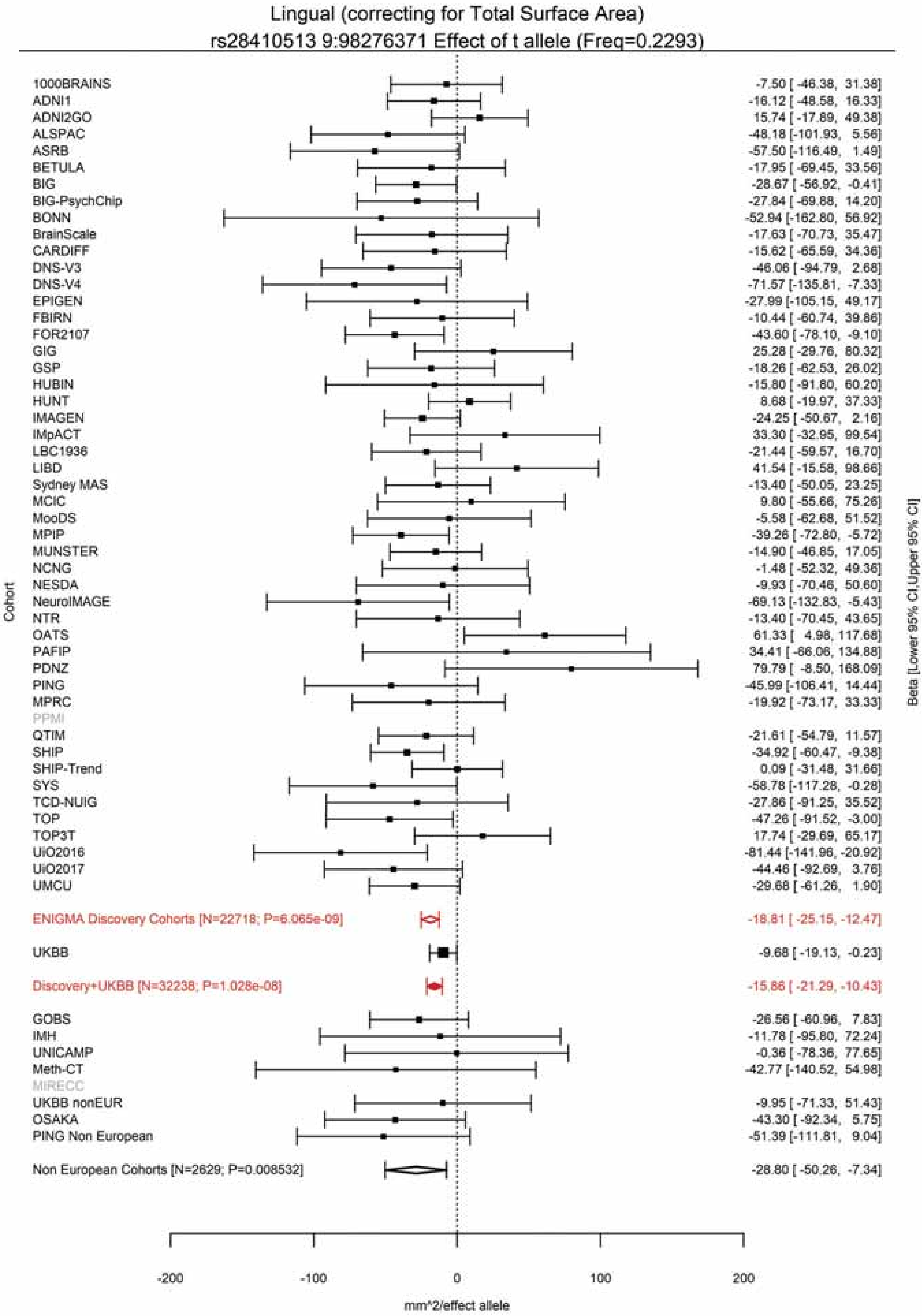
Lingual (correcting for Total Surface Area)
rs28410513 9:98276371 Effect of t allele (Freq=0.2293)
Cohort
Beta [Lower 95% CI,Upper 95% CI]
1000BRAINS -7.50 [ -46.38, 31.38]
ADNI1 -16.12 [ -48.58, 16.33]
ADNI2GO 15.74 [ -17.89, 49.38]
ALSPAC -48.18 [-101.93, 5.56]
ASRB -57.50 [-116.49, 1.49]
BETULA -17.95 [ -69.45, 33.56]
BIG -28.67 [ -56.92, -0.41]
BIG-PsychChip -27.84 [ -69.88, 14.20]
BONN -52.94 [-162.80, 56.92]
BrainScale -17.63 [ -70.73, 35.47]
CARDIFF -15.62 [ -65.59, 34.36]
DNS-V3 -46.06 [ -94.79, 2.68]
DNS-V4 -71.57 [-135.81, -7.33]
EPIGEN -27.99 [-105.15, 49.17]
FBIRN -10.44 [ -60.74, 39.86]
FOR2107 -43.60 [ -78.10, -9.10]
GIG 25.28 [ -29.76, 80.32]
GSP -18.26 [ -62.53, 26.02]
HUBIN -15.80 [ -91.80, 60.20]
HUNT 8.68 [ -19.97, 37.33]
IMAGEN -24.25 [ -50.67, 2.16]
IMpACT 33.30 [ -32.95, 99.54]
LBC1936 -21.44 [ -59.57, 16.70]
LIBD 41.54 [ -15.58, 98.66]
Sydney MAS -13.40 [ -50.05, 23.25]
MCIC 9.80 [ -55.66, 75.26]
MooDS -5.58 [ -62.68, 51.52]
MPIP -39.26 [ -72.80, -5.72]
MUNSTER -14.90 [ -46.85, 17.05]
NCNG -1.48 [ -52.32, 49.36]
NESDA -9.93 [ -70.46, 50.60]
NeuroIMAGE -69.13 [-132.83, -5.43]
NTR -13.40 [ -70.45, 43.65]
OATS 61.33 [ 4.98, 117.68]
PAFIP 34.41 [ -66.06, 134.88]
PDNZ 79.79 [ -8.50, 168.09]
PING -45.99 [-106.41, 14.44]
MPRC -19.92 [ -73.17, 33.33]
PPMI
QTIM -21.61 [ -54.79, 11.57]
SHIP -34.92 [ -60.47, -9.38]
SHIP-Trend 0.09 [ -31.48, 31.66]
SYS -58.78 [-117.28, -0.28]
TCD-NUIG -27.86 [ -91.25, 35.52]
TOP -47.26 [ -91.52, -3.00]
TOP3T 17.74 [ -29.69, 65.17]
UiO2016 -81.44 [-141.96, -20.92]
UiO2017 -44.46 [ -92.69, 3.76]
UMCU -29.68 [ -61.26, 1.90]
ENIGMA Discovery Cohorts [N=22718; P=6.065e-09] -18.81 [ -25.15, -12.47]
UKBB -9.68 [ -19.13, -0.23]
Discovery+UKBB [N=32238; P=1.028e-08] -15.86 [ -21.29, -10.43]
GOBS -26.56 [ -60.96, 7.83]
IMH -11.78 [ -95.80, 72.24]
UNICAMP -0.36 [ -78.36, 77.65]
Meth-CT -42.77 [-140.52, 54.98]
MIRECC
UKBB nonEUR -9.95 [ -71.33, 51.43]
OSAKA -43.30 [ -92.34, 5.75]
PING Non European -51.39 [-111.81, 9.04]
Non European Cohorts [N=2629; P=0.008532] -28.80 [ -50.26, -7.34]
-200
-100
0
100
200
mm^2/effect allele

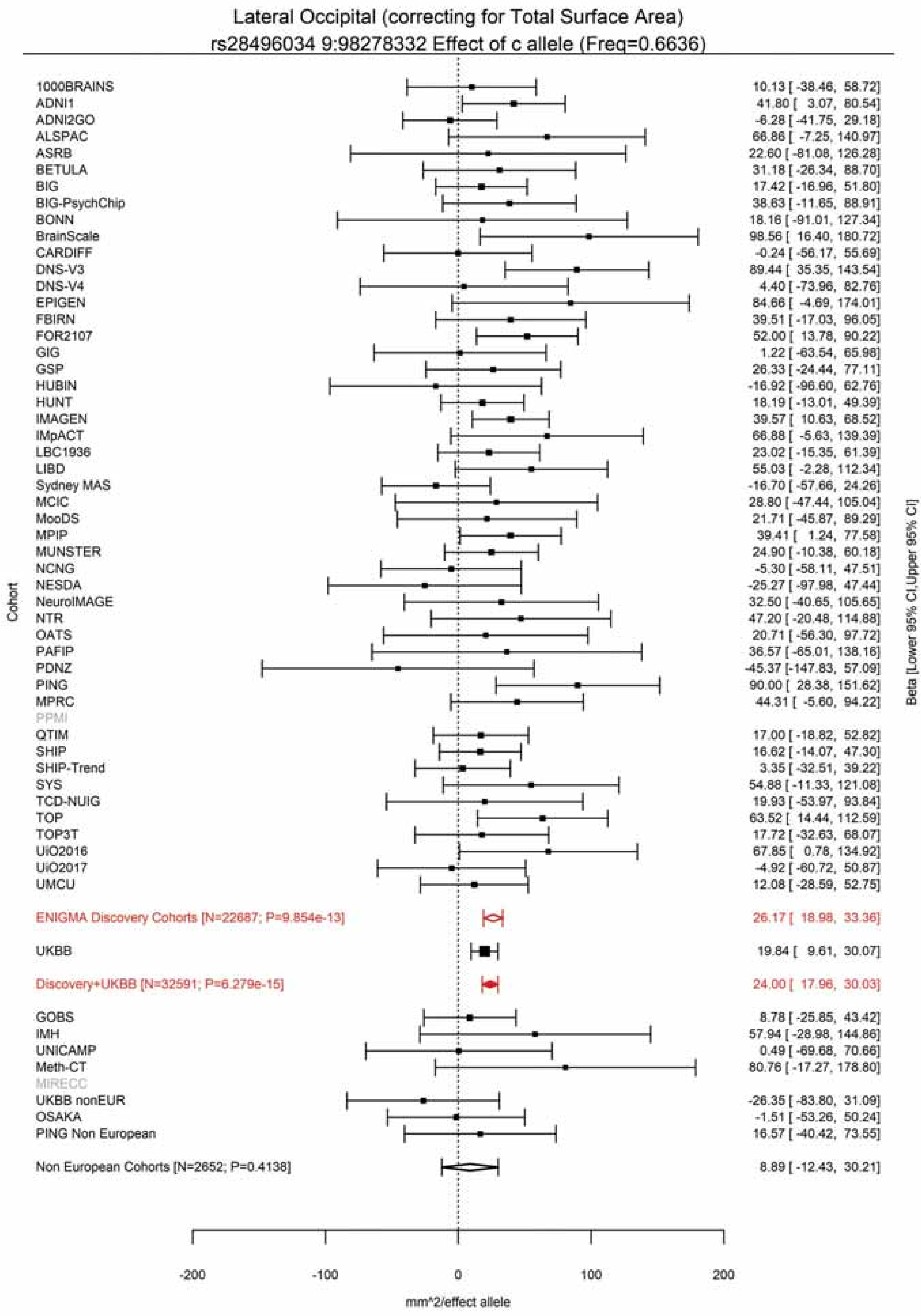

Lateral Occipital (correcting for Total Surface Area)
rs28496034 9:98278332 Effect of c allele (Freq=0.6636)
1000BRAINS 10.13 [ -38.46, 58.72]
ADNI1 41.80 [ 3.07, 80.54]
ADNI2GO -6.28 [ -41.75, 29.18]
ALSPAC 66.86 [ -7.25, 140.97]
ASRB 22.60 [ -81.08, 126.28]
BETULA 31.18 [ -26.34, 88.70]
BIG 17.42 [ -16.96, 51.80]
BIG-PsychChip 38.63 [ -11.65, 88.91]
BONN 18.16 [ -91.01, 127.34]
BrainScale 98.56 [ 16.40, 180.72]
CARDIFF -0.24 [ -56.17, 55.69]
DNS-V3 89.44 [ 35.35, 143.54]
DNS-V4 4.40 [ -73.96, 82.76]
EPIGEN 84.66 [ -4.69, 174.01]
FBIRN 39.51 [ -17.03, 96.05]
FOR2107 52.00 [ 13.78, 90.22]
GIG 1.22 [ -63.54, 65.98]
GSP 26.33 [ -24.44, 77.11]
HUBIN -16.92 [ -96.60, 62.76]
HUNT 18.19 [ -13.01, 49.39]
IMAGEN 39.57 [ 10.63, 68.52]
IMpACT 66.88 [ -5.63, 139.39]
LBC1936 23.02 [ -15.35, 61.39]
LIBD 55.03 [ -2.28, 112.34]
Sydney MAS -16.70 [ -57.66, 24.26]
MCIC 28.80 [ -47.44, 105.04]
MooDS 21.71 [ -45.87, 89.29]
MPIP 39.41 [ 1.24, 77.58]
MUNSTER 24.90 [ -10.38, 60.18]
NCNG -5.30 [ -58.11, 47.51]
NESDA -25.27 [ -97.98, 47.44]
NeuroIMAGE 32.50 [ -40.65, 105.65]
NTR 47.20 [ -20.48, 114.88]
OATS 20.71 [ -56.30, 97.72]
PAFIP 36.57 [ -65.01, 138.16]
PDNZ -45.37 [ -147.83, 57.09]
PING 90.00 [ 28.38, 151.62]
MPRC 44.31 [ -5.60, 94.22]
PPMI
QTIM 17.00 [ -18.82, 52.82]
SHIP 16.62 [ -14.07, 47.30]
SHIP-Trend 3.35 [ -32.51, 39.22]
SYS 54.88 [ -11.33, 121.08]
TCD-NUIG 19.93 [ -53.97, 93.84]
TOP 63.52 [ 14.44, 112.59]
TOP3T 17.72 [ -32.63, 68.07]
UiO2016 67.85 [ 0.78, 134.92]
UiO2017 -4.92 [ -60.72, 50.87]
UMCU 12.08 [ -28.59, 52.75]
ENIGMA Discovery Cohorts [N=22687; P=9.854e-13] 26.17 [ 18.98, 33.36]
UKBB 19.84 [ 9.61, 30.07]
Discovery+UKBB [N=32591; P=6.279e-15] 24.00 [ 17.96, 30.03]
GOBS 8.78 [ -25.85, 43.42]
IMH 57.94 [ -28.98, 144.86]
UNICAMP 0.49 [ -69.68, 70.66]
Meth-CT 80.76 [ -17.27, 178.80]
MIRECC
UKBB nonEUR -26.35 [ -83.80, 31.09]
OSAKA -1.51 [ -53.26, 50.24]
PING Non European 16.57 [ -40.42, 73.55]
Non European Cohorts [N=2652; P=0.4138] 8.89 [ -12.43, 30.21]
Cohort
Beta [Lower 95% CI,Upper 95% CI]
-200
-100
0
100
200
mm^2/effect allele

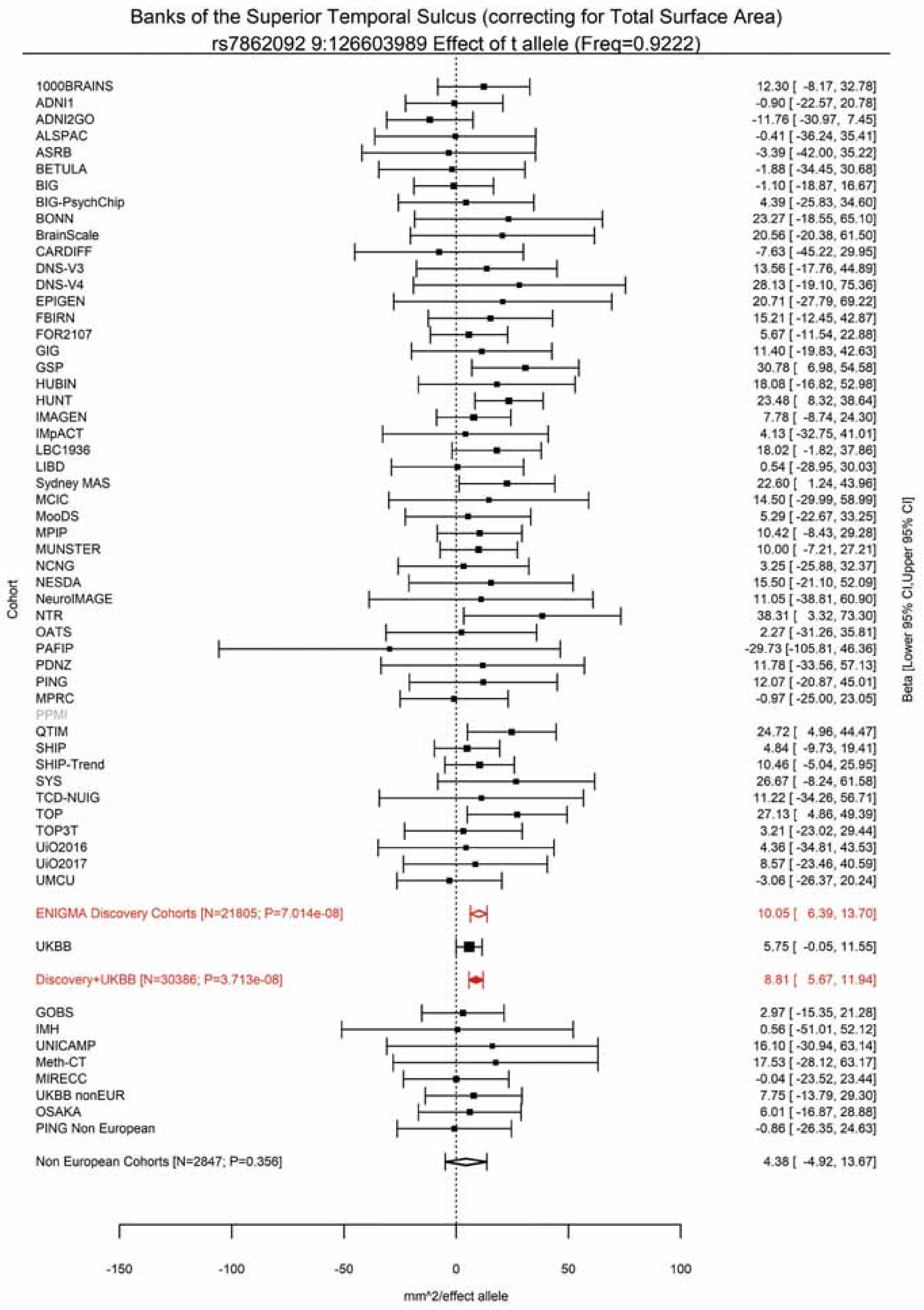

Banks of the Superior Temporal Sulcus (correcting for Total Surface Area)
rs7862092 9:126603989 Effect of t allele (Freq=0.9222)
Cohort
1000BRAINS 12.30 [ -8.17, 32.78]
ADNI1 -0.90 [ -22.57, 20.78]
ADNI2GO -11.76 [ -30.97, 7.45]
ALSPAC -0.41 [ -36.24, 35.41]
ASRB -3.39 [ -42.00, 35.22]
BETULA -1.88 [ -34.45, 30.68]
BIG -1.10 [ -18.87, 16.67]
BIG-PsychChip 4.39 [ -25.83, 34.60]
BONN 23.27 [ -18.55, 65.10]
BrainScale 20.56 [ -20.38, 61.50]
CARDIFF -7.63 [ -45.22, 29.95]
DNS-V3 13.56 [ -17.76, 44.89]
DNS-V4 28.13 [ -19.10, 75.36]
EPIGEN 20.71 [ -27.79, 69.22]
FBIRN 15.21 [ -12.45, 42.87]
FOR2107 5.67 [ -11.54, 22.88]
GIG 11.40 [ -19.83, 42.63]
GSP 30.78 [ 6.98, 54.58]
HUBIN 18.08 [ -16.82, 52.98]
HUNT 23.48 [ 8.32, 38.64]
IMAGEN 7.78 [ -8.74, 24.30]
IMpACT 4.13 [ -32.75, 41.01]
LBC1936 18.02 [ -1.82, 37.86]
LIBD 0.54 [ -28.95, 30.03]
Sydney MAS 22.60 [ 1.24, 43.96]
MCIC 14.50 [ -29.99, 58.99]
MooDS 5.29 [ -22.67, 33.25]
MPIP 10.42 [ -8.43, 29.28]
MUNSTER 10.00 [ -7.21, 27.21]
NCNG 3.25 [ -25.88, 32.37]
NESDA 15.50 [ -21.10, 52.09]
NeuroIMAGE 11.05 [ -38.81, 60.90]
NTR 38.31 [ 3.32, 73.30]
OATS 2.27 [ -31.26, 35.81]
PAFIP -29.73 [ -105.81, 46.36]
PDNZ 11.78 [ -33.56, 57.13]
PING 12.07 [ -20.87, 45.01]
MPRC -0.97 [ -25.00, 23.05]
PPMI
QTIM 24.72 [ 4.96, 44.47]
SHIP 4.84 [ -9.73, 19.41]
SHIP-Trend 10.46 [ -5.04, 25.95]
SYS 26.67 [ -8.24, 61.58]
TCD-NUIG 11.22 [ -34.26, 56.71]
TOP 27.13 [ 4.86, 49.39]
TOP3T 3.21 [ -23.02, 29.44]
UiO2016 4.36 [ -34.81, 43.53]
UiO2017 8.57 [ -23.46, 40.59]
UMCU -3.06 [ -26.37, 20.24]
ENIGMA Discovery Cohorts [N=21805; P=7.014e-08] 10.05 [ 6.39, 13.70]
UKBB 5.75 [ -0.05, 11.55]
Discovery+UKBB [N=30386; P=3.713e-08] 8.81 [ 5.67, 11.94]
GOBS 2.97 [ -15.35, 21.28]
IMH 0.56 [ -51.01, 52.12]
UNICAMP 16.10 [ -30.94, 63.14]
Meth-CT 17.53 [ -28.12, 63.17]
MIRECC -0.04 [ -23.52, 23.44]
UKBB nonEUR 7.75 [ -13.79, 29.30]
OSAKA 6.01 [ -16.87, 28.88]
PING Non European -0.86 [ -26.35, 24.63]
Non European Cohorts [N=2847; P=0.356] 4.38 [ -4.92, 13.67]
-150
-100
-50
0
50
100
mm^2/effect allele
Beta [Lower 95% CI,Upper 95% CI]

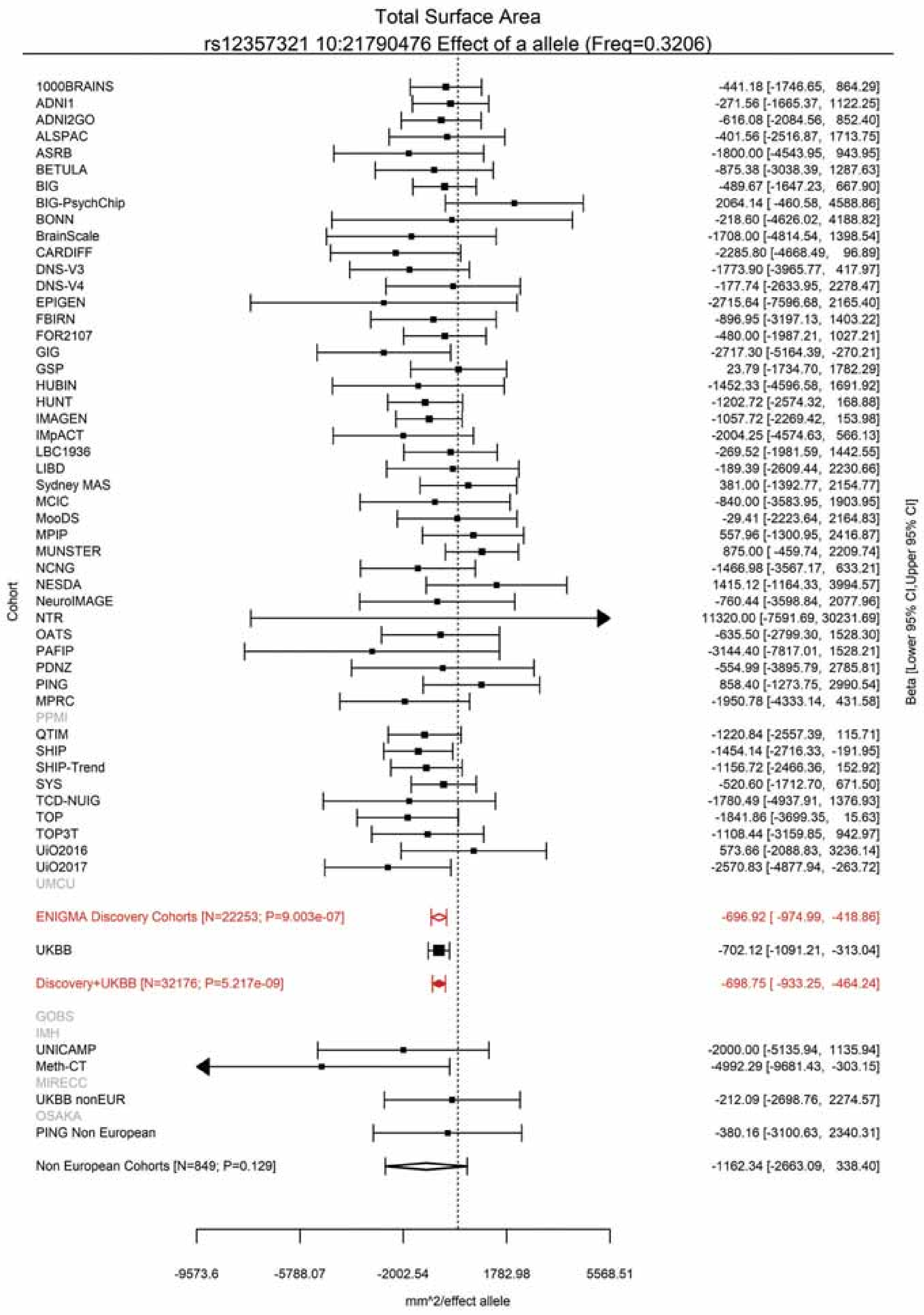
Total Surface Area
rs12357321 10:21790476 Effect of a allele (Freq=0.3206)
1000BRAINS -441.18 [-1746.65, 864.29]
ADNI1 -271.56 [-1665.37, 1122.25]
ADNI2GO -616.08 [-2084.56, 852.40]
ALSPAC -401.56 [-2516.87, 1713.75]
ASRB -1800.00 [-4543.95, 943.95]
BETULA -875.38 [-3038.39, 1287.63]
BIG -489.67 [-1647.23, 667.90]
BIG-PsychChip 2064.14 [-460.58, 4588.86]
BONN -218.60 [-4626.02, 4188.82]
BrainScale -1708.00 [-4814.54, 1398.54]
CARDIFF -2285.80 [-4668.49, 96.89]
DNS-V3 -1773.90 [-3965.77, 417.97]
DNS-V4 -177.74 [-2633.95, 2278.47]
EPIGEN -2715.64 [-7596.68, 2165.40]
FBIRN -896.95 [-3197.13, 1403.22]
FOR2107 -480.00 [-1987.21, 1027.21]
GIG -2717.30 [-5164.39, -270.21]
GSP 23.79 [-1734.70, 1782.29]
HUBIN -1452.33 [-4596.58, 1691.92]
HUNT -1202.72 [-2574.32, 168.88]
IMAGEN -1057.72 [-2269.42, 153.98]
IMpACT -2004.25 [-4574.63, 566.13]
LBC1936 -269.52 [-1981.59, 1442.55]
LIBD -189.39 [-2609.44, 2230.66]
Sydney MAS 381.00 [-1392.77, 2154.77]
MCIC -840.00 [-3583.95, 1903.95]
MooDS -29.41 [-2223.64, 2164.83]
MPIP 557.96 [-1300.95, 2416.87]
MUNSTER 875.00 [-459.74, 2209.74]
NCNG -1466.98 [-3567.17, 633.21]
NESDA 1415.12 [-1164.33, 3994.57]
NeuroIMAGE -760.44 [-3598.84, 2077.96]
NTR 11320.00 [-7591.69, 30231.69]
OATS -635.50 [-2799.30, 1528.30]
PAFIP -3144.40 [-7817.01, 1528.21]
PDNZ -554.99 [-3895.79, 2785.81]
PING 858.40 [-1273.75, 2990.54]
MPRC -1950.78 [-4333.14, 431.58]
PPMI
QTIM -1220.84 [-2557.39, 115.71]
SHIP -1454.14 [-2716.33, -191.95]
SHIP-Trend -1156.72 [-2466.36, 152.92]
SYS -520.60 [-1712.70, 671.50]
TCD-NUIG -1780.49 [-4937.91, 1376.93]
TOP -1841.86 [-3699.35, 15.63]
TOP3T -1108.44 [-3159.85, 942.97]
UiO2016 573.66 [-2088.83, 3236.14]
UiO2017 -2570.83 [-4877.94, -263.72]
UMCU
ENIGMA Discovery Cohorts [N=22253; P=9.003e-07] -696.92 [-974.99, -418.86]
UKBB -702.12 [-1091.21, -313.04]
Discovery+UKBB [N=32176; P=5.217e-09] -698.75 [-933.25, -464.24]
GOBS
IMH
UNICAMP -2000.00 [-5135.94, 1135.94]
Meth-CT -4992.29 [-9681.43, -303.15]
MIRECC
UKBB nonEUR -212.09 [-2698.76, 2274.57]
OSAKA
PING Non European -380.16 [-3100.63, 2340.31]
Non European Cohorts [N=849; P=0.129] -1162.34 [-2663.09, 338.40]
Cohort
Beta [Lower 95% CI,Upper 95% CI]
-9573.6
-5788.07
-2002.54
1782.98
5568.51
mm^2/effect allele

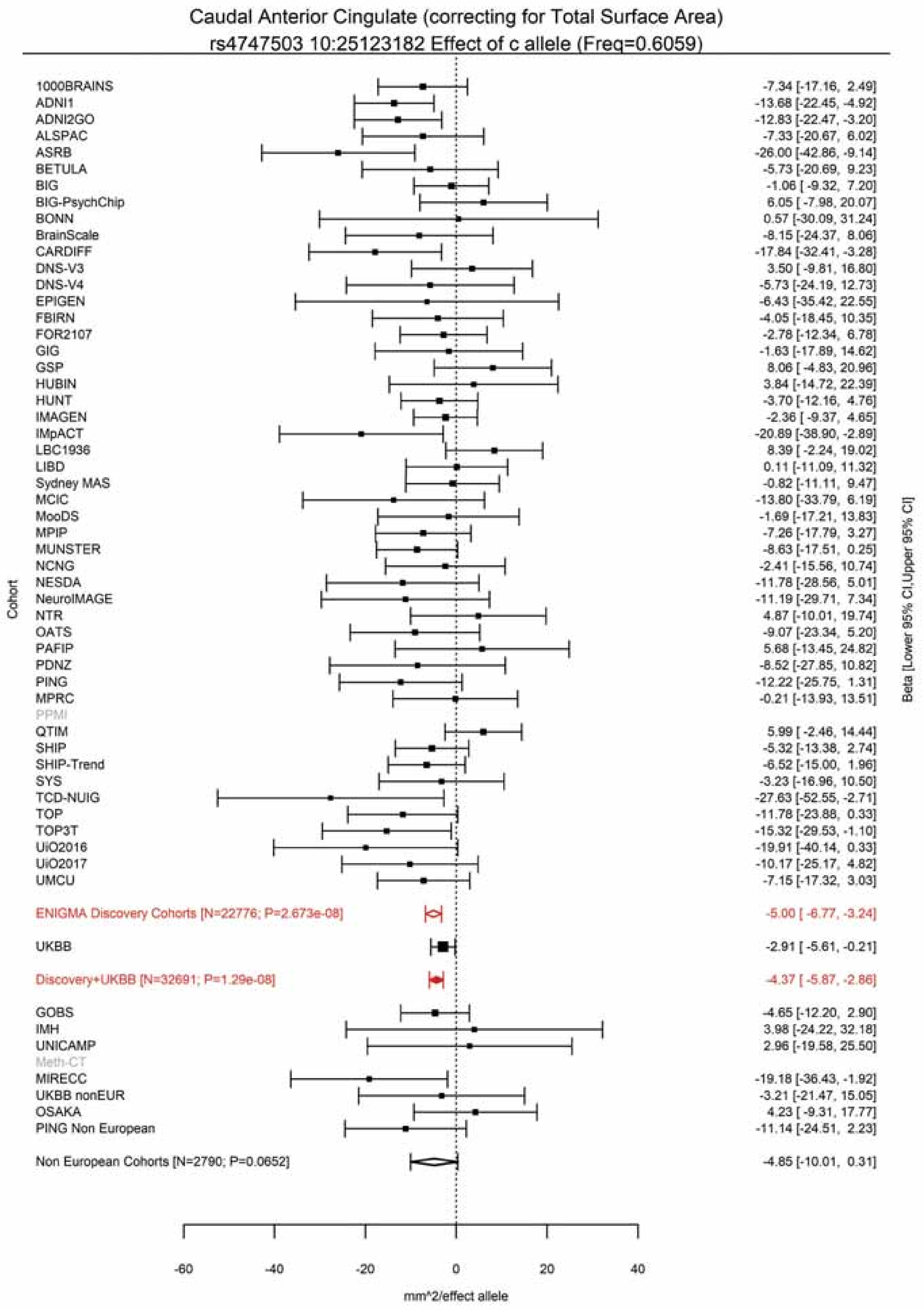
Caudal Anterior Cingulate (correcting for Total Surface Area)
rs4747503 10:25123182 Effect of c allele (Freq=0.6059)
Cohort
1000BRAINS -7.34 [-17.16, 2.49]
ADNI1 -13.68 [-22.45, -4.92]
ADNI2GO -12.83 [-22.47, -3.20]
ALSPAC -7.33 [-20.67, 6.02]
ASRB -26.00 [-42.86, -9.14]
BETULA -5.73 [-20.69, 9.23]
BIG -1.06 [ -9.32, 7.20]
BIG-PsychChip 6.05 [ -7.98, 20.07]
BONN 0.57 [-30.09, 31.24]
BrainScale -8.15 [-24.37, 8.06]
CARDIFF -17.84 [-32.41, -3.28]
DNS-V3 3.50 [ -9.81, 16.80]
DNS-V4 -5.73 [-24.19, 12.73]
EPIGEN -6.43 [-35.42, 22.55]
FBIRN -4.05 [-18.45, 10.35]
FOR2107 -2.78 [-12.34, 6.78]
GIG -1.63 [-17.89, 14.62]
GSP 8.06 [ -4.83, 20.96]
HUBIN 3.84 [-14.72, 22.39]
HUNT -3.70 [-12.16, 4.76]
IMAGEN -2.36 [ -9.37, 4.65]
IMpACT -20.89 [-38.90, -2.89]
LBC1936 8.39 [ -2.24, 19.02]
LIBD 0.11 [-11.09, 11.32]
Sydney MAS -0.82 [-11.11, 9.47]
MCIC -13.80 [-33.79, 6.19]
MooDS -1.69 [-17.21, 13.83]
MPIP -7.26 [-17.79, 3.27]
MUNSTER -8.63 [-17.51, 0.25]
NCNG -2.41 [-15.56, 10.74]
NESDA -11.78 [-28.56, 5.01]
NeuroIMAGE -11.19 [-29.71, 7.34]
NTR 4.87 [-10.01, 19.74]
OATS -9.07 [-23.34, 5.20]
PAFIP 5.68 [-13.45, 24.82]
PDNZ -8.52 [-27.85, 10.82]
PING -12.22 [-25.75, 1.31]
MPRC -0.21 [-13.93, 13.51]
PPMI
QTIM 5.99 [ -2.46, 14.44]
SHIP -5.32 [-13.38, 2.74]
SHIP-Trend -6.52 [-15.00, 1.96]
SYS -3.23 [-16.96, 10.50]
TCD-NUIG -27.63 [-52.55, -2.71]
TOP -11.78 [-23.88, 0.33]
TOP3T -15.32 [-29.53, -1.10]
UiO2016 -19.91 [-40.14, 0.33]
UiO2017 -10.17 [-25.17, 4.82]
UMCU -7.15 [-17.32, 3.03]
ENIGMA Discovery Cohorts [N=22776; P=2.673e-08] -5.00 [ -6.77, -3.24]
UKBB -2.91 [ -5.61, -0.21]
Discovery+UKBB [N=32691; P=1.29e-08] -4.37 [ -5.87, -2.86]
GOBS -4.65 [-12.20, 2.90]
IMH 3.98 [-24.22, 32.18]
UNICAMP 2.96 [-19.58, 25.50]
Meth-CT
MIRECC -19.18 [-36.43, -1.92]
UKBB nonEUR -3.21 [-21.47, 15.05]
OSAKA 4.23 [ -9.31, 17.77]
PING Non European -11.14 [-24.51, 2.23]
Non European Cohorts [N=2790; P=0.0652] -4.85 [-10.01, 0.31]
-60
-40
-20
0
20
40
mm^2/effect allele
Beta [Lower 95% CI,Upper 95% CI]

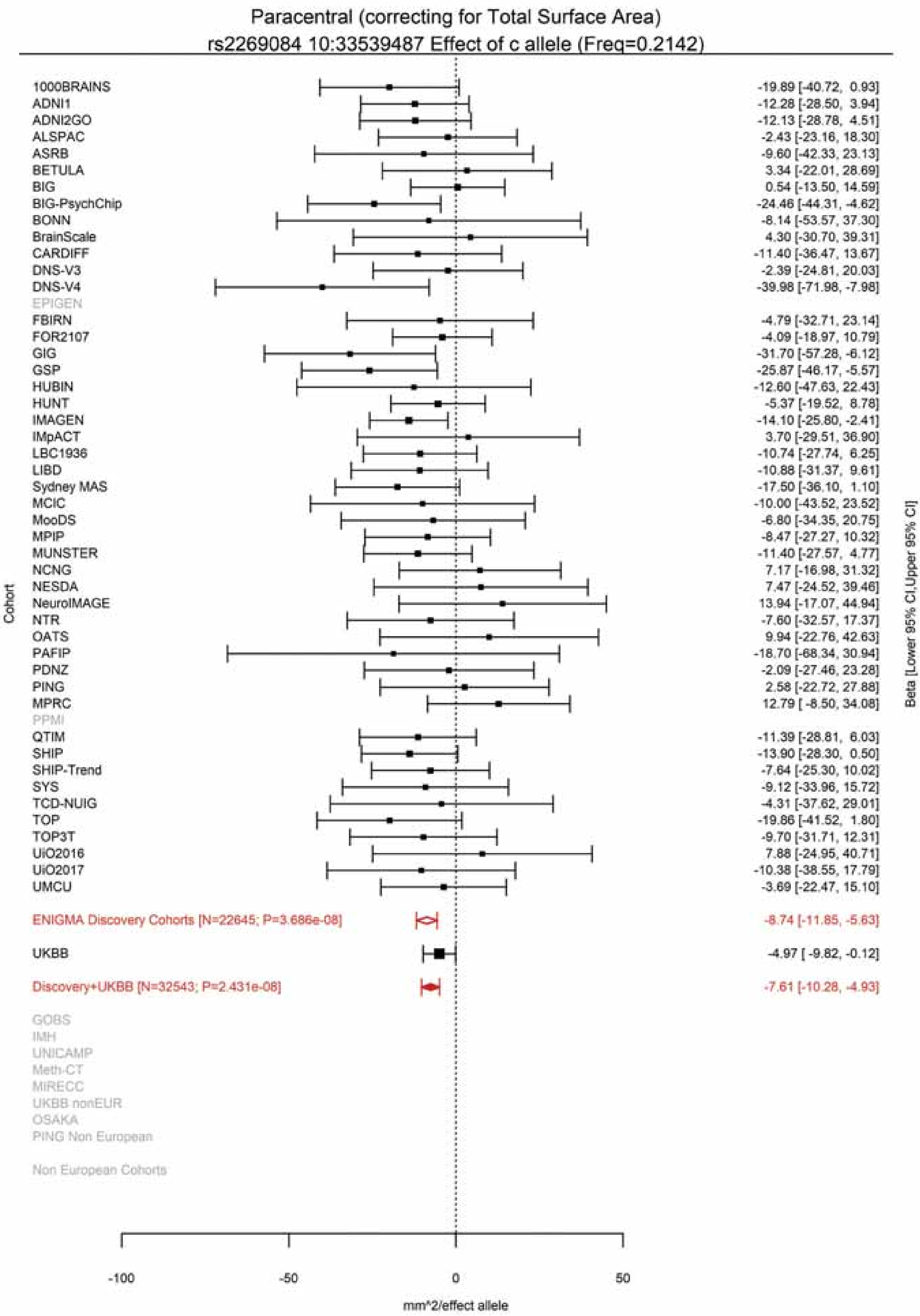
Paracentral (correcting for Total Surface Area)
rs2269084 10:33539487 Effect of c allele (Freq=0.2142)
Cohort
1000BRAINS -19.89 [-40.72, 0.93]
ADNI1 -12.28 [-28.50, 3.94]
ADNI2GO -12.13 [-28.78, 4.51]
ALSPAC -2.43 [-23.16, 18.30]
ASRB -9.60 [-42.33, 23.13]
BETULA 3.34 [-22.01, 28.69]
BIG 0.54 [-13.50, 14.59]
BIG-PsychChip -24.46 [-44.31, -4.62]
BONN -8.14 [-53.57, 37.30]
BrainScale 4.30 [-30.70, 39.31]
CARDIFF -11.40 [-36.47, 13.67]
DNS-V3 -2.39 [-24.81, 20.03]
DNS-V4 -39.98 [-71.98, -7.98]
EPIGEN
FBIRN -4.79 [-32.71, 23.14]
FOR2107 -4.09 [-18.97, 10.79]
GIG -31.70 [-57.28, -6.12]
GSP -25.87 [-46.17, -5.57]
HUBIN -12.60 [-47.63, 22.43]
HUNT -5.37 [-19.52, 8.78]
IMAGEN -14.10 [-25.80, -2.41]
IMpACT 3.70 [-29.51, 36.90]
LBC1936 -10.74 [-27.74, 6.25]
LIBD -10.88 [-31.37, 9.61]
Sydney MAS -17.50 [-36.10, 1.10]
MCIC -10.00 [-43.52, 23.52]
MooDS -6.80 [-34.35, 20.75]
MPIP -8.47 [-27.27, 10.32]
MUNSTER -11.40 [-27.57, 4.77]
NCNG 7.17 [-16.98, 31.32]
NESDA 7.47 [-24.52, 39.46]
NeuroIMAGE 13.94 [-17.07, 44.94]
NTR -7.60 [-32.57, 17.37]
OATS 9.94 [-22.76, 42.63]
PAFIP -18.70 [-68.34, 30.94]
PDNZ -2.09 [-27.46, 23.28]
PING 2.58 [-22.72, 27.88]
MPRC 12.79 [ -8.50, 34.08]
PPMI
QTIM -11.39 [-28.81, 6.03]
SHIP -13.90 [-28.30, 0.50]
SHIP-Trend -7.64 [-25.30, 10.02]
SYS -9.12 [-33.96, 15.72]
TCD-NUIG -4.31 [-37.62, 29.01]
TOP -19.86 [-41.52, 1.80]
TOP3T -9.70 [-31.71, 12.31]
UiO2016 7.88 [-24.95, 40.71]
UiO2017 -10.38 [-38.55, 17.79]
UMCU -3.69 [-22.47, 15.10]
ENIGMA Discovery Cohorts [N=22645; P=3.686e-08] -8.74 [-11.85, -5.63]
UKBB -4.97 [ -9.82, -0.12]
Discovery+UKBB [N=32543; P=2.431e-08] -7.61 [-10.28, -4.93]
GOBS
IMH
UNICAMP
Meth-CT
MIRECC
UKBB nonEUR
OSAKA
PING Non European
Non European Cohorts
Beta [Lower 95% CI,Upper 95% CI]
-100
-50
0
50
mm^2/effect allele

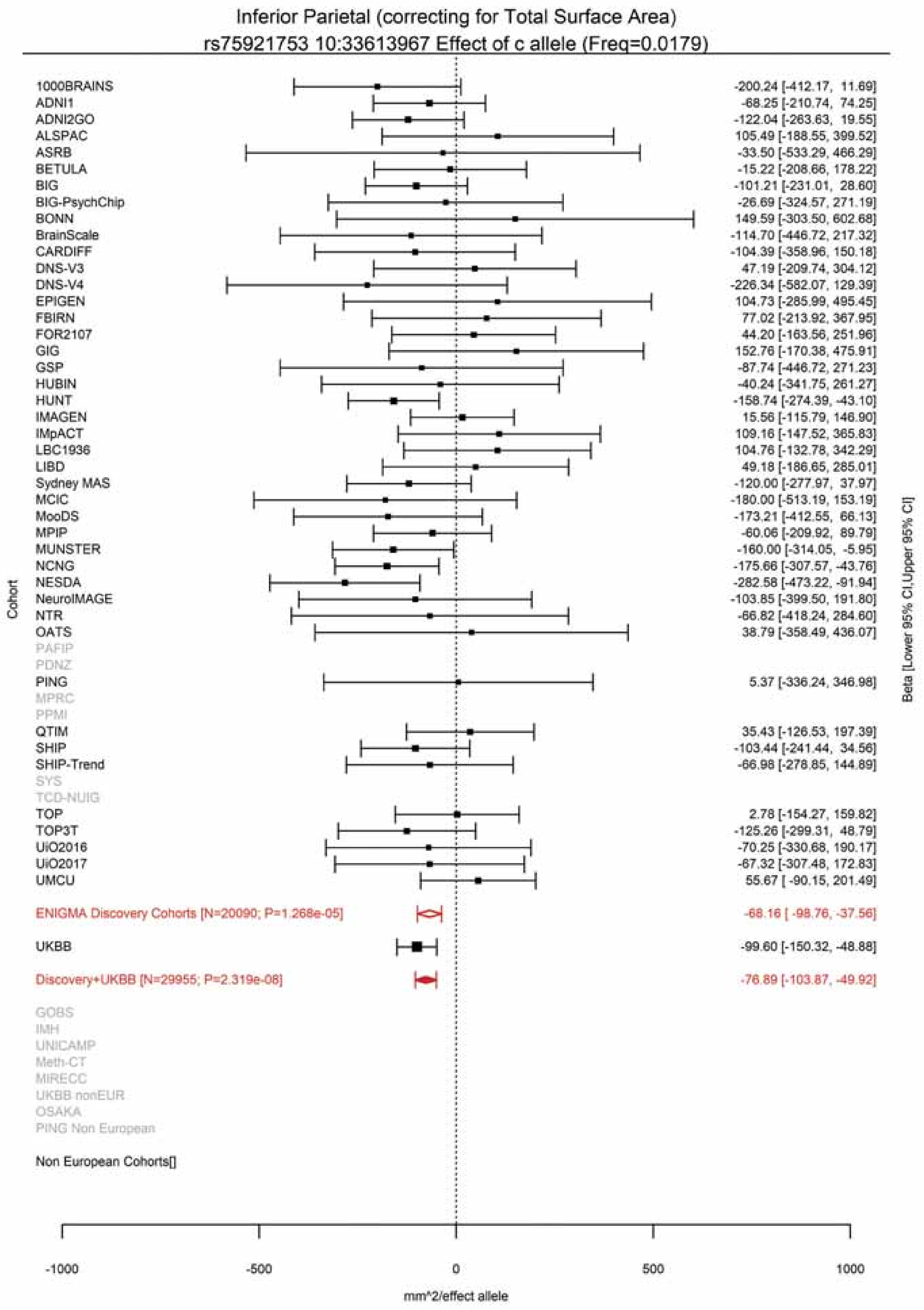

Inferior Parietal (correcting for Total Surface Area)
rs75921753 10:33613967 Effect of c allele (Freq=0.0179)
Cohort
Beta [Lower 95% CI,Upper 95% CI]
1000BRAINS -200.24 [-412.17, 11.69]
ADNI1 -68.25 [-210.74, 74.25]
ADNI2GO -122.04 [-263.63, 19.55]
ALSPAC 105.49 [-188.55, 399.52]
ASRB -33.50 [-533.29, 466.29]
BETULA -15.22 [-208.66, 178.22]
BIG -101.21 [-231.01, 28.60]
BIG-PsychChip -26.69 [-324.57, 271.19]
BONN 149.59 [-303.50, 602.68]
BrainScale -114.70 [-446.72, 217.32]
CARDIFF -104.39 [-358.96, 150.18]
DNS-V3 47.19 [-209.74, 304.12]
DNS-V4 -226.34 [-582.07, 129.39]
EPIGEN 104.73 [-285.99, 495.45]
FBIRN 77.02 [-213.92, 367.95]
FOR2107 44.20 [-163.56, 251.96]
GIG 152.76 [-170.38, 475.91]
GSP -87.74 [-446.72, 271.23]
HUBIN -40.24 [-341.75, 261.27]
HUNT -158.74 [-274.39, -43.10]
IMAGEN 15.56 [-115.79, 146.90]
IMpACT 109.16 [-147.52, 365.83]
LBC1936 104.76 [-132.78, 342.29]
LIBD 49.18 [-186.65, 285.01]
Sydney MAS -120.00 [-277.97, 37.97]
MCIC -180.00 [-513.19, 153.19]
MooDS -173.21 [-412.55, 66.13]
MPIP -60.06 [-209.92, 89.79]
MUNSTER -160.00 [-314.05, -5.95]
NCNG -175.66 [-307.57, -43.76]
NESDA -282.58 [-473.22, -91.94]
NeuroIMAGE -103.85 [-399.50, 191.80]
NTR -66.82 [-418.24, 284.60]
OATS 38.79 [-358.49, 436.07]
PAFIP
PDNZ
PING 5.37 [-336.24, 346.98]
MPRC
PPMI
QTIM 35.43 [-126.53, 197.39]
SHIP -103.44 [-241.44, 34.56]
SHIP-Trend -66.98 [-278.85, 144.89]
SYS
TCD-NUIG
TOP 2.78 [-154.27, 159.82]
TOP3T -125.26 [-299.31, 48.79]
UiO2016 -70.25 [-330.68, 190.17]
UiO2017 -67.32 [-307.48, 172.83]
UMCU 55.67 [ -90.15, 201.49]
ENIGMA Discovery Cohorts [N=20090; P=1.268e-05] -68.16 [ -98.76, -37.56]
UKBB -99.60 [-150.32, -48.88]
Discovery+UKBB [N=29955; P=2.319e-08] -76.89 [-103.87, -49.92]
GOBS
IMH
UNICAMP
Meth-CT
MIRECC
UKBB nonEUR
OSAKA
PING Non European
Non European Cohorts[]
-1000
-500
0
500
1000
mm^2/effect allele

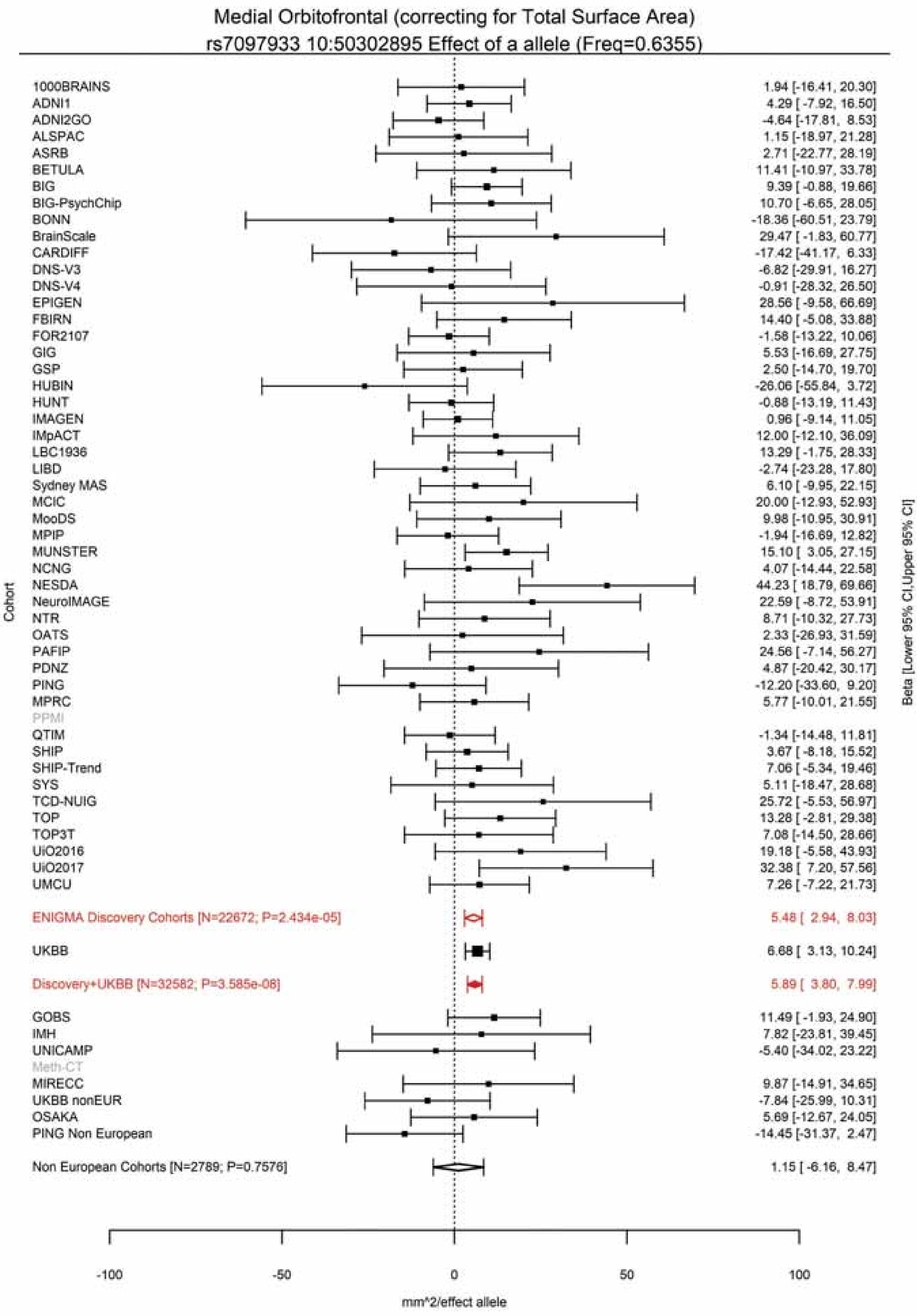

Medial Orbitofrontal (correcting for Total Surface Area)
rs7097933 10:50302895 Effect of a allele (Freq=0.6355)
Cohort
1000BRAINS 1.94 [-16.41, 20.30]
ADNI1 4.29 [ -7.92, 16.50]
ADNI2GO -4.64 [-17.81, 8.53]
ALSPAC 1.15 [-18.97, 21.28]
ASRB 2.71 [-22.77, 28.19]
BETULA 11.41 [-10.97, 33.78]
BIG 9.39 [ -0.88, 19.66]
BIG-PsychChip 10.70 [ -6.65, 28.05]
BONN -18.36 [-60.51, 23.79]
BrainScale 29.47 [ -1.83, 60.77]
CARDIFF -17.42 [-41.17, 6.33]
DNS-V3 -6.82 [-29.91, 16.27]
DNS-V4 -0.91 [-28.32, 26.50]
EPIGEN 28.56 [ -9.58, 66.69]
FBIRN 14.40 [ -5.08, 33.88]
FOR2107 -1.58 [-13.22, 10.06]
GIG 5.53 [-16.69, 27.75]
GSP 2.50 [-14.70, 19.70]
HUBIN -26.06 [-55.84, 3.72]
HUNT -0.88 [-13.19, 11.43]
IMAGEN 0.96 [ -9.14, 11.05]
IMpACT 12.00 [-12.10, 36.09]
LBC1936 13.29 [ -1.75, 28.33]
LIBD -2.74 [-23.28, 17.80]
Sydney MAS 6.10 [ -9.95, 22.15]
MCIC 20.00 [-12.93, 52.93]
MooDS 9.98 [-10.95, 30.91]
MPIP -1.94 [-16.69, 12.82]
MUNSTER 15.10 [ 3.05, 27.15]
NCNG 4.07 [-14.44, 22.58]
NESDA 44.23 [ 18.79, 69.66]
NeuroIMAGE 22.59 [ -8.72, 53.91]
NTR 8.71 [-10.32, 27.73]
OATS 2.33 [-26.93, 31.59]
PAFIP 24.56 [ -7.14, 56.27]
PDNZ 4.87 [-20.42, 30.17]
PING -12.20 [-33.60, 9.20]
MPRC 5.77 [-10.01, 21.55]
PPMI
QTIM -1.34 [-14.48, 11.81]
SHIP 3.67 [ -8.18, 15.52]
SHIP-Trend 7.06 [ -5.34, 19.46]
SYS 5.11 [-18.47, 28.68]
TCD-NUIG 25.72 [ -5.53, 56.97]
TOP 13.28 [ -2.81, 29.38]
TOP3T 7.08 [-14.50, 28.66]
UiO2016 19.18 [ -5.58, 43.93]
UiO2017 32.38 [ 7.20, 57.56]
UMCU 7.26 [ -7.22, 21.73]
ENIGMA Discovery Cohorts [N=22672; P=2.434e-05] 5.48 [ 2.94, 8.03]
UKBB 6.68 [ 3.13, 10.24]
Discovery+UKBB [N=32582; P=3.585e-08] 5.89 [ 3.80, 7.99]
GOBS 11.49 [ -1.93, 24.90]
IMH 7.82 [-23.81, 39.45]
UNICAMP -5.40 [-34.02, 23.22]
Meth-CT
MIRECC 9.87 [-14.91, 34.65]
UKBB nonEUR -7.84 [-25.99, 10.31]
OSAKA 5.69 [-12.67, 24.05]
PING Non European -14.45 [-31.37, 2.47]
Non European Cohorts [N=2789; P=0.7576] 1.15 [ -6.16, 8.47]
Beta [Lower 95% CI,Upper 95% CI]
-100
-50
0
50
100
mm^2/effect allele

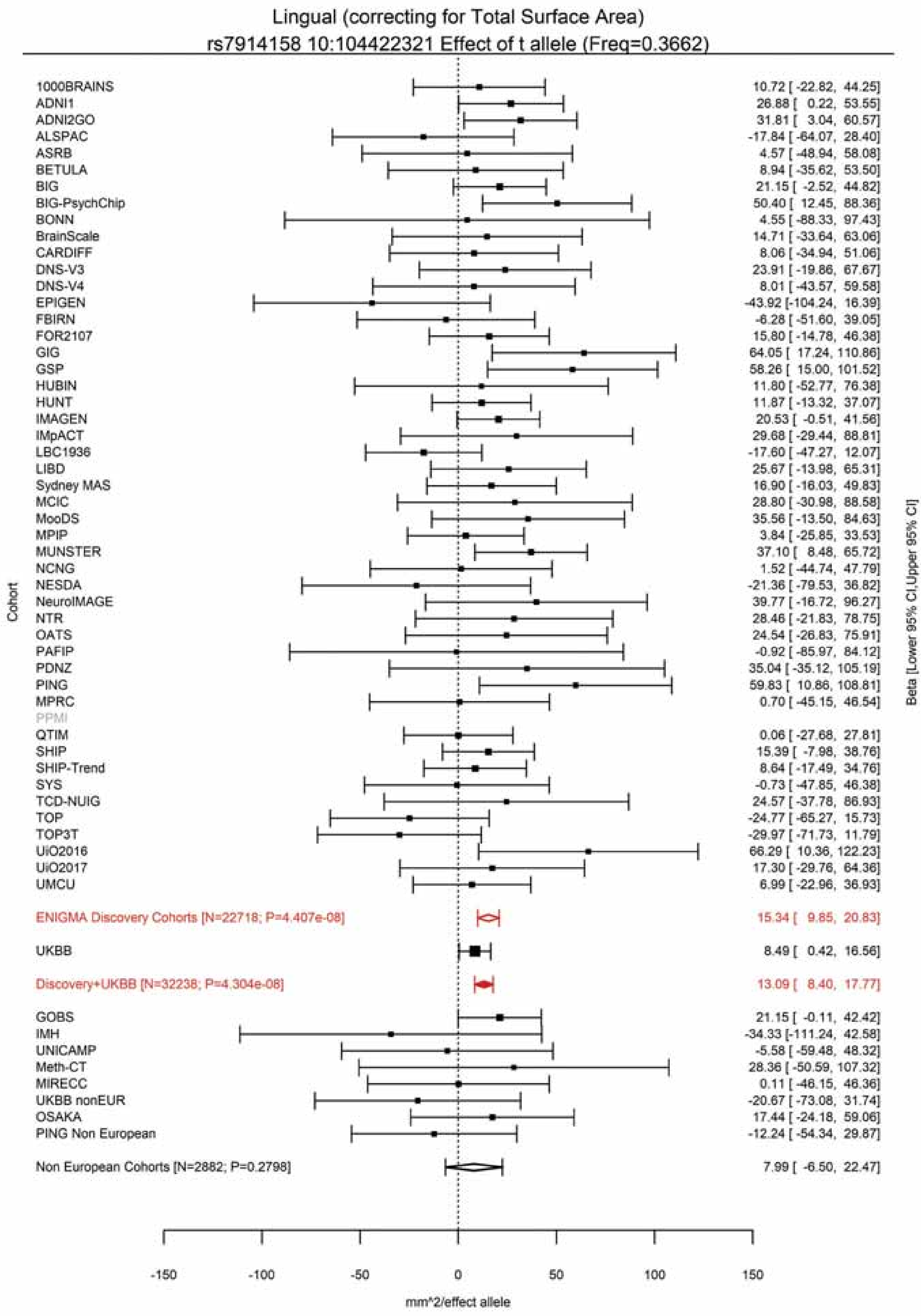

Lingual (correcting for Total Surface Area)
rs7914158 10:104422321 Effect of t allele (Freq=0.3662)
Cohort
Beta [Lower 95% CI,Upper 95% CI]
1000BRAINS 10.72 [ -22.82, 44.25]
ADNI1 26.88 [ 0.22, 53.55]
ADNI2GO 31.81 [ 3.04, 60.57]
ALSPAC -17.84 [ -64.07, 28.40]
ASRB 4.57 [ -48.94, 58.08]
BETULA 8.94 [ -35.62, 53.50]
BIG 21.15 [ -2.52, 44.82]
BIG-PsychChip 50.40 [ 12.45, 88.36]
BONN 4.55 [ -88.33, 97.43]
BrainScale 14.71 [ -33.64, 63.06]
CARDIFF 8.06 [ -34.94, 51.06]
DNS-V3 23.91 [ -19.86, 67.67]
DNS-V4 8.01 [ -43.57, 59.58]
EPIGEN -43.92 [-104.24, 16.39]
FBIRN -6.28 [ -51.60, 39.05]
FOR2107 15.80 [ -14.78, 46.38]
GIG 64.05 [ 17.24, 110.86]
GSP 58.26 [ 15.00, 101.52]
HUBIN 11.80 [ -52.77, 76.38]
HUNT 11.87 [ -13.32, 37.07]
IMAGEN 20.53 [ -0.51, 41.56]
IMpACT 29.68 [ -29.44, 88.81]
LBC1936 -17.60 [ -47.27, 12.07]
LIBD 25.67 [ -13.98, 65.31]
Sydney MAS 16.90 [ -16.03, 49.83]
MCIC 28.80 [ -30.98, 88.58]
MooDS 35.56 [ -13.50, 84.63]
MPIP 3.84 [ -25.85, 33.53]
MUNSTER 37.10 [ 8.48, 65.72]
NCNG 1.52 [ -44.74, 47.79]
NESDA -21.36 [ -79.53, 36.82]
NeuroIMAGE 39.77 [ -16.72, 96.27]
NTR 28.46 [ -21.83, 78.75]
OATS 24.54 [ -26.83, 75.91]
PAFIP -0.92 [ -85.97, 84.12]
PDNZ 35.04 [ -35.12, 105.19]
PING 59.83 [ 10.86, 108.81]
MPRC 0.70 [ -45.15, 46.54]
PPMI
QTIM 0.06 [ -27.68, 27.81]
SHIP 15.39 [ -7.98, 38.76]
SHIP-Trend 8.64 [ -17.49, 34.76]
SYS -0.73 [ -47.85, 46.38]
TCD-NUIG 24.57 [ -37.78, 86.93]
TOP -24.77 [ -65.27, 15.73]
TOP3T -29.97 [ -71.73, 11.79]
UiO2016 66.29 [ 10.36, 122.23]
UiO2017 17.30 [ -29.76, 64.36]
UMCU 6.99 [ -22.96, 36.93]
ENIGMA Discovery Cohorts [N=22718; P=4.407e-08] 15.34 [ 9.85, 20.83]
UKBB 8.49 [ 0.42, 16.56]
Discovery+UKBB [N=32238; P=4.304e-08] 13.09 [ 8.40, 17.77]
GOBS 21.15 [ -0.11, 42.42]
IMH -34.33 [-111.24, 42.58]
UNICAMP -5.58 [ -59.48, 48.32]
Meth-CT 28.36 [ -50.59, 107.32]
MIRECC 0.11 [ -46.15, 46.36]
UKBB nonEUR -20.67 [ -73.08, 31.74]
OSAKA 17.44 [ -24.18, 59.06]
PING Non European -12.24 [ -54.34, 29.87]
Non European Cohorts [N=2882; P=0.2798] 7.99 [ -6.50, 22.47]
-150
-100
-50
0
50
100
150
mm^2/effect allele

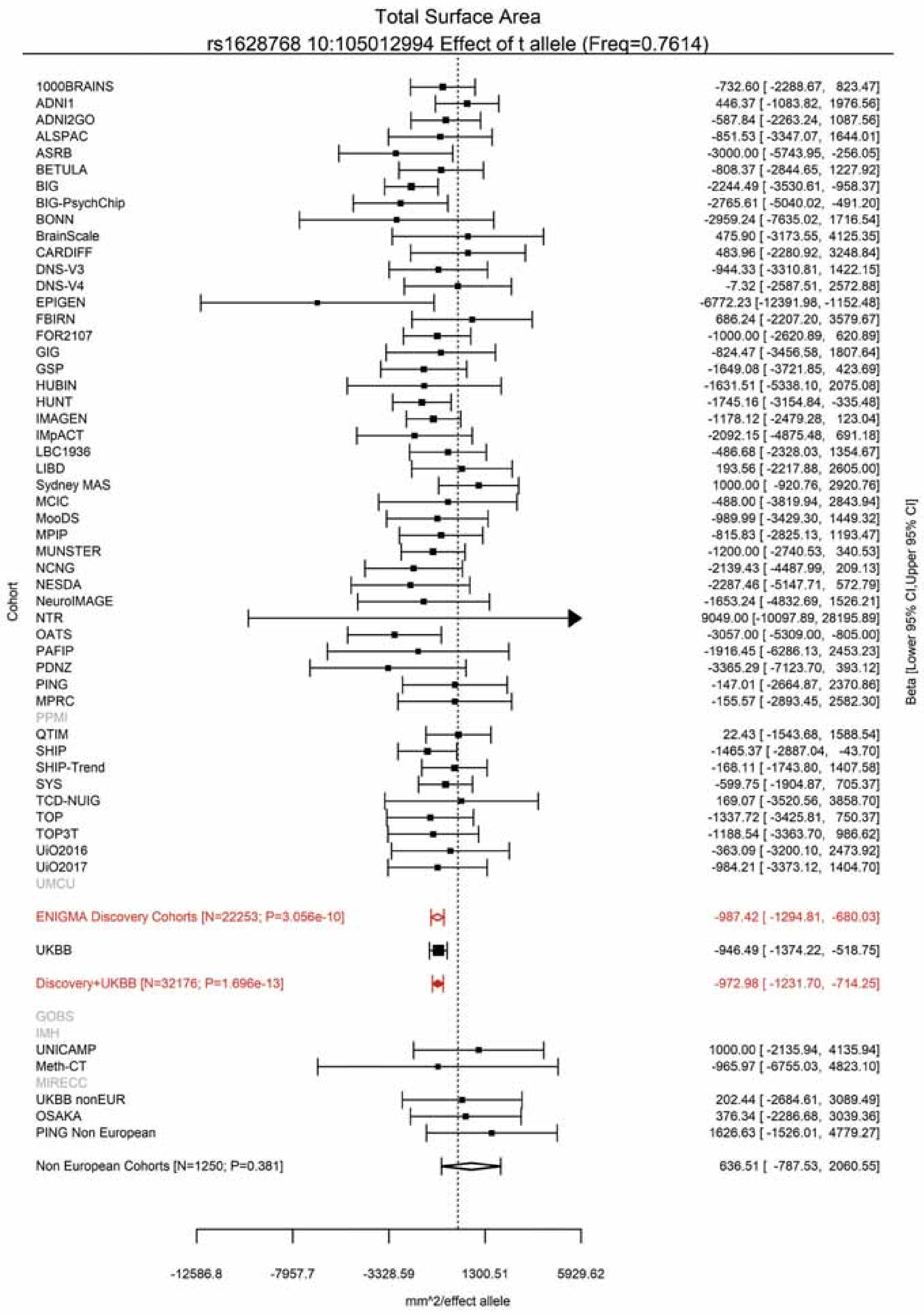
Total Surface Area
rs1628768 10:105012994 Effect of t allele (Freq=0.7614)
Cohort
1000BRAINS -732.60 [ -2288.67, 823.47]
ADNI1 446.37 [ -1083.82, 1976.56]
ADNI2GO -587.84 [ -2263.24, 1087.56]
ALSPAC -851.53 [ -3347.07, 1644.01]
ASRB -3000.00 [ -5743.95, -256.05]
BETULA -808.37 [ -2844.65, 1227.92]
BIG -2244.49 [ -3530.61, -958.37]
BIG-PsychChip -2765.61 [ -5040.02, -491.20]
BONN -2959.24 [ -7635.02, 1716.54]
BrainScale 475.90 [ -3173.55, 4125.35]
CARDIFF 483.96 [ -2280.92, 3248.84]
DNS-V3 -944.33 [ -3310.81, 1422.15]
DNS-V4 -7.32 [ -2587.51, 2572.88]
EPIGEN -6772.23 [-12391.98, -1152.48]
FBIRN 686.24 [ -2207.20, 3579.67]
FOR2107 -1000.00 [ -2620.89, 620.89]
GIG -824.47 [ -3456.58, 1807.64]
GSP -1649.08 [ -3721.85, 423.69]
HUBIN -1631.51 [ -5338.10, 2075.08]
HUNT -1745.16 [ -3154.84, -335.48]
IMAGEN -1178.12 [ -2479.28, 123.04]
IMpACT -2092.15 [ -4875.48, 691.18]
LBC1936 -486.68 [ -2328.03, 1354.67]
LIBD 193.56 [ -2217.88, 2605.00]
Sydney MAS 1000.00 [ -920.76, 2920.76]
MCIC -488.00 [ -3819.94, 2843.94]
MooDS -989.99 [ -3429.30, 1449.32]
MPIP -815.83 [ -2825.13, 1193.47]
MUNSTER -1200.00 [ -2740.53, 340.53]
NCNG -2139.43 [ -4487.99, 209.13]
NESDA -2287.46 [ -5147.71, 572.79]
NeuroIMAGE -1653.24 [ -4832.69, 1526.21]
NTR 9049.00 [-10097.89, 28195.89]
OATS -3057.00 [ -5309.00, -805.00]
PAFIP -1916.45 [ -6286.13, 2453.23]
PDNZ -3365.29 [ -7123.70, 393.12]
PING -147.01 [ -2664.87, 2370.86]
MPRC -155.57 [ -2893.45, 2582.30]
PPMI
QTIM 22.43 [ -1543.68, 1588.54]
SHIP -1465.37 [ -2887.04, -43.70]
SHIP-Trend -168.11 [ -1743.80, 1407.58]
SYS -599.75 [ -1904.87, 705.37]
TCD-NUIG 169.07 [ -3520.56, 3858.70]
TOP -1337.72 [ -3425.81, 750.37]
TOP3T -1188.54 [ -3363.70, 986.62]
UiO2016 -363.09 [ -3200.10, 2473.92]
UiO2017 -984.21 [ -3373.12, 1404.70]
UMCU
ENIGMA Discovery Cohorts [N=22253; P=3.056e-10] -987.42 [ -1294.81, -680.03]
UKBB -946.49 [ -1374.22, -518.75]
Discovery+UKBB [N=32176; P=1.696e-13] -972.98 [ -1231.70, -714.25]
GOBS
IMH
UNICAMP 1000.00 [ -2135.94, 4135.94]
Meth-CT -965.97 [ -6755.03, 4823.10]
MIRECC
UKBB nonEUR 202.44 [ -2684.61, 3089.49]
OSAKA 376.34 [ -2286.68, 3039.36]
PING Non European 1626.63 [ -1526.01, 4779.27]
Non European Cohorts [N=1250; P=0.381] 636.51 [ -787.53, 2060.55]
-12586.8
-7957.7
-3328.59
1300.51
5929.62
mm^2/effect allele
Beta [Lower 95% CI,Upper 95% CI]

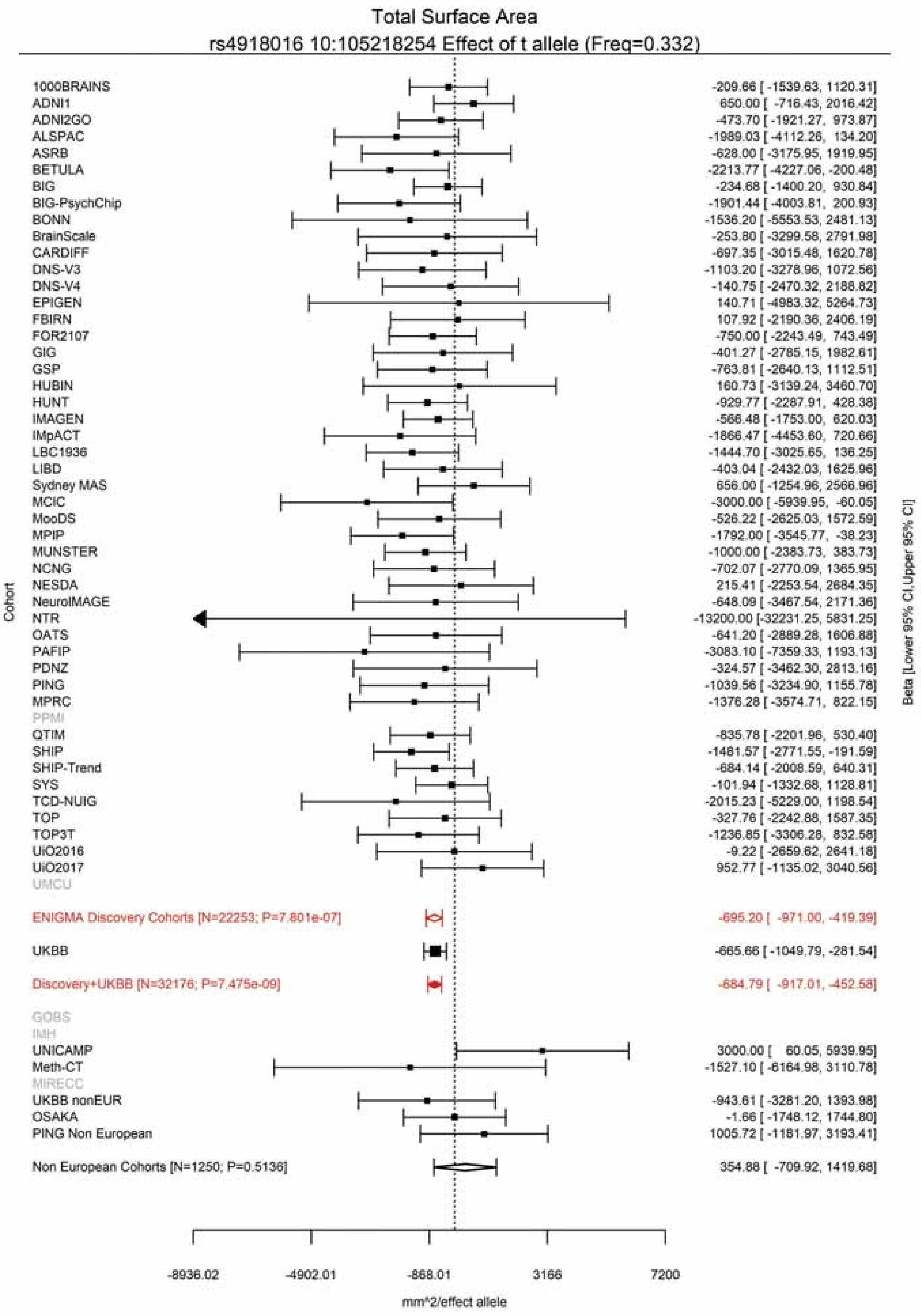

Total Surface Area
rs4918016 10:105218254 Effect of t allele (Freq=0.332)
Cohort
1000BRAINS -209.66 [ -1539.63, 1120.31]
ADNI1 650.00 [ -716.43, 2016.42]
ADNI2GO -473.70 [ -1921.27, 973.87]
ALSPAC -1989.03 [ -4112.26, 134.20]
ASRB -628.00 [ -3175.95, 1919.95]
BETULA -2213.77 [ -4227.06, -200.48]
BIG -234.68 [ -1400.20, 930.84]
BIG-PsychChip -1901.44 [ -4003.81, 200.93]
BONN -1536.20 [ -5553.53, 2481.13]
BrainScale -253.80 [ -3299.58, 2791.98]
CARDIFF -697.35 [ -3015.48, 1620.78]
DNS-V3 -1103.20 [ -3278.96, 1072.56]
DNS-V4 -140.75 [ -2470.32, 2188.82]
EPIGEN 140.71 [ -4983.32, 5264.73]
FBIRN 107.92 [ -2190.36, 2406.19]
FOR2107 -750.00 [ -2243.49, 743.49]
GIG -401.27 [ -2785.15, 1982.61]
GSP -763.81 [ -2640.13, 1112.51]
HUBIN 160.73 [ -3139.24, 3460.70]
HUNT -929.77 [ -2287.91, 428.38]
IMAGEN -566.48 [ -1753.00, 620.03]
IMpACT -1866.47 [ -4453.60, 720.66]
LBC1936 -1444.70 [ -3025.65, 136.25]
LIBD -403.04 [ -2432.03, 1625.96]
Sydney MAS 656.00 [ -1254.96, 2566.96]
MCIC -3000.00 [ -5939.95, -60.05]
MooDS -526.22 [ -2625.03, 1572.59]
MPIP -1792.00 [ -3545.77, -38.23]
MUNSTER -1000.00 [ -2383.73, 383.73]
NCNG -702.07 [ -2770.09, 1365.95]
NESDA 215.41 [ -2253.54, 2684.35]
NeuroIMAGE -648.09 [ -3467.54, 2171.36]
NTR -13200.00 [-32231.25, 5831.25]
OATS -641.20 [ -2889.28, 1606.88]
PAFIP -3083.10 [ -7359.33, 1193.13]
PDNZ -324.57 [ -3462.30, 2813.16]
PING -1039.56 [ -3234.90, 1155.78]
MPRC -1376.28 [ -3574.71, 822.15]
PPMI
QTIM -835.78 [ -2201.96, 530.40]
SHIP -1481.57 [ -2771.55, -191.59]
SHIP-Trend -684.14 [ -2008.59, 640.31]
SYS -101.94 [ -1332.68, 1128.81]
TCD-NUIG -2015.23 [ -5229.00, 1198.54]
TOP -327.76 [ -2242.88, 1587.35]
TOP3T -1236.85 [ -3306.28, 832.58]
UiO2016 -9.22 [ -2659.62, 2641.18]
UiO2017 952.77 [ -1135.02, 3040.56]
UMCU
ENIGMA Discovery Cohorts [N=22253; P=7.801e-07] -695.20 [ -971.00, -419.39]
UKBB -665.66 [ -1049.79, -281.54]
Discovery+UKBB [N=32176; P=7.475e-09] -684.79 [ -917.01, -452.58]
GOBS
IMH
UNICAMP 3000.00 [ 60.05, 5939.95]
Meth-CT -1527.10 [ -6164.98, 3110.78]
MIRECC
UKBB nonEUR -943.61 [ -3281.20, 1393.98]
OSAKA -1.66 [ -1748.12, 1744.80]
PING Non European 1005.72 [ -1181.97, 3193.41]
Non European Cohorts [N=1250; P=0.5136] 354.88 [ -709.92, 1419.68]
Beta [Lower 95% CI,Upper 95% CI]
-8936.02
-4902.01
-868.01
3166
7200
mm^2/effect allele

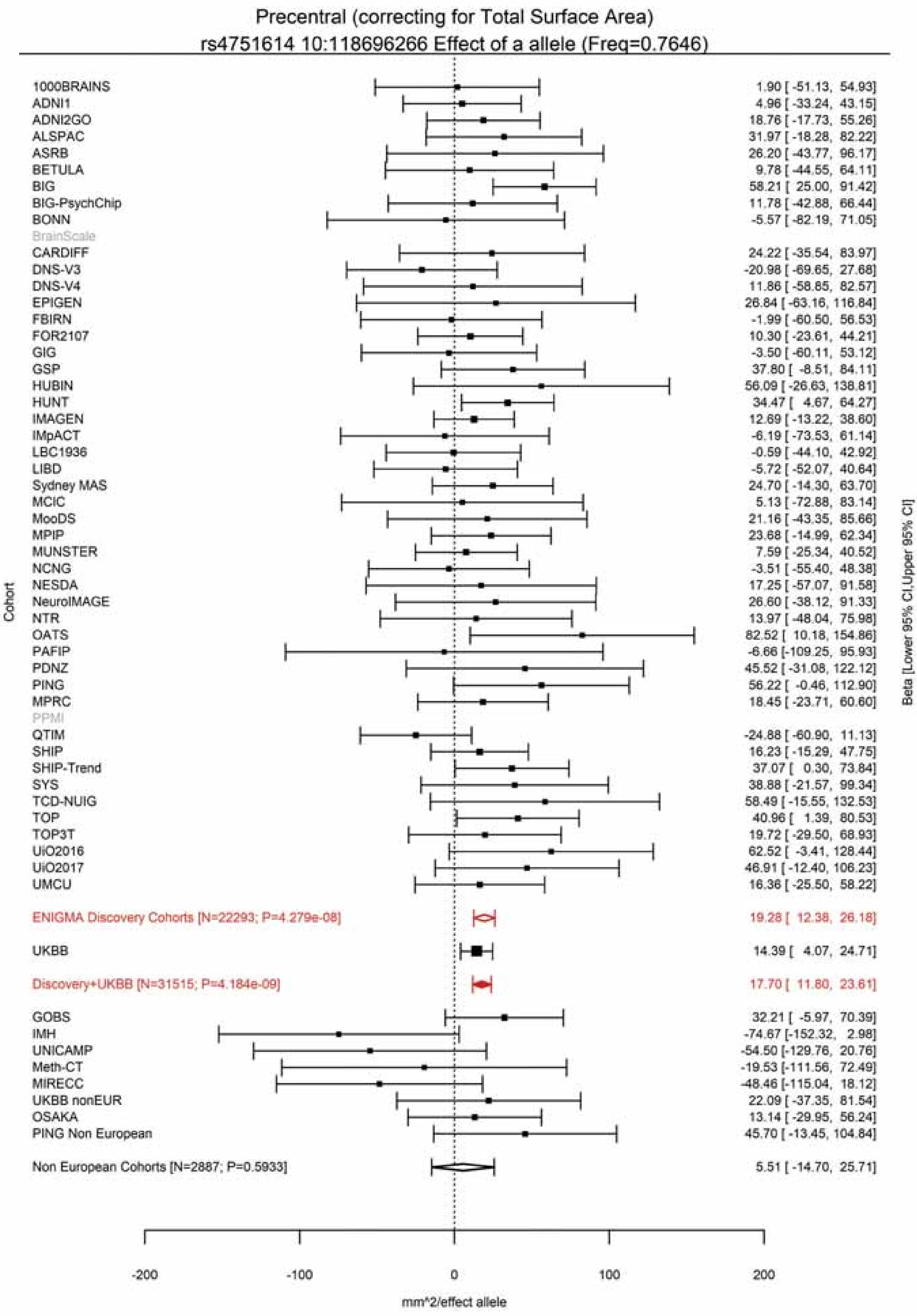
Precentral (correcting for Total Surface Area)
rs4751614 10:118696266 Effect of a allele (Freq=0.7646)
Cohort
1000BRAINS 1.90 [ -51.13, 54.93]
ADNI1 4.96 [ -33.24, 43.15]
ADNI2GO 18.76 [ -17.73, 55.26]
ALSPAC 31.97 [ -18.28, 82.22]
ASRB 26.20 [ -43.77, 96.17]
BETULA 9.78 [ -44.55, 64.11]
BIG 58.21 [ 25.00, 91.42]
BIG-PsychChip 11.78 [ -42.88, 66.44]
BONN -5.57 [ -82.19, 71.05]
BrainScale
CARDIFF 24.22 [ -35.54, 83.97]
DNS-V3 -20.98 [ -69.65, 27.68]
DNS-V4 11.86 [ -58.85, 82.57]
EPIGEN 26.84 [ -63.16, 116.84]
FBIRN -1.99 [ -60.50, 56.53]
FOR2107 10.30 [ -23.61, 44.21]
GIG -3.50 [ -60.11, 53.12]
GSP 37.80 [ -8.51, 84.11]
HUBIN 56.09 [ -26.63, 138.81]
HUNT 34.47 [ 4.67, 64.27]
IMAGEN 12.69 [ -13.22, 38.60]
IMpACT -6.19 [ -73.53, 61.14]
LBC1936 -0.59 [ -44.10, 42.92]
LIBD -5.72 [ -52.07, 40.64]
Sydney MAS 24.70 [ -14.30, 63.70]
MCIC 5.13 [ -72.88, 83.14]
MooDS 21.16 [ -43.35, 85.66]
MPIP 23.68 [ -14.99, 62.34]
MUNSTER 7.59 [ -25.34, 40.52]
NCNG -3.51 [ -55.40, 48.38]
NESDA 17.25 [ -57.07, 91.58]
NeuroIMAGE 26.60 [ -38.12, 91.33]
NTR 13.97 [ -48.04, 75.98]
OATS 82.52 [ 10.18, 154.86]
PAFIP -6.66 [-109.25, 95.93]
PDNZ 45.52 [ -31.08, 122.12]
PING 56.22 [ -0.46, 112.90]
MPRC 18.45 [ -23.71, 60.60]
PPMI
QTIM -24.88 [ -60.90, 11.13]
SHIP 16.23 [ -15.29, 47.75]
SHIP-Trend 37.07 [ 0.30, 73.84]
SYS 38.88 [ -21.57, 99.34]
TCD-NUIG 58.49 [ -15.55, 132.53]
TOP 40.96 [ 1.39, 80.53]
TOP3T 19.72 [ -29.50, 68.93]
UiO2016 62.52 [ -3.41, 128.44]
UiO2017 46.91 [ -12.40, 106.23]
UMCU 16.36 [ -25.50, 58.22]
ENIGMA Discovery Cohorts [N=22293; P=4.279e-08] 19.28 [ 12.38, 26.18]
UKBB 14.39 [ 4.07, 24.71]
Discovery+UKBB [N=31515; P=4.184e-09] 17.70 [ 11.80, 23.61]
GOBS 32.21 [ -5.97, 70.39]
IMH -74.67 [-152.32, 2.98]
UNICAMP -54.50 [-129.76, 20.76]
Meth-CT -19.53 [-111.56, 72.49]
MIRECC -48.46 [-115.04, 18.12]
UKBB nonEUR 22.09 [ -37.35, 81.54]
OSAKA 13.14 [ -29.95, 56.24]
PING Non European 45.70 [ -13.45, 104.84]
Non European Cohorts [N=2887; P=0.5933] 5.51 [ -14.70, 25.71]
-200
-100
0
100
200
mm^2/effect allele
Beta [Lower 95% CI,Upper 95% CI]

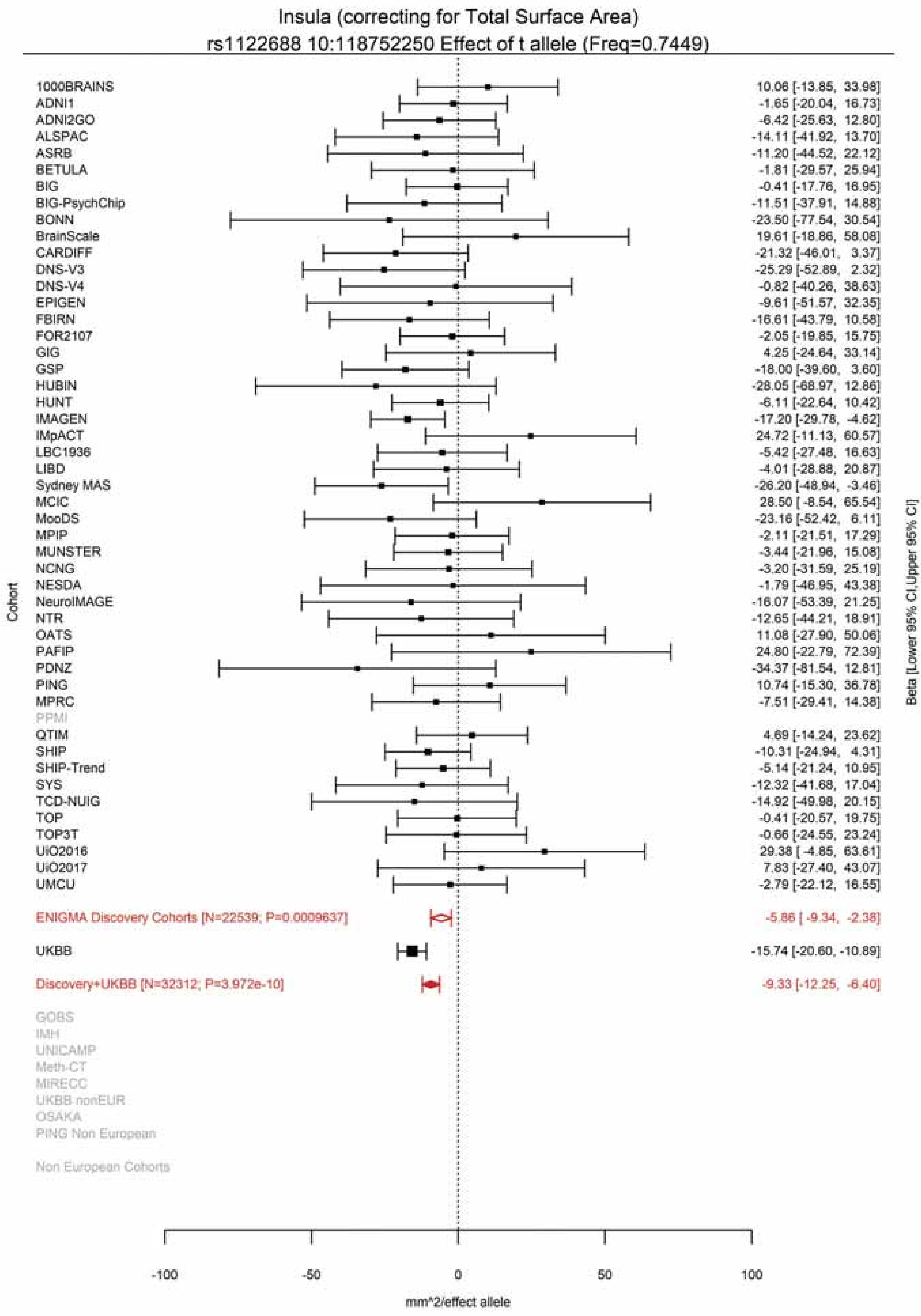
Insula (correcting for Total Surface Area)
rs1122688 10:118752250 Effect of t allele (Freq=0.7449)
Cohort
1000BRAINS 10.06 [-13.85, 33.98]
ADNI1 -1.65 [-20.04, 16.73]
ADNI2GO -6.42 [-25.63, 12.80]
ALSPAC -14.11 [-41.92, 13.70]
ASRB -11.20 [-44.52, 22.12]
BETULA -1.81 [-29.57, 25.94]
BIG -0.41 [-17.76, 16.95]
BIG-PsychChip -11.51 [-37.91, 14.88]
BONN -23.50 [-77.54, 30.54]
BrainScale 19.61 [-18.86, 58.08]
CARDIFF -21.32 [-46.01, 3.37]
DNS-V3 -25.29 [-52.89, 2.32]
DNS-V4 -0.82 [-40.26, 38.63]
EPIGEN -9.61 [-51.57, 32.35]
FBIRN -16.61 [-43.79, 10.58]
FOR2107 -2.05 [-19.85, 15.75]
GIG 4.25 [-24.64, 33.14]
GSP -18.00 [-39.60, 3.60]
HUBIN -28.05 [-68.97, 12.86]
HUNT -6.11 [-22.64, 10.42]
IMAGEN -17.20 [-29.78, -4.62]
IMpACT 24.72 [-11.13, 60.57]
LBC1936 -5.42 [-27.48, 16.63]
LIBD -4.01 [-28.88, 20.87]
Sydney MAS -26.20 [-48.94, -3.46]
MCIC 28.50 [ -8.54, 65.54]
MooDS -23.16 [-52.42, 6.11]
MPIP -2.11 [-21.51, 17.29]
MUNSTER -3.44 [-21.96, 15.08]
NCNG -3.20 [-31.59, 25.19]
NESDA -1.79 [-46.95, 43.38]
NeuroIMAGE -16.07 [-53.39, 21.25]
NTR -12.65 [-44.21, 18.91]
OATS 11.08 [-27.90, 50.06]
PAFIP 24.80 [-22.79, 72.39]
PDNZ -34.37 [-81.54, 12.81]
PING 10.74 [-15.30, 36.78]
MPRC -7.51 [-29.41, 14.38]
PPMI
QTIM 4.69 [-14.24, 23.62]
SHIP -10.31 [-24.94, 4.31]
SHIP-Trend -5.14 [-21.24, 10.95]
SYS -12.32 [-41.68, 17.04]
TCD-NUIG -14.92 [-49.98, 20.15]
TOP -0.41 [-20.57, 19.75]
TOP3T -0.66 [-24.55, 23.24]
UiO2016 29.38 [ -4.85, 63.61]
UiO2017 7.83 [-27.40, 43.07]
UMCU -2.79 [-22.12, 16.55]
ENIGMA Discovery Cohorts [N=22539; P=0.0009637] -5.86 [ -9.34, -2.38]
UKBB -15.74 [-20.60, -10.89]
Discovery+UKBB [N=32312; P=3.972e-10] -9.33 [-12.25, -6.40]
GOBS
IMH
UNICAMP
Meth-CT
MIRECC
UKBB nonEUR
OSAKA
PING Non European
Non European Cohorts
Beta [Lower 95% CI,Upper 95% CI]
-100
-50
0
50
100
mm^2/effect allele

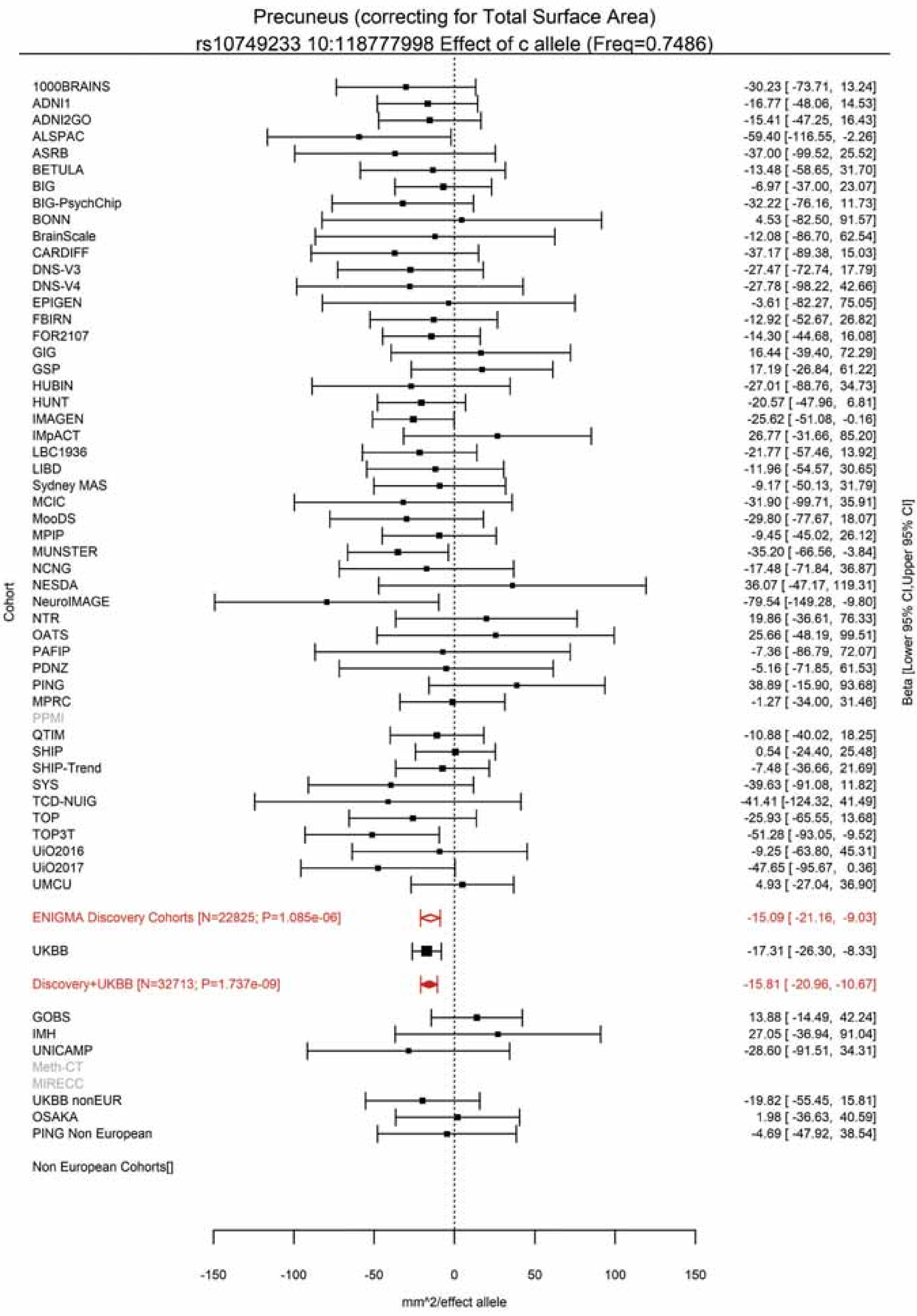

Precuneus (correcting for Total Surface Area)
rs10749233 10:118777998 Effect of c allele (Freq=0.7486)
Cohort
1000BRAINS -30.23 [ -73.71, 13.24]
ADNI1 -16.77 [ -48.06, 14.53]
ADNI2GO -15.41 [ -47.25, 16.43]
ALSPAC -59.40 [-116.55, -2.26]
ASRB -37.00 [ -99.52, 25.52]
BETULA -13.48 [ -58.65, 31.70]
BIG -6.97 [ -37.00, 23.07]
BIG-PsychChip -32.22 [ -76.16, 11.73]
BONN 4.53 [ -82.50, 91.57]
BrainScale -12.08 [ -86.70, 62.54]
CARDIFF -37.17 [ -89.38, 15.03]
DNS-V3 -27.47 [ -72.74, 17.79]
DNS-V4 -27.78 [ -98.22, 42.66]
EPIGEN -3.61 [ -82.27, 75.05]
FBIRN -12.92 [ -52.67, 26.82]
FOR2107 -14.30 [ -44.68, 16.08]
GIG 16.44 [ -39.40, 72.29]
GSP 17.19 [ -26.84, 61.22]
HUBIN -27.01 [ -88.76, 34.73]
HUNT -20.57 [ -47.96, 6.81]
IMAGEN -25.62 [ -51.08, -0.16]
IMpACT 26.77 [ -31.66, 85.20]
LBC1936 -21.77 [ -57.46, 13.92]
LIBD -11.96 [ -54.57, 30.65]
Sydney MAS -9.17 [ -50.13, 31.79]
MCIC -31.90 [ -99.71, 35.91]
MooDS -29.80 [ -77.67, 18.07]
MPIP -9.45 [ -45.02, 26.12]
MUNSTER -35.20 [ -66.56, -3.84]
NCNG -17.48 [ -71.84, 36.87]
NESDA 36.07 [ -47.17, 119.31]
NeuroIMAGE -79.54 [-149.28, -9.80]
NTR 19.86 [ -36.61, 76.33]
OATS 25.66 [ -48.19, 99.51]
PAFIP -7.36 [ -86.79, 72.07]
PDNZ -5.16 [ -71.85, 61.53]
PING 38.89 [ -15.90, 93.68]
MPRC -1.27 [ -34.00, 31.46]
PPMI
QTIM -10.88 [ -40.02, 18.25]
SHIP 0.54 [ -24.40, 25.48]
SHIP-Trend -7.48 [ -36.66, 21.69]
SYS -39.63 [ -91.08, 11.82]
TCD-NUIG -41.41 [-124.32, 41.49]
TOP -25.93 [ -65.55, 13.68]
TOP3T -51.28 [ -93.05, -9.52]
UiO2016 -9.25 [ -63.80, 45.31]
UiO2017 -47.65 [ -95.67, 0.36]
UMCU 4.93 [ -27.04, 36.90]
ENIGMA Discovery Cohorts [N=22825; P=1.085e-06] -15.09 [ -21.16, -9.03]
UKBB -17.31 [ -26.30, -8.33]
Discovery+UKBB [N=32713; P=1.737e-09] -15.81 [ -20.96, -10.67]
GOBS 13.88 [ -14.49, 42.24]
IMH 27.05 [ -36.94, 91.04]
UNICAMP -28.60 [ -91.51, 34.31]
Meth-CT
MIRECC
UKBB nonEUR -19.82 [ -55.45, 15.81]
OSAKA 1.98 [ -36.63, 40.59]
PING Non European -4.69 [ -47.92, 38.54]
Non European Cohorts[]
Beta [Lower 95% CI,Upper 95% CI]
-150
-100
-50
0
50
100
150
mm^2/effect allele

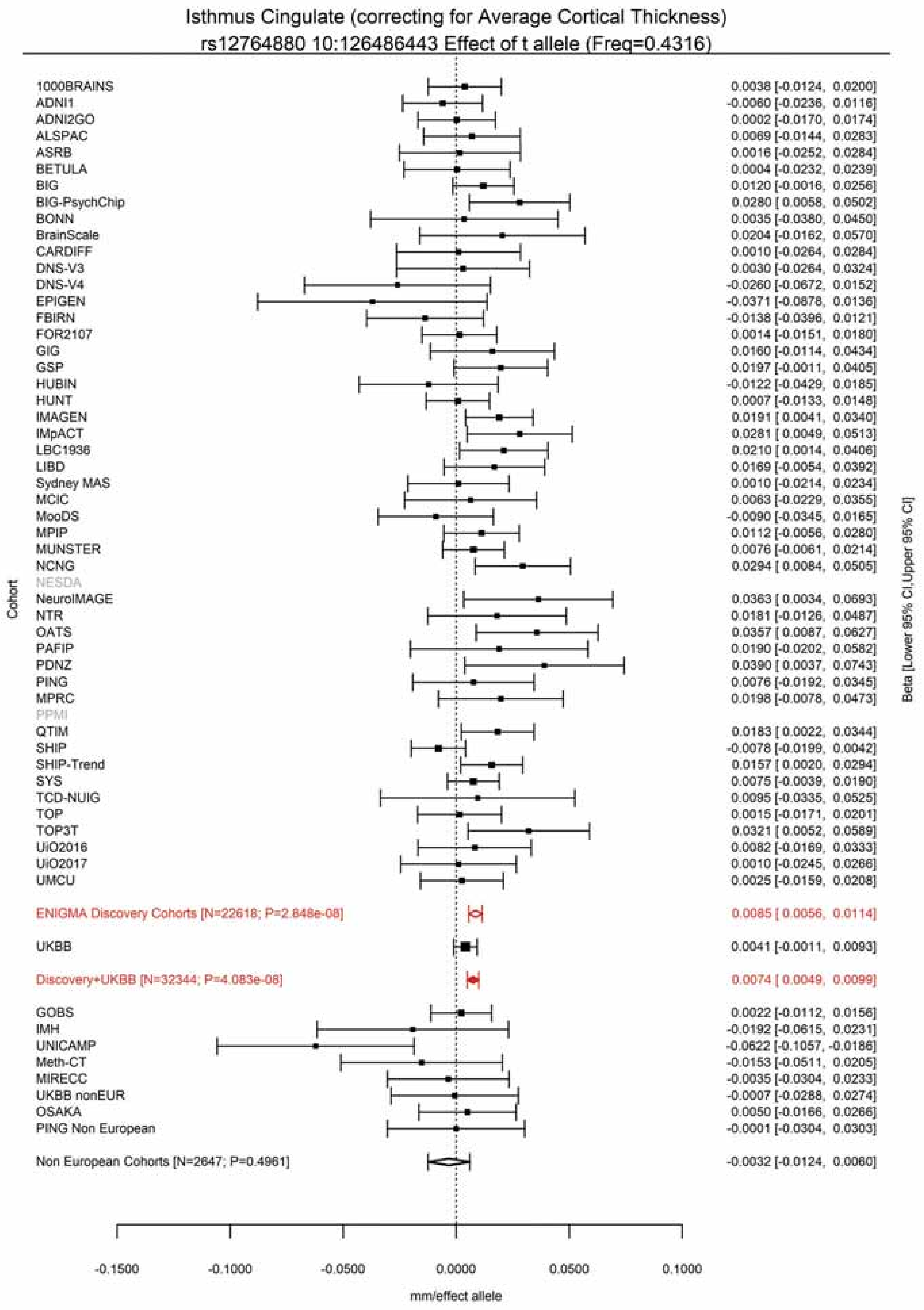

Isthmus Cingulate (correcting for Average Cortical Thickness)
rs12764880 10:126486443 Effect of t allele (Freq=0.4316)
Cohort
1000BRAINS 0.0038 [-0.0124, 0.0200]
ADNI1 -0.0060 [-0.0236, 0.0116]
ADNI2GO 0.0002 [-0.0170, 0.0174]
ALSPAC 0.0069 [-0.0144, 0.0283]
ASRB 0.0016 [-0.0252, 0.0284]
BETULA 0.0004 [-0.0232, 0.0239]
BIG 0.0120 [-0.0016, 0.0256]
BIG-PsychChip 0.0280 [ 0.0058, 0.0502]
BONN 0.0035 [-0.0380, 0.0450]
BrainScale 0.0204 [-0.0162, 0.0570]
CARDIFF 0.0010 [-0.0264, 0.0284]
DNS-V3 0.0030 [-0.0264, 0.0324]
DNS-V4 -0.0260 [-0.0672, 0.0152]
EPIGEN -0.0371 [-0.0878, 0.0136]
FBIRN -0.0138 [-0.0396, 0.0121]
FOR2107 0.0014 [-0.0151, 0.0180]
GIG 0.0160 [-0.0114, 0.0434]
GSP 0.0197 [-0.0011, 0.0405]
HUBIN -0.0122 [-0.0429, 0.0185]
HUNT 0.0007 [-0.0133, 0.0148]
IMAGEN 0.0191 [ 0.0041, 0.0340]
IMpACT 0.0281 [ 0.0049, 0.0513]
LBC1936 0.0210 [ 0.0014, 0.0406]
LIBD 0.0169 [-0.0054, 0.0392]
Sydney MAS 0.0010 [-0.0214, 0.0234]
MCIC 0.0063 [-0.0229, 0.0355]
MooDS -0.0090 [-0.0345, 0.0165]
MPIP 0.0112 [-0.0056, 0.0280]
MUNSTER 0.0076 [-0.0061, 0.0214]
NCNG 0.0294 [ 0.0084, 0.0505]
NESDA
NeuroIMAGE 0.0363 [ 0.0034, 0.0693]
NTR 0.0181 [-0.0126, 0.0487]
OATS 0.0357 [ 0.0087, 0.0627]
PAFIP 0.0190 [-0.0202, 0.0582]
PDNZ 0.0390 [ 0.0037, 0.0743]
PING 0.0076 [-0.0192, 0.0345]
MPRC 0.0198 [-0.0078, 0.0473]
PPMI
QTIM 0.0183 [ 0.0022, 0.0344]
SHIP -0.0078 [-0.0199, 0.0042]
SHIP-Trend 0.0157 [ 0.0020, 0.0294]
SYS 0.0075 [-0.0039, 0.0190]
TCD-NUIG 0.0095 [-0.0335, 0.0525]
TOP 0.0015 [-0.0171, 0.0201]
TOP3T 0.0321 [ 0.0052, 0.0589]
UiO2016 0.0082 [-0.0169, 0.0333]
UiO2017 0.0010 [-0.0245, 0.0266]
UMCU 0.0025 [-0.0159, 0.0208]
ENIGMA Discovery Cohorts [N=22618; P=2.848e-08] 0.0085 [ 0.0056, 0.0114]
UKBB 0.0041 [-0.0011, 0.0093]
Discovery+UKBB [N=32344; P=4.083e-08] 0.0074 [ 0.0049, 0.0099]
GOBS 0.0022 [-0.0112, 0.0156]
IMH -0.0192 [-0.0615, 0.0231]
UNICAMP -0.0622 [-0.1057, -0.0186]
Meth-CT -0.0153 [-0.0511, 0.0205]
MIRECC -0.0035 [-0.0304, 0.0233]
UKBB nonEUR -0.0007 [-0.0288, 0.0274]
OSAKA 0.0050 [-0.0166, 0.0266]
PING Non European -0.0001 [-0.0304, 0.0303]
Non European Cohorts [N=2647; P=0.4961] -0.0032 [-0.0124, 0.0060]
Beta [Lower 95% CI,Upper 95% CI]
-0.1500
-0.1000
-0.0500
0.0000
0.0500
0.1000
mm/effect allele

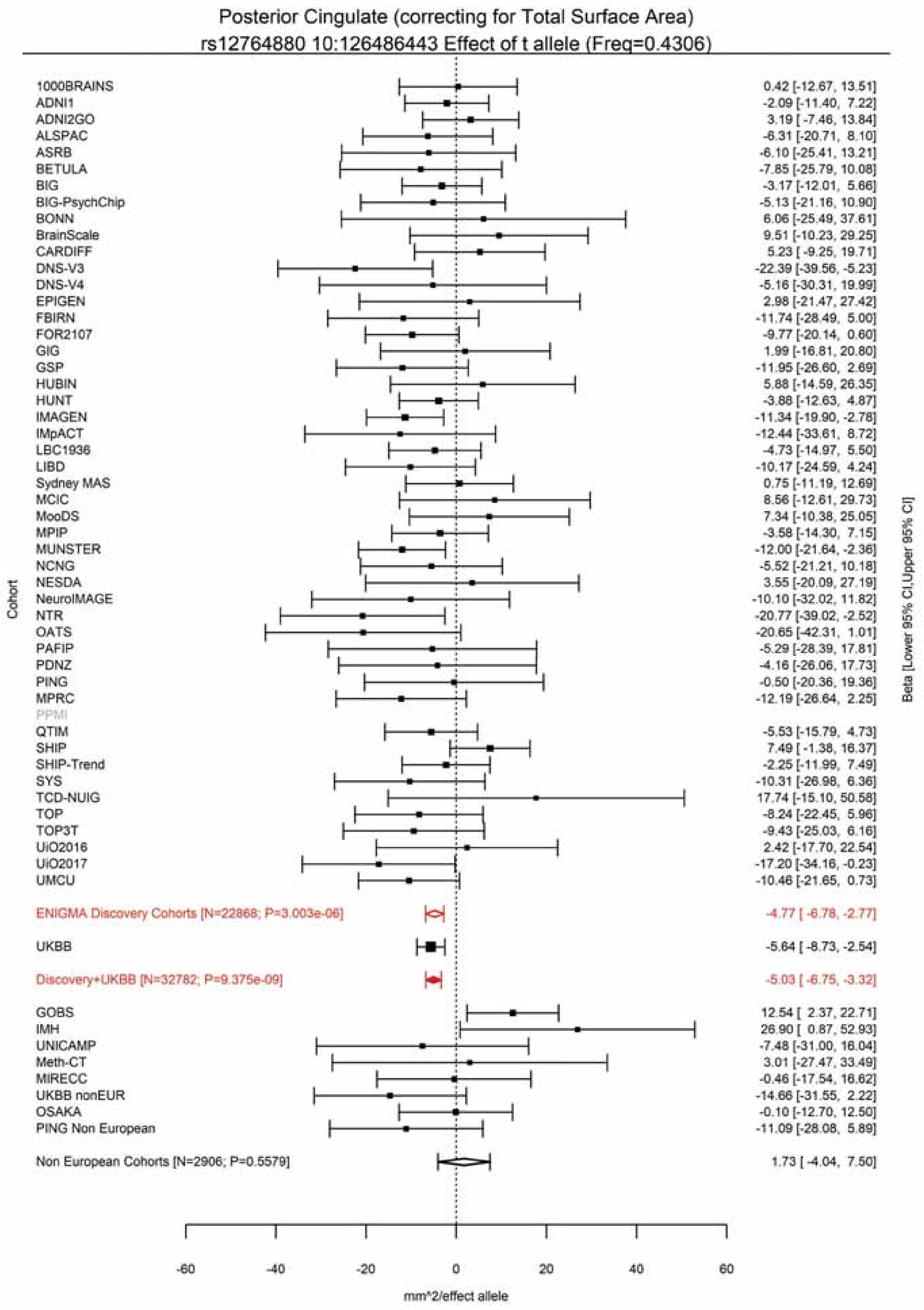
Posterior Cingulate (correcting for Total Surface Area)
rs12764880 10:126486443 Effect of t allele (Freq=0.4306)
Cohort
1000BRAINS 0.42 [-12.67, 13.51]
ADNI1 -2.09 [-11.40, 7.22]
ADNI2GO 3.19 [ -7.46, 13.84]
ALSPAC -6.31 [-20.71, 8.10]
ASRB -6.10 [-25.41, 13.21]
BETULA -7.85 [-25.79, 10.08]
BIG -3.17 [-12.01, 5.66]
BIG-PsychChip -5.13 [-21.16, 10.90]
BONN 6.06 [-25.49, 37.61]
BrainScale 9.51 [-10.23, 29.25]
CARDIFF 5.23 [ -9.25, 19.71]
DNS-V3 -22.39 [-39.56, -5.23]
DNS-V4 -5.16 [-30.31, 19.99]
EPIGEN 2.98 [-21.47, 27.42]
FBIRN -11.74 [-28.49, 5.00]
FOR2107 -9.77 [-20.14, 0.60]
GIG 1.99 [-16.81, 20.80]
GSP -11.95 [-26.60, 2.69]
HUBIN 5.88 [-14.59, 26.35]
HUNT -3.88 [-12.63, 4.87]
IMAGEN -11.34 [-19.90, -2.78]
IMpACT -12.44 [-33.61, 8.72]
LBC1936 -4.73 [-14.97, 5.50]
LIBD -10.17 [-24.59, 4.24]
Sydney MAS 0.75 [-11.19, 12.69]
MCIC 8.56 [-12.61, 29.73]
MooDS 7.34 [-10.38, 25.05]
MPIP -3.58 [-14.30, 7.15]
MUNSTER -12.00 [-21.64, -2.36]
NCNG -5.52 [-21.21, 10.18]
NESDA 3.55 [-20.09, 27.19]
NeuroIMAGE -10.10 [-32.02, 11.82]
NTR -20.77 [-39.02, -2.52]
OATS -20.65 [-42.31, 1.01]
PAFIP -5.29 [-28.39, 17.81]
PDNZ -4.16 [-26.06, 17.73]
PING -0.50 [-20.36, 19.36]
MPRC -12.19 [-26.64, 2.25]
PPMI
QTIM -5.53 [-15.79, 4.73]
SHIP 7.49 [ -1.38, 16.37]
SHIP-Trend -2.25 [-11.99, 7.49]
SYS -10.31 [-26.98, 6.36]
TCD-NUIG 17.74 [-15.10, 50.58]
TOP -8.24 [-22.45, 5.96]
TOP3T -9.43 [-25.03, 6.16]
UiO2016 2.42 [-17.70, 22.54]
UiO2017 -17.20 [-34.16, -0.23]
UMCU -10.46 [-21.65, 0.73]
ENIGMA Discovery Cohorts [N=22868; P=3.003e-06] -4.77 [ -6.78, -2.77]
UKBB -5.64 [ -8.73, -2.54]
Discovery+UKBB [N=32782; P=9.375e-09] -5.03 [ -6.75, -3.32]
GOBS 12.54 [ 2.37, 22.71]
IMH 26.90 [ 0.87, 52.93]
UNICAMP -7.48 [-31.00, 16.04]
Meth-CT 3.01 [-27.47, 33.49]
MIRECC -0.46 [-17.54, 16.62]
UKBB nonEUR -14.66 [-31.55, 2.22]
OSAKA -0.10 [-12.70, 12.50]
PING Non European -11.09 [-28.08, 5.89]
Non European Cohorts [N=2906; P=0.5579] 1.73 [ -4.04, 7.50]
Beta [Lower 95% CI,Upper 95% CI]
-60
-40
-20
0
20
40
60
mm^2/effect allele

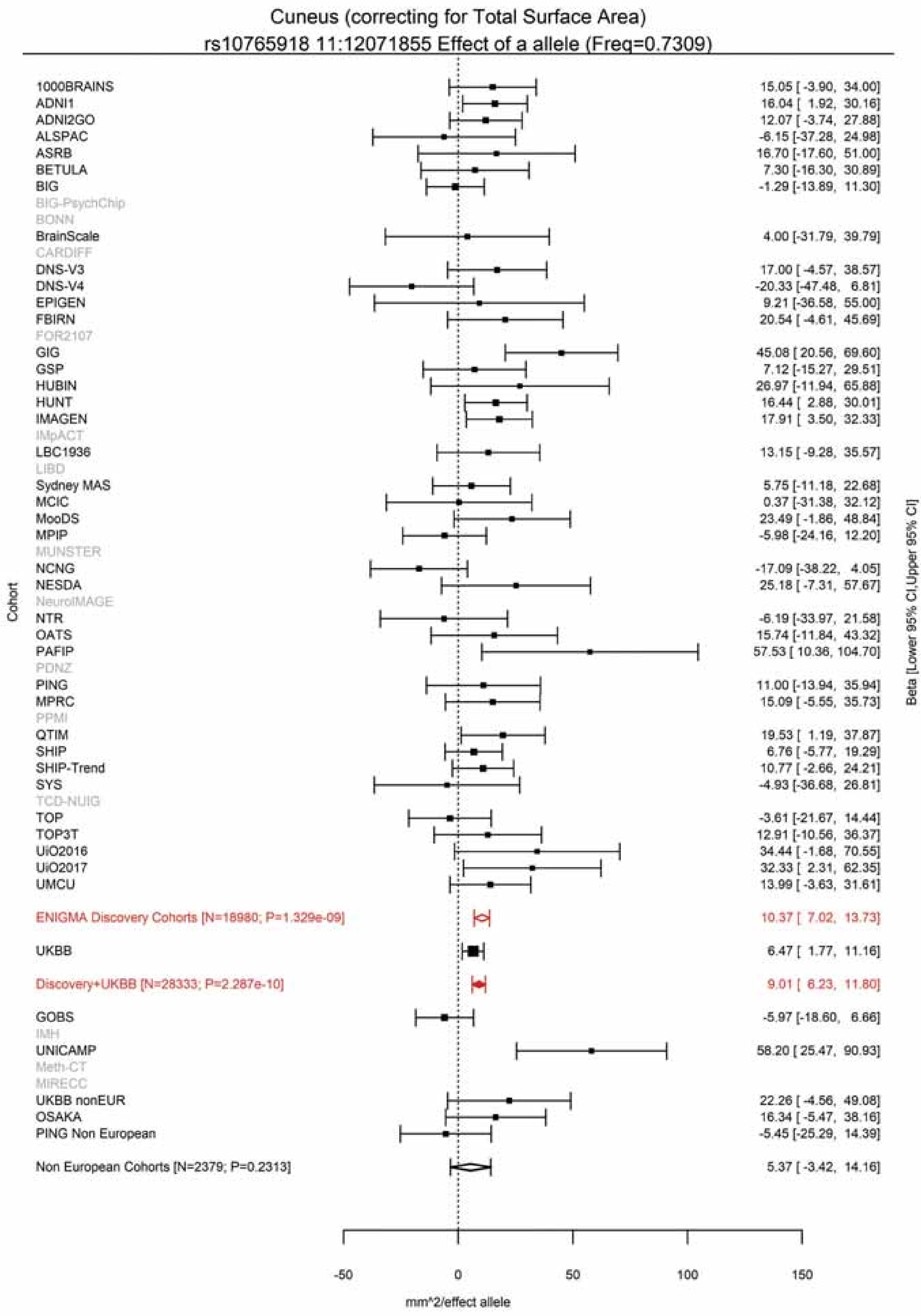
Cuneus (correcting for Total Surface Area)
rs10765918 11:12071855 Effect of a allele (Freq=0.7309)
1000BRAINS 15.05 [ -3.90, 34.00]
ADNI1 16.04 [ 1.92, 30.16]
ADNI2GO 12.07 [ -3.74, 27.88]
ALSPAC -6.15 [-37.28, 24.98]
ASRB 16.70 [-17.60, 51.00]
BETULA 7.30 [-16.30, 30.89]
BIG -1.29 [-13.89, 11.30]
BIG-PsychChip
BONN
BrainScale 4.00 [-31.79, 39.79]
CARDIFF
DNS-V3 17.00 [ -4.57, 38.57]
DNS-V4 -20.33 [-47.48, 6.81]
EPIGEN 9.21 [-36.58, 55.00]
FBIRN 20.54 [ -4.61, 45.69]
FOR2107
GIG 45.08 [ 20.56, 69.60]
GSP 7.12 [-15.27, 29.51]
HUBIN 26.97 [-11.94, 65.88]
HUNT 16.44 [ 2.88, 30.01]
IMAGEN 17.91 [ 3.50, 32.33]
IMpACT
LBC1936 13.15 [ -9.28, 35.57]
LIBD
Sydney MAS 5.75 [-11.18, 22.68]
MCIC 0.37 [-31.38, 32.12]
MooDS 23.49 [ -1.86, 48.84]
MPIP -5.98 [-24.16, 12.20]
MUNSTER
NCNG -17.09 [-38.22, 4.05]
NESDA 25.18 [ -7.31, 57.67]
NeuroIMAGE
NTR -6.19 [-33.97, 21.58]
OATS 15.74 [-11.84, 43.32]
PAFIP 57.53 [ 10.36, 104.70]
PDNZ
PING 11.00 [-13.94, 35.94]
MPRC 15.09 [ -5.55, 35.73]
PPMI
QTIM 19.53 [ 1.19, 37.87]
SHIP 6.76 [ -5.77, 19.29]
SHIP-Trend 10.77 [ -2.66, 24.21]
SYS -4.93 [-36.68, 26.81]
TCD-NUIG
TOP -3.61 [-21.67, 14.44]
TOP3T 12.91 [-10.56, 36.37]
UiO2016 34.44 [ -1.68, 70.55]
UiO2017 32.33 [ 2.31, 62.35]
UMCU 13.99 [ -3.63, 31.61]
ENIGMA Discovery Cohorts [N=18980; P=1.329e-09] 10.37 [ 7.02, 13.73]
UKBB 6.47 [ 1.77, 11.16]
Discovery+UKBB [N=28333; P=2.287e-10] 9.01 [ 6.23, 11.80]
GOBS -5.97 [-18.60, 6.66]
IMH
UNICAMP 58.20 [ 25.47, 90.93]
Meth-CT
MIRECC
UKBB nonEUR 22.26 [ -4.56, 49.08]
OSAKA 16.34 [ -5.47, 38.16]
PING Non European -5.45 [-25.29, 14.39]
Non European Cohorts [N=2379; P=0.2313] 5.37 [ -3.42, 14.16]
Cohort
Beta [Lower 95% CI,Upper 95% CI]
-50
0
50
100
150
mm^2/effect allele

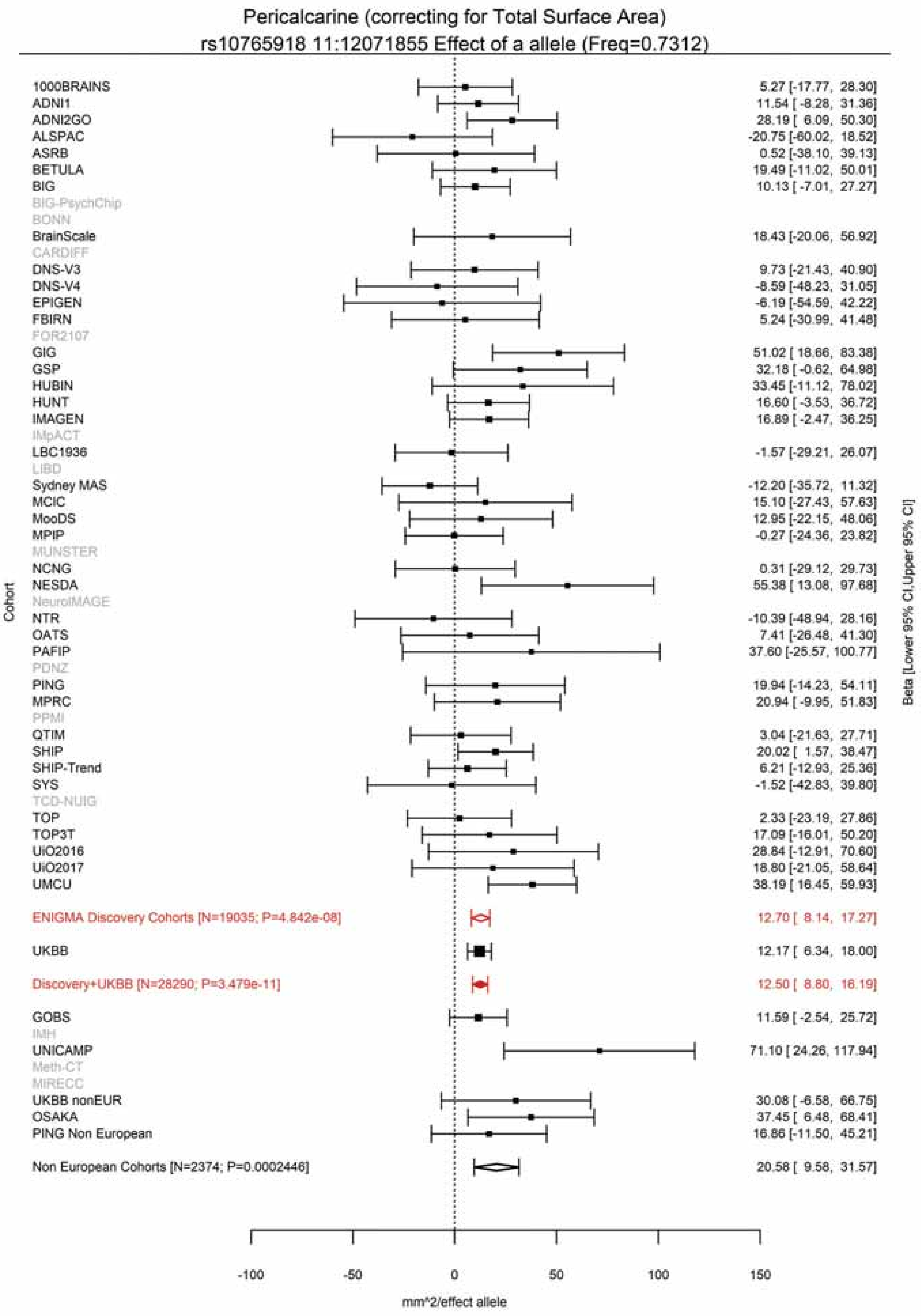
Pericalcarine (correcting for Total Surface Area)
rs10765918 11:12071855 Effect of a allele (Freq=0.7312)
1000BRAINS 5.27 [-17.77, 28.30]
ADNI1 11.54 [ -8.28, 31.36]
ADNI2GO 28.19 [ 6.09, 50.30]
ALSPAC -20.75 [-60.02, 18.52]
ASRB 0.52 [-38.10, 39.13]
BETULA 19.49 [-11.02, 50.01]
BIG 10.13 [ -7.01, 27.27]
BIG-PsychChip
BONN
BrainScale 18.43 [-20.06, 56.92]
CARDIFF
DNS-V3 9.73 [-21.43, 40.90]
DNS-V4 -8.59 [-48.23, 31.05]
EPIGEN -6.19 [-54.59, 42.22]
FBIRN 5.24 [-30.99, 41.48]
FOR2107
GIG 51.02 [ 18.66, 83.38]
GSP 32.18 [ -0.62, 64.98]
HUBIN 33.45 [-11.12, 78.02]
HUNT 16.60 [ -3.53, 36.72]
IMAGEN 16.89 [ -2.47, 36.25]
IMpACT
LBC1936 -1.57 [-29.21, 26.07]
LIBD
Sydney MAS -12.20 [-35.72, 11.32]
MCIC 15.10 [-27.43, 57.63]
MooDS 12.95 [-22.15, 48.06]
MPIP -0.27 [-24.36, 23.82]
MUNSTER
NCNG 0.31 [-29.12, 29.73]
NESDA 55.38 [ 13.08, 97.68]
NeuroIMAGE
NTR -10.39 [-48.94, 28.16]
OATS 7.41 [-26.48, 41.30]
PAFIP 37.60 [-25.57, 100.77]
PDNZ
PING 19.94 [-14.23, 54.11]
MPRC 20.94 [ -9.95, 51.83]
PPMI
QTIM 3.04 [-21.63, 27.71]
SHIP 20.02 [ 1.57, 38.47]
SHIP-Trend 6.21 [-12.93, 25.36]
SYS -1.52 [-42.83, 39.80]
TCD-NUIG
TOP 2.33 [-23.19, 27.86]
TOP3T 17.09 [-16.01, 50.20]
UiO2016 28.84 [-12.91, 70.60]
UiO2017 18.80 [-21.05, 58.64]
UMCU 38.19 [ 16.45, 59.93]
ENIGMA Discovery Cohorts [N=19035; P=4.842e-08] 12.70 [ 8.14, 17.27]
UKBB 12.17 [ 6.34, 18.00]
Discovery+UKBB [N=28290; P=3.479e-11] 12.50 [ 8.80, 16.19]
GOBS 11.59 [ -2.54, 25.72]
IMH
UNICAMP 71.10 [ 24.26, 117.94]
Meth-CT
MIRECC
UKBB nonEUR 30.08 [ -6.58, 66.75]
OSAKA 37.45 [ 6.48, 68.41]
PING Non European 16.86 [-11.50, 45.21]
Non European Cohorts [N=2374; P=0.0002446] 20.58 [ 9.58, 31.57]
Cohort
Beta [Lower 95% CI,Upper 95% CI]
-100
-50
0
50
100
150
mm^2/effect allele

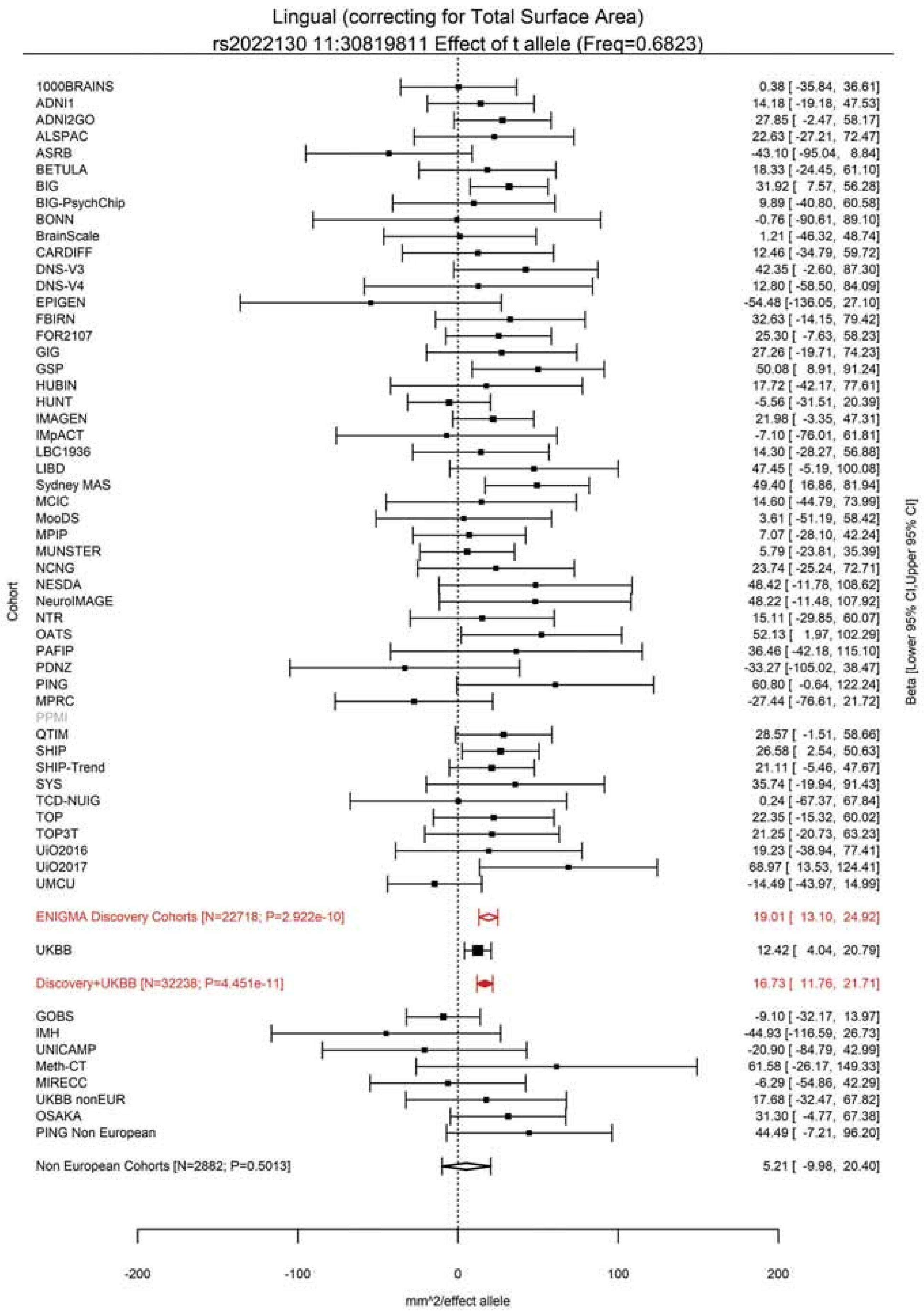

Lingual (correcting for Total Surface Area)
rs2022130 11:30819811 Effect of t allele (Freq=0.6823)
1000BRAINS 0.38 [ -35.84, 36.61]
ADNI1 14.18 [ -19.18, 47.53]
ADNI2GO 27.85 [ -2.47, 58.17]
ALSPAC 22.63 [ -27.21, 72.47]
ASRB -43.10 [ -95.04, 8.84]
BETULA 18.33 [ -24.45, 61.10]
BIG 31.92 [ 7.57, 56.28]
BIG-PsychChip 9.89 [ -40.80, 60.58]
BONN -0.76 [ -90.61, 89.10]
BrainScale 1.21 [ -46.32, 48.74]
CARDIFF 12.46 [ -34.79, 59.72]
DNS-V3 42.35 [ -2.60, 87.30]
DNS-V4 12.80 [ -58.50, 84.09]
EPIGEN -54.48 [-136.05, 27.10]
FBIRN 32.63 [ -14.15, 79.42]
FOR2107 25.30 [ -7.63, 58.23]
GIG 27.26 [ -19.71, 74.23]
GSP 50.08 [ 8.91, 91.24]
HUBIN 17.72 [ -42.17, 77.61]
HUNT -5.56 [ -31.51, 20.39]
IMAGEN 21.98 [ -3.35, 47.31]
IMpACT -7.10 [ -76.01, 61.81]
LBC1936 14.30 [ -28.27, 56.88]
LIBD 47.45 [ -5.19, 100.08]
Sydney MAS 49.40 [ 16.86, 81.94]
MCIC 14.60 [ -44.79, 73.99]
MooDS 3.61 [ -51.19, 58.42]
MPIP 7.07 [ -28.10, 42.24]
MUNSTER 5.79 [ -23.81, 35.39]
NCNG 23.74 [ -25.24, 72.71]
NESDA 48.42 [ -11.78, 108.62]
NeuroIMAGE 48.22 [ -11.48, 107.92]
NTR 15.11 [ -29.85, 60.07]
OATS 52.13 [ 1.97, 102.29]
PAFIP 36.46 [ -42.18, 115.10]
PDNZ -33.27 [-105.02, 38.47]
PING 60.80 [ -0.64, 122.24]
MPRC -27.44 [ -76.61, 21.72]
PPMI
QTIM 28.57 [ -1.51, 58.66]
SHIP 26.58 [ 2.54, 50.63]
SHIP-Trend 21.11 [ -5.46, 47.67]
SYS 35.74 [ -19.94, 91.43]
TCD-NUIG 0.24 [ -67.37, 67.84]
TOP 22.35 [ -15.32, 60.02]
TOP3T 21.25 [ -20.73, 63.23]
UiO2016 19.23 [ -38.94, 77.41]
UiO2017 68.97 [ 13.53, 124.41]
UMCU -14.49 [ -43.97, 14.99]
ENIGMA Discovery Cohorts [N=22718; P=2.922e-10] 19.01 [ 13.10, 24.92]
UKBB 12.42 [ 4.04, 20.79]
Discovery+UKBB [N=32238; P=4.451e-11] 16.73 [ 11.76, 21.71]
GOBS -9.10 [ -32.17, 13.97]
IMH -44.93 [-116.59, 26.73]
UNICAMP -20.90 [ -84.79, 42.99]
Meth-CT 61.58 [ -26.17, 149.33]
MIRECC -6.29 [ -54.86, 42.29]
UKBB nonEUR 17.68 [ -32.47, 67.82]
OSAKA 31.30 [ -4.77, 67.38]
PING Non European 44.49 [ -7.21, 96.20]
Non European Cohorts [N=2882; P=0.5013] 5.21 [ -9.98, 20.40]
Cohort
Beta [Lower 95% CI,Upper 95% CI]
-200
-100
0
100
200
mm^2/effect allele

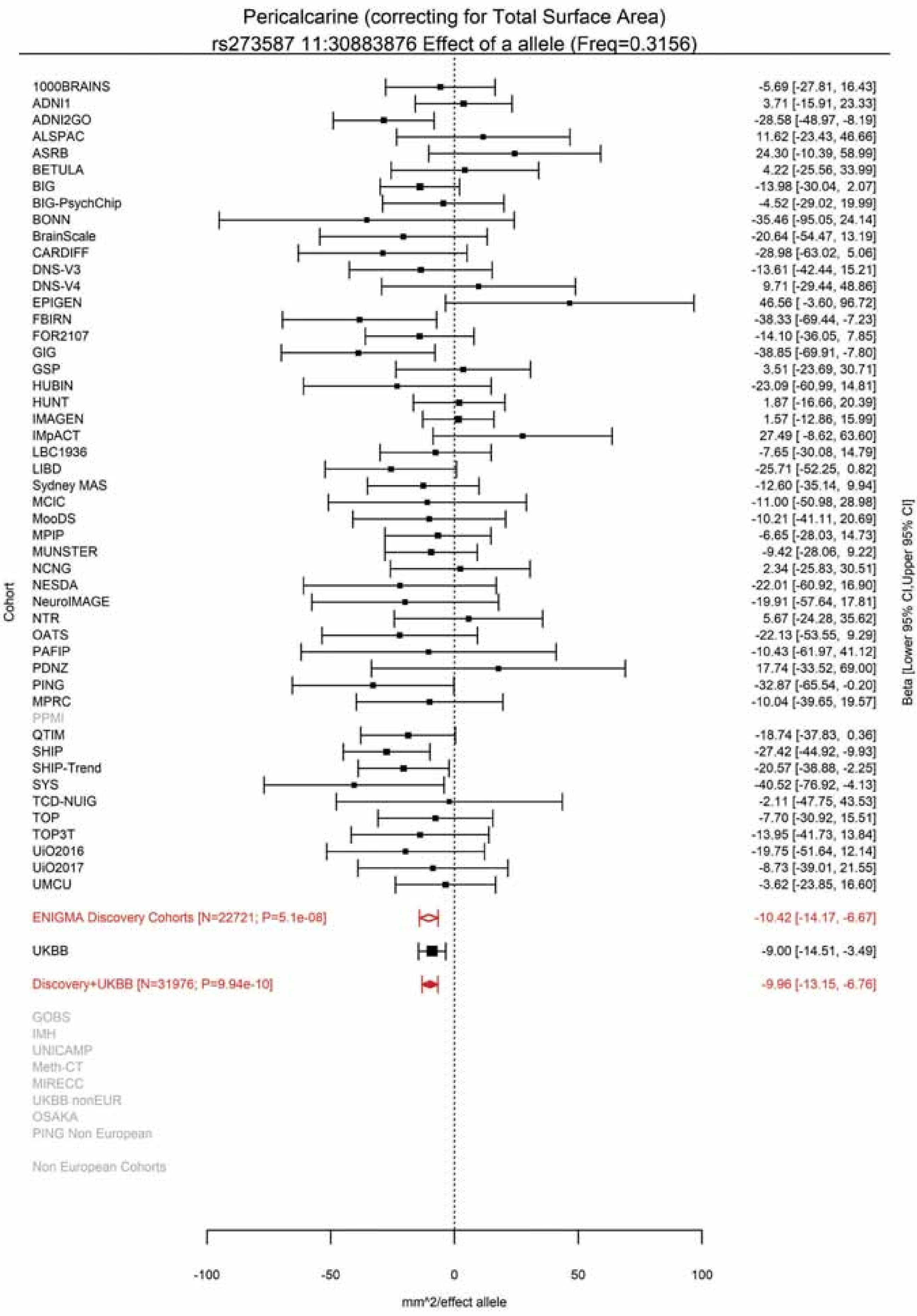

Pericalcarine (correcting for Total Surface Area)
rs273587 11:30883876 Effect of a allele (Freq=0.3156)
Cohort
Beta [Lower 95% CI,Upper 95% CI]
1000BRAINS -5.69 [-27.81, 16.43]
ADNI1 3.71 [-15.91, 23.33]
ADNI2GO -28.58 [-48.97, -8.19]
ALSPAC 11.62 [-23.43, 46.66]
ASRB 24.30 [-10.39, 58.99]
BETULA 4.22 [-25.56, 33.99]
BIG -13.98 [-30.04, 2.07]
BIG-PsychChip -4.52 [-29.02, 19.99]
BONN -35.46 [-95.05, 24.14]
BrainScale -20.64 [-54.47, 13.19]
CARDIFF -28.98 [-63.02, 5.06]
DNS-V3 -13.61 [-42.44, 15.21]
DNS-V4 9.71 [-29.44, 48.86]
EPIGEN 46.56 [ -3.60, 96.72]
FBIRN -38.33 [-69.44, -7.23]
FOR2107 -14.10 [-36.05, 7.85]
GIG -38.85 [-69.91, -7.80]
GSP 3.51 [-23.69, 30.71]
HUBIN -23.09 [-60.99, 14.81]
HUNT 1.87 [-16.66, 20.39]
IMAGEN 1.57 [-12.86, 15.99]
IMpACT 27.49 [ -8.62, 63.60]
LBC1936 -7.65 [-30.08, 14.79]
LIBD -25.71 [-52.25, 0.82]
Sydney MAS -12.60 [-35.14, 9.94]
MCIC -11.00 [-50.98, 28.98]
MooDS -10.21 [-41.11, 20.69]
MPIP -6.65 [-28.03, 14.73]
MUNSTER -9.42 [-28.06, 9.22]
NCNG 2.34 [-25.83, 30.51]
NESDA -22.01 [-60.92, 16.90]
NeuroIMAGE -19.91 [-57.64, 17.81]
NTR 5.67 [-24.28, 35.62]
OATS -22.13 [-53.55, 9.29]
PAFIP -10.43 [-61.97, 41.12]
PDNZ 17.74 [-33.52, 69.00]
PING -32.87 [-65.54, -0.20]
MPRC -10.04 [-39.65, 19.57]
PPMI
QTIM -18.74 [-37.83, 0.36]
SHIP -27.42 [-44.92, -9.93]
SHIP-Trend -20.57 [-38.88, -2.25]
SYS -40.52 [-76.92, -4.13]
TCD-NUIG -2.11 [-47.75, 43.53]
TOP -7.70 [-30.92, 15.51]
TOP3T -13.95 [-41.73, 13.84]
UiO2016 -19.75 [-51.64, 12.14]
UiO2017 -8.73 [-39.01, 21.55]
UMCU -3.62 [-23.85, 16.60]
ENIGMA Discovery Cohorts [N=22721; P=5.1e-08] -10.42 [-14.17, -6.67]
UKBB -9.00 [-14.51, -3.49]
Discovery+UKBB [N=31976; P=9.94e-10] -9.96 [-13.15, -6.76]
GOBS
IMH
UNICAMP
Meth-CT
MIRECC
UKBB nonEUR
OSAKA
PING Non European
Non European Cohorts
-100
-50
0
50
100
mm^2/effect allele

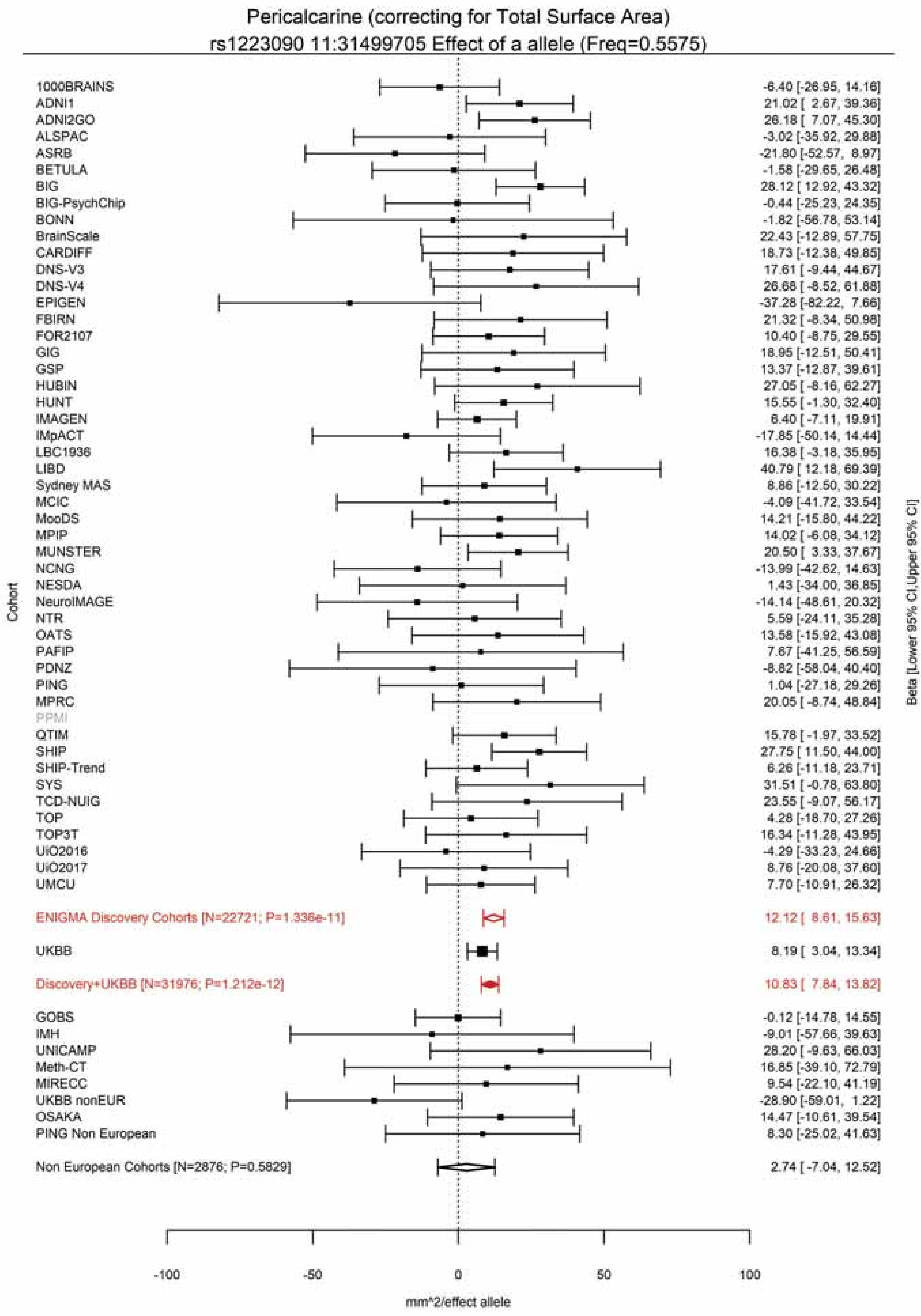

Pericalcarine (correcting for Total Surface Area)
rs1223090 11:31499705 Effect of a allele (Freq=0.5575)
Cohort
1000BRAINS -6.40 [-26.95, 14.16]
ADNI1 21.02 [ 2.67, 39.36]
ADNI2GO 26.18 [ 7.07, 45.30]
ALSPAC -3.02 [-35.92, 29.88]
ASRB -21.80 [-52.57, 8.97]
BETULA -1.58 [-29.65, 26.48]
BIG 28.12 [ 12.92, 43.32]
BIG-PsychChip -0.44 [-25.23, 24.35]
BONN -1.82 [-56.78, 53.14]
BrainScale 22.43 [-12.89, 57.75]
CARDIFF 18.73 [-12.38, 49.85]
DNS-V3 17.61 [ -9.44, 44.67]
DNS-V4 26.68 [ -8.52, 61.88]
EPIGEN -37.28 [-82.22, 7.66]
FBIRN 21.32 [ -8.34, 50.98]
FOR2107 10.40 [ -8.75, 29.55]
GIG 18.95 [-12.51, 50.41]
GSP 13.37 [-12.87, 39.61]
HUBIN 27.05 [ -8.16, 62.27]
HUNT 15.55 [ -1.30, 32.40]
IMAGEN 6.40 [ -7.11, 19.91]
IMpACT -17.85 [-50.14, 14.44]
LBC1936 16.38 [ -3.18, 35.95]
LIBD 40.79 [ 12.18, 69.39]
Sydney MAS 8.86 [-12.50, 30.22]
MCIC -4.09 [-41.72, 33.54]
MooDS 14.21 [-15.80, 44.22]
MPIP 14.02 [ -6.08, 34.12]
MUNSTER 20.50 [ 3.33, 37.67]
NCNG -13.99 [-42.62, 14.63]
NESDA 1.43 [-34.00, 36.85]
NeuroIMAGE -14.14 [-48.61, 20.32]
NTR 5.59 [-24.11, 35.28]
OATS 13.58 [-15.92, 43.08]
PAFIP 7.67 [-41.25, 56.59]
PDNZ -8.82 [-58.04, 40.40]
PING 1.04 [-27.18, 29.26]
MPRC 20.05 [ -8.74, 48.84]
PPMI
QTIM 15.78 [ -1.97, 33.52]
SHIP 27.75 [ 11.50, 44.00]
SHIP-Trend 6.26 [-11.18, 23.71]
SYS 31.51 [ -0.78, 63.80]
TCD-NUIG 23.55 [ -9.07, 56.17]
TOP 4.28 [-18.70, 27.26]
TOP3T 16.34 [-11.28, 43.95]
UiO2016 -4.29 [-33.23, 24.66]
UiO2017 8.76 [-20.08, 37.60]
UMCU 7.70 [-10.91, 26.32]
ENIGMA Discovery Cohorts [N=22721; P=1.336e-11] 12.12 [ 8.61, 15.63]
UKBB 8.19 [ 3.04, 13.34]
Discovery+UKBB [N=31976; P=1.212e-12] 10.83 [ 7.84, 13.82]
GOBS -0.12 [-14.78, 14.55]
IMH -9.01 [-57.66, 39.63]
UNICAMP 28.20 [ -9.63, 66.03]
Meth-CT 16.85 [-39.10, 72.79]
MIRECC 9.54 [-22.10, 41.19]
UKBB nonEUR -28.90 [-59.01, 1.22]
OSAKA 14.47 [-10.61, 39.54]
PING Non European 8.30 [-25.02, 41.63]
Non European Cohorts [N=2876; P=0.5829] 2.74 [ -7.04, 12.52]
Beta [Lower 95% CI,Upper 95% CI]
-100
-50
0
50
100
mm^2/effect allele

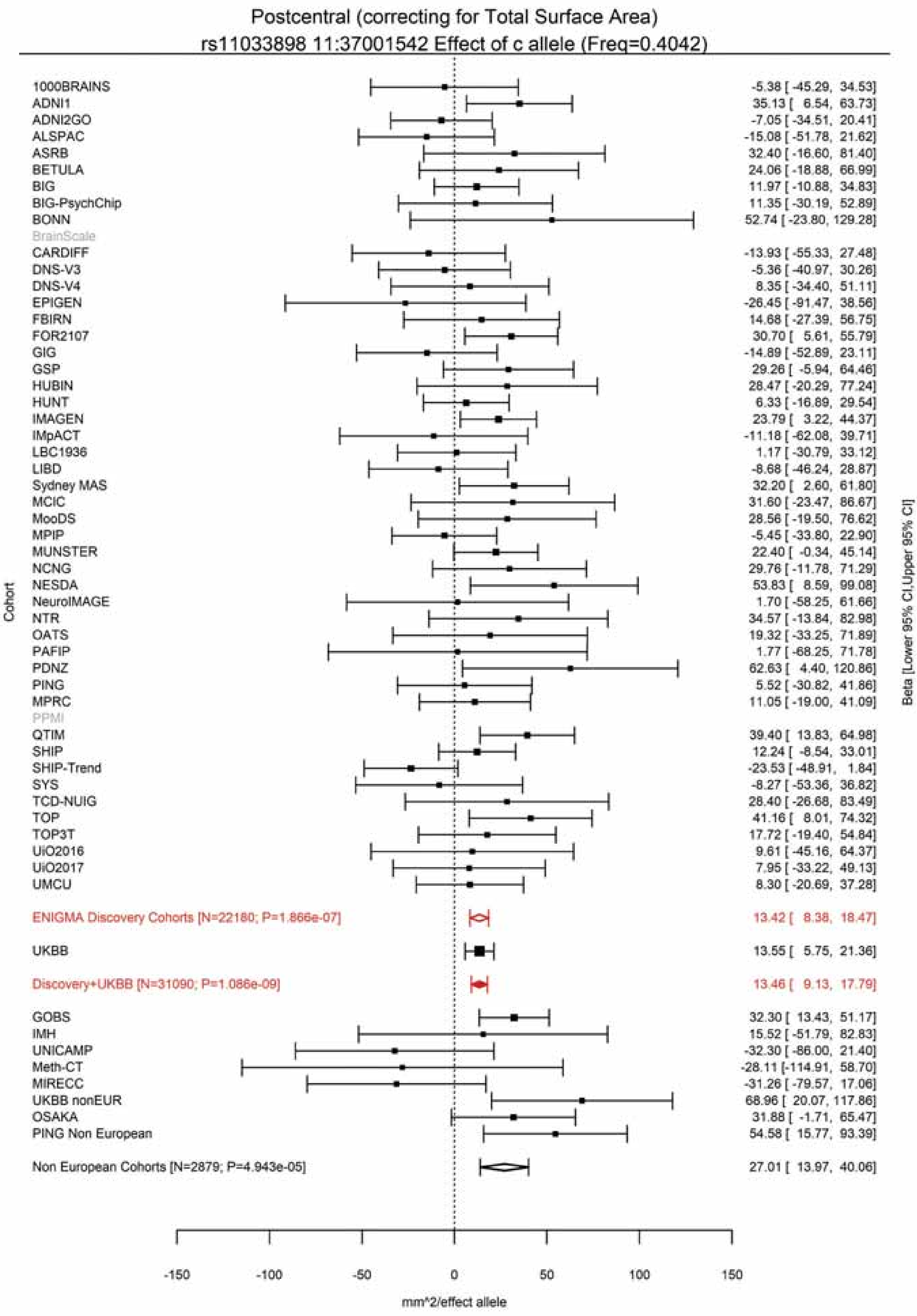
Postcentral (correcting for Total Surface Area)
rs11033898 11:37001542 Effect of c allele (Freq=0.4042)
1000BRAINS -5.38 [ -45.29, 34.53]
ADNI1 35.13 [ 6.54, 63.73]
ADNI2GO -7.05 [ -34.51, 20.41]
ALSPAC -15.08 [ -51.78, 21.62]
ASRB 32.40 [ -16.60, 81.40]
BETULA 24.06 [ -18.88, 66.99]
BIG 11.97 [ -10.88, 34.83]
BIG-PsychChip 11.35 [ -30.19, 52.89]
BONN 52.74 [ -23.80, 129.28]
BrainScale
CARDIFF -13.93 [ -55.33, 27.48]
DNS-V3 -5.36 [ -40.97, 30.26]
DNS-V4 8.35 [ -34.40, 51.11]
EPIGEN -26.45 [ -91.47, 38.56]
FBIRN 14.68 [ -27.39, 56.75]
FOR2107 30.70 [ 5.61, 55.79]
GIG -14.89 [ -52.89, 23.11]
GSP 29.26 [ -5.94, 64.46]
HUBIN 28.47 [ -20.29, 77.24]
HUNT 6.33 [ -16.89, 29.54]
IMAGEN 23.79 [ 3.22, 44.37]
IMpACT -11.18 [ -62.08, 39.71]
LBC1936 1.17 [ -30.79, 33.12]
LIBD -8.68 [ -46.24, 28.87]
Sydney MAS 32.20 [ 2.60, 61.80]
MCIC 31.60 [ -23.47, 86.67]
MooDS 28.56 [ -19.50, 76.62]
MPIP -5.45 [ -33.80, 22.90]
MUNSTER 22.40 [ -0.34, 45.14]
NCNG 29.76 [ -11.78, 71.29]
NESDA 53.83 [ 8.59, 99.08]
NeuroIMAGE 1.70 [ -58.25, 61.66]
NTR 34.57 [ -13.84, 82.98]
OATS 19.32 [ -33.25, 71.89]
PAFIP 1.77 [ -68.25, 71.78]
PDNZ 62.63 [ 4.40, 120.86]
PING 5.52 [ -30.82, 41.86]
MPRC 11.05 [ -19.00, 41.09]
PPMI
QTIM 39.40 [ 13.83, 64.98]
SHIP 12.24 [ -8.54, 33.01]
SHIP-Trend -23.53 [ -48.91, 1.84]
SYS -8.27 [ -53.36, 36.82]
TCD-NUIG 28.40 [ -26.68, 83.49]
TOP 41.16 [ 8.01, 74.32]
TOP3T 17.72 [ -19.40, 54.84]
UiO2016 9.61 [ -45.16, 64.37]
UiO2017 7.95 [ -33.22, 49.13]
UMCU 8.30 [ -20.69, 37.28]
ENIGMA Discovery Cohorts [N=22180; P=1.866e-07] 13.42 [ 8.38, 18.47]
UKBB 13.55 [ 5.75, 21.36]
Discovery+UKBB [N=31090; P=1.086e-09] 13.46 [ 9.13, 17.79]
GOBS 32.30 [ 13.43, 51.17]
IMH 15.52 [ -51.79, 82.83]
UNICAMP -32.30 [ -86.00, 21.40]
Meth-CT -28.11 [-114.91, 58.70]
MIRECC -31.26 [ -79.57, 17.06]
UKBB nonEUR 68.96 [ 20.07, 117.86]
OSAKA 31.88 [ -1.71, 65.47]
PING Non European 54.58 [ 15.77, 93.39]
Non European Cohorts [N=2879; P=4.943e-05] 27.01 [ 13.97, 40.06]
Cohort
Beta [Lower 95% CI,Upper 95% CI]
-150
-100
-50
0
50
100
150
mm^2/effect allele

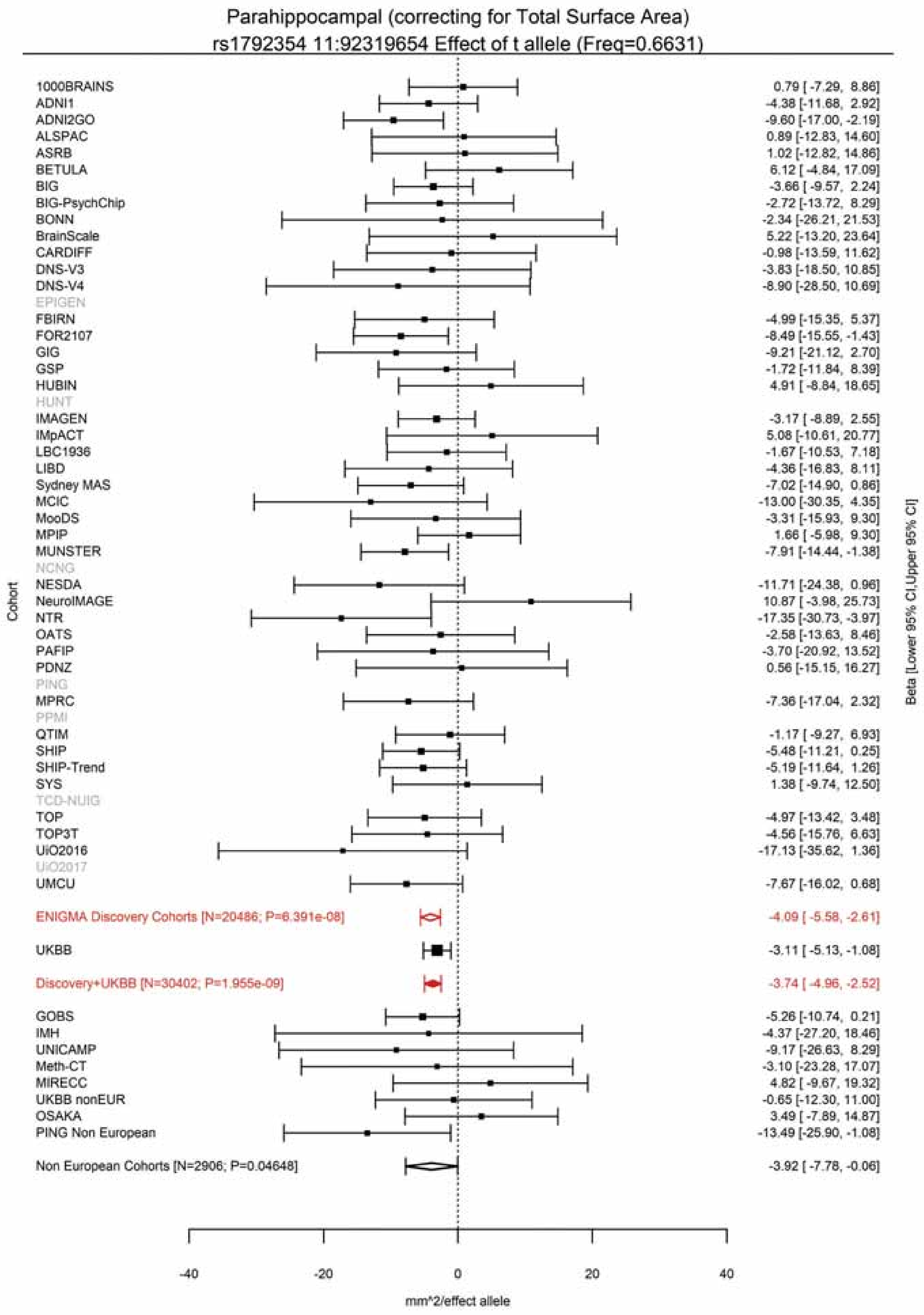

Parahippocampal (correcting for Total Surface Area)
rs1792354 11:92319654 Effect of t allele (Freq=0.6631)
Cohort
1000BRAINS 0.79 [ -7.29, 8.86]
ADNI1 -4.38 [-11.68, 2.92]
ADNI2GO -9.60 [-17.00, -2.19]
ALSPAC 0.89 [-12.83, 14.60]
ASRB 1.02 [-12.82, 14.86]
BETULA 6.12 [ -4.84, 17.09]
BIG -3.66 [ -9.57, 2.24]
BIG-PsychChip -2.72 [-13.72, 8.29]
BONN -2.34 [-26.21, 21.53]
BrainScale 5.22 [-13.20, 23.64]
CARDIFF -0.98 [-13.59, 11.62]
DNS-V3 -3.83 [-18.50, 10.85]
DNS-V4 -8.90 [-28.50, 10.69]
EPIGEN
FBIRN -4.99 [-15.35, 5.37]
FOR2107 -8.49 [-15.55, -1.43]
GIG -9.21 [-21.12, 2.70]
GSP -1.72 [-11.84, 8.39]
HUBIN 4.91 [ -8.84, 18.65]
HUNT
IMAGEN -3.17 [ -8.89, 2.55]
IMpACT 5.08 [-10.61, 20.77]
LBC1936 -1.67 [-10.53, 7.18]
LIBD -4.36 [-16.83, 8.11]
Sydney MAS -7.02 [-14.90, 0.86]
MCIC -13.00 [-30.35, 4.35]
MooDS -3.31 [-15.93, 9.30]
MPIP 1.66 [ -5.98, 9.30]
MUNSTER -7.91 [-14.44, -1.38]
NCNG
NESDA -11.71 [-24.38, 0.96]
NeuroIMAGE 10.87 [ -3.98, 25.73]
NTR -17.35 [-30.73, -3.97]
OATS -2.58 [-13.63, 8.46]
PAFIP -3.70 [-20.92, 13.52]
PDNZ 0.56 [-15.15, 16.27]
PING
MPRC -7.36 [-17.04, 2.32]
PPMI
QTIM -1.17 [ -9.27, 6.93]
SHIP -5.48 [-11.21, 0.25]
SHIP-Trend -5.19 [-11.64, 1.26]
SYS 1.38 [ -9.74, 12.50]
TCD-NUIG
TOP -4.97 [-13.42, 3.48]
TOP3T -4.56 [-15.76, 6.63]
UiO2016 -17.13 [-35.62, 1.36]
UiO2017
UMCU -7.67 [-16.02, 0.68]
ENIGMA Discovery Cohorts [N=20486; P=6.391e-08] -4.09 [ -5.58, -2.61]
UKBB -3.11 [ -5.13, -1.08]
Discovery+UKBB [N=30402; P=1.955e-09] -3.74 [ -4.96, -2.52]
GOBS -5.26 [-10.74, 0.21]
IMH -4.37 [-27.20, 18.46]
UNICAMP -9.17 [-26.63, 8.29]
Meth-CT -3.10 [-23.28, 17.07]
MIRECC 4.82 [ -9.67, 19.32]
UKBB nonEUR -0.65 [-12.30, 11.00]
OSAKA 3.49 [ -7.89, 14.87]
PING Non European -13.49 [-25.90, -1.08]
Non European Cohorts [N=2906; P=0.04648] -3.92 [ -7.78, -0.06]
Beta [Lower 95% CI,Upper 95% CI]
-40
-20
0
20
40
mm^2/effect allele

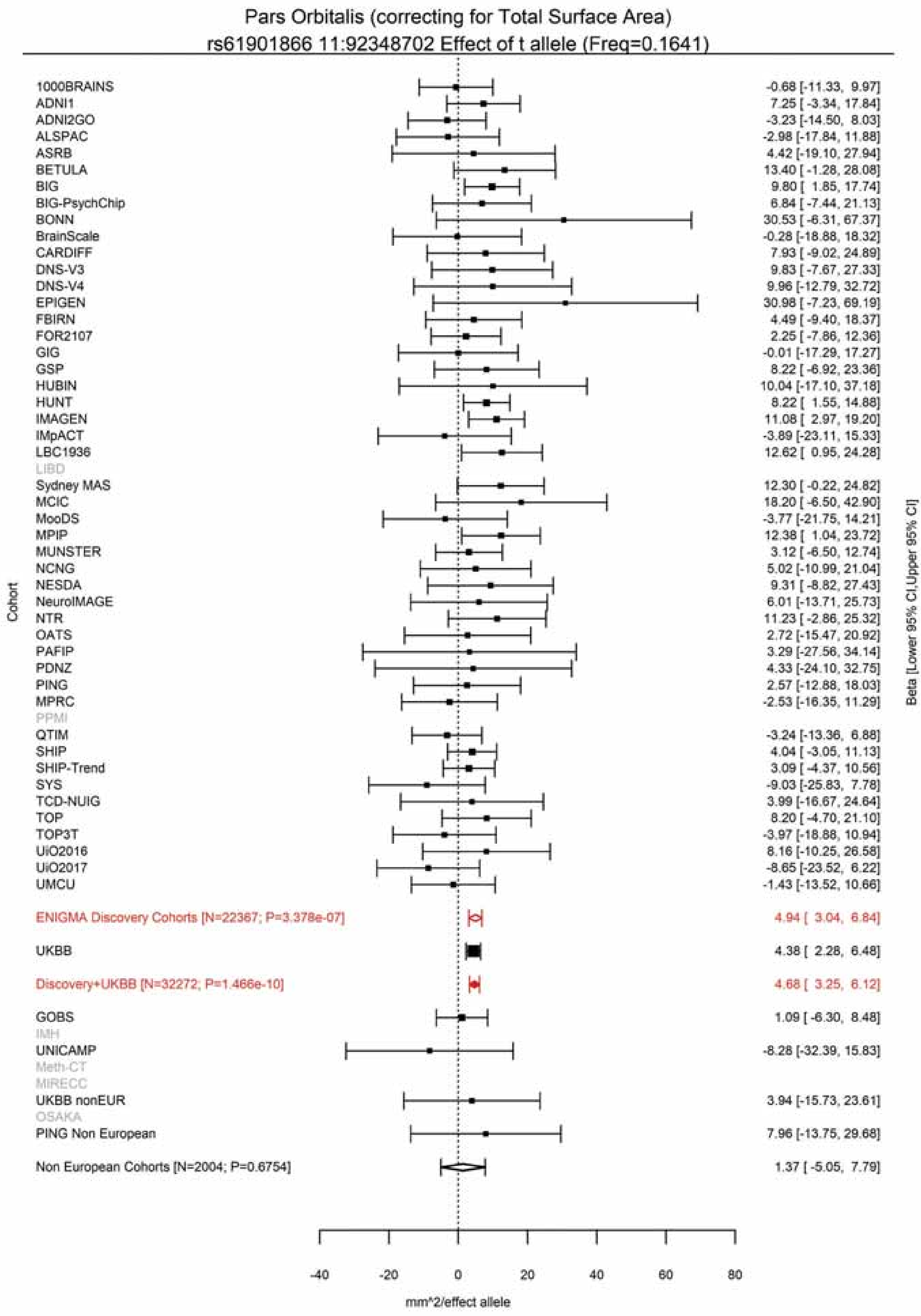
Pars Orbitalis (correcting for Total Surface Area)
rs61901866 11:92348702 Effect of t allele (Freq=0.1641)
1000BRAINS -0.68 [-11.33, 9.97]
ADNI1 7.25 [ -3.34, 17.84]
ADNI2GO -3.23 [-14.50, 8.03]
ALSPAC -2.98 [-17.84, 11.88]
ASRB 4.42 [-19.10, 27.94]
BETULA 13.40 [ -1.28, 28.08]
BIG 9.80 [ 1.85, 17.74]
BIG-PsychChip 6.84 [ -7.44, 21.13]
BONN 30.53 [ -6.31, 67.37]
BrainScale -0.28 [-18.88, 18.32]
CARDIFF 7.93 [ -9.02, 24.89]
DNS-V3 9.83 [ -7.67, 27.33]
DNS-V4 9.96 [-12.79, 32.72]
EPIGEN 30.98 [ -7.23, 69.19]
FBIRN 4.49 [ -9.40, 18.37]
FOR2107 2.25 [ -7.86, 12.36]
GIG -0.01 [-17.29, 17.27]
GSP 8.22 [ -6.92, 23.36]
HUBIN 10.04 [-17.10, 37.18]
HUNT 8.22 [ 1.55, 14.88]
IMAGEN 11.08 [ 2.97, 19.20]
IMpACT -3.89 [-23.11, 15.33]
LBC1936 12.62 [ 0.95, 24.28]
LIBD
Sydney MAS 12.30 [ -0.22, 24.82]
MCIC 18.20 [ -6.50, 42.90]
MooDS -3.77 [-21.75, 14.21]
MPIP 12.38 [ 1.04, 23.72]
MUNSTER 3.12 [ -6.50, 12.74]
NCNG 5.02 [-10.99, 21.04]
NESDA 9.31 [ -8.82, 27.43]
NeuroIMAGE 6.01 [-13.71, 25.73]
NTR 11.23 [ -2.86, 25.32]
OATS 2.72 [-15.47, 20.92]
PAFIP 3.29 [-27.56, 34.14]
PDNZ 4.33 [-24.10, 32.75]
PING 2.57 [-12.88, 18.03]
MPRC -2.53 [-16.35, 11.29]
PPMI
QTIM -3.24 [-13.36, 6.88]
SHIP 4.04 [ -3.05, 11.13]
SHIP-Trend 3.09 [ -4.37, 10.56]
SYS -9.03 [-25.83, 7.78]
TCD-NUIG 3.99 [-16.67, 24.64]
TOP 8.20 [ -4.70, 21.10]
TOP3T -3.97 [-18.88, 10.94]
UiO2016 8.16 [-10.25, 26.58]
UiO2017 -8.65 [-23.52, 6.22]
UMCU -1.43 [-13.52, 10.66]
ENIGMA Discovery Cohorts [N=22367; P=3.378e-07] 4.94 [ 3.04, 6.84]
UKBB 4.38 [ 2.28, 6.48]
Discovery+UKBB [N=32272; P=1.466e-10] 4.68 [ 3.25, 6.12]
GOBS 1.09 [ -6.30, 8.48]
IMH
UNICAMP -8.28 [-32.39, 15.83]
Meth-CT
MIRECC
UKBB nonEUR 3.94 [-15.73, 23.61]
OSAKA
PING Non European 7.96 [-13.75, 29.68]
Non European Cohorts [N=2004; P=0.6754] 1.37 [ -5.05, 7.79]
Cohort
Beta [Lower 95% CI,Upper 95% CI]
-40
-20
0
20
40
60
80
mm^2/effect allele

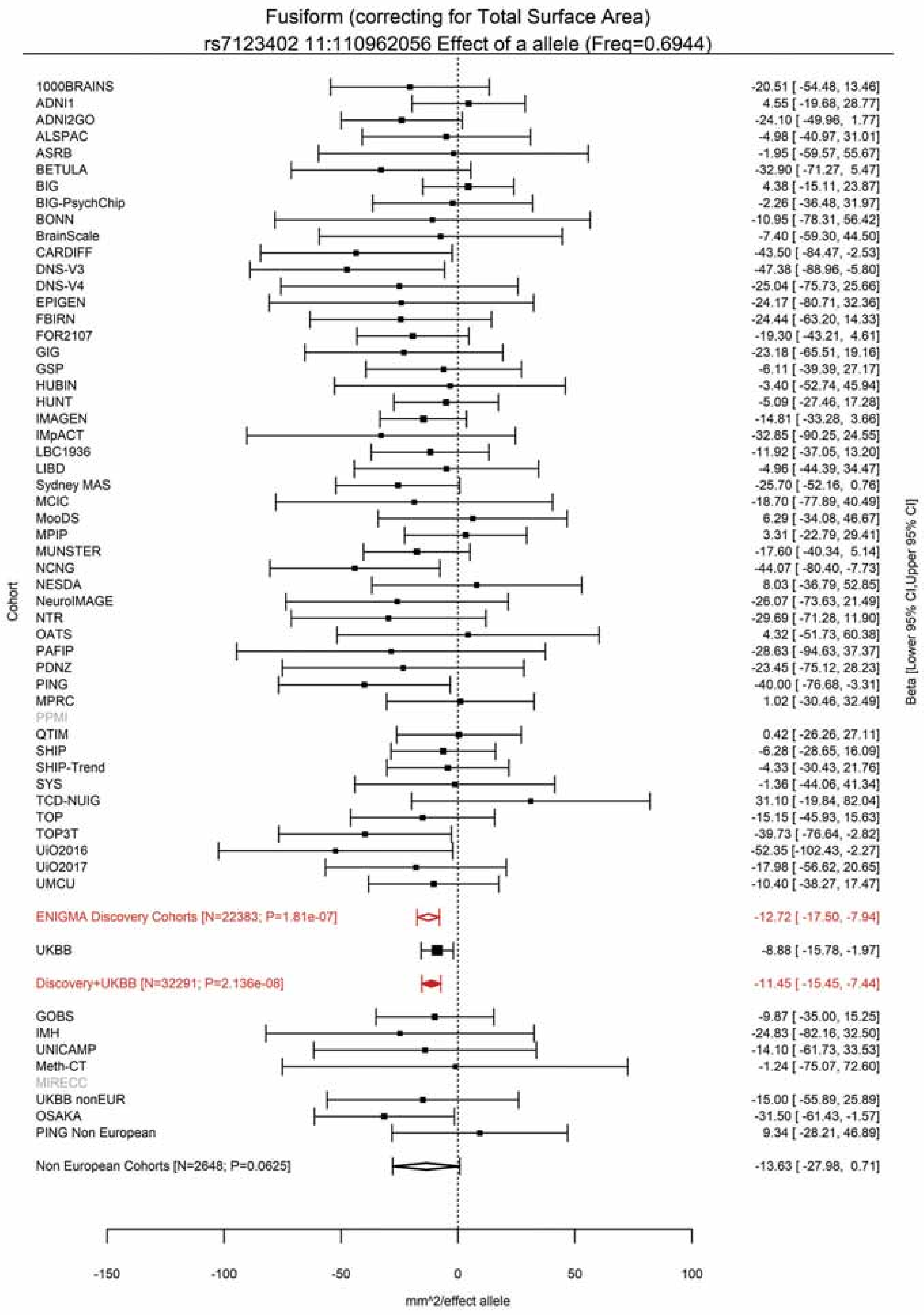
Fusiform (correcting for Total Surface Area)
rs7123402 11:110962056 Effect of a allele (Freq=0.6944)
Cohort
Beta [Lower 95% CI,Upper 95% CI]
1000BRAINS -20.51 [ -54.48, 13.46]
ADNI1 4.55 [ -19.68, 28.77]
ADNI2GO -24.10 [ -49.96, 1.77]
ALSPAC -4.98 [ -40.97, 31.01]
ASRB -1.95 [ -59.57, 55.67]
BETULA -32.90 [ -71.27, 5.47]
BIG 4.38 [ -15.11, 23.87]
BIG-PsychChip -2.26 [ -36.48, 31.97]
BONN -10.95 [ -78.31, 56.42]
BrainScale -7.40 [ -59.30, 44.50]
CARDIFF -43.50 [ -84.47, -2.53]
DNS-V3 -47.38 [ -88.96, -5.80]
DNS-V4 -25.04 [ -75.73, 25.66]
EPIGEN -24.17 [ -80.71, 32.36]
FBIRN -24.44 [ -63.20, 14.33]
FOR2107 -19.30 [ -43.21, 4.61]
GIG -23.18 [ -65.51, 19.16]
GSP -6.11 [ -39.39, 27.17]
HUBIN -3.40 [ -52.74, 45.94]
HUNT -5.09 [ -27.46, 17.28]
IMAGEN -14.81 [ -33.28, 3.66]
IMpACT -32.85 [ -90.25, 24.55]
LBC1936 -11.92 [ -37.05, 13.20]
LIBD -4.96 [ -44.39, 34.47]
Sydney MAS -25.70 [ -52.16, 0.76]
MCIC -18.70 [ -77.89, 40.49]
MooDS 6.29 [ -34.08, 46.67]
MPIP 3.31 [ -22.79, 29.41]
MUNSTER -17.60 [ -40.34, 5.14]
NCNG -44.07 [ -80.40, -7.73]
NESDA 8.03 [ -36.79, 52.85]
NeuroIMAGE -26.07 [ -73.63, 21.49]
NTR -29.69 [ -71.28, 11.90]
OATS 4.32 [ -51.73, 60.38]
PAFIP -28.63 [ -94.63, 37.37]
PDNZ -23.45 [ -75.12, 28.23]
PING -40.00 [ -76.68, -3.31]
MPRC 1.02 [ -30.46, 32.49]
PPMI
QTIM 0.42 [ -26.26, 27.11]
SHIP -6.28 [ -28.65, 16.09]
SHIP-Trend -4.33 [ -30.43, 21.76]
SYS -1.36 [ -44.06, 41.34]
TCD-NUIG 31.10 [ -19.84, 82.04]
TOP -15.15 [ -45.93, 15.63]
TOP3T -39.73 [ -76.64, -2.82]
UiO2016 -52.35 [-102.43, -2.27]
UiO2017 -17.98 [ -56.62, 20.65]
UMCU -10.40 [ -38.27, 17.47]
ENIGMA Discovery Cohorts [N=22383; P=1.81e-07] -12.72 [ -17.50, -7.94]
UKBB -8.88 [ -15.78, -1.97]
Discovery+UKBB [N=32291; P=2.136e-08] -11.45 [ -15.45, -7.44]
GOBS -9.87 [ -35.00, 15.25]
IMH -24.83 [ -82.16, 32.50]
UNICAMP -14.10 [ -61.73, 33.53]
Meth-CT -1.24 [ -75.07, 72.60]
MIRECC
UKBB nonEUR -15.00 [ -55.89, 25.89]
OSAKA -31.50 [ -61.43, -1.57]
PING Non European 9.34 [ -28.21, 46.89]
Non European Cohorts [N=2648; P=0.0625] -13.63 [ -27.98, 0.71]
-150
-100
-50
0
50
100
mm^2/effect allele

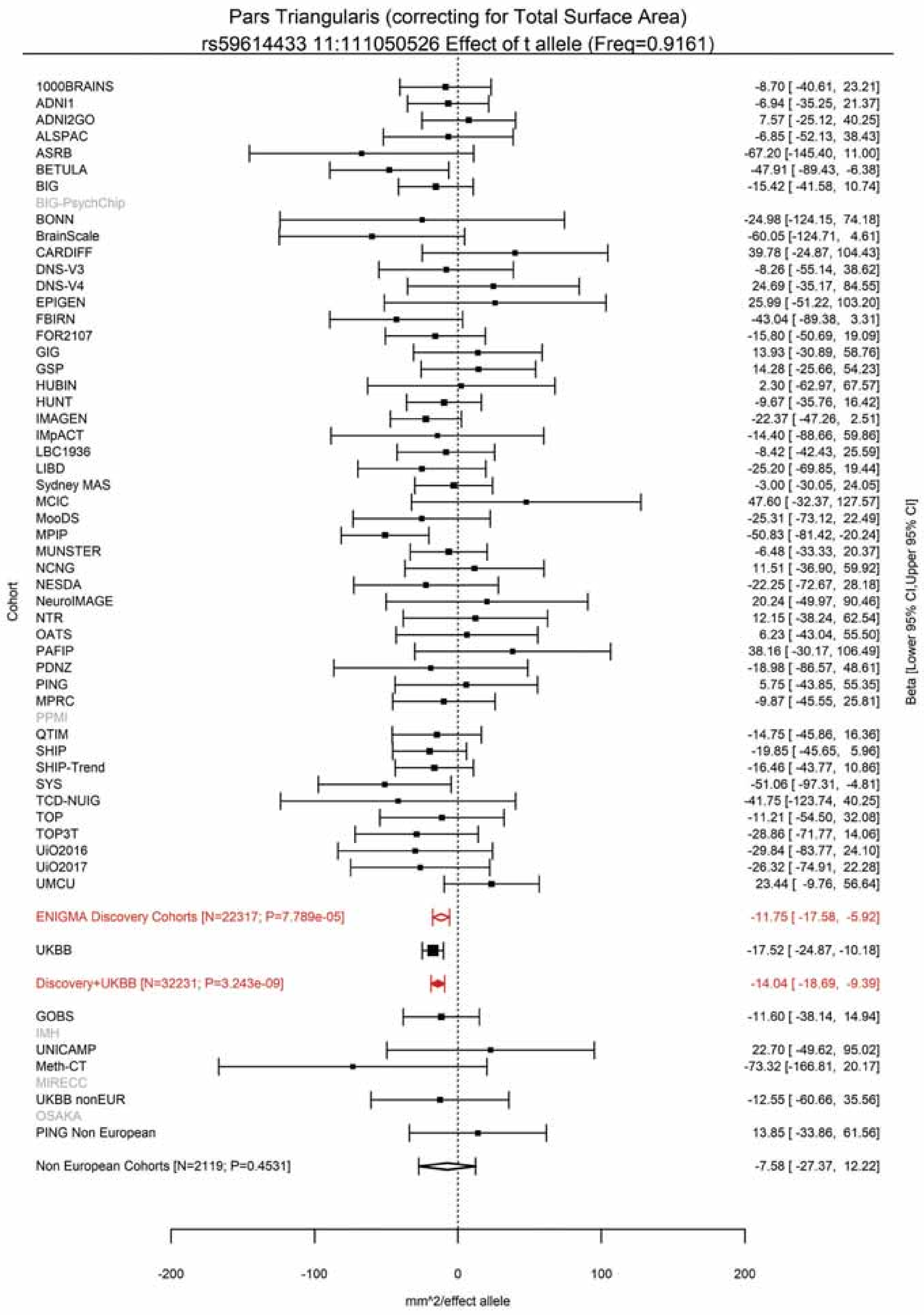

Pars Triangularis (correcting for Total Surface Area)
rs59614433 11:111050526 Effect of t allele (Freq=0.9161)
Cohort
1000BRAINS -8.70 [ -40.61, 23.21]
ADNI1 -6.94 [ -35.25, 21.37]
ADNI2GO 7.57 [ -25.12, 40.25]
ALSPAC -6.85 [ -52.13, 38.43]
ASRB -67.20 [-145.40, 11.00]
BETULA -47.91 [ -89.43, -6.38]
BIG -15.42 [ -41.58, 10.74]
BIG-PsychChip
BONN -24.98 [-124.15, 74.18]
BrainScale -60.05 [-124.71, 4.61]
CARDIFF 39.78 [ -24.87, 104.43]
DNS-V3 -8.26 [ -55.14, 38.62]
DNS-V4 24.69 [ -35.17, 84.55]
EPIGEN 25.99 [ -51.22, 103.20]
FBIRN -43.04 [ -89.38, 3.31]
FOR2107 -15.80 [ -50.69, 19.09]
GIG 13.93 [ -30.89, 58.76]
GSP 14.28 [ -25.66, 54.23]
HUBIN 2.30 [ -62.97, 67.57]
HUNT -9.67 [ -35.76, 16.42]
IMAGEN -22.37 [ -47.26, 2.51]
IMpACT -14.40 [ -88.66, 59.86]
LBC1936 -8.42 [ -42.43, 25.59]
LIBD -25.20 [ -69.85, 19.44]
Sydney MAS -3.00 [ -30.05, 24.05]
MCIC 47.60 [ -32.37, 127.57]
MooDS -25.31 [ -73.12, 22.49]
MPIP -50.83 [ -81.42, -20.24]
MUNSTER -6.48 [ -33.33, 20.37]
NCNG 11.51 [ -36.90, 59.92]
NESDA -22.25 [ -72.67, 28.18]
NeuroIMAGE 20.24 [ -49.97, 90.46]
NTR 12.15 [ -38.24, 62.54]
OATS 6.23 [ -43.04, 55.50]
PAFIP 38.16 [ -30.17, 106.49]
PDNZ -18.98 [ -86.57, 48.61]
PING 5.75 [ -43.85, 55.35]
MPRC -9.87 [ -45.55, 25.81]
PPMI
QTIM -14.75 [ -45.86, 16.36]
SHIP -19.85 [ -45.65, 5.96]
SHIP-Trend -16.46 [ -43.77, 10.86]
SYS -51.06 [ -97.31, -4.81]
TCD-NUIG -41.75 [-123.74, 40.25]
TOP -11.21 [ -54.50, 32.08]
TOP3T -28.86 [ -71.77, 14.06]
UiO2016 -29.84 [ -83.77, 24.10]
UiO2017 -26.32 [ -74.91, 22.28]
UMCU 23.44 [ -9.76, 56.64]
ENIGMA Discovery Cohorts [N=22317; P=7.789e-05] -11.75 [ -17.58, -5.92]
UKBB -17.52 [ -24.87, -10.18]
Discovery+UKBB [N=32231; P=3.243e-09] -14.04 [ -18.69, -9.39]
GOBS -11.60 [ -38.14, 14.94]
IMH
UNICAMP 22.70 [ -49.62, 95.02]
Meth-CT -73.32 [-166.81, 20.17]
MIRECC
UKBB nonEUR -12.55 [ -60.66, 35.56]
OSAKA
PING Non European 13.85 [ -33.86, 61.56]
Non European Cohorts [N=2119; P=0.4531] -7.58 [ -27.37, 12.22]
Beta [Lower 95% CI,Upper 95% CI]
-200
-100
0
100
200
mm^2/effect allele

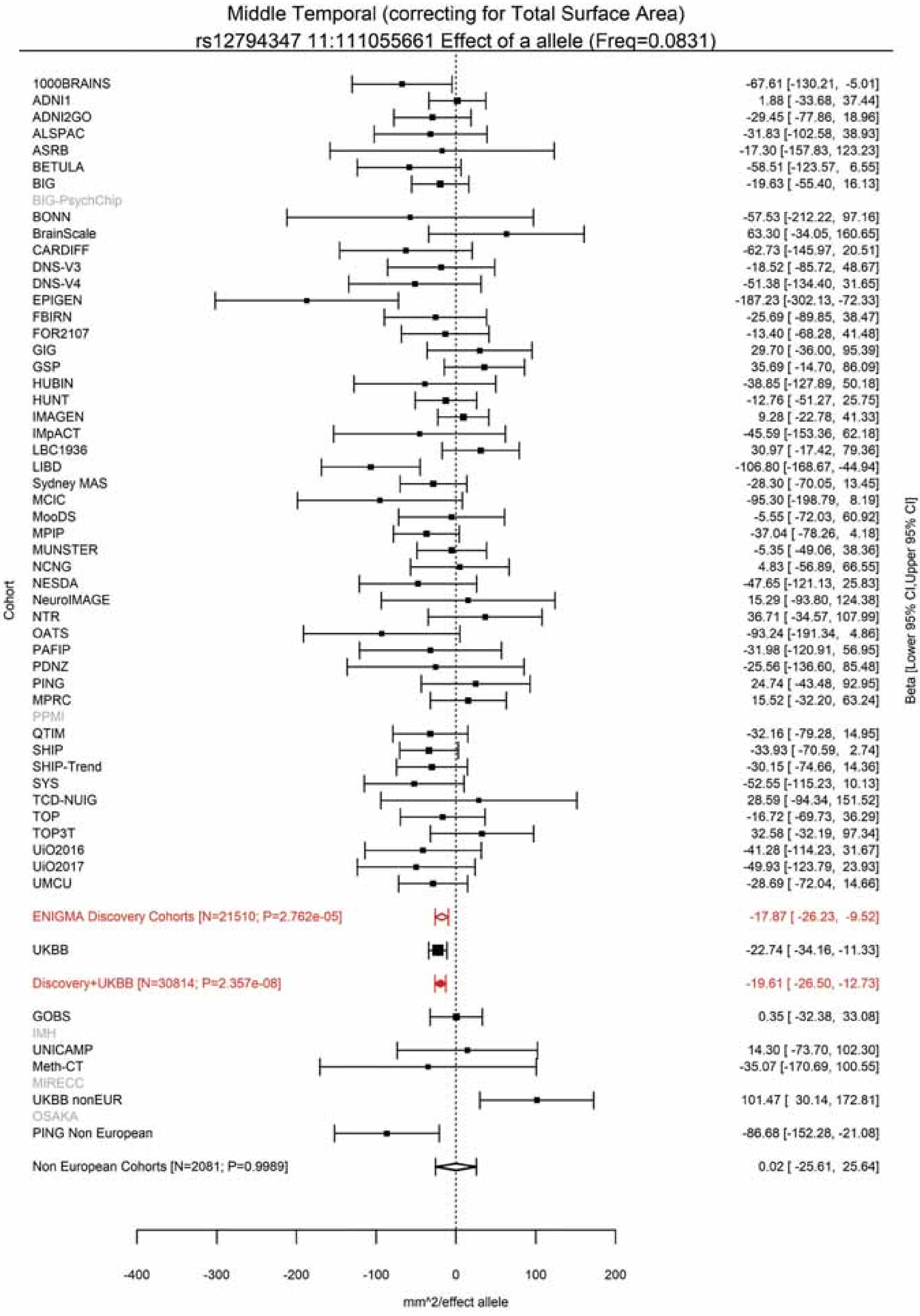
Middle Temporal (correcting for Total Surface Area)
rs12794347 11:111055661 Effect of a allele (Freq=0.0831)
1000BRAINS -67.61 [-130.21, -5.01]
ADNI1 1.88 [ -33.68, 37.44]
ADNI2GO -29.45 [ -77.86, 18.96]
ALSPAC -31.83 [-102.58, 38.93]
ASRB -17.30 [-157.83, 123.23]
BETULA -58.51 [-123.57, 6.55]
BIG -19.63 [ -55.40, 16.13]
BIG-PsychChip
BONN -57.53 [-212.22, 97.16]
BrainScale 63.30 [ -34.05, 160.65]
CARDIFF -62.73 [-145.97, 20.51]
DNS-V3 -18.52 [ -85.72, 48.67]
DNS-V4 -51.38 [-134.40, 31.65]
EPIGEN -187.23 [-302.13, -72.33]
FBIRN -25.69 [ -89.85, 38.47]
FOR2107 -13.40 [ -68.28, 41.48]
GIG 29.70 [ -36.00, 95.39]
GSP 35.69 [ -14.70, 86.09]
HUBIN -38.85 [-127.89, 50.18]
HUNT -12.76 [ -51.27, 25.75]
IMAGEN 9.28 [ -22.78, 41.33]
IMpACT -45.59 [-153.36, 62.18]
LBC1936 30.97 [ -17.42, 79.36]
LIBD -106.80 [-168.67, -44.94]
Sydney MAS -28.30 [ -70.05, 13.45]
MCIC -95.30 [-198.79, 8.19]
MooDS -5.55 [ -72.03, 60.92]
MPIP -37.04 [ -78.26, 4.18]
MUNSTER -5.35 [ -49.06, 38.36]
NCNG 4.83 [ -56.89, 66.55]
NESDA -47.65 [-121.13, 25.83]
NeuroIMAGE 15.29 [ -93.80, 124.38]
NTR 36.71 [ -34.57, 107.99]
OATS -93.24 [-191.34, 4.86]
PAFIP -31.98 [-120.91, 56.95]
PDNZ -25.56 [-136.60, 85.48]
PING 24.74 [ -43.48, 92.95]
MPRC 15.52 [ -32.20, 63.24]
PPMI
QTIM -32.16 [ -79.28, 14.95]
SHIP -33.93 [ -70.59, 2.74]
SHIP-Trend -30.15 [ -74.66, 14.36]
SYS -52.55 [-115.23, 10.13]
TCD-NUIG 28.59 [ -94.34, 151.52]
TOP -16.72 [ -69.73, 36.29]
TOP3T 32.58 [ -32.19, 97.34]
UiO2016 -41.28 [-114.23, 31.67]
UiO2017 -49.93 [-123.79, 23.93]
UMCU -28.69 [ -72.04, 14.66]
ENIGMA Discovery Cohorts [N=21510; P=2.762e-05] -17.87 [ -26.23, -9.52]
UKBB -22.74 [ -34.16, -11.33]
Discovery+UKBB [N=30814; P=2.357e-08] -19.61 [ -26.50, -12.73]
GOBS 0.35 [ -32.38, 33.08]
IMH
UNICAMP 14.30 [ -73.70, 102.30]
Meth-CT -35.07 [-170.69, 100.55]
MIRECC
UKBB nonEUR 101.47 [ 30.14, 172.81]
OSAKA
PING Non European -86.68 [-152.28, -21.08]
Non European Cohorts [N=2081; P=0.9989] 0.02 [ -25.61, 25.64]
Cohort
Beta [Lower 95% CI,Upper 95% CI]
-400
-300
-200
-100
0
100
200
mm^2/effect allele

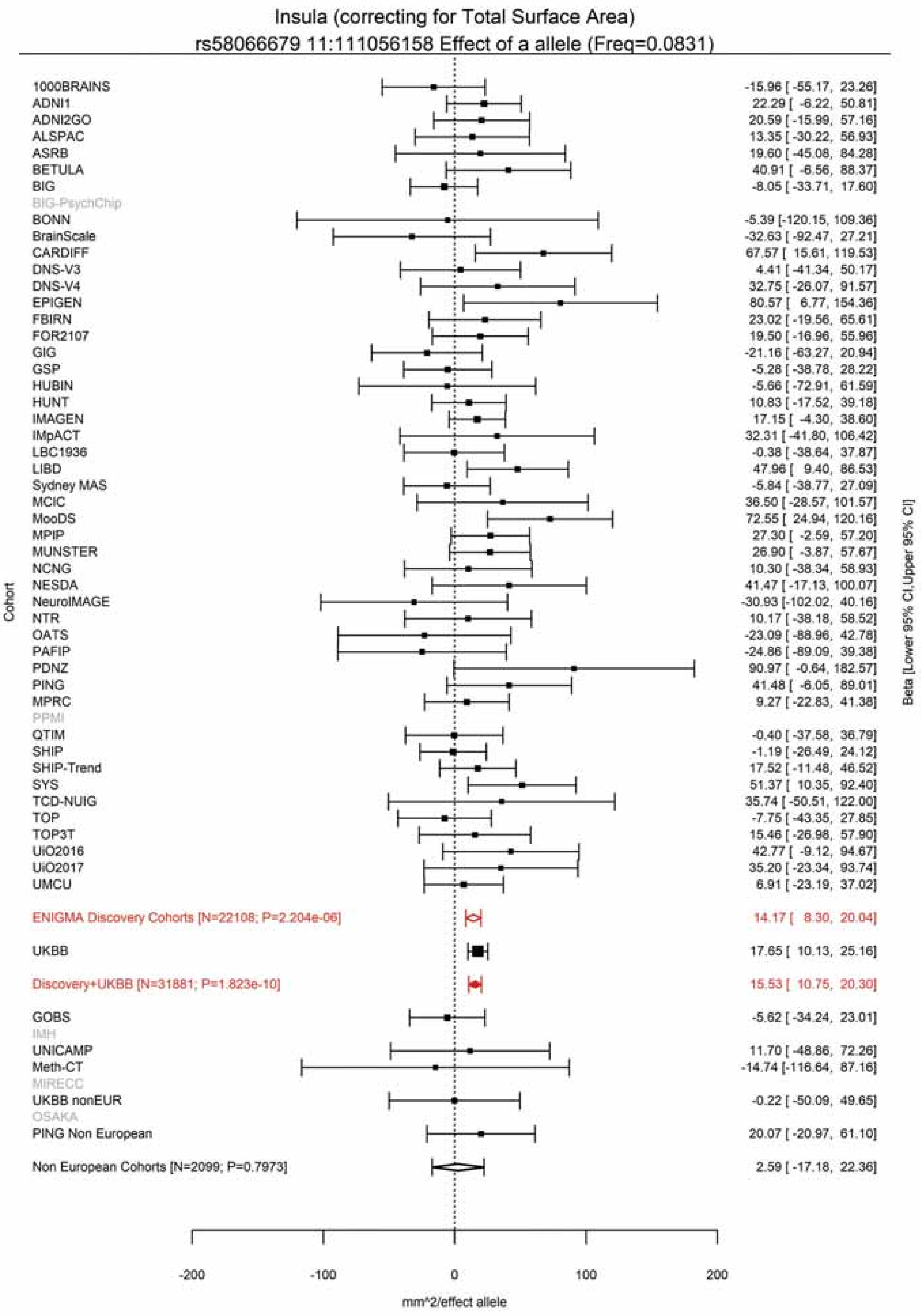

Insula (correcting for Total Surface Area)
rs58066679 11:111056158 Effect of a allele (Freq=0.0831)
1000BRAINS -15.96 [ -55.17, 23.26]
ADNI1 22.29 [ -6.22, 50.81]
ADNI2GO 20.59 [ -15.99, 57.16]
ALSPAC 13.35 [ -30.22, 56.93]
ASRB 19.60 [ -45.08, 84.28]
BETULA 40.91 [ -6.56, 88.37]
BIG -8.05 [ -33.71, 17.60]
BIG-PsychChip
BONN -5.39 [-120.15, 109.36]
BrainScale -32.63 [ -92.47, 27.21]
CARDIFF 67.57 [ 15.61, 119.53]
DNS-V3 4.41 [ -41.34, 50.17]
DNS-V4 32.75 [ -26.07, 91.57]
EPIGEN 80.57 [ 6.77, 154.36]
FBIRN 23.02 [ -19.56, 65.61]
FOR2107 19.50 [ -16.96, 55.96]
GIG -21.16 [ -63.27, 20.94]
GSP -5.28 [ -38.78, 28.22]
HUBIN -5.66 [ -72.91, 61.59]
HUNT 10.83 [ -17.52, 39.18]
IMAGEN 17.15 [ -4.30, 38.60]
IMpACT 32.31 [ -41.80, 106.42]
LBC1936 -0.38 [ -38.64, 37.87]
LIBD 47.96 [ 9.40, 86.53]
Sydney MAS -5.84 [ -38.77, 27.09]
MCIC 36.50 [ -28.57, 101.57]
MooDS 72.55 [ 24.94, 120.16]
MPIP 27.30 [ -2.59, 57.20]
MUNSTER 26.90 [ -3.87, 57.67]
NCNG 10.30 [ -38.34, 58.93]
NESDA 41.47 [ -17.13, 100.07]
NeuroIMAGE -30.93 [-102.02, 40.16]
NTR 10.17 [ -38.18, 58.52]
OATS -23.09 [ -88.96, 42.78]
PAFIP -24.86 [ -89.09, 39.38]
PDNZ 90.97 [ -0.64, 182.57]
PING 41.48 [ -6.05, 89.01]
MPRC 9.27 [ -22.83, 41.38]
PPMI
QTIM -0.40 [ -37.58, 36.79]
SHIP -1.19 [ -26.49, 24.12]
SHIP-Trend 17.52 [ -11.48, 46.52]
SYS 51.37 [ 10.35, 92.40]
TCD-NUIG 35.74 [ -50.51, 122.00]
TOP -7.75 [ -43.35, 27.85]
TOP3T 15.46 [ -26.98, 57.90]
UiO2016 42.77 [ -9.12, 94.67]
UiO2017 35.20 [ -23.34, 93.74]
UMCU 6.91 [ -23.19, 37.02]
ENIGMA Discovery Cohorts [N=22108; P=2.204e-06] 14.17 [ 8.30, 20.04]
UKBB 17.65 [ 10.13, 25.16]
Discovery+UKBB [N=31881; P=1.823e-10] 15.53 [ 10.75, 20.30]
GOBS -5.62 [ -34.24, 23.01]
IMH
UNICAMP 11.70 [ -48.86, 72.26]
Meth-CT -14.74 [-116.64, 87.16]
MIRECC
UKBB nonEUR -0.22 [ -50.09, 49.65]
OSAKA
PING Non European 20.07 [ -20.97, 61.10]
Non European Cohorts [N=2099; P=0.7973] 2.59 [ -17.18, 22.36]
Cohort
Beta [Lower 95% CI,Upper 95% CI]
-200
-100
0
100
200
mm^2/effect allele

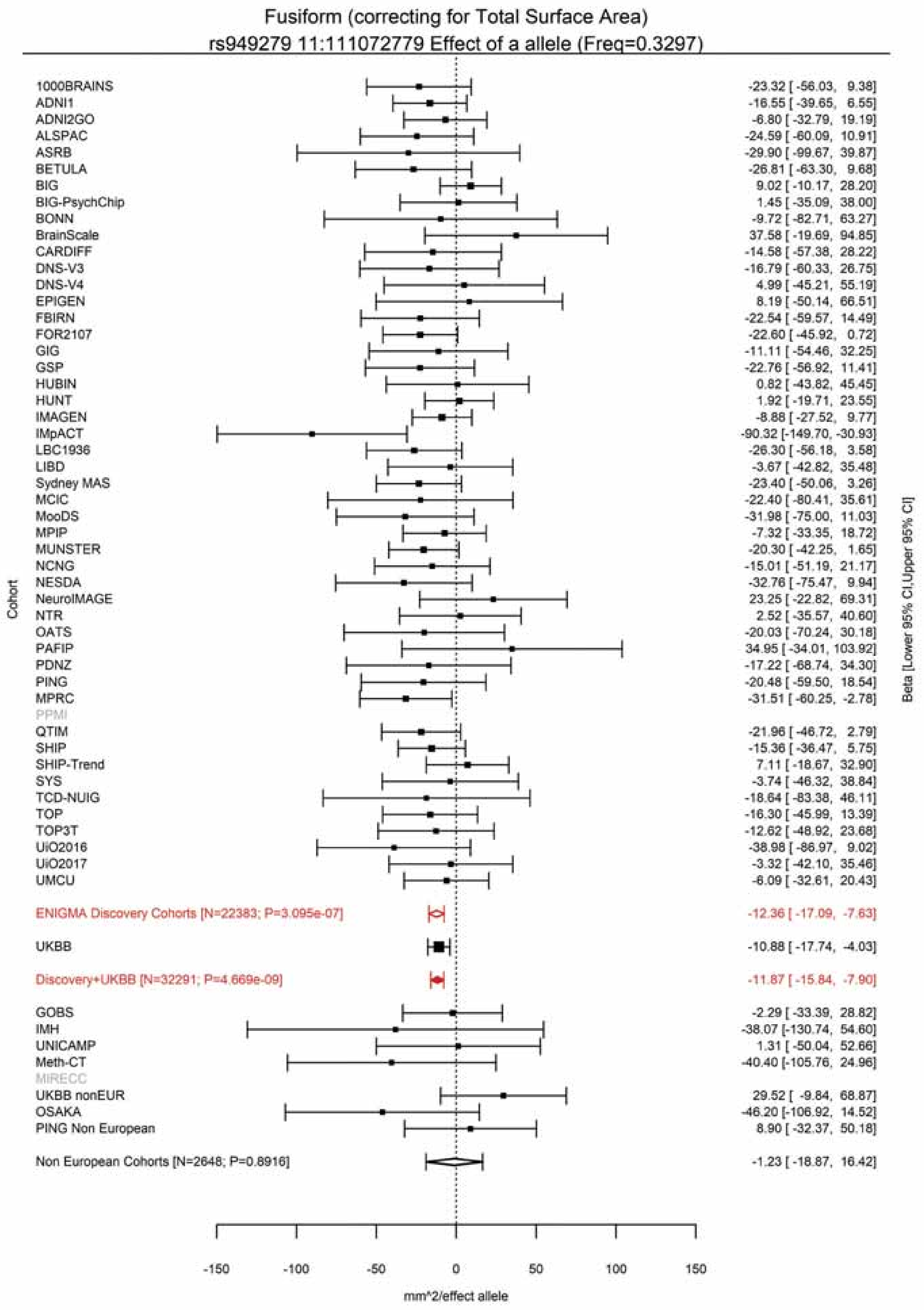
Fusiform (correcting for Total Surface Area)
rs949279 11:111072779 Effect of a allele (Freq=0.3297)
Cohort
1000BRAINS -23.32 [ -56.03, 9.38]
ADNI1 -16.55 [ -39.65, 6.55]
ADNI2GO -6.80 [ -32.79, 19.19]
ALSPAC -24.59 [ -60.09, 10.91]
ASRB -29.90 [ -99.67, 39.87]
BETULA -26.81 [ -63.30, 9.68]
BIG 9.02 [ -10.17, 28.20]
BIG-PsychChip 1.45 [ -35.09, 38.00]
BONN -9.72 [ -82.71, 63.27]
BrainScale 37.58 [ -19.69, 94.85]
CARDIFF -14.58 [ -57.38, 28.22]
DNS-V3 -16.79 [ -60.33, 26.75]
DNS-V4 4.99 [ -45.21, 55.19]
EPIGEN 8.19 [ -50.14, 66.51]
FBIRN -22.54 [ -59.57, 14.49]
FOR2107 -22.60 [ -45.92, 0.72]
GIG -11.11 [ -54.46, 32.25]
GSP -22.76 [ -56.92, 11.41]
HUBIN 0.82 [ -43.82, 45.45]
HUNT 1.92 [ -19.71, 23.55]
IMAGEN -8.88 [ -27.52, 9.77]
IMpACT -90.32 [-149.70, -30.93]
LBC1936 -26.30 [ -56.18, 3.58]
LIBD -3.67 [ -42.82, 35.48]
Sydney MAS -23.40 [ -50.06, 3.26]
MCIC -22.40 [ -80.41, 35.61]
MooDS -31.98 [ -75.00, 11.03]
MPIP -7.32 [ -33.35, 18.72]
MUNSTER -20.30 [ -42.25, 1.65]
NCNG -15.01 [ -51.19, 21.17]
NESDA -32.76 [ -75.47, 9.94]
NeuroIMAGE 23.25 [ -22.82, 69.31]
NTR 2.52 [ -35.57, 40.60]
OATS -20.03 [ -70.24, 30.18]
PAFIP 34.95 [ -34.01, 103.92]
PDNZ -17.22 [ -68.74, 34.30]
PING -20.48 [ -59.50, 18.54]
MPRC -31.51 [ -60.25, -2.78]
PPMI
QTIM -21.96 [ -46.72, 2.79]
SHIP -15.36 [ -36.47, 5.75]
SHIP-Trend 7.11 [ -18.67, 32.90]
SYS -3.74 [ -46.32, 38.84]
TCD-NUIG -18.64 [ -83.38, 46.11]
TOP -16.30 [ -45.99, 13.39]
TOP3T -12.62 [ -48.92, 23.68]
UiO2016 -38.98 [ -86.97, 9.02]
UiO2017 -3.32 [ -42.10, 35.46]
UMCU -6.09 [ -32.61, 20.43]
ENIGMA Discovery Cohorts [N=22383; P=3.095e-07] -12.36 [ -17.09, -7.63]
UKBB -10.88 [ -17.74, -4.03]
Discovery+UKBB [N=32291; P=4.669e-09] -11.87 [ -15.84, -7.90]
GOBS -2.29 [ -33.39, 28.82]
IMH -38.07 [-130.74, 54.60]
UNICAMP 1.31 [ -50.04, 52.66]
Meth-CT -40.40 [-105.76, 24.96]
MIRECC
UKBB nonEUR 29.52 [ -9.84, 68.87]
OSAKA -46.20 [-106.92, 14.52]
PING Non European 8.90 [ -32.37, 50.18]
Non European Cohorts [N=2648; P=0.8916] -1.23 [ -18.87, 16.42]
Beta [Lower 95% CI,Upper 95% CI]
-150
-100
-50
0
50
100
150
mm^2/effect allele

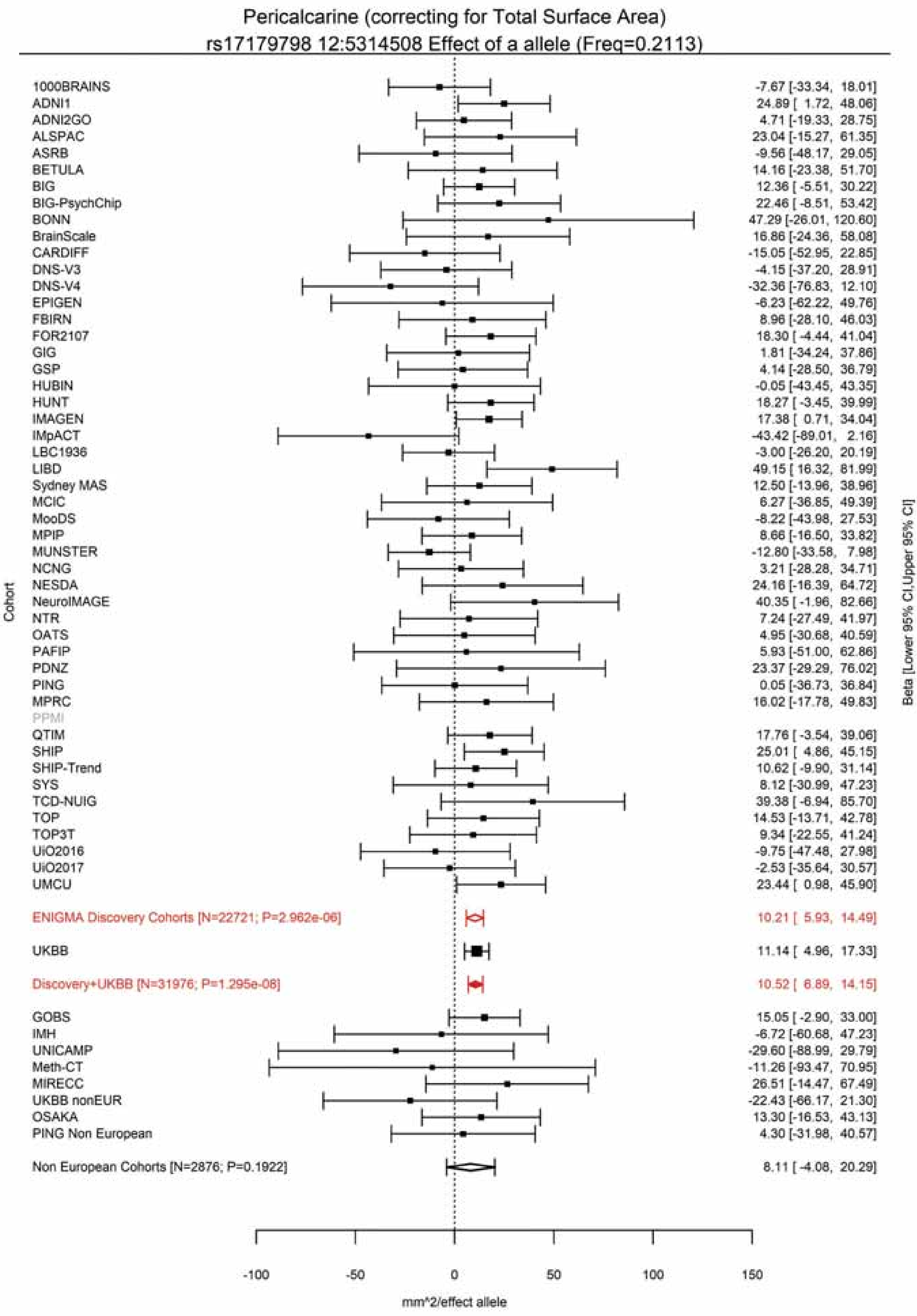
Pericalcarine (correcting for Total Surface Area)
rs17179798 12:5314508 Effect of a allele (Freq=0.2113)
Cohort
1000BRAINS -7.67 [-33.34, 18.01]
ADNI1 24.89 [ 1.72, 48.06]
ADNI2GO 4.71 [-19.33, 28.75]
ALSPAC 23.04 [-15.27, 61.35]
ASRB -9.56 [-48.17, 29.05]
BETULA 14.16 [-23.38, 51.70]
BIG 12.36 [ -5.51, 30.22]
BIG-PsychChip 22.46 [ -8.51, 53.42]
BONN 47.29 [-26.01, 120.60]
BrainScale 16.86 [-24.36, 58.08]
CARDIFF -15.05 [-52.95, 22.85]
DNS-V3 -4.15 [-37.20, 28.91]
DNS-V4 -32.36 [-76.83, 12.10]
EPIGEN -6.23 [-62.22, 49.76]
FBIRN 8.96 [-28.10, 46.03]
FOR2107 18.30 [ -4.44, 41.04]
GIG 1.81 [-34.24, 37.86]
GSP 4.14 [-28.50, 36.79]
HUBIN -0.05 [-43.45, 43.35]
HUNT 18.27 [ -3.45, 39.99]
IMAGEN 17.38 [ 0.71, 34.04]
IMpACT -43.42 [-89.01, 2.16]
LBC1936 -3.00 [-26.20, 20.19]
LIBD 49.15 [ 16.32, 81.99]
Sydney MAS 12.50 [-13.96, 38.96]
MCIC 6.27 [-36.85, 49.39]
MooDS -8.22 [-43.98, 27.53]
MPIP 8.66 [-16.50, 33.82]
MUNSTER -12.80 [-33.58, 7.98]
NCNG 3.21 [-28.28, 34.71]
NESDA 24.16 [-16.39, 64.72]
NeuroIMAGE 40.35 [ -1.96, 82.66]
NTR 7.24 [-27.49, 41.97]
OATS 4.95 [-30.68, 40.59]
PAFIP 5.93 [-51.00, 62.86]
PDNZ 23.37 [-29.29, 76.02]
PING 0.05 [-36.73, 36.84]
MPRC 16.02 [-17.78, 49.83]
PPMI
QTIM 17.76 [ -3.54, 39.06]
SHIP 25.01 [ 4.86, 45.15]
SHIP-Trend 10.62 [ -9.90, 31.14]
SYS 8.12 [-30.99, 47.23]
TCD-NUIG 39.38 [ -6.94, 85.70]
TOP 14.53 [-13.71, 42.78]
TOP3T 9.34 [-22.55, 41.24]
UiO2016 -9.75 [-47.48, 27.98]
UiO2017 -2.53 [-35.64, 30.57]
UMCU 23.44 [ 0.98, 45.90]
ENIGMA Discovery Cohorts [N=22721; P=2.962e-06] 10.21 [ 5.93, 14.49]
UKBB 11.14 [ 4.96, 17.33]
Discovery+UKBB [N=31976; P=1.295e-08] 10.52 [ 6.89, 14.15]
GOBS 15.05 [ -2.90, 33.00]
IMH -6.72 [-60.68, 47.23]
UNICAMP -29.60 [-88.99, 29.79]
Meth-CT -11.26 [-93.47, 70.95]
MIRECC 26.51 [-14.47, 67.49]
UKBB nonEUR -22.43 [-66.17, 21.30]
OSAKA 13.30 [-16.53, 43.13]
PING Non European 4.30 [-31.98, 40.57]
Non European Cohorts [N=2876; P=0.1922] 8.11 [ -4.08, 20.29]
Beta [Lower 95% CI,Upper 95% CI]
-100
-50
0
50
100
150
mm^2/effect allele

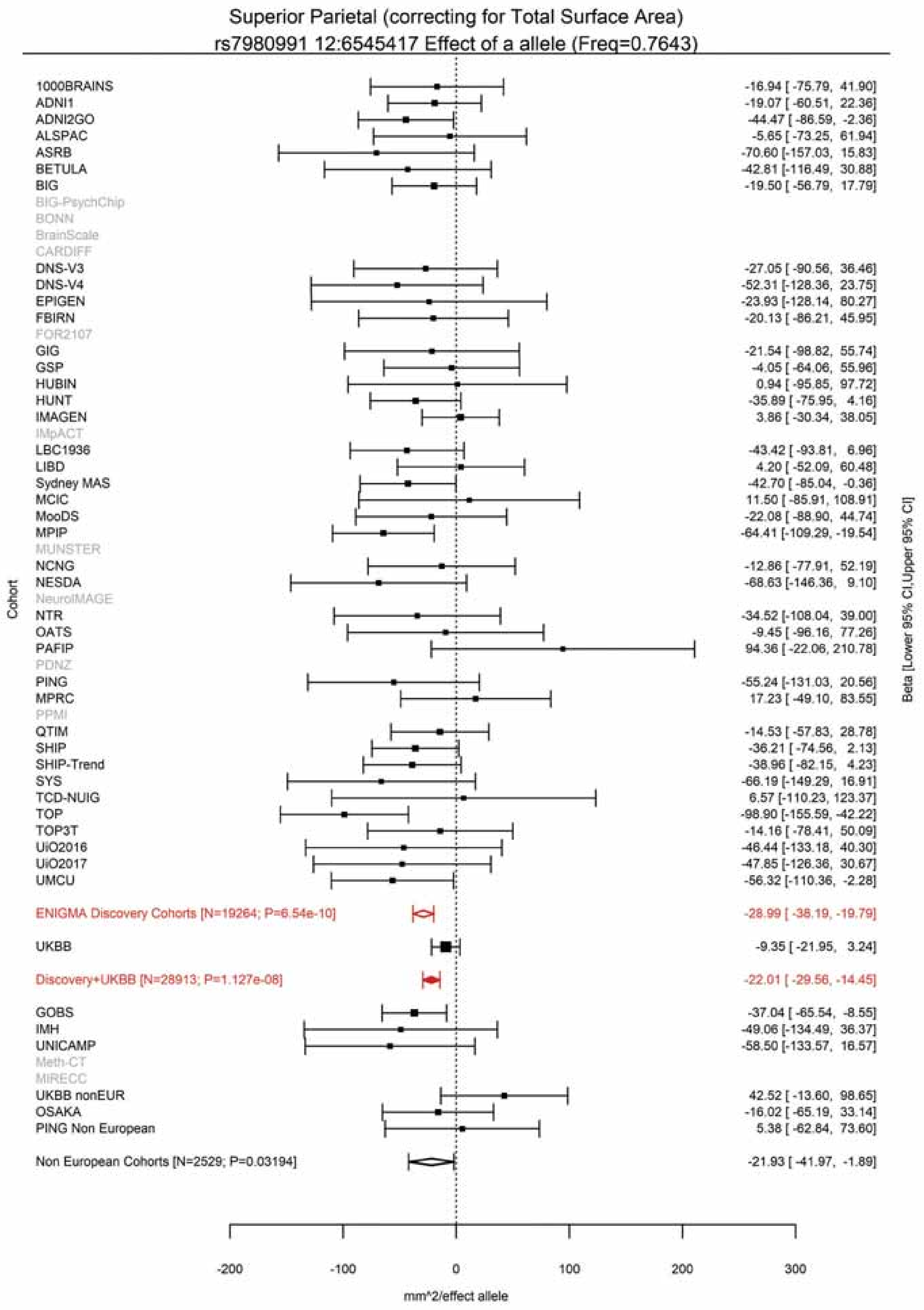

Superior Parietal (correcting for Total Surface Area)
rs7980991 12:6545417 Effect of a allele (Freq=0.7643)
1000BRAINS -16.94 [ -75.79, 41.90]
ADNI1 -19.07 [ -60.51, 22.36]
ADNI2GO -44.47 [ -86.59, -2.36]
ALSPAC -5.65 [ -73.25, 61.94]
ASRB -70.60 [-157.03, 15.83]
BETULA -42.81 [-116.49, 30.88]
BIG -19.50 [ -56.79, 17.79]
BIG-PsychChip
BONN
BrainScale
CARDIFF
DNS-V3 -27.05 [ -90.56, 36.46]
DNS-V4 -52.31 [-128.36, 23.75]
EPIGEN -23.93 [-128.14, 80.27]
FBIRN -20.13 [ -86.21, 45.95]
FOR2107
GIG -21.54 [ -98.82, 55.74]
GSP -4.05 [ -64.06, 55.96]
HUBIN 0.94 [ -95.85, 97.72]
HUNT -35.89 [ -75.95, 4.16]
IMAGEN 3.86 [ -30.34, 38.05]
IMpACT
LBC1936 -43.42 [ -93.81, 6.96]
LIBD 4.20 [ -52.09, 60.48]
Sydney MAS -42.70 [ -85.04, -0.36]
MCIC 11.50 [ -85.91, 108.91]
MooDS -22.08 [ -88.90, 44.74]
MPIP -64.41 [-109.29, -19.54]
MUNSTER
NCNG -12.86 [ -77.91, 52.19]
NESDA -68.63 [-146.36, 9.10]
NeuroIMAGE
NTR -34.52 [-108.04, 39.00]
OATS -9.45 [ -96.16, 77.26]
PAFIP 94.36 [ -22.06, 210.78]
PDNZ
PING -55.24 [-131.03, 20.56]
MPRC 17.23 [ -49.10, 83.55]
PPMI
QTIM -14.53 [ -57.83, 28.78]
SHIP -36.21 [ -74.56, 2.13]
SHIP-Trend -38.96 [ -82.15, 4.23]
SYS -66.19 [-149.29, 16.91]
TCD-NUIG 6.57 [-110.23, 123.37]
TOP -98.90 [-155.59, -42.22]
TOP3T -14.16 [ -78.41, 50.09]
UiO2016 -46.44 [-133.18, 40.30]
UiO2017 -47.85 [-126.36, 30.67]
UMCU -56.32 [-110.36, -2.28]
ENIGMA Discovery Cohorts [N=19264; P=6.54e-10] -28.99 [ -38.19, -19.79]
UKBB -9.35 [ -21.95, 3.24]
Discovery+UKBB [N=28913; P=1.127e-08] -22.01 [ -29.56, -14.45]
GOBS -37.04 [ -65.54, -8.55]
IMH -49.06 [-134.49, 36.37]
UNICAMP -58.50 [-133.57, 16.57]
Meth-CT
MIRECC
UKBB nonEUR 42.52 [ -13.60, 98.65]
OSAKA -16.02 [ -65.19, 33.14]
PING Non European 5.38 [ -62.84, 73.60]
Non European Cohorts [N=2529; P=0.03194] -21.93 [ -41.97, -1.89]
Cohort
Beta [Lower 95% CI,Upper 95% CI]
-200
-100
0
100
200
300
mm^2/effect allele

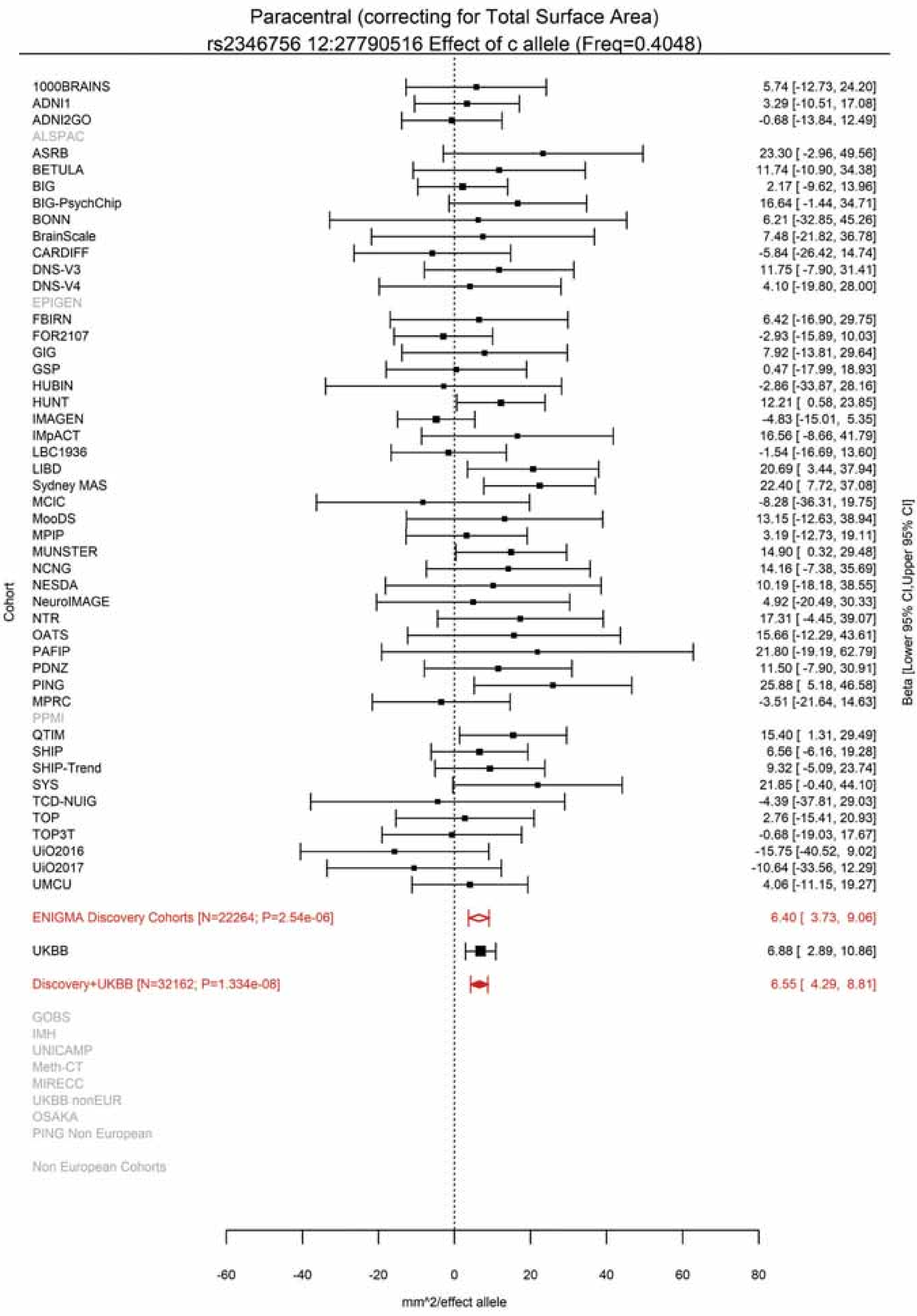

Paracentral (correcting for Total Surface Area)
rs2346756 12:27790516 Effect of c allele (Freq=0.4048)
1000BRAINS 5.74 [-12.73, 24.20]
ADNI1 3.29 [-10.51, 17.08]
ADNI2GO -0.68 [-13.84, 12.49]
ALSPAC
ASRB 23.30 [ -2.96, 49.56]
BETULA 11.74 [-10.90, 34.38]
BIG 2.17 [ -9.62, 13.96]
BIG-PsychChip 16.64 [ -1.44, 34.71]
BONN 6.21 [-32.85, 45.26]
BrainScale 7.48 [-21.82, 36.78]
CARDIFF -5.84 [-26.42, 14.74]
DNS-V3 11.75 [ -7.90, 31.41]
DNS-V4 4.10 [-19.80, 28.00]
EPIGEN
FBIRN 6.42 [-16.90, 29.75]
FOR2107 -2.93 [-15.89, 10.03]
GIG 7.92 [-13.81, 29.64]
GSP 0.47 [-17.99, 18.93]
HUBIN -2.86 [-33.87, 28.16]
HUNT 12.21 [ 0.58, 23.85]
IMAGEN -4.83 [-15.01, 5.35]
IMpACT 16.56 [ -8.66, 41.79]
LBC1936 -1.54 [-16.69, 13.60]
LIBD 20.69 [ 3.44, 37.94]
Sydney MAS 22.40 [ 7.72, 37.08]
MCIC -8.28 [-36.31, 19.75]
MooDS 13.15 [-12.63, 38.94]
MPIP 3.19 [-12.73, 19.11]
MUNSTER 14.90 [ 0.32, 29.48]
NCNG 14.16 [ -7.38, 35.69]
NESDA 10.19 [-18.18, 38.55]
NeuroIMAGE 4.92 [-20.49, 30.33]
NTR 17.31 [ -4.45, 39.07]
OATS 15.66 [-12.29, 43.61]
PAFIP 21.80 [-19.19, 62.79]
PDNZ 11.50 [ -7.90, 30.91]
PING 25.88 [ 5.18, 46.58]
MPRC -3.51 [-21.64, 14.63]
PPMI
QTIM 15.40 [ 1.31, 29.49]
SHIP 6.56 [ -6.16, 19.28]
SHIP-Trend 9.32 [ -5.09, 23.74]
SYS 21.85 [ -0.40, 44.10]
TCD-NUIG -4.39 [-37.81, 29.03]
TOP 2.76 [-15.41, 20.93]
TOP3T -0.68 [-19.03, 17.67]
UiO2016 -15.75 [-40.52, 9.02]
UiO2017 -10.64 [-33.56, 12.29]
UMCU 4.06 [-11.15, 19.27]
ENIGMA Discovery Cohorts [N=22264; P=2.54e-06] 6.40 [ 3.73, 9.06]
UKBB 6.88 [ 2.89, 10.86]
Discovery+UKBB [N=32162; P=1.334e-08] 6.55 [ 4.29, 8.81]
GOBS
IMH
UNICAMP
Meth-CT
MIRECC
UKBB nonEUR
OSAKA
PING Non European
Non European Cohorts
Cohort
Beta [Lower 95% CI,Upper 95% CI]
-60 -40 -20 0 20 40 60 80
mm^2/effect allele

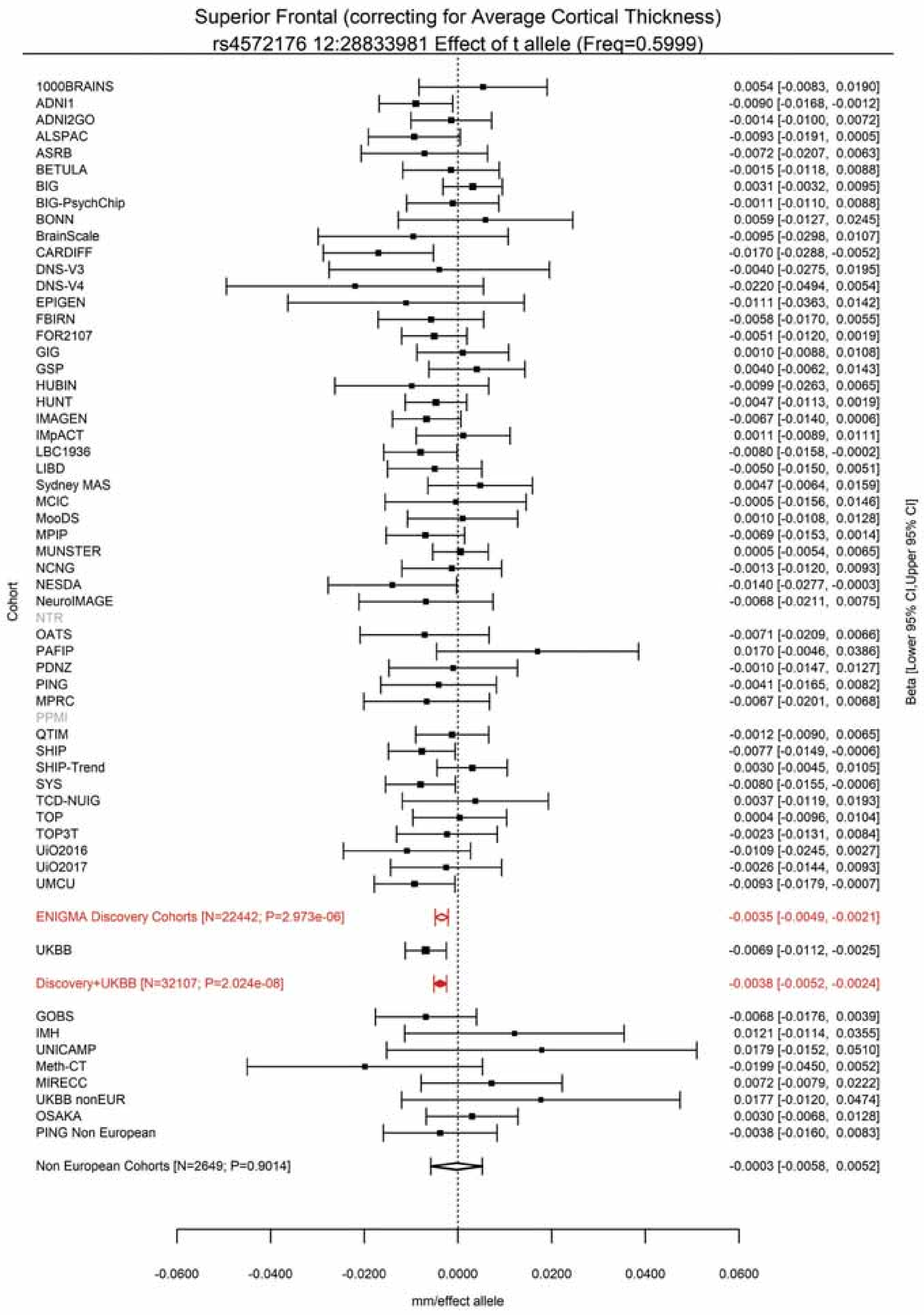

Superior Frontal (correcting for Average Cortical Thickness)
rs4572176 12:28833981 Effect of t allele (Freq=0.5999)
Cohort
Beta [Lower 95% CI,Upper 95% CI]
1000BRAINS 0.0054 [-0.0083, 0.0190]
ADNI1 -0.0090 [-0.0168, -0.0012]
ADNI2GO -0.0014 [-0.0100, 0.0072]
ALSPAC -0.0093 [-0.0191, 0.0005]
ASRB -0.0072 [-0.0207, 0.0063]
BETULA -0.0015 [-0.0118, 0.0088]
BIG 0.0031 [-0.0032, 0.0095]
BIG-PsychChip -0.0011 [-0.0110, 0.0088]
BONN 0.0059 [-0.0127, 0.0245]
BrainScale -0.0095 [-0.0298, 0.0107]
CARDIFF -0.0170 [-0.0288, -0.0052]
DNS-V3 -0.0040 [-0.0275, 0.0195]
DNS-V4 -0.0220 [-0.0494, 0.0054]
EPIGEN -0.0111 [-0.0363, 0.0142]
FBIRN -0.0058 [-0.0170, 0.0055]
FOR2107 -0.0051 [-0.0120, 0.0019]
GIG 0.0010 [-0.0088, 0.0108]
GSP 0.0040 [-0.0062, 0.0143]
HUBIN -0.0099 [-0.0263, 0.0065]
HUNT -0.0047 [-0.0113, 0.0019]
IMAGEN -0.0067 [-0.0140, 0.0006]
IMpACT 0.0011 [-0.0089, 0.0111]
LBC1936 -0.0080 [-0.0158, -0.0002]
LIBD -0.0050 [-0.0150, 0.0051]
Sydney MAS 0.0047 [-0.0064, 0.0159]
MCIC -0.0005 [-0.0156, 0.0146]
MooDS 0.0010 [-0.0108, 0.0128]
MPIP -0.0069 [-0.0153, 0.0014]
MUNSTER 0.0005 [-0.0054, 0.0065]
NCNG -0.0013 [-0.0120, 0.0093]
NESDA -0.0140 [-0.0277, -0.0003]
NeuroIMAGE -0.0068 [-0.0211, 0.0075]
NTR
OATS -0.0071 [-0.0209, 0.0066]
PAFIP 0.0170 [-0.0046, 0.0386]
PDNZ -0.0010 [-0.0147, 0.0127]
PING -0.0041 [-0.0165, 0.0082]
MPRC -0.0067 [-0.0201, 0.0068]
PPMI
QTIM -0.0012 [-0.0090, 0.0065]
SHIP -0.0077 [-0.0149, -0.0006]
SHIP-Trend 0.0030 [-0.0045, 0.0105]
SYS -0.0080 [-0.0155, -0.0006]
TCD-NUIG 0.0037 [-0.0119, 0.0193]
TOP 0.0004 [-0.0096, 0.0104]
TOP3T -0.0023 [-0.0131, 0.0084]
UiO2016 -0.0109 [-0.0245, 0.0027]
UiO2017 -0.0026 [-0.0144, 0.0093]
UMCU -0.0093 [-0.0179, -0.0007]
ENIGMA Discovery Cohorts [N=22442; P=2.973e-06] -0.0035 [-0.0049, -0.0021]
UKBB -0.0069 [-0.0112, -0.0025]
Discovery+UKBB [N=32107; P=2.024e-08] -0.0038 [-0.0052, -0.0024]
GOBS -0.0068 [-0.0176, 0.0039]
IMH 0.0121 [-0.0114, 0.0355]
UNICAMP 0.0179 [-0.0152, 0.0510]
Meth-CT -0.0199 [-0.0450, 0.0052]
MIRECC 0.0072 [-0.0079, 0.0222]
UKBB nonEUR 0.0177 [-0.0120, 0.0474]
OSAKA 0.0030 [-0.0068, 0.0128]
PING Non European -0.0038 [-0.0160, 0.0083]
Non European Cohorts [N=2649; P=0.9014] -0.0003 [-0.0058, 0.0052]
-0.0600
-0.0400
-0.0200
0.0000
0.0200
0.0400
0.0600
mm/effect allele

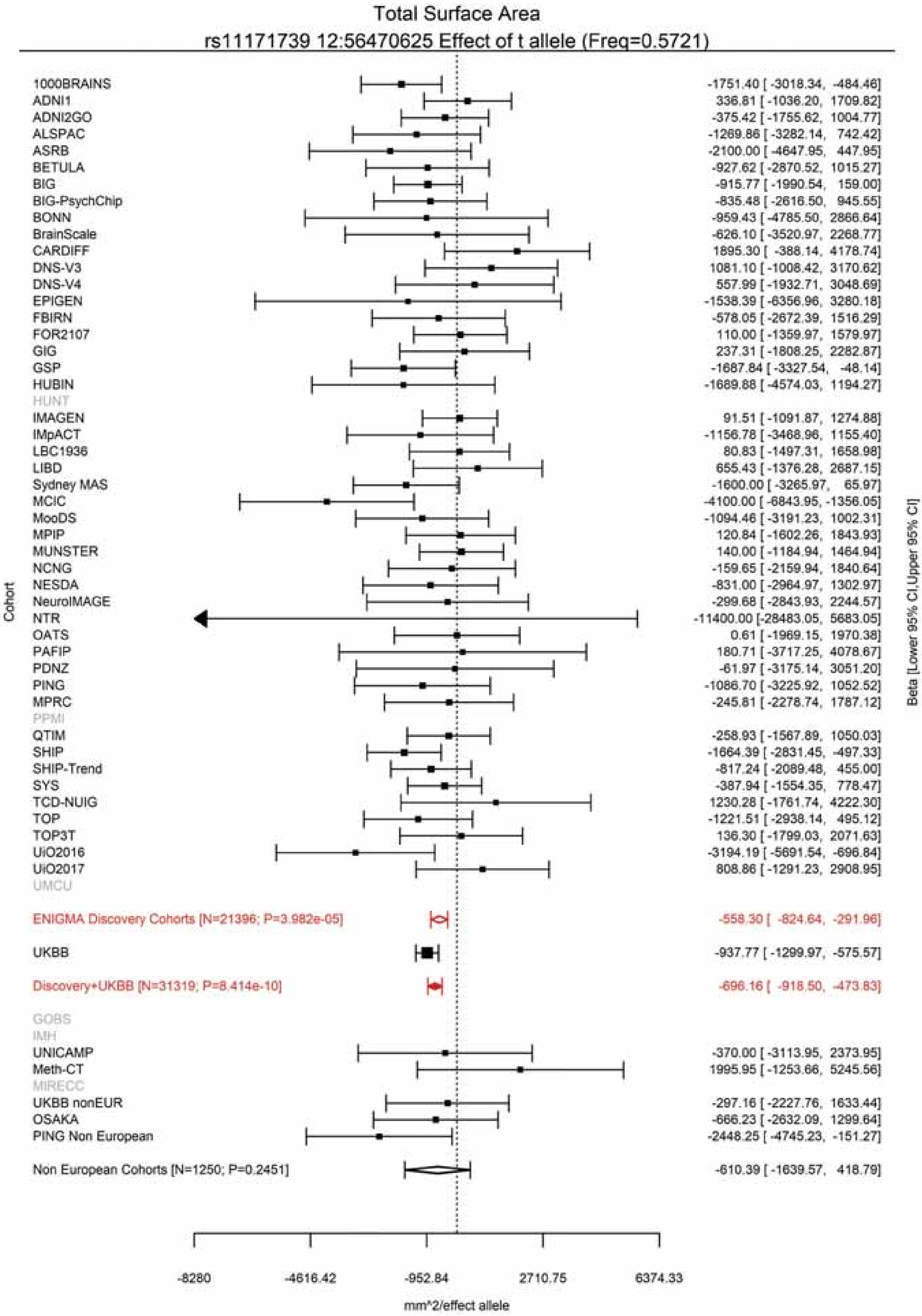
Total Surface Area
rs11171739 12:56470625 Effect of t allele (Freq=0.5721)
Cohort
1000BRAINS -1751.40 [ -3018.34, -484.46]
ADNI1 336.81 [ -1036.20, 1709.82]
ADNI2GO -375.42 [ -1755.62, 1004.77]
ALSPAC -1269.86 [ -3282.14, 742.42]
ASRB -2100.00 [ -4647.95, 447.95]
BETULA -927.62 [ -2870.52, 1015.27]
BIG -915.77 [ -1990.54, 159.00]
BIG-PsychChip -835.48 [ -2616.50, 945.55]
BONN -959.43 [ -4785.50, 2866.64]
BrainScale -626.10 [ -3520.97, 2268.77]
CARDIFF 1895.30 [ -388.14, 4178.74]
DNS-V3 1081.10 [ -1008.42, 3170.62]
DNS-V4 557.99 [ -1932.71, 3048.69]
EPIGEN -1538.39 [ -6356.96, 3280.18]
FBIRN -578.05 [ -2672.39, 1516.29]
FOR2107 110.00 [ -1359.97, 1579.97]
GIG 237.31 [ -1808.25, 2282.87]
GSP -1687.84 [ -3327.54, -48.14]
HUBIN -1669.88 [ -4574.03, 1194.27]
HUNT
IMAGEN 91.51 [ -1091.87, 1274.88]
IMpACT -1156.78 [ -3468.96, 1155.40]
LBC1936 80.83 [ -1497.31, 1658.98]
LIBD 655.43 [ -1376.28, 2687.15]
Sydney MAS -1600.00 [ -3265.97, 65.97]
MCIC -4100.00 [ -6843.95, -1356.05]
MooDS -1094.46 [ -3191.23, 1002.31]
MPIP 120.84 [ -1602.26, 1843.93]
MUNSTER 140.00 [ -1184.94, 1464.94]
NCNG -159.65 [ -2159.94, 1840.64]
NESDA -831.00 [ -2964.97, 1302.97]
NeuroIMAGE -299.68 [ -2843.93, 2244.57]
NTR -11400.00 [-28483.05, 5683.05]
OATS 0.61 [ -1969.15, 1970.38]
PAFIP 180.71 [ -3717.25, 4078.67]
PDNZ -61.97 [ -3175.14, 3051.20]
PING -1086.70 [ -3225.92, 1052.52]
MPRC -245.81 [ -2278.74, 1787.12]
PPMI
QTIM -258.93 [ -1567.89, 1050.03]
SHIP -1664.39 [ -2831.45, -497.33]
SHIP-Trend -817.24 [ -2089.48, 455.00]
SYS -387.94 [ -1554.35, 778.47]
TCD-NUIG 1230.28 [ -1761.74, 4222.30]
TOP -1221.51 [ -2938.14, 495.12]
TOP3T 136.30 [ -1799.03, 2071.63]
UiO2016 -3194.19 [ -5691.54, -696.84]
UiO2017 808.86 [ -1291.23, 2908.95]
UMCU
ENIGMA Discovery Cohorts [N=21396; P=3.982e-05] -558.30 [ -824.64, -291.96]
UKBB -937.77 [ -1299.97, -575.57]
Discovery+UKBB [N=31319; P=8.414e-10] -696.16 [ -918.50, -473.83]
GOBS
IMH
UNICAMP -370.00 [ -3113.95, 2373.95]
Meth-CT 1995.95 [ -1253.66, 5245.56]
MIRECC
UKBB nonEUR -297.16 [ -2227.76, 1633.44]
OSAKA -666.23 [ -2632.09, 1299.64]
PING Non European -2448.25 [ -4745.23, -151.27]
Non European Cohorts [N=1250; P=0.2451] -610.39 [ -1639.57, 418.79]
Beta [Lower 95% CI,Upper 95% CI]
-8280
-4616.42
-952.84
2710.75
6374.33
mm^2/effect allele

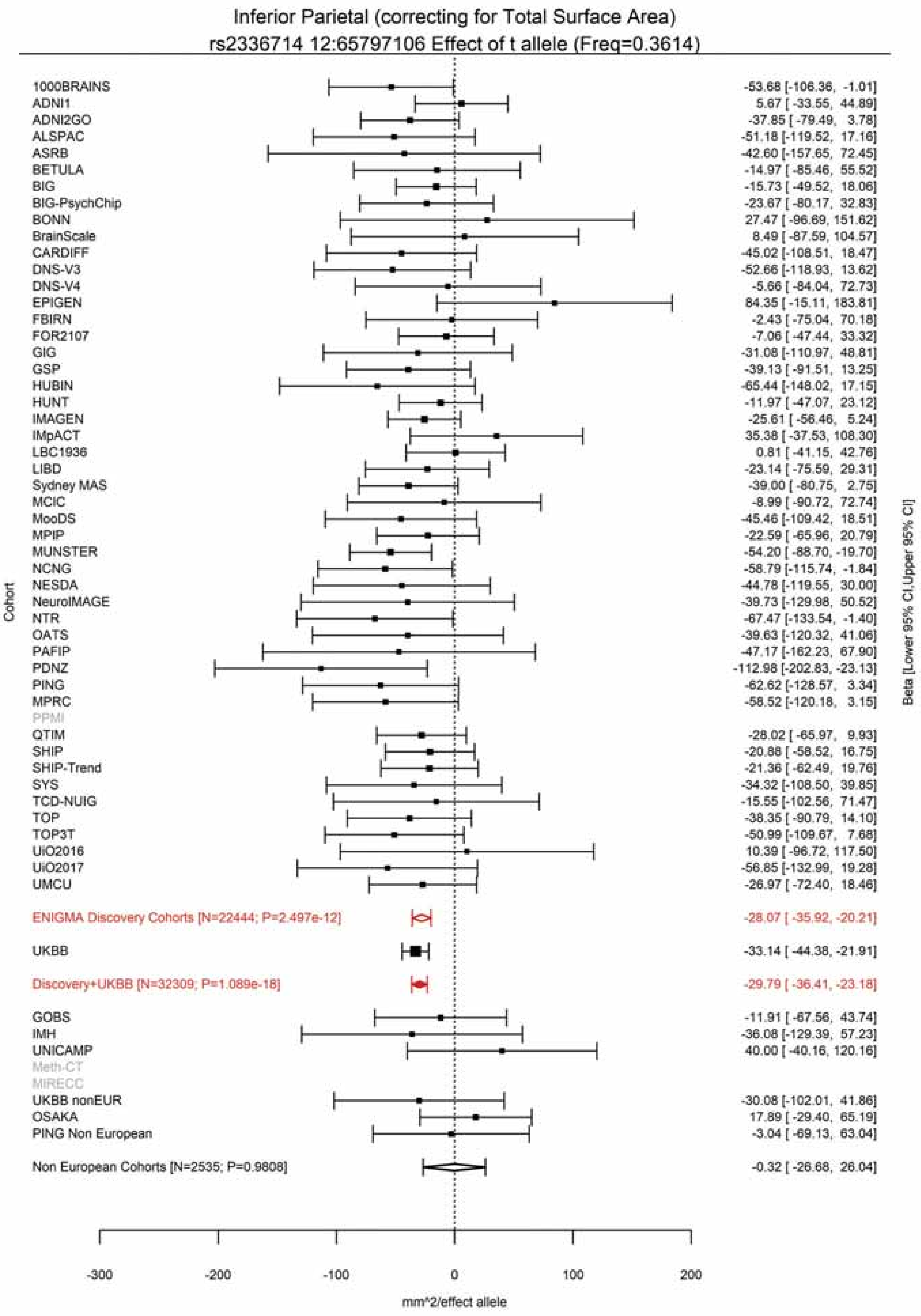

Inferior Parietal (correcting for Total Surface Area)
rs2336714 12:65797106 Effect of t allele (Freq=0.3614)
1000BRAINS -53.68 [-106.36, -1.01]
ADNI1 5.67 [ -33.55, 44.89]
ADNI2GO -37.85 [ -79.49, 3.78]
ALSPAC -51.18 [-119.52, 17.16]
ASRB -42.60 [-157.65, 72.45]
BETULA -14.97 [ -85.46, 55.52]
BIG -15.73 [ -49.52, 18.06]
BIG-PsychChip -23.67 [ -80.17, 32.83]
BONN 27.47 [ -96.69, 151.62]
BrainScale 8.49 [ -87.59, 104.57]
CARDIFF -45.02 [-108.51, 18.47]
DNS-V3 -52.66 [-118.93, 13.62]
DNS-V4 -5.66 [ -84.04, 72.73]
EPIGEN 84.35 [ -15.11, 183.81]
FBIRN -2.43 [ -75.04, 70.18]
FOR2107 -7.06 [ -47.44, 33.32]
GIG -31.08 [-110.97, 48.81]
GSP -39.13 [ -91.51, 13.25]
HUBIN -65.44 [-148.02, 17.15]
HUNT -11.97 [ -47.07, 23.12]
IMAGEN -25.61 [ -56.46, 5.24]
IMpACT 35.38 [ -37.53, 108.30]
LBC1936 0.81 [ -41.15, 42.76]
LIBD -23.14 [ -75.59, 29.31]
Sydney MAS -39.00 [ -80.75, 2.75]
MCIC -8.99 [ -90.72, 72.74]
MooDS -45.46 [-109.42, 18.51]
MPIP -22.59 [ -65.96, 20.79]
MUNSTER -54.20 [ -88.70, -19.70]
NCNG -58.79 [-115.74, -1.84]
NESDA -44.78 [-119.55, 30.00]
NeuroIMAGE -39.73 [-129.98, 50.52]
NTR -67.47 [-133.54, -1.40]
OATS -39.63 [-120.32, 41.06]
PAFIP -47.17 [-162.23, 67.90]
PDNZ -112.98 [-202.83, -23.13]
PING -62.62 [-128.57, 3.34]
MPRC -58.52 [-120.18, 3.15]
PPMI
QTIM -28.02 [ -65.97, 9.93]
SHIP -20.88 [ -58.52, 16.75]
SHIP-Trend -21.36 [ -62.49, 19.76]
SYS -34.32 [-108.50, 39.85]
TCD-NUIG -15.55 [-102.56, 71.47]
TOP -38.35 [ -90.79, 14.10]
TOP3T -50.99 [-109.67, 7.68]
UiO2016 10.39 [ -96.72, 117.50]
UiO2017 -56.85 [-132.99, 19.28]
UMCU -26.97 [ -72.40, 18.46]
ENIGMA Discovery Cohorts [N=22444; P=2.497e-12] -28.07 [ -35.92, -20.21]
UKBB -33.14 [ -44.38, -21.91]
Discovery+UKBB [N=32309; P=1.089e-18] -29.79 [ -36.41, -23.18]
GOBS -11.91 [ -67.56, 43.74]
IMH -36.08 [-129.39, 57.23]
UNICAMP 40.00 [ -40.16, 120.16]
Meth-CT
MIRECC
UKBB nonEUR -30.08 [-102.01, 41.86]
OSAKA 17.89 [ -29.40, 65.19]
PING Non European -3.04 [ -69.13, 63.04]
Non European Cohorts [N=2535; P=0.9808] -0.32 [ -26.68, 26.04]
Cohort
Beta [Lower 95% CI,Upper 95% CI]
-300
-200
-100
0
100
200
mm^2/effect allele

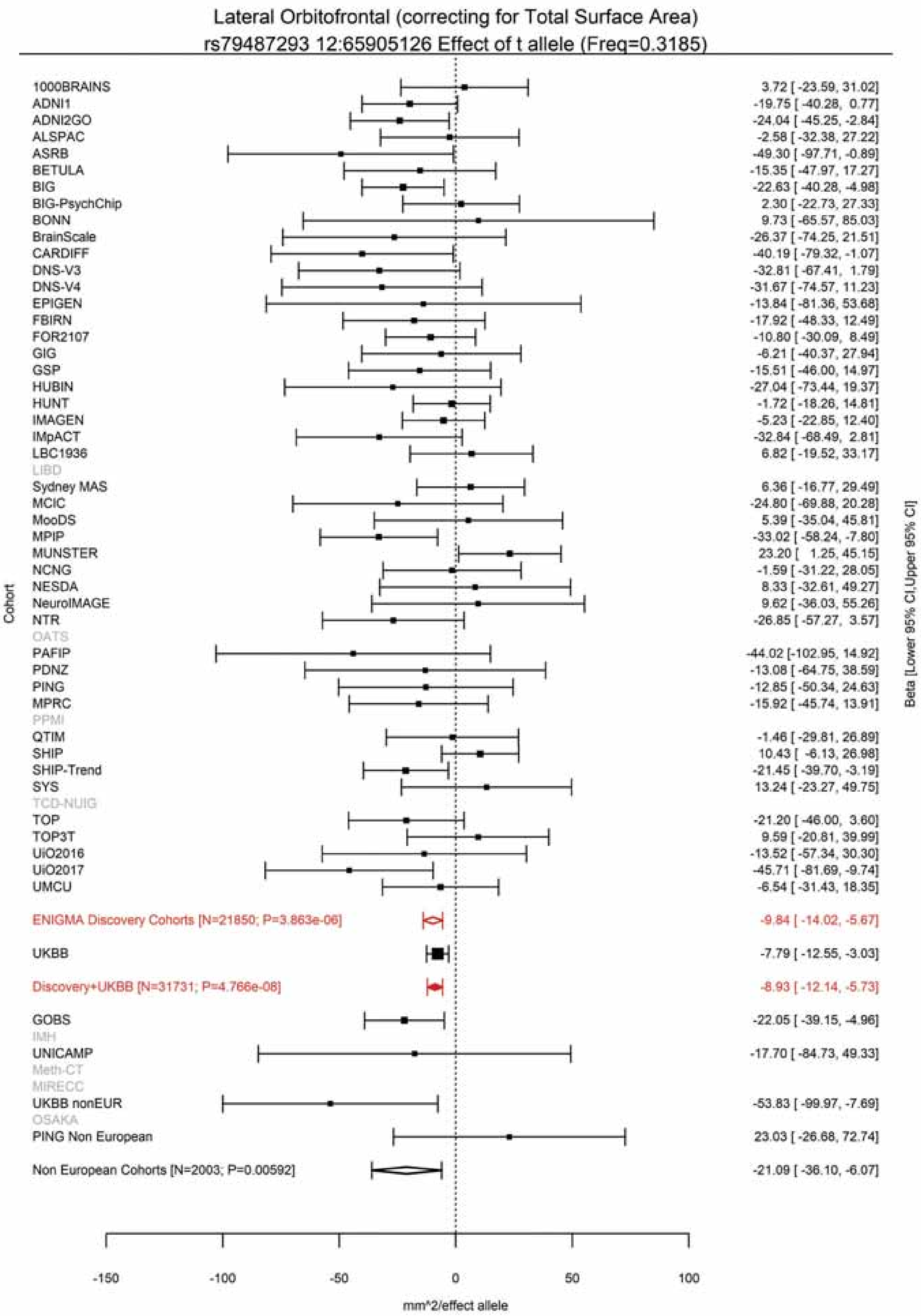
Lateral Orbitofrontal (correcting for Total Surface Area)
rs79487293 12:65905126 Effect of t allele (Freq=0.3185)
Cohort
Beta [Lower 95% CI,Upper 95% CI]
1000BRAINS 3.72 [ -23.59, 31.02]
ADNI1 -19.75 [ -40.28, 0.77]
ADNI2GO -24.04 [ -45.25, -2.84]
ALSPAC -2.58 [ -32.38, 27.22]
ASRB -49.30 [ -97.71, -0.89]
BETULA -15.35 [ -47.97, 17.27]
BIG -22.63 [ -40.28, -4.98]
BIG-PsychChip 2.30 [ -22.73, 27.33]
BONN 9.73 [ -65.57, 85.03]
BrainScale -26.37 [ -74.25, 21.51]
CARDIFF -40.19 [ -79.32, -1.07]
DNS-V3 -32.81 [ -67.41, 1.79]
DNS-V4 -31.67 [ -74.57, 11.23]
EPIGEN -13.84 [ -81.36, 53.68]
FBIRN -17.92 [ -48.33, 12.49]
FOR2107 -10.80 [ -30.09, 8.49]
GIG -6.21 [ -40.37, 27.94]
GSP -15.51 [ -46.00, 14.97]
HUBIN -27.04 [ -73.44, 19.37]
HUNT -1.72 [ -18.26, 14.81]
IMAGEN -5.23 [ -22.85, 12.40]
IMpACT -32.84 [ -68.49, 2.81]
LBC1936 6.82 [ -19.52, 33.17]
LIBD
Sydney MAS 6.36 [ -16.77, 29.49]
MCIC -24.80 [ -69.88, 20.28]
MooDS 5.39 [ -35.04, 45.81]
MPIP -33.02 [ -58.24, -7.80]
MUNSTER 23.20 [ 1.25, 45.15]
NCNG -1.59 [ -31.22, 28.05]
NESDA 8.33 [ -32.61, 49.27]
NeuroIMAGE 9.62 [ -36.03, 55.26]
NTR -26.85 [ -57.27, 3.57]
OATS
PAFIP -44.02 [-102.95, 14.92]
PDNZ -13.08 [ -64.75, 38.59]
PING -12.85 [ -50.34, 24.63]
MPRC -15.92 [ -45.74, 13.91]
PPMI
QTIM -1.46 [ -29.81, 26.89]
SHIP 10.43 [ -6.13, 26.98]
SHIP-Trend -21.45 [ -39.70, -3.19]
SYS 13.24 [ -23.27, 49.75]
TCD-NUIG
TOP -21.20 [ -46.00, 3.60]
TOP3T 9.59 [ -20.81, 39.99]
UiO2016 -13.52 [ -57.34, 30.30]
UiO2017 -45.71 [ -81.69, -9.74]
UMCU -6.54 [ -31.43, 18.35]
ENIGMA Discovery Cohorts [N=21850; P=3.863e-06] -9.84 [ -14.02, -5.67]
UKBB -7.79 [ -12.55, -3.03]
Discovery+UKBB [N=31731; P=4.766e-08] -8.93 [ -12.14, -5.73]
GOBS -22.05 [ -39.15, -4.96]
IMH
UNICAMP -17.70 [ -84.73, 49.33]
Meth-CT
MIRECC
UKBB nonEUR -53.83 [ -99.97, -7.69]
OSAKA
PING Non European 23.03 [ -26.68, 72.74]
Non European Cohorts [N=2003; P=0.00592] -21.09 [ -36.10, -6.07]
-150
-100
-50
0
50
100
mm^2/effect allele

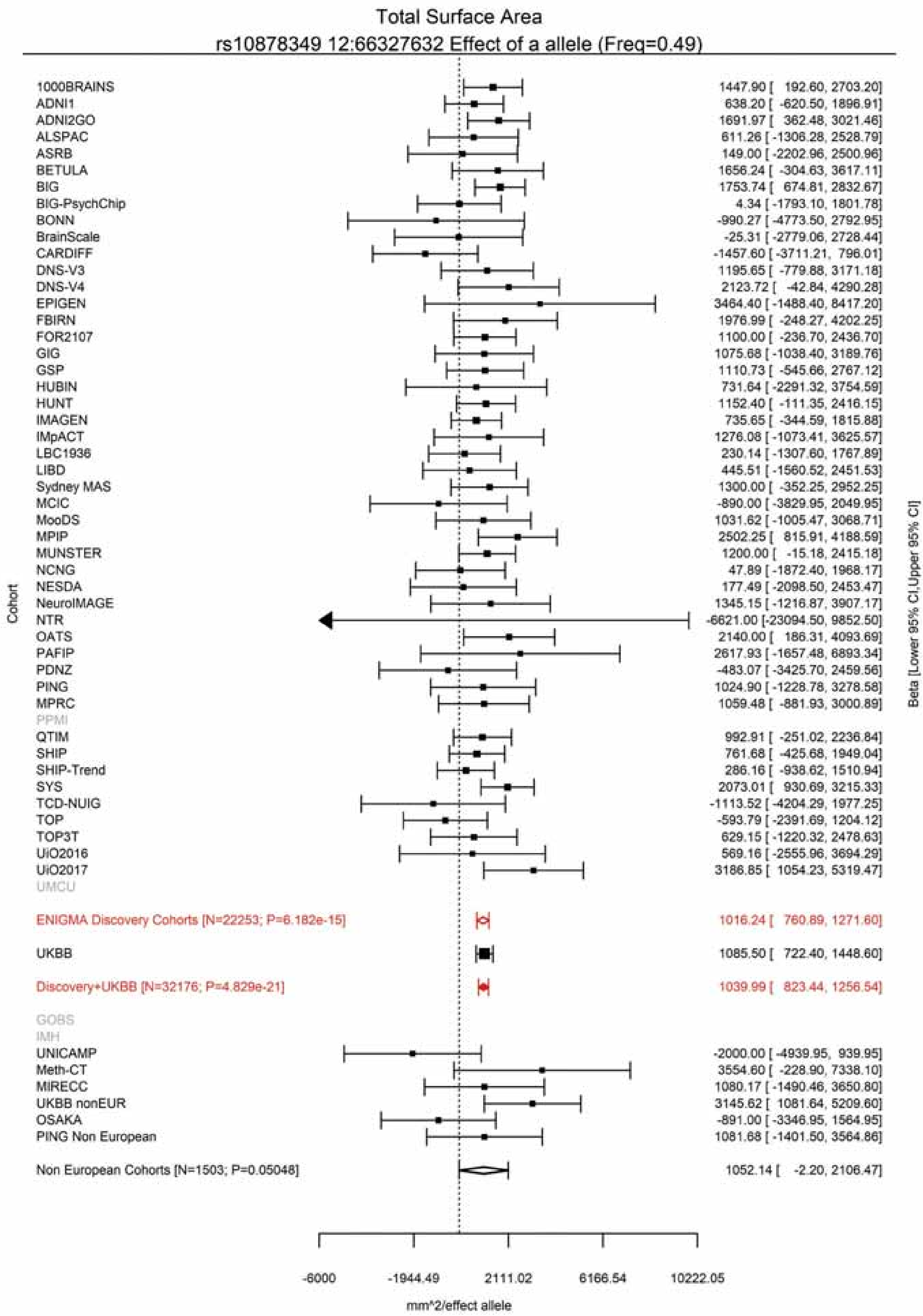
Total Surface Area
rs10878349 12:66327632 Effect of a allele (Freq=0.49)
Cohort
1000BRAINS 1447.90 [ 192.60, 2703.20]
ADNI1 638.20 [ -620.50, 1896.91]
ADNI2GO 1691.97 [ 362.48, 3021.46]
ALSPAC 611.26 [ -1306.28, 2528.79]
ASRB 149.00 [ -2202.96, 2500.96]
BETULA 1656.24 [ -304.63, 3617.11]
BIG 1753.74 [ 674.81, 2832.67]
BIG-PsychChip 4.34 [ -1793.10, 1801.78]
BONN -990.27 [ -4773.50, 2792.95]
BrainScale -25.31 [ -2779.06, 2728.44]
CARDIFF -1457.60 [ -3711.21, 796.01]
DNS-V3 1195.65 [ -779.88, 3171.18]
DNS-V4 2123.72 [ -42.84, 4290.28]
EPIGEN 3464.40 [ -1488.40, 8417.20]
FBIRN 1976.99 [ -248.27, 4202.25]
FOR2107 1100.00 [ -236.70, 2436.70]
GIG 1075.68 [ -1038.40, 3189.76]
GSP 1110.73 [ -545.66, 2767.12]
HUBIN 731.64 [ -2291.32, 3754.59]
HUNT 1152.40 [ -111.35, 2416.15]
IMAGEN 735.65 [ -344.59, 1815.88]
IMpACT 1276.08 [ -1073.41, 3625.57]
LBC1936 230.14 [ -1307.60, 1767.89]
LIBD 445.51 [ -1560.52, 2451.53]
Sydney MAS 1300.00 [ -352.25, 2952.25]
MCIC -890.00 [ -3829.95, 2049.95]
MooDS 1031.62 [ -1005.47, 3068.71]
MPIP 2502.25 [ 815.91, 4188.59]
MUNSTER 1200.00 [ -15.18, 2415.18]
NCNG 47.89 [ -1872.40, 1968.17]
NESDA 177.49 [ -2098.50, 2453.47]
NeuroIMAGE 1345.15 [ -1216.87, 3907.17]
NTR -6621.00 [-23094.50, 9852.50]
OATS 2140.00 [ 186.31, 4093.69]
PAFIP 2617.93 [ -1657.48, 6893.34]
PDNZ -483.07 [ -3425.70, 2459.56]
PING 1024.90 [ -1228.78, 3278.58]
MPRC 1059.48 [ -881.93, 3000.89]
PPMI
QTIM 992.91 [ -251.02, 2236.84]
SHIP 761.68 [ -425.68, 1949.04]
SHIP-Trend 286.16 [ -938.62, 1510.94]
SYS 2073.01 [ 930.69, 3215.33]
TCD-NUIG -1113.52 [ -4204.29, 1977.25]
TOP -593.79 [ -2391.69, 1204.12]
TOP3T 629.15 [ -1220.32, 2478.63]
UiO2016 569.16 [ -2555.96, 3694.29]
UiO2017 3186.85 [ 1054.23, 5319.47]
UMCU
ENIGMA Discovery Cohorts [N=22253; P=6.182e-15] 1016.24 [ 760.89, 1271.60]
UKBB 1085.50 [ 722.40, 1448.60]
Discovery+UKBB [N=32176; P=4.829e-21] 1039.99 [ 823.44, 1256.54]
GOBS
IMH
UNICAMP -2000.00 [ -4939.95, 939.95]
Meth-CT 3554.60 [ -228.90, 7338.10]
MIRECC 1080.17 [ -1490.46, 3650.80]
UKBB nonEUR 3145.62 [ 1081.64, 5209.60]
OSAKA -891.00 [ -3346.95, 1564.95]
PING Non European 1081.68 [ -1401.50, 3564.86]
Non European Cohorts [N=1503; P=0.05048] 1052.14 [ -2.20, 2106.47]
Beta [Lower 95% CI,Upper 95% CI]
-6000
-1944.49
2111.02
6166.54
10222.05
mm^2/effect allele

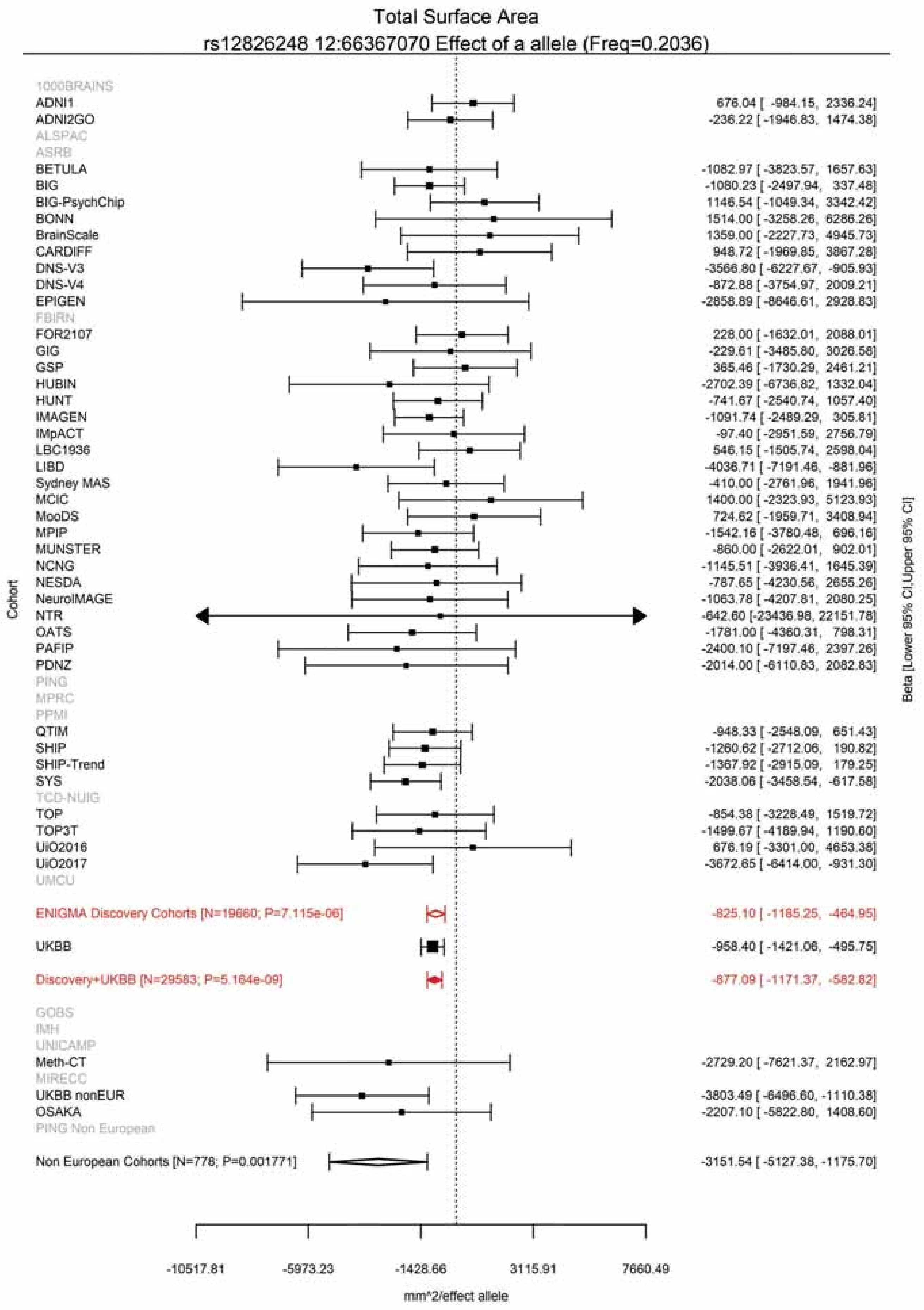
Total Surface Area
rs12826248 12:66367070 Effect of a allele (Freq=0.2036)
Cohort
1000BRAINS
ADNI1 676.04 [ -984.15, 2336.24]
ADNI2GO -236.22 [ -1946.83, 1474.38]
ALSPAC
ASRB
BETULA -1082.97 [ -3823.57, 1657.63]
BIG -1080.23 [ -2497.94, 337.48]
BIG-PsychChip 1146.54 [ -1049.34, 3342.42]
BONN 1514.00 [ -3258.26, 6286.26]
BrainScale 1359.00 [ -2227.73, 4945.73]
CARDIFF 948.72 [ -1969.85, 3867.28]
DNS-V3 -3566.80 [ -6227.67, -905.93]
DNS-V4 -872.88 [ -3754.97, 2009.21]
EPIGEN -2858.89 [ -8646.61, 2928.83]
FBIRN
FOR2107 228.00 [ -1632.01, 2088.01]
GIG -229.61 [ -3485.80, 3026.58]
GSP 365.46 [ -1730.29, 2461.21]
HUBIN -2702.39 [ -6736.82, 1332.04]
HUNT -741.67 [ -2540.74, 1057.40]
IMAGEN -1091.74 [ -2489.29, 305.81]
IMpACT -97.40 [ -2951.59, 2756.79]
LBC1936 546.15 [ -1505.74, 2598.04]
LIBD -4036.71 [ -7191.46, -881.96]
Sydney MAS -410.00 [ -2761.96, 1941.96]
MCIC 1400.00 [ -2323.93, 5123.93]
MooDS 724.62 [ -1959.71, 3408.94]
MPIP -1542.16 [ -3780.48, 696.16]
MUNSTER -860.00 [ -2622.01, 902.01]
NCNG -1145.51 [ -3936.41, 1645.39]
NESDA -787.65 [ -4230.56, 2655.26]
NeuroIMAGE -1063.78 [ -4207.81, 2080.25]
NTR -642.60 [-23436.98, 22151.78]
OATS -1781.00 [ -4360.31, 798.31]
PAFIP -2400.10 [ -7197.46, 2397.26]
PDNZ -2014.00 [ -6110.83, 2082.83]
PING
MPRC
PPMI
QTIM -948.33 [ -2548.09, 651.43]
SHIP -1260.62 [ -2712.06, 190.82]
SHIP-Trend -1367.92 [ -2915.09, 179.25]
SYS -2038.06 [ -3458.54, -617.58]
TCD-NUIG
TOP -854.38 [ -3228.49, 1519.72]
TOP3T -1499.67 [ -4189.94, 1190.60]
UiO2016 676.19 [ -3301.00, 4653.38]
UiO2017 -3672.65 [ -6414.00, -931.30]
UMCU
ENIGMA Discovery Cohorts [N=19660; P=7.115e-06] -825.10 [ -1185.25, -464.95]
UKBB -958.40 [ -1421.06, -495.75]
Discovery+UKBB [N=29583; P=5.164e-09] -877.09 [ -1171.37, -582.82]
GOBS
IMH
UNICAMP
Meth-CT -2729.20 [ -7621.37, 2162.97]
MIRECC
UKBB nonEUR -3803.49 [ -6496.60, -1110.38]
OSAKA -2207.10 [ -5822.80, 1408.60]
PING Non European
Non European Cohorts [N=778; P=0.001771] -3151.54 [ -5127.38, -1175.70]
Beta [Lower 95% CI,Upper 95% CI]
-10517.81
-5973.23
-1428.66
3115.91
7660.49
mm^2/effect allele

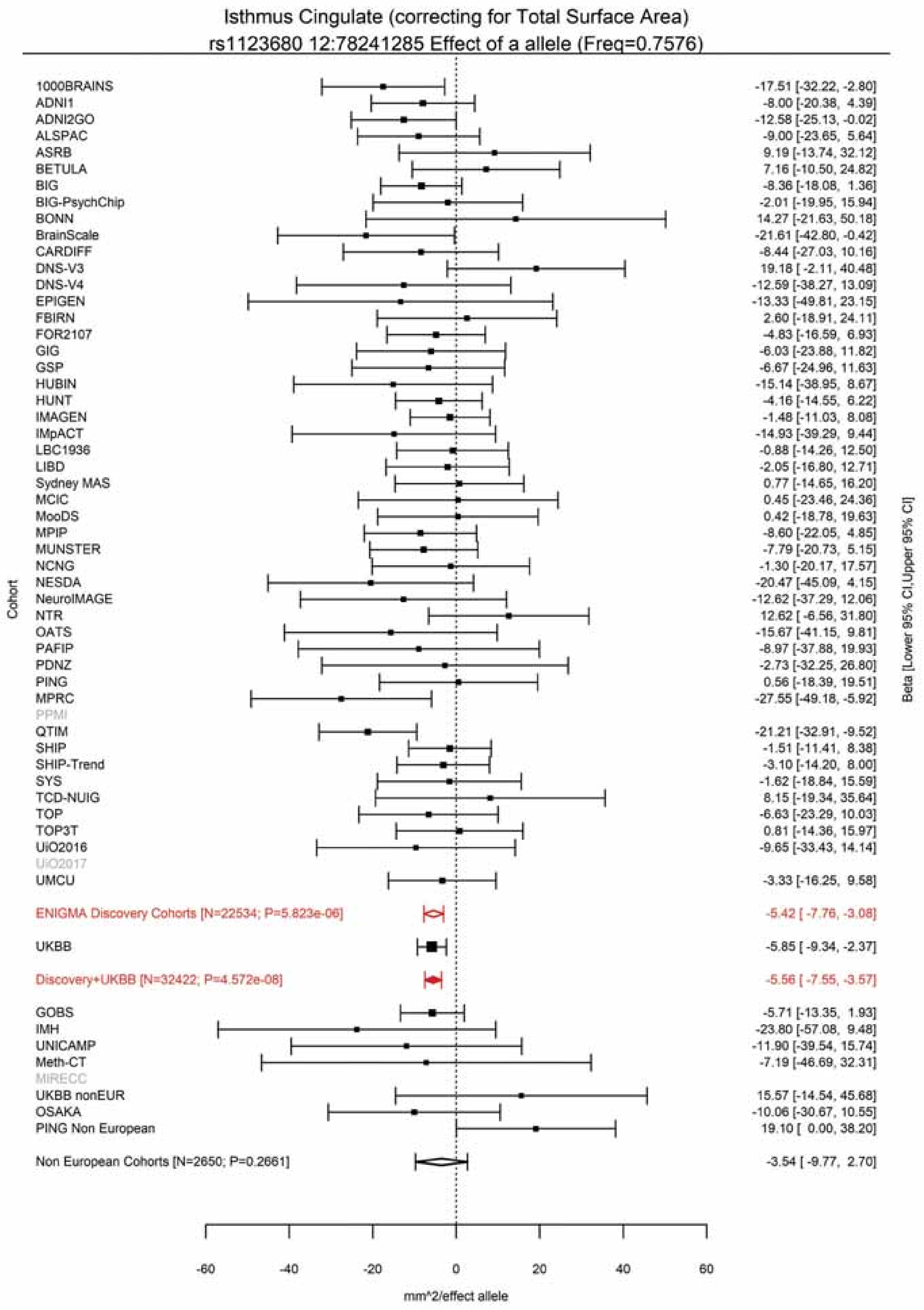
Isthmus Cingulate (correcting for Total Surface Area)
rs1123680 12:78241285 Effect of a allele (Freq=0.7576)
1000BRAINS -17.51 [-32.22, -2.80]
ADNI1 -8.00 [-20.38, 4.39]
ADNI2GO -12.58 [-25.13, -0.02]
ALSPAC -9.00 [-23.65, 5.64]
ASRB 9.19 [-13.74, 32.12]
BETULA 7.16 [-10.50, 24.82]
BIG -8.36 [-18.08, 1.36]
BIG-PsychChip -2.01 [-19.95, 15.94]
BONN 14.27 [-21.63, 50.18]
BrainScale -21.61 [-42.80, -0.42]
CARDIFF -8.44 [-27.03, 10.16]
DNS-V3 19.18 [-2.11, 40.48]
DNS-V4 -12.59 [-38.27, 13.09]
EPIGEN -13.33 [-49.81, 23.15]
FBIRN 2.60 [-18.91, 24.11]
FOR2107 -4.83 [-16.59, 6.93]
GIG -6.03 [-23.88, 11.82]
GSP -6.67 [-24.96, 11.63]
HUBIN -15.14 [-38.95, 8.67]
HUNT -4.16 [-14.55, 6.22]
IMAGEN -1.48 [-11.03, 8.08]
IMpACT -14.93 [-39.29, 9.44]
LBC1936 -0.88 [-14.26, 12.50]
LIBD -2.05 [-16.80, 12.71]
Sydney MAS 0.77 [-14.65, 16.20]
MCIC 0.45 [-23.46, 24.36]
MooDS 0.42 [-18.78, 19.63]
MPIP -8.60 [-22.05, 4.85]
MUNSTER -7.79 [-20.73, 5.15]
NCNG -1.30 [-20.17, 17.57]
NESDA -20.47 [-45.09, 4.15]
NeuroIMAGE -12.62 [-37.29, 12.06]
NTR 12.62 [-6.56, 31.80]
OATS -15.67 [-41.15, 9.81]
PAFIP -8.97 [-37.88, 19.93]
PDNZ -2.73 [-32.25, 26.80]
PING 0.56 [-18.39, 19.51]
MPRC -27.55 [-49.18, -5.92]
PPMI
QTIM -21.21 [-32.91, -9.52]
SHIP -1.51 [-11.41, 8.38]
SHIP-Trend -3.10 [-14.20, 8.00]
SYS -1.62 [-18.84, 15.59]
TCD-NUIG 8.15 [-19.34, 35.64]
TOP -6.63 [-23.29, 10.03]
TOP3T 0.81 [-14.36, 15.97]
UiO2016 -9.65 [-33.43, 14.14]
UiO2017
UMCU -3.33 [-16.25, 9.58]
ENIGMA Discovery Cohorts [N=22534; P=5.823e-06] -5.42 [-7.76, -3.08]
UKBB -5.85 [-9.34, -2.37]
Discovery+UKBB [N=32422; P=4.572e-08] -5.56 [-7.55, -3.57]
GOBS -5.71 [-13.35, 1.93]
IMH -23.80 [-57.08, 9.48]
UNICAMP -11.90 [-39.54, 15.74]
Meth-CT -7.19 [-46.69, 32.31]
MIRECC
UKBB nonEUR 15.57 [-14.54, 45.68]
OSAKA -10.06 [-30.67, 10.55]
PING Non European 19.10 [0.00, 38.20]
Non European Cohorts [N=2650; P=0.2661] -3.54 [-9.77, 2.70]
Cohort
Beta [Lower 95% CI,Upper 95% CI]
-60
-40
-20
0
20
40
60
mm^2/effect allele

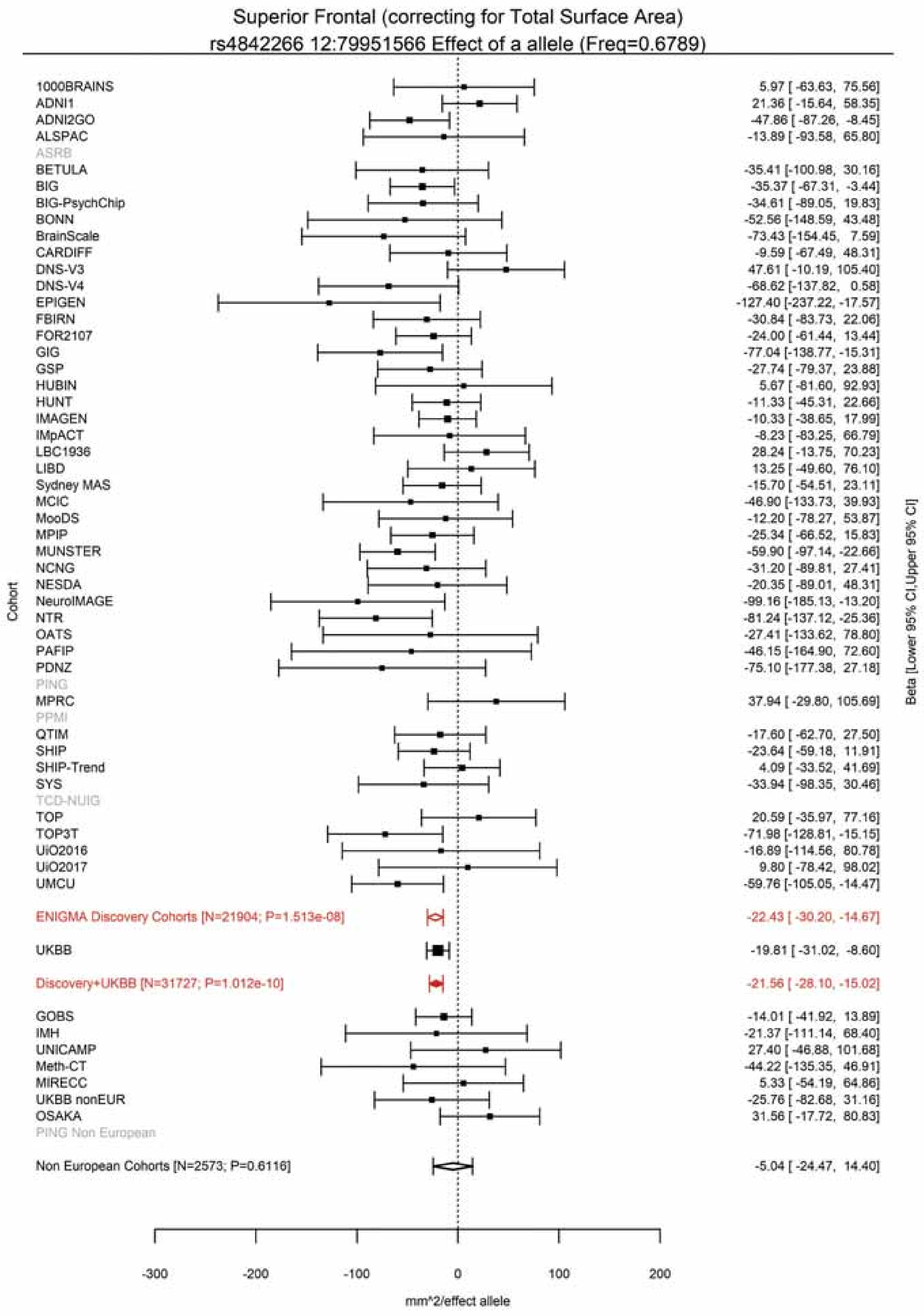
Superior Frontal (correcting for Total Surface Area)
rs4842266 12:79951566 Effect of a allele (Freq=0.6789)
Cohort
1000BRAINS 5.97 [ -63.63, 75.56]
ADNI1 21.36 [ -15.64, 58.35]
ADNI2GO -47.86 [ -87.26, -8.45]
ALSPAC -13.89 [ -93.58, 65.80]
ASRB
BETULA -35.41 [-100.98, 30.16]
BIG -35.37 [ -67.31, -3.44]
BIG-PsychChip -34.61 [ -89.05, 19.83]
BONN -52.56 [-148.59, 43.48]
BrainScale -73.43 [-154.45, 7.59]
CARDIFF -9.59 [ -67.49, 48.31]
DNS-V3 47.61 [ -10.19, 105.40]
DNS-V4 -68.62 [-137.82, 0.58]
EPIGEN -127.40 [-237.22, -17.57]
FBIRN -30.84 [ -83.73, 22.06]
FOR2107 -24.00 [ -61.44, 13.44]
GIG -77.04 [-138.77, -15.31]
GSP -27.74 [ -79.37, 23.88]
HUBIN 5.67 [ -81.60, 92.93]
HUNT -11.33 [ -45.31, 22.66]
IMAGEN -10.33 [ -38.65, 17.99]
IMpACT -8.23 [ -83.25, 66.79]
LBC1936 28.24 [ -13.75, 70.23]
LIBD 13.25 [ -49.60, 76.10]
Sydney MAS -15.70 [ -54.51, 23.11]
MCIC -46.90 [-133.73, 39.93]
MooDS -12.20 [ -78.27, 53.87]
MPIP -25.34 [ -66.52, 15.83]
MUNSTER -59.90 [ -97.14, -22.66]
NCNG -31.20 [ -89.81, 27.41]
NESDA -20.35 [ -89.01, 48.31]
NeuroIMAGE -99.16 [-185.13, -13.20]
NTR -81.24 [-137.12, -25.36]
OATS -27.41 [-133.62, 78.80]
PAFIP -46.15 [-164.90, 72.60]
PDNZ -75.10 [-177.38, 27.18]
PING
MPRC 37.94 [ -29.80, 105.69]
PPMI
QTIM -17.60 [ -62.70, 27.50]
SHIP -23.64 [ -59.18, 11.91]
SHIP-Trend 4.09 [ -33.52, 41.69]
SYS -33.94 [ -98.35, 30.46]
TCD-NUIG
TOP 20.59 [ -35.97, 77.16]
TOP3T -71.98 [-128.81, -15.15]
UiO2016 -16.89 [-114.56, 80.78]
UiO2017 9.80 [ -78.42, 98.02]
UMCU -59.76 [-105.05, -14.47]
ENIGMA Discovery Cohorts [N=21904; P=1.513e-08] -22.43 [ -30.20, -14.67]
UKBB -19.81 [ -31.02, -8.60]
Discovery+UKBB [N=31727; P=1.012e-10] -21.56 [ -28.10, -15.02]
GOBS -14.01 [ -41.92, 13.89]
IMH -21.37 [-111.14, 68.40]
UNICAMP 27.40 [ -46.88, 101.68]
Meth-CT -44.22 [-135.35, 46.91]
MIRECC 5.33 [ -54.19, 64.86]
UKBB nonEUR -25.76 [ -82.68, 31.16]
OSAKA 31.56 [ -17.72, 80.83]
PING Non European
Non European Cohorts [N=2573; P=0.6116] -5.04 [ -24.47, 14.40]
Beta [Lower 95% CI,Upper 95% CI]
-300
-200
-100
0
100
200
mm^2/effect allele

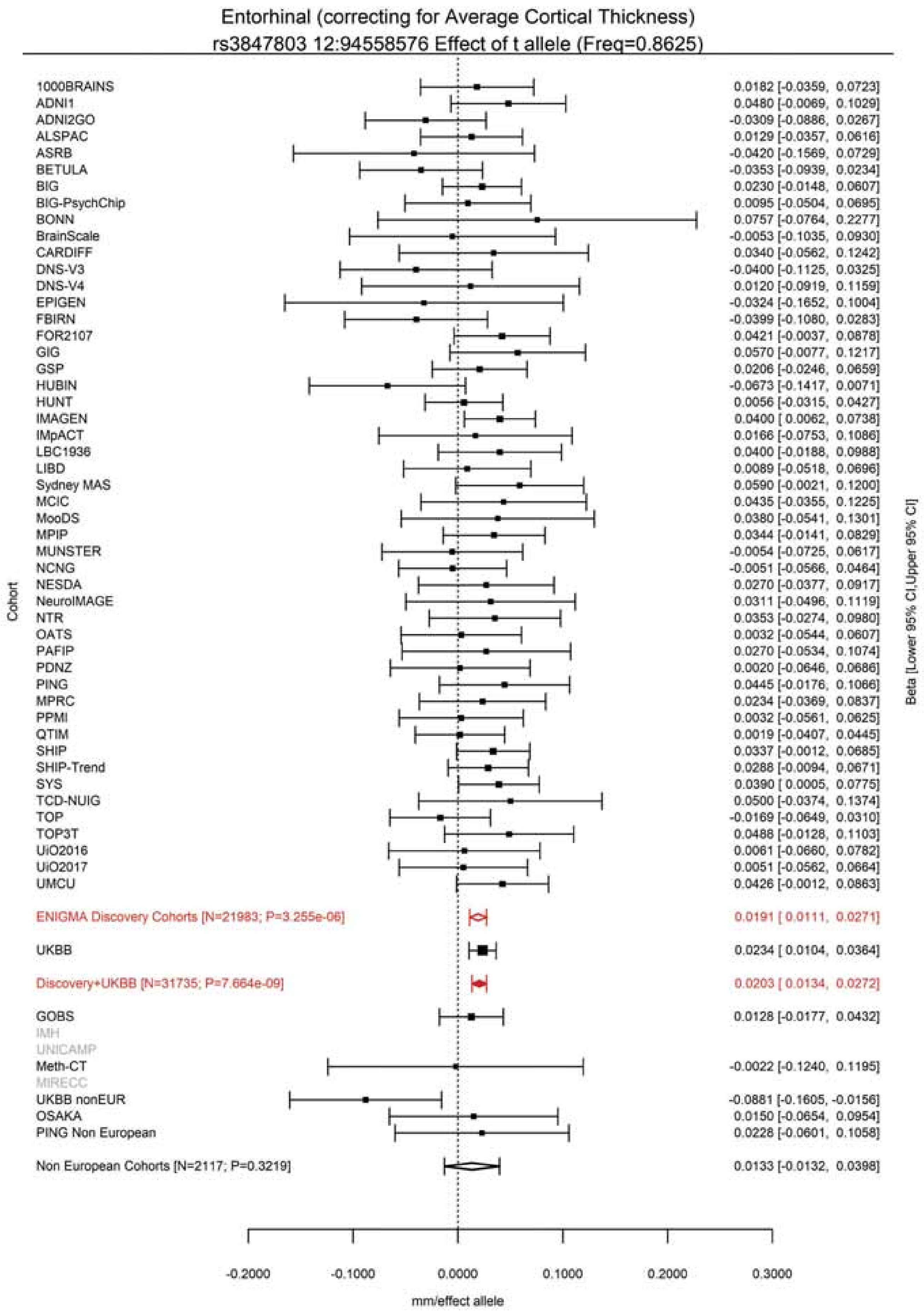
Entorhinal (correcting for Average Cortical Thickness)
rs3847803 12:94558576 Effect of t allele (Freq=0.8625)
Cohort
1000BRAINS 0.0182 [-0.0359, 0.0723]
ADNI1 0.0480 [-0.0069, 0.1029]
ADNI2GO -0.0309 [-0.0886, 0.0267]
ALSPAC 0.0129 [-0.0357, 0.0616]
ASRB -0.0420 [-0.1569, 0.0729]
BETULA -0.0353 [-0.0939, 0.0234]
BIG 0.0230 [-0.0148, 0.0607]
BIG-PsychChip 0.0095 [-0.0504, 0.0695]
BONN 0.0757 [-0.0764, 0.2277]
BrainScale -0.0053 [-0.1035, 0.0930]
CARDIFF 0.0340 [-0.0562, 0.1242]
DNS-V3 -0.0400 [-0.1125, 0.0325]
DNS-V4 0.0120 [-0.0919, 0.1159]
EPIGEN -0.0324 [-0.1652, 0.1004]
FBIRN -0.0399 [-0.1080, 0.0283]
FOR2107 0.0421 [-0.0037, 0.0878]
GIG 0.0570 [-0.0077, 0.1217]
GSP 0.0206 [-0.0246, 0.0659]
HUBIN -0.0673 [-0.1417, 0.0071]
HUNT 0.0056 [-0.0315, 0.0427]
IMAGEN 0.0400 [ 0.0062, 0.0738]
IMpACT 0.0166 [-0.0753, 0.1086]
LBC1936 0.0400 [-0.0188, 0.0988]
LIBD 0.0089 [-0.0518, 0.0696]
Sydney MAS 0.0590 [-0.0021, 0.1200]
MCIC 0.0435 [-0.0355, 0.1225]
MooDS 0.0380 [-0.0541, 0.1301]
MPIP 0.0344 [-0.0141, 0.0829]
MUNSTER -0.0054 [-0.0725, 0.0617]
NCNG -0.0051 [-0.0566, 0.0464]
NESDA 0.0270 [-0.0377, 0.0917]
NeuroIMAGE 0.0311 [-0.0496, 0.1119]
NTR 0.0353 [-0.0274, 0.0980]
OATS 0.0032 [-0.0544, 0.0607]
PAFIP 0.0270 [-0.0534, 0.1074]
PDNZ 0.0020 [-0.0646, 0.0686]
PING 0.0445 [-0.0176, 0.1066]
MPRC 0.0234 [-0.0369, 0.0837]
PPMI 0.0032 [-0.0561, 0.0625]
QTIM 0.0019 [-0.0407, 0.0445]
SHIP 0.0337 [-0.0012, 0.0685]
SHIP-Trend 0.0288 [-0.0094, 0.0671]
SYS 0.0390 [ 0.0005, 0.0775]
TCD-NUIG 0.0500 [-0.0374, 0.1374]
TOP -0.0169 [-0.0649, 0.0310]
TOP3T 0.0488 [-0.0128, 0.1103]
UiO2016 0.0061 [-0.0660, 0.0782]
UiO2017 0.0051 [-0.0562, 0.0664]
UMCU 0.0426 [-0.0012, 0.0863]
ENIGMA Discovery Cohorts [N=21983; P=3.255e-06] 0.0191 [ 0.0111, 0.0271]
UKBB 0.0234 [ 0.0104, 0.0364]
Discovery+UKBB [N=31735; P=7.664e-09] 0.0203 [ 0.0134, 0.0272]
GOBS 0.0128 [-0.0177, 0.0432]
IMH
UNICAMP
Meth-CT -0.0022 [-0.1240, 0.1195]
MIRECC
UKBB nonEUR -0.0881 [-0.1605, -0.0156]
OSAKA 0.0150 [-0.0654, 0.0954]
PING Non European 0.0228 [-0.0601, 0.1058]
Non European Cohorts [N=2117; P=0.3219] 0.0133 [-0.0132, 0.0398]
-0.2000
-0.1000
0.0000
0.1000
0.2000
0.3000
mm/effect allele
Beta [Lower 95% CI,Upper 95% CI]

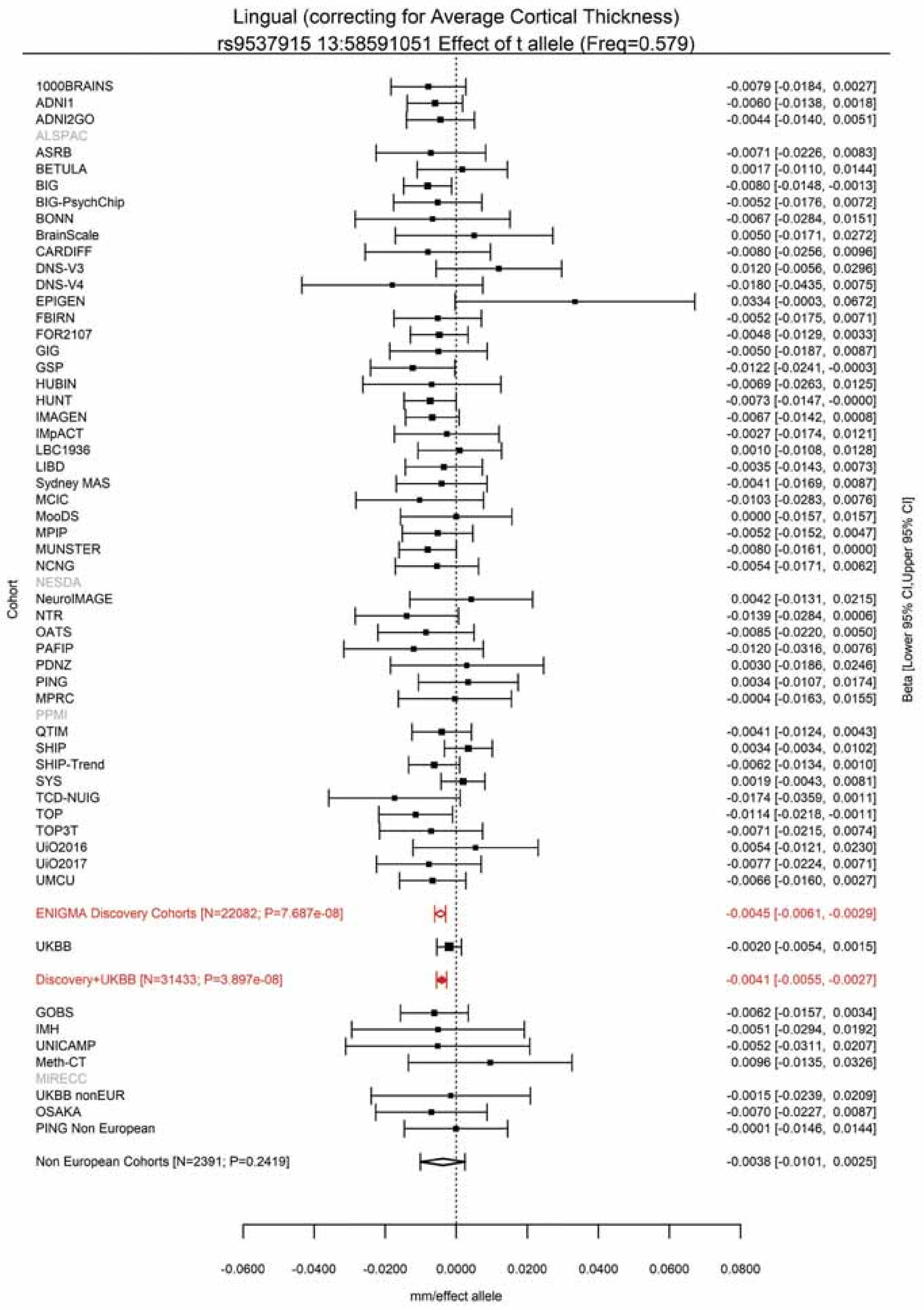

Lingual (correcting for Average Cortical Thickness)
rs9537915 13:58591051 Effect of t allele (Freq=0.579)
Cohort
1000BRAINS -0.0079 [-0.0184, 0.0027]
ADNI1 -0.0060 [-0.0138, 0.0018]
ADNI2GO -0.0044 [-0.0140, 0.0051]
ALSPAC
ASRB -0.0071 [-0.0226, 0.0083]
BETULA 0.0017 [-0.0110, 0.0144]
BIG -0.0080 [-0.0148, -0.0013]
BIG-PsychChip -0.0052 [-0.0176, 0.0072]
BONN -0.0067 [-0.0284, 0.0151]
BrainScale 0.0050 [-0.0171, 0.0272]
CARDIFF -0.0080 [-0.0256, 0.0096]
DNS-V3 0.0120 [-0.0056, 0.0296]
DNS-V4 -0.0180 [-0.0435, 0.0075]
EPIGEN 0.0334 [-0.0003, 0.0672]
FBIRN -0.0052 [-0.0175, 0.0071]
FOR2107 -0.0048 [-0.0129, 0.0033]
GIG -0.0050 [-0.0187, 0.0087]
GSP -0.0122 [-0.0241, -0.0003]
HUBIN -0.0069 [-0.0263, 0.0125]
HUNT -0.0073 [-0.0147, -0.0000]
IMAGEN -0.0067 [-0.0142, 0.0008]
IMpACT -0.0027 [-0.0174, 0.0121]
LBC1936 0.0010 [-0.0108, 0.0128]
LIBD -0.0035 [-0.0143, 0.0073]
Sydney MAS -0.0041 [-0.0169, 0.0087]
MCIC -0.0103 [-0.0283, 0.0076]
MooDS 0.0000 [-0.0157, 0.0157]
MPIP -0.0052 [-0.0152, 0.0047]
MUNSTER -0.0080 [-0.0161, 0.0000]
NCNG -0.0054 [-0.0171, 0.0062]
NESDA
NeuroIMAGE 0.0042 [-0.0131, 0.0215]
NTR -0.0139 [-0.0284, 0.0006]
OATS -0.0085 [-0.0220, 0.0050]
PAFIP -0.0120 [-0.0316, 0.0076]
PDNZ 0.0030 [-0.0186, 0.0246]
PING 0.0034 [-0.0107, 0.0174]
MPRC -0.0004 [-0.0163, 0.0155]
PPMI
QTIM -0.0041 [-0.0124, 0.0043]
SHIP 0.0034 [-0.0034, 0.0102]
SHIP-Trend -0.0062 [-0.0134, 0.0010]
SYS 0.0019 [-0.0043, 0.0081]
TCD-NUIG -0.0174 [-0.0359, 0.0011]
TOP -0.0114 [-0.0218, -0.0011]
TOP3T -0.0071 [-0.0215, 0.0074]
UiO2016 0.0054 [-0.0121, 0.0230]
UiO2017 -0.0077 [-0.0224, 0.0071]
UMCU -0.0066 [-0.0160, 0.0027]
ENIGMA Discovery Cohorts [N=22082; P=7.687e-08] -0.0045 [-0.0061, -0.0029]
UKBB -0.0020 [-0.0054, 0.0015]
Discovery+UKBB [N=31433; P=3.897e-08] -0.0041 [-0.0055, -0.0027]
GOBS -0.0062 [-0.0157, 0.0034]
IMH -0.0051 [-0.0294, 0.0192]
UNICAMP -0.0052 [-0.0311, 0.0207]
Meth-CT 0.0096 [-0.0135, 0.0326]
MIRECC
UKBB nonEUR -0.0015 [-0.0239, 0.0209]
OSAKA -0.0070 [-0.0227, 0.0087]
PING Non European -0.0001 [-0.0146, 0.0144]
Non European Cohorts [N=2391; P=0.2419] -0.0038 [-0.0101, 0.0025]
Beta [Lower 95% CI,Upper 95% CI]
-0.0600
-0.0400
-0.0200
0.0000
0.0200
0.0400
0.0600
0.0800
mm/effect allele

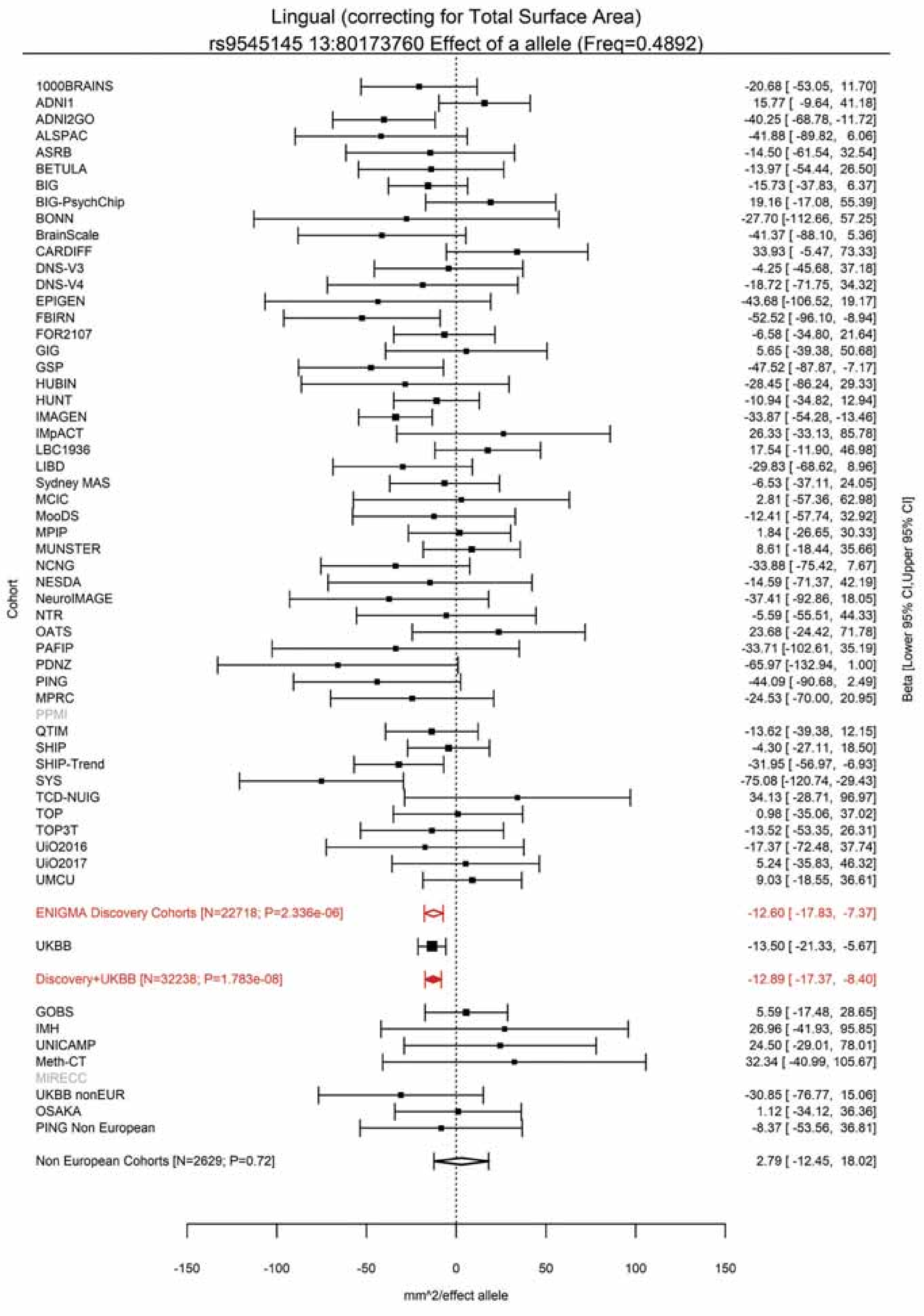
Lingual (correcting for Total Surface Area)
rs9545145 13:80173760 Effect of a allele (Freq=0.4892)
Cohort
1000BRAINS -20.68 [ -53.05, 11.70]
ADNI1 15.77 [ -9.64, 41.18]
ADNI2GO -40.25 [ -68.78, -11.72]
ALSPAC -41.88 [ -89.82, 6.06]
ASRB -14.50 [ -61.54, 32.54]
BETULA -13.97 [ -54.44, 26.50]
BIG -15.73 [ -37.83, 6.37]
BIG-PsychChip 19.16 [ -17.08, 55.39]
BONN -27.70 [-112.66, 57.25]
BrainScale -41.37 [ -88.10, 5.36]
CARDIFF 33.93 [ -5.47, 73.33]
DNS-V3 -4.25 [ -45.68, 37.18]
DNS-V4 -18.72 [ -71.75, 34.32]
EPIGEN -43.68 [-106.52, 19.17]
FBIRN -52.52 [ -96.10, -8.94]
FOR2107 -6.58 [ -34.80, 21.64]
GIG 5.65 [ -39.38, 50.68]
GSP -47.52 [ -87.87, -7.17]
HUBIN -28.45 [ -86.24, 29.33]
HUNT -10.94 [ -34.82, 12.94]
IMAGEN -33.87 [ -54.28, -13.46]
IMpACT 26.33 [ -33.13, 85.78]
LBC1936 17.54 [ -11.90, 46.98]
LIBD -29.83 [ -68.62, 8.96]
Sydney MAS -6.53 [ -37.11, 24.05]
MCIC 2.81 [ -57.36, 62.98]
MooDS -12.41 [ -57.74, 32.92]
MPIP 1.84 [ -26.65, 30.33]
MUNSTER 8.61 [ -18.44, 35.66]
NCNG -33.88 [ -75.42, 7.67]
NESDA -14.59 [ -71.37, 42.19]
NeuroIMAGE -37.41 [ -92.86, 18.05]
NTR -5.59 [ -55.51, 44.33]
OATS 23.68 [ -24.42, 71.78]
PAFIP -33.71 [-102.61, 35.19]
PDNZ -65.97 [-132.94, 1.00]
PING -44.09 [ -90.68, 2.49]
MPRC -24.53 [ -70.00, 20.95]
PPMI
QTIM -13.62 [ -39.38, 12.15]
SHIP -4.30 [ -27.11, 18.50]
SHIP-Trend -31.95 [ -56.97, -6.93]
SYS -75.08 [-120.74, -29.43]
TCD-NUIG 34.13 [ -28.71, 96.97]
TOP 0.98 [ -35.06, 37.02]
TOP3T -13.52 [ -53.35, 26.31]
UiO2016 -17.37 [ -72.48, 37.74]
UiO2017 5.24 [ -35.83, 46.32]
UMCU 9.03 [ -18.55, 36.61]
ENIGMA Discovery Cohorts [N=22718; P=2.336e-06] -12.60 [ -17.83, -7.37]
UKBB -13.50 [ -21.33, -5.67]
Discovery+UKBB [N=32238; P=1.783e-08] -12.89 [ -17.37, -8.40]
GOBS 5.59 [ -17.48, 28.65]
IMH 26.96 [ -41.93, 95.85]
UNICAMP 24.50 [ -29.01, 78.01]
Meth-CT 32.34 [ -40.99, 105.67]
MIRECC
UKBB nonEUR -30.85 [ -76.77, 15.06]
OSAKA 1.12 [ -34.12, 36.36]
PING Non European -8.37 [ -53.56, 36.81]
Non European Cohorts [N=2629; P=0.72] 2.79 [ -12.45, 18.02]
Beta [Lower 95% CI,Upper 95% CI]
-150
-100
-50
0
50
100
150
mm^2/effect allele

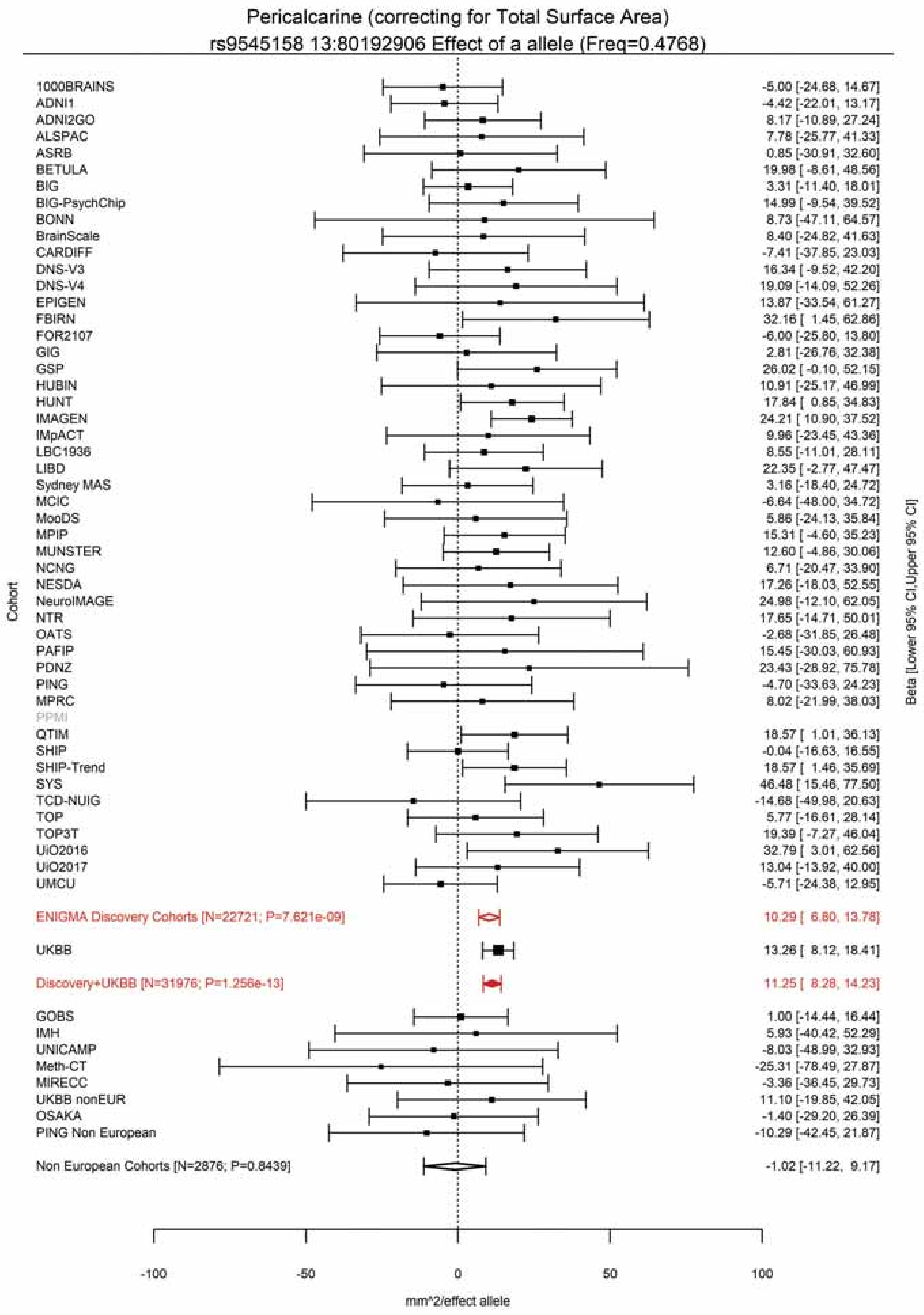
Pericalcarine (correcting for Total Surface Area)
rs9545158 13:80192906 Effect of a allele (Freq=0.4768)
Cohort
1000BRAINS -5.00 [-24.68, 14.67]
ADNI1 -4.42 [-22.01, 13.17]
ADNI2GO 8.17 [-10.89, 27.24]
ALSPAC 7.78 [-25.77, 41.33]
ASRB 0.85 [-30.91, 32.60]
BETULA 19.98 [ -8.61, 48.56]
BIG 3.31 [-11.40, 18.01]
BIG-PsychChip 14.99 [ -9.54, 39.52]
BONN 8.73 [-47.11, 64.57]
BrainScale 8.40 [-24.82, 41.63]
CARDIFF -7.41 [-37.85, 23.03]
DNS-V3 16.34 [ -9.52, 42.20]
DNS-V4 19.09 [-14.09, 52.26]
EPIGEN 13.87 [-33.54, 61.27]
FBIRN 32.16 [ 1.45, 62.86]
FOR2107 -6.00 [-25.80, 13.80]
GIG 2.81 [-26.76, 32.38]
GSP 26.02 [ -0.10, 52.15]
HUBIN 10.91 [-25.17, 46.99]
HUNT 17.84 [ 0.85, 34.83]
IMAGEN 24.21 [ 10.90, 37.52]
IMpACT 9.96 [-23.45, 43.36]
LBC1936 8.55 [-11.01, 28.11]
LIBD 22.35 [ -2.77, 47.47]
Sydney MAS 3.16 [-18.40, 24.72]
MCIC -6.64 [-48.00, 34.72]
MooDS 5.86 [-24.13, 35.84]
MPIP 15.31 [ -4.60, 35.23]
MUNSTER 12.60 [ -4.86, 30.06]
NCNG 6.71 [-20.47, 33.90]
NESDA 17.26 [-18.03, 52.55]
NeuroIMAGE 24.98 [-12.10, 62.05]
NTR 17.65 [-14.71, 50.01]
OATS -2.68 [-31.85, 26.48]
PAFIP 15.45 [-30.03, 60.93]
PDNZ 23.43 [-28.92, 75.78]
PING -4.70 [-33.63, 24.23]
MPRC 8.02 [-21.99, 38.03]
PPMI
QTIM 18.57 [ 1.01, 36.13]
SHIP -0.04 [-16.63, 16.55]
SHIP-Trend 18.57 [ 1.46, 35.69]
SYS 46.48 [ 15.46, 77.50]
TCD-NUIG -14.68 [-49.98, 20.63]
TOP 5.77 [-16.61, 28.14]
TOP3T 19.39 [ -7.27, 46.04]
UiO2016 32.79 [ 3.01, 62.56]
UiO2017 13.04 [-13.92, 40.00]
UMCU -5.71 [-24.38, 12.95]
ENIGMA Discovery Cohorts [N=22721; P=7.621e-09] 10.29 [ 6.80, 13.78]
UKBB 13.26 [ 8.12, 18.41]
Discovery+UKBB [N=31976; P=1.256e-13] 11.25 [ 8.28, 14.23]
GOBS 1.00 [-14.44, 16.44]
IMH 5.93 [-40.42, 52.29]
UNICAMP -8.03 [-48.99, 32.93]
Meth-CT -25.31 [-78.49, 27.87]
MIRECC -3.36 [-36.45, 29.73]
UKBB nonEUR 11.10 [-19.85, 42.05]
OSAKA -1.40 [-29.20, 26.39]
PING Non European -10.29 [-42.45, 21.87]
Non European Cohorts [N=2876; P=0.8439] -1.02 [-11.22, 9.17]
Beta [Lower 95% CI,Upper 95% CI]
-100
-50
0
50
100
mm^2/effect allele

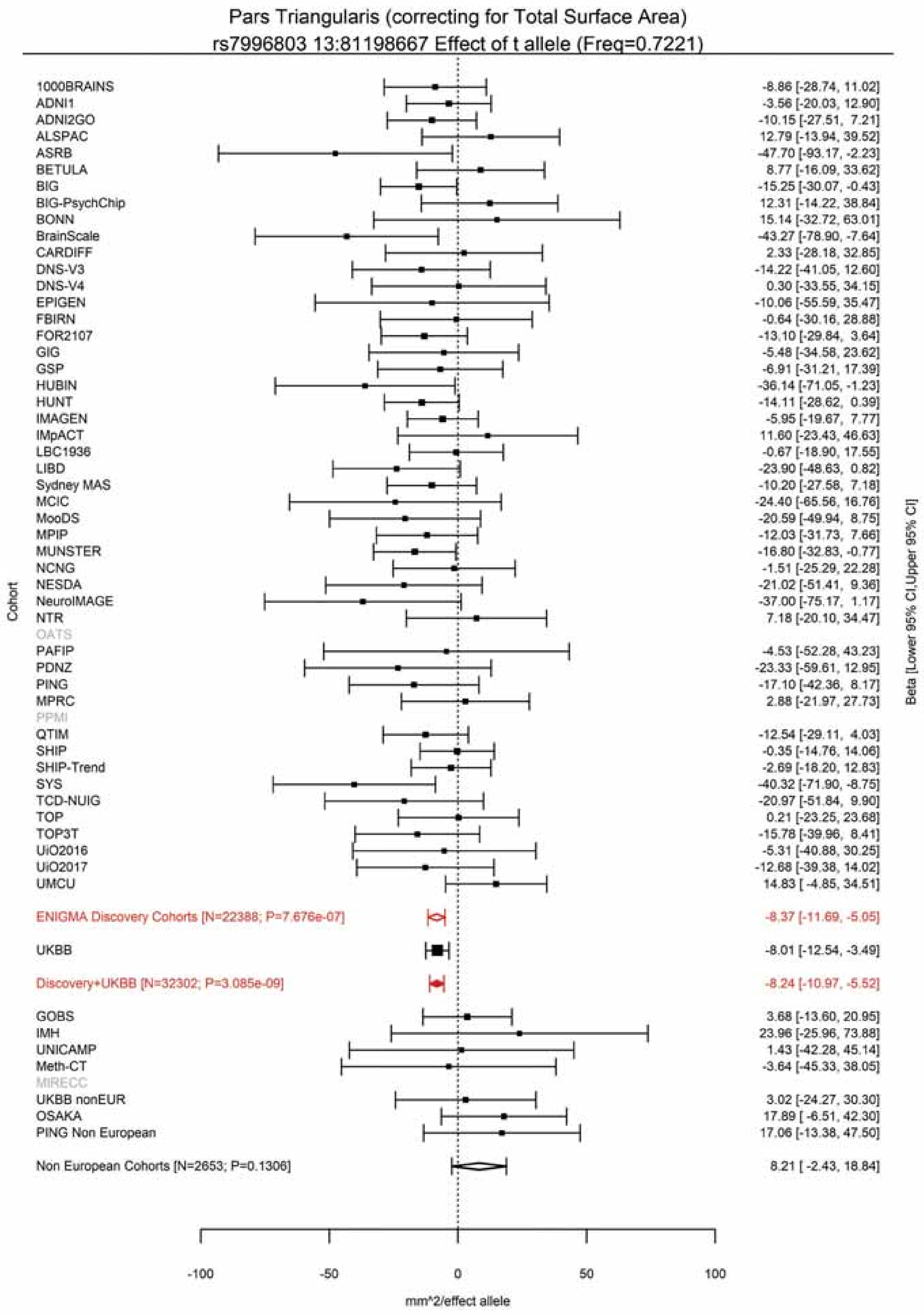
Pars Triangularis (correcting for Total Surface Area)
rs7996803 13:81198667 Effect of t allele (Freq=0.7221)
1000BRAINS -8.86 [-28.74, 11.02]
ADNI1 -3.56 [-20.03, 12.90]
ADNI2GO -10.15 [-27.51, 7.21]
ALSPAC 12.79 [-13.94, 39.52]
ASRB -47.70 [-93.17, -2.23]
BETULA 8.77 [-16.09, 33.62]
BIG -15.25 [-30.07, -0.43]
BIG-PsychChip 12.31 [-14.22, 38.84]
BONN 15.14 [-32.72, 63.01]
BrainScale -43.27 [-78.90, -7.64]
CARDIFF 2.33 [-28.18, 32.85]
DNS-V3 -14.22 [-41.05, 12.60]
DNS-V4 0.30 [-33.55, 34.15]
EPIGEN -10.06 [-55.59, 35.47]
FBIRN -0.64 [-30.16, 28.88]
FOR2107 -13.10 [-29.84, 3.64]
GIG -5.48 [-34.58, 23.62]
GSP -6.91 [-31.21, 17.39]
HUBIN -36.14 [-71.05, -1.23]
HUNT -14.11 [-28.62, 0.39]
IMAGEN -5.95 [-19.67, 7.77]
IMpACT 11.60 [-23.43, 46.63]
LBC1936 -0.67 [-18.90, 17.55]
LIBD -23.90 [-48.63, 0.82]
Sydney MAS -10.20 [-27.58, 7.18]
MCIC -24.40 [-65.56, 16.76]
MooDS -20.59 [-49.94, 8.75]
MPIP -12.03 [-31.73, 7.66]
MUNSTER -16.80 [-32.83, -0.77]
NCNG -1.51 [-25.29, 22.28]
NESDA -21.02 [-51.41, 9.36]
NeuroIMAGE -37.00 [-75.17, 1.17]
NTR 7.18 [-20.10, 34.47]
OATS
PAFIP -4.53 [-52.28, 43.23]
PDNZ -23.33 [-59.61, 12.95]
PING -17.10 [-42.36, 8.17]
MPRC 2.88 [-21.97, 27.73]
PPMI
QTIM -12.54 [-29.11, 4.03]
SHIP -0.35 [-14.76, 14.06]
SHIP-Trend -2.69 [-18.20, 12.83]
SYS -40.32 [-71.90, -8.75]
TCD-NUIG -20.97 [-51.84, 9.90]
TOP 0.21 [-23.25, 23.68]
TOP3T -15.78 [-39.96, 8.41]
UiO2016 -5.31 [-40.88, 30.25]
UiO2017 -12.68 [-39.38, 14.02]
UMCU 14.83 [ -4.85, 34.51]
ENIGMA Discovery Cohorts [N=22388; P=7.676e-07] -8.37 [-11.69, -5.05]
UKBB -8.01 [-12.54, -3.49]
Discovery+UKBB [N=32302; P=3.085e-09] -8.24 [-10.97, -5.52]
GOBS 3.68 [-13.60, 20.95]
IMH 23.96 [-25.96, 73.88]
UNICAMP 1.43 [-42.28, 45.14]
Meth-CT -3.64 [-45.33, 38.05]
MIRECC
UKBB nonEUR 3.02 [-24.27, 30.30]
OSAKA 17.89 [ -6.51, 42.30]
PING Non European 17.06 [-13.38, 47.50]
Non European Cohorts [N=2653; P=0.1306] 8.21 [ -2.43, 18.84]
Cohort
Beta [Lower 95% CI,Upper 95% CI]
-100
-50
0
50
100
mm^2/effect allele

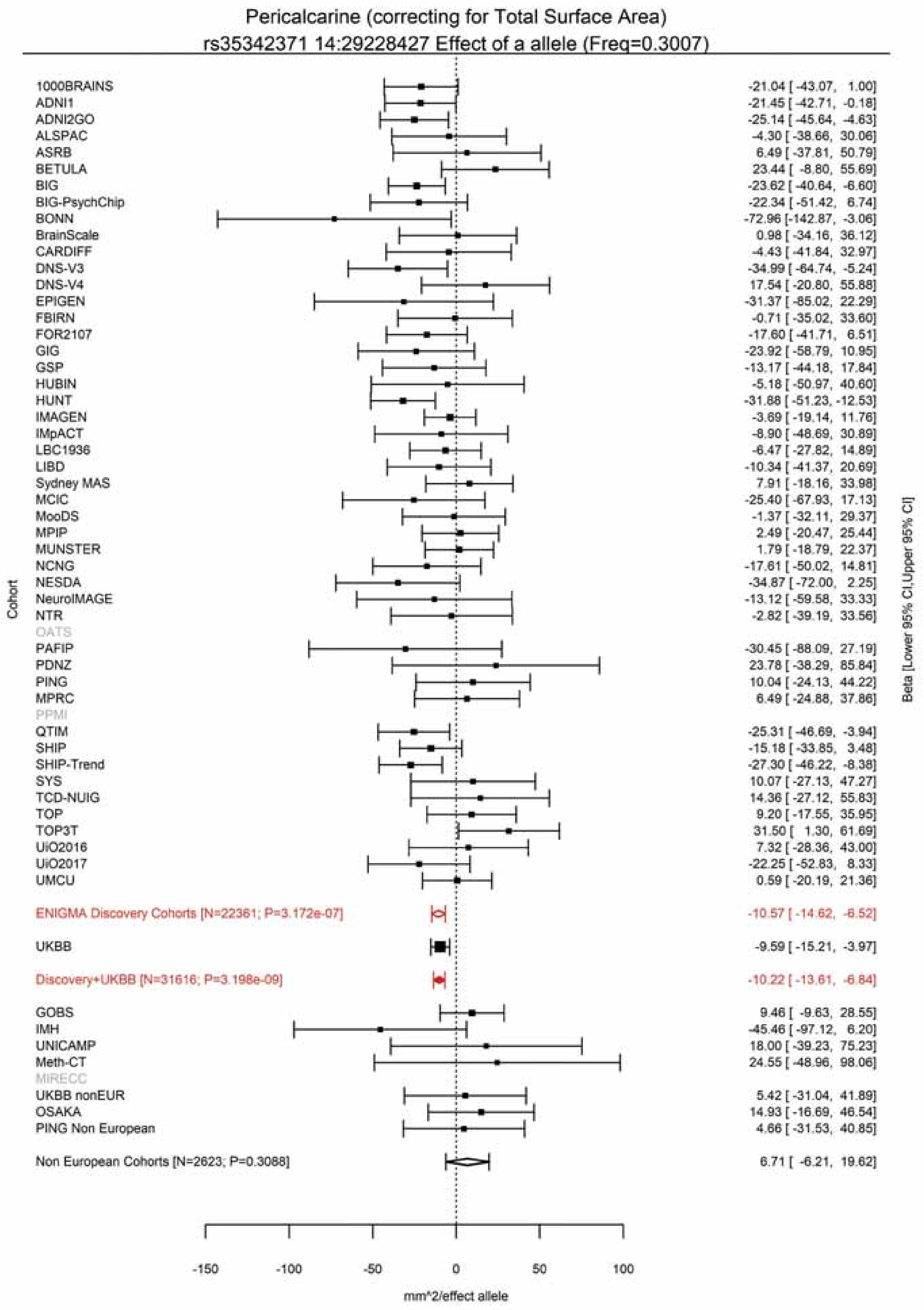

Pericalcarine (correcting for Total Surface Area)
rs35342371 14:29228427 Effect of a allele (Freq=0.3007)
Cohort
1000BRAINS -21.04 [ -43.07, 1.00]
ADNI1 -21.45 [ -42.71, -0.18]
ADNI2GO -25.14 [ -45.64, -4.63]
ALSPAC -4.30 [ -38.66, 30.06]
ASRB 6.49 [ -37.81, 50.79]
BETULA 23.44 [ -8.80, 55.69]
BIG -23.62 [ -40.64, -6.60]
BIG-PsychChip -22.34 [ -51.42, 6.74]
BONN -72.96 [-142.87, -3.06]
BrainScale 0.98 [ -34.16, 36.12]
CARDIFF -4.43 [ -41.84, 32.97]
DNS-V3 -34.99 [ -64.74, -5.24]
DNS-V4 17.54 [ -20.80, 55.88]
EPIGEN -31.37 [ -85.02, 22.29]
FBIRN -0.71 [ -35.02, 33.60]
FOR2107 -17.60 [ -41.71, 6.51]
GIG -23.92 [ -58.79, 10.95]
GSP -13.17 [ -44.18, 17.84]
HUBIN -5.18 [ -50.97, 40.60]
HUNT -31.88 [ -51.23, -12.53]
IMAGEN -3.69 [ -19.14, 11.76]
IMpACT -8.90 [ -48.69, 30.89]
LBC1936 -6.47 [ -27.82, 14.89]
LIBD -10.34 [ -41.37, 20.69]
Sydney MAS 7.91 [ -18.16, 33.98]
MCIC -25.40 [ -67.93, 17.13]
MooDS -1.37 [ -32.11, 29.37]
MPIP 2.49 [ -20.47, 25.44]
MUNSTER 1.79 [ -18.79, 22.37]
NCNG -17.61 [ -50.02, 14.81]
NESDA -34.87 [ -72.00, 2.25]
NeuroIMAGE -13.12 [ -59.58, 33.33]
NTR -2.82 [ -39.19, 33.56]
OATS
PAFIP -30.45 [ -88.09, 27.19]
PDNZ 23.78 [ -38.29, 85.84]
PING 10.04 [ -24.13, 44.22]
MPRC 6.49 [ -24.88, 37.86]
PPMI
QTIM -25.31 [ -46.69, -3.94]
SHIP -15.18 [ -33.85, 3.48]
SHIP-Trend -27.30 [ -46.22, -8.38]
SYS 10.07 [ -27.13, 47.27]
TCD-NUIG 14.36 [ -27.12, 55.83]
TOP 9.20 [ -17.55, 35.95]
TOP3T 31.50 [ 1.30, 61.69]
UiO2016 7.32 [ -28.36, 43.00]
UiO2017 -22.25 [ -52.83, 8.33]
UMCU 0.59 [ -20.19, 21.36]
ENIGMA Discovery Cohorts [N=22361; P=3.172e-07] -10.57 [ -14.62, -6.52]
UKBB -9.59 [ -15.21, -3.97]
Discovery+UKBB [N=31616; P=3.198e-09] -10.22 [ -13.61, -6.84]
GOBS 9.46 [ -9.63, 28.55]
IMH -45.46 [ -97.12, 6.20]
UNICAMP 18.00 [ -39.23, 75.23]
Meth-CT 24.55 [ -48.96, 98.06]
MIRECC
UKBB nonEUR 5.42 [ -31.04, 41.89]
OSAKA 14.93 [ -16.69, 46.54]
PING Non European 4.66 [ -31.53, 40.85]
Non European Cohorts [N=2623; P=0.3088] 6.71 [ -6.21, 19.62]
-150
-100
-50
0
50
100
mm^2/effect allele
Beta [Lower 95% CI,Upper 95% CI]

Lingual (correcting for Total Surface Area)
rs7148896 14:54316607 Effect of a allele (Freq=0.2877)

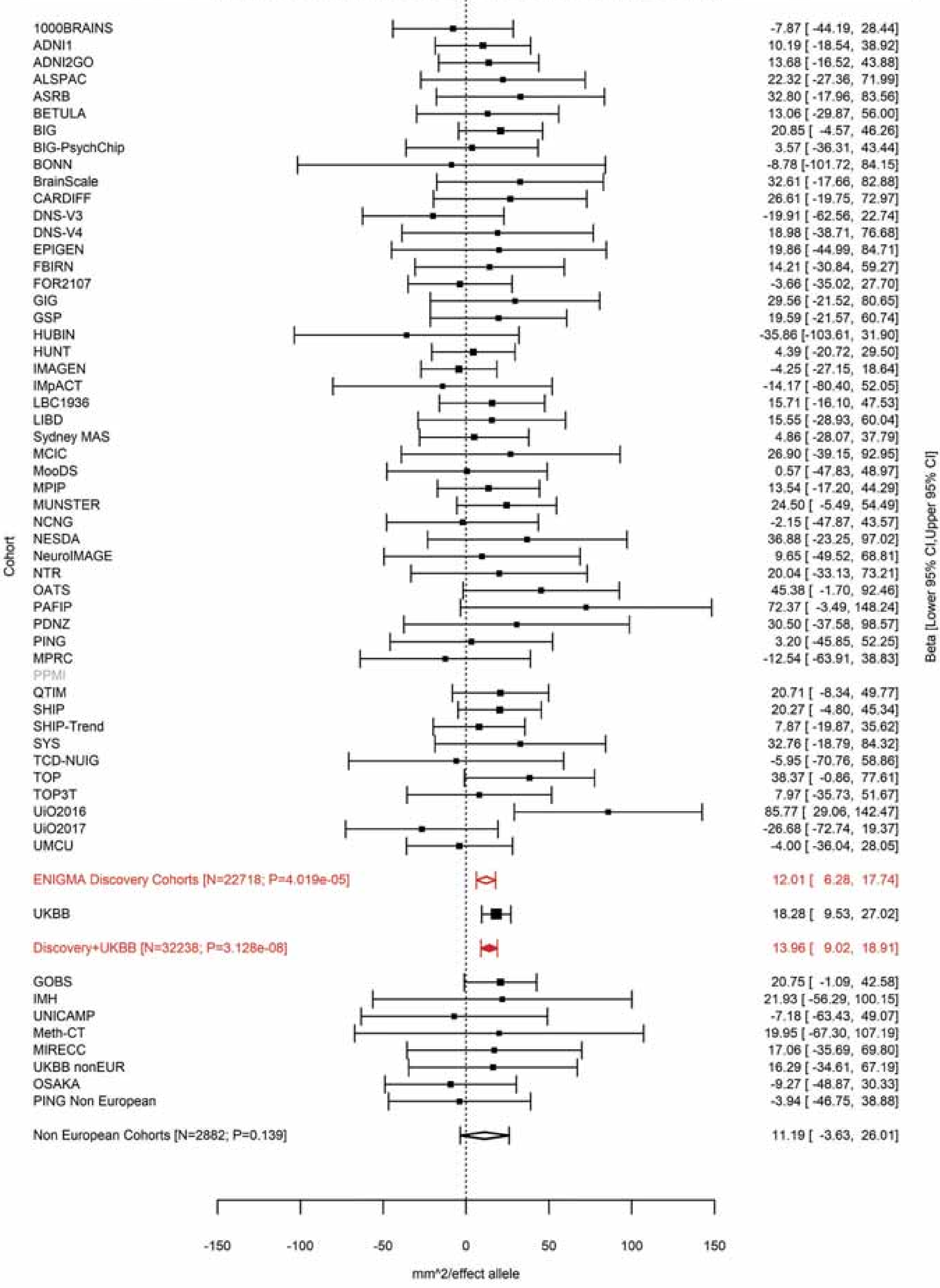

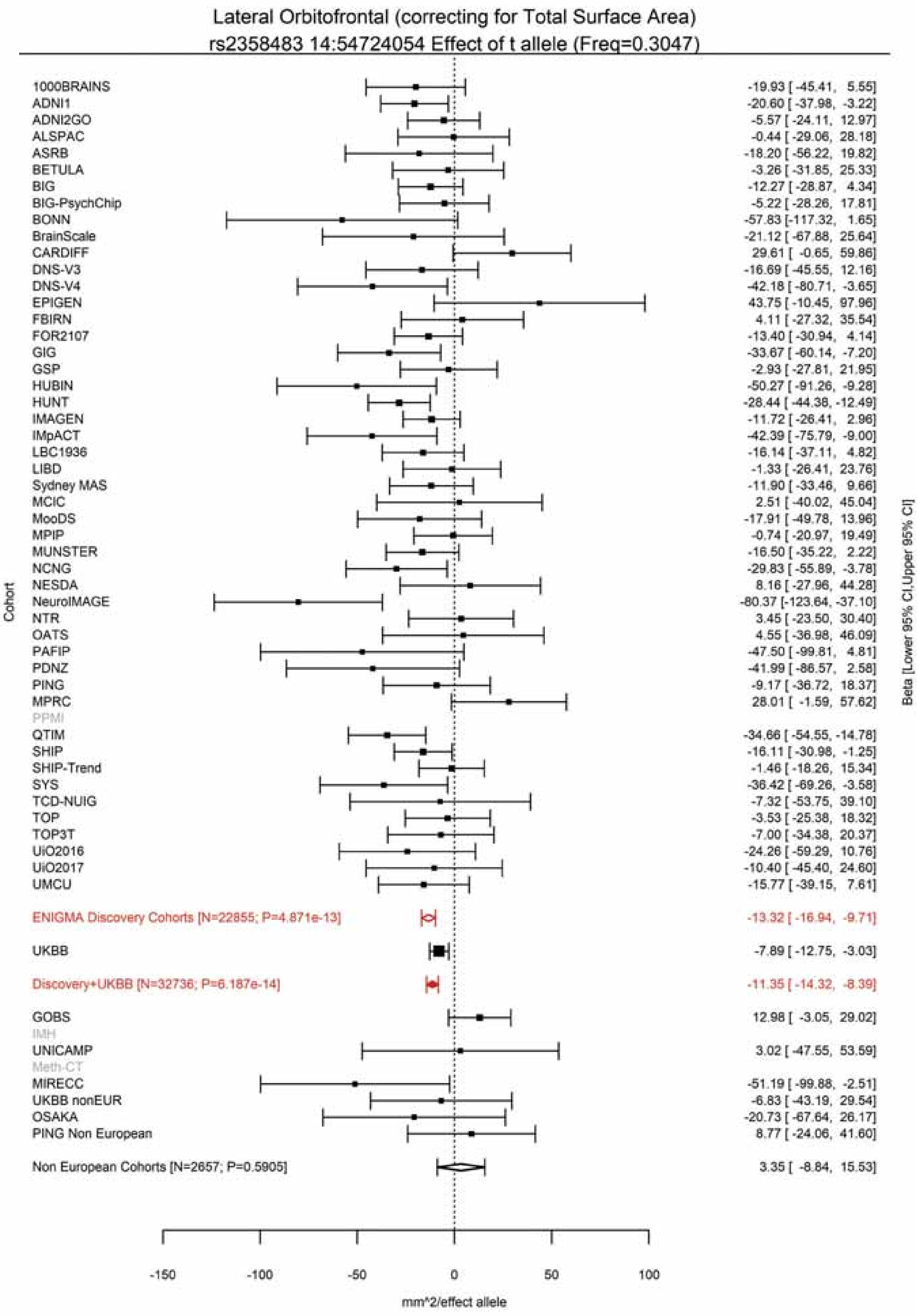

Lateral Orbitofrontal (correcting for Total Surface Area)
rs2358483 14:54724054 Effect of t allele (Freq=0.3047)
Cohort
1000BRAINS -19.93 [ -45.41, 5.55]
ADNI1 -20.60 [ -37.98, -3.22]
ADNI2GO -5.57 [ -24.11, 12.97]
ALSPAC -0.44 [ -29.06, 28.18]
ASRB -18.20 [ -56.22, 19.82]
BETULA -3.26 [ -31.85, 25.33]
BIG -12.27 [ -28.87, 4.34]
BIG-PsychChip -5.22 [ -28.26, 17.81]
BONN -57.83 [-117.32, 1.65]
BrainScale -21.12 [ -67.88, 25.64]
CARDIFF 29.61 [ -0.65, 59.86]
DNS-V3 -16.69 [ -45.55, 12.16]
DNS-V4 -42.18 [ -80.71, -3.65]
EPIGEN 43.75 [ -10.45, 97.96]
FBIRN 4.11 [ -27.32, 35.54]
FOR2107 -13.40 [ -30.94, 4.14]
GIG -33.67 [ -60.14, -7.20]
GSP -2.93 [ -27.81, 21.95]
HUBIN -50.27 [ -91.26, -9.28]
HUNT -28.44 [ -44.38, -12.49]
IMAGEN -11.72 [ -26.41, 2.96]
IMpACT -42.39 [ -75.79, -9.00]
LBC1936 -16.14 [ -37.11, 4.82]
LIBD -1.33 [ -26.41, 23.76]
Sydney MAS -11.90 [ -33.46, 9.66]
MCIC 2.51 [ -40.02, 45.04]
MooDS -17.91 [ -49.78, 13.96]
MPIP -0.74 [ -20.97, 19.49]
MUNSTER -16.50 [ -35.22, 2.22]
NCNG -29.83 [ -55.89, -3.78]
NESDA 8.16 [ -27.96, 44.28]
NeuroIMAGE -80.37 [-123.64, -37.10]
NTR 3.45 [ -23.50, 30.40]
OATS 4.55 [ -36.98, 46.09]
PAFIP -47.50 [ -99.81, 4.81]
PDNZ -41.99 [ -86.57, 2.58]
PING -9.17 [ -36.72, 18.37]
MPRC 28.01 [ -1.59, 57.62]
PPMI
QTIM -34.66 [ -54.55, -14.78]
SHIP -16.11 [ -30.98, -1.25]
SHIP-Trend -1.46 [ -18.26, 15.34]
SYS -36.42 [ -69.26, -3.58]
TCD-NUIG -7.32 [ -53.75, 39.10]
TOP -3.53 [ -25.38, 18.32]
TOP3T -7.00 [ -34.38, 20.37]
UiO2016 -24.26 [ -59.29, 10.76]
UiO2017 -10.40 [ -45.40, 24.60]
UMCU -15.77 [ -39.15, 7.61]
ENIGMA Discovery Cohorts [N=22855; P=4.871e-13] -13.32 [ -16.94, -9.71]
UKBB -7.89 [ -12.75, -3.03]
Discovery+UKBB [N=32736; P=6.187e-14] -11.35 [ -14.32, -8.39]
GOBS 12.98 [ -3.05, 29.02]
IMH
UNICAMP 3.02 [ -47.55, 53.59]
Meth-CT
MIRECC -51.19 [ -99.88, -2.51]
UKBB nonEUR -6.83 [ -43.19, 29.54]
OSAKA -20.73 [ -67.64, 26.17]
PING Non European 8.77 [ -24.06, 41.60]
Non European Cohorts [N=2657; P=0.5905] 3.35 [ -8.84, 15.53]
Beta [Lower 95% CI,Upper 95% CI]
-150
-100
-50
0
50
100
mm^2/effect allele

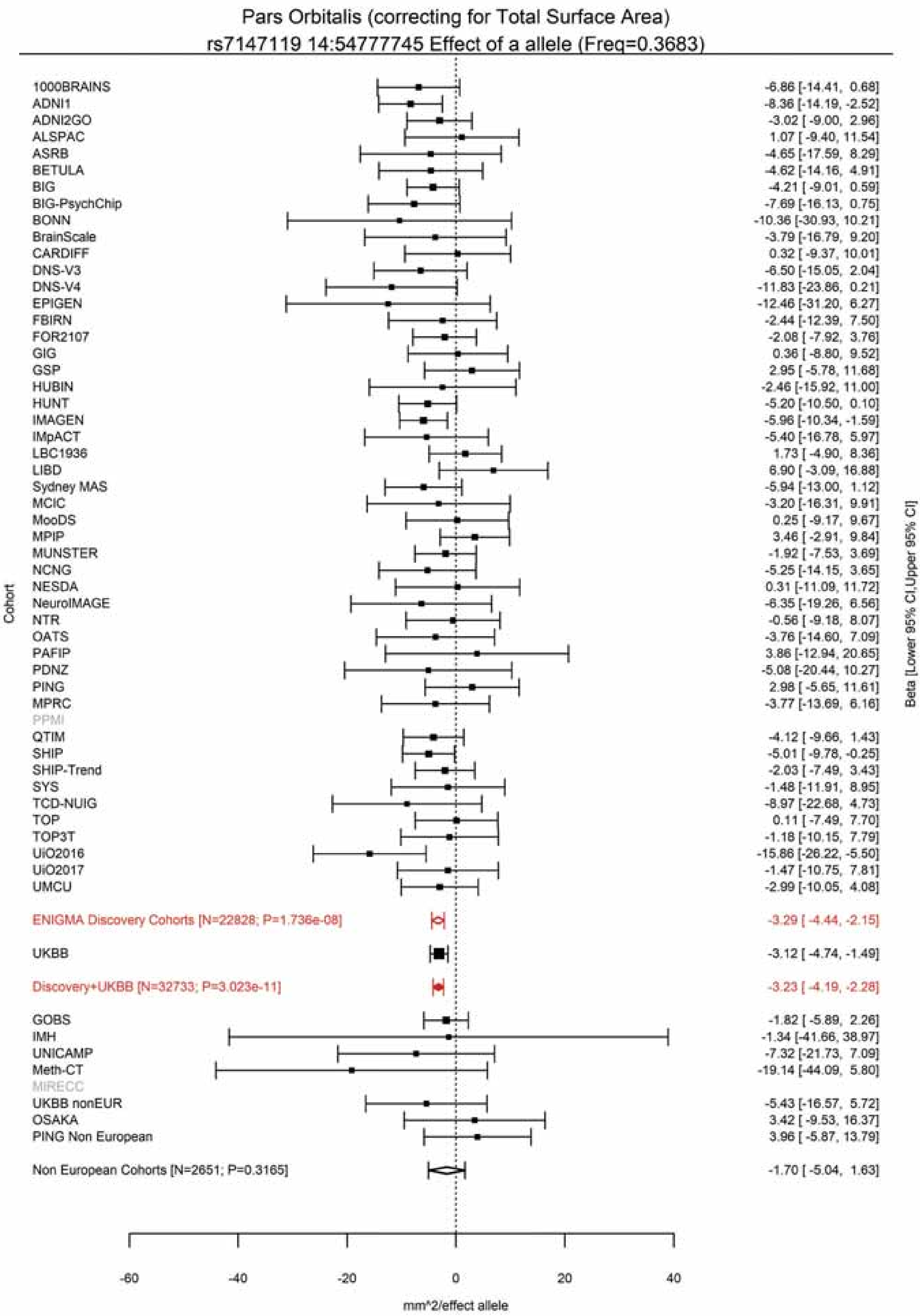

Pars Orbitalis (correcting for Total Surface Area)
rs7147119 14:54777745 Effect of a allele (Freq=0.3683)
Cohort
1000BRAINS -6.86 [-14.41, 0.68]
ADNI1 -8.36 [-14.19, -2.52]
ADNI2GO -3.02 [ -9.00, 2.96]
ALSPAC 1.07 [ -9.40, 11.54]
ASRB -4.65 [-17.59, 8.29]
BETULA -4.62 [-14.16, 4.91]
BIG -4.21 [ -9.01, 0.59]
BIG-PsychChip -7.69 [-16.13, 0.75]
BONN -10.36 [-30.93, 10.21]
BrainScale -3.79 [-16.79, 9.20]
CARDIFF 0.32 [ -9.37, 10.01]
DNS-V3 -6.50 [-15.05, 2.04]
DNS-V4 -11.83 [-23.86, 0.21]
EPIGEN -12.46 [-31.20, 6.27]
FBIRN -2.44 [-12.39, 7.50]
FOR2107 -2.08 [ -7.92, 3.76]
GIG 0.36 [ -8.80, 9.52]
GSP 2.95 [ -5.78, 11.68]
HUBIN -2.46 [-15.92, 11.00]
HUNT -5.20 [-10.50, 0.10]
IMAGEN -5.96 [-10.34, -1.59]
IMpACT -5.40 [-16.78, 5.97]
LBC1936 1.73 [ -4.90, 8.36]
LIBD 6.90 [ -3.09, 16.88]
Sydney MAS -5.94 [-13.00, 1.12]
MCIC -3.20 [-16.31, 9.91]
MooDS 0.25 [ -9.17, 9.67]
MPIP 3.46 [ -2.91, 9.84]
MUNSTER -1.92 [ -7.53, 3.69]
NCNG -5.25 [-14.15, 3.65]
NESDA 0.31 [-11.09, 11.72]
NeuroIMAGE -6.35 [-19.26, 6.56]
NTR -0.56 [ -9.18, 8.07]
OATS -3.76 [-14.60, 7.09]
PAFIP 3.86 [-12.94, 20.65]
PDNZ -5.08 [-20.44, 10.27]
PING 2.98 [ -5.65, 11.61]
MPRC -3.77 [-13.69, 6.16]
PPMI
QTIM -4.12 [ -9.66, 1.43]
SHIP -5.01 [ -9.78, -0.25]
SHIP-Trend -2.03 [ -7.49, 3.43]
SYS -1.48 [-11.91, 8.95]
TCD-NUIG -8.97 [-22.68, 4.73]
TOP 0.11 [ -7.49, 7.70]
TOP3T -1.18 [-10.15, 7.79]
UiO2016 -15.86 [-26.22, -5.50]
UiO2017 -1.47 [-10.75, 7.81]
UMCU -2.99 [-10.05, 4.08]
ENIGMA Discovery Cohorts [N=22828; P=1.736e-08] -3.29 [ -4.44, -2.15]
UKBB -3.12 [ -4.74, -1.49]
Discovery+UKBB [N=32733; P=3.023e-11] -3.23 [ -4.19, -2.28]
GOBS -1.82 [ -5.89, 2.26]
IMH -1.34 [-41.66, 38.97]
UNICAMP -7.32 [-21.73, 7.09]
Meth-CT -19.14 [-44.09, 5.80]
MIRECC
UKBB nonEUR -5.43 [-16.57, 5.72]
OSAKA 3.42 [ -9.53, 16.37]
PING Non European 3.96 [ -5.87, 13.79]
Non European Cohorts [N=2651; P=0.3165] -1.70 [ -5.04, 1.63]
Beta [Lower 95% CI,Upper 95% CI]
-60
-40
-20
0
20
40
mm^2/effect allele

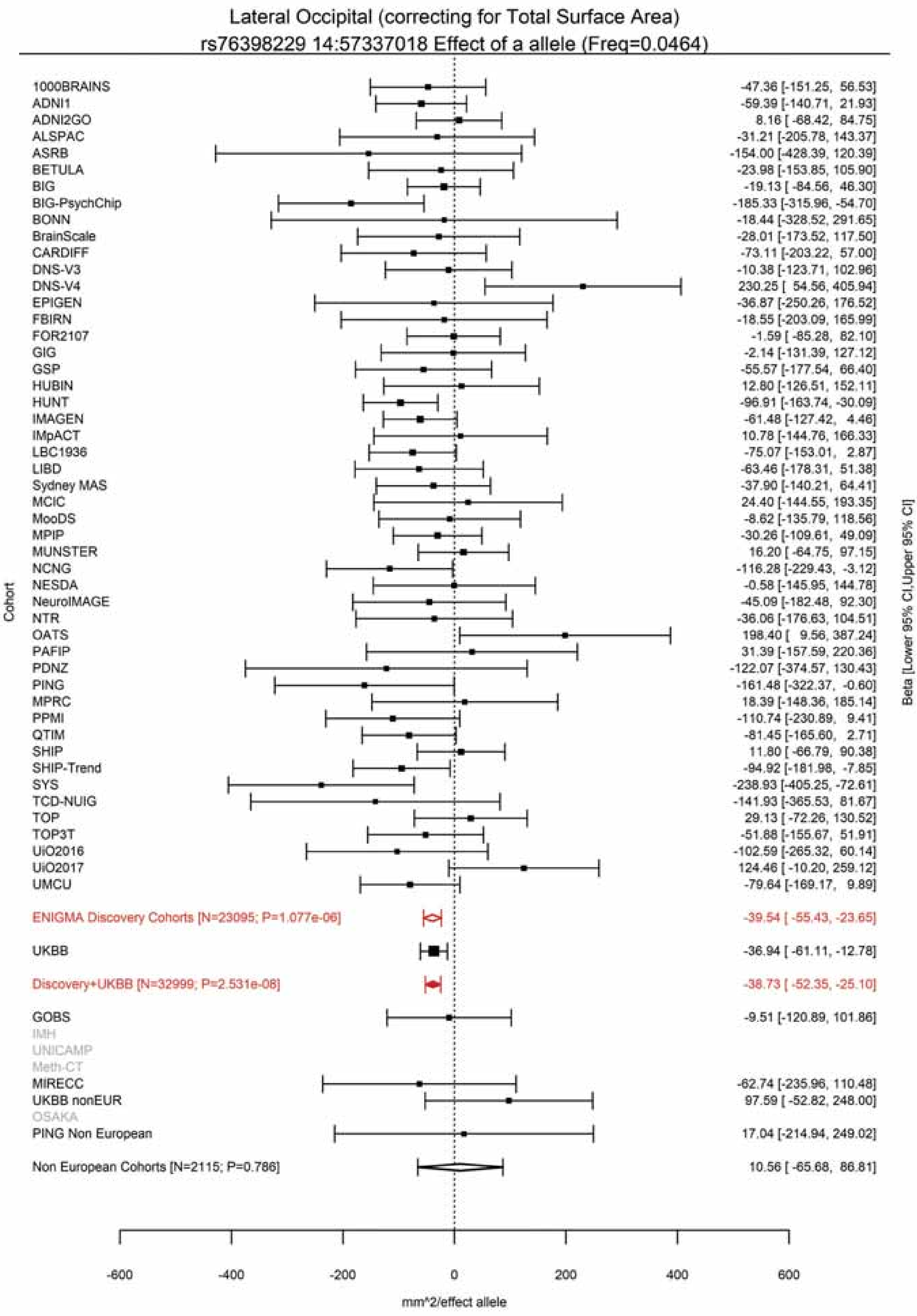
Lateral Occipital (correcting for Total Surface Area)
rs76398229 14:57337018 Effect of a allele (Freq=0.0464)
Cohort
1000BRAINS -47.36 [-151.25, 56.53]
ADNI1 -59.39 [-140.71, 21.93]
ADNI2GO 8.16 [ -68.42, 84.75]
ALSPAC -31.21 [-205.78, 143.37]
ASRB -154.00 [-428.39, 120.39]
BETULA -23.98 [-153.85, 105.90]
BIG -19.13 [ -84.56, 46.30]
BIG-PsychChip -185.33 [-315.96, -54.70]
BONN -18.44 [-328.52, 291.65]
BrainScale -28.01 [-173.52, 117.50]
CARDIFF -73.11 [-203.22, 57.00]
DNS-V3 -10.38 [-123.71, 102.96]
DNS-V4 230.25 [ 54.56, 405.94]
EPIGEN -36.87 [-250.26, 176.52]
FBIRN -18.55 [-203.09, 165.99]
FOR2107 -1.59 [ -85.28, 82.10]
GIG -2.14 [-131.39, 127.12]
GSP -55.57 [-177.54, 66.40]
HUBIN 12.80 [-126.51, 152.11]
HUNT -96.91 [-163.74, -30.09]
IMAGEN -61.48 [-127.42, 4.46]
IMpACT 10.78 [-144.76, 166.33]
LBC1936 -75.07 [-153.01, 2.87]
LIBD -63.46 [-178.31, 51.38]
Sydney MAS -37.90 [-140.21, 64.41]
MCIC 24.40 [-144.55, 193.35]
MooDS -8.62 [-135.79, 118.56]
MPIP -30.26 [-109.61, 49.09]
MUNSTER 16.20 [ -64.75, 97.15]
NCNG -116.28 [-229.43, -3.12]
NESDA -0.58 [-145.95, 144.78]
NeuroIMAGE -45.09 [-182.48, 92.30]
NTR -36.06 [-176.63, 104.51]
OATS 198.40 [ 9.56, 387.24]
PAFIP 31.39 [-157.59, 220.36]
PDNZ -122.07 [-374.57, 130.43]
PING -161.48 [-322.37, -0.60]
MPRC 18.39 [-148.36, 185.14]
PPMI -110.74 [-230.89, 9.41]
QTIM -81.45 [-165.60, 2.71]
SHIP 11.80 [ -66.79, 90.38]
SHIP-Trend -94.92 [-181.98, -7.85]
SYS -238.93 [-405.25, -72.61]
TCD-NUIG -141.93 [-365.53, 81.67]
TOP 29.13 [ -72.26, 130.52]
TOP3T -51.88 [-155.67, 51.91]
UiO2016 -102.59 [-265.32, 60.14]
UiO2017 124.46 [ -10.20, 259.12]
UMCU -79.64 [-169.17, 9.89]
ENIGMA Discovery Cohorts [N=23095; P=1.077e-06] -39.54 [ -55.43, -23.65]
UKBB -36.94 [ -61.11, -12.78]
Discovery+UKBB [N=32999; P=2.531e-08] -38.73 [ -52.35, -25.10]
GOBS -9.51 [-120.89, 101.86]
IMH
UNICAMP
Meth-CT
MIRECC -62.74 [-235.96, 110.48]
UKBB nonEUR 97.59 [ -52.82, 248.00]
OSAKA
PING Non European 17.04 [-214.94, 249.02]
Non European Cohorts [N=2115; P=0.786] 10.56 [ -65.68, 86.81]
Beta [Lower 95% CI,Upper 95% CI]
-600
-400
-200
0
200
400
600
mm^2/effect allele

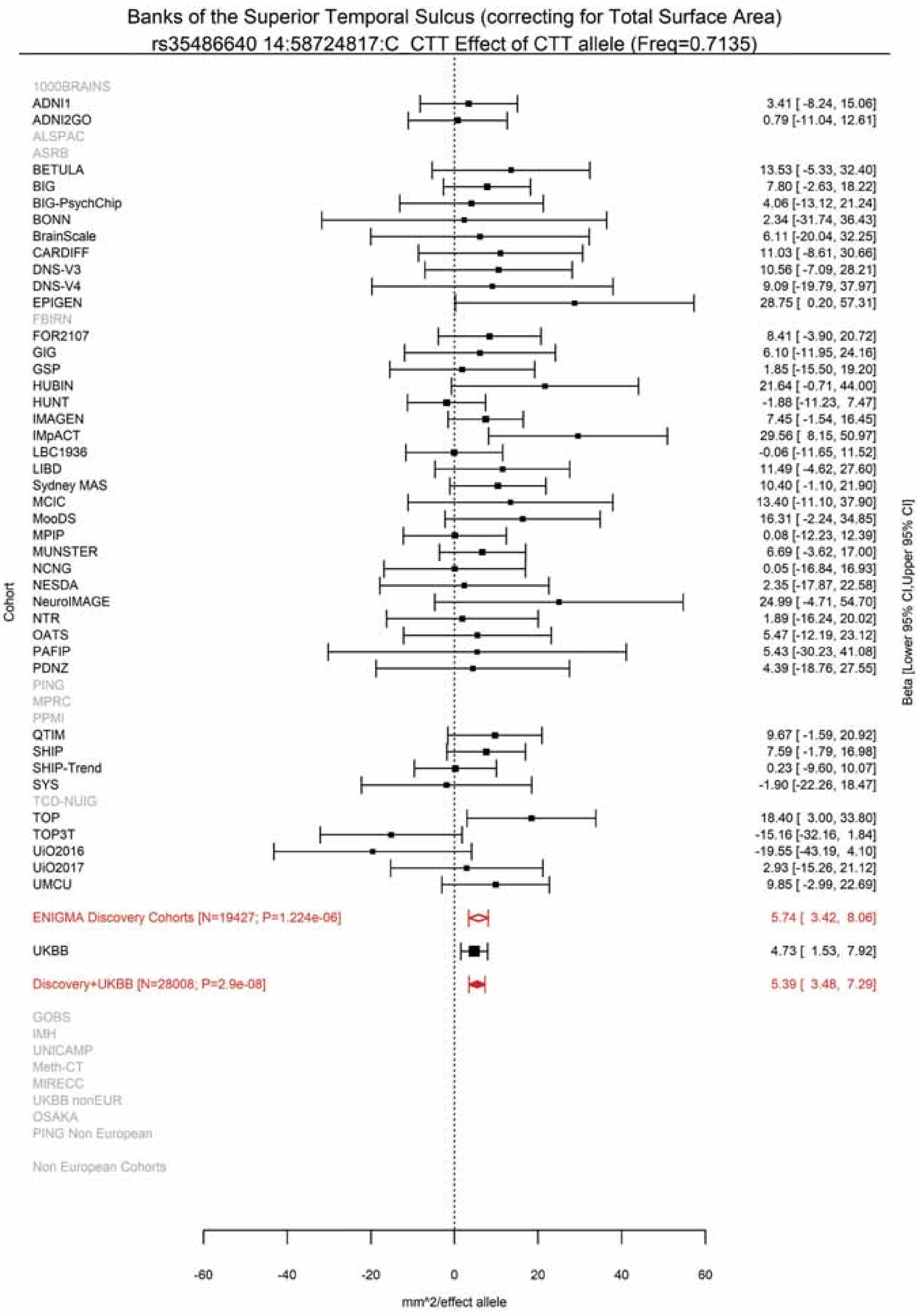

Banks of the Superior Temporal Sulcus (correcting for Total Surface Area)
rs35486640 14:58724817:C_CTT Effect of CTT allele (Freq=0.7135)
Cohort
1000BRAINS
ADNI1 3.41 [ -8.24, 15.06]
ADNI2GO 0.79 [-11.04, 12.61]
ALSPAC
ASRB
BETULA 13.53 [ -5.33, 32.40]
BIG 7.80 [ -2.63, 18.22]
BIG-PsychChip 4.06 [-13.12, 21.24]
BONN 2.34 [-31.74, 36.43]
BrainScale 6.11 [-20.04, 32.25]
CARDIFF 11.03 [ -8.61, 30.66]
DNS-V3 10.56 [ -7.09, 28.21]
DNS-V4 9.09 [-19.79, 37.97]
EPIGEN 28.75 [ 0.20, 57.31]
FBIRN
FOR2107 8.41 [ -3.90, 20.72]
GIG 6.10 [-11.95, 24.16]
GSP 1.85 [-15.50, 19.20]
HUBIN 21.64 [ -0.71, 44.00]
HUNT -1.88 [-11.23, 7.47]
IMAGEN 7.45 [ -1.54, 16.45]
IMpACT 29.56 [ 8.15, 50.97]
LBC1936 -0.06 [-11.65, 11.52]
LIBD 11.49 [ -4.62, 27.60]
Sydney MAS 10.40 [ -1.10, 21.90]
MCIC 13.40 [-11.10, 37.90]
MooDS 16.31 [ -2.24, 34.85]
MPIP 0.08 [-12.23, 12.39]
MUNSTER 6.69 [ -3.62, 17.00]
NCNG 0.05 [-16.84, 16.93]
NESDA 2.35 [-17.87, 22.58]
NeuroIMAGE 24.99 [ -4.71, 54.70]
NTR 1.89 [-16.24, 20.02]
OATS 5.47 [-12.19, 23.12]
PAFIP 5.43 [-30.23, 41.08]
PDNZ 4.39 [-18.76, 27.55]
PING
MPRC
PPMI
QTIM 9.67 [ -1.59, 20.92]
SHIP 7.59 [ -1.79, 16.98]
SHIP-Trend 0.23 [ -9.60, 10.07]
SYS -1.90 [-22.26, 18.47]
TCD-NUIG
TOP 18.40 [ 3.00, 33.80]
TOP3T -15.16 [-32.16, 1.84]
UiO2016 -19.55 [-43.19, 4.10]
UiO2017 2.93 [-15.26, 21.12]
UMCU 9.85 [ -2.99, 22.69]
ENIGMA Discovery Cohorts [N=19427; P=1.224e-06] 5.74 [ 3.42, 8.06]
UKBB 4.73 [ 1.53, 7.92]
Discovery+UKBB [N=28008; P=2.9e-08] 5.39 [ 3.48, 7.29]
GOBS
IMH
UNICAMP
Meth-CT
MIRECC
UKBB nonEUR
OSAKA
PING Non European
Non European Cohorts
-60
-40
-20
0
20
40
60
mm^2/effect allele
Beta [Lower 95% CI,Upper 95% CI]

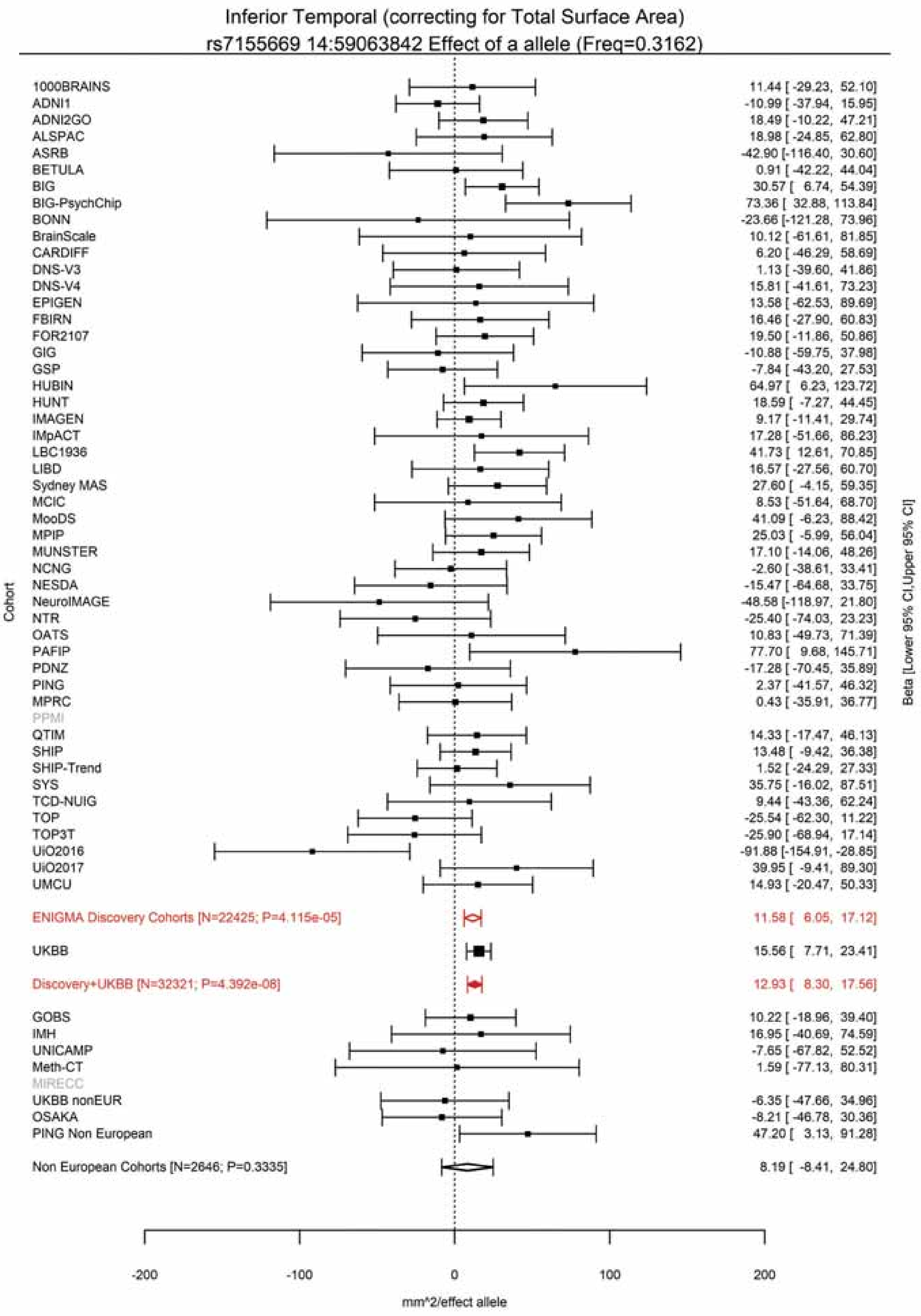

Inferior Temporal (correcting for Total Surface Area)
rs7155669 14:59063842 Effect of a allele (Freq=0.3162)
Cohort
Beta [Lower 95% CI,Upper 95% CI]
1000BRAINS 11.44 [ -29.23, 52.10]
ADNI1 -10.99 [ -37.94, 15.95]
ADNI2GO 18.49 [ -10.22, 47.21]
ALSPAC 18.98 [ -24.85, 62.80]
ASRB -42.90 [-116.40, 30.60]
BETULA 0.91 [ -42.22, 44.04]
BIG 30.57 [ 6.74, 54.39]
BIG-PsychChip 73.36 [ 32.88, 113.84]
BONN -23.66 [-121.28, 73.96]
BrainScale 10.12 [ -61.61, 81.85]
CARDIFF 6.20 [ -46.29, 58.69]
DNS-V3 1.13 [ -39.60, 41.86]
DNS-V4 15.81 [ -41.61, 73.23]
EPIGEN 13.58 [ -62.53, 89.69]
FBIRN 16.46 [ -27.90, 60.83]
FOR2107 19.50 [ -11.86, 50.86]
GIG -10.88 [ -59.75, 37.98]
GSP -7.84 [ -43.20, 27.53]
HUBIN 64.97 [ 6.23, 123.72]
HUNT 18.59 [ -7.27, 44.45]
IMAGEN 9.17 [ -11.41, 29.74]
IMpACT 17.28 [ -51.66, 86.23]
LBC1936 41.73 [ 12.61, 70.85]
LIBD 16.57 [ -27.56, 60.70]
Sydney MAS 27.60 [ -4.15, 59.35]
MCIC 8.53 [ -51.64, 68.70]
MooDS 41.09 [ -6.23, 88.42]
MPIP 25.03 [ -5.99, 56.04]
MUNSTER 17.10 [ -14.06, 48.26]
NCNG -2.60 [ -38.61, 33.41]
NESDA -15.47 [ -64.68, 33.75]
NeuroIMAGE -48.58 [-118.97, 21.80]
NTR -25.40 [ -74.03, 23.23]
OATS 10.83 [ -49.73, 71.39]
PAFIP 77.70 [ 9.68, 145.71]
PDNZ -17.28 [ -70.45, 35.89]
PING 2.37 [ -41.57, 46.32]
MPRC 0.43 [ -35.91, 36.77]
PPMI
QTIM 14.33 [ -17.47, 46.13]
SHIP 13.48 [ -9.42, 36.38]
SHIP-Trend 1.52 [ -24.29, 27.33]
SYS 35.75 [ -16.02, 87.51]
TCD-NUIG 9.44 [ -43.36, 62.24]
TOP -25.54 [ -62.30, 11.22]
TOP3T -25.90 [ -68.94, 17.14]
UiO2016 -91.88 [-154.91, -28.85]
UiO2017 39.95 [ -9.41, 89.30]
UMCU 14.93 [ -20.47, 50.33]
ENIGMA Discovery Cohorts [N=22425; P=4.115e-05] 11.58 [ 6.05, 17.12]
UKBB 15.56 [ 7.71, 23.41]
Discovery+UKBB [N=32321; P=4.392e-08] 12.93 [ 8.30, 17.56]
GOBS 10.22 [ -18.96, 39.40]
IMH 16.95 [ -40.69, 74.59]
UNICAMP -7.65 [ -67.82, 52.52]
Meth-CT 1.59 [ -77.13, 80.31]
MIRECC
UKBB nonEUR -6.35 [ -47.66, 34.96]
OSAKA -8.21 [ -46.78, 30.36]
PING Non European 47.20 [ 3.13, 91.28]
Non European Cohorts [N=2646; P=0.3335] 8.19 [ -8.41, 24.80]
-200
-100
0
100
200
mm^2/effect allele

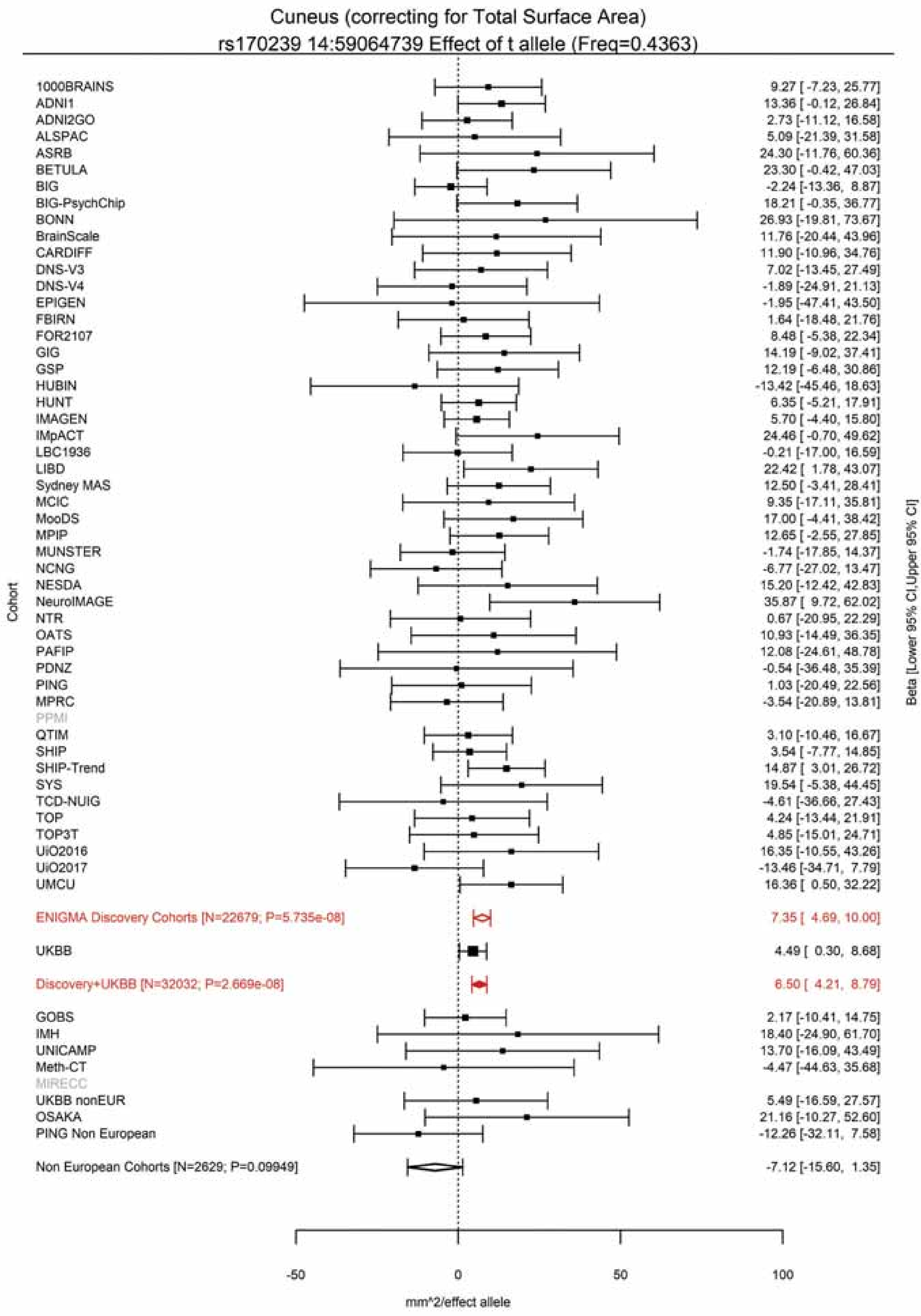

Cuneus (correcting for Total Surface Area)
rs170239 14:59064739 Effect of t allele (Freq=0.4363)
1000BRAINS 9.27 [ -7.23, 25.77]
ADNI1 13.36 [ -0.12, 26.84]
ADNI2GO 2.73 [-11.12, 16.58]
ALSPAC 5.09 [-21.39, 31.58]
ASRB 24.30 [-11.76, 60.36]
BETULA 23.30 [ -0.42, 47.03]
BIG -2.24 [-13.36, 8.87]
BIG-PsychChip 18.21 [ -0.35, 36.77]
BONN 26.93 [-19.81, 73.67]
BrainScale 11.76 [-20.44, 43.96]
CARDIFF 11.90 [-10.96, 34.76]
DNS-V3 7.02 [-13.45, 27.49]
DNS-V4 -1.89 [-24.91, 21.13]
EPIGEN -1.95 [-47.41, 43.50]
FBIRN 1.64 [-18.48, 21.76]
FOR2107 8.48 [ -5.38, 22.34]
GIG 14.19 [ -9.02, 37.41]
GSP 12.19 [ -6.48, 30.86]
HUBIN -13.42 [-45.46, 18.63]
HUNT 6.35 [ -5.21, 17.91]
IMAGEN 5.70 [ -4.40, 15.80]
IMpACT 24.46 [ -0.70, 49.62]
LBC1936 -0.21 [-17.00, 16.59]
LIBD 22.42 [ 1.78, 43.07]
Sydney MAS 12.50 [ -3.41, 28.41]
MCIC 9.35 [-17.11, 35.81]
MooDS 17.00 [ -4.41, 38.42]
MPIP 12.65 [ -2.55, 27.85]
MUNSTER -1.74 [-17.85, 14.37]
NCNG -6.77 [-27.02, 13.47]
NESDA 15.20 [-12.42, 42.83]
NeuroIMAGE 35.87 [ 9.72, 62.02]
NTR 0.67 [-20.95, 22.29]
OATS 10.93 [-14.49, 36.35]
PAFIP 12.08 [-24.61, 48.78]
PDNZ -0.54 [-36.48, 35.39]
PING 1.03 [-20.49, 22.56]
MPRC -3.54 [-20.89, 13.81]
PPMI
QTIM 3.10 [-10.46, 16.67]
SHIP 3.54 [ -7.77, 14.85]
SHIP-Trend 14.87 [ 3.01, 26.72]
SYS 19.54 [ -5.38, 44.45]
TCD-NUIG -4.61 [-36.66, 27.43]
TOP 4.24 [-13.44, 21.91]
TOP3T 4.85 [-15.01, 24.71]
UiO2016 16.35 [-10.55, 43.26]
UiO2017 -13.46 [-34.71, 7.79]
UMCU 16.36 [ 0.50, 32.22]
ENIGMA Discovery Cohorts [N=22679; P=5.735e-08] 7.35 [ 4.69, 10.00]
UKBB 4.49 [ 0.30, 8.68]
Discovery+UKBB [N=32032; P=2.669e-08] 6.50 [ 4.21, 8.79]
GOBS 2.17 [-10.41, 14.75]
IMH 18.40 [-24.90, 61.70]
UNICAMP 13.70 [-16.09, 43.49]
Meth-CT -4.47 [-44.63, 35.68]
MIRECC
UKBB nonEUR 5.49 [-16.59, 27.57]
OSAKA 21.16 [-10.27, 52.60]
PING Non European -12.26 [-32.11, 7.58]
Non European Cohorts [N=2629; P=0.09949] -7.12 [-15.60, 1.35]
Cohort
Beta [Lower 95% CI,Upper 95% CI]
-50
0
50
100
mm^2/effect allele

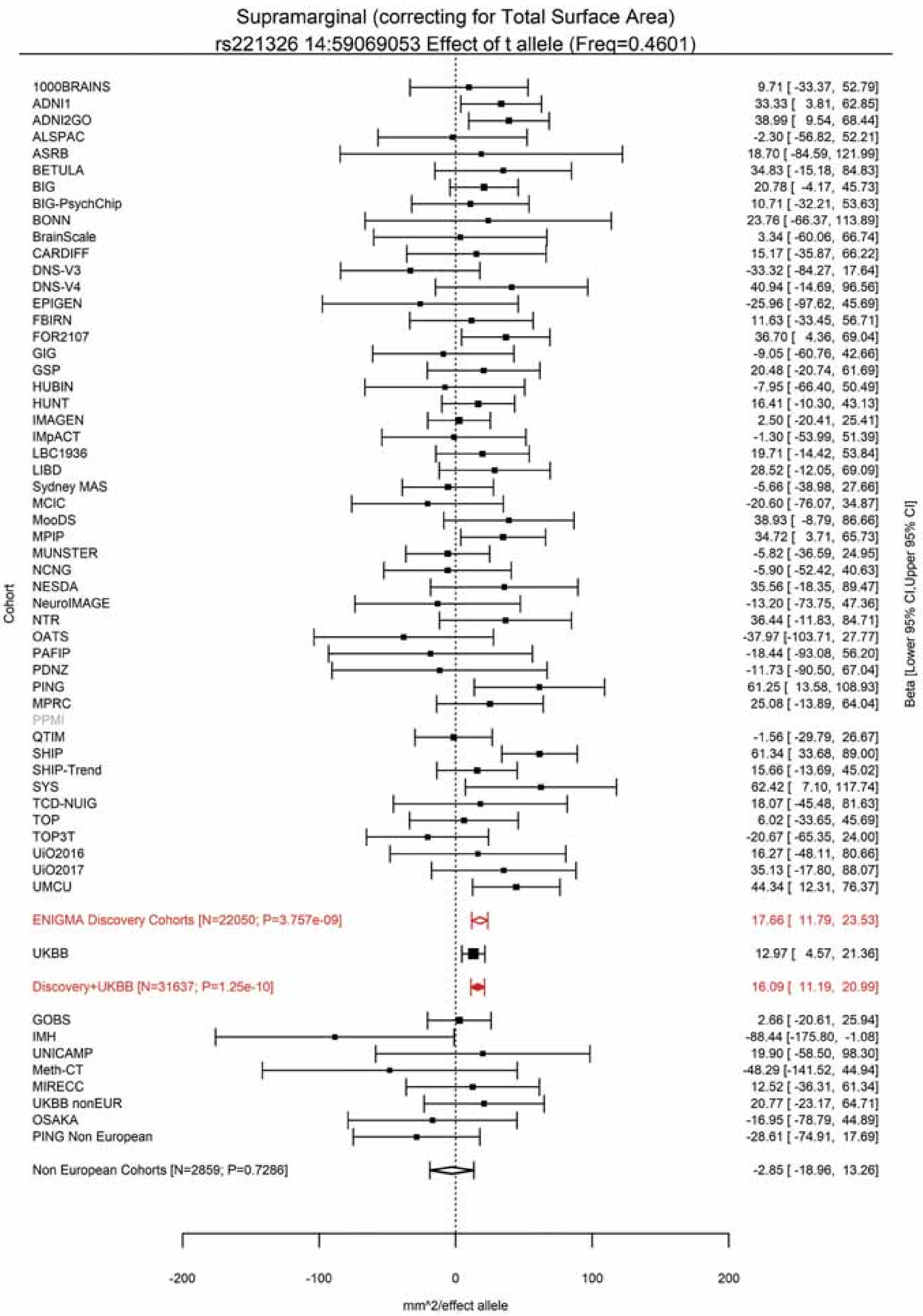
Supramarginal (correcting for Total Surface Area)
rs221326 14:59069053 Effect of t allele (Freq=0.4601)
Cohort
1000BRAINS 9.71 [ -33.37, 52.79]
ADNI1 33.33 [ 3.81, 62.85]
ADNI2GO 38.99 [ 9.54, 68.44]
ALSPAC -2.30 [ -56.82, 52.21]
ASRB 18.70 [ -84.59, 121.99]
BETULA 34.83 [ -15.18, 84.83]
BIG 20.78 [ -4.17, 45.73]
BIG-PsychChip 10.71 [ -32.21, 53.63]
BONN 23.76 [ -66.37, 113.89]
BrainScale 3.34 [ -60.06, 66.74]
CARDIFF 15.17 [ -35.87, 66.22]
DNS-V3 -33.32 [ -84.27, 17.64]
DNS-V4 40.94 [ -14.69, 96.56]
EPIGEN -25.96 [ -97.62, 45.69]
FBIRN 11.63 [ -33.45, 56.71]
FOR2107 36.70 [ 4.36, 69.04]
GIG -9.05 [ -60.76, 42.66]
GSP 20.48 [ -20.74, 61.69]
HUBIN -7.95 [ -66.40, 50.49]
HUNT 16.41 [ -10.30, 43.13]
IMAGEN 2.50 [ -20.41, 25.41]
IMpACT -1.30 [ -53.99, 51.39]
LBC1936 19.71 [ -14.42, 53.84]
LIBD 28.52 [ -12.05, 69.09]
Sydney MAS -5.66 [ -38.98, 27.66]
MCIC -20.60 [ -76.07, 34.87]
MooDS 38.93 [ -8.79, 86.66]
MPIP 34.72 [ 3.71, 65.73]
MUNSTER -5.82 [ -36.59, 24.95]
NCNG -5.90 [ -52.42, 40.63]
NESDA 35.56 [ -18.35, 89.47]
NeuroIMAGE -13.20 [ -73.75, 47.36]
NTR 36.44 [ -11.83, 84.71]
OATS -37.97 [-103.71, 27.77]
PAFIP -18.44 [ -93.08, 56.20]
PDNZ -11.73 [ -90.50, 67.04]
PING 61.25 [ 13.58, 108.93]
MPRC 25.08 [ -13.89, 64.04]
PPMI
QTIM -1.56 [ -29.79, 26.67]
SHIP 61.34 [ 33.68, 89.00]
SHIP-Trend 15.66 [ -13.69, 45.02]
SYS 62.42 [ 7.10, 117.74]
TCD-NUIG 18.07 [ -45.48, 81.63]
TOP 6.02 [ -33.65, 45.69]
TOP3T -20.67 [ -65.35, 24.00]
UiO2016 16.27 [ -48.11, 80.66]
UiO2017 35.13 [ -17.80, 88.07]
UMCU 44.34 [ 12.31, 76.37]
ENIGMA Discovery Cohorts [N=22050; P=3.757e-09] 17.66 [ 11.79, 23.53]
UKBB 12.97 [ 4.57, 21.36]
Discovery+UKBB [N=31637; P=1.25e-10] 16.09 [ 11.19, 20.99]
GOBS 2.66 [ -20.61, 25.94]
IMH -88.44 [-175.80, -1.08]
UNICAMP 19.90 [ -58.50, 98.30]
Meth-CT -48.29 [-141.52, 44.94]
MIRECC 12.52 [ -36.31, 61.34]
UKBB nonEUR 20.77 [ -23.17, 64.71]
OSAKA -16.95 [ -78.79, 44.89]
PING Non European -28.61 [ -74.91, 17.69]
Non European Cohorts [N=2859; P=0.7286] -2.85 [ -18.96, 13.26]
Beta [Lower 95% CI,Upper 95% CI]
-200
-100
0
100
200
mm^2/effect allele

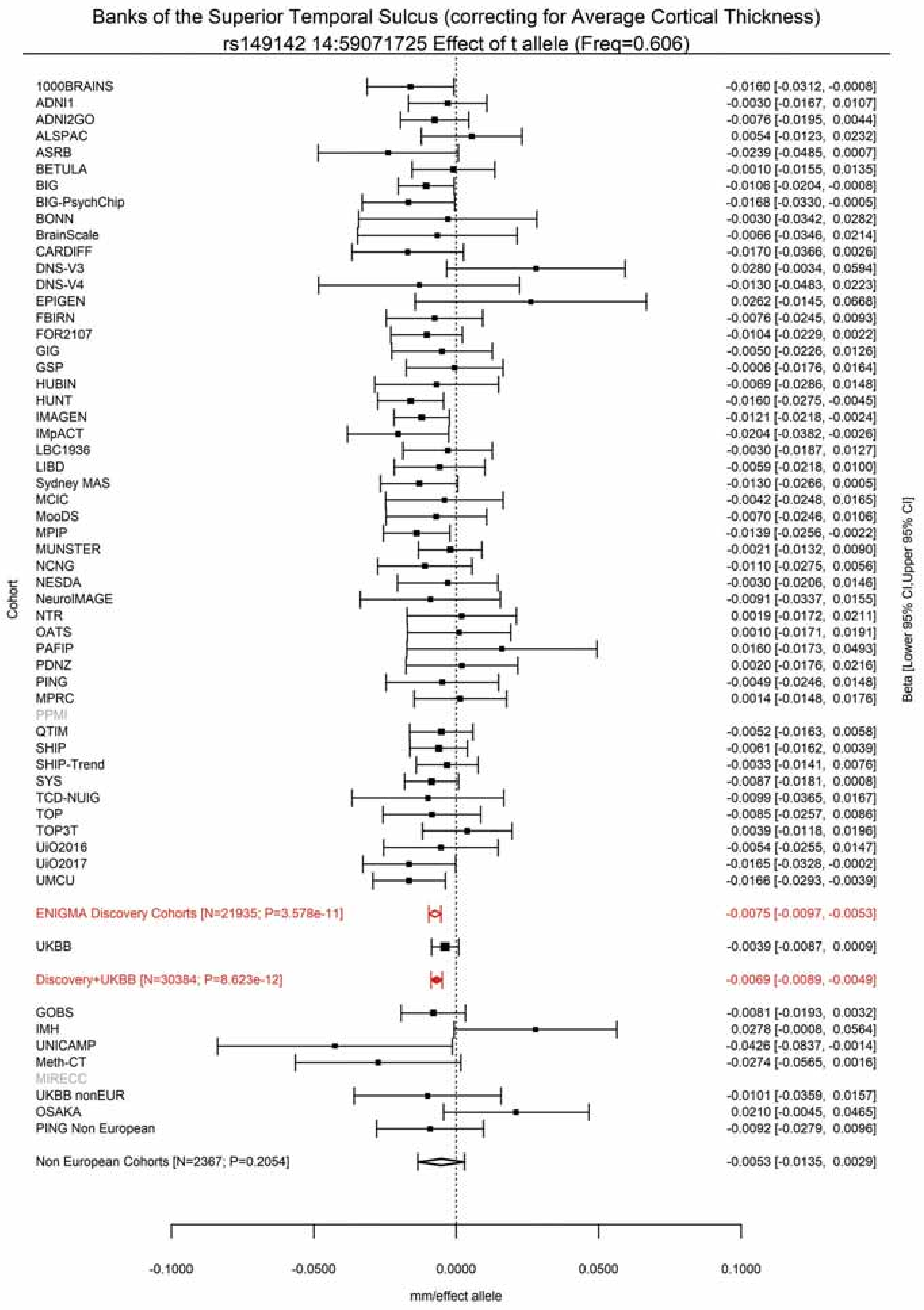

Banks of the Superior Temporal Sulcus (correcting for Average Cortical Thickness)
rs149142 14:59071725 Effect of t allele (Freq=0.606)
Cohort
1000BRAINS -0.0160 [-0.0312, -0.0008]
ADNI1 -0.0030 [-0.0167, 0.0107]
ADNI2GO -0.0076 [-0.0195, 0.0044]
ALSPAC 0.0054 [-0.0123, 0.0232]
ASRB -0.0239 [-0.0485, 0.0007]
BETULA -0.0010 [-0.0155, 0.0135]
BIG -0.0106 [-0.0204, -0.0008]
BIG-PsychChip -0.0168 [-0.0330, -0.0005]
BONN -0.0030 [-0.0342, 0.0282]
BrainScale -0.0066 [-0.0346, 0.0214]
CARDIFF -0.0170 [-0.0366, 0.0026]
DNS-V3 0.0280 [-0.0034, 0.0594]
DNS-V4 -0.0130 [-0.0483, 0.0223]
EPIGEN 0.0262 [-0.0145, 0.0668]
FBIRN -0.0076 [-0.0245, 0.0093]
FOR2107 -0.0104 [-0.0229, 0.0022]
GIG -0.0050 [-0.0226, 0.0126]
GSP -0.0006 [-0.0176, 0.0164]
HUBIN -0.0069 [-0.0286, 0.0148]
HUNT -0.0160 [-0.0275, -0.0045]
IMAGEN -0.0121 [-0.0218, -0.0024]
IMpACT -0.0204 [-0.0382, -0.0026]
LBC1936 -0.0030 [-0.0187, 0.0127]
LIBD -0.0059 [-0.0218, 0.0100]
Sydney MAS -0.0130 [-0.0266, 0.0005]
MCIC -0.0042 [-0.0248, 0.0165]
MooDS -0.0070 [-0.0246, 0.0106]
MPIP -0.0139 [-0.0256, -0.0022]
MUNSTER -0.0021 [-0.0132, 0.0090]
NCNG -0.0110 [-0.0275, 0.0056]
NESDA -0.0030 [-0.0206, 0.0146]
NeuroIMAGE -0.0091 [-0.0337, 0.0155]
NTR 0.0019 [-0.0172, 0.0211]
OATS 0.0010 [-0.0171, 0.0191]
PAFIP 0.0160 [-0.0173, 0.0493]
PDNZ 0.0020 [-0.0176, 0.0216]
PING -0.0049 [-0.0246, 0.0148]
MPRC 0.0014 [-0.0148, 0.0176]
PPMI
QTIM -0.0052 [-0.0163, 0.0058]
SHIP -0.0061 [-0.0162, 0.0039]
SHIP-Trend -0.0033 [-0.0141, 0.0076]
SYS -0.0087 [-0.0181, 0.0008]
TCD-NUIG -0.0099 [-0.0365, 0.0167]
TOP -0.0085 [-0.0257, 0.0086]
TOP3T 0.0039 [-0.0118, 0.0196]
UiO2016 -0.0054 [-0.0255, 0.0147]
UiO2017 -0.0165 [-0.0328, -0.0002]
UMCU -0.0166 [-0.0293, -0.0039]
ENIGMA Discovery Cohorts [N=21935; P=3.578e-11] -0.0075 [-0.0097, -0.0053]
UKBB -0.0039 [-0.0087, 0.0009]
Discovery+UKBB [N=30384; P=8.623e-12] -0.0069 [-0.0089, -0.0049]
GOBS -0.0081 [-0.0193, 0.0032]
IMH 0.0278 [-0.0008, 0.0564]
UNICAMP -0.0426 [-0.0837, -0.0014]
Meth-CT -0.0274 [-0.0565, 0.0016]
MIRECC
UKBB nonEUR -0.0101 [-0.0359, 0.0157]
OSAKA 0.0210 [-0.0045, 0.0465]
PING Non European -0.0092 [-0.0279, 0.0096]
Non European Cohorts [N=2367; P=0.2054] -0.0053 [-0.0135, 0.0029]
Beta [Lower 95% CI,Upper 95% CI]
-0.1000
-0.0500
0.0000
0.0500
0.1000
mm/effect allele

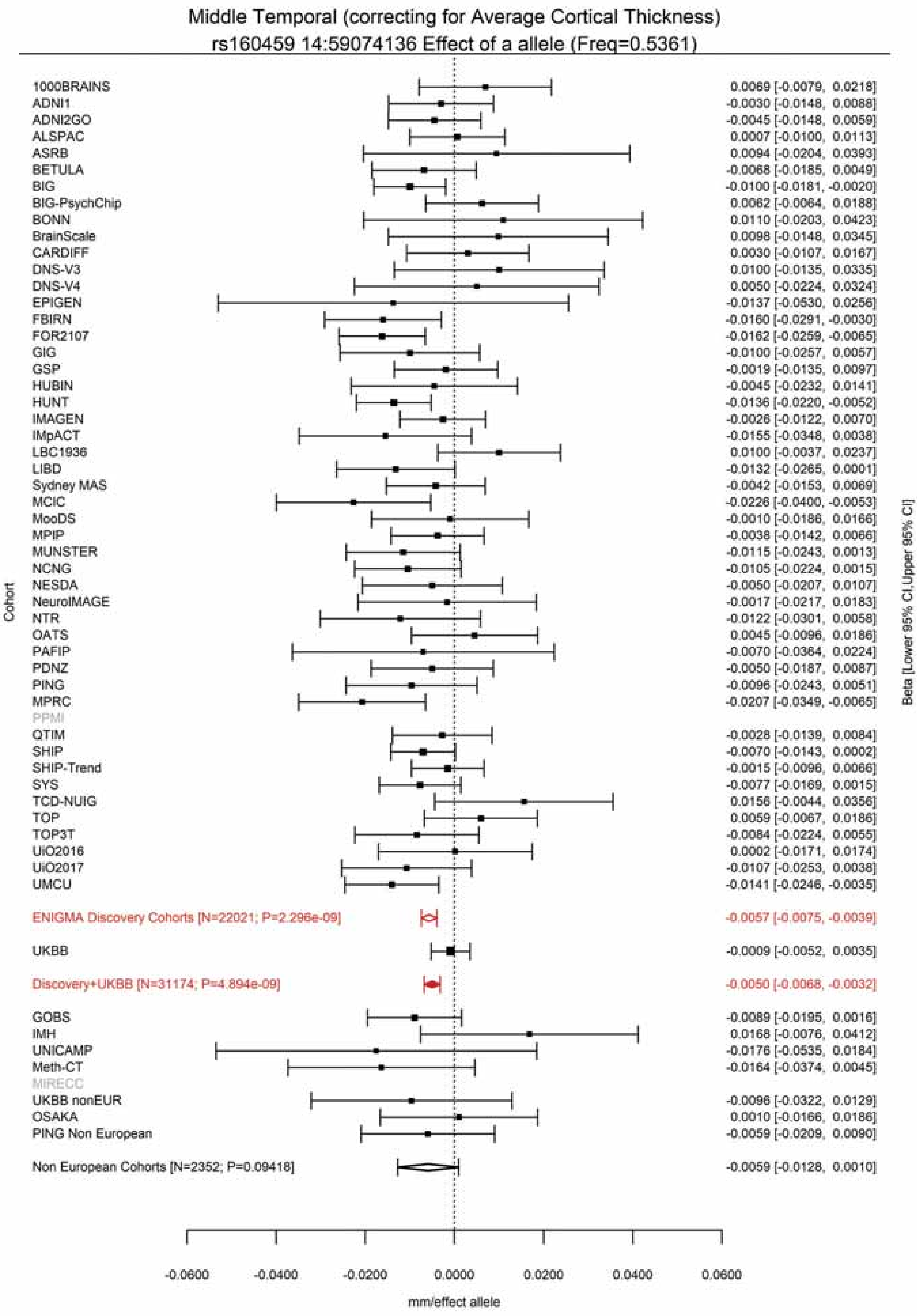

Middle Temporal (correcting for Average Cortical Thickness)
rs160459 14:59074136 Effect of a allele (Freq=0.5361)
Cohort
Beta [Lower 95% CI,Upper 95% CI]
1000BRAINS 0.0069 [-0.0079, 0.0218]
ADNI1 -0.0030 [-0.0148, 0.0088]
ADNI2GO -0.0045 [-0.0148, 0.0059]
ALSPAC 0.0007 [-0.0100, 0.0113]
ASRB 0.0094 [-0.0204, 0.0393]
BETULA -0.0068 [-0.0185, 0.0049]
BIG -0.0100 [-0.0181, -0.0020]
BIG-PsychChip 0.0062 [-0.0064, 0.0188]
BONN 0.0110 [-0.0203, 0.0423]
BrainScale 0.0098 [-0.0148, 0.0345]
CARDIFF 0.0030 [-0.0107, 0.0167]
DNS-V3 0.0100 [-0.0135, 0.0335]
DNS-V4 0.0050 [-0.0224, 0.0324]
EPIGEN -0.0137 [-0.0530, 0.0256]
FBIRN -0.0160 [-0.0291, -0.0030]
FOR2107 -0.0162 [-0.0259, -0.0065]
GIG -0.0100 [-0.0257, 0.0057]
GSP -0.0019 [-0.0135, 0.0097]
HUBIN -0.0045 [-0.0232, 0.0141]
HUNT -0.0136 [-0.0220, -0.0052]
IMAGEN -0.0026 [-0.0122, 0.0070]
IMpACT -0.0155 [-0.0348, 0.0038]
LBC1936 0.0100 [-0.0037, 0.0237]
LIBD -0.0132 [-0.0265, 0.0001]
Sydney MAS -0.0042 [-0.0153, 0.0069]
MCIC -0.0226 [-0.0400, -0.0053]
MooDS -0.0010 [-0.0186, 0.0166]
MPIP -0.0038 [-0.0142, 0.0066]
MUNSTER -0.0115 [-0.0243, 0.0013]
NCNG -0.0105 [-0.0224, 0.0015]
NESDA -0.0050 [-0.0207, 0.0107]
NeuroIMAGE -0.0017 [-0.0217, 0.0183]
NTR -0.0122 [-0.0301, 0.0058]
OATS 0.0045 [-0.0096, 0.0186]
PAFIP -0.0070 [-0.0364, 0.0224]
PDNZ -0.0050 [-0.0187, 0.0087]
PING -0.0096 [-0.0243, 0.0051]
MPRC -0.0207 [-0.0349, -0.0065]
PPMI
QTIM -0.0028 [-0.0139, 0.0084]
SHIP -0.0070 [-0.0143, 0.0002]
SHIP-Trend -0.0015 [-0.0096, 0.0066]
SYS -0.0077 [-0.0169, 0.0015]
TCD-NUIG 0.0156 [-0.0044, 0.0356]
TOP 0.0059 [-0.0067, 0.0186]
TOP3T -0.0084 [-0.0224, 0.0055]
UiO2016 0.0002 [-0.0171, 0.0174]
UiO2017 -0.0107 [-0.0253, 0.0038]
UMCU -0.0141 [-0.0246, -0.0035]
ENIGMA Discovery Cohorts [N=22021; P=2.296e-09] -0.0057 [-0.0075, -0.0039]
UKBB -0.0009 [-0.0052, 0.0035]
Discovery+UKBB [N=31174; P=4.894e-09] -0.0050 [-0.0068, -0.0032]
GOBS -0.0089 [-0.0195, 0.0016]
IMH 0.0168 [-0.0076, 0.0412]
UNICAMP -0.0176 [-0.0535, 0.0184]
Meth-CT -0.0164 [-0.0374, 0.0045]
MIRECC
UKBB nonEUR -0.0096 [-0.0322, 0.0129]
OSAKA 0.0010 [-0.0166, 0.0186]
PING Non European -0.0059 [-0.0209, 0.0090]
Non European Cohorts [N=2352; P=0.09418] -0.0059 [-0.0128, 0.0010]
-0.0600
-0.0400
-0.0200
0.0000
0.0200
0.0400
0.0600
mm/effect allele

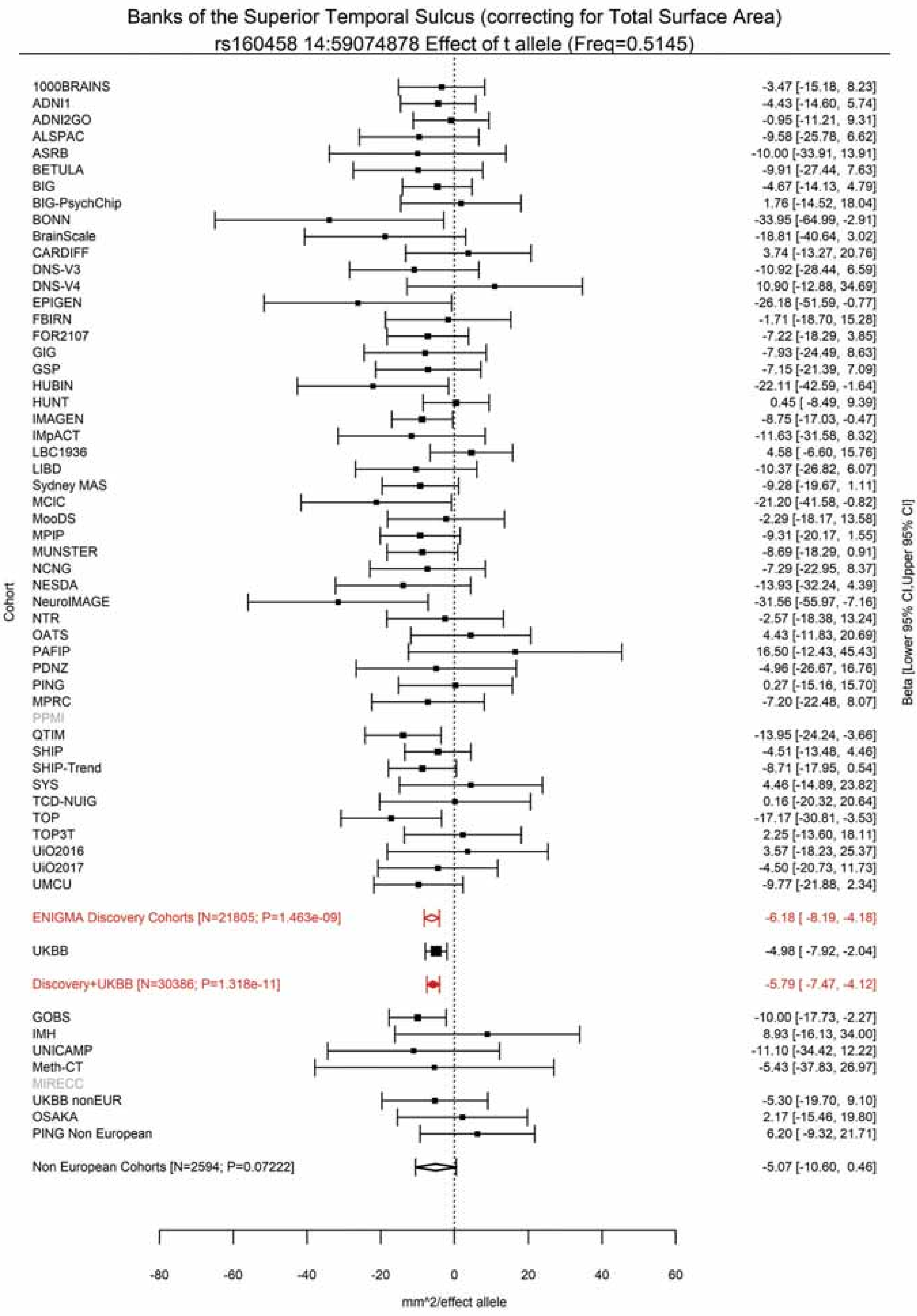

Banks of the Superior Temporal Sulcus (correcting for Total Surface Area)
rs160458 14:59074878 Effect of t allele (Freq=0.5145)
Cohort
1000BRAINS -3.47 [-15.18, 8.23]
ADNI1 -4.43 [-14.60, 5.74]
ADNI2GO -0.95 [-11.21, 9.31]
ALSPAC -9.58 [-25.78, 6.62]
ASRB -10.00 [-33.91, 13.91]
BETULA -9.91 [-27.44, 7.63]
BIG -4.67 [-14.13, 4.79]
BIG-PsychChip 1.76 [-14.52, 18.04]
BONN -33.95 [-64.99, -2.91]
BrainScale -18.81 [-40.64, 3.02]
CARDIFF 3.74 [-13.27, 20.76]
DNS-V3 -10.92 [-28.44, 6.59]
DNS-V4 10.90 [-12.88, 34.69]
EPIGEN -26.18 [-51.59, -0.77]
FBIRN -1.71 [-18.70, 15.28]
FOR2107 -7.22 [-18.29, 3.85]
GIG -7.93 [-24.49, 8.63]
GSP -7.15 [-21.39, 7.09]
HUBIN -22.11 [-42.59, -1.64]
HUNT 0.45 [-8.49, 9.39]
IMAGEN -8.75 [-17.03, -0.47]
IMpACT -11.63 [-31.58, 8.32]
LBC1936 4.58 [-6.60, 15.76]
LIBD -10.37 [-26.82, 6.07]
Sydney MAS -9.28 [-19.67, 1.11]
MCIC -21.20 [-41.58, -0.82]
MooDS -2.29 [-18.17, 13.58]
MPIP -9.31 [-20.17, 1.55]
MUNSTER -8.69 [-18.29, 0.91]
NCNG -7.29 [-22.95, 8.37]
NESDA -13.93 [-32.24, 4.39]
NeuroIMAGE -31.56 [-55.97, -7.16]
NTR -2.57 [-18.38, 13.24]
OATS 4.43 [-11.83, 20.69]
PAFIP 16.50 [-12.43, 45.43]
PDNZ -4.96 [-26.67, 16.76]
PING 0.27 [-15.16, 15.70]
MPRC -7.20 [-22.48, 8.07]
PPMI
QTIM -13.95 [-24.24, -3.66]
SHIP -4.51 [-13.48, 4.46]
SHIP-Trend -8.71 [-17.95, 0.54]
SYS 4.46 [-14.89, 23.82]
TCD-NUIG 0.16 [-20.32, 20.64]
TOP -17.17 [-30.81, -3.53]
TOP3T 2.25 [-13.60, 18.11]
UiO2016 3.57 [-18.23, 25.37]
UiO2017 -4.50 [-20.73, 11.73]
UMCU -9.77 [-21.88, 2.34]
ENIGMA Discovery Cohorts [N=21805; P=1.463e-09] -6.18 [-8.19, -4.18]
UKBB -4.98 [-7.92, -2.04]
Discovery+UKBB [N=30386; P=1.318e-11] -5.79 [-7.47, -4.12]
GOBS -10.00 [-17.73, -2.27]
IMH 8.93 [-16.13, 34.00]
UNICAMP -11.10 [-34.42, 12.22]
Meth-CT -5.43 [-37.83, 26.97]
MIRECC
UKBB nonEUR -5.30 [-19.70, 9.10]
OSAKA 2.17 [-15.46, 19.80]
PING Non European 6.20 [-9.32, 21.71]
Non European Cohorts [N=2594; P=0.07222] -5.07 [-10.60, 0.46]
-80 -60 -40 -20 0 20 40 60
mm^2/effect allele
Beta [Lower 95% CI,Upper 95% CI]

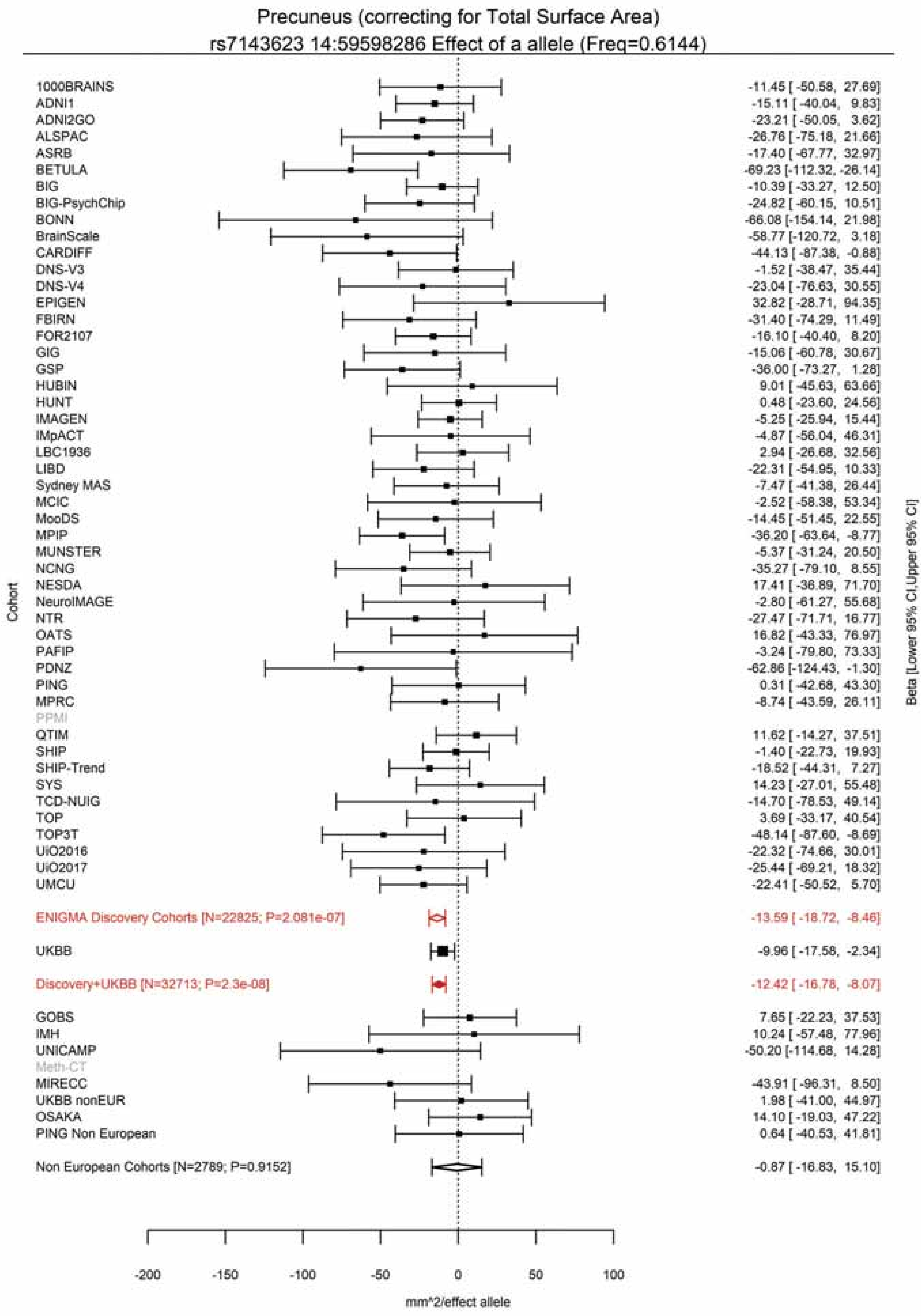

Precuneus (correcting for Total Surface Area)
rs7143623 14:59598286 Effect of a allele (Freq=0.6144)
1000BRAINS -11.45 [ -50.58, 27.69]
ADNI1 -15.11 [ -40.04, 9.83]
ADNI2GO -23.21 [ -50.05, 3.62]
ALSPAC -26.76 [ -75.18, 21.66]
ASRB -17.40 [ -67.77, 32.97]
BETULA -69.23 [-112.32, -26.14]
BIG -10.39 [ -33.27, 12.50]
BIG-PsychChip -24.82 [ -60.15, 10.51]
BONN -66.08 [-154.14, 21.98]
BrainScale -58.77 [-120.72, 3.18]
CARDIFF -44.13 [ -87.38, -0.88]
DNS-V3 -1.52 [ -38.47, 35.44]
DNS-V4 -23.04 [ -76.63, 30.55]
EPIGEN 32.82 [ -28.71, 94.35]
FBIRN -31.40 [ -74.29, 11.49]
FOR2107 -16.10 [ -40.40, 8.20]
GIG -15.06 [ -60.78, 30.67]
GSP -36.00 [ -73.27, 1.28]
HUBIN 9.01 [ -45.63, 63.66]
HUNT 0.48 [ -23.60, 24.56]
IMAGEN -5.25 [ -25.94, 15.44]
IMpACT -4.87 [ -56.04, 46.31]
LBC1936 2.94 [ -26.68, 32.56]
LIBD -22.31 [ -54.95, 10.33]
Sydney MAS -7.47 [ -41.38, 26.44]
MCIC -2.52 [ -58.38, 53.34]
MooDS -14.45 [ -51.45, 22.55]
MPIP -36.20 [ -63.64, -8.77]
MUNSTER -5.37 [ -31.24, 20.50]
NCNG -35.27 [ -79.10, 8.55]
NESDA 17.41 [ -36.89, 71.70]
NeuroIMAGE -2.80 [ -61.27, 55.68]
NTR -27.47 [ -71.71, 16.77]
OATS 16.82 [ -43.33, 76.97]
PAFIP -3.24 [ -79.80, 73.33]
PDNZ -62.86 [-124.43, -1.30]
PING 0.31 [ -42.68, 43.30]
MPRC -8.74 [ -43.59, 26.11]
PPMI
QTIM 11.62 [ -14.27, 37.51]
SHIP -1.40 [ -22.73, 19.93]
SHIP-Trend -18.52 [ -44.31, 7.27]
SYS 14.23 [ -27.01, 55.48]
TCD-NUIG -14.70 [ -78.53, 49.14]
TOP 3.69 [ -33.17, 40.54]
TOP3T -48.14 [ -87.60, -8.69]
UiO2016 -22.32 [ -74.66, 30.01]
UiO2017 -25.44 [ -69.21, 18.32]
UMCU -22.41 [ -50.52, 5.70]
ENIGMA Discovery Cohorts [N=22825; P=2.081e-07] -13.59 [ -18.72, -8.46]
UKBB -9.96 [ -17.58, -2.34]
Discovery+UKBB [N=32713; P=2.3e-08] -12.42 [ -16.78, -8.07]
GOBS 7.65 [ -22.23, 37.53]
IMH 10.24 [ -57.48, 77.96]
UNICAMP -50.20 [-114.68, 14.28]
Meth-CT
MIRECC -43.91 [ -96.31, 8.50]
UKBB nonEUR 1.98 [ -41.00, 44.97]
OSAKA 14.10 [ -19.03, 47.22]
PING Non European 0.64 [ -40.53, 41.81]
Non European Cohorts [N=2789; P=0.9152] -0.87 [ -16.83, 15.10]
Cohort
Beta [Lower 95% CI,Upper 95% CI]
-200 -150 -100 -50 0 50 100
mm^2/effect allele

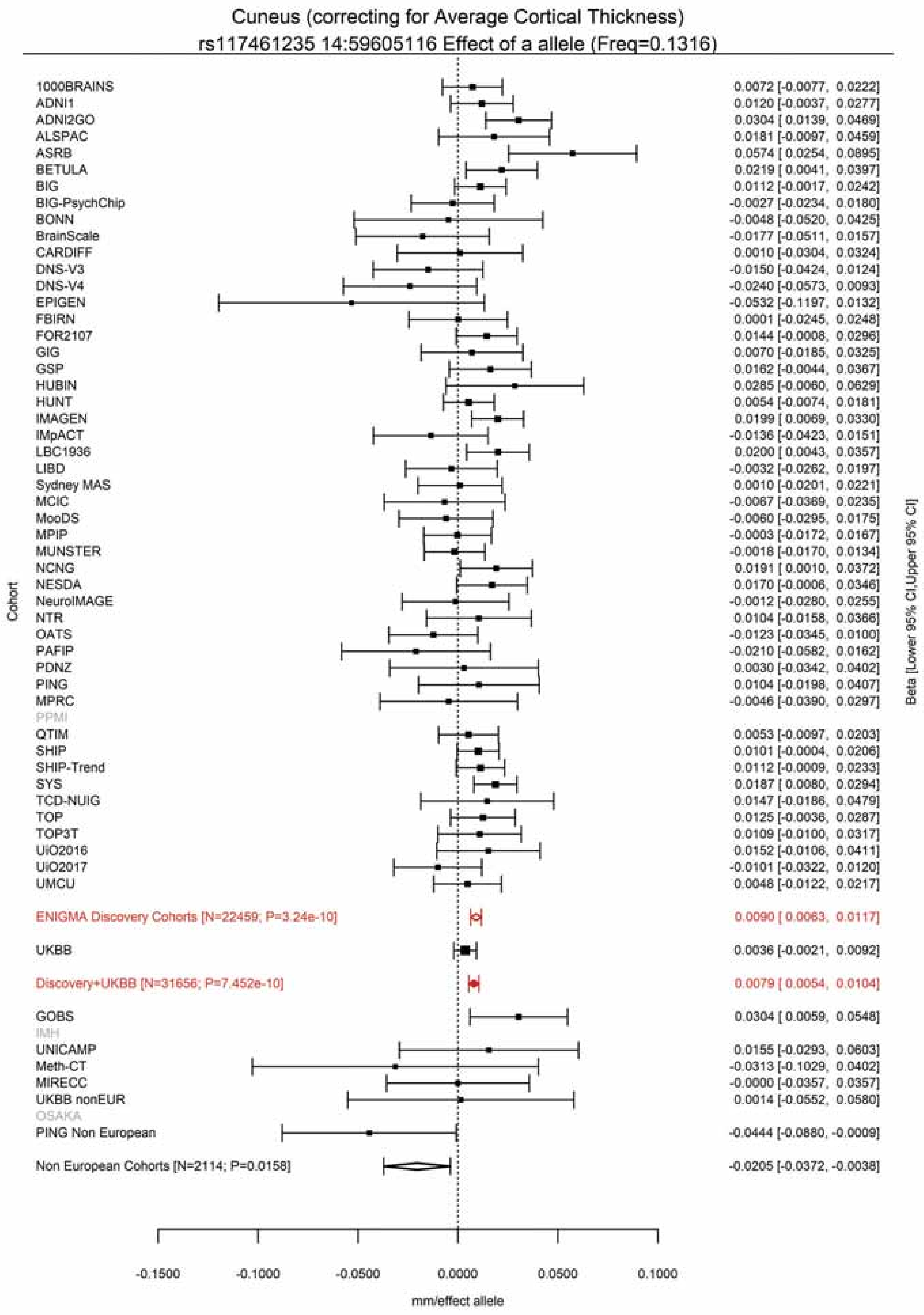
Cuneus (correcting for Average Cortical Thickness)
rs117461235 14:59605116 Effect of a allele (Freq=0.1316)
Cohort
1000BRAINS 0.0072 [-0.0077, 0.0222]
ADNI1 0.0120 [-0.0037, 0.0277]
ADNI2GO 0.0304 [ 0.0139, 0.0469]
ALSPAC 0.0181 [-0.0097, 0.0459]
ASRB 0.0574 [ 0.0254, 0.0895]
BETULA 0.0219 [ 0.0041, 0.0397]
BIG 0.0112 [-0.0017, 0.0242]
BIG-PsychChip -0.0027 [-0.0234, 0.0180]
BONN -0.0048 [-0.0520, 0.0425]
BrainScale -0.0177 [-0.0511, 0.0157]
CARDIFF 0.0010 [-0.0304, 0.0324]
DNS-V3 -0.0150 [-0.0424, 0.0124]
DNS-V4 -0.0240 [-0.0573, 0.0093]
EPIGEN -0.0532 [-0.1197, 0.0132]
FBIRN 0.0001 [-0.0245, 0.0248]
FOR2107 0.0144 [-0.0008, 0.0296]
GIG 0.0070 [-0.0185, 0.0325]
GSP 0.0162 [-0.0044, 0.0367]
HUBIN 0.0285 [-0.0060, 0.0629]
HUNT 0.0054 [-0.0074, 0.0181]
IMAGEN 0.0199 [ 0.0069, 0.0330]
IMpACT -0.0136 [-0.0423, 0.0151]
LBC1936 0.0200 [ 0.0043, 0.0357]
LIBD -0.0032 [-0.0262, 0.0197]
Sydney MAS 0.0010 [-0.0201, 0.0221]
MCIC -0.0067 [-0.0369, 0.0235]
MooDS -0.0060 [-0.0295, 0.0175]
MPIP -0.0003 [-0.0172, 0.0167]
MUNSTER -0.0018 [-0.0170, 0.0134]
NCNG 0.0191 [ 0.0010, 0.0372]
NESDA 0.0170 [-0.0006, 0.0346]
NeuroIMAGE -0.0012 [-0.0280, 0.0255]
NTR 0.0104 [-0.0158, 0.0366]
OATS -0.0123 [-0.0345, 0.0100]
PAFIP -0.0210 [-0.0582, 0.0162]
PDNZ 0.0030 [-0.0342, 0.0402]
PING 0.0104 [-0.0198, 0.0407]
MPRC -0.0046 [-0.0390, 0.0297]
PPMI
QTIM 0.0053 [-0.0097, 0.0203]
SHIP 0.0101 [-0.0004, 0.0206]
SHIP-Trend 0.0112 [-0.0009, 0.0233]
SYS 0.0187 [ 0.0080, 0.0294]
TCD-NUIG 0.0147 [-0.0186, 0.0479]
TOP 0.0125 [-0.0036, 0.0287]
TOP3T 0.0109 [-0.0100, 0.0317]
UiO2016 0.0152 [-0.0106, 0.0411]
UiO2017 -0.0101 [-0.0322, 0.0120]
UMCU 0.0048 [-0.0122, 0.0217]
ENIGMA Discovery Cohorts [N=22459; P=3.24e-10] 0.0090 [ 0.0063, 0.0117]
UKBB 0.0036 [-0.0021, 0.0092]
Discovery+UKBB [N=31656; P=7.452e-10] 0.0079 [ 0.0054, 0.0104]
GOBS 0.0304 [ 0.0059, 0.0548]
IMH
UNICAMP 0.0155 [-0.0293, 0.0603]
Meth-CT -0.0313 [-0.1029, 0.0402]
MIRECC -0.0000 [-0.0357, 0.0357]
UKBB nonEUR 0.0014 [-0.0552, 0.0580]
OSAKA
PING Non European -0.0444 [-0.0880, -0.0009]
Non European Cohorts [N=2114; P=0.0158] -0.0205 [-0.0372, -0.0038]
Beta [Lower 95% CI,Upper 95% CI]
-0.1500
-0.1000
-0.0500
0.0000
0.0500
0.1000
mm/effect allele

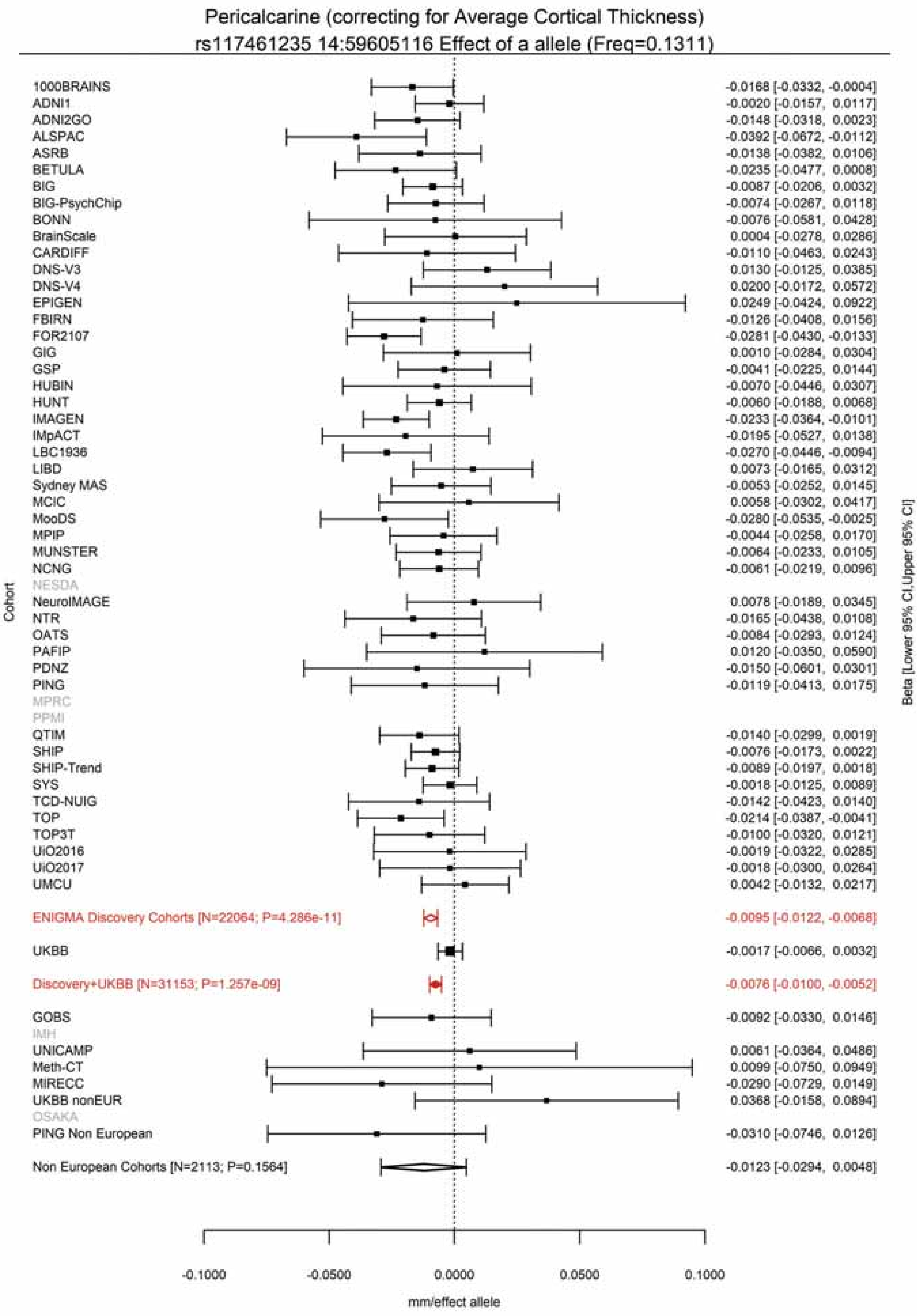

Pericalcarine (correcting for Average Cortical Thickness)
rs117461235 14:59605116 Effect of a allele (Freq=0.1311)
Cohort
1000BRAINS -0.0168 [-0.0332, -0.0004]
ADNI1 -0.0020 [-0.0157, 0.0117]
ADNI2GO -0.0148 [-0.0318, 0.0023]
ALSPAC -0.0392 [-0.0672, -0.0112]
ASRB -0.0138 [-0.0382, 0.0106]
BETULA -0.0235 [-0.0477, 0.0008]
BIG -0.0087 [-0.0206, 0.0032]
BIG-PsychChip -0.0074 [-0.0267, 0.0118]
BONN -0.0076 [-0.0581, 0.0428]
BrainScale 0.0004 [-0.0278, 0.0286]
CARDIFF -0.0110 [-0.0463, 0.0243]
DNS-V3 0.0130 [-0.0125, 0.0385]
DNS-V4 0.0200 [-0.0172, 0.0572]
EPIGEN 0.0249 [-0.0424, 0.0922]
FBIRN -0.0126 [-0.0408, 0.0156]
FOR2107 -0.0281 [-0.0430, -0.0133]
GIG 0.0010 [-0.0284, 0.0304]
GSP -0.0041 [-0.0225, 0.0144]
HUBIN -0.0070 [-0.0446, 0.0307]
HUNT -0.0060 [-0.0188, 0.0068]
IMAGEN -0.0233 [-0.0364, -0.0101]
IMpACT -0.0195 [-0.0527, 0.0138]
LBC1936 -0.0270 [-0.0446, -0.0094]
LIBD 0.0073 [-0.0165, 0.0312]
Sydney MAS -0.0053 [-0.0252, 0.0145]
MCIC 0.0058 [-0.0302, 0.0417]
MooDS -0.0280 [-0.0535, -0.0025]
MPIP -0.0044 [-0.0258, 0.0170]
MUNSTER -0.0064 [-0.0233, 0.0105]
NCNG -0.0061 [-0.0219, 0.0096]
NESDA
NeuroIMAGE 0.0078 [-0.0189, 0.0345]
NTR -0.0165 [-0.0438, 0.0108]
OATS -0.0084 [-0.0293, 0.0124]
PAFIP 0.0120 [-0.0350, 0.0590]
PDNZ -0.0150 [-0.0601, 0.0301]
PING -0.0119 [-0.0413, 0.0175]
MPRC
PPMI
QTIM -0.0140 [-0.0299, 0.0019]
SHIP -0.0076 [-0.0173, 0.0022]
SHIP-Trend -0.0089 [-0.0197, 0.0018]
SYS -0.0018 [-0.0125, 0.0089]
TCD-NUIG -0.0142 [-0.0423, 0.0140]
TOP -0.0214 [-0.0387, -0.0041]
TOP3T -0.0100 [-0.0320, 0.0121]
UiO2016 -0.0019 [-0.0322, 0.0285]
UiO2017 -0.0018 [-0.0300, 0.0264]
UMCU 0.0042 [-0.0132, 0.0217]
ENIGMA Discovery Cohorts [N=22064; P=4.286e-11] -0.0095 [-0.0122, -0.0068]
UKBB -0.0017 [-0.0066, 0.0032]
Discovery+UKBB [N=31153; P=1.257e-09] -0.0076 [-0.0100, -0.0052]
GOBS -0.0092 [-0.0330, 0.0146]
IMH
UNICAMP 0.0061 [-0.0364, 0.0486]
Meth-CT 0.0099 [-0.0750, 0.0949]
MIRECC -0.0290 [-0.0729, 0.0149]
UKBB nonEUR 0.0368 [-0.0158, 0.0894]
OSAKA
PING Non European -0.0310 [-0.0746, 0.0126]
Non European Cohorts [N=2113; P=0.1564] -0.0123 [-0.0294, 0.0048]
-0.1000
-0.0500
0.0000
0.0500
0.1000
mm/effect allele
Beta [Lower 95% CI,Upper 95% CI]

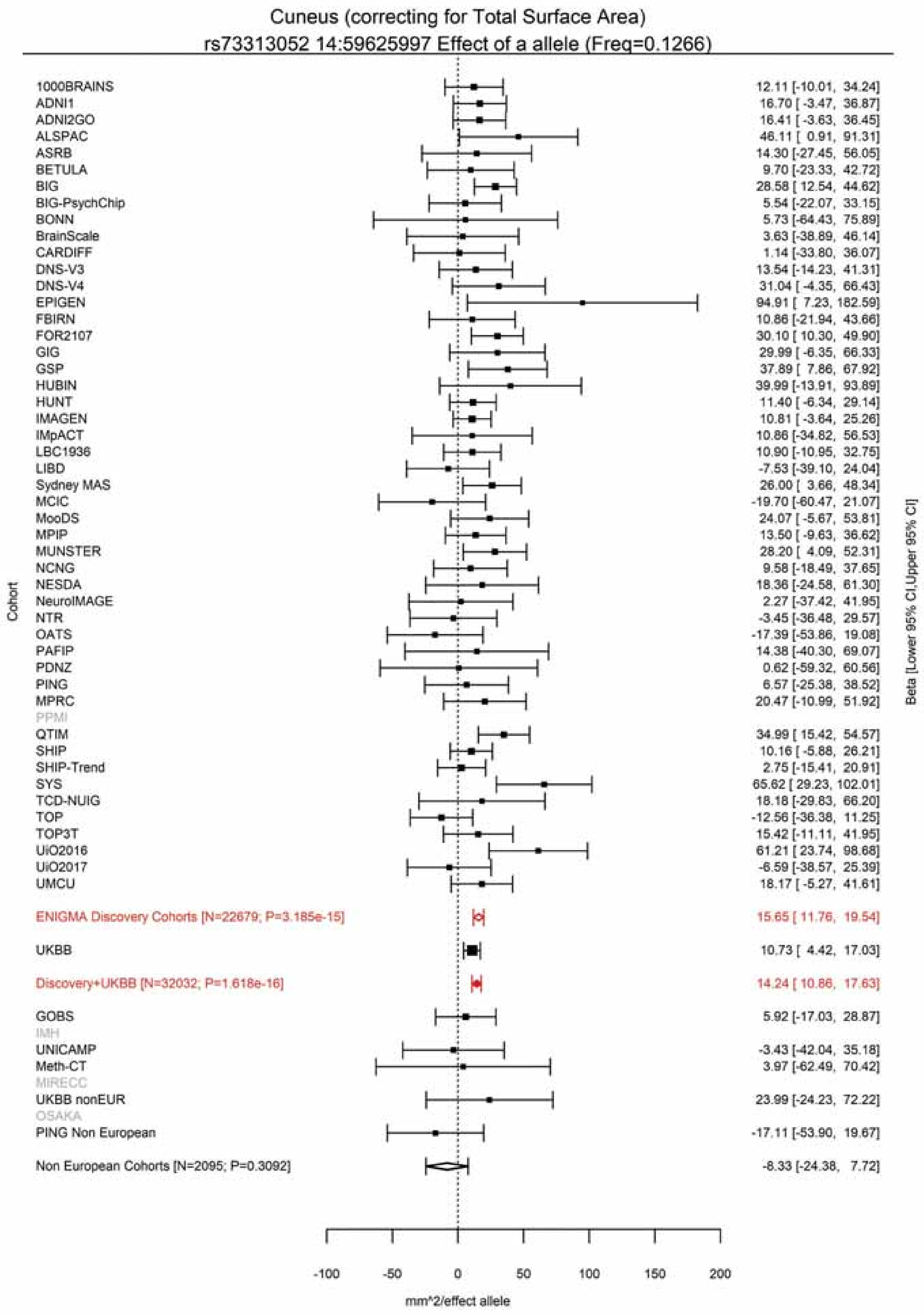
Cuneus (correcting for Total Surface Area)
rs73313052 14:59625997 Effect of a allele (Freq=0.1266)
Cohort
Beta [Lower 95% CI,Upper 95% CI]
1000BRAINS 12.11 [-10.01, 34.24]
ADNI1 16.70 [ -3.47, 36.87]
ADNI2GO 16.41 [ -3.63, 36.45]
ALSPAC 46.11 [ 0.91, 91.31]
ASRB 14.30 [-27.45, 56.05]
BETULA 9.70 [-23.33, 42.72]
BIG 28.58 [ 12.54, 44.62]
BIG-PsychChip 5.54 [-22.07, 33.15]
BONN 5.73 [-64.43, 75.89]
BrainScale 3.63 [-38.89, 46.14]
CARDIFF 1.14 [-33.80, 36.07]
DNS-V3 13.54 [-14.23, 41.31]
DNS-V4 31.04 [ -4.35, 66.43]
EPIGEN 94.91 [ 7.23, 182.59]
FBIRN 10.86 [-21.94, 43.66]
FOR2107 30.10 [ 10.30, 49.90]
GIG 29.99 [ -6.35, 66.33]
GSP 37.89 [ 7.86, 67.92]
HUBIN 39.99 [-13.91, 93.89]
HUNT 11.40 [ -6.34, 29.14]
IMAGEN 10.81 [ -3.64, 25.26]
IMpACT 10.86 [-34.82, 56.53]
LBC1936 10.90 [-10.95, 32.75]
LIBD -7.53 [-39.10, 24.04]
Sydney MAS 26.00 [ 3.66, 48.34]
MCIC -19.70 [-60.47, 21.07]
MooDS 24.07 [ -5.67, 53.81]
MPIP 13.50 [ -9.63, 36.62]
MUNSTER 28.20 [ 4.09, 52.31]
NCNG 9.58 [-18.49, 37.65]
NESDA 18.36 [-24.58, 61.30]
NeuroIMAGE 2.27 [-37.42, 41.95]
NTR -3.45 [-36.48, 29.57]
OATS -17.39 [-53.86, 19.08]
PAFIP 14.38 [-40.30, 69.07]
PDNZ 0.62 [-59.32, 60.56]
PING 6.57 [-25.38, 38.52]
MPRC 20.47 [-10.99, 51.92]
PPMI
QTIM 34.99 [ 15.42, 54.57]
SHIP 10.16 [ -5.88, 26.21]
SHIP-Trend 2.75 [-15.41, 20.91]
SYS 65.62 [ 29.23, 102.01]
TCD-NUIG 18.18 [-29.83, 66.20]
TOP -12.56 [-36.38, 11.25]
TOP3T 15.42 [-11.11, 41.95]
UiO2016 61.21 [ 23.74, 98.68]
UiO2017 -6.59 [-38.57, 25.39]
UMCU 18.17 [ -5.27, 41.61]
ENIGMA Discovery Cohorts [N=22679; P=3.185e-15] 15.65 [ 11.76, 19.54]
UKBB 10.73 [ 4.42, 17.03]
Discovery+UKBB [N=32032; P=1.618e-16] 14.24 [ 10.86, 17.63]
GOBS 5.92 [-17.03, 28.87]
IMH
UNICAMP -3.43 [-42.04, 35.18]
Meth-CT 3.97 [-62.49, 70.42]
MIRECC
UKBB nonEUR 23.99 [-24.23, 72.22]
OSAKA
PING Non European -17.11 [-53.90, 19.67]
Non European Cohorts [N=2095; P=0.3092] -8.33 [-24.38, 7.72]
-100
-50
0
50
100
150
200
mm^2/effect allele

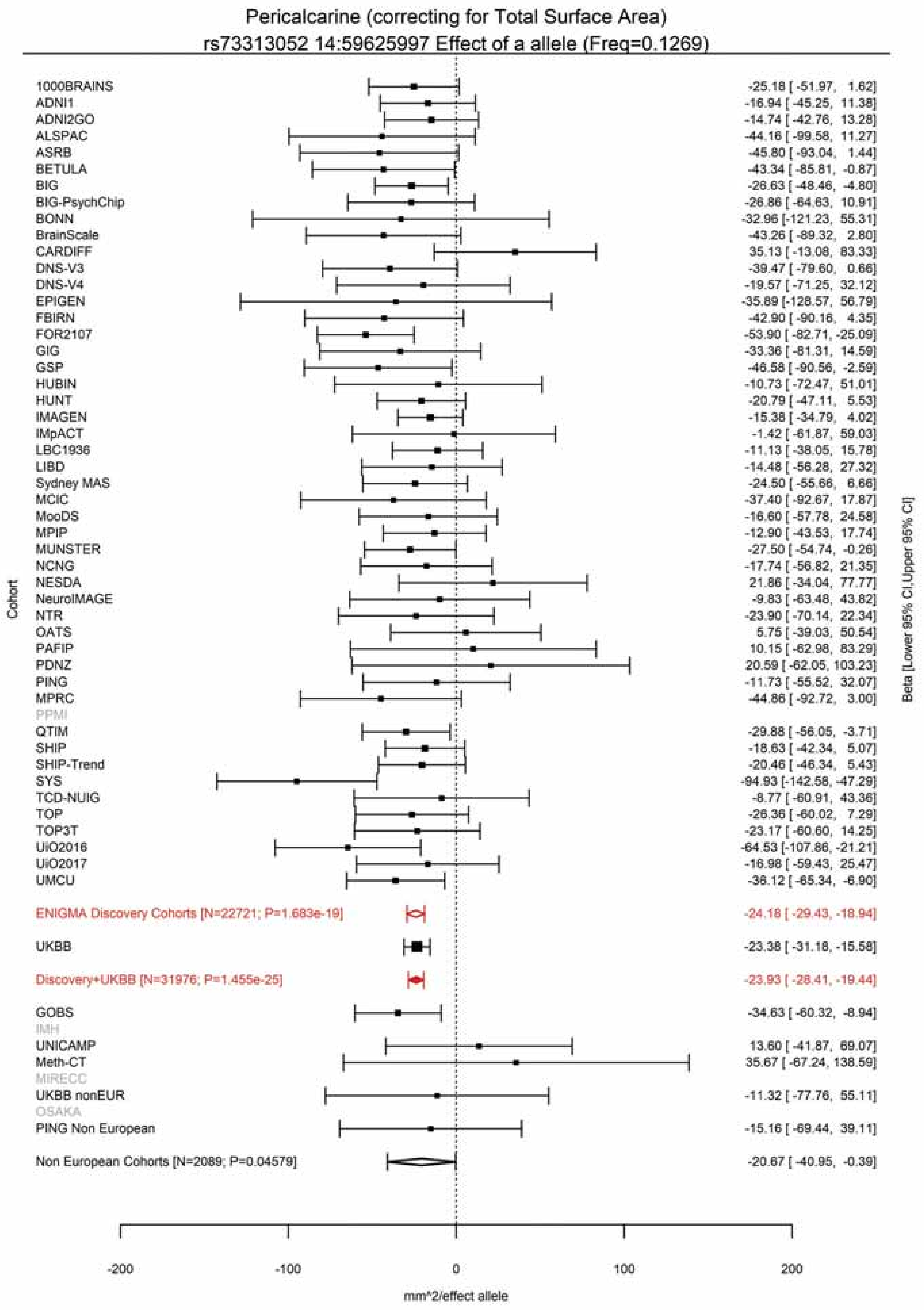
Pericalcarine (correcting for Total Surface Area)
rs73313052 14:59625997 Effect of a allele (Freq=0.1269)
Cohort
1000BRAINS -25.18 [ -51.97, 1.62]
ADNI1 -16.94 [ -45.25, 11.38]
ADNI2GO -14.74 [ -42.76, 13.28]
ALSPAC -44.16 [ -99.58, 11.27]
ASRB -45.80 [ -93.04, 1.44]
BETULA -43.34 [ -85.81, -0.87]
BIG -26.63 [ -48.46, -4.80]
BIG-PsychChip -26.86 [ -64.63, 10.91]
BONN -32.96 [-121.23, 55.31]
BrainScale -43.26 [ -89.32, 2.80]
CARDIFF 35.13 [ -13.08, 83.33]
DNS-V3 -39.47 [ -79.60, 0.66]
DNS-V4 -19.57 [ -71.25, 32.12]
EPIGEN -35.89 [-128.57, 56.79]
FBIRN -42.90 [ -90.16, 4.35]
FOR2107 -53.90 [ -82.71, -25.09]
GIG -33.36 [ -81.31, 14.59]
GSP -46.58 [ -90.56, -2.59]
HUBIN -10.73 [ -72.47, 51.01]
HUNT -20.79 [ -47.11, 5.53]
IMAGEN -15.38 [ -34.79, 4.02]
IMpACT -1.42 [ -61.87, 59.03]
LBC1936 -11.13 [ -38.05, 15.78]
LIBD -14.48 [ -56.28, 27.32]
Sydney MAS -24.50 [ -55.66, 6.66]
MCIC -37.40 [ -92.67, 17.87]
MooDS -16.60 [ -57.78, 24.58]
MPIP -12.90 [ -43.53, 17.74]
MUNSTER -27.50 [ -54.74, -0.26]
NCNG -17.74 [ -56.82, 21.35]
NESDA 21.86 [ -34.04, 77.77]
NeuroIMAGE -9.83 [ -63.48, 43.82]
NTR -23.90 [ -70.14, 22.34]
OATS 5.75 [ -39.03, 50.54]
PAFIP 10.15 [ -62.98, 83.29]
PDNZ 20.59 [ -62.05, 103.23]
PING -11.73 [ -55.52, 32.07]
MPRC -44.86 [ -92.72, 3.00]
PPMI
QTIM -29.88 [ -56.05, -3.71]
SHIP -18.63 [ -42.34, 5.07]
SHIP-Trend -20.46 [ -46.34, 5.43]
SYS -94.93 [-142.58, -47.29]
TCD-NUIG -8.77 [ -60.91, 43.36]
TOP -26.36 [ -60.02, 7.29]
TOP3T -23.17 [ -60.60, 14.25]
UiO2016 -64.53 [-107.86, -21.21]
UiO2017 -16.98 [ -59.43, 25.47]
UMCU -36.12 [ -65.34, -6.90]
ENIGMA Discovery Cohorts [N=22721; P=1.683e-19] -24.18 [ -29.43, -18.94]
UKBB -23.38 [ -31.18, -15.58]
Discovery+UKBB [N=31976; P=1.455e-25] -23.93 [ -28.41, -19.44]
GOBS -34.63 [ -60.32, -8.94]
IMH
UNICAMP 13.60 [ -41.87, 69.07]
Meth-CT 35.67 [ -67.24, 138.59]
MIRECC
UKBB nonEUR -11.32 [ -77.76, 55.11]
OSAKA
PING Non European -15.16 [ -69.44, 39.11]
Non European Cohorts [N=2089; P=0.04579] -20.67 [ -40.95, -0.39]
Beta [Lower 95% CI,Upper 95% CI]
-200
-100
0
100
200
mm^2/effect allele

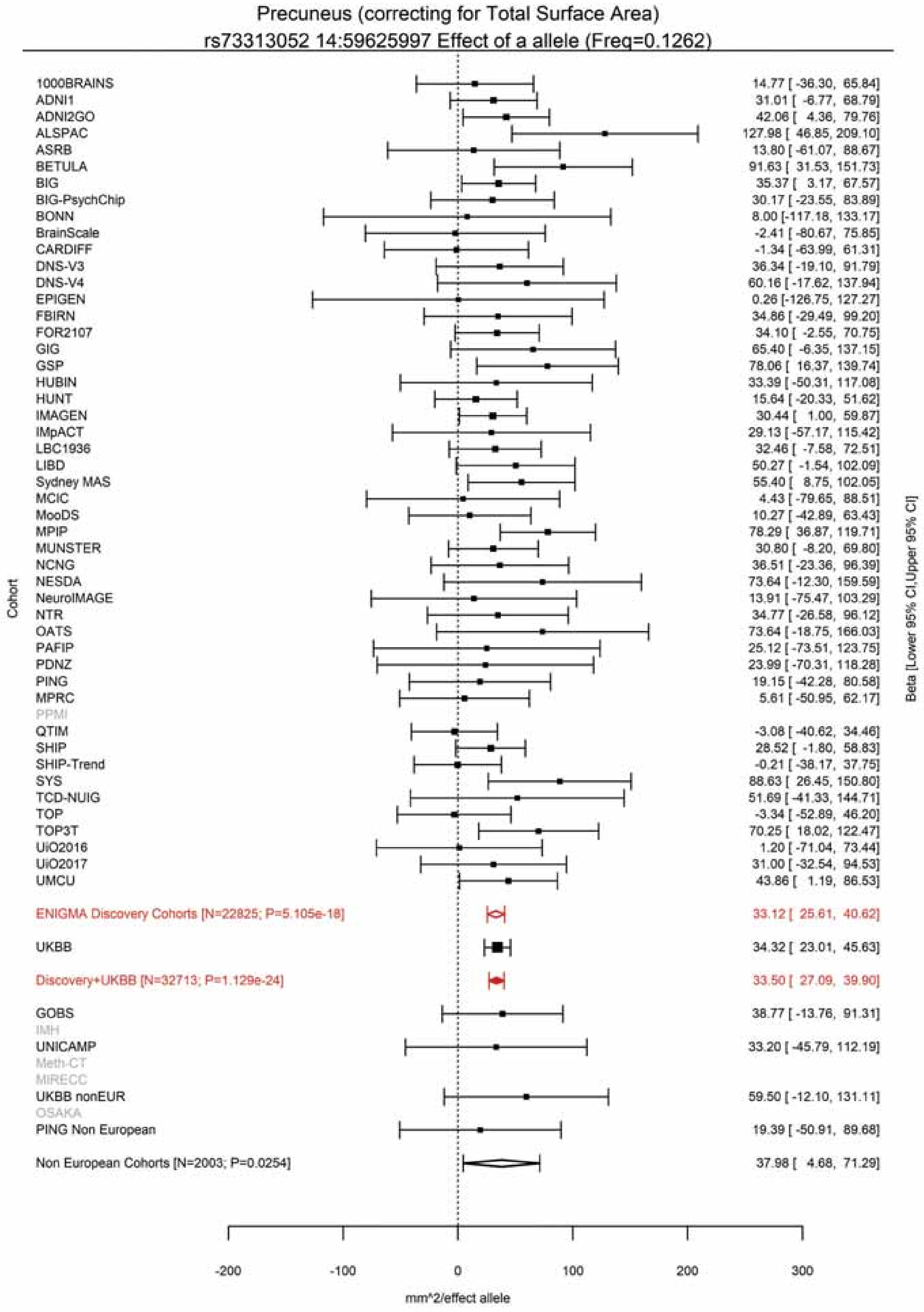

Precuneus (correcting for Total Surface Area)
rs73313052 14:59625997 Effect of a allele (Freq=0.1262)
Cohort
1000BRAINS 14.77 [ -36.30, 65.84]
ADNI1 31.01 [ -6.77, 68.79]
ADNI2GO 42.06 [ 4.36, 79.76]
ALSPAC 127.98 [ 46.85, 209.10]
ASRB 13.80 [ -61.07, 88.67]
BETULA 91.63 [ 31.53, 151.73]
BIG 35.37 [ 3.17, 67.57]
BIG-PsychChip 30.17 [ -23.55, 83.89]
BONN 8.00 [-117.18, 133.17]
BrainScale -2.41 [ -80.67, 75.85]
CARDIFF -1.34 [ -63.99, 61.31]
DNS-V3 36.34 [ -19.10, 91.79]
DNS-V4 60.16 [ -17.62, 137.94]
EPIGEN 0.26 [-126.75, 127.27]
FBIRN 34.86 [ -29.49, 99.20]
FOR2107 34.10 [ -2.55, 70.75]
GIG 65.40 [ -6.35, 137.15]
GSP 78.06 [ 16.37, 139.74]
HUBIN 33.39 [ -50.31, 117.08]
HUNT 15.64 [ -20.33, 51.62]
IMAGEN 30.44 [ 1.00, 59.87]
IMpACT 29.13 [ -57.17, 115.42]
LBC1936 32.46 [ -7.58, 72.51]
LIBD 50.27 [ -1.54, 102.09]
Sydney MAS 55.40 [ 8.75, 102.05]
MCIC 4.43 [ -79.65, 88.51]
MooDS 10.27 [ -42.89, 63.43]
MPIP 78.29 [ 36.87, 119.71]
MUNSTER 30.80 [ -8.20, 69.80]
NCNG 36.51 [ -23.36, 96.39]
NESDA 73.64 [ -12.30, 159.59]
NeuroIMAGE 13.91 [ -75.47, 103.29]
NTR 34.77 [ -26.58, 96.12]
OATS 73.64 [ -18.75, 166.03]
PAFIP 25.12 [ -73.51, 123.75]
PDNZ 23.99 [ -70.31, 118.28]
PING 19.15 [ -42.28, 80.58]
MPRC 5.61 [ -50.95, 62.17]
PPMI
QTIM -3.08 [ -40.62, 34.46]
SHIP 28.52 [ -1.80, 58.83]
SHIP-Trend -0.21 [ -38.17, 37.75]
SYS 88.63 [ 26.45, 150.80]
TCD-NUIG 51.69 [ -41.33, 144.71]
TOP -3.34 [ -52.89, 46.20]
TOP3T 70.25 [ 18.02, 122.47]
UiO2016 1.20 [ -71.04, 73.44]
UiO2017 31.00 [ -32.54, 94.53]
UMCU 43.86 [ 1.19, 86.53]
ENIGMA Discovery Cohorts [N=22825; P=5.105e-18] 33.12 [ 25.61, 40.62]
UKBB 34.32 [ 23.01, 45.63]
Discovery+UKBB [N=32713; P=1.129e-24] 33.50 [ 27.09, 39.90]
GOBS 38.77 [ -13.76, 91.31]
IMH
UNICAMP 33.20 [ -45.79, 112.19]
Meth-CT
MIRECC
UKBB nonEUR 59.50 [ -12.10, 131.11]
OSAKA
PING Non European 19.39 [ -50.91, 89.68]
Non European Cohorts [N=2003; P=0.0254] 37.98 [ 4.68, 71.29]
Beta [Lower 95% CI,Upper 95% CI]
-200
-100
0
100
200
300
mm^2/effect allele

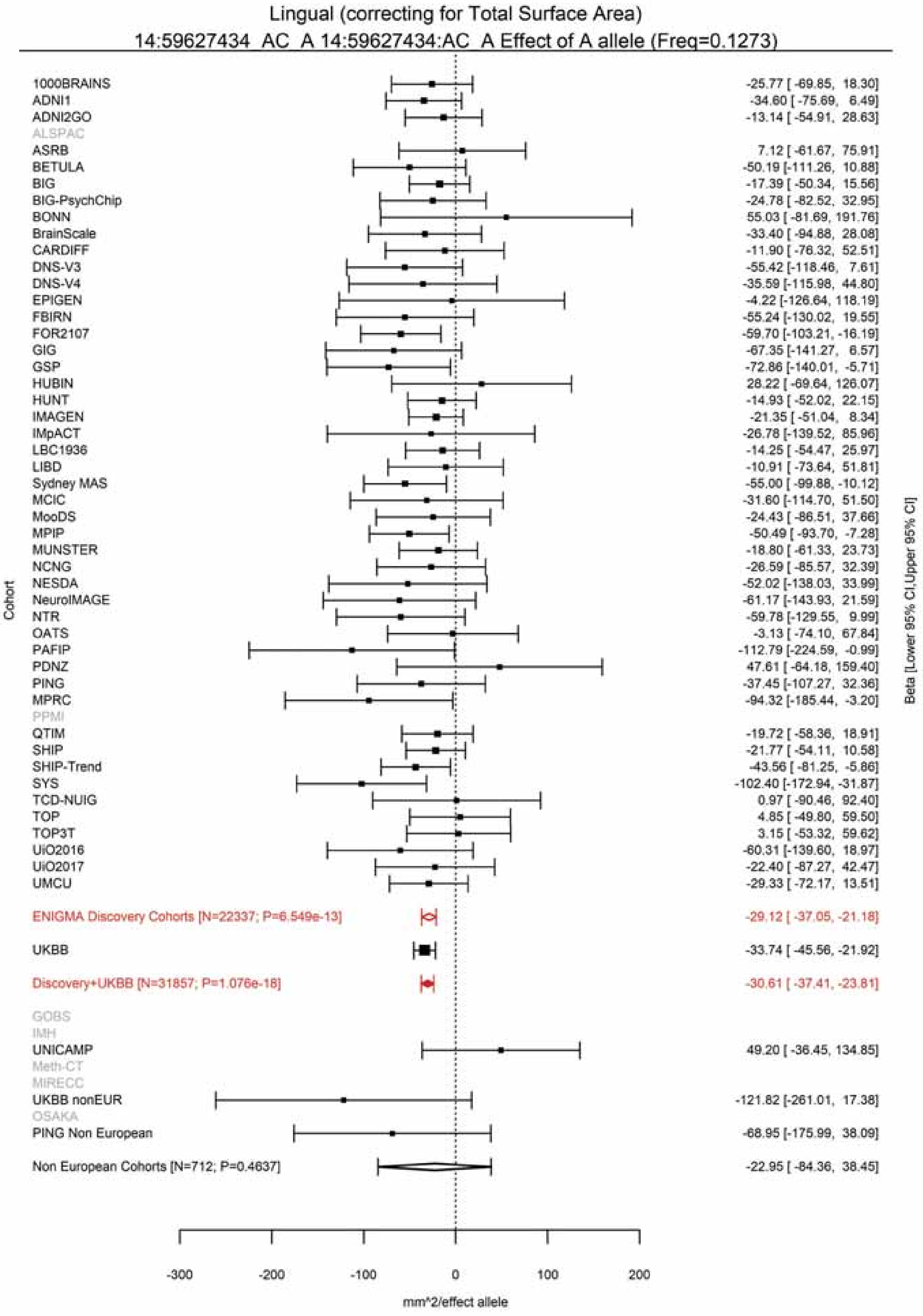

Lingual (correcting for Total Surface Area)
14:59627434_AC_A 14:59627434:AC_A Effect of A allele (Freq=0.1273)
1000BRAINS -25.77 [ -69.85, 18.30]
ADNI1 -34.60 [ -75.69, 6.49]
ADNI2GO -13.14 [ -54.91, 28.63]
ALSPAC
ASRB 7.12 [ -61.67, 75.91]
BETULA -50.19 [-111.26, 10.88]
BIG -17.39 [ -50.34, 15.56]
BIG-PsychChip -24.78 [ -82.52, 32.95]
BONN 55.03 [ -81.69, 191.76]
BrainScale -33.40 [ -94.88, 28.08]
CARDIFF -11.90 [ -76.32, 52.51]
DNS-V3 -55.42 [-118.46, 7.61]
DNS-V4 -35.59 [-115.98, 44.80]
EPIGEN -4.22 [-126.64, 118.19]
FBIRN -55.24 [-130.02, 19.55]
FOR2107 -59.70 [-103.21, -16.19]
GIG -67.35 [-141.27, 6.57]
GSP -72.86 [-140.01, -5.71]
HUBIN 28.22 [ -69.64, 126.07]
HUNT -14.93 [ -52.02, 22.15]
IMAGEN -21.35 [ -51.04, 8.34]
IMpACT -26.78 [-139.52, 85.96]
LBC1936 -14.25 [ -54.47, 25.97]
LIBD -10.91 [ -73.64, 51.81]
Sydney MAS -55.00 [ -99.88, -10.12]
MCIC -31.60 [-114.70, 51.50]
MooDS -24.43 [ -86.51, 37.66]
MPIP -50.49 [ -93.70, -7.28]
MUNSTER -18.80 [ -61.33, 23.73]
NCNG -26.59 [ -85.57, 32.39]
NESDA -52.02 [-138.03, 33.99]
NeuroIMAGE -61.17 [-143.93, 21.59]
NTR -59.78 [-129.55, 9.99]
OATS -3.13 [ -74.10, 67.84]
PAFIP -112.79 [-224.59, -0.99]
PDNZ 47.61 [ -64.18, 159.40]
PING -37.45 [-107.27, 32.36]
MPRC -94.32 [-185.44, -3.20]
PPMI
QTIM -19.72 [ -58.36, 18.91]
SHIP -21.77 [ -54.11, 10.58]
SHIP-Trend -43.56 [ -81.25, -5.86]
SYS -102.40 [-172.94, -31.87]
TCD-NUIG 0.97 [ -90.46, 92.40]
TOP 4.85 [ -49.80, 59.50]
TOP3T 3.15 [ -53.32, 59.62]
UiO2016 -60.31 [-139.60, 18.97]
UiO2017 -22.40 [ -87.27, 42.47]
UMCU -29.33 [ -72.17, 13.51]
ENIGMA Discovery Cohorts [N=22337; P=6.549e-13] -29.12 [ -37.05, -21.18]
UKBB -33.74 [ -45.56, -21.92]
Discovery+UKBB [N=31857; P=1.076e-18] -30.61 [ -37.41, -23.81]
GOBS
IMH
UNICAMP 49.20 [ -36.45, 134.85]
Meth-CT
MIRECC
UKBB nonEUR -121.82 [-261.01, 17.38]
OSAKA
PING Non European -68.95 [-175.99, 38.09]
Non European Cohorts [N=712; P=0.4637] -22.95 [ -84.36, 38.45]
Cohort
Beta [Lower 95% CI,Upper 95% CI]
-300
-200
-100
0
100
200
mm^2/effect allele

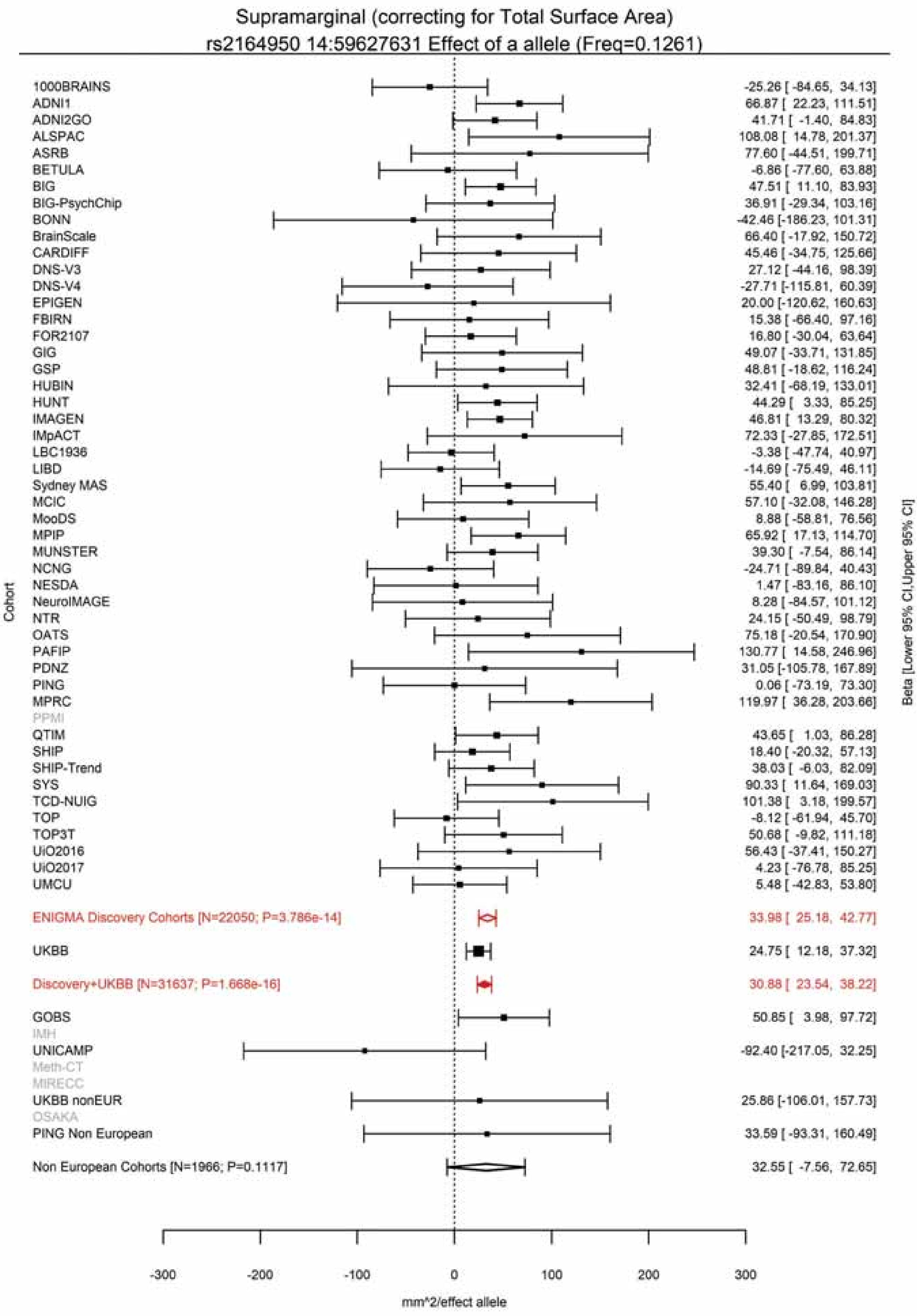

Supramarginal (correcting for Total Surface Area)
rs2164950 14:59627631 Effect of a allele (Freq=0.1261)
Cohort
1000BRAINS -25.26 [ -84.65, 34.13]
ADNI1 66.87 [ 22.23, 111.51]
ADNI2GO 41.71 [ -1.40, 84.83]
ALSPAC 108.08 [ 14.78, 201.37]
ASRB 77.60 [ -44.51, 199.71]
BETULA -6.86 [ -77.60, 63.88]
BIG 47.51 [ 11.10, 83.93]
BIG-PsychChip 36.91 [ -29.34, 103.16]
BONN -42.46 [-186.23, 101.31]
BrainScale 66.40 [ -17.92, 150.72]
CARDIFF 45.46 [ -34.75, 125.66]
DNS-V3 27.12 [ -44.16, 98.39]
DNS-V4 -27.71 [-115.81, 60.39]
EPIGEN 20.00 [-120.62, 160.63]
FBIRN 15.38 [ -66.40, 97.16]
FOR2107 16.80 [ -30.04, 63.64]
GIG 49.07 [ -33.71, 131.85]
GSP 48.81 [ -18.62, 116.24]
HUBIN 32.41 [ -68.19, 133.01]
HUNT 44.29 [ 3.33, 85.25]
IMAGEN 46.81 [ 13.29, 80.32]
IMpACT 72.33 [ -27.85, 172.51]
LBC1936 -3.38 [ -47.74, 40.97]
LIBD -14.69 [ -75.49, 46.11]
Sydney MAS 55.40 [ 6.99, 103.81]
MCIC 57.10 [ -32.08, 146.28]
MooDS 8.88 [ -58.81, 76.56]
MPIP 65.92 [ 17.13, 114.70]
MUNSTER 39.30 [ -7.54, 86.14]
NCNG -24.71 [ -89.84, 40.43]
NESDA 1.47 [ -83.16, 86.10]
NeuroIMAGE 8.28 [ -84.57, 101.12]
NTR 24.15 [ -50.49, 98.79]
OATS 75.18 [ -20.54, 170.90]
PAFIP 130.77 [ 14.58, 246.96]
PDNZ 31.05 [-105.78, 167.89]
PING 0.06 [ -73.19, 73.30]
MPRC 119.97 [ 36.28, 203.66]
PPMI
QTIM 43.65 [ 1.03, 86.28]
SHIP 18.40 [ -20.32, 57.13]
SHIP-Trend 38.03 [ -6.03, 82.09]
SYS 90.33 [ 11.64, 169.03]
TCD-NUIG 101.38 [ 3.18, 199.57]
TOP -8.12 [ -61.94, 45.70]
TOP3T 50.68 [ -9.82, 111.18]
UiO2016 56.43 [ -37.41, 150.27]
UiO2017 4.23 [ -76.78, 85.25]
UMCU 5.48 [ -42.83, 53.80]
ENIGMA Discovery Cohorts [N=22050; P=3.786e-14] 33.98 [ 25.18, 42.77]
UKBB 24.75 [ 12.18, 37.32]
Discovery+UKBB [N=31637; P=1.668e-16] 30.88 [ 23.54, 38.22]
GOBS 50.85 [ 3.98, 97.72]
IMH
UNICAMP -92.40 [-217.05, 32.25]
Meth-CT
MIRECC
UKBB nonEUR 25.86 [-106.01, 157.73]
OSAKA
PING Non European 33.59 [ -93.31, 160.49]
Non European Cohorts [N=1966; P=0.1117] 32.55 [ -7.56, 72.65]
-300
-200
-100
0
100
200
300
mm^2/effect allele
Beta [Lower 95% CI,Upper 95% CI]

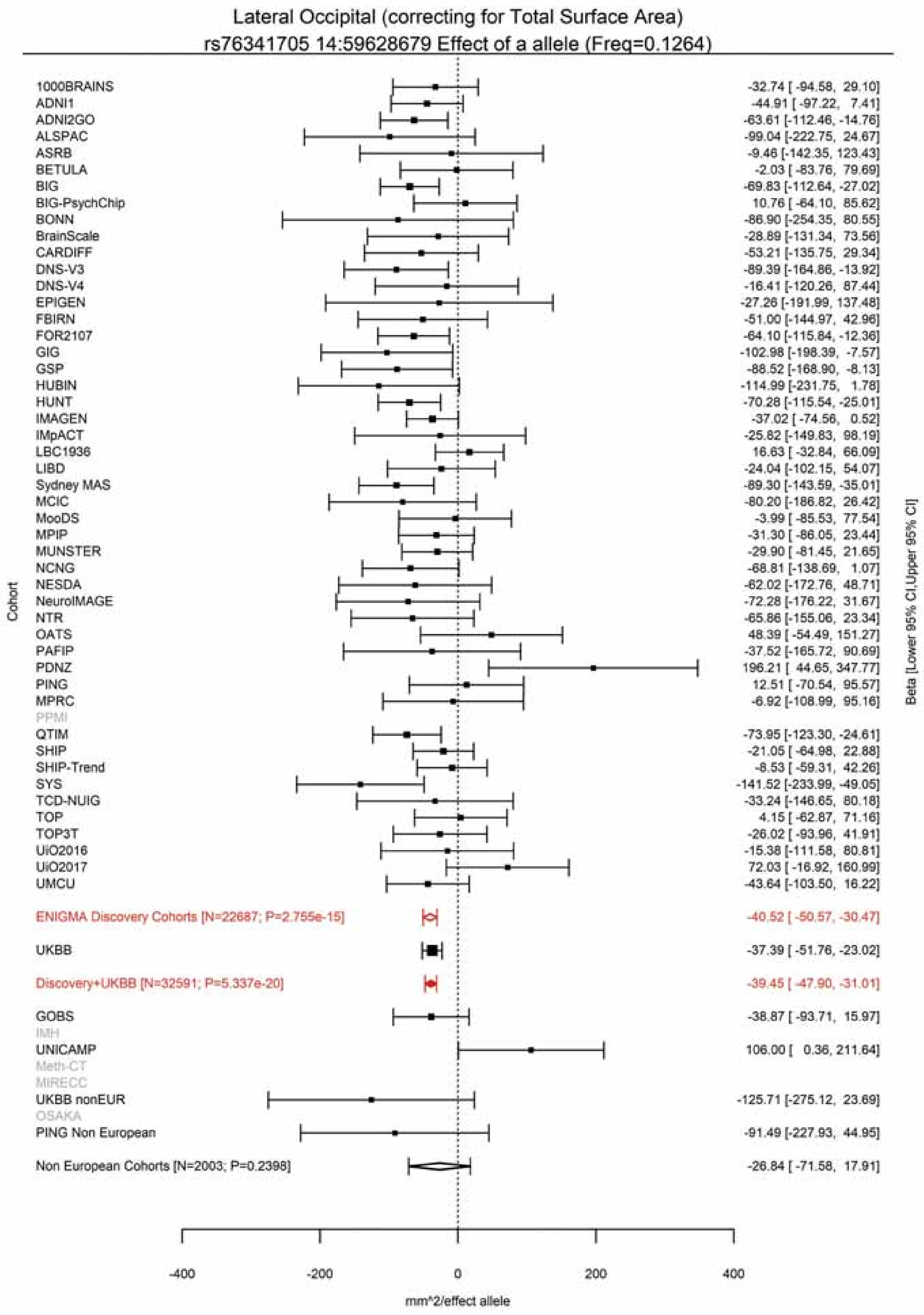
Lateral Occipital (correcting for Total Surface Area)
rs76341705 14:59628679 Effect of a allele (Freq=0.1264)
Cohort
Beta [Lower 95% CI,Upper 95% CI]
1000BRAINS -32.74 [ -94.58, 29.10]
ADNI1 -44.91 [ -97.22, 7.41]
ADNI2GO -63.61 [-112.46, -14.76]
ALSPAC -99.04 [-222.75, 24.67]
ASRB -9.46 [-142.35, 123.43]
BETULA -2.03 [ -83.76, 79.69]
BIG -69.83 [-112.64, -27.02]
BIG-PsychChip 10.76 [ -64.10, 85.62]
BONN -86.90 [-254.35, 80.55]
BrainScale -28.89 [-131.34, 73.56]
CARDIFF -53.21 [-135.75, 29.34]
DNS-V3 -89.39 [-164.86, -13.92]
DNS-V4 -16.41 [-120.26, 87.44]
EPIGEN -27.26 [-191.99, 137.48]
FBIRN -51.00 [-144.97, 42.96]
FOR2107 -64.10 [-115.84, -12.36]
GIG -102.98 [-198.39, -7.57]
GSP -88.52 [-168.90, -8.13]
HUBIN -114.99 [-231.75, 1.78]
HUNT -70.28 [-115.54, -25.01]
IMAGEN -37.02 [ -74.56, 0.52]
IMpACT -25.82 [-149.83, 98.19]
LBC1936 16.63 [ -32.84, 66.09]
LIBD -24.04 [-102.15, 54.07]
Sydney MAS -89.30 [-143.59, -35.01]
MCIC -80.20 [-186.82, 26.42]
MooDS -3.99 [ -85.53, 77.54]
MPIP -31.30 [ -86.05, 23.44]
MUNSTER -29.90 [ -81.45, 21.65]
NCNG -68.81 [-138.69, 1.07]
NESDA -62.02 [-172.76, 48.71]
NeuroIMAGE -72.28 [-176.22, 31.67]
NTR -65.86 [-155.06, 23.34]
OATS 48.39 [ -54.49, 151.27]
PAFIP -37.52 [-165.72, 90.69]
PDNZ 196.21 [ 44.65, 347.77]
PING 12.51 [ -70.54, 95.57]
MPRC -6.92 [-108.99, 95.16]
PPMI
QTIM -73.95 [-123.30, -24.61]
SHIP -21.05 [ -64.98, 22.88]
SHIP-Trend -8.53 [ -59.31, 42.26]
SYS -141.52 [-233.99, -49.05]
TCD-NUIG -33.24 [-146.65, 80.18]
TOP 4.15 [ -62.87, 71.16]
TOP3T -26.02 [ -93.96, 41.91]
UiO2016 -15.38 [-111.58, 80.81]
UiO2017 72.03 [ -16.92, 160.99]
UMCU -43.64 [-103.50, 16.22]
ENIGMA Discovery Cohorts [N=22687; P=2.755e-15] -40.52 [ -50.57, -30.47]
UKBB -37.39 [ -51.76, -23.02]
Discovery+UKBB [N=32591; P=5.337e-20] -39.45 [ -47.90, -31.01]
GOBS -38.87 [ -93.71, 15.97]
IMH
UNICAMP 106.00 [ 0.36, 211.64]
Meth-CT
MIRECC
UKBB nonEUR -125.71 [-275.12, 23.69]
OSAKA
PING Non European -91.49 [-227.93, 44.95]
Non European Cohorts [N=2003; P=0.2398] -26.84 [ -71.58, 17.91]
-400
-200
0
200
400
mm^2/effect allele

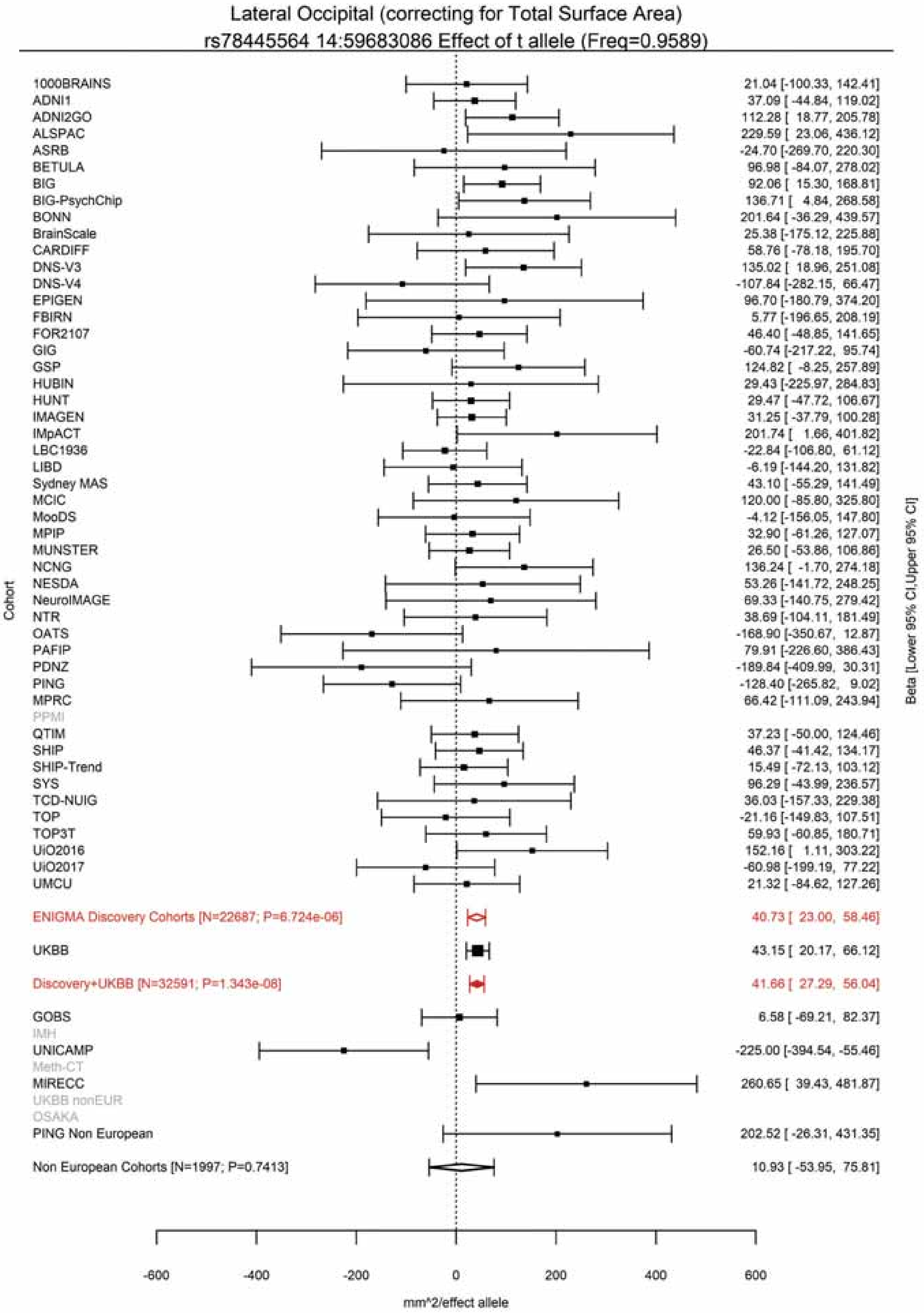

Lateral Occipital (correcting for Total Surface Area)
rs78445564 14:59683086 Effect of t allele (Freq=0.9589)
Cohort
1000BRAINS 21.04 [-100.33, 142.41]
ADNI1 37.09 [-44.84, 119.02]
ADNI2GO 112.28 [ 18.77, 205.78]
ALSPAC 229.59 [ 23.06, 436.12]
ASRB -24.70 [-269.70, 220.30]
BETULA 96.98 [-84.07, 278.02]
BIG 92.06 [ 15.30, 168.81]
BIG-PsychChip 136.71 [ 4.84, 268.58]
BONN 201.64 [-36.29, 439.57]
BrainScale 25.38 [-175.12, 225.88]
CARDIFF 58.76 [-78.18, 195.70]
DNS-V3 135.02 [ 18.96, 251.08]
DNS-V4 -107.84 [-282.15, 66.47]
EPIGEN 96.70 [-180.79, 374.20]
FBIRN 5.77 [-196.65, 208.19]
FOR2107 46.40 [-48.85, 141.65]
GIG -60.74 [-217.22, 95.74]
GSP 124.82 [ -8.25, 257.89]
HUBIN 29.43 [-225.97, 284.83]
HUNT 29.47 [-47.72, 106.67]
IMAGEN 31.25 [-37.79, 100.28]
IMpACT 201.74 [ 1.66, 401.82]
LBC1936 -22.84 [-106.80, 61.12]
LIBD -6.19 [-144.20, 131.82]
Sydney MAS 43.10 [-55.29, 141.49]
MCIC 120.00 [-85.80, 325.80]
MooDS -4.12 [-156.05, 147.80]
MPIP 32.90 [-61.26, 127.07]
MUNSTER 26.50 [-53.86, 106.86]
NCNG 136.24 [ -1.70, 274.18]
NESDA 53.26 [-141.72, 248.25]
NeuroIMAGE 69.33 [-140.75, 279.42]
NTR 38.69 [-104.11, 181.49]
OATS -168.90 [-350.67, 12.87]
PAFIP 79.91 [-226.60, 386.43]
PDNZ -189.84 [-409.99, 30.31]
PING -128.40 [-265.82, 9.02]
MPRC 66.42 [-111.09, 243.94]
PPMI
QTIM 37.23 [-50.00, 124.46]
SHIP 46.37 [-41.42, 134.17]
SHIP-Trend 15.49 [-72.13, 103.12]
SYS 96.29 [-43.99, 236.57]
TCD-NUIG 36.03 [-157.33, 229.38]
TOP -21.16 [-149.83, 107.51]
TOP3T 59.93 [-60.85, 180.71]
UiO2016 152.16 [ 1.11, 303.22]
UiO2017 -60.98 [-199.19, 77.22]
UMCU 21.32 [-84.62, 127.26]
ENIGMA Discovery Cohorts [N=22687; P=6.724e-06] 40.73 [ 23.00, 58.46]
UKBB 43.15 [ 20.17, 66.12]
Discovery+UKBB [N=32591; P=1.343e-08] 41.66 [ 27.29, 56.04]
GOBS 6.58 [-69.21, 82.37]
IMH
UNICAMP -225.00 [-394.54, -55.46]
Meth-CT
MIRECC 260.65 [ 39.43, 481.87]
UKBB nonEUR
OSAKA
PING Non European 202.52 [-26.31, 431.35]
Non European Cohorts [N=1997; P=0.7413] 10.93 [-53.95, 75.81]
Beta [Lower 95% CI,Upper 95% CI]
-600
-400
-200
0
200
400
600
mm^2/effect allele

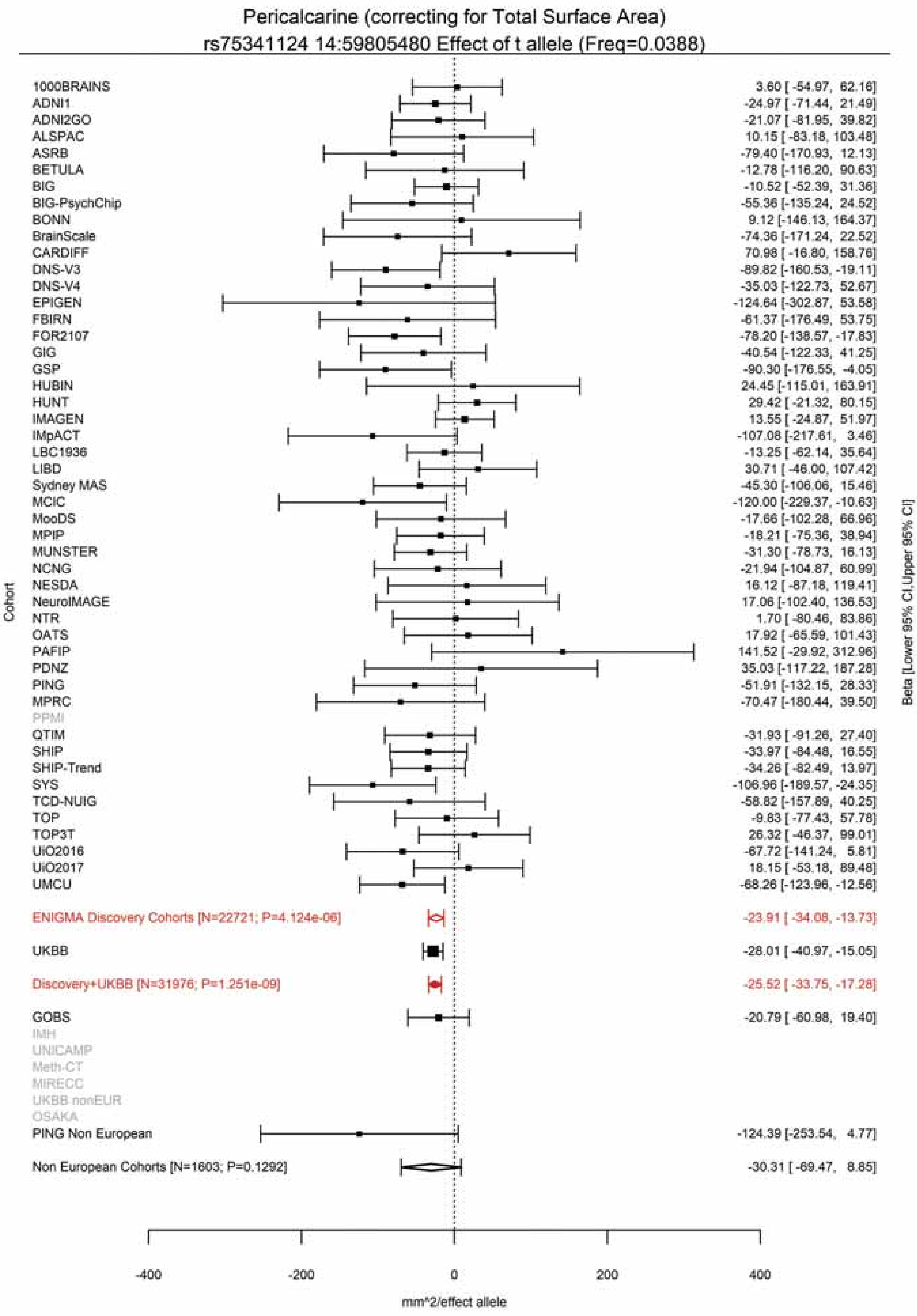

Pericalcarine (correcting for Total Surface Area)
rs75341124 14:59805480 Effect of t allele (Freq=0.0388)
1000BRAINS 3.60 [ -54.97, 62.16]
ADNI1 -24.97 [ -71.44, 21.49]
ADNI2GO -21.07 [ -81.95, 39.82]
ALSPAC 10.15 [ -83.18, 103.48]
ASRB -79.40 [-170.93, 12.13]
BETULA -12.78 [-116.20, 90.63]
BIG -10.52 [ -52.39, 31.36]
BIG-PsychChip -55.36 [-135.24, 24.52]
BONN 9.12 [-146.13, 164.37]
BrainScale -74.36 [-171.24, 22.52]
CARDIFF 70.98 [ -16.80, 158.76]
DNS-V3 -89.82 [-160.53, -19.11]
DNS-V4 -35.03 [-122.73, 52.67]
EPIGEN -124.64 [-302.87, 53.58]
FBIRN -61.37 [-176.49, 53.75]
FOR2107 -78.20 [-138.57, -17.83]
GIG -40.54 [-122.33, 41.25]
GSP -90.30 [-176.55, -4.05]
HUBIN 24.45 [-115.01, 163.91]
HUNT 29.42 [ -21.32, 80.15]
IMAGEN 13.55 [ -24.87, 51.97]
IMpACT -107.08 [-217.61, 3.46]
LBC1936 -13.25 [ -62.14, 35.64]
LIBD 30.71 [ -46.00, 107.42]
Sydney MAS -45.30 [-106.06, 15.46]
MCIC -120.00 [-229.37, -10.63]
MooDS -17.66 [-102.28, 66.96]
MPIP -18.21 [ -75.36, 38.94]
MUNSTER -31.30 [ -78.73, 16.13]
NCNG -21.94 [-104.87, 60.99]
NESDA 16.12 [ -87.18, 119.41]
NeuroIMAGE 17.06 [-102.40, 136.53]
NTR 1.70 [ -80.46, 83.86]
OATS 17.92 [ -65.59, 101.43]
PAFIP 141.52 [ -29.92, 312.96]
PDNZ 35.03 [-117.22, 187.28]
PING -51.91 [-132.15, 28.33]
MPRC -70.47 [-180.44, 39.50]
PPMI
QTIM -31.93 [ -91.26, 27.40]
SHIP -33.97 [ -84.48, 16.55]
SHIP-Trend -34.26 [ -82.49, 13.97]
SYS -106.96 [-189.57, -24.35]
TCD-NUIG -58.82 [-157.89, 40.25]
TOP -9.83 [ -77.43, 57.78]
TOP3T 26.32 [ -46.37, 99.01]
UiO2016 -67.72 [-141.24, 5.81]
UiO2017 18.15 [ -53.18, 89.48]
UMCU -68.26 [-123.96, -12.56]
ENIGMA Discovery Cohorts [N=22721; P=4.124e-06] -23.91 [ -34.08, -13.73]
UKBB -28.01 [ -40.97, -15.05]
Discovery+UKBB [N=31976; P=1.251e-09] -25.52 [ -33.75, -17.28]
GOBS -20.79 [ -60.98, 19.40]
IMH
UNICAMP
Meth-CT
MIRECC
UKBB nonEUR
OSAKA
PING Non European -124.39 [-253.54, 4.77]
Non European Cohorts [N=1603; P=0.1292] -30.31 [ -69.47, 8.85]
Cohort
Beta [Lower 95% CI,Upper 95% CI]
-400
-200
0
200
400
mm^2/effect allele

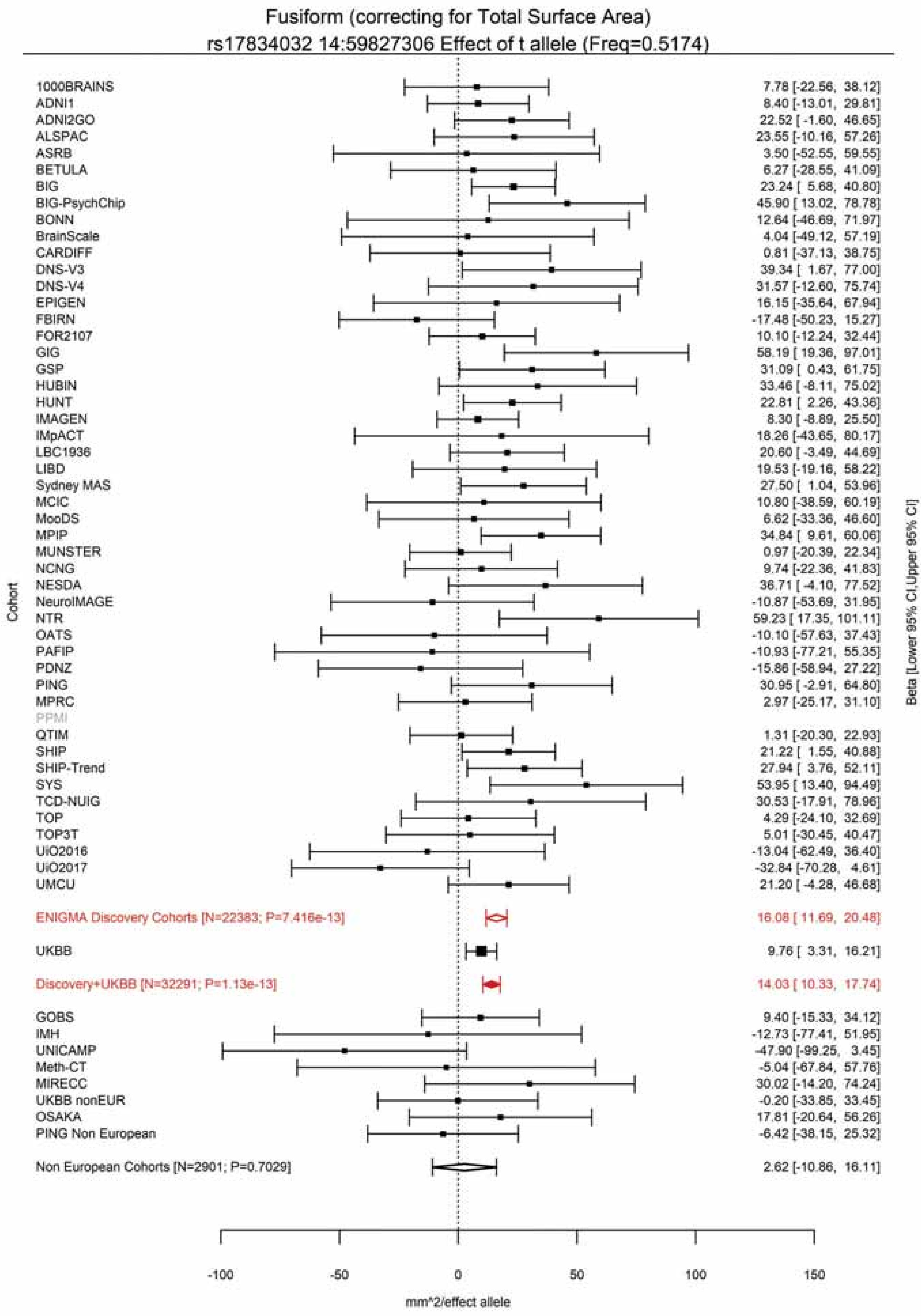
Fusiform (correcting for Total Surface Area)
rs17834032 14:59827306 Effect of t allele (Freq=0.5174)
1000BRAINS 7.78 [-22.56, 38.12]
ADNI1 8.40 [-13.01, 29.81]
ADNI2GO 22.52 [ -1.60, 46.65]
ALSPAC 23.55 [-10.16, 57.26]
ASRB 3.50 [-52.55, 59.55]
BETULA 6.27 [-28.55, 41.09]
BIG 23.24 [ 5.68, 40.80]
BIG-PsychChip 45.90 [ 13.02, 78.78]
BONN 12.64 [-46.69, 71.97]
BrainScale 4.04 [-49.12, 57.19]
CARDIFF 0.81 [-37.13, 38.75]
DNS-V3 39.34 [ 1.67, 77.00]
DNS-V4 31.57 [-12.60, 75.74]
EPIGEN 16.15 [-35.64, 67.94]
FBIRN -17.48 [-50.23, 15.27]
FOR2107 10.10 [-12.24, 32.44]
GIG 58.19 [ 19.36, 97.01]
GSP 31.09 [ 0.43, 61.75]
HUBIN 33.46 [ -8.11, 75.02]
HUNT 22.81 [ 2.26, 43.36]
IMAGEN 8.30 [ -8.89, 25.50]
IMpACT 18.26 [-43.65, 80.17]
LBC1936 20.60 [ -3.49, 44.69]
LIBD 19.53 [-19.16, 58.22]
Sydney MAS 27.50 [ 1.04, 53.96]
MCIC 10.80 [-38.59, 60.19]
MooDS 6.62 [-33.36, 46.60]
MPIP 34.84 [ 9.61, 60.06]
MUNSTER 0.97 [-20.39, 22.34]
NCNG 9.74 [-22.36, 41.83]
NESDA 36.71 [ -4.10, 77.52]
NeuroIMAGE -10.87 [-53.69, 31.95]
NTR 59.23 [ 17.35, 101.11]
OATS -10.10 [-57.63, 37.43]
PAFIP -10.93 [-77.21, 55.35]
PDNZ -15.86 [-58.94, 27.22]
PING 30.95 [ -2.91, 64.80]
MPRC 2.97 [-25.17, 31.10]
PPMI
QTIM 1.31 [-20.30, 22.93]
SHIP 21.22 [ 1.55, 40.88]
SHIP-Trend 27.94 [ 3.76, 52.11]
SYS 53.95 [ 13.40, 94.49]
TCD-NUIG 30.53 [-17.91, 78.96]
TOP 4.29 [-24.10, 32.69]
TOP3T 5.01 [-30.45, 40.47]
UiO2016 -13.04 [-62.49, 36.40]
UiO2017 -32.84 [-70.28, 4.61]
UMCU 21.20 [ -4.28, 46.68]
ENIGMA Discovery Cohorts [N=22383; P=7.416e-13] 16.08 [ 11.69, 20.48]
UKBB 9.76 [ 3.31, 16.21]
Discovery+UKBB [N=32291; P=1.13e-13] 14.03 [ 10.33, 17.74]
GOBS 9.40 [-15.33, 34.12]
IMH -12.73 [-77.41, 51.95]
UNICAMP -47.90 [-99.25, 3.45]
Meth-CT -5.04 [-67.84, 57.76]
MIRECC 30.02 [-14.20, 74.24]
UKBB nonEUR -0.20 [-33.85, 33.45]
OSAKA 17.81 [-20.64, 56.26]
PING Non European -6.42 [-38.15, 25.32]
Non European Cohorts [N=2901; P=0.7029] 2.62 [-10.86, 16.11]
Cohort
Beta [Lower 95% CI,Upper 95% CI]
-100
-50
0
50
100
150
mm^2/effect allele

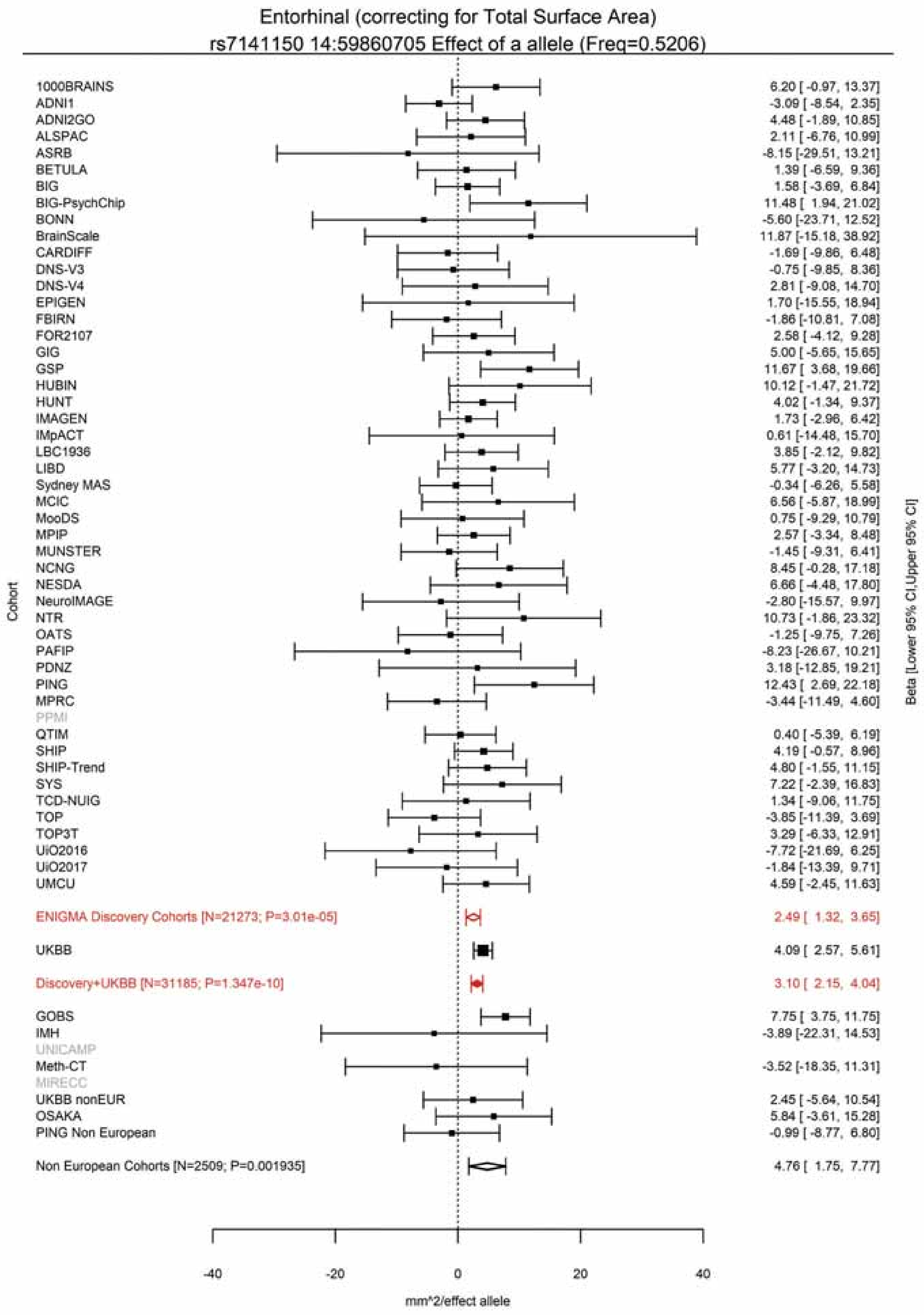
Entorhinal (correcting for Total Surface Area)
rs7141150 14:59860705 Effect of a allele (Freq=0.5206)
1000BRAINS 6.20 [ -0.97, 13.37]
ADNI1 -3.09 [ -8.54, 2.35]
ADNI2GO 4.48 [ -1.89, 10.85]
ALSPAC 2.11 [ -6.76, 10.99]
ASRB -8.15 [-29.51, 13.21]
BETULA 1.39 [ -6.59, 9.36]
BIG 1.58 [ -3.69, 6.84]
BIG-PsychChip 11.48 [ 1.94, 21.02]
BONN -5.60 [-23.71, 12.52]
BrainScale 11.87 [-15.18, 38.92]
CARDIFF -1.69 [ -9.86, 6.48]
DNS-V3 -0.75 [ -9.85, 8.36]
DNS-V4 2.81 [ -9.08, 14.70]
EPIGEN 1.70 [-15.55, 18.94]
FBIRN -1.86 [-10.81, 7.08]
FOR2107 2.58 [ -4.12, 9.28]
GIG 5.00 [ -5.65, 15.65]
GSP 11.67 [ 3.68, 19.66]
HUBIN 10.12 [ -1.47, 21.72]
HUNT 4.02 [ -1.34, 9.37]
IMAGEN 1.73 [ -2.96, 6.42]
IMpACT 0.61 [-14.48, 15.70]
LBC1936 3.85 [ -2.12, 9.82]
LIBD 5.77 [ -3.20, 14.73]
Sydney MAS -0.34 [ -6.26, 5.58]
MCIC 6.56 [ -5.87, 18.99]
MooDS 0.75 [ -9.29, 10.79]
MPIP 2.57 [ -3.34, 8.48]
MUNSTER -1.45 [ -9.31, 6.41]
NCNG 8.45 [ -0.28, 17.18]
NESDA 6.66 [ -4.48, 17.80]
NeuroIMAGE -2.80 [-15.57, 9.97]
NTR 10.73 [ -1.86, 23.32]
OATS -1.25 [ -9.75, 7.26]
PAFIP -8.23 [-26.67, 10.21]
PDNZ 3.18 [-12.85, 19.21]
PING 12.43 [ 2.69, 22.18]
MPRC -3.44 [-11.49, 4.60]
PPMI
QTIM 0.40 [ -5.39, 6.19]
SHIP 4.19 [ -0.57, 8.96]
SHIP-Trend 4.80 [ -1.55, 11.15]
SYS 7.22 [ -2.39, 16.83]
TCD-NUIG 1.34 [ -9.06, 11.75]
TOP -3.85 [-11.39, 3.69]
TOP3T 3.29 [ -6.33, 12.91]
UiO2016 -7.72 [-21.69, 6.25]
UiO2017 -1.84 [-13.39, 9.71]
UMCU 4.59 [ -2.45, 11.63]
ENIGMA Discovery Cohorts [N=21273; P=3.01e-05] 2.49 [ 1.32, 3.65]
UKBB 4.09 [ 2.57, 5.61]
Discovery+UKBB [N=31185; P=1.347e-10] 3.10 [ 2.15, 4.04]
GOBS 7.75 [ 3.75, 11.75]
IMH -3.89 [-22.31, 14.53]
UNICAMP
Meth-CT -3.52 [-18.35, 11.31]
MIRECC
UKBB nonEUR 2.45 [ -5.64, 10.54]
OSAKA 5.84 [ -3.61, 15.28]
PING Non European -0.99 [ -8.77, 6.80]
Non European Cohorts [N=2509; P=0.001935] 4.76 [ 1.75, 7.77]
Cohort
Beta [Lower 95% CI,Upper 95% CI]
-40
-20
0
20
40
mm^2/effect allele

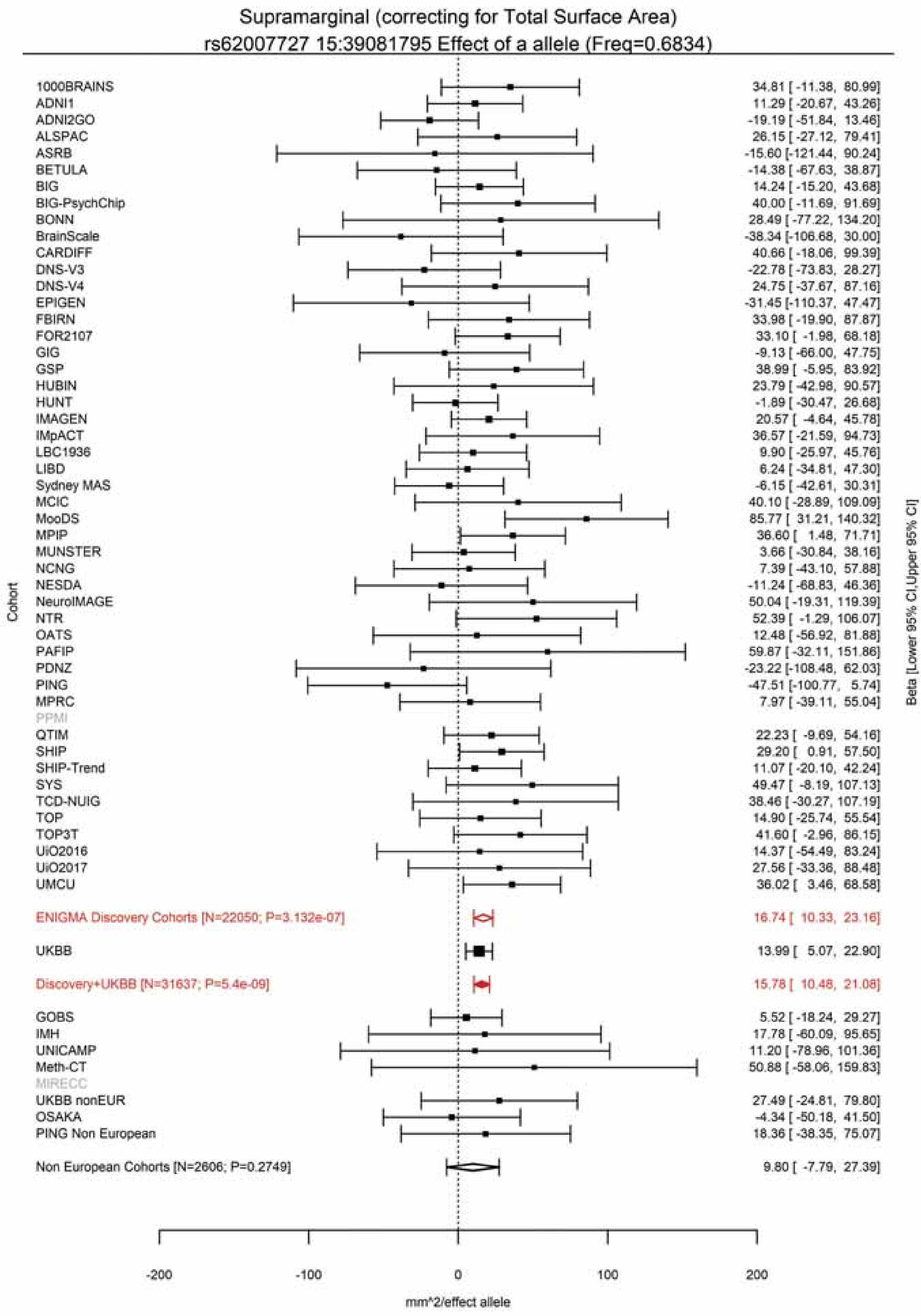
Supramarginal (correcting for Total Surface Area)
rs62007727 15:39081795 Effect of a allele (Freq=0.6834)
Cohort
1000BRAINS 34.81 [ -11.38, 80.99]
ADNI1 11.29 [ -20.67, 43.26]
ADNI2GO -19.19 [ -51.84, 13.46]
ALSPAC 26.15 [ -27.12, 79.41]
ASRB -15.60 [-121.44, 90.24]
BETULA -14.38 [ -67.63, 38.87]
BIG 14.24 [ -15.20, 43.68]
BIG-PsychChip 40.00 [ -11.69, 91.69]
BONN 28.49 [ -77.22, 134.20]
BrainScale -38.34 [-106.68, 30.00]
CARDIFF 40.66 [ -18.06, 99.39]
DNS-V3 -22.78 [ -73.83, 28.27]
DNS-V4 24.75 [ -37.67, 87.16]
EPIGEN -31.45 [-110.37, 47.47]
FBIRN 33.98 [ -19.90, 87.87]
FOR2107 33.10 [ -1.98, 68.18]
GIG -9.13 [ -66.00, 47.75]
GSP 38.99 [ -5.95, 83.92]
HUBIN 23.79 [ -42.98, 90.57]
HUNT -1.89 [ -30.47, 26.68]
IMAGEN 20.57 [ -4.64, 45.78]
IMpACT 36.57 [ -21.59, 94.73]
LBC1936 9.90 [ -25.97, 45.76]
LIBD 6.24 [ -34.81, 47.30]
Sydney MAS -6.15 [ -42.61, 30.31]
MCIC 40.10 [ -28.89, 109.09]
MooDS 85.77 [ 31.21, 140.32]
MPIP 36.60 [ 1.48, 71.71]
MUNSTER 3.66 [ -30.84, 38.16]
NCNG 7.39 [ -43.10, 57.88]
NESDA -11.24 [ -68.83, 46.36]
NeuroIMAGE 50.04 [ -19.31, 119.39]
NTR 52.39 [ -1.29, 106.07]
OATS 12.48 [ -56.92, 81.88]
PAFIP 59.87 [ -32.11, 151.86]
PDNZ -23.22 [-108.48, 62.03]
PING -47.51 [-100.77, 5.74]
MPRC 7.97 [ -39.11, 55.04]
PPMI
QTIM 22.23 [ -9.69, 54.16]
SHIP 29.20 [ 0.91, 57.50]
SHIP-Trend 11.07 [ -20.10, 42.24]
SYS 49.47 [ -8.19, 107.13]
TCD-NUIG 38.46 [ -30.27, 107.19]
TOP 14.90 [ -25.74, 55.54]
TOP3T 41.60 [ -2.96, 86.15]
UiO2016 14.37 [ -54.49, 83.24]
UiO2017 27.56 [ -33.36, 88.48]
UMCU 36.02 [ 3.46, 68.58]
ENIGMA Discovery Cohorts [N=22050; P=3.132e-07] 16.74 [ 10.33, 23.16]
UKBB 13.99 [ 5.07, 22.90]
Discovery+UKBB [N=31637; P=5.4e-09] 15.78 [ 10.48, 21.08]
GOBS 5.52 [ -18.24, 29.27]
IMH 17.78 [ -60.09, 95.65]
UNICAMP 11.20 [ -78.96, 101.36]
Meth-CT 50.88 [ -58.06, 159.83]
MIRECC
UKBB nonEUR 27.49 [ -24.81, 79.80]
OSAKA -4.34 [ -50.18, 41.50]
PING Non European 18.36 [ -38.35, 75.07]
Non European Cohorts [N=2606; P=0.2749] 9.80 [ -7.79, 27.39]
-200
-100
0
100
200
mm^2/effect allele
Beta [Lower 95% CI,Upper 95% CI]

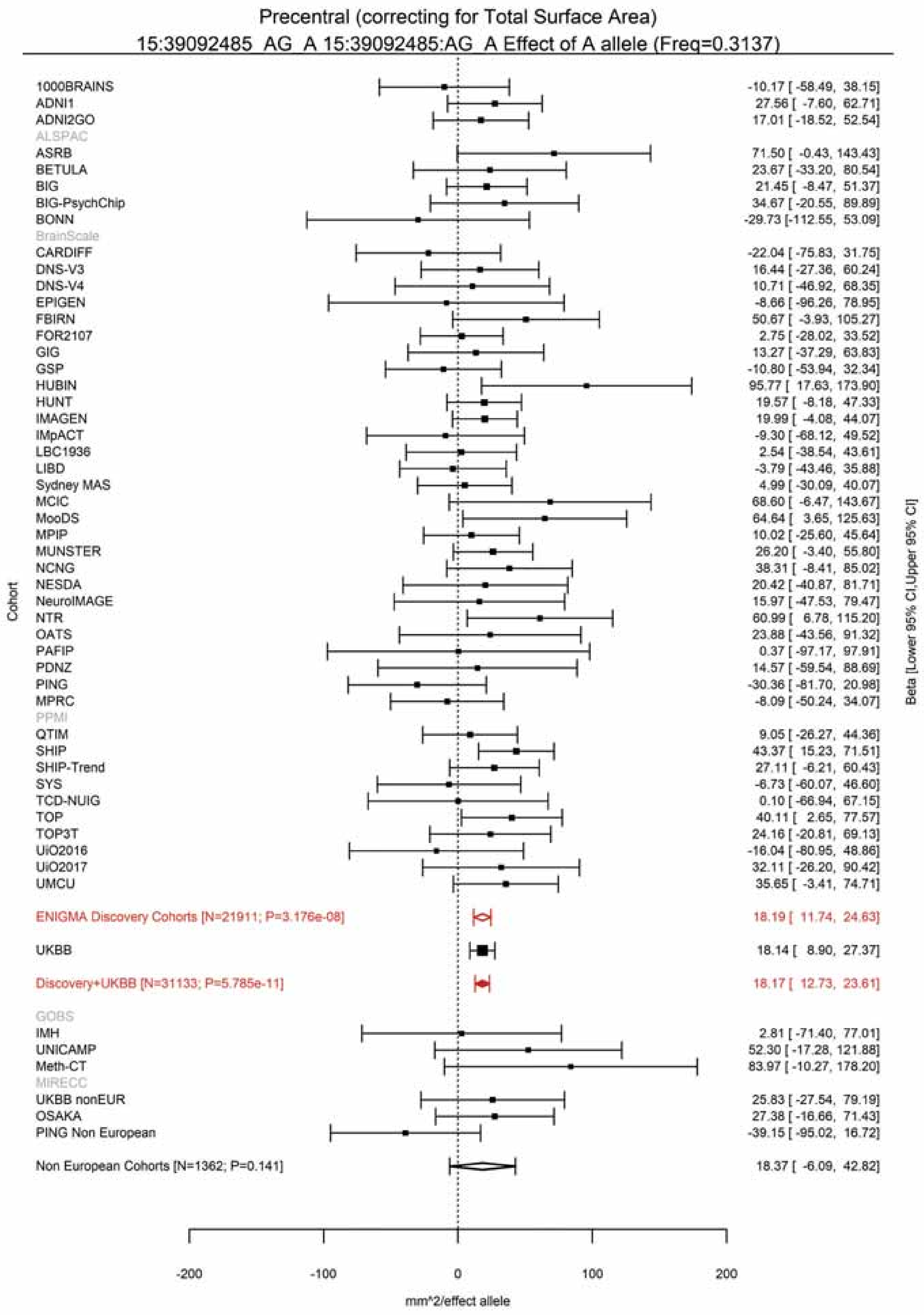
Precentral (correcting for Total Surface Area)
15:39092485_AG_A 15:39092485:AG_A Effect of A allele (Freq=0.3137)
1000BRAINS -10.17 [ -58.49, 38.15]
ADNI1 27.56 [ -7.60, 62.71]
ADNI2GO 17.01 [ -18.52, 52.54]
ALSPAC
ASRB 71.50 [ -0.43, 143.43]
BETULA 23.67 [ -33.20, 80.54]
BIG 21.45 [ -8.47, 51.37]
BIG-PsychChip 34.67 [ -20.55, 89.89]
BONN -29.73 [-112.55, 53.09]
BrainScale
CARDIFF -22.04 [ -75.83, 31.75]
DNS-V3 16.44 [ -27.36, 60.24]
DNS-V4 10.71 [ -46.92, 68.35]
EPIGEN -8.66 [ -96.26, 78.95]
FBIRN 50.67 [ -3.93, 105.27]
FOR2107 2.75 [ -28.02, 33.52]
GIG 13.27 [ -37.29, 63.83]
GSP -10.80 [ -53.94, 32.34]
HUBIN 95.77 [ 17.63, 173.90]
HUNT 19.57 [ -8.18, 47.33]
IMAGEN 19.99 [ -4.08, 44.07]
IMpACT -9.30 [ -68.12, 49.52]
LBC1936 2.54 [ -38.54, 43.61]
LIBD -3.79 [ -43.46, 35.88]
Sydney MAS 4.99 [ -30.09, 40.07]
MCIC 68.60 [ -6.47, 143.67]
MooDS 64.64 [ 3.65, 125.63]
MPIP 10.02 [ -25.60, 45.64]
MUNSTER 26.20 [ -3.40, 55.80]
NCNG 38.31 [ -8.41, 85.02]
NESDA 20.42 [ -40.87, 81.71]
NeuroIMAGE 15.97 [ -47.53, 79.47]
NTR 60.99 [ 6.78, 115.20]
OATS 23.88 [ -43.56, 91.32]
PAFIP 0.37 [ -97.17, 97.91]
PDNZ 14.57 [ -59.54, 88.69]
PING -30.36 [ -81.70, 20.98]
MPRC -8.09 [ -50.24, 34.07]
PPMI
QTIM 9.05 [ -26.27, 44.36]
SHIP 43.37 [ 15.23, 71.51]
SHIP-Trend 27.11 [ -6.21, 60.43]
SYS -6.73 [ -60.07, 46.60]
TCD-NUIG 0.10 [ -66.94, 67.15]
TOP 40.11 [ 2.65, 77.57]
TOP3T 24.16 [ -20.81, 69.13]
UiO2016 -16.04 [ -80.95, 48.86]
UiO2017 32.11 [ -26.20, 90.42]
UMCU 35.65 [ -3.41, 74.71]
ENIGMA Discovery Cohorts [N=21911; P=3.176e-08] 18.19 [ 11.74, 24.63]
UKBB 18.14 [ 8.90, 27.37]
Discovery+UKBB [N=31133; P=5.785e-11] 18.17 [ 12.73, 23.61]
GOBS
IMH 2.81 [ -71.40, 77.01]
UNICAMP 52.30 [ -17.28, 121.88]
Meth-CT 83.97 [ -10.27, 178.20]
MIRECC
UKBB nonEUR 25.83 [ -27.54, 79.19]
OSAKA 27.38 [ -16.66, 71.43]
PING Non European -39.15 [ -95.02, 16.72]
Non European Cohorts [N=1362; P=0.141] 18.37 [ -6.09, 42.82]
Cohort
Beta [Lower 95% CI,Upper 95% CI]
-200
-100
0
100
200
mm^2/effect allele

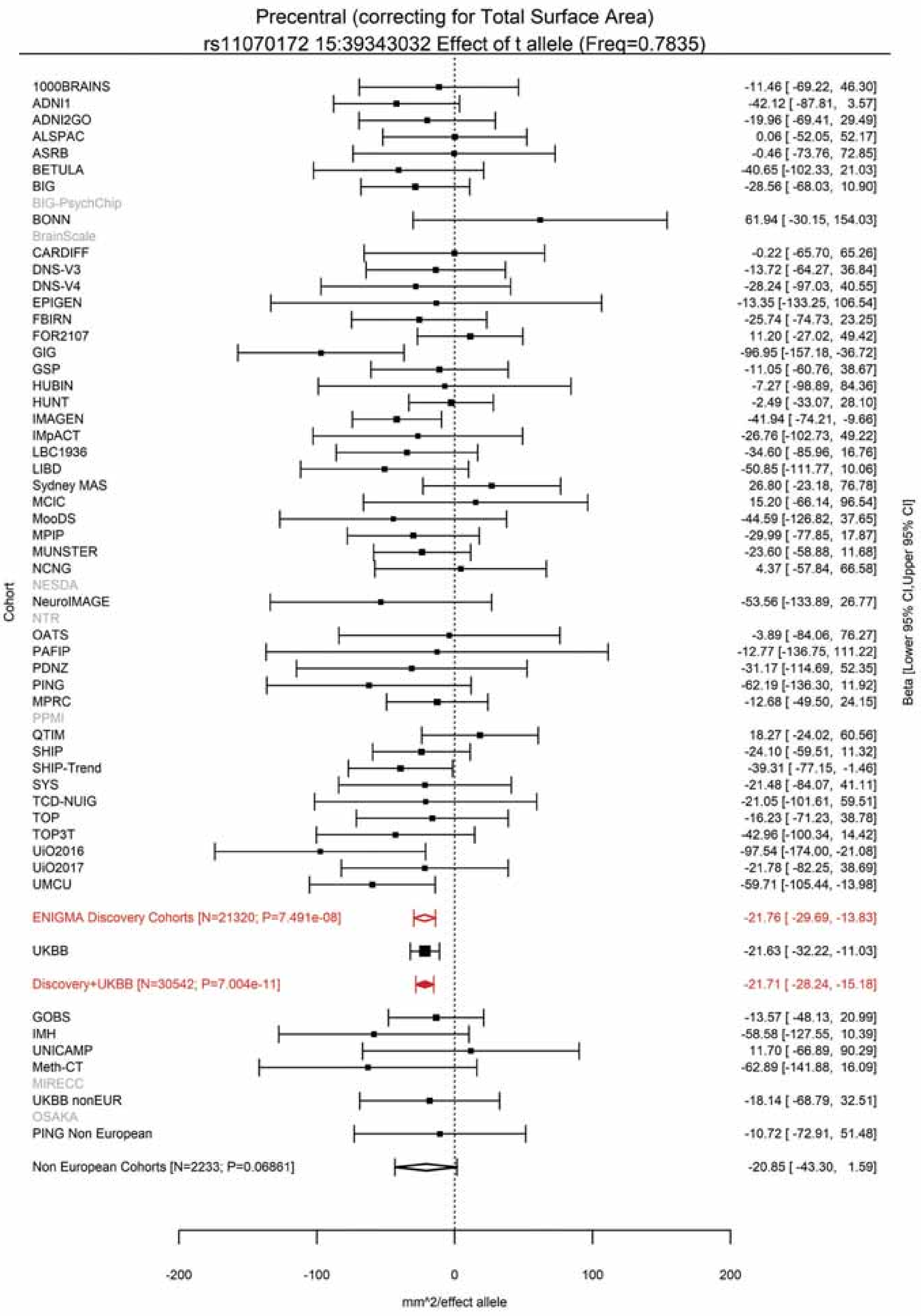
Precentral (correcting for Total Surface Area)
rs11070172 15:39343032 Effect of t allele (Freq=0.7835)
Cohort
Beta [Lower 95% CI,Upper 95% CI]
1000BRAINS -11.46 [ -69.22, 46.30]
ADNI1 -42.12 [ -87.81, 3.57]
ADNI2GO -19.96 [ -69.41, 29.49]
ALSPAC 0.06 [ -52.05, 52.17]
ASRB -0.46 [ -73.76, 72.85]
BETULA -40.65 [-102.33, 21.03]
BIG -28.56 [ -68.03, 10.90]
BIG-PsychChip
BONN 61.94 [ -30.15, 154.03]
BrainScale
CARDIFF -0.22 [ -65.70, 65.26]
DNS-V3 -13.72 [ -64.27, 36.84]
DNS-V4 -28.24 [ -97.03, 40.55]
EPIGEN -13.35 [-133.25, 106.54]
FBIRN -25.74 [ -74.73, 23.25]
FOR2107 11.20 [ -27.02, 49.42]
GIG -96.95 [-157.18, -36.72]
GSP -11.05 [ -60.76, 38.67]
HUBIN -7.27 [ -98.89, 84.36]
HUNT -2.49 [ -33.07, 28.10]
IMAGEN -41.94 [ -74.21, -9.66]
IMpACT -26.76 [-102.73, 49.22]
LBC1936 -34.60 [ -85.96, 16.76]
LIBD -50.85 [-111.77, 10.06]
Sydney MAS 26.80 [ -23.18, 76.78]
MCIC 15.20 [ -66.14, 96.54]
MooDS -44.59 [-126.82, 37.65]
MPIP -29.99 [ -77.85, 17.87]
MUNSTER -23.60 [ -58.88, 11.68]
NCNG 4.37 [ -57.84, 66.58]
NESDA
NeuroIMAGE -53.56 [-133.89, 26.77]
NTR
OATS -3.89 [ -84.06, 76.27]
PAFIP -12.77 [-136.75, 111.22]
PDNZ -31.17 [-114.69, 52.35]
PING -62.19 [-136.30, 11.92]
MPRC -12.68 [ -49.50, 24.15]
PPMI
QTIM 18.27 [ -24.02, 60.56]
SHIP -24.10 [ -59.51, 11.32]
SHIP-Trend -39.31 [ -77.15, -1.46]
SYS -21.48 [ -84.07, 41.11]
TCD-NUIG -21.05 [-101.61, 59.51]
TOP -16.23 [ -71.23, 38.78]
TOP3T -42.96 [-100.34, 14.42]
UiO2016 -97.54 [-174.00, -21.08]
UiO2017 -21.78 [ -82.25, 38.69]
UMCU -59.71 [-105.44, -13.98]
ENIGMA Discovery Cohorts [N=21320; P=7.491e-08] -21.76 [ -29.69, -13.83]
UKBB -21.63 [ -32.22, -11.03]
Discovery+UKBB [N=30542; P=7.004e-11] -21.71 [ -28.24, -15.18]
GOBS -13.57 [ -48.13, 20.99]
IMH -58.58 [-127.55, 10.39]
UNICAMP 11.70 [ -66.89, 90.29]
Meth-CT -62.89 [-141.88, 16.09]
MIRECC
UKBB nonEUR -18.14 [ -68.79, 32.51]
OSAKA
PING Non European -10.72 [ -72.91, 51.48]
Non European Cohorts [N=2233; P=0.06861] -20.85 [ -43.30, 1.59]
-200
-100
0
100
200
mm^2/effect allele

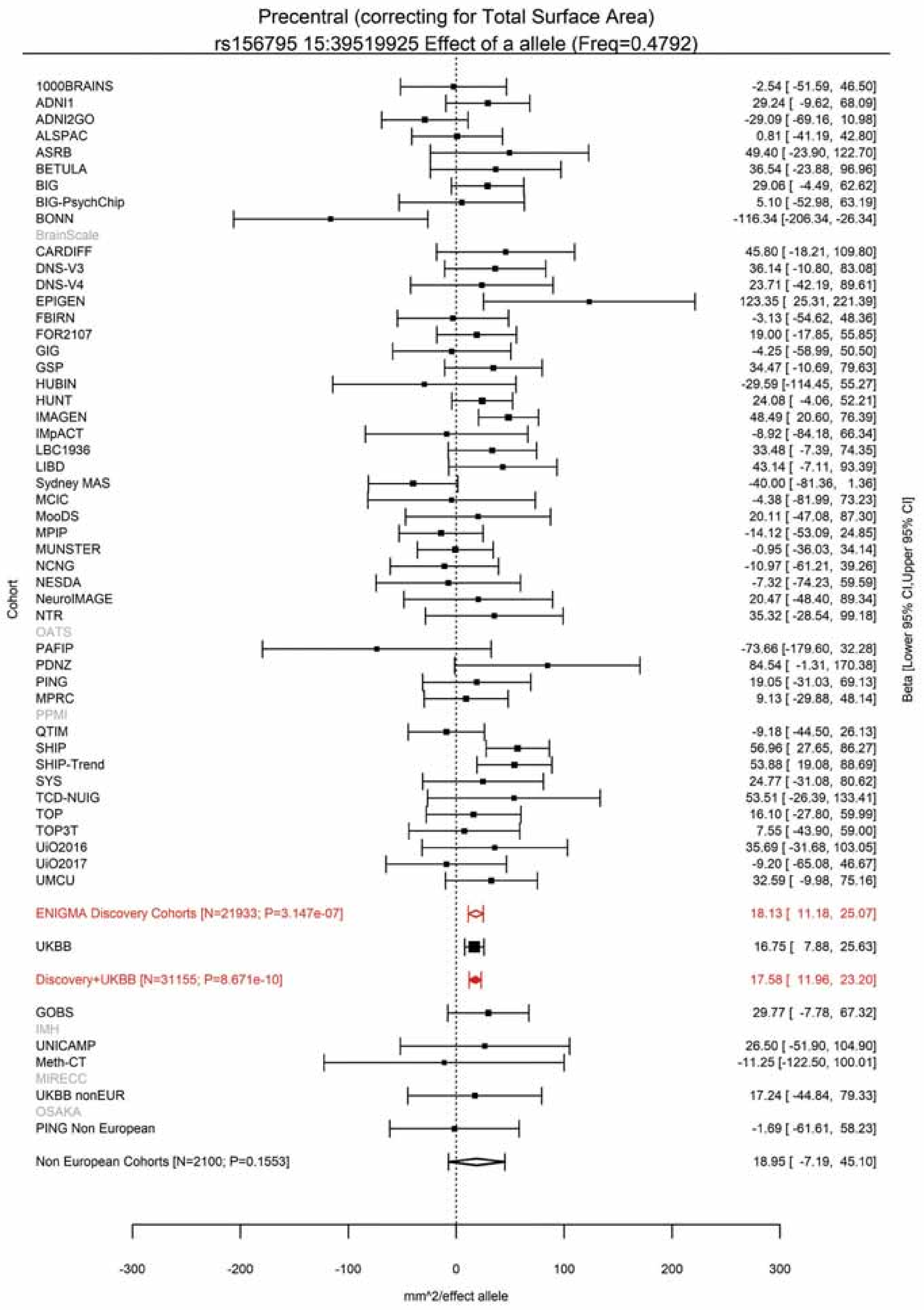
Precentral (correcting for Total Surface Area)
rs156795 15:39519925 Effect of a allele (Freq=0.4792)
Cohort
1000BRAINS -2.54 [ -51.59, 46.50]
ADNI1 29.24 [ -9.62, 68.09]
ADNI2GO -29.09 [ -69.16, 10.98]
ALSPAC 0.81 [ -41.19, 42.80]
ASRB 49.40 [ -23.90, 122.70]
BETULA 36.54 [ -23.88, 96.96]
BIG 29.06 [ -4.49, 62.62]
BIG-PsychChip 5.10 [ -52.98, 63.19]
BONN -116.34 [-206.34, -26.34]
BrainScale
CARDIFF 45.80 [ -18.21, 109.80]
DNS-V3 36.14 [ -10.80, 83.08]
DNS-V4 23.71 [ -42.19, 89.61]
EPIGEN 123.35 [ 25.31, 221.39]
FBIRN -3.13 [ -54.62, 48.36]
FOR2107 19.00 [ -17.85, 55.85]
GIG -4.25 [ -58.99, 50.50]
GSP 34.47 [ -10.69, 79.63]
HUBIN -29.59 [-114.45, 55.27]
HUNT 24.08 [ -4.06, 52.21]
IMAGEN 48.49 [ 20.60, 76.39]
IMpACT -8.92 [ -84.18, 66.34]
LBC1936 33.48 [ -7.39, 74.35]
LIBD 43.14 [ -7.11, 93.39]
Sydney MAS -40.00 [ -81.36, 1.36]
MCIC -4.38 [ -81.99, 73.23]
MooDS 20.11 [ -47.08, 87.30]
MPIP -14.12 [ -53.09, 24.85]
MUNSTER -0.95 [ -36.03, 34.14]
NCNG -10.97 [ -61.21, 39.26]
NESDA -7.32 [ -74.23, 59.59]
NeuroIMAGE 20.47 [ -48.40, 89.34]
NTR 35.32 [ -28.54, 99.18]
OATS
PAFIP -73.66 [-179.60, 32.28]
PDNZ 84.54 [ -1.31, 170.38]
PING 19.05 [ -31.03, 69.13]
MPRC 9.13 [ -29.88, 48.14]
PPMI
QTIM -9.18 [ -44.50, 26.13]
SHIP 56.96 [ 27.65, 86.27]
SHIP-Trend 53.88 [ 19.08, 88.69]
SYS 24.77 [ -31.08, 80.62]
TCD-NUIG 53.51 [ -26.39, 133.41]
TOP 16.10 [ -27.80, 59.99]
TOP3T 7.55 [ -43.90, 59.00]
UiO2016 35.69 [ -31.68, 103.05]
UiO2017 -9.20 [ -65.08, 46.67]
UMCU 32.59 [ -9.98, 75.16]
ENIGMA Discovery Cohorts [N=21933; P=3.147e-07] 18.13 [ 11.18, 25.07]
UKBB 16.75 [ 7.88, 25.63]
Discovery+UKBB [N=31155; P=8.671e-10] 17.58 [ 11.96, 23.20]
GOBS 29.77 [ -7.78, 67.32]
IMH
UNICAMP 26.50 [ -51.90, 104.90]
Meth-CT -11.25 [-122.50, 100.01]
MIRECC
UKBB nonEUR 17.24 [ -44.84, 79.33]
OSAKA
PING Non European -1.69 [ -61.61, 58.23]
Non European Cohorts [N=2100; P=0.1553] 18.95 [ -7.19, 45.10]
Beta [Lower 95% CI,Upper 95% CI]
-300
-200
-100
0
100
200
300
mm^2/effect allele

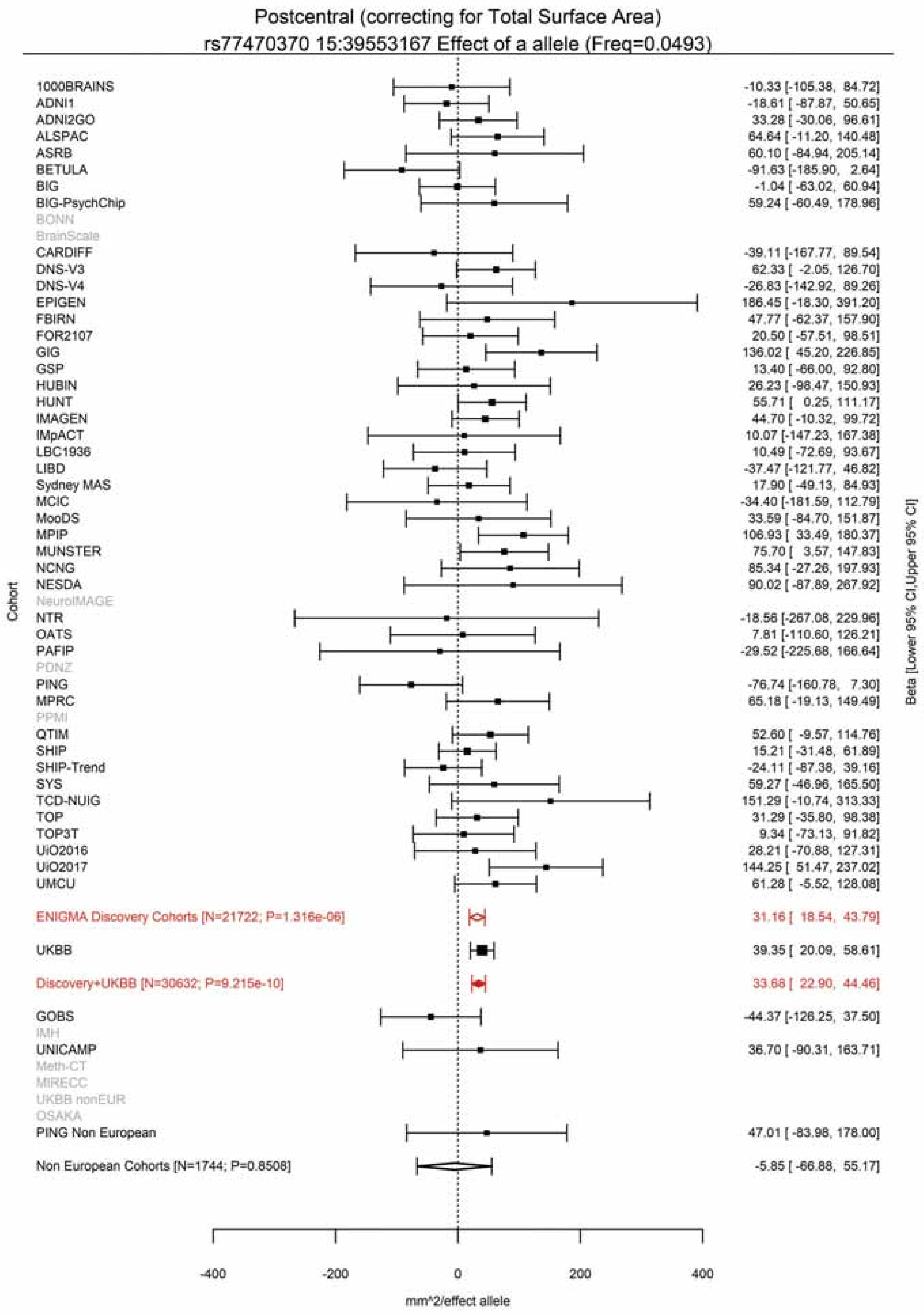
Postcentral (correcting for Total Surface Area)
rs77470370 15:39553167 Effect of a allele (Freq=0.0493)
1000BRAINS -10.33 [-105.38, 84.72]
ADNI1 -18.61 [ -87.87, 50.65]
ADNI2GO 33.28 [ -30.06, 96.61]
ALSPAC 64.64 [ -11.20, 140.48]
ASRB 60.10 [ -84.94, 205.14]
BETULA -91.63 [-185.90, 2.64]
BIG -1.04 [ -63.02, 60.94]
BIG-PsychChip 59.24 [ -60.49, 178.96]
BONN
BrainScale
CARDIFF -39.11 [-167.77, 89.54]
DNS-V3 62.33 [ -2.05, 126.70]
DNS-V4 -26.83 [-142.92, 89.26]
EPIGEN 186.45 [ -18.30, 391.20]
FBIRN 47.77 [ -62.37, 157.90]
FOR2107 20.50 [ -57.51, 98.51]
GIG 136.02 [ 45.20, 226.85]
GSP 13.40 [ -66.00, 92.80]
HUBIN 26.23 [ -98.47, 150.93]
HUNT 55.71 [ 0.25, 111.17]
IMAGEN 44.70 [ -10.32, 99.72]
IMpACT 10.07 [-147.23, 167.38]
LBC1936 10.49 [ -72.69, 93.67]
LIBD -37.47 [-121.77, 46.82]
Sydney MAS 17.90 [ -49.13, 84.93]
MCIC -34.40 [-181.59, 112.79]
MooDS 33.59 [ -84.70, 151.87]
MPIP 106.93 [ 33.49, 180.37]
MUNSTER 75.70 [ 3.57, 147.83]
NCNG 85.34 [ -27.26, 197.93]
NESDA 90.02 [ -87.89, 267.92]
NeuroIMAGE
NTR -18.56 [-267.08, 229.96]
OATS 7.81 [-110.60, 126.21]
PAFIP -29.52 [-225.68, 166.64]
PDNZ
PING -76.74 [-160.78, 7.30]
MPRC 65.18 [ -19.13, 149.49]
PPMI
QTIM 52.60 [ -9.57, 114.76]
SHIP 15.21 [ -31.48, 61.89]
SHIP-Trend -24.11 [ -87.38, 39.16]
SYS 59.27 [ -46.96, 165.50]
TCD-NUIG 151.29 [ -10.74, 313.33]
TOP 31.29 [ -35.80, 98.38]
TOP3T 9.34 [ -73.13, 91.82]
UiO2016 28.21 [ -70.88, 127.31]
UiO2017 144.25 [ 51.47, 237.02]
UMCU 61.28 [ -5.52, 128.08]
ENIGMA Discovery Cohorts [N=21722; P=1.316e-06] 31.16 [ 18.54, 43.79]
UKBB 39.35 [ 20.09, 58.61]
Discovery+UKBB [N=30632; P=9.215e-10] 33.68 [ 22.90, 44.46]
GOBS -44.37 [-126.25, 37.50]
IMH
UNICAMP 36.70 [ -90.31, 163.71]
Meth-CT
MIRECC
UKBB nonEUR
OSAKA
PING Non European 47.01 [ -83.98, 178.00]
Non European Cohorts [N=1744; P=0.8508] -5.85 [ -66.88, 55.17]
Cohort
Beta [Lower 95% CI,Upper 95% CI]
-400
-200
0
200
400
mm^2/effect allele

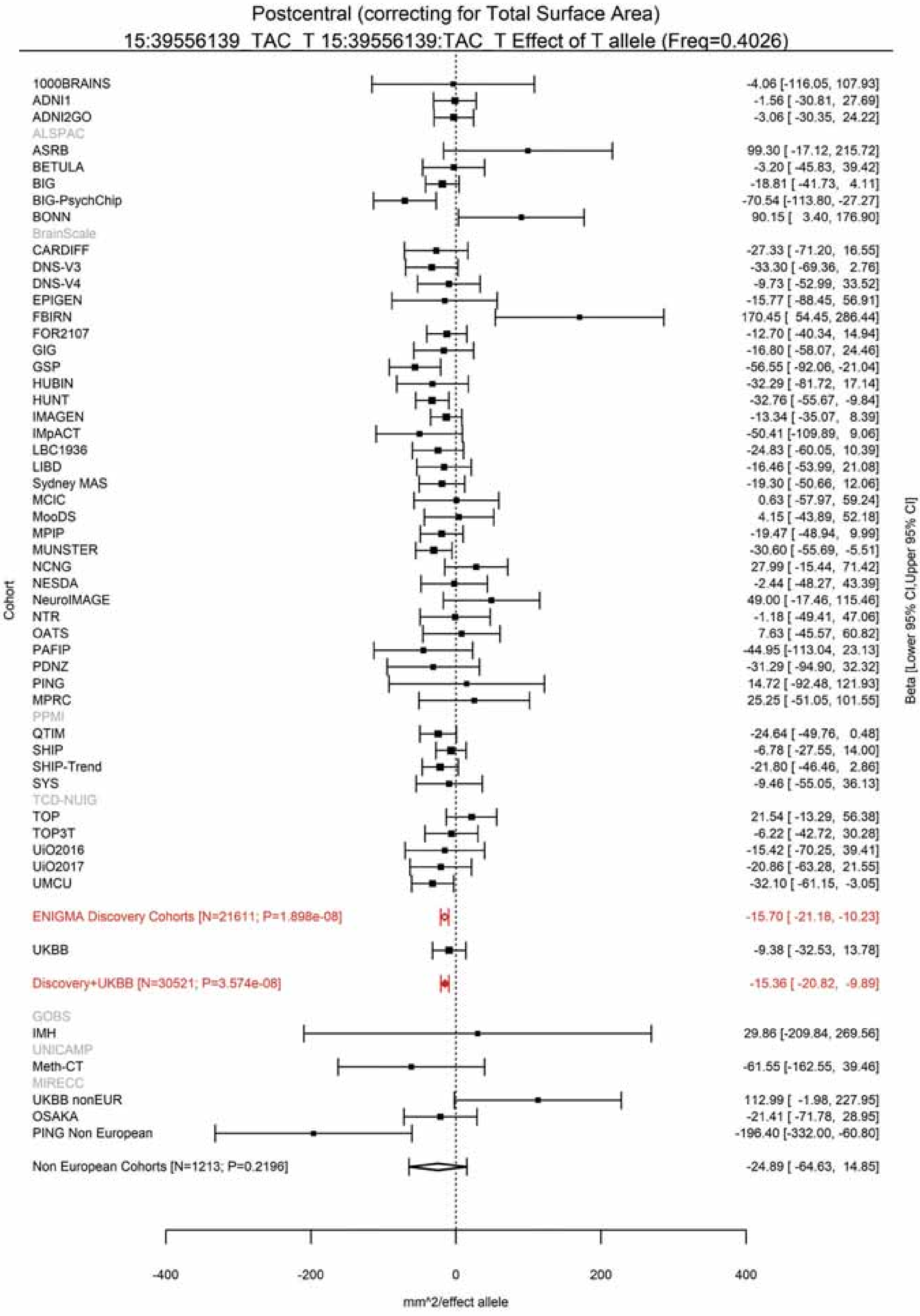
Postcentral (correcting for Total Surface Area)
15:39556139_TAC_T 15:39556139:TAC_T Effect of T allele (Freq=0.4026)
Cohort
1000BRAINS -4.06 [-116.05, 107.93]
ADNI1 -1.56 [ -30.81, 27.69]
ADNI2GO -3.06 [ -30.35, 24.22]
ALSPAC
ASRB 99.30 [ -17.12, 215.72]
BETULA -3.20 [ -45.83, 39.42]
BIG -18.81 [ -41.73, 4.11]
BIG-PsychChip -70.54 [-113.80, -27.27]
BONN 90.15 [ 3.40, 176.90]
BrainScale
CARDIFF -27.33 [ -71.20, 16.55]
DNS-V3 -33.30 [ -69.36, 2.76]
DNS-V4 -9.73 [ -52.99, 33.52]
EPIGEN -15.77 [ -88.45, 56.91]
FBIRN 170.45 [ 54.45, 286.44]
FOR2107 -12.70 [ -40.34, 14.94]
GIG -16.80 [ -58.07, 24.46]
GSP -56.55 [ -92.06, -21.04]
HUBIN -32.29 [ -81.72, 17.14]
HUNT -32.76 [ -55.67, -9.84]
IMAGEN -13.34 [ -35.07, 8.39]
IMpACT -50.41 [-109.89, 9.06]
LBC1936 -24.83 [ -60.05, 10.39]
LIBD -16.46 [ -53.99, 21.08]
Sydney MAS -19.30 [ -50.66, 12.06]
MCIC 0.63 [ -57.97, 59.24]
MooDS 4.15 [ -43.89, 52.18]
MPIP -19.47 [ -48.94, 9.99]
MUNSTER -30.60 [ -55.69, -5.51]
NCNG 27.99 [ -15.44, 71.42]
NESDA -2.44 [ -48.27, 43.39]
NeuroIMAGE 49.00 [ -17.46, 115.46]
NTR -1.18 [ -49.41, 47.06]
OATS 7.63 [ -45.57, 60.82]
PAFIP -44.95 [-113.04, 23.13]
PDNZ -31.29 [ -94.90, 32.32]
PING 14.72 [ -92.48, 121.93]
MPRC 25.25 [ -51.05, 101.55]
PPMI
QTIM -24.64 [ -49.76, 0.48]
SHIP -6.78 [ -27.55, 14.00]
SHIP-Trend -21.80 [ -46.46, 2.86]
SYS -9.46 [ -55.05, 36.13]
TCD-NUIG
TOP 21.54 [ -13.29, 56.38]
TOP3T -6.22 [ -42.72, 30.28]
UiO2016 -15.42 [ -70.25, 39.41]
UiO2017 -20.86 [ -63.28, 21.55]
UMCU -32.10 [ -61.15, -3.05]
ENIGMA Discovery Cohorts [N=21611; P=1.898e-08] -15.70 [ -21.18, -10.23]
UKBB -9.38 [ -32.53, 13.78]
Discovery+UKBB [N=30521; P=3.574e-08] -15.36 [ -20.82, -9.89]
GOBS
IMH 29.86 [-209.84, 269.56]
UNICAMP
Meth-CT -61.55 [-162.55, 39.46]
MIRECC
UKBB nonEUR 112.99 [ -1.98, 227.95]
OSAKA -21.41 [ -71.78, 28.95]
PING Non European -196.40 [-332.00, -60.80]
Non European Cohorts [N=1213; P=0.2196] -24.89 [ -64.63, 14.85]
Beta [Lower 95% CI,Upper 95% CI]
-400
-200
0
200
400
mm^2/effect allele

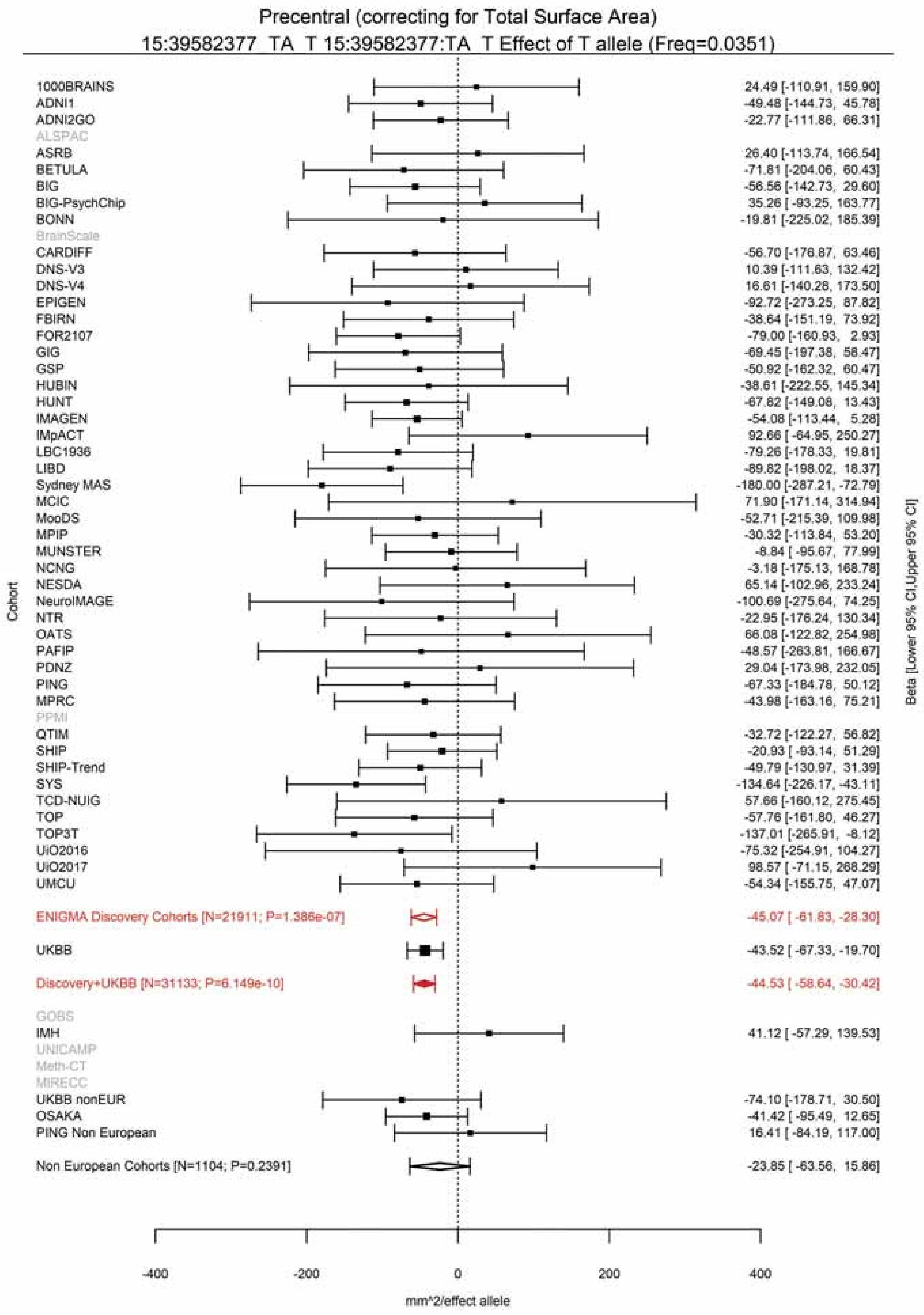
Precentral (correcting for Total Surface Area)
15:39582377_TA_T 15:39582377:TA_T Effect of T allele (Freq=0.0351)
Cohort
1000BRAINS 24.49 [-110.91, 159.90]
ADNI1 -49.48 [-144.73, 45.78]
ADNI2GO -22.77 [-111.86, 66.31]
ALSPAC
ASRB 26.40 [-113.74, 166.54]
BETULA -71.81 [-204.06, 60.43]
BIG -56.56 [-142.73, 29.60]
BIG-PsychChip 35.26 [ -93.25, 163.77]
BONN -19.81 [-225.02, 185.39]
BrainScale
CARDIFF -56.70 [-176.87, 63.46]
DNS-V3 10.39 [-111.63, 132.42]
DNS-V4 16.61 [-140.28, 173.50]
EPIGEN -92.72 [-273.25, 87.82]
FBIRN -38.64 [-151.19, 73.92]
FOR2107 -79.00 [-160.93, 2.93]
GIG -69.45 [-197.38, 58.47]
GSP -50.92 [-162.32, 60.47]
HUBIN -38.61 [-222.55, 145.34]
HUNT -67.82 [-149.08, 13.43]
IMAGEN -54.08 [-113.44, 5.28]
IMpACT 92.66 [ -64.95, 250.27]
LBC1936 -79.26 [-178.33, 19.81]
LIBD -89.82 [-198.02, 18.37]
Sydney MAS -180.00 [-287.21, -72.79]
MCIC 71.90 [-171.14, 314.94]
MooDS -52.71 [-215.39, 109.98]
MPIP -30.32 [-113.84, 53.20]
MUNSTER -8.84 [ -95.67, 77.99]
NCNG -3.18 [-175.13, 168.78]
NESDA 65.14 [-102.96, 233.24]
NeuroIMAGE -100.69 [-275.64, 74.25]
NTR -22.95 [-176.24, 130.34]
OATS 66.08 [-122.82, 254.98]
PAFIP -48.57 [-263.81, 166.67]
PDNZ 29.04 [-173.98, 232.05]
PING -67.33 [-184.78, 50.12]
MPRC -43.98 [-163.16, 75.21]
PPMI
QTIM -32.72 [-122.27, 56.82]
SHIP -20.93 [ -93.14, 51.29]
SHIP-Trend -49.79 [-130.97, 31.39]
SYS -134.64 [-226.17, -43.11]
TCD-NUIG 57.66 [-160.12, 275.45]
TOP -57.76 [-161.80, 46.27]
TOP3T -137.01 [-265.91, -8.12]
UiO2016 -75.32 [-254.91, 104.27]
UiO2017 98.57 [ -71.15, 268.29]
UMCU -54.34 [-155.75, 47.07]
ENIGMA Discovery Cohorts [N=21911; P=1.386e-07] -45.07 [ -61.83, -28.30]
UKBB -43.52 [ -67.33, -19.70]
Discovery+UKBB [N=31133; P=6.149e-10] -44.53 [ -58.64, -30.42]
GOBS
IMH 41.12 [ -57.29, 139.53]
UNICAMP
Meth-CT
MIRECC
UKBB nonEUR -74.10 [-178.71, 30.50]
OSAKA -41.42 [ -95.49, 12.65]
PING Non European 16.41 [ -84.19, 117.00]
Non European Cohorts [N=1104; P=0.2391] -23.85 [ -63.56, 15.86]
Beta [Lower 95% CI,Upper 95% CI]
-400
-200
0
200
400
mm^2/effect allele

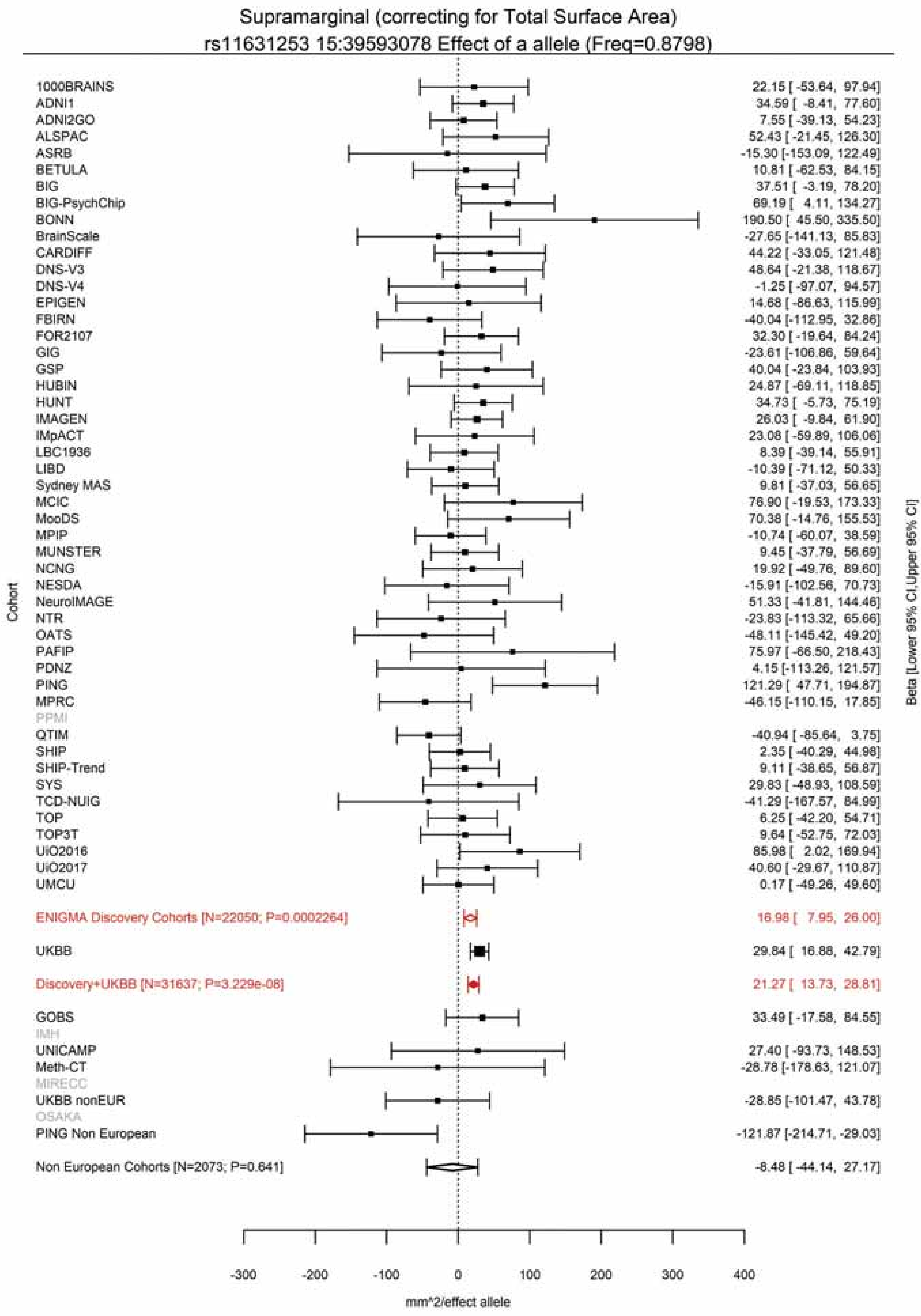
Supramarginal (correcting for Total Surface Area)
rs11631253 15:39593078 Effect of a allele (Freq=0.8798)
Cohort
1000BRAINS 22.15 [ -53.64, 97.94]
ADNI1 34.59 [ -8.41, 77.60]
ADNI2GO 7.55 [ -39.13, 54.23]
ALSPAC 52.43 [ -21.45, 126.30]
ASRB -15.30 [-153.09, 122.49]
BETULA 10.81 [ -62.53, 84.15]
BIG 37.51 [ -3.19, 78.20]
BIG-PsychChip 69.19 [ 4.11, 134.27]
BONN 190.50 [ 45.50, 335.50]
BrainScale -27.65 [-141.13, 85.83]
CARDIFF 44.22 [ -33.05, 121.48]
DNS-V3 48.64 [ -21.38, 118.67]
DNS-V4 -1.25 [ -97.07, 94.57]
EPIGEN 14.68 [ -86.63, 115.99]
FBIRN -40.04 [-112.95, 32.86]
FOR2107 32.30 [ -19.64, 84.24]
GIG -23.61 [-106.86, 59.64]
GSP 40.04 [ -23.84, 103.93]
HUBIN 24.87 [ -69.11, 118.85]
HUNT 34.73 [ -5.73, 75.19]
IMAGEN 26.03 [ -9.84, 61.90]
IMpACT 23.08 [ -59.89, 106.06]
LBC1936 8.39 [ -39.14, 55.91]
LIBD -10.39 [ -71.12, 50.33]
Sydney MAS 9.81 [ -37.03, 56.65]
MCIC 76.90 [ -19.53, 173.33]
MooDS 70.38 [ -14.76, 155.53]
MPIP -10.74 [ -60.07, 38.59]
MUNSTER 9.45 [ -37.79, 56.69]
NCNG 19.92 [ -49.76, 89.60]
NESDA -15.91 [-102.56, 70.73]
NeuroIMAGE 51.33 [ -41.81, 144.46]
NTR -23.83 [-113.32, 65.66]
OATS -48.11 [-145.42, 49.20]
PAFIP 75.97 [ -66.50, 218.43]
PDNZ 4.15 [-113.26, 121.57]
PING 121.29 [ 47.71, 194.87]
MPRC -46.15 [-110.15, 17.85]
PPMI
QTIM -40.94 [ -85.64, 3.75]
SHIP 2.35 [ -40.29, 44.98]
SHIP-Trend 9.11 [ -38.65, 56.87]
SYS 29.83 [ -48.93, 108.59]
TCD-NUIG -41.29 [-167.57, 84.99]
TOP 6.25 [ -42.20, 54.71]
TOP3T 9.64 [ -52.75, 72.03]
UiO2016 85.98 [ 2.02, 169.94]
UiO2017 40.60 [ -29.67, 110.87]
UMCU 0.17 [ -49.26, 49.60]
ENIGMA Discovery Cohorts [N=22050; P=0.0002264] 16.98 [ 7.95, 26.00]
UKBB 29.84 [ 16.88, 42.79]
Discovery+UKBB [N=31637; P=3.229e-08] 21.27 [ 13.73, 28.81]
GOBS 33.49 [ -17.58, 84.55]
IMH
UNICAMP 27.40 [ -93.73, 148.53]
Meth-CT -28.78 [-178.63, 121.07]
MIRECC
UKBB nonEUR -28.85 [-101.47, 43.78]
OSAKA
PING Non European -121.87 [-214.71, -29.03]
Non European Cohorts [N=2073; P=0.641] -8.48 [ -44.14, 27.17]
-300
-200
-100
0
100
200
300
400
mm^2/effect allele
Beta [Lower 95% CI,Upper 95% CI]

Postcentral (correcting for Average Cortical Thickness)
rs62002282 15:39595340 Effect of a allele (Freq=0.8786)

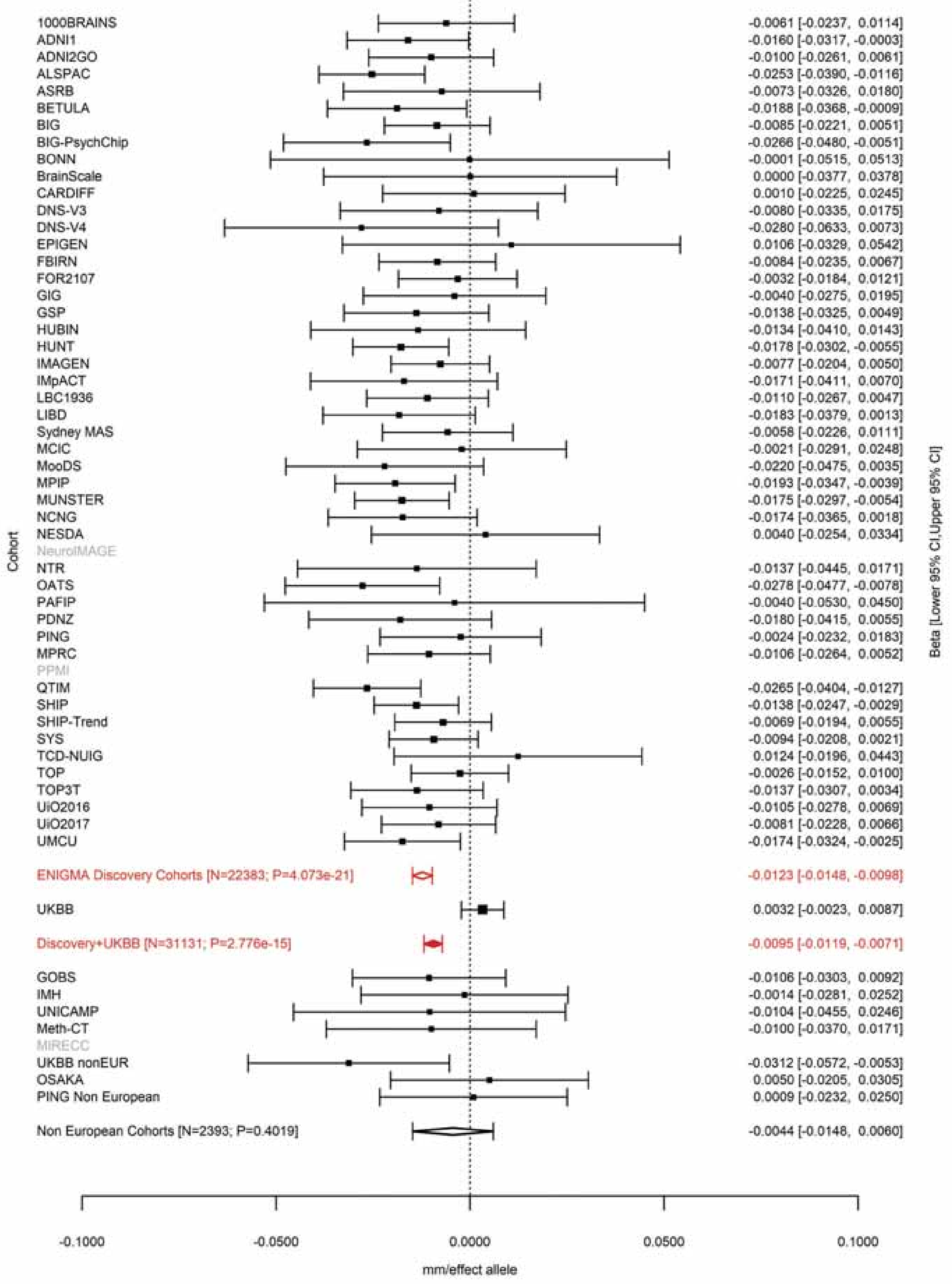

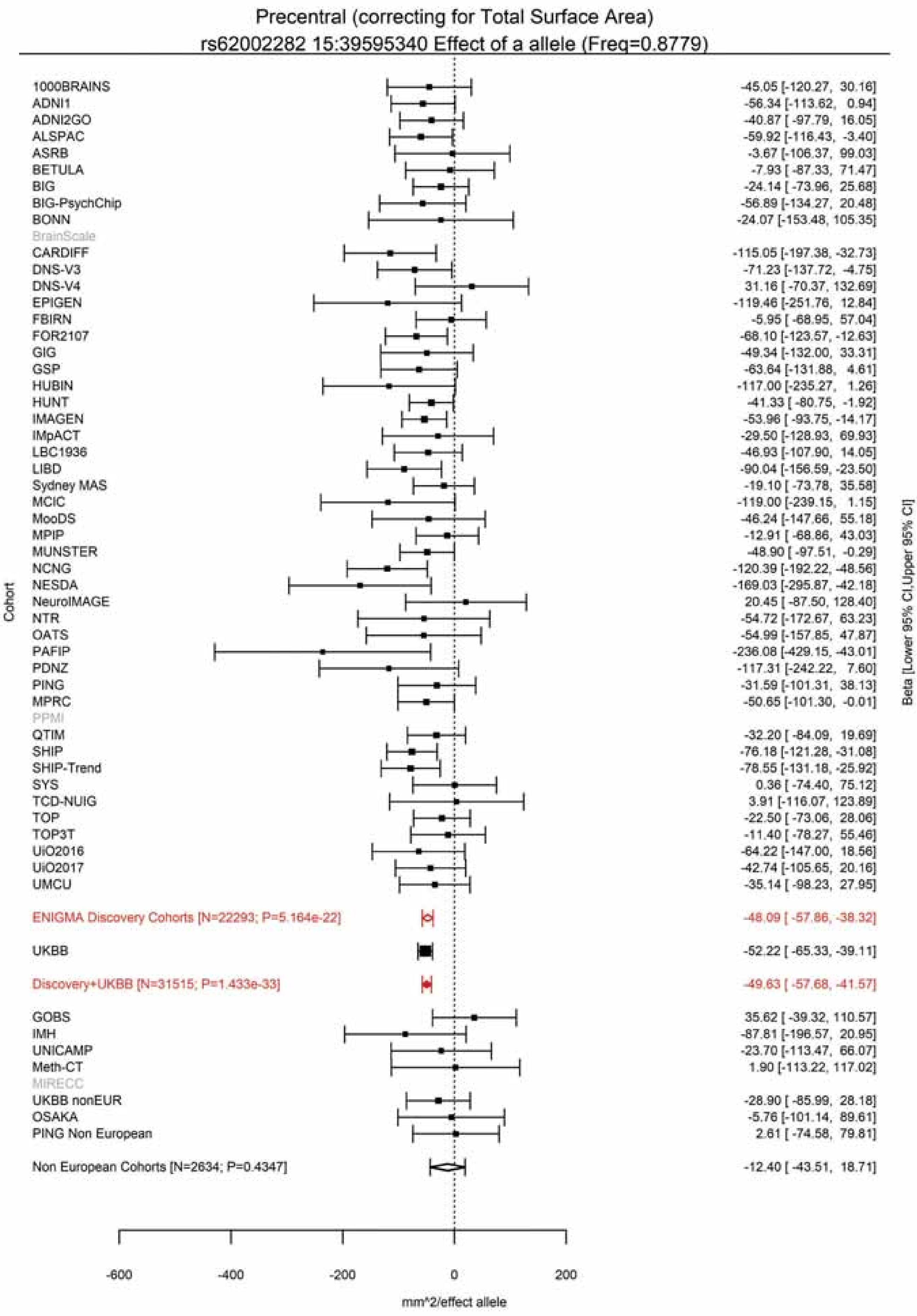
Precentral (correcting for Total Surface Area)
rs62002282 15:39595340 Effect of a allele (Freq=0.8779)
Cohort
1000BRAINS -45.05 [-120.27, 30.16]
ADNI1 -56.34 [-113.62, 0.94]
ADNI2GO -40.87 [ -97.79, 16.05]
ALSPAC -59.92 [-116.43, -3.40]
ASRB -3.67 [-106.37, 99.03]
BETULA -7.93 [ -87.33, 71.47]
BIG -24.14 [ -73.96, 25.68]
BIG-PsychChip -56.89 [-134.27, 20.48]
BONN -24.07 [-153.48, 105.35]
BrainScale
CARDIFF -115.05 [-197.38, -32.73]
DNS-V3 -71.23 [-137.72, -4.75]
DNS-V4 31.16 [ -70.37, 132.69]
EPIGEN -119.46 [-251.76, 12.84]
FBIRN -5.95 [ -68.95, 57.04]
FOR2107 -68.10 [-123.57, -12.63]
GIG -49.34 [-132.00, 33.31]
GSP -63.64 [-131.88, 4.61]
HUBIN -117.00 [-235.27, 1.26]
HUNT -41.33 [ -80.75, -1.92]
IMAGEN -53.96 [ -93.75, -14.17]
IMpACT -29.50 [-128.93, 69.93]
LBC1936 -46.93 [-107.90, 14.05]
LIBD -90.04 [-156.59, -23.50]
Sydney MAS -19.10 [ -73.78, 35.58]
MCIC -119.00 [-239.15, 1.15]
MooDS -46.24 [-147.66, 55.18]
MPIP -12.91 [ -68.86, 43.03]
MUNSTER -48.90 [ -97.51, -0.29]
NCNG -120.39 [-192.22, -48.56]
NESDA -169.03 [-295.87, -42.18]
NeuroIMAGE 20.45 [ -87.50, 128.40]
NTR -54.72 [-172.67, 63.23]
OATS -54.99 [-157.85, 47.87]
PAFIP -236.08 [-429.15, -43.01]
PDNZ -117.31 [-242.22, 7.60]
PING -31.59 [-101.31, 38.13]
MPRC -50.65 [-101.30, -0.01]
PPMI
QTIM -32.20 [ -84.09, 19.69]
SHIP -76.18 [-121.28, -31.08]
SHIP-Trend -78.55 [-131.18, -25.92]
SYS 0.36 [ -74.40, 75.12]
TCD-NUIG 3.91 [-116.07, 123.89]
TOP -22.50 [ -73.06, 28.06]
TOP3T -11.40 [ -78.27, 55.46]
UiO2016 -64.22 [-147.00, 18.56]
UiO2017 -42.74 [-105.65, 20.16]
UMCU -35.14 [ -98.23, 27.95]
ENIGMA Discovery Cohorts [N=22293; P=5.164e-22] -48.09 [ -57.86, -38.32]
UKBB -52.22 [ -65.33, -39.11]
Discovery+UKBB [N=31515; P=1.433e-33] -49.63 [ -57.68, -41.57]
GOBS 35.62 [ -39.32, 110.57]
IMH -87.81 [-196.57, 20.95]
UNICAMP -23.70 [-113.47, 66.07]
Meth-CT 1.90 [-113.22, 117.02]
MIRECC
UKBB nonEUR -28.90 [ -85.99, 28.18]
OSAKA -5.76 [-101.14, 89.61]
PING Non European 2.61 [ -74.58, 79.81]
Non European Cohorts [N=2634; P=0.4347] -12.40 [ -43.51, 18.71]
Beta [Lower 95% CI,Upper 95% CI]
-600
-400
-200
0
200
mm^2/effect allele

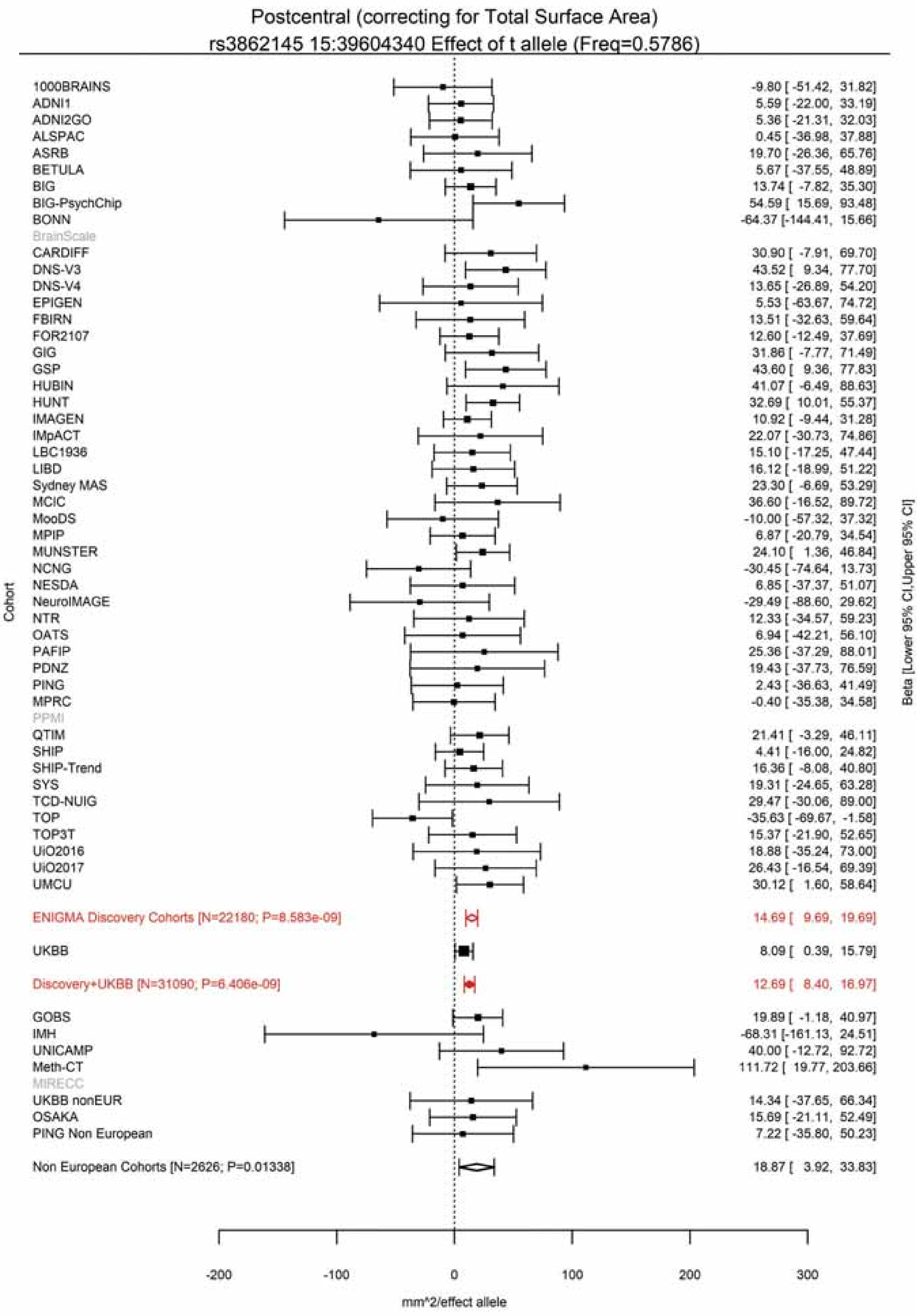
Postcentral (correcting for Total Surface Area)
rs3862145 15:39604340 Effect of t allele (Freq=0.5786)
1000BRAINS -9.80 [ -51.42, 31.82]
ADNI1 5.59 [ -22.00, 33.19]
ADNI2GO 5.36 [ -21.31, 32.03]
ALSPAC 0.45 [ -36.98, 37.88]
ASRB 19.70 [ -26.36, 65.76]
BETULA 5.67 [ -37.55, 48.89]
BIG 13.74 [ -7.82, 35.30]
BIG-PsychChip 54.59 [ 15.69, 93.48]
BONN -64.37 [-144.41, 15.66]
BrainScale
CARDIFF 30.90 [ -7.91, 69.70]
DNS-V3 43.52 [ 9.34, 77.70]
DNS-V4 13.65 [ -26.89, 54.20]
EPIGEN 5.53 [ -63.67, 74.72]
FBIRN 13.51 [ -32.63, 59.64]
FOR2107 12.60 [ -12.49, 37.69]
GIG 31.86 [ -7.77, 71.49]
GSP 43.60 [ 9.36, 77.83]
HUBIN 41.07 [ -6.49, 88.63]
HUNT 32.69 [ 10.01, 55.37]
IMAGEN 10.92 [ -9.44, 31.28]
IMpACT 22.07 [ -30.73, 74.86]
LBC1936 15.10 [ -17.25, 47.44]
LIBD 16.12 [ -18.99, 51.22]
Sydney MAS 23.30 [ -6.69, 53.29]
MCIC 36.60 [ -16.52, 89.72]
MooDS -10.00 [ -57.32, 37.32]
MPIP 6.87 [ -20.79, 34.54]
MUNSTER 24.10 [ 1.36, 46.84]
NCNG -30.45 [ -74.64, 13.73]
NESDA 6.85 [ -37.37, 51.07]
NeuroIMAGE -29.49 [ -88.60, 29.62]
NTR 12.33 [ -34.57, 59.23]
OATS 6.94 [ -42.21, 56.10]
PAFIP 25.36 [ -37.29, 88.01]
PDNZ 19.43 [ -37.73, 76.59]
PING 2.43 [ -36.63, 41.49]
MPRC -0.40 [ -35.38, 34.58]
PPMI
QTIM 21.41 [ -3.29, 46.11]
SHIP 4.41 [ -16.00, 24.82]
SHIP-Trend 16.36 [ -8.08, 40.80]
SYS 19.31 [ -24.65, 63.28]
TCD-NUIG 29.47 [ -30.06, 89.00]
TOP -35.63 [ -69.67, -1.58]
TOP3T 15.37 [ -21.90, 52.65]
UiO2016 18.88 [ -35.24, 73.00]
UiO2017 26.43 [ -16.54, 69.39]
UMCU 30.12 [ 1.60, 58.64]
ENIGMA Discovery Cohorts [N=22180; P=8.583e-09] 14.69 [ 9.69, 19.69]
UKBB 8.09 [ 0.39, 15.79]
Discovery+UKBB [N=31090; P=6.406e-09] 12.69 [ 8.40, 16.97]
GOBS 19.89 [ -1.18, 40.97]
IMH -68.31 [-161.13, 24.51]
UNICAMP 40.00 [ -12.72, 92.72]
Meth-CT 111.72 [ 19.77, 203.66]
MIRECC
UKBB nonEUR 14.34 [ -37.65, 66.34]
OSAKA 15.69 [ -21.11, 52.49]
PING Non European 7.22 [ -35.80, 50.23]
Non European Cohorts [N=2626; P=0.01338] 18.87 [ 3.92, 33.83]
Cohort
Beta [Lower 95% CI,Upper 95% CI]
-200
-100
0
100
200
300
mm^2/effect allele

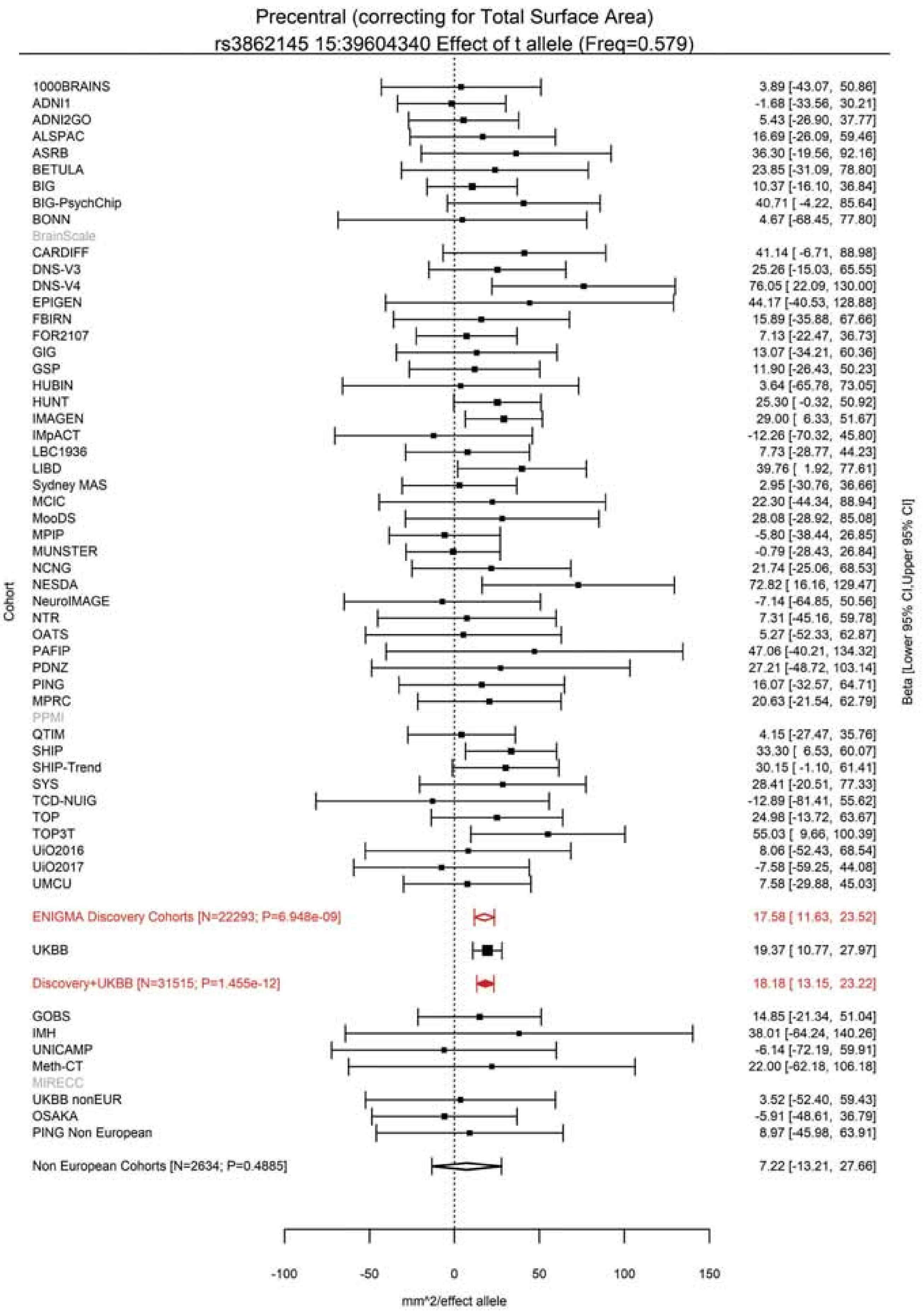
Precentral (correcting for Total Surface Area)
rs3862145 15:39604340 Effect of t allele (Freq=0.579)
Cohort
1000BRAINS 3.89 [-43.07, 50.86]
ADNI1 -1.68 [-33.56, 30.21]
ADNI2GO 5.43 [-26.90, 37.77]
ALSPAC 16.69 [-26.09, 59.46]
ASRB 36.30 [-19.56, 92.16]
BETULA 23.85 [-31.09, 78.80]
BIG 10.37 [-16.10, 36.84]
BIG-PsychChip 40.71 [ -4.22, 85.64]
BONN 4.67 [-68.45, 77.80]
BrainScale
CARDIFF 41.14 [ -6.71, 88.98]
DNS-V3 25.26 [-15.03, 65.55]
DNS-V4 76.05 [ 22.09, 130.00]
EPIGEN 44.17 [-40.53, 128.88]
FBIRN 15.89 [-35.88, 67.66]
FOR2107 7.13 [-22.47, 36.73]
GIG 13.07 [-34.21, 60.36]
GSP 11.90 [-26.43, 50.23]
HUBIN 3.64 [-65.78, 73.05]
HUNT 25.30 [ -0.32, 50.92]
IMAGEN 29.00 [ 6.33, 51.67]
IMpACT -12.26 [-70.32, 45.80]
LBC1936 7.73 [-28.77, 44.23]
LIBD 39.76 [ 1.92, 77.61]
Sydney MAS 2.95 [-30.76, 36.66]
MCIC 22.30 [-44.34, 88.94]
MooDS 28.08 [-28.92, 85.08]
MPIP -5.80 [-38.44, 26.85]
MUNSTER -0.79 [-28.43, 26.84]
NCNG 21.74 [-25.06, 68.53]
NESDA 72.82 [ 16.16, 129.47]
NeuroIMAGE -7.14 [-64.85, 50.56]
NTR 7.31 [-45.16, 59.78]
OATS 5.27 [-52.33, 62.87]
PAFIP 47.06 [-40.21, 134.32]
PDNZ 27.21 [-48.72, 103.14]
PING 16.07 [-32.57, 64.71]
MPRC 20.63 [-21.54, 62.79]
PPMI
QTIM 4.15 [-27.47, 35.76]
SHIP 33.30 [ 6.53, 60.07]
SHIP-Trend 30.15 [ -1.10, 61.41]
SYS 28.41 [-20.51, 77.33]
TCD-NUIG -12.89 [-81.41, 55.62]
TOP 24.98 [-13.72, 63.67]
TOP3T 55.03 [ 9.66, 100.39]
UiO2016 8.06 [-52.43, 68.54]
UiO2017 -7.58 [-59.25, 44.08]
UMCU 7.58 [-29.88, 45.03]
ENIGMA Discovery Cohorts [N=22293; P=6.948e-09] 17.58 [ 11.63, 23.52]
UKBB 19.37 [ 10.77, 27.97]
Discovery+UKBB [N=31515; P=1.455e-12] 18.18 [ 13.15, 23.22]
GOBS 14.85 [-21.34, 51.04]
IMH 38.01 [-64.24, 140.26]
UNICAMP -6.14 [-72.19, 59.91]
Meth-CT 22.00 [-62.18, 106.18]
MIRECC
UKBB nonEUR 3.52 [-52.40, 59.43]
OSAKA -5.91 [-48.61, 36.79]
PING Non European 8.97 [-45.98, 63.91]
Non European Cohorts [N=2634; P=0.4885] 7.22 [-13.21, 27.66]
-100 -50 0 50 100 150
mm^2/effect allele
Beta [Lower 95% CI,Upper 95% CI]

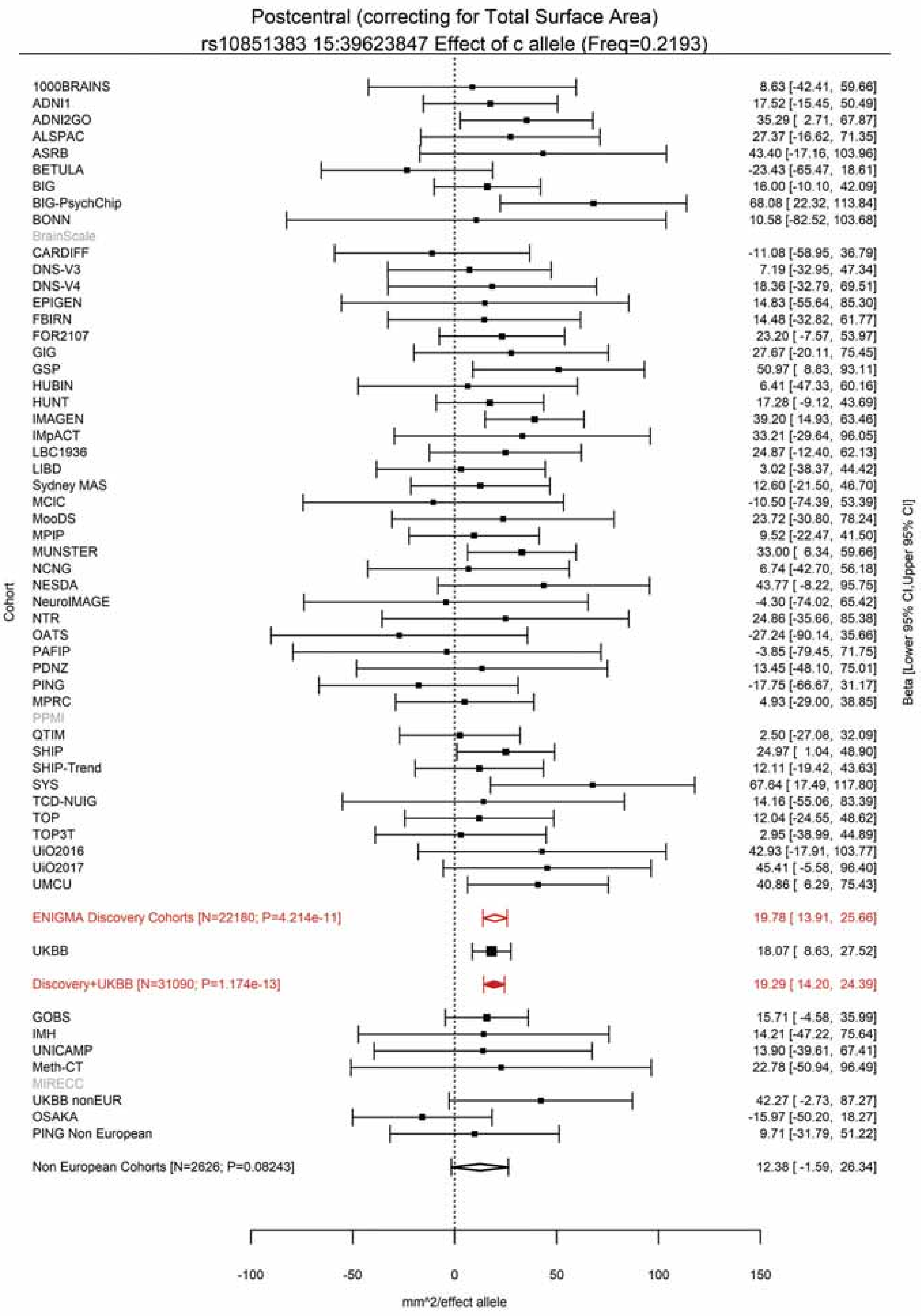

Postcentral (correcting for Total Surface Area)
rs10851383 15:39623847 Effect of c allele (Freq=0.2193)
Cohort
1000BRAINS 8.63 [-42.41, 59.66]
ADNI1 17.52 [-15.45, 50.49]
ADNI2GO 35.29 [ 2.71, 67.87]
ALSPAC 27.37 [-16.62, 71.35]
ASRB 43.40 [-17.16, 103.96]
BETULA -23.43 [-65.47, 18.61]
BIG 16.00 [-10.10, 42.09]
BIG-PsychChip 68.08 [ 22.32, 113.84]
BONN 10.58 [-82.52, 103.68]
BrainScale
CARDIFF -11.08 [-58.95, 36.79]
DNS-V3 7.19 [-32.95, 47.34]
DNS-V4 18.36 [-32.79, 69.51]
EPIGEN 14.83 [-55.64, 85.30]
FBIRN 14.48 [-32.82, 61.77]
FOR2107 23.20 [ -7.57, 53.97]
GIG 27.67 [-20.11, 75.45]
GSP 50.97 [ 8.83, 93.11]
HUBIN 6.41 [-47.33, 60.16]
HUNT 17.28 [ -9.12, 43.69]
IMAGEN 39.20 [ 14.93, 63.46]
IMpACT 33.21 [-29.64, 96.05]
LBC1936 24.87 [-12.40, 62.13]
LIBD 3.02 [-38.37, 44.42]
Sydney MAS 12.60 [-21.50, 46.70]
MCIC -10.50 [-74.39, 53.39]
MooDS 23.72 [-30.80, 78.24]
MPIP 9.52 [-22.47, 41.50]
MUNSTER 33.00 [ 6.34, 59.66]
NCNG 6.74 [-42.70, 56.18]
NESDA 43.77 [ -8.22, 95.75]
NeuroIMAGE -4.30 [-74.02, 65.42]
NTR 24.86 [-35.66, 85.38]
OATS -27.24 [-90.14, 35.66]
PAFIP -3.85 [-79.45, 71.75]
PDNZ 13.45 [-48.10, 75.01]
PING -17.75 [-66.67, 31.17]
MPRC 4.93 [-29.00, 38.85]
PPMI
QTIM 2.50 [-27.08, 32.09]
SHIP 24.97 [ 1.04, 48.90]
SHIP-Trend 12.11 [-19.42, 43.63]
SYS 67.64 [ 17.49, 117.80]
TCD-NUIG 14.16 [-55.06, 83.39]
TOP 12.04 [-24.55, 48.62]
TOP3T 2.95 [-38.99, 44.89]
UiO2016 42.93 [-17.91, 103.77]
UiO2017 45.41 [ -5.58, 96.40]
UMCU 40.86 [ 6.29, 75.43]
ENIGMA Discovery Cohorts [N=22180; P=4.214e-11] 19.78 [ 13.91, 25.66]
UKBB 18.07 [ 8.63, 27.52]
Discovery+UKBB [N=31090; P=1.174e-13] 19.29 [ 14.20, 24.39]
GOBS 15.71 [ -4.58, 35.99]
IMH 14.21 [-47.22, 75.64]
UNICAMP 13.90 [-39.61, 67.41]
Meth-CT 22.78 [-50.94, 96.49]
MIRECC
UKBB nonEUR 42.27 [ -2.73, 87.27]
OSAKA -15.97 [-50.20, 18.27]
PING Non European 9.71 [-31.79, 51.22]
Non European Cohorts [N=2626; P=0.08243] 12.38 [ -1.59, 26.34]
-100
-50
0
50
100
150
mm^2/effect allele
Beta [Lower 95% CI,Upper 95% CI]

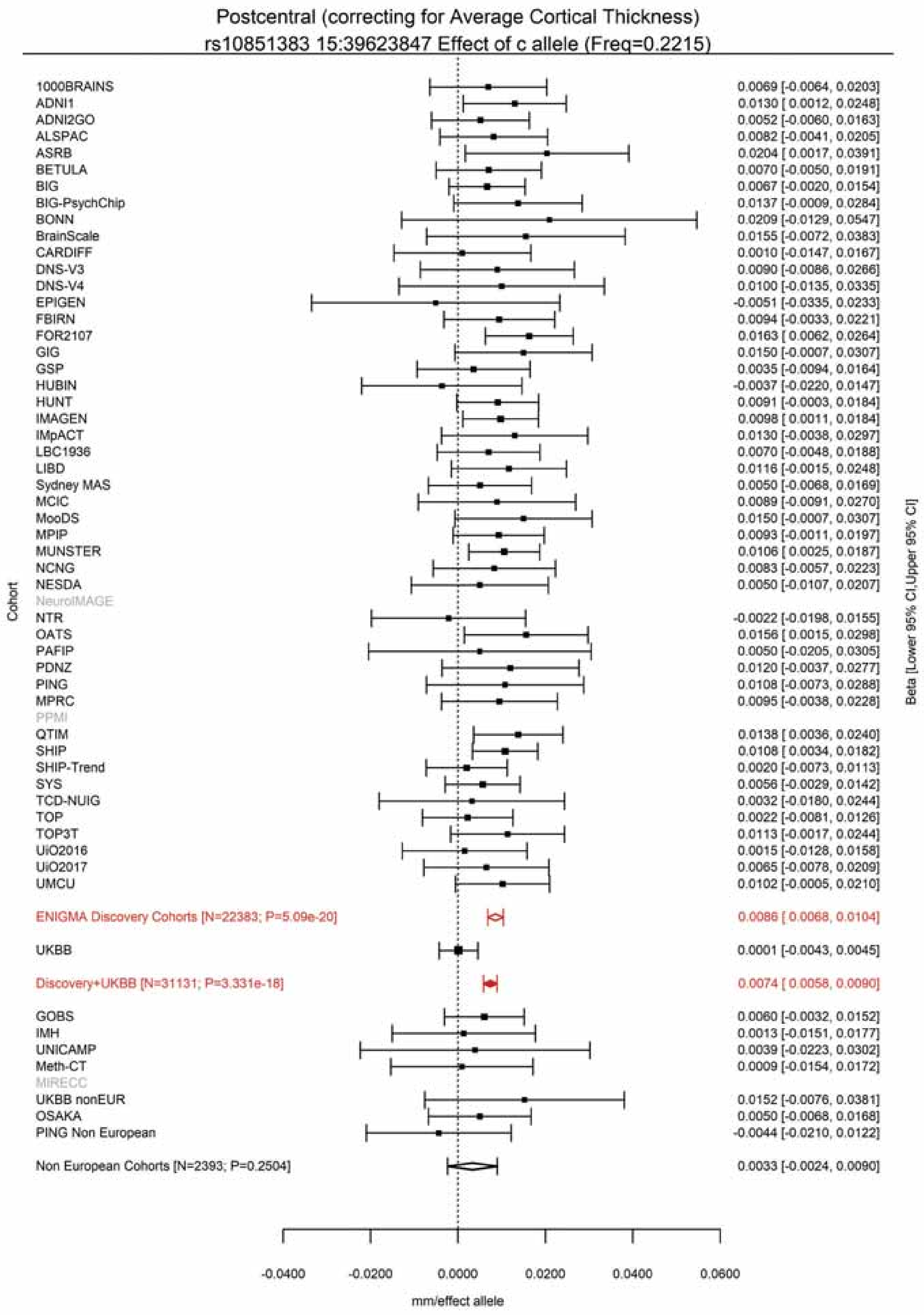
Postcentral (correcting for Average Cortical Thickness)
rs10851383 15:39623847 Effect of c allele (Freq=0.2215)
Cohort
Beta [Lower 95% CI,Upper 95% CI]
1000BRAINS 0.0069 [-0.0064, 0.0203]
ADNI1 0.0130 [ 0.0012, 0.0248]
ADNI2GO 0.0052 [-0.0060, 0.0163]
ALSPAC 0.0082 [-0.0041, 0.0205]
ASRB 0.0204 [ 0.0017, 0.0391]
BETULA 0.0070 [-0.0050, 0.0191]
BIG 0.0067 [-0.0020, 0.0154]
BIG-PsychChip 0.0137 [-0.0009, 0.0284]
BONN 0.0209 [-0.0129, 0.0547]
BrainScale 0.0155 [-0.0072, 0.0383]
CARDIFF 0.0010 [-0.0147, 0.0167]
DNS-V3 0.0090 [-0.0086, 0.0266]
DNS-V4 0.0100 [-0.0135, 0.0335]
EPIGEN -0.0051 [-0.0335, 0.0233]
FBIRN 0.0094 [-0.0033, 0.0221]
FOR2107 0.0163 [ 0.0062, 0.0264]
GIG 0.0150 [-0.0007, 0.0307]
GSP 0.0035 [-0.0094, 0.0164]
HUBIN -0.0037 [-0.0220, 0.0147]
HUNT 0.0091 [-0.0003, 0.0184]
IMAGEN 0.0098 [ 0.0011, 0.0184]
IMpACT 0.0130 [-0.0038, 0.0297]
LBC1936 0.0070 [-0.0048, 0.0188]
LIBD 0.0116 [-0.0015, 0.0248]
Sydney MAS 0.0050 [-0.0068, 0.0169]
MCIC 0.0089 [-0.0091, 0.0270]
MooDS 0.0150 [-0.0007, 0.0307]
MPIP 0.0093 [-0.0011, 0.0197]
MUNSTER 0.0106 [ 0.0025, 0.0187]
NCNG 0.0083 [-0.0057, 0.0223]
NESDA 0.0050 [-0.0107, 0.0207]
NeuroIMAGE
NTR -0.0022 [-0.0198, 0.0155]
OATS 0.0156 [ 0.0015, 0.0298]
PAFIP 0.0050 [-0.0205, 0.0305]
PDNZ 0.0120 [-0.0037, 0.0277]
PING 0.0108 [-0.0073, 0.0288]
MPRC 0.0095 [-0.0038, 0.0228]
PPMI
QTIM 0.0138 [ 0.0036, 0.0240]
SHIP 0.0108 [ 0.0034, 0.0182]
SHIP-Trend 0.0020 [-0.0073, 0.0113]
SYS 0.0056 [-0.0029, 0.0142]
TCD-NUIG 0.0032 [-0.0180, 0.0244]
TOP 0.0022 [-0.0081, 0.0126]
TOP3T 0.0113 [-0.0017, 0.0244]
UiO2016 0.0015 [-0.0128, 0.0158]
UiO2017 0.0065 [-0.0078, 0.0209]
UMCU 0.0102 [-0.0005, 0.0210]
ENIGMA Discovery Cohorts [N=22383; P=5.09e-20] 0.0086 [ 0.0068, 0.0104]
UKBB 0.0001 [-0.0043, 0.0045]
Discovery+UKBB [N=31131; P=3.331e-18] 0.0074 [ 0.0058, 0.0090]
GOBS 0.0060 [-0.0032, 0.0152]
IMH 0.0013 [-0.0151, 0.0177]
UNICAMP 0.0039 [-0.0223, 0.0302]
Meth-CT 0.0009 [-0.0154, 0.0172]
MIRECC
UKBB nonEUR 0.0152 [-0.0076, 0.0381]
OSAKA 0.0050 [-0.0068, 0.0168]
PING Non European -0.0044 [-0.0210, 0.0122]
Non European Cohorts [N=2393; P=0.2504] 0.0033 [-0.0024, 0.0090]
-0.0400
-0.0200
0.0000
0.0200
0.0400
0.0600
mm/effect allele

Pars Opercularis (correcting for Total Surface Area)
rs11070185 15:39625618 Effect of t allele (Freq=0.2186)

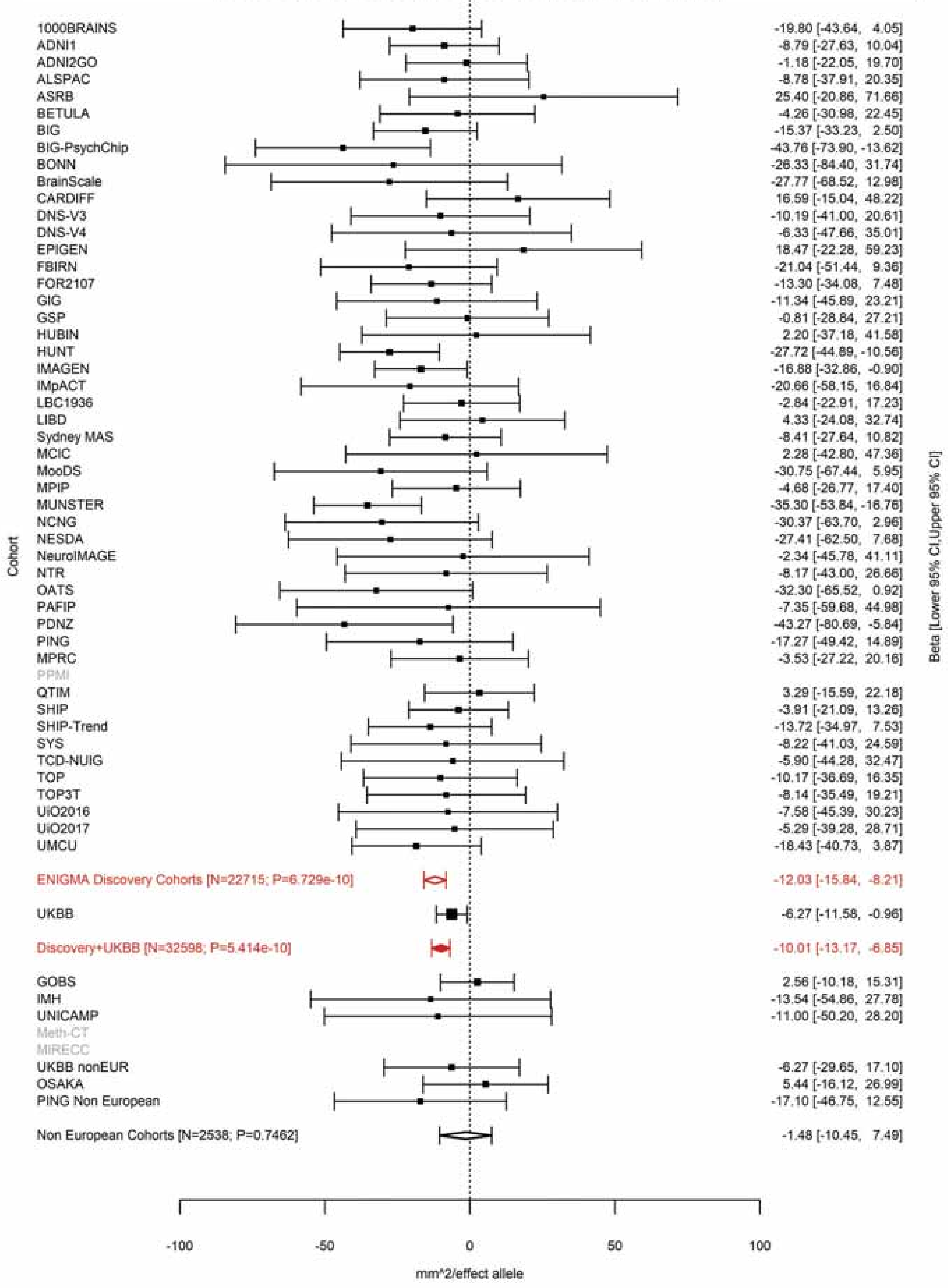

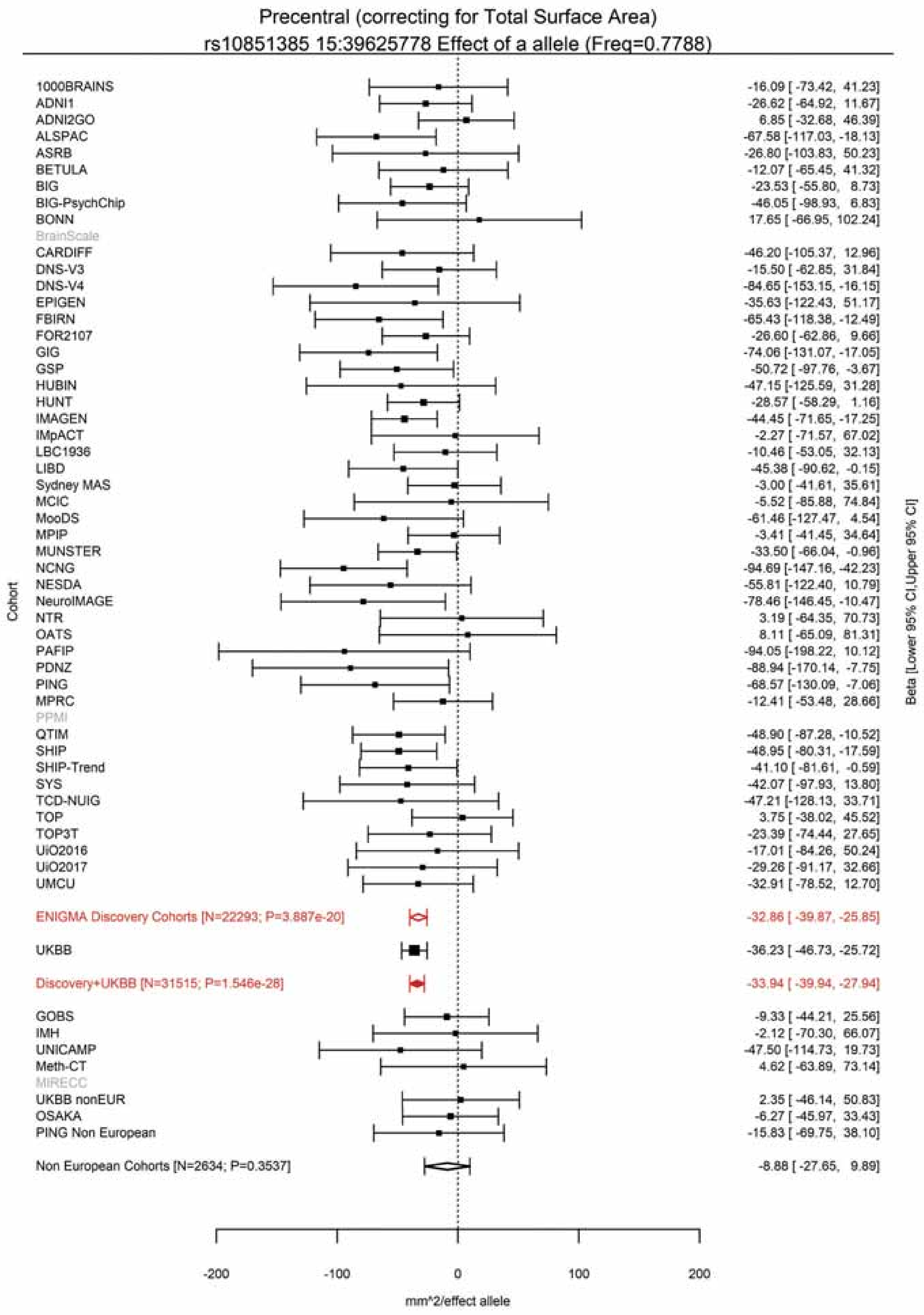
Precentral (correcting for Total Surface Area)
rs10851385 15:39625778 Effect of a allele (Freq=0.7788)
Cohort
1000BRAINS -16.09 [ -73.42, 41.23]
ADNI1 -26.62 [ -64.92, 11.67]
ADNI2GO 6.85 [ -32.68, 46.39]
ALSPAC -67.58 [-117.03, -18.13]
ASRB -26.80 [-103.83, 50.23]
BETULA -12.07 [ -65.45, 41.32]
BIG -23.53 [ -55.80, 8.73]
BIG-PsychChip -46.05 [ -98.93, 6.83]
BONN 17.65 [ -66.95, 102.24]
BrainScale
CARDIFF -46.20 [-105.37, 12.96]
DNS-V3 -15.50 [ -62.85, 31.84]
DNS-V4 -84.65 [-153.15, -16.15]
EPIGEN -35.63 [-122.43, 51.17]
FBIRN -65.43 [-118.38, -12.49]
FOR2107 -26.60 [ -62.86, 9.66]
GIG -74.06 [-131.07, -17.05]
GSP -50.72 [ -97.76, -3.67]
HUBIN -47.15 [-125.59, 31.28]
HUNT -28.57 [ -58.29, 1.16]
IMAGEN -44.45 [ -71.65, -17.25]
IMpACT -2.27 [ -71.57, 67.02]
LBC1936 -10.46 [ -53.05, 32.13]
LIBD -45.38 [ -90.62, -0.15]
Sydney MAS -3.00 [ -41.61, 35.61]
MCIC -5.52 [ -85.88, 74.84]
MooDS -61.46 [-127.47, 4.54]
MPIP -3.41 [ -41.45, 34.64]
MUNSTER -33.50 [ -66.04, -0.96]
NCNG -94.69 [-147.16, -42.23]
NESDA -55.81 [-122.40, 10.79]
NeuroIMAGE -78.46 [-146.45, -10.47]
NTR 3.19 [ -64.35, 70.73]
OATS 8.11 [ -65.09, 81.31]
PAFIP -94.05 [-198.22, 10.12]
PDNZ -88.94 [-170.14, -7.75]
PING -68.57 [-130.09, -7.06]
MPRC -12.41 [ -53.48, 28.66]
PPMI
QTIM -48.90 [ -87.28, -10.52]
SHIP -48.95 [ -80.31, -17.59]
SHIP-Trend -41.10 [ -81.61, -0.59]
SYS -42.07 [ -97.93, 13.80]
TCD-NUIG -47.21 [-128.13, 33.71]
TOP 3.75 [ -38.02, 45.52]
TOP3T -23.39 [ -74.44, 27.65]
UiO2016 -17.01 [ -84.26, 50.24]
UiO2017 -29.26 [ -91.17, 32.66]
UMCU -32.91 [ -78.52, 12.70]
ENIGMA Discovery Cohorts [N=22293; P=3.887e-20] -32.86 [ -39.87, -25.85]
UKBB -36.23 [ -46.73, -25.72]
Discovery+UKBB [N=31515; P=1.546e-28] -33.94 [ -39.94, -27.94]
GOBS -9.33 [ -44.21, 25.56]
IMH -2.12 [ -70.30, 66.07]
UNICAMP -47.50 [-114.73, 19.73]
Meth-CT 4.62 [ -63.89, 73.14]
MIRECC
UKBB nonEUR 2.35 [ -46.14, 50.83]
OSAKA -6.27 [ -45.97, 33.43]
PING Non European -15.83 [ -69.75, 38.10]
Non European Cohorts [N=2634; P=0.3537] -8.88 [ -27.65, 9.89]
Beta [Lower 95% CI,Upper 95% CI]
-200
-100
0
100
200
mm^2/effect allele

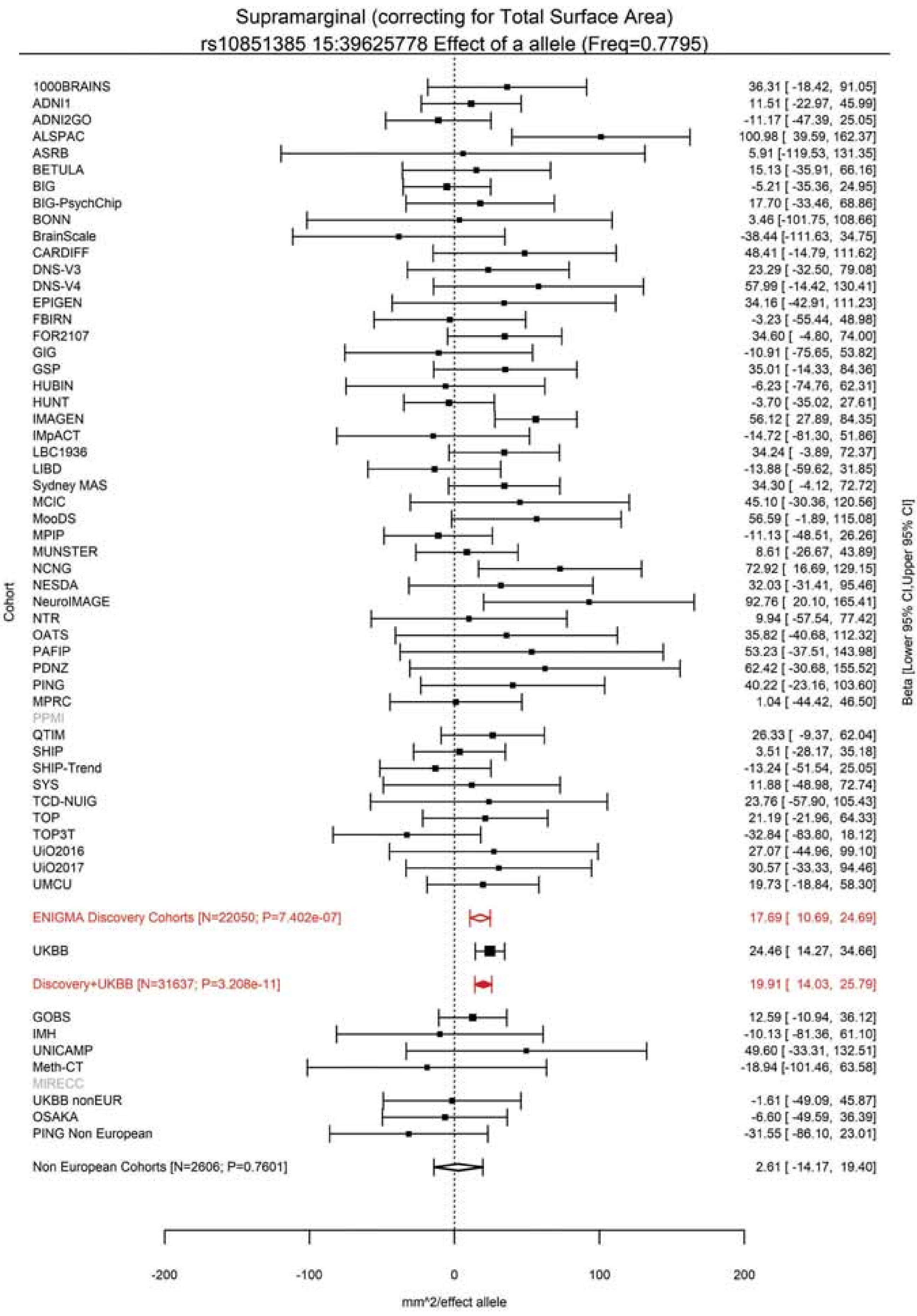

Supramarginal (correcting for Total Surface Area)
rs10851385 15:39625778 Effect of a allele (Freq=0.7795)
Cohort
1000BRAINS 36.31 [ -18.42, 91.05]
ADNI1 11.51 [ -22.97, 45.99]
ADNI2GO -11.17 [ -47.39, 25.05]
ALSPAC 100.98 [ 39.59, 162.37]
ASRB 5.91 [-119.53, 131.35]
BETULA 15.13 [ -35.91, 66.16]
BIG -5.21 [ -35.36, 24.95]
BIG-PsychChip 17.70 [ -33.46, 68.86]
BONN 3.46 [-101.75, 108.66]
BrainScale -38.44 [-111.63, 34.75]
CARDIFF 48.41 [ -14.79, 111.62]
DNS-V3 23.29 [ -32.50, 79.08]
DNS-V4 57.99 [ -14.42, 130.41]
EPIGEN 34.16 [ -42.91, 111.23]
FBIRN -3.23 [ -55.44, 48.98]
FOR2107 34.60 [ -4.80, 74.00]
GIG -10.91 [ -75.65, 53.82]
GSP 35.01 [ -14.33, 84.36]
HUBIN -6.23 [ -74.76, 62.31]
HUNT -3.70 [ -35.02, 27.61]
IMAGEN 56.12 [ 27.89, 84.35]
IMpACT -14.72 [ -81.30, 51.86]
LBC1936 34.24 [ -3.89, 72.37]
LIBD -13.88 [ -59.62, 31.85]
Sydney MAS 34.30 [ -4.12, 72.72]
MCIC 45.10 [ -30.36, 120.56]
MooDS 56.59 [ -1.89, 115.08]
MPIP -11.13 [ -48.51, 26.26]
MUNSTER 8.61 [ -26.67, 43.89]
NCNG 72.92 [ 16.69, 129.15]
NESDA 32.03 [ -31.41, 95.46]
NeuroIMAGE 92.76 [ 20.10, 165.41]
NTR 9.94 [ -57.54, 77.42]
OATS 35.82 [ -40.68, 112.32]
PAFIP 53.23 [ -37.51, 143.98]
PDNZ 62.42 [ -30.68, 155.52]
PING 40.22 [ -23.16, 103.60]
MPRC 1.04 [ -44.42, 46.50]
PPMI
QTIM 26.33 [ -9.37, 62.04]
SHIP 3.51 [ -28.17, 35.18]
SHIP-Trend -13.24 [ -51.54, 25.05]
SYS 11.88 [ -48.98, 72.74]
TCD-NUIG 23.76 [ -57.90, 105.43]
TOP 21.19 [ -21.96, 64.33]
TOP3T -32.84 [ -83.80, 18.12]
UiO2016 27.07 [ -44.96, 99.10]
UiO2017 30.57 [ -33.33, 94.46]
UMCU 19.73 [ -18.84, 58.30]
ENIGMA Discovery Cohorts [N=22050; P=7.402e-07] 17.69 [ 10.69, 24.69]
UKBB 24.46 [ 14.27, 34.66]
Discovery+UKBB [N=31637; P=3.208e-11] 19.91 [ 14.03, 25.79]
GOBS 12.59 [ -10.94, 36.12]
IMH -10.13 [ -81.36, 61.10]
UNICAMP 49.60 [ -33.31, 132.51]
Meth-CT -18.94 [-101.46, 63.58]
MIRECC
UKBB nonEUR -1.61 [ -49.09, 45.87]
OSAKA -6.60 [ -49.59, 36.39]
PING Non European -31.55 [ -86.10, 23.01]
Non European Cohorts [N=2606; P=0.7601] 2.61 [ -14.17, 19.40]
Beta [Lower 95% CI,Upper 95% CI]
-200
-100
0
100
200
mm^2/effect allele

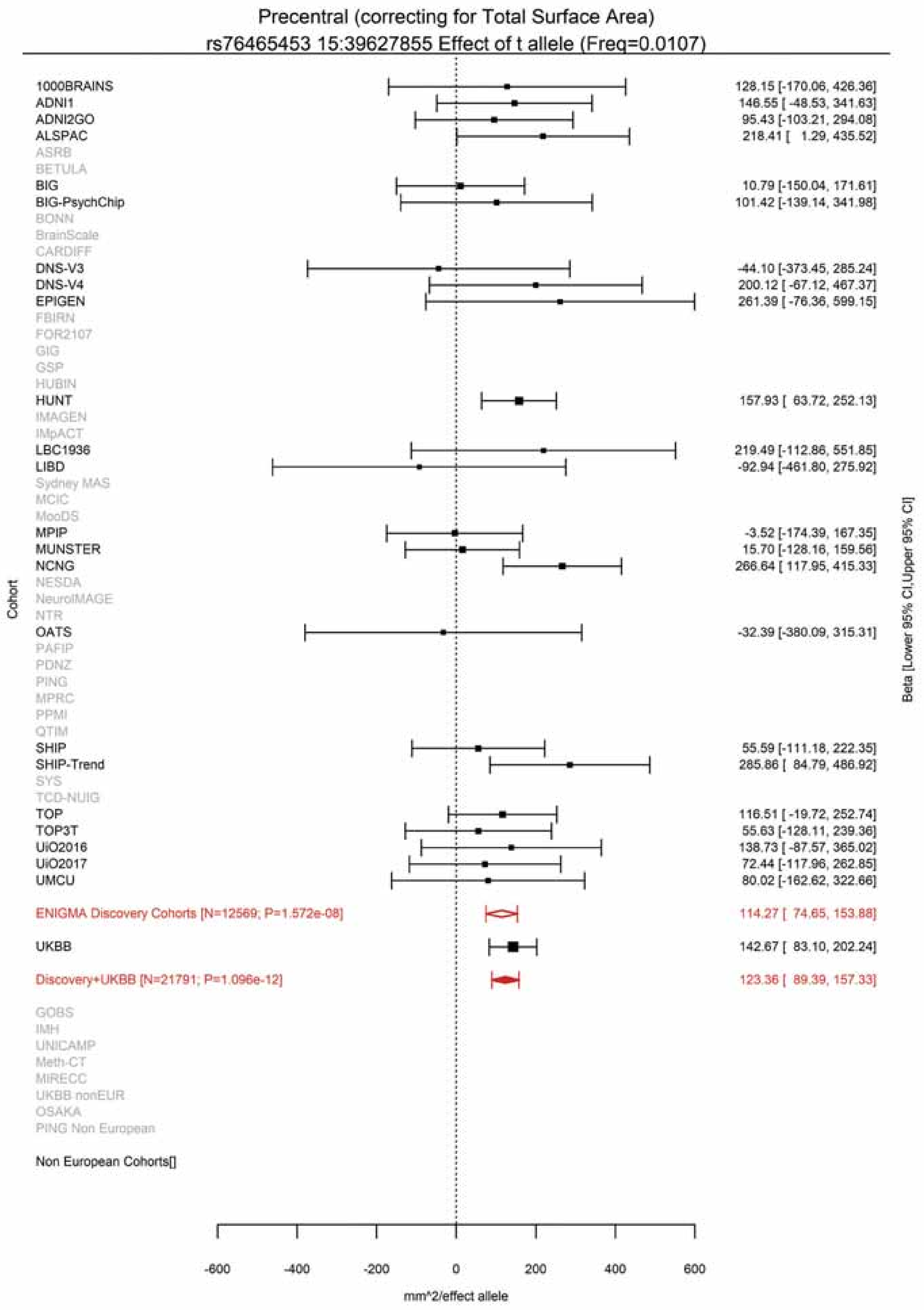

Precentral (correcting for Total Surface Area)
rs76465453 15:39627855 Effect of t allele (Freq=0.0107)
1000BRAINS 128.15 [-170.06, 426.36]
ADNI1 146.55 [ -48.53, 341.63]
ADNI2GO 95.43 [-103.21, 294.08]
ALSPAC 218.41 [ 1.29, 435.52]
ASRB
BETULA
BIG 10.79 [-150.04, 171.61]
BIG-PsychChip 101.42 [-139.14, 341.98]
BONN
BrainScale
CARDIFF
DNS-V3 -44.10 [-373.45, 285.24]
DNS-V4 200.12 [ -67.12, 467.37]
EPIGEN 261.39 [ -76.36, 599.15]
FBIRN
FOR2107
GIG
GSP
HUBIN
HUNT 157.93 [ 63.72, 252.13]
IMAGEN
IMpACT
LBC1936 219.49 [-112.86, 551.85]
LIBD -92.94 [-461.80, 275.92]
Sydney MAS
MCIC
MooDS
MPIP -3.52 [-174.39, 167.35]
MUNSTER 15.70 [-128.16, 159.56]
NCNG 266.64 [ 117.95, 415.33]
NESDA
NeuroIMAGE
NTR
OATS -32.39 [-380.09, 315.31]
PAFIP
PDNZ
PING
MPRC
PPMI
QTIM
SHIP 55.59 [-111.18, 222.35]
SHIP-Trend 285.86 [ 84.79, 486.92]
SYS
TCD-NUIG
TOP 116.51 [ -19.72, 252.74]
TOP3T 55.63 [-128.11, 239.36]
UiO2016 138.73 [ -87.57, 365.02]
UiO2017 72.44 [-117.96, 262.85]
UMCU 80.02 [-162.62, 322.66]
ENIGMA Discovery Cohorts [N=12569; P=1.572e-08] 114.27 [ 74.65, 153.88]
UKBB 142.67 [ 83.10, 202.24]
Discovery+UKBB [N=21791; P=1.096e-12] 123.36 [ 89.39, 157.33]
GOBS
IMH
UNICAMP
Meth-CT
MIRECC
UKBB nonEUR
OSAKA
PING Non European
Non European Cohorts[]
Cohort
Beta [Lower 95% CI,Upper 95% CI]
-600
-400
-200
0
200
400
600
mm^2/effect allele

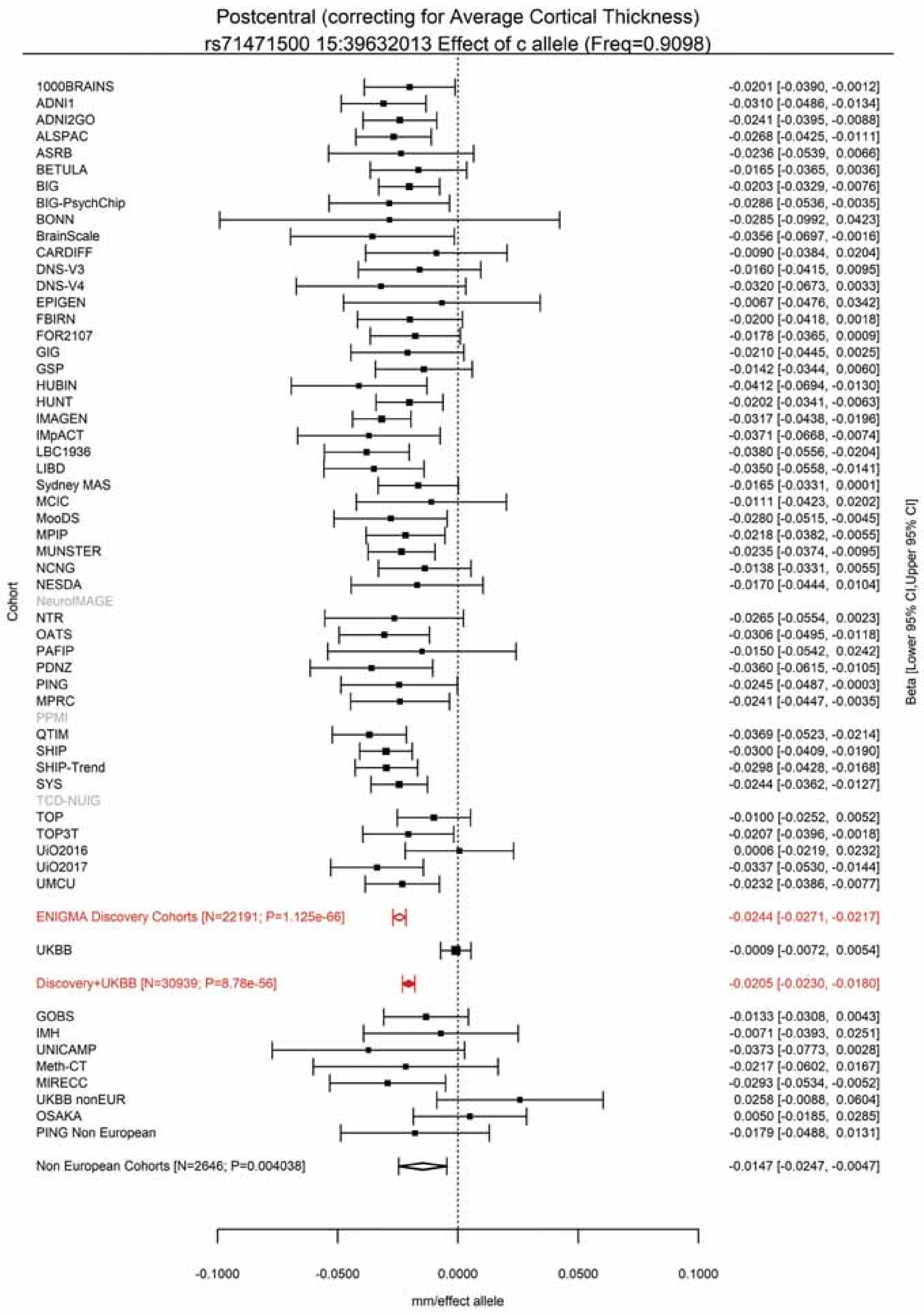

Postcentral (correcting for Average Cortical Thickness)
rs71471500 15:39632013 Effect of c allele (Freq=0.9098)
Cohort
1000BRAINS -0.0201 [-0.0390, -0.0012]
ADNI1 -0.0310 [-0.0486, -0.0134]
ADNI2GO -0.0241 [-0.0395, -0.0088]
ALSPAC -0.0268 [-0.0425, -0.0111]
ASRB -0.0236 [-0.0539, 0.0066]
BETULA -0.0165 [-0.0365, 0.0036]
BIG -0.0203 [-0.0329, -0.0076]
BIG-PsychChip -0.0286 [-0.0536, -0.0035]
BONN -0.0285 [-0.0992, 0.0423]
BrainScale -0.0356 [-0.0697, -0.0016]
CARDIFF -0.0090 [-0.0384, 0.0204]
DNS-V3 -0.0160 [-0.0415, 0.0095]
DNS-V4 -0.0320 [-0.0673, 0.0033]
EPIGEN -0.0067 [-0.0476, 0.0342]
FBIRN -0.0200 [-0.0418, 0.0018]
FOR2107 -0.0178 [-0.0365, 0.0009]
GIG -0.0210 [-0.0445, 0.0025]
GSP -0.0142 [-0.0344, 0.0060]
HUBIN -0.0412 [-0.0694, -0.0130]
HUNT -0.0202 [-0.0341, -0.0063]
IMAGEN -0.0317 [-0.0438, -0.0196]
IMpACT -0.0371 [-0.0668, -0.0074]
LBC1936 -0.0380 [-0.0556, -0.0204]
LIBD -0.0350 [-0.0558, -0.0141]
Sydney MAS -0.0165 [-0.0331, 0.0001]
MCIC -0.0111 [-0.0423, 0.0202]
MooDS -0.0280 [-0.0515, -0.0045]
MPIP -0.0218 [-0.0382, -0.0055]
MUNSTER -0.0235 [-0.0374, -0.0095]
NCNG -0.0138 [-0.0331, 0.0055]
NESDA -0.0170 [-0.0444, 0.0104]
NeuroIMAGE
NTR -0.0265 [-0.0554, 0.0023]
OATS -0.0306 [-0.0495, -0.0118]
PAFIP -0.0150 [-0.0542, 0.0242]
PDNZ -0.0360 [-0.0615, -0.0105]
PING -0.0245 [-0.0487, -0.0003]
MPRC -0.0241 [-0.0447, -0.0035]
PPMI
QTIM -0.0369 [-0.0523, -0.0214]
SHIP -0.0300 [-0.0409, -0.0190]
SHIP-Trend -0.0298 [-0.0428, -0.0168]
SYS -0.0244 [-0.0362, -0.0127]
TCD-NUIG
TOP -0.0100 [-0.0252, 0.0052]
TOP3T -0.0207 [-0.0396, -0.0018]
UiO2016 0.0006 [-0.0219, 0.0232]
UiO2017 -0.0337 [-0.0530, -0.0144]
UMCU -0.0232 [-0.0386, -0.0077]
ENIGMA Discovery Cohorts [N=22191; P=1.125e-66] -0.0244 [-0.0271, -0.0217]
UKBB -0.0009 [-0.0072, 0.0054]
Discovery+UKBB [N=30939; P=8.78e-56] -0.0205 [-0.0230, -0.0180]
GOBS -0.0133 [-0.0308, 0.0043]
IMH -0.0071 [-0.0393, 0.0251]
UNICAMP -0.0373 [-0.0773, 0.0028]
Meth-CT -0.0217 [-0.0602, 0.0167]
MIRECC -0.0293 [-0.0534, -0.0052]
UKBB nonEUR 0.0258 [-0.0088, 0.0604]
OSAKA 0.0050 [-0.0185, 0.0285]
PING Non European -0.0179 [-0.0488, 0.0131]
Non European Cohorts [N=2646; P=0.004038] -0.0147 [-0.0247, -0.0047]
Beta [Lower 95% CI,Upper 95% CI]
-0.1000
-0.0500
0.0000
0.0500
0.1000
mm/effect allele

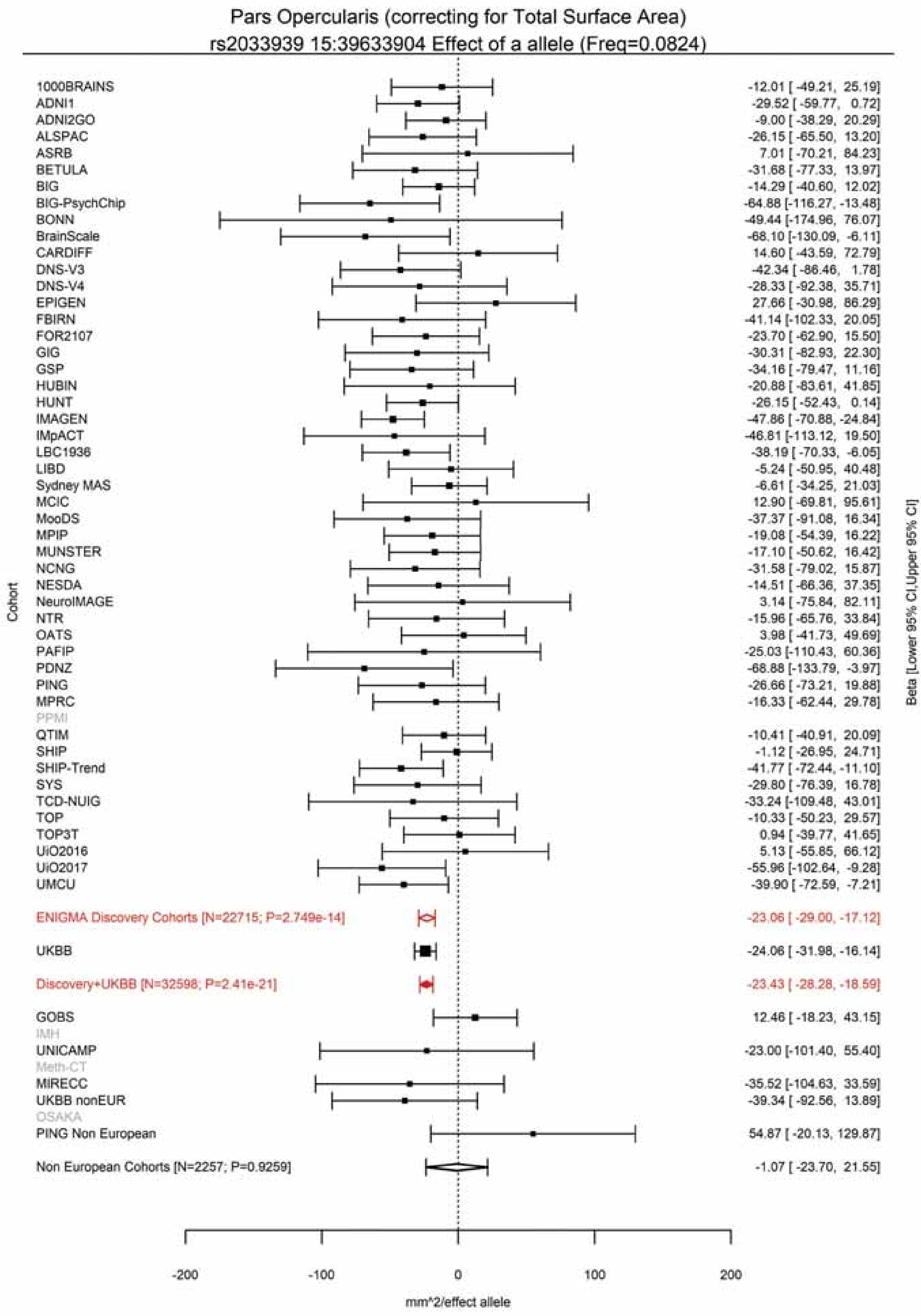

Pars Opercularis (correcting for Total Surface Area)
rs2033939 15:39633904 Effect of a allele (Freq=0.0824)
Cohort
1000BRAINS -12.01 [ -49.21, 25.19]
ADNI1 -29.52 [ -59.77, 0.72]
ADNI2GO -9.00 [ -38.29, 20.29]
ALSPAC -26.15 [ -65.50, 13.20]
ASRB 7.01 [ -70.21, 84.23]
BETULA -31.68 [ -77.33, 13.97]
BIG -14.29 [ -40.60, 12.02]
BIG-PsychChip -64.88 [-116.27, -13.48]
BONN -49.44 [-174.96, 76.07]
BrainScale -68.10 [-130.09, -6.11]
CARDIFF 14.60 [ -43.59, 72.79]
DNS-V3 -42.34 [ -86.46, 1.78]
DNS-V4 -28.33 [ -92.38, 35.71]
EPIGEN 27.66 [ -30.98, 86.29]
FBIRN -41.14 [-102.33, 20.05]
FOR2107 -23.70 [ -62.90, 15.50]
GIG -30.31 [ -82.93, 22.30]
GSP -34.16 [ -79.47, 11.16]
HUBIN -20.88 [ -83.61, 41.85]
HUNT -26.15 [ -52.43, 0.14]
IMAGEN -47.86 [ -70.88, -24.84]
IMpACT -46.81 [-113.12, 19.50]
LBC1936 -38.19 [ -70.33, -6.05]
LIBD -5.24 [ -50.95, 40.48]
Sydney MAS -6.61 [ -34.25, 21.03]
MCIC 12.90 [ -69.81, 95.61]
MooDS -37.37 [ -91.08, 16.34]
MPIP -19.08 [ -54.39, 16.22]
MUNSTER -17.10 [ -50.62, 16.42]
NCNG -31.58 [ -79.02, 15.87]
NESDA -14.51 [ -66.36, 37.35]
NeuroIMAGE 3.14 [ -75.84, 82.11]
NTR -15.96 [ -65.76, 33.84]
OATS 3.98 [ -41.73, 49.69]
PAFIP -25.03 [-110.43, 60.36]
PDNZ -68.88 [-133.79, -3.97]
PING -26.66 [ -73.21, 19.88]
MPRC -16.33 [ -62.44, 29.78]
PPMI
QTIM -10.41 [ -40.91, 20.09]
SHIP -1.12 [ -26.95, 24.71]
SHIP-Trend -41.77 [ -72.44, -11.10]
SYS -29.80 [ -76.39, 16.78]
TCD-NUIG -33.24 [-109.48, 43.01]
TOP -10.33 [ -50.23, 29.57]
TOP3T 0.94 [ -39.77, 41.65]
UiO2016 5.13 [ -55.85, 66.12]
UiO2017 -55.96 [-102.64, -9.28]
UMCU -39.90 [ -72.59, -7.21]
ENIGMA Discovery Cohorts [N=22715; P=2.749e-14] -23.06 [ -29.00, -17.12]
UKBB -24.06 [ -31.98, -16.14]
Discovery+UKBB [N=32598; P=2.41e-21] -23.43 [ -28.28, -18.59]
GOBS 12.46 [ -18.23, 43.15]
IMH
UNICAMP -23.00 [-101.40, 55.40]
Meth-CT
MIRECC -35.52 [-104.63, 33.59]
UKBB nonEUR -39.34 [ -92.56, 13.89]
OSAKA
PING Non European 54.87 [ -20.13, 129.87]
Non European Cohorts [N=2257; P=0.9259] -1.07 [ -23.70, 21.55]
Beta [Lower 95% CI,Upper 95% CI]
-200
-100
0
100
200
mm^2/effect allele

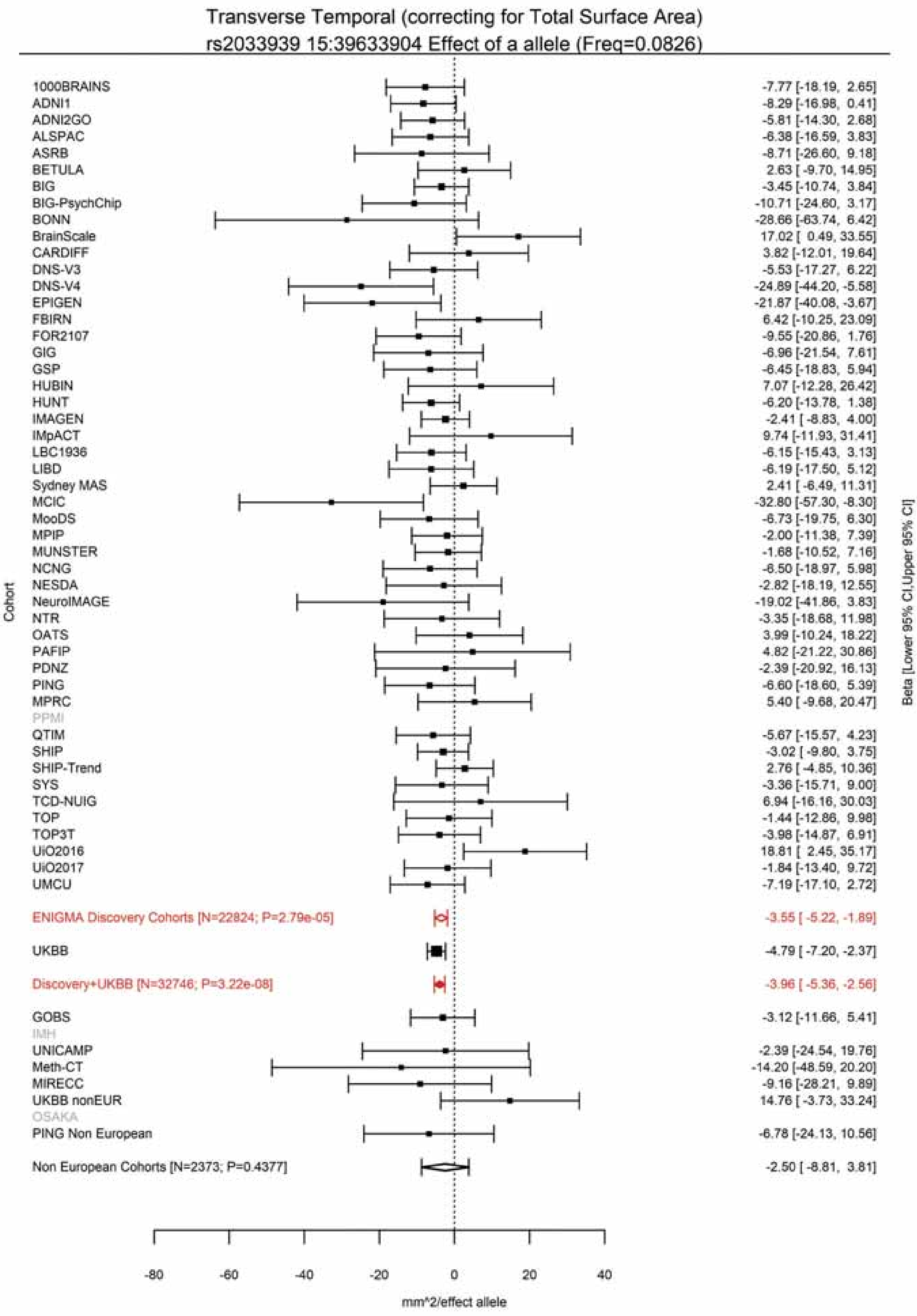
Transverse Temporal (correcting for Total Surface Area)
rs2033939 15:39633904 Effect of a allele (Freq=0.0826)
1000BRAINS -7.77 [-18.19, 2.65]
ADNI1 -8.29 [-16.98, 0.41]
ADNI2GO -5.81 [-14.30, 2.68]
ALSPAC -6.38 [-16.59, 3.83]
ASRB -8.71 [-26.60, 9.18]
BETULA 2.63 [-9.70, 14.95]
BIG -3.45 [-10.74, 3.84]
BIG-PsychChip -10.71 [-24.60, 3.17]
BONN -28.66 [-63.74, 6.42]
BrainScale 17.02 [0.49, 33.55]
CARDIFF 3.82 [-12.01, 19.64]
DNS-V3 -5.53 [-17.27, 6.22]
DNS-V4 -24.89 [-44.20, -5.58]
EPIGEN -21.87 [-40.08, -3.67]
FBIRN 6.42 [-10.25, 23.09]
FOR2107 -9.55 [-20.86, 1.76]
GIG -6.96 [-21.54, 7.61]
GSP -6.45 [-18.83, 5.94]
HUBIN 7.07 [-12.28, 26.42]
HUNT -6.20 [-13.78, 1.38]
IMAGEN -2.41 [-8.83, 4.00]
IMpACT 9.74 [-11.93, 31.41]
LBC1936 -6.15 [-15.43, 3.13]
LIBD -6.19 [-17.50, 5.12]
Sydney MAS 2.41 [-6.49, 11.31]
MCIC -32.80 [-57.30, -8.30]
MooDS -6.73 [-19.75, 6.30]
MPIP -2.00 [-11.38, 7.39]
MUNSTER -1.68 [-10.52, 7.16]
NCNG -6.50 [-18.97, 5.98]
NESDA -2.82 [-18.19, 12.55]
NeuroIMAGE -19.02 [-41.86, 3.83]
NTR -3.35 [-18.68, 11.98]
OATS 3.99 [-10.24, 18.22]
PAFIP 4.82 [-21.22, 30.86]
PDNZ -2.39 [-20.92, 16.13]
PING -6.60 [-18.60, 5.39]
MPRC 5.40 [-9.68, 20.47]
PPMI
QTIM -5.67 [-15.57, 4.23]
SHIP -3.02 [-9.80, 3.75]
SHIP-Trend 2.76 [-4.85, 10.36]
SYS -3.36 [-15.71, 9.00]
TCD-NUIG 6.94 [-16.16, 30.03]
TOP -1.44 [-12.86, 9.98]
TOP3T -3.98 [-14.87, 6.91]
UiO2016 18.81 [2.45, 35.17]
UiO2017 -1.84 [-13.40, 9.72]
UMCU -7.19 [-17.10, 2.72]
ENIGMA Discovery Cohorts [N=22824; P=2.79e-05] -3.55 [-5.22, -1.89]
UKBB -4.79 [-7.20, -2.37]
Discovery+UKBB [N=32746; P=3.22e-08] -3.96 [-5.36, -2.56]
GOBS -3.12 [-11.66, 5.41]
IMH
UNICAMP -2.39 [-24.54, 19.76]
Meth-CT -14.20 [-48.59, 20.20]
MIRECC -9.16 [-28.21, 9.89]
UKBB nonEUR 14.76 [-3.73, 33.24]
OSAKA
PING Non European -6.78 [-24.13, 10.56]
Non European Cohorts [N=2373; P=0.4377] -2.50 [-8.81, 3.81]
Cohort
Beta [Lower 95% CI,Upper 95% CI]
-80
-60
-40
-20
0
20
40
mm^2/effect allele

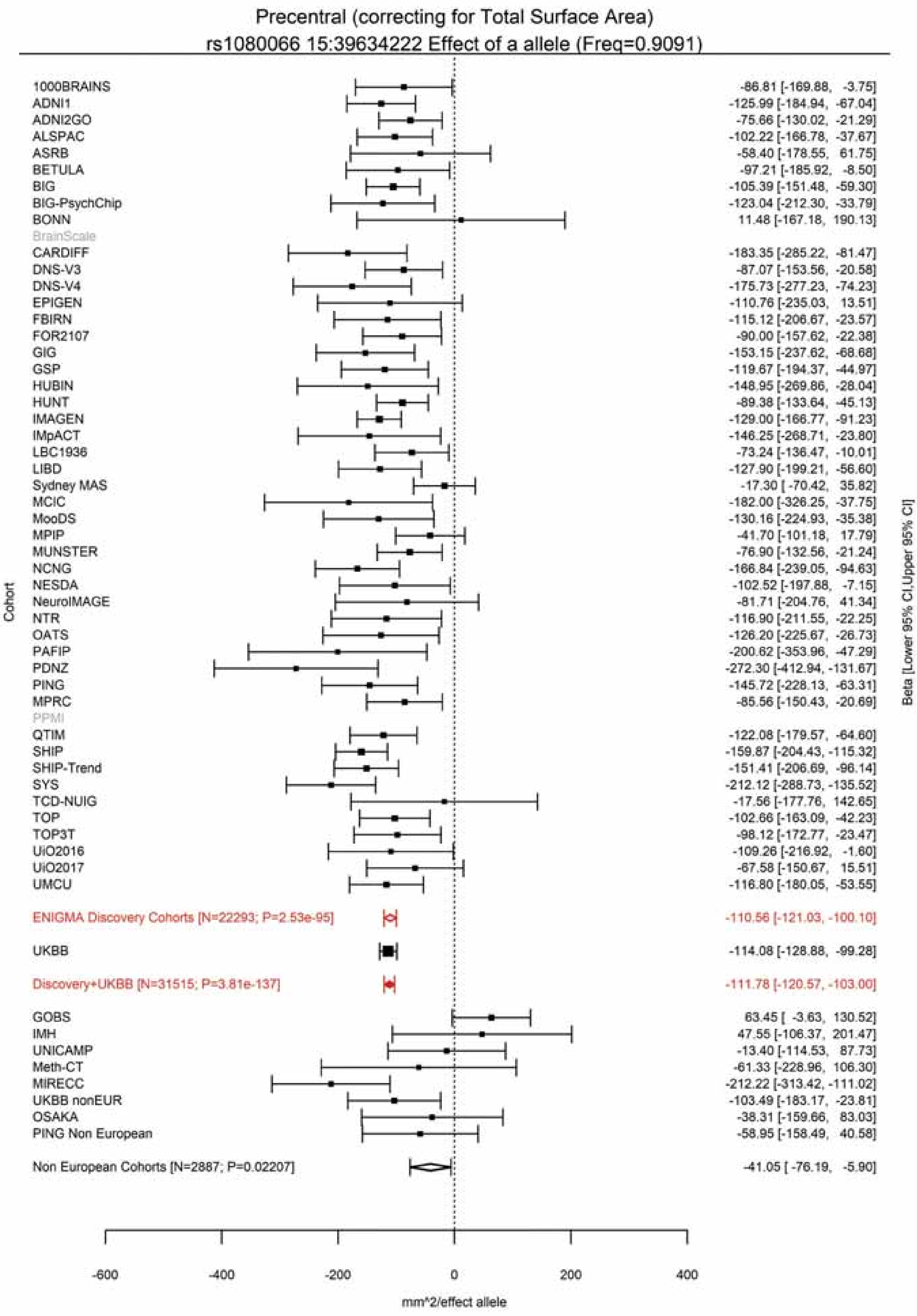
Precentral (correcting for Total Surface Area)
rs1080066 15:39634222 Effect of a allele (Freq=0.9091)
1000BRAINS -86.81 [-169.88, -3.75]
ADNI1 -125.99 [-184.94, -67.04]
ADNI2GO -75.66 [-130.02, -21.29]
ALSPAC -102.22 [-166.78, -37.67]
ASRB -58.40 [-178.55, 61.75]
BETULA -97.21 [-185.92, -8.50]
BIG -105.39 [-151.48, -59.30]
BIG-PsychChip -123.04 [-212.30, -33.79]
BONN 11.48 [-167.18, 190.13]
BrainScale
CARDIFF -183.35 [-285.22, -81.47]
DNS-V3 -87.07 [-153.56, -20.58]
DNS-V4 -175.73 [-277.23, -74.23]
EPIGEN -110.76 [-235.03, 13.51]
FBIRN -115.12 [-206.67, -23.57]
FOR2107 -90.00 [-157.62, -22.38]
GIG -153.15 [-237.62, -68.68]
GSP -119.67 [-194.37, -44.97]
HUBIN -148.95 [-269.86, -28.04]
HUNT -89.38 [-133.64, -45.13]
IMAGEN -129.00 [-166.77, -91.23]
IMpACT -146.25 [-268.71, -23.80]
LBC1936 -73.24 [-136.47, -10.01]
LIBD -127.90 [-199.21, -56.60]
Sydney MAS -17.30 [-70.42, 35.82]
MCIC -182.00 [-326.25, -37.75]
MooDS -130.16 [-224.93, -35.38]
MPIP -41.70 [-101.18, 17.79]
MUNSTER -76.90 [-132.56, -21.24]
NCNG -166.84 [-239.05, -94.63]
NESDA -102.52 [-197.88, -7.15]
NeuroIMAGE -81.71 [-204.76, 41.34]
NTR -116.90 [-211.55, -22.25]
OATS -126.20 [-225.67, -26.73]
PAFIP -200.62 [-353.96, -47.29]
PDNZ -272.30 [-412.94, -131.67]
PING -145.72 [-228.13, -63.31]
MPRC -85.56 [-150.43, -20.69]
PPMI
QTIM -122.08 [-179.57, -64.60]
SHIP -159.87 [-204.43, -115.32]
SHIP-Trend -151.41 [-206.69, -96.14]
SYS -212.12 [-288.73, -135.52]
TCD-NUIG -17.56 [-177.76, 142.65]
TOP -102.66 [-163.09, -42.23]
TOP3T -98.12 [-172.77, -23.47]
UiO2016 -109.26 [-216.92, -1.60]
UiO2017 -67.58 [-150.67, 15.51]
UMCU -116.80 [-180.05, -53.55]
ENIGMA Discovery Cohorts [N=22293; P=2.53e-95] -110.56 [-121.03, -100.10]
UKBB -114.08 [-128.88, -99.28]
Discovery+UKBB [N=31515; P=3.81e-137] -111.78 [-120.57, -103.00]
GOBS 63.45 [ -3.63, 130.52]
IMH 47.55 [-106.37, 201.47]
UNICAMP -13.40 [-114.53, 87.73]
Meth-CT -61.33 [-228.96, 106.30]
MIRECC -212.22 [-313.42, -111.02]
UKBB nonEUR -103.49 [-183.17, -23.81]
OSAKA -38.31 [-159.66, 83.03]
PING Non European -58.95 [-158.49, 40.58]
Non European Cohorts [N=2887; P=0.02207] -41.05 [ -76.19, -5.90]
Cohort
Beta [Lower 95% CI,Upper 95% CI]
-600
-400
-200
0
200
400
mm^2/effect allele

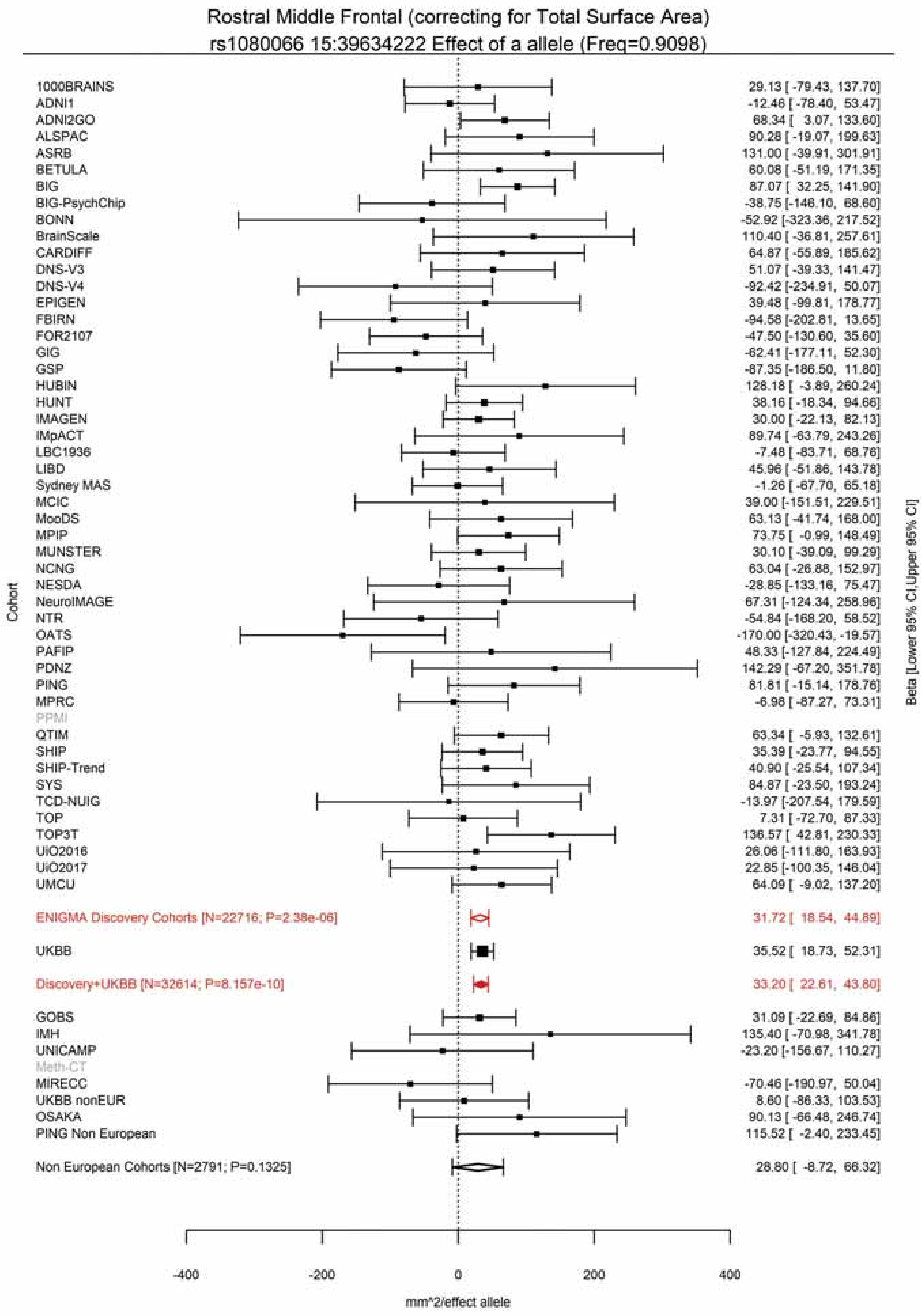

Rostral Middle Frontal (correcting for Total Surface Area)
rs1080066 15:39634222 Effect of a allele (Freq=0.9098)
1000BRAINS 29.13 [ -79.43, 137.70]
ADNI1 -12.46 [ -78.40, 53.47]
ADNI2GO 68.34 [ 3.07, 133.60]
ALSPAC 90.28 [ -19.07, 199.63]
ASRB 131.00 [ -39.91, 301.91]
BETULA 60.08 [ -51.19, 171.35]
BIG 87.07 [ 32.25, 141.90]
BIG-PsychChip -38.75 [-146.10, 68.60]
BONN -52.92 [-323.36, 217.52]
BrainScale 110.40 [ -36.81, 257.61]
CARDIFF 64.87 [ -55.89, 185.62]
DNS-V3 51.07 [ -39.33, 141.47]
DNS-V4 -92.42 [-234.91, 50.07]
EPIGEN 39.48 [ -99.81, 178.77]
FBIRN -94.58 [-202.81, 13.65]
FOR2107 -47.50 [-130.60, 35.60]
GIG -62.41 [-177.11, 52.30]
GSP -87.35 [-186.50, 11.80]
HUBIN 128.18 [ -3.89, 260.24]
HUNT 38.16 [ -18.34, 94.66]
IMAGEN 30.00 [ -22.13, 82.13]
IMpACT 89.74 [ -63.79, 243.26]
LBC1936 -7.48 [ -83.71, 68.76]
LIBD 45.96 [ -51.86, 143.78]
Sydney MAS -1.26 [ -67.70, 65.18]
MCIC 39.00 [-151.51, 229.51]
MooDS 63.13 [ -41.74, 168.00]
MPIP 73.75 [ -0.99, 148.49]
MUNSTER 30.10 [ -39.09, 99.29]
NCNG 63.04 [ -26.88, 152.97]
NESDA -28.85 [-133.16, 75.47]
NeuroIMAGE 67.31 [-124.34, 258.96]
NTR -54.84 [-168.20, 58.52]
OATS -170.00 [-320.43, -19.57]
PAFIP 48.33 [-127.84, 224.49]
PDNZ 142.29 [ -67.20, 351.78]
PING 81.81 [ -15.14, 178.76]
MPRC -6.98 [ -87.27, 73.31]
PPMI
QTIM 63.34 [ -5.93, 132.61]
SHIP 35.39 [ -23.77, 94.55]
SHIP-Trend 40.90 [ -25.54, 107.34]
SYS 84.87 [ -23.50, 193.24]
TCD-NUIG -13.97 [-207.54, 179.59]
TOP 7.31 [ -72.70, 87.33]
TOP3T 136.57 [ 42.81, 230.33]
UiO2016 26.06 [-111.80, 163.93]
UiO2017 22.85 [-100.35, 146.04]
UMCU 64.09 [ -9.02, 137.20]
ENIGMA Discovery Cohorts [N=22716; P=2.38e-06] 31.72 [ 18.54, 44.89]
UKBB 35.52 [ 18.73, 52.31]
Discovery+UKBB [N=32614; P=8.157e-10] 33.20 [ 22.61, 43.80]
GOBS 31.09 [ -22.69, 84.86]
IMH 135.40 [ -70.98, 341.78]
UNICAMP -23.20 [-156.67, 110.27]
Meth-CT
MIRECC -70.46 [-190.97, 50.04]
UKBB nonEUR 8.60 [ -86.33, 103.53]
OSAKA 90.13 [ -66.48, 246.74]
PING Non European 115.52 [ -2.40, 233.45]
Non European Cohorts [N=2791; P=0.1325] 28.80 [ -8.72, 66.32]
Cohort
Beta [Lower 95% CI,Upper 95% CI]
-400
-200
0
200
400
mm^2/effect allele

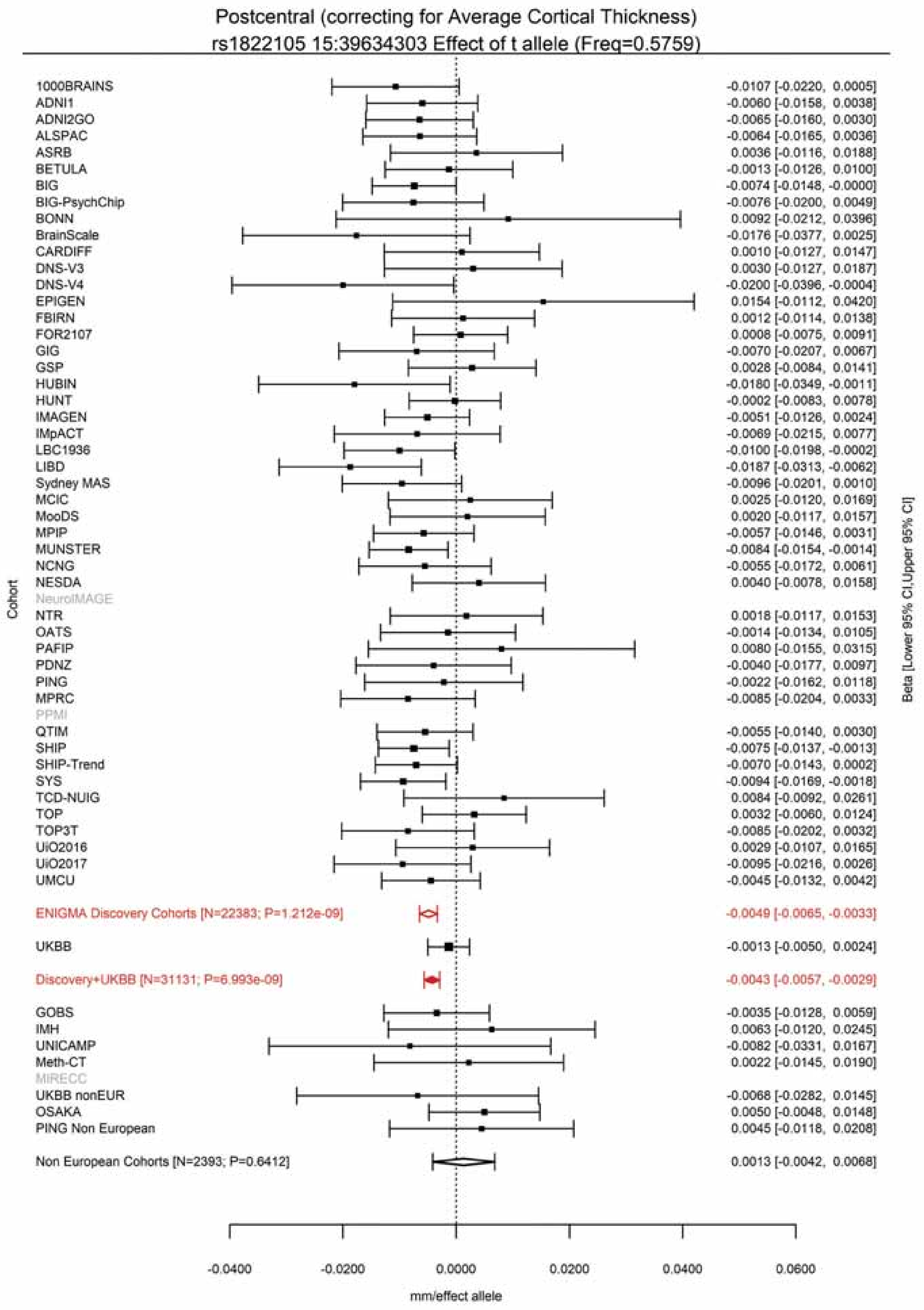

Postcentral (correcting for Average Cortical Thickness)
rs1822105 15:39634303 Effect of t allele (Freq=0.5759)
Cohort
1000BRAINS -0.0107 [-0.0220, 0.0005]
ADNI1 -0.0060 [-0.0158, 0.0038]
ADNI2GO -0.0065 [-0.0160, 0.0030]
ALSPAC -0.0064 [-0.0165, 0.0036]
ASRB 0.0036 [-0.0116, 0.0188]
BETULA -0.0013 [-0.0126, 0.0100]
BIG -0.0074 [-0.0148, -0.0000]
BIG-PsychChip -0.0076 [-0.0200, 0.0049]
BONN 0.0092 [-0.0212, 0.0396]
BrainScale -0.0176 [-0.0377, 0.0025]
CARDIFF 0.0010 [-0.0127, 0.0147]
DNS-V3 0.0030 [-0.0127, 0.0187]
DNS-V4 -0.0200 [-0.0396, -0.0004]
EPIGEN 0.0154 [-0.0112, 0.0420]
FBIRN 0.0012 [-0.0114, 0.0138]
FOR2107 0.0008 [-0.0075, 0.0091]
GIG -0.0070 [-0.0207, 0.0067]
GSP 0.0028 [-0.0084, 0.0141]
HUBIN -0.0180 [-0.0349, -0.0011]
HUNT -0.0002 [-0.0083, 0.0078]
IMAGEN -0.0051 [-0.0126, 0.0024]
IMpACT -0.0069 [-0.0215, 0.0077]
LBC1936 -0.0100 [-0.0198, -0.0002]
LIBD -0.0187 [-0.0313, -0.0062]
Sydney MAS -0.0096 [-0.0201, 0.0010]
MCIC 0.0025 [-0.0120, 0.0169]
MooDS 0.0020 [-0.0117, 0.0157]
MPIP -0.0057 [-0.0146, 0.0031]
MUNSTER -0.0084 [-0.0154, -0.0014]
NCNG -0.0055 [-0.0172, 0.0061]
NESDA 0.0040 [-0.0078, 0.0158]
NeuroIMAGE
NTR 0.0018 [-0.0117, 0.0153]
OATS -0.0014 [-0.0134, 0.0105]
PAFIP 0.0080 [-0.0155, 0.0315]
PDNZ -0.0040 [-0.0177, 0.0097]
PING -0.0022 [-0.0162, 0.0118]
MPRC -0.0085 [-0.0204, 0.0033]
PPMI
QTIM -0.0055 [-0.0140, 0.0030]
SHIP -0.0075 [-0.0137, -0.0013]
SHIP-Trend -0.0070 [-0.0143, 0.0002]
SYS -0.0094 [-0.0169, -0.0018]
TCD-NUIG 0.0084 [-0.0092, 0.0261]
TOP 0.0032 [-0.0060, 0.0124]
TOP3T -0.0085 [-0.0202, 0.0032]
UiO2016 0.0029 [-0.0107, 0.0165]
UiO2017 -0.0095 [-0.0216, 0.0026]
UMCU -0.0045 [-0.0132, 0.0042]
ENIGMA Discovery Cohorts [N=22383; P=1.212e-09] -0.0049 [-0.0065, -0.0033]
UKBB -0.0013 [-0.0050, 0.0024]
Discovery+UKBB [N=31131; P=6.993e-09] -0.0043 [-0.0057, -0.0029]
GOBS -0.0035 [-0.0128, 0.0059]
IMH 0.0063 [-0.0120, 0.0245]
UNICAMP -0.0082 [-0.0331, 0.0167]
Meth-CT 0.0022 [-0.0145, 0.0190]
MIRECC
UKBB nonEUR -0.0068 [-0.0282, 0.0145]
OSAKA 0.0050 [-0.0048, 0.0148]
PING Non European 0.0045 [-0.0118, 0.0208]
Non European Cohorts [N=2393; P=0.6412] 0.0013 [-0.0042, 0.0068]
Beta [Lower 95% CI,Upper 95% CI]
-0.0400
-0.0200
0.0000
0.0200
0.0400
0.0600
mm/effect allele

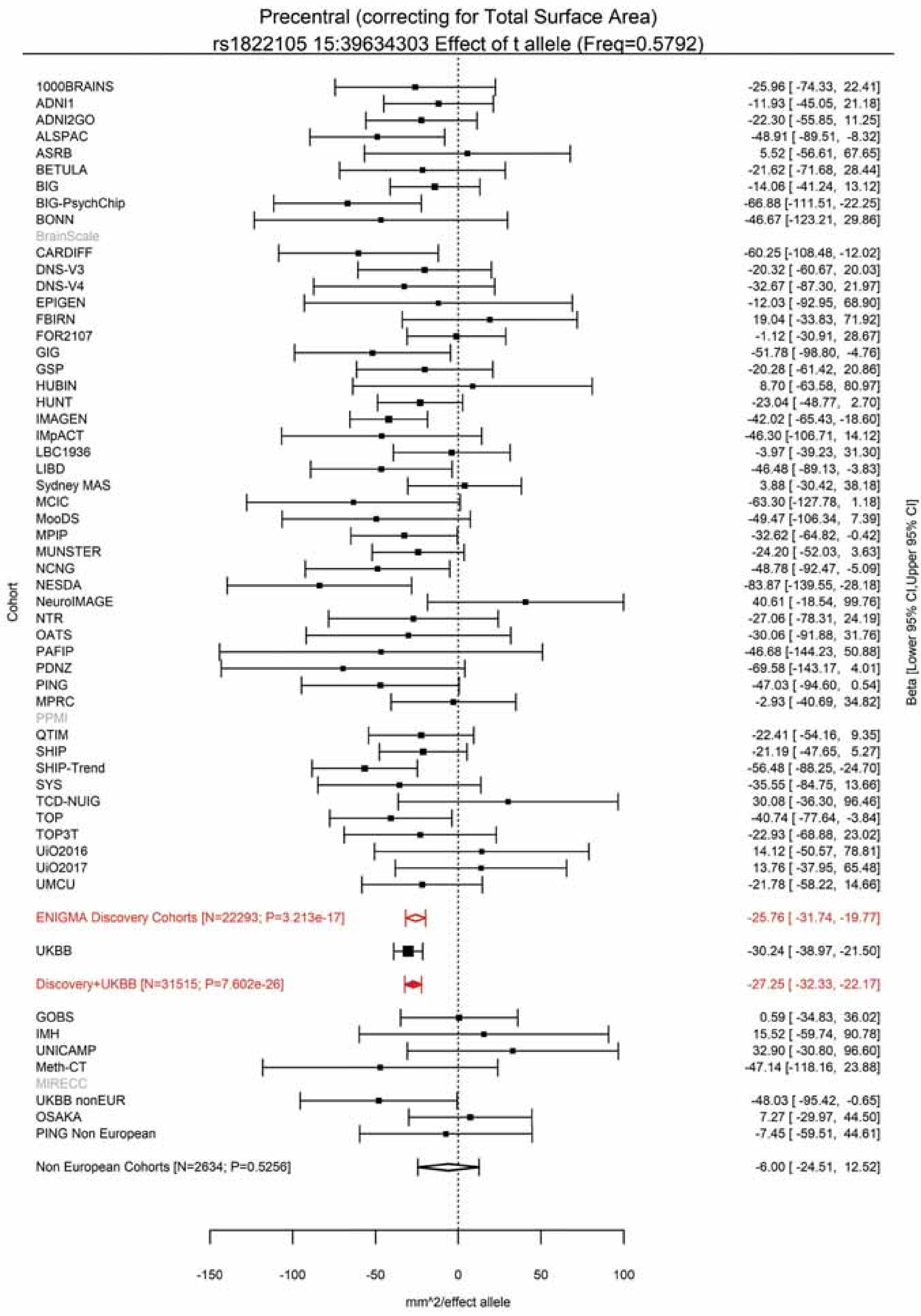
Precentral (correcting for Total Surface Area)
rs1822105 15:39634303 Effect of t allele (Freq=0.5792)
1000BRAINS -25.96 [ -74.33, 22.41]
ADNI1 -11.93 [ -45.05, 21.18]
ADNI2GO -22.30 [ -55.85, 11.25]
ALSPAC -48.91 [ -89.51, -8.32]
ASRB 5.52 [ -56.61, 67.65]
BETULA -21.62 [ -71.68, 28.44]
BIG -14.06 [ -41.24, 13.12]
BIG-PsychChip -66.88 [-111.51, -22.25]
BONN -46.67 [-123.21, 29.86]
BrainScale
CARDIFF -60.25 [-108.48, -12.02]
DNS-V3 -20.32 [ -60.67, 20.03]
DNS-V4 -32.67 [ -87.30, 21.97]
EPIGEN -12.03 [ -92.95, 68.90]
FBIRN 19.04 [ -33.83, 71.92]
FOR2107 -1.12 [ -30.91, 28.67]
GIG -51.78 [ -98.80, -4.76]
GSP -20.28 [ -61.42, 20.86]
HUBIN 8.70 [ -63.58, 80.97]
HUNT -23.04 [ -48.77, 2.70]
IMAGEN -42.02 [ -65.43, -18.60]
IMpACT -46.30 [-106.71, 14.12]
LBC1936 -3.97 [ -39.23, 31.30]
LIBD -46.48 [ -89.13, -3.83]
Sydney MAS 3.88 [ -30.42, 38.18]
MCIC -63.30 [-127.78, 1.18]
MooDS -49.47 [-106.34, 7.39]
MPIP -32.62 [ -64.82, -0.42]
MUNSTER -24.20 [ -52.03, 3.63]
NCNG -48.78 [ -92.47, -5.09]
NESDA -83.87 [-139.55, -28.18]
NeuroIMAGE 40.61 [ -18.54, 99.76]
NTR -27.06 [ -78.31, 24.19]
OATS -30.06 [ -91.88, 31.76]
PAFIP -46.68 [-144.23, 50.88]
PDNZ -69.58 [-143.17, 4.01]
PING -47.03 [ -94.60, 0.54]
MPRC -2.93 [ -40.69, 34.82]
PPMI
QTIM -22.41 [ -54.16, 9.35]
SHIP -21.19 [ -47.65, 5.27]
SHIP-Trend -56.48 [ -88.25, -24.70]
SYS -35.55 [ -84.75, 13.66]
TCD-NUIG 30.08 [ -36.30, 96.46]
TOP -40.74 [ -77.64, -3.84]
TOP3T -22.93 [ -68.88, 23.02]
UiO2016 14.12 [ -50.57, 78.81]
UiO2017 13.76 [ -37.95, 65.48]
UMCU -21.78 [ -58.22, 14.66]
ENIGMA Discovery Cohorts [N=22293; P=3.213e-17] -25.76 [ -31.74, -19.77]
UKBB -30.24 [ -38.97, -21.50]
Discovery+UKBB [N=31515; P=7.602e-26] -27.25 [ -32.33, -22.17]
GOBS 0.59 [ -34.83, 36.02]
IMH 15.52 [ -59.74, 90.78]
UNICAMP 32.90 [ -30.80, 96.60]
Meth-CT -47.14 [-118.16, 23.88]
MIRECC
UKBB nonEUR -48.03 [ -95.42, -0.65]
OSAKA 7.27 [ -29.97, 44.50]
PING Non European -7.45 [ -59.51, 44.61]
Non European Cohorts [N=2634; P=0.5256] -6.00 [ -24.51, 12.52]
Cohort
Beta [Lower 95% CI,Upper 95% CI]
-150
-100
-50
0
50
100
mm^2/effect allele

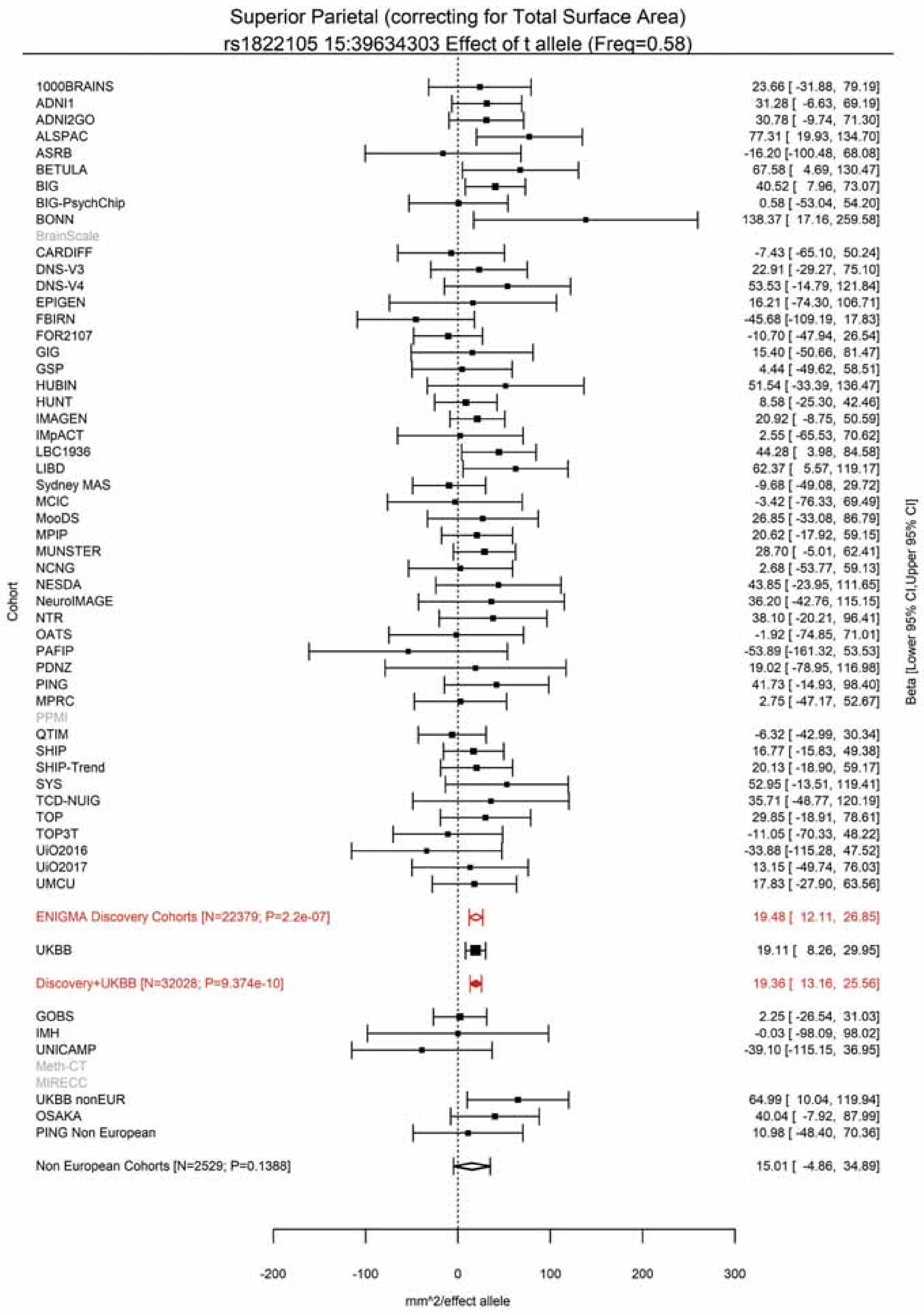
Superior Parietal (correcting for Total Surface Area)
rs1822105 15:39634303 Effect of t allele (Freq=0.58)
Cohort
1000BRAINS 23.66 [ -31.88, 79.19]
ADNI1 31.28 [ -6.63, 69.19]
ADNI2GO 30.78 [ -9.74, 71.30]
ALSPAC 77.31 [ 19.93, 134.70]
ASRB -16.20 [-100.48, 68.08]
BETULA 67.58 [ 4.69, 130.47]
BIG 40.52 [ 7.96, 73.07]
BIG-PsychChip 0.58 [ -53.04, 54.20]
BONN 138.37 [ 17.16, 259.58]
BrainScale
CARDIFF -7.43 [ -65.10, 50.24]
DNS-V3 22.91 [ -29.27, 75.10]
DNS-V4 53.53 [ -14.79, 121.84]
EPIGEN 16.21 [ -74.30, 106.71]
FBIRN -45.68 [-109.19, 17.83]
FOR2107 -10.70 [ -47.94, 26.54]
GIG 15.40 [ -50.66, 81.47]
GSP 4.44 [ -49.62, 58.51]
HUBIN 51.54 [ -33.39, 136.47]
HUNT 8.58 [ -25.30, 42.46]
IMAGEN 20.92 [ -8.75, 50.59]
IMpACT 2.55 [ -65.53, 70.62]
LBC1936 44.28 [ 3.98, 84.58]
LIBD 62.37 [ 5.57, 119.17]
Sydney MAS -9.68 [ -49.08, 29.72]
MCIC -3.42 [ -76.33, 69.49]
MooDS 26.85 [ -33.08, 86.79]
MPIP 20.62 [ -17.92, 59.15]
MUNSTER 28.70 [ -5.01, 62.41]
NCNG 2.68 [ -53.77, 59.13]
NESDA 43.85 [ -23.95, 111.65]
NeuroIMAGE 36.20 [ -42.76, 115.15]
NTR 38.10 [ -20.21, 96.41]
OATS -1.92 [ -74.85, 71.01]
PAFIP -53.89 [-161.32, 53.53]
PDNZ 19.02 [ -78.95, 116.98]
PING 41.73 [ -14.93, 98.40]
MPRC 2.75 [ -47.17, 52.67]
PPMI
QTIM -6.32 [ -42.99, 30.34]
SHIP 16.77 [ -15.83, 49.38]
SHIP-Trend 20.13 [ -18.90, 59.17]
SYS 52.95 [ -13.51, 119.41]
TCD-NUIG 35.71 [ -48.77, 120.19]
TOP 29.85 [ -18.91, 78.61]
TOP3T -11.05 [ -70.33, 48.22]
UiO2016 -33.88 [-115.28, 47.52]
UiO2017 13.15 [ -49.74, 76.03]
UMCU 17.83 [ -27.90, 63.56]
ENIGMA Discovery Cohorts [N=22379; P=2.2e-07] 19.48 [ 12.11, 26.85]
UKBB 19.11 [ 8.26, 29.95]
Discovery+UKBB [N=32028; P=9.374e-10] 19.36 [ 13.16, 25.56]
GOBS 2.25 [ -26.54, 31.03]
IMH -0.03 [ -98.09, 98.02]
UNICAMP -39.10 [-115.15, 36.95]
Meth-CT
MIRECC
UKBB nonEUR 64.99 [ 10.04, 119.94]
OSAKA 40.04 [ -7.92, 87.99]
PING Non European 10.98 [ -48.40, 70.36]
Non European Cohorts [N=2529; P=0.1388] 15.01 [ -4.86, 34.89]
Beta [Lower 95% CI,Upper 95% CI]
-200
-100
0
100
200
300
mm^2/effect allele

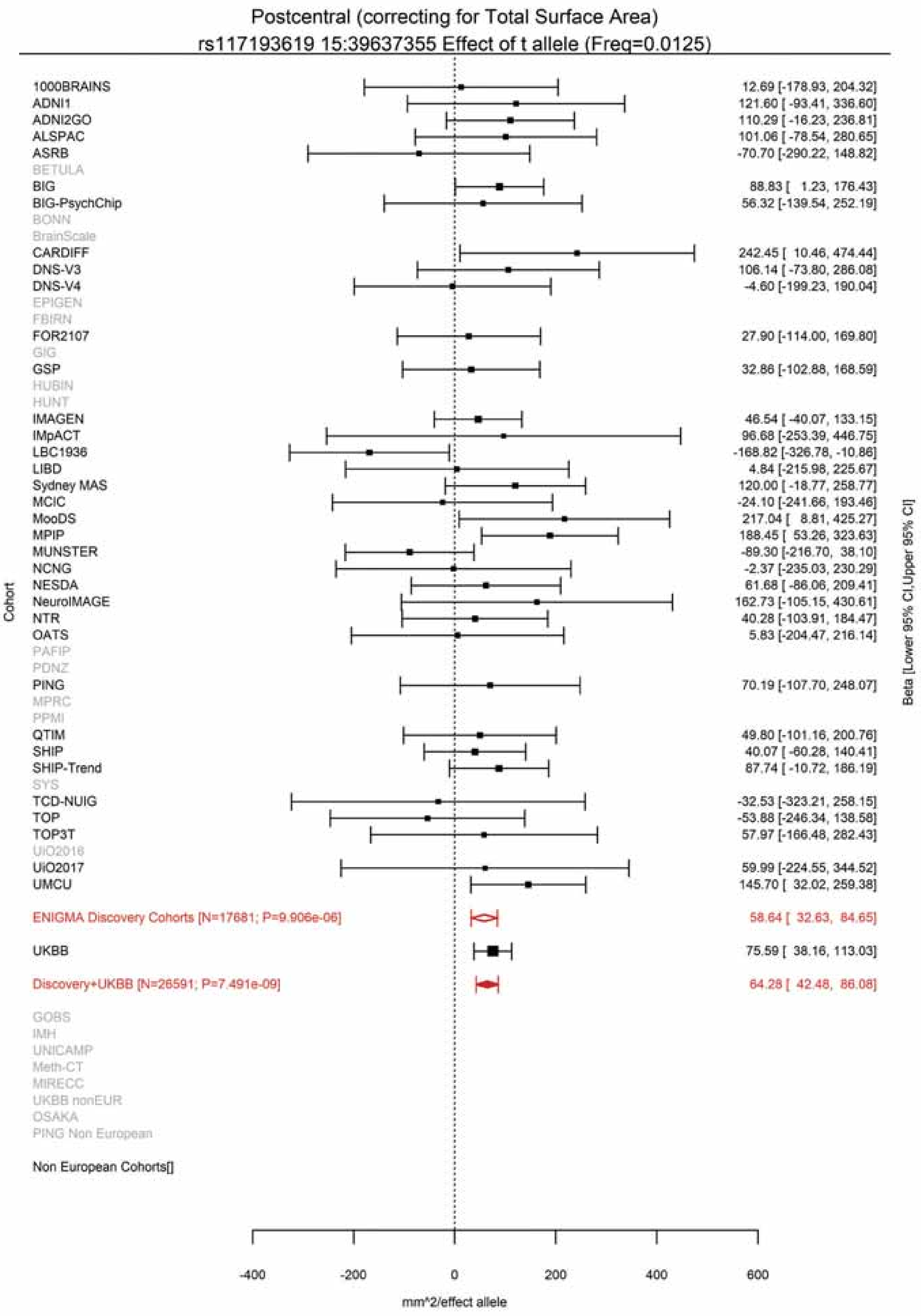
Postcentral (correcting for Total Surface Area)
rs117193619 15:39637355 Effect of t allele (Freq=0.0125)
1000BRAINS 12.69 [-178.93, 204.32]
ADNI1 121.60 [ -93.41, 336.60]
ADNI2GO 110.29 [ -16.23, 236.81]
ALSPAC 101.06 [ -78.54, 280.65]
ASRB -70.70 [-290.22, 148.82]
BETULA
BIG 88.83 [ 1.23, 176.43]
BIG-PsychChip 56.32 [-139.54, 252.19]
BONN
BrainScale
CARDIFF 242.45 [ 10.46, 474.44]
DNS-V3 106.14 [ -73.80, 286.08]
DNS-V4 -4.60 [-199.23, 190.04]
EPIGEN
FBIRN
FOR2107 27.90 [-114.00, 169.80]
GIG
GSP 32.86 [-102.88, 168.59]
HUBIN
HUNT
IMAGEN 46.54 [ -40.07, 133.15]
IMpACT 96.68 [-253.39, 446.75]
LBC1936 -168.82 [-326.78, -10.86]
LIBD 4.84 [-215.98, 225.67]
Sydney MAS 120.00 [ -18.77, 258.77]
MCIC -24.10 [-241.66, 193.46]
MooDS 217.04 [ 8.81, 425.27]
MPIP 188.45 [ 53.26, 323.63]
MUNSTER -89.30 [-216.70, 38.10]
NCNG -2.37 [-235.03, 230.29]
NESDA 61.68 [ -86.06, 209.41]
NeuroIMAGE 162.73 [-105.15, 430.61]
NTR 40.28 [-103.91, 184.47]
OATS 5.83 [-204.47, 216.14]
PAFIP
PDNZ
PING 70.19 [-107.70, 248.07]
MPRC
PPMI
QTIM 49.80 [-101.16, 200.76]
SHIP 40.07 [ -60.28, 140.41]
SHIP-Trend 87.74 [ -10.72, 186.19]
SYS
TCD-NUIG -32.53 [-323.21, 258.15]
TOP -53.88 [-246.34, 138.58]
TOP3T 57.97 [-166.48, 282.43]
UiO2016
UiO2017 59.99 [-224.55, 344.52]
UMCU 145.70 [ 32.02, 259.38]
ENIGMA Discovery Cohorts [N=17681; P=9.906e-06] 58.64 [ 32.63, 84.65]
UKBB 75.59 [ 38.16, 113.03]
Discovery+UKBB [N=26591; P=7.491e-09] 64.28 [ 42.48, 86.08]
GOBS
IMH
UNICAMP
Meth-CT
MIRECC
UKBB nonEUR
OSAKA
PING Non European
Non European Cohorts[]
Cohort
Beta [Lower 95% CI,Upper 95% CI]
-400
-200
0
200
400
600
mm^2/effect allele

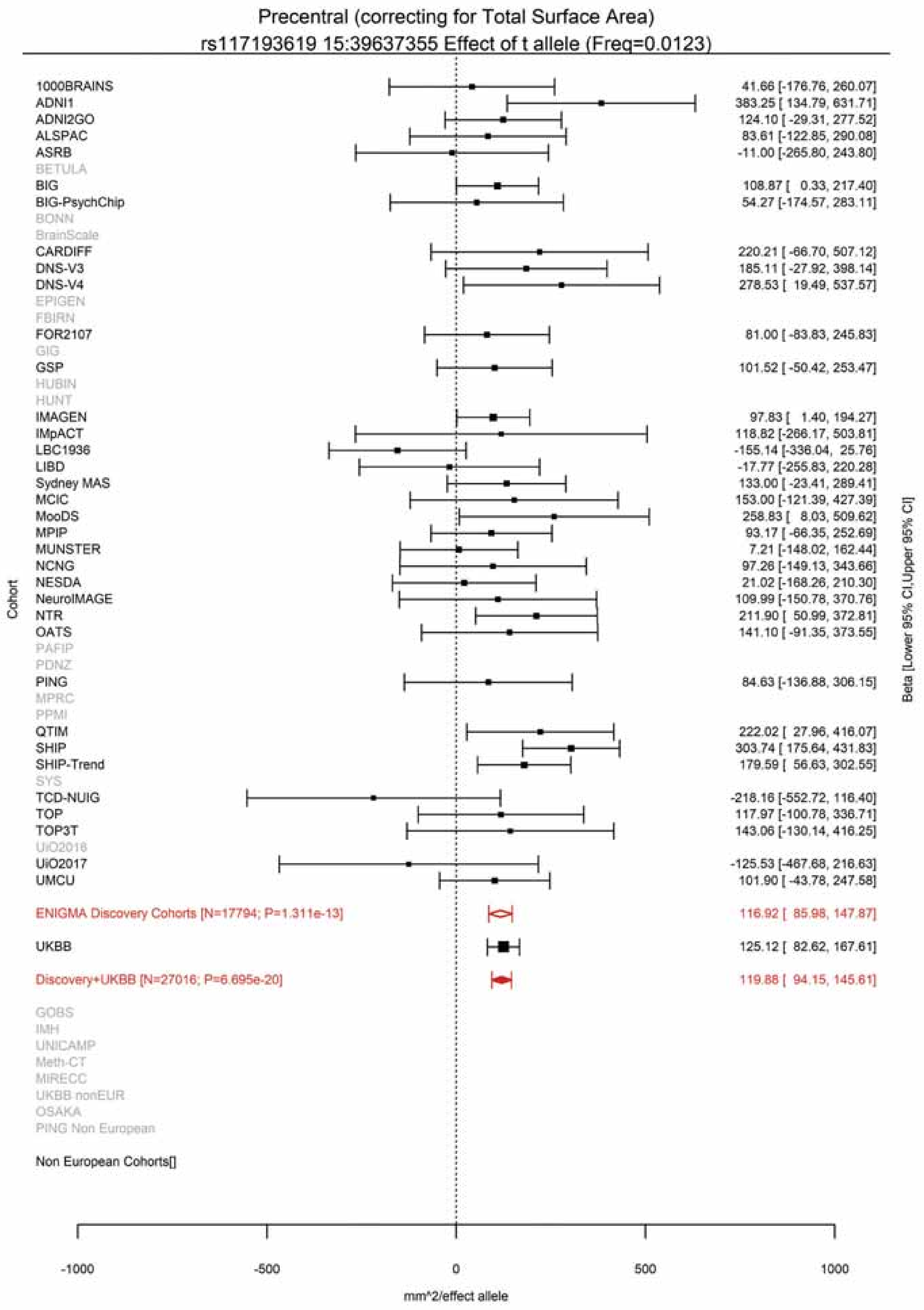
Precentral (correcting for Total Surface Area)
rs117193619 15:39637355 Effect of t allele (Freq=0.0123)
1000BRAINS 41.66 [-176.76, 260.07]
ADNI1 383.25 [ 134.79, 631.71]
ADNI2GO 124.10 [ -29.31, 277.52]
ALSPAC 83.61 [-122.85, 290.08]
ASRB -11.00 [-265.80, 243.80]
BETULA
BIG 108.87 [ 0.33, 217.40]
BIG-PsychChip 54.27 [-174.57, 283.11]
BONN
BrainScale
CARDIFF 220.21 [ -66.70, 507.12]
DNS-V3 185.11 [ -27.92, 398.14]
DNS-V4 278.53 [ 19.49, 537.57]
EPIGEN
FBIRN
FOR2107 81.00 [ -83.83, 245.83]
GIG
GSP 101.52 [ -50.42, 253.47]
HUBIN
HUNT
IMAGEN 97.83 [ 1.40, 194.27]
IMpACT 118.82 [-266.17, 503.81]
LBC1936 -155.14 [-336.04, 25.76]
LIBD -17.77 [-255.83, 220.28]
Sydney MAS 133.00 [ -23.41, 289.41]
MCIC 153.00 [-121.39, 427.39]
MooDS 258.83 [ 8.03, 509.62]
MPIP 93.17 [ -66.35, 252.69]
MUNSTER 7.21 [-148.02, 162.44]
NCNG 97.26 [-149.13, 343.66]
NESDA 21.02 [-168.26, 210.30]
NeuroIMAGE 109.99 [-150.78, 370.76]
NTR 211.90 [ 50.99, 372.81]
OATS 141.10 [ -91.35, 373.55]
PAFIP
PDNZ
PING 84.63 [-136.88, 306.15]
MPRC
PPMI
QTIM 222.02 [ 27.96, 416.07]
SHIP 303.74 [ 175.64, 431.83]
SHIP-Trend 179.59 [ 56.63, 302.55]
SYS
TCD-NUIG -218.16 [-552.72, 116.40]
TOP 117.97 [-100.78, 336.71]
TOP3T 143.06 [-130.14, 416.25]
UiO2016
UiO2017 -125.53 [-467.68, 216.63]
UMCU 101.90 [ -43.78, 247.58]
ENIGMA Discovery Cohorts [N=17794; P=1.311e-13] 116.92 [ 85.98, 147.87]
UKBB 125.12 [ 82.62, 167.61]
Discovery+UKBB [N=27016; P=6.695e-20] 119.88 [ 94.15, 145.61]
GOBS
IMH
UNICAMP
Meth-CT
MIRECC
UKBB nonEUR
OSAKA
PING Non European
Non European Cohorts[]
Cohort
Beta [Lower 95% CI,Upper 95% CI]
-1000
-500
0
500
1000
mm^2/effect allele

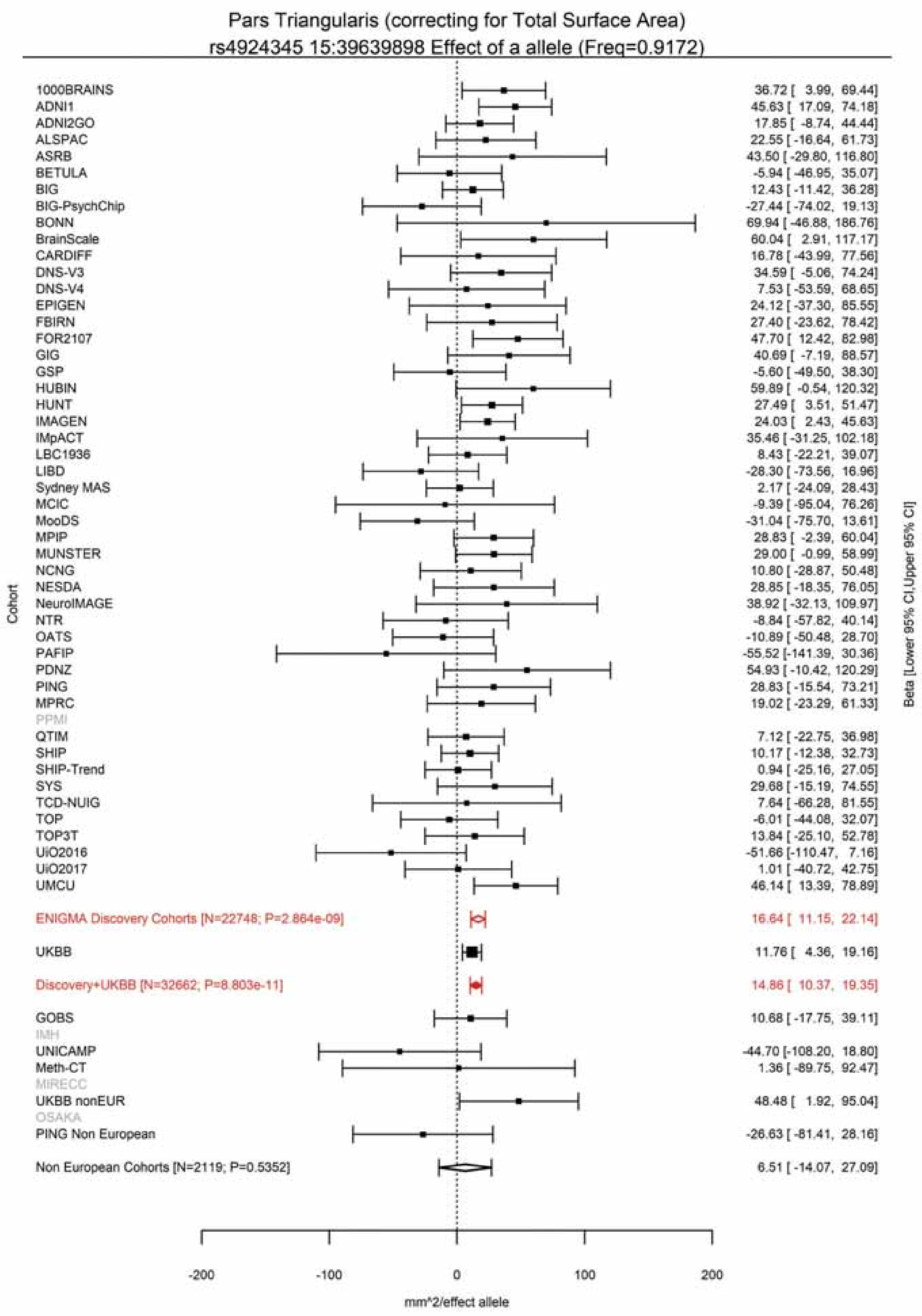
Pars Triangularis (correcting for Total Surface Area)
rs4924345 15:39639898 Effect of a allele (Freq=0.9172)
Cohort
Beta [Lower 95% CI,Upper 95% CI]
1000BRAINS 36.72 [ 3.99, 69.44]
ADNI1 45.63 [ 17.09, 74.18]
ADNI2GO 17.85 [ -8.74, 44.44]
ALSPAC 22.55 [ -16.64, 61.73]
ASRB 43.50 [ -29.80, 116.80]
BETULA -5.94 [ -46.95, 35.07]
BIG 12.43 [ -11.42, 36.28]
BIG-PsychChip -27.44 [ -74.02, 19.13]
BONN 69.94 [ -46.88, 186.76]
BrainScale 60.04 [ 2.91, 117.17]
CARDIFF 16.78 [ -43.99, 77.56]
DNS-V3 34.59 [ -5.06, 74.24]
DNS-V4 7.53 [ -53.59, 68.65]
EPIGEN 24.12 [ -37.30, 85.55]
FBIRN 27.40 [ -23.62, 78.42]
FOR2107 47.70 [ 12.42, 82.98]
GIG 40.69 [ -7.19, 88.57]
GSP -5.60 [ -49.50, 38.30]
HUBIN 59.89 [ -0.54, 120.32]
HUNT 27.49 [ 3.51, 51.47]
IMAGEN 24.03 [ 2.43, 45.63]
IMpACT 35.46 [ -31.25, 102.18]
LBC1936 8.43 [ -22.21, 39.07]
LIBD -28.30 [ -73.56, 16.96]
Sydney MAS 2.17 [ -24.09, 28.43]
MCIC -9.39 [ -95.04, 76.26]
MooDS -31.04 [ -75.70, 13.61]
MPIP 28.83 [ -2.39, 60.04]
MUNSTER 29.00 [ -0.99, 58.99]
NCNG 10.80 [ -28.87, 50.48]
NESDA 28.85 [ -18.35, 76.05]
NeuroIMAGE 38.92 [ -32.13, 109.97]
NTR -8.84 [ -57.82, 40.14]
OATS -10.89 [ -50.48, 28.70]
PAFIP -55.52 [-141.39, 30.36]
PDNZ 54.93 [ -10.42, 120.29]
PING 28.83 [ -15.54, 73.21]
MPRC 19.02 [ -23.29, 61.33]
PPMI
QTIM 7.12 [ -22.75, 36.98]
SHIP 10.17 [ -12.38, 32.73]
SHIP-Trend 0.94 [ -25.16, 27.05]
SYS 29.68 [ -15.19, 74.55]
TCD-NUIG 7.64 [ -66.28, 81.55]
TOP -6.01 [ -44.08, 32.07]
TOP3T 13.84 [ -25.10, 52.78]
UiO2016 -51.66 [-110.47, 7.16]
UiO2017 1.01 [ -40.72, 42.75]
UMCU 46.14 [ 13.39, 78.89]
ENIGMA Discovery Cohorts [N=22748; P=2.864e-09] 16.64 [ 11.15, 22.14]
UKBB 11.76 [ 4.36, 19.16]
Discovery+UKBB [N=32662; P=8.803e-11] 14.86 [ 10.37, 19.35]
GOBS 10.68 [ -17.75, 39.11]
IMH
UNICAMP -44.70 [-108.20, 18.80]
Meth-CT 1.36 [ -89.75, 92.47]
MIRECC
UKBB nonEUR 48.48 [ 1.92, 95.04]
OSAKA
PING Non European -26.63 [ -81.41, 28.16]
Non European Cohorts [N=2119; P=0.5352] 6.51 [ -14.07, 27.09]
-200
-100
0
100
200
mm^2/effect allele

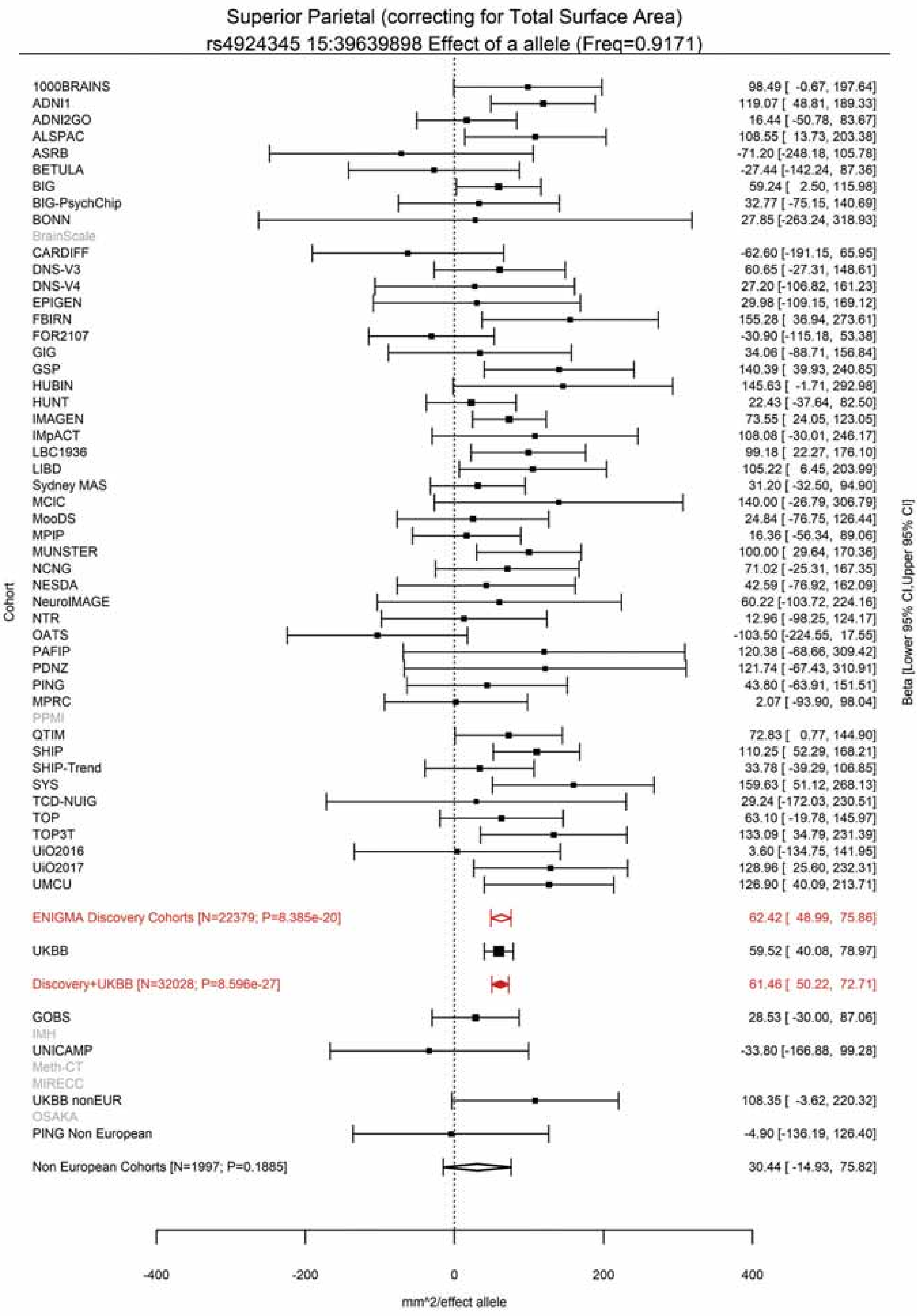

Superior Parietal (correcting for Total Surface Area)
rs4924345 15:39639898 Effect of a allele (Freq=0.9171)
Cohort
1000BRAINS 98.49 [ -0.67, 197.64]
ADNI1 119.07 [ 48.81, 189.33]
ADNI2GO 16.44 [ -50.78, 83.67]
ALSPAC 108.55 [ 13.73, 203.38]
ASRB -71.20 [-248.18, 105.78]
BETULA -27.44 [-142.24, 87.36]
BIG 59.24 [ 2.50, 115.98]
BIG-PsychChip 32.77 [ -75.15, 140.69]
BONN 27.85 [-263.24, 318.93]
BrainScale
CARDIFF -62.60 [-191.15, 65.95]
DNS-V3 60.65 [ -27.31, 148.61]
DNS-V4 27.20 [-106.82, 161.23]
EPIGEN 29.98 [-109.15, 169.12]
FBIRN 155.28 [ 36.94, 273.61]
FOR2107 -30.90 [-115.18, 53.38]
GIG 34.06 [ -88.71, 156.84]
GSP 140.39 [ 39.93, 240.85]
HUBIN 145.63 [ -1.71, 292.98]
HUNT 22.43 [ -37.64, 82.50]
IMAGEN 73.55 [ 24.05, 123.05]
IMpACT 108.08 [ -30.01, 246.17]
LBC1936 99.18 [ 22.27, 176.10]
LIBD 105.22 [ 6.45, 203.99]
Sydney MAS 31.20 [ -32.50, 94.90]
MCIC 140.00 [ -26.79, 306.79]
MooDS 24.84 [ -76.75, 126.44]
MPIP 16.36 [ -56.34, 89.06]
MUNSTER 100.00 [ 29.64, 170.36]
NCNG 71.02 [ -25.31, 167.35]
NESDA 42.59 [ -76.92, 162.09]
NeuroIMAGE 60.22 [-103.72, 224.16]
NTR 12.96 [ -98.25, 124.17]
OATS -103.50 [-224.55, 17.55]
PAFIP 120.38 [ -68.66, 309.42]
PDNZ 121.74 [ -67.43, 310.91]
PING 43.80 [ -63.91, 151.51]
MPRC 2.07 [ -93.90, 98.04]
PPMI
QTIM 72.83 [ 0.77, 144.90]
SHIP 110.25 [ 52.29, 168.21]
SHIP-Trend 33.78 [ -39.29, 106.85]
SYS 159.63 [ 51.12, 268.13]
TCD-NUIG 29.24 [-172.03, 230.51]
TOP 63.10 [ -19.78, 145.97]
TOP3T 133.09 [ 34.79, 231.39]
UiO2016 3.60 [-134.75, 141.95]
UiO2017 128.96 [ 25.60, 232.31]
UMCU 126.90 [ 40.09, 213.71]
ENIGMA Discovery Cohorts [N=22379; P=8.385e-20] 62.42 [ 48.99, 75.86]
UKBB 59.52 [ 40.08, 78.97]
Discovery+UKBB [N=32028; P=8.596e-27] 61.46 [ 50.22, 72.71]
GOBS 28.53 [ -30.00, 87.06]
IMH
UNICAMP -33.80 [-166.88, 99.28]
Meth-CT
MIRECC
UKBB nonEUR 108.35 [ -3.62, 220.32]
OSAKA
PING Non European -4.90 [-136.19, 126.40]
Non European Cohorts [N=1997; P=0.1885] 30.44 [ -14.93, 75.82]
-400
-200
0
200
400
mm^2/effect allele
Beta [Lower 95% CI,Upper 95% CI]

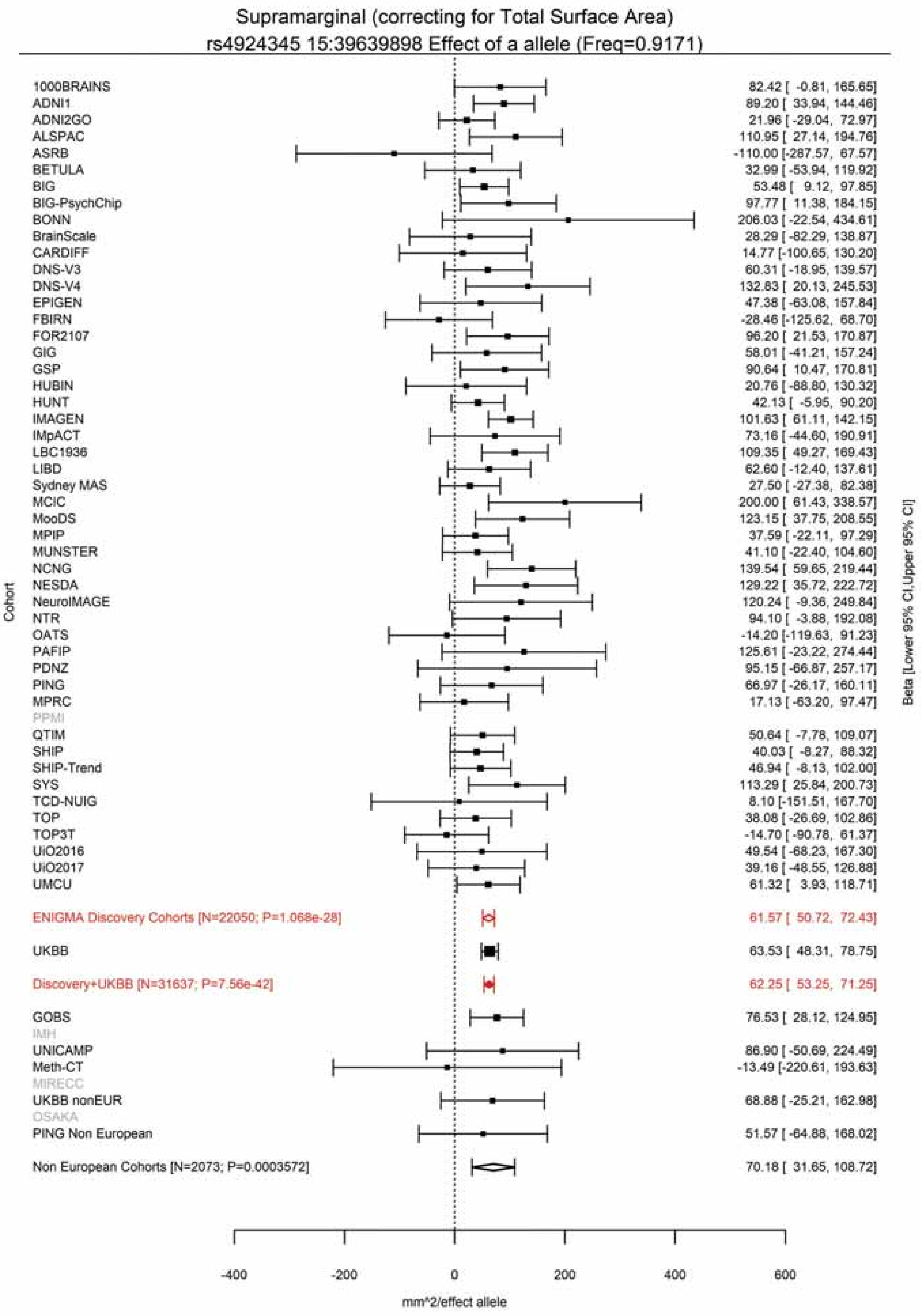
Supramarginal (correcting for Total Surface Area)
rs4924345 15:39639898 Effect of a allele (Freq=0.9171)
Cohort
1000BRAINS 82.42 [ -0.81, 165.65]
ADNI1 89.20 [ 33.94, 144.46]
ADNI2GO 21.96 [ -29.04, 72.97]
ALSPAC 110.95 [ 27.14, 194.76]
ASRB -110.00 [-287.57, 67.57]
BETULA 32.99 [ -53.94, 119.92]
BIG 53.48 [ 9.12, 97.85]
BIG-PsychChip 97.77 [ 11.38, 184.15]
BONN 206.03 [ -22.54, 434.61]
BrainScale 28.29 [ -82.29, 138.87]
CARDIFF 14.77 [-100.65, 130.20]
DNS-V3 60.31 [ -18.95, 139.57]
DNS-V4 132.83 [ 20.13, 245.53]
EPIGEN 47.38 [ -63.08, 157.84]
FBIRN -28.46 [-125.62, 68.70]
FOR2107 96.20 [ 21.53, 170.87]
GIG 58.01 [ -41.21, 157.24]
GSP 90.64 [ 10.47, 170.81]
HUBIN 20.76 [ -88.80, 130.32]
HUNT 42.13 [ -5.95, 90.20]
IMAGEN 101.63 [ 61.11, 142.15]
IMpACT 73.16 [ -44.60, 190.91]
LBC1936 109.35 [ 49.27, 169.43]
LIBD 62.60 [ -12.40, 137.61]
Sydney MAS 27.50 [ -27.38, 82.38]
MCIC 200.00 [ 61.43, 338.57]
MooDS 123.15 [ 37.75, 208.55]
MPIP 37.59 [ -22.11, 97.29]
MUNSTER 41.10 [ -22.40, 104.60]
NCNG 139.54 [ 59.65, 219.44]
NESDA 129.22 [ 35.72, 222.72]
NeuroIMAGE 120.24 [ -9.36, 249.84]
NTR 94.10 [ -3.88, 192.08]
OATS -14.20 [-119.63, 91.23]
PAFIP 125.61 [ -23.22, 274.44]
PDNZ 95.15 [ -66.87, 257.17]
PING 66.97 [ -26.17, 160.11]
MPRC 17.13 [ -63.20, 97.47]
PPMI
QTIM 50.64 [ -7.78, 109.07]
SHIP 40.03 [ -8.27, 88.32]
SHIP-Trend 46.94 [ -8.13, 102.00]
SYS 113.29 [ 25.84, 200.73]
TCD-NUIG 8.10 [-151.51, 167.70]
TOP 38.08 [ -26.69, 102.86]
TOP3T -14.70 [ -90.78, 61.37]
UiO2016 49.54 [ -68.23, 167.30]
UiO2017 39.16 [ -48.55, 126.88]
UMCU 61.32 [ 3.93, 118.71]
ENIGMA Discovery Cohorts [N=22050; P=1.068e-28] 61.57 [ 50.72, 72.43]
UKBB 63.53 [ 48.31, 78.75]
Discovery+UKBB [N=31637; P=7.56e-42] 62.25 [ 53.25, 71.25]
GOBS 76.53 [ 28.12, 124.95]
IMH
UNICAMP 86.90 [ -50.69, 224.49]
Meth-CT -13.49 [-220.61, 193.63]
MIRECC
UKBB nonEUR 68.88 [ -25.21, 162.98]
OSAKA
PING Non European 51.57 [ -64.88, 168.02]
Non European Cohorts [N=2073; P=0.0003572] 70.18 [ 31.65, 108.72]
Beta [Lower 95% CI,Upper 95% CI]
-400
-200
0
200
400
600
mm^2/effect allele

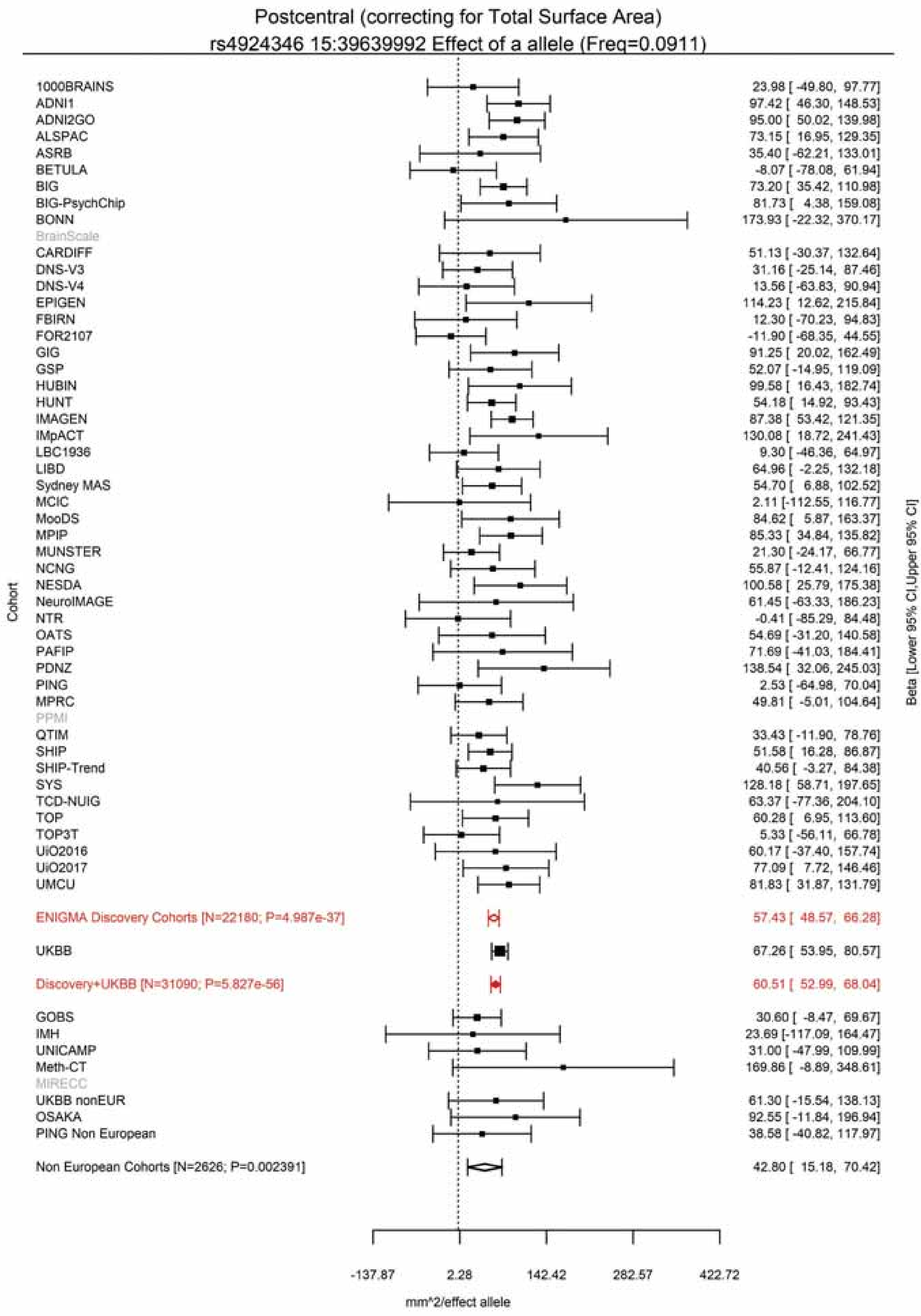
Postcentral (correcting for Total Surface Area)
rs4924346 15:39639992 Effect of a allele (Freq=0.0911)
1000BRAINS 23.98 [ -49.80, 97.77]
ADNI1 97.42 [ 46.30, 148.53]
ADNI2GO 95.00 [ 50.02, 139.98]
ALSPAC 73.15 [ 16.95, 129.35]
ASRB 35.40 [ -62.21, 133.01]
BETULA -8.07 [ -78.08, 61.94]
BIG 73.20 [ 35.42, 110.98]
BIG-PsychChip 81.73 [ 4.38, 159.08]
BONN 173.93 [ -22.32, 370.17]
BrainScale
CARDIFF 51.13 [ -30.37, 132.64]
DNS-V3 31.16 [ -25.14, 87.46]
DNS-V4 13.56 [ -63.83, 90.94]
EPIGEN 114.23 [ 12.62, 215.84]
FBIRN 12.30 [ -70.23, 94.83]
FOR2107 -11.90 [ -68.35, 44.55]
GIG 91.25 [ 20.02, 162.49]
GSP 52.07 [ -14.95, 119.09]
HUBIN 99.58 [ 16.43, 182.74]
HUNT 54.18 [ 14.92, 93.43]
IMAGEN 87.38 [ 53.42, 121.35]
IMpACT 130.08 [ 18.72, 241.43]
LBC1936 9.30 [ -46.36, 64.97]
LIBD 64.96 [ -2.25, 132.18]
Sydney MAS 54.70 [ 6.88, 102.52]
MCIC 2.11 [-112.55, 116.77]
MooDS 84.62 [ 5.87, 163.37]
MPIP 85.33 [ 34.84, 135.82]
MUNSTER 21.30 [ -24.17, 66.77]
NCNG 55.87 [ -12.41, 124.16]
NESDA 100.58 [ 25.79, 175.38]
NeuroIMAGE 61.45 [ -63.33, 186.23]
NTR -0.41 [ -85.29, 84.48]
OATS 54.69 [ -31.20, 140.58]
PAFIP 71.69 [ -41.03, 184.41]
PDNZ 138.54 [ 32.06, 245.03]
PING 2.53 [ -64.98, 70.04]
MPRC 49.81 [ -5.01, 104.64]
PPMI
QTIM 33.43 [ -11.90, 78.76]
SHIP 51.58 [ 16.28, 86.87]
SHIP-Trend 40.56 [ -3.27, 84.38]
SYS 128.18 [ 58.71, 197.65]
TCD-NUIG 63.37 [ -77.36, 204.10]
TOP 60.28 [ 6.95, 113.60]
TOP3T 5.33 [ -56.11, 66.78]
UiO2016 60.17 [ -37.40, 157.74]
UiO2017 77.09 [ 7.72, 146.46]
UMCU 81.83 [ 31.87, 131.79]
ENIGMA Discovery Cohorts [N=22180; P=4.987e-37] 57.43 [ 48.57, 66.28]
UKBB 67.26 [ 53.95, 80.57]
Discovery+UKBB [N=31090; P=5.827e-56] 60.51 [ 52.99, 68.04]
GOBS 30.60 [ -8.47, 69.67]
IMH 23.69 [-117.09, 164.47]
UNICAMP 31.00 [ -47.99, 109.99]
Meth-CT 169.86 [ -8.89, 348.61]
MIRECC
UKBB nonEUR 61.30 [ -15.54, 138.13]
OSAKA 92.55 [ -11.84, 196.94]
PING Non European 38.58 [ -40.82, 117.97]
Non European Cohorts [N=2626; P=0.002391] 42.80 [ 15.18, 70.42]
Cohort
Beta [Lower 95% CI,Upper 95% CI]
-137.87
2.28
142.42
282.57
422.72
mm^2/effect allele

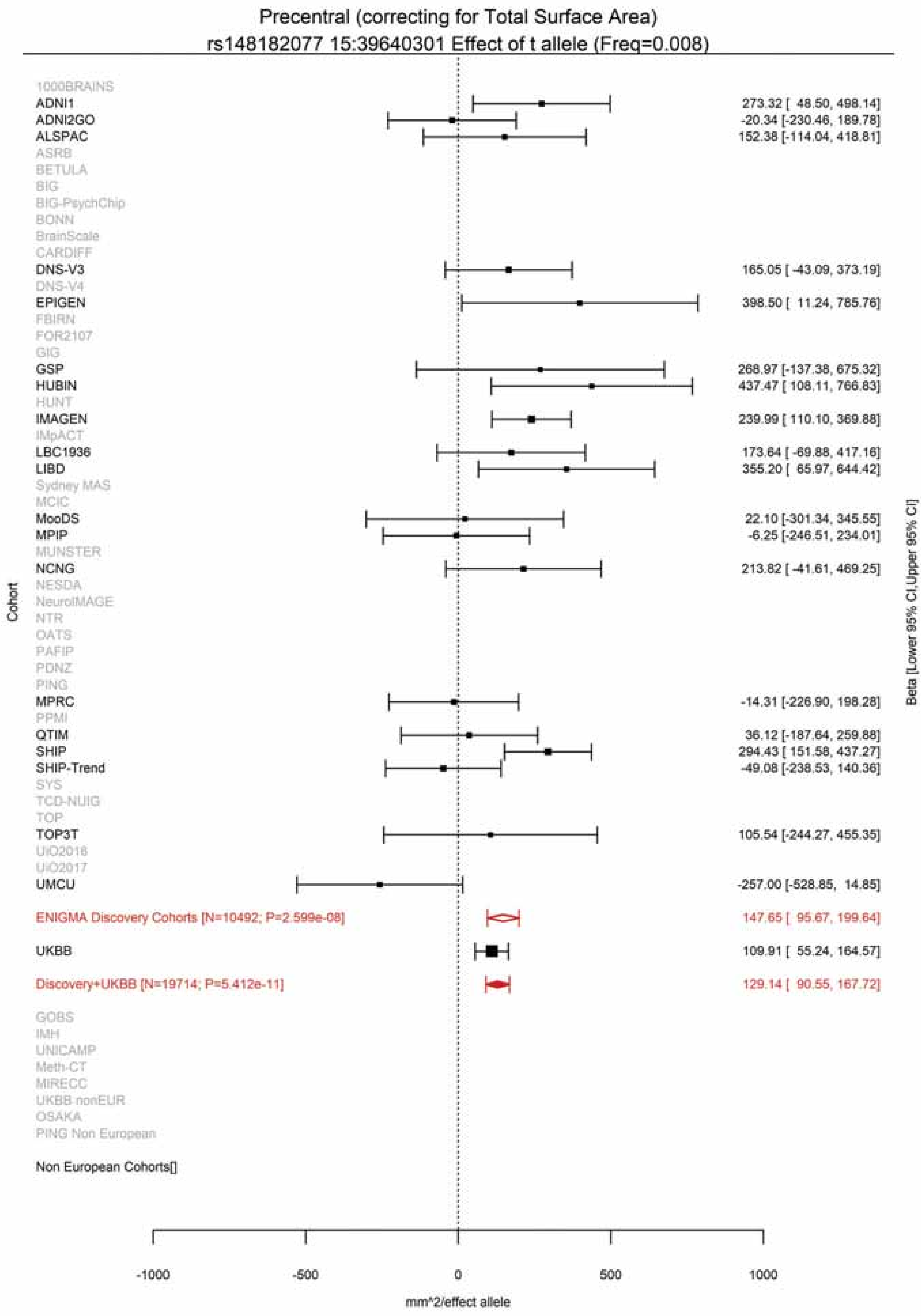

Precentral (correcting for Total Surface Area)
rs148182077 15:39640301 Effect of t allele (Freq=0.008)
Cohort
1000BRAINS
ADNI1 273.32 [ 48.50, 498.14]
ADNI2GO -20.34 [-230.46, 189.78]
ALSPAC 152.38 [-114.04, 418.81]
ASRB
BETULA
BIG
BIG-PsychChip
BONN
BrainScale
CARDIFF
DNS-V3 165.05 [ -43.09, 373.19]
DNS-V4
EPIGEN 398.50 [ 11.24, 785.76]
FBIRN
FOR2107
GIG
GSP 268.97 [-137.38, 675.32]
HUBIN 437.47 [ 108.11, 766.83]
HUNT
IMAGEN 239.99 [ 110.10, 369.88]
IMpACT
LBC1936 173.64 [ -69.88, 417.16]
LIBD 355.20 [ 65.97, 644.42]
Sydney MAS
MCIC
MooDS 22.10 [-301.34, 345.55]
MPIP -6.25 [-246.51, 234.01]
MUNSTER
NCNG 213.82 [ -41.61, 469.25]
NESDA
NeuroIMAGE
NTR
OATS
PAFIP
PDNZ
PING
MPRC -14.31 [-226.90, 198.28]
PPMI
QTIM 36.12 [-187.64, 259.88]
SHIP 294.43 [ 151.58, 437.27]
SHIP-Trend -49.08 [-238.53, 140.36]
SYS
TCD-NUIG
TOP
TOP3T 105.54 [-244.27, 455.35]
UiO2016
UiO2017
UMCU -257.00 [-528.85, 14.85]
ENIGMA Discovery Cohorts [N=10492; P=2.599e-08] 147.65 [ 95.67, 199.64]
UKBB 109.91 [ 55.24, 164.57]
Discovery+UKBB [N=19714; P=5.412e-11] 129.14 [ 90.55, 167.72]
GOBS
IMH
UNICAMP
Meth-CT
MIRECC
UKBB nonEUR
OSAKA
PING Non European
Non European Cohorts[]
Beta [Lower 95% CI,Upper 95% CI]
-1000
-500
0
500
1000
mm^2/effect allele

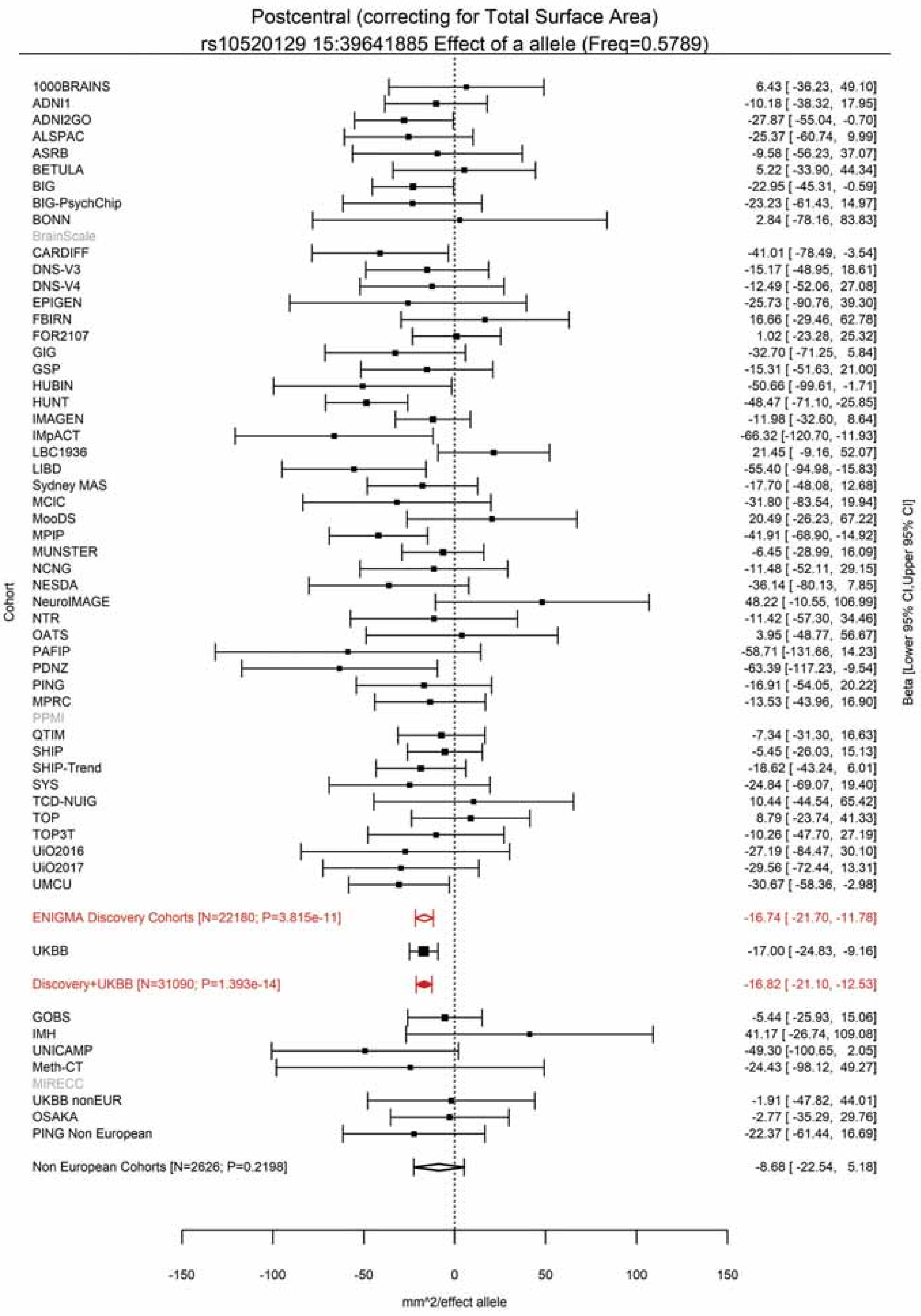
Postcentral (correcting for Total Surface Area)
rs10520129 15:39641885 Effect of a allele (Freq=0.5789)
Cohort
1000BRAINS 6.43 [ -36.23, 49.10]
ADNI1 -10.18 [ -38.32, 17.95]
ADNI2GO -27.87 [ -55.04, -0.70]
ALSPAC -25.37 [ -60.74, 9.99]
ASRB -9.58 [ -56.23, 37.07]
BETULA 5.22 [ -33.90, 44.34]
BIG -22.95 [ -45.31, -0.59]
BIG-PsychChip -23.23 [ -61.43, 14.97]
BONN 2.84 [ -78.16, 83.83]
BrainScale
CARDIFF -41.01 [ -78.49, -3.54]
DNS-V3 -15.17 [ -48.95, 18.61]
DNS-V4 -12.49 [ -52.06, 27.08]
EPIGEN -25.73 [ -90.76, 39.30]
FBIRN 16.66 [ -29.46, 62.78]
FOR2107 1.02 [ -23.28, 25.32]
GIG -32.70 [ -71.25, 5.84]
GSP -15.31 [ -51.63, 21.00]
HUBIN -50.66 [ -99.61, -1.71]
HUNT -48.47 [ -71.10, -25.85]
IMAGEN -11.98 [ -32.60, 8.64]
IMpACT -66.32 [-120.70, -11.93]
LBC1936 21.45 [ -9.16, 52.07]
LIBD -55.40 [ -94.98, -15.83]
Sydney MAS -17.70 [ -48.08, 12.68]
MCIC -31.80 [ -83.54, 19.94]
MooDS 20.49 [ -26.23, 67.22]
MPIP -41.91 [ -68.90, -14.92]
MUNSTER -6.45 [ -28.99, 16.09]
NCNG -11.48 [ -52.11, 29.15]
NESDA -36.14 [ -80.13, 7.85]
NeuroIMAGE 48.22 [ -10.55, 106.99]
NTR -11.42 [ -57.30, 34.46]
OATS 3.95 [ -48.77, 56.67]
PAFIP -58.71 [-131.66, 14.23]
PDNZ -63.39 [-117.23, -9.54]
PING -16.91 [ -54.05, 20.22]
MPRC -13.53 [ -43.96, 16.90]
PPMI
QTIM -7.34 [ -31.30, 16.63]
SHIP -5.45 [ -26.03, 15.13]
SHIP-Trend -18.62 [ -43.24, 6.01]
SYS -24.84 [ -69.07, 19.40]
TCD-NUIG 10.44 [ -44.54, 65.42]
TOP 8.79 [ -23.74, 41.33]
TOP3T -10.26 [ -47.70, 27.19]
UiO2016 -27.19 [ -84.47, 30.10]
UiO2017 -29.56 [ -72.44, 13.31]
UMCU -30.67 [ -58.36, -2.98]
ENIGMA Discovery Cohorts [N=22180; P=3.815e-11] -16.74 [ -21.70, -11.78]
UKBB -17.00 [ -24.83, -9.16]
Discovery+UKBB [N=31090; P=1.393e-14] -16.82 [ -21.10, -12.53]
GOBS -5.44 [ -25.93, 15.06]
IMH 41.17 [ -26.74, 109.08]
UNICAMP -49.30 [-100.65, 2.05]
Meth-CT -24.43 [ -98.12, 49.27]
MIRECC
UKBB nonEUR -1.91 [ -47.82, 44.01]
OSAKA -2.77 [ -35.29, 29.76]
PING Non European -22.37 [ -61.44, 16.69]
Non European Cohorts [N=2626; P=0.2198] -8.68 [ -22.54, 5.18]
Beta [Lower 95% CI,Upper 95% CI]
-150
-100
-50
0
50
100
150
mm^2/effect allele

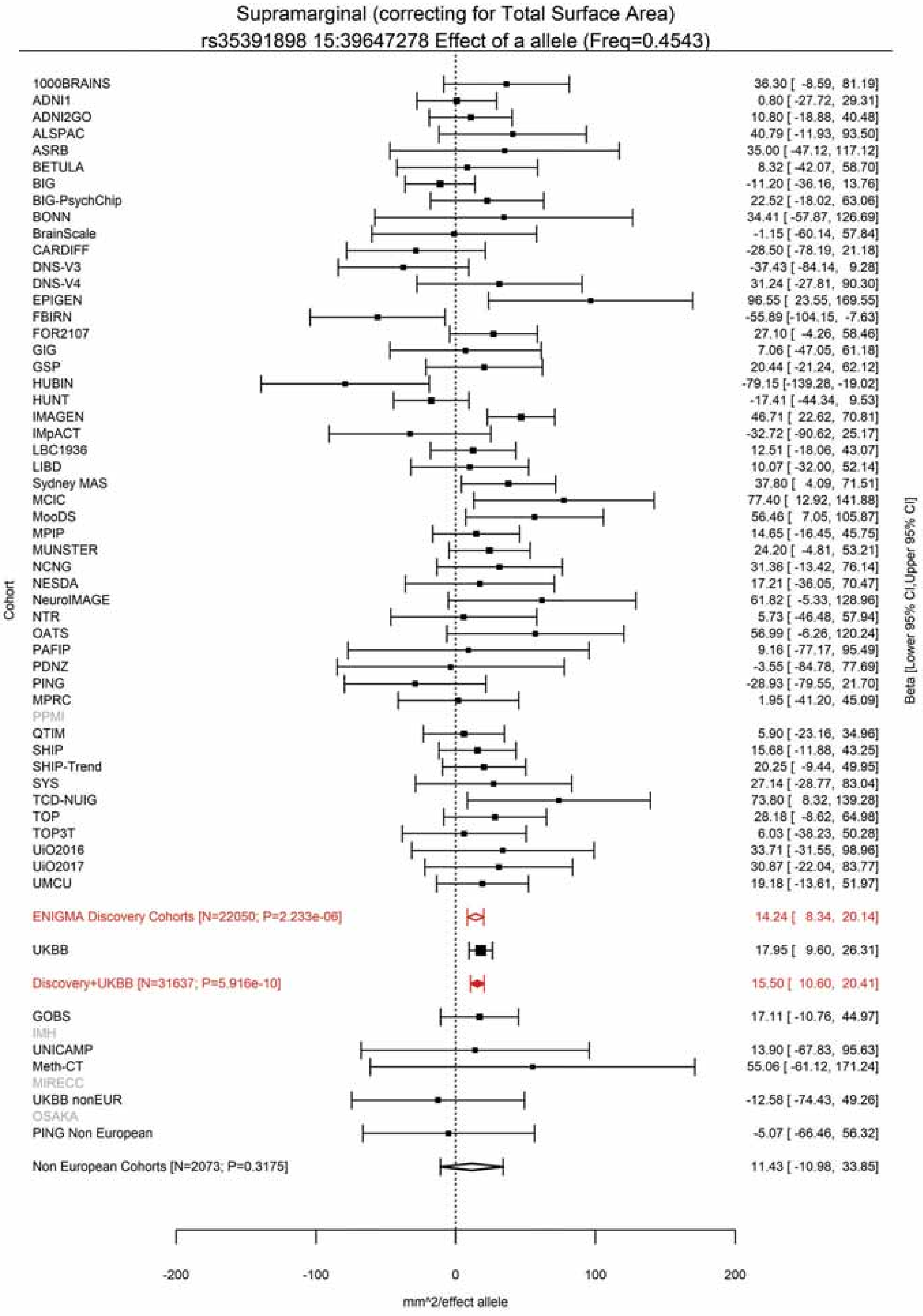

Supramarginal (correcting for Total Surface Area)
rs35391898 15:39647278 Effect of a allele (Freq=0.4543)
Cohort
1000BRAINS 36.30 [ -8.59, 81.19]
ADNI1 0.80 [ -27.72, 29.31]
ADNI2GO 10.80 [ -18.88, 40.48]
ALSPAC 40.79 [ -11.93, 93.50]
ASRB 35.00 [ -47.12, 117.12]
BETULA 8.32 [ -42.07, 58.70]
BIG -11.20 [ -36.16, 13.76]
BIG-PsychChip 22.52 [ -18.02, 63.06]
BONN 34.41 [ -57.87, 126.69]
BrainScale -1.15 [ -60.14, 57.84]
CARDIFF -28.50 [ -78.19, 21.18]
DNS-V3 -37.43 [ -84.14, 9.28]
DNS-V4 31.24 [ -27.81, 90.30]
EPIGEN 96.55 [ 23.55, 169.55]
FBIRN -55.89 [-104.15, -7.63]
FOR2107 27.10 [ -4.26, 58.46]
GIG 7.06 [ -47.05, 61.18]
GSP 20.44 [ -21.24, 62.12]
HUBIN -79.15 [-139.28, -19.02]
HUNT -17.41 [ -44.34, 9.53]
IMAGEN 46.71 [ 22.62, 70.81]
IMpACT -32.72 [ -90.62, 25.17]
LBC1936 12.51 [ -18.06, 43.07]
LIBD 10.07 [ -32.00, 52.14]
Sydney MAS 37.80 [ 4.09, 71.51]
MCIC 77.40 [ 12.92, 141.88]
MooDS 56.46 [ 7.05, 105.87]
MPIP 14.65 [ -16.45, 45.75]
MUNSTER 24.20 [ -4.81, 53.21]
NCNG 31.36 [ -13.42, 76.14]
NESDA 17.21 [ -36.05, 70.47]
NeuroIMAGE 61.82 [ -5.33, 128.96]
NTR 5.73 [ -46.48, 57.94]
OATS 56.99 [ -6.26, 120.24]
PAFIP 9.16 [ -77.17, 95.49]
PDNZ -3.55 [ -84.78, 77.69]
PING -28.93 [ -79.55, 21.70]
MPRC 1.95 [ -41.20, 45.09]
PPMI
QTIM 5.90 [ -23.16, 34.96]
SHIP 15.68 [ -11.88, 43.25]
SHIP-Trend 20.25 [ -9.44, 49.95]
SYS 27.14 [ -28.77, 83.04]
TCD-NUIG 73.80 [ 8.32, 139.28]
TOP 28.18 [ -8.62, 64.98]
TOP3T 6.03 [ -38.23, 50.28]
UiO2016 33.71 [ -31.55, 98.96]
UiO2017 30.87 [ -22.04, 83.77]
UMCU 19.18 [ -13.61, 51.97]
ENIGMA Discovery Cohorts [N=22050; P=2.233e-06] 14.24 [ 8.34, 20.14]
UKBB 17.95 [ 9.60, 26.31]
Discovery+UKBB [N=31637; P=5.916e-10] 15.50 [ 10.60, 20.41]
GOBS 17.11 [ -10.76, 44.97]
IMH
UNICAMP 13.90 [ -67.83, 95.63]
Meth-CT 55.06 [ -61.12, 171.24]
MIRECC
UKBB nonEUR -12.58 [ -74.43, 49.26]
OSAKA
PING Non European -5.07 [ -66.46, 56.32]
Non European Cohorts [N=2073; P=0.3175] 11.43 [ -10.98, 33.85]
Beta [Lower 95% CI,Upper 95% CI]
-200
-100
0
100
200
mm^2/effect allele

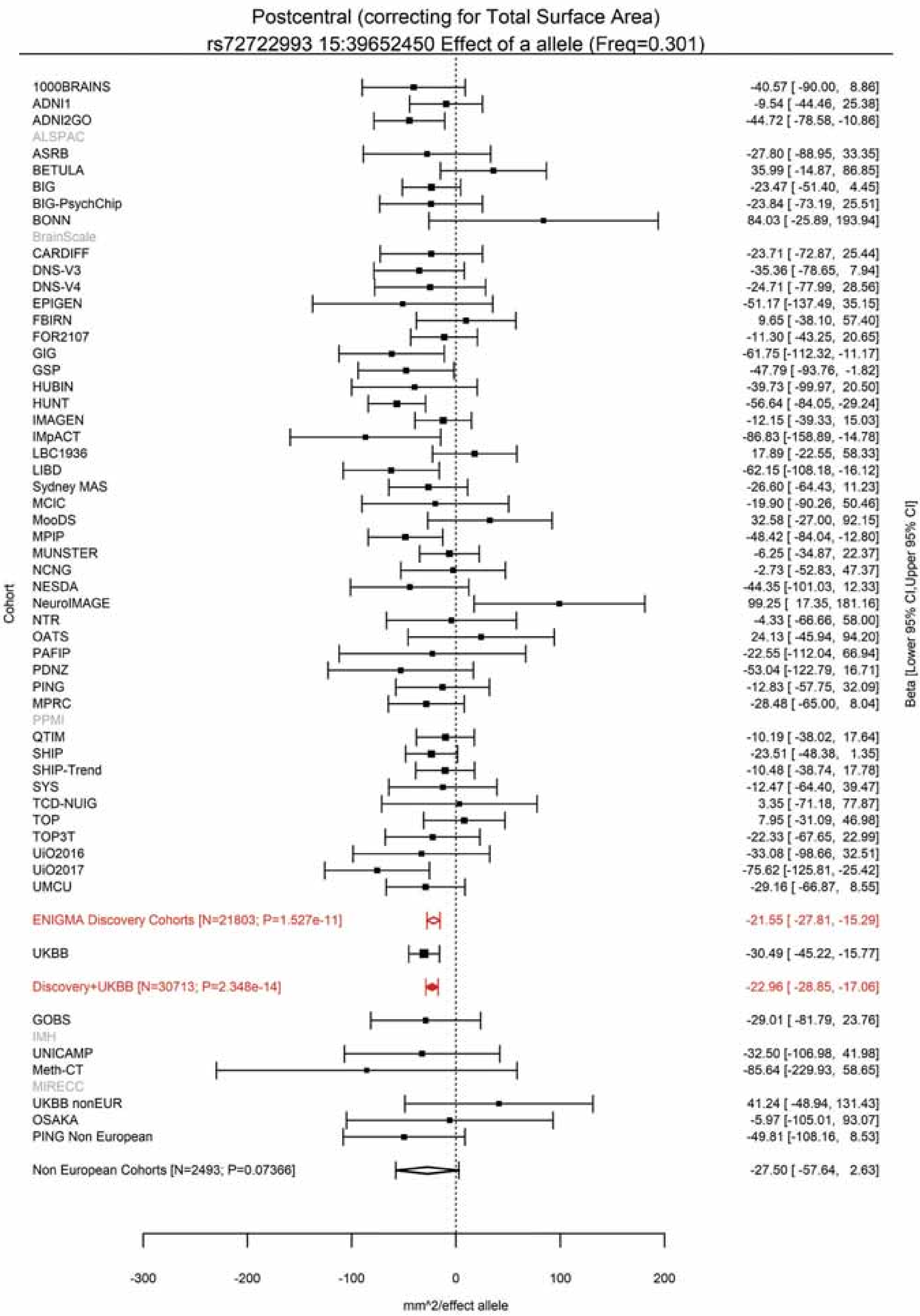

Postcentral (correcting for Total Surface Area)
rs72722993 15:39652450 Effect of a allele (Freq=0.301)
1000BRAINS -40.57 [ -90.00, 8.86]
ADNI1 -9.54 [ -44.46, 25.38]
ADNI2GO -44.72 [ -78.58, -10.86]
ALSPAC
ASRB -27.80 [ -88.95, 33.35]
BETULA 35.99 [ -14.87, 86.85]
BIG -23.47 [ -51.40, 4.45]
BIG-PsychChip -23.84 [ -73.19, 25.51]
BONN 84.03 [ -25.89, 193.94]
BrainScale
CARDIFF -23.71 [ -72.87, 25.44]
DNS-V3 -35.36 [ -78.65, 7.94]
DNS-V4 -24.71 [ -77.99, 28.56]
EPIGEN -51.17 [-137.49, 35.15]
FBIRN 9.65 [ -38.10, 57.40]
FOR2107 -11.30 [ -43.25, 20.65]
GIG -61.75 [-112.32, -11.17]
GSP -47.79 [ -93.76, -1.82]
HUBIN -39.73 [ -99.97, 20.50]
HUNT -56.64 [ -84.05, -29.24]
IMAGEN -12.15 [ -39.33, 15.03]
IMpACT -86.83 [-158.89, -14.78]
LBC1936 17.89 [ -22.55, 58.33]
LIBD -62.15 [-108.18, -16.12]
Sydney MAS -26.60 [ -64.43, 11.23]
MCIC -19.90 [ -90.26, 50.46]
MooDS 32.58 [ -27.00, 92.15]
MPIP -48.42 [ -84.04, -12.80]
MUNSTER -6.25 [ -34.87, 22.37]
NCNG -2.73 [ -52.83, 47.37]
NESDA -44.35 [-101.03, 12.33]
NeuroIMAGE 99.25 [ 17.35, 181.16]
NTR -4.33 [ -66.66, 58.00]
OATS 24.13 [ -45.94, 94.20]
PAFIP -22.55 [-112.04, 66.94]
PDNZ -53.04 [-122.79, 16.71]
PING -12.83 [ -57.75, 32.09]
MPRC -28.48 [ -65.00, 8.04]
PPMI
QTIM -10.19 [ -38.02, 17.64]
SHIP -23.51 [ -48.38, 1.35]
SHIP-Trend -10.48 [ -38.74, 17.78]
SYS -12.47 [ -64.40, 39.47]
TCD-NUIG 3.35 [ -71.18, 77.87]
TOP 7.95 [ -31.09, 46.98]
TOP3T -22.33 [ -67.65, 22.99]
UiO2016 -33.08 [ -98.66, 32.51]
UiO2017 -75.62 [-125.81, -25.42]
UMCU -29.16 [ -66.87, 8.55]
ENIGMA Discovery Cohorts [N=21803; P=1.527e-11] -21.55 [ -27.81, -15.29]
UKBB -30.49 [ -45.22, -15.77]
Discovery+UKBB [N=30713; P=2.348e-14] -22.96 [ -28.85, -17.06]
GOBS -29.01 [ -81.79, 23.76]
IMH
UNICAMP -32.50 [-106.98, 41.98]
Meth-CT -85.64 [-229.93, 58.65]
MIRECC
UKBB nonEUR 41.24 [ -48.94, 131.43]
OSAKA -5.97 [-105.01, 93.07]
PING Non European -49.81 [-108.16, 8.53]
Non European Cohorts [N=2493; P=0.07366] -27.50 [ -57.64, 2.63]
Cohort
Beta [Lower 95% CI,Upper 95% CI]
-300
-200
-100
0
100
200
mm^2/effect allele

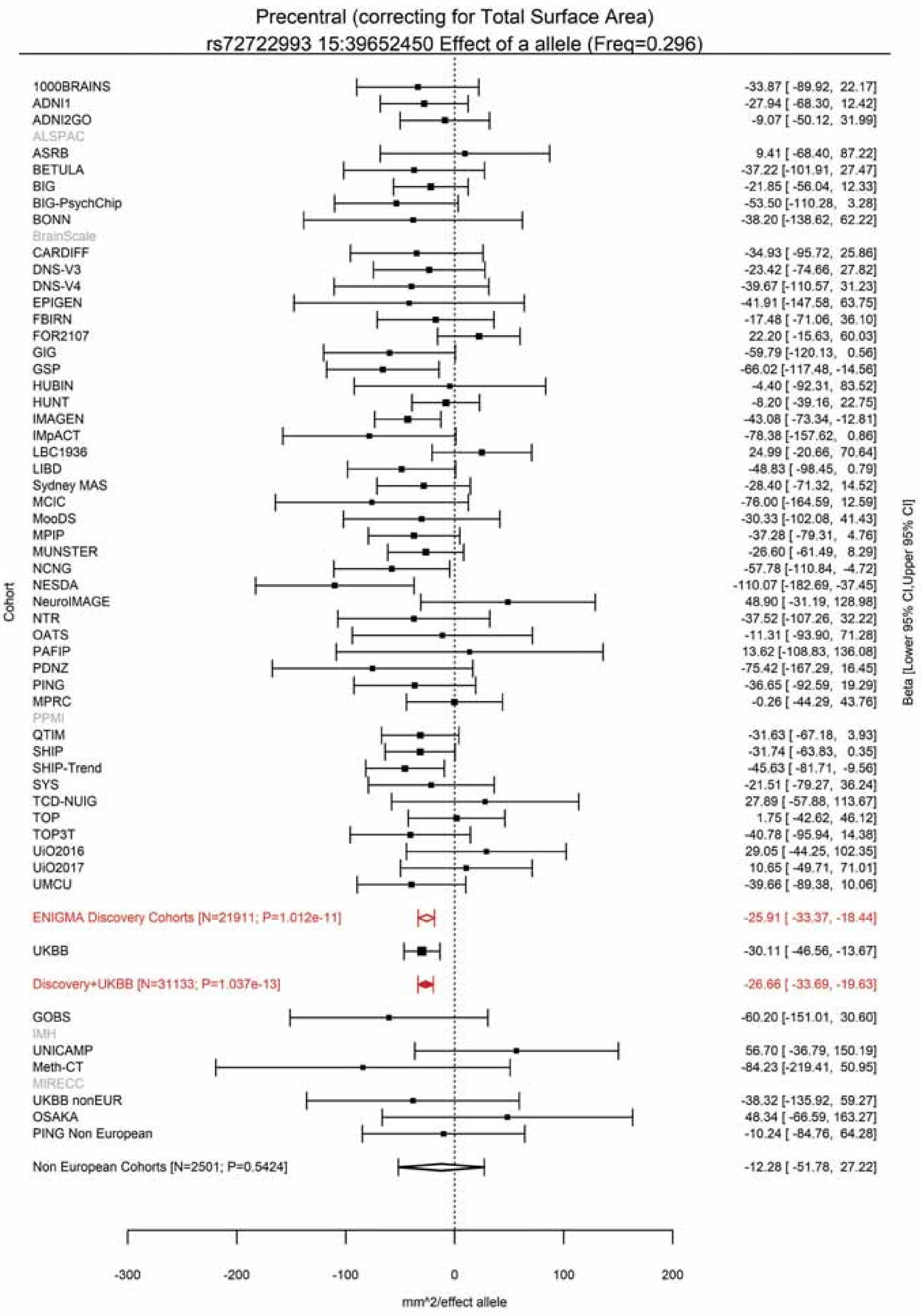

Precentral (correcting for Total Surface Area)
rs72722993 15:39652450 Effect of a allele (Freq=0.296)
Cohort
1000BRAINS -33.87 [ -89.92, 22.17]
ADNI1 -27.94 [ -68.30, 12.42]
ADNI2GO -9.07 [ -50.12, 31.99]
ALSPAC
ASRB 9.41 [ -68.40, 87.22]
BETULA -37.22 [-101.91, 27.47]
BIG -21.85 [ -56.04, 12.33]
BIG-PsychChip -53.50 [-110.28, 3.28]
BONN -38.20 [-138.62, 62.22]
BrainScale
CARDIFF -34.93 [ -95.72, 25.86]
DNS-V3 -23.42 [ -74.66, 27.82]
DNS-V4 -39.67 [-110.57, 31.23]
EPIGEN -41.91 [-147.58, 63.75]
FBIRN -17.48 [ -71.06, 36.10]
FOR2107 22.20 [ -15.63, 60.03]
GIG -59.79 [-120.13, 0.56]
GSP -66.02 [-117.48, -14.56]
HUBIN -4.40 [ -92.31, 83.52]
HUNT -8.20 [ -39.16, 22.75]
IMAGEN -43.08 [ -73.34, -12.81]
IMpACT -78.38 [-157.62, 0.86]
LBC1936 24.99 [ -20.66, 70.64]
LIBD -48.83 [ -98.45, 0.79]
Sydney MAS -28.40 [ -71.32, 14.52]
MCIC -76.00 [-164.59, 12.59]
MooDS -30.33 [-102.08, 41.43]
MPIP -37.28 [ -79.31, 4.76]
MUNSTER -26.60 [ -61.49, 8.29]
NCNG -57.78 [-110.84, -4.72]
NESDA -110.07 [-182.69, -37.45]
NeuroIMAGE 48.90 [ -31.19, 128.98]
NTR -37.52 [-107.26, 32.22]
OATS -11.31 [ -93.90, 71.28]
PAFIP 13.62 [-108.83, 136.08]
PDNZ -75.42 [-167.29, 16.45]
PING -36.65 [ -92.59, 19.29]
MPRC -0.26 [ -44.29, 43.76]
PPMI
QTIM -31.63 [ -67.18, 3.93]
SHIP -31.74 [ -63.83, 0.35]
SHIP-Trend -45.63 [ -81.71, -9.56]
SYS -21.51 [ -79.27, 36.24]
TCD-NUIG 27.89 [ -57.88, 113.67]
TOP 1.75 [ -42.62, 46.12]
TOP3T -40.78 [ -95.94, 14.38]
UiO2016 29.05 [ -44.25, 102.35]
UiO2017 10.65 [ -49.71, 71.01]
UMCU -39.66 [ -89.38, 10.06]
ENIGMA Discovery Cohorts [N=21911; P=1.012e-11] -25.91 [ -33.37, -18.44]
UKBB -30.11 [ -46.56, -13.67]
Discovery+UKBB [N=31133; P=1.037e-13] -26.66 [ -33.69, -19.63]
GOBS -60.20 [-151.01, 30.60]
IMH
UNICAMP 56.70 [ -36.79, 150.19]
Meth-CT -84.23 [-219.41, 50.95]
MIRECC
UKBB nonEUR -38.32 [-135.92, 59.27]
OSAKA 48.34 [ -66.59, 163.27]
PING Non European -10.24 [ -84.76, 64.28]
Non European Cohorts [N=2501; P=0.5424] -12.28 [ -51.78, 27.22]
-300
-200
-100
0
100
200
mm^2/effect allele
Beta [Lower 95% CI,Upper 95% CI]

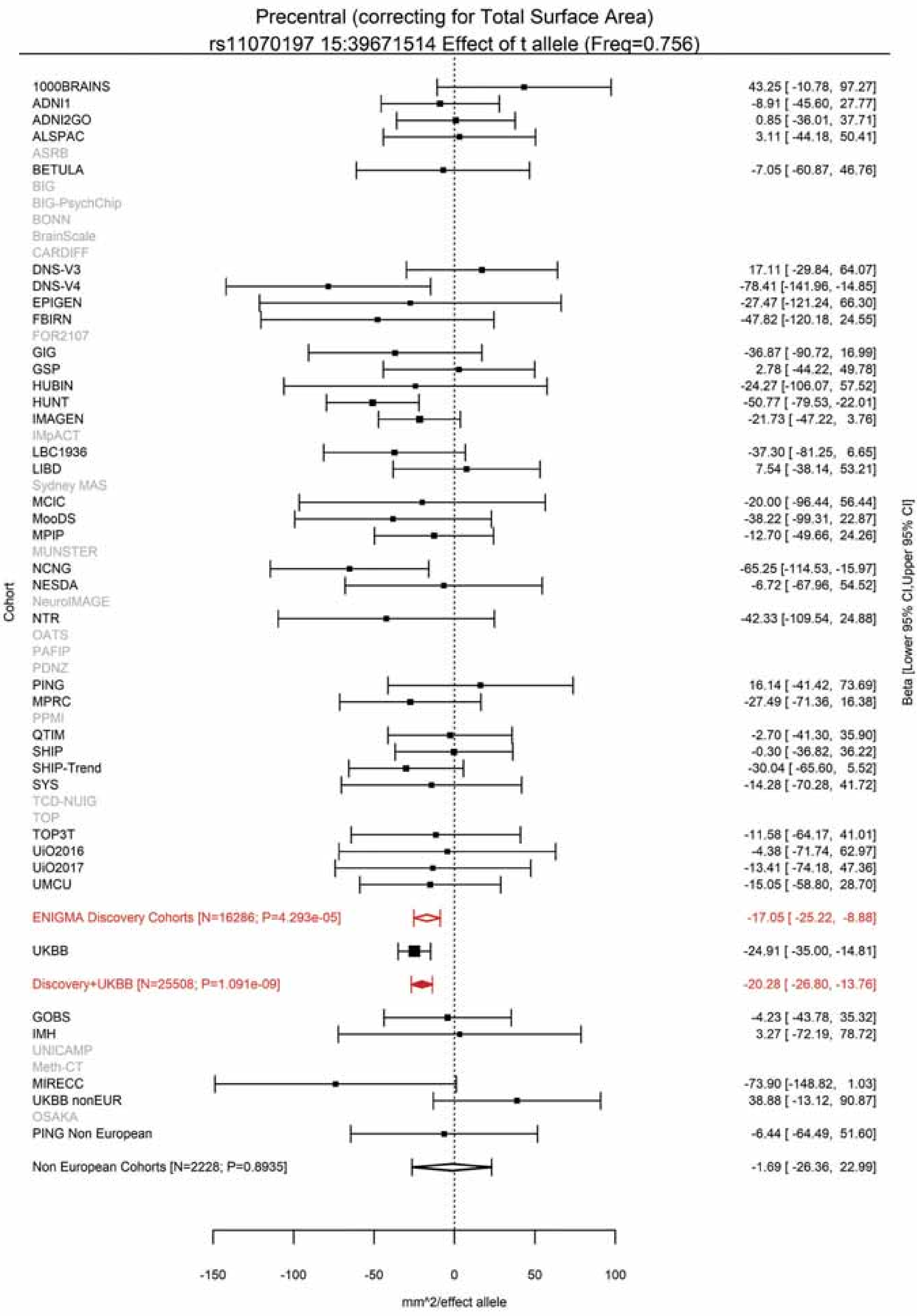

Precentral (correcting for Total Surface Area)
rs11070197 15:39671514 Effect of t allele (Freq=0.756)
1000BRAINS 43.25 [ -10.78, 97.27]
ADNI1 -8.91 [ -45.60, 27.77]
ADNI2GO 0.85 [ -36.01, 37.71]
ALSPAC 3.11 [ -44.18, 50.41]
ASRB
BETULA -7.05 [ -60.87, 46.76]
BIG
BIG-PsychChip
BONN
BrainScale
CARDIFF
DNS-V3 17.11 [ -29.84, 64.07]
DNS-V4 -78.41 [-141.96, -14.85]
EPIGEN -27.47 [-121.24, 66.30]
FBIRN -47.82 [-120.18, 24.55]
FOR2107
GIG -36.87 [ -90.72, 16.99]
GSP 2.78 [ -44.22, 49.78]
HUBIN -24.27 [-106.07, 57.52]
HUNT -50.77 [ -79.53, -22.01]
IMAGEN -21.73 [ -47.22, 3.76]
IMpACT
LBC1936 -37.30 [ -81.25, 6.65]
LIBD 7.54 [ -38.14, 53.21]
Sydney MAS
MCIC -20.00 [ -96.44, 56.44]
MooDS -38.22 [ -99.31, 22.87]
MPIP -12.70 [ -49.66, 24.26]
MUNSTER
NCNG -65.25 [-114.53, -15.97]
NESDA -6.72 [ -67.96, 54.52]
NeuroIMAGE
NTR -42.33 [-109.54, 24.88]
OATS
PAFIP
PDNZ
PING 16.14 [ -41.42, 73.69]
MPRC -27.49 [ -71.36, 16.38]
PPMI
QTIM -2.70 [ -41.30, 35.90]
SHIP -0.30 [ -36.82, 36.22]
SHIP-Trend -30.04 [ -65.60, 5.52]
SYS -14.28 [ -70.28, 41.72]
TCD-NUIG
TOP
TOP3T -11.58 [ -64.17, 41.01]
UiO2016 -4.38 [ -71.74, 62.97]
UiO2017 -13.41 [ -74.18, 47.36]
UMCU -15.05 [ -58.80, 28.70]
ENIGMA Discovery Cohorts [N=16286; P=4.293e-05] -17.05 [ -25.22, -8.88]
UKBB -24.91 [ -35.00, -14.81]
Discovery+UKBB [N=25508; P=1.091e-09] -20.28 [ -26.80, -13.76]
GOBS -4.23 [ -43.78, 35.32]
IMH 3.27 [ -72.19, 78.72]
UNICAMP
Meth-CT
MIRECC -73.90 [-148.82, 1.03]
UKBB nonEUR 38.88 [ -13.12, 90.87]
OSAKA
PING Non European -6.44 [ -64.49, 51.60]
Non European Cohorts [N=2228; P=0.8935] -1.69 [ -26.36, 22.99]
Cohort
Beta [Lower 95% CI,Upper 95% CI]
-150
-100
-50
0
50
100
mm^2/effect allele

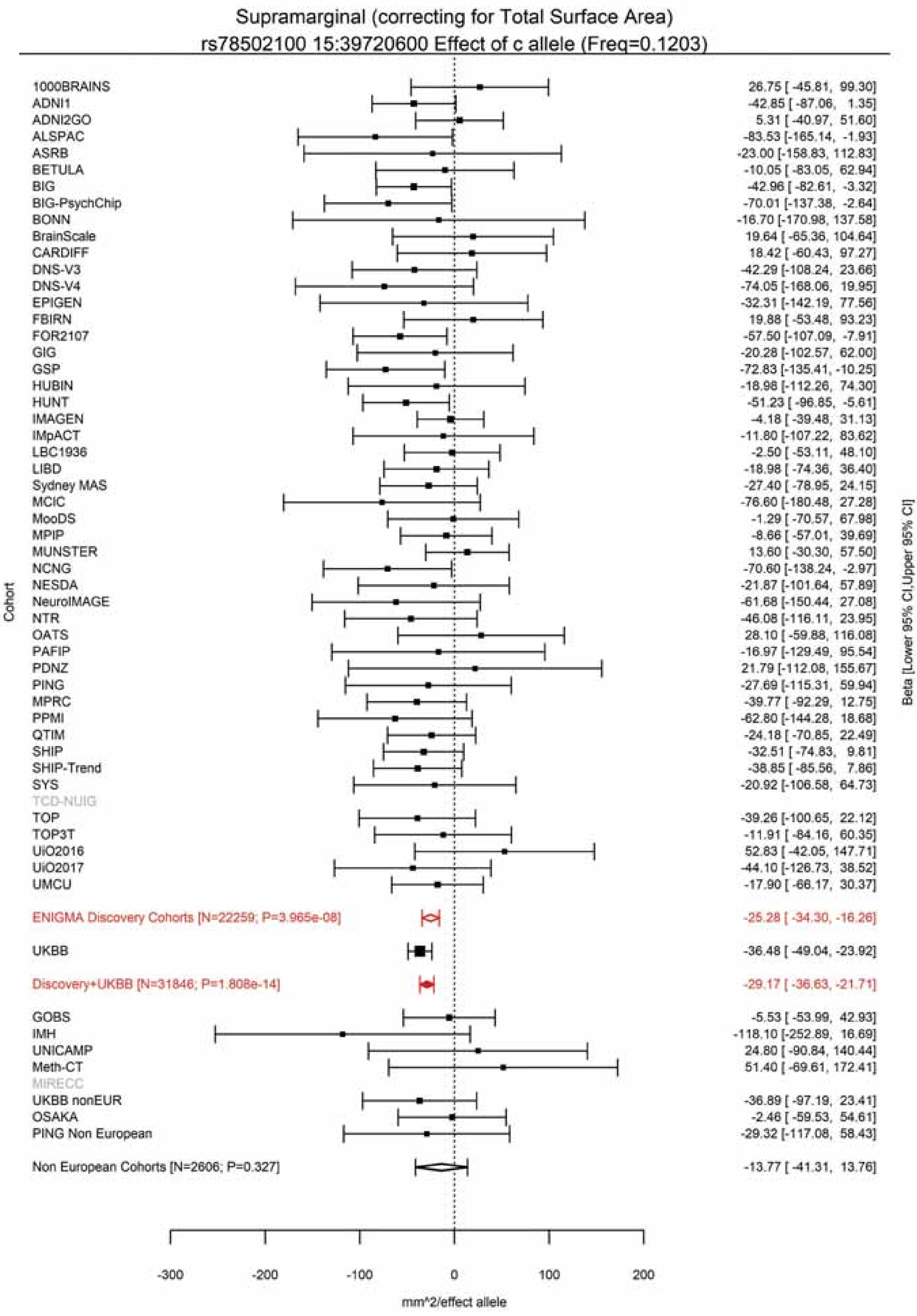

Supramarginal (correcting for Total Surface Area)
rs78502100 15:39720600 Effect of c allele (Freq=0.1203)
Cohort
1000BRAINS 26.75 [ -45.81, 99.30]
ADNI1 -42.85 [ -87.06, 1.35]
ADNI2GO 5.31 [ -40.97, 51.60]
ALSPAC -83.53 [-165.14, -1.93]
ASRB -23.00 [-158.83, 112.83]
BETULA -10.05 [ -83.05, 62.94]
BIG -42.96 [ -82.61, -3.32]
BIG-PsychChip -70.01 [-137.38, -2.64]
BONN -16.70 [-170.98, 137.58]
BrainScale 19.64 [ -65.36, 104.64]
CARDIFF 18.42 [ -60.43, 97.27]
DNS-V3 -42.29 [-108.24, 23.66]
DNS-V4 -74.05 [-168.06, 19.95]
EPIGEN -32.31 [-142.19, 77.56]
FBIRN 19.88 [ -53.48, 93.23]
FOR2107 -57.50 [-107.09, -7.91]
GIG -20.28 [-102.57, 62.00]
GSP -72.83 [-135.41, -10.25]
HUBIN -18.98 [-112.26, 74.30]
HUNT -51.23 [ -96.85, -5.61]
IMAGEN -4.18 [ -39.48, 31.13]
IMpACT -11.80 [-107.22, 83.62]
LBC1936 -2.50 [ -53.11, 48.10]
LIBD -18.98 [ -74.36, 36.40]
Sydney MAS -27.40 [ -78.95, 24.15]
MCIC -76.60 [-180.48, 27.28]
MooDS -1.29 [ -70.57, 67.98]
MPIP -8.66 [ -57.01, 39.69]
MUNSTER 13.60 [ -30.30, 57.50]
NCNG -70.60 [-138.24, -2.97]
NESDA -21.87 [-101.64, 57.89]
NeuroIMAGE -61.68 [-150.44, 27.08]
NTR -46.08 [-116.11, 23.95]
OATS 28.10 [ -59.88, 116.08]
PAFIP -16.97 [-129.49, 95.54]
PDNZ 21.79 [-112.08, 155.67]
PING -27.69 [-115.31, 59.94]
MPRC -39.77 [ -92.29, 12.75]
PPMI -62.80 [-144.28, 18.68]
QTIM -24.18 [ -70.85, 22.49]
SHIP -32.51 [ -74.83, 9.81]
SHIP-Trend -38.85 [ -85.56, 7.86]
SYS -20.92 [-106.58, 64.73]
TCD-NUIG
TOP -39.26 [-100.65, 22.12]
TOP3T -11.91 [ -84.16, 60.35]
UiO2016 52.83 [ -42.05, 147.71]
UiO2017 -44.10 [-126.73, 38.52]
UMCU -17.90 [ -66.17, 30.37]
ENIGMA Discovery Cohorts [N=22259; P=3.965e-08] -25.28 [ -34.30, -16.26]
UKBB -36.48 [ -49.04, -23.92]
Discovery+UKBB [N=31846; P=1.808e-14] -29.17 [ -36.63, -21.71]
GOBS -5.53 [ -53.99, 42.93]
IMH -118.10 [-252.89, 16.69]
UNICAMP 24.80 [ -90.84, 140.44]
Meth-CT 51.40 [ -69.61, 172.41]
MIRECC
UKBB nonEUR -36.89 [ -97.19, 23.41]
OSAKA -2.46 [ -59.53, 54.61]
PING Non European -29.32 [-117.08, 58.43]
Non European Cohorts [N=2606; P=0.327] -13.77 [ -41.31, 13.76]
-300
-200
-100
0
100
200
mm^2/effect allele
Beta [Lower 95% CI,Upper 95% CI]

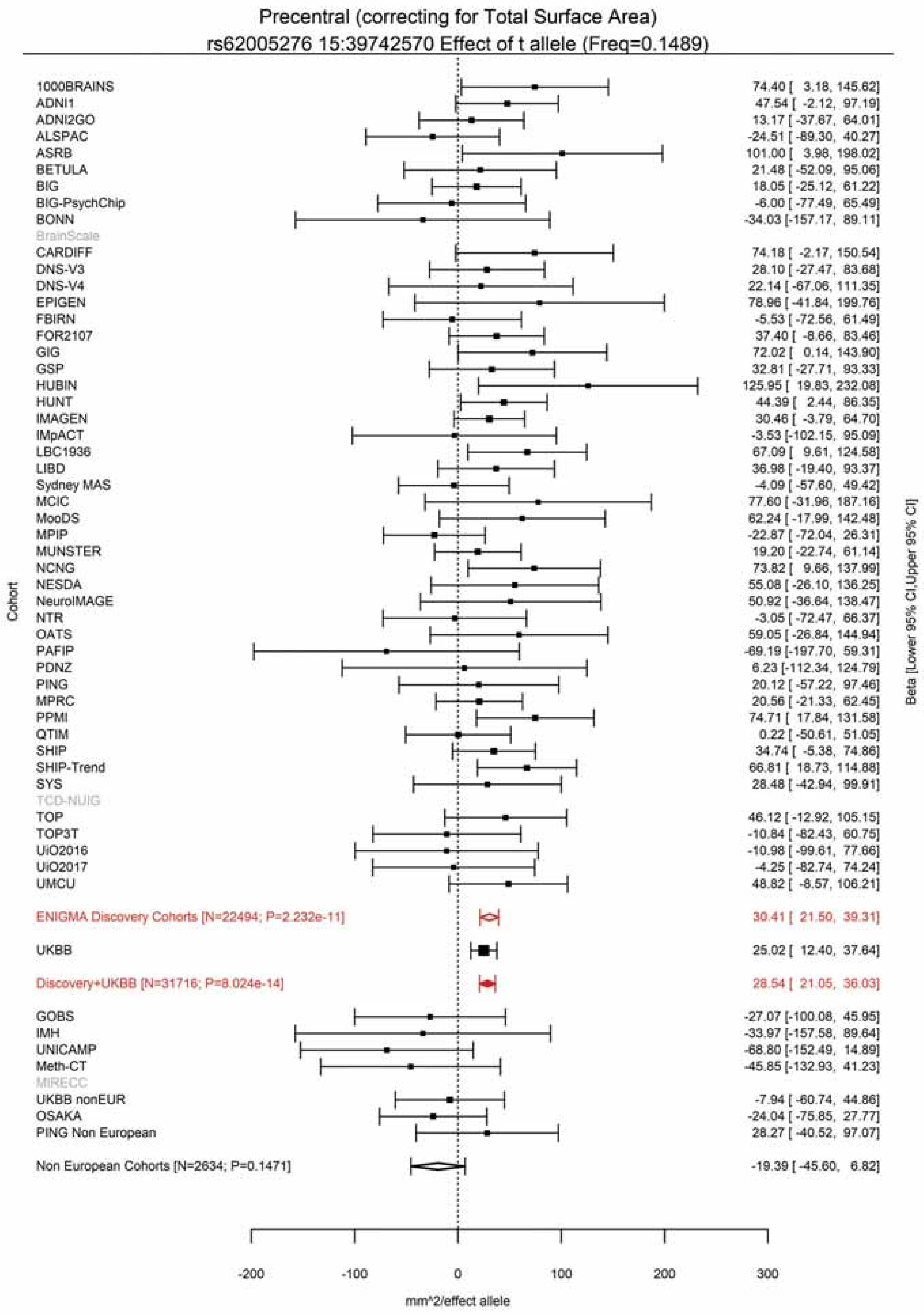
Precentral (correcting for Total Surface Area)
rs62005276 15:39742570 Effect of t allele (Freq=0.1489)
Cohort
1000BRAINS 74.40 [ 3.18, 145.62]
ADNI1 47.54 [ -2.12, 97.19]
ADNI2GO 13.17 [ -37.67, 64.01]
ALSPAC -24.51 [ -89.30, 40.27]
ASRB 101.00 [ 3.98, 198.02]
BETULA 21.48 [ -52.09, 95.06]
BIG 18.05 [ -25.12, 61.22]
BIG-PsychChip -6.00 [ -77.49, 65.49]
BONN -34.03 [-157.17, 89.11]
BrainScale
CARDIFF 74.18 [ -2.17, 150.54]
DNS-V3 28.10 [ -27.47, 83.68]
DNS-V4 22.14 [ -67.06, 111.35]
EPIGEN 78.96 [ -41.84, 199.76]
FBIRN -5.53 [ -72.56, 61.49]
FOR2107 37.40 [ -8.66, 83.46]
GIG 72.02 [ 0.14, 143.90]
GSP 32.81 [ -27.71, 93.33]
HUBIN 125.95 [ 19.83, 232.08]
HUNT 44.39 [ 2.44, 86.35]
IMAGEN 30.46 [ -3.79, 64.70]
IMpACT -3.53 [-102.15, 95.09]
LBC1936 67.09 [ 9.61, 124.58]
LIBD 36.98 [ -19.40, 93.37]
Sydney MAS -4.09 [ -57.60, 49.42]
MCIC 77.60 [ -31.96, 187.16]
MooDS 62.24 [ -17.99, 142.48]
MPIP -22.87 [ -72.04, 26.31]
MUNSTER 19.20 [ -22.74, 61.14]
NCNG 73.82 [ 9.66, 137.99]
NESDA 55.08 [ -26.10, 136.25]
NeuroIMAGE 50.92 [ -36.64, 138.47]
NTR -3.05 [ -72.47, 66.37]
OATS 59.05 [ -26.84, 144.94]
PAFIP -69.19 [-197.70, 59.31]
PDNZ 6.23 [-112.34, 124.79]
PING 20.12 [ -57.22, 97.46]
MPRC 20.56 [ -21.33, 62.45]
PPMI 74.71 [ 17.84, 131.58]
QTIM 0.22 [ -50.61, 51.05]
SHIP 34.74 [ -5.38, 74.86]
SHIP-Trend 66.81 [ 18.73, 114.88]
SYS 28.48 [ -42.94, 99.91]
TCD-NUIG
TOP 46.12 [ -12.92, 105.15]
TOP3T -10.84 [ -82.43, 60.75]
UiO2016 -10.98 [ -99.61, 77.66]
UiO2017 -4.25 [ -82.74, 74.24]
UMCU 48.82 [ -8.57, 106.21]
ENIGMA Discovery Cohorts [N=22494; P=2.232e-11] 30.41 [ 21.50, 39.31]
UKBB 25.02 [ 12.40, 37.64]
Discovery+UKBB [N=31716; P=8.024e-14] 28.54 [ 21.05, 36.03]
GOBS -27.07 [-100.08, 45.95]
IMH -33.97 [-157.58, 89.64]
UNICAMP -68.80 [-152.49, 14.89]
Meth-CT -45.85 [-132.93, 41.23]
MIRECC
UKBB nonEUR -7.94 [ -60.74, 44.86]
OSAKA -24.04 [ -75.85, 27.77]
PING Non European 28.27 [ -40.52, 97.07]
Non European Cohorts [N=2634; P=0.1471] -19.39 [ -45.60, 6.82]
-200
-100
0
100
200
300
mm^2/effect allele
Beta [Lower 95% CI,Upper 95% CI]

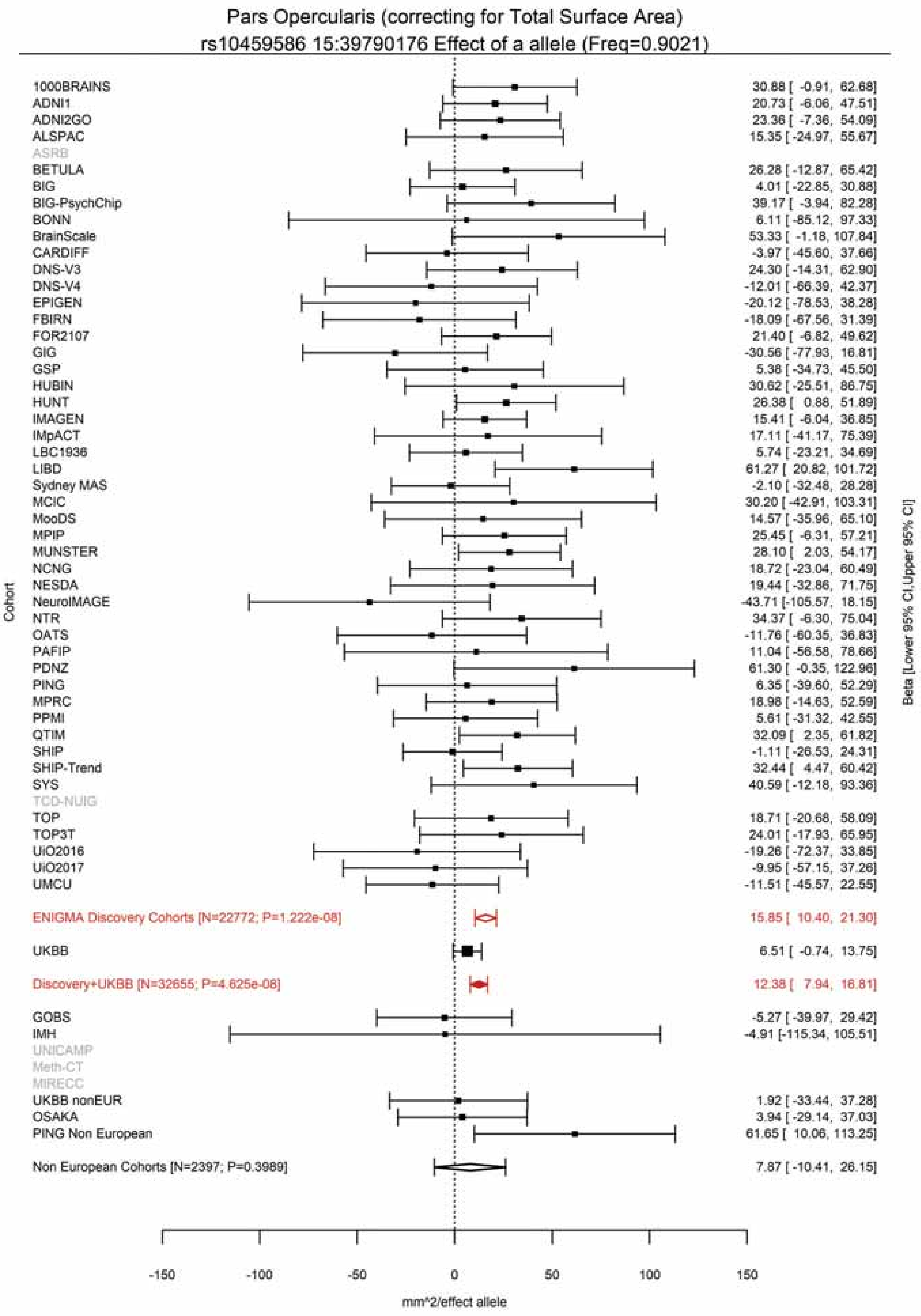

Pars Opercularis (correcting for Total Surface Area)
rs10459586 15:39790176 Effect of a allele (Freq=0.9021)
Cohort
1000BRAINS 30.88 [ -0.91, 62.68]
ADNI1 20.73 [ -6.06, 47.51]
ADNI2GO 23.36 [ -7.36, 54.09]
ALSPAC 15.35 [ -24.97, 55.67]
ASRB
BETULA 26.28 [ -12.87, 65.42]
BIG 4.01 [ -22.85, 30.88]
BIG-PsychChip 39.17 [ -3.94, 82.28]
BONN 6.11 [ -85.12, 97.33]
BrainScale 53.33 [ -1.18, 107.84]
CARDIFF -3.97 [ -45.60, 37.66]
DNS-V3 24.30 [ -14.31, 62.90]
DNS-V4 -12.01 [ -66.39, 42.37]
EPIGEN -20.12 [ -78.53, 38.28]
FBIRN -18.09 [ -67.56, 31.39]
FOR2107 21.40 [ -6.82, 49.62]
GIG -30.56 [ -77.93, 16.81]
GSP 5.38 [ -34.73, 45.50]
HUBIN 30.62 [ -25.51, 86.75]
HUNT 26.38 [ 0.88, 51.89]
IMAGEN 15.41 [ -6.04, 36.85]
IMpACT 17.11 [ -41.17, 75.39]
LBC1936 5.74 [ -23.21, 34.69]
LIBD 61.27 [ 20.82, 101.72]
Sydney MAS -2.10 [ -32.48, 28.28]
MCIC 30.20 [ -42.91, 103.31]
MooDS 14.57 [ -35.96, 65.10]
MPIP 25.45 [ -6.31, 57.21]
MUNSTER 28.10 [ 2.03, 54.17]
NCNG 18.72 [ -23.04, 60.49]
NESDA 19.44 [ -32.86, 71.75]
NeuroIMAGE -43.71 [-105.57, 18.15]
NTR 34.37 [ -6.30, 75.04]
OATS -11.76 [ -60.35, 36.83]
PAFIP 11.04 [ -56.58, 78.66]
PDNZ 61.30 [ -0.35, 122.96]
PING 6.35 [ -39.60, 52.29]
MPRC 18.98 [ -14.63, 52.59]
PPMI 5.61 [ -31.32, 42.55]
QTIM 32.09 [ 2.35, 61.82]
SHIP -1.11 [ -26.53, 24.31]
SHIP-Trend 32.44 [ 4.47, 60.42]
SYS 40.59 [ -12.18, 93.36]
TCD-NUIG
TOP 18.71 [ -20.68, 58.09]
TOP3T 24.01 [ -17.93, 65.95]
UiO2016 -19.26 [ -72.37, 33.85]
UiO2017 -9.95 [ -57.15, 37.26]
UMCU -11.51 [ -45.57, 22.55]
ENIGMA Discovery Cohorts [N=22772; P=1.222e-08] 15.85 [ 10.40, 21.30]
UKBB 6.51 [ -0.74, 13.75]
Discovery+UKBB [N=32655; P=4.625e-08] 12.38 [ 7.94, 16.81]
GOBS -5.27 [ -39.97, 29.42]
IMH -4.91 [-115.34, 105.51]
UNICAMP
Meth-CT
MIRECC
UKBB nonEUR 1.92 [ -33.44, 37.28]
OSAKA 3.94 [ -29.14, 37.03]
PING Non European 61.65 [ 10.06, 113.25]
Non European Cohorts [N=2397; P=0.3989] 7.87 [ -10.41, 26.15]
-150
-100
-50
0
50
100
150
mm^2/effect allele
Beta [Lower 95% CI,Upper 95% CI]

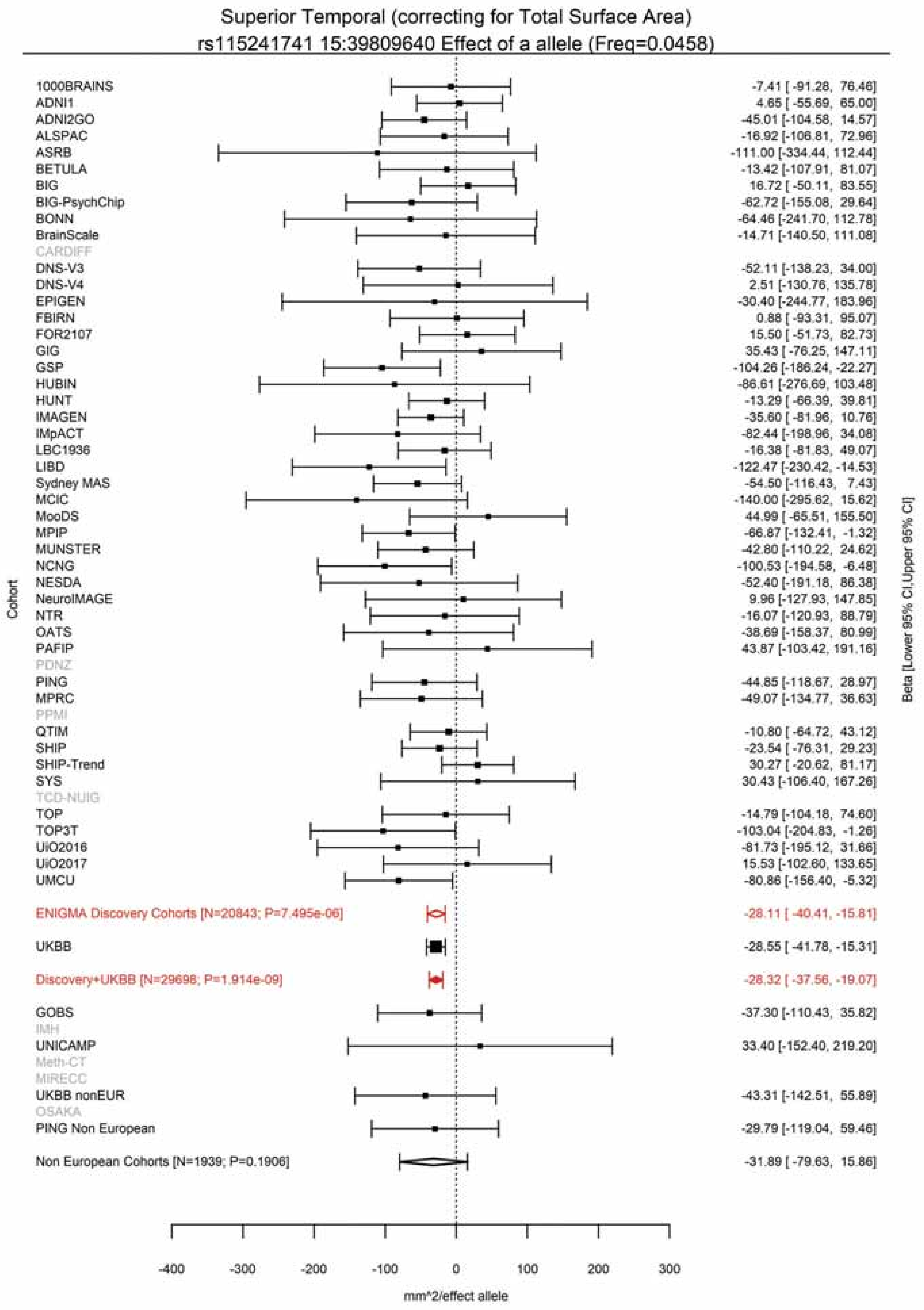
Superior Temporal (correcting for Total Surface Area)
rs115241741 15:39809640 Effect of a allele (Freq=0.0458)
Cohort
1000BRAINS -7.41 [ -91.28, 76.46]
ADNI1 4.65 [ -55.69, 65.00]
ADNI2GO -45.01 [-104.58, 14.57]
ALSPAC -16.92 [-106.81, 72.96]
ASRB -111.00 [-334.44, 112.44]
BETULA -13.42 [-107.91, 81.07]
BIG 16.72 [ -50.11, 83.55]
BIG-PsychChip -62.72 [-155.08, 29.64]
BONN -64.46 [-241.70, 112.78]
BrainScale -14.71 [-140.50, 111.08]
CARDIFF
DNS-V3 -52.11 [-138.23, 34.00]
DNS-V4 2.51 [-130.76, 135.78]
EPIGEN -30.40 [-244.77, 183.96]
FBIRN 0.88 [ -93.31, 95.07]
FOR2107 15.50 [ -51.73, 82.73]
GIG 35.43 [ -76.25, 147.11]
GSP -104.26 [-186.24, -22.27]
HUBIN -86.61 [-276.69, 103.48]
HUNT -13.29 [ -66.39, 39.81]
IMAGEN -35.60 [ -81.96, 10.76]
IMpACT -82.44 [-198.96, 34.08]
LBC1936 -16.38 [ -81.83, 49.07]
LIBD -122.47 [-230.42, -14.53]
Sydney MAS -54.50 [-116.43, 7.43]
MCIC -140.00 [-295.62, 15.62]
MooDS 44.99 [ -65.51, 155.50]
MPIP -66.87 [-132.41, -1.32]
MUNSTER -42.80 [-110.22, 24.62]
NCNG -100.53 [-194.58, -6.48]
NESDA -52.40 [-191.18, 86.38]
NeuroIMAGE 9.96 [-127.93, 147.85]
NTR -16.07 [-120.93, 88.79]
OATS -38.69 [-158.37, 80.99]
PAFIP 43.87 [-103.42, 191.16]
PDNZ
PING -44.85 [-118.67, 28.97]
MPRC -49.07 [-134.77, 36.63]
PPMI
QTIM -10.80 [ -64.72, 43.12]
SHIP -23.54 [ -76.31, 29.23]
SHIP-Trend 30.27 [ -20.62, 81.17]
SYS 30.43 [-106.40, 167.26]
TCD-NUIG
TOP -14.79 [-104.18, 74.60]
TOP3T -103.04 [-204.83, -1.26]
UiO2016 -81.73 [-195.12, 31.66]
UiO2017 15.53 [-102.60, 133.65]
UMCU -80.86 [-156.40, -5.32]
ENIGMA Discovery Cohorts [N=20843; P=7.495e-06] -28.11 [ -40.41, -15.81]
UKBB -28.55 [ -41.78, -15.31]
Discovery+UKBB [N=29698; P=1.914e-09] -28.32 [ -37.56, -19.07]
GOBS -37.30 [-110.43, 35.82]
IMH
UNICAMP 33.40 [-152.40, 219.20]
Meth-CT
MIRECC
UKBB nonEUR -43.31 [-142.51, 55.89]
OSAKA
PING Non European -29.79 [-119.04, 59.46]
Non European Cohorts [N=1939; P=0.1906] -31.89 [ -79.63, 15.86]
Beta [Lower 95% CI,Upper 95% CI]
-400
-300
-200
-100
0
100
200
300
mm^2/effect allele

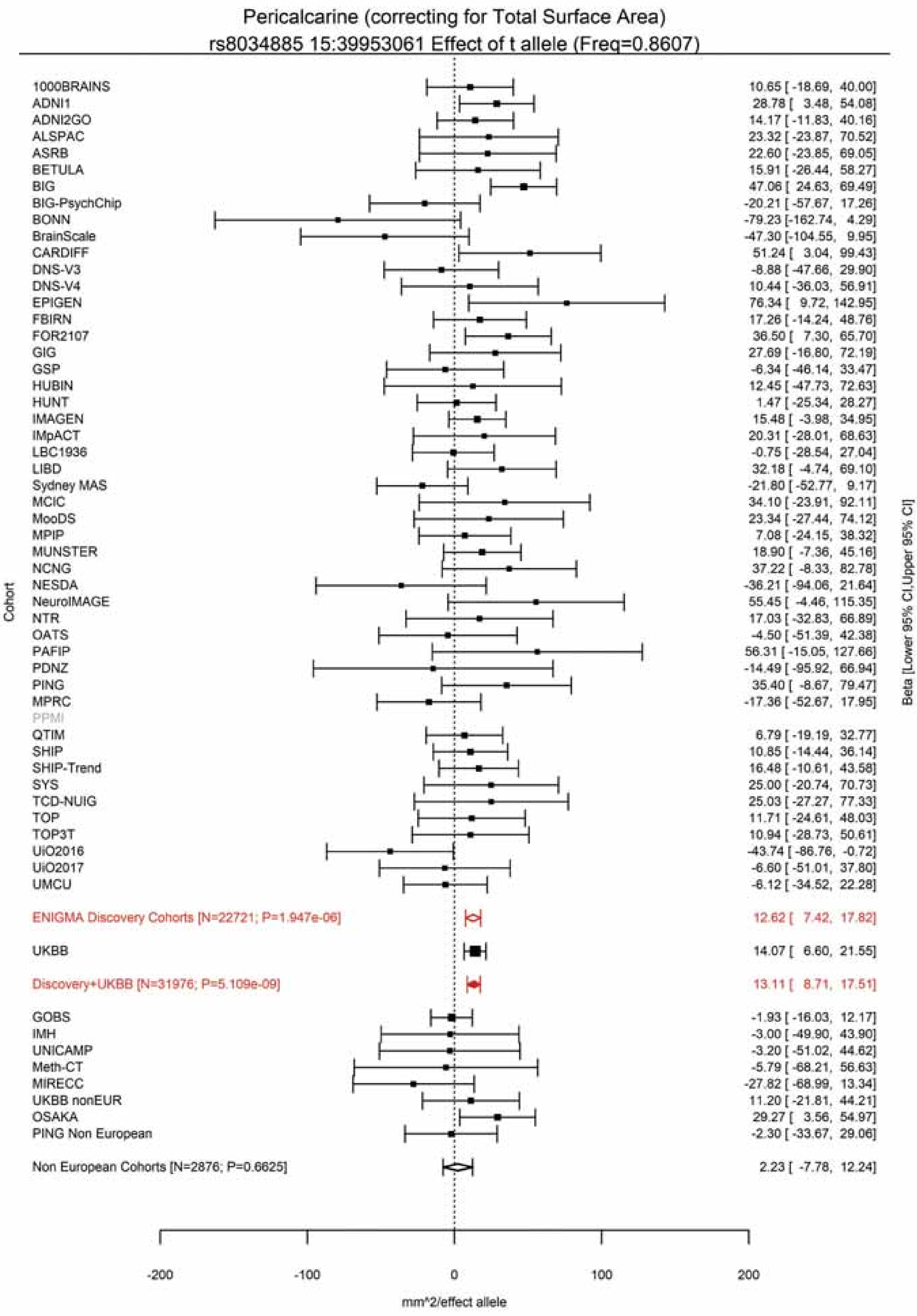

Pericalcarine (correcting for Total Surface Area)
rs8034885 15:39953061 Effect of t allele (Freq=0.8607)
Cohort
1000BRAINS 10.65 [ -18.69, 40.00]
ADNI1 28.78 [ 3.48, 54.08]
ADNI2GO 14.17 [ -11.83, 40.16]
ALSPAC 23.32 [ -23.87, 70.52]
ASRB 22.60 [ -23.85, 69.05]
BETULA 15.91 [ -26.44, 58.27]
BIG 47.06 [ 24.63, 69.49]
BIG-PsychChip -20.21 [ -57.67, 17.26]
BONN -79.23 [-162.74, 4.29]
BrainScale -47.30 [-104.55, 9.95]
CARDIFF 51.24 [ 3.04, 99.43]
DNS-V3 -8.88 [ -47.66, 29.90]
DNS-V4 10.44 [ -36.03, 56.91]
EPIGEN 76.34 [ 9.72, 142.95]
FBIRN 17.26 [ -14.24, 48.76]
FOR2107 36.50 [ 7.30, 65.70]
GIG 27.69 [ -16.80, 72.19]
GSP -6.34 [ -46.14, 33.47]
HUBIN 12.45 [ -47.73, 72.63]
HUNT 1.47 [ -25.34, 28.27]
IMAGEN 15.48 [ -3.98, 34.95]
IMpACT 20.31 [ -28.01, 68.63]
LBC1936 -0.75 [ -28.54, 27.04]
LIBD 32.18 [ -4.74, 69.10]
Sydney MAS -21.80 [ -52.77, 9.17]
MCIC 34.10 [ -23.91, 92.11]
MooDS 23.34 [ -27.44, 74.12]
MPIP 7.08 [ -24.15, 38.32]
MUNSTER 18.90 [ -7.36, 45.16]
NCNG 37.22 [ -8.33, 82.78]
NESDA -36.21 [ -94.06, 21.64]
NeuroIMAGE 55.45 [ -4.46, 115.35]
NTR 17.03 [ -32.83, 66.89]
OATS -4.50 [ -51.39, 42.38]
PAFIP 56.31 [ -15.05, 127.66]
PDNZ -14.49 [ -95.92, 66.94]
PING 35.40 [ -8.67, 79.47]
MPRC -17.36 [ -52.67, 17.95]
PPMI
QTIM 6.79 [ -19.19, 32.77]
SHIP 10.85 [ -14.44, 36.14]
SHIP-Trend 16.48 [ -10.61, 43.58]
SYS 25.00 [ -20.74, 70.73]
TCD-NUIG 25.03 [ -27.27, 77.33]
TOP 11.71 [ -24.61, 48.03]
TOP3T 10.94 [ -28.73, 50.61]
UiO2016 -43.74 [ -86.76, -0.72]
UiO2017 -6.60 [ -51.01, 37.80]
UMCU -6.12 [ -34.52, 22.28]
ENIGMA Discovery Cohorts [N=22721; P=1.947e-06] 12.62 [ 7.42, 17.82]
UKBB 14.07 [ 6.60, 21.55]
Discovery+UKBB [N=31976; P=5.109e-09] 13.11 [ 8.71, 17.51]
GOBS -1.93 [ -16.03, 12.17]
IMH -3.00 [ -49.90, 43.90]
UNICAMP -3.20 [ -51.02, 44.62]
Meth-CT -5.79 [ -68.21, 56.63]
MIRECC -27.82 [ -68.99, 13.34]
UKBB nonEUR 11.20 [ -21.81, 44.21]
OSAKA 29.27 [ 3.56, 54.97]
PING Non European -2.30 [ -33.67, 29.06]
Non European Cohorts [N=2876; P=0.6625] 2.23 [ -7.78, 12.24]
Beta [Lower 95% CI,Upper 95% CI]
-200
-100
0
100
200
mm^2/effect allele

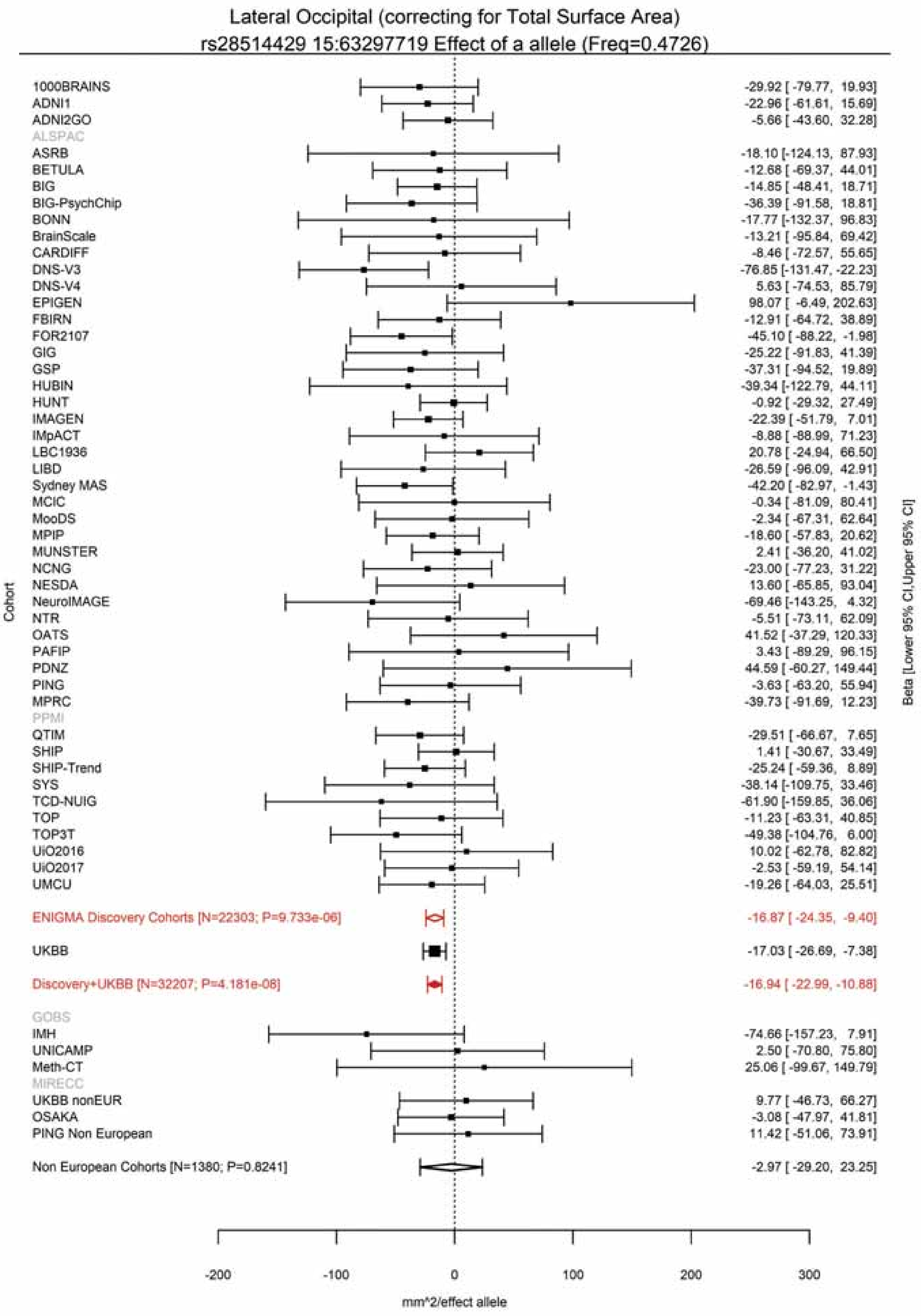
Lateral Occipital (correcting for Total Surface Area)
rs28514429 15:63297719 Effect of a allele (Freq=0.4726)
Cohort
1000BRAINS -29.92 [ -79.77, 19.93]
ADNI1 -22.96 [ -61.61, 15.69]
ADNI2GO -5.66 [ -43.60, 32.28]
ALSPAC
ASRB -18.10 [-124.13, 87.93]
BETULA -12.68 [ -69.37, 44.01]
BIG -14.85 [ -48.41, 18.71]
BIG-PsychChip -36.39 [ -91.58, 18.81]
BONN -17.77 [-132.37, 96.83]
BrainScale -13.21 [ -95.84, 69.42]
CARDIFF -8.46 [ -72.57, 55.65]
DNS-V3 -76.85 [-131.47, -22.23]
DNS-V4 5.63 [ -74.53, 85.79]
EPIGEN 98.07 [ -6.49, 202.63]
FBIRN -12.91 [ -64.72, 38.89]
FOR2107 -45.10 [ -88.22, -1.98]
GIG -25.22 [ -91.83, 41.39]
GSP -37.31 [ -94.52, 19.89]
HUBIN -39.34 [-122.79, 44.11]
HUNT -0.92 [ -29.32, 27.49]
IMAGEN -22.39 [ -51.79, 7.01]
IMpACT -8.88 [ -88.99, 71.23]
LBC1936 20.78 [ -24.94, 66.50]
LIBD -26.59 [ -96.09, 42.91]
Sydney MAS -42.20 [ -82.97, -1.43]
MCIC -0.34 [ -81.09, 80.41]
MooDS -2.34 [ -67.31, 62.64]
MPIP -18.60 [ -57.83, 20.62]
MUNSTER 2.41 [ -36.20, 41.02]
NCNG -23.00 [ -77.23, 31.22]
NESDA 13.60 [ -65.85, 93.04]
NeuroIMAGE -69.46 [-143.25, 4.32]
NTR -5.51 [ -73.11, 62.09]
OATS 41.52 [ -37.29, 120.33]
PAFIP 3.43 [ -89.29, 96.15]
PDNZ 44.59 [ -60.27, 149.44]
PING -3.63 [ -63.20, 55.94]
MPRC -39.73 [ -91.69, 12.23]
PPMI
QTIM -29.51 [ -66.67, 7.65]
SHIP 1.41 [ -30.67, 33.49]
SHIP-Trend -25.24 [ -59.36, 8.89]
SYS -38.14 [-109.75, 33.46]
TCD-NUIG -61.90 [-159.85, 36.06]
TOP -11.23 [ -63.31, 40.85]
TOP3T -49.38 [-104.76, 6.00]
UiO2016 10.02 [ -62.78, 82.82]
UiO2017 -2.53 [ -59.19, 54.14]
UMCU -19.26 [ -64.03, 25.51]
ENIGMA Discovery Cohorts [N=22303; P=9.733e-06] -16.87 [ -24.35, -9.40]
UKBB -17.03 [ -26.69, -7.38]
Discovery+UKBB [N=32207; P=4.181e-08] -16.94 [ -22.99, -10.88]
GOBS
IMH -74.66 [-157.23, 7.91]
UNICAMP 2.50 [ -70.80, 75.80]
Meth-CT 25.06 [ -99.67, 149.79]
MIRECC
UKBB nonEUR 9.77 [ -46.73, 66.27]
OSAKA -3.08 [ -47.97, 41.81]
PING Non European 11.42 [ -51.06, 73.91]
Non European Cohorts [N=1380; P=0.8241] -2.97 [ -29.20, 23.25]
Beta [Lower 95% CI,Upper 95% CI]
-200
-100
0
100
200
300
mm^2/effect allele

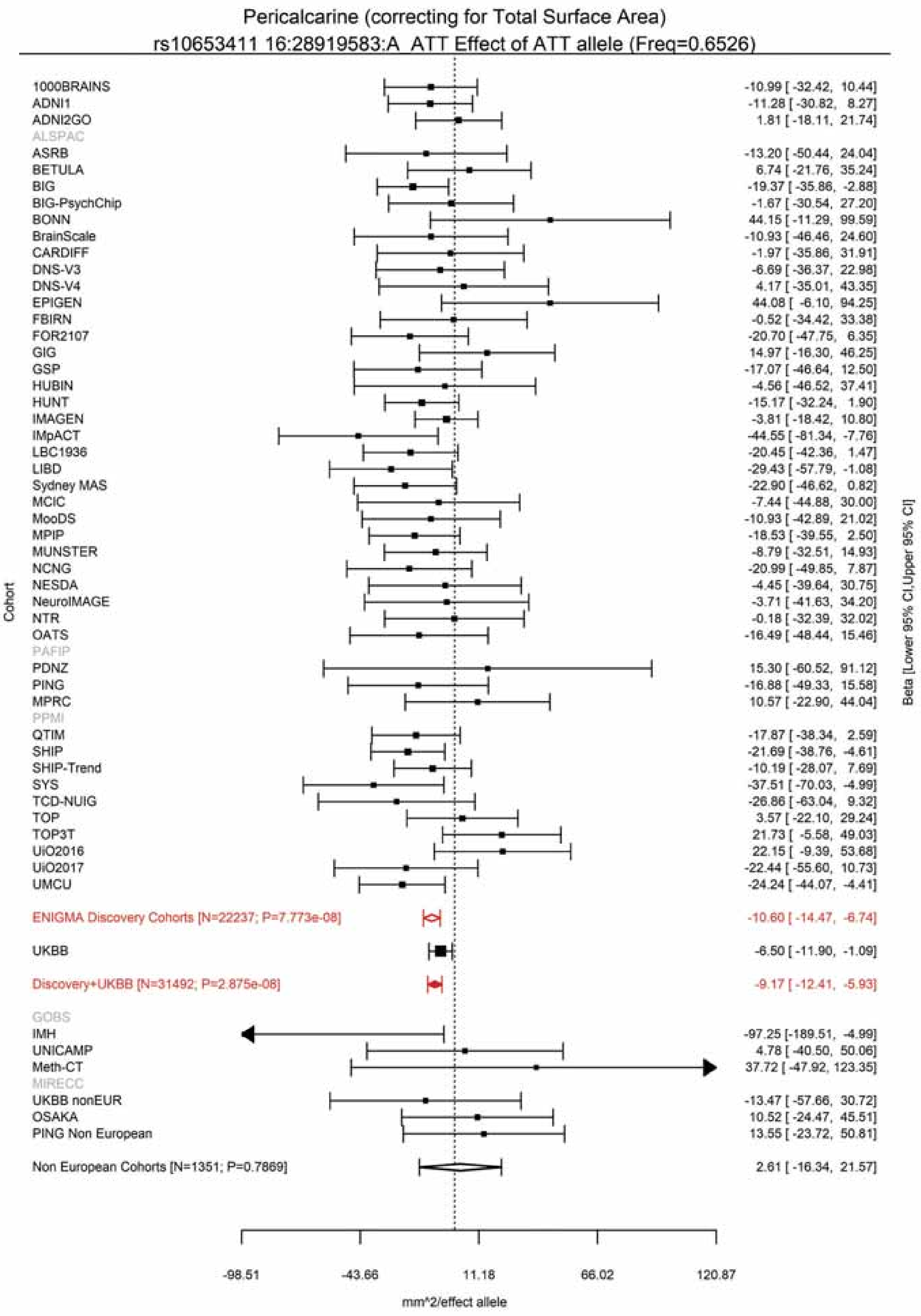
Pericalcarine (correcting for Total Surface Area)
rs10653411 16:28919583:A ATT Effect of ATT allele (Freq=0.6526)
Cohort
1000BRAINS -10.99 [ -32.42, 10.44]
ADNI1 -11.28 [ -30.82, 8.27]
ADNI2GO 1.81 [ -18.11, 21.74]
ALSPAC
ASRB -13.20 [ -50.44, 24.04]
BETULA 6.74 [ -21.76, 35.24]
BIG -19.37 [ -35.86, -2.88]
BIG-PsychChip -1.67 [ -30.54, 27.20]
BONN 44.15 [ -11.29, 99.59]
BrainScale -10.93 [ -46.46, 24.60]
CARDIFF -1.97 [ -35.86, 31.91]
DNS-V3 -6.69 [ -36.37, 22.98]
DNS-V4 4.17 [ -35.01, 43.35]
EPIGEN 44.08 [ -6.10, 94.25]
FBIRN -0.52 [ -34.42, 33.38]
FOR2107 -20.70 [ -47.75, 6.35]
GIG 14.97 [ -16.30, 46.25]
GSP -17.07 [ -46.64, 12.50]
HUBIN -4.56 [ -46.52, 37.41]
HUNT -15.17 [ -32.24, 1.90]
IMAGEN -3.81 [ -18.42, 10.80]
IMpACT -44.55 [ -81.34, -7.76]
LBC1936 -20.45 [ -42.36, 1.47]
LIBD -29.43 [ -57.79, -1.08]
Sydney MAS -22.90 [ -46.62, 0.82]
MCIC -7.44 [ -44.88, 30.00]
MooDS -10.93 [ -42.89, 21.02]
MPIP -18.53 [ -39.55, 2.50]
MUNSTER -8.79 [ -32.51, 14.93]
NCNG -20.99 [ -49.85, 7.87]
NESDA -4.45 [ -39.64, 30.75]
NeuroIMAGE -3.71 [ -41.63, 34.20]
NTR -0.18 [ -32.39, 32.02]
OATS -16.49 [ -48.44, 15.46]
PAFIP
PDNZ 15.30 [ -60.52, 91.12]
PING -16.88 [ -49.33, 15.58]
MPRC 10.57 [ -22.90, 44.04]
PPMI
QTIM -17.87 [ -38.34, 2.59]
SHIP -21.69 [ -38.76, -4.61]
SHIP-Trend -10.19 [ -28.07, 7.69]
SYS -37.51 [ -70.03, -4.99]
TCD-NUIG -26.86 [ -63.04, 9.32]
TOP 3.57 [ -22.10, 29.24]
TOP3T 21.73 [ -5.58, 49.03]
UiO2016 22.15 [ -9.39, 53.68]
UiO2017 -22.44 [ -55.60, 10.73]
UMCU -24.24 [ -44.07, -4.41]
ENIGMA Discovery Cohorts [N=22237; P=7.773e-08] -10.60 [ -14.47, -6.74]
UKBB -6.50 [ -11.90, -1.09]
Discovery+UKBB [N=31492; P=2.875e-08] -9.17 [ -12.41, -5.93]
GOBS
IMH -97.25 [-189.51, -4.99]
UNICAMP 4.78 [ -40.50, 50.06]
Meth-CT 37.72 [ -47.92, 123.35]
MIRECC
UKBB nonEUR -13.47 [ -57.66, 30.72]
OSAKA 10.52 [ -24.47, 45.51]
PING Non European 13.55 [ -23.72, 50.81]
Non European Cohorts [N=1351; P=0.7869] 2.61 [ -16.34, 21.57]
-98.51
-43.66
11.18
66.02
120.87
mm^2/effect allele
Beta [Lower 95% CI,Upper 95% CI]

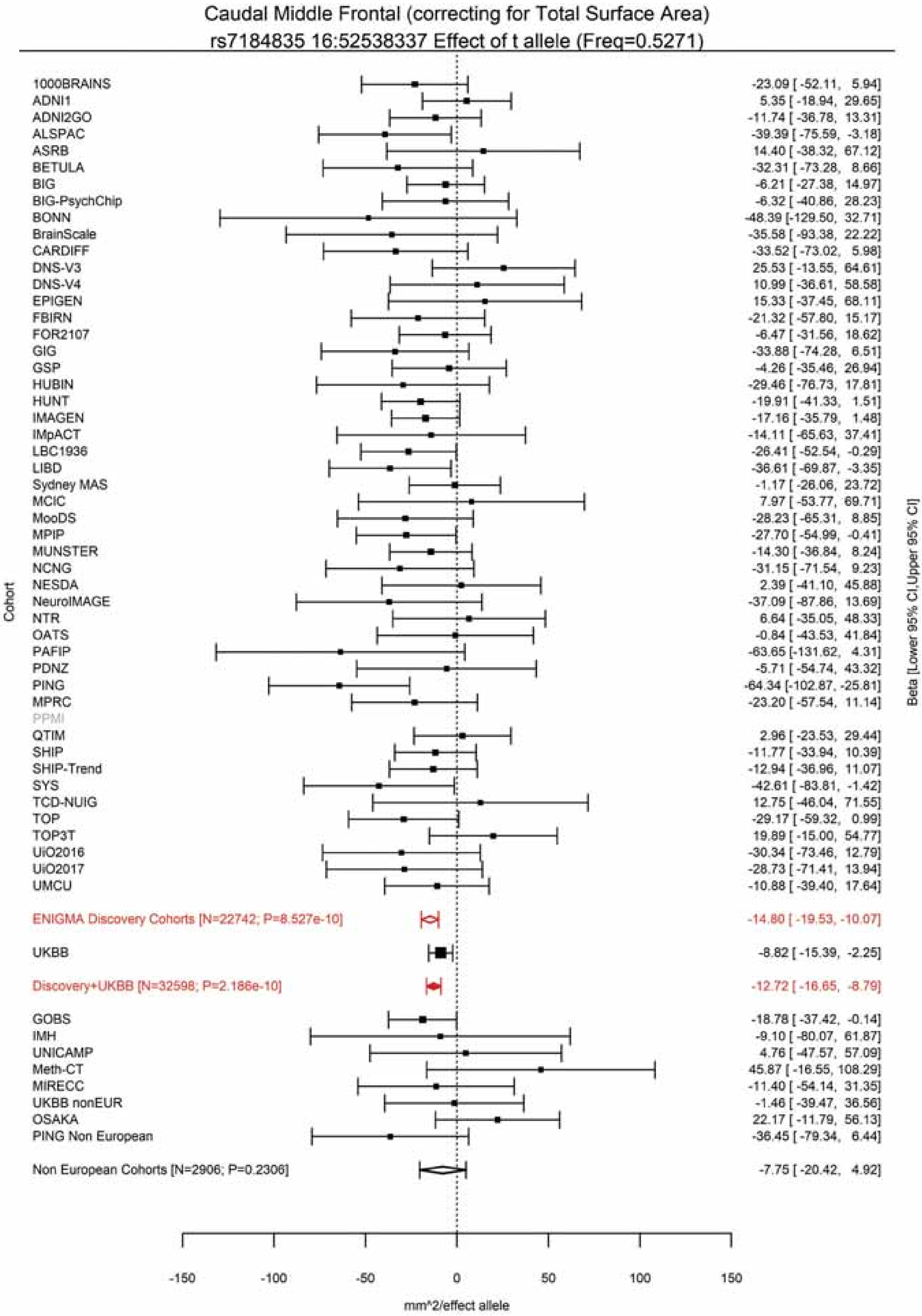
Caudal Middle Frontal (correcting for Total Surface Area)
rs7184835 16:52538337 Effect of t allele (Freq=0.5271)
Cohort
1000BRAINS -23.09 [ -52.11, 5.94]
ADNI1 5.35 [ -18.94, 29.65]
ADNI2GO -11.74 [ -36.78, 13.31]
ALSPAC -39.39 [ -75.59, -3.18]
ASRB 14.40 [ -38.32, 67.12]
BETULA -32.31 [ -73.28, 8.66]
BIG -6.21 [ -27.38, 14.97]
BIG-PsychChip -6.32 [ -40.86, 28.23]
BONN -48.39 [-129.50, 32.71]
BrainScale -35.58 [ -93.38, 22.22]
CARDIFF -33.52 [ -73.02, 5.98]
DNS-V3 25.53 [ -13.55, 64.61]
DNS-V4 10.99 [ -36.61, 58.58]
EPIGEN 15.33 [ -37.45, 68.11]
FBIRN -21.32 [ -57.80, 15.17]
FOR2107 -6.47 [ -31.56, 18.62]
GIG -33.88 [ -74.28, 6.51]
GSP -4.26 [ -35.46, 26.94]
HUBIN -29.46 [ -76.73, 17.81]
HUNT -19.91 [ -41.33, 1.51]
IMAGEN -17.16 [ -35.79, 1.48]
IMpACT -14.11 [ -65.63, 37.41]
LBC1936 -26.41 [ -52.54, -0.29]
LIBD -36.61 [ -69.87, -3.35]
Sydney MAS -1.17 [ -26.06, 23.72]
MCIC 7.97 [ -53.77, 69.71]
MooDS -28.23 [ -65.31, 8.85]
MPIP -27.70 [ -54.99, -0.41]
MUNSTER -14.30 [ -36.84, 8.24]
NCNG -31.15 [ -71.54, 9.23]
NESDA 2.39 [ -41.10, 45.88]
NeuroIMAGE -37.09 [ -87.86, 13.69]
NTR 6.64 [ -35.05, 48.33]
OATS -0.84 [ -43.53, 41.84]
PAFIP -63.65 [-131.62, 4.31]
PDNZ -5.71 [ -54.74, 43.32]
PING -64.34 [-102.87, -25.81]
MPRC -23.20 [ -57.54, 11.14]
PPMI
QTIM 2.96 [ -23.53, 29.44]
SHIP -11.77 [ -33.94, 10.39]
SHIP-Trend -12.94 [ -36.96, 11.07]
SYS -42.61 [ -83.81, -1.42]
TCD-NUIG 12.75 [ -46.04, 71.55]
TOP -29.17 [ -59.32, 0.99]
TOP3T 19.89 [ -15.00, 54.77]
UiO2016 -30.34 [ -73.46, 12.79]
UiO2017 -28.73 [ -71.41, 13.94]
UMCU -10.88 [ -39.40, 17.64]
ENIGMA Discovery Cohorts [N=22742; P=8.527e-10] -14.80 [ -19.53, -10.07]
UKBB -8.82 [ -15.39, -2.25]
Discovery+UKBB [N=32598; P=2.186e-10] -12.72 [ -16.65, -8.79]
GOBS -18.78 [ -37.42, -0.14]
IMH -9.10 [ -80.07, 61.87]
UNICAMP 4.76 [ -47.57, 57.09]
Meth-CT 45.87 [ -16.55, 108.29]
MIRECC -11.40 [ -54.14, 31.35]
UKBB nonEUR -1.46 [ -39.47, 36.56]
OSAKA 22.17 [ -11.79, 56.13]
PING Non European -36.45 [ -79.34, 6.44]
Non European Cohorts [N=2906; P=0.2306] -7.75 [ -20.42, 4.92]
Beta [Lower 95% CI,Upper 95% CI]
-150
-100
-50
0
50
100
150
mm^2/effect allele

Caudal Anterior Cingulate (correcting for Total Surface Area)
rs80241863 16:60827388 Effect of a allele (Freq=0.0127)

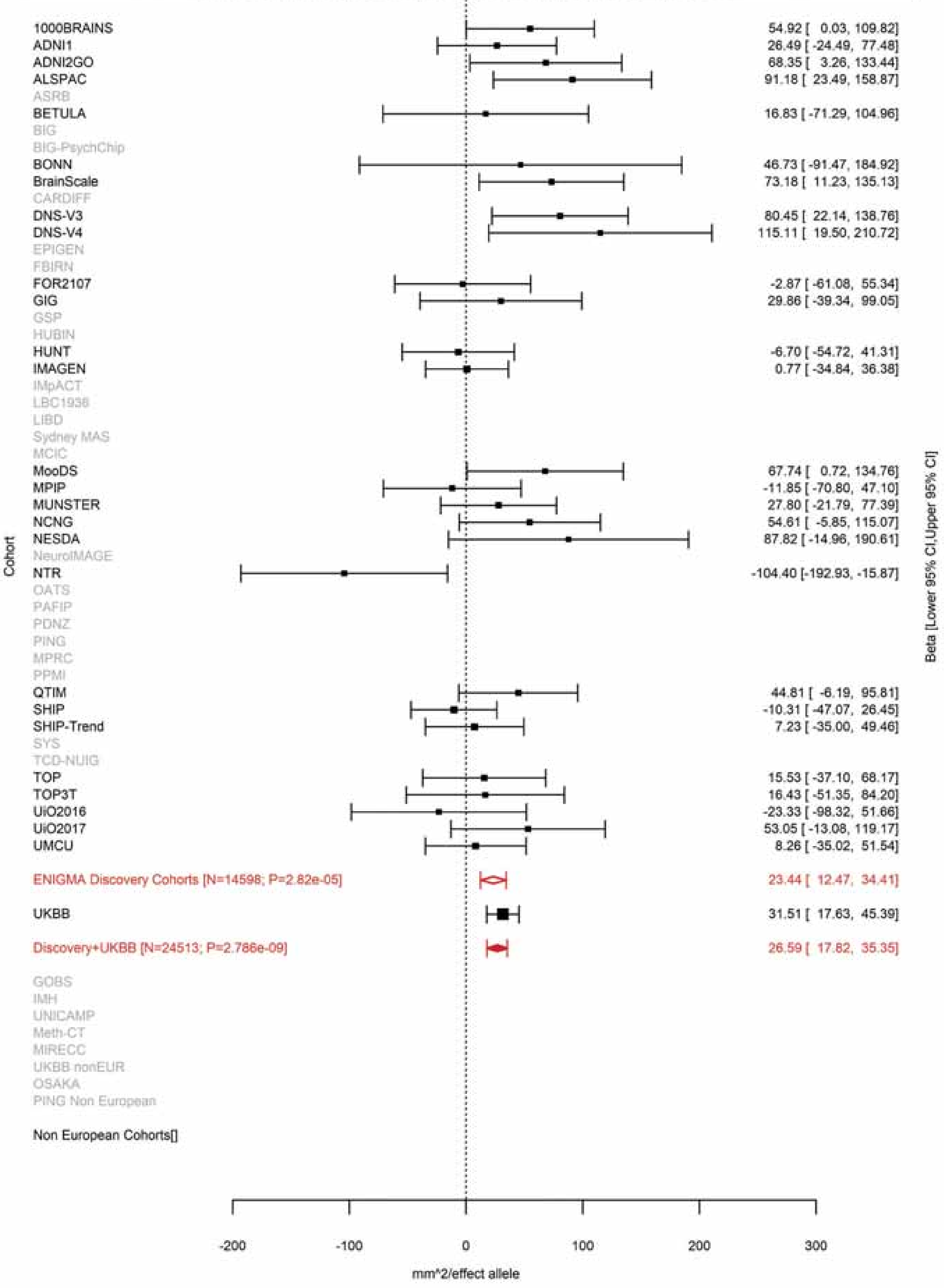

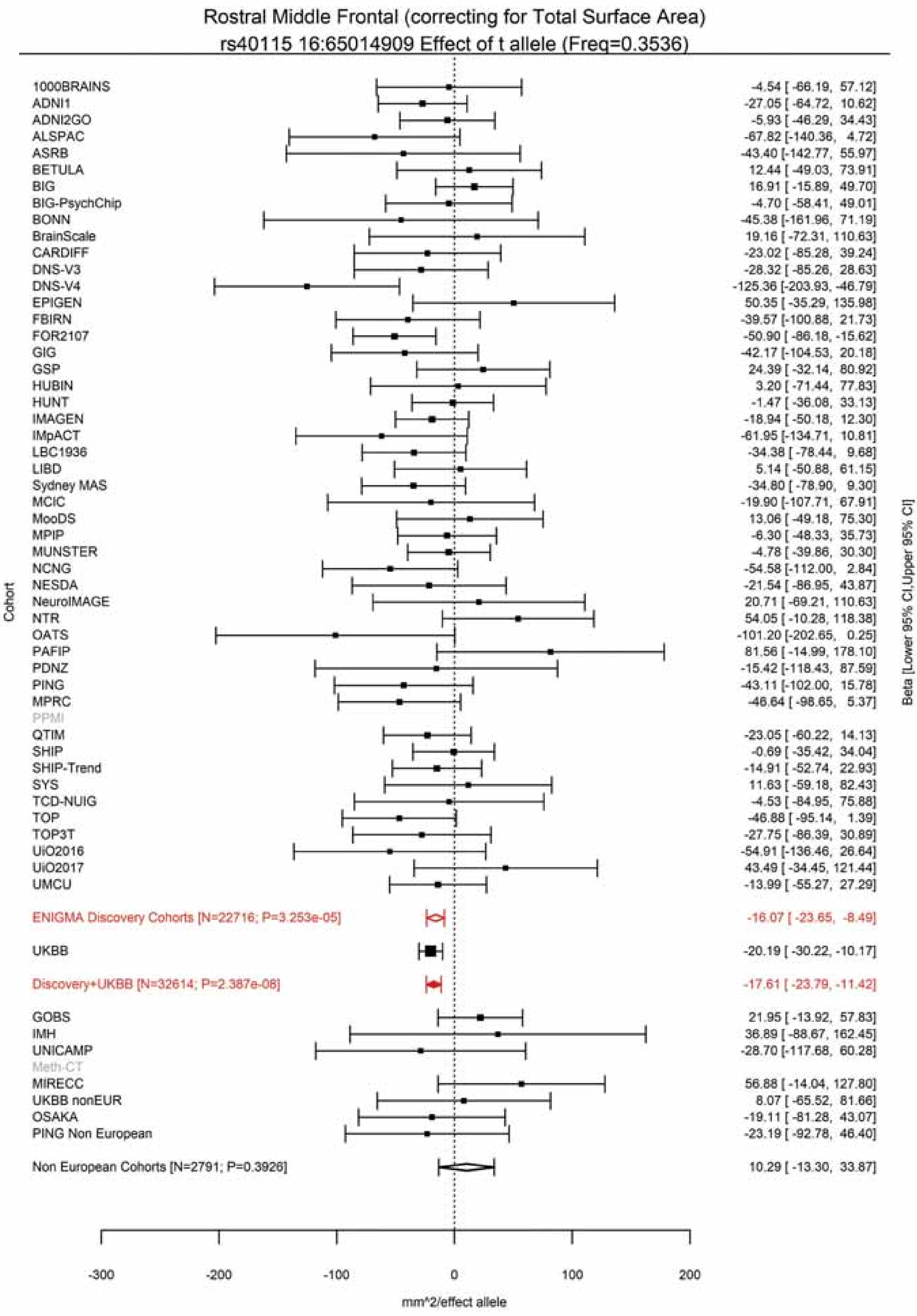

Rostral Middle Frontal (correcting for Total Surface Area)
rs40115 16:65014909 Effect of t allele (Freq=0.3536)
Cohort
1000BRAINS -4.54 [ -66.19, 57.12]
ADNI1 -27.05 [ -64.72, 10.62]
ADNI2GO -5.93 [ -46.29, 34.43]
ALSPAC -67.82 [-140.36, 4.72]
ASRB -43.40 [-142.77, 55.97]
BETULA 12.44 [ -49.03, 73.91]
BIG 16.91 [ -15.89, 49.70]
BIG-PsychChip -4.70 [ -58.41, 49.01]
BONN -45.38 [-161.96, 71.19]
BrainScale 19.16 [ -72.31, 110.63]
CARDIFF -23.02 [ -85.28, 39.24]
DNS-V3 -28.32 [ -85.26, 28.63]
DNS-V4 -125.36 [-203.93, -46.79]
EPIGEN 50.35 [ -35.29, 135.98]
FBIRN -39.57 [-100.88, 21.73]
FOR2107 -50.90 [ -86.18, -15.62]
GIG -42.17 [-104.53, 20.18]
GSP 24.39 [ -32.14, 80.92]
HUBIN 3.20 [ -71.44, 77.83]
HUNT -1.47 [ -36.08, 33.13]
IMAGEN -18.94 [ -50.18, 12.30]
IMpACT -61.95 [-134.71, 10.81]
LBC1936 -34.38 [ -78.44, 9.68]
LIBD 5.14 [ -50.88, 61.15]
Sydney MAS -34.80 [ -78.90, 9.30]
MCIC -19.90 [-107.71, 67.91]
MooDS 13.06 [ -49.18, 75.30]
MPIP -6.30 [ -48.33, 35.73]
MUNSTER -4.78 [ -39.86, 30.30]
NCNG -54.58 [-112.00, 2.84]
NESDA -21.54 [ -86.95, 43.87]
NeuroIMAGE 20.71 [ -69.21, 110.63]
NTR 54.05 [ -10.28, 118.38]
OATS -101.20 [-202.65, 0.25]
PAFIP 81.56 [ -14.99, 178.10]
PDNZ -15.42 [-118.43, 87.59]
PING -43.11 [-102.00, 15.78]
MPRC -46.64 [ -98.65, 5.37]
PPMI
QTIM -23.05 [ -60.22, 14.13]
SHIP -0.69 [ -35.42, 34.04]
SHIP-Trend -14.91 [ -52.74, 22.93]
SYS 11.63 [ -59.18, 82.43]
TCD-NUIG -4.53 [ -84.95, 75.88]
TOP -46.88 [ -95.14, 1.39]
TOP3T -27.75 [ -86.39, 30.89]
UiO2016 -54.91 [-136.46, 26.64]
UiO2017 43.49 [ -34.45, 121.44]
UMCU -13.99 [ -55.27, 27.29]
ENIGMA Discovery Cohorts [N=22716; P=3.253e-05] -16.07 [ -23.65, -8.49]
UKBB -20.19 [ -30.22, -10.17]
Discovery+UKBB [N=32614; P=2.387e-08] -17.61 [ -23.79, -11.42]
GOBS 21.95 [ -13.92, 57.83]
IMH 36.89 [ -88.67, 162.45]
UNICAMP -28.70 [-117.68, 60.28]
Meth-CT
MIRECC 56.88 [ -14.04, 127.80]
UKBB nonEUR 8.07 [ -65.52, 81.66]
OSAKA -19.11 [ -81.28, 43.07]
PING Non European -23.19 [ -92.78, 46.40]
Non European Cohorts [N=2791; P=0.3926] 10.29 [ -13.30, 33.87]
-300
-200
-100
0
100
200
mm^2/effect allele
Beta [Lower 95% CI,Upper 95% CI]

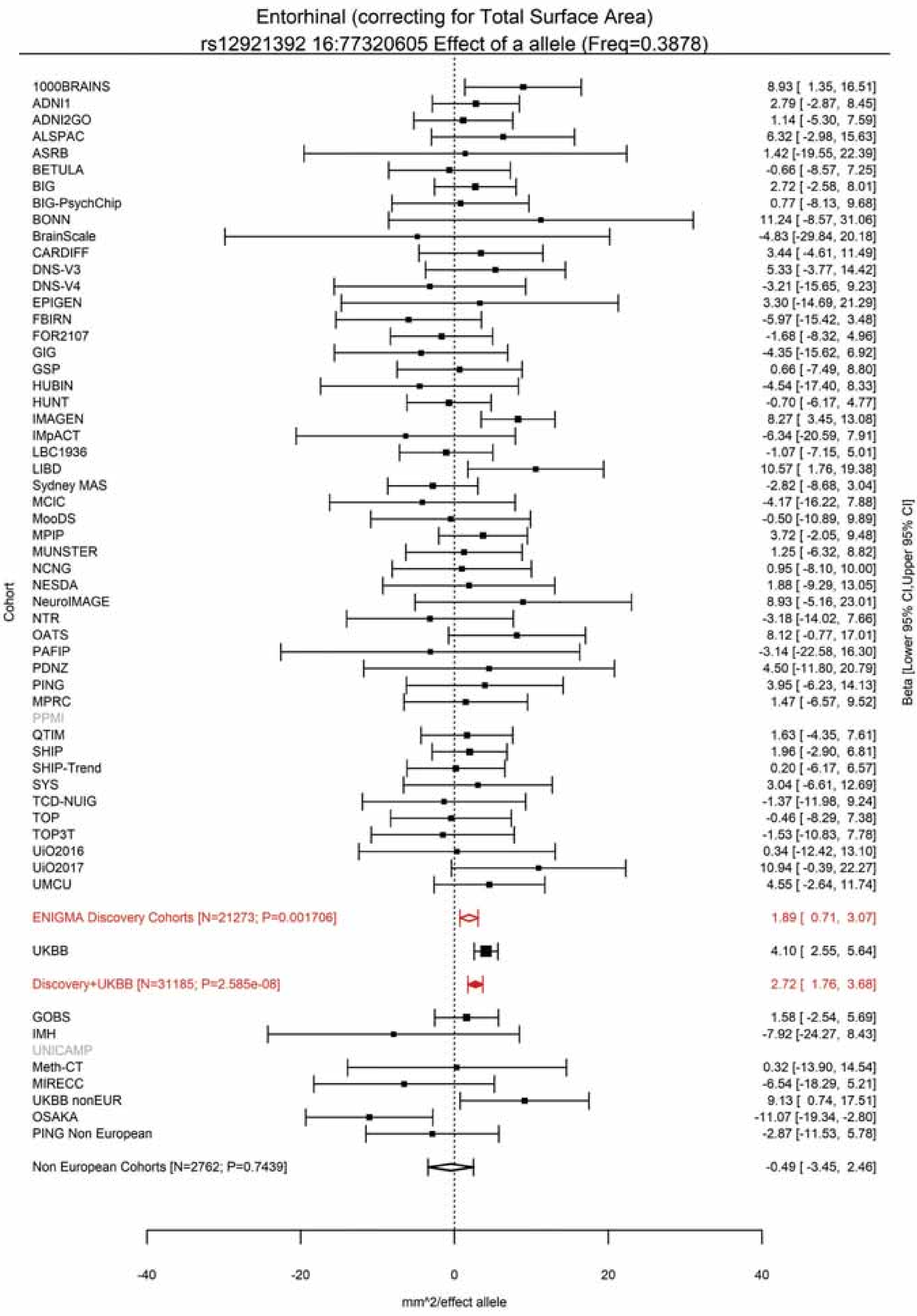

Entorhinal (correcting for Total Surface Area)
rs12921392 16:77320605 Effect of a allele (Freq=0.3878)
Cohort
Beta [Lower 95% CI,Upper 95% CI]
1000BRAINS 8.93 [ 1.35, 16.51]
ADNI1 2.79 [ -2.87, 8.45]
ADNI2GO 1.14 [ -5.30, 7.59]
ALSPAC 6.32 [ -2.98, 15.63]
ASRB 1.42 [-19.55, 22.39]
BETULA -0.66 [ -8.57, 7.25]
BIG 2.72 [ -2.58, 8.01]
BIG-PsychChip 0.77 [ -8.13, 9.68]
BONN 11.24 [ -8.57, 31.06]
BrainScale -4.83 [-29.84, 20.18]
CARDIFF 3.44 [ -4.61, 11.49]
DNS-V3 5.33 [ -3.77, 14.42]
DNS-V4 -3.21 [-15.65, 9.23]
EPIGEN 3.30 [-14.69, 21.29]
FBIRN -5.97 [-15.42, 3.48]
FOR2107 -1.68 [ -8.32, 4.96]
GIG -4.35 [-15.62, 6.92]
GSP 0.66 [ -7.49, 8.80]
HUBIN -4.54 [-17.40, 8.33]
HUNT -0.70 [ -6.17, 4.77]
IMAGEN 8.27 [ 3.45, 13.08]
IMpACT -6.34 [-20.59, 7.91]
LBC1936 -1.07 [ -7.15, 5.01]
LIBD 10.57 [ 1.76, 19.38]
Sydney MAS -2.82 [ -8.68, 3.04]
MCIC -4.17 [-16.22, 7.88]
MooDS -0.50 [-10.89, 9.89]
MPIP 3.72 [ -2.05, 9.48]
MUNSTER 1.25 [ -6.32, 8.82]
NCNG 0.95 [ -8.10, 10.00]
NESDA 1.88 [ -9.29, 13.05]
NeuroIMAGE 8.93 [ -5.16, 23.01]
NTR -3.18 [-14.02, 7.66]
OATS 8.12 [ -0.77, 17.01]
PAFIP -3.14 [-22.58, 16.30]
PDNZ 4.50 [-11.80, 20.79]
PING 3.95 [ -6.23, 14.13]
MPRC 1.47 [ -6.57, 9.52]
PPMI
QTIM 1.63 [ -4.35, 7.61]
SHIP 1.96 [ -2.90, 6.81]
SHIP-Trend 0.20 [ -6.17, 6.57]
SYS 3.04 [ -6.61, 12.69]
TCD-NUIG -1.37 [-11.98, 9.24]
TOP -0.46 [ -8.29, 7.38]
TOP3T -1.53 [-10.83, 7.78]
UiO2016 0.34 [-12.42, 13.10]
UiO2017 10.94 [ -0.39, 22.27]
UMCU 4.55 [ -2.64, 11.74]
ENIGMA Discovery Cohorts [N=21273; P=0.001706] 1.89 [ 0.71, 3.07]
UKBB 4.10 [ 2.55, 5.64]
Discovery+UKBB [N=31185; P=2.585e-08] 2.72 [ 1.76, 3.68]
GOBS 1.58 [ -2.54, 5.69]
IMH -7.92 [-24.27, 8.43]
UNICAMP
Meth-CT 0.32 [-13.90, 14.54]
MIRECC -6.54 [-18.29, 5.21]
UKBB nonEUR 9.13 [ 0.74, 17.51]
OSAKA -11.07 [-19.34, -2.80]
PING Non European -2.87 [-11.53, 5.78]
Non European Cohorts [N=2762; P=0.7439] -0.49 [ -3.45, 2.46]
-40
-20
0
20
40
mm^2/effect allele

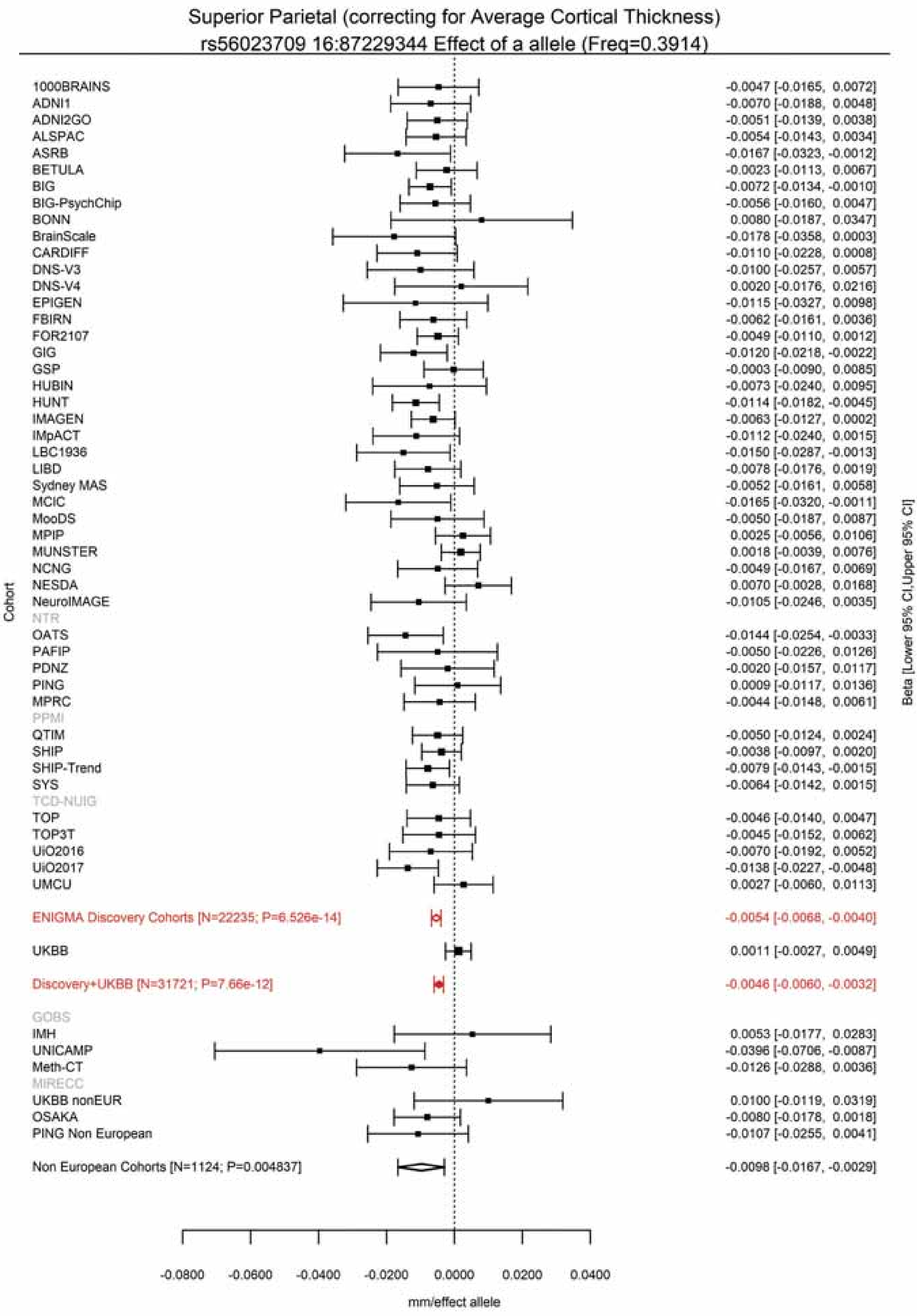

Superior Parietal (correcting for Average Cortical Thickness)
rs56023709 16:87229344 Effect of a allele (Freq=0.3914)
Cohort
1000BRAINS -0.0047 [-0.0165, 0.0072]
ADNI1 -0.0070 [-0.0188, 0.0048]
ADNI2GO -0.0051 [-0.0139, 0.0038]
ALSPAC -0.0054 [-0.0143, 0.0034]
ASRB -0.0167 [-0.0323, -0.0012]
BETULA -0.0023 [-0.0113, 0.0067]
BIG -0.0072 [-0.0134, -0.0010]
BIG-PsychChip -0.0056 [-0.0160, 0.0047]
BONN 0.0080 [-0.0187, 0.0347]
BrainScale -0.0178 [-0.0358, 0.0003]
CARDIFF -0.0110 [-0.0228, 0.0008]
DNS-V3 -0.0100 [-0.0257, 0.0057]
DNS-V4 0.0020 [-0.0176, 0.0216]
EPIGEN -0.0115 [-0.0327, 0.0098]
FBIRN -0.0062 [-0.0161, 0.0036]
FOR2107 -0.0049 [-0.0110, 0.0012]
GIG -0.0120 [-0.0218, -0.0022]
GSP -0.0003 [-0.0090, 0.0085]
HUBIN -0.0073 [-0.0240, 0.0095]
HUNT -0.0114 [-0.0182, -0.0045]
IMAGEN -0.0063 [-0.0127, 0.0002]
IMpACT -0.0112 [-0.0240, 0.0015]
LBC1936 -0.0150 [-0.0287, -0.0013]
LIBD -0.0078 [-0.0176, 0.0019]
Sydney MAS -0.0052 [-0.0161, 0.0058]
MCIC -0.0165 [-0.0320, -0.0011]
MooDS -0.0050 [-0.0187, 0.0087]
MPIP 0.0025 [-0.0056, 0.0106]
MUNSTER 0.0018 [-0.0039, 0.0076]
NCNG -0.0049 [-0.0167, 0.0069]
NESDA 0.0070 [-0.0028, 0.0168]
NeuroIMAGE -0.0105 [-0.0246, 0.0035]
NTR
OATS -0.0144 [-0.0254, -0.0033]
PAFIP -0.0050 [-0.0226, 0.0126]
PDNZ -0.0020 [-0.0157, 0.0117]
PING 0.0009 [-0.0117, 0.0136]
MPRC -0.0044 [-0.0148, 0.0061]
PPMI
QTIM -0.0050 [-0.0124, 0.0024]
SHIP -0.0038 [-0.0097, 0.0020]
SHIP-Trend -0.0079 [-0.0143, -0.0015]
SYS -0.0064 [-0.0142, 0.0015]
TCD-NUIG
TOP -0.0046 [-0.0140, 0.0047]
TOP3T -0.0045 [-0.0152, 0.0062]
UiO2016 -0.0070 [-0.0192, 0.0052]
UiO2017 -0.0138 [-0.0227, -0.0048]
UMCU 0.0027 [-0.0060, 0.0113]
ENIGMA Discovery Cohorts [N=22235; P=6.526e-14] -0.0054 [-0.0068, -0.0040]
UKBB 0.0011 [-0.0027, 0.0049]
Discovery+UKBB [N=31721; P=7.66e-12] -0.0046 [-0.0060, -0.0032]
GOBS
IMH 0.0053 [-0.0177, 0.0283]
UNICAMP -0.0396 [-0.0706, -0.0087]
Meth-CT -0.0126 [-0.0288, 0.0036]
MIRECC
UKBB nonEUR 0.0100 [-0.0119, 0.0319]
OSAKA -0.0080 [-0.0178, 0.0018]
PING Non European -0.0107 [-0.0255, 0.0041]
Non European Cohorts [N=1124; P=0.004837] -0.0098 [-0.0167, -0.0029]
Beta [Lower 95% CI,Upper 95% CI]
-0.0800
-0.0600
-0.0400
-0.0200
0.0000
0.0200
0.0400
mm/effect allele

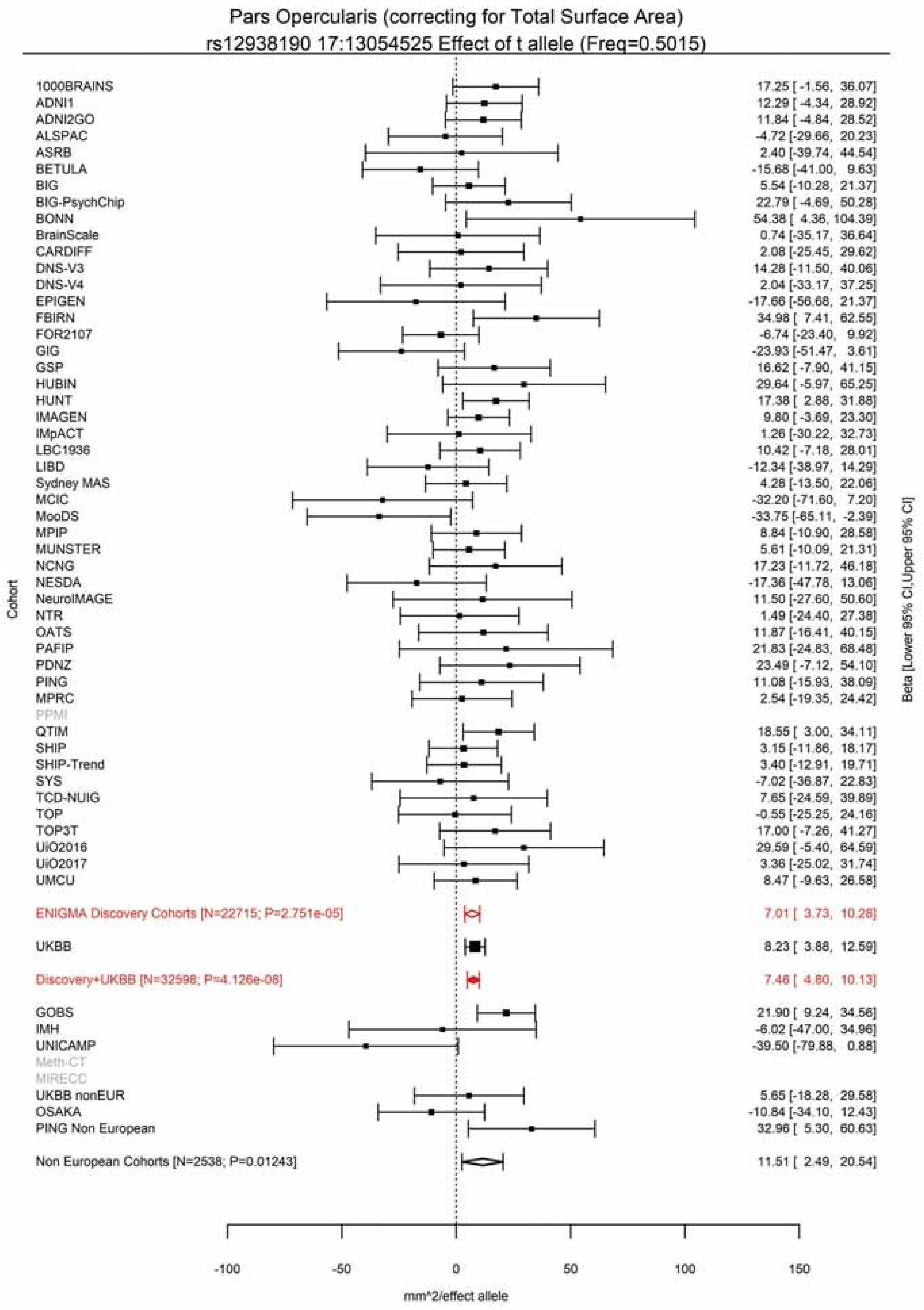
Pars Opercularis (correcting for Total Surface Area)
rs12938190 17:13054525 Effect of t allele (Freq=0.5015)
Cohort
1000BRAINS 17.25 [ -1.56, 36.07]
ADNI1 12.29 [ -4.34, 28.92]
ADNI2GO 11.84 [ -4.84, 28.52]
ALSPAC -4.72 [-29.66, 20.23]
ASRB 2.40 [-39.74, 44.54]
BETULA -15.68 [-41.00, 9.63]
BIG 5.54 [-10.28, 21.37]
BIG-PsychChip 22.79 [ -4.69, 50.28]
BONN 54.38 [ 4.36, 104.39]
BrainScale 0.74 [-35.17, 36.64]
CARDIFF 2.08 [-25.45, 29.62]
DNS-V3 14.28 [-11.50, 40.06]
DNS-V4 2.04 [-33.17, 37.25]
EPIGEN -17.66 [-56.68, 21.37]
FBIRN 34.98 [ 7.41, 62.55]
FOR2107 -6.74 [-23.40, 9.92]
GIG -23.93 [-51.47, 3.61]
GSP 16.62 [ -7.90, 41.15]
HUBIN 29.64 [ -5.97, 65.25]
HUNT 17.38 [ 2.88, 31.88]
IMAGEN 9.80 [ -3.69, 23.30]
IMpACT 1.26 [-30.22, 32.73]
LBC1936 10.42 [ -7.18, 28.01]
LIBD -12.34 [-38.97, 14.29]
Sydney MAS 4.28 [-13.50, 22.06]
MCIC -32.20 [-71.60, 7.20]
MooDS -33.75 [-65.11, -2.39]
MPIP 8.84 [-10.90, 28.58]
MUNSTER 5.61 [-10.09, 21.31]
NCNG 17.23 [-11.72, 46.18]
NESDA -17.36 [-47.78, 13.06]
NeuroIMAGE 11.50 [-27.60, 50.60]
NTR 1.49 [-24.40, 27.38]
OATS 11.87 [-16.41, 40.15]
PAFIP 21.83 [-24.83, 68.48]
PDNZ 23.49 [ -7.12, 54.10]
PING 11.08 [-15.93, 38.09]
MPRC 2.54 [-19.35, 24.42]
PPMI
QTIM 18.55 [ 3.00, 34.11]
SHIP 3.15 [-11.86, 18.17]
SHIP-Trend 3.40 [-12.91, 19.71]
SYS -7.02 [-36.87, 22.83]
TCD-NUIG 7.65 [-24.59, 39.89]
TOP -0.55 [-25.25, 24.16]
TOP3T 17.00 [ -7.26, 41.27]
UiO2016 29.59 [ -5.40, 64.59]
UiO2017 3.36 [-25.02, 31.74]
UMCU 8.47 [ -9.63, 26.58]
ENIGMA Discovery Cohorts [N=22715; P=2.751e-05] 7.01 [ 3.73, 10.28]
UKBB 8.23 [ 3.88, 12.59]
Discovery+UKBB [N=32598; P=4.126e-08] 7.46 [ 4.80, 10.13]
GOBS 21.90 [ 9.24, 34.56]
IMH -6.02 [-47.00, 34.96]
UNICAMP -39.50 [-79.88, 0.88]
Meth-CT
MIRECC
UKBB nonEUR 5.65 [-18.28, 29.58]
OSAKA -10.84 [-34.10, 12.43]
PING Non European 32.96 [ 5.30, 60.63]
Non European Cohorts [N=2538; P=0.01243] 11.51 [ 2.49, 20.54]
Beta [Lower 95% CI,Upper 95% CI]
-100
-50
0
50
100
150
mm^2/effect allele

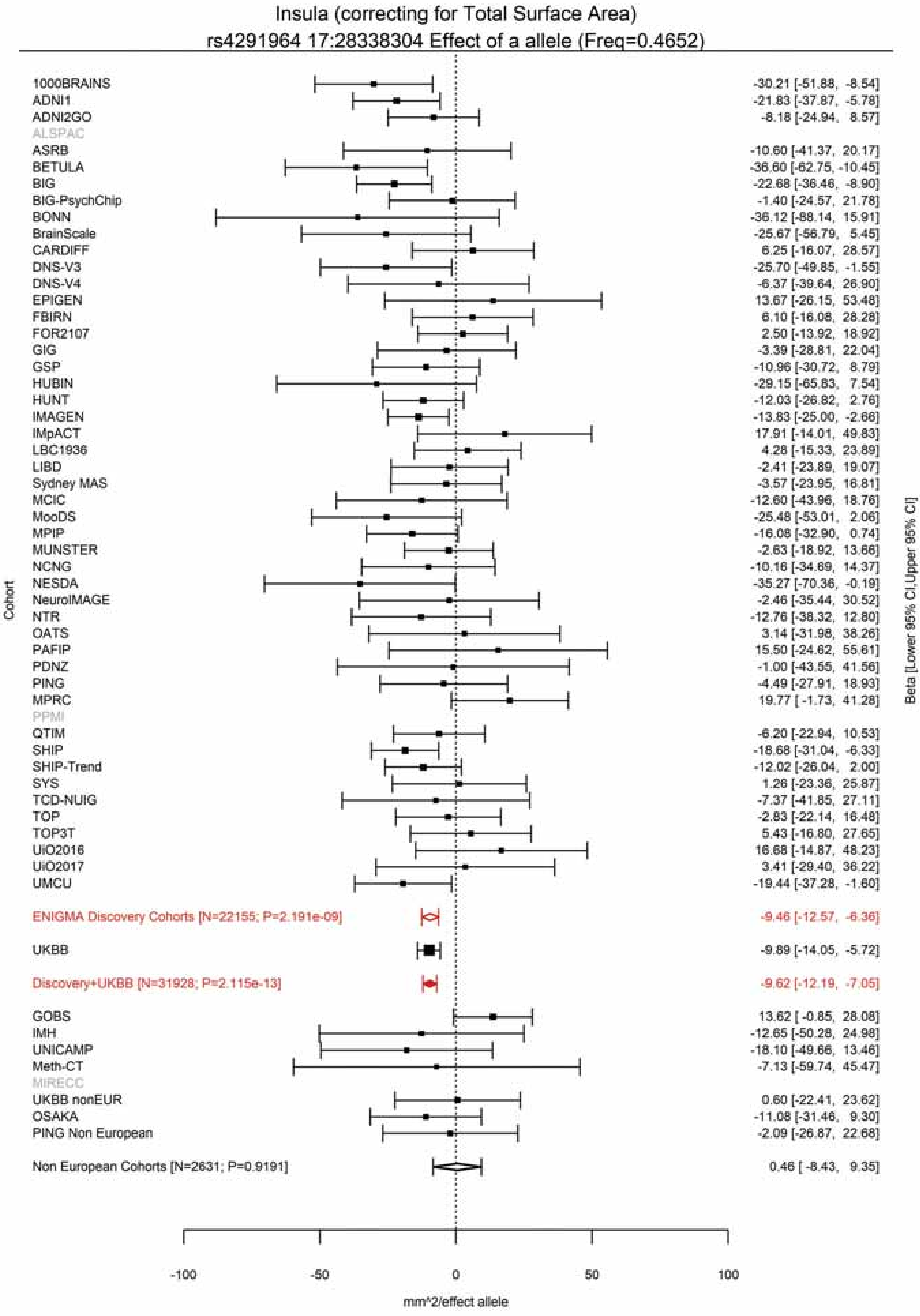

Insula (correcting for Total Surface Area)
rs4291964 17:28338304 Effect of a allele (Freq=0.4652)
Cohort
1000BRAINS -30.21 [-51.88, -8.54]
ADNI1 -21.83 [-37.87, -5.78]
ADNI2GO -8.18 [-24.94, 8.57]
ALSPAC
ASRB -10.60 [-41.37, 20.17]
BETULA -36.60 [-62.75, -10.45]
BIG -22.68 [-36.46, -8.90]
BIG-PsychChip -1.40 [-24.57, 21.78]
BONN -36.12 [-88.14, 15.91]
BrainScale -25.67 [-56.79, 5.45]
CARDIFF 6.25 [-16.07, 28.57]
DNS-V3 -25.70 [-49.85, -1.55]
DNS-V4 -6.37 [-39.64, 26.90]
EPIGEN 13.67 [-26.15, 53.48]
FBIRN 6.10 [-16.08, 28.28]
FOR2107 2.50 [-13.92, 18.92]
GIG -3.39 [-28.81, 22.04]
GSP -10.96 [-30.72, 8.79]
HUBIN -29.15 [-65.83, 7.54]
HUNT -12.03 [-26.82, 2.76]
IMAGEN -13.83 [-25.00, -2.66]
IMpACT 17.91 [-14.01, 49.83]
LBC1936 4.28 [-15.33, 23.89]
LIBD -2.41 [-23.89, 19.07]
Sydney MAS -3.57 [-23.95, 16.81]
MCIC -12.60 [-43.96, 18.76]
MooDS -25.48 [-53.01, 2.06]
MPIP -16.08 [-32.90, 0.74]
MUNSTER -2.63 [-18.92, 13.66]
NCNG -10.16 [-34.69, 14.37]
NESDA -35.27 [-70.36, -0.19]
NeuroIMAGE -2.46 [-35.44, 30.52]
NTR -12.76 [-38.32, 12.80]
OATS 3.14 [-31.98, 38.26]
PAFIP 15.50 [-24.62, 55.61]
PDNZ -1.00 [-43.55, 41.56]
PING -4.49 [-27.91, 18.93]
MPRC 19.77 [ -1.73, 41.28]
PPMI
QTIM -6.20 [-22.94, 10.53]
SHIP -18.68 [-31.04, -6.33]
SHIP-Trend -12.02 [-26.04, 2.00]
SYS 1.26 [-23.36, 25.87]
TCD-NUIG -7.37 [-41.85, 27.11]
TOP -2.83 [-22.14, 16.48]
TOP3T 5.43 [-16.80, 27.65]
UiO2016 16.68 [-14.87, 48.23]
UiO2017 3.41 [-29.40, 36.22]
UMCU -19.44 [-37.28, -1.60]
ENIGMA Discovery Cohorts [N=22155; P=2.191e-09] -9.46 [-12.57, -6.36]
UKBB -9.89 [-14.05, -5.72]
Discovery+UKBB [N=31928; P=2.115e-13] -9.62 [-12.19, -7.05]
GOBS 13.62 [ -0.85, 28.08]
IMH -12.65 [-50.28, 24.98]
UNICAMP -18.10 [-49.66, 13.46]
Meth-CT -7.13 [-59.74, 45.47]
MIRECC
UKBB nonEUR 0.60 [-22.41, 23.62]
OSAKA -11.08 [-31.46, 9.30]
PING Non European -2.09 [-26.87, 22.68]
Non European Cohorts [N=2631; P=0.9191] 0.46 [ -8.43, 9.35]
Beta [Lower 95% CI,Upper 95% CI]
-100
-50
0
50
100
mm^2/effect allele

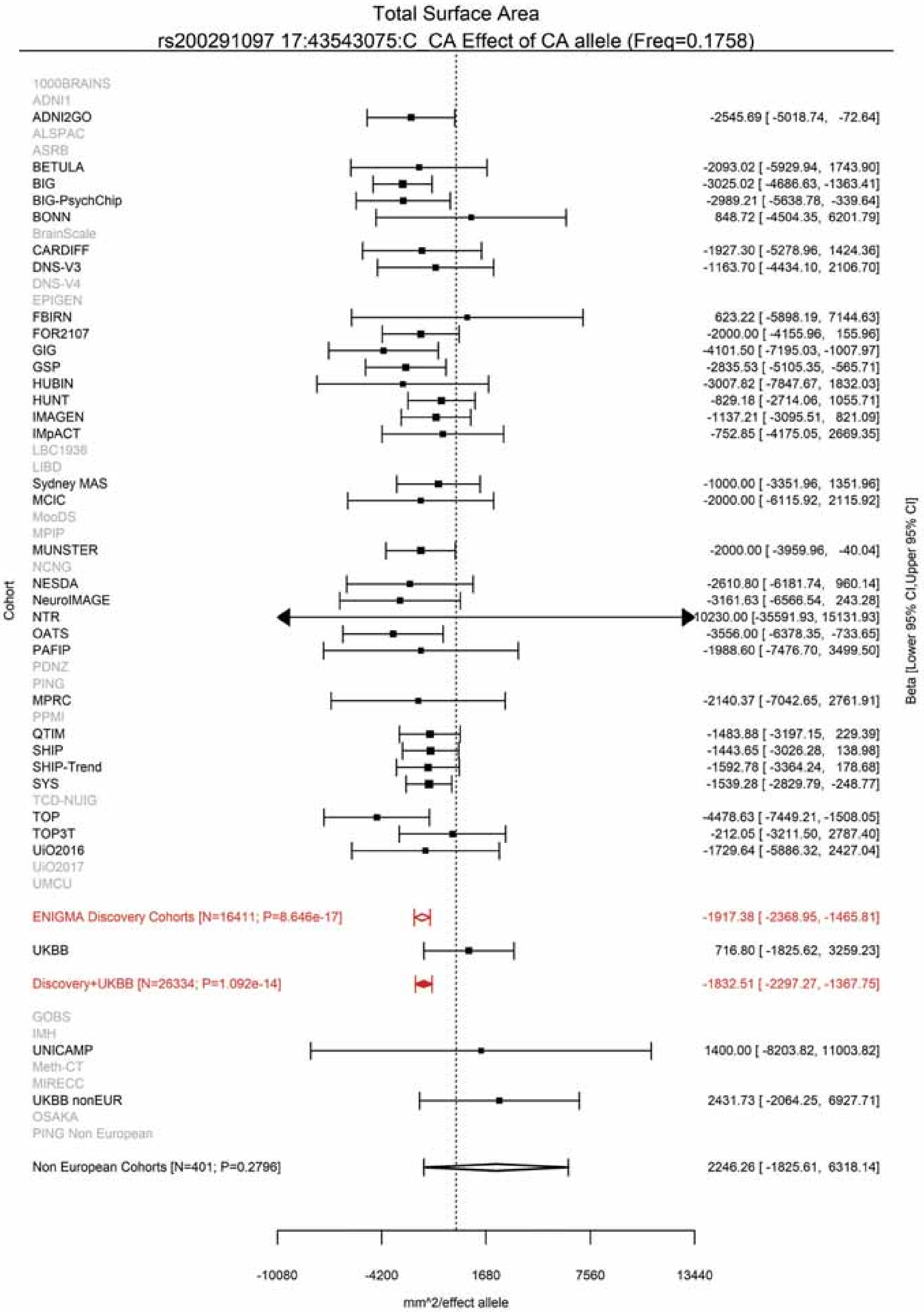

Total Surface Area
rs200291097 17:43543075:C_CA Effect of CA allele (Freq=0.1758)
Cohort
1000BRAINS
ADNI1
ADNI2GO -2545.69 [ -5018.74, -72.64]
ALSPAC
ASRB
BETULA -2093.02 [ -5929.94, 1743.90]
BIG -3025.02 [ -4686.63, -1363.41]
BIG-PsychChip -2989.21 [ -5638.78, -339.64]
BONN 848.72 [ -4504.35, 6201.79]
BrainScale
CARDIFF -1927.30 [ -5278.96, 1424.36]
DNS-V3 -1163.70 [ -4434.10, 2106.70]
DNS-V4
EPIGEN
FBIRN 623.22 [ -5898.19, 7144.63]
FOR2107 -2000.00 [ -4155.96, 155.96]
GIG -4101.50 [ -7195.03, -1007.97]
GSP -2835.53 [ -5105.35, -565.71]
HUBIN -3007.82 [ -7847.67, 1832.03]
HUNT -829.18 [ -2714.06, 1055.71]
IMAGEN -1137.21 [ -3095.51, 821.09]
IMpACT -752.85 [ -4175.05, 2669.35]
LBC1936
LIBD
Sydney MAS -1000.00 [ -3351.96, 1351.96]
MCIC -2000.00 [ -6115.92, 2115.92]
MooDS
MPIP
MUNSTER -2000.00 [ -3959.96, -40.04]
NCNG
NESDA -2610.80 [ -6181.74, 960.14]
NeuroIMAGE -3161.63 [ -6566.54, 243.28]
NTR -10230.00 [-35591.93, 15131.93]
OATS -3556.00 [ -6378.35, -733.65]
PAFIP -1988.60 [ -7476.70, 3499.50]
PDNZ
PING
MPRC -2140.37 [ -7042.65, 2761.91]
PPMI
QTIM -1483.88 [ -3197.15, 229.39]
SHIP -1443.65 [ -3026.28, 138.98]
SHIP-Trend -1592.78 [ -3364.24, 178.68]
SYS -1539.28 [ -2829.79, -248.77]
TCD-NUIG
TOP -4478.63 [ -7449.21, -1508.05]
TOP3T -212.05 [ -3211.50, 2787.40]
UiO2016 -1729.64 [ -5886.32, 2427.04]
UiO2017
UMCU
ENIGMA Discovery Cohorts [N=16411; P=8.646e-17] -1917.38 [ -2368.95, -1465.81]
UKBB 716.80 [ -1825.62, 3259.23]
Discovery+UKBB [N=26334; P=1.092e-14] -1832.51 [ -2297.27, -1367.75]
GOBS
IMH
UNICAMP 1400.00 [ -8203.82, 11003.82]
Meth-CT
MIRECC
UKBB nonEUR 2431.73 [ -2064.25, 6927.71]
OSAKA
PING Non European
Non European Cohorts [N=401; P=0.2796] 2246.26 [ -1825.61, 6318.14]
-10080
-4200
1680
7560
13440
mm^2/effect allele
Beta [Lower 95% CI,Upper 95% CI]

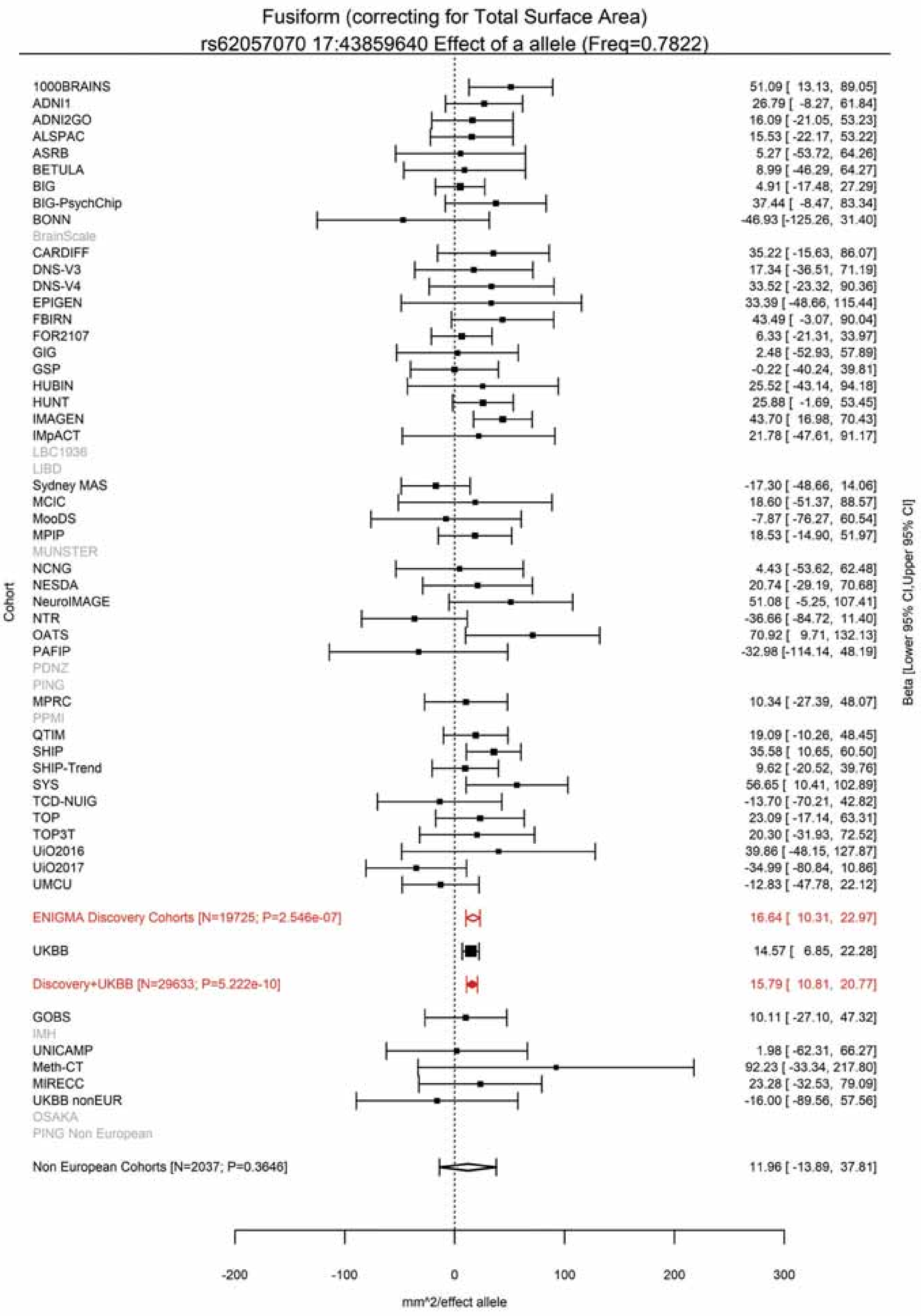

Fusiform (correcting for Total Surface Area)
rs62057070 17:43859640 Effect of a allele (Freq=0.7822)
1000BRAINS 51.09 [ 13.13, 89.05]
ADNI1 26.79 [ -8.27, 61.84]
ADNI2GO 16.09 [ -21.05, 53.23]
ALSPAC 15.53 [ -22.17, 53.22]
ASRB 5.27 [ -53.72, 64.26]
BETULA 8.99 [ -46.29, 64.27]
BIG 4.91 [ -17.48, 27.29]
BIG-PsychChip 37.44 [ -8.47, 83.34]
BONN -46.93 [-125.26, 31.40]
BrainScale
CARDIFF 35.22 [ -15.63, 86.07]
DNS-V3 17.34 [ -36.51, 71.19]
DNS-V4 33.52 [ -23.32, 90.36]
EPIGEN 33.39 [ -48.66, 115.44]
FBIRN 43.49 [ -3.07, 90.04]
FOR2107 6.33 [ -21.31, 33.97]
GIG 2.48 [ -52.93, 57.89]
GSP -0.22 [ -40.24, 39.81]
HUBIN 25.52 [ -43.14, 94.18]
HUNT 25.88 [ -1.69, 53.45]
IMAGEN 43.70 [ 16.98, 70.43]
IMpACT 21.78 [ -47.61, 91.17]
LBC1936
LIBD
Sydney MAS -17.30 [ -48.66, 14.06]
MCIC 18.60 [ -51.37, 88.57]
MooDS -7.87 [ -76.27, 60.54]
MPIP 18.53 [ -14.90, 51.97]
MUNSTER
NCNG 4.43 [ -53.62, 62.48]
NESDA 20.74 [ -29.19, 70.68]
NeuroIMAGE 51.08 [ -5.25, 107.41]
NTR -36.66 [ -84.72, 11.40]
OATS 70.92 [ 9.71, 132.13]
PAFIP -32.98 [-114.14, 48.19]
PDNZ
PING
MPRC 10.34 [ -27.39, 48.07]
PPMI
QTIM 19.09 [ -10.26, 48.45]
SHIP 35.58 [ 10.65, 60.50]
SHIP-Trend 9.62 [ -20.52, 39.76]
SYS 56.65 [ 10.41, 102.89]
TCD-NUIG -13.70 [ -70.21, 42.82]
TOP 23.09 [ -17.14, 63.31]
TOP3T 20.30 [ -31.93, 72.52]
UiO2016 39.86 [ -48.15, 127.87]
UiO2017 -34.99 [ -80.84, 10.86]
UMCU -12.83 [ -47.78, 22.12]
ENIGMA Discovery Cohorts [N=19725; P=2.546e-07] 16.64 [ 10.31, 22.97]
UKBB 14.57 [ 6.85, 22.28]
Discovery+UKBB [N=29633; P=5.222e-10] 15.79 [ 10.81, 20.77]
GOBS 10.11 [ -27.10, 47.32]
IMH
UNICAMP 1.98 [ -62.31, 66.27]
Meth-CT 92.23 [ -33.34, 217.80]
MIRECC 23.28 [ -32.53, 79.09]
UKBB nonEUR -16.00 [ -89.56, 57.56]
OSAKA
PING Non European
Non European Cohorts [N=2037; P=0.3646] 11.96 [ -13.89, 37.81]
Cohort
Beta [Lower 95% CI,Upper 95% CI]
-200
-100
0
100
200
300
mm^2/effect allele

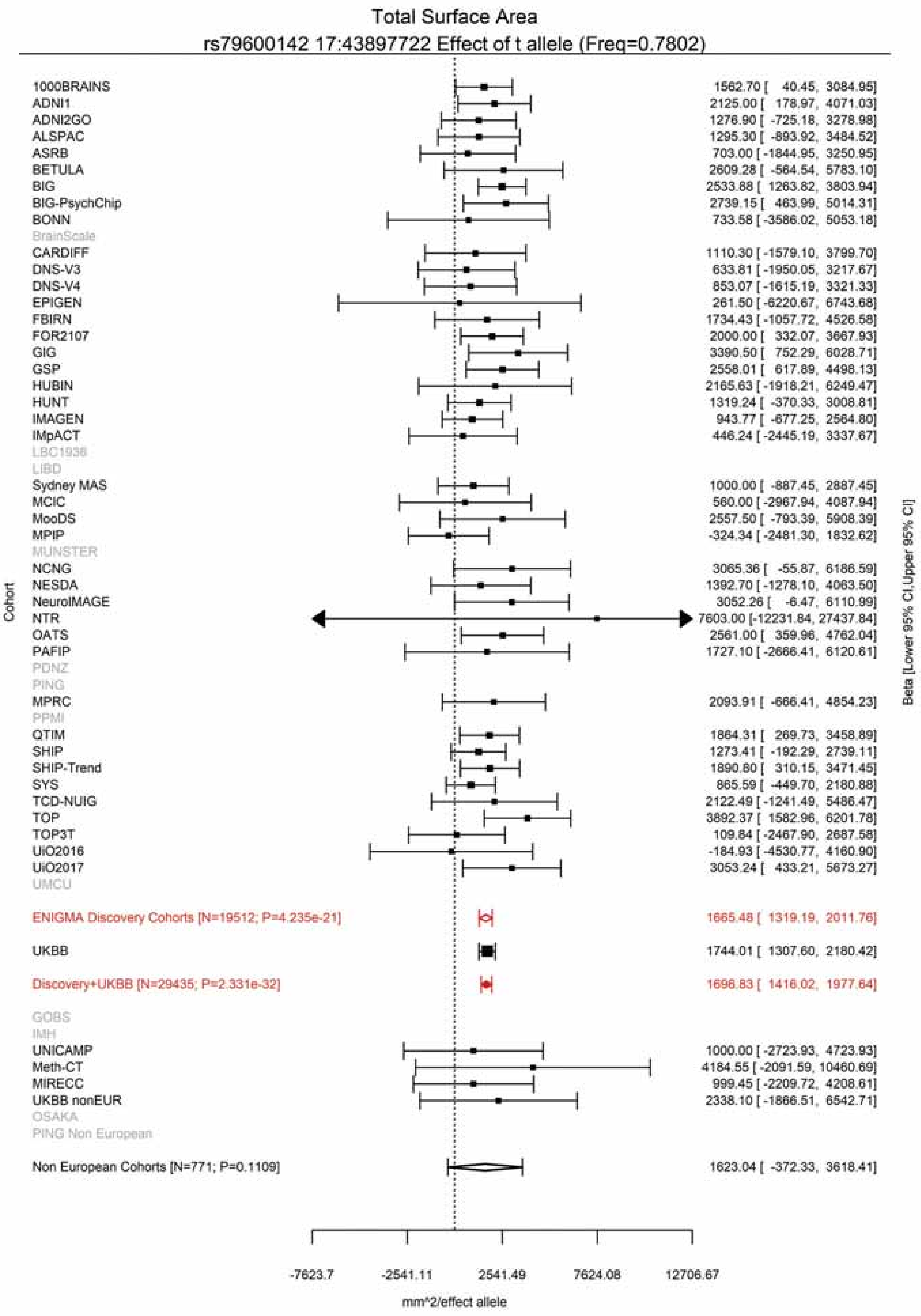
Total Surface Area
rs79600142 17:43897722 Effect of t allele (Freq=0.7802)
1000BRAINS 1562.70 [ 40.45, 3084.95]
ADNI1 2125.00 [ 178.97, 4071.03]
ADNI2GO 1276.90 [ -725.18, 3278.98]
ALSPAC 1295.30 [ -893.92, 3484.52]
ASRB 703.00 [ -1844.95, 3250.95]
BETULA 2609.28 [ -564.54, 5783.10]
BIG 2533.88 [ 1263.82, 3803.94]
BIG-PsychChip 2739.15 [ 463.99, 5014.31]
BONN 733.58 [ -3586.02, 5053.18]
BrainScale
CARDIFF 1110.30 [ -1579.10, 3799.70]
DNS-V3 633.81 [ -1950.05, 3217.67]
DNS-V4 853.07 [ -1615.19, 3321.33]
EPIGEN 261.50 [ -6220.67, 6743.68]
FBIRN 1734.43 [ -1057.72, 4526.58]
FOR2107 2000.00 [ 332.07, 3667.93]
GIG 3390.50 [ 752.29, 6028.71]
GSP 2558.01 [ 617.89, 4498.13]
HUBIN 2165.63 [ -1918.21, 6249.47]
HUNT 1319.24 [ -370.33, 3008.81]
IMAGEN 943.77 [ -677.25, 2564.80]
IMpACT 446.24 [ -2445.19, 3337.67]
LBC1936
LIBD
Sydney MAS 1000.00 [ -887.45, 2887.45]
MCIC 560.00 [ -2967.94, 4087.94]
MooDS 2557.50 [ -793.39, 5908.39]
MPIP -324.34 [ -2481.30, 1832.62]
MUNSTER
NCNG 3065.36 [ -55.87, 6186.59]
NESDA 1392.70 [ -1278.10, 4063.50]
NeuroIMAGE 3052.26 [ -6.47, 6110.99]
NTR 7603.00 [-12231.84, 27437.84]
OATS 2561.00 [ 359.96, 4762.04]
PAFIP 1727.10 [ -2666.41, 6120.61]
PDNZ
PING
MPRC 2093.91 [ -666.41, 4854.23]
PPMI
QTIM 1864.31 [ 269.73, 3458.89]
SHIP 1273.41 [ -192.29, 2739.11]
SHIP-Trend 1890.80 [ 310.15, 3471.45]
SYS 865.59 [ -449.70, 2180.88]
TCD-NUIG 2122.49 [ -1241.49, 5486.47]
TOP 3892.37 [ 1582.96, 6201.78]
TOP3T 109.84 [ -2467.90, 2687.58]
UiO2016 -184.93 [ -4530.77, 4160.90]
UiO2017 3053.24 [ 433.21, 5673.27]
UMCU
ENIGMA Discovery Cohorts [N=19512; P=4.235e-21] 1665.48 [ 1319.19, 2011.76]
UKBB 1744.01 [ 1307.60, 2180.42]
Discovery+UKBB [N=29435; P=2.331e-32] 1696.83 [ 1416.02, 1977.64]
GOBS
IMH
UNICAMP 1000.00 [ -2723.93, 4723.93]
Meth-CT 4184.55 [ -2091.59, 10460.69]
MIRECC 999.45 [ -2209.72, 4208.61]
UKBB nonEUR 2338.10 [ -1866.51, 6542.71]
OSAKA
PING Non European
Non European Cohorts [N=771; P=0.1109] 1623.04 [ -372.33, 3618.41]
Cohort
Beta [Lower 95% CI,Upper 95% CI]
-7623.7
-2541.11
2541.49
7624.08
12706.67
mm^2/effect allele

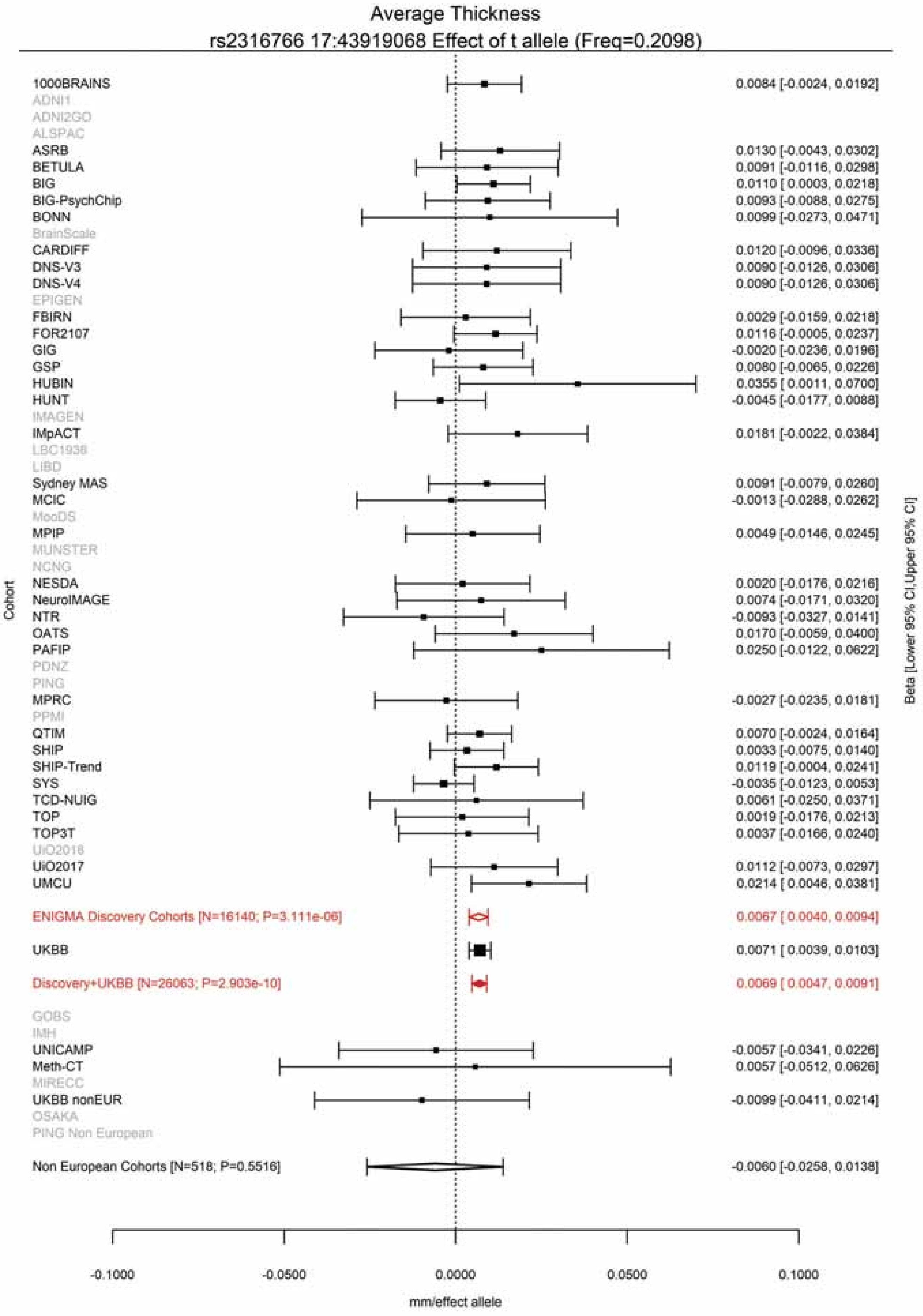
Average Thickness
rs2316766 17:43919068 Effect of t allele (Freq=0.2098)
Cohort
1000BRAINS 0.0084 [-0.0024, 0.0192]
ADNI1
ADNI2GO
ALSPAC
ASRB 0.0130 [-0.0043, 0.0302]
BETULA 0.0091 [-0.0116, 0.0298]
BIG 0.0110 [ 0.0003, 0.0218]
BIG-PsychChip 0.0093 [-0.0088, 0.0275]
BONN 0.0099 [-0.0273, 0.0471]
BrainScale
CARDIFF 0.0120 [-0.0096, 0.0336]
DNS-V3 0.0090 [-0.0126, 0.0306]
DNS-V4 0.0090 [-0.0126, 0.0306]
EPIGEN
FBIRN 0.0029 [-0.0159, 0.0218]
FOR2107 0.0116 [-0.0005, 0.0237]
GIG -0.0020 [-0.0236, 0.0196]
GSP 0.0080 [-0.0065, 0.0226]
HUBIN 0.0355 [ 0.0011, 0.0700]
HUNT -0.0045 [-0.0177, 0.0088]
IMAGEN
IMpACT 0.0181 [-0.0022, 0.0384]
LBC1936
LIBD
Sydney MAS 0.0091 [-0.0079, 0.0260]
MCIC -0.0013 [-0.0288, 0.0262]
MooDS
MPIP 0.0049 [-0.0146, 0.0245]
MUNSTER
NCNG
NESDA 0.0020 [-0.0176, 0.0216]
NeuroIMAGE 0.0074 [-0.0171, 0.0320]
NTR -0.0093 [-0.0327, 0.0141]
OATS 0.0170 [-0.0059, 0.0400]
PAFIP 0.0250 [-0.0122, 0.0622]
PDNZ
PING
MPRC -0.0027 [-0.0235, 0.0181]
PPMI
QTIM 0.0070 [-0.0024, 0.0164]
SHIP 0.0033 [-0.0075, 0.0140]
SHIP-Trend 0.0119 [-0.0004, 0.0241]
SYS -0.0035 [-0.0123, 0.0053]
TCD-NUIG 0.0061 [-0.0250, 0.0371]
TOP 0.0019 [-0.0176, 0.0213]
TOP3T 0.0037 [-0.0166, 0.0240]
UiO2016
UiO2017 0.0112 [-0.0073, 0.0297]
UMCU 0.0214 [ 0.0046, 0.0381]
ENIGMA Discovery Cohorts [N=16140; P=3.111e-06] 0.0067 [ 0.0040, 0.0094]
UKBB 0.0071 [ 0.0039, 0.0103]
Discovery+UKBB [N=26063; P=2.903e-10] 0.0069 [ 0.0047, 0.0091]
GOBS
IMH
UNICAMP -0.0057 [-0.0341, 0.0226]
Meth-CT 0.0057 [-0.0512, 0.0626]
MIRECC
UKBB nonEUR -0.0099 [-0.0411, 0.0214]
OSAKA
PING Non European
Non European Cohorts [N=518; P=0.5516] -0.0060 [-0.0258, 0.0138]
Beta [Lower 95% CI,Upper 95% CI]
-0.1000
-0.0500
0.0000
0.0500
0.1000
mm/effect allele

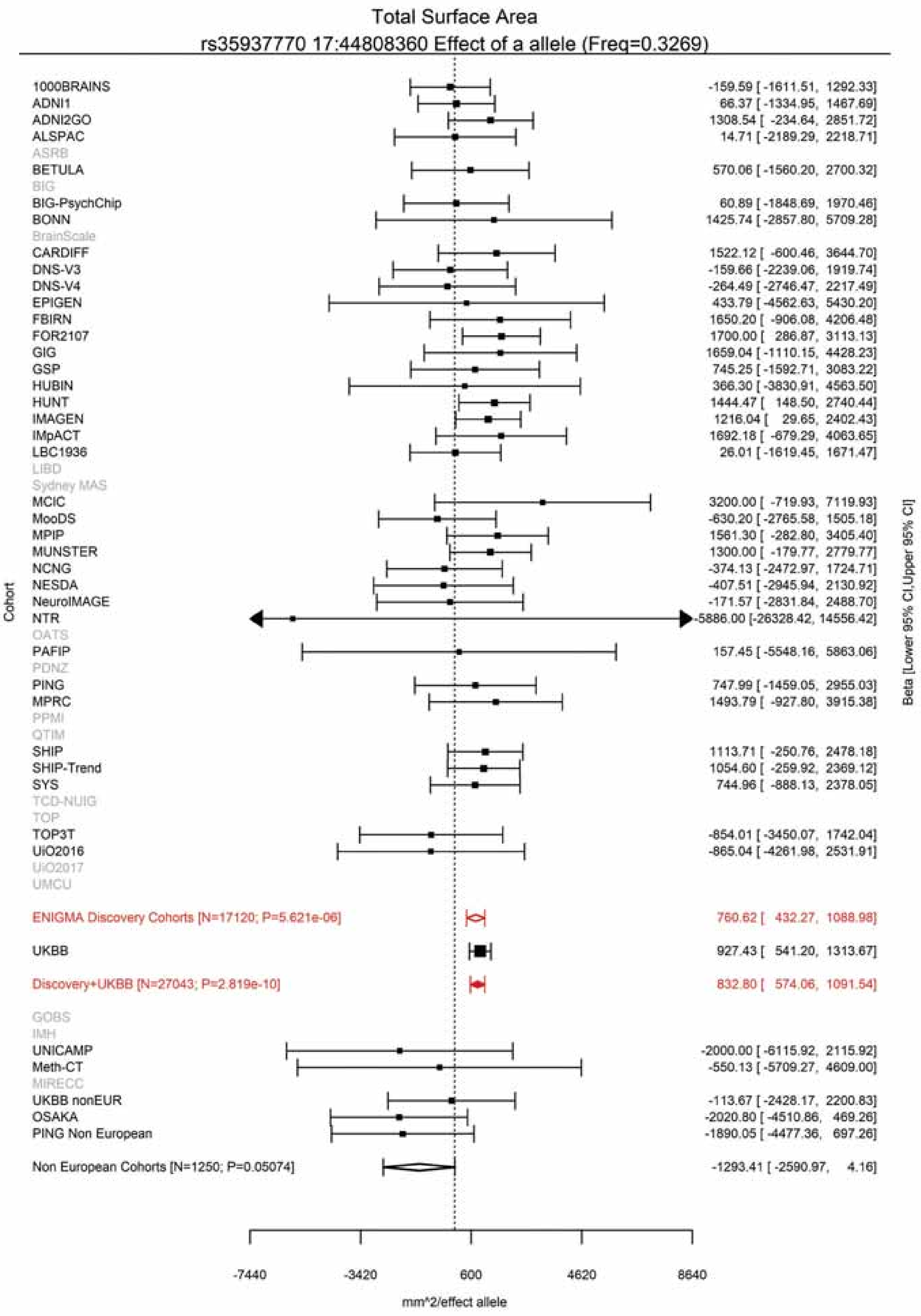
Total Surface Area
rs35937770 17:44808360 Effect of a allele (Freq=0.3269)
1000BRAINS -159.59 [ -1611.51, 1292.33]
ADNI1 66.37 [ -1334.95, 1467.69]
ADNI2GO 1308.54 [ -234.64, 2851.72]
ALSPAC 14.71 [ -2189.29, 2218.71]
ASRB
BETULA 570.06 [ -1560.20, 2700.32]
BIG
BIG-PsychChip 60.89 [ -1848.69, 1970.46]
BONN 1425.74 [ -2857.80, 5709.28]
BrainScale
CARDIFF 1522.12 [ -600.46, 3644.70]
DNS-V3 -159.66 [ -2239.06, 1919.74]
DNS-V4 -264.49 [ -2746.47, 2217.49]
EPIGEN 433.79 [ -4562.63, 5430.20]
FBIRN 1650.20 [ -906.08, 4206.48]
FOR2107 1700.00 [ 286.87, 3113.13]
GIG 1659.04 [ -1110.15, 4428.23]
GSP 745.25 [ -1592.71, 3083.22]
HUBIN 366.30 [ -3830.91, 4563.50]
HUNT 1444.47 [ 148.50, 2740.44]
IMAGEN 1216.04 [ 29.65, 2402.43]
IMpACT 1692.18 [ -679.29, 4063.65]
LBC1936 26.01 [ -1619.45, 1671.47]
LIBD
Sydney MAS
MCIC 3200.00 [ -719.93, 7119.93]
MooDS -630.20 [ -2765.58, 1505.18]
MPIP 1561.30 [ -282.80, 3405.40]
MUNSTER 1300.00 [ -179.77, 2779.77]
NCNG -374.13 [ -2472.97, 1724.71]
NESDA -407.51 [ -2945.94, 2130.92]
NeuroIMAGE -171.57 [ -2831.84, 2488.70]
NTR -5886.00 [-26328.42, 14556.42]
OATS
PAFIP 157.45 [ -5548.16, 5863.06]
PDNZ
PING 747.99 [ -1459.05, 2955.03]
MPRC 1493.79 [ -927.80, 3915.38]
PPMI
QTIM
SHIP 1113.71 [ -250.76, 2478.18]
SHIP-Trend 1054.60 [ -259.92, 2369.12]
SYS 744.96 [ -888.13, 2378.05]
TCD-NUIG
TOP
TOP3T -854.01 [ -3450.07, 1742.04]
UiO2016 -865.04 [ -4261.98, 2531.91]
UiO2017
UMCU
ENIGMA Discovery Cohorts [N=17120; P=5.621e-06] 760.62 [ 432.27, 1088.98]
UKBB 927.43 [ 541.20, 1313.67]
Discovery+UKBB [N=27043; P=2.819e-10] 832.80 [ 574.06, 1091.54]
GOBS
IMH
UNICAMP -2000.00 [ -6115.92, 2115.92]
Meth-CT -550.13 [ -5709.27, 4609.00]
MIRECC
UKBB nonEUR -113.67 [ -2428.17, 2200.83]
OSAKA -2020.80 [ -4510.86, 469.26]
PING Non European -1890.05 [ -4477.36, 697.26]
Non European Cohorts [N=1250; P=0.05074] -1293.41 [ -2590.97, 4.16]
Cohort
Beta [Lower 95% CI,Upper 95% CI]
-7440
-3420
600
4620
8640
mm^2/effect allele

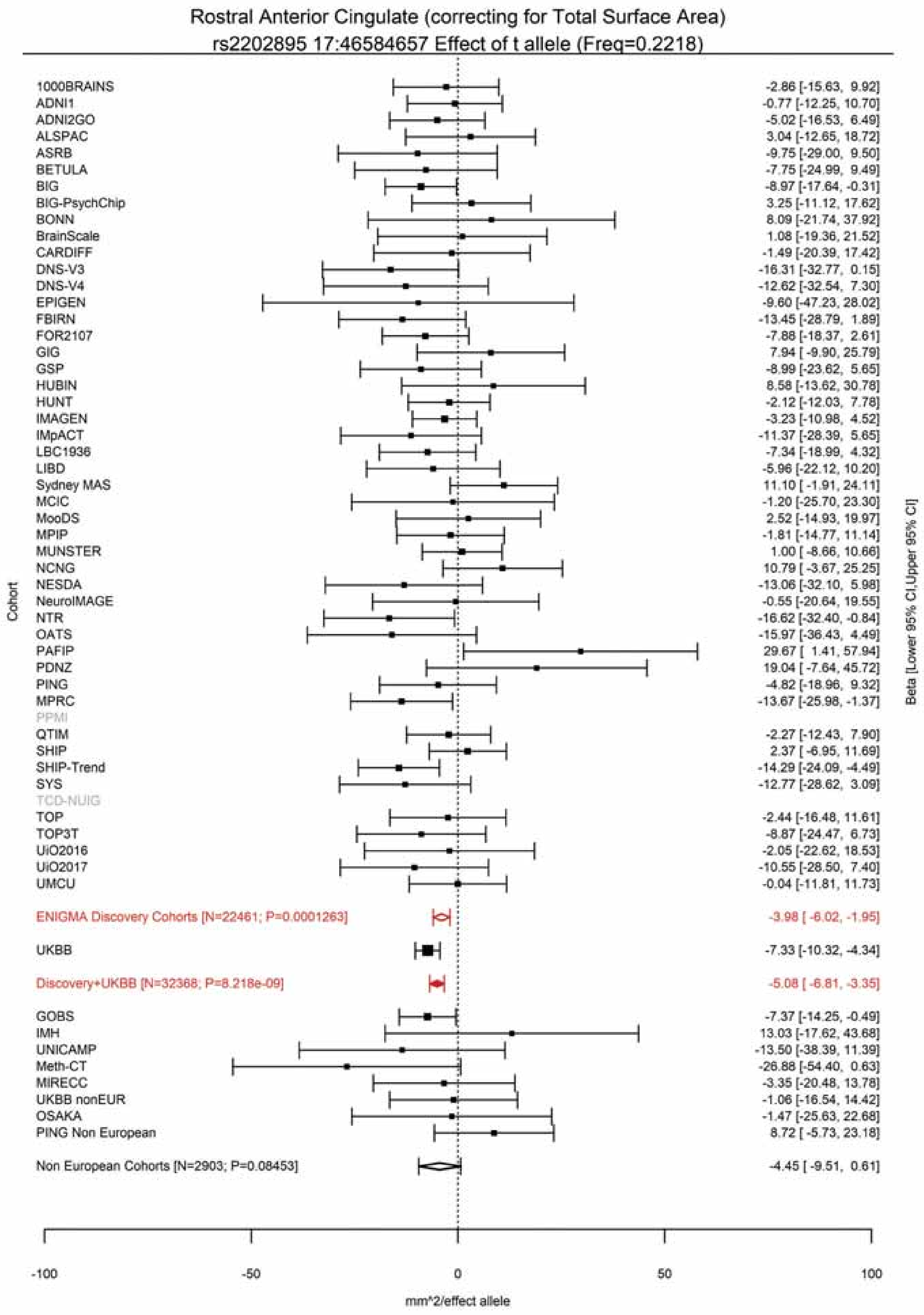
Rostral Anterior Cingulate (correcting for Total Surface Area)
rs2202895 17:46584657 Effect of t allele (Freq=0.2218)
Cohort
Beta [Lower 95% CI,Upper 95% CI]
1000BRAINS -2.86 [-15.63, 9.92]
ADNI1 -0.77 [-12.25, 10.70]
ADNI2GO -5.02 [-16.53, 6.49]
ALSPAC 3.04 [-12.65, 18.72]
ASRB -9.75 [-29.00, 9.50]
BETULA -7.75 [-24.99, 9.49]
BIG -8.97 [-17.64, -0.31]
BIG-PsychChip 3.25 [-11.12, 17.62]
BONN 8.09 [-21.74, 37.92]
BrainScale 1.08 [-19.36, 21.52]
CARDIFF -1.49 [-20.39, 17.42]
DNS-V3 -16.31 [-32.77, 0.15]
DNS-V4 -12.62 [-32.54, 7.30]
EPIGEN -9.60 [-47.23, 28.02]
FBIRN -13.45 [-28.79, 1.89]
FOR2107 -7.88 [-18.37, 2.61]
GIG 7.94 [-9.90, 25.79]
GSP -8.99 [-23.62, 5.65]
HUBIN 8.58 [-13.62, 30.78]
HUNT -2.12 [-12.03, 7.78]
IMAGEN -3.23 [-10.98, 4.52]
IMpACT -11.37 [-28.39, 5.65]
LBC1936 -7.34 [-18.99, 4.32]
LIBD -5.96 [-22.12, 10.20]
Sydney MAS 11.10 [-1.91, 24.11]
MCIC -1.20 [-25.70, 23.30]
MooDS 2.52 [-14.93, 19.97]
MPIP -1.81 [-14.77, 11.14]
MUNSTER 1.00 [-8.66, 10.66]
NCNG 10.79 [-3.67, 25.25]
NESDA -13.06 [-32.10, 5.98]
NeuroIMAGE -0.55 [-20.64, 19.55]
NTR -16.62 [-32.40, -0.84]
OATS -15.97 [-36.43, 4.49]
PAFIP 29.67 [1.41, 57.94]
PDNZ 19.04 [-7.64, 45.72]
PING -4.82 [-18.96, 9.32]
MPRC -13.67 [-25.98, -1.37]
PPMI
QTIM -2.27 [-12.43, 7.90]
SHIP 2.37 [-6.95, 11.69]
SHIP-Trend -14.29 [-24.09, -4.49]
SYS -12.77 [-28.62, 3.09]
TCD-NUIG
TOP -2.44 [-16.48, 11.61]
TOP3T -8.87 [-24.47, 6.73]
UiO2016 -2.05 [-22.62, 18.53]
UiO2017 -10.55 [-28.50, 7.40]
UMCU -0.04 [-11.81, 11.73]
ENIGMA Discovery Cohorts [N=22461; P=0.0001263] -3.98 [-6.02, -1.95]
UKBB -7.33 [-10.32, -4.34]
Discovery+UKBB [N=32368; P=8.218e-09] -5.08 [-6.81, -3.35]
GOBS -7.37 [-14.25, -0.49]
IMH 13.03 [-17.62, 43.68]
UNICAMP -13.50 [-38.39, 11.39]
Meth-CT -26.88 [-54.40, 0.63]
MIRECC -3.35 [-20.48, 13.78]
UKBB nonEUR -1.06 [-16.54, 14.42]
OSAKA -1.47 [-25.63, 22.68]
PING Non European 8.72 [-5.73, 23.18]
Non European Cohorts [N=2903; P=0.08453] -4.45 [-9.51, 0.61]
-100
-50
0
50
100
mm^2/effect allele

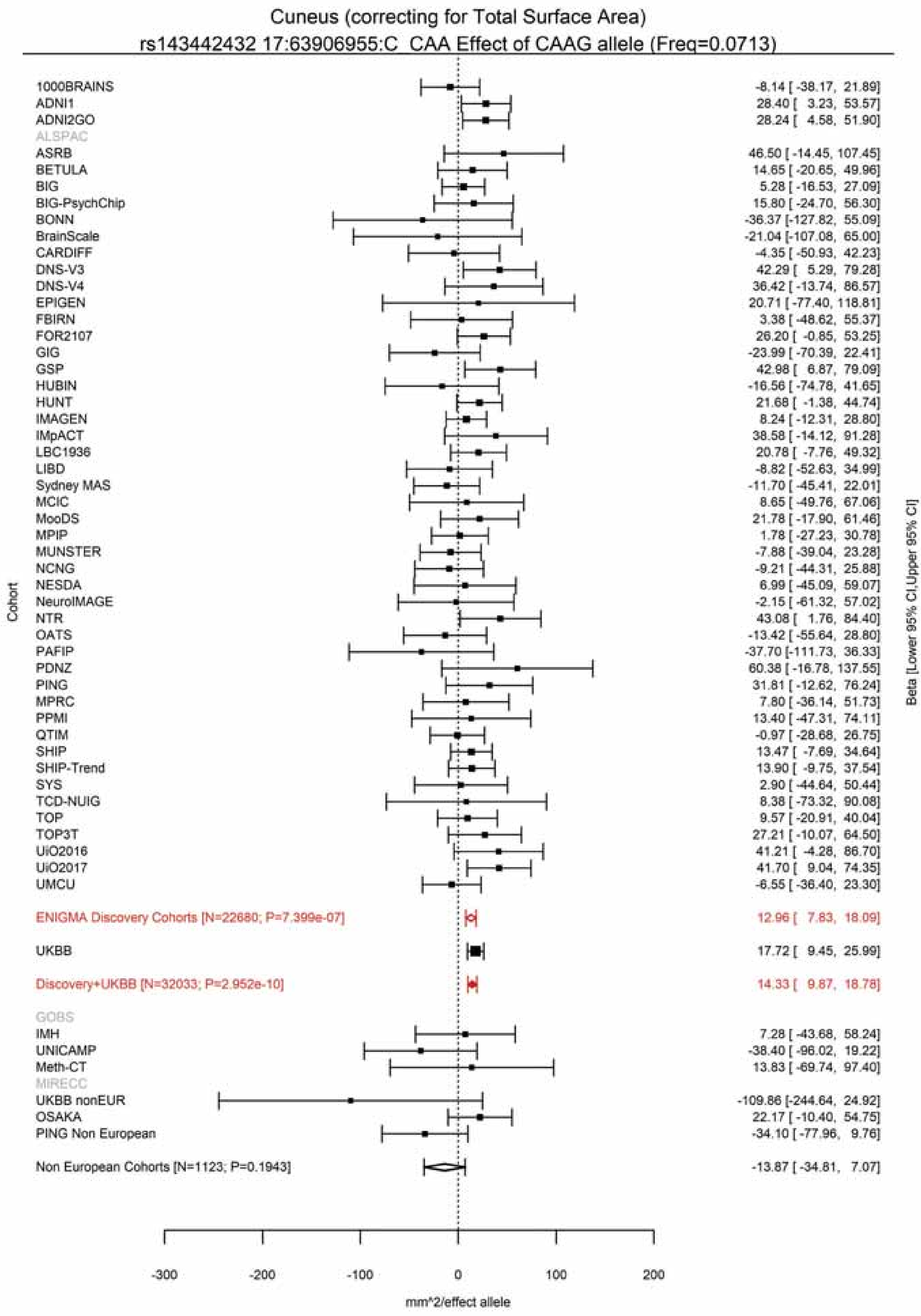

Cuneus (correcting for Total Surface Area)
rs143442432 17:63906955:C_CAA Effect of CAAG allele (Freq=0.0713)
Cohort
1000BRAINS -8.14 [ -38.17, 21.89]
ADNI1 28.40 [ 3.23, 53.57]
ADNI2GO 28.24 [ 4.58, 51.90]
ALSPAC
ASRB 46.50 [ -14.45, 107.45]
BETULA 14.65 [ -20.65, 49.96]
BIG 5.28 [ -16.53, 27.09]
BIG-PsychChip 15.80 [ -24.70, 56.30]
BONN -36.37 [-127.82, 55.09]
BrainScale -21.04 [-107.08, 65.00]
CARDIFF -4.35 [ -50.93, 42.23]
DNS-V3 42.29 [ 5.29, 79.28]
DNS-V4 36.42 [ -13.74, 86.57]
EPIGEN 20.71 [ -77.40, 118.81]
FBIRN 3.38 [ -48.62, 55.37]
FOR2107 26.20 [ -0.85, 53.25]
GIG -23.99 [ -70.39, 22.41]
GSP 42.98 [ 6.87, 79.09]
HUBIN -16.56 [ -74.78, 41.65]
HUNT 21.68 [ -1.38, 44.74]
IMAGEN 8.24 [ -12.31, 28.80]
IMpACT 38.58 [ -14.12, 91.28]
LBC1936 20.78 [ -7.76, 49.32]
LIBD -8.82 [ -52.63, 34.99]
Sydney MAS -11.70 [ -45.41, 22.01]
MCIC 8.65 [ -49.76, 67.06]
MooDS 21.78 [ -17.90, 61.46]
MPIP 1.78 [ -27.23, 30.78]
MUNSTER -7.88 [ -39.04, 23.28]
NCNG -9.21 [ -44.31, 25.88]
NESDA 6.99 [ -45.09, 59.07]
NeuroIMAGE -2.15 [ -61.32, 57.02]
NTR 43.08 [ 1.76, 84.40]
OATS -13.42 [ -55.64, 28.80]
PAFIP -37.70 [-111.73, 36.33]
PDNZ 60.38 [ -16.78, 137.55]
PING 31.81 [ -12.62, 76.24]
MPRC 7.80 [ -36.14, 51.73]
PPMI 13.40 [ -47.31, 74.11]
QTIM -0.97 [ -28.68, 26.75]
SHIP 13.47 [ -7.69, 34.64]
SHIP-Trend 13.90 [ -9.75, 37.54]
SYS 2.90 [ -44.64, 50.44]
TCD-NUIG 8.38 [ -73.32, 90.08]
TOP 9.57 [ -20.91, 40.04]
TOP3T 27.21 [ -10.07, 64.50]
UiO2016 41.21 [ -4.28, 86.70]
UiO2017 41.70 [ 9.04, 74.35]
UMCU -6.55 [ -36.40, 23.30]
ENIGMA Discovery Cohorts [N=22680; P=7.399e-07] 12.96 [ 7.83, 18.09]
UKBB 17.72 [ 9.45, 25.99]
Discovery+UKBB [N=32033; P=2.952e-10] 14.33 [ 9.87, 18.78]
GOBS
IMH 7.28 [ -43.68, 58.24]
UNICAMP -38.40 [ -96.02, 19.22]
Meth-CT 13.83 [ -69.74, 97.40]
MIRECC
UKBB nonEUR -109.86 [-244.64, 24.92]
OSAKA 22.17 [ -10.40, 54.75]
PING Non European -34.10 [ -77.96, 9.76]
Non European Cohorts [N=1123; P=0.1943] -13.87 [ -34.81, 7.07]
-300
-200
-100
0
100
200
mm^2/effect allele
Beta [Lower 95% CI,Upper 95% CI]

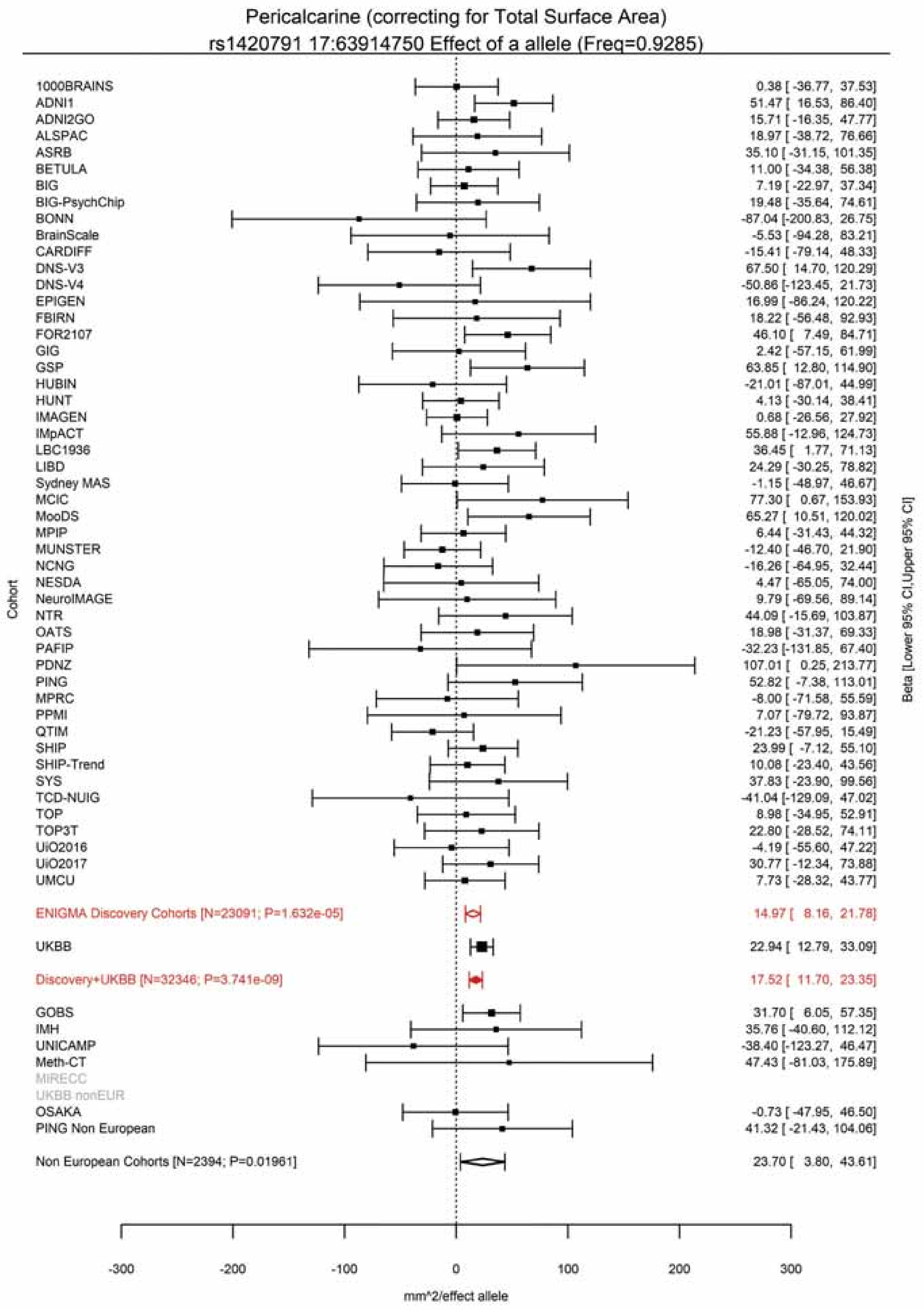

Pericalcarine (correcting for Total Surface Area)
rs1420791 17:63914750 Effect of a allele (Freq=0.9285)
Cohort
1000BRAINS 0.38 [ -36.77, 37.53]
ADNI1 51.47 [ 16.53, 86.40]
ADNI2GO 15.71 [ -16.35, 47.77]
ALSPAC 18.97 [ -38.72, 76.66]
ASRB 35.10 [ -31.15, 101.35]
BETULA 11.00 [ -34.38, 56.38]
BIG 7.19 [ -22.97, 37.34]
BIG-PsychChip 19.48 [ -35.64, 74.61]
BONN -87.04 [-200.83, 26.75]
BrainScale -5.53 [ -94.28, 83.21]
CARDIFF -15.41 [ -79.14, 48.33]
DNS-V3 67.50 [ 14.70, 120.29]
DNS-V4 -50.86 [-123.45, 21.73]
EPIGEN 16.99 [ -86.24, 120.22]
FBIRN 18.22 [ -56.48, 92.93]
FOR2107 46.10 [ 7.49, 84.71]
GIG 2.42 [ -57.15, 61.99]
GSP 63.85 [ 12.80, 114.90]
HUBIN -21.01 [ -87.01, 44.99]
HUNT 4.13 [ -30.14, 38.41]
IMAGEN 0.68 [ -26.56, 27.92]
IMpACT 55.88 [ -12.96, 124.73]
LBC1936 36.45 [ 1.77, 71.13]
LIBD 24.29 [ -30.25, 78.82]
Sydney MAS -1.15 [ -48.97, 46.67]
MCIC 77.30 [ 0.67, 153.93]
MooDS 65.27 [ 10.51, 120.02]
MPIP 6.44 [ -31.43, 44.32]
MUNSTER -12.40 [ -46.70, 21.90]
NCNG -16.26 [ -64.95, 32.44]
NESDA 4.47 [ -65.05, 74.00]
NeuroIMAGE 9.79 [ -69.56, 89.14]
NTR 44.09 [ -15.69, 103.87]
OATS 18.98 [ -31.37, 69.33]
PAFIP -32.23 [-131.85, 67.40]
PDNZ 107.01 [ 0.25, 213.77]
PING 52.82 [ -7.38, 113.01]
MPRC -8.00 [ -71.58, 55.59]
PPMI 7.07 [ -79.72, 93.87]
QTIM -21.23 [ -57.95, 15.49]
SHIP 23.99 [ -7.12, 55.10]
SHIP-Trend 10.08 [ -23.40, 43.56]
SYS 37.83 [ -23.90, 99.56]
TCD-NUIG -41.04 [-129.09, 47.02]
TOP 8.98 [ -34.95, 52.91]
TOP3T 22.80 [ -28.52, 74.11]
UiO2016 -4.19 [ -55.60, 47.22]
UiO2017 30.77 [ -12.34, 73.88]
UMCU 7.73 [ -28.32, 43.77]
ENIGMA Discovery Cohorts [N=23091; P=1.632e-05] 14.97 [ 8.16, 21.78]
UKBB 22.94 [ 12.79, 33.09]
Discovery+UKBB [N=32346; P=3.741e-09] 17.52 [ 11.70, 23.35]
GOBS 31.70 [ 6.05, 57.35]
IMH 35.76 [ -40.60, 112.12]
UNICAMP -38.40 [-123.27, 46.47]
Meth-CT 47.43 [ -81.03, 175.89]
MIRECC
UKBB nonEUR
OSAKA -0.73 [ -47.95, 46.50]
PING Non European 41.32 [ -21.43, 104.06]
Non European Cohorts [N=2394; P=0.01961] 23.70 [ 3.80, 43.61]
Beta [Lower 95% CI,Upper 95% CI]
-300
-200
-100
0
100
200
300
mm^2/effect allele

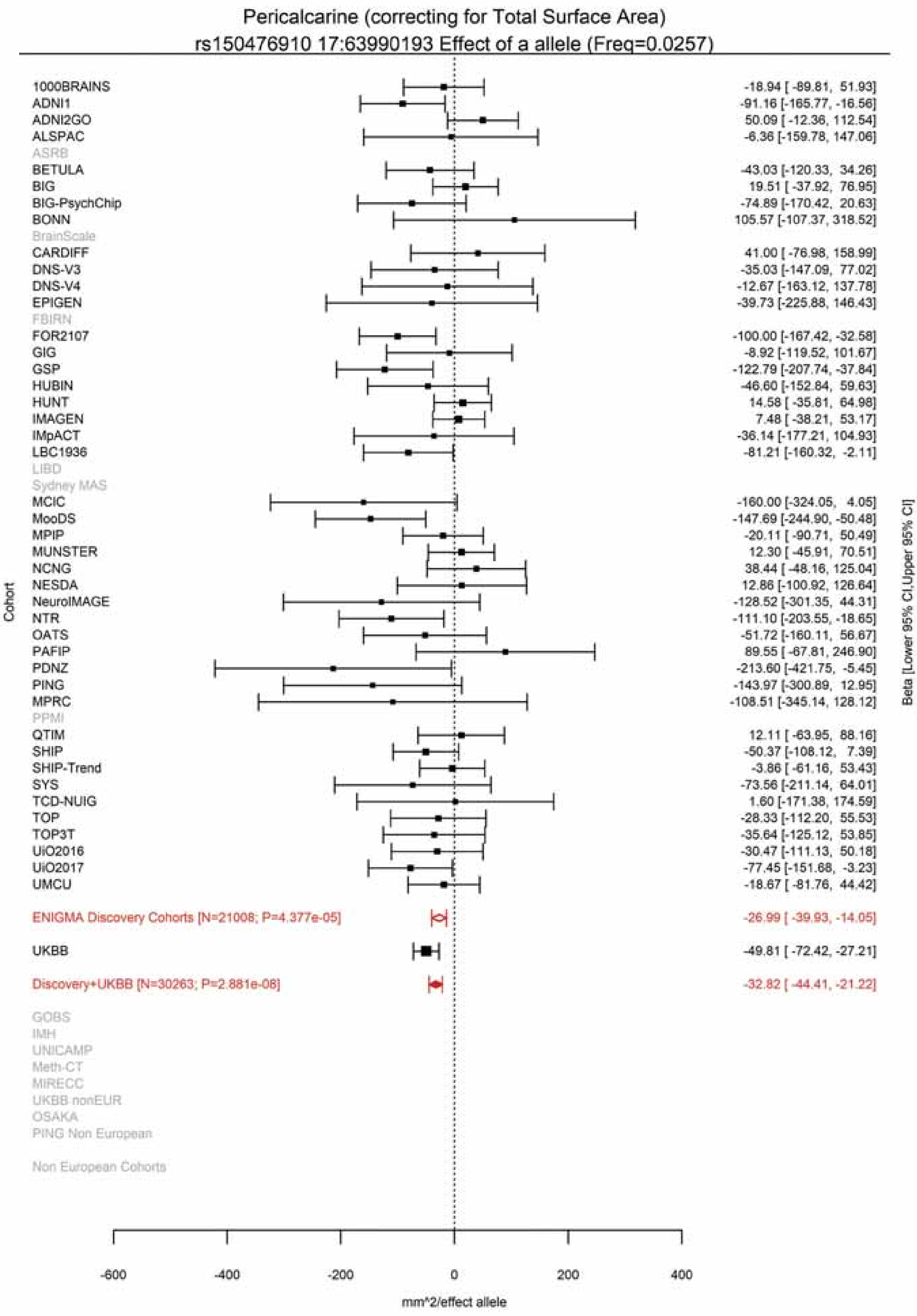
Pericalcarine (correcting for Total Surface Area)
rs150476910 17:63990193 Effect of a allele (Freq=0.0257)
1000BRAINS -18.94 [ -89.81, 51.93]
ADNI1 -91.16 [-165.77, -16.56]
ADNI2GO 50.09 [ -12.36, 112.54]
ALSPAC -6.36 [-159.78, 147.06]
ASRB
BETULA -43.03 [-120.33, 34.26]
BIG 19.51 [ -37.92, 76.95]
BIG-PsychChip -74.89 [-170.42, 20.63]
BONN 105.57 [-107.37, 318.52]
BrainScale
CARDIFF 41.00 [ -76.98, 158.99]
DNS-V3 -35.03 [-147.09, 77.02]
DNS-V4 -12.67 [-163.12, 137.78]
EPIGEN -39.73 [-225.88, 146.43]
FBIRN
FOR2107 -100.00 [-167.42, -32.58]
GIG -8.92 [-119.52, 101.67]
GSP -122.79 [-207.74, -37.84]
HUBIN -46.60 [-152.84, 59.63]
HUNT 14.58 [ -35.81, 64.98]
IMAGEN 7.48 [ -38.21, 53.17]
IMpACT -36.14 [-177.21, 104.93]
LBC1936 -81.21 [-160.32, -2.11]
LIBD
Sydney MAS
MCIC -160.00 [-324.05, 4.05]
MooDS -147.69 [-244.90, -50.48]
MPIP -20.11 [ -90.71, 50.49]
MUNSTER 12.30 [ -45.91, 70.51]
NCNG 38.44 [ -48.16, 125.04]
NESDA 12.86 [-100.92, 126.64]
NeuroIMAGE -128.52 [-301.35, 44.31]
NTR -111.10 [-203.55, -18.65]
OATS -51.72 [-160.11, 56.67]
PAFIP 89.55 [ -67.81, 246.90]
PDNZ -213.60 [-421.75, -5.45]
PING -143.97 [-300.89, 12.95]
MPRC -108.51 [-345.14, 128.12]
PPMI
QTIM 12.11 [ -63.95, 88.16]
SHIP -50.37 [-108.12, 7.39]
SHIP-Trend -3.86 [ -61.16, 53.43]
SYS -73.56 [-211.14, 64.01]
TCD-NUIG 1.60 [-171.38, 174.59]
TOP -28.33 [-112.20, 55.53]
TOP3T -35.64 [-125.12, 53.85]
UiO2016 -30.47 [-111.13, 50.18]
UiO2017 -77.45 [-151.68, -3.23]
UMCU -18.67 [ -81.76, 44.42]
ENIGMA Discovery Cohorts [N=21008; P=4.377e-05] -26.99 [ -39.93, -14.05]
UKBB -49.81 [ -72.42, -27.21]
Discovery+UKBB [N=30263; P=2.881e-08] -32.82 [ -44.41, -21.22]
GOBS
IMH
UNICAMP
Meth-CT
MIRECC
UKBB nonEUR
OSAKA
PING Non European
Non European Cohorts
Cohort
Beta [Lower 95% CI,Upper 95% CI]
-600
-400
-200
0
200
400
mm^2/effect allele

Superior Parietal (correcting for Total Surface Area)
rs17884482 19:12904400 Effect of t allele (Freq=0.0759)

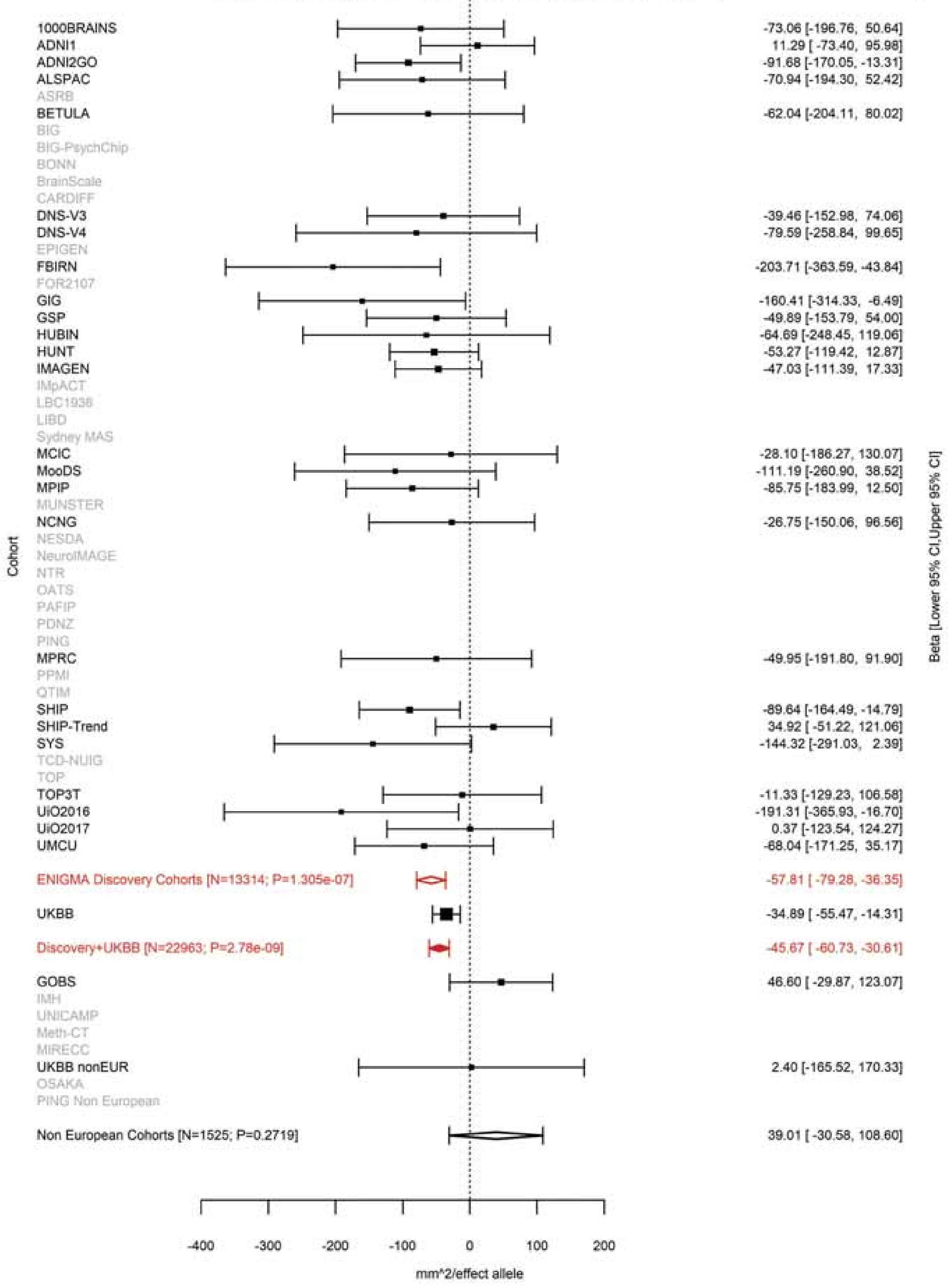

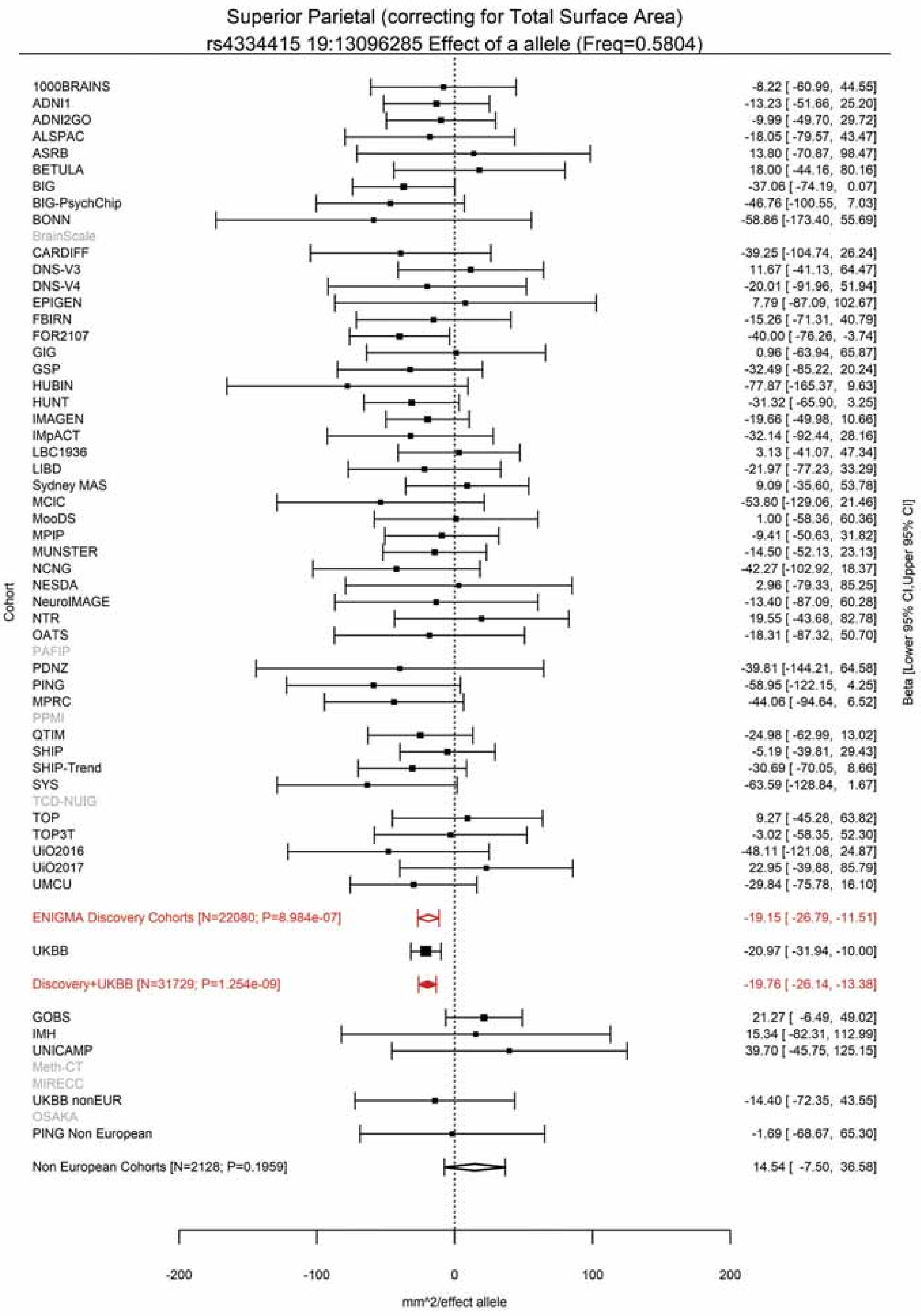
Superior Parietal (correcting for Total Surface Area)
rs4334415 19:13096285 Effect of a allele (Freq=0.5804)
Cohort
1000BRAINS -8.22 [ -60.99, 44.55]
ADNI1 -13.23 [ -51.66, 25.20]
ADNI2GO -9.99 [ -49.70, 29.72]
ALSPAC -18.05 [ -79.57, 43.47]
ASRB 13.80 [ -70.87, 98.47]
BETULA 18.00 [ -44.16, 80.16]
BIG -37.06 [ -74.19, 0.07]
BIG-PsychChip -46.76 [-100.55, 7.03]
BONN -58.86 [-173.40, 55.69]
BrainScale
CARDIFF -39.25 [-104.74, 26.24]
DNS-V3 11.67 [ -41.13, 64.47]
DNS-V4 -20.01 [ -91.96, 51.94]
EPIGEN 7.79 [ -87.09, 102.67]
FBIRN -15.26 [ -71.31, 40.79]
FOR2107 -40.00 [ -76.26, -3.74]
GIG 0.96 [ -63.94, 65.87]
GSP -32.49 [ -85.22, 20.24]
HUBIN -77.87 [-165.37, 9.63]
HUNT -31.32 [ -65.90, 3.25]
IMAGEN -19.66 [ -49.98, 10.66]
IMpACT -32.14 [ -92.44, 28.16]
LBC1936 3.13 [ -41.07, 47.34]
LIBD -21.97 [ -77.23, 33.29]
Sydney MAS 9.09 [ -35.60, 53.78]
MCIC -53.80 [-129.06, 21.46]
MooDS 1.00 [ -58.36, 60.36]
MPIP -9.41 [ -50.63, 31.82]
MUNSTER -14.50 [ -52.13, 23.13]
NCNG -42.27 [-102.92, 18.37]
NESDA 2.96 [ -79.33, 85.25]
NeuroIMAGE -13.40 [ -87.09, 60.28]
NTR 19.55 [ -43.68, 82.78]
OATS -18.31 [ -87.32, 50.70]
PAFIP
PDNZ -39.81 [-144.21, 64.58]
PING -58.95 [-122.15, 4.25]
MPRC -44.06 [ -94.64, 6.52]
PPMI
QTIM -24.98 [ -62.99, 13.02]
SHIP -5.19 [ -39.81, 29.43]
SHIP-Trend -30.69 [ -70.05, 8.66]
SYS -63.59 [-128.84, 1.67]
TCD-NUIG
TOP 9.27 [ -45.28, 63.82]
TOP3T -3.02 [ -58.35, 52.30]
UiO2016 -48.11 [-121.08, 24.87]
UiO2017 22.95 [ -39.88, 85.79]
UMCU -29.84 [ -75.78, 16.10]
ENIGMA Discovery Cohorts [N=22080; P=8.984e-07] -19.15 [ -26.79, -11.51]
UKBB -20.97 [ -31.94, -10.00]
Discovery+UKBB [N=31729; P=1.254e-09] -19.76 [ -26.14, -13.38]
GOBS 21.27 [ -6.49, 49.02]
IMH 15.34 [ -82.31, 112.99]
UNICAMP 39.70 [ -45.75, 125.15]
Meth-CT
MIRECC
UKBB nonEUR -14.40 [ -72.35, 43.55]
OSAKA
PING Non European -1.69 [ -68.67, 65.30]
Non European Cohorts [N=2128; P=0.1959] 14.54 [ -7.50, 36.58]
Beta [Lower 95% CI,Upper 95% CI]
-200
-100
0
100
200
mm^2/effect allele

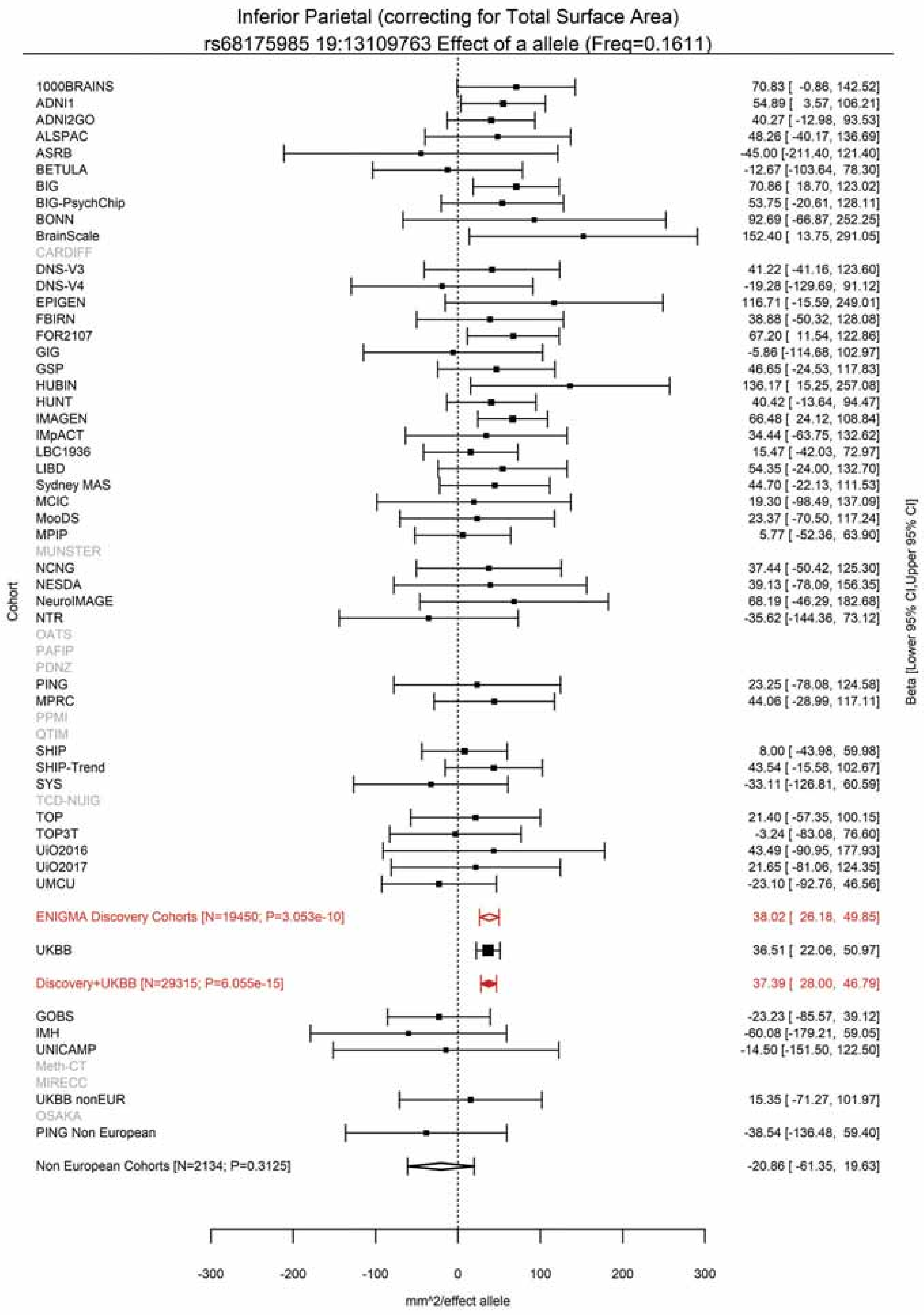

Inferior Parietal (correcting for Total Surface Area)
rs68175985 19:13109763 Effect of a allele (Freq=0.1611)
1000BRAINS 70.83 [ -0.86, 142.52]
ADNI1 54.89 [ 3.57, 106.21]
ADNI2GO 40.27 [ -12.98, 93.53]
ALSPAC 48.26 [ -40.17, 136.69]
ASRB -45.00 [-211.40, 121.40]
BETULA -12.67 [-103.64, 78.30]
BIG 70.86 [ 18.70, 123.02]
BIG-PsychChip 53.75 [ -20.61, 128.11]
BONN 92.69 [ -66.87, 252.25]
BrainScale 152.40 [ 13.75, 291.05]
CARDIFF
DNS-V3 41.22 [ -41.16, 123.60]
DNS-V4 -19.28 [-129.69, 91.12]
EPIGEN 116.71 [ -15.59, 249.01]
FBIRN 38.88 [ -50.32, 128.08]
FOR2107 67.20 [ 11.54, 122.86]
GIG -5.86 [-114.68, 102.97]
GSP 46.65 [ -24.53, 117.83]
HUBIN 136.17 [ 15.25, 257.08]
HUNT 40.42 [ -13.64, 94.47]
IMAGEN 66.48 [ 24.12, 108.84]
IMpACT 34.44 [ -63.75, 132.62]
LBC1936 15.47 [ -42.03, 72.97]
LIBD 54.35 [ -24.00, 132.70]
Sydney MAS 44.70 [ -22.13, 111.53]
MCIC 19.30 [ -98.49, 137.09]
MooDS 23.37 [ -70.50, 117.24]
MPIP 5.77 [ -52.36, 63.90]
MUNSTER
NCNG 37.44 [ -50.42, 125.30]
NESDA 39.13 [ -78.09, 156.35]
NeuroIMAGE 68.19 [ -46.29, 182.68]
NTR -35.62 [-144.36, 73.12]
OATS
PAFIP
PDNZ
PING 23.25 [ -78.08, 124.58]
MPRC 44.06 [ -28.99, 117.11]
PPMI
QTIM
SHIP 8.00 [ -43.98, 59.98]
SHIP-Trend 43.54 [ -15.58, 102.67]
SYS -33.11 [-126.81, 60.59]
TCD-NUIG
TOP 21.40 [ -57.35, 100.15]
TOP3T -3.24 [ -83.08, 76.60]
UiO2016 43.49 [ -90.95, 177.93]
UiO2017 21.65 [ -81.06, 124.35]
UMCU -23.10 [ -92.76, 46.56]
ENIGMA Discovery Cohorts [N=19450; P=3.053e-10] 38.02 [ 26.18, 49.85]
UKBB 36.51 [ 22.06, 50.97]
Discovery+UKBB [N=29315; P=6.055e-15] 37.39 [ 28.00, 46.79]
GOBS -23.23 [ -85.57, 39.12]
IMH -60.08 [-179.21, 59.05]
UNICAMP -14.50 [-151.50, 122.50]
Meth-CT
MIRECC
UKBB nonEUR 15.35 [ -71.27, 101.97]
OSAKA
PING Non European -38.54 [-136.48, 59.40]
Non European Cohorts [N=2134; P=0.3125] -20.86 [ -61.35, 19.63]
Cohort
Beta [Lower 95% CI,Upper 95% CI]
-300
-200
-100
0
100
200
300
mm^2/effect allele

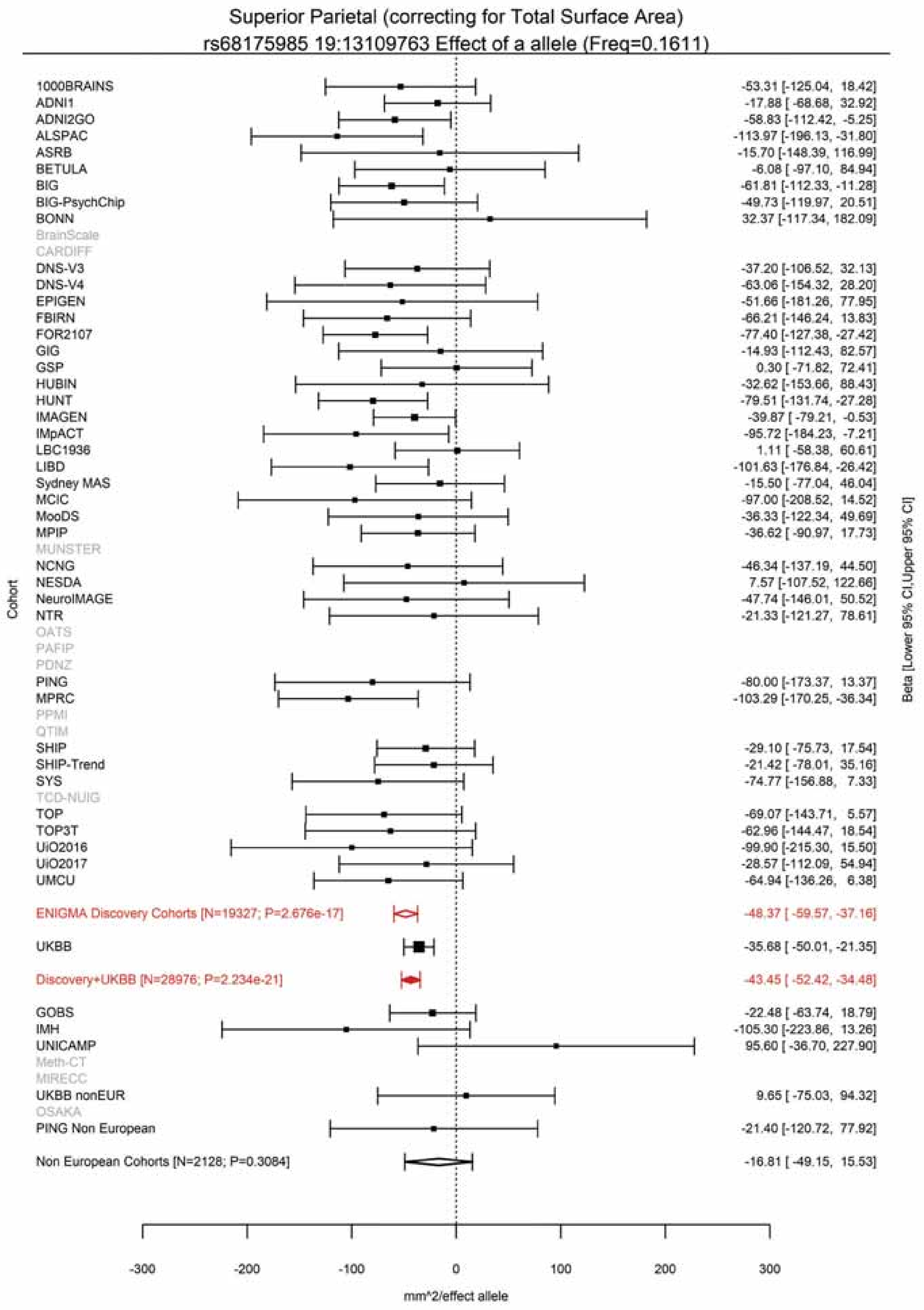

Superior Parietal (correcting for Total Surface Area)
rs68175985 19:13109763 Effect of a allele (Freq=0.1611)
Cohort
1000BRAINS -53.31 [-125.04, 18.42]
ADNI1 -17.88 [ -68.68, 32.92]
ADNI2GO -58.83 [-112.42, -5.25]
ALSPAC -113.97 [-196.13, -31.80]
ASRB -15.70 [-148.39, 116.99]
BETULA -6.08 [ -97.10, 84.94]
BIG -61.81 [-112.33, -11.28]
BIG-PsychChip -49.73 [-119.97, 20.51]
BONN 32.37 [-117.34, 182.09]
BrainScale
CARDIFF
DNS-V3 -37.20 [-106.52, 32.13]
DNS-V4 -63.06 [-154.32, 28.20]
EPIGEN -51.66 [-181.26, 77.95]
FBIRN -66.21 [-146.24, 13.83]
FOR2107 -77.40 [-127.38, -27.42]
GIG -14.93 [-112.43, 82.57]
GSP 0.30 [ -71.82, 72.41]
HUBIN -32.62 [-153.66, 88.43]
HUNT -79.51 [-131.74, -27.28]
IMAGEN -39.87 [ -79.21, -0.53]
IMpACT -95.72 [-184.23, -7.21]
LBC1936 1.11 [ -58.38, 60.61]
LIBD -101.63 [-176.84, -26.42]
Sydney MAS -15.50 [ -77.04, 46.04]
MCIC -97.00 [-208.52, 14.52]
MooDS -36.33 [-122.34, 49.69]
MPIP -36.62 [ -90.97, 17.73]
MUNSTER
NCNG -46.34 [-137.19, 44.50]
NESDA 7.57 [-107.52, 122.66]
NeuroIMAGE -47.74 [-146.01, 50.52]
NTR -21.33 [-121.27, 78.61]
OATS
PAFIP
PDNZ
PING -80.00 [-173.37, 13.37]
MPRC -103.29 [-170.25, -36.34]
PPMI
QTIM
SHIP -29.10 [ -75.73, 17.54]
SHIP-Trend -21.42 [ -78.01, 35.16]
SYS -74.77 [-156.88, 7.33]
TCD-NUIG
TOP -69.07 [-143.71, 5.57]
TOP3T -62.96 [-144.47, 18.54]
UiO2016 -99.90 [-215.30, 15.50]
UiO2017 -28.57 [-112.09, 54.94]
UMCU -64.94 [-136.26, 6.38]
ENIGMA Discovery Cohorts [N=19327; P=2.676e-17] -48.37 [ -59.57, -37.16]
UKBB -35.68 [ -50.01, -21.35]
Discovery+UKBB [N=28976; P=2.234e-21] -43.45 [ -52.42, -34.48]
GOBS -22.48 [ -63.74, 18.79]
IMH -105.30 [-223.86, 13.26]
UNICAMP 95.60 [ -36.70, 227.90]
Meth-CT
MIRECC
UKBB nonEUR 9.65 [ -75.03, 94.32]
OSAKA
PING Non European -21.40 [-120.72, 77.92]
Non European Cohorts [N=2128; P=0.3084] -16.81 [ -49.15, 15.53]
Beta [Lower 95% CI,Upper 95% CI]
-300 -200 -100 0 100 200 300
mm^2/effect allele

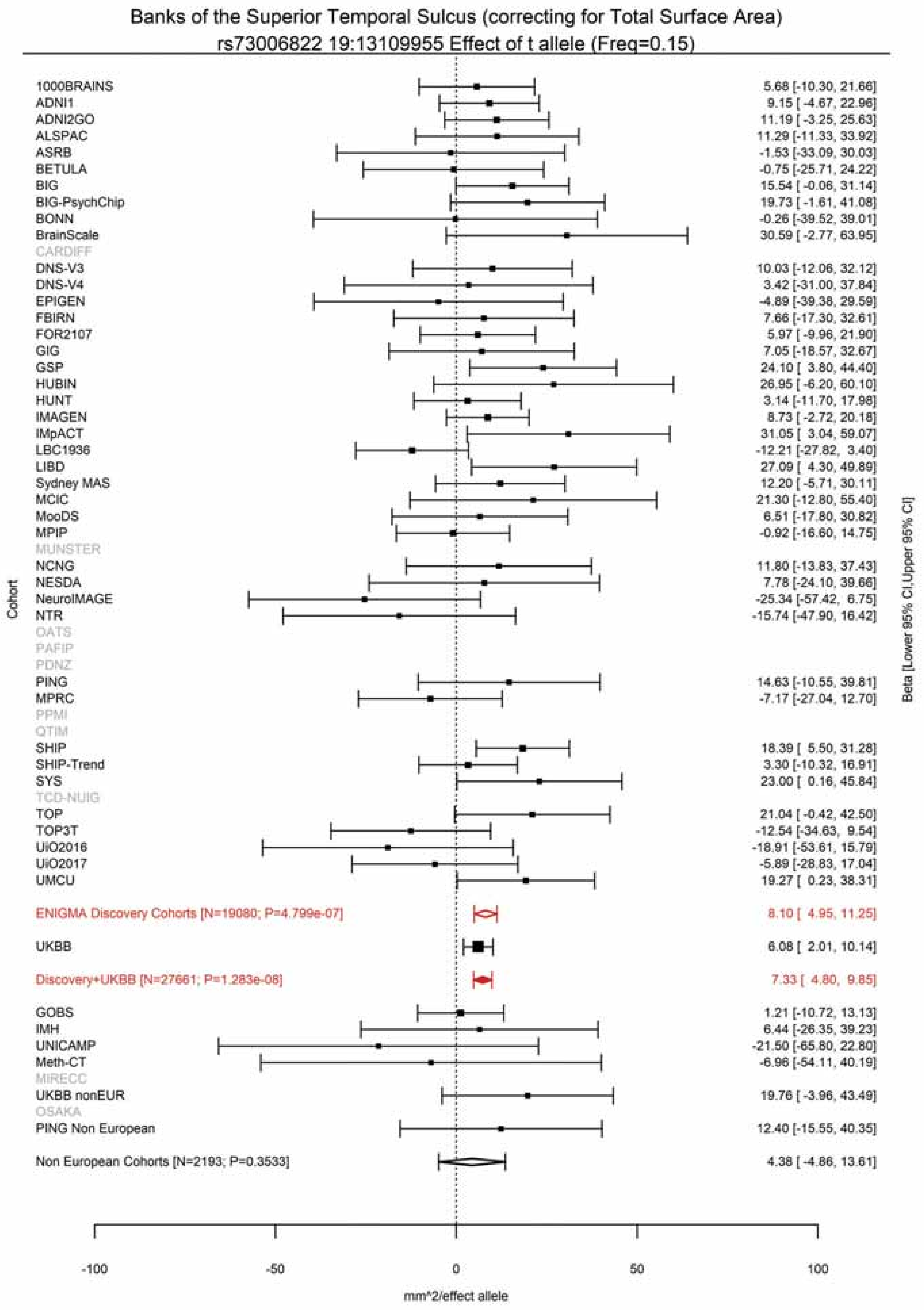
Banks of the Superior Temporal Sulcus (correcting for Total Surface Area)
rs73006822 19:13109955 Effect of t allele (Freq=0.15)
Cohort
Beta [Lower 95% CI,Upper 95% CI]
1000BRAINS 5.68 [-10.30, 21.66]
ADNI1 9.15 [ -4.67, 22.96]
ADNI2GO 11.19 [ -3.25, 25.63]
ALSPAC 11.29 [-11.33, 33.92]
ASRB -1.53 [-33.09, 30.03]
BETULA -0.75 [-25.71, 24.22]
BIG 15.54 [ -0.06, 31.14]
BIG-PsychChip 19.73 [ -1.61, 41.08]
BONN -0.26 [-39.52, 39.01]
BrainScale 30.59 [ -2.77, 63.95]
CARDIFF
DNS-V3 10.03 [-12.06, 32.12]
DNS-V4 3.42 [-31.00, 37.84]
EPIGEN -4.89 [-39.38, 29.59]
FBIRN 7.66 [-17.30, 32.61]
FOR2107 5.97 [ -9.96, 21.90]
GIG 7.05 [-18.57, 32.67]
GSP 24.10 [ 3.80, 44.40]
HUBIN 26.95 [ -6.20, 60.10]
HUNT 3.14 [-11.70, 17.98]
IMAGEN 8.73 [ -2.72, 20.18]
IMpACT 31.05 [ 3.04, 59.07]
LBC1936 -12.21 [-27.82, 3.40]
LIBD 27.09 [ 4.30, 49.89]
Sydney MAS 12.20 [ -5.71, 30.11]
MCIC 21.30 [-12.80, 55.40]
MooDS 6.51 [-17.80, 30.82]
MPIP -0.92 [-16.60, 14.75]
MUNSTER
NCNG 11.80 [-13.83, 37.43]
NESDA 7.78 [-24.10, 39.66]
NeuroIMAGE -25.34 [-57.42, 6.75]
NTR -15.74 [-47.90, 16.42]
OATS
PAFIP
PDNZ
PING 14.63 [-10.55, 39.81]
MPRC -7.17 [-27.04, 12.70]
PPMI
QTIM
SHIP 18.39 [ 5.50, 31.28]
SHIP-Trend 3.30 [-10.32, 16.91]
SYS 23.00 [ 0.16, 45.84]
TCD-NUIG
TOP 21.04 [ -0.42, 42.50]
TOP3T -12.54 [-34.63, 9.54]
UiO2016 -18.91 [-53.61, 15.79]
UiO2017 -5.89 [-28.83, 17.04]
UMCU 19.27 [ 0.23, 38.31]
ENIGMA Discovery Cohorts [N=19080; P=4.799e-07] 8.10 [ 4.95, 11.25]
UKBB 6.08 [ 2.01, 10.14]
Discovery+UKBB [N=27661; P=1.283e-08] 7.33 [ 4.80, 9.85]
GOBS 1.21 [-10.72, 13.13]
IMH 6.44 [-26.35, 39.23]
UNICAMP -21.50 [-65.80, 22.80]
Meth-CT -6.96 [-54.11, 40.19]
MIRECC
UKBB nonEUR 19.76 [ -3.96, 43.49]
OSAKA
PING Non European 12.40 [-15.55, 40.35]
Non European Cohorts [N=2193; P=0.3533] 4.38 [ -4.86, 13.61]
-100
-50
0
50
100
mm^2/effect allele

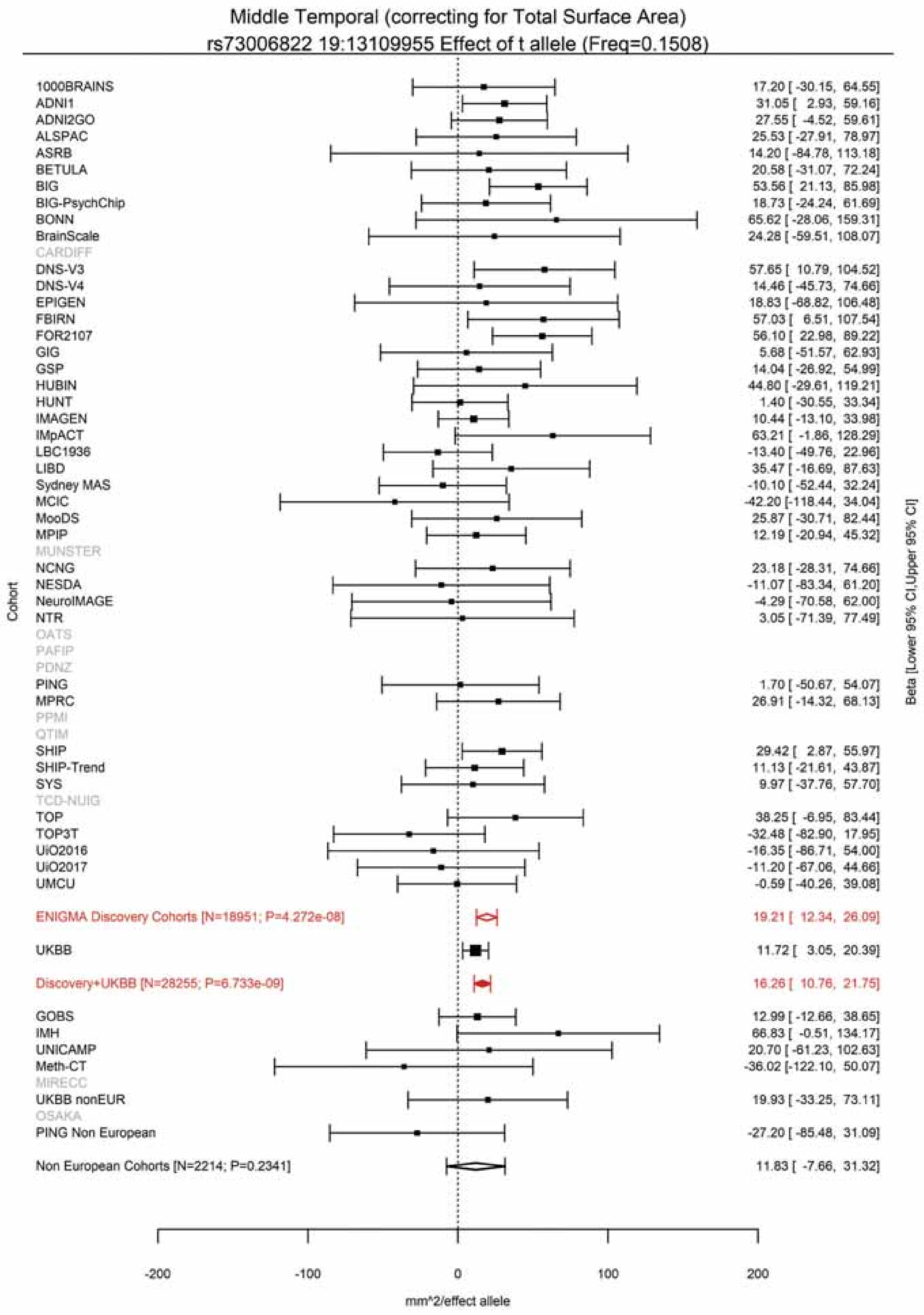
Middle Temporal (correcting for Total Surface Area)
rs73006822 19:13109955 Effect of t allele (Freq=0.1508)
1000BRAINS 17.20 [ -30.15, 64.55]
ADNI1 31.05 [ 2.93, 59.16]
ADNI2GO 27.55 [ -4.52, 59.61]
ALSPAC 25.53 [ -27.91, 78.97]
ASRB 14.20 [ -84.78, 113.18]
BETULA 20.58 [ -31.07, 72.24]
BIG 53.56 [ 21.13, 85.98]
BIG-PsychChip 18.73 [ -24.24, 61.69]
BONN 65.62 [ -28.06, 159.31]
BrainScale 24.28 [ -59.51, 108.07]
CARDIFF
DNS-V3 57.65 [ 10.79, 104.52]
DNS-V4 14.46 [ -45.73, 74.66]
EPIGEN 18.83 [ -68.82, 106.48]
FBIRN 57.03 [ 6.51, 107.54]
FOR2107 56.10 [ 22.98, 89.22]
GIG 5.68 [ -51.57, 62.93]
GSP 14.04 [ -26.92, 54.99]
HUBIN 44.80 [ -29.61, 119.21]
HUNT 1.40 [ -30.55, 33.34]
IMAGEN 10.44 [ -13.10, 33.98]
IMpACT 63.21 [ -1.86, 128.29]
LBC1936 -13.40 [ -49.76, 22.96]
LIBD 35.47 [ -16.69, 87.63]
Sydney MAS -10.10 [ -52.44, 32.24]
MCIC -42.20 [-118.44, 34.04]
MooDS 25.87 [ -30.71, 82.44]
MPIP 12.19 [ -20.94, 45.32]
MUNSTER
NCNG 23.18 [ -28.31, 74.66]
NESDA -11.07 [ -83.34, 61.20]
NeuroIMAGE -4.29 [ -70.58, 62.00]
NTR 3.05 [ -71.39, 77.49]
OATS
PAFIP
PDNZ
PING 1.70 [ -50.67, 54.07]
MPRC 26.91 [ -14.32, 68.13]
PPMI
QTIM
SHIP 29.42 [ 2.87, 55.97]
SHIP-Trend 11.13 [ -21.61, 43.87]
SYS 9.97 [ -37.76, 57.70]
TCD-NUIG
TOP 38.25 [ -6.95, 83.44]
TOP3T -32.48 [ -82.90, 17.95]
UiO2016 -16.35 [ -86.71, 54.00]
UiO2017 -11.20 [ -67.06, 44.66]
UMCU -0.59 [ -40.26, 39.08]
ENIGMA Discovery Cohorts [N=18951; P=4.272e-08] 19.21 [ 12.34, 26.09]
UKBB 11.72 [ 3.05, 20.39]
Discovery+UKBB [N=28255; P=6.733e-09] 16.26 [ 10.76, 21.75]
GOBS 12.99 [ -12.66, 38.65]
IMH 66.83 [ -0.51, 134.17]
UNICAMP 20.70 [ -61.23, 102.63]
Meth-CT -36.02 [-122.10, 50.07]
MIRECC
UKBB nonEUR 19.93 [ -33.25, 73.11]
OSAKA
PING Non European -27.20 [ -85.48, 31.09]
Non European Cohorts [N=2214; P=0.2341] 11.83 [ -7.66, 31.32]
Cohort
Beta [Lower 95% CI,Upper 95% CI]
-200
-100
0
100
200
mm^2/effect allele

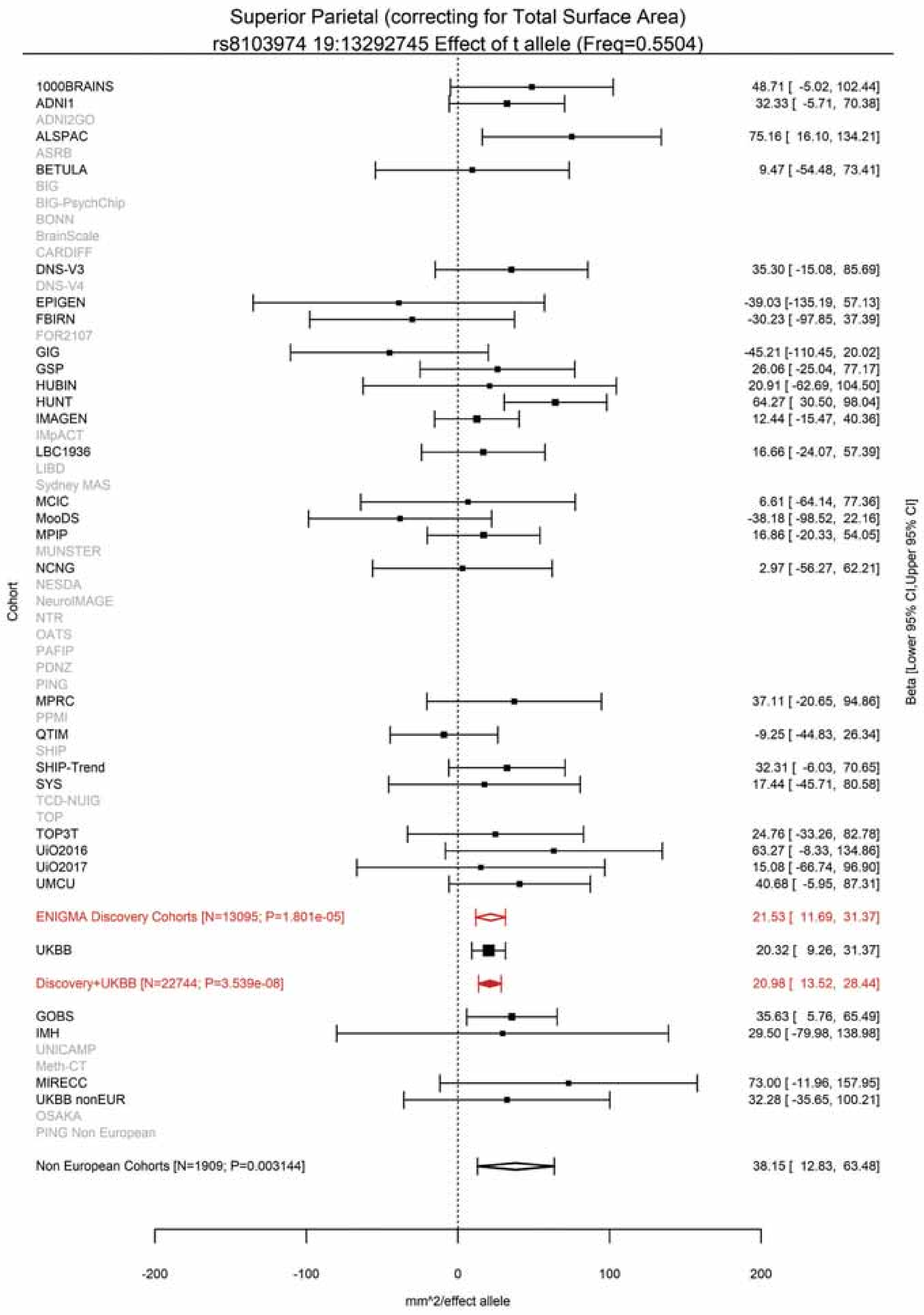
Superior Parietal (correcting for Total Surface Area)
rs8103974 19:13292745 Effect of t allele (Freq=0.5504)
1000BRAINS 48.71 [ -5.02, 102.44]
ADNI1 32.33 [ -5.71, 70.38]
ADNI2GO
ALSPAC 75.16 [ 16.10, 134.21]
ASRB
BETULA 9.47 [ -54.48, 73.41]
BIG
BIG-PsychChip
BONN
BrainScale
CARDIFF
DNS-V3 35.30 [ -15.08, 85.69]
DNS-V4
EPIGEN -39.03 [-135.19, 57.13]
FBIRN -30.23 [ -97.85, 37.39]
FOR2107
GIG -45.21 [-110.45, 20.02]
GSP 26.06 [ -25.04, 77.17]
HUBIN 20.91 [ -62.69, 104.50]
HUNT 64.27 [ 30.50, 98.04]
IMAGEN 12.44 [ -15.47, 40.36]
IMpACT
LBC1936 16.66 [ -24.07, 57.39]
LIBD
Sydney MAS
MCIC 6.61 [ -64.14, 77.36]
MooDS -38.18 [ -98.52, 22.16]
MPIP 16.86 [ -20.33, 54.05]
MUNSTER
NCNG 2.97 [ -56.27, 62.21]
NESDA
NeuroIMAGE
NTR
OATS
PAFIP
PDNZ
PING
MPRC 37.11 [ -20.65, 94.86]
PPMI
QTIM -9.25 [ -44.83, 26.34]
SHIP
SHIP-Trend 32.31 [ -6.03, 70.65]
SYS 17.44 [ -45.71, 80.58]
TCD-NUIG
TOP
TOP3T 24.76 [ -33.26, 82.78]
UiO2016 63.27 [ -8.33, 134.86]
UiO2017 15.08 [ -66.74, 96.90]
UMCU 40.68 [ -5.95, 87.31]
ENIGMA Discovery Cohorts [N=13095; P=1.801e-05] 21.53 [ 11.69, 31.37]
UKBB 20.32 [ 9.26, 31.37]
Discovery+UKBB [N=22744; P=3.539e-08] 20.98 [ 13.52, 28.44]
GOBS 35.63 [ 5.76, 65.49]
IMH 29.50 [ -79.98, 138.98]
UNICAMP
Meth-CT
MIRECC 73.00 [ -11.96, 157.95]
UKBB nonEUR 32.28 [ -35.65, 100.21]
OSAKA
PING Non European
Non European Cohorts [N=1909; P=0.003144] 38.15 [ 12.83, 63.48]
Cohort
Beta [Lower 95% CI,Upper 95% CI]
-200
-100
0
100
200
mm^2/effect allele

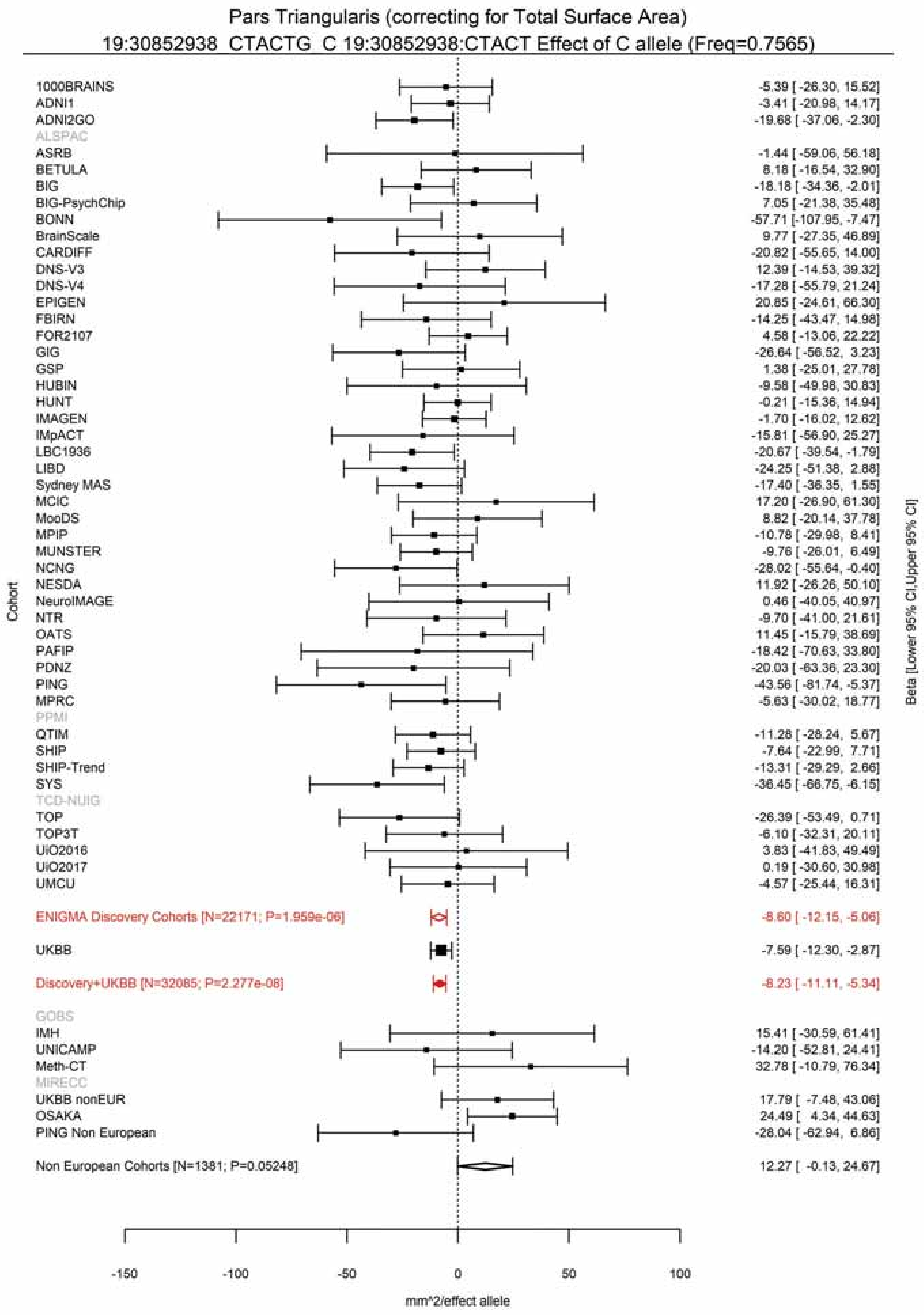
Pars Triangularis (correcting for Total Surface Area)
19:30852938_CTACTG_C 19:30852938:CTACT Effect of C allele (Freq=0.7565)
Cohort
1000BRAINS -5.39 [ -26.30, 15.52]
ADNI1 -3.41 [ -20.98, 14.17]
ADNI2GO -19.68 [ -37.06, -2.30]
ALSPAC
ASRB -1.44 [ -59.06, 56.18]
BETULA 8.18 [ -16.54, 32.90]
BIG -18.18 [ -34.36, -2.01]
BIG-PsychChip 7.05 [ -21.38, 35.48]
BONN -57.71 [-107.95, -7.47]
BrainScale 9.77 [ -27.35, 46.89]
CARDIFF -20.82 [ -55.65, 14.00]
DNS-V3 12.39 [ -14.53, 39.32]
DNS-V4 -17.28 [ -55.79, 21.24]
EPIGEN 20.85 [ -24.61, 66.30]
FBIRN -14.25 [ -43.47, 14.98]
FOR2107 4.58 [ -13.06, 22.22]
GIG -26.64 [ -56.52, 3.23]
GSP 1.38 [ -25.01, 27.78]
HUBIN -9.58 [ -49.98, 30.83]
HUNT -0.21 [ -15.36, 14.94]
IMAGEN -1.70 [ -16.02, 12.62]
IMpACT -15.81 [ -56.90, 25.27]
LBC1936 -20.67 [ -39.54, -1.79]
LIBD -24.25 [ -51.38, 2.88]
Sydney MAS -17.40 [ -36.35, 1.55]
MCIC 17.20 [ -26.90, 61.30]
MooDS 8.82 [ -20.14, 37.78]
MPIP -10.78 [ -29.98, 8.41]
MUNSTER -9.76 [ -26.01, 6.49]
NCNG -28.02 [ -55.64, -0.40]
NESDA 11.92 [ -26.26, 50.10]
NeuroIMAGE 0.46 [ -40.05, 40.97]
NTR -9.70 [ -41.00, 21.61]
OATS 11.45 [ -15.79, 38.69]
PAFIP -18.42 [ -70.63, 33.80]
PDNZ -20.03 [ -63.36, 23.30]
PING -43.56 [ -81.74, -5.37]
MPRC -5.63 [ -30.02, 18.77]
PPMI
QTIM -11.28 [ -28.24, 5.67]
SHIP -7.64 [ -22.99, 7.71]
SHIP-Trend -13.31 [ -29.29, 2.66]
SYS -36.45 [ -66.75, -6.15]
TCD-NUIG
TOP -26.39 [ -53.49, 0.71]
TOP3T -6.10 [ -32.31, 20.11]
UiO2016 3.83 [ -41.83, 49.49]
UiO2017 0.19 [ -30.60, 30.98]
UMCU -4.57 [ -25.44, 16.31]
ENIGMA Discovery Cohorts [N=22171; P=1.959e-06] -8.60 [ -12.15, -5.06]
UKBB -7.59 [ -12.30, -2.87]
Discovery+UKBB [N=32085; P=2.277e-08] -8.23 [ -11.11, -5.34]
GOBS
IMH 15.41 [ -30.59, 61.41]
UNICAMP -14.20 [ -52.81, 24.41]
Meth-CT 32.78 [ -10.79, 76.34]
MIRECC
UKBB nonEUR 17.79 [ -7.48, 43.06]
OSAKA 24.49 [ 4.34, 44.63]
PING Non European -28.04 [ -62.94, 6.86]
Non European Cohorts [N=1381; P=0.05248] 12.27 [ -0.13, 24.67]
-150
-100
-50
0
50
100
mm^2/effect allele
Beta [Lower 95% CI,Upper 95% CI]

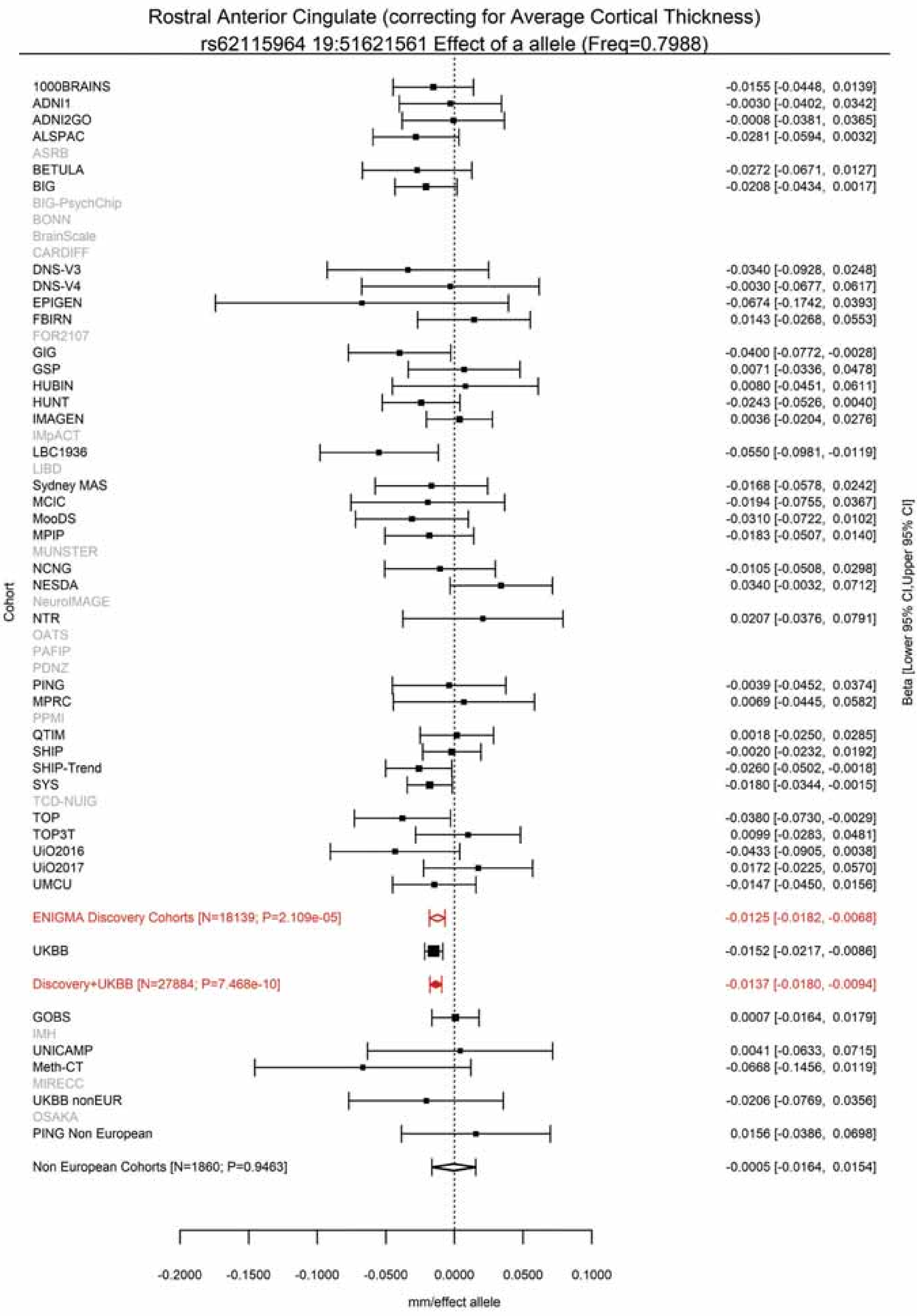
Rostral Anterior Cingulate (correcting for Average Cortical Thickness)
rs62115964 19:51621561 Effect of a allele (Freq=0.7988)
1000BRAINS -0.0155 [-0.0448, 0.0139]
ADNI1 -0.0030 [-0.0402, 0.0342]
ADNI2GO -0.0008 [-0.0381, 0.0365]
ALSPAC -0.0281 [-0.0594, 0.0032]
ASRB
BETULA -0.0272 [-0.0671, 0.0127]
BIG -0.0208 [-0.0434, 0.0017]
BIG-PsychChip
BONN
BrainScale
CARDIFF
DNS-V3 -0.0340 [-0.0928, 0.0248]
DNS-V4 -0.0030 [-0.0677, 0.0617]
EPIGEN -0.0674 [-0.1742, 0.0393]
FBIRN 0.0143 [-0.0268, 0.0553]
FOR2107
GIG -0.0400 [-0.0772, -0.0028]
GSP 0.0071 [-0.0336, 0.0478]
HUBIN 0.0080 [-0.0451, 0.0611]
HUNT -0.0243 [-0.0526, 0.0040]
IMAGEN 0.0036 [-0.0204, 0.0276]
IMpACT
LBC1936 -0.0550 [-0.0981, -0.0119]
LIBD
Sydney MAS -0.0168 [-0.0578, 0.0242]
MCIC -0.0194 [-0.0755, 0.0367]
MooDS -0.0310 [-0.0722, 0.0102]
MPIP -0.0183 [-0.0507, 0.0140]
MUNSTER
NCNG -0.0105 [-0.0508, 0.0298]
NESDA 0.0340 [-0.0032, 0.0712]
NeuroIMAGE
NTR 0.0207 [-0.0376, 0.0791]
OATS
PAFIP
PDNZ
PING -0.0039 [-0.0452, 0.0374]
MPRC 0.0069 [-0.0445, 0.0582]
PPMI
QTIM 0.0018 [-0.0250, 0.0285]
SHIP -0.0020 [-0.0232, 0.0192]
SHIP-Trend -0.0260 [-0.0502, -0.0018]
SYS -0.0180 [-0.0344, -0.0015]
TCD-NUIG
TOP -0.0380 [-0.0730, -0.0029]
TOP3T 0.0099 [-0.0283, 0.0481]
UiO2016 -0.0433 [-0.0905, 0.0038]
UiO2017 0.0172 [-0.0225, 0.0570]
UMCU -0.0147 [-0.0450, 0.0156]
ENIGMA Discovery Cohorts [N=18139; P=2.109e-05] -0.0125 [-0.0182, -0.0068]
UKBB -0.0152 [-0.0217, -0.0086]
Discovery+UKBB [N=27884; P=7.468e-10] -0.0137 [-0.0180, -0.0094]
GOBS 0.0007 [-0.0164, 0.0179]
IMH
UNICAMP 0.0041 [-0.0633, 0.0715]
Meth-CT -0.0668 [-0.1456, 0.0119]
MIRECC
UKBB nonEUR -0.0206 [-0.0769, 0.0356]
OSAKA
PING Non European 0.0156 [-0.0386, 0.0698]
Non European Cohorts [N=1860; P=0.9463] -0.0005 [-0.0164, 0.0154]
Cohort
Beta [Lower 95% CI,Upper 95% CI]
-0.2000 -0.1500 -0.1000 -0.0500 0.0000 0.0500 0.1000
mm/effect allele

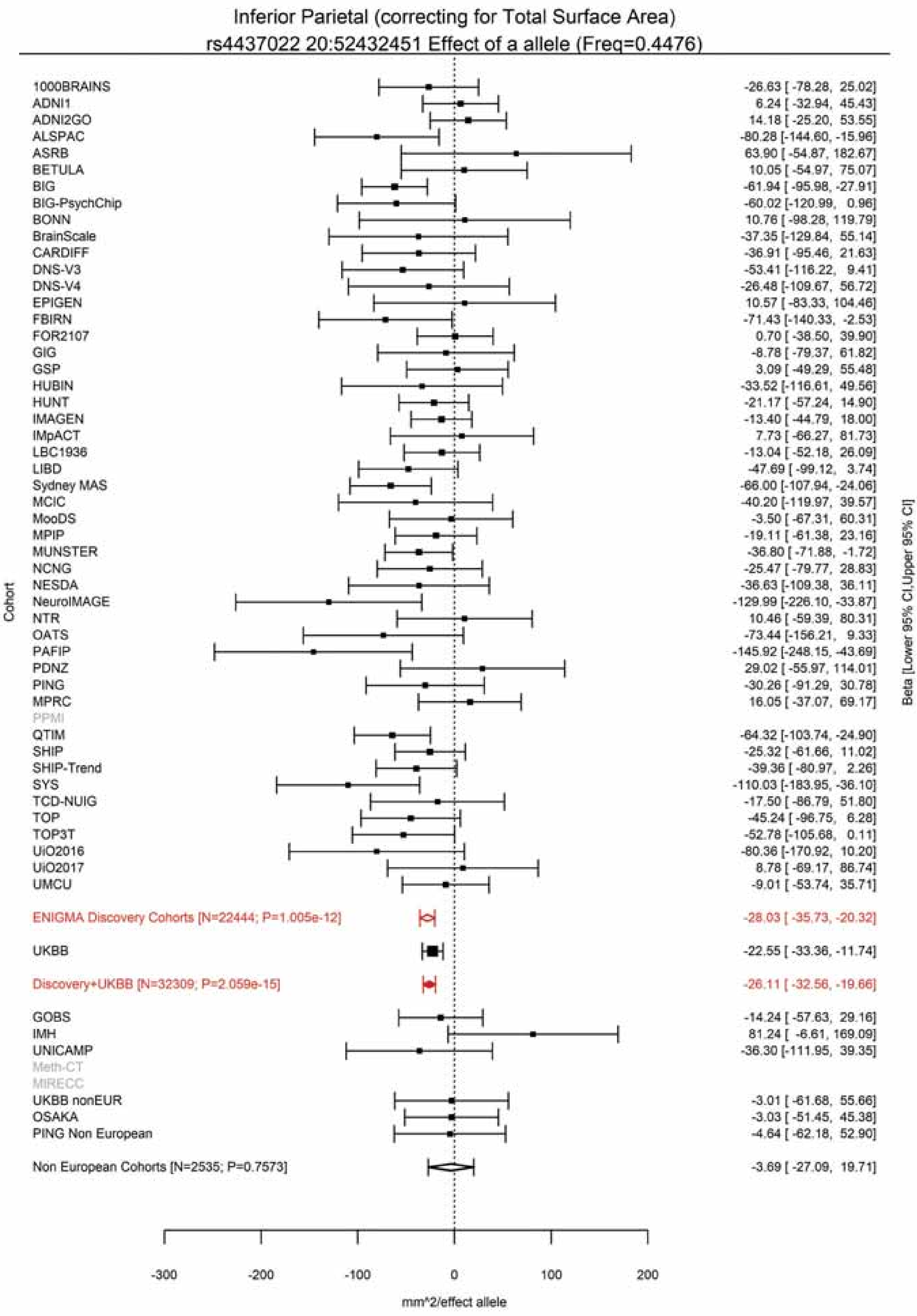
Inferior Parietal (correcting for Total Surface Area)
rs4437022 20:52432451 Effect of a allele (Freq=0.4476)
1000BRAINS -26.63 [ -78.28, 25.02]
ADNI1 6.24 [ -32.94, 45.43]
ADNI2GO 14.18 [ -25.20, 53.55]
ALSPAC -80.28 [-144.60, -15.96]
ASRB 63.90 [ -54.87, 182.67]
BETULA 10.05 [ -54.97, 75.07]
BIG -61.94 [ -95.98, -27.91]
BIG-PsychChip -60.02 [-120.99, 0.96]
BONN 10.76 [ -98.28, 119.79]
BrainScale -37.35 [-129.84, 55.14]
CARDIFF -36.91 [ -95.46, 21.63]
DNS-V3 -53.41 [-116.22, 9.41]
DNS-V4 -26.48 [-109.67, 56.72]
EPIGEN 10.57 [ -83.33, 104.46]
FBIRN -71.43 [-140.33, -2.53]
FOR2107 0.70 [ -38.50, 39.90]
GIG -8.78 [ -79.37, 61.82]
GSP 3.09 [ -49.29, 55.48]
HUBIN -33.52 [-116.61, 49.56]
HUNT -21.17 [ -57.24, 14.90]
IMAGEN -13.40 [ -44.79, 18.00]
IMpACT 7.73 [ -66.27, 81.73]
LBC1936 -13.04 [ -52.18, 26.09]
LIBD -47.69 [ -99.12, 3.74]
Sydney MAS -66.00 [-107.94, -24.06]
MCIC -40.20 [-119.97, 39.57]
MooDS -3.50 [ -67.31, 60.31]
MPIP -19.11 [ -61.38, 23.16]
MUNSTER -36.80 [ -71.88, -1.72]
NCNG -25.47 [ -79.77, 28.83]
NESDA -36.63 [-109.38, 36.11]
NeuroIMAGE -129.99 [-226.10, -33.87]
NTR 10.46 [ -59.39, 80.31]
OATS -73.44 [-156.21, 9.33]
PAFIP -145.92 [-248.15, -43.69]
PDNZ 29.02 [ -55.97, 114.01]
PING -30.26 [ -91.29, 30.78]
MPRC 16.05 [ -37.07, 69.17]
PPMI
QTIM -64.32 [-103.74, -24.90]
SHIP -25.32 [ -61.66, 11.02]
SHIP-Trend -39.36 [ -80.97, 2.26]
SYS -110.03 [-183.95, -36.10]
TCD-NUIG -17.50 [ -86.79, 51.80]
TOP -45.24 [ -96.75, 6.28]
TOP3T -52.78 [-105.68, 0.11]
UiO2016 -80.36 [-170.92, 10.20]
UiO2017 8.78 [ -69.17, 86.74]
UMCU -9.01 [ -53.74, 35.71]
ENIGMA Discovery Cohorts [N=22444; P=1.005e-12] -28.03 [ -35.73, -20.32]
UKBB -22.55 [ -33.36, -11.74]
Discovery+UKBB [N=32309; P=2.059e-15] -26.11 [ -32.56, -19.66]
GOBS -14.24 [ -57.63, 29.16]
IMH 81.24 [ -6.61, 169.09]
UNICAMP -36.30 [-111.95, 39.35]
Meth-CT
MIRECC
UKBB nonEUR -3.01 [ -61.68, 55.66]
OSAKA -3.03 [ -51.45, 45.38]
PING Non European -4.64 [ -62.18, 52.90]
Non European Cohorts [N=2535; P=0.7573] -3.69 [ -27.09, 19.71]
Cohort
Beta [Lower 95% CI,Upper 95% CI]
-300
-200
-100
0
100
200
mm^2/effect allele

Superior Parietal (correcting for Total Surface Area)
rs6022786 20:52447303 Effect of a allele (Freq=0.4125)

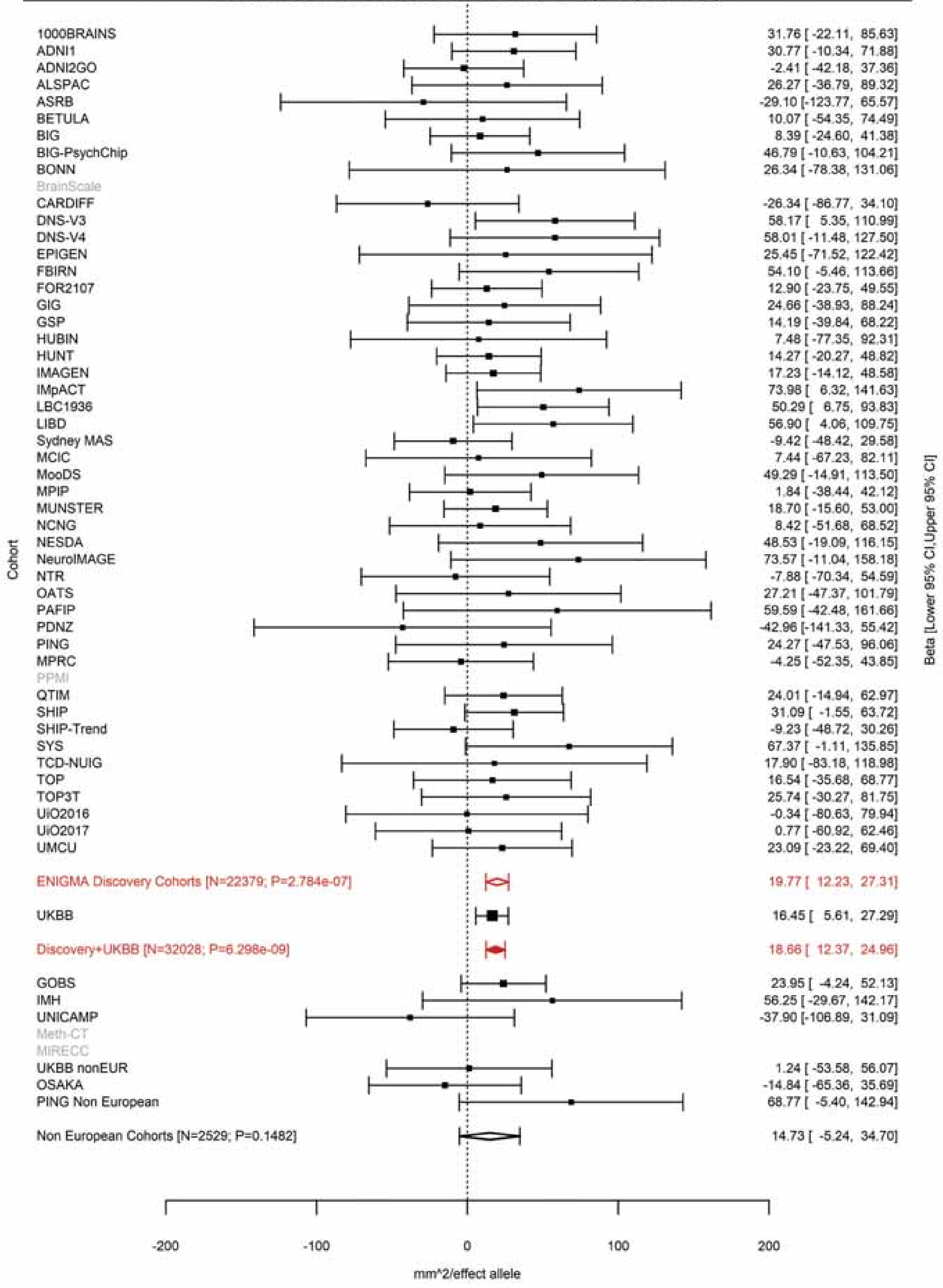

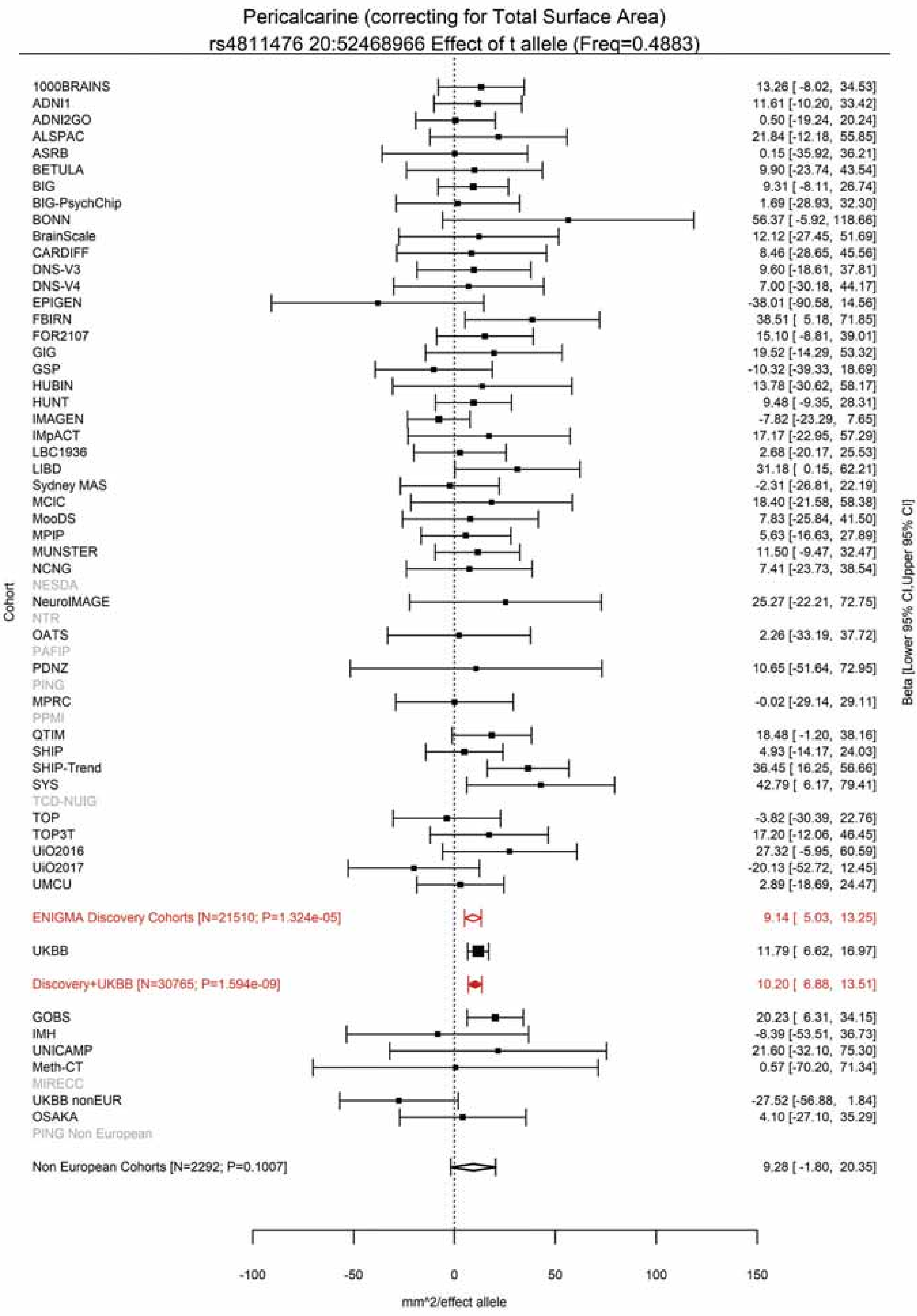
Pericalcarine (correcting for Total Surface Area)
rs4811476 20:52468966 Effect of t allele (Freq=0.4883)
Cohort
1000BRAINS 13.26 [ -8.02, 34.53]
ADNI1 11.61 [-10.20, 33.42]
ADNI2GO 0.50 [-19.24, 20.24]
ALSPAC 21.84 [-12.18, 55.85]
ASRB 0.15 [-35.92, 36.21]
BETULA 9.90 [-23.74, 43.54]
BIG 9.31 [ -8.11, 26.74]
BIG-PsychChip 1.69 [-28.93, 32.30]
BONN 56.37 [ -5.92, 118.66]
BrainScale 12.12 [-27.45, 51.69]
CARDIFF 8.46 [-28.65, 45.56]
DNS-V3 9.60 [-18.61, 37.81]
DNS-V4 7.00 [-30.18, 44.17]
EPIGEN -38.01 [-90.58, 14.56]
FBIRN 38.51 [ 5.18, 71.85]
FOR2107 15.10 [ -8.81, 39.01]
GIG 19.52 [-14.29, 53.32]
GSP -10.32 [-39.33, 18.69]
HUBIN 13.78 [-30.62, 58.17]
HUNT 9.48 [ -9.35, 28.31]
IMAGEN -7.82 [-23.29, 7.65]
IMpACT 17.17 [-22.95, 57.29]
LBC1936 2.68 [-20.17, 25.53]
LIBD 31.18 [ 0.15, 62.21]
Sydney MAS -2.31 [-26.81, 22.19]
MCIC 18.40 [-21.58, 58.38]
MooDS 7.83 [-25.84, 41.50]
MPIP 5.63 [-16.63, 27.89]
MUNSTER 11.50 [ -9.47, 32.47]
NCNG 7.41 [-23.73, 38.54]
NESDA
NeuroIMAGE 25.27 [-22.21, 72.75]
NTR
OATS 2.26 [-33.19, 37.72]
PAFIP
PDNZ 10.65 [-51.64, 72.95]
PING
MPRC -0.02 [-29.14, 29.11]
PPMI
QTIM 18.48 [ -1.20, 38.16]
SHIP 4.93 [-14.17, 24.03]
SHIP-Trend 36.45 [ 16.25, 56.66]
SYS 42.79 [ 6.17, 79.41]
TCD-NUIG
TOP -3.82 [-30.39, 22.76]
TOP3T 17.20 [-12.06, 46.45]
UiO2016 27.32 [ -5.95, 60.59]
UiO2017 -20.13 [-52.72, 12.45]
UMCU 2.89 [-18.69, 24.47]
ENIGMA Discovery Cohorts [N=21510; P=1.324e-05] 9.14 [ 5.03, 13.25]
UKBB 11.79 [ 6.62, 16.97]
Discovery+UKBB [N=30765; P=1.594e-09] 10.20 [ 6.88, 13.51]
GOBS 20.23 [ 6.31, 34.15]
IMH -8.39 [-53.51, 36.73]
UNICAMP 21.60 [-32.10, 75.30]
Meth-CT 0.57 [-70.20, 71.34]
MIRECC
UKBB nonEUR -27.52 [-56.88, 1.84]
OSAKA 4.10 [-27.10, 35.29]
PING Non European
Non European Cohorts [N=2292; P=0.1007] 9.28 [ -1.80, 20.35]
Beta [Lower 95% CI,Upper 95% CI]
-100
-50
0
50
100
150
mm^2/effect allele

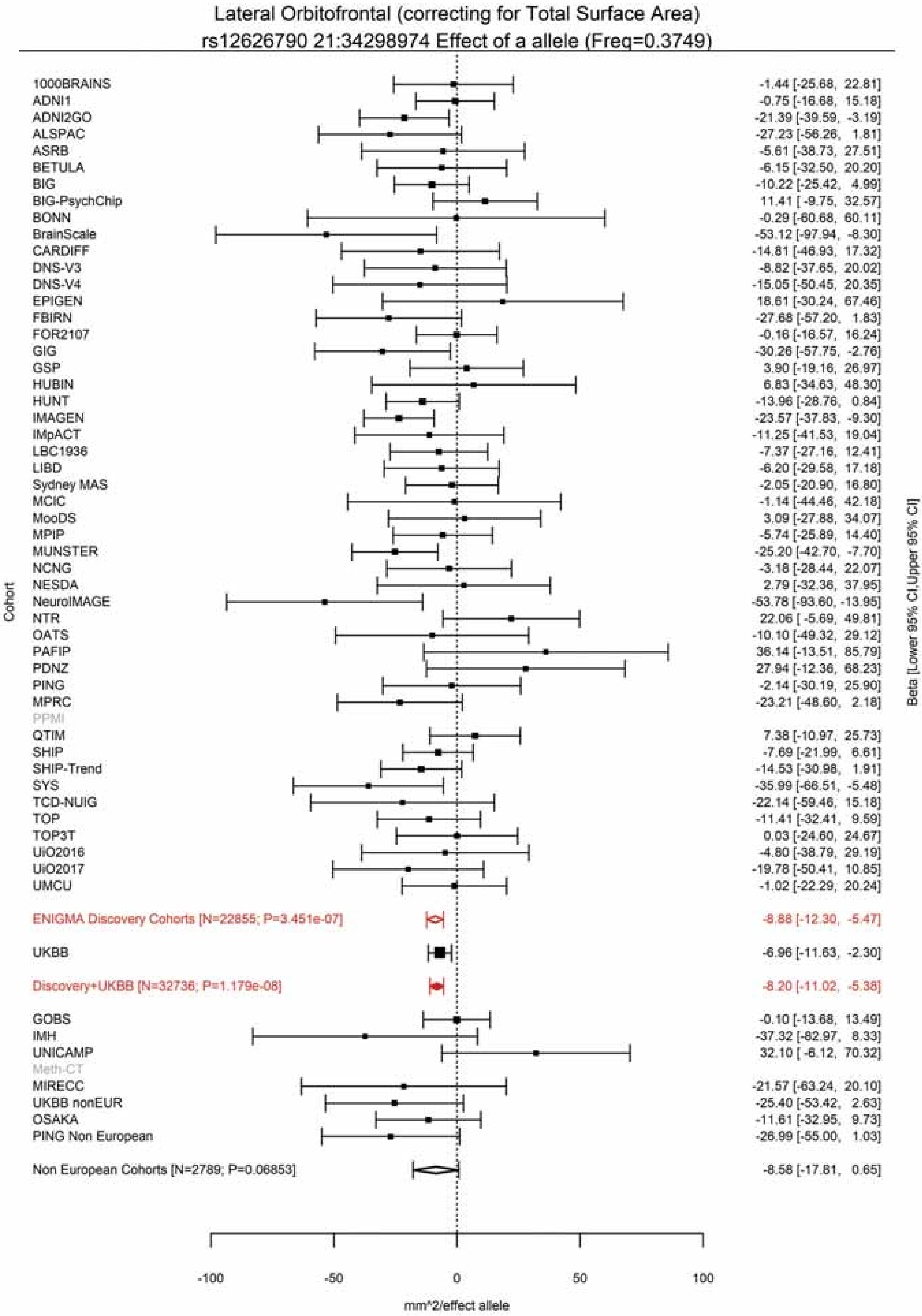
Lateral Orbitofrontal (correcting for Total Surface Area)
rs12626790 21:34298974 Effect of a allele (Freq=0.3749)
1000BRAINS -1.44 [-25.68, 22.81]
ADNI1 -0.75 [-16.68, 15.18]
ADNI2GO -21.39 [-39.59, -3.19]
ALSPAC -27.23 [-56.26, 1.81]
ASRB -5.61 [-38.73, 27.51]
BETULA -6.15 [-32.50, 20.20]
BIG -10.22 [-25.42, 4.99]
BIG-PsychChip 11.41 [ -9.75, 32.57]
BONN -0.29 [-60.68, 60.11]
BrainScale -53.12 [-97.94, -8.30]
CARDIFF -14.81 [-46.93, 17.32]
DNS-V3 -8.82 [-37.65, 20.02]
DNS-V4 -15.05 [-50.45, 20.35]
EPIGEN 18.61 [-30.24, 67.46]
FBIRN -27.68 [-57.20, 1.83]
FOR2107 -0.16 [-16.57, 16.24]
GIG -30.26 [-57.75, -2.76]
GSP 3.90 [-19.16, 26.97]
HUBIN 6.83 [-34.63, 48.30]
HUNT -13.96 [-28.76, 0.84]
IMAGEN -23.57 [-37.83, -9.30]
IMpACT -11.25 [-41.53, 19.04]
LBC1936 -7.37 [-27.16, 12.41]
LIBD -6.20 [-29.58, 17.18]
Sydney MAS -2.05 [-20.90, 16.80]
MCIC -1.14 [-44.46, 42.18]
MooDS 3.09 [-27.88, 34.07]
MPIP -5.74 [-25.89, 14.40]
MUNSTER -25.20 [-42.70, -7.70]
NCNG -3.18 [-28.44, 22.07]
NESDA 2.79 [-32.36, 37.95]
NeuroIMAGE -53.78 [-93.60, -13.95]
NTR 22.06 [ -5.69, 49.81]
OATS -10.10 [-49.32, 29.12]
PAFIP 36.14 [-13.51, 85.79]
PDNZ 27.94 [-12.36, 68.23]
PING -2.14 [-30.19, 25.90]
MPRC -23.21 [-48.60, 2.18]
PPMI
QTIM 7.38 [-10.97, 25.73]
SHIP -7.69 [-21.99, 6.61]
SHIP-Trend -14.53 [-30.98, 1.91]
SYS -35.99 [-66.51, -5.48]
TCD-NUIG -22.14 [-59.46, 15.18]
TOP -11.41 [-32.41, 9.59]
TOP3T 0.03 [-24.60, 24.67]
UiO2016 -4.80 [-38.79, 29.19]
UiO2017 -19.78 [-50.41, 10.85]
UMCU -1.02 [-22.29, 20.24]
ENIGMA Discovery Cohorts [N=22855; P=3.451e-07] -8.88 [-12.30, -5.47]
UKBB -6.96 [-11.63, -2.30]
Discovery+UKBB [N=32736; P=1.179e-08] -8.20 [-11.02, -5.38]
GOBS -0.10 [-13.68, 13.49]
IMH -37.32 [-82.97, 8.33]
UNICAMP 32.10 [ -6.12, 70.32]
Meth-CT
MIRECC -21.57 [-63.24, 20.10]
UKBB nonEUR -25.40 [-53.42, 2.63]
OSAKA -11.61 [-32.95, 9.73]
PING Non European -26.99 [-55.00, 1.03]
Non European Cohorts [N=2789; P=0.06853] -8.58 [-17.81, 0.65]
Cohort
Beta [Lower 95% CI,Upper 95% CI]
-100
-50
0
50
100
mm^2/effect allele

Postcentral (correcting for Average Cortical Thickness)
rs4823878 22:47200354 Effect of t allele (Freq=0.2889)

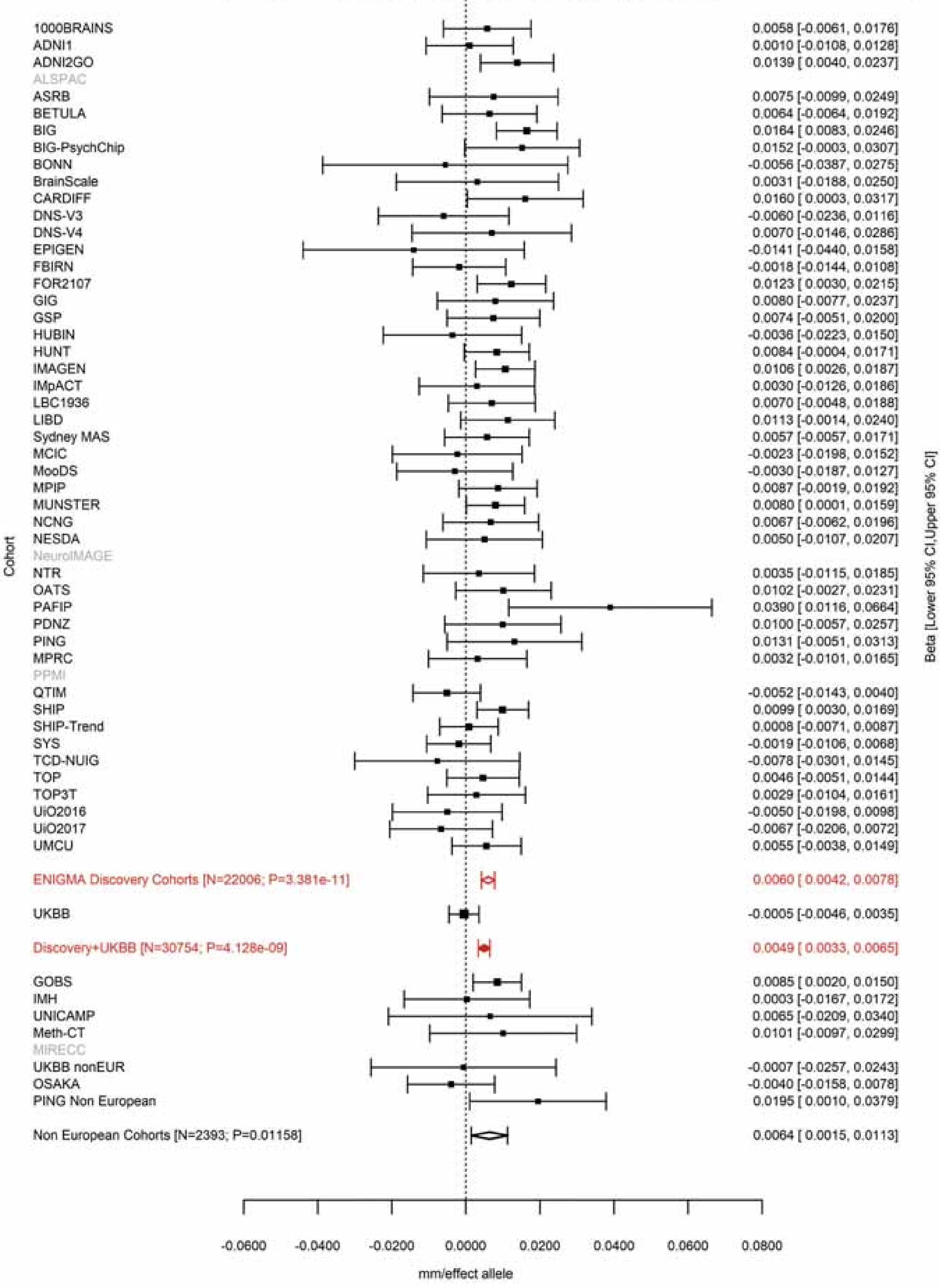
