## Supplemental_LocusZoom_Plots for "The genetic architecture of the human cerebral cortex"


| Locus # | Area | Trait | Marker | SNP | Page |
|---|---|---|---|---|---|
| 1 | Rostral Middle Frontal | SA | 1:17311882 | rs6682671 | 7 |
| 2 | Lingual | SA | 1:18962095 | rs1934057 | 8 |
| 3 | Superior Frontal | SA | 1:64235727 | rs4915928 | 9 |
| 4 | Posterior Cingulate | SA | 1:87633483:T_TGA | rs113198643 | 10 |
| 5 | Precuneus | SA | 1:87873700 | rs59373415 | 11 |
| 6 | Inferior Parietal | SA | 1:88216936 | rs305437 | 12 |
| 7 | Pericalcarine | SA | 1:88269969 | rs147753572 | 13 |
| 8 | Inferior Parietal | SA | 1:88423397 | rs1413536 | 14 |
| 9 | Lingual | SA | 1:93032583 | rs6603991 | 15 |
| 10 | Lingual | SA | 1:113239478 | rs2999158 | 16 |
| 11 | Pericalcarine | SA | 1:113239478 | rs2999158 | 17 |
| 12 | Pars Orbitalis | SA | 1:119762175 | rs72691108 | 18 |
| 13 | Lateral Orbitofrontal | SA | 1:119799099 | rs7529542 | 19 |
| 14 | Rostral Middle Frontal | TH | 1:247527273 | rs12058942 | 20 |
| 15 | Superior Parietal | SA | 2:4550112:CTG_C | rs769344141 | 21 |
| 16 | Superior Parietal | SA | 2:4550411 | rs688409 | 22 |
| 17 | Inferior Parietal | SA | 2:4554304 | rs639016 | 23 |
| 18 | Average Thickness | TH | 2:27148654:C_CT | rs11386753 | 24 |
| 19 | Precuneus | SA | 2:37076594 | rs62132521 | 25 |
| 20 | Precuneus | SA | 2:37079583 | rs62132522 | 26 |
| 21 | Isthmus Cingulate | SA | 2:37150793 | rs3770776 | 27 |
| 22 | Posterior Cingulate | SA | 2:37175444 | rs11695609 | 28 |
| 23 | Precuneus | SA | 2:37748367 | rs7559976 | 29 |
| 24 | Inferior Temporal | SA | 2:37899310 | rs9309013 | 30 |
| 25 | Superior Temporal | SA | 2:48277490 | rs7601767 | 31 |
| 26 | Lingual | TH | 2:49391013 | rs10495963 | 32 |
| 27 | Inferior Parietal | SA | 2:54615325 | rs79272390 | 33 |
| 28 | Superior Parietal | SA | 2:54615325 | rs79272390 | 34 |
| 29 | Pars Orbitalis | SA | 2:65971597 | rs2287283 | 35 |
| 30 | Pars Triangularis | SA | 2:65971789:C_CAG | rs142706617 | 36 |
| 31 | Caudal Middle Frontal | SA | 2:65974661:AG_A | rs770408932 | 37 |
| 32 | Average Thickness | TH | 2:98275354 | rs11692435 | 38 |
| 33 | Transverse Temporal | SA | 2:149940486 | rs2889657 | 39 |
| 34 | Superior Temporal | SA | 2:149994571 | rs13011264 | 40 |
| 35 | Superior Temporal | SA | 2:150007758 | rs389020 | 41 |
| 36 | Transverse Temporal | SA | 2:150037201 | rs11684511 | 42 |
| 37 | Caudal Anterior Cingulate | SA | 2:162822709 | rs13021985 | 43 |
| 38 | Lingual | SA | 2:162868858 | rs1014444 | 44 |
| 39 | Pericalcarine | SA | 2:162901327 | rs16822665 | 45 |
| 40 | Frontal Pole | SA | 2:188278203 | rs17464221 | 46 |
| 41 | Rostral Middle Frontal | SA | 2:188379320 | rs35612915 | 47 |
| 42 | Cuneus | SA | 2:204517033 | rs71427711 | 48 |
| 43 | Total Surface Area | SA | 3:28007315 | rs12630663 | 49 |
| 44 | Average Thickness | TH | 3:39489651 | rs533577 | 50 |
| 45 | Rostral Middle Frontal | TH | 3:55068255 | rs4955920 | 51 |
| 46 | Superior Parietal | SA | 3:71557865 | rs17718831 | 52 |
| 47 | Cuneus | SA | 3:104647183 | rs4895120 | 53 |
| 48 | Lingual | SA | 3:104662433 | rs13318870 | 54 |
| 49 | Lateral Occipital | SA | 3:104678133 | rs9863836 | 55 |
| 50 | Inferior Parietal | SA | 3:104724219 | rs9856782 | 56 |
| 51 | Pericalcarine | SA | 3:104724787 | rs971550 | 57 |
| 52 | Lingual | SA | 3:104735633 | rs28551708 | 58 |
| 53 | Lateral Occipital | SA | 3:118994959 | rs56007616 | 59 |
| 54 | Pericalcarine | SA | 3:118995379 | rs16829649 | 60 |
| 55 | Total Surface Area | SA | 3:141721762 | rs34464850 | 61 |
| 56 | Lingual | SA | 3:146993779 | rs61508189 | 62 |
| 57 | Pars Orbitalis | SA | 3:147093600 | rs1503738 | 63 |

| | | | | | |
|---|---|---|---|---|---|
| 58 | Postcentral | SA | 3:147105790 | rs2279830 | 64 |
| 59 | Pars Triangularis | SA | 3:147106319 | rs2279829 | 65 |
| 60 | Postcentral | SA | 3:147106319 | rs2279829 | 66 |
| 61 | Supramarginal | SA | 3:147106319 | rs2279829 | 67 |
| 62 | Pars Triangularis | SA | 3:147121751 | rs6766244 | 68 |
| 63 | Pars Triangularis | SA | 3:147171352 | rs9881533 | 69 |
| 64 | Pars Triangularis | SA | 3:147218844:CCAT_C | 3:147218844_CCAT_C | 70 |
| 65 | Postcentral | SA | 3:147227163 | rs78155705 | 71 |
| 66 | Lateral Occipital | SA | 3:177295215 | rs552305 | 72 |
| 67 | Precuneus | SA | 3:190657360 | rs905124 | 73 |
| 68 | Rostral Middle Frontal | SA | 3:193536820 | rs1165645 | 74 |
| 69 | Caudal Middle Frontal | SA | 3:193547618:T_TG | rs140045876 | 75 |
| 70 | Pericalcarine | SA | 4:980896 | rs11248061 | 76 |
| 71 | Lingual | SA | 4:1002448 | rs6812278 | 77 |
| 72 | Pericalcarine | SA | 4:1002448 | rs6812278 | 78 |
| 73 | Pericalcarine | SA | 4:19095652 | rs13115025 | 79 |
| 74 | Superior Parietal | SA | 4:53702875 | rs6554054 | 80 |
| 75 | Supramarginal | SA | 4:53738411 | rs2200225 | 81 |
| 76 | Temporal Pole | SA | 4:103112470 | rs6855246 | 82 |
| 77 | Total Surface Area | SA | 4:106009763 | rs2301718 | 83 |
| 78 | Superior Parietal | SA | 4:113416783 | rs6840242 | 84 |
| 79 | Average Thickness | TH | 4:121643239 | rs35021943 | 85 |
| 80 | Postcentral | SA | 4:127537734 | rs313135 | 86 |
| 81 | Pericalcarine | SA | 4:145284208 | rs7378179 | 87 |
| 82 | Fusiform | SA | 5:56082094 | rs10940512 | 88 |
| 83 | Lingual | SA | 5:60299840:C_CT | rs75625671 | 89 |
| 84 | Pericalcarine | SA | 5:60318973 | rs62367903 | 90 |
| 85 | Pericalcarine | SA | 5:60484660 | rs159540 | 91 |
| 86 | Posterior Cingulate | TH | 5:66144113 | rs12110247 | 92 |
| 87 | Posterior Cingulate | SA | 5:66155020 | rs72761270 | 93 |
| 88 | Total Surface Area | SA | 5:81092787 | rs386424 | 94 |
| 89 | Entorhinal | SA | 5:81882571 | rs4147321 | 95 |
| 90 | Parahippocampal | SA | 5:82350191 | rs27493 | 96 |
| 91 | Caudal Anterior Cingulate | SA | 5:82843609 | rs7728751 | 97 |
| 92 | Insula | SA | 5:82843609 | rs7728751 | 98 |
| 93 | Inferior Parietal | SA | 5:92090343 | rs34969 | 99 |
| 94 | Inferior Parietal | SA | 5:92134273 | rs27540 | 100 |
| 95 | Precuneus | SA | 5:92186429 | rs888814 | 101 |
| 96 | Superior Frontal | SA | 5:92187932 | rs17669337 | 102 |
| 97 | Superior Parietal | SA | 5:92797166 | rs115877304 | 103 |
| 98 | Superior Parietal | SA | 5:92948485 | rs114489117 | 104 |
| 99 | Precentral | SA | 5:92950673 | rs10064431 | 105 |
| 100 | Postcentral | SA | 5:93347836 | rs34322452 | 106 |
| 101 | Middle Temporal | SA | 5:93385558 | rs141834426 | 107 |
| 102 | Middle Temporal | SA | 5:93557702 | rs17376456 | 108 |
| 103 | Pericalcarine | SA | 5:124283820:T_TA | rs149998495 | 109 |
| 104 | Inferior Parietal | SA | 5:128971797 | rs62399042 | 110 |
| 105 | Caudal Middle Frontal | SA | 5:129018314 | rs30641 | 111 |
| 106 | Transverse Temporal | SA | 5:131341541 | rs7714191 | 112 |
| 107 | Total Surface Area | SA | 5:170778824 | rs7715167 | 113 |
| 108 | Caudal Middle Frontal | TH | 5:171927246 | rs11745941 | 114 |
| 109 | Lateral Orbitofrontal | SA | 6:7119134 | rs13208234 | 115 |
| 110 | Caudal Middle Frontal | SA | 6:92002653 | rs9345125 | 116 |
| 111 | Transverse Temporal | SA | 6:92009564 | rs4706391 | 117 |
| 112 | Precentral | SA | 6:92009596 | rs4706392 | 118 |
| 113 | Superior Parietal | SA | 6:92010245 | rs2144366 | 119 |
| 114 | Total Surface Area | SA | 6:108926496 | rs2802295 | 120 |
| 115 | Superior Frontal | SA | 6:126024301 | rs142301939 | 121 |
| 116 | Pars Opercularis | SA | 6:126070789 | rs7764016 | 122 |
| 117 | Total Surface Area | SA | 6:126412953 | rs11154343 | 123 |

| | | | | | |
|---|---|---|---|---|---|
| 118 | Cuneus | SA | 6:126613946 | rs76470478 | 124 |
| 119 | Pericalcarine | SA | 6:126666648 | rs117892760 | 125 |
| 120 | Lateral Orbitofrontal | SA | 6:126727908 | rs4897178 | 126 |
| 121 | Caudal Middle Frontal | SA | 6:126727930 | rs4897179 | 127 |
| 122 | Total Surface Area | SA | 6:126792095 | rs11759026 | 128 |
| 123 | Parahippocampal | SA | 6:126868567 | rs58321169 | 129 |
| 124 | Caudal Middle Frontal | SA | 6:126964510 | rs4273712 | 130 |
| 125 | Lateral Orbitofrontal | SA | 6:126995044 | rs1262478 | 131 |
| 126 | Total Surface Area | SA | 6:127000881 | rs74580701 | 132 |
| 127 | Total Surface Area | SA | 6:127025661 | rs13212044 | 133 |
| 128 | Lateral Occipital | SA | 6:127096181 | rs9401907 | 134 |
| 129 | Lingual | SA | 6:127096181 | rs9401907 | 135 |
| 130 | Pericalcarine | SA | 6:127096181 | rs9401907 | 136 |
| 131 | Total Surface Area | SA | 6:127204623 | rs9375477 | 137 |
| 132 | Pericalcarine | SA | 6:138865324 | rs4895532 | 138 |
| 133 | Precuneus | SA | 6:138868516 | rs9399245 | 139 |
| 134 | Rostral Anterior Cingulate | SA | 7:18737197 | rs1178101 | 140 |
| 135 | Cuneus | SA | 7:18881075 | rs12536836 | 141 |
| 136 | Pericalcarine | SA | 7:18883690 | rs6461386 | 142 |
| 137 | Lingual | SA | 7:18915666 | rs10237280 | 143 |
| 138 | Lateral Orbitofrontal | SA | 7:19520977 | rs73685918 | 144 |
| 139 | Lateral Orbitofrontal | SA | 7:19619484:T_TAG | rs371426453 | 145 |
| 140 | Lateral Orbitofrontal | SA | 7:19626622 | rs4721802 | 146 |
| 141 | Pars Triangularis | SA | 7:96202718 | rs10278627 | 147 |
| 142 | Pars Triangularis | SA | 7:96227720 | rs56290730 | 148 |
| 143 | Lingual | SA | 7:107237807 | rs7809950 | 149 |
| 144 | Superior Parietal | TH | 7:120721087 | rs73215353 | 150 |
| 145 | Superior Temporal | SA | 8:8605632 | rs4841029 | 151 |
| 146 | Lateral Orbitofrontal | SA | 8:8975103 | rs1822951 | 152 |
| 147 | Superior Temporal | SA | 8:9055772 | rs10094141 | 153 |
| 148 | Superior Temporal | SA | 8:9370559 | rs10100760 | 154 |
| 149 | Superior Temporal | SA | 8:9408271 | rs4840425 | 155 |
| 150 | Superior Temporal | SA | 8:10431422 | rs11250033 | 156 |
| 151 | Superior Temporal | SA | 8:10431486 | rs12548232 | 157 |
| 152 | Transverse Temporal | SA | 8:10943276 | rs2409691 | 158 |
| 153 | Superior Temporal | SA | 8:10962800 | rs1057626 | 159 |
| 154 | Superior Temporal | SA | 8:11109691 | rs2736373 | 160 |
| 155 | Pars Opercularis | SA | 8:71564667 | rs441890 | 161 |
| 156 | Inferior Parietal | SA | 8:93204400:T_TA | rs35576477 | 162 |
| 157 | Average Thickness | TH | 8:110585288 | rs7824177 | 163 |
| 158 | Rostral Middle Frontal | SA | 8:120596023 | rs10283100 | 164 |
| 159 | Postcentral | SA | 9:76157130 | rs11789773 | 165 |
| 160 | Superior Frontal | SA | 9:83480392 | rs76696867 | 166 |
| 161 | Pericalcarine | SA | 9:98257305 | rs28633576 | 167 |
| 162 | Lingual | SA | 9:98276371 | rs28410513 | 168 |
| 163 | Lateral Occipital | SA | 9:98278332 | rs28496034 | 169 |
| 164 | Banks of the Superior Temporal Sulcus | SA | 9:126603989 | rs7862092 | 170 |
| 165 | Total Surface Area | SA | 10:21790476 | rs12357321 | 171 |
| 166 | Caudal Anterior Cingulate | SA | 10:25123182 | rs4747503 | 172 |
| 167 | Paracentral | SA | 10:33539487 | rs2269084 | 173 |
| 168 | Inferior Parietal | SA | 10:33613967 | rs75921753 | 174 |
| 169 | Medial Orbitofrontal | SA | 10:50302895 | rs7097933 | 175 |
| 170 | Lingual | SA | 10:104422321 | rs7914158 | 176 |
| 171 | Total Surface Area | SA | 10:105012994 | rs1628768 | 177 |
| 172 | Total Surface Area | SA | 10:105218254 | rs4918016 | 178 |
| 173 | Precentral | SA | 10:118696266 | rs4751614 | 179 |
| 174 | Insula | SA | 10:118752250 | rs1122688 | 180 |
| 175 | Precuneus | SA | 10:118777998 | rs10749233 | 181 |
| 176 | Isthmus Cingulate | TH | 10:126486443 | rs12764880 | 182 |

| | | | | | |
|---|---|---|---|---|---|
| 177 | Posterior Cingulate | SA | 10:126486443 | rs12764880 | 183 |
| 178 | Cuneus | SA | 11:12071855 | rs10765918 | 184 |
| 179 | Pericalcarine | SA | 11:12071855 | rs10765918 | 185 |
| 180 | Lingual | SA | 11:30819811 | rs2022130 | 186 |
| 181 | Pericalcarine | SA | 11:30883876 | rs273587 | 187 |
| 182 | Pericalcarine | SA | 11:31499705 | rs1223090 | 188 |
| 183 | Postcentral | SA | 11:37001542 | rs11033898 | 189 |
| 184 | Parahippocampal | SA | 11:92319654 | rs1792354 | 190 |
| 185 | Pars Orbitalis | SA | 11:92348702 | rs61901866 | 191 |
| 186 | Fusiform | SA | 11:110962056 | rs7123402 | 192 |
| 187 | Pars Triangularis | SA | 11:111050526 | rs59614433 | 193 |
| 188 | Middle Temporal | SA | 11:111055661 | rs12794347 | 194 |
| 189 | Insula | SA | 11:111056158 | rs58066679 | 195 |
| 190 | Fusiform | SA | 11:111072779 | rs949279 | 196 |
| 191 | Pericalcarine | SA | 12:5314508 | rs17179798 | 197 |
| 192 | Superior Parietal | SA | 12:6545417 | rs7980991 | 198 |
| 193 | Paracentral | SA | 12:27790516 | rs2346756 | 199 |
| 194 | Superior Frontal | TH | 12:28833981 | rs4572176 | 200 |
| 195 | Total Surface Area | SA | 12:56470625 | rs11171739 | 201 |
| 196 | Inferior Parietal | SA | 12:65797106 | rs2336714 | 202 |
| 197 | Lateral Orbitofrontal | SA | 12:65905126 | rs79487293 | 203 |
| 198 | Total Surface Area | SA | 12:66327632 | rs10878349 | 204 |
| 199 | Total Surface Area | SA | 12:66367070 | rs12826248 | 205 |
| 200 | Isthmus Cingulate | SA | 12:78241285 | rs1123680 | 206 |
| 201 | Superior Frontal | SA | 12:79951566 | rs4842266 | 207 |
| 202 | Entorhinal | TH | 12:94558576 | rs3847803 | 208 |
| 203 | Lingual | TH | 13:58591051 | rs9537915 | 209 |
| 204 | Lingual | SA | 13:80173760 | rs9545145 | 210 |
| 205 | Pericalcarine | SA | 13:80192906 | rs9545158 | 211 |
| 206 | Pars Triangularis | SA | 13:81198667 | rs7996803 | 212 |
| 207 | Pericalcarine | SA | 14:29228427 | rs35342371 | 213 |
| 208 | Lingual | SA | 14:54316607 | rs7148896 | 214 |
| 209 | Lateral Orbitofrontal | SA | 14:54724054 | rs2358483 | 215 |
| 210 | Pars Orbitalis | SA | 14:54777745 | rs7147119 | 216 |
| 211 | Lateral Occipital | SA | 14:57337018 | rs76398229 | 217 |
| 212 | Banks of the Superior Temporal Sulcus | SA | 14:58724817:C_CTT | rs35486640 | 218 |
| 213 | Inferior Temporal | SA | 14:59063842 | rs7155669 | 219 |
| 214 | Cuneus | SA | 14:59064739 | rs170239 | 220 |
| 215 | Supramarginal | SA | 14:59069053 | rs221326 | 221 |
| 216 | Banks of the Superior Temporal Sulcus | TH | 14:59071725 | rs149142 | 222 |
| 217 | Middle Temporal | TH | 14:59074136 | rs160459 | 223 |
| 218 | Banks of the Superior Temporal Sulcus | SA | 14:59074878 | rs160458 | 224 |
| 219 | Precuneus | SA | 14:59598286 | rs7143623 | 225 |
| 220 | Cuneus | TH | 14:59605116 | rs117461235 | 226 |
| 221 | Pericalcarine | TH | 14:59605116 | rs117461235 | 227 |
| 222 | Cuneus | SA | 14:59625997 | rs73313052 | 228 |
| 223 | Pericalcarine | SA | 14:59625997 | rs73313052 | 229 |
| 224 | Precuneus | SA | 14:59625997 | rs73313052 | 230 |
| 225 | Lingual | SA | 14:59627434:AC_A | 14:59627434_AC_A | 231 |
| 226 | Supramarginal | SA | 14:59627631 | rs2164950 | 232 |
| 227 | Lateral Occipital | SA | 14:59628679 | rs76341705 | 233 |
| 228 | Lateral Occipital | SA | 14:59683086 | rs78445564 | 234 |
| 229 | Pericalcarine | SA | 14:59805480 | rs75341124 | 235 |
| 230 | Fusiform | SA | 14:59827306 | rs17834032 | 236 |
| 231 | Entorhinal | SA | 14:59860705 | rs7141150 | 237 |
| 232 | Supramarginal | SA | 15:39081795 | rs62007727 | 238 |
| 233 | Precentral | SA | 15:39092485:AG_A | 15:39092485_AG_A | 239 |

| | | | | | |
|---|---|---|---|---|---|
| 234 | Precentral | SA | 15:39343032 | rs11070172 | 240 |
| 235 | Precentral | SA | 15:39519925 | rs156795 | 241 |
| 236 | Postcentral | SA | 15:39553167 | rs77470370 | 242 |
| 237 | Postcentral | SA | 15:39556139:TAC_T | 15:39556139_TAC_T | 243 |
| 238 | Precentral | SA | 15:39582377:TA_T | 15:39582377_TA_T | 244 |
| 239 | Supramarginal | SA | 15:39593078 | rs11631253 | 245 |
| 240 | Postcentral | TH | 15:39595340 | rs62002282 | 246 |
| 241 | Precentral | SA | 15:39595340 | rs62002282 | 247 |
| 242 | Postcentral | SA | 15:39604340 | rs3862145 | 248 |
| 243 | Precentral | SA | 15:39604340 | rs3862145 | 249 |
| 244 | Postcentral | SA | 15:39623847 | rs10851383 | 250 |
| 245 | Postcentral | TH | 15:39623847 | rs10851383 | 251 |
| 246 | Pars Opercularis | SA | 15:39625618 | rs11070185 | 252 |
| 247 | Precentral | SA | 15:39625778 | rs10851385 | 253 |
| 248 | Supramarginal | SA | 15:39625778 | rs10851385 | 254 |
| 249 | Precentral | SA | 15:39627855 | rs76465453 | 255 |
| 250 | Postcentral | TH | 15:39632013 | rs71471500 | 256 |
| 251 | Pars Opercularis | SA | 15:39633904 | rs2033939 | 257 |
| 252 | Transverse Temporal | SA | 15:39633904 | rs2033939 | 258 |
| 253 | Precentral | SA | 15:39634222 | rs1080066 | 259 |
| 254 | Rostral Middle Frontal | SA | 15:39634222 | rs1080066 | 260 |
| 255 | Postcentral | TH | 15:39634303 | rs1822105 | 261 |
| 256 | Precentral | SA | 15:39634303 | rs1822105 | 262 |
| 257 | Superior Parietal | SA | 15:39634303 | rs1822105 | 263 |
| 258 | Postcentral | SA | 15:39637355 | rs117193619 | 264 |
| 259 | Precentral | SA | 15:39637355 | rs117193619 | 265 |
| 260 | Pars Triangularis | SA | 15:39639898 | rs4924345 | 266 |
| 261 | Superior Parietal | SA | 15:39639898 | rs4924345 | 267 |
| 262 | Supramarginal | SA | 15:39639898 | rs4924345 | 268 |
| 263 | Postcentral | SA | 15:39639992 | rs4924346 | 269 |
| 264 | Precentral | SA | 15:39640301 | rs148182077 | 270 |
| 265 | Postcentral | SA | 15:39641885 | rs10520129 | 271 |
| 266 | Supramarginal | SA | 15:39647278 | rs35391898 | 272 |
| 267 | Postcentral | SA | 15:39652450 | rs72722993 | 273 |
| 268 | Precentral | SA | 15:39652450 | rs72722993 | 274 |
| 269 | Precentral | SA | 15:39671514 | rs11070197 | 275 |
| 270 | Supramarginal | SA | 15:39720600 | rs78502100 | 276 |
| 271 | Precentral | SA | 15:39742570 | rs62005276 | 277 |
| 272 | Pars Opercularis | SA | 15:39790176 | rs10459586 | 278 |
| 273 | Superior Temporal | SA | 15:39809640 | rs115241741 | 279 |
| 274 | Pericalcarine | SA | 15:39953061 | rs8034885 | 280 |
| 275 | Lateral Occipital | SA | 15:63297719 | rs28514429 | 281 |
| 276 | Pericalcarine | SA | 16:28919583:A_ATT | rs10653411 | 282 |
| 277 | Caudal Middle Frontal | SA | 16:52538337 | rs7184835 | 283 |
| 278 | Caudal Anterior Cingulate | SA | 16:60827388 | rs80241863 | 284 |
| 279 | Rostral Middle Frontal | SA | 16:65014909 | rs40115 | 285 |
| 280 | Entorhinal | SA | 16:77320605 | rs12921392 | 286 |
| 281 | Superior Parietal | TH | 16:87229344 | rs56023709 | 287 |
| 282 | Pars Opercularis | SA | 17:13054525 | rs12938190 | 288 |
| 283 | Insula | SA | 17:28338304 | rs4291964 | 289 |
| 284 | Total Surface Area | SA | 17:43543075:C_CA | rs200291097 | 290 |
| 285 | Fusiform | SA | 17:43859640 | rs62057070 | 291 |
| 286 | Total Surface Area | SA | 17:43897722 | rs79600142 | 292 |
| 287 | Average Thickness | TH | 17:43919068 | rs2316766 | 293 |
| 288 | Total Surface Area | SA | 17:44808360 | rs35937770 | 294 |
| 289 | Rostral Anterior Cingulate | SA | 17:46584657 | rs2202895 | 295 |
| 290 | Cuneus | SA | 17:63906955:C_CAA | rs143442432 | 296 |
| 291 | Pericalcarine | SA | 17:63914750 | rs1420791 | 297 |
| 292 | Pericalcarine | SA | 17:63990193 | rs150476910 | 298 |
| 293 | Superior Parietal | SA | 19:12904400 | rs17884482 | 299 |

| | | | | | |
|---|---|---|---|---|---|
| 294 | Superior Parietal | SA | 19:13096285 | rs4334415 | 300 |
| 295 | Inferior Parietal | SA | 19:13109763 | rs68175985 | 301 |
| 296 | Superior Parietal | SA | 19:13109763 | rs68175985 | 302 |
| 297 | Banks of the Superior Temporal Sulcus | SA | 19:13109955 | rs73006822 | 303 |
| 298 | Middle Temporal | SA | 19:13109955 | rs73006822 | 304 |
| 299 | Superior Parietal | SA | 19:13292745 | rs8103974 | 305 |
| 300 | Pars Triangularis | SA | 19:30852938:CTACT_G_C | 19:30852938_CTACTG_C | 306 |
| 301 | Rostral Anterior Cingulate | TH | 19:51621561 | rs62115964 | 307 |
| 302 | Inferior Parietal | SA | 20:52432451 | rs4437022 | 308 |
| 303 | Superior Parietal | SA | 20:52447303 | rs6022786 | 309 |
| 304 | Pericalcarine | SA | 20:52468966 | rs4811476 | 310 |
| 305 | Lateral Orbitofrontal | SA | 21:34298974 | rs12626790 | 311 |
| 306 | Postcentral | TH | 22:47200354 | rs4823878 | 312 |

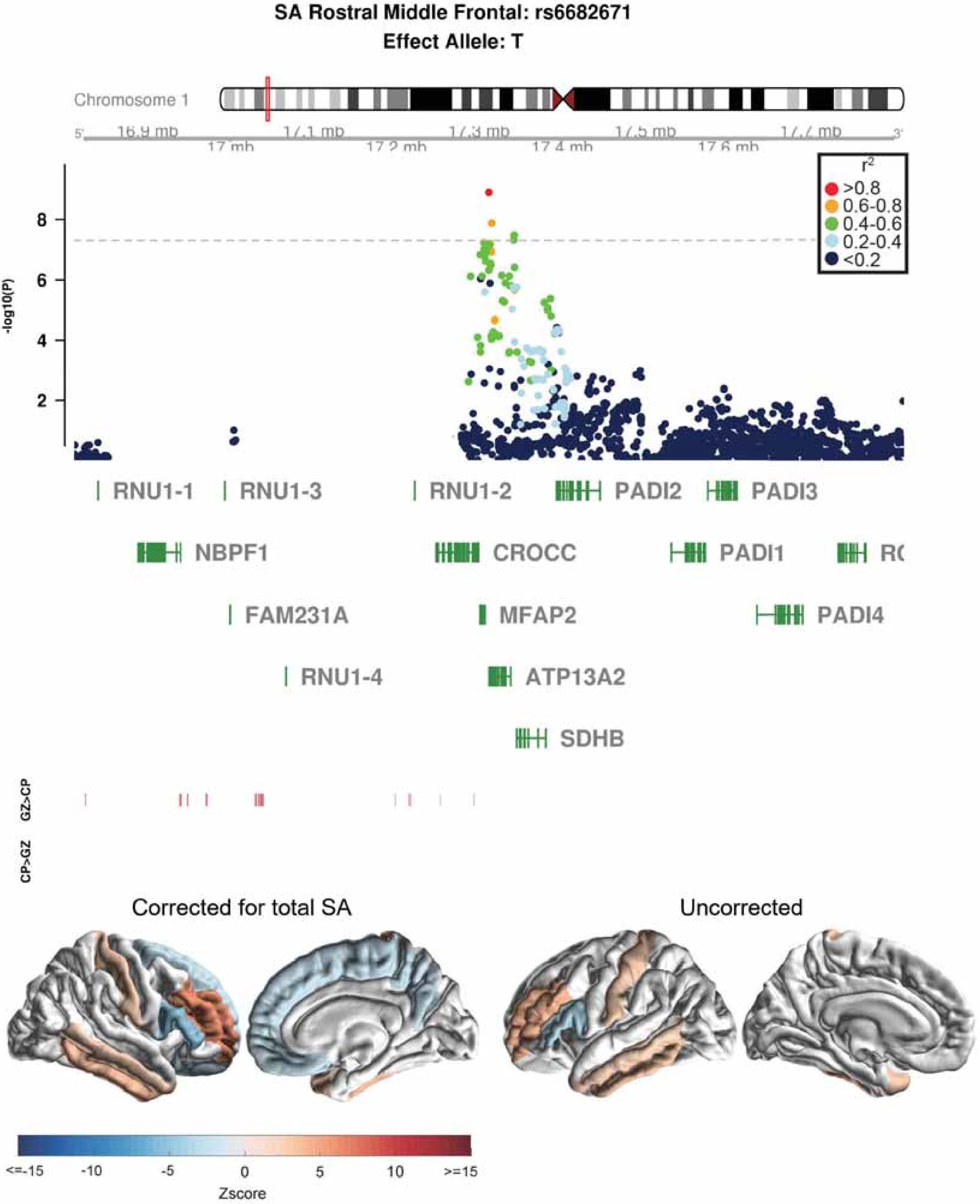

SA Rostral Middle Frontal: rs6682671
Effect Allele: T
Chromosome 1
16.9 mb
17 mb
17.1 mb
17.2 mb
17.3 mb
17.4 mb
17.5 mb
17.6 mb
17.7 mb
-log10(P)
r2
>0.8
0.6-0.8
0.4-0.6
0.2-0.4
<0.2
RNU1-1
RNU1-3
RNU1-2
PADI2
PADI3
NBPF1
CROCC
PADI1
FAM231A
MFAP2
PADI4
RNU1-4
ATP13A2
SDHB
GZ>CP
CP>GZ
Corrected for total SA
Uncorrected
<=-15
-10
-5
0
5
10
>=15
Zscore

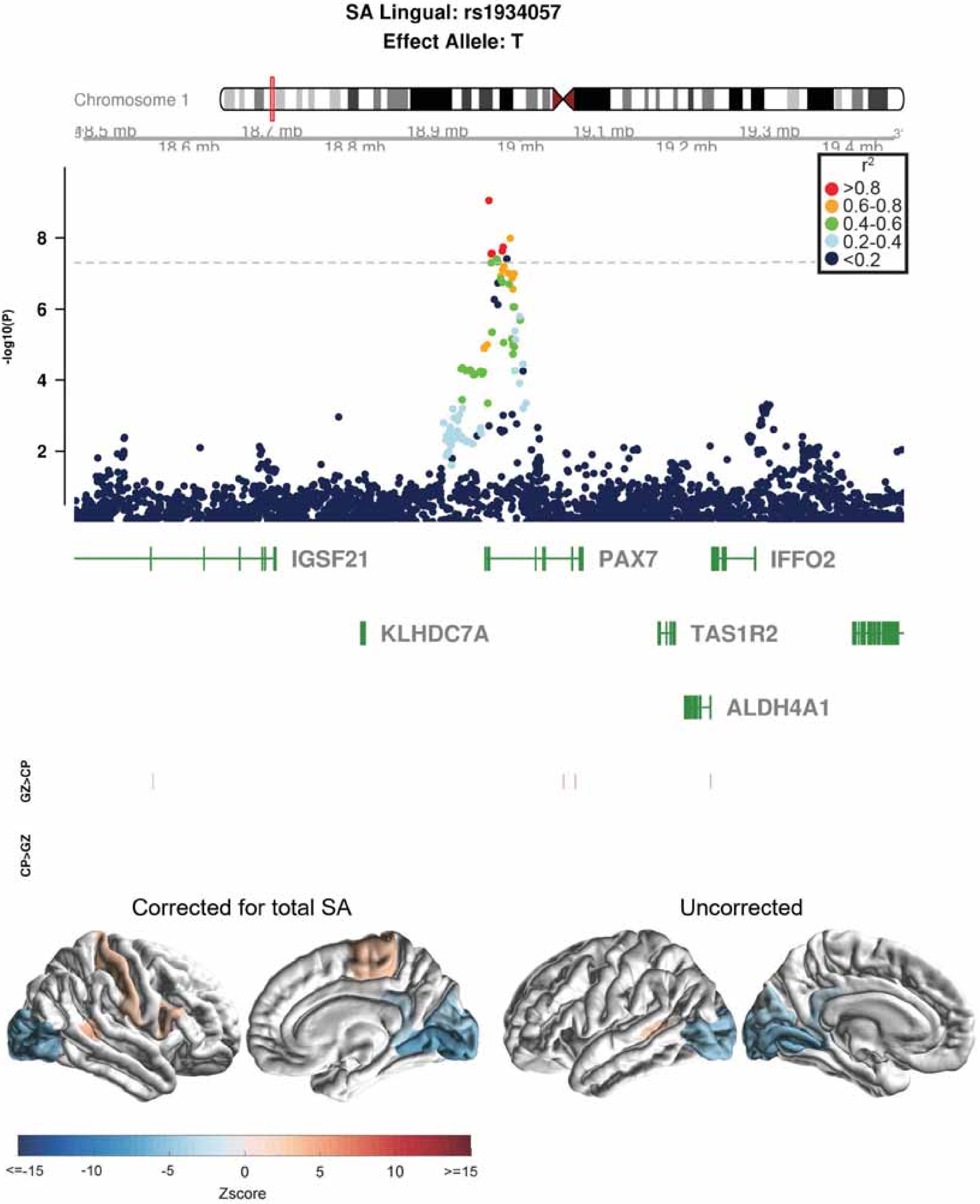
SA Lingual: rs1934057
Effect Allele: T
Chromosome 1
18.6 mb
18.7 mb
18.8 mb
18.9 mb
19 mb
19.1 mb
19.2 mb
19.3 mb
19.4 mb
r2
>0.8
0.6-0.8
0.4-0.6
0.2-0.4
<0.2
-log10(P)
IGSF21
PAX7
IFFO2
KLHDC7A
TAS1R2
ALDH4A1
GZ>CP
CP>GZ
Corrected for total SA
Uncorrected
<=-15
-10
-5
0
5
10
>=15
Zscore

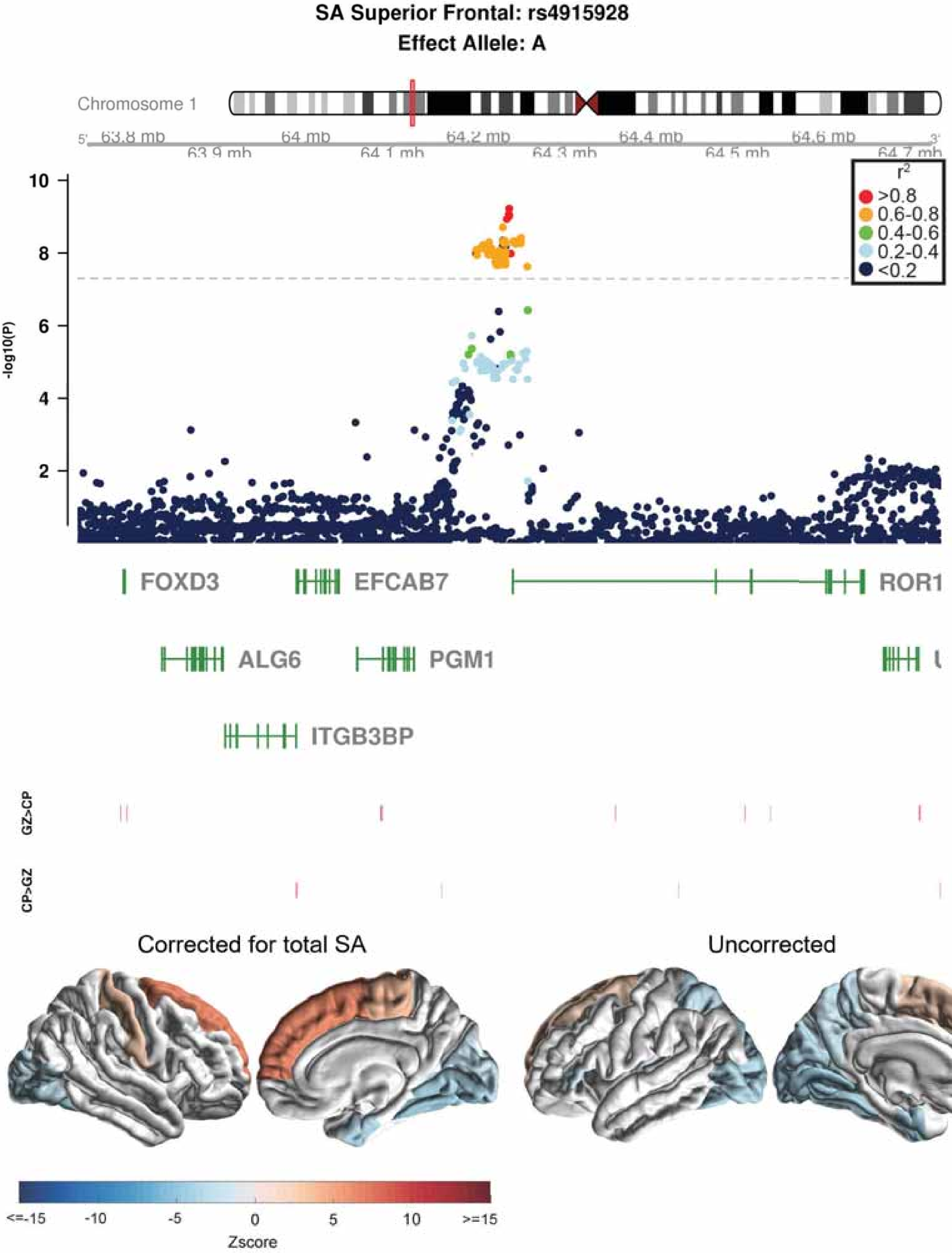
SA Superior Frontal: rs4915928
Effect Allele: A
Chromosome 1
63.8 mb
64 mb
64.2 mb
64.4 mb
64.6 mb
63.9 mb
64.1 mb
64.3 mb
64.5 mb
64.7 mb
-log10(P)
10
8
6
4
2
$r^2$
>0.8
0.6-0.8
0.4-0.6
0.2-0.4
<0.2
FOXD3
EFCAB7
ROR1
ALG6
PGM1
ITGB3BP
GZ>CP
CP>GZ
Corrected for total SA
Uncorrected
<=-15
-10
-5
0
5
10
>=15
Zscore

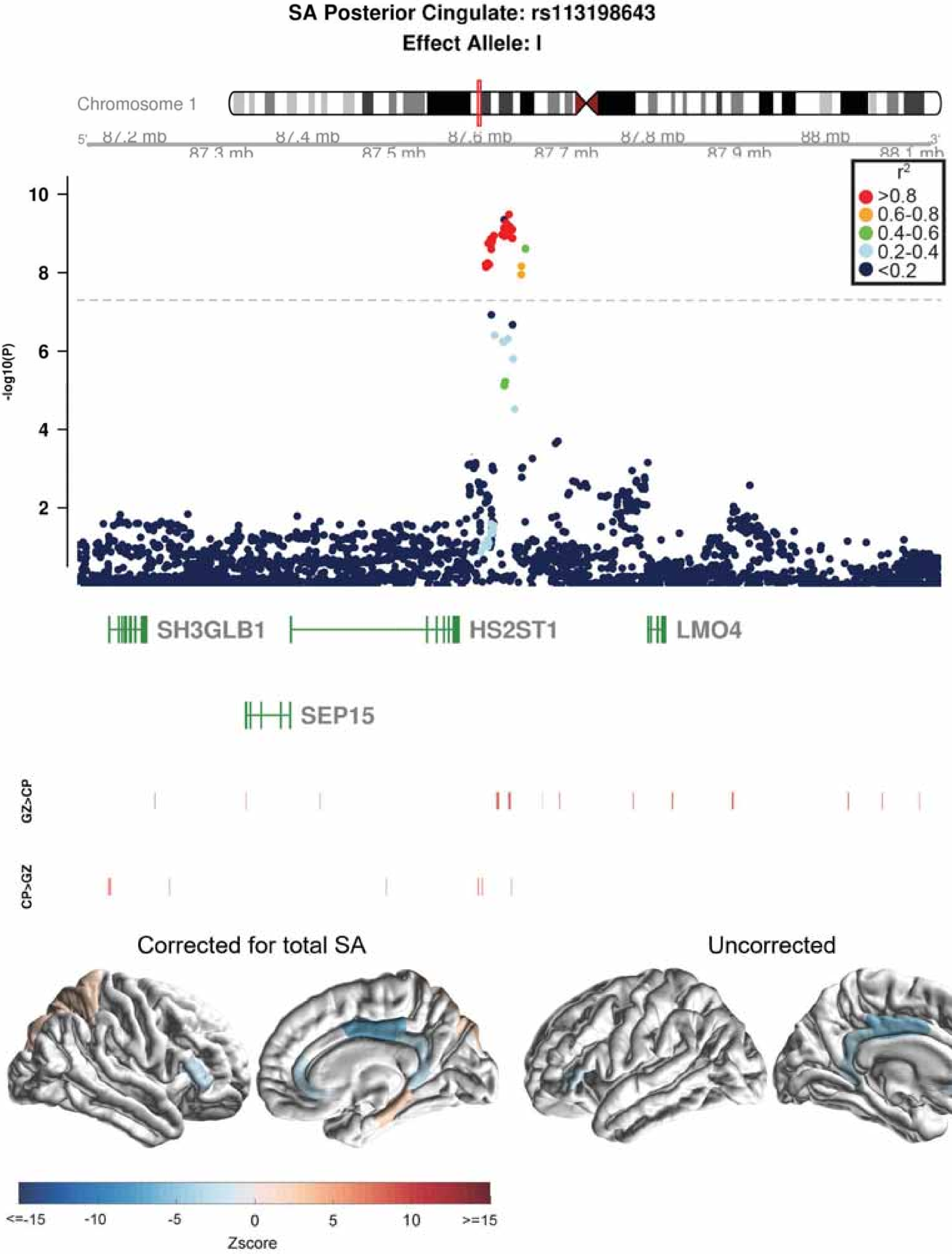

SA Posterior Cingulate: rs113198643
Effect Allele: I
Chromosome 1
5'
87.2 mb
87.3 mb
87.4 mb
87.5 mb
87.6 mb
87.7 mb
87.8 mb
87.9 mb
88 mb
88.1 mb
3'
r2
>0.8
0.6-0.8
0.4-0.6
0.2-0.4
<0.2
-log10(P)
10
8
6
4
2
SH3GLB1
HS2ST1
LMO4
SEP15
GZ>CP
CP>GZ
Corrected for total SA
Uncorrected
<=-15
-10
-5
0
5
10
>=15
Zscore

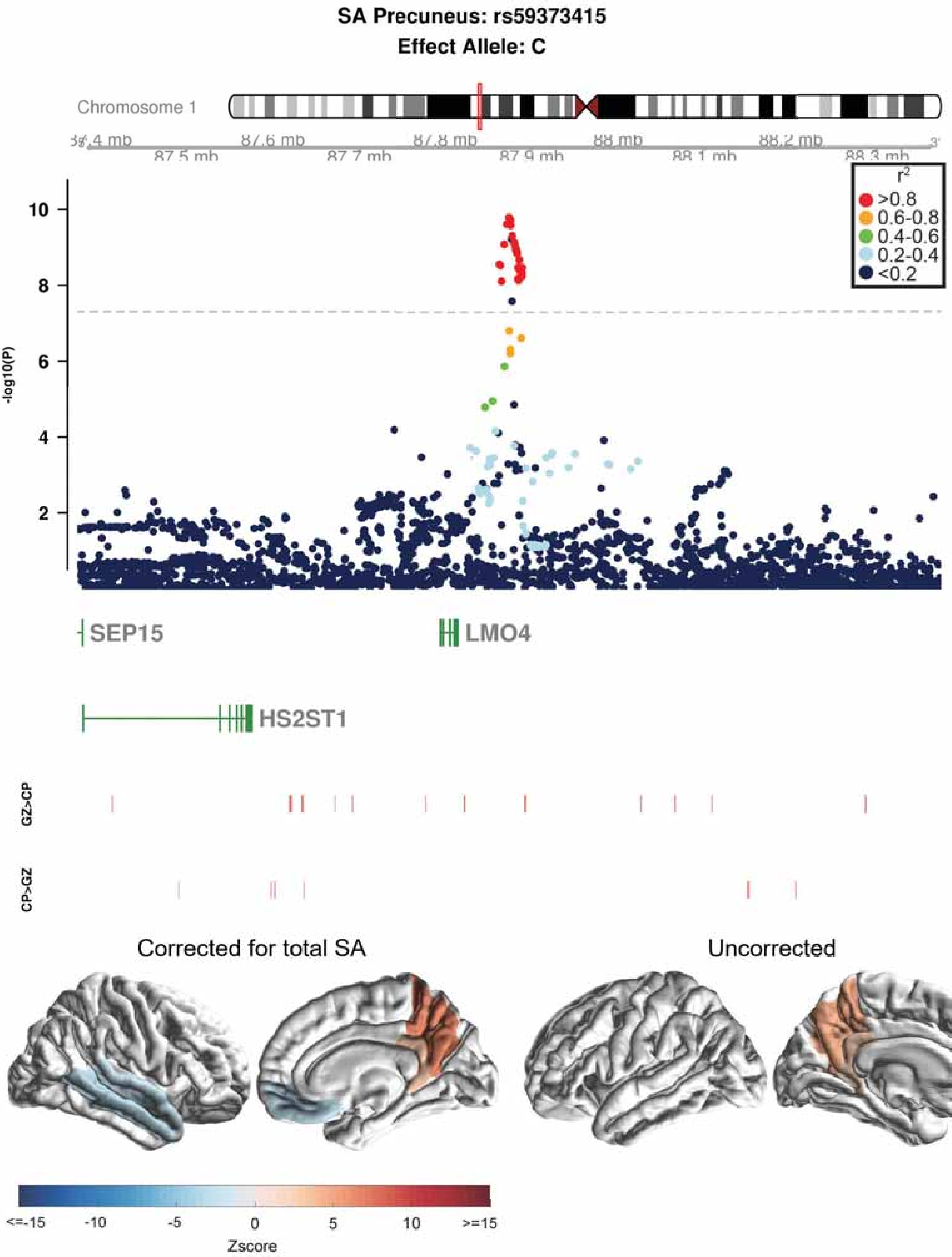

SA Precuneus: rs59373415
Effect Allele: C
Chromosome 1
87.5 mb
87.6 mb
87.7 mb
87.8 mb
87.9 mb
88 mb
88.1 mb
88.2 mb
88.3 mb
r2
>0.8
0.6-0.8
0.4-0.6
0.2-0.4
<0.2
-log10(P)
10
8
6
4
2
SEP15
LMO4
HS2ST1
GZ>CP
CP>GZ
Corrected for total SA
Uncorrected
<=-15
-10
-5
0
5
10
>=15
Zscore

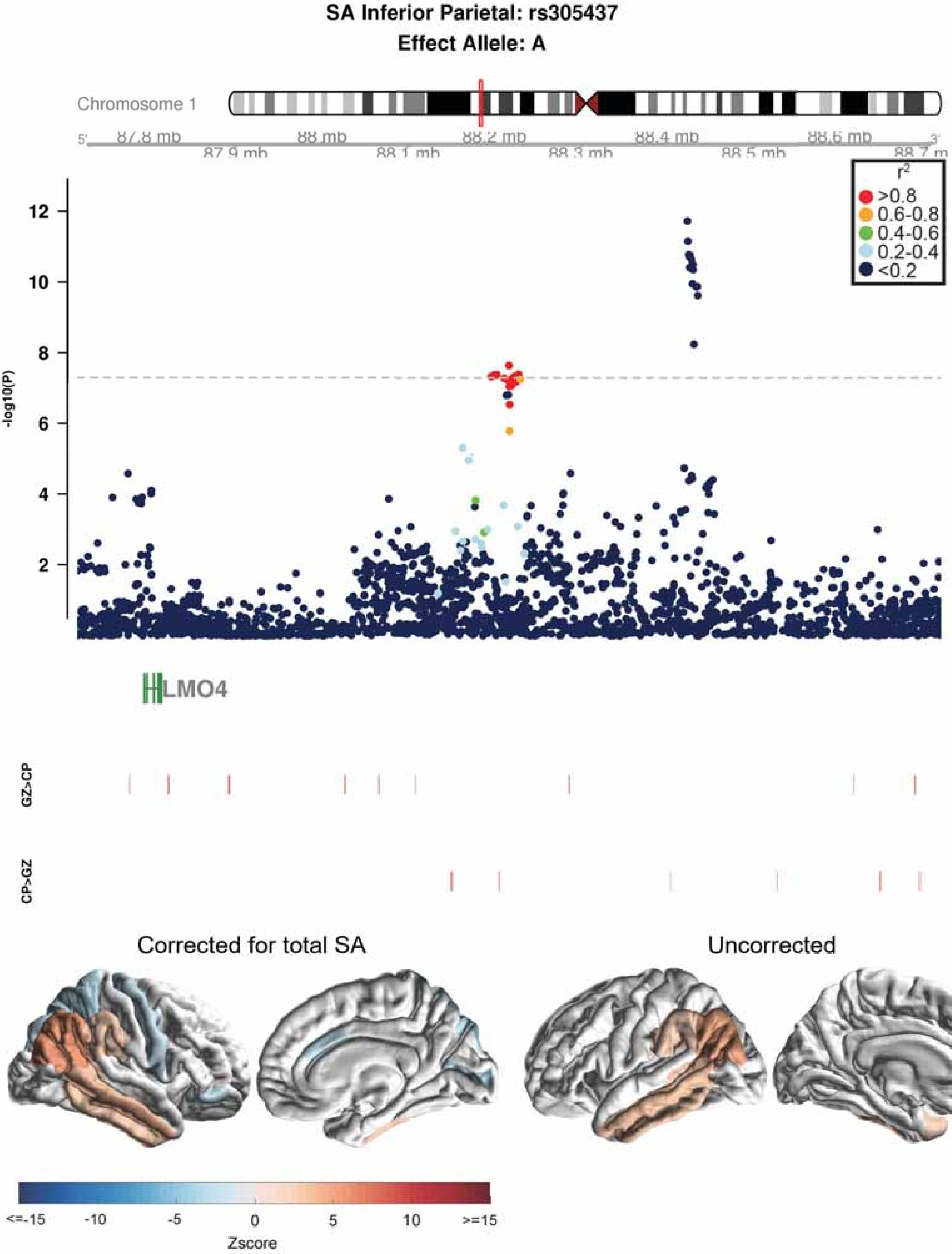
SA Inferior Parietal: rs305437
Effect Allele: A
Chromosome 1
87.8 mb
87.9 mb
88 mb
88.1 mb
88.2 mb
88.3 mb
88.4 mb
88.5 mb
88.6 mb
88.7 m
r2
>0.8
0.6-0.8
0.4-0.6
0.2-0.4
<0.2
-log10(P)
LMO4
GZ>CP
CP>GZ
Corrected for total SA
Uncorrected
<=-15
-10
-5
0
5
10
>=15
Zscore

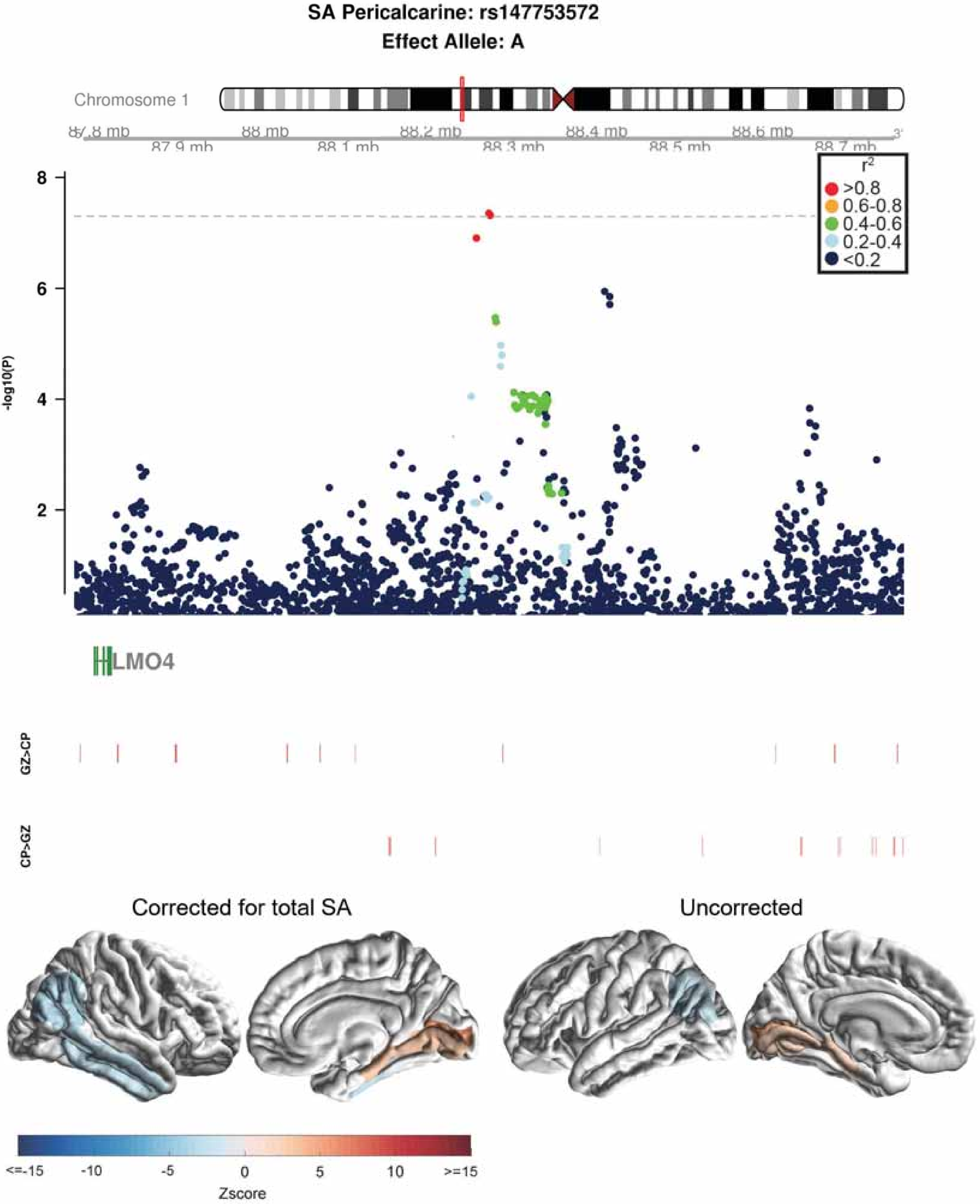
SA Pericalcarine: rs147753572
Effect Allele: A
Chromosome 1
87.8 mb
88 mb
88.2 mb
88.4 mb
88.6 mb
87.9 mb
88.1 mb
88.3 mb
88.5 mb
88.7 mb
r2
>0.8
0.6-0.8
0.4-0.6
0.2-0.4
<0.2
-log10(P)
8
6
4
2
LMO4
GZ>CP
CP>GZ
Corrected for total SA
Uncorrected
<=-15
-10
-5
0
5
10
>=15
Zscore

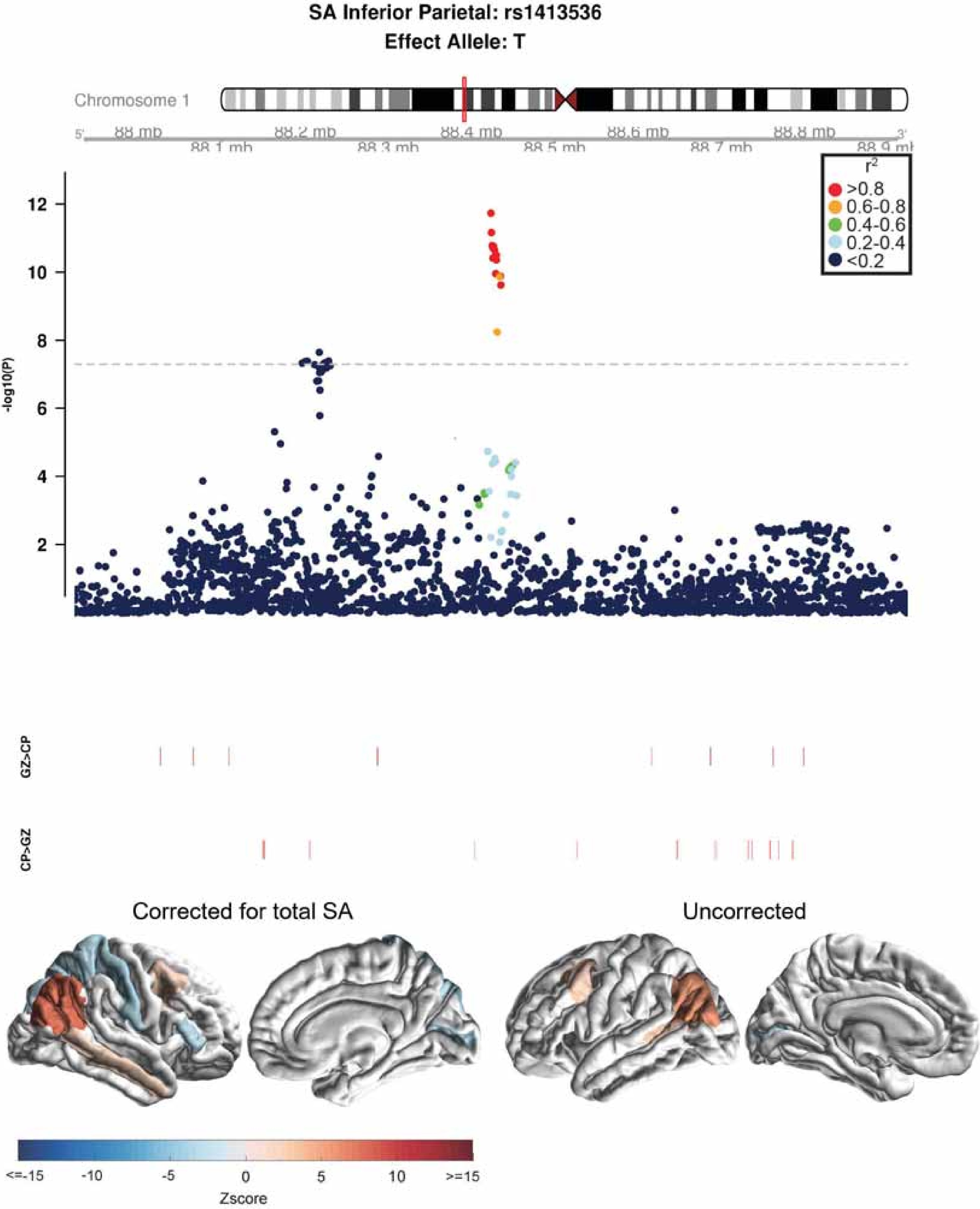
SA Inferior Parietal: rs1413536
Effect Allele: T
Chromosome 1
88 mb
88.1 mb
88.2 mb
88.3 mb
88.4 mb
88.5 mb
88.6 mb
88.7 mb
88.8 mb
r2
>0.8
0.6-0.8
0.4-0.6
0.2-0.4
<0.2
-log10(P)
12
10
8
6
4
2
GZ>CP
CP>GZ
Corrected for total SA
Uncorrected
<=-15
-10
-5
0
5
10
>=15
Zscore

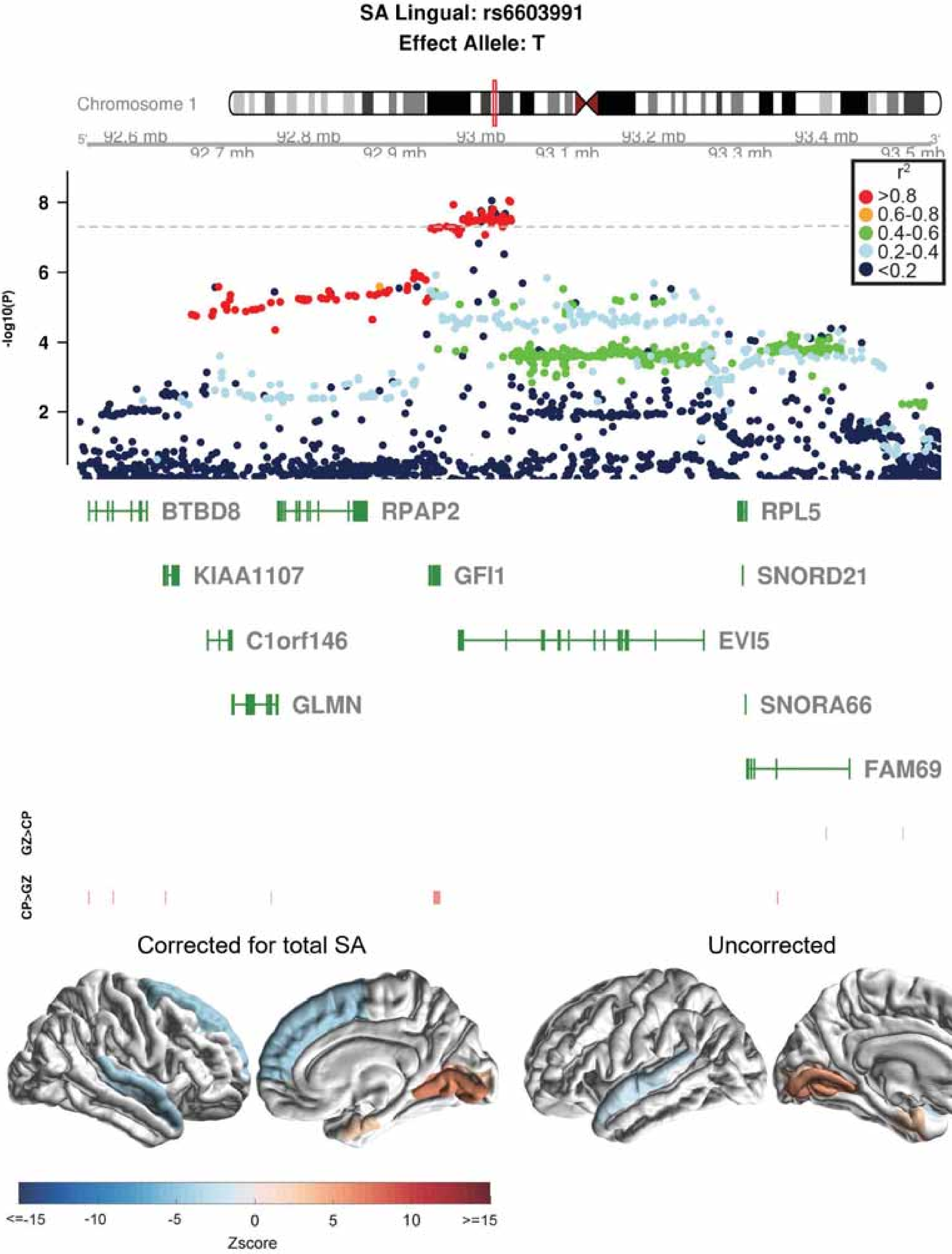

SA Lingual: rs6603991
Effect Allele: T
Chromosome 1
92.6 mb
92.7 mb
92.8 mb
92.9 mb
93 mb
93.1 mb
93.2 mb
93.3 mb
93.4 mb
93.5 mb
-log10(P)
r²
>0.8
0.6-0.8
0.4-0.6
0.2-0.4
<0.2
BTBD8
RPAP2
RPL5
KIAA1107
GFI1
SNORD21
C1orf146
EVI5
GLMN
SNORA66
FAM69
GZ>CP
CP>GZ
Corrected for total SA
Uncorrected
<=-15
-10
-5
0
5
10
>=15
Zscore

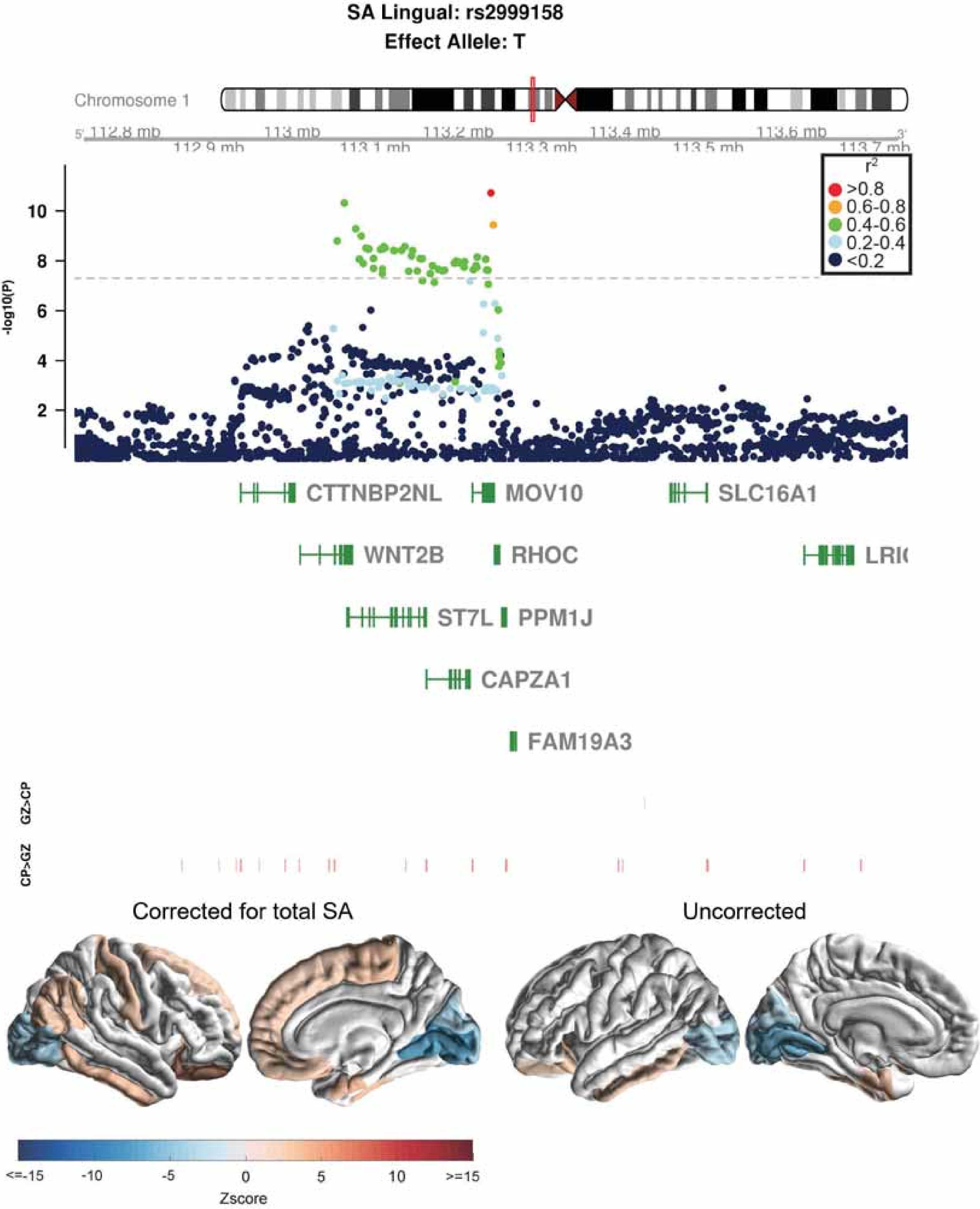
SA Lingual: rs2999158
Effect Allele: T
Chromosome 1
112.8 mb
112.9 mb
113 mb
113.1 mb
113.2 mb
113.3 mb
113.4 mb
113.5 mb
113.6 mb
113.7 mb
r²
>0.8
0.6-0.8
0.4-0.6
0.2-0.4
<0.2
-log10(P)
CTTNBP2NL
MOV10
SLC16A1
WNT2B
RHOC
ST7L
PPM1J
CAPZA1
FAM19A3
GZ>CP
CP>GZ
Corrected for total SA
Uncorrected
<=-15
-10
-5
0
5
10
>=15
Zscore

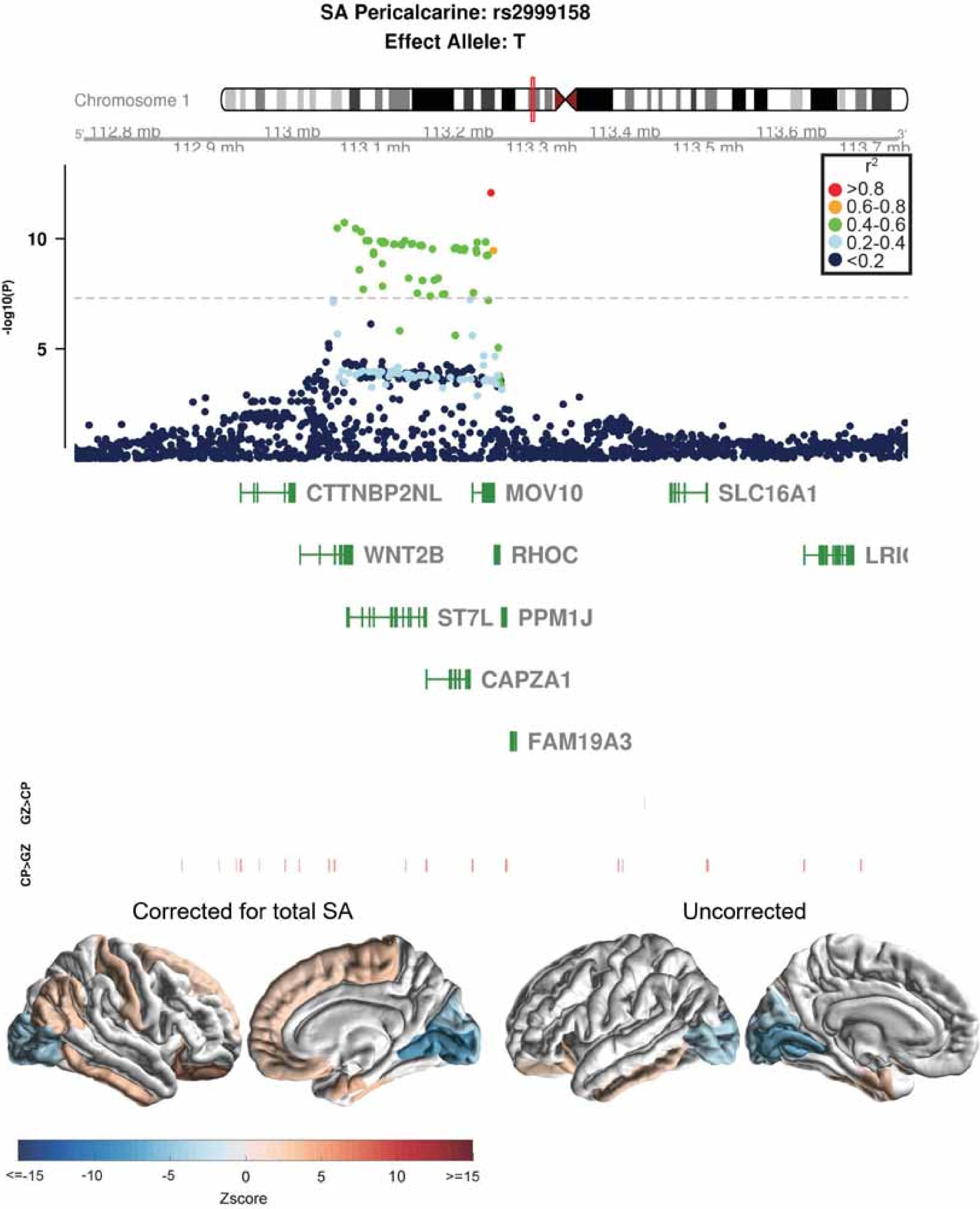
SA Pericalcarine: rs2999158
Effect Allele: T
Chromosome 1
112.8 mb
112.9 mb
113 mb
113.1 mb
113.2 mb
113.3 mb
113.4 mb
113.5 mb
113.6 mb
113.7 mb
r2
>0.8
0.6-0.8
0.4-0.6
0.2-0.4
<0.2
-log10(P)
10
5
CTTNBP2NL
MOV10
SLC16A1
WNT2B
RHOC
ST7L
PPM1J
CAPZA1
FAM19A3
GZ>CP
CP>GZ
Corrected for total SA
Uncorrected
<=-15
-10
-5
0
5
10
>=15
Zscore

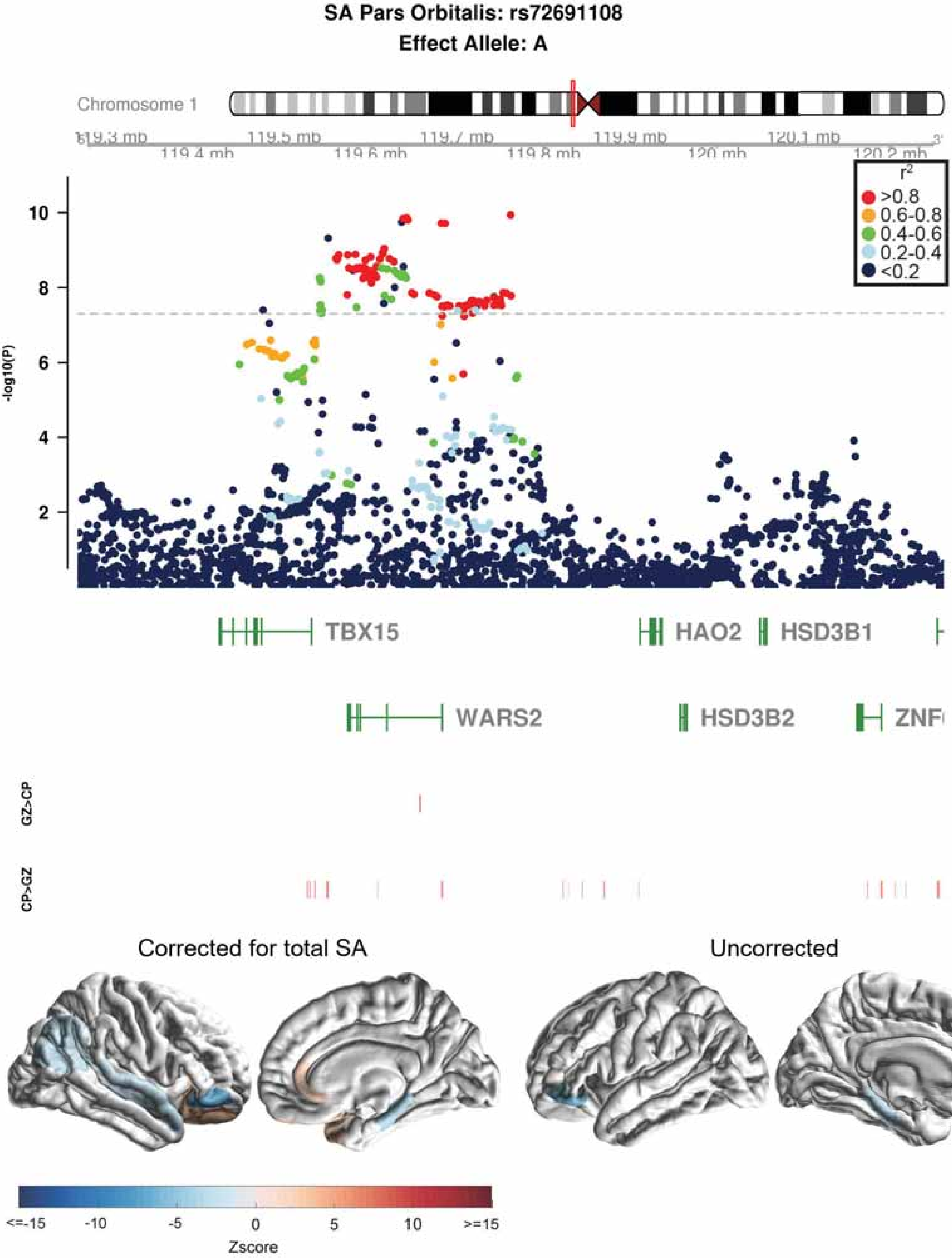

SA Pars Orbitalis: rs72691108
Effect Allele: A
Chromosome 1
119.3 mb
119.4 mb
119.5 mb
119.6 mb
119.7 mb
119.8 mb
119.9 mb
120 mb
120.1 mb
120.2 mb
r2
>0.8
0.6-0.8
0.4-0.6
0.2-0.4
<0.2
-log10(P)
10
8
6
4
2
TBX15
HAO2
HSD3B1
WARS2
HSD3B2
GZ>CP
CP>GZ
Corrected for total SA
Uncorrected
<=-15
-10
-5
0
5
10
>=15
Zscore

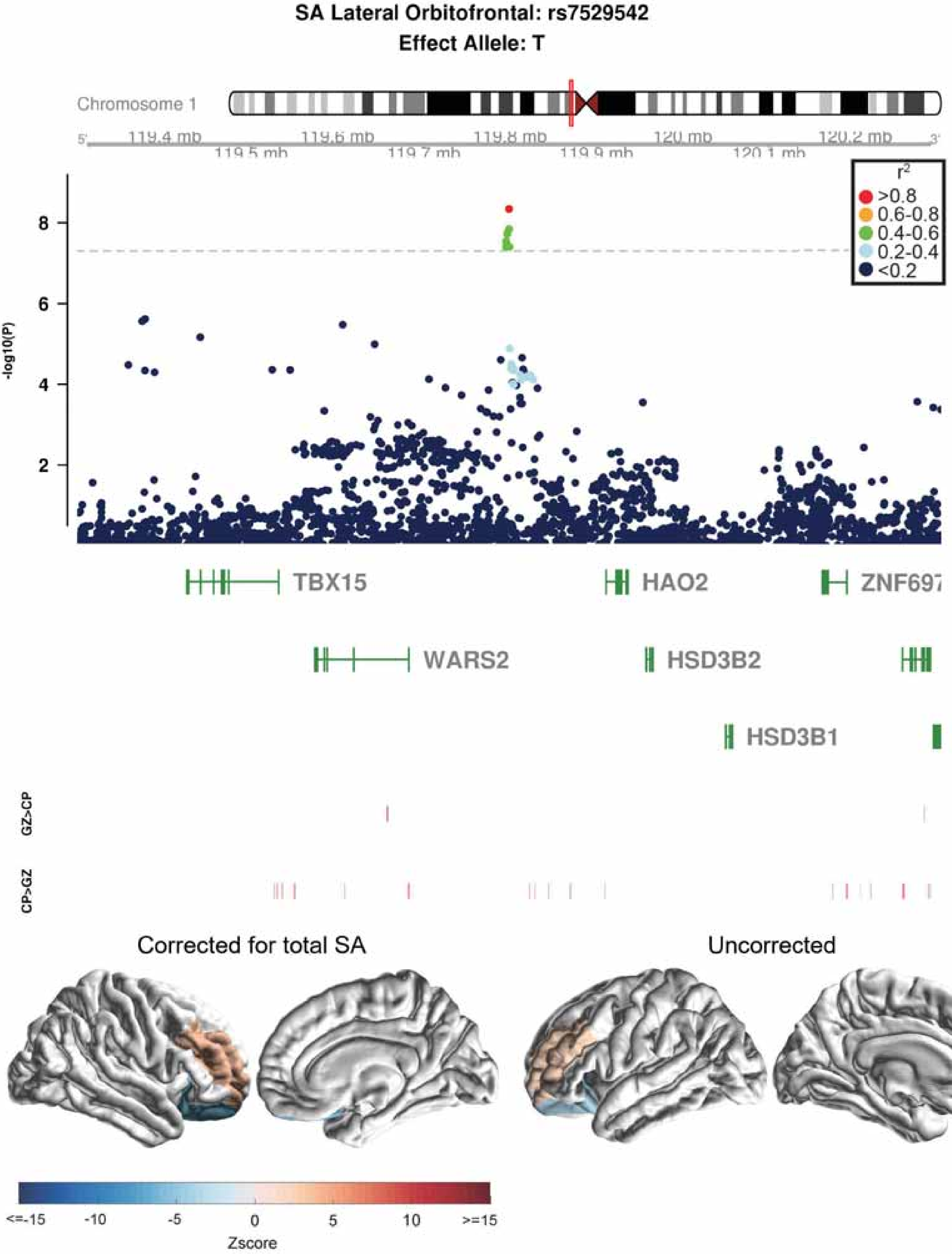
SA Lateral Orbitofrontal: rs7529542
Effect Allele: T
Chromosome 1
119.4 mb
119.5 mb
119.6 mb
119.7 mb
119.8 mb
119.9 mb
120 mb
120.1 mb
120.2 mb
r2
>0.8
0.6-0.8
0.4-0.6
0.2-0.4
<0.2
-log10(P)
TBX15
HAO2
ZNF697
WARS2
HSD3B2
HSD3B1
GZ>CP
CP>GZ
Corrected for total SA
Uncorrected
<=-15
-10
-5
0
5
10
>=15
Zscore

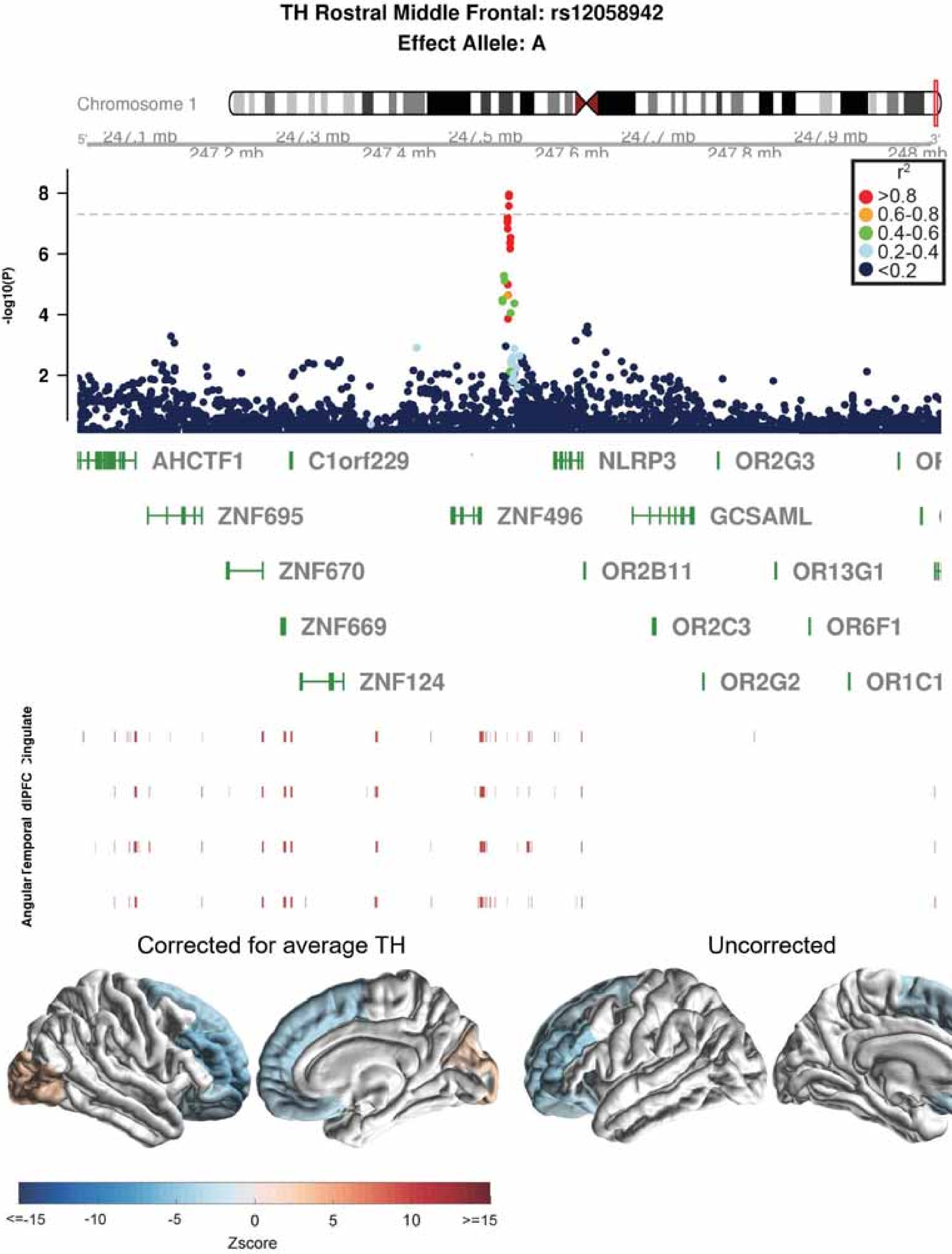
TH Rostral Middle Frontal: rs12058942
Effect Allele: A
Chromosome 1
247.1 mb
247.2 mb
247.3 mb
247.4 mb
247.5 mb
247.6 mb
247.7 mb
247.8 mb
247.9 mb
248 mb
-log10(P)
r2
>0.8
0.6-0.8
0.4-0.6
0.2-0.4
<0.2
AHCTF1
C1orf229
NLRP3
OR2G3
ZNF695
ZNF496
GCSAML
ZNF670
OR2B11
OR13G1
ZNF669
OR2C3
OR6F1
ZNF124
OR2G2
OR1C1
Angular
Temporal
dlPFC
Cingulate
Corrected for average TH
Uncorrected
<=-15
-10
-5
0
5
10
>=15
Zscore

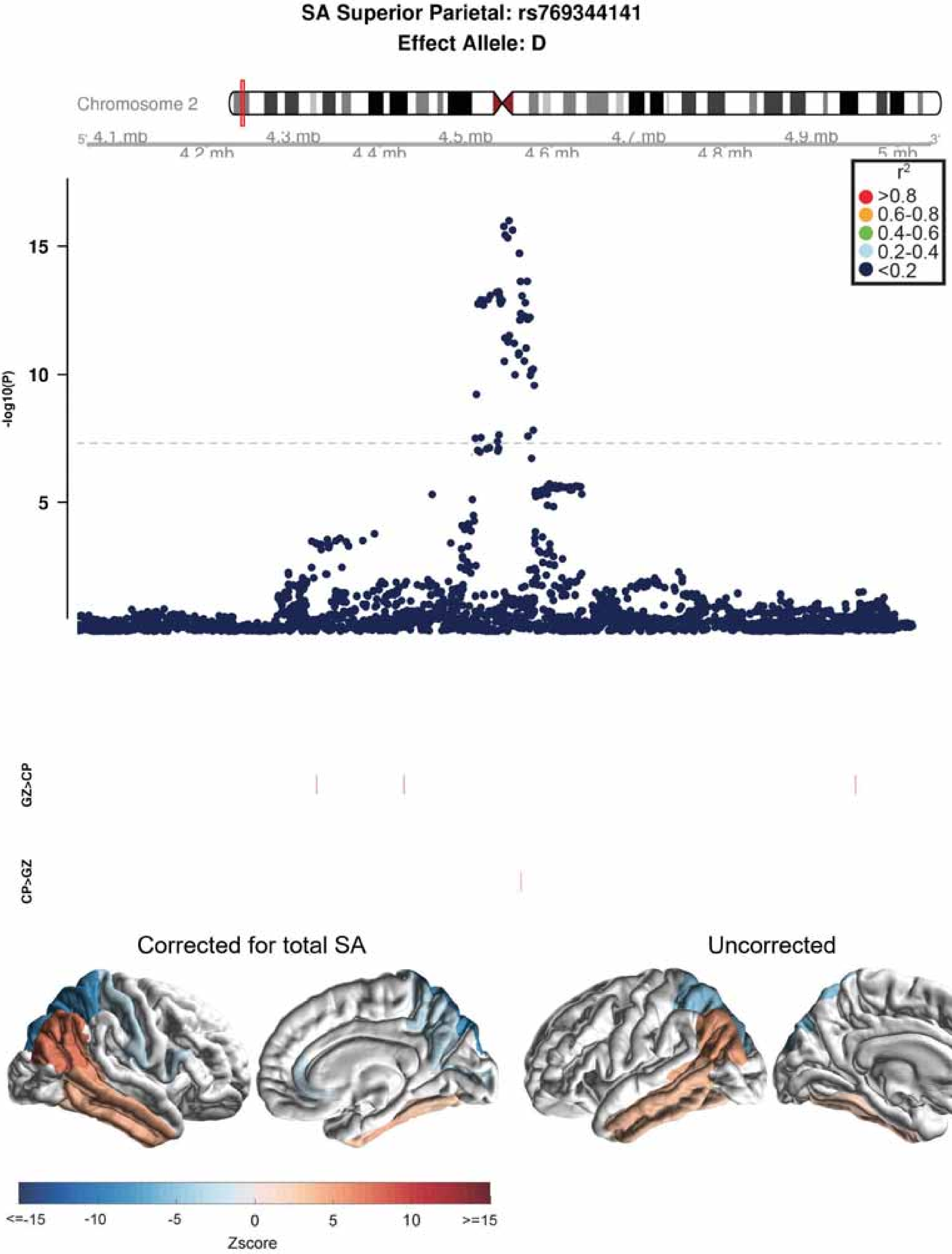

SA Superior Parietal: rs769344141
Effect Allele: D
Chromosome 2
4.1 mb
4.2 mb
4.3 mb
4.4 mb
4.5 mb
4.6 mb
4.7 mb
4.8 mb
4.9 mb
5 mb
r2
>0.8
0.6-0.8
0.4-0.6
0.2-0.4
<0.2
-log10(P)
15
10
5
GZ>CP
CP>GZ
Corrected for total SA
Uncorrected
<=-15
-10
-5
0
5
10
>=15
Zscore

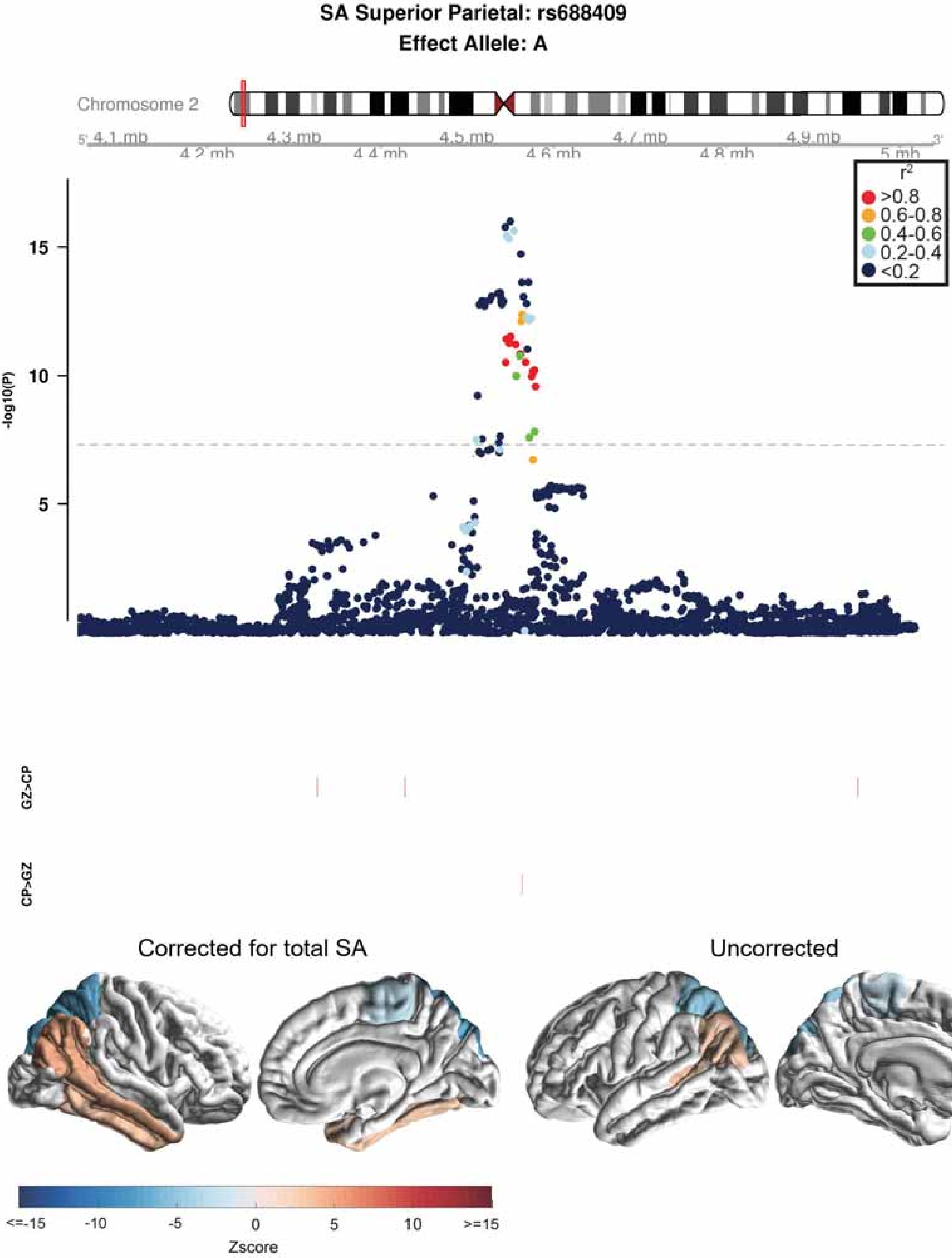

SA Superior Parietal: rs688409
Effect Allele: A
Chromosome 2
4.1 mb
4.2 mb
4.3 mb
4.4 mb
4.5 mb
4.6 mb
4.7 mb
4.8 mb
4.9 mb
5 mb
r2
>0.8
0.6-0.8
0.4-0.6
0.2-0.4
<0.2
-log10(P)
15
10
5
GZ>CP
CP>GZ
Corrected for total SA
Uncorrected
<=-15
-10
-5
0
5
10
>=15
Zscore

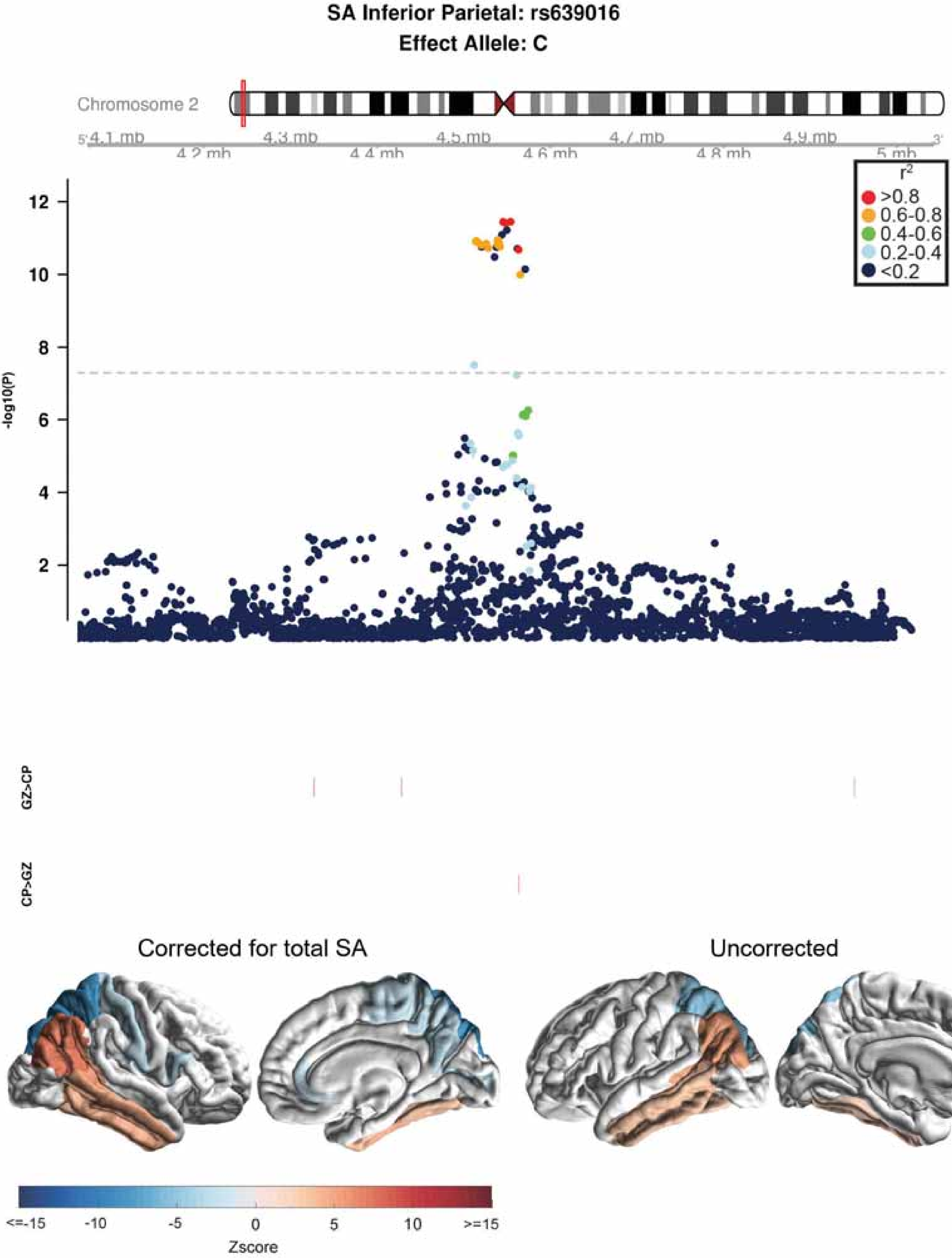

SA Inferior Parietal: rs639016
Effect Allele: C
Chromosome 2
4.1 mb
4.2 mb
4.3 mb
4.4 mb
4.5 mb
4.6 mb
4.7 mb
4.8 mb
4.9 mb
5 mb
r2
>0.8
0.6-0.8
0.4-0.6
0.2-0.4
<0.2
-log10(P)
GZ>CP
CP>GZ
Corrected for total SA
Uncorrected
<=-15
-10
-5
0
5
10
>=15
Zscore

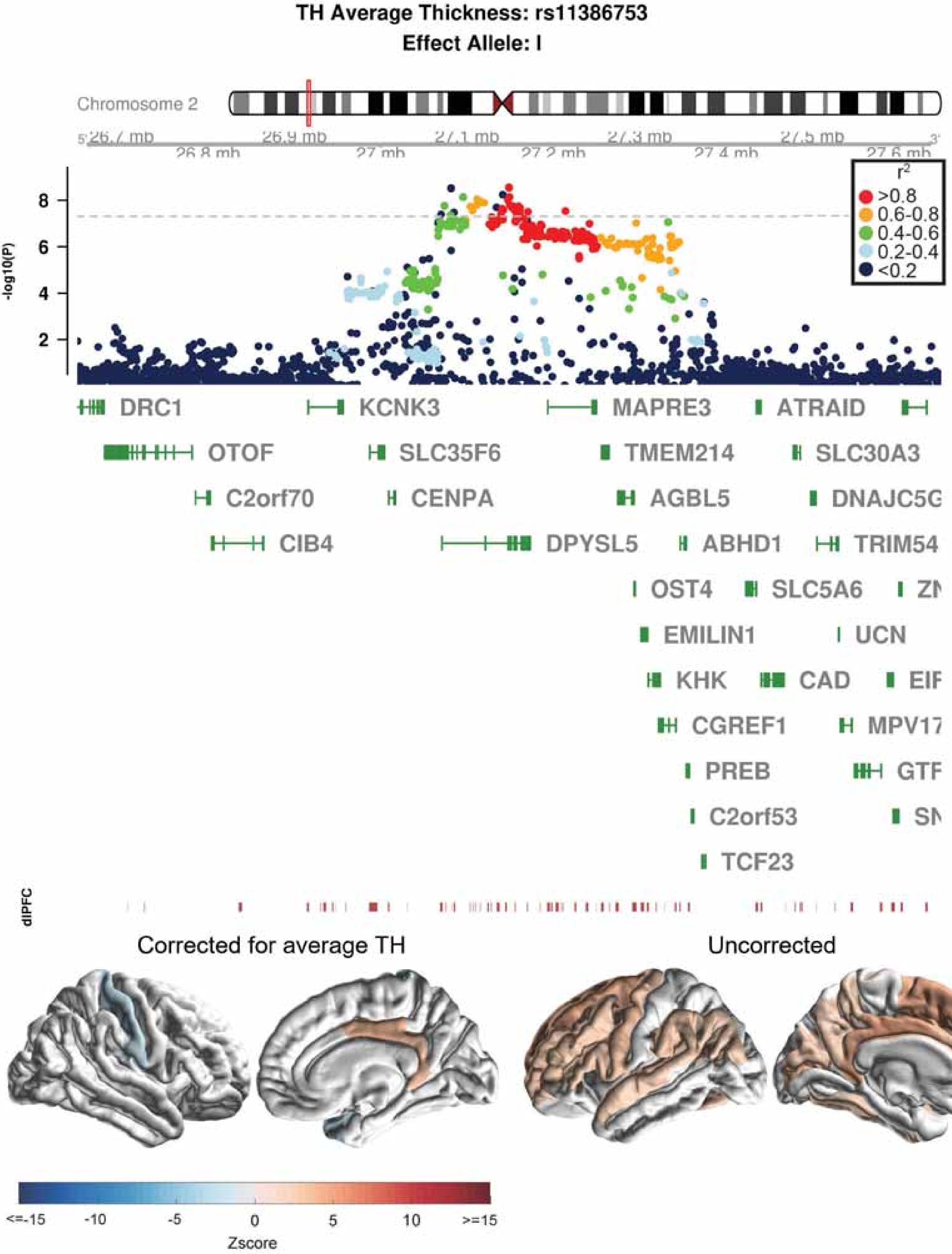
TH Average Thickness: rs11386753
Effect Allele: I
Chromosome 2
26.7 mb
26.8 mb
26.9 mb
27 mb
27.1 mb
27.2 mb
27.3 mb
27.4 mb
27.5 mb
27.6 mb
-log10(P)
r2
>0.8
0.6-0.8
0.4-0.6
0.2-0.4
<0.2
DRC1
KCNK3
MAPRE3
ATRAID
OTOF
SLC35F6
TMEM214
SLC30A3
C2orf70
CENPA
AGBL5
DNAJC5G
CIB4
DPYSL5
ABHD1
TRIM54
OST4
SLC5A6
EMILIN1
UCN
KHK
CAD
CGREF1
MPV17
PREB
C2orf53
TCF23
dlPFC
Corrected for average TH
Uncorrected
<=-15
-10
-5
0
5
10
>=15
Zscore

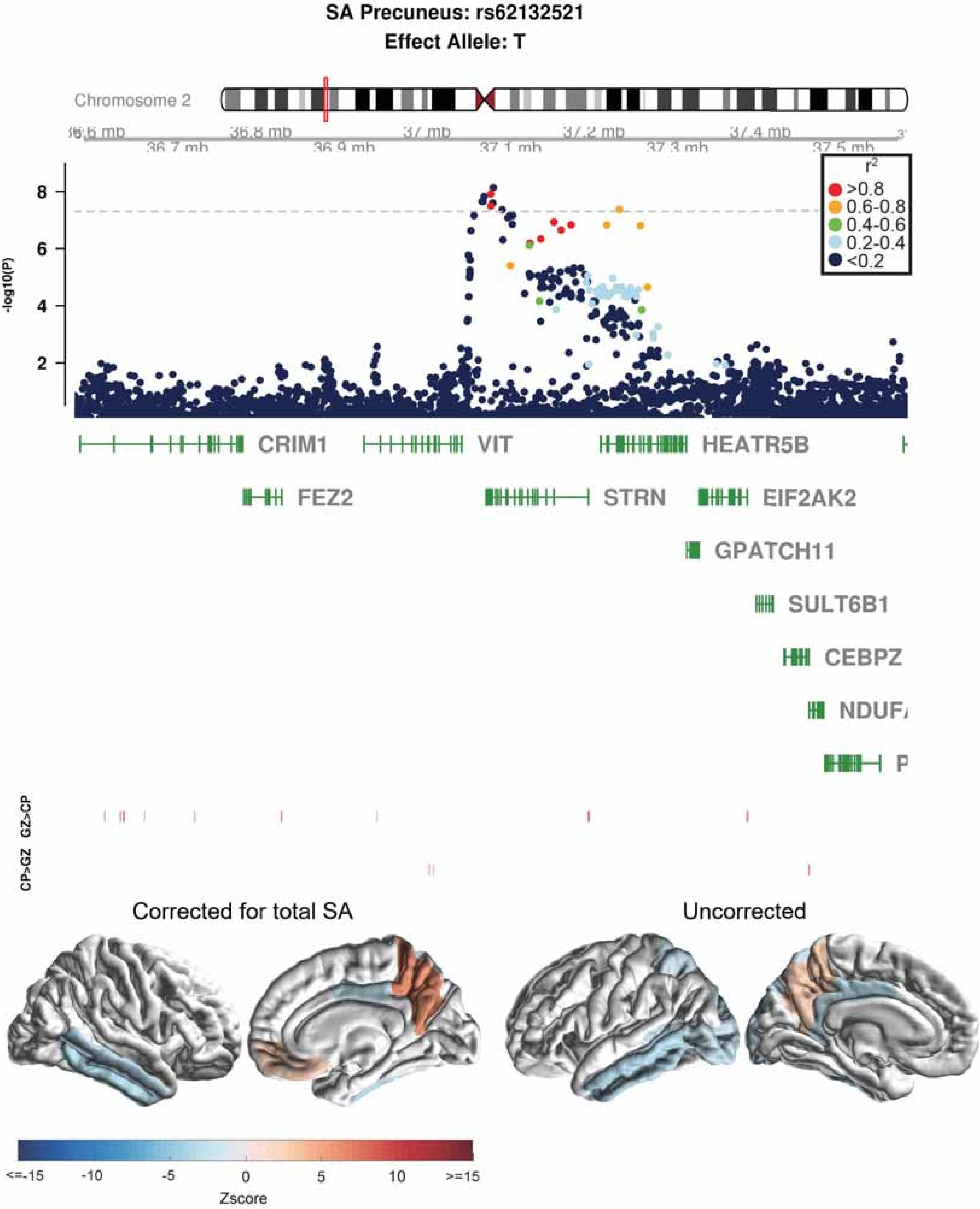
SA Precuneus: rs62132521
Effect Allele: T
Chromosome 2
36.8 mb
37 mb
37.2 mb
37.4 mb
36.7 mb
36.9 mb
37.1 mb
37.3 mb
37.5 mb
r²
>0.8
0.6-0.8
0.4-0.6
0.2-0.4
<0.2
-log10(P)
8
6
4
2
CRIM1
VIT
HEATR5B
FEZ2
STRN
EIF2AK2
GPATCH11
SULT6B1
CEBPZ
GZ>CP
CP>GZ
Corrected for total SA
Uncorrected
<=-15
-10
-5
0
5
10
>=15
Zscore

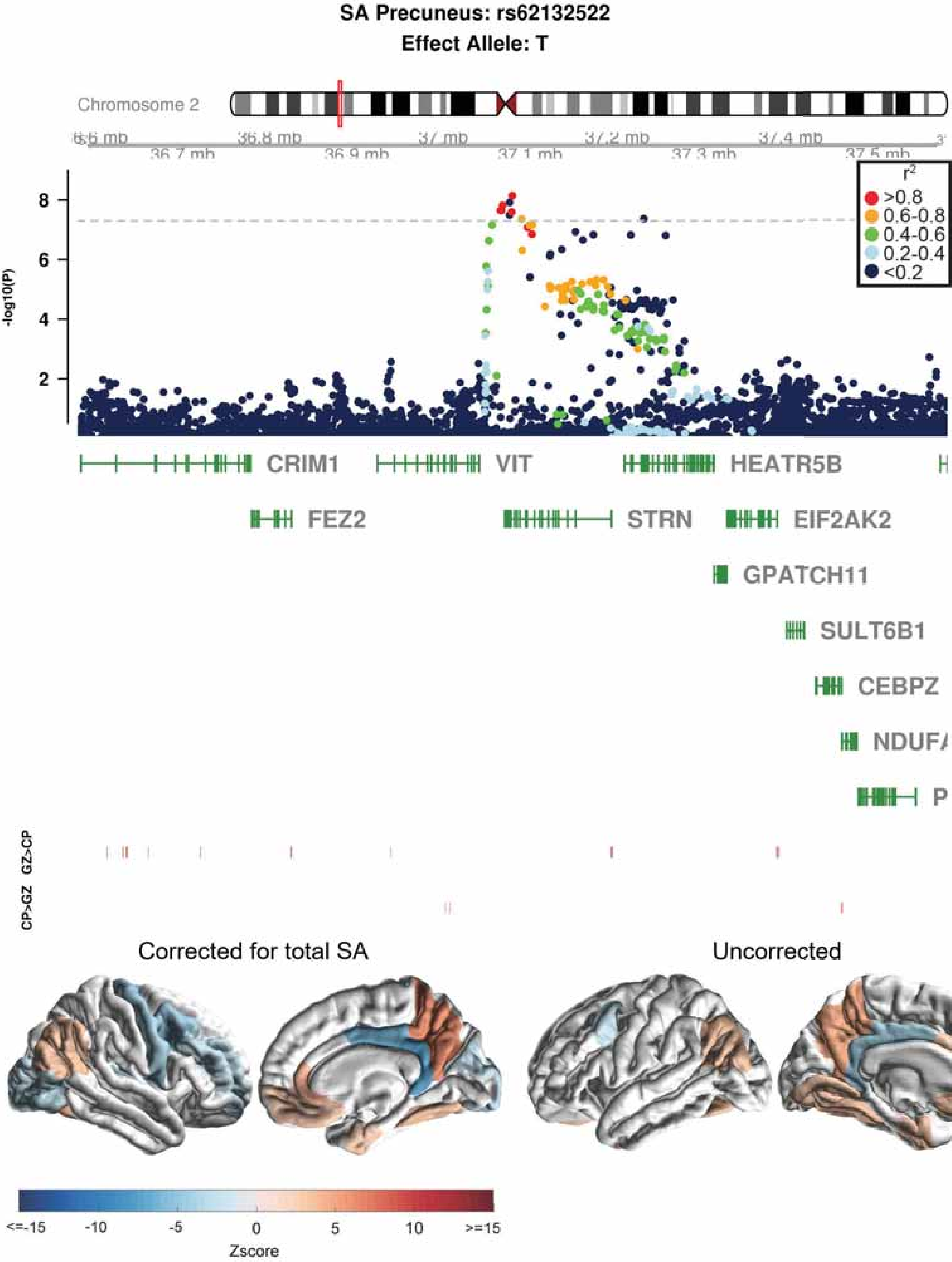

SA Precuneus: rs62132522
Effect Allele: T
Chromosome 2
36.8 mb
37 mb
37.2 mb
37.4 mb
36.7 mb
36.9 mb
37.1 mb
37.3 mb
37.5 mb
r2
>0.8
0.6-0.8
0.4-0.6
0.2-0.4
<0.2
-log10(P)
8
6
4
2
CRIM1
VIT
HEATR5B
FEZ2
STRN
EIF2AK2
GPATCH11
SULT6B1
CEBPZ
GZ>CP
CP>GZ
Corrected for total SA
Uncorrected
<=-15
-10
-5
0
5
10
>=15
Zscore

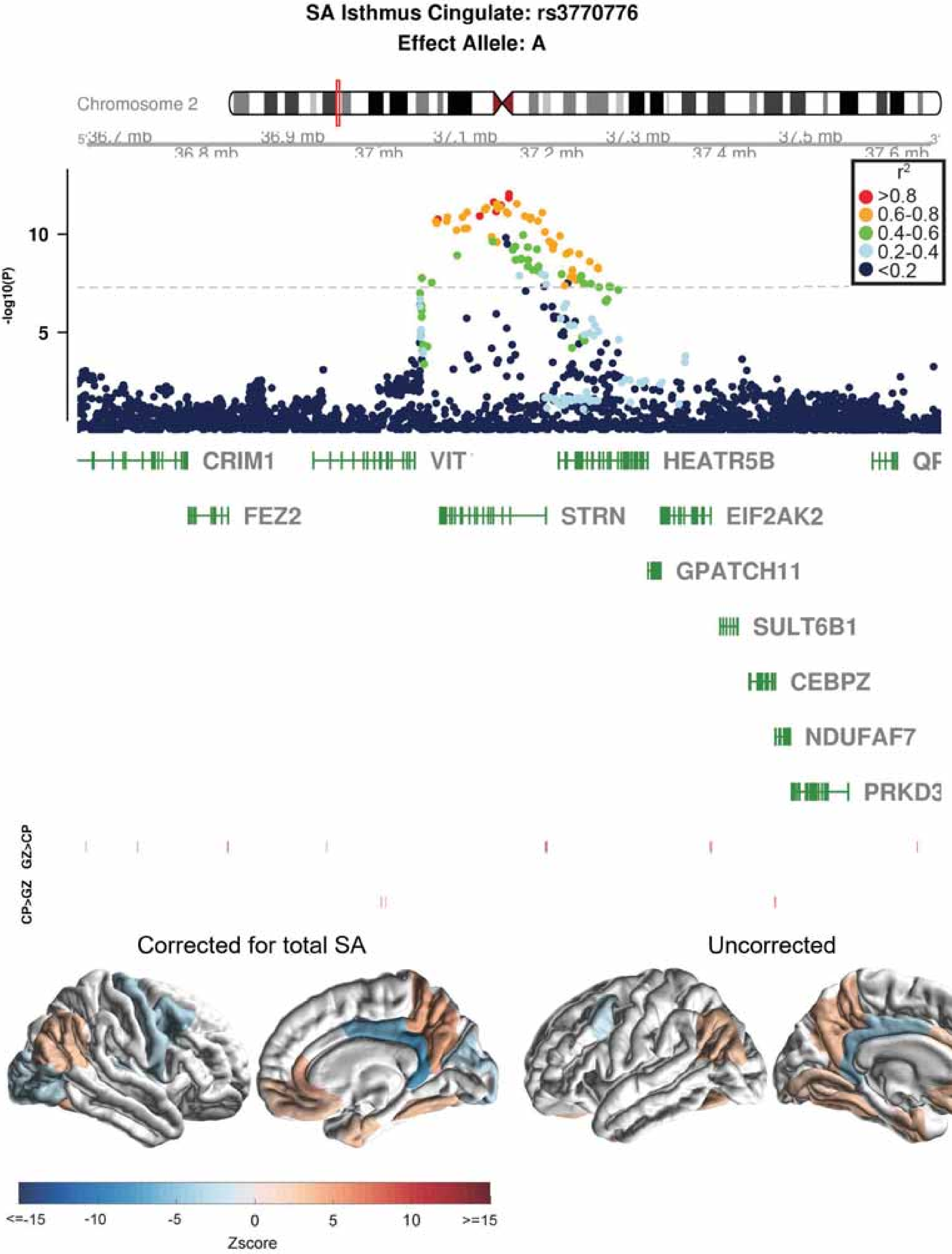

SA Isthmus Cingulate: rs3770776
Effect Allele: A
Chromosome 2
36.7 mb
36.8 mb
36.9 mb
37 mb
37.1 mb
37.2 mb
37.3 mb
37.4 mb
37.5 mb
37.6 mb
-log10(P)
10
5
$r^2$
>0.8
0.6-0.8
0.4-0.6
0.2-0.4
<0.2
CRIM1
VIT
HEATR5B
FEZ2
STRN
EIF2AK2
GPATCH11
SULT6B1
CEBPZ
NDUFAF7
PRKD3
GZ>CP
CP>GZ
Corrected for total SA
Uncorrected
<=-15
-10
-5
0
5
10
>=15
Zscore

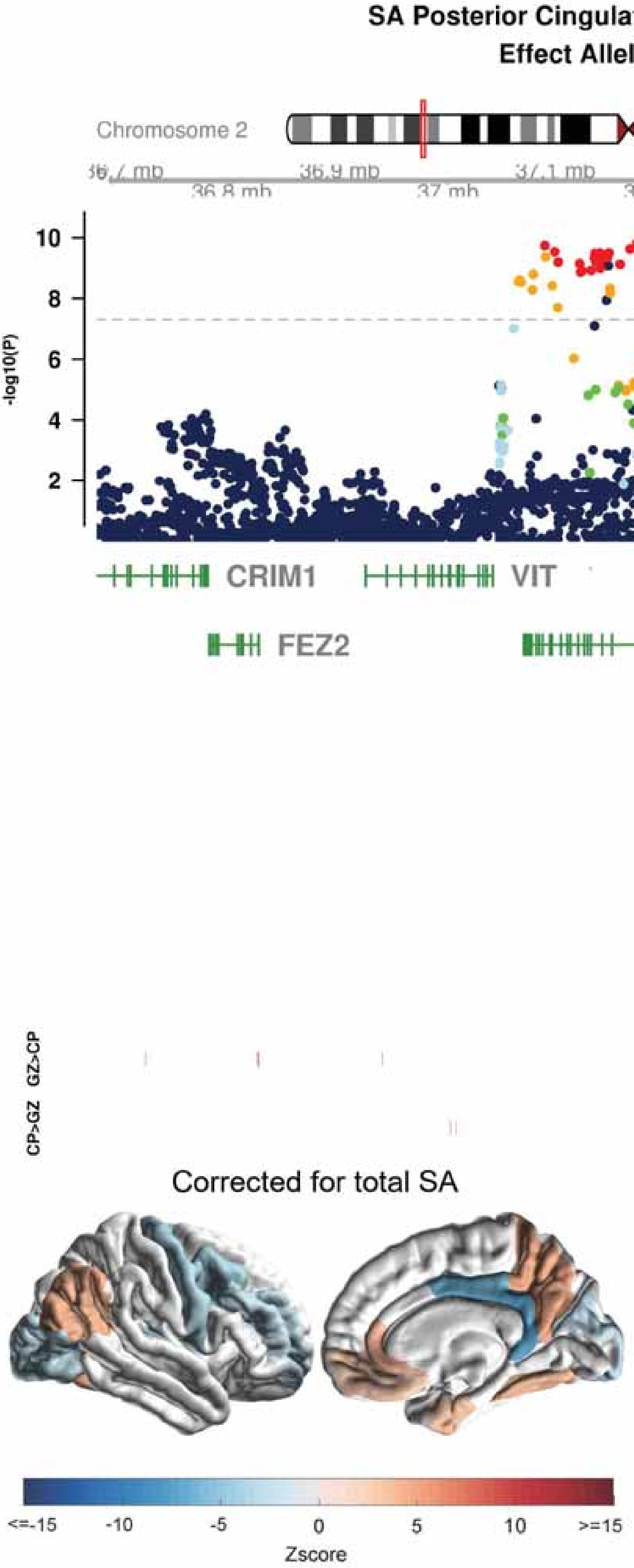

SA Posterior Cingulate: rs11695609
Effect Allele: T
Chromosome 2
36.7 mb
36.8 mb
36.9 mb
37 mb
37.1 mb
37.2 mb
37.3 mb
37.4 mb
37.5 mb
37.6 mb
-log10(P)
r2
>0.8
0.6-0.8
0.4-0.6
0.2-0.4
<0.2
CRIM1
VIT
HEATR5B
QPC
FEZ2
STRN
EIF2AK2
GPATCH11
SULT6B1
CEBPZ
NDUFAF7
PRKD3
GZ>CP
CP>GZ
Corrected for total SA
Uncorrected
<=-15
-10
-5
0
5
10
>=15
Zscore

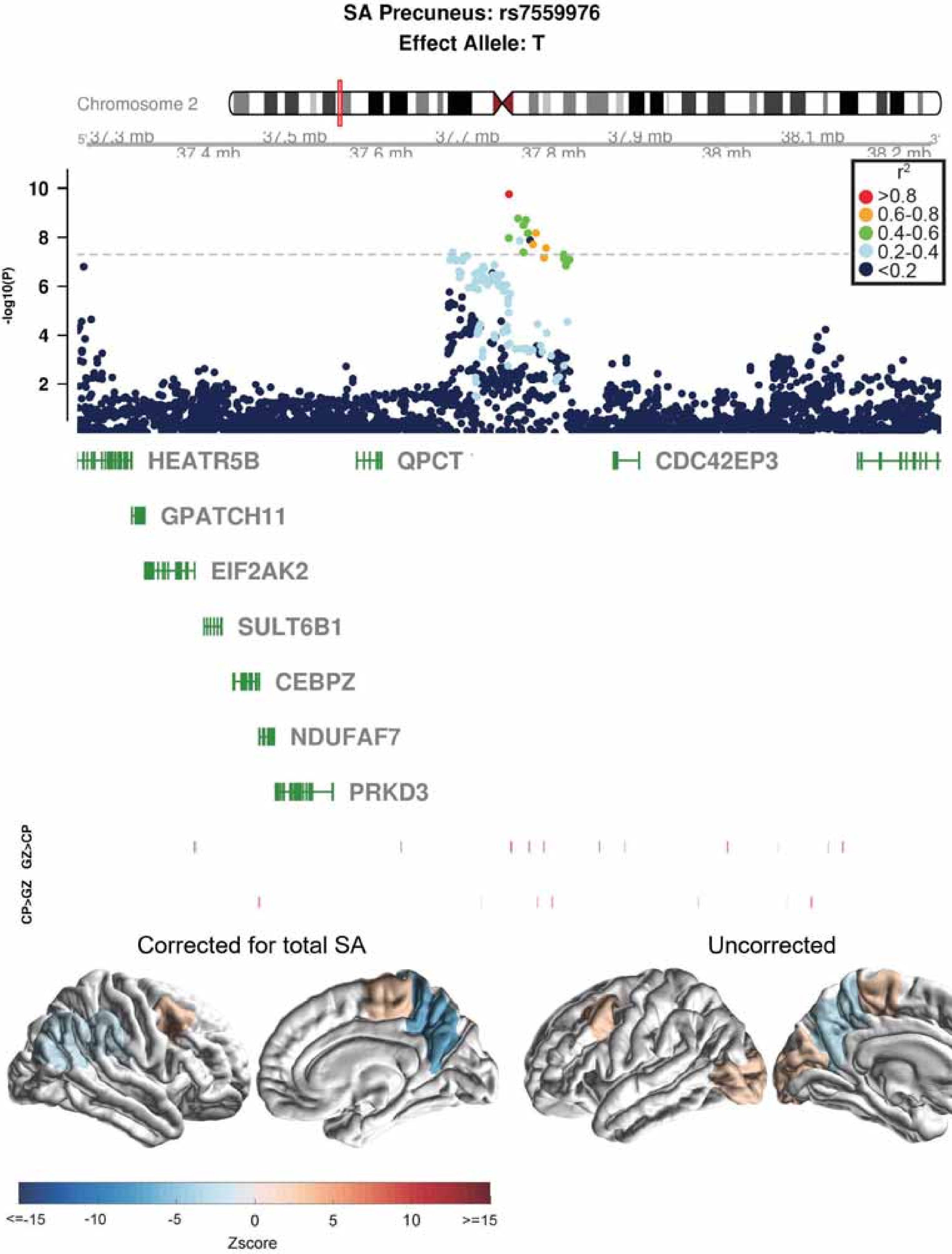

SA Precuneus: rs7559976
Effect Allele: T
Chromosome 2
37.3 mb
37.4 mb
37.5 mb
37.6 mb
37.7 mb
37.8 mb
37.9 mb
38 mb
38.1 mb
38.2 mb
-log10(P)
10
8
6
4
2
r2
>0.8
0.6-0.8
0.4-0.6
0.2-0.4
<0.2
HEATR5B
QPCT
CDC42EP3
GPATCH11
EIF2AK2
SULT6B1
CEBPZ
NDUFAF7
PRKD3
GZ>CP
CP>GZ
Corrected for total SA
Uncorrected
<=-15
-10
-5
0
5
10
>=15
Zscore

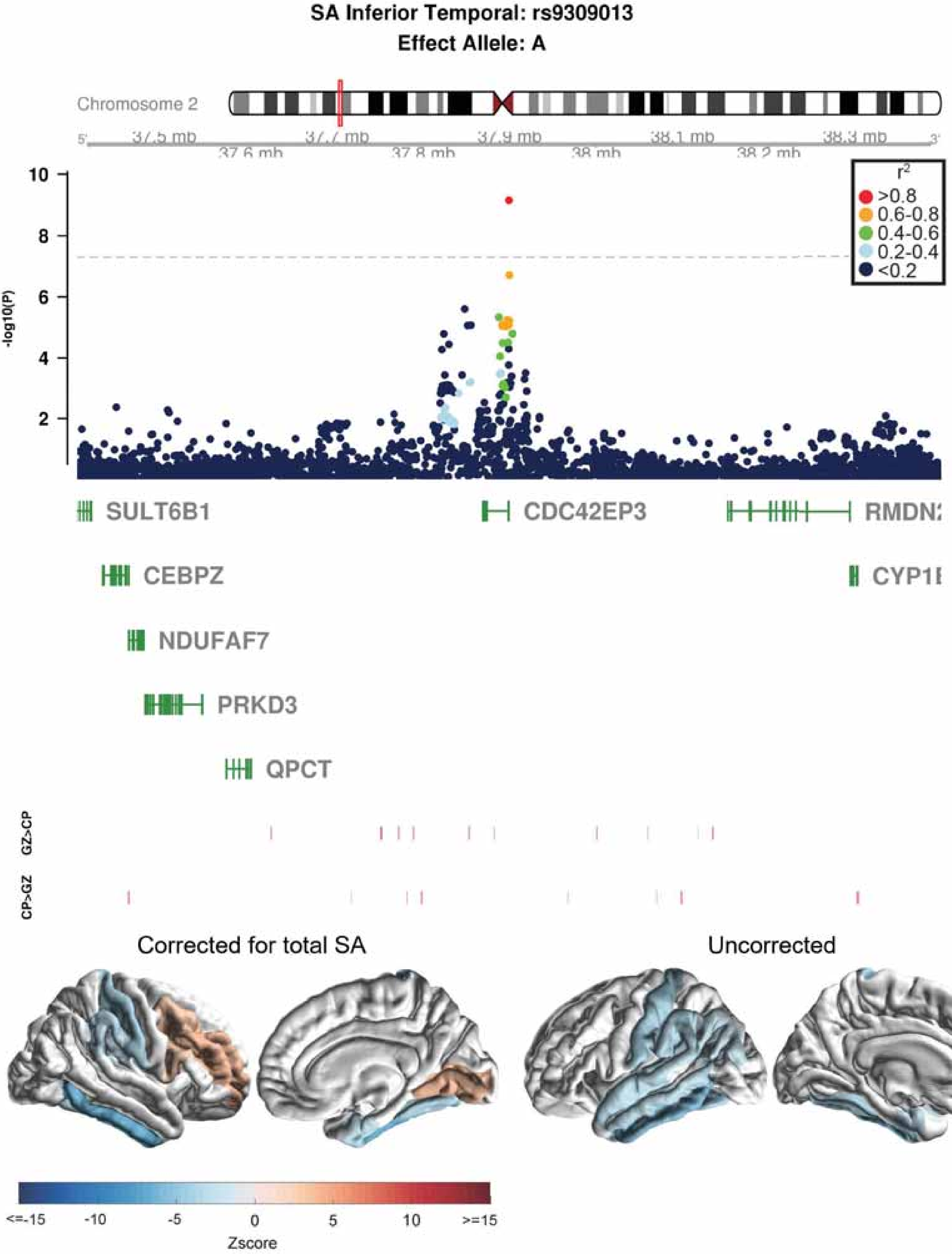

SA Inferior Temporal: rs9309013
Effect Allele: A
Chromosome 2
37.5 mb
37.6 mb
37.7 mb
37.8 mb
37.9 mb
38 mb
38.1 mb
38.2 mb
38.3 mb
-log10(P)
r2
>0.8
0.6-0.8
0.4-0.6
0.2-0.4
<0.2
SULT6B1
CDC42EP3
RMDN2
CEBPZ
CYP1B
NDUFAF7
PRKD3
QPCT
GZ>CP
CP>GZ
Corrected for total SA
Uncorrected
<=-15
-10
-5
0
5
10
>=15
Zscore

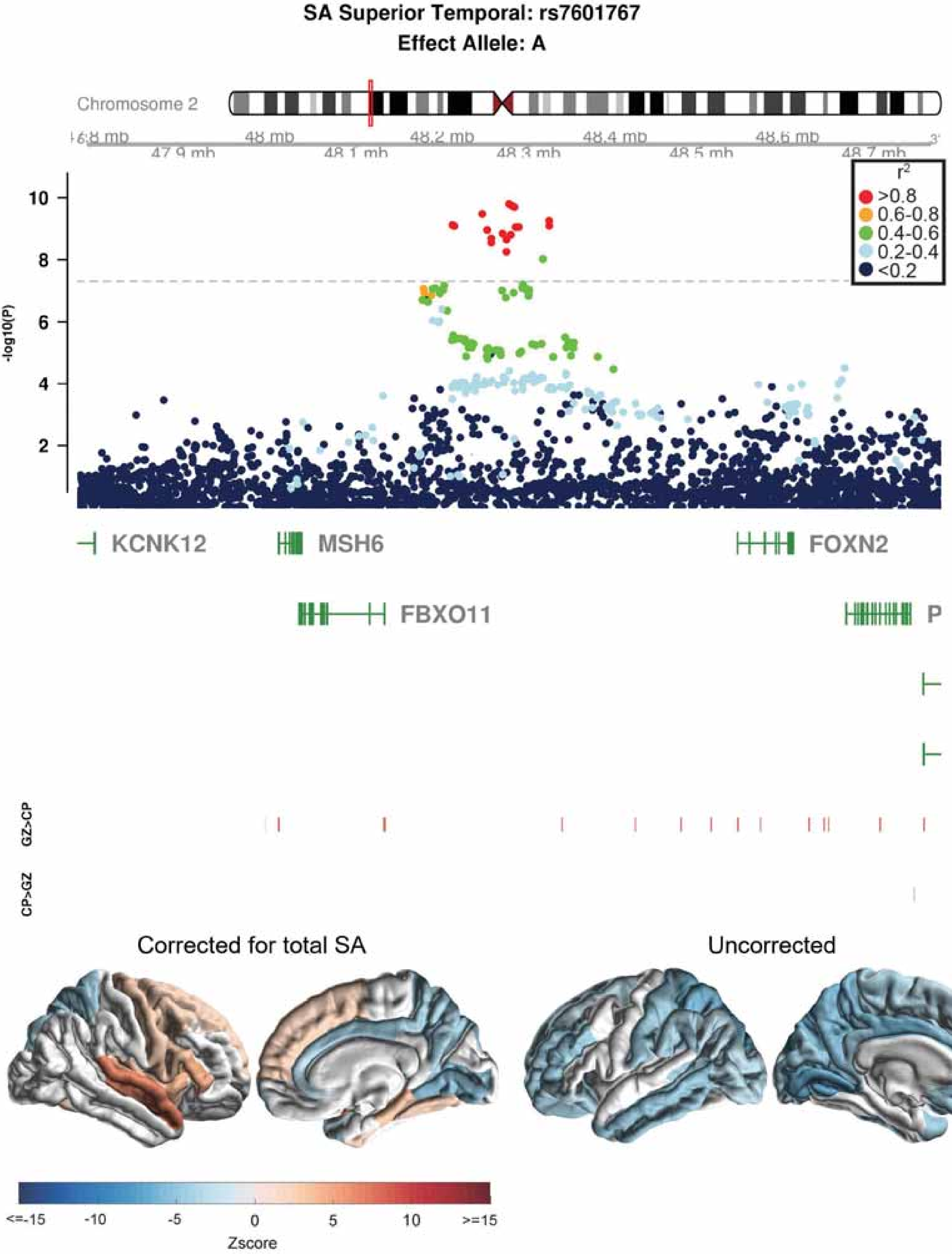

SA Superior Temporal: rs7601767
Effect Allele: A
Chromosome 2
48 mb
48.2 mb
48.4 mb
48.6 mb
47.9 mb
48.1 mb
48.3 mb
48.5 mb
48.7 mb
r2
>0.8
0.6-0.8
0.4-0.6
0.2-0.4
<0.2
-log10(P)
10
8
6
4
2
KCNK12
MSH6
FOXN2
FBXO11
P
GZ>CP
CP>GZ
Corrected for total SA
Uncorrected
<=-15
-10
-5
0
5
10
>=15
Zscore

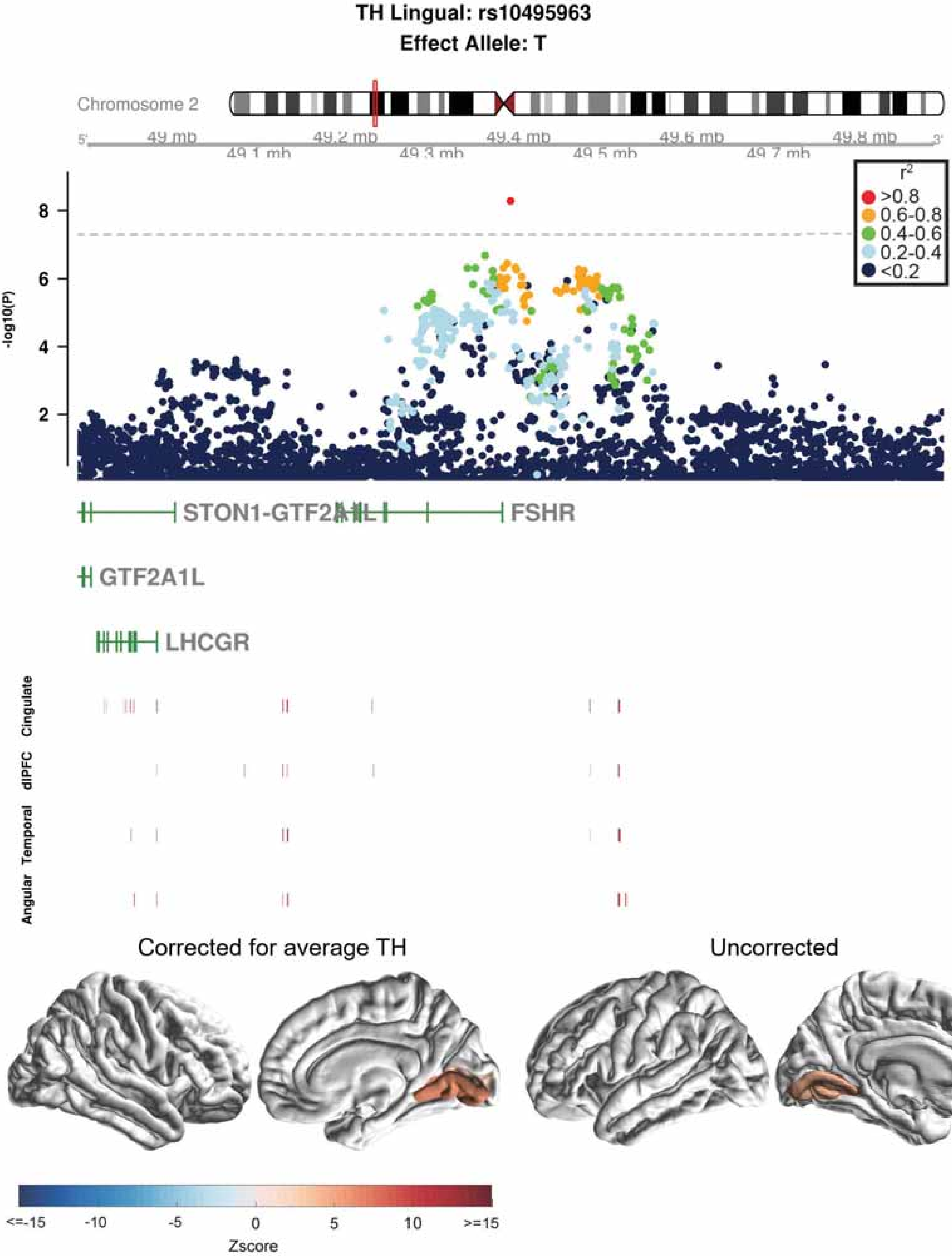

TH Lingual: rs10495963
Effect Allele: T
Chromosome 2
49 mb
49.1 mb
49.2 mb
49.3 mb
49.4 mb
49.5 mb
49.6 mb
49.7 mb
49.8 mb
r2
>0.8
0.6-0.8
0.4-0.6
0.2-0.4
<0.2
-log10(P)
STON1-GTF2A1L
FSHR
GTF2A1L
LHCGR
Cingulate
dlPFC
Temporal
Angular
Corrected for average TH
Uncorrected
<=-15
-10
-5
0
5
10
>=15
Zscore

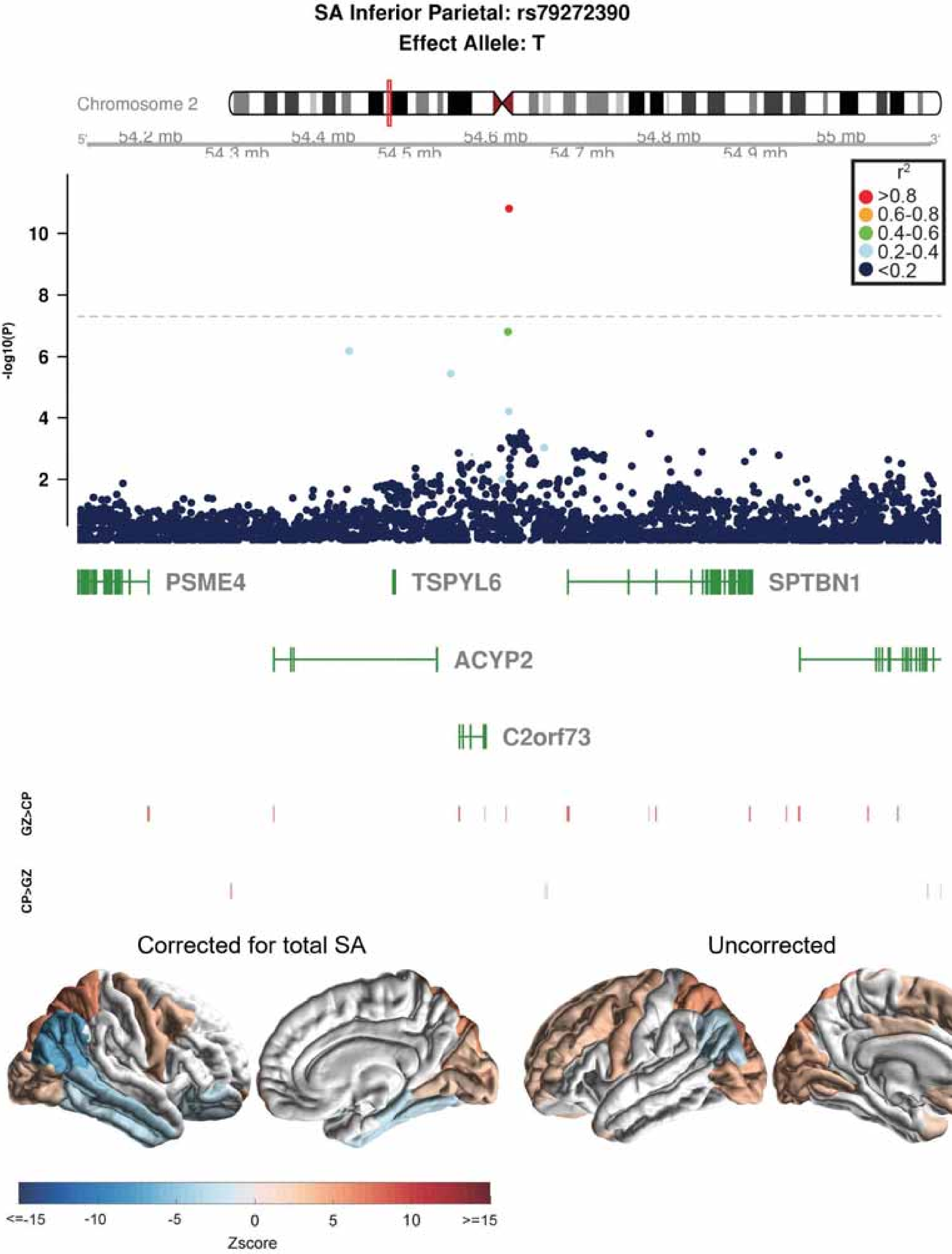
SA Inferior Parietal: rs79272390
Effect Allele: T
Chromosome 2
54.2 mb
54.3 mb
54.4 mb
54.5 mb
54.6 mb
54.7 mb
54.8 mb
54.9 mb
55 mb
r2
>0.8
0.6-0.8
0.4-0.6
0.2-0.4
<0.2
-log10(P)
PSME4
TSPYL6
SPTBN1
ACYP2
C2orf73
GZ>CP
CP>GZ
Corrected for total SA
Uncorrected
<=-15
-10
-5
0
5
10
>=15
Zscore

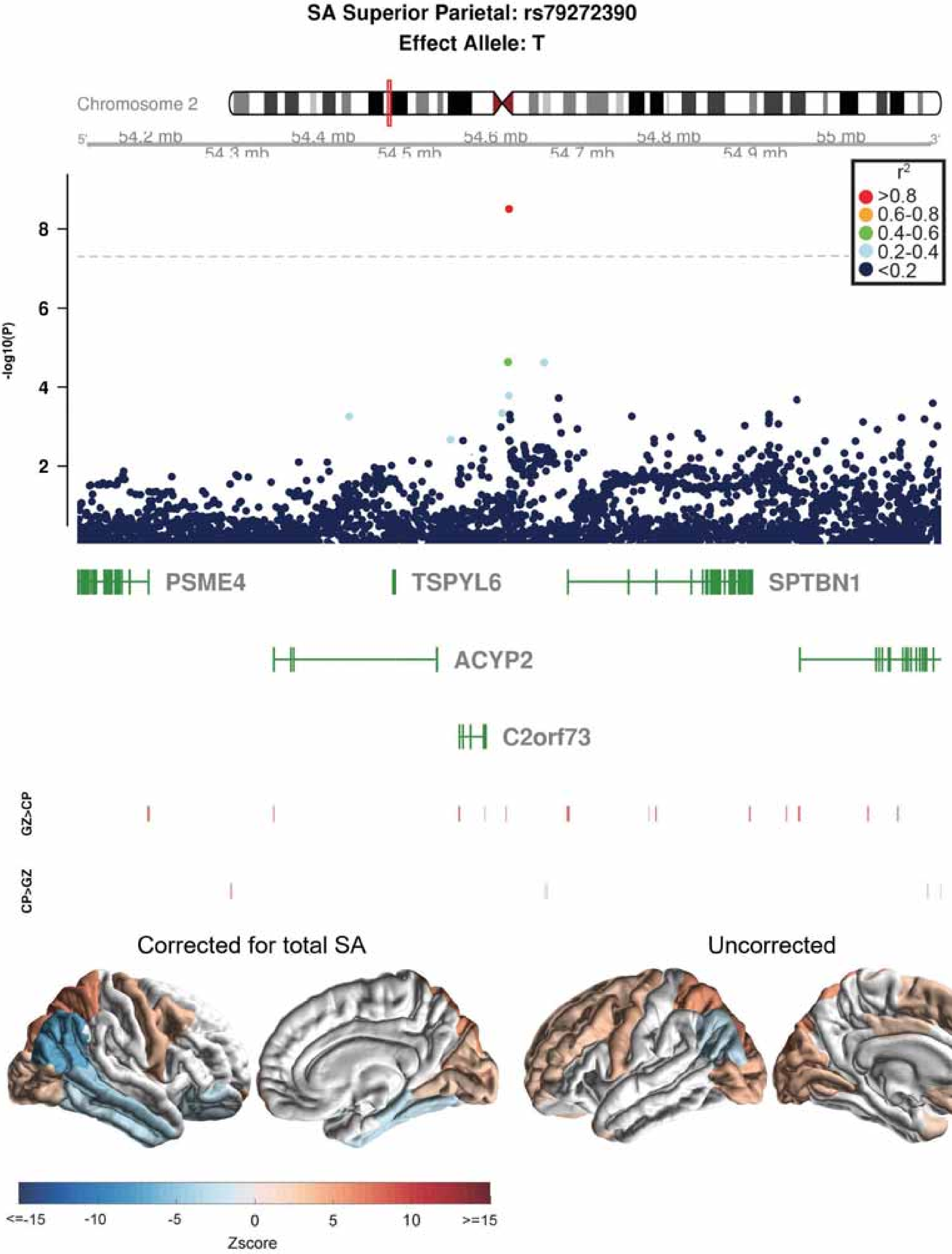
SA Superior Parietal: rs79272390
Effect Allele: T
Chromosome 2
54.2 mb
54.3 mb
54.4 mb
54.5 mb
54.6 mb
54.7 mb
54.8 mb
54.9 mb
55 mb
r2
>0.8
0.6-0.8
0.4-0.6
0.2-0.4
<0.2
-log10(P)
PSME4
TSPYL6
SPTBN1
ACYP2
C2orf73
GZ>CP
CP>GZ
Corrected for total SA
Uncorrected
<=-15
-10
-5
0
5
10
>=15
Zscore

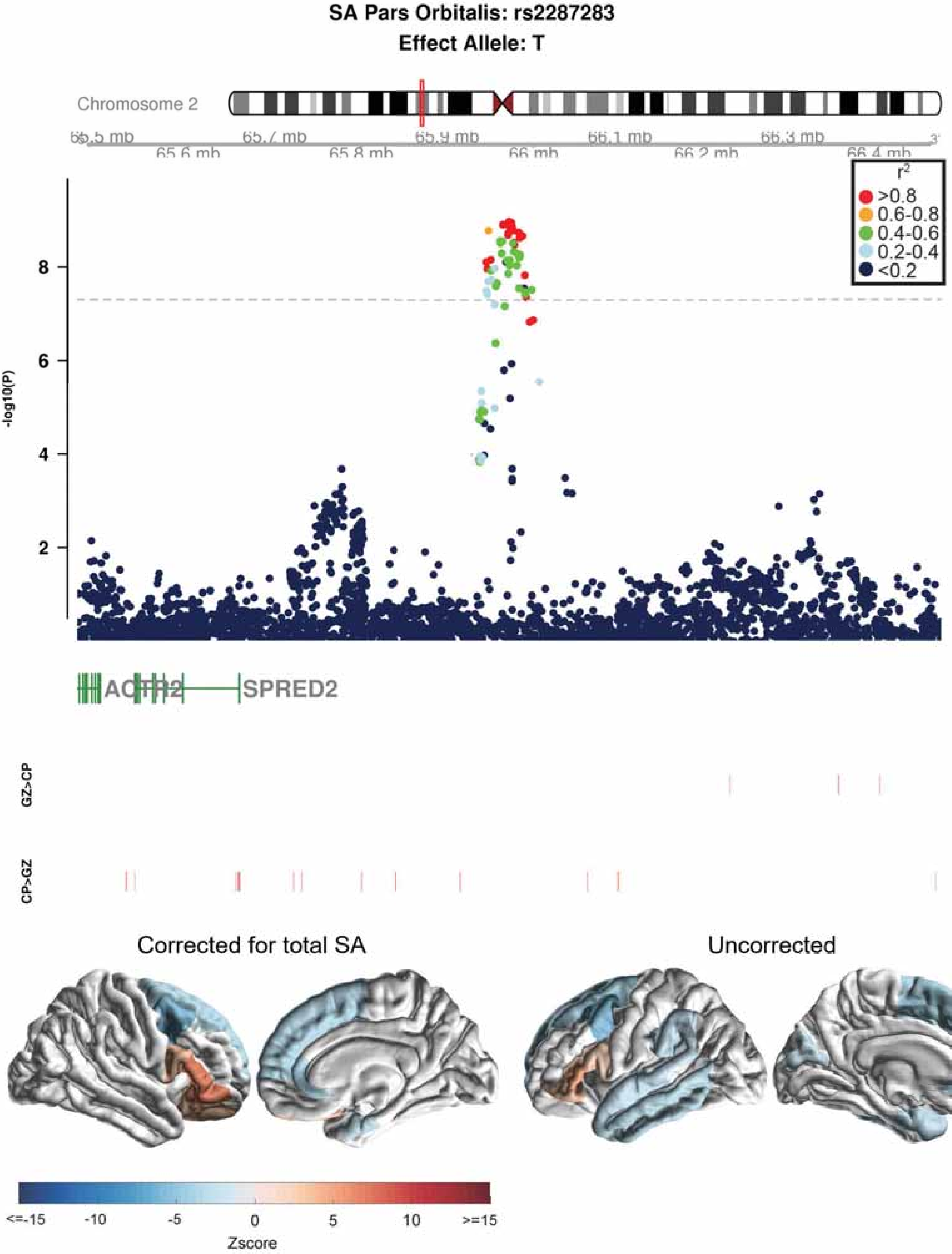

SA Pars Orbitalis: rs2287283
Effect Allele: T
Chromosome 2
65.5 mb
65.6 mb
65.7 mb
65.8 mb
65.9 mb
66 mb
66.1 mb
66.2 mb
66.3 mb
66.4 mb
r2
>0.8
0.6-0.8
0.4-0.6
0.2-0.4
<0.2
-log10(P)
2
4
6
8
SPRED2
GZ>CP
CP>GZ
Corrected for total SA
Uncorrected
<=-15
-10
-5
0
5
10
>=15
Zscore

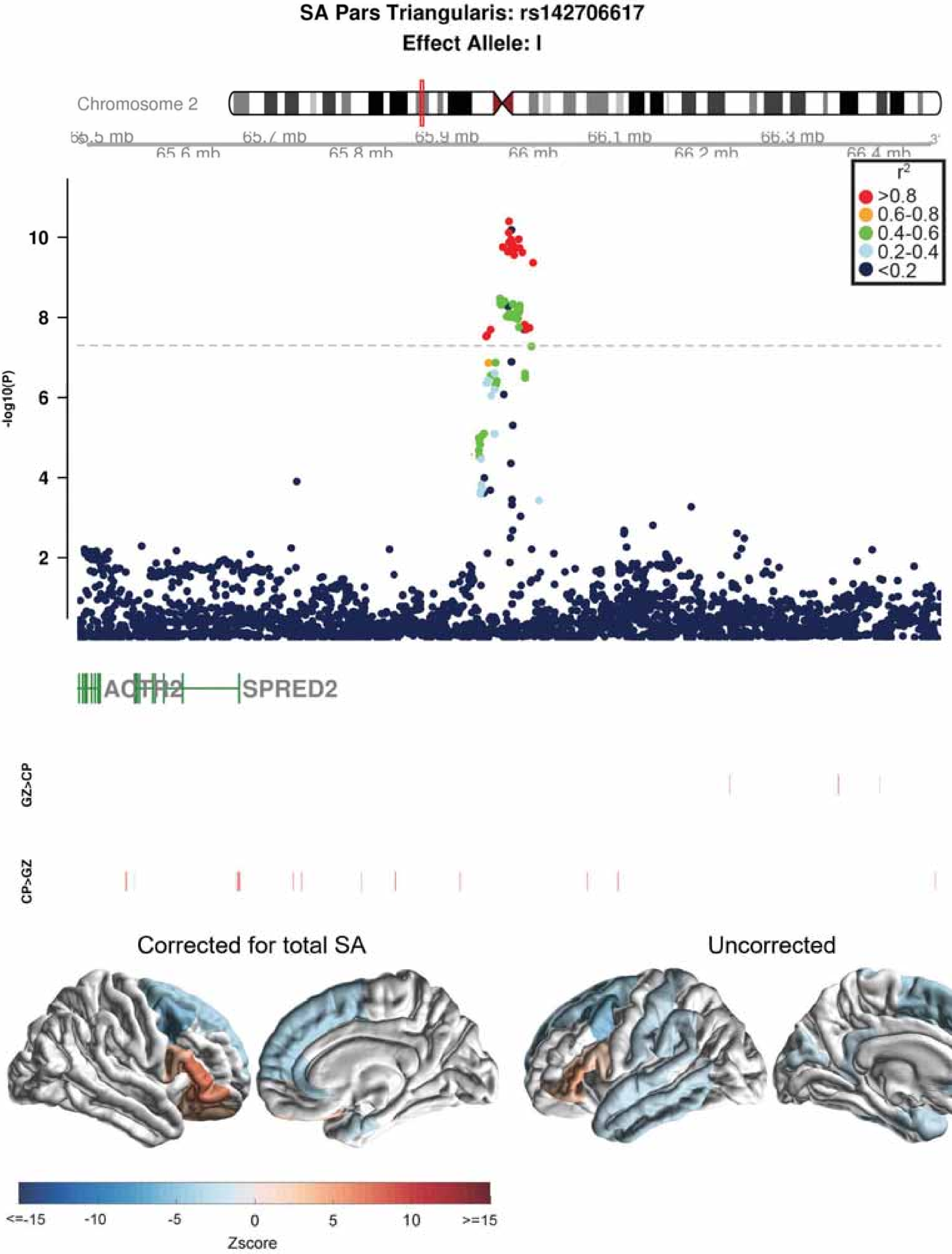
SA Pars Triangularis: rs142706617
Effect Allele: I
Chromosome 2
65.6 mb
65.7 mb
65.8 mb
65.9 mb
66 mb
66.1 mb
66.2 mb
66.3 mb
66.4 mb
r2
>0.8
0.6-0.8
0.4-0.6
0.2-0.4
<0.2
-log10(P)
10
8
6
4
2
ACTR2
SPRED2
GZ>CP
CP>GZ
Corrected for total SA
Uncorrected
<=-15
-10
-5
0
5
10
>=15
Zscore

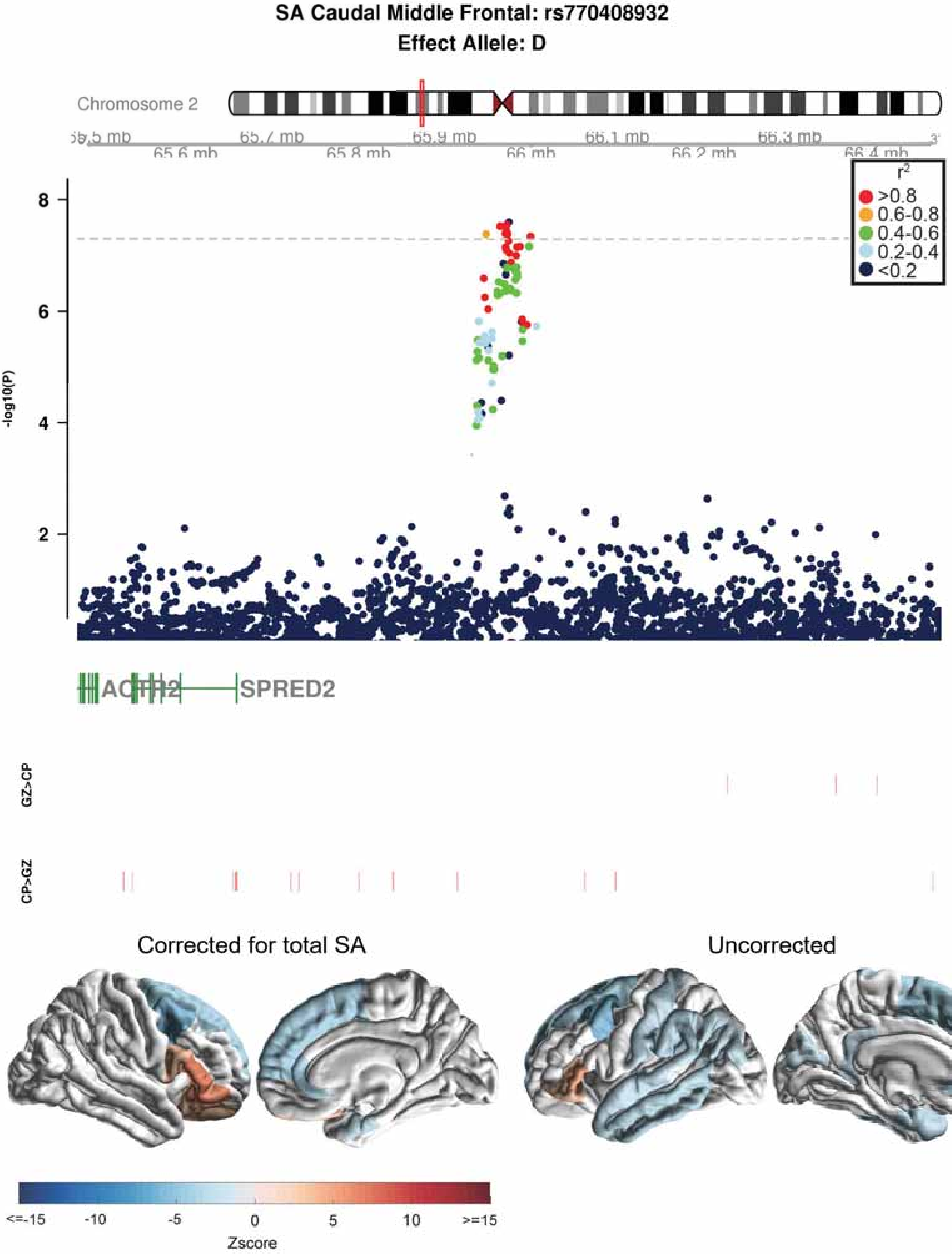
SA Caudal Middle Frontal: rs770408932
Effect Allele: D
Chromosome 2
65.7 mb
65.9 mb
66.1 mb
66.3 mb
65.6 mb
65.8 mb
66 mb
66.2 mb
66.4 mb
r2
>0.8
0.6-0.8
0.4-0.6
0.2-0.4
<0.2
-log10(P)
8
6
4
2
ACTR2
SPRED2
GZ>CP
CP>GZ
Corrected for total SA
Uncorrected
<=-15
-10
-5
0
5
10
>=15
Zscore

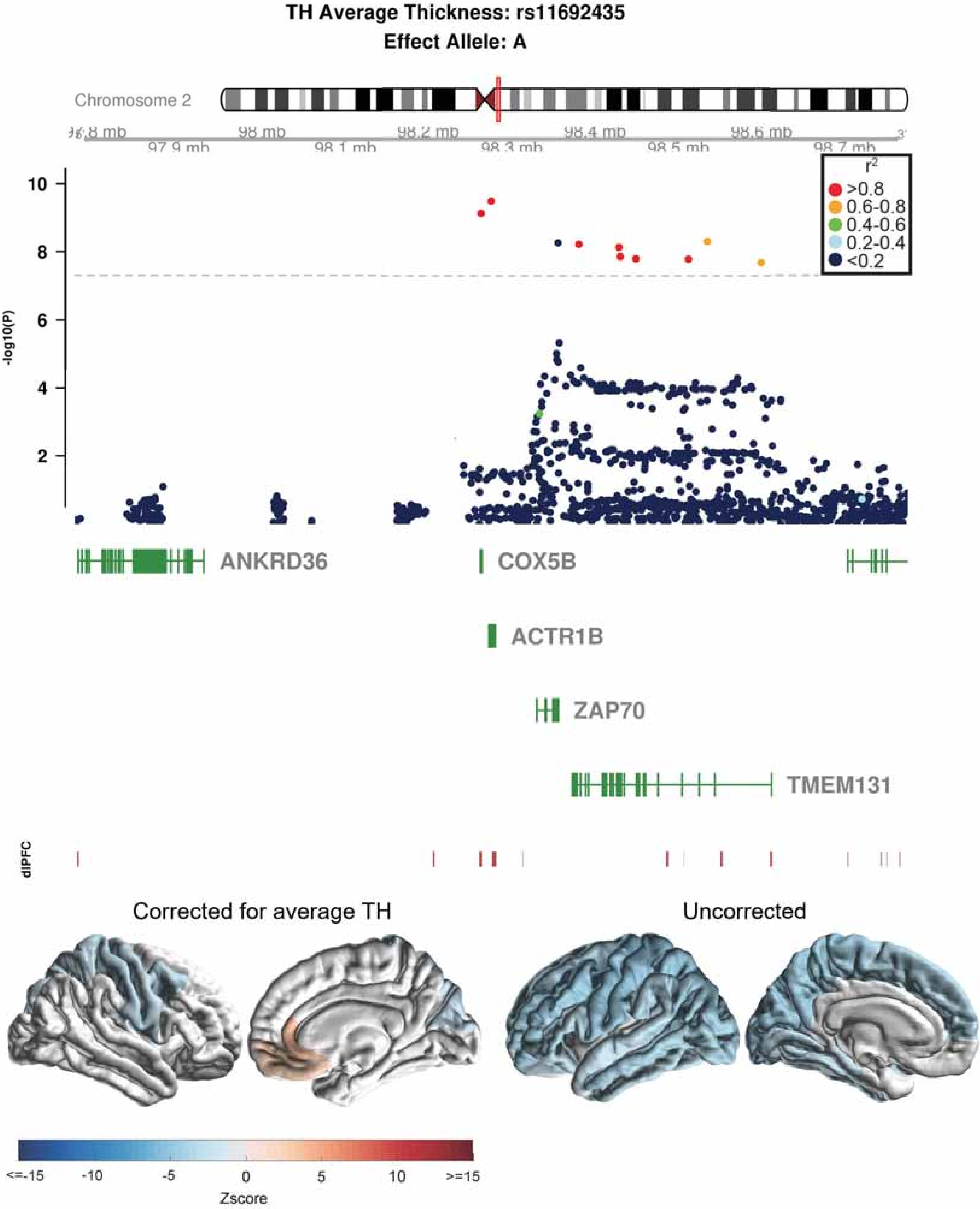
TH Average Thickness: rs11692435
Effect Allele: A
Chromosome 2
98 mb
98.2 mb
98.4 mb
98.6 mb
97.9 mb
98.1 mb
98.3 mb
98.5 mb
98.7 mb
-log10(P)
r2
>0.8
0.6-0.8
0.4-0.6
0.2-0.4
<0.2
ANKRD36
COX5B
ACTR1B
ZAP70
TMEM131
dlPFC
Corrected for average TH
Uncorrected
<=-15
-10
-5
0
5
10
>=15
Zscore

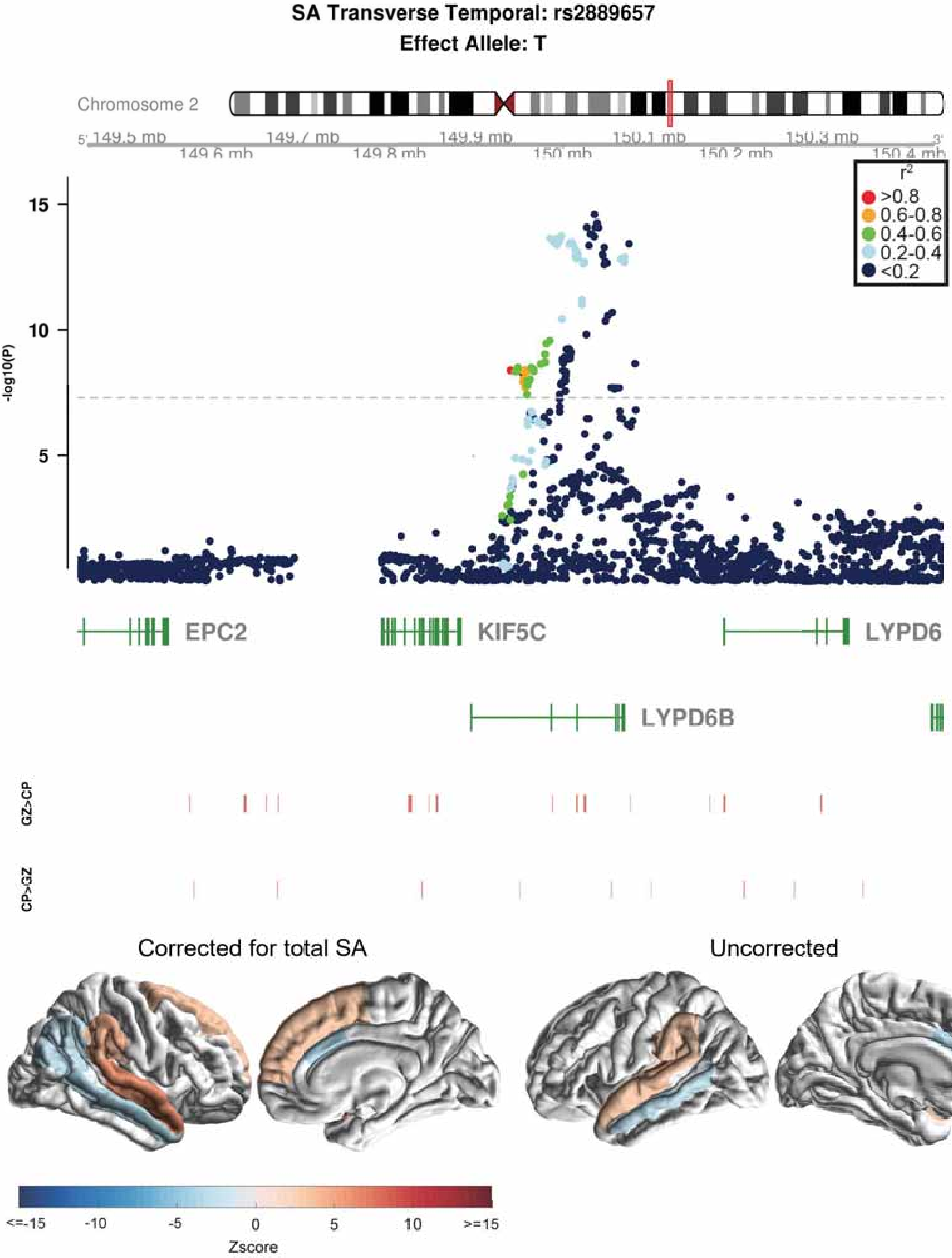

SA Transverse Temporal: rs2889657
Effect Allele: T
Chromosome 2
149.5 mb
149.6 mb
149.7 mb
149.8 mb
149.9 mb
150 mb
150.1 mb
150.2 mb
150.3 mb
150.4 mb
r2
>0.8
0.6-0.8
0.4-0.6
0.2-0.4
<0.2
-log10(P)
15
10
5
EPC2
KIF5C
LYPD6
LYPD6B
GZ>CP
CP>GZ
Corrected for total SA
Uncorrected
<=-15
-10
-5
0
5
10
>=15
Zscore

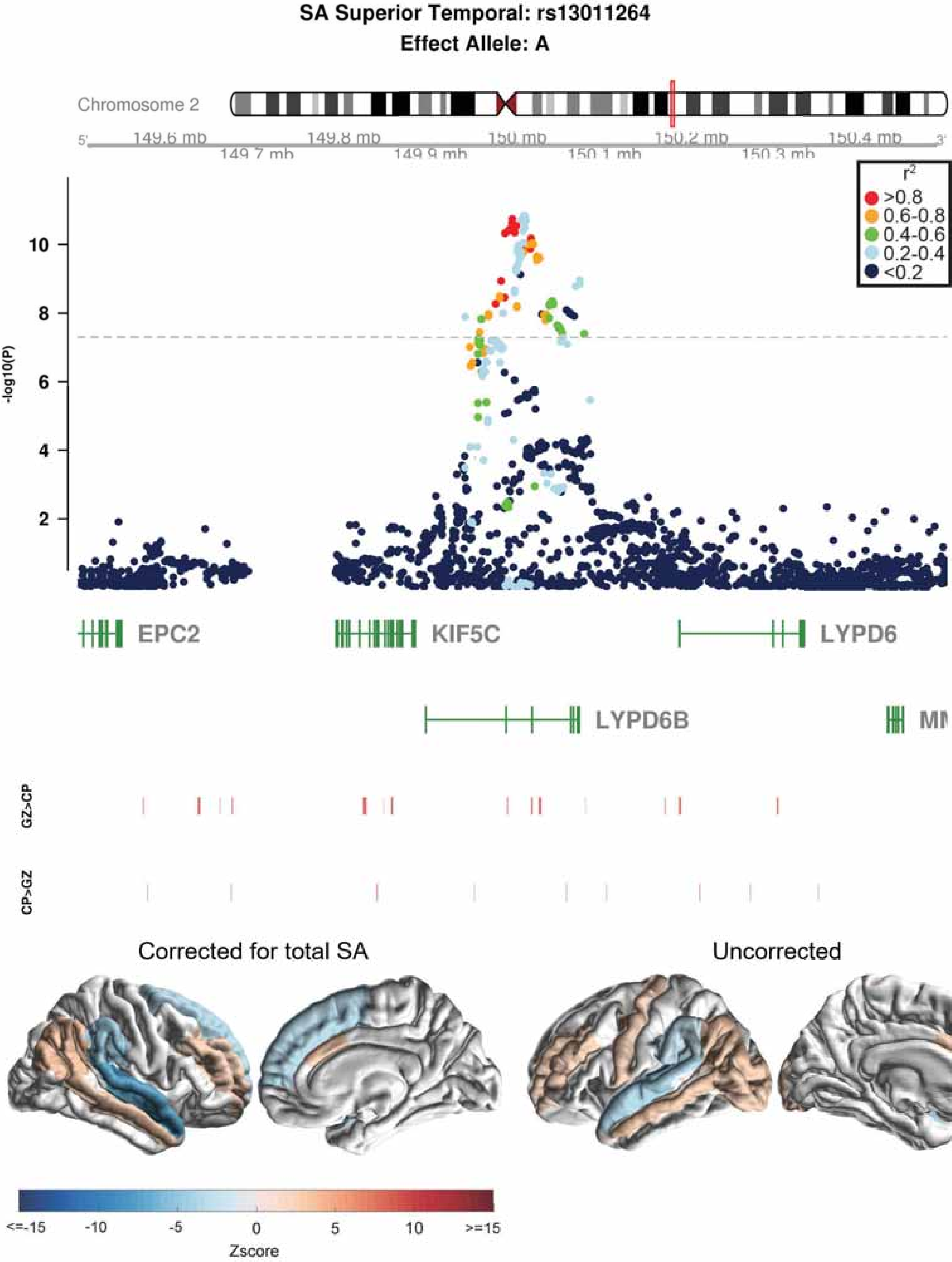
SA Superior Temporal: rs13011264
Effect Allele: A
Chromosome 2
149.6 mb
149.7 mb
149.8 mb
149.9 mb
150 mb
150.1 mb
150.2 mb
150.3 mb
150.4 mb
-log10(P)
r²
>0.8
0.6-0.8
0.4-0.6
0.2-0.4
<0.2
EPC2
KIF5C
LYPD6
LYPD6B
GZ>CP
CP>GZ
Corrected for total SA
Uncorrected
<=-15
-10
-5
0
5
10
>=15
Zscore

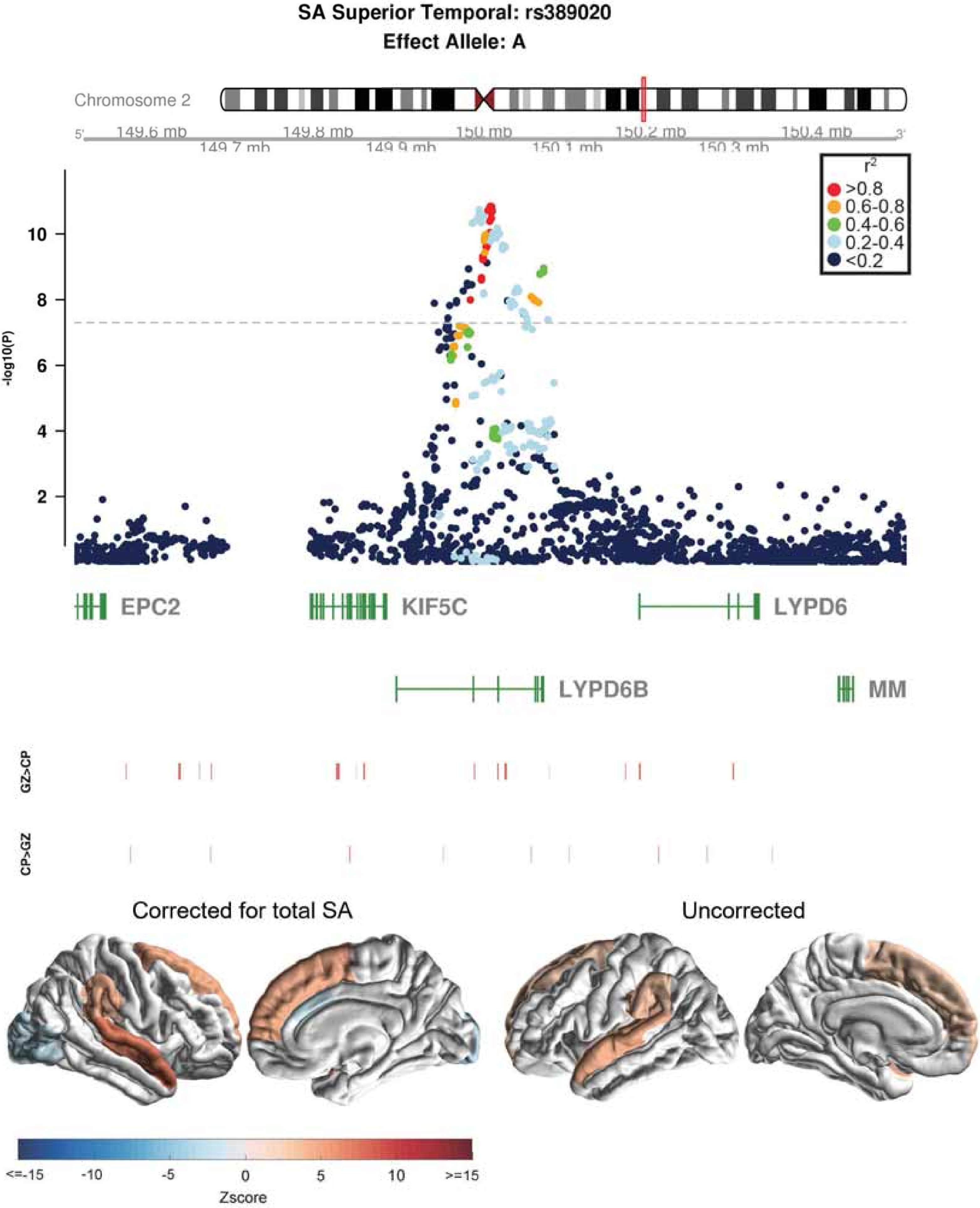
SA Superior Temporal: rs389020
Effect Allele: A
Chromosome 2
149.6 mb
149.7 mb
149.8 mb
149.9 mb
150 mb
150.1 mb
150.2 mb
150.3 mb
150.4 mb
r2
>0.8
0.6-0.8
0.4-0.6
0.2-0.4
<0.2
-log10(P)
EPC2
KIF5C
LYPD6
LYPD6B
MM
GZ>CP
CP>GZ
Corrected for total SA
Uncorrected
<=-15
-10
-5
0
5
10
>=15
Zscore

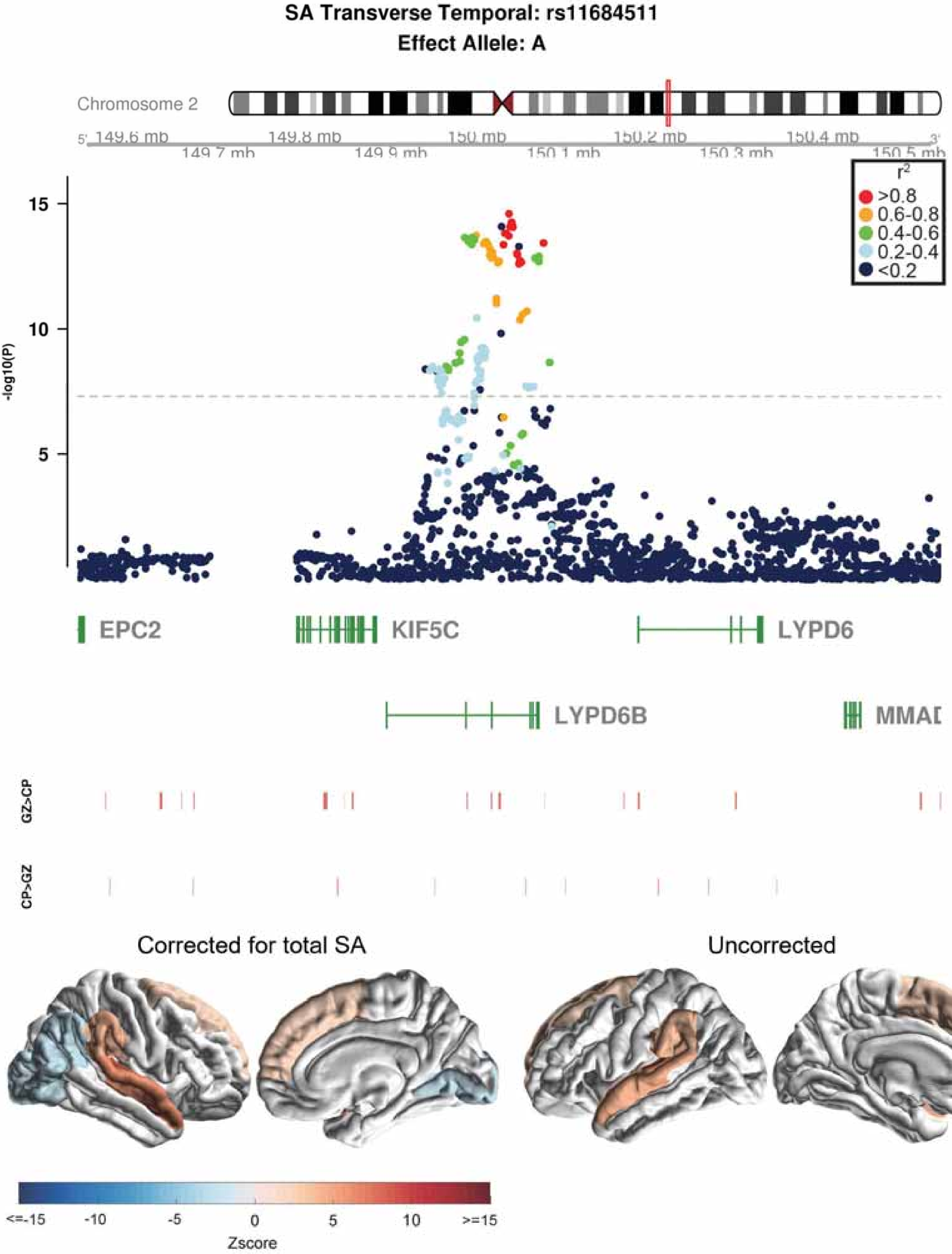
SA Transverse Temporal: rs11684511
Effect Allele: A
Chromosome 2
5' 149.6 mb
149.7 mb
149.8 mb
149.9 mb
150 mb
150.1 mb
150.2 mb
150.3 mb
150.4 mb
150.5 mb 3'
r2
>0.8
0.6-0.8
0.4-0.6
0.2-0.4
<0.2
-log10(P)
15
10
5
EPC2
KIF5C
LYPD6
LYPD6B
MMAD
GZ>CP
CP>GZ
Corrected for total SA
Uncorrected
<=-15
-10
-5
0
5
10
>=15
Zscore

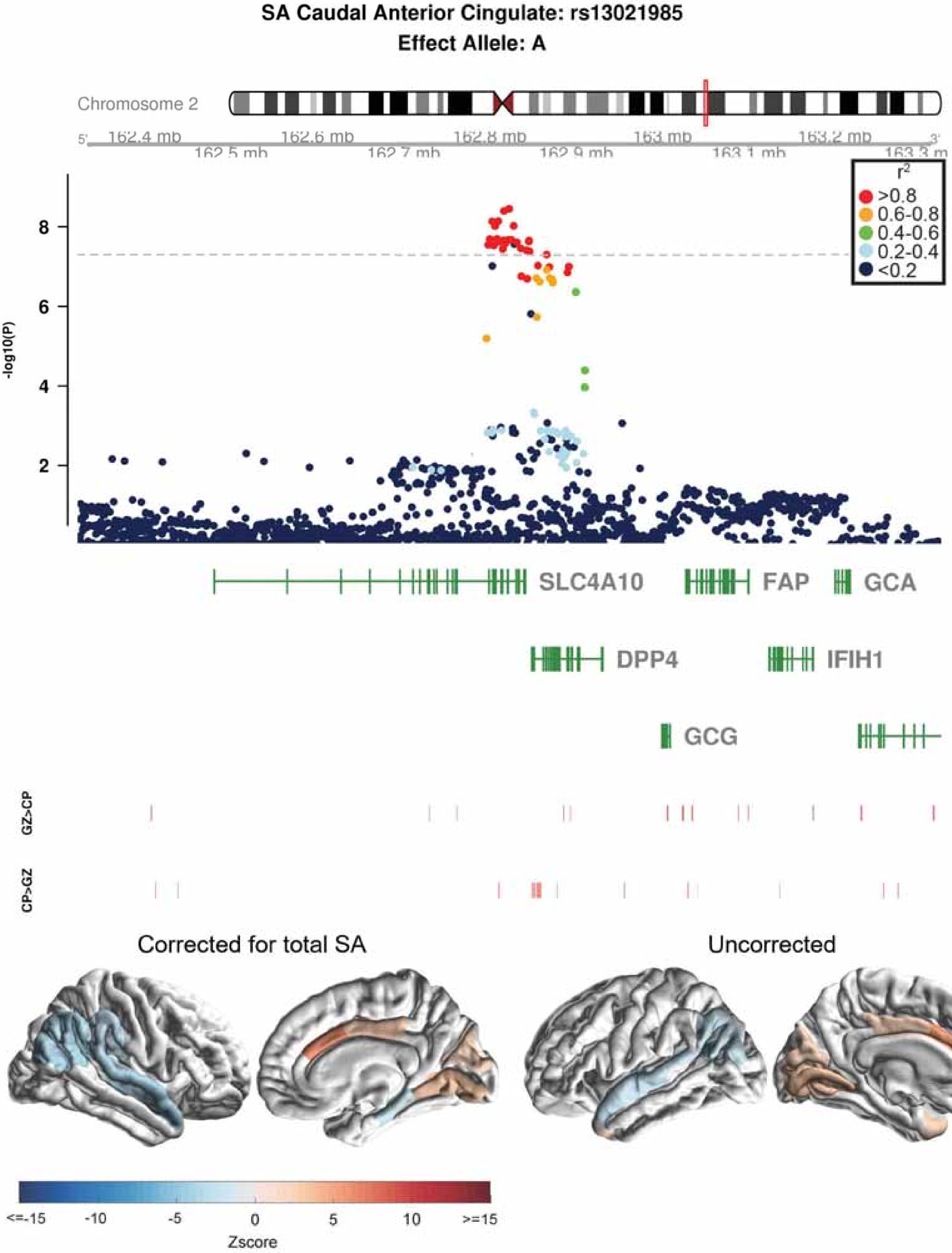
SA Caudal Anterior Cingulate: rs13021985
Effect Allele: A
Chromosome 2
162.4 mb
162.5 mb
162.6 mb
162.7 mb
162.8 mb
162.9 mb
163 mb
163.1 mb
163.2 mb
163.3 mb
-log10(P)
r²
>0.8
0.6-0.8
0.4-0.6
0.2-0.4
<0.2
SLC4A10
FAP
GCA
DPP4
IFIH1
GCG
GZ>CP
CP>GZ
Corrected for total SA
Uncorrected
<=-15
-10
-5
0
5
10
>=15
Zscore

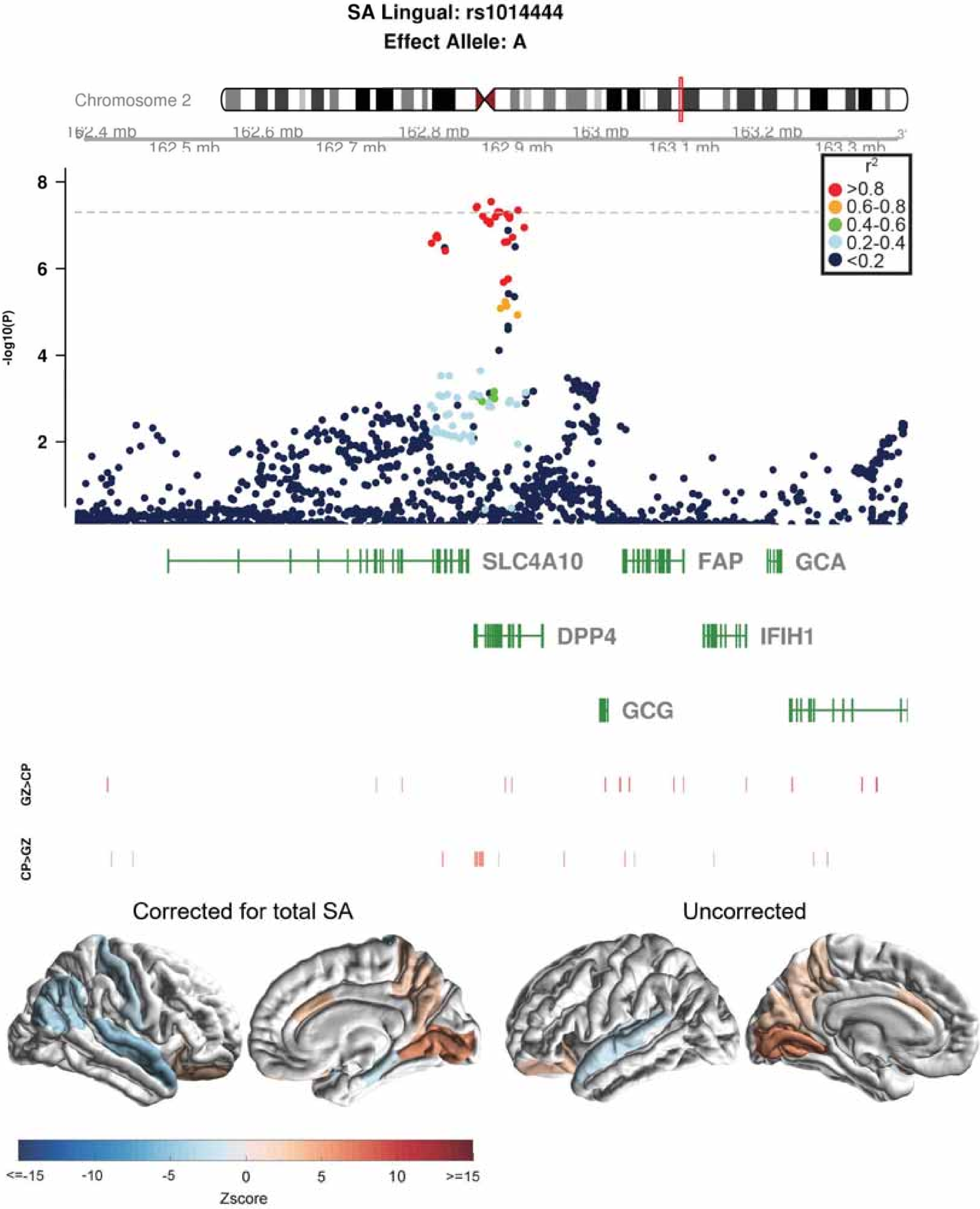

SA Lingual: rs1014444
Effect Allele: A
Chromosome 2
162.4 mb
162.5 mb
162.6 mb
162.7 mb
162.8 mb
162.9 mb
163 mb
163.1 mb
163.2 mb
163.3 mb
-log10(P)
8
6
4
2
r2
>0.8
0.6-0.8
0.4-0.6
0.2-0.4
<0.2
SLC4A10
FAP
GCA
DPP4
IFIH1
GCG
GZ>CP
CP>GZ
Corrected for total SA
Uncorrected
<=-15
-10
-5
0
5
10
>=15
Zscore

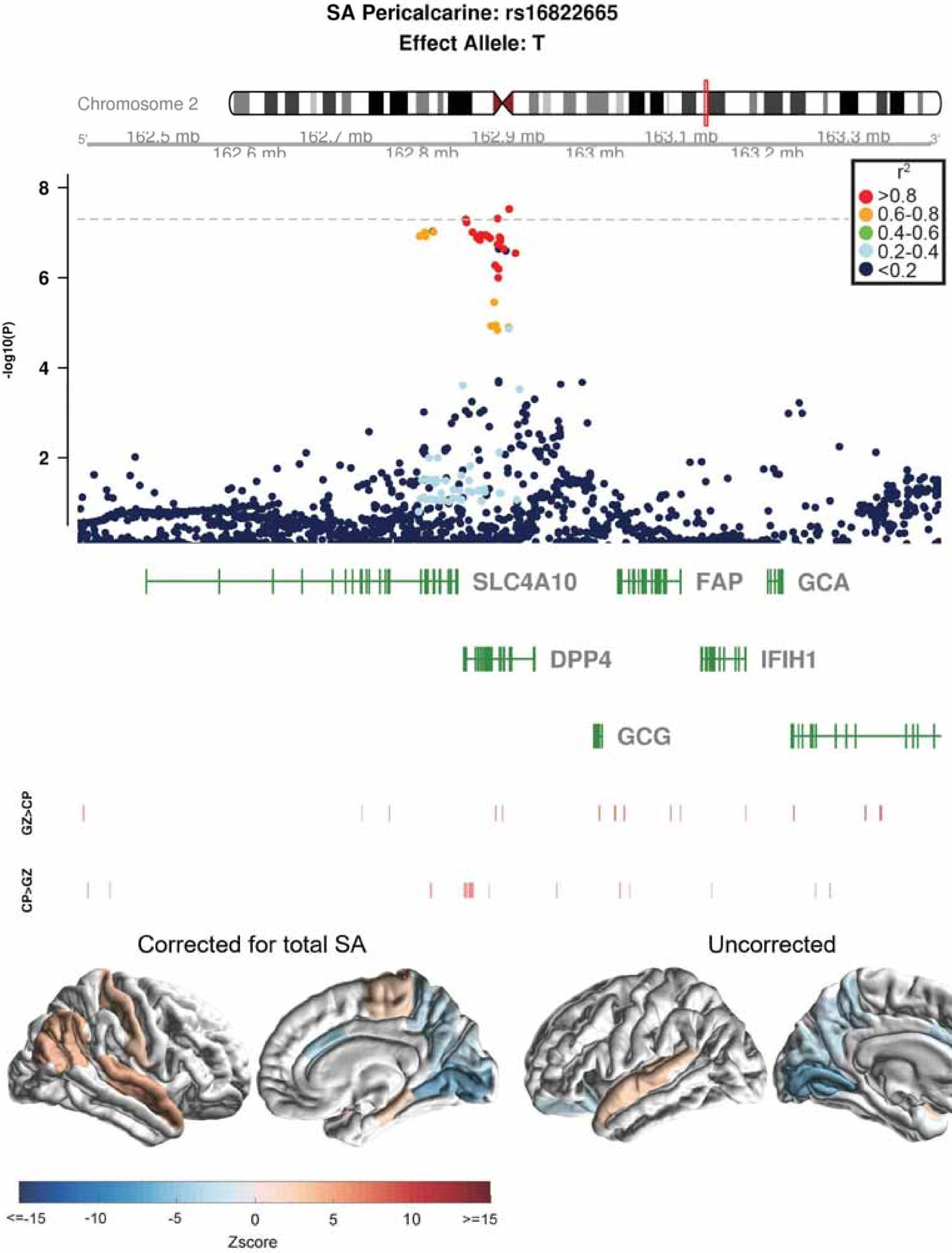
SA Pericalcarine: rs16822665
Effect Allele: T
Chromosome 2
162.5 mb
162.6 mb
162.7 mb
162.8 mb
162.9 mb
163 mb
163.1 mb
163.2 mb
163.3 mb
-log10(P)
r2
>0.8
0.6-0.8
0.4-0.6
0.2-0.4
<0.2
SLC4A10
FAP
GCA
DPP4
IFIH1
GCG
GZ>CP
CP>GZ
Corrected for total SA
Uncorrected
<=-15
-10
-5
0
5
10
>=15
Zscore

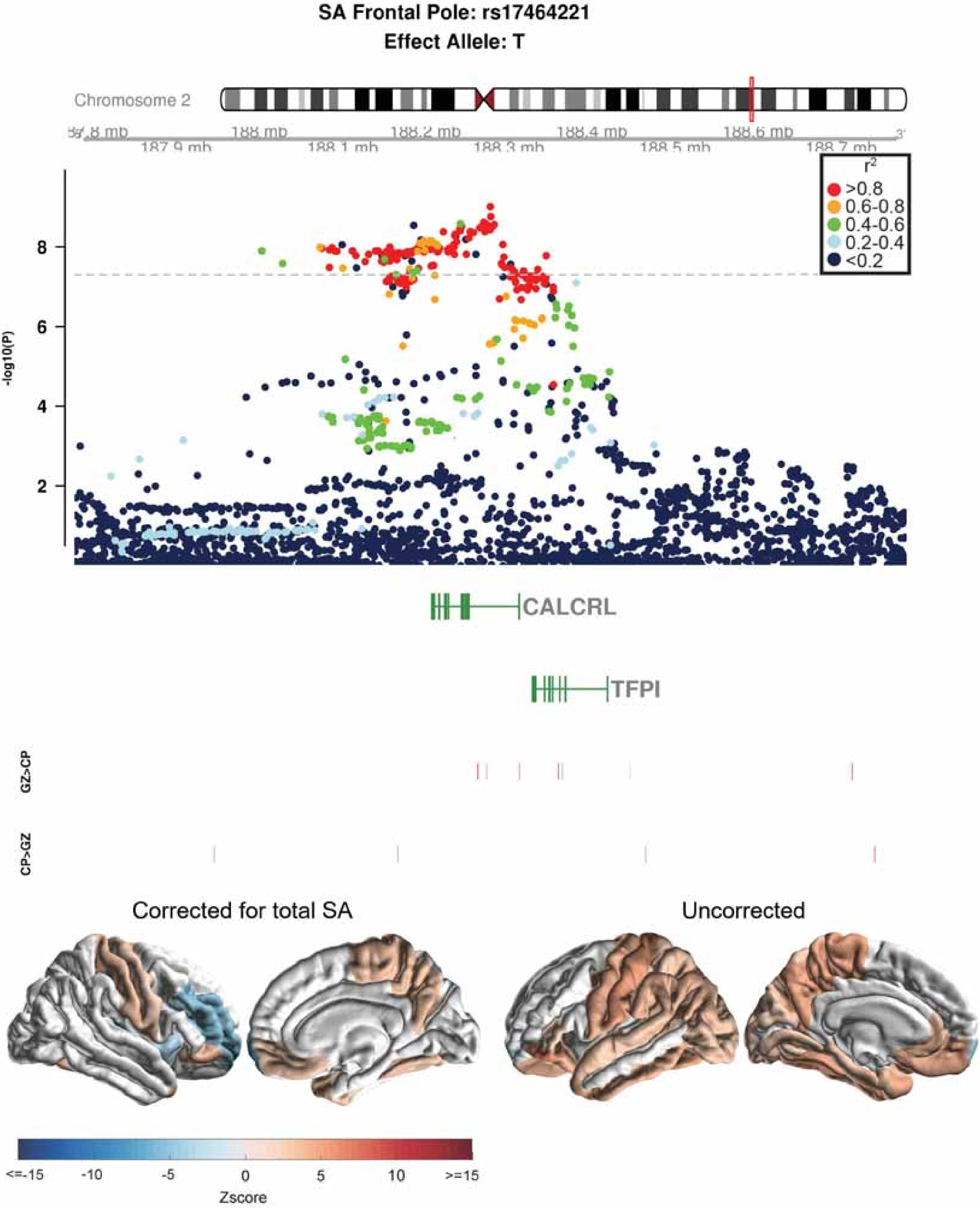
SA Frontal Pole: rs17464221
Effect Allele: T
Chromosome 2
188 mb
188.2 mb
188.4 mb
188.6 mb
187.9 mb
188.1 mb
188.3 mb
188.5 mb
188.7 mb
r2
>0.8
0.6-0.8
0.4-0.6
0.2-0.4
<0.2
-log10(P)
8
6
4
2
CALCRL
TFPI
GZ>CP
CP>GZ
Corrected for total SA
Uncorrected
<=-15
-10
-5
0
5
10
>=15
Zscore

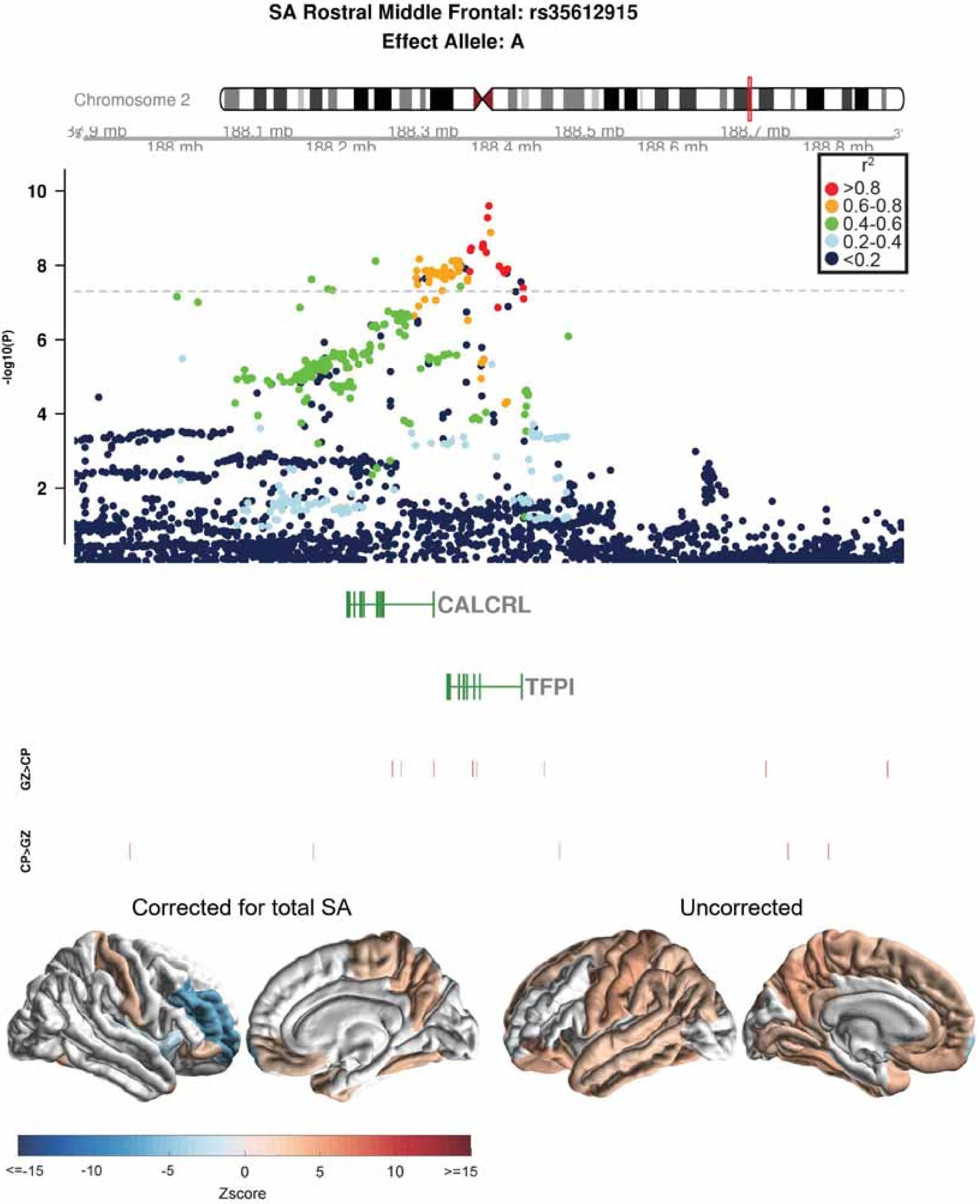

SA Rostral Middle Frontal: rs35612915
Effect Allele: A
Chromosome 2
188.1 mb
188.3 mb
188.5 mb
188.7 mb
188 mb
188.2 mb
188.4 mb
188.6 mb
188.8 mb
r2
>0.8
0.6-0.8
0.4-0.6
0.2-0.4
<0.2
-log10(P)
10
8
6
4
2
CALCRL
TFPI
GZ>CP
CP>GZ
Corrected for total SA
Uncorrected
<=-15
-10
-5
0
5
10
>=15
Zscore

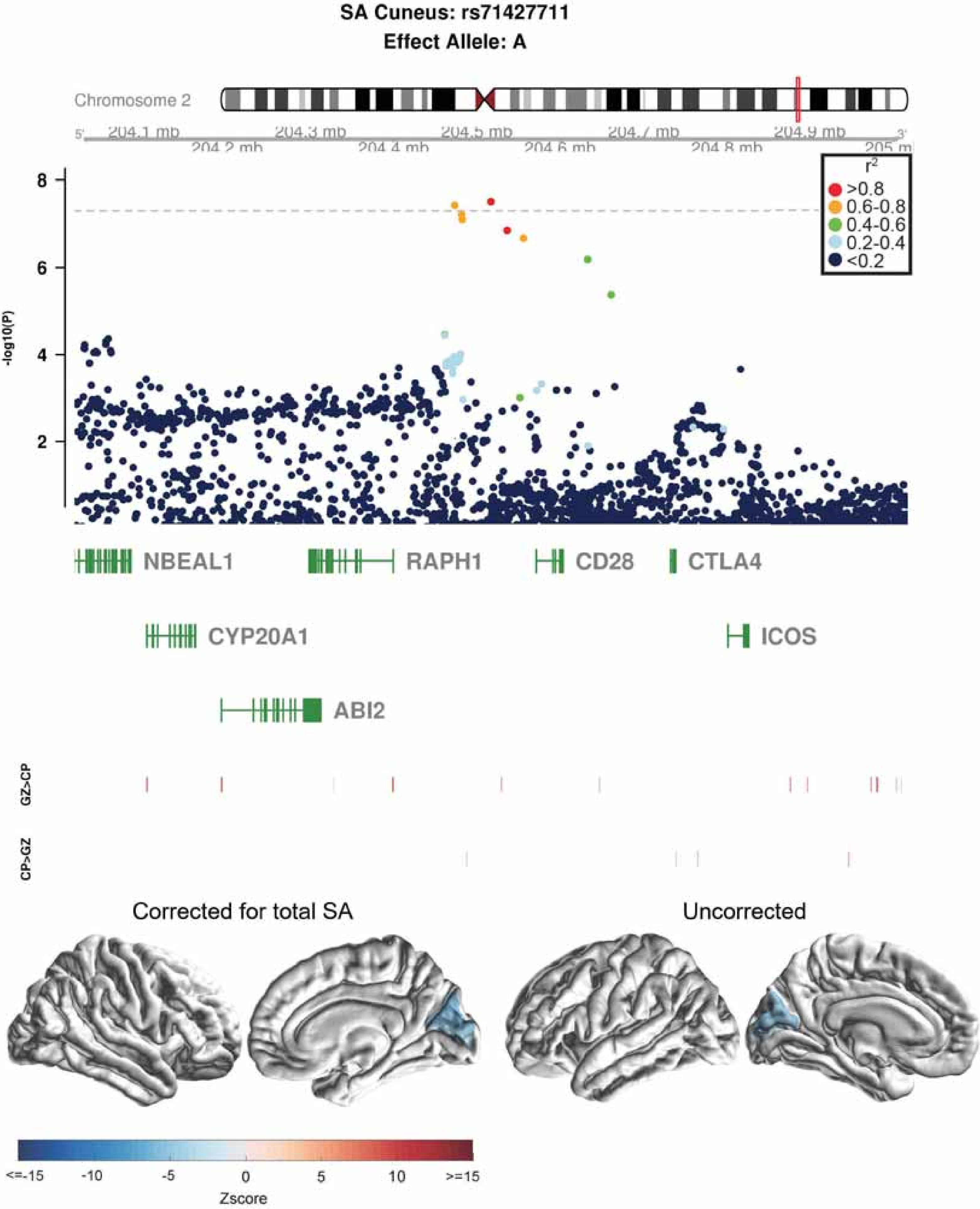
SA Cuneus: rs71427711
Effect Allele: A
Chromosome 2
204.1 mb
204.2 mb
204.3 mb
204.4 mb
204.5 mb
204.6 mb
204.7 mb
204.8 mb
204.9 mb
205 mb
-log10(P)
r2
>0.8
0.6-0.8
0.4-0.6
0.2-0.4
<0.2
NBEAL1
RAPH1
CD28
CTLA4
CYP20A1
ICOS
ABI2
GZ>CP
CP>GZ
Corrected for total SA
Uncorrected
<=-15
-10
-5
0
5
10
>=15
Zscore

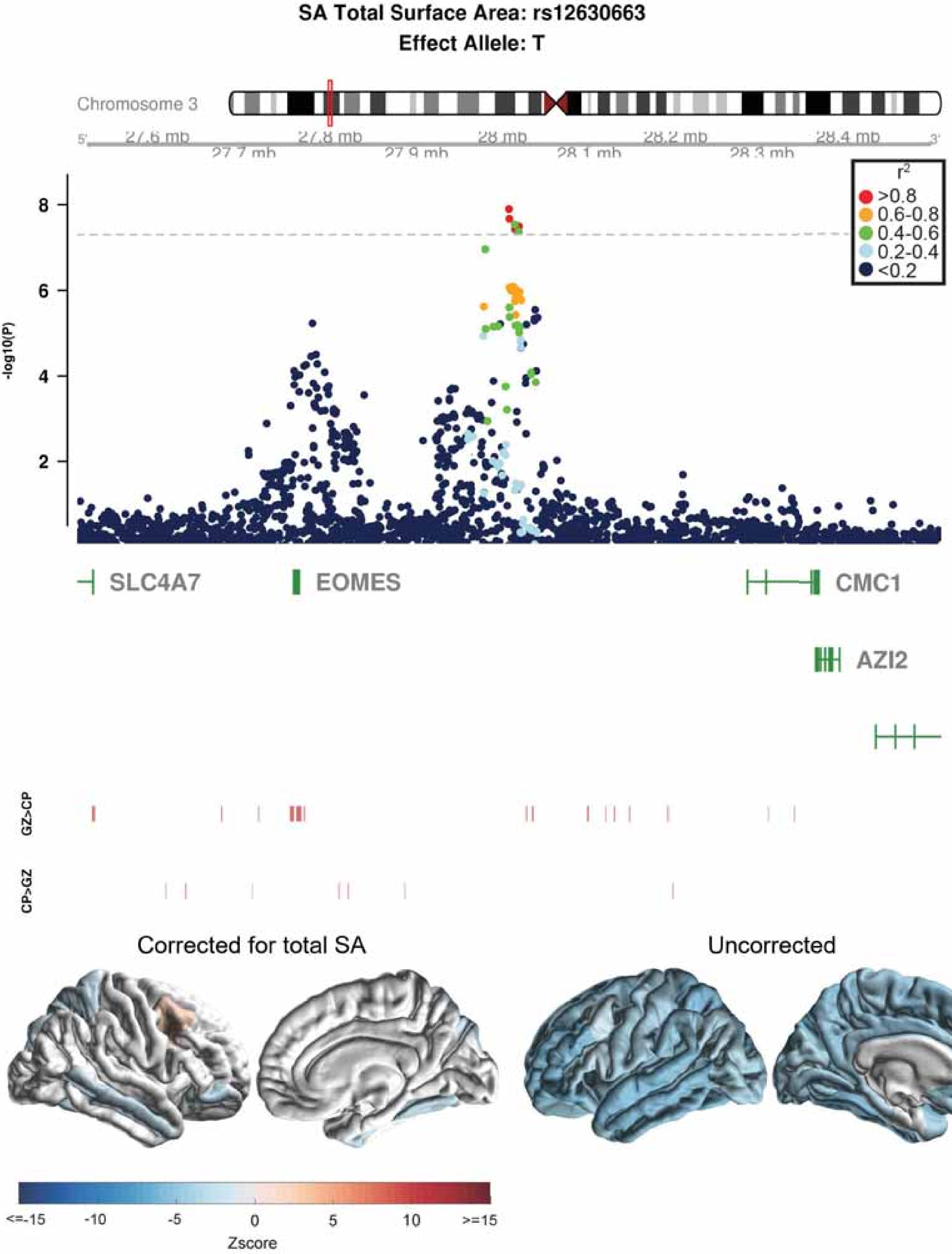

SA Total Surface Area: rs12630663
Effect Allele: T
Chromosome 3
27.6 mb
27.7 mb
27.8 mb
27.9 mb
28 mb
28.1 mb
28.2 mb
28.3 mb
28.4 mb
r2
>0.8
0.6-0.8
0.4-0.6
0.2-0.4
<0.2
-log10(P)
SLC4A7
EOMES
CMC1
AZI2
GZ>CP
CP>GZ
Corrected for total SA
Uncorrected
<=-15
-10
-5
0
5
10
>=15
Zscore

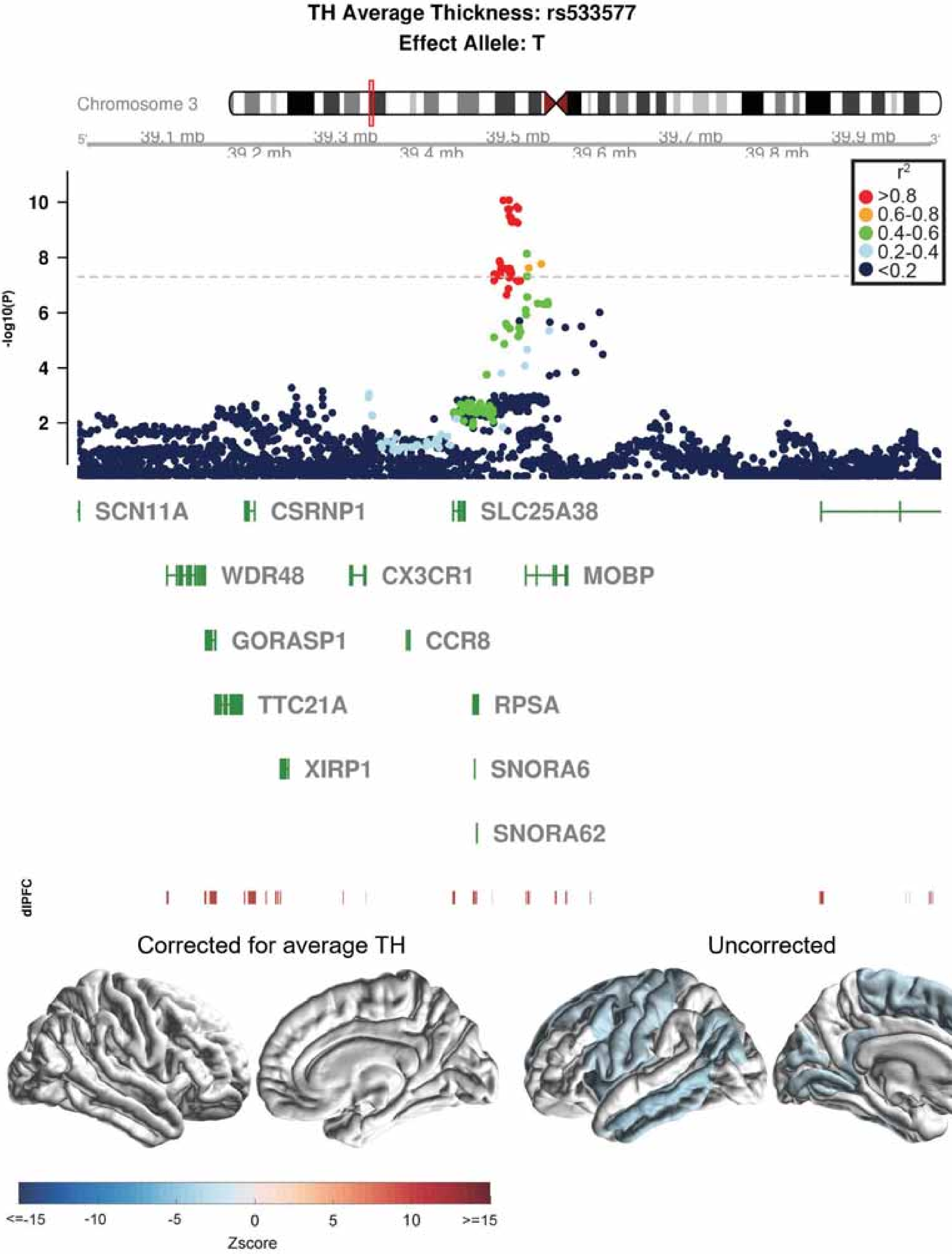

TH Average Thickness: rs533577
Effect Allele: T
Chromosome 3
39.1 mb
39.2 mb
39.3 mb
39.4 mb
39.5 mb
39.6 mb
39.7 mb
39.8 mb
39.9 mb
r2
>0.8
0.6-0.8
0.4-0.6
0.2-0.4
<0.2
-log10(P)
SCN11A
CSRNP1
SLC25A38
WDR48
CX3CR1
MOBP
GORASP1
CCR8
TTC21A
RPSA
XIRP1
SNORA6
SNORA62
dlPFC
Corrected for average TH
Uncorrected
<=-15
-10
-5
0
5
10
>=15
Zscore

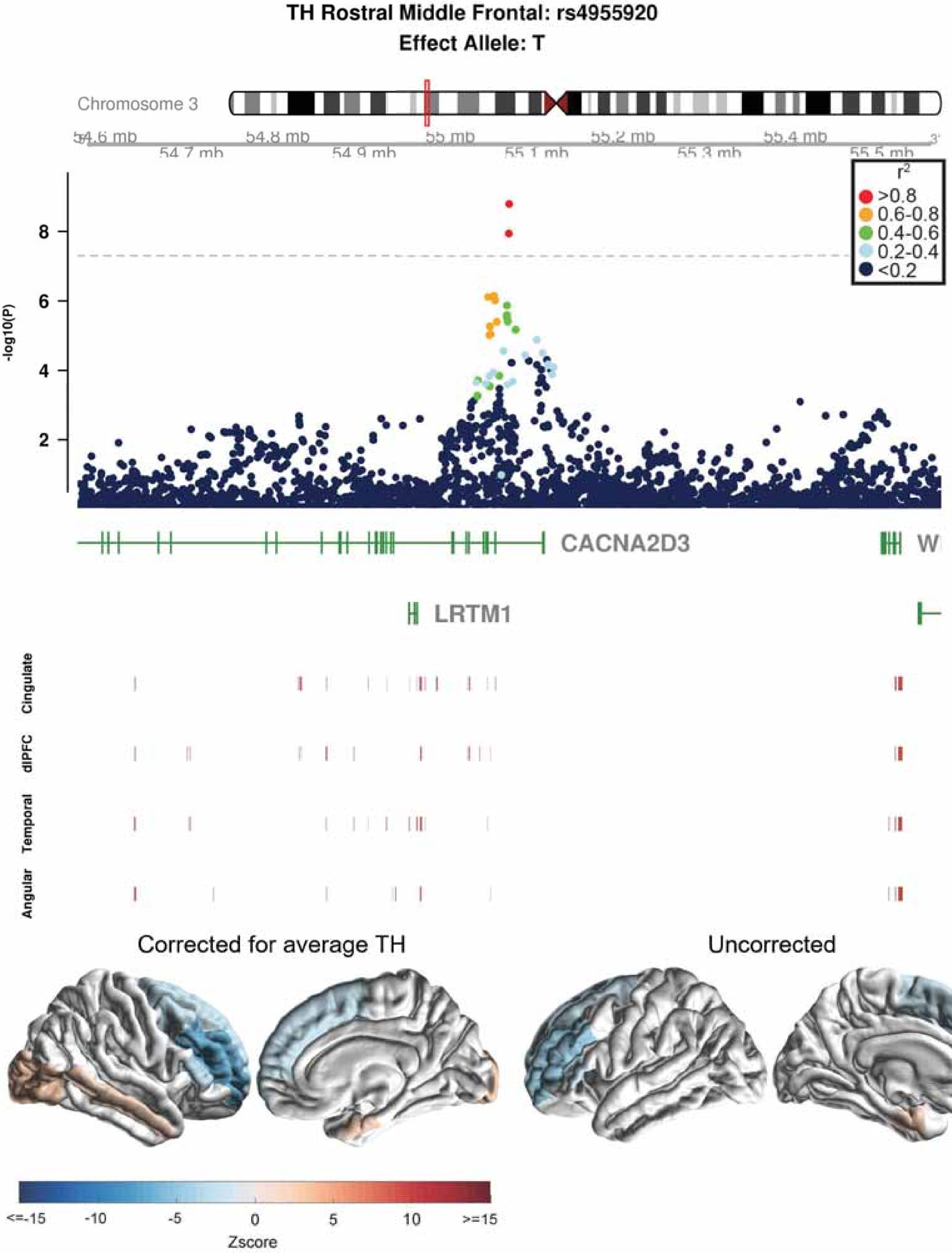

TH Rostral Middle Frontal: rs4955920
Effect Allele: T
Chromosome 3
54.6 mb
54.7 mb
54.8 mb
54.9 mb
55 mb
55.1 mb
55.2 mb
55.3 mb
55.4 mb
55.5 mb
-log10(P)
$r^2$
>0.8
0.6-0.8
0.4-0.6
0.2-0.4
<0.2
CACNA2D3
LRTM1
Cingulate
dlPFC
Temporal
Angular
Corrected for average TH
Uncorrected
<=-15
-10
-5
0
5
10
>=15
Zscore

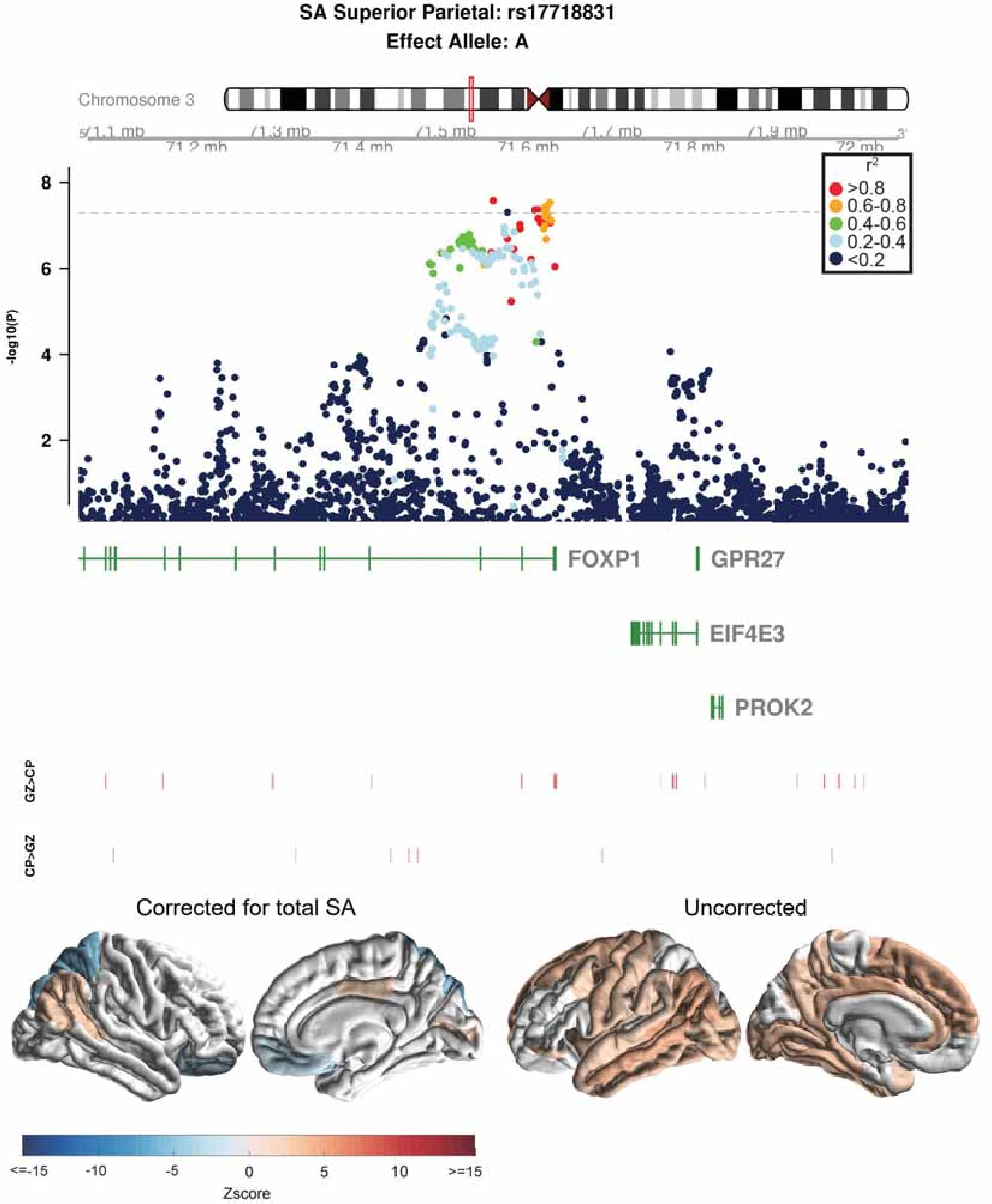

SA Superior Parietal: rs17718831
Effect Allele: A
Chromosome 3
71.1 mb
71.3 mb
71.5 mb
71.7 mb
71.9 mb
71.2 mb
71.4 mb
71.6 mb
71.8 mb
72 mb
-log10(P)
r2
>0.8
0.6-0.8
0.4-0.6
0.2-0.4
<0.2
FOXP1
GPR27
EIF4E3
PROK2
GZ>CP
CP>GZ
Corrected for total SA
Uncorrected
<=-15
-10
-5
0
5
10
>=15
Zscore

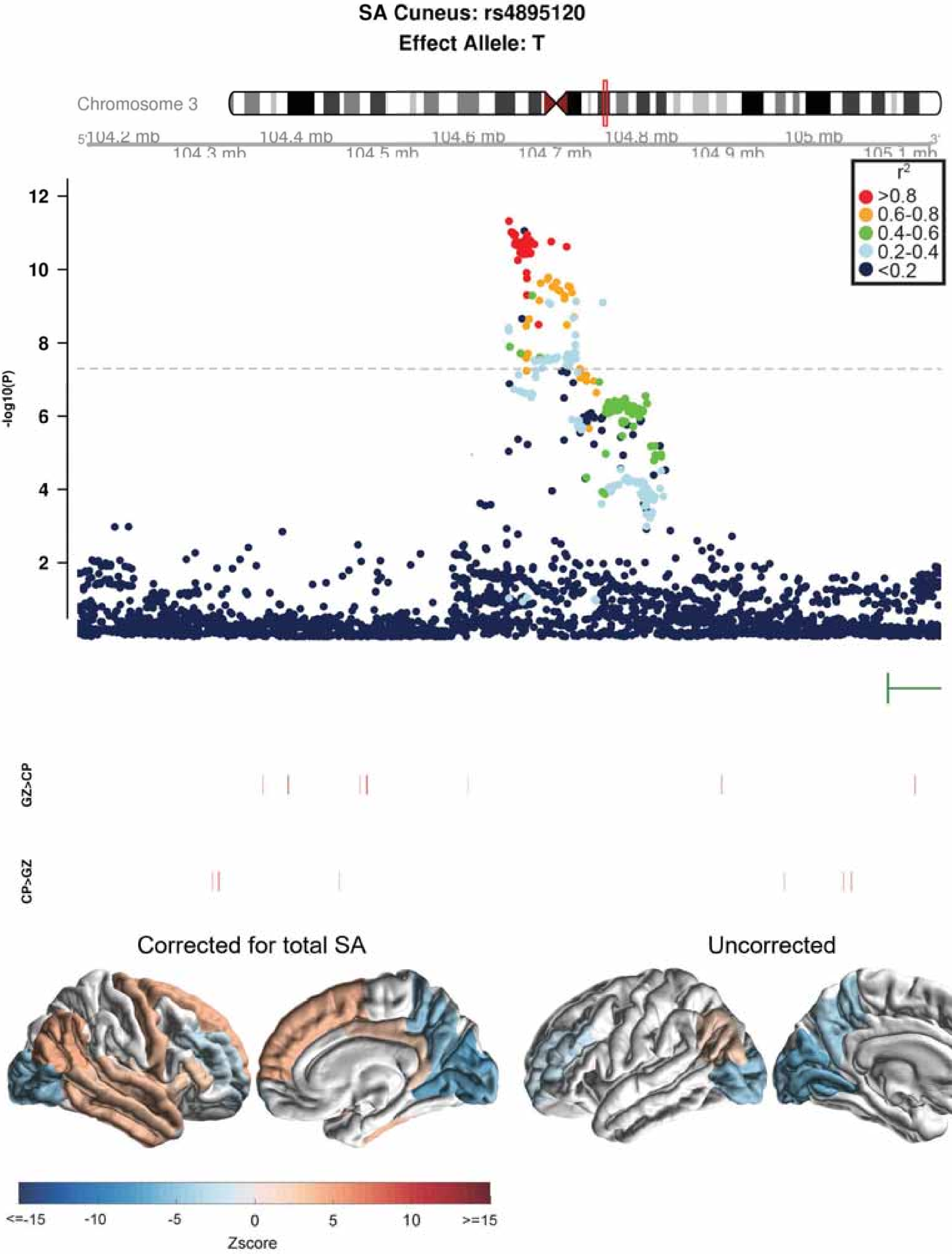
SA Cuneus: rs4895120
Effect Allele: T
Chromosome 3
104.2 mb
104.3 mb
104.4 mb
104.5 mb
104.6 mb
104.7 mb
104.8 mb
104.9 mb
105 mb
105.1 mb
-log10(P)
$r^2$
>0.8
0.6-0.8
0.4-0.6
0.2-0.4
<0.2
GZ>CP
CP>GZ
Corrected for total SA
Uncorrected
<=-15
-10
-5
0
5
10
>=15
Zscore

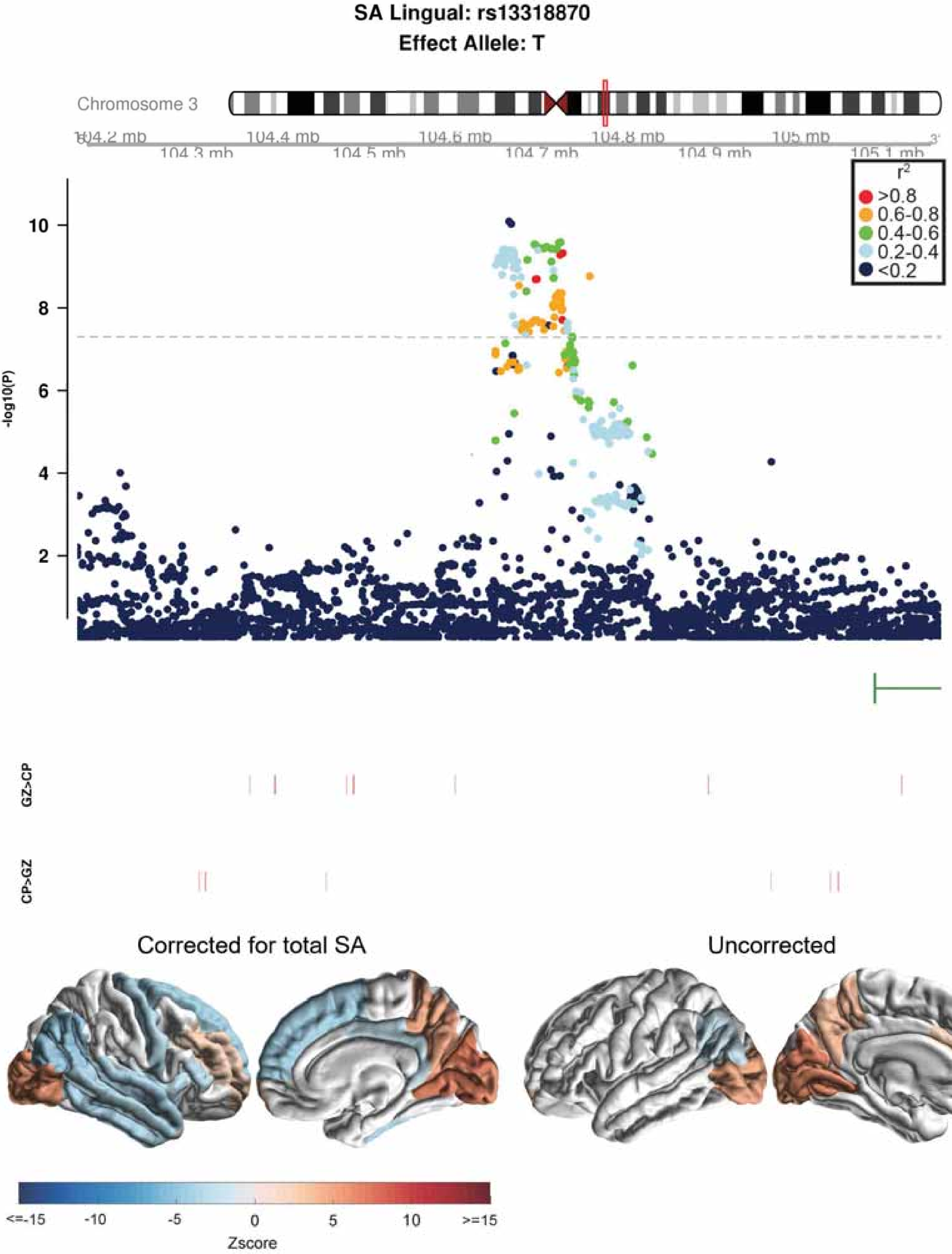

SA Lingual: rs13318870
Effect Allele: T
Chromosome 3
104.2 mb
104.3 mb
104.4 mb
104.5 mb
104.6 mb
104.7 mb
104.8 mb
104.9 mb
105 mb
105.1 mb
r²
>0.8
0.6-0.8
0.4-0.6
0.2-0.4
<0.2
-log10(P)
10
8
6
4
2
GZ>CP
CP>GZ
Corrected for total SA
Uncorrected
<=-15
-10
-5
0
5
10
>=15
Zscore

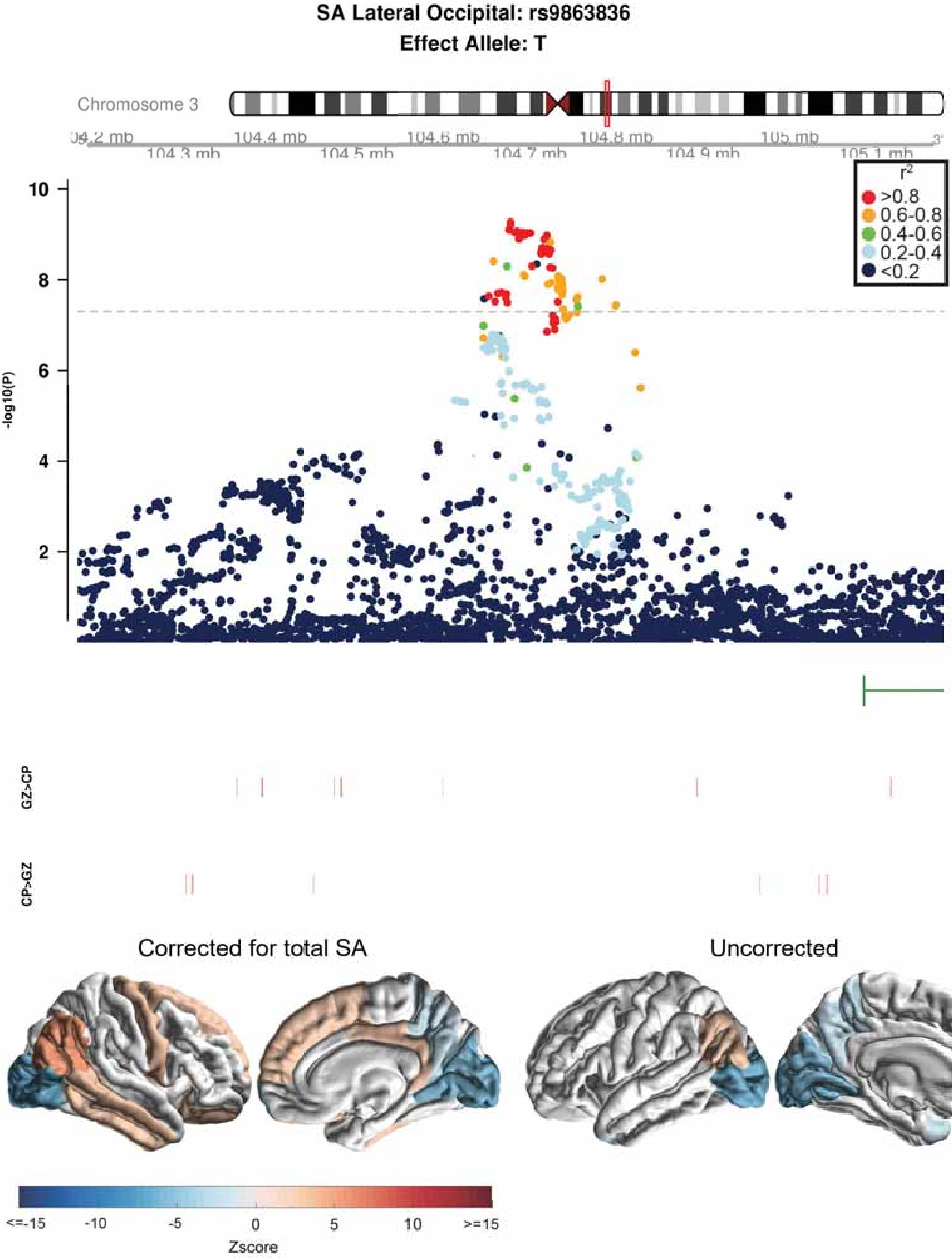

SA Lateral Occipital: rs9863836
Effect Allele: T
Chromosome 3
104.3 mb
104.4 mb
104.5 mb
104.6 mb
104.7 mb
104.8 mb
104.9 mb
105 mb
105.1 mb
-log10(P)
r2
>0.8
0.6-0.8
0.4-0.6
0.2-0.4
<0.2
GZ>CP
CP>GZ
Corrected for total SA
Uncorrected
<=-15
-10
-5
0
5
10
>=15
Zscore

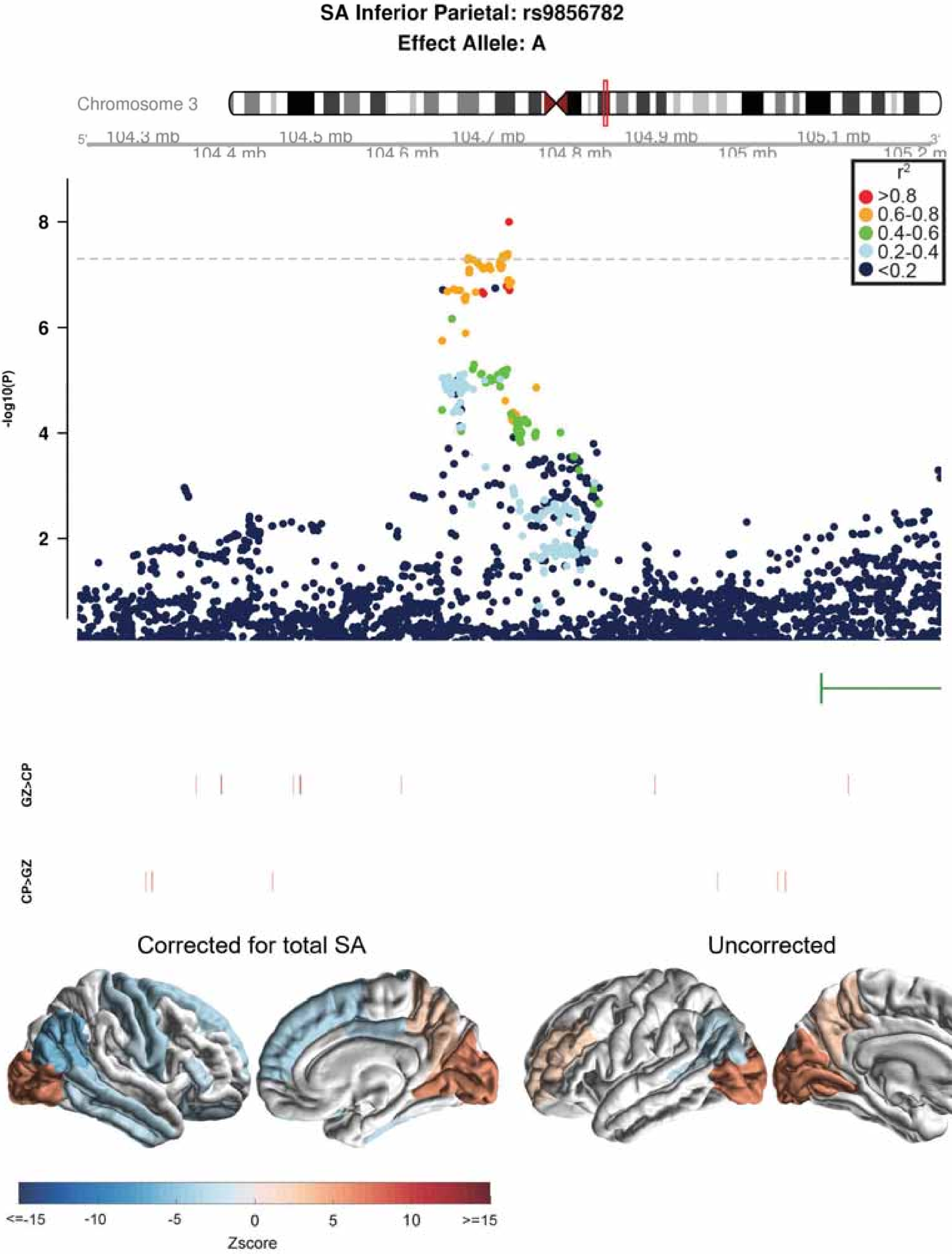
SA Inferior Parietal: rs9856782
Effect Allele: A
Chromosome 3
5'
104.3 mb
104.4 mb
104.5 mb
104.6 mb
104.7 mb
104.8 mb
104.9 mb
105 mb
105.1 mb
105.2 m
3'
$r^2$
>0.8
0.6-0.8
0.4-0.6
0.2-0.4
<0.2
-log10(P)
8
6
4
2
GZ>CP
CP>GZ
Corrected for total SA
Uncorrected
<=-15
-10
-5
0
5
10
>=15
Zscore

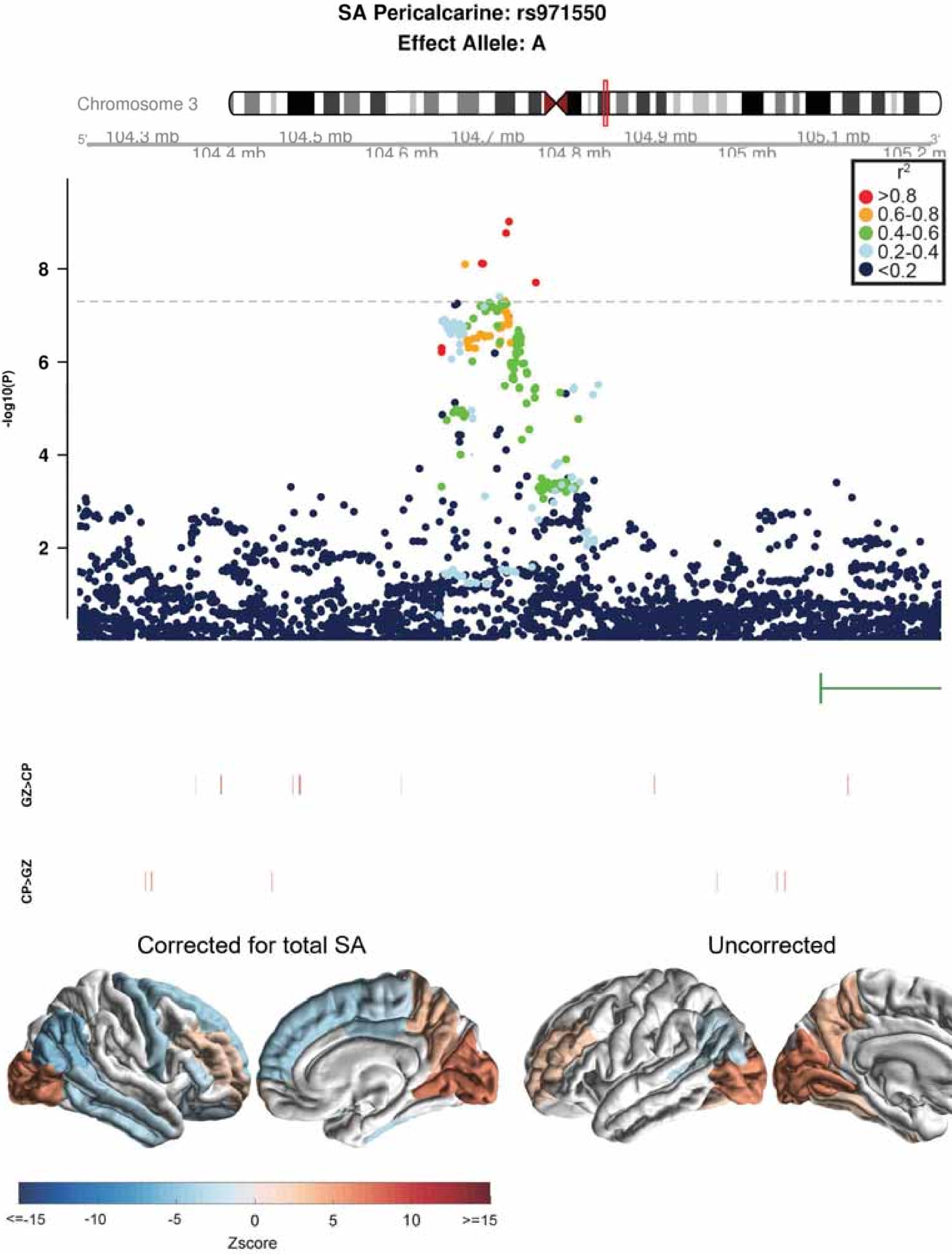

SA Pericalcarine: rs971550
Effect Allele: A
Chromosome 3
104.3 mb
104.4 mb
104.5 mb
104.6 mb
104.7 mb
104.8 mb
104.9 mb
105 mb
105.1 mb
105.2 mb
-log10(P)
2
4
6
8
$r^2$
>0.8
0.6-0.8
0.4-0.6
0.2-0.4
<0.2
GZ>CP
CP>GZ
Corrected for total SA
Uncorrected
<=-15
-10
-5
0
5
10
>=15
Zscore

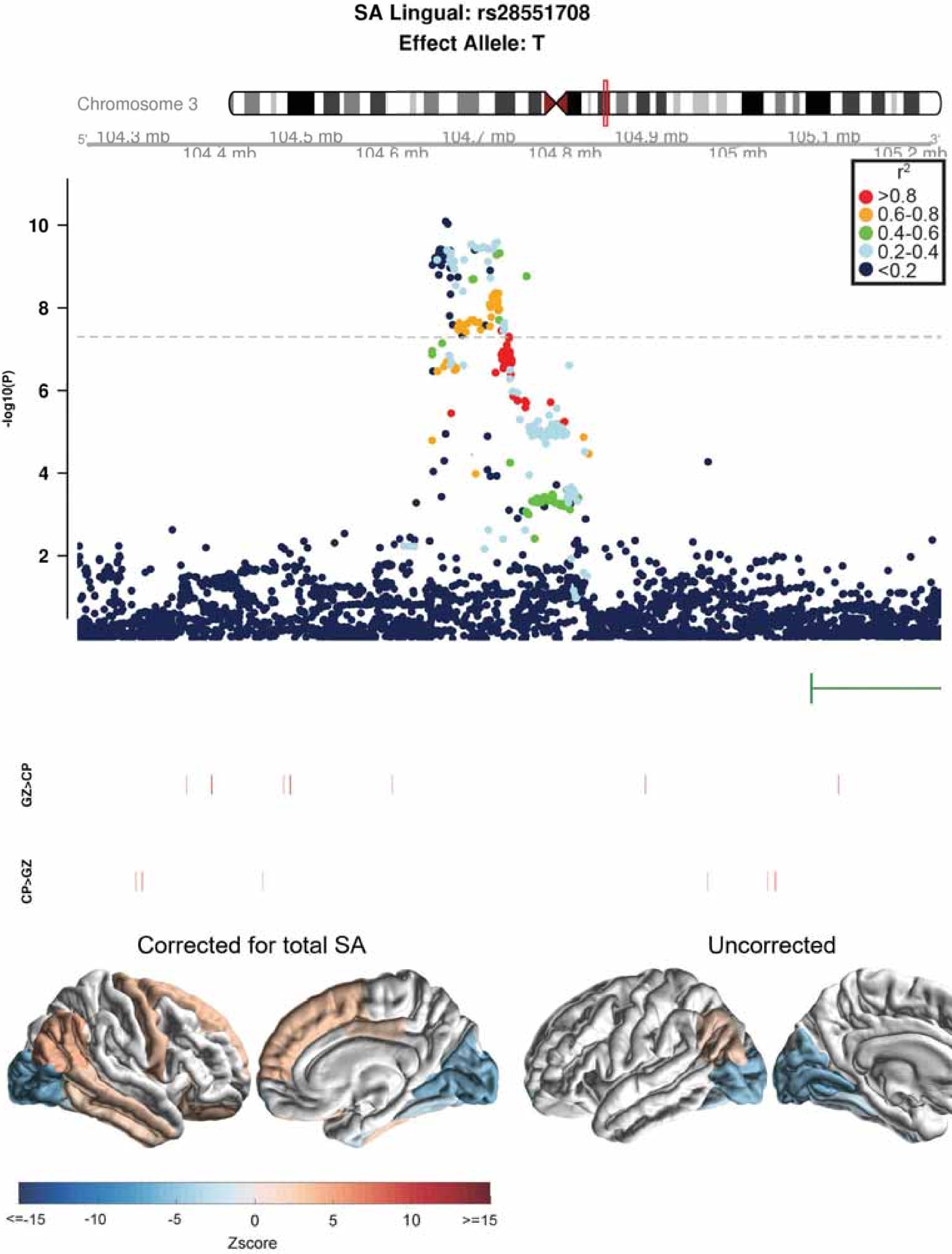

SA Lingual: rs28551708
Effect Allele: T
Chromosome 3
104.3 mb
104.4 mb
104.5 mb
104.6 mb
104.7 mb
104.8 mb
104.9 mb
105 mb
105.1 mb
105.2 mb
r2
>0.8
0.6-0.8
0.4-0.6
0.2-0.4
<0.2
-log10(P)
GZ>CP
CP>GZ
Corrected for total SA
Uncorrected
<=-15
-10
-5
0
5
10
>=15
Zscore

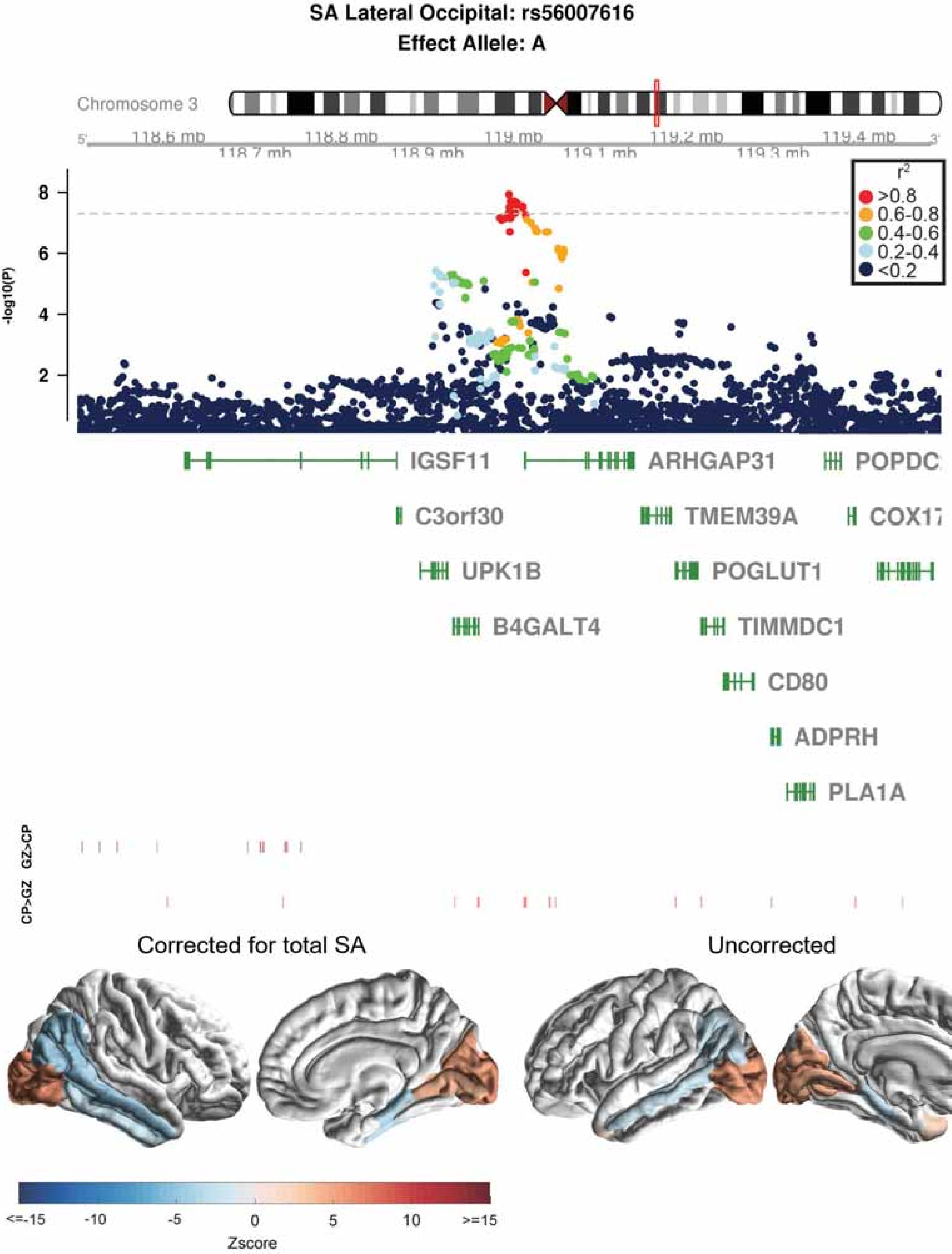

SA Lateral Occipital: rs56007616
Effect Allele: A
Chromosome 3
118.6 mb
118.7 mb
118.8 mb
118.9 mb
119 mb
119.1 mb
119.2 mb
119.3 mb
119.4 mb
-log10(P)
r2
>0.8
0.6-0.8
0.4-0.6
0.2-0.4
<0.2
IGSF11
ARHGAP31
POPDC
C3orf30
TMEM39A
COX17
UPK1B
POGLUT1
B4GALT4
TIMMDC1
CD80
ADPRH
PLA1A
GZ>CP
CP>GZ
Corrected for total SA
Uncorrected
<=-15
-10
-5
0
5
10
>=15
Zscore

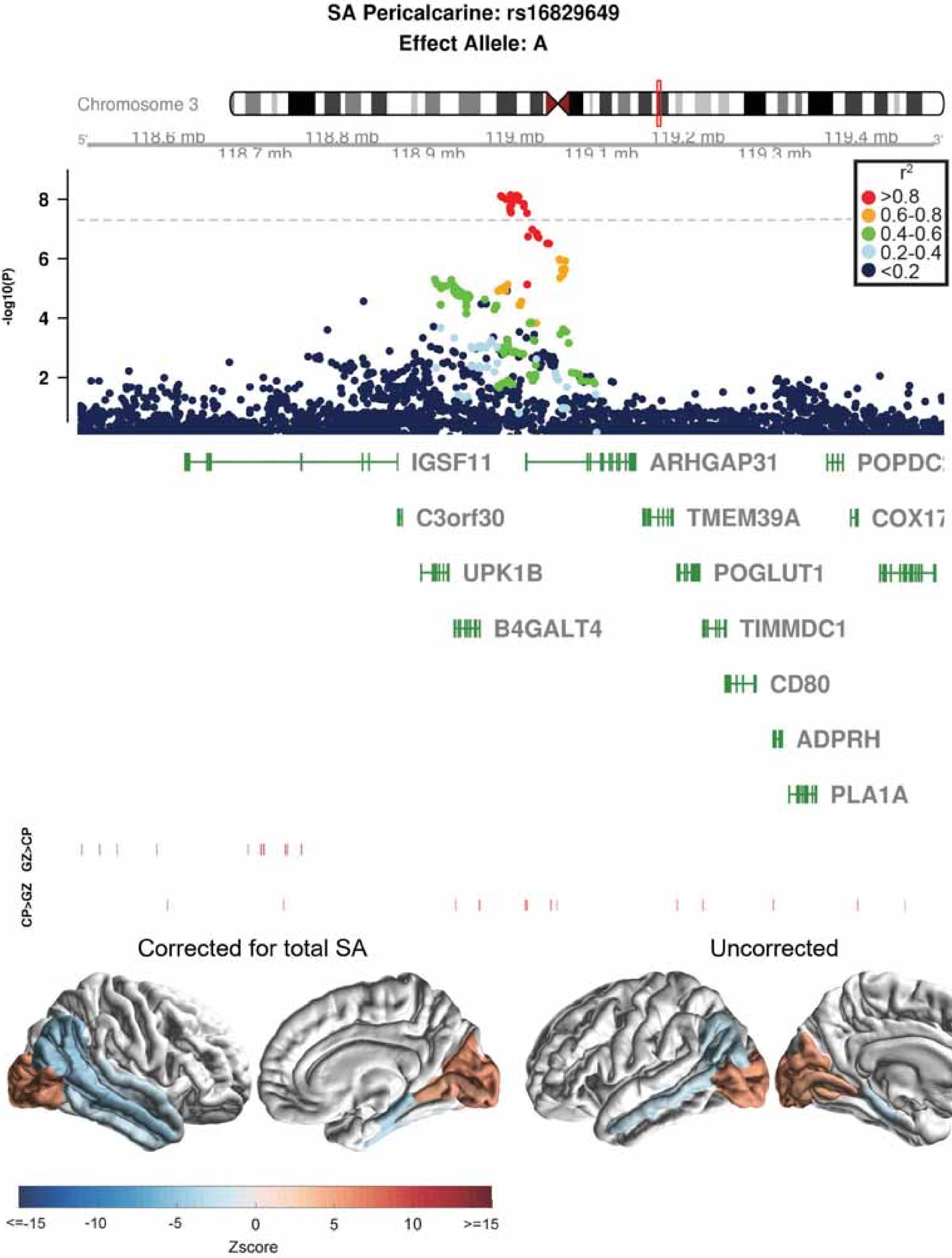

SA Pericalcarine: rs16829649
Effect Allele: A
Chromosome 3
118.6 mb
118.7 mb
118.8 mb
118.9 mb
119 mb
119.1 mb
119.2 mb
119.3 mb
119.4 mb
-log10(P)
r2
>0.8
0.6-0.8
0.4-0.6
0.2-0.4
<0.2
IGSF11
ARHGAP31
POPDC
C3orf30
TMEM39A
COX17
UPK1B
POGLUT1
B4GALT4
TIMMDC1
CD80
ADPRH
PLA1A
GZ>CP
CP>GZ
Corrected for total SA
Uncorrected
<=-15
-10
-5
0
5
10
>=15
Zscore

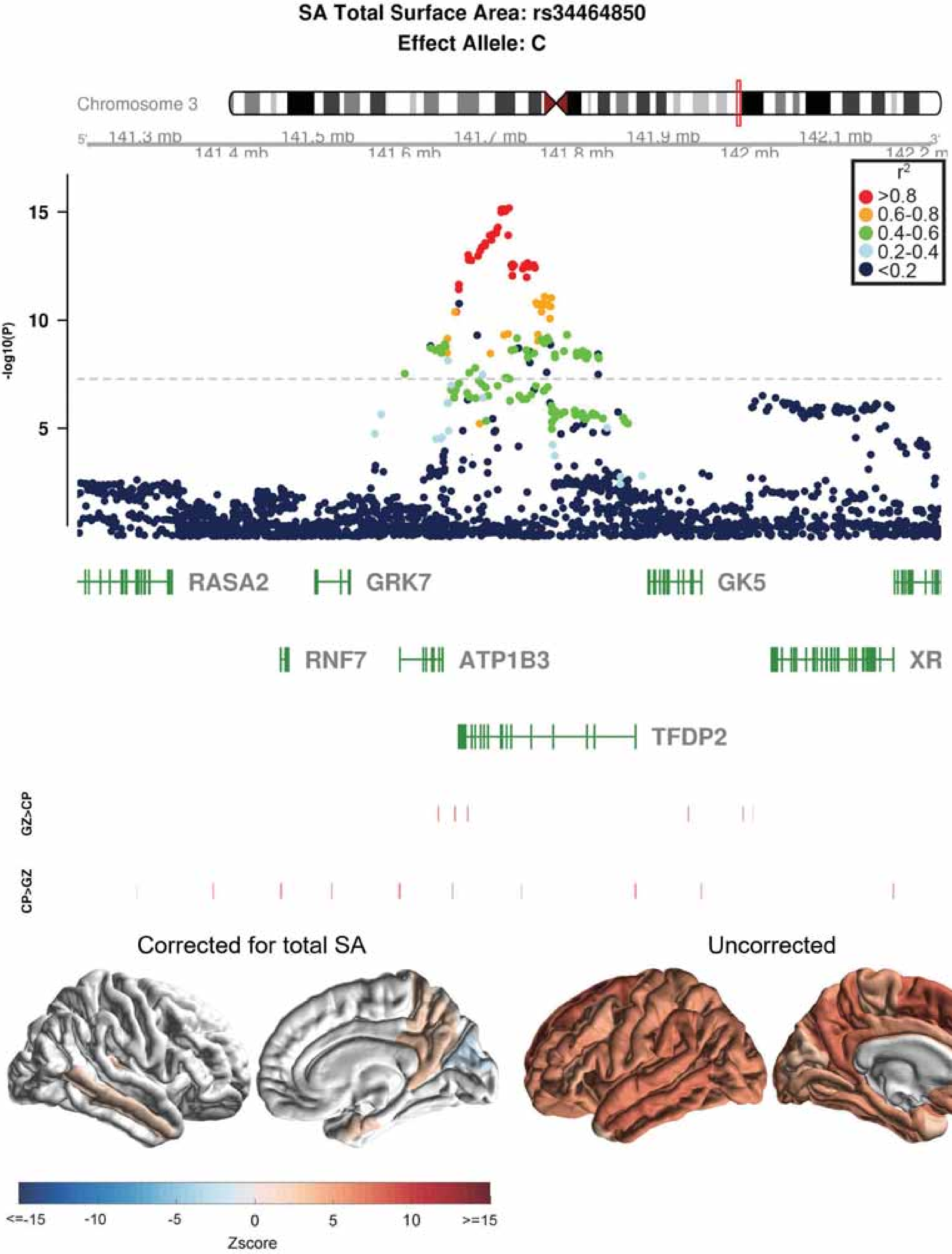

SA Total Surface Area: rs34464850
Effect Allele: C
Chromosome 3
141.3 mb
141.4 mb
141.5 mb
141.6 mb
141.7 mb
141.8 mb
141.9 mb
142 mb
142.1 mb
142.2 mb
$r^2$
>0.8
0.6-0.8
0.4-0.6
0.2-0.4
<0.2
-log10(P)
15
10
5
RASA2
GRK7
GK5
RNF7
ATP1B3
XR
TFDP2
GZ>CP
CP>GZ
Corrected for total SA
Uncorrected
<=-15
-10
-5
0
5
10
>=15
Zscore

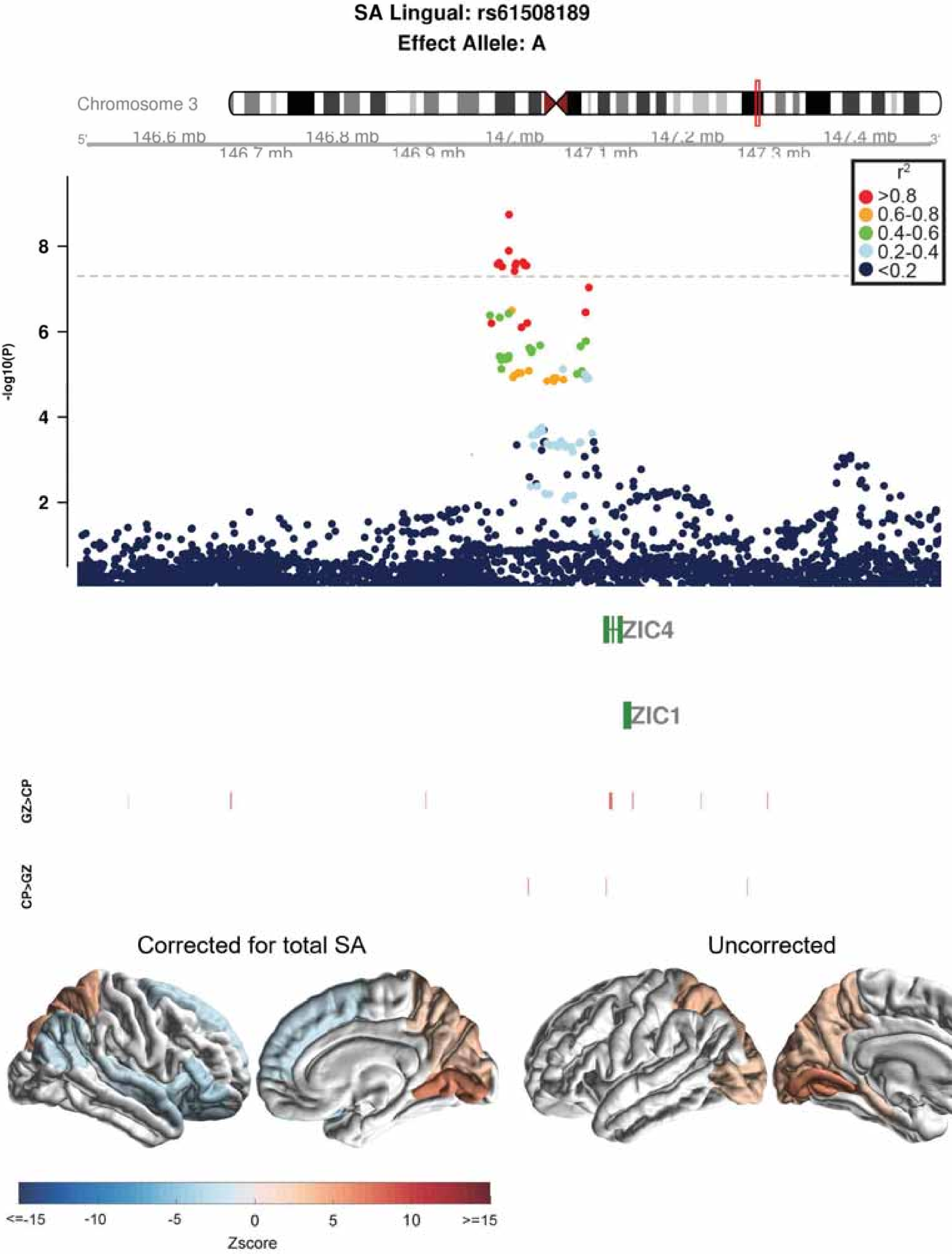
SA Lingual: rs61508189
Effect Allele: A
Chromosome 3
146.6 mb
146.7 mb
146.8 mb
146.9 mb
147 mb
147.1 mb
147.2 mb
147.3 mb
147.4 mb
r2
>0.8
0.6-0.8
0.4-0.6
0.2-0.4
<0.2
-log10(P)
ZIC4
ZIC1
GZ>CP
CP>GZ
Corrected for total SA
Uncorrected
<=-15
-10
-5
0
5
10
>=15
Zscore

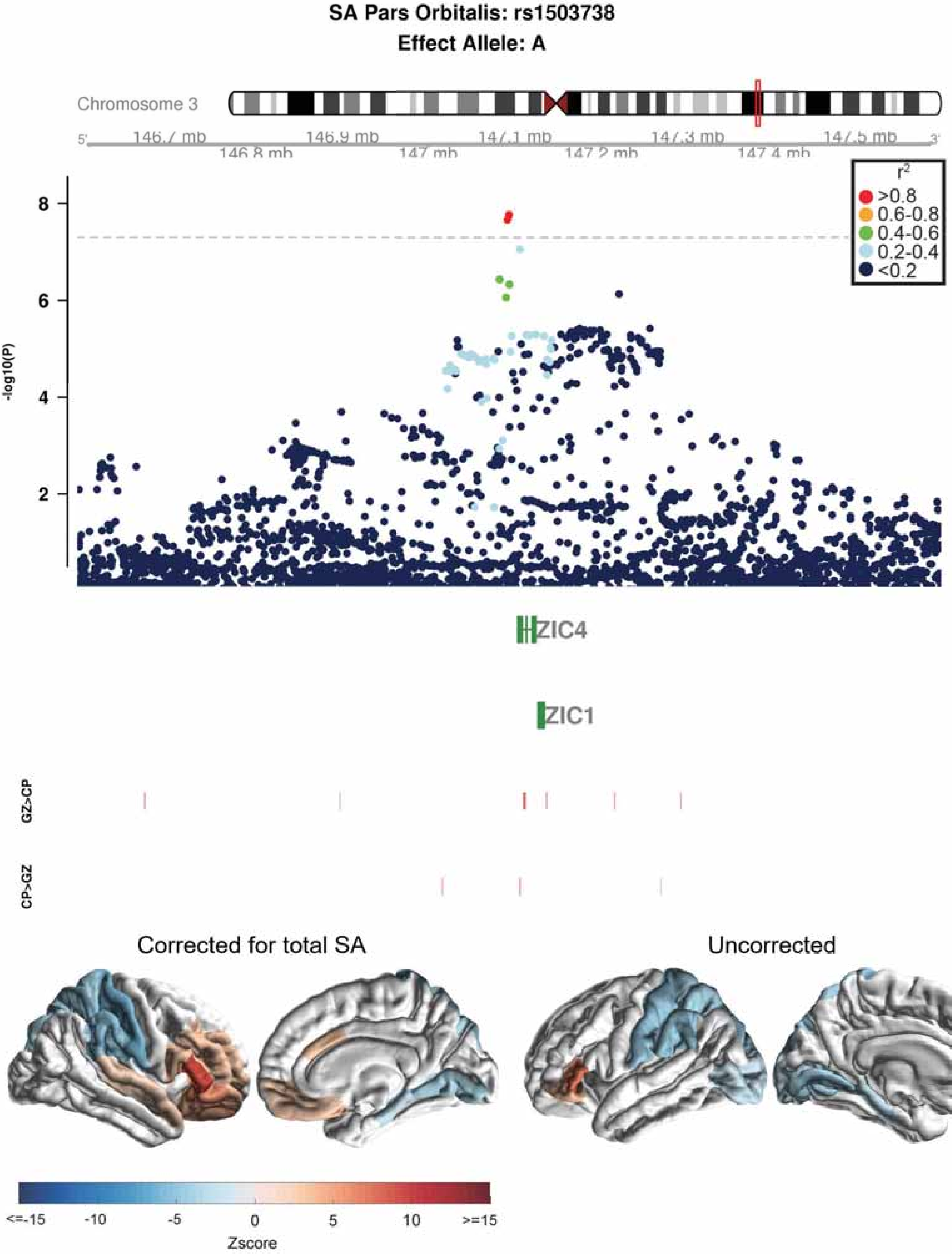

SA Pars Orbitalis: rs1503738
Effect Allele: A
Chromosome 3
146.8 mb
146.9 mb
147 mb
147.2 mb
147.3 mb
147.4 mb
r2
>0.8
0.6-0.8
0.4-0.6
0.2-0.4
<0.2
-log10(P)
8
6
4
2
ZIC4
ZIC1
GZ>CP
CP>GZ
Corrected for total SA
Uncorrected
<=-15
-10
-5
0
5
10
>=15
Zscore

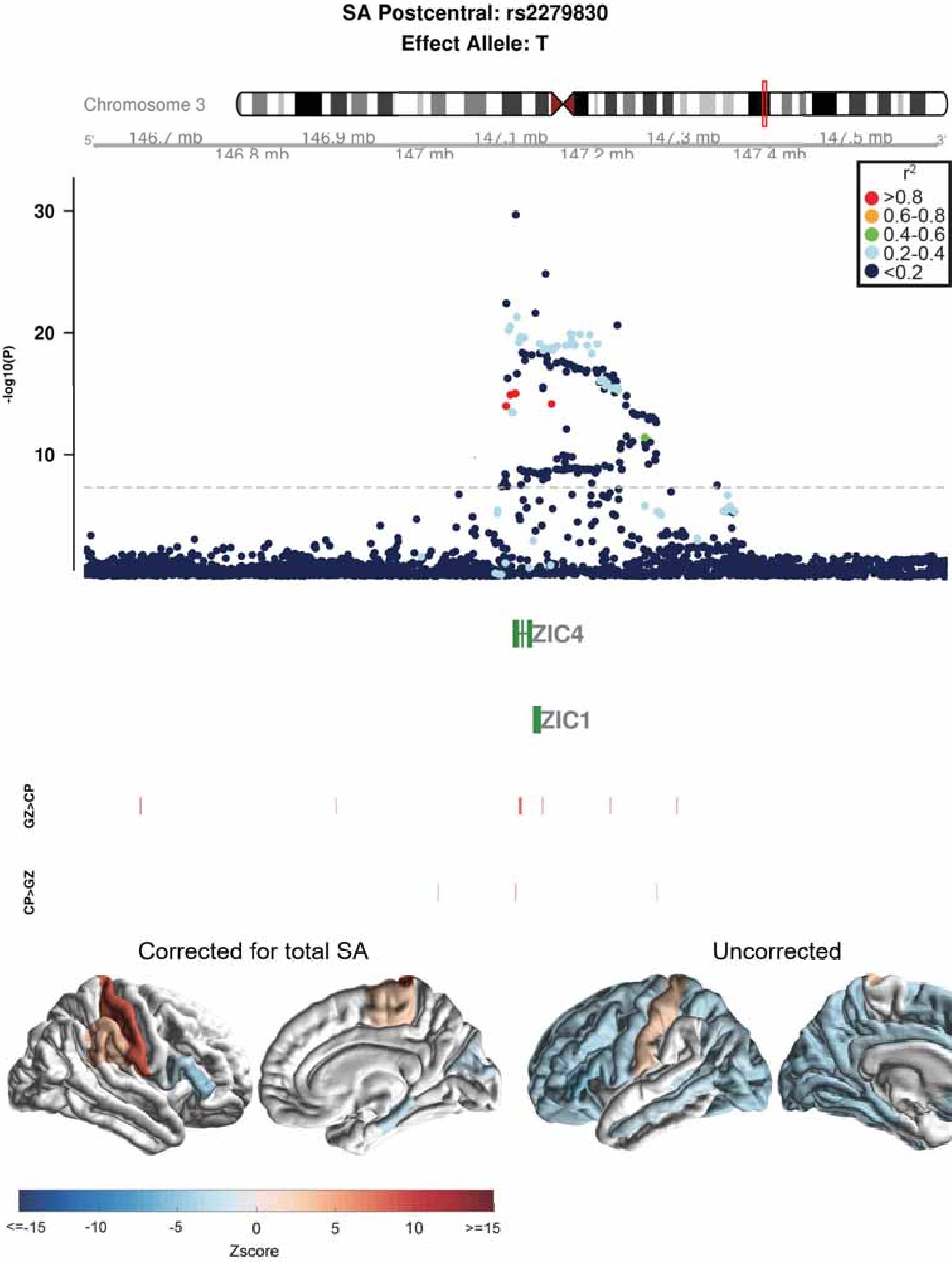
SA Postcentral: rs2279830
Effect Allele: T
Chromosome 3
146.7 mb
146.8 mb
146.9 mb
147 mb
147.1 mb
147.2 mb
147.3 mb
147.4 mb
147.5 mb
r2
>0.8
0.6-0.8
0.4-0.6
0.2-0.4
<0.2
-log10(P)
30
20
10
ZIC4
ZIC1
GZ>CP
CP>GZ
Corrected for total SA
Uncorrected
<=-15
-10
-5
0
5
10
>=15
Zscore

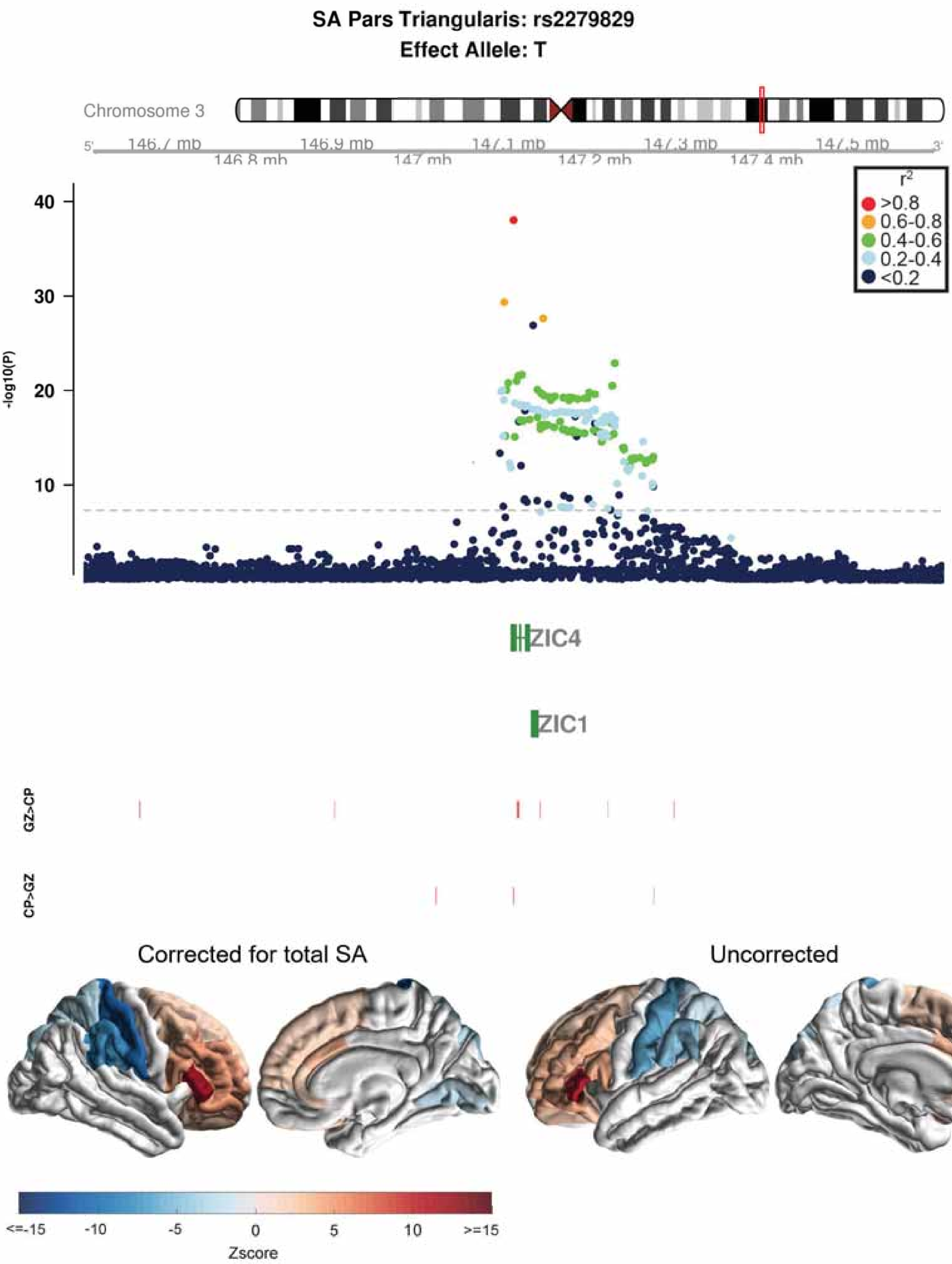
SA Pars Triangularis: rs2279829
Effect Allele: T
Chromosome 3
146.7 mb
146.8 mb
146.9 mb
147 mb
147.1 mb
147.2 mb
147.3 mb
147.4 mb
147.5 mb
$r^2$
>0.8
0.6-0.8
0.4-0.6
0.2-0.4
<0.2
-log10(P)
ZIC4
ZIC1
GZ>CP
CP>GZ
Corrected for total SA
Uncorrected
<=-15
-10
-5
0
5
10
>=15
Zscore

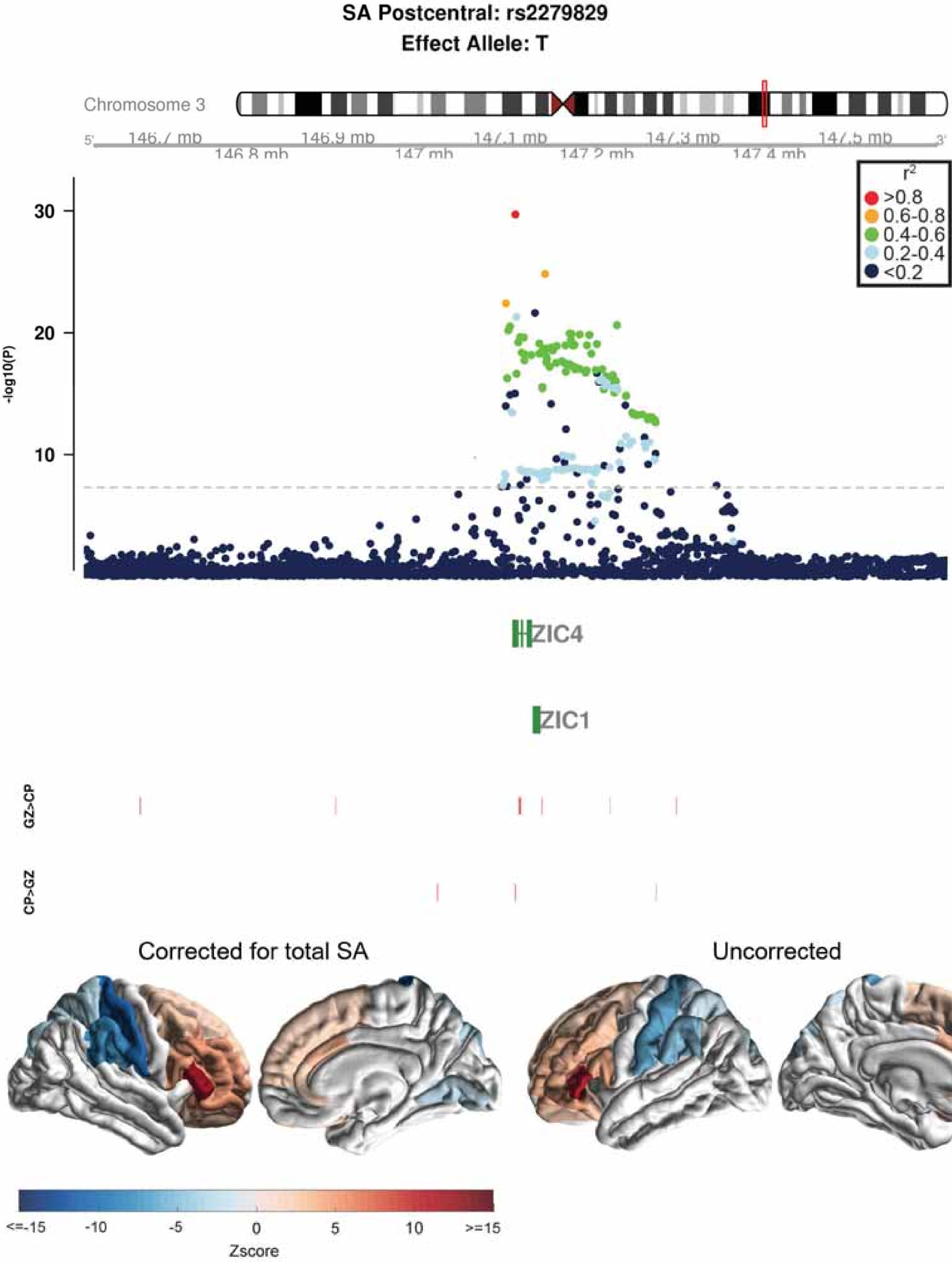

SA Postcentral: rs2279829
Effect Allele: T
Chromosome 3
146.7 mb
146.8 mb
146.9 mb
147 mb
147.1 mb
147.2 mb
147.3 mb
147.4 mb
147.5 mb
r2
>0.8
0.6-0.8
0.4-0.6
0.2-0.4
<0.2
-log10(P)
30
20
10
ZIC4
ZIC1
GZ>CP
CP>GZ
Corrected for total SA
Uncorrected
<=-15
-10
-5
0
5
10
>=15
Zscore

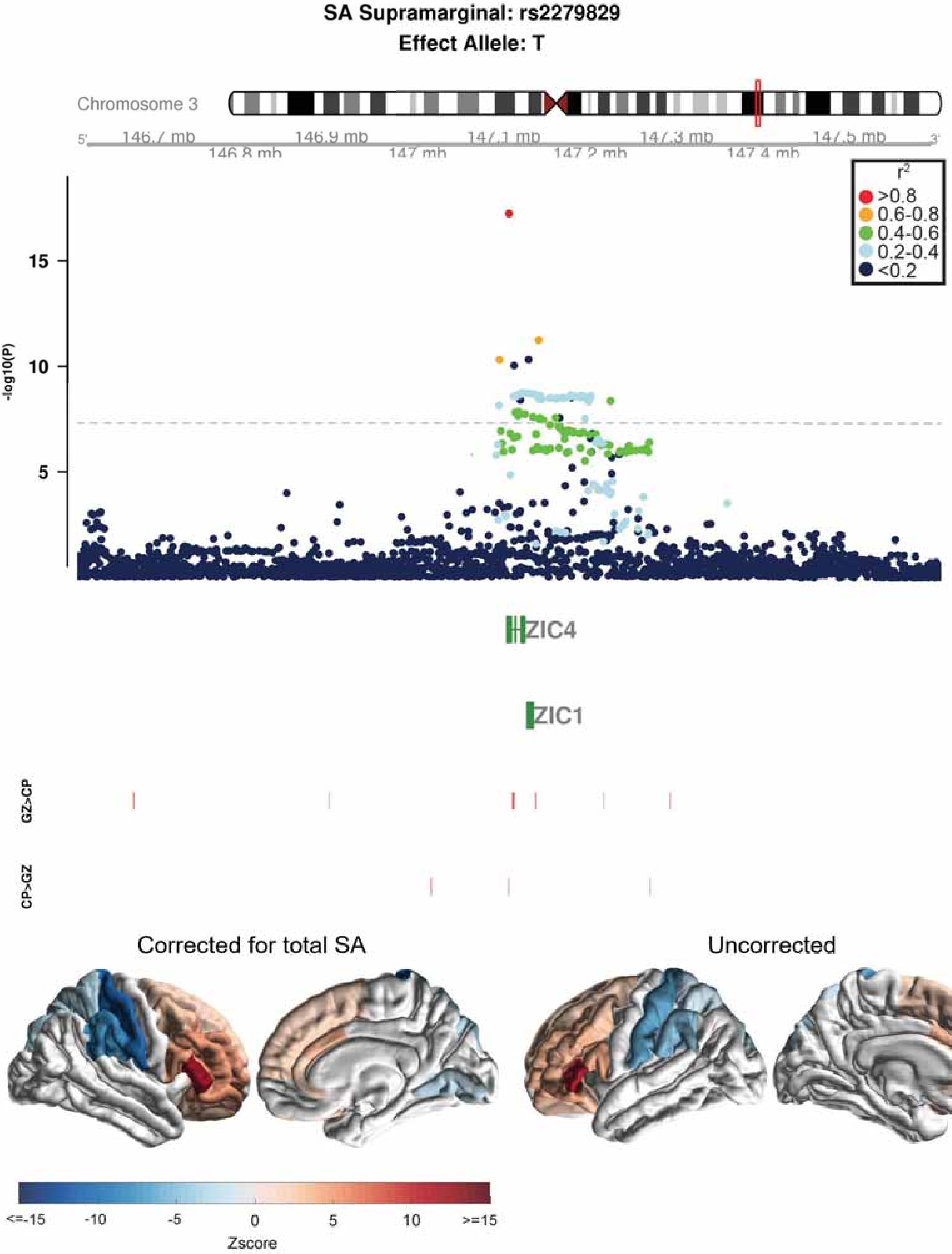

SA Supramarginal: rs2279829
Effect Allele: T
Chromosome 3
5'
146.7 mb
146.8 mb
146.9 mb
147 mb
147.1 mb
147.2 mb
147.3 mb
147.4 mb
147.5 mb
3'
$r^2$
>0.8
0.6-0.8
0.4-0.6
0.2-0.4
<0.2
-log10(P)
15
10
5
ZIC4
ZIC1
GZ>CP
CP>GZ
Corrected for total SA
Uncorrected
<=-15
-10
-5
0
5
10
>=15
Zscore

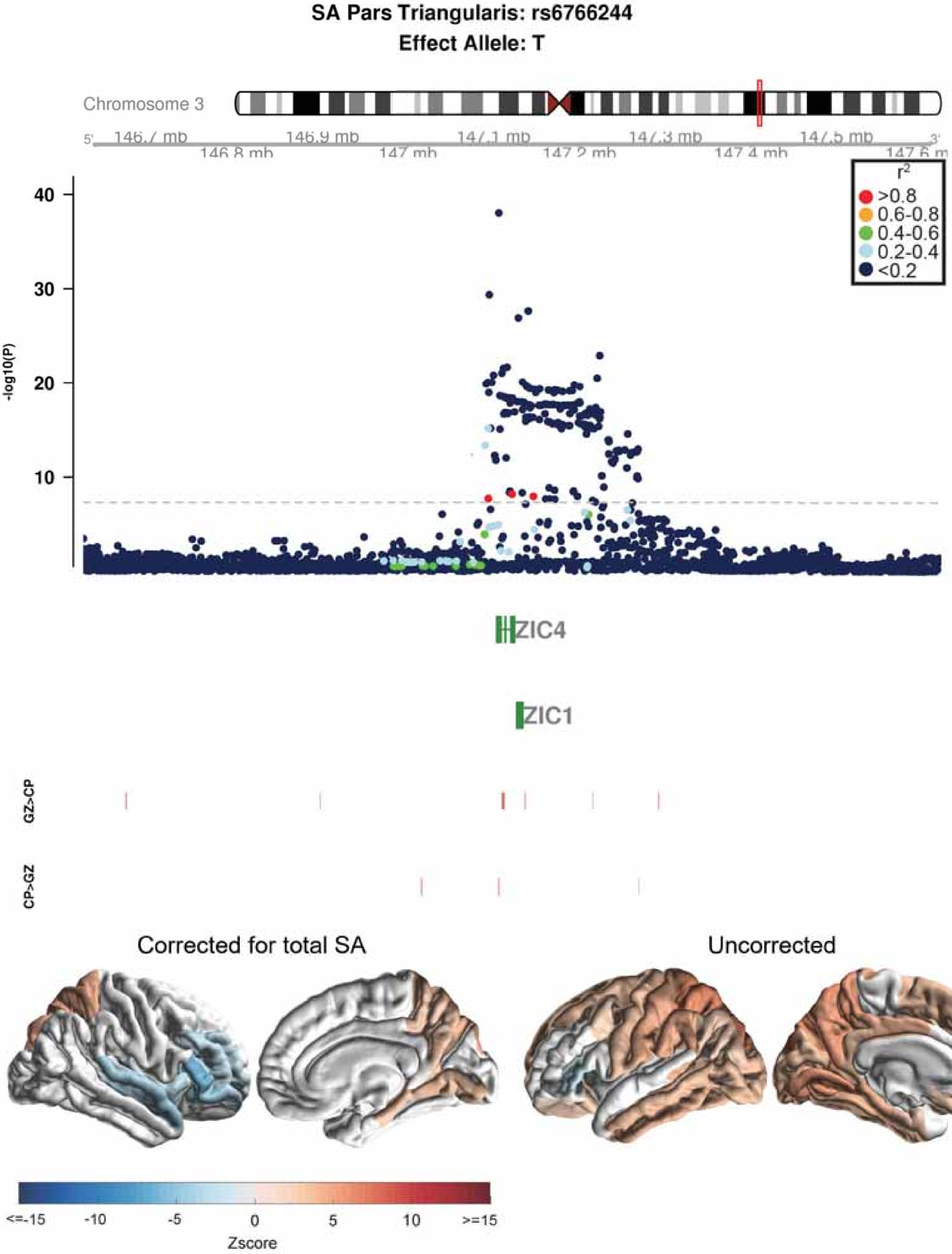

SA Pars Triangularis: rs6766244
Effect Allele: T
Chromosome 3
5'
146.7 mb
146.8 mb
146.9 mb
147 mb
147.1 mb
147.2 mb
147.3 mb
147.4 mb
147.5 mb
147.6 m
3'
-log10(P)
40
30
20
10
r2
>0.8
0.6-0.8
0.4-0.6
0.2-0.4
<0.2
ZIC4
ZIC1
GZ>CP
CP>GZ
Corrected for total SA
Uncorrected
<=-15
-10
-5
0
5
10
>=15
Zscore

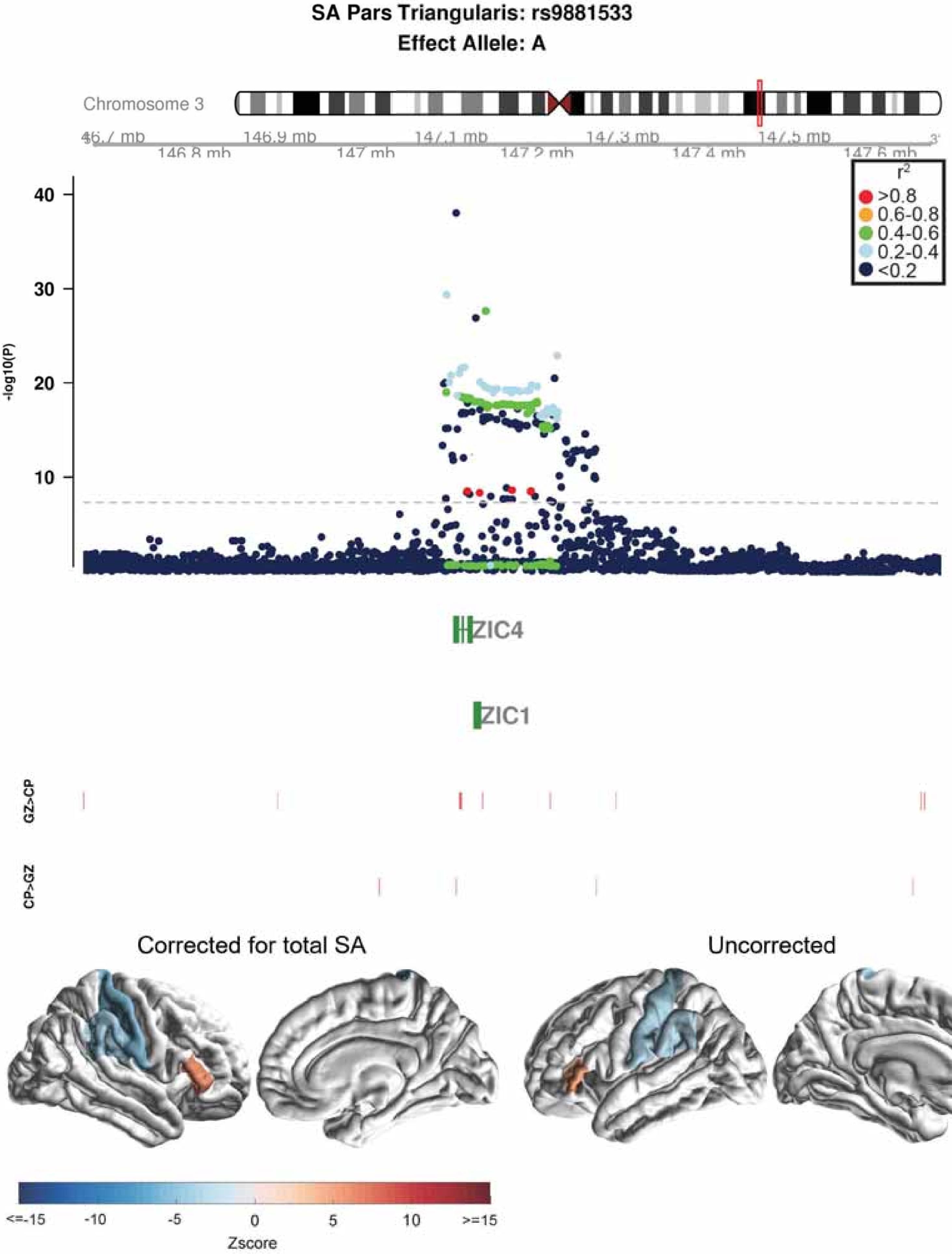

SA Pars Triangularis: rs9881533
Effect Allele: A
Chromosome 3
146.7 mb
146.8 mb
146.9 mb
147 mb
147.1 mb
147.2 mb
147.3 mb
147.4 mb
147.5 mb
147.6 mb
r2
>0.8
0.6-0.8
0.4-0.6
0.2-0.4
<0.2
-log10(P)
40
30
20
10
ZIC4
ZIC1
GZ>CP
CP>GZ
Corrected for total SA
Uncorrected
<=-15
-10
-5
0
5
10
>=15
Zscore

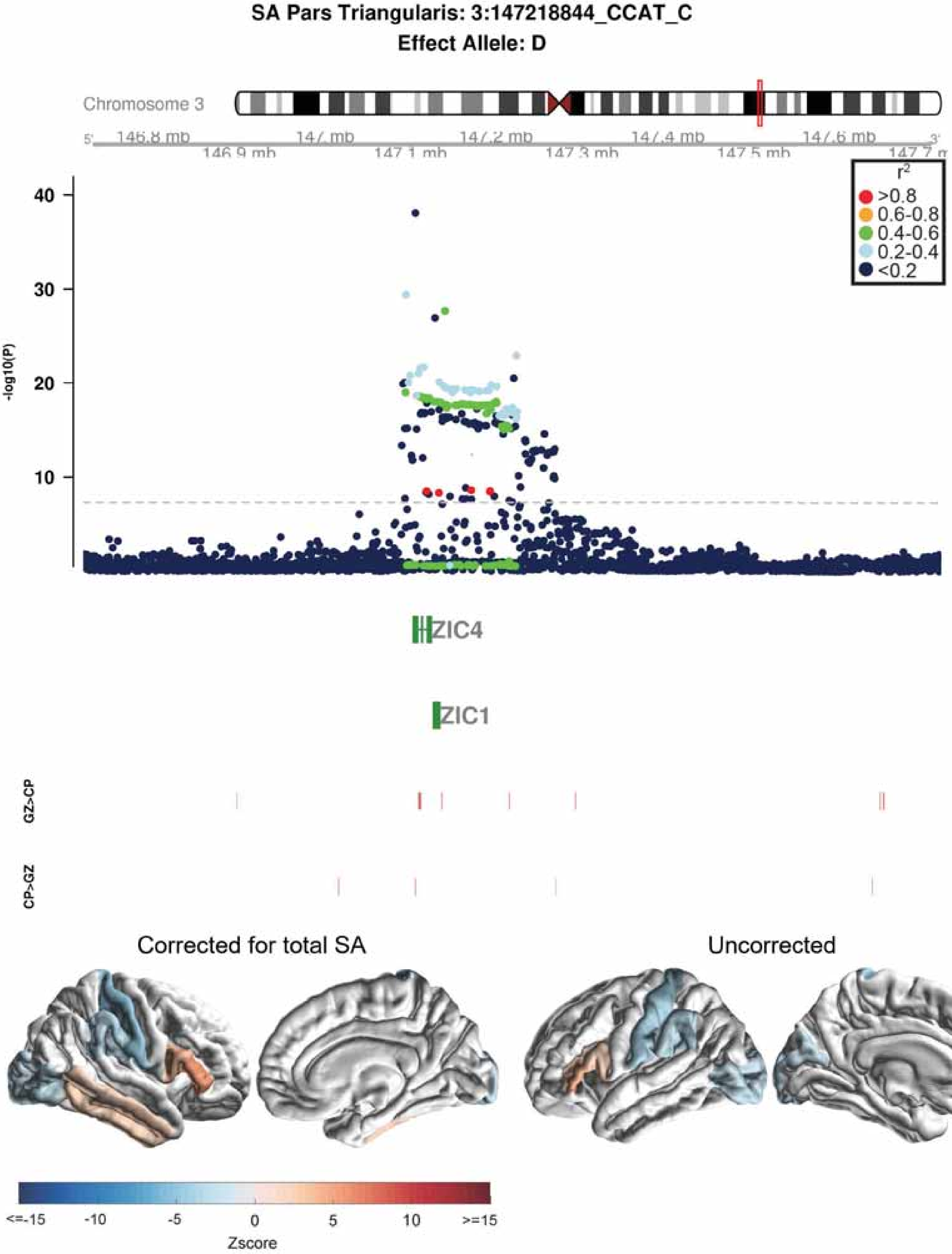
SA Pars Triangularis: 3:147218844_CCAT_C
Effect Allele: D
Chromosome 3
146.8 mb
146.9 mb
147 mb
147.1 mb
147.2 mb
147.3 mb
147.4 mb
147.5 mb
147.6 mb
r2
>0.8
0.6-0.8
0.4-0.6
0.2-0.4
<0.2
-log10(P)
40
30
20
10
ZIC4
ZIC1
GZ>CP
CP>GZ
Corrected for total SA
Uncorrected
<=-15
-10
-5
0
5
10
>=15
Zscore

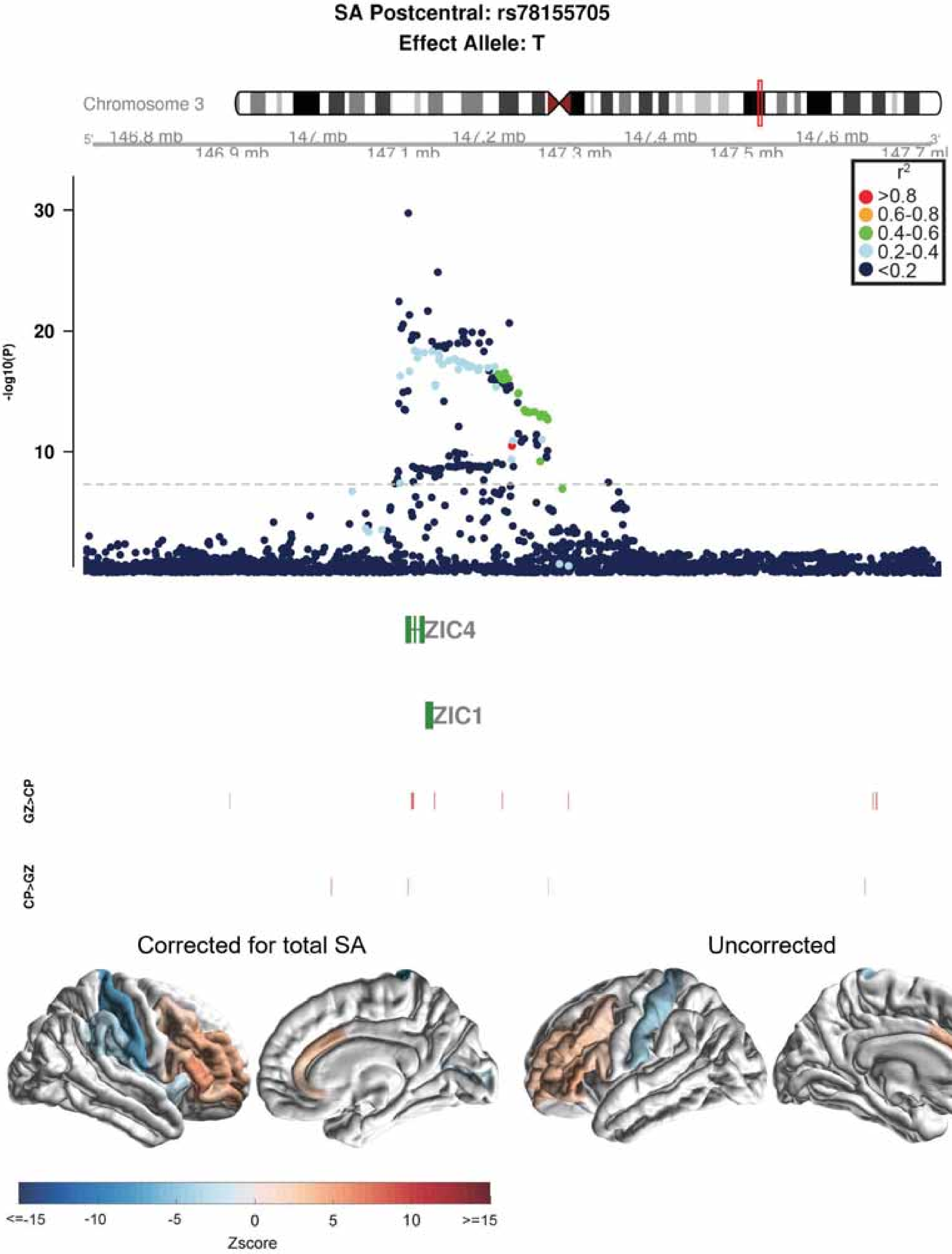
SA Postcentral: rs78155705
Effect Allele: T
Chromosome 3
146.8 mb
146.9 mb
147 mb
147.1 mb
147.2 mb
147.3 mb
147.4 mb
147.5 mb
147.6 mb
147.7 mb
r2
>0.8
0.6-0.8
0.4-0.6
0.2-0.4
<0.2
-log10(P)
30
20
10
ZIC4
ZIC1
GZ>CP
CP>GZ
Corrected for total SA
Uncorrected
<=-15
-10
-5
0
5
10
>=15
Zscore

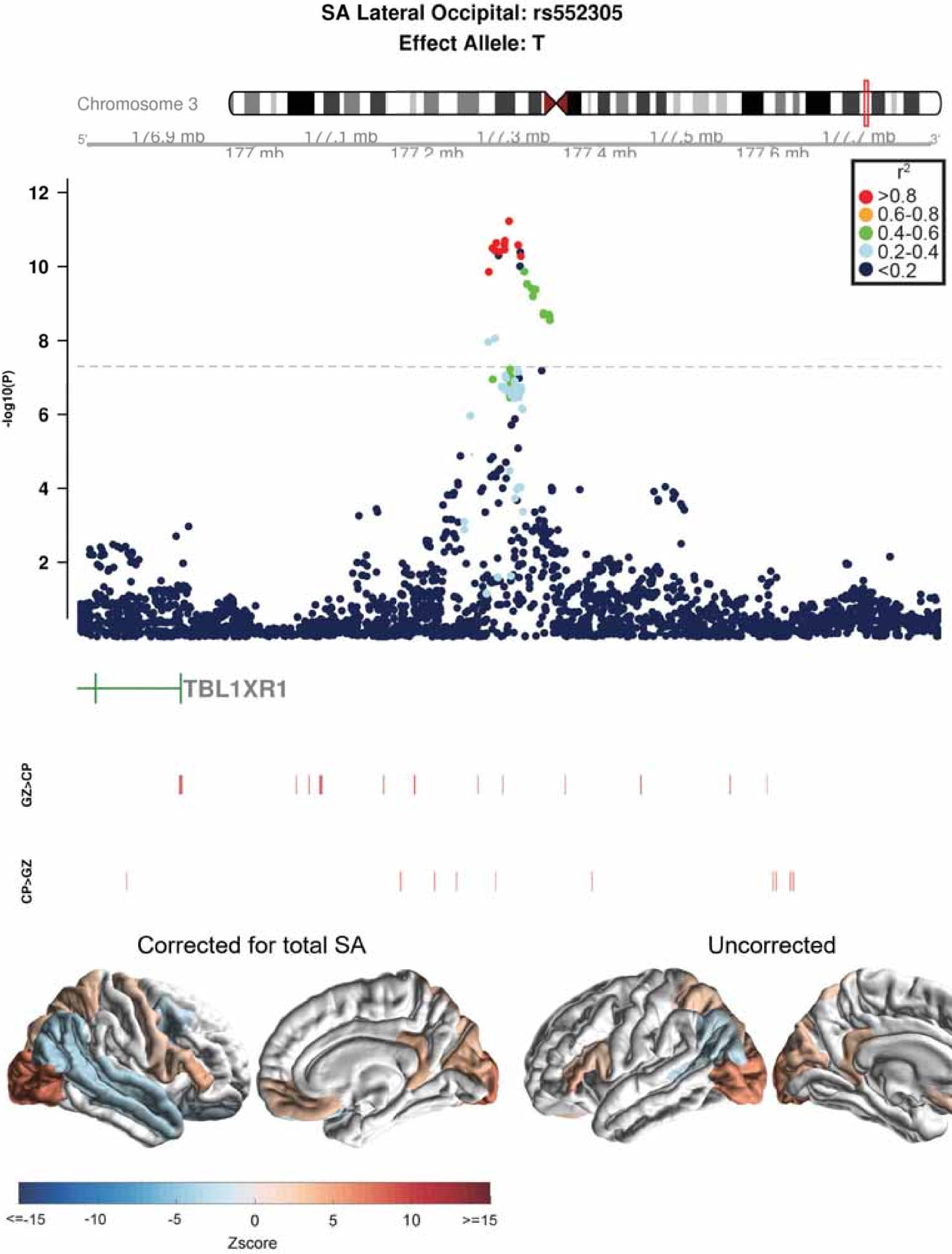
SA Lateral Occipital: rs552305
Effect Allele: T
Chromosome 3
177 mb
177.2 mb
177.4 mb
177.6 mb
r2
>0.8
0.6-0.8
0.4-0.6
0.2-0.4
<0.2
-log10(P)
TBL1XR1
GZ>CP
CP>GZ
Corrected for total SA
Uncorrected
<=-15
-10
-5
0
5
10
>=15
Zscore

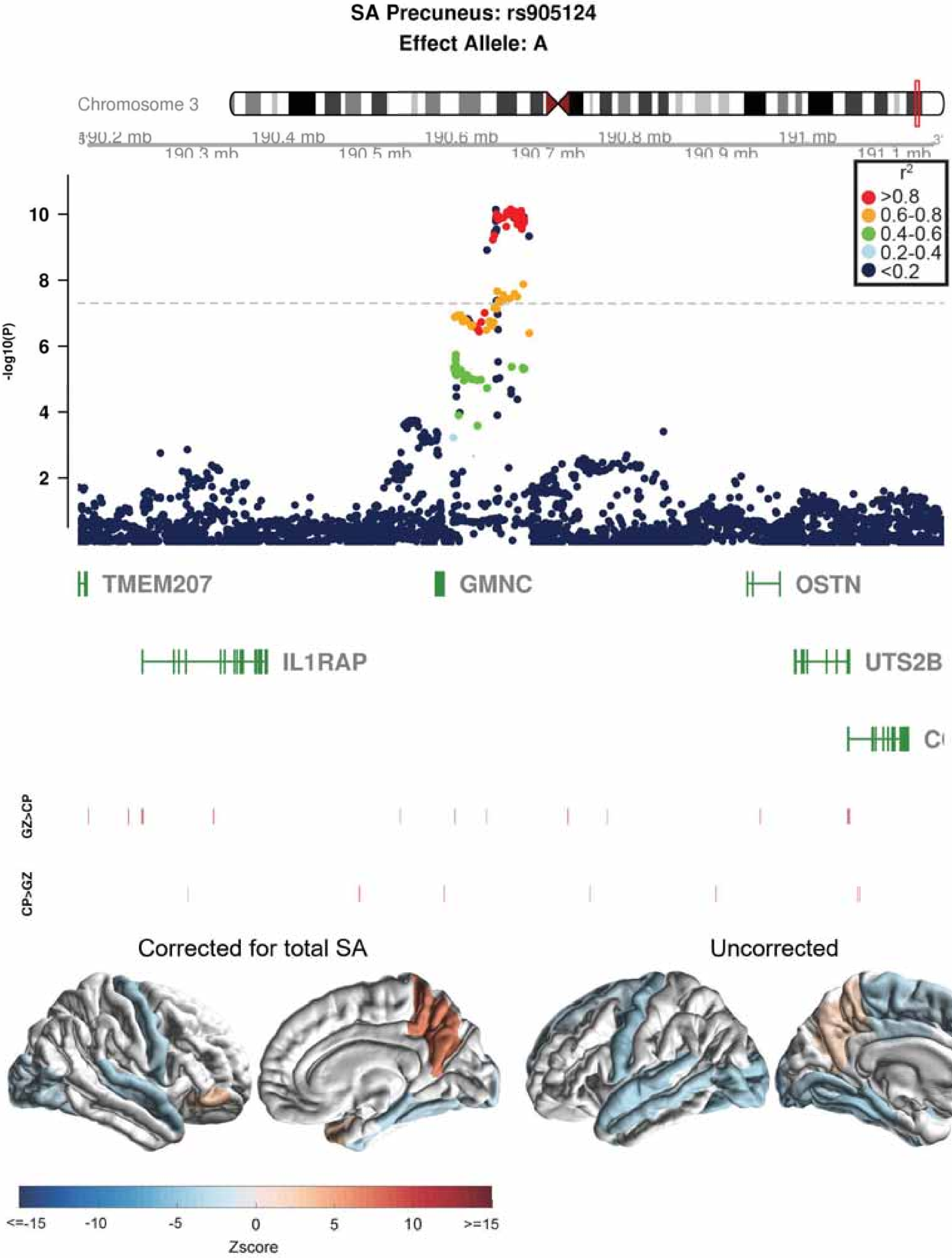

SA Precuneus: rs905124
Effect Allele: A
Chromosome 3
190.2 mb
190.3 mb
190.4 mb
190.5 mb
190.6 mb
190.7 mb
190.8 mb
190.9 mb
191 mb
191.1 mb
r2
>0.8
0.6-0.8
0.4-0.6
0.2-0.4
<0.2
-log10(P)
10
8
6
4
2
TMEM207
GMNC
OSTN
IL1RAP
UTS2B
GZ>CP
CP>GZ
Corrected for total SA
Uncorrected
<=-15
-10
-5
0
5
10
>=15
Zscore

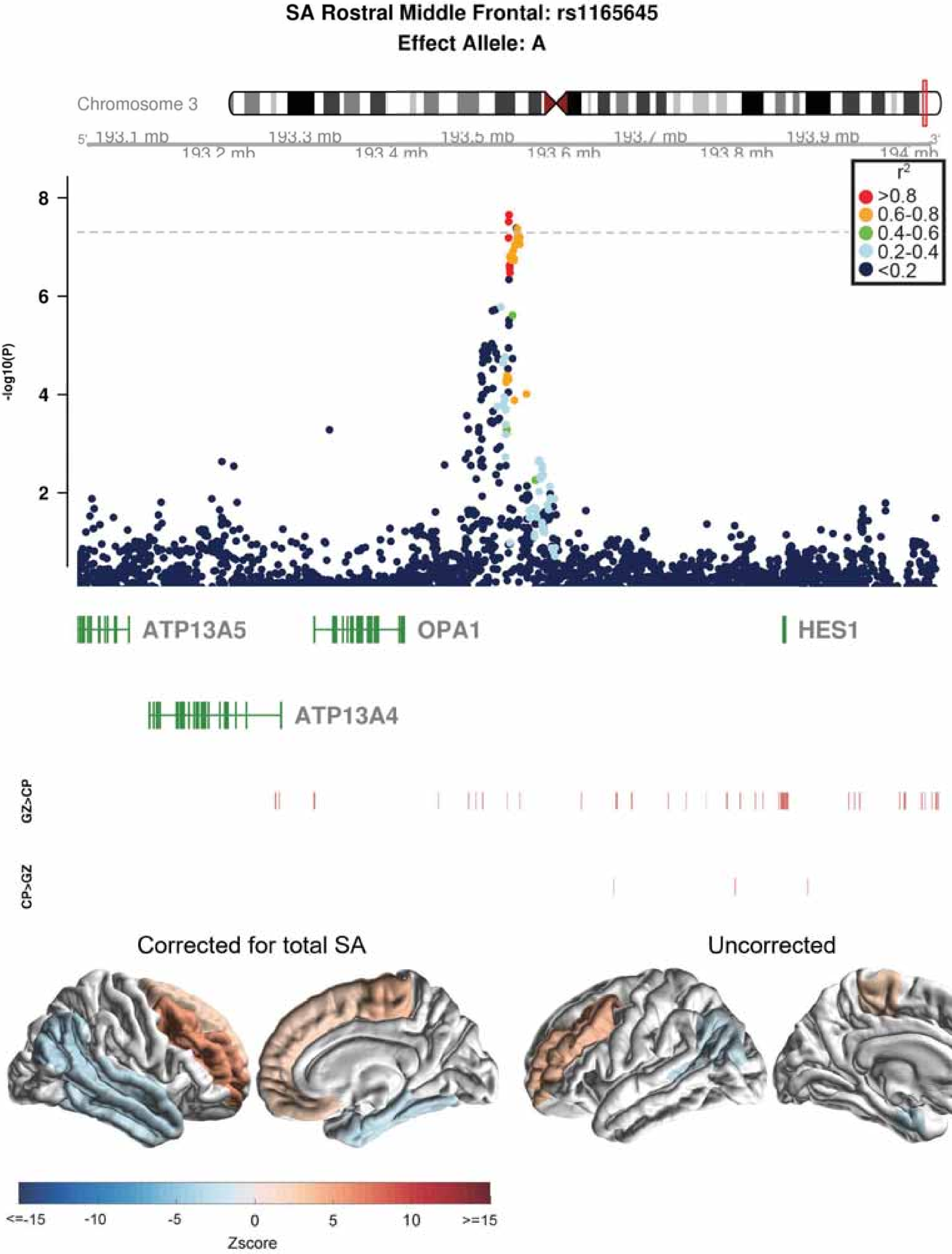

SA Rostral Middle Frontal: rs1165645
Effect Allele: A
Chromosome 3
193.1 mb
193.2 mb
193.3 mb
193.4 mb
193.5 mb
193.6 mb
193.7 mb
193.8 mb
193.9 mb
194 mb
r2
>0.8
0.6-0.8
0.4-0.6
0.2-0.4
<0.2
-log10(P)
ATP13A5
OPA1
HES1
ATP13A4
GZ>CP
CP>GZ
Corrected for total SA
Uncorrected
<=-15
-10
-5
0
5
10
>=15
Zscore

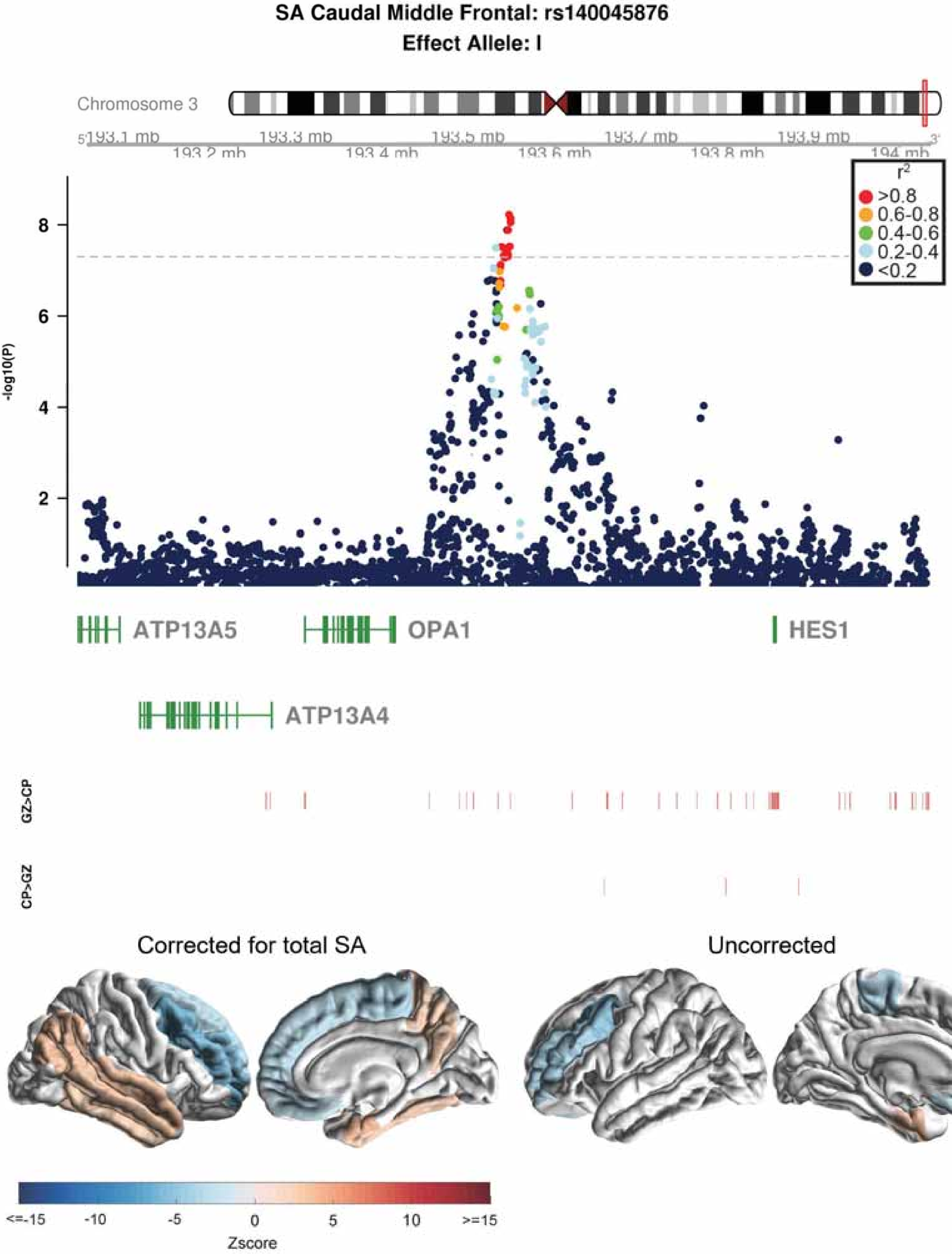
SA Caudal Middle Frontal: rs140045876
Effect Allele: I
Chromosome 3
193.1 mb
193.2 mb
193.3 mb
193.4 mb
193.5 mb
193.6 mb
193.7 mb
193.8 mb
193.9 mb
194 mb
-log10(P)
2
4
6
8
r2
>0.8
0.6-0.8
0.4-0.6
0.2-0.4
<0.2
ATP13A5
OPA1
HES1
ATP13A4
GZ>CP
CP>GZ
Corrected for total SA
Uncorrected
<=-15
-10
-5
0
5
10
>=15
Zscore

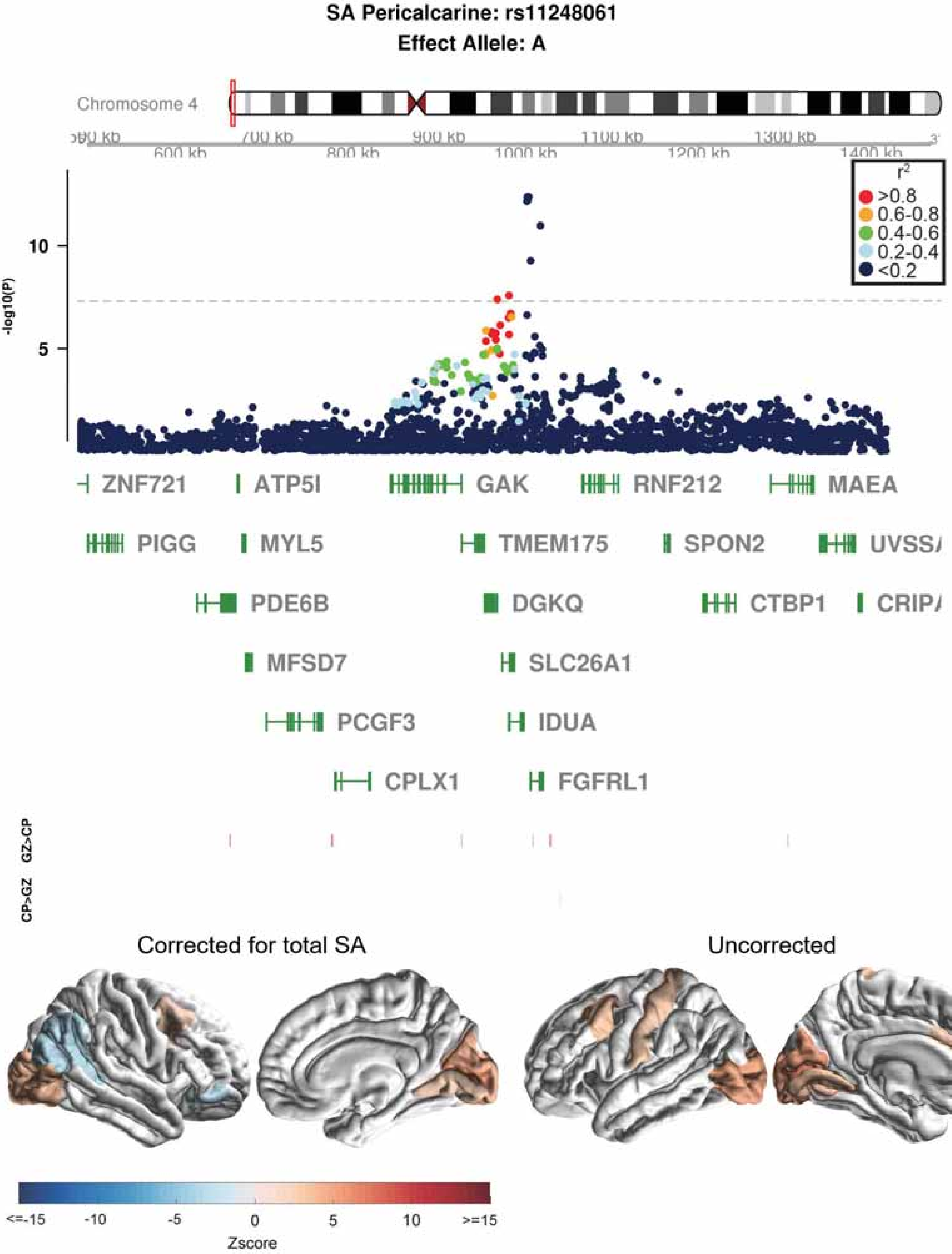
SA Pericalcarine: rs11248061
Effect Allele: A
Chromosome 4
600 kb
700 kb
800 kb
900 kb
1000 kb
1100 kb
1200 kb
1300 kb
1400 kb
r2
>0.8
0.6-0.8
0.4-0.6
0.2-0.4
<0.2
-log10(P)
10
5
ZNF721
ATP5I
GAK
RNF212
MAEA
PIGG
MYL5
TMEM175
SPON2
PDE6B
DGKQ
CTBP1
MFSD7
SLC26A1
PCGF3
IDUA
CPLX1
FGFRL1
GZ>CP
CP>GZ
Corrected for total SA
Uncorrected
<=-15
-10
-5
0
5
10
>=15
Zscore

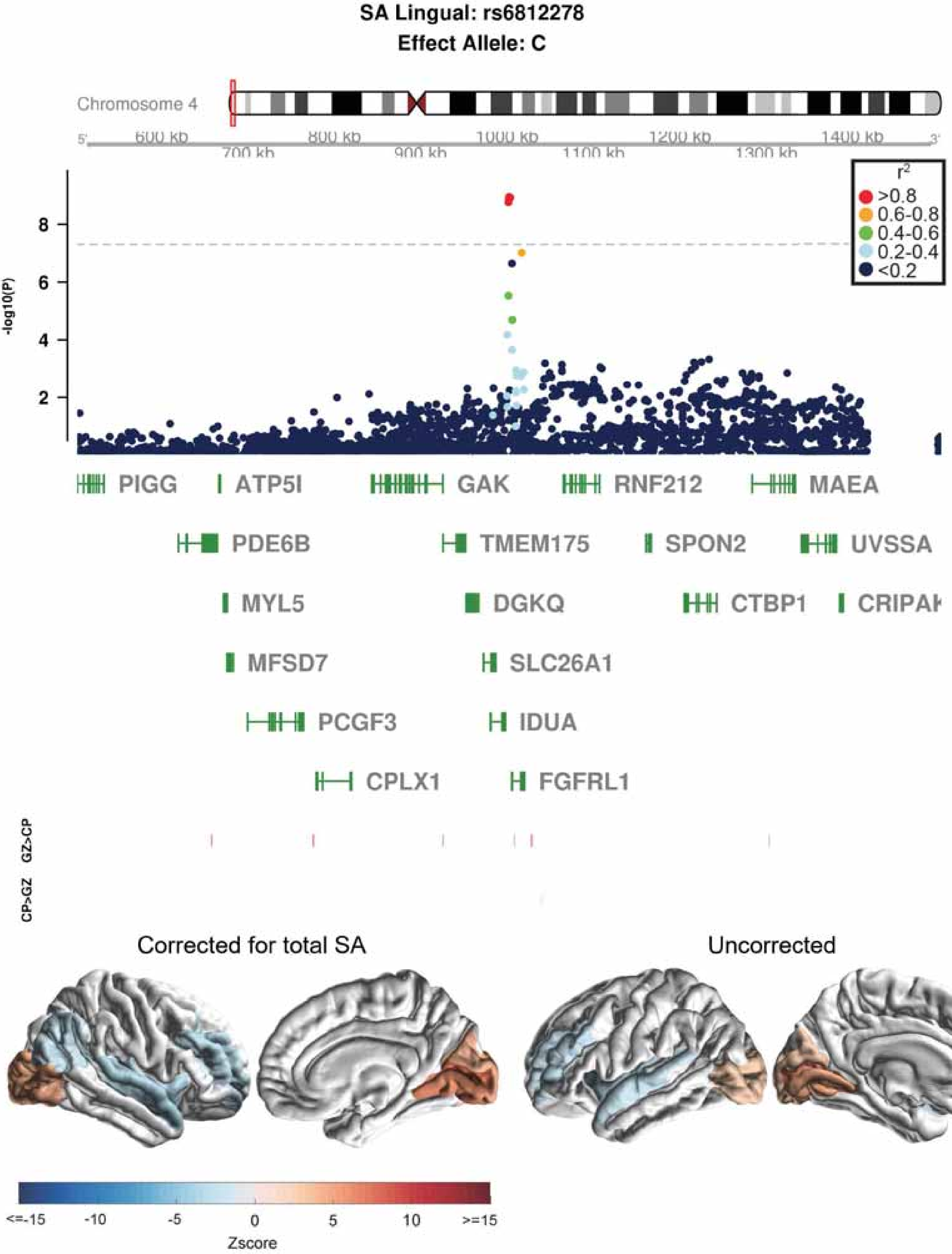

SA Lingual: rs6812278
Effect Allele: C
Chromosome 4
600 kb
700 kb
800 kb
900 kb
1000 kb
1100 kb
1200 kb
1300 kb
1400 kb
-log10(P)
2
4
6
8
r2
>0.8
0.6-0.8
0.4-0.6
0.2-0.4
<0.2
PIGG
ATP5I
GAK
RNF212
MAEA
PDE6B
TMEM175
SPON2
UVSSA
MYL5
DGKQ
CTBP1
CRIPAK
MFSD7
SLC26A1
PCGF3
IDUA
CPLX1
FGFRL1
GZ>CP
CP>GZ
Corrected for total SA
Uncorrected
<=-15
-10
-5
0
5
10
>=15
Zscore

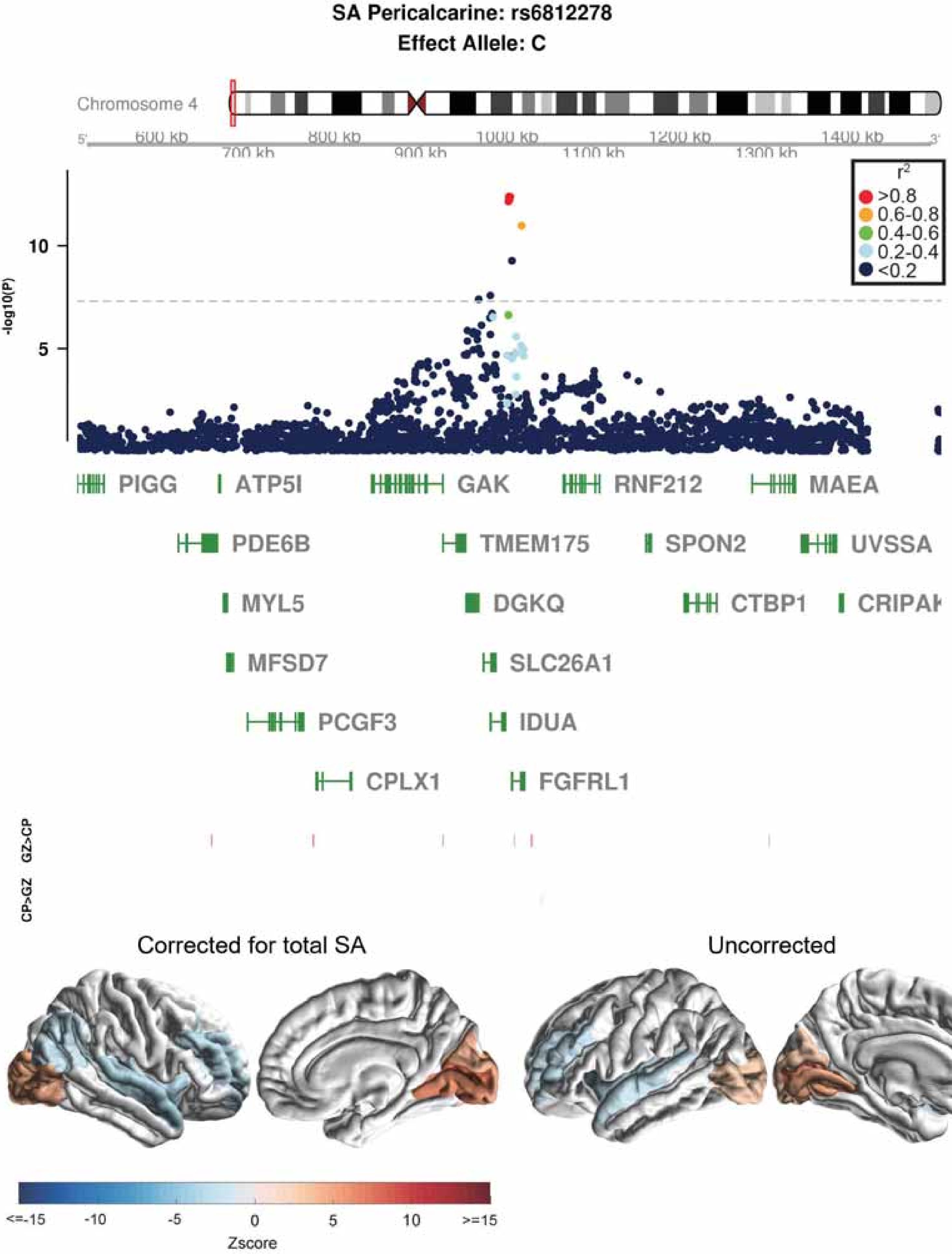

SA Pericalcarine: rs6812278
Effect Allele: C
Chromosome 4
600 kb
700 kb
800 kb
900 kb
1000 kb
1100 kb
1200 kb
1300 kb
1400 kb
-log10(P)
10
5
r2
>0.8
0.6-0.8
0.4-0.6
0.2-0.4
<0.2
PIGG
ATP5I
GAK
RNF212
MAEA
PDE6B
TMEM175
SPON2
UVSSA
MYL5
DGKQ
CTBP1
MFSD7
SLC26A1
PCGF3
IDUA
CPLX1
FGFRL1
GZ>CP
CP>GZ
Corrected for total SA
Uncorrected
<=-15
-10
-5
0
5
10
>=15
Zscore

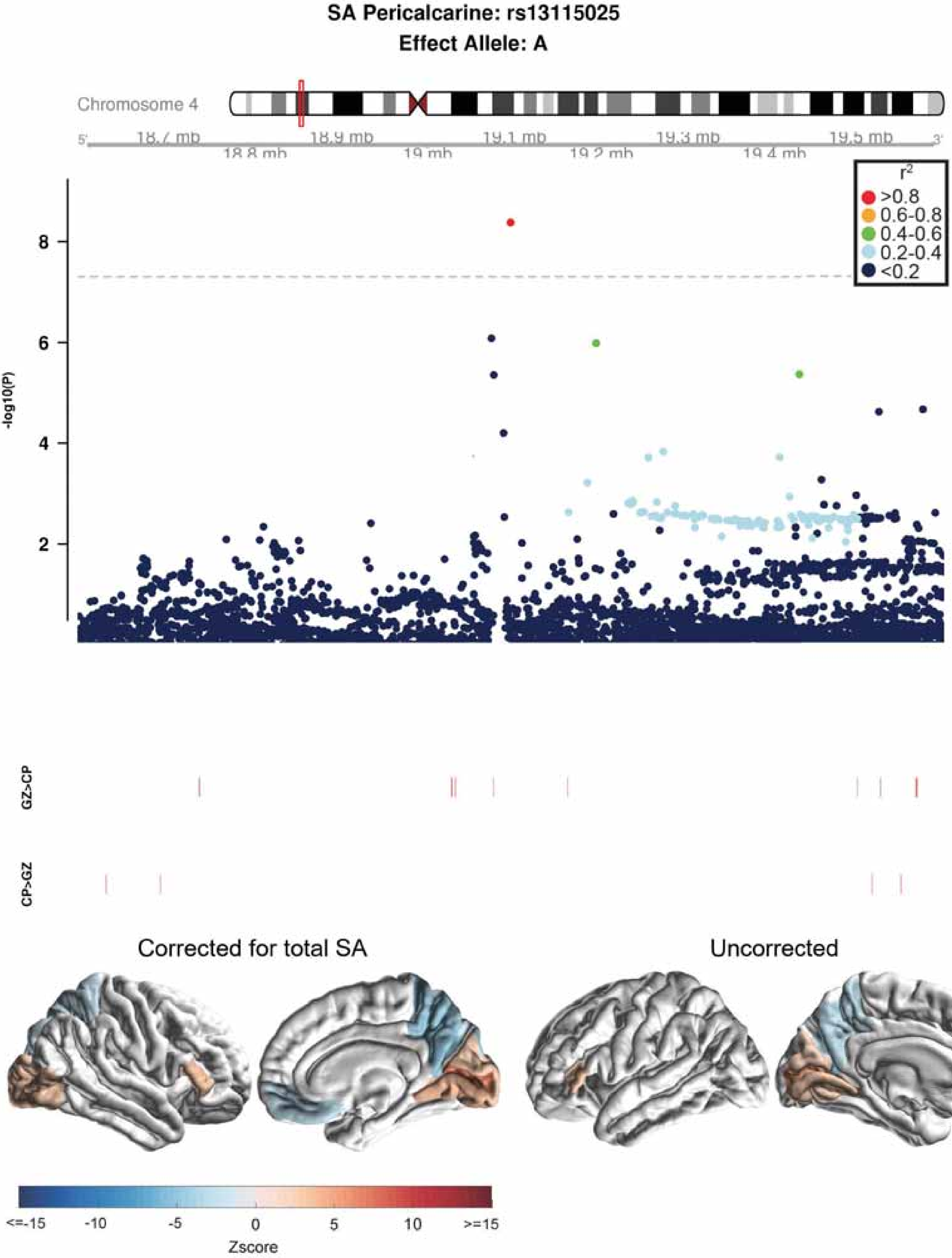
SA Pericalcarine: rs13115025
Effect Allele: A
Chromosome 4
18.7 mb
18.8 mb
18.9 mb
19 mb
19.1 mb
19.2 mb
19.3 mb
19.4 mb
19.5 mb
-log10(P)
r²
>0.8
0.6-0.8
0.4-0.6
0.2-0.4
<0.2
GZ>CP
CP>GZ
Corrected for total SA
Uncorrected
<=-15
-10
-5
0
5
10
>=15
Zscore

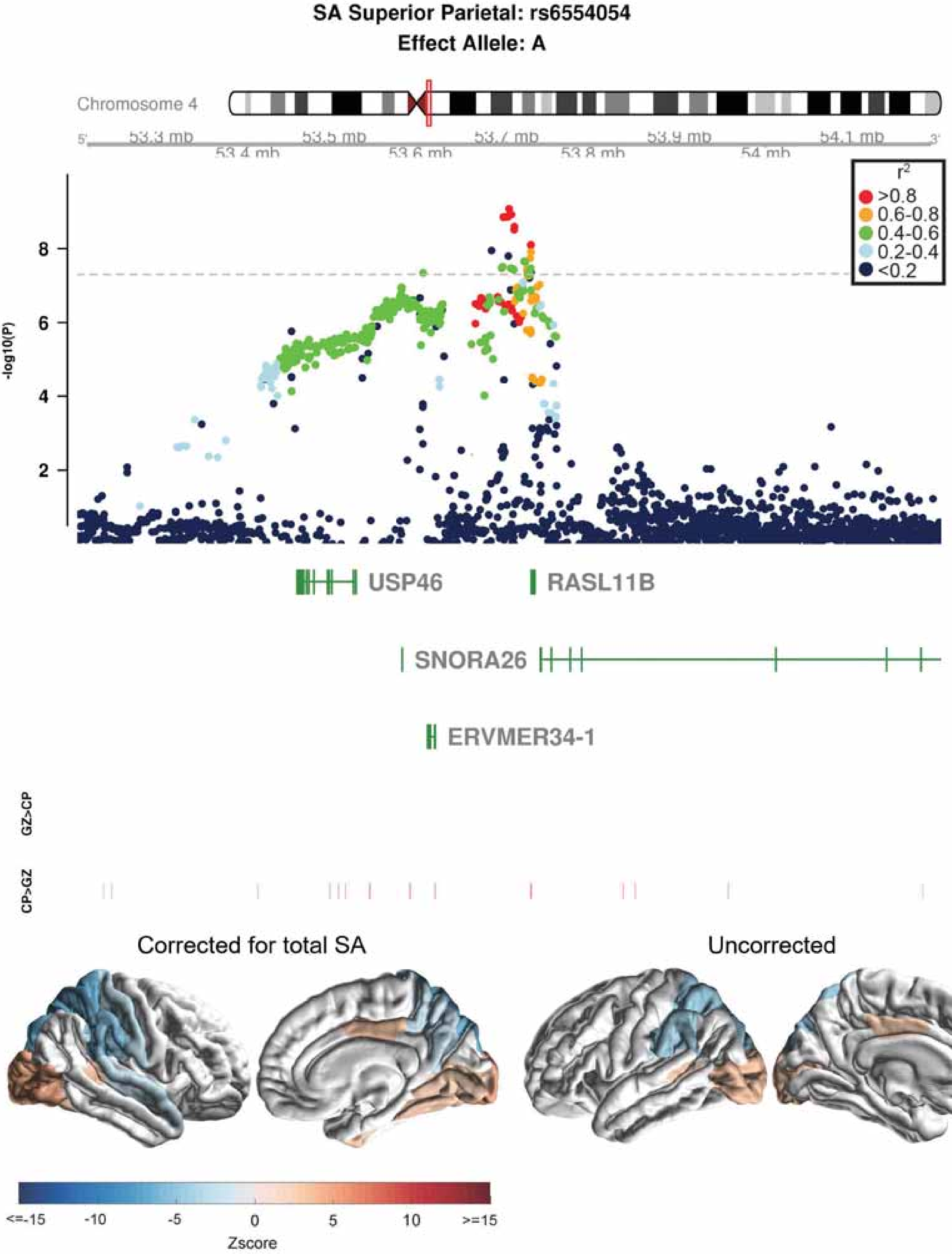
SA Superior Parietal: rs6554054
Effect Allele: A
Chromosome 4
53.3 mb
53.4 mb
53.5 mb
53.6 mb
53.7 mb
53.8 mb
53.9 mb
54 mb
54.1 mb
r2
>0.8
0.6-0.8
0.4-0.6
0.2-0.4
<0.2
-log10(P)
USP46
RASL11B
SNORA26
ERVMER34-1
GZ>CP
CP>GZ
Corrected for total SA
Uncorrected
<=-15
-10
-5
0
5
10
>=15
Zscore

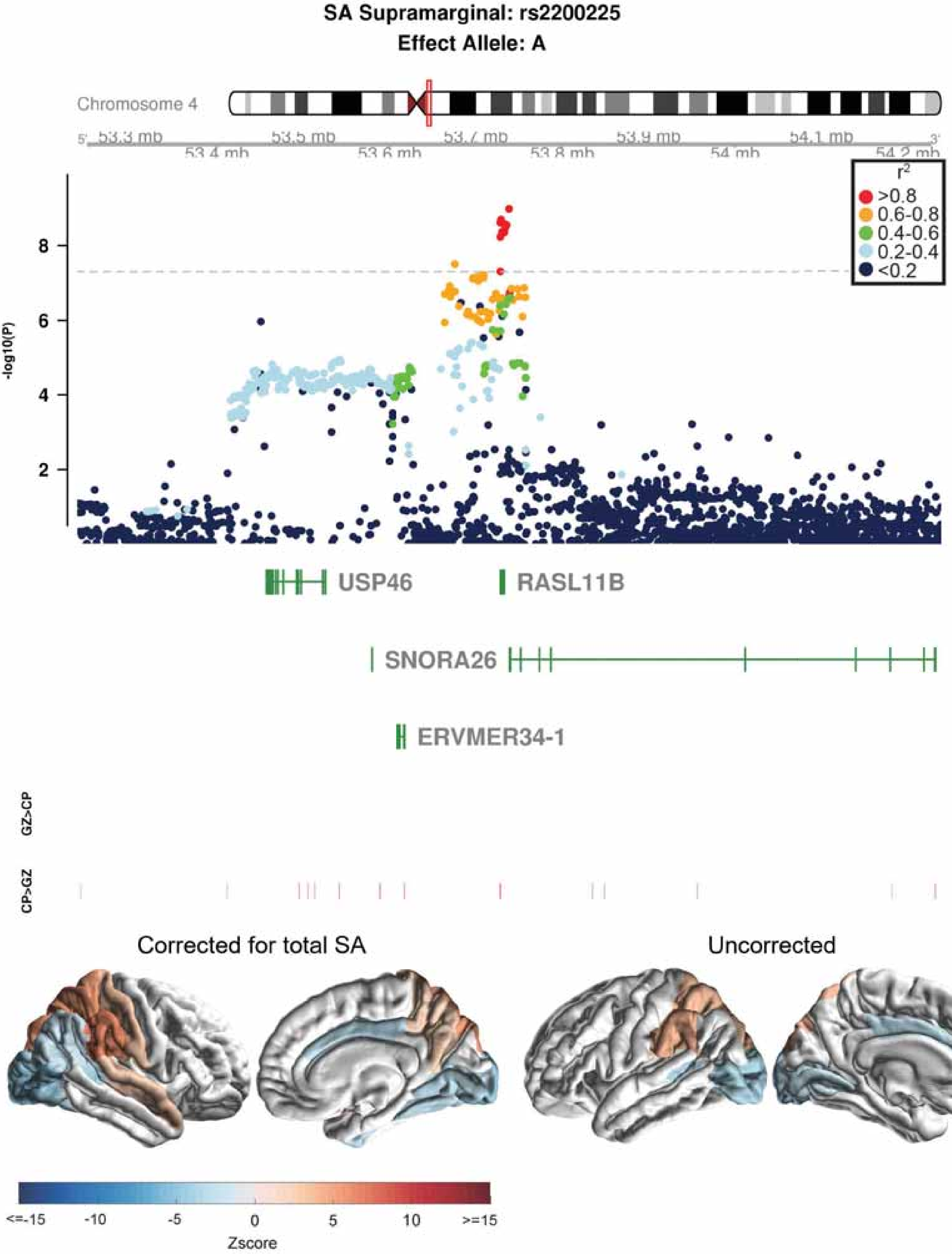
SA Supramarginal: rs2200225
Effect Allele: A
Chromosome 4
53.3 mb
53.4 mb
53.5 mb
53.6 mb
53.7 mb
53.8 mb
53.9 mb
54 mb
54.1 mb
54.2 mb
r2
>0.8
0.6-0.8
0.4-0.6
0.2-0.4
<0.2
-log10(P)
2
4
6
8
USP46
RASL11B
SNORA26
ERVMER34-1
GZ>CP
CP>GZ
Corrected for total SA
Uncorrected
<=-15
-10
-5
0
5
10
>=15
Zscore

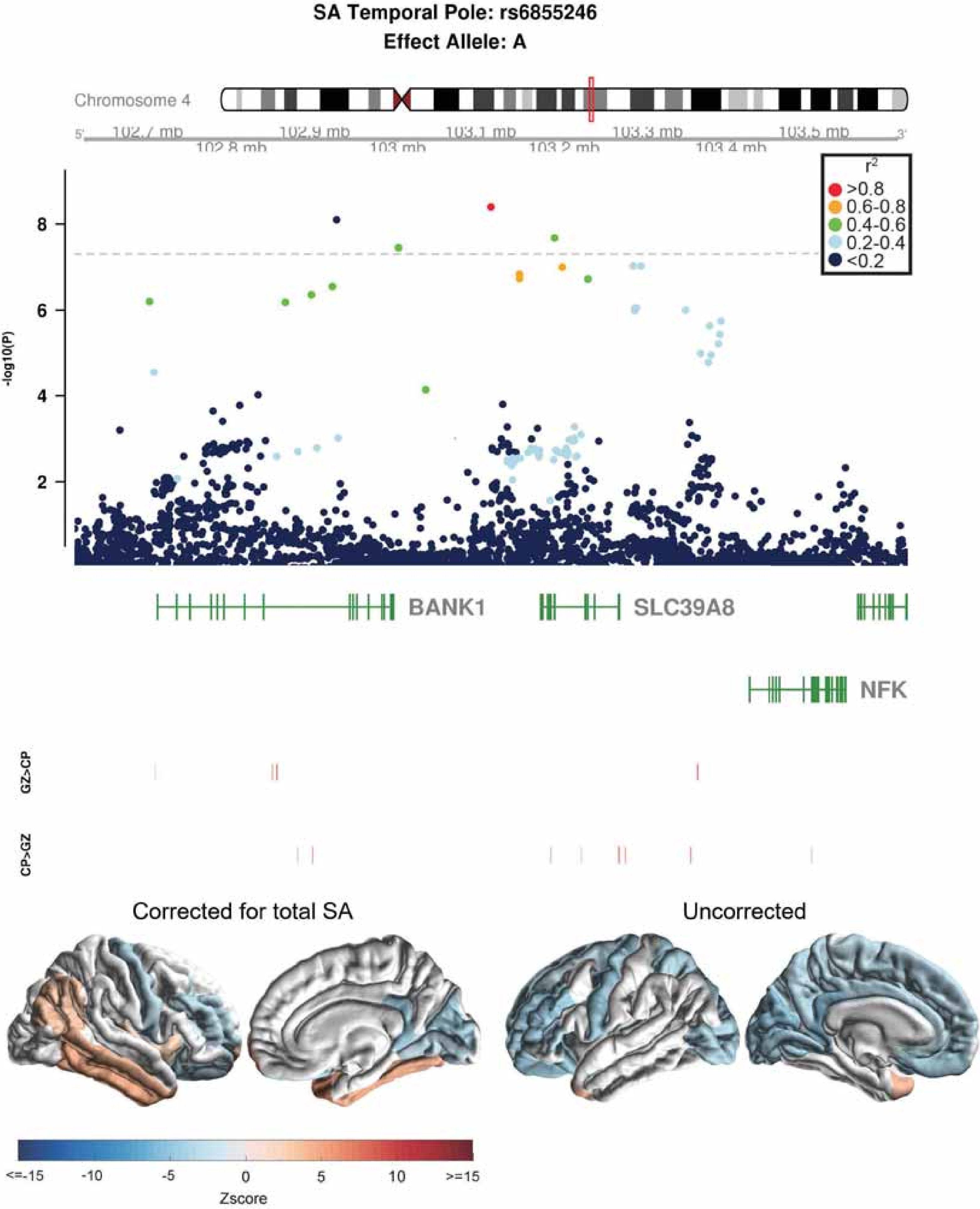
SA Temporal Pole: rs6855246
Effect Allele: A
Chromosome 4
102.7 mb
102.8 mb
102.9 mb
103 mb
103.1 mb
103.2 mb
103.3 mb
103.4 mb
103.5 mb
r2
>0.8
0.6-0.8
0.4-0.6
0.2-0.4
<0.2
-log10(P)
BANK1
SLC39A8
NFK
GZ>CP
CP>GZ
Corrected for total SA
Uncorrected
<=-15
-10
-5
0
5
10
>=15
Zscore

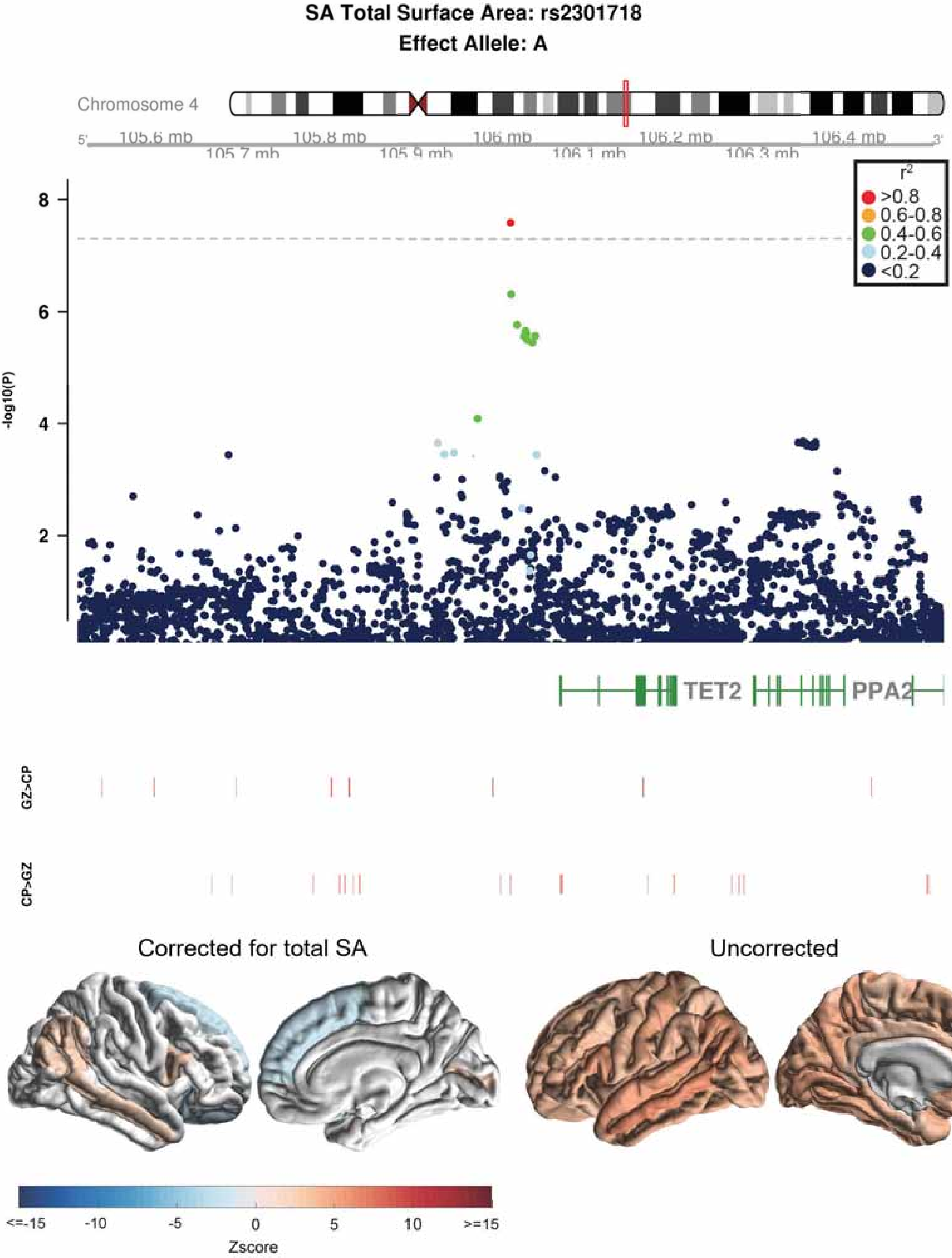
SA Total Surface Area: rs2301718
Effect Allele: A
Chromosome 4
5'
105.6 mb
105.7 mb
105.8 mb
105.9 mb
106 mb
106.1 mb
106.2 mb
106.3 mb
106.4 mb
3'
r2
>0.8
0.6-0.8
0.4-0.6
0.2-0.4
<0.2
-log10(P)
8
6
4
2
TET2
PPA2
GZ>CP
CP>GZ
Corrected for total SA
Uncorrected
<=-15
-10
-5
0
5
10
>=15
Zscore

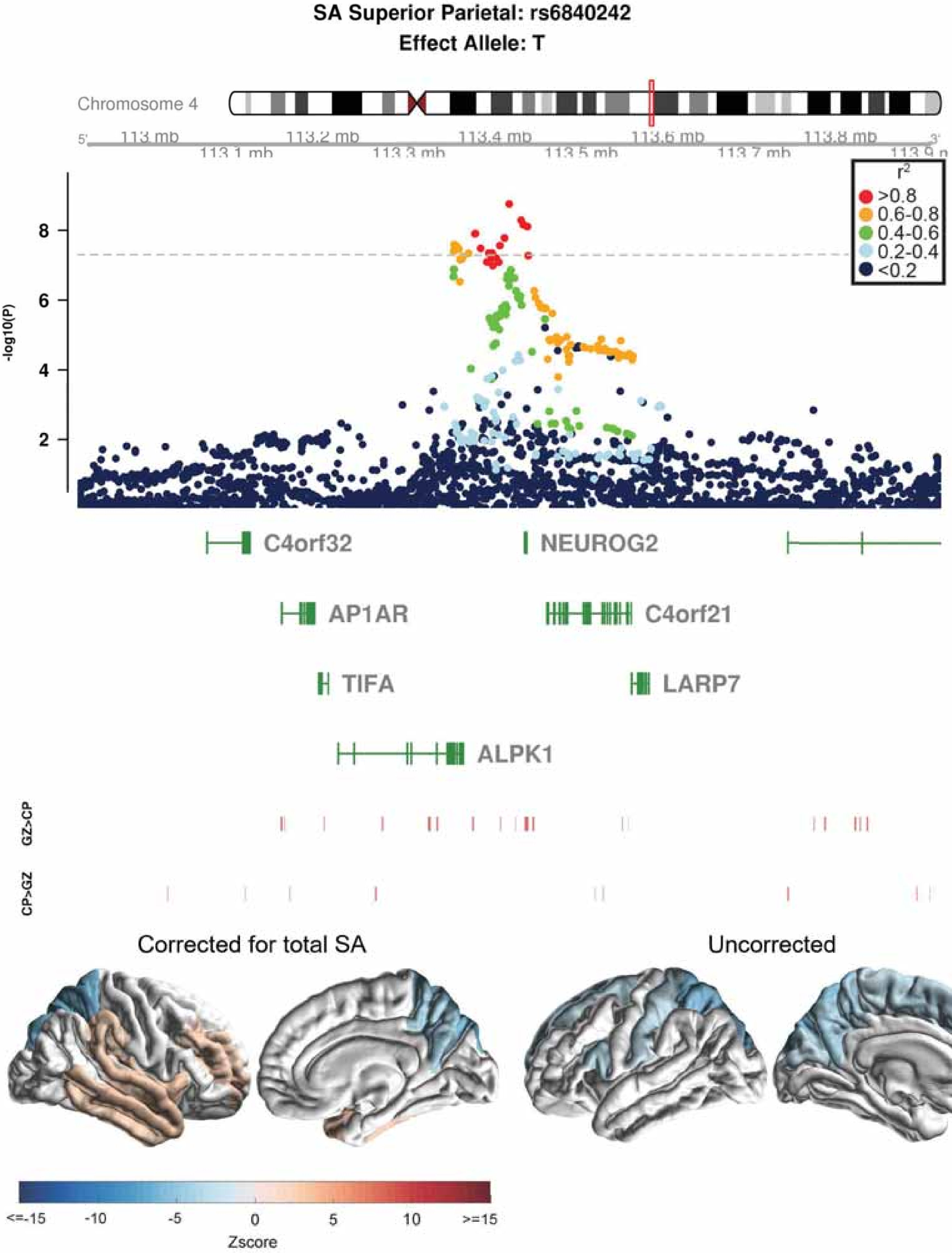

SA Superior Parietal: rs6840242
Effect Allele: T
Chromosome 4
113 mb
113.1 mb
113.2 mb
113.3 mb
113.4 mb
113.5 mb
113.6 mb
113.7 mb
113.8 mb
-log10(P)
r²
>0.8
0.6-0.8
0.4-0.6
0.2-0.4
<0.2
C4orf32
NEUROG2
AP1AR
C4orf21
TIFA
LARP7
ALPK1
GZ>CP
CP>GZ
Corrected for total SA
Uncorrected
<=-15
-10
-5
0
5
10
>=15
Zscore

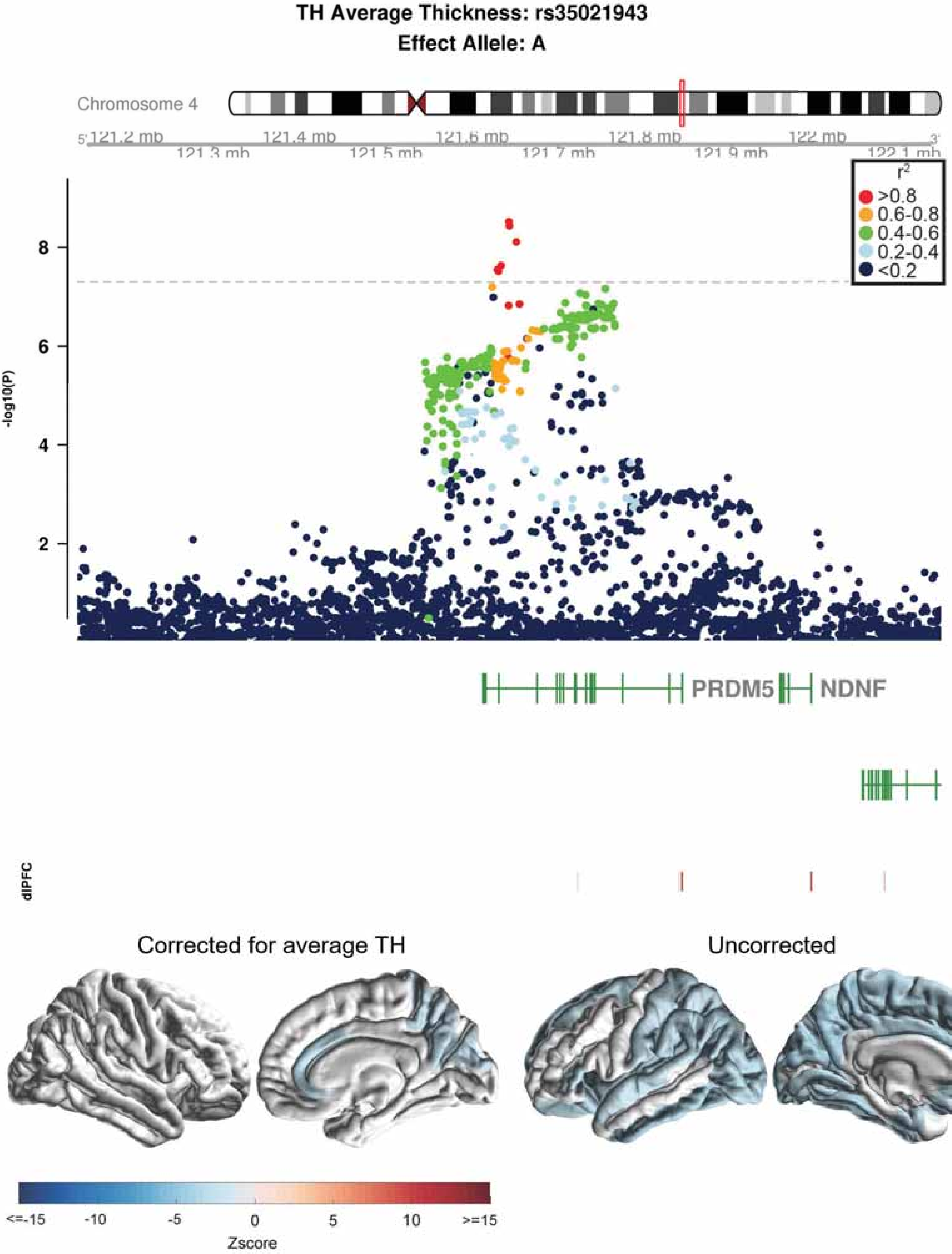
TH Average Thickness: rs35021943
Effect Allele: A
Chromosome 4
121.2 mb
121.3 mb
121.4 mb
121.5 mb
121.6 mb
121.7 mb
121.8 mb
121.9 mb
122 mb
122.1 mb
r2
>0.8
0.6-0.8
0.4-0.6
0.2-0.4
<0.2
-log10(P)
8
6
4
2
PRDM5
NDNF
dlPFC
Corrected for average TH
Uncorrected
<=-15
-10
-5
0
5
10
>=15
Zscore

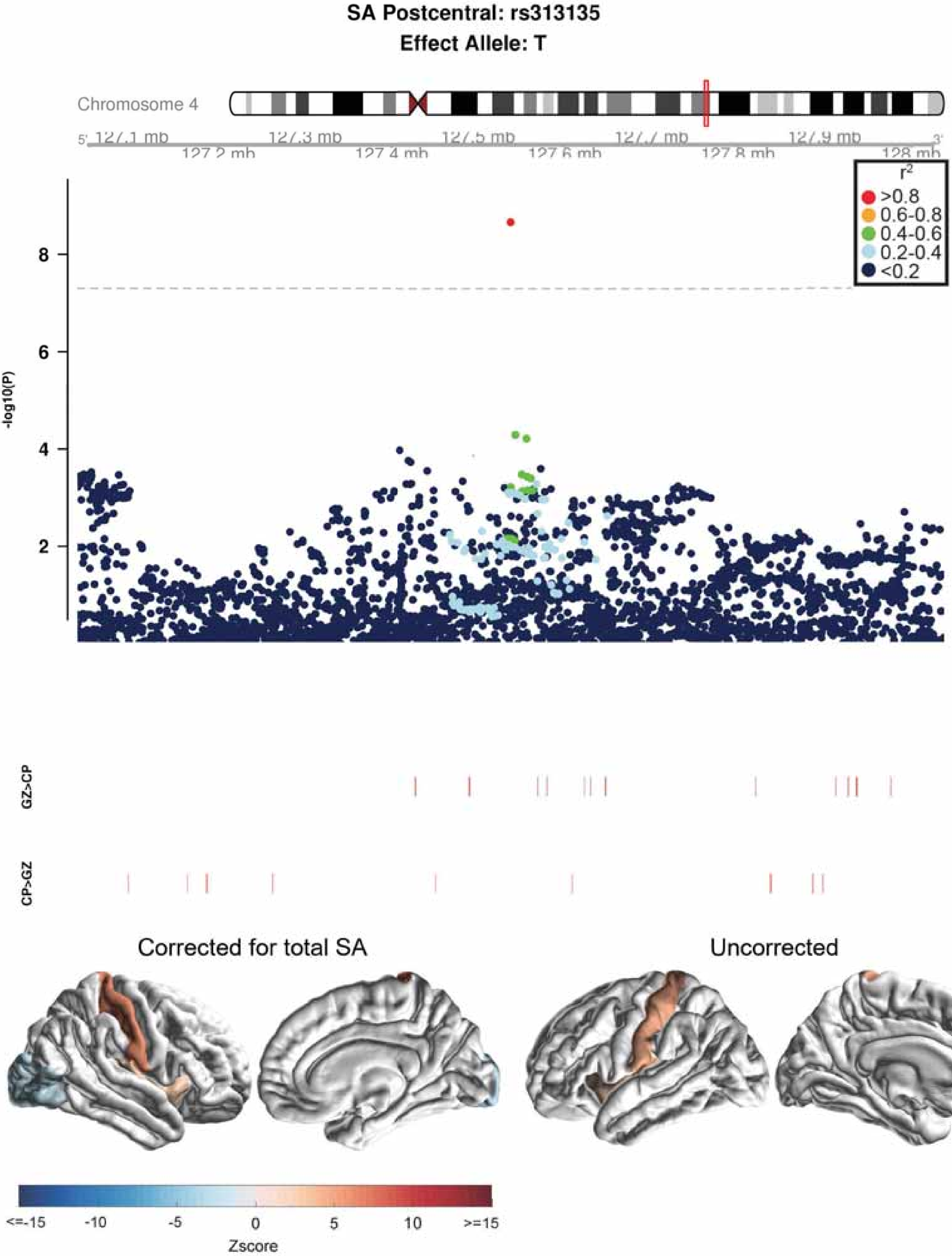

SA Postcentral: rs313135
Effect Allele: T
Chromosome 4
127.1 mb
127.2 mb
127.3 mb
127.4 mb
127.5 mb
127.6 mb
127.7 mb
127.8 mb
127.9 mb
128 mb
r2
>0.8
0.6-0.8
0.4-0.6
0.2-0.4
<0.2
-log10(P)
GZ>CP
CP>GZ
Corrected for total SA
Uncorrected
<=-15
-10
-5
0
5
10
>=15
Zscore

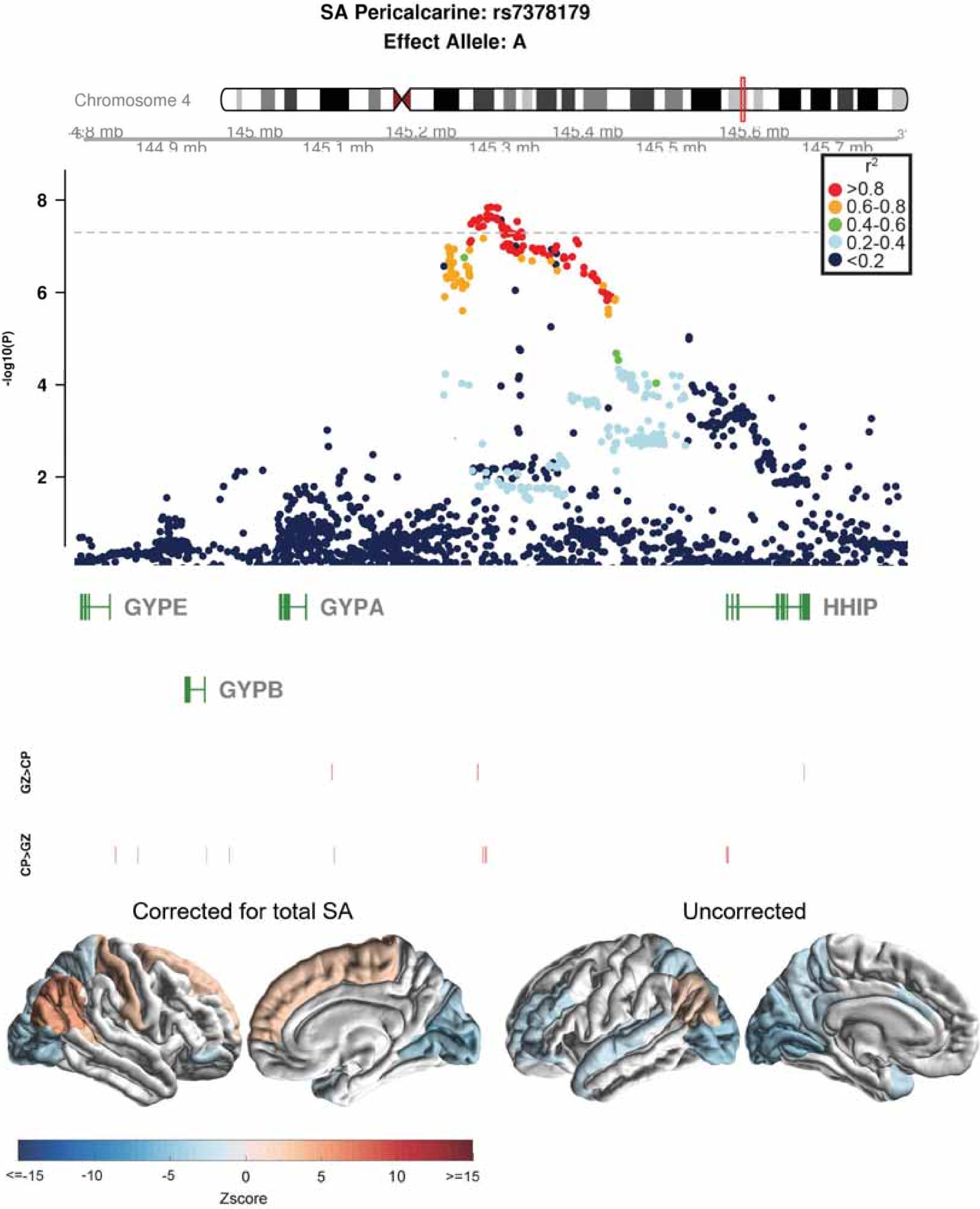
SA Pericalcarine: rs7378179
Effect Allele: A
Chromosome 4
145 mb
145.2 mb
145.4 mb
145.6 mb
144.9 mb
145.1 mb
145.3 mb
145.5 mb
145.7 mb
r²
>0.8
0.6-0.8
0.4-0.6
0.2-0.4
<0.2
-log10(P)
8
6
4
2
GYPE
GYPA
HHIP
GYPB
GZ>CP
CP>GZ
Corrected for total SA
Uncorrected
<=-15
-10
-5
0
5
10
>=15
Zscore

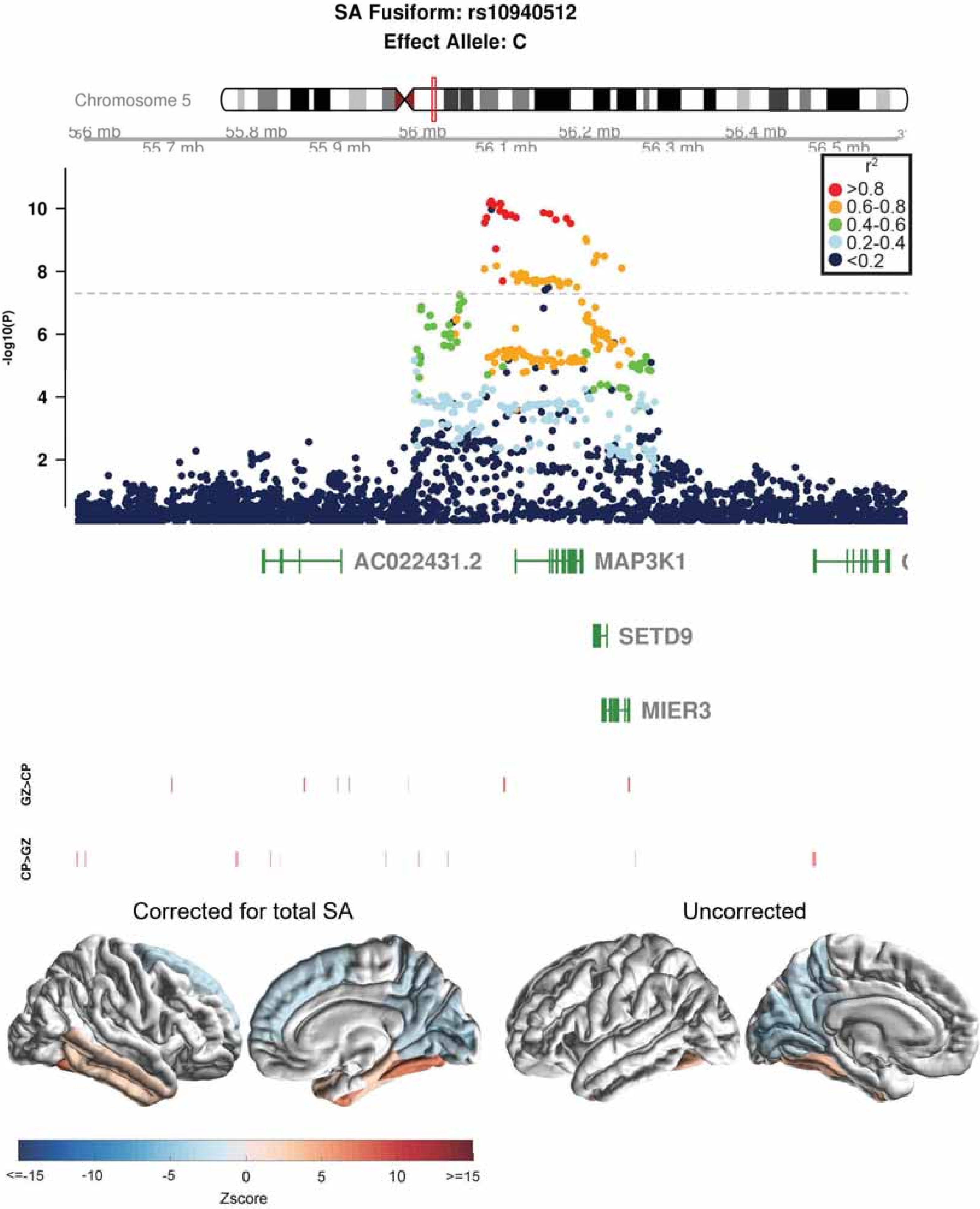

SA Fusiform: rs10940512
Effect Allele: C
Chromosome 5
55.6 mb
55.7 mb
55.8 mb
55.9 mb
56 mb
56.1 mb
56.2 mb
56.3 mb
56.4 mb
56.5 mb
-log10(P)
10
8
6
4
2
r2
>0.8
0.6-0.8
0.4-0.6
0.2-0.4
<0.2
AC022431.2
MAP3K1
SETD9
MIER3
GZ>CP
CP>GZ
Corrected for total SA
Uncorrected
<=-15
-10
-5
0
5
10
>=15
Zscore

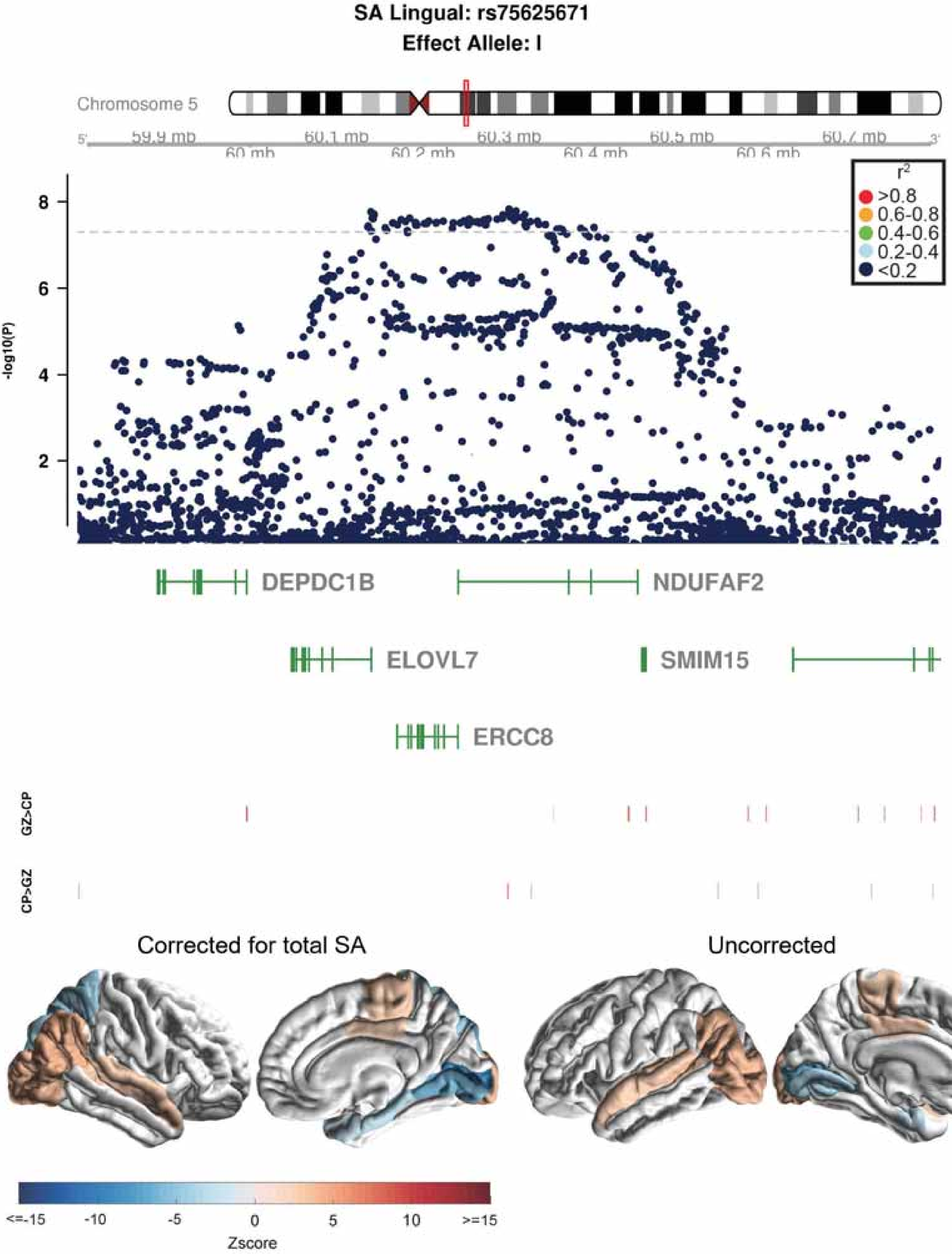

SA Lingual: rs75625671
Effect Allele: I
Chromosome 5
59.9 mb
60 mb
60.1 mb
60.2 mb
60.3 mb
60.4 mb
60.5 mb
60.6 mb
60.7 mb
r2
>0.8
0.6-0.8
0.4-0.6
0.2-0.4
<0.2
-log10(P)
8
6
4
2
DEPDC1B
NDUFAF2
ELOVL7
SMIM15
ERCC8
GZ>CP
CP>GZ
Corrected for total SA
Uncorrected
<=-15
-10
-5
0
5
10
>=15
Zscore

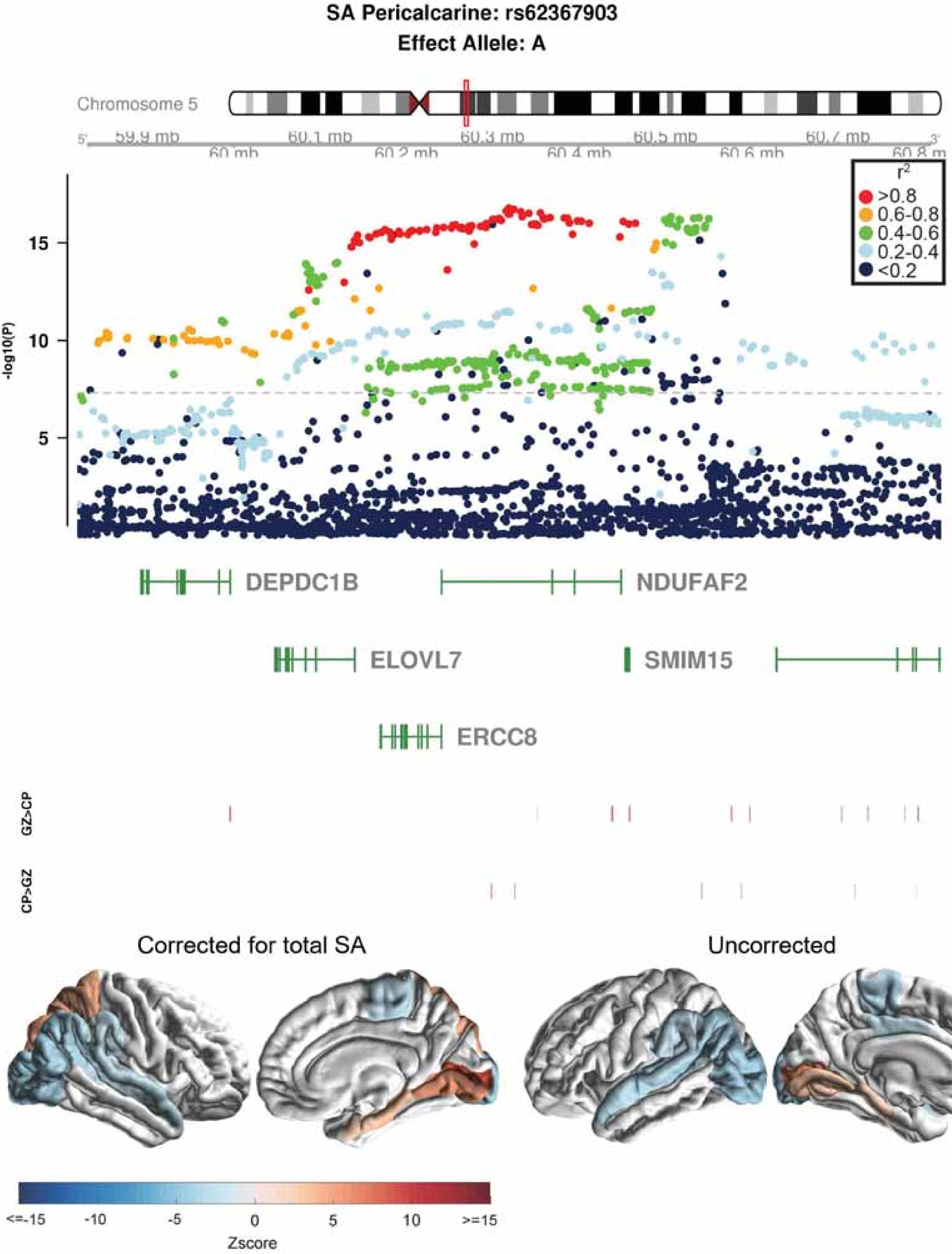

SA Pericalcarine: rs62367903
Effect Allele: A
Chromosome 5
59.9 mb
60 mb
60.1 mb
60.2 mb
60.3 mb
60.4 mb
60.5 mb
60.6 mb
60.7 mb
60.8 mb
-log10(P)
15
10
5
r2
>0.8
0.6-0.8
0.4-0.6
0.2-0.4
<0.2
DEPDC1B
NDUFAF2
ELOVL7
SMIM15
ERCC8
GZ>CP
CP>GZ
Corrected for total SA
Uncorrected
<=-15
-10
-5
0
5
10
>=15
Zscore

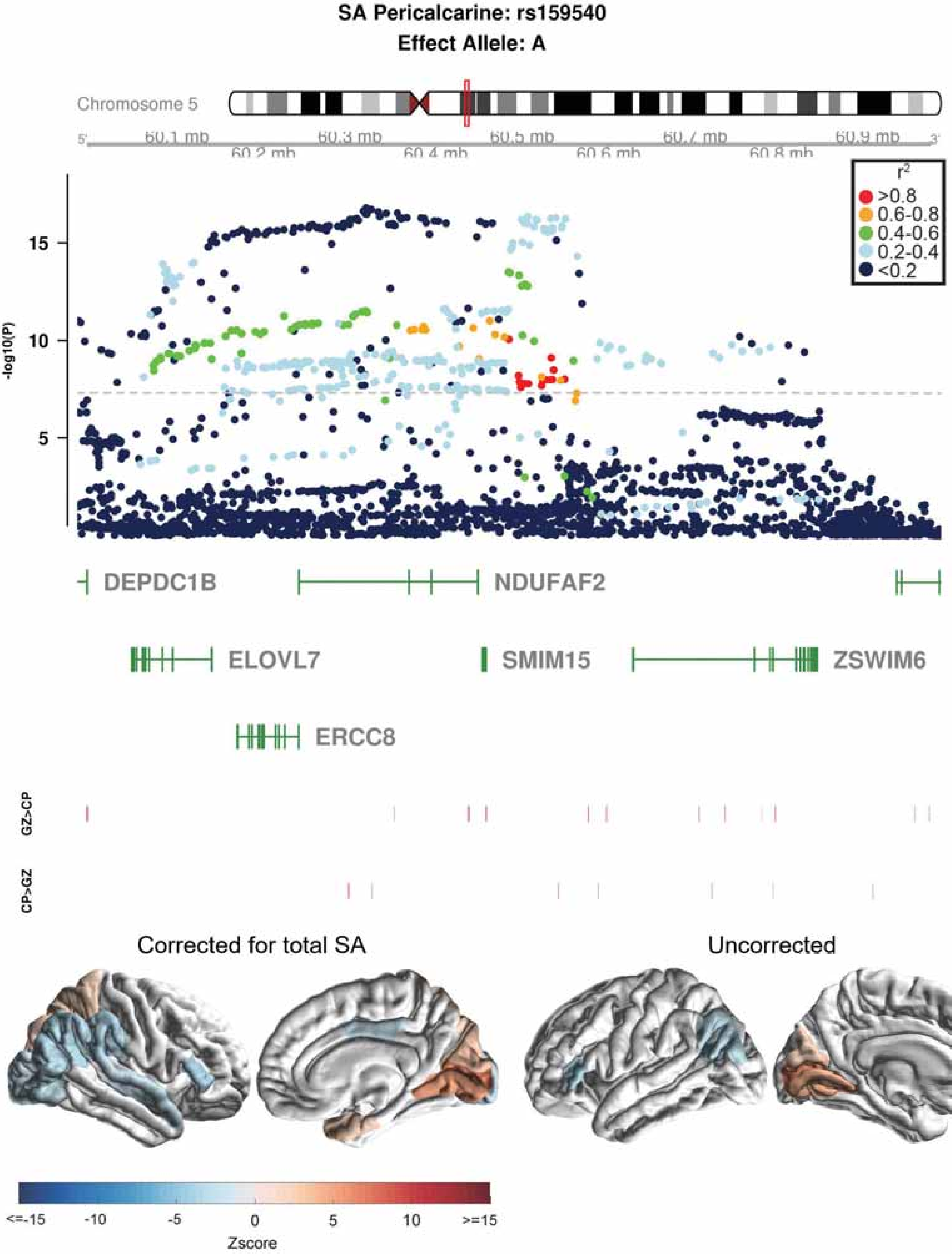

SA Pericalcarine: rs159540
Effect Allele: A
Chromosome 5
60.1 mb
60.2 mb
60.3 mb
60.4 mb
60.5 mb
60.6 mb
60.7 mb
60.8 mb
60.9 mb
r2
>0.8
0.6-0.8
0.4-0.6
0.2-0.4
<0.2
-log10(P)
15
10
5
DEPDC1B
NDUFAF2
ELOVL7
SMIM15
ZSWIM6
ERCC8
GZ>CP
CP>GZ
Corrected for total SA
Uncorrected
<=-15
-10
-5
0
5
10
>=15
Zscore

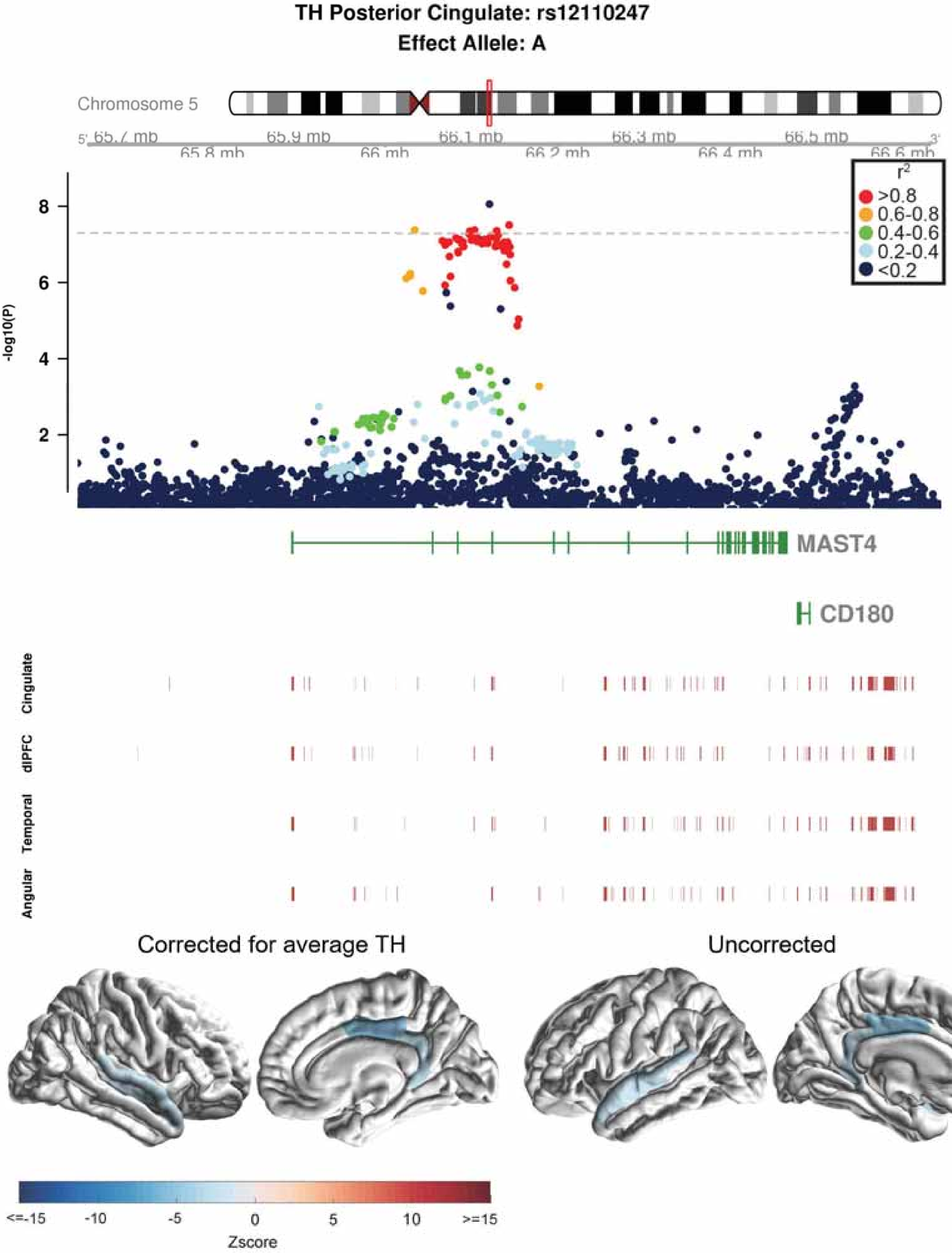

TH Posterior Cingulate: rs12110247
Effect Allele: A
Chromosome 5
65.7 mb
65.8 mb
65.9 mb
66 mb
66.1 mb
66.2 mb
66.3 mb
66.4 mb
66.5 mb
66.6 mb
r2
>0.8
0.6-0.8
0.4-0.6
0.2-0.4
<0.2
-log10(P)
8
6
4
2
MAST4
CD180
Cingulate
dlPFC
Temporal
Angular
Corrected for average TH
Uncorrected
<=-15
-10
-5
0
5
10
>=15
Zscore

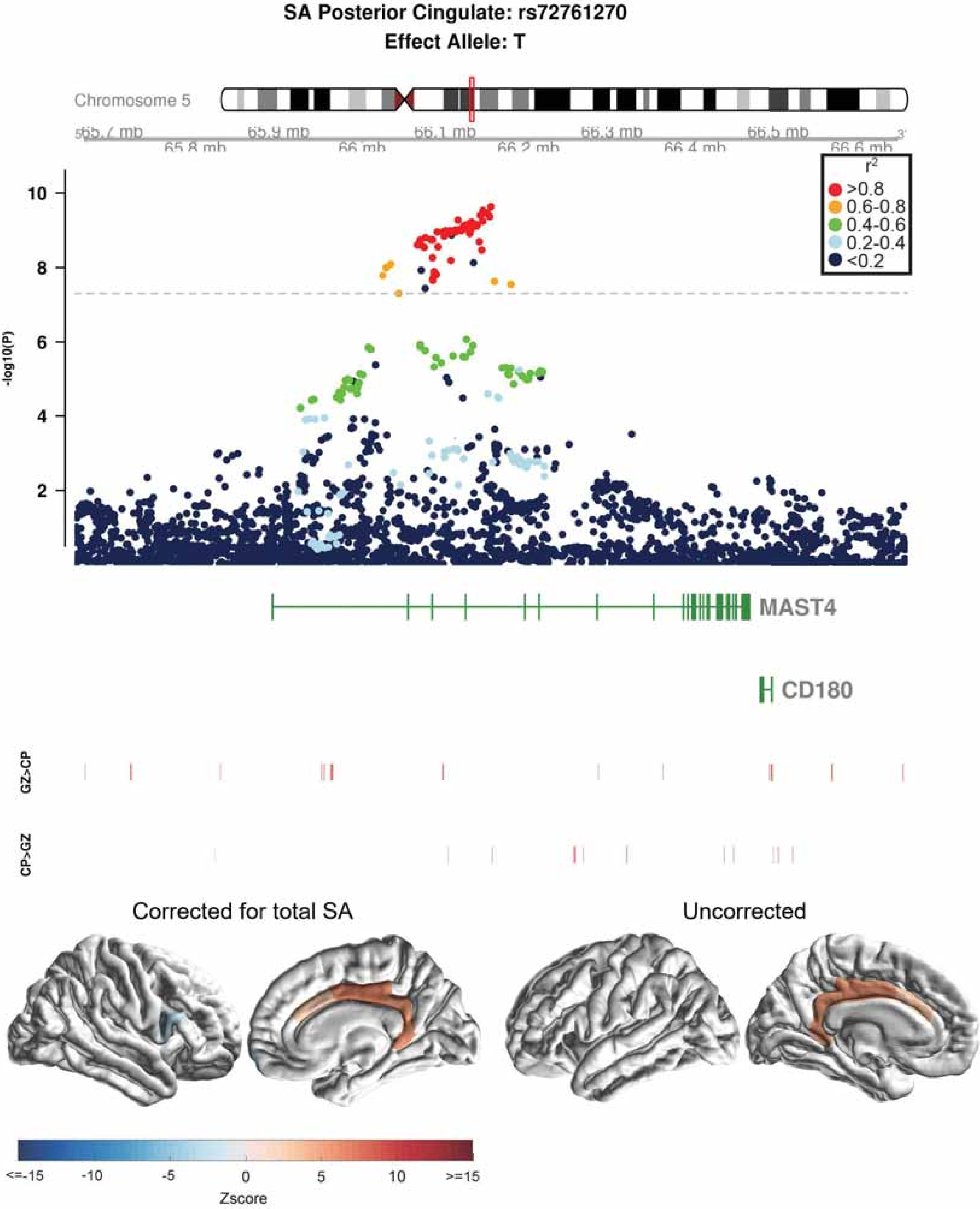

SA Posterior Cingulate: rs72761270
Effect Allele: T
Chromosome 5
65.7 mb
65.8 mb
65.9 mb
66 mb
66.1 mb
66.2 mb
66.3 mb
66.4 mb
66.5 mb
66.6 mb
r2
>0.8
0.6-0.8
0.4-0.6
0.2-0.4
<0.2
-log10(P)
10
8
6
4
2
MAST4
CD180
GZ>CP
CP>GZ
Corrected for total SA
Uncorrected
<=-15
-10
-5
0
5
10
>=15
Zscore

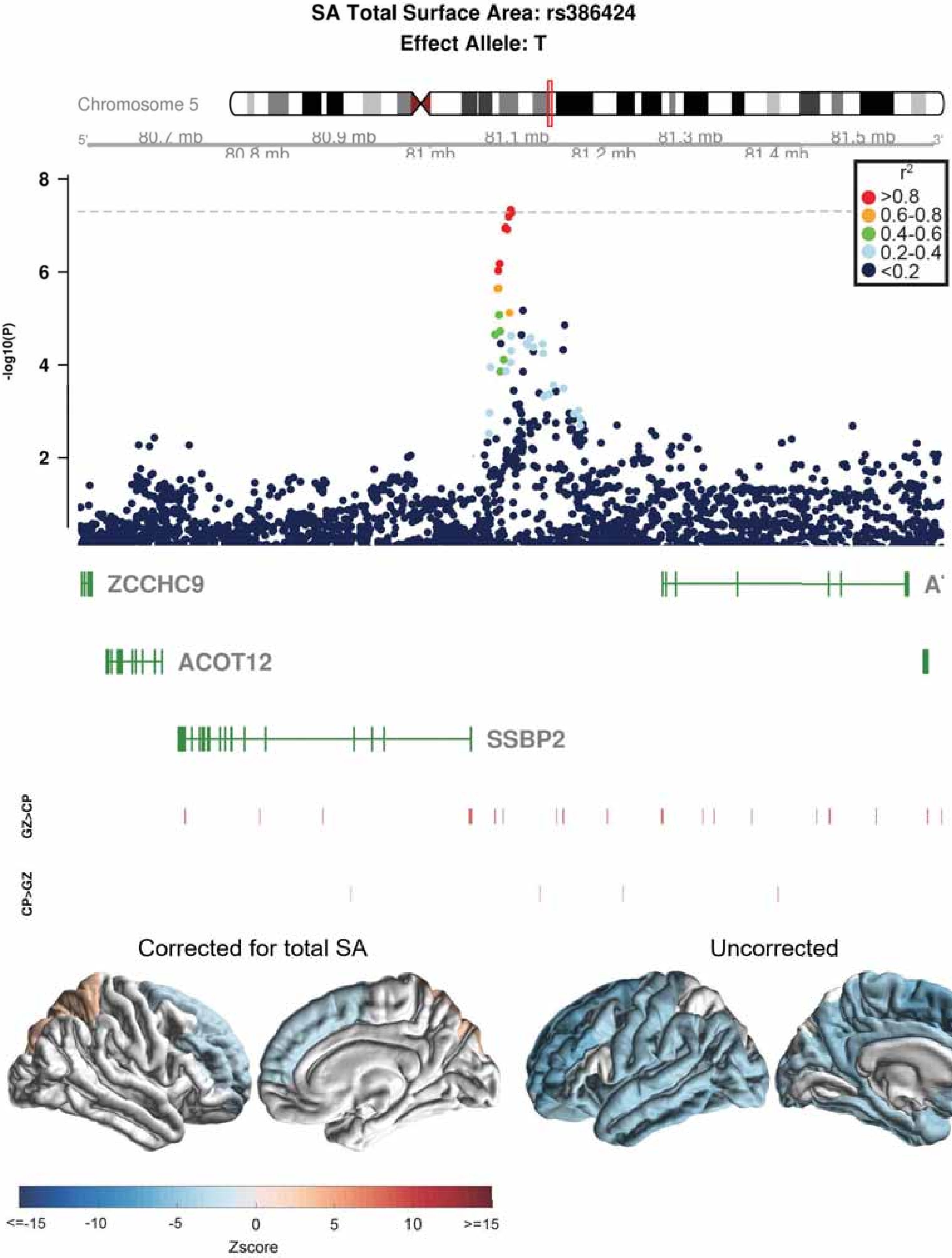
SA Total Surface Area: rs386424
Effect Allele: T
Chromosome 5
80.7 mb
80.8 mb
80.9 mb
81 mb
81.1 mb
81.2 mb
81.3 mb
81.4 mb
81.5 mb
-log10(P)
r2
>0.8
0.6-0.8
0.4-0.6
0.2-0.4
<0.2
ZCCHC9
ACOT12
SSBP2
GZ>CP
CP>GZ
Corrected for total SA
Uncorrected
<=-15
-10
-5
0
5
10
>=15
Zscore

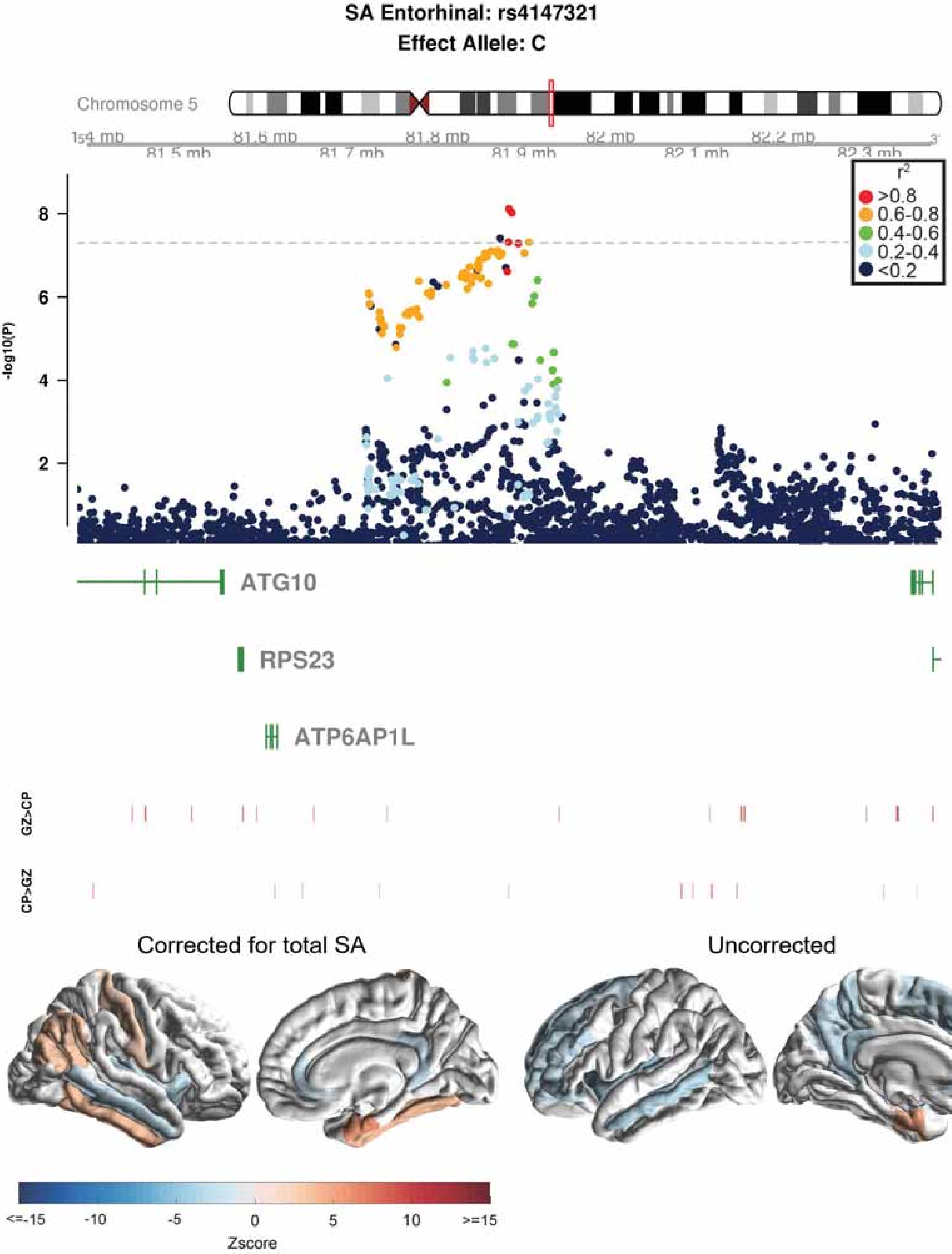

SA Entorhinal: rs4147321
Effect Allele: C
Chromosome 5
81.5 mb
81.6 mb
81.7 mb
81.8 mb
81.9 mb
82 mb
82.1 mb
82.2 mb
82.3 mb
-log10(P)
r²
>0.8
0.6-0.8
0.4-0.6
0.2-0.4
<0.2
ATG10
RPS23
ATP6AP1L
GZ>CP
CP>GZ
Corrected for total SA
Uncorrected
<=-15
-10
-5
0
5
10
>=15
Zscore

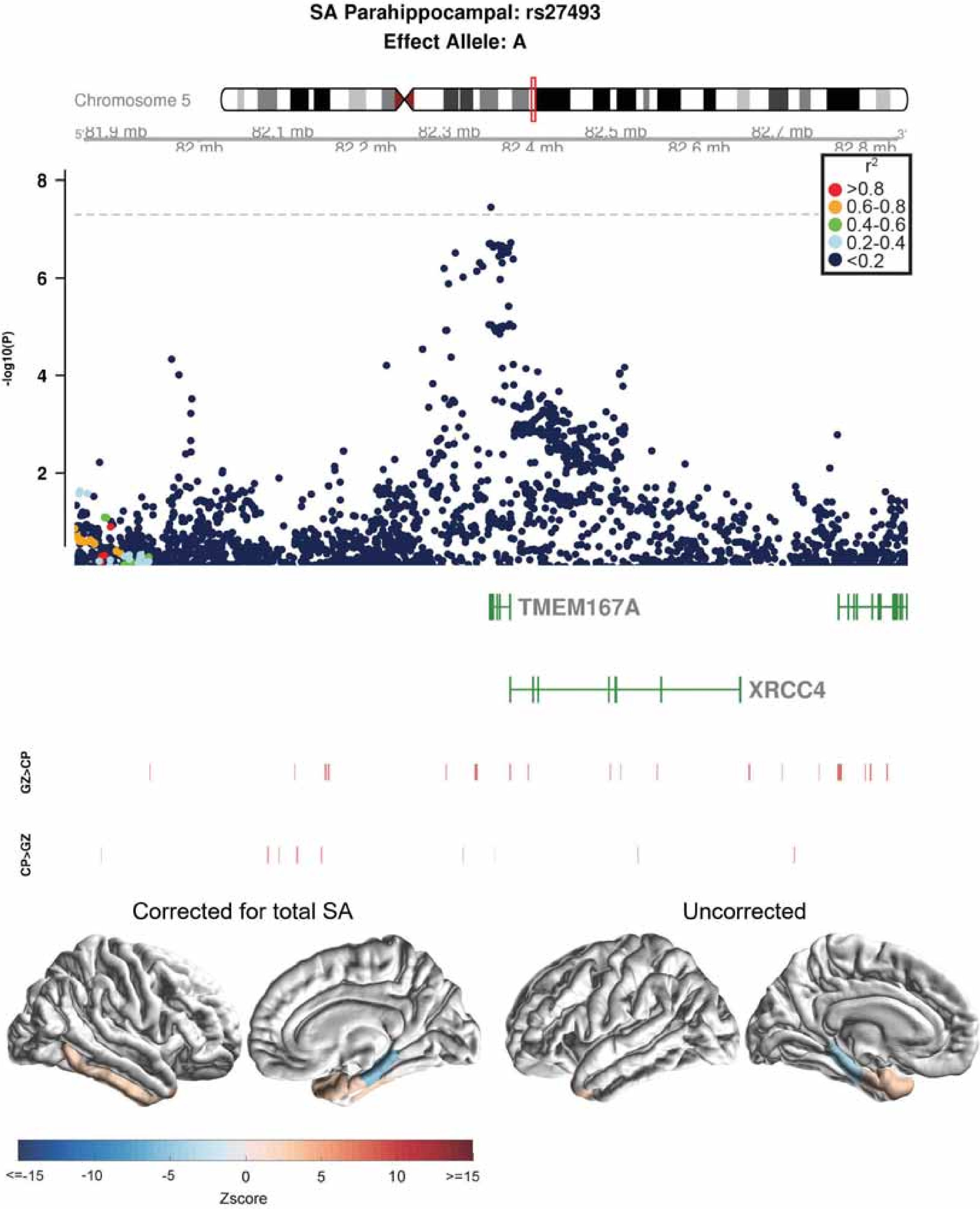
SA Parahippocampal: rs27493
Effect Allele: A
Chromosome 5
5' 81.9 mb
82 mb
82.1 mb
82.2 mb
82.3 mb
82.4 mb
82.5 mb
82.6 mb
82.7 mb
82.8 mb 3'
-log10(P)
8
6
4
2
r2
>0.8
0.6-0.8
0.4-0.6
0.2-0.4
<0.2
TMEM167A
XRCC4
GZ>CP
CP>GZ
Corrected for total SA
Uncorrected
<=-15
-10
-5
0
5
10
>=15
Zscore

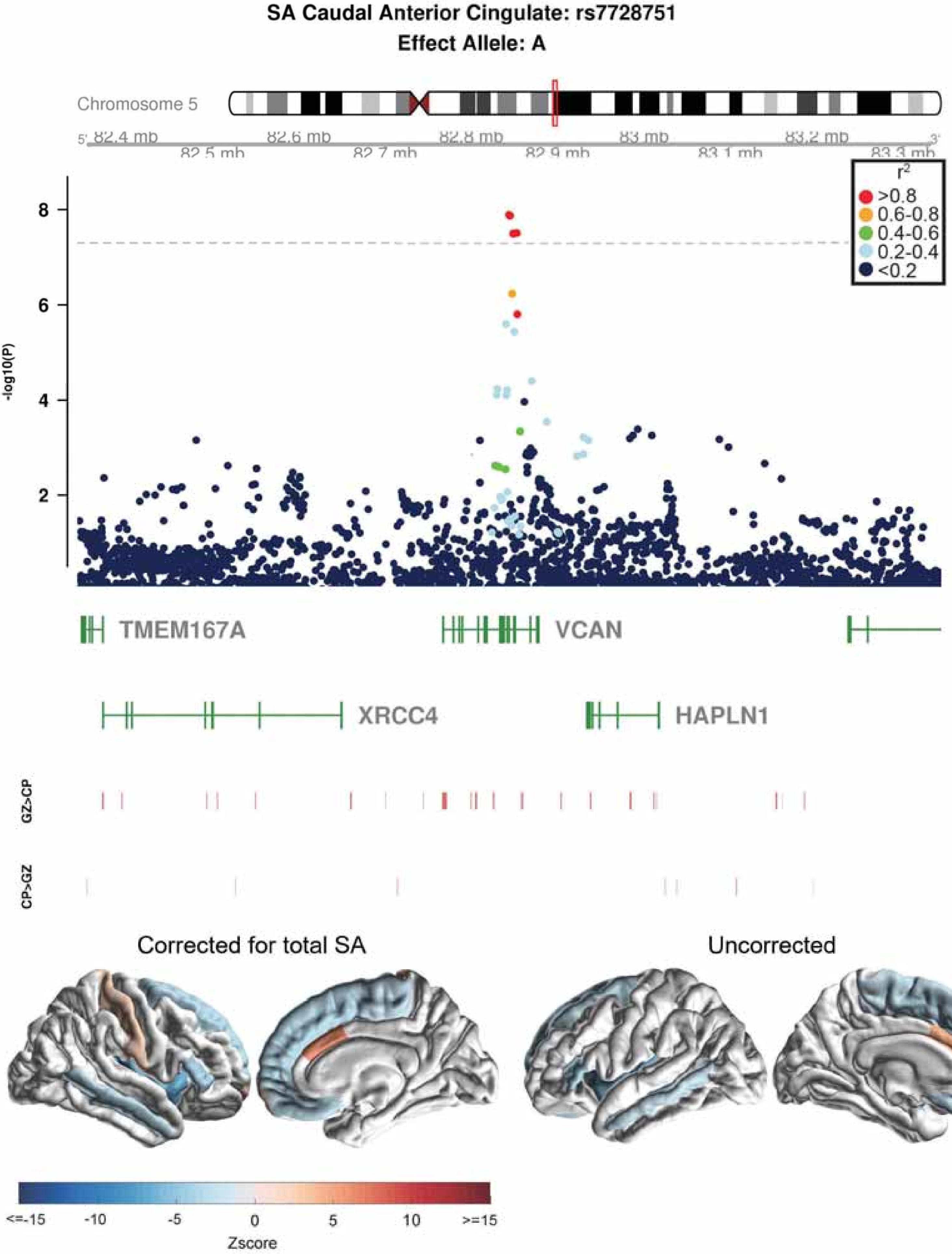
SA Caudal Anterior Cingulate: rs7728751
Effect Allele: A
Chromosome 5
82.4 mb
82.5 mb
82.6 mb
82.7 mb
82.8 mb
82.9 mb
83 mb
83.1 mb
83.2 mb
83.3 mb
5'
3'
r2
>0.8
0.6-0.8
0.4-0.6
0.2-0.4
<0.2
-log10(P)
8
6
4
2
TMEM167A
VCAN
XRCC4
HAPLN1
GZ>CP
CP>GZ
Corrected for total SA
Uncorrected
<=-15
-10
-5
0
5
10
>=15
Zscore

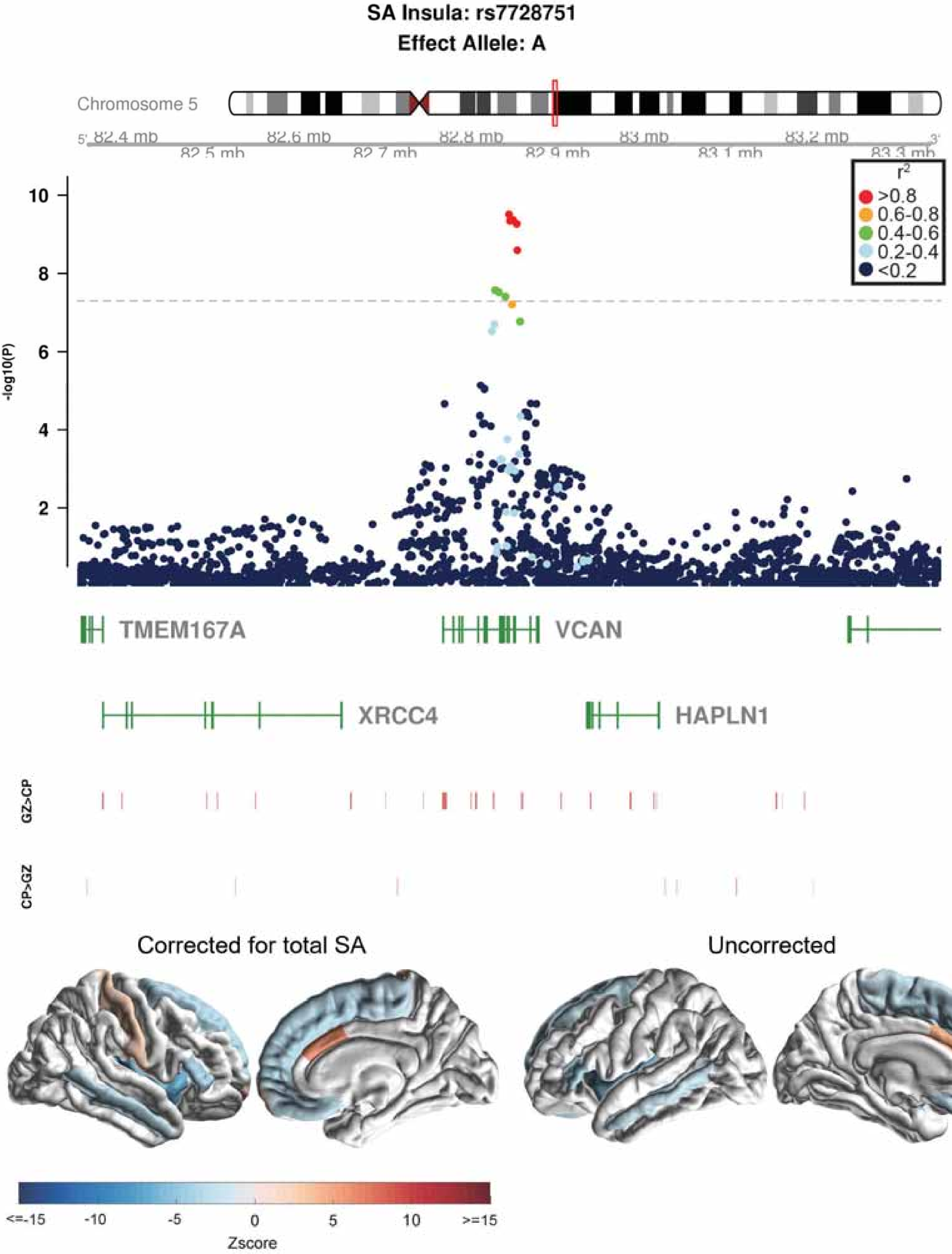
SA Insula: rs7728751
Effect Allele: A
Chromosome 5
82.4 mb
82.5 mb
82.6 mb
82.7 mb
82.8 mb
82.9 mb
83 mb
83.1 mb
83.2 mb
83.3 mb
-log10(P)
r2
>0.8
0.6-0.8
0.4-0.6
0.2-0.4
<0.2
TMEM167A
VCAN
XRCC4
HAPLN1
GZ>CP
CP>GZ
Corrected for total SA
Uncorrected
<=-15
-10
-5
0
5
10
>=15
Zscore

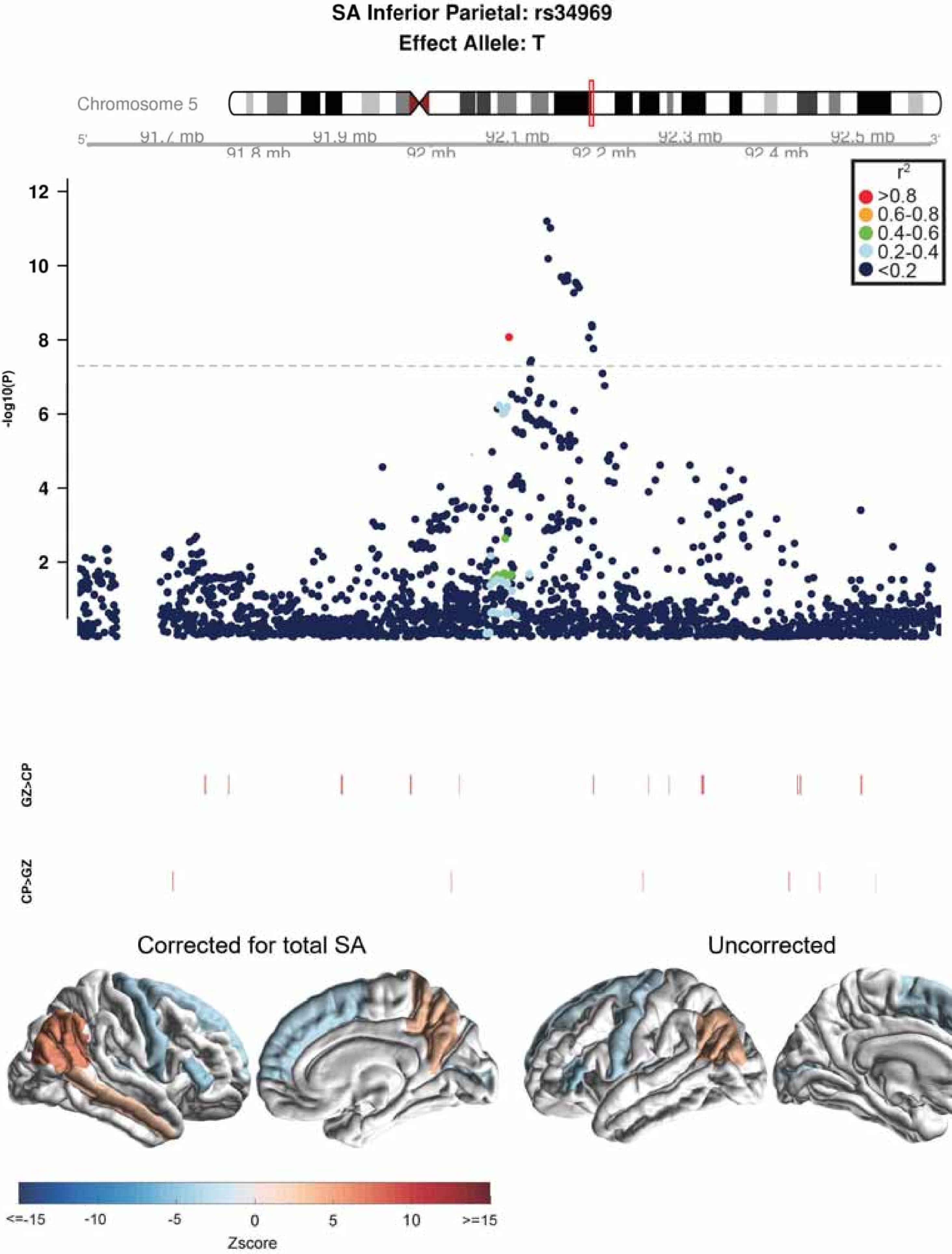

SA Inferior Parietal: rs34969
Effect Allele: T
Chromosome 5
5'
91.7 mb
91.8 mb
91.9 mb
92 mb
92.1 mb
92.2 mb
92.3 mb
92.4 mb
92.5 mb
3'
r2
>0.8
0.6-0.8
0.4-0.6
0.2-0.4
<0.2
-log10(P)
12
10
8
6
4
2
GZ>CP
CP>GZ
Corrected for total SA
Uncorrected
<=-15
-10
-5
0
5
10
>=15
Zscore

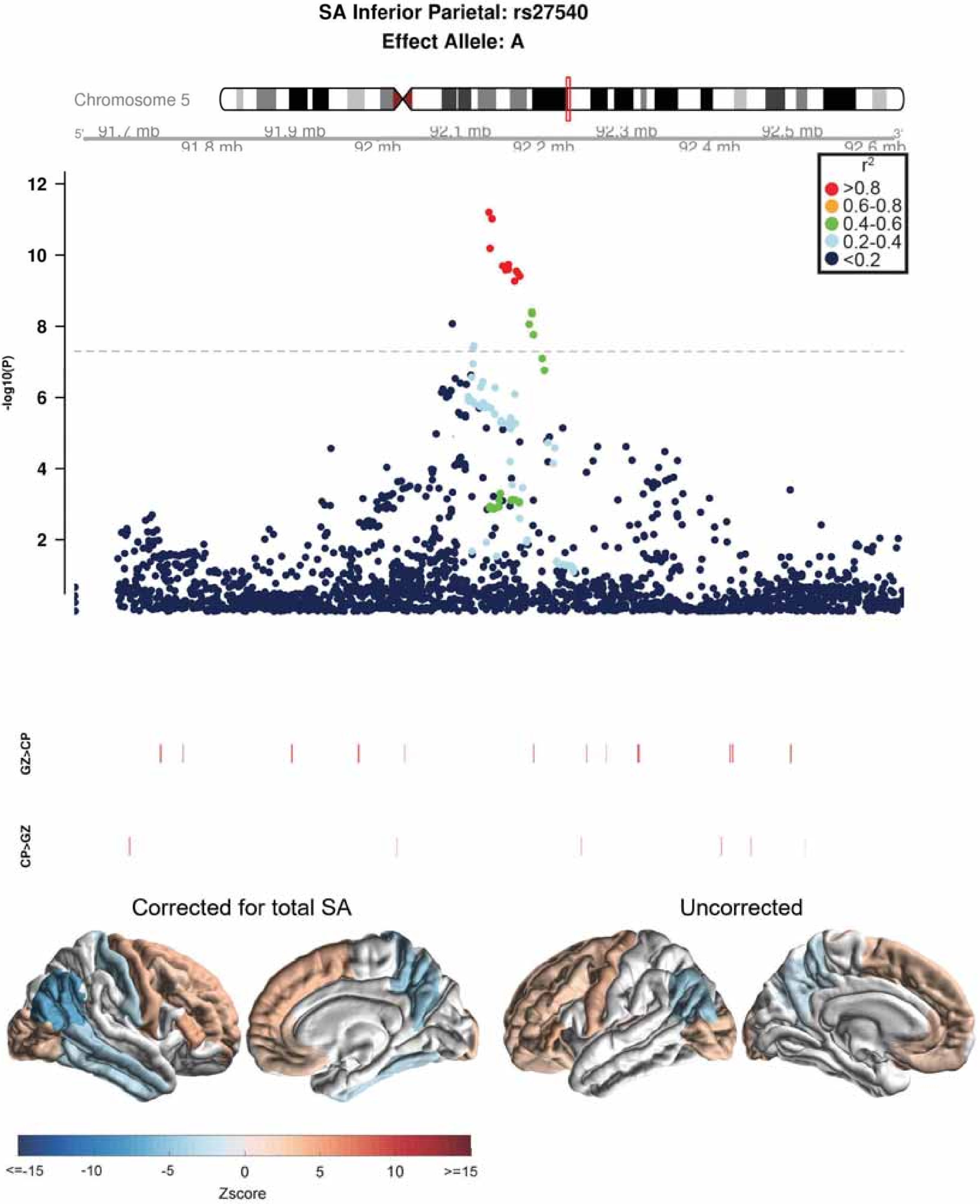
SA Inferior Parietal: rs27540
Effect Allele: A
Chromosome 5
5'
91.7 mb
91.9 mb
92.1 mb
92.3 mb
92.5 mb
3'
91.8 mb
92 mb
92.2 mb
92.4 mb
92.6 mb
$r^2$
>0.8
0.6-0.8
0.4-0.6
0.2-0.4
<0.2
-log10(P)
12
10
8
6
4
2
GZ>CP
CP>GZ
Corrected for total SA
Uncorrected
<=-15
-10
-5
0
5
10
>=15
Zscore

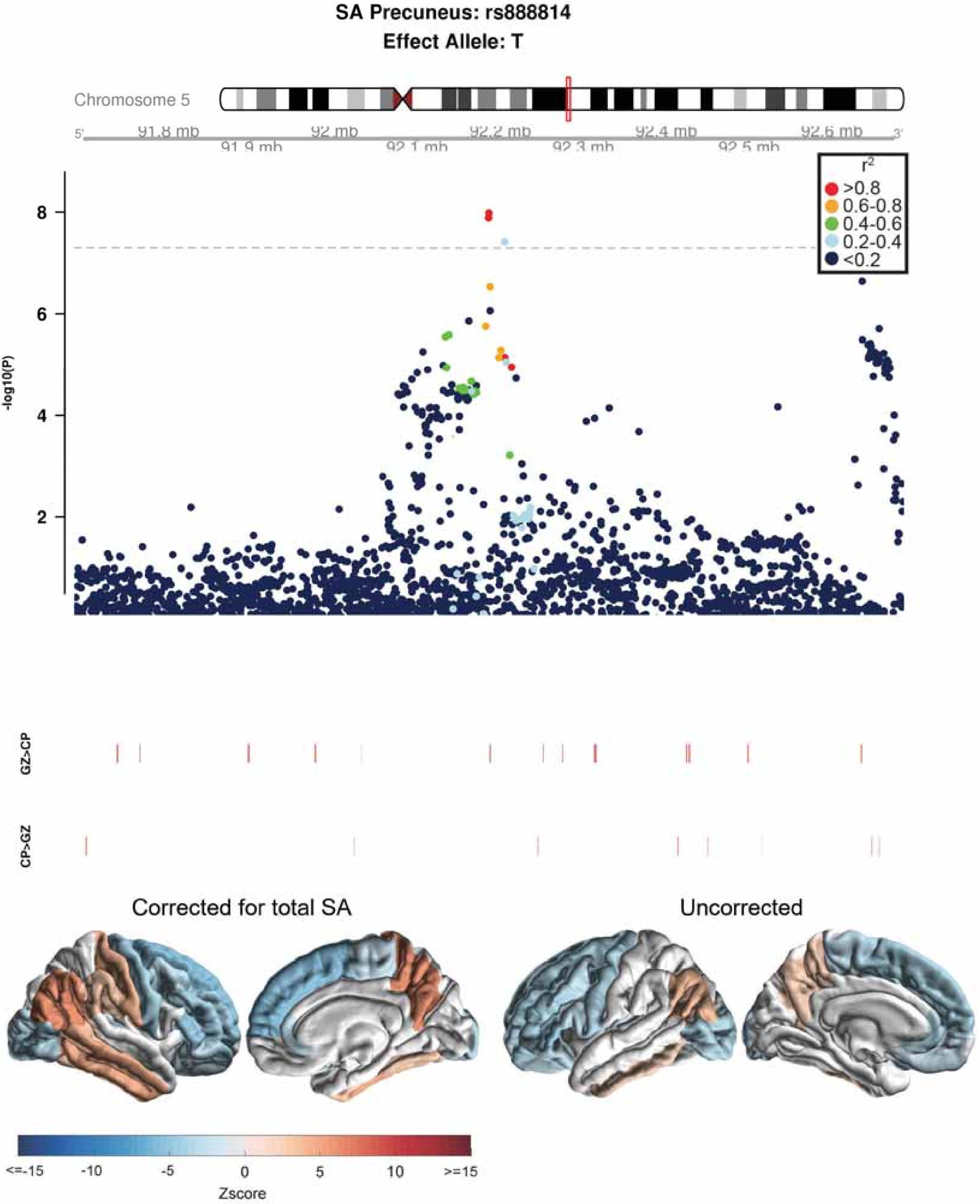
SA Precuneus: rs888814
Effect Allele: T
Chromosome 5
5'
91.8 mb
91.9 mb
92 mb
92.1 mb
92.2 mb
92.3 mb
92.4 mb
92.5 mb
92.6 mb
3'
r2
>0.8
0.6-0.8
0.4-0.6
0.2-0.4
<0.2
-log10(P)
8
6
4
2
GZ>CP
CP>GZ
Corrected for total SA
Uncorrected
<=-15
-10
-5
0
5
10
>=15
Zscore

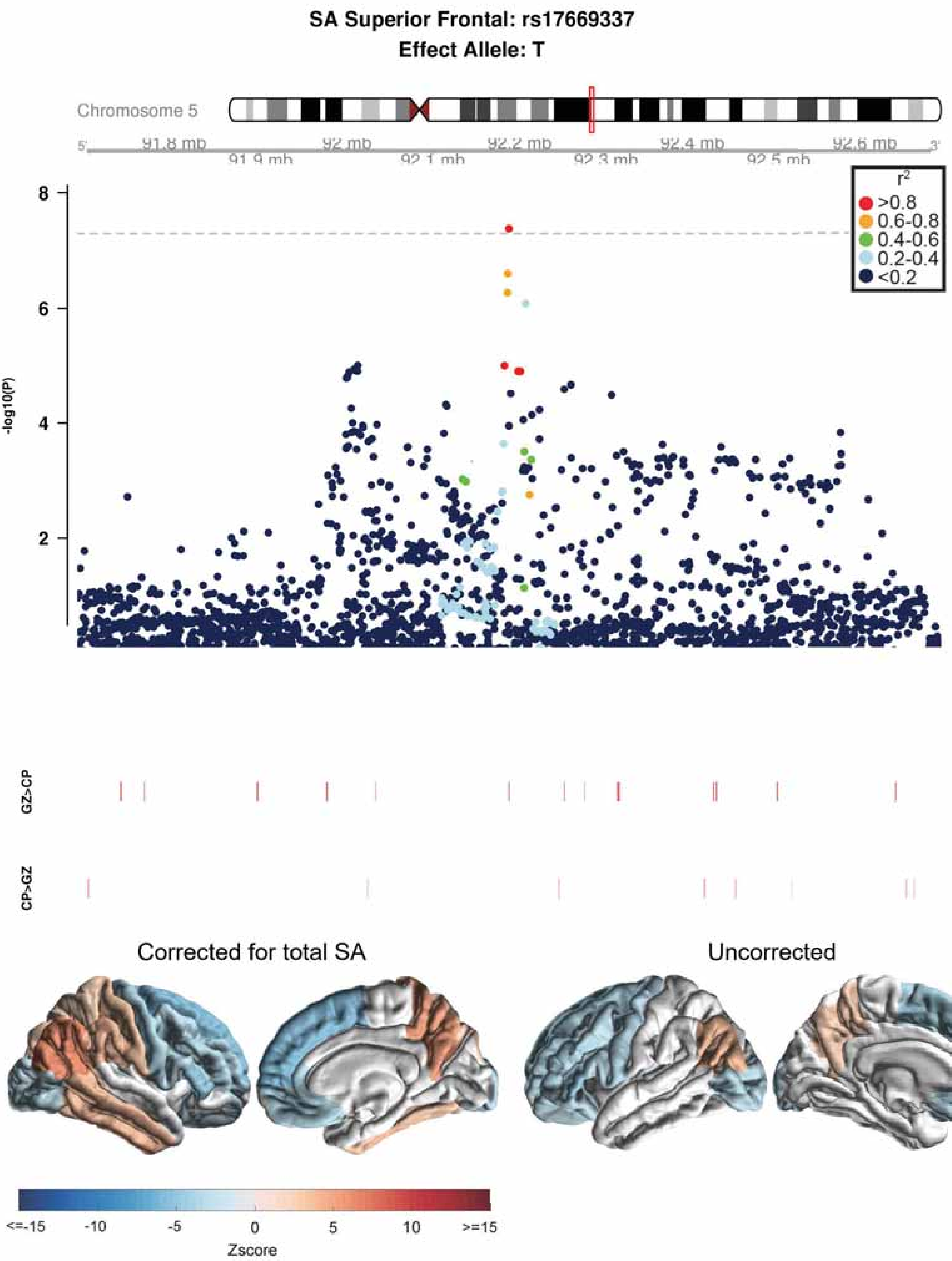
SA Superior Frontal: rs17669337
Effect Allele: T
Chromosome 5
91.8 mb
92 mb
92.2 mb
92.4 mb
92.6 mb
91.9 mb
92.1 mb
92.3 mb
92.5 mb
r2
>0.8
0.6-0.8
0.4-0.6
0.2-0.4
<0.2
-log10(P)
GZ>CP
CP>GZ
Corrected for total SA
Uncorrected
<=-15
-10
-5
0
5
10
>=15
Zscore

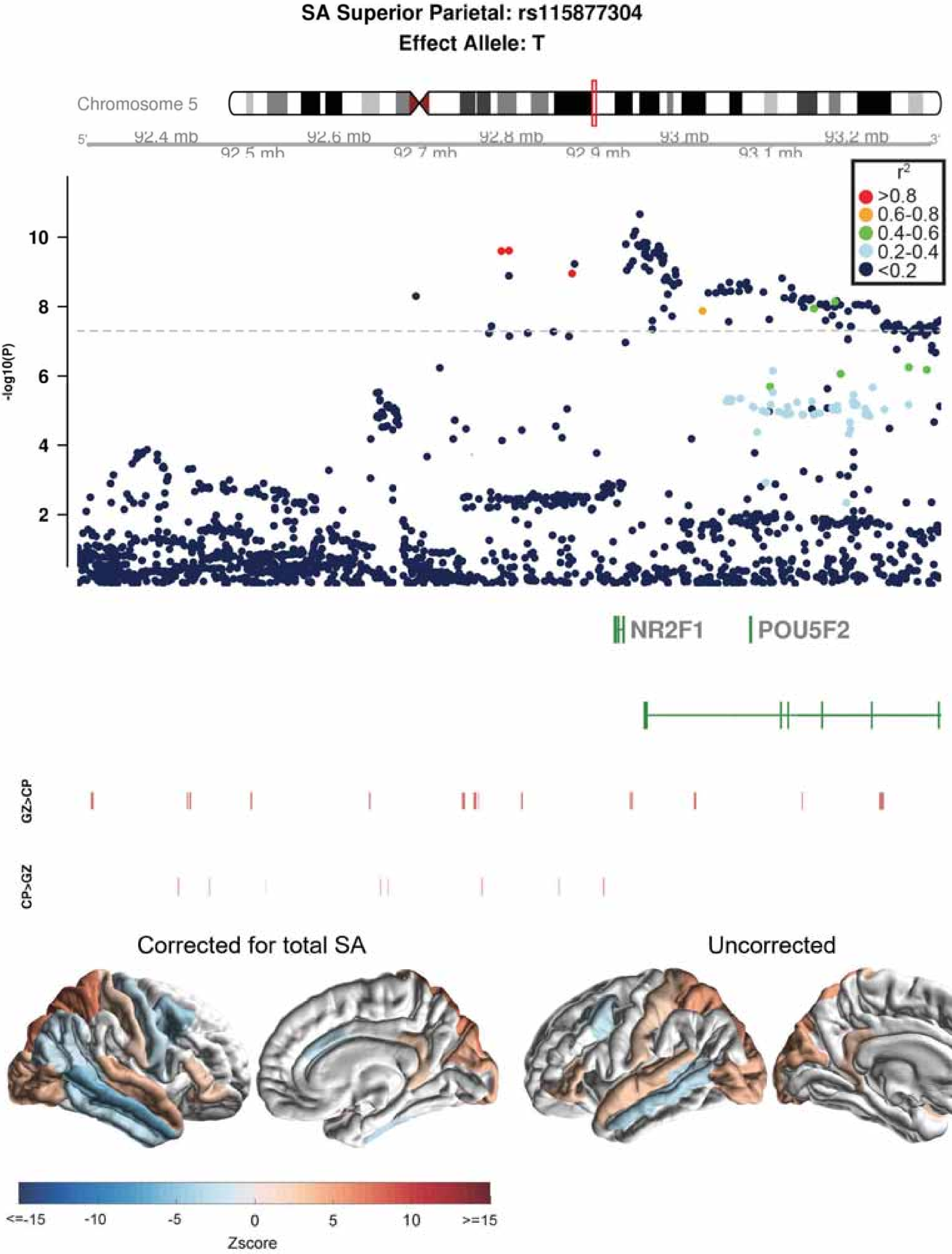
SA Superior Parietal: rs115877304
Effect Allele: T
Chromosome 5
92.4 mb
92.5 mb
92.6 mb
92.7 mb
92.8 mb
92.9 mb
93 mb
93.1 mb
93.2 mb
-log10(P)
r2
>0.8
0.6-0.8
0.4-0.6
0.2-0.4
<0.2
NR2F1
POU5F2
GZ>CP
CP>GZ
Corrected for total SA
Uncorrected
<=-15
-10
-5
0
5
10
>=15
Zscore

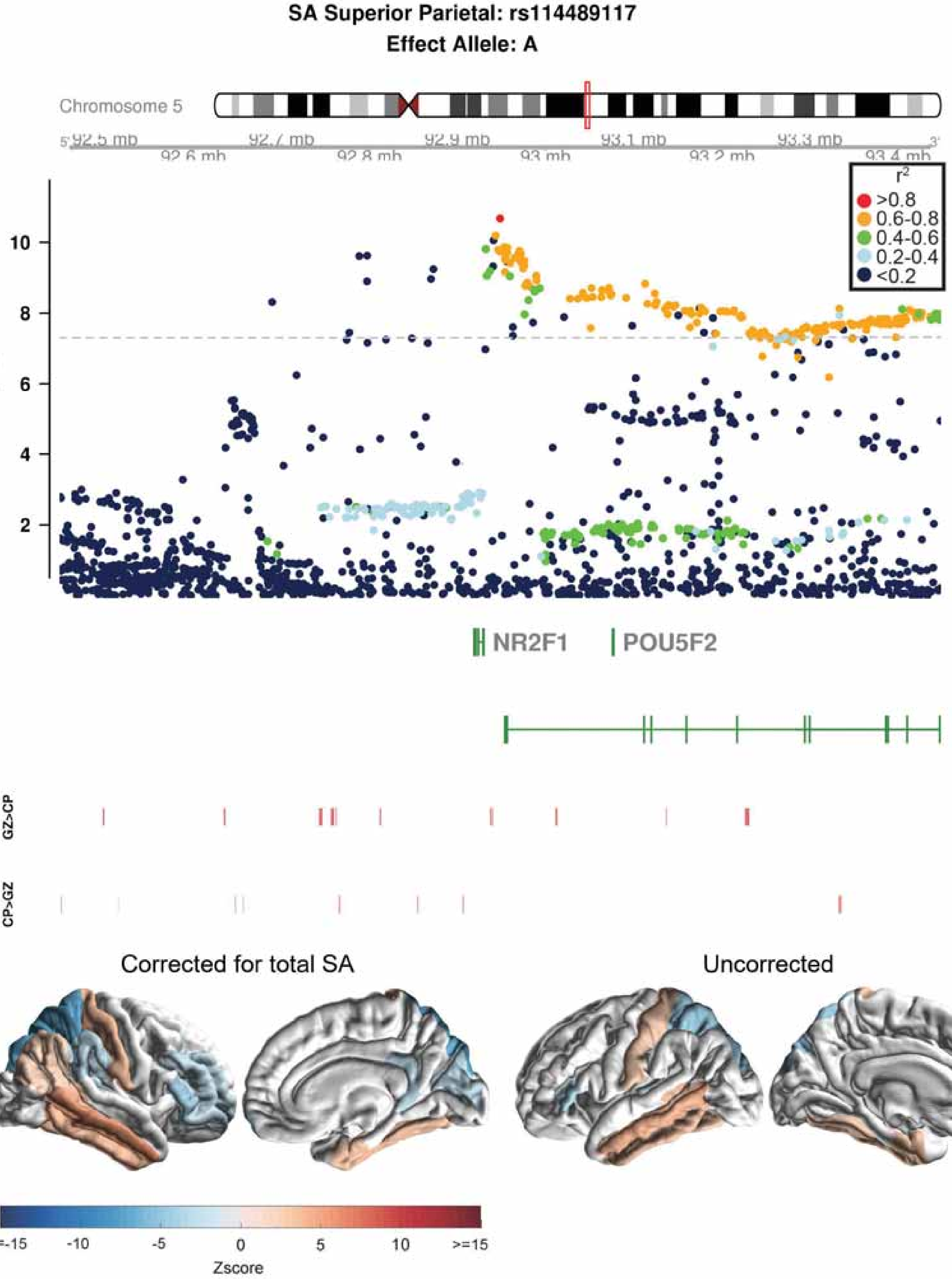

SA Superior Parietal: rs114489117
Effect Allele: A
Chromosome 5
92.5 mb
92.6 mb
92.7 mb
92.8 mb
92.9 mb
93 mb
93.1 mb
93.2 mb
93.3 mb
93.4 mb
r2
>0.8
0.6-0.8
0.4-0.6
0.2-0.4
<0.2
-log10(P)
10
8
6
4
2
NR2F1
POU5F2
GZ>CP
CP>GZ
Corrected for total SA
Uncorrected
<=-15
-10
-5
0
5
10
>=15
Zscore

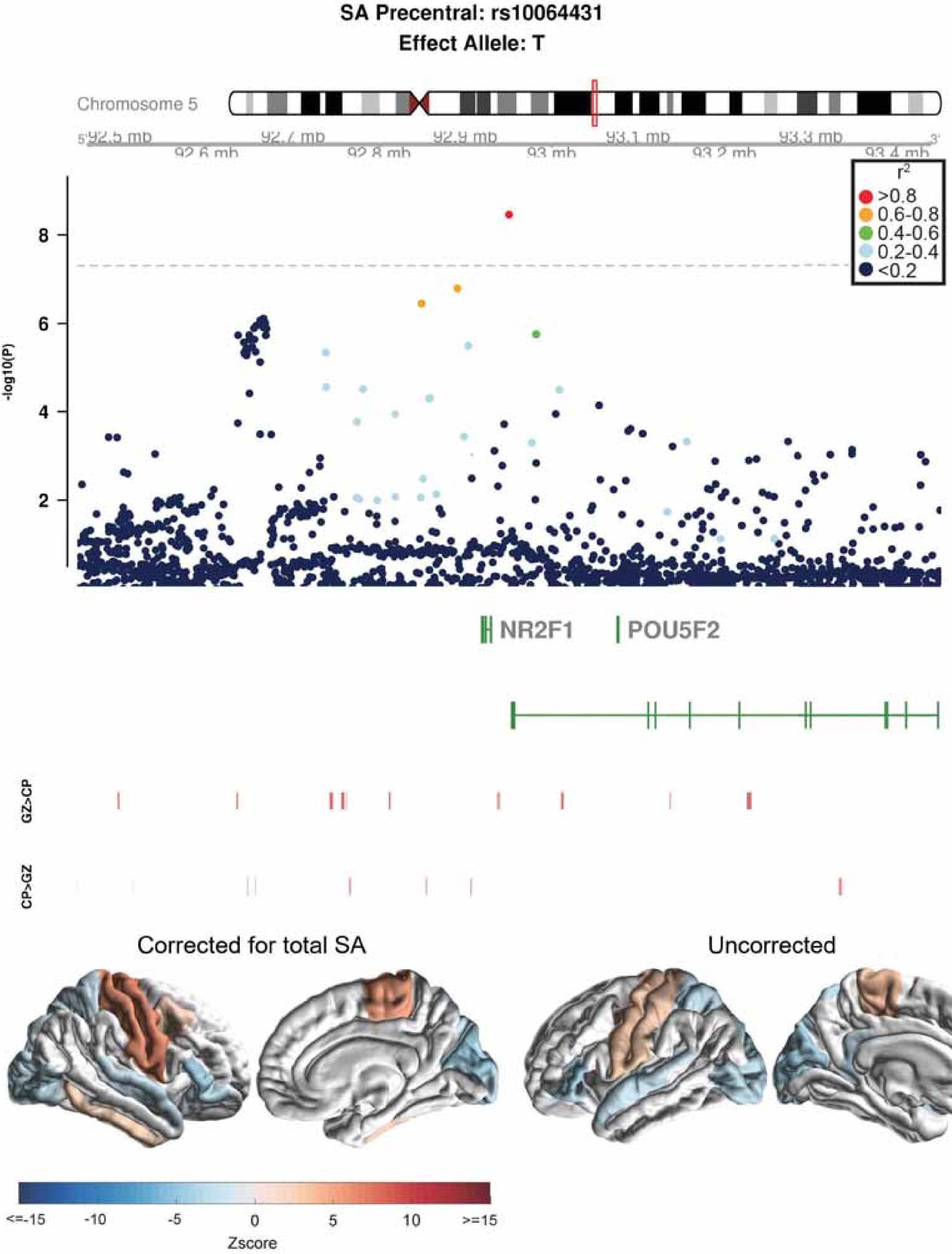

SA Precentral: rs10064431
Effect Allele: T
Chromosome 5
92.5 mb
92.6 mb
92.7 mb
92.8 mb
92.9 mb
93 mb
93.1 mb
93.2 mb
93.3 mb
93.4 mb
r2
>0.8
0.6-0.8
0.4-0.6
0.2-0.4
<0.2
-log10(P)
NR2F1
POU5F2
GZ>CP
CP>GZ
Corrected for total SA
Uncorrected
<=-15
-10
-5
0
5
10
>=15
Zscore

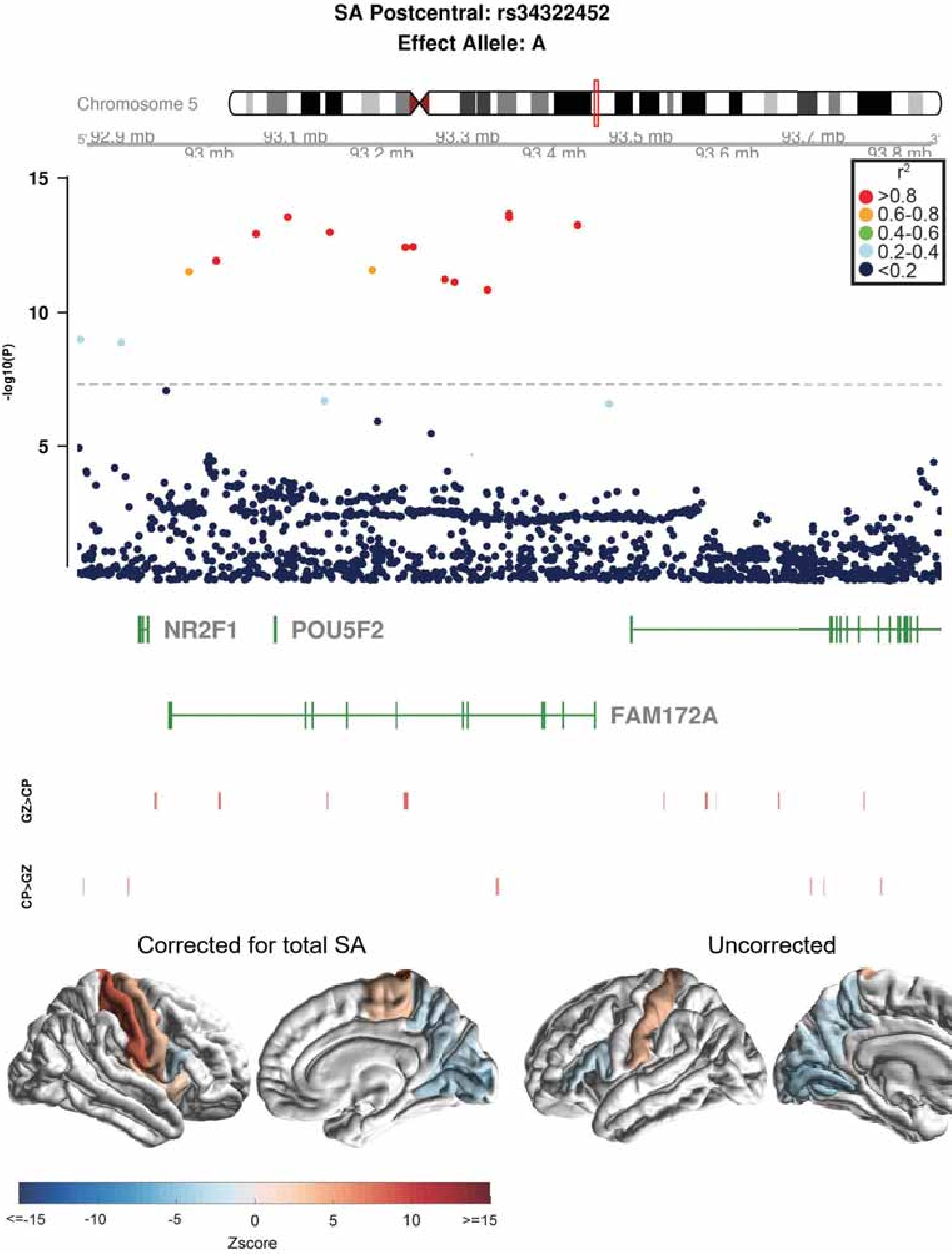
SA Postcentral: rs34322452
Effect Allele: A
Chromosome 5
92.9 mb
93 mb
93.1 mb
93.2 mb
93.3 mb
93.4 mb
93.5 mb
93.6 mb
93.7 mb
93.8 mb
-log10(P)
15
10
5
r2
>0.8
0.6-0.8
0.4-0.6
0.2-0.4
<0.2
NR2F1
POU5F2
FAM172A
GZ>CP
CP>GZ
Corrected for total SA
Uncorrected
<=-15
-10
-5
0
5
10
>=15
Zscore

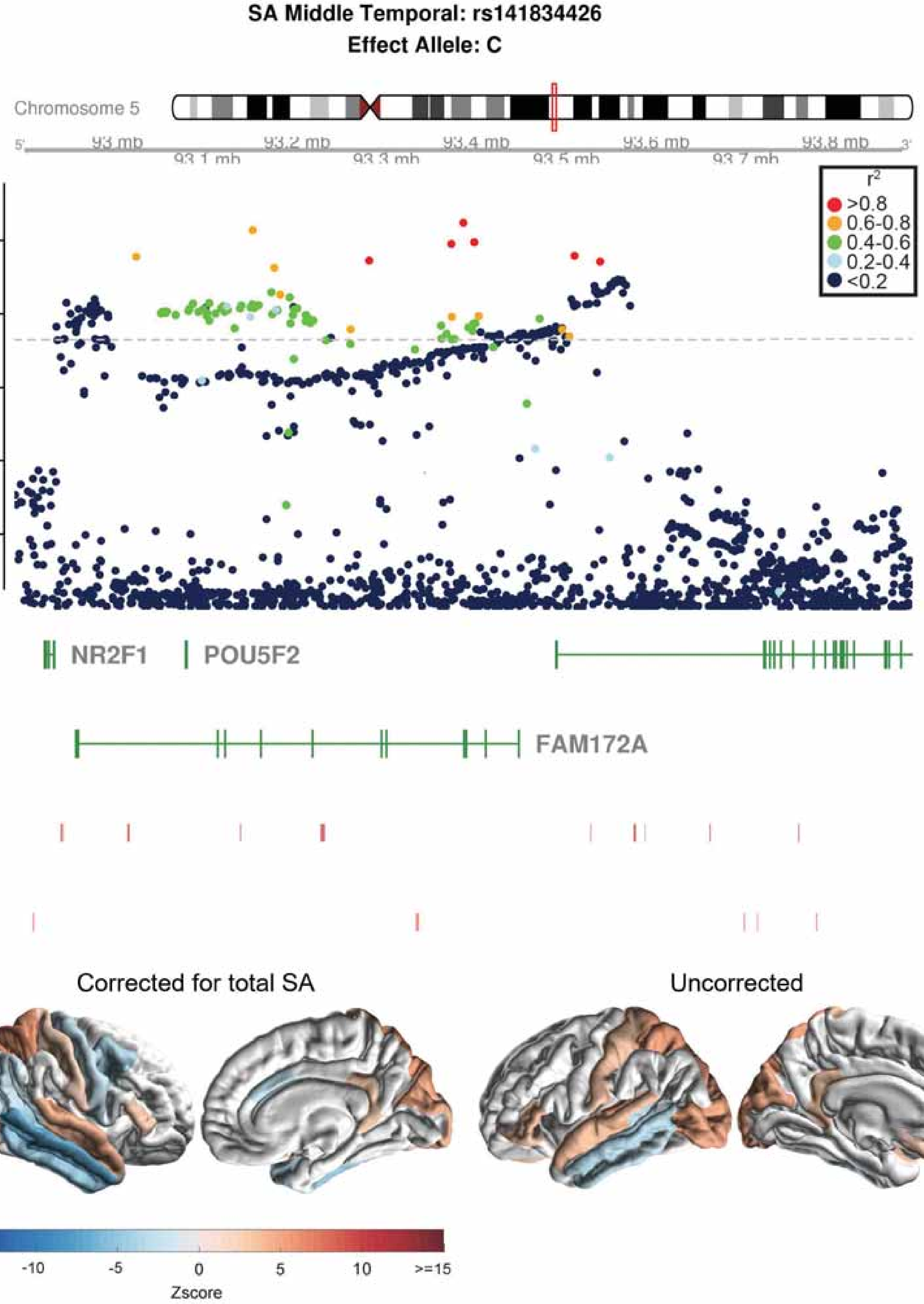

SA Middle Temporal: rs141834426
Effect Allele: C
Chromosome 5
93 mb
93.1 mb
93.2 mb
93.3 mb
93.4 mb
93.5 mb
93.6 mb
93.7 mb
93.8 mb
r2
>0.8
0.6-0.8
0.4-0.6
0.2-0.4
<0.2
-log10(P)
NR2F1
POU5F2
FAM172A
GZ>CP
CP>GZ
Corrected for total SA
Uncorrected
<=-15
-10
-5
0
5
10
>=15
Zscore

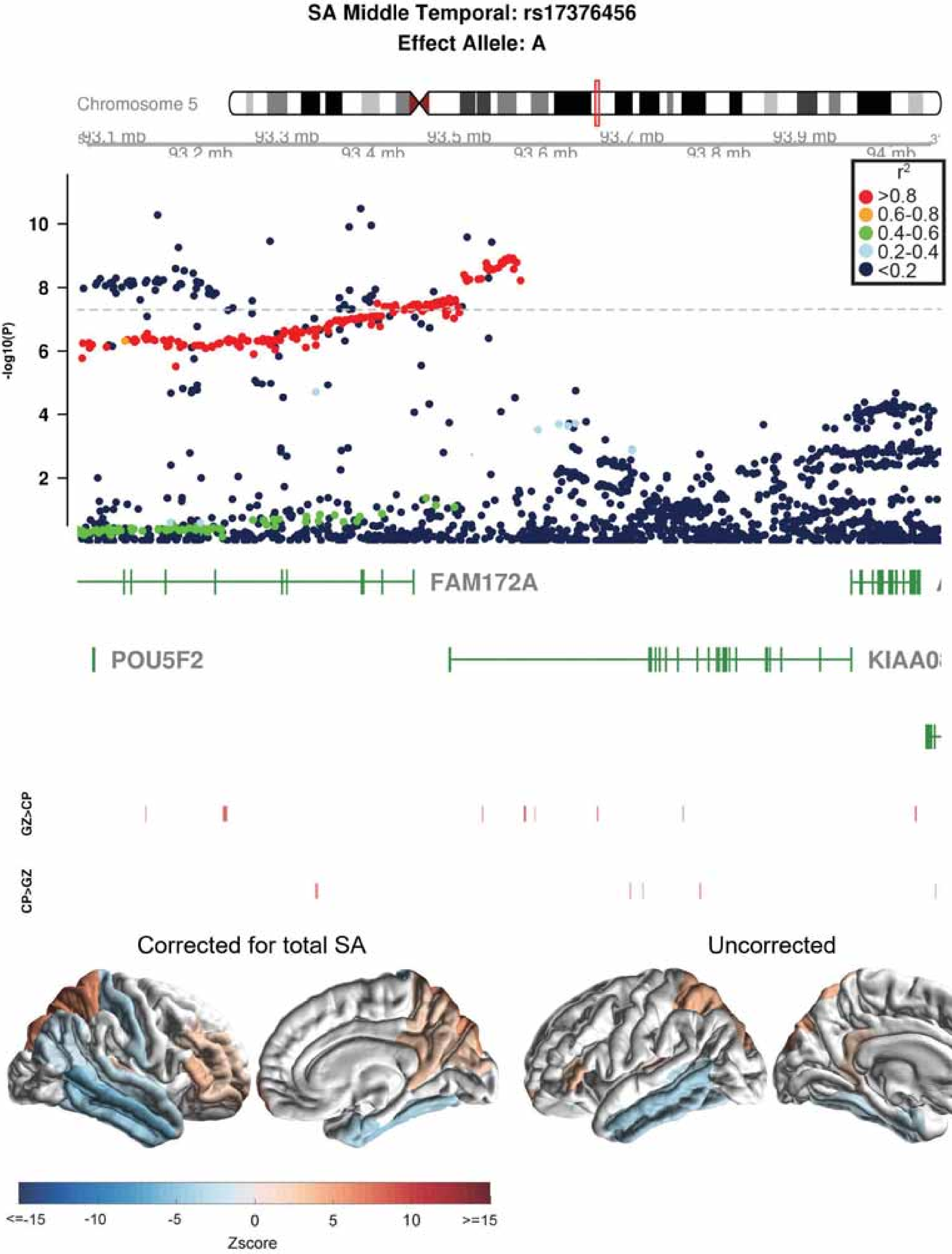
SA Middle Temporal: rs17376456
Effect Allele: A
Chromosome 5
93.1 mb
93.2 mb
93.3 mb
93.4 mb
93.5 mb
93.6 mb
93.7 mb
93.8 mb
93.9 mb
94 mb
r2
>0.8
0.6-0.8
0.4-0.6
0.2-0.4
<0.2
-log10(P)
10
8
6
4
2
FAM172A
POU5F2
GZ>CP
CP>GZ
Corrected for total SA
Uncorrected
<=-15
-10
-5
0
5
10
>=15
Zscore

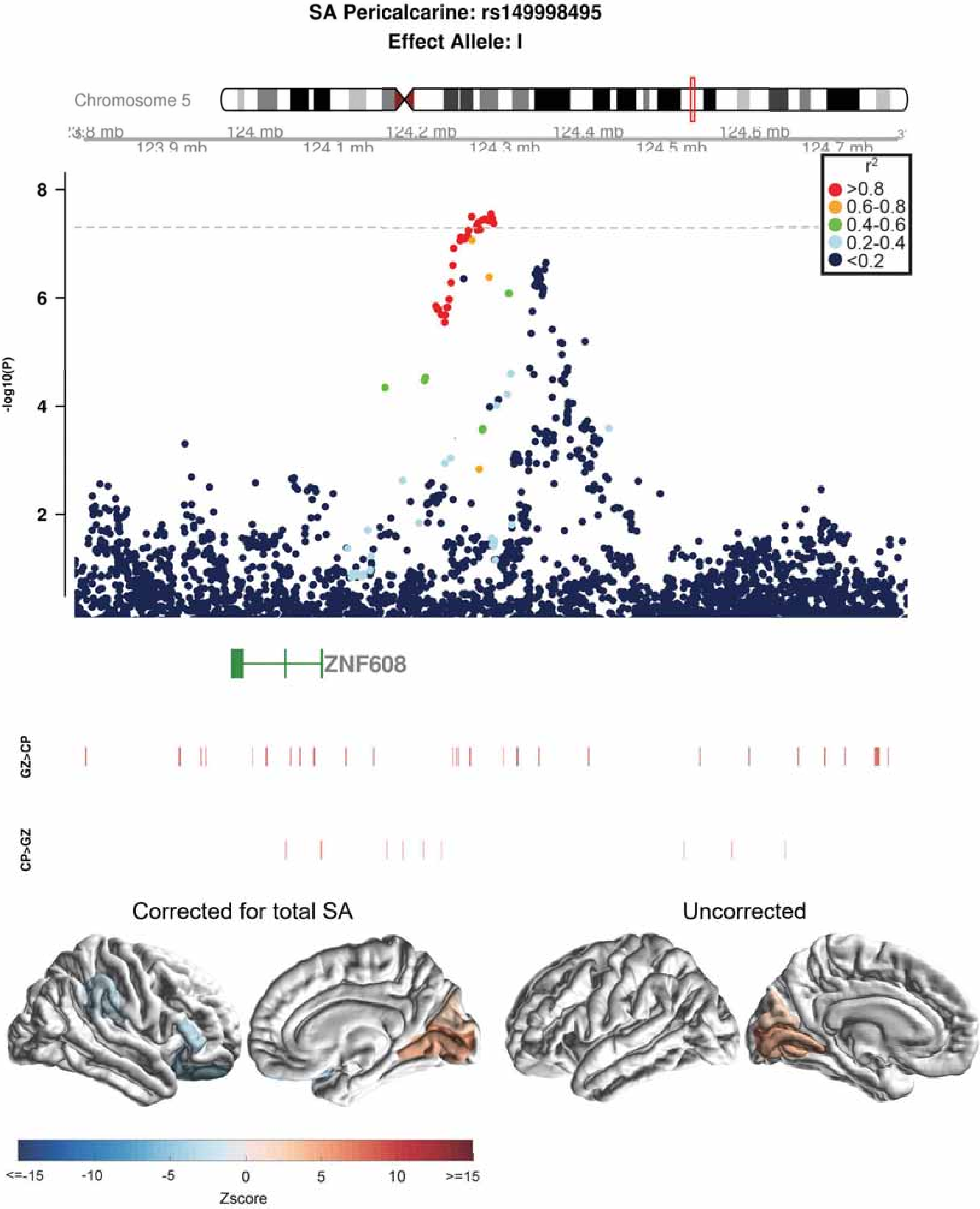

SA Pericalcarine: rs149998495
Effect Allele: I
Chromosome 5
124 mb
124.2 mb
124.4 mb
124.6 mb
123.9 mb
124.1 mb
124.3 mb
124.5 mb
124.7 mb
r2
>0.8
0.6-0.8
0.4-0.6
0.2-0.4
<0.2
8
6
4
2
-log10(P)
ZNF608
GZ>CP
CP>GZ
Corrected for total SA
Uncorrected
<=-15
-10
-5
0
5
10
>=15
Zscore

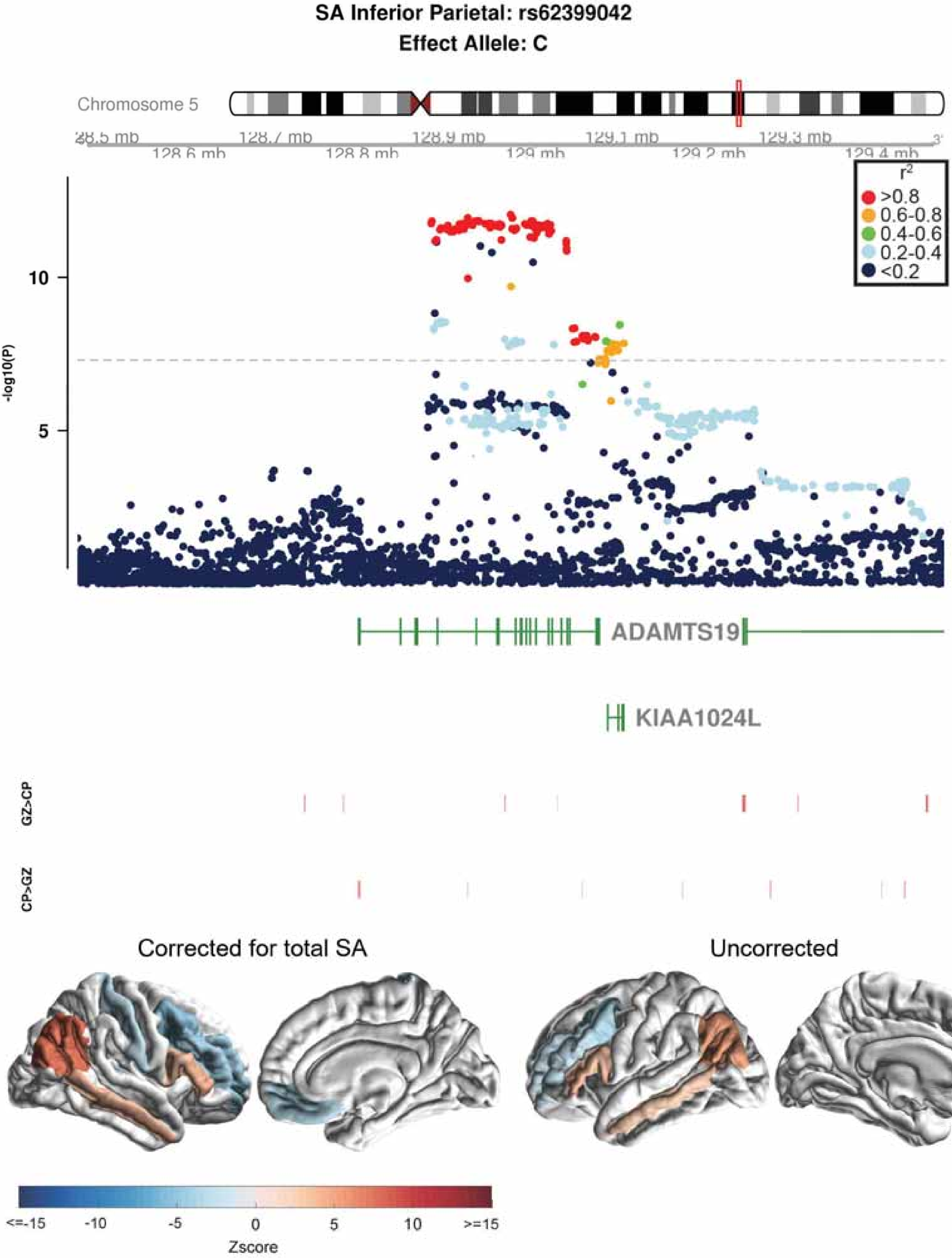
SA Inferior Parietal: rs62399042
Effect Allele: C
Chromosome 5
128.7 mb
128.9 mb
129.1 mb
129.3 mb
128.6 mb
128.8 mb
129 mb
129.2 mb
129.4 mb
r2
>0.8
0.6-0.8
0.4-0.6
0.2-0.4
<0.2
-log10(P)
10
5
ADAMTS19
KIAA1024L
GZ>CP
CP>GZ
Corrected for total SA
Uncorrected
<=-15
-10
-5
0
5
10
>=15
Zscore

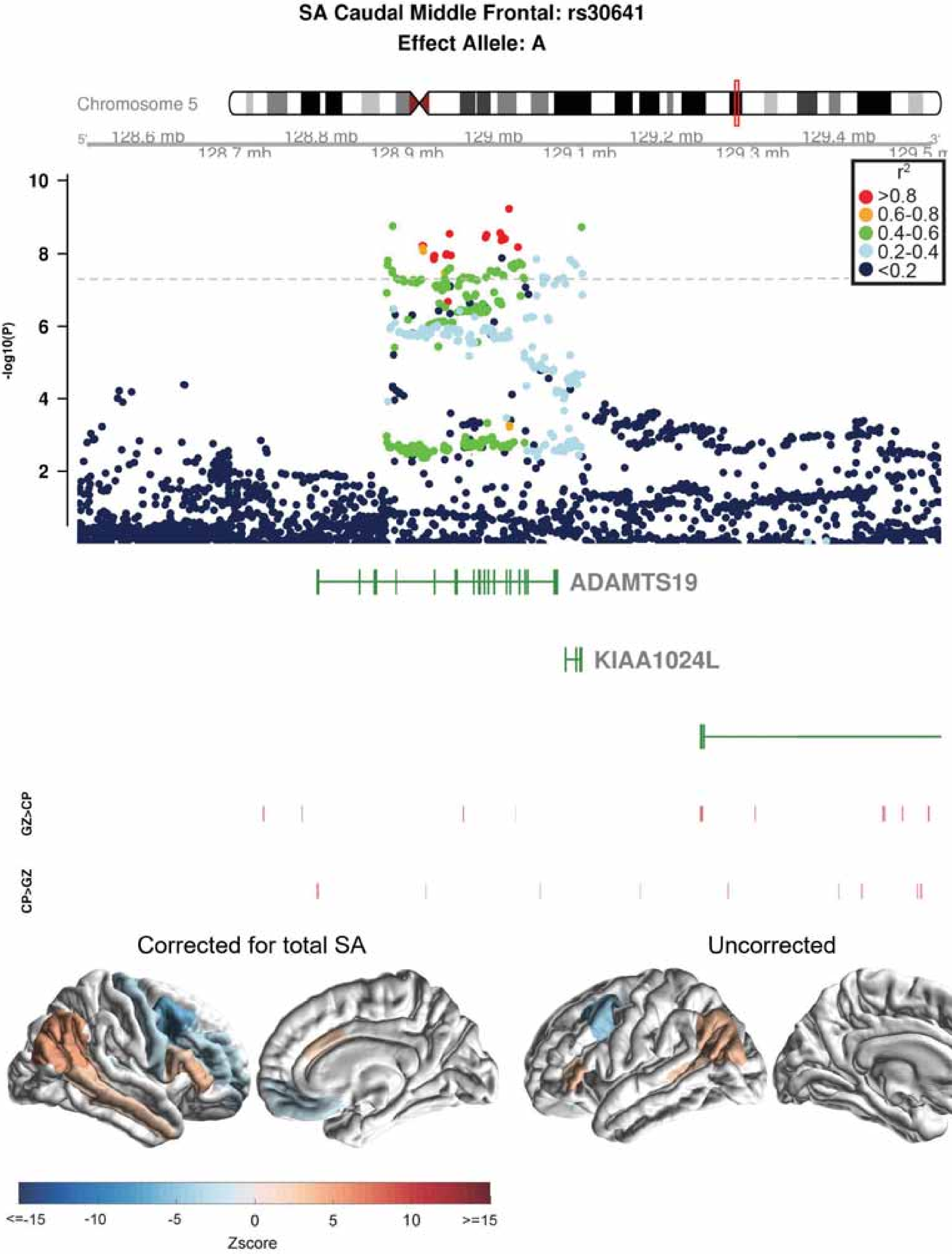
SA Caudal Middle Frontal: rs30641
Effect Allele: A
Chromosome 5
128.6 mb
128.7 mb
128.8 mb
128.9 mb
129 mb
129.1 mb
129.2 mb
129.3 mb
129.4 mb
-log10(P)
r2
>0.8
0.6-0.8
0.4-0.6
0.2-0.4
<0.2
ADAMTS19
KIAA1024L
GZ>CP
CP>GZ
Corrected for total SA
Uncorrected
<=-15
-10
-5
0
5
10
>=15
Zscore

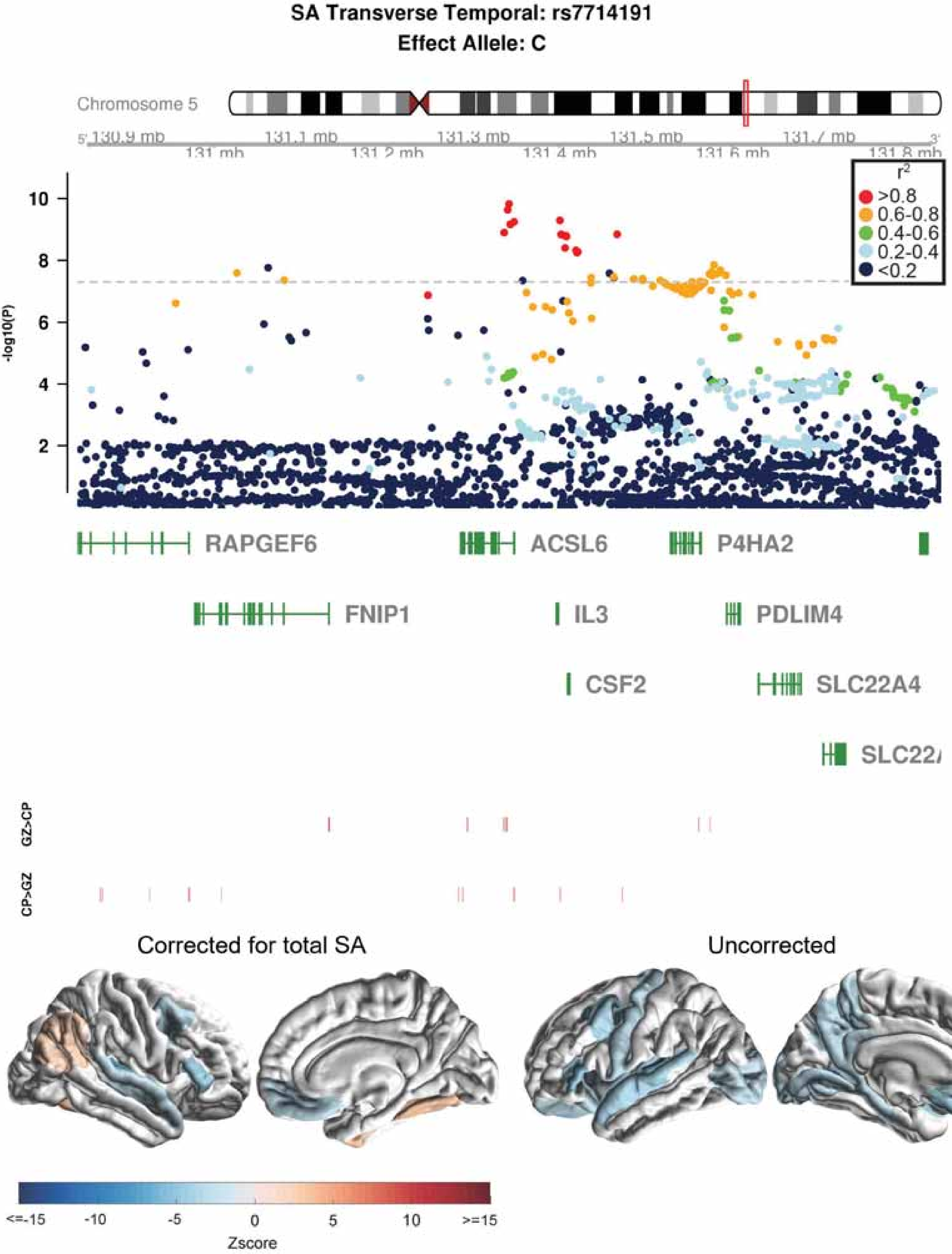

SA Transverse Temporal: rs7714191
Effect Allele: C
Chromosome 5
130.9 mb
131 mb
131.1 mb
131.2 mb
131.3 mb
131.4 mb
131.5 mb
131.6 mb
131.7 mb
131.8 mb
-log10(P)
r2
>0.8
0.6-0.8
0.4-0.6
0.2-0.4
<0.2
RAPGEF6
ACSL6
P4HA2
FNIP1
IL3
PDLIM4
CSF2
SLC22A4
SLC22/
GZ>CP
CP>GZ
Corrected for total SA
Uncorrected
<=-15
-10
-5
0
5
10
>=15
Zscore

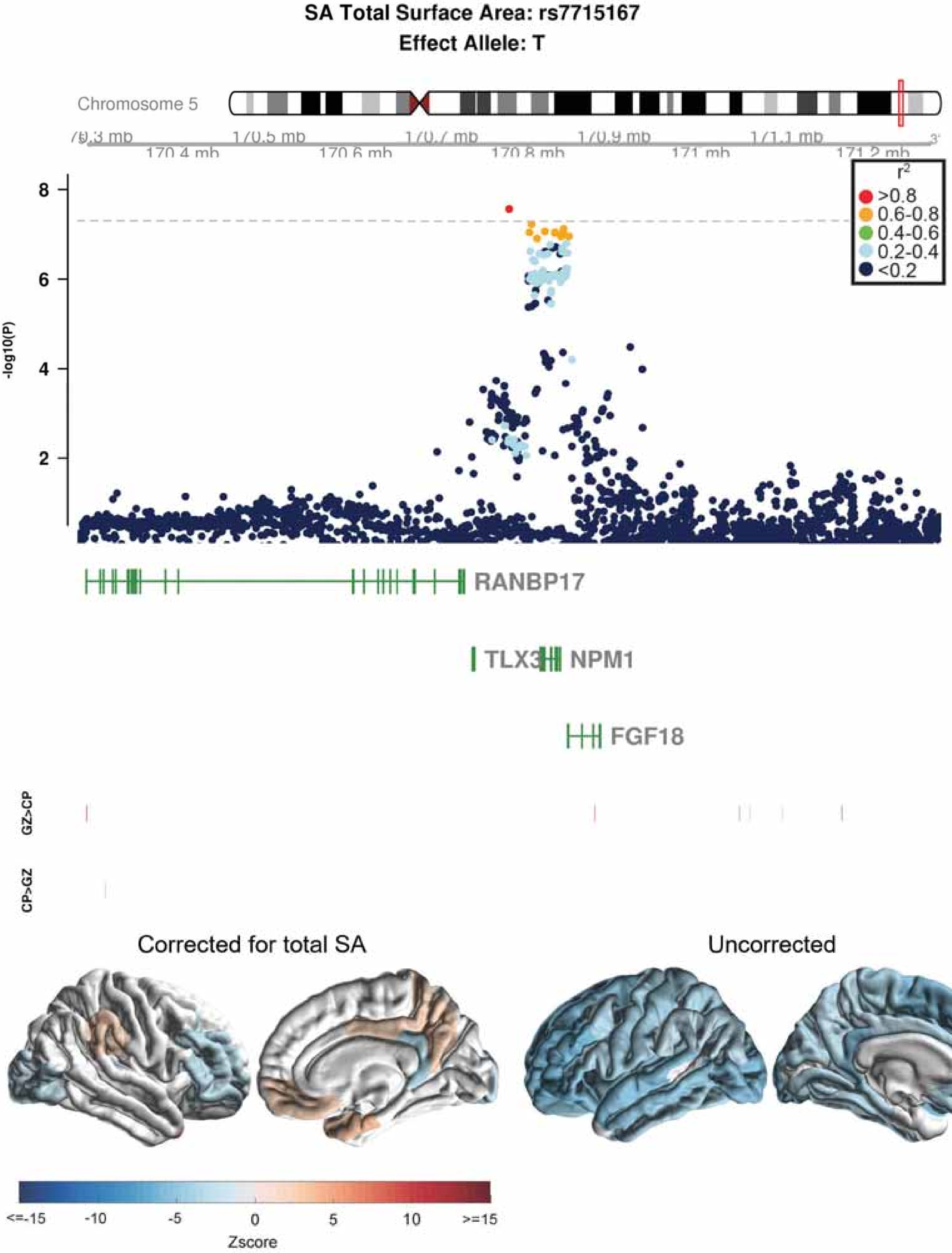
SA Total Surface Area: rs7715167
Effect Allele: T
Chromosome 5
-log10(P)
RANBP17
TLX3
NPM1
FGF18
GZ>CP
CP>GZ
Corrected for total SA
Uncorrected
Zscore

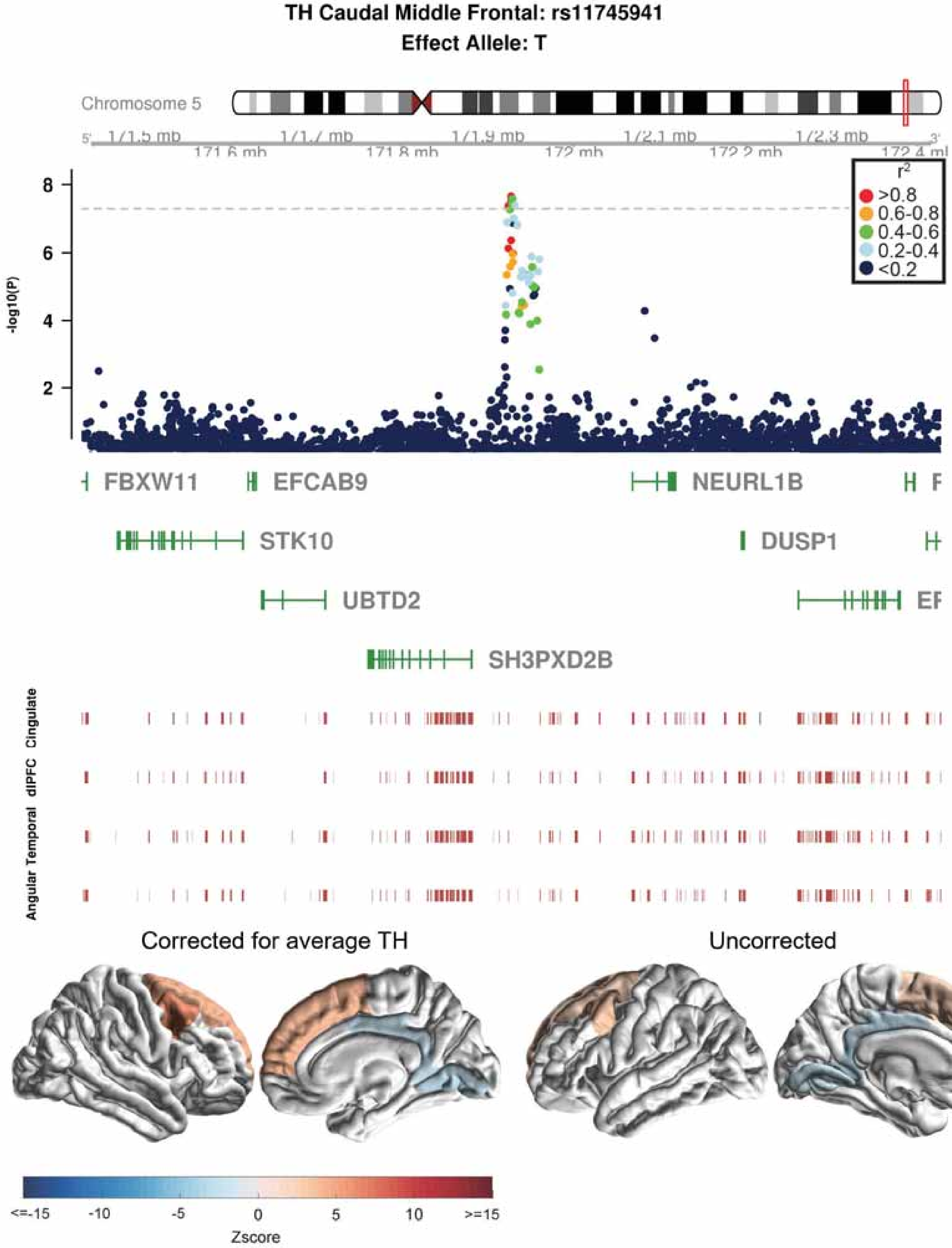
TH Caudal Middle Frontal: rs11745941
Effect Allele: T
Chromosome 5
171.6 mb
171.8 mb
172 mb
172.2 mb
-log10(P)
r2
>0.8
0.6-0.8
0.4-0.6
0.2-0.4
<0.2
FBXW11
EFCAB9
NEURL1B
STK10
DUSP1
UBTD2
SH3PXD2B
Cingulate
dlPFC
Temporal
Angular
Corrected for average TH
Uncorrected
Zscore
<=-15
-10
-5
0
5
10
>=15

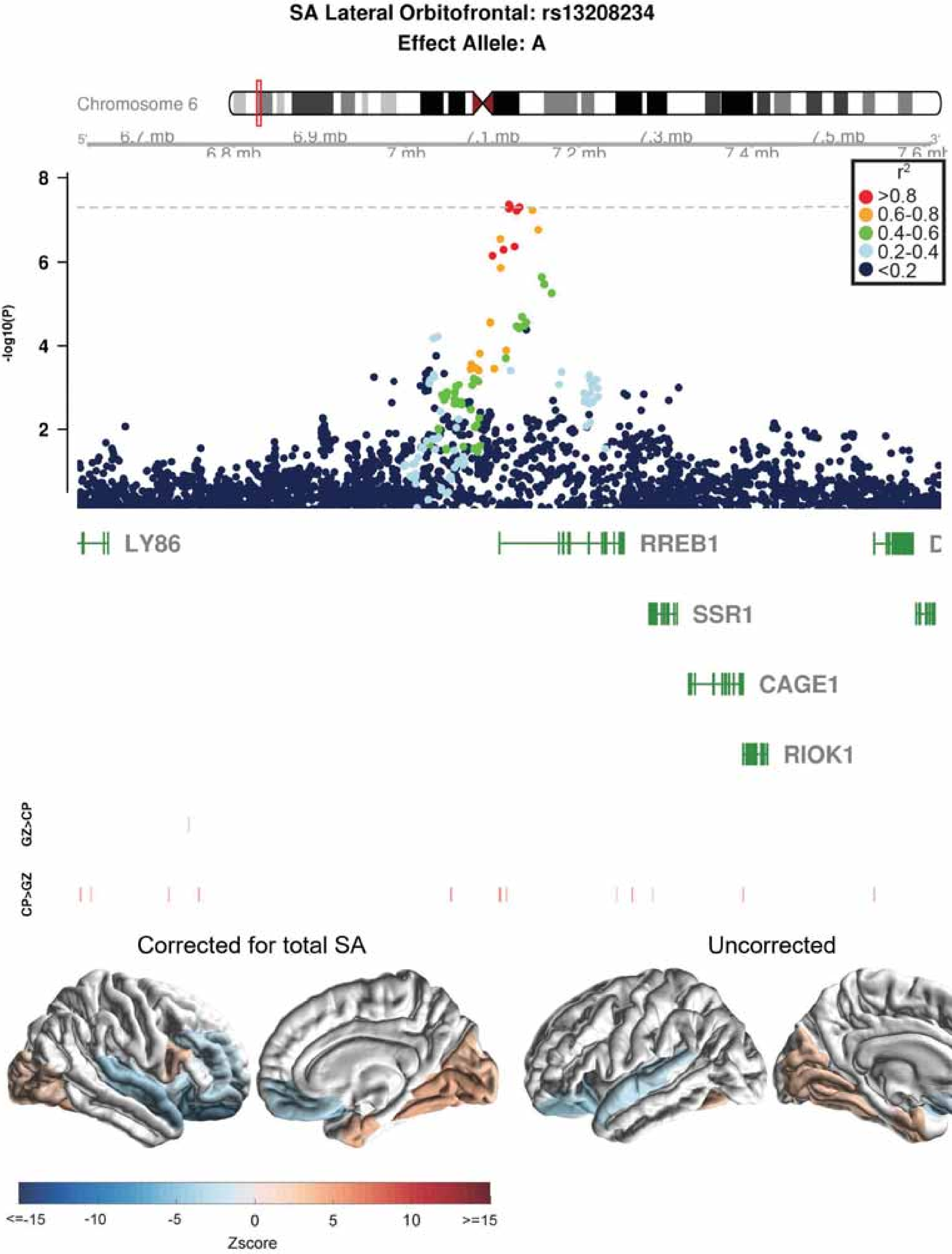
SA Lateral Orbitofrontal: rs13208234
Effect Allele: A
Chromosome 6
6.7 mb
6.8 mb
6.9 mb
7 mb
7.1 mb
7.2 mb
7.3 mb
7.4 mb
7.5 mb
7.6 mb
-log10(P)
r2
>0.8
0.6-0.8
0.4-0.6
0.2-0.4
<0.2
LY86
RREB1
SSR1
CAGE1
RIOK1
GZ>CP
CP>GZ
Corrected for total SA
Uncorrected
<=-15
-10
-5
0
5
10
>=15
Zscore

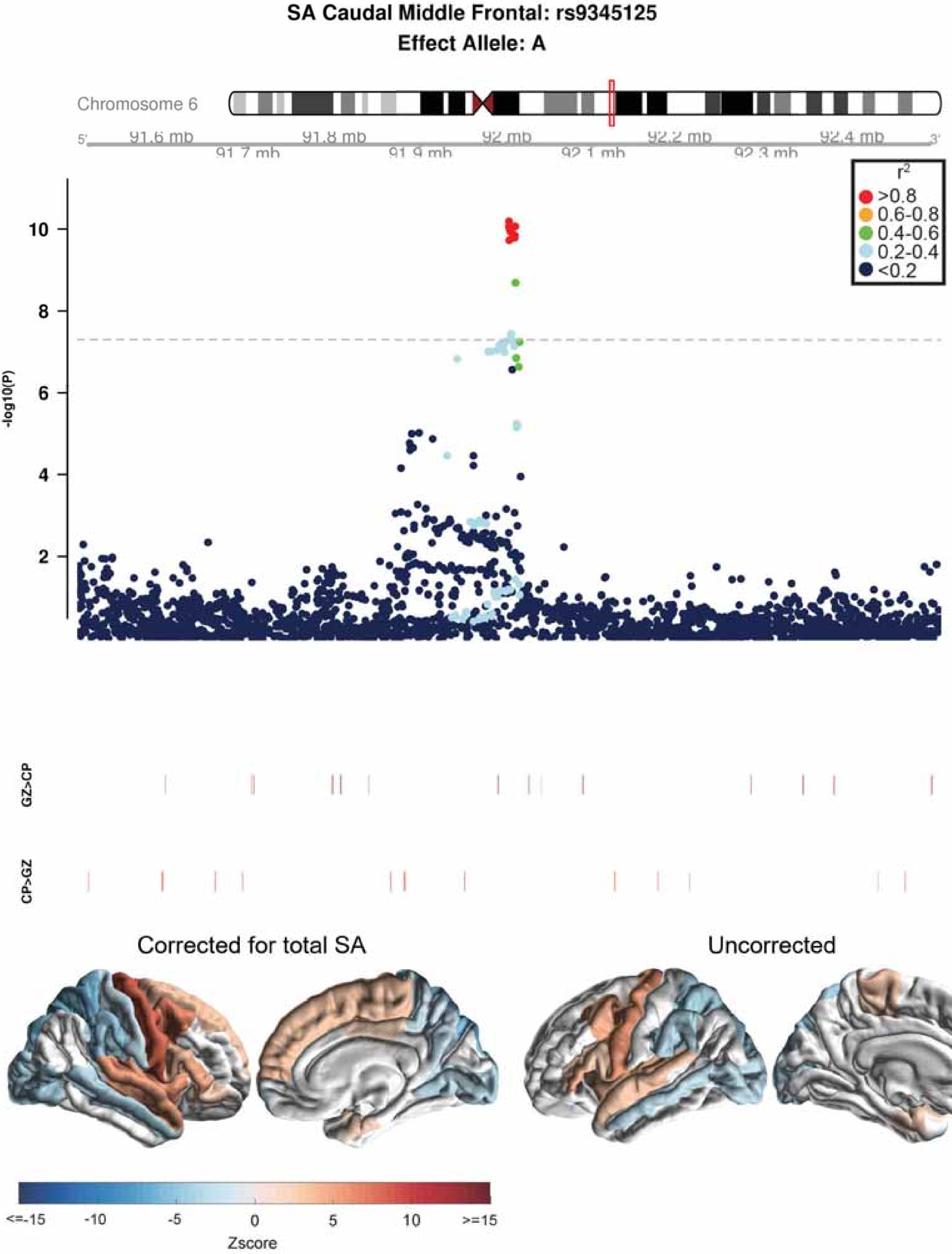
SA Caudal Middle Frontal: rs9345125
Effect Allele: A
Chromosome 6
91.6 mb
91.7 mb
91.8 mb
91.9 mb
92 mb
92.1 mb
92.2 mb
92.3 mb
92.4 mb
-log10(P)
$r^2$
>0.8
0.6-0.8
0.4-0.6
0.2-0.4
<0.2
GZ>CP
CP>GZ
Corrected for total SA
Uncorrected
<=-15
-10
-5
0
5
10
>=15
Zscore

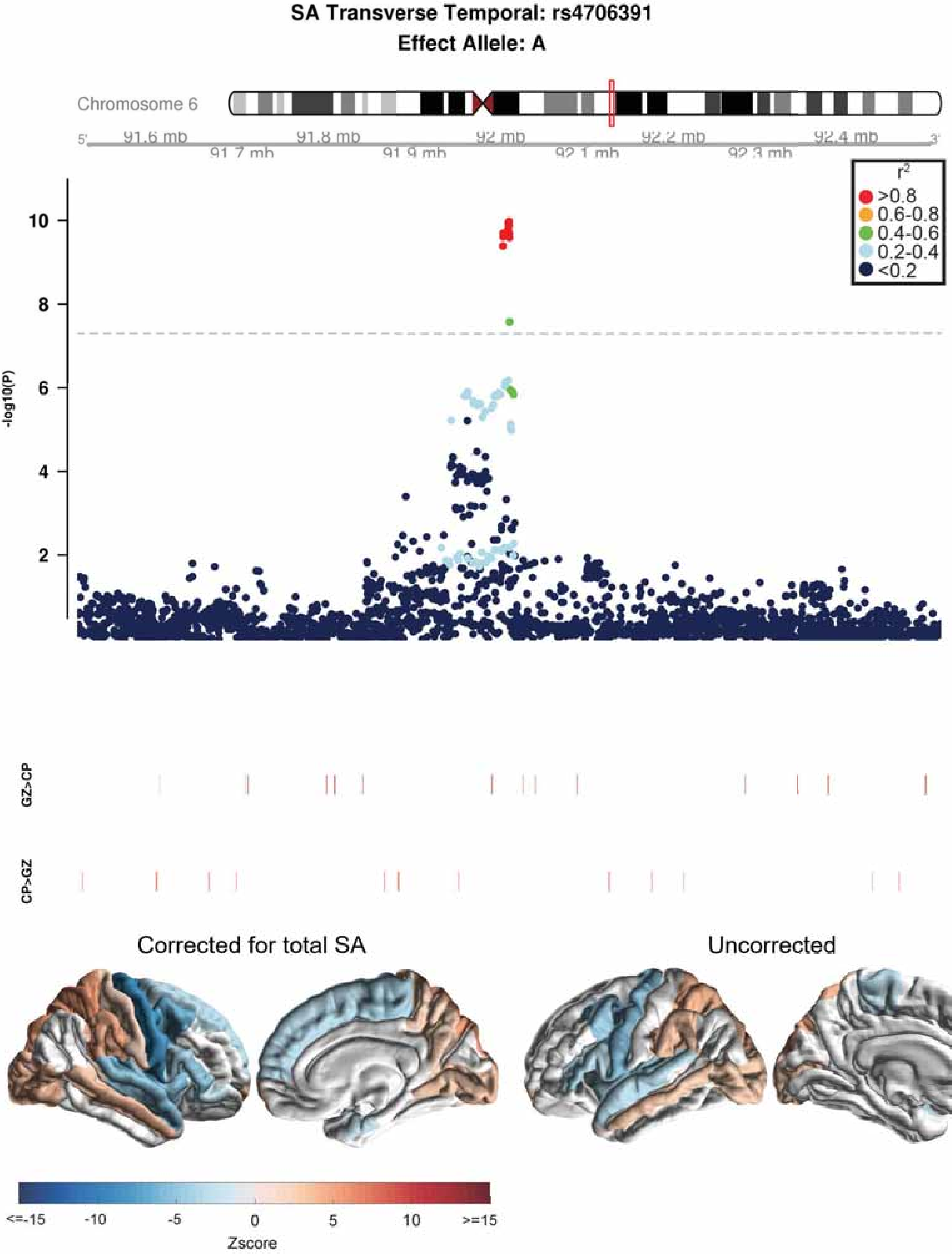
SA Transverse Temporal: rs4706391
Effect Allele: A
Chromosome 6
91.6 mb
91.7 mb
91.8 mb
91.9 mb
92 mb
92.1 mb
92.2 mb
92.3 mb
92.4 mb
-log10(P)
r2
>0.8
0.6-0.8
0.4-0.6
0.2-0.4
<0.2
GZ>CP
CP>GZ
Corrected for total SA
Uncorrected
<=-15
-10
-5
0
5
10
>=15
Zscore

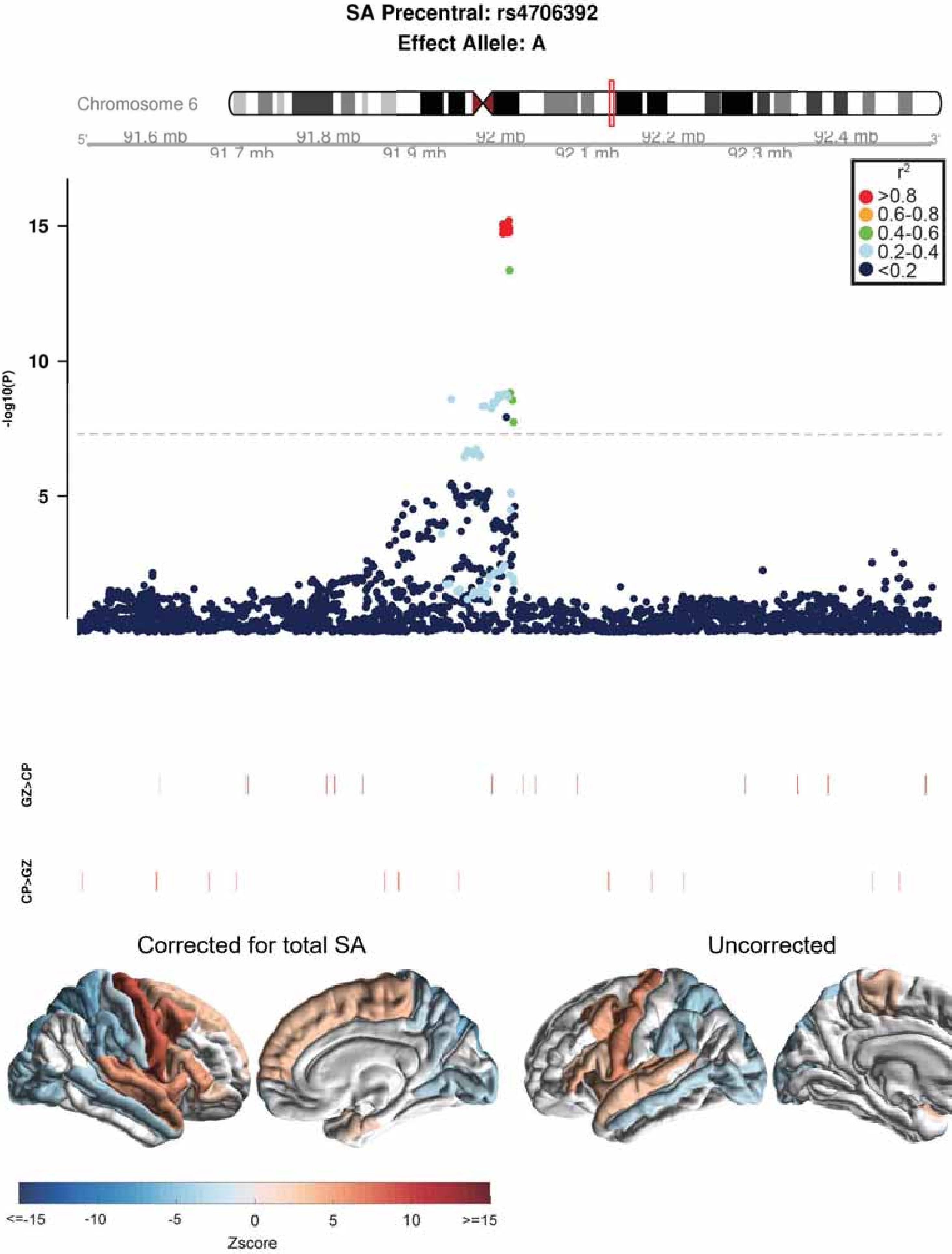

SA Precentral: rs4706392
Effect Allele: A
Chromosome 6
91.6 mb
91.7 mb
91.8 mb
91.9 mb
92 mb
92.1 mb
92.2 mb
92.3 mb
92.4 mb
r2
>0.8
0.6-0.8
0.4-0.6
0.2-0.4
<0.2
-log10(P)
15
10
5
GZ>CP
CP>GZ
Corrected for total SA
Uncorrected
<=-15
-10
-5
0
5
10
>=15
Zscore

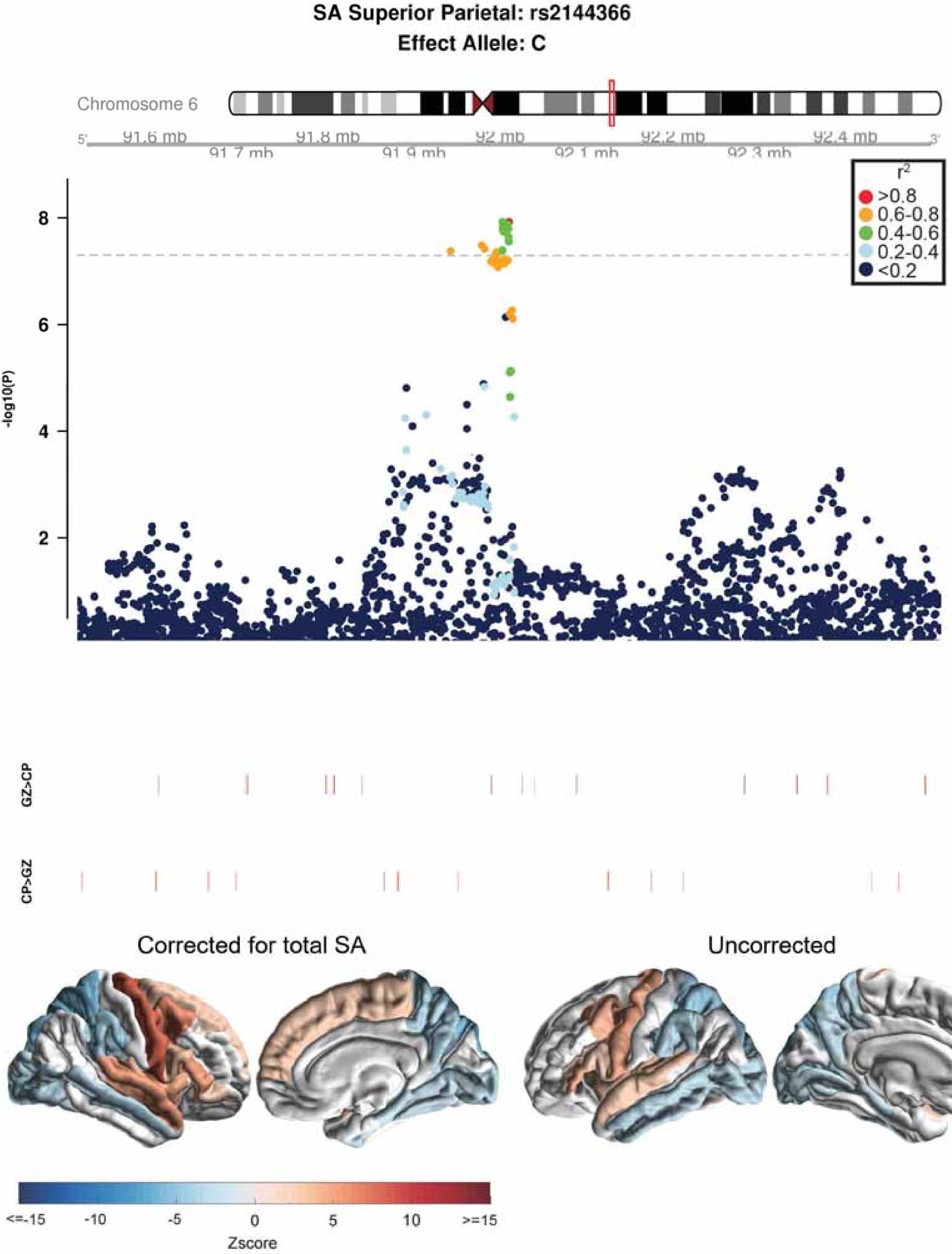

SA Superior Parietal: rs2144366
Effect Allele: C
Chromosome 6
91.6 mb
91.7 mb
91.8 mb
91.9 mb
92 mb
92.1 mb
92.2 mb
92.3 mb
92.4 mb
r2
>0.8
0.6-0.8
0.4-0.6
0.2-0.4
<0.2
-log10(P)
GZ>CP
CP>GZ
Corrected for total SA
Uncorrected
<=-15
-10
-5
0
5
10
>=15
Zscore

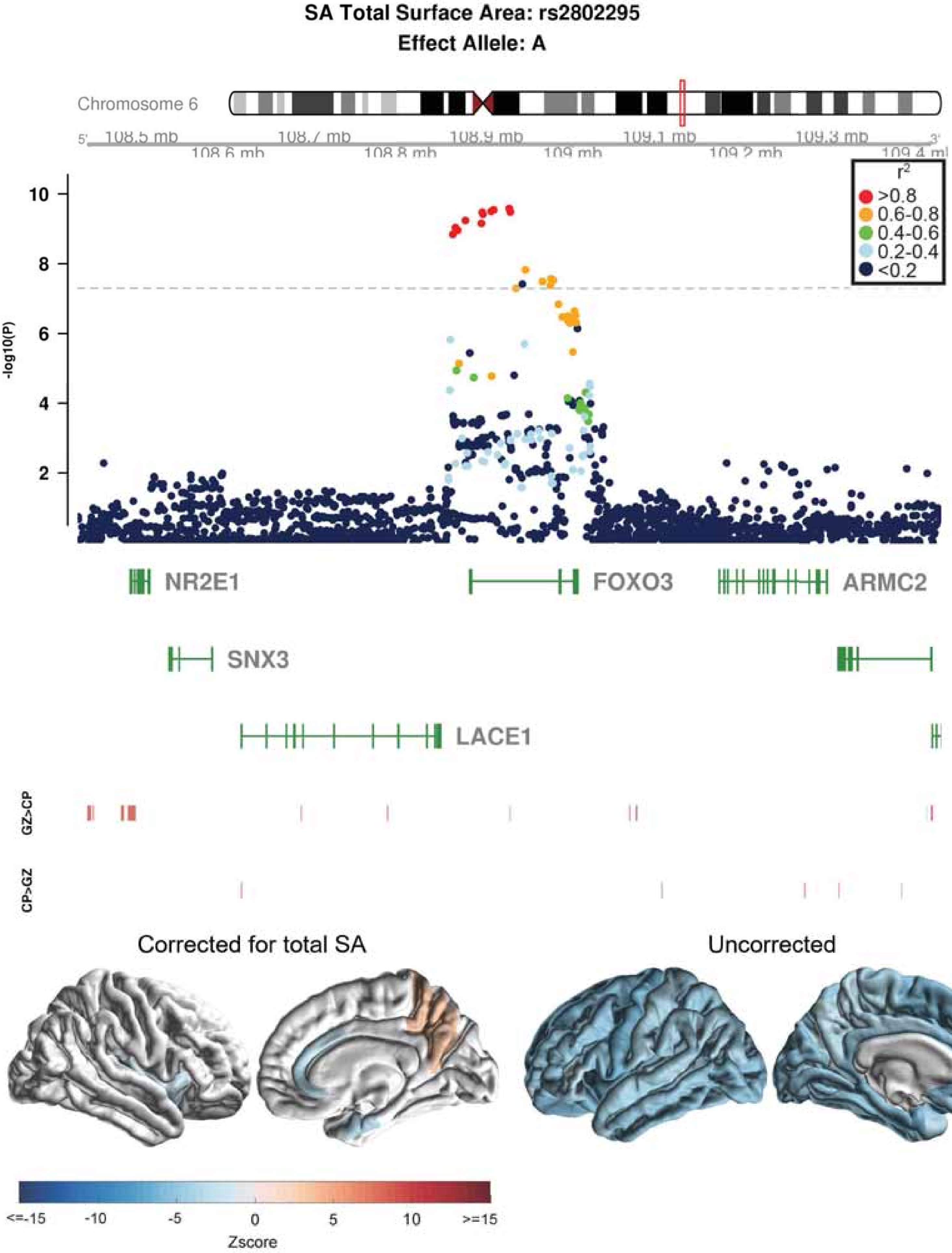
SA Total Surface Area: rs2802295
Effect Allele: A
Chromosome 6
5'
108.5 mb
108.6 mb
108.7 mb
108.8 mb
108.9 mb
109 mb
109.1 mb
109.2 mb
109.3 mb
109.4 mb
3'
-log10(P)
10
8
6
4
2
r²
>0.8
0.6-0.8
0.4-0.6
0.2-0.4
<0.2
NR2E1
FOXO3
ARMC2
SNX3
LACE1
GZ>CP
CP>GZ
Corrected for total SA
Uncorrected
<=-15
-10
-5
0
5
10
>=15
Zscore

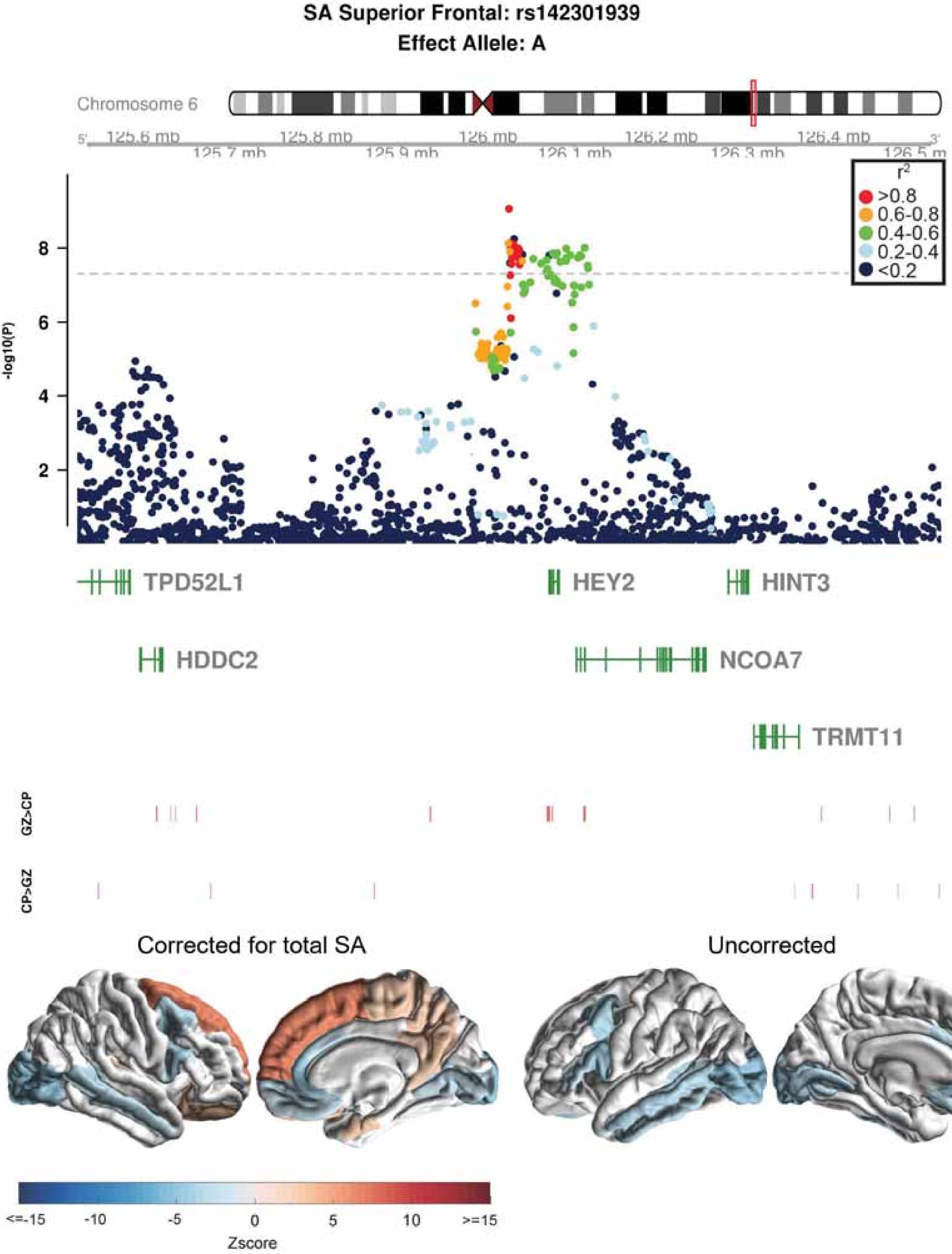
SA Superior Frontal: rs142301939
Effect Allele: A
Chromosome 6
125.6 mb
125.7 mb
125.8 mb
125.9 mb
126 mb
126.1 mb
126.2 mb
126.3 mb
126.4 mb
126.5 m
r2
>0.8
0.6-0.8
0.4-0.6
0.2-0.4
<0.2
-log10(P)
2
4
6
8
TPD52L1
HEY2
HINT3
HDDC2
NCOA7
TRMT11
GZ>CP
CP>GZ
Corrected for total SA
Uncorrected
<=-15
-10
-5
0
5
10
>=15
Zscore

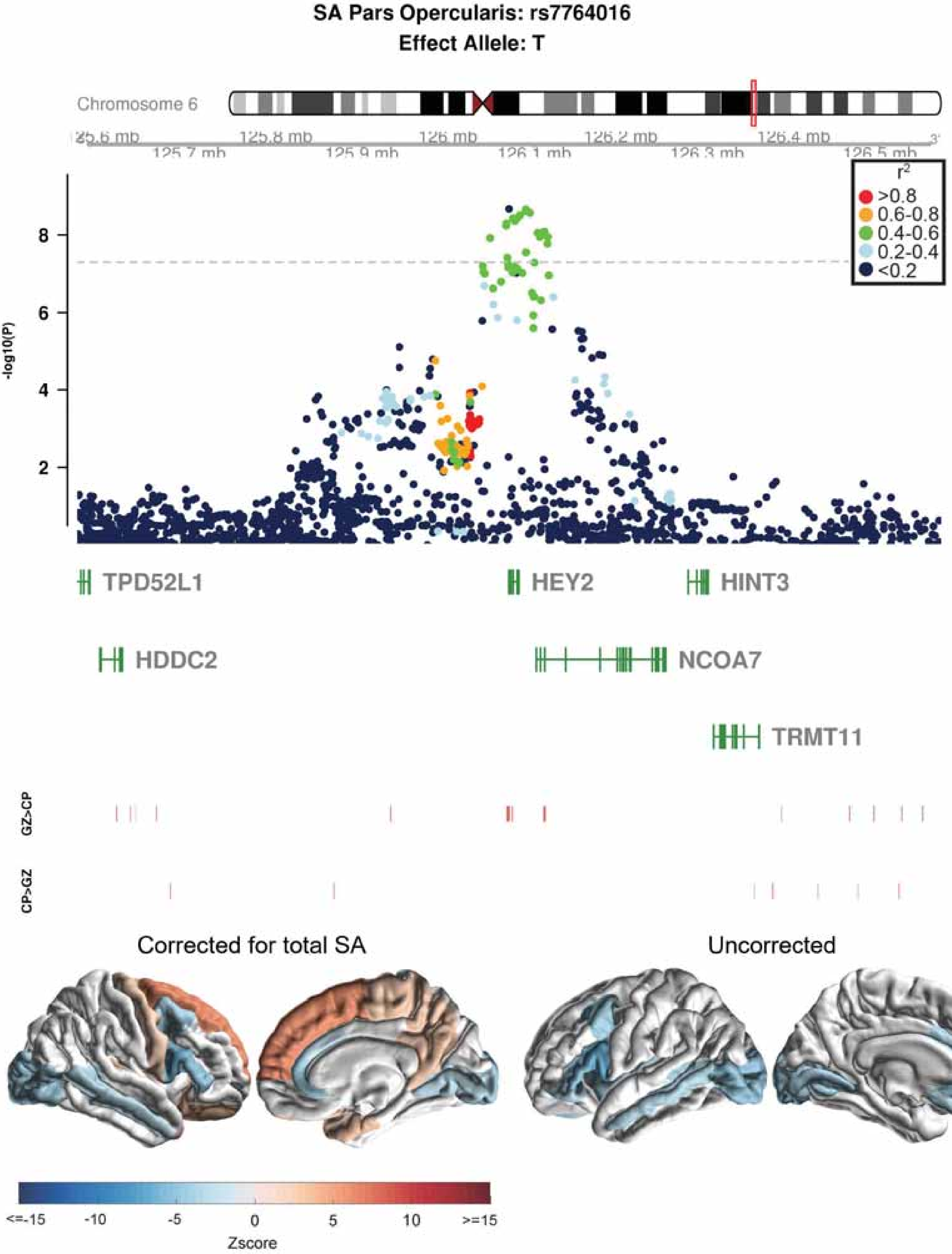
SA Pars Opercularis: rs7764016
Effect Allele: T
Chromosome 6
125.7 mb
125.8 mb
125.9 mb
126 mb
126.1 mb
126.2 mb
126.3 mb
126.4 mb
126.5 mb
-log10(P)
2
4
6
8
r2
>0.8
0.6-0.8
0.4-0.6
0.2-0.4
<0.2
TPD52L1
HEY2
HINT3
HDDC2
NCOA7
TRMT11
GZ>CP
CP>GZ
Corrected for total SA
Uncorrected
<=-15
-10
-5
0
5
10
>=15
Zscore

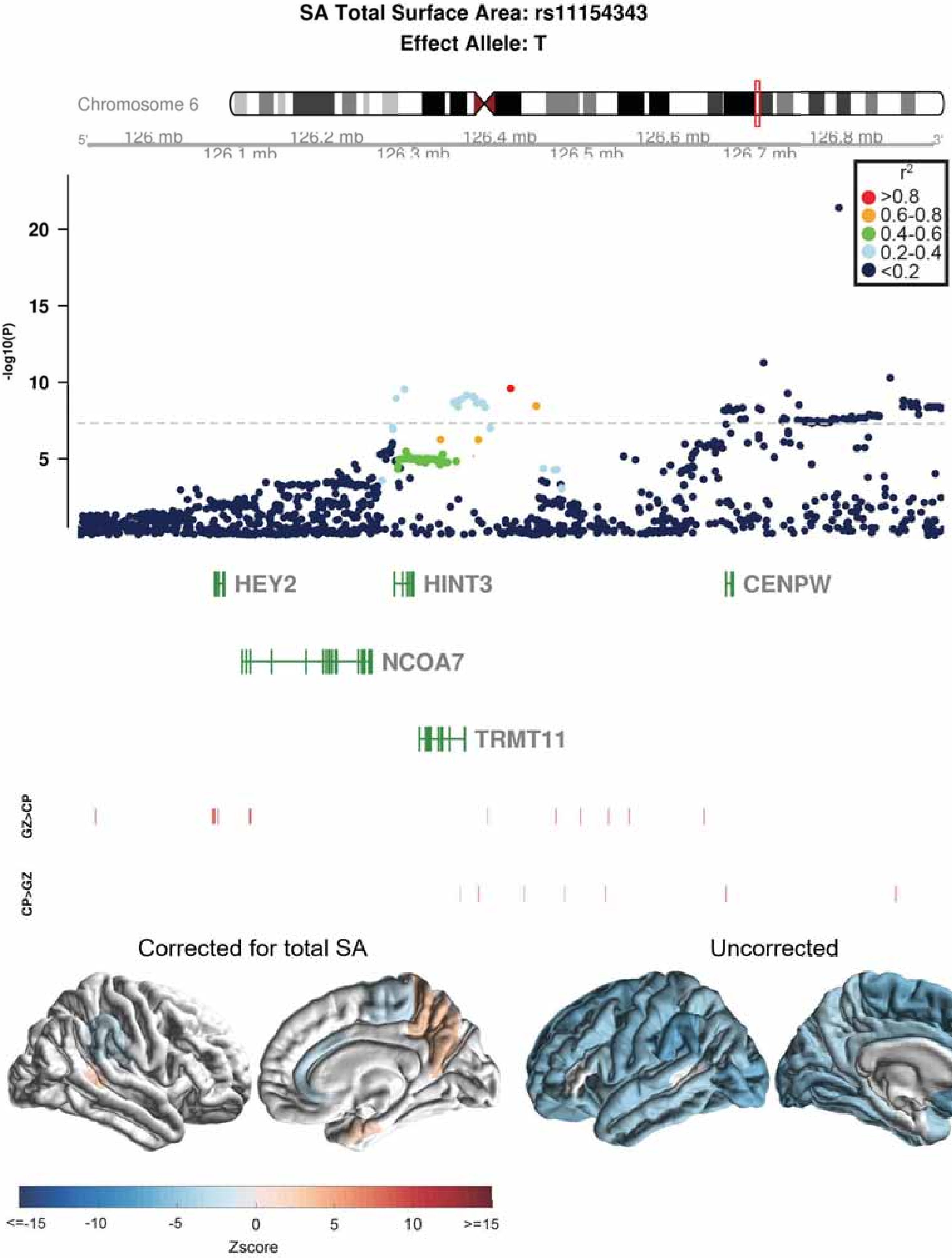

SA Total Surface Area: rs11154343
Effect Allele: T
Chromosome 6
126 mb
126.1 mb
126.2 mb
126.3 mb
126.4 mb
126.5 mb
126.6 mb
126.7 mb
126.8 mb
r2
>0.8
0.6-0.8
0.4-0.6
0.2-0.4
<0.2
-log10(P)
20
15
10
5
HEY2
HINT3
CENPW
NCOA7
TRMT11
GZ>CP
CP>GZ
Corrected for total SA
Uncorrected
<=-15
-10
-5
0
5
10
>=15
Zscore

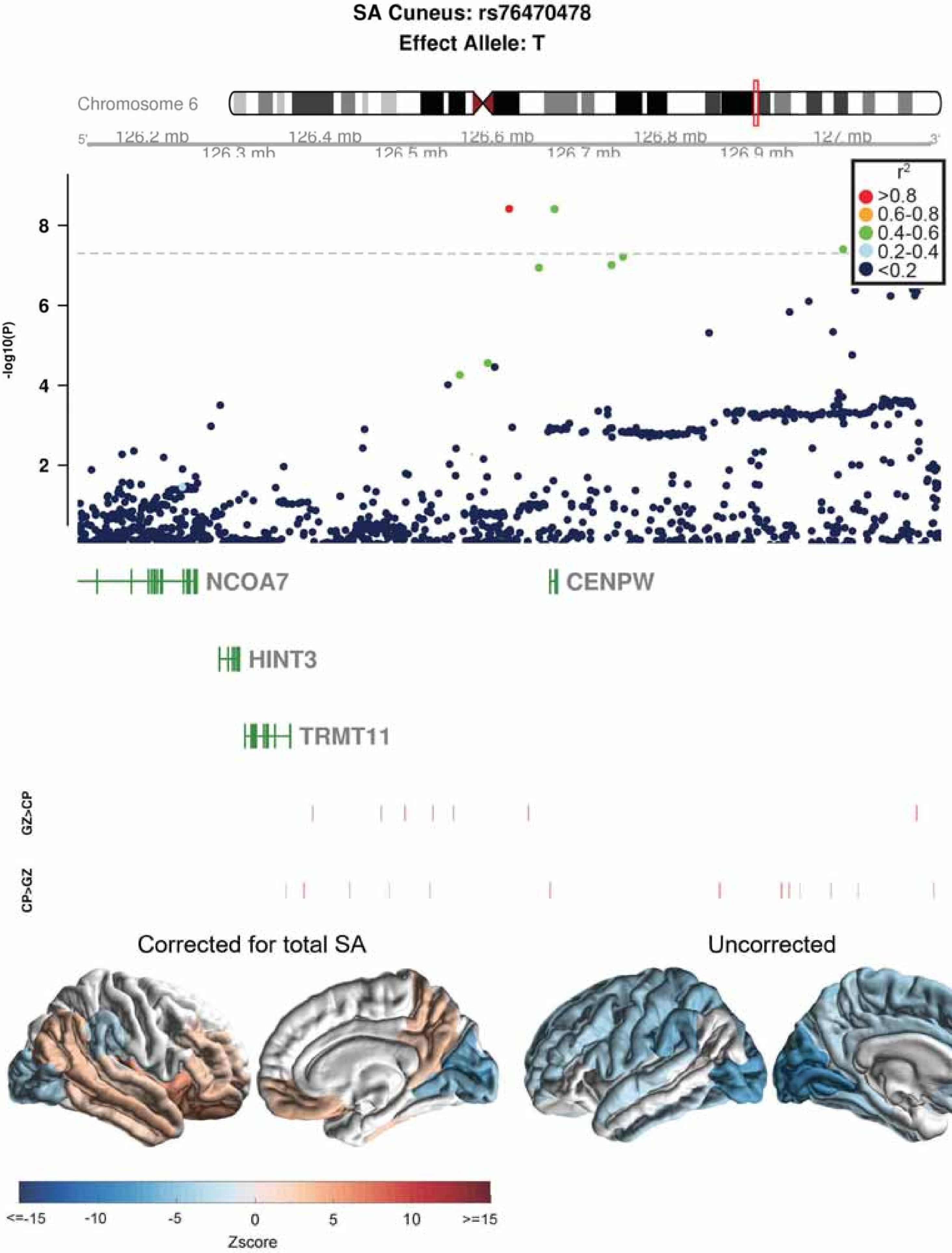
SA Cuneus: rs76470478
Effect Allele: T
Chromosome 6
126.2 mb
126.3 mb
126.4 mb
126.5 mb
126.6 mb
126.7 mb
126.8 mb
126.9 mb
127 mb
-log10(P)
$r^2$
>0.8
0.6-0.8
0.4-0.6
0.2-0.4
<0.2
NCOA7
CENPW
HINT3
TRMT11
GZ>CP
CP>GZ
Corrected for total SA
Uncorrected
<=-15
-10
-5
0
5
10
>=15
Zscore

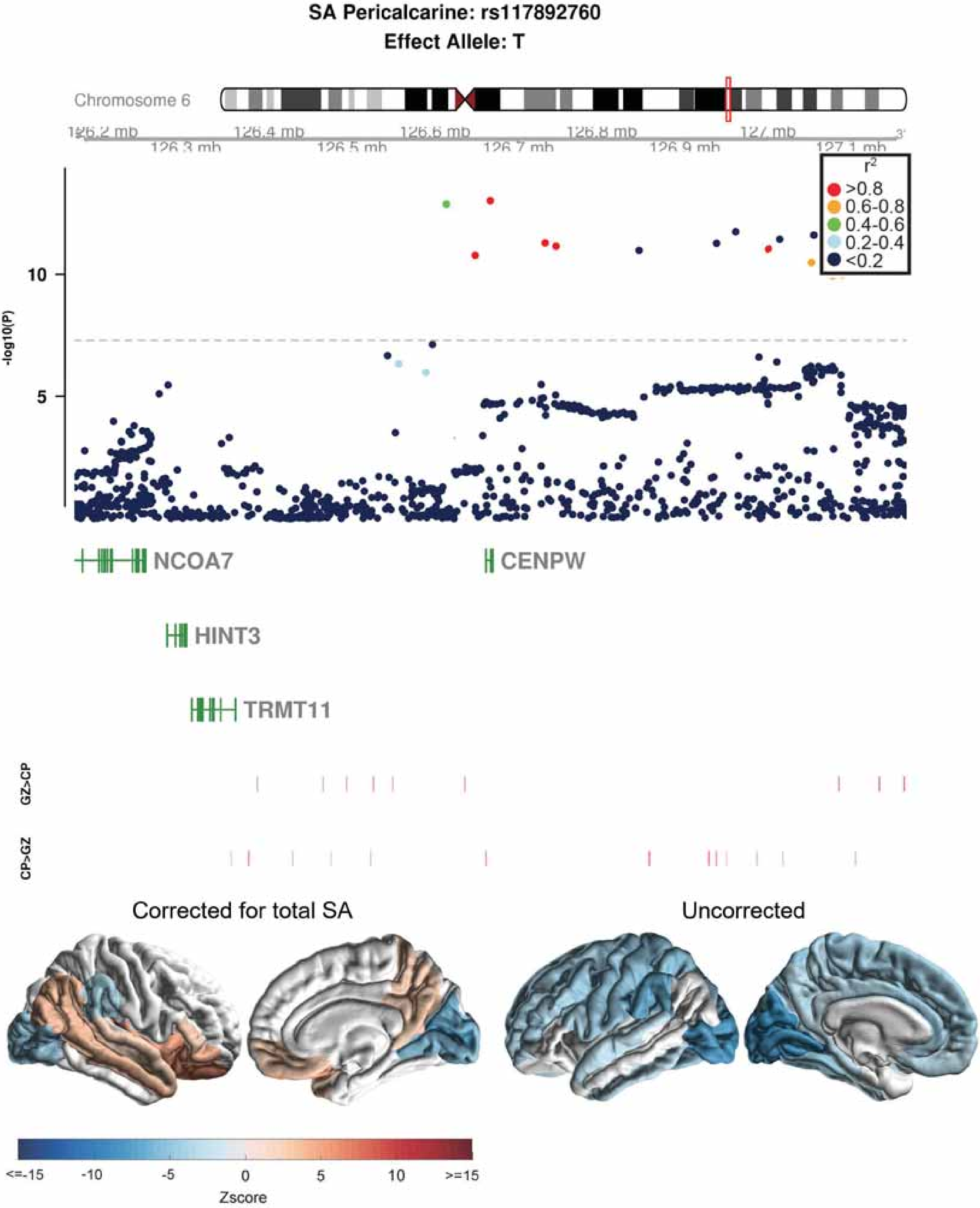
SA Pericalcarine: rs117892760
Effect Allele: T
Chromosome 6
126.2 mb
126.3 mb
126.4 mb
126.5 mb
126.6 mb
126.7 mb
126.8 mb
126.9 mb
127 mb
127.1 mb
3'
r2
>0.8
0.6-0.8
0.4-0.6
0.2-0.4
<0.2
-log10(P)
10
5
NCOA7
CENPW
HINT3
TRMT11
GZ>CP
CP>GZ
Corrected for total SA
Uncorrected
<=-15
-10
-5
0
5
10
>=15
Zscore

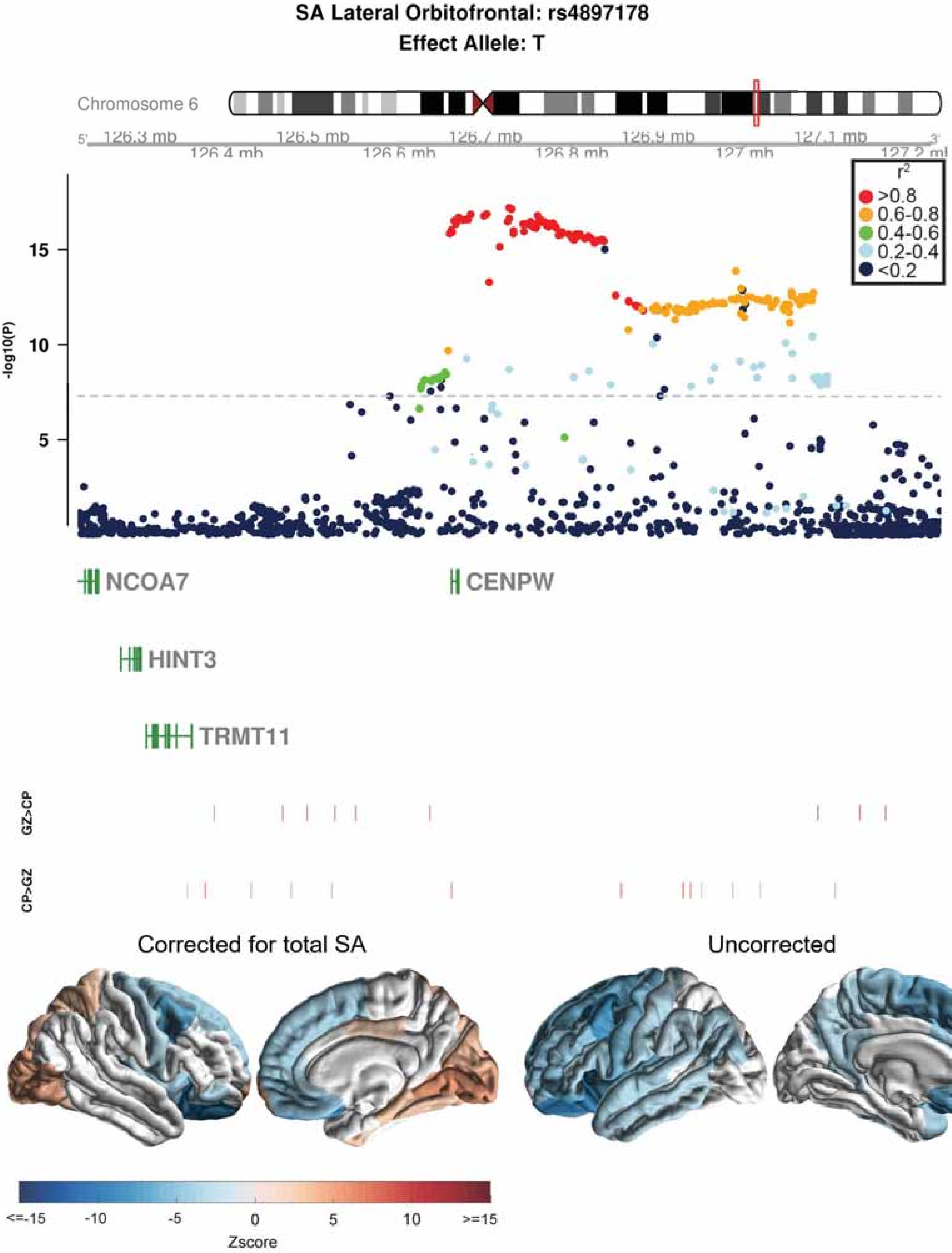

SA Lateral Orbitofrontal: rs4897178
Effect Allele: T
Chromosome 6
126.3 mb
126.4 mb
126.5 mb
126.6 mb
126.7 mb
126.8 mb
126.9 mb
127 mb
127.1 mb
-log10(P)
r2
>0.8
0.6-0.8
0.4-0.6
0.2-0.4
<0.2
NCOA7
CENPW
HINT3
TRMT11
GZ>CP
CP>GZ
Corrected for total SA
Uncorrected
<=-15
-10
-5
0
5
10
>=15
Zscore

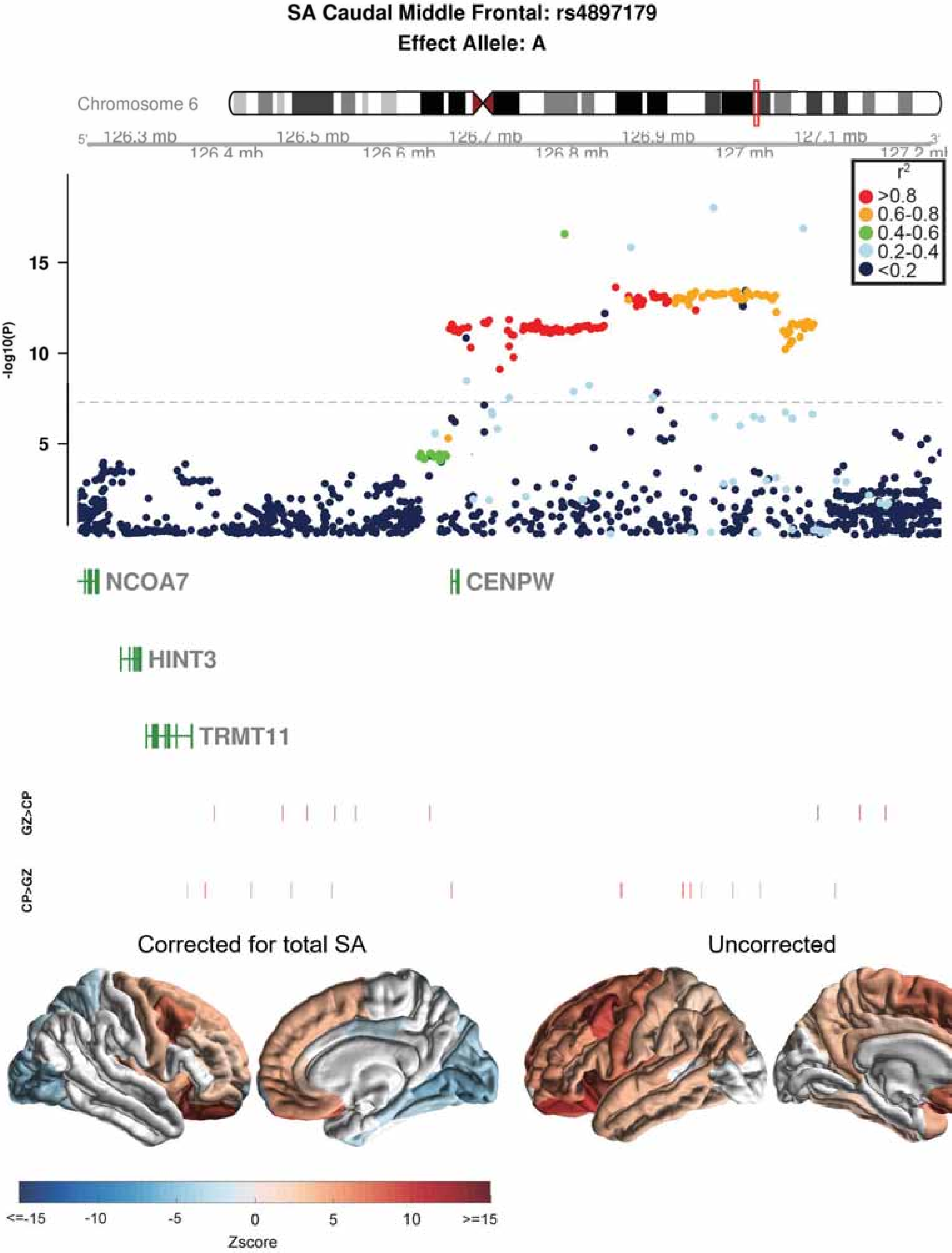
SA Caudal Middle Frontal: rs4897179
Effect Allele: A
Chromosome 6
126.3 mb
126.4 mb
126.5 mb
126.6 mb
126.7 mb
126.8 mb
126.9 mb
127 mb
127.1 mb
127.2 mb
r2
>0.8
0.6-0.8
0.4-0.6
0.2-0.4
<0.2
-log10(P)
15
10
5
NCOA7
CENPW
HINT3
TRMT11
GZ>CP
CP>GZ
Corrected for total SA
Uncorrected
<=-15
-10
-5
0
5
10
>=15
Zscore

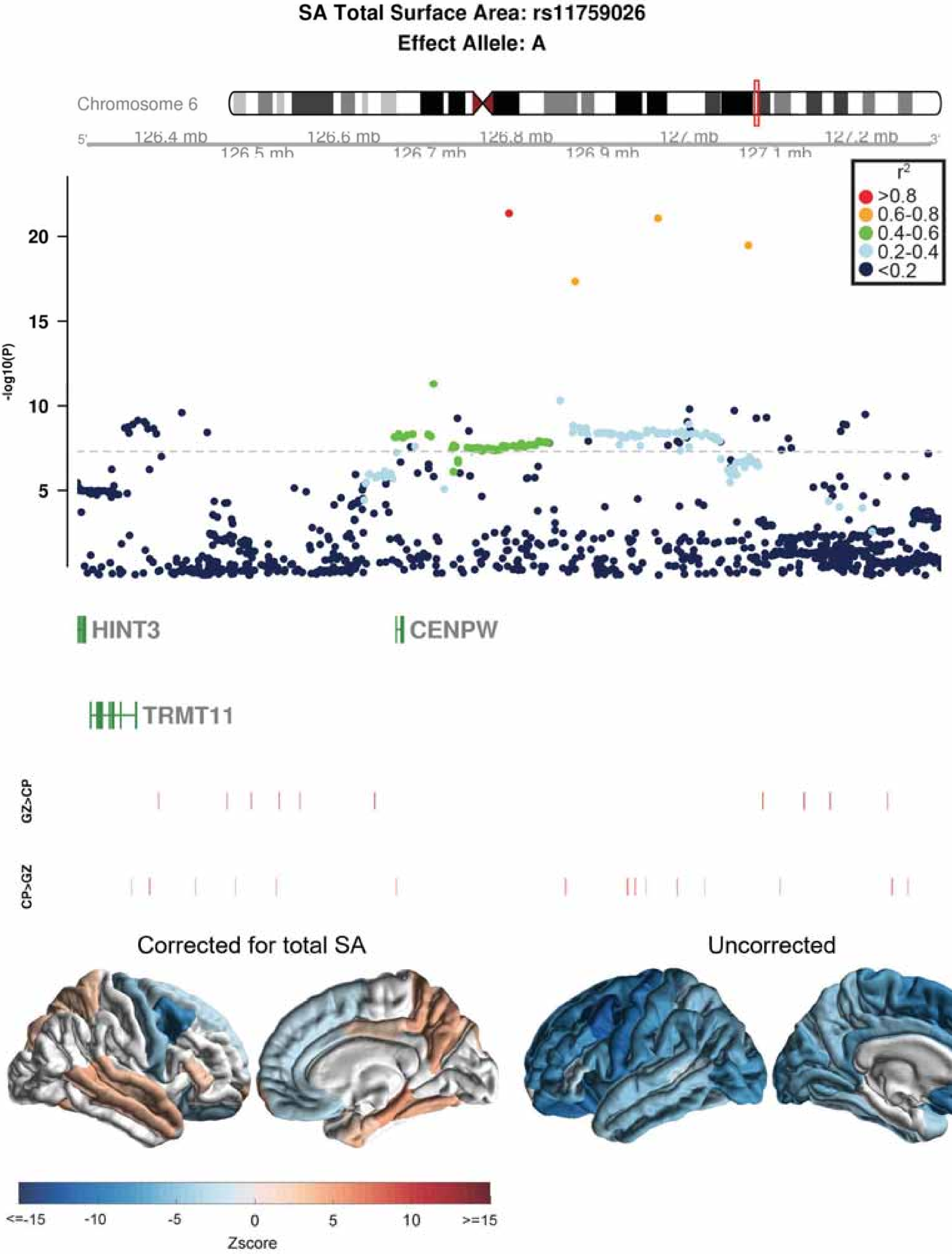

SA Total Surface Area: rs11759026
Effect Allele: A
Chromosome 6
5'
126.4 mb
126.5 mb
126.6 mb
126.7 mb
126.8 mb
126.9 mb
127 mb
127.1 mb
127.2 mb
3'
r2
>0.8
0.6-0.8
0.4-0.6
0.2-0.4
<0.2
-log10(P)
20
15
10
5
HINT3
CENPW
TRMT11
GZ>CP
CP>GZ
Corrected for total SA
Uncorrected
<=-15
-10
-5
0
5
10
>=15
Zscore

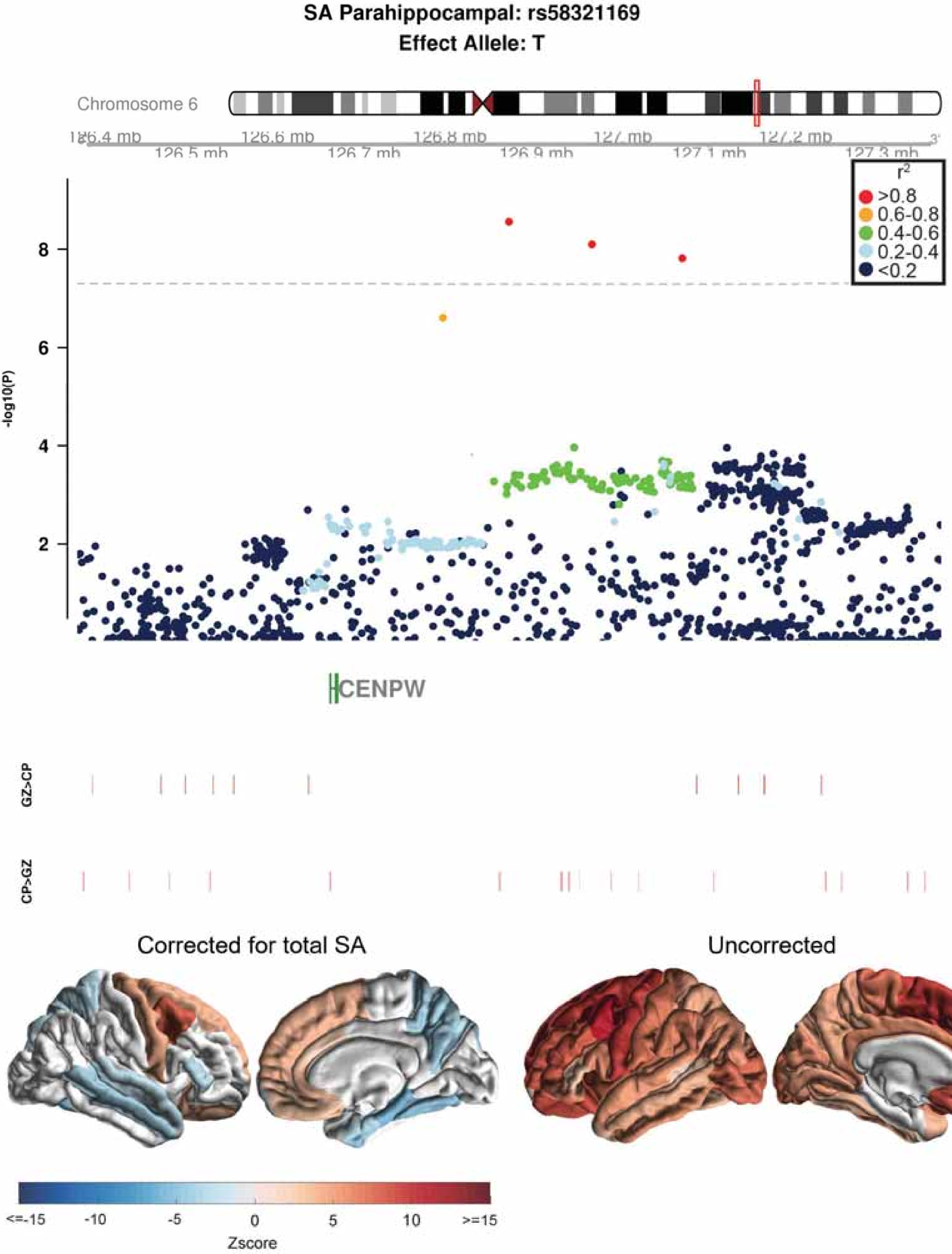
SA Parahippocampal: rs58321169
Effect Allele: T
Chromosome 6
126.4 mb
126.5 mb
126.6 mb
126.7 mb
126.8 mb
126.9 mb
127 mb
127.1 mb
127.2 mb
127.3 mb
r2
>0.8
0.6-0.8
0.4-0.6
0.2-0.4
<0.2
-log10(P)
2
4
6
8
CENPW
GZ>CP
CP>GZ
Corrected for total SA
Uncorrected
<=-15
-10
-5
0
5
10
>=15
Zscore

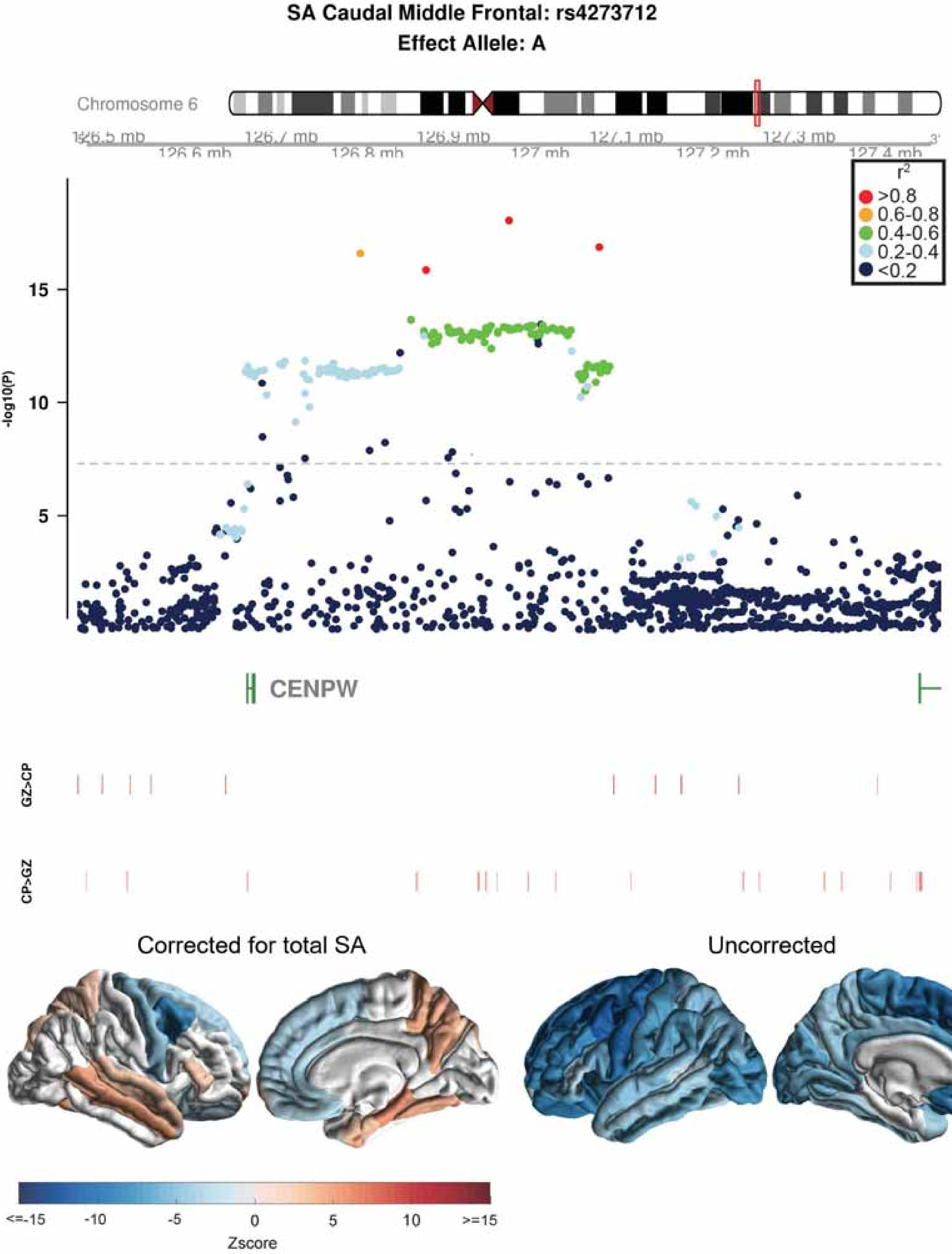

SA Caudal Middle Frontal: rs4273712
Effect Allele: A
Chromosome 6
126.5 mb
126.6 mb
126.7 mb
126.8 mb
126.9 mb
127 mb
127.1 mb
127.2 mb
127.3 mb
127.4 mb
r2
>0.8
0.6-0.8
0.4-0.6
0.2-0.4
<0.2
-log10(P)
15
10
5
CENPW
GZ>CP
CP>GZ
Corrected for total SA
Uncorrected
<=-15
-10
-5
0
5
10
>=15
Zscore

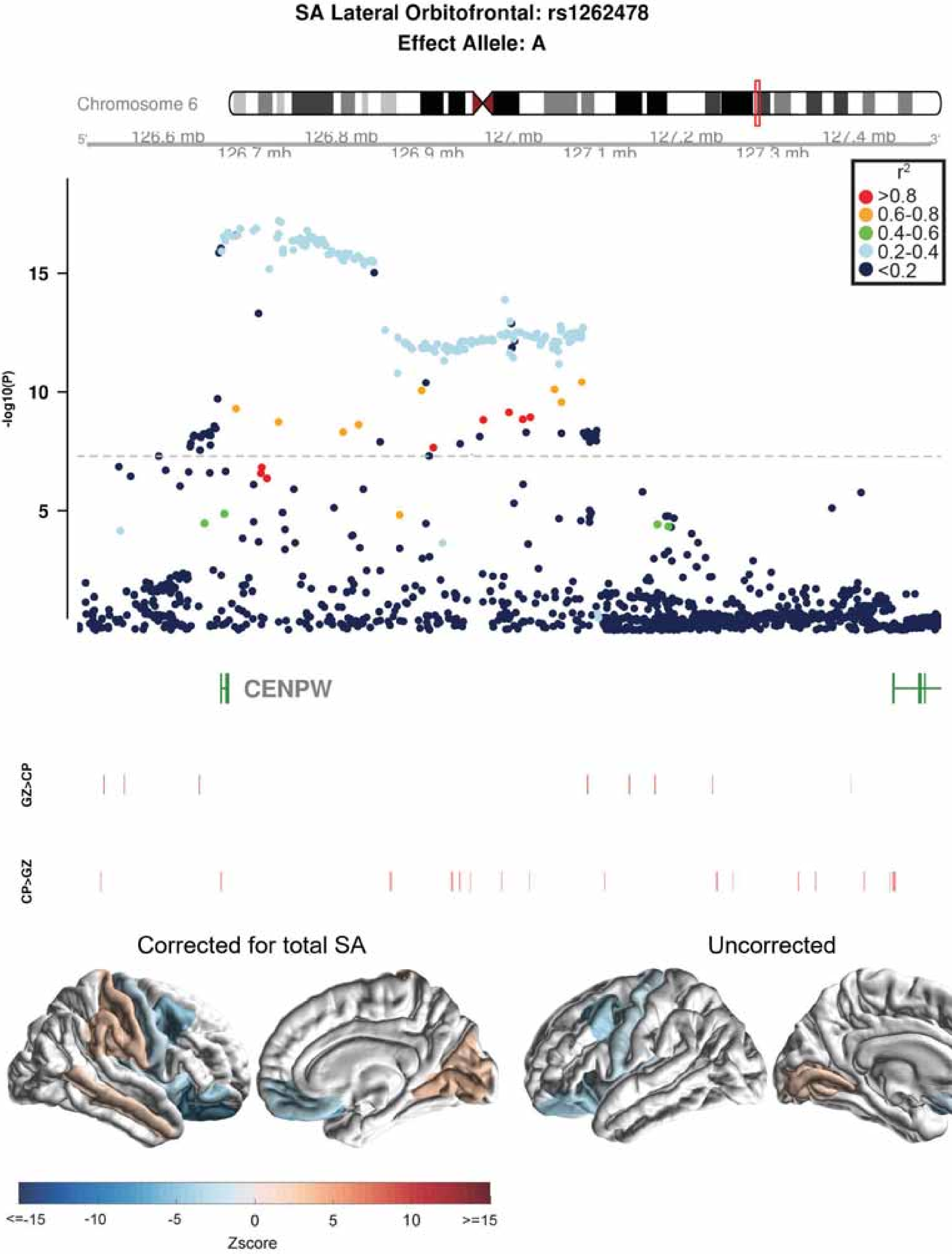
SA Lateral Orbitofrontal: rs1262478
Effect Allele: A
Chromosome 6
126.6 mb
126.7 mb
126.8 mb
126.9 mb
127 mb
127.1 mb
127.2 mb
127.3 mb
127.4 mb
r2
>0.8
0.6-0.8
0.4-0.6
0.2-0.4
<0.2
-log10(P)
15
10
5
CENPW
GZ>CP
CP>GZ
Corrected for total SA
Uncorrected
<=-15
-10
-5
0
5
10
>=15
Zscore

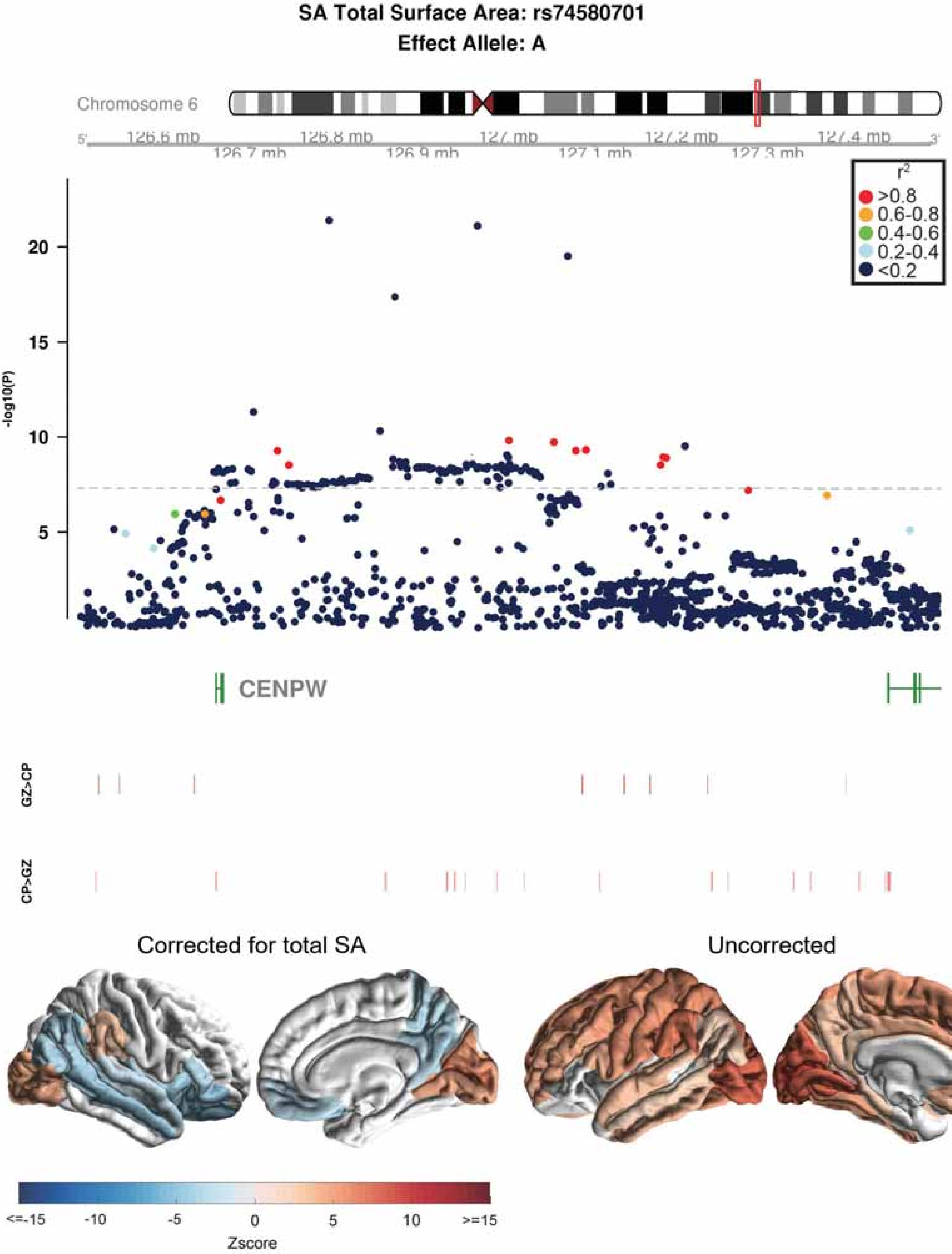
SA Total Surface Area: rs74580701
Effect Allele: A
Chromosome 6
126.6 mb
126.7 mb
126.8 mb
126.9 mb
127 mb
127.1 mb
127.2 mb
127.3 mb
127.4 mb
r2
>0.8
0.6-0.8
0.4-0.6
0.2-0.4
<0.2
-log10(P)
20
15
10
5
CENPW
GZ>CP
CP>GZ
Corrected for total SA
Uncorrected
<=-15
-10
-5
0
5
10
>=15
Zscore

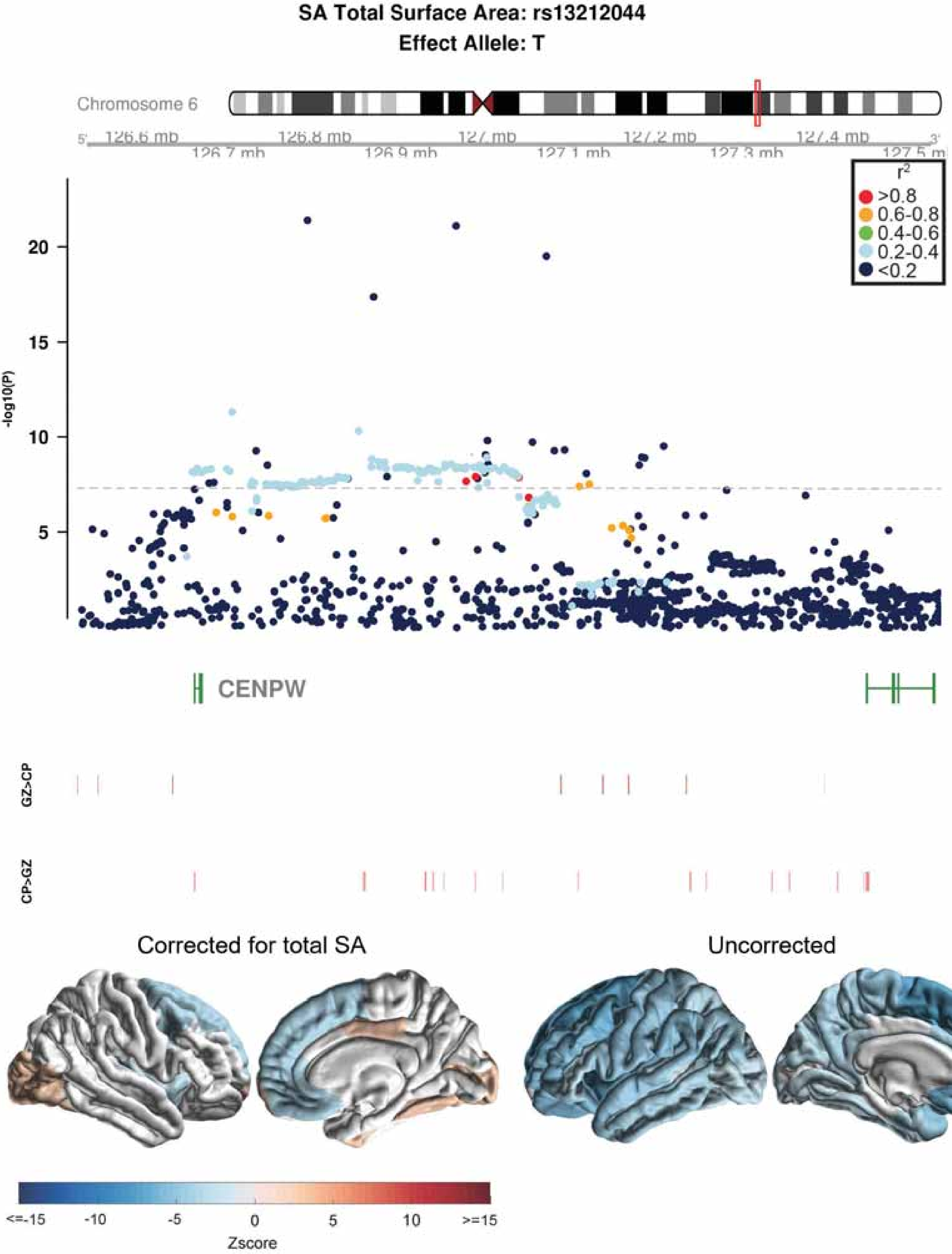

SA Total Surface Area: rs13212044
Effect Allele: T
Chromosome 6
126.6 mb
126.7 mb
126.8 mb
126.9 mb
127 mb
127.1 mb
127.2 mb
127.3 mb
127.4 mb
127.5 mb
r2
>0.8
0.6-0.8
0.4-0.6
0.2-0.4
<0.2
-log10(P)
20
15
10
5
CENPW
GZ>CP
CP>GZ
Corrected for total SA
Uncorrected
<=-15
-10
-5
0
5
10
>=15
Zscore

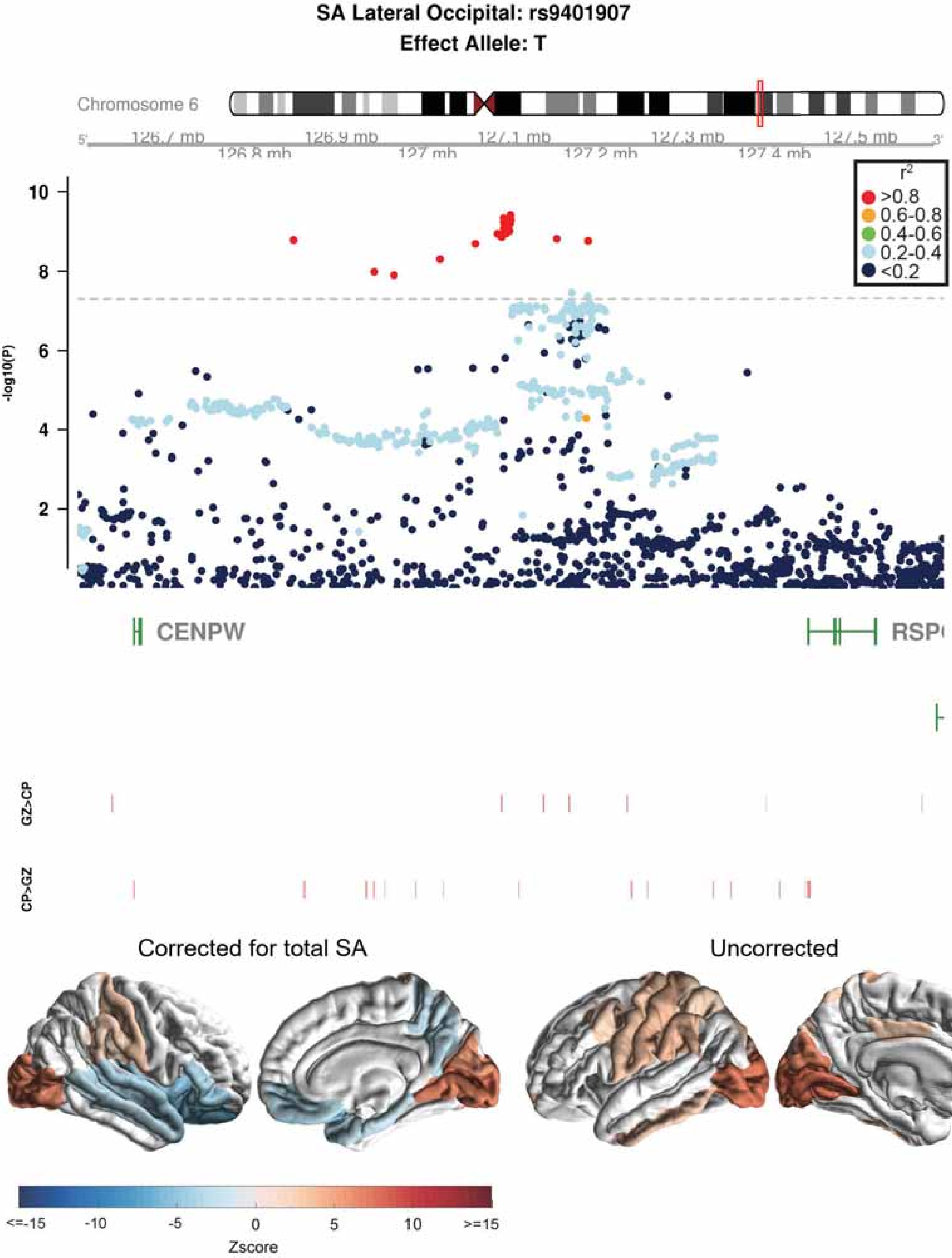
SA Lateral Occipital: rs9401907
Effect Allele: T
Chromosome 6
126.7 mb
126.8 mb
126.9 mb
127 mb
127.1 mb
127.2 mb
127.3 mb
127.4 mb
127.5 mb
r2
>0.8
0.6-0.8
0.4-0.6
0.2-0.4
<0.2
-log10(P)
CENPW
GZ>CP
CP>GZ
Corrected for total SA
Uncorrected
<=-15
-10
-5
0
5
10
>=15
Zscore

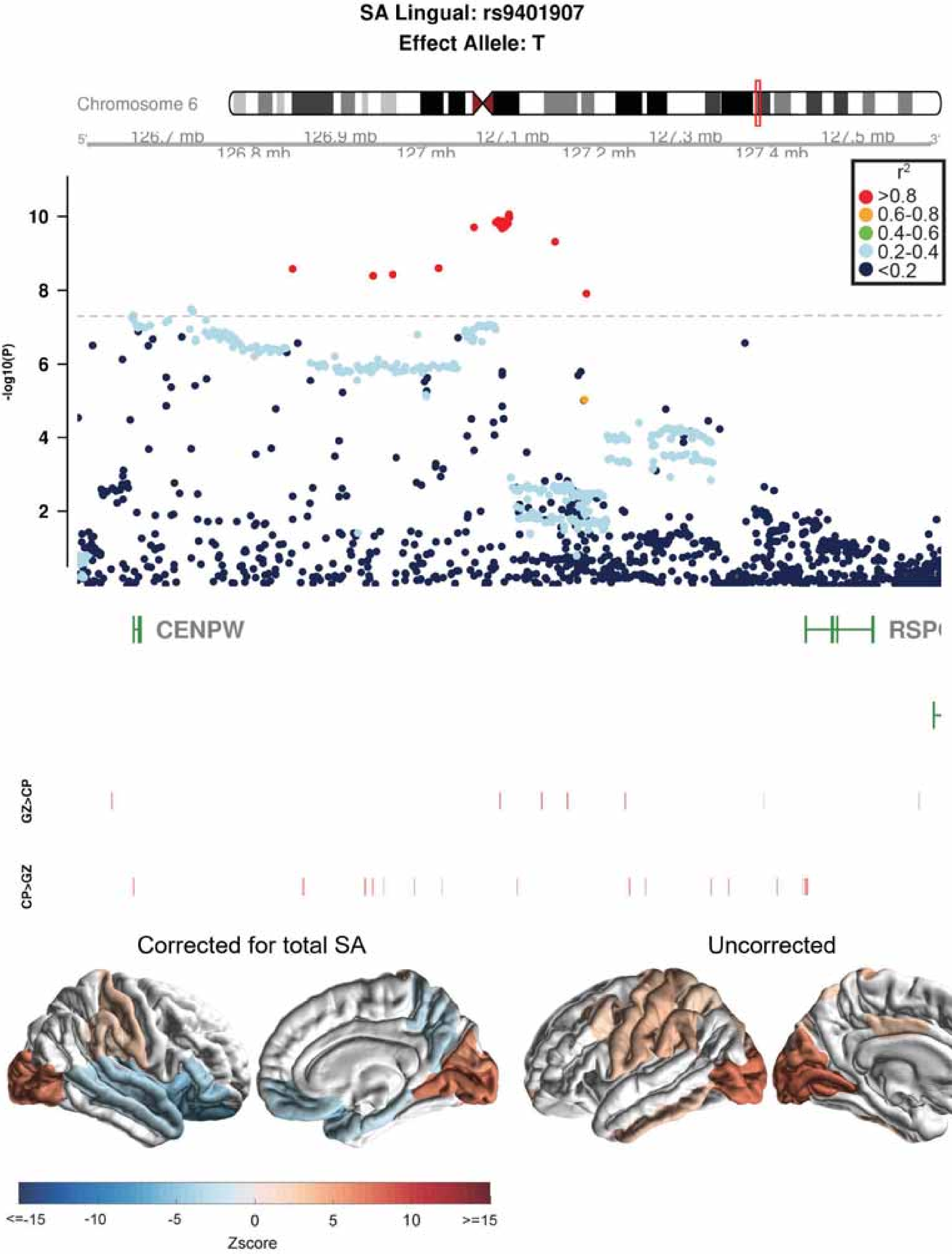
SA Lingual: rs9401907
Effect Allele: T
Chromosome 6
126.7 mb
126.8 mb
126.9 mb
127 mb
127.1 mb
127.2 mb
127.3 mb
127.4 mb
127.5 mb
r2
>0.8
0.6-0.8
0.4-0.6
0.2-0.4
<0.2
-log10(P)
10
8
6
4
2
CENPW
RSP
GZ>CP
CP>GZ
Corrected for total SA
Uncorrected
<=-15
-10
-5
0
5
10
>=15
Zscore

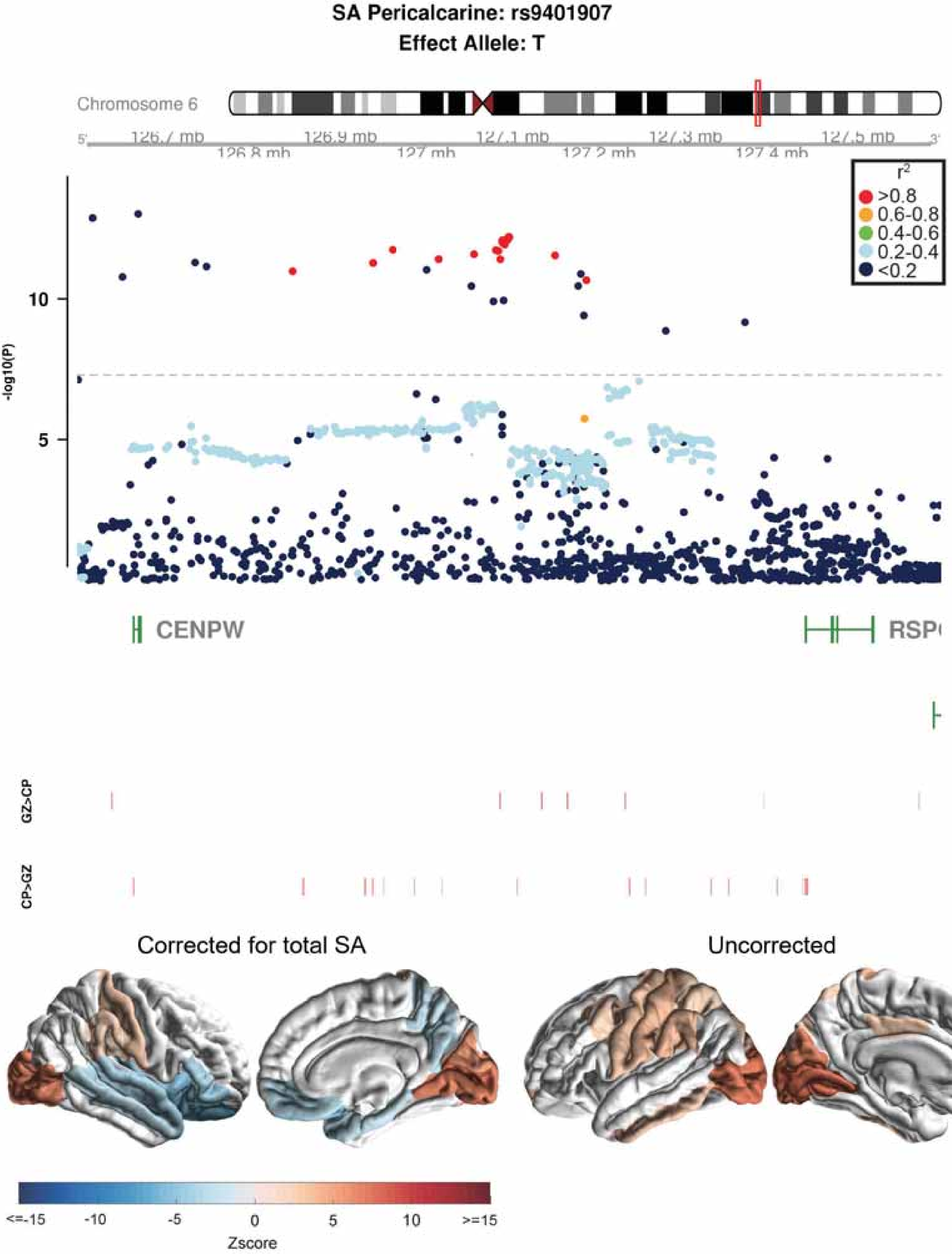
SA Pericalcarine: rs9401907
Effect Allele: T
Chromosome 6
126.7 mb
126.8 mb
126.9 mb
127 mb
127.1 mb
127.2 mb
127.3 mb
127.4 mb
127.5 mb
r2
>0.8
0.6-0.8
0.4-0.6
0.2-0.4
<0.2
-log10(P)
10
5
CENPW
RSP
GZ>CP
CP>GZ
Corrected for total SA
Uncorrected
<=-15
-10
-5
0
5
10
>=15
Zscore

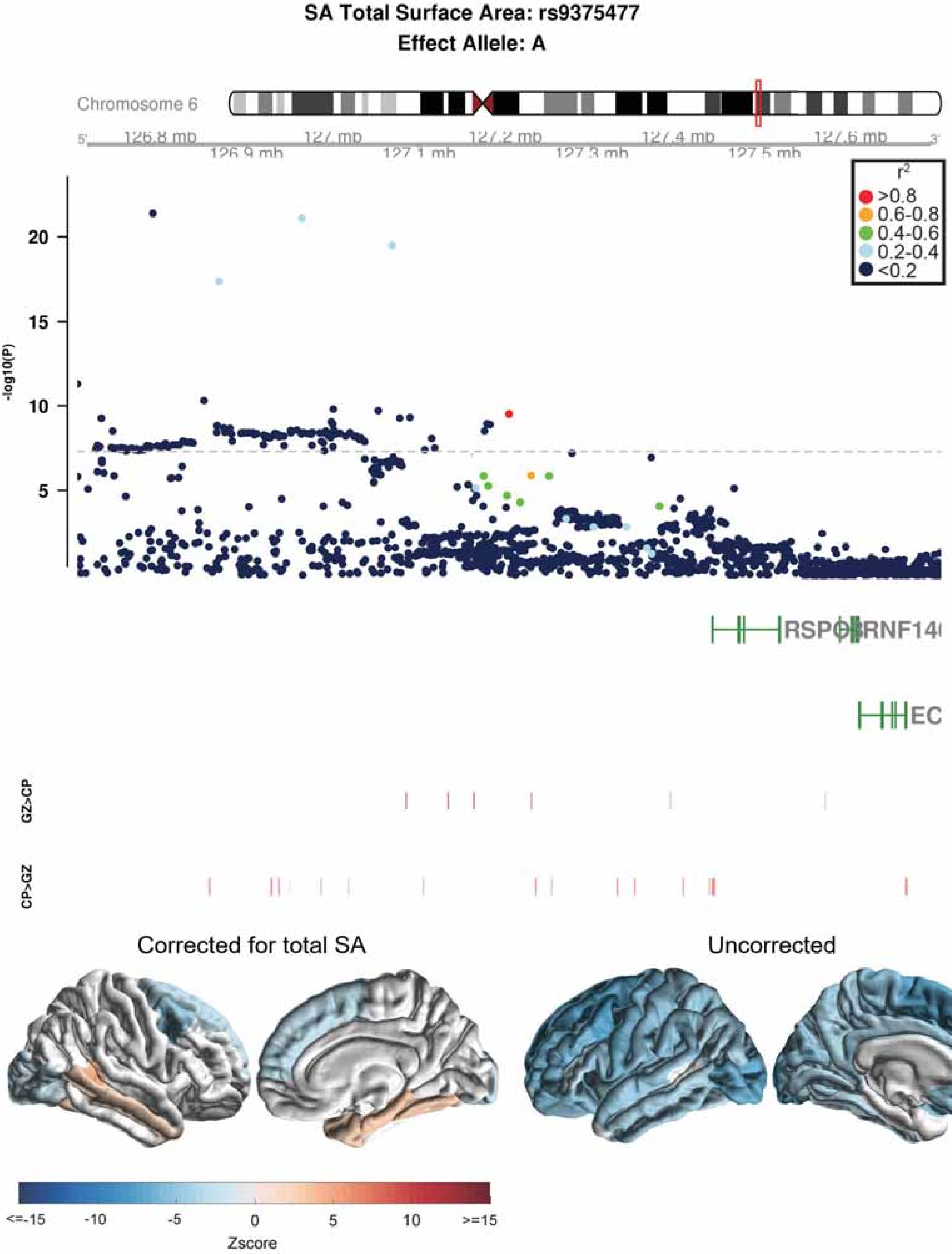

SA Total Surface Area: rs9375477
Effect Allele: A
Chromosome 6
126.8 mb
126.9 mb
127 mb
127.1 mb
127.2 mb
127.3 mb
127.4 mb
127.5 mb
127.6 mb
r2
>0.8
0.6-0.8
0.4-0.6
0.2-0.4
<0.2
-log10(P)
20
15
10
5
RSPO3
RNF146
EC
GZ>CP
CP>GZ
Corrected for total SA
Uncorrected
<=-15
-10
-5
0
5
10
>=15
Zscore

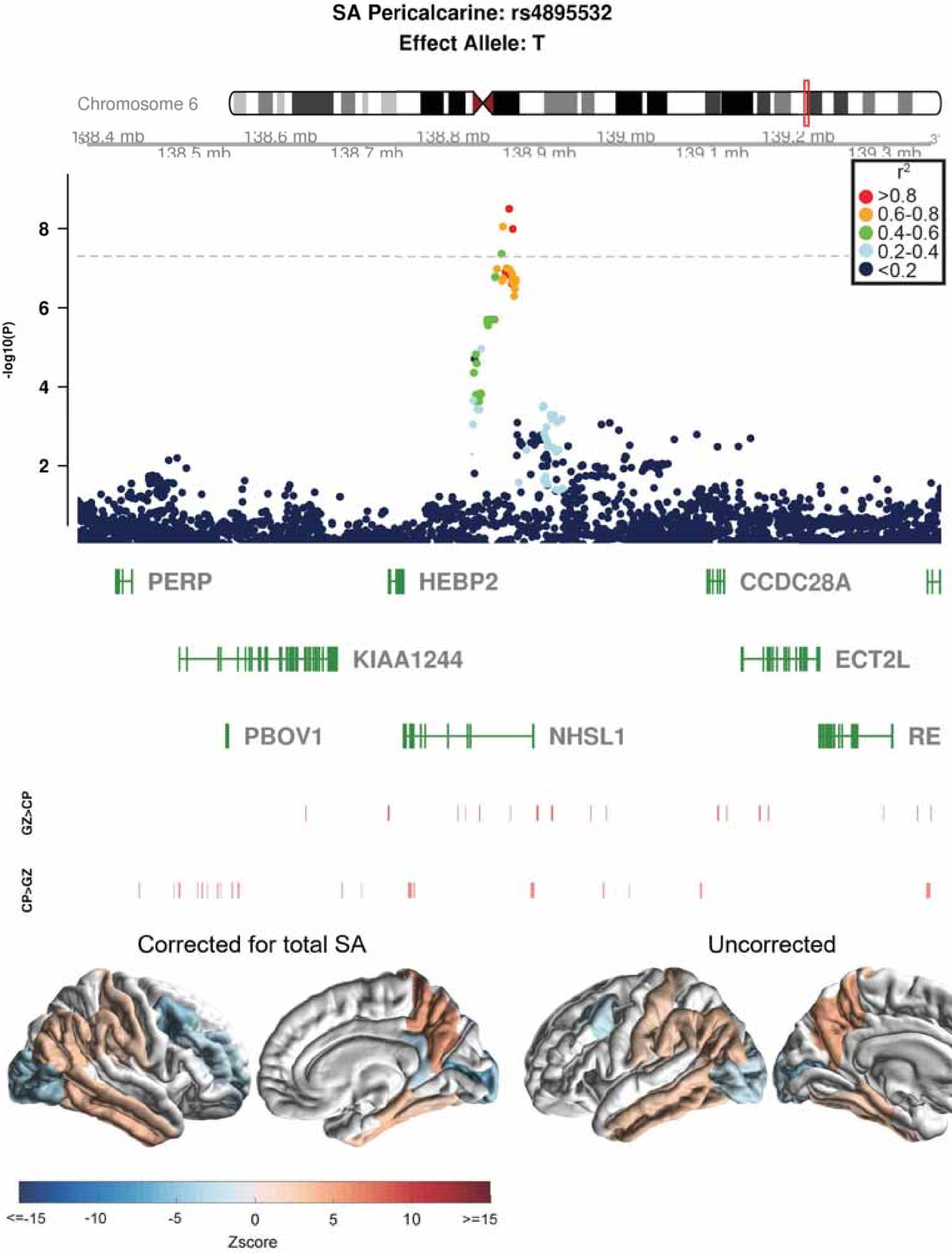
SA Pericalcarine: rs4895532
Effect Allele: T
Chromosome 6
138.4 mb
138.5 mb
138.6 mb
138.7 mb
138.8 mb
138.9 mb
139 mb
139.1 mb
139.2 mb
139.3 mb
-log10(P)
8
6
4
2
r2
>0.8
0.6-0.8
0.4-0.6
0.2-0.4
<0.2
PERP
HEBP2
CCDC28A
KIAA1244
ECT2L
PBOV1
NHSL1
RE
GZ>CP
CP>GZ
Corrected for total SA
Uncorrected
<=-15
-10
-5
0
5
10
>=15
Zscore

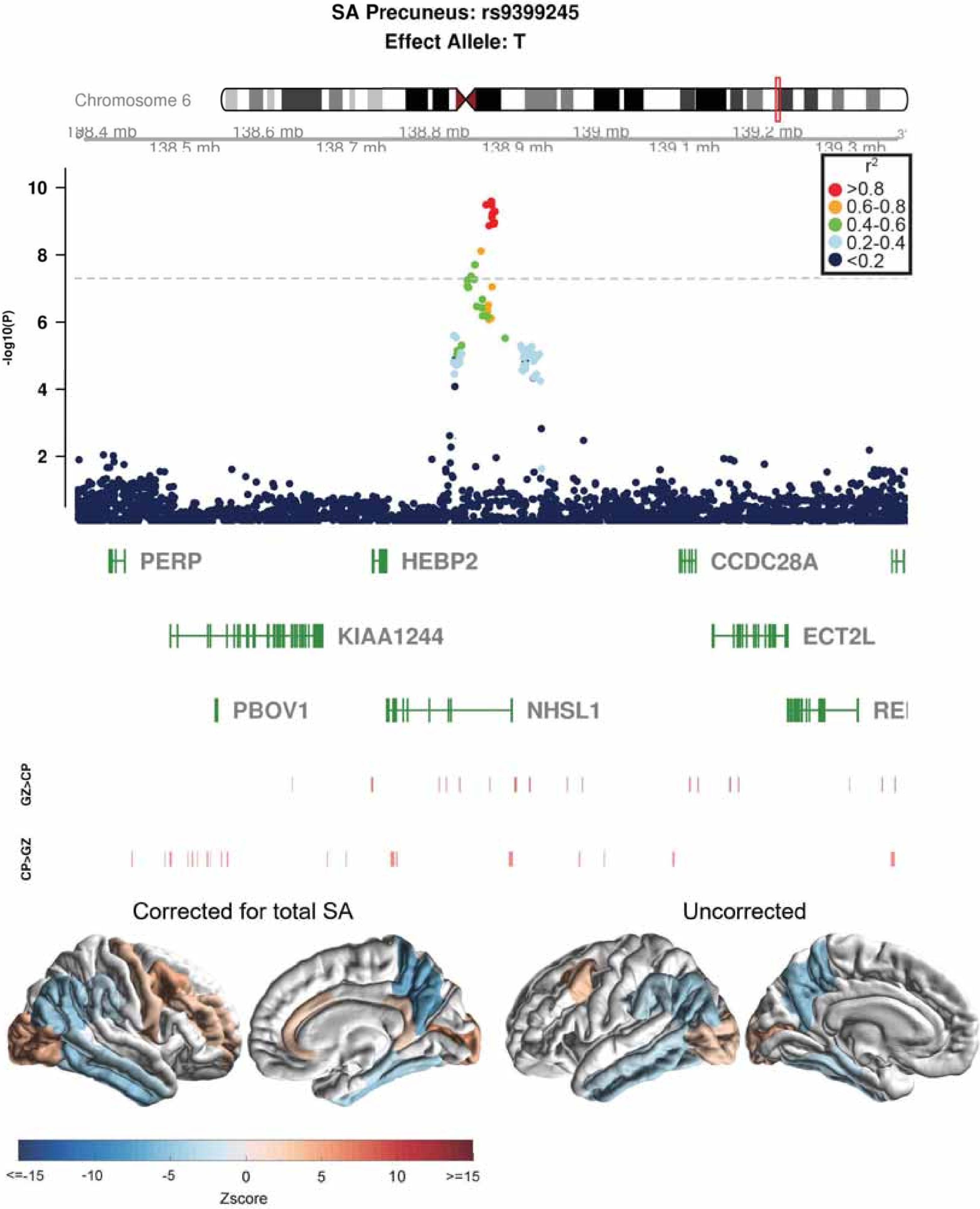
SA Precuneus: rs9399245
Effect Allele: T
Chromosome 6
138.4 mb
138.5 mb
138.6 mb
138.7 mb
138.8 mb
138.9 mb
139 mb
139.1 mb
139.2 mb
139.3 mb
-log10(P)
2
4
6
8
10
r2
>0.8
0.6-0.8
0.4-0.6
0.2-0.4
<0.2
PERP
HEBP2
CCDC28A
KIAA1244
ECT2L
PBOV1
NHSL1
GZ>CP
CP>GZ
Corrected for total SA
Uncorrected
<=-15
-10
-5
0
5
10
>=15
Zscore

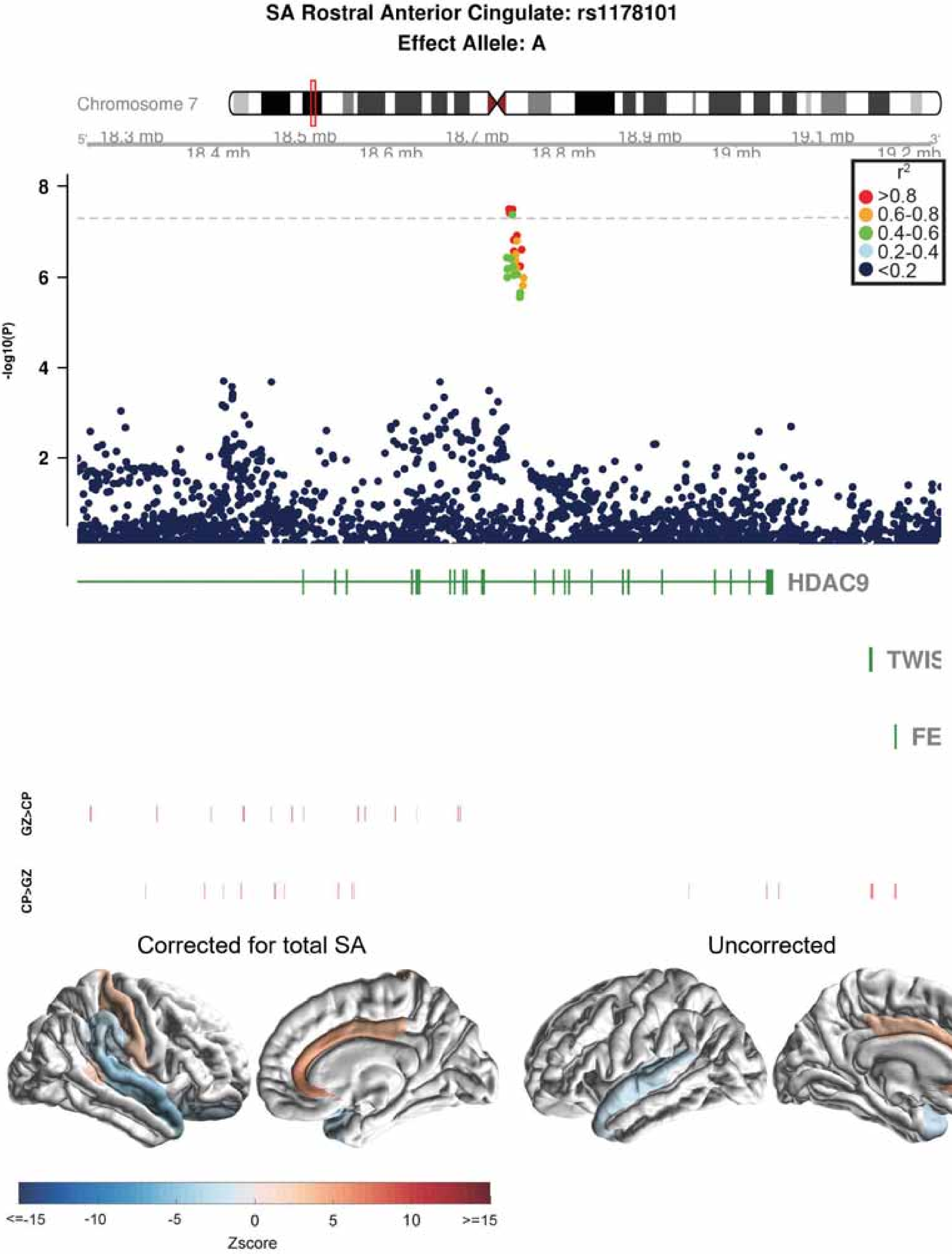

SA Rostral Anterior Cingulate: rs1178101
Effect Allele: A
Chromosome 7
18.3 mb
18.4 mb
18.5 mb
18.6 mb
18.7 mb
18.8 mb
18.9 mb
19 mb
19.1 mb
19.2 mb
r2
>0.8
0.6-0.8
0.4-0.6
0.2-0.4
<0.2
-log10(P)
8
6
4
2
HDAC9
TWIS
FE
GZ>CP
CP>GZ
Corrected for total SA
Uncorrected
<=-15
-10
-5
0
5
10
>=15
Zscore

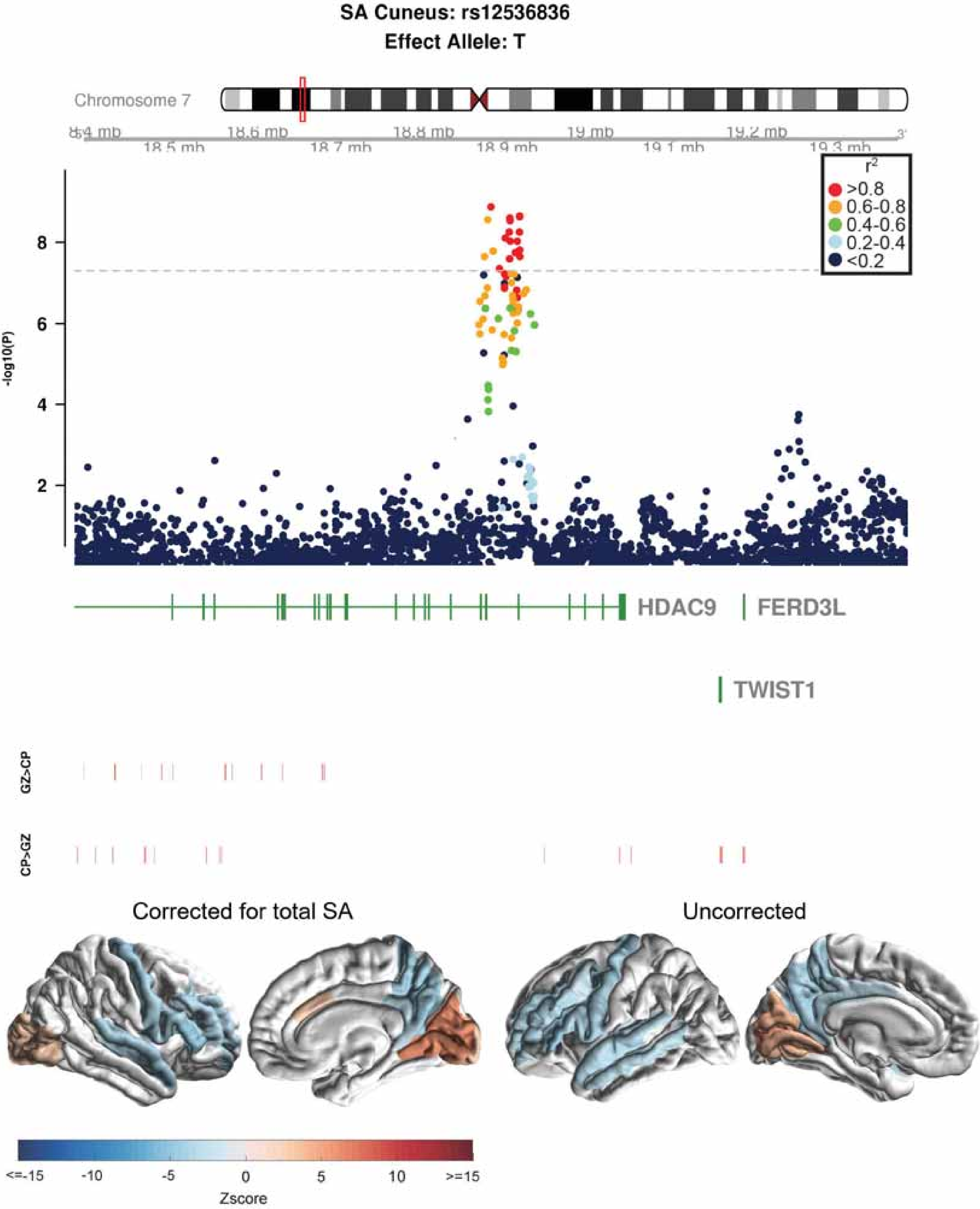

SA Cuneus: rs12536836
Effect Allele: T
Chromosome 7
18.5 mb
18.6 mb
18.7 mb
18.8 mb
18.9 mb
19 mb
19.1 mb
19.2 mb
19.3 mb
r2
>0.8
0.6-0.8
0.4-0.6
0.2-0.4
<0.2
-log10(P)
8
6
4
2
HDAC9
FERD3L
TWIST1
GZ>CP
CP>GZ
Corrected for total SA
Uncorrected
<=-15
-10
-5
0
5
10
>=15
Zscore

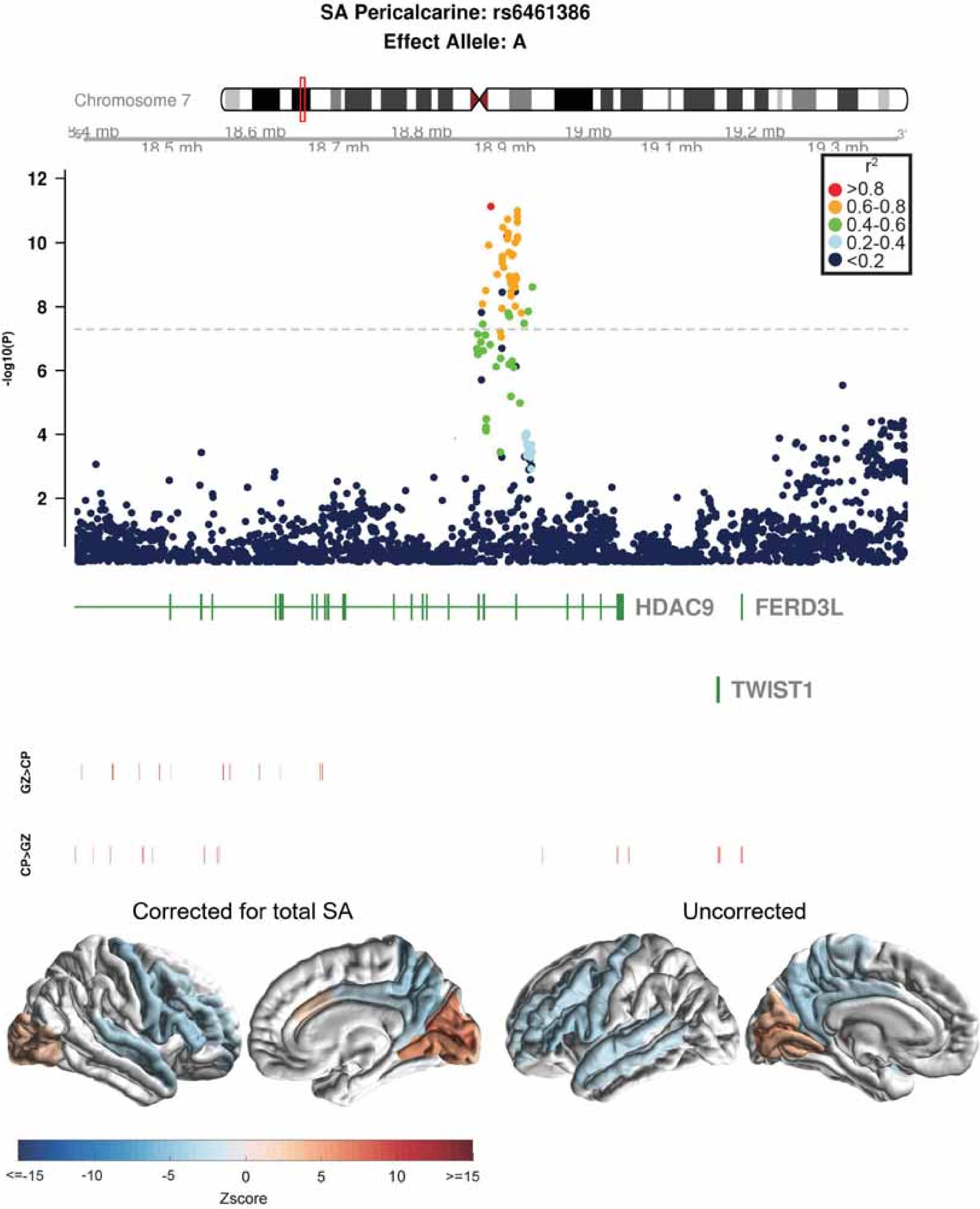

SA Pericalcarine: rs6461386
Effect Allele: A
Chromosome 7
18.4 mb
18.5 mb
18.6 mb
18.7 mb
18.8 mb
18.9 mb
19 mb
19.1 mb
19.2 mb
19.3 mb
r²
>0.8
0.6-0.8
0.4-0.6
0.2-0.4
<0.2
-log10(P)
12
10
8
6
4
2
HDAC9
FERD3L
TWIST1
GZ>CP
CP>GZ
Corrected for total SA
Uncorrected
<=-15
-10
-5
0
5
10
>=15
Zscore

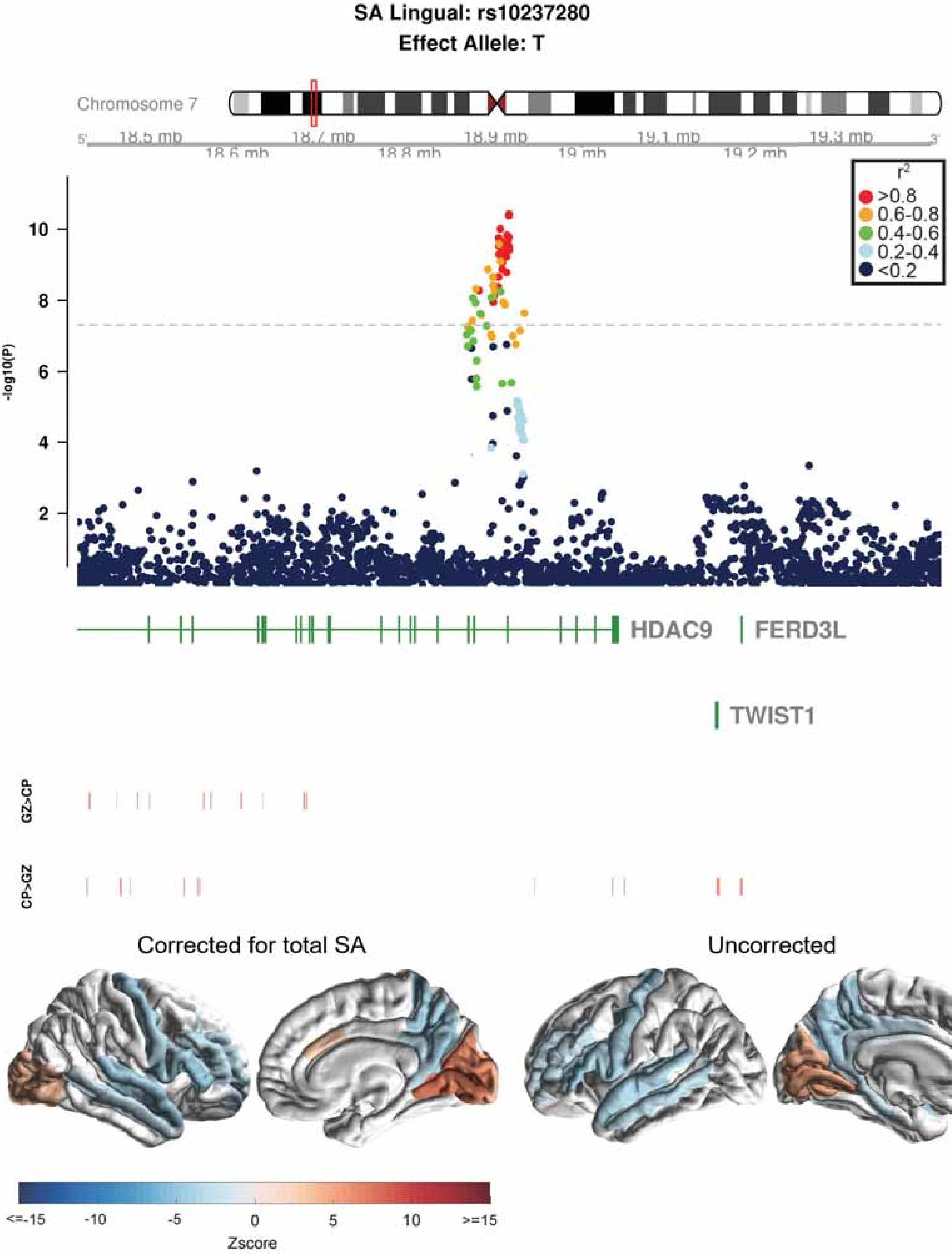

SA Lingual: rs10237280
Effect Allele: T
Chromosome 7
18.5 mb
18.6 mb
18.7 mb
18.8 mb
18.9 mb
19 mb
19.1 mb
19.2 mb
19.3 mb
-log10(P)
r2
>0.8
0.6-0.8
0.4-0.6
0.2-0.4
<0.2
HDAC9
FERD3L
TWIST1
GZ>CP
CP>GZ
Corrected for total SA
Uncorrected
<=-15
-10
-5
0
5
10
>=15
Zscore

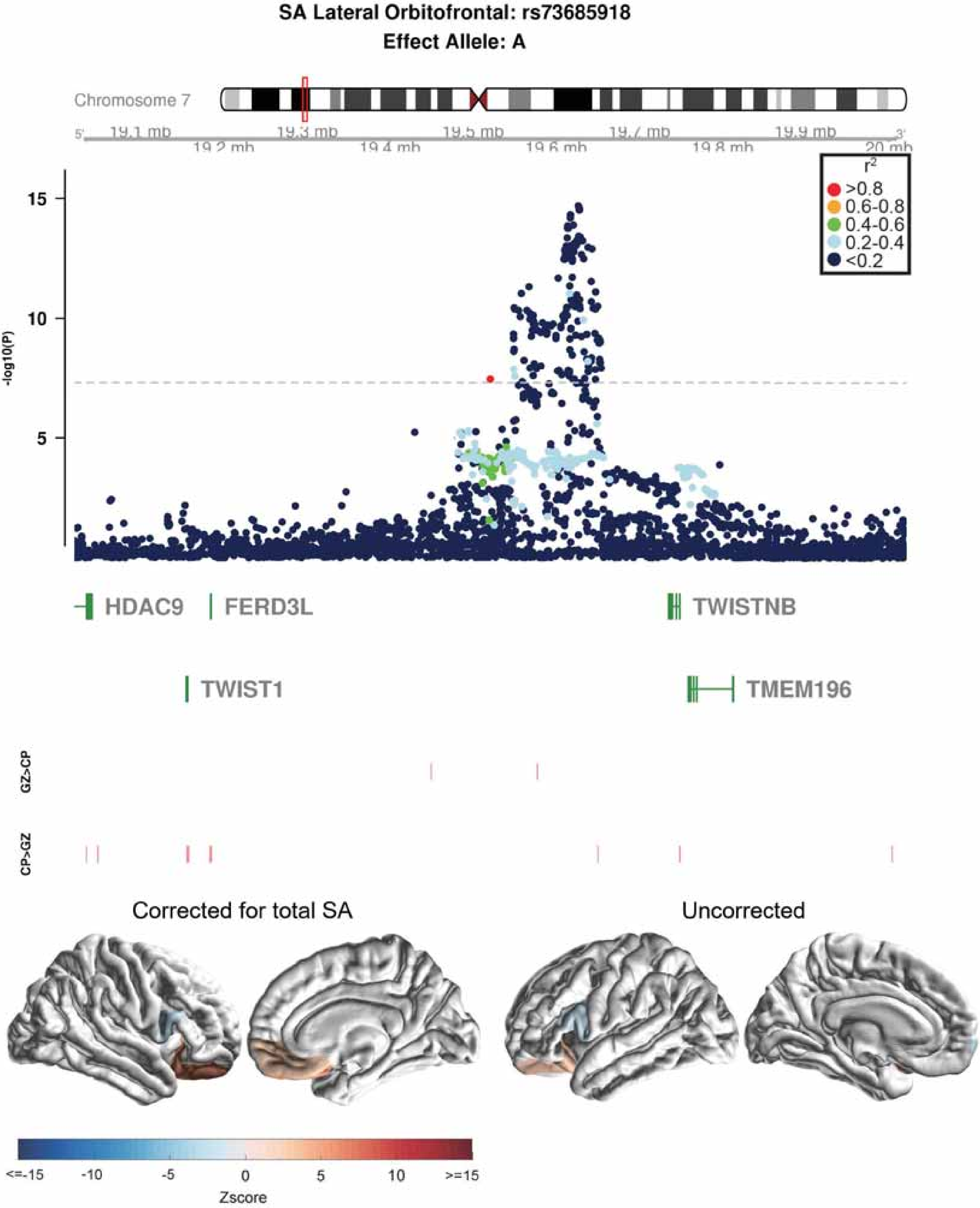

SA Lateral Orbitofrontal: rs73685918
Effect Allele: A
Chromosome 7
19.1 mb
19.2 mb
19.3 mb
19.4 mb
19.5 mb
19.6 mb
19.7 mb
19.8 mb
19.9 mb
20 mb
r2
>0.8
0.6-0.8
0.4-0.6
0.2-0.4
<0.2
-log10(P)
15
10
5
HDAC9
FERD3L
TWISTNB
TWIST1
TMEM196
GZ>CP
CP>GZ
Corrected for total SA
Uncorrected
<=-15
-10
-5
0
5
10
>=15
Zscore

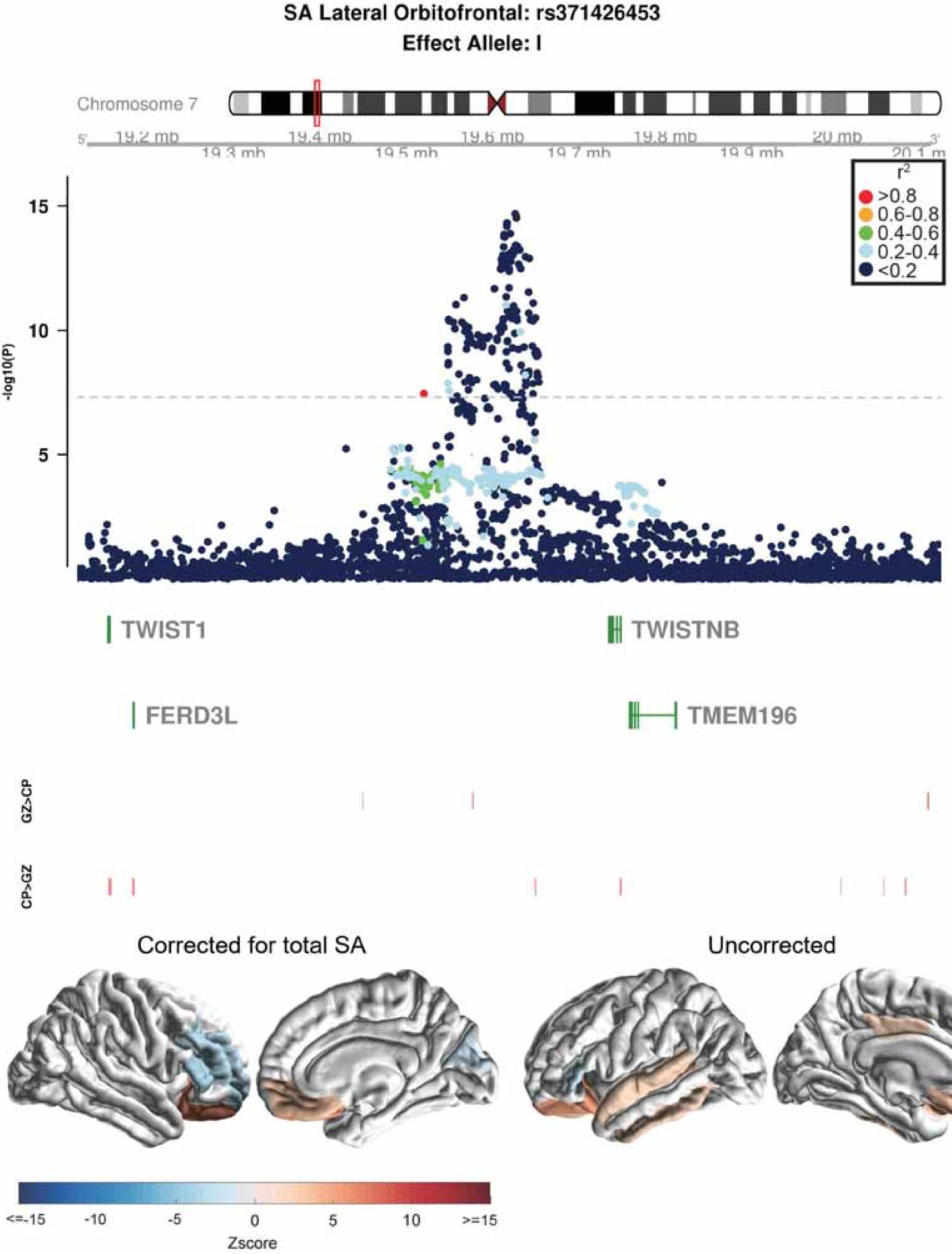

SA Lateral Orbitofrontal: rs371426453
Effect Allele: I
Chromosome 7
19.2 mb
19.3 mb
19.4 mb
19.5 mb
19.6 mb
19.7 mb
19.8 mb
19.9 mb
20 mb
20.1 m
-log10(P)
15
10
5
r2
>0.8
0.6-0.8
0.4-0.6
0.2-0.4
<0.2
TWIST1
TWISTNB
FERD3L
TMEM196
GZ>CP
CP>GZ
Corrected for total SA
Uncorrected
<=-15
-10
-5
0
5
10
>=15
Zscore

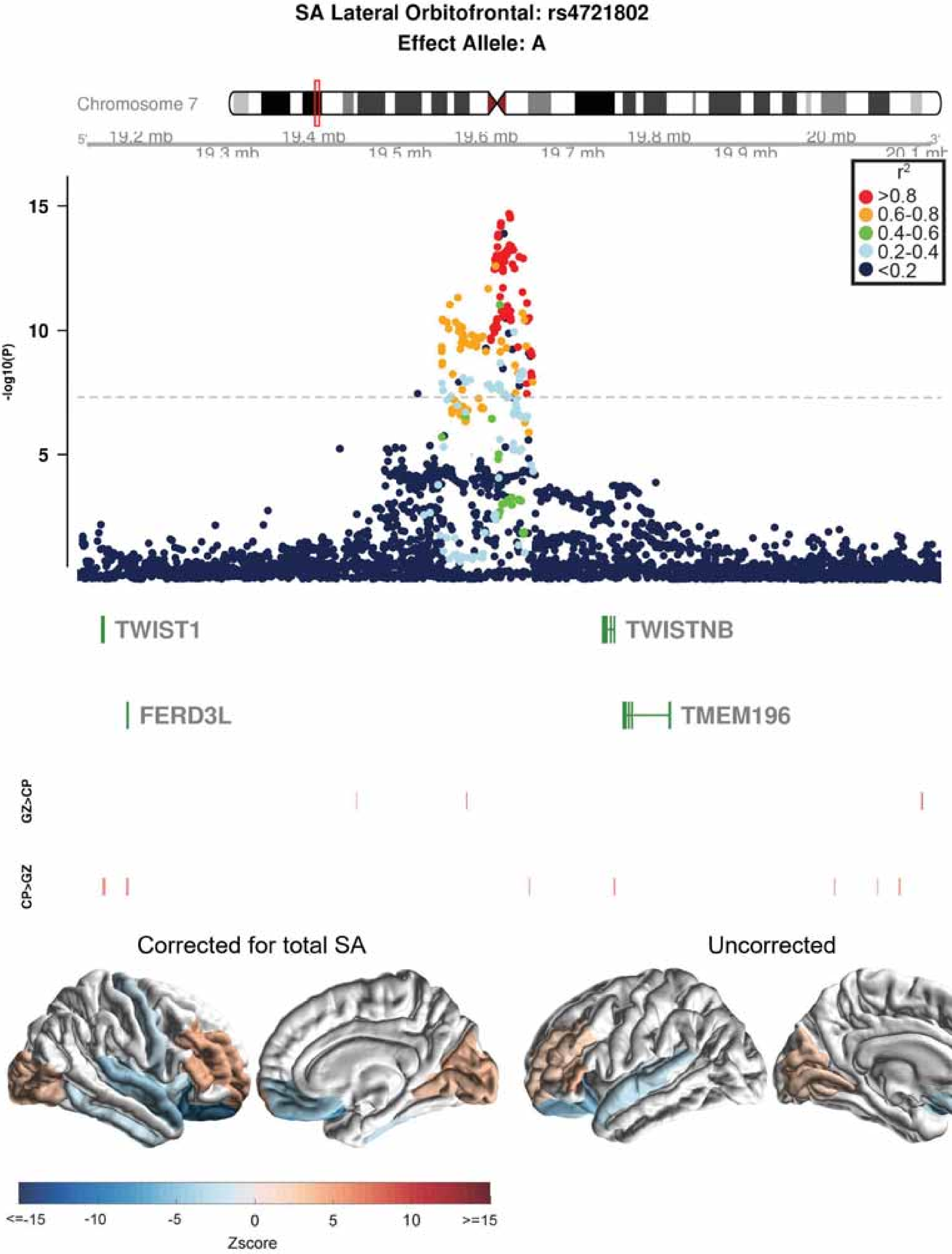

SA Lateral Orbitofrontal: rs4721802
Effect Allele: A
Chromosome 7
19.2 mb
19.3 mb
19.4 mb
19.5 mb
19.6 mb
19.7 mb
19.8 mb
19.9 mb
20 mb
20.1 mb
r2
>0.8
0.6-0.8
0.4-0.6
0.2-0.4
<0.2
-log10(P)
15
10
5
TWIST1
TWISTNB
FERD3L
TMEM196
GZ>CP
CP>GZ
Corrected for total SA
Uncorrected
<=-15
-10
-5
0
5
10
>=15
Zscore

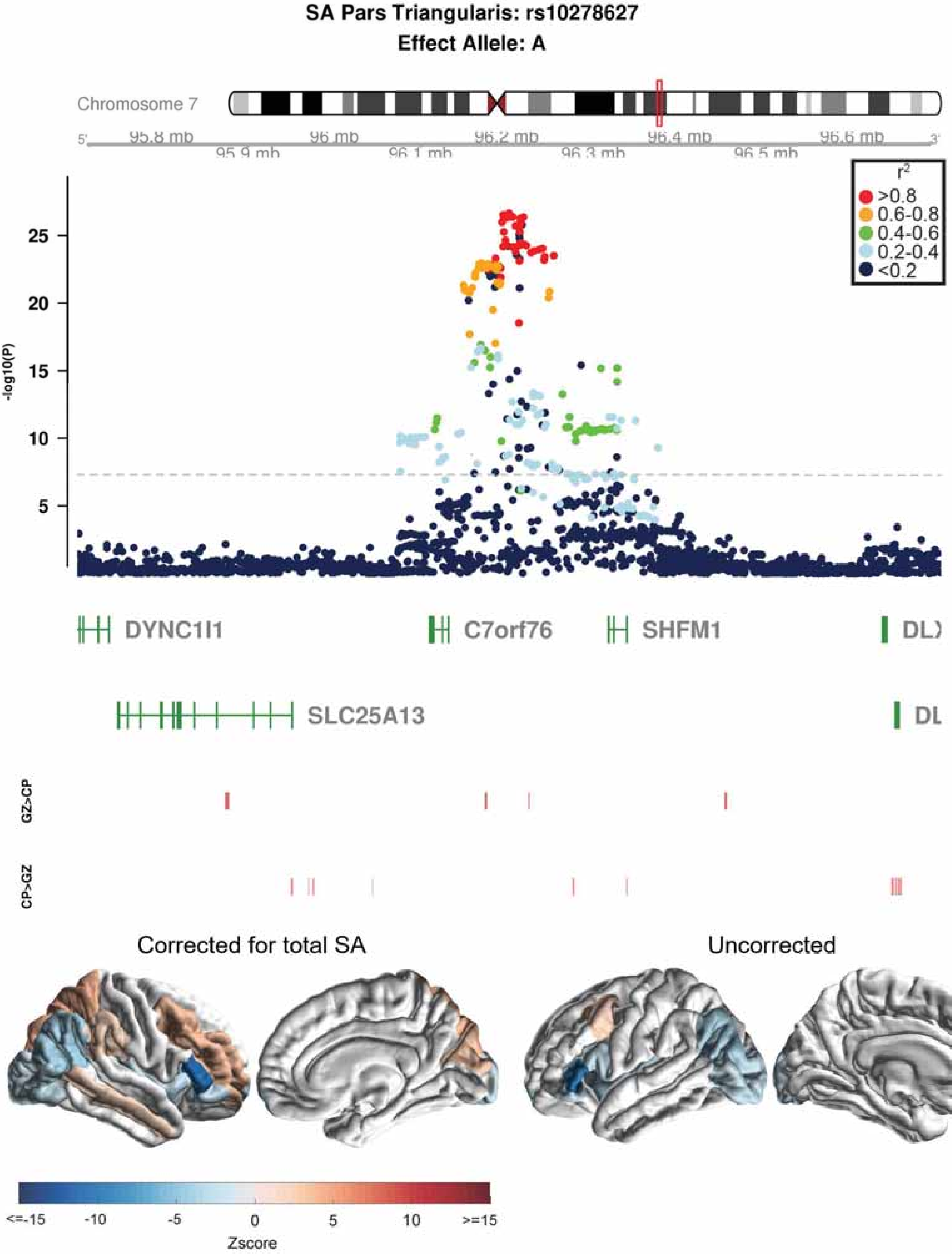
SA Pars Triangularis: rs10278627
Effect Allele: A
Chromosome 7
95.8 mb
95.9 mb
96 mb
96.1 mb
96.2 mb
96.3 mb
96.4 mb
96.5 mb
96.6 mb
5'
3'
r2
>0.8
0.6-0.8
0.4-0.6
0.2-0.4
<0.2
-log10(P)
25
20
15
10
5
DYNC1I1
C7orf76
SHFM1
SLC25A13
GZ>CP
CP>GZ
Corrected for total SA
Uncorrected
<=-15
-10
-5
0
5
10
>=15
Zscore

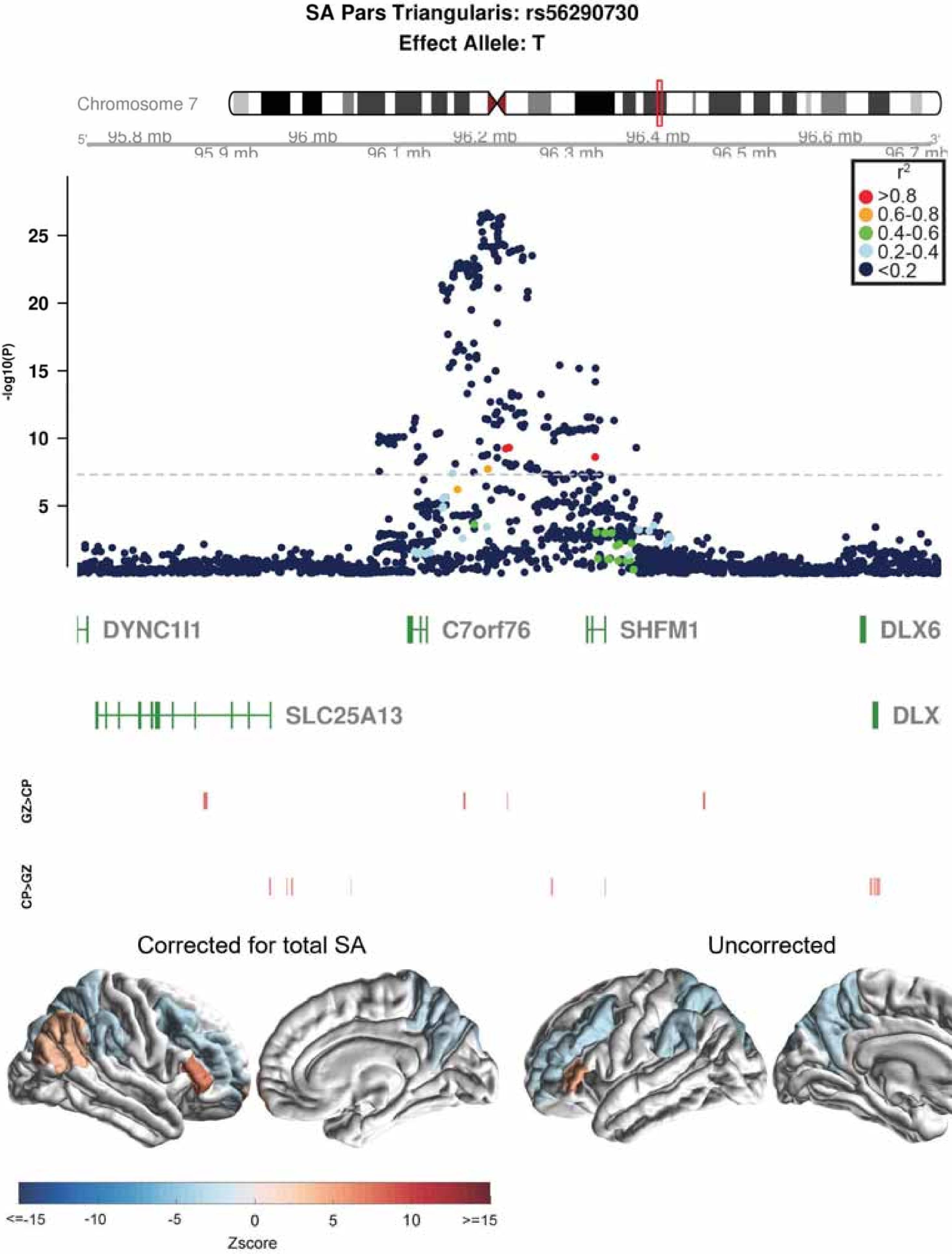
SA Pars Triangularis: rs56290730
Effect Allele: T
Chromosome 7
95.8 mb
95.9 mb
96 mb
96.1 mb
96.2 mb
96.3 mb
96.4 mb
96.5 mb
96.6 mb
96.7 mb
r2
>0.8
0.6-0.8
0.4-0.6
0.2-0.4
<0.2
-log10(P)
25
20
15
10
5
DYNC1I1
C7orf76
SHFM1
DLX6
SLC25A13
DLX
GZ>CP
CP>GZ
Corrected for total SA
Uncorrected
<=-15
-10
-5
0
5
10
>=15
Zscore

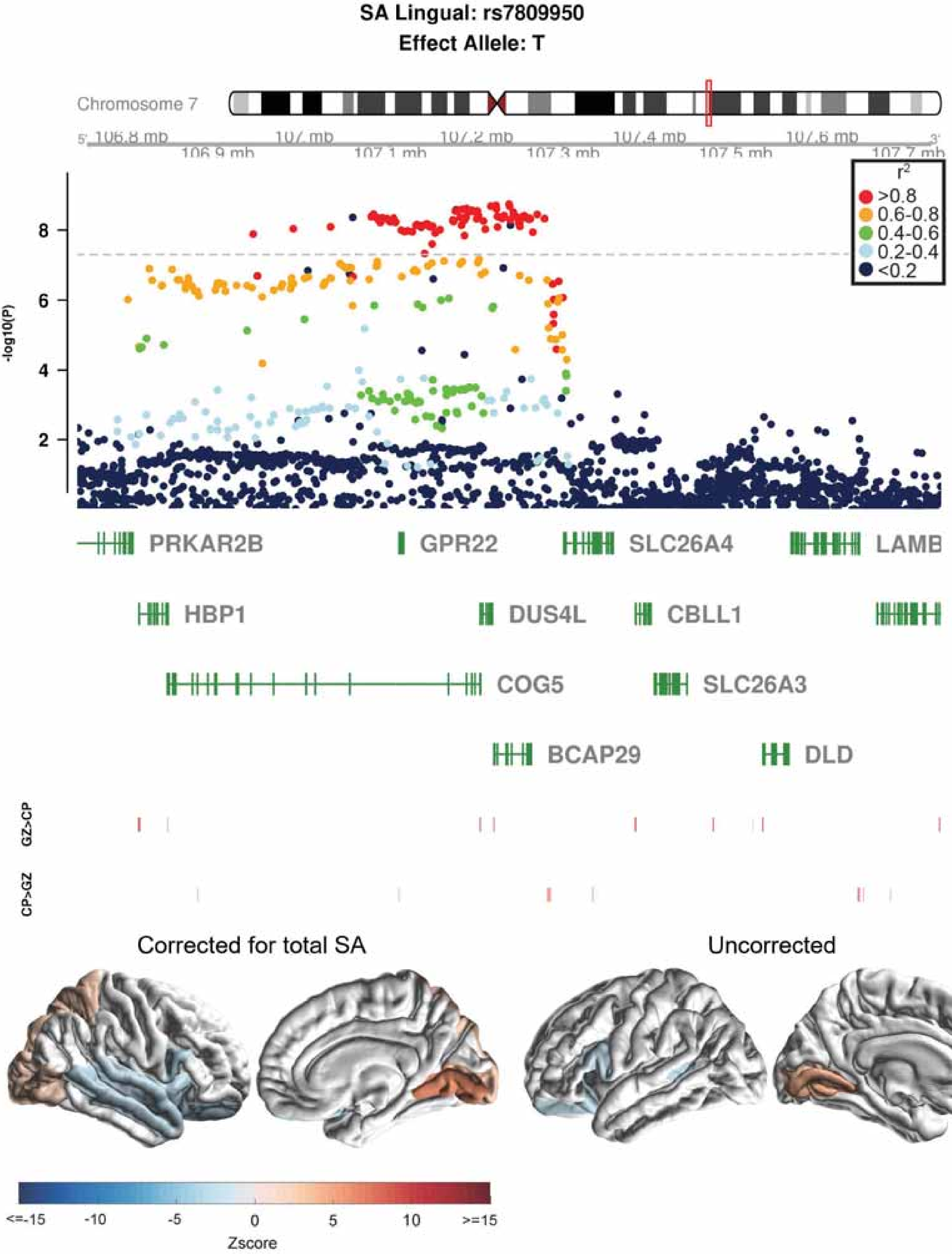

SA Lingual: rs7809950
Effect Allele: T
Chromosome 7
106.8 mb
107 mb
107.2 mb
107.4 mb
107.6 mb
106.9 mb
107.1 mb
107.3 mb
107.5 mb
107.7 mb
-log10(P)
8
6
4
2
r2
>0.8
0.6-0.8
0.4-0.6
0.2-0.4
<0.2
PRKAR2B
GPR22
SLC26A4
LAMB
HBP1
DUS4L
CBLL1
COG5
SLC26A3
BCAP29
DLD
GZ>CP
CP>GZ
Corrected for total SA
Uncorrected
<=-15
-10
-5
0
5
10
>=15
Zscore

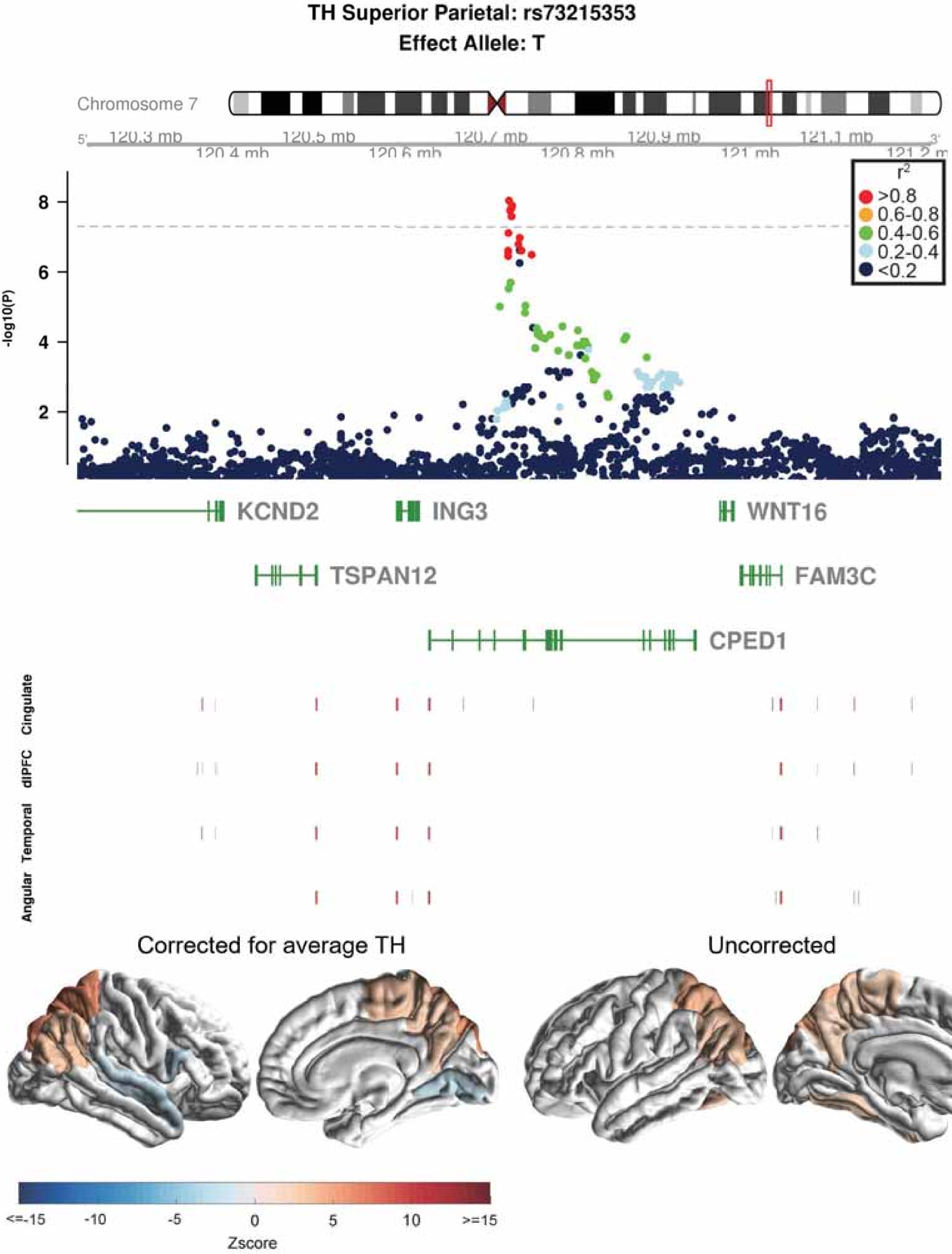

TH Superior Parietal: rs73215353
Effect Allele: T
Chromosome 7
120.3 mb
120.4 mb
120.5 mb
120.6 mb
120.7 mb
120.8 mb
120.9 mb
121 mb
121.1 mb
r2
>0.8
0.6-0.8
0.4-0.6
0.2-0.4
<0.2
-log10(P)
KCND2
ING3
WNT16
TSPAN12
FAM3C
CPED1
Cingulate
dlPFC
Temporal
Angular
Corrected for average TH
Uncorrected
<=-15
-10
-5
0
5
10
>=15
Zscore

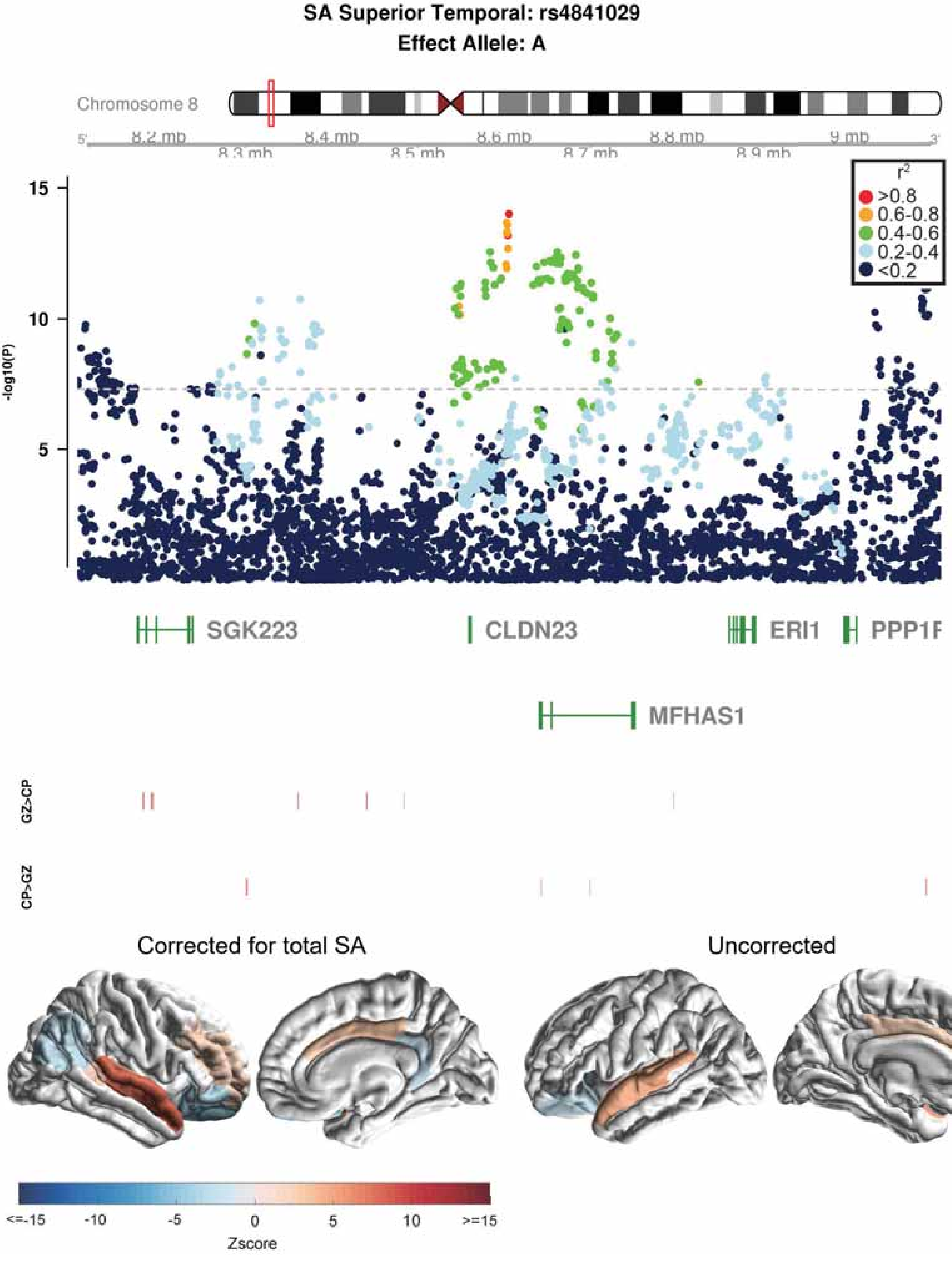

SA Superior Temporal: rs4841029
Effect Allele: A
Chromosome 8
5'
8.2 mb
8.3 mb
8.4 mb
8.5 mb
8.6 mb
8.7 mb
8.8 mb
8.9 mb
9 mb
3'
r2
>0.8
0.6-0.8
0.4-0.6
0.2-0.4
<0.2
15
10
5
-log10(P)
SGK223
CLDN23
ERI1
PPP1F
MFHAS1
GZ>CP
CP>GZ
Corrected for total SA
Uncorrected
<=-15
-10
-5
0
5
10
>=15
Zscore

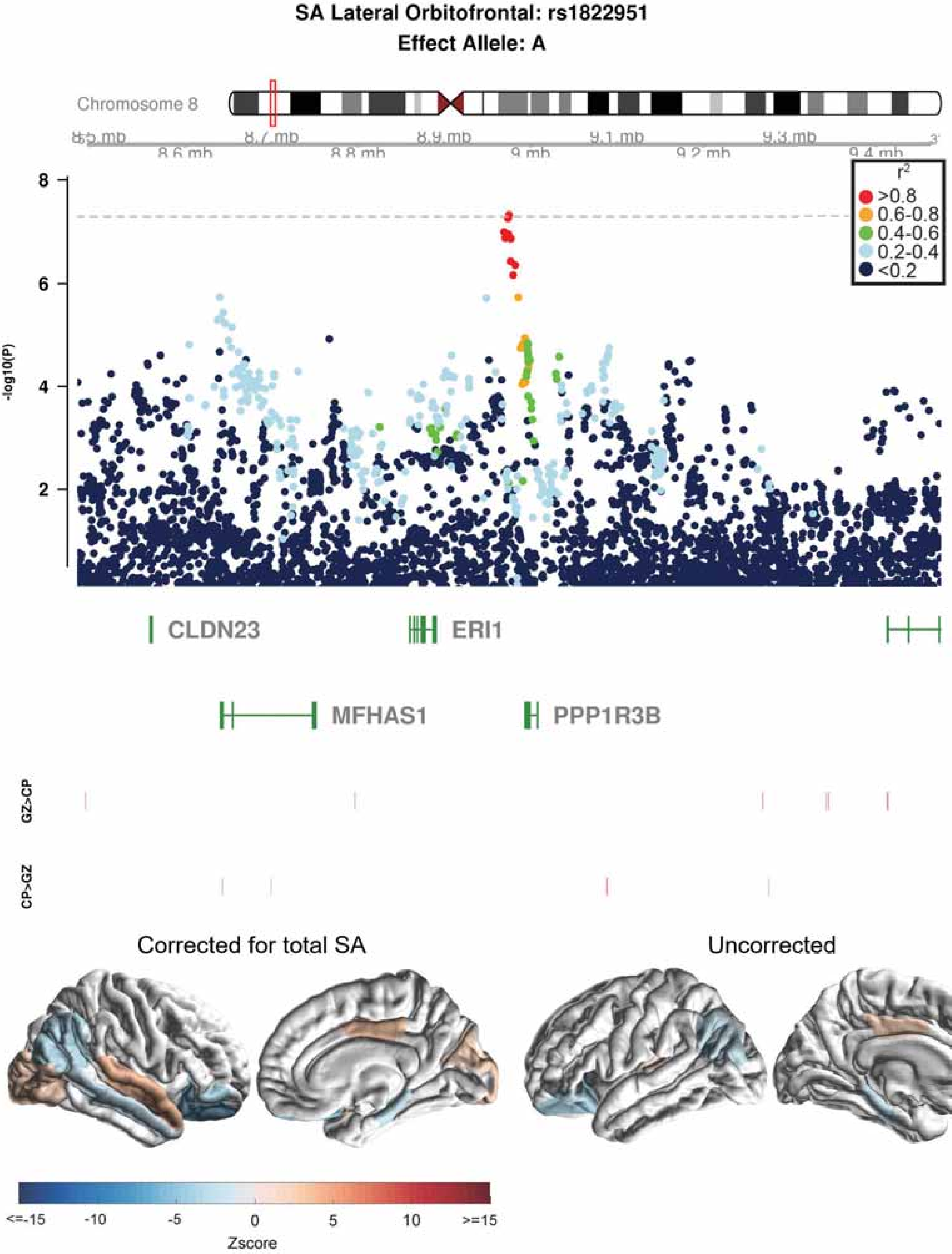

SA Lateral Orbitofrontal: rs1822951
Effect Allele: A
Chromosome 8
8.5 mb
8.6 mb
8.7 mb
8.8 mb
8.9 mb
9 mb
9.1 mb
9.2 mb
9.3 mb
9.4 mb
-log10(P)
8
6
4
2
$r^2$
>0.8
0.6-0.8
0.4-0.6
0.2-0.4
<0.2
CLDN23
ERI1
MFHAS1
PPP1R3B
GZ>CP
CP>GZ
Corrected for total SA
Uncorrected
<=-15
-10
-5
0
5
10
>=15
Zscore

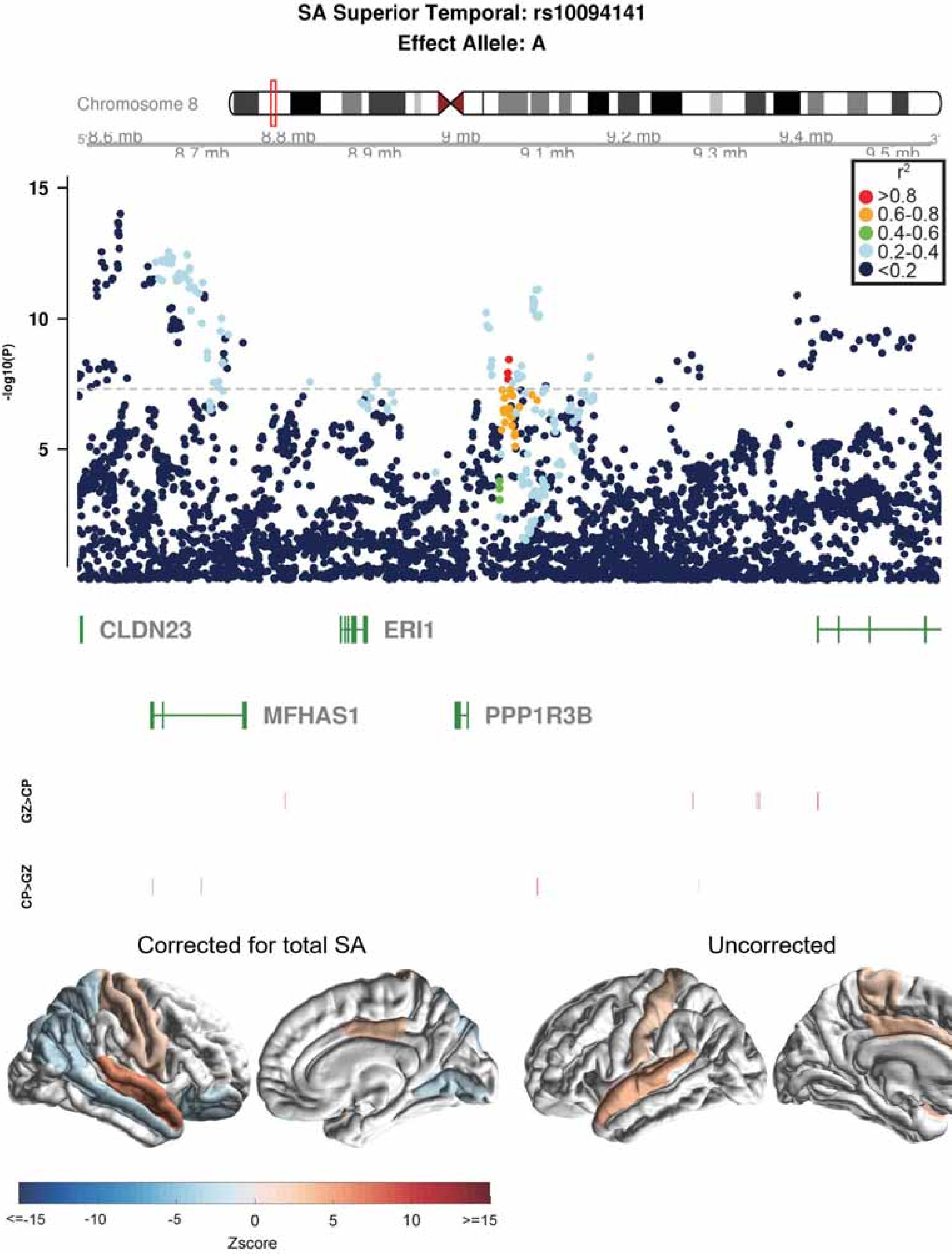
SA Superior Temporal: rs10094141
Effect Allele: A
Chromosome 8
8.6 mb
8.7 mb
8.8 mb
8.9 mb
9 mb
9.1 mb
9.2 mb
9.3 mb
9.4 mb
9.5 mb
r2
>0.8
0.6-0.8
0.4-0.6
0.2-0.4
<0.2
-log10(P)
15
10
5
CLDN23
ERI1
MFHAS1
PPP1R3B
GZ>CP
CP>GZ
Corrected for total SA
Uncorrected
<=-15
-10
-5
0
5
10
>=15
Zscore

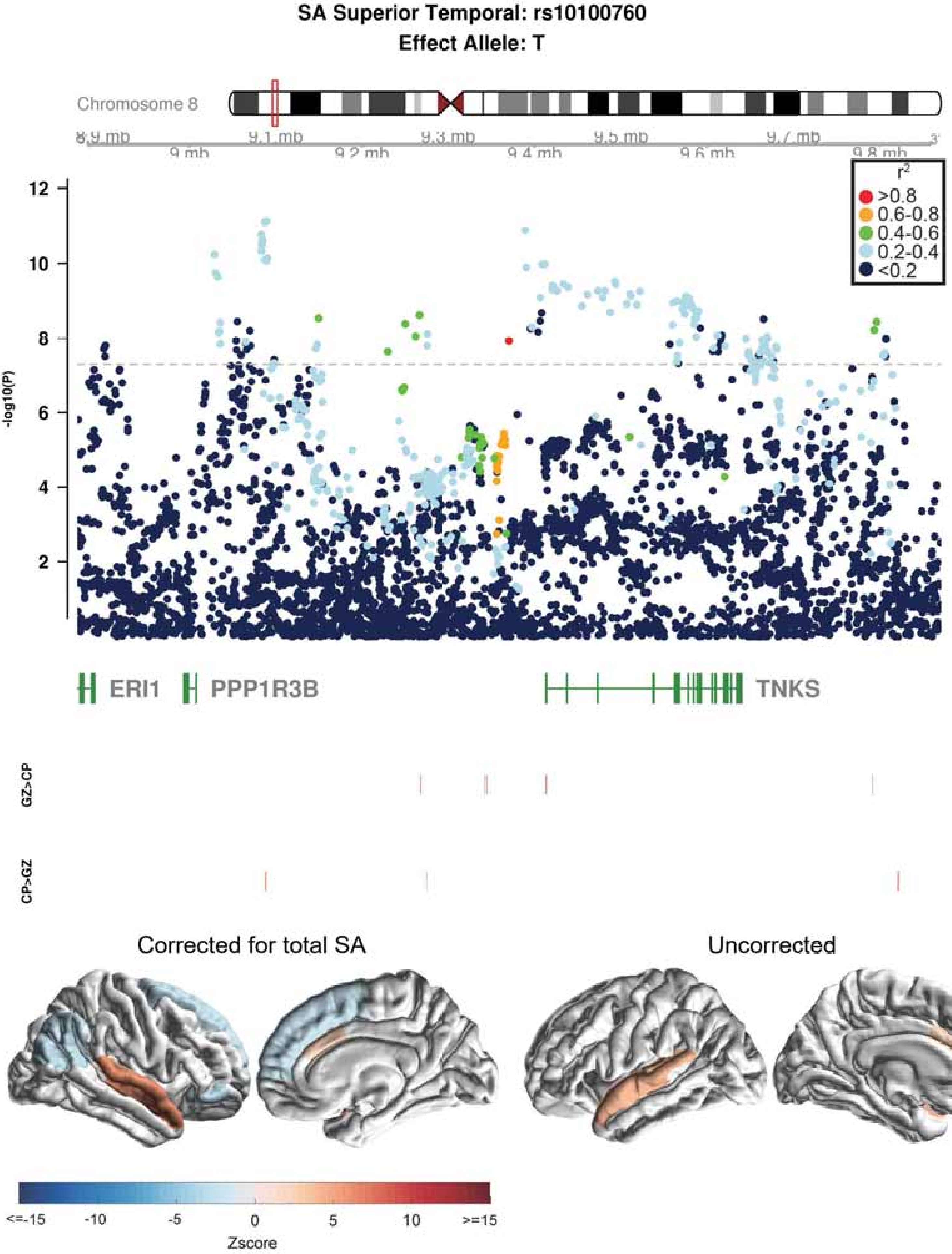
SA Superior Temporal: rs10100760
Effect Allele: T
Chromosome 8
8.9 mb
9 mb
9.1 mb
9.2 mb
9.3 mb
9.4 mb
9.5 mb
9.6 mb
9.7 mb
9.8 mb
r2
>0.8
0.6-0.8
0.4-0.6
0.2-0.4
<0.2
-log10(P)
12
10
8
6
4
2
ERI1
PPP1R3B
TNKS
GZ>CP
CP>GZ
Corrected for total SA
Uncorrected
<=-15
-10
-5
0
5
10
>=15
Zscore

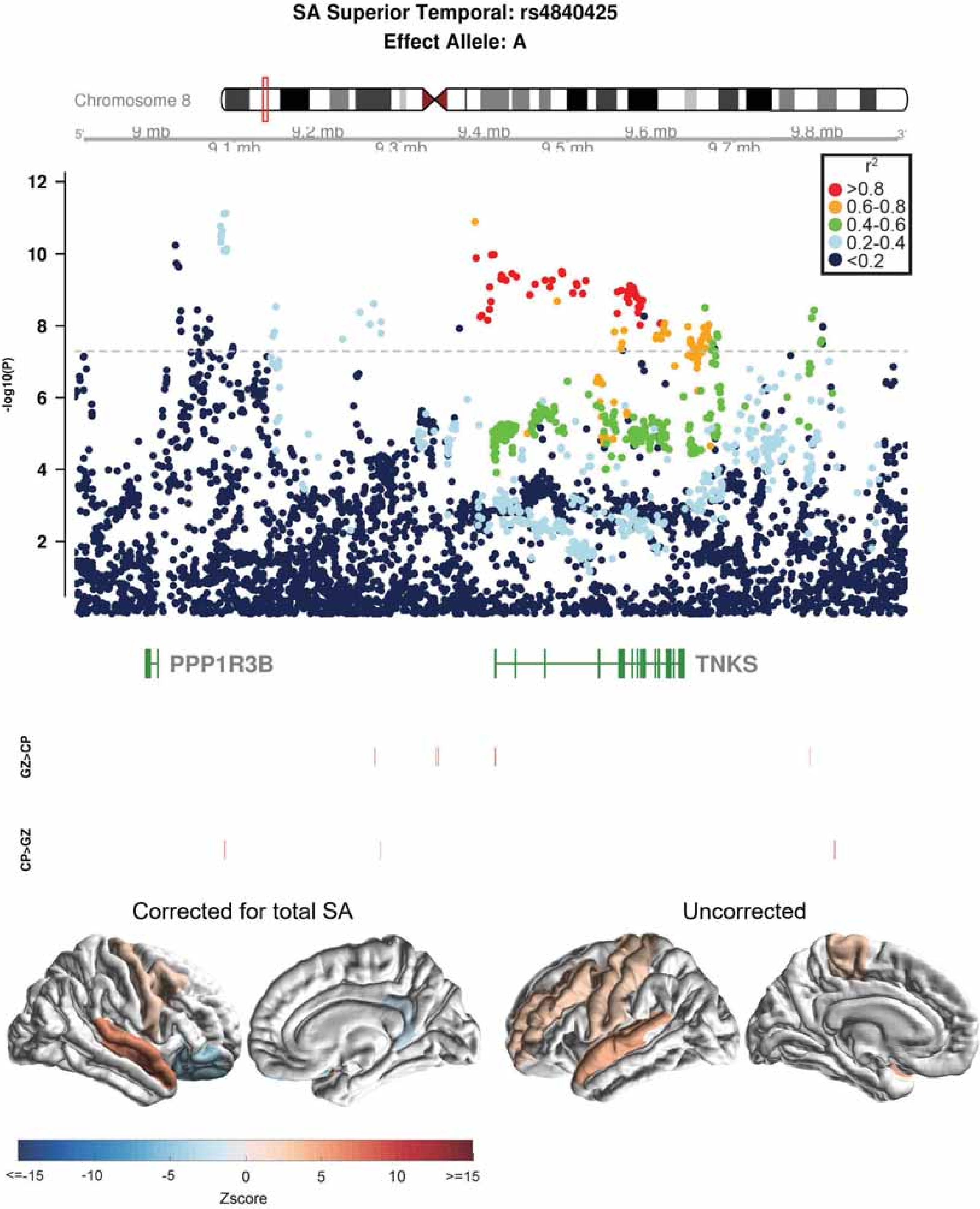
SA Superior Temporal: rs4840425
Effect Allele: A
Chromosome 8
9 mb
9.1 mb
9.2 mb
9.3 mb
9.4 mb
9.5 mb
9.6 mb
9.7 mb
9.8 mb
-log10(P)
r2
>0.8
0.6-0.8
0.4-0.6
0.2-0.4
<0.2
PPP1R3B
TNKS
GZ>CP
CP>GZ
Corrected for total SA
Uncorrected
<=-15
-10
-5
0
5
10
>=15
Zscore

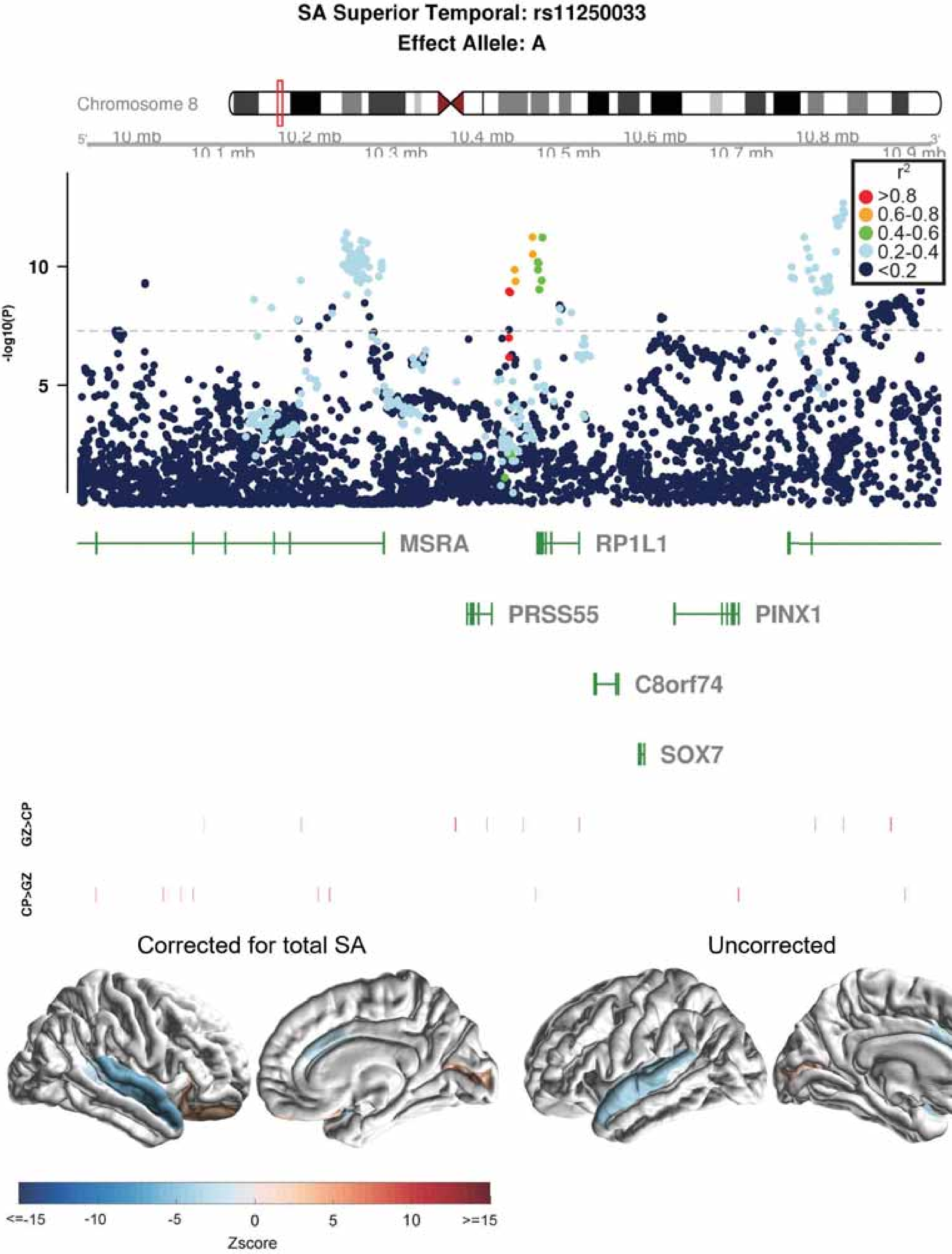

SA Superior Temporal: rs11250033
Effect Allele: A
Chromosome 8
10 mb
10.1 mb
10.2 mb
10.3 mb
10.4 mb
10.5 mb
10.6 mb
10.7 mb
10.8 mb
10.9 mb
-log10(P)
10
5
r2
>0.8
0.6-0.8
0.4-0.6
0.2-0.4
<0.2
MSRA
RP1L1
PRSS55
PINX1
C8orf74
SOX7
GZ>CP
CP>GZ
Corrected for total SA
Uncorrected
<=-15
-10
-5
0
5
10
>=15
Zscore

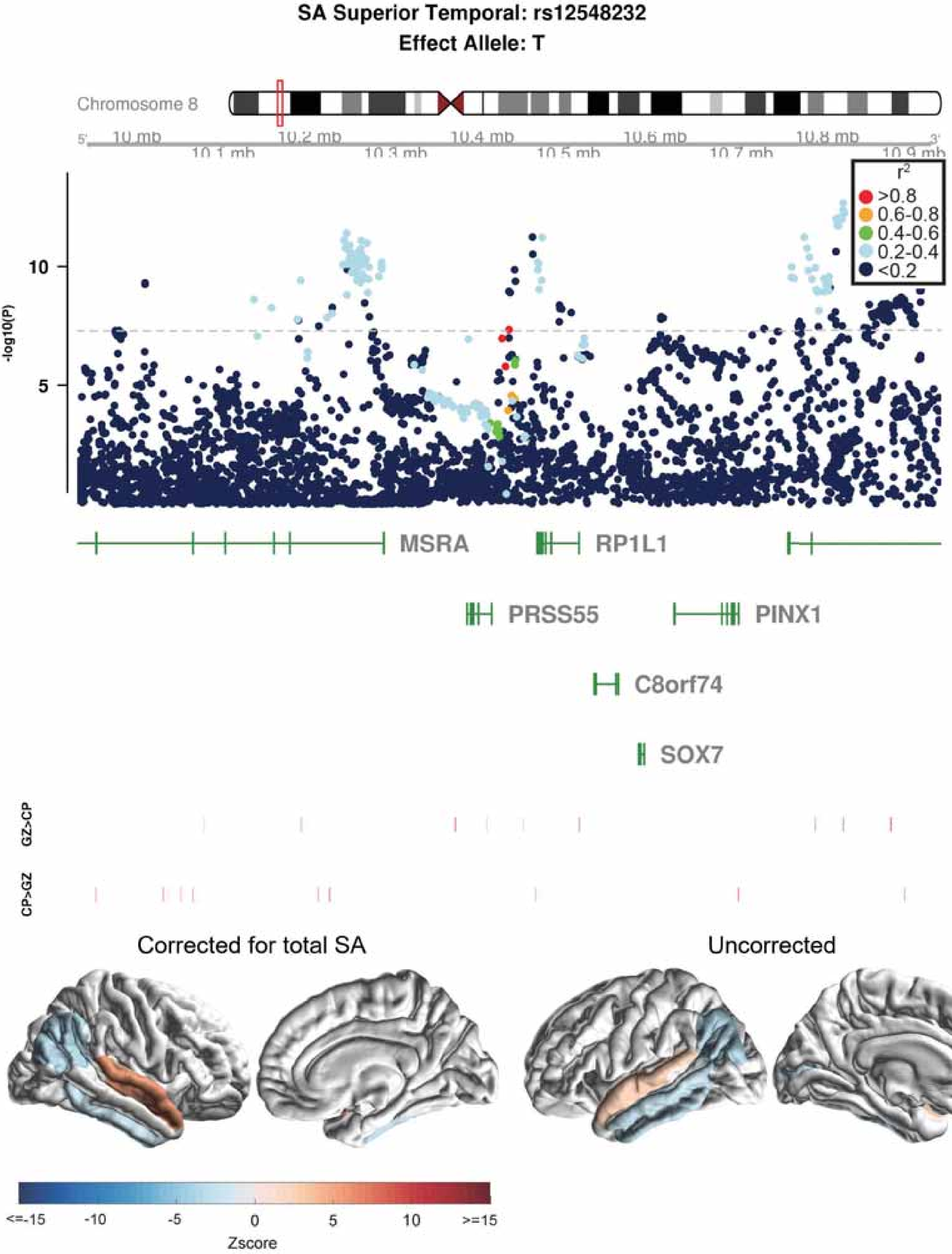

SA Superior Temporal: rs12548232
Effect Allele: T
Chromosome 8
10 mb
10.1 mb
10.2 mb
10.3 mb
10.4 mb
10.5 mb
10.6 mb
10.7 mb
10.8 mb
10.9 mb
-log10(P)
r2
>0.8
0.6-0.8
0.4-0.6
0.2-0.4
<0.2
MSRA
RP1L1
PRSS55
PINX1
C8orf74
SOX7
GZ>CP
CP>GZ
Corrected for total SA
Uncorrected
<=-15
-10
-5
0
5
10
>=15
Zscore

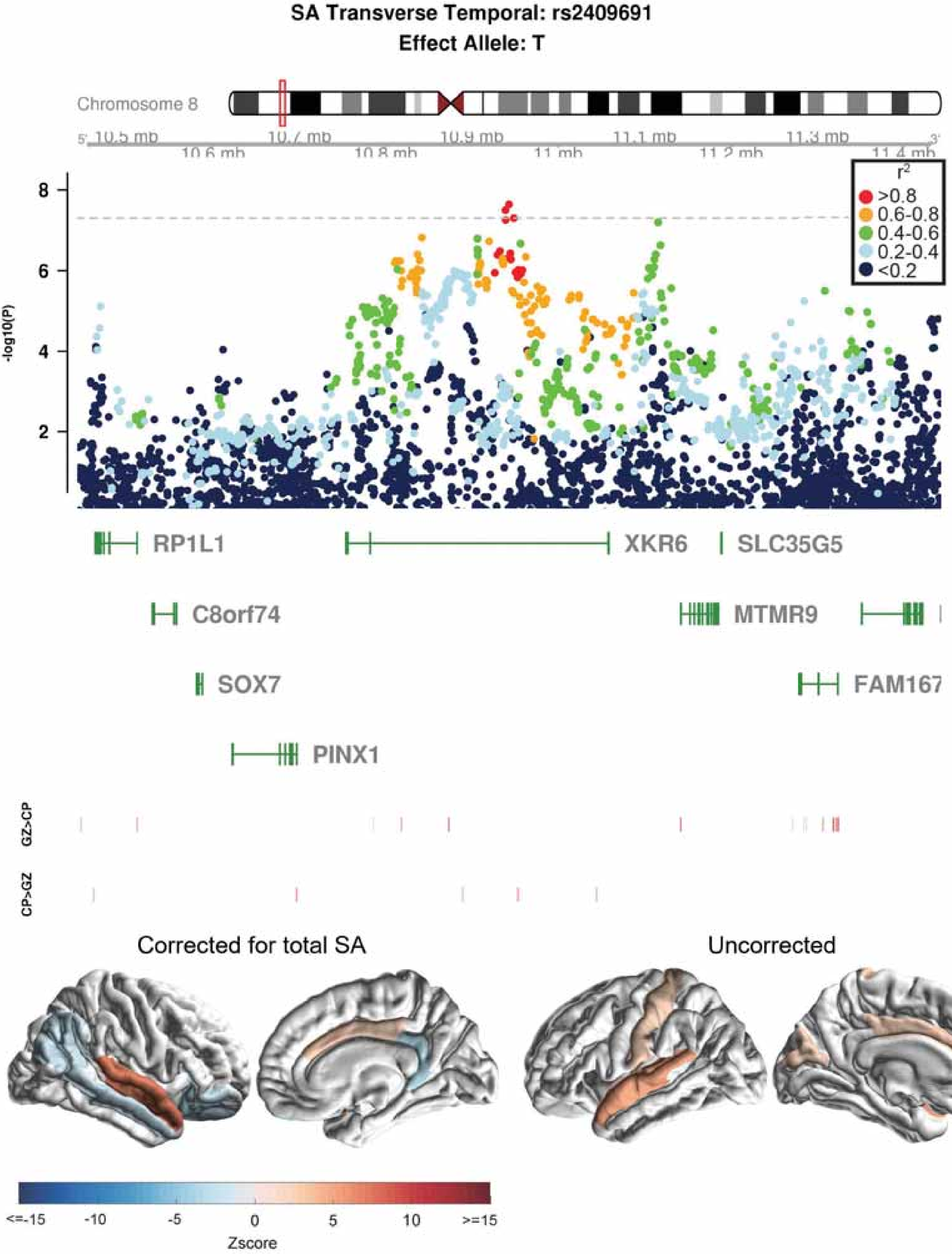

SA Transverse Temporal: rs2409691
Effect Allele: T
Chromosome 8
10.5 mb
10.6 mb
10.7 mb
10.8 mb
10.9 mb
11 mb
11.1 mb
11.2 mb
11.3 mb
11.4 mb
-log10(P)
r²
>0.8
0.6-0.8
0.4-0.6
0.2-0.4
<0.2
RP1L1
XKR6
SLC35G5
C8orf74
MTMR9
SOX7
FAM167
PINX1
GZ>CP
CP>GZ
Corrected for total SA
Uncorrected
<=-15
-10
-5
0
5
10
>=15
Zscore

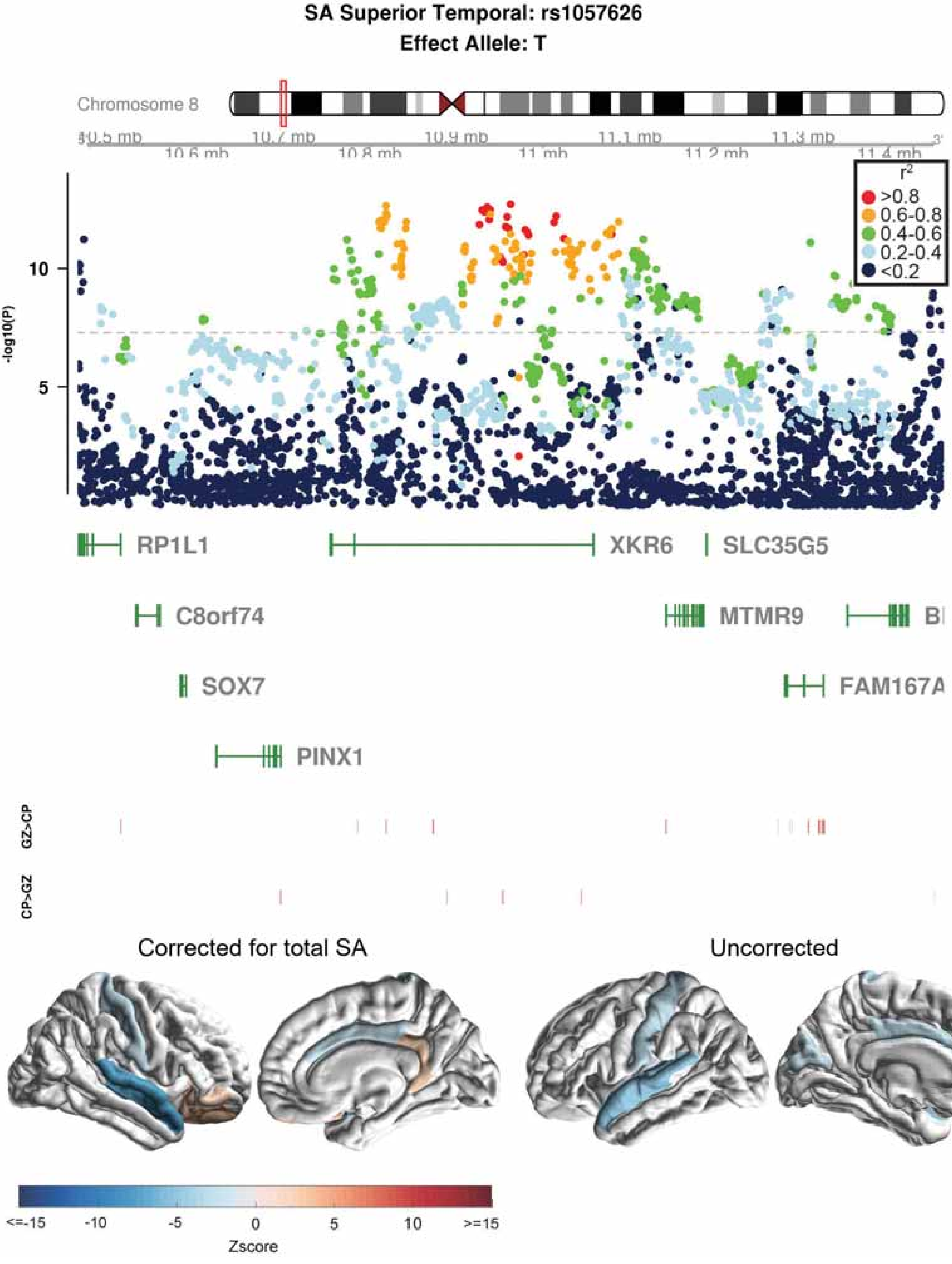
SA Superior Temporal: rs1057626
Effect Allele: T
Chromosome 8
10.5 mb
10.6 mb
10.7 mb
10.8 mb
10.9 mb
11 mb
11.1 mb
11.2 mb
11.3 mb
11.4 mb
r2
>0.8
0.6-0.8
0.4-0.6
0.2-0.4
<0.2
-log10(P)
10
5
RP1L1
XKR6
SLC35G5
C8orf74
MTMR9
SOX7
FAM167A
PINX1
GZ>CP
CP>GZ
Corrected for total SA
Uncorrected
<=-15
-10
-5
0
5
10
>=15
Zscore

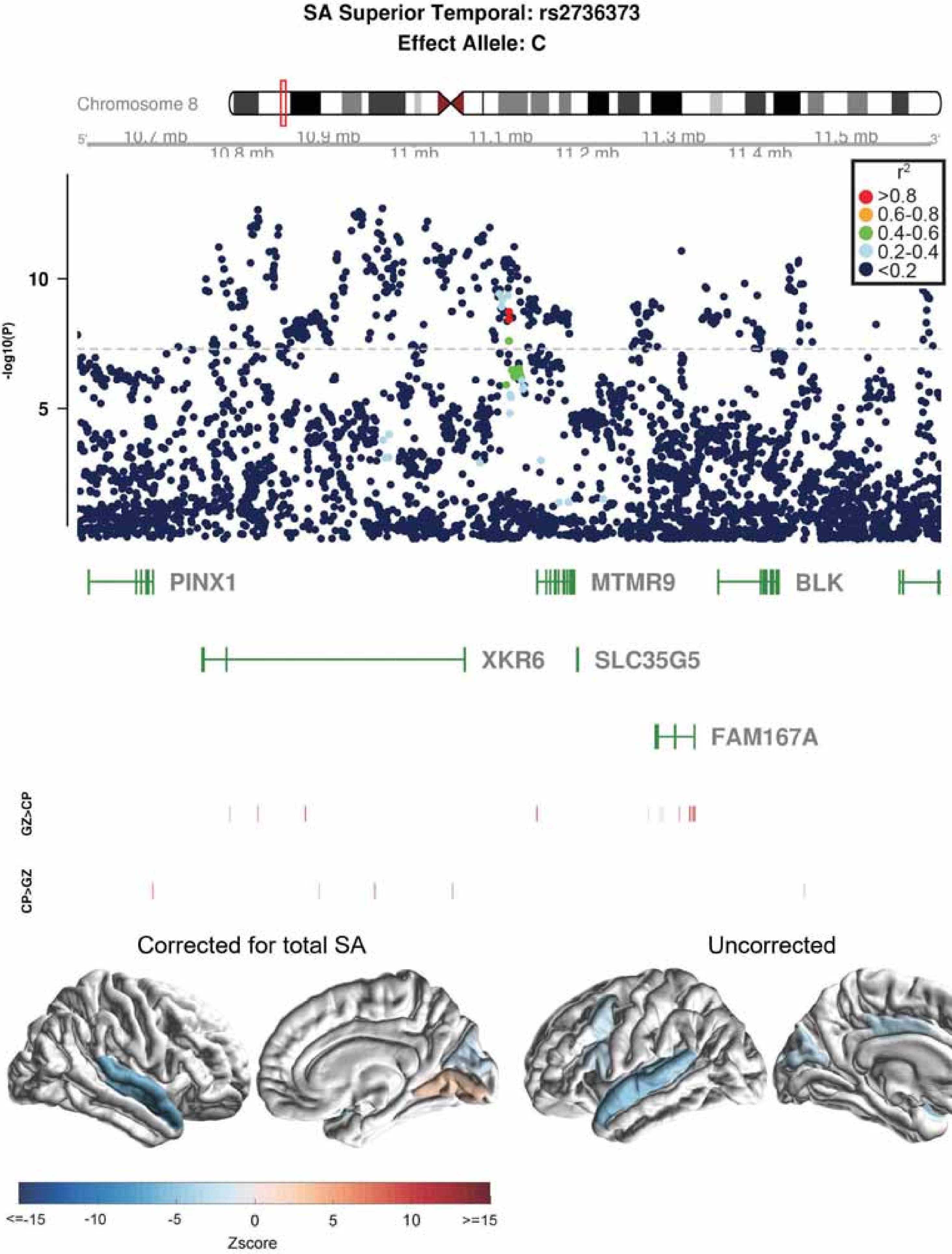

SA Superior Temporal: rs2736373
Effect Allele: C
Chromosome 8
10.7 mb
10.8 mb
10.9 mb
11 mb
11.1 mb
11.2 mb
11.3 mb
11.4 mb
11.5 mb
r2
>0.8
0.6-0.8
0.4-0.6
0.2-0.4
<0.2
-log10(P)
10
5
PINX1
MTMR9
BLK
XKR6
SLC35G5
FAM167A
GZ>CP
CP>GZ
Corrected for total SA
Uncorrected
<=-15
-10
-5
0
5
10
>=15
Zscore

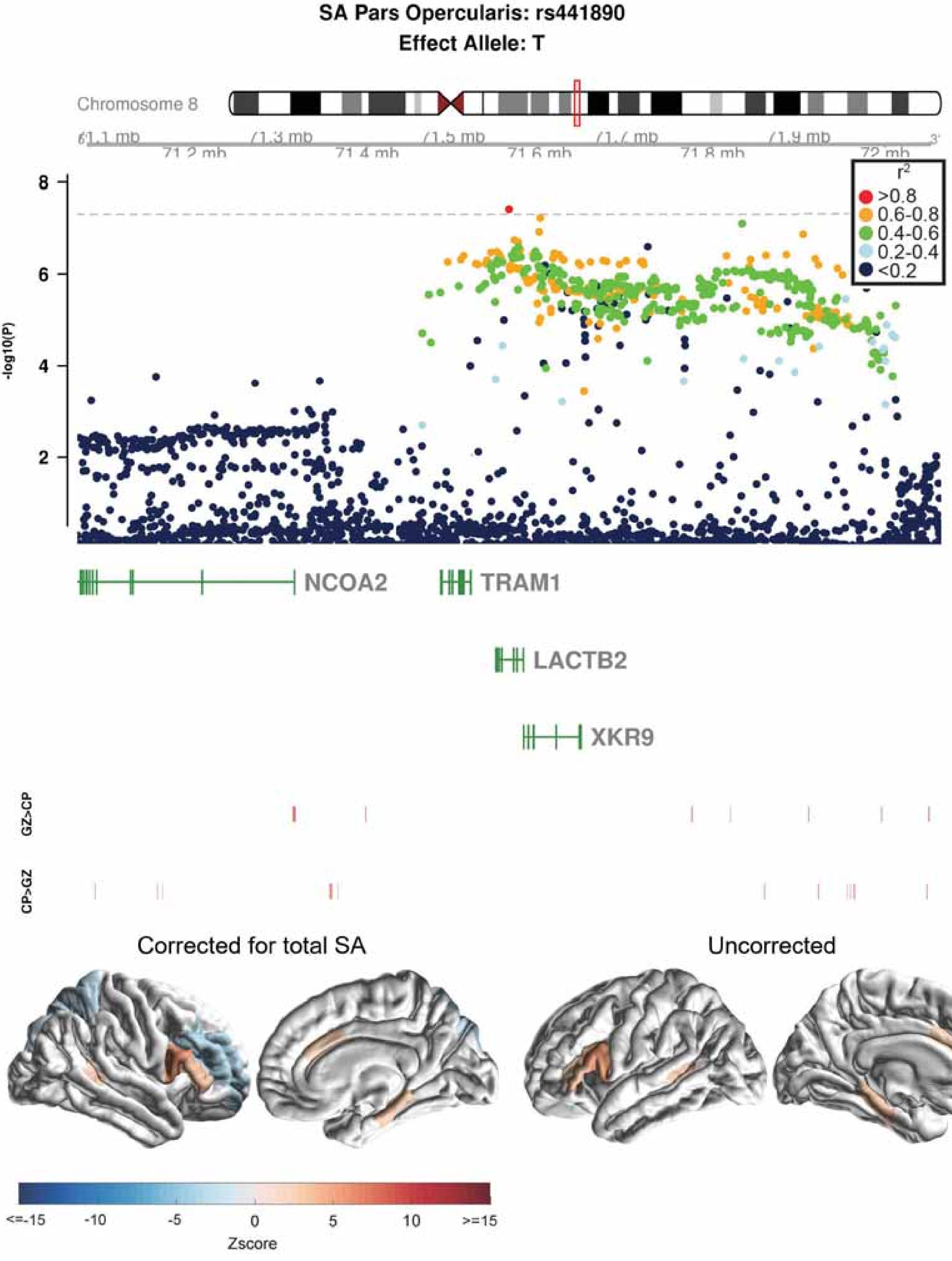

SA Pars Opercularis: rs441890
Effect Allele: T
Chromosome 8
71.2 mb
71.4 mb
71.6 mb
71.8 mb
72 mb
-log10(P)
8
6
4
2
r²
>0.8
0.6-0.8
0.4-0.6
0.2-0.4
<0.2
NCOA2
TRAM1
LACTB2
XKR9
GZ>CP
CP>GZ
Corrected for total SA
Uncorrected
<=-15
-10
-5
0
5
10
>=15
Zscore

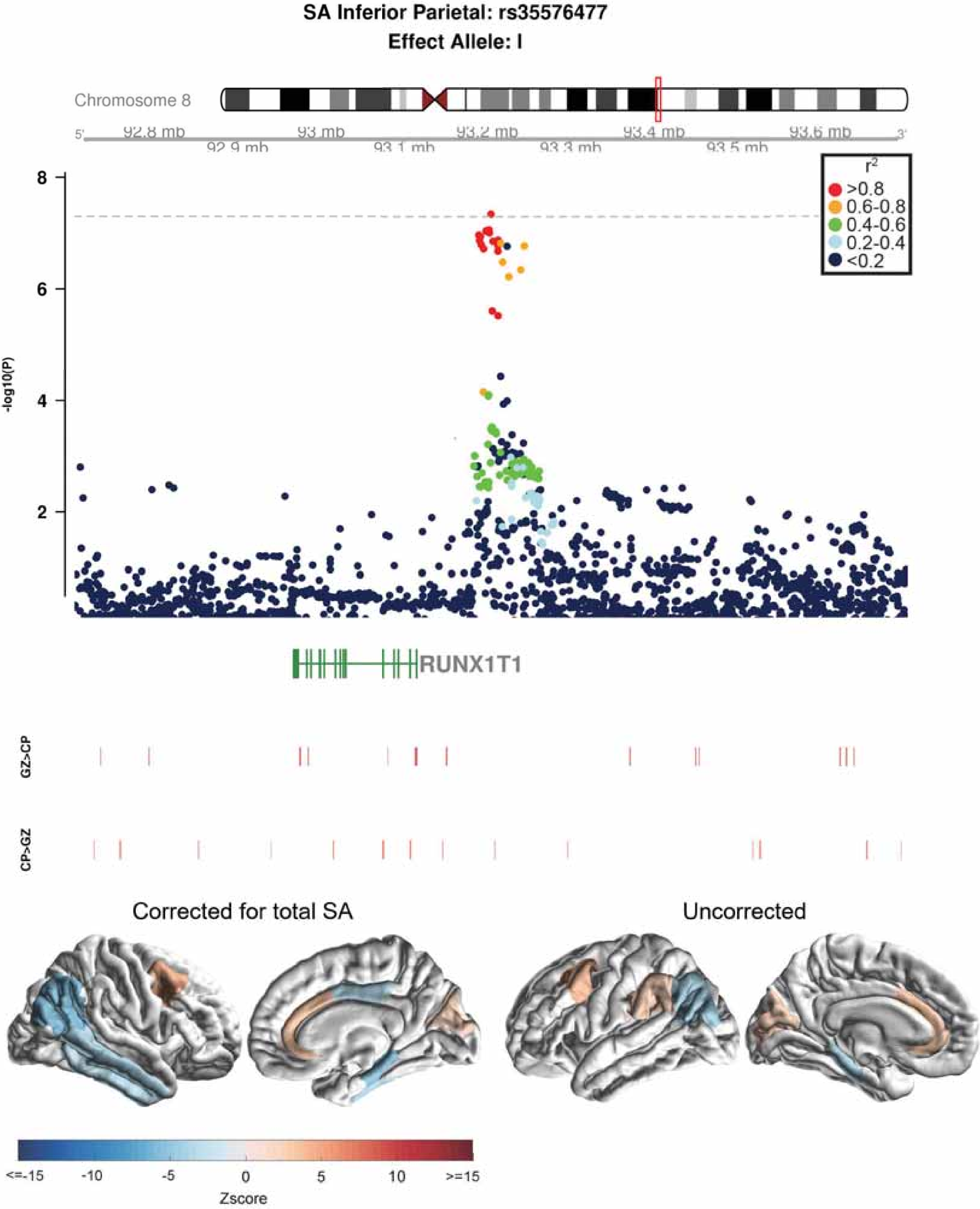

SA Inferior Parietal: rs35576477
Effect Allele: I
Chromosome 8
5'
92.8 mb
92.9 mb
93 mb
93.1 mb
93.2 mb
93.3 mb
93.4 mb
93.5 mb
93.6 mb
3'
r2
>0.8
0.6-0.8
0.4-0.6
0.2-0.4
<0.2
-log10(P)
8
6
4
2
RUNX1T1
GZ>CP
CP>GZ
Corrected for total SA
Uncorrected
<=-15
-10
-5
0
5
10
>=15
Zscore

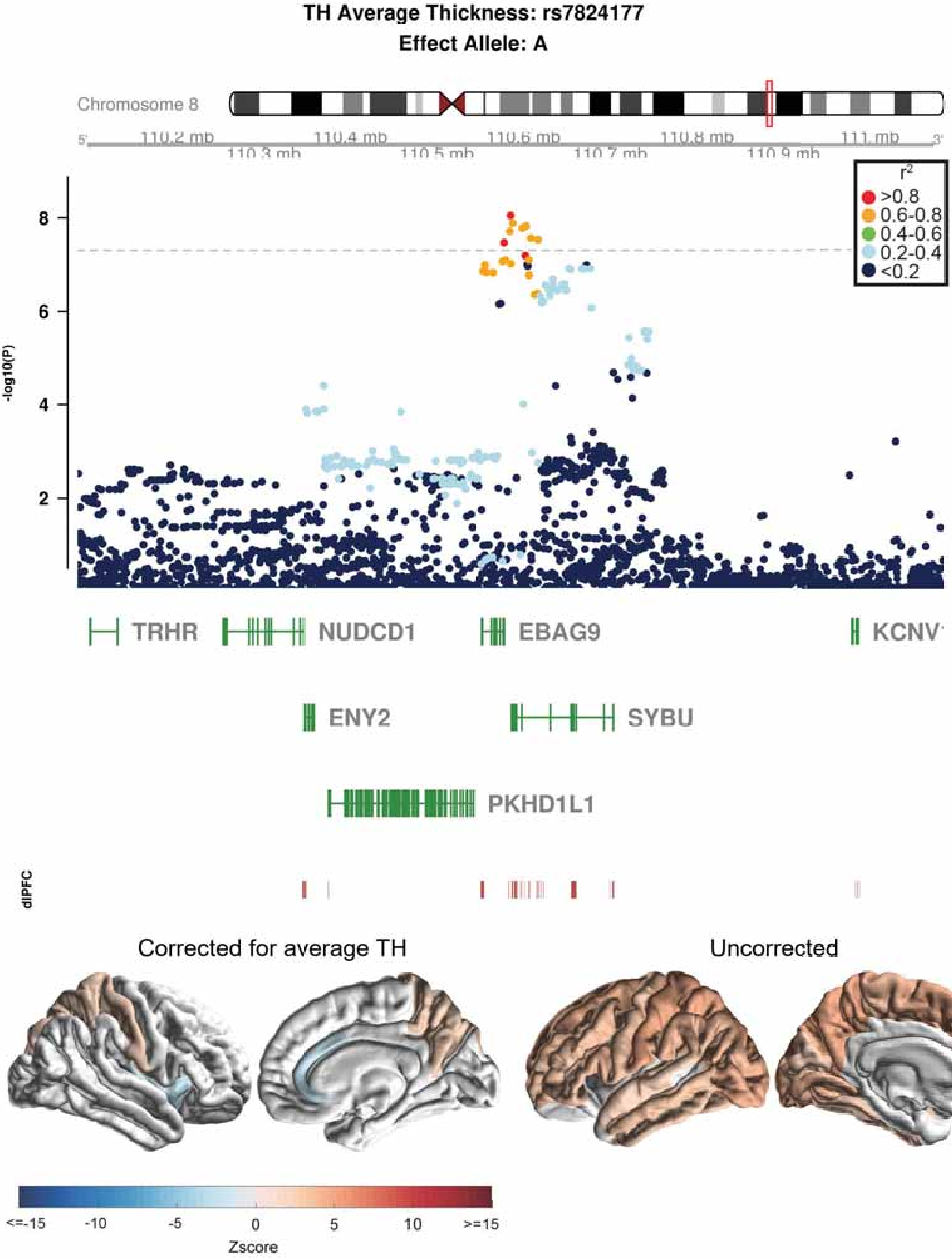
TH Average Thickness: rs7824177
Effect Allele: A
Chromosome 8
110.2 mb
110.3 mb
110.4 mb
110.5 mb
110.6 mb
110.7 mb
110.8 mb
110.9 mb
111 mb
-log10(P)
r2
>0.8
0.6-0.8
0.4-0.6
0.2-0.4
<0.2
TRHR
NUDCD1
EBAG9
KCNV
ENY2
SYBU
PKHD1L1
dlPFC
Corrected for average TH
Uncorrected
<=-15
-10
-5
0
5
10
>=15
Zscore

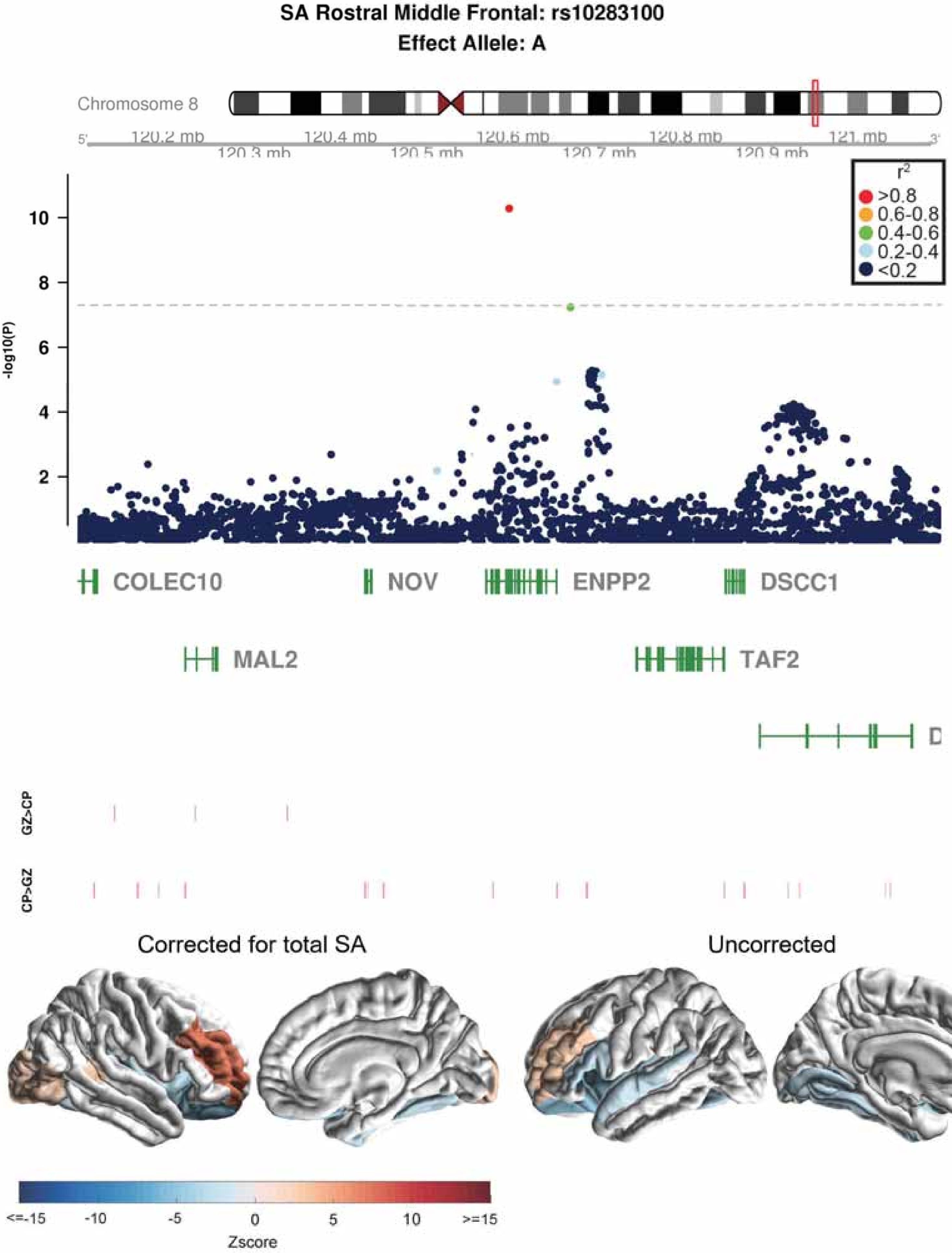
SA Rostral Middle Frontal: rs10283100
Effect Allele: A
Chromosome 8
120.2 mb
120.3 mb
120.4 mb
120.5 mb
120.6 mb
120.7 mb
120.8 mb
120.9 mb
121 mb
r2
>0.8
0.6-0.8
0.4-0.6
0.2-0.4
<0.2
-log10(P)
COLEC10
NOV
ENPP2
DSCC1
MAL2
TAF2
GZ>CP
CP>GZ
Corrected for total SA
Uncorrected
<=-15
-10
-5
0
5
10
>=15
Zscore

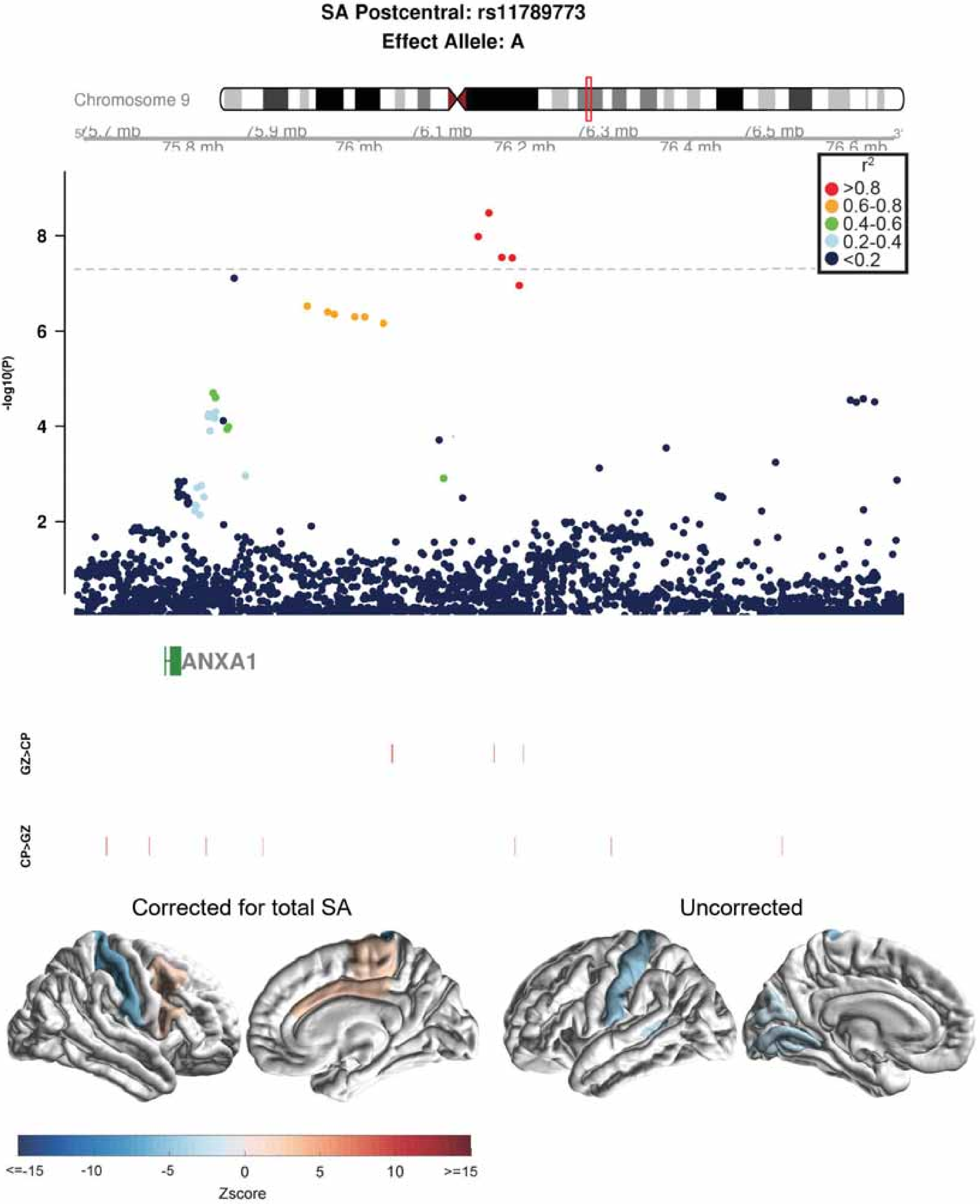
SA Postcentral: rs11789773
Effect Allele: A
Chromosome 9
75.7 mb
75.9 mb
76.1 mb
76.3 mb
76.5 mb
75.8 mb
76 mb
76.2 mb
76.4 mb
76.6 mb
r2
>0.8
0.6-0.8
0.4-0.6
0.2-0.4
<0.2
-log10(P)
8
6
4
2
ANXA1
GZ>CP
CP>GZ
Corrected for total SA
Uncorrected
<=-15
-10
-5
0
5
10
>=15
Zscore

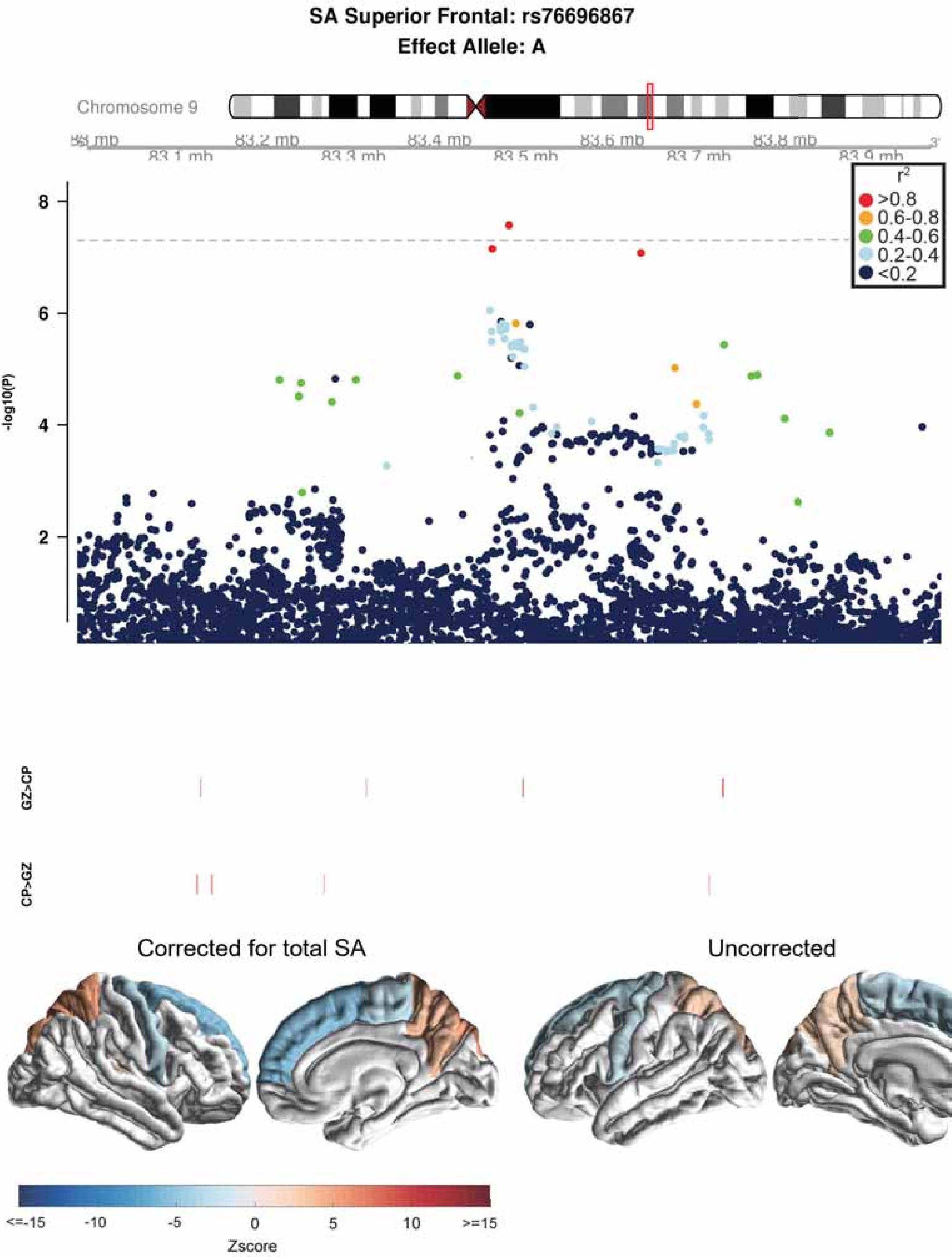
SA Superior Frontal: rs76696867
Effect Allele: A
Chromosome 9
83.1 mb
83.2 mb
83.3 mb
83.4 mb
83.5 mb
83.6 mb
83.7 mb
83.8 mb
83.9 mb
-log10(P)
8
6
4
2
r2
>0.8
0.6-0.8
0.4-0.6
0.2-0.4
<0.2
GZ>CP
CP>GZ
Corrected for total SA
Uncorrected
<=-15
-10
-5
0
5
10
>=15
Zscore

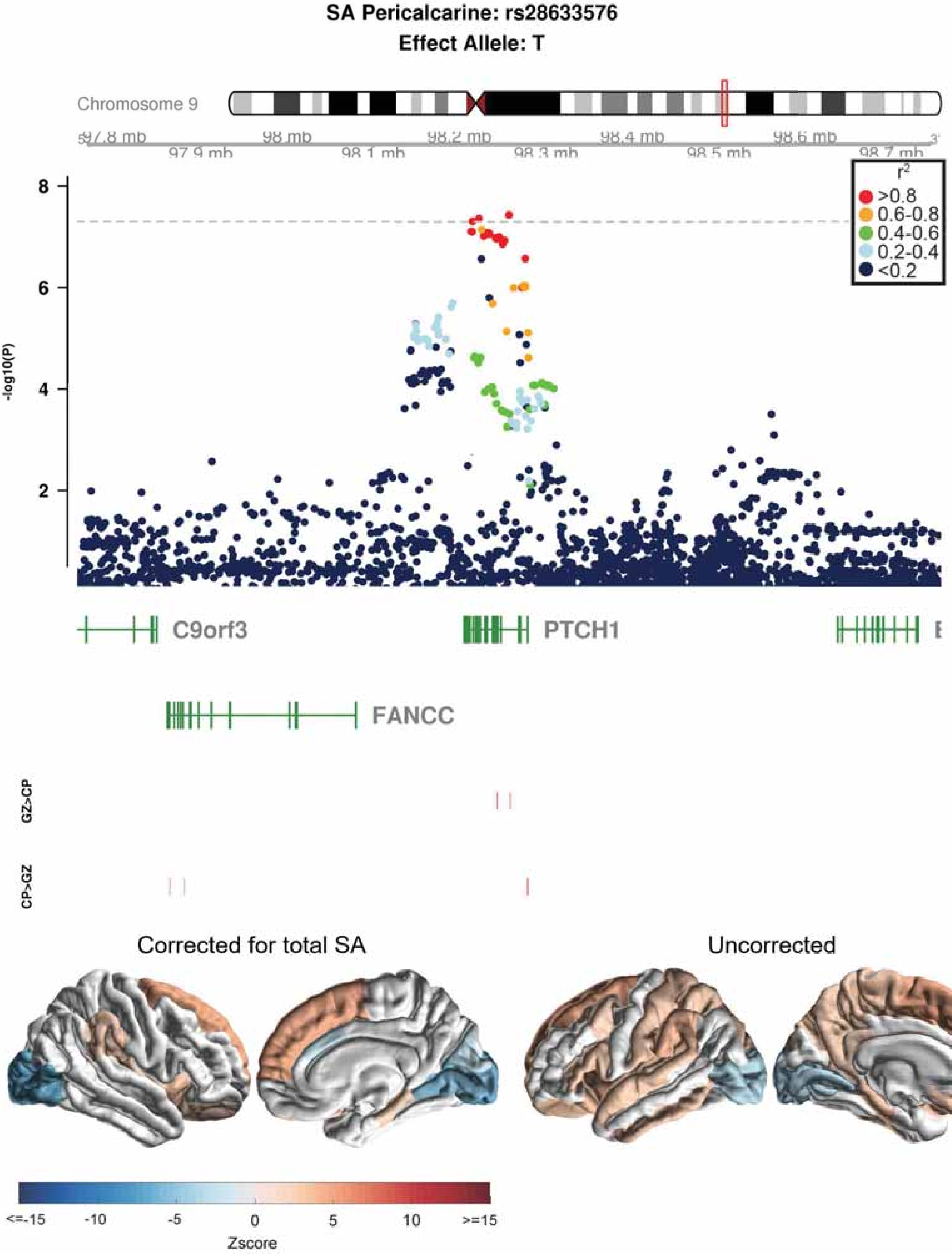

SA Pericalcarine: rs28633576
Effect Allele: T
Chromosome 9
97.8 mb
98 mb
98.2 mb
98.4 mb
98.6 mb
97.9 mb
98.1 mb
98.3 mb
98.5 mb
98.7 mb
r2
>0.8
0.6-0.8
0.4-0.6
0.2-0.4
<0.2
-log10(P)
8
6
4
2
C9orf3
PTCH1
FANCC
GZ>CP
CP>GZ
Corrected for total SA
Uncorrected
<=-15
-10
-5
0
5
10
>=15
Zscore

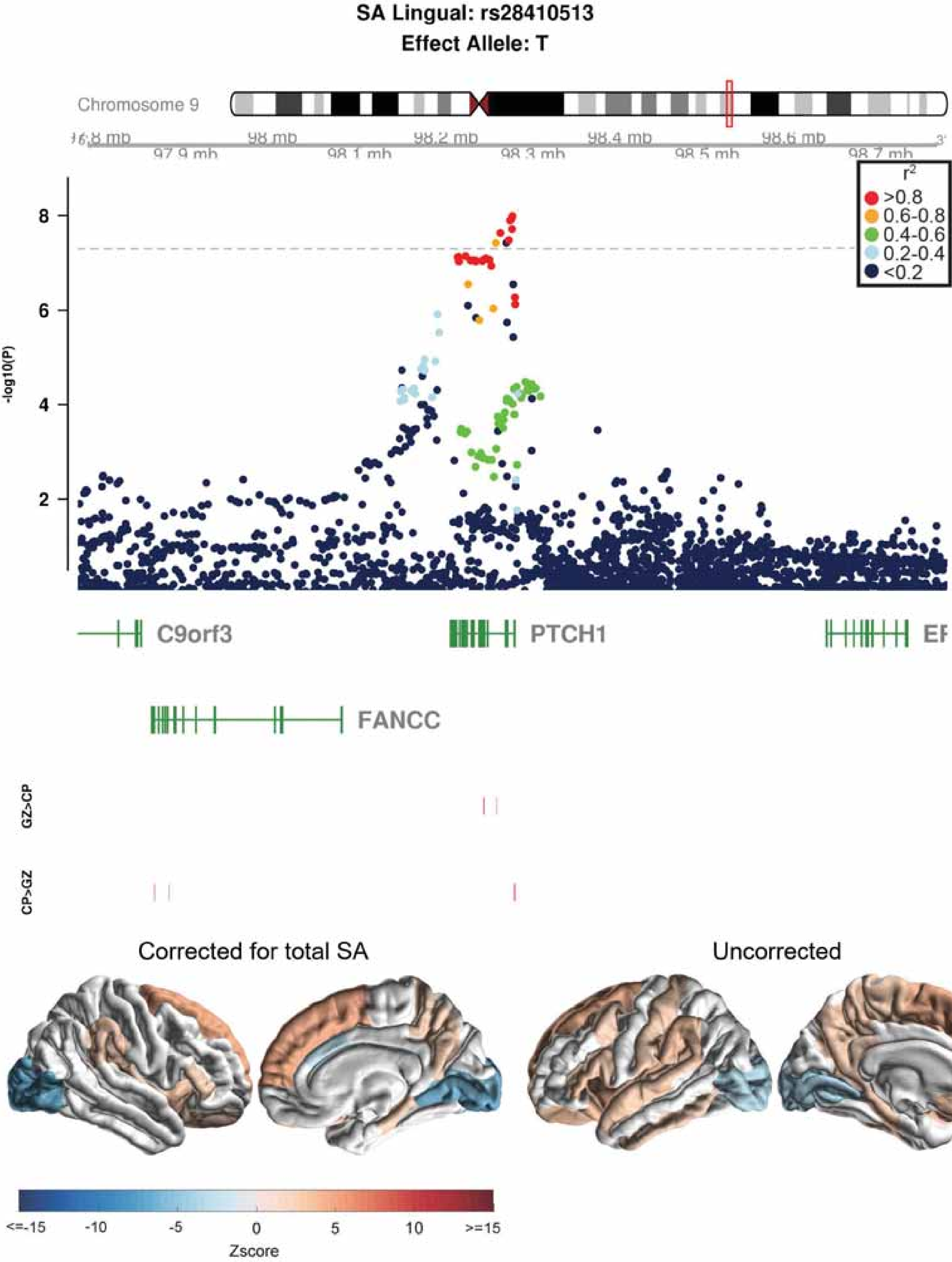

SA Lingual: rs28410513
Effect Allele: T
Chromosome 9
r2
>0.8
0.6-0.8
0.4-0.6
0.2-0.4
<0.2
-log10(P)
C9orf3
PTCH1
FANCC
GZ>CP
CP>GZ
Corrected for total SA
Uncorrected
Zscore

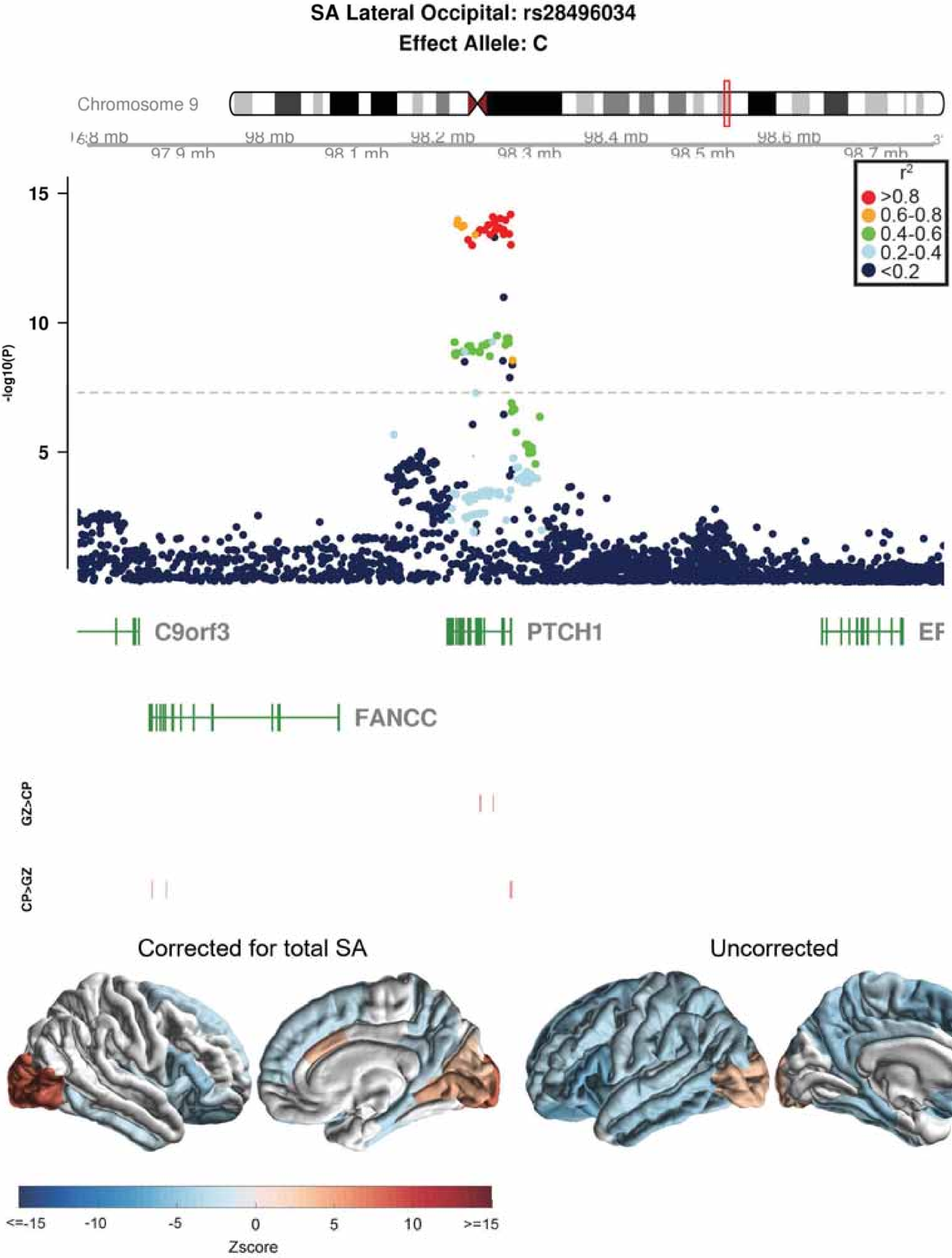
SA Lateral Occipital: rs28496034
Effect Allele: C
Chromosome 9
98 mb
98.2 mb
98.4 mb
98.6 mb
97.9 mb
98.1 mb
98.3 mb
98.5 mb
98.7 mb
r2
>0.8
0.6-0.8
0.4-0.6
0.2-0.4
<0.2
15
10
5
-log10(P)
C9orf3
PTCH1
FANCC
GZ>CP
CP>GZ
Corrected for total SA
Uncorrected
<=-15
-10
-5
0
5
10
>=15
Zscore

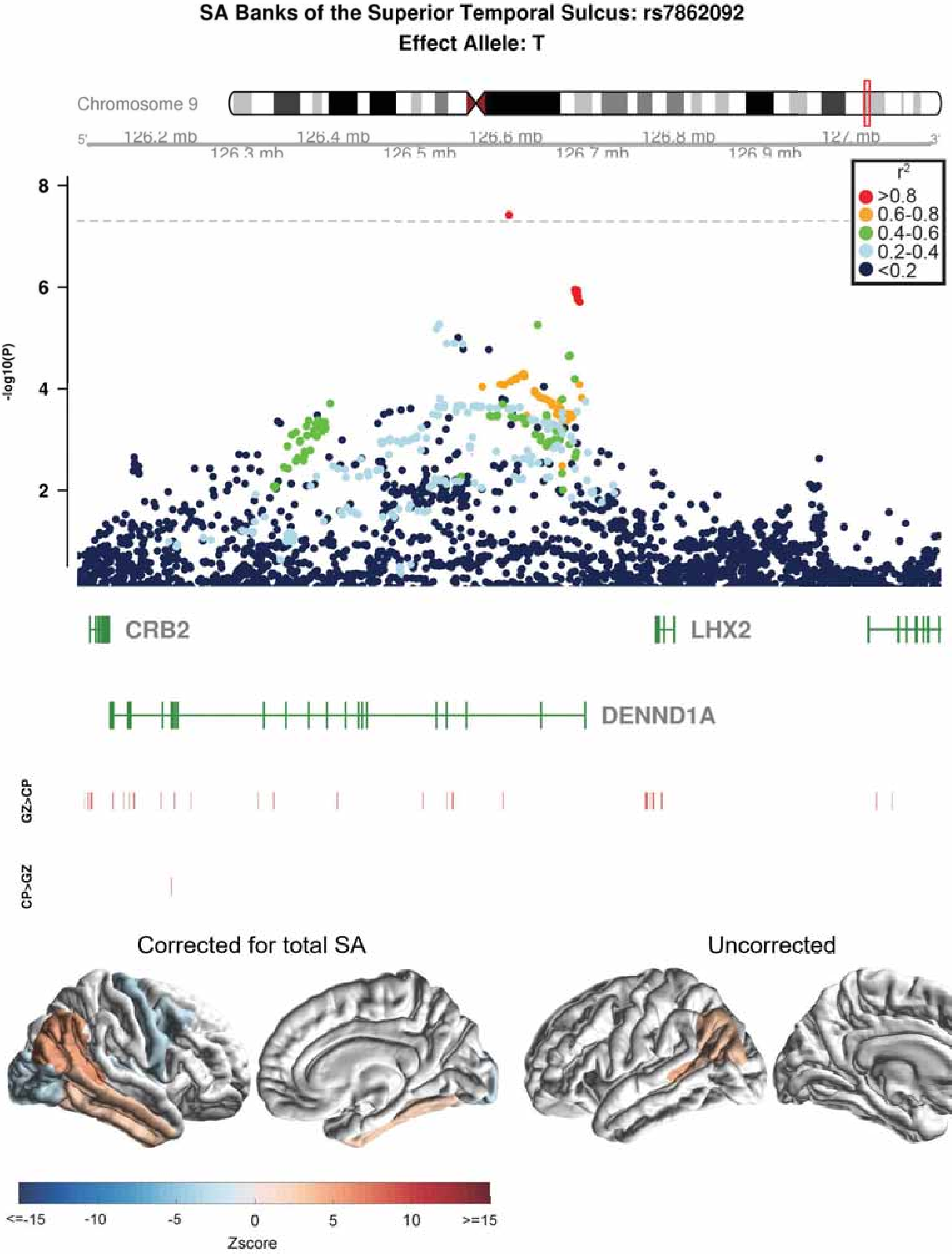
SA Banks of the Superior Temporal Sulcus: rs7862092
Effect Allele: T
Chromosome 9
126.2 mb
126.3 mb
126.4 mb
126.5 mb
126.6 mb
126.7 mb
126.8 mb
126.9 mb
127 mb
r2
>0.8
0.6-0.8
0.4-0.6
0.2-0.4
<0.2
-log10(P)
CRB2
LHX2
DENND1A
GZ>CP
CP>GZ
Corrected for total SA
Uncorrected
<=-15
-10
-5
0
5
10
>=15
Zscore

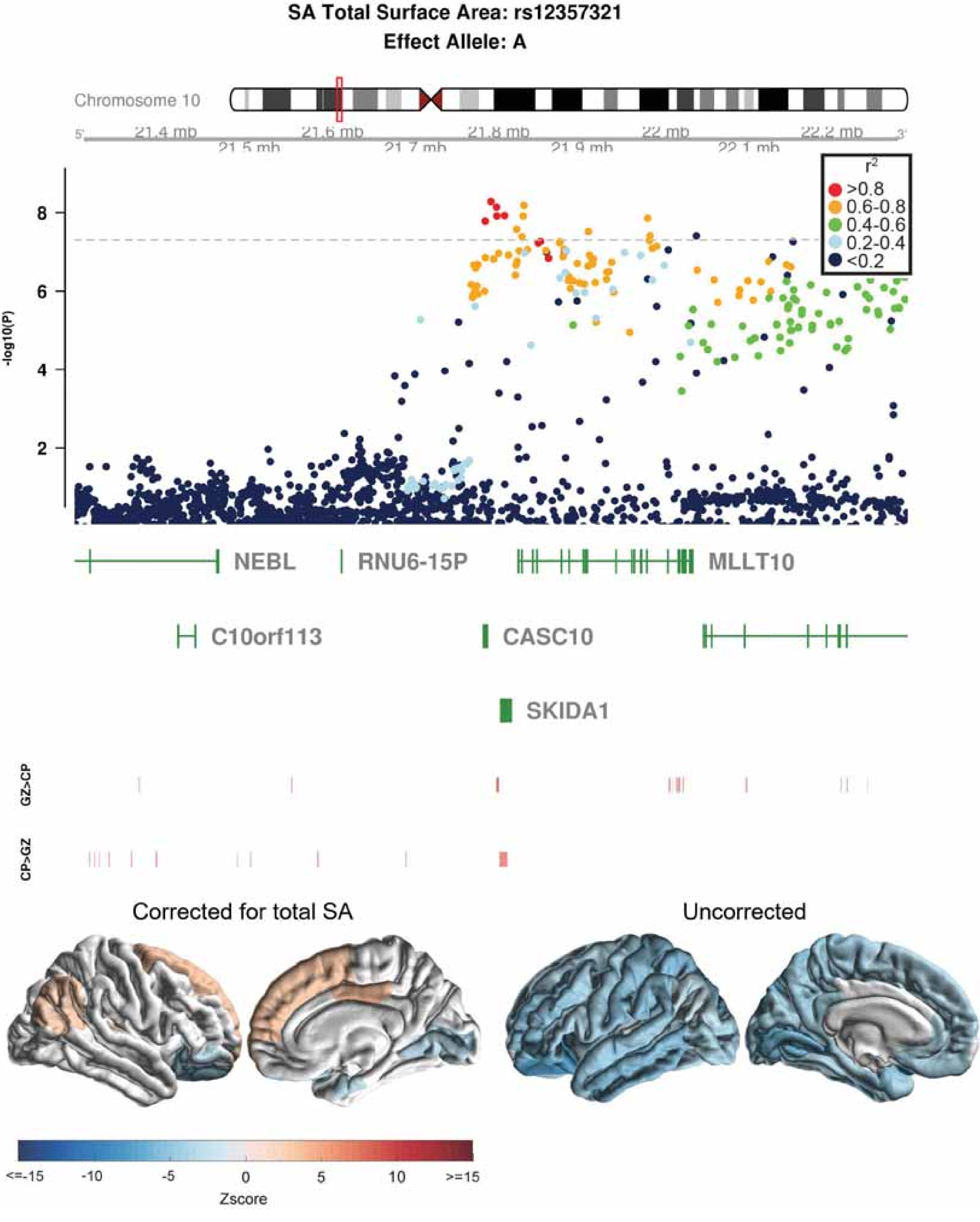

SA Total Surface Area: rs12357321
Effect Allele: A
Chromosome 10
21.4 mb
21.5 mb
21.6 mb
21.7 mb
21.8 mb
21.9 mb
22 mb
22.1 mb
22.2 mb
r2
>0.8
0.6-0.8
0.4-0.6
0.2-0.4
<0.2
-log10(P)
NEBL
RNU6-15P
MLLT10
C10orf113
CASC10
SKIDA1
GZ>CP
CP>GZ
Corrected for total SA
Uncorrected
<=-15
-10
-5
0
5
10
>=15
Zscore

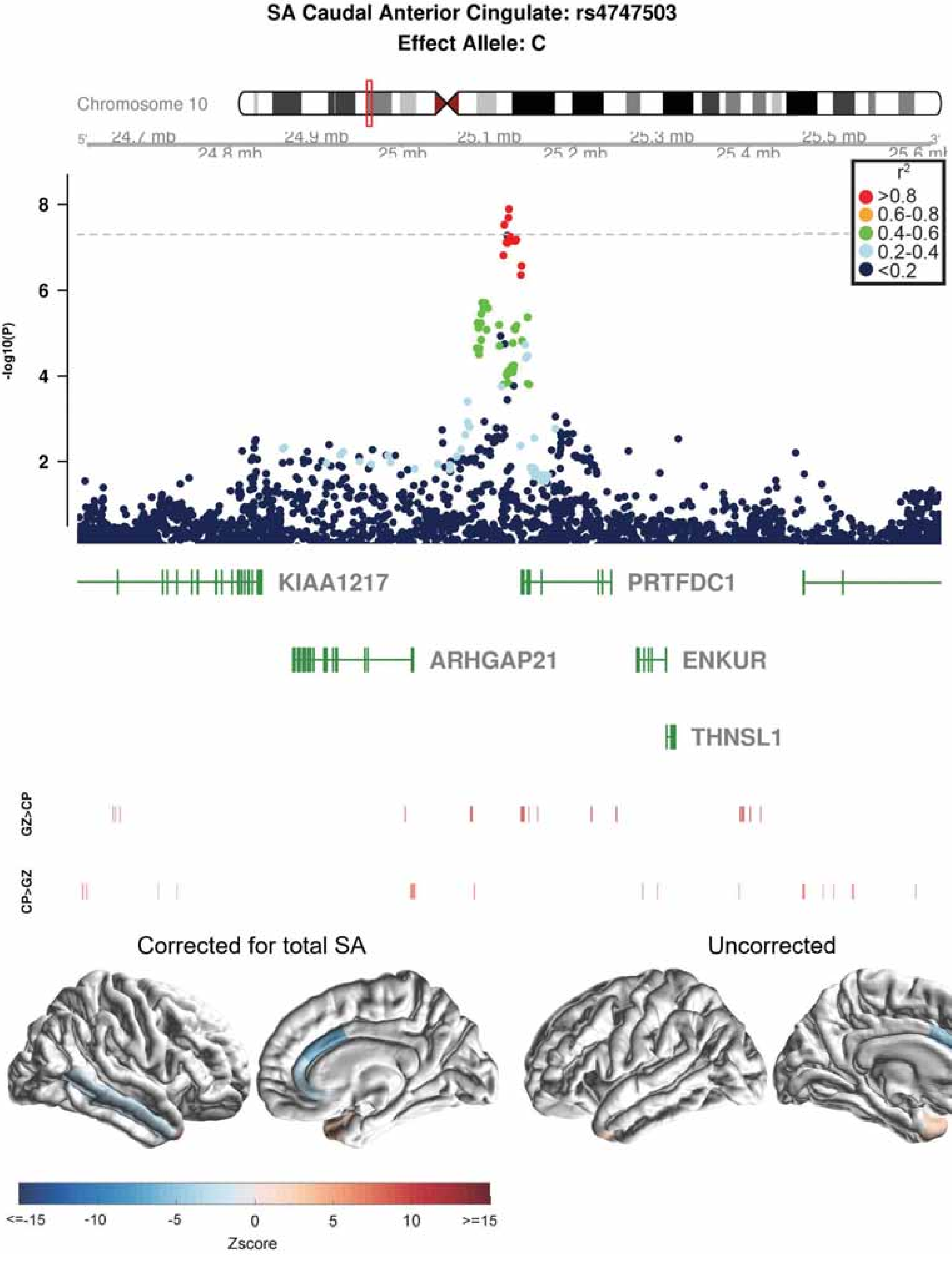
SA Caudal Anterior Cingulate: rs4747503
Effect Allele: C
Chromosome 10
24.7 mb
24.8 mb
24.9 mb
25 mb
25.1 mb
25.2 mb
25.3 mb
25.4 mb
25.5 mb
25.6 mb
r2
>0.8
0.6-0.8
0.4-0.6
0.2-0.4
<0.2
-log10(P)
8
6
4
2
KIAA1217
PRTFDC1
ARHGAP21
ENKUR
THNSL1
GZ>CP
CP>GZ
Corrected for total SA
Uncorrected
<=-15
-10
-5
0
5
10
>=15
Zscore

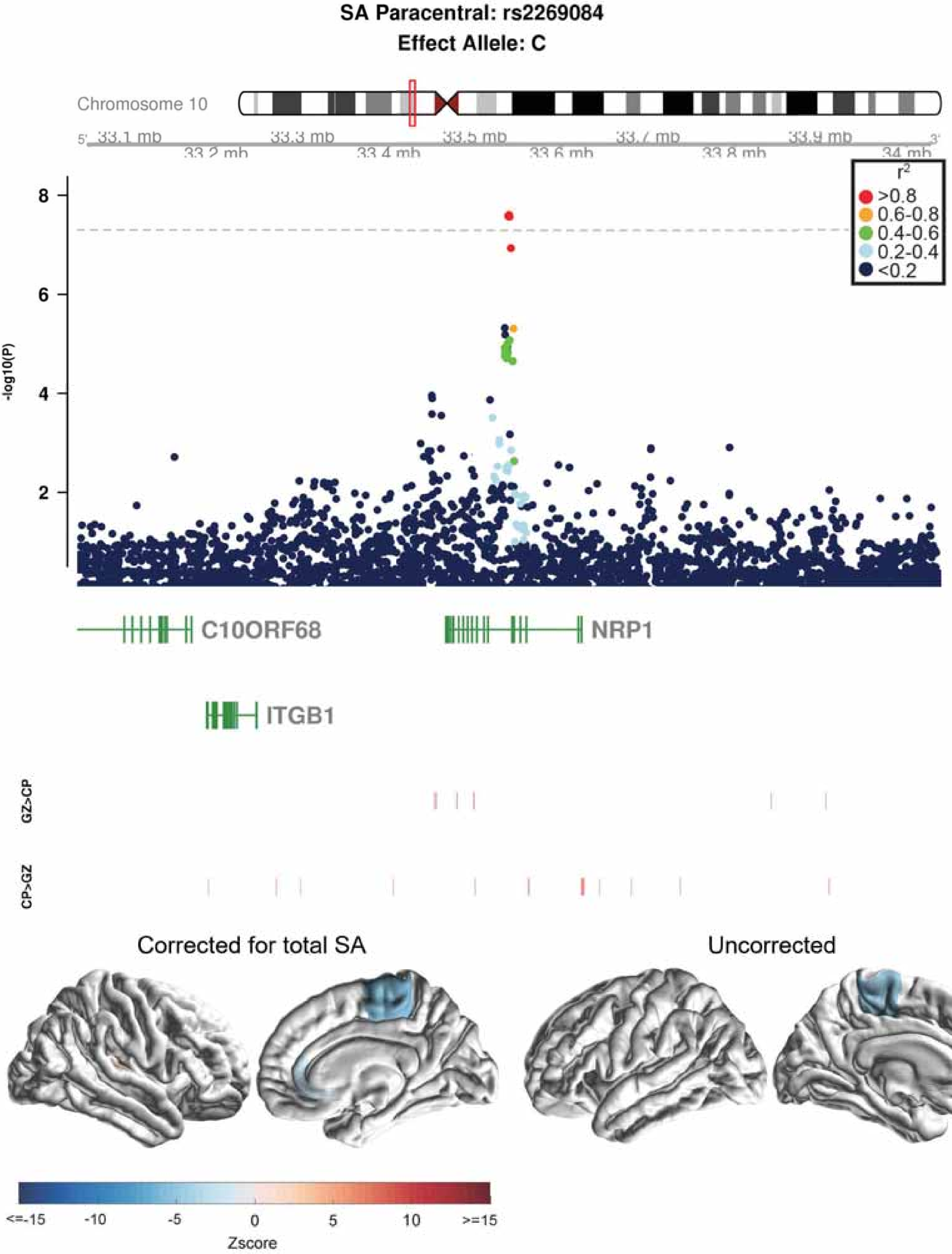

SA Paracentral: rs2269084
Effect Allele: C
Chromosome 10
5'
33.1 mb
33.2 mb
33.3 mb
33.4 mb
33.5 mb
33.6 mb
33.7 mb
33.8 mb
33.9 mb
34 mb
3'
r²
>0.8
0.6-0.8
0.4-0.6
0.2-0.4
<0.2
-log10(P)
8
6
4
2
C10ORF68
NRP1
ITGB1
GZ>CP
CP>GZ
Corrected for total SA
Uncorrected
<=-15
-10
-5
0
5
10
>=15
Zscore

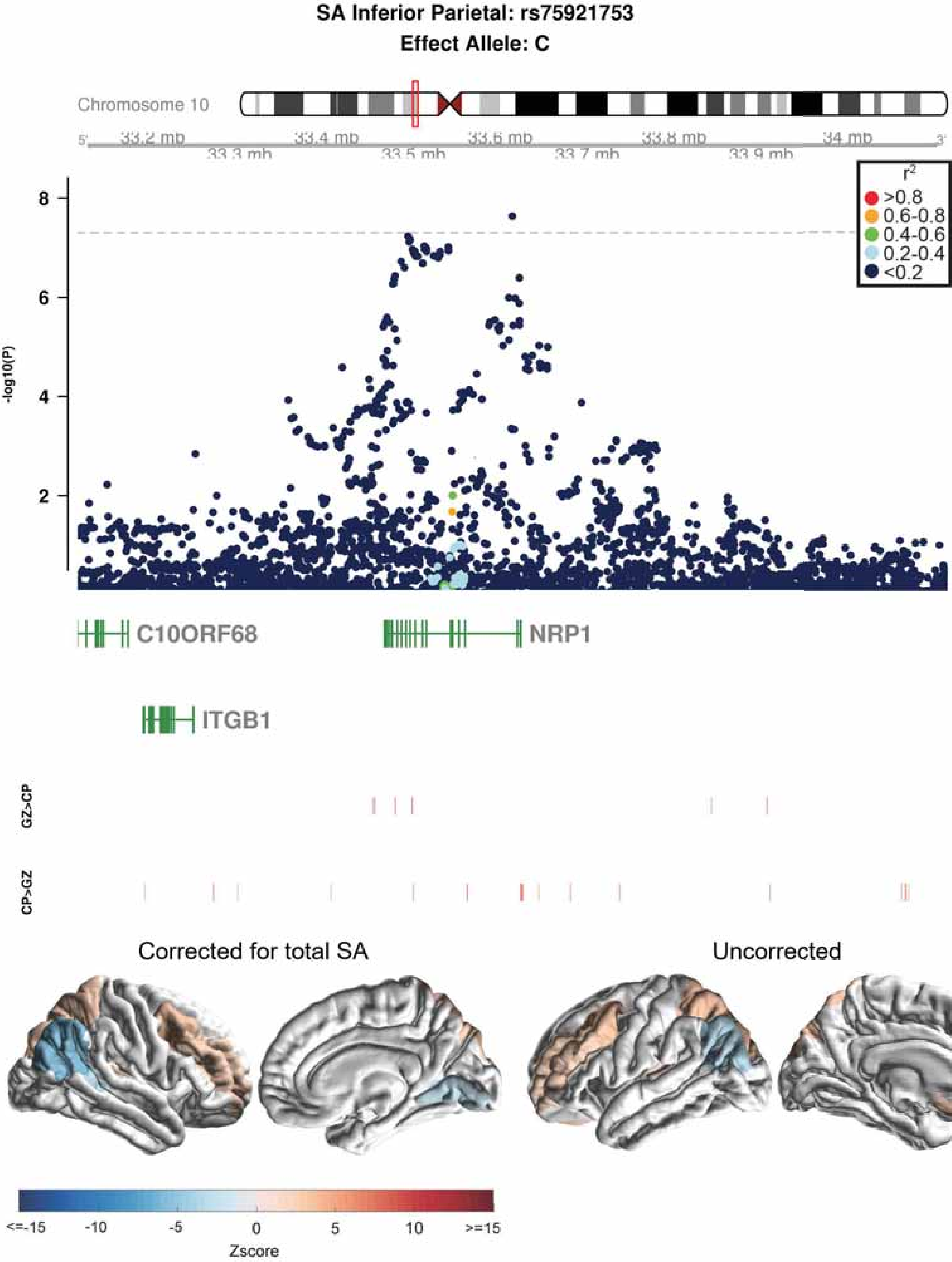
SA Inferior Parietal: rs75921753
Effect Allele: C
Chromosome 10
33.2 mb
33.3 mb
33.4 mb
33.5 mb
33.6 mb
33.7 mb
33.8 mb
33.9 mb
34 mb
-log10(P)
r2
>0.8
0.6-0.8
0.4-0.6
0.2-0.4
<0.2
C10ORF68
NRP1
ITGB1
GZ>CP
CP>GZ
Corrected for total SA
Uncorrected
<=-15
-10
-5
0
5
10
>=15
Zscore

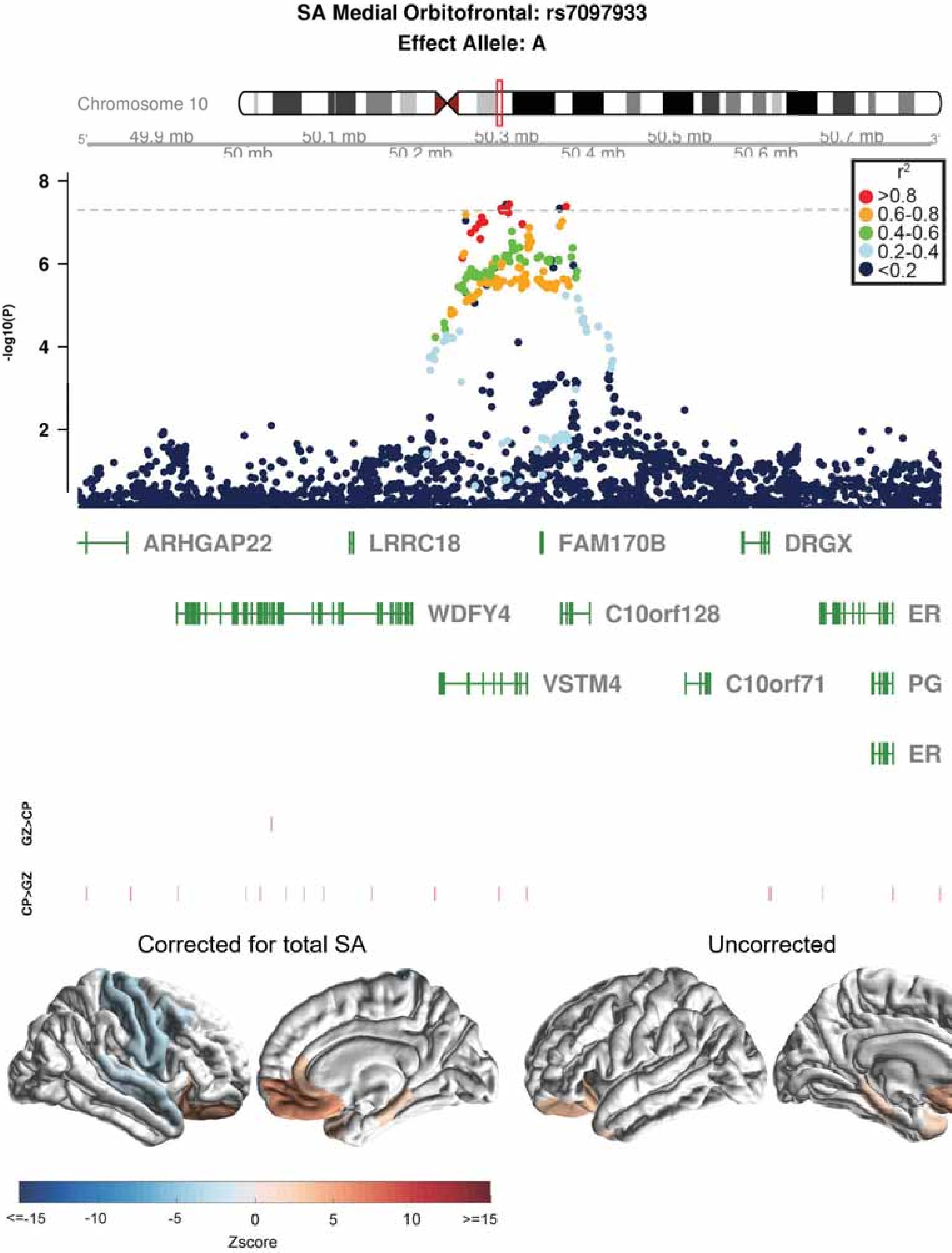

SA Medial Orbitofrontal: rs7097933
Effect Allele: A
Chromosome 10
49.9 mb
50 mb
50.1 mb
50.2 mb
50.3 mb
50.4 mb
50.5 mb
50.6 mb
50.7 mb
-log10(P)
r2
>0.8
0.6-0.8
0.4-0.6
0.2-0.4
<0.2
ARHGAP22
LRRC18
FAM170B
DRGX
WDFY4
C10orf128
ER
VSTM4
C10orf71
PG
ER
GZ>CP
CP>GZ
Corrected for total SA
Uncorrected
<=-15
-10
-5
0
5
10
>=15
Zscore

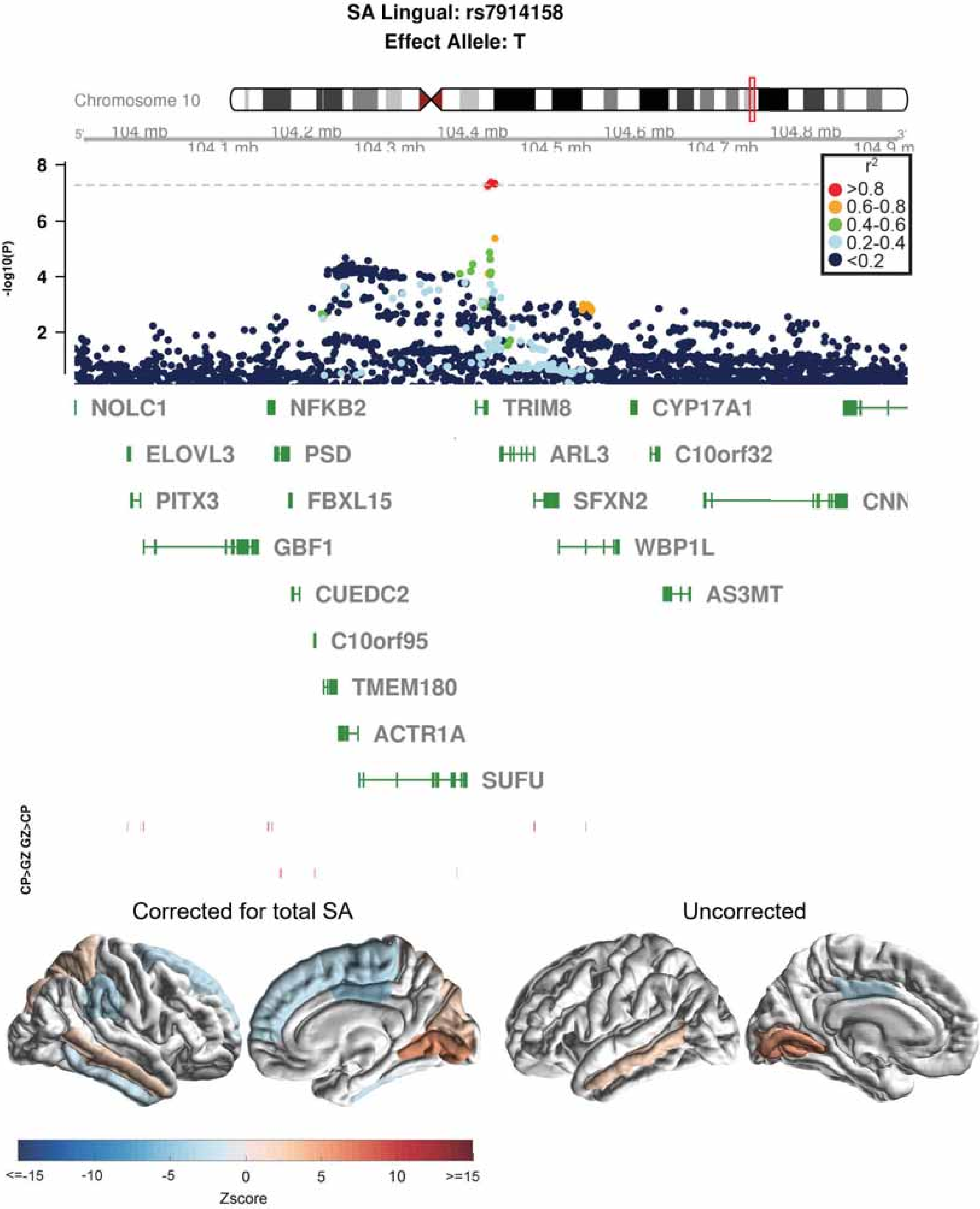
SA Lingual: rs7914158
Effect Allele: T
Chromosome 10
104 mb
104.1 mb
104.2 mb
104.3 mb
104.4 mb
104.5 mb
104.6 mb
104.7 mb
104.8 mb
-log10(P)
r2
>0.8
0.6-0.8
0.4-0.6
0.2-0.4
<0.2
NOLC1
NFKB2
TRIM8
CYP17A1
ELOVL3
PSD
ARL3
C10orf32
PITX3
FBXL15
SFXN2
GBF1
WBP1L
CUEDC2
AS3MT
C10orf95
TMEM180
ACTR1A
SUFU
CP>GZ GZ>CP
Corrected for total SA
Uncorrected
<=-15
-10
-5
0
5
10
>=15
Zscore

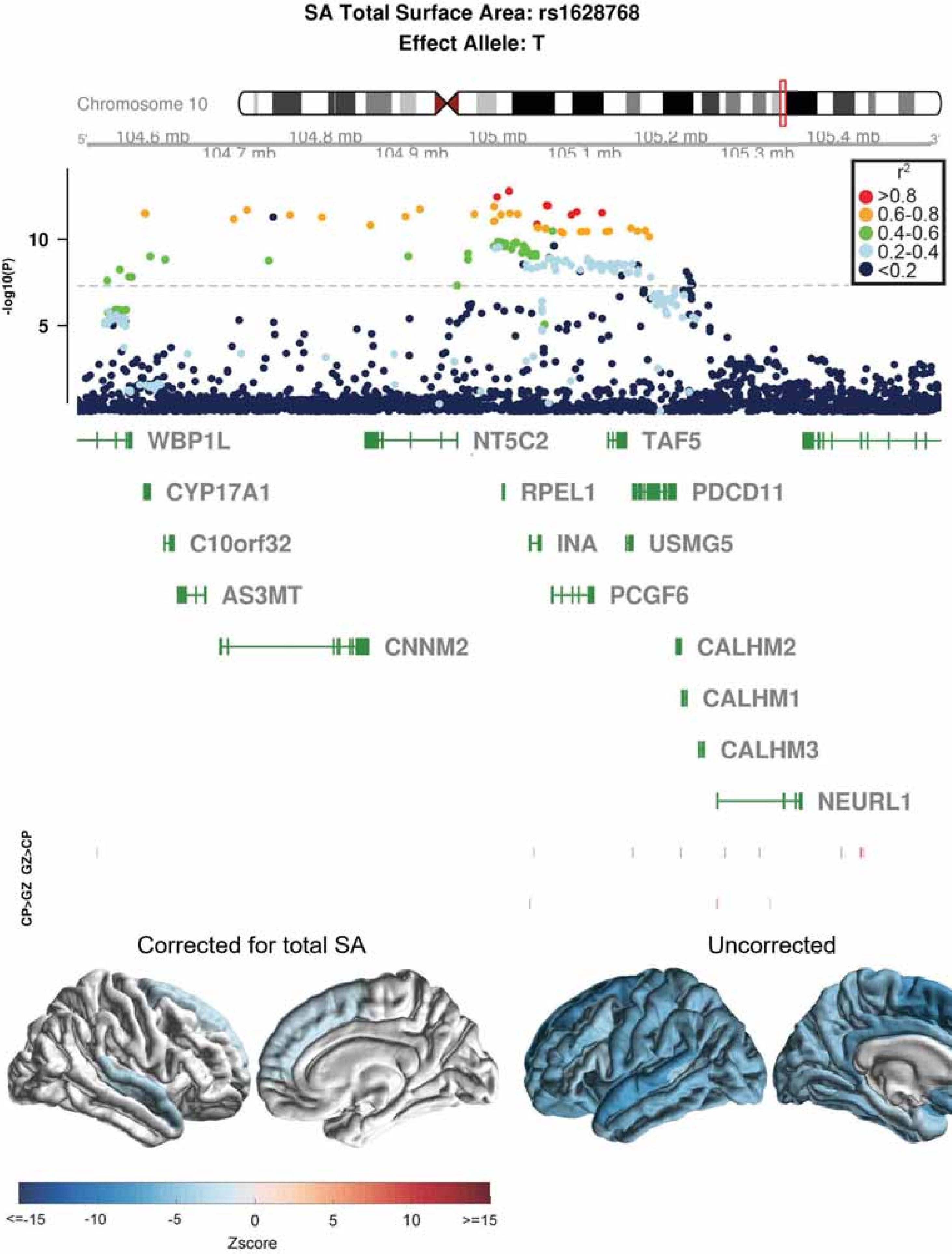
SA Total Surface Area: rs1628768
Effect Allele: T
Chromosome 10
104.6 mb
104.7 mb
104.8 mb
104.9 mb
105 mb
105.1 mb
105.2 mb
105.3 mb
105.4 mb
-log10(P)
10
5
r2
>0.8
0.6-0.8
0.4-0.6
0.2-0.4
<0.2
WBP1L
NT5C2
TAF5
CYP17A1
RPEL1
PDCD11
C10orf32
INA
USMG5
AS3MT
PCGF6
CNNM2
CALHM2
CALHM1
CALHM3
NEURL1
GZ>CP
CP>GZ
Corrected for total SA
Uncorrected
<=-15
-10
-5
0
5
10
>=15
Zscore

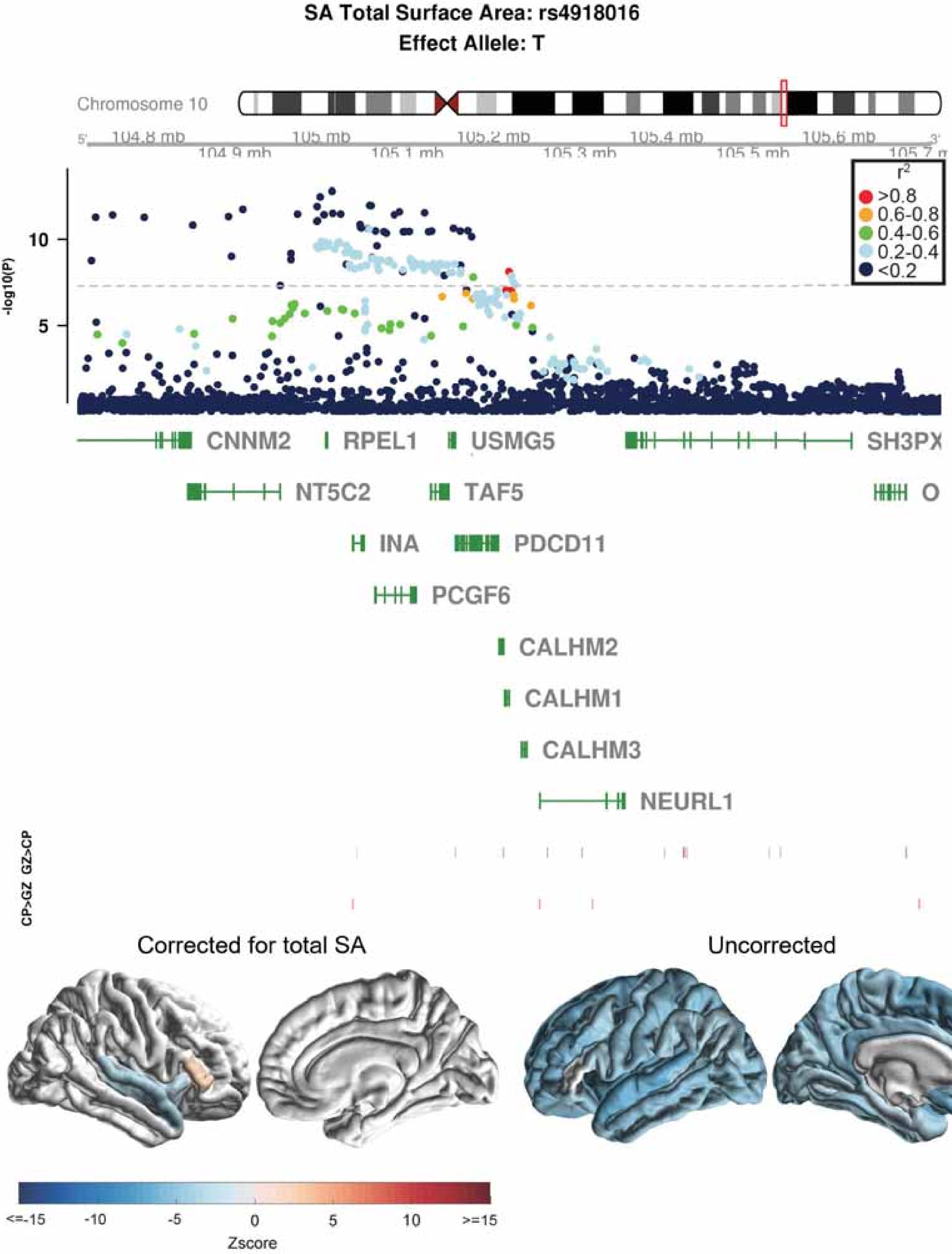

SA Total Surface Area: rs4918016
Effect Allele: T
Chromosome 10
104.8 mb
104.9 mb
105 mb
105.1 mb
105.2 mb
105.3 mb
105.4 mb
105.5 mb
105.6 mb
-log10(P)
10
5
r²
>0.8
0.6-0.8
0.4-0.6
0.2-0.4
<0.2
CNNM2
RPEL1
USMG5
SH3PX
NT5C2
TAF5
INA
PDCD11
PCGF6
CALHM2
CALHM1
CALHM3
NEURL1
GZ>CP
CP>GZ
Corrected for total SA
Uncorrected
<=-15
-10
-5
0
5
10
>=15
Zscore

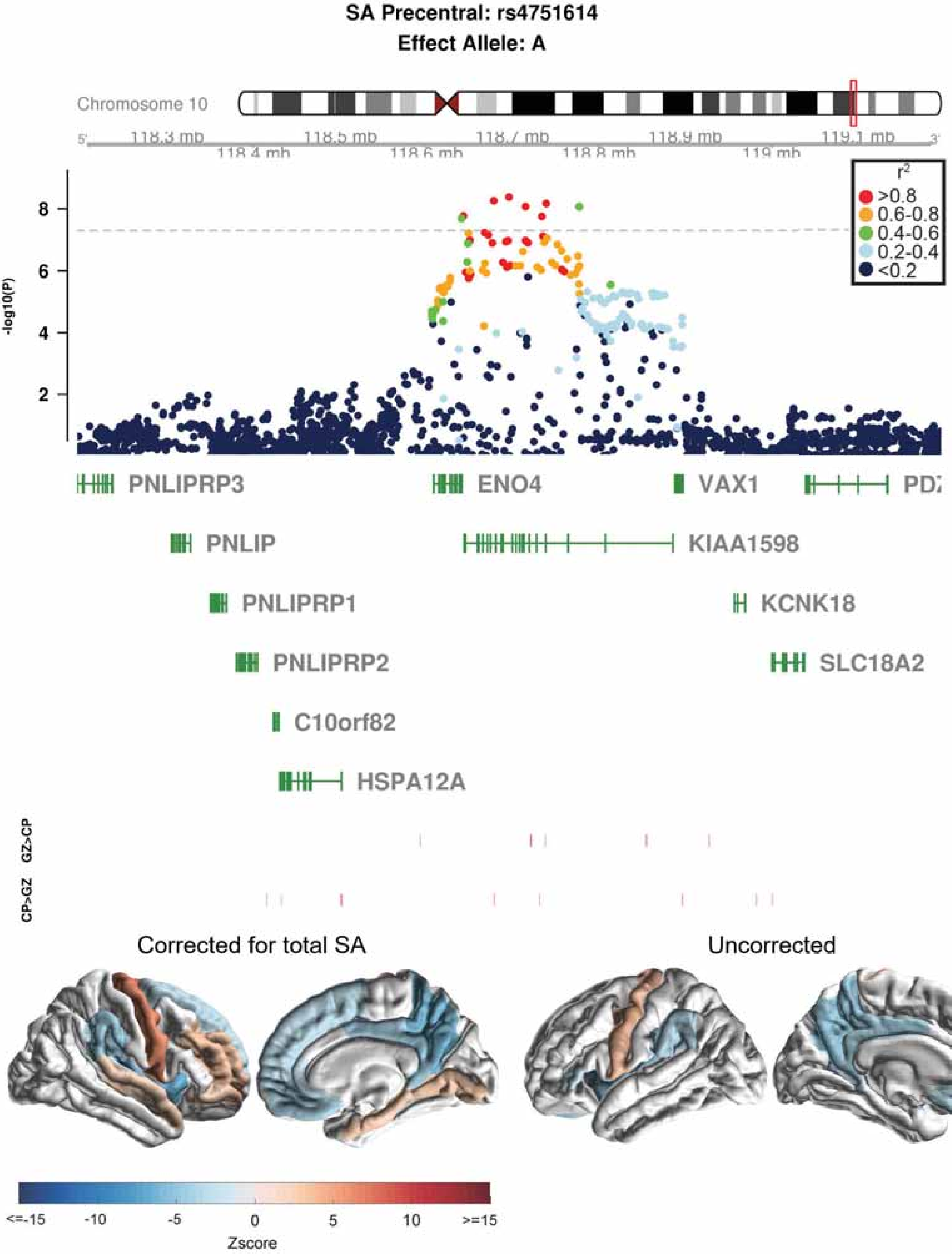
SA Precentral: rs4751614
Effect Allele: A
Chromosome 10
118.3 mb
118.4 mb
118.5 mb
118.6 mb
118.7 mb
118.8 mb
118.9 mb
119 mb
119.1 mb
-log10(P)
r²
>0.8
0.6-0.8
0.4-0.6
0.2-0.4
<0.2
PNLIPRP3
ENO4
VAX1
PNLIP
KIAA1598
PNLIPRP1
KCNK18
PNLIPRP2
SLC18A2
C10orf82
HSPA12A
GZ>CP
CP>GZ
Corrected for total SA
Uncorrected
<=-15
-10
-5
0
5
10
>=15
Zscore

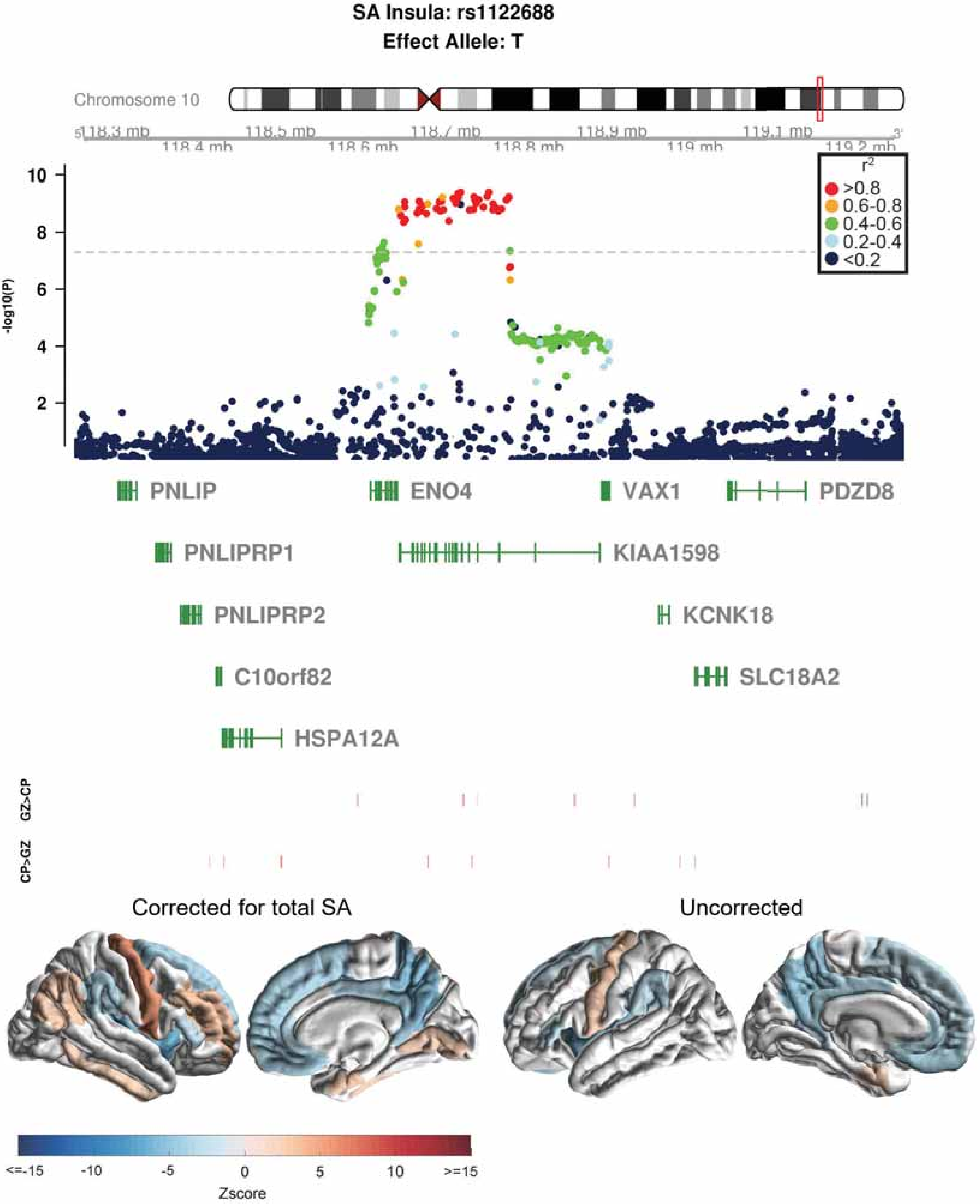
SA Insula: rs1122688
Effect Allele: T
Chromosome 10
118.3 mb
118.4 mb
118.5 mb
118.6 mb
118.7 mb
118.8 mb
118.9 mb
119 mb
119.1 mb
119.2 mb
-log10(P)
r2
>0.8
0.6-0.8
0.4-0.6
0.2-0.4
<0.2
PNLIP
ENO4
VAX1
PDZD8
PNLIPRP1
KIAA1598
PNLIPRP2
KCNK18
C10orf82
SLC18A2
HSPA12A
GZ>CP
CP>GZ
Corrected for total SA
Uncorrected
<=-15
-10
-5
0
5
10
>=15
Zscore

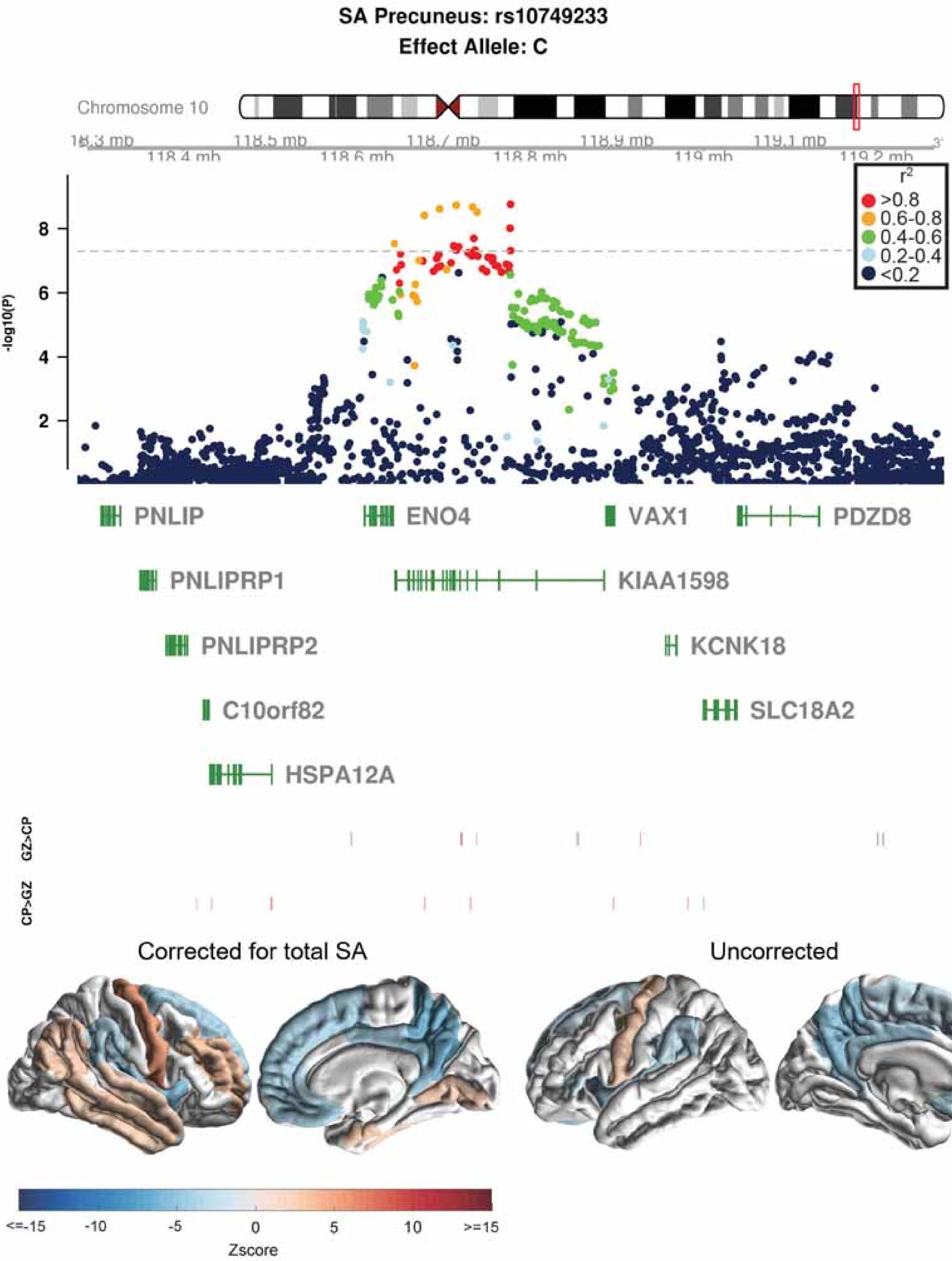

SA Precuneus: rs10749233
Effect Allele: C
Chromosome 10
118.5 mb
118.7 mb
118.9 mb
119.1 mb
118.4 mb
118.6 mb
118.8 mb
119 mb
119.2 mb
-log10(P)
8
6
4
2
r2
>0.8
0.6-0.8
0.4-0.6
0.2-0.4
<0.2
PNLIP
ENO4
VAX1
PDZD8
PNLIPRP1
KIAA1598
PNLIPRP2
KCNK18
C10orf82
SLC18A2
HSPA12A
GZ>CP
CP>GZ
Corrected for total SA
Uncorrected
<=-15
-10
-5
0
5
10
>=15
Zscore

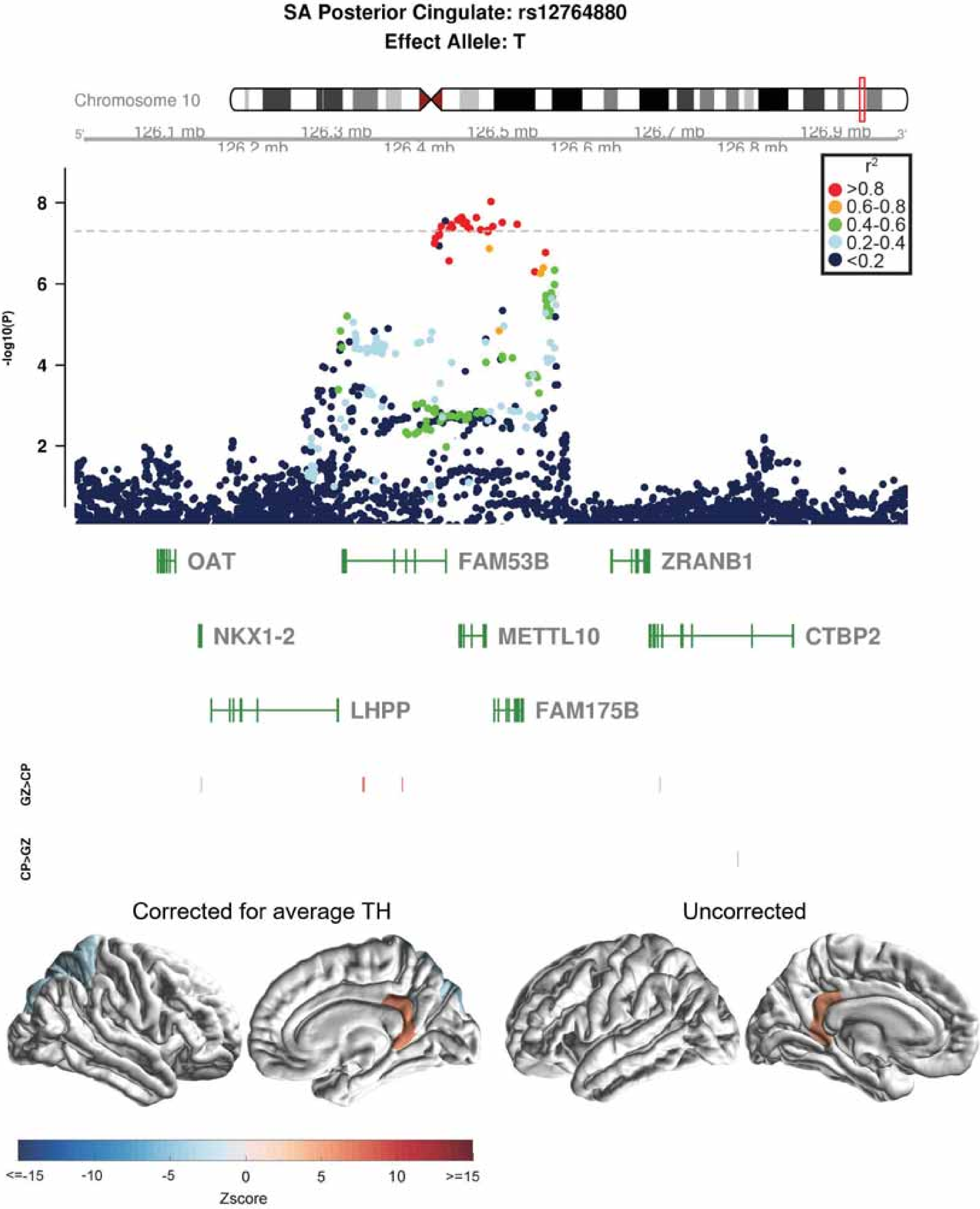

SA Posterior Cingulate: rs12764880
Effect Allele: T
Chromosome 10
126.1 mb
126.2 mb
126.3 mb
126.4 mb
126.5 mb
126.6 mb
126.7 mb
126.8 mb
126.9 mb
r2
>0.8
0.6-0.8
0.4-0.6
0.2-0.4
<0.2
-log10(P)
OAT
FAM53B
ZRANB1
NKX1-2
METTL10
CTBP2
LHPP
FAM175B
GZ>CP
CP>GZ
Corrected for average TH
Uncorrected
<=-15
-10
-5
0
5
10
>=15
Zscore

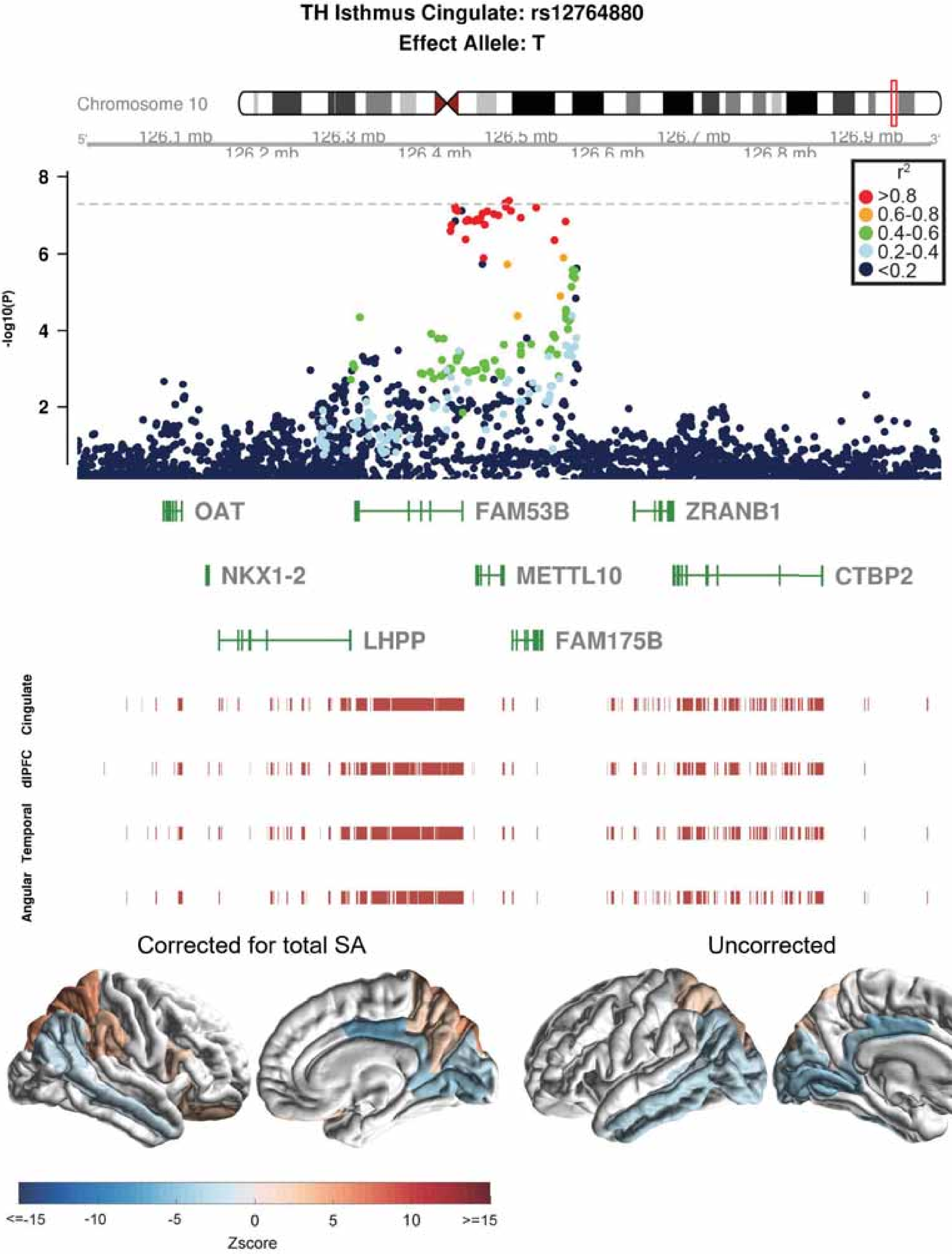

TH Isthmus Cingulate: rs12764880
Effect Allele: T
Chromosome 10
126.1 mb
126.2 mb
126.3 mb
126.4 mb
126.5 mb
126.6 mb
126.7 mb
126.8 mb
126.9 mb
-log10(P)
r2
>0.8
0.6-0.8
0.4-0.6
0.2-0.4
<0.2
OAT
FAM53B
ZRANB1
NKX1-2
METTL10
CTBP2
LHPP
FAM175B
Cingulate
dlPFC
Temporal
Angular
Corrected for total SA
Uncorrected
<=-15
-10
-5
0
5
10
>=15
Zscore

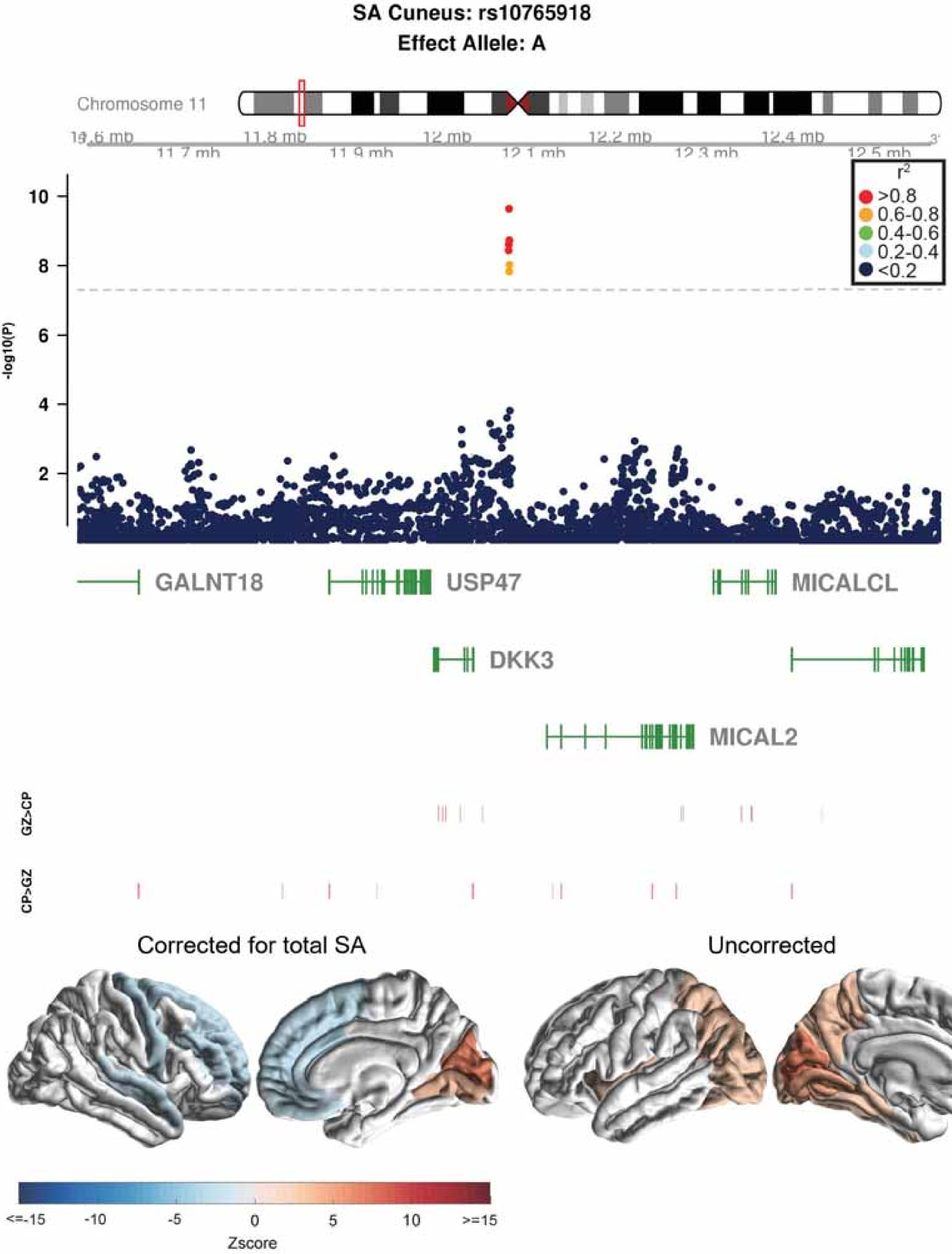

SA Cuneus: rs10765918
Effect Allele: A
Chromosome 11
11.8 mb
11.9 mb
12 mb
12.1 mb
12.2 mb
12.3 mb
12.4 mb
12.5 mb
-log10(P)
r2
>0.8
0.6-0.8
0.4-0.6
0.2-0.4
<0.2
GALNT18
USP47
MICALCL
DKK3
MICAL2
GZ>CP
CP>GZ
Corrected for total SA
Uncorrected
<=-15
-10
-5
0
5
10
>=15
Zscore

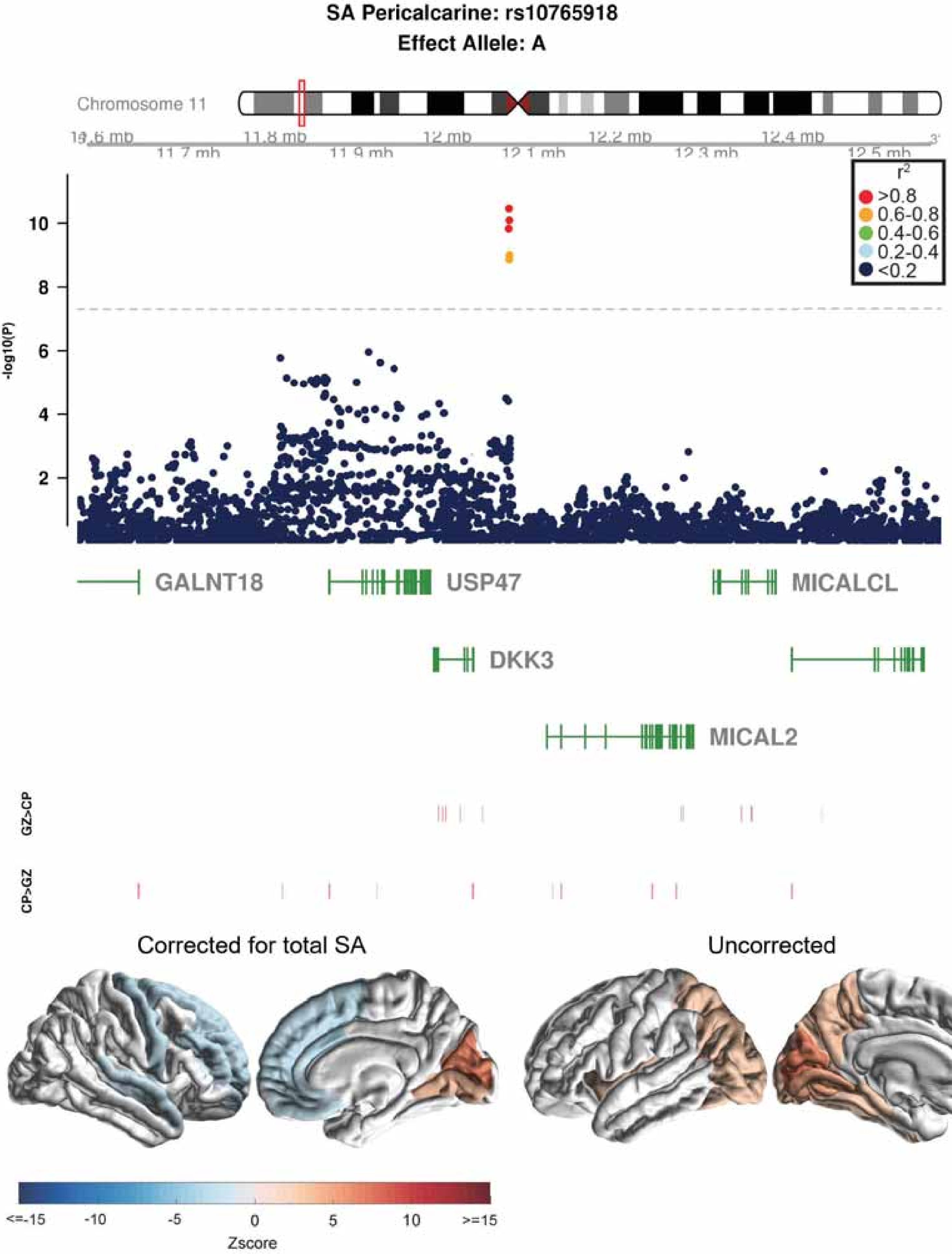
SA Pericalcarine: rs10765918
Effect Allele: A
Chromosome 11
11.7 mb
11.8 mb
11.9 mb
12 mb
12.1 mb
12.2 mb
12.3 mb
12.4 mb
12.5 mb
r²
>0.8
0.6-0.8
0.4-0.6
0.2-0.4
<0.2
-log10(P)
10
8
6
4
2
GALNT18
USP47
MICALCL
DKK3
MICAL2
GZ>CP
CP>GZ
Corrected for total SA
Uncorrected
<=-15
-10
-5
0
5
10
>=15
Zscore

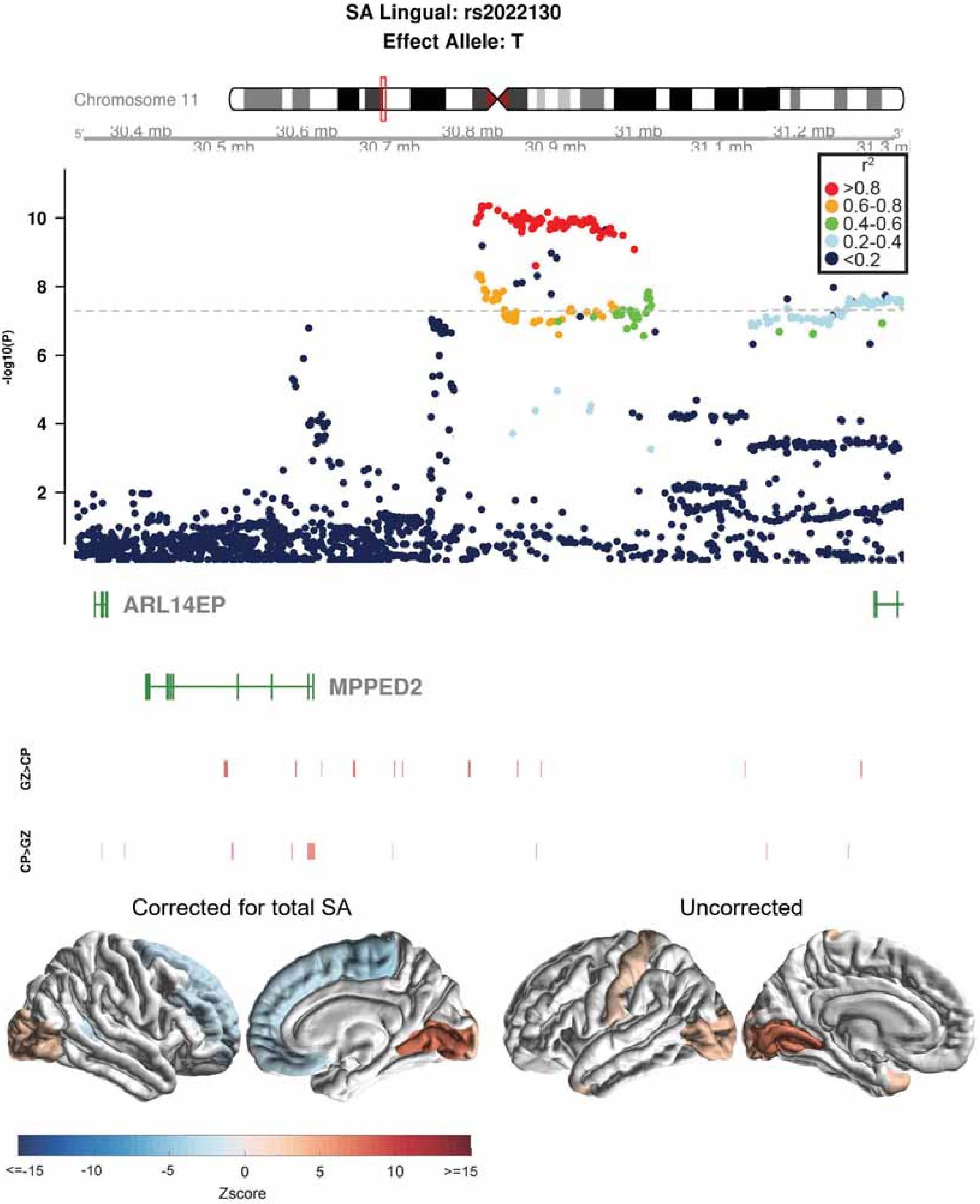
SA Lingual: rs2022130
Effect Allele: T
Chromosome 11
30.4 mb
30.5 mb
30.6 mb
30.7 mb
30.8 mb
30.9 mb
31 mb
31.1 mb
31.2 mb
31.3 mb
r2
>0.8
0.6-0.8
0.4-0.6
0.2-0.4
<0.2
-log10(P)
ARL14EP
MPPED2
GZ>CP
CP>GZ
Corrected for total SA
Uncorrected
<=-15
-10
-5
0
5
10
>=15
Zscore

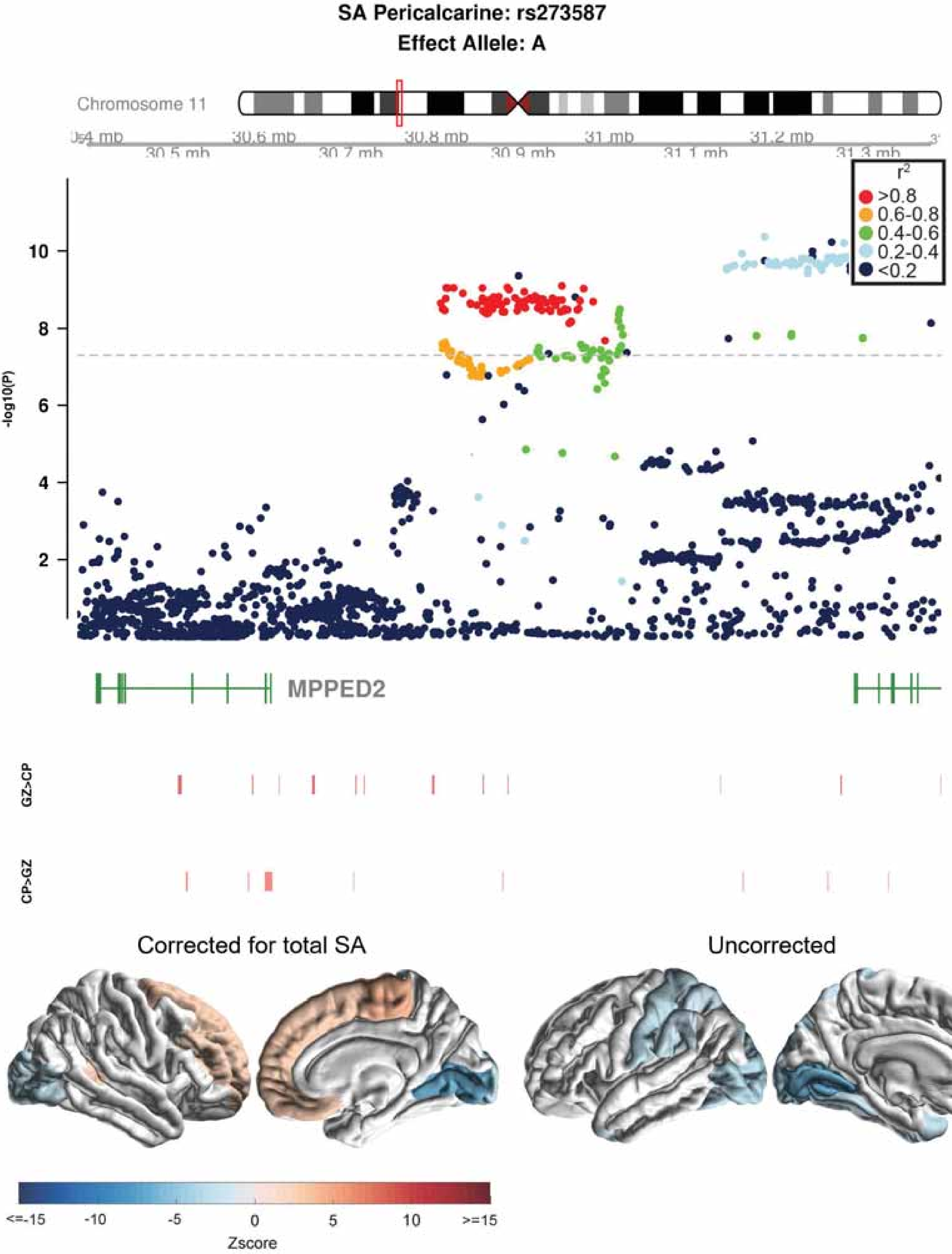

SA Pericalcarine: rs273587
Effect Allele: A
Chromosome 11
30.4 mb
30.5 mb
30.6 mb
30.7 mb
30.8 mb
30.9 mb
31 mb
31.1 mb
31.2 mb
31.3 mb
r2
>0.8
0.6-0.8
0.4-0.6
0.2-0.4
<0.2
-log10(P)
2
4
6
8
10
MPPED2
GZ>CP
CP>GZ
Corrected for total SA
Uncorrected
<=-15
-10
-5
0
5
10
>=15
Zscore

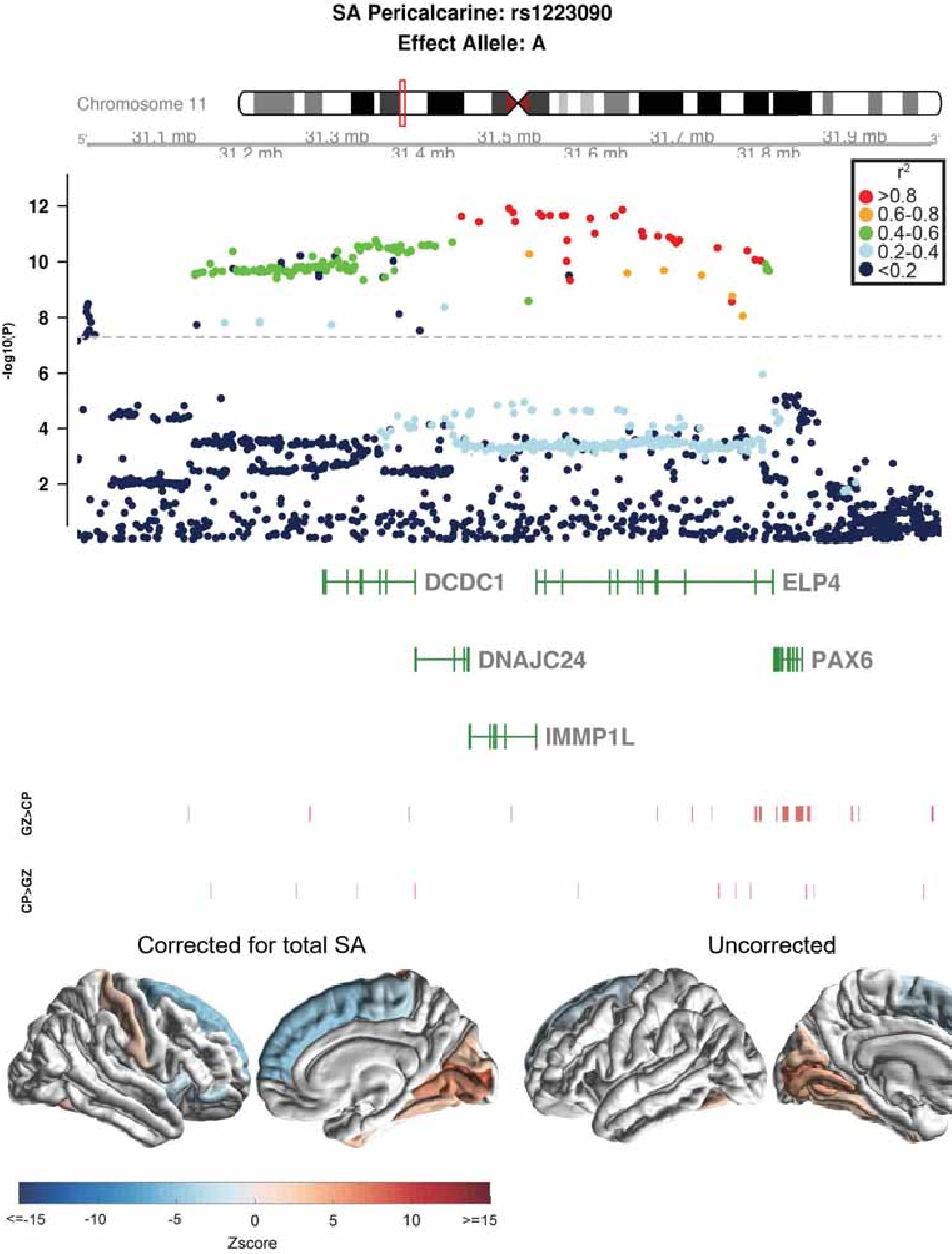

SA Pericalcarine: rs1223090
Effect Allele: A
Chromosome 11
31.1 mb
31.2 mb
31.3 mb
31.4 mb
31.5 mb
31.6 mb
31.7 mb
31.8 mb
31.9 mb
-log10(P)
r2
>0.8
0.6-0.8
0.4-0.6
0.2-0.4
<0.2
DCDC1
ELP4
DNAJC24
PAX6
IMMP1L
GZ>CP
CP>GZ
Corrected for total SA
Uncorrected
<=-15
-10
-5
0
5
10
>=15
Zscore

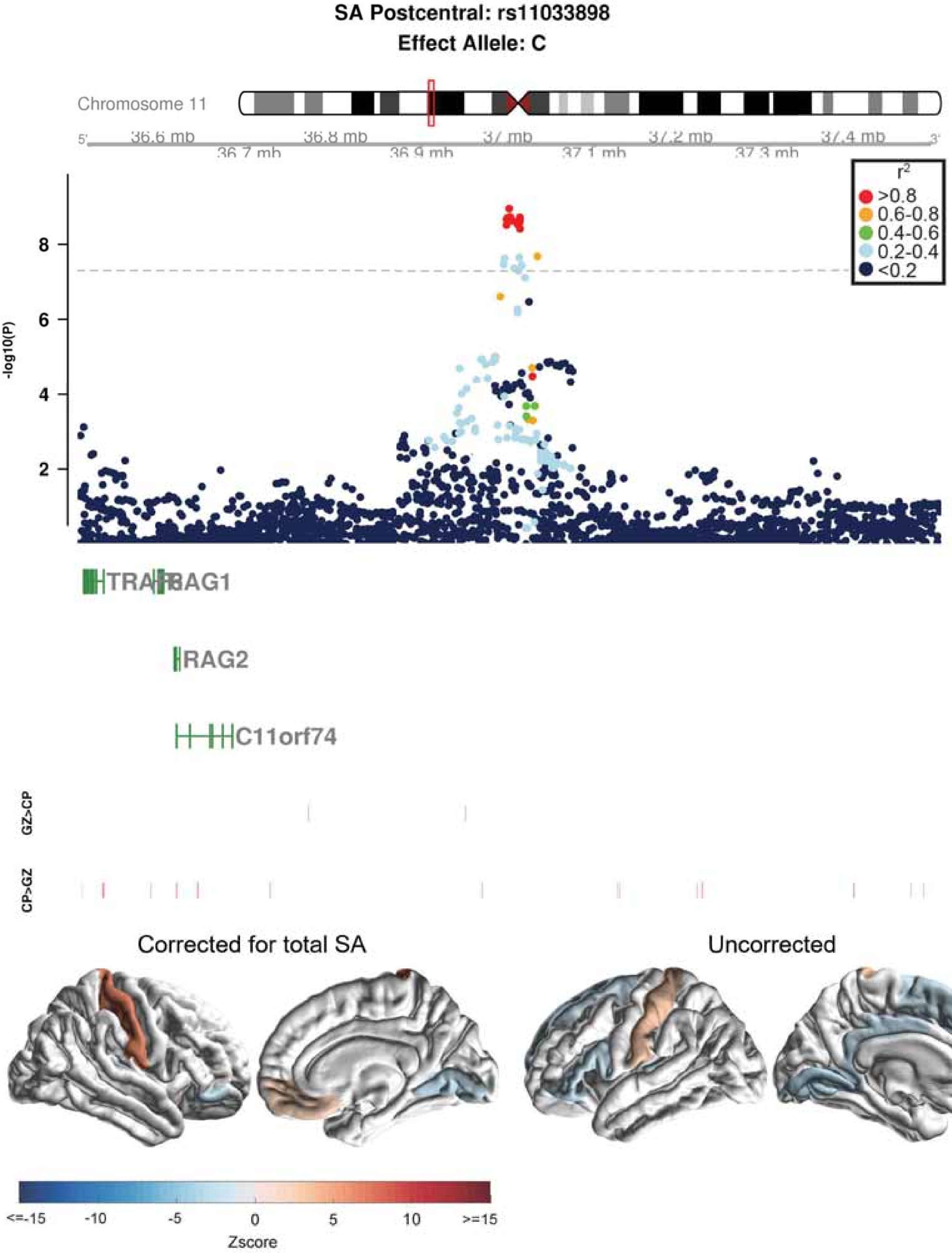

SA Postcentral: rs11033898
Effect Allele: C
Chromosome 11
36.6 mb
36.7 mb
36.8 mb
36.9 mb
37 mb
37.1 mb
37.2 mb
37.3 mb
37.4 mb
-log10(P)
r2
>0.8
0.6-0.8
0.4-0.6
0.2-0.4
<0.2
RAG1
RAG2
C11orf74
GZ>CP
CP>GZ
Corrected for total SA
Uncorrected
<=-15
-10
-5
0
5
10
>=15
Zscore

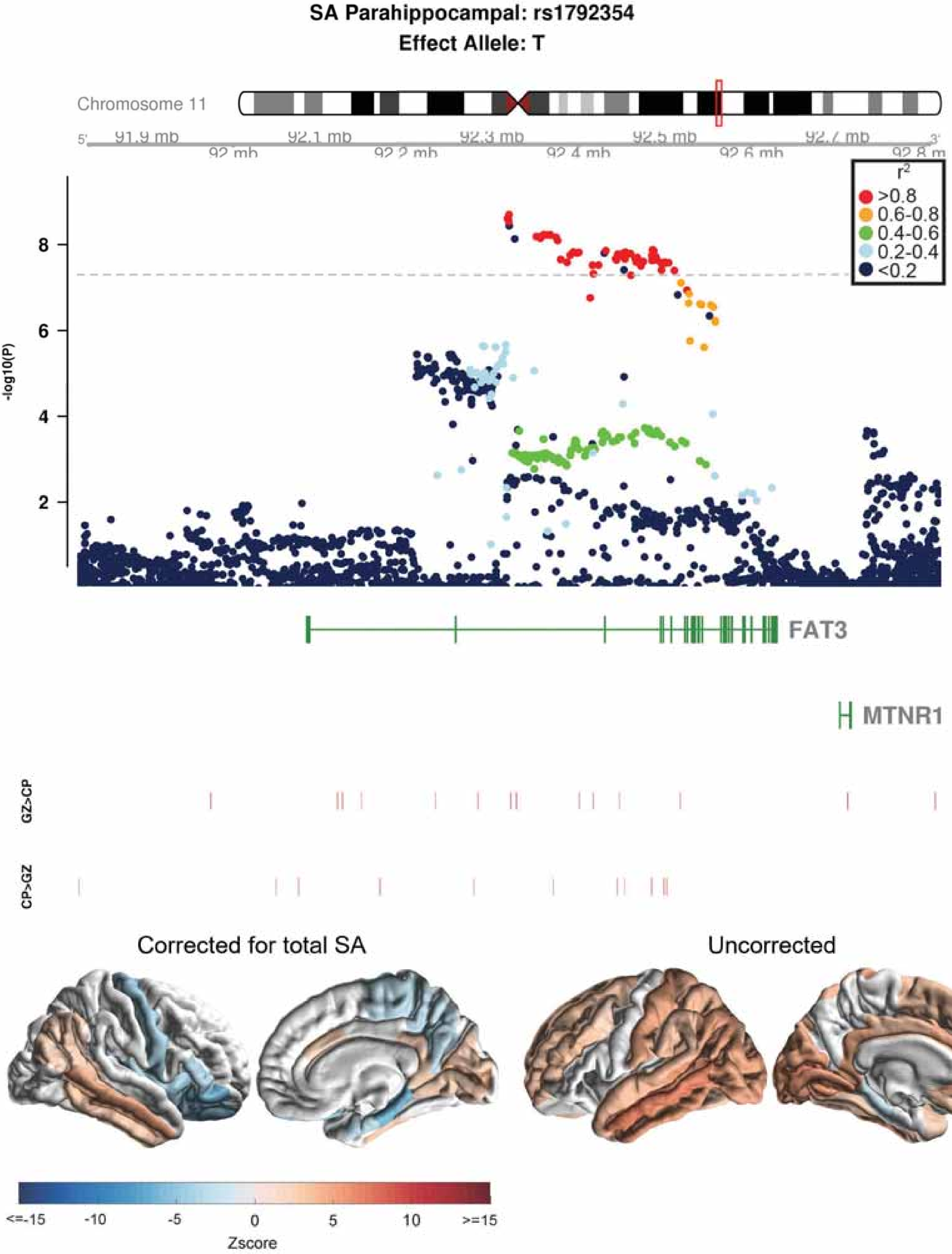

SA Parahippocampal: rs1792354
Effect Allele: T
Chromosome 11
91.9 mb
92.1 mb
92.3 mb
92.5 mb
92.7 mb
92 mb
92.2 mb
92.4 mb
92.6 mb
92.8 m
r2
>0.8
0.6-0.8
0.4-0.6
0.2-0.4
<0.2
-log10(P)
FAT3
MTNR1
GZ>CP
CP>GZ
Corrected for total SA
Uncorrected
<=-15
-10
-5
0
5
10
>=15
Zscore

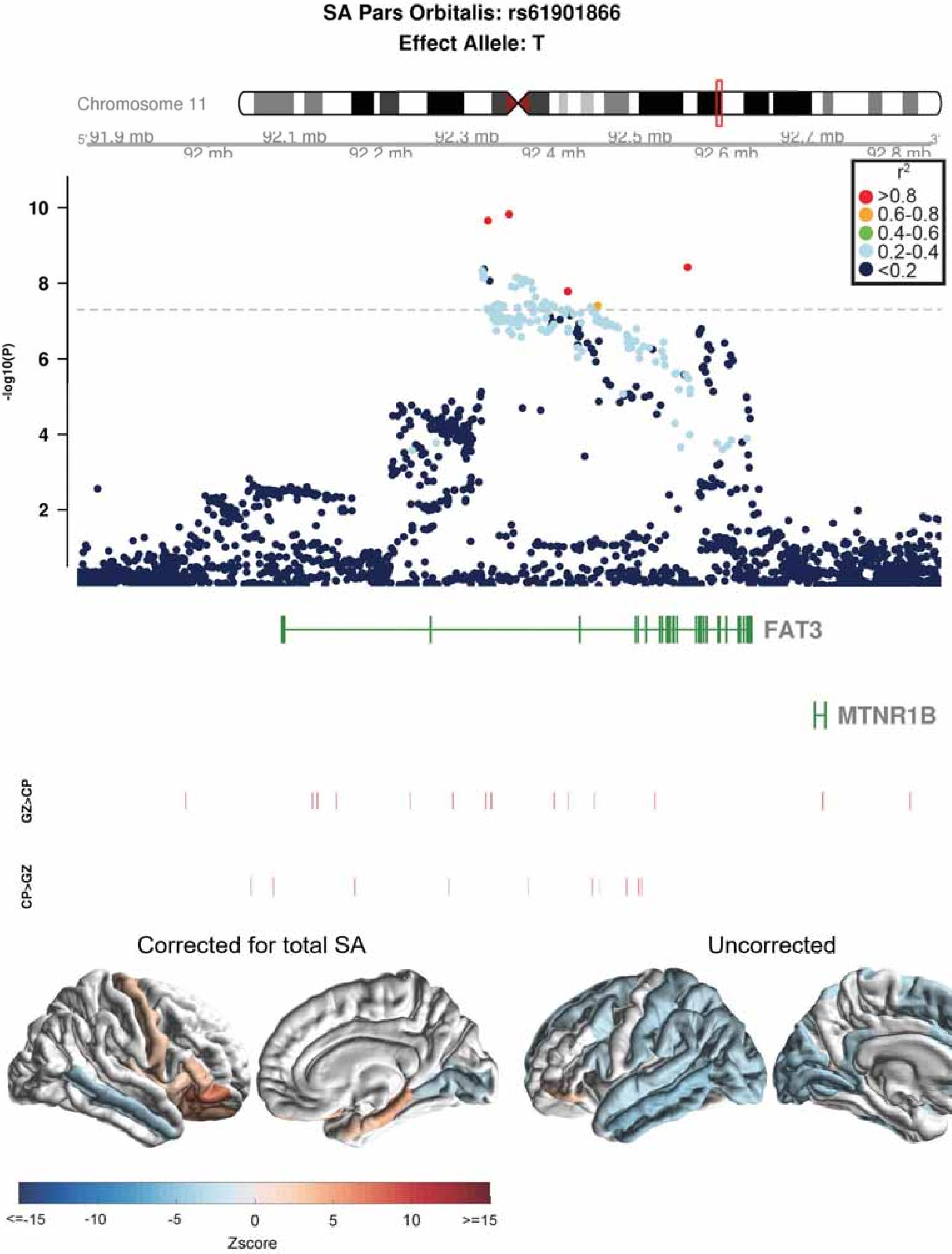

SA Pars Orbitalis: rs61901866
Effect Allele: T
Chromosome 11
91.9 mb
92.1 mb
92.3 mb
92.5 mb
92.7 mb
92 mb
92.2 mb
92.4 mb
92.6 mb
92.8 mb
r2
>0.8
0.6-0.8
0.4-0.6
0.2-0.4
<0.2
-log10(P)
FAT3
MTNR1B
GZ>CP
CP>GZ
Corrected for total SA
Uncorrected
<=-15
-10
-5
0
5
10
>=15
Zscore

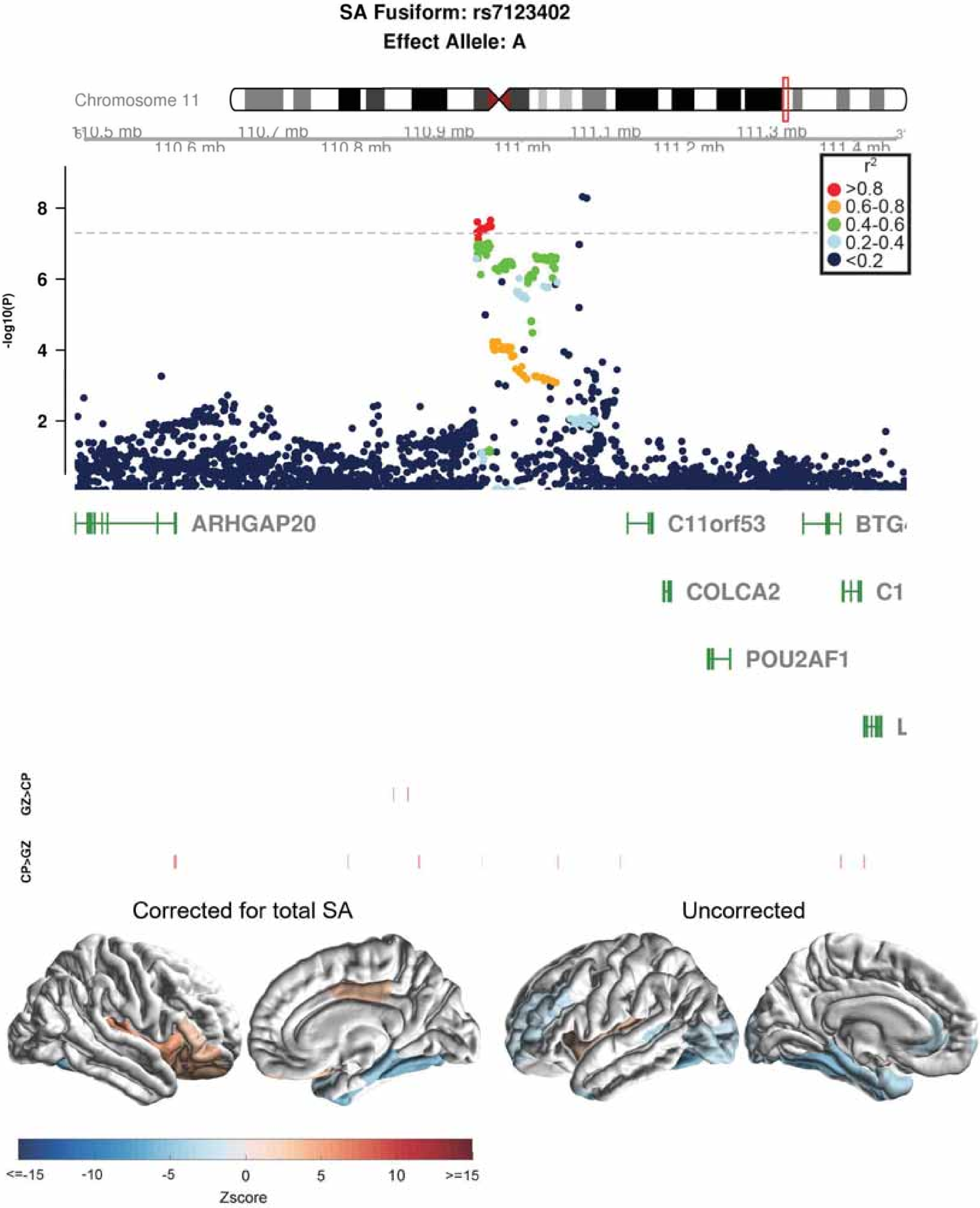

SA Fusiform: rs7123402
Effect Allele: A
Chromosome 11
110.5 mb
110.6 mb
110.7 mb
110.8 mb
110.9 mb
111 mb
111.1 mb
111.2 mb
111.3 mb
111.4 mb
-log10(P)
8
6
4
2
r2
>0.8
0.6-0.8
0.4-0.6
0.2-0.4
<0.2
ARHGAP20
C11orf53
BTG
COLCA2
C1
POU2AF1
L
GZ>CP
CP>GZ
Corrected for total SA
Uncorrected
<=-15
-10
-5
0
5
10
>=15
Zscore

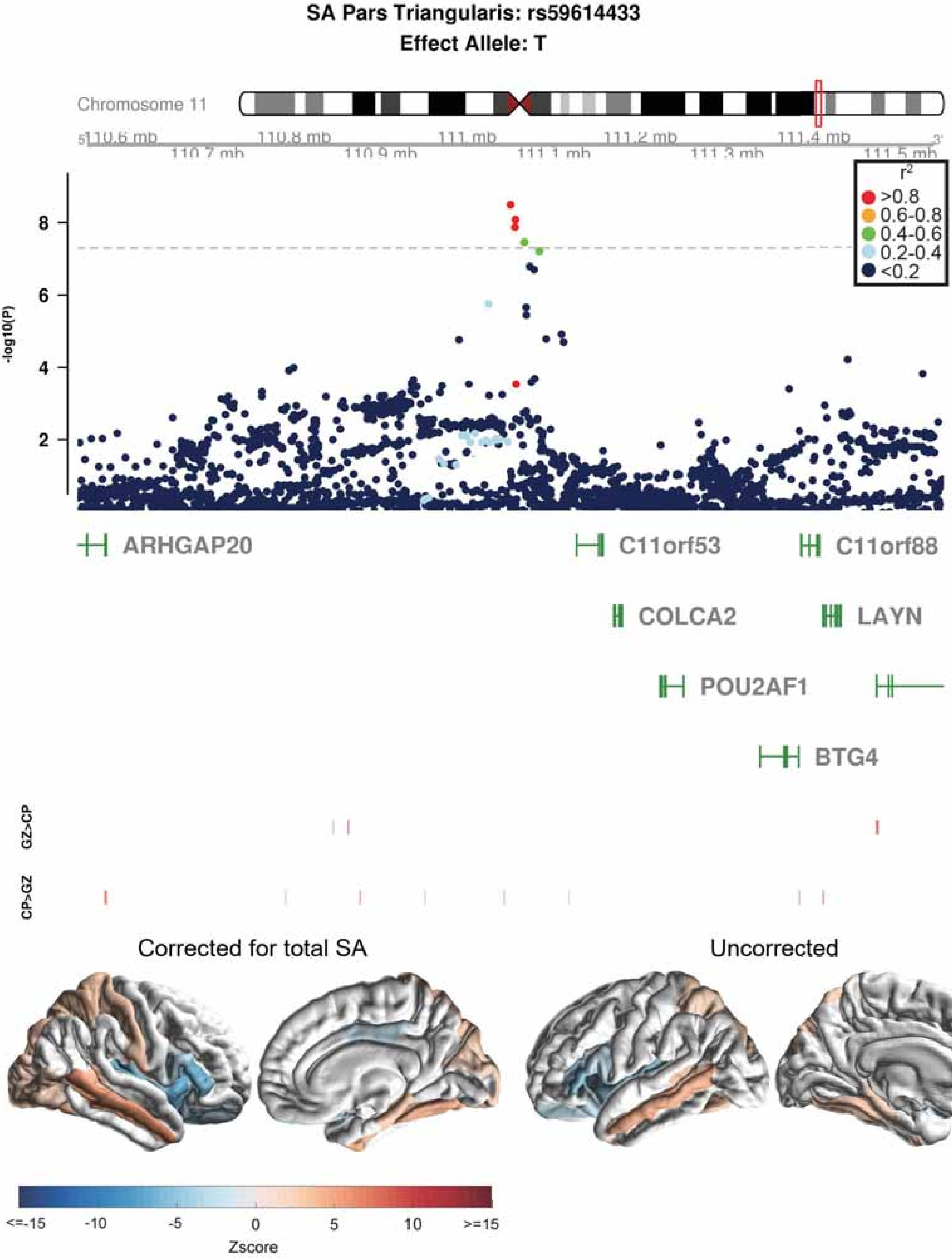

SA Pars Triangularis: rs59614433
Effect Allele: T
Chromosome 11
110.6 mb
110.7 mb
110.8 mb
110.9 mb
111 mb
111.1 mb
111.2 mb
111.3 mb
111.4 mb
111.5 mb
-log10(P)
r2
>0.8
0.6-0.8
0.4-0.6
0.2-0.4
<0.2
ARHGAP20
C11orf53
C11orf88
COLCA2
LAYN
POU2AF1
BTG4
GZ>CP
CP>GZ
Corrected for total SA
Uncorrected
<=-15
-10
-5
0
5
10
>=15
Zscore

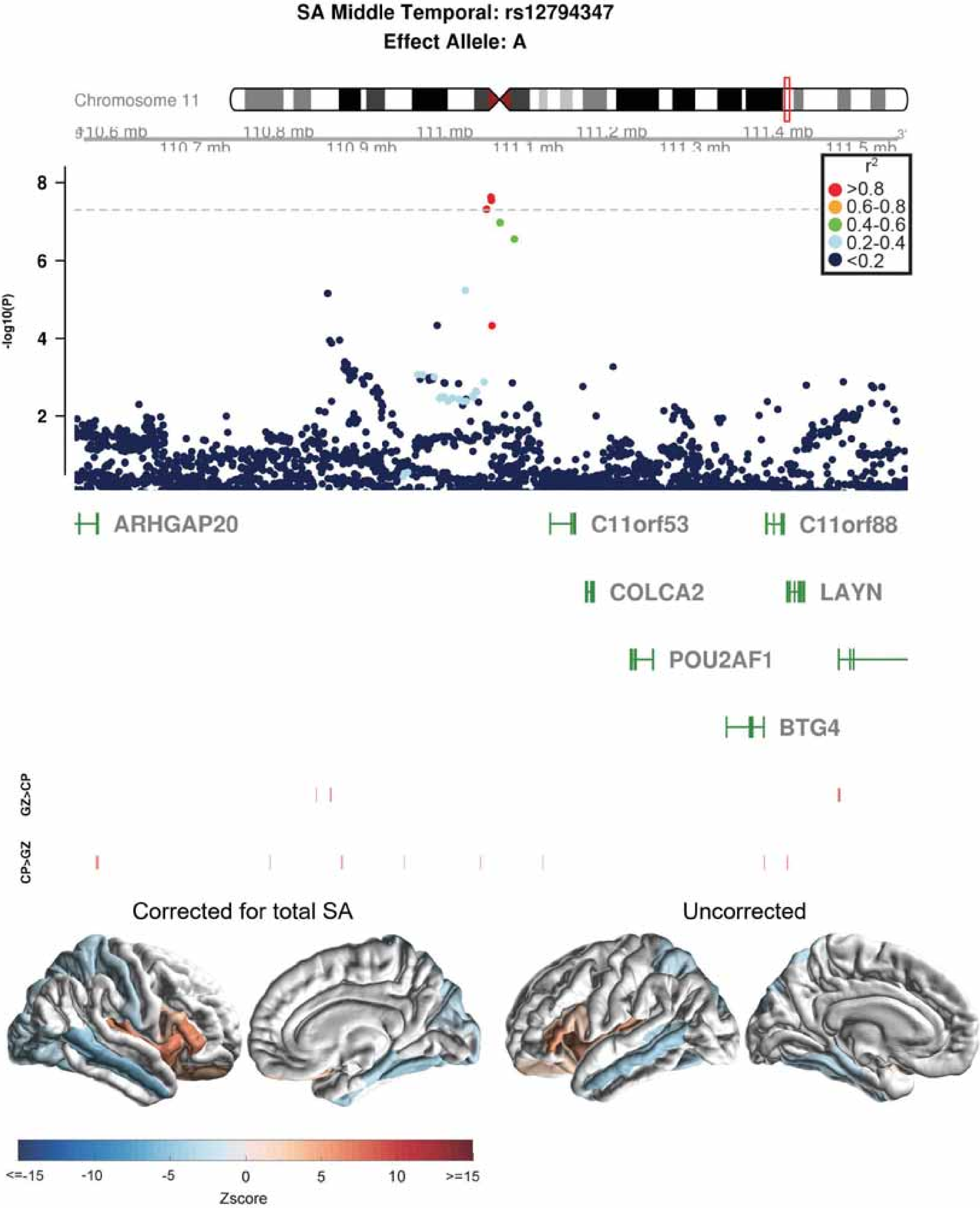
SA Middle Temporal: rs12794347
Effect Allele: A
Chromosome 11
110.6 mb
110.7 mb
110.8 mb
110.9 mb
111 mb
111.1 mb
111.2 mb
111.3 mb
111.4 mb
111.5 mb
r2
>0.8
0.6-0.8
0.4-0.6
0.2-0.4
<0.2
-log10(P)
ARHGAP20
C11orf53
C11orf88
COLCA2
LAYN
POU2AF1
BTG4
GZ>CP
CP>GZ
Corrected for total SA
Uncorrected
<=-15
-10
-5
0
5
10
>=15
Zscore

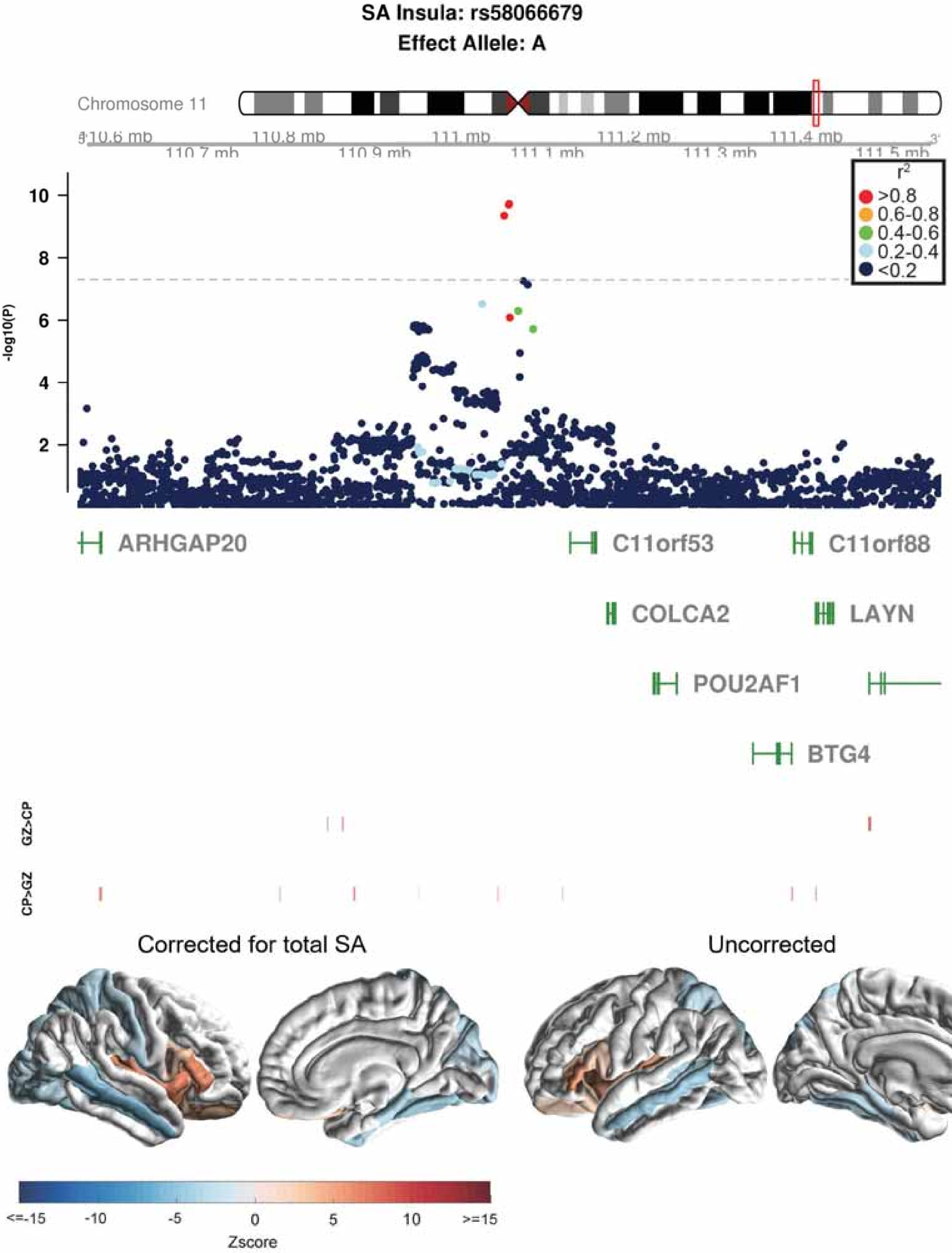
SA Insula: rs58066679
Effect Allele: A
Chromosome 11
110.6 mb
110.7 mb
110.8 mb
110.9 mb
111 mb
111.1 mb
111.2 mb
111.3 mb
111.4 mb
111.5 mb
-log10(P)
r2
>0.8
0.6-0.8
0.4-0.6
0.2-0.4
<0.2
ARHGAP20
C11orf53
C11orf88
COLCA2
LAYN
POU2AF1
BTG4
GZ>CP
CP>GZ
Corrected for total SA
Uncorrected
<=-15
-10
-5
0
5
10
>=15
Zscore

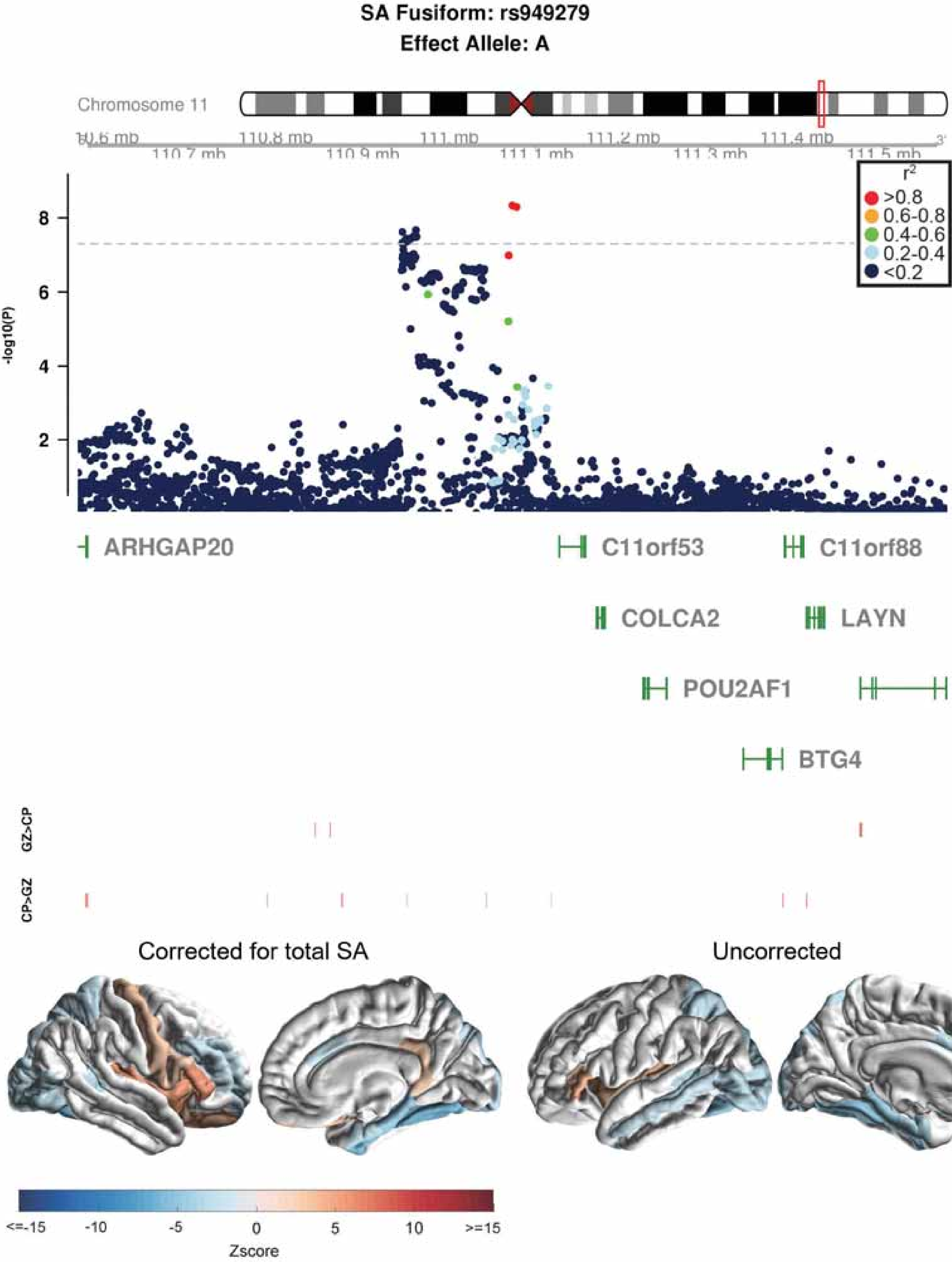

SA Fusiform: rs949279
Effect Allele: A
Chromosome 11
110.6 mb
110.7 mb
110.8 mb
110.9 mb
111 mb
111.1 mb
111.2 mb
111.3 mb
111.4 mb
111.5 mb
-log10(P)
8
6
4
2
r2
>0.8
0.6-0.8
0.4-0.6
0.2-0.4
<0.2
ARHGAP20
C11orf53
C11orf88
COLCA2
LAYN
POU2AF1
BTG4
GZ>CP
CP>GZ
Corrected for total SA
Uncorrected
<=-15
-10
-5
0
5
10
>=15
Zscore

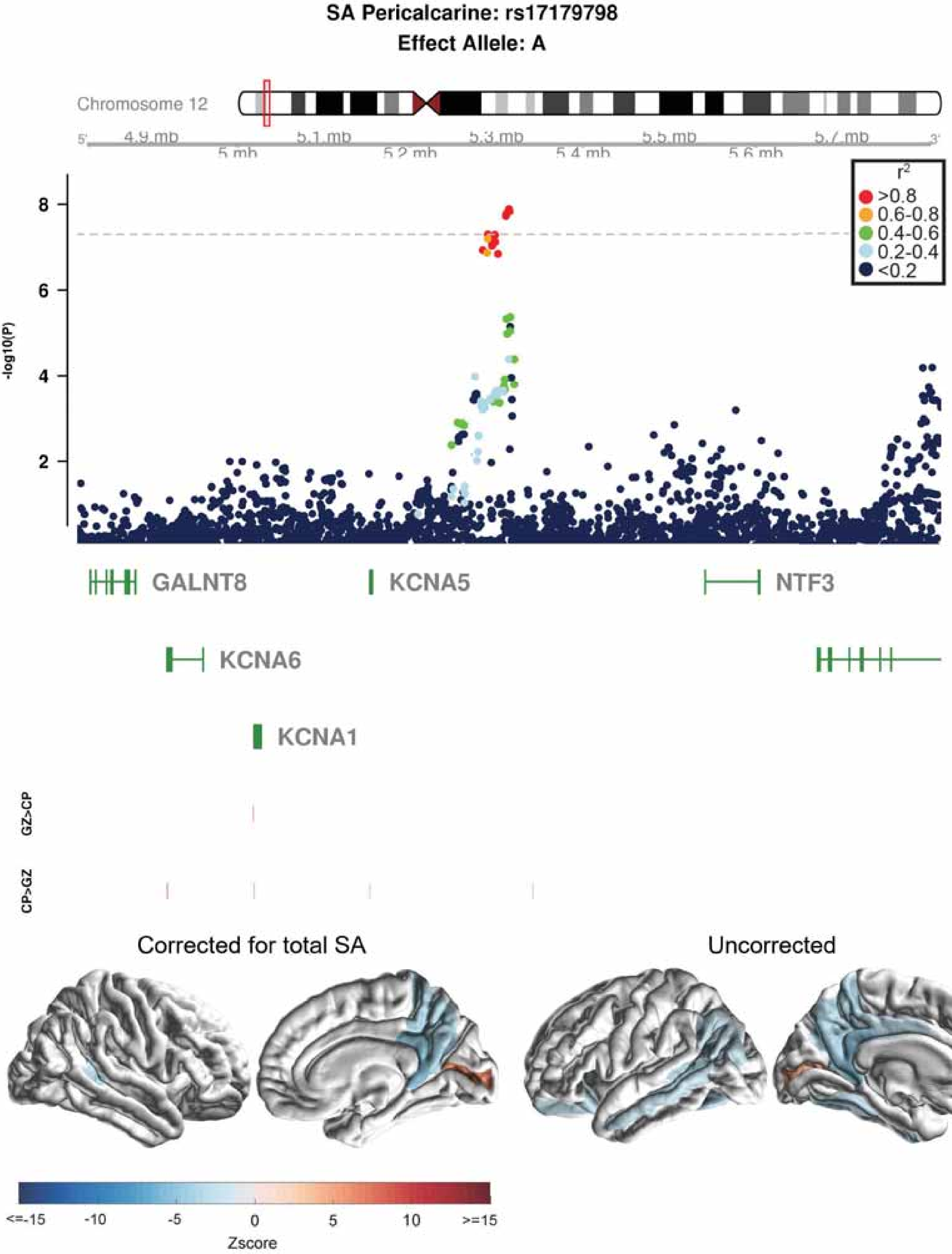
SA Pericalcarine: rs17179798
Effect Allele: A
Chromosome 12
4.9 mb
5 mb
5.1 mb
5.2 mb
5.3 mb
5.4 mb
5.5 mb
5.6 mb
5.7 mb
-log10(P)
r2
>0.8
0.6-0.8
0.4-0.6
0.2-0.4
<0.2
GALNT8
KCNA5
NTF3
KCNA6
KCNA1
GZ>CP
CP>GZ
Corrected for total SA
Uncorrected
Zscore

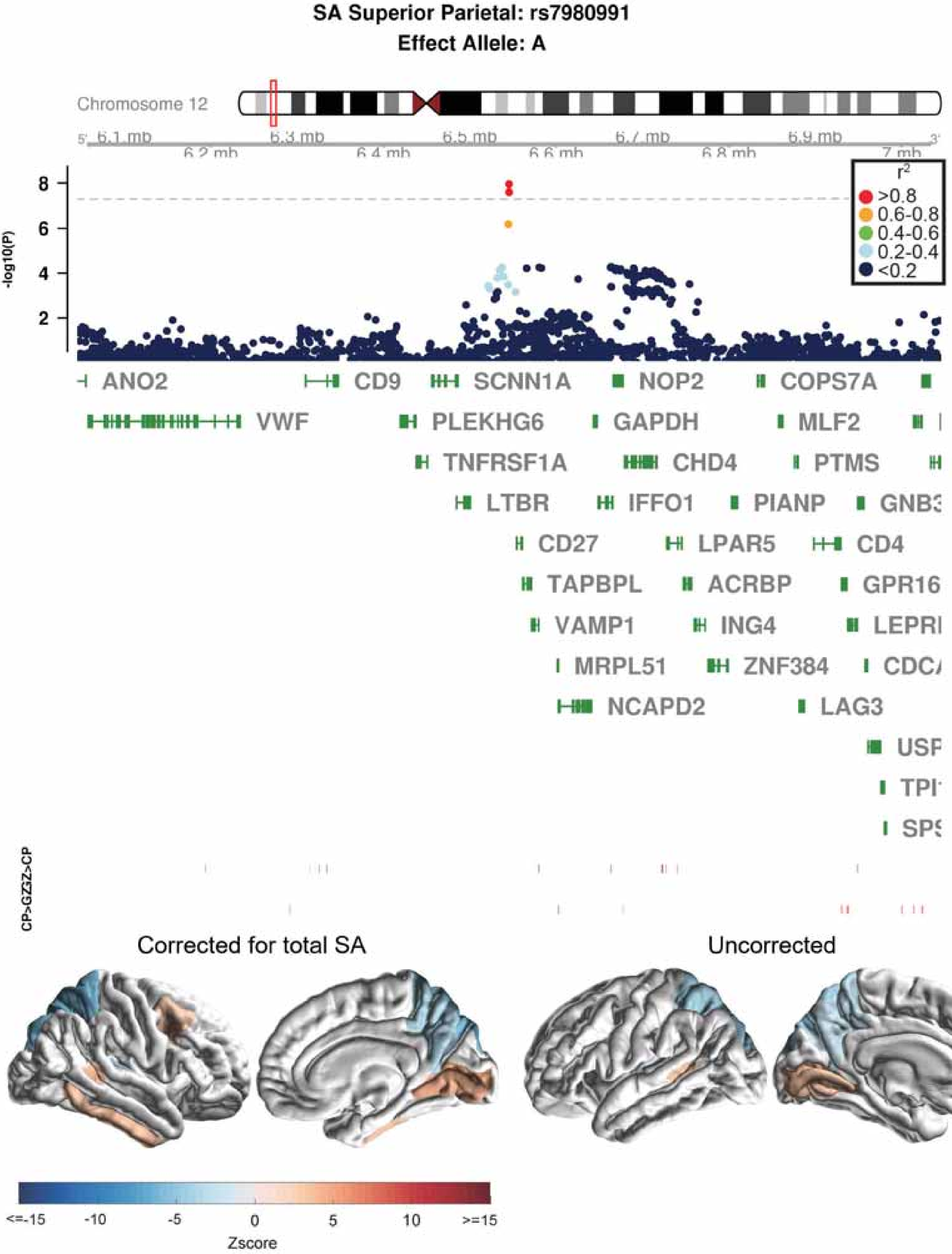
SA Superior Parietal: rs7980991
Effect Allele: A
Chromosome 12
6.1 mb
6.2 mb
6.3 mb
6.4 mb
6.5 mb
6.6 mb
6.7 mb
6.8 mb
6.9 mb
7 mb
-log10(P)
$r^2$
>0.8
0.6-0.8
0.4-0.6
0.2-0.4
<0.2
ANO2
CD9
SCNN1A
NOP2
COPS7A
VWF
PLEKHG6
GAPDH
MLF2
TNFRSF1A
CHD4
PTMS
LTBR
IFFO1
PIANP
CD27
LPAR5
CD4
TAPBPL
ACRBP
VAMP1
ING4
MRPL51
ZNF384
NCAPD2
LAG3
CP>GZ
GZ>CP
Corrected for total SA
Uncorrected
<=-15
-10
-5
0
5
10
>=15
Zscore

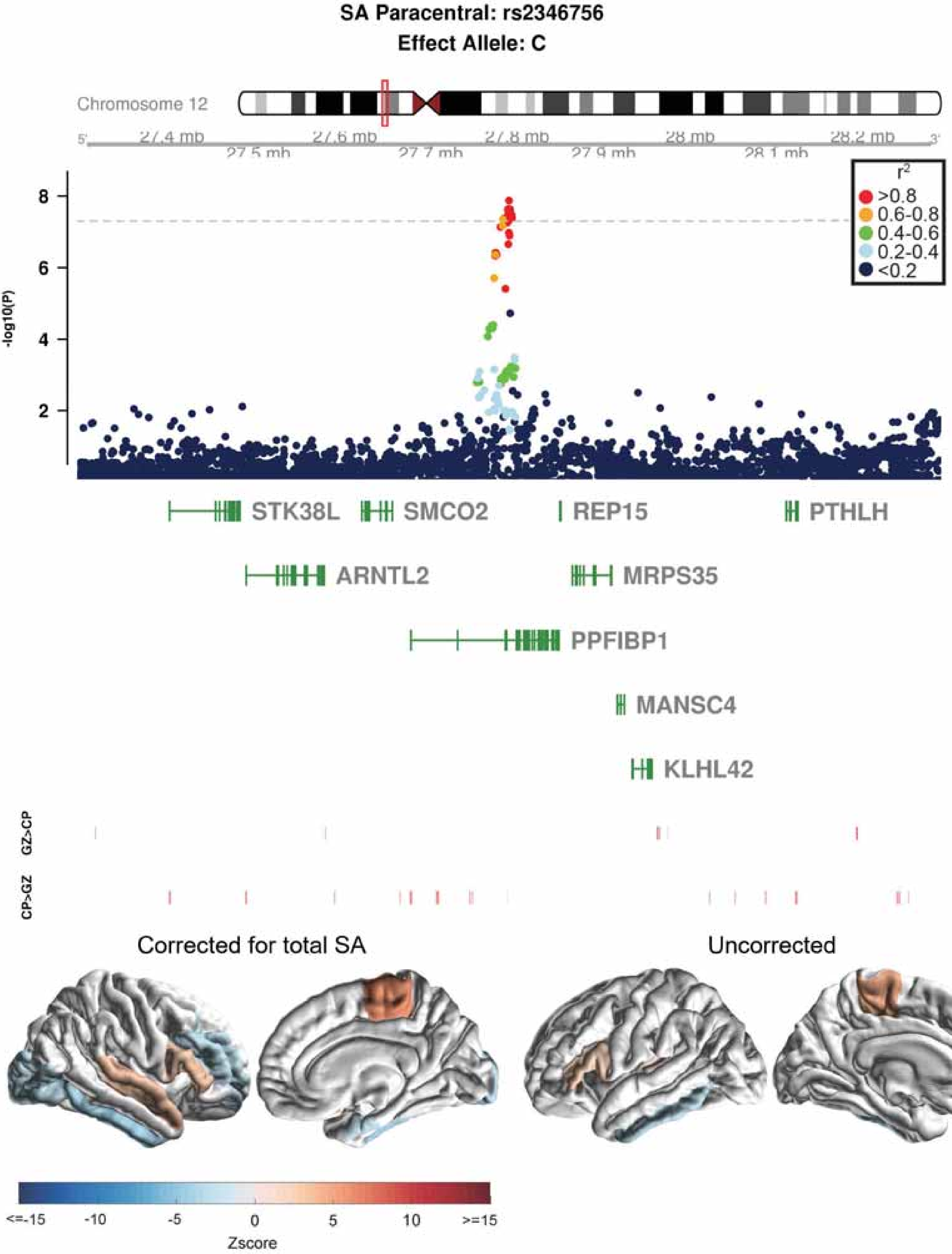

SA Paracentral: rs2346756
Effect Allele: C
Chromosome 12
27.4 mb
27.5 mb
27.6 mb
27.7 mb
27.8 mb
27.9 mb
28 mb
28.1 mb
28.2 mb
-log10(P)
r2
>0.8
0.6-0.8
0.4-0.6
0.2-0.4
<0.2
STK38L
SMCO2
REP15
PTHLH
ARNTL2
MRPS35
PPFIBP1
MANSC4
KLHL42
GZ>CP
CP>GZ
Corrected for total SA
Uncorrected
<=-15
-10
-5
0
5
10
>=15
Zscore

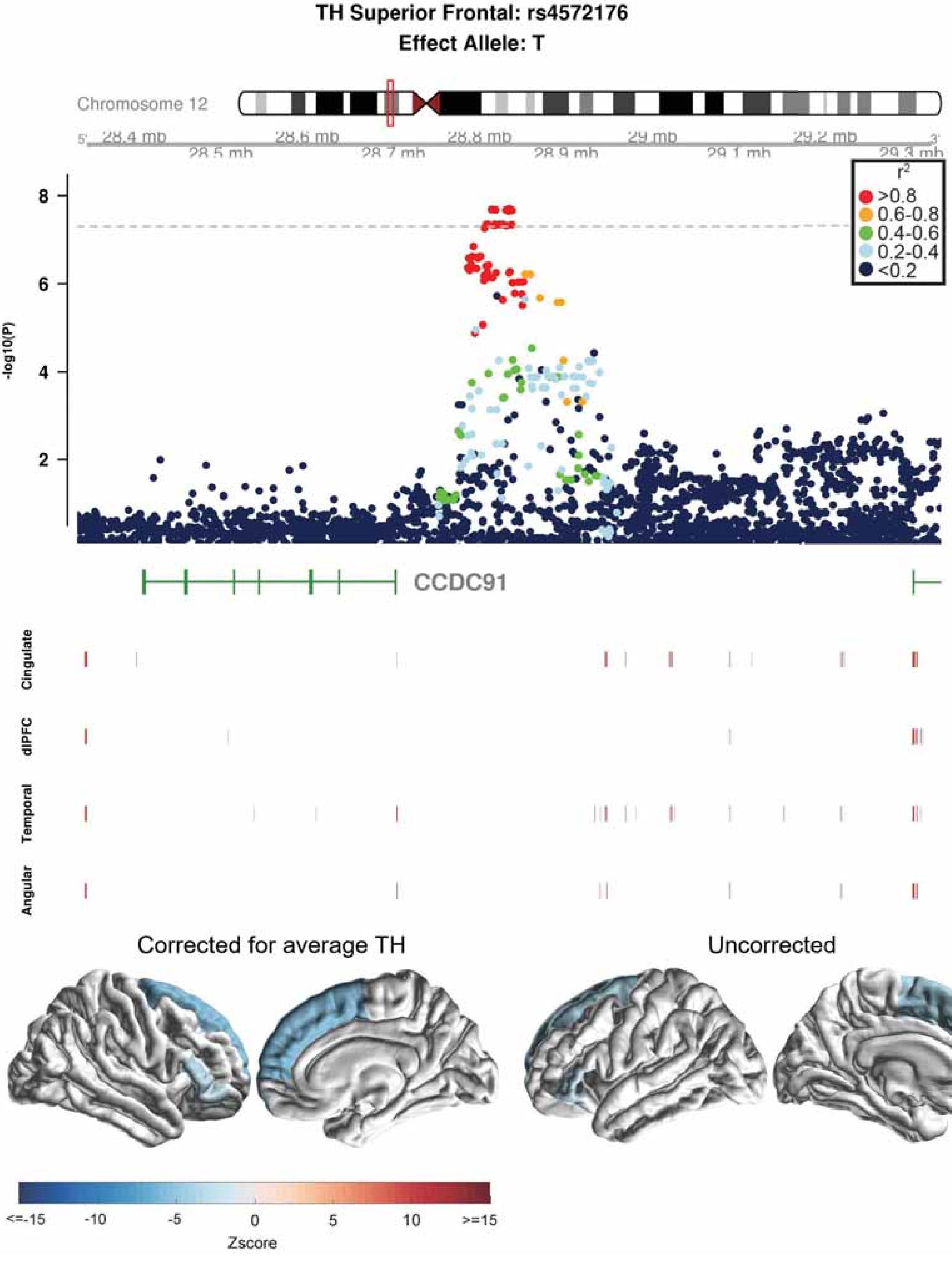

TH Superior Frontal: rs4572176
Effect Allele: T
Chromosome 12
28.4 mb
28.5 mb
28.6 mb
28.7 mb
28.8 mb
28.9 mb
29 mb
29.1 mb
29.2 mb
29.3 mb
r2
>0.8
0.6-0.8
0.4-0.6
0.2-0.4
<0.2
-log10(P)
CCDC91
Cingulate
dlPFC
Temporal
Angular
Corrected for average TH
Uncorrected
<=-15
-10
-5
0
5
10
>=15
Zscore

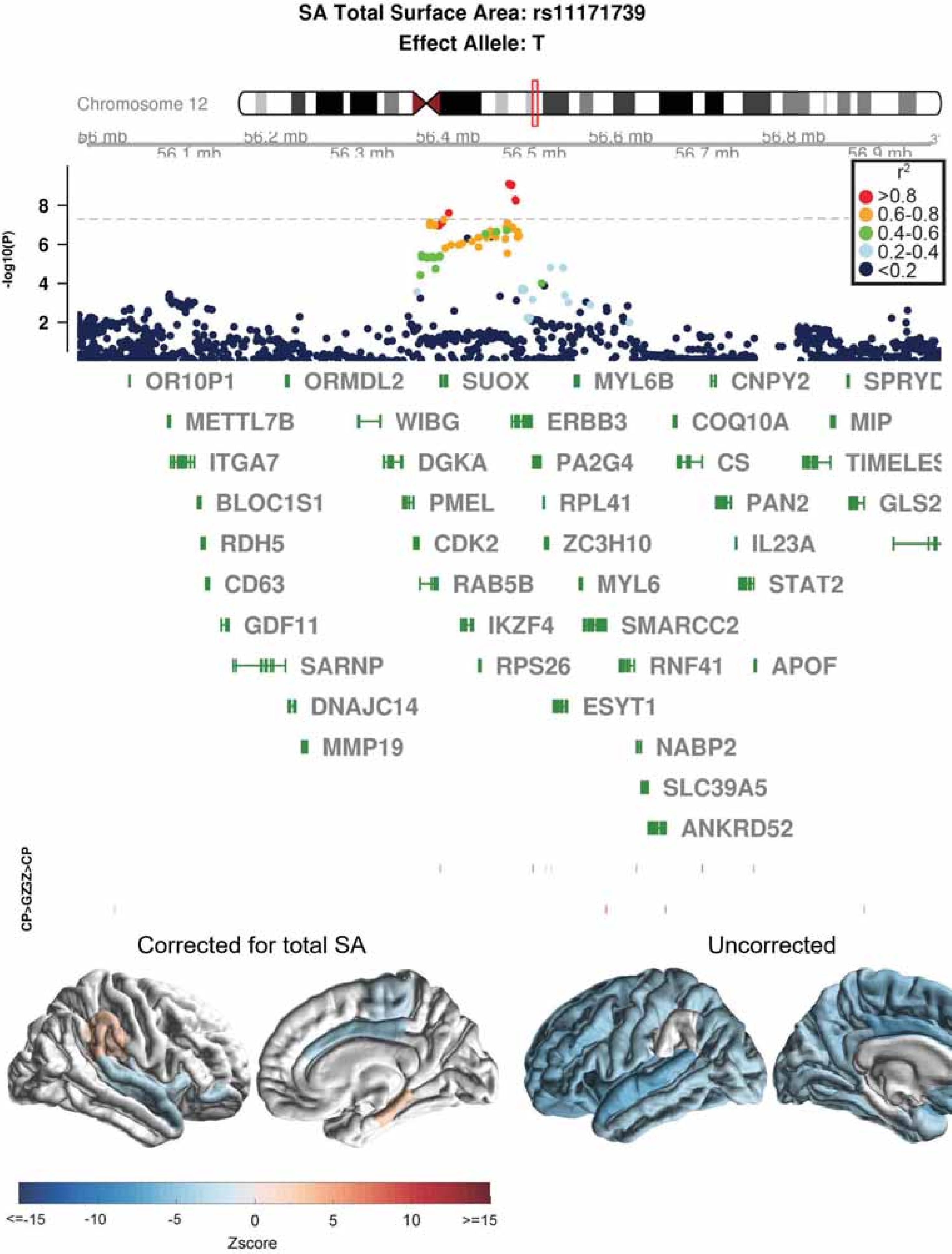

SA Total Surface Area: rs11171739
Effect Allele: T
Chromosome 12
56.1 mb
56.2 mb
56.3 mb
56.4 mb
56.5 mb
56.6 mb
56.7 mb
56.8 mb
56.9 mb
-log10(P)
2
4
6
8
r²
>0.8
0.6-0.8
0.4-0.6
0.2-0.4
<0.2
OR10P1
ORMDL2
SUOX
MYL6B
CNPY2
METTL7B
WIBG
ERBB3
COQ10A
MIP
ITGA7
DGKA
PA2G4
CS
BLOC1S1
PMEL
RPL41
PAN2
GLS2
RDH5
CDK2
ZC3H10
IL23A
CD63
RAB5B
MYL6
STAT2
GDF11
IKZF4
SMARCC2
SARNP
RPS26
RNF41
APOF
DNAJC14
ESYT1
MMP19
NABP2
SLC39A5
ANKRD52
CP>GZ
Z>CP
Corrected for total SA
Uncorrected
<=-15
-10
-5
0
5
10
>=15
Zscore

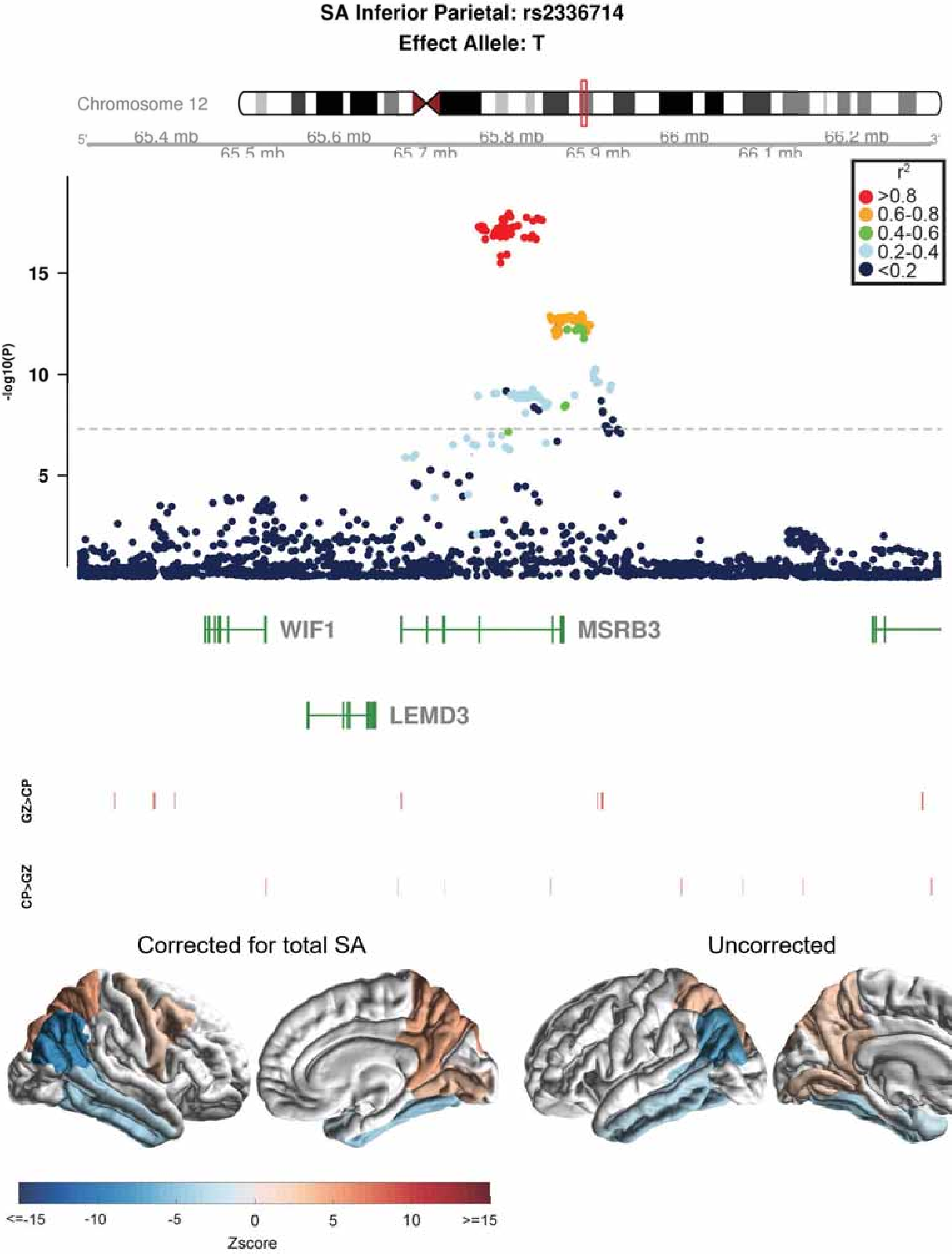
SA Inferior Parietal: rs2336714
Effect Allele: T
Chromosome 12
65.4 mb
65.5 mb
65.6 mb
65.7 mb
65.8 mb
65.9 mb
66 mb
66.1 mb
66.2 mb
-log10(P)
r2
>0.8
0.6-0.8
0.4-0.6
0.2-0.4
<0.2
WIF1
MSRB3
LEMD3
GZ>CP
CP>GZ
Corrected for total SA
Uncorrected
Zscore
<=-15
-10
-5
0
5
10
>=15

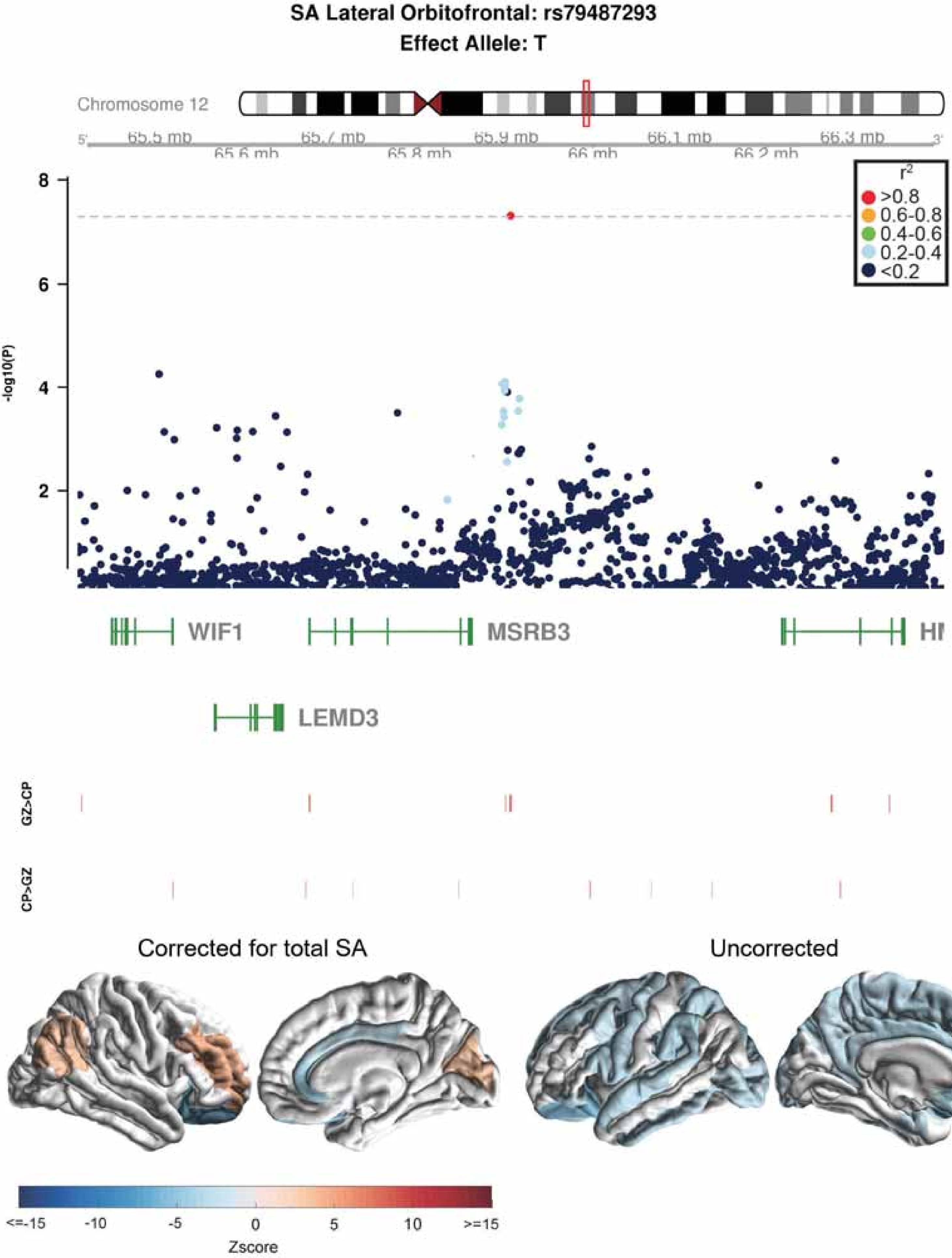

SA Lateral Orbitofrontal: rs79487293
Effect Allele: T
Chromosome 12
65.5 mb
65.6 mb
65.7 mb
65.8 mb
65.9 mb
66 mb
66.1 mb
66.2 mb
66.3 mb
5'
3'
r2
>0.8
0.6-0.8
0.4-0.6
0.2-0.4
<0.2
-log10(P)
8
6
4
2
WIF1
MSRB3
LEMD3
GZ>CP
CP>GZ
Corrected for total SA
Uncorrected
<=-15
-10
-5
0
5
10
>=15
Zscore

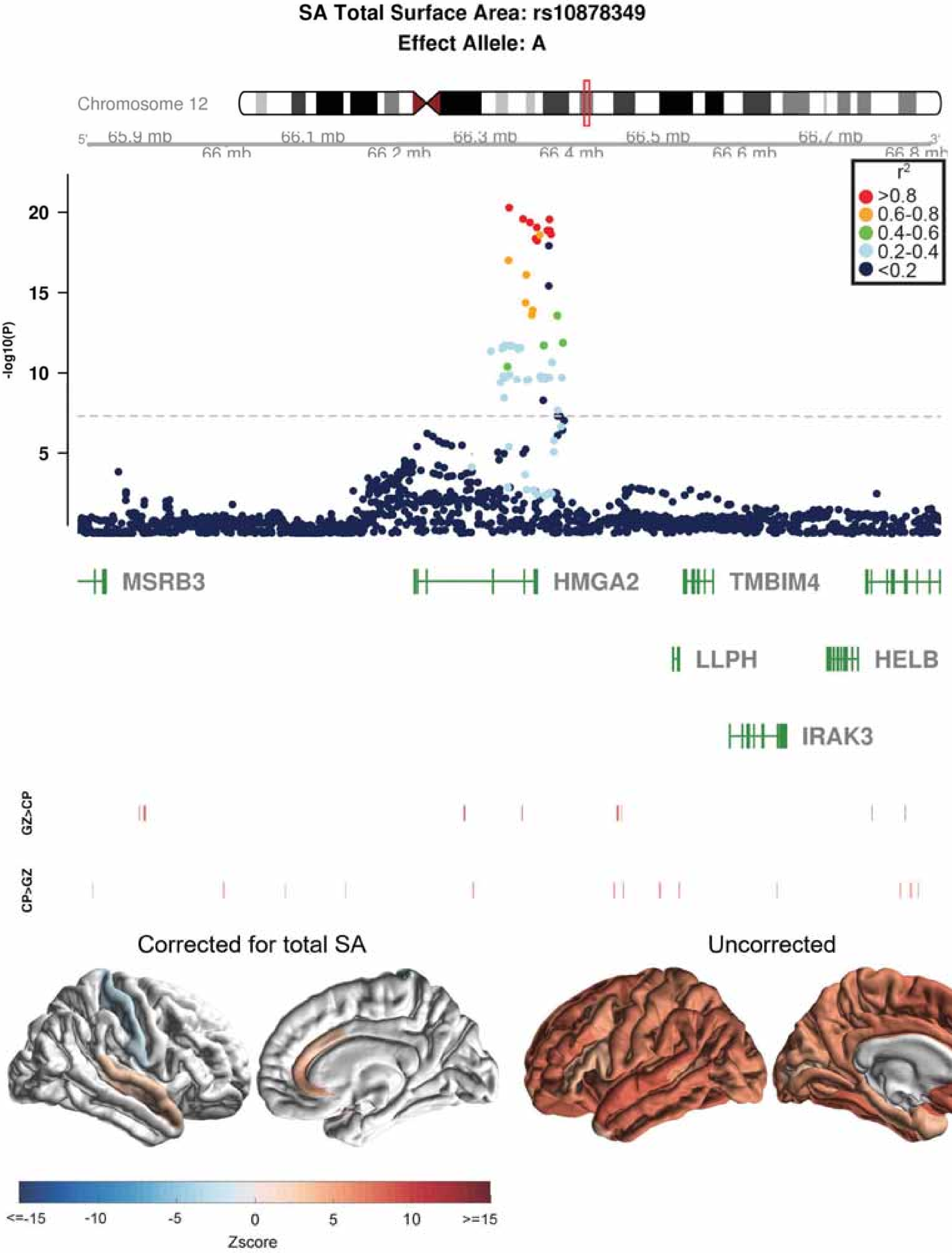

SA Total Surface Area: rs10878349
Effect Allele: A
Chromosome 12
5'
65.9 mb
66 mb
66.1 mb
66.2 mb
66.3 mb
66.4 mb
66.5 mb
66.6 mb
66.7 mb
66.8 mb
3'
r2
>0.8
0.6-0.8
0.4-0.6
0.2-0.4
<0.2
-log10(P)
20
15
10
5
MSRB3
HMGA2
TMBIM4
LLPH
HELB
IRAK3
GZ>CP
CP>GZ
Corrected for total SA
Uncorrected
<=-15
-10
-5
0
5
10
>=15
Zscore

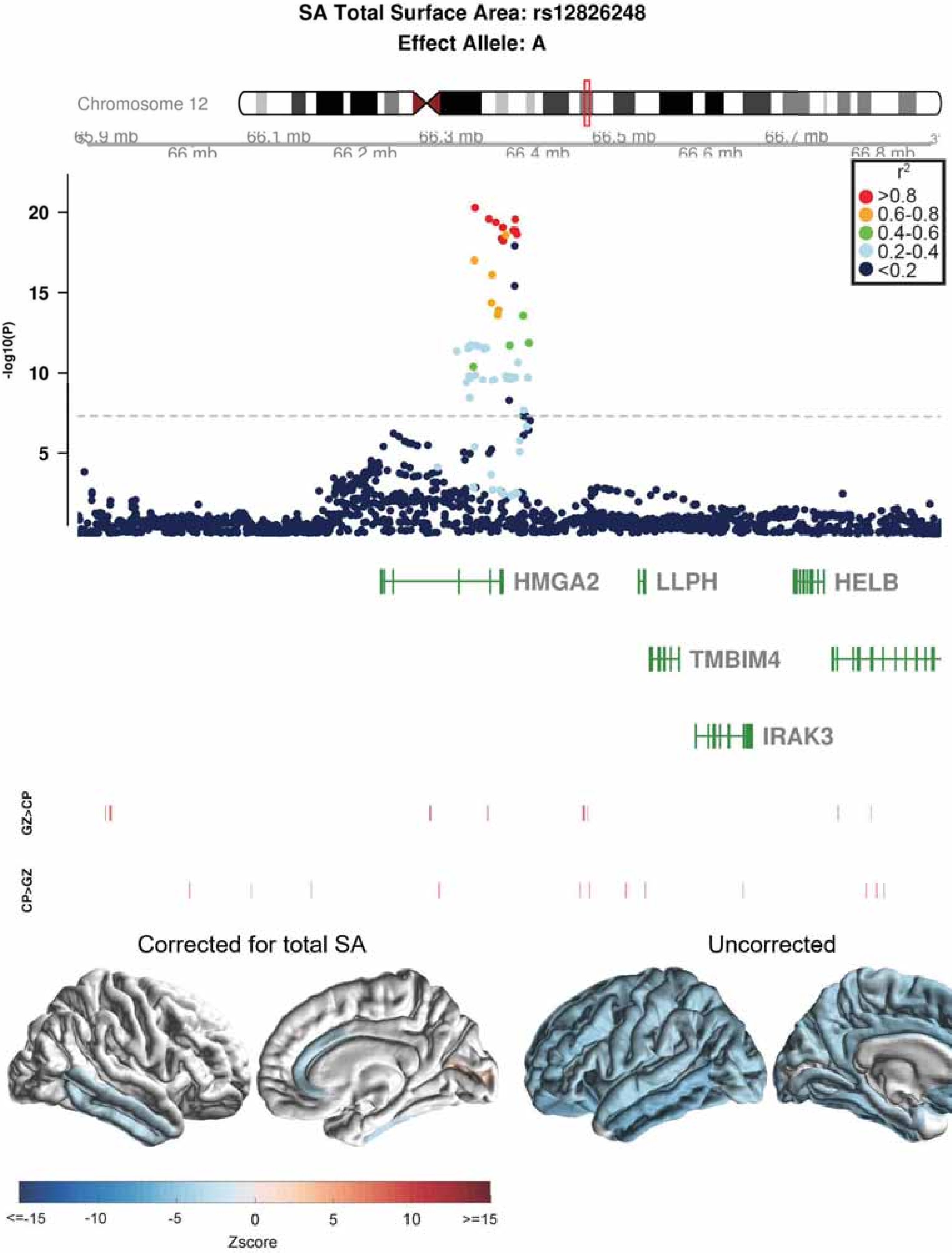

SA Total Surface Area: rs12826248
Effect Allele: A
Chromosome 12
65.9 mb
66 mb
66.1 mb
66.2 mb
66.3 mb
66.4 mb
66.5 mb
66.6 mb
66.7 mb
66.8 mb
r2
>0.8
0.6-0.8
0.4-0.6
0.2-0.4
<0.2
-log10(P)
20
15
10
5
HMGA2
LLPH
HELB
TMBIM4
IRAK3
GZ>CP
CP>GZ
Corrected for total SA
Uncorrected
<=-15
-10
-5
0
5
10
>=15
Zscore

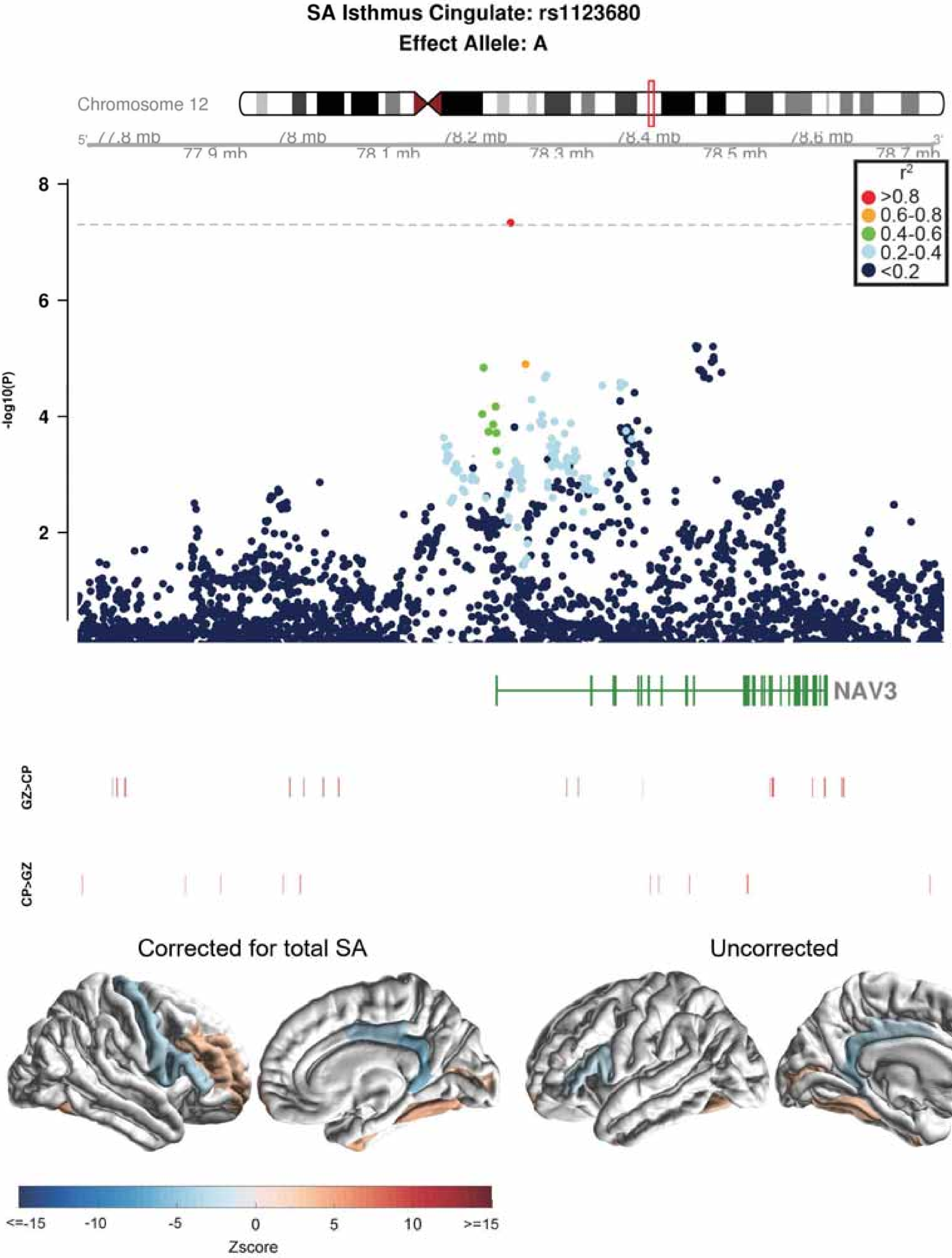

SA Isthmus Cingulate: rs1123680
Effect Allele: A
Chromosome 12
77.8 mb
77.9 mb
78 mb
78.1 mb
78.2 mb
78.3 mb
78.4 mb
78.5 mb
78.6 mb
78.7 mb
-log10(P)
r²
>0.8
0.6-0.8
0.4-0.6
0.2-0.4
<0.2
NAV3
GZ>CP
CP>GZ
Corrected for total SA
Uncorrected
Zscore

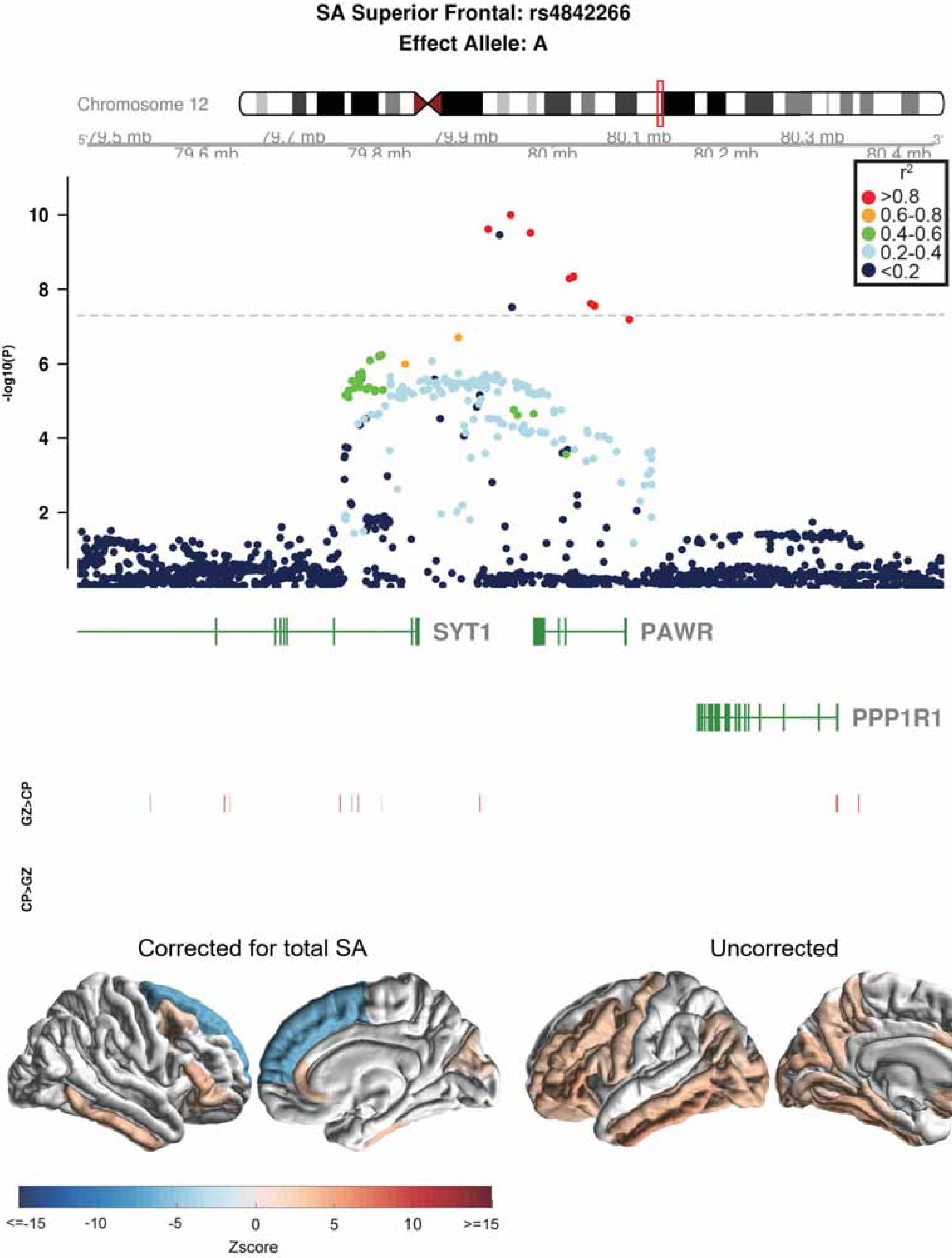

SA Superior Frontal: rs4842266
Effect Allele: A
Chromosome 12
79.5 mb
79.6 mb
79.7 mb
79.8 mb
79.9 mb
80 mb
80.1 mb
80.2 mb
80.3 mb
80.4 mb
r2
>0.8
0.6-0.8
0.4-0.6
0.2-0.4
<0.2
-log10(P)
10
8
6
4
2
SYT1
PAWR
PPP1R1
GZ>CP
CP>GZ
Corrected for total SA
Uncorrected
<=-15
-10
-5
0
5
10
>=15
Zscore

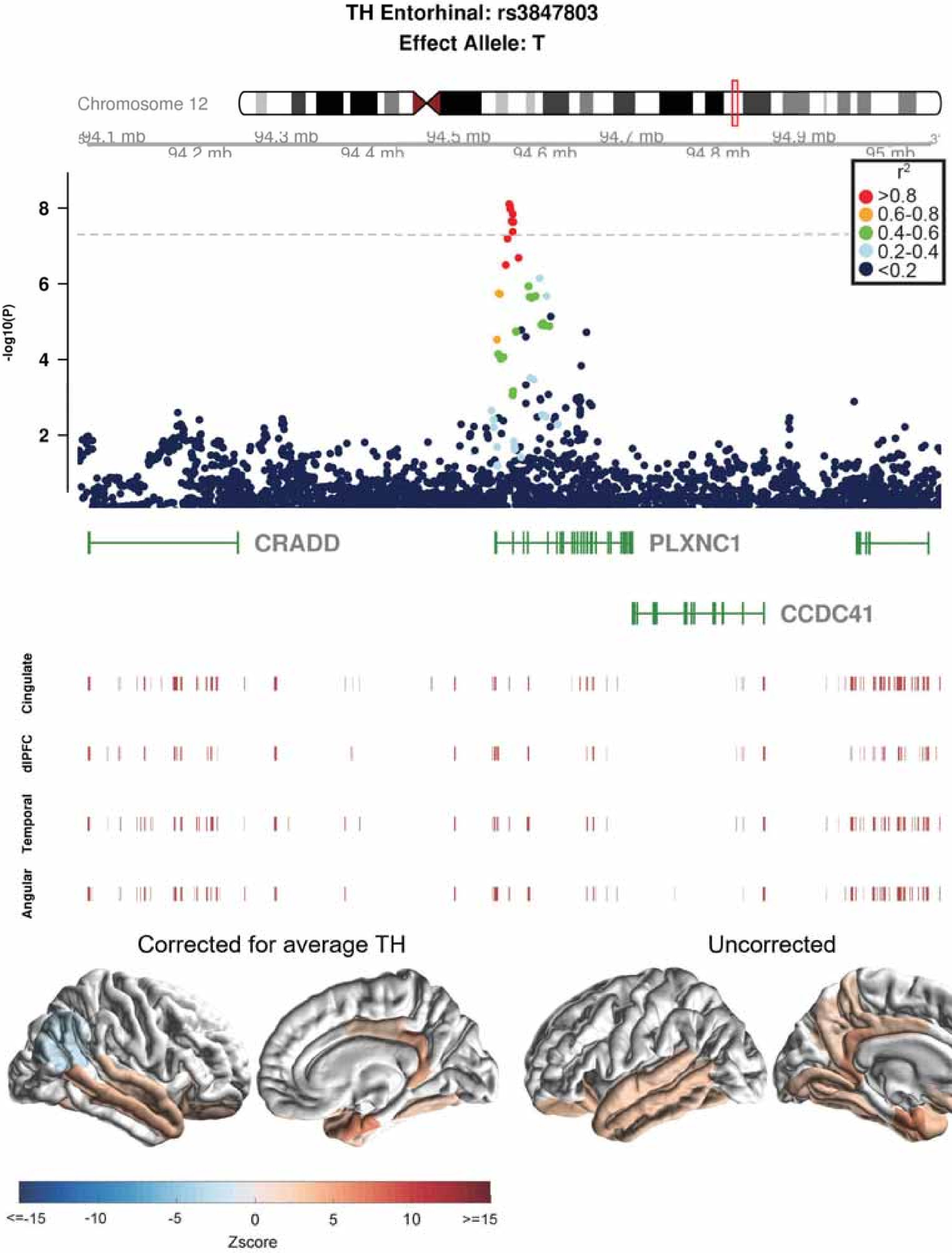
TH Entorhinal: rs3847803
Effect Allele: T
Chromosome 12
94.1 mb
94.2 mb
94.3 mb
94.4 mb
94.5 mb
94.6 mb
94.7 mb
94.8 mb
94.9 mb
95 mb
-log10(P)
8
6
4
2
$r^2$
>0.8
0.6-0.8
0.4-0.6
0.2-0.4
<0.2
CRADD
PLXNC1
CCDC41
Cingulate
dlPFC
Temporal
Angular
Corrected for average TH
Uncorrected
<=-15
-10
-5
0
5
10
>=15
Zscore

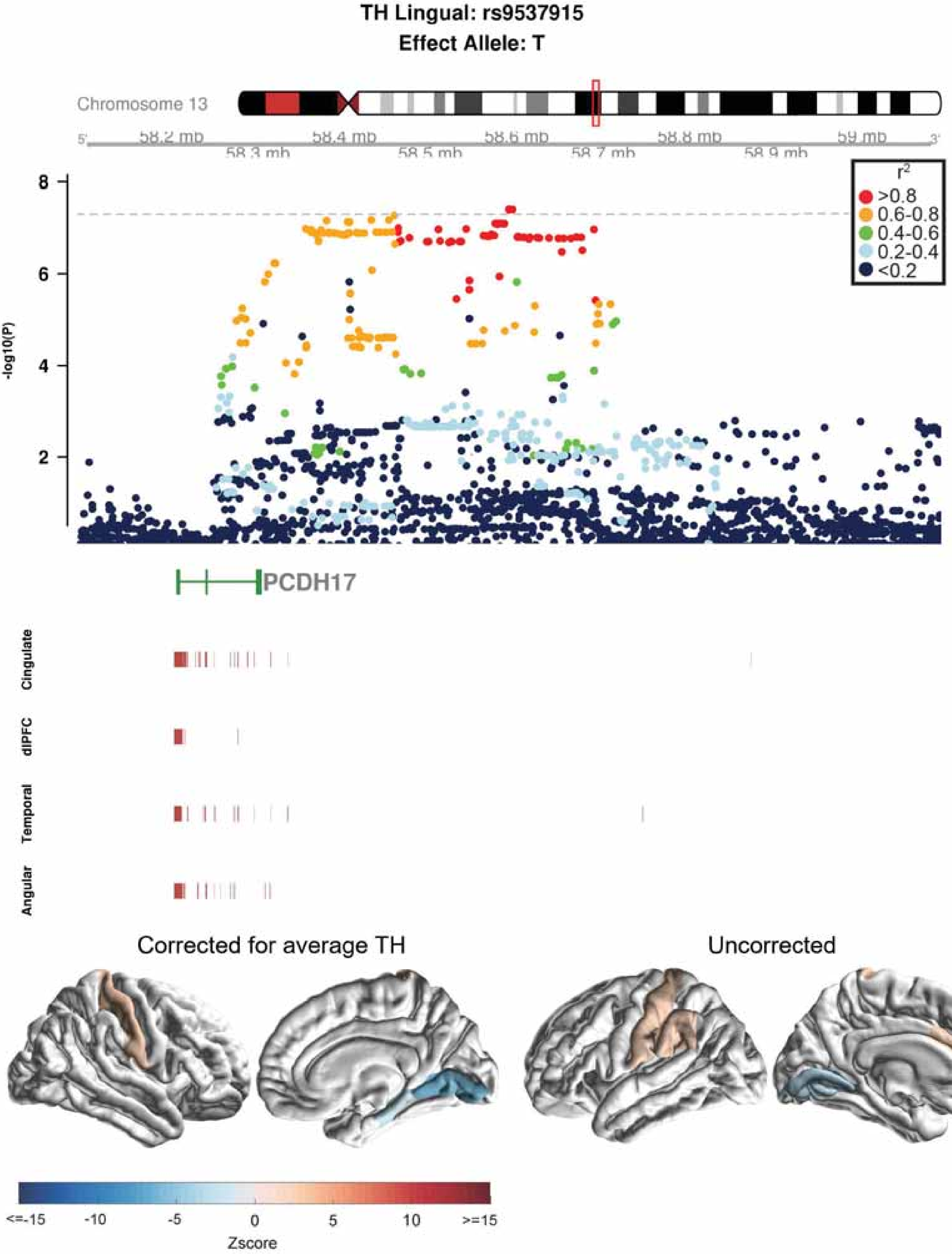

TH Lingual: rs9537915
Effect Allele: T
Chromosome 13
58.2 mb
58.3 mb
58.4 mb
58.5 mb
58.6 mb
58.7 mb
58.8 mb
58.9 mb
59 mb
-log10(P)
r²
>0.8
0.6-0.8
0.4-0.6
0.2-0.4
<0.2
PCDH17
Cingulate
dlPFC
Temporal
Angular
Corrected for average TH
Uncorrected
<=-15
-10
-5
0
5
10
>=15
Zscore

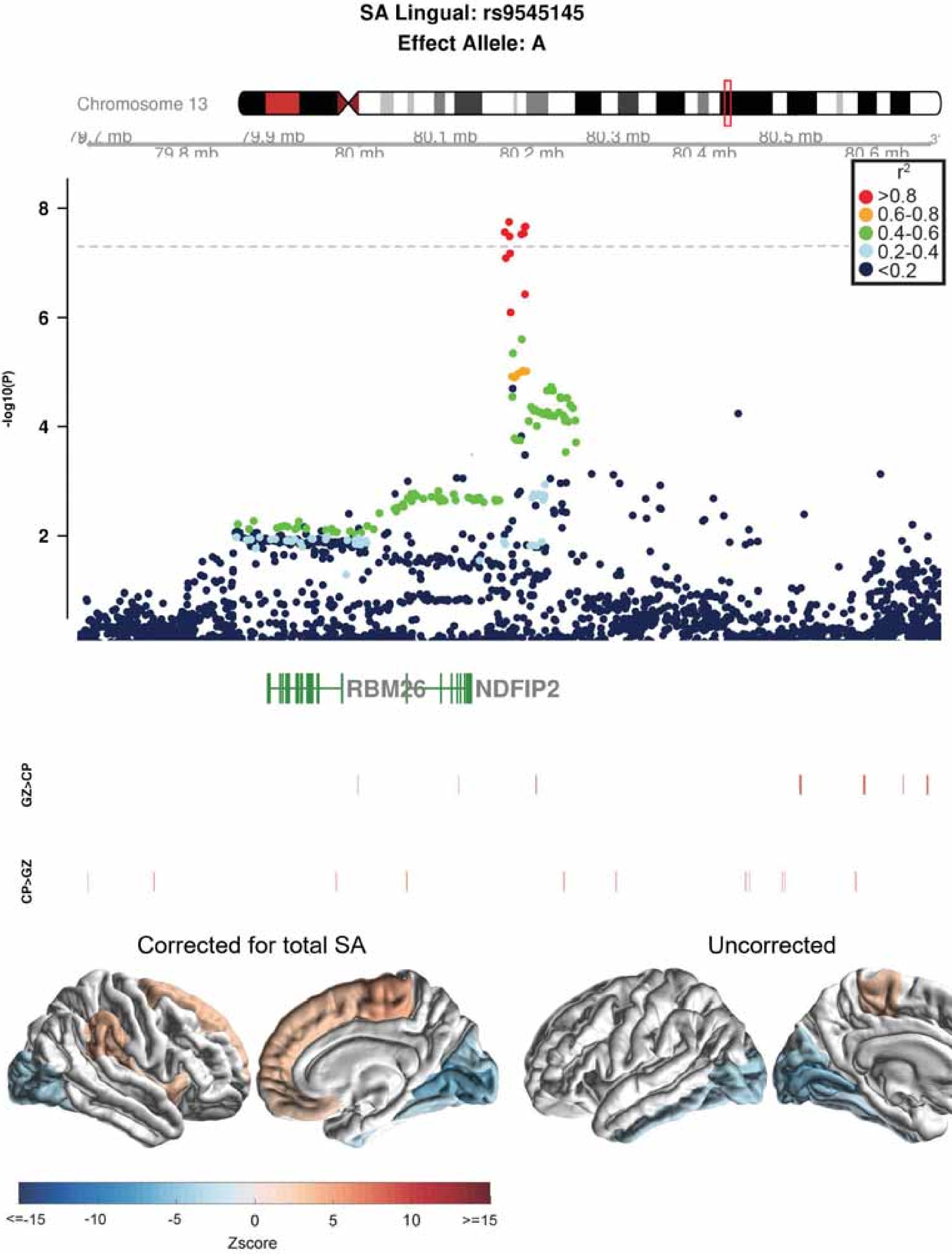

SA Lingual: rs9545145
Effect Allele: A
Chromosome 13
79.7 mb
79.8 mb
79.9 mb
80 mb
80.1 mb
80.2 mb
80.3 mb
80.4 mb
80.5 mb
80.6 mb
r2
>0.8
0.6-0.8
0.4-0.6
0.2-0.4
<0.2
-log10(P)
RBM26
NDFIP2
GZ>CP
CP>GZ
Corrected for total SA
Uncorrected
<=-15
-10
-5
0
5
10
>=15
Zscore

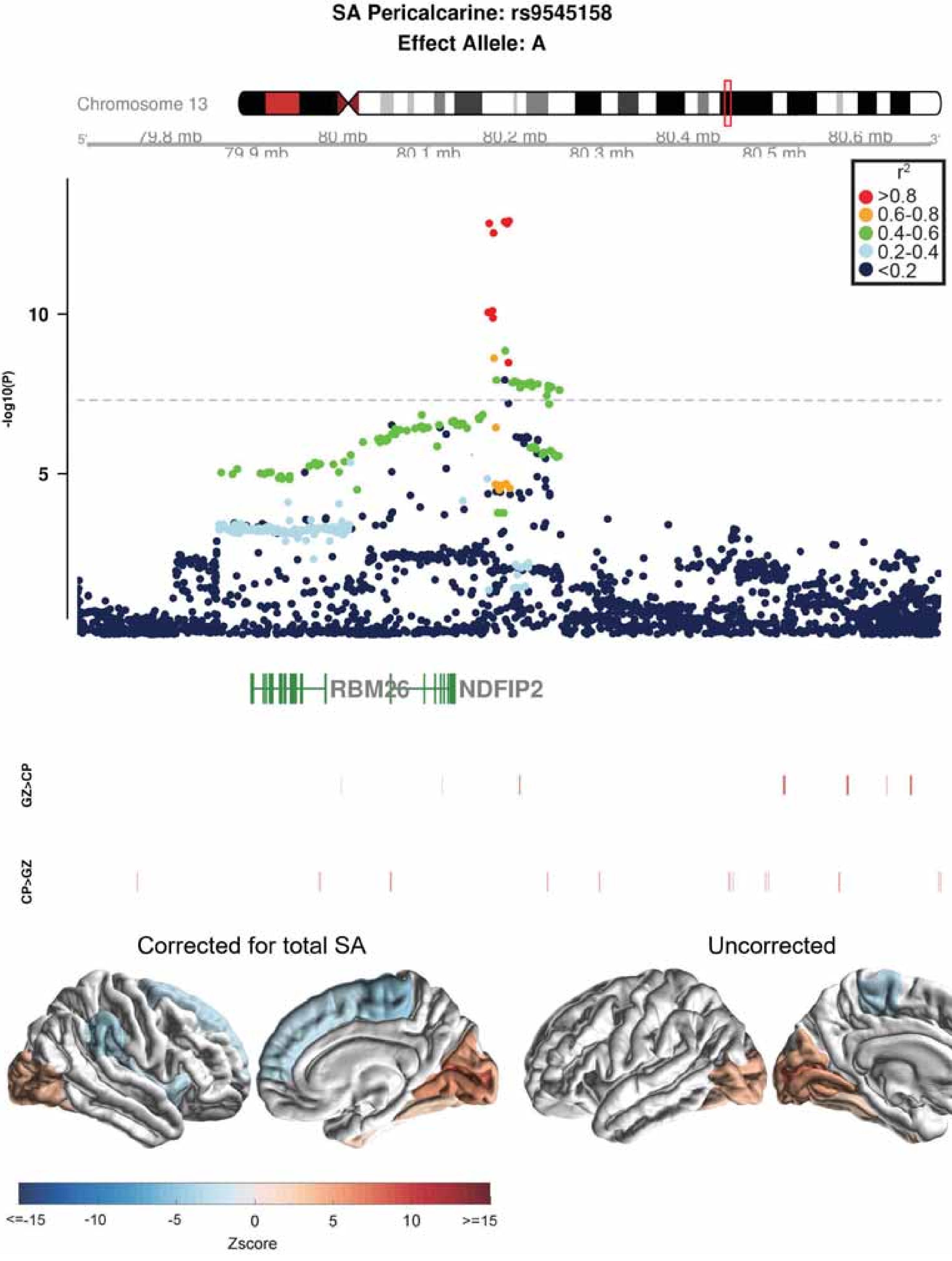

SA Pericalcarine: rs9545158
Effect Allele: A
Chromosome 13
79.8 mb
80 mb
80.2 mb
80.4 mb
80.6 mb
79.9 mb
80.1 mb
80.3 mb
80.5 mb
r2
>0.8
0.6-0.8
0.4-0.6
0.2-0.4
<0.2
-log10(P)
10
5
RBM26
NDFIP2
GZ>CP
CP>GZ
Corrected for total SA
Uncorrected
<=-15
-10
-5
0
5
10
>=15
Zscore

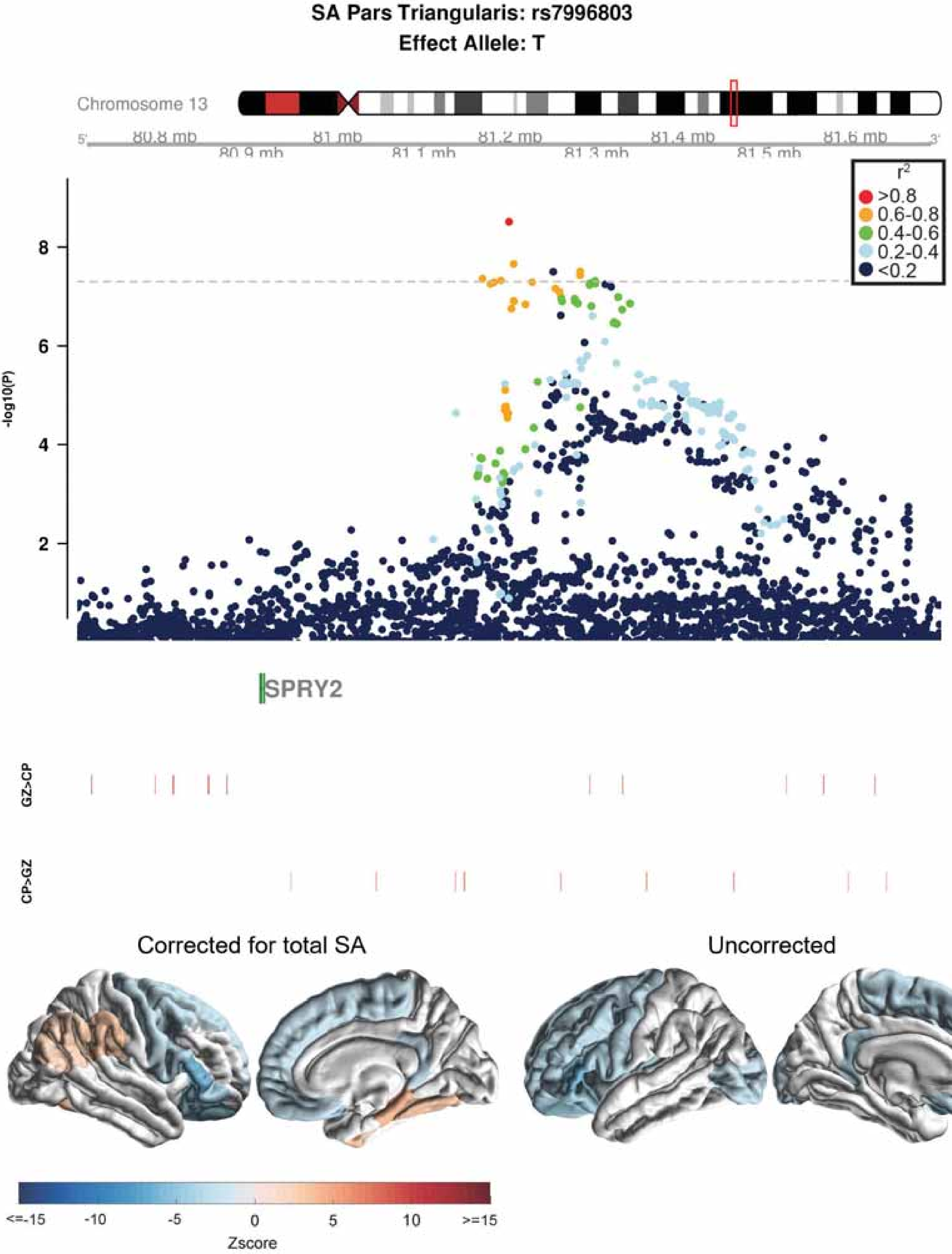
SA Pars Triangularis: rs7996803
Effect Allele: T
Chromosome 13
80.8 mb
80.9 mb
81 mb
81.1 mb
81.2 mb
81.3 mb
81.4 mb
81.5 mb
81.6 mb
r2
>0.8
0.6-0.8
0.4-0.6
0.2-0.4
<0.2
-log10(P)
8
6
4
2
SPRY2
GZ>CP
CP>GZ
Corrected for total SA
Uncorrected
<=-15
-10
-5
0
5
10
>=15
Zscore

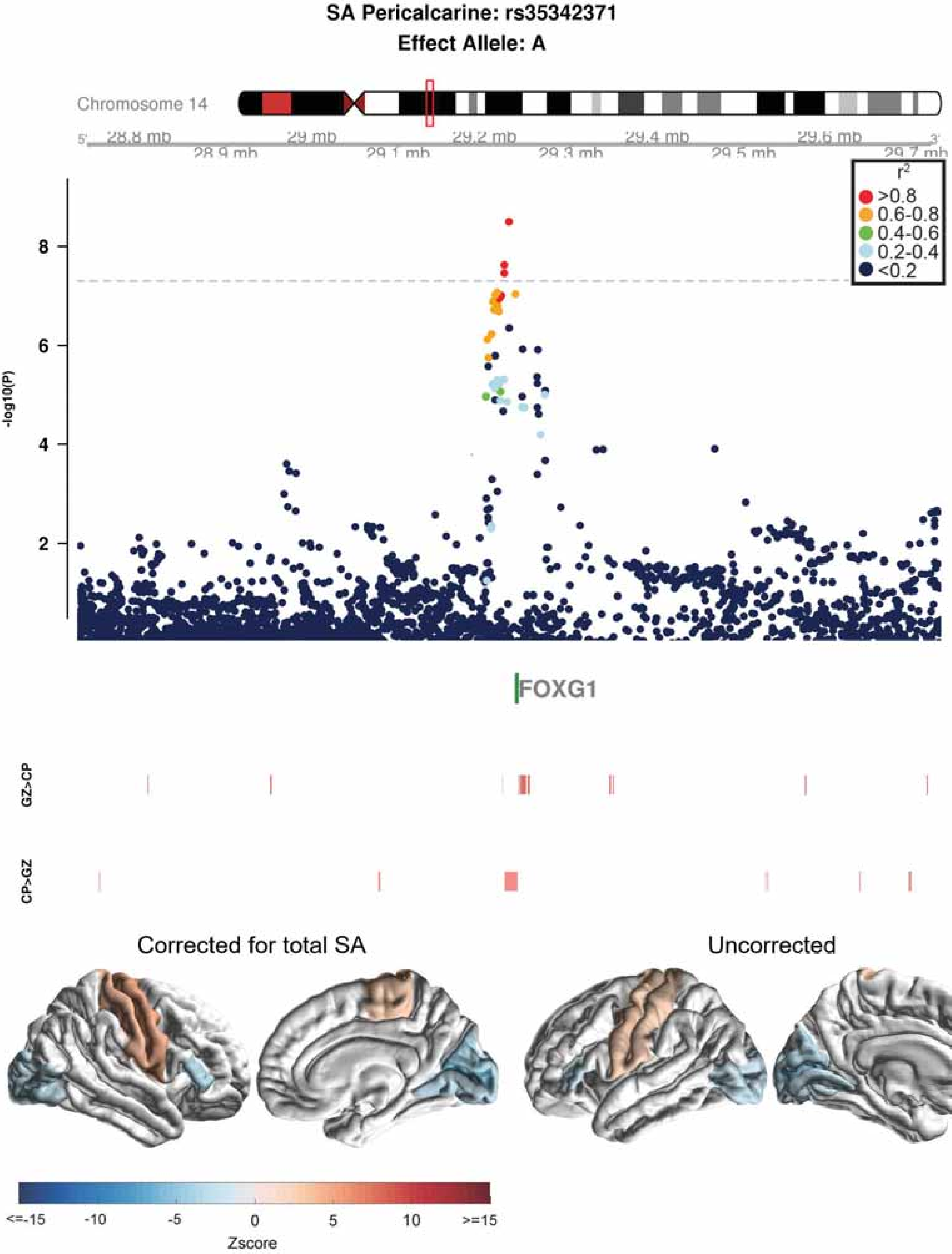

SA Pericalcarine: rs35342371
Effect Allele: A
Chromosome 14
28.8 mb
28.9 mb
29 mb
29.1 mb
29.2 mb
29.3 mb
29.4 mb
29.5 mb
29.6 mb
29.7 mb
r2
>0.8
0.6-0.8
0.4-0.6
0.2-0.4
<0.2
-log10(P)
2
4
6
8
FOXG1
GZ>CP
CP>GZ
Corrected for total SA
Uncorrected
<=-15
-10
-5
0
5
10
>=15
Zscore

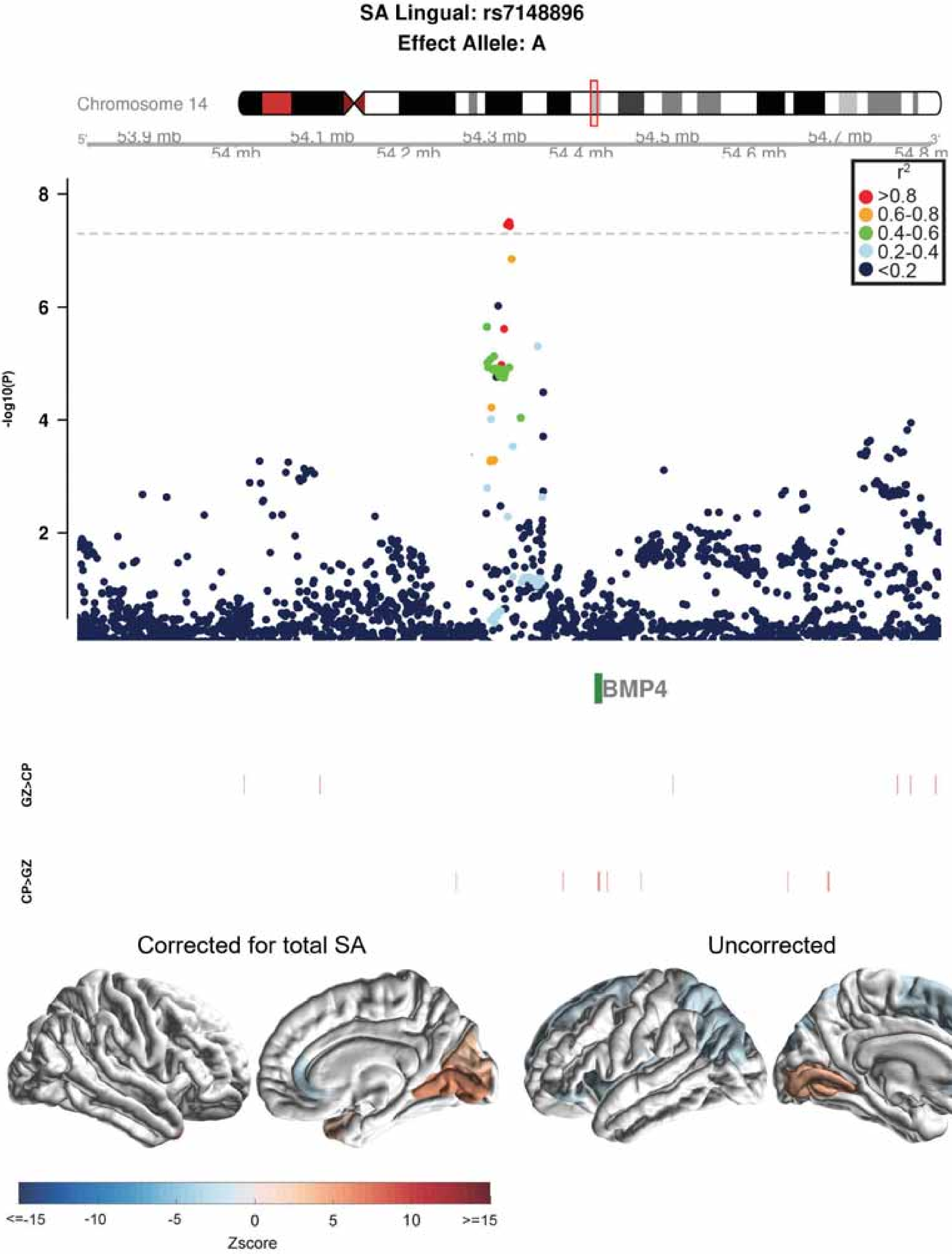
SA Lingual: rs7148896
Effect Allele: A
Chromosome 14
53.9 mb
54 mb
54.1 mb
54.2 mb
54.3 mb
54.4 mb
54.5 mb
54.6 mb
54.7 mb
54.8 m
r2
>0.8
0.6-0.8
0.4-0.6
0.2-0.4
<0.2
-log10(P)
8
6
4
2
BMP4
GZ>CP
CP>GZ
Corrected for total SA
Uncorrected
<=-15
-10
-5
0
5
10
>=15
Zscore

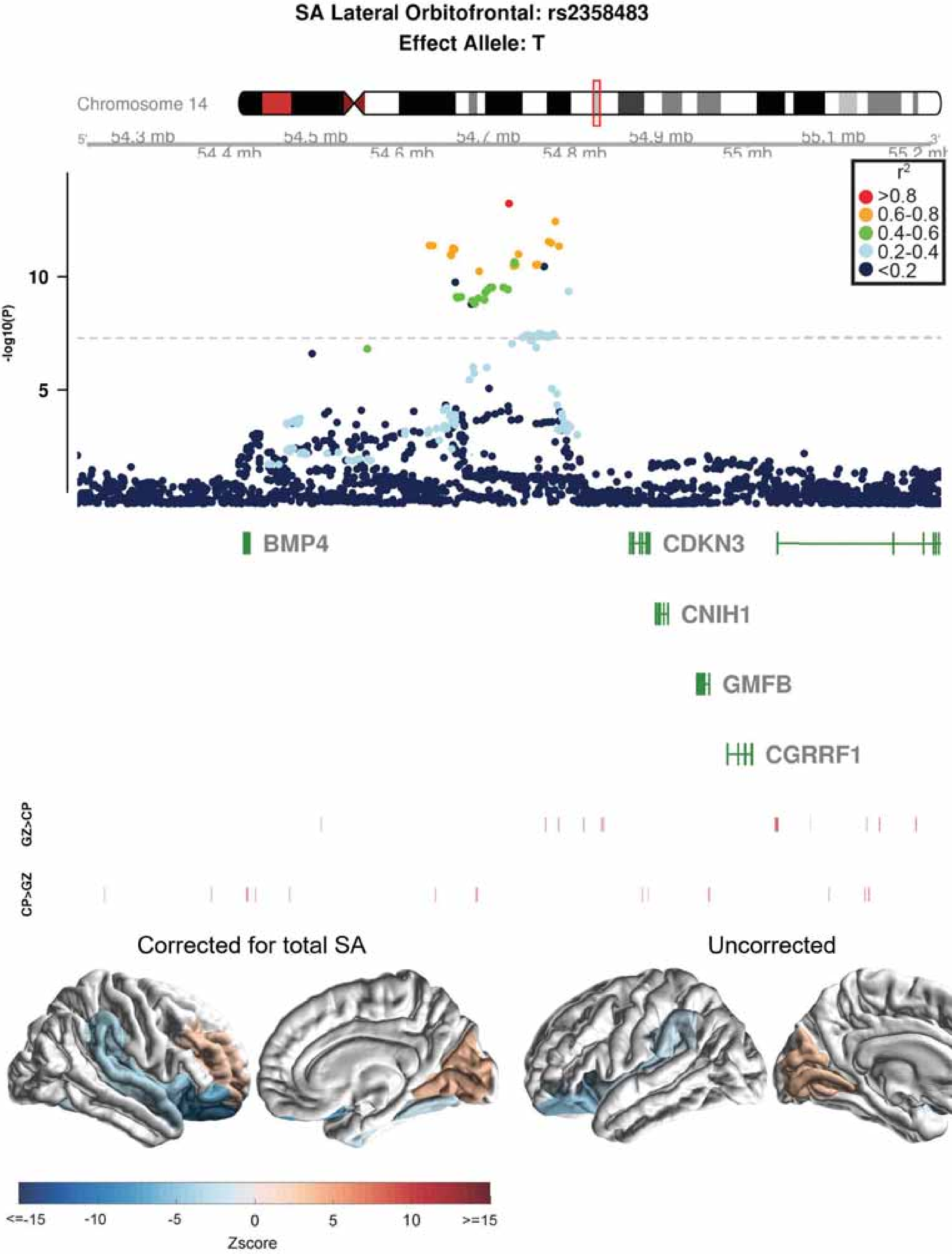
SA Lateral Orbitofrontal: rs2358483
Effect Allele: T
Chromosome 14
54.3 mb
54.4 mb
54.5 mb
54.6 mb
54.7 mb
54.8 mb
54.9 mb
55 mb
55.1 mb
-log10(P)
10
5
r2
>0.8
0.6-0.8
0.4-0.6
0.2-0.4
<0.2
BMP4
CDKN3
CNIH1
GMFB
CGRRF1
GZ>CP
CP>GZ
Corrected for total SA
Uncorrected
<=-15
-10
-5
0
5
10
>=15
Zscore

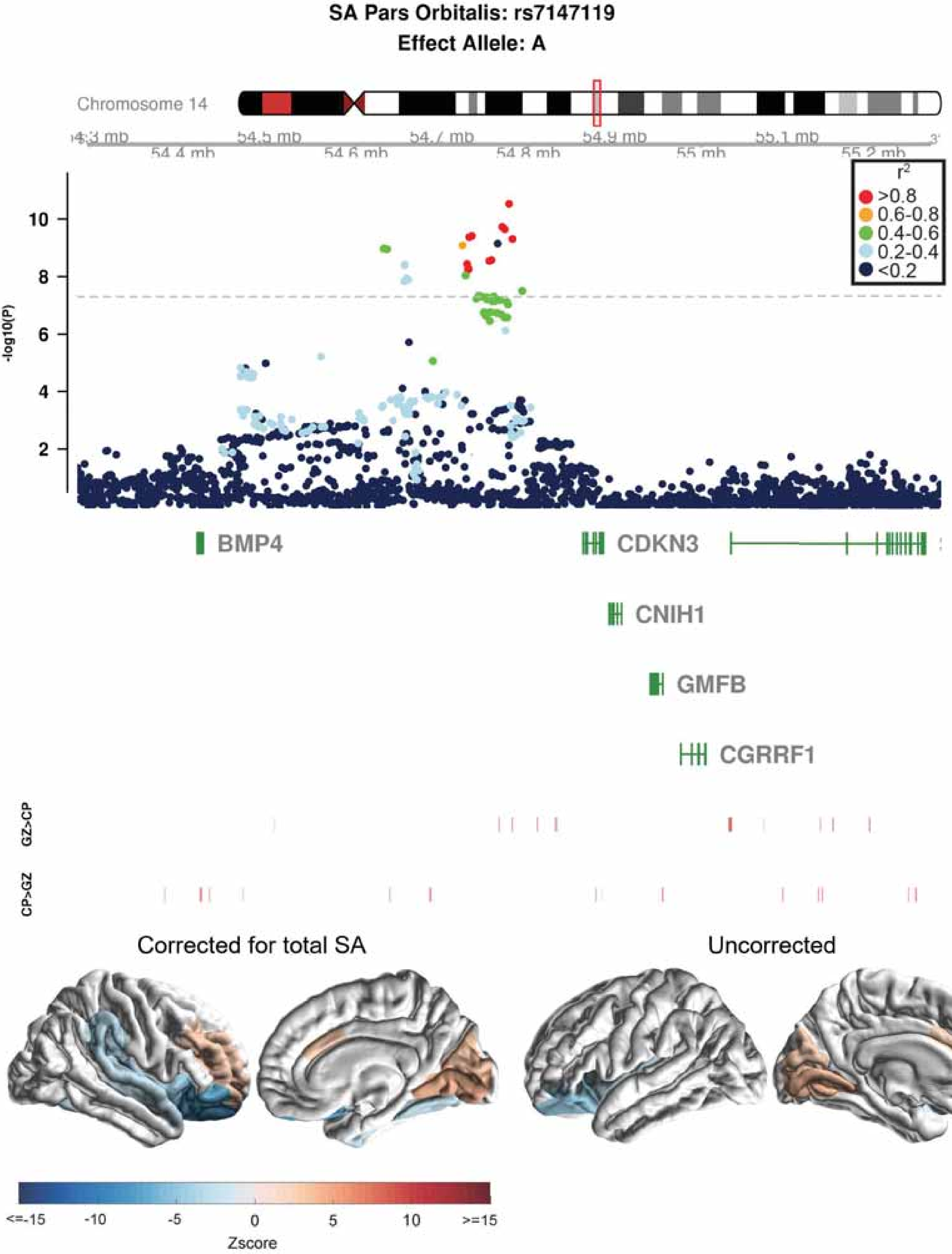
SA Pars Orbitalis: rs7147119
Effect Allele: A
Chromosome 14
54.4 mb
54.5 mb
54.6 mb
54.7 mb
54.8 mb
54.9 mb
55 mb
55.1 mb
55.2 mb
r2
>0.8
0.6-0.8
0.4-0.6
0.2-0.4
<0.2
-log10(P)
10
8
6
4
2
BMP4
CDKN3
CNIH1
GMFB
CGRRF1
GZ>CP
CP>GZ
Corrected for total SA
Uncorrected
<=-15
-10
-5
0
5
10
>=15
Zscore

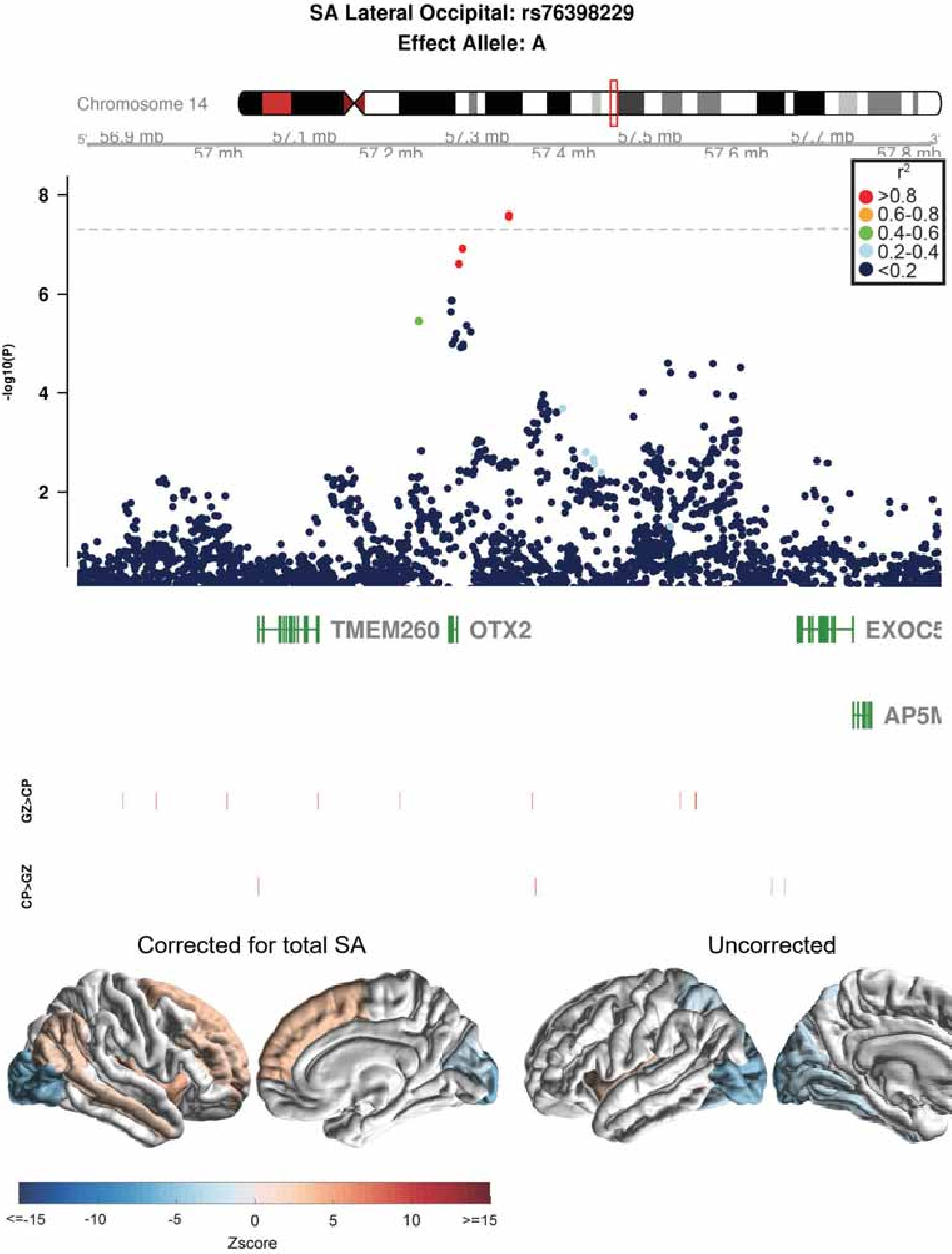
SA Lateral Occipital: rs76398229
Effect Allele: A
Chromosome 14
56.9 mb
57 mb
57.1 mb
57.2 mb
57.3 mb
57.4 mb
57.5 mb
57.6 mb
57.7 mb
57.8 mb
r2
>0.8
0.6-0.8
0.4-0.6
0.2-0.4
<0.2
-log10(P)
8
6
4
2
TMEM260
OTX2
EXOC5
AP5M
GZ>CP
CP>GZ
Corrected for total SA
Uncorrected
<=-15
-10
-5
0
5
10
>=15
Zscore

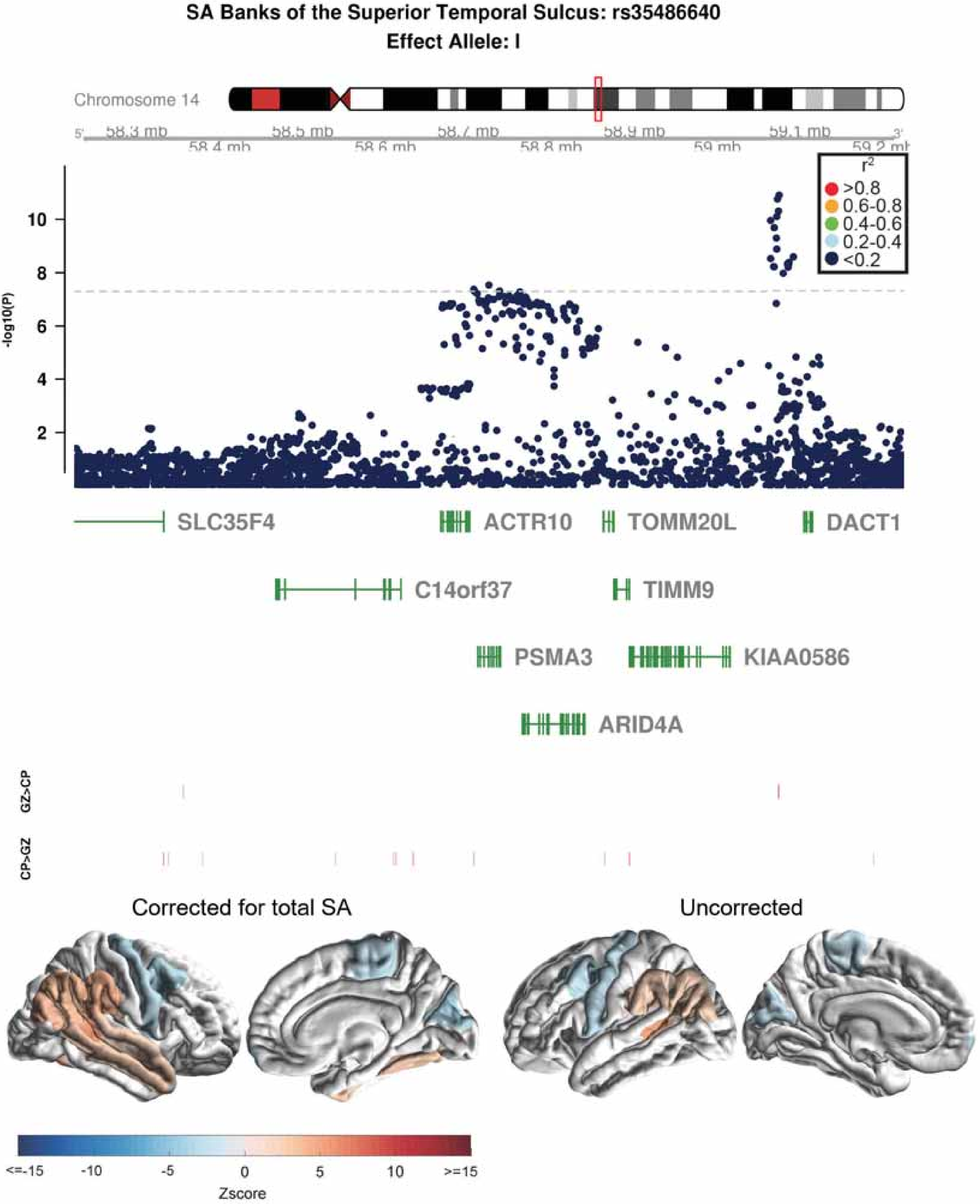
SA Banks of the Superior Temporal Sulcus: rs35486640
Effect Allele: I
Chromosome 14
58.3 mb
58.4 mb
58.5 mb
58.6 mb
58.7 mb
58.8 mb
58.9 mb
59 mb
59.1 mb
59.2 mb
r2
>0.8
0.6-0.8
0.4-0.6
0.2-0.4
<0.2
-log10(P)
SLC35F4
ACTR10
TOMM20L
DACT1
C14orf37
TIMM9
PSMA3
KIAA0586
ARID4A
GZ>CP
CP>GZ
Corrected for total SA
Uncorrected
<=-15
-10
-5
0
5
10
>=15
Zscore

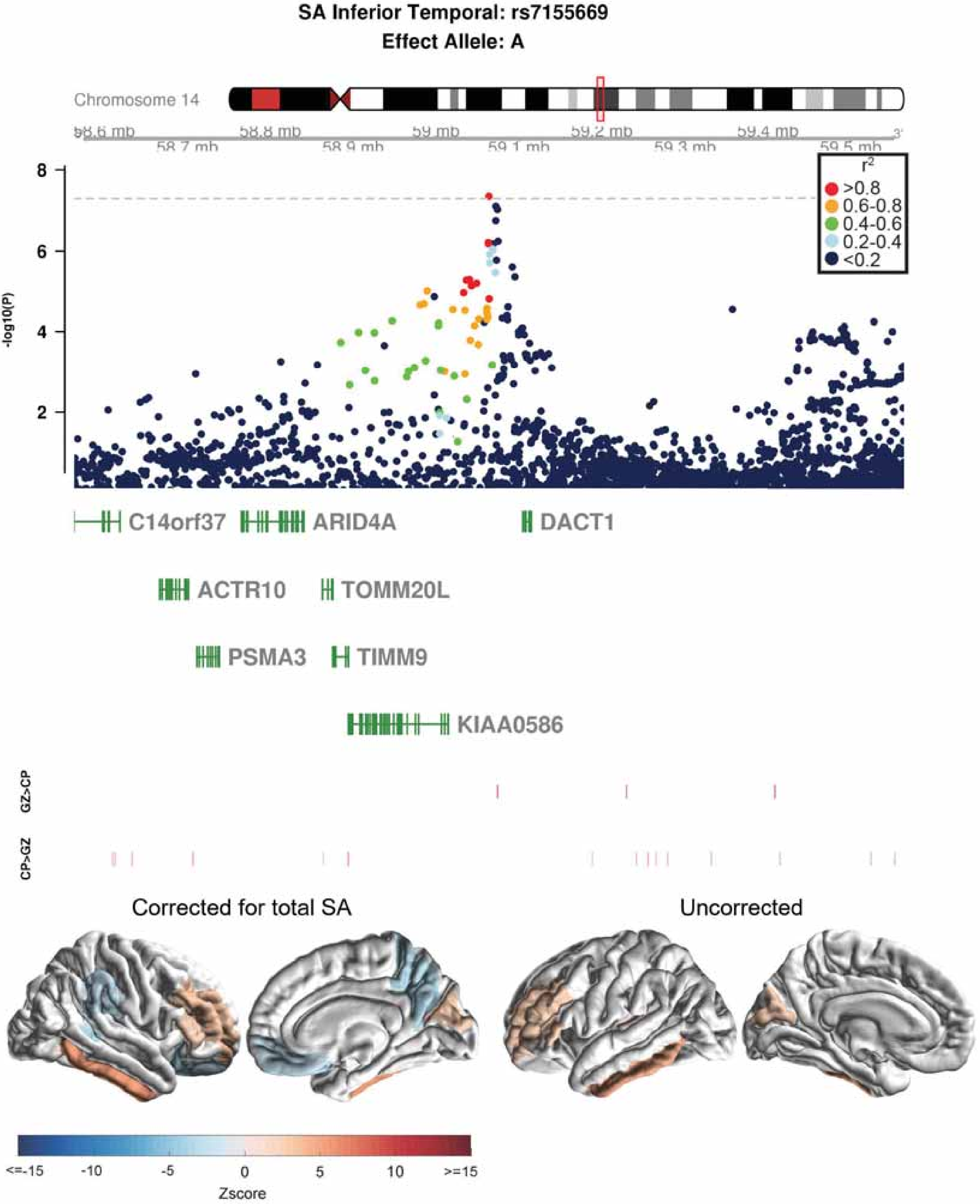
SA Inferior Temporal: rs7155669
Effect Allele: A
Chromosome 14
58.6 mb
58.7 mb
58.8 mb
58.9 mb
59 mb
59.1 mb
59.2 mb
59.3 mb
59.4 mb
59.5 mb
-log10(P)
r2
>0.8
0.6-0.8
0.4-0.6
0.2-0.4
<0.2
C14orf37
ARID4A
DACT1
ACTR10
TOMM20L
PSMA3
TIMM9
KIAA0586
GZ>CP
CP>GZ
Corrected for total SA
Uncorrected
<=-15
-10
-5
0
5
10
>=15
Zscore

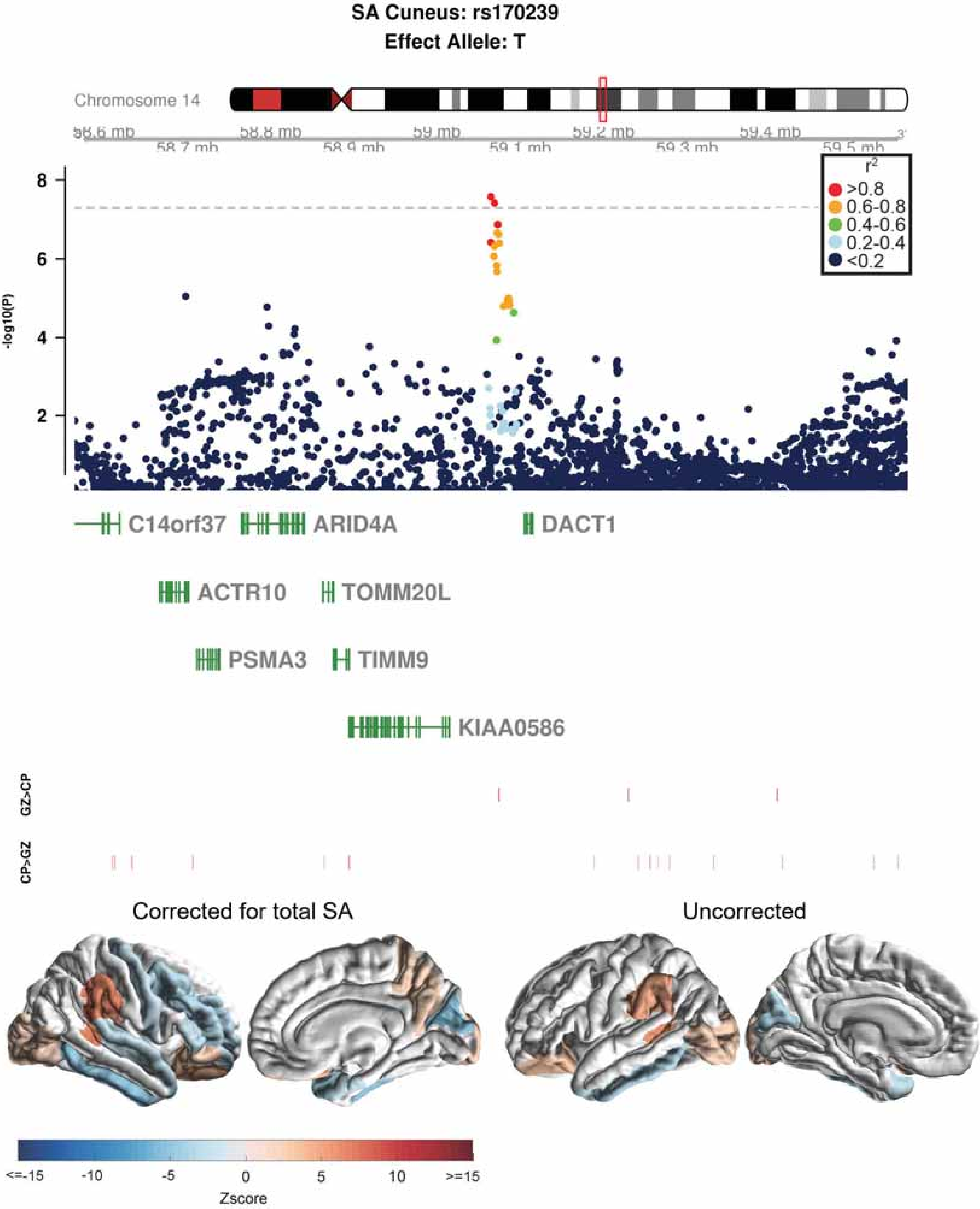

SA Cuneus: rs170239
Effect Allele: T
Chromosome 14
58.6 mb
58.7 mb
58.8 mb
58.9 mb
59 mb
59.1 mb
59.2 mb
59.3 mb
59.4 mb
59.5 mb
-log10(P)
r2
>0.8
0.6-0.8
0.4-0.6
0.2-0.4
<0.2
C14orf37
ARID4A
DACT1
ACTR10
TOMM20L
PSMA3
TIMM9
KIAA0586
GZ>CP
CP>GZ
Corrected for total SA
Uncorrected
<=-15
-10
-5
0
5
10
>=15
Zscore

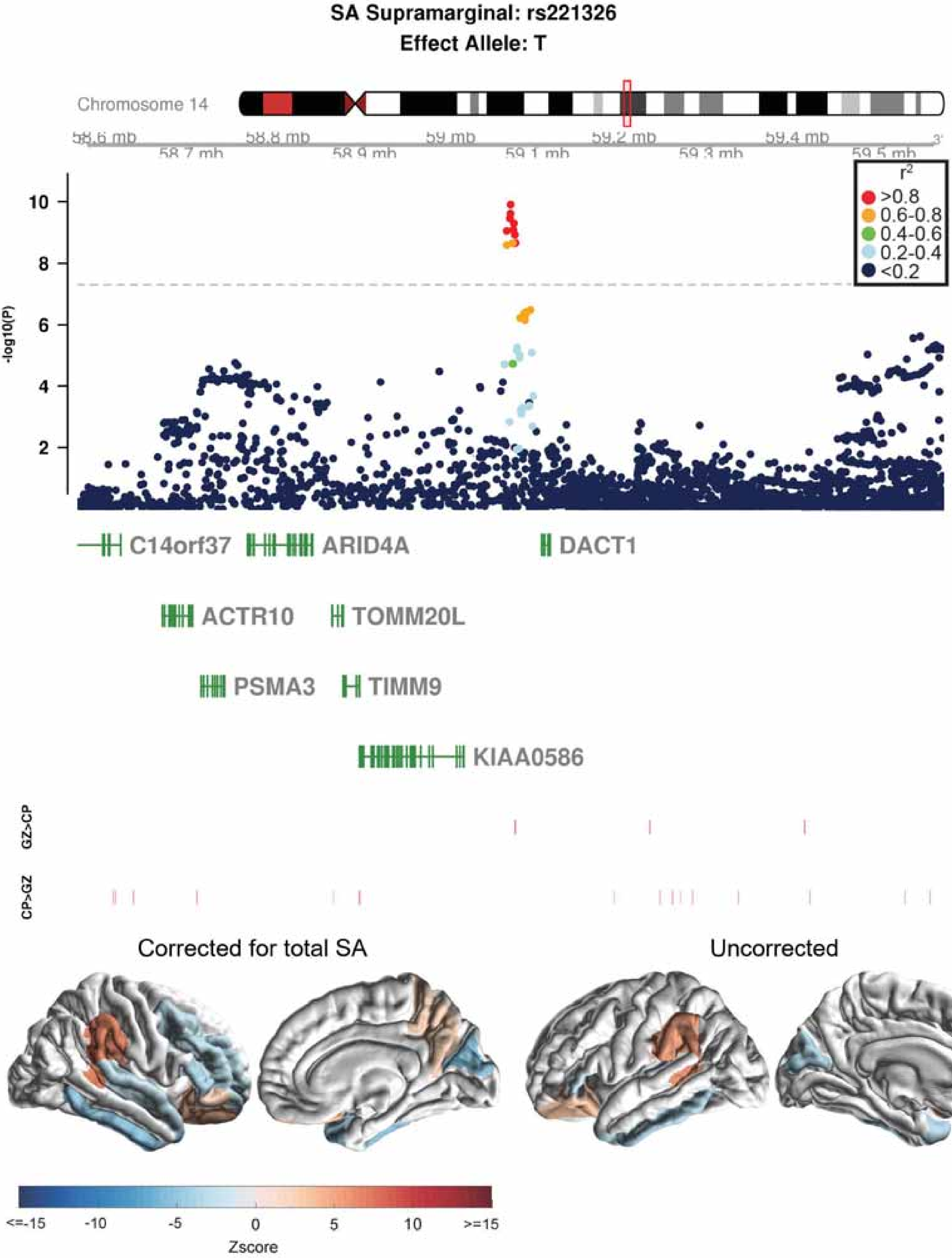

SA Supramarginal: rs221326
Effect Allele: T
Chromosome 14
58.6 mb
58.7 mb
58.8 mb
58.9 mb
59 mb
59.1 mb
59.2 mb
59.3 mb
59.4 mb
59.5 mb
-log10(P)
r2
>0.8
0.6-0.8
0.4-0.6
0.2-0.4
<0.2
C14orf37
ARID4A
DACT1
ACTR10
TOMM20L
PSMA3
TIMM9
KIAA0586
GZ>CP
CP>GZ
Corrected for total SA
Uncorrected
<=-15
-10
-5
0
5
10
>=15
Zscore

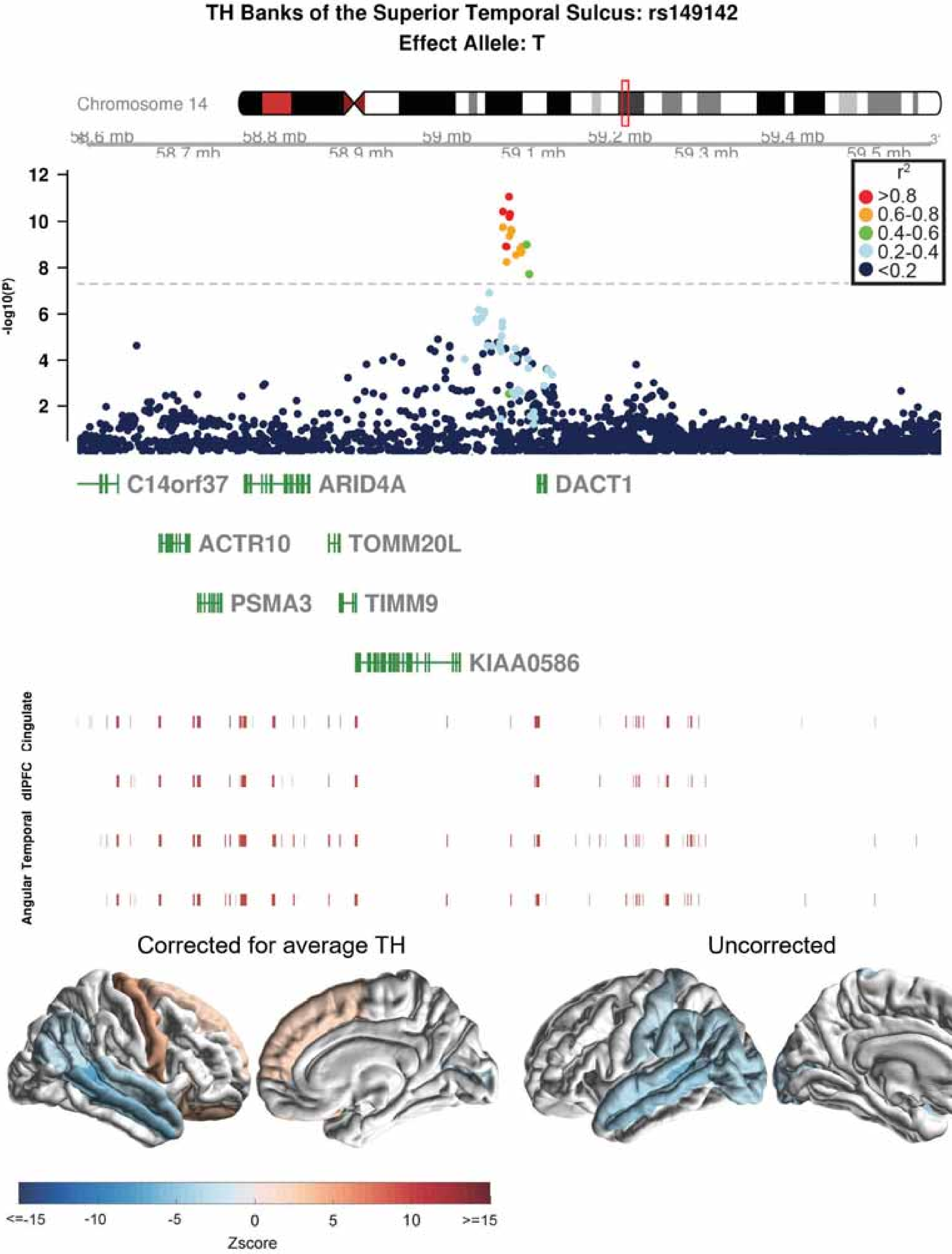
TH Banks of the Superior Temporal Sulcus: rs149142
Effect Allele: T
Chromosome 14
58.6 mb
58.7 mb
58.8 mb
58.9 mb
59 mb
59.1 mb
59.2 mb
59.3 mb
59.4 mb
59.5 mb
-log10(P)
r2
>0.8
0.6-0.8
0.4-0.6
0.2-0.4
<0.2
C14orf37
ARID4A
DACT1
ACTR10
TOMM20L
PSMA3
TIMM9
KIAA0586
Cingulate
dlPFC
Temporal
Angular
Corrected for average TH
Uncorrected
<=-15
-10
-5
0
5
10
>=15
Zscore

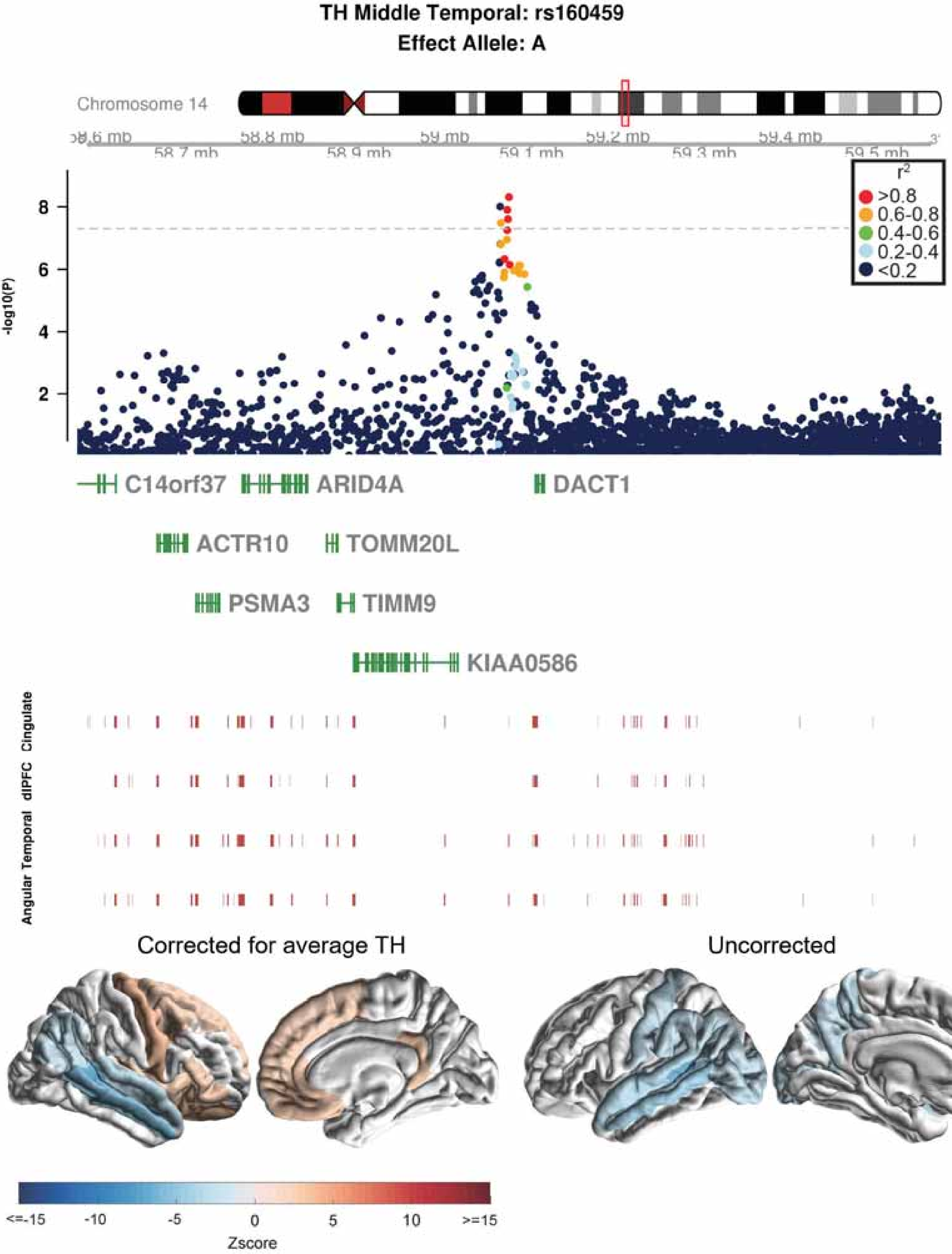

TH Middle Temporal: rs160459
Effect Allele: A
Chromosome 14
58.6 mb
58.7 mb
58.8 mb
58.9 mb
59 mb
59.1 mb
59.2 mb
59.3 mb
59.4 mb
59.5 mb
-log10(P)
r2
>0.8
0.6-0.8
0.4-0.6
0.2-0.4
<0.2
C14orf37
ARID4A
DACT1
ACTR10
TOMM20L
PSMA3
TIMM9
KIAA0586
Cingulate
dlPFC
Temporal
Angular
Corrected for average TH
Uncorrected
<=-15
-10
-5
0
5
10
>=15
Zscore

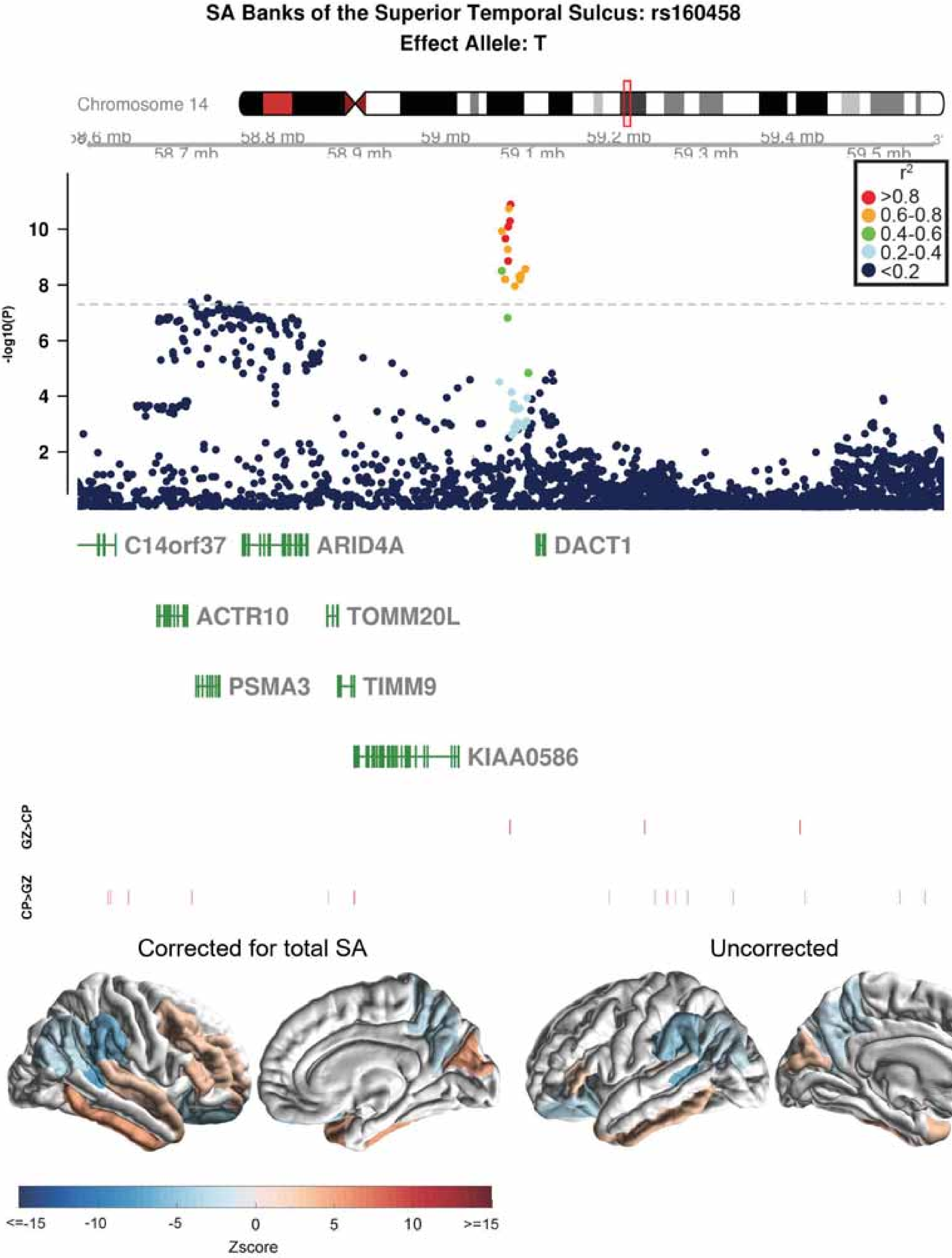
SA Banks of the Superior Temporal Sulcus: rs160458
Effect Allele: T
Chromosome 14
58.8 mb
59 mb
59.2 mb
59.4 mb
58.7 mb
58.9 mb
59.1 mb
59.3 mb
59.5 mb
r2
>0.8
0.6-0.8
0.4-0.6
0.2-0.4
<0.2
-log10(P)
2
4
6
8
10
C14orf37
ARID4A
DACT1
ACTR10
TOMM20L
PSMA3
TIMM9
KIAA0586
GZ>CP
CP>GZ
Corrected for total SA
Uncorrected
<=-15
-10
-5
0
5
10
>=15
Zscore

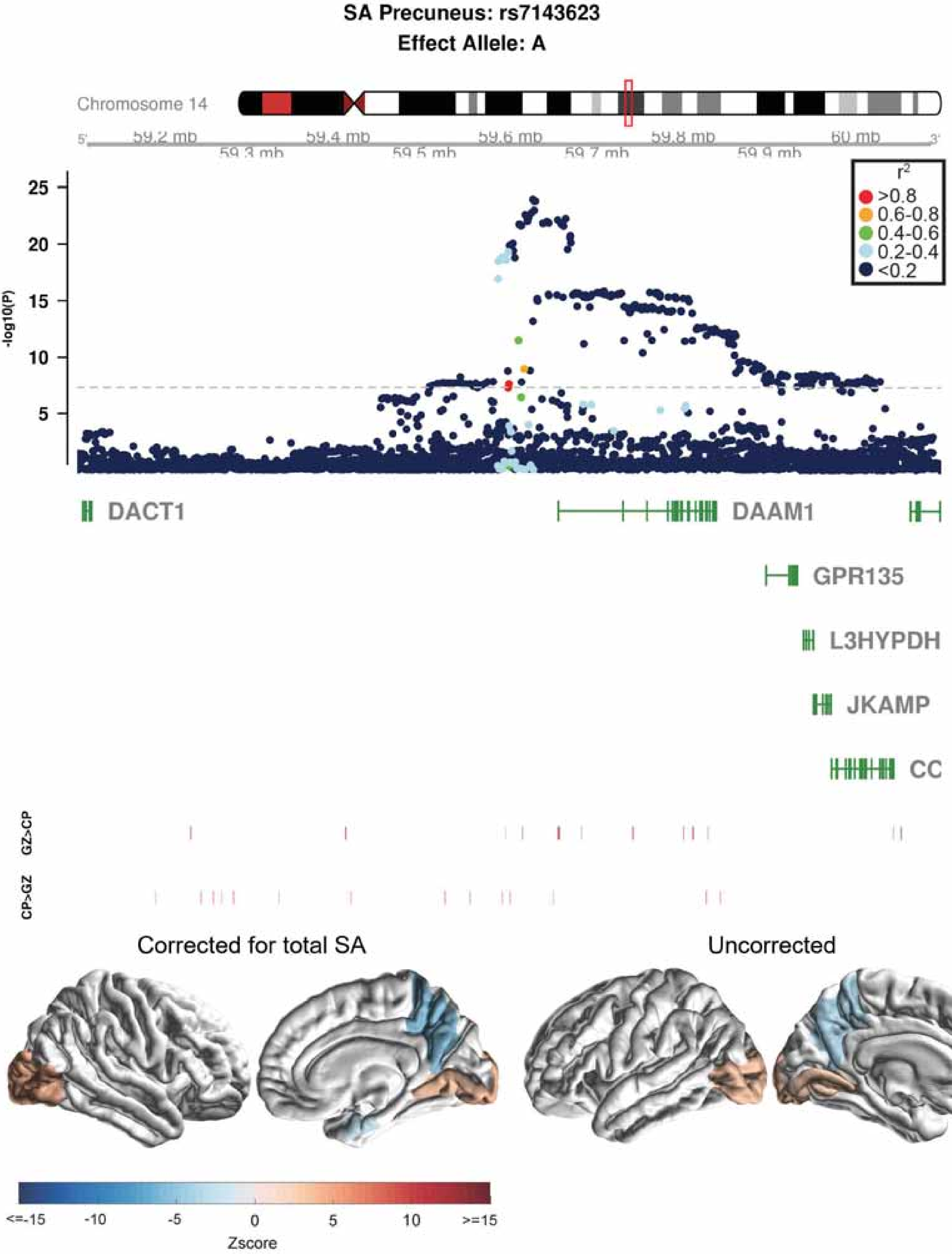
SA Precuneus: rs7143623
Effect Allele: A
Chromosome 14
59.2 mb
59.3 mb
59.4 mb
59.5 mb
59.6 mb
59.7 mb
59.8 mb
59.9 mb
60 mb
-log10(P)
r2
>0.8
0.6-0.8
0.4-0.6
0.2-0.4
<0.2
DACT1
DAAM1
GPR135
L3HYPDH
JKAMP
CC
GZ>CP
CP>GZ
Corrected for total SA
Uncorrected
<=-15
-10
-5
0
5
10
>=15
Zscore

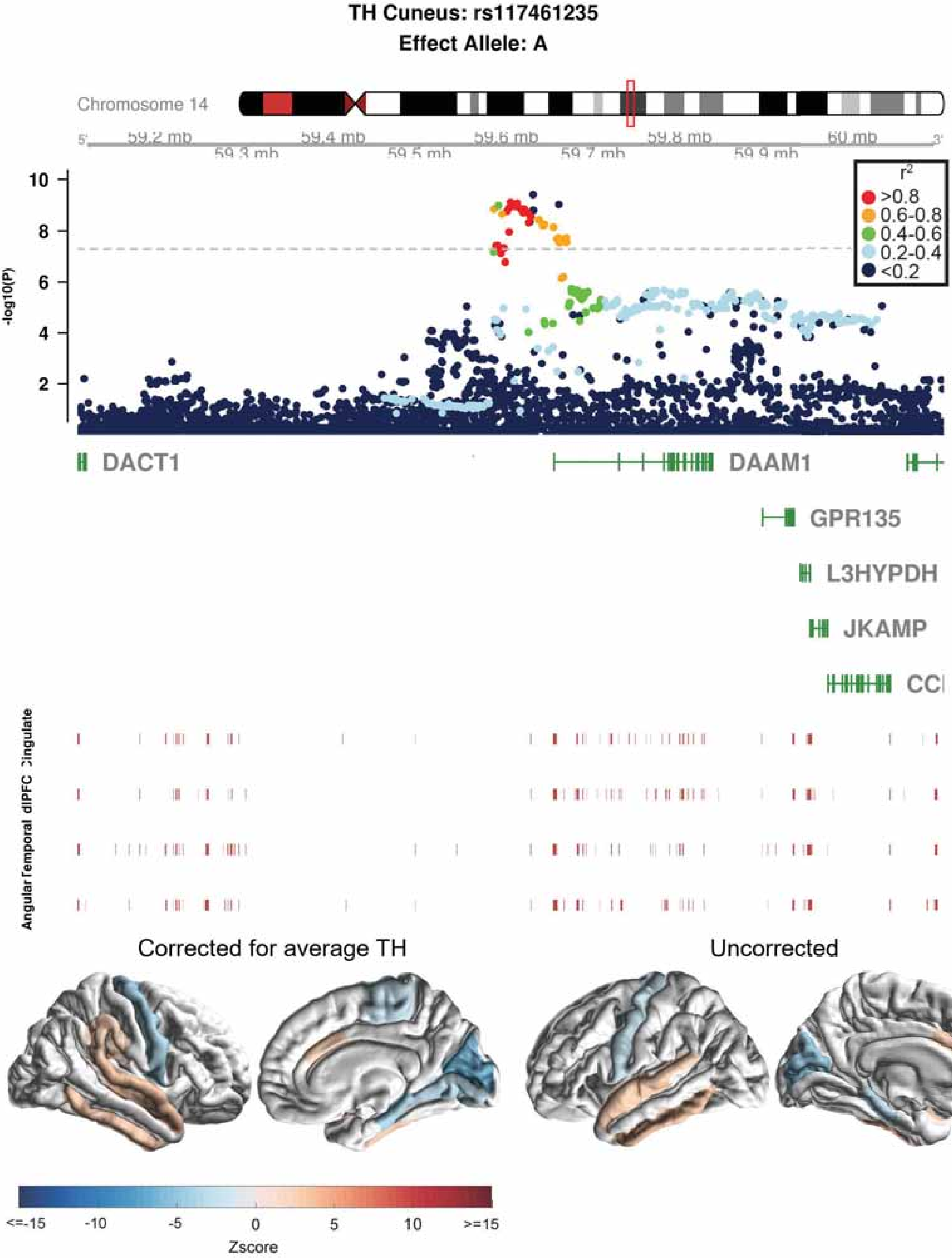
TH Cuneus: rs117461235
Effect Allele: A
Chromosome 14
59.2 mb
59.3 mb
59.4 mb
59.5 mb
59.6 mb
59.7 mb
59.8 mb
59.9 mb
60 mb
-log10(P)
10
8
6
4
2
r2
>0.8
0.6-0.8
0.4-0.6
0.2-0.4
<0.2
DACT1
DAAM1
GPR135
L3HYPDH
JKAMP
Angular
dlPFC
Corrected for average TH
Uncorrected
<=-15
-10
-5
0
5
10
>=15
Zscore

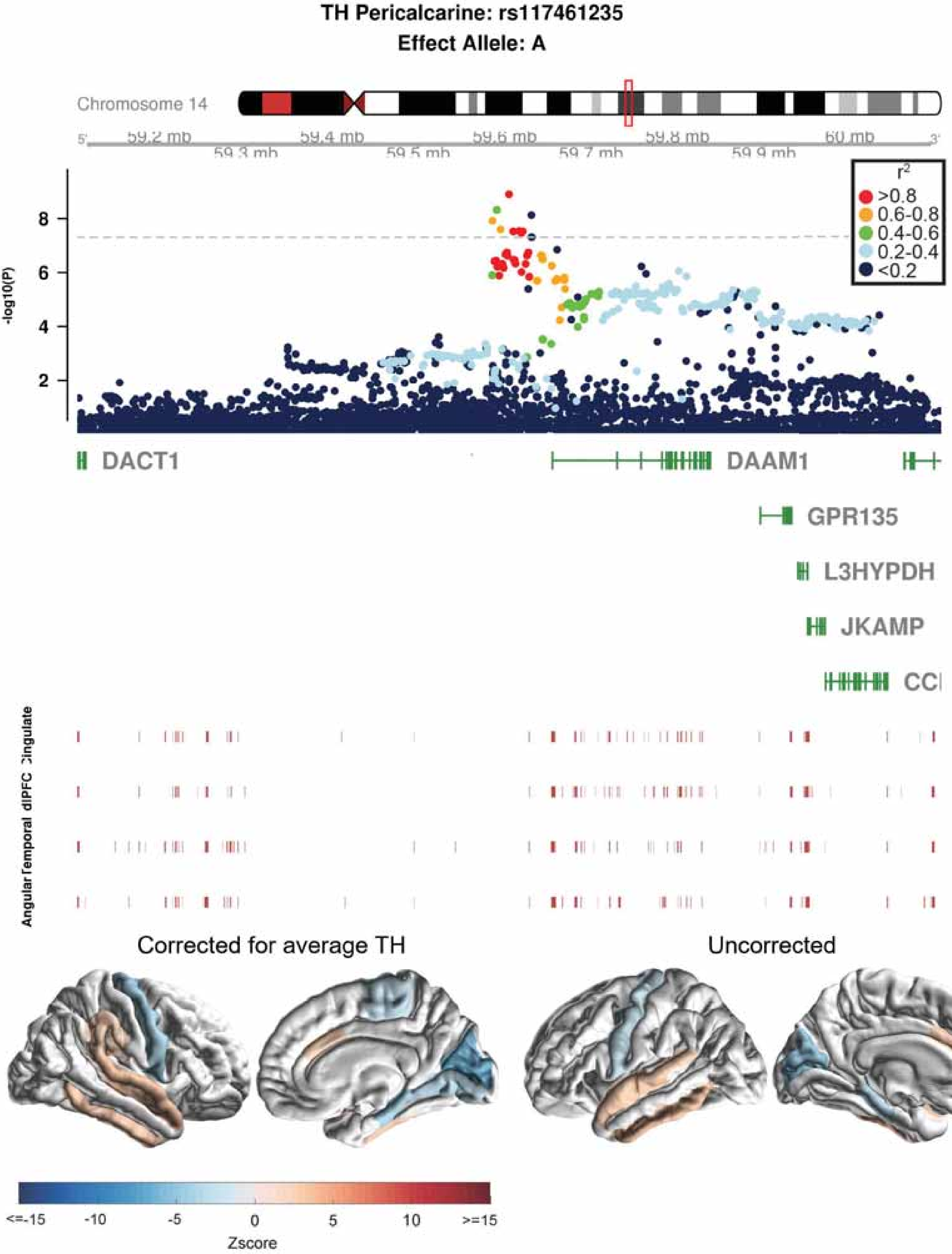

TH Pericalcarine: rs117461235
Effect Allele: A
Chromosome 14
59.2 mb
59.3 mb
59.4 mb
59.5 mb
59.6 mb
59.7 mb
59.8 mb
59.9 mb
60 mb
r²
>0.8
0.6-0.8
0.4-0.6
0.2-0.4
<0.2
-log10(P)
8
6
4
2
DACT1
DAAM1
GPR135
L3HYPDH
JKAMP
CCI
Angular
Temporal
dlPFC
Cingulate
Corrected for average TH
Uncorrected
<=-15
-10
-5
0
5
10
>=15
Zscore

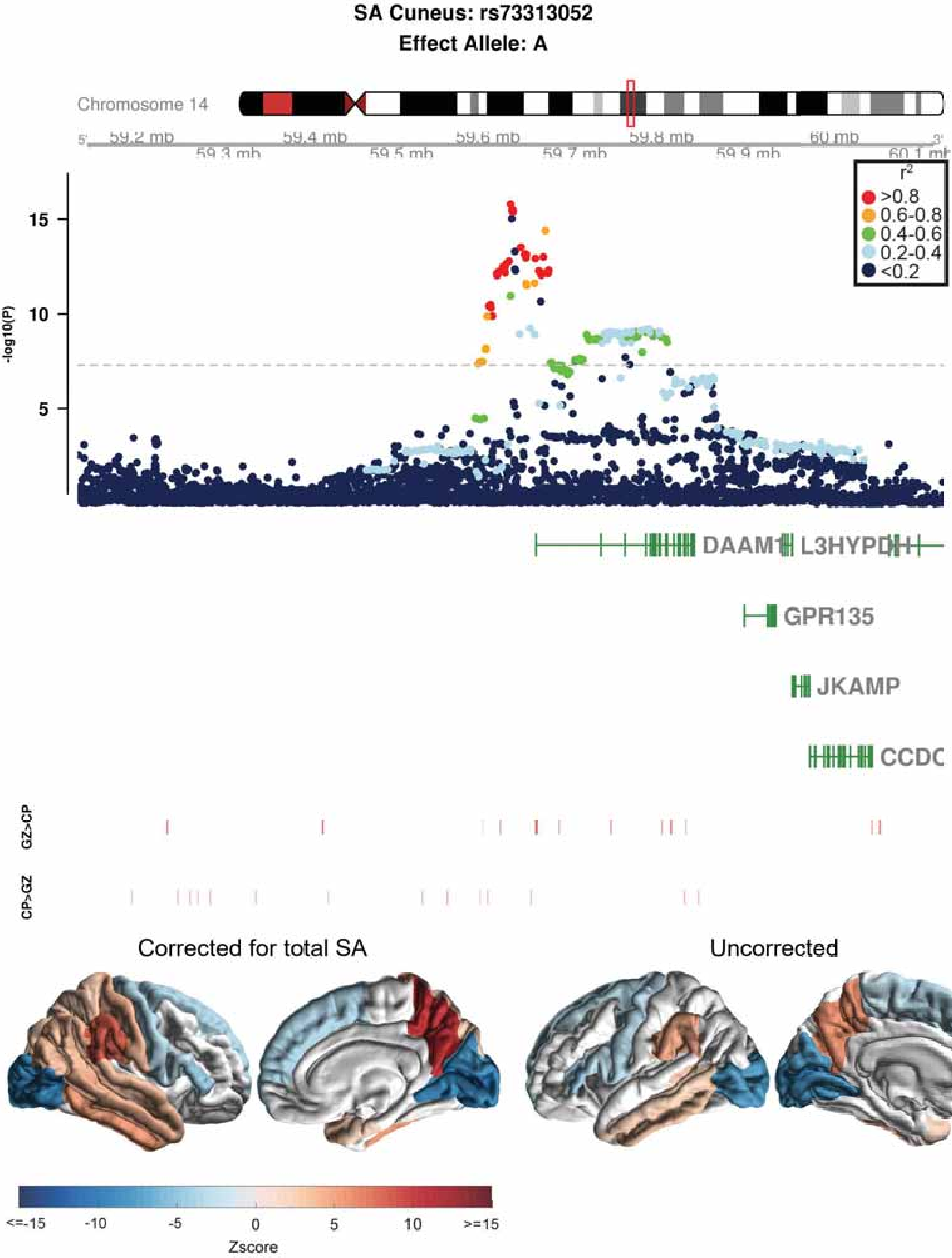
SA Cuneus: rs73313052
Effect Allele: A
Chromosome 14
59.2 mb
59.3 mb
59.4 mb
59.5 mb
59.6 mb
59.7 mb
59.8 mb
59.9 mb
60 mb
60.1 mb
r2
>0.8
0.6-0.8
0.4-0.6
0.2-0.4
<0.2
-log10(P)
15
10
5
DAAM1
L3HYPDH
GPR135
JKAMP
CCDC
GZ>CP
CP>GZ
Corrected for total SA
Uncorrected
<=-15
-10
-5
0
5
10
>=15
Zscore

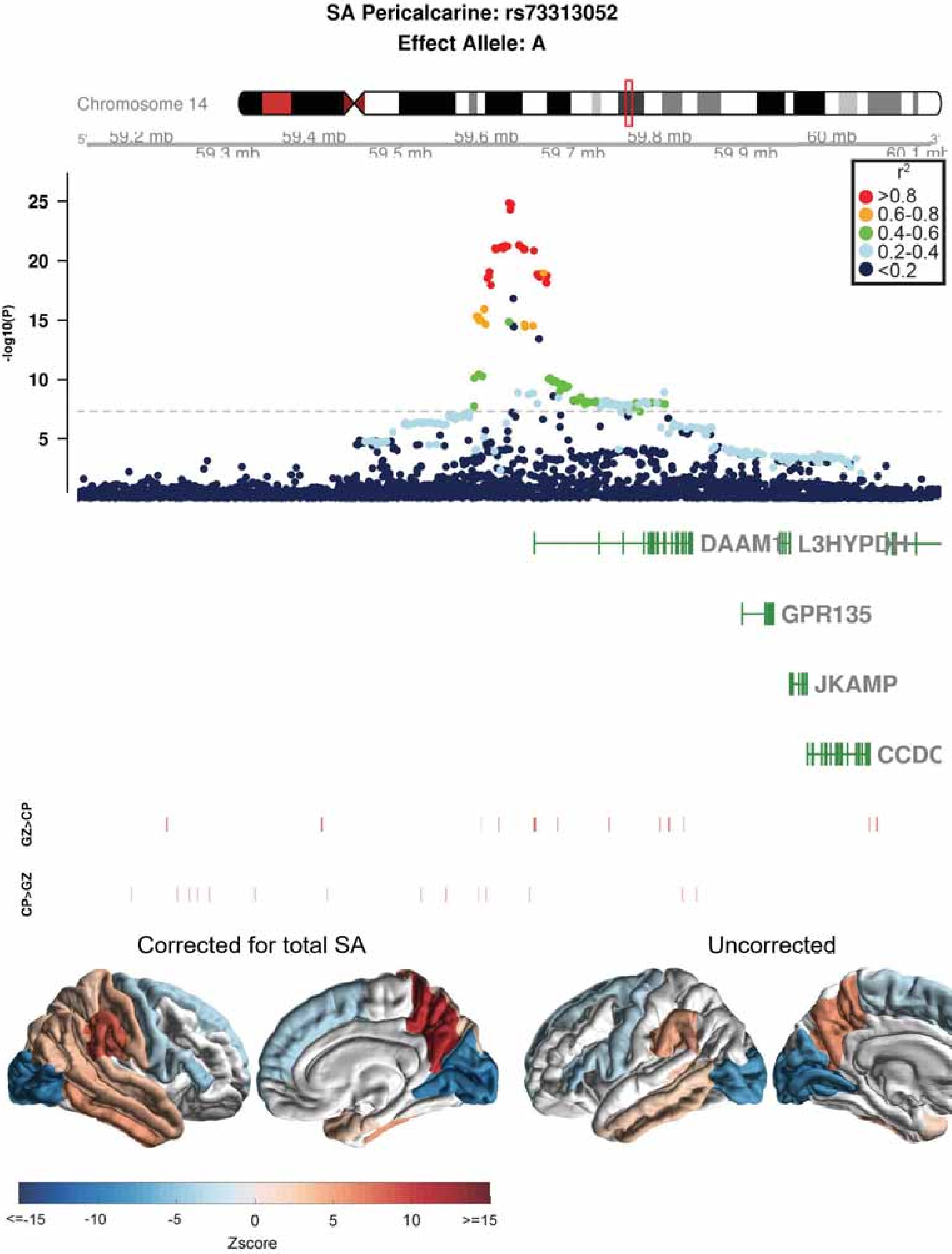

SA Pericalcarine: rs73313052
Effect Allele: A
Chromosome 14
59.2 mb
59.3 mb
59.4 mb
59.5 mb
59.6 mb
59.7 mb
59.8 mb
59.9 mb
60 mb
60.1 mb
-log10(P)
25
20
15
10
5
$r^2$
>0.8
0.6-0.8
0.4-0.6
0.2-0.4
<0.2
DAAM1
L3HYPDH
GPR135
JKAMP
CCDC
GZ>CP
CP>GZ
Corrected for total SA
Uncorrected
<=-15
-10
-5
0
5
10
>=15
Zscore

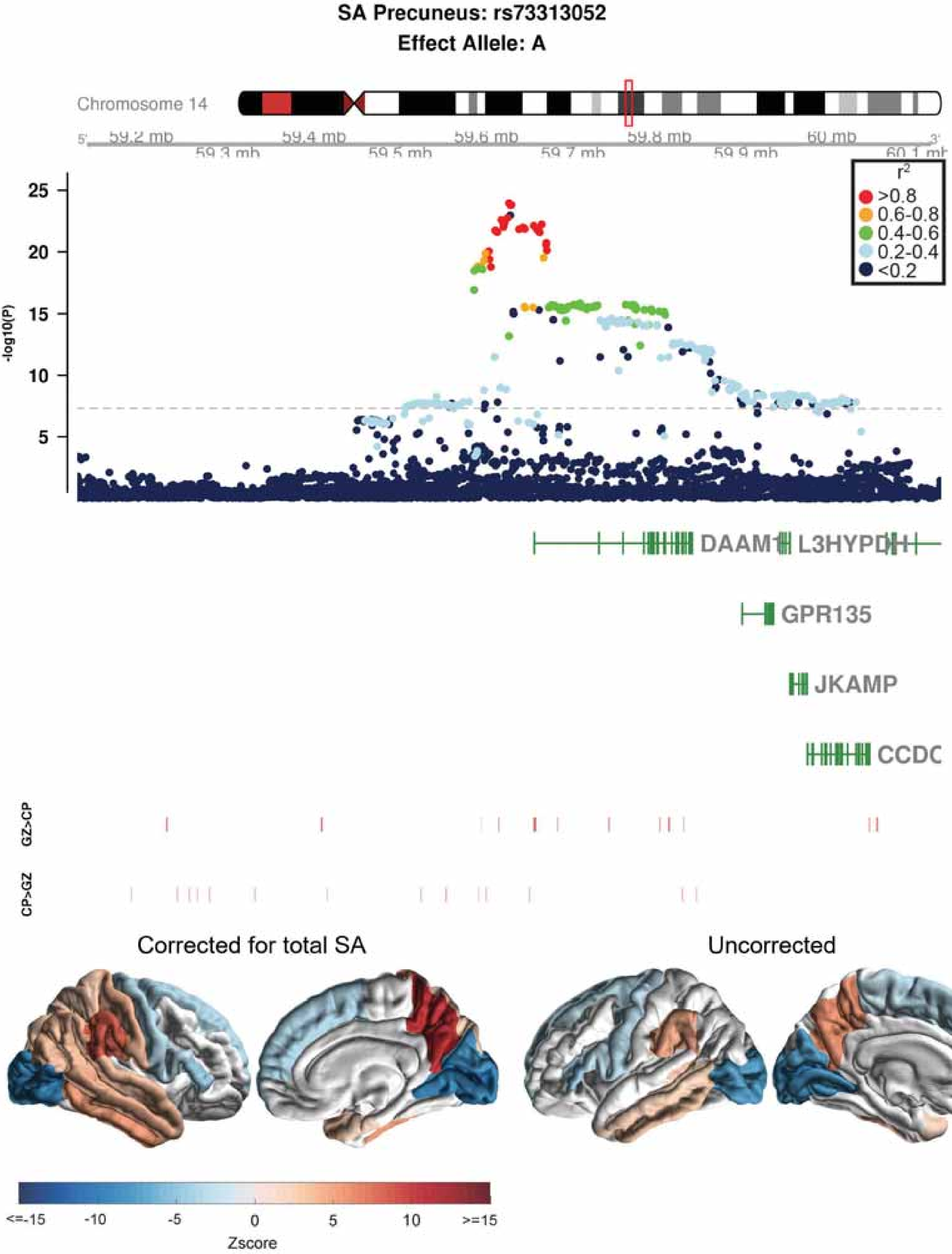
SA Precuneus: rs73313052
Effect Allele: A
Chromosome 14
59.2 mb
59.3 mb
59.4 mb
59.5 mb
59.6 mb
59.7 mb
59.8 mb
59.9 mb
60 mb
60.1 mb
-log10(P)
25
20
15
10
5
r2
>0.8
0.6-0.8
0.4-0.6
0.2-0.4
<0.2
DAAM1
L3HYPDH
GPR135
JKAMP
CCDC
GZ>CP
CP>GZ
Corrected for total SA
Uncorrected
<=-15
-10
-5
0
5
10
>=15
Zscore

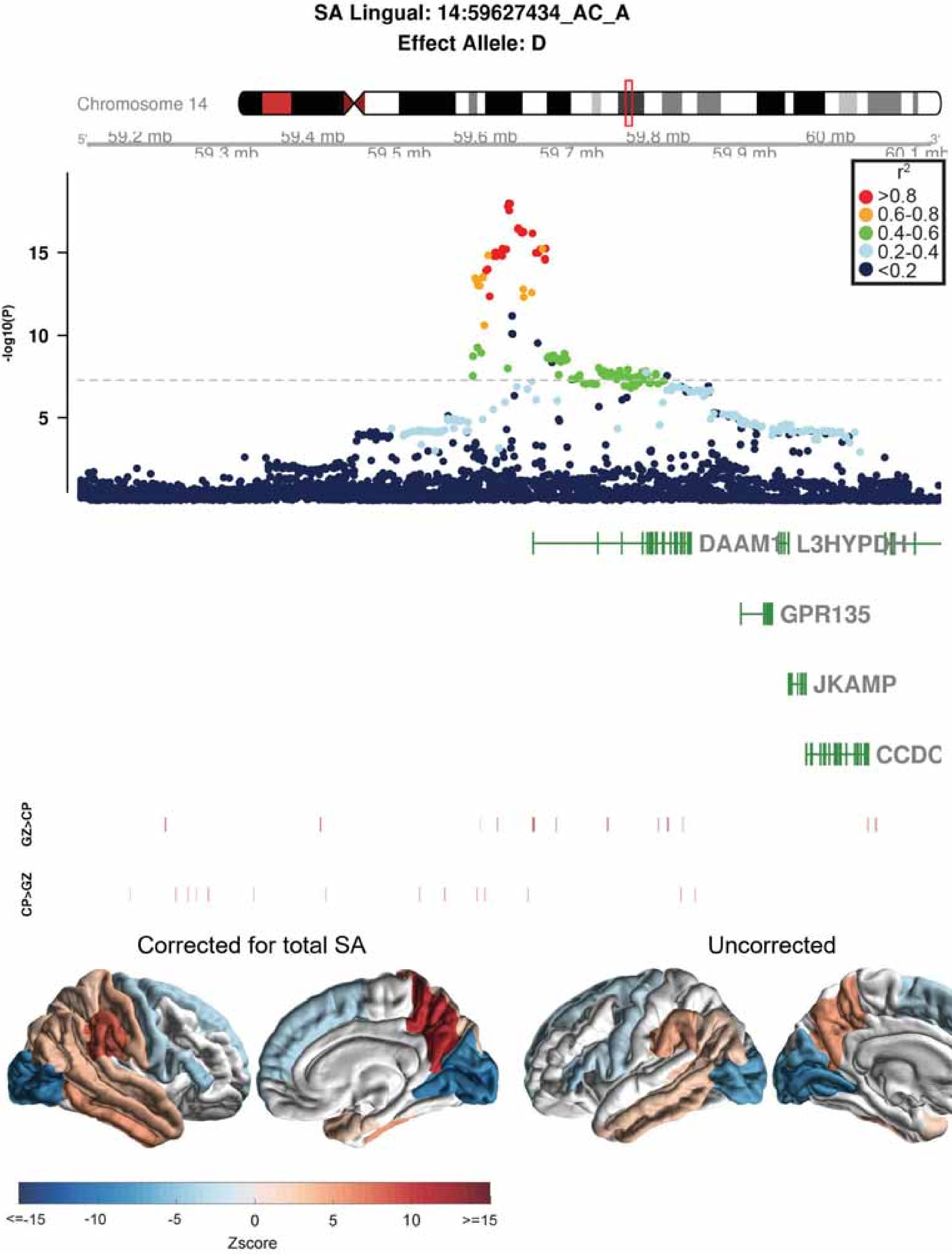

SA Lingual: 14:59627434_AC_A
Effect Allele: D
Chromosome 14
59.2 mb
59.3 mb
59.4 mb
59.5 mb
59.6 mb
59.7 mb
59.8 mb
59.9 mb
60 mb
60.1 mb
r2
>0.8
0.6-0.8
0.4-0.6
0.2-0.4
<0.2
-log10(P)
15
10
5
DAAM1
L3HYPDH
GPR135
JKAMP
CCDC
GZ>CP
CP>GZ
Corrected for total SA
Uncorrected
<=-15
-10
-5
0
5
10
>=15
Zscore

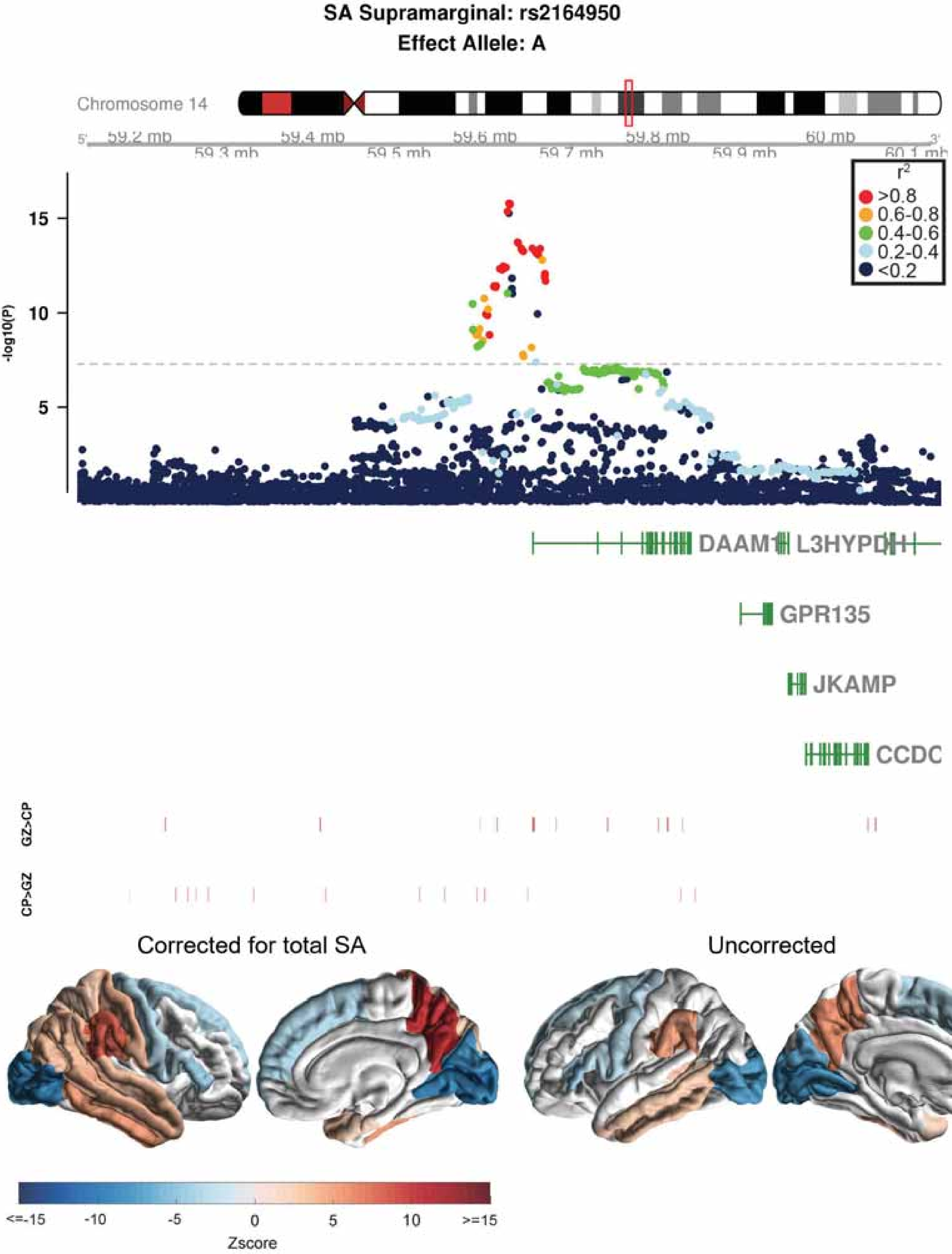

SA Supramarginal: rs2164950
Effect Allele: A
Chromosome 14
59.2 mb
59.3 mb
59.4 mb
59.5 mb
59.6 mb
59.7 mb
59.8 mb
59.9 mb
60 mb
60.1 mb
r2
>0.8
0.6-0.8
0.4-0.6
0.2-0.4
<0.2
-log10(P)
15
10
5
DAAM1
L3HYPDH
GPR135
JKAMP
CCDC
GZ>CP
CP>GZ
Corrected for total SA
Uncorrected
<=-15
-10
-5
0
5
10
>=15
Zscore

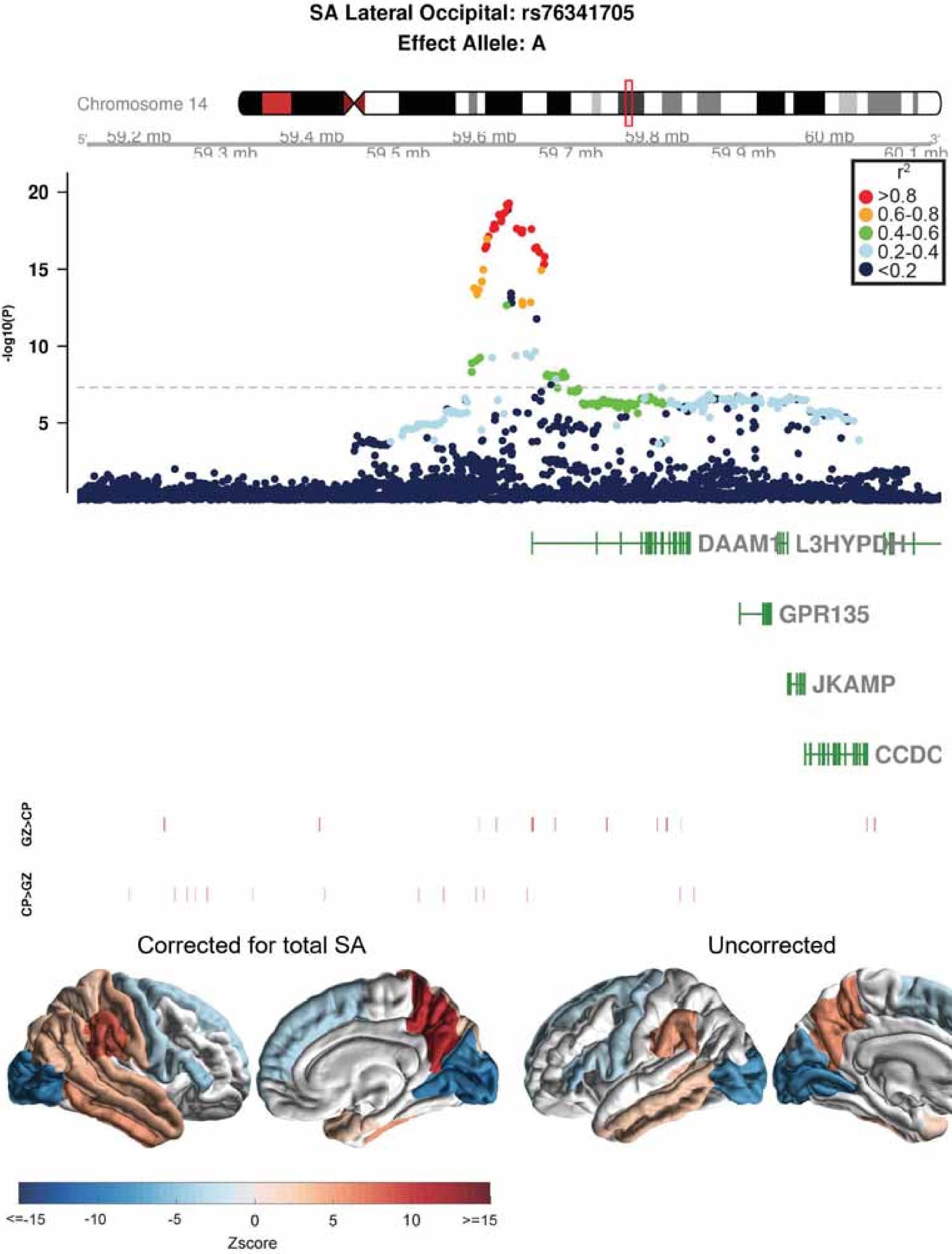

SA Lateral Occipital: rs76341705
Effect Allele: A
Chromosome 14
59.2 mb
59.3 mb
59.4 mb
59.5 mb
59.6 mb
59.7 mb
59.8 mb
59.9 mb
60 mb
60.1 mb
-log10(P)
r2
>0.8
0.6-0.8
0.4-0.6
0.2-0.4
<0.2
DAAM1
L3HYPDH
GPR135
JKAMP
CCDC
GZ>CP
CP>GZ
Corrected for total SA
Uncorrected
Zscore

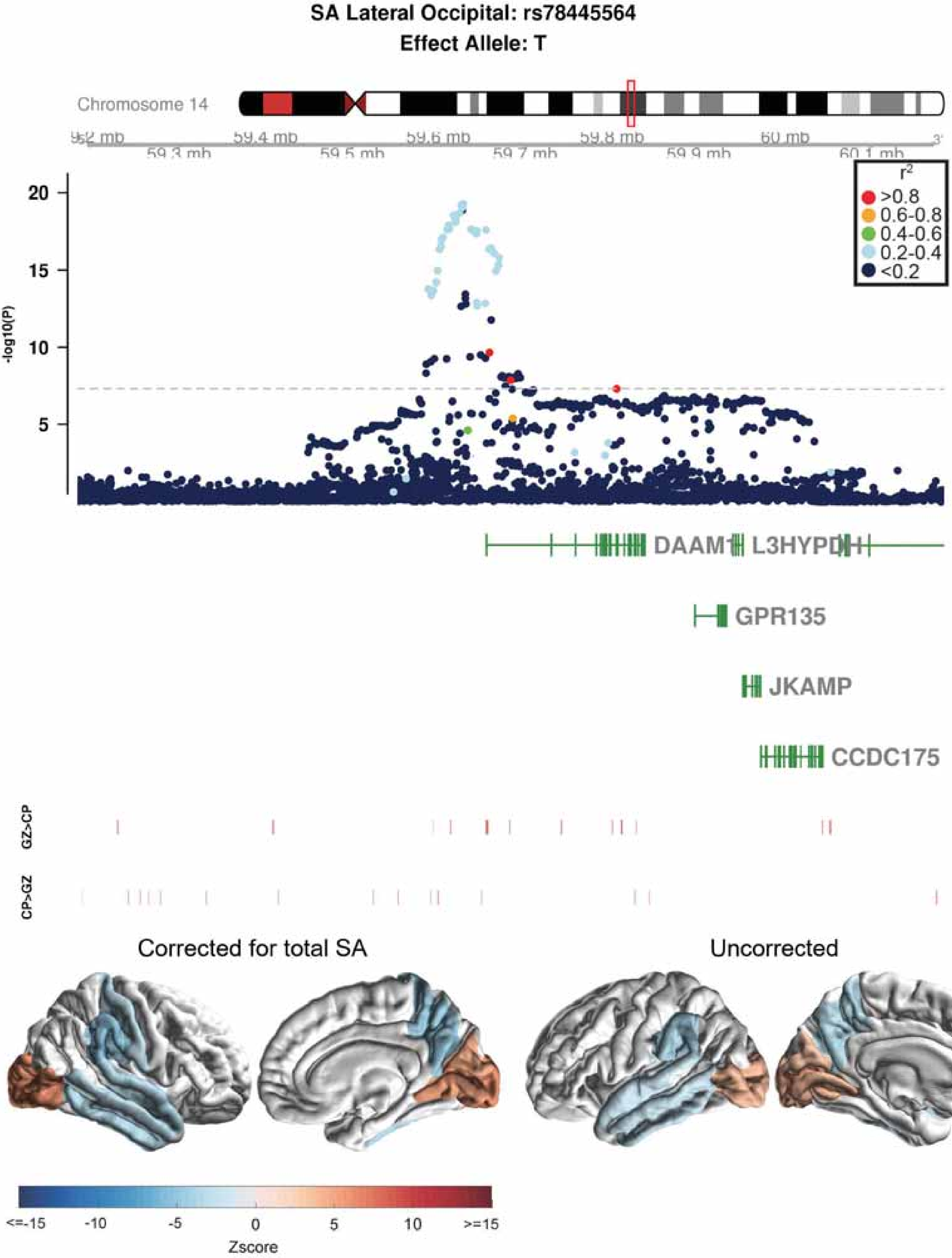

SA Lateral Occipital: rs78445564
Effect Allele: T
Chromosome 14
59.2 mb
59.3 mb
59.4 mb
59.5 mb
59.6 mb
59.7 mb
59.8 mb
59.9 mb
60 mb
60.1 mb
r2
>0.8
0.6-0.8
0.4-0.6
0.2-0.4
<0.2
-log10(P)
20
15
10
5
DAAM1
L3HYPDH
GPR135
JKAMP
CCDC175
GZ>CP
CP>GZ
Corrected for total SA
Uncorrected
<=-15
-10
-5
0
5
10
>=15
Zscore

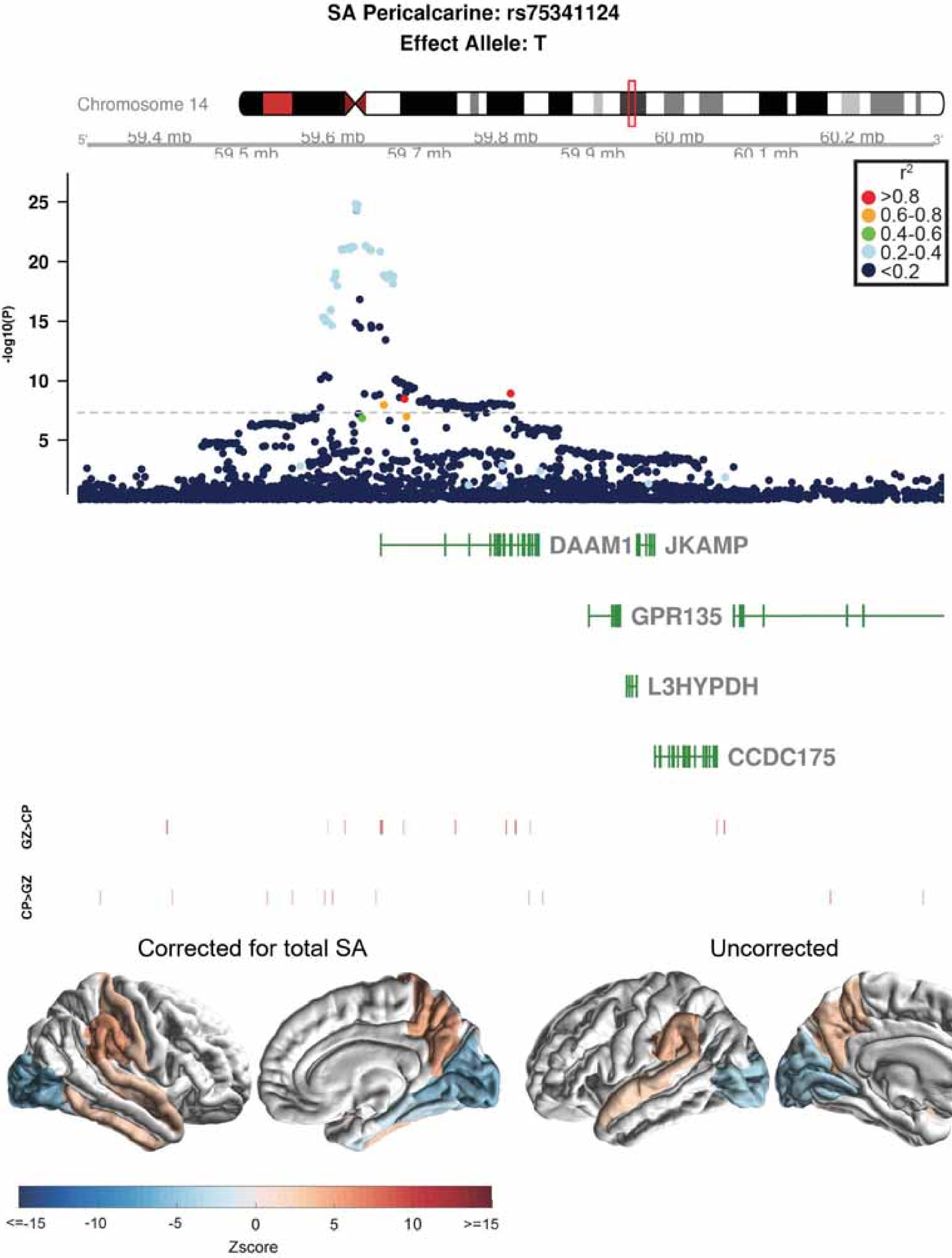

SA Pericalcarine: rs75341124
Effect Allele: T
Chromosome 14
59.4 mb
59.5 mb
59.6 mb
59.7 mb
59.8 mb
59.9 mb
60 mb
60.1 mb
60.2 mb
-log10(P)
$r^2$
>0.8
0.6-0.8
0.4-0.6
0.2-0.4
<0.2
DAAM1
JKAMP
GPR135
L3HYPDH
CCDC175
GZ>CP
CP>GZ
Corrected for total SA
Uncorrected
<=-15
-10
-5
0
5
10
>=15
Zscore

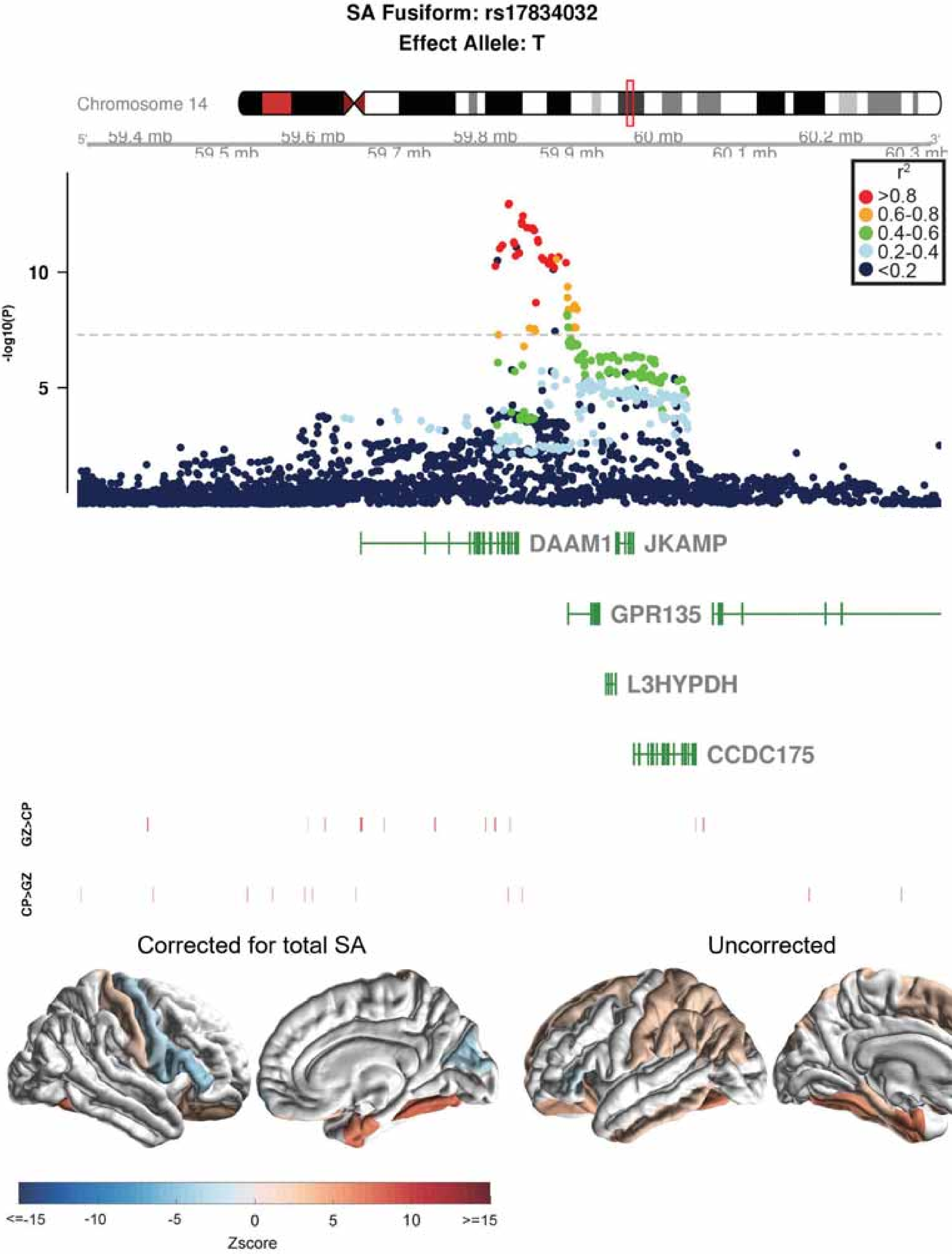
SA Fusiform: rs17834032
Effect Allele: T
Chromosome 14
59.4 mb
59.5 mb
59.6 mb
59.7 mb
59.8 mb
59.9 mb
60 mb
60.1 mb
60.2 mb
60.3 mb
-log10(P)
10
5
r2
>0.8
0.6-0.8
0.4-0.6
0.2-0.4
<0.2
DAAM1
JKAMP
GPR135
L3HYPDH
CCDC175
GZ>CP
CP>GZ
Corrected for total SA
Uncorrected
<=-15
-10
-5
0
5
10
>=15
Zscore

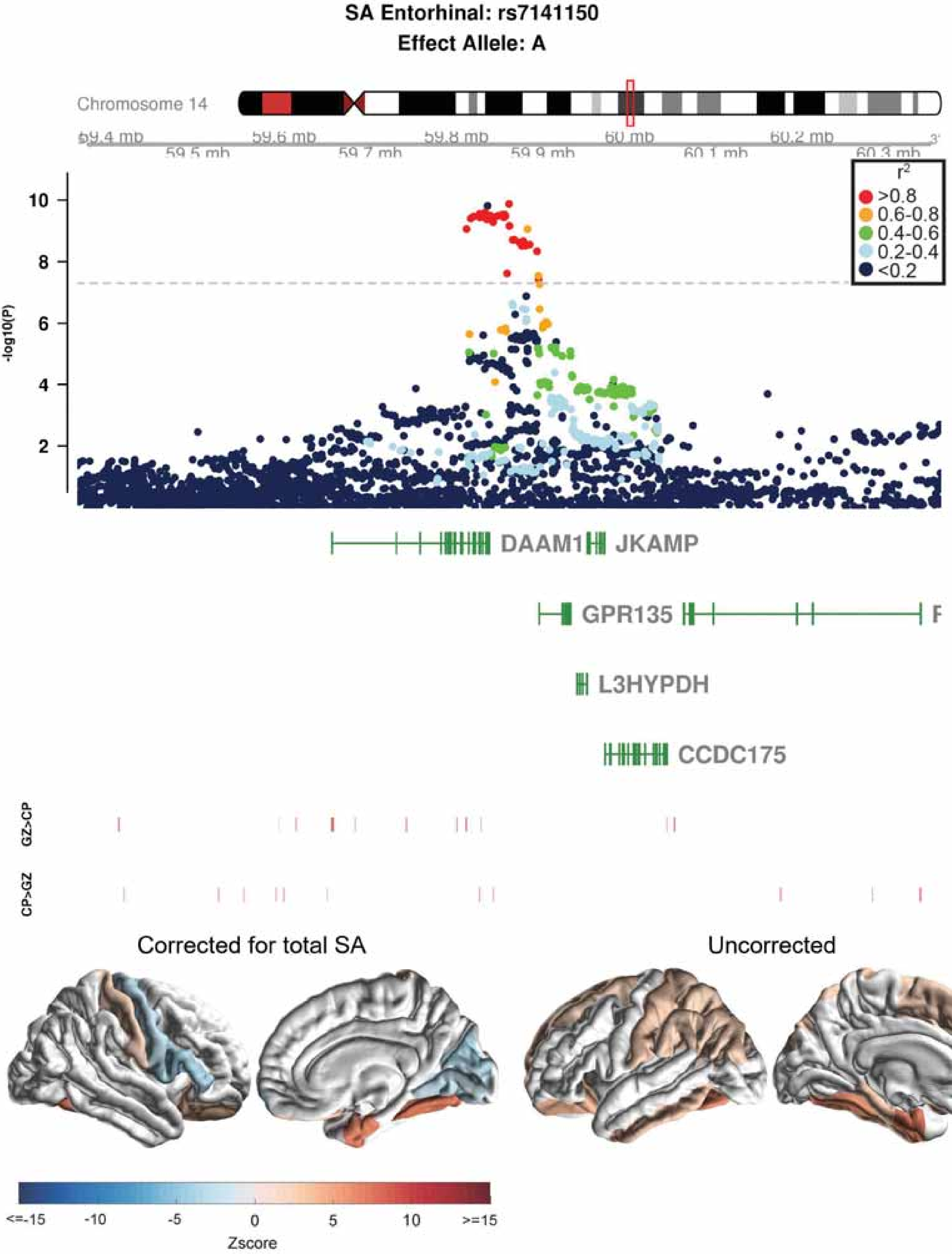

SA Entorhinal: rs7141150
Effect Allele: A
Chromosome 14
59.4 mb
59.5 mb
59.6 mb
59.7 mb
59.8 mb
59.9 mb
60 mb
60.1 mb
60.2 mb
60.3 mb
-log10(P)
10
8
6
4
2
r2
>0.8
0.6-0.8
0.4-0.6
0.2-0.4
<0.2
DAAM1
JKAMP
GPR135
L3HYPDH
CCDC175
GZ>CP
CP>GZ
Corrected for total SA
Uncorrected
<=-15
-10
-5
0
5
10
>=15
Zscore

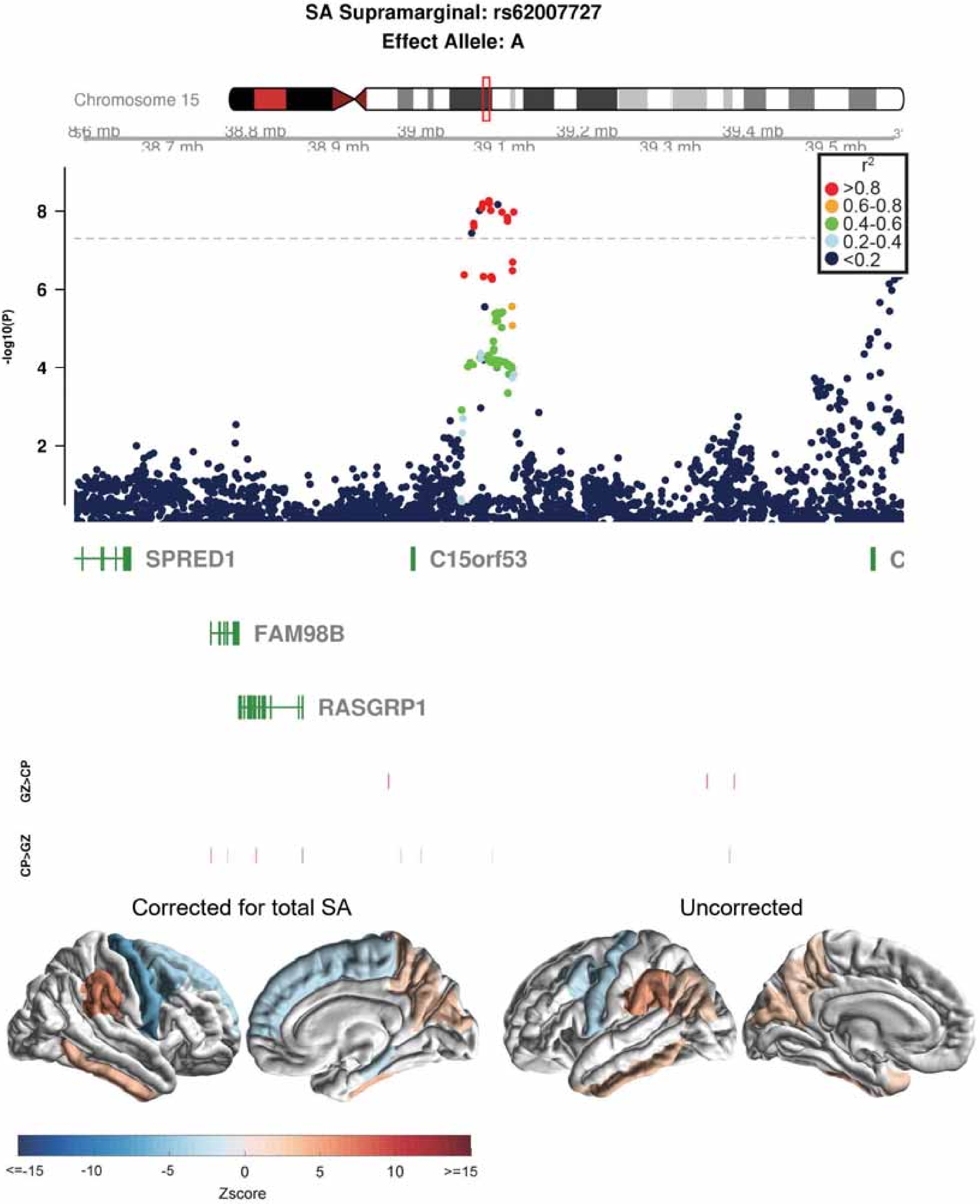
SA Supramarginal: rs62007727
Effect Allele: A
Chromosome 15
38.6 mb
38.7 mb
38.8 mb
38.9 mb
39 mb
39.1 mb
39.2 mb
39.3 mb
39.4 mb
39.5 mb
r2
>0.8
0.6-0.8
0.4-0.6
0.2-0.4
<0.2
-log10(P)
8
6
4
2
SPRED1
C15orf53
FAM98B
RASGRP1
GZ>CP
CP>GZ
Corrected for total SA
Uncorrected
<=-15
-10
-5
0
5
10
>=15
Zscore

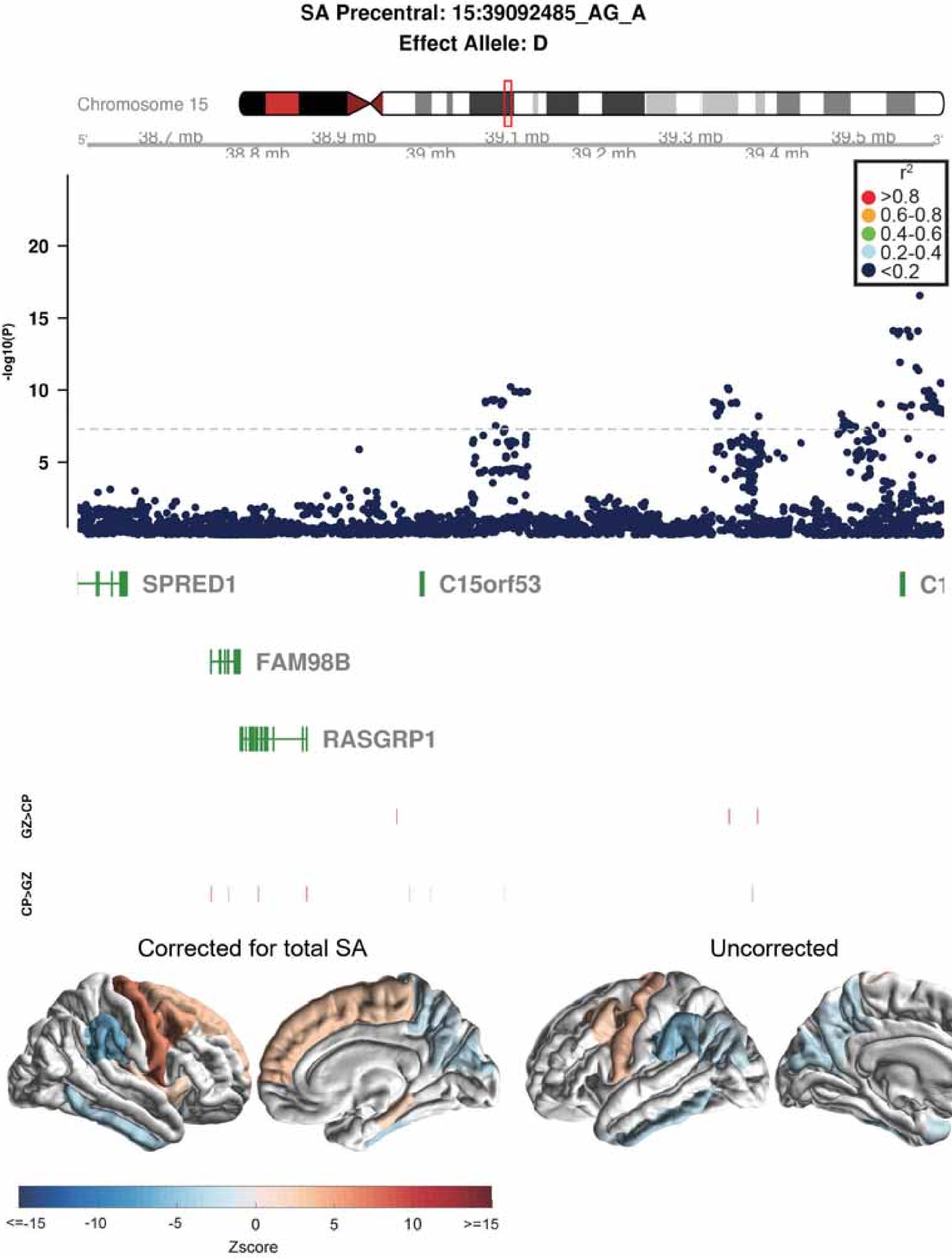

SA Precentral: 15:39092485_AG_A
Effect Allele: D
Chromosome 15
38.7 mb
38.8 mb
38.9 mb
39 mb
39.1 mb
39.2 mb
39.3 mb
39.4 mb
39.5 mb
-log10(P)
20
15
10
5
r2
>0.8
0.6-0.8
0.4-0.6
0.2-0.4
<0.2
SPRED1
C15orf53
FAM98B
RASGRP1
GZ>CP
CP>GZ
Corrected for total SA
Uncorrected
<=-15
-10
-5
0
5
10
>=15
Zscore

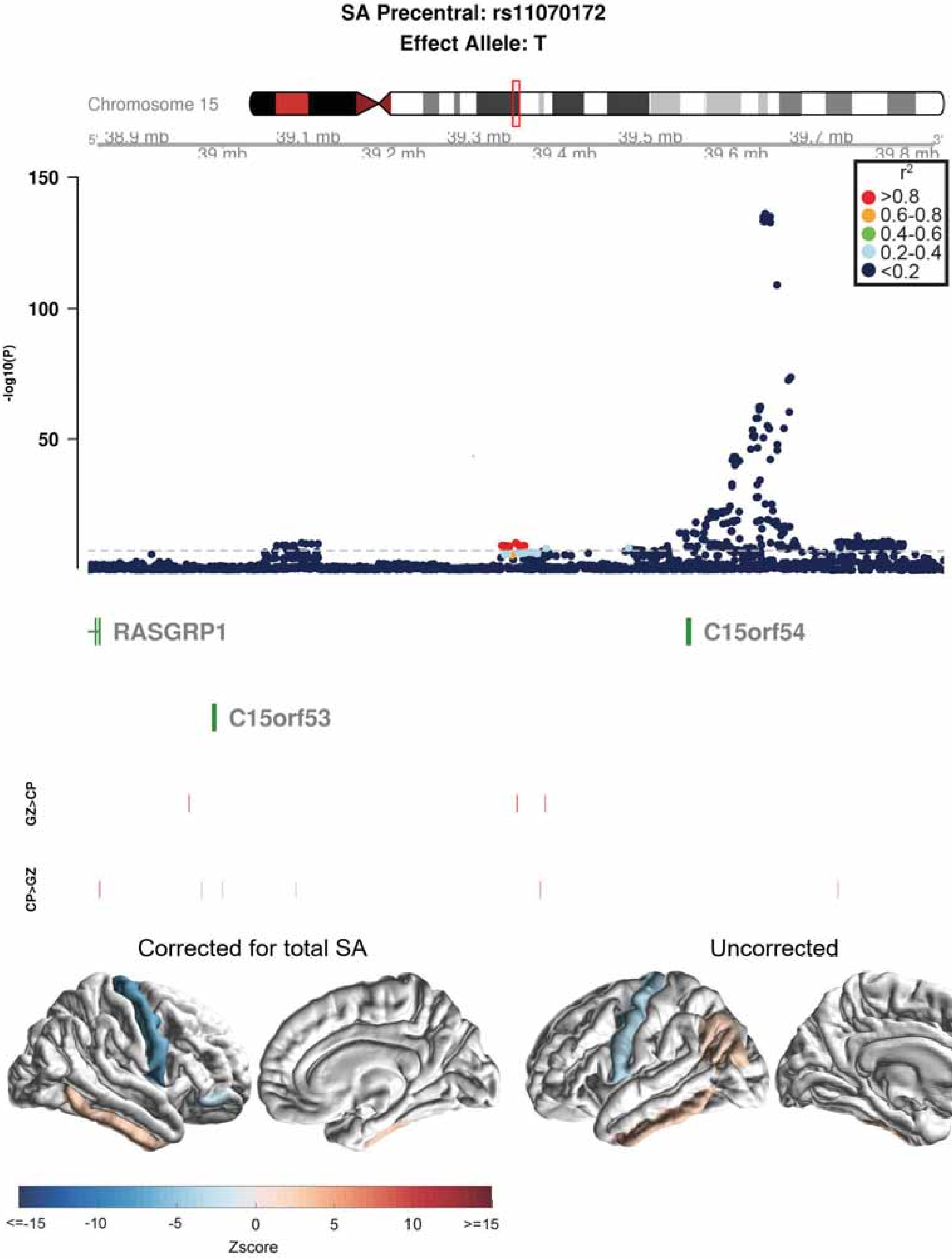

SA Precentral: rs11070172
Effect Allele: T
Chromosome 15
38.9 mb
39 mb
39.1 mb
39.2 mb
39.3 mb
39.4 mb
39.5 mb
39.6 mb
39.7 mb
39.8 mb
-log10(P)
150
100
50
r2
>0.8
0.6-0.8
0.4-0.6
0.2-0.4
<0.2
RASGRP1
C15orf54
C15orf53
GZ>CP
CP>GZ
Corrected for total SA
Uncorrected
<=-15
-10
-5
0
5
10
>=15
Zscore

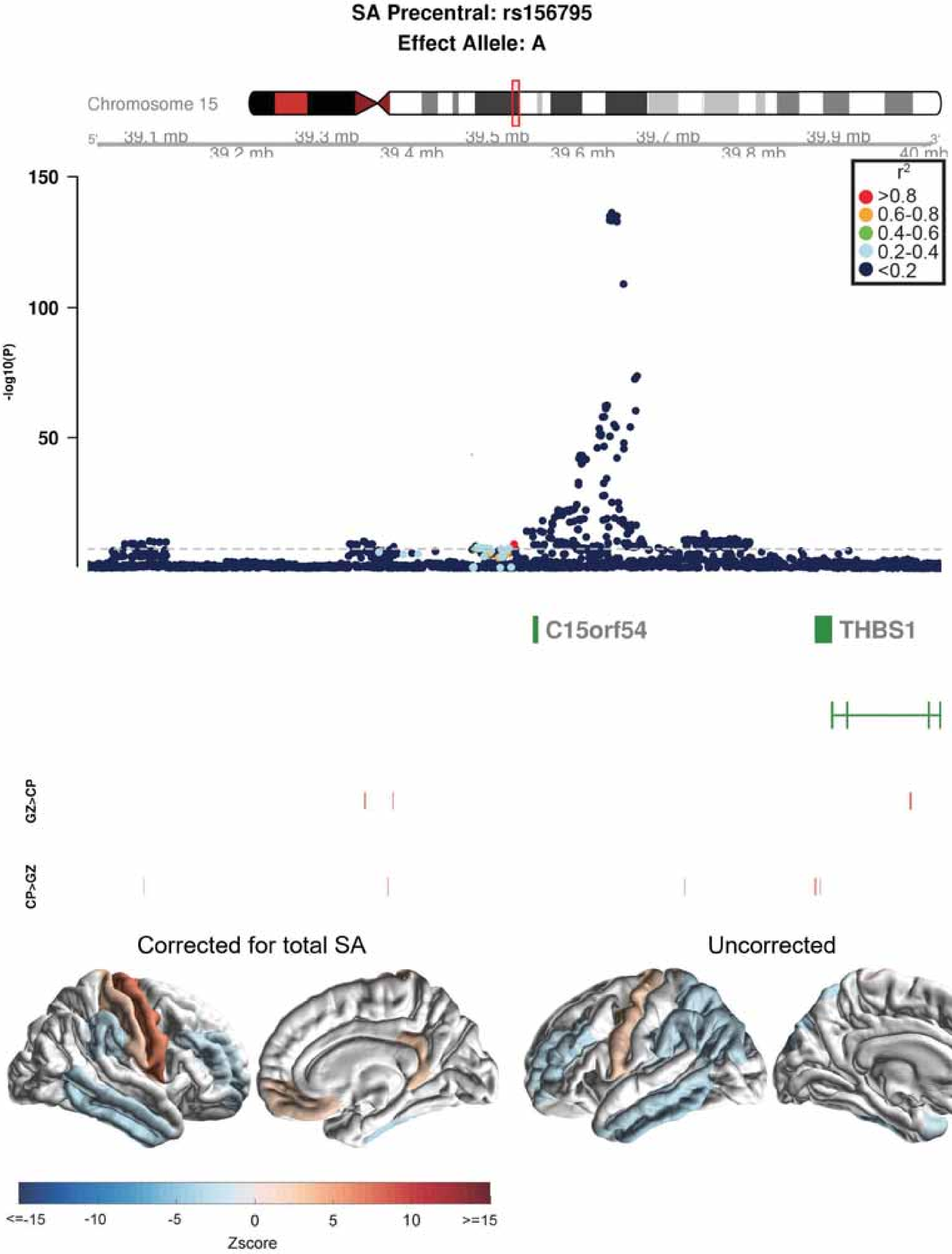

SA Precentral: rs156795
Effect Allele: A
Chromosome 15
5'
39.1 mb
39.2 mb
39.3 mb
39.4 mb
39.5 mb
39.6 mb
39.7 mb
39.8 mb
39.9 mb
40 mb
3'
150
100
50
-log10(P)
r2
>0.8
0.6-0.8
0.4-0.6
0.2-0.4
<0.2
C15orf54
THBS1
GZ>CP
CP>GZ
Corrected for total SA
Uncorrected
<=-15
-10
-5
0
5
10
>=15
Zscore

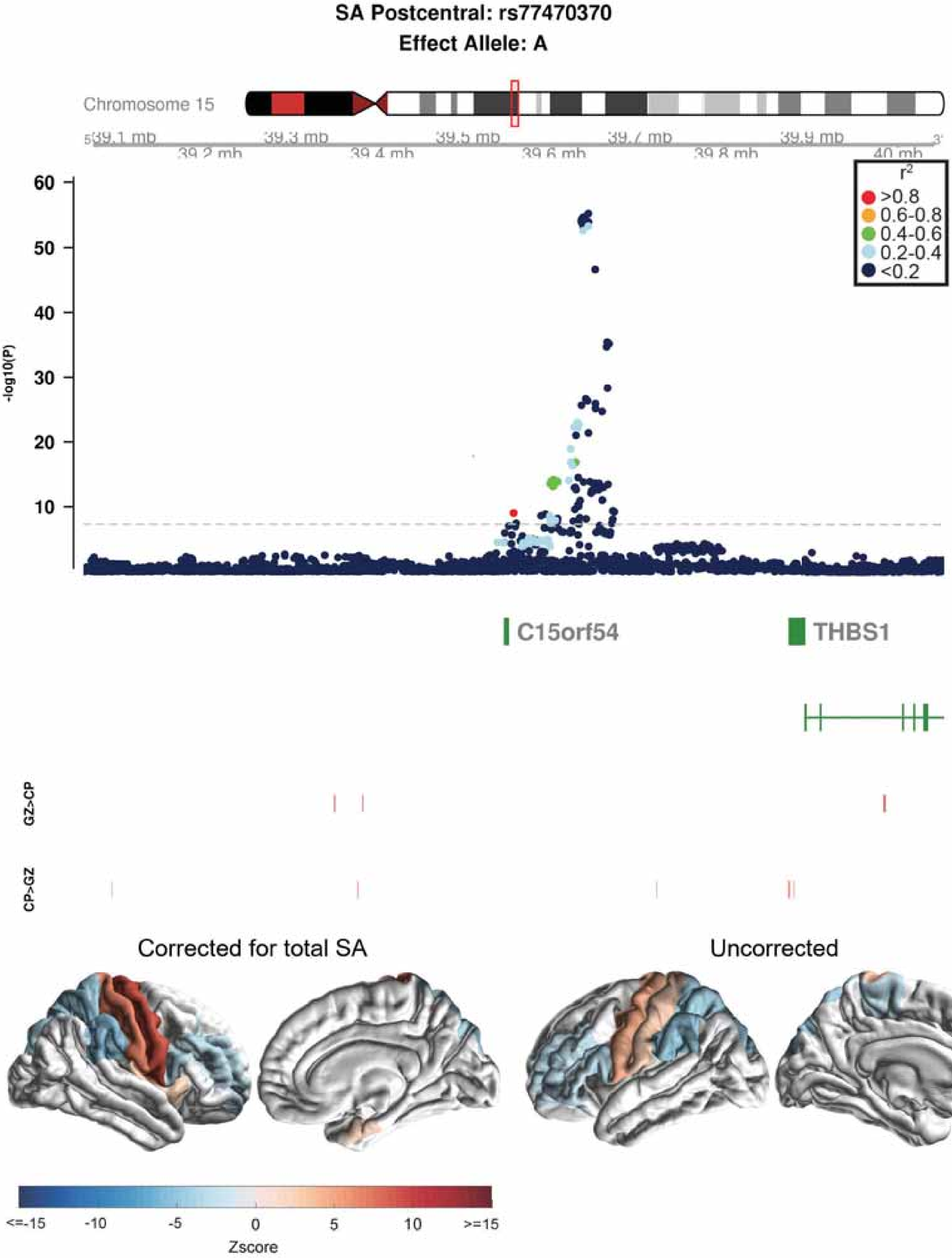

SA Postcentral: rs77470370
Effect Allele: A
Chromosome 15
39.1 mb
39.2 mb
39.3 mb
39.4 mb
39.5 mb
39.6 mb
39.7 mb
39.8 mb
39.9 mb
40 mb
r2
>0.8
0.6-0.8
0.4-0.6
0.2-0.4
<0.2
-log10(P)
60
50
40
30
20
10
C15orf54
THBS1
GZ>CP
CP>GZ
Corrected for total SA
Uncorrected
<=-15
-10
-5
0
5
10
>=15
Zscore

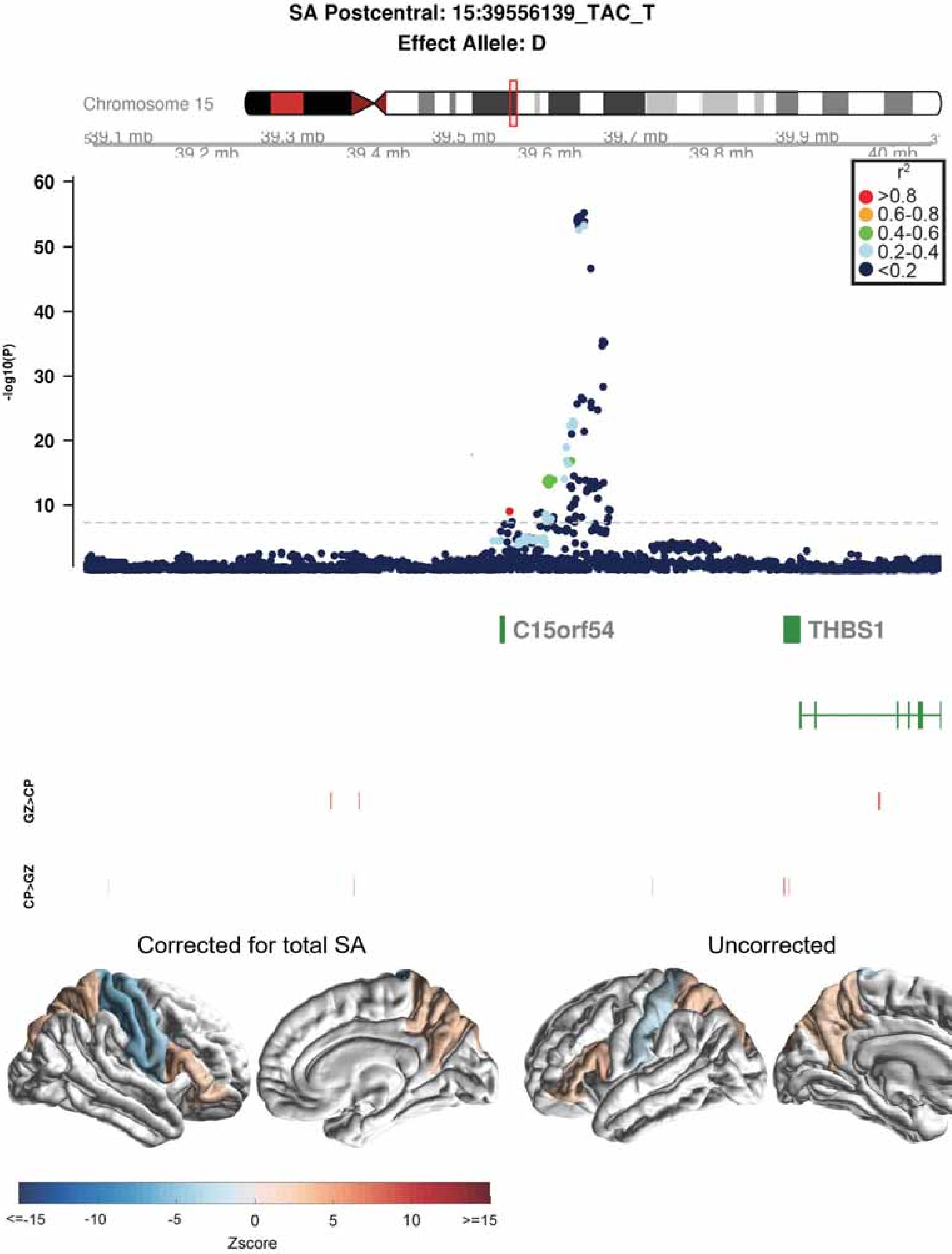
SA Postcentral: 15:39556139_TAC_T
Effect Allele: D
Chromosome 15
39.1 mb
39.2 mb
39.3 mb
39.4 mb
39.5 mb
39.6 mb
39.7 mb
39.8 mb
39.9 mb
40 mb
-log10(P)
r2
>0.8
0.6-0.8
0.4-0.6
0.2-0.4
<0.2
C15orf54
THBS1
GZ>CP
CP>GZ
Corrected for total SA
Uncorrected
<=-15
-10
-5
0
5
10
>=15
Zscore

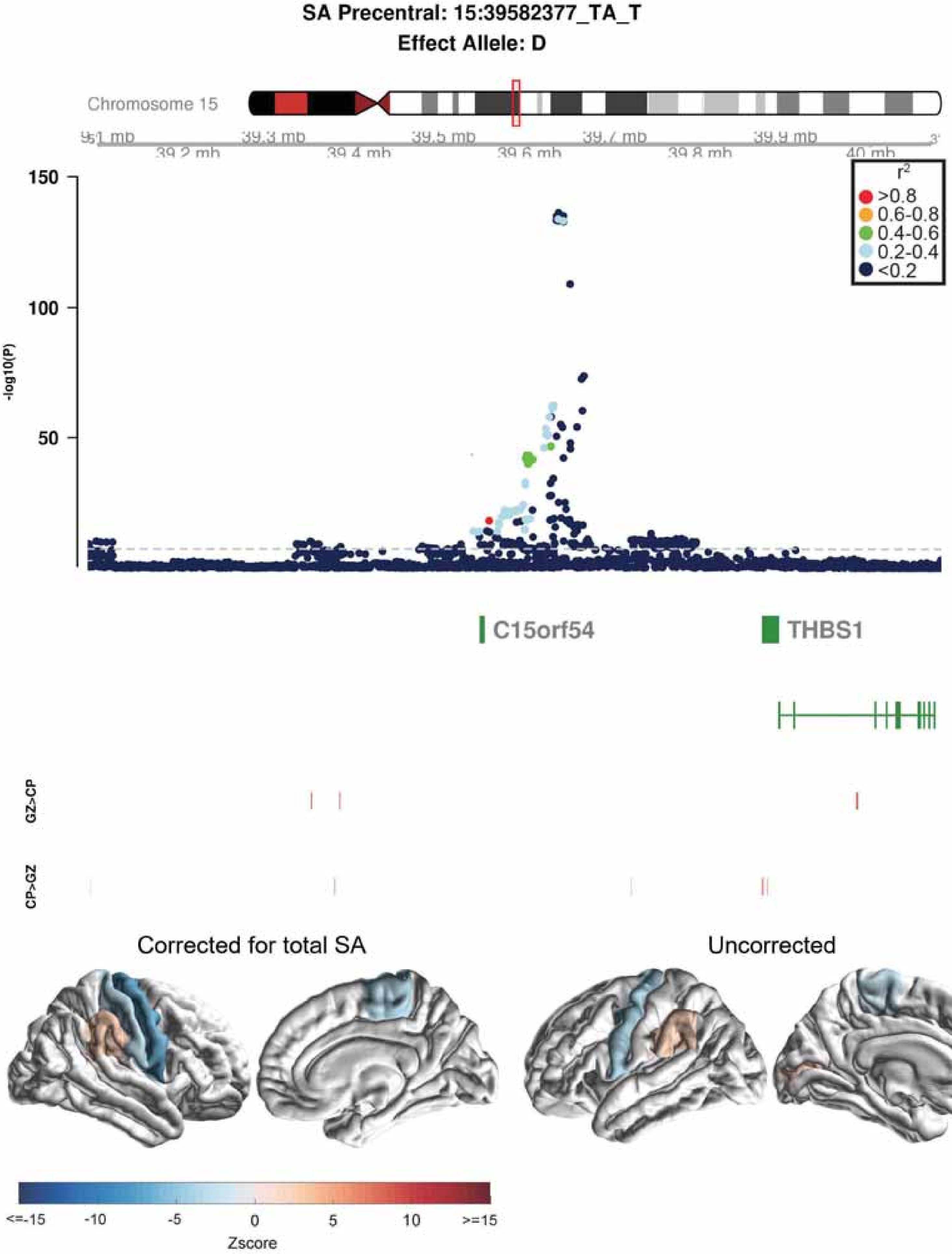
SA Precentral: 15:39582377_TA_T
Effect Allele: D
Chromosome 15
39.2 mb
39.3 mb
39.4 mb
39.5 mb
39.6 mb
39.7 mb
39.8 mb
39.9 mb
40 mb
150
100
50
-log10(P)
r2
>0.8
0.6-0.8
0.4-0.6
0.2-0.4
<0.2
C15orf54
THBS1
GZ>CP
CP>GZ
Corrected for total SA
Uncorrected
<=-15
-10
-5
0
5
10
>=15
Zscore

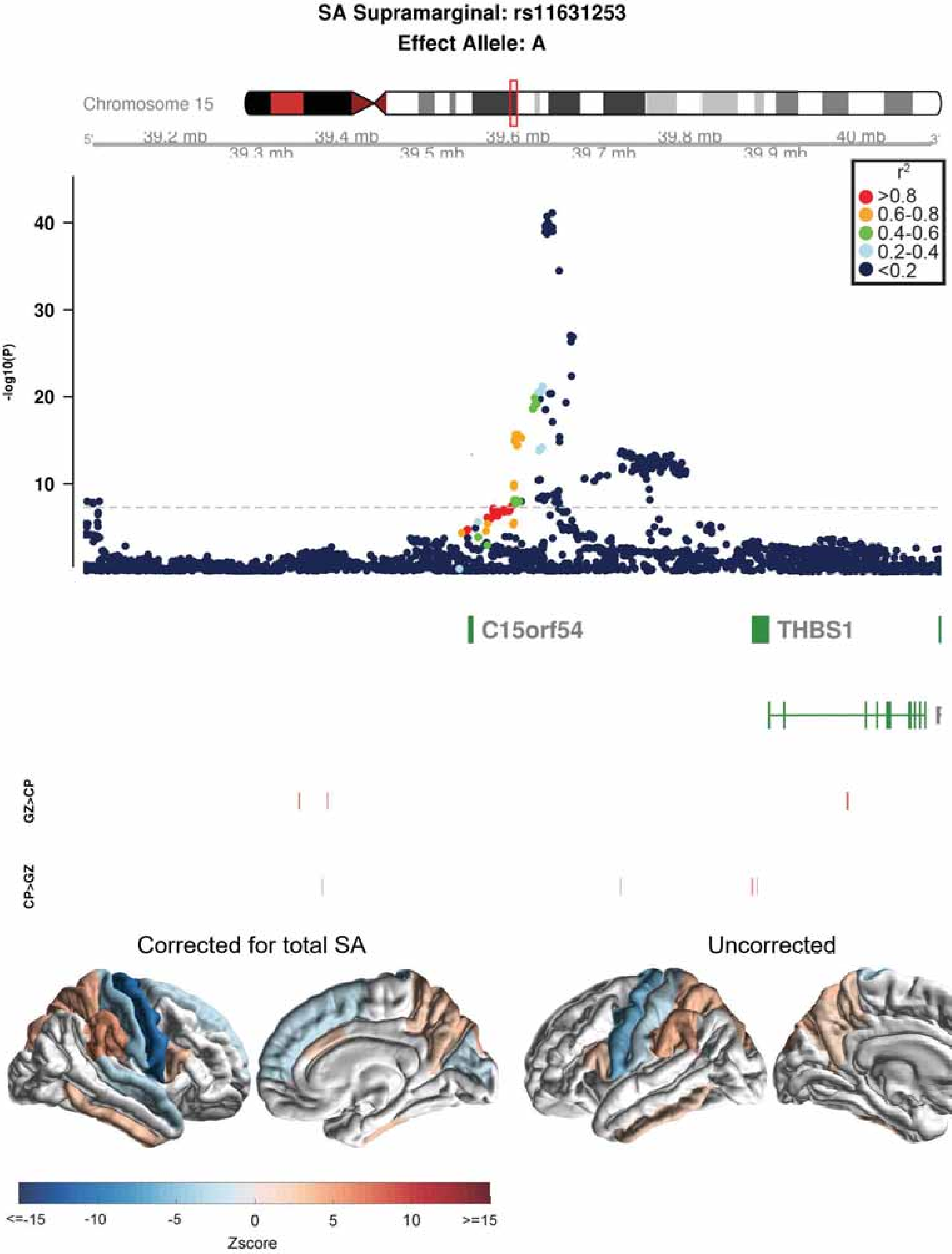

SA Supramarginal: rs11631253
Effect Allele: A
Chromosome 15
39.2 mb
39.3 mb
39.4 mb
39.5 mb
39.6 mb
39.7 mb
39.8 mb
39.9 mb
40 mb
-log10(P)
40
30
20
10
r2
>0.8
0.6-0.8
0.4-0.6
0.2-0.4
<0.2
C15orf54
THBS1
GZ>CP
CP>GZ
Corrected for total SA
Uncorrected
<=-15
-10
-5
0
5
10
>=15
Zscore

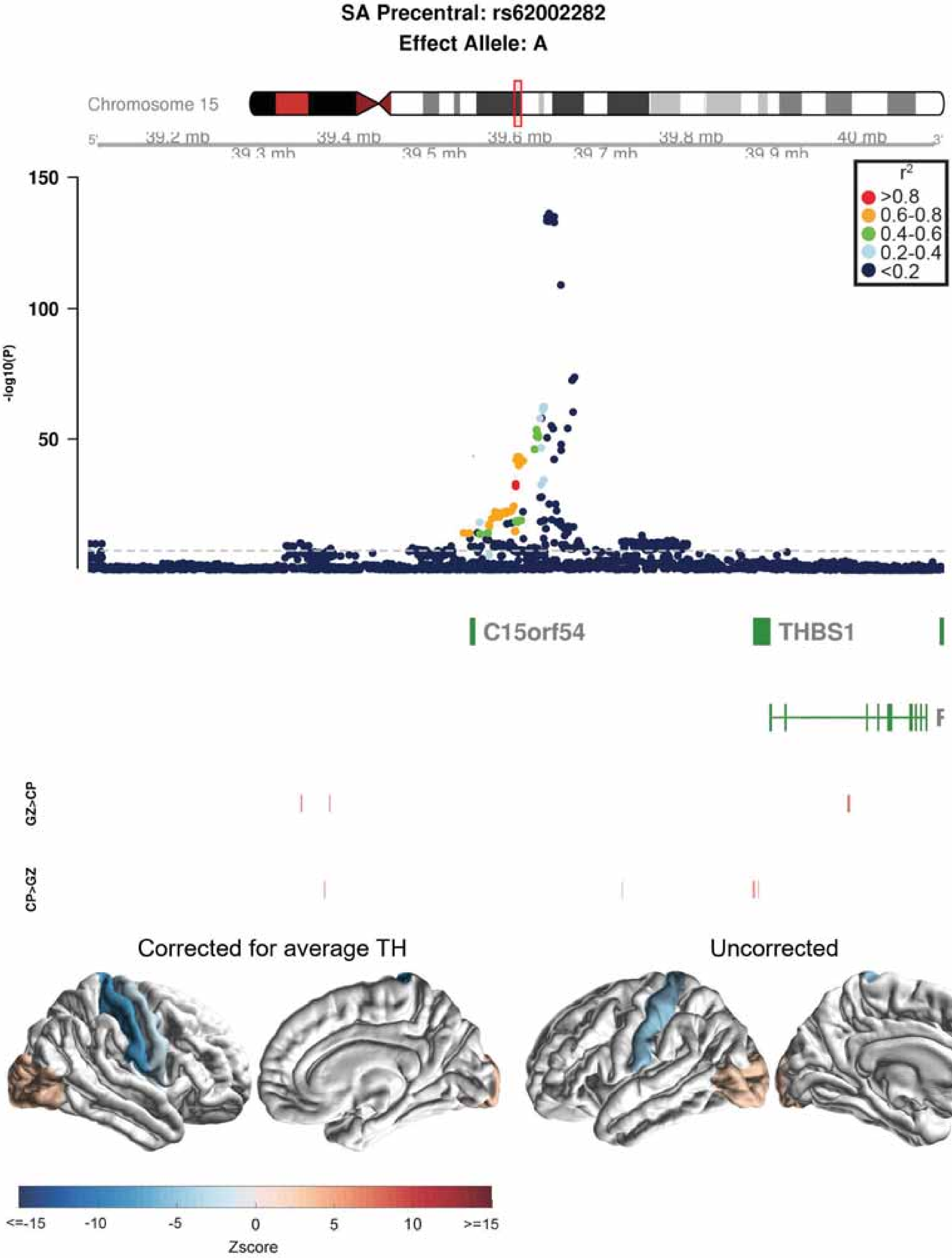
SA Precentral: rs62002282
Effect Allele: A
Chromosome 15
39.2 mb
39.3 mb
39.4 mb
39.5 mb
39.6 mb
39.7 mb
39.8 mb
39.9 mb
40 mb
150
100
50
-log10(P)
r2
>0.8
0.6-0.8
0.4-0.6
0.2-0.4
<0.2
C15orf54
THBS1
GZ>CP
CP>GZ
Corrected for average TH
Uncorrected
<=-15
-10
-5
0
5
10
>=15
Zscore

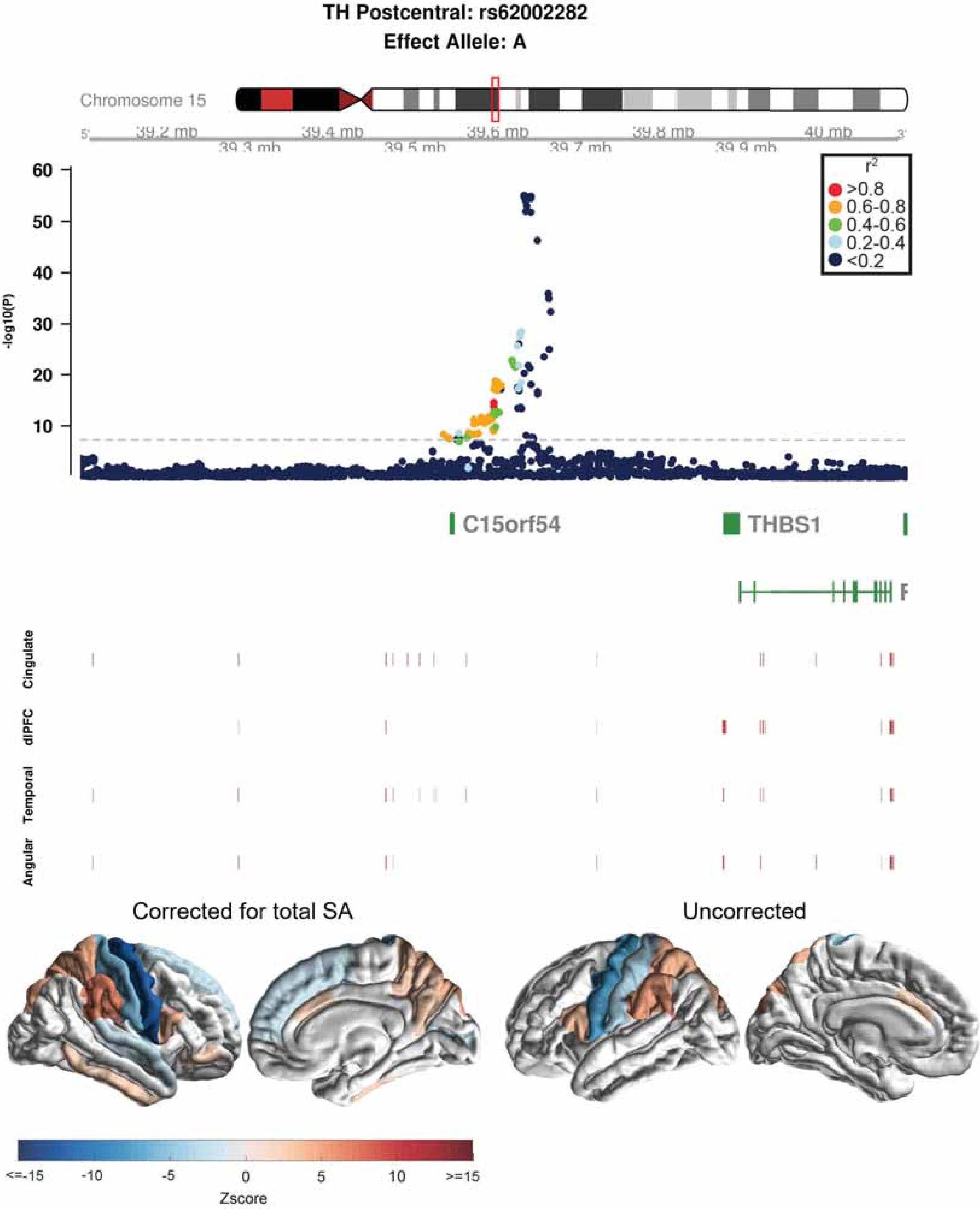
TH Postcentral: rs62002282
Effect Allele: A
Chromosome 15
39.2 mb
39.3 mb
39.4 mb
39.5 mb
39.6 mb
39.7 mb
39.8 mb
39.9 mb
40 mb
-log10(P)
r²
>0.8
0.6-0.8
0.4-0.6
0.2-0.4
<0.2
C15orf54
THBS1
Cingulate
dlPFC
Temporal
Angular
Corrected for total SA
Uncorrected
<=-15
-10
-5
0
5
10
>=15
Zscore

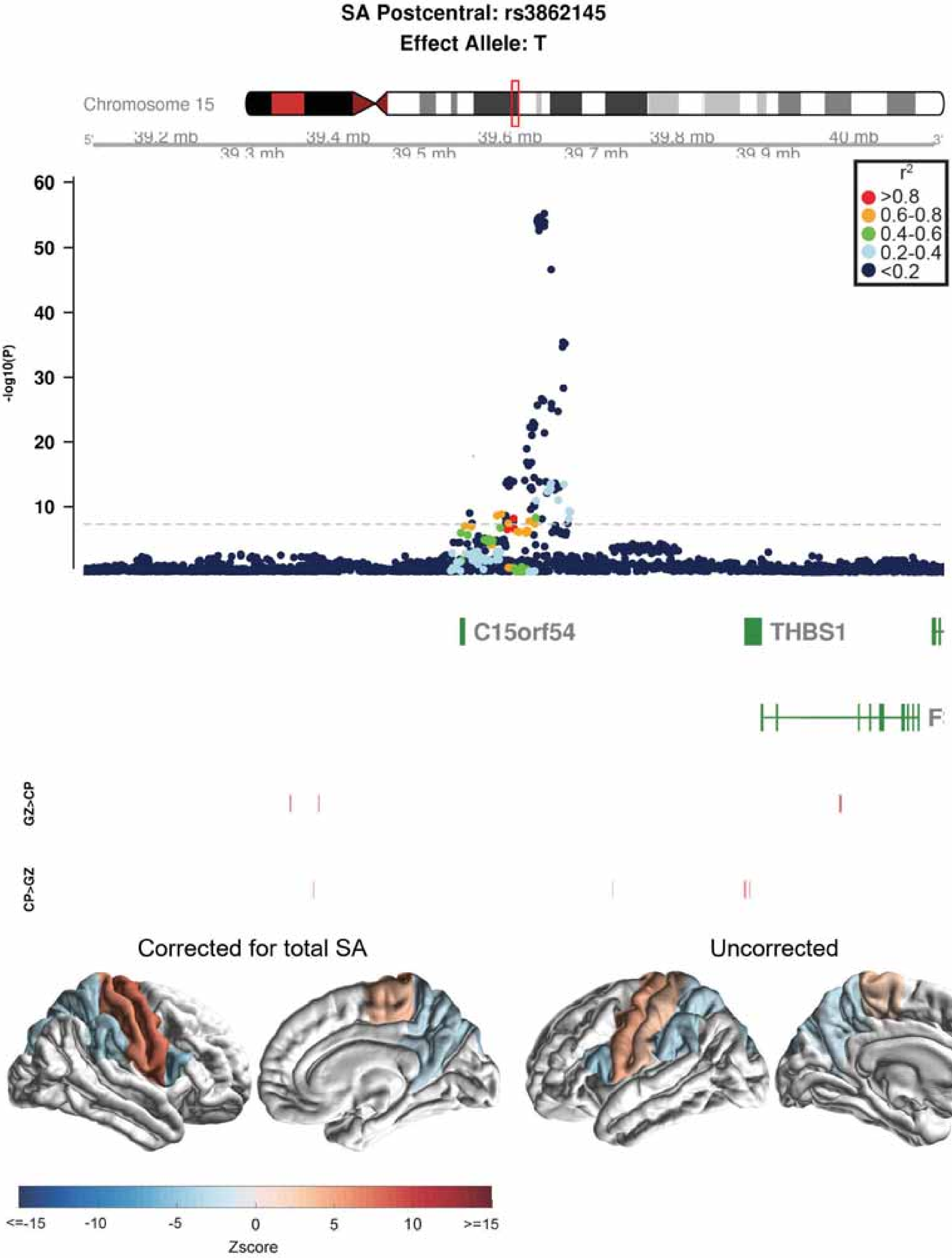
SA Postcentral: rs3862145
Effect Allele: T
Chromosome 15
39.2 mb
39.3 mb
39.4 mb
39.5 mb
39.6 mb
39.7 mb
39.8 mb
39.9 mb
40 mb
-log10(P)
r2
>0.8
0.6-0.8
0.4-0.6
0.2-0.4
<0.2
C15orf54
THBS1
GZ>CP
CP>GZ
Corrected for total SA
Uncorrected
<=-15
-10
-5
0
5
10
>=15
Zscore

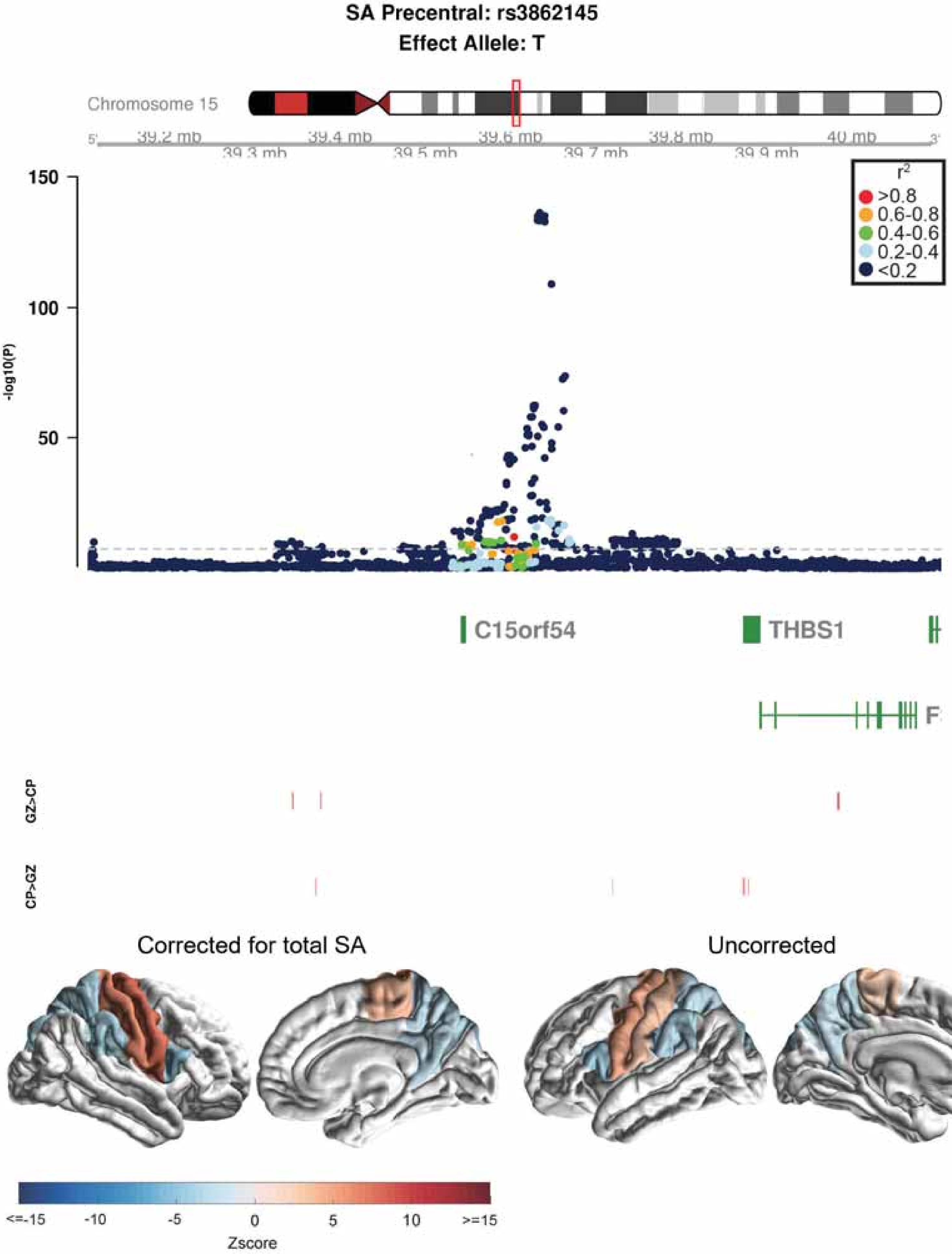
SA Precentral: rs3862145
Effect Allele: T
Chromosome 15
39.2 mb
39.3 mb
39.4 mb
39.5 mb
39.6 mb
39.7 mb
39.8 mb
39.9 mb
40 mb
-log10(P)
150
100
50
r2
>0.8
0.6-0.8
0.4-0.6
0.2-0.4
<0.2
C15orf54
THBS1
GZ>CP
CP>GZ
Corrected for total SA
Uncorrected
<=-15
-10
-5
0
5
10
>=15
Zscore

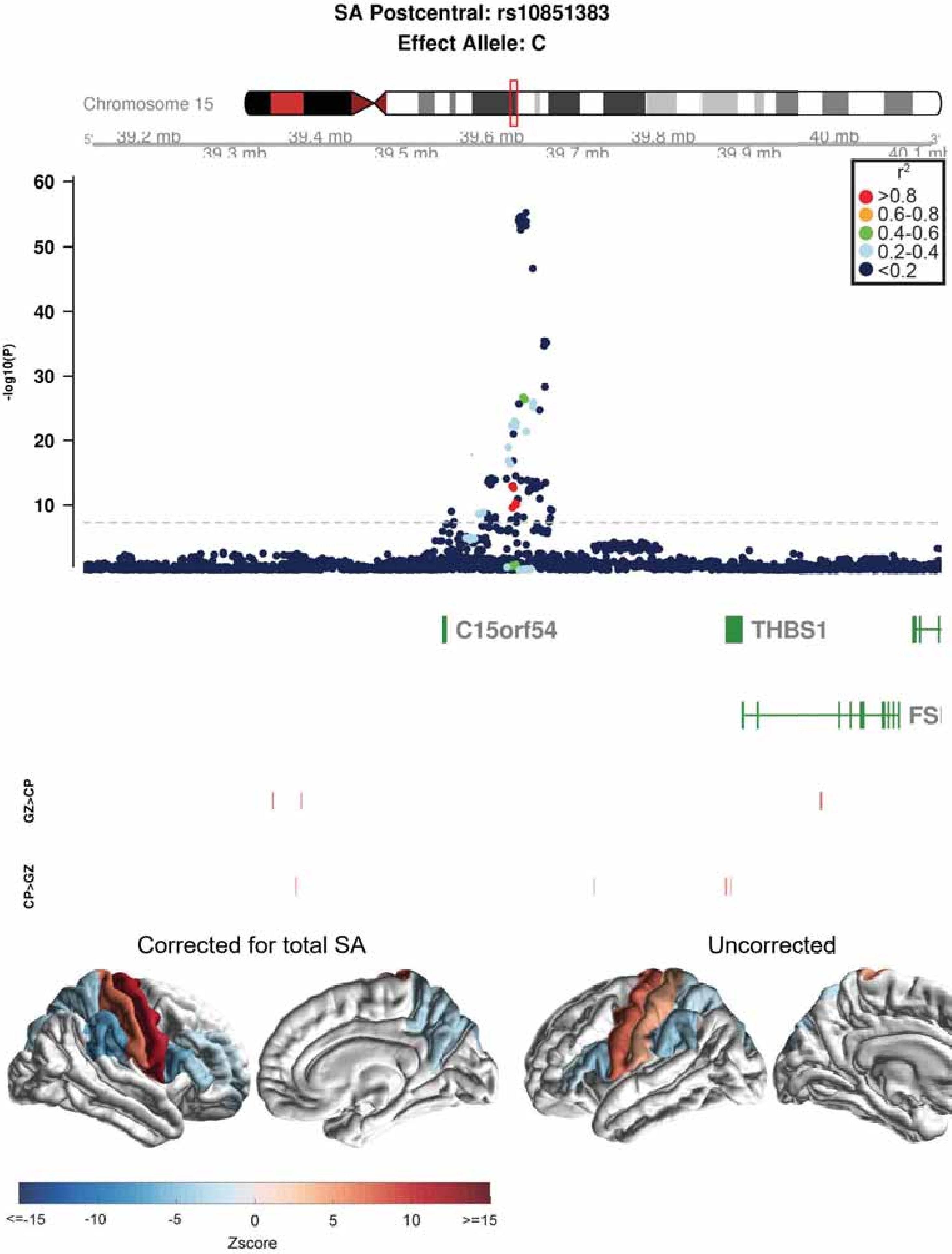

SA Postcentral: rs10851383
Effect Allele: C
Chromosome 15
39.2 mb
39.3 mb
39.4 mb
39.5 mb
39.6 mb
39.7 mb
39.8 mb
39.9 mb
40 mb
40.1 mb
r2
>0.8
0.6-0.8
0.4-0.6
0.2-0.4
<0.2
-log10(P)
60
50
40
30
20
10
C15orf54
THBS1
GZ>CP
CP>GZ
Corrected for total SA
Uncorrected
<=-15
-10
-5
0
5
10
>=15
Zscore

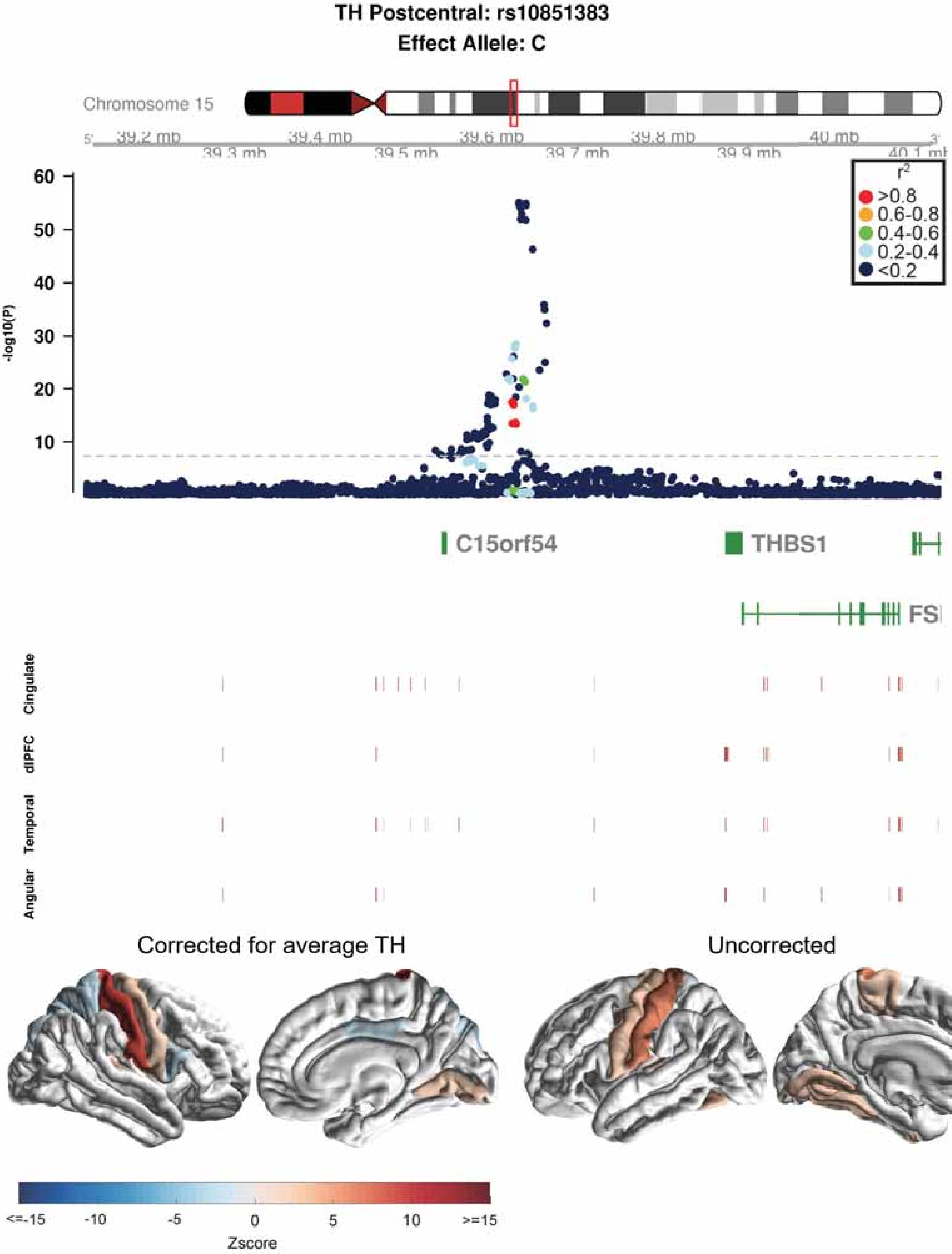

TH Postcentral: rs10851383
Effect Allele: C
Chromosome 15
39.2 mb
39.3 mb
39.4 mb
39.5 mb
39.6 mb
39.7 mb
39.8 mb
39.9 mb
40 mb
-log10(P)
60
50
40
30
20
10
r2
>0.8
0.6-0.8
0.4-0.6
0.2-0.4
<0.2
C15orf54
THBS1
Cingulate
dlPFC
Temporal
Angular
Corrected for average TH
Uncorrected
<=-15
-10
-5
0
5
10
>=15
Zscore

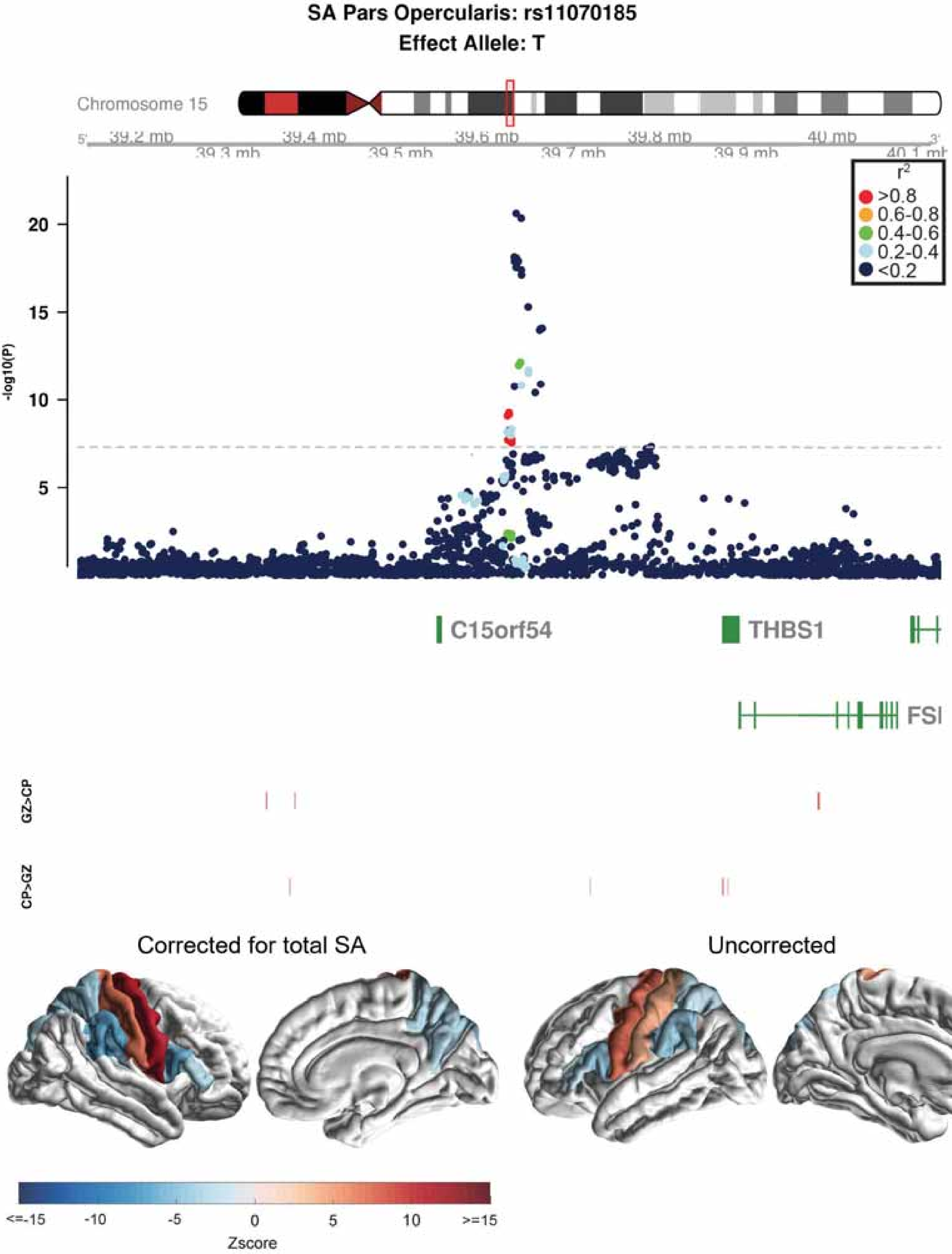

SA Pars Opercularis: rs11070185
Effect Allele: T
Chromosome 15
39.2 mb
39.3 mb
39.4 mb
39.5 mb
39.6 mb
39.7 mb
39.8 mb
39.9 mb
40 mb
-log10(P)
20
15
10
5
r2
>0.8
0.6-0.8
0.4-0.6
0.2-0.4
<0.2
C15orf54
THBS1
GZ>CP
CP>GZ
Corrected for total SA
Uncorrected
<=-15
-10
-5
0
5
10
>=15
Zscore

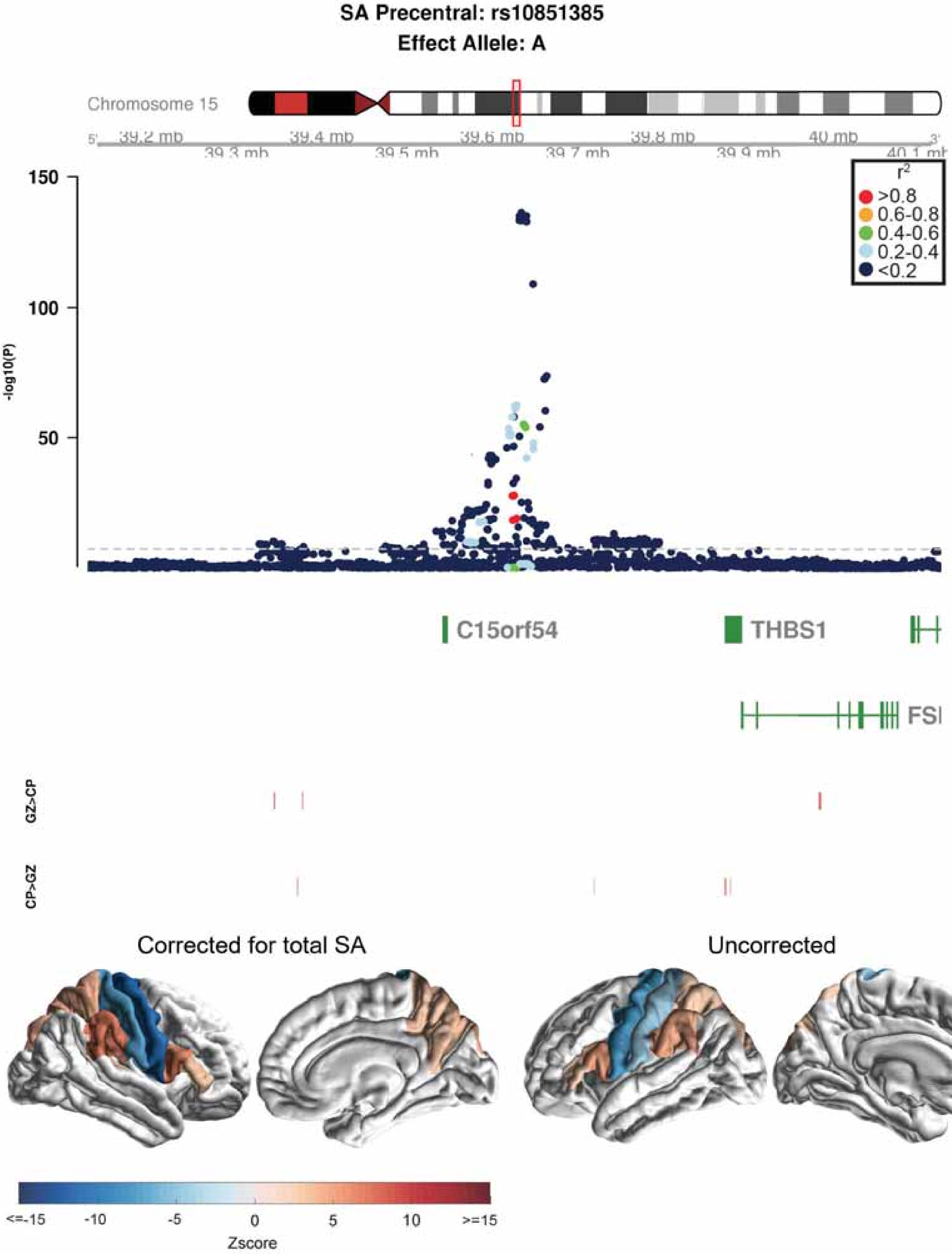

SA Precentral: rs10851385
Effect Allele: A
Chromosome 15
39.2 mb
39.3 mb
39.4 mb
39.5 mb
39.6 mb
39.7 mb
39.8 mb
39.9 mb
40 mb
40.1 mb
150
100
50
-log10(P)
r2
>0.8
0.6-0.8
0.4-0.6
0.2-0.4
<0.2
C15orf54
THBS1
FSI
GZ>CP
CP>GZ
Corrected for total SA
Uncorrected
<=-15
-10
-5
0
5
10
>=15
Zscore

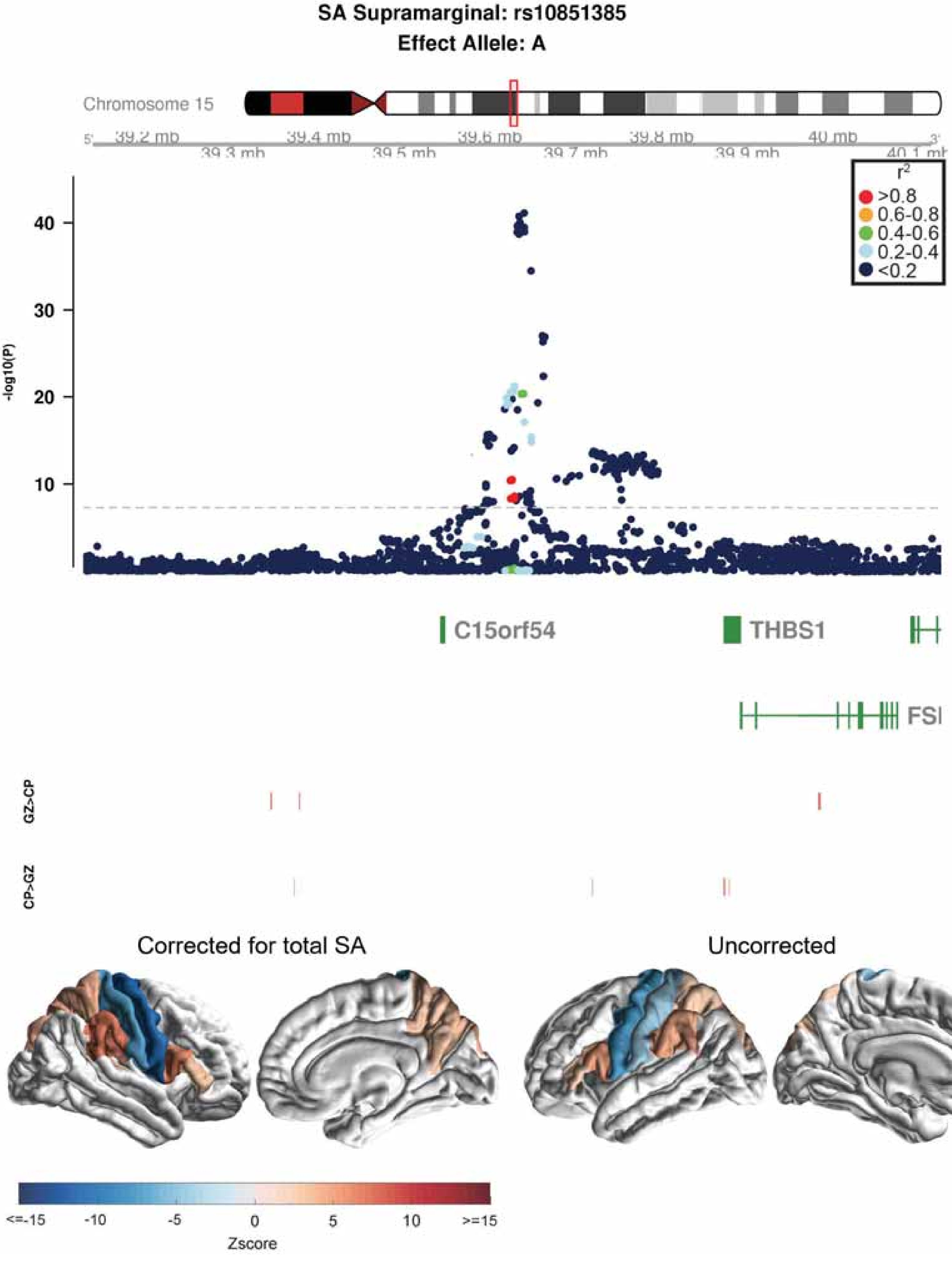

SA Supramarginal: rs10851385
Effect Allele: A
Chromosome 15
39.2 mb
39.3 mb
39.4 mb
39.5 mb
39.6 mb
39.7 mb
39.8 mb
39.9 mb
40 mb
40.1 mb
r2
>0.8
0.6-0.8
0.4-0.6
0.2-0.4
<0.2
-log10(P)
10
20
30
40
C15orf54
THBS1
FSI
GZ>CP
CP>GZ
Corrected for total SA
Uncorrected
<=-15
-10
-5
0
5
10
>=15
Zscore

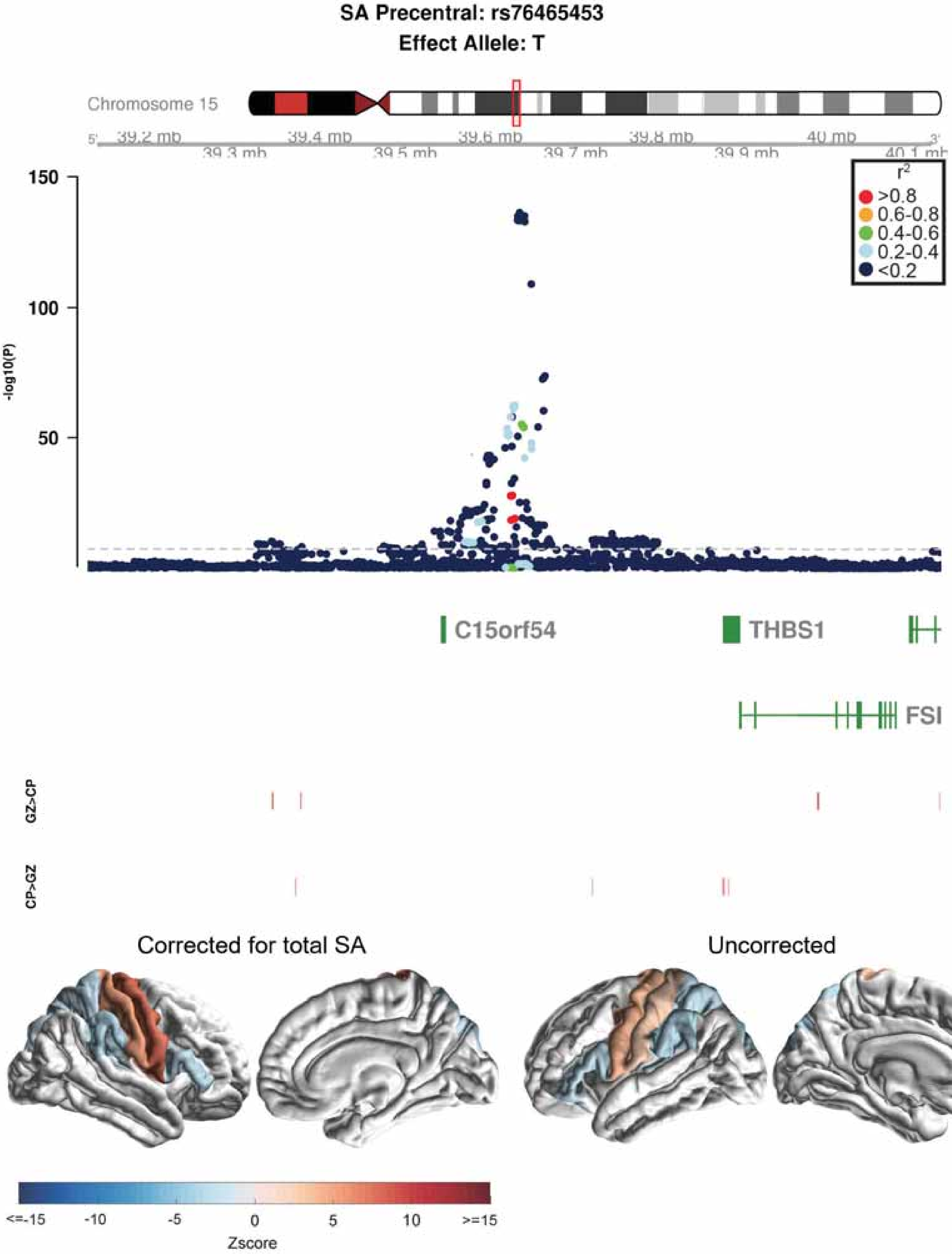

SA Precentral: rs76465453
Effect Allele: T
Chromosome 15
39.2 mb
39.3 mb
39.4 mb
39.5 mb
39.6 mb
39.7 mb
39.8 mb
39.9 mb
40 mb
40.1 mb
150
100
50
-log10(P)
r2
>0.8
0.6-0.8
0.4-0.6
0.2-0.4
<0.2
C15orf54
THBS1
FSI
GZ>CP
CP>GZ
Corrected for total SA
Uncorrected
<=-15
-10
-5
0
5
10
>=15
Zscore

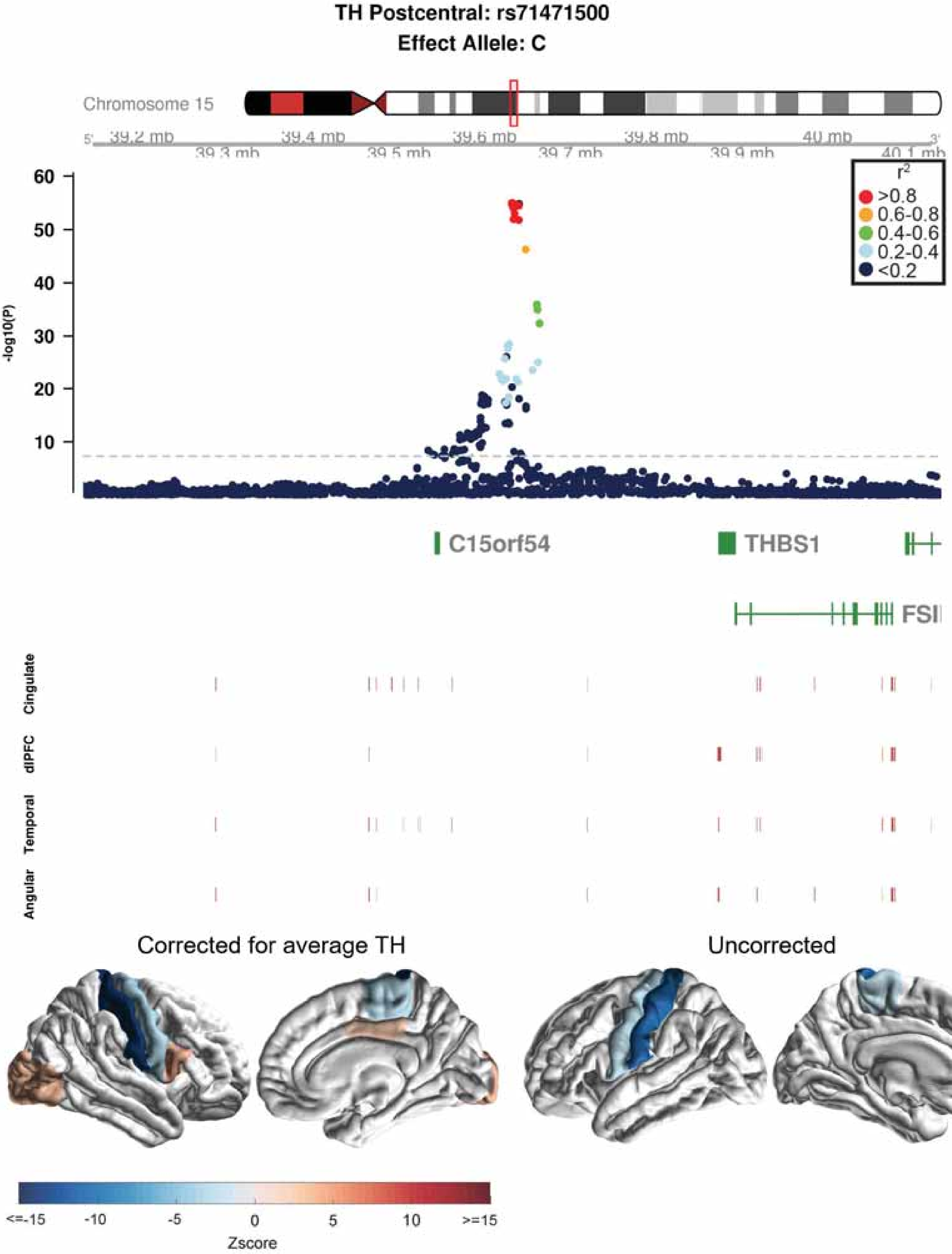

TH Postcentral: rs71471500
Effect Allele: C
Chromosome 15
39.2 mb
39.3 mb
39.4 mb
39.5 mb
39.6 mb
39.7 mb
39.8 mb
39.9 mb
40 mb
40.1 mb
-log10(P)
60
50
40
30
20
10
r2
>0.8
0.6-0.8
0.4-0.6
0.2-0.4
<0.2
C15orf54
THBS1
FSI
Cingulate
dlPFC
Temporal
Angular
Corrected for average TH
Uncorrected
<=-15
-10
-5
0
5
10
>=15
Zscore

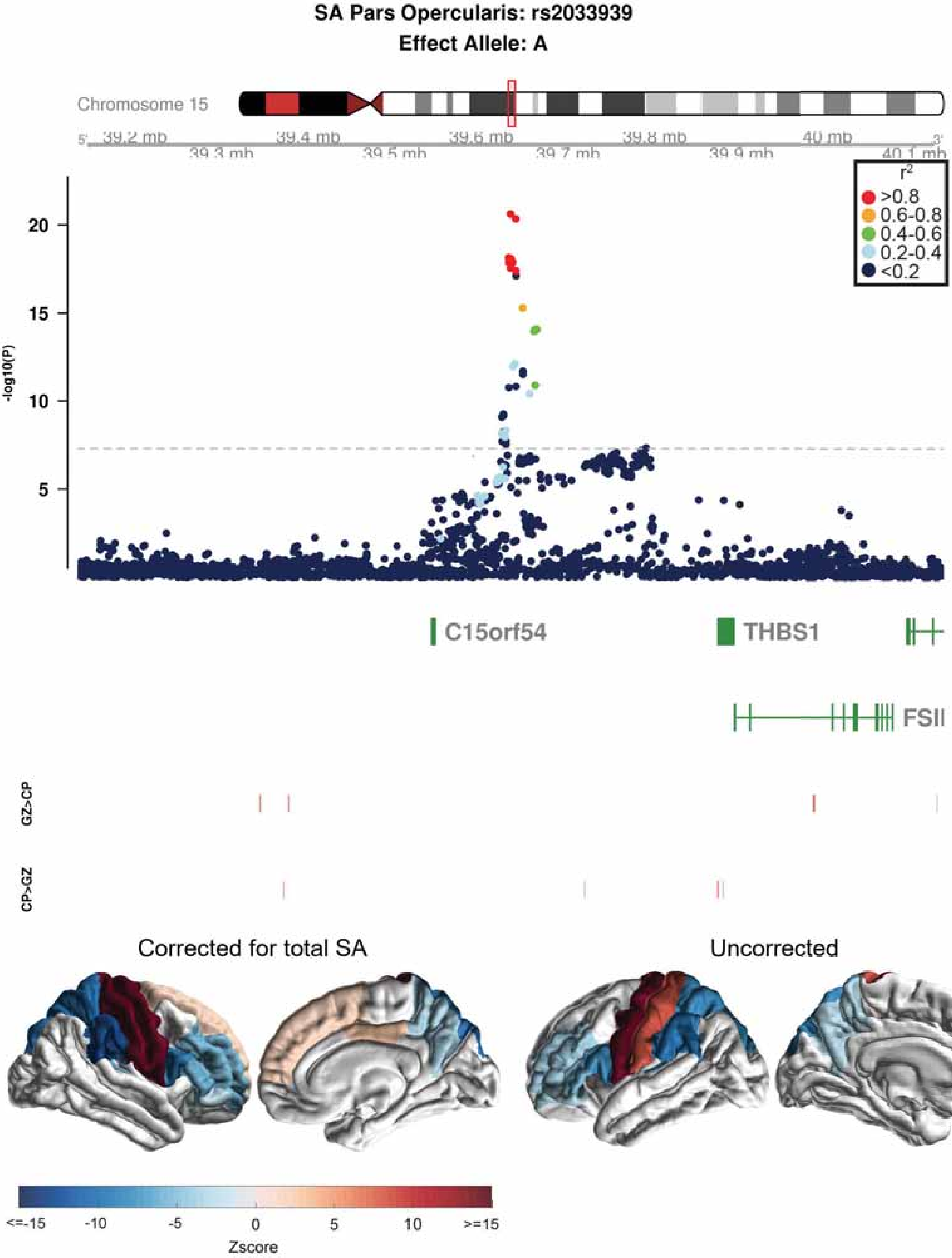
SA Pars Opercularis: rs2033939
Effect Allele: A
Chromosome 15
39.2 mb
39.3 mb
39.4 mb
39.5 mb
39.6 mb
39.7 mb
39.8 mb
39.9 mb
40 mb
40.1 mb
r2
>0.8
0.6-0.8
0.4-0.6
0.2-0.4
<0.2
-log10(P)
20
15
10
5
C15orf54
THBS1
FSIP1
GZ>CP
CP>GZ
Corrected for total SA
Uncorrected
<=-15
-10
-5
0
5
10
>=15
Zscore

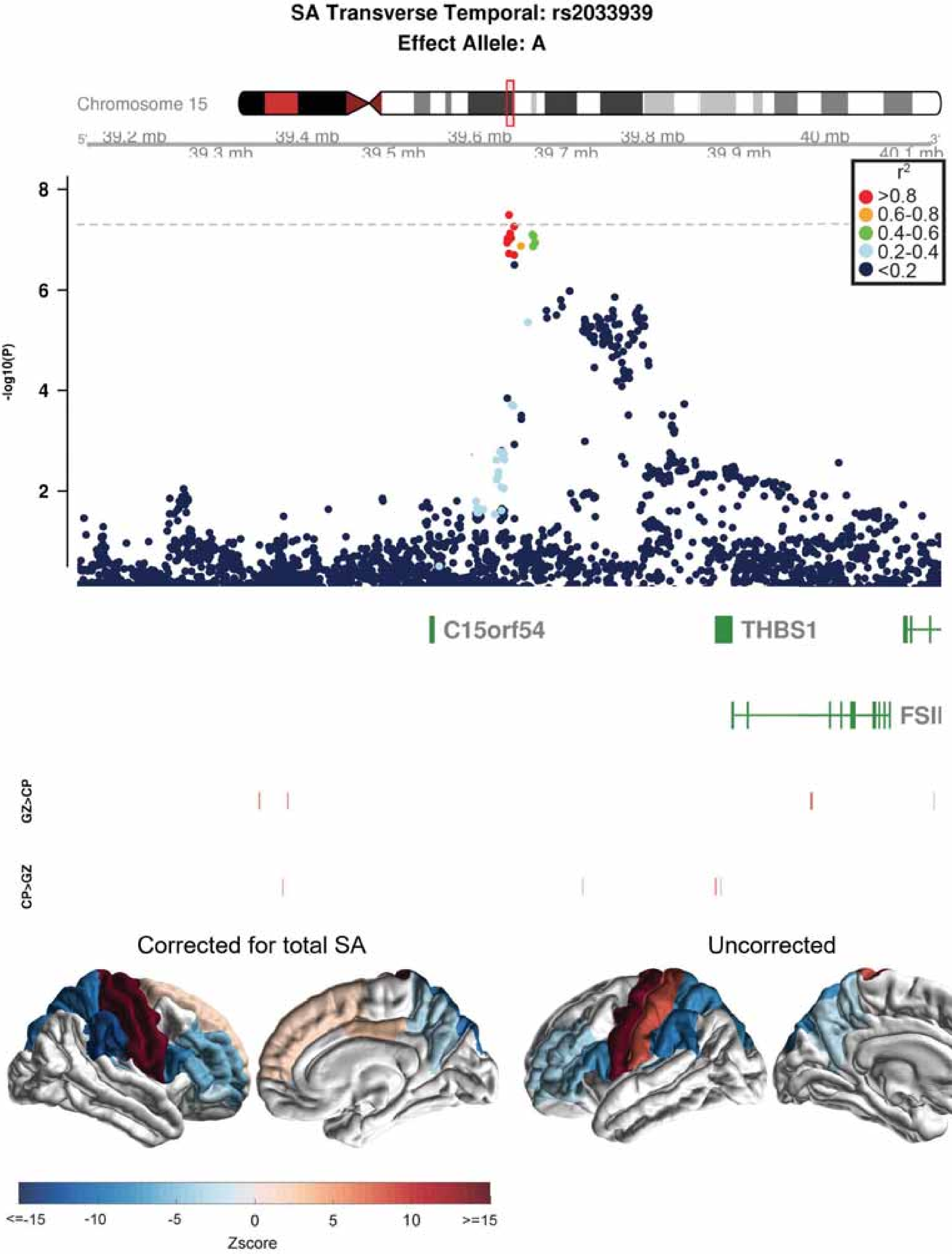
SA Transverse Temporal: rs2033939
Effect Allele: A
Chromosome 15
39.2 mb
39.3 mb
39.4 mb
39.5 mb
39.6 mb
39.7 mb
39.8 mb
39.9 mb
40 mb
40.1 mb
r2
>0.8
0.6-0.8
0.4-0.6
0.2-0.4
<0.2
-log10(P)
8
6
4
2
C15orf54
THBS1
FSIP1
GZ>CP
CP>GZ
Corrected for total SA
Uncorrected
<=-15
-10
-5
0
5
10
>=15
Zscore

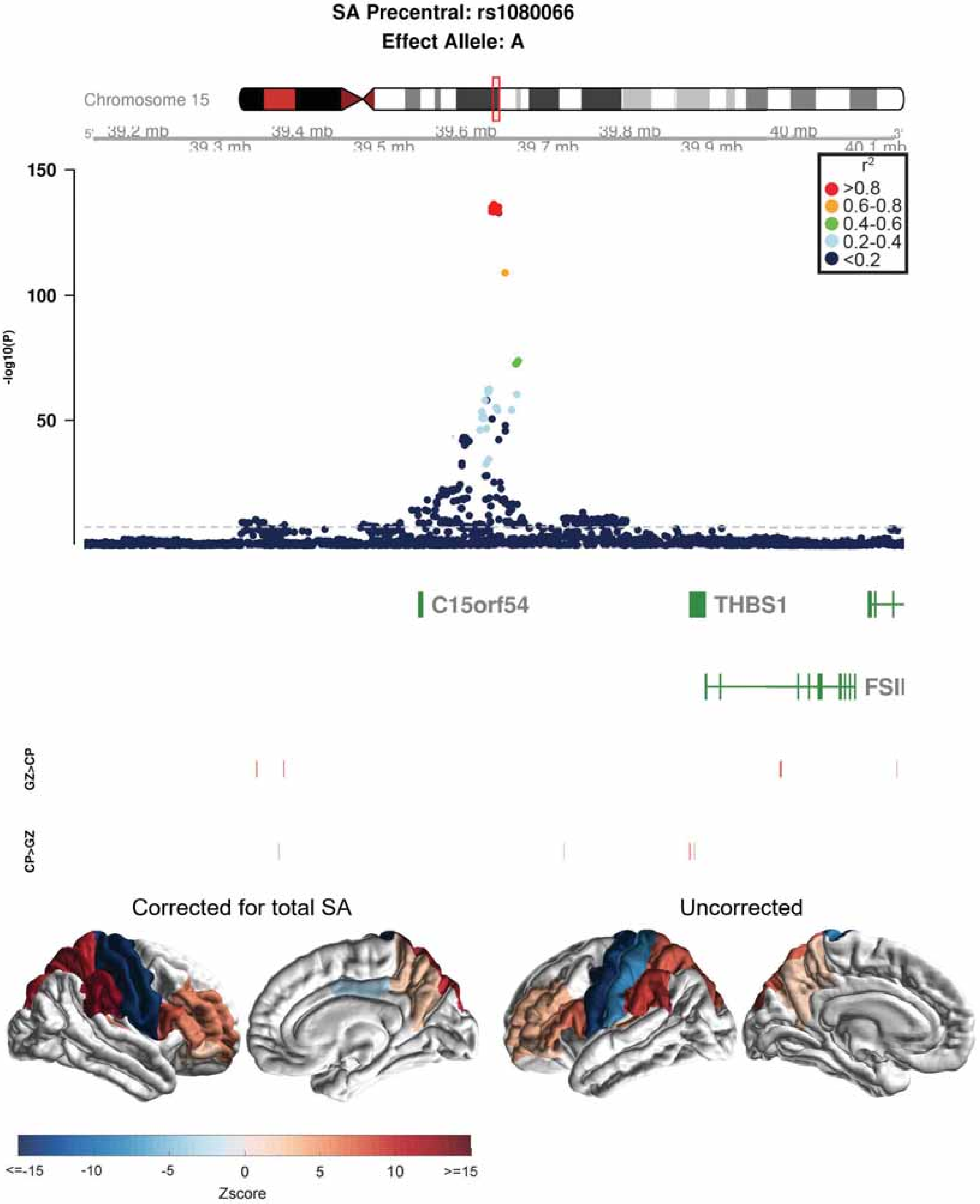

SA Precentral: rs1080066
Effect Allele: A
Chromosome 15
39.2 mb
39.3 mb
39.4 mb
39.5 mb
39.6 mb
39.7 mb
39.8 mb
39.9 mb
40 mb
40.1 mb
150
100
50
-log10(P)
r2
>0.8
0.6-0.8
0.4-0.6
0.2-0.4
<0.2
C15orf54
THBS1
FSII
GZ>CP
CP>GZ
Corrected for total SA
Uncorrected
<=-15
-10
-5
0
5
10
>=15
Zscore

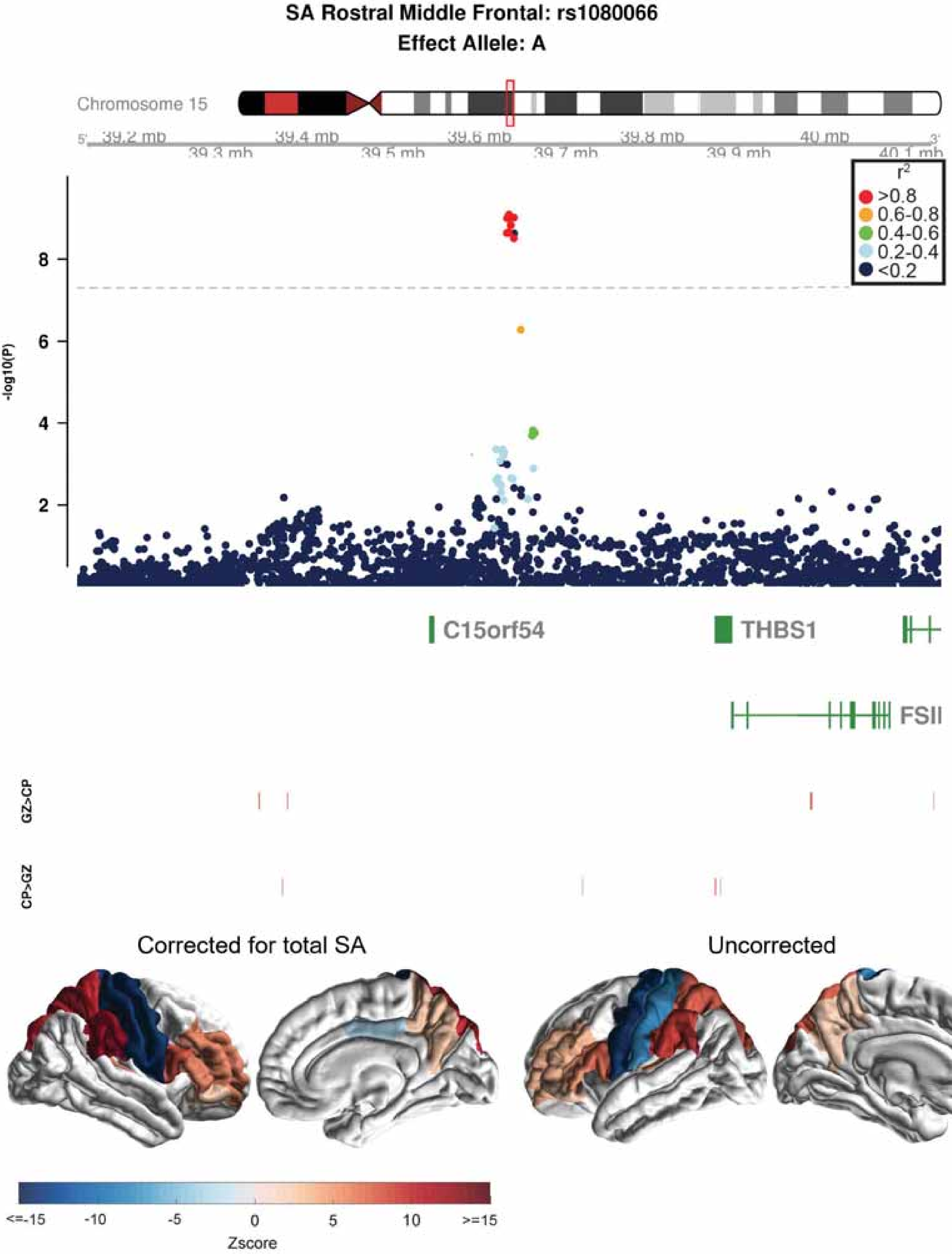
SA Rostral Middle Frontal: rs1080066
Effect Allele: A
Chromosome 15
39.2 mb
39.3 mb
39.4 mb
39.5 mb
39.6 mb
39.7 mb
39.8 mb
39.9 mb
40 mb
40.1 mb
r2
>0.8
0.6-0.8
0.4-0.6
0.2-0.4
<0.2
-log10(P)
C15orf54
THBS1
FSIP1
GZ>CP
CP>GZ
Corrected for total SA
Uncorrected
<=-15
-10
-5
0
5
10
>=15
Zscore

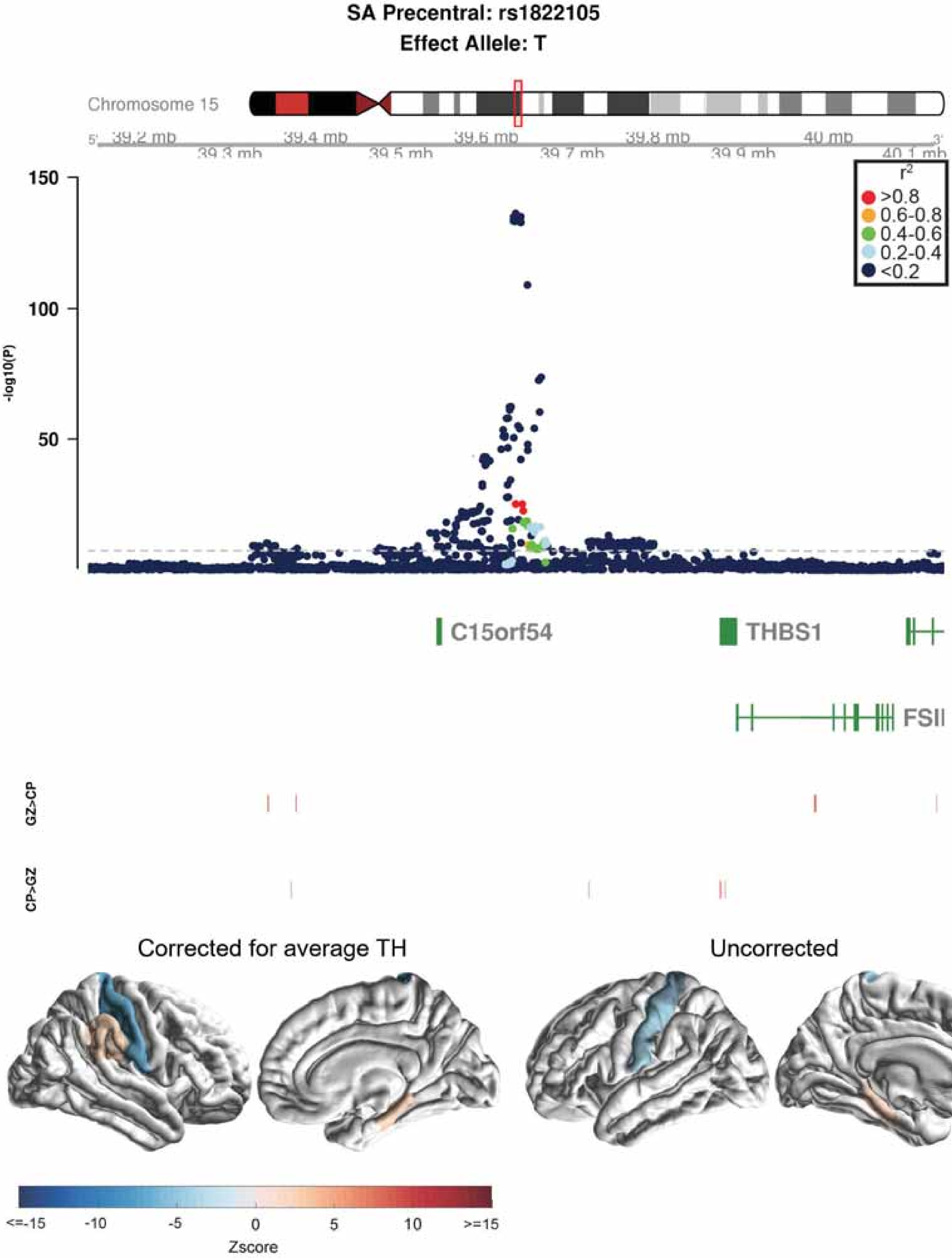
SA Precentral: rs1822105
Effect Allele: T
Chromosome 15
39.2 mb
39.3 mb
39.4 mb
39.5 mb
39.6 mb
39.7 mb
39.8 mb
39.9 mb
40 mb
40.1 mb
r²
>0.8
0.6-0.8
0.4-0.6
0.2-0.4
<0.2
150
100
50
-log10(P)
C15orf54
THBS1
FSIP1
GZ>CP
CP>GZ
Corrected for average TH
Uncorrected
<=-15
-10
-5
0
5
10
>=15
Zscore

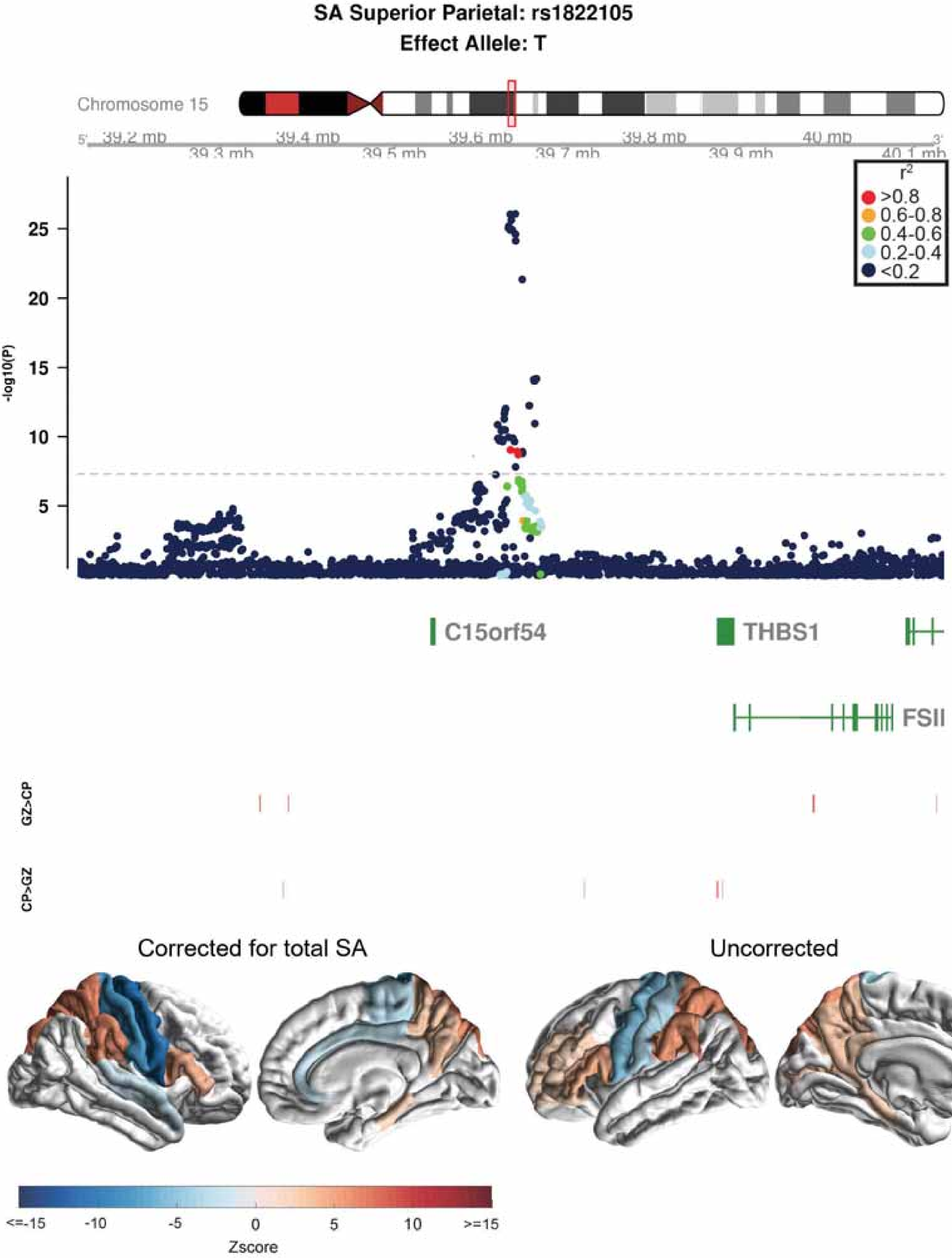

SA Superior Parietal: rs1822105
Effect Allele: T
Chromosome 15
39.2 mb
39.3 mb
39.4 mb
39.5 mb
39.6 mb
39.7 mb
39.8 mb
39.9 mb
40 mb
40.1 mb
r2
>0.8
0.6-0.8
0.4-0.6
0.2-0.4
<0.2
-log10(P)
25
20
15
10
5
C15orf54
THBS1
FSIP1
GZ>CP
CP>GZ
Corrected for total SA
Uncorrected
<=-15
-10
-5
0
5
10
>=15
Zscore

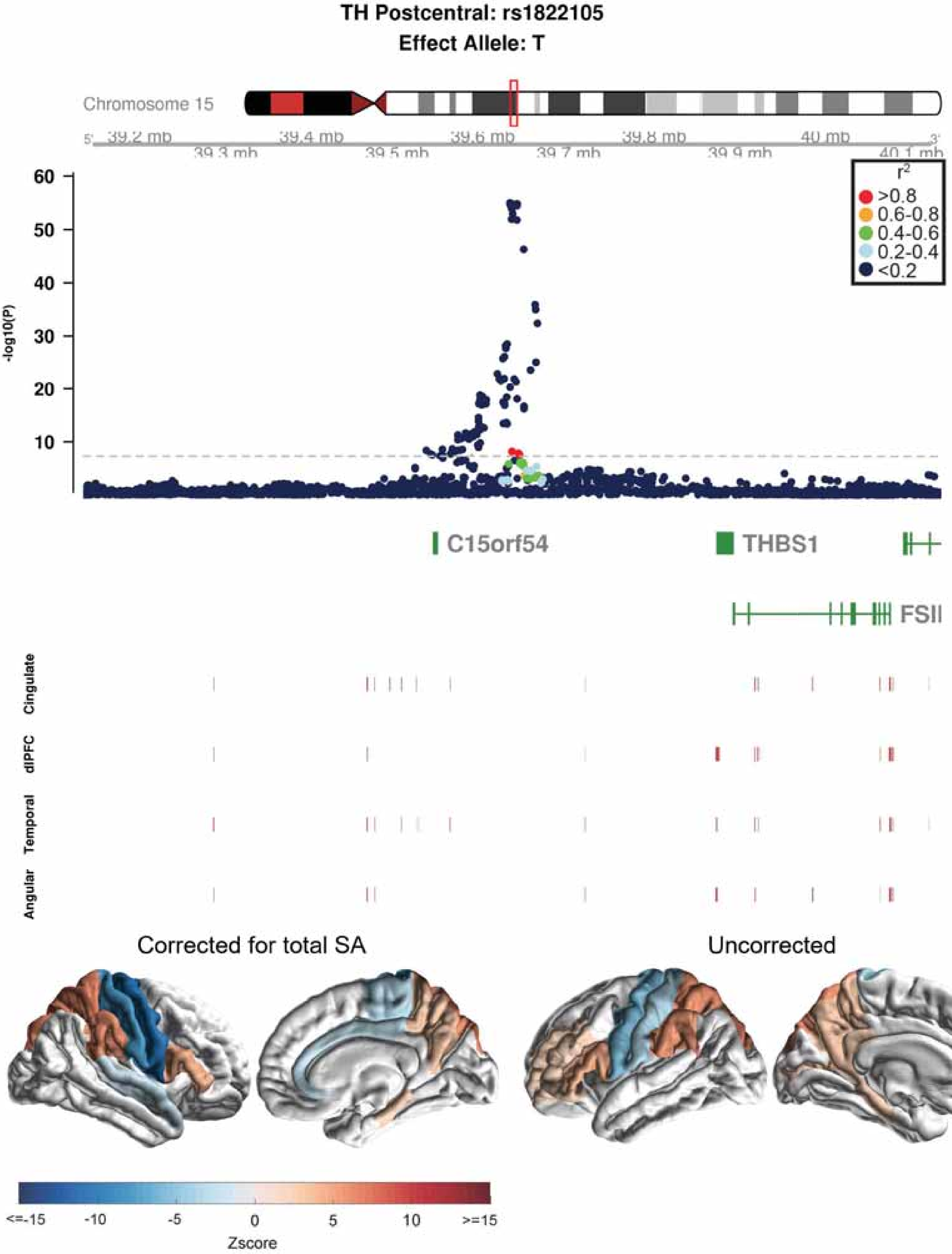
TH Postcentral: rs1822105
Effect Allele: T
Chromosome 15
39.2 mb
39.3 mb
39.4 mb
39.5 mb
39.6 mb
39.7 mb
39.8 mb
39.9 mb
40 mb
40.1 mb
-log10(P)
60
50
40
30
20
10
$r^2$
>0.8
0.6-0.8
0.4-0.6
0.2-0.4
<0.2
C15orf54
THBS1
FSIP1
Cingulate
dlPFC
Temporal
Angular
Corrected for total SA
Uncorrected
<=-15
-10
-5
0
5
10
>=15
Zscore

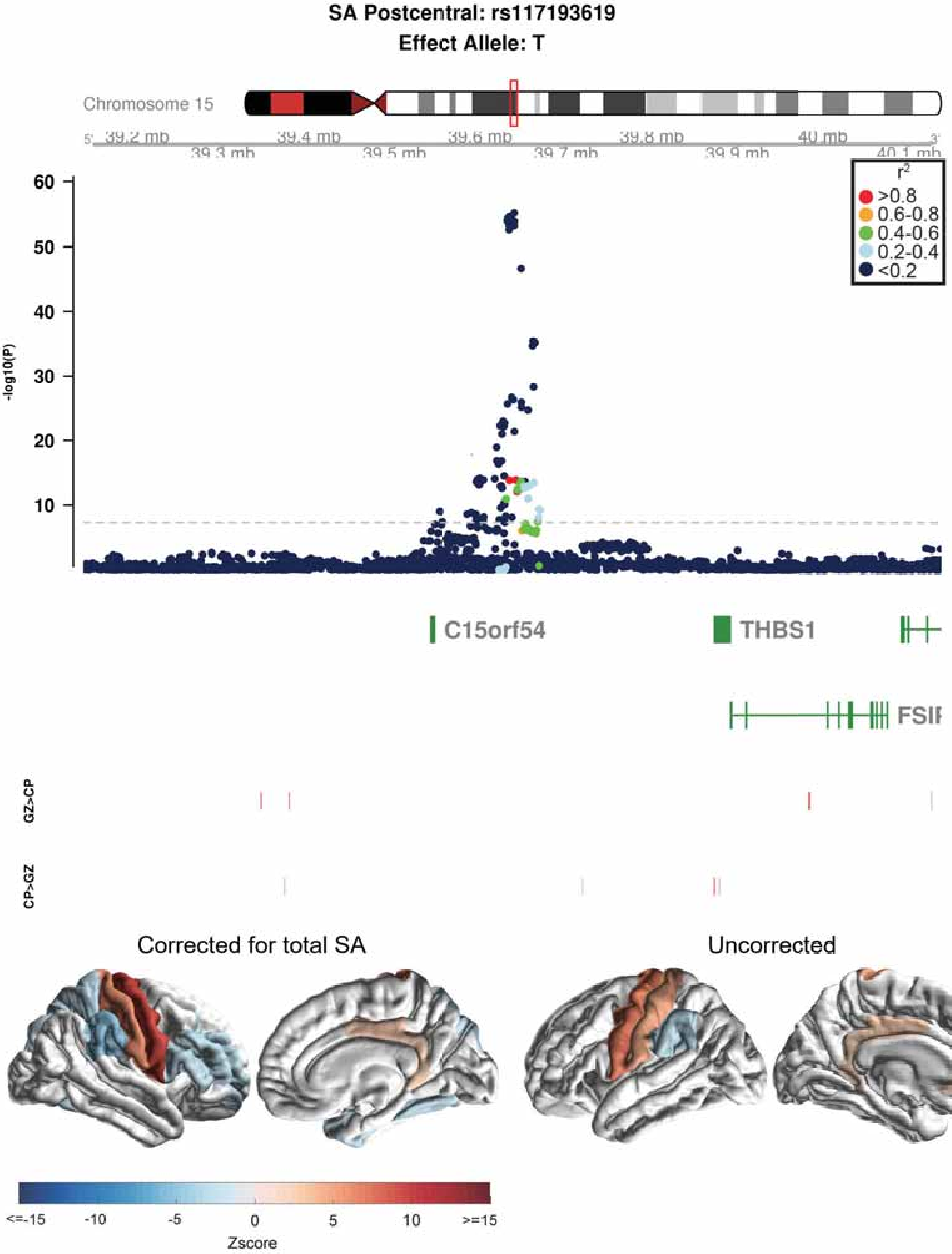
SA Postcentral: rs117193619
Effect Allele: T
Chromosome 15
39.2 mb
39.3 mb
39.4 mb
39.5 mb
39.6 mb
39.7 mb
39.8 mb
39.9 mb
40 mb
40.1 mb
5'
3'
r2
>0.8
0.6-0.8
0.4-0.6
0.2-0.4
<0.2
-log10(P)
60
50
40
30
20
10
C15orf54
THBS1
FSIP
GZ>CP
CP>GZ
Corrected for total SA
Uncorrected
<=-15
-10
-5
0
5
10
>=15
Zscore

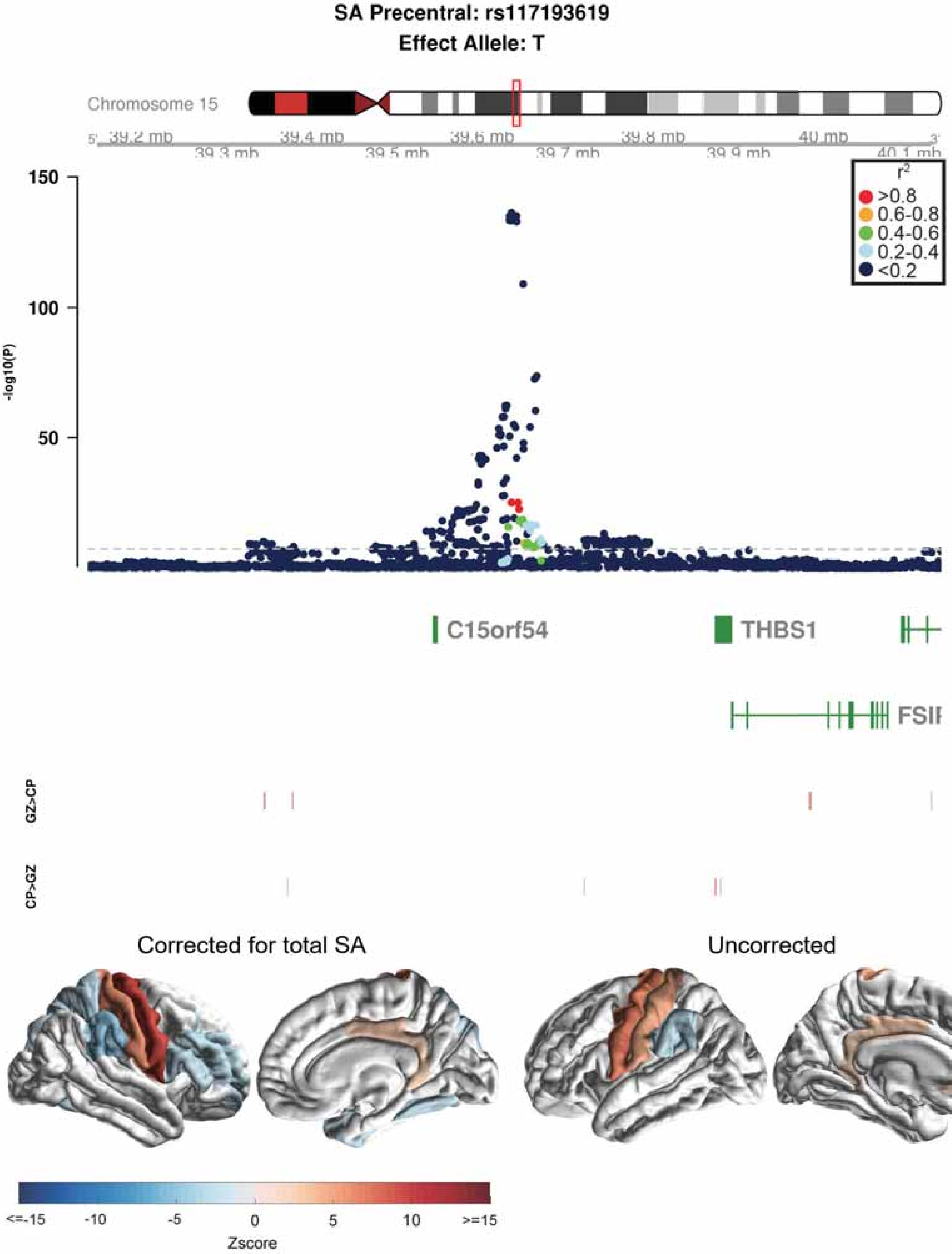
SA Precentral: rs117193619
Effect Allele: T
Chromosome 15
39.2 mb
39.3 mb
39.4 mb
39.5 mb
39.6 mb
39.7 mb
39.8 mb
39.9 mb
40 mb
40.1 mb
-log10(P)
150
100
50
r2
>0.8
0.6-0.8
0.4-0.6
0.2-0.4
<0.2
C15orf54
THBS1
FSII
GZ>CP
CP>GZ
Corrected for total SA
Uncorrected
<=-15
-10
-5
0
5
10
>=15
Zscore

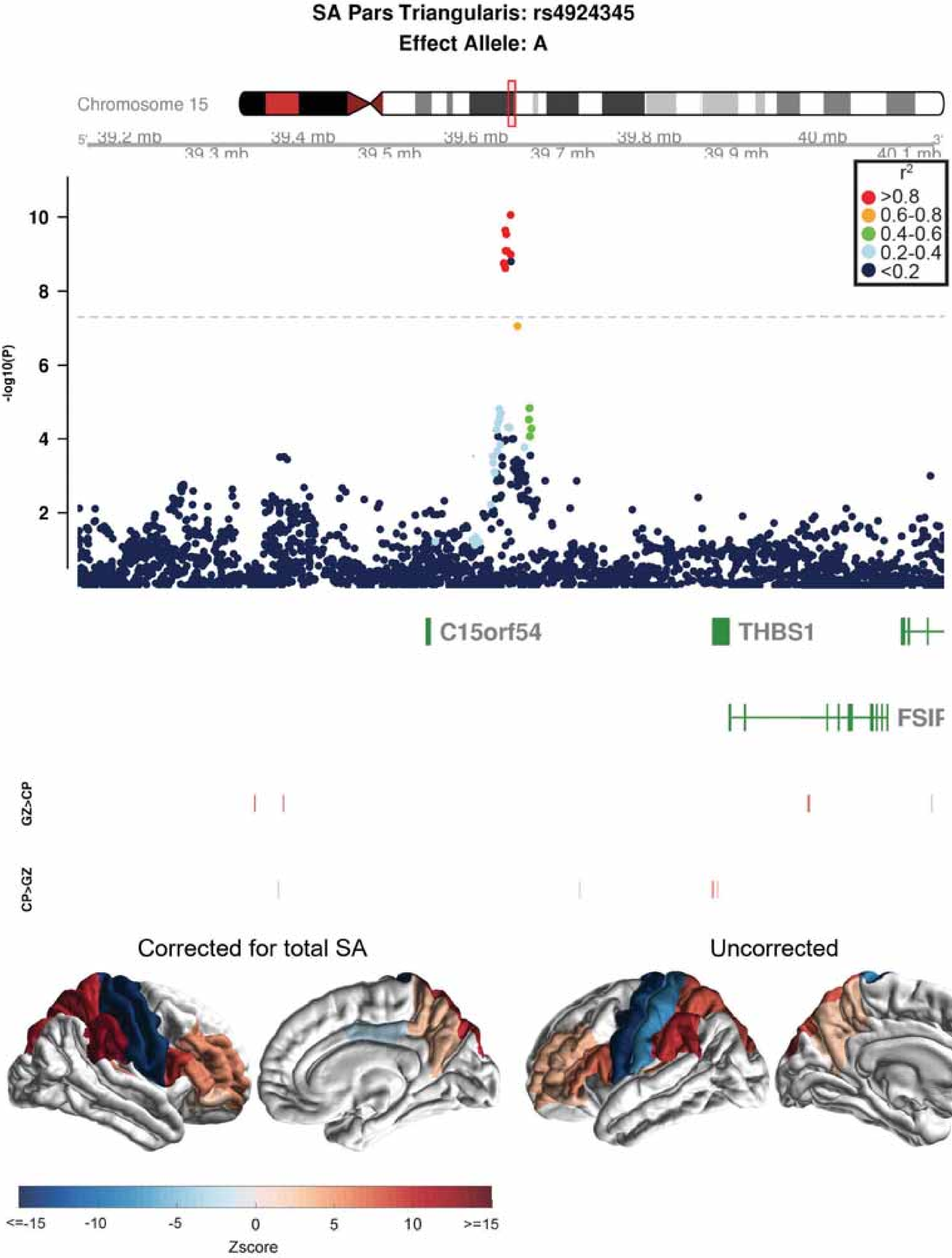
SA Pars Triangularis: rs4924345
Effect Allele: A
Chromosome 15
39.2 mb
39.3 mb
39.4 mb
39.5 mb
39.6 mb
39.7 mb
39.8 mb
39.9 mb
40 mb
40.1 mb
r2
>0.8
0.6-0.8
0.4-0.6
0.2-0.4
<0.2
-log10(P)
10
8
6
4
2
C15orf54
THBS1
FSIP
GZ>CP
CP>GZ
Corrected for total SA
Uncorrected
<=-15
-10
-5
0
5
10
>=15
Zscore

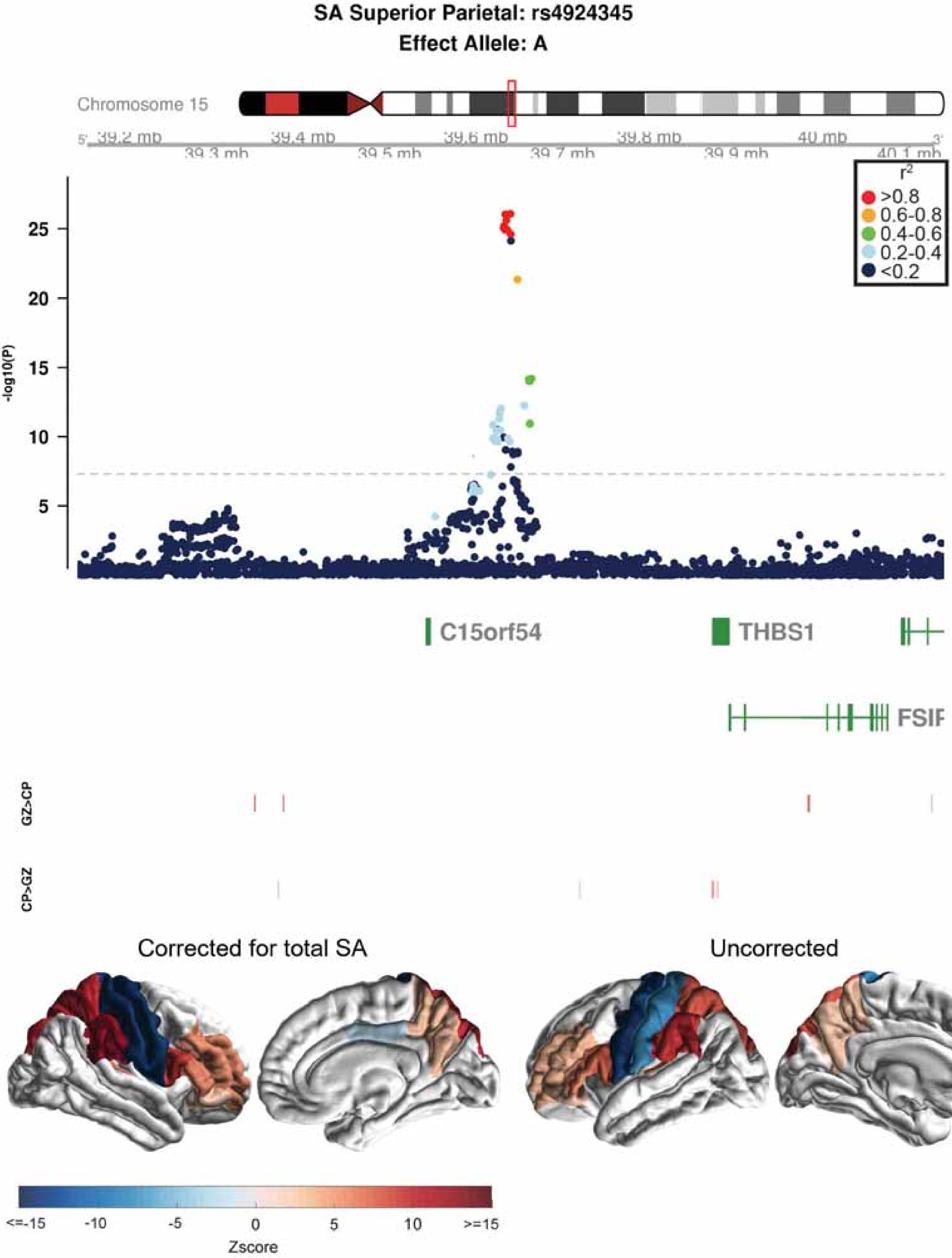
SA Superior Parietal: rs4924345
Effect Allele: A
Chromosome 15
39.2 mb
39.3 mb
39.4 mb
39.5 mb
39.6 mb
39.7 mb
39.8 mb
39.9 mb
40 mb
40.1 mb
r2
>0.8
0.6-0.8
0.4-0.6
0.2-0.4
<0.2
-log10(P)
25
20
15
10
5
C15orf54
THBS1
FSIP
GZ>CP
CP>GZ
Corrected for total SA
Uncorrected
<=-15
-10
-5
0
5
10
>=15
Zscore

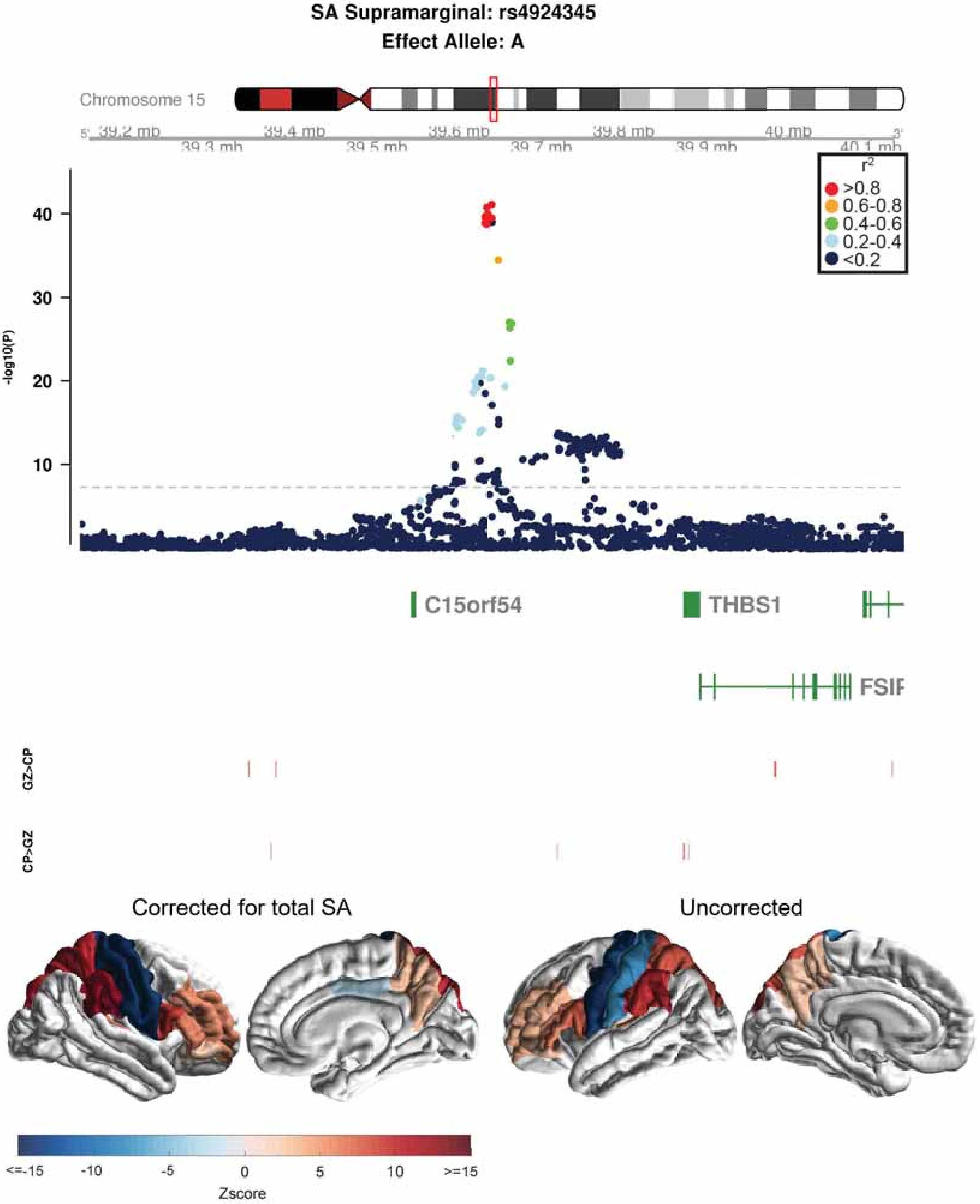

SA Supramarginal: rs4924345
Effect Allele: A
Chromosome 15
39.2 mb
39.3 mb
39.4 mb
39.5 mb
39.6 mb
39.7 mb
39.8 mb
39.9 mb
40 mb
40.1 mb
r2
>0.8
0.6-0.8
0.4-0.6
0.2-0.4
<0.2
-log10(P)
40
30
20
10
C15orf54
THBS1
FSIP
GZ>CP
CP>GZ
Corrected for total SA
Uncorrected
<=-15
-10
-5
0
5
10
>=15
Zscore

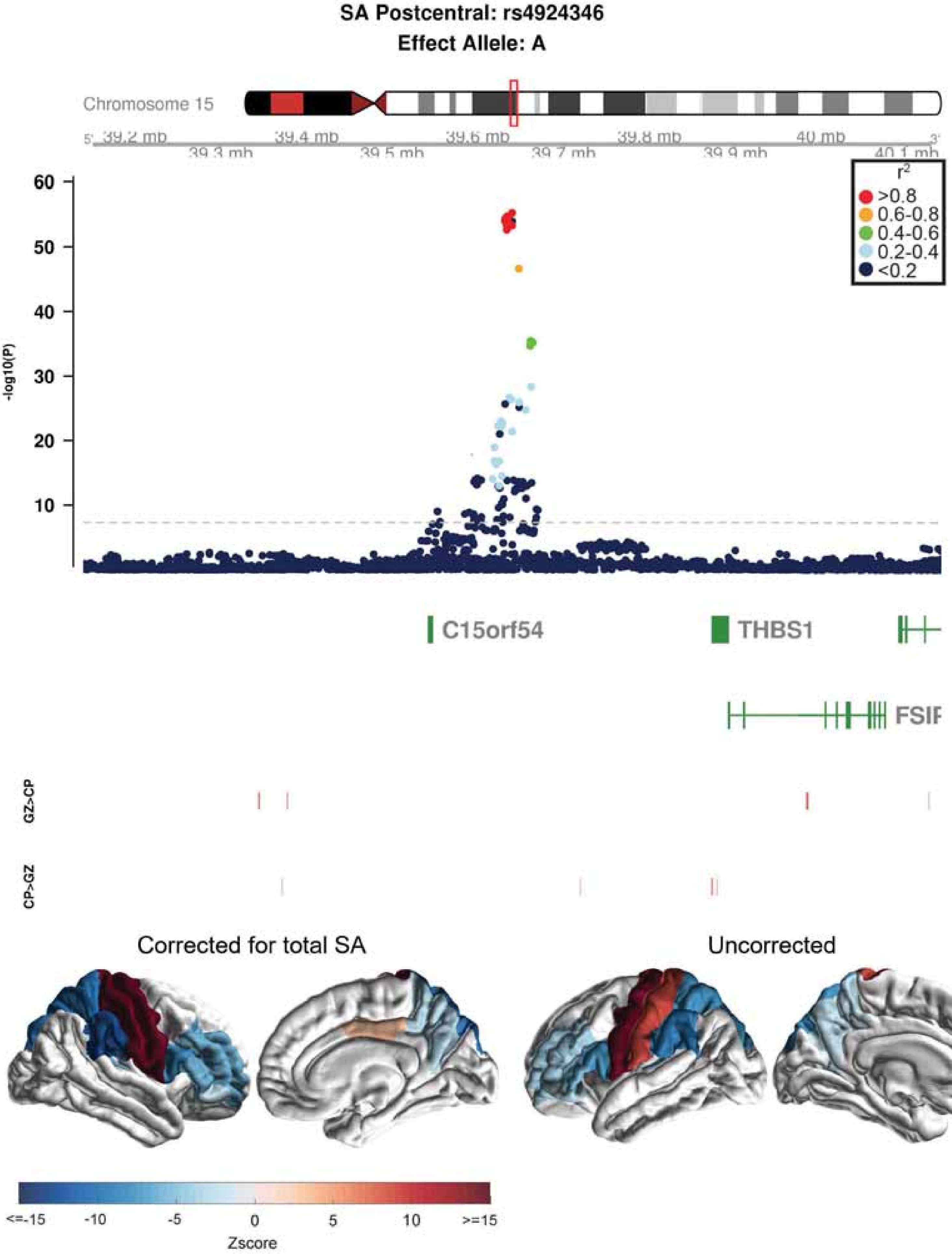

SA Postcentral: rs4924346
Effect Allele: A
Chromosome 15
39.2 mb
39.3 mb
39.4 mb
39.5 mb
39.6 mb
39.7 mb
39.8 mb
39.9 mb
40 mb
40.1 mb
r2
>0.8
0.6-0.8
0.4-0.6
0.2-0.4
<0.2
-log10(P)
C15orf54
THBS1
FSIP
GZ>CP
CP>GZ
Corrected for total SA
Uncorrected
<=-15
-10
-5
0
5
10
>=15
Zscore

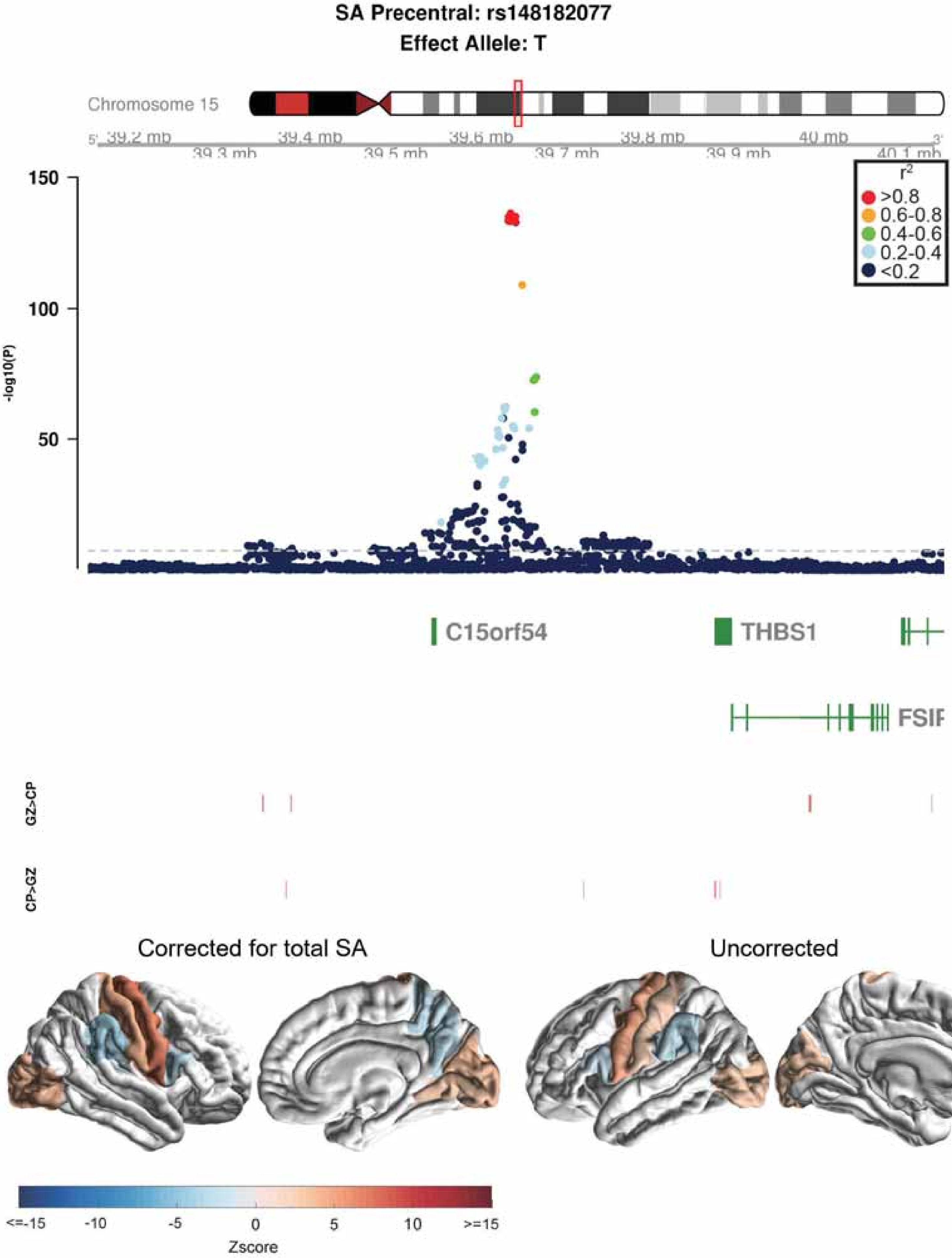
SA Precentral: rs148182077
Effect Allele: T
Chromosome 15
39.2 mb
39.3 mb
39.4 mb
39.5 mb
39.6 mb
39.7 mb
39.8 mb
39.9 mb
40 mb
40.1 mb
150
100
50
-log10(P)
r2
>0.8
0.6-0.8
0.4-0.6
0.2-0.4
<0.2
C15orf54
THBS1
FSIP
GZ>CP
CP>GZ
Corrected for total SA
Uncorrected
<=-15
-10
-5
0
5
10
>=15
Zscore

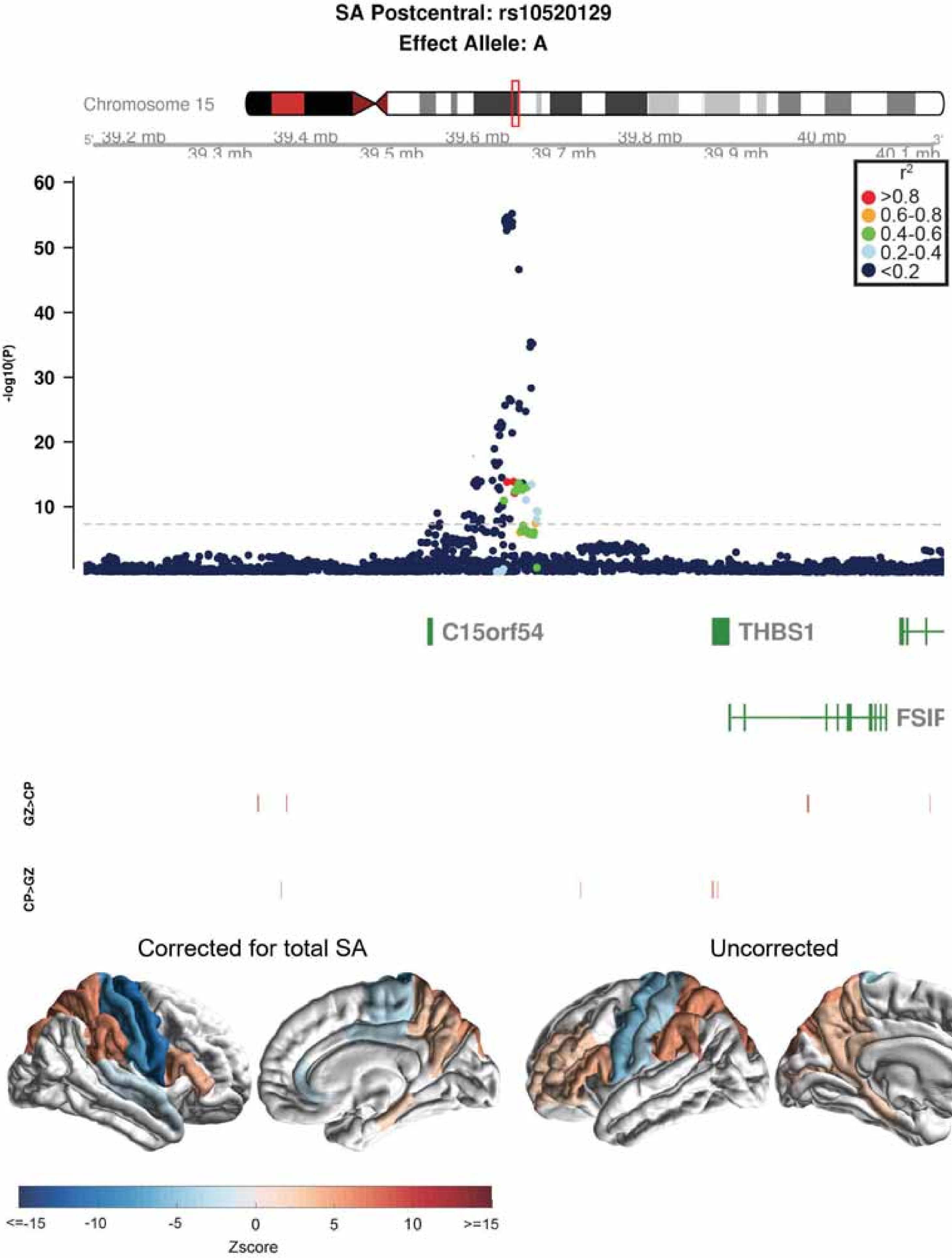

SA Postcentral: rs10520129
Effect Allele: A
Chromosome 15
39.2 mb
39.3 mb
39.4 mb
39.5 mb
39.6 mb
39.7 mb
39.8 mb
39.9 mb
40 mb
40.1 mb
-log10(P)
60
50
40
30
20
10
r2
>0.8
0.6-0.8
0.4-0.6
0.2-0.4
<0.2
C15orf54
THBS1
FSIP
GZ>CP
CP>GZ
Corrected for total SA
Uncorrected
<=-15
-10
-5
0
5
10
>=15
Zscore

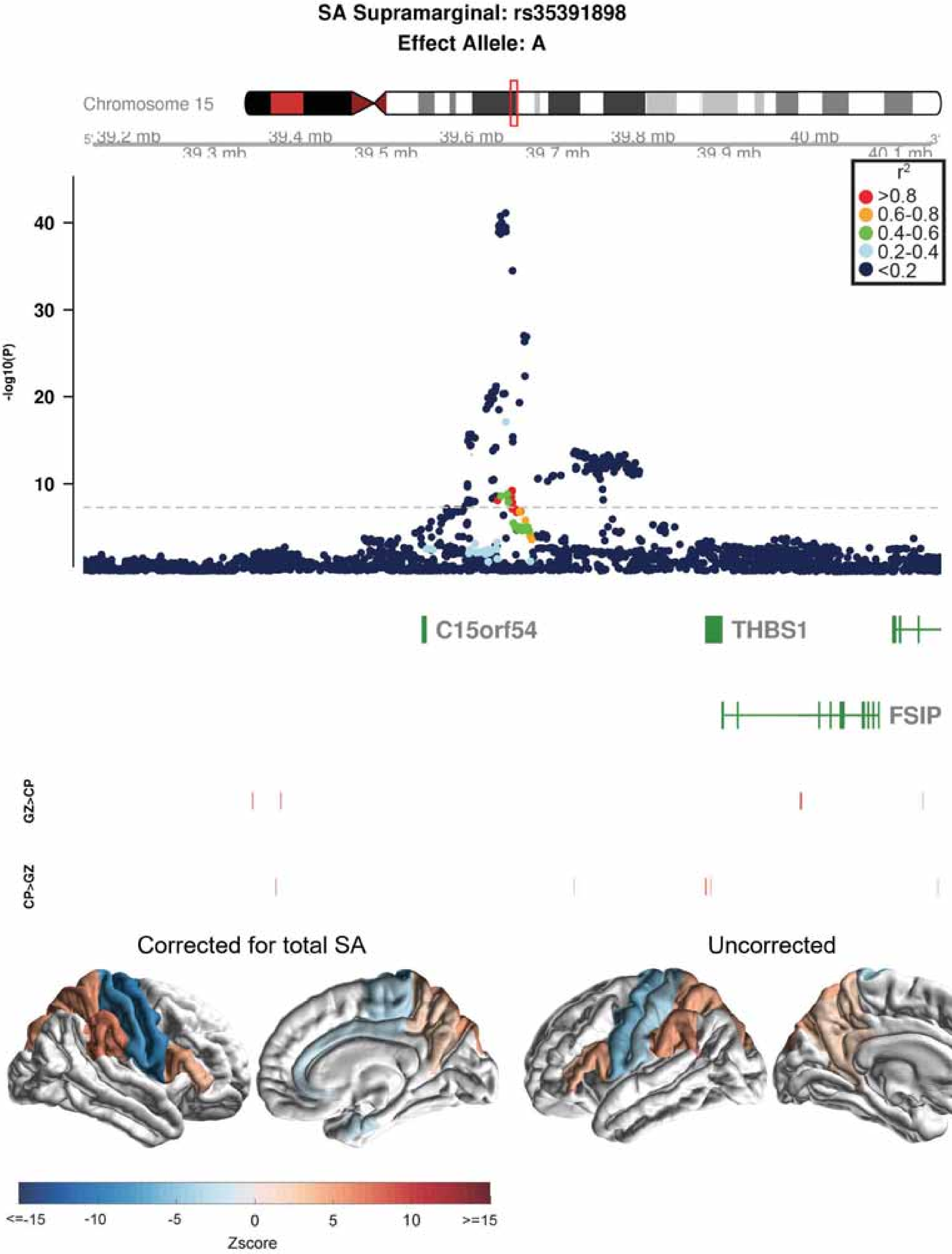

SA Supramarginal: rs35391898
Effect Allele: A
Chromosome 15
39.2 mb
39.3 mb
39.4 mb
39.5 mb
39.6 mb
39.7 mb
39.8 mb
39.9 mb
40 mb
40.1 mb
r2
>0.8
0.6-0.8
0.4-0.6
0.2-0.4
<0.2
-log10(P)
40
30
20
10
C15orf54
THBS1
FSIP
GZ>CP
CP>GZ
Corrected for total SA
Uncorrected
<=-15
-10
-5
0
5
10
>=15
Zscore

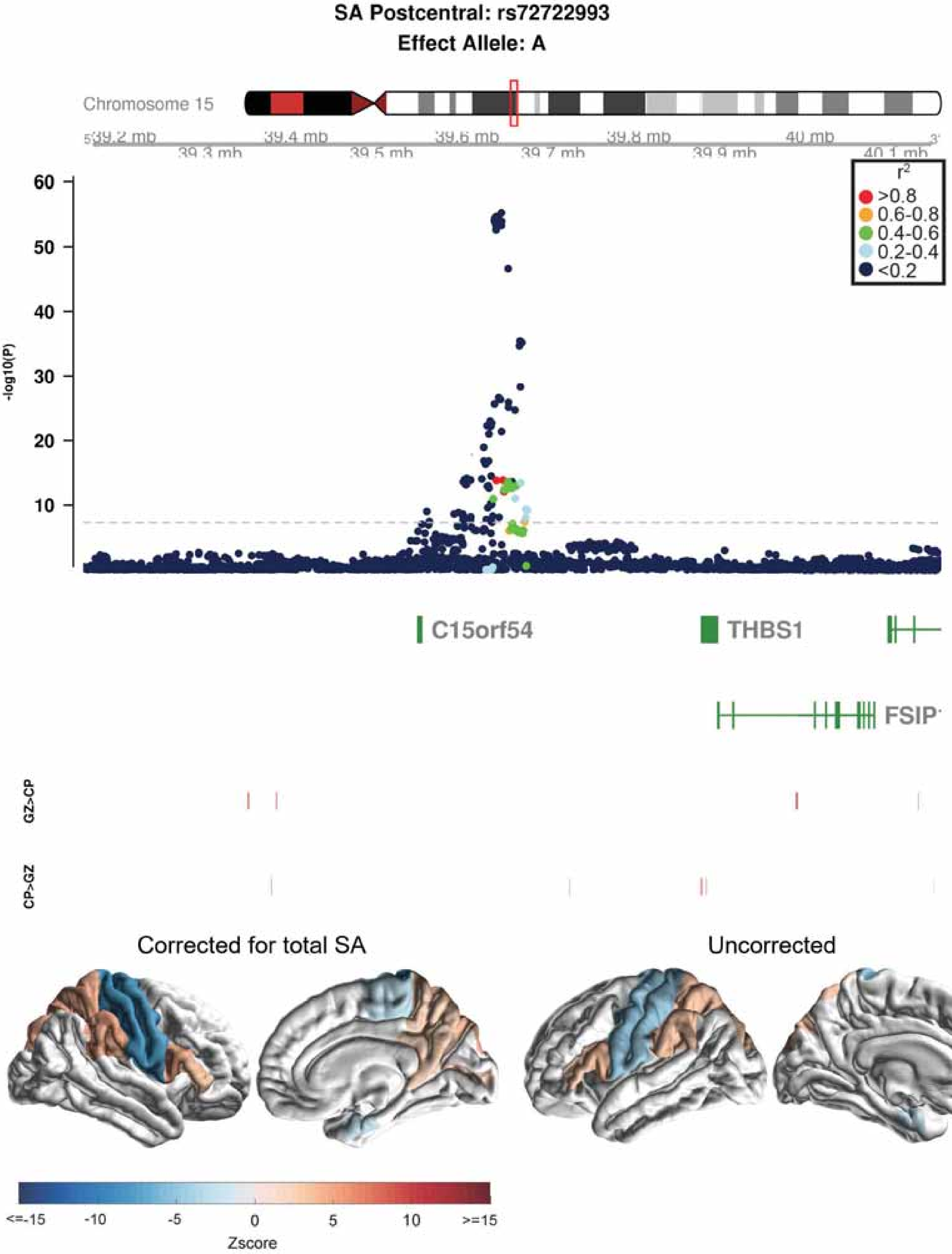
SA Postcentral: rs72722993
Effect Allele: A
Chromosome 15
39.2 mb
39.3 mb
39.4 mb
39.5 mb
39.6 mb
39.7 mb
39.8 mb
39.9 mb
40 mb
40.1 mb
-log10(P)
r2
>0.8
0.6-0.8
0.4-0.6
0.2-0.4
<0.2
C15orf54
THBS1
FSIP
GZ>CP
CP>GZ
Corrected for total SA
Uncorrected
<=-15
-10
-5
0
5
10
>=15
Zscore

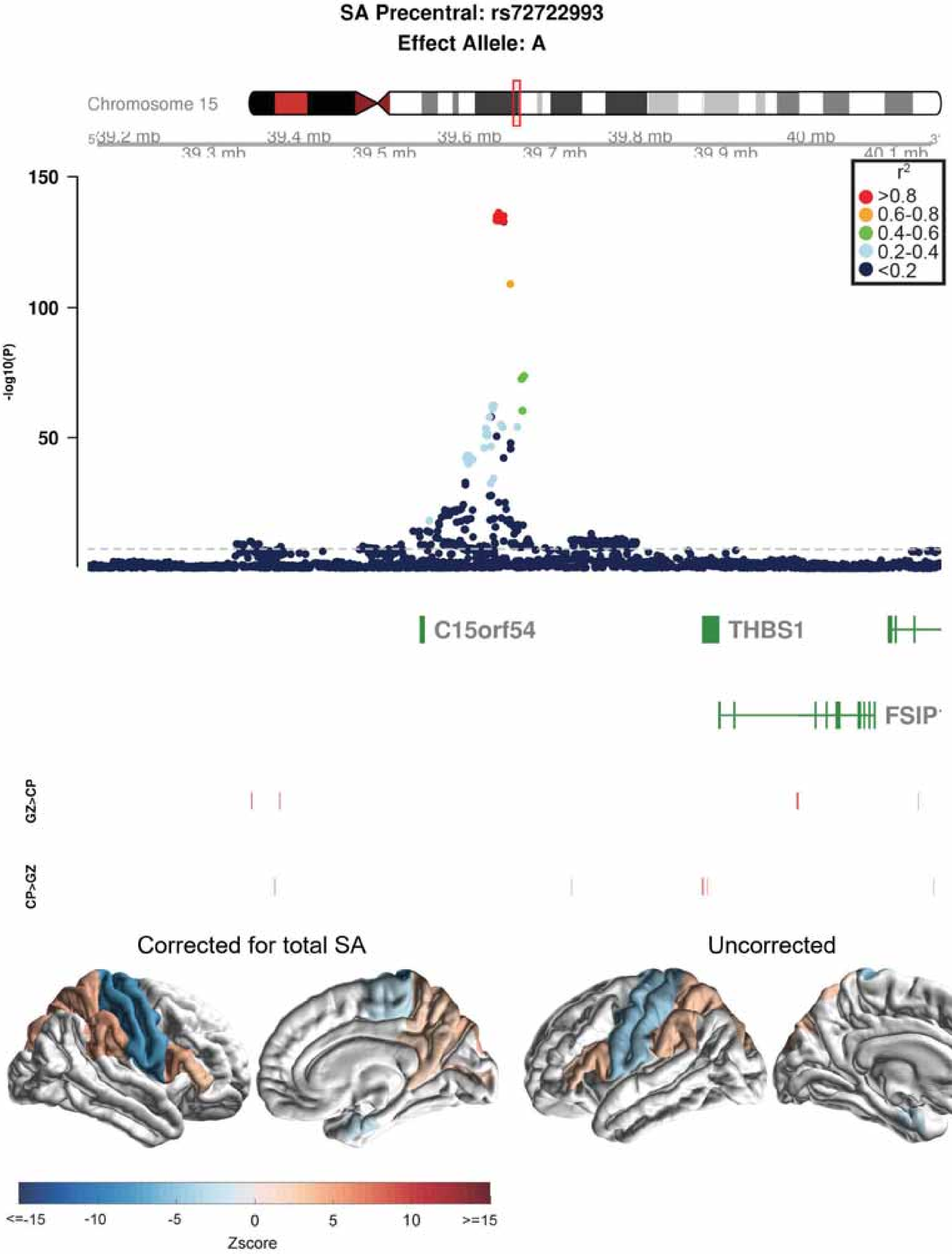
SA Precentral: rs72722993
Effect Allele: A
Chromosome 15
39.2 mb
39.3 mb
39.4 mb
39.5 mb
39.6 mb
39.7 mb
39.8 mb
39.9 mb
40 mb
40.1 mb
-log10(P)
150
100
50
r2
>0.8
0.6-0.8
0.4-0.6
0.2-0.4
<0.2
C15orf54
THBS1
FSIP
GZ>CP
CP>GZ
Corrected for total SA
Uncorrected
<=-15
-10
-5
0
5
10
>=15
Zscore

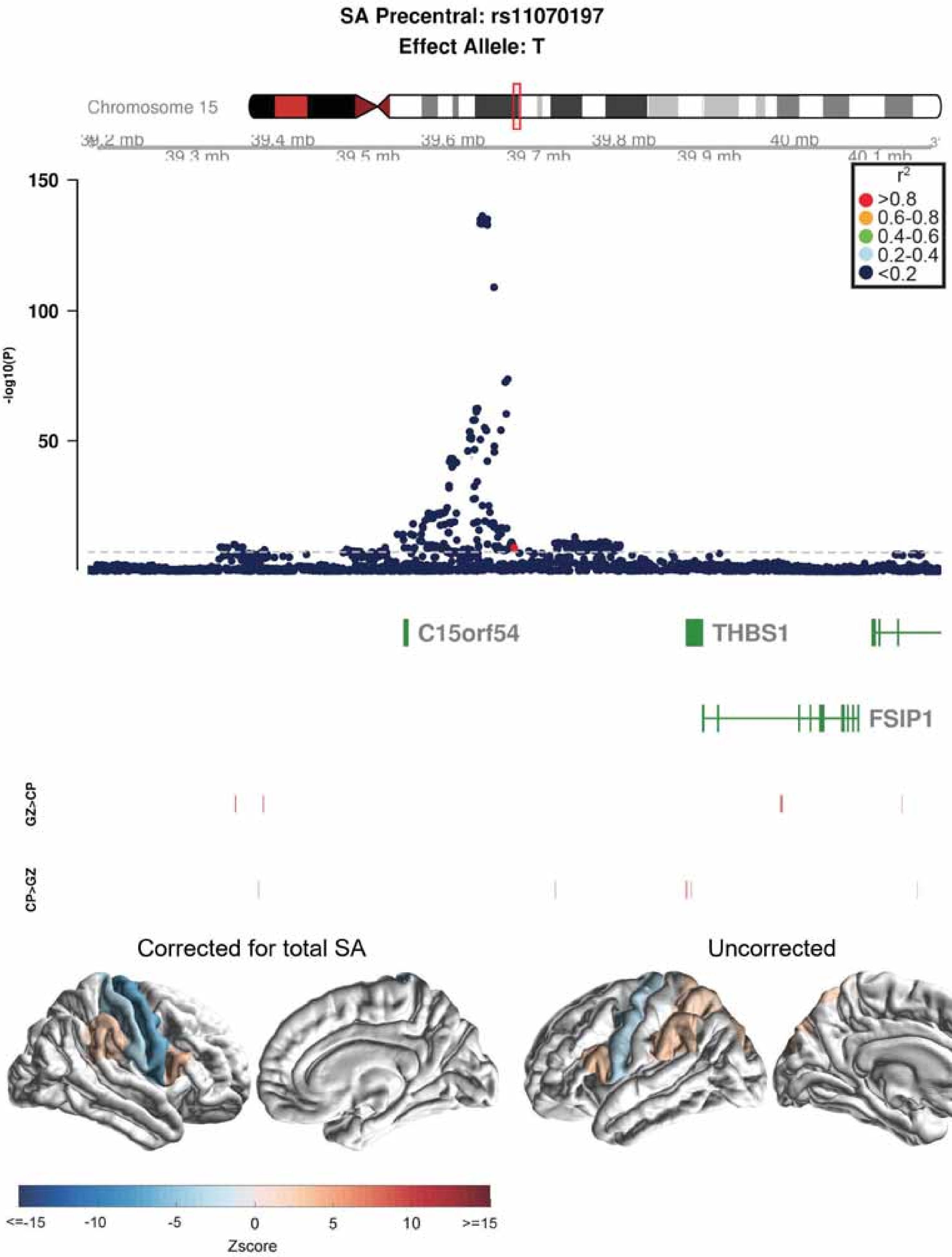

SA Precentral: rs11070197
Effect Allele: T
Chromosome 15
39.2 mb
39.3 mb
39.4 mb
39.5 mb
39.6 mb
39.7 mb
39.8 mb
39.9 mb
40 mb
40.1 mb
150
100
50
-log10(P)
$r^2$
>0.8
0.6-0.8
0.4-0.6
0.2-0.4
<0.2
C15orf54
THBS1
FSIP1
GZ>CP
CP>GZ
Corrected for total SA
Uncorrected
<=-15
-10
-5
0
5
10
>=15
Zscore

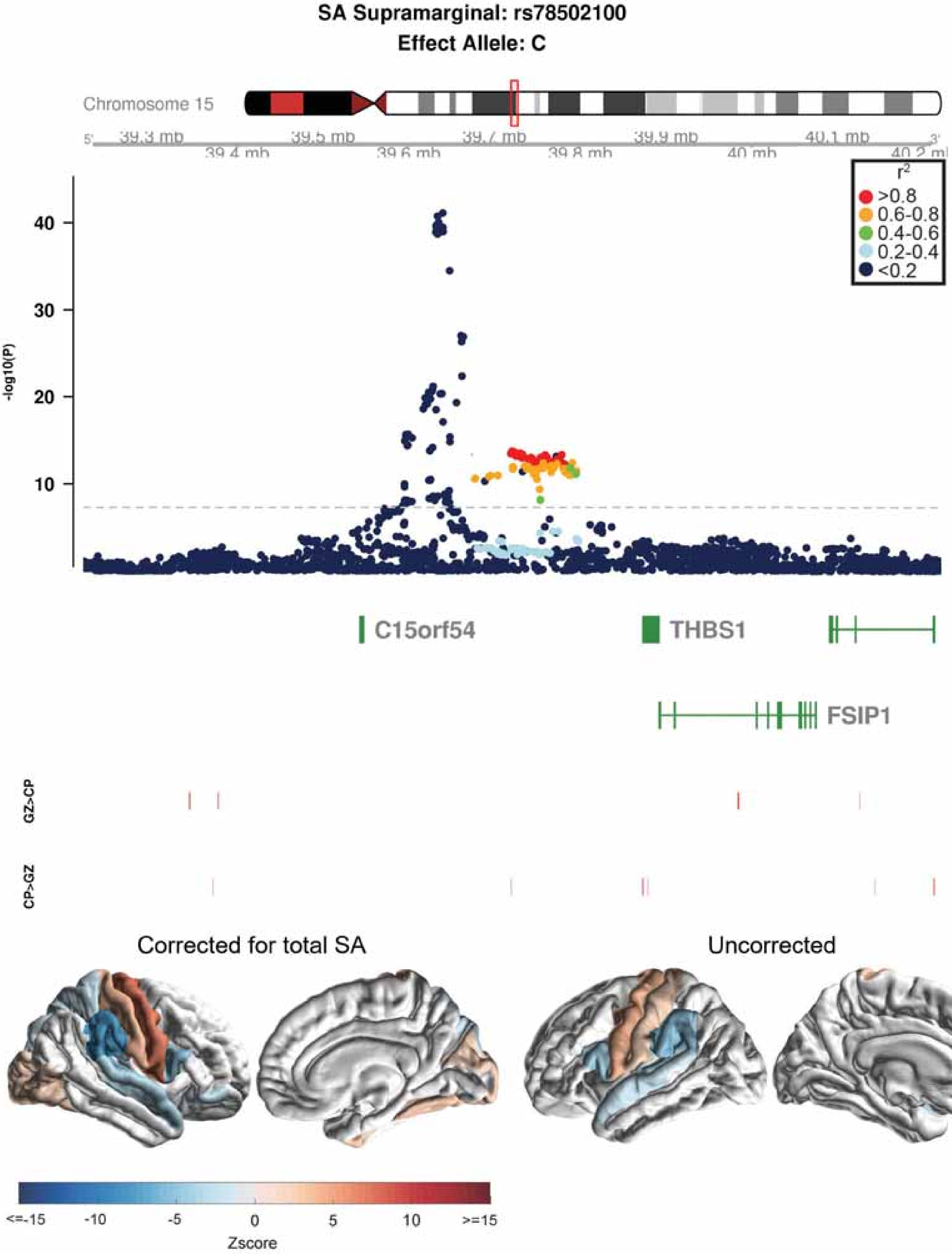
SA Supramarginal: rs78502100
Effect Allele: C
Chromosome 15
39.3 mb
39.4 mb
39.5 mb
39.6 mb
39.7 mb
39.8 mb
39.9 mb
40 mb
40.1 mb
40.2 mb
r2
>0.8
0.6-0.8
0.4-0.6
0.2-0.4
<0.2
-log10(P)
40
30
20
10
C15orf54
THBS1
FSIP1
GZ>CP
CP>GZ
Corrected for total SA
Uncorrected
<=-15
-10
-5
0
5
10
>=15
Zscore

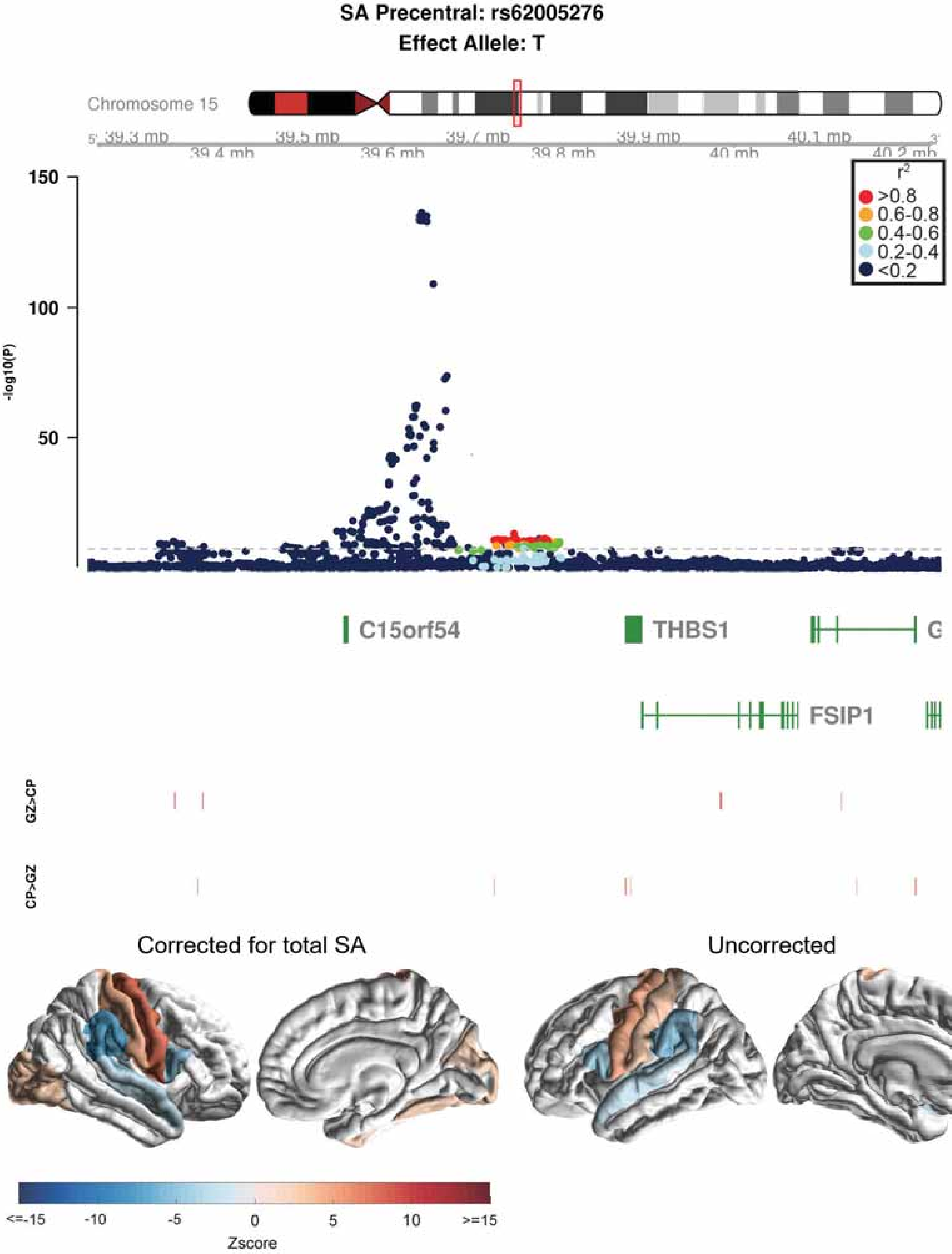
SA Precentral: rs62005276
Effect Allele: T
Chromosome 15
39.3 mb
39.4 mb
39.5 mb
39.6 mb
39.7 mb
39.8 mb
39.9 mb
40 mb
40.1 mb
40.2 mb
r2
>0.8
0.6-0.8
0.4-0.6
0.2-0.4
<0.2
150
100
50
-log10(P)
C15orf54
THBS1
FSIP1
GZ>CP
CP>GZ
Corrected for total SA
Uncorrected
<=-15
-10
-5
0
5
10
>=15
Zscore

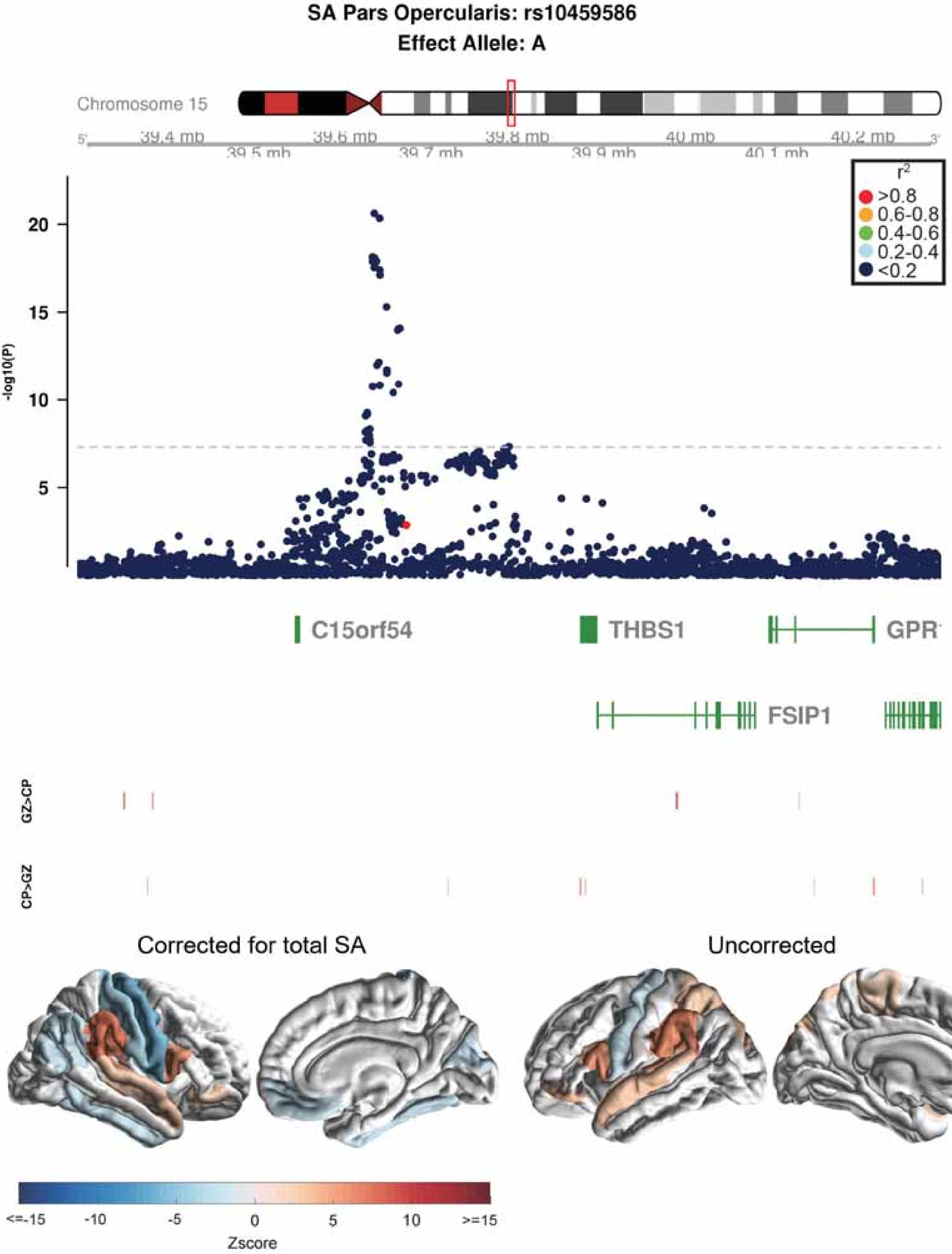
SA Pars Opercularis: rs10459586
Effect Allele: A
Chromosome 15
39.4 mb
39.5 mb
39.6 mb
39.7 mb
39.8 mb
39.9 mb
40 mb
40.1 mb
40.2 mb
-log10(P)
20
15
10
5
r2
>0.8
0.6-0.8
0.4-0.6
0.2-0.4
<0.2
C15orf54
THBS1
GPR
FSIP1
GZ>CP
CP>GZ
Corrected for total SA
Uncorrected
<=-15
-10
-5
0
5
10
>=15
Zscore

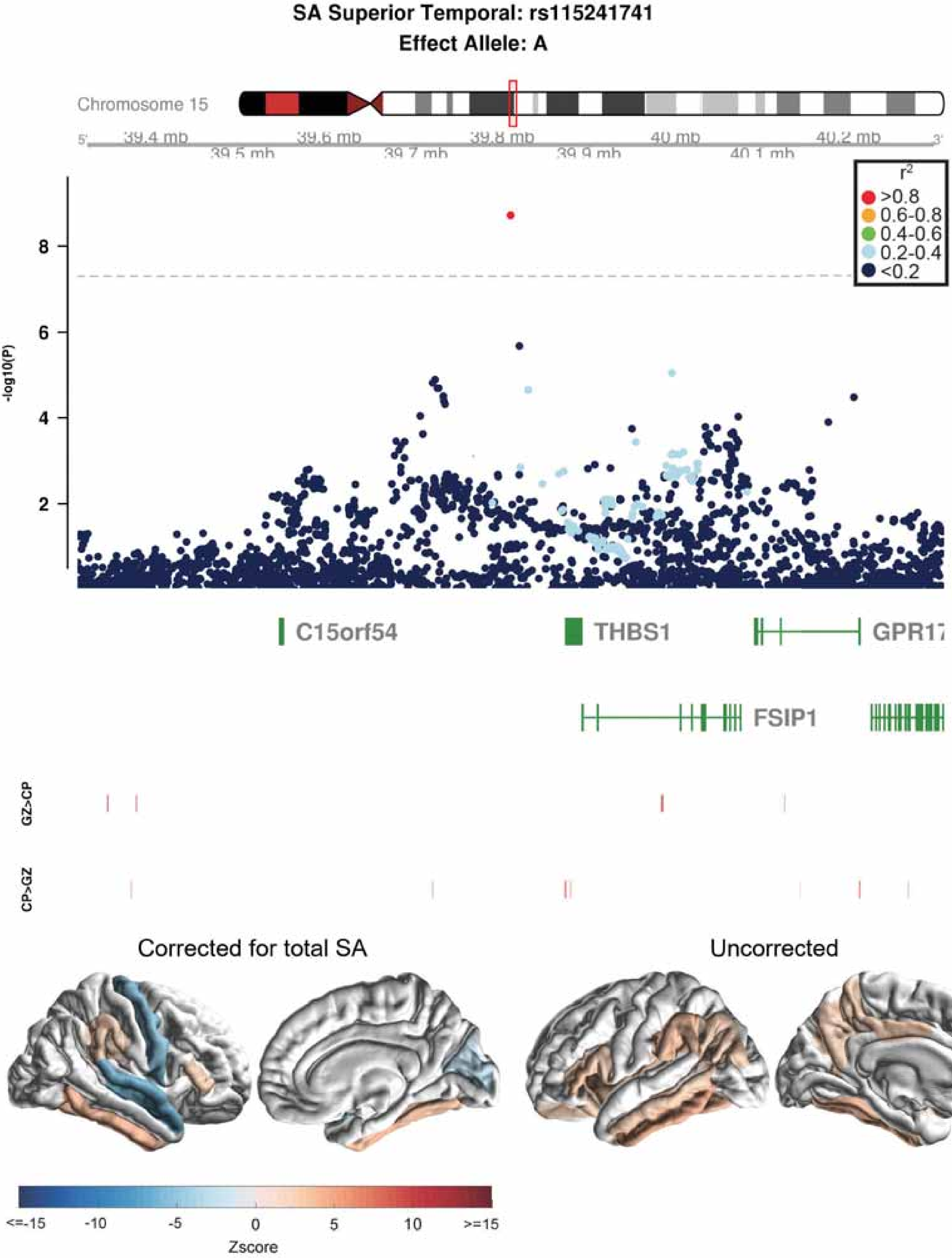
SA Superior Temporal: rs115241741
Effect Allele: A
Chromosome 15
39.4 mb
39.5 mb
39.6 mb
39.7 mb
39.8 mb
39.9 mb
40 mb
40.1 mb
40.2 mb
r²
>0.8
0.6-0.8
0.4-0.6
0.2-0.4
<0.2
-log10(P)
8
6
4
2
C15orf54
THBS1
GPR17
FSIP1
GZ>CP
CP>GZ
Corrected for total SA
Uncorrected
<=-15
-10
-5
0
5
10
>=15
Zscore

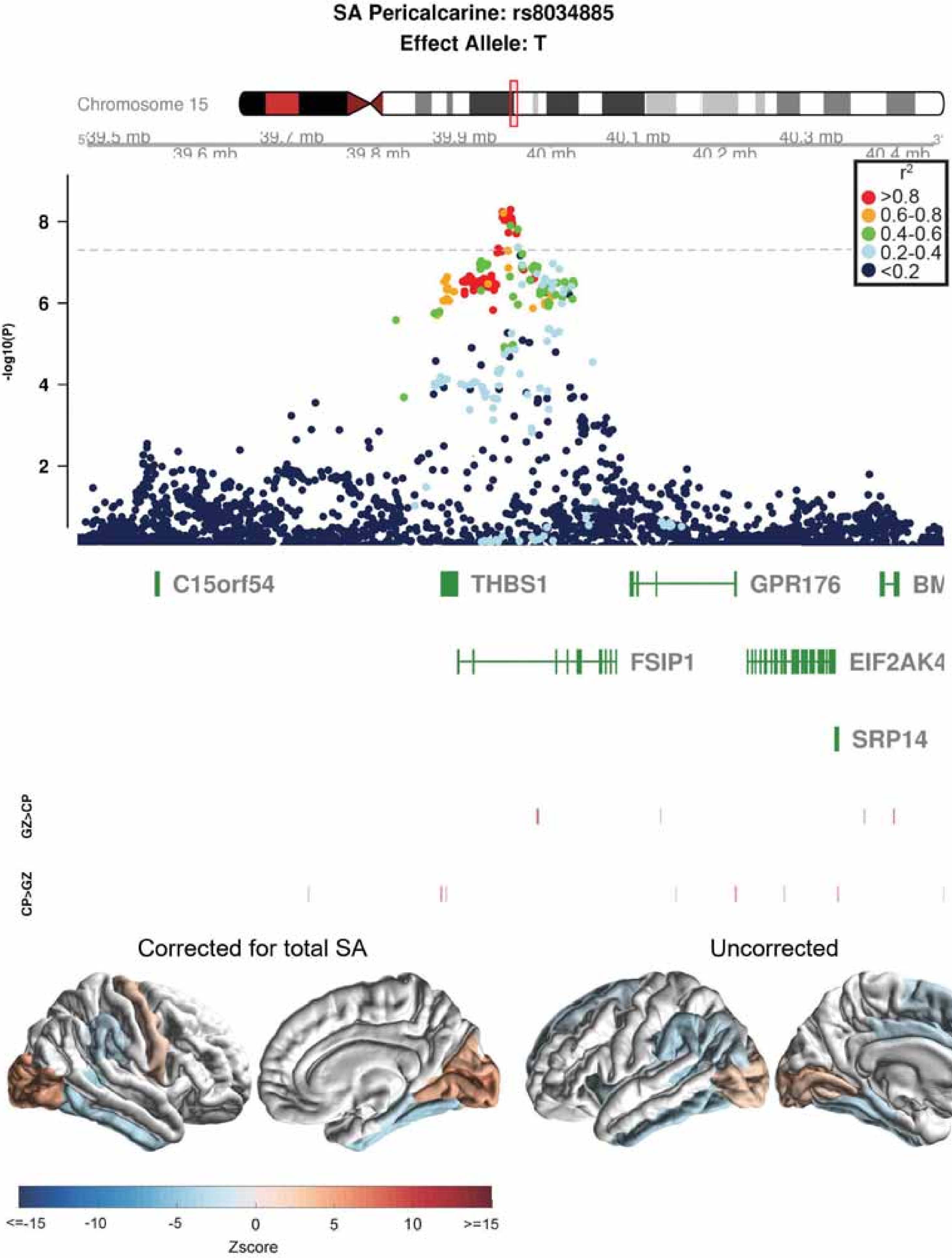
SA Pericalcarine: rs8034885
Effect Allele: T
Chromosome 15
39.5 mb
39.6 mb
39.7 mb
39.8 mb
39.9 mb
40 mb
40.1 mb
40.2 mb
40.3 mb
40.4 mb
r2
>0.8
0.6-0.8
0.4-0.6
0.2-0.4
<0.2
-log10(P)
2
4
6
8
C15orf54
THBS1
GPR176
FSIP1
EIF2AK4
SRP14
GZ>CP
CP>GZ
Corrected for total SA
Uncorrected
<=-15
-10
-5
0
5
10
>=15
Zscore

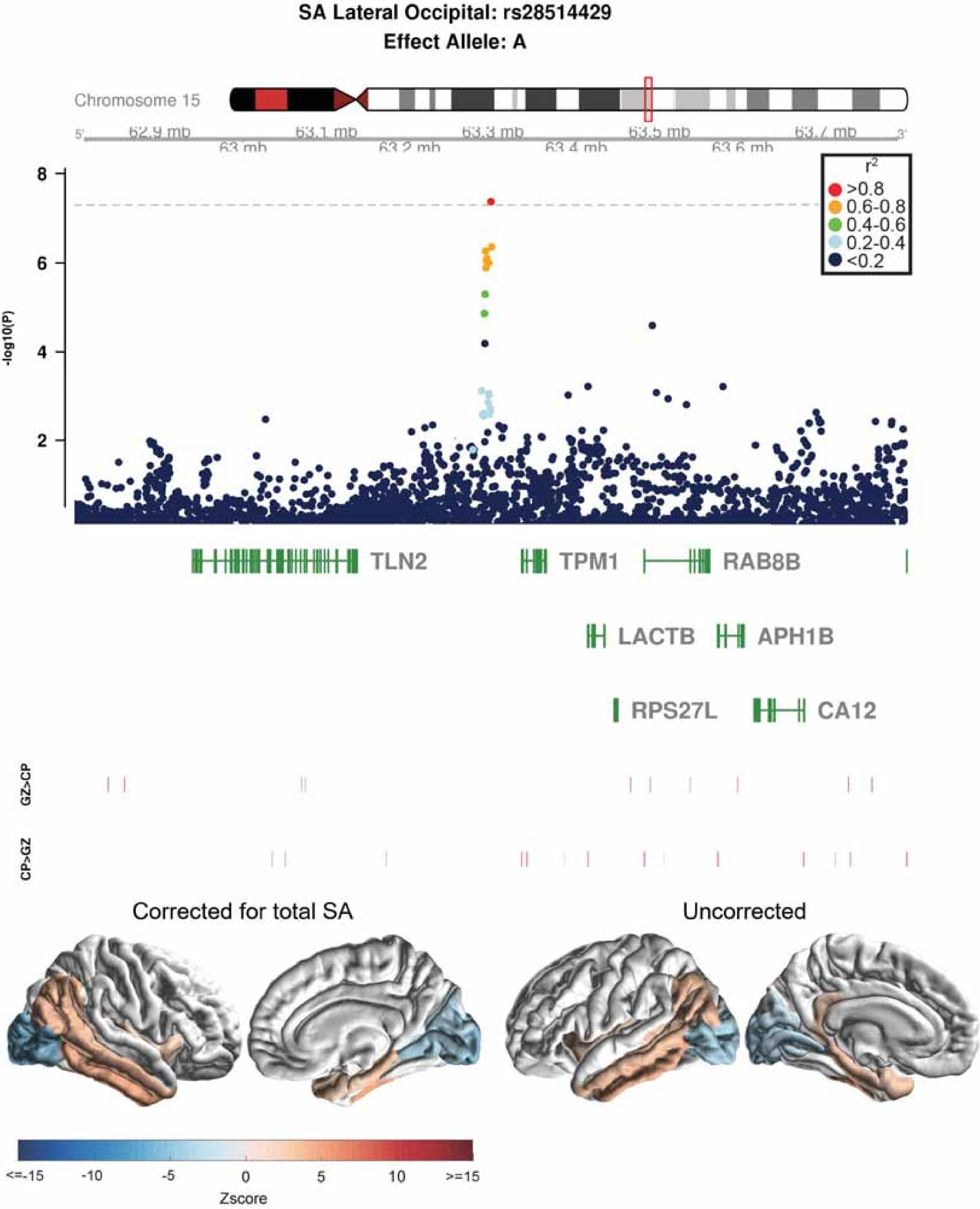

SA Lateral Occipital: rs28514429
Effect Allele: A
Chromosome 15
62.9 mb
63 mb
63.1 mb
63.2 mb
63.3 mb
63.4 mb
63.5 mb
63.6 mb
63.7 mb
-log10(P)
r2
>0.8
0.6-0.8
0.4-0.6
0.2-0.4
<0.2
TLN2
TPM1
RAB8B
LACTB
APH1B
RPS27L
CA12
GZ>CP
CP>GZ
Corrected for total SA
Uncorrected
<=-15
-10
-5
0
5
10
>=15
Zscore

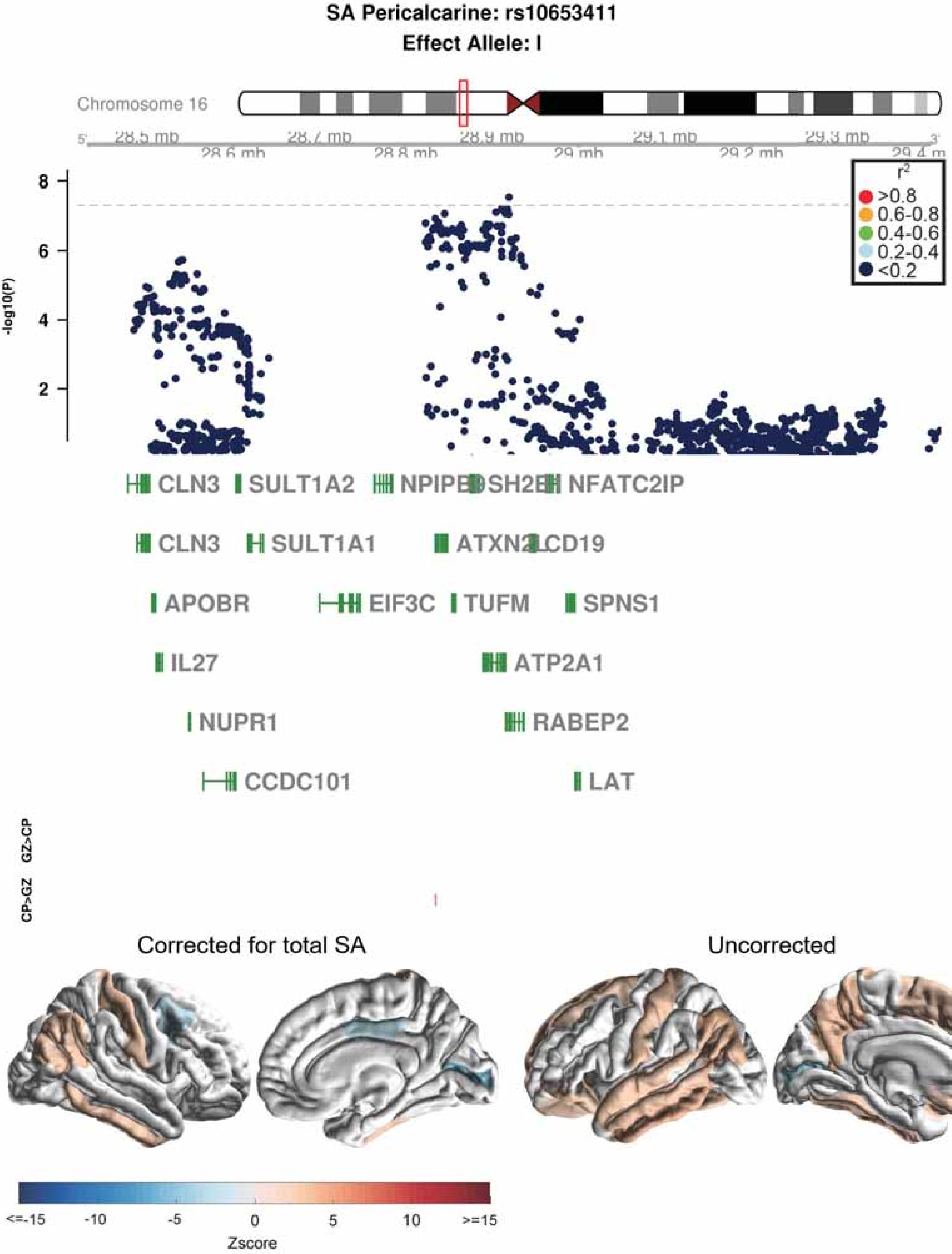
SA Pericalcarine: rs10653411
Effect Allele: I
Chromosome 16
28.5 mb
28.6 mb
28.7 mb
28.8 mb
28.9 mb
29 mb
29.1 mb
29.2 mb
29.3 mb
29.4 m
-log10(P)
r2
>0.8
0.6-0.8
0.4-0.6
0.2-0.4
<0.2
CLN3
SULT1A2
NPIPB
SH2B
NFATC2IP
CLN3
SULT1A1
ATXN2L
CD19
APOBR
EIF3C
TUFM
SPNS1
IL27
ATP2A1
NUPR1
RABEP2
CCDC101
LAT
GZ>CP
CP>GZ
Corrected for total SA
Uncorrected
<=-15
-10
-5
0
5
10
>=15
Zscore

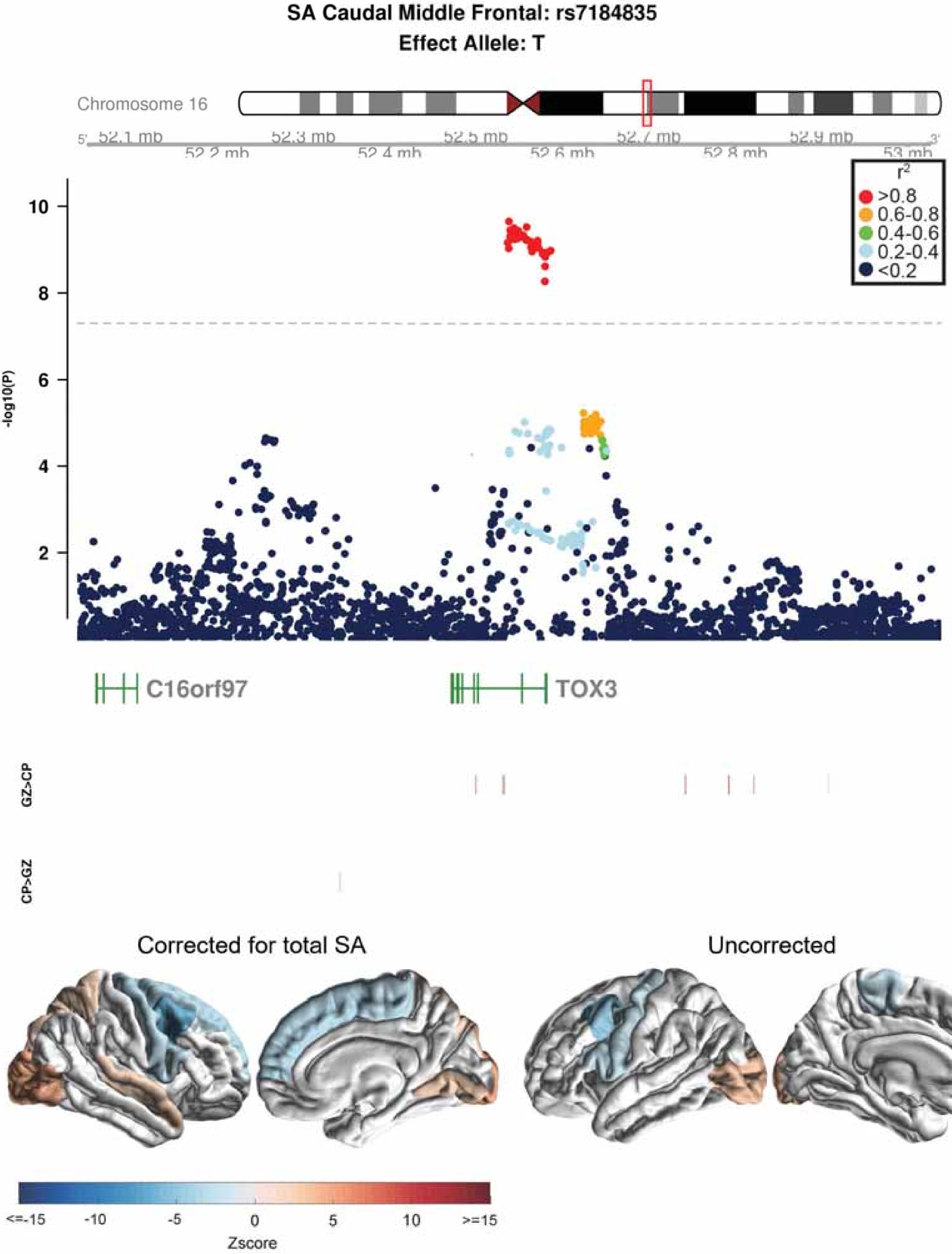

SA Caudal Middle Frontal: rs7184835
Effect Allele: T
Chromosome 16
5'
52.1 mb
52.2 mb
52.3 mb
52.4 mb
52.5 mb
52.6 mb
52.7 mb
52.8 mb
52.9 mb
53 mb
3'
$r^2$
>0.8
0.6-0.8
0.4-0.6
0.2-0.4
<0.2
-log10(P)
10
8
6
4
2
C16orf97
TOX3
GZ>CP
CP>GZ
Corrected for total SA
Uncorrected
<=-15
-10
-5
0
5
10
>=15
Zscore

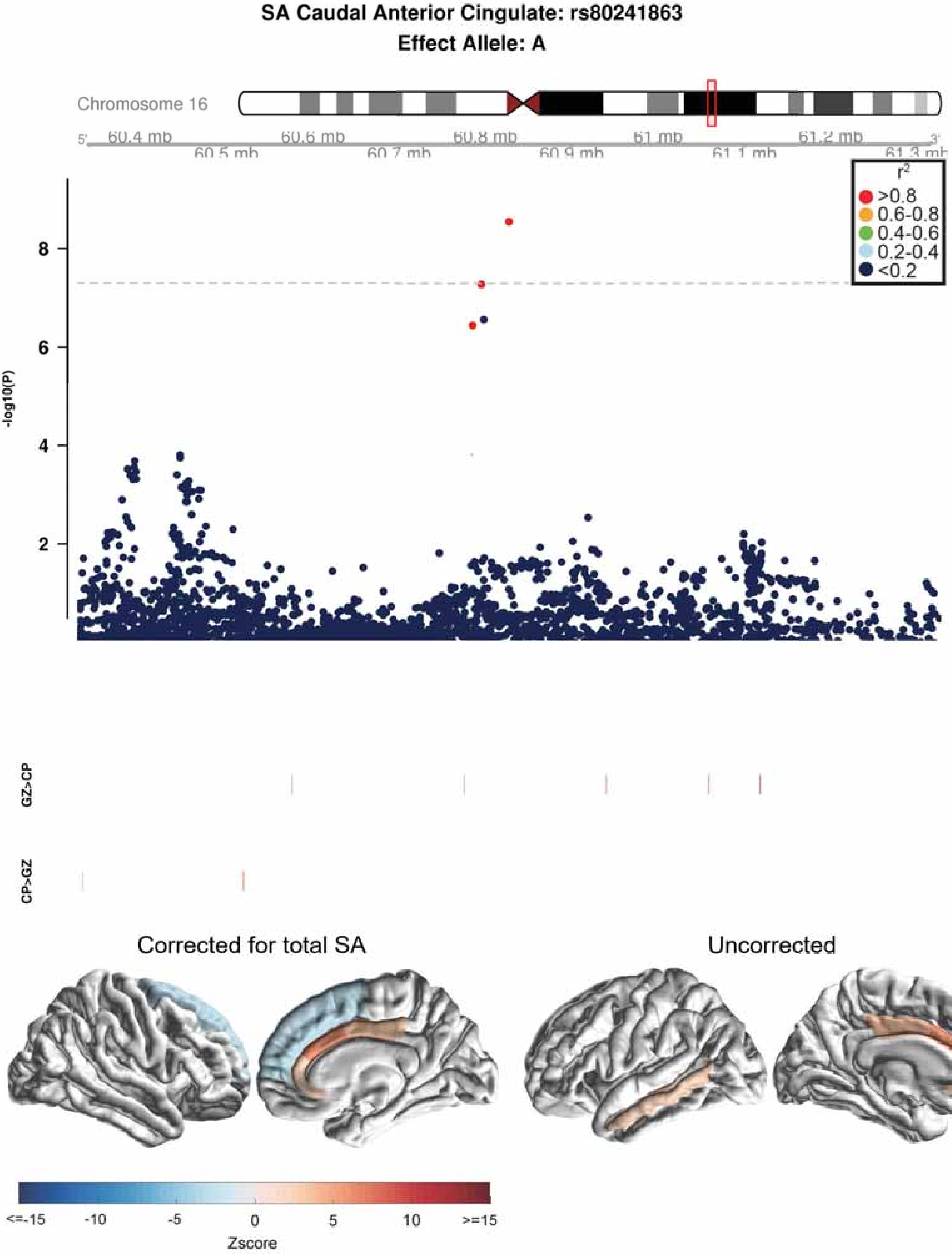
SA Caudal Anterior Cingulate: rs80241863
Effect Allele: A
Chromosome 16
5'
60.4 mb
60.5 mb
60.6 mb
60.7 mb
60.8 mb
60.9 mb
61 mb
61.1 mb
61.2 mb
61.3 mb
3'
r2
>0.8
0.6-0.8
0.4-0.6
0.2-0.4
<0.2
-log10(P)
8
6
4
2
GZ>CP
CP>GZ
Corrected for total SA
Uncorrected
<=-15
-10
-5
0
5
10
>=15
Zscore

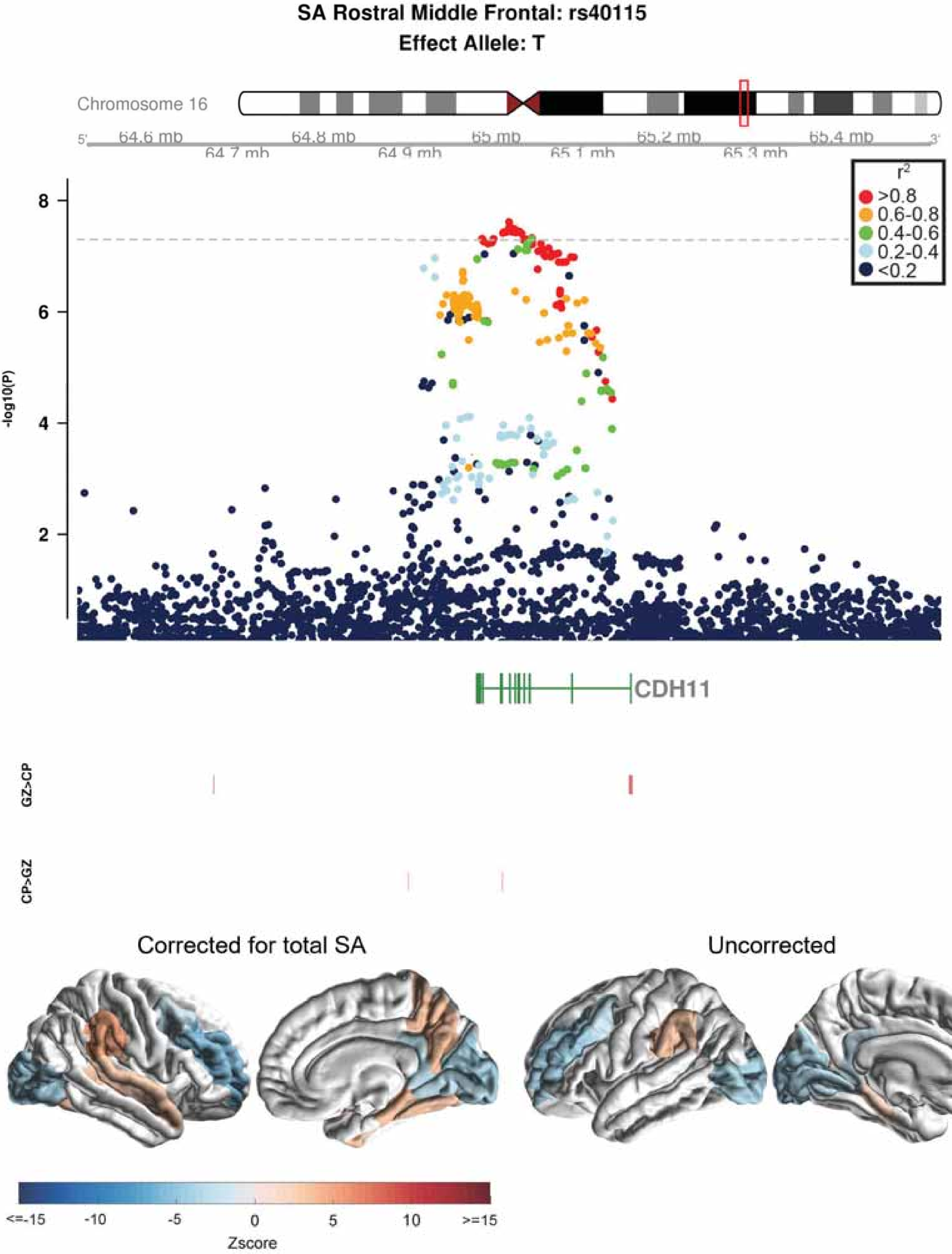
SA Rostral Middle Frontal: rs40115
Effect Allele: T
Chromosome 16
5'
64.6 mb
64.7 mb
64.8 mb
64.9 mb
65 mb
65.1 mb
65.2 mb
65.3 mb
65.4 mb
3'
r²
>0.8
0.6-0.8
0.4-0.6
0.2-0.4
<0.2
-log10(P)
8
6
4
2
CDH11
GZ>CP
CP>GZ
Corrected for total SA
Uncorrected
<=-15
-10
-5
0
5
10
>=15
Zscore

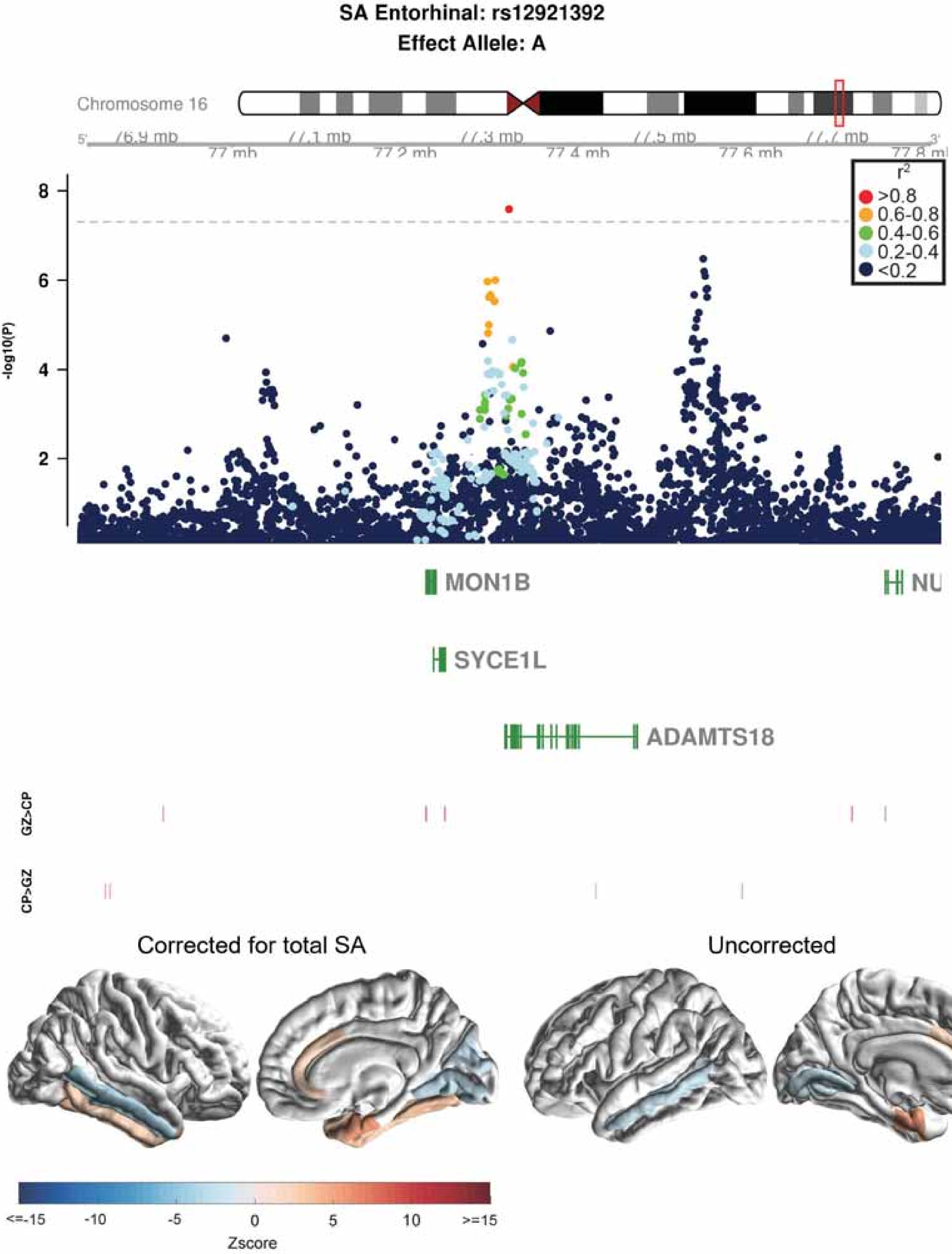

SA Entorhinal: rs12921392
Effect Allele: A
Chromosome 16
5'
3'
76.9 mb
77 mb
77.1 mb
77.2 mb
77.3 mb
77.4 mb
77.5 mb
77.6 mb
77.7 mb
77.8 mb
$r^2$
>0.8
0.6-0.8
0.4-0.6
0.2-0.4
<0.2
-log10(P)
8
6
4
2
MON1B
NU
SYCE1L
ADAMTS18
GZ>CP
CP>GZ
Corrected for total SA
Uncorrected
<=-15
-10
-5
0
5
10
>=15
Zscore

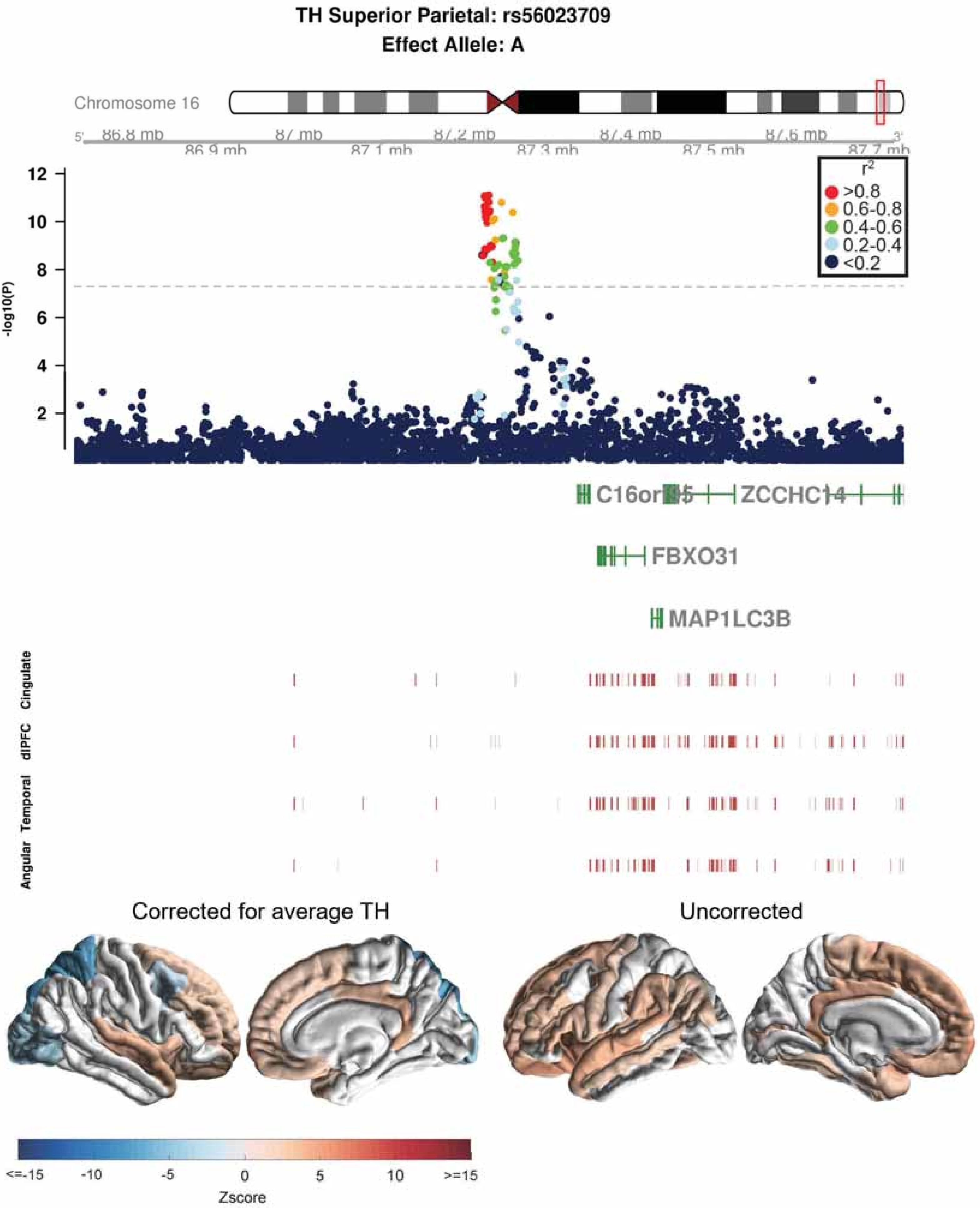

TH Superior Parietal: rs56023709
Effect Allele: A
Chromosome 16
86.8 mb
86.9 mb
87 mb
87.1 mb
87.2 mb
87.3 mb
87.4 mb
87.5 mb
87.6 mb
87.7 mb
-log10(P)
r2
>0.8
0.6-0.8
0.4-0.6
0.2-0.4
<0.2
C16orf95
ZCCHC14
FBXO31
MAP1LC3B
Cingulate
dlPFC
Temporal
Angular
Corrected for average TH
Uncorrected
<=-15
-10
-5
0
5
10
>=15
Zscore

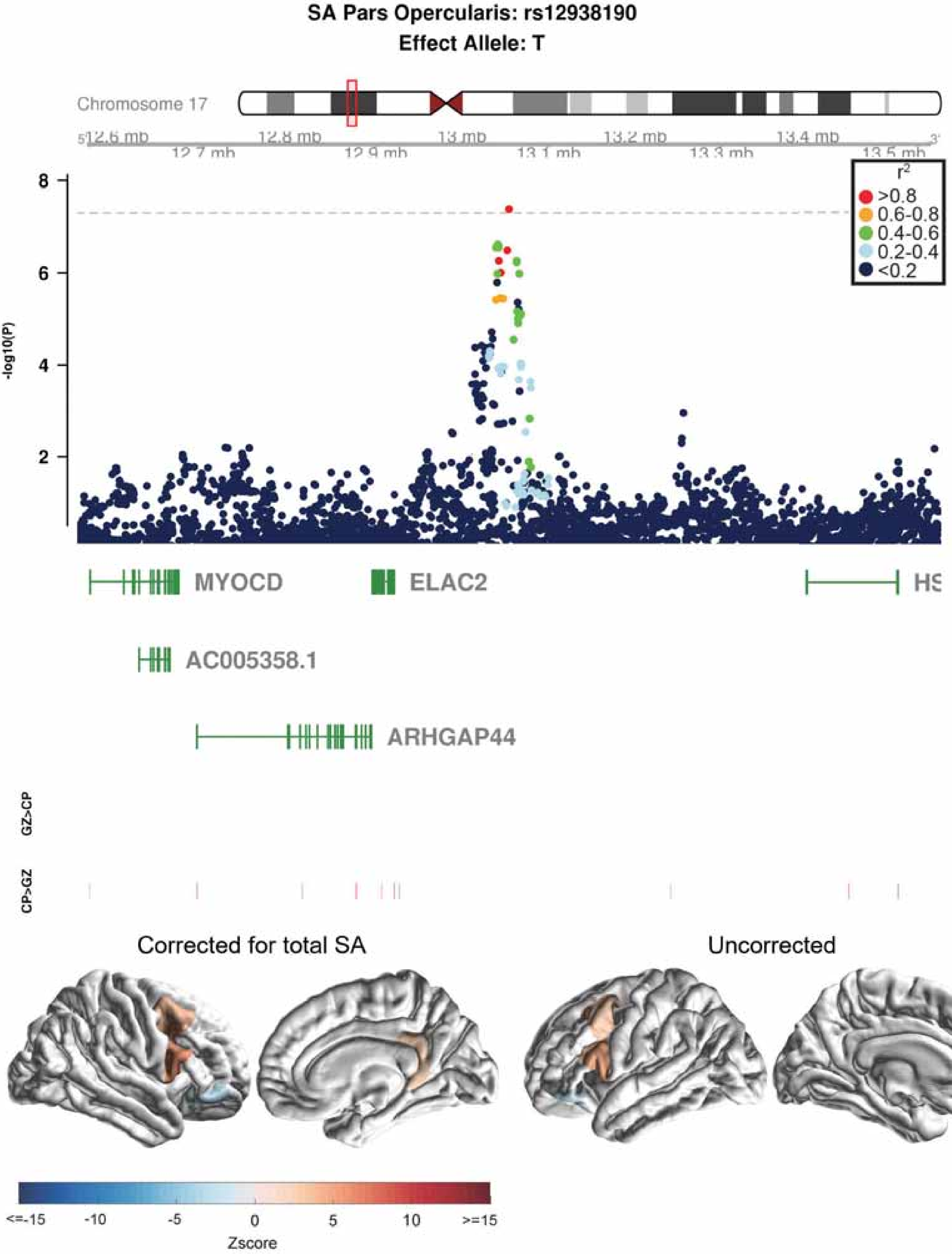
SA Pars Opercularis: rs12938190
Effect Allele: T
Chromosome 17
12.6 mb
12.7 mb
12.8 mb
12.9 mb
13 mb
13.1 mb
13.2 mb
13.3 mb
13.4 mb
13.5 mb
-log10(P)
8
6
4
2
r2
>0.8
0.6-0.8
0.4-0.6
0.2-0.4
<0.2
MYOCD
ELAC2
AC005358.1
ARHGAP44
GZ>CP
CP>GZ
Corrected for total SA
Uncorrected
<=-15
-10
-5
0
5
10
>=15
Zscore

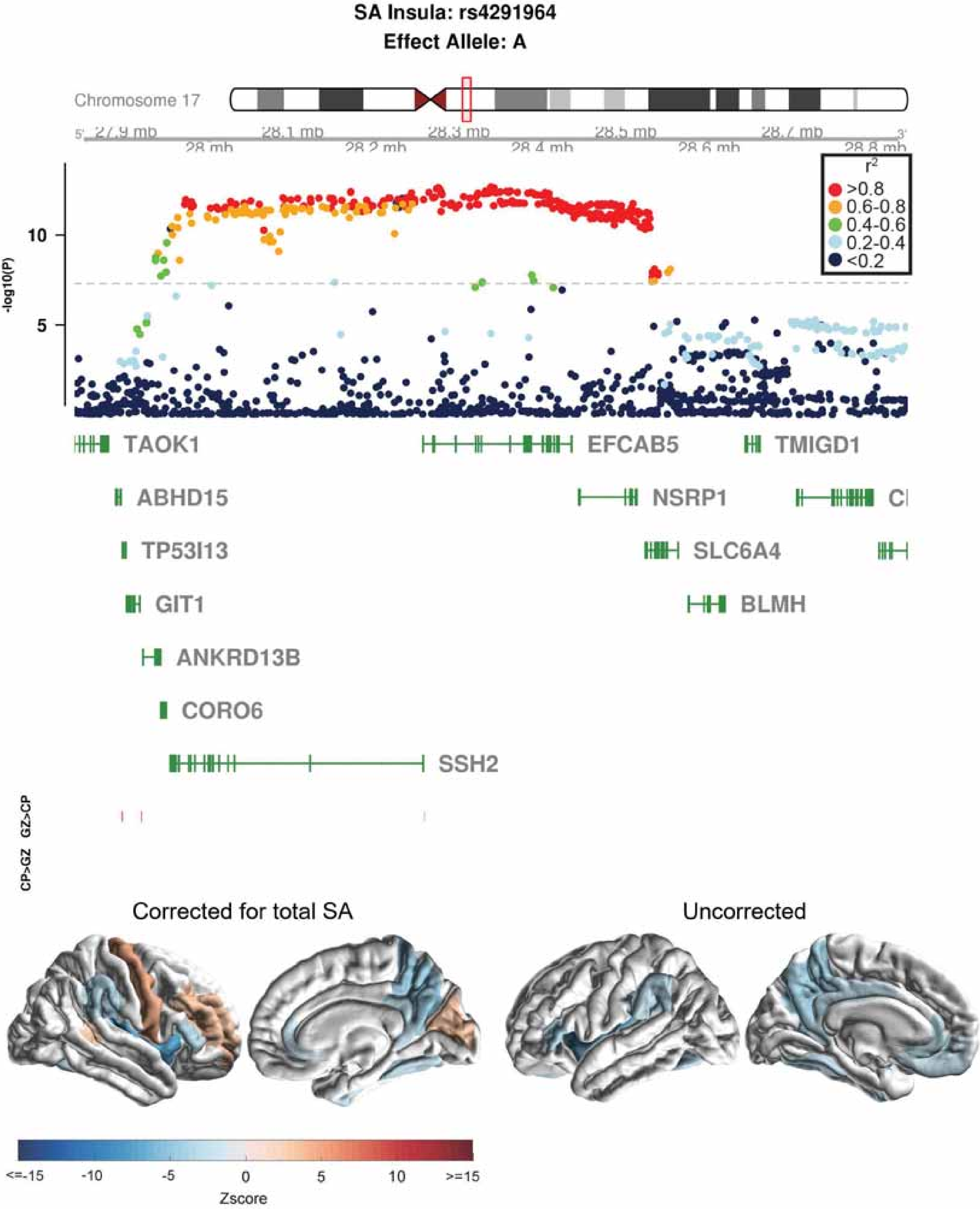
SA Insula: rs4291964
Effect Allele: A
Chromosome 17
27.9 mb
28 mb
28.1 mb
28.2 mb
28.3 mb
28.4 mb
28.5 mb
28.6 mb
28.7 mb
28.8 mb
-log10(P)
10
5
r2
>0.8
0.6-0.8
0.4-0.6
0.2-0.4
<0.2
TAOK1
EFCAB5
TMIGD1
ABHD15
NSRP1
TP53I13
SLC6A4
GIT1
BLMH
ANKRD13B
CORO6
SSH2
GZ>CP
CP>GZ
Corrected for total SA
Uncorrected
<=-15
-10
-5
0
5
10
>=15
Zscore

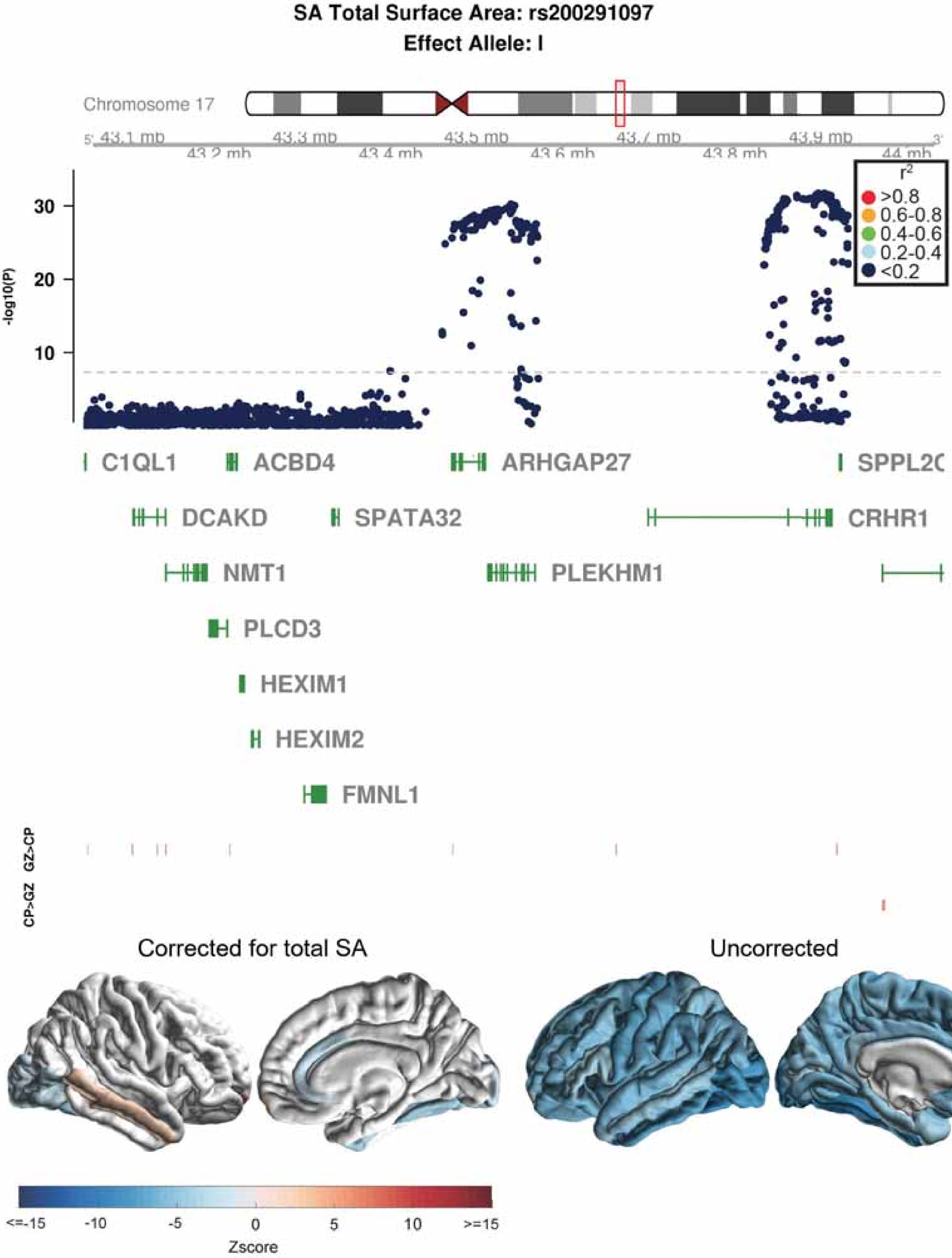
SA Total Surface Area: rs200291097
Effect Allele: I
Chromosome 17
43.1 mb
43.2 mb
43.3 mb
43.4 mb
43.5 mb
43.6 mb
43.7 mb
43.8 mb
43.9 mb
44 mb
-log10(P)
30
20
10
r2
>0.8
0.6-0.8
0.4-0.6
0.2-0.4
<0.2
C1QL1
ACBD4
ARHGAP27
SPPL2C
DCAKD
SPATA32
CRHR1
NMT1
PLEKHM1
PLCD3
HEXIM1
HEXIM2
FMNL1
GZ>CP
CP>GZ
Corrected for total SA
Uncorrected
<=-15
-10
-5
0
5
10
>=15
Zscore

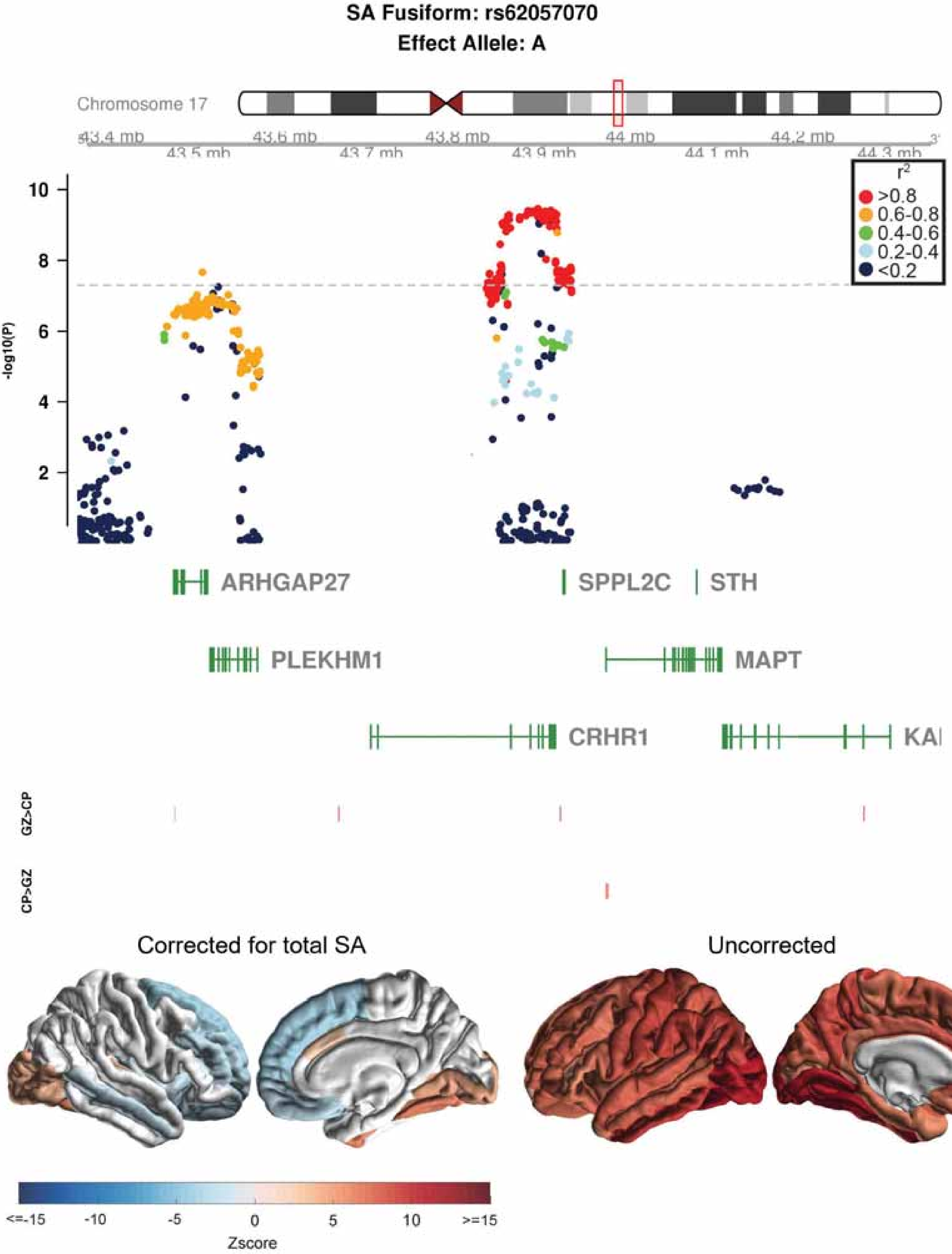

SA Fusiform: rs62057070
Effect Allele: A
Chromosome 17
43.4 mb
43.5 mb
43.6 mb
43.7 mb
43.8 mb
43.9 mb
44 mb
44.1 mb
44.2 mb
44.3 mb
-log10(P)
r2
>0.8
0.6-0.8
0.4-0.6
0.2-0.4
<0.2
ARHGAP27
SPPL2C
STH
PLEKHM1
MAPT
CRHR1
KA
GZ>CP
CP>GZ
Corrected for total SA
Uncorrected
<=-15
-10
-5
0
5
10
>=15
Zscore

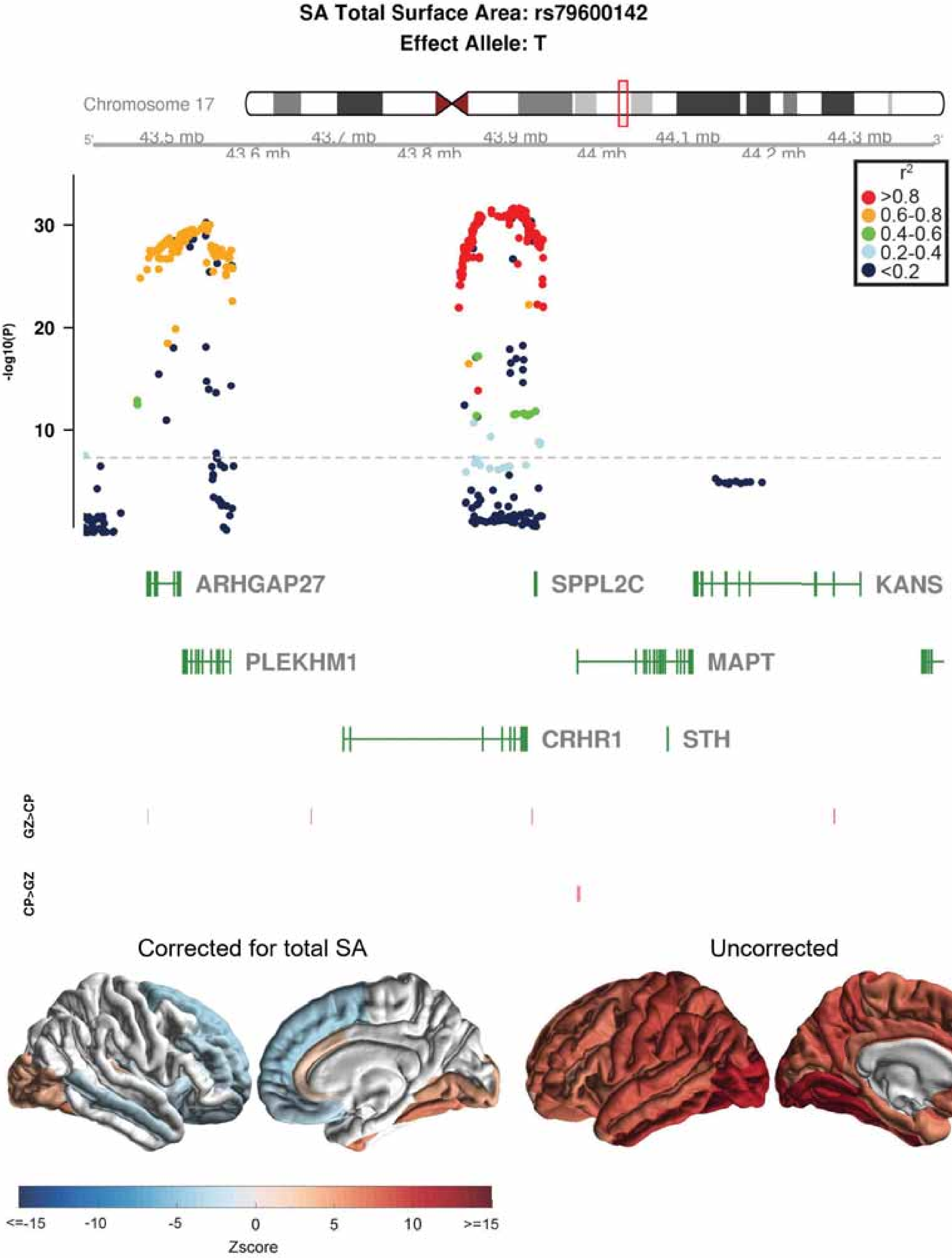

SA Total Surface Area: rs79600142
Effect Allele: T
Chromosome 17
43.5 mb
43.6 mb
43.7 mb
43.8 mb
43.9 mb
44 mb
44.1 mb
44.2 mb
44.3 mb
r2
>0.8
0.6-0.8
0.4-0.6
0.2-0.4
<0.2
-log10(P)
30
20
10
ARHGAP27
SPPL2C
KANS
PLEKHM1
MAPT
CRHR1
STH
GZ>CP
CP>GZ
Corrected for total SA
Uncorrected
<=-15
-10
-5
0
5
10
>=15
Zscore

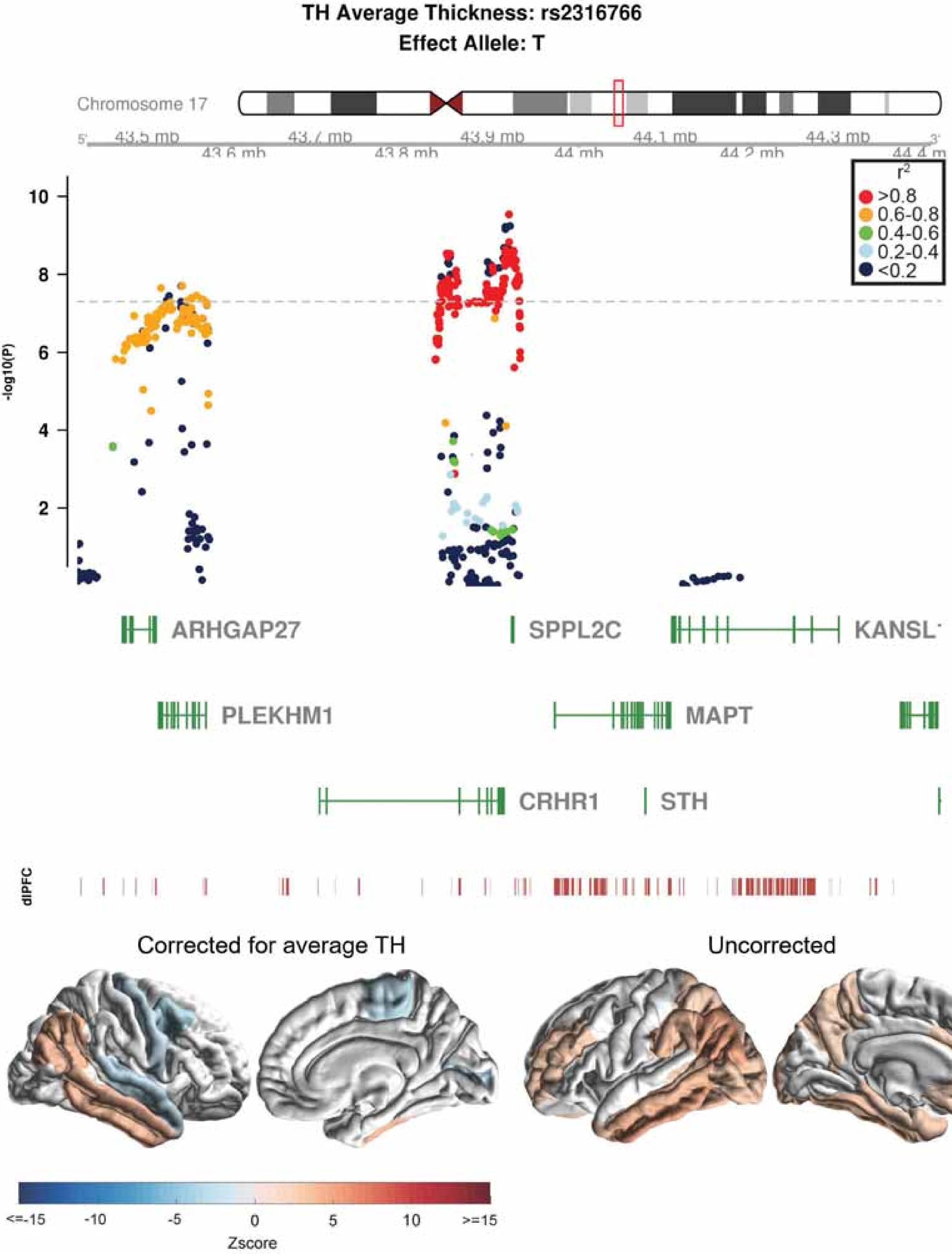

TH Average Thickness: rs2316766
Effect Allele: T
Chromosome 17
43.5 mb
43.6 mb
43.7 mb
43.8 mb
43.9 mb
44 mb
44.1 mb
44.2 mb
44.3 mb
r2
>0.8
0.6-0.8
0.4-0.6
0.2-0.4
<0.2
-log10(P)
10
8
6
4
2
ARHGAP27
SPPL2C
KANSL
PLEKHM1
MAPT
CRHR1
STH
dlPFC
Corrected for average TH
Uncorrected
<=-15
-10
-5
0
5
10
>=15
Zscore

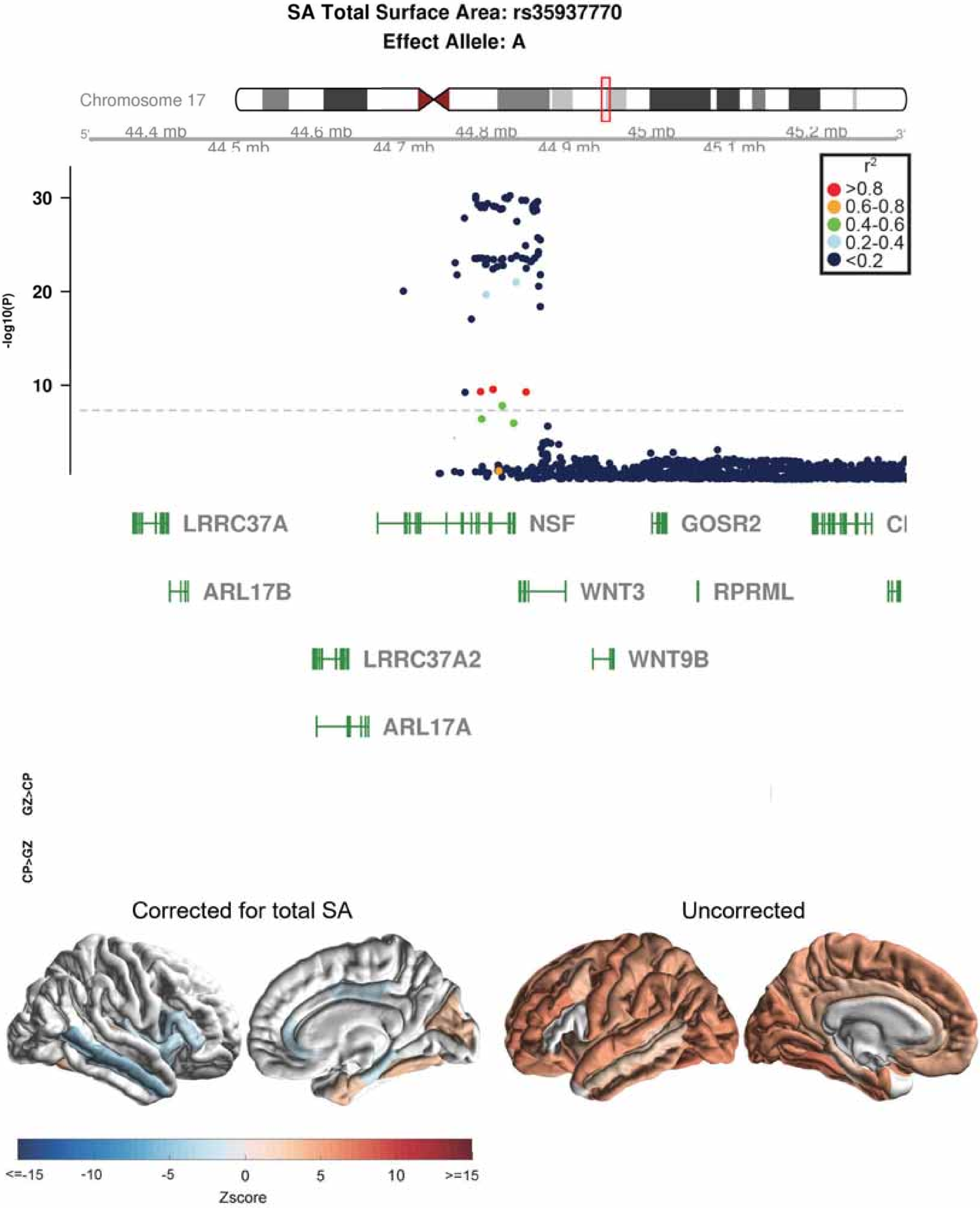
SA Total Surface Area: rs35937770
Effect Allele: A
Chromosome 17
44.4 mb
44.5 mb
44.6 mb
44.7 mb
44.8 mb
44.9 mb
45 mb
45.1 mb
45.2 mb
-log10(P)
30
20
10
r2
>0.8
0.6-0.8
0.4-0.6
0.2-0.4
<0.2
LRRC37A
NSF
GOSR2
ARL17B
WNT3
RPRML
LRRC37A2
WNT9B
ARL17A
GZ>CP
CP>GZ
Corrected for total SA
Uncorrected
<=-15
-10
-5
0
5
10
>=15
Zscore

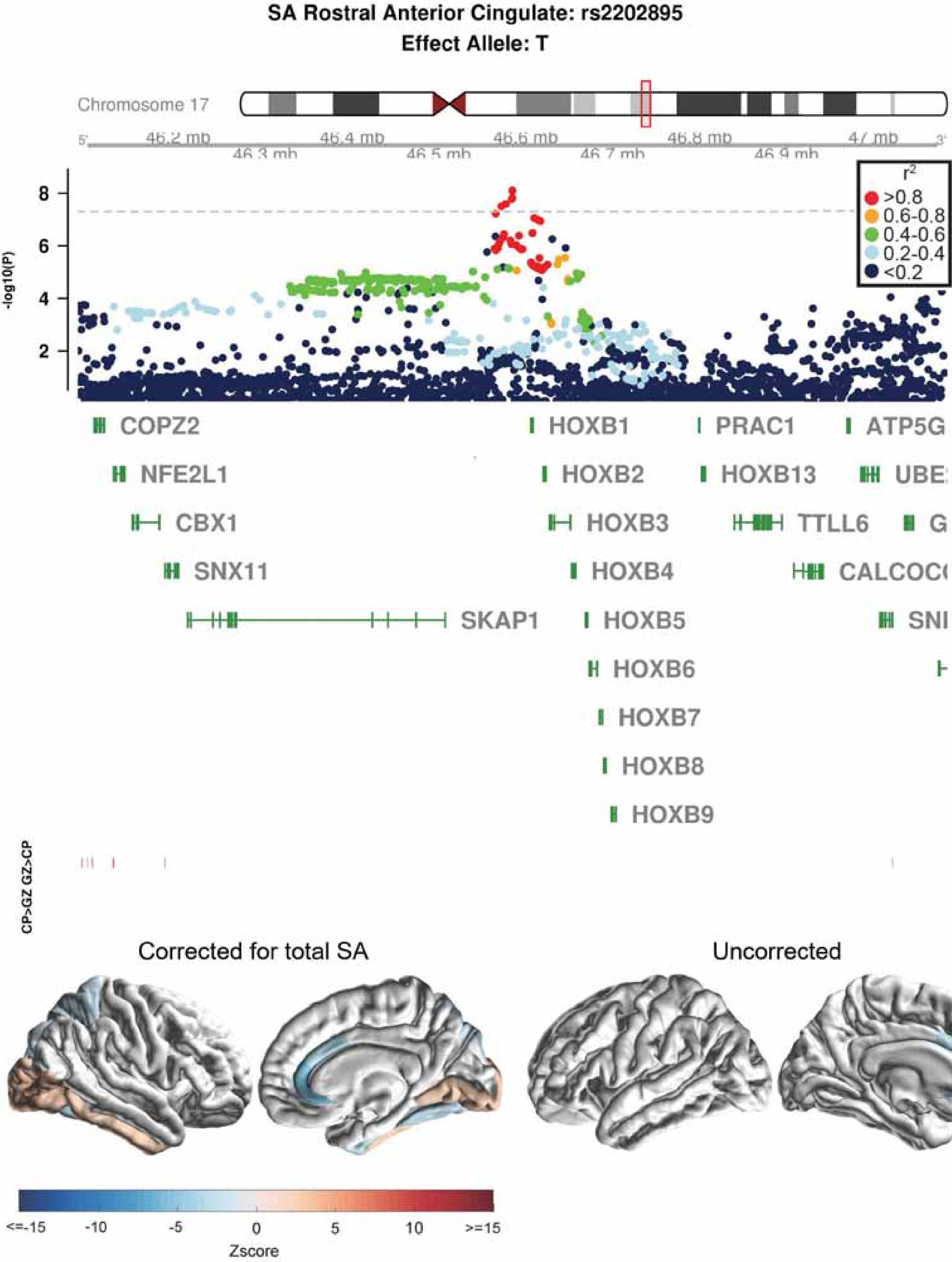

SA Rostral Anterior Cingulate: rs2202895
Effect Allele: T
Chromosome 17
46.2 mb
46.3 mb
46.4 mb
46.5 mb
46.6 mb
46.7 mb
46.8 mb
46.9 mb
47 mb
-log10(P)
r2
>0.8
0.6-0.8
0.4-0.6
0.2-0.4
<0.2
COPZ2
NFE2L1
CBX1
SNX11
SKAP1
HOXB1
HOXB2
HOXB3
HOXB4
HOXB5
HOXB6
HOXB7
HOXB8
HOXB9
PRAC1
HOXB13
TTLL6
CALCOC
ATP5G
UBE
G
SNI
CP>GZ GZ>CP
Corrected for total SA
Uncorrected
<=-15
-10
-5
0
5
10
>=15
Zscore

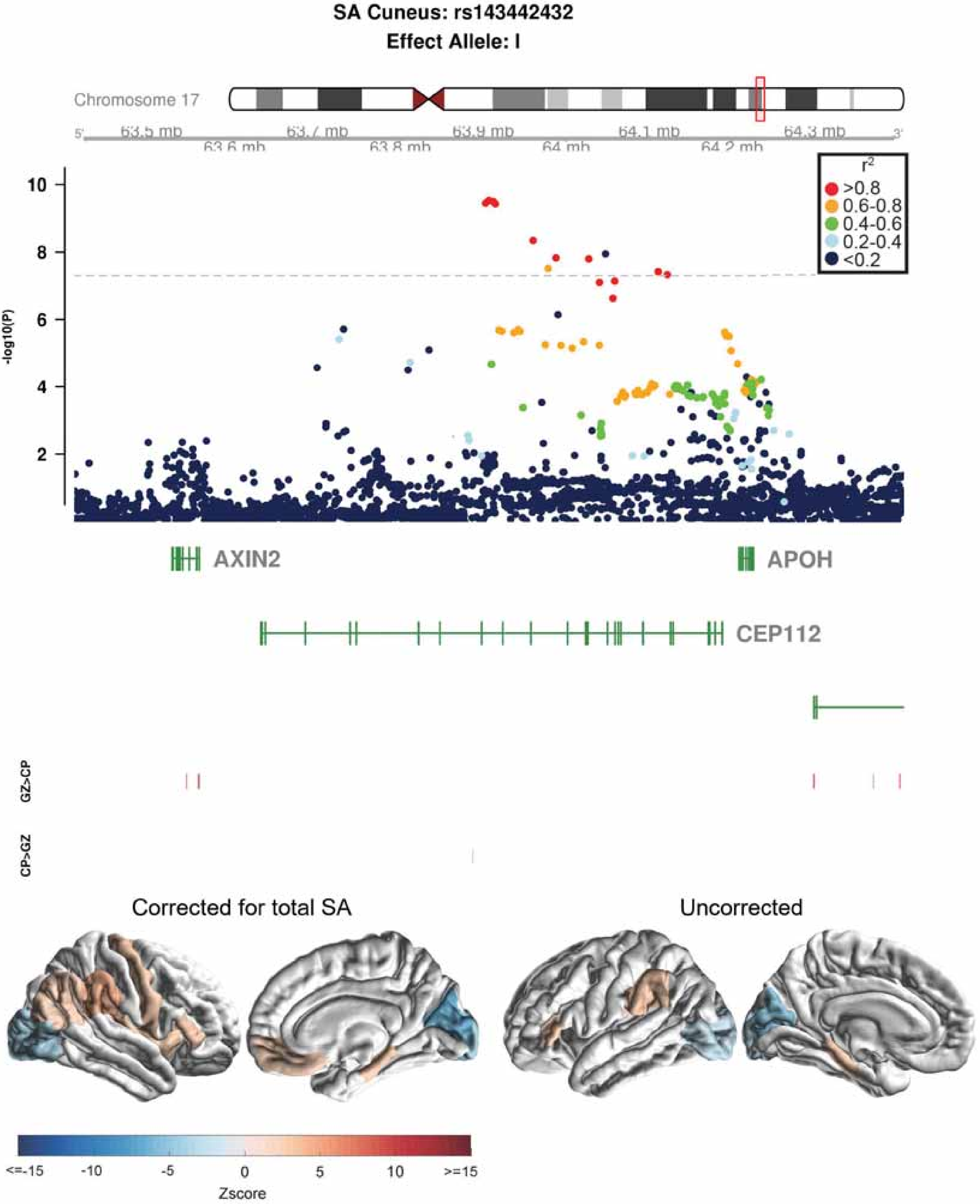

SA Cuneus: rs143442432
Effect Allele: I
Chromosome 17
63.5 mb
63.6 mb
63.7 mb
63.8 mb
63.9 mb
64 mb
64.1 mb
64.2 mb
64.3 mb
r2
>0.8
0.6-0.8
0.4-0.6
0.2-0.4
<0.2
-log10(P)
AXIN2
APOH
CEP112
GZ>CP
CP>GZ
Corrected for total SA
Uncorrected
<=-15
-10
-5
0
5
10
>=15
Zscore

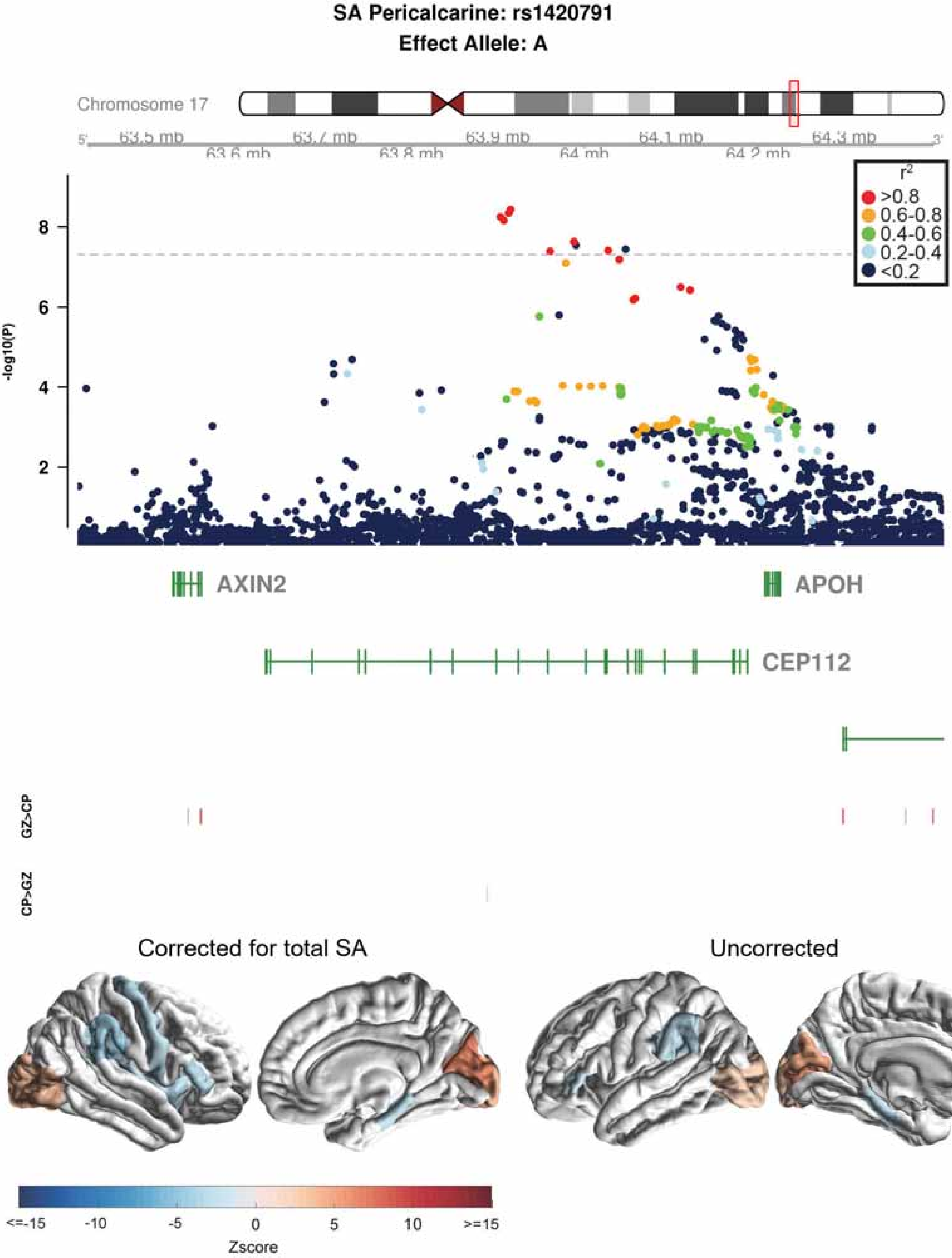
SA Pericalcarine: rs1420791
Effect Allele: A
Chromosome 17
63.5 mb
63.6 mb
63.7 mb
63.8 mb
63.9 mb
64 mb
64.1 mb
64.2 mb
64.3 mb
r2
>0.8
0.6-0.8
0.4-0.6
0.2-0.4
<0.2
-log10(P)
AXIN2
APOH
CEP112
GZ>CP
CP>GZ
Corrected for total SA
Uncorrected
<=-15
-10
-5
0
5
10
>=15
Zscore

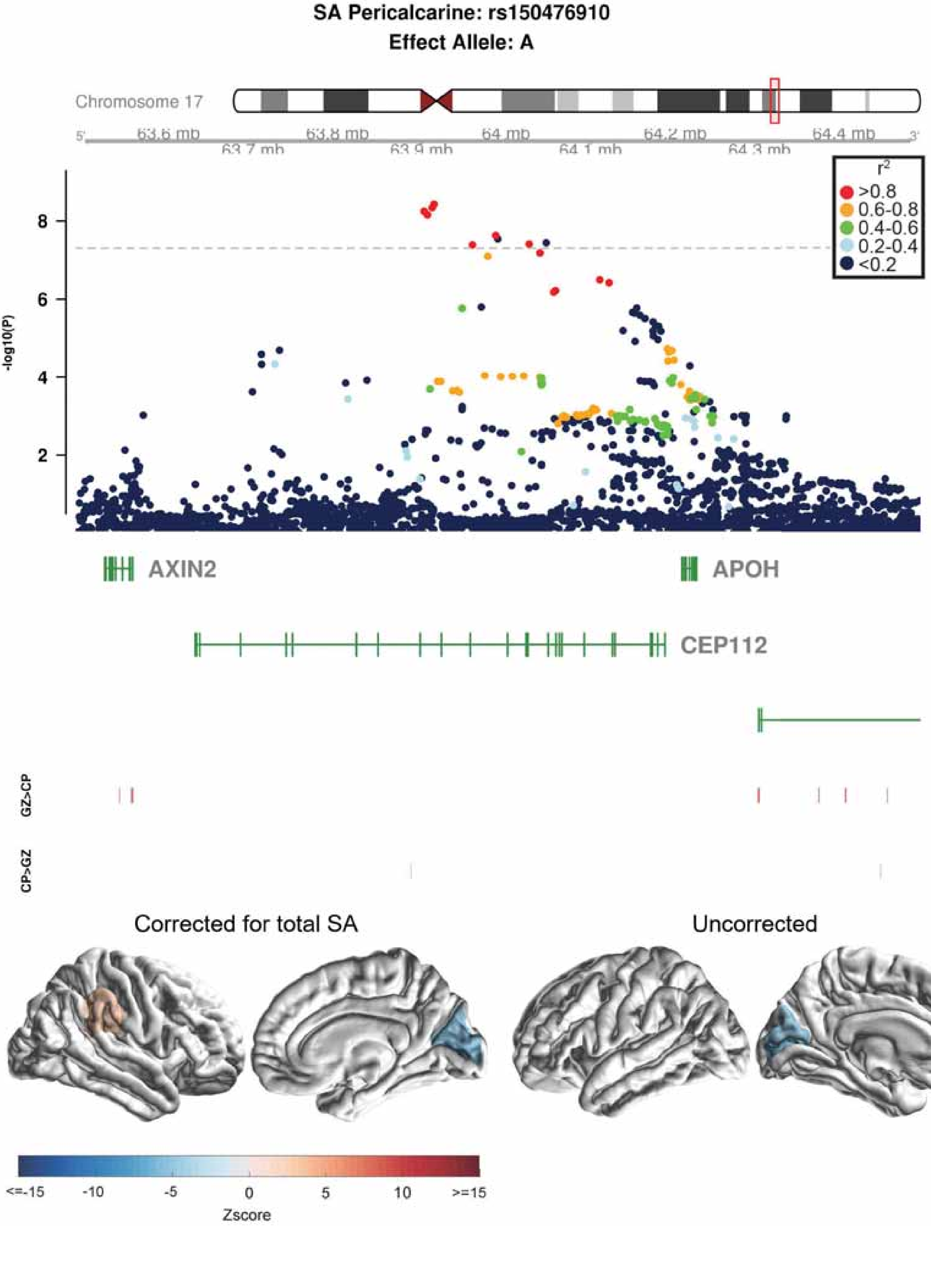

SA Pericalcarine: rs150476910
Effect Allele: A
Chromosome 17
63.6 mb
63.7 mb
63.8 mb
63.9 mb
64 mb
64.1 mb
64.2 mb
64.3 mb
64.4 mb
-log10(P)
r2
>0.8
0.6-0.8
0.4-0.6
0.2-0.4
<0.2
AXIN2
APOH
CEP112
GZ>CP
CP>GZ
Corrected for total SA
Uncorrected
<=-15
-10
-5
0
5
10
>=15
Zscore

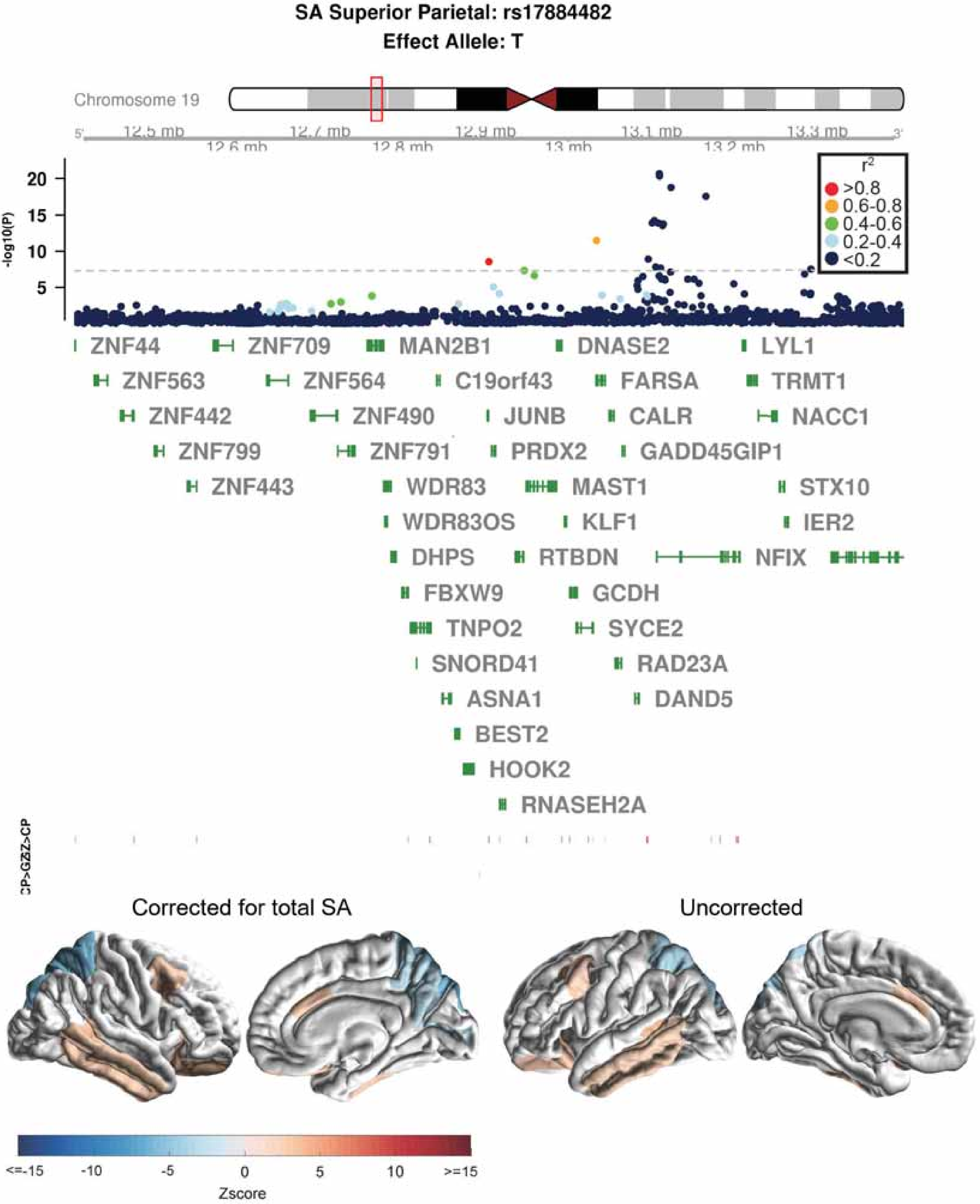
SA Superior Parietal: rs17884482
Effect Allele: T
Chromosome 19
12.5 mb
12.6 mb
12.7 mb
12.8 mb
12.9 mb
13 mb
13.1 mb
13.2 mb
13.3 mb
-log10(P)
20
15
10
5
r2
>0.8
0.6-0.8
0.4-0.6
0.2-0.4
<0.2
ZNF44
ZNF709
MAN2B1
DNASE2
LYL1
ZNF563
ZNF564
C19orf43
FARSA
TRMT1
ZNF442
ZNF490
JUNB
CALR
NACC1
ZNF799
ZNF791
PRDX2
GADD45GIP1
ZNF443
WDR83
MAST1
STX10
WDR83OS
KLF1
IER2
DHPS
RTBDN
NFIX
FBXW9
GCDH
TNPO2
SYCE2
SNORD41
RAD23A
ASNA1
DAND5
BEST2
HOOK2
RNASEH2A
Corrected for total SA
Uncorrected
<=-15
-10
-5
0
5
10
>=15
Zscore

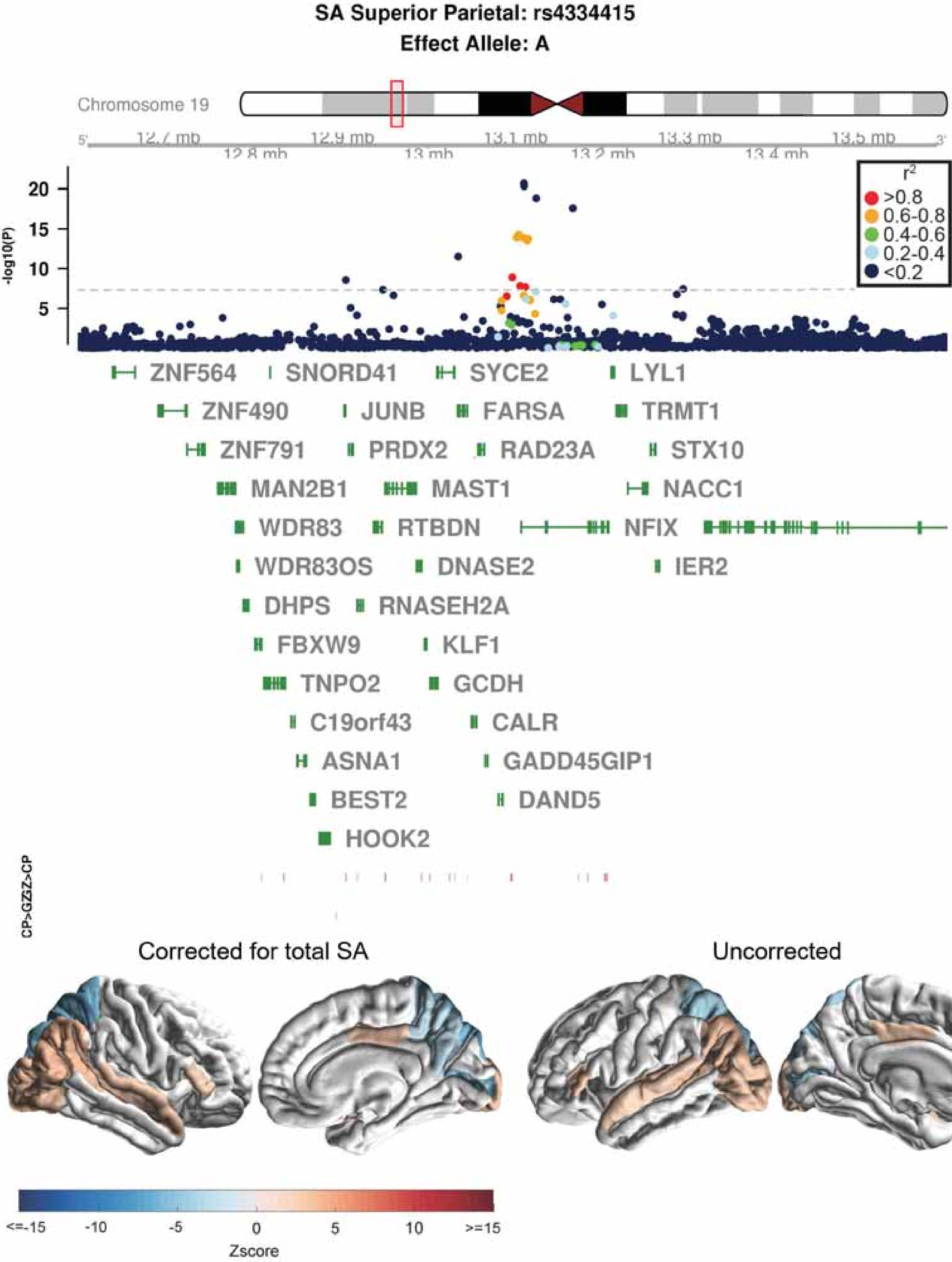

SA Superior Parietal: rs4334415
Effect Allele: A
Chromosome 19
12.7 mb
12.8 mb
12.9 mb
13 mb
13.1 mb
13.2 mb
13.3 mb
13.4 mb
13.5 mb
-log10(P)
r2
>0.8
0.6-0.8
0.4-0.6
0.2-0.4
<0.2
ZNF564
SNORD41
SYCE2
LYL1
ZNF490
JUNB
FARSA
TRMT1
ZNF791
PRDX2
RAD23A
STX10
MAN2B1
MAST1
NACC1
WDR83
RTBDN
NFIX
WDR83OS
DNASE2
IER2
DHPS
RNASEH2A
FBXW9
KLF1
TNPO2
GCDH
C19orf43
CALR
ASNA1
GADD45GIP1
BEST2
DAND5
HOOK2
CP>GZ
GZ>CP
Corrected for total SA
Uncorrected
<=-15
-10
-5
0
5
10
>=15
Zscore

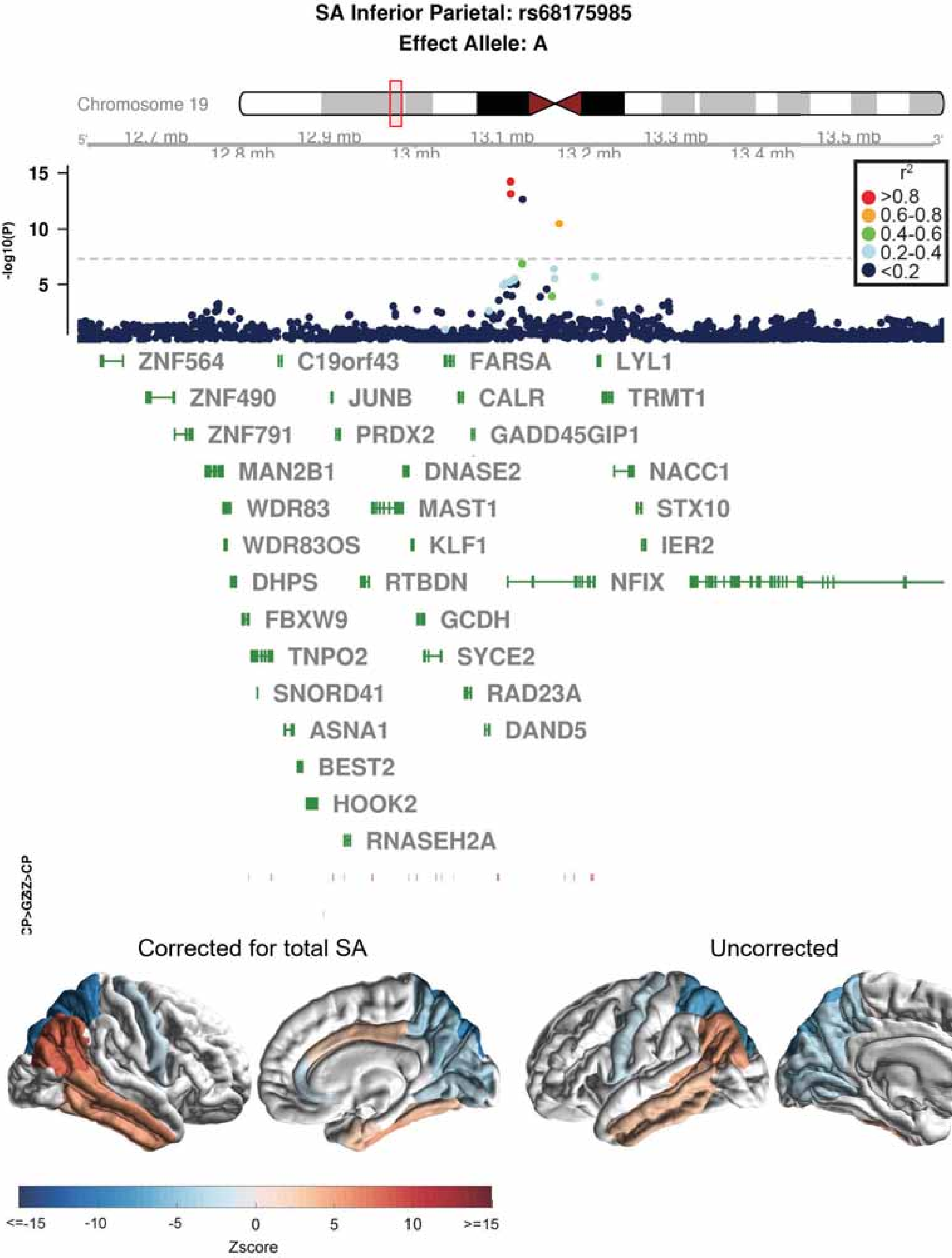

SA Inferior Parietal: rs68175985
Effect Allele: A
Chromosome 19
12.7 mb
12.8 mb
12.9 mb
13 mb
13.1 mb
13.2 mb
13.3 mb
13.4 mb
13.5 mb
-log10(P)
15
10
5
r2
>0.8
0.6-0.8
0.4-0.6
0.2-0.4
<0.2
ZNF564
C19orf43
FARSA
LYL1
ZNF490
JUNB
CALR
TRMT1
ZNF791
PRDX2
GADD45GIP1
MAN2B1
DNASE2
NACC1
WDR83
MAST1
STX10
WDR83OS
KLF1
IER2
DHPS
RTBDN
NFIX
FBXW9
GCDH
TNPO2
SYCE2
SNORD41
RAD23A
ASNA1
DAND5
BEST2
HOOK2
RNASEH2A
CP>GZ
Z>CP
Corrected for total SA
Uncorrected
<=-15
-10
-5
0
5
10
>=15
Zscore

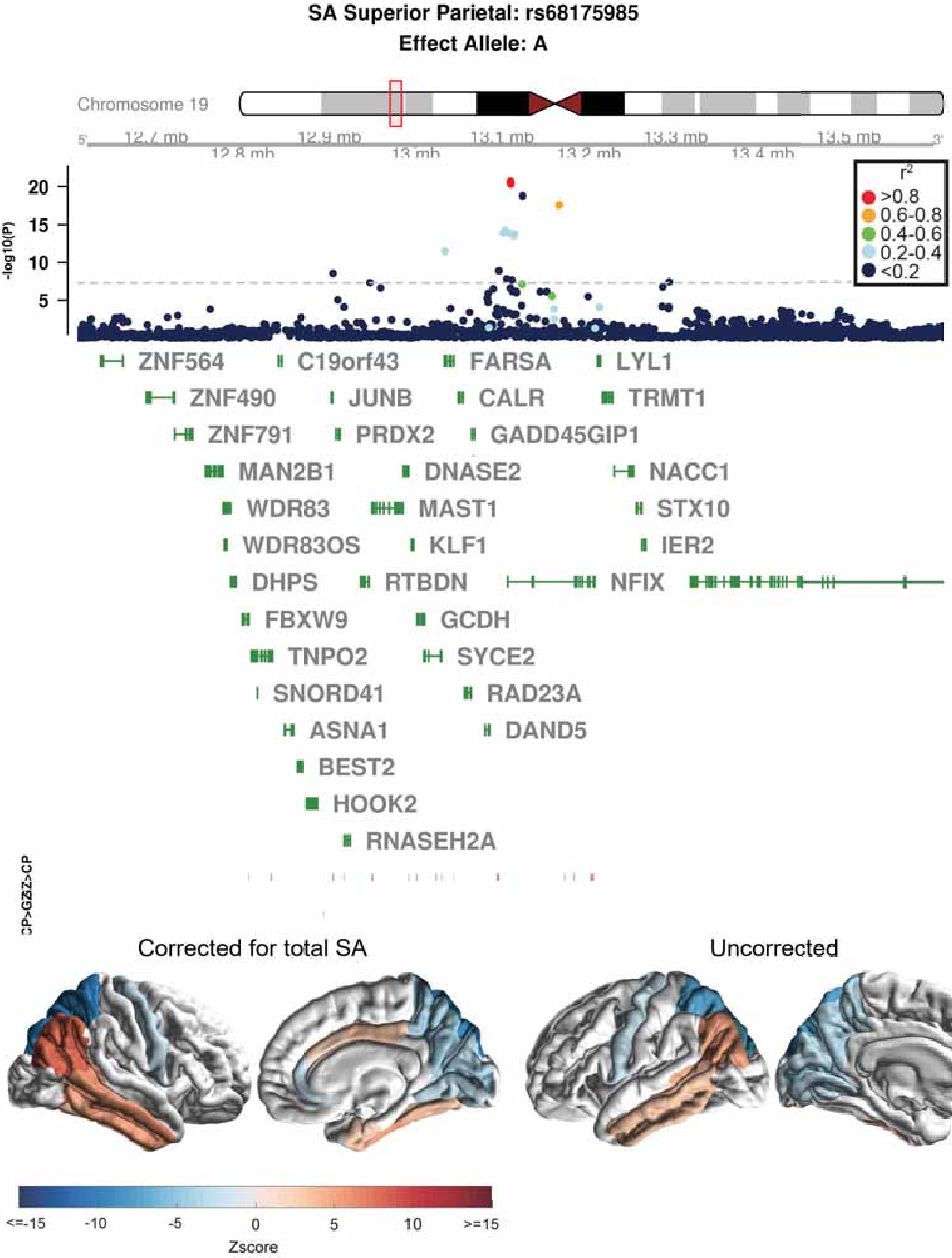
SA Superior Parietal: rs68175985
Effect Allele: A
Chromosome 19
12.7 mb
12.8 mb
12.9 mb
13 mb
13.1 mb
13.2 mb
13.3 mb
13.4 mb
13.5 mb
-log10(P)
20
15
10
5
r2
>0.8
0.6-0.8
0.4-0.6
0.2-0.4
<0.2
ZNF564
C19orf43
FARSA
LYL1
ZNF490
JUNB
CALR
TRMT1
ZNF791
PRDX2
GADD45GIP1
MAN2B1
DNASE2
NACC1
WDR83
MAST1
STX10
WDR83OS
KLF1
IER2
DHPS
RTBDN
NFIX
FBXW9
GCDH
TNPO2
SYCE2
SNORD41
RAD23A
ASNA1
DAND5
BEST2
HOOK2
RNASEH2A
CP>GZ
GZ>CP
Corrected for total SA
Uncorrected
<=-15
-10
-5
0
5
10
>=15
Zscore

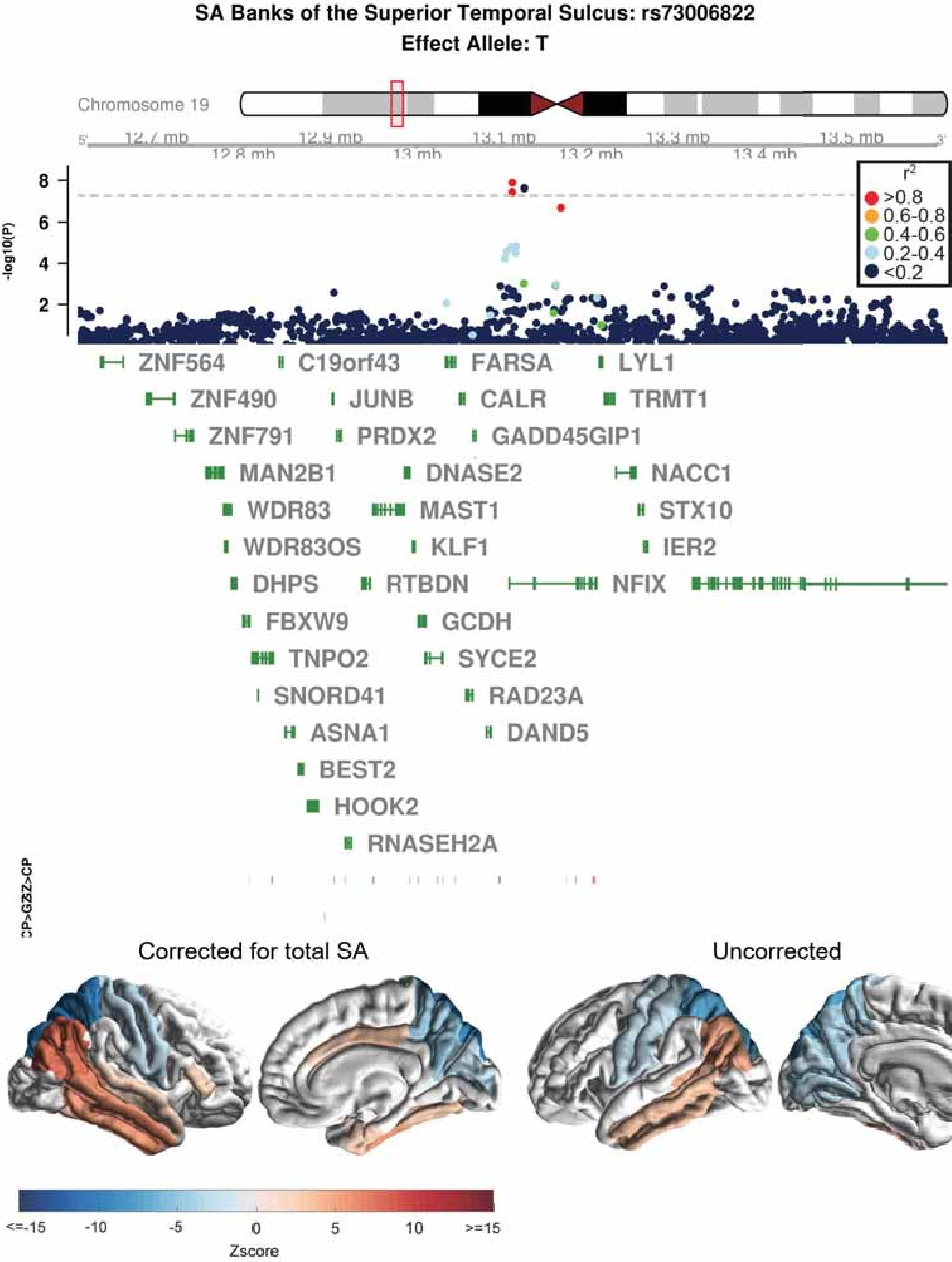
SA Banks of the Superior Temporal Sulcus: rs73006822
Effect Allele: T
Chromosome 19
12.7 mb
12.8 mb
12.9 mb
13 mb
13.1 mb
13.2 mb
13.3 mb
13.4 mb
13.5 mb
-log10(P)
8
6
4
2
r2
>0.8
0.6-0.8
0.4-0.6
0.2-0.4
<0.2
ZNF564
C19orf43
FARSA
LYL1
ZNF490
JUNB
CALR
TRMT1
ZNF791
PRDX2
GADD45GIP1
MAN2B1
DNASE2
NACC1
WDR83
MAST1
STX10
WDR83OS
KLF1
IER2
DHPS
RTBDN
NFIX
FBXW9
GCDH
TNPO2
SYCE2
SNORD41
RAD23A
ASNA1
DAND5
BEST2
HOOK2
RNASEH2A
Corrected for total SA
Uncorrected
<=-15
-10
-5
0
5
10
>=15
Zscore

**SA Middle Temporal: rs73006822**
**Effect Allele: T**

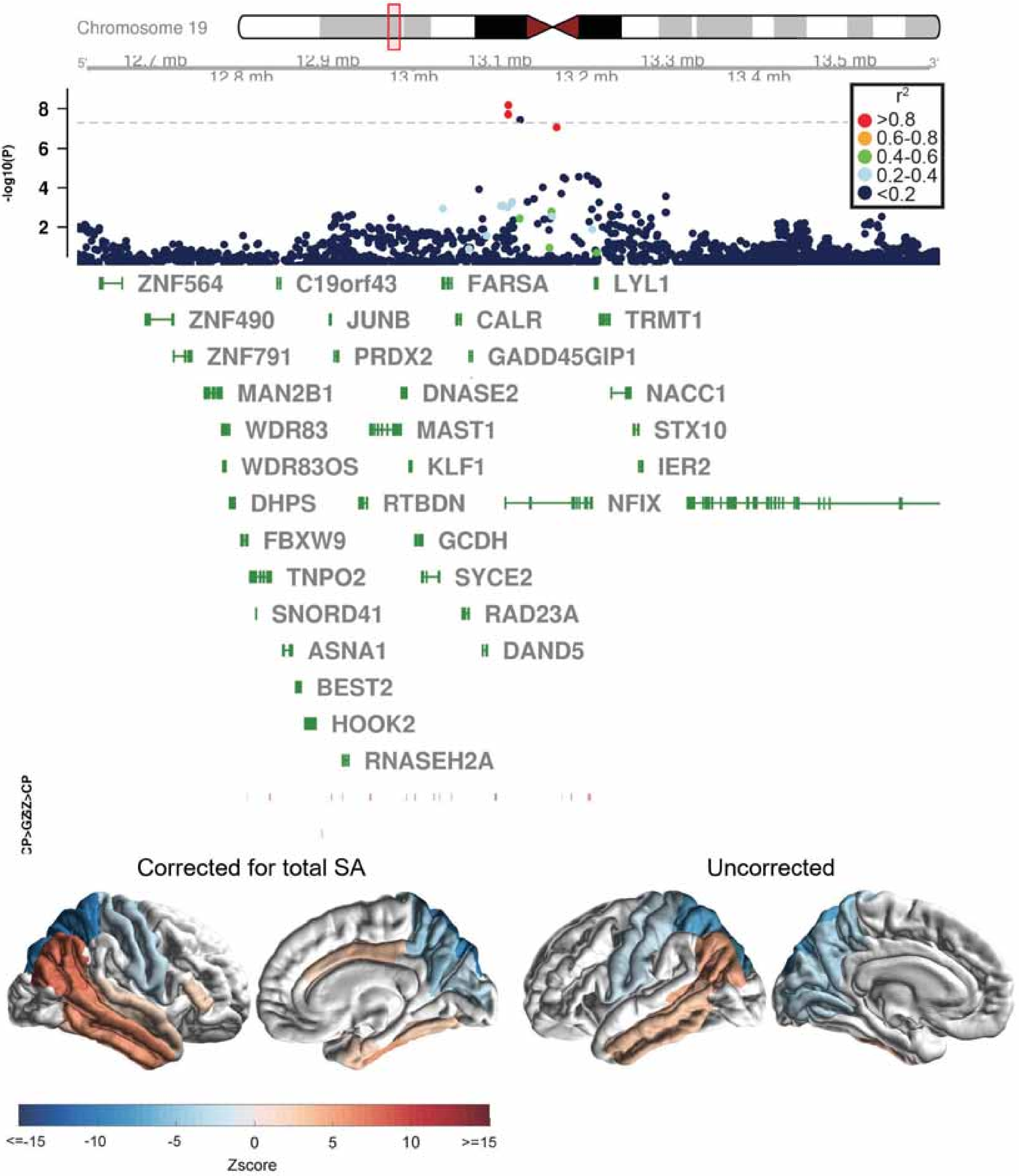

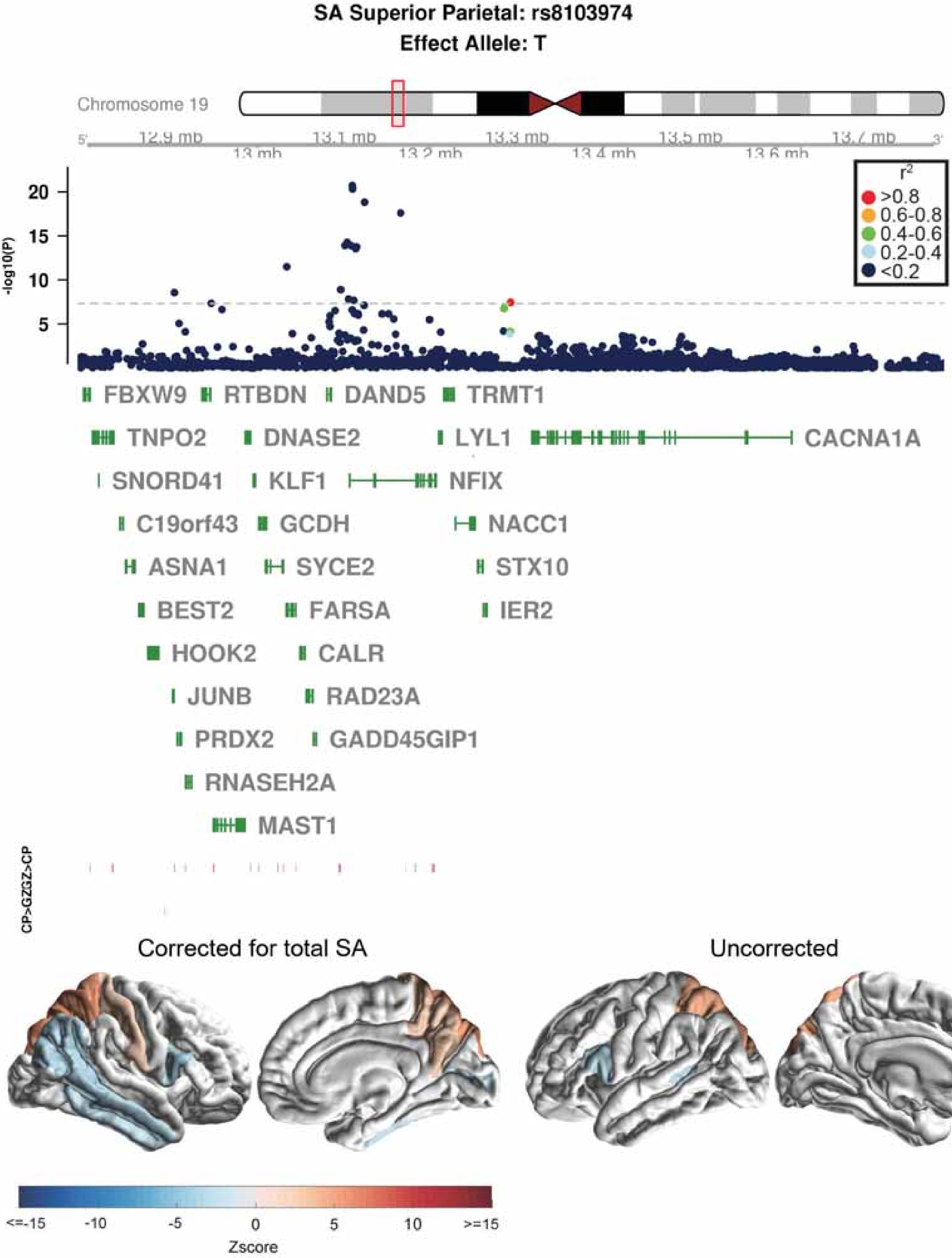
SA Superior Parietal: rs8103974
Effect Allele: T
Chromosome 19
5'
12.9 mb
13 mb
13.1 mb
13.2 mb
13.3 mb
13.4 mb
13.5 mb
13.6 mb
13.7 mb
3'
-log10(P)
20
15
10
5
$r^2$
>0.8
0.6-0.8
0.4-0.6
0.2-0.4
<0.2
FBXW9
RTBDN
DAND5
TRMT1
TNPO2
DNASE2
LYL1
CACNA1A
SNORD41
KLF1
NFIX
C19orf43
GCDH
NACC1
ASNA1
SYCE2
STX10
BEST2
FARSA
IER2
HOOK2
CALR
JUNB
RAD23A
PRDX2
GADD45GIP1
RNASEH2A
MAST1
CP>GZGZ>CP
Corrected for total SA
Uncorrected
<=-15
-10
-5
0
5
10
>=15
Zscore

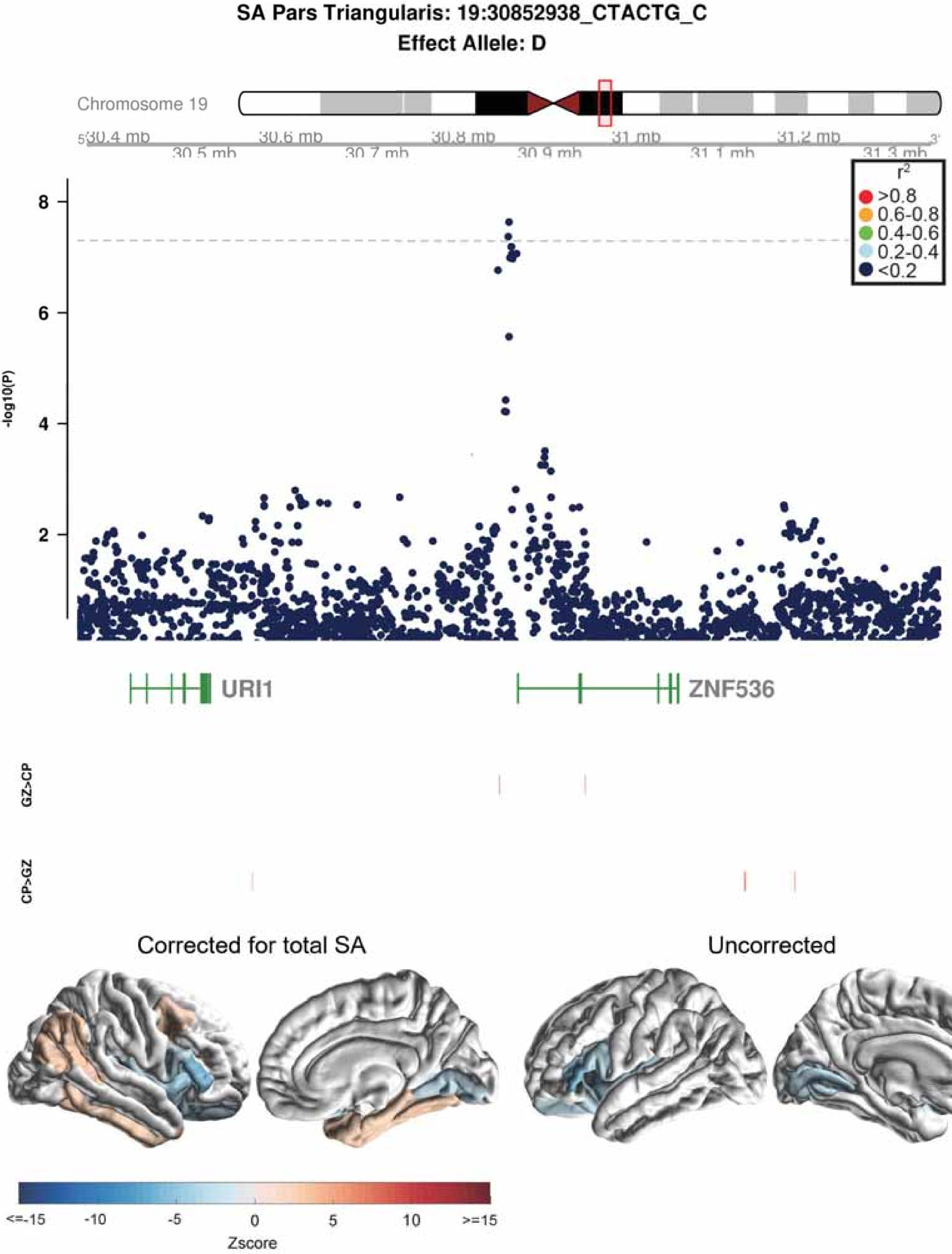

SA Pars Triangularis: 19:30852938_CTACTG_C
Effect Allele: D
Chromosome 19
30.4 mb
30.5 mb
30.6 mb
30.7 mb
30.8 mb
30.9 mb
31 mb
31.1 mb
31.2 mb
31.3 mb
r2
>0.8
0.6-0.8
0.4-0.6
0.2-0.4
<0.2
-log10(P)
URI1
ZNF536
GZ>CP
CP>GZ
Corrected for total SA
Uncorrected
<=-15
-10
-5
0
5
10
>=15
Zscore

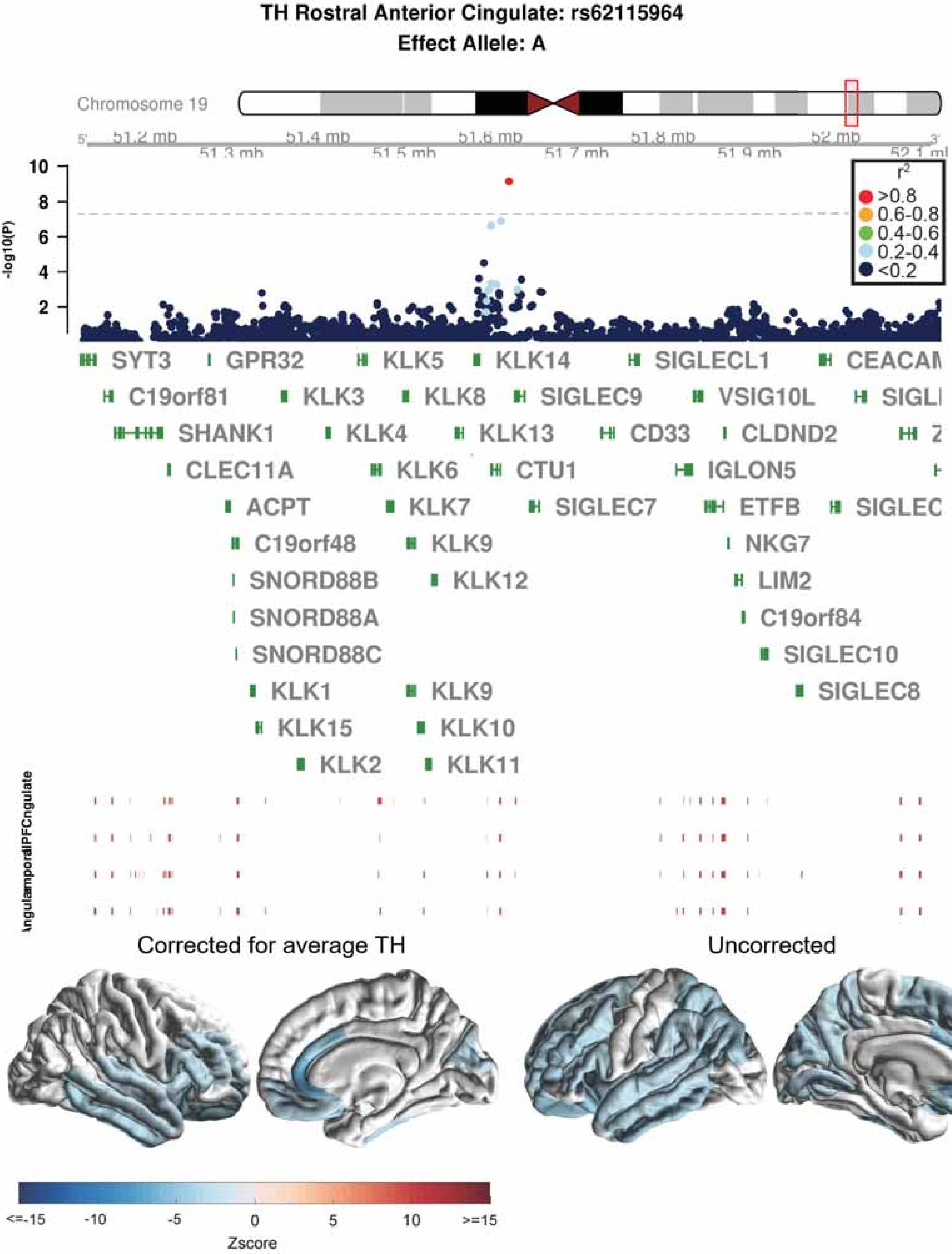
TH Rostral Anterior Cingulate: rs62115964
Effect Allele: A
Chromosome 19
51.2 mb
51.3 mb
51.4 mb
51.5 mb
51.6 mb
51.7 mb
51.8 mb
51.9 mb
52 mb
-log10(P)
r2
>0.8
0.6-0.8
0.4-0.6
0.2-0.4
<0.2
SYT3
GPR32
KLK5
KLK14
SIGLECL1
C19orf81
KLK3
KLK8
SIGLEC9
VSIG10L
SHANK1
KLK4
KLK13
CD33
CLDND2
CLEC11A
KLK6
CTU1
IGLON5
ACPT
KLK7
SIGLEC7
ETFB
C19orf48
KLK9
NKG7
SNORD88B
KLK12
LIM2
SNORD88A
C19orf84
SNORD88C
SIGLEC10
KLK1
KLK9
SIGLEC8
KLK15
KLK10
KLK2
KLK11
Corrected for average TH
Uncorrected
<=-15
-10
-5
0
5
10
>=15
Zscore

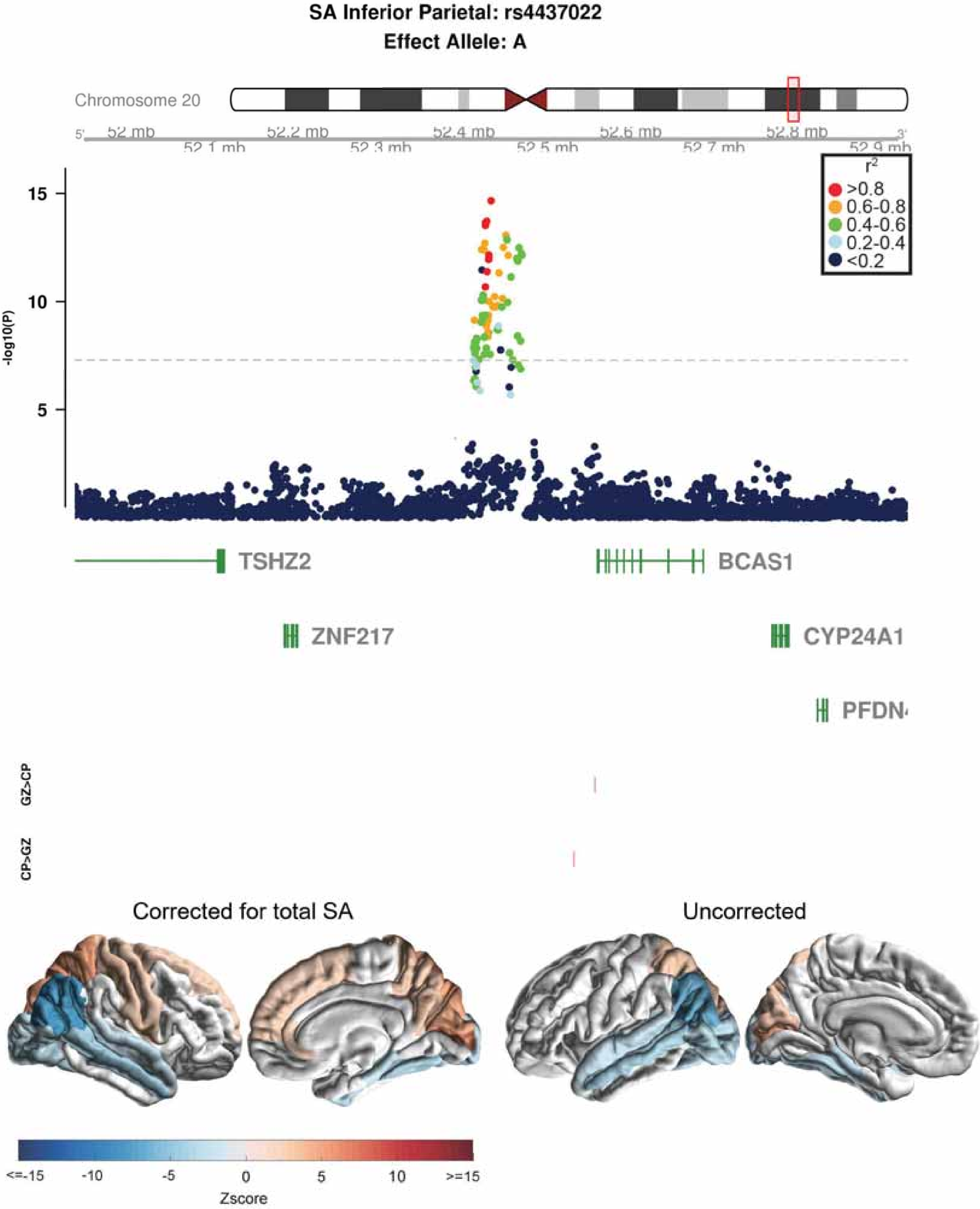
SA Inferior Parietal: rs4437022
Effect Allele: A
Chromosome 20
52 mb
52.1 mb
52.2 mb
52.3 mb
52.4 mb
52.5 mb
52.6 mb
52.7 mb
52.8 mb
52.9 mb
r2
>0.8
0.6-0.8
0.4-0.6
0.2-0.4
<0.2
-log10(P)
15
10
5
TSHZ2
BCAS1
ZNF217
CYP24A1
PFDN
GZ>CP
CP>GZ
Corrected for total SA
Uncorrected
<=-15
-10
-5
0
5
10
>=15
Zscore

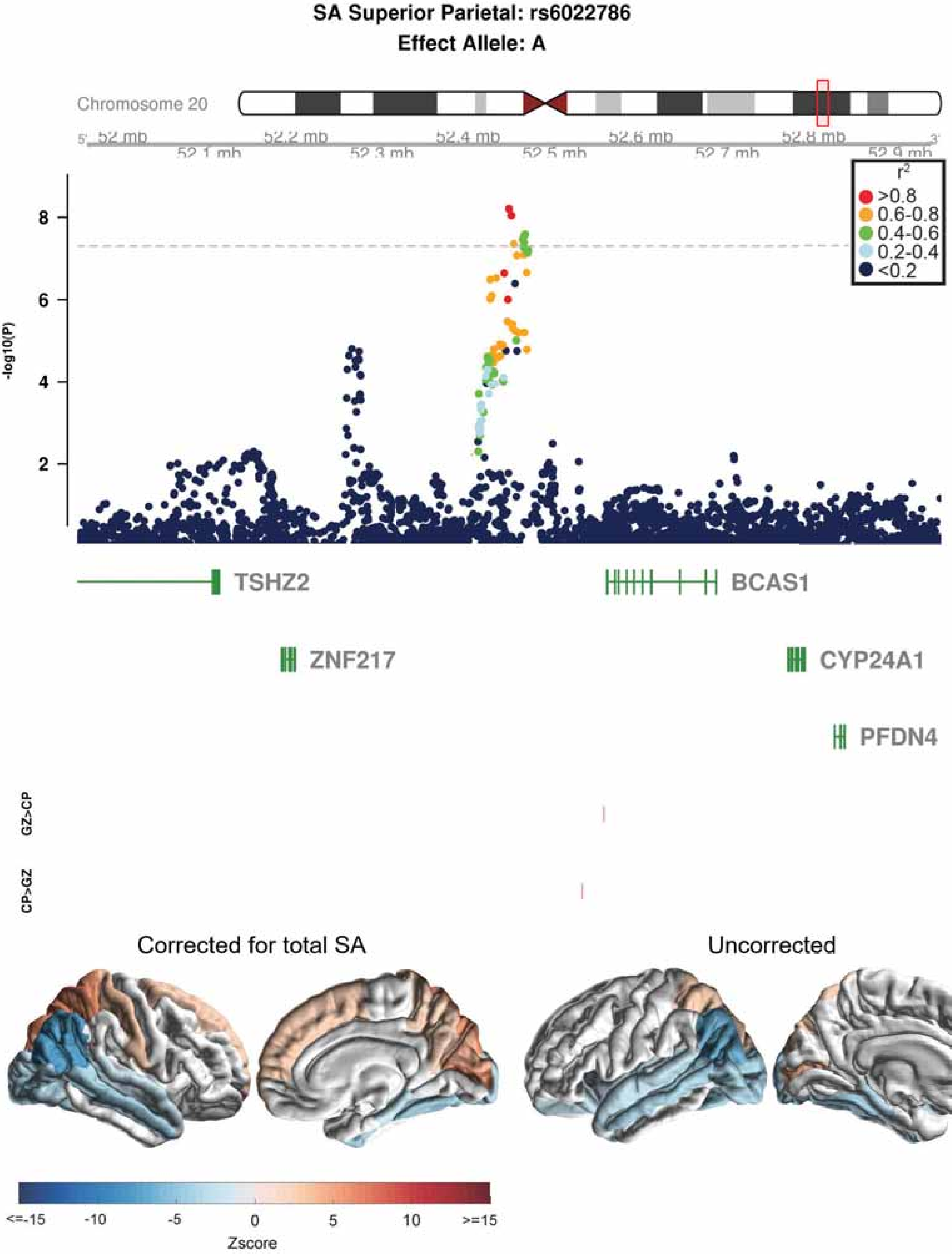

SA Superior Parietal: rs6022786
Effect Allele: A
Chromosome 20
52 mb
52.1 mb
52.2 mb
52.3 mb
52.4 mb
52.5 mb
52.6 mb
52.7 mb
52.8 mb
52.9 mb
5'
3'
-log10(P)
2
4
6
8
r2
>0.8
0.6-0.8
0.4-0.6
0.2-0.4
<0.2
TSHZ2
BCAS1
ZNF217
CYP24A1
PFDN4
GZ>CP
CP>GZ
Corrected for total SA
Uncorrected
<=-15
-10
-5
0
5
10
>=15
Zscore

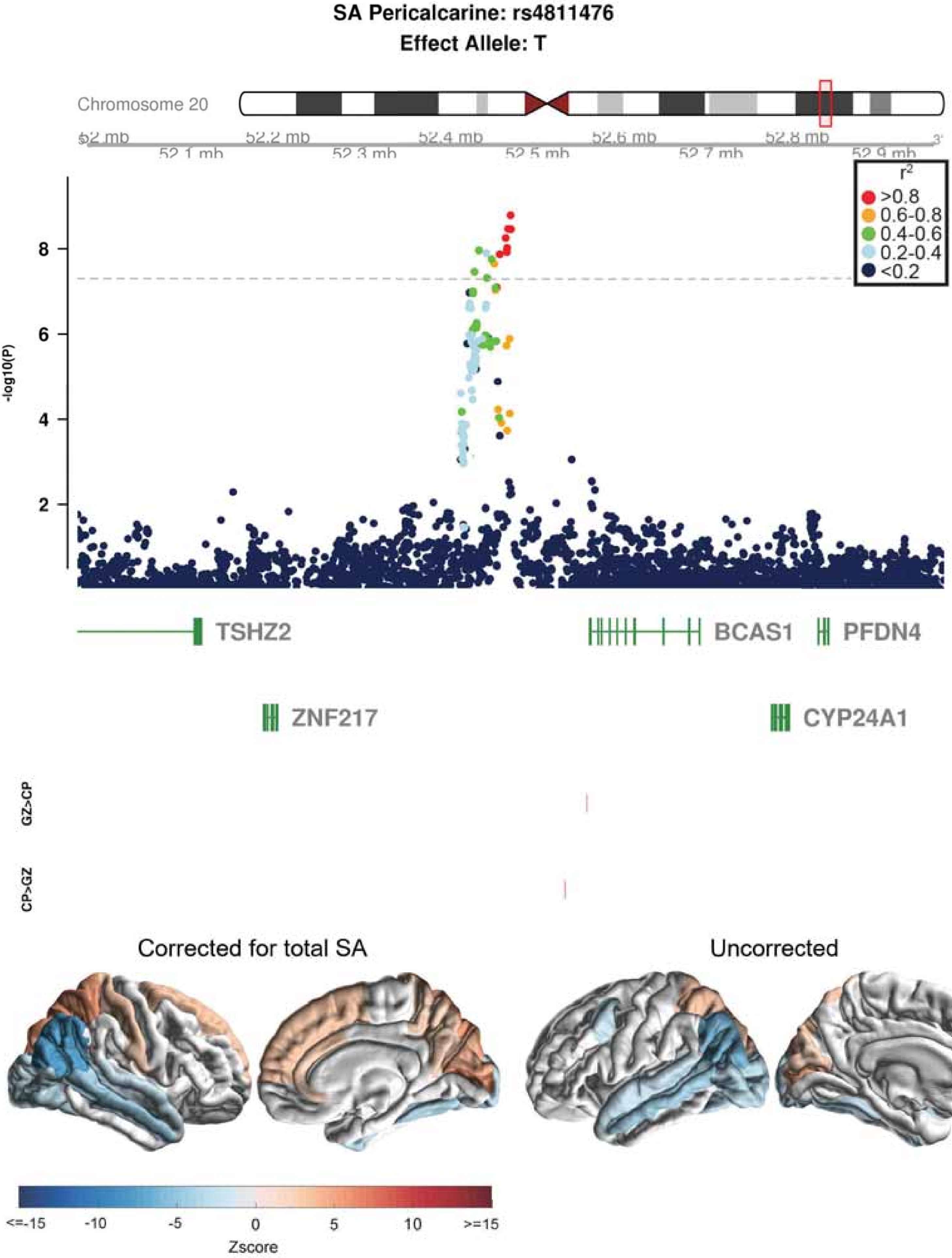
SA Pericalcarine: rs4811476
Effect Allele: T
Chromosome 20
52 mb
52.1 mb
52.2 mb
52.3 mb
52.4 mb
52.5 mb
52.6 mb
52.7 mb
52.8 mb
52.9 mb
r2
>0.8
0.6-0.8
0.4-0.6
0.2-0.4
<0.2
-log10(P)
TSHZ2
BCAS1
PFDN4
ZNF217
CYP24A1
GZ>CP
CP>GZ
Corrected for total SA
Uncorrected
<=-15
-10
-5
0
5
10
>=15
Zscore

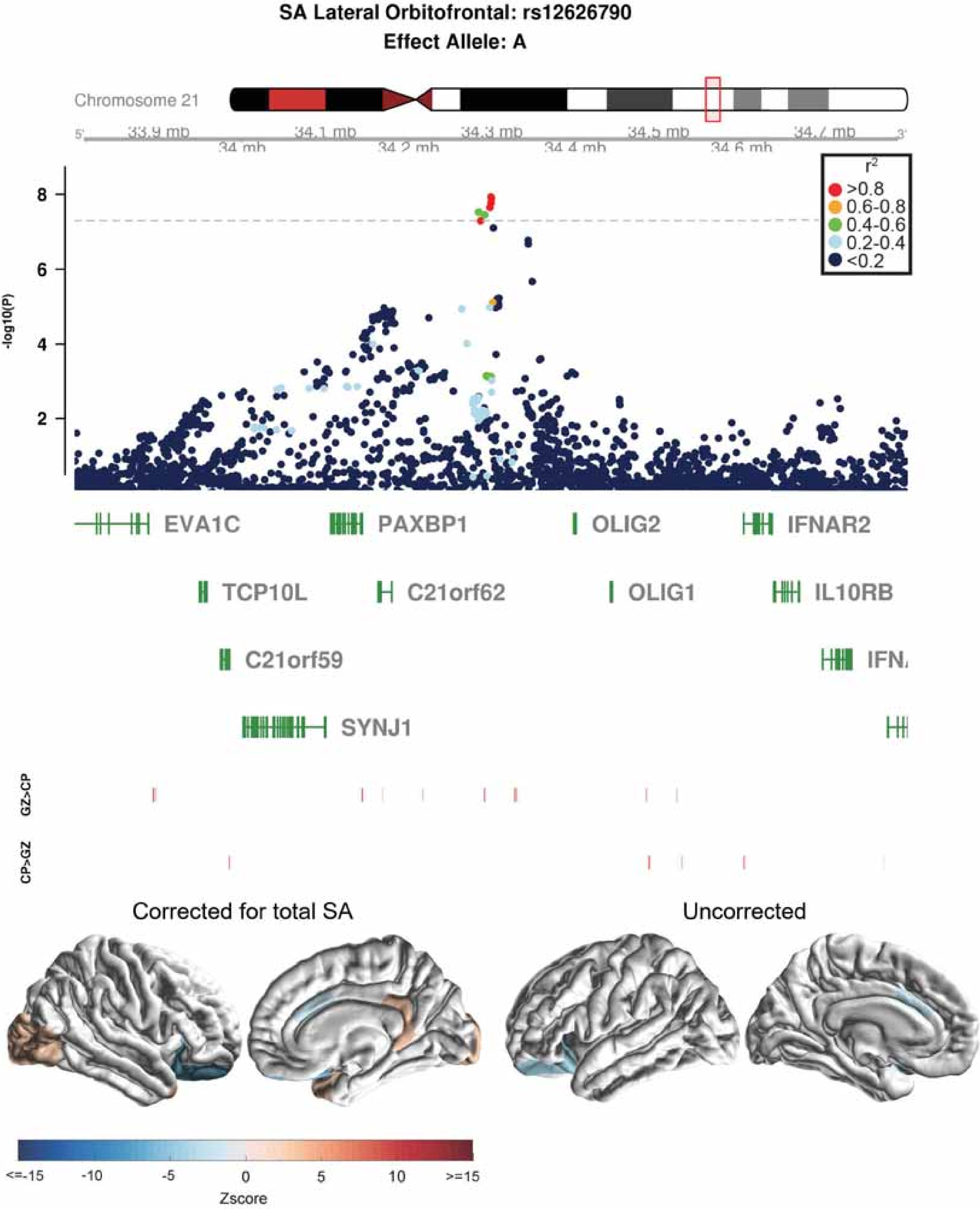

SA Lateral Orbitofrontal: rs12626790
Effect Allele: A
Chromosome 21
33.9 mb
34 mb
34.1 mb
34.2 mb
34.3 mb
34.4 mb
34.5 mb
34.6 mb
34.7 mb
-log10(P)
r2
>0.8
0.6-0.8
0.4-0.6
0.2-0.4
<0.2
EVA1C
PAXBP1
OLIG2
IFNAR2
TCP10L
C21orf62
OLIG1
IL10RB
C21orf59
SYNJ1
GZ>CP
CP>GZ
Corrected for total SA
Uncorrected
<=-15
-10
-5
0
5
10
>=15
Zscore

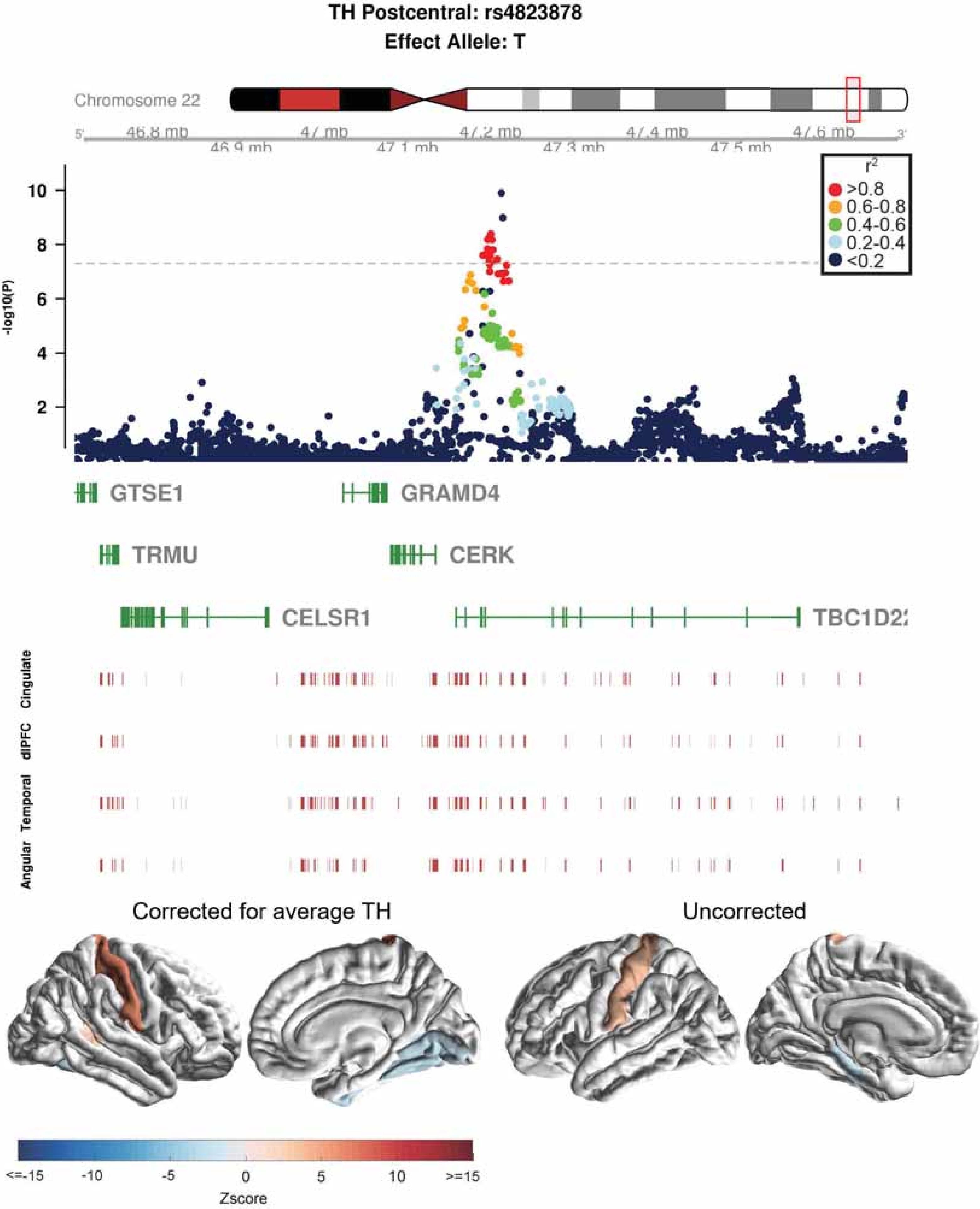

TH Postcentral: rs4823878
Effect Allele: T
Chromosome 22
46.8 mb
46.9 mb
47 mb
47.1 mb
47.2 mb
47.3 mb
47.4 mb
47.5 mb
47.6 mb
r2
>0.8
0.6-0.8
0.4-0.6
0.2-0.4
<0.2
-log10(P)
10
8
6
4
2
GTSE1
GRAMD4
TRMU
CERK
CELSR1
Cingulate
dlPFC
Temporal
Angular
Corrected for average TH
Uncorrected
<=-15
-10
-5
0
5
10
>=15
Zscore
